## Supplementary Table 2 for "MAFB drives differentiation by permitting WT1 binding to podocyte specific promoters"

### ChIP-Seq analysis for MAFB and WT1

| | |
|---|---|
| Worksheet 0 | Summary |
| Worksheet 1 | MAFB peaks identified by IDR analysis |
| Worksheet 2 | GREAT analysis for MAFB peaks |
| Worksheet 3 | WT1 peaks identified by IDR analysis |
| Worksheet 4 | GREAT analysis for WT1 peaks |
| Worksheet 5 | Regions that overlap between WT1 and MAFB |
| Worksheet 6 | GREAT analysis of overlapping regions |

| MAFB peaks | | | | | | | | | | | | | | | H3K4me3 | | | | | | H3K4me1 | | | | | |
|---|---|---|---|---|---|---|---|---|---|---|---|---|---|---|---|---|---|---|---|---|---|---|---|---|---|---|
| Chrom | Start | End | IDR score | annotation | geneChr | geneStart | geneEnd | geneLength | geneStrand | transcriptId | distanceToTSS | geneName | geneId | podocytes specific | Chrom | Start | End | -LOG10(pval | fold | -LOG10(q-va | Chrom | Start | End | -LOG10(pval | fold | -LOG10(q-value) |
| chr2 | 25452963 | 25453154 | 587 | Downstream (1-2kb) | 2 | 25456838 | 25458676 | 1839 | 1 | ENSMUST00 | -3684 | Clic3 | ENSMUSG0000001 | true | . | . | . | . | . | . | chr2 | 25446553 | 25456355 | 6.09426 | 4.46882 | 3.92693 |
| chr6 | 70073928 | 70074050 | 587 | Promoter (<=1kb) | 6 | 70074024 | 70074584 | 561 | 2 | ENSMUST00 | 534 | Igkv6-32 | ENSMUSG0000007 | false | . | . | . | . | . | . | chr6 | 70072263 | 70075068 | 3.76337 | 3.46811 | 1.98668 |
| chr5 | 35233245 | 35233420 | 114 | Distal Intergenic | 5 | 35278566 | 35281761 | 3196 | 1 | ENSMUST00 | -45146 | Adra2c | ENSMUSG0000004 | false | . | . | . | . | . | . | chr5 | 35229105 | 35239186 | 4.86991 | 4.0349 | 2.9021 |
| chr8 | 104886805 | 104887096 | 529 | Distal Intergenic | 8 | 104867488 | 104874080 | 6593 | 1 | ENSMUST00 | 19318 | Ces2d-ps | ENSMUSG0000003 | false | . | . | . | . | . | . | chr8 | 104884771 | 104889614 | 5.64618 | 4.36566 | 3.53177 |
| chr8 | 104867314 | 104867666 | 519 | Promoter (<=1kb) | 8 | 104867488 | 104874080 | 6593 | 1 | ENSMUST00 | 0 | Ces2d-ps | ENSMUSG0000003 | false | . | . | . | . | . | . | chr8 | 104864625 | 104870535 | 4.84736 | 4.01234 | 2.88046 |
| chr14 | 57056404 | 57056672 | 588 | Promoter (1-2kb) | 14 | 57035608 | 57058030 | 22423 | 2 | ENSMUST00 | 1358 | Gja3 | ENSMUSG0000004 | false | chr14 | 57056396 | 57058454 | 32.01914 | 97.37792 | 93.87154 | chr14 | 57053149 | 57056474 | 4.05021 | 3.5021 | 2.24184 |
| chr7 | 30459561 | 30459728 | 555 | Promoter (<=1kb) | 7 | 30459725 | 30487223 | 27499 | 1 | ENSMUST00 | 0 | Nphs1 | ENSMUSG0000000 | true | chr7 | 30459332 | 30463950 | 22.44583 | 74.56482 | 71.50171 | . | . | . | . | . | . |
| chr16 | 45110613 | 45110834 | 594 | Intron (ENSMUST00000099498.3/ENSMUSG00 | 16 | 45094053 | 45127924 | 33872 | 1 | ENSMUST00 | 16561 | Ccdc80 | ENSMUSG0000002 | false | . | . | . | . | . | . | chr16 | 45109400 | 45112859 | 4.13036 | 3.69823 | 2.28418 |
| chr1 | 156309319 | 156309613 | 591 | Promoter (1-2kb) | 1 | 156310727 | 156328035 | 17309 | 1 | ENSMUST00 | -1114 | Nphs2 | ENSMUSG0000002 | true | chr1 | 156309732 | 156310571 | 8.92274 | 20.60519 | 18.07254 | . | . | . | . | . | . |
| chr15 | 84696178 | 84696412 | 543 | Intron (ENSMUST00000171460.2/ENSMUSG00 | 15 | 84681002 | 84703672 | 22671 | 1 | ENSMUST00 | 15177 | Prr5 | ENSMUSG0000003 | false | . | . | . | . | . | . | chr15 | 84682178 | 84700461 | 6.48581 | 4.70787 | 4.24565 |
| chr4 | 62699353 | 62699588 | 114 | Promoter (<=1kb) | 4 | 62699466 | 62703018 | 3553 | 1 | ENSMUST00 | 0 | Rgs3 | ENSMUSG0000005 | false | . | . | . | . | . | . | chr4 | 62694793 | 62699426 | 7.42731 | 5.01341 | 4.9919 |
| chr17 | 49477431 | 49477568 | 114 | Intron (ENSMUST00000057610.6/ENSMUSG00 | 17 | 49452284 | 49455435 | 3152 | 1 | ENSMUST00 | 25148 | Mocs1 | ENSMUSG0000006 | false | . | . | . | . | . | . | chr17 | 49468167 | 49480126 | 6.43614 | 4.71108 | 4.19031 |
| chr8 | 61008825 | 61009076 | 534 | Intron (ENSMUST00000120689.2/ENSMUSG00 | 8 | 60993241 | 61030600 | 37360 | 1 | ENSMUST00 | 15585 | Nek1 | ENSMUSG0000003 | true | . | . | . | . | . | . | chr8 | 61007494 | 61010706 | 4.67783 | 3.84507 | 2.76552 |
| chr5 | 19180868 | 19181188 | 510 | Distal Intergenic | 5 | 19202368 | 19226555 | 24188 | 2 | ENSMUST00 | 45367 | 4921504A21 | ENSMUSG0000009 | true | . | . | . | . | . | . | chr5 | 19177836 | 19184145 | 4.31014 | 3.73966 | 2.43836 |
| chrX | 166385063 | 166385172 | 604 | Intron (ENSMUST00000112235.2/ENSMUSG00 | 20 | 166344842 | 166373287 | 28446 | 1 | ENSMUST00 | 40222 | Gpm6b | ENSMUSG0000003 | false | . | . | . | . | . | . | . | . | . | . | . | . |
| chr9 | 60637104 | 60637360 | 550 | Intron (ENSMUST00000150060.2/ENSMUSG00 | 9 | 60610270 | 60649710 | 39441 | 2 | ENSMUST00 | 12350 | Lrrc49 | ENSMUSG0000004 | true | . | . | . | . | . | . | chr9 | 60623926 | 60640440 | 4.81608 | 3.88284 | 2.88797 |
| chr2 | 32617181 | 32617356 | 574 | Intron (ENSMUST00000140983.2/ENSMUSG00 | 2 | 32621758 | 32635058 | 13301 | 1 | ENSMUST00 | -4402 | Ak1 | ENSMUSG0000002 | false | . | . | . | . | . | . | chr2 | 32605009 | 32639275 | 6.86759 | 4.92928 | 4.53618 |
| chr6 | 18543915 | 18544306 | 615 | Distal Intergenic | 6 | 18366547 | 18514869 | 148323 | 2 | ENSMUST00 | -29047 | Cttnbp2 | ENSMUSG0000000 | true | . | . | . | . | . | . | chr6 | 18539839 | 18548637 | 4.5475 | 3.89202 | 2.64481 |
| chr14 | 70160831 | 70161028 | 596 | Promoter (1-2kb) | 14 | 70154405 | 70159502 | 5098 | 2 | ENSMUST00 | -1330 | 9930012K11 | ENSMUSG0000004 | false | . | . | . | . | . | . | chr14 | 70156516 | 70175803 | 7.12683 | 5 | 4.75261 |
| chr8 | 121280477 | 121280768 | 424 | Intron (ENSMUST00000127664.1/ENSMUSG00 | 8 | 121256747 | 121356836 | 100090 | 2 | ENSMUST00 | 76068 | Gm26815 | ENSMUSG0000009 | false | . | . | . | . | . | . | chr8 | 121277448 | 121284034 | 5.99549 | 4.54477 | 3.8074 |
| chr6 | 140433286 | 140433494 | 114 | Intron (ENSMUST00000087622.3/ENSMUSG00 | 6 | 140424099 | 140594906 | 170808 | 1 | ENSMUST00 | 9188 | Plekha5 | ENSMUSG0000003 | true | . | . | . | . | . | . | chr6 | 140425415 | 140441290 | 5.09666 | 4.0883 | 3.09824 |
| chr11 | 32383789 | 32384143 | 662 | Intron (ENSMUST00000038753.5/ENSMUSG00 | 11 | 32347840 | 32390300 | 42461 | 1 | ENSMUST00 | 35950 | Sh3pxd2b | ENSMUSG0000004 | false | . | . | . | . | . | . | chr11 | 32383288 | 32386252 | 5.8032 | 4.27592 | 3.68113 |
| chr10 | 86340102 | 86340227 | 619 | Intron (ENSMUST00000020234.8/ENSMUSG00 | 10 | 86302854 | 86343981 | 41128 | 1 | ENSMUST00 | 37249 | Timp3 | ENSMUSG0000002 | false | . | . | . | . | . | . | chr10 | 86303145 | 86347283 | 7.80266 | 5.33584 | 5.30789 |
| chr10 | 22868403 | 22868624 | 588 | Distal Intergenic | 10 | 22817279 | 22820128 | 2850 | 2 | ENSMUST00 | -48276 | Tcf21 | ENSMUSG0000004 | true | . | . | . | . | . | . | chr10 | 22866220 | 22874539 | 4.6225 | 3.87067 | 2.69769 |
| chr9 | 63611068 | 63611246 | 571 | Intron (ENSMUST00000041551.7/ENSMUSG00 | 9 | 63602655 | 63641889 | 39235 | 1 | ENSMUST00 | 8414 | Aagab | ENSMUSG0000003 | false | . | . | . | . | . | . | chr9 | 63603357 | 63616669 | 6.44347 | 4.69808 | 4.19638 |
| chr10 | 122029860 | 122030078 | 574 | Intron (ENSMUST00000020322.6/ENSMUSG00 | 10 | 121780991 | 122047308 | 266318 | 2 | ENSMUST00 | 17230 | Srgap1 | ENSMUSG0000002 | true | . | . | . | . | . | . | chr10 | 121945243 | 122036389 | 5.74397 | 4.3951 | 3.62974 |
| chr1 | 153543060 | 153543319 | 114 | Intron (ENSMUST00000041874.8/ENSMUSG00 | 1 | 153548422 | 153549711 | 1290 | 2 | ENSMUST00 | 6392 | Npl | ENSMUSG0000004 | false | . | . | . | . | . | . | chr1 | 153539148 | 153550339 | 5.26804 | 4.15937 | 3.23842 |
| chr2 | 32907336 | 32907628 | 506 | Intron (ENSMUST00000028135.9/ENSMUSG00 | 2 | 32910908 | 32912587 | 1680 | 1 | ENSMUST00 | -3280 | Fam129b | ENSMUSG0000002 | false | . | . | . | . | . | . | chr2 | 32877160 | 32918190 | 7.77045 | 5.19056 | 5.2936 |
| chr1 | 152799996 | 152800343 | 593 | Promoter (<=1kb) | 1 | 152800194 | 152836991 | 36798 | 1 | ENSMUST00 | 0 | Ncf2 | ENSMUSG0000002 | true | chr1 | 152800193 | 152801982 | 14.44046 | 33.88361 | 31.37022 | chr1 | 152788034 | 152800060 | 6.38067 | 4.71623 | 4.13727 |
| chr17 | 5591633 | 5591828 | 609 | Intron (ENSMUST00000089185.4/ENSMUSG00 | 17 | 5492600 | 5753891 | 261292 | 1 | ENSMUST00 | 99034 | Zdhhc14 | ENSMUSG0000003 | true | . | . | . | . | . | . | chr17 | 5587797 | 5599687 | 5.93548 | 4.48203 | 3.77591 |
| chr17 | 5612424 | 5612702 | 578 | Intron (ENSMUST00000089185.4/ENSMUSG00 | 17 | 5492600 | 5753891 | 261292 | 1 | ENSMUST00 | 119825 | Zdhhc14 | ENSMUSG0000003 | true | . | . | . | . | . | . | chr17 | 5611281 | 5615799 | 4.75775 | 3.90015 | 2.81034 |
| chr5 | 24724939 | 24725267 | 589 | Intron (ENSMUST00000068693.6/ENSMUSG00 | 5 | 24712265 | 24730680 | 18416 | 2 | ENSMUST00 | 5413 | Wdr86 | ENSMUSG0000005 | false | . | . | . | . | . | . | chr5 | 24723425 | 24734625 | 5.82193 | 4.37932 | 3.69553 |
| chr6 | 116006861 | 116007131 | 566 | Distal Intergenic | 6 | 115954811 | 115995005 | 40195 | 2 | ENSMUST00 | -11857 | Plxnd1 | ENSMUSG0000003 | false | . | . | . | . | . | . | chr6 | 116004192 | 116008178 | 6.32655 | 4.46684 | 4.10455 |
| chr5 | 64337025 | 64337412 | 114 | Intron (ENSMUST00000043893.7/ENSMUSG00 | 5 | 64349945 | 64350589 | 645 | 1 | ENSMUST00 | -12533 | Tbc1d1 | ENSMUSG0000002 | false | . | . | . | . | . | . | chr5 | 64333951 | 64340633 | 6.27533 | 4.60721 | 4.05113 |
| chr6 | 52233552 | 52233733 | 114 | Promoter (<=1kb) | 6 | 52231199 | 52233324 | 2126 | 2 | ENSMUST00 | -229 | Hoxa10 | ENSMUSG0000000 | false | chr6 | 52227601 | 52235632 | 41.69514 | 140.65352 | 135.74945 | . | . | . | . | . | . |
| chr16 | 33872134 | 33872340 | 593 | Intron (ENSMUST00000115028.4/ENSMUSG00 | 16 | 33885026 | 33910585 | 25560 | 1 | ENSMUST00 | -12686 | Itgb5 | ENSMUSG0000002 | true | . | . | . | . | . | . | chr16 | 33868993 | 33898226 | 6.80171 | 4.94523 | 4.47886 |
| chr16 | 58514787 | 58515018 | 538 | Intron (ENSMUST00000137035.2/ENSMUSG00 | 16 | 58502861 | 58507898 | 5038 | 2 | ENSMUST00 | -6890 | St3gal6 | ENSMUSG0000002 | true | . | . | . | . | . | . | chr16 | 58496830 | 58520680 | 5.70067 | 4.33546 | 3.59278 |
| chr11 | 69880992 | 69881443 | 399 | Promoter (<=1kb) | 11 | 69878261 | 69881406 | 3146 | 2 | ENSMUST00 | 0 | Kctd11 | ENSMUSG0000004 | false | chr11 | 69879287 | 69881992 | 16.78262 | 53.63071 | 50.86883 | . | . | . | . | . | . |
| chr1 | 127306458 | 127306781 | 636 | Promoter (<=1kb) | 1 | 127306762 | 127483704 | 176943 | 1 | ENSMUST00 | 0 | Mgat5 | ENSMUSG0000003 | true | chr1 | 127306652 | 127307946 | 25.3802 | 72.59858 | 69.5683 | . | . | . | . | . | . |
| chr2 | 27758427 | 27758649 | 114 | Intron (ENSMUST00000077257.6/ENSMUSG00 | 2 | 27740201 | 27748671 | 8471 | 1 | ENSMUST00 | 18227 | Rxra | ENSMUSG0000001 | false | . | . | . | . | . | . | chr2 | 27756216 | 27760543 | 7.96442 | 5.34041 | 5.43403 |
| chr11 | 8494100 | 8494421 | 628 | Intron (ENSMUST00000020695.7/ENSMUSG00 | 11 | 8451070 | 8468675 | 17606 | 2 | ENSMUST00 | -25426 | Tns3 | ENSMUSG0000002 | false | chr11 | 8494386 | 8495188 | 5.6323 | 9.47728 | 7.31216 | chr11 | 8476638 | 8500780 | 7.03559 | 4.95227 | 4.67872 |
| chr4 | 66450124 | 66450325 | 583 | Distal Intergenic | 4 | 65380803 | 66404611 | 1023809 | 2 | ENSMUST00 | -45514 | Astn2 | ENSMUSG0000002 | true | . | . | . | . | . | . | chr4 | 66449477 | 66451280 | 4.51968 | 3.80583 | 2.65848 |
| chr9 | 57831471 | 57831807 | 641 | Promoter (2-3kb) | 9 | 57790513 | 57834236 | 43724 | 2 | ENSMUST00 | 2429 | Arid3b | ENSMUSG0000000 | false | . | . | . | . | . | . | chr9 | 57827202 | 57836152 | 5.62883 | 4.39478 | 3.50932 |
| chr18 | 61894653 | 61894888 | 611 | Intron (ENSMUST00000049378.9/ENSMUSG00 | 18 | 61799395 | 61911824 | 112430 | 2 | ENSMUST00 | 16936 | Ablim3 | ENSMUSG0000003 | false | . | . | . | . | . | . | chr18 | 61877692 | 61898809 | 5.45644 | 4.15279 | 3.39554 |
| chr4 | 123282580 | 123282748 | 541 | Promoter (<=1kb) | 4 | 123282825 | 123298925 | 16101 | 1 | ENSMUST00 | -77 | Pabpc4 | ENSMUSG0000001 | false | chr4 | 123282579 | 123284220 | 24.32174 | 95.77222 | 92.30251 | . | . | . | . | . | . |
| chr8 | 70418957 | 70419091 | 590 | Intron (ENSMUST00000076615.5/ENSMUSG00 | 8 | 70406127 | 70431285 | 25159 | 2 | ENSMUST00 | 12194 | Crtc1 | ENSMUSG0000000 | false | . | . | . | . | . | . | chr8 | 70415799 | 70434143 | 6.2581 | 4.55457 | 4.06384 |
| chr6 | 132463691 | 132463814 | 805 | Distal Intergenic | 6 | 132361041 | 132364134 | 3094 | 2 | ENSMUST00 | -99558 | Gm8882 | ENSMUSG0000003 | false | chr6 | 132463619 | 132463896 | 29.41006 | 85.09721 | 81.85307 | . | . | . | . | . | . |
| chr1 | 44951203 | 44951314 | 790 | Distal Intergenic | 1 | 44924149 | 44924893 | 745 | 2 | ENSMUST00 | -26311 | RP23-231B1 | ENSMUSG0000010 | false | chr1 | 44951123 | 44951386 | 16.66573 | 41.91806 | 39.30453 | . | . | . | . | . | . |
| chr7 | 98060011 | 98060119 | 773 | Intron (ENSMUST00000107128.2/ENSMUSG00 | 7 | 98062643 | 98065202 | 2560 | 2 | ENSMUST00 | 5083 | Myo7a | ENSMUSG0000003 | false | . | . | . | . | . | . | . | . | . | . | . | . |
| chr9 | 19575415 | 19575525 | 760 | Distal Intergenic | 9 | 19566435 | 19567382 | 948 | 2 | ENSMUST00 | -8034 | Olfr854 | ENSMUSG0000009 | false | chr9 | 19575345 | 19575603 | 9.94526 | 25.28477 | 22.8772 | . | . | . | . | . | . |
| chr6 | 125876898 | 125877197 | 749 | Intron (ENSMUST00000159111.2/ENSMUSG00 | 6 | 125884415 | 125884524 | 110 | 1 | ENSMUST00 | -7218 | Gm24947 | ENSMUSG0000008 | false | chr6 | 125876838 | 125877186 | 9.38717 | 20.36066 | 18.02005 | . | . | . | . | . | . |
| chr10 | 37263702 | 37263818 | 740 | Distal Intergenic | 10 | 37337486 | 37379652 | 42167 | 1 | ENSMUST00 | -73668 | Gm26535 | ENSMUSG0000009 | false | . | . | . | . | . | . | . | . | . | . | . | . |
| chr18 | 45056018 | 45056124 | 732 | Distal Intergenic | 18 | 45089326 | 45090314 | 989 | 2 | ENSMUST00 | 34190 | Gm26928 | ENSMUSG0000009 | false | chr18 | 45055938 | 45056205 | 12.46556 | 27.94533 | 25.5059 | . | . | . | . | . | . |
| chr17 | 56199988 | 56200099 | 726 | Intron (ENSMUST00000038794.4/ENSMUSG00 | 17 | 56175744 | 56183920 | 8177 | 2 | ENSMUST00 | -16069 | D17Wsu104 | ENSMUSG0000001 | false | chr17 | 56199916 | 56200182 | 21.65864 | 58.36299 | 55.53803 | . | . | . | . | . | . |
| chr17 | 4626163 | 4626599 | 719 | Distal Intergenic | 17 | 4678020 | 4678126 | 107 | 1 | ENSMUST00 | -51421 | Gm23186 | ENSMUSG0000009 | false | chr17 | 4626080 | 4626682 | 79.22768 | 354.44785 | 346.18814 | . | . | . | . | . | . |
| chr7 | 127345551 | 127345781 | 714 | Promoter (<=1kb) | 7 | 127342797 | 127345314 | 2518 | 2 | ENSMUST00 | -238 | Zfp768 | ENSMUSG0000004 | false | chr7 | 127344643 | 127346973 | 28.22734 | 101.53345 | 97.92473 | . | . | . | . | . | . |
| chr2 | 60402751 | 60402850 | 709 | Distal Intergenic | 2 | 60411525 | 60412579 | 1055 | 1 | ENSMUST00 | -8675 | Gm13580 | ENSMUSG0000008 | false | . | . | . | . | . | . | . | . | . | . | . | . |
| chr7 | 19344961 | 19345201 | 703 | Promoter (<=1kb) | 7 | 19345101 | 19355474 | 10374 | 1 | ENSMUST00 | 0 | Ercc1 | ENSMUSG0000000 | false | chr7 | 19344647 | 19345778 | 16.59601 | 56.40767 | 53.61239 | chr7 | 19340662 | 19346039 | 5.44993 | 4.1911 | 3.38706 |
| chr16 | 72804355 | 72804447 | 699 | Intron (ENSMUST00000023600.7/ENSMUSG00 | 16 | 72663149 | 73046095 | 382947 | 1 | ENSMUST00 | 141207 | Robo1 | ENSMUSG0000002 | false | . | . | . | . | . | . | . | . | . | . | . | . |
| chr11 | 73047705 | 73047930 | 695 | Promoter (<=1kb) | 11 | 73047783 | 73080217 | 32435 | 1 | ENSMUST00 | 0 | 1200014J11 | ENSMUSG0000002 | false | chr11 | 73047761 | 73048865 | 24.12502 | 101.02714 | 97.43493 | . | . | . | . | . | . |
| chr2 | 84826804 | 84827122 | 692 | Promoter (<=1kb) | 2 | 84826997 | 84830213 | 3217 | 1 | ENSMUST00 | 0 | Timm10 | ENSMUSG0000002 | false | chr2 | 84826391 | 84827862 | 37.27983 | 118.82364 | 114.72062 | chr2 | 84826306 | 84828168 | 5.68195 | 4.30482 | 3.60799 |
| chr10 | 22731264 | 22731554 | 684 | Promoter (<=1kb) | 10 | 22703879 | 22731447 | 27569 | 2 | ENSMUST00 | 0 | Tbpl1 | ENSMUSG0000007 | false | chr10 | 22730228 | 22732192 | 39.29051 | 134.21745 | 129.56854 | . | . | . | . | . | . |
| chr12 | 83631963 | 83632324 | 681 | Promoter (<=1kb) | 12 | 83632252 | 83683123 | 50872 | 1 | ENSMUST00 | 0 | Rbm25 | ENSMUSG0000001 | false | chr12 | 83632162 | 83633479 | 32.16725 | 95.65015 | 92.18271 | . | . | . | . | . | . |
| chr7 | 126975957 | 126976203 | 678 | Promoter (<=1kb) | 7 | 126972625 | 126976098 | 3474 | 2 | ENSMUST00 | 0 | D830044I16 | ENSMUSG0000009 | false | chr7 | 126976071 | 126977438 | 37.60551 | 160.01111 | 154.36346 | . | . | . | . | . | . |
| chr11 | 120484506 | 120484778 | 673 | Promoter (<=1kb) | 11 | 120484613 | 120489065 | 4453 | 1 | ENSMUST00 | 0 | Mrpl12 | ENSMUSG0000003 | false | chr11 | 120484051 | 120485543 | 42.33011 | 146.25195 | 141.10883 | chr11 | 120483856 | 120486834 | 4.26692 | 3.47603 | 2.4501 |
| chr11 | 50174584 | 50174902 | 671 | Promoter (<=1kb) | 11 | 50174849 | 50200115 | 25267 | 1 | ENSMUST00 | 0 | 3010026O09 | ENSMUSG0000002 | false | chr11 | 50174812 | 50175473 | 19.72513 | 62.50151 | 59.62064 | chr11 | 50174421 | 50189350 | 5.97902 | 4.44839 | 3.83251 |
| chr8 | 66312617 | 66312718 | 666 | Intron (ENSMUST00000152320.2/ENSMUSG00 | 8 | 66386294 | 66470421 | 84128 | 1 | ENSMUST00 | -73576 | March1 | ENSMUSG0000003 | false | . | . | . | . | . | . | . | . | . | . | . | . |
| chr16 | 37011510 | 37011866 | 664 | Promoter (<=1kb) | 16 | 37011786 | 37094992 | 83207 | 1 | ENSMUST00 | 0 | Polq | ENSMUSG0000003 | false | chr16 | 37011163 | 37012278 | 15.11128 | 52.55099 | 49.80313 | chr16 | 37011043 | 37012490 | 5.42283 | 4.05299 | 3.38571 |
| chr1 | 163994634 | 163994981 | 660 | Promoter (<=1kb) | 1 | 163994889 | 163997243 | 2355 | 1 | ENSMUST00 | 0 | Mettl18 | ENSMUSG0000004 | false | chr1 | 163994206 | 163995669 | 25.99992 | 84.80917 | 81.57088 | chr1 | 163991574 | 163996008 | 5.65475 | 4.33948 | 3.57649 |
| chr1 | 90603270 | 90603508 | 658 | Promoter (<=1kb) | 1 | 90603476 | 90613124 | 9649 | 1 | ENSMUST00 | 0 | Cops8 | ENSMUSG0000003 | false | chr1 | 90603053 | 90604104 | 41.42819 | 139.07397 | 134.22679 | . | . | . | . | . | . |
| chr15 | 75616398 | 75616734 | 656 | Promoter (<=1kb) | 15 | 75616679 | 75625300 | 8622 | 1 | ENSMUST00 | 0 | Zfp41 | ENSMUSG0000004 | false | chr15 | 75615707 | 75617294 | 17.71499 | 57.83842 | 55.02271 | chr15 | 75615615 | 75617342 | 5.21478 | 4.18872 | 3.17278 |
| chr9 | 66945885 | 66946285 | 654 | Promoter (<=1kb) | 9 | 66946086 | 66949512 | 3427 | 1 | ENSMUST00 | 0 | Rps27l | ENSMUSG0000003 | false | chr9 | 66945369 | 66946939 | 33.73597 | 126.30199 | 121.9427 | . | . | . | . | . | . |
| chr15 | 102325998 | 102326225 | 653 | Promoter (<=1kb) | 15 | 102326116 | 102331873 | 5758 | 1 | ENSMUST00 | 0 | Pfdn5 | ENSMUSG0000000 | false | chr15 | 102325753 | 102327357 | 28.48738 | 97.07422 | 93.57343 | . | . | . | . | . | . |
| chr6 | 70116960 | 70117160 | 651 | Promoter (<=1kb) | 6 | 70117061 | 70117617 | 557 | 2 | ENSMUST00 | 457 | Igkv8-30 | ENSMUSG0000007 | false | . | . | . | . | . | . | chr6 | 70115225 | 70118243 | 3.61941 | 3.39411 | 1.85895 |
| chr1 | 52844877 | 52845014 | 649 | Promoter (<=1kb) | 1 | 52844929 | 52848693 | 3765 | 1 | ENSMUST00 | 0 | Hibch | ENSMUSG0000004 | false | chr1 | 52844968 | 52845904 | 18.77433 | 54.14365 | 51.37372 | chr1 | 52844108 | 52846226 | 4.34574 | 3.73717 | 2.50184 |
| chr5 | 131519713 | 131519810 | 648 | Intron (ENSMUST00000161226.3/ENSMUSG00 | 5 | 131476742 | 131534397 | 57656 | 2 | ENSMUST00 | 14587 | Auts2 | ENSMUSG0000002 | true | . | . | . | . | . | . | . | . | . | . | . | . |
| chr15 | 5617871 | 5618057 | 646 | Distal Intergenic | 15 | 5864306 | 5864419 | 114 | 1 | ENSMUST00 | -246249 | Gm27760 | ENSMUSG0000009 | false | . | . | . | . | . | . | chr15 | 5617091 | 5619625 | 4.3034 | 3.71772 | 2.48127 |
| chr7 | 122101808 | 122102039 | 643 | Promoter (<=1kb) | 7 | 122085403 | 122101886 | 16484 | 2 | ENSMUST00 | 0 | Ndufab1 | ENSMUSG0000003 | false | chr7 | 122100946 | 122102271 | 34.36326 | 108.53439 | 104.74065 | chr7 | 122098290 | 122102367 | 5.71206 | 4.36131 | 3.60933 |
| chr1 | 93478233 | 93478572 | 642 | Promoter (<=1kb) | 1 | 93440688 | 93478377 | 37690 | 2 | ENSMUST00 | 0 | Hdlbp | ENSMUSG0000003 | false | chr1 | 93477516 | 93480357 | 31.43325 | 99.35611 | 95.79983 | . | . | . | . | . | . |
| chr1 | 143777101 | 143777309 | 639 | Promoter (<=1kb) | 1 | 143777272 | 143807305 | 30034 | 1 | ENSMUST00 | 0 | Uchl5 | ENSMUSG0000001 | false | chr1 | 143776236 | 143778292 | 27.97104 | 107.99458 | 104.2211 | . | . | . | . | . | . |
| chr5 | 52782967 | 52783129 | 638 | Promoter (<=1kb) | 5 | 52783055 | 52823571 | 40517 | 1 | ENSMUST00 | 0 | Zcchc4 | ENSMUSG0000002 | false | chr5 | 52782729 | 52783710 | 36.60697 | 119.77473 | 115.64114 | . | . | . | . | . | . |
| chr14 | 107075290 | 107075385 | 637 | Distal Intergenic | 14 | 106766030 | 106766156 | 127 | 1 | ENSMUST00 | 309261 | Gm23437 | ENSMUSG0000007 | false | . | . | . | . | . | . | . | . | . | . | . | . |
| chr6 | 6578531 | 6578847 | 635 | Promoter (<=1kb) | 6 | 6557294 | 6578658 | 21365 | 2 | ENSMUST00 | 0 | Shfm1 | ENSMUSG0000004 | false | chr6 | 6577340 | 6579136 | 30.03893 | 101.89226 | 98.2715 | . | . | . | . | . | . |
| chr11 | 25100071 | 25100227 | 632 | Distal Intergenic | 11 | 25232626 | 25367133 | 134508 | 1 | ENSMUST00 | -132399 | 4933427E13 | ENSMUSG0000003 | false | chr11 | 25099986 | 25100286 | 15.80552 | 59.41874 | 56.58204 | . | . | . | . | . | . |
| chr7 | 24904196 | 24904530 | 631 | Promoter (<=1kb) | 7 | 24904358 | 24926589 | 22232 | 1 | ENSMUST00 | 0 | Arhgef1 | ENSMUSG0000004 | false | chr7 | 24902634 | 24904205 | 26.65955 | 86.95457 | 83.67183 | . | . | . | . | . | . |
| chr1 | 93211537 | 93211806 | 630 | Distal Intergenic | 1 | 93215899 | 93231072 | 15174 | 1 | ENSMUST00 | -4093 | E030010N08 | ENSMUSG0000008 | false | . | . | . | . | . | . | chr1 | 93210088 | 93211973 | 5.50157 | 4.34556 | 3.43814 |
| chr19 | 8880841 | 8881036 | 629 | Promoter (<=1kb) | 19 | 8880016 | 8880933 | 918 | 2 | ENSMUST00 | 0 | AI462493 | ENSMUSG0000007 | false | chr19 | 8879994 | 8881359 | 28.70854 | 85.79402 | 82.53136 | chr19 | 8878949 | 8881392 | 4.78404 | 4.03421 | 2.82388 |
| chr19 | 38836434 | 38836633 | 627 | Promoter (<=1kb) | 19 | 38836579 | 38919923 | 83345 | 1 | ENSMUST00 | 0 | Tbc1d12 | ENSMUSG0000004 | false | chr19 | 38836126 | 38837723 | 33.26678 | 112.81776 | 108.90401 | . | . | . | . | . | . |
| chr17 | 36958376 | 36958657 | 626 | Promoter (<=1kb) | 17 | 36958592 | 36965622 | 7031 | 1 | ENSMUST00 | 0 | Znrd1as | ENSMUSG0000003 | false | chr17 | 36957679 | 36959243 | 19.17126 | 69.02763 | 66.05415 | chr17 | 36956323 | 36959716 | 5.61048 | 4.33555 | 3.51521 |
| chr5 | 145204386 | 145204586 | 625 | Promoter (<=1kb) | 5 | 145204562 | 145221750 | 17189 | 1 | ENSMUST00 | 0 | Zkscan5 | ENSMUSG0000005 | false | chr5 | 145203613 | 145205310 | 22.19715 | 59.28558 | 56.44936 | chr5 | 145203828 | 145206444 | 3.93342 | 3.32478 | 2.15132 |
| chr9 | 79759727 | 79759918 | 625 | Promoter (<=1kb) | 9 | 79755241 | 79759853 | 4613 | 2 | ENSMUST00 | 0 | Cox7a2 | ENSMUSG0000003 | false | chr9 | 79758867 | 79760573 | 31.07114 | 95.73475 | 92.26586 | . | . | . | . | . | . |
| chr15 | 89196450 | 89196671 | 624 | Promoter (<=1kb) | 15 | 89182217 | 89196474 | 14258 | 2 | ENSMUST00 | 0 | Dennd6b | ENSMUSG0000001 | false | chr15 | 89195732 | 89197334 | 31.96831 | 119.80809 | 115.67081 | . | . | . | . | . | . |
| chr1 | 133025356 | 133025518 | 622 | Promoter (<=1kb) | 1 | 132986220 | 133025416 | 39197 | 2 | ENSMUST00 | 0 | Mdm4 | ENSMUSG0000005 | false | chr1 | 133024460 | 133026209 | 28.90751 | 104.81161 | 101.11721 | . | . | . | . | . | . |
| chr13 | 113418276 | 113418463 | 621 | Distal Intergenic | 13 | 113403088 | 113403188 | 101 | 1 | ENSMUST00 | 15189 | Gm22385 | ENSMUSG0000008 | false | . | . | . | . | . | . | chr13 | 113417937 | 113419056 | 4.53797 | 3.83939 | 2.66852 |
| chr13 | 37531848 | 37532120 | 620 | Distal Intergenic | 13 | 37508940 | 37512074 | 3135 | 1 | ENSMUST00 | 22909 | RP23-21N15 | ENSMUSG0000010 | false | . | . | . | . | . | . | chr13 | 37526947 | 37535595 | 4.83754 | 3.94 | 2.87988 |
| chr17 | 3114769 | 3115224 | 619 | Promoter (<=1kb) | 17 | 3114972 | 3198855 | 83884 | 1 | ENSMUST00 | 0 | Scaf8 | ENSMUSG0000004 | false | chr17 | 3113719 | 3116459 | 36.98618 | 117.25745 | 113.20857 | . | . | . | . | . | . |
| chr7 | 40192452 | 40192673 | 618 | Promoter (1-2kb) | 7 | 40190935 | 40191324 | 390 | 1 | ENSMUST00 | 1518 | Gm6851 | ENSMUSG0000005 | false | . | . | . | . | . | . | . | . | . | . | . | . |
| chr6 | 124965452 | 124965632 | 617 | Promoter (<=1kb) | 6 | 124958413 | 124965538 | 7126 | 2 | ENSMUST00 | 0 | Cops7a | ENSMUSG0000003 | false | chr6 | 124964714 | 124966415 | 31.91637 | 110.38313 | 106.53261 | chr6 | 124964520 | 124966898 | 6.53504 | 4.78964 | 4.24247 |
| chr2 | 158361473 | 158361698 | 616 | Promoter (<=1kb) | 2 | 158355307 | 158361510 | 6204 | 2 | ENSMUST00 | 0 | 9430008C03 | ENSMUSG0000008 | false | chr2 | 158360817 | 158361635 | 27.82288 | 84.25431 | 81.02234 | chr2 | 158360808 | 158362259 | 4.72392 | 3.99093 | 2.78378 |
| chr16 | 52000169 | 52000446 | 615 | Distal Intergenic | 16 | 52031549 | 52208047 | 176499 | 1 | ENSMUST00 | -31103 | Cblb | ENSMUSG0000002 | true | . | . | . | . | . | . | chr16 | 51999695 | 52002797 | 4.10448 | 3.61196 | 2.26807 |
| chr5 | 81519487 | 81519765 | 614 | Intron (ENSMUST00000132375.2/ENSMUSG00 | 5 | 81645989 | 81688522 | 42534 | 1 | ENSMUST00 | -126224 | Lphn3 | ENSMUSG0000003 | false | chr5 | 81519672 | 81520026 | 5.15049 | 9.68248 | 7.51107 | chr5 | 81518468 | 81520895 | 3.83839 | 3.51106 | 2.04204 |
| chr12 | 94281002 | 94281214 | 613 | Distal Intergenic | 12 | 93927004 | 93929102 | 2099 | 2 | ENSMUST00 | -351901 | Gm9726 | ENSMUSG0000009 | false | . | . | . | . | . | . | chr12 | 94279611 | 94284656 | 3.79808 | 3.5006 | 2.0053 |
| chr13 | 42242122 | 42242453 | 613 | Distal Intergenic | 13 | 42301270 | 42307989 | 6720 | 1 | ENSMUST00 | -58817 | Edn1 | ENSMUSG0000002 | false | . | . | . | . | . | . | chr13 | 42235582 | 42245410 | 5.3203 | 4.22032 | 3.27886 |
| chr16 | 50786294 | 50786471 | 612 | Promoter (2-3kb) | 16 | 50783478 | 50784065 | 588 | 1 | ENSMUST00 | 2817 | 1700116B05 | ENSMUSG0000009 | false | . | . | . | . | . | . | chr16 | 50781927 | 50788895 | 5.4458 | 4.28026 | 3.36614 |
| chr7 | 126475337 | 126475506 | 611 | Promoter (<=1kb) | 7 | 126466994 | 126475130 | 8137 | 2 | ENSMUST00 | -208 | Sh2b1 | ENSMUSG0000003 | false | chr7 | 126474394 | 126476021 | 33.52805 | 106.05383 | 102.33234 | chr7 | 126472698 | 126475728 | 4.32439 | 3.73622 | 2.44657 |
| chr2 | 4652028 | 4652235 | 610 | Promoter (<=1kb) | 2 | 4622058 | 4652086 | 30029 | 2 | ENSMUST00 | 0 | Prpf18 | ENSMUSG0000003 | false | chr2 | 4651042 | 4652227 | 27.24215 | 82.29425 | 79.10038 | . | . | . | . | . | . |
| chr4 | 126321637 | 126321782 | 610 | Promoter (<=1kb) | 4 | 126316047 | 126321703 | 5657 | 2 | ENSMUST00 | 0 | Adprhl2 | ENSMUSG0000004 | false | chr4 | 126320878 | 126322123 | 37.54867 | 136.59039 | 131.84451 | . | . | . | . | . | . |
| chr1 | 143503859 | 143504022 | 608 | Distal Intergenic | 1 | 143633851 | 143636577 | 2727 | 2 | ENSMUST00 | 132555 | Cdc73 | ENSMUSG0000002 | false | . | . | . | . | . | . | . | . | . | . | . | . |
| chr15 | 78877311 | 78877582 | 608 | Promoter (<=1kb) | 15 | 78877190 | 78894585 | 17396 | 1 | ENSMUST00 | 122 | Gga1 | ENSMUSG0000003 | false | chr15 | 78876271 | 78878467 | 27.78601 | 91.88203 | 88.49705 | . | . | . | . | . | . |
| chr16 | 32877560 | 32877843 | 607 | Promoter (<=1kb) | 16 | 32877781 | 32908875 | 31095 | 1 | ENSMUST00 | 0 | Fyttd1 | ENSMUSG0000002 | false | chr16 | 32877139 | 32879191 | 27.85705 | 92.98368 | 89.57738 | . | . | . | . | . | . |
| chr2 | 50110219 | 50110384 | 607 | Intron (ENSMUST00000112712.4/ENSMUSG00 | 2 | 50128369 | 50193563 | 65195 | 1 | ENSMUST00 | -17985 | Lypd6 | ENSMUSG0000005 | false | . | . | . | . | . | . | chr2 | 50109251 | 50111092 | 3.6132 | 3.35253 | 1.86513 |
| chr15 | 76899829 | 76900051 | 606 | Promoter (<=1kb) | 15 | 76899910 | 76901305 | 1396 | 1 | ENSMUST00 | 0 | Commd5 | ENSMUSG0000005 | false | chr15 | 76899239 | 76900771 | 22.83944 | 61.22662 | 58.36318 | chr15 | 76898961 | 76901093 | 6.34124 | 4.67101 | 4.14957 |
| chr2 | 180171369 | 180171526 | 606 | Promoter (<=1kb) | 2 | 180171485 | 180176286 | 4802 | 1 | ENSMUST00 | 0 | Adrm1 | ENSMUSG0000003 | false | chr2 | 180171482 | 180172920 | 31.01414 | 97.6227 | 94.10706 | chr2 | 180171070 | 180174457 | 6.60325 | 4.53247 | 4.35711 |
| chr10 | 62216777 | 62216860 | 605 | Intron (ENSMUST00000047883.9/ENSMUSG00 | 10 | 62185396 | 62231218 | 45823 | 2 | ENSMUST00 | 14358 | Tspan15 | ENSMUSG0000003 | false | . | . | . | . | . | . | chr10 | 62185685 | 62229768 | 8.28034 | 5.46771 | 5.70297 |
| chr9 | 95954518 | 95954861 | 605 | Promoter (<=1kb) | 9 | 95954760 | 96053027 | 98268 | 1 | ENSMUST00 | 0 | Xrn1 | ENSMUSG0000003 | false | chr9 | 95954057 | 95955654 | 32.63288 | 132.66988 | 128.08362 | . | . | . | . | . | . |
| chr10 | 43540914 | 43541106 | 604 | Promoter (<=1kb) | 10 | 43525121 | 43540994 | 15874 | 2 | ENSMUST00 | 0 | 1700021F05 | ENSMUSG0000001 | false | chr10 | 43540096 | 43541068 | 37.09717 | 122.76616 | 118.52905 | chr10 | 43536760 | 43541464 | 4.73951 | 3.86652 | 2.82674 |
| chr17 | 83288188 | 83288417 | 604 | Promoter (<=1kb) | 17 | 83288546 | 83289028 | 483 | 1 | ENSMUST00 | -129 | Rpl21-ps7 | ENSMUSG0000009 | false | . | . | . | . | . | . | chr17 | 83287263 | 83289778 | 7.57114 | 5.04054 | 5.14387 |
| chr11 | 82871267 | 82871429 | 603 | Promoter (<=1kb) | 11 | 82803822 | 82871210 | 67389 | 2 | ENSMUST00 | -58 | Rffl | ENSMUSG0000002 | false | chr11 | 82870035 | 82871276 | 36.78915 | 128.67926 | 124.23671 | chr11 | 82869837 | 82871608 | 3.13383 | 3.0708 | 1.52114 |
| chr9 | 32672456 | 32672608 | 603 | Intron (ENSMUST00000183879.1/ENSMUSG00 | 9 | 32696022 | 32757820 | 61799 | 1 | ENSMUST00 | -23414 | Ets1 | ENSMUSG0000003 | true | chr9 | 32671954 | 32672786 | 8.53631 | 15.91019 | 13.6344 | chr9 | 32666505 | 32679281 | 4.67115 | 3.90561 | 2.75084 |
| chr6 | 39810927 | 39811028 | 602 | Promoter (<=1kb) | 6 | 39801804 | 39810988 | 9185 | 2 | ENSMUST00 | 0 | Mrps33 | ENSMUSG0000002 | false | chr6 | 39810232 | 39811219 | 18.02578 | 59.25625 | 56.42029 | . | . | . | . | . | . |
| chr1 | 39762143 | 39762248 | 601 | Distal Intergenic | 1 | 39665306 | 39720997 | 55692 | 2 | ENSMUST00 | -41147 | Rfx8 | ENSMUSG0000005 | false | chr1 | 39761731 | 39762300 | 12.68821 | 30.86953 | 28.39186 | chr1 | 39759438 | 39762551 | 4.33134 | 3.78958 | 2.45114 |
| chr7 | 48925264 | 48925509 | 601 | Intron (ENSMUST00000184336.1/ENSMUSG00 | 7 | 48908716 | 48963824 | 55109 | 1 | ENSMUST00 | 16549 | Nav2 | ENSMUSG0000005 | false | . | . | . | . | . | . | chr7 | 48924797 | 48926851 | 7.38669 | 5.17388 | 4.96231 |
| chr8 | 107031162 | 107031395 | 601 | Promoter (<=1kb) | 8 | 107031222 | 107045755 | 14534 | 1 | ENSMUST00 | 0 | Vps4a | ENSMUSG0000003 | false | chr8 | 107030699 | 107032235 | 37.92415 | 138.37384 | 133.55817 | . | . | . | . | . | . |
| chr19 | 45304688 | 45304861 | 600 | Promoter (1-2kb) | 19 | 45300463 | 45303034 | 2572 | 2 | ENSMUST00 | -1655 | RP23-188J8. | ENSMUSG0000010 | false | . | . | . | . | . | . | chr19 | 45300529 | 45315943 | 6.02178 | 4.57633 | 3.84298 |
| chr6 | 114921773 | 114922125 | 600 | Promoter (<=1kb) | 6 | 114861424 | 114921821 | 60398 | 2 | ENSMUST00 | 0 | Vgll4 | ENSMUSG0000003 | true | chr6 | 114917117 | 114921811 | 29.60582 | 98.11335 | 94.5914 | chr6 | 114922002 | 114923292 | 3.84635 | 3.53185 | 2.04366 |
| chr1 | 36352475 | 36352590 | 599 | Intron (ENSMUST00000185912.1/ENSMUSG00 | 1 | 36348682 | 36348758 | 77 | 1 | ENSMUST00 | -3718 | Mir6898 | ENSMUSG0000009 | false | . | . | . | . | . | . | chr1 | 36350640 | 36368234 | 5.46208 | 4.18784 | 3.39932 |
| chr16 | 25549707 | 25550005 | 599 | Distal Intergenic | 16 | 25683763 | 25821318 | 137556 | 1 | ENSMUST00 | -133758 | Trp63 | ENSMUSG0000002 | false | . | . | . | . | . | . | . | . | . | . | . | . |
| chr1 | 125760316 | 125760420 | 598 | Intron (ENSMUST00000027581.6/ENSMUSG00 | 1 | 125840003 | 125873173 | 33171 | 1 | ENSMUST00 | -79583 | Gpr39 | ENSMUSG0000002 | false | . | . | . | . | . | . | chr1 | 125759485 | 125760922 | 3.8444 | 3.47019 | 2.06578 |
| chr2 | 120093774 | 120093938 | 598 | Intron (ENSMUST00000028755.7/ENSMUSG00 | 2 | 120076214 | 120085772 | 9559 | 2 | ENSMUST00 | -8003 | Sptbn5 | ENSMUSG0000007 | false | . | . | . | . | . | . | chr2 | 120090578 | 120107323 | 5.14183 | 4.07311 | 3.14279 |
| chr16 | 13671818 | 13671967 | 597 | Promoter (<=1kb) | 16 | 13671858 | 13703612 | 31755 | 1 | ENSMUST00 | 0 | Bfar | ENSMUSG0000002 | false | chr16 | 13671401 | 13672923 | 43.50657 | 151.72289 | 146.37334 | . | . | . | . | . | . |
| chr2 | 106974307 | 106974523 | 597 | Promoter (<=1kb) | 2 | 106962527 | 106974424 | 11898 | 2 | ENSMUST00 | 0 | Arl14ep | ENSMUSG0000002 | false | chr2 | 106973606 | 106974761 | 26.6505 | 78.45867 | 75.33303 | chr2 | 106973065 | 106974784 | 4.27265 | 3.80927 | 2.39771 |
| chr6 | 70215864 | 70216062 | 597 | Promoter (<=1kb) | 6 | 70215433 | 70215957 | 525 | 1 | ENSMUST00 | 432 | Igkv6-25 | ENSMUSG0000009 | false | . | . | . | . | . | . | chr6 | 70213474 | 70219351 | 5.14256 | 4.19463 | 3.11118 |
| chr11 | 51967522 | 51967753 | 596 | Promoter (<=1kb) | 11 | 51967661 | 51977334 | 9674 | 1 | ENSMUST00 | 0 | Cdkn2aipnl | ENSMUSG0000002 | false | chr11 | 51967584 | 51968496 | 22.2526 | 73.30839 | 70.26766 | chr11 | 51967324 | 51968537 | 5.98126 | 4.53235 | 3.82674 |
| chr2 | 157048528 | 157048715 | 596 | Intron (ENSMUST00000069098.6/ENSMUSG00 | 2 | 157037050 | 157040239 | 3190 | 2 | ENSMUST00 | -8290 | Soga1 | ENSMUSG0000005 | false | . | . | . | . | . | . | chr2 | 157047505 | 157049845 | 3.42377 | 3.07497 | 1.74981 |
| chr16 | 74093141 | 74093367 | 595 | Intron (ENSMUST00000117785.2/ENSMUSG00 | 16 | 74058899 | 74059033 | 135 | 2 | ENSMUST00 | -34109 | Gm22163 | ENSMUSG0000008 | false | . | . | . | . | . | . | chr16 | 74090760 | 74094201 | 4.77226 | 3.99214 | 2.80493 |
| chr15 | 99701985 | 99702359 | 594 | Promoter (<=1kb) | 15 | 99702287 | 99713995 | 11709 | 1 | ENSMUST00 | 0 | Smarcd1 | ENSMUSG0000002 | false | chr15 | 99701792 | 99702839 | 37.30079 | 124.19571 | 119.91465 | chr15 | 99700422 | 99705143 | 3.90355 | 3.54268 | 2.10055 |
| chr13 | 119428834 | 119428999 | 593 | Promoter (<=1kb) | 13 | 119428822 | 119458218 | 29297 | 1 | ENSMUST00 | 0 | Paip1 | ENSMUSG0000002 | false | chr13 | 119428264 | 119430336 | 31.16539 | 106.97112 | 103.21663 | . | . | . | . | . | . |
| chr3 | 132835187 | 132835388 | 592 | Intron (ENSMUST00000169172.2/ENSMUSG00 | 3 | 132787225 | 132787335 | 111 | 2 | ENSMUST00 | -47853 | Gm22322 | ENSMUSG0000008 | false | . | . | . | . | . | . | . | . | . | . | . | . |
| chr7 | 46033564 | 46033740 | 592 | Promoter (<=1kb) | 7 | 46033698 | 46084212 | 50515 | 1 | ENSMUST00 | 0 | Nomo1 | ENSMUSG0000003 | false | chr7 | 46033517 | 46034659 | 31.66066 | 102.77801 | 99.14294 | . | . | . | . | . | . |
| chr4 | 43025791 | 43025895 | 591 | Promoter (<=1kb) | 4 | 43017635 | 43025819 | 8185 | 2 | ENSMUST00 | 0 | Pigo | ENSMUSG0000002 | false | chr4 | 43025093 | 43025900 | 16.59601 | 56.40767 | 53.61239 | . | . | . | . | . | . |
| chr6 | 70240248 | 70240516 | 591 | Distal Intergenic | 6 | 70260409 | 70260933 | 525 | 2 | ENSMUST00 | 20417 | Igkv6-23 | ENSMUSG0000009 | false | . | . | . | . | . | . | chr6 | 70239615 | 70241403 | 4.75318 | 3.82127 | 2.83041 |
| chr10 | 84409025 | 84409126 | 590 | Intron (ENSMUST00000020220.9/ENSMUSG00 | 10 | 84375296 | 84392388 | 17093 | 2 | ENSMUST00 | -16638 | Nuak1 | ENSMUSG0000002 | false | . | . | . | . | . | . | chr10 | 84408525 | 84412493 | 5.17253 | 4.02084 | 3.19694 |
| chr15 | 36496744 | 36496906 | 590 | Promoter (<=1kb) | 15 | 36477668 | 36496791 | 19124 | 2 | ENSMUST00 | 0 | Ankrd46 | ENSMUSG0000004 | false | chr15 | 36495983 | 36497122 | 27.18312 | 111.36749 | 107.49936 | chr15 | 36487947 | 36503116 | 6.4169 | 4.70464 | 4.17261 |
| chr1 | 45795193 | 45795574 | 589 | Promoter (<=1kb) | 1 | 45795449 | 45823604 | 28156 | 1 | ENSMUST00 | 0 | Wdr75 | ENSMUSG0000002 | false | chr1 | 45794798 | 45796261 | 37.58501 | 119.35227 | 115.22867 | . | . | . | . | . | . |
| chr12 | 111516901 | 111517035 | 589 | Distal Intergenic | 12 | 111538101 | 111546753 | 8653 | 1 | ENSMUST00 | -21066 | Eif5 | ENSMUSG0000002 | false | . | . | . | . | . | . | chr12 | 111513836 | 111518852 | 5.17158 | 4.07062 | 3.1695 |
| chr4 | 149860376 | 149860597 | 588 | Distal Intergenic | 4 | 149817732 | 149817840 | 109 | 1 | ENSMUST00 | 42645 | n-R5s193 | ENSMUSG0000006 | false | . | . | . | . | . | . | chr4 | 149859847 | 149861376 | 5.80362 | 4.28726 | 3.6996 |
| chr2 | 28467915 | 28468169 | 587 | Promoter (<=1kb) | 2 | 28468066 | 28471178 | 3113 | 1 | ENSMUST00 | 0 | Mrps2 | ENSMUSG0000003 | false | chr2 | 28467819 | 28468909 | 37.97791 | 128.62614 | 124.18539 | chr2 | 28467586 | 28468947 | 4.87461 | 3.96608 | 2.94727 |
| chr10 | 68445931 | 68446189 | 586 | Intron (ENSMUST00000020103.3/ENSMUSG00 | 10 | 68436280 | 68525767 | 89488 | 2 | ENSMUST00 | 79578 | 1700040L02 | ENSMUSG0000001 | false | . | . | . | . | . | . | chr10 | 68445235 | 68447123 | 5.83517 | 4.52043 | 3.68283 |
| chr11 | 100619593 | 100619867 | 586 | Promoter (<=1kb) | 11 | 100590894 | 100619597 | 28704 | 2 | ENSMUST00 | 0 | Dnajc7 | ENSMUSG0000001 | false | chr11 | 100618769 | 100620432 | 29.28796 | 98.53321 | 94.99628 | . | . | . | . | . | . |
| chr12 | 69410848 | 69411030 | 586 | Distal Intergenic | 12 | 69371835 | 69372458 | 624 | 2 | ENSMUST00 | -38391 | Gm9887 | ENSMUSG0000005 | false | . | . | . | . | . | . | chr12 | 69405873 | 69415994 | 4.13026 | 3.54336 | 2.31242 |
| chr14 | 14120899 | 14121074 | 585 | Promoter (<=1kb) | 14 | 14112184 | 14120984 | 8801 | 2 | ENSMUST00 | 0 | Psmd6 | ENSMUSG0000002 | false | chr14 | 14119872 | 14121052 | 41.83324 | 144.2525 | 139.19977 | chr14 | 14119398 | 14121211 | 5.34325 | 4.16111 | 3.32385 |
| chr2 | 153241358 | 153241613 | 585 | Promoter (<=1kb) | 2 | 153241533 | 153270247 | 28715 | 1 | ENSMUST00 | 0 | Pofut1 | ENSMUSG0000004 | true | chr2 | 153239613 | 153242107 | 24.32174 | 95.77222 | 92.30251 | . | . | . | . | . | . |
| chr14 | 45507174 | 45507316 | 584 | Intron (ENSMUST00000045905.7/ENSMUSG00 | 14 | 45504672 | 45529786 | 25115 | 2 | ENSMUST00 | 22470 | Fermt2 | ENSMUSG0000003 | true | . | . | . | . | . | . | chr14 | 45489964 | 45528839 | 6.12493 | 4.54024 | 3.93691 |
| chr4 | 150784817 | 150785023 | 584 | Distal Intergenic | 4 | 150825733 | 150839575 | 13843 | 1 | ENSMUST00 | -40710 | Gm13049 | ENSMUSG0000008 | false | . | . | . | . | . | . | chr4 | 150784510 | 150786637 | 4.42467 | 3.8393 | 2.5188 |
| chr8 | 128544599 | 128544764 | 584 | Distal Intergenic | 8 | 128650997 | 128651103 | 107 | 1 | ENSMUST00 | -106233 | Gm26002 | ENSMUSG0000007 | false | . | . | . | . | . | . | chr8 | 128519464 | 128548431 | 5.17648 | 4.1349 | 3.15487 |
| chr18 | 76088011 | 76088375 | 583 | Intron (ENSMUST00000058997.8/ENSMUSG00 | 18 | 76059458 | 76146443 | 86986 | 1 | ENSMUST00 | 28554 | Zbtb7c | ENSMUSG0000004 | true | . | . | . | . | . | . | chr18 | 76061445 | 76132293 | 6.70015 | 4.79833 | 4.4158 |
| chr5 | 23787583 | 23787752 | 583 | Promoter (<=1kb) | 5 | 23787711 | 23790003 | 2293 | 1 | ENSMUST00 | 0 | Rint1 | ENSMUSG0000002 | false | chr5 | 23787616 | 23788615 | 39.52876 | 141.27539 | 136.34271 | chr5 | 23787360 | 23789287 | 5.95812 | 4.527 | 3.83932 |
| chr13 | 104743015 | 104743170 | 582 | Intron (ENSMUST00000022228.7/ENSMUSG00 | 13 | 104792484 | 104838360 | 45877 | 1 | ENSMUST00 | -49314 | Srek1ip1 | ENSMUSG0000002 | false | . | . | . | . | . | . | chr13 | 104740457 | 104744715 | 5.11238 | 4.12621 | 3.09044 |
| chr17 | 5189837 | 5190082 | 582 | Intron (ENSMUST00000092723.5/ENSMUSG00 | 17 | 5110781 | 5112119 | 1339 | 2 | ENSMUST00 | -77719 | Gm15599 | ENSMUSG0000008 | false | chr17 | 5189499 | 5190676 | 15.70458 | 43.19576 | 40.56621 | chr17 | 5188721 | 5199127 | 4.36014 | 3.6534 | 2.51291 |
| chr6 | 121146971 | 121147218 | 582 | Distal Intergenic | 6 | 121183667 | 121196185 | 12519 | 1 | ENSMUST00 | -36449 | Pex26 | ENSMUSG0000006 | false | . | . | . | . | . | . | chr6 | 121143778 | 121148146 | 5.47647 | 4.19044 | 3.41087 |
| chr17 | 47368720 | 47368938 | 581 | Promoter (<=1kb) | 17 | 47368887 | 47378679 | 9793 | 1 | ENSMUST00 | 0 | Mrps10 | ENSMUSG0000003 | false | chr17 | 47368416 | 47369691 | 28.1615 | 93.53628 | 90.11485 | . | . | . | . | . | . |
| chr2 | 180701986 | 180702122 | 581 | Promoter (<=1kb) | 2 | 180685024 | 180701992 | 16969 | 2 | ENSMUST00 | 0 | Dido1 | ENSMUSG0000003 | false | chr2 | 180700691 | 180702845 | 31.25561 | 98.0451 | 94.52408 | . | . | . | . | . | . |
| chr8 | 61665126 | 61665289 | 581 | Intron (ENSMUST00000121785.2/ENSMUSG00 | 8 | 61513248 | 61591142 | 77895 | 2 | ENSMUST00 | -73985 | Palld | ENSMUSG0000005 | true | . | . | . | . | . | . | chr8 | 61662521 | 61666943 | 4.86592 | 3.97418 | 2.91383 |
| chr11 | 97437928 | 97438272 | 580 | Promoter (1-2kb) | 11 | 97436285 | 97444228 | 7944 | 1 | ENSMUST00 | 1644 | Arhgap23 | ENSMUSG0000004 | true | . | . | . | . | . | . | chr11 | 97431804 | 97440135 | 9.81823 | 6.18842 | 6.94298 |
| chr18 | 9958043 | 9958217 | 580 | Promoter (<=1kb) | 18 | 9958180 | 9995484 | 37305 | 1 | ENSMUST00 | 0 | Thoc1 | ENSMUSG0000002 | false | chr18 | 9957507 | 9958955 | 23.46333 | 91.19009 | 87.82124 | . | . | . | . | . | . |
| chr5 | 136953155 | 136953329 | 580 | Promoter (<=1kb) | 5 | 136953275 | 136966232 | 12958 | 1 | ENSMUST00 | 0 | Fis1 | ENSMUSG0000001 | false | chr5 | 136952722 | 136954222 | 22.63194 | 64.29881 | 61.39256 | . | . | . | . | . | . |
| chr11 | 121794597 | 121794898 | 579 | Distal Intergenic | 11 | 121803107 | 121810319 | 7213 | 1 | ENSMUST00 | -8209 | Gm12589 | ENSMUSG0000008 | false | . | . | . | . | . | . | chr11 | 121787526 | 121808717 | 6.67832 | 4.85129 | 4.38811 |
| chr4 | 132212035 | 132212466 | 579 | Promoter (<=1kb) | 4 | 132184912 | 132212303 | 27392 | 2 | ENSMUST00 | 0 | Ythdf2 | ENSMUSG0000004 | false | chr4 | 132210818 | 132212608 | 36.04346 | 118.78738 | 114.68494 | . | . | . | . | . | . |
| chr9 | 58541951 | 58542185 | 579 | Promoter (1-2kb) | 9 | 58524300 | 58540941 | 16642 | 2 | ENSMUST00 | -1011 | Cd276 | ENSMUSG0000003 | false | . | . | . | . | . | . | chr9 | 58541298 | 58542496 | 5.43317 | 4.11368 | 3.40583 |
| chr11 | 49203385 | 49203688 | 578 | Promoter (<=1kb) | 11 | 49203485 | 49213636 | 10152 | 1 | ENSMUST00 | 0 | Zfp62 | ENSMUSG0000004 | false | chr11 | 49202015 | 49204776 | 21.75525 | 76.8157 | 73.7174 | . | . | . | . | . | . |
| chr5 | 103629555 | 103629691 | 578 | Promoter (<=1kb) | 5 | 103605711 | 103629403 | 23693 | 2 | ENSMUST00 | -153 | Slc10a6 | ENSMUSG0000002 | false | . | . | . | . | . | . | chr5 | 103627004 | 103630959 | 5.1666 | 4.08182 | 3.15645 |
| chr12 | 101913011 | 101913233 | 577 | Promoter (<=1kb) | 12 | 101886220 | 101913123 | 26904 | 2 | ENSMUST00 | 0 | Trip11 | ENSMUSG0000002 | false | chr12 | 101911905 | 101913759 | 29.58769 | 108.11006 | 104.32592 | . | . | . | . | . | . |
| chr17 | 46052201 | 46052365 | 577 | Distal Intergenic | 17 | 46016996 | 46032377 | 15382 | 2 | ENSMUST00 | -19825 | Vegfa | ENSMUSG0000002 | true | . | . | . | . | . | . | chr17 | 46048792 | 46077235 | 6.11576 | 4.54762 | 3.92959 |
| chr14 | 50785440 | 50785674 | 576 | Promoter (<=1kb) | 14 | 50765415 | 50785518 | 20104 | 2 | ENSMUST00 | 0 | Ttc5 | ENSMUSG0000000 | false | chr14 | 50784823 | 50785681 | 31.62822 | 118.12274 | 114.03909 | chr14 | 50783693 | 50785856 | 3.59414 | 3.28458 | 1.88913 |
| chr7 | 110615259 | 110615685 | 576 | Promoter (<=1kb) | 7 | 110308013 | 110614922 | 306910 | 2 | ENSMUST00 | -338 | Sbf2 | ENSMUSG0000003 | true | chr7 | 110613377 | 110615752 | 31.19317 | 93.97161 | 90.54182 | . | . | . | . | . | . |
| chr8 | 70610242 | 70610426 | 576 | Promoter (1-2kb) | 8 | 70597491 | 70608314 | 10824 | 2 | ENSMUST00 | -1929 | Ssbp4 | ENSMUSG0000007 | false | chr8 | 70610311 | 70610820 | 10.04446 | 20.40655 | 18.06574 | chr8 | 70604127 | 70612475 | 6.23684 | 4.68532 | 4.01553 |
| chr15 | 74709224 | 74709596 | 575 | Promoter (<=1kb) | 15 | 74702301 | 74709535 | 7235 | 2 | ENSMUST00 | 0 | Jrk | ENSMUSG0000004 | false | chr15 | 74708602 | 74709818 | 29.58769 | 108.11006 | 104.32592 | chr15 | 74708335 | 74710396 | 4.89568 | 4.03818 | 2.92069 |
| chr19 | 5366591 | 5366985 | 575 | Promoter (<=1kb) | 19 | 5366813 | 5371511 | 4699 | 1 | ENSMUST00 | 0 | Eif1ad | ENSMUSG0000002 | false | chr19 | 5366022 | 5367764 | 27.5664 | 95.37588 | 91.91415 | . | . | . | . | . | . |
| chr4 | 115709082 | 115709344 | 575 | Promoter (1-2kb) | 4 | 115707251 | 115708303 | 1053 | 1 | ENSMUST00 | 1832 | Gm12849 | ENSMUSG0000008 | false | . | . | . | . | . | . | chr4 | 115707353 | 115710474 | 3.80958 | 3.35728 | 2.063 |
| chr1 | 90442898 | 90443107 | 574 | Distal Intergenic | 1 | 90453394 | 90454501 | 1108 | 2 | ENSMUST00 | 11394 | RP24-154L11 | ENSMUSG0000010 | false | . | . | . | . | . | . | chr1 | 90438990 | 90445431 | 5.52456 | 4.20032 | 3.45273 |
| chr14 | 73509915 | 73510070 | 573 | Promoter (<=1kb) | 14 | 73510049 | 73518545 | 8497 | 1 | ENSMUST00 | 0 | Med4 | ENSMUSG0000002 | false | chr14 | 73509938 | 73510748 | 29.66345 | 100.21 | 96.63225 | chr14 | 73509549 | 73512613 | 4.53431 | 3.79887 | 2.62423 |
| chr5 | 140777154 | 140777360 | 573 | Intron (ENSMUST00000000153.4/ENSMUSG00 | 5 | 140749283 | 140761439 | 12157 | 1 | ENSMUST00 | 27872 | Amz1 | ENSMUSG0000005 | false | . | . | . | . | . | . | chr5 | 140761728 | 140779292 | 6.36667 | 4.61767 | 4.14055 |
| chr8 | 91522387 | 91522476 | 573 | Intron (ENSMUST00000069718.8/ENSMUSG00 | 8 | 91467685 | 91666946 | 199262 | 1 | ENSMUST00 | 54703 | Fto | ENSMUSG0000005 | false | . | . | . | . | . | . | chr8 | 91521033 | 91528017 | 5.83853 | 4.42897 | 3.70928 |
| chr4 | 136602688 | 136602954 | 572 | Promoter (<=1kb) | 4 | 136550540 | 136602723 | 52184 | 2 | ENSMUST00 | 0 | Kdm1a | ENSMUSG0000003 | false | chr4 | 136601489 | 136602829 | 37.62274 | 127.9987 | 123.57949 | . | . | . | . | . | . |
| chr5 | 3543748 | 3543968 | 572 | Promoter (<=1kb) | 5 | 3543844 | 3569810 | 25967 | 1 | ENSMUST00 | 0 | Fam133b | ENSMUSG0000005 | false | chr5 | 3543740 | 3544744 | 23.32969 | 96.49238 | 93.00913 | . | . | . | . | . | . |
| chr9 | 58253058 | 58253243 | 571 | Promoter (<=1kb) | 9 | 58253164 | 58262523 | 9360 | 1 | ENSMUST00 | 0 | Stoml1 | ENSMUSG0000003 | false | chr9 | 58253191 | 58253958 | 42.22279 | 143.81401 | 138.77701 | chr9 | 58252556 | 58255789 | 4.55946 | 3.87216 | 2.64575 |
| chr18 | 77713949 | 77714118 | 570 | Promoter (<=1kb) | 18 | 77633281 | 77714010 | 80730 | 2 | ENSMUST00 | 0 | 8030462N17 | ENSMUSG0000004 | false | chr18 | 77712518 | 77714741 | 34.68901 | 133.45216 | 128.82967 | . | . | . | . | . | . |
| chr2 | 33606011 | 33606128 | 570 | Intron (ENSMUST00000041730.5/ENSMUSG00 | 2 | 33567156 | 33632283 | 65128 | 2 | ENSMUST00 | 26155 | Lmx1b | ENSMUSG0000003 | true | . | . | . | . | . | . | chr2 | 33565260 | 33638297 | 6.78481 | 4.81457 | 4.48235 |
| chr7 | 25316829 | 25316988 | 570 | Promoter (<=1kb) | 7 | 25317164 | 25365917 | 48754 | 1 | ENSMUST00 | -176 | Megf8 | ENSMUSG0000004 | false | chr7 | 25316780 | 25318001 | 32.44658 | 100.11784 | 96.54536 | chr7 | 25314793 | 25322508 | 5.81353 | 4.33708 | 3.6977 |
| chr3 | 116221785 | 116222049 | 569 | Distal Intergenic | 3 | 116249654 | 116253484 | 3831 | 2 | ENSMUST00 | 31435 | Gpr88 | ENSMUSG0000006 | false | . | . | . | . | . | . | . | . | . | . | . | . |
| chr5 | 32863614 | 32863801 | 569 | Promoter (<=1kb) | 5 | 32863701 | 32887048 | 23348 | 1 | ENSMUST00 | 0 | Depdc5 | ENSMUSG0000003 | false | chr5 | 32863170 | 32864374 | 20.32388 | 57.71489 | 54.90119 | . | . | . | . | . | . |
| chr6 | 10186376 | 10186468 | 569 | Distal Intergenic | 6 | 10172022 | 10172099 | 78 | 2 | ENSMUST00 | -14278 | Gm24104 | ENSMUSG0000009 | false | . | . | . | . | . | . | . | . | . | . | . | . |
| chr1 | 179546272 | 179546468 | 568 | Promoter (<=1kb) | 1 | 179546370 | 179627478 | 81109 | 1 | ENSMUST00 | 0 | Cnst | ENSMUSG0000003 | false | chr1 | 179545202 | 179547773 | 26.61085 | 108.21047 | 104.42454 | . | . | . | . | . | . |
| chr17 | 56036707 | 56036892 | 568 | Promoter (<=1kb) | 17 | 56016753 | 56036637 | 19885 | 2 | ENSMUST00 | -71 | Sh3gl1 | ENSMUSG0000000 | false | . | . | . | . | . | . | . | . | . | . | . | . |
| chr17 | 47695026 | 47695191 | 567 | Promoter (<=1kb) | 17 | 47695198 | 47704286 | 9089 | 1 | ENSMUST00 | -7 | Frs3 | ENSMUSG0000002 | false | chr17 | 47694413 | 47695873 | 29.73511 | 88.5301 | 85.22105 | . | . | . | . | . | . |
| chr5 | 103606423 | 103606665 | 567 | 3' UTR | 5 | 103602587 | 103602879 | 293 | 1 | ENSMUST00 | 3837 | Gm23085 | ENSMUSG0000008 | false | . | . | . | . | . | . | chr5 | 103602894 | 103607755 | 4.47885 | 3.74726 | 2.61032 |
| chr13 | 21530864 | 21531210 | 566 | Promoter (<=1kb) | 13 | 21513222 | 21531120 | 17899 | 2 | ENSMUST00 | 0 | Zkscan8 | ENSMUSG0000006 | false | chr13 | 21530204 | 21531455 | 41.44478 | 148.06744 | 142.8609 | . | . | . | . | . | . |
| chr6 | 72743833 | 72744169 | 566 | Intron (ENSMUST00000069536.6/ENSMUSG00 | 6 | 72632976 | 72788598 | 155623 | 2 | ENSMUST00 | 44429 | Tcf7l1 | ENSMUSG0000005 | false | . | . | . | . | . | . | chr6 | 72739919 | 72749221 | 7.65916 | 5.17359 | 5.21352 |
| chr1 | 74587986 | 74588370 | 565 | Promoter (<=1kb) | 1 | 74565127 | 74588028 | 22902 | 2 | ENSMUST00 | 0 | Zfp142 | ENSMUSG0000002 | false | chr1 | 74588268 | 74588770 | 17.64787 | 49.73517 | 47.02245 | . | . | . | . | . | . |
| chr12 | 84617357 | 84617600 | 565 | Promoter (<=1kb) | 12 | 84602531 | 84617466 | 14936 | 2 | ENSMUST00 | 0 | Abcd4 | ENSMUSG0000002 | false | chr12 | 84616765 | 84618108 | 27.20707 | 96.65539 | 93.16586 | . | . | . | . | . | . |
| chr5 | 135806733 | 135806973 | 564 | Promoter (<=1kb) | 5 | 135806890 | 135841953 | 35064 | 1 | ENSMUST00 | 0 | Srrm3 | ENSMUSG0000003 | false | . | . | . | . | . | . | . | . | . | . | . | . |
| chr9 | 62608582 | 62608670 | 564 | Distal Intergenic | 9 | 62677826 | 62783982 | 106157 | 1 | ENSMUST00 | -69156 | Itga11 | ENSMUSG0000003 | false | . | . | . | . | . | . | . | . | . | . | . | . |
| chrX | 99216819 | 99217086 | 564 | Distal Intergenic | 20 | 99138131 | 99147230 | 9100 | 1 | ENSMUST00 | 78689 | Efnb1 | ENSMUSG0000003 | false | chrX | 99216303 | 99217267 | 11.94783 | 25.26529 | 22.85895 | . | . | . | . | . | . |
| chr11 | 58103937 | 58104177 | 563 | Promoter (<=1kb) | 11 | 58104153 | 58118594 | 14442 | 1 | ENSMUST00 | 0 | Cnot8 | ENSMUSG0000002 | false | chr11 | 58104039 | 58105114 | 38.13901 | 127.82562 | 123.40861 | . | . | . | . | . | . |
| chr19 | 44562665 | 44562884 | 563 | Promoter (<=1kb) | 19 | 44562850 | 44563952 | 1103 | 1 | ENSMUST00 | 0 | Hif1an | ENSMUSG0000003 | false | chr19 | 44562423 | 44563784 | 28.95052 | 83.36136 | 80.14891 | chr19 | 44562321 | 44564336 | 5.10712 | 4.10999 | 3.13372 |
| chr11 | 63144302 | 63144490 | 562 | Intron (ENSMUST00000108702.2/ENSMUSG00 | 11 | 63133068 | 63158566 | 25499 | 1 | ENSMUST00 | 11235 | Pmp22 | ENSMUSG0000001 | false | . | . | . | . | . | . | . | . | . | . | . | . |
| chr7 | 120635114 | 120635253 | 562 | Promoter (<=1kb) | 7 | 120635176 | 120659478 | 24303 | 1 | ENSMUST00 | 0 | Uqcrc2 | ENSMUSG0000003 | false | chr7 | 120634318 | 120636330 | 39.37288 | 136.56555 | 131.82602 | . | . | . | . | . | . |
| chr1 | 172082577 | 172082822 | 561 | Promoter (<=1kb) | 1 | 172066013 | 172082795 | 16783 | 2 | ENSMUST00 | 0 | Ncstn | ENSMUSG0000003 | false | chr1 | 172081970 | 172083760 | 30.77379 | 96.20782 | 92.72798 | . | . | . | . | . | . |
| chr13 | 45897687 | 45897917 | 561 | Intron (ENSMUST00000091628.4/ENSMUSG00 | 13 | 45549758 | 45964991 | 415234 | 2 | ENSMUST00 | 67074 | Atxn1 | ENSMUSG0000004 | false | . | . | . | . | . | . | chr13 | 45897011 | 45898862 | 3.09204 | 2.93398 | 1.47583 |
| chr2 | 122575492 | 122575750 | 561 | Distal Intergenic | 2 | 122595489 | 122598534 | 3046 | 2 | ENSMUST00 | 22784 | Gatm | ENSMUSG0000002 | false | . | . | . | . | . | . | chr2 | 122574487 | 122577662 | 5.6028 | 4.4306 | 3.49482 |

| | | | | | | | | | | | | | | | | | | | | | | | | | | |
|---|---|---|---|---|---|---|---|---|---|---|---|---|---|---|---|---|---|---|---|---|---|---|---|---|---|---|
| chr15 | 80132973 | 80133203 | 560 | Promoter (<=1kb) | 15 | 80133154 | 80161701 | 28548 | 1 | ENSMUST00 | 0 | Tab1 | ENSMUSG0000002 | false | chr15 | 80132695 | 80134176 | 24.78212 | 78.86476 | 75.73294 | chr15 | 80132452 | 80135076 | 5.60141 | 4.37527 | 3.50624 |
| chr3 | 96727405 | 96727638 | 560 | Promoter (<=1kb) | 3 | 96727611 | 96791155 | 63545 | 1 | ENSMUST00 | 0 | Rnf115 | ENSMUSG0000002 | false | chr3 | 96726884 | 96728769 | 34.74108 | 114.39095 | 110.42815 | . | . | . | . | . | . |
| chr11 | 76763533 | 76763713 | 559 | Promoter (<=1kb) | 11 | 76726602 | 76763579 | 36978 | 2 | ENSMUST00 | 0 | Gosr1 | ENSMUSG0000001 | false | chr11 | 76762717 | 76763684 | 29.3377 | 90.74815 | 87.38457 | . | . | . | . | . | . |
| chr11 | 48816999 | 48817429 | 558 | Promoter (<=1kb) | 11 | 48806404 | 48817353 | 10950 | 2 | ENSMUST00 | 0 | Trim41 | ENSMUSG0000004 | false | chr11 | 48815920 | 48818558 | 26.65955 | 86.95457 | 83.67183 | . | . | . | . | . | . |
| chr7 | 112680121 | 112680275 | 558 | Promoter (<=1kb) | 7 | 112613468 | 112680058 | 66591 | 2 | ENSMUST00 | -64 | 2310014F06 | ENSMUSG0000008 | false | chr7 | 112678205 | 112681471 | 36.76257 | 113.72284 | 109.77938 | . | . | . | . | . | . |
| chr5 | 143112967 | 143113208 | 557 | Promoter (<=1kb) | 5 | 142990893 | 143113020 | 122128 | 2 | ENSMUST00 | 0 | Rnf216 | ENSMUSG0000004 | false | chr5 | 143111903 | 143113458 | 29.34276 | 99.62727 | 96.06711 | chr5 | 143100232 | 143113412 | 4.82716 | 3.92416 | 2.88218 |
| chr16 | 33380600 | 33380874 | 556 | Promoter (<=1kb) | 16 | 33380775 | 33503903 | 123129 | 1 | ENSMUST00 | 0 | Zfp148 | ENSMUSG0000002 | false | chr16 | 33380162 | 33382282 | 22.99841 | 82.87405 | 79.66897 | . | . | . | . | . | . |
| chr2 | 26132995 | 26133189 | 556 | Distal Intergenic | 2 | 26138546 | 26139543 | 998 | 1 | ENSMUST00 | -5357 | 1810012K08 | ENSMUSG0000008 | false | chr2 | 26132583 | 26133600 | 9.51244 | 19.51414 | 17.18665 | chr2 | 26129136 | 26135084 | 6.01114 | 4.43692 | 3.85388 |
| chr18 | 10617701 | 10617842 | 554 | Promoter (<=1kb) | 18 | 10617796 | 10628230 | 10435 | 1 | ENSMUST00 | 0 | Snrpd1 | ENSMUSG0000002 | false | chr18 | 10617431 | 10618746 | 36.76141 | 116.86796 | 112.83147 | chr18 | 10617345 | 10619627 | 3.67087 | 3.44448 | 1.89704 |
| chr5 | 114707683 | 114707852 | 554 | Promoter (<=1kb) | 5 | 114707760 | 114722327 | 14568 | 1 | ENSMUST00 | 0 | Tchp | ENSMUSG0000002 | false | chr5 | 114707703 | 114708580 | 28.53699 | 95.19626 | 91.7378 | chr5 | 114707363 | 114708929 | 7.03659 | 5.04842 | 4.64939 |
| chr7 | 45717885 | 45718181 | 552 | Promoter (<=1kb) | 7 | 45709467 | 45718002 | 8536 | 2 | ENSMUST00 | 0 | Sphk2 | ENSMUSG0000005 | false | chr7 | 45717180 | 45719165 | 29.60582 | 98.11335 | 94.5914 | . | . | . | . | . | . |
| chr7 | 105736409 | 105736735 | 551 | Promoter (<=1kb) | 7 | 105736592 | 105742925 | 6334 | 1 | ENSMUST00 | 0 | Ilk | ENSMUSG0000003 | false | chr7 | 105735728 | 105738289 | 44.26889 | 153.06256 | 147.66058 | . | . | . | . | . | . |
| chr7 | 127636984 | 127637190 | 550 | Distal Intergenic | 7 | 127608756 | 127615797 | 7042 | 2 | ENSMUST00 | -21188 | Zfp629 | ENSMUSG0000004 | false | chr7 | 127636649 | 127637757 | 14.64398 | 38.39655 | 35.82735 | chr7 | 127636481 | 127637911 | 5.47361 | 4.28826 | 3.42456 |
| chr4 | 116651641 | 116651773 | 549 | Promoter (<=1kb) | 4 | 116636510 | 116651680 | 15171 | 2 | ENSMUST00 | 0 | Akr1a1 | ENSMUSG0000002 | false | chr4 | 116650295 | 116652083 | 33.7247 | 108.45113 | 104.65779 | . | . | . | . | . | . |
| chr17 | 66419507 | 66419717 | 548 | Intron (ENSMUST00000086693.6/ENSMUSG00 | 17 | 66413631 | 66413984 | 354 | 2 | ENSMUST00 | -5524 | Gm4705 | ENSMUSG0000009 | false | . | . | . | . | . | . | chr17 | 66401889 | 66429482 | 6.16935 | 4.5489 | 3.97689 |
| chr18 | 33463794 | 33464200 | 548 | Promoter (<=1kb) | 18 | 33437019 | 33464029 | 27011 | 2 | ENSMUST00 | 0 | Nrep | ENSMUSG0000004 | false | chr18 | 33462104 | 33464772 | 29.3377 | 90.74815 | 87.38457 | . | . | . | . | . | . |
| chr6 | 83775754 | 83775969 | 547 | Promoter (<=1kb) | 6 | 83768885 | 83775813 | 6929 | 2 | ENSMUST00 | 0 | Tex261 | ENSMUSG0000001 | false | chr6 | 83774759 | 83776337 | 32.94367 | 134.34938 | 129.69266 | chr6 | 83774182 | 83776885 | 4.59104 | 3.72868 | 2.70779 |
| chr1 | 118388998 | 118389476 | 546 | Promoter (<=1kb) | 1 | 118389058 | 118609432 | 220375 | 1 | ENSMUST00 | 0 | Clasp1 | ENSMUSG0000006 | false | chr1 | 118387964 | 118390405 | 29.66345 | 100.21 | 96.63225 | . | . | . | . | . | . |
| chr11 | 54744293 | 54744501 | 545 | Promoter (2-3kb) | 11 | 54746357 | 54746824 | 468 | 2 | ENSMUST00 | 2323 | Gm12229 | ENSMUSG0000008 | false | . | . | . | . | . | . | chr11 | 54722256 | 54750513 | 5.8032 | 4.39 | 3.67574 |
| chr1 | 170885405 | 170885547 | 544 | Promoter (<=1kb) | 1 | 170873498 | 170885457 | 11960 | 2 | ENSMUST00 | 0 | Dusp12 | ENSMUSG0000002 | false | chr1 | 170884514 | 170885772 | 43.01277 | 150.85193 | 145.53938 | . | . | . | . | . | . |
| chr18 | 53744488 | 53744814 | 544 | Promoter (<=1kb) | 18 | 53681724 | 53744547 | 62824 | 2 | ENSMUST00 | 0 | Cep120 | ENSMUSG0000004 | false | chr18 | 53743447 | 53745668 | 31.14379 | 104.89771 | 101.20105 | . | . | . | . | . | . |
| chr4 | 116732409 | 116732670 | 542 | Intron (ENSMUST00000124628.2/ENSMUSG00 | 4 | 116725899 | 116725989 | 91 | 1 | ENSMUST00 | 6511 | Gm27688 | ENSMUSG0000009 | false | . | . | . | . | . | . | chr4 | 116731176 | 116741002 | 4.98596 | 4.04488 | 2.98621 |
| chr7 | 132852423 | 132852864 | 541 | Promoter (<=1kb) | 7 | 132827457 | 132852673 | 25217 | 2 | ENSMUST00 | 0 | Mettl10 | ENSMUSG0000003 | false | chr7 | 132851815 | 132852798 | 34.84858 | 109.37691 | 105.56304 | chr7 | 132850885 | 132853191 | 4.1659 | 3.50231 | 2.36259 |
| chr2 | 16683068 | 16683158 | 539 | Intron (ENSMUST00000028081.7/ENSMUSG00 | 2 | 16638696 | 16751690 | 112995 | 1 | ENSMUST00 | 44373 | Plxdc2 | ENSMUSG0000002 | false | . | . | . | . | . | . | . | . | . | . | . | . |
| chr7 | 16618636 | 16618859 | 538 | Promoter (<=1kb) | 7 | 16618627 | 16619832 | 1206 | 2 | ENSMUST00 | 973 | RP24-490A2 | ENSMUSG0000010 | false | . | . | . | . | . | . | chr7 | 16615607 | 16621870 | 5.3619 | 4.10589 | 3.33445 |
| chr15 | 79670718 | 79670915 | 537 | Promoter (<=1kb) | 15 | 79670861 | 79672862 | 2002 | 1 | ENSMUST00 | 0 | Tomm22 | ENSMUSG0000002 | false | chr15 | 79670476 | 79671513 | 41.84328 | 149.90991 | 144.6302 | . | . | . | . | . | . |
| chr11 | 100622806 | 100622968 | 536 | Promoter (<=1kb) | 11 | 100622876 | 100627607 | 4732 | 1 | ENSMUST00 | 0 | Nkiras2 | ENSMUSG0000001 | false | chr11 | 100622779 | 100623594 | 18.2599 | 77.14165 | 74.04132 | . | . | . | . | . | . |
| chr18 | 13787507 | 13787638 | 535 | Intron (ENSMUST00000025288.7/ENSMUSG00 | 18 | 13687013 | 13972733 | 285721 | 2 | ENSMUST00 | 185095 | Zfp521 | ENSMUSG0000002 | true | chr18 | 13786645 | 13787751 | 6.86732 | 15.30181 | 13.03456 | . | . | . | . | . | . |
| chr6 | 22356026 | 22356271 | 535 | Promoter (<=1kb) | 6 | 22306520 | 22356166 | 49647 | 2 | ENSMUST00 | 0 | Fam3c | ENSMUSG0000002 | false | chr6 | 22355441 | 22356405 | 28.45235 | 93.08366 | 89.67602 | chr6 | 22353838 | 22356671 | 3.43704 | 3.28121 | 1.74509 |
| chr2 | 85037052 | 85037216 | 533 | Promoter (<=1kb) | 2 | 85037234 | 85047109 | 9876 | 1 | ENSMUST00 | -18 | Ssrp1 | ENSMUSG0000002 | false | chr2 | 85036527 | 85038036 | 34.95629 | 115.81918 | 111.80935 | . | . | . | . | . | . |
| chr13 | 21362679 | 21362925 | 532 | Promoter (<=1kb) | 13 | 21362827 | 21369824 | 6998 | 1 | ENSMUST00 | 0 | Zscan12 | ENSMUSG0000003 | false | chr13 | 21362385 | 21363516 | 25.17393 | 93.69764 | 90.27328 | chr13 | 21362419 | 21364810 | 3.63587 | 3.27994 | 1.90693 |
| chr5 | 34170814 | 34171018 | 532 | Promoter (1-2kb) | 5 | 34007198 | 34169448 | 162251 | 2 | ENSMUST00 | -1367 | Poln | ENSMUSG0000004 | false | . | . | . | . | . | . | chr5 | 34168693 | 34173571 | 5.15082 | 4.13032 | 3.13292 |
| chr13 | 110160128 | 110160458 | 530 | Intron (ENSMUST00000167824.1/ENSMUSG00 | 13 | 110054187 | 110280206 | 226020 | 2 | ENSMUST00 | 119748 | Rab3c | ENSMUSG0000002 | false | . | . | . | . | . | . | . | . | . | . | . | . |
| chr18 | 23989467 | 23989713 | 528 | Promoter (<=1kb) | 18 | 23989632 | 24005376 | 15745 | 1 | ENSMUST00 | 0 | Zfp35 | ENSMUSG0000006 | false | chr18 | 23989012 | 23990303 | 21.04643 | 56.48158 | 53.68221 | chr18 | 23985209 | 23991248 | 3.708 | 3.41132 | 1.98244 |
| chr2 | 129625322 | 129625446 | 528 | Intron (ENSMUST00000161620.2/ENSMUSG00 | 2 | 129620685 | 129628044 | 7360 | 1 | ENSMUST00 | 4638 | Sirpa | ENSMUSG0000003 | true | . | . | . | . | . | . | chr2 | 129621436 | 129635375 | 5.74226 | 4.37432 | 3.64032 |
| chr11 | 116490285 | 116490725 | 527 | Promoter (<=1kb) | 11 | 116470845 | 116490314 | 19470 | 2 | ENSMUST00 | 0 | Prpsap1 | ENSMUSG0000001 | false | chr11 | 116489203 | 116491230 | 35.98183 | 119.73972 | 115.60696 | . | . | . | . | . | . |
| chr17 | 35841193 | 35841427 | 526 | Promoter (<=1kb) | 17 | 35841515 | 35859670 | 18156 | 1 | ENSMUST00 | -88 | Mdc1 | ENSMUSG0000006 | false | chr17 | 35840826 | 35841315 | 12.99241 | 30.10301 | 27.6364 | . | . | . | . | . | . |
| chr6 | 106800069 | 106800186 | 525 | Promoter (<=1kb) | 6 | 106780201 | 106800074 | 19874 | 2 | ENSMUST00 | 0 | Crbn | ENSMUSG0000000 | false | chr6 | 106799187 | 106800432 | 34.3275 | 112.61823 | 108.70669 | chr6 | 106798218 | 106800458 | 4.81515 | 3.86877 | 2.89302 |
| chr18 | 35815583 | 35815702 | 523 | Distal Intergenic | 18 | 35829818 | 35861687 | 31870 | 1 | ENSMUST00 | -14116 | Cxxc5 | ENSMUSG0000004 | false | . | . | . | . | . | . | chr18 | 35812838 | 35818671 | 7.77207 | 5.31683 | 5.31473 |
| chr13 | 21453687 | 21453841 | 522 | Promoter (<=1kb) | 13 | 21442175 | 21453727 | 11553 | 2 | ENSMUST00 | 0 | Zscan26 | ENSMUSG0000002 | false | chr13 | 21452567 | 21453886 | 16.30988 | 55.04095 | 52.25988 | chr13 | 21451174 | 21454345 | 4.35522 | 3.70388 | 2.51477 |
| chr1 | 13589819 | 13590108 | 521 | Promoter (<=1kb) | 1 | 13564698 | 13589910 | 25213 | 2 | ENSMUST00 | 0 | Tram1 | ENSMUSG0000002 | false | chr1 | 13588480 | 13590440 | 31.85236 | 101.09525 | 97.496 | . | . | . | . | . | . |
| chr14 | 55722194 | 55722352 | 520 | Promoter (<=1kb) | 14 | 55715873 | 55722198 | 6326 | 2 | ENSMUST00 | 0 | Rabggta | ENSMUSG0000004 | false | chr14 | 55721408 | 55722416 | 37.48268 | 121.30223 | 117.11643 | . | . | . | . | . | . |
| chr15 | 10485877 | 10486221 | 518 | Promoter (<=1kb) | 15 | 10474779 | 10485947 | 11169 | 2 | ENSMUST00 | 0 | Brix1 | ENSMUSG0000002 | false | chr15 | 10485017 | 10486692 | 24.86315 | 92.1349 | 88.74321 | . | . | . | . | . | . |
| chr9 | 32657244 | 32657445 | 517 | Intron (ENSMUST00000183879.1/ENSMUSG00 | 9 | 32636239 | 32651476 | 15238 | 1 | ENSMUST00 | 21006 | Ets1 | ENSMUSG0000003 | true | . | . | . | . | . | . | chr9 | 32656242 | 32658110 | 6.87778 | 4.79195 | 4.58763 |
| chr8 | 107056689 | 107056853 | 516 | Promoter (<=1kb) | 8 | 107046289 | 107056689 | 10401 | 2 | ENSMUST00 | -1 | Cog8 | ENSMUSG0000003 | false | chr8 | 107055713 | 107057716 | 40.65603 | 138.83176 | 133.99156 | . | . | . | . | . | . |
| chr8 | 83566615 | 83566905 | 515 | Promoter (<=1kb) | 8 | 83566671 | 83571626 | 4956 | 1 | ENSMUST00 | 0 | Ndufb7 | ENSMUSG0000003 | false | chr8 | 83566640 | 83567815 | 39.54136 | 129.19044 | 124.73553 | . | . | . | . | . | . |
| chr3 | 125228376 | 125228470 | 513 | Distal Intergenic | 3 | 125404094 | 125686644 | 282551 | 1 | ENSMUST00 | -175624 | Ndst4 | ENSMUSG0000002 | false | chr3 | 125228295 | 125228542 | 6.00779 | 10.46335 | 8.27668 | . | . | . | . | . | . |
| chrX | 7721836 | 7721987 | 512 | Promoter (<=1kb) | 20 | 7721973 | 7726292 | 4320 | 1 | ENSMUST00 | 0 | Wdr45 | ENSMUSG0000003 | false | chrX | 7721487 | 7722729 | 12.8669 | 28.00426 | 25.56298 | . | . | . | . | . | . |
| chr8 | 88362120 | 88362280 | 509 | Promoter (<=1kb) | 8 | 88331059 | 88362194 | 31136 | 2 | ENSMUST00 | 0 | Brd7 | ENSMUSG0000003 | false | chr8 | 88360901 | 88363027 | 43.4858 | 141.53311 | 136.59532 | . | . | . | . | . | . |
| chr11 | 59506166 | 59506345 | 508 | Promoter (<=1kb) | 11 | 59499544 | 59506220 | 6677 | 2 | ENSMUST00 | 0 | Zkscan17 | ENSMUSG0000002 | false | chr11 | 59505222 | 59507156 | 33.04283 | 115.53907 | 111.53828 | . | . | . | . | . | . |
| chr4 | 133369566 | 133369851 | 507 | Promoter (<=1kb) | 4 | 133369706 | 133423702 | 53997 | 1 | ENSMUST00 | 0 | Slc9a1 | ENSMUSG0000002 | false | chr4 | 133369618 | 133371516 | 31.91637 | 110.38313 | 106.53261 | chr4 | 133369126 | 133372459 | 4.80288 | 3.86439 | 2.88403 |
| chr12 | 69361226 | 69361491 | 505 | Promoter (<=1kb) | 12 | 69361182 | 69361481 | 300 | 2 | ENSMUST00 | 0 | Gm26191 | ENSMUSG0000009 | false | chr12 | 69361102 | 69361383 | 13.18045 | 29.78312 | 27.32097 | . | . | . | . | . | . |
| chr17 | 46782629 | 46782849 | 504 | Promoter (<=1kb) | 17 | 46773907 | 46782656 | 8750 | 2 | ENSMUST00 | 0 | Rpl7l1 | ENSMUSG0000006 | false | chr17 | 46781856 | 46783237 | 23.29623 | 62.85914 | 59.97368 | . | . | . | . | . | . |
| chr17 | 28518117 | 28518304 | 503 | Promoter (<=1kb) | 17 | 28399105 | 28517524 | 118420 | 2 | ENSMUST00 | -594 | Fkbp5 | ENSMUSG0000002 | false | chr17 | 28517555 | 28518704 | 14.29631 | 42.77194 | 40.14703 | chr17 | 28514373 | 28519487 | 5.73469 | 4.2575 | 3.64316 |
| chr11 | 23067735 | 23067981 | 501 | Promoter (<=1kb) | 11 | 23067998 | 23075574 | 7577 | 1 | ENSMUST00 | -17 | Gm12059 | ENSMUSG0000008 | false | chr11 | 23067253 | 23068481 | 27.34985 | 78.73917 | 75.61149 | chr11 | 23067141 | 23068590 | 4.89236 | 3.95874 | 2.96174 |
| chr18 | 34776841 | 34776979 | 500 | Promoter (<=1kb) | 18 | 34777008 | 34839370 | 62363 | 1 | ENSMUST00 | -29 | Kdm3b | ENSMUSG0000003 | false | chr18 | 34776612 | 34778537 | 35.62435 | 116.99458 | 112.94973 | . | . | . | . | . | . |
| chr8 | 87933878 | 87934027 | 499 | Intron (ENSMUST00000109655.3/ENSMUSG00 | 8 | 87783316 | 87944052 | 160737 | 2 | ENSMUST00 | 10025 | Zfp423 | ENSMUSG0000004 | true | . | . | . | . | . | . | chr8 | 87909819 | 87941142 | 6.06882 | 4.48607 | 3.90276 |
| chr4 | 34687161 | 34687402 | 498 | Promoter (<=1kb) | 4 | 34674979 | 34687255 | 12277 | 2 | ENSMUST00 | 0 | Slc35a1 | ENSMUSG0000002 | false | chr4 | 34686600 | 34687906 | 28.9509 | 90.07051 | 86.72839 | . | . | . | . | . | . |
| chr19 | 47280914 | 47281270 | 497 | Intron (ENSMUST00000081619.3/ENSMUSG00 | 19 | 47273519 | 47273589 | 71 | 2 | ENSMUST00 | -7326 | Mir6995 | ENSMUSG0000009 | false | . | . | . | . | . | . | chr19 | 47274861 | 47283196 | 8.38768 | 5.49381 | 5.82214 |
| chr18 | 53793661 | 53793792 | 495 | Distal Intergenic | 18 | 53681724 | 53744547 | 62824 | 2 | ENSMUST00 | -49115 | Cep120 | ENSMUSG0000004 | false | . | . | . | . | . | . | chr18 | 53792269 | 53794132 | 4.9428 | 4.11433 | 2.94935 |
| chr11 | 96065249 | 96065582 | 494 | Promoter (<=1kb) | 11 | 96047432 | 96065388 | 17957 | 2 | ENSMUST00 | 0 | Ube2z | ENSMUSG0000001 | false | chr11 | 96064220 | 96065900 | 22.47347 | 97.37944 | 93.87287 | . | . | . | . | . | . |
| chr6 | 114883288 | 114883427 | 493 | Intron (ENSMUST00000032459.8/ENSMUSG00 | 6 | 114860628 | 114875107 | 14480 | 2 | ENSMUST00 | -8182 | Vgll4 | ENSMUSG0000003 | true | . | . | . | . | . | . | chr6 | 114853844 | 114919520 | 8.05355 | 5.36011 | 5.51688 |
| chr11 | 87422928 | 87423122 | 492 | Promoter (<=1kb) | 11 | 87422867 | 87423030 | 164 | 1 | ENSMUST00 | 62 | Rnu1a1 | ENSMUSG0000009 | false | . | . | . | . | . | . | . | . | . | . | . | . |
| chr15 | 101283827 | 101284114 | 489 | Promoter (<=1kb) | 15 | 101284301 | 101290934 | 6634 | 1 | ENSMUST00 | -187 | Atg101 | ENSMUSG0000003 | false | chr15 | 101283352 | 101285088 | 25.48472 | 95.2657 | 91.8051 | . | . | . | . | . | . |
| chr11 | 83298916 | 83299018 | 486 | Promoter (<=1kb) | 11 | 83294642 | 83298977 | 4336 | 2 | ENSMUST00 | 0 | Pex12 | ENSMUSG0000001 | false | chr11 | 83298225 | 83299602 | 30.87849 | 93.42764 | 90.01563 | chr11 | 83297206 | 83299946 | 4.20388 | 3.69202 | 2.34489 |
| chr15 | 27525344 | 27525649 | 483 | Intron (ENSMUST00000022875.6/ENSMUSG00 | 15 | 27571449 | 27593163 | 21715 | 1 | ENSMUST00 | -45800 | Ank | ENSMUSG0000002 | false | . | . | . | . | . | . | . | . | . | . | . | . |
| chr18 | 69344284 | 69344560 | 482 | Promoter (<=1kb) | 18 | 69344492 | 69565506 | 221015 | 1 | ENSMUST00 | 0 | Tcf4 | ENSMUSG0000005 | false | chr18 | 69343931 | 69345637 | 23.09106 | 69.49222 | 66.50756 | chr18 | 69343638 | 69345358 | 3.2489 | 3.19933 | 1.55274 |
| chr9 | 88553226 | 88553309 | 478 | Promoter (<=1kb) | 9 | 88553236 | 88553525 | 290 | 1 | ENSMUST00 | 0 | Gm26461 | ENSMUSG0000009 | false | chr9 | 88553144 | 88553395 | 4.97263 | 8.68936 | 6.53912 | . | . | . | . | . | . |
| chr13 | 21506323 | 21506671 | 477 | Promoter (1-2kb) | 13 | 21504382 | 21505056 | 675 | 2 | ENSMUST00 | -1268 | Zfp389 | ENSMUSG0000008 | false | chr13 | 21505925 | 21506836 | 28.36989 | 85.2065 | 81.96005 | chr13 | 21505803 | 21506932 | 3.87999 | 3.3988 | 2.12773 |
| chr12 | 84996103 | 84996337 | 475 | Promoter (<=1kb) | 12 | 84996321 | 85070492 | 74172 | 1 | ENSMUST00 | 0 | Ylpm1 | ENSMUSG0000002 | false | chr12 | 84996250 | 84997380 | 39.12038 | 124.14072 | 119.86815 | . | . | . | . | . | . |
| chr8 | 82939866 | 82940258 | 473 | Intron (ENSMUST00000078525.5/ENSMUSG00 | 8 | 82863356 | 83091271 | 227916 | 1 | ENSMUST00 | 76511 | Rnf150 | ENSMUSG0000004 | true | . | . | . | . | . | . | chr8 | 82936463 | 82941511 | 3.87884 | 3.42569 | 2.13119 |
| chr16 | 26049406 | 26049978 | 472 | Intron (ENSMUST00000039990.5/ENSMUSG00 | 16 | 26031810 | 26040870 | 9061 | 2 | ENSMUST00 | -8537 | AU015336 | ENSMUSG0000009 | false | . | . | . | . | . | . | chr16 | 26035252 | 26056185 | 6.03158 | 4.46161 | 3.87334 |
| chr14 | 60264972 | 60265292 | 468 | Promoter (<=1kb) | 14 | 60265205 | 60302370 | 37166 | 1 | ENSMUST00 | 0 | Mtmr6 | ENSMUSG0000002 | false | chr14 | 60265179 | 60266233 | 37.7199 | 126.00829 | 121.65485 | chr14 | 60264849 | 60267973 | 4.58611 | 3.73389 | 2.70745 |
| chr10 | 119065504 | 119065739 | 464 | Distal Intergenic | 10 | 119092905 | 119103107 | 10203 | 1 | ENSMUST00 | -27166 | RP23-360L4. | ENSMUSG0000010 | false | . | . | . | . | . | . | chr10 | 119061401 | 119066309 | 4.92666 | 3.97999 | 2.96336 |
| chr3 | 75930732 | 75931033 | 462 | Intron (ENSMUST00000117242.2/ENSMUSG00 | 3 | 75909856 | 75956650 | 46795 | 2 | ENSMUST00 | 25617 | Golim4 | ENSMUSG0000003 | true | . | . | . | . | . | . | chr3 | 75908736 | 75932827 | 4.90077 | 3.96047 | 2.93621 |
| chr17 | 86618553 | 86618846 | 460 | Intron (ENSMUST00000097275.3/ENSMUSG00 | 17 | 86703298 | 86703407 | 110 | 2 | ENSMUST00 | 84561 | Gm22235 | ENSMUSG0000008 | false | . | . | . | . | . | . | chr17 | 86616853 | 86621360 | 5.8599 | 4.38553 | 3.73139 |
| chr14 | 69732573 | 69732865 | 458 | Promoter (<=1kb) | 14 | 69732784 | 69745869 | 13086 | 1 | ENSMUST00 | 0 | 4930480K23 | ENSMUSG0000009 | false | chr14 | 69731620 | 69733869 | 41.18432 | 138.64781 | 133.82259 | . | . | . | . | . | . |
| chr19 | 10952070 | 10952268 | 456 | Promoter (2-3kb) | 19 | 10941481 | 10949266 | 7786 | 2 | ENSMUST00 | -2805 | Ccdc86 | ENSMUSG0000002 | false | . | . | . | . | . | . | chr19 | 10946315 | 10954190 | 5.98357 | 4.50647 | 3.81648 |
| chr11 | 31828057 | 31828264 | 452 | Distal Intergenic | 11 | 31800688 | 31824524 | 23837 | 2 | ENSMUST00 | -3534 | D630024D03 | ENSMUSG0000008 | false | . | . | . | . | . | . | chr11 | 31821641 | 31832598 | 5.80216 | 4.3489 | 3.6737 |
| chr7 | 127615224 | 127615428 | 450 | Promoter (<=1kb) | 7 | 127607031 | 127615272 | 8242 | 2 | ENSMUST00 | 0 | Zfp629 | ENSMUSG0000004 | false | chr7 | 127614375 | 127616118 | 24.60788 | 97.30984 | 93.80507 | . | . | . | . | . | . |
| chr5 | 92834538 | 92834783 | 448 | Intron (ENSMUST00000113055.3/ENSMUSG00 | 5 | 92804220 | 92809488 | 5269 | 2 | ENSMUST00 | -25051 | 4932430I15R | ENSMUSG0000009 | false | . | . | . | . | . | . | chr5 | 92813516 | 92845987 | 5.7053 | 4.3919 | 3.59815 |
| chr5 | 136888530 | 136888697 | 445 | Distal Intergenic | 5 | 136757489 | 136883209 | 125721 | 2 | ENSMUST00 | -5322 | Col26a1 | ENSMUSG0000000 | false | . | . | . | . | . | . | chr5 | 136885321 | 136889075 | 4.822 | 3.84455 | 2.90361 |
| chr9 | 120577249 | 120577403 | 443 | Promoter (<=1kb) | 9 | 120577346 | 120578072 | 727 | 1 | ENSMUST00 | 0 | 5830454E08 | ENSMUSG0000005 | false | chr9 | 120576838 | 120577979 | 22.37683 | 79.83291 | 76.68538 | chr9 | 120576533 | 120578356 | 5.20142 | 4.23175 | 3.16252 |
| chr4 | 151933650 | 151933767 | 439 | Promoter (<=1kb) | 4 | 151933691 | 151982137 | 48447 | 1 | ENSMUST00 | 0 | Dnajc11 | ENSMUSG0000003 | false | chr4 | 151933681 | 151934863 | 34.92026 | 124.22972 | 119.94103 | . | . | . | . | . | . |
| chr15 | 38470642 | 38470826 | 436 | Distal Intergenic | 15 | 38490473 | 38490547 | 75 | 2 | ENSMUST00 | 19721 | Mir6951 | ENSMUSG0000009 | false | chr15 | 38470059 | 38471389 | 13.87847 | 36.33116 | 33.78668 | chr15 | 38468227 | 38474123 | 4.55881 | 3.82285 | 2.65383 |
| chr1 | 23264325 | 23264602 | 434 | Distal Intergenic | 1 | 23272264 | 23272345 | 82 | 1 | ENSMUST00 | -7662 | Mir30a | ENSMUSG0000006 | false | . | . | . | . | . | . | chr1 | 23258890 | 23271405 | 4.63528 | 3.84316 | 2.71286 |
| chr3 | 51916199 | 51916588 | 427 | Intron (ENSMUST00000121440.2/ENSMUSG00 | 3 | 51828912 | 51829028 | 117 | 1 | ENSMUST00 | 87288 | n-R5s196 | ENSMUSG0000006 | false | . | . | . | . | . | . | chr3 | 51904229 | 51919287 | 6.09838 | 4.57081 | 3.90639 |
| chr3 | 89859020 | 89859208 | 412 | Distal Intergenic | 3 | 89831370 | 89858834 | 27465 | 1 | ENSMUST00 | 27651 | She | ENSMUSG0000004 | false | . | . | . | . | . | . | chr3 | 89858315 | 89886122 | 6.15651 | 4.57192 | 3.96514 |
| chr14 | 30951184 | 30951474 | 408 | Promoter (<=1kb) | 14 | 30951486 | 30987439 | 35954 | 1 | ENSMUST00 | -12 | Nek4 | ENSMUSG0000002 | false | chr14 | 30950835 | 30952082 | 29.63113 | 94.2009 | 90.76292 | chr14 | 30950690 | 30952434 | 5.14147 | 4.20193 | 3.1205 |
| chr15 | 102470199 | 102470541 | 403 | Promoter (<=1kb) | 15 | 102470632 | 102500059 | 29428 | 1 | ENSMUST00 | -91 | Pcbp2 | ENSMUSG0000005 | false | chr15 | 102470489 | 102471870 | 22.78592 | 76.10319 | 73.01487 | chr15 | 102469905 | 102470225 | 4.3255 | 3.84212 | 2.44764 |
| chr9 | 44772783 | 44773036 | 391 | Promoter (<=1kb) | 9 | 44772935 | 44792714 | 19780 | 1 | ENSMUST00 | 0 | Ift46 | ENSMUSG0000000 | false | chr9 | 44772876 | 44773729 | 25.98482 | 77.30067 | 74.19341 | chr9 | 44772488 | 44774750 | 6.44521 | 4.83499 | 4.18512 |
| chr10 | 81402067 | 81402224 | 387 | Promoter (1-2kb) | 10 | 81399196 | 81400554 | 1359 | 2 | ENSMUST00 | -1514 | Nfic | ENSMUSG0000005 | false | chr10 | 81401906 | 81402504 | 5.97392 | 9.35129 | 7.18948 | chr10 | 81392649 | 81426205 | 8.72751 | 5.63067 | 6.0679 |
| chr1 | 133610190 | 133610410 | 114 | Promoter (<=1kb) | 1 | 133603871 | 133610291 | 6421 | 2 | ENSMUST00 | 0 | Snrpe | ENSMUSG0000009 | false | chr1 | 133609373 | 133610411 | 25.61729 | 89.79346 | 86.4557 | . | . | . | . | . | . |
| chr1 | 44584883 | 44585045 | 114 | Intron (ENSMUST00000074525.4/ENSMUSG00 | 1 | 44552597 | 44708632 | 156036 | 1 | ENSMUST00 | 32287 | Gulp1 | ENSMUSG0000005 | true | . | . | . | . | . | . | chr1 | 44583660 | 44586527 | 5.26874 | 4.13444 | 3.26857 |
| chr10 | 84389019 | 84389164 | 114 | Intron (ENSMUST00000020220.9/ENSMUSG00 | 10 | 84375296 | 84392388 | 17093 | 2 | ENSMUST00 | 3224 | Nuak1 | ENSMUSG0000002 | false | . | . | . | . | . | . | chr10 | 84386221 | 84399643 | 5.21453 | 4.13999 | 3.18346 |
| chr11 | 116274077 | 116274196 | 114 | Promoter (<=1kb) | 11 | 116245166 | 116274135 | 28970 | 2 | ENSMUST00 | 0 | Srp68 | ENSMUSG0000002 | false | chr11 | 116273124 | 116274837 | 26.68262 | 76.66692 | 73.57355 | . | . | . | . | . | . |
| chr11 | 76945517 | 76945801 | 114 | Promoter (<=1kb) | 11 | 76945729 | 76946758 | 1030 | 1 | ENSMUST00 | 0 | Blmh | ENSMUSG0000002 | false | chr11 | 76945142 | 76946667 | 29.48955 | 94.93548 | 91.48817 | chr11 | 76944160 | 76947151 | 4.10471 | 3.58068 | 2.28813 |
| chr11 | 101732810 | 101732934 | 114 | Promoter (<=1kb) | 11 | 101732919 | 101735696 | 2778 | 1 | ENSMUST00 | 0 | Dhx8 | ENSMUSG0000003 | false | chr11 | 101732893 | 101733889 | 22.10065 | 59.96099 | 57.11441 | chr11 | 101727843 | 101734081 | 5.39683 | 4.28202 | 3.36272 |
| chr11 | 117076637 | 117076801 | 114 | Promoter (<=1kb) | 11 | 117076790 | 117077457 | 668 | 1 | ENSMUST00 | 0 | 2810008D09 | ENSMUSG0000008 | false | chr11 | 117076313 | 117077570 | 39.81545 | 135.14195 | 130.45284 | chr11 | 117076150 | 117083562 | 5.19932 | 3.9166 | 3.20454 |
| chr12 | 85608219 | 85608403 | 114 | Exon (ENSMUST00000177587.2/ENSMUSG000 | 12 | 85600018 | 85639878 | 39861 | 1 | ENSMUST00 | 8202 | Jdp2 | ENSMUSG0000003 | true | . | . | . | . | . | . | chr12 | 85599922 | 85615323 | 5.04005 | 3.93431 | 3.0761 |
| chr16 | 87454878 | 87455063 | 114 | Promoter (<=1kb) | 16 | 87454979 | 87483310 | 28332 | 1 | ENSMUST00 | 0 | Usp16 | ENSMUSG0000002 | false | chr16 | 87454436 | 87456031 | 38.65522 | 135.28891 | 130.59163 | . | . | . | . | . | . |
| chr16 | 52013175 | 52013380 | 114 | Distal Intergenic | 16 | 52031549 | 52208047 | 176499 | 1 | ENSMUST00 | -18169 | Cblb | ENSMUSG0000002 | true | . | . | . | . | . | . | chr16 | 52009607 | 52016621 | 4.28782 | 3.76463 | 2.41076 |
| chr17 | 5849457 | 5849782 | 114 | Intron (ENSMUST00000002436.9/ENSMUSG00 | 17 | 5841328 | 5931033 | 89706 | 1 | ENSMUST00 | 8130 | Snx9 | ENSMUSG0000002 | false | chr17 | 5849210 | 5849793 | 9.38717 | 20.36066 | 18.02005 | chr17 | 5842208 | 5850409 | 5.53242 | 4.3591 | 3.4424 |
| chr17 | 45563839 | 45563943 | 114 | Promoter (<=1kb) | 17 | 45563964 | 45567669 | 3706 | 1 | ENSMUST00 | -21 | Slc35b2 | ENSMUSG0000003 | false | chr17 | 45563840 | 45564964 | 39.39634 | 133.30592 | 128.68834 | chr17 | 45563577 | 45571950 | 4.52176 | 3.76389 | 2.64295 |
| chr17 | 45931243 | 45931494 | 114 | Distal Intergenic | 17 | 46018712 | 46021372 | 2661 | 2 | ENSMUST00 | 89878 | Vegfa | ENSMUSG0000002 | true | . | . | . | . | . | . | chr17 | 45928605 | 45934413 | 4.63767 | 3.72884 | 2.75421 |
| chr17 | 56613321 | 56613451 | 114 | Promoter (<=1kb) | 17 | 56613395 | 56614243 | 849 | 1 | ENSMUST00 | 0 | Rpl36 | ENSMUSG0000005 | false | chr17 | 56613301 | 56614568 | 20.88808 | 77.73511 | 74.61916 | chr17 | 56608775 | 56618499 | 4.98769 | 4.03202 | 3.02403 |
| chr17 | 35331725 | 35331848 | 114 | Distal Intergenic | 17 | 35335909 | 35336292 | 384 | 1 | ENSMUST00 | -4061 | Gm8741 | ENSMUSG0000009 | false | . | . | . | . | . | . | . | . | . | . | . | . |
| chr17 | 43554623 | 43554811 | 114 | Distal Intergenic | 17 | 43534174 | 43543639 | 9466 | 2 | ENSMUST00 | -10985 | Ankrd66 | ENSMUSG0000009 | false | . | . | . | . | . | . | chr17 | 43554410 | 43555332 | 4.40255 | 3.73011 | 2.56571 |
| chr18 | 38277860 | 38278048 | 114 | Exon (ENSMUST00000025311.5/ENSMUSG000 | 18 | 38267092 | 38284401 | 17310 | 2 | ENSMUST00 | 6353 | Pcdh12 | ENSMUSG0000002 | false | . | . | . | . | . | . | chr18 | 38274249 | 38279359 | 5.63879 | 4.38394 | 3.54879 |
| chr18 | 5441180 | 5441328 | 114 | Distal Intergenic | 18 | 5518415 | 5521109 | 2695 | 2 | ENSMUST00 | 79781 | Gm10125 | ENSMUSG0000006 | false | . | . | . | . | . | . | . | . | . | . | . | . |
| chr18 | 11973814 | 11974028 | 114 | Distal Intergenic | 18 | 11840361 | 11840438 | 78 | 2 | ENSMUST00 | -133377 | Mir1901 | ENSMUSG0000008 | false | . | . | . | . | . | . | chr18 | 11971839 | 11979509 | 5.50442 | 4.29031 | 3.42052 |
| chr19 | 10605523 | 10605623 | 114 | Promoter (<=1kb) | 19 | 10605625 | 10629813 | 24189 | 1 | ENSMUST00 | -2 | Ddb1 | ENSMUSG0000002 | false | chr19 | 10603045 | 10606602 | 41.22328 | 133.19821 | 128.59248 | . | . | . | . | . | . |
| chr19 | 31531514 | 31531642 | 114 | Intron (ENSMUST00000065067.7/ENSMUSG00 | 19 | 30568547 | 31664370 | 1095824 | 2 | ENSMUST00 | 132728 | Prkg1 | ENSMUSG0000005 | true | . | . | . | . | . | . | chr19 | 31525771 | 31533375 | 4.9636 | 4.09548 | 2.97296 |
| chr2 | 128128065 | 128128159 | 114 | Promoter (<=1kb) | 2 | 128128131 | 128128737 | 607 | 1 | ENSMUST00 | 0 | Bcl2l11 | ENSMUSG0000002 | false | chr2 | 128125321 | 128128418 | 32.64849 | 123.19247 | 118.94151 | . | . | . | . | . | . |
| chr2 | 69080874 | 69081057 | 114 | Intron (ENSMUST00000028426.3/ENSMUSG00 | 2 | 69135800 | 69189330 | 53531 | 1 | ENSMUST00 | -54743 | Nostrin | ENSMUSG0000003 | false | . | . | . | . | . | . | chr2 | 69076857 | 69090376 | 5.44132 | 4.24372 | 3.37539 |
| chr2 | 33166087 | 33166290 | 114 | Promoter (1-2kb) | 2 | 33137636 | 33168222 | 30587 | 2 | ENSMUST00 | 1932 | Ralgps1 | ENSMUSG0000003 | false | . | . | . | . | . | . | chr2 | 33154669 | 33171007 | 7.40171 | 5.13174 | 4.97591 |
| chr2 | 167421572 | 167421945 | 114 | Promoter (<=1kb) | 2 | 167421712 | 167477000 | 55289 | 1 | ENSMUST00 | 0 | Slc9a8 | ENSMUSG0000003 | false | chr2 | 167420983 | 167422465 | 24.86315 | 92.1349 | 88.74321 | chr2 | 167420660 | 167422853 | 4.22416 | 3.64759 | 2.38613 |
| chr2 | 29696451 | 29696681 | 114 | Intron (ENSMUST00000095087.5/ENSMUSG00 | 2 | 29702482 | 29740363 | 37882 | 1 | ENSMUST00 | -5801 | Rapgef1 | ENSMUSG0000003 | false | . | . | . | . | . | . | chr2 | 29690471 | 29698739 | 5.92345 | 4.4721 | 3.78221 |
| chr2 | 164460907 | 164461079 | 114 | Promoter (<=1kb) | 2 | 164460971 | 164465508 | 4538 | 1 | ENSMUST00 | 0 | Sys1 | ENSMUSG0000004 | false | chr2 | 164460919 | 164462087 | 31.43241 | 100.35837 | 96.77719 | . | . | . | . | . | . |
| chr2 | 159270560 | 159270753 | 114 | Distal Intergenic | 2 | 159139361 | 159139492 | 132 | 1 | ENSMUST00 | 131200 | Gm25090 | ENSMUSG0000007 | false | . | . | . | . | . | . | chr2 | 159269480 | 159284862 | 4.11414 | 3.61003 | 2.27932 |
| chr4 | 43031349 | 43031515 | 114 | Promoter (<=1kb) | 4 | 43028229 | 43031456 | 3228 | 2 | ENSMUST00 | 0 | Stoml2 | ENSMUSG0000002 | false | chr4 | 43030580 | 43031848 | 29.85632 | 88.73936 | 85.41974 | chr4 | 43030228 | 43032060 | 5.27107 | 4.15375 | 3.2536 |
| chr4 | 124714649 | 124714867 | 114 | Promoter (<=1kb) | 4 | 124714776 | 124732460 | 17685 | 1 | ENSMUST00 | 0 | Sf3a3 | ENSMUSG0000002 | false | chr4 | 124714395 | 124715568 | 26.69297 | 77.60641 | 74.495 | chr4 | 124714358 | 124715736 | 4.08613 | 3.28607 | 2.28504 |
| chr4 | 141078342 | 141078459 | 114 | Promoter (<=1kb) | 4 | 141066512 | 141078357 | 11846 | 2 | ENSMUST00 | 0 | Necap2 | ENSMUSG0000002 | false | chr4 | 141077406 | 141078403 | 36.66269 | 126.29414 | 121.93508 | chr4 | 141073554 | 141078823 | 5.95773 | 4.50816 | 3.80002 |
| chr4 | 106650851 | 106650998 | 114 | Promoter (<=1kb) | 4 | 106651069 | 106653539 | 2471 | 1 | ENSMUST00 | -71 | Pars2 | ENSMUSG0000004 | false | . | . | . | . | . | . | chr4 | 106649637 | 106652500 | 5.3602 | 4.00044 | 3.33642 |
| chr5 | 138155648 | 138155866 | 114 | Promoter (<=1kb) | 5 | 138139702 | 138155744 | 16043 | 2 | ENSMUST00 | 0 | Zfp113 | ENSMUSG0000003 | false | chr5 | 138154912 | 138155829 | 27.03504 | 88.5911 | 85.27629 | chr5 | 138154312 | 138156292 | 4.80915 | 3.98217 | 2.86575 |
| chr5 | 119325825 | 119326141 | 114 | Distal Intergenic | 5 | 119297245 | 119297361 | 117 | 1 | ENSMUST00 | 28581 | n-R5s175 | ENSMUSG0000008 | false | . | . | . | . | . | . | . | . | . | . | . | . |
| chr6 | 47835601 | 47835731 | 114 | Promoter (<=1kb) | 6 | 47835661 | 47843647 | 7987 | 1 | ENSMUST00 | 0 | Zfp398 | ENSMUSG0000006 | false | chr6 | 47835563 | 47836836 | 41.84113 | 139.79433 | 134.92708 | chr6 | 47835331 | 47838254 | 3.87408 | 3.48275 | 2.12306 |
| chr6 | 70386500 | 70386896 | 114 | Promoter (<=1kb) | 6 | 70386672 | 70387238 | 567 | 2 | ENSMUST00 | 342 | Igkv8-16 | ENSMUSG0000007 | false | . | . | . | . | . | . | chr6 | 70385956 | 70388311 | 6.31575 | 4.65551 | 4.07275 |
| chr7 | 130547255 | 130547442 | 114 | Promoter (<=1kb) | 7 | 130532523 | 130547381 | 14859 | 2 | ENSMUST00 | 0 | Nsmce4a | ENSMUSG0000004 | false | chr7 | 130546352 | 130547429 | 28.91247 | 96.86193 | 93.36482 | . | . | . | . | . | . |
| chr8 | 126853071 | 126853291 | 114 | Distal Intergenic | 8 | 126944847 | 126944980 | 134 | 2 | ENSMUST00 | 91689 | Gm26397 | ENSMUSG0000006 | false | . | . | . | . | . | . | chr8 | 126847776 | 126853210 | 4.5877 | 3.85968 | 2.679 |
| chr8 | 121241958 | 121242162 | 114 | Intron (ENSMUST00000127664.1/ENSMUSG00 | 8 | 121198714 | 121201835 | 3122 | 2 | ENSMUST00 | -40124 | 5033426O07 | ENSMUSG0000009 | false | . | . | . | . | . | . | chr8 | 121236124 | 121247254 | 7.60766 | 5.2125 | 5.14118 |
| chr9 | 14641847 | 14642063 | 114 | Distal Intergenic | 9 | 14593444 | 14615483 | 22040 | 2 | ENSMUST00 | -26365 | Amotl1 | ENSMUSG0000001 | false | . | . | . | . | . | . | . | . | . | . | . | . |
| chr9 | 71674521 | 71674693 | 114 | Intron (ENSMUST00000072899.3/ENSMUSG00 | 9 | 71632633 | 71648350 | 15718 | 2 | ENSMUST00 | -26172 | Cgnl1 | ENSMUSG0000003 | false | . | . | . | . | . | . | chr9 | 71672716 | 71681830 | 5.44209 | 4.19703 | 3.39991 |
| chr9 | 57765142 | 57765292 | 114 | Promoter (<=1kb) | 9 | 57750711 | 57765860 | 15150 | 2 | ENSMUST00 | 568 | Clk3 | ENSMUSG0000003 | false | chr9 | 57763942 | 57766341 | 37.17318 | 134.81126 | 130.13722 | . | . | . | . | . | . |
| chr9 | 65032281 | 65032426 | 114 | Promoter (<=1kb) | 9 | 65032458 | 65082650 | 50193 | 1 | ENSMUST00 | -32 | Dpp8 | ENSMUSG0000003 | false | chr9 | 65032063 | 65033438 | 35.98183 | 119.73972 | 115.60696 | chr9 | 65031928 | 65034604 | 4.72792 | 3.93879 | 2.82826 |
| chr9 | 122865929 | 122866047 | 114 | Distal Intergenic | 9 | 122851914 | 122862128 | 10215 | 2 | ENSMUST00 | -3802 | Zfp445 | ENSMUSG0000004 | false | chr9 | 122864928 | 122866263 | 22.5292 | 69.37365 | 66.39268 | . | . | . | . | . | . |
| chr9 | 32646915 | 32647076 | 114 | Intron (ENSMUST00000183879.1/ENSMUSG00 | 9 | 32636239 | 32651476 | 15238 | 1 | ENSMUST00 | 10677 | Ets1 | ENSMUSG0000003 | true | . | . | . | . | . | . | chr9 | 32645640 | 32647917 | 6.06489 | 4.58743 | 3.88733 |

| Ontology | # Term Name | Binom Ran | Binom Raw P-Value | Binom FDR Q-Val | Binom Fold | Binom Obs | Binom Reg | Hyper Ran | Hyper FDR | Hyper Fold | Hyper Obs | Hyper Tota | Hyper Gene Set Coverage |
|---|---|---|---|---|---|---|---|---|---|---|---|---|---|
| GO Biological processes | **cell-matrix adhesion** | 18 | 5.85829E-06 | 0.00426028 | 4.381677 | 14 | 4.03% | 36 | 0.000564 | 5.116914 | 13 | 107 | 2.56% |

The test set of 347 genomic regions picked 508 (2%) of all 21,395 genes.
*GO Biological Process* has 13,090 terms covering 17,925 (84%) of all 21,395 genes, and 1,163,819 term - gene associations.
13,090 ontology terms (100%) were tested using an annotation count range of [1, Inf].

| Ontology | # Term Name | Binom Ran | Binom Raw P-Value | Binom FDR Q-Val | Binom Fold | Binom Obs | Binom Reg | Hyper Ran | Hyper FDR | Hyper Fold | Hyper Obs | Hyper Tota | Hyper Gene Set Coverage |
|---|---|---|---|---|---|---|---|---|---|---|---|---|---|
| GO Cellular Component | **integral component of organelle membrane** | 17 | 1.02214E-05 | 0.001018535 | 4.1671 | 14 | 4.03% | 22 | 0.002786 | 3.617337 | 14 | 163 | 2.76% |
| | **intracellular ribonucleoprotein complex** | 18 | 1.4706E-05 | 0.001383993 | 2.13601 | 37 | 10.66% | 26 | 0.019372 | 1.837615 | 37 | 848 | 7.28% |
| | **ribonucleoprotein complex** | 19 | 1.50341E-05 | 0.001340411 | 2.133812 | 37 | 10.66% | 27 | 0.019514 | 1.833291 | 37 | 850 | 7.28% |
| | **intrinsic component of organelle membrane** | 20 | 1.74273E-05 | 0.001476088 | 3.968736 | 14 | 4.03% | 24 | 0.005551 | 3.369291 | 14 | 175 | 2.76% |
| | **podosome** | 24 | 0.000155077 | 0.010945816 | 7.699724 | 6 | 1.73% | 23 | 0.005694 | 8.151511 | 6 | 31 | 1.18% |

The test set of 347 genomic regions picked 508 genes (2%) of all 21,395 genes.
*GO Cellular Component has 1,* 1,694 ontology terms were tested (100%) using an annotation count range of [1, 1000].
GREAT version 4.0.4
Species assembly: mm10
Association rule: Basal+extension: 5000 bp upstream, 1000 bp downstream, 1000000 bp max extension, curated regulatory domains included

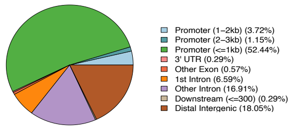

**MAFB peaks distribution**

| Ontology | # Term Name | Binom Ran | Binom Raw P-Value | Binom FDR Q-Val | Binom Fold | Binom Obs | Binom Reg | Hyper Ran | Hyper FDR | Hyper Fold | Hyper Obs | Hyper Tota | Hyper Gene Set Coverage |
|---|---|---|---|---|---|---|---|---|---|---|---|---|---|
| | **abnormal mesangial matrix morphology** | 1 | 1.30693E-06 | 0.012513855 | 6.62086 | 11 | 3.17% | 3 | 0.001076 | 8.099258 | 10 | 52 | 1.97% |
| | **dilated capillary** | 2 | 1.70015E-06 | 0.008139487 | 12.81292 | 7 | 2.02% | 100 | 0.043617 | 10.52904 | 4 | 16 | 0.79% |
| | **abnormal podocyte slit diaphragm morphology** | 4 | 3.17048E-06 | 0.007589325 | 11.64359 | 7 | 2.02% | 4 | 0.001064 | 18.04978 | 6 | 14 | 1.18% |
| | **expanded mesangial matrix** | 6 | 6.73046E-06 | 0.010740694 | 6.226018 | 10 | 2.88% | 12 | 0.00188 | 7.580906 | 9 | 50 | 1.77% |
| | **podocyte foot process effacement** | 7 | 1.17467E-05 | 0.016067821 | 5.833323 | 10 | 2.88% | 10 | 0.001307 | 8.064793 | 9 | 47 | 1.77% |
| | **abnormal renal glomerulus basement membrane morphology** | 8 | 1.46608E-05 | 0.017547133 | 5.682886 | 10 | 2.88% | 16 | 0.00322 | 6.891732 | 9 | 55 | 1.77% |
| | **abnormal podocyte slit junction morphology** | 9 | 1.54222E-05 | 0.016407475 | 9.100778 | 7 | 2.02% | 9 | 0.001211 | 15.79355 | 6 | 16 | 1.18% |
| | **abnormal podocyte foot process morphology** | 10 | 4.11219E-05 | 0.039374219 | 4.556228 | 11 | 3.17% | 13 | 0.002132 | 6.479406 | 10 | 65 | 1.97% |
| | **abnormal glomerular endothelium fenestra morphology** | 11 | 4.25316E-05 | 0.037021859 | 9.774284 | 6 | 1.73% | 14 | 0.002077 | 19.1437 | 5 | 11 | 0.98% |
| | **abnormal capillary morphology** | 12 | 4.756E-05 | 0.037948893 | 3.420542 | 15 | 4.32% | 18 | 0.0053 | 4.679571 | 12 | 108 | 2.36% |

The test set of 347 genomic regions picked 508 (2%) of all 21,395 genes.
*Mouse Phenotype has 9,575 terms covering 9,654 (45%) of all 21,395 genes, and 705,265 term - gene associations.*
9,575 ontology terms (100%) were tested using an annotation count range of [1, Inf].

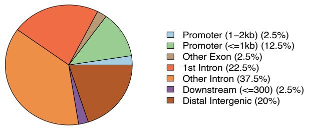

**MAFB peaks distribution (podocytes specific)**

| * podocyte foot process effacement - associated genes |
|---|
| Nphs1 (-80) |
| Vegfa (-19914) |
| Vegfa (+101000) |
| Nphs2 (-1261) |
| Ilk (-20) |
| Lmx1b (+34441) |
| Tcf21 (-48340) |
| Ehd4 (+60750) |
| Itgb1 (-140972) |

| Chrom | Start | End | IDR score | annotation | geneChr | geneStart | geneEnd | geneLength | geneStrand | transcriptId | distanceToTS | geneName | geneId | Podocyte specific | Chrom | Start | End | -LOG10(pvalue) | fold | -LOG10(q-value) | Chrom | Start | End | -LOG10(pvalue) | fold | -LOG10(q-value) |
|---|---|---|---|---|---|---|---|---|---|---|---|---|---|---|---|---|---|---|---|---|---|---|---|---|---|---|
| chr13 | 5861060 | 5861990 | 1000 | Promoter (<=1kb) | 13 | 5861489 | 5870393 | 8905 | 1 | ENSMUST00 | 0 | Klf6 | ENSMUSG0C | false | chr13 | 5860614 | 5861246 | 24.30896 | 73.44389 | 70.40032 | . | . | . | . | . | . |
| chr9 | 57520873 | 57521610 | 1000 | Promoter (<=1kb) | 9 | 57521232 | 57532424 | 11193 | 1 | ENSMUST00 | 0 | Cox5a | ENSMUSG0C | false | chr9 | 57520719 | 57522380 | 36.27325 | 129.91212 | 125.42401 | chr9 | 57520538 | 57520919 | 3.81619 | 3.5197 | 2.04033 |
| chrX | 7762631 | 7762972 | 1000 | Promoter (<=1kb) | 20 | 7762658 | 7775202 | 12545 | 1 | ENSMUST00 | 0 | Tfe3 | ENSMUSG0C | false | chrX | 7762650 | 7764496 | 22.51707 | 59.83231 | 56.98677 | . | . | . | . | . | . |
| chr11 | 108920155 | 108921351 | 1000 | Promoter (<=1kb) | 11 | 108920349 | 108950783 | 30435 | 1 | ENSMUST00 | 0 | Axin2 | ENSMUSG0C | false | chr11 | 108920064 | 108922921 | 42.74922 | 148.11885 | 142.90198 | . | . | . | . | . | . |
| chr5 | 140803995 | 140804323 | 1000 | Intron (ENSMUST00000000153.4 | 5 | 140758408 | 140830431 | 72024 | 2 | ENSMUST00 | 26108 | Gna12 | ENSMUSG0C | false | . | . | . | . | . | . | chr5 | 140787727 | 140831108 | 6.24285 | 4.56075 | 4.04432 |
| chr5 | 140827918 | 140828241 | 1000 | Promoter (2-3kb) | 5 | 140758408 | 140830431 | 72024 | 2 | ENSMUST00 | 2190 | Gna12 | ENSMUSG0C | false | . | . | . | . | . | . | chr5 | 140787727 | 140831108 | 6.24285 | 4.56075 | 4.04432 |
| chr9 | 50659260 | 50660142 | 1000 | Promoter (<=1kb) | 9 | 50634633 | 50659780 | 25148 | 2 | ENSMUST00 | 0 | Dlat | ENSMUSG0C | false | chr9 | 50658630 | 50660762 | 35.20524 | 115.20684 | 111.21615 | . | . | . | . | . | . |
| chr5 | 143015756 | 143016898 | 1000 | Promoter (<=1kb) | 7 | 143017409 | 143018903 | 1495 | 1 | ENSMUST00 | -511 | Tspan32 | ENSMUSG0C | false | . | . | . | . | . | . | chr7 | 143004528 | 143022688 | 7.71992 | 5.26439 | 5.23131 |
| chr5 | 45530709 | 45531260 | 1000 | Intron (ENSMUST00000000260.7 | 13 | 45538809 | 45542550 | 3742 | 1 | ENSMUST00 | -7549 | Gmpr | ENSMUSG0C | false | . | . | . | . | . | . | chr13 | 45522834 | 45531762 | 4.69665 | 3.835 | 2.77414 |
| chr11 | 89060434 | 89061052 | 1000 | Promoter (<=1kb) | 11 | 89037582 | 89060748 | 23167 | 2 | ENSMUST00 | 0 | Dgke | ENSMUSG0C | false | chr11 | 89059680 | 89061154 | 36.9999 | 115.1778 | 111.19392 | . | . | . | . | . | . |
| chr11 | 74831681 | 74832522 | 1000 | Promoter (<=1kb) | 11 | 74830920 | 74845725 | 14806 | 1 | ENSMUST00 | 762 | Mnt | ENSMUSG0C | false | chr11 | 74832439 | 74833147 | 8.8423 | 19.93079 | 17.59637 | . | . | . | . | . | . |
| chr11 | 74829932 | 74831447 | 1000 | Promoter (<=1kb) | 11 | 74830920 | 74845725 | 14806 | 1 | ENSMUST00 | 0 | Mnt | ENSMUSG0C | false | chr11 | 74829633 | 74830333 | 18.54859 | 48.96143 | 46.25894 | chr11 | 74826002 | 74829950 | 5.69928 | 4.35746 | 3.60487 |
| chr16 | 18392377 | 18393458 | 1000 | Promoter (<=1kb) | 16 | 18392588 | 18407076 | 14489 | 1 | ENSMUST00 | 0 | Arvcf | ENSMUSG0C | true | chr16 | 18392488 | 18395490 | 30.94804 | 114.76607 | 110.79114 | chr16 | 18376326 | 18392535 | 8.53722 | 5.66709 | 5.90216 |
| chr5 | 18396151 | 18396482 | 1000 | Promoter (1-2kb) | 16 | 18397724 | 18399004 | 1281 | 1 | ENSMUST00 | -1242 | Arvcf | ENSMUSG0C | true | . | . | . | . | . | . | chr16 | 18394817 | 18413117 | 4.98764 | 3.91476 | 3.01618 |
| chr5 | 152636365 | 152637073 | 1000 | Distal Intergenic | 2 | 152632967 | 152635198 | 2232 | 1 | ENSMUST00 | 3399 | Rem1 | ENSMUSG0C | false | . | . | . | . | . | . | chr2 | 152636211 | 152636888 | 3.09468 | 3.02977 | 1.49453 |
| chr11 | 6546354 | 6547277 | 1000 | Promoter (<=1kb) | 11 | 6546887 | 6596744 | 49858 | 1 | ENSMUST00 | 0 | Ccm2 | ENSMUSG0C | false | chr11 | 6546029 | 6548154 | 25.90858 | 83.69981 | 80.48042 | . | . | . | . | . | . |
| chr5 | 18344207 | 18344546 | 1000 | Distal Intergenic | 6 | 18367526 | 18382860 | 15335 | 2 | ENSMUST00 | 38314 | Cttnbp2 | ENSMUSG0C | true | . | . | . | . | . | . | chr6 | 18344340 | 18345282 | 2.96416 | 2.87223 | 1.37558 |
| chr5 | 18543809 | 18544254 | 1000 | Distal Intergenic | 6 | 18366547 | 18514869 | 148323 | 2 | ENSMUST00 | -28941 | Cttnbp2 | ENSMUSG0C | true | . | . | . | . | . | . | chr6 | 18539839 | 18548637 | 4.5475 | 3.89202 | 2.64481 |
| chr18 | 24205160 | 24206144 | 1000 | Promoter (<=1kb) | 18 | 24205344 | 24286818 | 81475 | 1 | ENSMUST00 | 0 | Galnt1 | ENSMUSG0C | false | chr18 | 24205060 | 24207222 | 35.27562 | 110.11647 | 106.27654 | . | . | . | . | . | . |
| chr15 | 80013731 | 80015194 | 1000 | Promoter (<=1kb) | 15 | 79995900 | 80014808 | 18909 | 2 | ENSMUST00 | 0 | Pdgfb | ENSMUSG0C | false | chr15 | 80012006 | 80015223 | 33.97523 | 103.77077 | 100.10409 | chr15 | 80009666 | 80015472 | 5.02785 | 4.1066 | 3.0255 |
| chr5 | 101182251 | 101182695 | 1000 | Intron (ENSMUST00000000544.1 | 15 | 101174067 | 101213679 | 39613 | 1 | ENSMUST00 | 8185 | Acvr1b | ENSMUSG0C | false | . | . | . | . | . | . | chr15 | 101174616 | 101193068 | 5.45533 | 4.23935 | 3.38802 |
| chr15 | 101173526 | 101174376 | 1000 | Promoter (<=1kb) | 15 | 101174067 | 101213679 | 39613 | 1 | ENSMUST00 | 0 | Acvr1b | ENSMUSG0C | false | chr15 | 101173412 | 101176072 | 37.34335 | 127.50397 | 123.10989 | . | . | . | . | . | . |
| chr5 | 99978763 | 99979539 | 1000 | Promoter (<=1kb) | 5 | 99958426 | 99978938 | 20513 | 2 | ENSMUST00 | 0 | Hnrnpd | ENSMUSG0C | false | chr5 | 99979164 | 99979864 | 23.80619 | 80.75649 | 77.58801 | . | . | . | . | . | . |
| chr5 | 99977880 | 99978580 | 1000 | Promoter (<=1kb) | 5 | 99962283 | 99978624 | 16342 | 2 | ENSMUST00 | 44 | Hnrnpd | ENSMUSG0C | false | chr5 | 99976691 | 99979009 | 35.32033 | 108.13151 | 104.34708 | . | . | . | . | . | . |
| chr17 | 6601504 | 6601915 | 1000 | Promoter (<=1kb) | 17 | 6601671 | 6609679 | 8009 | 1 | ENSMUST00 | 0 | Dynlt1c | ENSMUSG0C | false | chr17 | 6601259 | 6601680 | 14.6238 | 41.22485 | 38.6191 | . | . | . | . | . | . |
| chr5 | 77847587 | 77848005 | 1000 | Promoter (1-2kb) | 11 | 77849451 | 77853454 | 4004 | 1 | ENSMUST00 | -1446 | Myo18a | ENSMUSG0C | false | . | . | . | . | . | . | chr11 | 77846749 | 77848431 | 6.09013 | 4.22878 | 3.94777 |
| chr5 | 77779322 | 77779646 | 1000 | Promoter (2-3kb) | 11 | 77777316 | 77864848 | 87533 | 1 | ENSMUST00 | 2007 | Myo18a | ENSMUSG0C | false | . | . | . | . | . | . | . | . | . | . | . | . |
| chr11 | 77802196 | 77802983 | 1000 | Promoter (<=1kb) | 11 | 77801375 | 77865056 | 63682 | 1 | ENSMUST00 | 822 | Myo18a | ENSMUSG0C | false | chr11 | 77802210 | 77803345 | 11.2646 | 26.49366 | 24.07236 | chr11 | 77799596 | 77817107 | 5.11096 | 4.06241 | 3.11534 |
| chr6 | 83033925 | 83034303 | 1000 | Promoter (<=1kb) | 6 | 83034173 | 83037555 | 3383 | 1 | ENSMUST00 | 0 | Loxl3 | ENSMUSG0C | false | chr6 | 83032551 | 83034569 | 30.73327 | 90.24956 | 86.89949 | chr6 | 83033860 | 83034780 | 4.38398 | 3.86548 | 2.48332 |
| chr17 | 53566857 | 53567825 | 1000 | Promoter (<=1kb) | 17 | 53566861 | 53624491 | 57631 | 1 | ENSMUST00 | 0 | Kat2b | ENSMUSG0C | false | chr17 | 53566162 | 53568773 | 27.88725 | 99.90214 | 96.33501 | . | . | . | . | . | . |
| chr10 | 128695913 | 128696706 | 1000 | Promoter (<=1kb) | 10 | 128677193 | 128696268 | 19076 | 2 | ENSMUST00 | 0 | Rab5b | ENSMUSG0C | false | chr10 | 128695458 | 128696358 | 27.04123 | 118.00974 | 113.93439 | chr10 | 128696587 | 128697833 | 3.41017 | 3.27945 | 1.68197 |
| chr5 | 12643920 | 12644361 | 1000 | Intron (ENSMUST00000164774.2 | 8 | 12614279 | 12637982 | 23704 | 2 | ENSMUST00 | -5939 | Tubgcp3 | ENSMUSG0C | false | . | . | . | . | . | . | chr8 | 12641450 | 12645514 | 4.8959 | 3.94853 | 2.96492 |
| chr11 | 52283008 | 52283555 | 1000 | Promoter (<=1kb) | 11 | 52253236 | 52283014 | 29779 | 2 | ENSMUST00 | 0 | Tcf7 | ENSMUSG0C | false | chr11 | 52282532 | 52283722 | 33.55804 | 103.05054 | 99.40515 | chr11 | 52282605 | 52284796 | 6.38136 | 4.60082 | 4.19702 |
| chr5 | 89613795 | 89614270 | 1000 | Intron (ENSMUST00000000811.7 | 3 | 89662518 | 89672072 | 9555 | 1 | ENSMUST00 | -48248 | Kcnn3 | ENSMUSG0C | false | . | . | . | . | . | . | chr3 | 89612218 | 89615065 | 5.00601 | 4.1185 | 3.01634 |
| chr11 | 85139558 | 85140441 | 1000 | Promoter (<=1kb) | 11 | 84984442 | 85140038 | 155597 | 2 | ENSMUST00 | 0 | Usp32 | ENSMUSG0C | false | chr11 | 85138661 | 85140048 | 33.08631 | 99.21803 | 95.66874 | . | . | . | . | . | . |
| chr5 | 123973503 | 123974408 | 1000 | Promoter (<=1kb) | 5 | 123973628 | 124003211 | 29584 | 1 | ENSMUST00 | 0 | Hip1r | ENSMUSG0C | false | chr5 | 123973277 | 123975236 | 31.50646 | 96.49045 | 93.00759 | chr5 | 123973252 | 123973542 | 4.29604 | 3.82118 | 2.43654 |
| chr5 | 123985107 | 123985505 | 1000 | Intron (ENSMUST00000000939.9 | 5 | 123980254 | 123989984 | 9731 | 1 | ENSMUST00 | 4854 | Hip1r | ENSMUSG0C | false | . | . | . | . | . | . | chr5 | 123974676 | 123993001 | 6.443 | 4.63774 | 4.21224 |
| chr15 | 75678645 | 75678980 | 1000 | Promoter (<=1kb) | 15 | 75657035 | 75678800 | 21766 | 2 | ENSMUST00 | 0 | Top1mt | ENSMUSG0C | true | chr15 | 75677956 | 75683746 | 14.91789 | 45.43439 | 42.77634 | chr15 | 75674100 | 75679277 | 5.51092 | 4.23146 | 3.4513 |
| chr6 | 52233382 | 52234275 | 1000 | Promoter (<=1kb) | 6 | 52231199 | 52233324 | 2126 | 2 | ENSMUST00 | -59 | Hoxa10 | ENSMUSG0C | false | chr6 | 52227601 | 52235632 | 41.69514 | 140.65352 | 135.74945 | . | . | . | . | . | . |
| chr6 | 52191705 | 52192177 | 1000 | Promoter (<=1kb) | 6 | 52189671 | 52191753 | 2083 | 2 | ENSMUST00 | 0 | Hoxa4 | ENSMUSG0C | false | chr6 | 52191910 | 52192593 | 18.95815 | 63.55374 | 60.65727 | chr6 | 52188874 | 52193165 | 4.2446 | 3.64299 | 2.41132 |
| chr3 | 130060170 | 130060989 | 1000 | Promoter (<=1kb) | 3 | 129982759 | 130060907 | 78149 | 2 | ENSMUST00 | 0 | Sec24b | ENSMUSG0C | false | chr3 | 130059441 | 130061742 | 35.78724 | 111.00061 | 107.13931 | . | . | . | . | . | . |
| chr11 | 51650708 | 51651220 | 1000 | Promoter (<=1kb) | 11 | 51643063 | 51650842 | 7780 | 2 | ENSMUST00 | 0 | N4bp3 | ENSMUSG0C | false | chr11 | 51649934 | 51651437 | 34.00537 | 102.80987 | 99.17244 | chr11 | 51646060 | 51651285 | 3.79144 | 3.32194 | 2.04478 |
| chr5 | 136517992 | 136518541 | 1000 | Promoter (2-3kb) | 4 | 136521395 | 136546394 | 25000 | 1 | ENSMUST00 | -2854 | Luzp1 | ENSMUSG0C | false | . | . | . | . | . | . | chr4 | 136493307 | 136526569 | 5.23798 | 4.16966 | 3.21541 |
| chr11 | 78511998 | 78512482 | 1000 | Promoter (<=1kb) | 11 | 78512193 | 78522736 | 10544 | 1 | ENSMUST00 | 0 | Poldip2 | ENSMUSG0C | false | chr11 | 78511525 | 78512886 | 30.33064 | 129.04645 | 124.59567 | chr11 | 78511174 | 78513341 | 4.49595 | 3.91671 | 2.56963 |
| chr1 | 36511538 | 36512071 | 1000 | Promoter (<=1kb) | 1 | 36511867 | 36528237 | 16371 | 1 | ENSMUST00 | 0 | Cnnm3 | ENSMUSG0C | false | chr1 | 36511402 | 36513251 | 37.50825 | 119.21938 | 115.10198 | . | . | . | . | . | . |
| chr1 | 36445167 | 36445561 | 1000 | Promoter (<=1kb) | 1 | 36422065 | 36445271 | 23207 | 2 | ENSMUST00 | 0 | Lman2l | ENSMUSG0C | false | chr1 | 36444527 | 36445300 | 38.58512 | 125.36365 | 121.04008 | chr1 | 36444391 | 36445814 | 4.73369 | 3.87037 | 2.79314 |
| chr5 | 47916951 | 47917285 | 1000 | Intron (ENSMUST00000001202.9 | 20 | 47912467 | 47915582 | 3116 | 1 | ENSMUST00 | 4485 | Ocrl | ENSMUSG0C | false | . | . | . | . | . | . | . | . | . | . | . | . |
| chrX | 47912265 | 47912759 | 1000 | Promoter (<=1kb) | 20 | 47912387 | 47965868 | 53482 | 1 | ENSMUST00 | 0 | Ocrl | ENSMUSG0C | false | chrX | 47912348 | 47913327 | 26.65286 | 74.78688 | 71.72144 | . | . | . | . | . | . |
| chr12 | 100199913 | 100200951 | 1000 | Promoter (<=1kb) | 12 | 100200144 | 100209806 | 9663 | 1 | ENSMUST00 | 0 | Calm1 | ENSMUSG0C | false | chr12 | 100198919 | 100201212 | 21.48562 | 91.50835 | 88.13527 | chr12 | 100200702 | 100205277 | 4.60381 | 3.89787 | 2.67349 |
| chr10 | 78351929 | 78352793 | 1000 | Promoter (<=1kb) | 10 | 78269178 | 78352442 | 83265 | 2 | ENSMUST00 | 0 | Agpat3 | ENSMUSG0C | false | chr10 | 78351249 | 78352952 | 26.1063 | 98.41757 | 94.88632 | chr10 | 78348875 | 78353922 | 4.98596 | 4.08669 | 2.99507 |
| chr7 | 31150867 | 31151234 | 1000 | Promoter (<=1kb) | 7 | 31130127 | 31151012 | 20886 | 2 | ENSMUST00 | 0 | Gramd1a | ENSMUSG0C | false | chr7 | 31150227 | 31151795 | 24.0356 | 94.23969 | 90.80066 | . | . | . | . | . | . |
| chr15 | 102406134 | 102407152 | 1000 | Promoter (<=1kb) | 15 | 102406143 | 102408235 | 2093 | 1 | ENSMUST00 | 0 | Sp1 | ENSMUSG0C | false | chr15 | 102406202 | 102407269 | 21.92219 | 60.49891 | 57.64639 | chr15 | 102407054 | 102421808 | 4.47126 | 3.73085 | 2.59402 |
| chr15 | 102246919 | 102248258 | 1000 | Promoter (<=1kb) | 15 | 102234940 | 102246500 | 11561 | 2 | ENSMUST00 | -420 | Rarg | ENSMUSG0C | false | chr15 | 102244836 | 102248267 | 30.17592 | 94.17345 | 90.73757 | chr15 | 102241789 | 102251883 | 5.05731 | 4.11666 | 3.0557 |
| chr15 | 102257041 | 102257829 | 1000 | Promoter (<=1kb) | 15 | 102234938 | 102257517 | 22580 | 2 | ENSMUST00 | 0 | Rarg | ENSMUSG0C | false | chr15 | 102256411 | 102258508 | 31.97522 | 95.31979 | 91.85828 | chr15 | 102255556 | 102259358 | 6.54281 | 4.71118 | 4.27807 |
| chr11 | 74925591 | 74926388 | 1000 | Promoter (<=1kb) | 11 | 74906359 | 74925798 | 19440 | 2 | ENSMUST00 | 0 | Srr | ENSMUSG0C | false | chr11 | 74924900 | 74926913 | 33.41831 | 117.26761 | 113.21259 | . | . | . | . | . | . |
| chr5 | 129285919 | 129286361 | 1000 | Promoter (1-2kb) | 4 | 129287617 | 129308559 | 20943 | 1 | ENSMUST00 | -1256 | Sync | ENSMUSG0C | false | . | . | . | . | . | . | chr4 | 129284848 | 129289902 | 6.10204 | 4.63545 | 3.89161 |
| chr5 | 129147535 | 129148037 | 1000 | 3' UTR | 4 | 129136999 | 129144593 | 7595 | 1 | ENSMUST00 | 10537 | Fndc5 | ENSMUSG0C | false | . | . | . | . | . | . | chr4 | 129141119 | 129150343 | 5.31606 | 4.18214 | 3.25946 |
| chr3 | 88141881 | 88142831 | 1000 | Promoter (<=1kb) | 3 | 88142372 | 88172086 | 29715 | 1 | ENSMUST00 | 0 | Mef2d | ENSMUSG0C | false | chr3 | 88141285 | 88144231 | 33.92091 | 108.79443 | 104.99046 | chr3 | 88136354 | 88141917 | 6.47517 | 4.6068 | 4.24981 |
| chr3 | 88138367 | 88138989 | 1000 | Distal Intergenic | 3 | 88142372 | 88172086 | 29715 | 1 | ENSMUST00 | -3383 | Mef2d | ENSMUSG0C | false | chr3 | 88138472 | 88139892 | 22.69969 | 67.91293 | 64.95291 | chr3 | 88136354 | 88141917 | 6.47517 | 4.6068 | 4.24981 |
| chr10 | 77111217 | 77111530 | 1000 | Promoter (2-3kb) | 10 | 77052181 | 77113676 | 61496 | 2 | ENSMUST00 | 2146 | Col18a1 | ENSMUSG0C | false | chr10 | 77110823 | 77114286 | 25.79081 | 75.126 | 72.05579 | chr10 | 77096587 | 77112087 | 5.00583 | 4.02872 | 3.02421 |
| chr11 | 97186994 | 97188585 | 1000 | Promoter (<=1kb) | 11 | 97159714 | 97187881 | 28168 | 2 | ENSMUST00 | 0 | Kpnb1 | ENSMUSG0C | false | chr11 | 97186741 | 97188025 | 22.31878 | 85.15456 | 81.90936 | chr11 | 97186077 | 97187057 | 5.69029 | 4.43525 | 3.55435 |
| chr11 | 97279902 | 97280840 | 1000 | Promoter (<=1kb) | 11 | 97205842 | 97280486 | 74645 | 2 | ENSMUST00 | 0 | Npepps | ENSMUSG0C | false | chr11 | 97279259 | 97281276 | 33.7938 | 119.00099 | 114.88904 | . | . | . | . | . | . |
| chr5 | 101965551 | 101966008 | 1000 | Promoter (1-2kb) | 11 | 101962458 | 101967015 | 4558 | 2 | ENSMUST00 | 1007 | Sost | ENSMUSG0C | false | . | . | . | . | . | . | chr11 | 101962847 | 101969318 | 4.37367 | 3.70571 | 2.49923 |
| chr11 | 95075196 | 95075709 | 1000 | Promoter (1-2kb) | 11 | 95044474 | 95076714 | 32241 | 2 | ENSMUST00 | 1005 | Itga3 | ENSMUSG0C | true | chr11 | 95073478 | 95078046 | 40.23456 | 136.98251 | 132.21786 | . | . | . | . | . | . |
| chr11 | 95076461 | 95077325 | 1000 | Promoter (<=1kb) | 11 | 95044474 | 95076714 | 32241 | 2 | ENSMUST00 | 0 | Itga3 | ENSMUSG0C | true | chr11 | 95073478 | 95078046 | 40.23456 | 136.98251 | 132.21786 | . | . | . | . | . | . |
| chr11 | 95071222 | 95071598 | 1000 | Promoter (2-3kb) | 11 | 95044474 | 95069198 | 24725 | 2 | ENSMUST00 | -2025 | Itga3 | ENSMUSG0C | true | chr11 | 95071021 | 95073215 | 8.80182 | 17.30588 | 15.00915 | chr11 | 95064128 | 95074961 | 7.24424 | 5.01152 | 4.86589 |
| chr5 | 95080733 | 95081160 | 1000 | Distal Intergenic | 11 | 95044474 | 95076801 | 32328 | 2 | ENSMUST00 | -3933 | Itga3 | ENSMUSG0C | true | . | . | . | . | . | . | chr11 | 95077539 | 95082670 | 7.743 | 5.23471 | 5.25962 |
| chr5 | 100391016 | 100391662 | 1000 | Intron (ENSMUST00000001592.9 | 11 | 100383404 | 100386770 | 3367 | 2 | ENSMUST00 | -4247 | Jup | ENSMUSG0C | false | . | . | . | . | . | . | chr11 | 100384756 | 100396489 | 9.22555 | 5.81405 | 6.48761 |
| chr11 | 100387900 | 100388478 | 1000 | Promoter (1-2kb) | 11 | 100383404 | 100386770 | 3367 | 2 | ENSMUST00 | -1131 | Jup | ENSMUSG0C | false | chr11 | 100388449 | 100388773 | 4.7098 | 6.74815 | 4.64672 | chr11 | 100384756 | 100396489 | 9.22555 | 5.81405 | 6.48761 |
| chr11 | 100396898 | 100397479 | 1000 | Promoter (<=1kb) | 11 | 100370619 | 100397749 | 27131 | 2 | ENSMUST00 | 270 | Jup | ENSMUSG0C | false | chr11 | 100394893 | 100397863 | 34.54477 | 122.48211 | 118.25522 | . | . | . | . | . | . |
| chr11 | 100395216 | 100395690 | 1000 | Promoter (2-3kb) | 11 | 100370619 | 100397749 | 27131 | 2 | ENSMUST00 | 2059 | Jup | ENSMUSG0C | false | chr11 | 100394893 | 100397863 | 34.54477 | 122.48211 | 118.25522 | chr11 | 100384756 | 100396489 | 9.22555 | 5.81405 | 6.48761 |
| chr17 | 26561264 | 26561729 | 1000 | Promoter (<=1kb) | 17 | 26561489 | 26656931 | 95443 | 1 | ENSMUST00 | 0 | Ergic1 | ENSMUSG0C | false | chr17 | 26561044 | 26562480 | 21.75525 | 76.8157 | 73.7174 | . | . | . | . | . | . |
| chr6 | 113306403 | 113307313 | 1000 | Promoter (<=1kb) | 6 | 113307181 | 113324857 | 17677 | 1 | ENSMUST00 | 0 | Brpf1 | ENSMUSG0C | false | chr6 | 113306728 | 113308496 | 20.57128 | 91.19001 | 87.82124 | . | . | . | . | . | . |
| chr15 | 102999084 | 102999733 | 1000 | Promoter (<=1kb) | 15 | 103000529 | 103010006 | 9478 | 1 | ENSMUST00 | -796 | Hoxc6 | ENSMUSG0C | false | chr15 | 102997422 | 103001867 | 28.24444 | 85.94312 | 82.67722 | . | . | . | . | . | . |
| chr15 | 103005448 | 103005744 | 1000 | Intron (ENSMUST00000174869.1 | 15 | 103009573 | 103011881 | 2309 | 1 | ENSMUST00 | -3829 | Hoxc6 | ENSMUSG0C | false | chr15 | 103005091 | 103006170 | 8.48197 | 18.2582 | 15.9485 | chr15 | 103001250 | 103018640 | 5.43629 | 4.24027 | 3.36324 |
| chr5 | 148552354 | 148553046 | 1000 | Promoter (<=1kb) | 5 | 148504631 | 148552788 | 48158 | 2 | ENSMUST00 | 0 | Ubl3 | ENSMUSG0C | false | chr5 | 148550262 | 148553141 | 24.82645 | 85.46523 | 82.21117 | . | . | . | . | . | . |
| chr5 | 136967615 | 136967929 | 1000 | Promoter (<=1kb) | 5 | 136967869 | 136975844 | 7976 | 1 | ENSMUST00 | 0 | Cldn15 | ENSMUSG0C | false | chr5 | 136967757 | 136968173 | 5.47834 | 9.2208 | 7.06269 | chr5 | 136953761 | 136969776 | 6.09241 | 4.43302 | 3.92021 |
| chr5 | 101056840 | 101057323 | 1000 | Intron (ENSMUST00000103110.4 | 11 | 101070012 | 101077672 | 7661 | 1 | ENSMUST00 | -12689 | Naglu | ENSMUSG0C | false | . | . | . | . | . | . | chr11 | 101054317 | 101067609 | 9.20102 | 5.81671 | 6.45648 |
| chr5 | 101060248 | 101060663 | 1000 | Intron (ENSMUST00000103110.4 | 11 | 101070012 | 101077672 | 7661 | 1 | ENSMUST00 | -9349 | Naglu | ENSMUSG0C | false | . | . | . | . | . | . | chr11 | 101054317 | 101067609 | 9.20102 | 5.81671 | 6.45648 |
| chr6 | 29735301 | 29736431 | 1000 | Promoter (<=1kb) | 6 | 29735694 | 29761365 | 25672 | 1 | ENSMUST00 | 0 | Smo | ENSMUSG0C | true | chr6 | 29735189 | 29737881 | 36.78915 | 128.67926 | 124.23671 | . | . | . | . | . | . |
| chr5 | 145831884 | 145832750 | 1000 | Intron (ENSMUST00000110005.2 | 2 | 145822289 | 145845087 | 22799 | 1 | ENSMUST00 | 9596 | Rin2 | ENSMUSG0C | true | . | . | . | . | . | . | chr2 | 145827498 | 145846856 | 5.11522 | 4.05923 | 3.13101 |
| chr5 | 74647490 | 74647861 | 1000 | Distal Intergenic | 2 | 74655616 | 74659419 | 3804 | 2 | ENSMUST00 | 11558 | Evx2 | ENSMUSG0C | false | . | . | . | . | . | . | chr2 | 74647095 | 74649142 | 4.60439 | 3.91436 | 2.6796 |
| chr9 | 25252000 | 25253232 | 1000 | Promoter (<=1kb) | 9 | 25252439 | 25308571 | 56133 | 1 | ENSMUST00 | 0 | Sep-07 | ENSMUSG0C | false | chr9 | 25251253 | 25253817 | 39.05061 | 143.74966 | 138.71384 | . | . | . | . | . | . |
| chr2 | 31953562 | 31954051 | 1000 | Intron (ENSMUST00000001920.7 | 2 | 31950517 | 31971338 | 20822 | 1 | ENSMUST00 | 3046 | Aif1l | ENSMUSG0C | false | chr2 | 31949970 | 31953731 | 33.10969 | 106.34647 | 102.61562 | chr2 | 31951476 | 31965598 | 6.45762 | 4.68343 | 4.21645 |
| chr5 | 74866481 | 74867115 | 1000 | Distal Intergenic | 17 | 75005568 | 75391767 | 386200 | 1 | ENSMUST00 | -138453 | Ltbp1 | ENSMUSG0C | false | . | . | . | . | . | . | chr17 | 74864637 | 74871267 | 6.04854 | 4.62257 | 3.85767 |
| chr11 | 21370455 | 21371689 | 1000 | Promoter (<=1kb) | 11 | 21321138 | 21370477 | 49340 | 2 | ENSMUST00 | 0 | Ugp2 | ENSMUSG0C | false | chr11 | 21369063 | 21371817 | 43.65555 | 141.82639 | 136.87535 | chr11 | 21371475 | 21372797 | 4.93412 | 3.98363 | 3.0008 |
| chr8 | 84799424 | 84800109 | 1000 | Promoter (<=1kb) | 8 | 84713795 | 84800024 | 86230 | 2 | ENSMUST00 | 0 | Nfix | ENSMUSG0C | false | chr8 | 84799139 | 84800864 | 25.4311 | 72.68642 | 69.65319 | chr8 | 84798889 | 84799757 | 3.73782 | 3.43156 | 1.97468 |
| chr5 | 20644782 | 20645130 | 1000 | Exon (ENSMUST00000115375.2/ | 20 | 20658326 | 20683179 | 24854 | 1 | ENSMUST00 | -13196 | Uba1 | ENSMUSG0C | false | . | . | . | . | . | . | . | . | . | . | . | . |
| chr5 | 125385455 | 125386079 | 1000 | Distal Intergenic | 8 | 125418063 | 125492710 | 74648 | 2 | ENSMUST00 | 106631 | Sipa1l2 | ENSMUSG0C | false | . | . | . | . | . | . | chr8 | 125384422 | 125392083 | 5.56972 | 4.2897 | 3.4878 |
| chr8 | 125569558 | 125570153 | 1000 | Distal Intergenic | 8 | 125418063 | 125492710 | 74648 | 2 | ENSMUST00 | -76849 | Sipa1l2 | ENSMUSG0C | false | chr8 | 125569988 | 125570633 | 28.31422 | 85.10979 | 81.86543 | . | . | . | . | . | . |
| chr5 | 125532860 | 125533211 | 1000 | Distal Intergenic | 8 | 125418063 | 125492710 | 74648 | 2 | ENSMUST00 | -40151 | Sipa1l2 | ENSMUSG0C | false | . | . | . | . | . | . | chr8 | 125526488 | 125537403 | 5.56127 | 4.26582 | 3.48444 |
| chrX | 73824799 | 73825162 | 1000 | Promoter (<=1kb) | 20 | 73793359 | 73824969 | 31611 | 2 | ENSMUST00 | 0 | Pdzd4 | ENSMUSG0C | false | chrX | 73824175 | 73825239 | 16.7644 | 42.82802 | 40.20103 | . | . | . | . | . | . |
| chr9 | 44880994 | 44881914 | 1000 | Promoter (<=1kb) | 9 | 44803355 | 44881274 | 77920 | 2 | ENSMUST00 | 0 | Kmt2a | ENSMUSG0C | false | chr9 | 44879791 | 44881351 | 27.97104 | 107.99458 | 104.2211 | . | . | . | . | . | . |
| chr7 | 16309339 | 16309758 | 1000 | Promoter (<=1kb) | 7 | 16309616 | 16318205 | 8590 | 1 | ENSMUST00 | 0 | Bbc3 | ENSMUSG0C | false | chr7 | 16309242 | 16310716 | 29.86959 | 86.83979 | 83.56236 | chr7 | 16309137 | 16315495 | 5.75225 | 4.42325 | 3.62489 |
| chr7 | 16299526 | 16300006 | 1000 | Distal Intergenic | 7 | 16309616 | 16318205 | 8590 | 1 | ENSMUST00 | -9610 | Bbc3 | ENSMUSG0C | false | chr7 | 16299552 | 16300220 | 10.05864 | 20.96754 | 18.61967 | chr7 | 16299190 | 16300473 | 6.90451 | 4.95101 | 4.53691 |
| chr7 | 16309961 | 16310556 | 1000 | Promoter (<=1kb) | 7 | 16310455 | 16313738 | 3284 | 1 | ENSMUST00 | 0 | Bbc3 | ENSMUSG0C | false | chr7 | 16309242 | 16310716 | 29.86959 | 86.83979 | 83.56236 | chr7 | 16309137 | 16315495 | 5.75225 | 4.42325 | 3.62489 |
| chr2 | 91070216 | 91070629 | 1000 | Promoter (<=1kb) | 2 | 91061791 | 91070229 | 8439 | 2 | ENSMUST00 | 0 | Slc39a13 | ENSMUSG0C | false | chr2 | 91069362 | 91071119 | 31.06412 | 92.76744 | 89.36313 | . | . | . | . | . | . |
| chr5 | 27790946 | 27791723 | 1000 | Promoter (<=1kb) | 5 | 27740667 | 27791339 | 50673 | 2 | ENSMUST00 | 0 | Paxip1 | ENSMUSG0C | false | chr5 | 27790221 | 27792376 | 30.35223 | 94.4818 | 91.04041 | . | . | . | . | . | . |
| chr6 | 71439818 | 71440871 | 1000 | Promoter (<=1kb) | 6 | 71388634 | 71440637 | 52004 | 2 | ENSMUST00 | 0 | Rmnd5a | ENSMUSG0C | false | chr6 | 71439112 | 71440634 | 25.79552 | 96.83904 | 93.34669 | . | . | . | . | . | . |
| chr5 | 71428325 | 71428705 | 1000 | Intron (ENSMUST00000002292.9 | 6 | 71388634 | 71440637 | 52004 | 2 | ENSMUST00 | 11932 | Rmnd5a | ENSMUSG0C | false | . | . | . | . | . | . | chr6 | 71413520 | 71434591 | 5.21097 | 4.11837 | 3.19483 |
| chr3 | 104817472 | 104818821 | 1000 | Promoter (<=1kb) | 3 | 104794836 | 104818242 | 23407 | 2 | ENSMUST00 | 0 | Mov10 | ENSMUSG0C | false | chr3 | 104817345 | 104818906 | 33.04283 | 115.53907 | 111.53828 | chr3 | 104815706 | 104817623 | 3.1219 | 3.13828 | 1.52588 |
| chr17 | 28349983 | 28351119 | 1000 | Promoter (<=1kb) | 17 | 28331671 | 28350346 | 18676 | 2 | ENSMUST00 | 0 | Tead3 | ENSMUSG0C | false | chr17 | 28349620 | 28351838 | 25.84672 | 90.2271 | 86.88143 | chr17 | 28348863 | 28350136 | 3.74746 | 3.51912 | 1.95823 |
| chr5 | 28253287 | 28253613 | 1000 | Intron (ENSMUST00000002320.9 | 17 | 28271708 | 28273163 | 1456 | 1 | ENSMUST00 | -18095 | Ppard | ENSMUSG0C | false | . | . | . | . | . | . | chr17 | 28251377 | 28258644 | 5.73452 | 4.39083 | 3.62025 |
| chr5 | 5868920 | 5869407 | 1000 | Intron (ENSMUST00000002436.9 | 17 | 5841328 | 5931033 | 89706 | 1 | ENSMUST00 | 27593 | Snx9 | ENSMUSG0C | false | . | . | . | . | . | . | chr17 | 5867189 | 5879266 | 5.9622 | 4.44173 | 3.80171 |
| chr17 | 5840997 | 5842081 | 1000 | Promoter (<=1kb) | 17 | 5841328 | 5931033 | 89706 | 1 | ENSMUST00 | 0 | Snx9 | ENSMUSG0C | false | chr17 | 5840733 | 5842592 | 25.50663 | 88.63404 | 85.31516 | . | . | . | . | . | . |
| chr8 | 71381775 | 71382534 | 1000 | Promoter (<=1kb) | 8 | 71374123 | 71381906 | 7784 | 2 | ENSMUST00 | 0 | Nr2f6 | ENSMUSG0C | false | chr8 | 71380023 | 71382831 | 37.92415 | 138.37384 | 133.55817 | chr8 | 71378170 | 71383957 | 4.97224 | 3.98404 | 3.01306 |
| chr6 | 39725076 | 39725715 | 1000 | Promoter (<=1kb) | 6 | 39603237 | 39725463 | 122227 | 2 | ENSMUST00 | 0 | Braf | ENSMUSG0C | false | chr6 | 39724148 | 39725926 | 34.54477 | 122.48211 | 118.25522 | . | . | . | . | . | . |
| chr17 | 24638714 | 24639854 | 1000 | 3' UTR | 17 | 24639282 | 24644957 | 5676 | 2 | ENSMUST00 | 5103 | Slc9a3r2 | ENSMUSG0C | false | chr17 | 24639673 | 24640299 | 7.1665 | 12.77688 | 10.5477 | chr17 | 24630730 | 24649025 | 8.70874 | 5.59711 | 6.05272 |
| chr17 | 24649493 | 24650551 | 1000 | Promoter (<=1kb) | 17 | 24639282 | 24650305 | 11024 | 2 | ENSMUST00 | 0 | Slc9a3r2 | ENSMUSG0C | false | chr17 | 24648245 | 24650753 | 34.54477 | 122.48211 | 118.25522 | chr17 | 24650464 | 24654716 | 4.48905 | 3.89646 | 2.57513 |
| chr17 | 24643959 | 24644650 | 1000 | Promoter (<=1kb) | 17 | 24639282 | 24644957 | 5676 | 2 | ENSMUST00 | 307 | Slc9a3r2 | ENSMUSG0C | false | chr17 | 24642925 | 24645980 | 11.05033 | 23.18759 | 20.81059 | chr17 | 24630730 | 24649025 | 8.70874 | 5.59711 | 6.05272 |
| chr7 | 25788105 | 25788929 | 1000 | Promoter (<=1kb) | 7 | 25757273 | 25788687 | 31415 | 2 | ENSMUST00 | 0 | Axl | ENSMUSG0C | false | chr7 | 25787969 | 25789058 | 19.75988 | 50.25065 | 47.53236 | chr7 | 25785754 | 25789387 | 5.83807 | 4.47654 | 3.68809 |
| chr5 | 25764239 | 25764665 | 1000 | Promoter (<=1kb) | 7 | 25761369 | 25764413 | 3045 | 2 | ENSMUST00 | 0 | Axl | ENSMUSG0C | false | . | . | . | . | . | . | chr7 | 25764498 | 25766367 | 3.02714 | 3.03523 | 1.43374 |
| chr5 | 25787583 | 25787867 | 1000 | Promoter (<=1kb) | 7 | 25785939 | 25787572 | 1634 | 2 | ENSMUST00 | -12 | Axl | ENSMUSG0C | false | . | . | . | . | . | . | chr7 | 25785754 | 25789387 | 5.83807 | 4.47654 | 3.68809 |
| chr7 | 25781291 | 25782374 | 1000 | Intron (ENSMUST00000002677.5 | 7 | 25778680 | 25787066 | 8387 | 2 | ENSMUST00 | 4692 | Axl | ENSMUSG0C | false | chr7 | 25781465 | 25782062 | 14.5262 | 33.35895 | 30.8513 | chr7 | 25779352 | 25783967 | 6.79659 | 4.90679 | 4.46063 |
| chr7 | 25689070 | 25689564 | 1000 | Promoter (<=1kb) | 7 | 25688927 | 25697213 | 8287 | 1 | ENSMUST00 | 144 | Tgfb1 | ENSMUSG0C | false | chr7 | 25686448 | 25689783 | 37.30079 | 124.19571 | 119.91465 | chr7 | 25686345 | 25690723 | 4.57586 | 3.80181 | 2.69028 |
| chr7 | 25686667 | 25687540 | 1000 | Promoter (<=1kb) | 7 | 25687002 | 25705077 | 18076 | 1 | ENSMUST00 | 0 | Tgfb1 | ENSMUSG0C | false | chr7 | 25686448 | 25689783 | 37.30079 | 124.19571 | 119.91465 | chr7 | 25686345 | 25690723 | 4.57586 | 3.80181 | 2.69028 |
| chr12 | 50648806 | 50649570 | 1000 | Promoter (<=1kb) | 12 | 50341231 | 50649223 | 307993 | 2 | ENSMUST00 | 0 | Prkd1 | ENSMUSG0C | false | chr12 | 50647570 | 50649686 | 30.41442 | 103.5799 | 99.9162 | chr12 | 50645591 | 50649902 | 3.80439 | 3.39333 | 2.04575 |
| chr16 | 38452455 | 38453156 | 1000 | Promoter (<=1kb) | 16 | 38445399 | 38452689 | 7291 | 2 | ENSMUST00 | 0 | Adprh | ENSMUSG0C | false | chr16 | 38452104 | 38453454 | 26.18681 | 91.82585 | 88.44535 | chr16 | 38451617 | 38453366 | 3.45903 | 3.29779 | 1.72186 |
| chr1 | 52499849 | 52501040 | 1000 | Promoter (<=1kb) | 1 | 52457294 | 52500300 | 43007 | 2 | ENSMUST00 | 0 | Nab1 | ENSMUSG0C | true | chr1 | 52498999 | 52501178 | 29.22523 | 92.5029 | 89.10189 | chr1 | 52500792 | 52502775 | 4.05106 | 3.61019 | 2.24027 |
| chr8 | 83741115 | 83741542 | 1000 | Promoter (<=1kb) | 8 | 83723251 | 83741326 | 18076 | 2 | ENSMUST00 | 0 | Cd97 | ENSMUSG0C | false | chr8 | 83740029 | 83742098 | 22.15371 | 67.81625 | 64.85734 | . | . | . | . | . | . |
| chr8 | 4238602 | 4239298 | 1000 | Promoter (<=1kb) | 8 | 4238740 | 4247896 | 9157 | 1 | ENSMUST00 | 0 | Map2k7 | ENSMUSG0C | false | chr8 | 4237910 | 4239927 | 29.35304 | 95.67908 | 92.21078 | . | . | . | . | . | . |
| chr7 | 19769036 | 19769367 | 1000 | Promoter (1-2kb) | 7 | 19763673 | 19770495 | 6823 | 2 | ENSMUST00 | 1128 | Bcam | ENSMUSG0C | false | chr7 | 19768283 | 19771284 | 29.58769 | 108.11006 | 104.32592 | chr7 | 19764082 | 19769494 | 5.42164 | 4.13482 | 3.3954 |
| chr5 | 19756279 | 19756983 | 1000 | Promoter (<=1kb) | 7 | 19756557 | 19756951 | 395 | 2 | ENSMUST00 | 0 | Bcam | ENSMUSG0C | false | . | . | . | . | . | . | chr7 | 19755936 | 19759036 | 6.05107 | 4.29254 | 3.90932 |
| chr7 | 19629156 | 19629707 | 1000 | Promoter (<=1kb) | 7 | 19606217 | 19629438 | 23222 | 2 | ENSMUST00 | 0 | Relb | ENSMUSG0C | false | chr7 | 19628045 | 19629963 | 39.02359 | 127.20569 | 122.81842 | . | . | . | . | . | . |
| chr12 | 31949176 | 31949980 | 1000 | Promoter (<=1kb) | 12 | 31932833 | 31949608 | 16776 | 2 | ENSMUST00 | 0 | Hbp1 | ENSMUSG0C | false | chr12 | 31948692 | 31950317 | 42.96212 | 142.86137 | 137.8616 | . | . | . | . | . | . |
| chr4 | 55532294 | 55532976 | 1000 | Promoter (<=1kb) | 4 | 55527143 | 55532466 | 5324 | 2 | ENSMUST00 | 0 | Klf4 | ENSMUSG0C | false | chr4 | 55530022 | 55533908 | 24.6666 | 67.80661 | 64.85043 | chr4 | 55530100 | 55532342 | 5.76405 | 4.432 | 3.65261 |
| chr4 | 133967148 | 133967708 | 1000 | Promoter (<=1kb) | 4 | 133964990 | 133967441 | 2452 | 2 | ENSMUST00 | 0 | Hmgn2 | ENSMUSG0C | false | chr4 | 133966850 | 133968680 | 36.12563 | 122.12295 | 117.91594 | . | . | . | . | . | . |
| chr10 | 80115298 | 80116106 | 1000 | Promoter (<=1kb) | 10 | 80115803 | 80130469 | 14667 | 1 | ENSMUST00 | 0 | Stk11 | ENSMUSG0C | false | chr10 | 80114764 | 80115501 | 12.92336 | 34.59913 | 32.07519 | . | . | . | . | . | . |
| chr5 | 80141214 | 80141624 | 1000 | Promoter (<=1kb) | 10 | 80141457 | 80145813 | 4357 | 1 | ENSMUST00 | 0 | Atp5d | ENSMUSG0C | false | . | . | . | . | . | . | chr10 | 80140751 | 80148113 | 6.54123 | 4.83987 | 4.26386 |
| chr7 | 17027673 | 17028207 | 1000 | Promoter (<=1kb) | 7 | 17004640 | 17027916 | 23277 | 2 | ENSMUST00 | 0 | Ppp5c | ENSMUSG0C | false | chr7 | 17026986 | 17028377 | 42.72282 | 144.68874 | 139.61232 | chr7 | 17023136 | 17028530 | 4.02124 | 3.53869 | 2.21297 |
| chr11 | 98202815 | 98203530 | 1000 | Promoter (<=1kb) | 11 | 98203059 | 98251564 | 48506 | 1 | ENSMUST00 | 0 | Cdk12 | ENSMUSG0C | false | chr11 | 98201924 | 98205269 | 34.00884 | 130.01486 | 125.52227 | . | . | . | . | . | . |
| chr7 | 25395847 | 25396222 | 1000 | Promoter (<=1kb) | 7 | 25379527 | 25395987 | 16461 | 2 | ENSMUST00 | 0 | Lipe | ENSMUSG0C | false | chr7 | 25394610 | 25396560 | 28.08037 | 85.65662 | 82.39679 | . | . | . | . | . | . |
| chr1 | 39535573 | 39535957 | 1000 | Promoter (<=1kb) | 1 | 39535802 | 39546873 | 11072 | 1 | ENSMUST00 | 0 | Cnot11 | ENSMUSG0C | false | chr1 | 39535231 | 39536703 | 32.98857 | 124.89146 | 120.57959 | chr1 | 39534965 | 39537974 | 4.86004 | 4.012 | 2.91035 |
| chr6 | 122819621 | 122820321 | 1000 | Promoter (<=1kb) | 6 | 122819914 | 122845366 | 25453 | 1 | ENSMUST00 | 0 | Foxj2 | ENSMUSG0C | false | chr6 | 122819193 | 122821504 | 22.03264 | 83.65932 | 80.44393 | chr6 | 122818679 | 122828548 | 6.07039 | 4.57354 | 3.88889 |
| chr6 | 122810715 | 122811710 | 1000 | Downstream (1-2kb) | 6 | 122819914 | 122845366 | 25453 | 1 | ENSMUST00 | -8204 | Foxj2 | ENSMUSG0C | false | chr6 | 122810365 | 122810935 | 15.64407 | 45.32187 | 42.668 | . | . | . | . | . | . |
| chr6 | 121109700 | 121110638 | 1000 | Distal Intergenic | 6 | 121007241 | 121081609 | 74369 | 2 | ENSMUST00 | -28092 | Mical3 | ENSMUSG0C | false | chr6 | 121108833 | 121110689 | 33.45073 | 115.2328 | 111.2408 | chr6 | 121110555 | 121114977 | 5.58267 | 4.27018 | 3.52272 |
| chr5 | 121118045 | 121118708 | 1000 | Distal Intergenic | 6 | 121007241 | 121081609 | 74369 | 2 | ENSMUST00 | -36437 | Mical3 | ENSMUSG0C | false | . | . | . | . | . | . | chr6 | 121116518 | 121123608 | 4.83566 | 3.92225 | 2.89449 |
| chr5 | 45000417 | 45001072 | 1000 | Promoter (<=1kb) | 7 | 45000501 | 45001257 | 757 | 1 | ENSMUST00 | 0 | Irf3 | ENSMUSG0C | false | . | . | . | . | . | . | chr7 | 44998207 | 45002477 | 7.02584 | 4.93652 | 4.70726 |
| chr19 | 60890014 | 60890569 | 1000 | Promoter (<=1kb) | 19 | 60889749 | 61092553 | 202805 | 1 | ENSMUST00 | 266 | Grk5 | ENSMUSG0C | true | chr19 | 60889254 | 60892116 | 41.53183 | 140.36792 | 135.47412 | . | . | . | . | . | . |
| chr5 | 70632843 | 70633195 | 1000 | Intron (ENSMUST00000133983.2 | 17 | 70561787 | 70816093 | 254307 | 1 | ENSMUST00 | 71057 | Dlgap1 | ENSMUSG0C | false | . | . | . | . | . | . | chr17 | 70632093 | 70633548 | 3.26882 | 3.13755 | 1.6048 |
| chr4 | 3938220 | 3939155 | 1000 | Promoter (<=1kb) | 4 | 3900996 | 3938382 | 37387 | 2 | ENSMUST00 | 0 | Plag1 | ENSMUSG0C | false | chr4 | 3936695 | 3938445 | 34.36702 | 111.64683 | 107.76208 | . | . | . | . | . | . |
| chr9 | 21238619 | 21239482 | 1000 | Promoter (<=1kb) | 9 | 21229730 | 21239332 | 9603 | 2 | ENSMUST00 | 0 | Keap1 | ENSMUSG0C | false | chr9 | 21237217 | 21239638 | 34.78613 | 113.42423 | 109.48744 | . | . | . | . | . | . |
| chr5 | 111273208 | 111273724 | 1000 | Intron (ENSMUST00000038739.4 | 8 | 111157898 | 111259216 | 101319 | 2 | ENSMUST00 | -13993 | Glg1 | ENSMUSG0C | false | . | . | . | . | . | . | chr8 | 111272059 | 111276120 | 4.19776 | 3.69472 | 2.34274 |
| chr10 | 80656396 | 80657102 | 1000 | Promoter (<=1kb) | 10 | 80642617 | 80656494 | 13878 | 2 | ENSMUST00 | 0 | Btbd2 | ENSMUSG0C | false | chr10 | 80655644 | 80656595 | 25.90858 | 83.69981 | 80.48042 | chr10 | 80654838 | 80661529 | 4.56231 | 3.86979 | 2.64938 |
| chr5 | 98632161 | 98632545 | 1000 | Promoter (<=1kb) | 15 | 98632220 | 98644530 | 12311 | 1 | ENSMUST00 | 0 | Cacnb3 | ENSMUSG0C | false | . | . | . | . | . | . | chr15 | 98631220 | 98635579 | 5.30417 | 4.16229 | 3.25613 |
| chr15 | 98634479 | 98635342 | 1000 | Promoter (<=1kb) | 15 | 98634764 | 98644530 | 9767 | 1 | ENSMUST00 | 0 | Cacnb3 | ENSMUSG0C | false | chr15 | 98634150 | 98635460 | 24.55236 | 90.57755 | 87.22021 | chr15 | 98631220 | 98635579 | 5.30417 | 4.16229 | 3.25613 |
| chr5 | 108904029 | 108904440 | 1000 | Intron (ENSMUST00000106650.3 | 4 | 108890290 | 108941188 | 50899 | 1 | ENSMUST00 | 13740 | Rab3b | ENSMUSG0C | true | . | . | . | . | . | . | chr4 | 108891993 | 108908067 | 6.16979 | 4.57777 | 3.97619 |
| chr5 | 108894839 | 108895485 | 1000 | Intron (ENSMUST00000106650.3 | 4 | 108890290 | 108941188 | 50899 | 1 | ENSMUST00 | 4550 | Rab3b | ENSMUSG0C | true | . | . | . | . | . | . | chr4 | 108891993 | 108908067 | 6.16979 | 4.57777 | 3.97619 |
| chr5 | 108902382 | 108902773 | 1000 | Intron (ENSMUST00000106650.3 | 4 | 108890290 | 108941188 | 50899 | 1 | ENSMUST00 | 12093 | Rab3b | ENSMUSG0C | true | . | . | . | . | . | . | chr4 | 108891993 | 108908067 | 6.16979 | 4.57777 | 3.97619 |
| chr5 | 108905828 | 108906255 | 1000 | Intron (ENSMUST00000106650.3 | 4 | 108890290 | 108941188 | 50899 | 1 | ENSMUST00 | 15539 | Rab3b | ENSMUSG0C | true | . | . | . | . | . | . | chr4 | 108891993 | 108908067 | 6.16979 | 4.57777 | 3.97619 |
| chr6 | 29211946 | 29212491 | 1000 | Promoter (<=1kb) | 6 | 29200437 | 29212249 | 11813 | 2 | ENSMUST00 | 0 | Impdh1 | ENSMUSG0C | false | chr6 | 29211215 | 29212626 | 26.69297 | 77.60641 | 74.495 | chr6 | 29210289 | 29212833 | 3.63297 | 3.34578 | 1.93407 |
| chr17 | 35699934 | 35701083 | 1000 | Promoter (<=1kb) | 17 | 35681567 | 35700610 | 19044 | 2 | ENSMUST00 | 0 | Ddr1 | ENSMUSG0C | false | chr17 | 35699302 | 35701573 | 21.75525 | 76.8157 | 73.7174 | chr17 | 35696430 | 35706756 | 5.35332 | 4.19448 | 3.30476 |
| chr5 | 35721776 | 35722151 | 1000 | Distal Intergenic | 17 | 35690059 | 35704621 | 14563 | 2 | ENSMUST00 | -17156 | Ddr1 | ENSMUSG0C | false | . | . | . | . | . | . | chr17 | 35719181 | 35723461 | 7.09118 | 5.05512 | 4.74957 |
| chr5 | 35720273 | 35720914 | 1000 | Distal Intergenic | 17 | 35690059 | 35704621 | 14563 | 2 | ENSMUST00 | -15653 | Ddr1 | ENSMUSG0C | false | . | . | . | . | . | . | chr17 | 35719181 | 35723461 | 7.09118 | 5.05512 | 4.74957 |
| chr7 | 19319929 | 19320332 | 1000 | Distal Intergenic | 7 | 19302721 | 19310045 | 7325 | 2 | ENSMUST00 | -9885 | Fosb | ENSMUSG0C | false | chr7 | 19319673 | 19320088 | 13.78596 | 30.80405 | 28.32676 | chr7 | 19319605 | 19322418 | 5.10098 | 4.09515 | 3.1244 |
| chr7 | 19326448 | 19326820 | 1000 | Distal Intergenic | 7 | 19302721 | 19310045 | 7325 | 2 | ENSMUST00 | -16404 | Fosb | ENSMUSG0C | false | chr7 | 19326495 | 19327541 | 9.28568 | 20.18442 | 17.84641 | chr7 | 19326443 | 19328062 | 5.82901 | 4.29291 | 3.72172 |
| chr5 | 46661612 | 46661997 | 1000 | Distal Intergenic | 19 | 46668087 | 46669513 | 1427 | 2 | ENSMUST00 | 7516 | Cyp17a1 | ENSMUSG0C | false | . | . | . | . | . | . | chr19 | 46659100 | 46667582 | 5.73382 | 4.29769 | 3.62886 |
| chr5 | 70417973 | 70418393 | 1000 | Intron (ENSMUST00000076615.5 | 8 | 70406127 | 70431285 | 25159 | 2 | ENSMUST00 | 12892 | Crtc1 | ENSMUSG0C | false | . | . | . | . | . | . | chr8 | 70415799 | 70434143 | 6.2581 | 4.55457 | 4.06384 |
| chr8 | 70418823 | 70419275 | 1000 | Intron (ENSMUST00000076615.5 | 8 | 70406127 | 70431285 | 25159 | 2 | ENSMUST00 | 12010 | Crtc1 | ENSMUSG0C | false | chr8 | 70419225 | 70419465 | 5.08786 | 7.92609 | 5.7945 | chr8 | 70415799 | 70434143 | 6.2581 | 4.55457 | 4.06384 |
| chr8 | 70439176 | 70439855 | 1000 | Promoter (<=1kb) | 8 | 70382355 | 70439579 | 57225 | 2 | ENSMUST00 | 0 | Crtc1 | ENSMUSG0C | false | chr8 | 70438562 | 70440049 | 22.67078 | 69.62844 | 66.64306 | chr8 | 70435195 | 70440006 | 5.15203 | 3.96331 | 3.13378 |
| chr2 | 112492814 | 112493536 | 1000 | Promoter (<=1kb) | 2 | 112492964 | 112634573 | 141610 | 1 | ENSMUST00 | 0 | Aven | ENSMUSG0C | false | chr2 | 112492346 | 112494140 | 30.77379 | 96.20782 | 92.72798 | . | . | . | . | . | . |
| chr5 | 121323226 | 121323644 | 1000 | Promoter (<=1kb) | 1 | 121307237 | 121323191 | 15955 | 2 | ENSMUST00 | -36 | Insig2 | ENSMUSG0C | false | . | . | . | . | . | . | chr1 | 121323223 | 121330049 | 4.29763 | 3.74518 | 2.43067 |
| chr1 | 121327540 | 121328265 | 1000 | Promoter (<=1kb) | 1 | 121304353 | 121328024 | 23672 | 2 | ENSMUST00 | 0 | Insig2 | ENSMUSG0C | false | chr1 | 121326874 | 121328605 | 36.04672 | 129.50026 | 125.02953 | chr1 | 121323223 | 121330049 | 4.29763 | 3.74518 | 2.43067 |
| chr5 | 116371680 | 116372029 | 1000 | Intron (ENSMUST00000106484.2 | 4 | 116322434 | 116405986 | 83553 | 2 | ENSMUST00 | 33957 | Mast2 | ENSMUSG0C | false | . | . | . | . | . | . | chr4 | 116370781 | 116372701 | 4.10369 | 3.57203 | 2.30662 |
| chr5 | 116411413 | 116411828 | 1000 | Intron (ENSMUST00000106484.2 | 4 | 116332944 | 116406807 | 73864 | 2 | ENSMUST00 | -4607 | Mast2 | ENSMUSG0C | false | . | . | . | . | . | . | chr4 | 116409347 | 116414289 | 5.31339 | 4.1986 | 3.25664 |
| chr11 | 90249088 | 90249800 | 1000 | Promoter (<=1kb) | 11 | 90249456 | 90268465 | 19010 | 1 | ENSMUST00 | 0 | Mmd | ENSMUSG0C | false | chr11 | 90249309 | 90250643 | 26.03857 | 105.07202 | 101.37238 | chr11 | 90249797 | 90256670 | 5.12591 | 4.07412 | 3.12837 |
| chr11 | 90417765 | 90418315 | 1000 | Distal Intergenic | 11 | 90336536 | 90390895 | 54360 | 2 | ENSMUST00 | -26871 | Hlf | ENSMUSG0C | false | chr11 | 90417641 | 90418483 | 11.60586 | 25.27969 | 22.87289 | chr11 | 90416888 | 90418766 | 5.13082 | 4.09616 | 3.16842 |
| chr16 | 36071301 | 36072016 | 1000 | Promoter (<=1kb) | 16 | 36043844 | 36071515 | 27672 | 2 | ENSMUST00 | 0 | Fam162a | ENSMUSG0C | false | chr16 | 36070577 | 36072663 | 32.25957 | 103.83389 | 100.16508 | . | . | . | . | . | . |
| chr5 | 91430308 | 91430736 | 1000 | Distal Intergenic | 13 | 91460283 | 91688866 | 228584 | 1 | ENSMUST00 | -29547 | Ssbp2 | ENSMUSG0C | true | . | . | . | . | . | . | chr13 | 91426307 | 91431680 | 4.15997 | 3.65164 | 2.33694 |
| chr5 | 91626861 | 91627231 | 1000 | Intron (ENSMUST00000156547.2 | 13 | 91670437 | 91691432 | 20996 | 1 | ENSMUST00 | -43206 | Ssbp2 | ENSMUSG0C | true | . | . | . | . | . | . | chr13 | 91624801 | 91629586 | 3.91658 | 3.53471 | 2.11541 |
| chr13 | 91460335 | 91462151 | 1000 | Promoter (<=1kb) | 13 | 91461050 | 91703429 | 242380 | 1 | ENSMUST00 | 0 | Ssbp2 | ENSMUSG0C | true | chr13 | 91460982 | 91462809 | 23.93078 | 87.47927 | 84.18233 | chr13 | 91458688 | 91460345 | 4.45753 | 3.74015 | 2.57891 |
| chr5 | 91448842 | 91449250 | 1000 | Distal Intergenic | 13 | 91460283 | 91688866 | 228584 | 1 | ENSMUST00 | -11033 | Ssbp2 | ENSMUSG0C | true | . | . | . | . | . | . | chr13 | 91448747 | 91450867 | 5.64427 | 4.25992 | 3.57976 |
| chr5 | 158347819 | 158348223 | 1000 | Intron (ENSMUST00000004133.8 | 1 | 158245271 | 158356260 | 110990 | 2 | ENSMUST00 | 8037 | Brinp2 | ENSMUSG0C | false | . | . | . | . | . | . | chr1 | 158347048 | 158348746 | 3.89142 | 3.60489 | 2.07461 |
| chr11 | 100970494 | 100970981 | 1000 | Promoter (<=1kb) | 11 | 100956733 | 100970887 | 14155 | 2 | ENSMUST00 | 0 | Ptrf | ENSMUSG0C | false | chr11 | 100968093 | 100971128 | 34.92026 | 124.22972 | 119.94103 | . | . | . | . | . | . |
| chr11 | 100968900 | 100969984 | 1000 | Promoter (<=1kb) | 11 | 100964407 | 100970551 | 6145 | 2 | ENSMUST00 | 567 | Ptrf | ENSMUSG0C | false | chr11 | 100968093 | 100971128 | 34.92026 | 124.22972 | 119.94103 | . | . | . | . | . | . |
| chr19 | 5689426 | 5690339 | 1000 | Promoter (<=1kb) | 19 | 5689131 | 5702862 | 13732 | 1 | ENSMUST00 | 296 | Map3k11 | ENSMUSG0C | false | chr19 | 5687926 | 5692029 | 32.27334 | 102.84337 | 99.20039 | . | . | . | . | . | . |
| chr7 | 27605600 | 27605981 | 1000 | Promoter (<=1kb) | 7 | 27605797 | 27628869 | 23073 | 1 | ENSMUST00 | 0 | Akt2 | ENSMUSG0C | false | chr7 | 27605855 | 27606384 | 8.76716 | 17.74342 | 15.43897 | chr7 | 27592472 | 27610994 | 7.14155 | 5.01543 | 4.77185 |
| chr5 | 27602909 | 27603238 | 1000 | Promoter (2-3kb) | 7 | 27605797 | 27628869 | 23073 | 1 | ENSMUST00 | -2559 | Akt2 | ENSMUSG0C | false | . | . | . | . | . | . | chr7 | 27592472 | 27610994 | 7.14155 | 5.01543 | 4.77185 |
| chr5 | 27607463 | 27607972 | 1000 | Promoter (<=1kb) | 7 | 27607748 | 27633246 | 25499 | 1 | ENSMUST00 | 0 | Akt2 | ENSMUSG0C | false | . | . | . | . | . | . | chr7 | 27592472 | 27610994 | 7.14155 | 5.01543 | 4.77185 |
| chr16 | 4642089 | 4642736 | 1000 | Promoter (2-3kb) | 16 | 4639989 | 4687292 | 47304 | 1 | ENSMUST00 | 2101 | Dnaja3 | ENSMUSG0C | false | chr16 | 4642600 | 4643276 | 9.44616 | 20.46285 | 18.12126 | chr16 | 4642609 | 4661657 | 7.56188 | 5.16905 | 5.12366 |
| chr16 | 4789532 | 4790129 | 1000 | Promoter (<=1kb) | 16 | 4765461 | 4789990 | 24530 | 2 | ENSMUST00 | 0 | Cdip1 | ENSMUSG0C | false | chr16 | 4789130 | 4790637 | 21.17422 | 79.20749 | 76.06625 | . | . | . | . | . | . |
| chr2 | 72285539 | 72286120 | 1000 | Promoter (<=1kb) | 2 | 72285637 | 72442610 | 156974 | 1 | ENSMUST00 | 0 | Zak | ENSMUSG0C | false | chr2 | 72285363 | 72286768 | 34.61343 | 111.03911 | 107.17758 | . | . | . | . | . | . |
| chr5 | 33223786 | 33224405 | 1000 | Intron (ENSMUST00000004208.5 | 2 | 33216098 | 33230124 | 14027 | 1 | ENSMUST00 | 7689 | Angptl2 | ENSMUSG0C | false | . | . | . | . | . | . | chr2 | 33217442 | 33227221 | 7.68866 | 5.14665 | 5.24566 |
| chr6 | 124755930 | 124757123 | 1000 | Promoter (<=1kb) | 6 | 124739189 | 124756021 | 16833 | 2 | ENSMUST00 | 0 | Atn1 | ENSMUSG0C | false | chr6 | 124755579 | 124757546 | 29.25995 | 87.70835 | 84.40896 | chr6 | 124757014 | 124759681 | 6.18502 | 4.6504 | 3.98377 |
| chr1 | 80339908 | 80341255 | 1000 | Promoter (<=1kb) | 1 | 80264923 | 80340480 | 75558 | 2 | ENSMUST00 | 0 | Cul3 | ENSMUSG0C | false | chr1 | 80338787 | 80341787 | 27.03504 | 88.5911 | 85.27629 | . | . | . | . | . | . |
| chr7 | 4776823 | 4777394 | 1000 | Promoter (<=1kb) | 7 | 4773055 | 4778141 | 5087 | 2 | ENSMUST00 | 747 | Il11 | ENSMUSG0C | false | chr7 | 4776455 | 4777620 | 26.65286 | 74.78688 | 71.72144 | chr7 | 4776575 | 4777871 | 3.74963 | 3.37318 | 1.99283 |
| chr5 | 136796856 | 136797222 | 1000 | Intron (ENSMUST00000057497.7 | 5 | 136757663 | 136773046 | 15384 | 2 | ENSMUST00 | -23811 | Col26a1 | ENSMUSG0C | false | . | . | . | . | . | . | . | . | . | . | . | . |
| chr5 | 136886869 | 136887498 | 1000 | Distal Intergenic | 5 | 136757489 | 136883209 | 125721 | 2 | ENSMUST00 | -3661 | Col26a1 | ENSMUSG0C | false | . | . | . | . | . | . | chr5 | 136885321 | 136889075 | 4.822 | 3.84455 | 2.90361 |
| chr1 | 119504225 | 119505124 | 1000 | Promoter (<=1kb) | 1 | 119470305 | 119504794 | 34490 | 2 | ENSMUST00 | 0 | Ralb | ENSMUSG0C | false | chr1 | 119503795 | 119504823 | 22.37683 | 79.83291 | 76.68538 | chr1 | 119503231 | 119506162 | 4.18156 | 3.70473 | 2.32105 |
| chr7 | 97082629 | 97083178 | 1000 | Promoter (<=1kb) | 7 | 97081751 | 97308940 | 227190 | 1 | ENSMUST00 | 879 | Gab2 | ENSMUSG0C | true | chr7 | 97081213 | 97083356 | 37.54867 | 136.59039 | 131.84451 | chr7 | 97082937 | 97085999 | 4.33316 | 3.67721 | 2.47772 |
| chr5 | 97129007 | 97129324 | 1000 | Intron (ENSMUST00000004622.5 | 7 | 97081751 | 97308940 | 227190 | 1 | ENSMUST00 | 47257 | Gab2 | ENSMUSG0C | true | . | . | . | . | . | . | chr7 | 97125901 | 97131259 | 4.77256 | 3.97515 | 2.81568 |
| chr5 | 113908038 | 113909013 | 1000 | Promoter (<=1kb) | 5 | 113842436 | 113908705 | 66270 | 2 | ENSMUST00 | 0 | Coro1c | ENSMUSG0C | false | chr5 | 113907486 | 113909403 | 41.69731 | 136.21887 | 131.48979 | . | . | . | . | . | . |
| chr8 | 3394753 | 3395475 | 1000 | Promoter (1-2kb) | 8 | 3393073 | 3429639 | 36567 | 1 | ENSMUST00 | 1681 | Arhgef18 | ENSMUSG0C | true | chr8 | 3392521 | 3397432 | 33.45073 | 115.2328 | 111.2408 | chr8 | 3394895 | 3435896 | 6.08039 | 4.58876 | 3.89109 |
| chr8 | 3400581 | 3400979 | 1000 | Intron (ENSMUST00000131773.2 | 8 | 3393073 | 3429639 | 36567 | 1 | ENSMUST00 | 7509 | Arhgef18 | ENSMUSG0C | true | chr8 | 3400828 | 3402931 | 12.40753 | 28.46989 | 26.02371 | chr8 | 3394895 | 3435896 | 6.08039 | 4.58876 | 3.89109 |

| | | | | | | | | | | | | | | | | | | | | | | | | | | |
|---|---|---|---|---|---|---|---|---|---|---|---|---|---|---|---|---|---|---|---|---|---|---|---|---|---|---|
| chr5 | 3431018 | 3431477 | 1000 | Intron (ENSMUST00000131773.2 | 8 | 3437011 | 3456600 | 19590 | 1 | ENSMUST00 | -5534 | Arhgef18 | ENSMUSG0C | true | . | . | . | . | . | . | chr8 | 3394895 | 3435896 | 6.08039 | 4.58876 | 3.89109 |
| chr3 | 142881187 | 142882502 | 1000 | Promoter (<=1kb) | 3 | 142790902 | 142882004 | 91103 | 2 | ENSMUST00 | 0 | Pkn2 | ENSMUSG0C | false | chr3 | 142880321 | 142883208 | 30.90292 | 132.31535 | 127.74332 | . | . | . | . | . | . |
| chr5 | 57831347 | 57831930 | 1000 | Promoter (2-3kb) | 9 | 57790513 | 57834236 | 43724 | 2 | ENSMUST00 | 2306 | Arid3b | ENSMUSG0C | false | . | . | . | . | . | . | chr9 | 57827202 | 57836152 | 5.62883 | 4.39478 | 3.50932 |
| chr9 | 57836529 | 57837023 | 1000 | Promoter (<=1kb) | 9 | 57790353 | 57836706 | 46354 | 2 | ENSMUST00 | 0 | Arid3b | ENSMUSG0C | false | chr9 | 57835728 | 57836991 | 37.30079 | 124.19571 | 119.91465 | . | . | . | . | . | . |
| chr5 | 137045843 | 137046287 | 1000 | Promoter (<=1kb) | 5 | 137034993 | 137046135 | 11143 | 2 | ENSMUST00 | 0 | Ap1s1 | ENSMUSG0C | false | chr5 | 137045243 | 137046377 | 24.70746 | 75.05766 | 71.98973 | chr5 | 137044809 | 137046626 | 3.62664 | 3.44087 | 1.85822 |
| chr3 | 87972219 | 87972877 | 1000 | Promoter (1-2kb) | 3 | 87971129 | 87980451 | 9323 | 1 | ENSMUST00 | 1091 | Nes | ENSMUSG0C | false | chr3 | 87970233 | 87975133 | 30.04451 | 127.41808 | 123.02824 | . | . | . | . | . | . |
| chr5 | 87982714 | 87983616 | 1000 | Distal Intergenic | 3 | 87971129 | 87980451 | 9323 | 1 | ENSMUST00 | 11586 | Nes | ENSMUSG0C | false | . | . | . | . | . | . | chr3 | 87973300 | 87988519 | 8.41459 | 5.52768 | 5.81173 |
| chr3 | 87970854 | 87971894 | 1000 | Promoter (<=1kb) | 3 | 87971078 | 87973391 | 2314 | 1 | ENSMUST00 | 0 | Nes | ENSMUSG0C | false | chr3 | 87970233 | 87975133 | 30.04451 | 127.41808 | 123.02824 | . | . | . | . | . | . |
| chr10 | 81167566 | 81168035 | 1000 | Promoter (<=1kb) | 10 | 81153266 | 81167923 | 14658 | 2 | ENSMUST00 | 0 | Pias4 | ENSMUSG0C | false | chr10 | 81166899 | 81167857 | 29.28796 | 98.53321 | 94.99628 | chr10 | 81159044 | 81168639 | 4.74134 | 3.95933 | 2.79323 |
| chr5 | 5463935 | 5464243 | 1000 | Distal Intergenic | 17 | 5432295 | 5440259 | 7965 | 2 | ENSMUST00 | -23677 | Tmem242 | ENSMUSG0C | false | . | . | . | . | . | . | chr17 | 5463779 | 5464999 | 3.17784 | 3.09065 | 1.53401 |
| chr5 | 136019837 | 136020360 | 1000 | Promoter (<=1kb) | 5 | 136019558 | 136032864 | 13307 | 1 | ENSMUST00 | 280 | Dtx2 | ENSMUSG0C | false | . | . | . | . | . | . | chr5 | 136014251 | 136022977 | 5.92015 | 4.36442 | 3.79999 |
| chr5 | 136014873 | 136015346 | 1000 | Intron (ENSMUST00000111142.3 | 5 | 136019558 | 136032864 | 13307 | 1 | ENSMUST00 | -4212 | Dtx2 | ENSMUSG0C | false | . | . | . | . | . | . | chr5 | 136014251 | 136022977 | 5.92015 | 4.36442 | 3.79999 |
| chr6 | 51469871 | 51471324 | 1000 | Promoter (<=1kb) | 6 | 51460434 | 51469894 | 9461 | 2 | ENSMUST00 | 0 | Hnrnpa2b1 | ENSMUSG0C | false | chr6 | 51467769 | 51469887 | 37.34335 | 127.50397 | 123.10989 | chr6 | 51471223 | 51473362 | 3.56396 | 3.31503 | 1.85003 |
| chr2 | 102658667 | 102659114 | 1000 | Promoter (<=1kb) | 2 | 102658659 | 102790784 | 132126 | 1 | ENSMUST00 | 9 | Slc1a2 | ENSMUSG0C | false | chr2 | 102658165 | 102659605 | 12.30394 | 36.76666 | 34.21749 | chr2 | 102657794 | 102659588 | 4.74898 | 3.79819 | 2.83875 |
| chr5 | 118441576 | 118441927 | 1000 | Promoter (2-3kb) | 2 | 118438842 | 118452290 | 13449 | 1 | ENSMUST00 | 2735 | Eif2ak4 | ENSMUSG0C | false | . | . | . | . | . | . | chr2 | 118441009 | 118442438 | 4.30326 | 3.5596 | 2.4599 |
| chr15 | 66968257 | 66969004 | 1000 | Promoter (<=1kb) | 15 | 66943136 | 66967940 | 24805 | 2 | ENSMUST00 | -318 | Ndrg1 | ENSMUSG0C | false | chr15 | 66968167 | 66970335 | 29.58769 | 108.11006 | 104.32592 | chr15 | 66959859 | 66970307 | 5.98016 | 4.46549 | 3.81257 |
| chr5 | 66962257 | 66962650 | 1000 | Intron (ENSMUST00000005256.7 | 15 | 66943079 | 66967925 | 24847 | 2 | ENSMUST00 | 5275 | Ndrg1 | ENSMUSG0C | false | . | . | . | . | . | . | chr15 | 66959859 | 66970307 | 5.98016 | 4.46549 | 3.81257 |
| chr15 | 66969569 | 66970237 | 1000 | Promoter (<=1kb) | 15 | 66929321 | 66969640 | 40320 | 2 | ENSMUST00 | 0 | Ndrg1 | ENSMUSG0C | false | chr15 | 66968167 | 66970335 | 29.58769 | 108.11006 | 104.32592 | chr15 | 66959859 | 66970307 | 5.98016 | 4.46549 | 3.81257 |
| chr11 | 69758182 | 69759060 | 1000 | Promoter (<=1kb) | 11 | 69733997 | 69758637 | 24641 | 2 | ENSMUST00 | 0 | Polr2a | ENSMUSG0C | false | chr11 | 69758740 | 69759758 | 16.32424 | 48.10018 | 45.4107 | . | . | . | . | . | . |
| chr6 | 54594954 | 54595551 | 1000 | Promoter (<=1kb) | 6 | 54595111 | 54645839 | 50729 | 1 | ENSMUST00 | 0 | Plekha8 | ENSMUSG0C | false | chr6 | 54594686 | 54596059 | 33.5288 | 108.10788 | 104.32592 | chr6 | 54590683 | 54597216 | 5.00553 | 4.10098 | 3.01554 |
| chr13 | 58215143 | 58215863 | 1000 | Promoter (<=1kb) | 13 | 58176156 | 58215653 | 39498 | 2 | ENSMUST00 | 0 | Ubqln1 | ENSMUSG0C | false | chr13 | 58214535 | 58216489 | 42.33011 | 146.25195 | 141.10883 | . | . | . | . | . | . |
| chr5 | 55168923 | 55169534 | 1000 | Distal Intergenic | 13 | 55156117 | 55156904 | 788 | 1 | ENSMUST00 | 12807 | Fgfr4 | ENSMUSG0C | false | . | . | . | . | . | . | chr13 | 55161802 | 55178663 | 6.64938 | 4.78575 | 4.37276 |
| chr5 | 77882549 | 77883219 | 1000 | Distal Intergenic | 15 | 77915047 | 77915700 | 654 | 2 | ENSMUST00 | 32481 | Txn2 | ENSMUSG0C | false | . | . | . | . | . | . | chr15 | 77880695 | 77893977 | 6.24651 | 4.66367 | 4.02172 |
| chr5 | 77886473 | 77886962 | 1000 | Distal Intergenic | 15 | 77915047 | 77915700 | 654 | 2 | ENSMUST00 | 28738 | Txn2 | ENSMUSG0C | false | . | . | . | . | . | . | chr15 | 77880695 | 77893977 | 6.24651 | 4.66367 | 4.02172 |
| chr5 | 77920533 | 77921120 | 1000 | Intron (ENSMUST00000173631.2 | 15 | 77915047 | 77915700 | 654 | 2 | ENSMUST00 | -4834 | Txn2 | ENSMUSG0C | false | . | . | . | . | . | . | chr15 | 77919576 | 77921027 | 3.49679 | 3.35195 | 1.76156 |
| chr17 | 88064468 | 88066595 | 1000 | Promoter (<=1kb) | 17 | 87990859 | 88065028 | 74170 | 2 | ENSMUST00 | 0 | Fbxo11 | ENSMUSG0C | false | chr17 | 88063293 | 88066770 | 36.04346 | 118.78738 | 114.68494 | . | . | . | . | . | . |
| chr5 | 135149118 | 135149867 | 1000 | Promoter (<=1kb) | 5 | 135149657 | 135163873 | 14217 | 1 | ENSMUST00 | 0 | Tbl2 | ENSMUSG0C | false | chr5 | 135148518 | 135149494 | 22.78188 | 63.69425 | 60.796 | . | . | . | . | . | . |
| chr13 | 13438235 | 13439232 | 1000 | Promoter (<=1kb) | 13 | 13437602 | 13512275 | 74674 | 1 | ENSMUST00 | 634 | Nid1 | ENSMUSG0C | false | chr13 | 13438186 | 13439077 | 16.88215 | 57.7818 | 54.96725 | chr13 | 13437021 | 13441430 | 4.37956 | 3.67113 | 2.54828 |
| chr13 | 13437256 | 13437790 | 1000 | Promoter (<=1kb) | 13 | 13437602 | 13512275 | 74674 | 1 | ENSMUST00 | 0 | Nid1 | ENSMUSG0C | false | chr13 | 13437424 | 13437964 | 14.58106 | 49.93175 | 47.21736 | chr13 | 13437021 | 13441430 | 4.37956 | 3.67113 | 2.54828 |
| chr5 | 75089335 | 75089767 | 1000 | Distal Intergenic | 8 | 75093591 | 75100596 | 7006 | 1 | ENSMUST00 | -3824 | Hmox1 | ENSMUSG0C | false | . | . | . | . | . | . | chr8 | 75087664 | 75090479 | 5.67279 | 4.43567 | 3.54346 |
| chr11 | 59664868 | 59665414 | 1000 | Promoter (2-3kb) | 11 | 59662507 | 59780854 | 118348 | 1 | ENSMUST00 | 2362 | Mprip | ENSMUSG0C | false | chr11 | 59661712 | 59664872 | 16.78262 | 53.63071 | 50.86883 | chr11 | 59663434 | 59690186 | 5.94708 | 4.45384 | 3.79525 |
| chr5 | 59670915 | 59671298 | 1000 | Intron (ENSMUST00000138234.1 | 11 | 59662507 | 59780854 | 118348 | 1 | ENSMUST00 | 8409 | Mprip | ENSMUSG0C | false | . | . | . | . | . | . | chr11 | 59663434 | 59690186 | 5.94708 | 4.45384 | 3.79525 |
| chr5 | 59707982 | 59708657 | 1000 | Intron (ENSMUST00000138234.1 | 11 | 59727681 | 59776602 | 48922 | 1 | ENSMUST00 | -19024 | Mprip | ENSMUSG0C | false | . | . | . | . | . | . | chr11 | 59704908 | 59728229 | 5.43634 | 4.16695 | 3.39654 |
| chr7 | 25281789 | 25282940 | 1000 | Promoter (<=1kb) | 7 | 25282179 | 25294145 | 11967 | 1 | ENSMUST00 | 0 | Cic | ENSMUSG0C | false | chr7 | 25281477 | 25282003 | 8.26071 | 16.87976 | 14.58833 | chr7 | 25269284 | 25281823 | 5.87867 | 4.36785 | 3.75349 |
| chr8 | 83972370 | 83973307 | 1000 | Promoter (<=1kb) | 8 | 83972978 | 83996445 | 23468 | 1 | ENSMUST00 | 0 | Prkaca | ENSMUSG0C | false | chr8 | 83972900 | 83974112 | 30.94804 | 114.76607 | 110.79114 | . | . | . | . | . | . |
| chr7 | 67950733 | 67952753 | 1000 | Promoter (<=1kb) | 7 | 67952859 | 68226780 | 273922 | 1 | ENSMUST00 | -106 | Igf1r | ENSMUSG0C | false | chr7 | 67950399 | 67954090 | 38.55812 | 129.64772 | 125.16568 | chr7 | 67950130 | 67950802 | 3.26575 | 3.17849 | 1.56514 |
| chr5 | 68058198 | 68058510 | 1000 | Intron (ENSMUST00000005671.8 | 7 | 67952859 | 68226780 | 273922 | 1 | ENSMUST00 | 105340 | Igf1r | ENSMUSG0C | false | . | . | . | . | . | . | chr7 | 68056560 | 68067870 | 6.6301 | 4.72849 | 4.37098 |
| chr5 | 110988038 | 110988671 | 1000 | Distal Intergenic | 7 | 110868270 | 110982461 | 114192 | 2 | ENSMUST00 | -5578 | Mrvi1 | ENSMUSG0C | false | . | . | . | . | . | . | chr7 | 110982186 | 110991297 | 5.7251 | 4.32045 | 3.62307 |
| chr5 | 110983640 | 110984029 | 1000 | Promoter (1-2kb) | 7 | 110868270 | 110982461 | 114192 | 2 | ENSMUST00 | -1180 | Mrvi1 | ENSMUSG0C | false | . | . | . | . | . | . | chr7 | 110982186 | 110991297 | 5.7251 | 4.32045 | 3.62307 |
| chr7 | 80993132 | 80994157 | 1000 | Promoter (<=1kb) | 7 | 80993681 | 81045164 | 51484 | 1 | ENSMUST00 | 0 | Zfp592 | ENSMUSG0C | false | chr7 | 80992503 | 80994889 | 39.66568 | 134.87851 | 130.20418 | . | . | . | . | . | . |
| chr5 | 54789412 | 54790070 | 1000 | Intron (ENSMUST00000005798.8 | 12 | 54746349 | 54795703 | 49355 | 2 | ENSMUST00 | 5633 | Snx6 | ENSMUSG0C | false | . | . | . | . | . | . | chr12 | 54786094 | 54794667 | 4.88503 | 3.75256 | 2.94543 |
| chr5 | 54805930 | 54806416 | 1000 | Distal Intergenic | 12 | 54746349 | 54795703 | 49355 | 2 | ENSMUST00 | -10228 | Snx6 | ENSMUSG0C | false | . | . | . | . | . | . | chr12 | 54804705 | 54806959 | 4.74004 | 3.7775 | 2.82865 |
| chr1 | 171222475 | 171222831 | 1000 | Promoter (<=1kb) | 1 | 171217802 | 171222514 | 4713 | 2 | ENSMUST00 | 0 | Tomm40l | ENSMUSG0C | false | chr1 | 171222739 | 171223191 | 14.66885 | 35.63733 | 33.10225 | . | . | . | . | . | . |
| chr10 | 128303090 | 128303778 | 1000 | Promoter (<=1kb) | 10 | 128303335 | 128321358 | 18024 | 1 | ENSMUST00 | 0 | Pan2 | ENSMUSG0C | false | chr10 | 128302721 | 128304467 | 22.32483 | 64.62918 | 61.71925 | . | . | . | . | . | . |
| chr10 | 128337575 | 128338249 | 1000 | Promoter (<=1kb) | 10 | 128337832 | 128362479 | 24648 | 1 | ENSMUST00 | 0 | Cs | ENSMUSG0C | false | chr10 | 128337679 | 128338932 | 34.81759 | 111.39526 | 107.52535 | . | . | . | . | . | . |
| chr5 | 110709633 | 110710026 | 1000 | Distal Intergenic | 7 | 110768206 | 110811919 | 43714 | 1 | ENSMUST00 | -58180 | Ampd3 | ENSMUSG0C | false | . | . | . | . | . | . | . | . | . | . | . | . |
| chr8 | 105635955 | 105637026 | 1000 | Promoter (<=1kb) | 8 | 105636568 | 105681647 | 45080 | 1 | ENSMUST00 | 0 | Ctcf | ENSMUSG0C | false | chr8 | 105635762 | 105638088 | 33.69596 | 103.2888 | 99.6348 | . | . | . | . | . | . |
| chr5 | 165837235 | 165838270 | 1000 | Intron (ENSMUST00000161971.2 | 1 | 165858216 | 165861555 | 3340 | 1 | ENSMUST00 | -19946 | Cd247 | ENSMUSG0C | false | . | . | . | . | . | . | chr1 | 165833325 | 165839586 | 4.76919 | 3.8939 | 2.83602 |
| chr2 | 122765073 | 122765569 | 1000 | Promoter (<=1kb) | 2 | 122765237 | 122809569 | 44333 | 1 | ENSMUST00 | 0 | Sqrdl | ENSMUSG0C | false | chr2 | 122764897 | 122766045 | 28.08619 | 80.9371 | 77.76572 | chr2 | 122764587 | 122766206 | 3.04729 | 3.04371 | 1.42073 |
| chr3 | 138489036 | 138489782 | 1000 | Promoter (<=1kb) | 3 | 138458960 | 138489382 | 30423 | 2 | ENSMUST00 | 0 | Metap1 | ENSMUSG0C | false | chr3 | 138488534 | 138490299 | 37.30082 | 116.74746 | 112.71554 | . | . | . | . | . | . |
| chr1 | 13372308 | 13374686 | 1000 | Promoter (<=1kb) | 1 | 13139105 | 13372439 | 233335 | 2 | ENSMUST00 | 0 | Ncoa2 | ENSMUSG0C | false | chr1 | 13371872 | 13374991 | 29.92778 | 109.76675 | 105.9399 | chr1 | 13374358 | 13375928 | 3.38584 | 3.27552 | 1.71874 |
| chr6 | 92091270 | 92092496 | 1000 | Promoter (<=1kb) | 6 | 92091390 | 92173057 | 81668 | 1 | ENSMUST00 | 0 | Nr2c2 | ENSMUSG0C | false | chr6 | 92090752 | 92093151 | 37.50639 | 125.63292 | 121.30651 | . | . | . | . | . | . |
| chr16 | 21210798 | 21211346 | 1000 | Promoter (<=1kb) | 16 | 21211095 | 21222967 | 11873 | 1 | ENSMUST00 | 0 | Ephb3 | ENSMUSG0C | false | chr16 | 21210692 | 21211697 | 12.99241 | 30.10301 | 27.6364 | chr16 | 21210154 | 21211833 | 6.28753 | 4.67942 | 4.06639 |
| chr16 | 3872151 | 3872598 | 1000 | Promoter (<=1kb) | 16 | 3872375 | 3900811 | 28437 | 1 | ENSMUST00 | 0 | Naa60 | ENSMUSG0C | false | chr16 | 3871717 | 3873084 | 26.57451 | 76.48061 | 73.3894 | chr16 | 3870833 | 3873104 | 4.39932 | 3.74019 | 2.56894 |
| chr5 | 4285712 | 4286124 | 1000 | Promoter (2-3kb) | 19 | 4279466 | 4283137 | 3672 | 2 | ENSMUST00 | -2576 | Ankrd13d | ENSMUSG0C | false | . | . | . | . | . | . | chr19 | 4284895 | 4286987 | 6.47429 | 4.47141 | 4.25312 |
| chr11 | 96075745 | 96076198 | 1000 | Promoter (<=1kb) | 11 | 96068852 | 96075670 | 6819 | 2 | ENSMUST00 | -76 | Atp5g1 | ENSMUSG0C | false | chr11 | 96076096 | 96076468 | 17.85986 | 47.76698 | 45.0813 | . | . | . | . | . | . |
| chr11 | 75630669 | 75631064 | 1000 | Promoter (<=1kb) | 11 | 75630988 | 75648871 | 17884 | 1 | ENSMUST00 | 0 | Inpp5k | ENSMUSG0C | false | chr11 | 75630258 | 75632213 | 25.57207 | 77.50074 | 74.3909 | . | . | . | . | . | . |
| chr5 | 131169091 | 131169433 | 1000 | Intron (ENSMUST00000124096.2 | 7 | 131174402 | 131250945 | 76544 | 1 | ENSMUST00 | -4969 | 5430419D17 | ENSMUSG0C | false | . | . | . | . | . | . | chr7 | 131168859 | 131170525 | 3.42235 | 3.26063 | 1.71507 |
| chr5 | 130960003 | 130960321 | 1000 | Intron (ENSMUST00000150717.2 | 7 | 130943489 | 130985660 | 42172 | 1 | ENSMUST00 | 16515 | Htra1 | ENSMUSG0C | false | . | . | . | . | . | . | chr7 | 130956209 | 130988327 | 6.29742 | 4.64648 | 4.07297 |
| chr5 | 130964657 | 130965068 | 1000 | Intron (ENSMUST00000150717.2 | 7 | 130981670 | 130982443 | 774 | 1 | ENSMUST00 | -16602 | Htra1 | ENSMUSG0C | false | . | . | . | . | . | . | chr7 | 130956209 | 130988327 | 6.29742 | 4.64648 | 4.07297 |
| chr5 | 130982362 | 130982802 | 1000 | Promoter (<=1kb) | 7 | 130982730 | 130985396 | 2667 | 1 | ENSMUST00 | 0 | Htra1 | ENSMUSG0C | false | . | . | . | . | . | . | chr7 | 130956209 | 130988327 | 6.29742 | 4.64648 | 4.07297 |
| chr8 | 69088434 | 69088921 | 1000 | Promoter (<=1kb) | 8 | 69088646 | 69113711 | 25066 | 1 | ENSMUST00 | 0 | Atp6v1b2 | ENSMUSG0C | false | chr8 | 69088367 | 69089628 | 35.15467 | 116.16882 | 112.14974 | chr8 | 69088166 | 69088510 | 3.77816 | 3.53524 | 2.00135 |
| chr1 | 74284607 | 74285238 | 1000 | Promoter (<=1kb) | 1 | 74279840 | 74284738 | 4899 | 2 | ENSMUST00 | 0 | Aamp | ENSMUSG0C | false | chr1 | 74283791 | 74284772 | 31.24818 | 92.1069 | 88.71768 | . | . | . | . | . | . |
| chr1 | 74303734 | 74304390 | 1000 | Promoter (<=1kb) | 1 | 74288251 | 74304351 | 16101 | 2 | ENSMUST00 | 0 | Tmbim1 | ENSMUSG0C | false | chr1 | 74302782 | 74304983 | 33.62711 | 105.2041 | 101.50168 | . | . | . | . | . | . |
| chr5 | 74249179 | 74249937 | 1000 | Intron (ENSMUST00000113820.3 | 1 | 74236550 | 74267988 | 31439 | 1 | ENSMUST00 | 12630 | Arpc2 | ENSMUSG0C | false | . | . | . | . | . | . | chr1 | 74237722 | 74252941 | 5.81759 | 4.46841 | 3.67748 |
| chr5 | 74240445 | 74240764 | 1000 | Intron (ENSMUST00000113820.3 | 1 | 74236550 | 74267988 | 31439 | 1 | ENSMUST00 | 3896 | Arpc2 | ENSMUSG0C | false | . | . | . | . | . | . | chr1 | 74237722 | 74252941 | 5.81759 | 4.46841 | 3.67748 |
| chr10 | 75667232 | 75668210 | 1000 | Intron (ENSMUST00000001712.6 | 10 | 75636619 | 75644008 | 7390 | 2 | ENSMUST00 | -23225 | Susd2 | ENSMUSG0C | false | chr10 | 75666959 | 75668360 | 30.28657 | 98.33401 | 94.80609 | chr10 | 75666654 | 75667316 | 4.433 | 3.80176 | 2.58418 |
| chr5 | 75677942 | 75678248 | 1000 | Intron (ENSMUST00000001712.6 | 10 | 75636619 | 75644008 | 7390 | 2 | ENSMUST00 | -33935 | Susd2 | ENSMUSG0C | false | . | . | . | . | . | . | chr10 | 75675682 | 75679496 | 5.67318 | 4.21243 | 3.60466 |
| chr5 | 75587905 | 75588803 | 1000 | Promoter (<=1kb) | 10 | 75589340 | 75617200 | 27861 | 1 | ENSMUST00 | -537 | Ggt5 | ENSMUSG0C | false | . | . | . | . | . | . | chr10 | 75585895 | 75593802 | 6.23311 | 4.61744 | 4.02118 |
| chr12 | 113142500 | 113142964 | 1000 | Promoter (2-3kb) | 12 | 113140236 | 113145504 | 5269 | 1 | ENSMUST00 | 2265 | Crip2 | ENSMUSG0C | false | chr12 | 113141417 | 113143358 | 10.95946 | 24.18275 | 21.79051 | chr12 | 113139363 | 113149705 | 6.62613 | 4.83812 | 4.33223 |
| chr12 | 113146459 | 113147033 | 1000 | Distal Intergenic | 12 | 113152012 | 113153877 | 1866 | 1 | ENSMUST00 | -4979 | Crip1 | ENSMUSG0C | false | chr12 | 113146275 | 113147394 | 12.01557 | 29.04976 | 26.5957 | chr12 | 113139363 | 113149705 | 6.62613 | 4.83812 | 4.33223 |
| chr8 | 122677841 | 122678906 | 1000 | Promoter (<=1kb) | 8 | 122629463 | 122678072 | 48610 | 2 | ENSMUST00 | 0 | Cbfa2t3 | ENSMUSG0C | false | chr8 | 122676506 | 122679041 | 35.62435 | 116.99458 | 112.94973 | chr8 | 122678840 | 122679099 | 4.38206 | 3.81687 | 2.54679 |
| chr5 | 122661066 | 122661603 | 1000 | Intron (ENSMUST00000127664.1 | 8 | 122639018 | 122646466 | 7449 | 2 | ENSMUST00 | -14601 | Cbfa2t3 | ENSMUSG0C | false | . | . | . | . | . | . | chr8 | 122648562 | 122676943 | 7.87697 | 5.30313 | 5.37575 |
| chr5 | 122665247 | 122665947 | 1000 | Intron (ENSMUST00000127664.1 | 8 | 122629463 | 122678072 | 48610 | 2 | ENSMUST00 | 12125 | Cbfa2t3 | ENSMUSG0C | false | . | . | . | . | . | . | chr8 | 122648562 | 122676943 | 7.87697 | 5.30313 | 5.37575 |
| chr5 | 122655750 | 122656686 | 1000 | Intron (ENSMUST00000127664.1 | 8 | 122639018 | 122646466 | 7449 | 2 | ENSMUST00 | -9285 | Cbfa2t3 | ENSMUSG0C | false | . | . | . | . | . | . | chr8 | 122648562 | 122676943 | 7.87697 | 5.30313 | 5.37575 |
| chr15 | 85205820 | 85206145 | 1000 | Promoter (<=1kb) | 15 | 85205949 | 85286535 | 80587 | 1 | ENSMUST00 | 0 | Fbln1 | ENSMUSG0C | false | chr15 | 85205804 | 85206354 | 13.0727 | 32.19727 | 29.70434 | chr15 | 85205970 | 85207117 | 4.68094 | 3.81377 | 2.77898 |
| chr5 | 171377889 | 171378329 | 1000 | Intron (ENSMUST00000111286.3 | 1 | 171370355 | 171388285 | 17931 | 1 | ENSMUST00 | 7535 | Pvrl4 | ENSMUSG0C | false | . | . | . | . | . | . | chr1 | 171369697 | 171385331 | 5.48452 | 4.28914 | 3.41912 |
| chr5 | 47228621 | 47229349 | 1000 | Promoter (<=1kb) | 19 | 47228843 | 47259440 | 30598 | 1 | ENSMUST00 | 0 | Neurl1a | ENSMUSG0C | false | . | . | . | . | . | . | chr19 | 47227865 | 47230866 | 5.65458 | 4.14361 | 3.57668 |
| chr5 | 141330045 | 141330347 | 1000 | Distal Intergenic | 4 | 141316913 | 141320901 | 3989 | 1 | ENSMUST00 | 13133 | Epha2 | ENSMUSG0C | false | . | . | . | . | . | . | chr4 | 141328684 | 141330883 | 5.01172 | 4.05296 | 3.02662 |
| chr5 | 167712142 | 167712485 | 1000 | Intron (ENSMUST00000141923.1 | 2 | 167691181 | 167697801 | 6621 | 1 | ENSMUST00 | 20962 | A530013C23 | ENSMUSG0C | false | . | . | . | . | . | . | chr2 | 167711295 | 167715576 | 5.46164 | 4.25954 | 3.37957 |
| chr2 | 71872886 | 71873682 | 1000 | Promoter (<=1kb) | 2 | 71873224 | 71903858 | 30635 | 1 | ENSMUST00 | 0 | Pdk1 | ENSMUSG0C | false | chr2 | 71872647 | 71874146 | 32.27147 | 102.84008 | 99.19724 | chr2 | 71872214 | 71874436 | 4.5115 | 3.86061 | 2.62983 |
| chr10 | 79840565 | 79841543 | 1000 | Distal Intergenic | 10 | 79854427 | 79864432 | 10006 | 1 | ENSMUST00 | -12884 | Ptbp1 | ENSMUSG0C | false | chr10 | 79841181 | 79841634 | 13.42322 | 30.19304 | 27.72564 | chr10 | 79831825 | 79846178 | 5.85392 | 4.4421 | 3.72163 |
| chr10 | 79853480 | 79853964 | 1000 | Promoter (<=1kb) | 10 | 79854427 | 79864432 | 10006 | 1 | ENSMUST00 | -463 | Ptbp1 | ENSMUSG0C | false | chr10 | 79852880 | 79854274 | 16.95696 | 43.908 | 41.27272 | chr10 | 79849905 | 79854106 | 7.60217 | 5.2407 | 5.14883 |
| chr10 | 79854166 | 79855429 | 1000 | Promoter (<=1kb) | 10 | 79854427 | 79864432 | 10006 | 1 | ENSMUST00 | 0 | Ptbp1 | ENSMUSG0C | false | chr10 | 79852880 | 79854274 | 16.95696 | 43.908 | 41.27272 | . | . | . | . | . | . |
| chr14 | 30715246 | 30716636 | 1000 | Promoter (<=1kb) | 14 | 30715615 | 30822713 | 107099 | 1 | ENSMUST00 | 0 | Sfmbt1 | ENSMUSG0C | false | chr14 | 30714921 | 30717340 | 30.08562 | 102.9821 | 99.33765 | chr14 | 30714740 | 30715478 | 4.38377 | 3.77971 | 2.50702 |
| chr5 | 30968368 | 30969003 | 1000 | Promoter (<=1kb) | 5 | 30968677 | 30977018 | 8342 | 1 | ENSMUST00 | 0 | Tcf23 | ENSMUSG0C | false | . | . | . | . | . | . | chr5 | 30968096 | 30970822 | 6.06707 | 4.59041 | 3.91406 |
| chr7 | 30459355 | 30459800 | 1000 | Promoter (<=1kb) | 7 | 30459725 | 30487223 | 27499 | 1 | ENSMUST00 | 0 | Nphs1 | ENSMUSG0C | true | chr7 | 30459332 | 30463950 | 22.44583 | 74.56482 | 71.50171 | . | . | . | . | . | . |
| chr5 | 30477699 | 30478433 | 1000 | Promoter (<=1kb) | 7 | 30477865 | 30486881 | 9017 | 1 | ENSMUST00 | 0 | Nphs1 | ENSMUSG0C | true | . | . | . | . | . | . | chr7 | 30473123 | 30482465 | 5.32795 | 4.21182 | 3.28024 |
| chr5 | 30441457 | 30441970 | 1000 | Exon (ENSMUST00000006828.7/ | 7 | 30434981 | 30445582 | 10602 | 2 | ENSMUST00 | 3612 | Aplp1 | ENSMUSG0C | false | . | . | . | . | . | . | chr7 | 30438998 | 30446114 | 5.82673 | 4.30727 | 3.72584 |
| chr7 | 30445157 | 30445797 | 1000 | Promoter (<=1kb) | 7 | 30434981 | 30445582 | 10602 | 2 | ENSMUST00 | 0 | Aplp1 | ENSMUSG0C | false | chr7 | 30444157 | 30445874 | 38.03245 | 121.19012 | 117.00504 | chr7 | 30438998 | 30446114 | 5.82673 | 4.30727 | 3.72584 |
| chr5 | 108600044 | 108600456 | 1000 | Promoter (2-3kb) | 9 | 108578826 | 108597600 | 18775 | 2 | ENSMUST00 | -2445 | P4htm | ENSMUSG0C | false | . | . | . | . | . | . | chr9 | 108596599 | 108602820 | 4.71757 | 3.8751 | 2.78944 |
| chr17 | 31564377 | 31565513 | 1000 | Promoter (<=1kb) | 17 | 31564749 | 31605481 | 40733 | 1 | ENSMUST00 | 0 | Pknox1 | ENSMUSG0C | false | chr17 | 31563690 | 31564428 | 19.72434 | 50.19016 | 47.47384 | chr17 | 31565498 | 31568376 | 3.44226 | 3.0595 | 1.77677 |
| chr10 | 127063196 | 127063663 | 1000 | Promoter (<=1kb) | 10 | 127063534 | 127067920 | 4387 | 1 | ENSMUST00 | 0 | Cdk4 | ENSMUSG0C | false | chr10 | 127063046 | 127063233 | 4.55837 | 6.25963 | 4.17328 | chr10 | 127062912 | 127063203 | 5.32116 | 4.30419 | 3.27153 |
| chr18 | 6240574 | 6240970 | 1000 | Promoter (<=1kb) | 18 | 6214569 | 6241470 | 26902 | 2 | ENSMUST00 | 500 | Kif5b | ENSMUSG0C | true | chr18 | 6239314 | 6242188 | 31.91637 | 110.38313 | 106.53261 | . | . | . | . | . | . |
| chr2 | 166154630 | 166155752 | 1000 | Promoter (<=1kb) | 2 | 166073089 | 166155663 | 82575 | 2 | ENSMUST00 | 0 | Sulf2 | ENSMUSG0C | false | chr2 | 166153906 | 166155964 | 23.28017 | 72.50986 | 69.48009 | chr2 | 166153340 | 166156360 | 4.42045 | 3.79293 | 2.51697 |
| chr5 | 166115732 | 166116570 | 1000 | Intron (ENSMUST00000109249.3 | 2 | 166074815 | 166078119 | 3305 | 2 | ENSMUST00 | -37614 | Sulf2 | ENSMUSG0C | false | . | . | . | . | . | . | chr2 | 166111370 | 166117038 | 4.57128 | 3.82866 | 2.65452 |
| chr2 | 166091757 | 166092253 | 1000 | Intron (ENSMUST00000109249.3 | 2 | 166074815 | 166078119 | 3305 | 2 | ENSMUST00 | -13639 | Sulf2 | ENSMUSG0C | false | chr2 | 166092157 | 166093106 | 5.905 | 11.42563 | 9.22457 | chr2 | 166085159 | 166097943 | 5.73745 | 4.40556 | 3.61152 |
| chr5 | 101221278 | 101221681 | 1000 | Intron (ENSMUST00000107284.2 | 11 | 101197389 | 101226426 | 29038 | 2 | ENSMUST00 | 4745 | Ezh1 | ENSMUSG0C | false | . | . | . | . | . | . | chr11 | 101218378 | 101226786 | 4.71132 | 3.90034 | 2.78728 |
| chr5 | 101189379 | 101190263 | 1000 | 3' UTR | 11 | 101193275 | 101194422 | 1148 | 2 | ENSMUST00 | 4159 | Ezh1 | ENSMUSG0C | false | . | . | . | . | . | . | chr11 | 101187692 | 101193026 | 5.10079 | 3.9423 | 3.13132 |
| chr11 | 101226078 | 101226632 | 1000 | Promoter (<=1kb) | 11 | 101191115 | 101226463 | 35349 | 2 | ENSMUST00 | 0 | Ezh1 | ENSMUSG0C | false | chr11 | 101225447 | 101226781 | 39.32978 | 125.57745 | 121.25191 | chr11 | 101218378 | 101226786 | 4.71132 | 3.90034 | 2.78728 |
| chr17 | 34933975 | 34934434 | 1000 | Promoter (2-3kb) | 17 | 34931863 | 34932765 | 903 | 1 | ENSMUST00 | 2113 | Neu1 | ENSMUSG0C | false | chr17 | 34933721 | 34934805 | 13.7551 | 32.05274 | 29.56132 | chr17 | 34930771 | 34935446 | 5.8546 | 4.41264 | 3.721 |
| chr5 | 3647246 | 3648054 | 1000 | Promoter (<=1kb) | 5 | 3639961 | 3647934 | 7974 | 2 | ENSMUST00 | 0 | Gatad1 | ENSMUSG0C | false | chr5 | 3646616 | 3648093 | 39.1345 | 126.3175 | 121.95772 | . | . | . | . | . | . |
| chr17 | 20945229 | 20945813 | 1000 | Promoter (<=1kb) | 17 | 20945311 | 20965916 | 20606 | 1 | ENSMUST00 | 0 | Ppp2r1a | ENSMUSG0C | false | chr17 | 20944553 | 20945245 | 15.08796 | 37.05233 | 34.49837 | . | . | . | . | . | . |
| chr5 | 71497898 | 71498344 | 1000 | 3' UTR | 8 | 71489459 | 71491411 | 1953 | 1 | ENSMUST00 | 8440 | Gtpbp3 | ENSMUSG0C | false | . | . | . | . | . | . | chr8 | 71495272 | 71500046 | 3.90309 | 3.28772 | 2.14928 |
| chr4 | 47352850 | 47353907 | 1000 | Promoter (<=1kb) | 4 | 47353222 | 47414931 | 61710 | 1 | ENSMUST00 | 0 | Tgfbr1 | ENSMUSG0C | false | chr4 | 47352092 | 47354375 | 44.11508 | 143.73827 | 138.70853 | chr4 | 47353809 | 47354428 | 4.27066 | 3.68633 | 2.401 |
| chr13 | 93303312 | 93304335 | 1000 | Promoter (<=1kb) | 13 | 93303757 | 93386744 | 82988 | 1 | ENSMUST00 | 0 | Homer1 | ENSMUSG0C | false | chr13 | 93303134 | 93306100 | 31.28813 | 116.44205 | 112.41405 | . | . | . | . | . | . |
| chr11 | 87404182 | 87404731 | 1000 | Promoter (<=1kb) | 11 | 87376645 | 87404523 | 27879 | 2 | ENSMUST00 | 0 | Rad51c | ENSMUSG0C | false | chr11 | 87403994 | 87405030 | 24.03115 | 75.67385 | 72.59326 | chr11 | 87404056 | 87405008 | 4.6104 | 3.65619 | 2.71132 |
| chr9 | 108306180 | 108307005 | 1000 | Promoter (<=1kb) | 9 | 108306205 | 108337939 | 31735 | 1 | ENSMUST00 | 0 | Rhoa | ENSMUSG0C | false | chr9 | 108306306 | 108308038 | 39.80159 | 147.3544 | 142.17488 | . | . | . | . | . | . |
| chr14 | 25457363 | 25458541 | 1000 | Promoter (<=1kb) | 14 | 25459185 | 25538492 | 79308 | 1 | ENSMUST00 | -644 | Zmiz1 | ENSMUSG0C | false | chr14 | 25457181 | 25461653 | 31.54088 | 108.67455 | 104.87239 | chr14 | 25456064 | 25458833 | 4.92503 | 4.03101 | 2.94873 |
| chr5 | 45151307 | 45151771 | 1000 | Promoter (2-3kb) | 7 | 45147883 | 45149179 | 1297 | 2 | ENSMUST00 | -2129 | Aldh16a1 | ENSMUSG0C | false | . | . | . | . | . | . | chr7 | 45148391 | 45152553 | 6.06164 | 4.57874 | 3.88674 |
| chr7 | 45053968 | 45054583 | 1000 | Promoter (<=1kb) | 7 | 45053895 | 45054880 | 986 | 2 | ENSMUST00 | 297 | Prrg2 | ENSMUSG0C | false | chr7 | 45052080 | 45054223 | 37.68163 | 117.40437 | 113.3432 | chr7 | 45053821 | 45055150 | 3.76211 | 3.51314 | 1.99435 |
| chr4 | 133753394 | 133754696 | 1000 | Promoter (<=1kb) | 4 | 133679008 | 133753611 | 74604 | 2 | ENSMUST00 | 0 | Arid1a | ENSMUSG0C | false | chr4 | 133751327 | 133754985 | 28.90751 | 104.81161 | 101.11721 | chr4 | 133754644 | 133758088 | 8.04793 | 5.46477 | 5.4798 |
| chrX | 10717133 | 10717561 | 1000 | Promoter (<=1kb) | 20 | 10717390 | 10719690 | 2301 | 1 | ENSMUST00 | 0 | Mid1ip1 | ENSMUSG0C | false | chrX | 10717005 | 10719476 | 18.84082 | 47.13427 | 44.45482 | . | . | . | . | . | . |
| chr7 | 27486675 | 27487698 | 1000 | Promoter (<=1kb) | 7 | 27486910 | 27490316 | 3407 | 1 | ENSMUST00 | 0 | Sertad1 | ENSMUSG0C | false | chr7 | 27485942 | 27486823 | 16.43109 | 40.78362 | 38.18623 | . | . | . | . | . | . |
| chr7 | 27490297 | 27491066 | 1000 | Promoter (2-3kb) | 7 | 27487375 | 27489507 | 2133 | 1 | ENSMUST00 | 2923 | Sertad1 | ENSMUSG0C | false | chr7 | 27486916 | 27490677 | 29.02802 | 87.30634 | 84.01743 | chr7 | 27488220 | 27512085 | 6.36425 | 4.68802 | 4.12603 |
| chr7 | 27481430 | 27481972 | 1000 | Distal Intergenic | 7 | 27486910 | 27490316 | 3407 | 1 | ENSMUST00 | -4938 | Sertad1 | ENSMUSG0C | false | chr7 | 27481298 | 27482075 | 8.58007 | 16.45236 | 14.16747 | chr7 | 27480885 | 27484597 | 5.28079 | 4.2597 | 3.27298 |
| chr10 | 93310209 | 93311377 | 1000 | Promoter (<=1kb) | 10 | 93247414 | 93311135 | 63722 | 2 | ENSMUST00 | 0 | Elk3 | ENSMUSG0C | false | chr10 | 93309491 | 93311606 | 23.28017 | 72.50986 | 69.48009 | chr10 | 93308789 | 93310554 | 3.32746 | 3.20035 | 1.64244 |
| chr9 | 25151132 | 25152175 | 1000 | Promoter (<=1kb) | 9 | 25108130 | 25151781 | 43652 | 2 | ENSMUST00 | 0 | Herpud2 | ENSMUSG0C | true | chr9 | 25149881 | 25152473 | 25.75243 | 103.50989 | 99.85224 | . | . | . | . | . | . |
| chr8 | 105860185 | 105860924 | 1000 | Promoter (<=1kb) | 8 | 105860634 | 105880402 | 19769 | 1 | ENSMUST00 | 0 | Nutf2 | ENSMUSG0C | false | chr8 | 105859948 | 105861575 | 31.96831 | 119.80809 | 115.67081 | chr8 | 105854235 | 105860201 | 4.58143 | 3.86089 | 2.66378 |
| chr1 | 152766181 | 152766915 | 1000 | Promoter (<=1kb) | 1 | 152766542 | 152775597 | 9056 | 1 | ENSMUST00 | 0 | Arpc5 | ENSMUSG0C | false | chr1 | 152765791 | 152768258 | 42.74922 | 148.11885 | 142.90198 | . | . | . | . | . | . |
| chr4 | 82505738 | 82506906 | 1000 | Promoter (<=1kb) | 4 | 82290173 | 82505770 | 215598 | 2 | ENSMUST00 | 0 | Nfib | ENSMUSG0C | false | chr4 | 82504132 | 82507030 | 22.41791 | 60.5052 | 57.65255 | chr4 | 82504215 | 82507496 | 4.80875 | 3.91795 | 2.86519 |
| chr3 | 88553480 | 88554503 | 1000 | Promoter (<=1kb) | 3 | 88553716 | 88569725 | 16010 | 1 | ENSMUST00 | 0 | Ubqln4 | ENSMUSG0C | false | chr3 | 88553099 | 88554546 | 32.67316 | 105.57997 | 101.8659 | chr3 | 88554421 | 88555584 | 3.94379 | 3.6433 | 2.15459 |
| chr5 | 6756834 | 6757127 | 1000 | Distal Intergenic | 16 | 6809222 | 7409813 | 600592 | 1 | ENSMUST00 | -52095 | Rbfox1 | ENSMUSG0C | false | . | . | . | . | . | . | . | . | . | . | . | . |
| chr3 | 100685221 | 100686291 | 1000 | Promoter (<=1kb) | 3 | 100562208 | 100685503 | 123296 | 2 | ENSMUST00 | 0 | Man1a2 | ENSMUSG0C | true | chr3 | 100680989 | 100686556 | 30.41442 | 103.5799 | 99.9162 | chr3 | 100686276 | 100688595 | 4.03041 | 3.5226 | 2.23762 |
| chr11 | 102218599 | 102219243 | 1000 | Promoter (<=1kb) | 11 | 102195749 | 102218929 | 23181 | 2 | ENSMUST00 | 0 | Hdac5 | ENSMUSG0C | false | chr11 | 102218423 | 102218882 | 10.54274 | 25.8059 | 23.39399 | chr11 | 102213449 | 102231135 | 7.77054 | 5.31127 | 5.26338 |
| chr13 | 17943667 | 17944574 | 1000 | Promoter (<=1kb) | 13 | 17880575 | 17944239 | 63665 | 2 | ENSMUST00 | 0 | Rala | ENSMUSG0C | false | chr13 | 17943115 | 17944986 | 35.88896 | 115.35191 | 111.35841 | . | . | . | . | . | . |
| chr2 | 172939999 | 172940898 | 1000 | Promoter (<=1kb) | 2 | 172868012 | 172940321 | 72310 | 2 | ENSMUST00 | 0 | Bmp7 | ENSMUSG0C | false | chr2 | 172938608 | 172940993 | 30.78865 | 90.34473 | 86.99165 | chr2 | 172924796 | 172941177 | 5.52242 | 4.25198 | 3.44947 |
| chr5 | 172932229 | 172932570 | 1000 | Intron (ENSMUST00000009143.7 | 2 | 172916118 | 172940092 | 23975 | 2 | ENSMUST00 | 7522 | Bmp7 | ENSMUSG0C | false | . | . | . | . | . | . | chr2 | 172924796 | 172941177 | 5.52242 | 4.25198 | 3.44947 |
| chr5 | 172931279 | 172931862 | 1000 | Intron (ENSMUST00000009143.7 | 2 | 172916118 | 172940092 | 23975 | 2 | ENSMUST00 | 8230 | Bmp7 | ENSMUSG0C | false | . | . | . | . | . | . | chr2 | 172924796 | 172941177 | 5.52242 | 4.25198 | 3.44947 |
| chr5 | 115299872 | 115300403 | 1000 | Promoter (<=1kb) | 5 | 115298431 | 115299435 | 1005 | 2 | ENSMUST00 | -438 | Dynll1 | ENSMUSG0C | false | chr5 | 115299554 | 115300971 | 33.47463 | 112.14942 | 108.24687 | . | . | . | . | . | . |
| chr5 | 79391479 | 79392135 | 1000 | Intron (ENSMUST00000178522.1 | 15 | 79360684 | 79402919 | 42236 | 2 | ENSMUST00 | 10784 | Tmem184b | ENSMUSG0C | false | . | . | . | . | . | . | chr15 | 79389106 | 79402713 | 7.35715 | 5.09621 | 4.94899 |
| chr5 | 79398174 | 79398500 | 1000 | Intron (ENSMUST00000178522.1 | 15 | 79360684 | 79402919 | 42236 | 2 | ENSMUST00 | 4419 | Tmem184b | ENSMUSG0C | false | . | . | . | . | . | . | chr15 | 79389106 | 79402713 | 7.35715 | 5.09621 | 4.94899 |
| chr5 | 4800663 | 4800981 | 1000 | Intron (ENSMUST00000109910.3 | 11 | 4794431 | 4815240 | 20810 | 2 | ENSMUST00 | 14259 | Nf2 | ENSMUSG0C | false | . | . | . | . | . | . | chr11 | 4797551 | 4801222 | 4.03147 | 3.59013 | 2.22825 |
| chr5 | 48572720 | 48573026 | 1000 | Promoter (<=1kb) | 6 | 48569698 | 48572670 | 2973 | 2 | ENSMUST00 | -51 | Rarres2 | ENSMUSG0C | false | . | . | . | . | . | . | chr6 | 48572590 | 48573536 | 4.40813 | 3.65652 | 2.53247 |
| chr5 | 77591217 | 77591715 | 1000 | Promoter (<=1kb) | 10 | 77592337 | 77597130 | 4794 | 1 | ENSMUST00 | -622 | Pttg1ip | ENSMUSG0C | false | . | . | . | . | . | . | chr10 | 77580829 | 77594032 | 8.74443 | 5.72701 | 6.06549 |
| chr10 | 77583914 | 77584436 | 1000 | Promoter (<=1kb) | 10 | 77583945 | 77592999 | 9055 | 1 | ENSMUST00 | 0 | Pttg1ip | ENSMUSG0C | false | chr10 | 77581692 | 77584244 | 38.6007 | 121.10697 | 116.92481 | chr10 | 77580829 | 77594032 | 8.74443 | 5.72701 | 6.06549 |
| chr5 | 98919971 | 98920291 | 1000 | Intron (ENSMUST00000109401.2 | 13 | 98839019 | 98926384 | 87366 | 2 | ENSMUST00 | 6093 | Tnpo1 | ENSMUSG0C | false | . | . | . | . | . | . | chr13 | 98907937 | 98925350 | 5.77769 | 4.39128 | 3.65662 |
| chr2 | 32712651 | 32713745 | 1000 | Promoter (<=1kb) | 2 | 32707438 | 32712784 | 5347 | 2 | ENSMUST00 | 0 | Cdk9 | ENSMUSG0C | false | chr2 | 32711568 | 32713700 | 30.94804 | 114.76607 | 110.79114 | . | . | . | . | . | . |
| chr5 | 13357804 | 13358551 | 1000 | Promoter (<=1kb) | 16 | 13358421 | 13417529 | 59109 | 1 | ENSMUST00 | 0 | Mkl2 | ENSMUSG0C | false | . | . | . | . | . | . | chr16 | 13356427 | 13361066 | 5.99793 | 4.55412 | 3.81793 |
| chr5 | 27247192 | 27247540 | 1000 | Promoter (<=1kb) | 2 | 27188393 | 27247266 | 58874 | 2 | ENSMUST00 | 0 | Sardh | ENSMUSG0C | false | . | . | . | . | . | . | chr2 | 27244364 | 27247885 | 5.39346 | 4.1779 | 3.34031 |
| chr2 | 27426481 | 27427303 | 1000 | Promoter (<=1kb) | 2 | 27262104 | 27426912 | 164809 | 2 | ENSMUST00 | 0 | Vav2 | ENSMUSG0C | false | chr2 | 27425224 | 27427512 | 33.03282 | 126.03245 | 121.67878 | chr2 | 27424994 | 27428278 | 5.51507 | 4.36394 | 3.41315 |
| chr7 | 31041980 | 31042314 | 1000 | Promoter (<=1kb) | 7 | 31032730 | 31042118 | 9389 | 2 | ENSMUST00 | 0 | Fxyd5 | ENSMUSG0C | false | chr7 | 31041678 | 31042331 | 6.75876 | 12.51203 | 10.2897 | chr7 | 31041819 | 31042862 | 3.72006 | 3.46317 | 1.94342 |
| chr9 | 113930824 | 113931346 | 1000 | Promoter (<=1kb) | 9 | 113930934 | 113977199 | 46266 | 1 | ENSMUST00 | 0 | Ubp1 | ENSMUSG0C | false | chr9 | 113930277 | 113930872 | 17.64787 | 49.73517 | 47.02245 | . | . | . | . | . | . |
| chr5 | 132333673 | 132334132 | 1000 | Distal Intergenic | 1 | 132327596 | 132330450 | 2855 | 1 | ENSMUST00 | 6078 | Nuak2 | ENSMUSG0C | false | . | . | . | . | . | . | chr1 | 132317064 | 132339308 | 6.74463 | 4.86917 | 4.43585 |
| chr5 | 132319258 | 132319616 | 1000 | Intron (ENSMUST00000072177.8 | 1 | 132316165 | 132317736 | 1572 | 1 | ENSMUST00 | 3094 | Nuak2 | ENSMUSG0C | false | . | . | . | . | . | . | chr1 | 132317064 | 132339308 | 6.74463 | 4.86917 | 4.43585 |
| chr1 | 106759614 | 106760106 | 1000 | Promoter (<=1kb) | 1 | 106720459 | 106759727 | 39269 | 2 | ENSMUST00 | 0 | Kdsr | ENSMUSG0C | false | chr1 | 106758646 | 106760410 | 25.79552 | 96.83904 | 93.34669 | . | . | . | . | . | . |
| chr9 | 107399508 | 107400344 | 1000 | Promoter (<=1kb) | 9 | 107399612 | 107529346 | 129735 | 1 | ENSMUST00 | 0 | Cacna2d2 | ENSMUSG0C | false | chr9 | 107399489 | 107400750 | 28.36989 | 85.2065 | 81.96005 | chr9 | 107397435 | 107400901 | 4.18586 | 3.66016 | 2.37838 |
| chr9 | 107377963 | 107379078 | 1000 | Distal Intergenic | 9 | 107399612 | 107529346 | 129735 | 1 | ENSMUST00 | -20534 | Cacna2d2 | ENSMUSG0C | false | chr9 | 107377927 | 107378950 | 7.82203 | 16.59716 | 14.30959 | chr9 | 107377204 | 107379940 | 5.16612 | 3.84758 | 3.18046 |
| chr5 | 107397707 | 107398067 | 1000 | Promoter (1-2kb) | 9 | 107399612 | 107529346 | 129735 | 1 | ENSMUST00 | -1545 | Cacna2d2 | ENSMUSG0C | false | . | . | . | . | . | . | chr9 | 107397435 | 107400901 | 4.18586 | 3.66016 | 2.37838 |
| chr11 | 94497997 | 94499167 | 1000 | Promoter (<=1kb) | 11 | 94496002 | 94498250 | 2249 | 2 | ENSMUST00 | 0 | Epn3 | ENSMUSG0C | false | chr11 | 94495773 | 94500513 | 28.91859 | 89.0447 | 85.71584 | . | . | . | . | . | . |
| chr11 | 94496852 | 94497246 | 1000 | Promoter (1-2kb) | 11 | 94496002 | 94498250 | 2249 | 2 | ENSMUST00 | 1004 | Epn3 | ENSMUSG0C | false | chr11 | 94495773 | 94500513 | 28.91859 | 89.0447 | 85.71584 | chr11 | 94489158 | 94497346 | 7.88949 | 5.284 | 5.39442 |
| chr8 | 123332516 | 123333072 | 1000 | Promoter (<=1kb) | 8 | 123332713 | 123369515 | 36803 | 1 | ENSMUST00 | 0 | Spire2 | ENSMUSG0C | false | chr8 | 123332553 | 123333171 | 11.9418 | 26.44599 | 24.02746 | chr8 | 123332449 | 123334319 | 4.77198 | 4.02173 | 2.81056 |
| chr7 | 47050437 | 47051136 | 1000 | Promoter (<=1kb) | 7 | 47050640 | 47054776 | 4137 | 1 | ENSMUST00 | 0 | Tmem86a | ENSMUSG0C | false | chr7 | 47050135 | 47051334 | 45.49427 | 162.13052 | 156.40686 | chr7 | 47049975 | 47052123 | 4.32289 | 3.71355 | 2.49836 |
| chr1 | 133908359 | 133908994 | 1000 | Promoter (<=1kb) | 1 | 133897200 | 133907999 | 10800 | 2 | ENSMUST00 | -361 | Optc | ENSMUSG0C | false | chr1 | 133908664 | 133908930 | 7.27821 | 12.55215 | 10.32949 | chr1 | 133907821 | 133909507 | 5.20087 | 4.24008 | 3.15894 |
| chr4 | 109676200 | 109677100 | 1000 | Promoter (<=1kb) | 4 | 109676588 | 109963960 | 287373 | 1 | ENSMUST00 | 0 | Faf1 | ENSMUSG0C | false | chr4 | 109675990 | 109677632 | 27.78601 | 91.88203 | 88.49705 | . | . | . | . | . | . |
| chr12 | 111487141 | 111487459 | 1000 | Promoter (1-2kb) | 12 | 111484611 | 111485823 | 1213 | 2 | ENSMUST00 | -1319 | Gm266 | ENSMUSG0C | false | chr12 | 111484468 | 111487284 | 26.28407 | 85.32412 | 82.07336 | chr12 | 111482224 | 111493728 | 5.00155 | 3.95404 | 3.01745 |
| chr5 | 111488328 | 111488790 | 1000 | Promoter (2-3kb) | 12 | 111484611 | 111485823 | 1213 | 2 | ENSMUST00 | -2506 | Gm266 | ENSMUSG0C | false | . | . | . | . | . | . | chr12 | 111482224 | 111493728 | 5.00155 | 3.95404 | 3.01745 |
| chr5 | 180266529 | 180266967 | 1000 | Distal Intergenic | 1 | 180245638 | 180263438 | 17801 | 2 | ENSMUST00 | -3092 | Psen2 | ENSMUSG0C | false | . | . | . | . | . | . | chr1 | 180265074 | 180268890 | 5.67353 | 4.29274 | 3.59167 |
| chr19 | 10183301 | 10183713 | 1000 | Promoter (<=1kb) | 19 | 10182888 | 10196870 | 13983 | 1 | ENSMUST00 | 414 | Fads1 | ENSMUSG0C | false | chr19 | 10182386 | 10184244 | 28.56742 | 103.16996 | 99.51957 | chr19 | 10183538 | 10184863 | 4.33066 | 3.83076 | 2.45135 |
| chr5 | 101994163 | 101994585 | 1000 | Distal Intergenic | 11 | 101987056 | 101992264 | 5209 | 1 | ENSMUST00 | 7108 | 1700006E09 | ENSMUSG0C | false | . | . | . | . | . | . | chr11 | 101993115 | 102001120 | 5.75336 | 4.46092 | 3.63249 |
| chr5 | 101997938 | 101998541 | 1000 | Downstream (1-2kb) | 11 | 101987056 | 101992264 | 5209 | 1 | ENSMUST00 | 10883 | 1700006E09 | ENSMUSG0C | false | . | . | . | . | . | . | chr11 | 101993115 | 102001120 | 5.75336 | 4.46092 | 3.63249 |
| chr12 | 113000279 | 113001299 | 1000 | Promoter (<=1kb) | 12 | 112959862 | 113000621 | 40760 | 2 | ENSMUST00 | 0 | Brf1 | ENSMUSG0C | false | chr12 | 112999391 | 113000799 | 35.88896 | 115.35191 | 111.35841 | chr12 | 113001174 | 113002260 | 6.39041 | 4.74356 | 4.16935 |
| chr12 | 17544475 | 17545086 | 1000 | Promoter (<=1kb) | 12 | 17544873 | 17551502 | 6630 | 1 | ENSMUST00 | 0 | Odc1 | ENSMUSG0C | false | chr12 | 17544060 | 17544701 | 30.67071 | 98.01645 | 94.49575 | . | . | . | . | . | . |
| chr11 | 77493055 | 77494061 | 1000 | Promoter (<=1kb) | 11 | 77493562 | 77507786 | 14225 | 1 | ENSMUST00 | 0 | Git1 | ENSMUSG0C | false | chr11 | 77493384 | 77494525 | 30.37608 | 91.57915 | 88.20459 | chr11 | 77488828 | 77499060 | 5.56381 | 4.29511 | 3.47349 |
| chr9 | 69989227 | 69989811 | 1000 | Promoter (<=1kb) | 9 | 69989466 | 70008317 | 18852 | 1 | ENSMUST00 | 0 | Bnip2 | ENSMUSG0C | false | chr9 | 69988988 | 69989351 | 7.76973 | 17.46802 | 15.16681 | . | . | . | . | . | . |
| chr17 | 46282451 | 46283221 | 1000 | Promoter (<=1kb) | 17 | 46257851 | 46283026 | 25176 | 2 | ENSMUST00 | 0 | Tjap1 | ENSMUSG0C | true | chr17 | 46281444 | 46283018 | 26.02615 | 74.62016 | 71.5567 | chr17 | 46283145 | 46283539 | 6.55648 | 4.89539 | 4.26555 |
| chr8 | 83899116 | 83900657 | 1000 | Promoter (<=1kb) | 8 | 83900105 | 83941954 | 41850 | 1 | ENSMUST00 | 0 | Lphn1 | ENSMUSG0C | false | chr8 | 83899124 | 83901392 | 23.65566 | 74.08845 | 71.03342 | chr8 | 83898878 | 83902314 | 4.8258 | 3.94283 | 2.85241 |
| chr11 | 95990910 | 95991347 | 1000 | Intron (ENSMUST00000013559.2 | 11 | 95973979 | 95982611 | 8633 | 2 | ENSMUST00 | -8300 | Igf2bp1 | ENSMUSG0C | false | chr11 | 95990773 | 95991742 | 15.77044 | 42.5731 | 39.95169 | chr11 | 95987920 | 95991850 | 5.55484 | 4.35809 | 3.47278 |
| chr5 | 170339636 | 170340118 | 1000 | Distal Intergenic | 2 | 170347931 | 170349740 | 1810 | 2 | ENSMUST00 | 9622 | Bcas1 | ENSMUSG0C | false | . | . | . | . | . | . | chr2 | 170335805 | 170341326 | 4.65119 | 3.71249 | 2.7524 |
| chr5 | 71317160 | 71317525 | 1000 | Distal Intergenic | 9 | 71215789 | 71296243 | 80455 | 1 | ENSMUST00 | 101372 | Aldh1a2 | ENSMUSG0C | false | . | . | . | . | . | . | chr9 | 71316102 | 71318710 | 5.61234 | 4.3594 | 3.50702 |
| chr15 | 36531932 | 36532423 | 1000 | Intron (ENSMUST00000013755.6 | 15 | 36537508 | 36545687 | 8180 | 2 | ENSMUST00 | 13264 | Snx31 | ENSMUSG0C | false | chr15 | 36531830 | 36532522 | 11.4883 | 23.92001 | 21.53215 | chr15 | 36532272 | 36534656 | 6.80791 | 4.93214 | 4.4797 |
| chr5 | 31080166 | 31080573 | 1000 | Distal Intergenic | 5 | 31075813 | 31076664 | 852 | 1 | ENSMUST00 | 4354 | Cad | ENSMUSG0C | false | . | . | . | . | . | . | chr5 | 31078705 | 31083714 | 4.9121 | 3.90464 | 2.97754 |
| chr5 | 58419078 | 58419385 | 1000 | Promoter (1-2kb) | 11 | 58421111 | 58426024 | 4914 | 1 | ENSMUST00 | -1726 | 1810065E05 | ENSMUSG0C | false | . | . | . | . | . | . | chr11 | 58418007 | 58419412 | 4.33494 | 3.62905 | 2.50215 |
| chr19 | 32757334 | 32758159 | 1000 | Promoter (<=1kb) | 19 | 32757497 | 32826160 | 68664 | 1 | ENSMUST00 | 0 | Pten | ENSMUSG0C | false | chr19 | 32755517 | 32759967 | 38.34636 | 121.73327 | 117.53308 | . | . | . | . | . | . |
| chr17 | 34898215 | 34899171 | 1000 | Promoter (<=1kb) | 17 | 34898469 | 34914051 | 15583 | 1 | ENSMUST00 | 0 | Ehmt2 | ENSMUSG0C | false | chr17 | 34897900 | 34900008 | 27.89552 | 88.209 | 84.90351 | . | . | . | . | . | . |
| chr8 | 26119075 | 26119900 | 1000 | Promoter (<=1kb) | 8 | 26119368 | 26143871 | 24504 | 1 | ENSMUST00 | 0 | Rnf170 | ENSMUSG0C | false | chr8 | 26118269 | 26120274 | 44.25352 | 156.47305 | 150.94255 | . | . | . | . | . | . |
| chr7 | 30291429 | 30291862 | 1000 | Promoter (<=1kb) | 7 | 30291672 | 30308367 | 16696 | 1 | ENSMUST00 | 0 | Clip3 | ENSMUSG0C | false | chr7 | 30291531 | 30291939 | 5.35832 | 8.05648 | 5.92315 | chr7 | 30291104 | 30291757 | 4.17375 | 3.73964 | 2.30783 |
| chr16 | 97851260 | 97852136 | 1000 | Promoter (<=1kb) | 16 | 97791520 | 97851850 | 60331 | 2 | ENSMUST00 | 0 | Prdm15 | ENSMUSG0C | false | chr16 | 97850536 | 97852480 | 37.91852 | 123.12976 | 118.88376 | . | . | . | . | . | . |
| chr2 | 119547451 | 119548013 | 1000 | Promoter (<=1kb) | 2 | 119547697 | 119587027 | 39331 | 1 | ENSMUST00 | 0 | Chp1 | ENSMUSG0C | false | chr2 | 119547092 | 119548523 | 22.37683 | 79.83291 | 76.68538 | . | . | . | . | . | . |
| chr5 | 114667501 | 114667839 | 1000 | Downstream (1-2kb) | 5 | 114622152 | 114658421 | 36270 | 2 | ENSMUST00 | -9081 | Trpv4 | ENSMUSG0C | false | chr5 | 114667261 | 114667584 | 8.11812 | 15.68191 | 13.40967 | chr5 | 114666775 | 114668238 | 4.11799 | 3.63419 | 2.28025 |
| chr5 | 114639484 | 114639826 | 1000 | Intron (ENSMUST00000071968.3 | 5 | 114622157 | 114636261 | 14105 | 2 | ENSMUST00 | -3224 | Trpv4 | ENSMUSG0C | false | . | . | . | . | . | . | chr5 | 114639039 | 114640025 | 6.47212 | 4.77674 | 4.20762 |
| chr13 | 58113115 | 58113966 | 1000 | Promoter (<=1kb) | 13 | 58004221 | 58113592 | 109372 | 2 | ENSMUST00 | 0 | Klhl3 | ENSMUSG0C | false | chr13 | 58112415 | 58113896 | 24.50127 | 70.18111 | 67.18839 | chr13 | 58113704 | 58114637 | 3.99499 | 3.54384 | 2.20119 |
| chr5 | 100584278 | 100584782 | 1000 | Promoter (<=1kb) | 11 | 100582818 | 100584786 | 1969 | 2 | ENSMUST00 | 4 | Dnajc7 | ENSMUSG0C | false | . | . | . | . | . | . | chr11 | 100581886 | 100586248 | 5.02584 | 3.69183 | 3.06436 |
| chr11 | 100619068 | 100619801 | 1000 | Promoter (<=1kb) | 11 | 100590887 | 100619576 | 28690 | 2 | ENSMUST00 | 0 | Dnajc7 | ENSMUSG0C | false | chr11 | 100618769 | 100620432 | 29.28796 | 98.53321 | 94.99628 | . | . | . | . | . | . |
| chr16 | 4594055 | 4595182 | 1000 | Promoter (<=1kb) | 16 | 4594713 | 4615957 | 21245 | 1 | ENSMUST00 | 0 | Glis2 | ENSMUSG0C | false | chr16 | 4593688 | 4596406 | 25.15761 | 80.47005 | 77.31023 | chr16 | 4591906 | 4609359 | 7.50331 | 5.19962 | 5.05636 |
| chr16 | 4586467 | 4587108 | 1000 | Distal Intergenic | 16 | 4594713 | 4615957 | 21245 | 1 | ENSMUST00 | -7605 | Glis2 | ENSMUSG0C | false | chr16 | 4586876 | 4587231 | 11.00893 | 22.55865 | 20.19038 | chr16 | 4585831 | 4590326 | 6.32379 | 4.67695 | 4.08213 |
| chr16 | 4595614 | 4595975 | 1000 | Promoter (<=1kb) | 16 | 4594713 | 4615957 | 21245 | 1 | ENSMUST00 | 902 | Glis2 | ENSMUSG0C | false | chr16 | 4593688 | 4596406 | 25.15761 | 80.47005 | 77.31023 | chr16 | 4591906 | 4609359 | 7.50331 | 5.19962 | 5.05636 |
| chr2 | 128698590 | 128698992 | 1000 | Promoter (<=1kb) | 2 | 128698956 | 128802894 | 103939 | 1 | ENSMUST00 | 0 | Mertk | ENSMUSG0C | true | chr2 | 128697969 | 128703570 | 36.46257 | 120.58523 | 116.42456 | . | . | . | . | . | . |
| chr17 | 12318424 | 12318959 | 1000 | Promoter (<=1kb) | 17 | 12227621 | 12318660 | 91040 | 2 | ENSMUST00 | 0 | Map3k4 | ENSMUSG0C | false | chr17 | 12317546 | 12319360 | 45.68299 | 161.30347 | 155.60802 | . | . | . | . | . | . |
| chr8 | 122550738 | 122551722 | 1000 | Promoter (<=1kb) | 8 | 122481698 | 122551329 | 69632 | 2 | ENSMUST00 | 0 | Piezo1 | ENSMUSG0C | false | chr8 | 122550054 | 122551803 | 32.25957 | 103.83389 | 100.16508 | chr8 | 122535817 | 122552157 | 5.24095 | 4.06992 | 3.22847 |
| chr14 | 31831253 | 31831621 | 1000 | Promoter (<=1kb) | 14 | 31700015 | 31830415 | 130401 | 2 | ENSMUST00 | -839 | Ankrd28 | ENSMUSG0C | false | chr14 | 31829329 | 31831818 | 34.38402 | 121.12925 | 116.94628 | chr14 | 31831266 | 31832071 | 2.73408 | 2.73826 | 1.19769 |
| chr14 | 31829951 | 31831025 | 1000 | Promoter (<=1kb) | 14 | 31700015 | 31830415 | 130401 | 2 | ENSMUST00 | 0 | Ankrd28 | ENSMUSG0C | false | chr14 | 31829329 | 31831818 | 34.38402 | 121.12925 | 116.94628 | . | . | . | . | . | . |
| chr5 | 83482504 | 83482971 | 1000 | Intron (ENSMUST00000014698.4 | 6 | 83481213 | 83487010 | 5798 | 2 | ENSMUST00 | 4039 | Dguok | ENSMUSG0C | false | . | . | . | . | . | . | chr6 | 83480571 | 83484057 | 6.0858 | 4.49292 | 3.91508 |
| chr4 | 151861179 | 151862748 | 1000 | Promoter (<=1kb) | 4 | 150917322 | 151861669 | 944348 | 2 | ENSMUST00 | 0 | Camta1 | ENSMUSG0C | false | chr4 | 151860740 | 151861826 | 27.29387 | 78.64282 | 75.51551 | chr4 | 151860052 | 151861191 | 3.62661 | 3.39286 | 1.91577 |
| chr5 | 151181325 | 151181814 | 1000 | Intron (ENSMUST00000169423.3 | 4 | 151086039 | 151139584 | 53546 | 2 | ENSMUST00 | -41742 | Camta1 | ENSMUSG0C | false | . | . | . | . | . | . | chr4 | 151162762 | 151183610 | 6.13162 | 4.48987 | 3.9543 |

| | | | | | | | | | | | | | | | | | | | | | | | | | | |
|---|---|---|---|---|---|---|---|---|---|---|---|---|---|---|---|---|---|---|---|---|---|---|---|---|---|---|
| chr5 | 151180098 | 151180605 | 1000 | Intron (ENSMUST00000169423.3 | 4 | 151086039 | 151139584 | 53546 | 2 | ENSMUST00 | -40515 | Camta1 | ENSMUSG0C | false | . | . | . | . | . | . | chr4 | 151162762 | 151183610 | 6.13162 | 4.48987 | 3.9543 |
| chr6 | 52164923 | 52165360 | 1000 | Promoter (<=1kb) | 6 | 52162511 | 52164831 | 2321 | 2 | ENSMUST00 | -93 | Hoxa2 | ENSMUSG0C | false | chr6 | 52164166 | 52165356 | 8.01187 | 19.41161 | 17.08515 | chr6 | 52159328 | 52167512 | 5.47105 | 4.07894 | 3.42628 |
| chr17 | 15375740 | 15376696 | 1000 | Promoter (<=1kb) | 17 | 15367354 | 15375976 | 8623 | 2 | ENSMUST00 | 0 | Dll1 | ENSMUSG0C | false | chr17 | 15374183 | 15377041 | 23.4661 | 79.19912 | 76.05865 | chr17 | 15371856 | 15381968 | 5.90641 | 4.43925 | 3.75812 |
| chr8 | 105346621 | 105347229 | 1000 | Promoter (<=1kb) | 8 | 105329163 | 105347909 | 18747 | 2 | ENSMUST00 | 680 | Fhod1 | ENSMUSG0C | false | chr8 | 105346518 | 105349557 | 29.3377 | 90.74815 | 87.38457 | chr8 | 105344468 | 105347416 | 3.64916 | 3.3804 | 1.9168 |
| chr5 | 69089075 | 69089428 | 1000 | Distal Intergenic | 14 | 69029289 | 69041401 | 12113 | 1 | ENSMUST00 | 59787 | Stc1 | ENSMUSG0C | false | . | . | . | . | . | . | . | . | . | . | . | . |
| chr5 | 105460721 | 105461104 | 1000 | Exon (ENSMUST00000109355.3/ | 8 | 105467683 | 105468662 | 980 | 2 | ENSMUST00 | 7558 | Tppp3 | ENSMUSG0C | false | . | . | . | . | . | . | chr8 | 105459144 | 105464255 | 4.46543 | 3.81004 | 2.56294 |
| chr5 | 32610880 | 32611735 | 1000 | Promoter (<=1kb) | 5 | 32611171 | 32687066 | 75896 | 1 | ENSMUST00 | 0 | Yes1 | ENSMUSG0C | false | chr5 | 32610381 | 32612364 | 41.49189 | 142.53105 | 137.54099 | . | . | . | . | . | . |
| chr5 | 32458740 | 32459599 | 1000 | Promoter (<=1kb) | 5 | 32458974 | 32495523 | 36550 | 1 | ENSMUST00 | 0 | Ppp1cb | ENSMUSG0C | false | chr5 | 32458307 | 32458851 | 12.75224 | 30.33398 | 27.86505 | . | . | . | . | . | . |
| chr6 | 88198471 | 88198834 | 1000 | Promoter (<=1kb) | 6 | 88198664 | 88207030 | 8367 | 1 | ENSMUST00 | 0 | Gata2 | ENSMUSG0C | false | chr6 | 88196796 | 88199942 | 17.60262 | 45.78123 | 43.11721 | chr6 | 88193546 | 88201175 | 5.2385 | 4.1386 | 3.21754 |
| chr2 | 25557612 | 25558048 | 1000 | Promoter (<=1kb) | 2 | 25557847 | 25562082 | 4236 | 1 | ENSMUST00 | 0 | Edf1 | ENSMUSG0C | false | chr2 | 25557062 | 25558826 | 28.56742 | 103.16996 | 99.51957 | . | . | . | . | . | . |
| chr5 | 25452632 | 25453208 | 1000 | Downstream (1-2kb) | 2 | 25456838 | 25458676 | 1839 | 1 | ENSMUST00 | -3630 | Clic3 | ENSMUSG0C | true | . | . | . | . | . | . | chr2 | 25446553 | 25456355 | 6.09426 | 4.46882 | 3.92693 |
| chr17 | 25188076 | 25188774 | 1000 | Promoter (<=1kb) | 17 | 25188397 | 25234443 | 46047 | 1 | ENSMUST00 | 0 | Unkl | ENSMUSG0C | false | chr17 | 25187913 | 25189669 | 20.34014 | 89.76241 | 86.42476 | . | . | . | . | . | . |
| chr5 | 66325361 | 66325788 | 1000 | Intron (ENSMUST00000015277.8 | 7 | 66341500 | 66343265 | 1766 | 2 | ENSMUST00 | 17477 | Lrrk1 | ENSMUSG0C | false | . | . | . | . | . | . | chr7 | 66320849 | 66326654 | 5.84047 | 4.37327 | 3.7133 |
| chr5 | 66446844 | 66447230 | 1000 | Distal Intergenic | 7 | 66390892 | 66427517 | 36626 | 2 | ENSMUST00 | -19328 | Aldh1a3 | ENSMUSG0C | false | . | . | . | . | . | . | chr7 | 66438246 | 66451395 | 6.06663 | 4.59192 | 3.88434 |
| chr12 | 80260125 | 80260872 | 1000 | Promoter (<=1kb) | 12 | 80167542 | 80260371 | 92830 | 2 | ENSMUST00 | 0 | Actn1 | ENSMUSG0C | true | chr12 | 80258838 | 80261410 | 36.31995 | 116.10173 | 112.08785 | . | . | . | . | . | . |
| chr7 | 28808607 | 28809327 | 1000 | Promoter (<=1kb) | 7 | 28808806 | 28816960 | 8155 | 1 | ENSMUST00 | 0 | Hnrnpl | ENSMUSG0C | false | chr7 | 28808053 | 28810551 | 34.4409 | 118.06123 | 113.98219 | . | . | . | . | . | . |
| chr5 | 46095303 | 46095637 | 1000 | Distal Intergenic | 19 | 46075863 | 46085528 | 9666 | 1 | ENSMUST00 | 19441 | Nolc1 | ENSMUSG0C | false | . | . | . | . | . | . | chr19 | 46095155 | 46096272 | 3.38118 | 3.30696 | 1.69023 |
| chrX | 71364533 | 71364921 | 1000 | Promoter (<=1kb) | 20 | 71364760 | 71388756 | 23997 | 1 | ENSMUST00 | 0 | Mtmr1 | ENSMUSG0C | false | chrX | 71364259 | 71365317 | 19.75988 | 50.25065 | 47.53236 | . | . | . | . | . | . |
| chr10 | 8885647 | 8886216 | 1000 | Promoter (<=1kb) | 10 | 8722219 | 8886070 | 163852 | 2 | ENSMUST00 | 0 | Sash1 | ENSMUSG0C | false | chr10 | 8885006 | 8886830 | 27.49702 | 86.54257 | 83.27029 | . | . | . | . | . | . |
| chr9 | 65293909 | 65294487 | 1000 | Promoter (<=1kb) | 9 | 65294260 | 65330657 | 36398 | 1 | ENSMUST00 | 0 | Clpx | ENSMUSG0C | false | chr9 | 65293303 | 65295315 | 39.4261 | 145.54997 | 140.44281 | . | . | . | . | . | . |
| chr10 | 13964310 | 13965264 | 1000 | Promoter (<=1kb) | 10 | 13966075 | 14150589 | 184515 | 1 | ENSMUST00 | -811 | Hivep2 | ENSMUSG0C | true | chr10 | 13964189 | 13968566 | 36.09911 | 114.6679 | 110.69713 | . | . | . | . | . | . |
| chr10 | 13965413 | 13967658 | 1000 | Promoter (<=1kb) | 10 | 13966075 | 14150589 | 184515 | 1 | ENSMUST00 | 0 | Hivep2 | ENSMUSG0C | true | chr10 | 13964189 | 13968566 | 36.09911 | 114.6679 | 110.69713 | . | . | . | . | . | . |
| chr5 | 114821641 | 114822198 | 1000 | Promoter (2-3kb) | 13 | 114818905 | 114825230 | 6326 | 1 | ENSMUST00 | 2737 | Mocs2 | ENSMUSG0C | false | . | . | . | . | . | . | chr13 | 114817110 | 114824321 | 4.75411 | 3.77476 | 2.83645 |
| chr5 | 114863423 | 114863904 | 1000 | Intron (ENSMUST00000056117.8 | 13 | 114818905 | 114825230 | 6326 | 1 | ENSMUST00 | 44519 | Mocs2 | ENSMUSG0C | false | . | . | . | . | . | . | chr13 | 114861630 | 114865097 | 4.17427 | 3.69893 | 2.32429 |
| chr17 | 46383313 | 46384009 | 1000 | Promoter (<=1kb) | 17 | 46383731 | 46414814 | 31084 | 1 | ENSMUST00 | 0 | Zfp318 | ENSMUSG0C | false | chr17 | 46383068 | 46384821 | 27.54716 | 98.2761 | 94.74957 | chr17 | 46382906 | 46383319 | 4.23108 | 3.76413 | 2.35299 |
| chr17 | 46552832 | 46553463 | 1000 | Promoter (2-3kb) | 17 | 46546839 | 46556162 | 9324 | 2 | ENSMUST00 | 2699 | Srf | ENSMUSG0C | false | chr17 | 46552796 | 46553732 | 12.92336 | 34.59913 | 32.07519 | chr17 | 46552021 | 46555146 | 5.82865 | 4.25137 | 3.72202 |
| chr5 | 180214416 | 180215183 | 1000 | Intron (ENSMUST00000015791.5 | 2 | 180176373 | 180225859 | 49487 | 2 | ENSMUST00 | 10676 | Lama5 | ENSMUSG0C | true | . | . | . | . | . | . | chr2 | 180202228 | 180224817 | 7.70036 | 5.22812 | 5.23282 |
| chr2 | 180227156 | 180227987 | 1000 | Promoter (1-2kb) | 2 | 180176373 | 180225859 | 49487 | 2 | ENSMUST00 | -1298 | Lama5 | ENSMUSG0C | true | chr2 | 180227200 | 180228692 | 9.76265 | 21.55577 | 19.19873 | chr2 | 180225908 | 180236447 | 7.12665 | 5.06061 | 4.74583 |
| chr5 | 5693653 | 5694511 | 1000 | Promoter (<=1kb) | 5 | 5665954 | 5694060 | 28107 | 2 | ENSMUST00 | 0 | Steap2 | ENSMUSG0C | false | chr5 | 5693212 | 5694665 | 28.71633 | 95.52087 | 92.05702 | chr5 | 5692022 | 5693664 | 3.8443 | 3.49796 | 2.10494 |
| chr5 | 95307090 | 95307453 | 1000 | Promoter (<=1kb) | 3 | 95296096 | 95307176 | 11081 | 2 | ENSMUST00 | 0 | Anxa9 | ENSMUSG0C | false | . | . | . | . | . | . | chr3 | 95305900 | 95308095 | 5.28343 | 4.07767 | 3.26216 |
| chr7 | 84409121 | 84410288 | 1000 | Promoter (<=1kb) | 7 | 84246281 | 84409959 | 163679 | 2 | ENSMUST00 | 0 | Arnt2 | ENSMUSG0C | false | chr7 | 84409363 | 84410707 | 27.24215 | 82.29425 | 79.10038 | . | . | . | . | . | . |
| chr3 | 95995556 | 95996281 | 1000 | Promoter (<=1kb) | 3 | 95988429 | 95996001 | 7573 | 2 | ENSMUST00 | 0 | Plekho1 | ENSMUSG0C | true | chr3 | 95992078 | 95997273 | 32.06657 | 104.51031 | 100.83098 | . | . | . | . | . | . |
| chr5 | 96040067 | 96040418 | 1000 | Intron (ENSMUST00000015891.5 | 3 | 96056887 | 96057719 | 833 | 2 | ENSMUST00 | 17301 | Vps45 | ENSMUSG0C | false | . | . | . | . | . | . | chr3 | 96034741 | 96043819 | 4.83962 | 3.81336 | 2.91218 |
| chr5 | 95966118 | 95966497 | 1000 | Distal Intergenic | 3 | 95944204 | 95945523 | 1320 | 1 | ENSMUST00 | 21915 | Anp32e | ENSMUSG0C | false | . | . | . | . | . | . | chr3 | 95966255 | 95967083 | 6.73943 | 4.75384 | 4.45244 |
| chr5 | 95957497 | 95957851 | 1000 | Distal Intergenic | 3 | 95944204 | 95945523 | 1320 | 1 | ENSMUST00 | 13294 | Anp32e | ENSMUSG0C | false | . | . | . | . | . | . | chr3 | 95956444 | 95958338 | 3.56994 | 3.33746 | 1.87248 |
| chr3 | 95893744 | 95894758 | 1000 | Promoter (<=1kb) | 3 | 95893969 | 95896758 | 2790 | 1 | ENSMUST00 | 0 | Aph1a | ENSMUSG0C | false | chr3 | 95893407 | 95895298 | 23.7035 | 64.41534 | 61.50821 | . | . | . | . | . | . |
| chr10 | 7955347 | 7957014 | 1000 | Promoter (<=1kb) | 10 | 7905653 | 7956230 | 50578 | 2 | ENSMUST00 | 0 | Tab2 | ENSMUSG0C | true | chr10 | 7954510 | 7957213 | 36.91095 | 117.12714 | 113.07945 | . | . | . | . | . | . |
| chr5 | 137557658 | 137558027 | 1000 | Intron (ENSMUST00000058210.7 | 6 | 137477668 | 137571017 | 93350 | 2 | ENSMUST00 | 12990 | Eps8 | ENSMUSG0C | false | . | . | . | . | . | . | chr6 | 137557484 | 137559802 | 2.90855 | 2.92957 | 1.30626 |
| chr11 | 50210031 | 50210815 | 1000 | Promoter (<=1kb) | 11 | 50199366 | 50210791 | 11426 | 2 | ENSMUST00 | 0 | Sqstm1 | ENSMUSG0C | false | chr11 | 50208222 | 50211414 | 38.97723 | 131.47449 | 126.92584 | . | . | . | . | . | . |
| chr2 | 75703446 | 75705162 | 1000 | Promoter (<=1kb) | 2 | 75675513 | 75704641 | 29129 | 2 | ENSMUST00 | 0 | Nfe2l2 | ENSMUSG0C | false | chr2 | 75702478 | 75705502 | 45.52735 | 154.11624 | 148.67334 | . | . | . | . | . | . |
| chr5 | 27774167 | 27774777 | 1000 | Distal Intergenic | 2 | 27740201 | 27748671 | 8471 | 1 | ENSMUST00 | 33967 | Rxra | ENSMUSG0C | false | . | . | . | . | . | . | chr2 | 27773884 | 27775146 | 4.63578 | 3.76664 | 2.75434 |
| chr5 | 27745856 | 27746551 | 1000 | Intron (ENSMUST00000077257.6 | 2 | 27740201 | 27748671 | 8471 | 1 | ENSMUST00 | 5656 | Rxra | ENSMUSG0C | false | . | . | . | . | . | . | chr2 | 27742587 | 27752199 | 6.62442 | 4.72897 | 4.35907 |
| chr5 | 27758371 | 27759014 | 1000 | Intron (ENSMUST00000077257.6 | 2 | 27740201 | 27748671 | 8471 | 1 | ENSMUST00 | 18171 | Rxra | ENSMUSG0C | false | . | . | . | . | . | . | chr2 | 27756216 | 27760543 | 7.96442 | 5.34041 | 5.43403 |
| chr2 | 27676013 | 27677213 | 1000 | Promoter (<=1kb) | 2 | 27676440 | 27741898 | 65459 | 1 | ENSMUST00 | 0 | Rxra | ENSMUSG0C | false | chr2 | 27676111 | 27677129 | 29.4107 | 87.008 | 83.72497 | chr2 | 27675717 | 27676874 | 4.76733 | 3.9469 | 2.80838 |
| chr5 | 95675396 | 95675796 | 1000 | Promoter (1-2kb) | 3 | 95676750 | 95677151 | 402 | 2 | ENSMUST00 | 1355 | Adamtsl4 | ENSMUSG0C | false | . | . | . | . | . | . | chr3 | 95660803 | 95677602 | 6.84699 | 4.91959 | 4.52508 |
| chr5 | 87249573 | 87250027 | 1000 | Downstream (<1kb) | 3 | 87250758 | 87263599 | 12842 | 2 | ENSMUST00 | 13572 | Fcrls | ENSMUSG0C | false | . | . | . | . | . | . | chr3 | 87249427 | 87250553 | 4.83816 | 4.04781 | 2.86717 |
| chr5 | 116472319 | 116472719 | 1000 | Promoter (<=1kb) | 11 | 116471376 | 116473243 | 1868 | 2 | ENSMUST00 | 524 | Prpsap1 | ENSMUSG0C | false | . | . | . | . | . | . | chr11 | 116471953 | 116473758 | 6.45738 | 4.60369 | 4.25195 |
| chr2 | 143914642 | 143915148 | 1000 | Promoter (<=1kb) | 2 | 143915320 | 143943324 | 28005 | 1 | ENSMUST00 | -172 | Dstn | ENSMUSG0C | false | chr2 | 143914317 | 143916573 | 31.85236 | 101.09525 | 97.496 | . | . | . | . | . | . |
| chr13 | 56135228 | 56135664 | 1000 | Promoter (<=1kb) | 13 | 56073619 | 56135631 | 62013 | 2 | ENSMUST00 | 0 | H2afy | ENSMUSG0C | false | chr13 | 56134430 | 56136012 | 34.00884 | 130.01486 | 125.52227 | chr13 | 56132562 | 56136152 | 4.04555 | 3.54219 | 2.25967 |
| chr5 | 56088498 | 56088902 | 1000 | Promoter (1-2kb) | 13 | 56074359 | 56090058 | 15700 | 2 | ENSMUST00 | 1156 | H2afy | ENSMUSG0C | false | . | . | . | . | . | . | chr13 | 56086171 | 56089366 | 3.97432 | 3.50427 | 2.19372 |
| chr1 | 177796197 | 177796850 | 1000 | Promoter (<=1kb) | 1 | 177762962 | 177796511 | 33550 | 2 | ENSMUST00 | 0 | Adss | ENSMUSG0C | false | chr1 | 177794901 | 177796574 | 32.98857 | 124.89146 | 120.57959 | . | . | . | . | . | . |
| chr13 | 112927272 | 112927936 | 1000 | Promoter (<=1kb) | 13 | 112867780 | 112927380 | 59601 | 2 | ENSMUST00 | 0 | Skiv2l2 | ENSMUSG0C | false | chr13 | 112926380 | 112928598 | 22.03264 | 83.65932 | 80.44393 | . | . | . | . | . | . |
| chrX | 36329921 | 36330281 | 1000 | Promoter (<=1kb) | 20 | 36329216 | 36341754 | 12539 | 1 | ENSMUST00 | 706 | Lonrf3 | ENSMUSG0C | false | chrX | 36327630 | 36330397 | 18.84082 | 47.13427 | 44.45482 | . | . | . | . | . | . |
| chr1 | 192854883 | 192856208 | 1000 | Promoter (<=1kb) | 1 | 192844431 | 192855752 | 11322 | 2 | ENSMUST00 | 0 | Sertad4 | ENSMUSG0C | false | chr1 | 192854608 | 192856186 | 17.10421 | 70.31819 | 67.3235 | chr1 | 192854282 | 192856232 | 4.07264 | 3.45109 | 2.27548 |
| chr2 | 105126170 | 105127048 | 1000 | Promoter (<=1kb) | 2 | 105126529 | 105173612 | 47084 | 1 | ENSMUST00 | 0 | Wt1 | ENSMUSG0C | true | chr2 | 105122726 | 105134202 | 33.52805 | 106.05383 | 102.33234 | . | . | . | . | . | . |
| chr13 | 29983861 | 29985030 | 1000 | Promoter (<=1kb) | 13 | 29918784 | 29984219 | 65436 | 2 | ENSMUST00 | 0 | E2f3 | ENSMUSG0C | true | chr13 | 29983304 | 29986029 | 23.74947 | 92.7123 | 89.30923 | chr13 | 29984693 | 29985427 | 3.50471 | 3.39001 | 1.76259 |
| chr5 | 146884877 | 146885286 | 1000 | Distal Intergenic | 6 | 146888487 | 147032025 | 143539 | 1 | ENSMUST00 | -3201 | Ppfibp1 | ENSMUSG0C | false | . | . | . | . | . | . | chr6 | 146884620 | 146892279 | 5.32906 | 4.13735 | 3.32548 |
| chr6 | 146888276 | 146889005 | 1000 | Promoter (<=1kb) | 6 | 146888487 | 147032025 | 143539 | 1 | ENSMUST00 | 0 | Ppfibp1 | ENSMUSG0C | false | chr6 | 146887899 | 146889364 | 33.1106 | 101.26704 | 97.66567 | chr6 | 146884620 | 146892279 | 5.32906 | 4.13735 | 3.32548 |
| chr5 | 194927700 | 194928256 | 1000 | Distal Intergenic | 1 | 194938821 | 194961292 | 22472 | 1 | ENSMUST00 | -10565 | Cd34 | ENSMUSG0C | false | . | . | . | . | . | . | . | . | . | . | . | . |
| chr5 | 146907472 | 146907980 | 1000 | Distal Intergenic | 5 | 146948657 | 146955614 | 6958 | 1 | ENSMUST00 | -40677 | Gtf3a | ENSMUSG0C | false | . | . | . | . | . | . | chr5 | 146903456 | 146909712 | 6.67519 | 4.71367 | 4.3822 |
| chr15 | 77970635 | 77971270 | 1000 | Promoter (<=1kb) | 15 | 77958998 | 77970824 | 11827 | 2 | ENSMUST00 | 0 | Eif3d | ENSMUSG0C | false | chr15 | 77969917 | 77971584 | 19.74353 | 71.90538 | 68.88615 | . | . | . | . | . | . |
| chr11 | 116023868 | 116024386 | 1000 | Promoter (<=1kb) | 11 | 116023509 | 116024473 | 965 | 2 | ENSMUST00 | 87 | H3f3b | ENSMUSG0C | false | chr11 | 116023135 | 116024573 | 35.28817 | 116.40383 | 112.37991 | . | . | . | . | . | . |
| chr5 | 83471494 | 83472046 | 1000 | Distal Intergenic | 15 | 83375607 | 83464606 | 89000 | 2 | ENSMUST00 | -6889 | Pacsin2 | ENSMUSG0C | false | . | . | . | . | . | . | chr15 | 83468832 | 83472485 | 7.51947 | 5.23752 | 5.06228 |
| chr5 | 83440675 | 83441087 | 1000 | Intron (ENSMUST00000171436.2 | 15 | 83375607 | 83432850 | 57244 | 2 | ENSMUST00 | -7826 | Pacsin2 | ENSMUSG0C | false | . | . | . | . | . | . | chr15 | 83417027 | 83463739 | 6.18372 | 4.57623 | 3.99178 |
| chr1 | 12718064 | 12718820 | 1000 | Promoter (<=1kb) | 1 | 12718459 | 12822916 | 104458 | 1 | ENSMUST00 | 0 | Sulf1 | ENSMUSG0C | true | chr1 | 12718393 | 12719676 | 8.39879 | 16.15083 | 13.87107 | chr1 | 12710824 | 12724742 | 4.87179 | 3.95158 | 2.92304 |
| chr5 | 12715405 | 12715843 | 1000 | Promoter (2-3kb) | 1 | 12718459 | 12822916 | 104458 | 1 | ENSMUST00 | -2616 | Sulf1 | ENSMUSG0C | true | . | . | . | . | . | . | chr1 | 12710824 | 12724742 | 4.87179 | 3.95158 | 2.92304 |
| chr5 | 12697849 | 12698330 | 1000 | Intron (ENSMUST00000177608.2 | 1 | 12692532 | 12852489 | 159958 | 1 | ENSMUST00 | 5318 | Sulf1 | ENSMUSG0C | true | . | . | . | . | . | . | chr1 | 12693290 | 12699363 | 4.74812 | 3.91894 | 2.8045 |
| chr2 | 162931343 | 162932316 | 1000 | Promoter (<=1kb) | 2 | 162931528 | 162937121 | 5594 | 1 | ENSMUST00 | 0 | Srsf6 | ENSMUSG0C | false | chr2 | 162930504 | 162932824 | 42.33011 | 146.25195 | 141.10883 | . | . | . | . | . | . |
| chr5 | 164420820 | 164421350 | 1000 | Downstream (2-3kb) | 2 | 164389393 | 164405160 | 15768 | 2 | ENSMUST00 | -15661 | Matn4 | ENSMUSG0C | false | . | . | . | . | . | . | chr2 | 164415904 | 164437244 | 5.93196 | 4.473 | 3.77292 |
| chr2 | 164432404 | 164433241 | 1000 | Intron (ENSMUST00000017153.3 | 2 | 164424247 | 164443186 | 18940 | 2 | ENSMUST00 | 9945 | Sdc4 | ENSMUSG0C | true | chr2 | 164431717 | 164432497 | 10.37607 | 20.95961 | 18.61343 | chr2 | 164415904 | 164437244 | 5.93196 | 4.473 | 3.77292 |
| chr2 | 164436940 | 164437380 | 1000 | Intron (ENSMUST00000017153.3 | 2 | 164424247 | 164443186 | 18940 | 2 | ENSMUST00 | 5806 | Sdc4 | ENSMUSG0C | true | chr2 | 164432671 | 164438188 | 15.62409 | 36.57069 | 34.02363 | chr2 | 164415904 | 164437244 | 5.93196 | 4.473 | 3.77292 |
| chr2 | 164438310 | 164438795 | 1000 | Intron (ENSMUST00000017153.3 | 2 | 164424247 | 164443186 | 18940 | 2 | ENSMUST00 | 4391 | Sdc4 | ENSMUSG0C | true | chr2 | 164438320 | 164443697 | 20.02967 | 73.35366 | 70.31202 | . | . | . | . | . | . |
| chr2 | 164442965 | 164443527 | 1000 | Promoter (<=1kb) | 2 | 164424247 | 164443186 | 18940 | 2 | ENSMUST00 | 0 | Sdc4 | ENSMUSG0C | true | chr2 | 164438320 | 164443697 | 20.02967 | 73.35366 | 70.31202 | chr2 | 164443397 | 164445791 | 4.18218 | 3.75066 | 2.31744 |
| chrX | 36111824 | 36112265 | 1000 | Promoter (<=1kb) | 20 | 36112110 | 36171259 | 59150 | 1 | ENSMUST00 | 0 | Il13ra1 | ENSMUSG0C | false | chrX | 36111740 | 36113074 | 25.27427 | 69.73028 | 66.74294 | . | . | . | . | . | . |
| chr2 | 51146952 | 51148141 | 1000 | Promoter (<=1kb) | 2 | 51137069 | 51149094 | 12026 | 2 | ENSMUST00 | 953 | Rnd3 | ENSMUSG0C | false | chr2 | 51146410 | 51149200 | 38.09013 | 122.35755 | 118.14518 | . | . | . | . | . | . |
| chr11 | 101170280 | 101171671 | 1000 | Promoter (<=1kb) | 11 | 101170523 | 101178316 | 7794 | 1 | ENSMUST00 | 0 | Cntnap1 | ENSMUSG0C | false | chr11 | 101169063 | 101171430 | 33.5288 | 108.10788 | 104.32592 | . | . | . | . | . | . |
| chr11 | 98708079 | 98709813 | 1000 | Promoter (<=1kb) | 11 | 98707266 | 98708003 | 738 | 2 | ENSMUST00 | -77 | Med24 | ENSMUSG0C | false | chr11 | 98709428 | 98710246 | 6.46168 | 12.41373 | 10.19419 | chr11 | 98700043 | 98711662 | 7.93936 | 5.23478 | 5.43738 |
| chr11 | 98053122 | 98053687 | 1000 | Promoter (<=1kb) | 11 | 98036623 | 98053462 | 16840 | 2 | ENSMUST00 | 0 | Stac2 | ENSMUSG0C | false | chr11 | 98051992 | 98053609 | 18.51447 | 52.87213 | 50.12019 | chr11 | 98051686 | 98053723 | 4.08379 | 3.44401 | 2.28578 |
| chr11 | 118428995 | 118429574 | 1000 | Promoter (<=1kb) | 11 | 118428499 | 118449963 | 21465 | 1 | ENSMUST00 | 497 | C1qtnf1 | ENSMUSG0C | false | chr11 | 118427431 | 118430359 | 22.74714 | 65.36804 | 62.44672 | chr11 | 118411596 | 118448618 | 7.23691 | 5.03811 | 4.84329 |
| chr11 | 118433416 | 118433933 | 1000 | Promoter (<=1kb) | 11 | 118433853 | 118449963 | 16111 | 1 | ENSMUST00 | 0 | C1qtnf1 | ENSMUSG0C | false | chr11 | 118433766 | 118434600 | 11.73508 | 26.0954 | 23.67916 | chr11 | 118411596 | 118448618 | 7.23691 | 5.03811 | 4.84329 |
| chr5 | 118439059 | 118439726 | 1000 | Intron (ENSMUST00000124861.1 | 11 | 118443471 | 118443850 | 380 | 1 | ENSMUST00 | -3745 | C1qtnf1 | ENSMUSG0C | false | . | . | . | . | . | . | chr11 | 118411596 | 118448618 | 7.23691 | 5.03811 | 4.84329 |
| chr5 | 118437807 | 118438320 | 1000 | Intron (ENSMUST00000124861.1 | 11 | 118433853 | 118449963 | 16111 | 1 | ENSMUST00 | 3955 | C1qtnf1 | ENSMUSG0C | false | . | . | . | . | . | . | chr11 | 118411596 | 118448618 | 7.23691 | 5.03811 | 4.84329 |
| chr11 | 118430169 | 118430587 | 1000 | Promoter (1-2kb) | 11 | 118428499 | 118449963 | 21465 | 1 | ENSMUST00 | 1671 | C1qtnf1 | ENSMUSG0C | false | chr11 | 118430506 | 118432556 | 10.50614 | 21.17611 | 18.82507 | chr11 | 118411596 | 118448618 | 7.23691 | 5.03811 | 4.84329 |
| chr11 | 118354989 | 118355759 | 1000 | Promoter (<=1kb) | 11 | 118301069 | 118355506 | 54438 | 2 | ENSMUST00 | 0 | Timp2 | ENSMUSG0C | false | chr11 | 118353231 | 118355882 | 34.66032 | 110.08408 | 106.24455 | chr11 | 118355503 | 118356375 | 9.06481 | 5.7183 | 6.30018 |
| chr5 | 16644735 | 16645055 | 1000 | Distal Intergenic | 14 | 16430842 | 16575472 | 144631 | 2 | ENSMUST00 | -69264 | Rarb | ENSMUSG0C | false | . | . | . | . | . | . | chr14 | 16644680 | 16646107 | 3.23926 | 3.08978 | 1.60785 |
| chr5 | 16677123 | 16677466 | 1000 | Distal Intergenic | 14 | 16430842 | 16575472 | 144631 | 2 | ENSMUST00 | -101652 | Rarb | ENSMUSG0C | false | . | . | . | . | . | . | chr14 | 16676569 | 16680764 | 4.30938 | 3.66931 | 2.43952 |
| chr5 | 99043153 | 99043510 | 1000 | Promoter (<=1kb) | 11 | 99043597 | 99051920 | 8324 | 1 | ENSMUST00 | -87 | Igfbp4 | ENSMUSG0C | false | . | . | . | . | . | . | chr11 | 99042650 | 99044634 | 3.6257 | 3.37225 | 1.87388 |
| chr11 | 79992873 | 79993640 | 1000 | Promoter (<=1kb) | 11 | 79993106 | 80034122 | 41017 | 1 | ENSMUST00 | 0 | Suz12 | ENSMUSG0C | false | chr11 | 79992377 | 79994546 | 39.66383 | 132.67865 | 128.09186 | . | . | . | . | . | . |
| chr5 | 80049996 | 80050409 | 1000 | Intron (ENSMUST00000103233.4 | 11 | 80056217 | 80057899 | 1683 | 2 | ENSMUST00 | 7490 | Crlf3 | ENSMUSG0C | true | . | . | . | . | . | . | chr11 | 80046430 | 80057838 | 5.61843 | 4.37638 | 3.5175 |
| chr5 | 80056223 | 80056553 | 1000 | Promoter (1-2kb) | 11 | 80056217 | 80057899 | 1683 | 2 | ENSMUST00 | 1346 | Crlf3 | ENSMUSG0C | true | . | . | . | . | . | . | chr11 | 80046430 | 80057838 | 5.61843 | 4.37638 | 3.5175 |
| chr5 | 76447781 | 76448200 | 1000 | Intron (ENSMUST00000094012.5 | 11 | 76423048 | 76437220 | 14173 | 2 | ENSMUST00 | -10562 | Abr | ENSMUSG0C | false | . | . | . | . | . | . | chr11 | 76424298 | 76452713 | 6.90719 | 4.90323 | 4.58627 |
| chr5 | 76468314 | 76468626 | 1000 | Promoter (<=1kb) | 11 | 76416751 | 76468465 | 51715 | 2 | ENSMUST00 | 0 | Abr | ENSMUSG0C | false | . | . | . | . | . | . | chr11 | 76460441 | 76470304 | 6.56354 | 4.63779 | 4.31962 |
| chr11 | 76437044 | 76437544 | 1000 | Promoter (<=1kb) | 11 | 76423048 | 76437220 | 14173 | 2 | ENSMUST00 | 0 | Abr | ENSMUSG0C | false | chr11 | 76434132 | 76437894 | 35.38397 | 108.24114 | 104.45254 | chr11 | 76424298 | 76452713 | 6.90719 | 4.90323 | 4.58627 |
| chr2 | 165323920 | 165324273 | 1000 | Promoter (2-3kb) | 2 | 165289155 | 165326479 | 37325 | 2 | ENSMUST00 | 2206 | Elmo2 | ENSMUSG0C | false | chr2 | 165323665 | 165324081 | 5.10133 | 8.58512 | 6.43618 | chr2 | 165307563 | 165325045 | 8.91187 | 5.755 | 6.22321 |
| chr5 | 165314173 | 165314751 | 1000 | Promoter (<=1kb) | 2 | 165298279 | 165314306 | 16028 | 2 | ENSMUST00 | 0 | Elmo2 | ENSMUSG0C | false | . | . | . | . | . | . | chr2 | 165307563 | 165325045 | 8.91187 | 5.755 | 6.22321 |
| chr5 | 165308644 | 165309103 | 1000 | Promoter (2-3kb) | 2 | 165304694 | 165311642 | 6949 | 2 | ENSMUST00 | 2539 | Elmo2 | ENSMUSG0C | false | . | . | . | . | . | . | chr2 | 165307563 | 165325045 | 8.91187 | 5.755 | 6.22321 |
| chr5 | 80294881 | 80295764 | 1000 | Distal Intergenic | 11 | 80300912 | 80355955 | 55044 | 1 | ENSMUST00 | -5148 | Rhbdl3 | ENSMUSG0C | true | . | . | . | . | . | . | chr11 | 80292230 | 80297765 | 6.78559 | 4.84623 | 4.46696 |
| chr11 | 80300514 | 80301342 | 1000 | Promoter (<=1kb) | 11 | 80300912 | 80355955 | 55044 | 1 | ENSMUST00 | 0 | Rhbdl3 | ENSMUSG0C | true | chr11 | 80300381 | 80302828 | 37.51873 | 124.57824 | 120.27934 | chr11 | 80298848 | 80300705 | 5.14473 | 4.22038 | 3.11169 |
| chr5 | 80305087 | 80305510 | 1000 | Intron (ENSMUST00000017836.7 | 11 | 80300912 | 80355955 | 55044 | 1 | ENSMUST00 | 4176 | Rhbdl3 | ENSMUSG0C | true | . | . | . | . | . | . | chr11 | 80302459 | 80307589 | 4.65342 | 3.84878 | 2.73157 |
| chr5 | 164511214 | 164511635 | 1000 | Distal Intergenic | 2 | 164499916 | 164501639 | 1724 | 1 | ENSMUST00 | 11299 | Pigt | ENSMUSG0C | false | . | . | . | . | . | . | chr2 | 164509177 | 164517266 | 5.6289 | 4.32047 | 3.52645 |
| chr11 | 101772407 | 101772838 | 1000 | Promoter (<=1kb) | 11 | 101770354 | 101772181 | 1828 | 2 | ENSMUST00 | -227 | Etv4 | ENSMUSG0C | false | chr11 | 101772347 | 101773047 | 12.99241 | 30.10301 | 27.6364 | chr11 | 101771997 | 101773182 | 4.23046 | 3.65955 | 2.38585 |
| chr2 | 164485964 | 164487211 | 1000 | Promoter (<=1kb) | 2 | 164486106 | 164490767 | 4662 | 1 | ENSMUST00 | 0 | Dbndd2 | ENSMUSG0C | false | chr2 | 164485834 | 164487505 | 36.17194 | 114.79427 | 110.81874 | chr2 | 164485179 | 164487759 | 5.96839 | 4.54196 | 3.82394 |
| chr11 | 75651380 | 75652104 | 1000 | Promoter (<=1kb) | 11 | 75651510 | 75673910 | 22401 | 1 | ENSMUST00 | 0 | Myo1c | ENSMUSG0C | false | chr11 | 75651407 | 75657466 | 25.17393 | 93.69764 | 90.27328 | . | . | . | . | . | . |
| chr11 | 75653786 | 75654407 | 1000 | Promoter (1-2kb) | 11 | 75655873 | 75673908 | 18036 | 1 | ENSMUST00 | -1466 | Myo1c | ENSMUSG0C | false | chr11 | 75651407 | 75657466 | 25.17393 | 93.69764 | 90.27328 | chr11 | 75652271 | 75655636 | 6.31319 | 4.65216 | 4.07889 |
| chr11 | 101119577 | 101120170 | 1000 | Promoter (<=1kb) | 11 | 101096322 | 101119843 | 23522 | 2 | ENSMUST00 | 0 | Fam134c | ENSMUSG0C | false | chr11 | 101118885 | 101119919 | 32.62678 | 97.43105 | 93.92233 | . | . | . | . | . | . |
| chr17 | 53478767 | 53479847 | 1000 | Promoter (<=1kb) | 17 | 53479234 | 53507671 | 28438 | 1 | ENSMUST00 | 0 | Rab5a | ENSMUSG0C | false | chr17 | 53478468 | 53481041 | 31.85236 | 101.09525 | 97.496 | . | . | . | . | . | . |
| chr5 | 110537834 | 110538175 | 1000 | Intron (ENSMUST00000084985.4 | 12 | 110485739 | 110583061 | 97323 | 1 | ENSMUST00 | 52096 | Ppp2r5c | ENSMUSG0C | false | . | . | . | . | . | . | chr12 | 110530824 | 110543309 | 5.16932 | 4.1173 | 3.16417 |
| chr5 | 167317918 | 167318247 | 1000 | Intron (ENSMUST00000109221.3 | 2 | 167307610 | 167313900 | 6291 | 2 | ENSMUST00 | -4019 | B4galt5 | ENSMUSG0C | false | . | . | . | . | . | . | chr2 | 167313423 | 167319941 | 5.07451 | 3.89375 | 3.10085 |
| chr5 | 167250689 | 167251070 | 1000 | Distal Intergenic | 2 | 167202907 | 167240604 | 37698 | 2 | ENSMUST00 | -10086 | Ptgis | ENSMUSG0C | false | . | . | . | . | . | . | chr2 | 167248720 | 167256438 | 5.64516 | 4.34556 | 3.54025 |
| chr5 | 143638489 | 143638829 | 1000 | Promoter (2-3kb) | 5 | 143636351 | 143692554 | 56204 | 1 | ENSMUST00 | 2139 | Cyth3 | ENSMUSG0C | false | . | . | . | . | . | . | chr5 | 143628431 | 143643057 | 5.82289 | 4.44189 | 3.69355 |
| chr5 | 83164865 | 83165373 | 1000 | Intron (ENSMUST00000018186.1 | 15 | 83160385 | 83170177 | 9793 | 2 | ENSMUST00 | 4804 | Cyb5r3 | ENSMUSG0C | false | . | . | . | . | . | . | chr15 | 83162089 | 83173300 | 7.98675 | 5.40074 | 5.45307 |
| chr15 | 83170008 | 83170395 | 1000 | Promoter (<=1kb) | 15 | 83160385 | 83170177 | 9793 | 2 | ENSMUST00 | 0 | Cyb5r3 | ENSMUSG0C | false | chr15 | 83169965 | 83170528 | 5.81673 | 9.0994 | 6.94297 | chr15 | 83162089 | 83173300 | 7.98675 | 5.40074 | 5.45307 |
| chr5 | 118559843 | 118561464 | 1000 | Promoter (<=1kb) | 5 | 118560719 | 118765435 | 204717 | 1 | ENSMUST00 | 0 | Med13l | ENSMUSG0C | false | chr5 | 118559542 | 118561980 | 28.96508 | 85.28318 | 82.03651 | chr5 | 118559570 | 118559870 | 2.83457 | 2.83494 | 1.27261 |
| chr5 | 139796893 | 139797205 | 1000 | Promoter (<=1kb) | 5 | 139797514 | 139802331 | 4818 | 1 | ENSMUST00 | -309 | Mafk | ENSMUSG0C | false | chr5 | 139796675 | 139797064 | 13.83063 | 32.8443 | 30.34371 | chr5 | 139792309 | 139799489 | 7.14368 | 5.03259 | 4.78629 |
| chr5 | 143841090 | 143841550 | 1000 | Distal Intergenic | 1 | 143794717 | 143806941 | 12225 | 1 | ENSMUST00 | 46374 | Uchl5 | ENSMUSG0C | false | . | . | . | . | . | . | chr1 | 143836631 | 143841216 | 5.72203 | 4.37792 | 3.60808 |
| chr11 | 70654448 | 70655267 | 1000 | Promoter (<=1kb) | 11 | 70651850 | 70654644 | 2795 | 2 | ENSMUST00 | 0 | Pfn1 | ENSMUSG0C | false | chr11 | 70652869 | 70656609 | 41.07278 | 140.67711 | 135.76277 | . | . | . | . | . | . |
| chr2 | 163994742 | 163995283 | 1000 | Promoter (<=1kb) | 2 | 163994960 | 164018588 | 23629 | 1 | ENSMUST00 | 0 | Ywhab | ENSMUSG0C | false | chr2 | 163994559 | 163996605 | 32.3084 | 121.498 | 117.30614 | chr2 | 163994551 | 164011391 | 5.67263 | 4.35304 | 3.57315 |
| chr5 | 79058242 | 79058734 | 1000 | Promoter (<=1kb) | 11 | 79045183 | 79057627 | 12445 | 2 | ENSMUST00 | -616 | Ksr1 | ENSMUSG0C | true | . | . | . | . | . | . | chr11 | 79045596 | 79144720 | 7.09579 | 4.93486 | 4.74337 |
| chr5 | 79116640 | 79117154 | 1000 | Intron (ENSMUST00000018478.5 | 11 | 79013440 | 79146407 | 132968 | 2 | ENSMUST00 | 29253 | Ksr1 | ENSMUSG0C | true | . | . | . | . | . | . | chr11 | 79045596 | 79144720 | 7.09579 | 4.93486 | 4.74337 |
| chr5 | 79063509 | 79063981 | 1000 | Intron (ENSMUST00000018478.5 | 11 | 79044920 | 79059875 | 14956 | 2 | ENSMUST00 | -3635 | Ksr1 | ENSMUSG0C | true | . | . | . | . | . | . | chr11 | 79045596 | 79144720 | 7.09579 | 4.93486 | 4.74337 |
| chr5 | 79107818 | 79108194 | 1000 | Intron (ENSMUST00000018478.5 | 11 | 79013440 | 79146407 | 132968 | 2 | ENSMUST00 | 38213 | Ksr1 | ENSMUSG0C | true | . | . | . | . | . | . | chr11 | 79045596 | 79144720 | 7.09579 | 4.93486 | 4.74337 |
| chr5 | 79081281 | 79081714 | 1000 | Intron (ENSMUST00000018478.5 | 11 | 79044920 | 79059875 | 14956 | 2 | ENSMUST00 | -21407 | Ksr1 | ENSMUSG0C | true | . | . | . | . | . | . | chr11 | 79045596 | 79144720 | 7.09579 | 4.93486 | 4.74337 |
| chr5 | 79065714 | 79066124 | 1000 | Intron (ENSMUST00000018478.5 | 11 | 79044920 | 79059875 | 14956 | 2 | ENSMUST00 | -5840 | Ksr1 | ENSMUSG0C | true | . | . | . | . | . | . | chr11 | 79045596 | 79144720 | 7.09579 | 4.93486 | 4.74337 |
| chr11 | 65807011 | 65807614 | 1000 | Promoter (<=1kb) | 11 | 65807175 | 65829238 | 22064 | 1 | ENSMUST00 | 0 | Zkscan6 | ENSMUSG0C | false | chr11 | 65806297 | 65808006 | 24.48636 | 83.88964 | 80.66612 | . | . | . | . | . | . |
| chr11 | 106919992 | 106921007 | 1000 | Promoter (<=1kb) | 11 | 106820066 | 106920715 | 100650 | 2 | ENSMUST00 | 0 | Smurf2 | ENSMUSG0C | false | chr11 | 106918841 | 106920785 | 26.86698 | 95.04005 | 91.5885 | . | . | . | . | . | . |
| chr11 | 106787507 | 106789437 | 1000 | Promoter (<=1kb) | 11 | 106789252 | 106818860 | 29609 | 1 | ENSMUST00 | 0 | Cep95 | ENSMUSG0C | false | chr11 | 106786098 | 106790164 | 37.82652 | 124.04217 | 119.77048 | . | . | . | . | . | . |
| chr11 | 88067239 | 88069381 | 1000 | Promoter (<=1kb) | 11 | 88068279 | 88084729 | 16451 | 1 | ENSMUST00 | 0 | Vezf1 | ENSMUSG0C | false | chr11 | 88067386 | 88069722 | 35.29575 | 125.98199 | 121.63858 | chr11 | 88066600 | 88067607 | 5.08706 | 4.07747 | 3.08682 |
| chr5 | 88180968 | 88181577 | 1000 | Promoter (<=1kb) | 11 | 88182303 | 88183280 | 978 | 1 | ENSMUST00 | -726 | Cuedc1 | ENSMUSG0C | true | . | . | . | . | . | . | chr11 | 88146778 | 88183132 | 6.97604 | 4.89368 | 4.62779 |
| chr11 | 53519593 | 53520345 | 1000 | Promoter (<=1kb) | 11 | 53519746 | 53544095 | 24350 | 1 | ENSMUST00 | 0 | Sep-08 | ENSMUSG0C | false | chr11 | 53519402 | 53521185 | 40.11361 | 128.01187 | 123.59241 | chr11 | 53519096 | 53521911 | 4.4349 | 3.84194 | 2.57204 |
| chr5 | 104213590 | 104213960 | 1000 | Distal Intergenic | 11 | 104231390 | 104332090 | 100701 | 1 | ENSMUST00 | -17430 | Mapt | ENSMUSG0C | true | . | . | . | . | . | . | chr11 | 104212269 | 104220562 | 4.58875 | 3.79375 | 2.6827 |
| chr11 | 104441315 | 104442740 | 1000 | Promoter (<=1kb) | 11 | 104333229 | 104442291 | 109063 | 2 | ENSMUST00 | 0 | Kansl1 | ENSMUSG0C | false | chr11 | 104440325 | 104443479 | 24.60788 | 97.30984 | 93.80507 | . | . | . | . | . | . |
| chr5 | 86962037 | 86962621 | 1000 | Intron (ENSMUST00000018571.4 | 11 | 86936425 | 86993707 | 57283 | 2 | ENSMUST00 | 31086 | Ypel2 | ENSMUSG0C | false | . | . | . | . | . | . | chr11 | 86954328 | 86964828 | 6.34543 | 4.58398 | 4.13993 |
| chr11 | 69368155 | 69369646 | 1000 | Promoter (<=1kb) | 11 | 69343273 | 69369406 | 26134 | 2 | ENSMUST00 | 0 | Chd3 | ENSMUSG0C | false | chr11 | 69366750 | 69369724 | 26.28407 | 85.32412 | 82.07336 | chr11 | 69364078 | 69368657 | 6.32118 | 4.46781 | 4.09186 |
| chr11 | 69366400 | 69367003 | 1000 | Promoter (1-2kb) | 11 | 69345136 | 69364779 | 19644 | 2 | ENSMUST00 | -1622 | Chd3 | ENSMUSG0C | false | chr11 | 69366750 | 69369724 | 26.28407 | 85.32412 | 82.07336 | chr11 | 69364078 | 69368657 | 6.32118 | 4.46781 | 4.09186 |
| chr5 | 69364912 | 69365341 | 1000 | Promoter (<=1kb) | 11 | 69345136 | 69364779 | 19644 | 2 | ENSMUST00 | -134 | Chd3 | ENSMUSG0C | false | . | . | . | . | . | . | chr11 | 69364078 | 69368657 | 6.32118 | 4.46781 | 4.09186 |
| chr11 | 69415280 | 69415615 | 1000 | Promoter (1-2kb) | 11 | 69398508 | 69413675 | 15168 | 2 | ENSMUST00 | -1606 | Kdm6b | ENSMUSG0C | false | chr11 | 69415361 | 69416249 | 20.21941 | 51.82391 | 49.08421 | chr11 | 69410823 | 69417315 | 7.23301 | 5.12321 | 4.82613 |
| chr11 | 69415810 | 69416583 | 1000 | Promoter (2-3kb) | 11 | 69398508 | 69413675 | 15168 | 2 | ENSMUST00 | -2136 | Kdm6b | ENSMUSG0C | false | chr11 | 69415361 | 69416249 | 20.21941 | 51.82391 | 49.08421 | chr11 | 69410823 | 69417315 | 7.23301 | 5.12321 | 4.82613 |
| chr11 | 69409440 | 69410230 | 1000 | Intron (ENSMUST00000094077.4 | 11 | 69398508 | 69413675 | 15168 | 2 | ENSMUST00 | 3445 | Kdm6b | ENSMUSG0C | false | chr11 | 69408142 | 69412281 | 27.88725 | 99.90214 | 96.33501 | chr11 | 69407007 | 69409461 | 6.0002 | 4.43125 | 3.83149 |
| chr11 | 69417496 | 69418326 | 1000 | Downstream (2-3kb) | 11 | 69398508 | 69413675 | 15168 | 2 | ENSMUST00 | -3822 | Kdm6b | ENSMUSG0C | false | chr11 | 69416361 | 69418915 | 33.54585 | 101.01101 | 97.41918 | . | . | . | . | . | . |
| chr11 | 69418556 | 69419150 | 1000 | Downstream (1-2kb) | 11 | 69398508 | 69413675 | 15168 | 2 | ENSMUST00 | -4882 | Kdm6b | ENSMUSG0C | false | chr11 | 69416361 | 69418915 | 33.54585 | 101.01101 | 97.41918 | . | . | . | . | . | . |
| chr11 | 62456944 | 62457899 | 1000 | Promoter (<=1kb) | 11 | 62316426 | 62457332 | 140907 | 2 | ENSMUST00 | 0 | Ncor1 | ENSMUSG0C | false | chr11 | 62455956 | 62459233 | 26.32471 | 106.6389 | 102.89503 | . | . | . | . | . | . |
| chr11 | 87126808 | 87127524 | 1000 | Promoter (<=1kb) | 11 | 87127077 | 87220683 | 93607 | 2 | ENSMUST00 | 0 | Trim37 | ENSMUSG0C | false | chr11 | 87126333 | 87127988 | 43.50226 | 149.44124 | 144.17897 | chr11 | 87126325 | 87129601 | 3.63904 | 3.33705 | 1.89687 |
| chr11 | 69996774 | 69997574 | 1000 | Promoter (<=1kb) | 11 | 69996941 | 69998496 | 1556 | 1 | ENSMUST00 | 0 | Phf23 | ENSMUSG0C | false | chr11 | 69995731 | 69997499 | 38.91574 | 128.10115 | 123.6785 | chr11 | 69997404 | 69998177 | 3.98266 | 3.55094 | 2.20032 |
| chr5 | 55414186 | 55414716 | 1000 | Intron (ENSMUST00000108858.2 | 11 | 55394500 | 55419898 | 25399 | 2 | ENSMUST00 | 5182 | Sparc | ENSMUSG0C | true | . | . | . | . | . | . | chr11 | 55405757 | 55417699 | 6.44715 | 4.69413 | 4.21932 |
| chr11 | 96849685 | 96850294 | 1000 | Promoter (<=1kb) | 11 | 96849870 | 96851590 | 1721 | 1 | ENSMUST00 | 0 | Copz2 | ENSMUSG0C | false | chr11 | 96849555 | 96850735 | 27.24215 | 82.29425 | 79.10038 | chr11 | 96848128 | 96851397 | 5.18652 | 4.07181 | 3.20166 |
| chr11 | 102407350 | 102407928 | 1000 | Promoter (<=1kb) | 11 | 102402985 | 102407489 | 4505 | 2 | ENSMUST00 | 0 | Slc25a39 | ENSMUSG0C | false | chr11 | 102406444 | 102408601 | 32.64849 | 123.19247 | 118.94151 | chr11 | 102407921 | 102409731 | 4.81374 | 4.01023 | 2.85015 |
| chr5 | 96995104 | 96995700 | 1000 | Distal Intergenic | 11 | 96954219 | 96982959 | 28741 | 2 | ENSMUST00 | -12146 | Sp2 | ENSMUSG0C | false | . | . | . | . | . | . | chr11 | 96984955 | 96998855 | 6.28932 | 4.5897 | 4.08329 |
| chr11 | 96977143 | 96977818 | 1000 | Promoter (<=1kb) | 11 | 96953340 | 96977688 | 24349 | 2 | ENSMUST00 | 0 | Sp2 | ENSMUSG0C | false | chr11 | 96976345 | 96977744 | 37.54867 | 136.59039 | 131.84451 | . | . | . | . | . | . |
| chr11 | 96997208 | 96997784 | 1000 | Distal Intergenic | 11 | 96954219 | 96982959 | 28741 | 2 | ENSMUST00 | -14250 | Sp2 | ENSMUSG0C | false | chr11 | 96997117 | 96997561 | 6.28665 | 10.55282 | 8.36284 | chr11 | 96984955 | 96998855 | 6.28932 | 4.5897 | 4.08329 |
| chr5 | 110555746 | 110556407 | 1000 | Intron (ENSMUST00000084985.4 | 12 | 110601452 | 110666945 | 65494 | 1 | ENSMUST00 | -45045 | Dync1h1 | ENSMUSG0C | false | . | . | . | . | . | . | chr12 | 110549007 | 110557689 | 5.99039 | 4.4389 | 3.83972 |
| chr11 | 68852486 | 68853395 | 1000 | Promoter (<=1kb) | 11 | 68821434 | 68853135 | 31702 | 2 | ENSMUST00 | 0 | Ndel1 | ENSMUSG0C | false | chr11 | 68851734 | 68853148 | 35.99199 | 121.88605 | 117.68387 | . | . | . | . | . | . |
| chr5 | 69777927 | 69778287 | 1000 | Promoter (2-3kb) | 11 | 69775406 | 69779673 | 4268 | 1 | ENSMUST00 | 2522 | Zbtb4 | ENSMUSG0C | false | . | . | . | . | . | . | chr11 | 69771591 | 69779554 | 5.27322 | 3.95146 | 3.25945 |
| chr11 | 69695489 | 69696187 | 1000 | Promoter (<=1kb) | 11 | 69682577 | 69696098 | 13522 | 2 | ENSMUST00 | 0 | Tnfsf12Tnfsf | ENSMUSG0C | false | chr11 | 69695084 | 69696451 | 30.37608 | 91.57915 | 88.20459 | chr11 | 69688439 | 69697396 | 5.01103 | 4.04632 | 3.01977 |
| chr5 | 142470984 | 142471585 | 1000 | Promoter (<=1kb) | 7 | 142471843 | 142494599 | 22757 | 1 | ENSMUST00 | -258 | Lsp1 | ENSMUSG0C | false | . | . | . | . | . | . | chr7 | 142469601 | 142472837 | 7.10498 | 5.05047 | 4.74174 |
| chr19 | 42163887 | 42164525 | 1000 | Promoter (<=1kb) | 19 | 42163951 | 42177726 | 13776 | 1 | ENSMUST00 | 0 | Zfyve27 | ENSMUSG0C | false | chr19 | 42163549 | 42163994 | 4.31067 | 6.37555 | 4.28397 | chr19 | 42160246 | 42168006 | 5.40781 | 4.29476 | 3.33041 |
| chr19 | 42128825 | 42129492 | 1000 | Promoter (<=1kb) | 19 | 42123273 | 42129059 | 5787 | 2 | ENSMUST00 | 0 | Avpi1 | ENSMUSG0C | false | chr19 | 42128190 | 42129561 | 25.50663 | 88.63404 | 85.31516 | chr19 | 42127135 | 42130834 | 4.67193 | 3.96929 | 2.72193 |
| chr11 | 35838352 | 35838617 | 1000 | 3' UTR | 11 | 35808381 | 35834506 | 26126 | 2 | ENSMUST00 | -3847 | Rars | ENSMUSG0C | false | chr11 | 35838072 | 35838821 | 10.5671 | 23.5101 | 21.12713 | chr11 | 35837903 | 35838819 | 3.9286 | 3.48188 | 2.11713 |
| chr5 | 115400842 | 115401268 | 1000 | Promoter (2-3kb) | 11 | 115403752 | 115412674 | 8923 | 1 | ENSMUST00 | -2484 | Ict1 | ENSMUSG0C | false | . | . | . | . | . | . | chr11 | 115400497 | 115401635 | 5.00385 | 3.84016 | 3.04939 |
| chr5 | 97302243 | 97302727 | 1000 | Distal Intergenic | 11 | 97315716 | 97329920 | 14205 | 1 | ENSMUST00 | -12989 | Mrpl45 | ENSMUSG0C | false | . | . | . | . | . | . | chr11 | 97301032 | 97303320 | 6.18902 | 4.68642 | 4.0451 |
| chr5 | 77025471 | 77025846 | 1000 | Promoter (2-3kb) | 15 | 77015489 | 77022787 | 7299 | 2 | ENSMUST00 | -2685 | Mb | ENSMUSG0C | false | . | . | . | . | . | . | chr15 | 77025092 | 77026286 | 4.80928 | 3.8186 | 2.89726 |
| chr5 | 54115345 | 54115734 | 1000 | Promoter (2-3kb) | 11 | 54113203 | 54114274 | 1072 | 1 | ENSMUST00 | 2143 | P4ha2 | ENSMUSG0C | false | . | . | . | . | . | . | chr11 | 54110812 | 54117439 | 8.16018 | 5.41308 | 5.62178 |
| chr5 | 54115844 | 54116337 | 1000 | Promoter (2-3kb) | 11 | 54113203 | 54114274 | 1072 | 1 | ENSMUST00 | 2642 | P4ha2 | ENSMUSG0C | false | . | . | . | . | . | . | chr11 | 54110812 | 54117439 | 8.16018 | 5.41308 | 5.62178 |
| chr7 | 99535392 | 99535965 | 1000 | Promoter (<=1kb) | 7 | 99535466 | 99606771 | 71306 | 1 | ENSMUST00 | 0 | Arrb1 | ENSMUSG0C | false | chr7 | 99535128 | 99537920 | 25.99992 | 84.80917 | 81.57088 | chr7 | 99534502 | 99535527 | 5.81443 | 4.51927 | 3.65705 |
| chr5 | 99570780 | 99571090 | 1000 | Promoter (1-2kb) | 7 | 99573064 | 99588128 | 15065 | 1 | ENSMUST00 | -1974 | Arrb1 | ENSMUSG0C | false | . | . | . | . | . | . | chr7 | 99536482 | 99576026 | 7.32182 | 4.95837 | 4.9327 |
| chr5 | 99528511 | 99528884 | 1000 | Distal Intergenic | 7 | 99535466 | 99606771 | 71306 | 1 | ENSMUST00 | -6582 | Arrb1 | ENSMUSG0C | false | . | . | . | . | . | . | chr7 | 99522439 | 99531882 | 5.37849 | 4.25246 | 3.31489 |
| chr5 | 99529451 | 99529820 | 1000 | Distal Intergenic | 7 | 99535466 | 99606771 | 71306 | 1 | ENSMUST00 | -5646 | Arrb1 | ENSMUSG0C | false | . | . | . | . | . | . | chr7 | 99522439 | 99531882 | 5.37849 | 4.25246 | 3.31489 |
| chr11 | 60933547 | 60933918 | 1000 | Promoter (1-2kb) | 11 | 60932066 | 60952794 | 20729 | 1 | ENSMUST00 | 1482 | Map2k3 | ENSMUSG0C | false | chr11 | 60933399 | 60934322 | 6.12159 | 11.42045 | 9.22029 | chr11 | 60932782 | 60942948 | 7.29531 | 5.06712 | 4.89209 |
| chr5 | 33018476 | 33019338 | 1000 | Promoter (<=1kb) | 5 | 33018816 | 33027966 | 9151 | 1 | ENSMUST00 | 0 | Ywhah | ENSMUSG0C | false | chr5 | 33018136 | 33018705 | 23.5743 | 68.56921 | 65.60024 | . | . | . | . | . | . |
| chr5 | 54573253 | 54573694 | 1000 | Distal Intergenic | 6 | 54505890 | 54566489 | 60600 | 2 | ENSMUST00 | -6765 | Scrn1 | ENSMUSG0C | false | . | . | . | . | . | . | chr6 | 54572187 | 54576142 | 4.48629 | 3.68427 | 2.62741 |
| chr13 | 24800679 | 24801915 | 1000 | Promoter (<=1kb) | 13 | 24801657 | 24814013 | 12357 | 1 | ENSMUST00 | 0 | BC005537 | ENSMUSG0C | false | chr13 | 24799815 | 24800731 | 21.02725 | 63.18829 | 60.29588 | . | . | . | . | . | . |
| chr8 | 70593884 | 70594548 | 1000 | Promoter (<=1kb) | 8 | 70594481 | 70597288 | 2808 | 1 | ENSMUST00 | 0 | Isyna1 | ENSMUSG0C | false | chr8 | 70593499 | 70595422 | 22.78592 | 76.10319 | 73.01487 | chr8 | 70592285 | 70603072 | 6.42958 | 4.72399 | 4.17224 |
| chr5 | 70593104 | 70593488 | 1000 | Promoter (<=1kb) | 8 | 70594481 | 70597288 | 2808 | 1 | ENSMUST00 | -993 | Isyna1 | ENSMUSG0C | false | . | . | . | . | . | . | chr8 | 70592285 | 70603072 | 6.42958 | 4.72399 | 4.17224 |
| chr5 | 78119614 | 78120106 | 1000 | Promoter (<=1kb) | 15 | 77991748 | 78120028 | 128281 | 2 | ENSMUST00 | 0 | Cacng2 | ENSMUSG0C | false | . | . | . | . | . | . | chr15 | 78119433 | 78120204 | 3.1749 | 2.91547 | 1.56306 |
| chr5 | 33657852 | 33658577 | 1000 | Promoter (<=1kb) | 5 | 33653216 | 33657977 | 4762 | 2 | ENSMUST00 | 0 | Tmem129 | ENSMUSG0C | false | chr5 | 33656992 | 33659141 | 23.12601 | 77.64799 | 74.53377 | . | . | . | . | . | . |
| chr17 | 24695981 | 24696482 | 1000 | Promoter (<=1kb) | 17 | 24696234 | 24700529 | 4296 | 1 | ENSMUST00 | 0 | Noxo1 | ENSMUSG0C | false | chr17 | 24695460 | 24696926 | 28.91247 | 96.86193 | 93.36482 | chr17 | 24695110 | 24697166 | 4.05217 | 3.62049 | 2.27412 |
| chr5 | 16917813 | 16918453 | 1000 | Promoter (<=1kb) | 7 | 16915937 | 16917203 | 1267 | 2 | ENSMUST00 | -611 | Calm3 | ENSMUSG0C | false | . | . | . | . | . | . | chr7 | 16913262 | 16922976 | 7.95697 | 5.3563 | 5.42533 |
| chr8 | 83652293 | 83652808 | 1000 | Promoter (<=1kb) | 8 | 83652678 | 83664788 | 12111 | 1 | ENSMUST00 | 0 | Gipc1 | ENSMUSG0C | false | chr8 | 83651941 | 83653788 | 35.38397 | 108.24114 | 104.45254 | . | . | . | . | . | . |
| chr5 | 83666274 | 83666957 | 1000 | Promoter (<=1kb) | 8 | 83666833 | 83671268 | 4436 | 1 | ENSMUST00 | 0 | Ptger1 | ENSMUSG0C | false | . | . | . | . | . | . | chr8 | 83664333 | 83669382 | 6.72627 | 4.88407 | 4.41607 |
| chr18 | 47368576 | 47369298 | 1000 | Promoter (<=1kb) | 18 | 47245254 | 47368830 | 123577 | 2 | ENSMUST00 | 0 | Sema6a | ENSMUSG0C | false | chr18 | 47366217 | 47370796 | 17.00442 | 50.91425 | 48.18763 | chr18 | 47366285 | 47371323 | 4.47824 | 3.76553 | 2.58752 |
| chr1 | 177257106 | 177258848 | 1000 | Promoter (<=1kb) | 1 | 177022116 | 177258203 | 236088 | 2 | ENSMUST00 | 0 | Akt3 | ENSMUSG0C | true | chr1 | 177256972 | 177259264 | 37.21373 | 117.65135 | 113.58522 | . | . | . | . | . | . |
| chr5 | 72434995 | 72435268 | 1000 | Promoter (<=1kb) | 8 | 72424183 | 72435156 | 10974 | 2 | ENSMUST00 | 0 | Calr3 | ENSMUSG0C | false | . | . | . | . | . | . | chr8 | 72434698 | 72435801 | 6.41426 | 4.80533 | 4.15198 |
| chr5 | 3633845 | 3634505 | 1000 | Distal Intergenic | 10 | 3540288 | 3552074 | 11787 | 1 | ENSMUST00 | 93558 | Iyd | ENSMUSG0C | false | . | . | . | . | . | . | chr10 | 3633305 | 3635178 | 5.97277 | 4.61135 | 3.79128 |
| chr10 | 4432048 | 4432659 | 1000 | Promoter (<=1kb) | 10 | 4401915 | 4432388 | 30474 | 2 | ENSMUST00 | 0 | Rmnd1 | ENSMUSG0C | false | chr10 | 4431665 | 4432359 | 21.40274 | 64.72344 | 61.81032 | chr10 | 4422045 | 4437373 | 5.83291 | 4.40627 | 3.70343 |
| chr10 | 5823762 | 5824487 | 1000 | Promoter (<=1kb) | 10 | 5811887 | 5823910 | 12024 | 2 | ENSMUST00 | 0 | Mtrf1l | ENSMUSG0C | false | chr10 | 5823125 | 5825026 | 23.4661 | 79.19912 | 76.05865 | . | . | . | . | . | . |
| chr10 | 34483213 | 34483600 | 1000 | Promoter (<=1kb) | 10 | 34483400 | 34611226 | 127827 | 1 | ENSMUST00 | 0 | Frk | ENSMUSG0C | false | chr10 | 34483224 | 34485079 | 21.59801 | 56.60135 | 53.80005 | chr10 | 34482834 | 34483449 | 3.14215 | 3.11292 | 1.52688 |
| chr10 | 30842266 | 30843233 | 1000 | Promoter (<=1kb) | 10 | 30832359 | 30842801 | 10443 | 2 | ENSMUST00 | 0 | Hey2 | ENSMUSG0C | false | chr10 | 30841853 | 30844572 | 23.80619 | 80.75649 | 77.58801 | chr10 | 30843121 | 30844566 | 4.49887 | 3.80495 | 2.63395 |
| chr10 | 7662604 | 7663750 | 1000 | Promoter (<=1kb) | 10 | 7629373 | 7663264 | 33892 | 2 | ENSMUST00 | 0 | Pcmt1 | ENSMUSG0C | false | chr10 | 7662375 | 7663966 | 31.77438 | 112.18448 | 108.28101 | chr10 | 7663688 | 7664269 | 4.57359 | 3.90425 | 2.64889 |
| chr10 | 43540562 | 43541201 | 1000 | Promoter (<=1kb) | 10 | 43525121 | 43540994 | 15874 | 2 | ENSMUST00 | 0 | 1700021F05I | ENSMUSG0C | false | chr10 | 43540096 | 43541068 | 37.09717 | 122.76616 | 118.52905 | chr10 | 43536760 | 43541464 | 4.73951 | 3.86652 | 2.82674 |
| chr10 | 42761335 | 42761758 | 1000 | Promoter (<=1kb) | 10 | 42761496 | 42832514 | 71019 | 1 | ENSMUST00 | 0 | Sec63 | ENSMUSG0C | false | chr10 | 42761070 | 42761389 | 11.15939 | 31.88065 | 29.39184 | . | . | . | . | . | . |
| chr5 | 13107157 | 13107649 | 1000 | Intron (ENSMUST00000123135.2 | 10 | 13113465 | 13127341 | 13877 | 1 | ENSMUST00 | -5816 | Plagl1 | ENSMUSG0C | true | . | . | . | . | . | . | chr10 | 13102931 | 13117612 | 5.30894 | 4.20106 | 3.26917 |
| chr10 | 12868033 | 12868554 | 1000 | Distal Intergenic | 10 | 12382188 | 12861735 | 479548 | 2 | ENSMUST00 | -6299 | Utrn | ENSMUSG0C | true | chr10 | 12866850 | 12869855 | 26.89699 | 109.78668 | 105.95901 | . | . | . | . | . | . |
| chr10 | 12868924 | 12869499 | 1000 | Distal Intergenic | 10 | 12382188 | 12861735 | 479548 | 2 | ENSMUST00 | -7190 | Utrn | ENSMUSG0C | true | chr10 | 12866850 | 12869855 | 26.89699 | 109.78668 | 105.95901 | . | . | . | . | . | . |
| chr10 | 39731180 | 39732894 | 1000 | Promoter (<=1kb) | 10 | 39732118 | 39875211 | 143094 | 1 | ENSMUST00 | 0 | Rev3l | ENSMUSG0C | false | chr10 | 39730769 | 39733802 | 22.79947 | 93.49365 | 90.07696 | . | . | . | . | . | . |
| chr5 | 39387541 | 39387852 | 1000 | Intron (ENSMUST00000148152.2 | 10 | 39369803 | 39564423 | 194621 | 1 | ENSMUST00 | 17739 | Fyn | ENSMUSG0C | true | . | . | . | . | . | . | chr10 | 39371015 | 39390933 | 5.77617 | 4.40934 | 3.64788 |
| chr5 | 39378239 | 39378742 | 1000 | Intron (ENSMUST00000148152.2 | 10 | 39369803 | 39564423 | 194621 | 1 | ENSMUST00 | 8437 | Fyn | ENSMUSG0C | true | . | . | . | . | . | . | chr10 | 39371015 | 39390933 | 5.77617 | 4.40934 | 3.64788 |

| | | | | | | | | | | | | | | | | | | | | | | | | | | |
|---|---|---|---|---|---|---|---|---|---|---|---|---|---|---|---|---|---|---|---|---|---|---|---|---|---|---|
| chr10 | 39369600 | 39370988 | 1000 | Promoter (<=1kb) | 10 | 39369764 | 39565381 | 195618 | 1 | ENSMUST00 | 0 | Fyn | ENSMUSG0C | true | chr10 | 39369298 | 39371780 | 26.17789 | 120.04698 | 115.90433 | . | . | . | . | . | . |
| chr5 | 39459660 | 39460207 | 1000 | Intron (ENSMUST00000148152.2 | 10 | 39454915 | 39564355 | 109441 | 1 | ENSMUST00 | 4746 | Fyn | ENSMUSG0C | true | . | . | . | . | . | . | chr10 | 39448300 | 39473913 | 6.46111 | 4.64933 | 4.22352 |
| chr5 | 39487435 | 39487744 | 1000 | Intron (ENSMUST00000148152.2 | 10 | 39511739 | 39564422 | 52684 | 1 | ENSMUST00 | -23995 | Fyn | ENSMUSG0C | true | . | . | . | . | . | . | chr10 | 39483418 | 39514959 | 6.19365 | 4.56605 | 3.99401 |
| chr5 | 39495195 | 39495587 | 1000 | Intron (ENSMUST00000148152.2 | 10 | 39511739 | 39564422 | 52684 | 1 | ENSMUST00 | -16152 | Fyn | ENSMUSG0C | true | . | . | . | . | . | . | chr10 | 39483418 | 39514959 | 6.19365 | 4.56605 | 3.99401 |
| chr5 | 39463164 | 39463554 | 1000 | Intron (ENSMUST00000148152.2 | 10 | 39454915 | 39564355 | 109441 | 1 | ENSMUST00 | 8250 | Fyn | ENSMUSG0C | true | . | . | . | . | . | . | chr10 | 39448300 | 39473913 | 6.46111 | 4.64933 | 4.22352 |
| chr5 | 18072855 | 18073238 | 1000 | Intron (ENSMUST00000154718.2 | 10 | 18056004 | 18125144 | 69141 | 1 | ENSMUST00 | 16852 | Reps1 | ENSMUSG0C | false | . | . | . | . | . | . | chr10 | 18064965 | 18077164 | 5.38699 | 4.24746 | 3.33301 |
| chr5 | 18070137 | 18070523 | 1000 | Intron (ENSMUST00000154718.2 | 10 | 18056004 | 18125144 | 69141 | 1 | ENSMUST00 | 14134 | Reps1 | ENSMUSG0C | false | . | . | . | . | . | . | chr10 | 18064965 | 18077164 | 5.38699 | 4.24746 | 3.33301 |
| chr10 | 18055369 | 18056208 | 1000 | Promoter (<=1kb) | 10 | 18055861 | 18125130 | 69270 | 1 | ENSMUST00 | 0 | Reps1 | ENSMUSG0C | false | chr10 | 18055122 | 18057311 | 35.94236 | 112.30728 | 108.40293 | . | . | . | . | . | . |
| chr5 | 53752574 | 53752977 | 1000 | Promoter (1-2kb) | 10 | 53696997 | 53750909 | 53913 | 2 | ENSMUST00 | -1666 | Fam184a | ENSMUSG0C | true | . | . | . | . | . | . | chr10 | 53751243 | 53753430 | 4.93021 | 4.02583 | 2.95304 |
| chr10 | 53596543 | 53597126 | 1000 | Promoter (<=1kb) | 10 | 53596961 | 53609213 | 12253 | 1 | ENSMUST00 | 0 | Asf1a | ENSMUSG0C | false | chr10 | 53596886 | 53598164 | 37.19513 | 119.7376 | 115.60696 | . | . | . | . | . | . |
| chr10 | 52381849 | 52382426 | 1000 | Promoter (<=1kb) | 10 | 52337024 | 52382124 | 45101 | 2 | ENSMUST00 | 0 | Gopc | ENSMUSG0C | false | chr10 | 52380966 | 52382996 | 24.89402 | 98.85246 | 95.31046 | . | . | . | . | . | . |
| chr10 | 14705230 | 14705734 | 1000 | Promoter (<=1kb) | 10 | 14705591 | 14766771 | 61181 | 1 | ENSMUST00 | 0 | Nmbr | ENSMUSG0C | false | chr10 | 14704375 | 14705862 | 37.4597 | 129.87831 | 125.39164 | . | . | . | . | . | . |
| chr10 | 67096281 | 67097300 | 1000 | Promoter (<=1kb) | 10 | 67009189 | 67096988 | 87800 | 2 | ENSMUST00 | 0 | Reep3 | ENSMUSG0C | false | chr10 | 67095916 | 67097940 | 31.16539 | 106.97112 | 103.21663 | chr10 | 67097202 | 67100674 | 5.16813 | 4.17858 | 3.14766 |
| chr10 | 108161711 | 108162883 | 1000 | Promoter (<=1kb) | 10 | 108162400 | 108277575 | 115176 | 1 | ENSMUST00 | 0 | Ppp1r12a | ENSMUSG0C | false | chr10 | 108161189 | 108163933 | 32.33361 | 140.55707 | 135.65718 | . | . | . | . | . | . |
| chr10 | 59322955 | 59323925 | 1000 | Promoter (<=1kb) | 10 | 59323296 | 59373304 | 50009 | 1 | ENSMUST00 | 0 | P4ha1 | ENSMUSG0C | false | chr10 | 59322839 | 59324728 | 37.44653 | 124.45157 | 120.15771 | chr10 | 59322422 | 59323063 | 4.46606 | 3.96523 | 2.55107 |
| chr10 | 58394156 | 58394553 | 1000 | Promoter (<=1kb) | 10 | 58394373 | 58421783 | 27411 | 1 | ENSMUST00 | 0 | Lims1 | ENSMUSG0C | false | chr10 | 58394347 | 58395666 | 19.69817 | 53.33889 | 50.58166 | chr10 | 58389168 | 58394233 | 6.02364 | 4.49468 | 3.84 |
| chr10 | 98913880 | 98915772 | 1000 | Promoter (<=1kb) | 10 | 98915152 | 99026143 | 110992 | 1 | ENSMUST00 | 0 | Atp2b1 | ENSMUSG0C | true | chr10 | 98913573 | 98916645 | 37.17318 | 134.81126 | 130.13722 | . | . | . | . | . | . |
| chr5 | 68355044 | 68355580 | 1000 | Distal Intergenic | 10 | 68095593 | 68278726 | 183134 | 2 | ENSMUST00 | -76319 | Arid5b | ENSMUSG0C | false | . | . | . | . | . | . | chr10 | 68354519 | 68357142 | 3.97382 | 3.56536 | 2.14982 |
| chr10 | 68276308 | 68276918 | 1000 | Promoter (1-2kb) | 10 | 68095593 | 68278726 | 183134 | 2 | ENSMUST00 | 1808 | Arid5b | ENSMUSG0C | false | chr10 | 68275857 | 68280673 | 31.01414 | 97.6227 | 94.10706 | chr10 | 68274867 | 68276489 | 4.06595 | 3.59413 | 2.27098 |
| chr10 | 89744754 | 89745106 | 1000 | Promoter (<=1kb) | 10 | 89744991 | 89819869 | 74879 | 1 | ENSMUST00 | 0 | Uhrf1bp1l | ENSMUSG0C | false | chr10 | 89744559 | 89746078 | 32.3084 | 121.498 | 117.30614 | chr10 | 89744229 | 89744781 | 3.4804 | 3.33557 | 1.74551 |
| chr5 | 99201789 | 99202202 | 1000 | Distal Intergenic | 10 | 99192892 | 99196850 | 3959 | 1 | ENSMUST00 | 8898 | Poc1b | ENSMUSG0C | false | . | . | . | . | . | . | chr10 | 99199695 | 99205008 | 4.87834 | 4.03183 | 2.90004 |
| chr5 | 99171876 | 99172531 | 1000 | Promoter (<=1kb) | 10 | 99171284 | 99197028 | 25745 | 1 | ENSMUST00 | 593 | Poc1b | ENSMUSG0C | false | . | . | . | . | . | . | chr10 | 99168497 | 99173864 | 6.60781 | 4.81227 | 4.31674 |
| chr10 | 99262643 | 99263304 | 1000 | Promoter (<=1kb) | 10 | 99263231 | 99267488 | 4258 | 1 | ENSMUST00 | 0 | Dusp6 | ENSMUSG0C | false | chr10 | 99261081 | 99265157 | 20.41348 | 80.25478 | 77.09972 | . | . | . | . | . | . |
| chr10 | 99261443 | 99262086 | 1000 | Promoter (1-2kb) | 10 | 99263231 | 99267488 | 4258 | 1 | ENSMUST00 | -1145 | Dusp6 | ENSMUSG0C | false | chr10 | 99261081 | 99265157 | 20.41348 | 80.25478 | 77.09972 | chr10 | 99260424 | 99261473 | 3.26993 | 2.94736 | 1.61653 |
| chr10 | 91181477 | 91181995 | 1000 | Distal Intergenic | 10 | 91160417 | 91171582 | 11166 | 2 | ENSMUST00 | -9896 | Tmpo | ENSMUSG0C | false | chr10 | 91180987 | 91183079 | 27.08035 | 80.13548 | 76.98148 | . | . | . | . | . | . |
| chr10 | 91170621 | 91171947 | 1000 | Promoter (<=1kb) | 10 | 91147571 | 91171582 | 24012 | 2 | ENSMUST00 | 0 | Tmpo | ENSMUSG0C | false | chr10 | 91170021 | 91172319 | 32.64849 | 123.19247 | 118.94151 | . | . | . | . | . | . |
| chr5 | 21921499 | 21921886 | 1000 | Intron (ENSMUST00000120509.2 | 10 | 21928470 | 21990402 | 61933 | 1 | ENSMUST00 | -6584 | Sgk1 | ENSMUSG0C | false | . | . | . | . | . | . | . | . | . | . | . | . |
| chr5 | 25381944 | 25382277 | 1000 | Intron (ENSMUST00000053748.9 | 10 | 25359798 | 25523519 | 163722 | 1 | ENSMUST00 | 22147 | Epb4.1l2 | ENSMUSG0C | false | . | . | . | . | . | . | chr10 | 25381636 | 25382706 | 3.02522 | 2.95865 | 1.42049 |
| chr10 | 25359260 | 25360464 | 1000 | Promoter (<=1kb) | 10 | 25359798 | 25523519 | 163722 | 1 | ENSMUST00 | 0 | Epb4.1l2 | ENSMUSG0C | false | chr10 | 25359196 | 25361334 | 35.84351 | 110.06287 | 106.22365 | . | . | . | . | . | . |
| chr5 | 20906897 | 20907468 | 1000 | Distal Intergenic | 10 | 20952547 | 21080428 | 127882 | 1 | ENSMUST00 | -45079 | Ahi1 | ENSMUSG0C | false | . | . | . | . | . | . | chr10 | 20906355 | 20912462 | 3.89411 | 3.4388 | 2.10651 |
| chr10 | 20906028 | 20906462 | 1000 | Distal Intergenic | 10 | 20952547 | 21080428 | 127882 | 1 | ENSMUST00 | -46085 | Ahi1 | ENSMUSG0C | false | chr10 | 20905969 | 20906735 | 8.07187 | 15.14287 | 12.87755 | chr10 | 20906355 | 20912462 | 3.89411 | 3.4388 | 2.10651 |
| chr10 | 20952039 | 20952994 | 1000 | Promoter (<=1kb) | 10 | 20952547 | 21080428 | 127882 | 1 | ENSMUST00 | 0 | Ahi1 | ENSMUSG0C | false | chr10 | 20951930 | 20953388 | 30.03893 | 101.89226 | 98.2715 | . | . | . | . | . | . |
| chr5 | 19698943 | 19699550 | 1000 | Distal Intergenic | 10 | 19712587 | 19760053 | 47467 | 1 | ENSMUST00 | -13037 | Il20ra | ENSMUSG0C | false | . | . | . | . | . | . | chr10 | 19698197 | 19700877 | 4.00105 | 3.62539 | 2.17033 |
| chr10 | 93160501 | 93161768 | 1000 | Promoter (<=1kb) | 10 | 93160876 | 93241342 | 80467 | 1 | ENSMUST00 | 0 | Cdk17 | ENSMUSG0C | false | chr10 | 93160225 | 93162546 | 40.13043 | 161.53474 | 155.83125 | . | . | . | . | . | . |
| chr5 | 94466380 | 94466783 | 1000 | Intron (ENSMUST00000132743.1 | 10 | 94514857 | 94590956 | 76100 | 1 | ENSMUST00 | -48074 | Tmcc3 | ENSMUSG0C | false | . | . | . | . | . | . | chr10 | 94462174 | 94467854 | 4.65377 | 3.90587 | 2.72223 |
| chr10 | 94514572 | 94515459 | 1000 | Promoter (<=1kb) | 10 | 94514857 | 94590956 | 76100 | 1 | ENSMUST00 | 0 | Tmcc3 | ENSMUSG0C | false | chr10 | 94514294 | 94516659 | 32.48404 | 109.34737 | 105.53431 | . | . | . | . | . | . |
| chr10 | 94688516 | 94689064 | 1000 | Promoter (<=1kb) | 10 | 94688790 | 94790336 | 101547 | 1 | ENSMUST00 | 0 | Cep83 | ENSMUSG0C | false | chr10 | 94687776 | 94689607 | 40.62631 | 137.67056 | 132.88402 | . | . | . | . | . | . |
| chr10 | 95415633 | 95416339 | 1000 | Promoter (<=1kb) | 10 | 95411490 | 95416212 | 4723 | 2 | ENSMUST00 | 0 | Socs2 | ENSMUSG0C | false | chr10 | 95414179 | 95418016 | 35.49308 | 110.49252 | 106.63927 | chr10 | 95413852 | 95418311 | 4.09983 | 3.68592 | 2.25149 |
| chr10 | 95416427 | 95417320 | 1000 | Promoter (<=1kb) | 10 | 95411490 | 95416857 | 5368 | 2 | ENSMUST00 | 0 | Socs2 | ENSMUSG0C | false | chr10 | 95414179 | 95418016 | 35.49308 | 110.49252 | 106.63927 | chr10 | 95413852 | 95418311 | 4.09983 | 3.68592 | 2.25149 |
| chr10 | 84440284 | 84441213 | 1000 | Promoter (<=1kb) | 10 | 84370905 | 84440597 | 69693 | 2 | ENSMUST00 | 0 | Nuak1 | ENSMUSG0C | false | chr10 | 84439467 | 84441707 | 23.61999 | 85.9385 | 82.67356 | chr10 | 84437726 | 84441929 | 5.36859 | 4.15154 | 3.34486 |
| chr5 | 84383270 | 84383860 | 1000 | Intron (ENSMUST00000020220.9 | 10 | 84375296 | 84392388 | 17093 | 2 | ENSMUST00 | 8528 | Nuak1 | ENSMUSG0C | false | . | . | . | . | . | . | chr10 | 84383011 | 84385039 | 5.40407 | 4.29151 | 3.33393 |
| chr10 | 84576260 | 84577558 | 1000 | Promoter (<=1kb) | 10 | 84576626 | 84614359 | 37734 | 1 | ENSMUST00 | 0 | Tcp11l2 | ENSMUSG0C | false | chr10 | 84575899 | 84577895 | 39.05061 | 143.74966 | 138.71384 | . | . | . | . | . | . |
| chr5 | 84713369 | 84713745 | 1000 | Exon (ENSMUST00000077175.5/ | 10 | 84756062 | 84906144 | 150083 | 1 | ENSMUST00 | -42317 | Rfx4 | ENSMUSG0C | false | . | . | . | . | . | . | chr10 | 84710435 | 84714947 | 4.41 | 3.78387 | 2.54057 |
| chr5 | 85482014 | 85482339 | 1000 | Intron (ENSMUST00000105307.2 | 10 | 85386827 | 85640594 | 253768 | 1 | ENSMUST00 | 95188 | Btbd11 | ENSMUSG0C | false | . | . | . | . | . | . | chr10 | 85480415 | 85482646 | 3.19147 | 3.13817 | 1.53939 |
| chr5 | 86341564 | 86341971 | 1000 | Intron (ENSMUST00000020234.8 | 10 | 86302854 | 86343981 | 41128 | 1 | ENSMUST00 | 38711 | Timp3 | ENSMUSG0C | false | . | . | . | . | . | . | chr10 | 86303145 | 86347283 | 7.80266 | 5.33584 | 5.30789 |
| chr5 | 86339936 | 86340518 | 1000 | Intron (ENSMUST00000020234.8 | 10 | 86302854 | 86343981 | 41128 | 1 | ENSMUST00 | 37083 | Timp3 | ENSMUSG0C | false | . | . | . | . | . | . | chr10 | 86303145 | 86347283 | 7.80266 | 5.33584 | 5.30789 |
| chr10 | 63338193 | 63339505 | 1000 | Promoter (<=1kb) | 10 | 63319007 | 63339032 | 20026 | 2 | ENSMUST00 | 0 | Sirt1 | ENSMUSG0C | false | chr10 | 63336957 | 63339812 | 34.20045 | 107.22294 | 103.46436 | chr10 | 63339368 | 63339804 | 5.71072 | 4.38972 | 3.59547 |
| chr10 | 63243588 | 63244280 | 1000 | Promoter (<=1kb) | 10 | 63243797 | 63317881 | 74085 | 1 | ENSMUST00 | 0 | Herc4 | ENSMUSG0C | false | chr10 | 63243089 | 63243701 | 16.95696 | 43.908 | 41.27272 | . | . | . | . | . | . |
| chr5 | 63156893 | 63157433 | 1000 | Intron (ENSMUST00000095580.2 | 10 | 63115795 | 63203952 | 88158 | 2 | ENSMUST00 | 46519 | Mypn | ENSMUSG0C | false | . | . | . | . | . | . | chr10 | 63155333 | 63159268 | 5.31639 | 4.17941 | 3.28492 |
| chr10 | 62650926 | 62651769 | 1000 | Promoter (<=1kb) | 10 | 62616023 | 62651218 | 35196 | 2 | ENSMUST00 | 0 | Ddx50 | ENSMUSG0C | false | chr10 | 62650076 | 62651299 | 37.17318 | 134.81126 | 130.13722 | . | . | . | . | . | . |
| chr10 | 61782979 | 61783714 | 1000 | Promoter (<=1kb) | 10 | 61738665 | 61784147 | 45483 | 2 | ENSMUST00 | 433 | H2afy2 | ENSMUSG0C | false | chr10 | 61782647 | 61784301 | 28.56742 | 103.16996 | 99.51957 | . | . | . | . | . | . |
| chr10 | 61648338 | 61648886 | 1000 | Promoter (<=1kb) | 10 | 61648552 | 61674168 | 25617 | 1 | ENSMUST00 | 0 | Ppa1 | ENSMUSG0C | false | chr10 | 61647907 | 61649648 | 35.2897 | 111.17733 | 107.31322 | chr10 | 61647817 | 61649792 | 4.45275 | 3.87032 | 2.54756 |
| chr10 | 61383062 | 61383744 | 1000 | Promoter (<=1kb) | 10 | 61319657 | 61383523 | 63867 | 2 | ENSMUST00 | 0 | Pald1 | ENSMUSG0C | false | chr10 | 61382020 | 61383994 | 31.64465 | 94.75051 | 91.30649 | chr10 | 61368996 | 61384383 | 5.02563 | 4.04497 | 3.06054 |
| chr5 | 125713167 | 125713512 | 1000 | Distal Intergenic | 10 | 125966219 | 126015359 | 49141 | 1 | ENSMUST00 | -252707 | Lrig3 | ENSMUSG0C | false | . | . | . | . | . | . | . | . | . | . | . | . |
| chr5 | 59974039 | 59974511 | 1000 | Distal Intergenic | 10 | 59988404 | 59990383 | 1980 | 2 | ENSMUST00 | 15872 | Anapc16 | ENSMUSG0C | false | . | . | . | . | . | . | chr10 | 59970305 | 59981054 | 7.15674 | 4.95725 | 4.79406 |
| chr5 | 59919418 | 59920052 | 1000 | Distal Intergenic | 10 | 59949675 | 59951770 | 2096 | 2 | ENSMUST00 | 31718 | Ddit4 | ENSMUSG0C | false | . | . | . | . | . | . | chr10 | 59918664 | 59922070 | 6.15988 | 4.45838 | 4.00029 |
| chr10 | 59879364 | 59879985 | 1000 | Promoter (<=1kb) | 10 | 59879556 | 59899302 | 19747 | 1 | ENSMUST00 | 0 | Dnajb12 | ENSMUSG0C | false | chr10 | 59878813 | 59880788 | 22.79947 | 93.49365 | 90.07696 | . | . | . | . | . | . |
| chr10 | 119239200 | 119240420 | 1000 | Promoter (<=1kb) | 10 | 119199255 | 119240055 | 40801 | 2 | ENSMUST00 | 0 | Cand1 | ENSMUSG0C | false | chr10 | 119238719 | 119241052 | 38.25672 | 133.48087 | 128.85442 | chr10 | 119237624 | 119239358 | 3.35864 | 3.16661 | 1.68671 |
| chr5 | 121955683 | 121956388 | 1000 | Intron (ENSMUST00000020322.6 | 10 | 121780991 | 122047308 | 266318 | 2 | ENSMUST00 | 90920 | Srgap1 | ENSMUSG0C | true | . | . | . | . | . | . | chr10 | 121945243 | 122036389 | 5.74397 | 4.3951 | 3.62974 |
| chr5 | 121904136 | 121904620 | 1000 | Intron (ENSMUST00000020322.6 | 10 | 121806640 | 121823133 | 16494 | 2 | ENSMUST00 | -81004 | Srgap1 | ENSMUSG0C | true | . | . | . | . | . | . | chr10 | 121883772 | 121938386 | 5.42616 | 4.28464 | 3.35663 |
| chr5 | 121872963 | 121873663 | 1000 | Intron (ENSMUST00000020322.6 | 10 | 121806640 | 121823133 | 16494 | 2 | ENSMUST00 | -49831 | Srgap1 | ENSMUSG0C | true | . | . | . | . | . | . | chr10 | 121869425 | 121882678 | 5.26441 | 4.14787 | 3.23352 |
| chr5 | 121921053 | 121921501 | 1000 | Intron (ENSMUST00000020322.6 | 10 | 121806640 | 121823133 | 16494 | 2 | ENSMUST00 | -97921 | Srgap1 | ENSMUSG0C | true | . | . | . | . | . | . | chr10 | 121883772 | 121938386 | 5.42616 | 4.28464 | 3.35663 |
| chr11 | 21238576 | 21239531 | 1000 | Promoter (<=1kb) | 11 | 21239281 | 21321122 | 81842 | 1 | ENSMUST00 | 0 | Vps54 | ENSMUSG0C | false | chr11 | 21238334 | 21240491 | 39.02359 | 127.20569 | 122.81842 | . | . | . | . | . | . |
| chr11 | 21091002 | 21092082 | 1000 | Promoter (<=1kb) | 11 | 21091291 | 21143005 | 51715 | 1 | ENSMUST00 | 0 | Peli1 | ENSMUSG0C | false | chr11 | 21091018 | 21093123 | 36.04346 | 118.78738 | 114.68494 | . | . | . | . | . | . |
| chr10 | 115384393 | 115385941 | 1000 | Promoter (<=1kb) | 10 | 115369966 | 115384435 | 14470 | 2 | ENSMUST00 | 0 | Thap2 | ENSMUSG0C | false | chr10 | 115383165 | 115387078 | 36.02846 | 130.54782 | 126.03754 | . | . | . | . | . | . |
| chr5 | 20038760 | 20039217 | 1000 | Distal Intergenic | 11 | 20078686 | 20112867 | 34182 | 2 | ENSMUST00 | 73650 | Actr2 | ENSMUSG0C | false | . | . | . | . | . | . | chr11 | 20033774 | 20044254 | 5.74342 | 4.41024 | 3.62353 |
| chr10 | 116580960 | 116582366 | 1000 | Promoter (<=1kb) | 10 | 116485161 | 116581511 | 96351 | 2 | ENSMUST00 | 0 | Cnot2 | ENSMUSG0C | false | chr10 | 116580190 | 116583053 | 31.70051 | 127.6562 | 123.25787 | . | . | . | . | . | . |
| chr5 | 117006174 | 117006518 | 1000 | Intron (ENSMUST00000020378.4 | 10 | 116986314 | 117025040 | 38727 | 1 | ENSMUST00 | 19861 | Best3 | ENSMUSG0C | false | . | . | . | . | . | . | chr10 | 117005657 | 117006889 | 3.13058 | 3.03532 | 1.48733 |
| chr10 | 117224111 | 117224668 | 1000 | Promoter (<=1kb) | 10 | 117215142 | 117224507 | 9366 | 2 | ENSMUST00 | 0 | Yeats4 | ENSMUSG0C | false | chr10 | 117223735 | 117225006 | 37.63617 | 126.94061 | 122.55849 | chr10 | 117222875 | 117225088 | 5.7615 | 4.41212 | 3.66971 |
| chr11 | 12037088 | 12037807 | 1000 | Promoter (<=1kb) | 11 | 11930508 | 12037401 | 106894 | 2 | ENSMUST00 | 0 | Grb10 | ENSMUSG0C | false | chr11 | 12035826 | 12038123 | 26.00477 | 73.67448 | 70.62826 | chr11 | 12034418 | 12038326 | 3.27373 | 3.08101 | 1.6369 |
| chr10 | 117629227 | 117629622 | 1000 | Promoter (<=1kb) | 10 | 117629500 | 117635373 | 5874 | 1 | ENSMUST00 | 0 | Cpm | ENSMUSG0C | false | chr10 | 117628811 | 117630031 | 22.30831 | 66.34106 | 63.40397 | chr10 | 117628514 | 117629932 | 4.4395 | 3.80427 | 2.55824 |
| chr5 | 117717654 | 117717998 | 1000 | Distal Intergenic | 10 | 117688888 | 117710758 | 21871 | 2 | ENSMUST00 | -6897 | Mdm2 | ENSMUSG0C | false | . | . | . | . | . | . | chr10 | 117716080 | 117718805 | 4.59649 | 3.91279 | 2.67762 |
| chr5 | 80729482 | 80729778 | 1000 | Intron (ENSMUST00000020420.7 | 10 | 80706978 | 80742211 | 35234 | 2 | ENSMUST00 | 12433 | Ap3d1 | ENSMUSG0C | false | . | . | . | . | . | . | chr10 | 80728990 | 80731555 | 6.89288 | 4.80117 | 4.60479 |
| chr10 | 111506047 | 111506618 | 1000 | Promoter (<=1kb) | 10 | 111506286 | 111508645 | 2360 | 1 | ENSMUST00 | 0 | Phlda1 | ENSMUSG0C | false | chr10 | 111505983 | 111508558 | 27.78601 | 91.88203 | 88.49705 | chr10 | 111505291 | 111506262 | 3.63613 | 3.38327 | 1.88233 |
| chr5 | 111543665 | 111544232 | 1000 | Distal Intergenic | 10 | 111507696 | 111508165 | 470 | 1 | ENSMUST00 | 35970 | Phlda1 | ENSMUSG0C | false | . | . | . | . | . | . | chr10 | 111541392 | 111545493 | 5.88782 | 4.51568 | 3.73516 |
| chr10 | 118103102 | 118103702 | 1000 | Distal Intergenic | 10 | 118141787 | 118168997 | 27211 | 1 | ENSMUST00 | -38085 | Mdm1 | ENSMUSG0C | false | chr10 | 118103047 | 118103667 | 10.13814 | 22.76719 | 20.39335 | chr10 | 118103500 | 118103806 | 5.28107 | 4.12922 | 3.26788 |
| chr5 | 121055456 | 121056288 | 1000 | Intron (ENSMUST00000020439.5 | 10 | 121034024 | 121100645 | 66622 | 1 | ENSMUST00 | 21433 | Wif1 | ENSMUSG0C | false | . | . | . | . | . | . | chr10 | 121053259 | 121060302 | 4.44657 | 3.81891 | 2.55183 |
| chr5 | 144999986 | 145000403 | 1000 | Intron (ENSMUST00000036579.8 | 4 | 144974926 | 144983961 | 9036 | 2 | ENSMUST00 | -16026 | Vps13d | ENSMUSG0C | false | . | . | . | . | . | . | chr4 | 144995438 | 145002495 | 5.44672 | 4.21292 | 3.37905 |
| chr5 | 144960337 | 144960874 | 1000 | Distal Intergenic | 4 | 144972623 | 144974683 | 2061 | 2 | ENSMUST00 | 13809 | Vps13d | ENSMUSG0C | false | . | . | . | . | . | . | chr4 | 144960361 | 144961325 | 3.23854 | 3.18414 | 1.54142 |
| chr10 | 81350233 | 81350822 | 1000 | Promoter (<=1kb) | 10 | 81346048 | 81350480 | 4433 | 2 | ENSMUST00 | 0 | Hmg20b | ENSMUSG0C | false | chr10 | 81349311 | 81350520 | 43.01277 | 150.85193 | 145.53938 | chr10 | 81339953 | 81351319 | 5.28841 | 4.20398 | 3.24758 |
| chr10 | 82698812 | 82699450 | 1000 | Promoter (<=1kb) | 10 | 82699007 | 82742428 | 43422 | 1 | ENSMUST00 | 0 | Hcfc2 | ENSMUSG0C | false | chr10 | 82698621 | 82700018 | 27.66025 | 106.38611 | 102.65278 | chr10 | 82697328 | 82698818 | 5.29331 | 4.15672 | 3.28985 |
| chr10 | 83359850 | 83360248 | 1000 | Promoter (<=1kb) | 10 | 83360221 | 83368835 | 8615 | 1 | ENSMUST00 | 0 | D10Wsu102e | ENSMUSG0C | false | chr10 | 83359694 | 83360997 | 29.22133 | 96.43108 | 92.94852 | chr10 | 83357992 | 83362008 | 4.45361 | 3.80665 | 2.55649 |
| chr9 | 106170513 | 106171141 | 1000 | Promoter (<=1kb) | 9 | 106170928 | 106191136 | 20209 | 1 | ENSMUST00 | 0 | Wdr82 | ENSMUSG0C | false | chr9 | 106170267 | 106173420 | 30.60524 | 96.90557 | 93.40668 | chr9 | 106169574 | 106170598 | 3.76934 | 3.45845 | 2.03549 |
| chr11 | 32221210 | 32222432 | 1000 | Promoter (<=1kb) | 11 | 32209585 | 32222300 | 12716 | 2 | ENSMUST00 | 0 | Rhbdf1 | ENSMUSG0C | false | chr11 | 32221136 | 32222863 | 33.13715 | 105.37189 | 101.66248 | chr11 | 32220195 | 32227427 | 6.51865 | 4.73291 | 4.25781 |
| chr11 | 32267156 | 32267847 | 1000 | Promoter (<=1kb) | 11 | 32225628 | 32267689 | 42062 | 2 | ENSMUST00 | 0 | Nprl3 | ENSMUSG0C | false | chr11 | 32266754 | 32267959 | 26.52689 | 93.43019 | 90.01563 | chr11 | 32265771 | 32268130 | 4.22439 | 3.59981 | 2.41508 |
| chr11 | 30267045 | 30268574 | 1000 | Promoter (<=1kb) | 11 | 30099395 | 30268169 | 168775 | 2 | ENSMUST00 | 0 | Sptbn1 | ENSMUSG0C | false | chr11 | 30266216 | 30269045 | 34.68901 | 133.45216 | 128.82967 | . | . | . | . | . | . |
| chr11 | 49243662 | 49244891 | 1000 | Promoter (<=1kb) | 11 | 49244191 | 49263022 | 18832 | 1 | ENSMUST00 | 0 | Mgat1 | ENSMUSG0C | false | chr11 | 49243444 | 49244480 | 15.68842 | 38.79056 | 36.21557 | chr11 | 49244640 | 49253728 | 6.81123 | 4.86749 | 4.50404 |
| chr11 | 49248597 | 49249040 | 1000 | Promoter (<=1kb) | 11 | 49248957 | 49251637 | 2681 | 1 | ENSMUST00 | 0 | Mgat1 | ENSMUSG0C | false | chr11 | 49249000 | 49249511 | 8.85273 | 17.39095 | 15.09274 | chr11 | 49244640 | 49253728 | 6.81123 | 4.86749 | 4.50404 |
| chr11 | 51605856 | 51606457 | 1000 | Promoter (<=1kb) | 11 | 51600100 | 51606847 | 6748 | 2 | ENSMUST00 | 390 | Hnrnpab | ENSMUSG0C | false | chr11 | 51605527 | 51607486 | 35.02911 | 135.1772 | 130.48503 | . | . | . | . | . | . |
| chr11 | 53300334 | 53300961 | 1000 | Promoter (<=1kb) | 11 | 53259814 | 53300457 | 40644 | 2 | ENSMUST00 | 0 | Hspa4 | ENSMUSG0C | false | chr11 | 53298936 | 53301316 | 37.74981 | 149.12509 | 143.87827 | . | . | . | . | . | . |
| chr11 | 49712295 | 49713070 | 1000 | Promoter (<=1kb) | 11 | 49671503 | 49712723 | 41221 | 2 | ENSMUST00 | 0 | Cnot6 | ENSMUSG0C | false | chr11 | 49711694 | 49713635 | 32.66734 | 113.81544 | 109.86345 | . | . | . | . | . | . |
| chr5 | 50176954 | 50177758 | 1000 | Promoter (2-3kb) | 11 | 50174907 | 50197024 | 22118 | 1 | ENSMUST00 | 2048 | 3010026O09 | ENSMUSG0C | false | . | . | . | . | . | . | chr11 | 50174421 | 50189350 | 5.97902 | 4.44839 | 3.83251 |
| chr5 | 50184831 | 50185244 | 1000 | Intron (ENSMUST00000020647.4 | 11 | 50174907 | 50197024 | 22118 | 1 | ENSMUST00 | 9925 | 3010026O09 | ENSMUSG0C | false | . | . | . | . | . | . | chr11 | 50174421 | 50189350 | 5.97902 | 4.44839 | 3.83251 |
| chr5 | 50178383 | 50178794 | 1000 | Intron (ENSMUST00000020647.4 | 11 | 50174907 | 50197024 | 22118 | 1 | ENSMUST00 | 3477 | 3010026O09 | ENSMUSG0C | false | . | . | . | . | . | . | chr11 | 50174421 | 50189350 | 5.97902 | 4.44839 | 3.83251 |
| chr5 | 51852112 | 51852856 | 1000 | Intron (ENSMUST00000020655.8 | 11 | 51842301 | 51857125 | 14825 | 2 | ENSMUST00 | 4269 | Jade2 | ENSMUSG0C | false | . | . | . | . | . | . | chr11 | 51849911 | 51856404 | 5.82151 | 4.48521 | 3.68239 |
| chr11 | 51855537 | 51857556 | 1000 | Promoter (<=1kb) | 11 | 51813460 | 51857396 | 43937 | 2 | ENSMUST00 | 0 | Jade2 | ENSMUSG0C | false | chr11 | 51855275 | 51859109 | 33.55804 | 103.05054 | 99.40515 | chr11 | 51849911 | 51856404 | 5.82151 | 4.48521 | 3.68239 |
| chr5 | 5243732 | 5244681 | 1000 | Exon (ENSMUST00000020662.9/ | 11 | 5204833 | 5238448 | 33616 | 2 | ENSMUST00 | -5285 | Kremen1 | ENSMUSG0C | true | . | . | . | . | . | . | chr11 | 5235885 | 5260441 | 6.3098 | 4.58865 | 4.10038 |
| chr5 | 5263285 | 5263655 | 1000 | Promoter (1-2kb) | 11 | 5191552 | 5261558 | 70007 | 2 | ENSMUST00 | -1728 | Kremen1 | ENSMUSG0C | true | . | . | . | . | . | . | chr11 | 5262124 | 5273545 | 5.78378 | 4.43575 | 3.65277 |
| chr5 | 5267279 | 5267669 | 1000 | Distal Intergenic | 11 | 5191552 | 5261558 | 70007 | 2 | ENSMUST00 | -5722 | Kremen1 | ENSMUSG0C | true | . | . | . | . | . | . | chr11 | 5262124 | 5273545 | 5.78378 | 4.43575 | 3.65277 |
| chr11 | 52360564 | 52361903 | 1000 | Promoter (<=1kb) | 11 | 52360860 | 52389397 | 28538 | 1 | ENSMUST00 | 0 | Vdac1 | ENSMUSG0C | false | chr11 | 52360043 | 52362840 | 43.41453 | 148.15424 | 142.93486 | . | . | . | . | . | . |
| chr11 | 8468114 | 8468912 | 1000 | Promoter (<=1kb) | 11 | 8451070 | 8468675 | 17606 | 2 | ENSMUST00 | 0 | Tns3 | ENSMUSG0C | false | chr11 | 8467453 | 8469249 | 6.91491 | 17.20307 | 14.90901 | chr11 | 8438250 | 8474887 | 7.10839 | 4.95354 | 4.75045 |
| chr11 | 8624386 | 8624996 | 1000 | Intron (ENSMUST00000020695.7 | 11 | 8431652 | 8664535 | 232884 | 2 | ENSMUST00 | 39539 | Tns3 | ENSMUSG0C | false | chr11 | 8623718 | 8624944 | 19.77295 | 52.66102 | 49.91236 | chr11 | 8620899 | 8637313 | 4.64963 | 3.85973 | 2.73583 |
| chr5 | 8701460 | 8701959 | 1000 | Distal Intergenic | 11 | 8431652 | 8664535 | 232884 | 2 | ENSMUST00 | -36926 | Tns3 | ENSMUSG0C | false | . | . | . | . | . | . | chr11 | 8700767 | 8702823 | 3.98942 | 3.51087 | 2.20444 |
| chr11 | 8493988 | 8494521 | 1000 | Intron (ENSMUST00000020695.7 | 11 | 8451070 | 8468675 | 17606 | 2 | ENSMUST00 | -25314 | Tns3 | ENSMUSG0C | false | chr11 | 8494386 | 8495188 | 5.6323 | 9.47728 | 7.31216 | chr11 | 8476638 | 8500780 | 7.03559 | 4.95227 | 4.67872 |
| chr5 | 8484491 | 8484956 | 1000 | Intron (ENSMUST00000020695.7 | 11 | 8451070 | 8468675 | 17606 | 2 | ENSMUST00 | -15817 | Tns3 | ENSMUSG0C | false | . | . | . | . | . | . | chr11 | 8476638 | 8500780 | 7.03559 | 4.95227 | 4.67872 |
| chr11 | 8663909 | 8664763 | 1000 | Promoter (<=1kb) | 11 | 8431652 | 8664535 | 232884 | 2 | ENSMUST00 | 0 | Tns3 | ENSMUSG0C | false | chr11 | 8663227 | 8664945 | 27.54716 | 98.2761 | 94.74957 | . | . | . | . | . | . |
| chr11 | 8662664 | 8662993 | 1000 | Promoter (1-2kb) | 11 | 8431652 | 8664535 | 232884 | 2 | ENSMUST00 | 1542 | Tns3 | ENSMUSG0C | false | chr11 | 8662432 | 8662983 | 6.82388 | 11.81097 | 9.59978 | chr11 | 8643123 | 8662898 | 5.56545 | 4.3124 | 3.47336 |
| chr5 | 8706121 | 8707316 | 1000 | Distal Intergenic | 11 | 8431652 | 8664535 | 232884 | 2 | ENSMUST00 | -41587 | Tns3 | ENSMUSG0C | false | . | . | . | . | . | . | chr11 | 8705433 | 8709145 | 4.66499 | 3.73072 | 2.75954 |
| chr11 | 8495029 | 8495435 | 1000 | Intron (ENSMUST00000020695.7 | 11 | 8451070 | 8468675 | 17606 | 2 | ENSMUST00 | -26355 | Tns3 | ENSMUSG0C | false | chr11 | 8494386 | 8495188 | 5.6323 | 9.47728 | 7.31216 | chr11 | 8476638 | 8500780 | 7.03559 | 4.95227 | 4.67872 |
| chr5 | 8496590 | 8497011 | 1000 | Intron (ENSMUST00000020695.7 | 11 | 8451070 | 8468675 | 17606 | 2 | ENSMUST00 | -27916 | Tns3 | ENSMUSG0C | false | . | . | . | . | . | . | chr11 | 8476638 | 8500780 | 7.03559 | 4.95227 | 4.67872 |
| chr1 | 134063045 | 134063773 | 1000 | Distal Intergenic | 1 | 134075170 | 134079120 | 3951 | 2 | ENSMUST00 | 15347 | Btg2 | ENSMUSG0C | false | chr1 | 134062917 | 134063765 | 7.39463 | 13.15164 | 10.92004 | chr1 | 134058485 | 134067448 | 6.65638 | 4.82565 | 4.35972 |
| chr1 | 134079049 | 134079667 | 1000 | Promoter (<=1kb) | 1 | 134075170 | 134079120 | 3951 | 2 | ENSMUST00 | 0 | Btg2 | ENSMUSG0C | false | chr1 | 134075527 | 134079138 | 22.90469 | 70.93823 | 67.93287 | chr1 | 134079534 | 134084201 | 5.79764 | 4.37454 | 3.66417 |
| chr1 | 134092866 | 134093447 | 1000 | Distal Intergenic | 1 | 134075170 | 134079120 | 3951 | 2 | ENSMUST00 | -13747 | Btg2 | ENSMUSG0C | false | chr1 | 134092728 | 134092987 | 8.73111 | 16.23101 | 13.94854 | chr1 | 134091960 | 134094329 | 5.68815 | 4.30744 | 3.58143 |
| chr1 | 134077684 | 134078465 | 1000 | Promoter (<=1kb) | 1 | 134075170 | 134079120 | 3951 | 2 | ENSMUST00 | 655 | Btg2 | ENSMUSG0C | false | chr1 | 134075527 | 134079138 | 22.90469 | 70.93823 | 67.93287 | . | . | . | . | . | . |
| chr5 | 7230065 | 7230411 | 1000 | Distal Intergenic | 11 | 7207686 | 7213923 | 6238 | 2 | ENSMUST00 | -16143 | Igfbp3 | ENSMUSG0C | false | . | . | . | . | . | . | chr11 | 7229857 | 7231164 | 3.13128 | 2.88332 | 1.50848 |
| chr5 | 6999357 | 6999736 | 1000 | Distal Intergenic | 11 | 7063489 | 7178506 | 115018 | 1 | ENSMUST00 | -63753 | Adcy1 | ENSMUSG0C | true | . | . | . | . | . | . | chr11 | 6996705 | 7001264 | 4.35399 | 3.75461 | 2.48212 |
| chr5 | 7129741 | 7130164 | 1000 | Intron (ENSMUST00000020706.4 | 11 | 7159882 | 7172879 | 12998 | 1 | ENSMUST00 | -29718 | Adcy1 | ENSMUSG0C | true | . | . | . | . | . | . | chr11 | 7111251 | 7145535 | 5.23684 | 4.17059 | 3.20426 |
| chr5 | 7156354 | 7156809 | 1000 | Intron (ENSMUST00000020706.4 | 11 | 7159882 | 7172879 | 12998 | 1 | ENSMUST00 | -3073 | Adcy1 | ENSMUSG0C | true | . | . | . | . | . | . | chr11 | 7147428 | 7161440 | 5.80144 | 4.4368 | 3.66324 |
| chr5 | 7135244 | 7135637 | 1000 | Intron (ENSMUST00000020706.4 | 11 | 7159882 | 7172879 | 12998 | 1 | ENSMUST00 | -24245 | Adcy1 | ENSMUSG0C | true | . | . | . | . | . | . | chr11 | 7111251 | 7145535 | 5.23684 | 4.17059 | 3.20426 |
| chr5 | 7134362 | 7134721 | 1000 | Intron (ENSMUST00000020706.4 | 11 | 7159882 | 7172879 | 12998 | 1 | ENSMUST00 | -25161 | Adcy1 | ENSMUSG0C | true | . | . | . | . | . | . | chr11 | 7111251 | 7145535 | 5.23684 | 4.17059 | 3.20426 |
| chr5 | 7149443 | 7149822 | 1000 | Exon (ENSMUST00000020706.4/ | 11 | 7159882 | 7172879 | 12998 | 1 | ENSMUST00 | -10060 | Adcy1 | ENSMUSG0C | true | . | . | . | . | . | . | chr11 | 7147428 | 7161440 | 5.80144 | 4.4368 | 3.66324 |
| chr5 | 3567848 | 3568515 | 1000 | Distal Intergenic | 11 | 3532460 | 3540612 | 8153 | 2 | ENSMUST00 | -27237 | Smtn | ENSMUSG0C | false | . | . | . | . | . | . | chr11 | 3563541 | 3569418 | 5.36923 | 4.11866 | 3.32794 |
| chr5 | 3573059 | 3573546 | 1000 | Distal Intergenic | 11 | 3532460 | 3540612 | 8153 | 2 | ENSMUST00 | -32448 | Smtn | ENSMUSG0C | false | . | . | . | . | . | . | chr11 | 3571707 | 3576040 | 6.68511 | 4.75587 | 4.39333 |
| chr5 | 5892330 | 5892785 | 1000 | Distal Intergenic | 11 | 5896637 | 5898782 | 2146 | 2 | ENSMUST00 | 5997 | Myl7 | ENSMUSG0C | false | . | . | . | . | . | . | chr11 | 5886065 | 5902928 | 6.65142 | 4.85213 | 4.35765 |
| chr11 | 5862753 | 5863246 | 1000 | Promoter (<=1kb) | 11 | 5861947 | 5872088 | 10142 | 1 | ENSMUST00 | 807 | Aebp1 | ENSMUSG0C | false | chr11 | 5861239 | 5863555 | 28.12373 | 86.68959 | 83.41467 | chr11 | 5859631 | 5868965 | 5.43399 | 4.25115 | 3.37271 |
| chr11 | 5861581 | 5862249 | 1000 | Promoter (<=1kb) | 11 | 5861947 | 5872088 | 10142 | 1 | ENSMUST00 | 0 | Aebp1 | ENSMUSG0C | false | chr11 | 5861239 | 5863555 | 28.12373 | 86.68959 | 83.41467 | chr11 | 5859631 | 5868965 | 5.43399 | 4.25115 | 3.37271 |
| chr11 | 87986846 | 87987937 | 1000 | Promoter (<=1kb) | 11 | 87979525 | 87987533 | 8009 | 2 | ENSMUST00 | 0 | Dynll2 | ENSMUSG0C | false | chr11 | 87986201 | 87988020 | 37.7199 | 126.00829 | 121.65485 | . | . | . | . | . | . |
| chr5 | 87991843 | 87992318 | 1000 | Distal Intergenic | 11 | 87979525 | 87987533 | 8009 | 2 | ENSMUST00 | -4311 | Dynll2 | ENSMUSG0C | false | . | . | . | . | . | . | chr11 | 87991111 | 87992330 | 4.60228 | 3.97233 | 2.69142 |
| chr11 | 57801450 | 57802439 | 1000 | Promoter (<=1kb) | 11 | 57801637 | 57810217 | 8581 | 1 | ENSMUST00 | 0 | Sap30l | ENSMUSG0C | false | chr11 | 57800608 | 57802887 | 26.86698 | 95.04005 | 91.5885 | chr11 | 57802356 | 57805403 | 5.69306 | 4.43094 | 3.56353 |
| chr5 | 57635295 | 57635612 | 1000 | Intron (ENSMUST00000142486.1 | 11 | 57645442 | 57787514 | 142073 | 1 | ENSMUST00 | -9830 | Galnt10 | ENSMUSG0C | true | . | . | . | . | . | . | chr11 | 57634136 | 57636581 | 6.50675 | 4.79828 | 4.24501 |
| chr11 | 85310659 | 85311503 | 1000 | Promoter (<=1kb) | 11 | 85311244 | 85347066 | 35823 | 1 | ENSMUST00 | 0 | Ppm1d | ENSMUSG0C | false | chr11 | 85310340 | 85313068 | 36.46257 | 120.58523 | 116.42456 | . | . | . | . | . | . |
| chr11 | 84869701 | 84870291 | 1000 | Promoter (<=1kb) | 11 | 84832806 | 84870202 | 37397 | 2 | ENSMUST00 | 0 | Ggnbp2 | ENSMUSG0C | false | chr11 | 84868751 | 84871497 | 31.96831 | 119.80809 | 115.67081 | . | . | . | . | . | . |
| chr11 | 84870386 | 84871084 | 1000 | Promoter (<=1kb) | 11 | 84832361 | 84870738 | 38378 | 2 | ENSMUST00 | 0 | Ggnbp2 | ENSMUSG0C | false | chr11 | 84868751 | 84871497 | 31.96831 | 119.80809 | 115.67081 | . | . | . | . | . | . |
| chr5 | 90603144 | 90603615 | 1000 | 3' UTR | 11 | 90619271 | 90630917 | 11647 | 2 | ENSMUST00 | 27302 | Stxbp4 | ENSMUSG0C | false | . | . | . | . | . | . | chr11 | 90603231 | 90604698 | 3.73443 | 3.30488 | 1.99053 |
| chr5 | 65142075 | 65142589 | 1000 | Intron (ENSMUST00000047463.9 | 11 | 65151910 | 65159890 | 7981 | 2 | ENSMUST00 | 17301 | 1700086D15 | ENSMUSG0C | false | . | . | . | . | . | . | chr11 | 65138738 | 65143476 | 5.25256 | 4.17623 | 3.25476 |
| chr5 | 89901006 | 89901660 | 1000 | Distal Intergenic | 11 | 89982665 | 90002894 | 20230 | 2 | ENSMUST00 | 101234 | Pctp | ENSMUSG0C | false | . | . | . | . | . | . | chr11 | 89896221 | 89903570 | 4.59506 | 3.93873 | 2.65513 |
| chr12 | 33147518 | 33147884 | 1000 | Promoter (<=1kb) | 12 | 33147693 | 33251146 | 103454 | 1 | ENSMUST00 | 0 | Atxn7l1 | ENSMUSG0C | false | chr12 | 33146768 | 33148505 | 30.4989 | 91.7916 | 88.41422 | . | . | . | . | . | . |
| chr12 | 33303369 | 33303850 | 1000 | Promoter (<=1kb) | 12 | 33302515 | 33368273 | 65759 | 1 | ENSMUST00 | 855 | Atxn7l1 | ENSMUSG0C | false | chr12 | 33302526 | 33303732 | 21.79225 | 59.43074 | 56.59393 | chr12 | 33303260 | 33304234 | 4.40756 | 3.79132 | 2.55531 |
| chr12 | 32953729 | 32954819 | 1000 | Promoter (<=1kb) | 12 | 32953891 | 32979050 | 25160 | 1 | ENSMUST00 | 0 | Sypl | ENSMUSG0C | false | chr12 | 32953340 | 32955472 | 34.36702 | 111.64683 | 107.76208 | . | . | . | . | . | . |
| chr5 | 17250971 | 17251602 | 1000 | Distal Intergenic | 12 | 17266545 | 17281810 | 15266 | 1 | ENSMUST00 | -14943 | Pdia6 | ENSMUSG0C | false | . | . | . | . | . | . | chr12 | 17248844 | 17258774 | 6.51242 | 4.76263 | 4.24509 |
| chr5 | 32767946 | 32768900 | 1000 | Distal Intergenic | 12 | 32820335 | 32853369 | 33035 | 1 | ENSMUST00 | -51435 | Nampt | ENSMUSG0C | true | . | . | . | . | . | . | chr12 | 32765525 | 32771065 | 5.20276 | 4.14872 | 3.21917 |
| chr12 | 32820001 | 32820923 | 1000 | Promoter (<=1kb) | 12 | 32820335 | 32853369 | 33035 | 1 | ENSMUST00 | 0 | Nampt | ENSMUSG0C | true | chr12 | 32819564 | 32820103 | 13.94362 | 38.54297 | 35.97016 | . | . | . | . | . | . |
| chr5 | 32717557 | 32717973 | 1000 | Distal Intergenic | 12 | 32820335 | 32853369 | 33035 | 1 | ENSMUST00 | -102362 | Nampt | ENSMUSG0C | true | . | . | . | . | . | . | chr12 | 32713700 | 32718759 | 5.2438 | 4.25503 | 3.20864 |
| chr5 | 32616202 | 32616627 | 1000 | Distal Intergenic | 12 | 32820335 | 32853369 | 33035 | 1 | ENSMUST00 | -203708 | Nampt | ENSMUSG0C | true | . | . | . | . | . | . | chr12 | 32615385 | 32619245 | 4.32667 | 3.73703 | 2.45364 |
| chr5 | 32716192 | 32716591 | 1000 | Distal Intergenic | 12 | 32820335 | 32853369 | 33035 | 1 | ENSMUST00 | -103744 | Nampt | ENSMUSG0C | true | . | . | . | . | . | . | chr12 | 32713700 | 32718759 | 5.2438 | 4.25503 | 3.20864 |
| chr5 | 32621846 | 32622400 | 1000 | Distal Intergenic | 12 | 32820335 | 32853369 | 33035 | 1 | ENSMUST00 | -197935 | Nampt | ENSMUSG0C | true | . | . | . | . | . | . | chr12 | 32620273 | 32624861 | 4.21551 | 3.67106 | 2.3608 |
| chr12 | 8673934 | 8674855 | 1000 | Promoter (<=1kb) | 12 | 8674134 | 8751454 | 77321 | 1 | ENSMUST00 | 0 | Pum2 | ENSMUSG0C | false | chr12 | 8673672 | 8675883 | 37.44409 | 118.04973 | 113.97176 | . | . | . | . | . | . |
| chr12 | 15816188 | 15817203 | 1000 | Promoter (<=1kb) | 12 | 15791727 | 15816785 | 25059 | 2 | ENSMUST00 | 0 | Trib2 | ENSMUSG0C | true | chr12 | 15810380 | 15817601 | 30.94804 | 114.76607 | 110.79114 | chr12 | 15817199 | 15817789 | 3.96419 | 3.64607 | 2.13854 |
| chr5 | 8394849 | 8395327 | 1000 | Distal Intergenic | 12 | 8313433 | 8343824 | 30392 | 1 | ENSMUST00 | 81417 | Hs1bp3 | ENSMUSG0C | false | . | . | . | . | . | . | chr12 | 8393920 | 8398878 | 5.80676 | 4.22852 | 3.69621 |
| chr12 | 11265746 | 11266658 | 1000 | Promoter (<=1kb) | 12 | 11265886 | 11319785 | 53900 | 1 | ENSMUST00 | 0 | Smc6 | ENSMUSG0C | false | chr12 | 11265266 | 11267287 | 30.14656 | 119.38538 | 115.2599 | . | . | . | . | . | . |
| chr5 | 8042738 | 8043086 | 1000 | Distal Intergenic | 12 | 8012359 | 8016835 | 4477 | 1 | ENSMUST00 | 30380 | Apob | ENSMUSG0C | false | . | . | . | . | . | . | chr12 | 8040794 | 8044474 | 3.85644 | 3.34118 | 2.10912 |
| chr11 | 109425753 | 109426341 | 1000 | Promoter (<=1kb) | 11 | 109425946 | 109438148 | 12203 | 1 | ENSMUST00 | 0 | Amz2 | ENSMUSG0C | false | chr11 | 109425224 | 109425788 | 18.02173 | 47.27326 | 44.5908 | chr11 | 109425269 | 109427542 | 3.52176 | 3.33391 | 1.79135 |
| chr11 | 109362481 | 109363051 | 1000 | Promoter (<=1kb) | 11 | 109362831 | 109380475 | 17645 | 1 | ENSMUST00 | 0 | Gna13 | ENSMUSG0C | false | chr11 | 109361906 | 109362668 | 24.45608 | 69.21248 | 66.23546 | . | . | . | . | . | . |
| chr11 | 109650845 | 109651696 | 1000 | Promoter (<=1kb) | 11 | 109650921 | 109669656 | 18736 | 1 | ENSMUST00 | 0 | Prkar1a | ENSMUSG0C | false | chr11 | 109649925 | 109652561 | 31.28813 | 116.44205 | 112.41405 | . | . | . | . | . | . |
| chr5 | 10136155 | 10136508 | 1000 | Distal Intergenic | 12 | 10369973 | 10390170 | 20198 | 1 | ENSMUST00 | -233465 | Nt5c1b | ENSMUSG0C | false | . | . | . | . | . | . | chr12 | 10133274 | 10136926 | 4.13946 | 3.60598 | 2.34383 |
| chr12 | 5375123 | 5376060 | 1000 | Promoter (<=1kb) | 12 | 5077472 | 5375682 | 298211 | 2 | ENSMUST00 | 0 | Klhl29 | ENSMUSG0C | true | chr12 | 5373404 | 5376248 | 26.18681 | 91.82585 | 88.44535 | chr12 | 5372978 | 5376323 | 3.31501 | 3.17659 | 1.67432 |
| chr5 | 28726292 | 28726637 | 1000 | Intron (ENSMUST00000168129.3 | 12 | 28690629 | 28750394 | 59766 | 2 | ENSMUST00 | 23757 | Trappc12 | ENSMUSG0C | false | . | . | . | . | . | . | chr12 | 28725582 | 28727685 | 5.32815 | 4.19638 | 3.32814 |
| chr12 | 24651037 | 24652248 | 1000 | Promoter (<=1kb) | 12 | 24651371 | 24662774 | 11404 | 1 | ENSMUST00 | 0 | Klf11 | ENSMUSG0C | false | chr12 | 24650599 | 24652468 | 35.02911 | 135.1772 | 130.48503 | . | . | . | . | . | . |
| chr12 | 24593031 | 24593658 | 1000 | Intron (ENSMUST00000085553.6 | 12 | 24580749 | 24617391 | 36643 | 1 | ENSMUST00 | 12283 | Grhl1 | ENSMUSG0C | false | chr12 | 24592715 | 24593894 | 15.37719 | 36.84684 | 34.29707 | chr12 | 24590560 | 24593996 | 5.62 | 4.35318 | 3.56592 |
| chr12 | 24571927 | 24572484 | 1000 | Promoter (<=1kb) | 12 | 24572287 | 24617387 | 45101 | 1 | ENSMUST00 | 0 | Grhl1 | ENSMUSG0C | false | chr12 | 24571767 | 24572102 | 13.50614 | 30.33286 | 27.86432 | chr12 | 24571597 | 24576131 | 4.40813 | 3.73639 | 2.52624 |
| chr12 | 3806391 | 3808802 | 1000 | Promoter (<=1kb) | 12 | 3806498 | 3867484 | 60987 | 1 | ENSMUST00 | 0 | Dnmt3a | ENSMUSG0C | false | chr12 | 3806158 | 3809012 | 29.81577 | 89.63673 | 86.30187 | chr12 | 3808261 | 3828410 | 6.1255 | 4.55618 | 3.94004 |
| chr5 | 3907418 | 3907763 | 1000 | Promoter (<=1kb) | 12 | 3906816 | 3908547 | 1732 | 1 | ENSMUST00 | 603 | Dnmt3a | ENSMUSG0C | false | . | . | . | . | . | . | chr12 | 3906604 | 3908070 | 5.84142 | 4.21456 | 3.72158 |
| chr12 | 3309500 | 3310399 | 1000 | Promoter (<=1kb) | 12 | 3247430 | 3309969 | 62540 | 2 | ENSMUST00 | 0 | Rab10 | ENSMUSG0C | false | chr12 | 3307840 | 3310000 | 20.49778 | 85.71664 | 82.45538 | . | . | . | . | . | . |
| chr11 | 105292122 | 105293385 | 1000 | Promoter (<=1kb) | 11 | 105292643 | 105326891 | 34249 | 1 | ENSMUST00 | 0 | Mrc2 | ENSMUSG0C | true | chr11 | 105291956 | 105293037 | 37.73948 | 119.61958 | 115.49072 | chr11 | 105290918 | 105292533 | 6.34946 | 4.68425 | 4.11073 |
| chr11 | 82870992 | 82871448 | 1000 | Promoter (<=1kb) | 11 | 82803822 | 82871210 | 67389 | 2 | ENSMUST00 | 0 | Rffl | ENSMUSG0C | false | chr11 | 82870035 | 82871276 | 36.78915 | 128.67926 | 124.23671 | chr11 | 82869837 | 82871608 | 3.13383 | 3.0708 | 1.52114 |
| chr11 | 106084187 | 106085640 | 1000 | Promoter (<=1kb) | 11 | 106084613 | 106155446 | 70834 | 1 | ENSMUST00 | 0 | Map3k3 | ENSMUSG0C | false | chr11 | 106083606 | 106084248 | 24.38567 | 80.91491 | 77.74369 | . | . | . | . | . | . |
| chr5 | 82339925 | 82340556 | 1000 | Distal Intergenic | 11 | 82176657 | 82305690 | 129034 | 2 | ENSMUST00 | -34236 | Ccl1 | ENSMUSG0C | false | . | . | . | . | . | . | chr11 | 82339249 | 82341416 | 3.96002 | 3.37992 | 2.18187 |
| chr5 | 81185427 | 81186038 | 1000 | Intron (ENSMUST00000066197.6 | 11 | 80880169 | 81153321 | 273153 | 2 | ENSMUST00 | -32107 | Asic2 | ENSMUSG0C | true | . | . | . | . | . | . | chr11 | 81181914 | 81191363 | 4.49237 | 3.66069 | 2.62262 |
| chr11 | 115164209 | 115164966 | 1000 | Promoter (<=1kb) | 11 | 115163341 | 115181181 | 17841 | 1 | ENSMUST00 | 869 | Slc9a3r1 | ENSMUSG0C | false | chr11 | 115162609 | 115165876 | 34.68901 | 133.45216 | 128.82967 | . | . | . | . | . | . |
| chr11 | 115535609 | 115536537 | 1000 | Promoter (<=1kb) | 11 | 115523102 | 115535817 | 12716 | 2 | ENSMUST00 | 0 | Sumo2 | ENSMUSG0C | false | chr11 | 115534905 | 115536788 | 28.22734 | 101.53345 | 97.92473 | . | . | . | . | . | . |
| chr11 | 74637960 | 74638913 | 1000 | 3' UTR | 11 | 74649495 | 74656551 | 7057 | 1 | ENSMUST00 | -10582 | Cluh | ENSMUSG0C | false | chr11 | 74637560 | 74638919 | 27.78601 | 91.88203 | 88.49705 | chr11 | 74637263 | 74639420 | 5.74907 | 4.37305 | 3.62361 |
| chr11 | 74649122 | 74649614 | 1000 | Promoter (<=1kb) | 11 | 74649495 | 74656551 | 7057 | 1 | ENSMUST00 | 0 | Cluh | ENSMUSG0C | false | chr11 | 74648841 | 74650947 | 35.67123 | 127.73884 | 123.33056 | chr11 | 74648886 | 74653397 | 4.70613 | 3.95834 | 2.75929 |
| chr11 | 74723623 | 74724678 | 1000 | Promoter (<=1kb) | 11 | 74673949 | 74724335 | 50387 | 2 | ENSMUST00 | 0 | Pafah1b1 | ENSMUSG0C | false | chr11 | 74722916 | 74725014 | 35.02911 | 135.1772 | 130.48503 | . | . | . | . | . | . |
| chr5 | 115779185 | 115779584 | 1000 | Intron (ENSMUST00000093912.5 | 11 | 115775756 | 115799033 | 23278 | 1 | ENSMUST00 | 3430 | 2310067B10 | ENSMUSG0C | false | . | . | . | . | . | . | chr11 | 115773727 | 115781059 | 5.21271 | 4.08187 | 3.23295 |
| chr5 | 115904993 | 115905450 | 1000 | Promoter (1-2kb) | 11 | 115896809 | 115907253 | 10445 | 2 | ENSMUST00 | 1803 | Recql5 | ENSMUSG0C | false | . | . | . | . | . | . | chr11 | 115899613 | 115906730 | 7.08174 | 4.8342 | 4.74864 |
| chr5 | 115832546 | 115833219 | 1000 | Promoter (<=1kb) | 11 | 115831968 | 115844883 | 12916 | 1 | ENSMUST00 | 579 | Llgl2 | ENSMUSG0C | false | . | . | . | . | . | . | chr11 | 115830477 | 115834921 | 5.14143 | 3.93185 | 3.14914 |
| chr11 | 72607003 | 72608617 | 1000 | Promoter (<=1kb) | 11 | 72607283 | 72686481 | 79199 | 1 | ENSMUST00 | 0 | Ube2g1 | ENSMUSG0C | false | chr11 | 72606702 | 72608877 | 36.38946 | 142.11884 | 137.14919 | . | . | . | . | . | . |
| chr5 | 116621150 | 116621715 | 1000 | Promoter (2-3kb) | 11 | 116598165 | 116624200 | 26036 | 2 | ENSMUST00 | 2485 | Rhbdf2 | ENSMUSG0C | false | . | . | . | . | . | . | chr11 | 116608011 | 116628763 | 6.7697 | 4.86532 | 4.46452 |
| chr5 | 116615730 | 116616167 | 1000 | Intron (ENSMUST00000103029.4 | 11 | 116598165 | 116624200 | 26036 | 2 | ENSMUST00 | 8033 | Rhbdf2 | ENSMUSG0C | false | . | . | . | . | . | . | chr11 | 116608011 | 116628763 | 6.7697 | 4.86532 | 4.46452 |
| chr5 | 116630572 | 116630922 | 1000 | Distal Intergenic | 11 | 116598169 | 116627019 | 28851 | 2 | ENSMUST00 | -3554 | Rhbdf2 | ENSMUSG0C | false | . | . | . | . | . | . | . | . | . | . | . | . |
| chr11 | 116827338 | 116828005 | 1000 | Promoter (<=1kb) | 11 | 116807913 | 116828003 | 20091 | 2 | ENSMUST00 | 0 | Mxra7 | ENSMUSG0C | false | chr11 | 116827118 | 116828605 | 34.90036 | 108.43528 | 104.64344 | chr11 | 116810954 | 116829199 | 5.76656 | 4.27097 | 3.65606 |
| chr5 | 117120551 | 117121565 | 1000 | Intron (ENSMUST00000127090.2 | 11 | 117115252 | 117128763 | 13512 | 1 | ENSMUST00 | 5300 | Sec14l1 | ENSMUSG0C | false | . | . | . | . | . | . | chr11 | 117116362 | 117143393 | 7.76107 | 5.12248 | 5.29175 |
| chr5 | 70608692 | 70609153 | 1000 | Promoter (<=1kb) | 11 | 70609738 | 70610709 | 972 | 1 | ENSMUST00 | -585 | Mink1 | ENSMUSG0C | false | . | . | . | . | . | . | chr11 | 70607550 | 70610065 | 6.43934 | 4.60183 | 4.22641 |
| chr11 | 70562196 | 70564145 | 1000 | Promoter (<=1kb) | 11 | 70562881 | 70614482 | 51602 | 1 | ENSMUST00 | 0 | Mink1 | ENSMUSG0C | false | chr11 | 70562192 | 70564153 | 43.65555 | 141.82639 | 136.87535 | chr11 | 70561696 | 70562582 | 6.04011 | 4.18219 | 3.90045 |
| chr5 | 76343282 | 76343663 | 1000 | Intron (ENSMUST00000021204.3 | 11 | 76358803 | 76399068 | 40266 | 2 | ENSMUST00 | 55405 | Nxn | ENSMUSG0C | true | . | . | . | . | . | . | chr11 | 76341422 | 76344449 | 6.16599 | 4.61127 | 3.96773 |
| chr11 | 75732629 | 75733741 | 1000 | Promoter (<=1kb) | 11 | 75732869 | 75765845 | 32977 | 1 | ENSMUST00 | 0 | Ywhae | ENSMUSG0C | false | chr11 | 75732399 | 75734879 | 22.96739 | 100.34364 | 96.76433 | . | . | . | . | . | . |
| chr11 | 94321423 | 94322278 | 1000 | Promoter (<=1kb) | 11 | 94287890 | 94321958 | 34069 | 2 | ENSMUST00 | 0 | Luc7l3 | ENSMUSG0C | false | chr11 | 94320406 | 94322646 | 41.67902 | 156.4369 | 150.90947 | . | . | . | . | . | . |
| chr11 | 94327758 | 94328640 | 1000 | Promoter (<=1kb) | 11 | 94328001 | 94341841 | 13841 | 1 | ENSMUST00 | 0 | Ankrd40 | ENSMUSG0C | false | chr11 | 94327395 | 94329063 | 29.37071 | 89.8338 | 86.49576 | . | . | . | . | . | . |
| chr5 | 94470335 | 94470861 | 1000 | Promoter (2-3kb) | 11 | 94426952 | 94473641 | 46690 | 2 | ENSMUST00 | 2780 | Cacna1g | ENSMUSG0C | false | . | . | . | . | . | . | chr11 | 94468980 | 94476124 | 5.80314 | 4.36554 | 3.67462 |
| chr5 | 97031992 | 97032483 | 1000 | Promoter (1-2kb) | 11 | 97030134 | 97032276 | 2143 | 1 | ENSMUST00 | 1859 | Scrn2 | ENSMUSG0C | false | . | . | . | . | . | . | chr11 | 97029231 | 97033997 | 7.09985 | 5.06752 | 4.72888 |
| chr11 | 98148959 | 98150709 | 1000 | Promoter (<=1kb) | 11 | 98082556 | 98149612 | 67057 | 2 | ENSMUST00 | 0 | Fbxl20 | ENSMUSG0C | false | chr11 | 98148294 | 98150721 | 36.42221 | 131.26616 | 126.73016 | chr11 | 98150567 | 98150859 | 4.50841 | 3.90899 | 2.5838 |
| chr5 | 98139169 | 98139491 | 1000 | Intron (ENSMUST00000103143.4 | 11 | 98091091 | 98149606 | 58516 | 2 | ENSMUST00 | 10115 | Fbxl20 | ENSMUSG0C | false | . | . | . | . | . | . | chr11 | 98131727 | 98148623 | 5.44202 | 4.28943 | 3.37216 |
| chr8 | 70865380 | 70866023 | 1000 | Promoter (2-3kb) | 8 | 70863127 | 70865428 | 2302 | 1 | ENSMUST00 | 2254 | A230052G05 | ENSMUSG0C | false | chr8 | 70864890 | 70866158 | 27.41053 | 90.23362 | 86.88379 | chr8 | 70862668 | 70866754 | 4.93202 | 3.84805 | 2.97579 |
| chr5 | 98776837 | 98777216 | 1000 | Promoter (1-2kb) | 11 | 98767932 | 98775333 | 7402 | 2 | ENSMUST00 | -1505 | Nr1d1 | ENSMUSG0C | false | . | . | . | . | . | . | chr11 | 98775631 | 98778417 | 8.10463 | 5.49842 | 5.54643 |
| chr5 | 98781933 | 98782471 | 1000 | Distal Intergenic | 11 | 98767932 | 98775333 | 7402 | 2 | ENSMUST00 | -6601 | Nr1d1 | ENSMUSG0C | false | . | . | . | . | . | . | chr11 | 98780022 | 98794752 | 7.02549 | 5.01464 | 4.66092 |

| | | | | | | | | | | | | | | | | | | | | | | | | | | |
|---|---|---|---|---|---|---|---|---|---|---|---|---|---|---|---|---|---|---|---|---|---|---|---|---|---|---|
| chr5 | 98765543 | 98766315 | 1000 | Intron (ENSMUST00000064187.6 | 11 | 98769347 | 98770426 | 1080 | 2 | ENSMUST00 | 4111 | Nr1d1 | ENSMUSG0C | false | . | . | . | . | . | . | chr11 | 98764539 | 98774391 | 7.88503 | 5.36472 | 5.3871 |
| chr11 | 98773610 | 98774224 | 1000 | Promoter (1-2kb) | 11 | 98767932 | 98775333 | 7402 | 2 | ENSMUST00 | 1109 | Nr1d1 | ENSMUSG0C | false | chr11 | 98773371 | 98773844 | 11.73508 | 26.0954 | 23.67916 | chr11 | 98764539 | 98774391 | 7.88503 | 5.36472 | 5.3871 |
| chr11 | 69096777 | 69097439 | 1000 | Promoter (1-2kb) | 11 | 69098948 | 69109960 | 11013 | 1 | ENSMUST00 | -1509 | Per1 | ENSMUSG0C | true | chr11 | 69097223 | 69098719 | 11.28697 | 24.15705 | 21.76529 | . | . | . | . | . | . |
| chr5 | 69087009 | 69087640 | 1000 | Promoter (<=1kb) | 11 | 69088490 | 69092384 | 3895 | 1 | ENSMUST00 | -850 | Vamp2 | ENSMUSG0C | false | . | . | . | . | . | . | chr11 | 69084778 | 69088340 | 5.05808 | 4.13119 | 3.04009 |
| chr11 | 100712168 | 100712606 | 1000 | Promoter (<=1kb) | 11 | 100704746 | 100712173 | 7428 | 2 | ENSMUST00 | 0 | Kat2a | ENSMUSG0C | false | chr11 | 100711979 | 100712568 | 29.64488 | 92.26185 | 88.869 | chr11 | 100711238 | 100714526 | 3.77343 | 3.43721 | 2.02212 |
| chr11 | 102316493 | 102318504 | 1000 | Promoter (<=1kb) | 11 | 102305924 | 102316680 | 10757 | 2 | ENSMUST00 | 0 | Ubtf | ENSMUSG0C | false | chr11 | 102316341 | 102321029 | 40.78419 | 137.94737 | 133.15321 | chr11 | 102314909 | 102316738 | 4.43369 | 3.72652 | 2.57056 |
| chr11 | 103697639 | 103698124 | 1000 | Promoter (<=1kb) | 11 | 103676849 | 103697898 | 21050 | 2 | ENSMUST00 | 0 | Gosr2 | ENSMUSG0C | false | chr11 | 103696863 | 103698327 | 32.09266 | 107.61995 | 103.85494 | . | . | . | . | . | . |
| chr12 | 65073359 | 65074124 | 1000 | Promoter (<=1kb) | 12 | 65062436 | 65073944 | 11509 | 2 | ENSMUST00 | 0 | Fkbp3 | ENSMUSG0C | false | chr12 | 65073017 | 65076367 | 39.97929 | 126.69925 | 122.32521 | . | . | . | . | . | . |
| chr5 | 51708808 | 51709241 | 1000 | Intron (ENSMUST00000021338.8 | 12 | 51608541 | 51691914 | 83374 | 2 | ENSMUST00 | -16895 | Strn3 | ENSMUSG0C | false | . | . | . | . | . | . | chr12 | 51707758 | 51710706 | 5.09134 | 3.89181 | 3.11469 |
| chr12 | 51690830 | 51692450 | 1000 | Promoter (<=1kb) | 12 | 51608541 | 51691914 | 83374 | 2 | ENSMUST00 | 0 | Strn3 | ENSMUSG0C | false | chr12 | 51690328 | 51693137 | 43.58744 | 151.86528 | 146.50356 | . | . | . | . | . | . |
| chr5 | 91703502 | 91703931 | 1000 | Exon (ENSMUST00000111609.2/ | 12 | 91703413 | 91713974 | 10562 | 2 | ENSMUST00 | 10043 | Ston2 | ENSMUSG0C | false | . | . | . | . | . | . | chr12 | 91702783 | 91704861 | 4.12339 | 3.50428 | 2.33392 |
| chr12 | 91588987 | 91590622 | 1000 | Promoter (<=1kb) | 12 | 91555916 | 91590009 | 34094 | 2 | ENSMUST00 | 0 | Gtf2a1 | ENSMUSG0C | false | chr12 | 91587910 | 91591057 | 26.41709 | 100.00125 | 96.43182 | . | . | . | . | . | . |
| chr12 | 69296474 | 69297062 | 1000 | Promoter (<=1kb) | 12 | 69296681 | 69301117 | 4437 | 1 | ENSMUST00 | 0 | Klhdc2 | ENSMUSG0C | false | chr12 | 69296264 | 69297928 | 40.24926 | 142.56519 | 137.57257 | chr12 | 69295864 | 69296490 | 3.57998 | 3.3568 | 1.88887 |
| chr5 | 98108560 | 98109314 | 1000 | Distal Intergenic | 12 | 98202304 | 98259459 | 57156 | 2 | ENSMUST00 | 150145 | Galc | ENSMUSG0C | false | . | . | . | . | . | . | chr12 | 98105830 | 98109674 | 4.79379 | 4.05834 | 2.82062 |
| chr5 | 98187430 | 98187753 | 1000 | Distal Intergenic | 12 | 98202304 | 98259459 | 57156 | 2 | ENSMUST00 | 71706 | Galc | ENSMUSG0C | false | . | . | . | . | . | . | chr12 | 98186645 | 98190709 | 3.56295 | 3.33099 | 1.87083 |
| chr5 | 98106403 | 98106983 | 1000 | Distal Intergenic | 12 | 98202304 | 98259459 | 57156 | 2 | ENSMUST00 | 152476 | Galc | ENSMUSG0C | false | . | . | . | . | . | . | chr12 | 98105830 | 98109674 | 4.79379 | 4.05834 | 2.82062 |
| chr12 | 98736917 | 98737978 | 1000 | Promoter (<=1kb) | 12 | 98676741 | 98737405 | 60665 | 2 | ENSMUST00 | 0 | Ptpn21 | ENSMUSG0C | false | chr12 | 98736301 | 98738314 | 30.35223 | 94.4818 | 91.04041 | chr12 | 98737864 | 98738792 | 4.80463 | 4.02113 | 2.85834 |
| chr5 | 98755753 | 98756097 | 1000 | Intron (ENSMUST00000057000.1 | 12 | 98746968 | 98787774 | 40807 | 1 | ENSMUST00 | 8786 | Zc3h14 | ENSMUSG0C | false | . | . | . | . | . | . | chr12 | 98747766 | 98757789 | 5.14082 | 4.11124 | 3.1444 |
| chr12 | 98746789 | 98747514 | 1000 | Promoter (<=1kb) | 12 | 98746968 | 98787774 | 40807 | 1 | ENSMUST00 | 0 | Zc3h14 | ENSMUSG0C | false | chr12 | 98746611 | 98748039 | 33.09821 | 104.28432 | 100.60858 | . | . | . | . | . | . |
| chr5 | 55443098 | 55443948 | 1000 | Distal Intergenic | 12 | 55407383 | 55412287 | 4905 | 1 | ENSMUST00 | 35716 | Psma6 | ENSMUSG0C | false | . | . | . | . | . | . | chr12 | 55437151 | 55453280 | 7.38007 | 5.14554 | 4.94833 |
| chr5 | 55446570 | 55447264 | 1000 | Distal Intergenic | 12 | 55407383 | 55412287 | 4905 | 1 | ENSMUST00 | 39188 | Psma6 | ENSMUSG0C | false | . | . | . | . | . | . | chr12 | 55437151 | 55453280 | 7.38007 | 5.14554 | 4.94833 |
| chr5 | 87093202 | 87093922 | 1000 | Distal Intergenic | 12 | 87098025 | 87102350 | 4326 | 2 | ENSMUST00 | 8428 | Ngb | ENSMUSG0C | false | . | . | . | . | . | . | chr12 | 87088240 | 87097968 | 6.4867 | 4.74287 | 4.23052 |
| chr12 | 46818717 | 46819648 | 1000 | Promoter (<=1kb) | 12 | 46694517 | 46818775 | 124259 | 2 | ENSMUST00 | 0 | Nova1 | ENSMUSG0C | false | chr12 | 46817868 | 46820117 | 31.63878 | 95.72812 | 92.25946 | . | . | . | . | . | . |
| chr5 | 75520527 | 75521085 | 1000 | Intron (ENSMUST00000021447.7 | 12 | 75450883 | 75596200 | 145318 | 2 | ENSMUST00 | 75115 | Ppp2r5e | ENSMUSG0C | false | . | . | . | . | . | . | chr12 | 75515237 | 75524877 | 4.10046 | 3.54733 | 2.29131 |
| chr12 | 77238578 | 77239366 | 1000 | Promoter (<=1kb) | 12 | 77239065 | 77476338 | 237274 | 1 | ENSMUST00 | 0 | Fut8 | ENSMUSG0C | false | chr12 | 77237711 | 77240432 | 34.68901 | 133.45216 | 128.82967 | . | . | . | . | . | . |
| chr5 | 77261419 | 77261770 | 1000 | Intron (ENSMUST00000171770.3 | 12 | 77239065 | 77476338 | 237274 | 1 | ENSMUST00 | 22355 | Fut8 | ENSMUSG0C | false | . | . | . | . | . | . | chr12 | 77259417 | 77265718 | 4.06173 | 3.53787 | 2.26097 |
| chr12 | 69986743 | 69987450 | 1000 | Promoter (<=1kb) | 12 | 69965012 | 69987002 | 21991 | 2 | ENSMUST00 | 0 | Sav1 | ENSMUSG0C | false | chr12 | 69986329 | 69987632 | 32.29185 | 112.09676 | 108.19604 | . | . | . | . | . | . |
| chr12 | 70111090 | 70112260 | 1000 | Promoter (<=1kb) | 12 | 70018740 | 70111925 | 93186 | 2 | ENSMUST00 | 0 | Nin | ENSMUSG0C | false | chr12 | 70110700 | 70112394 | 38.54033 | 140.59962 | 135.69896 | chr12 | 70112138 | 70112836 | 3.41059 | 3.29764 | 1.70724 |
| chr5 | 70522533 | 70522932 | 1000 | Distal Intergenic | 12 | 70459841 | 70460576 | 736 | 1 | ENSMUST00 | 62693 | Tmx1 | ENSMUSG0C | false | . | . | . | . | . | . | chr12 | 70521815 | 70523173 | 2.7558 | 2.82423 | 1.19401 |
| chr12 | 72760857 | 72761972 | 1000 | Promoter (<=1kb) | 12 | 72761211 | 72794940 | 33730 | 1 | ENSMUST00 | 0 | Ppm1a | ENSMUSG0C | false | chr12 | 72760286 | 72762586 | 40.89836 | 130.45236 | 125.94646 | . | . | . | . | . | . |
| chr5 | 104855610 | 104855980 | 1000 | Intron (ENSMUST00000109936.1 | 12 | 104763114 | 104865076 | 101963 | 2 | ENSMUST00 | 9096 | Clmn | ENSMUSG0C | false | . | . | . | . | . | . | chr12 | 104848512 | 104856935 | 5.59388 | 4.3553 | 3.507 |
| chr5 | 104837194 | 104837527 | 1000 | Intron (ENSMUST00000109936.1 | 12 | 104763114 | 104865076 | 101963 | 2 | ENSMUST00 | 27549 | Clmn | ENSMUSG0C | false | . | . | . | . | . | . | chr12 | 104833203 | 104841822 | 7.03146 | 5.08004 | 4.65729 |
| chr12 | 73584543 | 73584934 | 1000 | Promoter (<=1kb) | 12 | 73584797 | 73778185 | 193389 | 1 | ENSMUST00 | 0 | Prkch | ENSMUSG0C | false | chr12 | 73584164 | 73586065 | 27.82288 | 84.25431 | 81.02234 | chr12 | 73584217 | 73585953 | 6.08961 | 4.55846 | 3.88338 |
| chr5 | 73892554 | 73892905 | 1000 | Distal Intergenic | 12 | 73901375 | 73947530 | 46156 | 1 | ENSMUST00 | -8470 | Hif1a | ENSMUSG0C | false | . | . | . | . | . | . | chr12 | 73891785 | 73893600 | 4.77056 | 3.96621 | 2.84275 |
| chr12 | 73907504 | 73908747 | 1000 | Promoter (<=1kb) | 12 | 73907904 | 73933076 | 25173 | 1 | ENSMUST00 | 0 | Hif1a | ENSMUSG0C | false | chr12 | 73907239 | 73909233 | 47.35943 | 168.9256 | 162.98769 | . | . | . | . | . | . |
| chr12 | 105784205 | 105785319 | 1000 | Promoter (<=1kb) | 12 | 105784694 | 105838807 | 54114 | 1 | ENSMUST00 | 0 | Papola | ENSMUSG0C | false | chr12 | 105783997 | 105786120 | 29.75681 | 92.45776 | 89.05831 | chr12 | 105783996 | 105784224 | 3.47175 | 3.37411 | 1.7243 |
| chr12 | 78748531 | 78749530 | 1000 | Promoter (<=1kb) | 12 | 78748947 | 78840711 | 91765 | 1 | ENSMUST00 | 0 | Mpp5 | ENSMUSG0C | true | chr12 | 78747954 | 78748798 | 24.86315 | 92.1349 | 88.74321 | . | . | . | . | . | . |
| chr5 | 80120457 | 80120837 | 1000 | Exon (ENSMUST00000180643.1/ | 12 | 80107760 | 80113013 | 5254 | 2 | ENSMUST00 | -7445 | Zfp36l1 | ENSMUSG0C | false | . | . | . | . | . | . | chr12 | 80116083 | 80133218 | 5.88241 | 4.4251 | 3.73792 |
| chr12 | 80114909 | 80115950 | 1000 | Promoter (1-2kb) | 12 | 80107760 | 80113013 | 5254 | 2 | ENSMUST00 | -1897 | Zfp36l1 | ENSMUSG0C | false | chr12 | 80113160 | 80116456 | 21.91337 | 68.25864 | 65.29428 | . | . | . | . | . | . |
| chr12 | 80112859 | 80113979 | 1000 | Promoter (<=1kb) | 12 | 80107760 | 80113013 | 5254 | 2 | ENSMUST00 | 0 | Zfp36l1 | ENSMUSG0C | false | chr12 | 80109381 | 80113014 | 31.85236 | 101.09525 | 97.496 | chr12 | 80113722 | 80114749 | 4.8306 | 4.07928 | 2.85837 |
| chr12 | 80790277 | 80791245 | 1000 | Promoter (<=1kb) | 12 | 80790532 | 80880832 | 90301 | 1 | ENSMUST00 | 0 | 4933426M11 | ENSMUSG0C | false | chr12 | 80789969 | 80791597 | 33.7938 | 119.00099 | 114.88904 | . | . | . | . | . | . |
| chr5 | 80824214 | 80824596 | 1000 | Intron (ENSMUST00000068519.5 | 12 | 80790532 | 80880832 | 90301 | 1 | ENSMUST00 | 33683 | 4933426M11 | ENSMUSG0C | false | . | . | . | . | . | . | chr12 | 80813773 | 80841711 | 4.74421 | 3.89481 | 2.82027 |
| chr12 | 81859855 | 81860993 | 1000 | Promoter (<=1kb) | 12 | 81860030 | 82000924 | 140895 | 1 | ENSMUST00 | 0 | Pcnx | ENSMUSG0C | false | chr12 | 81859409 | 81861509 | 37.88559 | 139.40556 | 134.54842 | . | . | . | . | . | . |
| chr5 | 113032600 | 113033099 | 1000 | Intron (ENSMUST00000084891.4 | 12 | 113014508 | 113074401 | 59894 | 1 | ENSMUST00 | 18093 | Pacs2 | ENSMUSG0C | true | . | . | . | . | . | . | chr12 | 113022909 | 113045363 | 6.33099 | 4.57761 | 4.12561 |
| chr12 | 113098357 | 113098782 | 1000 | Promoter (<=1kb) | 12 | 113098401 | 113136669 | 38269 | 1 | ENSMUST00 | 0 | Mta1 | ENSMUSG0C | false | chr12 | 113097661 | 113099119 | 34.40292 | 104.50784 | 100.82993 | chr12 | 113097010 | 113101275 | 4.60631 | 3.85511 | 2.73345 |
| chr13 | 9763887 | 9765065 | 1000 | Promoter (<=1kb) | 13 | 9684836 | 9764423 | 79588 | 2 | ENSMUST00 | 0 | Zmynd11 | ENSMUSG0C | false | chr13 | 9763025 | 9765435 | 35.13211 | 109.86813 | 106.03947 | . | . | . | . | . | . |
| chr12 | 116280671 | 116281502 | 1000 | Promoter (<=1kb) | 12 | 116281222 | 116373096 | 91875 | 1 | ENSMUST00 | 0 | Esyt2 | ENSMUSG0C | false | chr12 | 116280328 | 116282437 | 41.31276 | 141.09892 | 136.17311 | . | . | . | . | . | . |
| chr12 | 100724770 | 100725513 | 1000 | Promoter (<=1kb) | 12 | 100549778 | 100725028 | 175251 | 2 | ENSMUST00 | 0 | Rps6ka5 | ENSMUSG0C | false | chr12 | 100723877 | 100725193 | 31.1999 | 108.05416 | 104.27962 | . | . | . | . | . | . |
| chr5 | 102416387 | 102416805 | 1000 | Intron (ENSMUST00000110020.2 | 12 | 102394084 | 102423769 | 29686 | 2 | ENSMUST00 | 6964 | Lgmn | ENSMUSG0C | false | . | . | . | . | . | . | chr12 | 102416275 | 102416904 | 3.26986 | 3.05119 | 1.61933 |
| chr5 | 102532827 | 102533115 | 1000 | Distal Intergenic | 12 | 102554969 | 102565027 | 10059 | 1 | ENSMUST00 | -21854 | Chga | ENSMUSG0C | false | . | . | . | . | . | . | chr12 | 102532272 | 102536644 | 5.12842 | 3.99596 | 3.13602 |
| chr5 | 102535817 | 102536210 | 1000 | Distal Intergenic | 12 | 102554969 | 102565027 | 10059 | 1 | ENSMUST00 | -18759 | Chga | ENSMUSG0C | false | . | . | . | . | . | . | chr12 | 102532272 | 102536644 | 5.12842 | 3.99596 | 3.13602 |
| chr12 | 103531989 | 103532555 | 1000 | Promoter (<=1kb) | 12 | 103532283 | 103613831 | 81549 | 1 | ENSMUST00 | 0 | Ppp4r4 | ENSMUSG0C | false | chr12 | 103531957 | 103533229 | 31.01414 | 97.6227 | 94.10706 | chr12 | 103531586 | 103533339 | 4.21042 | 3.59621 | 2.40654 |
| chr7 | 36697175 | 36698436 | 1000 | Promoter (<=1kb) | 7 | 36698118 | 36773553 | 75436 | 1 | ENSMUST00 | 0 | Tshz3 | ENSMUSG0C | false | chr7 | 36695587 | 36698932 | 23.93078 | 87.47927 | 84.18233 | chr7 | 36694422 | 36697367 | 4.16665 | 3.7399 | 2.31095 |
| chr13 | 3537831 | 3538834 | 1000 | Promoter (<=1kb) | 13 | 3538075 | 3566261 | 28187 | 1 | ENSMUST00 | 0 | Gdi2 | ENSMUSG0C | false | chr13 | 3537313 | 3537923 | 25.56571 | 75.64988 | 72.57267 | . | . | . | . | . | . |
| chr12 | 83486624 | 83488038 | 1000 | Promoter (<=1kb) | 12 | 83213751 | 83487716 | 273966 | 2 | ENSMUST00 | 0 | Dpf3 | ENSMUSG0C | false | chr12 | 83485304 | 83489226 | 31.85236 | 101.09525 | 97.496 | . | . | . | . | . | . |
| chr5 | 83785816 | 83786308 | 1000 | Intron (ENSMUST00000121733.2 | 12 | 83763662 | 83781132 | 17471 | 1 | ENSMUST00 | 22155 | Papln | ENSMUSG0C | false | . | . | . | . | . | . | chr12 | 83784767 | 83787118 | 5.56589 | 4.30947 | 3.51946 |
| chr12 | 85454963 | 85455716 | 1000 | Distal Intergenic | 12 | 85473890 | 85477273 | 3384 | 1 | ENSMUST00 | -18174 | Fos | ENSMUSG0C | false | chr12 | 85454794 | 85455155 | 7.50973 | 14.65382 | 12.39649 | chr12 | 85454295 | 85457445 | 6.11803 | 4.61014 | 3.93027 |
| chr12 | 85471930 | 85472465 | 1000 | Promoter (1-2kb) | 12 | 85473890 | 85477273 | 3384 | 1 | ENSMUST00 | -1425 | Fos | ENSMUSG0C | false | chr12 | 85471798 | 85472829 | 14.80502 | 36.56548 | 34.02003 | chr12 | 85471270 | 85476604 | 4.14697 | 3.66678 | 2.30093 |
| chr12 | 85473315 | 85473683 | 1000 | Promoter (<=1kb) | 12 | 85473890 | 85477273 | 3384 | 1 | ENSMUST00 | -207 | Fos | ENSMUSG0C | false | chr12 | 85472966 | 85475150 | 24.03115 | 75.67385 | 72.59326 | chr12 | 85471270 | 85476604 | 4.14697 | 3.66678 | 2.30093 |
| chr12 | 85485893 | 85486460 | 1000 | Distal Intergenic | 12 | 85474911 | 85476063 | 1153 | 1 | ENSMUST00 | 10983 | Fos | ENSMUSG0C | false | chr12 | 85485608 | 85487883 | 27.24215 | 82.29425 | 79.10038 | chr12 | 85485423 | 85488221 | 5.47077 | 4.36176 | 3.37833 |
| chr5 | 86691133 | 86691582 | 1000 | Intron (ENSMUST00000021681.3 | 12 | 86678700 | 86692091 | 13392 | 1 | ENSMUST00 | 12434 | Vash1 | ENSMUSG0C | false | . | . | . | . | . | . | chr12 | 86685220 | 86694692 | 6.20303 | 4.49357 | 4.02263 |
| chr5 | 108223277 | 108223843 | 1000 | Intron (ENSMUST00000136175.1 | 12 | 108179738 | 108203359 | 23622 | 1 | ENSMUST00 | 43540 | Ccnk | ENSMUSG0C | false | . | . | . | . | . | . | chr12 | 108216919 | 108227928 | 7.31121 | 5.08617 | 4.91072 |
| chr5 | 108621954 | 108622295 | 1000 | Intron (ENSMUST00000021689.7 | 12 | 108635753 | 108688513 | 52761 | 1 | ENSMUST00 | -13458 | Evl | ENSMUSG0C | false | . | . | . | . | . | . | chr12 | 108621324 | 108622893 | 4.57161 | 3.80617 | 2.67109 |
| chr12 | 108793960 | 108794738 | 1000 | Promoter (<=1kb) | 12 | 108792973 | 108816632 | 23660 | 1 | ENSMUST00 | 988 | Yy1 | ENSMUSG0C | false | chr12 | 108792910 | 108795425 | 33.7938 | 119.00099 | 114.88904 | . | . | . | . | . | . |
| chr5 | 108816615 | 108816979 | 1000 | 3' UTR | 12 | 108806239 | 108815564 | 9326 | 1 | ENSMUST00 | 10377 | Yy1 | ENSMUSG0C | false | . | . | . | . | . | . | chr12 | 108816381 | 108817783 | 6.34861 | 4.65931 | 4.16546 |
| chr5 | 63152753 | 63153305 | 1000 | Intron (ENSMUST00000054963.9 | 14 | 63145153 | 63177793 | 32641 | 2 | ENSMUST00 | 24488 | Fdft1 | ENSMUSG0C | false | . | . | . | . | . | . | chr14 | 63148622 | 63154226 | 5.17179 | 4.06034 | 3.1827 |
| chr14 | 63163868 | 63165231 | 1000 | Exon (ENSMUST00000054963.9/ | 14 | 63145153 | 63177793 | 32641 | 2 | ENSMUST00 | 12562 | Fdft1 | ENSMUSG0C | false | chr14 | 63164409 | 63165221 | 18.80236 | 49.39901 | 46.69355 | chr14 | 63159286 | 63169501 | 4.41178 | 3.64285 | 2.55932 |
| chr5 | 111446294 | 111446841 | 1000 | Promoter (<=1kb) | 12 | 111446084 | 111453825 | 7742 | 1 | ENSMUST00 | 211 | Tnfaip2 | ENSMUSG0C | false | . | . | . | . | . | . | chr12 | 111443489 | 111451985 | 9.04148 | 5.70305 | 6.33624 |
| chr5 | 111516750 | 111517207 | 1000 | Distal Intergenic | 12 | 111538101 | 111546753 | 8653 | 1 | ENSMUST00 | -20894 | Eif5 | ENSMUSG0C | false | . | . | . | . | . | . | chr12 | 111513836 | 111518852 | 5.17158 | 4.07062 | 3.1695 |
| chr12 | 111514813 | 111515178 | 1000 | Distal Intergenic | 12 | 111538101 | 111546753 | 8653 | 1 | ENSMUST00 | -22923 | Eif5 | ENSMUSG0C | false | chr12 | 111514819 | 111515302 | 7.45394 | 13.24889 | 11.01597 | chr12 | 111513836 | 111518852 | 5.17158 | 4.07062 | 3.1695 |
| chr5 | 111884657 | 111885167 | 1000 | Intron (ENSMUST00000054815.8 | 12 | 111866013 | 111908040 | 42028 | 2 | ENSMUST00 | 22873 | Ppp1r13b | ENSMUSG0C | false | . | . | . | . | . | . | chr12 | 111880166 | 111903803 | 5.80743 | 4.40294 | 3.67276 |
| chr12 | 111907452 | 111908511 | 1000 | Promoter (<=1kb) | 12 | 111828458 | 111908055 | 79598 | 2 | ENSMUST00 | 0 | Ppp1r13b | ENSMUSG0C | false | chr12 | 111907130 | 111908809 | 30.59681 | 95.89824 | 92.42339 | chr12 | 111908371 | 111908912 | 4.76823 | 3.99405 | 2.82466 |
| chr5 | 111816276 | 111816894 | 1000 | Promoter (2-3kb) | 12 | 111814170 | 111828388 | 14219 | 1 | ENSMUST00 | 2107 | Zfyve21 | ENSMUSG0C | false | . | . | . | . | . | . | chr12 | 111813203 | 111823116 | 5.80855 | 4.38461 | 3.68777 |
| chr5 | 111780377 | 111780788 | 1000 | Promoter (1-2kb) | 12 | 111781947 | 111807844 | 25898 | 1 | ENSMUST00 | -1159 | Klc1 | ENSMUSG0C | false | . | . | . | . | . | . | chr12 | 111776671 | 111782879 | 5.48019 | 4.12221 | 3.43546 |
| chr5 | 111794111 | 111794521 | 1000 | Promoter (<=1kb) | 12 | 111795352 | 111806726 | 11375 | 1 | ENSMUST00 | -831 | Klc1 | ENSMUSG0C | false | . | . | . | . | . | . | chr12 | 111793212 | 111795551 | 3.92102 | 3.25803 | 2.15046 |
| chr5 | 111938541 | 111939169 | 1000 | Distal Intergenic | 12 | 111961376 | 111966977 | 5602 | 2 | ENSMUST00 | 27808 | 2010107E04I | ENSMUSG0C | false | . | . | . | . | . | . | chr12 | 111931379 | 111939617 | 5.47845 | 4.18739 | 3.42754 |
| chr13 | 14062972 | 14064358 | 1000 | Promoter (<=1kb) | 13 | 14052449 | 14063401 | 10953 | 2 | ENSMUST00 | 0 | Ggps1 | ENSMUSG0C | false | chr13 | 14062233 | 14063503 | 23.09106 | 69.49222 | 66.50756 | . | . | . | . | . | . |
| chr13 | 13393322 | 13393881 | 1000 | Promoter (<=1kb) | 13 | 13357623 | 13393624 | 36002 | 2 | ENSMUST00 | 0 | Gpr137b | ENSMUSG0C | false | chr13 | 13392538 | 13394560 | 25.29353 | 74.26213 | 71.20484 | chr13 | 13391893 | 13394867 | 5.33543 | 4.08284 | 3.32348 |
| chr5 | 24964696 | 24964991 | 1000 | Intron (ENSMUST00000021773.6 | 13 | 24955854 | 24962818 | 6965 | 1 | ENSMUST00 | 8843 | Gpld1 | ENSMUSG0C | false | . | . | . | . | . | . | chr13 | 24963888 | 24966111 | 4.23688 | 3.62664 | 2.42177 |
| chr5 | 41457343 | 41457953 | 1000 | Intron (ENSMUST00000021794.7 | 13 | 41309916 | 41487320 | 177405 | 2 | ENSMUST00 | 29367 | Nedd9 | ENSMUSG0C | false | . | . | . | . | . | . | chr13 | 41453612 | 41462201 | 4.95064 | 3.98019 | 2.98994 |
| chr13 | 42102931 | 42103337 | 1000 | Intron (ENSMUST00000060148.5 | 13 | 42052021 | 42185026 | 133006 | 1 | ENSMUST00 | 50911 | Hivep1 | ENSMUSG0C | false | chr13 | 42103197 | 42103826 | 5.905 | 11.42563 | 9.22457 | chr13 | 42098974 | 42105815 | 5.30539 | 4.14893 | 3.26488 |
| chr13 | 42051021 | 42052993 | 1000 | Promoter (<=1kb) | 13 | 42052021 | 42185026 | 133006 | 1 | ENSMUST00 | 0 | Hivep1 | ENSMUSG0C | false | chr13 | 42050686 | 42051679 | 18.39885 | 52.66533 | 49.91441 | . | . | . | . | . | . |
| chr5 | 42242086 | 42242587 | 1000 | Distal Intergenic | 13 | 42301270 | 42307989 | 6720 | 1 | ENSMUST00 | -58683 | Edn1 | ENSMUSG0C | false | . | . | . | . | . | . | chr13 | 42235582 | 42245410 | 5.3203 | 4.22032 | 3.27886 |
| chr13 | 43559753 | 43560530 | 1000 | Promoter (<=1kb) | 13 | 43538412 | 43560191 | 21780 | 2 | ENSMUST00 | 0 | Mcur1 | ENSMUSG0C | false | chr13 | 43558962 | 43561028 | 18.68094 | 84.05138 | 80.82243 | . | . | . | . | . | . |
| chr13 | 46501782 | 46502253 | 1000 | Promoter (<=1kb) | 13 | 46501848 | 46649642 | 147795 | 1 | ENSMUST00 | 0 | Cap2 | ENSMUSG0C | false | chr13 | 46501688 | 46502622 | 15.85026 | 49.49464 | 46.78799 | chr13 | 46501933 | 46502609 | 3.92604 | 3.57635 | 2.18225 |
| chr13 | 46727094 | 46728168 | 1000 | Promoter (<=1kb) | 13 | 46679905 | 46727940 | 48036 | 2 | ENSMUST00 | 0 | Nup153 | ENSMUSG0C | false | chr13 | 46726280 | 46728669 | 36.67561 | 120.95901 | 116.78313 | . | . | . | . | . | . |
| chr13 | 47104964 | 47105384 | 1000 | Promoter (<=1kb) | 13 | 47098155 | 47105830 | 7676 | 2 | ENSMUST00 | 446 | Dek | ENSMUSG0C | false | chr13 | 47104757 | 47106907 | 41.73432 | 142.95721 | 137.9541 | chr13 | 47103120 | 47105043 | 3.49727 | 3.23513 | 1.78199 |
| chr13 | 47105560 | 47106479 | 1000 | Promoter (<=1kb) | 13 | 47084782 | 47106190 | 21409 | 2 | ENSMUST00 | 0 | Dek | ENSMUSG0C | false | chr13 | 47104757 | 47106907 | 41.73432 | 142.95721 | 137.9541 | . | . | . | . | . | . |
| chr13 | 48227686 | 48228172 | 1000 | Intron (ENSMUST00000180921.1 | 13 | 48261427 | 48264036 | 2610 | 1 | ENSMUST00 | -33255 | Id4 | ENSMUSG0C | false | chr13 | 48228002 | 48228409 | 10.10971 | 19.98882 | 17.65231 | chr13 | 48226993 | 48228600 | 5.9694 | 4.56095 | 3.77735 |
| chr13 | 48260667 | 48261268 | 1000 | Promoter (<=1kb) | 13 | 48261427 | 48264036 | 2610 | 1 | ENSMUST00 | -159 | Id4 | ENSMUSG0C | false | chr13 | 48260400 | 48260882 | 9.09833 | 17.80034 | 15.4957 | . | . | . | . | . | . |
| chr5 | 33018800 | 33019093 | 1000 | Distal Intergenic | 13 | 33027416 | 33041884 | 14469 | 1 | ENSMUST00 | -8323 | Serpinb9b | ENSMUSG0C | false | . | . | . | . | . | . | chr13 | 33018470 | 33019502 | 3.86877 | 3.21976 | 2.11243 |
| chr13 | 34652097 | 34653393 | 1000 | Promoter (<=1kb) | 13 | 34627840 | 34652671 | 24832 | 2 | ENSMUST00 | 0 | Pxdc1 | ENSMUSG0C | true | chr13 | 34650232 | 34653634 | 30.94804 | 114.76607 | 110.79114 | . | . | . | . | . | . |
| chr13 | 34875410 | 34876015 | 1000 | Promoter (<=1kb) | 13 | 34875494 | 34902877 | 27384 | 1 | ENSMUST00 | 0 | Prpf4b | ENSMUSG0C | false | chr13 | 34873812 | 34876956 | 44.11508 | 143.73827 | 138.70853 | . | . | . | . | . | . |
| chr13 | 36117181 | 36117964 | 1000 | Promoter (<=1kb) | 13 | 36117411 | 36537592 | 420182 | 1 | ENSMUST00 | 0 | Fars2 | ENSMUSG0C | false | chr13 | 36116667 | 36118632 | 26.65955 | 86.95457 | 83.67183 | . | . | . | . | . | . |
| chr13 | 51846105 | 51846736 | 1000 | Promoter (<=1kb) | 13 | 51846678 | 51848468 | 1791 | 1 | ENSMUST00 | 0 | Gadd45g | ENSMUSG0C | false | chr13 | 51845444 | 51849264 | 29.81577 | 89.63673 | 86.30187 | chr13 | 51845166 | 51850273 | 5.3175 | 4.22752 | 3.26442 |
| chr5 | 63256693 | 63257240 | 1000 | Intron (ENSMUST00000091560.5 | 13 | 63282142 | 63326096 | 43955 | 1 | ENSMUST00 | -24902 | 2010111I01R | ENSMUSG0C | false | . | . | . | . | . | . | chr13 | 63252739 | 63272327 | 6.37956 | 4.73666 | 4.1425 |
| chr13 | 63239803 | 63240170 | 1000 | Exon (ENSMUST00000091560.5/ | 13 | 63282142 | 63326096 | 43955 | 1 | ENSMUST00 | -41972 | 2010111I01R | ENSMUSG0C | false | chr13 | 63239341 | 63244158 | 33.94791 | 109.87469 | 106.04007 | . | . | . | . | . | . |
| chr5 | 63023739 | 63024092 | 1000 | Intron (ENSMUST00000091560.5 | 13 | 63015167 | 63302284 | 287118 | 1 | ENSMUST00 | 8573 | 2010111I01R | ENSMUSG0C | false | . | . | . | . | . | . | chr13 | 63022988 | 63024656 | 3.8164 | 3.49736 | 2.0232 |
| chr13 | 63252394 | 63252671 | 1000 | Intron (ENSMUST00000091560.5 | 13 | 63282142 | 63326096 | 43955 | 1 | ENSMUST00 | -29471 | 2010111I01R | ENSMUSG0C | false | chr13 | 63249311 | 63254013 | 20.9555 | 57.98476 | 55.16582 | . | . | . | . | . | . |
| chr5 | 63154975 | 63155383 | 1000 | Intron (ENSMUST00000091560.5 | 13 | 63282142 | 63326096 | 43955 | 1 | ENSMUST00 | -126759 | 2010111I01R | ENSMUSG0C | false | . | . | . | . | . | . | chr13 | 63154418 | 63156276 | 3.9579 | 3.54325 | 2.15684 |
| chr5 | 63496872 | 63497424 | 1000 | Promoter (<=1kb) | 13 | 63347512 | 63497278 | 149767 | 2 | ENSMUST00 | 0 | Fancc | ENSMUSG0C | false | . | . | . | . | . | . | chr13 | 63494672 | 63498953 | 4.92111 | 4.0798 | 2.93352 |
| chr13 | 63563001 | 63565453 | 1000 | Promoter (<=1kb) | 13 | 63511533 | 63565520 | 53988 | 2 | ENSMUST00 | 67 | Ptch1 | ENSMUSG0C | false | chr13 | 63562847 | 63568718 | 33.69019 | 111.49343 | 107.61708 | . | . | . | . | . | . |
| chr5 | 53361417 | 53361690 | 1000 | Intron (ENSMUST00000021920.6 | 13 | 53332716 | 53377365 | 44650 | 2 | ENSMUST00 | 15675 | Sptlc1 | ENSMUSG0C | false | . | . | . | . | . | . | chr13 | 53361252 | 53362641 | 4.45514 | 3.75044 | 2.60525 |
| chr5 | 63957206 | 63957525 | 1000 | Distal Intergenic | 13 | 63872455 | 63900302 | 27848 | 1 | ENSMUST00 | 84752 | Ercc6l2 | ENSMUSG0C | false | . | . | . | . | . | . | chr13 | 63956458 | 63957971 | 4.92241 | 3.98838 | 2.98086 |
| chr5 | 64400761 | 64401468 | 1000 | Distal Intergenic | 13 | 64363214 | 64370306 | 7093 | 2 | ENSMUST00 | -30456 | Ctsl | ENSMUSG0C | false | . | . | . | . | . | . | chr13 | 64399856 | 64401068 | 5.60494 | 4.19581 | 3.53693 |
| chr13 | 55211028 | 55212974 | 1000 | Promoter (1-2kb) | 13 | 55209782 | 55318325 | 108544 | 1 | ENSMUST00 | 1247 | Nsd1 | ENSMUSG0C | true | chr13 | 55210377 | 55213952 | 20.60195 | 76.26862 | 73.17842 | chr13 | 55209838 | 55211508 | 4.00761 | 3.6192 | 2.1939 |
| chr5 | 55742283 | 55742735 | 1000 | Intron (ENSMUST00000021958.5 | 13 | 55727368 | 55776830 | 49463 | 1 | ENSMUST00 | 14916 | Pcbd2 | ENSMUSG0C | false | . | . | . | . | . | . | chr13 | 55739452 | 55743934 | 5.40467 | 4.31375 | 3.33541 |
| chr13 | 55727171 | 55727763 | 1000 | Promoter (<=1kb) | 13 | 55727368 | 55776828 | 49461 | 1 | ENSMUST00 | 0 | Pcbd2 | ENSMUSG0C | false | chr13 | 55726885 | 55728189 | 23.12601 | 77.64799 | 74.53377 | . | . | . | . | . | . |
| chr13 | 58401398 | 58403243 | 1000 | Promoter (<=1kb) | 13 | 58391239 | 58402464 | 11226 | 2 | ENSMUST00 | 0 | Hnrnpk | ENSMUSG0C | false | chr13 | 58400187 | 58403692 | 36.42221 | 131.26616 | 126.73016 | . | . | . | . | . | . |
| chr13 | 58273543 | 58274877 | 1000 | Promoter (<=1kb) | 13 | 58233351 | 58274188 | 40838 | 2 | ENSMUST00 | 0 | Gkap1 | ENSMUSG0C | false | chr13 | 58273089 | 58275023 | 36.88899 | 121.33285 | 117.14651 | chr13 | 58274588 | 58275274 | 2.76589 | 2.88043 | 1.21076 |
| chr5 | 59646475 | 59647250 | 1000 | Intron (ENSMUST00000022039.5 | 13 | 59632725 | 59634639 | 1915 | 1 | ENSMUST00 | 13751 | Naa35 | ENSMUSG0C | false | . | . | . | . | . | . | chr13 | 59644933 | 59657508 | 6.35164 | 4.69646 | 4.11411 |
| chr13 | 59675288 | 59676022 | 1000 | Promoter (<=1kb) | 13 | 59634996 | 59675784 | 40789 | 2 | ENSMUST00 | 0 | Golm1 | ENSMUSG0C | false | chr13 | 59674664 | 59676081 | 19.38504 | 61.02572 | 58.1656 | chr13 | 59671152 | 59676286 | 4.54264 | 3.87482 | 2.62246 |
| chr13 | 59556816 | 59557517 | 1000 | Promoter (<=1kb) | 13 | 59449538 | 59557289 | 107752 | 2 | ENSMUST00 | 0 | Agtpbp1 | ENSMUSG0C | false | chr13 | 59556269 | 59557388 | 36.46257 | 120.58523 | 116.42456 | . | . | . | . | . | . |
| chr13 | 60601749 | 60602310 | 1000 | Promoter (<=1kb) | 13 | 60601947 | 60763185 | 161239 | 1 | ENSMUST00 | 0 | Dapk1 | ENSMUSG0C | false | chr13 | 60601515 | 60602641 | 33.50033 | 109.08844 | 105.27855 | chr13 | 60601514 | 60603209 | 3.2068 | 3.14798 | 1.56191 |
| chr13 | 75943383 | 75944166 | 1000 | Promoter (<=1kb) | 13 | 75869537 | 75943867 | 74331 | 2 | ENSMUST00 | 0 | Rhobtb3 | ENSMUSG0C | false | chr13 | 75942785 | 75944428 | 27.34946 | 104.78247 | 101.09169 | chr13 | 75944085 | 75944844 | 4.84221 | 3.9859 | 2.88848 |
| chr13 | 69533526 | 69535101 | 1000 | Promoter (<=1kb) | 13 | 69533746 | 69615468 | 81723 | 1 | ENSMUST00 | 0 | Nsun2 | ENSMUSG0C | false | chr13 | 69532246 | 69535657 | 30.14656 | 119.38538 | 115.2599 | chr13 | 69530175 | 69533610 | 4.29829 | 3.51675 | 2.45714 |
| chr13 | 100650913 | 100651719 | 1000 | Promoter (<=1kb) | 13 | 100617165 | 100651059 | 33895 | 2 | ENSMUST00 | 0 | Rad17 | ENSMUSG0C | false | chr13 | 100650333 | 100652423 | 37.7199 | 126.00829 | 121.65485 | . | . | . | . | . | . |
| chr13 | 100616539 | 100617064 | 1000 | Promoter (<=1kb) | 13 | 100595957 | 100616944 | 20988 | 2 | ENSMUST00 | 0 | Marveld2 | ENSMUSG0C | false | chr13 | 100615532 | 100617288 | 22.68762 | 81.35053 | 78.1749 | chr13 | 100614708 | 100617785 | 3.56775 | 3.31396 | 1.86847 |
| chr13 | 98015948 | 98016635 | 1000 | Intron (ENSMUST00000109426.1 | 13 | 97898595 | 98206165 | 307571 | 2 | ENSMUST00 | 189530 | Arhgef28 | ENSMUSG0C | false | chr13 | 98015595 | 98016881 | 15.62409 | 36.57069 | 34.02363 | . | . | . | . | . | . |
| chr13 | 96542457 | 96543471 | 1000 | Promoter (<=1kb) | 13 | 96480689 | 96542485 | 61797 | 2 | ENSMUST00 | 0 | Polk | ENSMUSG0C | false | chr13 | 96541915 | 96544264 | 32.3084 | 121.498 | 117.30614 | . | . | . | . | . | . |
| chr13 | 95891612 | 95892139 | 1000 | Promoter (<=1kb) | 13 | 95627177 | 95891922 | 264746 | 2 | ENSMUST00 | 0 | Iqgap2 | ENSMUSG0C | true | chr13 | 95890627 | 95892671 | 39.90052 | 130.90422 | 126.38404 | . | . | . | . | . | . |
| chr5 | 94263939 | 94264419 | 1000 | Promoter (<=1kb) | 13 | 94225023 | 94265146 | 40124 | 2 | ENSMUST00 | 727 | Scamp1 | ENSMUSG0C | false | . | . | . | . | . | . | chr13 | 94255339 | 94269967 | 4.75211 | 3.92563 | 2.82403 |
| chr13 | 107021735 | 107022613 | 1000 | Promoter (<=1kb) | 13 | 106958996 | 107022118 | 63123 | 2 | ENSMUST00 | 0 | Kif2a | ENSMUSG0C | false | chr13 | 107021328 | 107023109 | 29.74442 | 97.36826 | 93.86232 | chr13 | 107019988 | 107021739 | 3.31265 | 3.19165 | 1.67098 |
| chr13 | 109902838 | 109903317 | 1000 | Promoter (<=1kb) | 13 | 109903102 | 109951387 | 48286 | 1 | ENSMUST00 | 0 | Pde4d | ENSMUSG0C | false | chr13 | 109902335 | 109904319 | 23.12601 | 77.64799 | 74.53377 | chr13 | 109899147 | 109906052 | 4.63584 | 3.90735 | 2.69774 |
| chr5 | 109822357 | 109823227 | 1000 | Intron (ENSMUST00000122041.2 | 13 | 109757221 | 109761612 | 4392 | 1 | ENSMUST00 | 65137 | Pde4d | ENSMUSG0C | false | . | . | . | . | . | . | chr13 | 109818398 | 109826347 | 5.304 | 4.21765 | 3.25083 |
| chr5 | 109652097 | 109652472 | 1000 | Intron (ENSMUST00000122041.2 | 13 | 109632780 | 109951364 | 318585 | 1 | ENSMUST00 | 19318 | Pde4d | ENSMUSG0C | false | . | . | . | . | . | . | chr13 | 109651021 | 109654287 | 4.95044 | 4.04334 | 2.9771 |
| chr5 | 92642121 | 92643384 | 1000 | Intron (ENSMUST00000049488.7 | 13 | 92611138 | 92711944 | 100807 | 1 | ENSMUST00 | 30984 | Serinc5 | ENSMUSG0C | false | . | . | . | . | . | . | . | . | . | . | . | . |
| chr13 | 103920104 | 103921854 | 1000 | Promoter (<=1kb) | 13 | 103818787 | 103920514 | 101728 | 2 | ENSMUST00 | 0 | Erbb2ip | ENSMUSG0C | false | chr13 | 103918287 | 103922058 | 26.18681 | 91.82585 | 88.44535 | chr13 | 103921648 | 103922087 | 3.44311 | 3.29355 | 1.70184 |
| chr13 | 104228400 | 104228941 | 1000 | Promoter (<=1kb) | 13 | 104205121 | 104228844 | 23724 | 2 | ENSMUST00 | 0 | Ppwd1 | ENSMUSG0C | false | chr13 | 104227893 | 104229418 | 30.91854 | 102.46954 | 98.8397 | . | . | . | . | . | . |
| chr14 | 14702706 | 14703704 | 1000 | Promoter (<=1kb) | 14 | 14703025 | 14799943 | 96919 | 1 | ENSMUST00 | 0 | Slc4a7 | ENSMUSG0C | false | chr14 | 14702478 | 14704027 | 31.16539 | 106.97112 | 103.21663 | . | . | . | . | . | . |
| chr14 | 14012339 | 14013596 | 1000 | Promoter (<=1kb) | 14 | 14012491 | 14107296 | 94806 | 1 | ENSMUST00 | 0 | Atxn7 | ENSMUSG0C | false | chr14 | 14011969 | 14013852 | 27.88725 | 99.90214 | 96.33501 | chr14 | 14013494 | 14015042 | 5.34563 | 4.26062 | 3.31381 |
| chr5 | 12111805 | 12112145 | 1000 | Intron (ENSMUST00000022264.7 | 14 | 12091160 | 12122132 | 30973 | 1 | ENSMUST00 | 20646 | Ptprg | ENSMUSG0C | false | . | . | . | . | . | . | chr14 | 12109629 | 12114213 | 5.04991 | 4.17996 | 3.04465 |
| chr5 | 8164486 | 8164830 | 1000 | 3' UTR | 14 | 8165991 | 8172992 | 7002 | 2 | ENSMUST00 | 8162 | Pdhb | ENSMUSG0C | false | . | . | . | . | . | . | chr14 | 8163574 | 8164817 | 3.75761 | 3.51599 | 1.96326 |
| chr14 | 8213757 | 8214504 | 1000 | Promoter (<=1kb) | 14 | 8214148 | 8223567 | 9420 | 1 | ENSMUST00 | 0 | Kctd6 | ENSMUSG0C | false | chr14 | 8213589 | 8215726 | 30.43737 | 91.68518 | 88.31002 | chr14 | 8213679 | 8216950 | 4.45532 | 3.83392 | 2.56051 |
| chr13 | 111807804 | 111809897 | 1000 | Promoter (<=1kb) | 13 | 111746428 | 111808993 | 62566 | 2 | ENSMUST00 | 0 | Map3k1 | ENSMUSG0C | false | chr13 | 111806971 | 111810084 | 41.81742 | 134.22485 | 129.57179 | . | . | . | . | . | . |
| chr14 | 21499539 | 21500284 | 1000 | Promoter (<=1kb) | 14 | 21499770 | 21672071 | 172302 | 1 | ENSMUST00 | 0 | Kat6b | ENSMUSG0C | false | chr14 | 21499033 | 21500514 | 27.83657 | 122.73275 | 118.49821 | chr14 | 21498879 | 21499657 | 3.94281 | 3.5563 | 2.11848 |
| chr14 | 21830944 | 21831687 | 1000 | Promoter (<=1kb) | 14 | 21831269 | 21845879 | 14611 | 1 | ENSMUST00 | 0 | Vdac2 | ENSMUSG0C | false | chr14 | 21830875 | 21833127 | 30.94804 | 114.76607 | 110.79114 | chr14 | 21820134 | 21830991 | 6.22936 | 4.53212 | 4.04608 |
| chr5 | 21823827 | 21824256 | 1000 | Distal Intergenic | 14 | 21831269 | 21845879 | 14611 | 1 | ENSMUST00 | -7013 | Vdac2 | ENSMUSG0C | false | . | . | . | . | . | . | chr14 | 21820134 | 21830991 | 6.22936 | 4.53212 | 4.04608 |
| chr5 | 21822814 | 21823347 | 1000 | Distal Intergenic | 14 | 21831269 | 21845879 | 14611 | 1 | ENSMUST00 | -7922 | Vdac2 | ENSMUSG0C | false | . | . | . | . | . | . | chr14 | 21820134 | 21830991 | 6.22936 | 4.53212 | 4.04608 |
| chr14 | 18330968 | 18332116 | 1000 | Promoter (<=1kb) | 14 | 18282722 | 18331859 | 49138 | 2 | ENSMUST00 | 0 | Ube2e1 | ENSMUSG0C | false | chr14 | 18330705 | 18332441 | 21.7465 | 82.16968 | 78.97962 | chr14 | 18326214 | 18331038 | 3.75249 | 3.44576 | 2.01182 |
| chr14 | 18238213 | 18239592 | 1000 | Promoter (<=1kb) | 14 | 18204054 | 18239127 | 35074 | 2 | ENSMUST00 | 0 | Nr1d2 | ENSMUSG0C | false | chr14 | 18237831 | 18239850 | 37.42379 | 126.5659 | 122.20106 | . | . | . | . | . | . |
| chr5 | 24205252 | 24205898 | 1000 | Intron (ENSMUST00000073687.7 | 14 | 24133953 | 24177789 | 43837 | 2 | ENSMUST00 | -27464 | Dlg5 | ENSMUSG0C | true | . | . | . | . | . | . | chr14 | 24191579 | 24244546 | 6.62149 | 4.76248 | 4.34728 |
| chr14 | 24245241 | 24246173 | 1000 | Promoter (<=1kb) | 14 | 24133953 | 24245920 | 111968 | 2 | ENSMUST00 | 0 | Dlg5 | ENSMUSG0C | true | chr14 | 24244111 | 24246301 | 38.14117 | 119.25307 | 115.13219 | chr14 | 24246016 | 24248610 | 4.17386 | 3.6478 | 2.3352 |
| chr14 | 34310230 | 34310954 | 1000 | Promoter (<=1kb) | 14 | 34310727 | 34345144 | 34418 | 1 | ENSMUST00 | 0 | Glud1 | ENSMUSG0C | false | chr14 | 34310717 | 34312662 | 35.98183 | 119.73972 | 115.60696 | . | . | . | . | . | . |
| chr5 | 34444934 | 34445295 | 1000 | Intron (ENSMUST00000049005.9 | 14 | 34414486 | 34460556 | 46071 | 2 | ENSMUST00 | 15261 | Bmpr1a | ENSMUSG0C | true | . | . | . | . | . | . | chr14 | 34439321 | 34450550 | 4.3338 | 3.66849 | 2.47835 |
| chr14 | 34501951 | 34503228 | 1000 | Promoter (<=1kb) | 14 | 34410734 | 34502687 | 91954 | 2 | ENSMUST00 | 0 | Bmpr1a | ENSMUSG0C | true | chr14 | 34500567 | 34503606 | 29.52499 | 116.10789 | 112.09218 | . | . | . | . | . | . |
| chr5 | 34532529 | 34532869 | 1000 | Intron (ENSMUST00000090040.5 | 14 | 34511619 | 34522801 | 11183 | 2 | ENSMUST00 | -9729 | 9230112D13 | ENSMUSG0C | false | . | . | . | . | . | . | chr14 | 34531588 | 34534123 | 5.19143 | 4.23736 | 3.15886 |
| chr5 | 19746196 | 19746580 | 1000 | Distal Intergenic | 14 | 19751257 | 19811787 | 60531 | 1 | ENSMUST00 | -4677 | Nid2 | ENSMUSG0C | false | . | . | . | . | . | . | chr14 | 19744839 | 19746690 | 5.58286 | 4.31065 | 3.52346 |
| chr14 | 20545780 | 20546734 | 1000 | Promoter (<=1kb) | 14 | 20499364 | 20546573 | 47210 | 2 | ENSMUST00 | 0 | Ppp3cb | ENSMUSG0C | false | chr14 | 20544831 | 20546595 | 34.50201 | 115.01646 | 111.03505 | . | . | . | . | . | . |
| chr5 | 20768571 | 20768954 | 1000 | Intron (ENSMUST00000071816.6 | 14 | 20734875 | 20794088 | 59214 | 2 | ENSMUST00 | 25134 | Camk2g | ENSMUSG0C | false | . | . | . | . | . | . | chr14 | 20768338 | 20769428 | 5.6471 | 4.31539 | 3.57275 |
| chr14 | 20793769 | 20794335 | 1000 | Promoter (<=1kb) | 14 | 20734875 | 20794088 | 59214 | 2 | ENSMUST00 | 0 | Camk2g | ENSMUSG0C | false | chr14 | 20793027 | 20794619 | 33.45292 | 101.85796 | 98.2438 | chr14 | 20790572 | 20794626 | 4.91791 | 4.02378 | 2.98507 |
| chr14 | 20928890 | 20929471 | 1000 | Promoter (<=1kb) | 14 | 20929433 | 21033673 | 104241 | 1 | ENSMUST00 | 0 | Vcl | ENSMUSG0C | false | chr14 | 20928550 | 20929186 | 22.91325 | 62.20349 | 59.32608 | . | . | . | . | . | . |
| chr14 | 45219175 | 45219856 | 1000 | Promoter (<=1kb) | 14 | 45134448 | 45219394 | 84947 | 2 | ENSMUST00 | 0 | Txndc16 | ENSMUSG0C | false | chr14 | 45219632 | 45221251 | 28.22734 | 101.53345 | 97.92473 | . | . | . | . | . | . |
| chr14 | 46882467 | 46883230 | 1000 | Promoter (<=1kb) | 14 | 46882854 | 47103150 | 220297 | 1 | ENSMUST00 | 0 | Samd4 | ENSMUSG0C | true | chr14 | 46882071 | 46885731 | 34.78613 | 113.42423 | 109.48744 | chr14 | 46881114 | 46887653 | 4.95941 | 4.04007 | 2.96585 |
| chr5 | 46999401 | 47000066 | 1000 | Promoter (1-2kb) | 14 | 47001340 | 47003357 | 2018 | 1 | ENSMUST00 | -1274 | Samd4 | ENSMUSG0C | true | . | . | . | . | . | . | chr14 | 46999778 | 47001724 | 4.91477 | 3.97818 | 2.95137 |
| chr14 | 46883394 | 46884335 | 1000 | Promoter (<=1kb) | 14 | 46883506 | 46884646 | 1141 | 1 | ENSMUST00 | 0 | Samd4 | ENSMUSG0C | true | chr14 | 46882071 | 46885731 | 34.78613 | 113.42423 | 109.48744 | chr14 | 46881114 | 46887653 | 4.95941 | 4.04007 | 2.96585 |
| chr14 | 47649082 | 47649979 | 1000 | Promoter (<=1kb) | 14 | 47649311 | 47736557 | 87247 | 1 | ENSMUST00 | 0 | Ktn1 | ENSMUSG0C | false | chr14 | 47648782 | 47650473 | 28.56742 | 103.16996 | 99.51957 | . | . | . | . | . | . |
| chr5 | 47888976 | 47889472 | 1000 | Distal Intergenic | 14 | 47729659 | 47736561 | 6903 | 1 | ENSMUST00 | 159318 | Ktn1 | ENSMUSG0C | false | . | . | . | . | . | . | chr14 | 47887039 | 47893837 | 4.36041 | 3.66154 | 2.50717 |
| chr5 | 47857312 | 47857881 | 1000 | Distal Intergenic | 14 | 47729659 | 47736561 | 6903 | 1 | ENSMUST00 | 127654 | Ktn1 | ENSMUSG0C | false | . | . | . | . | . | . | chr14 | 47856634 | 47859992 | 4.68948 | 3.93507 | 2.78767 |
| chr5 | 47823178 | 47823662 | 1000 | Distal Intergenic | 14 | 47729659 | 47736561 | 6903 | 1 | ENSMUST00 | 93520 | Ktn1 | ENSMUSG0C | false | . | . | . | . | . | . | chr14 | 47821002 | 47826875 | 4.48487 | 3.87377 | 2.58413 |
| chr5 | 47890241 | 47890742 | 1000 | Distal Intergenic | 14 | 47729659 | 47736561 | 6903 | 1 | ENSMUST00 | 160583 | Ktn1 | ENSMUSG0C | false | . | . | . | . | . | . | chr14 | 47887039 | 47893837 | 4.36041 | 3.66154 | 2.50717 |
| chr14 | 48120566 | 48121229 | 1000 | Promoter (<=1kb) | 14 | 48120869 | 48260883 | 140015 | 1 | ENSMUST00 | 0 | Peli2 | ENSMUSG0C | false | chr14 | 48119853 | 48122326 | 38.55812 | 129.64772 | 125.16568 | . | . | . | . | . | . |
| chr5 | 26357276 | 26357640 | 1000 | Promoter (<=1kb) | 14 | 26357547 | 26400456 | 42910 | 1 | ENSMUST00 | 0 | 4933413J09R | ENSMUSG0C | false | . | . | . | . | . | . | . | . | . | . | . | . |
| chr5 | 31400957 | 31401451 | 1000 | Intron (ENSMUST00000140002.2 | 14 | 31377595 | 31407385 | 29791 | 2 | ENSMUST00 | 5934 | Sh3bp5 | ENSMUSG0C | false | . | . | . | . | . | . | chr14 | 31400398 | 31416000 | 4.95975 | 4.04832 | 2.97568 |
| chr5 | 31402511 | 31402871 | 1000 | Intron (ENSMUST00000140002.2 | 14 | 31377595 | 31407385 | 29791 | 2 | ENSMUST00 | 4514 | Sh3bp5 | ENSMUSG0C | false | . | . | . | . | . | . | chr14 | 31400398 | 31416000 | 4.95975 | 4.04832 | 2.97568 |
| chr14 | 31336513 | 31337111 | 1000 | Promoter (<=1kb) | 14 | 31336638 | 31371984 | 35347 | 1 | ENSMUST00 | 0 | Capn7 | ENSMUSG0C | false | chr14 | 31336155 | 31337880 | 45.12022 | 156.84895 | 151.30879 | . | . | . | . | . | . |
| chr5 | 31226767 | 31227479 | 1000 | Intron (ENSMUST00000090180.2 | 14 | 31217860 | 31230350 | 12491 | 1 | ENSMUST00 | 8908 | Sema3g | ENSMUSG0C | true | . | . | . | . | . | . | chr14 | 31219988 | 31235200 | 8.317 | 5.47646 | 5.7231 |
| chr5 | 31229378 | 31229802 | 1000 | 3' UTR | 14 | 31217860 | 31230350 | 12491 | 1 | ENSMUST00 | 11519 | Sema3g | ENSMUSG0C | true | . | . | . | . | . | . | chr14 | 31219988 | 31235200 | 8.317 | 5.47646 | 5.7231 |
| chr14 | 31223800 | 31224237 | 1000 | Intron (ENSMUST00000090180.2 | 14 | 31217860 | 31230350 | 12491 | 1 | ENSMUST00 | 5941 | Sema3g | ENSMUSG0C | true | chr14 | 31220307 | 31223809 | 14.70503 | 33.66069 | 31.14909 | chr14 | 31219988 | 31235200 | 8.317 | 5.47646 | 5.7231 |
| chr14 | 31194882 | 31195558 | 1000 | Promoter (2-3kb) | 14 | 31177501 | 31192619 | 15119 | 2 | ENSMUST00 | -2264 | Nisch | ENSMUSG0C | false | chr14 | 31194886 | 31195731 | 6.48596 | 10.87719 | 8.68403 | chr14 | 31192553 | 31200159 | 6.28843 | 4.53257 | 4.09199 |
| chr14 | 61439252 | 61440293 | 1000 | Promoter (<=1kb) | 14 | 61365186 | 61439947 | 74762 | 2 | ENSMUST00 | 0 | Kpna3 | ENSMUSG0C | false | chr14 | 61438917 | 61440050 | 38.58512 | 125.36365 | 121.04008 | . | . | . | . | . | . |
| chr5 | 33412143 | 33412440 | 1000 | Promoter (1-2kb) | 14 | 33382170 | 33410994 | 28825 | 2 | ENSMUST00 | -1150 | Mapk8 | ENSMUSG0C | false | . | . | . | . | . | . | chr14 | 33411253 | 33416467 | 4.59246 | 3.88709 | 2.67806 |
| chr14 | 33446655 | 33447450 | 1000 | Promoter (<=1kb) | 14 | 33377898 | 33447158 | 69261 | 2 | ENSMUST00 | 0 | Mapk8 | ENSMUSG0C | false | chr14 | 33447382 | 33447720 | 7.54398 | 13.82373 | 11.57928 | . | . | . | . | . | . |
| chr14 | 56885691 | 56887501 | 1000 | Promoter (<=1kb) | 14 | 56887795 | 56962579 | 74785 | 1 | ENSMUST00 | -294 | Zmym2 | ENSMUSG0C | false | chr14 | 56885267 | 56886353 | 32.63288 | 132.66988 | 128.08362 | chr14 | 56887483 | 56890847 | 4.61055 | 3.79377 | 2.70445 |
| chr5 | 57736967 | 57737475 | 1000 | Promoter (2-3kb) | 14 | 57722189 | 57734300 | 12112 | 2 | ENSMUST00 | -2668 | Lats2 | ENSMUSG0C | false | . | . | . | . | . | . | chr14 | 57704250 | 57744175 | 6.91785 | 4.80068 | 4.61004 |
| chr14 | 57745797 | 57746571 | 1000 | Promoter (<=1kb) | 14 | 57642077 | 57746103 | 104027 | 2 | ENSMUST00 | 0 | Lats2 | ENSMUSG0C | false | chr14 | 57743310 | 57746899 | 34.76187 | 122.87653 | 118.63768 | . | . | . | . | . | . |
| chr5 | 57804235 | 57804620 | 1000 | 3' UTR | 14 | 57798638 | 57802335 | 3698 | 1 | ENSMUST00 | 5598 | Sap18 | ENSMUSG0C | false | . | . | . | . | . | . | chr14 | 57799380 | 57808560 | 5.47838 | 4.2987 | 3.40457 |
| chr14 | 58071898 | 58072522 | 1000 | Promoter (<=1kb) | 14 | 58072686 | 58110174 | 37489 | 1 | ENSMUST00 | -164 | Fgf9 | ENSMUSG0C | false | chr14 | 58071755 | 58073331 | 21.02725 | 63.18829 | 60.29588 | chr14 | 58069813 | 58076920 | 5.9191 | 4.53596 | 3.79262 |
| chr14 | 65097640 | 65098366 | 1000 | Promoter (<=1kb) | 14 | 65052060 | 65098106 | 46047 | 2 | ENSMUST00 | 0 | Extl3 | ENSMUSG0C | false | chr14 | 65097178 | 65098671 | 30.45704 | 104.66754 | 100.97924 | chr14 | 65096235 | 65101381 | 5.61541 | 4.33316 | 3.53079 |
| chr5 | 65098882 | 65099464 | 1000 | Promoter (<=1kb) | 14 | 65052060 | 65098106 | 46047 | 2 | ENSMUST00 | -777 | Extl3 | ENSMUSG0C | false | . | . | . | . | . | . | chr14 | 65096235 | 65101381 | 5.61541 | 4.33316 | 3.53079 |
| chr5 | 65113456 | 65113852 | 1000 | Distal Intergenic | 14 | 65052060 | 65098106 | 46047 | 2 | ENSMUST00 | -15351 | Extl3 | ENSMUSG0C | false | . | . | . | . | . | . | chr14 | 65111316 | 65115414 | 5.33089 | 4.19802 | 3.31303 |
| chr14 | 59440611 | 59441155 | 1000 | Promoter (<=1kb) | 14 | 59440981 | 59548903 | 107923 | 1 | ENSMUST00 | 0 | Cab39l | ENSMUSG0C | false | chr14 | 59439678 | 59441779 | 18.33657 | 60.6815 | 57.82544 | . | . | . | . | . | . |
| chr14 | 60265083 | 60265571 | 1000 | Promoter (<=1kb) | 14 | 60265205 | 60302370 | 37166 | 1 | ENSMUST00 | 0 | Mtmr6 | ENSMUSG0C | false | chr14 | 60265179 | 60266233 | 37.7199 | 126.00829 | 121.65485 | chr14 | 60264849 | 60267973 | 4.58611 | 3.73389 | 2.70745 |
| chr14 | 60634515 | 60634886 | 1000 | Promoter (<=1kb) | 14 | 60634729 | 60764556 | 129828 | 1 | ENSMUST00 | 0 | Spata13 | ENSMUSG0C | true | chr14 | 60634125 | 60635526 | 18.72985 | 51.651 | 48.91581 | chr14 | 60633986 | 60644204 | 5.19538 | 4.099 | 3.1763 |
| chr14 | 74731997 | 74732368 | 1000 | Promoter (<=1kb) | 14 | 74732297 | 74750765 | 18469 | 1 | ENSMUST00 | 0 | Esd | ENSMUSG0C | false | chr14 | 74731818 | 74733133 | 29.2476 | 106.45832 | 102.71833 | . | . | . | . | . | . |
| chr5 | 74762719 | 74763143 | 1000 | Intron (ENSMUST00000088970.5 | 14 | 74738506 | 74749755 | 11250 | 1 | ENSMUST00 | 24214 | Esd | ENSMUSG0C | false | . | . | . | . | . | . | chr14 | 74762567 | 74765258 | 4.8918 | 3.93148 | 2.96478 |
| chr14 | 75284049 | 75284678 | 1000 | Promoter (<=1kb) | 14 | 75284373 | 75344426 | 60054 | 1 | ENSMUST00 | 0 | Zc3h13 | ENSMUSG0C | false | chr14 | 75283583 | 75285322 | 30.26787 | 111.42836 | 107.5545 | . | . | . | . | . | . |
| chr14 | 76504121 | 76504708 | 1000 | Promoter (<=1kb) | 14 | 76504214 | 76507763 | 3550 | 1 | ENSMUST00 | 0 | Tsc22d1 | ENSMUSG0C | false | chr14 | 76503316 | 76505898 | 28.17892 | 90.64638 | 87.2878 | . | . | . | . | . | . |
| chr14 | 76505713 | 76506425 | 1000 | Promoter (<=1kb) | 14 | 76505098 | 76507002 | 1905 | 1 | ENSMUST00 | 616 | Tsc22d1 | ENSMUSG0C | false | chr14 | 76506045 | 76507699 | 17.34451 | 52.33425 | 49.58816 | . | . | . | . | . | . |
| chr14 | 76413988 | 76416165 | 1000 | Promoter (<=1kb) | 14 | 76414961 | 76507762 | 92802 | 1 | ENSMUST00 | 0 | Tsc22d1 | ENSMUSG0C | false | chr14 | 76413767 | 76416715 | 31.43241 | 100.35837 | 96.77719 | . | . | . | . | . | . |
| chr14 | 78536421 | 78537175 | 1000 | Promoter (<=1kb) | 14 | 78492246 | 78536808 | 44563 | 2 | ENSMUST00 | 0 | Akap11 | ENSMUSG0C | true | chr14 | 78535987 | 78537459 | 33.41831 | 117.26761 | 113.21259 | chr14 | 78522737 | 78539051 | 5.22391 | 4.1452 | 3.19494 |
| chr14 | 79480626 | 79482098 | 1000 | Promoter (<=1kb) | 14 | 79459937 | 79481268 | 21332 | 2 | ENSMUST00 | 0 | Wbp4 | ENSMUSG0C | false | chr14 | 79480316 | 79483475 | 31.4752 | 135.60028 | 130.8949 | . | . | . | . | . | . |
| chr14 | 66868458 | 66868930 | 1000 | Promoter (<=1kb) | 14 | 66802864 | 66868688 | 65825 | 2 | ENSMUST00 | 0 | Dpysl2 | ENSMUSG0C | true | chr14 | 66866432 | 66869441 | 25.50663 | 88.63404 | 85.31516 | . | . | . | . | . | . |
| chr14 | 67008374 | 67009444 | 1000 | Promoter (<=1kb) | 14 | 66985239 | 67008856 | 23618 | 2 | ENSMUST00 | 0 | Bnip3l | ENSMUSG0C | false | chr14 | 67007547 | 67009614 | 30.78991 | 105.27288 | 101.56436 | chr14 | 67009407 | 67010049 | 4.13178 | 3.70175 | 2.29871 |
| chr5 | 70170479 | 70171064 | 1000 | Promoter (<=1kb) | 14 | 70171512 | 70171986 | 475 | 2 | ENSMUST00 | 922 | Pdlim2 | ENSMUSG0C | true | . | . | . | . | . | . | chr14 | 70156516 | 70175803 | 7.12683 | 5 | 4.75261 |

| | | | | | | | | | | | | | | | | | | | | | | | | | | |
|---|---|---|---|---|---|---|---|---|---|---|---|---|---|---|---|---|---|---|---|---|---|---|---|---|---|---|
| chr14 | 70206079 | 70206959 | 1000 | Promoter (<=1kb) | 14 | 70180468 | 70207637 | 27170 | 2 | ENSMUST00 | 678 | Sorbs3 | ENSMUSG0C | false | chr14 | 70205110 | 70207222 | 19.02488 | 51.36755 | 48.63442 | chr14 | 70202314 | 70210660 | 7.58237 | 5.24911 | 5.1119 |
| chr14 | 70766275 | 70767044 | 1000 | Promoter (<=1kb) | 14 | 70654246 | 70766628 | 112383 | 2 | ENSMUST00 | 0 | Xpo7 | ENSMUSG0C | false | chr14 | 70765491 | 70767927 | 35.29575 | 125.98199 | 121.63858 | . | . | . | . | . | . |
| chr14 | 73325263 | 73326259 | 1000 | Promoter (<=1kb) | 14 | 73195502 | 73325822 | 130321 | 2 | ENSMUST00 | 0 | Rb1 | ENSMUSG0C | false | chr14 | 73324628 | 73326699 | 39.39101 | 131.10754 | 126.58253 | . | . | . | . | . | . |
| chr14 | 73142847 | 73143828 | 1000 | Promoter (<=1kb) | 14 | 73142863 | 73207843 | 64981 | 1 | ENSMUST00 | 0 | Rcbtb2 | ENSMUSG0C | false | chr14 | 73142491 | 73144603 | 28.8526 | 98.7319 | 95.19158 | chr14 | 73142369 | 73142959 | 3.43994 | 3.28792 | 1.70762 |
| chr5 | 73388572 | 73389039 | 1000 | Distal Intergenic | 14 | 73362231 | 73385271 | 23041 | 2 | ENSMUST00 | -3302 | Itm2b | ENSMUSG0C | false | . | . | . | . | . | . | chr14 | 73385739 | 73391282 | 5.53227 | 4.41051 | 3.43569 |
| chr5 | 73447979 | 73448485 | 1000 | Distal Intergenic | 14 | 73510049 | 73518545 | 8497 | 1 | ENSMUST00 | -61564 | Med4 | ENSMUSG0C | false | . | . | . | . | . | . | chr14 | 73441241 | 73453651 | 4.9946 | 4.03042 | 3.01369 |
| chr5 | 73453178 | 73453587 | 1000 | Distal Intergenic | 14 | 73510049 | 73518545 | 8497 | 1 | ENSMUST00 | -56462 | Med4 | ENSMUSG0C | false | . | . | . | . | . | . | chr14 | 73441241 | 73453651 | 4.9946 | 4.03042 | 3.01369 |
| chr14 | 101653738 | 101654499 | 1000 | Promoter (<=1kb) | 14 | 101653967 | 101696124 | 42158 | 1 | ENSMUST00 | 0 | Uchl3 | ENSMUSG0C | false | chr14 | 101653775 | 101654926 | 45.08533 | 152.20454 | 146.8354 | chr14 | 101652886 | 101655028 | 3.72742 | 3.5118 | 1.94849 |
| chr14 | 105896698 | 105898107 | 1000 | Promoter (<=1kb) | 14 | 105891949 | 105896819 | 4871 | 2 | ENSMUST00 | 0 | Spry2 | ENSMUSG0C | false | chr14 | 105894686 | 105899120 | 22.31878 | 85.15456 | 81.90936 | . | . | . | . | . | . |
| chr14 | 105483320 | 105483706 | 1000 | Distal Intergenic | 14 | 105498788 | 105505622 | 6835 | 1 | ENSMUST00 | -15082 | 4930449E01I | ENSMUSG0C | false | chr14 | 105483079 | 105483514 | 8.80131 | 17.30502 | 15.00832 | chr14 | 105483114 | 105484006 | 4.03816 | 3.55321 | 2.22355 |
| chr14 | 105176577 | 105177448 | 1000 | Promoter (<=1kb) | 14 | 105114965 | 105176860 | 61896 | 2 | ENSMUST00 | 0 | Rbm26 | ENSMUSG0C | false | chr14 | 105176084 | 105177955 | 37.6931 | 122.73721 | 118.50226 | . | . | . | . | . | . |
| chr5 | 103485680 | 103486030 | 1000 | Distal Intergenic | 14 | 103513341 | 103613346 | 100006 | 1 | ENSMUST00 | -27311 | Scel | ENSMUSG0C | true | . | . | . | . | . | . | chr14 | 103483842 | 103487218 | 5.85521 | 4.42685 | 3.71922 |
| chr5 | 103471070 | 103471498 | 1000 | Distal Intergenic | 14 | 103513341 | 103613346 | 100006 | 1 | ENSMUST00 | -41843 | Scel | ENSMUSG0C | true | . | . | . | . | . | . | chr14 | 103469233 | 103476308 | 5.58573 | 4.30613 | 3.50205 |
| chr5 | 103474689 | 103475103 | 1000 | Distal Intergenic | 14 | 103513341 | 103613346 | 100006 | 1 | ENSMUST00 | -38238 | Scel | ENSMUSG0C | true | . | . | . | . | . | . | chr14 | 103469233 | 103476308 | 5.58573 | 4.30613 | 3.50205 |
| chr14 | 103098811 | 103099986 | 1000 | Promoter (<=1kb) | 14 | 103080239 | 103099554 | 19316 | 2 | ENSMUST00 | 0 | Fbxl3 | ENSMUSG0C | false | chr14 | 103098163 | 103100145 | 33.5288 | 108.10788 | 104.32592 | . | . | . | . | . | . |
| chr15 | 8443442 | 8444860 | 1000 | Promoter (<=1kb) | 15 | 8291225 | 8444463 | 153239 | 2 | ENSMUST00 | 0 | Nipbl | ENSMUSG0C | false | chr15 | 8442340 | 8446554 | 43.84918 | 145.52415 | 140.42462 | . | . | . | . | . | . |
| chr5 | 6909661 | 6910134 | 1000 | Distal Intergenic | 15 | 6815037 | 6874969 | 59933 | 2 | ENSMUST00 | -34693 | Osmr | ENSMUSG0C | false | . | . | . | . | . | . | chr15 | 6908820 | 6910699 | 4.2014 | 3.36418 | 2.38171 |
| chr15 | 6386395 | 6387022 | 1000 | Promoter (<=1kb) | 15 | 6386598 | 6440705 | 54108 | 1 | ENSMUST00 | 0 | Dab2 | ENSMUSG0C | false | chr15 | 6386203 | 6388628 | 26.19333 | 73.09374 | 70.054 | chr15 | 6383263 | 6386496 | 4.13234 | 3.62074 | 2.30439 |
| chr14 | 54575508 | 54576296 | 1000 | Promoter (1-2kb) | 14 | 54567469 | 54577661 | 10193 | 2 | ENSMUST00 | 1365 | Ajuba | ENSMUSG0C | false | chr14 | 54574109 | 54578573 | 28.90341 | 112.84859 | 108.93253 | chr14 | 54568405 | 54575512 | 5.48983 | 4.2726 | 3.41523 |
| chr14 | 54572633 | 54573290 | 1000 | Intron (ENSMUST00000054487.8 | 14 | 54567469 | 54577661 | 10193 | 2 | ENSMUST00 | 4371 | Ajuba | ENSMUSG0C | false | chr14 | 54572758 | 54573818 | 11.05033 | 23.18759 | 20.81059 | chr14 | 54568405 | 54575512 | 5.48983 | 4.2726 | 3.41523 |
| chr5 | 54777832 | 54778280 | 1000 | Intron (ENSMUST00000022787.6 | 14 | 54722216 | 54781886 | 59671 | 2 | ENSMUST00 | 3606 | Slc7a8 | ENSMUSG0C | false | . | . | . | . | . | . | . | . | . | . | . | . |
| chr5 | 4790252 | 4790724 | 1000 | Intron (ENSMUST00000162585.2 | 15 | 4765945 | 4791164 | 25220 | 1 | ENSMUST00 | 24308 | C6 | ENSMUSG0C | false | . | . | . | . | . | . | chr15 | 4787398 | 4791363 | 4.57556 | 3.88581 | 2.65656 |
| chr15 | 3979157 | 3979847 | 1000 | Promoter (<=1kb) | 15 | 3965445 | 3979573 | 14129 | 2 | ENSMUST00 | 0 | Fbxo4 | ENSMUSG0C | true | chr15 | 3978820 | 3979702 | 24.53253 | 73.83858 | 70.79035 | chr15 | 3979789 | 3980961 | 6.07489 | 4.70467 | 3.88525 |
| chr5 | 54897239 | 54897854 | 1000 | Promoter (2-3kb) | 14 | 54894833 | 54895890 | 1058 | 1 | ENSMUST00 | 2407 | Pabpn1 | ENSMUSG0C | false | . | . | . | . | . | . | chr14 | 54894784 | 54900877 | 6.84642 | 4.93575 | 4.51393 |
| chr15 | 12549264 | 12550304 | 1000 | Promoter (<=1kb) | 15 | 12407282 | 12549278 | 141997 | 2 | ENSMUST00 | 0 | Pdzd2 | ENSMUSG0C | false | chr15 | 12548931 | 12550332 | 24.50502 | 69.29691 | 66.31957 | chr15 | 12548770 | 12551102 | 4.41991 | 3.85423 | 2.51643 |
| chr15 | 12321280 | 12322300 | 1000 | Promoter (<=1kb) | 15 | 12321496 | 12351265 | 29770 | 1 | ENSMUST00 | 0 | Golph3 | ENSMUSG0C | true | chr15 | 12320620 | 12323549 | 33.04283 | 115.53907 | 111.53828 | . | . | . | . | . | . |
| chr15 | 12117318 | 12118837 | 1000 | Promoter (<=1kb) | 15 | 12117831 | 12185683 | 67853 | 1 | ENSMUST00 | 0 | Zfr | ENSMUSG0C | false | chr15 | 12117175 | 12119324 | 36.04346 | 118.78738 | 114.68494 | chr15 | 12116945 | 12117384 | 5.50418 | 4.45757 | 3.42236 |
| chr14 | 54926427 | 54927099 | 1000 | Promoter (<=1kb) | 14 | 54916543 | 54926788 | 10246 | 2 | ENSMUST00 | 0 | Efs | ENSMUSG0C | false | chr14 | 54925688 | 54927212 | 28.49099 | 81.63235 | 78.4519 | chr14 | 54925510 | 54927179 | 4.46987 | 3.77551 | 2.5809 |
| chr14 | 55014528 | 55015030 | 1000 | Promoter (<=1kb) | 14 | 55015454 | 55024135 | 8682 | 1 | ENSMUST00 | -424 | Ngdn | ENSMUSG0C | false | chr14 | 55014352 | 55015181 | 22.31642 | 65.48328 | 62.55884 | chr14 | 55013838 | 55014602 | 3.90442 | 3.64245 | 2.08615 |
| chr15 | 11794611 | 11795056 | 1000 | Distal Intergenic | 15 | 11839896 | 11905674 | 65779 | 2 | ENSMUST00 | 110618 | Npr3 | ENSMUSG0C | true | . | . | . | . | . | . | chr15 | 11792640 | 11797553 | 4.61736 | 3.73311 | 2.73454 |
| chr15 | 11803691 | 11804005 | 1000 | Distal Intergenic | 15 | 11839896 | 11905674 | 65779 | 2 | ENSMUST00 | 101669 | Npr3 | ENSMUSG0C | true | . | . | . | . | . | . | chr15 | 11801200 | 11804147 | 3.48727 | 3.14603 | 1.82901 |
| chr15 | 11862743 | 11863574 | 1000 | Intron (ENSMUST00000066529.3 | 15 | 11839896 | 11905674 | 65779 | 2 | ENSMUST00 | 42100 | Npr3 | ENSMUSG0C | true | . | . | . | . | . | . | chr15 | 11851382 | 11874908 | 5.13432 | 4.1471 | 3.11197 |
| chr14 | 55560335 | 55560964 | 1000 | Promoter (<=1kb) | 14 | 55560482 | 55561099 | 618 | 1 | ENSMUST00 | 0 | Dcaf11 | ENSMUSG0C | false | chr14 | 55559308 | 55561525 | 31.37882 | 97.26304 | 93.75943 | . | . | . | . | . | . |
| chr14 | 55584943 | 55585502 | 1000 | Promoter (<=1kb) | 14 | 55581516 | 55584946 | 3431 | 2 | ENSMUST00 | 0 | Emc9 | ENSMUSG0C | false | chr14 | 55584681 | 55585616 | 22.06604 | 78.32127 | 75.1984 | chr14 | 55584461 | 55586283 | 3.75718 | 3.36177 | 2.02066 |
| chr15 | 32515870 | 32516313 | 1000 | Intron (ENSMUST00000067458.6 | 15 | 32244813 | 32696341 | 451529 | 1 | ENSMUST00 | 271058 | Sema5a | ENSMUSG0C | true | . | . | . | . | . | . | chr15 | 32499761 | 32540421 | 4.87542 | 4.01677 | 2.90332 |
| chr15 | 32265566 | 32265957 | 1000 | Intron (ENSMUST00000067458.6 | 15 | 32244813 | 32696341 | 451529 | 1 | ENSMUST00 | 20754 | Sema5a | ENSMUSG0C | true | . | . | . | . | . | . | chr15 | 32254606 | 32269959 | 6.09798 | 4.55757 | 3.9265 |
| chr15 | 32533582 | 32533966 | 1000 | Intron (ENSMUST00000067458.6 | 15 | 32244813 | 32696341 | 451529 | 1 | ENSMUST00 | 288770 | Sema5a | ENSMUSG0C | true | . | . | . | . | . | . | chr15 | 32499761 | 32540421 | 4.87542 | 4.01677 | 2.90332 |
| chr15 | 31568612 | 31569015 | 1000 | Promoter (<=1kb) | 15 | 31568791 | 31588259 | 19469 | 1 | ENSMUST00 | 0 | Cmbl | ENSMUSG0C | false | chr15 | 31568362 | 31568899 | 6.36264 | 14.31869 | 12.06804 | chr15 | 31562637 | 31570270 | 5.23895 | 4.15286 | 3.24218 |
| chr15 | 31322702 | 31323369 | 1000 | Intron (ENSMUST00000110410.3 | 15 | 31293186 | 31367532 | 74347 | 2 | ENSMUST00 | 44163 | Ankrd33b | ENSMUSG0C | false | . | . | . | . | . | . | chr15 | 31315773 | 31357869 | 5.77982 | 4.40135 | 3.65864 |
| chr15 | 31381886 | 31382470 | 1000 | Distal Intergenic | 15 | 31291479 | 31367759 | 76281 | 2 | ENSMUST00 | -14128 | Ankrd33b | ENSMUSG0C | false | . | . | . | . | . | . | chr15 | 31381071 | 31383131 | 5.98082 | 4.61734 | 3.80958 |
| chr15 | 31375222 | 31375623 | 1000 | Distal Intergenic | 15 | 31291479 | 31367759 | 76281 | 2 | ENSMUST00 | -7464 | Ankrd33b | ENSMUSG0C | false | . | . | . | . | . | . | chr15 | 31372886 | 31376371 | 5.15987 | 4.04366 | 3.16706 |
| chr15 | 31392664 | 31393141 | 1000 | Distal Intergenic | 15 | 31291479 | 31367759 | 76281 | 2 | ENSMUST00 | -24906 | Ankrd33b | ENSMUSG0C | false | . | . | . | . | . | . | chr15 | 31391477 | 31394264 | 4.8383 | 4.01609 | 2.88704 |
| chr9 | 63187276 | 63187683 | 1000 | Intron (ENSMUST00000034920.9 | 9 | 63138170 | 63148961 | 10792 | 2 | ENSMUST00 | -38316 | Skor1 | ENSMUSG0C | false | . | . | . | . | . | . | chr9 | 63186693 | 63188542 | 5.98817 | 4.5808 | 3.86015 |
| chr15 | 10691414 | 10691875 | 1000 | Intron (ENSMUST00000090339.4 | 15 | 10568979 | 10713540 | 144562 | 2 | ENSMUST00 | 21665 | Rai14 | ENSMUSG0C | true | . | . | . | . | . | . | chr15 | 10658484 | 10698797 | 6.0561 | 4.48411 | 3.88941 |
| chr15 | 10648390 | 10648771 | 1000 | Intron (ENSMUST00000090339.4 | 15 | 10568979 | 10713540 | 144562 | 2 | ENSMUST00 | 64769 | Rai14 | ENSMUSG0C | true | . | . | . | . | . | . | chr15 | 10640772 | 10650356 | 5.68862 | 4.34425 | 3.58498 |
| chr15 | 10714420 | 10715183 | 1000 | Promoter (<=1kb) | 15 | 10568979 | 10714631 | 145653 | 2 | ENSMUST00 | 0 | Rai14 | ENSMUSG0C | true | chr15 | 10712738 | 10715620 | 38.0703 | 123.3938 | 119.13641 | chr15 | 10714952 | 10715950 | 4.43224 | 3.7607 | 2.59681 |
| chr15 | 10663757 | 10664535 | 1000 | Intron (ENSMUST00000090339.4 | 15 | 10568979 | 10713540 | 144562 | 2 | ENSMUST00 | 49005 | Rai14 | ENSMUSG0C | true | . | . | . | . | . | . | chr15 | 10658484 | 10698797 | 6.0561 | 4.48411 | 3.88941 |
| chr15 | 10481779 | 10482078 | 1000 | Intron (ENSMUST00000022855.6 | 15 | 10478767 | 10485385 | 6619 | 2 | ENSMUST00 | 3307 | Brix1 | ENSMUSG0C | false | . | . | . | . | . | . | . | . | . | . | . | . |
| chr15 | 9071069 | 9071617 | 1000 | Promoter (<=1kb) | 15 | 9071260 | 9110487 | 39228 | 1 | ENSMUST00 | 0 | Nadk2 | ENSMUSG0C | false | chr15 | 9070331 | 9072400 | 37.86004 | 121.9582 | 117.75514 | . | . | . | . | . | . |
| chr15 | 32778556 | 32779143 | 1000 | Distal Intergenic | 15 | 32920723 | 33034730 | 114008 | 1 | ENSMUST00 | -141580 | Sdc2 | ENSMUSG0C | false | chr15 | 32778424 | 32779830 | 17.6846 | 53.76268 | 50.99834 | . | . | . | . | . | . |
| chr15 | 32920459 | 32921134 | 1000 | Promoter (<=1kb) | 15 | 32920723 | 33034730 | 114008 | 1 | ENSMUST00 | 0 | Sdc2 | ENSMUSG0C | false | chr15 | 32920480 | 32923199 | 20.32388 | 57.71489 | 54.90119 | . | . | . | . | . | . |
| chr15 | 27884719 | 27885067 | 1000 | Intron (ENSMUST00000090247.5 | 15 | 27730652 | 28025848 | 295197 | 2 | ENSMUST00 | 140781 | Trio | ENSMUSG0C | false | . | . | . | . | . | . | chr15 | 27883064 | 27885475 | 6.45121 | 4.7154 | 4.23282 |
| chr15 | 25872118 | 25872582 | 1000 | Intron (ENSMUST00000022881.8 | 15 | 25843264 | 25973687 | 130424 | 1 | ENSMUST00 | 28855 | Fam134b | ENSMUSG0C | false | . | . | . | . | . | . | chr15 | 25850428 | 25881315 | 5.59878 | 4.34742 | 3.49714 |
| chr15 | 25876687 | 25877190 | 1000 | Intron (ENSMUST00000022881.8 | 15 | 25843264 | 25973687 | 130424 | 1 | ENSMUST00 | 33424 | Fam134b | ENSMUSG0C | false | . | . | . | . | . | . | chr15 | 25850428 | 25881315 | 5.59878 | 4.34742 | 3.49714 |
| chr15 | 25914192 | 25914510 | 1000 | Intron (ENSMUST00000022881.8 | 15 | 25940887 | 25972340 | 31454 | 1 | ENSMUST00 | -26377 | Fam134b | ENSMUSG0C | false | . | . | . | . | . | . | chr15 | 25908288 | 25917989 | 5.38375 | 4.21727 | 3.3349 |
| chr15 | 25622246 | 25623395 | 1000 | Promoter (<=1kb) | 15 | 25622525 | 25813673 | 191149 | 1 | ENSMUST00 | 0 | Myo10 | ENSMUSG0C | false | chr15 | 25621457 | 25622305 | 13.41152 | 31.46699 | 28.98322 | chr15 | 25623293 | 25647336 | 5.36713 | 4.20892 | 3.31906 |
| chr15 | 36282365 | 36283420 | 1000 | Promoter (<=1kb) | 15 | 36239934 | 36283147 | 43214 | 2 | ENSMUST00 | 0 | Rnf19a | ENSMUSG0C | false | chr15 | 36281706 | 36283694 | 33.50033 | 109.08844 | 105.27855 | . | . | . | . | . | . |
| chr15 | 36608876 | 36609758 | 1000 | Promoter (<=1kb) | 15 | 36595661 | 36608973 | 13313 | 2 | ENSMUST00 | 0 | Pabpc1 | ENSMUSG0C | false | chr15 | 36607571 | 36609023 | 36.83603 | 116.99731 | 112.9521 | . | . | . | . | . | . |
| chr15 | 36792372 | 36794687 | 1000 | Promoter (<=1kb) | 15 | 36770770 | 36794538 | 23769 | 2 | ENSMUST00 | 0 | Ywhaz | ENSMUSG0C | false | chr15 | 36790381 | 36794895 | 37.74981 | 149.12509 | 143.87827 | . | . | . | . | . | . |
| chr15 | 41788540 | 41789551 | 1000 | Promoter (<=1kb) | 15 | 41789034 | 41861045 | 72012 | 1 | ENSMUST00 | 0 | Oxr1 | ENSMUSG0C | false | chr15 | 41788058 | 41790113 | 30.78991 | 105.27288 | 101.56436 | . | . | . | . | . | . |
| chr15 | 51798761 | 51799335 | 1000 | Exon (ENSMUST00000022925.8/ | 15 | 51786558 | 51865481 | 78924 | 2 | ENSMUST00 | 66146 | Eif3h | ENSMUSG0C | false | . | . | . | . | . | . | chr15 | 51798313 | 51799851 | 3.80716 | 3.3001 | 2.07143 |
| chr15 | 51987198 | 51987584 | 1000 | Intron (ENSMUST00000022927.9 | 15 | 51962605 | 51991760 | 29156 | 2 | ENSMUST00 | 4176 | Rad21 | ENSMUSG0C | true | . | . | . | . | . | . | chr15 | 51973657 | 51989901 | 5.03068 | 4.09477 | 3.03288 |
| chr15 | 35155527 | 35156077 | 1000 | Promoter (<=1kb) | 15 | 34875496 | 35155806 | 280311 | 2 | ENSMUST00 | 0 | Stk3 | ENSMUSG0C | false | chr15 | 35154948 | 35155944 | 36.16867 | 113.74171 | 109.7979 | . | . | . | . | . | . |
| chr15 | 68929052 | 68929420 | 1000 | Promoter (<=1kb) | 15 | 68928420 | 69093512 | 165093 | 1 | ENSMUST00 | 633 | Khdrbs3 | ENSMUSG0C | false | chr15 | 68927803 | 68930333 | 25.15761 | 80.47005 | 77.31023 | chr15 | 68929353 | 68934612 | 3.53741 | 3.34464 | 1.80709 |
| chr15 | 68258315 | 68259316 | 1000 | Promoter (<=1kb) | 15 | 68083766 | 68258856 | 175091 | 2 | ENSMUST00 | 0 | Zfat | ENSMUSG0C | false | chr15 | 68257894 | 68259920 | 25.57207 | 77.50074 | 74.3909 | . | . | . | . | . | . |
| chr15 | 44705779 | 44706102 | 1000 | Intron (ENSMUST00000110267.2 | 15 | 44671859 | 44752458 | 80600 | 2 | ENSMUST00 | 46356 | Sybu | ENSMUSG0C | false | . | . | . | . | . | . | chr15 | 44705107 | 44706651 | 4.49728 | 3.70565 | 2.63764 |
| chr15 | 59018996 | 59019358 | 1000 | Intron (ENSMUST00000080371.6 | 15 | 58941244 | 59082026 | 140783 | 2 | ENSMUST00 | 62668 | Mtss1 | ENSMUSG0C | true | . | . | . | . | . | . | chr15 | 58965186 | 59078715 | 7.47325 | 5.12649 | 5.04432 |
| chr15 | 59081918 | 59082362 | 1000 | Promoter (<=1kb) | 15 | 58941244 | 59082026 | 140783 | 2 | ENSMUST00 | 0 | Mtss1 | ENSMUSG0C | true | chr15 | 59075869 | 59083046 | 40.64776 | 144.39505 | 139.32506 | chr15 | 59082267 | 59100762 | 5.7793 | 4.47653 | 3.63859 |
| chr15 | 59003861 | 59004216 | 1000 | Intron (ENSMUST00000080371.6 | 15 | 58933810 | 58939489 | 5680 | 1 | ENSMUST00 | 70052 | Ndufb9 | ENSMUSG0C | false | . | . | . | . | . | . | chr15 | 58965186 | 59078715 | 7.47325 | 5.12649 | 5.04432 |
| chr15 | 64043194 | 64043566 | 1000 | Intron (ENSMUST00000063838.5 | 15 | 63929097 | 64060448 | 131352 | 2 | ENSMUST00 | 16882 | Fam49b | ENSMUSG0C | false | . | . | . | . | . | . | chr15 | 64041925 | 64044151 | 5.14293 | 3.93163 | 3.15621 |
| chr15 | 63997699 | 63998365 | 1000 | Promoter (<=1kb) | 15 | 63930944 | 63997969 | 67026 | 2 | ENSMUST00 | 0 | Fam49b | ENSMUSG0C | false | chr15 | 63997162 | 63998791 | 34.52358 | 106.75686 | 103.01088 | . | . | . | . | . | . |
| chr15 | 85735119 | 85735787 | 1000 | Promoter (<=1kb) | 15 | 85735564 | 85802163 | 66600 | 1 | ENSMUST00 | 0 | Ppara | ENSMUSG0C | false | chr15 | 85734614 | 85736405 | 22.88266 | 105.72421 | 102.00799 | chr15 | 85734520 | 85736392 | 4.95624 | 3.96493 | 3.01306 |
| chr15 | 81802287 | 81802884 | 1000 | Promoter (<=1kb) | 15 | 81802421 | 81826861 | 24441 | 1 | ENSMUST00 | 0 | Tef | ENSMUSG0C | false | chr15 | 81802642 | 81804673 | 20.25921 | 52.6909 | 49.93975 | chr15 | 81800930 | 81810375 | 6.79861 | 4.85005 | 4.49384 |
| chr15 | 81804600 | 81804921 | 1000 | Promoter (1-2kb) | 15 | 81802826 | 81805911 | 3086 | 1 | ENSMUST00 | 1775 | Tef | ENSMUSG0C | false | chr15 | 81802642 | 81804673 | 20.25921 | 52.6909 | 49.93975 | chr15 | 81800930 | 81810375 | 6.79861 | 4.85005 | 4.49384 |
| chr15 | 81810624 | 81811891 | 1000 | Promoter (<=1kb) | 15 | 81811046 | 81821256 | 10211 | 1 | ENSMUST00 | 0 | Tef | ENSMUSG0C | false | chr15 | 81810073 | 81812169 | 29.06085 | 90.26336 | 86.91321 | chr15 | 81811644 | 81815593 | 6.40149 | 4.7636 | 4.14853 |
| chr15 | 81399427 | 81400198 | 1000 | Promoter (<=1kb) | 15 | 81400138 | 81456807 | 56670 | 1 | ENSMUST00 | 0 | Xpnpep3 | ENSMUSG0C | false | chr15 | 81398585 | 81400981 | 38.20206 | 124.69703 | 120.39601 | . | . | . | . | . | . |
| chr15 | 80366872 | 80367233 | 1000 | Intron (ENSMUST00000160424.2 | 15 | 80389328 | 80390917 | 1590 | 1 | ENSMUST00 | -22095 | Cacna1i | ENSMUSG0C | false | . | . | . | . | . | . | chr15 | 80365842 | 80367398 | 3.76725 | 3.49438 | 1.97906 |
| chr15 | 79774147 | 79774613 | 1000 | Promoter (<=1kb) | 15 | 79761453 | 79774383 | 12931 | 2 | ENSMUST00 | 0 | Dnal4 | ENSMUSG0C | false | chr15 | 79773737 | 79775005 | 32.79732 | 102.7449 | 99.11129 | chr15 | 79773369 | 79775294 | 3.90809 | 3.4885 | 2.14969 |
| chr15 | 79687181 | 79688353 | 1000 | Promoter (<=1kb) | 15 | 79674250 | 79687872 | 13623 | 2 | ENSMUST00 | 0 | Josd1 | ENSMUSG0C | false | chr15 | 79686797 | 79688582 | 34.59177 | 109.96477 | 106.12952 | chr15 | 79680355 | 79687261 | 5.9641 | 4.46501 | 3.81779 |
| chr15 | 79436700 | 79436987 | 1000 | Promoter (2-3kb) | 15 | 79426399 | 79439018 | 12620 | 2 | ENSMUST00 | 2031 | Csnk1e | ENSMUSG0C | false | . | . | . | . | . | . | chr15 | 79428565 | 79442760 | 6.13618 | 4.60978 | 3.93942 |
| chr15 | 78899547 | 78900088 | 1000 | Promoter (<=1kb) | 15 | 78899667 | 78919032 | 19366 | 1 | ENSMUST00 | 0 | Sh3bp1 | ENSMUSG0C | false | chr15 | 78899411 | 78900583 | 27.18262 | 78.45122 | 75.32643 | chr15 | 78899068 | 78901639 | 5.59654 | 4.37707 | 3.5321 |
| chr15 | 77764276 | 77764869 | 1000 | Promoter (2-3kb) | 15 | 77765328 | 77767190 | 1863 | 2 | ENSMUST00 | 2321 | Myh9 | ENSMUSG0C | true | . | . | . | . | . | . | chr15 | 77763188 | 77768392 | 3.94204 | 3.48635 | 2.15878 |
| chr15 | 77784001 | 77784851 | 1000 | Intron (ENSMUST00000016771.7 | 15 | 77765328 | 77767190 | 1863 | 2 | ENSMUST00 | -16812 | Myh9 | ENSMUSG0C | true | . | . | . | . | . | . | chr15 | 77781877 | 77840076 | 8.49833 | 5.61339 | 5.87304 |
| chr15 | 77792176 | 77792594 | 1000 | 3' UTR | 15 | 77794652 | 77813231 | 18580 | 2 | ENSMUST00 | 20637 | Myh9 | ENSMUSG0C | true | . | . | . | . | . | . | chr15 | 77781877 | 77840076 | 8.49833 | 5.61339 | 5.87304 |
| chr15 | 77842021 | 77842891 | 1000 | Promoter (<=1kb) | 15 | 77760587 | 77842175 | 81589 | 2 | ENSMUST00 | 0 | Myh9 | ENSMUSG0C | true | chr15 | 77837742 | 77842188 | 46.1021 | 163.20312 | 157.4384 | . | . | . | . | . | . |
| chr15 | 77798198 | 77799228 | 1000 | Intron (ENSMUST00000016771.7 | 15 | 77794652 | 77813231 | 18580 | 2 | ENSMUST00 | 14003 | Myh9 | ENSMUSG0C | true | . | . | . | . | . | . | chr15 | 77781877 | 77840076 | 8.49833 | 5.61339 | 5.87304 |
| chr15 | 95520562 | 95520912 | 1000 | Intron (ENSMUST00000166170.2 | 15 | 95219933 | 95528252 | 308320 | 2 | ENSMUST00 | 7340 | Nell2 | ENSMUSG0C | true | . | . | . | . | . | . | chr15 | 95520023 | 95520886 | 3.22516 | 3.16037 | 1.60283 |
| chr15 | 95492194 | 95492594 | 1000 | Intron (ENSMUST00000166170.2 | 15 | 95219933 | 95528252 | 308320 | 2 | ENSMUST00 | 35658 | Nell2 | ENSMUSG0C | true | . | . | . | . | . | . | chr15 | 95489294 | 95492902 | 4.20527 | 3.69546 | 2.35743 |
| chr15 | 96698720 | 96699948 | 1000 | Promoter (<=1kb) | 15 | 96687394 | 96699698 | 12305 | 2 | ENSMUST00 | 0 | Slc38a2 | ENSMUSG0C | false | chr15 | 96697901 | 96700652 | 37.70819 | 121.69438 | 117.49508 | . | . | . | . | . | . |
| chr15 | 82147035 | 82148012 | 1000 | Promoter (<=1kb) | 15 | 82147266 | 82205376 | 58111 | 1 | ENSMUST00 | 0 | Srebf2 | ENSMUSG0C | false | chr15 | 82146565 | 82148559 | 40.89836 | 130.45236 | 125.94646 | chr15 | 82147972 | 82148752 | 3.38648 | 3.27132 | 1.70407 |
| chr15 | 97000094 | 97000547 | 1000 | Intron (ENSMUST00000023101.4 | 15 | 96996765 | 97020322 | 23558 | 2 | ENSMUST00 | 19775 | Slc38a4 | ENSMUSG0C | false | . | . | . | . | . | . | chr15 | 96998486 | 97005550 | 6.1246 | 4.57017 | 3.93131 |
| chr15 | 97677990 | 97678466 | 1000 | Exon (ENSMUST00000023104.5/ | 15 | 97675105 | 97705822 | 30718 | 2 | ENSMUST00 | 27356 | Rpap3 | ENSMUSG0C | false | . | . | . | . | . | . | chr15 | 97677139 | 97678904 | 5.63414 | 4.40595 | 3.53672 |
| chr15 | 97695862 | 97696330 | 1000 | Intron (ENSMUST00000023104.5 | 15 | 97675105 | 97705822 | 30718 | 2 | ENSMUST00 | 9492 | Rpap3 | ENSMUSG0C | false | . | . | . | . | . | . | chr15 | 97693179 | 97698449 | 6.12751 | 4.61178 | 3.94234 |
| chr15 | 97680299 | 97680674 | 1000 | Exon (ENSMUST00000023104.5/ | 15 | 97675105 | 97705822 | 30718 | 2 | ENSMUST00 | 25148 | Rpap3 | ENSMUSG0C | false | . | . | . | . | . | . | chr15 | 97679921 | 97681129 | 3.98552 | 3.40561 | 2.19195 |
| chr15 | 97721157 | 97721616 | 1000 | Promoter (<=1kb) | 15 | 97711019 | 97721150 | 10132 | 2 | ENSMUST00 | -8 | Endou | ENSMUSG0C | false | . | . | . | . | . | . | chr15 | 97716040 | 97732730 | 6.4081 | 4.59676 | 4.19736 |
| chr15 | 97729985 | 97730773 | 1000 | Promoter (<=1kb) | 15 | 97711019 | 97731405 | 20387 | 2 | ENSMUST00 | 632 | Endou | ENSMUSG0C | false | . | . | . | . | . | . | chr15 | 97716040 | 97732730 | 6.4081 | 4.59676 | 4.19736 |
| chr15 | 97762800 | 97763187 | 1000 | Promoter (<=1kb) | 15 | 97759364 | 97761879 | 2516 | 2 | ENSMUST00 | -922 | Rapgef3 | ENSMUSG0C | false | . | . | . | . | . | . | chr15 | 97747182 | 97765592 | 5.8605 | 4.39118 | 3.71878 |
| chr15 | 97766848 | 97768215 | 1000 | Promoter (<=1kb) | 15 | 97744770 | 97767846 | 23077 | 2 | ENSMUST00 | 0 | Rapgef3 | ENSMUSG0C | false | chr15 | 97764504 | 97768671 | 37.30079 | 124.19571 | 119.91465 | chr15 | 97768171 | 97769108 | 5.04582 | 4.14049 | 3.03535 |
| chr15 | 97832135 | 97832758 | 1000 | Promoter (<=1kb) | 15 | 97792681 | 97831767 | 39087 | 2 | ENSMUST00 | -369 | Hdac7 | ENSMUSG0C | false | chr15 | 97830319 | 97832239 | 27.11239 | 76.48743 | 73.39479 | chr15 | 97819806 | 97833530 | 5.44469 | 4.22025 | 3.37668 |
| chr15 | 97854506 | 97854961 | 1000 | 3' UTR | 15 | 97857537 | 97859698 | 2162 | 2 | ENSMUST00 | 4737 | Vdr | ENSMUSG0C | false | . | . | . | . | . | . | chr15 | 97854125 | 97855146 | 4.67403 | 3.93512 | 2.78433 |
| chr15 | 97907259 | 97908412 | 1000 | Promoter (<=1kb) | 15 | 97854425 | 97908310 | 53886 | 2 | ENSMUST00 | 0 | Vdr | ENSMUSG0C | false | chr15 | 97907246 | 97908866 | 21.37461 | 59.55079 | 56.71094 | chr15 | 97902462 | 97908754 | 5.09257 | 3.97616 | 3.09637 |
| chr16 | 10992879 | 10993424 | 1000 | Promoter (<=1kb) | 16 | 10959275 | 10993121 | 33847 | 2 | ENSMUST00 | 0 | Litaf | ENSMUSG0C | false | chr16 | 10991534 | 10993175 | 37.7199 | 126.00829 | 121.65485 | chr16 | 10993368 | 10994055 | 4.72976 | 3.96363 | 2.80521 |
| chr16 | 10313721 | 10314106 | 1000 | Promoter (<=1kb) | 16 | 10281749 | 10313968 | 32220 | 2 | ENSMUST00 | 0 | Emp2 | ENSMUSG0C | false | chr16 | 10313103 | 10314401 | 32.62678 | 97.43105 | 93.92233 | chr16 | 10310537 | 10314468 | 4.78089 | 3.80422 | 2.85259 |
| chr16 | 8829195 | 8830266 | 1000 | Promoter (<=1kb) | 16 | 8830100 | 8858916 | 28817 | 1 | ENSMUST00 | 0 | 1810013L24f | ENSMUSG0C | false | chr16 | 8828855 | 8829800 | 23.05105 | 65.89761 | 62.96862 | chr16 | 8828890 | 8829363 | 5.23197 | 4.26298 | 3.18058 |
| chr16 | 26354740 | 26355097 | 1000 | Downstream (1-2kb) | 16 | 26356642 | 26371841 | 15200 | 2 | ENSMUST00 | 16744 | Cldn1 | ENSMUSG0C | false | . | . | . | . | . | . | chr16 | 26354195 | 26360022 | 4.63445 | 3.93945 | 2.69823 |
| chr16 | 4213059 | 4214216 | 1000 | Promoter (<=1kb) | 16 | 4081328 | 4213407 | 132080 | 2 | ENSMUST00 | 0 | Crebbp | ENSMUSG0C | false | chr16 | 4212030 | 4214851 | 31.70772 | 93.8754 | 90.44607 | . | . | . | . | . | . |
| chr16 | 30083552 | 30083924 | 1000 | Distal Intergenic | 16 | 30066338 | 30067319 | 982 | 1 | ENSMUST00 | 17215 | Hes1 | ENSMUSG0C | false | . | . | . | . | . | . | chr16 | 30080758 | 30087022 | 7.36233 | 5.17806 | 4.92775 |
| chr16 | 30068383 | 30069063 | 1000 | Promoter (2-3kb) | 16 | 30066338 | 30067319 | 982 | 1 | ENSMUST00 | 2046 | Hes1 | ENSMUSG0C | false | . | . | . | . | . | . | chr16 | 30067043 | 30072068 | 6.09156 | 4.6344 | 3.88906 |
| chr16 | 30102524 | 30103509 | 1000 | Distal Intergenic | 16 | 30066338 | 30067319 | 982 | 1 | ENSMUST00 | 36187 | Hes1 | ENSMUSG0C | false | . | . | . | . | . | . | chr16 | 30101885 | 30104898 | 5.41046 | 4.30607 | 3.34416 |
| chr16 | 30063084 | 30064581 | 1000 | Promoter (<=1kb) | 16 | 30064384 | 30065569 | 1186 | 1 | ENSMUST00 | 0 | Hes1 | ENSMUSG0C | false | chr16 | 30063120 | 30067742 | 24.1047 | 77.63509 | 74.52327 | chr16 | 30059278 | 30065013 | 4.58644 | 3.91861 | 2.6533 |
| chr16 | 30069959 | 30071041 | 1000 | Distal Intergenic | 16 | 30066338 | 30067319 | 982 | 1 | ENSMUST00 | 3622 | Hes1 | ENSMUSG0C | false | chr16 | 30070559 | 30071103 | 8.65254 | 17.05594 | 14.76351 | chr16 | 30067043 | 30072068 | 6.09156 | 4.6344 | 3.88906 |
| chr15 | 76315380 | 76315695 | 1000 | Intron (ENSMUST00000165279.1 | 15 | 76300954 | 76307389 | 6436 | 2 | ENSMUST00 | -7992 | Oplah | ENSMUSG0C | false | . | . | . | . | . | . | chr15 | 76313569 | 76316064 | 4.30644 | 3.73169 | 2.43513 |
| chr15 | 76206124 | 76206585 | 1000 | Promoter (<=1kb) | 15 | 76170974 | 76206321 | 35348 | 2 | ENSMUST00 | 0 | Plec | ENSMUSG0C | false | chr15 | 76205354 | 76207876 | 14.73538 | 34.38367 | 31.86286 | chr15 | 76192493 | 76209331 | 6.98012 | 4.91238 | 4.64105 |
| chr15 | 76229243 | 76229869 | 1000 | Promoter (<=1kb) | 15 | 76170975 | 76229497 | 58523 | 2 | ENSMUST00 | 0 | Plec | ENSMUSG0C | false | chr15 | 76229079 | 76230123 | 17.64787 | 49.73517 | 47.02245 | chr15 | 76227744 | 76232701 | 5.05766 | 4.09409 | 3.05283 |
| chr15 | 76208119 | 76208871 | 1000 | Promoter (<=1kb) | 15 | 76170974 | 76208253 | 37280 | 2 | ENSMUST00 | 0 | Plec | ENSMUSG0C | false | chr15 | 76208303 | 76208821 | 4.50687 | 8.15551 | 6.02026 | chr15 | 76192493 | 76209331 | 6.98012 | 4.91238 | 4.64105 |
| chr15 | 76206707 | 76207797 | 1000 | Promoter (<=1kb) | 15 | 76170976 | 76206322 | 35347 | 2 | ENSMUST00 | -386 | Plec | ENSMUSG0C | false | chr15 | 76205354 | 76207876 | 14.73538 | 34.38367 | 31.86286 | chr15 | 76192493 | 76209331 | 6.98012 | 4.91238 | 4.64105 |
| chr15 | 76199336 | 76200117 | 1000 | Promoter (<=1kb) | 15 | 76170974 | 76199835 | 28862 | 2 | ENSMUST00 | 0 | Plec | ENSMUSG0C | false | chr15 | 76197865 | 76200595 | 17.16829 | 59.16322 | 56.32936 | chr15 | 76192493 | 76209331 | 6.98012 | 4.91238 | 4.64105 |
| chr15 | 75873374 | 75873771 | 1000 | Distal Intergenic | 15 | 75862339 | 75867404 | 5066 | 1 | ENSMUST00 | 11036 | Gsdmd | ENSMUSG0C | false | . | . | . | . | . | . | chr15 | 75872819 | 75873778 | 3.75432 | 3.44183 | 2.03141 |
| chr15 | 75862112 | 75862460 | 1000 | Promoter (<=1kb) | 15 | 75862339 | 75867404 | 5066 | 1 | ENSMUST00 | 0 | Gsdmd | ENSMUSG0C | false | chr15 | 75861839 | 75862989 | 18.02336 | 51.19578 | 48.46415 | chr15 | 75860973 | 75863138 | 4.66669 | 4.00112 | 2.71589 |
| chr15 | 75720724 | 75721064 | 1000 | Distal Intergenic | 15 | 75704394 | 75714225 | 9832 | 1 | ENSMUST00 | 16331 | Rhpn1 | ENSMUSG0C | true | . | . | . | . | . | . | chr15 | 75708646 | 75721294 | 7.3449 | 5.11049 | 4.92697 |
| chr15 | 75696868 | 75697376 | 1000 | Distal Intergenic | 15 | 75704280 | 75714408 | 10129 | 1 | ENSMUST00 | -6904 | Rhpn1 | ENSMUSG0C | true | . | . | . | . | . | . | chr15 | 75682738 | 75703169 | 5.10271 | 4.14364 | 3.08563 |
| chr15 | 73422854 | 73423361 | 1000 | Promoter (<=1kb) | 15 | 73205105 | 73423191 | 218087 | 2 | ENSMUST00 | 0 | Ptk2 | ENSMUSG0C | false | chr15 | 73421126 | 73424166 | 30.41442 | 103.5799 | 99.9162 | . | . | . | . | . | . |
| chr15 | 89140011 | 89140971 | 1000 | Promoter (<=1kb) | 15 | 89130585 | 89140703 | 10119 | 2 | ENSMUST00 | 0 | Mapk12 | ENSMUSG0C | false | chr15 | 89139588 | 89141123 | 28.03146 | 79.91027 | 76.76085 | chr15 | 89138742 | 89141795 | 4.26771 | 3.7871 | 2.38616 |
| chr15 | 93335783 | 93337204 | 1000 | Promoter (<=1kb) | 15 | 93284278 | 93336808 | 52531 | 2 | ENSMUST00 | 0 | Yaf2 | ENSMUSG0C | true | chr15 | 93335351 | 93337833 | 28.90341 | 112.84859 | 108.93253 | . | . | . | . | . | . |
| chr15 | 93398074 | 93398562 | 1000 | Promoter (<=1kb) | 15 | 93386097 | 93398301 | 12205 | 2 | ENSMUST00 | 0 | Zcrb1 | ENSMUSG0C | false | chr15 | 93397435 | 93399514 | 25.16654 | 87.04672 | 83.76057 | . | . | . | . | . | . |
| chr16 | 52452834 | 52453386 | 1000 | Promoter (<=1kb) | 16 | 52305264 | 52452654 | 147391 | 2 | ENSMUST00 | -181 | Alcam | ENSMUSG0C | true | chr16 | 52450004 | 52454370 | 27.24215 | 82.29425 | 79.10038 | . | . | . | . | . | . |
| chr16 | 52060878 | 52061302 | 1000 | Intron (ENSMUST00000114471.1 | 16 | 52031549 | 52208047 | 176499 | 1 | ENSMUST00 | 29330 | Cblb | ENSMUSG0C | true | . | . | . | . | . | . | chr16 | 52047691 | 52073501 | 5.26548 | 4.17513 | 3.22843 |
| chr16 | 52000101 | 52000544 | 1000 | Distal Intergenic | 16 | 52031549 | 52208047 | 176499 | 1 | ENSMUST00 | -31005 | Cblb | ENSMUSG0C | true | . | . | . | . | . | . | chr16 | 51999695 | 52002797 | 4.10448 | 3.61196 | 2.26807 |
| chr16 | 52032056 | 52032639 | 1000 | Promoter (<=1kb) | 16 | 52031549 | 52208047 | 176499 | 1 | ENSMUST00 | 508 | Cblb | ENSMUSG0C | true | chr16 | 52030893 | 52033595 | 36.7977 | 133.03651 | 128.43381 | . | . | . | . | . | . |
| chr16 | 50429508 | 50430285 | 1000 | Promoter (<=1kb) | 16 | 50224847 | 50429912 | 205066 | 2 | ENSMUST00 | 0 | Bbx | ENSMUSG0C | false | chr16 | 50429049 | 50432934 | 35.20524 | 115.20684 | 111.21615 | chr16 | 50428891 | 50429605 | 5.03387 | 4.10776 | 3.06722 |
| chr16 | 50364263 | 50364855 | 1000 | Intron (ENSMUST00000138166.2 | 16 | 50197159 | 50330993 | 133835 | 2 | ENSMUST00 | -33271 | Bbx | ENSMUSG0C | false | chr16 | 50363548 | 50364971 | 14.74478 | 37.16129 | 34.60659 | chr16 | 50360653 | 50367128 | 4.05902 | 3.45869 | 2.25848 |
| chr16 | 46496537 | 46497748 | 1000 | Promoter (<=1kb) | 16 | 46387706 | 46496967 | 109262 | 2 | ENSMUST00 | 0 | Pvrl3 | ENSMUSG0C | false | chr16 | 46495554 | 46498310 | 33.87313 | 113.90014 | 109.94483 | chr16 | 46494791 | 46496542 | 3.91473 | 3.51485 | 2.15353 |
| chr16 | 14180725 | 14181157 | 1000 | Promoter (2-3kb) | 16 | 14183573 | 14190520 | 6948 | 1 | ENSMUST00 | -2416 | Nde1 | ENSMUSG0C | false | . | . | . | . | . | . | chr16 | 14174692 | 14185475 | 6.74955 | 4.90863 | 4.43818 |
| chr16 | 13902843 | 13903490 | 1000 | Promoter (<=1kb) | 16 | 13903161 | 13949619 | 46459 | 1 | ENSMUST00 | 0 | Mpv17l | ENSMUSG0C | false | chr16 | 13902166 | 13904013 | 37.17318 | 134.81126 | 130.13722 | . | . | . | . | . | . |
| chr16 | 70313858 | 70314769 | 1000 | Promoter (<=1kb) | 16 | 70313949 | 70569716 | 255768 | 1 | ENSMUST00 | 0 | Gbe1 | ENSMUSG0C | false | chr16 | 70313499 | 70315103 | 32.47248 | 106.24805 | 102.52307 | . | . | . | . | . | . |
| chr16 | 43547310 | 43548258 | 1000 | Intron (ENSMUST00000132881.2 | 16 | 43510308 | 43619137 | 108830 | 1 | ENSMUST00 | 37003 | Zbtb20 | ENSMUSG0C | false | . | . | . | . | . | . | chr16 | 43543186 | 43567712 | 5.70766 | 4.3604 | 3.60355 |
| chr16 | 42874846 | 42876566 | 1000 | Promoter (<=1kb) | 16 | 42875881 | 43463175 | 587295 | 1 | ENSMUST00 | 0 | Zbtb20 | ENSMUSG0C | false | chr16 | 42875791 | 42876932 | 27.34889 | 81.53893 | 78.36224 | . | . | . | . | . | . |
| chr16 | 8738678 | 8739147 | 1000 | Promoter (<=1kb) | 16 | 8689600 | 8738420 | 48821 | 2 | ENSMUST00 | -259 | Usp7 | ENSMUSG0C | false | chr16 | 8736868 | 8739534 | 23.17719 | 89.67316 | 86.33752 | . | . | . | . | . | . |
| chr16 | 58523348 | 58523850 | 1000 | Promoter (<=1kb) | 16 | 58468125 | 58523426 | 55302 | 2 | ENSMUST00 | 0 | St3gal6 | ENSMUSG0C | true | chr16 | 58520006 | 58524671 | 32.66734 | 113.81544 | 109.86345 | . | . | . | . | . | . |
| chr16 | 58524018 | 58524421 | 1000 | Promoter (<=1kb) | 16 | 58469753 | 58524243 | 54491 | 2 | ENSMUST00 | 0 | St3gal6 | ENSMUSG0C | true | chr16 | 58520006 | 58524671 | 32.66734 | 113.81544 | 109.86345 | chr16 | 58524145 | 58524764 | 3.42104 | 3.29537 | 1.69247 |
| chr16 | 58514614 | 58515115 | 1000 | Intron (ENSMUST00000137035.2 | 16 | 58502861 | 58507898 | 5038 | 2 | ENSMUST00 | -6717 | St3gal6 | ENSMUSG0C | true | chr16 | 58512380 | 58514683 | 5.72276 | 11.47427 | 9.27082 | chr16 | 58496830 | 58520680 | 5.70067 | 4.33546 | 3.59278 |
| chr16 | 57639506 | 57639828 | 1000 | Intron (ENSMUST00000089332.4 | 16 | 57302199 | 57606825 | 304627 | 2 | ENSMUST00 | -32682 | Cmss1 | ENSMUSG0C | false | . | . | . | . | . | . | chr16 | 57639073 | 57640235 | 2.96267 | 2.94599 | 1.37041 |
| chr16 | 81200439 | 81200902 | 1000 | Promoter (<=1kb) | 16 | 81200697 | 81597805 | 397109 | 1 | ENSMUST00 | 0 | Ncam2 | ENSMUSG0C | false | chr16 | 81200246 | 81201154 | 18.98792 | 49.7182 | 47.00772 | chr16 | 81200172 | 81201202 | 3.25647 | 3.0767 | 1.61268 |
| chr16 | 17405178 | 17406212 | 1000 | Promoter (<=1kb) | 16 | 17406003 | 17430826 | 24824 | 1 | ENSMUST00 | 0 | Snap29 | ENSMUSG0C | false | chr16 | 17404953 | 17406958 | 21.47392 | 86.08567 | 82.81886 | chr16 | 17404145 | 17405257 | 5.32313 | 4.11147 | 3.28882 |
| chr16 | 31663728 | 31664474 | 1000 | Promoter (<=1kb) | 16 | 31663909 | 31871846 | 207938 | 1 | ENSMUST00 | 0 | Dlg1 | ENSMUSG0C | false | chr16 | 31663033 | 31664931 | 42.5703 | 137.72931 | 132.9411 | . | . | . | . | . | . |
| chr16 | 32078902 | 32079489 | 1000 | Promoter (<=1kb) | 16 | 32016290 | 32079342 | 63053 | 2 | ENSMUST00 | 0 | Pak2 | ENSMUSG0C | false | chr16 | 32077855 | 32079646 | 33.06624 | 113.49754 | 109.55873 | . | . | . | . | . | . |
| chr16 | 16557694 | 16558070 | 1000 | Promoter (1-2kb) | 16 | 16472266 | 16559990 | 87725 | 2 | ENSMUST00 | 1920 | Fgd4 | ENSMUSG0C | false | . | . | . | . | . | . | chr16 | 16553683 | 16561166 | 3.95114 | 3.54564 | 2.14289 |
| chr16 | 16358412 | 16359097 | 1000 | Promoter (<=1kb) | 16 | 16312235 | 16359030 | 46796 | 2 | ENSMUST00 | 0 | Dnm1l | ENSMUSG0C | false | chr16 | 16357981 | 16359179 | 34.78613 | 113.42423 | 109.48744 | chr16 | 16357313 | 16359368 | 4.87697 | 4.10997 | 2.90737 |
| chr16 | 32672895 | 32673336 | 1000 | Promoter (<=1kb) | 16 | 32673154 | 32679466 | 6313 | 1 | ENSMUST00 | 0 | Tnk2 | ENSMUSG0C | false | . | . | . | . | . | . | chr16 | 32672269 | 32673547 | 5.40226 | 4.37666 | 3.33168 |
| chr16 | 32643791 | 32644636 | 1000 | Promoter (<=1kb) | 16 | 32643874 | 32668279 | 24406 | 1 | ENSMUST00 | 0 | Tnk2 | ENSMUSG0C | false | chr16 | 32643583 | 32644999 | 29.41006 | 85.09721 | 81.85307 | chr16 | 32643570 | 32646072 | 3.40142 | 3.12364 | 1.73465 |
| chr16 | 32660539 | 32661358 | 1000 | Promoter (1-2kb) | 16 | 32658925 | 32668263 | 9339 | 1 | ENSMUST00 | 1615 | Tnk2 | ENSMUSG0C | false | . | . | . | . | . | . | chr16 | 32656126 | 32665456 | 6.5689 | 4.78559 | 4.29525 |
| chr16 | 32656759 | 32657109 | 1000 | Promoter (<=1kb) | 16 | 32657023 | 32668271 | 11249 | 1 | ENSMUST00 | 0 | Tnk2 | ENSMUSG0C | false | . | . | . | . | . | . | chr16 | 32656126 | 32665456 | 6.5689 | 4.78559 | 4.29525 |
| chr16 | 32877309 | 32878253 | 1000 | Promoter (<=1kb) | 16 | 32877500 | 32891409 | 13910 | 1 | ENSMUST00 | 0 | Fyttd1 | ENSMUSG0C | false | chr16 | 32877139 | 32879191 | 27.85705 | 92.98368 | 89.57738 | . | . | . | . | . | . |
| chr16 | 32939386 | 32939944 | 1000 | Intron (ENSMUST00000023491.7 | 16 | 32954698 | 32961734 | 7037 | 1 | ENSMUST00 | -14754 | Lrch3 | ENSMUSG0C | false | . | . | . | . | . | . | chr16 | 32934520 | 32940949 | 5.24063 | 4.24028 | 3.19657 |
| chr16 | 33178670 | 33179181 | 1000 | Distal Intergenic | 16 | 33185071 | 33243311 | 58241 | 1 | ENSMUST00 | -5890 | Osbpl11 | ENSMUSG0C | false | chr16 | 33178310 | 33178964 | 5.86754 | 9.51579 | 7.34688 | chr16 | 33177267 | 33180707 | 3.93566 | 3.39129 | 2.15789 |
| chr16 | 33380577 | 33381582 | 1000 | Promoter (<=1kb) | 16 | 33380775 | 33503903 | 123129 | 1 | ENSMUST00 | 0 | Zfp148 | ENSMUSG0C | false | chr16 | 33380162 | 33382282 | 22.99841 | 82.87405 | 79.66897 | . | . | . | . | . | . |
| chr16 | 33449276 | 33449679 | 1000 | Intron (ENSMUST00000165418.2 | 16 | 33400445 | 33498643 | 98199 | 1 | ENSMUST00 | 48832 | Zfp148 | ENSMUSG0C | false | . | . | . | . | . | . | chr16 | 33446425 | 33454527 | 4.54295 | 3.87525 | 2.63424 |
| chr16 | 38088648 | 38090147 | 1000 | Promoter (<=1kb) | 16 | 38089001 | 38246084 | 157084 | 1 | ENSMUST00 | 0 | Gsk3b | ENSMUSG0C | false | chr16 | 38088967 | 38093101 | 44.26367 | 154.19713 | 148.75002 | . | . | . | . | . | . |
| chr16 | 37814778 | 37815144 | 1000 | Promoter (<=1kb) | 16 | 37815706 | 37834426 | 18721 | 1 | ENSMUST00 | -562 | Fstl1 | ENSMUSG0C | false | . | . | . | . | . | . | chr16 | 37812310 | 37815411 | 5.39793 | 4.26551 | 3.3534 |
| chr16 | 33891523 | 33893022 | 1000 | Promoter (<=1kb) | 16 | 33891947 | 33899402 | 7456 | 1 | ENSMUST00 | 0 | Itgb5 | ENSMUSG0C | true | . | . | . | . | . | . | chr16 | 33868993 | 33898226 | 6.80171 | 4.94523 | 4.47886 |
| chr16 | 33922093 | 33922472 | 1000 | Promoter (<=1kb) | 16 | 33922486 | 33944148 | 21663 | 1 | ENSMUST00 | -14 | Itgb5 | ENSMUSG0C | true | . | . | . | . | . | . | chr16 | 33911227 | 33952498 | 7.40288 | 5.1092 | 4.9863 |
| chr16 | 33871925 | 33872475 | 1000 | Intron (ENSMUST00000115028.4 | 16 | 33885026 | 33910585 | 25560 | 1 | ENSMUST00 | -12551 | Itgb5 | ENSMUSG0C | true | . | . | . | . | . | . | chr16 | 33868993 | 33898226 | 6.80171 | 4.94523 | 4.47886 |
| chr16 | 33832635 | 33833477 | 1000 | Promoter (2-3kb) | 16 | 33829677 | 33949153 | 119477 | 1 | ENSMUST00 | 2959 | Itgb5 | ENSMUSG0C | true | chr16 | 33828900 | 33834465 | 38.14117 | 119.25307 | 115.13219 | chr16 | 33832776 | 33846769 | 5.19312 | 4.13271 | 3.17439 |
| chr16 | 33941219 | 33941749 | 1000 | Intron (ENSMUST00000115028.4 | 16 | 33922486 | 33944148 | 21663 | 1 | ENSMUST00 | 18734 | Itgb5 | ENSMUSG0C | true | . | . | . | . | . | . | chr16 | 33911227 | 33952498 | 7.40288 | 5.1092 | 4.9863 |
| chr16 | 33937545 | 33938039 | 1000 | Intron (ENSMUST00000115028.4 | 16 | 33922486 | 33944148 | 21663 | 1 | ENSMUST00 | 15060 | Itgb5 | ENSMUSG0C | true | . | . | . | . | . | . | chr16 | 33911227 | 33952498 | 7.40288 | 5.1092 | 4.9863 |
| chr16 | 37617716 | 37618149 | 1000 | Intron (ENSMUST00000160847.1 | 16 | 37621849 | 37632020 | 10172 | 1 | ENSMUST00 | -3700 | Hgd | ENSMUSG0C | false | . | . | . | . | . | . | chr16 | 37614959 | 37618754 | 4.96558 | 4.04706 | 2.98024 |
| chr16 | 35178577 | 35179017 | 1000 | Intron (ENSMUST00000114913.1 | 16 | 35154877 | 35305737 | 150861 | 1 | ENSMUST00 | 23701 | Adcy5 | ENSMUSG0C | true | . | . | . | . | . | . | chr16 | 35176357 | 35187367 | 5.48942 | 4.28454 | 3.41207 |
| chr16 | 35154605 | 35155474 | 1000 | Promoter (<=1kb) | 16 | 35154877 | 35305737 | 150861 | 1 | ENSMUST00 | 0 | Adcy5 | ENSMUSG0C | true | chr16 | 35154215 | 35157004 | 38.68979 | 124.4694 | 120.17514 | chr16 | 35153837 | 35158793 | 4.33176 | 3.75595 | 2.45108 |
| chr16 | 20702544 | 20702961 | 1000 | 3' UTR | 16 | 20707706 | 20708734 | 1029 | 2 | ENSMUST00 | 5773 | Clcn2 | ENSMUSG0C | false | . | . | . | . | . | . | chr16 | 20701677 | 20703464 | 7.05743 | 4.96928 | 4.74708 |
| chr16 | 20715919 | 20717241 | 1000 | Promoter (<=1kb) | 16 | 20702964 | 20716117 | 13154 | 2 | ENSMUST00 | 0 | Clcn2 | ENSMUSG0C | false | chr16 | 20715653 | 20718713 | 17.53424 | 68.85774 | 65.88623 | chr16 | 20714689 | 20719050 | 5.50285 | 4.33961 | 3.41363 |
| chr16 | 78576553 | 78577136 | 1000 | Promoter (<=1kb) | 16 | 78544012 | 78576657 | 32646 | 2 | ENSMUST00 | 0 | D16Ertd472e | ENSMUSG0C | false | chr16 | 78575560 | 78577511 | 39.06023 | 122.966 | 118.72274 | . | . | . | . | . | . |
| chr16 | 77013467 | 77014440 | 1000 | Promoter (<=1kb) | 16 | 77014069 | 77116779 | 102711 | 1 | ENSMUST00 | 0 | Usp25 | ENSMUSG0C | false | chr16 | 77013187 | 77016626 | 34.36702 | 111.64683 | 107.76208 | . | . | . | . | . | . |
| chr16 | 23233461 | 23233853 | 1000 | Intron (ENSMUST00000023601.8 | 16 | 23226021 | 23360350 | 134330 | 1 | ENSMUST00 | 7441 | St6gal1 | ENSMUSG0C | true | . | . | . | . | . | . | chr16 | 23227288 | 23275612 | 5.51887 | 4.26961 | 3.44483 |
| chr16 | 23358686 | 23359256 | 1000 | Promoter (<=1kb) | 16 | 23358316 | 23360348 | 2033 | 1 | ENSMUST00 | 371 | St6gal1 | ENSMUSG0C | true | . | . | . | . | . | . | chr16 | 23355278 | 23361898 | 5.18328 | 4.158 | 3.17046 |
| chr16 | 23240591 | 23241221 | 1000 | Intron (ENSMUST00000023601.8 | 16 | 23226021 | 23360350 | 134330 | 1 | ENSMUST00 | 14571 | St6gal1 | ENSMUSG0C | true | . | . | . | . | . | . | chr16 | 23227288 | 23275612 | 5.51887 | 4.26961 | 3.44483 |
| chr16 | 23256673 | 23257150 | 1000 | 5' UTR | 16 | 23269629 | 23321179 | 51551 | 1 | ENSMUST00 | -12479 | St6gal1 | ENSMUSG0C | true | . | . | . | . | . | . | chr16 | 23227288 | 23275612 | 5.51887 | 4.26961 | 3.44483 |
| chr16 | 23239597 | 23240163 | 1000 | Intron (ENSMUST00000023601.8 | 16 | 23226021 | 23360350 | 134330 | 1 | ENSMUST00 | 13577 | St6gal1 | ENSMUSG0C | true | . | . | . | . | . | . | chr16 | 23227288 | 23275612 | 5.51887 | 4.26961 | 3.44483 |
| chr16 | 23232612 | 23233118 | 1000 | Intron (ENSMUST00000023601.8 | 16 | 23226021 | 23360350 | 134330 | 1 | ENSMUST00 | 6592 | St6gal1 | ENSMUSG0C | true | . | . | . | . | . | . | chr16 | 23227288 | 23275612 | 5.51887 | 4.26961 | 3.44483 |
| chr16 | 23266232 | 23266686 | 1000 | Promoter (2-3kb) | 16 | 23269629 | 23321179 | 51551 | 1 | ENSMUST00 | -2943 | St6gal1 | ENSMUSG0C | true | . | . | . | . | . | . | chr16 | 23227288 | 23275612 | 5.51887 | 4.26961 | 3.44483 |
| chr16 | 85800891 | 85801662 | 1000 | Promoter (<=1kb) | 16 | 85795476 | 85801481 | 6006 | 2 | ENSMUST00 | 0 | Adamts1 | ENSMUSG0C | false | chr16 | 85800671 | 85803715 | 26.05581 | 78.34985 | 75.22605 | chr16 | 85800084 | 85805307 | 4.30005 | 3.70427 | 2.42091 |
| chr16 | 86252630 | 86253014 | 1000 | Distal Intergenic | 16 | 85858170 | 85901125 | 42956 | 2 | ENSMUST00 | -351506 | Adamts5 | ENSMUSG0C | false | . | . | . | . | . | . | . | . | . | . | . | . |
| chr16 | 95701862 | 95702872 | 1000 | Promoter (<=1kb) | 16 | 95702075 | 95721051 | 18977 | 1 | ENSMUST00 | 0 | Ets2 | ENSMUSG0C | false | chr16 | 95701830 | 95704226 | 35.74757 | 115.10552 | 111.12301 | chr16 | 95702821 | 95710369 | 4.85927 | 3.9975 | 2.89152 |
| chr16 | 95703870 | 95704281 | 1000 | Promoter (1-2kb) | 16 | 95702500 | 95712564 | 10065 | 1 | ENSMUST00 | 1371 | Ets2 | ENSMUSG0C | false | chr16 | 95701830 | 95704226 | 35.74757 | 115.10552 | 111.12301 | chr16 | 95702821 | 95710369 | 4.85927 | 3.9975 | 2.89152 |
| chr16 | 94568991 | 94571065 | 1000 | Promoter (<=1kb) | 16 | 94570010 | 94695067 | 125058 | 1 | ENSMUST00 | 0 | Dyrk1a | ENSMUSG0C | false | chr16 | 94568436 | 94571673 | 30.75929 | 103.19345 | 99.54015 | . | . | . | . | . | . |
| chr16 | 94526391 | 94527189 | 1000 | Promoter (<=1kb) | 16 | 94497783 | 94526830 | 29048 | 2 | ENSMUST00 | 0 | Dscr3 | ENSMUSG0C | false | chr16 | 94525785 | 94527653 | 28.08037 | 85.65662 | 82.39679 | . | . | . | . | . | . |
| chr16 | 35982966 | 35984092 | 1000 | Promoter (<=1kb) | 16 | 35983315 | 36037131 | 53817 | 1 | ENSMUST00 | 0 | Kpna1 | ENSMUSG0C | false | chr16 | 35982507 | 35984385 | 31.14594 | 106.93581 | 103.18726 | . | . | . | . | . | . |
| chr16 | 62872557 | 62873079 | 1000 | Intron (ENSMUST00000023629.8 | 16 | 62854307 | 62929346 | 75040 | 1 | ENSMUST00 | 18251 | Pros1 | ENSMUSG0C | false | . | . | . | . | . | . | chr16 | 62866329 | 62875410 | 4.59164 | 3.90036 | 2.66788 |
| chr16 | 96081448 | 96083026 | 1000 | Promoter (<=1kb) | 16 | 95992092 | 96082428 | 90337 | 2 | ENSMUST00 | 0 | Brwd1 | ENSMUSG0C | false | chr16 | 96081146 | 96083699 | 52.79799 | 184.38852 | 178.14023 | chr16 | 96078872 | 96081487 | 4.99734 | 4.08815 | 3.03594 |
| chr16 | 94370146 | 94371603 | 1000 | Promoter (<=1kb) | 16 | 94358763 | 94370289 | 11527 | 2 | ENSMUST00 | 0 | Pigp | ENSMUSG0C | false | chr16 | 94369721 | 94372158 | 31.62754 | 97.69553 | 94.17948 | . | . | . | . | . | . |
| chr16 | 92458308 | 92458780 | 1000 | Intron (ENSMUST00000060005.9 | 16 | 92391953 | 92466146 | 74194 | 2 | ENSMUST00 | 7366 | Rcan1 | ENSMUSG0C | false | . | . | . | . | . | . | chr16 | 92457736 | 92459012 | 5.42695 | 4.38556 | 3.34182 |
| chr16 | 91728950 | 91729758 | 1000 | Promoter (<=1kb) | 16 | 91729281 | 91870210 | 140930 | 1 | ENSMUST00 | 0 | Itsn1 | ENSMUSG0C | false | chr16 | 91729283 | 91730791 | 25.84672 | 90.2271 | 86.88143 | . | . | . | . | . | . |
| chr16 | 91546812 | 91547415 | 1000 | Promoter (<=1kb) | 16 | 91547072 | 91565169 | 18098 | 1 | ENSMUST00 | 0 | Ifngr2 | ENSMUSG0C | false | chr16 | 91546579 | 91548069 | 34.68901 | 133.45216 | 128.82967 | . | . | . | . | . | . |
| chr16 | 91010848 | 91011719 | 1000 | Promoter (<=1kb) | 16 | 90936092 | 91011308 | 75217 | 2 | ENSMUST00 | 0 | Synj1 | ENSMUSG0C | false | chr16 | 91009983 | 91011976 | 43.18097 | 163.77325 | 157.99008 | chr16 | 91011602 | 91012168 | 4.8253 | 4.11011 | 2.8497 |
| chr16 | 90242950 | 90243282 | 1000 | Exon (ENSMUST00000163419.2/ | 16 | 90220742 | 90226322 | 5581 | 1 | ENSMUST00 | 22209 | Sod1 | ENSMUSG0C | false | . | . | . | . | . | . | chr16 | 90242093 | 90244300 | 3.91131 | 3.56718 | 2.11434 |
| chr1 | 182019597 | 182020473 | 1000 | Promoter (<=1kb) | 1 | 181904445 | 182019980 | 115536 | 2 | ENSMUST00 | 0 | Enah | ENSMUSG0C | false | chr1 | 182018483 | 182020835 | 33.32351 | 103.65904 | 99.99326 | chr1 | 182020444 | 182022061 | 3.77107 | 3.51478 | 2.0277 |

| | | | | | | | | | | | | | | | | | | | | | | | | | | |
|---|---|---|---|---|---|---|---|---|---|---|---|---|---|---|---|---|---|---|---|---|---|---|---|---|---|---|
| chr15 | 98793336 | 98793671 | 1000 | 3' UTR | 15 | 98789857 | 98793837 | 3981 | 1 | ENSMUST00 | 3480 | Wnt1 | ENSMUSG0C | false | . | . | . | . | . | . | chr15 | 98789240 | 98794934 | 6.19157 | 4.38059 | 4.01653 |
| chr15 | 98933799 | 98934223 | 1000 | Promoter (<=1kb) | 15 | 98932776 | 98934438 | 1663 | 2 | ENSMUST00 | 215 | Tuba1b | ENSMUSG0C | false | chr15 | 98933157 | 98935385 | 28.03146 | 79.91027 | 76.76085 | . | . | . | . | . | . |
| chr15 | 99332886 | 99333246 | 1000 | Promoter (2-3kb) | 15 | 99327844 | 99335727 | 7884 | 2 | ENSMUST00 | 2481 | Fmnl3 | ENSMUSG0C | false | . | . | . | . | . | . | chr15 | 99329528 | 99333786 | 5.15197 | 4.14725 | 3.19381 |
| chr15 | 99373294 | 99373633 | 1000 | Promoter (2-3kb) | 15 | 99317225 | 99370482 | 53258 | 2 | ENSMUST00 | -2813 | Fmnl3 | ENSMUSG0C | false | . | . | . | . | . | . | . | . | . | . | . | . |
| chr15 | 99702025 | 99702671 | 1000 | Promoter (<=1kb) | 15 | 99702287 | 99713995 | 11709 | 1 | ENSMUST00 | 0 | Smarcd1 | ENSMUSG0C | false | chr15 | 99701792 | 99702839 | 37.30079 | 124.19571 | 119.91465 | chr15 | 99700422 | 99705143 | 3.90355 | 3.54268 | 2.10055 |
| chr15 | 99716880 | 99717179 | 1000 | Promoter (<=1kb) | 15 | 99717515 | 99725005 | 7491 | 1 | ENSMUST00 | -336 | Gpd1 | ENSMUSG0C | false | . | . | . | . | . | . | chr15 | 99714873 | 99718983 | 6.00883 | 4.38121 | 3.85462 |
| chr15 | 99772234 | 99773318 | 1000 | Promoter (<=1kb) | 15 | 99734881 | 99772462 | 37582 | 2 | ENSMUST00 | 0 | Cers5 | ENSMUSG0C | false | chr15 | 99771336 | 99772882 | 32.27147 | 102.84008 | 99.19724 | chr15 | 99772584 | 99773527 | 3.978 | 3.62458 | 2.1751 |
| chr15 | 99787359 | 99787991 | 1000 | Intron (ENSMUST00000109024.3 | 15 | 99781239 | 99783973 | 2735 | 2 | ENSMUST00 | -3387 | Lima1 | ENSMUSG0C | false | . | . | . | . | . | . | chr15 | 99784837 | 99793304 | 6.22394 | 4.62904 | 4.01375 |
| chr15 | 99972651 | 99973411 | 1000 | Promoter (<=1kb) | 15 | 99972780 | 100016351 | 43572 | 1 | ENSMUST00 | 0 | Larp4 | ENSMUSG0C | false | chr15 | 99971934 | 99974376 | 45.03415 | 147.57545 | 142.38377 | . | . | . | . | . | . |
| chr15 | 100227673 | 100228487 | 1000 | Promoter (<=1kb) | 15 | 100227819 | 100261244 | 33426 | 1 | ENSMUST00 | 0 | Atf1 | ENSMUSG0C | false | chr15 | 100227015 | 100229191 | 43.22482 | 143.31757 | 138.30418 | . | . | . | . | . | . |
| chr15 | 101266384 | 101267101 | 1000 | Promoter (<=1kb) | 15 | 101266846 | 101274792 | 7947 | 1 | ENSMUST00 | 0 | Nr4a1 | ENSMUSG0C | false | chr15 | 101266077 | 101268132 | 17.60262 | 45.78123 | 43.11721 | chr15 | 101265606 | 101266484 | 7.74147 | 5.47315 | 5.24075 |
| chr15 | 101246023 | 101246728 | 1000 | Distal Intergenic | 15 | 101266846 | 101274792 | 7947 | 1 | ENSMUST00 | -20118 | Nr4a1 | ENSMUSG0C | false | chr15 | 101245840 | 101246344 | 8.63619 | 18.02194 | 15.71366 | chr15 | 101245518 | 101249264 | 7.5155 | 5.18344 | 5.06701 |
| chr15 | 101253875 | 101254539 | 1000 | Distal Intergenic | 15 | 101266846 | 101274792 | 7947 | 1 | ENSMUST00 | -12307 | Nr4a1 | ENSMUSG0C | false | chr15 | 101253655 | 101254885 | 17.71499 | 57.83842 | 55.02271 | chr15 | 101253463 | 101256526 | 5.2177 | 4.17424 | 3.18119 |
| chr15 | 102516647 | 102517284 | 1000 | Promoter (<=1kb) | 15 | 102497649 | 102516836 | 19188 | 2 | ENSMUST00 | 0 | Map3k12 | ENSMUSG0C | false | chr15 | 102516392 | 102517958 | 23.16618 | 65.22538 | 62.30607 | . | . | . | . | . | . |
| chr15 | 102746607 | 102747471 | 1000 | Distal Intergenic | 15 | 102706779 | 102722178 | 15400 | 2 | ENSMUST00 | -24430 | Calcoco1 | ENSMUSG0C | false | chr15 | 102746412 | 102747635 | 19.52753 | 50.64327 | 47.92091 | chr15 | 102746212 | 102747761 | 5.68938 | 4.39528 | 3.57972 |
| chr10 | 52417132 | 52417700 | 1000 | Promoter (<=1kb) | 10 | 52417547 | 52440192 | 22646 | 1 | ENSMUST00 | 0 | Nus1 | ENSMUSG0C | false | chr10 | 52416804 | 52418699 | 39.33916 | 127.75399 | 123.34521 | chr10 | 52416544 | 52417173 | 5.20649 | 4.12755 | 3.21533 |
| chr4 | 123749787 | 123750730 | 1000 | Promoter (<=1kb) | 4 | 123734559 | 123750345 | 15787 | 2 | ENSMUST00 | 0 | Akirin1 | ENSMUSG0C | false | chr4 | 123749157 | 123750996 | 34.78613 | 113.42423 | 109.48744 | . | . | . | . | . | . |
| chr3 | 51219716 | 51220127 | 1000 | Distal Intergenic | 3 | 51224447 | 51251644 | 27198 | 1 | ENSMUST00 | -4320 | Ccrn4l | ENSMUSG0C | false | . | . | . | . | . | . | chr3 | 51212148 | 51224324 | 4.29558 | 3.75668 | 2.42602 |
| chr16 | 14361266 | 14361829 | 1000 | Promoter (<=1kb) | 16 | 14361558 | 14473902 | 112345 | 1 | ENSMUST00 | 0 | Abcc1 | ENSMUSG0C | false | chr16 | 14361174 | 14362412 | 35.84351 | 110.06287 | 106.22365 | chr16 | 14358851 | 14365335 | 4.28413 | 3.68925 | 2.44342 |
| chrX | 56731353 | 56731834 | 1000 | Promoter (<=1kb) | 20 | 56731787 | 56793346 | 61560 | 1 | ENSMUST00 | 0 | Fhl1 | ENSMUSG0C | false | chrX | 56730978 | 56731656 | 15.16456 | 35.10921 | 32.57929 | . | . | . | . | . | . |
| chr5 | 134577485 | 134577979 | 1000 | Distal Intergenic | 5 | 134582690 | 134598328 | 15639 | 1 | ENSMUST00 | -4711 | Rfc2 | ENSMUSG0C | false | . | . | . | . | . | . | chr5 | 134575355 | 134583628 | 5.43561 | 4.21716 | 3.39276 |
| chr5 | 134571715 | 134572324 | 1000 | Distal Intergenic | 5 | 134582690 | 134598328 | 15639 | 1 | ENSMUST00 | -10366 | Rfc2 | ENSMUSG0C | false | chr5 | 134571532 | 134572669 | 18.19667 | 46.80325 | 44.13025 | . | . | . | . | . | . |
| chr5 | 134580970 | 134581742 | 1000 | Promoter (<=1kb) | 5 | 134582690 | 134598328 | 15639 | 1 | ENSMUST00 | -948 | Rfc2 | ENSMUSG0C | false | . | . | . | . | . | . | chr5 | 134575355 | 134583628 | 5.43561 | 4.21716 | 3.39276 |
| chr10 | 79704230 | 79704570 | 1000 | Promoter (<=1kb) | 10 | 79704491 | 79711969 | 7479 | 1 | ENSMUST00 | 0 | Bsg | ENSMUSG0C | false | chr10 | 79704012 | 79705228 | 43.19602 | 139.91907 | 135.04468 | chr10 | 79703702 | 79710218 | 6.35568 | 4.66584 | 4.11889 |
| chr14 | 20165303 | 20165953 | 1000 | Intron (ENSMUST00000024011.8 | 14 | 20140058 | 20181782 | 41725 | 2 | ENSMUST00 | 15829 | Kcnk5 | ENSMUSG0C | false | . | . | . | . | . | . | chr14 | 20164261 | 20167568 | 5.88344 | 4.44836 | 3.78459 |
| chr17 | 47688831 | 47689246 | 1000 | Promoter (<=1kb) | 17 | 47689030 | 47699551 | 10522 | 1 | ENSMUST00 | 0 | Frs3 | ENSMUSG0C | false | chr17 | 47687260 | 47689501 | 34.78613 | 113.42423 | 109.48744 | chr17 | 47686192 | 47689517 | 5.23678 | 4.09637 | 3.20214 |
| chr8 | 119651453 | 119651805 | 1000 | Distal Intergenic | 8 | 119666523 | 119688222 | 21700 | 1 | ENSMUST00 | -14718 | Wfdc1 | ENSMUSG0C | false | . | . | . | . | . | . | chr8 | 119649334 | 119652098 | 3.6622 | 3.27726 | 1.93865 |
| chr8 | 119674059 | 119674630 | 1000 | Intron (ENSMUST00000024107.5 | 8 | 119666523 | 119688222 | 21700 | 1 | ENSMUST00 | 7537 | Wfdc1 | ENSMUSG0C | false | . | . | . | . | . | . | chr8 | 119673252 | 119675209 | 4.95381 | 4.00765 | 3.02498 |
| chr5 | 24451613 | 24452395 | 1000 | Promoter (<=1kb) | 5 | 24452177 | 24502047 | 49871 | 1 | ENSMUST00 | 0 | Agap3 | ENSMUSG0C | false | chr5 | 24451349 | 24453343 | 26.32003 | 79.74272 | 76.59713 | . | . | . | . | . | . |
| chr2 | 180724545 | 180725449 | 1000 | Promoter (<=1kb) | 2 | 180725263 | 180742280 | 17018 | 1 | ENSMUST00 | 0 | Slc17a9 | ENSMUSG0C | false | chr2 | 180724765 | 180725766 | 11.56301 | 29.49975 | 27.03959 | chr2 | 180723317 | 180727178 | 5.35965 | 4.251 | 3.32587 |
| chr14 | 55828743 | 55829270 | 1000 | Exon (ENSMUST00000024179.5/ | 14 | 55824795 | 55833943 | 9149 | 1 | ENSMUST00 | 3949 | Nfatc4 | ENSMUSG0C | false | . | . | . | . | . | . | chr14 | 55828948 | 55829541 | 3.00088 | 2.93752 | 1.37143 |
| chr17 | 66479228 | 66479623 | 1000 | Distal Intergenic | 17 | 66497490 | 66500319 | 2830 | 2 | ENSMUST00 | 20696 | Rab12 | ENSMUSG0C | false | . | . | . | . | . | . | chr17 | 66479003 | 66481332 | 5.23585 | 4.25884 | 3.18608 |
| chr17 | 66480321 | 66480789 | 1000 | Distal Intergenic | 17 | 66497490 | 66500319 | 2830 | 2 | ENSMUST00 | 19530 | Rab12 | ENSMUSG0C | false | . | . | . | . | . | . | chr17 | 66479003 | 66481332 | 5.23585 | 4.25884 | 3.18608 |
| chr4 | 155962129 | 155962649 | 1000 | Promoter (<=1kb) | 4 | 155962318 | 155966629 | 4312 | 1 | ENSMUST00 | 0 | Fam132a | ENSMUSG0C | false | chr4 | 155961847 | 155963148 | 24.30838 | 70.74035 | 67.73997 | chr4 | 155959571 | 155964198 | 4.05326 | 3.65443 | 2.23161 |
| chr11 | 69117857 | 69118341 | 1000 | Promoter (2-3kb) | 11 | 69120404 | 69124055 | 3652 | 1 | ENSMUST00 | -2063 | Hes7 | ENSMUSG0C | false | chr11 | 69117442 | 69119156 | 30.22771 | 102.23431 | 98.60725 | . | . | . | . | . | . |
| chr17 | 3314098 | 3314499 | 1000 | Distal Intergenic | 17 | 3326573 | 3519397 | 192825 | 1 | ENSMUST00 | -12074 | Tiam2 | ENSMUSG0C | false | chr17 | 3314035 | 3314882 | 23.26691 | 63.66726 | 60.76967 | chr17 | 3314042 | 3314917 | 3.346 | 3.29615 | 1.64963 |
| chr17 | 5940746 | 5941876 | 1000 | Promoter (<=1kb) | 17 | 5941280 | 6032793 | 91514 | 1 | ENSMUST00 | 0 | Synj2 | ENSMUSG0C | false | chr17 | 5940860 | 5941188 | 8.80131 | 17.30502 | 15.00832 | chr17 | 5940308 | 5943466 | 5.84544 | 4.5265 | 3.68746 |
| chr17 | 6948208 | 6948912 | 1000 | Promoter (<=1kb) | 17 | 6904716 | 6948356 | 43641 | 2 | ENSMUST00 | 0 | Rsph3b | ENSMUSG0C | false | chr17 | 6947623 | 6949113 | 25.6494 | 102.3303 | 98.70287 | chr17 | 6947432 | 6949497 | 5.35243 | 3.95142 | 3.35222 |
| chr17 | 12169256 | 12169754 | 1000 | Intron (ENSMUST00000164143.2 | 17 | 12151772 | 12219079 | 67308 | 1 | ENSMUST00 | 17485 | Agpat4 | ENSMUSG0C | true | . | . | . | . | . | . | chr17 | 12154205 | 12176743 | 5.8126 | 4.38993 | 3.68034 |
| chr17 | 12758031 | 12758422 | 1000 | Intron (ENSMUST00000160932.1 | 17 | 12748653 | 12751412 | 2760 | 2 | ENSMUST00 | -6620 | Igf2r | ENSMUSG0C | false | . | . | . | . | . | . | chr17 | 12755029 | 12763409 | 5.11764 | 4.11122 | 3.109 |
| chr17 | 17624398 | 17625660 | 1000 | Promoter (<=1kb) | 17 | 17527723 | 17624489 | 96767 | 2 | ENSMUST00 | 0 | Lnpep | ENSMUSG0C | false | chr17 | 17622986 | 17626193 | 32.3084 | 121.498 | 117.30614 | . | . | . | . | . | . |
| chr17 | 17592926 | 17593256 | 1000 | Intron (ENSMUST00000041047.2 | 17 | 17527723 | 17624489 | 96767 | 2 | ENSMUST00 | 31233 | Lnpep | ENSMUSG0C | false | . | . | . | . | . | . | chr17 | 17590519 | 17594385 | 3.80339 | 3.55532 | 2.01239 |
| chr17 | 15704505 | 15705758 | 1000 | Promoter (<=1kb) | 17 | 15704967 | 15772610 | 67644 | 1 | ENSMUST00 | 0 | Chd1 | ENSMUSG0C | false | chr17 | 15703965 | 15706505 | 38.04976 | 128.75285 | 124.30675 | . | . | . | . | . | . |
| chr17 | 8524562 | 8525518 | 1000 | Promoter (<=1kb) | 17 | 8525372 | 8981671 | 456300 | 1 | ENSMUST00 | 0 | Pde10a | ENSMUSG0C | true | chr17 | 8524255 | 8525207 | 15.50707 | 38.47921 | 35.908 | chr17 | 8523521 | 8524629 | 6.19063 | 4.41558 | 4.00719 |
| chr17 | 8487672 | 8488242 | 1000 | Distal Intergenic | 17 | 8525372 | 8981671 | 456300 | 1 | ENSMUST00 | -37130 | Pde10a | ENSMUSG0C | true | . | . | . | . | . | . | chr17 | 8485107 | 8491054 | 6.22769 | 4.62859 | 4.02201 |
| chr17 | 8800501 | 8802036 | 1000 | Promoter (<=1kb) | 17 | 8801693 | 8899189 | 97497 | 1 | ENSMUST00 | 0 | Pde10a | ENSMUSG0C | true | chr17 | 8800574 | 8803185 | 23.32542 | 76.20397 | 73.11496 | chr17 | 8800007 | 8800858 | 3.76827 | 3.53857 | 2.01089 |
| chr17 | 8508052 | 8508847 | 1000 | Distal Intergenic | 17 | 8525372 | 8981671 | 456300 | 1 | ENSMUST00 | -16525 | Pde10a | ENSMUSG0C | true | . | . | . | . | . | . | chr17 | 8504525 | 8510234 | 7.21779 | 5.10275 | 4.81866 |
| chr17 | 23677191 | 23677691 | 1000 | Promoter (<=1kb) | 17 | 23675447 | 23677449 | 2003 | 2 | ENSMUST00 | 0 | Tnfrsf12a | ENSMUSG0C | false | chr17 | 23676112 | 23677653 | 26.28407 | 85.32412 | 82.07336 | chr17 | 23673161 | 23678044 | 6.00185 | 4.61543 | 3.82218 |
| chr17 | 42975426 | 42975826 | 1000 | Distal Intergenic | 17 | 43016555 | 43089188 | 72634 | 1 | ENSMUST00 | -40729 | Tnfrsf21 | ENSMUSG0C | false | . | . | . | . | . | . | chr17 | 42974956 | 42976182 | 4.63592 | 3.77553 | 2.73822 |
| chr17 | 51178692 | 51179688 | 1000 | Promoter (<=1kb) | 17 | 50733127 | 51179349 | 446223 | 2 | ENSMUST00 | 0 | Tbc1d5 | ENSMUSG0C | false | chr17 | 51177960 | 51179858 | 32.27147 | 102.84008 | 99.19724 | . | . | . | . | . | . |
| chr17 | 51832404 | 51833300 | 1000 | Promoter (<=1kb) | 17 | 51736187 | 51832672 | 96486 | 2 | ENSMUST00 | 0 | Satb1 | ENSMUSG0C | false | chr17 | 51831671 | 51833605 | 15.90661 | 56.5361 | 53.73628 | chr17 | 51831453 | 51834265 | 5.04811 | 4.05581 | 3.04873 |
| chr17 | 51812671 | 51813723 | 1000 | Promoter (<=1kb) | 17 | 51739476 | 51812680 | 73205 | 2 | ENSMUST00 | 0 | Satb1 | ENSMUSG0C | false | chr17 | 51811751 | 51813952 | 31.80866 | 99.00968 | 95.46461 | chr17 | 51812712 | 51813956 | 3.12583 | 3.09326 | 1.51685 |
| chr17 | 45685650 | 45686489 | 1000 | Promoter (<=1kb) | 17 | 45686322 | 45698495 | 12174 | 1 | ENSMUST00 | 0 | Mrpl14 | ENSMUSG0C | false | chr17 | 45685260 | 45687208 | 40.58316 | 133.18515 | 128.57994 | chr17 | 45684856 | 45687918 | 4.08585 | 3.36391 | 2.29225 |
| chr17 | 45595286 | 45595971 | 1000 | Promoter (<=1kb) | 17 | 45585203 | 45595570 | 10368 | 2 | ENSMUST00 | 0 | Slc29a1 | ENSMUSG0C | false | chr17 | 45594883 | 45595981 | 28.49099 | 81.63235 | 78.4519 | chr17 | 45591091 | 45600246 | 5.33186 | 4.18113 | 3.28168 |
| chr17 | 45596470 | 45597100 | 1000 | Promoter (<=1kb) | 17 | 45590112 | 45597066 | 6955 | 2 | ENSMUST00 | 0 | Slc29a1 | ENSMUSG0C | false | . | . | . | . | . | . | chr17 | 45591091 | 45600246 | 5.33186 | 4.18113 | 3.28168 |
| chr17 | 45573700 | 45574492 | 1000 | Promoter (<=1kb) | 17 | 45571018 | 45573271 | 2254 | 2 | ENSMUST00 | -430 | Hsp90ab1 | ENSMUSG0C | false | chr17 | 45571510 | 45574755 | 48.52219 | 165.10075 | 159.2771 | . | . | . | . | . | . |
| chr17 | 45572367 | 45573544 | 1000 | Promoter (<=1kb) | 17 | 45567775 | 45573271 | 5497 | 2 | ENSMUST00 | 0 | Hsp90ab1 | ENSMUSG0C | false | chr17 | 45571510 | 45574755 | 48.52219 | 165.10075 | 159.2771 | . | . | . | . | . | . |
| chr17 | 46063187 | 46064203 | 1000 | Distal Intergenic | 17 | 46016996 | 46032377 | 15382 | 2 | ENSMUST00 | -30811 | Vegfa | ENSMUSG0C | true | . | . | . | . | . | . | chr17 | 46048792 | 46077235 | 6.11576 | 4.54762 | 3.92959 |
| chr17 | 46030476 | 46031377 | 1000 | Promoter (<=1kb) | 17 | 46018831 | 46031356 | 12526 | 2 | ENSMUST00 | 0 | Vegfa | ENSMUSG0C | true | chr17 | 46023564 | 46033809 | 32.66734 | 113.81544 | 109.86345 | . | . | . | . | . | . |
| chr17 | 46051930 | 46052506 | 1000 | Distal Intergenic | 17 | 46016996 | 46032377 | 15382 | 2 | ENSMUST00 | -19554 | Vegfa | ENSMUSG0C | true | . | . | . | . | . | . | chr17 | 46048792 | 46077235 | 6.11576 | 4.54762 | 3.92959 |
| chr17 | 46049547 | 46050047 | 1000 | Distal Intergenic | 17 | 46016996 | 46032377 | 15382 | 2 | ENSMUST00 | -17171 | Vegfa | ENSMUSG0C | true | . | . | . | . | . | . | chr17 | 46048792 | 46077235 | 6.11576 | 4.54762 | 3.92959 |
| chr17 | 46040603 | 46040979 | 1000 | Distal Intergenic | 17 | 46016996 | 46032377 | 15382 | 2 | ENSMUST00 | -8227 | Vegfa | ENSMUSG0C | true | . | . | . | . | . | . | chr17 | 46033010 | 46047864 | 6.11349 | 4.63753 | 3.9166 |
| chr17 | 44238949 | 44239847 | 1000 | Intron (ENSMUST00000024755.5 | 17 | 44188572 | 44280168 | 91597 | 1 | ENSMUST00 | 50378 | Clic5 | ENSMUSG0C | true | . | . | . | . | . | . | chr17 | 44197784 | 44272727 | 5.95006 | 4.47222 | 3.79255 |
| chr17 | 44216926 | 44217571 | 1000 | Intron (ENSMUST00000024755.5 | 17 | 44188572 | 44280168 | 91597 | 1 | ENSMUST00 | 28355 | Clic5 | ENSMUSG0C | true | . | . | . | . | . | . | chr17 | 44197784 | 44272727 | 5.95006 | 4.47222 | 3.79255 |
| chr17 | 44197265 | 44197697 | 1000 | Intron (ENSMUST00000024755.5 | 17 | 44188572 | 44280168 | 91597 | 1 | ENSMUST00 | 8694 | Clic5 | ENSMUSG0C | true | . | . | . | . | . | . | . | . | . | . | . | . |
| chr17 | 63498789 | 63500641 | 1000 | Promoter (<=1kb) | 17 | 63057452 | 63500017 | 442566 | 2 | ENSMUST00 | 0 | Fbxl17 | ENSMUSG0C | false | chr17 | 63498560 | 63500998 | 43.83784 | 156.88544 | 151.34334 | chr17 | 63495586 | 63498947 | 4.56477 | 3.87714 | 2.69804 |
| chr17 | 46129801 | 46130116 | 1000 | Intron (ENSMUST00000148581.2 | 17 | 46125458 | 46128793 | 3336 | 1 | ENSMUST00 | 4344 | Mrps18a | ENSMUSG0C | false | . | . | . | . | . | . | chr17 | 46129283 | 46132591 | 5.69216 | 4.32564 | 3.62564 |
| chr17 | 46117686 | 46118028 | 1000 | Exon (ENSMUST00000024763.4/ | 17 | 46122803 | 46128870 | 6068 | 1 | ENSMUST00 | -4775 | Mrps18a | ENSMUSG0C | false | . | . | . | . | . | . | chr17 | 46110303 | 46120605 | 4.89899 | 3.8479 | 2.96109 |
| chr17 | 46629028 | 46630331 | 1000 | Promoter (<=1kb) | 17 | 46564451 | 46629504 | 65054 | 2 | ENSMUST00 | 0 | Ptk7 | ENSMUSG0C | false | chr17 | 46628274 | 46630084 | 29.66345 | 100.21 | 96.63225 | chr17 | 46605355 | 46631878 | 5.60661 | 4.26306 | 3.52431 |
| chr17 | 46619227 | 46619669 | 1000 | Intron (ENSMUST00000181301.1 | 17 | 46564451 | 46629504 | 65054 | 2 | ENSMUST00 | 9835 | Ptk7 | ENSMUSG0C | false | . | . | . | . | . | . | chr17 | 46605355 | 46631878 | 5.60661 | 4.26306 | 3.52431 |
| chr17 | 46610229 | 46610519 | 1000 | Intron (ENSMUST00000181301.1 | 17 | 46564451 | 46629504 | 65054 | 2 | ENSMUST00 | 18985 | Ptk7 | ENSMUSG0C | false | . | . | . | . | . | . | chr17 | 46605355 | 46631878 | 5.60661 | 4.26306 | 3.52431 |
| chr17 | 47737430 | 47738493 | 1000 | Promoter (<=1kb) | 17 | 47737600 | 47786084 | 48485 | 1 | ENSMUST00 | 0 | Tfeb | ENSMUSG0C | false | chr17 | 47737000 | 47739098 | 25.17393 | 93.69764 | 90.27328 | chr17 | 47738161 | 47742214 | 6.43047 | 4.64153 | 4.19494 |
| chr17 | 47923756 | 47925281 | 1000 | Promoter (<=1kb) | 17 | 47867133 | 47924645 | 57513 | 2 | ENSMUST00 | 0 | Foxp4 | ENSMUSG0C | false | chr17 | 47923048 | 47925384 | 42.73649 | 138.01634 | 133.21471 | chr17 | 47921415 | 47923951 | 4.36195 | 3.56297 | 2.51175 |
| chr17 | 49738301 | 49738874 | 1000 | Promoter (<=1kb) | 17 | 49738753 | 49755390 | 16638 | 1 | ENSMUST00 | 0 | Kif6 | ENSMUSG0C | false | . | . | . | . | . | . | chr17 | 49737398 | 49739673 | 3.63017 | 3.31092 | 1.88958 |
| chr17 | 32283908 | 32284853 | 1000 | Promoter (<=1kb) | 17 | 32196274 | 32284097 | 87824 | 2 | ENSMUST00 | 0 | Brd4 | ENSMUSG0C | false | chr17 | 32283246 | 32285341 | 38.14666 | 123.52654 | 119.2661 | . | . | . | . | . | . |
| chr17 | 29341837 | 29342307 | 1000 | Promoter (<=1kb) | 17 | 29340792 | 29342930 | 2139 | 2 | ENSMUST00 | 623 | Mtch1 | ENSMUSG0C | false | . | . | . | . | . | . | chr17 | 29338838 | 29346864 | 7.3536 | 5.16659 | 4.92198 |
| chr17 | 29490203 | 29491169 | 1000 | Promoter (<=1kb) | 17 | 29490812 | 29496111 | 5300 | 1 | ENSMUST00 | 0 | Pim1 | ENSMUSG0C | false | chr17 | 29489687 | 29490498 | 17.34451 | 52.33425 | 49.58816 | . | . | . | . | . | . |
| chr17 | 29436965 | 29437827 | 1000 | Distal Intergenic | 17 | 29490812 | 29496111 | 5300 | 1 | ENSMUST00 | -52985 | Pim1 | ENSMUSG0C | false | chr17 | 29436910 | 29437547 | 8.29801 | 20.47831 | 18.13586 | chr17 | 29436890 | 29442166 | 6.63806 | 4.90552 | 4.35203 |
| chr17 | 29710544 | 29710901 | 1000 | Intron (ENSMUST00000137079.2 | 17 | 29699999 | 29716963 | 16965 | 2 | ENSMUST00 | 6062 | Ccdc167 | ENSMUSG0C | false | . | . | . | . | . | . | chr17 | 29708581 | 29713798 | 5.89693 | 4.35848 | 3.78064 |
| chr17 | 30609320 | 30609945 | 1000 | Promoter (2-3kb) | 17 | 30592866 | 30612659 | 19794 | 2 | ENSMUST00 | 2714 | Glo1 | ENSMUSG0C | false | . | . | . | . | . | . | chr17 | 30607119 | 30613115 | 5.78656 | 4.46694 | 3.65304 |
| chr17 | 30888707 | 30889077 | 1000 | Distal Intergenic | 17 | 30901867 | 30936510 | 34644 | 1 | ENSMUST00 | -12790 | Glp1r | ENSMUSG0C | false | . | . | . | . | . | . | chr17 | 30887881 | 30890561 | 4.19003 | 3.65743 | 2.3543 |
| chr17 | 31746422 | 31747081 | 1000 | Distal Intergenic | 17 | 31677933 | 31681722 | 3790 | 1 | ENSMUST00 | 68490 | Cryaa | ENSMUSG0C | false | . | . | . | . | . | . | chr17 | 31746063 | 31748899 | 6.3812 | 4.77923 | 4.18039 |
| chr17 | 67928300 | 67928859 | 1000 | Intron (ENSMUST00000024840.6 | 17 | 67845110 | 67950940 | 105831 | 2 | ENSMUST00 | 22081 | Arhgap28 | ENSMUSG0C | true | . | . | . | . | . | . | chr17 | 67915917 | 67949162 | 5.38115 | 4.23981 | 3.32205 |
| chr17 | 67995516 | 67995951 | 1000 | Intron (ENSMUST00000024840.6 | 17 | 67842713 | 68004120 | 161408 | 2 | ENSMUST00 | 8169 | Arhgap28 | ENSMUSG0C | true | . | . | . | . | . | . | chr17 | 67993413 | 68002591 | 4.56742 | 3.83999 | 2.64221 |
| chr17 | 67988873 | 67989618 | 1000 | Intron (ENSMUST00000024840.6 | 17 | 67842713 | 68004120 | 161408 | 2 | ENSMUST00 | 14502 | Arhgap28 | ENSMUSG0C | true | . | . | . | . | . | . | chr17 | 67966988 | 67992447 | 5.42867 | 4.20296 | 3.37432 |
| chr17 | 68022775 | 68023175 | 1000 | Distal Intergenic | 17 | 67842713 | 68004120 | 161408 | 2 | ENSMUST00 | -18656 | Arhgap28 | ENSMUSG0C | true | . | . | . | . | . | . | chr17 | 68022712 | 68025663 | 3.74105 | 3.42105 | 1.96936 |
| chr17 | 67931019 | 67931787 | 1000 | Intron (ENSMUST00000024840.6 | 17 | 67845110 | 67950940 | 105831 | 2 | ENSMUST00 | 19153 | Arhgap28 | ENSMUSG0C | true | . | . | . | . | . | . | chr17 | 67915917 | 67949162 | 5.38115 | 4.23981 | 3.32205 |
| chr17 | 71001837 | 71003010 | 1000 | Promoter (<=1kb) | 17 | 70993656 | 71002554 | 8899 | 2 | ENSMUST00 | 0 | Myl12a | ENSMUSG0C | false | chr17 | 71000626 | 71002523 | 25.18546 | 107.13963 | 103.3826 | chr17 | 71002949 | 71004324 | 5.31425 | 4.22635 | 3.25541 |
| chr17 | 32388981 | 32389637 | 1000 | Promoter (<=1kb) | 17 | 32354061 | 32389391 | 35331 | 2 | ENSMUST00 | 0 | Wiz | ENSMUSG0C | false | chr17 | 32388391 | 32389436 | 41.21593 | 136.4875 | 131.75212 | chr17 | 32387254 | 32389899 | 4.13531 | 3.70276 | 2.29084 |
| chr17 | 71183272 | 71184096 | 1000 | Promoter (<=1kb) | 17 | 71183556 | 71232306 | 48751 | 1 | ENSMUST00 | 0 | Lpin2 | ENSMUSG0C | false | chr17 | 71182601 | 71187348 | 37.59121 | 124.70534 | 120.4041 | . | . | . | . | . | . |
| chr17 | 71223725 | 71224063 | 1000 | Intron (ENSMUST00000180743.1 | 17 | 71231433 | 71238085 | 6653 | 1 | ENSMUST00 | -7370 | Lpin2 | ENSMUSG0C | false | . | . | . | . | . | . | chr17 | 71186127 | 71229089 | 5.51375 | 4.29183 | 3.43736 |
| chr17 | 71168679 | 71169213 | 1000 | Intron (ENSMUST00000180743.1 | 17 | 71182560 | 71235119 | 52560 | 1 | ENSMUST00 | -13347 | Lpin2 | ENSMUSG0C | false | . | . | . | . | . | . | chr17 | 71157703 | 71180230 | 5.23179 | 4.17367 | 3.19377 |
| chr17 | 71161636 | 71162046 | 1000 | Intron (ENSMUST00000180743.1 | 17 | 71182560 | 71235119 | 52560 | 1 | ENSMUST00 | -20514 | Lpin2 | ENSMUSG0C | false | . | . | . | . | . | . | chr17 | 71157703 | 71180230 | 5.23179 | 4.17367 | 3.19377 |
| chr17 | 71125553 | 71126036 | 1000 | Intron (ENSMUST00000180743.1 | 17 | 71182560 | 71235119 | 52560 | 1 | ENSMUST00 | -56524 | Lpin2 | ENSMUSG0C | false | . | . | . | . | . | . | chr17 | 71120006 | 71131073 | 6.06535 | 4.44566 | 3.90237 |
| chr17 | 73821516 | 73821881 | 1000 | Intron (ENSMUST00000024860.7 | 17 | 73804841 | 73832093 | 27253 | 1 | ENSMUST00 | 16676 | Ehd3 | ENSMUSG0C | false | . | . | . | . | . | . | chr17 | 73820868 | 73822953 | 3.78029 | 3.50156 | 2.03372 |
| chr17 | 74338576 | 74339360 | 1000 | Promoter (<=1kb) | 17 | 74338195 | 74391113 | 52127 | 1 | ENSMUST00 | 0 | Spast | ENSMUSG0C | false | chr17 | 74338195 | 74340082 | 22.26925 | 90.51499 | 87.16024 | . | . | . | . | . | . |
| chr17 | 78199678 | 78201689 | 1000 | Promoter (<=1kb) | 17 | 78200248 | 78376592 | 176345 | 1 | ENSMUST00 | 0 | Crim1 | ENSMUSG0C | true | chr17 | 78199554 | 78200068 | 22.2725 | 61.10292 | 58.24237 | . | . | . | . | . | . |
| chr17 | 78736174 | 78736967 | 1000 | Promoter (<=1kb) | 17 | 78649913 | 78737196 | 87284 | 2 | ENSMUST00 | 229 | Strn | ENSMUSG0C | false | chr17 | 78735419 | 78737563 | 40.53359 | 134.1972 | 129.55179 | . | . | . | . | . | . |
| chr17 | 64331336 | 64332432 | 1000 | Promoter (<=1kb) | 17 | 64281005 | 64331916 | 50912 | 2 | ENSMUST00 | 0 | Pja2 | ENSMUSG0C | false | chr17 | 64330658 | 64331974 | 40.91335 | 133.75845 | 129.12437 | . | . | . | . | . | . |
| chr17 | 64600525 | 64601622 | 1000 | Promoter (<=1kb) | 17 | 64600736 | 64755110 | 154375 | 1 | ENSMUST00 | 0 | Man2a1 | ENSMUSG0C | false | chr17 | 64600157 | 64602660 | 29.2476 | 106.45832 | 102.71833 | . | . | . | . | . | . |
| chr17 | 79771699 | 79772218 | 1000 | Distal Intergenic | 17 | 79706953 | 79715041 | 8089 | 2 | ENSMUST00 | -56659 | Cyp1b1 | ENSMUSG0C | false | . | . | . | . | . | . | chr17 | 79771691 | 79772641 | 3.97581 | 3.24085 | 2.19552 |
| chr17 | 79693265 | 79693637 | 1000 | Distal Intergenic | 17 | 79706953 | 79715041 | 8089 | 2 | ENSMUST00 | 21404 | Cyp1b1 | ENSMUSG0C | false | . | . | . | . | . | . | chr17 | 79691085 | 79694381 | 4.42587 | 3.85913 | 2.52129 |
| chr17 | 79757518 | 79758418 | 1000 | Distal Intergenic | 17 | 79706953 | 79715041 | 8089 | 2 | ENSMUST00 | -42478 | Cyp1b1 | ENSMUSG0C | false | . | . | . | . | . | . | chr17 | 79757395 | 79759261 | 6.31224 | 4.7144 | 4.13614 |
| chr17 | 79728234 | 79728832 | 1000 | Distal Intergenic | 17 | 79706953 | 79715041 | 8089 | 2 | ENSMUST00 | -13194 | Cyp1b1 | ENSMUSG0C | false | . | . | . | . | . | . | chr17 | 79726503 | 79733779 | 5.74212 | 4.41435 | 3.61439 |
| chr17 | 79715496 | 79716239 | 1000 | Promoter (<=1kb) | 17 | 79706953 | 79715041 | 8089 | 2 | ENSMUST00 | -456 | Cyp1b1 | ENSMUSG0C | false | chr17 | 79712319 | 79716277 | 16.88344 | 48.35773 | 45.66254 | chr17 | 79710186 | 79716400 | 4.02257 | 3.60551 | 2.19921 |
| chr17 | 65612610 | 65614351 | 1000 | Promoter (<=1kb) | 17 | 65580056 | 65613555 | 33500 | 2 | ENSMUST00 | 0 | Vapa | ENSMUSG0C | false | chr17 | 65612263 | 65614741 | 35.76354 | 117.23875 | 113.19022 | chr17 | 65594200 | 65612648 | 4.71795 | 3.88153 | 2.80257 |
| chr17 | 80061246 | 80062538 | 1000 | Promoter (<=1kb) | 17 | 80029487 | 80062227 | 32741 | 2 | ENSMUST00 | 0 | Hnrnpll | ENSMUSG0C | false | chr17 | 80060963 | 80063060 | 39.4261 | 145.54997 | 140.44281 | . | . | . | . | . | . |
| chr17 | 66111426 | 66111999 | 1000 | Promoter (<=1kb) | 17 | 66111546 | 66120503 | 8958 | 1 | ENSMUST00 | 0 | Wash | ENSMUSG0C | false | chr17 | 66111173 | 66112438 | 32.64849 | 123.19247 | 118.94151 | . | . | . | . | . | . |
| chr17 | 24141019 | 24141852 | 1000 | Promoter (<=1kb) | 17 | 24073680 | 24141595 | 67916 | 2 | ENSMUST00 | 0 | Pdpk1 | ENSMUSG0C | false | chr17 | 24140408 | 24142347 | 31.16539 | 106.97112 | 103.21663 | . | . | . | . | . | . |
| chr17 | 86732114 | 86732410 | 1000 | Distal Intergenic | 17 | 86753907 | 86833410 | 79504 | 1 | ENSMUST00 | -21497 | Epas1 | ENSMUSG0C | false | . | . | . | . | . | . | chr17 | 86731427 | 86732847 | 5.50348 | 4.2254 | 3.45485 |
| chr17 | 86769805 | 86770355 | 1000 | Intron (ENSMUST00000024954.9 | 17 | 86753907 | 86833410 | 79504 | 1 | ENSMUST00 | 15899 | Epas1 | ENSMUSG0C | false | . | . | . | . | . | . | chr17 | 86767047 | 86773493 | 5.37356 | 4.22421 | 3.31931 |
| chr17 | 86753086 | 86754287 | 1000 | Promoter (<=1kb) | 17 | 86753907 | 86833410 | 79504 | 1 | ENSMUST00 | 0 | Epas1 | ENSMUSG0C | false | chr17 | 86752839 | 86756072 | 28.88092 | 86.09254 | 82.82554 | . | . | . | . | . | . |
| chr17 | 86785250 | 86785615 | 1000 | Intron (ENSMUST00000024954.9 | 17 | 86753907 | 86833410 | 79504 | 1 | ENSMUST00 | 31344 | Epas1 | ENSMUSG0C | false | . | . | . | . | . | . | chr17 | 86779326 | 86788064 | 5.8917 | 4.48312 | 3.73481 |
| chr17 | 86962861 | 86963501 | 1000 | Promoter (<=1kb) | 17 | 86963082 | 87000069 | 36988 | 1 | ENSMUST00 | 0 | Rhoq | ENSMUSG0C | false | chr17 | 86962102 | 86964034 | 24.24157 | 89.02564 | 85.70093 | . | . | . | . | . | . |
| chr17 | 24849746 | 24850494 | 1000 | Promoter (<=1kb) | 17 | 24850490 | 24864450 | 13961 | 1 | ENSMUST00 | 0 | Hagh | ENSMUSG0C | false | chr17 | 24850465 | 24851465 | 27.97104 | 107.99458 | 104.2211 | chr17 | 24848532 | 24849812 | 5.30561 | 4.14175 | 3.29074 |
| chr17 | 25083466 | 25084014 | 1000 | Promoter (2-3kb) | 17 | 25057702 | 25081181 | 23480 | 2 | ENSMUST00 | -2286 | Tmem204 | ENSMUSG0C | false | chr17 | 25081688 | 25083565 | 15.01947 | 39.77812 | 37.19162 | chr17 | 25077267 | 25086277 | 7.00235 | 5.00822 | 4.65933 |
| chr17 | 25015299 | 25016173 | 1000 | Promoter (<=1kb) | 17 | 25016091 | 25092801 | 76711 | 1 | ENSMUST00 | 0 | Ift140 | ENSMUSG0C | false | chr17 | 25013319 | 25015876 | 45.1915 | 151.24933 | 145.91733 | . | . | . | . | . | . |
| chr17 | 25555778 | 25556993 | 1000 | Distal Intergenic | 17 | 25567988 | 25570214 | 2227 | 2 | ENSMUST00 | 13221 | Sox8 | ENSMUSG0C | false | chr17 | 25556206 | 25556515 | 4.35105 | 6.99241 | 4.88445 | chr17 | 25553801 | 25557994 | 6.34028 | 4.52881 | 4.128 |
| chr17 | 26138411 | 26139137 | 1000 | Promoter (<=1kb) | 17 | 26138688 | 26195811 | 57124 | 1 | ENSMUST00 | 0 | Axin1 | ENSMUSG0C | false | chr17 | 26138184 | 26139766 | 23.61999 | 85.9385 | 82.67356 | chr17 | 26138075 | 26143392 | 4.01058 | 3.46276 | 2.21164 |
| chr17 | 26505997 | 26506371 | 1000 | Promoter (1-2kb) | 17 | 26506516 | 26507723 | 1208 | 2 | ENSMUST00 | 1352 | Dusp1 | ENSMUSG0C | false | chr17 | 26505964 | 26508522 | 39.08731 | 125.158 | 120.83892 | . | . | . | . | . | . |
| chr17 | 26932899 | 26933525 | 1000 | Promoter (<=1kb) | 17 | 26933127 | 26937890 | 4764 | 1 | ENSMUST00 | 0 | Phf1 | ENSMUSG0C | false | chr17 | 26932571 | 26934332 | 41.111 | 131.90991 | 127.34579 | chr17 | 26931803 | 26941775 | 5.54742 | 4.33953 | 3.45882 |
| chr17 | 56830729 | 56831434 | 1000 | Promoter (<=1kb) | 17 | 56775897 | 56831008 | 55112 | 2 | ENSMUST00 | 0 | Rfx2 | ENSMUSG0C | false | chr17 | 56829772 | 56831129 | 26.19333 | 73.09374 | 70.054 | chr17 | 56829026 | 56831906 | 3.81685 | 3.41159 | 2.04998 |
| chr17 | 27622693 | 27623137 | 1000 | Promoter (<=1kb) | 17 | 27580574 | 27622964 | 42391 | 2 | ENSMUST00 | 0 | Nudt3 | ENSMUSG0C | false | chr17 | 27622163 | 27624145 | 39.98587 | 128.8752 | 124.42683 | . | . | . | . | . | . |
| chr17 | 27909076 | 27910266 | 1000 | Promoter (<=1kb) | 17 | 27901122 | 27909242 | 8121 | 2 | ENSMUST00 | 0 | Taf11 | ENSMUSG0C | false | chr17 | 27906960 | 27910842 | 24.39013 | 102.54827 | 98.91666 | chr17 | 27910040 | 27912863 | 6.51503 | 4.71773 | 4.25399 |
| chr17 | 28517495 | 28518459 | 1000 | Promoter (<=1kb) | 17 | 28399097 | 28517524 | 118428 | 2 | ENSMUST00 | 0 | Fkbp5 | ENSMUSG0C | false | chr17 | 28517555 | 28518704 | 14.29631 | 42.77194 | 40.14703 | chr17 | 28514373 | 28519487 | 5.73469 | 4.2575 | 3.64316 |
| chr18 | 4352745 | 4353647 | 1000 | Promoter (<=1kb) | 18 | 4331327 | 4352953 | 21627 | 2 | ENSMUST00 | 0 | Map3k8 | ENSMUSG0C | false | chr18 | 4352435 | 4354157 | 31.84599 | 102.09332 | 98.46812 | chr18 | 4346232 | 4355520 | 4.92123 | 3.89467 | 2.98176 |
| chr17 | 80479430 | 80480813 | 1000 | Promoter (<=1kb) | 17 | 80393752 | 80480453 | 86702 | 2 | ENSMUST00 | 0 | Sos1 | ENSMUSG0C | false | chr17 | 80478875 | 80481064 | 37.22259 | 125.13314 | 120.81451 | chr17 | 80480707 | 80483852 | 4.9619 | 4.13598 | 2.96023 |
| chr17 | 80482824 | 80483283 | 1000 | Promoter (2-3kb) | 17 | 80393752 | 80480453 | 86702 | 2 | ENSMUST00 | -2372 | Sos1 | ENSMUSG0C | false | chr17 | 80482568 | 80483679 | 21.13848 | 54.99947 | 52.22001 | chr17 | 80480707 | 80483852 | 4.9619 | 4.13598 | 2.96023 |
| chr17 | 80706613 | 80707496 | 1000 | Intron (ENSMUST00000025089.7 | 17 | 80580512 | 80728093 | 147582 | 2 | ENSMUST00 | 20597 | Map4k3 | ENSMUSG0C | false | . | . | . | . | . | . | chr17 | 80689291 | 80712683 | 5.99153 | 4.56609 | 3.82474 |
| chr17 | 80727480 | 80728306 | 1000 | Promoter (<=1kb) | 17 | 80580512 | 80728093 | 147582 | 2 | ENSMUST00 | 0 | Map4k3 | ENSMUSG0C | false | chr17 | 80726713 | 80728878 | 38.77008 | 154.42084 | 148.9682 | . | . | . | . | . | . |
| chr17 | 80944405 | 80944738 | 1000 | Promoter (<=1kb) | 17 | 80944632 | 81001816 | 57185 | 1 | ENSMUST00 | 0 | Tmem178 | ENSMUSG0C | false | chr17 | 80944259 | 80945584 | 29.22958 | 88.61999 | 85.30408 | . | . | . | . | . | . |
| chr17 | 83181611 | 83181942 | 1000 | Distal Intergenic | 17 | 83215292 | 83225070 | 9779 | 1 | ENSMUST00 | -33350 | Pkdcc | ENSMUSG0C | false | . | . | . | . | . | . | chr17 | 83180936 | 83184029 | 5.7038 | 4.50903 | 3.57371 |
| chr17 | 83214118 | 83215839 | 1000 | Promoter (<=1kb) | 17 | 83215292 | 83225070 | 9779 | 1 | ENSMUST00 | 0 | Pkdcc | ENSMUSG0C | false | chr17 | 83213509 | 83220055 | 37.87741 | 120.9216 | 116.74998 | chr17 | 83212533 | 83214208 | 5.39581 | 4.15799 | 3.38073 |
| chr17 | 83197238 | 83197969 | 1000 | Distal Intergenic | 17 | 83215292 | 83225070 | 9779 | 1 | ENSMUST00 | -17323 | Pkdcc | ENSMUSG0C | false | . | . | . | . | . | . | chr17 | 83193803 | 83205137 | 6.15255 | 4.63808 | 3.94752 |
| chr18 | 10725360 | 10726731 | 1000 | Promoter (<=1kb) | 18 | 10725548 | 10818578 | 93031 | 1 | ENSMUST00 | 0 | Mib1 | ENSMUSG0C | false | chr18 | 10725216 | 10727065 | 36.7977 | 133.03651 | 128.43381 | . | . | . | . | . | . |
| chr17 | 26916666 | 26917086 | 1000 | Promoter (<=1kb) | 17 | 26917091 | 26932579 | 15489 | 1 | ENSMUST00 | -5 | Kifc5b | ENSMUSG0C | false | chr17 | 26917006 | 26917751 | 31.06164 | 91.78615 | 88.409 | chr17 | 26916484 | 26917975 | 4.83942 | 3.95995 | 2.87158 |
| chr18 | 16809098 | 16809666 | 1000 | Promoter (<=1kb) | 18 | 16588877 | 16809246 | 220370 | 2 | ENSMUST00 | 0 | Cdh2 | ENSMUSG0C | false | chr18 | 16806869 | 16809934 | 28.91859 | 89.0447 | 85.71584 | chr18 | 16806883 | 16809951 | 5.69424 | 4.22686 | 3.60459 |
| chr18 | 21001169 | 21002284 | 1000 | Promoter (<=1kb) | 18 | 21001341 | 21028223 | 26883 | 1 | ENSMUST00 | 0 | Rnf138 | ENSMUSG0C | false | chr18 | 21001005 | 21001722 | 27.88725 | 99.90214 | 96.33501 | . | . | . | . | . | . |
| chr17 | 34031570 | 34032541 | 1000 | Promoter (<=1kb) | 17 | 34028267 | 34031690 | 3424 | 2 | ENSMUST00 | 0 | Slc39a7 | ENSMUSG0C | false | chr17 | 34030328 | 34032845 | 34.00537 | 102.80987 | 99.17244 | . | . | . | . | . | . |
| chr17 | 34120135 | 34120703 | 1000 | Promoter (<=1kb) | 17 | 34112030 | 34120411 | 8382 | 2 | ENSMUST00 | 0 | Brd2 | ENSMUSG0C | false | chr17 | 34119478 | 34123860 | 39.82221 | 128.59171 | 124.15125 | . | . | . | . | . | . |
| chr17 | 34120893 | 34121939 | 1000 | Promoter (<=1kb) | 17 | 34112023 | 34121747 | 9725 | 2 | ENSMUST00 | 0 | Brd2 | ENSMUSG0C | false | chr17 | 34119478 | 34123860 | 39.82221 | 128.59171 | 124.15125 | . | . | . | . | . | . |
| chr18 | 34954268 | 34954991 | 1000 | Promoter (<=1kb) | 18 | 34937414 | 34954357 | 16944 | 2 | ENSMUST00 | 0 | Hspa9 | ENSMUSG0C | false | chr18 | 34953189 | 34955577 | 43.43732 | 157.3205 | 151.76292 | . | . | . | . | . | . |
| chr18 | 32162779 | 32163673 | 1000 | Promoter (<=1kb) | 18 | 32163089 | 32236751 | 73663 | 1 | ENSMUST00 | 0 | Map3k2 | ENSMUSG0C | false | chr18 | 32163027 | 32164275 | 38.60941 | 130.82727 | 126.30969 | . | . | . | . | . | . |
| chr17 | 35164427 | 35165242 | 1000 | Promoter (<=1kb) | 17 | 35149076 | 35164897 | 15822 | 2 | ENSMUST00 | 0 | Prrc2a | ENSMUSG0C | false | chr17 | 35165145 | 35166016 | 20.11728 | 54.87862 | 52.10172 | . | . | . | . | . | . |
| chr18 | 31930934 | 31931520 | 1000 | Promoter (<=1kb) | 18 | 31931507 | 31958619 | 27113 | 1 | ENSMUST00 | 0 | Lims2 | ENSMUSG0C | false | chr18 | 31930574 | 31933785 | 25.16654 | 87.04672 | 83.76057 | chr18 | 31918598 | 31931008 | 4.89771 | 3.99887 | 2.93822 |
| chr18 | 13892935 | 13893279 | 1000 | Intron (ENSMUST00000025288.7 | 18 | 13687013 | 13972733 | 285721 | 2 | ENSMUST00 | 79454 | Zfp521 | ENSMUSG0C | true | . | . | . | . | . | . | chr18 | 13884925 | 13898147 | 5.50345 | 4.32012 | 3.42059 |
| chr18 | 13972173 | 13973130 | 1000 | Promoter (<=1kb) | 18 | 13687013 | 13972733 | 285721 | 2 | ENSMUST00 | 0 | Zfp521 | ENSMUSG0C | true | chr18 | 13969413 | 13973642 | 27.20707 | 96.65539 | 93.16586 | . | . | . | . | . | . |
| chr18 | 38419884 | 38420490 | 1000 | Promoter (<=1kb) | 18 | 38418975 | 38464399 | 45425 | 1 | ENSMUST00 | 910 | Ndfip1 | ENSMUSG0C | false | chr18 | 38418542 | 38420768 | 41.33371 | 134.48721 | 129.82906 | . | . | . | . | . | . |
| chr18 | 39489192 | 39491732 | 1000 | Promoter (<=1kb) | 18 | 39410545 | 39490674 | 80130 | 2 | ENSMUST00 | 0 | Nr3c1 | ENSMUSG0C | false | chr18 | 39488384 | 39492046 | 43.01277 | 150.85193 | 145.53938 | . | . | . | . | . | . |
| chr18 | 38276511 | 38276879 | 1000 | Intron (ENSMUST00000025311.5 | 18 | 38267092 | 38284401 | 17310 | 2 | ENSMUST00 | 7522 | Pcdh12 | ENSMUSG0C | false | . | . | . | . | . | . | chr18 | 38274249 | 38279359 | 5.63879 | 4.38394 | 3.54879 |
| chr18 | 37934860 | 37935838 | 1000 | Promoter (<=1kb) | 18 | 37843601 | 37935476 | 91876 | 2 | ENSMUST00 | 0 | Diap1 | ENSMUSG0C | false | chr18 | 37934343 | 37935812 | 23.5948 | 97.99914 | 94.47849 | . | . | . | . | . | . |
| chr18 | 37877754 | 37878145 | 1000 | Intron (ENSMUST00000115634.2 | 18 | 37896157 | 37897942 | 1786 | 2 | ENSMUST00 | 19797 | Diap1 | ENSMUSG0C | false | . | . | . | . | . | . | chr18 | 37877014 | 37880631 | 4.24513 | 3.6811 | 2.39992 |
| chr17 | 37045759 | 37046711 | 1000 | Promoter (<=1kb) | 17 | 37045966 | 37075067 | 29102 | 1 | ENSMUST00 | 0 | Gabbr1 | ENSMUSG0C | false | chr17 | 37045570 | 37046946 | 28.49099 | 81.63235 | 78.4519 | chr17 | 37046495 | 37047737 | 4.67942 | 3.98203 | 2.75765 |
| chr18 | 36515594 | 36516123 | 1000 | Promoter (<=1kb) | 18 | 36504929 | 36515805 | 10877 | 2 | ENSMUST00 | 0 | Hbegf | ENSMUSG0C | false | chr18 | 36514845 | 36516217 | 22.5292 | 69.37365 | 66.39268 | chr18 | 36515027 | 36516769 | 5.99343 | 4.63012 | 3.80678 |
| chr18 | 70564288 | 70564728 | 1000 | Distal Intergenic | 18 | 70568189 | 70626131 | 57943 | 1 | ENSMUST00 | -3461 | Mbd2 | ENSMUSG0C | false | . | . | . | . | . | . | chr18 | 70563875 | 70566315 | 4.39151 | 3.83268 | 2.50973 |
| chr18 | 70568029 | 70568748 | 1000 | Promoter (<=1kb) | 18 | 70568189 | 70626131 | 57943 | 1 | ENSMUST00 | 0 | Mbd2 | ENSMUSG0C | false | chr18 | 70567903 | 70569818 | 33.41831 | 117.26761 | 113.21259 | . | . | . | . | . | . |
| chr18 | 73703152 | 73704114 | 1000 | Promoter (<=1kb) | 18 | 73639009 | 73703780 | 64772 | 2 | ENSMUST00 | 0 | Smad4 | ENSMUSG0C | false | chr18 | 73702684 | 73704403 | 31.97681 | 101.31316 | 97.71113 | . | . | . | . | . | . |
| chr18 | 67497840 | 67498349 | 1000 | Intron (ENSMUST00000115050.2 | 18 | 67464849 | 67480581 | 15733 | 1 | ENSMUST00 | 32992 | Slmo1 | ENSMUSG0C | false | . | . | . | . | . | . | chr18 | 67497228 | 67503404 | 5.50174 | 4.28432 | 3.4286 |
| chr18 | 52831885 | 52832241 | 1000 | Intron (ENSMUST00000115410.3 | 18 | 52837964 | 52915586 | 77623 | 1 | ENSMUST00 | -5723 | Sncaip | ENSMUSG0C | true | . | . | . | . | . | . | chr18 | 52827941 | 52852025 | 4.46224 | 3.78219 | 2.57547 |
| chr18 | 52959886 | 52960327 | 1000 | Distal Intergenic | 18 | 52838010 | 52915312 | 77303 | 1 | ENSMUST00 | 121877 | Sncaip | ENSMUSG0C | true | . | . | . | . | . | . | chr18 | 52959247 | 52960346 | 3.19542 | 3.04355 | 1.57202 |
| chr18 | 68059097 | 68059659 | 1000 | Intron (ENSMUST00000063775.3 | 18 | 67933257 | 68255549 | 322293 | 1 | ENSMUST00 | 125841 | Ldlrad4 | ENSMUSG0C | false | chr18 | 68058798 | 68059342 | 11.02877 | 22.59171 | 20.22099 | chr18 | 68057251 | 68060499 | 3.88822 | 3.53367 | 2.0861 |
| chr18 | 79109661 | 79110425 | 1000 | Promoter (<=1kb) | 18 | 78750380 | 79109391 | 359012 | 2 | ENSMUST00 | -271 | Setbp1 | ENSMUSG0C | false | chr18 | 79108575 | 79110521 | 27.41053 | 90.23362 | 86.88379 | chr18 | 79108135 | 79110642 | 3.71667 | 3.46258 | 1.9533 |
| chr18 | 80933858 | 80934210 | 1000 | Promoter (<=1kb) | 18 | 80734143 | 80934058 | 199916 | 2 | ENSMUST00 | 0 | Atp9b | ENSMUSG0C | false | chr18 | 80933292 | 80934494 | 28.90751 | 104.81161 | 101.11721 | . | . | . | . | . | . |
| chr18 | 61335885 | 61336342 | 1000 | Intron (ENSMUST00000075299.7 | 18 | 61275023 | 61289924 | 14902 | 1 | ENSMUST00 | 60863 | Pde6a | ENSMUSG0C | false | . | . | . | . | . | . | chr18 | 61333315 | 61342581 | 5.33079 | 4.15853 | 3.28921 |
| chr18 | 61554855 | 61555447 | 1000 | Promoter (<=1kb) | 18 | 61555274 | 61590061 | 34788 | 1 | ENSMUST00 | 0 | Csnk1a1 | ENSMUSG0C | true | chr18 | 61554545 | 61555087 | 17.6846 | 53.76268 | 50.99834 | chr18 | 61553845 | 61554903 | 4.01397 | 3.66734 | 2.1757 |
| chr18 | 63691936 | 63692564 | 1000 | Promoter (<=1kb) | 18 | 63662801 | 63692359 | 29559 | 2 | ENSMUST00 | 0 | Txnl1 | ENSMUSG0C | false | chr18 | 63691141 | 63692990 | 42.12499 | 142.52214 | 137.54099 | . | . | . | . | . | . |
| chr18 | 64887080 | 64888532 | 1000 | Promoter (<=1kb) | 18 | 64887756 | 65217826 | 330071 | 1 | ENSMUST00 | 0 | Nedd4l | ENSMUSG0C | false | chr18 | 64886807 | 64889297 | 39.4261 | 145.54997 | 140.44281 | . | . | . | . | . | . |
| chr18 | 56707378 | 56707994 | 1000 | Promoter (<=1kb) | 18 | 56707813 | 56753424 | 45612 | 1 | ENSMUST00 | 0 | Lmnb1 | ENSMUSG0C | false | chr18 | 56707031 | 56709440 | 34.17132 | 121.80145 | 117.60049 | . | . | . | . | . | . |
| chr18 | 57354510 | 57355305 | 1000 | Promoter (<=1kb) | 18 | 57354733 | 57392719 | 37987 | 1 | ENSMUST00 | 0 | Prrc1 | ENSMUSG0C | false | chr18 | 57354231 | 57355888 | 21.48562 | 91.50835 | 88.13527 | . | . | . | . | . | . |
| chr18 | 57262387 | 57262836 | 1000 | Intron (ENSMUST00000075770.7 | 18 | 57354733 | 57392719 | 37987 | 1 | ENSMUST00 | -91897 | Prrc1 | ENSMUSG0C | false | . | . | . | . | . | . | chr18 | 57261733 | 57263619 | 4.28578 | 3.81059 | 2.44812 |
| chr18 | 57877993 | 57878844 | 1000 | Promoter (<=1kb) | 18 | 57878678 | 57946821 | 68144 | 1 | ENSMUST00 | 0 | Slc12a2 | ENSMUSG0C | false | chr18 | 57877616 | 57879893 | 32.27147 | 102.84008 | 99.19724 | . | . | . | . | . | . |
| chr18 | 60514168 | 60514417 | 1000 | Distal Intergenic | 18 | 60526221 | 60558762 | 32542 | 1 | ENSMUST00 | -11804 | Dctn4 | ENSMUSG0C | false | . | . | . | . | . | . | chr18 | 60514211 | 60514501 | 4.31225 | 3.74521 | 2.47249 |
| chr18 | 60746887 | 60747246 | 1000 | Distal Intergenic | 18 | 60774510 | 60778546 | 4037 | 1 | ENSMUST00 | -27264 | Rps14 | ENSMUSG0C | false | chr18 | 60747065 | 60749034 | 29.53108 | 87.21563 | 83.92796 | chr18 | 60742488 | 60750880 | 5.84853 | 4.46265 | 3.69893 |
| chr18 | 60744601 | 60745032 | 1000 | Distal Intergenic | 18 | 60774510 | 60778546 | 4037 | 1 | ENSMUST00 | -29478 | Rps14 | ENSMUSG0C | false | . | . | . | . | . | . | chr18 | 60742488 | 60750880 | 5.84853 | 4.46265 | 3.69893 |
| chr18 | 61041300 | 61041720 | 1000 | Distal Intergenic | 18 | 61045150 | 61085061 | 39912 | 1 | ENSMUST00 | -3430 | Pdgfrb | ENSMUSG0C | false | . | . | . | . | . | . | chr18 | 61040114 | 61041700 | 3.78605 | 3.33004 | 2.0456 |
| chr18 | 61059525 | 61059963 | 1000 | Intron (ENSMUST00000025522.5 | 18 | 61045200 | 61085056 | 39857 | 1 | ENSMUST00 | 14326 | Pdgfrb | ENSMUSG0C | false | . | . | . | . | . | . | chr18 | 61054129 | 61061450 | 6.93727 | 4.97103 | 4.59192 |
| chr18 | 61058356 | 61058894 | 1000 | Intron (ENSMUST00000025522.5 | 18 | 61045200 | 61085056 | 39857 | 1 | ENSMUST00 | 13157 | Pdgfrb | ENSMUSG0C | false | . | . | . | . | . | . | chr18 | 61054129 | 61061450 | 6.93727 | 4.97103 | 4.59192 |
| chr18 | 61057106 | 61057586 | 1000 | Intron (ENSMUST00000025522.5 | 18 | 61045200 | 61085056 | 39857 | 1 | ENSMUST00 | 11907 | Pdgfrb | ENSMUSG0C | false | . | . | . | . | . | . | chr18 | 61054129 | 61061450 | 6.93727 | 4.97103 | 4.59192 |
| chr18 | 61054320 | 61055039 | 1000 | Intron (ENSMUST00000025522.5 | 18 | 61045200 | 61085056 | 39857 | 1 | ENSMUST00 | 9121 | Pdgfrb | ENSMUSG0C | false | . | . | . | . | . | . | chr18 | 61054129 | 61061450 | 6.93727 | 4.97103 | 4.59192 |
| chr19 | 16132249 | 16133629 | 1000 | Promoter (<=1kb) | 19 | 16132831 | 16387463 | 254633 | 1 | ENSMUST00 | 0 | Gnaq | ENSMUSG0C | false | chr19 | 16131695 | 16134440 | 35.66819 | 113.91924 | 109.96352 | . | . | . | . | . | . |
| chr19 | 14508777 | 14509155 | 1000 | Intron (ENSMUST00000052011.8 | 19 | 14448072 | 14597983 | 149912 | 2 | ENSMUST00 | 88828 | Tle4 | ENSMUSG0C | false | . | . | . | . | . | . | chr19 | 14496852 | 14509645 | 5.46229 | 4.19323 | 3.40627 |
| chr19 | 14499991 | 14500582 | 1000 | Intron (ENSMUST00000052011.8 | 19 | 14448072 | 14597983 | 149912 | 2 | ENSMUST00 | 97401 | Tle4 | ENSMUSG0C | false | . | . | . | . | . | . | chr19 | 14496852 | 14509645 | 5.46229 | 4.19323 | 3.40627 |
| chr19 | 10003737 | 10004149 | 1000 | Promoter (2-3kb) | 19 | 10001669 | 10035586 | 33918 | 1 | ENSMUST00 | 2069 | Rab3il1 | ENSMUSG0C | false | . | . | . | . | . | . | chr19 | 10002861 | 10004845 | 5.28195 | 4.22648 | 3.25231 |
| chr19 | 10041187 | 10041888 | 1000 | Promoter (<=1kb) | 19 | 10041548 | 10059671 | 18124 | 1 | ENSMUST00 | 0 | Fads3 | ENSMUSG0C | false | chr19 | 10041037 | 10042840 | 43.67043 | 145.21399 | 140.12416 | chr19 | 10036253 | 10050591 | 5.31313 | 4.14637 | 3.29774 |
| chr19 | 12698128 | 12698525 | 1000 | Promoter (2-3kb) | 19 | 12695814 | 12719117 | 23304 | 1 | ENSMUST00 | 2315 | Keg1 | ENSMUSG0C | false | . | . | . | . | . | . | chr19 | 12696240 | 12700177 | 3.98585 | 3.46598 | 2.18713 |
| chr19 | 12794634 | 12795937 | 1000 | Promoter (<=1kb) | 19 | 12763660 | 12796126 | 32467 | 2 | ENSMUST00 | 189 | Zfp91 | ENSMUSG0C | false | chr19 | 12793084 | 12795521 | 45.20664 | 150.13857 | 144.8454 | . | . | . | . | . | . |
| chr19 | 17837149 | 17837945 | 1000 | Promoter (<=1kb) | 19 | 17432832 | 17837632 | 404801 | 2 | ENSMUST00 | 0 | Pcsk5 | ENSMUSG0C | true | chr19 | 17835376 | 17838194 | 27.88725 | 99.90214 | 96.33501 | . | . | . | . | . | . |
| chr19 | 17821307 | 17821698 | 1000 | Intron (ENSMUST00000025618.9 | 19 | 17432832 | 17837632 | 404801 | 2 | ENSMUST00 | 15934 | Pcsk5 | ENSMUSG0C | true | . | . | . | . | . | . | chr19 | 17818630 | 17824830 | 5.42481 | 4.09227 | 3.38732 |
| chr19 | 18670663 | 18671485 | 1000 | Promoter (<=1kb) | 19 | 18670780 | 18704788 | 34009 | 1 | ENSMUST00 | 0 | 2410127L17F | ENSMUSG0C | false | chr19 | 18670839 | 18672337 | 21.95811 | 99.8556 | 96.2922 | . | . | . | . | . | . |
| chr19 | 18652554 | 18652874 | 1000 | Distal Intergenic | 19 | 18670780 | 18704788 | 34009 | 1 | ENSMUST00 | -17906 | 2410127L17F | ENSMUSG0C | false | . | . | . | . | . | . | chr19 | 18651328 | 18653764 | 3.85964 | 3.54663 | 2.06466 |
| chr19 | 21271763 | 21273149 | 1000 | Promoter (<=1kb) | 19 | 21272278 | 21282289 | 10012 | 1 | ENSMUST00 | 0 | Zfand5 | ENSMUSG0C | false | chr19 | 21271332 | 21275517 | 33.89795 | 105.67399 | 101.95845 | . | . | . | . | . | . |
| chr19 | 21778208 | 21779318 | 1000 | Promoter (<=1kb) | 19 | 21778342 | 21858327 | 79986 | 1 | ENSMUST00 | 0 | Tmem2 | ENSMUSG0C | false | chr19 | 21777956 | 21779995 | 36.53154 | 123.91184 | 119.64436 | . | . | . | . | . | . |
| chr19 | 7165010 | 7166369 | 1000 | Intron (ENSMUST00000040261.5 | 19 | 7199540 | 7200464 | 925 | 2 | ENSMUST00 | 34095 | Otub1 | ENSMUSG0C | false | chr19 | 7164595 | 7166503 | 20.9437 | 83.15932 | 79.95254 | chr19 | 7163724 | 7167388 | 4.90862 | 4.06292 | 2.93319 |
| chr19 | 29016116 | 29016495 | 1000 | 3' UTR | 19 | 29025901 | 29027244 | 1344 | 2 | ENSMUST00 | 10749 | Ak3 | ENSMUSG0C | false | . | . | . | . | . | . | chr19 | 29014856 | 29017424 | 4.87891 | 4.07337 | 2.94904 |
| chr19 | 29251121 | 29252051 | 1000 | Promoter (<=1kb) | 19 | 29251828 | 29313080 | 61253 | 1 | ENSMUST00 | 0 | Jak2 | ENSMUSG0C | false | chr19 | 29250740 | 29253200 | 30.94804 | 114.76607 | 110.79114 | chr19 | 29249752 | 29251228 | 3.49627 | 3.347 | 1.76395 |

| | | | | | | | | | | | | | | | | | | | | | | | | | | |
|---|---|---|---|---|---|---|---|---|---|---|---|---|---|---|---|---|---|---|---|---|---|---|---|---|---|---|
| chr19 | 36208433 | 36208731 | 1000 | Distal Intergenic | 19 | 36111965 | 36119844 | 7880 | 2 | ENSMUST00 | -88590 | Ankrd1 | ENSMUSG0C | false | | | | | | | chr19 | 36207242 | 36208897 | 6.42505 | 4.7324 | 4.18181 |
| chr19 | 36348053 | 36349033 | 1000 | Distal Intergenic | 19 | 36379067 | 36456192 | 77126 | 1 | ENSMUST00 | -30034 | Pcgf5 | ENSMUSG0C | false | chr19 | 36347832 | 36349860 | 35.37319 | 123.98246 | 119.71299 | | | | | | |
| chr19 | 36789835 | 36790182 | 1000 | Distal Intergenic | 19 | 36834232 | 36893477 | 59246 | 1 | ENSMUST00 | -44050 | Tnks2 | ENSMUSG0C | false | | | | | | | chr19 | 36789412 | 36790251 | 4.11996 | 3.71536 | 2.29227 |
| chr19 | 24173702 | 24174584 | 1000 | Intron (ENSMUST00000099558.4 | 19 | 24094523 | 24225026 | 130504 | 2 | ENSMUST00 | 50442 | Tjp2 | ENSMUSG0C | false | chr19 | 24173413 | 24175288 | 24.82645 | 85.46523 | 82.21117 | | | | | | |
| chr19 | 24242209 | 24242561 | 1000 | Distal Intergenic | 19 | 24094523 | 24225026 | 130504 | 2 | ENSMUST00 | -17184 | Tjp2 | ENSMUSG0C | false | | | | | | | chr19 | 24241966 | 24242745 | 3.24905 | 3.18611 | 1.61578 |
| chr19 | 5871242 | 5871614 | 1000 | Promoter (1-2kb) | 19 | 5865184 | 5873430 | 8247 | 2 | ENSMUST00 | 1816 | Frmd8 | ENSMUSG0C | false | | | | | | | chr19 | 5869707 | 5876358 | 6.4359 | 4.66821 | 4.19169 |
| chr19 | 3260237 | 3260604 | 1000 | Downstream (<1kb) | 19 | 3265364 | 3267716 | 2353 | 2 | ENSMUST00 | 7112 | Ighmbp2 | ENSMUSG0C | false | | | | | | | chr19 | 3259601 | 3263900 | 6.21113 | 4.29911 | 4.03454 |
| chr19 | 4150842 | 4151211 | 1000 | Promoter (<=1kb) | 19 | 4151370 | 4153444 | 2075 | 1 | ENSMUST00 | -159 | Coro1b | ENSMUSG0C | false | | | | | | | chr19 | 4147701 | 4152092 | 7.29509 | 4.99142 | 4.89155 |
| chr19 | 3851560 | 3852669 | 1000 | Promoter (<=1kb) | 19 | 3851773 | 3894365 | 42593 | 1 | ENSMUST00 | 0 | Chka | ENSMUSG0C | false | chr19 | 3851080 | 3852947 | 28.56742 | 103.16996 | 99.51957 | chr19 | 3852602 | 3853434 | 3.36619 | 3.2596 | 1.66762 |
| chr19 | 4231599 | 4232141 | 1000 | Promoter (<=1kb) | 19 | 4231899 | 4233631 | 1733 | 1 | ENSMUST00 | 0 | Pold4 | ENSMUSG0C | false | chr19 | 4231890 | 4232260 | 20.56672 | 54.83671 | 52.06188 | chr19 | 4228207 | 4233506 | 8.46056 | 5.68198 | 5.82099 |
| chr19 | 4230404 | 4230772 | 1000 | Promoter (1-2kb) | 19 | 4231899 | 4233631 | 1733 | 1 | ENSMUST00 | -1127 | Pold4 | ENSMUSG0C | false | | | | | | | chr19 | 4228207 | 4233506 | 8.46056 | 5.68198 | 5.82099 |
| chr19 | 5087851 | 5088367 | 1000 | Promoter (<=1kb) | 19 | 5088538 | 5092879 | 4342 | 1 | ENSMUST00 | -171 | Yif1a | ENSMUSG0C | false | chr19 | 5087724 | 5089304 | 32.03847 | 101.42104 | 97.81718 | chr19 | 5087381 | 5089846 | 7.12625 | 5.02773 | 4.78191 |
| chr19 | 4625175 | 4625777 | 1000 | Promoter (<=1kb) | 19 | 4622551 | 4625617 | 3067 | 2 | ENSMUST00 | 0 | Rce1 | ENSMUSG0C | false | chr19 | 4624885 | 4626451 | 35.15467 | 116.16882 | 112.14974 | | | | | | |
| chr19 | 23834380 | 23834710 | 1000 | Intron (ENSMUST00000025830.7 | 19 | 23758876 | 23949597 | 190722 | 1 | ENSMUST00 | 75505 | Apba1 | ENSMUSG0C | false | | | | | | | chr19 | 23833702 | 23834965 | 3.95657 | 3.54905 | 2.15815 |
| chr19 | 3323083 | 3324108 | 1000 | Promoter (<=1kb) | 19 | 3323301 | 3385732 | 62432 | 1 | ENSMUST00 | 0 | Cpt1a | ENSMUSG0C | false | chr19 | 3321491 | 3326282 | 34.67015 | 117.41676 | 113.35385 | | | | | | |
| chr19 | 3324981 | 3325724 | 1000 | Promoter (1-2kb) | 19 | 3323301 | 3385732 | 62432 | 1 | ENSMUST00 | 1681 | Cpt1a | ENSMUSG0C | false | chr19 | 3321491 | 3326282 | 34.67015 | 117.41676 | 113.35385 | | | | | | |
| chr19 | 3641879 | 3642196 | 1000 | Intron (ENSMUST00000025856.1 | 19 | 3584836 | 3615792 | 30957 | 2 | ENSMUST00 | -26088 | Lrp5 | ENSMUSG0C | false | | | | | | | chr19 | 3635207 | 3643434 | 7.206 | 4.9739 | 4.82746 |
| chr19 | 3638488 | 3639084 | 1000 | Intron (ENSMUST00000025856.1 | 19 | 3584836 | 3615792 | 30957 | 2 | ENSMUST00 | -22697 | Lrp5 | ENSMUSG0C | false | | | | | | | chr19 | 3635207 | 3643434 | 7.206 | 4.9739 | 4.82746 |
| chr19 | 26604238 | 26605316 | 1000 | Promoter (<=1kb) | 19 | 26605115 | 26778321 | 173207 | 1 | ENSMUST00 | 0 | Smarca2 | ENSMUSG0C | false | chr19 | 26603929 | 26608671 | 23.80619 | 80.75649 | 77.58801 | chr19 | 26603658 | 26604251 | 3.60276 | 3.40366 | 1.84422 |
| chr19 | 26606235 | 26608129 | 1000 | Promoter (<=1kb) | 19 | 26607270 | 26778273 | 171004 | 1 | ENSMUST00 | 0 | Smarca2 | ENSMUSG0C | false | chr19 | 26603929 | 26608671 | 23.80619 | 80.75649 | 77.58801 | chr19 | 26607920 | 26623329 | 6.40254 | 4.73837 | 4.15669 |
| chr19 | 26612865 | 26613365 | 1000 | Intron (ENSMUST00000025862.8 | 19 | 26607270 | 26778273 | 171004 | 1 | ENSMUST00 | 5596 | Smarca2 | ENSMUSG0C | false | | | | | | | chr19 | 26607920 | 26623329 | 6.40254 | 4.73837 | 4.15669 |
| chr19 | 5741682 | 5742383 | 1000 | Promoter (<=1kb) | 19 | 5740904 | 5758532 | 17629 | 1 | ENSMUST00 | 779 | Ltbp3 | ENSMUSG0C | false | chr19 | 5741479 | 5742777 | 34.46491 | 104.61457 | 100.92722 | chr19 | 5741392 | 5744229 | 5.31289 | 4.14221 | 3.30114 |
| chr19 | 6015009 | 6015952 | 1000 | Promoter (<=1kb) | 19 | 5988546 | 6015247 | 26702 | 2 | ENSMUST00 | 0 | Capn1 | ENSMUSG0C | false | chr19 | 6014421 | 6016110 | 26.04686 | 75.56942 | 72.49508 | chr19 | 6013845 | 6016206 | 3.97113 | 3.3261 | 2.19189 |
| chr19 | 6349320 | 6349638 | 1000 | Promoter (1-2kb) | 19 | 6350909 | 6351580 | 672 | 1 | ENSMUST00 | -1271 | Map4k2 | ENSMUSG0C | false | | | | | | | chr19 | 6348258 | 6350133 | 4.06996 | 3.65609 | 2.27574 |
| chr19 | 6363305 | 6364546 | 1000 | Promoter (<=1kb) | 19 | 6363690 | 6378030 | 14341 | 1 | ENSMUST00 | 0 | Sf1 | ENSMUSG0C | false | chr19 | 6363840 | 6364939 | 39.80159 | 147.3544 | 142.17488 | | | | | | |
| chr19 | 6909393 | 6909850 | 1000 | Promoter (<=1kb) | 19 | 6906846 | 6909617 | 2772 | 2 | ENSMUST00 | 0 | Prdx5 | ENSMUSG0C | false | chr19 | 6908992 | 6910951 | 29.114 | 86.49567 | 83.2249 | | | | | | |
| chr19 | 53902936 | 53904068 | 1000 | Promoter (<=1kb) | 19 | 53903351 | 53929861 | 26511 | 1 | ENSMUST00 | 0 | Pdcd4 | ENSMUSG0C | false | chr19 | 53902548 | 53904667 | 41.07278 | 140.67711 | 135.76277 | | | | | | |
| chr19 | 53943847 | 53945487 | 1000 | Promoter (<=1kb) | 19 | 53944306 | 54033268 | 88963 | 1 | ENSMUST00 | 0 | Shoc2 | ENSMUSG0C | false | chr19 | 53943307 | 53944593 | 22.78592 | 76.10319 | 73.01487 | | | | | | |
| chr19 | 55096645 | 55097004 | 1000 | Promoter (2-3kb) | 19 | 55069734 | 55099451 | 29718 | 2 | ENSMUST00 | 2447 | Gpam | ENSMUSG0C | false | | | | | | | chr19 | 55091677 | 55104004 | 5.06737 | 4.08692 | 3.0685 |
| chr19 | 55098485 | 55098957 | 1000 | Promoter (<=1kb) | 19 | 55069734 | 55099451 | 29718 | 2 | ENSMUST00 | 494 | Gpam | ENSMUSG0C | false | chr19 | 55098118 | 55100331 | 30.33064 | 129.04645 | 124.59567 | chr19 | 55091677 | 55104004 | 5.06737 | 4.08692 | 3.0685 |
| chr19 | 55147504 | 55147939 | 1000 | Distal Intergenic | 19 | 55180733 | 55189665 | 8933 | 1 | ENSMUST00 | -32794 | Tectb | ENSMUSG0C | false | | | | | | | chr19 | 55146941 | 55148779 | 3.36787 | 2.98527 | 1.70172 |
| chr19 | 55158636 | 55159115 | 1000 | Distal Intergenic | 19 | 55180733 | 55189665 | 8933 | 1 | ENSMUST00 | -21618 | Tectb | ENSMUSG0C | false | | | | | | | chr19 | 55157727 | 55160983 | 4.8336 | 4.06861 | 2.86323 |
| chr19 | 55750245 | 55750564 | 1000 | Intron (ENSMUST00000061496.1 | 19 | 55742845 | 55910681 | 167837 | 1 | ENSMUST00 | 7401 | Tcf7l2 | ENSMUSG0C | false | | | | | | | chr19 | 55744268 | 55760177 | 6.20416 | 4.6193 | 3.99773 |
| chr19 | 55758201 | 55758458 | 1000 | Intron (ENSMUST00000061496.1 | 19 | 55742845 | 55910681 | 167837 | 1 | ENSMUST00 | 15357 | Tcf7l2 | ENSMUSG0C | false | | | | | | | chr19 | 55744268 | 55760177 | 6.20416 | 4.6193 | 3.99773 |
| chr19 | 55741393 | 55741901 | 1000 | Promoter (<=1kb) | 19 | 55741810 | 55933654 | 191845 | 1 | ENSMUST00 | 0 | Tcf7l2 | ENSMUSG0C | false | chr19 | 55741009 | 55745168 | 33.32866 | 126.59486 | 122.2257 | | | | | | |
| chr19 | 60873501 | 60873952 | 1000 | Promoter (<=1kb) | 19 | 60864051 | 60874556 | 10506 | 2 | ENSMUST00 | 604 | Prdx3 | ENSMUSG0C | false | | | | | | | chr19 | 60870881 | 60875062 | 4.28747 | 3.68047 | 2.4328 |
| chr19 | 38504555 | 38505144 | 1000 | Intron (ENSMUST00000181994.1 | 19 | 38524197 | 38785030 | 260834 | 1 | ENSMUST00 | -19053 | Plce1 | ENSMUSG0C | true | | | | | | | chr19 | 38495787 | 38511149 | 5.1247 | 4.07357 | 3.12523 |
| chr19 | 38521426 | 38521868 | 1000 | Promoter (2-3kb) | 19 | 38524197 | 38785030 | 260834 | 1 | ENSMUST00 | -2329 | Plce1 | ENSMUSG0C | true | | | | | | | chr19 | 38517349 | 38541346 | 5.17509 | 4.14931 | 3.15371 |
| chr19 | 38522114 | 38522632 | 1000 | Promoter (1-2kb) | 19 | 38524197 | 38785030 | 260834 | 1 | ENSMUST00 | -1565 | Plce1 | ENSMUSG0C | true | | | | | | | chr19 | 38517349 | 38541346 | 5.17509 | 4.14931 | 3.15371 |
| chr19 | 38480715 | 38481525 | 1000 | Promoter (<=1kb) | 19 | 38481109 | 38524066 | 42958 | 1 | ENSMUST00 | 0 | Plce1 | ENSMUSG0C | true | chr19 | 38480630 | 38483282 | 36.42221 | 131.26616 | 126.73016 | chr19 | 38475288 | 38480960 | 4.89146 | 3.88312 | 2.9558 |
| chr19 | 38528408 | 38528953 | 1000 | Intron (ENSMUST00000182481.2 | 19 | 38524209 | 38782795 | 258587 | 1 | ENSMUST00 | 4200 | Plce1 | ENSMUSG0C | true | chr19 | 38527399 | 38528412 | 6.15186 | 10.70359 | 8.51182 | chr19 | 38517349 | 38541346 | 5.17509 | 4.14931 | 3.15371 |
| chr19 | 38496519 | 38497021 | 1000 | Intron (ENSMUST00000181994.1 | 19 | 38481121 | 38783430 | 302310 | 1 | ENSMUST00 | 15399 | Plce1 | ENSMUSG0C | true | | | | | | | chr19 | 38495787 | 38511149 | 5.1247 | 4.07357 | 3.12523 |
| chr19 | 41263420 | 41264321 | 1000 | Promoter (<=1kb) | 19 | 41210842 | 41264004 | 53163 | 2 | ENSMUST00 | 0 | Tm9sf3 | ENSMUSG0C | false | chr19 | 41262676 | 41264809 | 29.66345 | 100.21 | 96.63225 | | | | | | |
| chr19 | 41482145 | 41483523 | 1000 | Promoter (<=1kb) | 19 | 41482645 | 41562246 | 79602 | 1 | ENSMUST00 | 0 | Lcor | ENSMUSG0C | false | chr19 | 41481135 | 41484006 | 27.41053 | 90.23362 | 86.88379 | | | | | | |
| chr19 | 53412191 | 53412622 | 1000 | Distal Intergenic | 19 | 53379214 | 53390573 | 11360 | 2 | ENSMUST00 | -21619 | Smndc1 | ENSMUSG0C | false | | | | | | | chr19 | 53409099 | 53414773 | 6.07728 | 4.55041 | 3.90583 |
| chr19 | 53329267 | 53329732 | 1000 | Promoter (<=1kb) | 19 | 53329413 | 53375810 | 46398 | 1 | ENSMUST00 | 0 | Mxi1 | ENSMUSG0C | false | chr19 | 53329381 | 53330780 | 33.94791 | 109.87469 | 106.04007 | | | | | | |
| chr19 | 53312595 | 53314379 | 1000 | Promoter (2-3kb) | 19 | 53310506 | 53375810 | 65305 | 1 | ENSMUST00 | 2090 | Mxi1 | ENSMUSG0C | false | chr19 | 53310158 | 53314217 | 35.4076 | 109.31139 | 105.49952 | | | | | | |
| chr19 | 53141316 | 53143328 | 1000 | Promoter (<=1kb) | 19 | 53142756 | 53247399 | 104644 | 1 | ENSMUST00 | 0 | Add3 | ENSMUSG0C | false | chr19 | 53140463 | 53142585 | 18.59898 | 66.17558 | 63.2431 | chr19 | 53140644 | 53142162 | 3.94157 | 3.62917 | 2.12504 |
| chr19 | 53052102 | 53052734 | 1000 | Distal Intergenic | 19 | 52991183 | 53040214 | 49032 | 2 | ENSMUST00 | -11889 | Xpnpep1 | ENSMUSG0C | false | | | | | | | chr19 | 53051279 | 53053121 | 5.22639 | 4.00985 | 3.22206 |
| chr19 | 53040295 | 53040692 | 1000 | Promoter (<=1kb) | 19 | 52991183 | 53040214 | 49032 | 2 | ENSMUST00 | -82 | Xpnpep1 | ENSMUSG0C | false | chr19 | 53040211 | 53041383 | 6.46168 | 12.41373 | 10.19419 | chr19 | 53037126 | 53042423 | 5.46211 | 4.19899 | 3.43522 |
| chr19 | 46501140 | 46502489 | 1000 | Promoter (<=1kb) | 19 | 46501648 | 46516444 | 14797 | 1 | ENSMUST00 | 0 | Trim8 | ENSMUSG0C | false | chr19 | 46500135 | 46506359 | 34.21972 | 118.71816 | 114.6235 | | | | | | |
| chr19 | 46503464 | 46504634 | 1000 | Promoter (1-2kb) | 19 | 46501648 | 46516444 | 14797 | 1 | ENSMUST00 | 1817 | Trim8 | ENSMUSG0C | false | chr19 | 46500135 | 46506359 | 34.21972 | 118.71816 | 114.6235 | | | | | | |
| chr19 | 46572897 | 46573439 | 1000 | Promoter (<=1kb) | 19 | 46531109 | 46573085 | 41977 | 2 | ENSMUST00 | 0 | Arl3 | ENSMUSG0C | false | chr19 | 46572370 | 46573955 | 25.53309 | 82.08177 | 78.89293 | chr19 | 46570703 | 46574043 | 4.04911 | 3.61752 | 2.23627 |
| chr19 | 46864623 | 46865145 | 1000 | Intron (ENSMUST00000077666.4 | 19 | 46891509 | 46898669 | 7161 | 2 | ENSMUST00 | 33524 | Nt5c2 | ENSMUSG0C | false | | | | | | | chr19 | 46861574 | 46866144 | 4.90893 | 4.07981 | 2.93026 |
| chr19 | 46961713 | 46962710 | 1000 | Promoter (<=1kb) | 19 | 46886831 | 46962351 | 75521 | 2 | ENSMUST00 | 0 | Nt5c2 | ENSMUSG0C | false | chr19 | 46960757 | 46963137 | 30.03893 | 101.89226 | 98.2715 | | | | | | |
| chr19 | 47563758 | 47564167 | 1000 | Distal Intergenic | 19 | 47579678 | 47645246 | 65569 | 1 | ENSMUST00 | -15511 | Sik | ENSMUSG0C | false | | | | | | | | | | | | |
| chr19 | 56396926 | 56397543 | 1000 | Promoter (<=1kb) | 19 | 56397129 | 56442344 | 45216 | 1 | ENSMUST00 | 0 | Casp7 | ENSMUSG0C | false | chr19 | 56396578 | 56397904 | 31.01414 | 97.6227 | 94.10706 | chr19 | 56395639 | 56398457 | 4.88044 | 4.04486 | 2.91368 |
| chr19 | 56976928 | 56977428 | 1000 | Intron (ENSMUST00000118800.2 | 19 | 56930098 | 57001046 | 70949 | 2 | ENSMUST00 | 23618 | Afap1l2 | ENSMUSG0C | false | | | | | | | chr19 | 56976918 | 56977864 | 2.98594 | 3.03906 | 1.39572 |
| chr19 | 57098433 | 57098909 | 1000 | Intron (ENSMUST00000079360.6 | 19 | 57036479 | 57118913 | 82435 | 2 | ENSMUST00 | 20004 | Ablim1 | ENSMUSG0C | false | | | | | | | chr19 | 57097552 | 57100265 | 4.71946 | 3.8426 | 2.80811 |
| chr19 | 57197243 | 57197663 | 1000 | Promoter (<=1kb) | 19 | 57033274 | 57197631 | 164358 | 2 | ENSMUST00 | 0 | Ablim1 | ENSMUSG0C | false | chr19 | 57197103 | 57197837 | 17.19387 | 43.56631 | 40.93357 | chr19 | 57196729 | 57198433 | 5.38507 | 4.32428 | 3.3283 |
| chr19 | 58833589 | 58834020 | 1000 | Intron (ENSMUST00000066285.4 | 19 | 58795751 | 58860984 | 65234 | 2 | ENSMUST00 | 26964 | Hspa12a | ENSMUSG0C | true | | | | | | | chr19 | 58814047 | 58859817 | 7.46659 | 5.12677 | 5.03638 |
| chr19 | 58838346 | 58838187 | 1000 | Intron (ENSMUST00000066285.4 | 19 | 58795751 | 58860984 | 65234 | 2 | ENSMUST00 | 21797 | Hspa12a | ENSMUSG0C | true | | | | | | | chr19 | 58814047 | 58859817 | 7.46659 | 5.12677 | 5.03638 |
| chr19 | 58860798 | 58861234 | 1000 | Promoter (<=1kb) | 19 | 58795751 | 58860984 | 65234 | 2 | ENSMUST00 | 0 | Hspa12a | ENSMUSG0C | true | chr19 | 58859113 | 58861512 | 26.1063 | 98.41757 | 94.88632 | | | | | | |
| chr7 | 81839744 | 81840158 | 1000 | Distal Intergenic | 7 | 81792075 | 81829431 | 37357 | 2 | ENSMUST00 | -10314 | Btbd1 | ENSMUSG0C | false | | | | | | | chr7 | 81839624 | 81842577 | 4.6832 | 3.90829 | 2.75525 |
| chr7 | 81933605 | 81934656 | 1000 | Promoter (<=1kb) | 7 | 81881251 | 81934473 | 53223 | 2 | ENSMUST00 | 0 | Hdgfrp3 | ENSMUSG0C | false | chr7 | 81932850 | 81934959 | 25.84672 | 90.2271 | 86.88143 | chr7 | 81934595 | 81934980 | 3.43958 | 3.29178 | 1.70597 |
| chr11 | 120572596 | 120573536 | 1000 | Promoter (<=1kb) | 11 | 120560298 | 120573253 | 12956 | 2 | ENSMUST00 | 0 | P4hb | ENSMUSG0C | false | chr11 | 120573364 | 120574265 | 28.22734 | 101.53345 | 97.92473 | | | | | | |
| chr7 | 141876099 | 141876617 | 1000 | Intron (ENSMUST00000055819.7 | 7 | 141874813 | 141902415 | 27603 | 2 | ENSMUST00 | 25798 | Tollip | ENSMUSG0C | false | | | | | | | chr7 | 141873818 | 141880865 | 5.51196 | 4.11411 | 3.46302 |
| chr7 | 142035063 | 142035512 | 1000 | Intron (ENSMUST00000084418.2 | 7 | 142008553 | 142061034 | 52482 | 2 | ENSMUST00 | 25522 | Mob2 | ENSMUSG0C | true | | | | | | | chr7 | 142029403 | 142051829 | 5.68259 | 4.36057 | 3.56764 |
| chr19 | 41980721 | 41982505 | 1000 | Promoter (<=1kb) | 19 | 41941086 | 41981114 | 40029 | 2 | ENSMUST00 | 0 | Mms19 | ENSMUSG0C | false | chr19 | 41982037 | 41982830 | 34.43156 | 129.74289 | 125.25954 | | | | | | |
| chr11 | 120950819 | 120951211 | 1000 | Promoter (<=1kb) | 11 | 120951138 | 120954212 | 3075 | 1 | ENSMUST00 | 0 | Slc16a3 | ENSMUSG0C | false | chr11 | 120950486 | 120950962 | 6.80124 | 12.58378 | 10.36022 | chr11 | 120948533 | 120951419 | 5.55747 | 4.1386 | 3.50262 |
| chr11 | 120990194 | 120991276 | 1000 | Promoter (<=1kb) | 11 | 120961749 | 120991092 | 29344 | 2 | ENSMUST00 | 0 | Csnk1d | ENSMUSG0C | false | chr11 | 120989882 | 120991796 | 26.812 | 131.41335 | 126.87386 | | | | | | |
| chr19 | 43699151 | 43699626 | 1000 | Intron (ENSMUST00000081079.5 | 19 | 43689840 | 43722574 | 32735 | 1 | ENSMUST00 | 9312 | Entpd7 | ENSMUSG0C | false | | | | | | | chr19 | 43690389 | 43702907 | 5.7357 | 4.40542 | 3.61051 |
| chr19 | 44069566 | 44070030 | 1000 | Promoter (<=1kb) | 19 | 44034944 | 44069775 | 34832 | 2 | ENSMUST00 | 0 | Erlin1 | ENSMUSG0C | false | chr19 | 44068955 | 44070157 | 26.72788 | 101.58999 | 97.97913 | | | | | | |
| chr19 | 44248003 | 44248463 | 1000 | Promoter (<=1kb) | 19 | 44203288 | 44244016 | 40729 | 1 | ENSMUST00 | 0 | Scd3 | ENSMUSG0C | false | chr19 | 44247327 | 44248258 | 13.89301 | 35.6666 | 33.12951 | chr19 | 44245947 | 44250507 | 5.28952 | 3.98409 | 3.27786 |
| chr19 | 44293505 | 44294179 | 1000 | Promoter (<=1kb) | 19 | 44293676 | 44306864 | 13189 | 1 | ENSMUST00 | 0 | Scd2 | ENSMUSG0C | false | chr19 | 44293182 | 44296580 | 35.73082 | 112.98314 | 109.06473 | chr19 | 44291679 | 44293667 | 6.41134 | 4.6985 | 4.20915 |
| chr19 | 44266777 | 44267506 | 1000 | Distal Intergenic | 19 | 44293676 | 44306864 | 13189 | 1 | ENSMUST00 | -26170 | Scd2 | ENSMUSG0C | false | chr19 | 44266714 | 44267515 | 7.9533 | 20.3031 | 17.96454 | chr19 | 44266096 | 44268813 | 5.62181 | 4.43946 | 3.49853 |
| chr19 | 44283853 | 44285032 | 1000 | Promoter (<=1kb) | 19 | 44293676 | 44306864 | 13189 | 1 | ENSMUST00 | -8644 | Scd2 | ENSMUSG0C | false | chr19 | 44284311 | 44285094 | 16.58925 | 40.3276 | 37.73373 | chr19 | 44280498 | 44288140 | 5.36112 | 4.2053 | 3.32931 |
| chr19 | 45079390 | 45080048 | 1000 | Distal Intergenic | 19 | 45076139 | 45079289 | 3151 | 1 | ENSMUST00 | 3252 | Kazald1 | ENSMUSG0C | false | chr19 | 45079650 | 45080851 | 8.87028 | 22.65516 | 20.28383 | chr19 | 45074094 | 45081464 | 5.80167 | 4.45472 | 3.66688 |
| chr19 | 45363433 | 45364187 | 1000 | Promoter (<=1kb) | 19 | 45363734 | 45533341 | 169608 | 1 | ENSMUST00 | 0 | Btrc | ENSMUSG0C | true | chr19 | 45363722 | 45365113 | 33.94791 | 109.87469 | 106.04007 | | | | | | |
| chr19 | 45355590 | 45356045 | 1000 | Distal Intergenic | 19 | 45363734 | 45533341 | 169608 | 1 | ENSMUST00 | -7689 | Btrc | ENSMUSG0C | true | | | | | | | | | | | | |
| chr19 | 45529276 | 45529635 | 1000 | 3' UTR | 19 | 45552275 | 45560531 | 8257 | 2 | ENSMUST00 | 30896 | Poll | ENSMUSG0C | false | | | | | | | chr19 | 45528687 | 45529829 | 3.80698 | 3.27472 | 2.06001 |
| chr19 | 45783146 | 45783966 | 1000 | Promoter (<=1kb) | 19 | 45750261 | 45783520 | 33260 | 2 | ENSMUST00 | 0 | Mgea5 | ENSMUSG0C | false | chr19 | 45781513 | 45783689 | 39.11016 | 156.19371 | 150.67332 | | | | | | |
| chr19 | 46039228 | 46040114 | 1000 | Promoter (<=1kb) | 19 | 46032594 | 46039621 | 7028 | 2 | ENSMUST00 | 0 | Ldb1 | ENSMUSG0C | false | chr19 | 46038333 | 46040333 | 26.84644 | 76.94898 | 73.84959 | chr19 | 46036064 | 46044344 | 7.61565 | 5.16237 | 5.15378 |
| chr19 | 46357724 | 46358214 | 1000 | Promoter (<=1kb) | 19 | 46356939 | 46367978 | 11040 | 1 | ENSMUST00 | 786 | Tmem180 | ENSMUSG0C | false | chr19 | 46356699 | 46358297 | 28.48018 | 82.55319 | 79.35308 | chr19 | 46356343 | 46359337 | 3.73595 | 3.20855 | 2.00706 |
| chr19 | 46453649 | 46454422 | 1000 | Intron (ENSMUST00000039922.7 | 19 | 46457906 | 46475634 | 17729 | 1 | ENSMUST00 | -3484 | Sufu | ENSMUSG0C | false | | | | | | | chr19 | 46452134 | 46463645 | 7.09488 | 4.77154 | 4.75738 |
| chr9 | 59539387 | 59540280 | 1000 | Promoter (<=1kb) | 9 | 59539667 | 59565105 | 25439 | 1 | ENSMUST00 | 0 | Hexa | ENSMUSG0C | false | chr9 | 59539139 | 59540326 | 31.43325 | 99.35611 | 95.79983 | chr9 | 59539082 | 59541022 | 3.68846 | 3.29982 | 1.97715 |
| chr9 | 59617060 | 59617653 | 1000 | Promoter (<=1kb) | 9 | 59617284 | 59650290 | 33007 | 1 | ENSMUST00 | 0 | Parp6 | ENSMUSG0C | false | chr9 | 59616681 | 59618061 | 42.09286 | 140.23274 | 135.34074 | | | | | | |
| chr14 | 7839201 | 7839508 | 1000 | Intron (ENSMUST00000052678.8 | 14 | 7817957 | 7951588 | 133632 | 1 | ENSMUST00 | 21245 | Flnb | ENSMUSG0C | false | | | | | | | chr14 | 7838162 | 7841540 | 4.72856 | 3.91446 | 2.79241 |
| chrX | 155278044 | 155278373 | 1000 | Intron (ENSMUST00000026324.9 | 20 | 155293843 | 155296913 | 3071 | 1 | ENSMUST00 | -15470 | Acot9 | ENSMUSG0C | false | | | | | | | | | | | | |
| chr2 | 90580017 | 90581078 | 1000 | Promoter (<=1kb) | 2 | 90432889 | 90580647 | 147759 | 2 | ENSMUST00 | 0 | Ptprj | ENSMUSG0C | false | chr2 | 90579154 | 90581189 | 29.58769 | 108.11006 | 104.32592 | chr2 | 90580890 | 90584657 | 5.85999 | 4.40403 | 3.72491 |
| chr2 | 90543024 | 90543585 | 1000 | Intron (ENSMUST00000168621.2 | 2 | 90432889 | 90580647 | 147759 | 2 | ENSMUST00 | 37062 | Ptprj | ENSMUSG0C | false | | | | | | | chr2 | 90529589 | 90563488 | 5.67436 | 4.3718 | 3.56584 |
| chr8 | 121950363 | 121950909 | 1000 | Promoter (<=1kb) | 8 | 121950492 | 122026177 | 75686 | 1 | ENSMUST00 | 0 | Banp | ENSMUSG0C | false | chr8 | 121950115 | 121951651 | 26.65955 | 86.95457 | 83.67183 | | | | | | |
| chr12 | 119238552 | 119239088 | 1000 | Promoter (<=1kb) | 12 | 119158022 | 119238302 | 80781 | 2 | ENSMUST00 | 0 | Itgb8 | ENSMUSG0C | false | chr12 | 119236888 | 119239435 | 23.00904 | 72.91611 | 69.8813 | | | | | | |
| chr12 | 119230486 | 119230834 | 1000 | Intron (ENSMUST00000026360.8 | 12 | 119201463 | 119238770 | 37308 | 2 | ENSMUST00 | 7936 | Itgb8 | ENSMUSG0C | false | | | | | | | chr12 | 119229504 | 119231936 | 4.37975 | 3.80807 | 2.50326 |
| chr12 | 118300746 | 118302558 | 1000 | Promoter (<=1kb) | 12 | 118234933 | 118301440 | 66508 | 2 | ENSMUST00 | 0 | Sp4 | ENSMUSG0C | false | chr12 | 118300522 | 118302892 | 46.18486 | 178.62439 | 172.4859 | | | | | | |
| chr7 | 59228404 | 59229606 | 1000 | Promoter (<=1kb) | 7 | 59228752 | 59306727 | 77976 | 1 | ENSMUST00 | 0 | Ube3a | ENSMUSG0C | false | chr7 | 59227911 | 59230196 | 24.92035 | 105.60457 | 101.88994 | chr7 | 59229503 | 59237758 | 4.50519 | 3.85138 | 2.59643 |
| chr5 | 130171582 | 130172214 | 1000 | Promoter (<=1kb) | 5 | 130171798 | 130214342 | 42545 | 1 | ENSMUST00 | 0 | Rabgef1 | ENSMUSG0C | false | chr5 | 130171056 | 130173066 | 30.60796 | 113.09483 | 109.17188 | | | | | | |
| chr10 | 128909594 | 128910453 | 1000 | Promoter (<=1kb) | 10 | 128909866 | 128912816 | 2951 | 1 | ENSMUST00 | 0 | Cd63 | ENSMUSG0C | false | chr10 | 128909451 | 128910702 | 26.41709 | 100.00125 | 96.43182 | chr10 | 128909324 | 128911668 | 3.28049 | 3.18712 | 1.60449 |
| chr10 | 128805302 | 128805795 | 1000 | Promoter (<=1kb) | 10 | 128805676 | 128819446 | 13771 | 1 | ENSMUST00 | 0 | Dnajc14 | ENSMUSG0C | false | chr10 | 128803556 | 128806228 | 31.43325 | 99.35611 | 95.79983 | | | | | | |
| chr10 | 128626646 | 128627035 | 1000 | Promoter (<=1kb) | 10 | 128624530 | 128626506 | 1977 | 2 | ENSMUST00 | -141 | Rps26 | ENSMUSG0C | false | chr10 | 128625655 | 128626818 | 29.09105 | 93.24493 | 89.8345 | | | | | | |
| chr10 | 128458988 | 128459823 | 1000 | Promoter (<=1kb) | 10 | 128459236 | 128490173 | 30938 | 1 | ENSMUST00 | 0 | Smarcc2 | ENSMUSG0C | false | chr10 | 128458440 | 128460755 | 40.9965 | 143.89526 | 138.84959 | | | | | | |
| chr11 | 119942184 | 119943158 | 1000 | Promoter (<=1kb) | 11 | 119942763 | 120002621 | 59859 | 1 | ENSMUST00 | 0 | Baiap2 | ENSMUSG0C | false | chr11 | 119942167 | 119942813 | 28.71633 | 95.52087 | 92.05702 | chr11 | 119937453 | 119942377 | 6.8796 | 4.95571 | 4.54178 |
| chr11 | 120378532 | 120378960 | 1000 | Promoter (<=1kb) | 11 | 120369055 | 120378735 | 9681 | 2 | ENSMUST00 | 0 | 2310003H01 | ENSMUSG0C | false | chr11 | 120377964 | 120379202 | 27.20707 | 96.65539 | 93.16586 | chr11 | 120377430 | 120379478 | 5.12109 | 3.90641 | 3.12801 |
| chr10 | 127665571 | 127666054 | 1000 | Promoter (<=1kb) | 10 | 127664925 | 127665481 | 557 | 2 | ENSMUST00 | -91 | Nab2 | ENSMUSG0C | false | chr10 | 127662238 | 127666785 | 29.66345 | 100.21 | 96.63225 | | | | | | |
| chr10 | 127288443 | 127289495 | 1000 | Promoter (<=1kb) | 10 | 127281956 | 127288890 | 6935 | 2 | ENSMUST00 | 0 | Mbd6 | ENSMUSG0C | false | chr10 | 127287777 | 127290398 | 27.94004 | 83.5095 | 80.29546 | | | | | | |
| chr10 | 126900821 | 126901483 | 1000 | Promoter (<=1kb) | 10 | 126868430 | 126901371 | 32942 | 2 | ENSMUST00 | 0 | Xrcc6bp1 | ENSMUSG0C | false | chr10 | 126900716 | 126901734 | 31.51432 | 94.52584 | 91.08421 | chr10 | 126900637 | 126902095 | 3.89305 | 3.3404 | 2.14343 |
| chr10 | 126928151 | 126928766 | 1000 | Distal Intergenic | 10 | 126887249 | 126901371 | 14123 | 2 | ENSMUST00 | -26781 | Xrcc6bp1 | ENSMUSG0C | false | chr10 | 126927626 | 126929244 | 27.66126 | 83.97217 | 80.74496 | chr10 | 126926132 | 126932821 | 6.41453 | 4.71647 | 4.17073 |
| chr13 | 119428710 | 119429621 | 1000 | Promoter (<=1kb) | 13 | 119428922 | 119458218 | 29297 | 1 | ENSMUST00 | 0 | Paip1 | ENSMUSG0C | false | chr13 | 119428264 | 119430336 | 31.16539 | 106.97112 | 103.21663 | | | | | | |
| chr13 | 119409345 | 119409680 | 1000 | Promoter (<=1kb) | 13 | 119393472 | 119408997 | 15526 | 2 | ENSMUST00 | -349 | Nnt | ENSMUSG0C | false | chr13 | 119408593 | 119409890 | 31.43241 | 100.35837 | 96.77719 | | | | | | |
| chr7 | 141467292 | 141468008 | 1000 | Promoter (<=1kb) | 7 | 141467362 | 141471479 | 4118 | 1 | ENSMUST00 | 0 | Cd151 | ENSMUSG0C | false | chr7 | 141466983 | 141469576 | 34.00537 | 102.80987 | 99.17244 | chr7 | 141456108 | 141486560 | 6.62038 | 4.76228 | 4.33989 |
| chr7 | 141476181 | 141477214 | 1000 | Promoter (<=1kb) | 7 | 141476400 | 141493421 | 17022 | 1 | ENSMUST00 | 0 | Tspan4 | ENSMUSG0C | false | chr7 | 141477003 | 141478311 | 10.0704 | 19.92358 | 17.59046 | chr7 | 141456108 | 141486560 | 6.62038 | 4.76228 | 4.33989 |
| chr5 | 130029174 | 130029713 | 1000 | Promoter (<=1kb) | 5 | 130029290 | 130060789 | 31500 | 1 | ENSMUST00 | 0 | Crcp | ENSMUSG0C | false | chr5 | 130028838 | 130030023 | 31.16539 | 106.97112 | 103.21663 | chr5 | 130028227 | 130031394 | 5.013 | 3.98377 | 3.05368 |
| chr14 | 121052872 | 121053331 | 1000 | Intron (ENSMUST00000135010.2 | 14 | 121035564 | 121283744 | 248181 | 1 | ENSMUST00 | 17309 | Farp1 | ENSMUSG0C | false | | | | | | | chr14 | 121049104 | 121054112 | 4.4634 | 3.79705 | 2.61369 |
| chr14 | 121035420 | 121036978 | 1000 | Promoter (<=1kb) | 14 | 121035564 | 121283744 | 248181 | 1 | ENSMUST00 | 0 | Farp1 | ENSMUSG0C | false | chr14 | 121034923 | 121037682 | 40.93993 | 131.61429 | 127.06391 | chr14 | 121036588 | 121040222 | 3.78661 | 3.39253 | 2.03321 |
| chr14 | 121596534 | 121596908 | 1000 | Intron (ENSMUST00000040700.7 | 14 | 121495621 | 121505254 | 45634 | 2 | ENSMUST00 | -91281 | Slc15a1 | ENSMUSG0C | false | | | | | | | chr14 | 121591562 | 121597928 | 4.46674 | 3.7725 | 2.5655 |
| chr14 | 121797517 | 121798119 | 1000 | Promoter (<=1kb) | 14 | 121542039 | 121797734 | 255696 | 2 | ENSMUST00 | 0 | Dock9 | ENSMUSG0C | false | chr14 | 121797106 | 121798657 | 30.78991 | 105.27288 | 101.56436 | chr14 | 121796689 | 121797551 | 5.26302 | 4.11317 | 3.26617 |
| chr11 | 117654439 | 117655106 | 1000 | Promoter (<=1kb) | 11 | 117654573 | 117763428 | 108856 | 1 | ENSMUST00 | 0 | Tnrc6c | ENSMUSG0C | false | chr11 | 117653596 | 117656293 | 35.34781 | 110.24136 | 106.39787 | | | | | | |
| chr11 | 117779336 | 117780341 | 1000 | Promoter (<=1kb) | 11 | 117765988 | 117779648 | 13661 | 2 | ENSMUST00 | 0 | Tmc6 | ENSMUSG0C | false | chr11 | 117778803 | 117781710 | 25.50448 | 78.30592 | 75.18496 | chr11 | 117776172 | 117784117 | 7.16057 | 5.04978 | 4.77595 |
| chr11 | 118421407 | 118421962 | 1000 | Promoter (2-3kb) | 11 | 118406289 | 118419086 | 12798 | 2 | ENSMUST00 | -2322 | Cant1 | ENSMUSG0C | false | | | | | | | chr11 | 118411596 | 118448618 | 7.23691 | 5.03811 | 4.84329 |
| chr11 | 119040616 | 119041336 | 1000 | Promoter (<=1kb) | 11 | 119036305 | 119040969 | 4665 | 2 | ENSMUST00 | 0 | Cbx8 | ENSMUSG0C | false | chr11 | 119040284 | 119040989 | 32.16725 | 95.65015 | 92.18271 | chr11 | 119034192 | 119045356 | 5.8015 | 4.39225 | 3.68327 |
| chr11 | 119794706 | 119795060 | 1000 | Intron (ENSMUST00000125583.2 | 11 | 119798773 | 119846955 | 48183 | 1 | ENSMUST00 | -3713 | Rptor | ENSMUSG0C | false | | | | | | | chr11 | 119792283 | 119799048 | 4.99119 | 4.00086 | 3.05034 |
| chr11 | 119683268 | 119683633 | 1000 | Intron (ENSMUST00000125583.2 | 11 | 119655944 | 119865810 | 209867 | 1 | ENSMUST00 | 27325 | Rptor | ENSMUSG0C | false | | | | | | | chr11 | 119680454 | 119686890 | 5.62726 | 4.17162 | 3.55435 |
| chr6 | 31563483 | 31564167 | 1000 | Promoter (<=1kb) | 6 | 31519488 | 31563981 | 44494 | 2 | ENSMUST00 | 0 | Podxl | ENSMUSG0C | true | chr6 | 31557739 | 31563996 | 41.04627 | 146.22916 | 141.09622 | | | | | | |
| chr16 | 87698743 | 87699383 | 1000 | Promoter (<=1kb) | 16 | 87698945 | 87733346 | 34402 | 1 | ENSMUST00 | 0 | Bach1 | ENSMUSG0C | false | chr16 | 87698632 | 87700430 | 33.54585 | 101.01101 | 97.41918 | | | | | | |
| chr16 | 87454653 | 87455249 | 1000 | Promoter (<=1kb) | 16 | 87454703 | 87483517 | 28815 | 1 | ENSMUST00 | 0 | Usp16 | ENSMUSG0C | false | chr16 | 87454436 | 87456031 | 38.65522 | 135.28891 | 130.59163 | | | | | | |
| chr9 | 109098919 | 109099473 | 1000 | Promoter (1-2kb) | 9 | 109093844 | 109097796 | 3953 | 2 | ENSMUST00 | -1124 | Gm7628 | ENSMUSG0C | false | chr9 | 109094798 | 109099441 | 38.03245 | 121.19012 | 117.00504 | chr9 | 109096591 | 109107997 | 6.47539 | 4.76075 | 4.21763 |
| chr9 | 109038316 | 109038862 | 1000 | Promoter (<=1kb) | 9 | 109038456 | 109059577 | 21122 | 1 | ENSMUST00 | 0 | Shisa5 | ENSMUSG0C | false | chr9 | 109037832 | 109039925 | 20.74539 | 66.97287 | 64.02668 | chr9 | 109037836 | 109041243 | 5.05646 | 4.05192 | 3.04189 |
| chr17 | 25808776 | 25809456 | 1000 | Promoter (<=1kb) | 17 | 25809085 | 25821244 | 12160 | 1 | ENSMUST00 | 0 | Fbxl16 | ENSMUSG0C | false | chr17 | 25808581 | 25809868 | 23.65566 | 74.08845 | 71.03342 | chr17 | 25808169 | 25810198 | 5.88694 | 4.51487 | 3.72021 |
| chr17 | 25813426 | 25813820 | 1000 | Intron (ENSMUST00000180868.1 | 17 | 25809085 | 25821244 | 12160 | 1 | ENSMUST00 | 4342 | Fbxl16 | ENSMUSG0C | false | | | | | | | chr17 | 25812733 | 25814026 | 4.42382 | 3.77758 | 2.55243 |
| chr5 | 30154733 | 30155500 | 1000 | Promoter (<=1kb) | 5 | 30118301 | 30155162 | 36862 | 2 | ENSMUST00 | 0 | Hadha | ENSMUSG0C | false | chr5 | 30154081 | 30156417 | 32.98857 | 124.89146 | 120.57959 | | | | | | |
| chr3 | 40744951 | 40745791 | 1000 | Promoter (<=1kb) | 3 | 40709382 | 40781598 | 36217 | 1 | ENSMUST00 | 0 | Hspa4l | ENSMUSG0C | false | chr3 | 40744587 | 40746633 | 37.54867 | 136.59039 | 131.88451 | | | | | | |
| chr3 | 41563259 | 41563891 | 1000 | Promoter (<=1kb) | 3 | 41563383 | 41616862 | 53480 | 1 | ENSMUST00 | 0 | Jade1 | ENSMUSG0C | false | chr3 | 41563120 | 41566467 | 31.85236 | 101.09525 | 97.496 | | | | | | |
| chr3 | 41564629 | 41566356 | 1000 | Promoter (<=1kb) | 3 | 41564880 | 41616862 | 51983 | 1 | ENSMUST00 | 0 | Jade1 | ENSMUSG0C | false | chr3 | 41563120 | 41566467 | 31.85236 | 101.09525 | 97.496 | chr3 | 41566113 | 41566734 | 4.13105 | 3.71639 | 2.27618 |
| chr3 | 41582268 | 41582679 | 1000 | Intron (ENSMUST00000126865.5 | 3 | 41564880 | 41616862 | 51983 | 1 | ENSMUST00 | 17389 | Jade1 | ENSMUSG0C | false | | | | | | | chr3 | 41578092 | 41584442 | 3.98127 | 3.43002 | 2.20558 |
| chr2 | 9935988 | 9936282 | 1000 | Intron (ENSMUST00000026888.5 | 2 | 9915291 | 9941751 | 26461 | 2 | ENSMUST00 | 5469 | Taf3 | ENSMUSG0C | false | | | | | | | chr2 | 9935380 | 9936307 | 4.45243 | 3.8765 | 2.54041 |
| chr7 | 74065461 | 74065893 | 1000 | Distal Intergenic | 7 | 73939119 | 74013682 | 74564 | 2 | ENSMUST00 | -51780 | St8sia2 | ENSMUSG0C | false | | | | | | | chr7 | 74055877 | 74067021 | 4.21683 | 3.65298 | 2.3751 |
| chr7 | 74554077 | 74554696 | 1000 | Promoter (<=1kb) | 7 | 74275419 | 74554633 | 279215 | 2 | ENSMUST00 | 0 | Slco3a1 | ENSMUSG0C | false | chr7 | 74553987 | 74555444 | 20.88808 | 77.73511 | 74.61916 | chr7 | 74553891 | 74555730 | 4.58187 | 3.89898 | 2.67227 |
| chr11 | 120498783 | 120499278 | 1000 | Promoter (1-2kb) | 11 | 120497725 | 120498377 | 553 | 1 | ENSMUST00 | 1059 | Slc25a10 | ENSMUSG0C | false | | | | | | | chr11 | 120498222 | 120499875 | 8.23022 | 5.66564 | 5.6364 |
| chr8 | 128685278 | 128686042 | 1000 | Promoter (<=1kb) | 8 | 128685654 | 128733200 | 47547 | 1 | ENSMUST00 | 0 | Itgb1 | ENSMUSG0C | true | chr8 | 128684852 | 128687886 | 35.51585 | 112.61008 | 108.7009 | | | | | | |
| chr8 | 127400535 | 127400875 | 1000 | 3' UTR | 8 | 127405445 | 127433141 | 27697 | 1 | ENSMUST00 | -4570 | Pard3 | ENSMUSG0C | false | | | | | | | chr8 | 127399292 | 127402981 | 4.28842 | 3.73916 | 2.48402 |
| chr8 | 127229507 | 127229926 | 1000 | Intron (ENSMUST00000162309.2 | 8 | 127299585 | 127402577 | 102993 | 1 | ENSMUST00 | -69659 | Pard3 | ENSMUSG0C | false | | | | | | | chr8 | 127217291 | 127236676 | 4.80205 | 3.9295 | 2.85412 |
| chr8 | 127063617 | 127064762 | 1000 | Promoter (<=1kb) | 8 | 127063893 | 127611970 | 548078 | 1 | ENSMUST00 | 0 | Pard3 | ENSMUSG0C | false | chr8 | 127063243 | 127065529 | 31.01414 | 97.6227 | 94.10706 | | | | | | |
| chr8 | 127080373 | 127080772 | 1000 | Intron (ENSMUST00000162309.2 | 8 | 127073240 | 127426684 | 353445 | 1 | ENSMUST00 | 7134 | Pard3 | ENSMUSG0C | false | | | | | | | chr8 | 127076670 | 127082769 | 4.56434 | 3.69108 | 2.68332 |
| chr6 | 47877363 | 47878061 | 1000 | Promoter (<=1kb) | 6 | 47877656 | 47892813 | 15158 | 1 | ENSMUST00 | 0 | Zfp282 | ENSMUSG0C | false | chr6 | 47877134 | 47878849 | 28.53699 | 95.19626 | 91.7378 | | | | | | |
| chr6 | 47812998 | 47813884 | 1000 | Promoter (<=1kb) | 6 | 47796142 | 47813152 | 17371 | 2 | ENSMUST00 | 0 | Pdia4 | ENSMUSG0C | false | chr6 | 47812239 | 47814216 | 31.19258 | 92.98883 | 89.58222 | | | | | | |
| chr5 | 139252176 | 139253095 | 1000 | Promoter (<=1kb) | 5 | 139252324 | 139270051 | 17728 | 1 | ENSMUST00 | 0 | Get4 | ENSMUSG0C | false | chr5 | 139251817 | 139253823 | 20.20131 | 69.382 | 66.39919 | | | | | | |
| chrX | 42067958 | 42068831 | 1000 | Promoter (<=1kb) | 20 | 42068398 | 42109656 | 41259 | 1 | ENSMUST00 | 0 | Xiap | ENSMUSG0C | false | chrX | 42067931 | 42068500 | 14.70503 | 33.66069 | 31.14909 | | | | | | |
| chr13 | 54574926 | 54575476 | 1000 | Promoter (<=1kb) | 13 | 54575015 | 54581128 | 6114 | 1 | ENSMUST00 | 0 | Arl10 | ENSMUSG0C | false | chr13 | 54574538 | 54576156 | 29.2476 | 106.45832 | 102.71833 | | | | | | |
| chr13 | 54661360 | 54661655 | 1000 | 3' UTR | 13 | 54651441 | 54655546 | 4106 | 1 | ENSMUST00 | 9920 | Faf2 | ENSMUSG0C | false | | | | | | | chr13 | 54660184 | 54663353 | 5.27729 | 4.15331 | 3.28111 |
| chr18 | 75368621 | 75369558 | 1000 | Promoter (<=1kb) | 18 | 75368883 | 75395479 | 26597 | 1 | ENSMUST00 | 0 | Smad7 | ENSMUSG0C | false | chr18 | 75366329 | 75373500 | 23.3092 | 84.4034 | 81.16875 | chr18 | 75369277 | 75392875 | 5.74724 | 4.44386 | 3.62103 |
| chr18 | 75370316 | 75370955 | 1000 | Promoter (<=1kb) | 18 | 75369976 | 75394365 | 24390 | 1 | ENSMUST00 | 341 | Smad7 | ENSMUSG0C | false | chr18 | 75366329 | 75373500 | 23.3092 | 84.4034 | 81.16875 | chr18 | 75369277 | 75392875 | 5.74724 | 4.44386 | 3.62103 |
| chr18 | 75366417 | 75368032 | 1000 | Promoter (<=1kb) | 18 | 75367529 | 75395935 | 28407 | 1 | ENSMUST00 | 0 | Smad7 | ENSMUSG0C | false | chr18 | 75366329 | 75373500 | 23.3092 | 84.4034 | 81.16875 | | | | | | |
| chr9 | 4309116 | 4309846 | 1000 | Promoter (<=1kb) | 9 | 4309743 | 4331721 | 21979 | 1 | ENSMUST00 | 0 | Kbtbd3 | ENSMUSG0C | false | chr9 | 4308674 | 4310563 | 29.66345 | 100.21 | 96.63225 | | | | | | |
| chr1 | 9700188 | 9700986 | 1000 | Promoter (<=1kb) | 1 | 9667415 | 9700209 | 32795 | 2 | ENSMUST00 | 0 | Mybl1 | ENSMUSG0C | false | chr1 | 9699653 | 9701230 | 25.84672 | 90.2271 | 86.88143 | | | | | | |
| chr1 | 9797726 | 9798692 | 1000 | Promoter (<=1kb) | 1 | 9798107 | 9900843 | 102737 | 1 | ENSMUST00 | 0 | Sgk3 | ENSMUSG0C | false | chr1 | 9797518 | 9799601 | 35.84351 | 110.06287 | 106.22365 | | | | | | |
| chr1 | 16104292 | 16106037 | 1000 | Promoter (<=1kb) | 1 | 16105774 | 16133734 | 27961 | 1 | ENSMUST00 | 0 | Rdh10 | ENSMUSG0C | false | chr1 | 16103566 | 16106735 | 36.88168 | 122.38801 | 118.16834 | | | | | | |
| chr1 | 64120851 | 64121997 | 1000 | Promoter (<=1kb) | 1 | 64029447 | 64121491 | 92045 | 2 | ENSMUST00 | 0 | Klf7 | ENSMUSG0C | true | chr1 | 64119053 | 64121494 | 21.42557 | 69.98924 | 66.99868 | | | | | | |
| chr1 | 64107435 | 64108033 | 1000 | Intron (ENSMUST00000114086.2 | 1 | 64041926 | 64121482 | 79557 | 2 | ENSMUST00 | 13449 | Klf7 | ENSMUSG0C | true | | | | | | | chr1 | 64074337 | 64110188 | 4.96577 | 4.0468 | 2.98015 |
| chr1 | 63534305 | 63534794 | 1000 | Promoter (2-3kb) | 1 | 63536967 | 63540473 | 3507 | 1 | ENSMUST00 | -2173 | Adam23 | ENSMUSG0C | false | | | | | | | chr1 | 63531930 | 63535491 | 5.14638 | 4.11455 | 3.17286 |
| chr1 | 62703003 | 62703786 | 1000 | Promoter (<=1kb) | 1 | 62703285 | 62739932 | 36648 | 1 | ENSMUST00 | 0 | Nrp2 | ENSMUSG0C | false | chr1 | 62702681 | 62704220 | 30.99688 | 91.67474 | 88.2998 | chr1 | 62702750 | 62703833 | 4.14697 | 3.70229 | 2.29286 |
| chr1 | 55087695 | 55088453 | 1000 | Promoter (<=1kb) | 1 | 55077835 | 55088024 | 10190 | 2 | ENSMUST00 | 0 | Hspd1 | ENSMUSG0C | false | chr1 | 55086793 | 55088783 | 35.24744 | 113.18649 | 109.26083 | | | | | | |
| chr1 | 55052314 | 55052682 | 1000 | Promoter (<=1kb) | 1 | 55052770 | 55071953 | 19184 | 1 | ENSMUST00 | -88 | Coq10b | ENSMUSG0C | false | chr1 | 55052067 | 55054172 | 31.62822 | 118.12274 | 114.03909 | chr1 | 55051595 | 55052397 | 3.46365 | 3.21627 | 1.77134 |
| chr1 | 55052931 | 55053427 | 1000 | Promoter (<=1kb) | 1 | 55052869 | 55054093 | 1225 | 1 | ENSMUST00 | 63 | Coq10b | ENSMUSG0C | false | chr1 | 55052067 | 55054172 | 31.62822 | 118.12274 | 114.03909 | | | | | | |
| chr1 | 46852705 | 46854238 | 1000 | Promoter (<=1kb) | 1 | 46807544 | 46854048 | 46503 | 2 | ENSMUST00 | 0 | Slc39a10 | ENSMUSG0C | false | chr1 | 46853640 | 46854715 | 23.88927 | 69.11851 | 66.14216 | | | | | | |
| chr1 | 46823861 | 46824177 | 1000 | Intron (ENSMUST00000027131.5 | 1 | 46817602 | 46835899 | 18298 | 2 | ENSMUST00 | 11722 | Slc39a10 | ENSMUSG0C | false | | | | | | | chr1 | 46822688 | 46825673 | 5.15565 | 4.16126 | 3.12928 |
| chr1 | 69686728 | 69687903 | 1000 | Promoter (<=1kb) | 1 | 69538582 | 69687245 | 148664 | 2 | ENSMUST00 | 0 | Ikzf2 | ENSMUSG0C | false | chr1 | 69685182 | 69688560 | 36.31549 | 112.95179 | 109.03448 | | | | | | |
| chr1 | 67043960 | 67044327 | 1000 | Distal Intergenic | 1 | 67000517 | 67038872 | 38356 | 2 | ENSMUST00 | -5089 | Lancl1 | ENSMUSG0C | false | | | | | | | | | | | | |
| chr1 | 66862655 | 66863592 | 1000 | Promoter (<=1kb) | 1 | 66830839 | 66863277 | 32439 | 2 | ENSMUST00 | 0 | Acadl | ENSMUSG0C | false | chr1 | 66862165 | 66864406 | 20.109 | 88.33971 | 85.03291 | chr1 | 66861739 | 66862707 | 4.46152 | 3.72984 | 2.58968 |
| chr1 | 66816351 | 66817878 | 1000 | Promoter (<=1kb) | 1 | 66719363 | 66817546 | 98184 | 2 | ENSMUST00 | 0 | Kansl1l | ENSMUSG0C | false | chr1 | 66815744 | 66818353 | 39.77934 | 136.18091 | 131.45303 | | | | | | |
| chr1 | 60097972 | 60098785 | 1000 | Promoter (<=1kb) | 1 | 60098247 | 60151032 | 52786 | 1 | ENSMUST00 | 0 | Carf | ENSMUSG0C | false | chr1 | 60097058 | 60099158 | 34.78613 | 113.42423 | 109.48744 | | | | | | |
| chr1 | 58422999 | 58423371 | 1000 | Promoter (<=1kb) | 1 | 58410189 | 58424066 | 13878 | 2 | ENSMUST00 | 695 | Clk1 | ENSMUSG0C | false | chr1 | 58421831 | 58424072 | 32.94367 | 134.34938 | 129.69266 | | | | | | |
| chr1 | 39900423 | 39901478 | 1000 | Promoter (<=1kb) | 1 | 39900913 | 40026310 | 125398 | 1 | ENSMUST00 | 0 | Map4k4 | ENSMUSG0C | false | chr1 | 39900215 | 39903023 | 36.76257 | 113.72284 | 109.77938 | | | | | | |
| chr1 | 39877651 | 39878303 | 1000 | Distal Intergenic | 1 | 39900913 | 40026310 | 125398 | 1 | ENSMUST00 | -22610 | Map4k4 | ENSMUSG0C | false | | | | | | | chr1 | 39877583 | 39877980 | 4.97772 | 3.67671 | 3.01848 |
| chr1 | 39192636 | 39193153 | 1000 | Promoter (<=1kb) | 1 | 39193731 | 39287615 | 93885 | 1 | ENSMUST00 | -578 | Npas2 | ENSMUSG0C | false | chr1 | 39192448 | 39192740 | 7.7331 | 13.70582 | 11.46404 | chr1 | 39191875 | 39193145 | 5.19027 | 4.18822 | 3.14326 |
| chr1 | 39193340 | 39194311 | 1000 | Promoter (<=1kb) | 1 | 39193731 | 39287615 | 93885 | 1 | ENSMUST00 | 0 | Npas2 | ENSMUSG0C | false | chr1 | 39192818 | 39194810 | 34.27164 | 115.65593 | 111.648 | | | | | | |
| chr1 | 38129410 | 38130122 | 1000 | Promoter (<=1kb) | 1 | 38052787 | 38129662 | 76876 | 2 | ENSMUST00 | 0 | Rev1 | ENSMUSG0C | false | chr1 | 38128784 | 38130448 | 28.56742 | 103.16996 | 99.51957 | | | | | | |
| chr1 | 37719465 | 37720155 | 1000 | Promoter (<=1kb) | 1 | 37611677 | 37720085 | 108409 | 2 | ENSMUST00 | 0 | 2010300C02 | ENSMUSG0C | false | chr1 | 37719170 | 37720553 | 28.90751 | 104.81161 | 101.11721 | chr1 | 37718376 | 37720815 | 3.24524 | 3.17821 | 1.55126 |
| chr1 | 36920114 | 36920565 | 1000 | Intron (ENSMUST00000027290.6 | 1 | 36792194 | 36939527 | 147334 | 2 | ENSMUST00 | 18962 | Tmem131 | ENSMUSG0C | false | | | | | | | chr1 | 36912955 | 36935642 | 6.00762 | 4.46937 | 3.851 |
| chr1 | 36934068 | 36934394 | 1000 | Intron (ENSMUST00000027290.6 | 1 | 36792194 | 36939527 | 147334 | 2 | ENSMUST00 | 5133 | Tmem131 | ENSMUSG0C | false | | | | | | | chr1 | 36912955 | 36935642 | 6.00762 | 4.46937 | 3.851 |
| chr1 | 36555925 | 36558445 | 1000 | Promoter (<=1kb) | 1 | 36548642 | 36558381 | 9740 | 2 | ENSMUST00 | 0 | Sema4c | ENSMUSG0C | false | chr1 | 36555717 | 36559135 | 31.54088 | 108.67455 | 104.87239 | chr1 | 36553805 | 36557197 | 5.79041 | 4.37481 | 3.71177 |
| chr1 | 34132907 | 34133312 | 1000 | Intron (ENSMUST00000182513.2 | 1 | 34121476 | 34164237 | 42762 | 1 | ENSMUST00 | 11432 | Dst | ENSMUSG0C | true | | | | | | | chr1 | 34126917 | 34135308 | 5.76734 | 4.2451 | 3.65458 |
| chr1 | 34114259 | 34114731 | 1000 | Intron (ENSMUST00000182513.2 | 1 | 34121476 | 34164237 | 42762 | 1 | ENSMUST00 | -6745 | Dst | ENSMUSG0C | true | | | | | | | chr1 | 34109258 | 34115357 | 5.04006 | 4.12901 | 3.03082 |
| chr1 | 34011087 | 34012281 | 1000 | Promoter (<=1kb) | 1 | 34012822 | 34308661 | 296840 | 1 | ENSMUST00 | 0 | Dst | ENSMUSG0C | true | chr1 | 34010872 | 34012641 | 32.67761 | 108.66347 | 104.86716 | | | | | | |
| chr1 | 34089064 | 34089486 | 1000 | Promoter (<=1kb) | 1 | 34089292 | 34119153 | 29862 | 1 | ENSMUST00 | 0 | Dst | ENSMUSG0C | true | | | | | | | chr1 | 34088296 | 34092016 | 4.09824 | 3.46506 | 2.30058 |
| chr1 | 23382843 | 23383614 | 1000 | Promoter (<=1kb) | 1 | 23366424 | 23383175 | 16752 | 2 | ENSMUST00 | 0 | Ogfrl1 | ENSMUSG0C | false | chr1 | 23382281 | 23383443 | 25.17393 | 93.69764 | 90.27328 | chr1 | 23381904 | 23384022 | 4.77311 | 3.97076 | 2.81045 |
| chr1 | 74794442 | 74794787 | 1000 | Promoter (<=1kb) | 1 | 74793363 | 74803495 | 10133 | 1 | ENSMUST00 | 1080 | Wnt10a | ENSMUSG0C | false | chr1 | 74794379 | 74795025 | 7.35251 | 12.67314 | 10.44663 | chr1 | 74794410 | 74795190 | 4.23631 | 3.3073 | 2.42063 |
| chr1 | 74391246 | 74392373 | 1000 | Promoter (<=1kb) | 1 | 74391509 | 74397285 | 5777 | 1 | ENSMUST00 | 0 | Ctdsp1 | ENSMUSG0C | false | chr1 | 74390907 | 74393162 | 38.14117 | 119.25307 | 115.13219 | | | | | | |
| chr1 | 74386282 | 74387358 | 1000 | Promoter (2-3kb) | 1 | 74383600 | 74385197 | 1598 | 1 | ENSMUST00 | 2683 | Slc11a1 | ENSMUSG0C | false | chr1 | 74386524 | 74387500 | 22.32483 | 64.62918 | 61.71925 | chr1 | 74386378 | 74391163 | 4.3995 | 3.8026 | 2.50604 |
| chr1 | 75381865 | 75382501 | 1000 | Promoter (<=1kb) | 1 | 75382114 | 75388345 | 6232 | 1 | ENSMUST00 | 0 | Speg | ENSMUSG0C | true | chr1 | 75381728 | 75383103 | 23.65566 | 74.08845 | 71.03342 | chr1 | 75381009 | 75384668 | 4.13538 | 3.53379 | 2.3139 |
| chr1 | 75384301 | 75385674 | 1000 | Promoter (<=1kb) | 1 | 75384700 | 75387948 | 3249 | 1 | ENSMUST00 | 0 | Speg | ENSMUSG0C | true | chr1 | 75384399 | 75387322 | 24.11245 | 66.85627 | 63.91418 | chr1 | 75381009 | 75384668 | 4.13538 | 3.53379 | 2.3139 |
| chr1 | 75387168 | 75388026 | 1000 | Promoter (1-2kb) | 1 | 75385610 | 75432304 | 46695 | 1 | ENSMUST00 | 1559 | Speg | ENSMUSG0C | true | chr1 | 75384399 | 75387322 | 24.11245 | 66.85627 | 63.91418 | chr1 | 75386249 | 75400648 | 7.0728 | 5.02664 | 4.70473 |
| chr1 | 75355969 | 75356870 | 1000 | Promoter (<=1kb) | 1 | 75360329 | 75368579 | 8251 | 1 | ENSMUST00 | 0 | Des | ENSMUSG0C | false | | | | | | | chr1 | 75355622 | 75362384 | 5.80475 | 4.47779 | 3.65868 |
| chr1 | 86111801 | 86112089 | 1000 | Promoter (<=1kb) | 1 | 86099026 | 86111970 | 12945 | 2 | ENSMUST00 | 0 | Htr2b | ENSMUSG0C | false | | | | | | | chr1 | 86110542 | 86112721 | 4.74905 | 3.94947 | 2.79786 |
| chr1 | 86526133 | 86527918 | 1000 | Promoter (<=1kb) | 1 | 86526726 | 86530712 | 3987 | 1 | ENSMUST00 | 0 | Ptma | ENSMUSG0C | false | chr1 | 86525464 | 86529269 | 41.35843 | 143.41548 | 138.39708 | | | | | | |
| chr1 | 86565239 | 86565803 | 1000 | Intron (ENSMUST00000027444.9 | 1 | 86543520 | 86582517 | 38998 | 2 | ENSMUST00 | 16714 | Pde6d | ENSMUSG0C | false | | | | | | | chr1 | 86564962 | 86566749 | 5.23729 | 4.09068 | 3.22251 |

| | | | | | | | | | | | | | | | | | | | | | | | | | | |
|---|---|---|---|---|---|---|---|---|---|---|---|---|---|---|---|---|---|---|---|---|---|---|---|---|---|---|
| chr1 | 93478131 | 93479270 | 1000 | Promoter (<=1kb) | 1 | 93478964 | 93510260 | 31297 | 1 | ENSMUST00 | 0 | Sep-02 | ENSMUSG0C | false | chr1 | 93477516 | 93480357 | 31.43325 | 99.35611 | 95.79983 | | | | | | |
| chr1 | 93685369 | 93685731 | 1000 | Promoter (<=1kb) | 1 | 93685660 | 93695764 | 10105 | 1 | ENSMUST00 | 0 | Bok | ENSMUSG0C | false | chr1 | 93685234 | 93686792 | 23.13664 | 67.80279 | 64.84689 | chr1 | 93685377 | 93687202 | 3.48118 | 3.17875 | 1.7988 |
| chr1 | 88403375 | 88403794 | 1000 | Distal Intergenic | 1 | 88406961 | 88426438 | 19478 | 1 | ENSMUST00 | -3167 | Spp2 | ENSMUSG0C | false | | | | | | | chr1 | 88399071 | 88408669 | 4.86164 | 3.89722 | 2.91633 |
| chr1 | 91052851 | 91053538 | 1000 | Promoter (<=1kb) | 1 | 91053444 | 91128944 | 75501 | 1 | ENSMUST00 | 0 | Lrrfip1 | ENSMUSG0C | true | chr1 | 91051607 | 91058713 | 31.85236 | 101.09525 | 97.496 | | | | | | |
| chr1 | 91077761 | 91078291 | 1000 | Intron (ENSMUST00000189505.1 | 1 | 91053586 | 91128287 | 74702 | 1 | ENSMUST00 | 24176 | Lrrfip1 | ENSMUSG0C | true | chr1 | 91078153 | 91078453 | 6.20072 | 10.05428 | 7.87816 | chr1 | 91055222 | 91105818 | 7.56473 | 5.21453 | 5.11657 |
| chr1 | 91094184 | 91094686 | 1000 | Intron (ENSMUST00000189505.1 | 1 | 91111214 | 91116921 | 5708 | 1 | ENSMUST00 | -16530 | Lrrfip1 | ENSMUSG0C | true | | | | | | | chr1 | 91055222 | 91105818 | 7.56473 | 5.21453 | 5.11657 |
| chr1 | 91061019 | 91061369 | 1000 | Intron (ENSMUST00000189505.1 | 1 | 91053586 | 91128287 | 74702 | 1 | ENSMUST00 | 7434 | Lrrfip1 | ENSMUSG0C | true | | | | | | | chr1 | 91055222 | 91105818 | 7.56473 | 5.21453 | 5.11657 |
| chr1 | 91059879 | 91060745 | 1000 | Intron (ENSMUST00000189505.1 | 1 | 91053586 | 91128287 | 74702 | 1 | ENSMUST00 | 6294 | Lrrfip1 | ENSMUSG0C | true | | | | | | | chr1 | 91055222 | 91105818 | 7.56473 | 5.21453 | 5.11657 |
| chr1 | 92179571 | 92181197 | 1000 | Promoter (<=1kb) | 1 | 91928779 | 92180341 | 251563 | 2 | ENSMUST00 | 0 | Hdac4 | ENSMUSG0C | false | chr1 | 92180566 | 92181428 | 27.03504 | 88.5911 | 85.27629 | chr1 | 92177024 | 92179782 | 6.0152 | 4.45666 | 3.84635 |
| chr8 | 14873710 | 14874016 | 1000 | Distal Intergenic | 8 | 14881335 | 14894716 | 13382 | 1 | ENSMUST00 | -7319 | Cln8 | ENSMUSG0C | false | | | | | | | | | | | | |
| chr1 | 97765767 | 97766243 | 1000 | Intron (ENSMUST00000042509.7 | 1 | 97770172 | 97792969 | 22798 | 1 | ENSMUST00 | -3929 | Gin1 | ENSMUSG0C | false | | | | | | | chr1 | 97759244 | 97769003 | 4.59138 | 3.86323 | 2.67265 |
| chr1 | 97976852 | 97977438 | 1000 | Promoter (<=1kb) | 1 | 97821100 | 97977255 | 156156 | 2 | ENSMUST00 | 0 | Pam | ENSMUSG0C | false | chr1 | 97976586 | 97977623 | 11.92996 | 37.32891 | 34.77106 | | | | | | |
| chr1 | 125434840 | 125435974 | 1000 | Promoter (<=1kb) | 1 | 125392905 | 125435727 | 42823 | 2 | ENSMUST00 | 0 | Actr3 | ENSMUSG0C | false | chr1 | 125435824 | 125436528 | 8.8423 | 19.93079 | 17.59637 | | | | | | |
| chr1 | 125592950 | 125593229 | 1000 | Intron (ENSMUST00000027580.5 | 1 | 125584513 | 125595148 | 10636 | 1 | ENSMUST00 | 8438 | Slc35f5 | ENSMUSG0C | false | | | | | | | chr1 | 125591834 | 125593588 | 3.02054 | 2.97214 | 1.41847 |
| chr1 | 125819166 | 125819555 | 1000 | Intron (ENSMUST00000027581.6 | 1 | 125840003 | 125873173 | 33171 | 1 | ENSMUST00 | -20448 | Gpr39 | ENSMUSG0C | false | | | | | | | chr1 | 125818051 | 125819930 | 3.07955 | 3.05828 | 1.48317 |
| chr1 | 127697801 | 127698132 | 1000 | Distal Intergenic | 1 | 127490898 | 127679548 | 188651 | 2 | ENSMUST00 | -18254 | Tmem163 | ENSMUSG0C | false | | | | | | | | | | | | |
| chr1 | 127812725 | 127813198 | 1000 | Downstream (2-3kb) | 1 | 127796571 | 127808061 | 11491 | 1 | ENSMUST00 | 16155 | Ccnt2 | ENSMUSG0C | false | | | | | | | | | | | | |
| chr1 | 143503701 | 143504111 | 1000 | Distal Intergenic | 1 | 143633851 | 143636577 | 2727 | 2 | ENSMUST00 | 132466 | Cdc73 | ENSMUSG0C | false | chr1 | 143503168 | 143503845 | 4.29207 | 7.17162 | 5.05983 | | | | | | |
| chr1 | 119624470 | 119624837 | 1000 | Intron (ENSMUST00000052404.7 | 1 | 119617655 | 119634000 | 16346 | 2 | ENSMUST00 | 9163 | Epb4.1l5 | ENSMUSG0C | false | | | | | | | chr1 | 119619960 | 119646490 | 4.72133 | 3.93355 | 2.78285 |
| chr1 | 119626780 | 119627203 | 1000 | Intron (ENSMUST00000052404.7 | 1 | 119617655 | 119634000 | 16346 | 2 | ENSMUST00 | 6797 | Epb4.1l5 | ENSMUSG0C | false | | | | | | | chr1 | 119619960 | 119646490 | 4.72133 | 3.93355 | 2.78285 |
| chr1 | 119836964 | 119837833 | 1000 | Promoter (<=1kb) | 1 | 119652467 | 119837613 | 185147 | 2 | ENSMUST00 | 0 | Ptpn4 | ENSMUSG0C | false | chr1 | 119836203 | 119837955 | 38.7176 | 123.44448 | 119.18625 | | | | | | |
| chr1 | 120176183 | 120176663 | 1000 | Intron (ENSMUST00000027637.7 | 1 | 120160480 | 120173566 | 13087 | 1 | ENSMUST00 | 15704 | 3110009E18F | ENSMUSG0C | false | | | | | | | chr1 | 120176280 | 120176602 | 3.25137 | 3.00897 | 1.59696 |
| chr1 | 120226082 | 120226457 | 1000 | 3' UTR | 1 | 120190757 | 120265255 | 74499 | 2 | ENSMUST00 | 38798 | Steap3 | ENSMUSG0C | false | | | | | | | chr1 | 120220892 | 120232141 | 6.47152 | 4.73253 | 4.21633 |
| chr1 | 120223592 | 120223947 | 1000 | Intron (ENSMUST00000140490.2 | 1 | 120190757 | 120265255 | 74499 | 2 | ENSMUST00 | 41308 | Steap3 | ENSMUSG0C | false | | | | | | | chr1 | 120220892 | 120232141 | 6.47152 | 4.73253 | 4.21633 |
| chr1 | 130455205 | 130455497 | 1000 | Promoter (2-3kb) | 1 | 130446661 | 130452416 | 5756 | 2 | ENSMUST00 | -2790 | Cd55 | ENSMUSG0C | false | | | | | | | chr1 | 130454416 | 130457212 | 4.01862 | 3.43866 | 2.23507 |
| chr1 | 136415152 | 136415844 | 1000 | Promoter (<=1kb) | 1 | 136415271 | 136417792 | 2522 | 1 | ENSMUST00 | 0 | Ddx59 | ENSMUSG0C | false | chr1 | 136414780 | 136415964 | 17.28728 | 67.4904 | 64.54075 | | | | | | |
| chr1 | 136011644 | 136012333 | 1000 | Intron (ENSMUST00000165125.2 | 1 | 136008219 | 136035030 | 26812 | 1 | ENSMUST00 | 3426 | Tmem9 | ENSMUSG0C | false | | | | | | | chr1 | 136010191 | 136020472 | 4.34132 | 3.68886 | 2.47402 |
| chr1 | 135739297 | 135739781 | 1000 | 5' UTR | 1 | 135729147 | 135752232 | 23086 | 1 | ENSMUST00 | 10151 | Csrp1 | ENSMUSG0C | false | | | | | | | chr1 | 135739364 | 135741558 | 3.96261 | 3.64046 | 2.14039 |
| chr1 | 135146555 | 135147627 | 1000 | Promoter (<=1kb) | 1 | 135146824 | 135156269 | 9446 | 1 | ENSMUST00 | 0 | Arl8a | ENSMUSG0C | false | chr1 | 135146032 | 135147917 | 42.26237 | 140.52768 | 135.6286 | | | | | | |
| chr1 | 132476908 | 132477691 | 1000 | Promoter (<=1kb) | 1 | 132477365 | 132504490 | 27126 | 1 | ENSMUST00 | 0 | Rbbp5 | ENSMUSG0C | false | chr1 | 132476575 | 132478169 | 32.69058 | 104.59052 | 100.90593 | | | | | | |
| chr1 | 133712378 | 133713281 | 1000 | Intron (ENSMUST00000143567.2 | 1 | 133716323 | 133717929 | 1607 | 2 | ENSMUST00 | 4648 | Atp2b4 | ENSMUSG0C | false | | | | | | | chr1 | 133710712 | 133713642 | 4.44558 | 3.60857 | 2.58097 |
| chr1 | 133695939 | 133696401 | 1000 | Distal Intergenic | 1 | 133701639 | 133701881 | 243 | 2 | ENSMUST00 | 5480 | Atp2b4 | ENSMUSG0C | false | | | | | | | chr1 | 133689861 | 133697286 | 7.10741 | 4.96538 | 4.74647 |
| chr1 | 156035919 | 156036760 | 1000 | Promoter (<=1kb) | 1 | 156004599 | 156036480 | 31882 | 2 | ENSMUST00 | 0 | Tor1aip1 | ENSMUSG0C | false | chr1 | 156034962 | 156037742 | 30.60796 | 113.09483 | 109.17188 | | | | | | |
| chr1 | 155416780 | 155417866 | 1000 | Promoter (<=1kb) | 1 | 155275701 | 155417309 | 141609 | 2 | ENSMUST00 | 0 | Xpr1 | ENSMUSG0C | false | chr1 | 155416315 | 155417702 | 39.81545 | 135.14195 | 130.45284 | | | | | | |
| chr1 | 155158512 | 155159506 | 1000 | Promoter (<=1kb) | 1 | 155158703 | 155203517 | 44815 | 1 | ENSMUST00 | 0 | Stx6 | ENSMUSG0C | false | chr1 | 155158626 | 155159615 | 31.10056 | 111.96791 | 108.07632 | chr1 | 155159461 | 155160615 | 4.58695 | 3.87053 | 2.69672 |
| chr1 | 153266906 | 153267299 | 1000 | Intron (ENSMUST00000027752.9 | 1 | 153248617 | 153250461 | 1845 | 2 | ENSMUST00 | -16446 | Lamc1 | ENSMUSG0C | false | | | | | | | chr1 | 153262519 | 153267853 | 5.74044 | 4.34534 | 3.65009 |
| chr1 | 152809684 | 152810297 | 1000 | Promoter (<=1kb) | 1 | 152810890 | 152824291 | 13402 | 1 | ENSMUST00 | -593 | Ncf2 | ENSMUSG0C | true | | | | | | | chr1 | 152801274 | 152821507 | 4.847 | 3.96927 | 2.88719 |
| chr1 | 152799911 | 152800889 | 1000 | Promoter (<=1kb) | 1 | 152800194 | 152836991 | 36798 | 1 | ENSMUST00 | 0 | Ncf2 | ENSMUSG0C | true | chr1 | 152800193 | 152801982 | 14.44046 | 33.88361 | 31.37022 | chr1 | 152788034 | 152800060 | 6.38067 | 4.71623 | 4.13727 |
| chr1 | 152641243 | 152641635 | 1000 | Intron (ENSMUST00000111859.2 | 1 | 152516951 | 152625217 | 108267 | 2 | ENSMUST00 | -16027 | Rgl1 | ENSMUSG0C | false | | | | | | | chr1 | 152640105 | 152641821 | 5.71815 | 4.15119 | 3.61493 |
| chr1 | 152609555 | 152609980 | 1000 | Intron (ENSMUST00000111859.2 | 1 | 152557543 | 152625029 | 67487 | 2 | ENSMUST00 | 15049 | Rgl1 | ENSMUSG0C | false | | | | | | | chr1 | 152606016 | 152624201 | 4.72366 | 3.90653 | 2.80197 |
| chr1 | 151709022 | 151709339 | 1000 | Exon (ENSMUST00000148810.2/ | 1 | 151677167 | 151719342 | 42176 | 1 | ENSMUST00 | 31856 | Fam129a | ENSMUSG0C | false | | | | | | | chr1 | 151708542 | 151711678 | 5.58747 | 4.37812 | 3.48937 |
| chr1 | 151571183 | 151571614 | 1000 | Promoter (<=1kb) | 1 | 151571186 | 151644898 | 73713 | 1 | ENSMUST00 | 0 | Fam129a | ENSMUSG0C | false | chr1 | 151571107 | 151571892 | 22.31642 | 65.48328 | 62.55884 | chr1 | 151570798 | 151571960 | 3.79403 | 3.46119 | 2.05566 |
| chr1 | 151500162 | 151501254 | 1000 | Promoter (<=1kb) | 1 | 151469405 | 151500811 | 31407 | 2 | ENSMUST00 | 0 | Rnf2 | ENSMUSG0C | false | chr1 | 151499350 | 151501598 | 39.06023 | 122.966 | 118.72274 | | | | | | |
| chr1 | 179959375 | 179960635 | 1000 | Promoter (<=1kb) | 1 | 179960472 | 180163156 | 202685 | 1 | ENSMUST00 | 0 | Cdc42bpa | ENSMUSG0C | false | chr1 | 179959117 | 179962379 | 36.31549 | 112.95179 | 109.03448 | chr1 | 179958717 | 179959481 | 3.12091 | 3.05174 | 1.44918 |
| chr1 | 178405628 | 178406718 | 1000 | Promoter (<=1kb) | 1 | 178405881 | 178483247 | 77367 | 1 | ENSMUST00 | 0 | Efcab2 | ENSMUSG0C | false | chr1 | 178405272 | 178406881 | 32.66734 | 113.81544 | 109.86345 | | | | | | |
| chr1 | 180568760 | 180569291 | 1000 | Promoter (<=1kb) | 1 | 180568955 | 180601253 | 32299 | 1 | ENSMUST00 | 0 | Parp1 | ENSMUSG0C | false | chr1 | 180568491 | 180569887 | 28.90751 | 104.81161 | 101.11721 | | | | | | |
| chr1 | 178318093 | 178319211 | 1000 | Promoter (<=1kb) | 1 | 178319130 | 178322335 | 3206 | 1 | ENSMUST00 | 0 | Cox20 | ENSMUSG0C | false | chr1 | 178317405 | 178320432 | 27.92142 | 85.37866 | 82.12785 | | | | | | |
| chr1 | 182408781 | 182409582 | 1000 | Promoter (<=1kb) | 1 | 182409172 | 182462432 | 53261 | 1 | ENSMUST00 | 0 | Trp53bp2 | ENSMUSG0C | false | chr1 | 182408150 | 182410996 | 46.39859 | 154.48299 | 149.02768 | | | | | | |
| chr1 | 182460847 | 182461291 | 1000 | Intron (ENSMUST00000117245.1 | 1 | 182409172 | 182462432 | 53261 | 1 | ENSMUST00 | 51676 | Trp53bp2 | ENSMUSG0C | false | | | | | | | chr1 | 182456749 | 182463620 | 6.24432 | 4.67121 | 4.01623 |
| chr1 | 181143901 | 181144493 | 1000 | Promoter (<=1kb) | 1 | 181093423 | 181144157 | 50735 | 2 | ENSMUST00 | 0 | Nvl | ENSMUSG0C | false | chr1 | 181143562 | 181145514 | 30.41442 | 103.5799 | 99.9162 | | | | | | |
| chr1 | 172436679 | 172437251 | 1000 | Distal Intergenic | 1 | 172475358 | 172478575 | 3218 | 1 | ENSMUST00 | -38107 | Slamf9 | ENSMUSG0C | false | | | | | | | chr1 | 172428189 | 172437787 | 6.0765 | 4.59609 | 3.89569 |
| chr1 | 172169769 | 172170168 | 1000 | Intron (ENSMUST00000074144.5 | 1 | 172148015 | 172196393 | 48379 | 1 | ENSMUST00 | 21755 | Dcaf8 | ENSMUSG0C | false | | | | | | | chr1 | 172167155 | 172175493 | 5.04442 | 4.07617 | 3.05469 |
| chr1 | 172147769 | 172148183 | 1000 | Promoter (<=1kb) | 1 | 172148015 | 172196393 | 48379 | 1 | ENSMUST00 | 0 | Dcaf8 | ENSMUSG0C | false | chr1 | 172147589 | 172149840 | 28.36679 | 125.90208 | 121.5666 | | | | | | |
| chr1 | 167269950 | 167270287 | 1000 | Intron (ENSMUST00000027839.8 | 1 | 167226090 | 167285127 | 59038 | 1 | ENSMUST00 | 14840 | Uck2 | ENSMUSG0C | true | | | | | | | chr1 | 167266805 | 167272027 | 6.87977 | 4.85774 | 4.55209 |
| chr1 | 167284235 | 167285378 | 1000 | Promoter (<=1kb) | 1 | 167226090 | 167285127 | 59038 | 2 | ENSMUST00 | 0 | Uck2 | ENSMUSG0C | true | chr1 | 167283561 | 167285860 | 36.53154 | 123.91184 | 119.64436 | chr1 | 167285263 | 167287397 | 6.041 | 4.2974 | 3.88441 |
| chr1 | 166001578 | 166003627 | 1000 | Promoter (<=1kb) | 1 | 165875457 | 166002609 | 127153 | 2 | ENSMUST00 | 0 | Pou2f1 | ENSMUSG0C | false | chr1 | 166002909 | 166003759 | 35.70928 | 138.63979 | 133.81488 | chr1 | 165989861 | 166001740 | 5.21788 | 4.20224 | 3.18435 |
| chr1 | 166000727 | 166001491 | 1000 | Promoter (1-2kb) | 1 | 165875457 | 166002609 | 127153 | 2 | ENSMUST00 | 1118 | Pou2f1 | ENSMUSG0C | false | chr1 | 166000437 | 166002688 | 31.91637 | 110.38313 | 106.53261 | chr1 | 165989861 | 166001740 | 5.21788 | 4.20224 | 3.18435 |
| chr1 | 165536126 | 165536654 | 1000 | Intron (ENSMUST00000111440.2 | 1 | 165543104 | 165551306 | 8203 | 1 | ENSMUST00 | -6450 | Adcy10 | ENSMUSG0C | false | | | | | | | chr1 | 165535220 | 165538794 | 4.5506 | 3.7509 | 2.65503 |
| chr1 | 156938848 | 156939857 | 1000 | Promoter (<=1kb) | 1 | 156804166 | 156939602 | 135437 | 2 | ENSMUST00 | 0 | Ralgps2 | ENSMUSG0C | false | chr1 | 156938566 | 156940198 | 31.43325 | 99.35611 | 95.79983 | chr1 | 156939741 | 156940217 | 2.83583 | 2.86695 | 1.26405 |
| chr1 | 156558561 | 156559677 | 1000 | Promoter (<=1kb) | 1 | 156558786 | 156625376 | 66591 | 1 | ENSMUST00 | 0 | Abl2 | ENSMUSG0C | false | chr1 | 156558296 | 156559937 | 29.28796 | 98.53321 | 94.99628 | | | | | | |
| chr1 | 156309145 | 156309691 | 1000 | Promoter (1-2kb) | 1 | 156310727 | 156328035 | 17309 | 1 | ENSMUST00 | -1036 | Nphs2 | ENSMUSG0C | true | | | | | | | chr1 | 156307023 | 156309151 | 5.03822 | 4.14188 | 3.02784 |
| chr1 | 189883386 | 189883697 | 1000 | Promoter (1-2kb) | 1 | 189880642 | 189882372 | 1731 | 2 | ENSMUST00 | -1015 | Smyd2 | ENSMUSG0C | false | | | | | | | chr1 | 189882207 | 189883993 | 4.01143 | 3.29808 | 2.23338 |
| chr1 | 189921791 | 189922622 | 1000 | Promoter (<=1kb) | 1 | 189880492 | 189922363 | 41872 | 2 | ENSMUST00 | 0 | Smyd2 | ENSMUSG0C | false | chr1 | 189921639 | 189922858 | 30.84885 | 99.33006 | 95.77879 | chr1 | 189921022 | 189923343 | 4.90167 | 3.9307 | 2.96327 |
| chr1 | 189750207 | 189750775 | 1000 | Intron (ENSMUST00000097442.3 | 1 | 189728268 | 189876689 | 148422 | 1 | ENSMUST00 | 21940 | Ptpn14 | ENSMUSG0C | true | | | | | | | chr1 | 189730267 | 189810107 | 6.93269 | 4.87829 | 4.60521 |
| chr1 | 189726105 | 189726518 | 1000 | Promoter (1-2kb) | 1 | 189728268 | 189876689 | 148422 | 1 | ENSMUST00 | -1750 | Ptpn14 | ENSMUSG0C | true | | | | | | | chr1 | 189725312 | 189727709 | 6.43393 | 4.82858 | 4.16176 |
| chr1 | 189860935 | 189861400 | 1000 | Promoter (1-2kb) | 1 | 189862659 | 189863433 | 775 | 1 | ENSMUST00 | -1259 | Ptpn14 | ENSMUSG0C | true | | | | | | | chr1 | 189860050 | 189861575 | 5.8146 | 4.278 | 3.71424 |
| chr1 | 189727342 | 189728719 | 1000 | Promoter (<=1kb) | 1 | 189728268 | 189876689 | 148422 | 1 | ENSMUST00 | 0 | Ptpn14 | ENSMUSG0C | true | chr1 | 189727043 | 189728064 | 11.4883 | 23.92001 | 21.53215 | chr1 | 189725312 | 189727709 | 6.43393 | 4.82858 | 4.16176 |
| chr1 | 189778356 | 189778734 | 1000 | Intron (ENSMUST00000097442.3 | 1 | 189786633 | 189876695 | 90063 | 1 | ENSMUST00 | -7899 | Ptpn14 | ENSMUSG0C | true | | | | | | | chr1 | 189730267 | 189810107 | 6.93269 | 4.87829 | 4.60521 |
| chr1 | 185287112 | 185287497 | 1000 | Intron (ENSMUST00000027929.4 | 1 | 185285569 | 185288282 | 2714 | 1 | ENSMUST00 | 785 | Iars2 | ENSMUSG0C | false | | | | | | | chr1 | 185286918 | 185287503 | 3.65081 | 3.37335 | 1.88391 |
| chr1 | 184995199 | 184995533 | 1000 | Intron (ENSMUST00000027929.4 | 1 | 184886443 | 184999549 | 103107 | 2 | ENSMUST00 | 4016 | Mark1 | ENSMUSG0C | false | | | | | | | chr1 | 184994929 | 184996042 | 4.2827 | 3.70322 | 2.42268 |
| chr1 | 191717793 | 191718901 | 1000 | Promoter (<=1kb) | 1 | 191717834 | 191719398 | 1565 | 1 | ENSMUST00 | 0 | Lpgat1 | ENSMUSG0C | false | chr1 | 191717578 | 191719447 | 33.65816 | 114.56059 | 110.59126 | | | | | | |
| chr1 | 191396148 | 191397494 | 1000 | Promoter (<=1kb) | 1 | 191351981 | 191397041 | 45061 | 2 | ENSMUST00 | 0 | Ppp2r5a | ENSMUSG0C | false | chr1 | 191395099 | 191398253 | 42.9543 | 141.72932 | 136.78545 | | | | | | |
| chr1 | 191465184 | 191465460 | 1000 | Distal Intergenic | 1 | 191351981 | 191397041 | 45061 | 2 | ENSMUST00 | -68144 | Ppp2r5a | ENSMUSG0C | false | | | | | | | chr1 | 191457318 | 191471879 | 4.86599 | 4.03205 | 2.90022 |
| chr1 | 191470599 | 191470955 | 1000 | Distal Intergenic | 1 | 191351981 | 191397041 | 45061 | 2 | ENSMUST00 | -73559 | Ppp2r5a | ENSMUSG0C | false | | | | | | | chr1 | 191457318 | 191471879 | 4.86599 | 4.03205 | 2.90022 |
| chr1 | 191183258 | 191183794 | 1000 | Promoter (<=1kb) | 1 | 191170296 | 191183340 | 13045 | 2 | ENSMUST00 | 0 | Atf3 | ENSMUSG0C | false | chr1 | 191182292 | 191186167 | 31.08358 | 101.75523 | 98.14187 | | | | | | |
| chr1 | 190928418 | 190929551 | 1000 | Promoter (<=1kb) | 1 | 190928492 | 190938435 | 9944 | 1 | ENSMUST00 | 0 | Angel2 | ENSMUSG0C | false | chr1 | 190927800 | 190929765 | 33.31514 | 101.62023 | 98.00909 | chr1 | 190929480 | 190930832 | 3.95744 | 3.35513 | 2.18307 |
| chr1 | 194795305 | 194795756 | 1000 | Intron (ENSMUST00000027952.6 | 1 | 194804710 | 194814227 | 9518 | 1 | ENSMUST00 | -8954 | Plxna2 | ENSMUSG0C | false | | | | | | | chr1 | 194792408 | 194796164 | 5.30049 | 4.0042 | 3.277 |
| chr2 | 4559444 | 4560105 | 1000 | Promoter (<=1kb) | 2 | 4559754 | 4597989 | 38236 | 1 | ENSMUST00 | 0 | Frmd4a | ENSMUSG0C | false | chr2 | 4558843 | 4560357 | 23.75302 | 64.50008 | 61.59195 | | | | | | |
| chr2 | 4881473 | 4882603 | 1000 | Promoter (<=1kb) | 2 | 4881564 | 4910557 | 28994 | 1 | ENSMUST00 | 0 | Sephs1 | ENSMUSG0C | false | chr2 | 4880757 | 4883313 | 40.19742 | 128.15663 | 123.73233 | | | | | | |
| chr2 | 5053069 | 5053375 | 1000 | Exon (ENSMUST00000114996.2/ | 2 | 5020644 | 5063938 | 43295 | 2 | ENSMUST00 | 10563 | Optn | ENSMUSG0C | true | | | | | | | chr2 | 5041736 | 5062961 | 5.08721 | 4.02119 | 3.09837 |
| chr1 | 167802653 | 167803413 | 1000 | Intron (ENSMUST00000111377.2 | 1 | 167689557 | 167848741 | 159185 | 1 | ENSMUST00 | 113097 | Lmx1a | ENSMUSG0C | false | chr1 | 167802629 | 167803394 | 12.24318 | 32.0257 | 29.53485 | chr1 | 167802693 | 167807845 | 5.01097 | 4.10905 | 3.01062 |
| chr1 | 162570362 | 162571014 | 1000 | Promoter (<=1kb) | 1 | 162570515 | 162599084 | 28570 | 1 | ENSMUST00 | 0 | Vamp4 | ENSMUSG0C | true | chr1 | 162569974 | 162571928 | 32.61975 | 142.21701 | 137.24615 | | | | | | |
| chr1 | 161413310 | 161413913 | 1000 | Intron (ENSMUST00000028024.4 | 1 | 161395409 | 161418410 | 23002 | 1 | ENSMUST00 | 17902 | Tnfsf4 | ENSMUSG0C | false | | | | | | | chr1 | 161413203 | 161414372 | 3.22922 | 3.14182 | 1.60538 |
| chr2 | 13544126 | 13544806 | 1000 | Promoter (<=1kb) | 2 | 13509014 | 13544664 | 35651 | 2 | ENSMUST00 | 0 | Trdmt1 | ENSMUSG0C | false | chr2 | 13543589 | 13545024 | 26.27418 | 74.13734 | 71.08211 | chr2 | 13543514 | 13545903 | 3.77504 | 3.46504 | 1.9846 |
| chr2 | 13372022 | 13372610 | 1000 | Intron (ENSMUST00000091436.5 | 2 | 13445021 | 13455624 | 10604 | 2 | ENSMUST00 | 83014 | Cubn | ENSMUSG0C | false | | | | | | | chr2 | 13370643 | 13373711 | 6.22319 | 4.67348 | 4.04606 |
| chr2 | 19037522 | 19038054 | 1000 | Distal Intergenic | 2 | 18998343 | 18999801 | 1459 | 1 | ENSMUST00 | 39180 | 4930426L09F | ENSMUSG0C | false | | | | | | | | | | | | |
| chr2 | 18913080 | 18913391 | 1000 | Intron (ENSMUST00000006912.6 | 2 | 18847497 | 18906274 | 58778 | 2 | ENSMUST00 | -6807 | Pip4k2a | ENSMUSG0C | false | | | | | | | chr2 | 18912756 | 18913227 | 3.23007 | 3.05553 | 1.58796 |
| chr2 | 18676194 | 18677846 | 1000 | Promoter (<=1kb) | 2 | 18677018 | 18686629 | 9612 | 1 | ENSMUST00 | 0 | Bmi1 | ENSMUSG0C | false | chr2 | 18676027 | 18678781 | 33.42125 | 102.81408 | 99.17643 | chr2 | 18675837 | 18676333 | 4.13276 | 3.37412 | 2.32646 |
| chr2 | 18680797 | 18681336 | 1000 | Promoter (<=1kb) | 2 | 18680994 | 18683030 | 2037 | 1 | ENSMUST00 | 0 | Bmi1 | ENSMUSG0C | false | chr2 | 18680675 | 18681355 | 10.47775 | 22.22469 | 19.86031 | chr2 | 18678073 | 18681251 | 3.54817 | 3.2383 | 1.84176 |
| chr2 | 18064076 | 18065686 | 1000 | Promoter (<=1kb) | 2 | 18064394 | 18122348 | 57955 | 1 | ENSMUST00 | 0 | Mllt10 | ENSMUSG0C | false | chr2 | 18063641 | 18065839 | 37.36699 | 117.91644 | 113.84495 | chr2 | 18065601 | 18068867 | 3.87188 | 3.32856 | 2.1109 |
| chr2 | 16738225 | 16738684 | 1000 | Intron (ENSMUST00000028081.7 | 2 | 16638696 | 16751690 | 112995 | 1 | ENSMUST00 | 99530 | Plxdc2 | ENSMUSG0C | false | | | | | | | chr2 | 16737358 | 16740367 | 4.50316 | 3.92059 | 2.58461 |
| chr2 | 16450914 | 16451560 | 1000 | Intron (ENSMUST00000028081.7 | 2 | 16357117 | 16751857 | 394741 | 1 | ENSMUST00 | 93798 | Plxdc2 | ENSMUSG0C | false | | | | | | | chr2 | 16441603 | 16456820 | 4.67386 | 3.87664 | 2.74338 |
| chr2 | 16355879 | 16356754 | 1000 | Promoter (<=1kb) | 2 | 16356304 | 16755839 | 399536 | 1 | ENSMUST00 | 0 | Plxdc2 | ENSMUSG0C | false | chr2 | 16355582 | 16356803 | 33.72937 | 104.36225 | 100.686 | chr2 | 16355500 | 16356137 | 4.39573 | 3.77814 | 2.50243 |
| chr2 | 38647484 | 38648142 | 1000 | Distal Intergenic | 2 | 38588036 | 38644087 | 56052 | 2 | ENSMUST00 | -3398 | Psmb7 | ENSMUSG0C | false | | | | | | | chr2 | 38641933 | 38648528 | 6.14313 | 4.52664 | 3.9704 |
| chr2 | 39007876 | 39008398 | 1000 | Promoter (<=1kb) | 2 | 39008076 | 39015877 | 7802 | 1 | ENSMUST00 | 0 | Arpc5l | ENSMUSG0C | false | chr2 | 39007436 | 39009098 | 36.59948 | 124.03223 | 119.76238 | | | | | | |
| chr2 | 48948959 | 48950240 | 1000 | Promoter (<=1kb) | 2 | 48902824 | 48949276 | 46453 | 2 | ENSMUST00 | 0 | Orc4 | ENSMUSG0C | false | chr2 | 48948386 | 48950830 | 38.55812 | 129.64772 | 125.15668 | | | | | | |
| chr1 | 60409216 | 60410372 | 1000 | Promoter (<=1kb) | 1 | 60409619 | 60481158 | 71540 | 1 | ENSMUST00 | 0 | Abi2 | ENSMUSG0C | false | chr1 | 60409123 | 60410773 | 34.49972 | 105.69308 | 101.97717 | chr1 | 60408911 | 60409266 | 2.95527 | 2.89808 | 1.35513 |
| chr2 | 32906947 | 32907675 | 1000 | Intron (ENSMUST00000028135.9 | 2 | 32910908 | 32912587 | 1680 | 1 | ENSMUST00 | -3233 | Fam129b | ENSMUSG0C | false | | | | | | | chr2 | 32877160 | 32918190 | 7.77045 | 5.19056 | 5.2936 |
| chr2 | 32914927 | 32915372 | 1000 | Intron (ENSMUST00000028135.9 | 2 | 32914711 | 32921024 | 6314 | 1 | ENSMUST00 | 217 | Fam129b | ENSMUSG0C | false | | | | | | | chr2 | 32877160 | 32918190 | 7.77045 | 5.19056 | 5.2936 |
| chr2 | 32856131 | 32856666 | 1000 | Distal Intergenic | 2 | 32787602 | 32847239 | 59638 | 2 | ENSMUST00 | -8893 | Stxbp1 | ENSMUSG0C | true | | | | | | | chr2 | 32854131 | 32859041 | 4.82363 | 3.96907 | 2.86865 |
| chr2 | 32846808 | 32847509 | 1000 | Promoter (<=1kb) | 2 | 32787602 | 32847239 | 59638 | 2 | ENSMUST00 | 0 | Stxbp1 | ENSMUSG0C | true | chr2 | 32845892 | 32847353 | 20.14837 | 78.81087 | 75.68276 | chr2 | 32847436 | 32849706 | 4.90633 | 4.08733 | 2.91164 |
| chr2 | 29525292 | 29525644 | 1000 | Distal Intergenic | 2 | 29504871 | 29524681 | 19811 | 1 | ENSMUST00 | 20422 | Med27 | ENSMUSG0C | false | | | | | | | chr2 | 29524461 | 29527515 | 4.02551 | 3.40888 | 2.23655 |
| chr2 | 28840034 | 28840773 | 1000 | Promoter (<=1kb) | 2 | 28840406 | 28847511 | 7106 | 1 | ENSMUST00 | 0 | Ddx31 | ENSMUSG0C | false | chr2 | 28839265 | 28841606 | 36.73285 | 125.34267 | 121.02122 | | | | | | |
| chr2 | 32606775 | 32607981 | 1000 | Promoter (<=1kb) | 2 | 32606961 | 32620804 | 13844 | 1 | ENSMUST00 | 0 | St6galnac6 | ENSMUSG0C | true | chr2 | 32606837 | 32611447 | 41.10367 | 138.50676 | 133.686 | chr2 | 32605009 | 32639275 | 6.86759 | 4.92928 | 4.53618 |
| chr2 | 32608422 | 32609098 | 1000 | Promoter (<=1kb) | 2 | 32608464 | 32620804 | 12341 | 1 | ENSMUST00 | 0 | St6galnac6 | ENSMUSG0C | true | chr2 | 32606837 | 32611447 | 41.10367 | 138.50676 | 133.686 | chr2 | 32605009 | 32639275 | 6.86759 | 4.92928 | 4.53618 |
| chr2 | 32616975 | 32617472 | 1000 | Intron (ENSMUST00000140983.2 | 2 | 32621758 | 32635058 | 13301 | 1 | ENSMUST00 | -4286 | Ak1 | ENSMUSG0C | false | | | | | | | chr2 | 32605009 | 32639275 | 6.86759 | 4.92928 | 4.53618 |
| chr2 | 32637791 | 32638348 | 1000 | Intron (ENSMUST00000127603.1 | 2 | 32631167 | 32646805 | 15639 | 1 | ENSMUST00 | 6625 | Ak1 | ENSMUSG0C | false | | | | | | | chr2 | 32605009 | 32639275 | 6.86759 | 4.92928 | 4.53618 |
| chr2 | 32625208 | 32625584 | 1000 | Promoter (<=1kb) | 2 | 32625437 | 32635054 | 9618 | 1 | ENSMUST00 | 0 | Ak1 | ENSMUSG0C | false | chr2 | 32625128 | 32627071 | 9.13756 | 26.51177 | 24.09012 | chr2 | 32605009 | 32639275 | 6.86759 | 4.92928 | 4.53618 |
| chr2 | 57237835 | 57238584 | 1000 | Promoter (<=1kb) | 2 | 57238297 | 57370719 | 132423 | 1 | ENSMUST00 | 0 | Gpd2 | ENSMUSG0C | false | chr2 | 57237553 | 57239193 | 29.88806 | 96.63187 | 93.14587 | | | | | | |
| chr2 | 28205478 | 28205817 | 1000 | Promoter (<=1kb) | 2 | 28205659 | 28214425 | 8767 | 1 | ENSMUST00 | 0 | Olfm1 | ENSMUSG0C | false | chr2 | 28205247 | 28206399 | 21.37461 | 59.55079 | 56.71094 | chr2 | 28204455 | 28206564 | 4.38036 | 3.63785 | 2.53503 |
| chr2 | 58516807 | 58517245 | 1000 | Promoter (<=1kb) | 2 | 58446434 | 58516272 | 69839 | 2 | ENSMUST00 | -536 | Acvr1 | ENSMUSG0C | false | | | | | | | chr2 | 58513168 | 58518712 | 4.38962 | 3.79661 | 2.50474 |
| chr2 | 58566552 | 58567789 | 1000 | Promoter (<=1kb) | 2 | 58446441 | 58566826 | 120386 | 2 | ENSMUST00 | 0 | Acvr1 | ENSMUSG0C | false | chr2 | 58566062 | 58567838 | 35.20524 | 115.20684 | 111.21615 | chr2 | 58564379 | 58566673 | 4.43859 | 3.87739 | 2.53336 |
| chr2 | 31771093 | 31771510 | 1000 | Intron (ENSMUST00000142554.2 | 2 | 31760173 | 31779024 | 18852 | 1 | ENSMUST00 | 10921 | Abl1 | ENSMUSG0C | false | | | | | | | chr2 | 31760789 | 31776841 | 6.98405 | 4.98586 | 4.62933 |
| chr2 | 31749237 | 31749696 | 1000 | Intron (ENSMUST00000142554.2 | 2 | 31759943 | 31804227 | 44285 | 1 | ENSMUST00 | -10247 | Abl1 | ENSMUSG0C | false | | | | | | | chr2 | 31740620 | 31752695 | 6.24045 | 4.61237 | 4.04053 |
| chr2 | 30981872 | 30982516 | 1000 | Promoter (<=1kb) | 2 | 30972833 | 30981941 | 9109 | 2 | ENSMUST00 | 0 | BC005624 | ENSMUSG0C | false | chr2 | 30981257 | 30983331 | 26.32471 | 106.6389 | 102.89503 | chr2 | 30980611 | 30987237 | 6.32613 | 4.73432 | 4.08648 |
| chr2 | 31025411 | 31025956 | 1000 | Downstream (<1kb) | 2 | 31019819 | 31021381 | 1563 | 1 | ENSMUST00 | 5593 | Usp20 | ENSMUSG0C | false | | | | | | | chr2 | 31024280 | 31028152 | 6.17587 | 4.51305 | 4.02148 |
| chr2 | 30390464 | 30390916 | 1000 | Promoter (1-2kb) | 2 | 30392254 | 30400529 | 8276 | 1 | ENSMUST00 | -1338 | Dolpp1 | ENSMUSG0C | false | | | | | | | chr2 | 30389556 | 30392102 | 6.42071 | 4.83408 | 4.15444 |
| chr2 | 35258252 | 35258881 | 1000 | Distal Intergenic | 2 | 35279663 | 35293567 | 13905 | 1 | ENSMUST00 | -20782 | Gsn | ENSMUSG0C | true | chr2 | 35255869 | 35258912 | 37.27983 | 118.82364 | 114.72062 | chr2 | 35257475 | 35289539 | 6.87726 | 4.86078 | 4.55899 |
| chr2 | 35272518 | 35272843 | 1000 | Distal Intergenic | 2 | 35279663 | 35293567 | 13905 | 1 | ENSMUST00 | -6820 | Gsn | ENSMUSG0C | true | | | | | | | chr2 | 35257475 | 35289539 | 6.87726 | 4.86078 | 4.55899 |
| chr2 | 35270425 | 35270947 | 1000 | Distal Intergenic | 2 | 35279663 | 35293567 | 13905 | 1 | ENSMUST00 | -8716 | Gsn | ENSMUSG0C | true | | | | | | | chr2 | 35257475 | 35289539 | 6.87726 | 4.86078 | 4.55899 |
| chr2 | 35302515 | 35303003 | 1000 | Exon (ENSMUST00000028239.6/ | 2 | 35282651 | 35284828 | 2178 | 1 | ENSMUST00 | 19865 | Gsn | ENSMUSG0C | true | | | | | | | chr2 | 35298059 | 35305401 | 6.1769 | 4.48478 | 4.00912 |
| chr2 | 35256166 | 35256538 | 1000 | Distal Intergenic | 2 | 35279663 | 35293567 | 13905 | 1 | ENSMUST00 | -23125 | Gsn | ENSMUSG0C | true | chr2 | 35255869 | 35258912 | 37.27983 | 118.82364 | 114.72062 | | | | | | |
| chr2 | 35271771 | 35272172 | 1000 | Distal Intergenic | 2 | 35279663 | 35293567 | 13905 | 1 | ENSMUST00 | -7491 | Gsn | ENSMUSG0C | true | | | | | | | chr2 | 35257475 | 35289539 | 6.87726 | 4.86078 | 4.55899 |
| chr2 | 35621819 | 35622209 | 1000 | Promoter (<=1kb) | 2 | 35621981 | 35730992 | 109012 | 1 | ENSMUST00 | 0 | Dab2ip | ENSMUSG0C | false | chr2 | 35621712 | 35622768 | 21.74415 | 57.67707 | 54.86403 | chr2 | 35621446 | 35624679 | 3.46766 | 3.27166 | 1.7613 |
| chr2 | 26122361 | 26123080 | 1000 | Promoter (<=1kb) | 2 | 26055535 | 26122818 | 67284 | 2 | ENSMUST00 | 0 | Nacc2 | ENSMUSG0C | false | chr2 | 26121900 | 26123466 | 25.50663 | 88.63404 | 85.31516 | chr2 | 26113529 | 26124879 | 5.32832 | 4.16544 | 3.29828 |
| chr2 | 25546566 | 25547045 | 1000 | Promoter (<=1kb) | 2 | 25517982 | 25546940 | 28959 | 2 | ENSMUST00 | 0 | Traf2 | ENSMUSG0C | false | chr2 | 25546078 | 25547023 | 26.22066 | 76.7888 | 73.6919 | chr2 | 25544324 | 25547714 | 3.93492 | 3.49381 | 2.14495 |
| chr2 | 25428451 | 25428892 | 1000 | Promoter (<=1kb) | 2 | 25428703 | 25448540 | 19838 | 1 | ENSMUST00 | 0 | Abca2 | ENSMUSG0C | false | chr2 | 25428578 | 25430241 | 29.28796 | 98.53321 | 94.99628 | chr2 | 25427945 | 25432516 | 4.59668 | 3.73174 | 2.70916 |
| chr2 | 60962022 | 60963672 | 1000 | Promoter (<=1kb) | 2 | 60750193 | 60963192 | 213000 | 2 | ENSMUST00 | 0 | Rbms1 | ENSMUSG0C | true | chr2 | 60961189 | 60963342 | 36.17514 | 115.85 | 111.8395 | | | | | | |
| chr2 | 60893012 | 60893542 | 1000 | Intron (ENSMUST00000028347.7 | 2 | 60842297 | 60889030 | 46734 | 2 | ENSMUST00 | -3983 | Rbms1 | ENSMUSG0C | true | | | | | | | chr2 | 60890567 | 60932725 | 5.60936 | 4.28041 | 3.52606 |
| chr2 | 60913289 | 60913735 | 1000 | Intron (ENSMUST00000028347.7 | 2 | 60842297 | 60889030 | 46734 | 2 | ENSMUST00 | -24260 | Rbms1 | ENSMUSG0C | true | | | | | | | chr2 | 60890567 | 60932725 | 5.60936 | 4.28041 | 3.52606 |
| chr2 | 60880971 | 60881834 | 1000 | Promoter (<=1kb) | 2 | 60751953 | 60881438 | 129486 | 2 | ENSMUST00 | 0 | Rbms1 | ENSMUSG0C | true | chr2 | 60879546 | 60881897 | 39.78054 | 131.78781 | 127.22884 | chr2 | 60881615 | 60885197 | 3.88249 | 3.34299 | 2.11842 |
| chr2 | 24416729 | 24417538 | 1000 | Distal Intergenic | 2 | 24421394 | 24423082 | 1689 | 1 | ENSMUST00 | 5544 | Pax8 | ENSMUSG0C | false | | | | | | | chr2 | 24416123 | 24419481 | 7.36874 | 5.15599 | 4.92704 |
| chr2 | 60209269 | 60210557 | 1000 | Promoter (<=1kb) | 2 | 60209887 | 60249211 | 39325 | 1 | ENSMUST00 | 0 | | 39142 | ENSMUSG0C false | chr2 | 60208946 | 60211053 | 35.8325 | 117.35963 | 113.30182 | | | | | | |
| chr2 | 59908944 | 59909310 | 1000 | Intron (ENSMUST00000112550.2 | 2 | 59903185 | 59912536 | 9352 | 2 | ENSMUST00 | 3226 | Baz2b | ENSMUSG0C | false | | | | | | | chr2 | 59908018 | 59910001 | 5.38891 | 4.20374 | 3.35822 |
| chr2 | 59908486 | 59908851 | 1000 | Intron (ENSMUST00000112550.2 | 2 | 59903185 | 59912536 | 9352 | 2 | ENSMUST00 | 3685 | Baz2b | ENSMUSG0C | false | | | | | | | chr2 | 59908018 | 59910001 | 5.38891 | 4.20374 | 3.35822 |
| chr2 | 60124605 | 60125689 | 1000 | Promoter (<=1kb) | 2 | 60004549 | 60125592 | 121044 | 2 | ENSMUST00 | 0 | Baz2b | ENSMUSG0C | false | chr2 | 60123837 | 60126567 | 34.90036 | 108.43528 | 104.64344 | | | | | | |
| chr2 | 80580639 | 80581763 | 1000 | Promoter (<=1kb) | 2 | 80500512 | 80581264 | 80753 | 2 | ENSMUST00 | 0 | Nckap1 | ENSMUSG0C | false | chr2 | 80579951 | 80581855 | 33.7938 | 119.00099 | 114.88904 | | | | | | |
| chr2 | 80315323 | 80315900 | 1000 | Promoter (<=1kb) | 2 | 80315466 | 80324788 | 9323 | 1 | ENSMUST00 | 0 | Dnajc10 | ENSMUSG0C | false | chr2 | 80314850 | 80316280 | 35.8325 | 117.35963 | 113.30182 | | | | | | |
| chr2 | 79634761 | 79636236 | 1000 | Promoter (<=1kb) | 2 | 79635352 | 79672966 | 37615 | 1 | ENSMUST00 | 0 | Ssfa2 | ENSMUSG0C | false | chr2 | 79635281 | 79637748 | 36.04672 | 129.50026 | 125.02953 | chr2 | 79634542 | 79634780 | 3.2652 | 3.25085 | 1.56529 |
| chr2 | 79601660 | 79602307 | 1000 | Distal Intergenic | 2 | 79635352 | 79672966 | 37615 | 1 | ENSMUST00 | -33045 | Ssfa2 | ENSMUSG0C | false | | | | | | | chr2 | 79597578 | 79605221 | 5.34068 | 4.26617 | 3.27267 |
| chr2 | 78868541 | 78869968 | 1000 | Promoter (<=1kb) | 2 | 78869047 | 78921293 | 52247 | 1 | ENSMUST00 | 0 | Ube2e3 | ENSMUSG0C | false | chr2 | 78868289 | 78870641 | 32.77523 | 100.68713 | 97.10149 | | | | | | |
| chr2 | 77926565 | 77926855 | 1000 | Promoter (2-3kb) | 2 | 77895766 | 77924528 | 28763 | 2 | ENSMUST00 | -2038 | Cwc22 | ENSMUSG0C | false | | | | | | | chr2 | 77925120 | 77928489 | 4.62512 | 3.88231 | 2.76109 |
| chr2 | 71146286 | 71146719 | 1000 | Intron (ENSMUST00000111831.2 | 2 | 71117923 | 71142926 | 25004 | 2 | ENSMUST00 | 28364 | Cybrd1 | ENSMUSG0C | false | | | | | | | chr2 | 71145769 | 71148129 | 4.63881 | 3.72682 | 2.74889 |
| chr2 | 77746494 | 77746843 | 1000 | Intron (ENSMUST00000111831.2 | 2 | 77410634 | 77719739 | 309106 | 2 | ENSMUST00 | -26756 | Zfp385b | ENSMUSG0C | true | | | | | | | chr2 | 77739513 | 77748524 | 4.386 | 3.76738 | 2.50742 |
| chr2 | 77777813 | 77778315 | 1000 | Intron (ENSMUST00000111831.2 | 2 | 77411100 | 77816816 | 405717 | 2 | ENSMUST00 | 38501 | Zfp385b | ENSMUSG0C | true | | | | | | | chr2 | 77767132 | 77782197 | 4.901 | 4.03491 | 2.92527 |
| chr2 | 77815775 | 77817266 | 1000 | Promoter (<=1kb) | 2 | 77411100 | 77816816 | 405717 | 2 | ENSMUST00 | 0 | Zfp385b | ENSMUSG0C | true | chr2 | 77814761 | 77817865 | 29.66345 | 100.21 | 96.63225 | | | | | | |
| chr2 | 68471045 | 68472433 | 1000 | Promoter (<=1kb) | 2 | 68210445 | 68472268 | 261824 | 2 | ENSMUST00 | 0 | Stk39 | ENSMUSG0C | false | chr2 | 68470868 | 68472728 | 28.24444 | 85.94312 | 82.67722 | chr2 | 68470804 | 68471527 | 5.5833 | 4.47434 | 3.47381 |
| chr2 | 68294576 | 68295069 | 1000 | Intron (ENSMUST00000102715.3 | 2 | 68211740 | 68251356 | 39617 | 2 | ENSMUST00 | -43221 | Stk39 | ENSMUSG0C | false | | | | | | | chr2 | 68294057 | 68295252 | 3.15572 | 3.10029 | 1.47759 |
| chr2 | 68873743 | 68874424 | 1000 | Intron (ENSMUST00000028426.3 | 2 | 68861586 | 69108644 | 247059 | 1 | ENSMUST00 | 12158 | Cers6 | ENSMUSG0C | true | | | | | | | chr2 | 68863137 | 68874811 | 5.56542 | 4.31322 | 3.46908 |
| chr2 | 68871308 | 68871923 | 1000 | Intron (ENSMUST00000028426.3 | 2 | 68861586 | 69108644 | 247059 | 1 | ENSMUST00 | 9723 | Cers6 | ENSMUSG0C | true | | | | | | | chr2 | 68863137 | 68874811 | 5.56542 | 4.31322 | 3.46908 |
| chr2 | 83723851 | 83724987 | 1000 | Promoter (<=1kb) | 2 | 83724397 | 83756797 | 32401 | 1 | ENSMUST00 | 0 | Itgav | ENSMUSG0C | false | chr2 | 83723513 | 83725691 | 36.83125 | 118.04518 | 113.96772 | chr2 | 83722849 | 83723926 | 6.87961 | 4.53604 | 4.58852 |
| chr2 | 74877230 | 74877684 | 1000 | 3' UTR | 2 | 74868981 | 74876504 | 7524 | 1 | ENSMUST00 | 8250 | Mtx2 | ENSMUSG0C | false | | | | | | | chr2 | 74876682 | 74877448 | 4.1845 | 3.64109 | 2.36705 |
| chr2 | 74704623 | 74706129 | 1000 | Promoter (<=1kb) | 2 | 74704905 | 74707933 | 3029 | 1 | ENSMUST00 | 0 | Hoxd8 | ENSMUSG0C | false | chr2 | 74704033 | 74706369 | 28.93225 | 84.27598 | 81.04385 | chr2 | 74706083 | 74707555 | 3.44102 | 3.31626 | 1.70848 |
| chr2 | 72978653 | 72980547 | 1000 | Promoter (<=1kb) | 2 | 72936427 | 72980446 | 44020 | 2 | ENSMUST00 | 0 | Sp3 | ENSMUSG0C | false | chr2 | 72977775 | 72981250 | 47.79134 | 159.19145 | 153.57301 | | | | | | |
| chr2 | 105003593 | 105003992 | 1000 | Promoter (2-3kb) | 2 | 104999659 | 105006823 | 7165 | 2 | ENSMUST00 | 2831 | Eif3m | ENSMUSG0C | false | | | | | | | chr2 | 104988432 | 105014256 | 4.21481 | 3.66036 | 2.36618 |
| chr2 | 104711692 | 104712534 | 1000 | Promoter (<=1kb) | 2 | 104678017 | 104712156 | 34140 | 2 | ENSMUST00 | 0 | Tcp11l1 | ENSMUSG0C | false | chr2 | 104711578 | 104712863 | 21.26469 | 95.5017 | 92.03861 | chr2 | 104710641 | 104713231 | 3.28802 | 3.23988 | 1.61394 |
| chr2 | 104493174 | 104494709 | 1000 | Promoter (<=1kb) | 2 | 104426481 | 104494446 | 67966 | 2 | ENSMUST00 | 0 | Hipk3 | ENSMUSG0C | false | chr2 | 104492244 | 104495149 | 37.91348 | 118.86035 | 114.75604 | | | | | | |
| chr2 | 104027503 | 104028001 | 1000 | Promoter (<=1kb) | 2 | 104027721 | 104063240 | 35520 | 1 | ENSMUST00 | 0 | Fbxo3 | ENSMUSG0C | false | chr2 | 104027441 | 104029300 | 40.51682 | 136.37238 | 131.6409 | chr2 | 104027008 | 104032574 | 4.26272 | 3.69895 | 2.38834 |
| chr2 | 103463791 | 103464273 | 1000 | Intron (ENSMUST00000028610.4 | 2 | 103448738 | 103449823 | 1086 | 1 | ENSMUST00 | 15054 | Elf5 | ENSMUSG0C | false | | | | | | | chr2 | 103460288 | 103464996 | 6.02312 | 4.48461 | 3.8784 |
| chr2 | 102552103 | 102552571 | 1000 | Promoter (1-2kb) | 2 | 102550149 | 102635648 | 85500 | 1 | ENSMUST00 | 1955 | Pamr1 | ENSMUSG0C | false | | | | | | | chr2 | 102549829 | 102553160 | 4.40894 | 3.75042 | 2.53025 |
| chr2 | 102400412 | 102401428 | 1000 | Promoter (<=1kb) | 2 | 102300119 | 102400900 | 100782 | 2 | ENSMUST00 | 0 | Trim44 | ENSMUSG0C | false | chr2 | 102399957 | 102401748 | 23.12601 | 77.64799 | 74.53377 | chr2 | 102396641 | 102401748 | 4.00385 | 3.54495 | 2.18924 |
| chr2 | 122575383 | 122575807 | 1000 | Distal Intergenic | 2 | 122595489 | 122598534 | 3046 | 2 | ENSMUST00 | 22727 | Gatm | ENSMUSG0C | false | | | | | | | chr2 | 122574487 | 122577662 | 5.6028 | 4.4306 | 3.49482 |
| chr2 | 124609578 | 124610936 | 1000 | Promoter (<=1kb) | 2 | 124610106 | 124662885 | 52780 | 1 | ENSMUST00 | 0 | Sema6d | ENSMUSG0C | false | chr2 | 124609423 | 124611639 | 34.90432 | 112.58753 | 108.68004 | chr2 | 124609497 | 124609794 | 3.38489 | 3.25996 | 1.7292 |
| chr2 | 125247095 | 125247676 | 1000 | Promoter (<=1kb) | 2 | 125247190 | 125257748 | 10559 | 1 | ENSMUST00 | 0 | Dut | ENSMUSG0C | false | chr2 | 125246816 | 125248232 | 15.90661 | 56.5361 | 53.73628 | chr2 | 125241248 | 125248606 | 4.13809 | 3.66552 | 2.3035 |
| chr2 | 116063719 | 116064328 | 1000 | Promoter (<=1kb) | 2 | 116049084 | 116064468 | 15385 | 2 | ENSMUST00 | 140 | Meis2 | ENSMUSG0C | false | chr2 | 116060041 | 116064637 | 29.3377 | 90.74815 | 87.38457 | | | | | | |
| chr2 | 93462381 | 93463316 | 1000 | Promoter (<=1kb) | 2 | 93419111 | 93462411 | 43301 | 2 | ENSMUST00 | 0 | Cd82 | ENSMUSG0C | false | chr2 | 93462618 | 93463003 | 10.33961 | 21.99109 | 19.62971 | chr2 | 93434899 | 93464965 | 6.68053 | 4.76412 | 4.39602 |
| chr2 | 92599340 | 92600402 | 1000 | Promoter (<=1kb) | 2 | 92599707 | 92615250 | 15544 | 1 | ENSMUST00 | 0 | Chst1 | ENSMUSG0C | true | chr2 | 92599547 | 92603771 | 22.31878 | 85.15456 | 81.90936 | chr2 | 92588753 | 92599426 | 3.68402 | 3.42737 | 1.91817 |
| chr2 | 91651716 | 91652106 | 1000 | Promoter (<=1kb) | 2 | 91651747 | 91668328 | 16582 | 1 | ENSMUST00 | 0 | Arhgap1 | ENSMUSG0C | false | chr2 | 91648776 | 91651859 | 28.06786 | 91.41865 | 88.0463 | chr2 | 91650770 | 91658145 | 7.66222 | 5.14416 | 5.21273 |
| chr2 | 136678480 | 136679384 | 1000 | Promoter (<=1kb) | 2 | 136713453 | 136782428 | 68976 | 1 | ENSMUST00 | -34069 | Snap25 | ENSMUSG0C | false | chr2 | 136678336 | 136680011 | 21.76566 | 71.50774 | 68.49327 | chr2 | 136677840 | 136679972 | 5.06206 | 4.18367 | 3.04713 |
| chr2 | 137114273 | 137115123 | 1000 | Promoter (<=1kb) | 2 | 137081456 | 137116644 | 35189 | 2 | ENSMUST00 | 1521 | Jag1 | ENSMUSG0C | false | chr2 | 137114213 | 137117889 | 37.86728 | 126.26704 | 121.91066 | chr2 | 137113547 | 137116039 | 3.34662 | 3.2517 | 1.63481 |
| chr2 | 120563693 | 120564027 | 1000 | Promoter (<=1kb) | 2 | 120506820 | 120563807 | 56988 | 2 | ENSMUST00 | 0 | Zfp106 | ENSMUSG0C | false | chr2 | 120562969 | 120563923 | 24.69612 | 110.86208 | 107.00209 | chr2 | 120556472 | 120564946 | 4.79451 | 3.96218 | 2.8439 |
| chr2 | 120124112 | 120124558 | 1000 | Intron (ENSMUST00000028755.7 | 2 | 120102220 | 120129725 | 27506 | 1 | ENSMUST00 | 5167 | Ehd4 | ENSMUSG0C | false | | | | | | | chr2 | 120119955 | 120135886 | 7.40237 | 5.05905 | 4.97897 |
| chr2 | 119799208 | 119800177 | 1000 | Promoter (<=1kb) | 2 | 119799239 | 119805650 | 6412 | 1 | ENSMUST00 | 0 | Tyro3 | ENSMUSG0C | false | chr2 | 119799062 | 119801370 | 28.53699 | 95.19626 | 91.7378 | chr2 | 119798123 | 119811353 | 5.76878 | 4.44113 | 3.64054 |
| chr2 | 119802741 | 119803332 | 1000 | Promoter (<=1kb) | 2 | 119803087 | 119807990 | 4904 | 1 | ENSMUST00 | 0 | Tyro3 | ENSMUSG0C | false | | | | | | | chr2 | 119798123 | 119811353 | 5.76878 | 4.44113 | 3.64054 |
| chr2 | 119803576 | 119804933 | 1000 | Promoter (<=1kb) | 2 | 119803087 | 119807990 | 4904 | 1 | ENSMUST00 | 490 | Tyro3 | ENSMUSG0C | false | chr2 | 119804550 | 119805857 | 6.83737 | 14.36054 | 12.10883 | chr2 | 119798123 | 119811353 | 5.76878 | 4.44113 | 3.64054 |
| chr2 | 130450038 | 130450883 | 1000 | Promoter (<=1kb) | 2 | 130450278 | 130554026 | 103749 | 1 | ENSMUST00 | 0 | Ptpra | ENSMUSG0C | false | chr2 | 130449526 | 130451893 | 28.91859 | 89.0447 | 85.75584 | chr2 | 130450791 | 130459150 | 5.02255 | 3.97532 | 3.06464 |
| chr2 | 130839813 | 130840447 | 1000 | Promoter (<=1kb) | 2 | 130706200 | 130840134 | 133935 | 2 | ENSMUST00 | 0 | 4930402H24 | ENSMUSG0C | false | chr2 | 130838998 | 130840815 | 30.45735 | 121.03082 | 116.85291 | | | | | | |
| chr2 | 130906238 | 130906979 | 1000 | Promoter (<=1kb) | 2 | 130812866 | 130906406 | 93541 | 2 | ENSMUST00 | 0 | 4930402H24 | ENSMUSG0C | false | chr2 | 130905723 | 130907477 | 34.0961 | 113.25239 | 109.32404 | | | | | | |
| chr2 | 130832684 | 130832991 | 1000 | Intron (ENSMUST00000044766.5 | 2 | 130709130 | 130839687 | 130558 | 2 | ENSMUST00 | 6696 | 4930402H24 | ENSMUSG0C | false | | | | | | | | | | | | |
| chr2 | 131491670 | 131492364 | 1000 | Promoter (<=1kb) | 2 | 131491764 | 131525183 | 33420 | 1 | ENSMUST00 | 0 | Smox | ENSMUSG0C | false | chr2 | 131491475 | 131492673 | 23.85991 | 99.51073 | 95.95212 | chr2 | 131491358 | 131493017 | 5.14431 | 4.1824 | 3.12649 |
| chr2 | 132253188 | 132253540 | 1000 | Promoter (<=1kb) | 2 | 132249162 | 132253314 | 4153 | 2 | ENSMUST00 | 0 | Pcna | ENSMUSG0C | false | chr2 | 132252245 | 132253796 | 29.2142 | 114.47596 | 110.50954 | | | | | | |
| chr2 | 117120506 | 117121863 | 1000 | Promoter (<=1kb) | 2 | 117121374 | 117182279 | 60906 | 1 | ENSMUST00 | 0 | Spred1 | ENSMUSG0C | false | chr2 | 117119728 | 117123489 | 34.36702 | 111.64683 | 107.76208 | | | | | | |
| chr2 | 126674867 | 126676085 | 1000 | Promoter (<=1kb) | 2 | 126627442 | 126675568 | 48127 | 2 | ENSMUST00 | 0 | Gabpb1 | ENSMUSG0C | false | chr2 | 126673657 | 126676639 | 33.69293 | 107.36673 | 103.60477 | chr2 | 126671908 | 126674879 | 4.26535 | 3.74639 | 2.42629 |
| chr2 | 126875752 | 126876446 | 1000 | Promoter (<=1kb) | 2 | 126791565 | 126876230 | 84666 | 2 | ENSMUST00 | 0 | Trpm7 | ENSMUSG0C | false | chr2 | 126874723 | 126876927 | 30.03893 | 101.89226 | 98.2715 | | | | | | |
| chr2 | 126932585 | 126933320 | 1000 | Promoter (<=1kb) | 2 | 126890391 | 126933235 | 42845 | 2 | ENSMUST00 | 0 | Sppl2a | ENSMUSG0C | false | chr2 | 126932250 | 126933908 | 27.57193 | 78.19525 | 75.07539 | chr2 | 126932016 | 126934758 | 3.88106 | 3.56702 | 2.07437 |
| chr2 | 127335843 | 127336456 | 1000 | Promoter (<=1kb) | 2 | 127336159 | 127338376 | 2218 | 1 | ENSMUST00 | 0 | Dusp2 | ENSMUSG0C | false | chr2 | 127335645 | 127336933 | 23.47016 | 67.50422 | 64.55198 | chr2 | 127335502 | 127336851 | 3.68203 | 3.46992 | 1.95461 |
| chr2 | 128127204 | 128128032 | 1000 | Promoter (<=1kb) | 2 | 128127575 | 128130145 | 2571 | 1 | ENSMUST00 | 0 | Bcl2l11 | ENSMUSG0C | false | chr2 | 128125321 | 128128418 | 32.64849 | 123.19247 | 118.94151 | | | | | | |
| chr2 | 144270340 | 144271112 | 1000 | Promoter (<=1kb) | 2 | 144250123 | 144270572 | 20450 | 2 | ENSMUST00 | 0 | Snx5 | ENSMUSG0C | false | chr2 | 144269317 | 144271392 | 35.41093 | 115.56754 | 111.56183 | | | | | | |
| chr2 | 148677200 | 148678061 | 1000 | Promoter (2-3kb) | 2 | 148681023 | 148692949 | 11927 | 1 | ENSMUST00 | -2962 | Gzf1 | ENSMUSG0C | false | chr2 | 148677699 | 148678246 | 12.2252 | 27.53752 | 25.10386 | chr2 | 148677241 | 148680520 | 4.09823 | 3.61342 | 2.31085 |
| chr2 | 154544104 | 154544394 | 1000 | Promoter (2-3kb) | 2 | 154545045 | 154546794 | 1750 | 2 | ENSMUST00 | 2400 | Necab3 | ENSMUSG0C | false | | | | | | | chr2 | 154543547 | 154545182 | 5.9 | 4.54717 | 3.74586 |
| chr2 | 154566623 | 154566973 | 1000 | Promoter (2-3kb) | 2 | 154559634 | 154569720 | 10087 | 2 | ENSMUST00 | 2747 | E2f1 | ENSMUSG0C | false | | | | | | | chr2 | 154555502 | 154571286 | 6.03478 | 4.51051 | 3.86945 |
| chr3 | 9004214 | 9004823 | 1000 | Promoter (<=1kb) | 3 | 8928626 | 9004478 | 75853 | 2 | ENSMUST00 | 0 | Tpd52 | ENSMUSG0C | false | chr3 | 9002549 | 9004984 | 35.29575 | 125.98199 | 121.63858 | chr3 | 9004804 | 9008983 | 6.88132 | 5.02165 | 4.53704 |
| chr3 | 9834059 | 9834970 | 1000 | Promoter (<=1kb) | 3 | 9687479 | 9833679 | 146201 | 2 | ENSMUST00 | -381 | Pag1 | ENSMUSG0C | true | chr3 | 9830234 | 9834375 | 30.45735 | 121.03082 | 116.85291 | chr3 | 9833871 | 9836075 | 5.07786 | 4.14961 | 3.07524 |
| chr3 | 9885137 | 9885504 | 1000 | Distal Intergenic | 3 | 9687479 | 9833679 | 146201 | 2 | ENSMUST00 | -51459 | Pag1 | ENSMUSG0C | true | | | | | | | chr3 | 9883934 | 9888608 | 5.45831 | 4.15879 | 3.42513 |
| chr2 | 173173624 | 173174508 | 1000 | Distal Intergenic | 2 | 173153082 | 173159271 | 6190 | 1 | ENSMUST00 | 20543 | Pck1 | ENSMUSG0C | false | chr2 | 173173553 | 173174191 | 9.38717 | 20.36066 | 18.02005 | chr2 | 173171318 | 173175002 | 5.00733 | 4.01065 | 3.02514 |

| | | | | | | | | | | | | | | | | | | | | | | | | | | |
|---|---|---|---|---|---|---|---|---|---|---|---|---|---|---|---|---|---|---|---|---|---|---|---|---|---|---|
| chr2 | 173459242 | 173459721 | 1000 | Distal Intergenic | 2 | 173522586 | 173528501 | 5916 | 1 | ENSMUST00 | -62865 | 1700021F07I | ENSMUSG0( | false | . | . | . | . | . | . | chr2 | 173456952 | 173460566 | 4.97635 | 3.87678 | 3.01934 |
| chr2 | 174075767 | 174076400 | 1000 | Promoter (<=1kb) | 2 | 174076308 | 174092433 | 16126 | 1 | ENSMUST00 | 0 | Stx16 | ENSMUSG0( | false | chr2 | 174075923 | 174077510 | 33.5288 | 108.10788 | 104.32592 | chr2 | 174073295 | 174092361 | 5.14014 | 4.03613 | 3.13424 |
| chr2 | 174329536 | 174330882 | 1000 | Promoter (<=1kb) | 2 | 174330006 | 174346744 | 16739 | 1 | ENSMUST00 | 0 | Gnas | ENSMUSG0( | true | chr2 | 174326582 | 174332166 | 45.47067 | 151.73354 | 146.38181 | chr2 | 174328392 | 174329735 | 3.14359 | 3.05236 | 1.50739 |
| chr2 | 174342794 | 174343472 | 1000 | Promoter (1-2kb) | 2 | 174341704 | 174345409 | 3706 | 1 | ENSMUST00 | 1091 | Gnas | ENSMUSG0( | true | . | . | . | . | . | . | chr2 | 174331748 | 174347496 | 6.56852 | 4.7785 | 4.29829 |
| chr3 | 9943270 | 9943831 | 1000 | Distal Intergenic | 3 | 10012548 | 10016607 | 4060 | 1 | ENSMUST00 | -68717 | Fabp5 | ENSMUSG0( | false | . | . | . | . | . | . | chr3 | 9939293 | 9947650 | 5.5458 | 4.38502 | 3.43991 |
| chr2 | 167942244 | 167942577 | 1000 | Intron (ENSMUST00000147210.2 | 2 | 167951048 | 167965067 | 14020 | 1 | ENSMUST00 | -8471 | Ptpn1 | ENSMUSG0( | false | . | . | . | . | . | . | chr2 | 167933270 | 167963827 | 6.4817 | 4.58258 | 4.24812 |
| chr2 | 167931553 | 167932542 | 1000 | Promoter (<=1kb) | 2 | 167932057 | 167974737 | 42681 | 1 | ENSMUST00 | 0 | Ptpn1 | ENSMUSG0( | false | chr2 | 167931436 | 167933867 | 38.36185 | 128.21687 | 123.78947 | chr2 | 167931002 | 167932040 | 5.09214 | 3.9953 | 3.13461 |
| chr3 | 14663997 | 14664433 | 1000 | Distal Intergenic | 3 | 14641727 | 14663002 | 21276 | 1 | ENSMUST00 | 22271 | Car13 | ENSMUSG0( | false | . | . | . | . | . | . | chr3 | 14662383 | 14667691 | 4.57202 | 3.81155 | 2.66936 |
| chr2 | 181457894 | 181458180 | 1000 | Promoter (1-2kb) | 2 | 181424228 | 181459379 | 35152 | 2 | ENSMUST00 | 1199 | Zbtb46 | ENSMUSG0( | false | chr2 | 181457277 | 181460692 | 38.47989 | 123.03305 | 118.78763 | chr2 | 181430299 | 181458286 | 5.18444 | 4.09234 | 3.18307 |
| chr2 | 181458536 | 181459770 | 1000 | Promoter (<=1kb) | 2 | 181387762 | 181459426 | 71665 | 2 | ENSMUST00 | 0 | Zbtb46 | ENSMUSG0( | false | chr2 | 181457277 | 181460692 | 38.47989 | 123.03305 | 118.78763 | . | . | . | . | . | . |
| chr2 | 181460283 | 181460826 | 1000 | Promoter (<=1kb) | 2 | 181421554 | 181459426 | 37873 | 2 | ENSMUST00 | -858 | Zbtb46 | ENSMUSG0( | false | chr2 | 181457277 | 181460692 | 38.47989 | 123.03305 | 118.78763 | chr2 | 181459802 | 181464590 | 6.32717 | 4.72232 | 4.08879 |
| chr2 | 154790570 | 154792226 | 1000 | Promoter (<=1kb) | 2 | 154791096 | 154867245 | 76150 | 2 | ENSMUST00 | 0 | Raly | ENSMUSG0( | false | chr2 | 154790229 | 154790835 | 11.64009 | 27.76484 | 25.32713 | . | . | . | . | . | . |
| chr2 | 156179991 | 156180744 | 1000 | Promoter (<=1kb) | 2 | 156147239 | 156180238 | 33000 | 2 | ENSMUST00 | 0 | Rbm39 | ENSMUSG0( | false | chr2 | 156180424 | 156181016 | 14.34627 | 35.7714 | 33.23358 | chr2 | 156180702 | 156190707 | 4.70851 | 3.89185 | 2.76608 |
| chr2 | 156474991 | 156476051 | 1000 | Promoter (<=1kb) | 2 | 156475550 | 156499674 | 24125 | 1 | ENSMUST00 | 0 | Epb4.1l1 | ENSMUSG0( | false | chr2 | 156474878 | 156476371 | 19.12836 | 53.15374 | 50.39941 | chr2 | 156474015 | 156476347 | 4.92019 | 3.95587 | 2.95967 |
| chr3 | 21935818 | 21936313 | 1000 | Distal Intergenic | 3 | 22076652 | 22216594 | 139943 | 1 | ENSMUST00 | -140339 | Tbl1xr1 | ENSMUSG0( | false | chr3 | 21935258 | 21936108 | 16.08362 | 38.04472 | 35.47989 | chr3 | 21933908 | 21938060 | 4.69098 | 3.94511 | 2.74663 |
| chr3 | 22075847 | 22077416 | 1000 | Promoter (<=1kb) | 3 | 22076652 | 22216594 | 139943 | 1 | ENSMUST00 | 0 | Tbl1xr1 | ENSMUSG0( | false | chr3 | 22075480 | 22078230 | 23.74947 | 92.7123 | 89.30923 | chr3 | 22074990 | 22075923 | 4.72547 | 4.02511 | 2.78047 |
| chr2 | 157423952 | 157424660 | 1000 | Promoter (<=1kb) | 2 | 157424116 | 157455707 | 31592 | 1 | ENSMUST00 | 0 | Src | ENSMUSG0( | false | chr2 | 157423815 | 157424803 | 22.08709 | 62.48929 | 59.61029 | chr2 | 157422849 | 157424900 | 4.13667 | 3.69664 | 2.28375 |
| chr2 | 158794627 | 158795080 | 1000 | Promoter (<=1kb) | 2 | 158794807 | 158858214 | 63408 | 1 | ENSMUST00 | 0 | Dhx35 | ENSMUSG0( | false | chr2 | 158794749 | 158795388 | 31.997 | 117.77848 | 113.70881 | chr2 | 158794093 | 158795592 | 4.92207 | 3.90097 | 2.97224 |
| chr3 | 31094028 | 31095935 | 1000 | Promoter (<=1kb) | 3 | 31095058 | 31122577 | 27520 | 1 | ENSMUST00 | 0 | Skil | ENSMUSG0( | false | chr3 | 31093380 | 31096252 | 27.34946 | 104.78247 | 101.09169 | . | . | . | . | . | . |
| chr3 | 32365205 | 32365932 | 1000 | Promoter (<=1kb) | 3 | 32334796 | 32365620 | 30825 | 2 | ENSMUST00 | 0 | Zmat3 | ENSMUSG0( | false | chr3 | 32364866 | 32366223 | 34.00537 | 102.80987 | 99.17244 | chr3 | 32364539 | 32366564 | 4.95499 | 4.10713 | 3.02244 |
| chr3 | 32396875 | 32397895 | 1000 | Promoter (<=1kb) | 3 | 32397671 | 32466107 | 68437 | 1 | ENSMUST00 | 0 | Pik3ca | ENSMUSG0( | false | chr3 | 32395974 | 32398506 | 30.09291 | 96.00345 | 92.52599 | . | . | . | . | . | . |
| chr3 | 32510374 | 32510966 | 1000 | Distal Intergenic | 3 | 32515295 | 32520685 | 5391 | 1 | ENSMUST00 | -4329 | Zfp639 | ENSMUSG0( | false | chr3 | 32509946 | 32511591 | 26.28407 | 85.32412 | 82.07336 | . | . | . | . | . | . |
| chr3 | 32529342 | 32529883 | 1000 | Promoter (<=1kb) | 3 | 32529465 | 32579239 | 49775 | 1 | ENSMUST00 | 0 | Mfn1 | ENSMUSG0( | false | chr3 | 32529073 | 32530325 | 21.7465 | 82.16968 | 78.97962 | . | . | . | . | . | . |
| chr3 | 34019604 | 34020312 | 1000 | Promoter (<=1kb) | 3 | 34020079 | 34069342 | 49264 | 1 | ENSMUST00 | 0 | Fxr1 | ENSMUSG0( | false | chr3 | 34019437 | 34021363 | 40.65367 | 138.82761 | 133.98773 | . | . | . | . | . | . |
| chr3 | 30418627 | 30418981 | 1000 | Intron (ENSMUST00000173495.2 | 3 | 30236569 | 30509487 | 272919 | 2 | ENSMUST00 | 90506 | Mecom | ENSMUSG0( | false | . | . | . | . | . | . | chr3 | 30418279 | 30419052 | 4.28244 | 3.76159 | 2.40702 |
| chr3 | 28658422 | 28658700 | 1000 | Intron (ENSMUST00000160934.2 | 3 | 28650027 | 28665945 | 15919 | 1 | ENSMUST00 | 8396 | Tnik | ENSMUSG0( | false | . | . | . | . | . | . | . | . | . | . | . | . |
| chr3 | 27988086 | 27988511 | 1000 | Intron (ENSMUST00000120834.2 | 3 | 27984314 | 28133234 | 148921 | 1 | ENSMUST00 | 3773 | Pld1 | ENSMUSG0( | true | . | . | . | . | . | . | chr3 | 27985243 | 27995015 | 5.4487 | 4.25198 | 3.39101 |
| chr3 | 51483742 | 51484341 | 1000 | Promoter (<=1kb) | 3 | 51483966 | 51495905 | 11940 | 1 | ENSMUST00 | 0 | Rab33b | ENSMUSG0( | false | chr3 | 51483418 | 51485063 | 34.56782 | 111.9988 | 108.10535 | chr3 | 51483072 | 51486962 | 3.68755 | 3.45943 | 1.91118 |
| chr3 | 60500566 | 60502033 | 1000 | Promoter (<=1kb) | 3 | 60501252 | 60629748 | 128497 | 1 | ENSMUST00 | 0 | Mbnl1 | ENSMUSG0( | false | chr3 | 60500555 | 60503503 | 30.26787 | 111.42836 | 107.5545 | chr3 | 60499106 | 60500728 | 4.14588 | 3.59843 | 2.32696 |
| chr3 | 68508753 | 68509101 | 1000 | Intron (ENSMUST00000029346.7 | 3 | 68494208 | 68626463 | 132256 | 1 | ENSMUST00 | 14546 | Schip1 | ENSMUSG0( | false | . | . | . | . | . | . | chr3 | 68505326 | 68513825 | 4.36634 | 3.64257 | 2.51287 |
| chr3 | 68587694 | 68588241 | 1000 | Intron (ENSMUST00000029346.7 | 3 | 68584179 | 68626464 | 42286 | 1 | ENSMUST00 | 3516 | Schip1 | ENSMUSG0( | false | . | . | . | . | . | . | chr3 | 68574597 | 68620372 | 5.61397 | 4.36647 | 3.51459 |
| chr3 | 68576441 | 68576957 | 1000 | Intron (ENSMUST00000029346.7 | 3 | 68572245 | 68626464 | 54220 | 1 | ENSMUST00 | 4197 | Schip1 | ENSMUSG0( | false | . | . | . | . | . | . | chr3 | 68574597 | 68620372 | 5.61397 | 4.36647 | 3.51459 |
| chr3 | 69126380 | 69127385 | 1000 | Promoter (<=1kb) | 3 | 69067149 | 69127113 | 59965 | 2 | ENSMUST00 | 0 | Kpna4 | ENSMUSG0( | false | chr3 | 69125615 | 69127992 | 34.86761 | 110.44456 | 106.5922 | . | . | . | . | . | . |
| chr3 | 69316516 | 69316952 | 1000 | Promoter (<=1kb) | 3 | 69316861 | 69560802 | 243942 | 1 | ENSMUST00 | 0 | Ppm1l | ENSMUSG0( | false | chr3 | 69315733 | 69318906 | 39.61801 | 127.15503 | 122.77008 | . | . | . | . | . | . |
| chr3 | 57282696 | 57283249 | 1000 | Downstream (2-3kb) | 3 | 57285611 | 57294889 | 9279 | 2 | ENSMUST00 | 11640 | Tm4sf1 | ENSMUSG0( | true | . | . | . | . | . | . | chr3 | 57280868 | 57284151 | 4.72748 | 3.92953 | 2.78228 |
| chr3 | 57293864 | 57294323 | 1000 | Promoter (<=1kb) | 3 | 57285611 | 57294889 | 9279 | 2 | ENSMUST00 | 566 | Tm4sf1 | ENSMUSG0( | true | chr3 | 57290579 | 57295802 | 25.29353 | 74.26213 | 71.20484 | . | . | . | . | . | . |
| chr3 | 57575402 | 57576825 | 1000 | Promoter (<=1kb) | 3 | 57455649 | 57575828 | 120180 | 2 | ENSMUST00 | 0 | Wwtr1 | ENSMUSG0( | false | chr3 | 57573515 | 57576914 | 34.45293 | 106.63475 | 102.89172 | chr3 | 57576560 | 57577215 | 4.66715 | 3.98072 | 2.72344 |
| chr3 | 63295247 | 63296326 | 1000 | Promoter (<=1kb) | 3 | 63295872 | 63382231 | 86360 | 1 | ENSMUST00 | 0 | Mme | ENSMUSG0( | false | chr3 | 63295209 | 63297475 | 22.99841 | 82.87405 | 79.66897 | . | . | . | . | . | . |
| chr3 | 63229189 | 63229549 | 1000 | Distal Intergenic | 3 | 63295872 | 63382231 | 86360 | 1 | ENSMUST00 | -66323 | Mme | ENSMUSG0( | false | . | . | . | . | . | . | chr3 | 63228800 | 63230717 | 4.15744 | 3.69041 | 2.31423 |
| chr3 | 63325655 | 63325957 | 1000 | Intron (ENSMUST00000029400.1 | 3 | 63295872 | 63382231 | 86360 | 1 | ENSMUST00 | 29784 | Mme | ENSMUSG0( | false | . | . | . | . | . | . | chr3 | 63319672 | 63327568 | 4.32678 | 3.74428 | 2.45114 |
| chr3 | 65957407 | 65958713 | 1000 | Promoter (<=1kb) | 3 | 65946151 | 65958162 | 12012 | 2 | ENSMUST00 | 0 | Ccnl1 | ENSMUSG0( | false | chr3 | 65956813 | 65959201 | 23.70827 | 104.82465 | 101.13002 | . | . | . | . | . | . |
| chr3 | 65974140 | 65974972 | 1000 | Distal Intergenic | 3 | 65946151 | 65958249 | 12099 | 2 | ENSMUST00 | -15892 | Ccnl1 | ENSMUSG0( | false | chr3 | 65973958 | 65975550 | 17.4042 | 56.42813 | 53.62976 | . | . | . | . | . | . |
| chr3 | 65871113 | 65871645 | 1000 | Distal Intergenic | 3 | 65946782 | 65947998 | 1217 | 2 | ENSMUST00 | 76353 | Ccnl1 | ENSMUSG0( | false | . | . | . | . | . | . | chr3 | 65871081 | 65872232 | 5.62746 | 4.08851 | 3.55332 |
| chr3 | 66981021 | 66981691 | 1000 | Promoter (<=1kb) | 3 | 66971727 | 66981318 | 9592 | 2 | ENSMUST00 | 0 | Shox2 | ENSMUSG0( | false | chr3 | 66980702 | 66981759 | 17.10022 | 42.66158 | 40.03807 | chr3 | 66980768 | 66983184 | 5.10016 | 4.15926 | 3.08667 |
| chr3 | 101109789 | 101110566 | 1000 | Promoter (<=1kb) | 3 | 101040232 | 101110278 | 70047 | 2 | ENSMUST00 | 0 | Ptgfrn | ENSMUSG0( | false | chr3 | 101109427 | 101110666 | 30.2678 | 92.36906 | 88.97462 | chr3 | 101108916 | 101110639 | 6.14603 | 4.39569 | 3.96977 |
| chr3 | 98267123 | 98267649 | 1000 | Distal Intergenic | 3 | 98280435 | 98310732 | 30298 | 1 | ENSMUST00 | -12786 | Hmgcs2 | ENSMUSG0( | false | . | . | . | . | . | . | chr3 | 98266143 | 98268440 | 4.28966 | 3.8256 | 2.41395 |
| chr3 | 98042497 | 98043137 | 1000 | Intron (ENSMUST00000079812.6 | 3 | 98013538 | 98150367 | 136830 | 1 | ENSMUST00 | 28960 | Notch2 | ENSMUSG0( | false | chr3 | 98042299 | 98042889 | 7.88522 | 15.75676 | 13.48224 | chr3 | 98014795 | 98071300 | 5.94878 | 4.48977 | 3.79636 |
| chr3 | 98013278 | 98013700 | 1000 | Promoter (<=1kb) | 3 | 98013538 | 98150367 | 136830 | 1 | ENSMUST00 | 0 | Notch2 | ENSMUSG0( | false | chr3 | 98012900 | 98014942 | 35.22365 | 121.58154 | 117.38489 | . | . | . | . | . | . |
| chr3 | 93520799 | 93521367 | 1000 | Promoter (<=1kb) | 3 | 93520488 | 93526287 | 5800 | 1 | ENSMUST00 | 312 | S100a11 | ENSMUSG0( | false | chr3 | 93520371 | 93523048 | 29.97852 | 88.95019 | 85.62817 | chr3 | 93520128 | 93524973 | 5.6407 | 4.38912 | 3.52992 |
| chr3 | 93537145 | 93537669 | 1000 | Distal Intergenic | 3 | 93520488 | 93526287 | 5800 | 1 | ENSMUST00 | 16658 | S100a11 | ENSMUSG0( | false | . | . | . | . | . | . | chr3 | 93536763 | 93538864 | 4.17413 | 3.64148 | 2.3708 |
| chr3 | 90465660 | 90466551 | 1000 | Promoter (<=1kb) | 3 | 90450591 | 90465866 | 15276 | 2 | ENSMUST00 | 0 | Npr1 | ENSMUSG0( | false | chr3 | 90462429 | 90466691 | 46.53382 | 157.01303 | 151.46753 | . | . | . | . | . | . |
| chr3 | 90432750 | 90433941 | 1000 | Promoter (<=1kb) | 3 | 90391380 | 90433636 | 42257 | 2 | ENSMUST00 | 0 | Ints3 | ENSMUSG0( | false | chr3 | 90432562 | 90433609 | 20.06521 | 63.98471 | 61.08233 | chr3 | 90424548 | 90432958 | 5.60023 | 4.26885 | 3.5145 |
| chr3 | 90051704 | 90052944 | 1000 | Promoter (<=1kb) | 3 | 90052814 | 90063341 | 10528 | 1 | ENSMUST00 | 0 | 4933434E20I | ENSMUSG0( | false | chr3 | 90052818 | 90053990 | 29.66345 | 100.21 | 96.63225 | . | . | . | . | . | . |
| chr3 | 89876222 | 89876717 | 1000 | Intron (ENSMUST00000029559.6 | 3 | 89869324 | 89913162 | 43839 | 2 | ENSMUST00 | 36445 | Il6ra | ENSMUSG0( | true | . | . | . | . | . | . | chr3 | 89858315 | 89886122 | 6.15651 | 4.57192 | 3.96514 |
| chr3 | 89875005 | 89875518 | 1000 | Intron (ENSMUST00000029559.6 | 3 | 89869324 | 89913162 | 43839 | 2 | ENSMUST00 | 37644 | Il6ra | ENSMUSG0( | true | . | . | . | . | . | . | chr3 | 89858315 | 89886122 | 6.15651 | 4.57192 | 3.96514 |
| chr3 | 89872255 | 89873073 | 1000 | Intron (ENSMUST00000029559.6 | 3 | 89869324 | 89913162 | 43839 | 2 | ENSMUST00 | 40089 | Il6ra | ENSMUSG0( | true | . | . | . | . | . | . | chr3 | 89858315 | 89886122 | 6.15651 | 4.57192 | 3.96514 |
| chr3 | 129161670 | 129162051 | 1000 | Distal Intergenic | 3 | 129199878 | 129219590 | 19713 | 1 | ENSMUST00 | -37827 | Pitx2 | ENSMUSG0( | false | . | . | . | . | . | . | chr3 | 129161397 | 129162289 | 3.00052 | 2.97734 | 1.39987 |
| chr3 | 129321514 | 129321988 | 1000 | Intron (ENSMUST00000029658.8 | 3 | 129270277 | 129332264 | 61988 | 2 | ENSMUST00 | 10276 | Enpep | ENSMUSG0( | true | . | . | . | . | . | . | chr3 | 129303627 | 129327892 | 5.35019 | 4.2164 | 3.29511 |
| chr3 | 129396939 | 129397725 | 1000 | Distal Intergenic | 3 | 129298594 | 129332720 | 34127 | 2 | ENSMUST00 | -64220 | Enpep | ENSMUSG0( | true | . | . | . | . | . | . | chr3 | 129395845 | 129398980 | 5.387 | 4.2788 | 3.37761 |
| chr3 | 129317317 | 129317668 | 1000 | Intron (ENSMUST00000029658.8 | 3 | 129276764 | 129309034 | 32271 | 2 | ENSMUST00 | -8284 | Enpep | ENSMUSG0( | true | . | . | . | . | . | . | chr3 | 129303627 | 129327892 | 5.35019 | 4.2164 | 3.29511 |
| chr3 | 132716092 | 132716398 | 1000 | Intron (ENSMUST00000029664.6 | 3 | 132684144 | 132838506 | 154363 | 1 | ENSMUST00 | 31949 | Tbck | ENSMUSG0( | false | . | . | . | . | . | . | chr3 | 132713886 | 132717995 | 3.93357 | 3.5311 | 2.15077 |
| chr3 | 152210069 | 152211241 | 1000 | Promoter (<=1kb) | 3 | 152210458 | 152236830 | 26373 | 1 | ENSMUST00 | 0 | Fubp1 | ENSMUSG0( | false | chr3 | 152209361 | 152212419 | 43.58744 | 151.86528 | 146.50356 | . | . | . | . | . | . |
| chr3 | 89393573 | 89395217 | 1000 | Promoter (<=1kb) | 3 | 89381138 | 89393678 | 12541 | 2 | ENSMUST00 | 0 | Zbtb7b | ENSMUSG0( | false | chr3 | 89391380 | 89396724 | 30.32912 | 88.589 | 85.27629 | . | . | . | . | . | . |
| chr3 | 88949847 | 88951147 | 1000 | Promoter (<=1kb) | 3 | 88950622 | 89079375 | 128754 | 1 | ENSMUST00 | 0 | Ash1l | ENSMUSG0( | false | chr3 | 88949487 | 88952169 | 27.34946 | 104.78247 | 101.09169 | chr3 | 88947692 | 88949850 | 3.96427 | 3.55235 | 2.16164 |
| chr3 | 88636438 | 88636844 | 1000 | Promoter (2-3kb) | 3 | 88634323 | 88635076 | 754 | 1 | ENSMUST00 | 2116 | Arhgef2 | ENSMUSG0( | false | . | . | . | . | . | . | chr3 | 88636079 | 88636914 | 3.23425 | 3.10046 | 1.58581 |
| chr3 | 88509869 | 88510431 | 1000 | Promoter (<=1kb) | 3 | 88502753 | 88509956 | 7204 | 2 | ENSMUST00 | 0 | Lmna | ENSMUSG0( | false | chr3 | 88509595 | 88510627 | 13.73236 | 30.71385 | 28.24041 | chr3 | 88505561 | 88513944 | 4.89659 | 4.0178 | 2.9339 |
| chr3 | 88501921 | 88502365 | 1000 | Promoter (<=1kb) | 3 | 88483205 | 88503307 | 20103 | 2 | ENSMUST00 | 942 | Lmna | ENSMUSG0( | false | chr3 | 88501420 | 88503310 | 37.81545 | 144.80544 | 139.72641 | chr3 | 88487350 | 88502372 | 7.5952 | 5.18047 | 5.13989 |
| chr3 | 88492890 | 88493346 | 1000 | Promoter (<=1kb) | 3 | 88483206 | 88493312 | 10107 | 2 | ENSMUST00 | 0 | Lmna | ENSMUSG0( | false | chr3 | 88492514 | 88493067 | 16.66573 | 41.91806 | 39.30453 | chr3 | 88487350 | 88502372 | 7.5952 | 5.18047 | 5.13989 |
| chr3 | 86862046 | 86862566 | 1000 | Intron (ENSMUST00000029719.8 | 3 | 86786150 | 86920884 | 134735 | 2 | ENSMUST00 | 58318 | Dclk2 | ENSMUSG0( | false | . | . | . | . | . | . | chr3 | 86860113 | 86863731 | 5.14551 | 4.12467 | 3.13623 |
| chr3 | 86223439 | 86225054 | 1000 | Promoter (<=1kb) | 3 | 86224690 | 86782693 | 558004 | 1 | ENSMUST00 | 0 | Lrba | ENSMUSG0( | false | chr3 | 86223104 | 86225747 | 28.32768 | 117.73571 | 113.66808 | . | . | . | . | . | . |
| chr3 | 86056779 | 86057049 | 1000 | Intron (ENSMUST00000182969.2 | 3 | 86070915 | 86130526 | 59612 | 1 | ENSMUST00 | -13866 | Sh3d19 | ENSMUSG0( | false | . | . | . | . | . | . | chr3 | 86053225 | 86060898 | 5.02373 | 4.03933 | 3.02522 |
| chr3 | 85970754 | 85971593 | 1000 | Promoter (<=1kb) | 3 | 85971109 | 86083538 | 112430 | 1 | ENSMUST00 | 0 | Sh3d19 | ENSMUSG0( | false | chr3 | 85970752 | 85972621 | 36.31701 | 113.99837 | 110.04225 | . | . | . | . | . | . |
| chr3 | 84814208 | 84815110 | 1000 | Promoter (<=1kb) | 3 | 84815268 | 84979198 | 163931 | 1 | ENSMUST00 | -158 | Fbxw7 | ENSMUSG0( | false | chr3 | 84813640 | 84816188 | 32.64849 | 123.19247 | 118.94151 | . | . | . | . | . | . |
| chr3 | 97158330 | 97159104 | 1000 | Promoter (<=1kb) | 3 | 97158777 | 97176574 | 17798 | 1 | ENSMUST00 | 0 | Acp6 | ENSMUSG0( | false | chr3 | 97158037 | 97159684 | 24.78212 | 78.86476 | 75.73294 | . | . | . | . | . | . |
| chr3 | 96696761 | 96697600 | 1000 | Promoter (<=1kb) | 3 | 96697076 | 96699692 | 2617 | 1 | ENSMUST00 | 0 | Pias3 | ENSMUSG0( | false | chr3 | 96696049 | 96697777 | 31.07893 | 124.33488 | 120.0426 | chr3 | 96697530 | 96699524 | 3.78952 | 3.48552 | 2.01252 |
| chr3 | 95818541 | 95819949 | 1000 | Promoter (<=1kb) | 3 | 95759873 | 95818953 | 59081 | 2 | ENSMUST00 | 0 | Rprd2 | ENSMUSG0( | false | chr3 | 95819497 | 95820267 | 16.33881 | 42.83351 | 40.20603 | chr3 | 95819928 | 95821126 | 3.14049 | 3.04093 | 1.51995 |
| chr3 | 122439036 | 122439442 | 1000 | Intron (ENSMUST00000029766.4 | 3 | 122419780 | 122530180 | 110401 | 1 | ENSMUST00 | 19257 | Bcar3 | ENSMUSG0( | true | . | . | . | . | . | . | chr3 | 122430877 | 122440688 | 4.67283 | 3.8714 | 2.75199 |
| chr3 | 122419561 | 122420802 | 1000 | Promoter (<=1kb) | 3 | 122419780 | 122530180 | 110401 | 1 | ENSMUST00 | 0 | Bcar3 | ENSMUSG0( | true | chr3 | 122418997 | 122422602 | 20.02967 | 73.35366 | 70.31202 | . | . | . | . | . | . |
| chr3 | 122245271 | 122245959 | 1000 | Promoter (<=1kb) | 3 | 122245557 | 122269984 | 24428 | 1 | ENSMUST00 | 0 | Gclm | ENSMUSG0( | false | chr3 | 122245528 | 122246629 | 34.11942 | 109.14127 | 105.33086 | chr3 | 122244749 | 122248146 | 3.39548 | 3.18231 | 1.70436 |
| chr3 | 122072997 | 122073517 | 1000 | Promoter (<=1kb) | 3 | 122074456 | 122087931 | 13476 | 1 | ENSMUST00 | -939 | Abca4 | ENSMUSG0( | false | . | . | . | . | . | . | chr3 | 122072522 | 122075417 | 4.27961 | 3.54856 | 2.44708 |
| chr3 | 122048012 | 122048792 | 1000 | Promoter (1-2kb) | 3 | 122046577 | 122097742 | 51166 | 1 | ENSMUST00 | 1436 | Abca4 | ENSMUSG0( | false | . | . | . | . | . | . | chr3 | 122045994 | 122049731 | 5.3457 | 4.16097 | 3.31733 |
| chr3 | 121814712 | 121815674 | 1000 | Promoter (<=1kb) | 3 | 121758909 | 121815302 | 56394 | 2 | ENSMUST00 | 0 | Abcd3 | ENSMUSG0( | false | chr3 | 121814226 | 121815863 | 36.3185 | 115.04841 | 111.06631 | . | . | . | . | . | . |
| chr3 | 119782438 | 119783674 | 1000 | Promoter (<=1kb) | 3 | 119718742 | 119783388 | 64647 | 2 | ENSMUST00 | 0 | Ptbp2 | ENSMUSG0( | false | chr3 | 119782091 | 119783812 | 33.41831 | 117.26761 | 113.21259 | . | . | . | . | . | . |
| chr3 | 94499873 | 94500305 | 1000 | 3' UTR | 3 | 94464985 | 94480341 | 15357 | 2 | ENSMUST00 | -19533 | Riiad1 | ENSMUSG0( | false | . | . | . | . | . | . | chr3 | 94490465 | 94502279 | 6.08744 | 4.57653 | 3.90418 |
| chr3 | 94495828 | 94496217 | 1000 | Downstream (1-2kb) | 3 | 94464985 | 94480341 | 15357 | 2 | ENSMUST00 | -15488 | Riiad1 | ENSMUSG0( | false | . | . | . | . | . | . | chr3 | 94490465 | 94502279 | 6.08744 | 4.57653 | 3.90418 |
| chr3 | 94309909 | 94310313 | 1000 | Promoter (<=1kb) | 3 | 94310089 | 94332540 | 22452 | 1 | ENSMUST00 | 0 | Them4 | ENSMUSG0( | false | chr3 | 94309394 | 94310950 | 18.31284 | 64.75944 | 61.84419 | . | . | . | . | . | . |
| chr3 | 138742059 | 138743185 | 1000 | Promoter (<=1kb) | 3 | 138742195 | 138904433 | 162239 | 1 | ENSMUST00 | 0 | Tspan5 | ENSMUSG0( | true | chr3 | 138741744 | 138743902 | 34.54477 | 122.48211 | 118.25522 | . | . | . | . | . | . |
| chr3 | 138654949 | 138655275 | 1000 | Distal Intergenic | 3 | 138742195 | 138904433 | 162239 | 1 | ENSMUST00 | -86920 | Tspan5 | ENSMUSG0( | true | . | . | . | . | . | . | chr3 | 138654963 | 138655653 | 4.12312 | 3.47781 | 2.30438 |
| chr3 | 138735834 | 138736167 | 1000 | Distal Intergenic | 3 | 138742195 | 138904433 | 162239 | 1 | ENSMUST00 | -6028 | Tspan5 | ENSMUSG0( | true | . | . | . | . | . | . | chr3 | 138735674 | 138736247 | 4.7574 | 3.97877 | 2.83378 |
| chr3 | 136669542 | 136671917 | 1000 | Promoter (<=1kb) | 3 | 136670124 | 136797838 | 127715 | 1 | ENSMUST00 | 0 | Ppp3ca | ENSMUSG0( | false | chr3 | 136669097 | 136672052 | 32.68175 | 111.76729 | 107.88066 | chr3 | 136671767 | 136676247 | 4.09701 | 3.47833 | 2.2891 |
| chr3 | 135603087 | 135603474 | 1000 | Promoter (<=1kb) | 3 | 135584745 | 135603796 | 19052 | 2 | ENSMUST00 | 322 | Nfkb1 | ENSMUSG0( | false | . | . | . | . | . | . | chr3 | 135601632 | 135607878 | 6.30926 | 4.60341 | 4.09794 |
| chr3 | 135681365 | 135681741 | 1000 | Intron (ENSMUST00000029812.8 | 3 | 135584655 | 135691546 | 106892 | 2 | ENSMUST00 | 9805 | Nfkb1 | ENSMUSG0( | false | . | . | . | . | . | . | chr3 | 135672933 | 135683932 | 5.7582 | 4.40397 | 3.63963 |
| chr3 | 135690820 | 135692126 | 1000 | Promoter (<=1kb) | 3 | 135584655 | 135691546 | 106892 | 2 | ENSMUST00 | 0 | Nfkb1 | ENSMUSG0( | false | chr3 | 135689869 | 135692324 | 30.41442 | 103.5799 | 99.9162 | chr3 | 135691923 | 135692687 | 4.36783 | 3.8163 | 2.48079 |
| chr3 | 135423010 | 135423702 | 1000 | Promoter (<=1kb) | 3 | 135410012 | 135423407 | 13396 | 2 | ENSMUST00 | 0 | Cisd2 | ENSMUSG0( | false | chr3 | 135422276 | 135424307 | 33.76198 | 105.43819 | 101.727 | . | . | . | . | . | . |
| chr3 | 135249035 | 135249563 | 1000 | Intron (ENSMUST00000062893.7 | 3 | 135281221 | 135304426 | 23206 | 1 | ENSMUST00 | -31658 | Bdh2 | ENSMUSG0( | false | . | . | . | . | . | . | . | . | . | . | . | . |
| chr3 | 159839486 | 159839867 | 1000 | Promoter (<=1kb) | 3 | 159839737 | 159935833 | 96097 | 1 | ENSMUST00 | 0 | Wls | ENSMUSG0( | false | chr3 | 159839398 | 159841336 | 24.71438 | 74.15884 | 71.1036 | chr3 | 159839066 | 159839528 | 3.02329 | 3.00889 | 1.37263 |
| chr3 | 148916167 | 148916636 | 1000 | Intron (ENSMUST00000106128.2 | 3 | 148815586 | 148954635 | 139050 | 2 | ENSMUST00 | 37999 | Lphn2 | ENSMUSG0( | false | . | . | . | . | . | . | chr3 | 148911765 | 148919652 | 4.92564 | 4.02806 | 2.96113 |
| chr3 | 148988634 | 148990683 | 1000 | Promoter (<=1kb) | 3 | 148888941 | 148989316 | 100376 | 2 | ENSMUST00 | 0 | Lphn2 | ENSMUSG0( | false | chr3 | 148986735 | 148991143 | 40.70288 | 146.73875 | 141.58145 | . | . | . | . | . | . |
| chr3 | 148914974 | 148915259 | 1000 | Intron (ENSMUST00000106128.2 | 3 | 148815586 | 148954635 | 139050 | 2 | ENSMUST00 | 39376 | Lphn2 | ENSMUSG0( | false | . | . | . | . | . | . | chr3 | 148911765 | 148919652 | 4.92564 | 4.02806 | 2.96113 |
| chr3 | 146499452 | 146500264 | 1000 | Promoter (<=1kb) | 3 | 146457203 | 146499753 | 42551 | 2 | ENSMUST00 | 0 | Spata1 | ENSMUSG0( | false | chr3 | 146498729 | 146501117 | 24.0356 | 94.23969 | 90.80066 | . | . | . | . | . | . |
| chr3 | 145923894 | 145925091 | 1000 | Promoter (<=1kb) | 3 | 145924378 | 145934283 | 9906 | 1 | ENSMUST00 | 0 | Bcl10 | ENSMUSG0( | false | chr3 | 145923480 | 145924025 | 14.6238 | 41.22485 | 38.6191 | chr3 | 145925046 | 145931258 | 5.71227 | 4.41121 | 3.5893 |
| chr3 | 145775309 | 145775620 | 1000 | Intron (ENSMUST00000029845.9 | 3 | 145760748 | 145893266 | 132519 | 1 | ENSMUST00 | 14562 | Ddah1 | ENSMUSG0( | false | . | . | . | . | . | . | chr3 | 145775024 | 145776036 | 4.48767 | 3.77975 | 2.59191 |
| chr3 | 145613316 | 145613624 | 1000 | Distal Intergenic | 3 | 145646971 | 145649985 | 3015 | 2 | ENSMUST00 | 36361 | Cyr61 | ENSMUSG0( | false | . | . | . | . | . | . | chr3 | 145612815 | 145615799 | 6.92855 | 4.96671 | 4.59673 |
| chr4 | 9642102 | 9642432 | 1000 | Promoter (1-2kb) | 4 | 9575267 | 9643717 | 68451 | 2 | ENSMUST00 | 1285 | Asph | ENSMUSG0( | false | . | . | . | . | . | . | chr4 | 9640579 | 9643792 | 4.15327 | 3.71367 | 2.30625 |
| chr4 | 14863808 | 14864623 | 1000 | Promoter (<=1kb) | 4 | 14864076 | 14915176 | 51101 | 1 | ENSMUST00 | 0 | Tmem55a | ENSMUSG0( | false | chr4 | 14863572 | 14865346 | 22.68762 | 81.35053 | 78.1749 | . | . | . | . | . | . |
| chr4 | 3835502 | 3835925 | 1000 | Promoter (<=1kb) | 4 | 3831334 | 3835665 | 4332 | 2 | ENSMUST00 | 0 | Rps20 | ENSMUSG0( | false | chr4 | 3834635 | 3836092 | 33.38818 | 124.5882 | 120.28912 | . | . | . | . | . | . |
| chr4 | 6453384 | 6454423 | 1000 | Promoter (<=1kb) | 4 | 6396207 | 6454271 | 58065 | 2 | ENSMUST00 | 0 | Nsmaf | ENSMUSG0( | false | chr4 | 6452287 | 6454754 | 30.32912 | 88.589 | 85.27629 | chr4 | 6454380 | 6454869 | 4.25157 | 3.73291 | 2.38375 |
| chr3 | 144201871 | 144202787 | 1000 | Promoter (<=1kb) | 3 | 144188530 | 144202317 | 13788 | 2 | ENSMUST00 | 0 | Lmo4 | ENSMUSG0( | false | chr3 | 144201264 | 144203047 | 28.46397 | 86.32589 | 83.05701 | chr3 | 144195322 | 144201886 | 4.23824 | 3.66035 | 2.38273 |
| chr3 | 144204648 | 144206019 | 1000 | Promoter (<=1kb) | 3 | 144188530 | 144205220 | 16691 | 2 | ENSMUST00 | 0 | Lmo4 | ENSMUSG0( | false | chr3 | 144203799 | 144206219 | 33.76535 | 103.4086 | 99.75368 | . | . | . | . | . | . |
| chr3 | 144198513 | 144199163 | 1000 | Intron (ENSMUST00000121796.2 | 3 | 144201623 | 144202300 | 678 | 2 | ENSMUST00 | 3137 | Lmo4 | ENSMUSG0( | false | chr3 | 144198116 | 144199381 | 27.41053 | 90.23362 | 86.88379 | chr3 | 144195322 | 144201886 | 4.23824 | 3.66035 | 2.38273 |
| chr4 | 34687060 | 34687591 | 1000 | Promoter (<=1kb) | 4 | 34663257 | 34687438 | 24182 | 2 | ENSMUST00 | 0 | Slc35a1 | ENSMUSG0( | false | chr4 | 34686600 | 34687906 | 28.9509 | 90.07051 | 86.72839 | . | . | . | . | . | . |
| chr4 | 46450776 | 46452544 | 1000 | Promoter (<=1kb) | 4 | 46450902 | 46472657 | 21756 | 1 | ENSMUST00 | 0 | Anp32b | ENSMUSG0( | false | chr4 | 46450413 | 46452821 | 36.76257 | 113.72284 | 109.77938 | chr4 | 46452472 | 46460187 | 5.16199 | 4.03176 | 3.18287 |
| chr4 | 46590264 | 46590596 | 1000 | Intron (ENSMUST00000107756.2 | 4 | 46543001 | 46601891 | 58891 | 2 | ENSMUST00 | 11295 | Coro2a | ENSMUSG0( | false | . | . | . | . | . | . | chr4 | 46590375 | 46590606 | 3.88865 | 3.66898 | 2.0726 |
| chr4 | 47227755 | 47228207 | 1000 | Intron (ENSMUST00000082303.7 | 4 | 47233267 | 47245512 | 12246 | 1 | ENSMUST00 | -5060 | Col15a1 | ENSMUSG0( | false | . | . | . | . | . | . | chr4 | 47224612 | 47229766 | 8.01781 | 5.39367 | 5.47329 |
| chr4 | 47225973 | 47226400 | 1000 | Intron (ENSMUST00000082303.7 | 4 | 47233267 | 47245512 | 12246 | 1 | ENSMUST00 | -6867 | Col15a1 | ENSMUSG0( | false | . | . | . | . | . | . | chr4 | 47224612 | 47229766 | 8.01781 | 5.39367 | 5.47329 |
| chr4 | 66450072 | 66450460 | 1000 | Distal Intergenic | 4 | 65380803 | 66404611 | 1023809 | 2 | ENSMUST00 | -45462 | Astn2 | ENSMUSG0( | true | . | . | . | . | . | . | chr4 | 66449477 | 66451280 | 4.51968 | 3.80583 | 2.65848 |
| chr4 | 59188885 | 59189770 | 1000 | Promoter (<=1kb) | 4 | 59189257 | 59222833 | 33577 | 1 | ENSMUST00 | 0 | Ugcg | ENSMUSG0( | false | chr4 | 59188732 | 59190495 | 16.43683 | 59.22888 | 56.39454 | . | . | . | . | . | . |
| chr4 | 59558411 | 59559065 | 1000 | Distal Intergenic | 4 | 59471868 | 59549364 | 77497 | 2 | ENSMUST00 | -9048 | Ptbp3 | ENSMUSG0( | false | . | . | . | . | . | . | chr4 | 59557291 | 59561763 | 5.13074 | 4.04635 | 3.12464 |
| chr4 | 59548526 | 59549649 | 1000 | Promoter (<=1kb) | 4 | 59471868 | 59549364 | 77497 | 2 | ENSMUST00 | 0 | Ptbp3 | ENSMUSG0( | false | chr4 | 59549506 | 59550194 | 26.78979 | 82.44355 | 79.24416 | . | . | . | . | . | . |
| chr4 | 59805602 | 59806252 | 1000 | Promoter (<=1kb) | 4 | 59805840 | 59904737 | 98898 | 1 | ENSMUST00 | 0 | Snx30 | ENSMUSG0( | false | chr4 | 59805261 | 59807196 | 42.09286 | 140.23274 | 135.34074 | . | . | . | . | . | . |
| chr4 | 62515458 | 62515941 | 1000 | Intron (ENSMUST00000137448.1 | 4 | 62509169 | 62519881 | 10713 | 2 | ENSMUST00 | 3940 | Alad | ENSMUSG0( | false | chr4 | 62515264 | 62516219 | 8.2976 | 15.51635 | 13.24704 | chr4 | 62514012 | 62519352 | 6.75426 | 4.89888 | 4.45045 |
| chr4 | 76135730 | 76136037 | 1000 | Promoter (1-2kb) | 4 | 76133158 | 76134143 | 986 | 2 | ENSMUST00 | -1588 | Ptprd | ENSMUSG0( | true | . | . | . | . | . | . | chr4 | 76135127 | 76136689 | 4.49958 | 3.8553 | 2.6296 |
| chr4 | 76449482 | 76450605 | 1000 | Promoter (<=1kb) | 4 | 75941238 | 76449980 | 508743 | 2 | ENSMUST00 | 0 | Ptprd | ENSMUSG0( | true | chr4 | 76447761 | 76450636 | 29.52499 | 116.10789 | 112.09218 | . | . | . | . | . | . |
| chr4 | 40721974 | 40723365 | 1000 | Promoter (<=1kb) | 4 | 40722150 | 40723795 | 1646 | 1 | ENSMUST00 | 0 | Dnaja1 | ENSMUSG0( | false | chr4 | 40721673 | 40724055 | 32.90589 | 143.88075 | 138.83896 | chr4 | 40721031 | 40722036 | 4.9544 | 3.94642 | 2.99762 |
| chr4 | 53439976 | 53440988 | 1000 | Promoter (<=1kb) | 4 | 53440413 | 53550200 | 109788 | 1 | ENSMUST00 | 0 | Slc44a1 | ENSMUSG0( | false | chr4 | 53439450 | 53441541 | 36.53934 | 116.48274 | 112.45372 | chr4 | 53440872 | 53442103 | 3.17801 | 3.10931 | 1.49868 |
| chr4 | 40874115 | 40874780 | 1000 | Distal Intergenic | 4 | 40809154 | 40854005 | 44852 | 2 | ENSMUST00 | -20111 | B4galt1 | ENSMUSG0( | true | . | . | . | . | . | . | chr4 | 40872190 | 40879418 | 5.9612 | 4.47527 | 3.8083 |
| chr4 | 40970705 | 40971482 | 1000 | Promoter (<=1kb) | 4 | 40970906 | 41012608 | 41703 | 1 | ENSMUST00 | 0 | Nfx1 | ENSMUSG0( | false | chr4 | 40970221 | 40972014 | 36.9999 | 115.1778 | 111.19392 | . | . | . | . | . | . |
| chr4 | 56802148 | 56802819 | 1000 | Promoter (<=1kb) | 4 | 56749680 | 56802331 | 52652 | 2 | ENSMUST00 | 0 | Ikbkap | ENSMUSG0( | false | chr4 | 56801586 | 56803102 | 28.04154 | 116.13699 | 112.1206 | chr4 | 56802808 | 56803650 | 3.21231 | 3.10793 | 1.57206 |
| chr4 | 57143350 | 57143851 | 1000 | Promoter (<=1kb) | 4 | 56991972 | 57143437 | 151466 | 2 | ENSMUST00 | 0 | Epb4.1l4b | ENSMUSG0( | false | chr4 | 57142712 | 57143905 | 21.92336 | 59.65632 | 56.81357 | chr4 | 57141647 | 57143994 | 5.43058 | 4.265 | 3.39554 |
| chr4 | 41387801 | 41388162 | 1000 | Exon (ENSMUST00000108060.4/ | 4 | 41371725 | 41388692 | 16968 | 1 | ENSMUST00 | 16077 | Ubap1 | ENSMUSG0( | false | . | . | . | . | . | . | chr4 | 41383957 | 41390580 | 5.50843 | 4.28488 | 3.4386 |
| chr4 | 41566170 | 41566588 | 1000 | Promoter (2-3kb) | 4 | 41517691 | 41569509 | 51819 | 2 | ENSMUST00 | 2921 | Fam219a | ENSMUSG0( | false | . | . | . | . | . | . | chr4 | 41563319 | 41570177 | 5.29575 | 4.11611 | 3.25298 |
| chr4 | 41639853 | 41640448 | 1000 | Promoter (<=1kb) | 4 | 41638144 | 41640324 | 2181 | 2 | ENSMUST00 | 0 | Enho | ENSMUSG0( | false | chr4 | 41639574 | 41640212 | 19.43846 | 52.88828 | 50.13554 | chr4 | 41638609 | 41640955 | 4.6365 | 3.86589 | 2.73563 |
| chr4 | 43442857 | 43443628 | 1000 | Promoter (<=1kb) | 4 | 43443321 | 43447794 | 4474 | 1 | ENSMUST00 | 0 | Tesk1 | ENSMUSG0( | false | chr4 | 43442217 | 43444119 | 26.86698 | 95.04005 | 91.5885 | . | . | . | . | . | . |
| chr4 | 43492745 | 43493399 | 1000 | Promoter (<=1kb) | 4 | 43492900 | 43494205 | 1306 | 1 | ENSMUST00 | 0 | Ccdc107 | ENSMUSG0( | false | chr4 | 43492637 | 43494474 | 33.62833 | 117.64893 | 113.58318 | . | . | . | . | . | . |
| chr4 | 83485726 | 83486851 | 1000 | Promoter (<=1kb) | 4 | 83455680 | 83486459 | 30780 | 2 | ENSMUST00 | 0 | Psip1 | ENSMUSG0( | false | chr4 | 83485372 | 83487207 | 26.28407 | 85.32412 | 82.07336 | . | . | . | . | . | . |
| chr4 | 87933704 | 87934163 | 1000 | Intron (ENSMUST00000078090.6 | 4 | 87832781 | 87874542 | 41762 | 2 | ENSMUST00 | -59163 | Mllt3 | ENSMUSG0( | false | . | . | . | . | . | . | chr4 | 87933223 | 87936886 | 6.33341 | 4.75754 | 4.09872 |
| chr4 | 87965787 | 87966310 | 1000 | Intron (ENSMUST00000078090.6 | 4 | 87837419 | 88032813 | 195395 | 2 | ENSMUST00 | 66503 | Mllt3 | ENSMUSG0( | false | . | . | . | . | . | . | chr4 | 87965174 | 87967170 | 4.92813 | 4.04539 | 2.99716 |
| chr4 | 105230224 | 105230580 | 1000 | Intron (ENSMUST00000064139.7 | 4 | 105157347 | 105232764 | 75418 | 1 | ENSMUST00 | 72878 | Ppap2b | ENSMUSG0( | false | . | . | . | . | . | . | chr4 | 105228748 | 105233860 | 4.36829 | 3.65429 | 2.53169 |
| chr4 | 105157087 | 105158531 | 1000 | Promoter (<=1kb) | 4 | 105157347 | 105232764 | 75418 | 1 | ENSMUST00 | 0 | Ppap2b | ENSMUSG0( | false | chr4 | 105156812 | 105160389 | 33.08674 | 107.33151 | 103.57004 | . | . | . | . | . | . |
| chr4 | 105088880 | 105089215 | 1000 | Intron (ENSMUST00000030243.7 | 4 | 105029874 | 105109890 | 80017 | 2 | ENSMUST00 | 20675 | Prkaa2 | ENSMUSG0( | false | . | . | . | . | . | . | chr4 | 105086489 | 105092977 | 3.96927 | 3.56321 | 2.15859 |
| chr4 | 105109272 | 105110169 | 1000 | Promoter (<=1kb) | 4 | 105029874 | 105109890 | 80017 | 2 | ENSMUST00 | 0 | Prkaa2 | ENSMUSG0( | false | chr4 | 105108502 | 105110562 | 33.18295 | 113.70766 | 109.76701 | . | . | . | . | . | . |
| chr4 | 103189106 | 103189585 | 1000 | Intron (ENSMUST00000150285.2 | 4 | 103210553 | 103214672 | 4120 | 2 | ENSMUST00 | 25087 | Slc35d1 | ENSMUSG0( | false | . | . | . | . | . | . | chr4 | 103187114 | 103191723 | 6.26381 | 4.72526 | 4.03761 |
| chr4 | 103114128 | 103114990 | 1000 | Promoter (<=1kb) | 4 | 103114390 | 103115904 | 1515 | 1 | ENSMUST00 | 0 | Mier1 | ENSMUSG0( | false | chr4 | 103113428 | 103115228 | 28.09372 | 84.72628 | 81.48907 | . | . | . | . | . | . |
| chr4 | 102759216 | 102759867 | 1000 | Promoter (<=1kb) | 4 | 102759222 | 102883347 | 124126 | 1 | ENSMUST00 | 0 | Sgip1 | ENSMUSG0( | true | chr4 | 102759017 | 102763519 | 24.30838 | 70.74035 | 67.73997 | chr4 | 102757338 | 102759387 | 5.4839 | 4.36075 | 3.39518 |
| chr4 | 102511507 | 102511915 | 1000 | Intron (ENSMUST00000106911.2 | 4 | 102470397 | 102488046 | 17650 | 1 | ENSMUST00 | 41111 | Pde4b | ENSMUSG0( | true | chr4 | 102511137 | 102512052 | 7.14186 | 14.46535 | 12.21151 | chr4 | 102502963 | 102513725 | 5.24222 | 4.19698 | 3.21121 |
| chr4 | 102510940 | 102511268 | 1000 | Intron (ENSMUST00000106911.2 | 4 | 102470397 | 102488046 | 17650 | 1 | ENSMUST00 | 40544 | Pde4b | ENSMUSG0( | true | chr4 | 102511137 | 102512052 | 7.14186 | 14.46535 | 12.21151 | chr4 | 102502963 | 102513725 | 5.24222 | 4.19698 | 3.21121 |
| chr4 | 102569922 | 102570680 | 1000 | Promoter (<=1kb) | 4 | 102570095 | 102605847 | 35753 | 1 | ENSMUST00 | 0 | Pde4b | ENSMUSG0( | true | chr4 | 102568208 | 102575841 | 28.36989 | 85.2065 | 81.96005 | . | . | . | . | . | . |
| chr4 | 101146797 | 101147322 | 1000 | Intron (ENSMUST00000038463.9 | 4 | 101154249 | 101155119 | 871 | 2 | ENSMUST00 | 7797 | Jak1 | ENSMUSG0( | false | . | . | . | . | . | . | chr4 | 101141547 | 101149273 | 5.88936 | 4.49025 | 3.75587 |
| chr4 | 101164187 | 101164499 | 1000 | Exon (ENSMUST00000102781.4/ | 4 | 101154249 | 101155119 | 871 | 2 | ENSMUST00 | -9069 | Jak1 | ENSMUSG0( | false | . | . | . | . | . | . | chr4 | 101163490 | 101165968 | 6.91547 | 5.06184 | 4.56003 |
| chr4 | 101182534 | 101183028 | 1000 | Promoter (2-3kb) | 4 | 101179591 | 101185425 | 5835 | 2 | ENSMUST00 | 2397 | Jak1 | ENSMUSG0( | false | . | . | . | . | . | . | chr4 | 101181182 | 101183583 | 3.81139 | 3.3875 | 2.07097 |
| chr4 | 118123108 | 118123481 | 1000 | Intron (ENSMUST00000030263.3 | 4 | 117932160 | 118134882 | 202723 | 2 | ENSMUST00 | 11401 | St3gal3 | ENSMUSG0( | true | . | . | . | . | . | . | chr4 | 118119621 | 118134187 | 5.32651 | 4.16451 | 3.29068 |
| chr4 | 118015791 | 118016171 | 1000 | Intron (ENSMUST00000030263.3 | 4 | 117940354 | 118031882 | 91529 | 2 | ENSMUST00 | 15711 | St3gal3 | ENSMUSG0( | true | . | . | . | . | . | . | chr4 | 118013880 | 118023443 | 5.10307 | 3.90399 | 3.12768 |
| chr4 | 118103109 | 118103499 | 1000 | Intron (ENSMUST00000030263.3 | 4 | 117932160 | 118134882 | 202723 | 2 | ENSMUST00 | 31383 | St3gal3 | ENSMUSG0( | true | . | . | . | . | . | . | chr4 | 118092492 | 118118436 | 4.65273 | 3.87829 | 2.74128 |
| chr4 | 117835855 | 117836793 | 1000 | Promoter (<=1kb) | 4 | 117835438 | 117857171 | 21734 | 1 | ENSMUST00 | 418 | Slc6a9 | ENSMUSG0( | false | chr4 | 117835153 | 117836940 | 26.1063 | 98.41757 | 94.88632 | . | . | . | . | . | . |
| chr4 | 117835053 | 117835531 | 1000 | Promoter (<=1kb) | 4 | 117835333 | 117875196 | 39864 | 1 | ENSMUST00 | 0 | Slc6a9 | ENSMUSG0( | false | chr4 | 117835153 | 117836940 | 26.1063 | 98.41757 | 94.88632 | chr4 | 117834903 | 117835570 | 4.36839 | 3.73728 | 2.48824 |
| chr4 | 111432295 | 111432648 | 1000 | Promoter (<=1kb) | 4 | 111433197 | 111460296 | 27100 | 1 | ENSMUST00 | -549 | Bend5 | ENSMUSG0( | false | . | . | . | . | . | . | chr4 | 111431432 | 111434570 | 5.44711 | 4.23984 | 3.42447 |
| chr4 | 109666475 | 109667193 | 1000 | Promoter (<=1kb) | 4 | 109660876 | 109667189 | 6314 | 2 | ENSMUST00 | 0 | Cdkn2c | ENSMUSG0( | false | chr4 | 109664951 | 109667518 | 26.52689 | 93.43019 | 90.01563 | . | . | . | . | . | . |
| chr4 | 109280098 | 109280905 | 1000 | Promoter (<=1kb) | 4 | 109280268 | 109387817 | 107550 | 1 | ENSMUST00 | 0 | Eps15 | ENSMUSG0( | false | chr4 | 109279957 | 109281494 | 39.93851 | 132.06328 | 127.49693 | chr4 | 109279037 | 109285662 | 5.18324 | 4.14445 | 3.14817 |
| chr4 | 99120253 | 99121165 | 1000 | Promoter (<=1kb) | 4 | 98936665 | 99120915 | 184251 | 2 | ENSMUST00 | 0 | Dock7 | ENSMUSG0( | false | chr4 | 99120070 | 99121403 | 30.11778 | 90.15907 | 86.81646 | chr4 | 99115814 | 99120323 | 4.02327 | 3.53714 | 2.20238 |
| chr4 | 109229770 | 109230109 | 1000 | Distal Intergenic | 4 | 109234485 | 109254251 | 19767 | 1 | ENSMUST00 | -4376 | Calr4 | ENSMUSG0( | false | . | . | . | . | . | . | chr4 | 109229533 | 109230353 | 4.08542 | 3.36327 | 2.28724 |
| chr4 | 109103400 | 109104007 | 1000 | Intron (ENSMUST00000084366.7 | 4 | 109087410 | 109108031 | 20622 | 2 | ENSMUST00 | 4024 | Osbpl9 | ENSMUSG0( | true | . | . | . | . | . | . | chr4 | 109078531 | 109116893 | 5.3508 | 4.12801 | 3.31157 |
| chr4 | 109117752 | 109118367 | 1000 | Promoter (<=1kb) | 4 | 109061145 | 109118022 | 56878 | 2 | ENSMUST00 | 0 | Osbpl9 | ENSMUSG0( | true | chr4 | 109116279 | 109118694 | 14.97413 | 40.41832 | 37.82371 | chr4 | 109118252 | 109153965 | 5.18833 | 4.10127 | 3.17184 |
| chr4 | 109149105 | 109149617 | 1000 | Intron (ENSMUST00000160774.2 | 4 | 109062087 | 109156610 | 94524 | 2 | ENSMUST00 | 6993 | Osbpl9 | ENSMUSG0( | true | . | . | . | . | . | . | chr4 | 109118252 | 109153965 | 5.18833 | 4.10127 | 3.17184 |
| chr4 | 98923008 | 98924363 | 1000 | Promoter (<=1kb) | 4 | 98923810 | 98935543 | 11734 | 1 | ENSMUST00 | 0 | Usp1 | ENSMUSG0( | false | chr4 | 98922058 | 98924971 | 37.63617 | 126.94061 | 122.55849 | . | . | . | . | . | . |
| chr4 | 97774912 | 97775449 | 1000 | Promoter (2-3kb) | 4 | 97777608 | 98118871 | 341264 | 1 | ENSMUST00 | -2159 | Nfia | ENSMUSG0( | false | chr4 | 97774995 | 97776568 | 13.83063 | 32.8443 | 30.34371 | chr4 | 97773491 | 97777053 | 4.35712 | 3.65762 | 2.50478 |
| chr4 | 97772072 | 97772605 | 1000 | Promoter (<=1kb) | 4 | 97772734 | 98000466 | 227733 | 1 | ENSMUST00 | -129 | Nfia | ENSMUSG0( | false | chr4 | 97771503 | 97773573 | 21.79372 | 61.12524 | 58.26245 | . | . | . | . | . | . |
| chr4 | 97776665 | 97778991 | 1000 | Promoter (<=1kb) | 4 | 97777608 | 98118871 | 341264 | 1 | ENSMUST00 | 0 | Nfia | ENSMUSG0( | false | chr4 | 97776660 | 97779285 | 35.45657 | 113.55091 | 109.61168 | chr4 | 97773491 | 97777053 | 4.35712 | 3.65762 | 2.50478 |
| chr4 | 130663038 | 130664569 | 1000 | Promoter (<=1kb) | 4 | 130663321 | 130781564 | 118244 | 1 | ENSMUST00 | 0 | Pum1 | ENSMUSG0( | false | chr4 | 130662855 | 130664831 | 30.78991 | 105.27288 | 101.56436 | . | . | . | . | . | . |
| chr4 | 143283923 | 143284533 | 1000 | Intron (ENSMUST00000030317.8 | 4 | 143268346 | 143299463 | 31118 | 2 | ENSMUST00 | 14930 | Pdpn | ENSMUSG0( | true | . | . | . | . | . | . | chr4 | 143277480 | 143296542 | 5.36947 | 4.22359 | 3.31454 |
| chr4 | 143281278 | 143281659 | 1000 | Intron (ENSMUST00000030317.8 | 4 | 143268346 | 143299463 | 31118 | 2 | ENSMUST00 | 17804 | Pdpn | ENSMUSG0( | true | . | . | . | . | . | . | chr4 | 143277480 | 143296542 | 5.36947 | 4.22359 | 3.31454 |
| chr4 | 107801876 | 107802402 | 1000 | Promoter (<=1kb) | 4 | 107802261 | 107874167 | 71907 | 1 | ENSMUST00 | 0 | Lrp8 | ENSMUSG0( | false | chr4 | 107801651 | 107804211 | 19.04495 | 59.55748 | 56.71581 | chr4 | 107801652 | 107804349 | 5.50588 | 4.26202 | 3.44614 |
| chr4 | 107253068 | 107254055 | 1000 | Promoter (<=1kb) | 4 | 107233514 | 107253532 | 20019 | 2 | ENSMUST00 | 0 | Lrrc42 | ENSMUSG0( | false | chr4 | 107252934 | 107254526 | 24.60788 | 97.30984 | 93.80507 | . | . | . | . | . | . |
| chr4 | 120747295 | 120747708 | 1000 | Promoter (<=1kb) | 4 | 120696138 | 120747248 | 51111 | 2 | ENSMUST00 | -48 | Kcnq4 | ENSMUSG0( | false | chr4 | 120746669 | 120747856 | 30.17592 | 94.17345 | 90.73757 | chr4 | 120746793 | 120748407 | 3.65921 | 3.45413 | 1.89786 |
| chr4 | 119959346 | 119959786 | 1000 | Intron (ENSMUST00000143837.1 | 4 | 120094489 | 120134557 | 40069 | 1 | ENSMUST00 | -134703 | Hivep3 | ENSMUSG0( | false | . | . | . | . | . | . | chr4 | 119959265 | 119960702 | 4.05547 | 3.56823 | 2.26748 |
| chr4 | 120161007 | 120161461 | 1000 | Promoter (<=1kb) | 4 | 120161206 | 120167360 | 6155 | 1 | ENSMUST00 | 0 | Edn2 | ENSMUSG0( | false | . | . | . | . | . | . | chr4 | 120159047 | 120164442 | 5.41379 | 4.28743 | 3.34632 |
| chr4 | 120172089 | 120172437 | 1000 | Distal Intergenic | 4 | 120161206 | 120167360 | 6155 | 1 | ENSMUST00 | 10884 | Edn2 | ENSMUSG0( | false | . | . | . | . | . | . | chr4 | 120167621 | 120173542 | 4.94324 | 4.02027 | 3.0083 |
| chr4 | 120148666 | 120149007 | 1000 | Distal Intergenic | 4 | 120161206 | 120167360 | 6155 | 1 | ENSMUST00 | -12199 | Edn2 | ENSMUSG0( | false | . | . | . | . | . | . | chr4 | 120148364 | 120152992 | 6.6025 | 4.82084 | 4.30672 |
| chr4 | 119244403 | 119244887 | 1000 | Promoter (<=1kb) | 4 | 119245020 | 119248133 | 3114 | 1 | ENSMUST00 | -133 | Lepre1 | ENSMUSG0( | false | . | . | . | . | . | . | chr4 | 119237720 | 119248571 | 6.98267 | 4.77088 | 4.67122 |
| chr4 | 119108608 | 119109477 | 1000 | Promoter (<=1kb) | 4 | 119108711 | 119137330 | 28620 | 1 | ENSMUST00 | 0 | Slc2a1 | ENSMUSG0( | false | chr4 | 119108327 | 119109801 | 33.94791 | 109.87469 | 106.04007 | . | . | . | . | . | . |
| chr4 | 123411544 | 123412567 | 1000 | Intron (ENSMUST00000097897.5 | 4 | 123397275 | 123407515 | 10241 | 2 | ENSMUST00 | -4030 | Macf1 | ENSMUSG0( | true | chr4 | 123410776 | 123412702 | 38.13901 | 127.82562 | 123.40861 | chr4 | 123412273 | 123429396 | 5.99663 | 4.50351 | 3.82082 |
| chr4 | 123680847 | 123681256 | 1000 | Intron (ENSMUST00000097897.5 | 4 | 123349633 | 123684360 | 334728 | 2 | ENSMUST00 | 3104 | Macf1 | ENSMUSG0( | true | . | . | . | . | . | . | chr4 | 123680314 | 123684036 | 6.59939 | 4.7744 | 4.34308 |
| chr4 | 123663939 | 123665183 | 1000 | Promoter (<=1kb) | 4 | 123456349 | 123664752 | 208404 | 2 | ENSMUST00 | 0 | Macf1 | ENSMUSG0( | true | chr4 | 123662851 | 123665633 | 27.48563 | 91.33697 | 87.96664 | . | . | . | . | . | . |
| chr4 | 123005081 | 123005552 | 1000 | Distal Intergenic | 4 | 122999339 | 123000517 | 1179 | 1 | ENSMUST00 | 5743 | Mycl | ENSMUSG0( | false | . | . | . | . | . | . | chr4 | 123001757 | 123006115 | 4.15155 | 3.63106 | 2.34596 |
| chr4 | 122955270 | 122955719 | 1000 | Intron (ENSMUST00000152308.2 | 4 | 122956732 | 122959990 | 3259 | 2 | ENSMUST00 | 4271 | Mfsd2a | ENSMUSG0( | false | . | . | . | . | . | . | chr4 | 122954640 | 122956079 | 6.54975 | 4.80752 | 4.27091 |
| chr4 | 122885798 | 122886256 | 1000 | Promoter (<=1kb) | 4 | 122859047 | 122885911 | 26865 | 2 | ENSMUST00 | 0 | Cap1 | ENSMUSG0( | false | chr4 | 122884994 | 122885972 | 43.75897 | 147.62399 | 142.42812 | . | . | . | . | . | . |
| chr4 | 135949810 | 135950277 | 1000 | Promoter (1-2kb) | 4 | 135948530 | 135958749 | 10220 | 1 | ENSMUST00 | 1281 | Hmgcl | ENSMUSG0( | false | . | . | . | . | . | . | chr4 | 135947277 | 135951324 | 7.33391 | 5.10337 | 4.93063 |
| chr4 | 135918955 | 135919336 | 1000 | Promoter (1-2kb) | 4 | 135920735 | 135940311 | 19577 | 1 | ENSMUST00 | -1399 | Fuca1 | ENSMUSG0( | false | . | . | . | . | . | . | chr4 | 135918448 | 135922378 | 4.72722 | 3.88867 | 2.82458 |
| chr4 | 135876606 | 135877056 | 1000 | Promoter (2-3kb) | 4 | 135870918 | 135873850 | 2933 | 2 | ENSMUST00 | -2757 | Pnrc2 | ENSMUSG0( | false | chr4 | 135876153 | 135877006 | 10.96168 | 23.03899 | 20.66252 | chr4 | 135874210 | 135879466 | 7.31253 | 5.17261 | 4.90109 |
| chr4 | 135855718 | 135856719 | 1000 | Promoter (<=1kb) | 4 | 135855747 | 135865815 | 10069 | 1 | ENSMUST00 | 0 | Srsf10 | ENSMUSG0( | false | chr4 | 135855184 | 135857243 | 28.22734 | 101.53345 | 97.92473 | . | . | . | . | . | . |
| chr4 | 117299956 | 117300277 | 1000 | Promoter (<=1kb) | 4 | 117285889 | 117300028 | 14140 | 2 | ENSMUST00 | 0 | Rnf220 | ENSMUSG0( | false | . | . | . | . | . | . | chr4 | 117299438 | 117304815 | 5.89906 | 4.2418 | 3.7845 |
| chr4 | 117484664 | 117485264 | 1000 | Intron (ENSMUST00000030439.9 | 4 | 117490235 | 117495011 | 4777 | 2 | ENSMUST00 | 9747 | Rnf220 | ENSMUSG0( | false | chr4 | 117483689 | 117485136 | 21.02725 | 63.18829 | 60.29588 | chr4 | 117483621 | 117485642 | 4.64697 | 3.79413 | 2.74242 |
| chr4 | 116220798 | 116222015 | 1000 | Promoter (<=1kb) | 4 | 116221618 | 116259576 | 37959 | 1 | ENSMUST00 | 0 | Pik3r3 | ENSMUSG0( | false | chr4 | 116221357 | 116223082 | 27.62344 | 81.07606 | 77.90333 | . | . | . | . | . | . |
| chr4 | 116160687 | 116161129 | 1000 | Downstream (<1kb) | 4 | 116157392 | 116159025 | 1634 | 1 | ENSMUST00 | 3296 | Pomgnt1 | ENSMUSG0( | false | . | . | . | . | . | . | chr4 | 116160310 | 116163250 | 3.88843 | 3.42834 | 2.1395 |
| chr4 | 115827815 | 115828663 | 1000 | Promoter (<=1kb) | 4 | 115828092 | 115836185 | 8094 | 1 | ENSMUST00 | 0 | Mob3c | ENSMUSG0( | false | chr4 | 115827538 | 115828949 | 25.71568 | 110.22335 | 106.38094 | chr4 | 115827101 | 115830708 | 3.84654 | 3.3576 | 2.09622 |
| chr4 | 115784745 | 115785190 | 1000 | Promoter (<=1kb) | 4 | 115784812 | 115798006 | 13195 | 1 | ENSMUST00 | 0 | Atpaf1 | ENSMUSG0( | false | chr4 | 115784604 | 115785862 | 38.16103 | 126.78209 | 122.40619 | chr4 | 115784203 | 115787303 | 4.48855 | 3.86239 | 2.62516 |

| | | | | | | | | | | | | | | | | | | | | | | | | | | |
|---|---|---|---|---|---|---|---|---|---|---|---|---|---|---|---|---|---|---|---|---|---|---|---|---|---|---|
| chr4 | 139250095 | 139250476 | 1000 | Intron (ENSMUST00000102508.4 | 4 | 139233383 | 139291783 | 58401 | 1 | ENSMUST00 | 16713 | Capzb | ENSMUSG0C | false | . | . | . | . | . | . | chr4 | 139241332 | 139253338 | 6.42884 | 4.70135 | 4.1813 |
| chr4 | 138216906 | 138217593 | 1000 | Promoter (<=1kb) | 4 | 138217127 | 138229757 | 12631 | 1 | ENSMUST00 | 0 | Hp1bp3 | ENSMUSG0C | false | chr4 | 138215850 | 138218184 | 36.7977 | 133.03651 | 128.43381 | . | . | . | . | . | . |
| chr4 | 137984228 | 137984603 | 1000 | Distal Intergenic | 4 | 137993022 | 138105419 | 112398 | 1 | ENSMUST00 | -8419 | Eif4g3 | ENSMUSG0C | false | . | . | . | . | . | . | chr4 | 137981682 | 137986058 | 4.74012 | 3.90891 | 2.83992 |
| chr4 | 137485199 | 137485679 | 1000 | Intron (ENSMUST00000030547.9 | 4 | 137468803 | 137570629 | 101827 | 1 | ENSMUST00 | 16397 | Hspg2 | ENSMUSG0C | false | . | . | . | . | . | . | chr4 | 137484181 | 137487113 | 4.37241 | 3.84074 | 2.48161 |
| chr5 | 21055270 | 21056477 | 1000 | Promoter (<=1kb) | 5 | 20986645 | 21055911 | 69267 | 2 | ENSMUST00 | 0 | Ptpn12 | ENSMUSG0C | true | chr5 | 21053536 | 21057043 | 37.92415 | 138.37384 | 133.55817 | . | . | . | . | . | . |
| chr4 | 129108529 | 129108927 | 1000 | Promoter (1-2kb) | 4 | 129106869 | 129110544 | 3676 | 1 | ENSMUST00 | 1661 | Tmem54 | ENSMUSG0C | false | . | . | . | . | . | . | chr4 | 129106734 | 129114119 | 7.73893 | 5.31745 | 5.25102 |
| chr4 | 129096593 | 129096995 | 1000 | Distal Intergenic | 4 | 129105548 | 129111626 | 6079 | 1 | ENSMUST00 | -8553 | Tmem54 | ENSMUSG0C | false | . | . | . | . | . | . | chr4 | 129095284 | 129100764 | 4.19173 | 3.68477 | 2.35145 |
| chr4 | 129819714 | 129820954 | 1000 | Promoter (<=1kb) | 4 | 129820221 | 129849978 | 29758 | 1 | ENSMUST00 | 0 | Ptp4a2 | ENSMUSG0C | false | chr4 | 129819570 | 129821757 | 26.65955 | 86.95457 | 83.67183 | chr4 | 129818600 | 129819771 | 5.74365 | 4.5042 | 3.61498 |
| chr4 | 129742084 | 129742893 | 1000 | Promoter (<=1kb) | 4 | 129703164 | 129742297 | 39134 | 2 | ENSMUST00 | 0 | Khdrbs1 | ENSMUSG0C | false | chr4 | 129740674 | 129743306 | 43.83784 | 156.88544 | 151.34334 | . | . | . | . | . | . |
| chr4 | 128734121 | 128734565 | 1000 | Intron (ENSMUST00000030588.7 | 4 | 128727982 | 128747132 | 19151 | 1 | ENSMUST00 | 6140 | Phc2 | ENSMUSG0C | false | chr4 | 128733817 | 128734160 | 10.15658 | 21.13245 | 18.78179 | chr4 | 128728570 | 128744451 | 8.13086 | 5.39142 | 5.59738 |
| chr4 | 128727278 | 128728149 | 1000 | Promoter (<=1kb) | 4 | 128727581 | 128752881 | 25301 | 1 | ENSMUST00 | 0 | Phc2 | ENSMUSG0C | false | chr4 | 128727024 | 128730437 | 31.25623 | 96.05631 | 92.57859 | . | . | . | . | . | . |
| chr4 | 129542582 | 129543028 | 1000 | Promoter (<=1kb) | 4 | 129516104 | 129542713 | 26610 | 2 | ENSMUST00 | 0 | Hdac1 | ENSMUSG0C | false | chr4 | 129541578 | 129543395 | 28.41342 | 89.1249 | 85.7944 | . | . | . | . | . | . |
| chr4 | 129378640 | 129379058 | 1000 | Promoter (<=1kb) | 4 | 129353628 | 129378116 | 24489 | 2 | ENSMUST00 | -525 | Zbtb8a | ENSMUSG0C | true | . | . | . | . | . | . | chr4 | 129378069 | 129387044 | 4.79047 | 3.93835 | 2.85504 |
| chr4 | 135352808 | 135353769 | 1000 | Promoter (<=1kb) | 4 | 135320484 | 135353321 | 32838 | 2 | ENSMUST00 | 0 | Srrm1 | ENSMUSG0C | false | chr4 | 135352017 | 135353251 | 36.14217 | 127.51552 | 123.12013 | . | . | . | . | . | . |
| chr4 | 127907220 | 127907453 | 1000 | Distal Intergenic | 4 | 127927592 | 127970951 | 43360 | 2 | ENSMUST00 | 63498 | CK137956 | ENSMUSG0C | false | . | . | . | . | . | . | chr4 | 127903416 | 127908019 | 5.64655 | 4.40601 | 3.54849 |
| chr4 | 134798805 | 134799165 | 1000 | Distal Intergenic | 4 | 134802759 | 134811213 | 8455 | 2 | ENSMUST00 | 12048 | Tmem57 | ENSMUSG0C | false | . | . | . | . | . | . | chr4 | 134795977 | 134799487 | 3.57784 | 3.26114 | 1.86282 |
| chr4 | 126753294 | 126754134 | 1000 | Promoter (<=1kb) | 4 | 126753544 | 126791894 | 38351 | 1 | ENSMUST00 | 0 | AU040320 | ENSMUSG0C | false | chr4 | 126752441 | 126754692 | 30.78991 | 105.27288 | 101.56436 | . | . | . | . | . | . |
| chr4 | 134370491 | 134371082 | 1000 | Promoter (1-2kb) | 4 | 134356372 | 134372558 | 16187 | 2 | ENSMUST00 | 1476 | Extl1 | ENSMUSG0C | false | chr4 | 134370510 | 134372917 | 25.7338 | 71.40817 | 68.39619 | chr4 | 134370182 | 134373591 | 4.152 | 3.65801 | 2.34914 |
| chr4 | 126259555 | 126260189 | 1000 | Promoter (2-3kb) | 4 | 126262325 | 126275883 | 13559 | 1 | ENSMUST00 | -2136 | Trappc3 | ENSMUSG0C | false | . | . | . | . | . | . | chr4 | 126258644 | 126263639 | 8.15369 | 5.4381 | 5.58566 |
| chr4 | 126227179 | 126227666 | 1000 | Distal Intergenic | 4 | 126232963 | 126233548 | 586 | 2 | ENSMUST00 | 5882 | Map7d1 | ENSMUSG0C | false | chr4 | 126227181 | 126227508 | 4.57821 | 7.98325 | 5.85079 | chr4 | 126227116 | 126227984 | 4.92277 | 4.07484 | 2.94025 |
| chr4 | 133378239 | 133378748 | 1000 | Intron (ENSMUST00000030669.7 | 4 | 133370644 | 133412136 | 41493 | 1 | ENSMUST00 | 7596 | Slc9a1 | ENSMUSG0C | false | . | . | . | . | . | . | chr4 | 133376103 | 133381987 | 6.34167 | 4.47881 | 4.14398 |
| chr4 | 133391232 | 133391572 | 1000 | Intron (ENSMUST00000030669.7 | 4 | 133370644 | 133412136 | 41493 | 1 | ENSMUST00 | 20589 | Slc9a1 | ENSMUSG0C | false | . | . | . | . | . | . | chr4 | 133388640 | 133392585 | 5.44586 | 4.13417 | 3.40204 |
| chr4 | 133377482 | 133377834 | 1000 | Intron (ENSMUST00000030669.7 | 4 | 133370644 | 133412136 | 41493 | 1 | ENSMUST00 | 6839 | Slc9a1 | ENSMUSG0C | false | . | . | . | . | . | . | chr4 | 133376103 | 133381987 | 6.34167 | 4.47881 | 4.14398 |
| chr4 | 125114648 | 125115134 | 1000 | Distal Intergenic | 4 | 125107976 | 125109428 | 1453 | 1 | ENSMUST00 | 6673 | Meaf6 | ENSMUSG0C | false | . | . | . | . | . | . | chr4 | 125106469 | 125127334 | 7.64816 | 5.22391 | 5.18478 |
| chr4 | 133117569 | 133118122 | 1000 | Distal Intergenic | 4 | 133130505 | 133199756 | 69252 | 1 | ENSMUST00 | -12383 | Wasf2 | ENSMUSG0C | true | . | . | . | . | . | . | chr4 | 133116131 | 133119516 | 5.27243 | 4.1108 | 3.23852 |
| chr4 | 124850058 | 124850913 | 1000 | Promoter (<=1kb) | 4 | 124850679 | 124855242 | 4564 | 1 | ENSMUST00 | 0 | Yrdc | ENSMUSG0C | false | chr4 | 124849422 | 124852147 | 42.27695 | 136.11826 | 131.39047 | . | . | . | . | . | . |
| chr4 | 132261057 | 132261724 | 1000 | Promoter (<=1kb) | 4 | 132222946 | 132261521 | 38576 | 2 | ENSMUST00 | 0 | Gmeb1 | ENSMUSG0C | false | chr4 | 132260423 | 132262171 | 33.15098 | 102.34641 | 98.71844 | . | . | . | . | . | . |
| chr4 | 124708622 | 124709276 | 1000 | Distal Intergenic | 4 | 124714776 | 124732460 | 17685 | 1 | ENSMUST00 | -5500 | Sf3a3 | ENSMUSG0C | false | chr4 | 124708582 | 124709363 | 11.90309 | 30.75663 | 28.28125 | chr4 | 124707919 | 124709324 | 4.20088 | 3.66847 | 2.34474 |
| chr4 | 132048531 | 132049388 | 1000 | Promoter (<=1kb) | 4 | 131923429 | 132049078 | 125650 | 2 | ENSMUST00 | 0 | Epb4.1 | ENSMUSG0C | false | chr4 | 132047947 | 132049501 | 33.06624 | 113.49754 | 109.55873 | . | . | . | . | . | . |
| chr4 | 131873361 | 131874240 | 1000 | Promoter (<=1kb) | 4 | 131873617 | 131901706 | 28090 | 1 | ENSMUST00 | 0 | Srsf4 | ENSMUSG0C | false | chr4 | 131872840 | 131874986 | 41.76281 | 135.22986 | 130.53618 | . | . | . | . | . | . |
| chr4 | 141793323 | 141794019 | 1000 | Promoter (<=1kb) | 4 | 141793612 | 141798086 | 4475 | 1 | ENSMUST00 | 0 | Casp9 | ENSMUSG0C | false | chr4 | 141793602 | 141794545 | 30.4989 | 91.7916 | 88.41422 | . | . | . | . | . | . |
| chr4 | 152185276 | 152186246 | 1000 | Promoter (<=1kb) | 4 | 152185765 | 152217796 | 32032 | 1 | ENSMUST00 | 0 | Acot7 | ENSMUSG0C | false | chr4 | 152185346 | 152186529 | 6.80177 | 13.42905 | 11.19262 | chr4 | 152184507 | 152187757 | 6.3131 | 4.6237 | 4.09891 |
| chr4 | 152153609 | 152154036 | 1000 | Promoter (1-2kb) | 4 | 152120336 | 152152371 | 32036 | 2 | ENSMUST00 | -1239 | Espn | ENSMUSG0C | false | chr4 | 152153765 | 152154225 | 7.96309 | 14.9624 | 12.70017 | chr4 | 152151222 | 152154727 | 5.03763 | 4.15389 | 3.04161 |
| chr4 | 152130741 | 152131244 | 1000 | Promoter (<=1kb) | 4 | 152128142 | 152131230 | 3089 | 2 | ENSMUST00 | 0 | Espn | ENSMUSG0C | false | chr4 | 152130570 | 152131958 | 14.28371 | 39.87881 | 37.29034 | chr4 | 152130436 | 152132154 | 8.52612 | 5.76366 | 5.8734 |
| chr5 | 24907750 | 24909024 | 1000 | Promoter (<=1kb) | 5 | 24862748 | 24908469 | 45722 | 2 | ENSMUST00 | 0 | Prkag2 | ENSMUSG0C | false | chr5 | 24907736 | 24909252 | 19.50994 | 80.00812 | 76.85559 | . | . | . | . | . | . |
| chr5 | 24842175 | 24843332 | 1000 | Promoter (<=1kb) | 5 | 24802823 | 24842624 | 39802 | 2 | ENSMUST00 | 0 | Rheb | ENSMUSG0C | false | chr5 | 24840760 | 24842707 | 34.74108 | 114.39095 | 110.42815 | . | . | . | . | . | . |
| chr5 | 24604604 | 24605206 | 1000 | Promoter (2-3kb) | 5 | 24592622 | 24602012 | 9391 | 2 | ENSMUST00 | -2593 | Smarcd3 | ENSMUSG0C | false | chr5 | 24604312 | 24605347 | 25.34239 | 74.34718 | 71.28911 | chr5 | 24601136 | 24606675 | 7.02969 | 4.92989 | 4.67941 |
| chr5 | 24444961 | 24445410 | 1000 | Promoter (<=1kb) | 5 | 24438440 | 24445194 | 6755 | 2 | ENSMUST00 | 0 | Fastk | ENSMUSG0C | false | chr5 | 24444466 | 24445612 | 23.61999 | 85.9385 | 82.67356 | chr5 | 24443601 | 24448395 | 3.70015 | 3.3356 | 1.98478 |
| chr4 | 150863013 | 150863381 | 1000 | Intron (ENSMUST00000073600.3 | 4 | 150855073 | 150868887 | 13815 | 1 | ENSMUST00 | 7941 | Errfi1 | ENSMUSG0C | false | . | . | . | . | . | . | chr4 | 150855582 | 150865541 | 4.69628 | 3.88452 | 2.77676 |
| chr4 | 149108915 | 149109265 | 1000 | Promoter (1-2kb) | 4 | 149107769 | 149108234 | 466 | 1 | ENSMUST00 | 1147 | Dffa | ENSMUSG0C | false | . | . | . | . | . | . | chr4 | 149108669 | 149110047 | 3.5294 | 3.35161 | 1.77934 |
| chr4 | 148900226 | 148901708 | 1000 | Promoter (<=1kb) | 4 | 148901149 | 148932896 | 31748 | 1 | ENSMUST00 | 0 | Casz1 | ENSMUSG0C | true | chr4 | 148900254 | 148902977 | 36.76141 | 116.86796 | 112.83147 | chr4 | 148899659 | 148900708 | 3.28238 | 3.21283 | 1.63769 |
| chr4 | 148923765 | 148924413 | 1000 | Intron (ENSMUST00000094464.4 | 4 | 148933114 | 148940511 | 7398 | 1 | ENSMUST00 | -8701 | Casz1 | ENSMUSG0C | true | . | . | . | . | . | . | chr4 | 148901819 | 148933146 | 7.00771 | 4.91323 | 4.66351 |
| chr4 | 149517959 | 149518525 | 1000 | Promoter (<=1kb) | 4 | 149518236 | 149566437 | 48202 | 2 | ENSMUST00 | 0 | Ctnnbip1 | ENSMUSG0C | false | chr4 | 149518122 | 149519059 | 28.50037 | 84.48055 | 81.24493 | chr4 | 149518326 | 149523161 | 4.73706 | 3.91161 | 2.81733 |
| chr4 | 148517760 | 148518664 | 1000 | Intron (ENSMUST00000103221.4 | 4 | 148495183 | 148500460 | 5278 | 2 | ENSMUST00 | -17301 | Angptl7 | ENSMUSG0C | false | . | . | . | . | . | . | chr4 | 148507077 | 148521698 | 5.44417 | 4.08041 | 3.40789 |
| chr4 | 153968393 | 153968983 | 1000 | Intron (ENSMUST00000030893.2 | 4 | 153969130 | 153975051 | 5922 | 2 | ENSMUST00 | 6068 | Dffb | ENSMUSG0C | false | . | . | . | . | . | . | chr4 | 153965866 | 153969508 | 5.82524 | 4.34627 | 3.6948 |
| chr4 | 154293811 | 154294453 | 1000 | Intron (ENSMUST00000144145.2 | 4 | 154278486 | 154299892 | 21407 | 2 | ENSMUST00 | 5439 | Arhgef16 | ENSMUSG0C | true | chr4 | 154294295 | 154295715 | 10.25605 | 26.45831 | 24.03902 | chr4 | 154277148 | 154295397 | 6.63389 | 4.70774 | 4.36575 |
| chr4 | 154279494 | 154280158 | 1000 | Promoter (2-3kb) | 4 | 154278486 | 154282499 | 4014 | 2 | ENSMUST00 | 2341 | Arhgef16 | ENSMUSG0C | true | . | . | . | . | . | . | chr4 | 154277148 | 154295397 | 6.63389 | 4.70774 | 4.36575 |
| chr4 | 155150472 | 155151115 | 1000 | Downstream (2-3kb) | 4 | 155154075 | 155188296 | 34222 | 2 | ENSMUST00 | 37181 | Ski | ENSMUSG0C | false | chr4 | 155150178 | 155151714 | 24.46993 | 79.22128 | 76.07978 | chr4 | 155144041 | 155153706 | 6.53431 | 4.75254 | 4.26776 |
| chr4 | 155178247 | 155178627 | 1000 | Intron (ENSMUST00000129314.1 | 4 | 155154075 | 155188296 | 34222 | 2 | ENSMUST00 | 9669 | Ski | ENSMUSG0C | false | . | . | . | . | . | . | chr4 | 155159960 | 155221990 | 6.99931 | 4.87167 | 4.66225 |
| chr4 | 155280337 | 155280956 | 1000 | Intron (ENSMUST00000103178.5 | 4 | 155271133 | 155273042 | 1910 | 2 | ENSMUST00 | -7296 | Prkcz | ENSMUSG0C | false | . | . | . | . | . | . | chr4 | 155279978 | 155286227 | 8.23924 | 5.53895 | 5.64266 |
| chr4 | 155282155 | 155282934 | 1000 | Intron (ENSMUST00000103178.5 | 4 | 155271133 | 155273042 | 1910 | 2 | ENSMUST00 | -9114 | Prkcz | ENSMUSG0C | false | . | . | . | . | . | . | chr4 | 155279978 | 155286227 | 8.23924 | 5.53895 | 5.64266 |
| chr4 | 155662650 | 155662989 | 1000 | Intron (ENSMUST00000151843.2 | 4 | 155656943 | 155659583 | 2641 | 2 | ENSMUST00 | -3068 | Mib2 | ENSMUSG0C | false | . | . | . | . | . | . | chr4 | 155660616 | 155668643 | 5.1671 | 4.02817 | 3.14947 |
| chr4 | 155651564 | 155652039 | 1000 | Promoter (<=1kb) | 4 | 155650657 | 155652025 | 1369 | 2 | ENSMUST00 | 0 | Mmp23 | ENSMUSG0C | false | chr4 | 155651243 | 155655658 | 26.60493 | 74.70471 | 71.64091 | chr4 | 155648087 | 155656351 | 6.88407 | 4.99201 | 4.54195 |
| chr4 | 155563211 | 155564102 | 1000 | Promoter (<=1kb) | 4 | 155563700 | 155591001 | 27302 | 1 | ENSMUST00 | 0 | Nadk | ENSMUSG0C | false | chr4 | 155562706 | 155563361 | 15.10403 | 36.38161 | 33.83695 | . | . | . | . | . | . |
| chr5 | 50058606 | 50059280 | 1000 | Promoter (<=1kb) | 5 | 49959951 | 50058996 | 99046 | 2 | ENSMUST00 | 0 | Gpr125 | ENSMUSG0C | false | chr5 | 50057827 | 50059574 | 31.56046 | 98.57728 | 95.04015 | chr5 | 50056734 | 50059965 | 3.90271 | 3.57355 | 2.104 |
| chr5 | 45102581 | 45102998 | 1000 | Distal Intergenic | 5 | 45155011 | 45169564 | 14554 | 2 | ENSMUST00 | 66566 | 4930435H24 | ENSMUSG0C | false | . | . | . | . | . | . | chr5 | 45101119 | 45103933 | 4.41648 | 3.65968 | 2.56686 |
| chr5 | 34573555 | 34574114 | 1000 | Promoter (<=1kb) | 5 | 34573664 | 34632308 | 58645 | 1 | ENSMUST00 | 0 | Add1 | ENSMUSG0C | false | chr5 | 34573033 | 34575112 | 30.78991 | 105.27288 | 101.56436 | . | . | . | . | . | . |
| chr5 | 32151224 | 32151723 | 1000 | Intron (ENSMUST00000031017.9 | 5 | 32136472 | 32157831 | 21360 | 1 | ENSMUST00 | 14753 | Fosl2 | ENSMUSG0C | false | . | . | . | . | . | . | chr5 | 32137682 | 32152653 | 5.37733 | 4.11983 | 3.34655 |
| chr5 | 32132913 | 32134087 | 1000 | Promoter (2-3kb) | 5 | 32136472 | 32157831 | 21360 | 1 | ENSMUST00 | -2385 | Fosl2 | ENSMUSG0C | false | chr5 | 32132837 | 32134178 | 20.96848 | 54.70951 | 51.93532 | chr5 | 32134025 | 32134768 | 4.75985 | 3.72224 | 2.84628 |
| chr5 | 32135016 | 32136973 | 1000 | Promoter (<=1kb) | 5 | 32136472 | 32157831 | 21360 | 1 | ENSMUST00 | 0 | Fosl2 | ENSMUSG0C | false | chr5 | 32134342 | 32139172 | 29.71419 | 101.30199 | 97.70005 | . | . | . | . | . | . |
| chr5 | 31564760 | 31565481 | 1000 | Distal Intergenic | 5 | 31543594 | 31554038 | 10445 | 1 | ENSMUST00 | 21167 | Slc4a1ap | ENSMUSG0C | false | . | . | . | . | . | . | chr5 | 31563326 | 31569220 | 5.93745 | 4.50421 | 3.77782 |
| chr5 | 31240592 | 31241499 | 1000 | Promoter (<=1kb) | 5 | 31240864 | 31251560 | 10697 | 1 | ENSMUST00 | 0 | Nrbp1 | ENSMUSG0C | false | chr5 | 31240267 | 31242202 | 30.14656 | 119.38538 | 115.2599 | . | . | . | . | . | . |
| chr5 | 73373092 | 73373551 | 1000 | Distal Intergenic | 5 | 73406076 | 73453435 | 47360 | 1 | ENSMUST00 | -32525 | Cwh43 | ENSMUSG0C | false | . | . | . | . | . | . | chr5 | 73373175 | 73374464 | 4.83905 | 4.00804 | 2.87246 |
| chr5 | 30913129 | 30913783 | 1000 | Promoter (<=1kb) | 5 | 30913402 | 30921278 | 7877 | 1 | ENSMUST00 | 0 | Emilin1 | ENSMUSG0C | false | chr5 | 30913426 | 30914740 | 13.78596 | 30.80405 | 28.32676 | chr5 | 30904966 | 30918702 | 6.19394 | 4.49227 | 4.02489 |
| chr5 | 30711505 | 30712013 | 1000 | Promoter (<=1kb) | 5 | 30711564 | 30799375 | 87812 | 1 | ENSMUST00 | 0 | Dpysl5 | ENSMUSG0C | false | chr5 | 30711168 | 30713618 | 23.43614 | 63.09834 | 60.20888 | chr5 | 30709944 | 30713870 | 5.07954 | 4.08117 | 3.07289 |
| chr5 | 30768484 | 30768920 | 1000 | Intron (ENSMUST00000114729.2 | 5 | 30778130 | 30786934 | 8805 | 1 | ENSMUST00 | -9210 | Dpysl5 | ENSMUSG0C | false | . | . | . | . | . | . | chr5 | 30767822 | 30769676 | 5.68014 | 4.25879 | 3.61281 |
| chr5 | 64287265 | 64287588 | 1000 | Intron (ENSMUST00000043893.7 | 5 | 64298530 | 64350919 | 52390 | 1 | ENSMUST00 | -10942 | Tbc1d1 | ENSMUSG0C | false | . | . | . | . | . | . | chr5 | 64273894 | 64301350 | 6.13726 | 4.51007 | 3.95856 |
| chr5 | 64367405 | 64367715 | 1000 | Distal Intergenic | 5 | 64349945 | 64350589 | 645 | 1 | ENSMUST00 | 17461 | Tbc1d1 | ENSMUSG0C | false | . | . | . | . | . | . | chr5 | 64363933 | 64368552 | 6.57121 | 4.78957 | 4.29438 |
| chr5 | 64292588 | 64292982 | 1000 | Intron (ENSMUST00000043893.7 | 5 | 64298530 | 64350919 | 52390 | 1 | ENSMUST00 | -5548 | Tbc1d1 | ENSMUSG0C | false | . | . | . | . | . | . | chr5 | 64273894 | 64301350 | 6.13726 | 4.51007 | 3.95856 |
| chr5 | 64291625 | 64291985 | 1000 | Intron (ENSMUST00000043893.7 | 5 | 64298530 | 64350919 | 52390 | 1 | ENSMUST00 | -6545 | Tbc1d1 | ENSMUSG0C | false | . | . | . | . | . | . | chr5 | 64273894 | 64301350 | 6.13726 | 4.51007 | 3.95856 |
| chr5 | 64296262 | 64296842 | 1000 | Promoter (1-2kb) | 5 | 64298530 | 64350919 | 52390 | 1 | ENSMUST00 | -1688 | Tbc1d1 | ENSMUSG0C | false | . | . | . | . | . | . | chr5 | 64273894 | 64301350 | 6.13726 | 4.51007 | 3.95856 |
| chr5 | 64298330 | 64298791 | 1000 | Promoter (<=1kb) | 5 | 64298530 | 64350919 | 52390 | 1 | ENSMUST00 | 0 | Tbc1d1 | ENSMUSG0C | false | . | . | . | . | . | . | chr5 | 64273894 | 64301350 | 6.13726 | 4.51007 | 3.95856 |
| chr5 | 64159819 | 64160485 | 1000 | Promoter (<=1kb) | 5 | 64159821 | 64351478 | 191658 | 1 | ENSMUST00 | 0 | Tbc1d1 | ENSMUSG0C | false | chr5 | 64159212 | 64161446 | 28.90617 | 85.18155 | 81.93615 | . | . | . | . | . | . |
| chr5 | 64230049 | 64230729 | 1000 | Promoter (<=1kb) | 5 | 64230299 | 64351482 | 121184 | 1 | ENSMUST00 | 0 | Tbc1d1 | ENSMUSG0C | false | chr5 | 64229858 | 64231190 | 32.25254 | 99.78123 | 96.21911 | . | . | . | . | . | . |
| chr5 | 64811128 | 64812457 | 1000 | Promoter (<=1kb) | 5 | 64812339 | 64829482 | 17144 | 1 | ENSMUST00 | 0 | Klf3 | ENSMUSG0C | false | chr5 | 64811664 | 64813679 | 12.8669 | 28.00426 | 25.56298 | chr5 | 64804440 | 64817688 | 5.05462 | 4.05864 | 3.05357 |
| chr5 | 64802678 | 64804291 | 1000 | Promoter (<=1kb) | 5 | 64803523 | 64830129 | 26607 | 1 | ENSMUST00 | 0 | Klf3 | ENSMUSG0C | false | chr5 | 64802420 | 64806226 | 29.58769 | 108.11006 | 104.32592 | chr5 | 64802418 | 64802923 | 4.84437 | 4.09155 | 2.86093 |
| chr5 | 64969537 | 64970310 | 1000 | Promoter (<=1kb) | 5 | 64970071 | 65041874 | 71804 | 1 | ENSMUST00 | 0 | Fam114a1 | ENSMUSG0C | false | chr5 | 64969293 | 64970870 | 35.57682 | 111.67485 | 107.78959 | chr5 | 64969030 | 64969627 | 3.61567 | 3.37093 | 1.90625 |
| chr5 | 65022002 | 65022385 | 1000 | Intron (ENSMUST00000031080.9 | 5 | 64972425 | 65005891 | 33467 | 1 | ENSMUST00 | 49578 | Fam114a1 | ENSMUSG0C | false | . | . | . | . | . | . | chr5 | 65019418 | 65022572 | 7.26089 | 5.21682 | 4.84068 |
| chr5 | 65697639 | 65698707 | 1000 | Promoter (<=1kb) | 5 | 65615260 | 65697856 | 82597 | 2 | ENSMUST00 | 0 | Pds5a | ENSMUSG0C | false | chr5 | 65696596 | 65698919 | 35.06079 | 109.74464 | 105.9221 | . | . | . | . | . | . |
| chr5 | 66609361 | 66609746 | 1000 | Intron (ENSMUST00000159786.2 | 5 | 66451982 | 66618640 | 166659 | 2 | ENSMUST00 | 8894 | Apbb2 | ENSMUSG0C | false | . | . | . | . | . | . | chr5 | 66558047 | 66617728 | 6.24773 | 4.56399 | 4.04918 |
| chr5 | 66581586 | 66581948 | 1000 | Intron (ENSMUST00000159786.2 | 5 | 66452267 | 66546957 | 94691 | 2 | ENSMUST00 | -34630 | Apbb2 | ENSMUSG0C | false | . | . | . | . | . | . | chr5 | 66558047 | 66617728 | 6.24773 | 4.56399 | 4.04918 |
| chr5 | 66549680 | 66550074 | 1000 | Promoter (2-3kb) | 5 | 66452267 | 66546957 | 94691 | 2 | ENSMUST00 | -2724 | Apbb2 | ENSMUSG0C | false | . | . | . | . | . | . | chr5 | 66502348 | 66557053 | 5.30285 | 4.18237 | 3.2769 |
| chr5 | 66617869 | 66619011 | 1000 | Promoter (<=1kb) | 5 | 66298861 | 66618828 | 319968 | 2 | ENSMUST00 | 0 | Apbb2 | ENSMUSG0C | false | chr5 | 66617073 | 66618806 | 29.18609 | 122.55783 | 118.32927 | chr5 | 66618990 | 66621424 | 6.79745 | 5.02559 | 4.46868 |
| chr5 | 66340154 | 66340492 | 1000 | Promoter (2-3kb) | 5 | 66301888 | 66337665 | 35778 | 2 | ENSMUST00 | -2490 | Apbb2 | ENSMUSG0C | false | . | . | . | . | . | . | chr5 | 66334637 | 66346684 | 5.95013 | 4.37258 | 3.81282 |
| chr5 | 66537811 | 66538199 | 1000 | Intron (ENSMUST00000159786.2 | 5 | 66452267 | 66546957 | 94691 | 2 | ENSMUST00 | 8758 | Apbb2 | ENSMUSG0C | false | . | . | . | . | . | . | chr5 | 66502348 | 66557053 | 5.30285 | 4.18237 | 3.2769 |
| chr5 | 66436694 | 66437200 | 1000 | Intron (ENSMUST00000159786.2 | 5 | 66364974 | 66451629 | 86656 | 2 | ENSMUST00 | 14429 | Apbb2 | ENSMUSG0C | false | . | . | . | . | . | . | chr5 | 66427186 | 66442420 | 5.04827 | 4.07617 | 3.04663 |
| chr5 | 66496966 | 66497362 | 1000 | Intron (ENSMUST00000159786.2 | 5 | 66451995 | 66514830 | 62836 | 2 | ENSMUST00 | 17468 | Apbb2 | ENSMUSG0C | false | . | . | . | . | . | . | chr5 | 66486679 | 66499653 | 5.75258 | 4.40582 | 3.62359 |
| chr5 | 71656769 | 71657113 | 1000 | Promoter (1-2kb) | 5 | 71569734 | 71658308 | 88575 | 2 | ENSMUST00 | 1195 | Gabra4 | ENSMUSG0C | true | chr5 | 71656804 | 71658689 | 20.74539 | 66.97287 | 64.02668 | chr5 | 71654292 | 71656931 | 4.51213 | 3.7492 | 2.62395 |
| chr5 | 75043907 | 75044909 | 1000 | Promoter (<=1kb) | 5 | 75001026 | 75044774 | 43749 | 2 | ENSMUST00 | 0 | Chic2 | ENSMUSG0C | false | chr5 | 75043434 | 75045043 | 24.39013 | 102.54827 | 98.91666 | . | . | . | . | . | . |
| chr5 | 76303696 | 76305225 | 1000 | Promoter (<=1kb) | 5 | 76212177 | 76304548 | 92372 | 2 | ENSMUST00 | 0 | Clock | ENSMUSG0C | false | chr5 | 76303304 | 76305577 | 36.46257 | 120.58523 | 116.42456 | . | . | . | . | . | . |
| chr5 | 77265170 | 77266835 | 1000 | Promoter (<=1kb) | 5 | 77265491 | 77286432 | 20942 | 1 | ENSMUST00 | 0 | Rest | ENSMUSG0C | false | chr5 | 77265048 | 77267299 | 34.36702 | 111.64683 | 107.76208 | . | . | . | . | . | . |
| chr5 | 77357502 | 77358698 | 1000 | Intron (ENSMUST00000046746.6 | 5 | 77340368 | 77342936 | 2569 | 1 | ENSMUST00 | 17135 | Polr2b | ENSMUSG0C | false | chr5 | 77357532 | 77358996 | 8.63619 | 18.02194 | 15.71366 | chr5 | 77353137 | 77404361 | 6.51707 | 4.75542 | 4.25399 |
| chr5 | 77369350 | 77369768 | 1000 | Intron (ENSMUST00000046746.6 | 5 | 77340368 | 77342936 | 2569 | 1 | ENSMUST00 | 28983 | Polr2b | ENSMUSG0C | false | . | . | . | . | . | . | chr5 | 77353137 | 77404361 | 6.51707 | 4.75542 | 4.25399 |
| chr5 | 86065485 | 86065880 | 1000 | Promoter (<=1kb) | 5 | 86012043 | 86065583 | 53541 | 2 | ENSMUST00 | 0 | Cenpc1 | ENSMUSG0C | false | chr5 | 86065747 | 86067028 | 16.08362 | 38.04472 | 35.47989 | . | . | . | . | . | . |
| chr5 | 108268515 | 108269059 | 1000 | Promoter (<=1kb) | 5 | 108268897 | 108280526 | 11630 | 1 | ENSMUST00 | 0 | Dr1 | ENSMUSG0C | false | chr5 | 108268150 | 108270837 | 37.7199 | 126.00829 | 121.65485 | chr5 | 108267921 | 108274932 | 4.1791 | 3.66976 | 2.3326 |
| chr5 | 107359509 | 107360012 | 1000 | Promoter (<=1kb) | 5 | 107359837 | 107386361 | 26525 | 1 | ENSMUST00 | 0 | Brdt | ENSMUSG0C | false | chr5 | 107359351 | 107360408 | 18.44084 | 48.77526 | 46.0739 | . | . | . | . | . | . |
| chr5 | 107046346 | 107047085 | 1000 | Distal Intergenic | 5 | 106975238 | 106979323 | 4086 | 1 | ENSMUST00 | 71109 | Cdc7 | ENSMUSG0C | false | . | . | . | . | . | . | chr5 | 107045311 | 107052433 | 4.95844 | 4.05869 | 2.97671 |
| chr5 | 107288862 | 107289887 | 1000 | Promoter (<=1kb) | 5 | 107106570 | 107289629 | 183060 | 2 | ENSMUST00 | 0 | Tgfbr3 | ENSMUSG0C | true | chr5 | 107287431 | 107289736 | 27.66025 | 106.38611 | 102.65278 | chr5 | 107289838 | 107290141 | 5.04736 | 3.94977 | 3.06601 |
| chr5 | 107167691 | 107168260 | 1000 | Intron (ENSMUST00000031224.9 | 5 | 107109281 | 107192367 | 83087 | 2 | ENSMUST00 | 24107 | Tgfbr3 | ENSMUSG0C | true | . | . | . | . | . | . | chr5 | 107143848 | 107172978 | 6.44265 | 4.73295 | 4.19318 |
| chr5 | 107157379 | 107157840 | 1000 | Intron (ENSMUST00000031224.9 | 5 | 107118197 | 107132923 | 14727 | 2 | ENSMUST00 | -24457 | Tgfbr3 | ENSMUSG0C | true | . | . | . | . | . | . | chr5 | 107143848 | 107172978 | 6.44265 | 4.73295 | 4.19318 |
| chr5 | 107148271 | 107148770 | 1000 | Intron (ENSMUST00000031224.9 | 5 | 107118197 | 107132923 | 14727 | 2 | ENSMUST00 | -15349 | Tgfbr3 | ENSMUSG0C | true | . | . | . | . | . | . | chr5 | 107143848 | 107172978 | 6.44265 | 4.73295 | 4.19318 |
| chr5 | 107087367 | 107087836 | 1000 | Distal Intergenic | 5 | 107118197 | 107132923 | 14727 | 2 | ENSMUST00 | 45087 | Tgfbr3 | ENSMUSG0C | true | . | . | . | . | . | . | chr5 | 107086709 | 107090182 | 5.8703 | 4.45482 | 3.73103 |
| chr5 | 107088208 | 107088637 | 1000 | Distal Intergenic | 5 | 107118197 | 107132923 | 14727 | 2 | ENSMUST00 | 44286 | Tgfbr3 | ENSMUSG0C | true | . | . | . | . | . | . | chr5 | 107086709 | 107090182 | 5.8703 | 4.45482 | 3.73103 |
| chr5 | 103691933 | 103692482 | 1000 | Promoter (<=1kb) | 5 | 103692374 | 103816428 | 124055 | 1 | ENSMUST00 | 0 | Aff1 | ENSMUSG0C | false | chr5 | 103691558 | 103694062 | 29.66345 | 100.21 | 96.63225 | . | . | . | . | . | . |
| chr5 | 101736975 | 101737326 | 1000 | Distal Intergenic | 5 | 101765130 | 101823858 | 58729 | 1 | ENSMUST00 | -27804 | Cds1 | ENSMUSG0C | false | . | . | . | . | . | . | chr5 | 101736512 | 101737520 | 4.93064 | 4.15598 | 2.9383 |
| chr5 | 101764735 | 101765752 | 1000 | Promoter (<=1kb) | 5 | 101765130 | 101823858 | 58729 | 1 | ENSMUST00 | 0 | Cds1 | ENSMUSG0C | false | chr5 | 101764380 | 101766811 | 35.62435 | 116.99458 | 112.94973 | chr5 | 101764333 | 101764738 | 3.38512 | 3.27657 | 1.69634 |
| chr5 | 99252616 | 99253239 | 1000 | Promoter (<=1kb) | 5 | 99217426 | 99252927 | 35502 | 2 | ENSMUST00 | 0 | Rasgef1b | ENSMUSG0C | false | chr5 | 99251538 | 99253454 | 33.41831 | 117.26761 | 113.21259 | chr5 | 99250794 | 99254240 | 4.50812 | 3.84986 | 2.59187 |
| chr5 | 98030170 | 98031343 | 1000 | Promoter (<=1kb) | 5 | 97882783 | 98030733 | 147951 | 2 | ENSMUST00 | 0 | Antxr2 | ENSMUSG0C | false | chr5 | 98030106 | 98031628 | 31.84599 | 102.09332 | 98.46812 | chr5 | 98030062 | 98031838 | 3.81485 | 3.46729 | 2.05899 |
| chr5 | 117359937 | 117361013 | 1000 | Promoter (<=1kb) | 5 | 117361022 | 117370922 | 9901 | 1 | ENSMUST00 | -9 | Wsb2 | ENSMUSG0C | false | chr5 | 117360938 | 117362519 | 8.01187 | 19.41161 | 17.08515 | chr5 | 117358665 | 117372620 | 6.58746 | 4.86077 | 4.29355 |
| chr5 | 117356398 | 117357630 | 1000 | Promoter (<=1kb) | 5 | 117357304 | 117378601 | 21298 | 1 | ENSMUST00 | 0 | Wsb2 | ENSMUSG0C | false | chr5 | 117356598 | 117359461 | 46.57093 | 159.38457 | 153.76053 | chr5 | 117355308 | 117356903 | 3.84951 | 3.55203 | 2.04497 |
| chr5 | 88797253 | 88797542 | 1000 | Distal Intergenic | 5 | 88764996 | 88783281 | 18286 | 1 | ENSMUST00 | 32258 | Dck | ENSMUSG0C | false | . | . | . | . | . | . | . | . | . | . | . | . |
| chr5 | 90443883 | 90444278 | 1000 | Distal Intergenic | 5 | 90460889 | 90476603 | 15715 | 1 | ENSMUST00 | -16611 | Alb | ENSMUSG0C | false | . | . | . | . | . | . | chr5 | 90441825 | 90444677 | 4.46758 | 3.87924 | 2.57299 |
| chr5 | 92931308 | 92931871 | 1000 | Intron (ENSMUST00000113055.3 | 5 | 92925407 | 92940455 | 15049 | 1 | ENSMUST00 | 5902 | Shroom3 | ENSMUSG0C | true | . | . | . | . | . | . | chr5 | 92909495 | 92934912 | 5.22466 | 4.15662 | 3.20252 |
| chr5 | 92856201 | 92856577 | 1000 | Intron (ENSMUST00000113055.3 | 5 | 92897993 | 92965308 | 67316 | 1 | ENSMUST00 | -41416 | Shroom3 | ENSMUSG0C | true | chr5 | 92856185 | 92858092 | 12.98899 | 30.74239 | 28.2684 | chr5 | 92847315 | 92878106 | 5.13148 | 4.10707 | 3.12438 |
| chr5 | 93205389 | 93206803 | 1000 | Promoter (<=1kb) | 5 | 93206518 | 93213475 | 6958 | 1 | ENSMUST00 | 0 | 2010109A12 | ENSMUSG0C | false | chr5 | 93205257 | 93207249 | 43.23485 | 145.58217 | 140.47385 | chr5 | 93204579 | 93205443 | 4.51128 | 3.85893 | 2.63295 |
| chr5 | 91962504 | 91963171 | 1000 | Promoter (<=1kb) | 5 | 91948904 | 91962556 | 13653 | 2 | ENSMUST00 | 0 | Rchy1 | ENSMUSG0C | false | chr5 | 91961707 | 91963799 | 27.88725 | 99.90214 | 96.33501 | . | . | . | . | . | . |
| chr5 | 124248988 | 124250380 | 1000 | Promoter (<=1kb) | 5 | 124118690 | 124249760 | 131071 | 2 | ENSMUST00 | 0 | Pitpnm2 | ENSMUSG0C | false | chr5 | 124248582 | 124250472 | 21.92106 | 61.34632 | 58.48102 | chr5 | 124246347 | 124251396 | 3.7821 | 3.43635 | 2.01379 |
| chr5 | 92137605 | 92138188 | 1000 | Promoter (<=1kb) | 5 | 92137938 | 92202794 | 64857 | 1 | ENSMUST00 | 0 | Uso1 | ENSMUSG0C | false | chr5 | 92137086 | 92138804 | 40.33899 | 132.76065 | 128.17128 | . | . | . | . | . | . |
| chr5 | 129416522 | 129416920 | 1000 | Distal Intergenic | 5 | 129501231 | 129571383 | 70153 | 1 | ENSMUST00 | -84311 | Sfswap | ENSMUSG0C | false | . | . | . | . | . | . | chr5 | 129405729 | 129418273 | 5.29114 | 4.12018 | 3.25854 |
| chr5 | 129531701 | 129532056 | 1000 | Intron (ENSMUST00000053737.7 | 5 | 129501231 | 129571383 | 70153 | 1 | ENSMUST00 | 30471 | Sfswap | ENSMUSG0C | false | . | . | . | . | . | . | chr5 | 129526676 | 129541751 | 6.43735 | 4.65853 | 4.2033 |
| chr5 | 129540680 | 129540976 | 1000 | Intron (ENSMUST00000053737.7 | 5 | 129501231 | 129571383 | 70153 | 1 | ENSMUST00 | 39450 | Sfswap | ENSMUSG0C | false | . | . | . | . | . | . | chr5 | 129526676 | 129541751 | 6.43735 | 4.65853 | 4.2033 |
| chr5 | 129538857 | 129539229 | 1000 | Intron (ENSMUST00000053737.7 | 5 | 129501231 | 129571383 | 70153 | 1 | ENSMUST00 | 37627 | Sfswap | ENSMUSG0C | false | . | . | . | . | . | . | chr5 | 129526676 | 129541751 | 6.43735 | 4.65853 | 4.2033 |
| chr5 | 129353648 | 129354071 | 1000 | Distal Intergenic | 5 | 129501231 | 129571383 | 70153 | 1 | ENSMUST00 | -147160 | Sfswap | ENSMUSG0C | false | . | . | . | . | . | . | chr5 | 129352556 | 129353896 | 4.94267 | 3.66921 | 3.00192 |
| chr5 | 123116143 | 123116428 | 1000 | Exon (ENSMUST00000067505.9/ | 5 | 123103044 | 123132396 | 29353 | 2 | ENSMUST00 | 15968 | Rhof | ENSMUSG0C | false | . | . | . | . | . | . | chr5 | 123115071 | 123118332 | 7.00575 | 5.0104 | 4.65686 |
| chr7 | 100706197 | 100707030 | 1000 | Promoter (<=1kb) | 7 | 100706651 | 100841313 | 134663 | 1 | ENSMUST00 | 0 | Fam168a | ENSMUSG0C | false | chr7 | 100705978 | 100707534 | 37.68163 | 117.40437 | 113.3432 | . | . | . | . | . | . |
| chr5 | 122501545 | 122502691 | 1000 | Promoter (<=1kb) | 5 | 122442702 | 122502225 | 59524 | 2 | ENSMUST00 | 0 | Atp2a2 | ENSMUSG0C | false | chr5 | 122500881 | 122503112 | 30.60556 | 103.92632 | 100.25623 | . | . | . | . | . | . |
| chr5 | 122707346 | 122708043 | 1000 | Promoter (<=1kb) | 5 | 122707584 | 122729044 | 21461 | 1 | ENSMUST00 | 0 | P2rx4 | ENSMUSG0C | false | chr5 | 122707005 | 122708548 | 27.1957 | 84.10433 | 80.87383 | . | . | . | . | . | . |
| chr5 | 122820771 | 122821614 | 1000 | Promoter (<=1kb) | 5 | 122787469 | 122821339 | 33871 | 2 | ENSMUST00 | 0 | Anapc5 | ENSMUSG0C | false | chr5 | 122820560 | 122821729 | 28.90751 | 104.81161 | 101.11721 | chr5 | 122820039 | 122820855 | 6.64478 | 4.88675 | 4.4071 |
| chr5 | 122899628 | 122900742 | 1000 | Promoter (<=1kb) | 5 | 122870668 | 122900355 | 29688 | 2 | ENSMUST00 | 0 | Kdm2b | ENSMUSG0C | false | chr5 | 122899282 | 122901119 | 35.17274 | 107.8772 | 104.10775 | chr5 | 122897461 | 122899669 | 3.71793 | 3.42206 | 1.94972 |
| chr5 | 125177487 | 125178378 | 1000 | Promoter (<=1kb) | 5 | 125017162 | 125179053 | 161892 | 2 | ENSMUST00 | 675 | Ncor2 | ENSMUSG0C | false | chr5 | 125177014 | 125179760 | 33.54585 | 101.01101 | 97.41918 | chr5 | 125124978 | 125178451 | 6.70656 | 4.76008 | 4.42049 |
| chr5 | 125178701 | 125179440 | 1000 | Promoter (<=1kb) | 5 | 125017153 | 125179214 | 162062 | 2 | ENSMUST00 | 0 | Ncor2 | ENSMUSG0C | false | chr5 | 125177014 | 125179760 | 33.54585 | 101.01101 | 97.41918 | chr5 | 125179405 | 125180638 | 5.22729 | 4.30919 | 3.18758 |
| chr5 | 125175331 | 125175665 | 1000 | Intron (ENSMUST00000111402.3 | 5 | 125017162 | 125179053 | 161892 | 2 | ENSMUST00 | 3388 | Ncor2 | ENSMUSG0C | false | chr5 | 125175332 | 125175506 | 5.79018 | 9.39037 | 7.2276 | chr5 | 125124978 | 125178451 | 6.70656 | 4.76008 | 4.42049 |
| chr5 | 125040755 | 125041143 | 1000 | Promoter (1-2kb) | 5 | 125017153 | 125042812 | 25660 | 2 | ENSMUST00 | 1669 | Ncor2 | ENSMUSG0C | false | . | . | . | . | . | . | chr5 | 125038618 | 125048555 | 7.1223 | 4.95277 | 4.74471 |
| chr5 | 125046278 | 125046704 | 1000 | Intron (ENSMUST00000111402.3 | 5 | 125017153 | 125042812 | 25660 | 2 | ENSMUST00 | -3467 | Ncor2 | ENSMUSG0C | false | . | . | . | . | . | . | chr5 | 125038618 | 125048555 | 7.1223 | 4.95277 | 4.74471 |
| chr5 | 125058070 | 125058739 | 1000 | Promoter (<=1kb) | 5 | 125030800 | 125058400 | 27601 | 2 | ENSMUST00 | 0 | Ncor2 | ENSMUSG0C | false | chr5 | 125055981 | 125058881 | 22.19715 | 59.28558 | 56.44936 | chr5 | 125050765 | 125060996 | 6.95919 | 4.79882 | 4.64527 |
| chr5 | 125465608 | 125466039 | 1000 | Distal Intergenic | 5 | 125475814 | 125517412 | 41599 | 1 | ENSMUST00 | -9775 | Aacs | ENSMUSG0C | false | chr5 | 125465513 | 125465775 | 6.83737 | 14.36054 | 12.10883 | . | . | . | . | . | . |
| chr5 | 110770489 | 110771273 | 1000 | Promoter (<=1kb) | 5 | 110664373 | 110770527 | 106155 | 2 | ENSMUST00 | 0 | Ep400 | ENSMUSG0C | false | chr5 | 110769828 | 110770769 | 36.42221 | 131.26616 | 126.73016 | chr5 | 110771111 | 110773942 | 2.95676 | 2.94983 | 1.37662 |
| chr5 | 110807987 | 110808598 | 1000 | Promoter (1-2kb) | 5 | 110784488 | 110810097 | 25610 | 2 | ENSMUST00 | 1499 | Ulk1 | ENSMUSG0C | false | chr5 | 110807512 | 110810111 | 35.89541 | 128.14636 | 123.72295 | chr5 | 110797106 | 110809391 | 6.92459 | 4.77628 | 4.6086 |
| chr5 | 115631663 | 115632472 | 1000 | Promoter (<=1kb) | 5 | 115631929 | 115647736 | 15808 | 1 | ENSMUST00 | 0 | Rab35 | ENSMUSG0C | false | chr5 | 115631117 | 115632856 | 34.54275 | 123.54154 | 119.28033 | . | . | . | . | . | . |
| chr5 | 115521504 | 115522112 | 1000 | Promoter (1-2kb) | 5 | 115520305 | 115555982 | 35678 | 1 | ENSMUST00 | 1200 | Pxn | ENSMUSG0C | false | . | . | . | . | . | . | chr5 | 115507551 | 115550291 | 7.76238 | 5.17462 | 5.28263 |
| chr5 | 115531213 | 115532103 | 1000 | Intron (ENSMUST00000067268.9 | 5 | 115542914 | 115546880 | 3967 | 1 | ENSMUST00 | -10811 | Pxn | ENSMUSG0C | false | . | . | . | . | . | . | chr5 | 115507551 | 115550291 | 7.76238 | 5.17462 | 5.28263 |
| chr5 | 115340966 | 115341375 | 1000 | Promoter (<=1kb) | 5 | 115341225 | 115343569 | 2345 | 1 | ENSMUST00 | 0 | Triap1 | ENSMUSG0C | false | chr5 | 115340384 | 115342249 | 38.27806 | 124.82941 | 120.52397 | chr5 | 115340029 | 115342966 | 4.05856 | 3.60449 | 2.24123 |
| chr5 | 115319300 | 115319854 | 1000 | Distal Intergenic | 5 | 115327177 | 115333080 | 5904 | 1 | ENSMUST00 | -7323 | Srsf9 | ENSMUSG0C | false | chr5 | 115319420 | 115320102 | 7.88522 | 15.75676 | 13.48224 | chr5 | 115316050 | 115320647 | 6.90815 | 4.93568 | 4.56444 |
| chr5 | 115326887 | 115327640 | 1000 | Promoter (<=1kb) | 5 | 115327177 | 115333080 | 5904 | 1 | ENSMUST00 | 0 | Srsf9 | ENSMUSG0C | false | chr5 | 115326237 | 115328553 | 34.00549 | 104.83981 | 101.14426 | . | . | . | . | . | . |
| chr5 | 115010978 | 115011491 | 1000 | Promoter (<=1kb) | 5 | 115011137 | 115098790 | 87654 | 1 | ENSMUST00 | 0 | Sppl3 | ENSMUSG0C | false | chr5 | 115010549 | 115012400 | 30.28657 | 98.33401 | 94.80609 | . | . | . | . | . | . |
| chr6 | 13601652 | 13602023 | 1000 | Intron (ENSMUST00000031554.3 | 6 | 13603382 | 13607748 | 4367 | 2 | ENSMUST00 | 5725 | Tmem168 | ENSMUSG0C | false | . | . | . | . | . | . | chr6 | 13600188 | 13602491 | 4.08541 | 3.57122 | 2.29589 |
| chr5 | 140610177 | 140611766 | 1000 | Promoter (2-3kb) | 5 | 140607341 | 140615543 | 8203 | 1 | ENSMUST00 | 2837 | Lfng | ENSMUSG0C | false | chr5 | 140610419 | 140611796 | 11.32191 | 27.20368 | 24.77393 | chr5 | 140608605 | 140612857 | 6.78704 | 4.88371 | 4.48462 |
| chr6 | 13069636 | 13070121 | 1000 | Promoter (<=1kb) | 6 | 13069759 | 13075094 | 5336 | 1 | ENSMUST00 | 0 | Tmem106b | ENSMUSG0C | false | chr6 | 13069130 | 13070822 | 39.05061 | 143.74966 | 138.71384 | . | . | . | . | . | . |
| chr5 | 114435603 | 114436023 | 1000 | Intron (ENSMUST00000031560.8 | 5 | 114433278 | 114443982 | 10705 | 2 | ENSMUST00 | 7959 | Mmab | ENSMUSG0C | false | . | . | . | . | . | . | chr5 | 114433544 | 114437611 | 6.67384 | 4.89909 | 4.39621 |
| chr5 | 142905722 | 142906283 | 1000 | Promoter (<=1kb) | 5 | 142904351 | 142905821 | 1471 | 2 | ENSMUST00 | 0 | Actb | ENSMUSG0C | false | chr5 | 142902951 | 142907768 | 40.65367 | 138.82761 | 133.98773 | . | . | . | . | . | . |
| chr5 | 145114049 | 145114482 | 1000 | Promoter (<=1kb) | 5 | 145114215 | 145129699 | 15485 | 1 | ENSMUST00 | 0 | Arpc1b | ENSMUSG0C | false | chr5 | 145113817 | 145115268 | 30.59503 | 95.89513 | 92.42037 | chr5 | 145113465 | 145118752 | 4.56678 | 3.84193 | 2.6654 |
| chr5 | 146230853 | 146231759 | 1000 | Promoter (<=1kb) | 5 | 146231230 | 146302874 | 71645 | 1 | ENSMUST00 | 0 | Cdk8 | ENSMUSG0C | false | chr5 | 146230632 | 146232287 | 34.46491 | 104.61457 | 100.92722 | chr5 | 146231694 | 146233828 | 5.7749 | 4.43344 | 3.63628 |
| chr6 | 8509220 | 8510468 | 1000 | Promoter (<=1kb) | 6 | 8509600 | 8597548 | 87949 | 1 | ENSMUST00 | 0 | Glcci1 | ENSMUSG0C | false | chr6 | 8508635 | 8513604 | 37.63617 | 126.94061 | 122.55849 | . | . | . | . | . | . |
| chr6 | 8511371 | 8511794 | 1000 | Promoter (<=1kb) | 6 | 8511227 | 8579647 | 68421 | 1 | ENSMUST00 | 145 | Glcci1 | ENSMUSG0C | false | chr6 | 8508635 | 8513604 | 37.63617 | 126.94061 | 122.55849 | . | . | . | . | . | . |
| chr5 | 146794467 | 146795288 | 1000 | Promoter (<=1kb) | 5 | 146734809 | 146795006 | 60198 | 2 | ENSMUST00 | 0 | Usp12 | ENSMUSG0C | false | chr5 | 146793690 | 146794900 | 18.27513 | 72.99583 | 69.9605 | . | . | . | . | . | . |
| chr5 | 146844728 | 146845236 | 1000 | Promoter (<=1kb) | 5 | 146845071 | 146847726 | 2656 | 1 | ENSMUST00 | 0 | Rasl11a | ENSMUSG0C | true | chr5 | 146844279 | 146848368 | 25.50663 | 88.63404 | 85.31516 | . | . | . | . | . | . |
| chr5 | 146841260 | 146841842 | 1000 | Distal Intergenic | 5 | 146845071 | 146847726 | 2656 | 1 | ENSMUST00 | -3229 | Rasl11a | ENSMUSG0C | true | . | . | . | . | . | . | chr5 | 146833824 | 146844691 | 4.65777 | 3.84315 | 2.73401 |
| chr5 | 147429698 | 147431219 | 1000 | Promoter (<=1kb) | 5 | 147430161 | 147548502 | 118342 | 1 | ENSMUST00 | 0 | Pan3 | ENSMUSG0C | false | chr5 | 147429490 | 147431889 | 33.65816 | 114.56059 | 110.59126 | chr5 | 147429109 | 147429817 | 3.80699 | 3.46191 | 2.01948 |
| chr5 | 147860225 | 147860844 | 1000 | Promoter (<=1kb) | 5 | 147860628 | 147875778 | 15151 | 1 | ENSMUST00 | 0 | Pomp | ENSMUSG0C | false | chr5 | 147859955 | 147861658 | 38.67513 | 141.95352 | 136.99454 | . | . | . | . | . | . |
| chr6 | 4505120 | 4505849 | 1000 | Promoter (<=1kb) | 6 | 4505493 | 4541543 | 36051 | 1 | ENSMUST00 | 0 | Col1a2 | ENSMUSG0C | false | chr6 | 4505135 | 4506838 | 11.90309 | 30.75663 | 28.28125 | chr6 | 4502270 | 4505461 | 5.06239 | 4.16142 | 3.04655 |
| chr6 | 22355772 | 22356389 | 1000 | Promoter (<=1kb) | 6 | 22306520 | 22356166 | 49647 | 2 | ENSMUST00 | 0 | Fam3c | ENSMUSG0C | false | chr6 | 22355441 | 22356405 | 28.45235 | 93.08366 | 89.67602 | chr6 | 22353838 | 22356671 | 3.43704 | 3.28121 | 1.74509 |
| chr5 | 132248549 | 132248848 | 1000 | Intron (ENSMUST00000161226.3 | 5 | 132030909 | 132050409 | 19501 | 2 | ENSMUST00 | -198141 | Auts2 | ENSMUSG0C | true | . | . | . | . | . | . | chr5 | 132247616 | 132251716 | 6.17277 | 4.49794 | 3.986 |
| chr5 | 131855328 | 131855831 | 1000 | Intron (ENSMUST00000161226.3 | 5 | 131439080 | 132050337 | 611258 | 2 | ENSMUST00 | 194506 | Auts2 | ENSMUSG0C | true | . | . | . | . | . | . | chr5 | 131850429 | 131856702 | 4.35797 | 3.75025 | 2.5053 |
| chr5 | 131537966 | 131539021 | 1000 | Promoter (<=1kb) | 5 | 131439298 | 131538687 | 99390 | 2 | ENSMUST00 | 0 | Auts2 | ENSMUSG0C | true | chr5 | 131536784 | 131539135 | 30.78865 | 90.34473 | 86.99165 | . | . | . | . | . | . |
| chr5 | 131881888 | 131882309 | 1000 | Intron (ENSMUST00000161226.3 | 5 | 131439080 | 132050337 | 611258 | 2 | ENSMUST00 | 168028 | Auts2 | ENSMUSG0C | true | . | . | . | . | . | . | chr5 | 131878259 | 131884481 | 4.40984 | 3.61316 | 2.56303 |
| chr5 | 132542956 | 132543922 | 1000 | Promoter (<=1kb) | 5 | 131437333 | 132543344 | 1106012 | 2 | ENSMUST00 | 0 | Auts2 | ENSMUSG0C | true | chr5 | 132541111 | 132544655 | 32.62678 | 97.43105 | 93.92233 | . | . | . | . | . | . |
| chr5 | 134772524 | 134772973 | 1000 | Distal Intergenic | 5 | 134703781 | 134747241 | 43461 | 2 | ENSMUST00 | -25284 | Eln | ENSMUSG0C | false | . | . | . | . | . | . | chr5 | 134770712 | 134774141 | 6.42644 | 4.70561 | 4.20353 |
| chr6 | 24619238 | 24619849 | 1000 | Exon (ENSMUST00000031695.9/ | 6 | 24597771 | 24605414 | 7644 | 1 | ENSMUST00 | 21468 | Lmod2 | ENSMUSG0C | false | . | . | . | . | . | . | chr6 | 24617540 | 24624235 | 4.20396 | 3.68982 | 2.35048 |
| chr6 | 24664230 | 24665294 | 1000 | Promoter (<=1kb) | 6 | 24613805 | 24665009 | 51205 | 2 | ENSMUST00 | 0 | Wasl | ENSMUSG0C | true | chr6 | 24662156 | 24665827 | 35.95432 | 114.41653 | 110.45183 | . | . | . | . | . | . |
| chr6 | 24667094 | 24667403 | 1000 | Promoter (2-3kb) | 6 | 24613805 | 24665009 | 51205 | 2 | ENSMUST00 | -2086 | Wasl | ENSMUSG0C | true | . | . | . | . | . | . | chr6 | 24665487 | 24668704 | 4.27254 | 3.70959 | 2.40612 |
| chr6 | 24530644 | 24531218 | 1000 | Promoter (2-3kb) | 6 | 24528144 | 24570984 | 42841 | 1 | ENSMUST00 | 2501 | Asb15 | ENSMUSG0C | true | . | . | . | . | . | . | chr6 | 24526474 | 24534302 | 5.36778 | 4.25265 | 3.31628 |
| chr6 | 24564142 | 24564494 | 1000 | Exon (ENSMUST00000031696.4/ | 6 | 24556072 | 24573164 | 17093 | 1 | ENSMUST00 | 8071 | Asb15 | ENSMUSG0C | true | . | . | . | . | . | . | chr6 | 24562559 | 24572536 | 4.84657 | 3.96593 | 2.89537 |
| chr6 | 47453887 | 47455031 | 1000 | Promoter (<=1kb) | 6 | 47453904 | 47523766 | 69863 | 1 | ENSMUST00 | 0 | Cul1 | ENSMUSG0C | false | chr6 | 47453518 | 47455734 | 34.16929 | 120.7392 | 116.5701 | . | . | . | . | . | . |
| chr5 | 135962447 | 135962935 | 1000 | Promoter (<=1kb) | 5 | 135960224 | 135962327 | 2104 | 2 | ENSMUST00 | -121 | Srcrb4d | ENSMUSG0C | false | chr5 | 135962842 | 135963512 | 30.28657 | 98.33401 | 94.80609 | chr5 | 135961199 | 135963745 | 5.89265 | 4.45768 | 3.73349 |
| chr5 | 136556562 | 136557050 | 1000 | Intron (ENSMUST00000176216.3 | 5 | 136274486 | 136565432 | 290947 | 2 | ENSMUST00 | 8382 | Cux1 | ENSMUSG0C | false | . | . | . | . | . | . | chr5 | 136548415 | 136565609 | 5.81364 | 4.39336 | 3.67644 |
| chr5 | 136522926 | 136523505 | 1000 | Intron (ENSMUST00000176216.3 | 5 | 136274486 | 136565432 | 290947 | 2 | ENSMUST00 | 41927 | Cux1 | ENSMUSG0C | false | . | . | . | . | . | . | chr5 | 136497915 | 136525985 | 5.56595 | 4.30446 | 3.49012 |
| chr5 | 136565919 | 136566772 | 1000 | Promoter (<=1kb) | 5 | 136274455 | 136566351 | 291897 | 2 | ENSMUST00 | 0 | Cux1 | ENSMUSG0C | false | chr5 | 136565019 | 136569262 | 36.46257 | 120.58523 | 116.42456 | . | . | . | . | . | . |
| chr5 | 137509985 | 137510999 | 1000 | Intron (ENSMUST00000111038.2 | 5 | 137501439 | 137502518 | 1080 | 2 | ENSMUST00 | -7468 | Pop7 | ENSMUSG0C | false | chr5 | 137509828 | 137511544 | 33.15098 | 102.34641 | 98.71844 | . | . | . | . | . | . |
| chr5 | 137611255 | 137611588 | 1000 | Promoter (<=1kb) | 5 | 137605103 | 137611408 | 6306 | 2 | ENSMUST00 | 0 | Pcolce | ENSMUSG0C | false | chr5 | 137610176 | 137611493 | 12.58308 | 29.40204 | 26.94344 | chr5 | 137608490 | 137613729 | 4.66714 | 3.91261 | 2.74881 |
| chr5 | 137745666 | 137746438 | 1000 | Promoter (<=1kb) | 5 | 137745730 | 137768450 | 22721 | 1 | ENSMUST00 | 0 | Tsc22d4 | ENSMUSG0C | false | chr5 | 137745433 | 137747808 | 26.28407 | 85.32412 | 82.07336 | chr5 | 137745353 | 137749156 | 5.54799 | 4.16841 | 3.47549 |
| chr5 | 137786152 | 137786882 | 1000 | Promoter (<=1kb) | 5 | 137781906 | 137786663 | 4758 | 2 | ENSMUST00 | 0 | Mepce | ENSMUSG0C | false | chr5 | 137785173 | 137786774 | 32.29185 | 112.09676 | 108.19604 | . | . | . | . | . | . |
| chr6 | 5725524 | 5726060 | 1000 | Promoter (<=1kb) | 6 | 5725639 | 6028039 | 302401 | 1 | ENSMUST00 | 0 | Dync1i1 | ENSMUSG0C | true | chr6 | 5725457 | 5726859 | 30.08928 | 92.05882 | 88.67145 | chr6 | 5725530 | 5728662 | 3.95916 | 3.55866 | 2.15556 |
| chr6 | 29767908 | 29768805 | 1000 | Promoter (<=1kb) | 6 | 29768011 | 29768727 | 717 | 1 | ENSMUST00 | 0 | Ahcyl2 | ENSMUSG0C | false | chr6 | 29768392 | 29769584 | 32.67316 | 105.57997 | 101.8659 | . | . | . | . | . | . |
| chr6 | 52640096 | 52640755 | 1000 | Promoter (<=1kb) | 6 | 52546228 | 52640389 | 94162 | 2 | ENSMUST00 | 0 | Hibadh | ENSMUSG0C | false | chr6 | 52639334 | 52640742 | 32.25957 | 103.83389 | 100.16508 | chr6 | 52640607 | 52641951 | 2.91105 | 2.91297 | 1.34931 |
| chr6 | 56831838 | 56832359 | 1000 | Promoter (<=1kb) | 6 | 56832059 | 56879358 | 47300 | 1 | ENSMUST00 | 0 | Fkbp9 | ENSMUSG0C | false | chr6 | 56831538 | 56832862 | 24.99264 | 71.03027 | 68.02251 | . | . | . | . | . | . |
| chr6 | 38554645 | 38555093 | 1000 | Promoter (<=1kb) | 6 | 38555723 | 38589286 | 33564 | 1 | ENSMUST00 | -630 | Luc7l2 | ENSMUSG0C | false | . | . | . | . | . | . | chr6 | 38552660 | 38556442 | 4.75764 | 4.02374 | 2.79597 |
| chr6 | 38550625 | 38552362 | 1000 | Promoter (<=1kb) | 6 | 38551334 | 38605040 | 53707 | 1 | ENSMUST00 | 0 | Luc7l2 | ENSMUSG0C | false | chr6 | 38550233 | 38551299 | 33.09821 | 104.28432 | 100.60858 | . | . | . | . | . | . |
| chr6 | 37870420 | 37871318 | 1000 | Promoter (<=1kb) | 6 | 37870811 | 37921505 | 50695 | 1 | ENSMUST00 | 0 | Trim24 | ENSMUSG0C | false | chr6 | 37870693 | 37872083 | 30.26787 | 111.42836 | 107.5545 | . | . | . | . | . | . |
| chr6 | 42349436 | 42350368 | 1000 | Promoter (<=1kb) | 6 | 42349828 | 42358392 | 8565 | 1 | ENSMUST00 | 0 | Zyx | ENSMUSG0C | false | chr6 | 42348851 | 42351089 | 28.90751 | 104.81161 | 101.11721 | . | . | . | . | . | . |
| chr6 | 42264674 | 42265394 | 1000 | Promoter (<=1kb) | 6 | 42264985 | 42282508 | 17524 | 1 | ENSMUST00 | 0 | Casp2 | ENSMUSG0C | false | chr6 | 42264469 | 42266101 | 26.52689 | 93.43019 | 90.01563 | . | . | . | . | . | . |
| chr6 | 38917412 | 38917769 | 1000 | Promoter (1-2kb) | 6 | 38918980 | 39084585 | 165606 | 1 | ENSMUST00 | -1211 | Tbxas1 | ENSMUSG0C | false | . | . | . | . | . | . | chr6 | 38910891 | 38920200 | 4.83904 | 3.9552 | 2.88392 |
| chr6 | 38919149 | 38919505 | 1000 | Promoter (<=1kb) | 6 | 38919024 | 39084585 | 165562 | 1 | ENSMUST00 | 126 | Tbxas1 | ENSMUSG0C | false | . | . | . | . | . | . | chr6 | 38910891 | 38920200 | 4.83904 | 3.9552 | 2.88392 |
| chr6 | 38875340 | 38876867 | 1000 | Promoter (<=1kb) | 6 | 38875404 | 38952973 | 77570 | 1 | ENSMUST00 | 0 | Tbxas1 | ENSMUSG0C | false | chr6 | 38874171 | 38877070 | 40.55112 | 132.03412 | 127.46851 | . | . | . | . | . | . |
| chr6 | 86785150 | 86785607 | 1000 | Intron (ENSMUST00000001187.9 | 6 | 86736840 | 86793584 | 56745 | 2 | ENSMUST00 | 7977 | Anxa4 | ENSMUSG0C | true | . | . | . | . | . | . | chr6 | 86761092 | 86794291 | 5.26625 | 4.09177 | 3.24957 |
| chr6 | 86222444 | 86222793 | 1000 | Intron (ENSMUST00000032066.9 | 6 | 86195251 | 86275449 | 80199 | 1 | ENSMUST00 | 27194 | Tgfa | ENSMUSG0C | false | . | . | . | . | . | . | chr6 | 86221760 | 86228479 | 4.74973 | 3.9282 | 2.84464 |

| | | | | | | | | | | | | | | | | | | | | | | | | | | |
|---|---|---|---|---|---|---|---|---|---|---|---|---|---|---|---|---|---|---|---|---|---|---|---|---|---|---|
| chr6 | 86195001 | 86195736 | 1000 | Promoter (<=1kb) | 6 | 86195251 | 86275449 | 80199 | 1 | ENSMUST00 | 0 | Tgfa | ENSMUSG0( | false | chr6 | 86194626 | 86196458 | 21.46036 | 80.68572 | 77.52129 | chr6 | 86193178 | 86199547 | 4.62388 | 3.88572 | 2.72166 |
| chr6 | 86086801 | 86087231 | 1000 | Exon (ENSMUST00000032069.5/ | 6 | 86078084 | 86119555 | 41472 | 1 | ENSMUST00 | 8718 | Add2 | ENSMUSG0( | false | . | . | . | . | . | . | . | . | . | . | . | . |
| chr6 | 86048401 | 86048914 | 1000 | Distal Intergenic | 6 | 86078084 | 86119555 | 41472 | 1 | ENSMUST00 | -29170 | Add2 | ENSMUSG0( | false | . | . | . | . | . | . | chr6 | 86047475 | 86050613 | 5.36426 | 3.9853 | 3.33876 |
| chr6 | 86074815 | 86075285 | 1000 | Promoter (2-3kb) | 6 | 86078084 | 86119555 | 41472 | 1 | ENSMUST00 | -2799 | Add2 | ENSMUSG0( | false | . | . | . | . | . | . | . | . | . | . | . | . |
| chr6 | 85973363 | 85973721 | 1000 | Distal Intergenic | 6 | 85954962 | 85961667 | 6706 | 2 | ENSMUST00 | -11697 | Dusp11 | ENSMUSG0( | false | . | . | . | . | . | . | chr6 | 85972591 | 85974188 | 5.53372 | 4.31869 | 3.45925 |
| chr6 | 85451037 | 85452090 | 1000 | Promoter (<=1kb) | 6 | 85451505 | 85468477 | 16973 | 1 | ENSMUST00 | 0 | Cct7 | ENSMUSG0( | false | chr6 | 85450706 | 85452985 | 39.97575 | 133.22411 | 128.61581 | chr6 | 85450883 | 85451159 | 3.46257 | 3.34771 | 1.78304 |
| chr6 | 119478951 | 119480187 | 1000 | Promoter (<=1kb) | 6 | 119479668 | 119482630 | 2963 | 1 | ENSMUST00 | 0 | Fbxl14 | ENSMUSG0( | false | chr6 | 119478571 | 119481610 | 42.04218 | 136.8161 | 132.06688 | chr6 | 119480182 | 119485346 | 5.55867 | 4.12801 | 3.49284 |
| chr6 | 92850235 | 92850765 | 1000 | Intron (ENSMUST00000113438.2 | 6 | 92793010 | 92860145 | 67136 | 2 | ENSMUST00 | 9380 | Adamts9 | ENSMUSG0( | false | . | . | . | . | . | . | chr6 | 92848230 | 92851329 | 4.41535 | 3.76664 | 2.52236 |
| chr6 | 94699745 | 94700542 | 1000 | Promoter (<=1kb) | 6 | 94604529 | 94700158 | 95630 | 2 | ENSMUST00 | 0 | Lrig1 | ENSMUSG0( | false | chr6 | 94698891 | 94701598 | 25.17393 | 93.69764 | 90.27328 | . | . | . | . | . | . |
| chr6 | 97205462 | 97205840 | 1000 | Promoter (<=1kb) | 6 | 97183670 | 97205560 | 21891 | 2 | ENSMUST00 | 0 | Uba3 | ENSMUSG0( | false | chr6 | 97204495 | 97205669 | 33.91391 | 110.8507 | 106.99079 | chr6 | 97203508 | 97206081 | 4.08611 | 3.56641 | 2.27121 |
| chr6 | 99295457 | 99295808 | 1000 | Intron (ENSMUST00000113322.3 | 6 | 99161940 | 99266532 | 104593 | 2 | ENSMUST00 | -28926 | Foxp1 | ENSMUSG0( | false | . | . | . | . | . | . | chr6 | 99295015 | 99296233 | 4.87422 | 3.9142 | 2.92936 |
| chr6 | 99439600 | 99440361 | 1000 | Intron (ENSMUST00000176565.2 | 6 | 99308377 | 99436396 | 128020 | 2 | ENSMUST00 | -3205 | Foxp1 | ENSMUSG0( | false | . | . | . | . | . | . | chr6 | 99437543 | 99441621 | 6.52464 | 4.67579 | 4.28601 |
| chr6 | 99094568 | 99095039 | 1000 | Promoter (1-2kb) | 6 | 99009933 | 99096205 | 86273 | 2 | ENSMUST00 | 1166 | Foxp1 | ENSMUSG0( | false | chr6 | 99094864 | 99096288 | 7.48194 | 15.52168 | 13.25109 | chr6 | 99089995 | 99102481 | 4.84431 | 4.00533 | 2.88029 |
| chr6 | 99520400 | 99523675 | 1000 | Promoter (<=1kb) | 6 | 98929189 | 99520997 | 591809 | 2 | ENSMUST00 | 0 | Foxp1 | ENSMUSG0( | false | chr6 | 99520242 | 99523905 | 29.33348 | 103.60687 | 99.9429 | chr6 | 99518709 | 99520624 | 3.86078 | 3.5582 | 2.05513 |
| chr6 | 88841853 | 88842743 | 1000 | Promoter (<=1kb) | 6 | 88835915 | 88841935 | 6021 | 2 | ENSMUST00 | 0 | Abtb1 | ENSMUSG0( | false | chr6 | 88841141 | 88842056 | 29.28796 | 98.53321 | 94.99628 | . | . | . | . | . | . |
| chr6 | 88784846 | 88785294 | 1000 | Intron (ENSMUST00000113585.3 | 6 | 88835915 | 88841935 | 6021 | 2 | ENSMUST00 | 56641 | Abtb1 | ENSMUSG0( | false | . | . | . | . | . | . | chr6 | 88783365 | 88788752 | 6.6838 | 4.87557 | 4.38323 |
| chr6 | 88816970 | 88817280 | 1000 | Intron (ENSMUST00000113585.3 | 6 | 88835915 | 88841935 | 6021 | 2 | ENSMUST00 | 24655 | Abtb1 | ENSMUSG0( | false | . | . | . | . | . | . | . | . | . | . | . | . |
| chr6 | 89342039 | 89342533 | 1000 | Exon (ENSMUST00000163139.2/ | 6 | 89319015 | 89357717 | 38703 | 2 | ENSMUST00 | 15184 | Plxna1 | ENSMUSG0( | false | . | . | . | . | . | . | chr6 | 89339950 | 89361705 | 7.61354 | 5.20417 | 5.1492 |
| chr6 | 89344876 | 89345486 | 1000 | Intron (ENSMUST00000163139.2 | 6 | 89319015 | 89357717 | 38703 | 2 | ENSMUST00 | 12231 | Plxna1 | ENSMUSG0( | false | chr6 | 89345210 | 89345510 | 6.47316 | 10.85638 | 8.66333 | chr6 | 89339950 | 89361705 | 7.61354 | 5.20417 | 5.1492 |
| chr6 | 89351455 | 89352040 | 1000 | Intron (ENSMUST00000163139.2 | 6 | 89319015 | 89357717 | 38703 | 2 | ENSMUST00 | 5677 | Plxna1 | ENSMUSG0( | false | . | . | . | . | . | . | chr6 | 89339950 | 89361705 | 7.61354 | 5.20417 | 5.1492 |
| chr6 | 90462152 | 90462800 | 1000 | Promoter (<=1kb) | 6 | 90462576 | 90475238 | 12663 | 1 | ENSMUST00 | 0 | Klf15 | ENSMUSG0( | false | chr6 | 90462123 | 90463911 | 27.3492 | 80.60155 | 77.44116 | chr6 | 90462103 | 90464387 | 3.24764 | 3.18149 | 1.56356 |
| chr6 | 90984089 | 90984507 | 1000 | Distal Intergenic | 6 | 91013317 | 91017429 | 4113 | 2 | ENSMUST00 | 32922 | Nup210 | ENSMUSG0( | false | . | . | . | . | . | . | chr6 | 90982644 | 90988187 | 5.21445 | 4.07964 | 3.20504 |
| chr6 | 91675089 | 91675645 | 1000 | Distal Intergenic | 6 | 91684067 | 91759063 | 74997 | 1 | ENSMUST00 | -8422 | Slc6a6 | ENSMUSG0( | false | . | . | . | . | . | . | chr6 | 91674008 | 91677067 | 5.15817 | 4.19556 | 3.1256 |
| chr6 | 91650184 | 91650573 | 1000 | Distal Intergenic | 6 | 91684067 | 91759063 | 74997 | 1 | ENSMUST00 | -33494 | Slc6a6 | ENSMUSG0( | false | . | . | . | . | . | . | chr6 | 91646829 | 91651203 | 4.70686 | 3.97322 | 2.75882 |
| chr6 | 108212545 | 108213452 | 1000 | Promoter (<=1kb) | 6 | 108213118 | 108551109 | 337992 | 1 | ENSMUST00 | 0 | Itpr1 | ENSMUSG0( | false | chr6 | 108212049 | 108214484 | 34.95504 | 111.6348 | 107.75649 | . | . | . | . | . | . |
| chr6 | 108659217 | 108660866 | 1000 | Promoter (<=1kb) | 6 | 108660629 | 108666925 | 6297 | 1 | ENSMUST00 | 0 | Bhlhe40 | ENSMUSG0( | false | chr6 | 108658527 | 108661954 | 18.36477 | 56.64417 | 53.84045 | chr6 | 108660861 | 108663170 | 4.0955 | 3.59916 | 2.27642 |
| chr6 | 108840482 | 108840881 | 1000 | Intron (ENSMUST00000089162.3 | 6 | 108828641 | 108859356 | 30716 | 1 | ENSMUST00 | 11842 | Edem1 | ENSMUSG0( | false | . | . | . | . | . | . | chr6 | 108829425 | 108843436 | 5.60264 | 4.30322 | 3.51143 |
| chr6 | 108838606 | 108839051 | 1000 | Intron (ENSMUST00000089162.3 | 6 | 108828641 | 108859356 | 30716 | 1 | ENSMUST00 | 9966 | Edem1 | ENSMUSG0( | false | . | . | . | . | . | . | chr6 | 108829425 | 108843436 | 5.60264 | 4.30322 | 3.51143 |
| chr6 | 118173850 | 118174425 | 1000 | Exon (ENSMUST00000032201.5/ | 6 | 118151748 | 118197744 | 45997 | 2 | ENSMUST00 | 23319 | Ret | ENSMUSG0( | false | . | . | . | . | . | . | chr6 | 118174093 | 118174403 | 2.84832 | 2.84921 | 1.25968 |
| chr6 | 124930788 | 124931837 | 1000 | Promoter (<=1kb) | 6 | 124931388 | 124936138 | 4751 | 1 | ENSMUST00 | 0 | Mlf2 | ENSMUSG0( | false | chr6 | 124931319 | 124932573 | 26.86698 | 95.04005 | 91.5885 | chr6 | 124929040 | 124930795 | 6.34457 | 4.75855 | 4.11572 |
| chr6 | 124916988 | 124918354 | 1000 | Promoter (<=1kb) | 6 | 124913681 | 124917946 | 4266 | 2 | ENSMUST00 | 0 | Ptms | ENSMUSG0( | false | chr6 | 124915360 | 124918752 | 39.97929 | 126.69925 | 122.32521 | . | . | . | . | . | . |
| chr6 | 116005596 | 116006031 | 1000 | Distal Intergenic | 6 | 115954811 | 115995005 | 40195 | 2 | ENSMUST00 | -10592 | Plxnd1 | ENSMUSG0( | false | . | . | . | . | . | . | chr6 | 116004192 | 116008178 | 6.32655 | 4.46684 | 4.10455 |
| chr6 | 115962688 | 115962996 | 1000 | Exon (ENSMUST00000015511.9/ | 6 | 115955800 | 115956364 | 565 | 2 | ENSMUST00 | -6325 | Plxnd1 | ENSMUSG0( | false | . | . | . | . | . | . | chr6 | 115962317 | 115964032 | 5.23294 | 4.22926 | 3.1832 |
| chr6 | 116006706 | 116007185 | 1000 | Distal Intergenic | 6 | 115954811 | 115995005 | 40195 | 2 | ENSMUST00 | -11702 | Plxnd1 | ENSMUSG0( | false | . | . | . | . | . | . | chr6 | 116004192 | 116008178 | 6.32655 | 4.46684 | 4.10455 |
| chr6 | 116192817 | 116193729 | 1000 | Promoter (<=1kb) | 6 | 116018618 | 116193486 | 174869 | 2 | ENSMUST00 | 0 | Tmcc1 | ENSMUSG0( | false | chr6 | 116192457 | 116194100 | 20.49778 | 85.71664 | 82.45538 | . | . | . | . | . | . |
| chr6 | 119562435 | 119562945 | 1000 | Distal Intergenic | 6 | 119432531 | 119544347 | 111817 | 2 | ENSMUST00 | -18089 | Wnt5b | ENSMUSG0( | false | . | . | . | . | . | . | chr6 | 119561849 | 119572810 | 6.78174 | 4.90966 | 4.46221 |
| chr6 | 119566713 | 119567076 | 1000 | Distal Intergenic | 6 | 119432531 | 119544347 | 111817 | 2 | ENSMUST00 | -22367 | Wnt5b | ENSMUSG0( | false | . | . | . | . | . | . | chr6 | 119561849 | 119572810 | 6.78174 | 4.90966 | 4.46221 |
| chr6 | 119565404 | 119566216 | 1000 | Distal Intergenic | 6 | 119432531 | 119544347 | 111817 | 2 | ENSMUST00 | -21058 | Wnt5b | ENSMUSG0( | false | . | . | . | . | . | . | chr6 | 119561849 | 119572810 | 6.78174 | 4.90966 | 4.46221 |
| chr6 | 119543831 | 119544587 | 1000 | Promoter (<=1kb) | 6 | 119432531 | 119544347 | 111817 | 2 | ENSMUST00 | 0 | Wnt5b | ENSMUSG0( | false | chr6 | 119543362 | 119544750 | 23.4661 | 79.19912 | 76.05865 | chr6 | 119543230 | 119544710 | 3.3312 | 3.21596 | 1.64135 |
| chr6 | 131387915 | 131388635 | 1000 | Promoter (<=1kb) | 6 | 131364858 | 131388450 | 23593 | 2 | ENSMUST00 | 0 | Ybx3 | ENSMUSG0( | false | chr6 | 131387180 | 131389119 | 22.78592 | 76.10319 | 73.01487 | . | . | . | . | . | . |
| chr6 | 134050433 | 134050801 | 1000 | Intron (ENSMUST00000081028.7 | 6 | 134035969 | 134248284 | 212316 | 1 | ENSMUST00 | 14465 | Etv6 | ENSMUSG0( | false | . | . | . | . | . | . | chr6 | 134037014 | 134053440 | 5.9149 | 4.52781 | 3.75986 |
| chr6 | 134034952 | 134036203 | 1000 | Promoter (<=1kb) | 6 | 134035700 | 134270158 | 234459 | 1 | ENSMUST00 | 0 | Etv6 | ENSMUSG0( | false | chr6 | 134034454 | 134036769 | 27.34946 | 104.78247 | 101.09169 | . | . | . | . | . | . |
| chr6 | 134043586 | 134044016 | 1000 | Intron (ENSMUST00000081028.7 | 6 | 134035969 | 134248284 | 212316 | 1 | ENSMUST00 | 7618 | Etv6 | ENSMUSG0( | false | . | . | . | . | . | . | chr6 | 134037014 | 134053440 | 5.9149 | 4.52781 | 3.75986 |
| chr6 | 134566358 | 134567430 | 1000 | Promoter (<=1kb) | 6 | 134446476 | 134566913 | 120438 | 2 | ENSMUST00 | 0 | Lrp6 | ENSMUSG0( | false | chr6 | 134565101 | 134567582 | 36.69013 | 115.69205 | 111.68276 | . | . | . | . | . | . |
| chr6 | 134489748 | 134490115 | 1000 | Intron (ENSMUST00000032322.9 | 6 | 134450483 | 134471293 | 20811 | 2 | ENSMUST00 | -18456 | Lrp6 | ENSMUSG0( | false | . | . | . | . | . | . | chr6 | 134486406 | 134495584 | 4.53888 | 3.83472 | 2.64166 |
| chr6 | 134560979 | 134561310 | 1000 | Intron (ENSMUST00000032322.9 | 6 | 134446476 | 134566913 | 120438 | 2 | ENSMUST00 | 5603 | Lrp6 | ENSMUSG0( | false | . | . | . | . | . | . | chr6 | 134553711 | 134565300 | 4.91825 | 4.02409 | 2.93578 |
| chr6 | 137444569 | 137444866 | 1000 | Intron (ENSMUST00000077115.7 | 6 | 137410721 | 137464629 | 53909 | 1 | ENSMUST00 | 33849 | Ptpro | ENSMUSG0( | true | . | . | . | . | . | . | chr6 | 137442771 | 137450357 | 4.6569 | 3.76236 | 2.76369 |
| chr6 | 137247282 | 137247668 | 1000 | Distal Intergenic | 6 | 137252299 | 137464629 | 212331 | 1 | ENSMUST00 | -4631 | Ptpro | ENSMUSG0( | true | . | . | . | . | . | . | chr6 | 137244902 | 137251836 | 5.39138 | 4.30457 | 3.32314 |
| chr6 | 137287635 | 137288191 | 1000 | Intron (ENSMUST00000077115.7 | 6 | 137252299 | 137464629 | 212331 | 1 | ENSMUST00 | 35337 | Ptpro | ENSMUSG0( | true | . | . | . | . | . | . | chr6 | 137275709 | 137305131 | 5.12249 | 4.15603 | 3.10285 |
| chr6 | 137280610 | 137281059 | 1000 | Intron (ENSMUST00000077115.7 | 6 | 137252299 | 137464629 | 212331 | 1 | ENSMUST00 | 28312 | Ptpro | ENSMUSG0( | true | . | . | . | . | . | . | chr6 | 137275709 | 137305131 | 5.12249 | 4.15603 | 3.10285 |
| chr6 | 137318341 | 137318794 | 1000 | Intron (ENSMUST00000077115.7 | 6 | 137252299 | 137464629 | 212331 | 1 | ENSMUST00 | 66043 | Ptpro | ENSMUSG0( | true | . | . | . | . | . | . | chr6 | 137312428 | 137331531 | 4.39206 | 3.74363 | 2.51481 |
| chr6 | 140433089 | 140433545 | 1000 | Intron (ENSMUST00000087622.3 | 6 | 140424099 | 140594906 | 170808 | 1 | ENSMUST00 | 8991 | Plekha5 | ENSMUSG0( | true | . | . | . | . | . | . | chr6 | 140425415 | 140441290 | 5.09666 | 4.0883 | 3.09824 |
| chr6 | 140423683 | 140424500 | 1000 | Promoter (<=1kb) | 6 | 140424099 | 140594906 | 170808 | 1 | ENSMUST00 | 0 | Plekha5 | ENSMUSG0( | true | chr6 | 140423533 | 140426226 | 36.12563 | 122.12295 | 117.91594 | . | . | . | . | . | . |
| chr6 | 140623120 | 140624313 | 1000 | Promoter (<=1kb) | 6 | 140623499 | 140652826 | 29328 | 1 | ENSMUST00 | 0 | Aebp2 | ENSMUSG0( | false | chr6 | 140622736 | 140625497 | 34.4409 | 118.06123 | 113.98219 | . | . | . | . | . | . |
| chr6 | 141434254 | 141434630 | 1000 | Intron (ENSMUST00000043259.8 | 6 | 141524386 | 141569720 | 45335 | 1 | ENSMUST00 | -89756 | Slco1c1 | ENSMUSG0( | false | . | . | . | . | . | . | chr6 | 141422605 | 141443938 | 4.72585 | 3.94437 | 2.77999 |
| chr6 | 141397079 | 141397414 | 1000 | Intron (ENSMUST00000043259.8 | 6 | 141524386 | 141569720 | 45335 | 1 | ENSMUST00 | -126972 | Slco1c1 | ENSMUSG0( | false | . | . | . | . | . | . | chr6 | 141387816 | 141400666 | 4.81448 | 4.05067 | 2.84093 |
| chr6 | 142506354 | 142506902 | 1000 | Promoter (<=1kb) | 6 | 142495568 | 142507805 | 12238 | 2 | ENSMUST00 | 903 | Ldhb | ENSMUSG0( | false | . | . | . | . | . | . | chr6 | 142505920 | 142508471 | 4.32485 | 3.68555 | 2.45254 |
| chr6 | 145866037 | 145866981 | 1000 | Promoter (<=1kb) | 6 | 145858243 | 145865558 | 7316 | 2 | ENSMUST00 | -480 | Bhlhe41 | ENSMUSG0( | false | chr6 | 145864843 | 145866819 | 26.13664 | 77.56537 | 74.45414 | chr6 | 145861038 | 145867407 | 3.647 | 3.29023 | 1.93146 |
| chr6 | 145746242 | 145747179 | 1000 | Promoter (<=1kb) | 6 | 145746748 | 145821079 | 74332 | 1 | ENSMUST00 | 0 | Rassf8 | ENSMUSG0( | true | chr6 | 145745878 | 145749136 | 38.25672 | 133.48087 | 128.85442 | . | . | . | . | . | . |
| chr6 | 145781959 | 145782404 | 1000 | Intron (ENSMUST00000111704.2 | 6 | 145808383 | 145817584 | 9202 | 1 | ENSMUST00 | -25979 | Rassf8 | ENSMUSG0( | true | . | . | . | . | . | . | chr6 | 145748529 | 145783801 | 6.70363 | 4.81765 | 4.41144 |
| chr6 | 113046012 | 113046658 | 1000 | Promoter (<=1kb) | 6 | 113046225 | 113068273 | 22049 | 1 | ENSMUST00 | 0 | Thumpd3 | ENSMUSG0( | false | chr6 | 113045546 | 113047223 | 37.49531 | 120.259 | 116.11364 | chr6 | 113045130 | 113048455 | 4.06407 | 3.62223 | 2.26594 |
| chr6 | 145250032 | 145250679 | 1000 | Promoter (<=1kb) | 6 | 145216699 | 145250239 | 33541 | 2 | ENSMUST00 | 0 | Kras | ENSMUSG0( | false | chr6 | 145249108 | 145251136 | 31.16539 | 106.97112 | 103.21663 | . | . | . | . | . | . |
| chr6 | 142756573 | 142757082 | 1000 | Promoter (<=1kb) | 6 | 142756686 | 142775714 | 19029 | 1 | ENSMUST00 | 0 | Cmas | ENSMUSG0( | false | chr6 | 142756393 | 142757636 | 30.41442 | 103.5799 | 99.9162 | chr6 | 142752999 | 142758406 | 4.17228 | 3.60399 | 2.35749 |
| chr6 | 146302661 | 146303066 | 1000 | Intron (ENSMUST00000053273.9 | 6 | 146290486 | 146294284 | 3799 | 2 | ENSMUST00 | -8378 | Itpr2 | ENSMUSG0( | false | . | . | . | . | . | . | chr6 | 146300604 | 146305187 | 5.0928 | 4.02763 | 3.09253 |
| chr6 | 146642223 | 146642591 | 1000 | Promoter (<=1kb) | 6 | 146642547 | 146650602 | 8056 | 1 | ENSMUST00 | 0 | Med21 | ENSMUSG0( | false | chr6 | 146642099 | 146643520 | 32.51422 | 118.74544 | 114.65017 | . | . | . | . | . | . |
| chr6 | 149101153 | 149101824 | 1000 | Promoter (<=1kb) | 6 | 148988071 | 149101680 | 113610 | 2 | ENSMUST00 | 0 | Dennd5b | ENSMUSG0( | false | chr6 | 149100192 | 149102240 | 24.60788 | 97.30984 | 93.80507 | . | . | . | . | . | . |
| chr6 | 114858685 | 114859047 | 1000 | Promoter (2-3kb) | 6 | 114856151 | 114859222 | 3072 | 1 | ENSMUST00 | 2535 | Atg7 | ENSMUSG0( | false | . | . | . | . | . | . | chr6 | 114853844 | 114919520 | 8.05355 | 5.36011 | 5.51688 |
| chr6 | 114874629 | 114875597 | 1000 | Promoter (<=1kb) | 6 | 114860628 | 114875107 | 14480 | 2 | ENSMUST00 | 0 | Vgll4 | ENSMUSG0( | true | chr6 | 114873653 | 114875291 | 9.18239 | 21.07545 | 18.72512 | chr6 | 114853844 | 114919520 | 8.05355 | 5.36011 | 5.51688 |
| chr6 | 114882435 | 114882948 | 1000 | Intron (ENSMUST00000032459.8 | 6 | 114860628 | 114875107 | 14480 | 2 | ENSMUST00 | -7329 | Vgll4 | ENSMUSG0( | true | . | . | . | . | . | . | chr6 | 114853844 | 114919520 | 8.05355 | 5.36011 | 5.51688 |
| chr6 | 114968767 | 114969302 | 1000 | Promoter (<=1kb) | 6 | 114862321 | 114969180 | 106860 | 2 | ENSMUST00 | 0 | Vgll4 | ENSMUSG0( | true | chr6 | 114967501 | 114970246 | 26.65955 | 86.95457 | 83.67183 | chr6 | 114966882 | 114970611 | 4.21249 | 3.6549 | 2.39033 |
| chr6 | 114873936 | 114874444 | 1000 | Promoter (<=1kb) | 6 | 114862191 | 114875020 | 12830 | 2 | ENSMUST00 | 576 | Vgll4 | ENSMUSG0( | true | chr6 | 114873653 | 114875291 | 9.18239 | 21.07545 | 18.72512 | chr6 | 114853844 | 114919520 | 8.05355 | 5.36011 | 5.51688 |
| chr6 | 114919548 | 114920039 | 1000 | Promoter (1-2kb) | 6 | 114861424 | 114921821 | 60398 | 2 | ENSMUST00 | 1782 | Vgll4 | ENSMUSG0( | true | chr6 | 114917117 | 114921811 | 29.60582 | 98.11335 | 94.5914 | . | . | . | . | . | . |
| chr6 | 114921632 | 114922233 | 1000 | Promoter (<=1kb) | 6 | 114861424 | 114921821 | 60398 | 2 | ENSMUST00 | 0 | Vgll4 | ENSMUSG0( | true | chr6 | 114917117 | 114921811 | 29.60582 | 98.11335 | 94.5914 | chr6 | 114922002 | 114923292 | 3.84635 | 3.53185 | 2.04366 |
| chr6 | 125313580 | 125314108 | 1000 | Promoter (<=1kb) | 6 | 125306571 | 125313885 | 7315 | 2 | ENSMUST00 | 0 | Ltbr | ENSMUSG0( | false | chr6 | 125312582 | 125314359 | 37.09717 | 122.76616 | 118.52905 | chr6 | 125312075 | 125314545 | 4.01201 | 3.55317 | 2.20327 |
| chr6 | 125349363 | 125350070 | 1000 | Promoter (<=1kb) | 6 | 125349451 | 125357316 | 7866 | 1 | ENSMUST00 | 0 | Tnfrsf1a | ENSMUSG0( | false | chr6 | 125349048 | 125350835 | 29.69128 | 86.53342 | 83.26198 | chr6 | 125348821 | 125350036 | 3.90265 | 3.35579 | 2.12777 |
| chr6 | 128044736 | 128045117 | 1000 | Intron (ENSMUST00000112173.2 | 6 | 127965747 | 128034592 | 68846 | 2 | ENSMUST00 | -10145 | Tspan9 | ENSMUSG0( | false | . | . | . | . | . | . | chr6 | 128044193 | 128045661 | 3.65656 | 3.43096 | 1.88672 |
| chr6 | 128124120 | 128124895 | 1000 | Promoter (<=1kb) | 6 | 127967166 | 128124316 | 157151 | 2 | ENSMUST00 | 0 | Tspan9 | ENSMUSG0( | false | . | . | . | . | . | . | . | . | . | . | . | . |
| chr6 | 128300110 | 128301068 | 1000 | Promoter (<=1kb) | 6 | 128227143 | 128300813 | 73671 | 2 | ENSMUST00 | 0 | Tead4 | ENSMUSG0( | false | chr6 | 128299157 | 128301648 | 37.17739 | 121.83733 | 117.63558 | chr6 | 128300936 | 128302171 | 4.08937 | 3.62369 | 2.292 |
| chr6 | 128275528 | 128275935 | 1000 | Promoter (<=1kb) | 6 | 128224288 | 128275577 | 51290 | 2 | ENSMUST00 | 0 | Tead4 | ENSMUSG0( | false | . | . | . | . | . | . | chr6 | 128273173 | 128278366 | 5.75281 | 4.34192 | 3.66756 |
| chr7 | 19269656 | 19270406 | 1000 | Promoter (1-2kb) | 7 | 19256929 | 19271817 | 14889 | 2 | ENSMUST00 | 1411 | Vasp | ENSMUSG0( | false | chr7 | 19269289 | 19272839 | 36.83125 | 118.04518 | 113.96772 | chr7 | 19265603 | 19271238 | 5.52106 | 4.28447 | 3.4309 |
| chr7 | 19082959 | 19083507 | 1000 | Promoter (<=1kb) | 7 | 19083849 | 19093821 | 9973 | 1 | ENSMUST00 | -342 | Dmpk | ENSMUSG0( | false | . | . | . | . | . | . | chr7 | 19075255 | 19089763 | 7.24488 | 4.98539 | 4.86301 |
| chr7 | 19078618 | 19079015 | 1000 | Promoter (<=1kb) | 7 | 19078026 | 19082502 | 4477 | 1 | ENSMUST00 | 593 | Dmwd | ENSMUSG0( | false | . | . | . | . | . | . | chr7 | 19075255 | 19089763 | 7.24488 | 4.98539 | 4.86301 |
| chr7 | 35663981 | 35664370 | 1000 | Distal Intergenic | 7 | 35641982 | 35647521 | 5540 | 2 | ENSMUST00 | -16461 | Pdcd5 | ENSMUSG0( | false | . | . | . | . | . | . | chr7 | 35663371 | 35664343 | 3.81569 | 3.47534 | 2.0737 |
| chr7 | 5060992 | 5061707 | 1000 | Promoter (<=1kb) | 7 | 5062143 | 5079945 | 17803 | 1 | ENSMUST00 | -436 | U2af2 | ENSMUSG0( | false | chr7 | 5060226 | 5063489 | 33.42125 | 102.81408 | 99.17643 | . | . | . | . | . | . |
| chr8 | 68062067 | 68062700 | 1000 | Promoter (<=1kb) | 8 | 67790470 | 68062286 | 271817 | 2 | ENSMUST00 | 0 | Psd3 | ENSMUSG0( | true | chr8 | 68061707 | 68062981 | 32.02775 | 102.41263 | 98.78432 | . | . | . | . | . | . |
| chr8 | 67948520 | 67948921 | 1000 | Promoter (2-3kb) | 8 | 67903648 | 67951879 | 48232 | 2 | ENSMUST00 | 2958 | Psd3 | ENSMUSG0( | true | chr8 | 67947543 | 67950750 | 20.65177 | 61.66082 | 58.78991 | chr8 | 67935983 | 67949268 | 5.35768 | 4.23511 | 3.29751 |
| chr7 | 35333796 | 35334422 | 1000 | Promoter (<=1kb) | 7 | 35334170 | 35392289 | 58120 | 1 | ENSMUST00 | 0 | Rhpn2 | ENSMUSG0( | false | chr7 | 35333177 | 35333959 | 25.15761 | 80.47005 | 77.31023 | . | . | . | . | . | . |
| chr7 | 34653905 | 34655823 | 1000 | Promoter (<=1kb) | 7 | 34639014 | 34654398 | 15385 | 2 | ENSMUST00 | 0 | Kctd15 | ENSMUSG0( | false | chr7 | 34652958 | 34655928 | 33.10969 | 106.34647 | 102.61562 | chr7 | 34652678 | 34654142 | 3.66219 | 3.34771 | 1.95131 |
| chr7 | 34618773 | 34619112 | 1000 | Distal Intergenic | 7 | 34639015 | 34652837 | 13823 | 2 | ENSMUST00 | 33725 | Kctd15 | ENSMUSG0( | false | . | . | . | . | . | . | chr7 | 34617606 | 34621108 | 5.39621 | 4.21255 | 3.3617 |
| chr7 | 65902388 | 65903066 | 1000 | Intron (ENSMUST00000176199.2 | 7 | 65862432 | 66050340 | 187909 | 1 | ENSMUST00 | 39957 | Pcsk6 | ENSMUSG0( | true | . | . | . | . | . | . | chr7 | 65898106 | 65906640 | 5.15164 | 4.00869 | 3.16921 |
| chr7 | 65393621 | 65394149 | 1000 | Distal Intergenic | 7 | 65296165 | 65371239 | 75075 | 2 | ENSMUST00 | -22383 | Tjp1 | ENSMUSG0( | true | . | . | . | . | . | . | chr7 | 65392078 | 65396169 | 7.03857 | 5.03568 | 4.68832 |
| chr7 | 65413976 | 65414828 | 1000 | Distal Intergenic | 7 | 65296165 | 65371239 | 75075 | 2 | ENSMUST00 | -42738 | Tjp1 | ENSMUSG0( | true | . | . | . | . | . | . | chr7 | 65409463 | 65433128 | 6.06932 | 4.54697 | 3.8847 |
| chr7 | 65415295 | 65415644 | 1000 | Distal Intergenic | 7 | 65296165 | 65371239 | 75075 | 2 | ENSMUST00 | -44057 | Tjp1 | ENSMUSG0( | true | . | . | . | . | . | . | chr7 | 65409463 | 65433128 | 6.06932 | 4.54697 | 3.8847 |
| chr7 | 65370389 | 65371480 | 1000 | Promoter (<=1kb) | 7 | 65296165 | 65371239 | 75075 | 2 | ENSMUST00 | 0 | Tjp1 | ENSMUSG0( | true | chr7 | 65366546 | 65371948 | 37.39668 | 122.22031 | 118.01075 | . | . | . | . | . | . |
| chr7 | 65378752 | 65379754 | 1000 | Distal Intergenic | 7 | 65296165 | 65371239 | 75075 | 2 | ENSMUST00 | -7514 | Tjp1 | ENSMUSG0( | true | . | . | . | . | . | . | chr7 | 65376995 | 65383642 | 5.04739 | 4.06847 | 3.06175 |
| chr7 | 80688162 | 80689191 | 1000 | Promoter (<=1kb) | 7 | 80586627 | 80688877 | 102251 | 2 | ENSMUST00 | 0 | Crtc3 | ENSMUSG0( | false | chr7 | 80687626 | 80689456 | 30.90191 | 98.42355 | 94.89227 | . | . | . | . | . | . |
| chr7 | 80554350 | 80554931 | 1000 | Distal Intergenic | 7 | 80454993 | 80535119 | 80127 | 2 | ENSMUST00 | -19232 | Blm | ENSMUSG0( | false | . | . | . | . | . | . | chr7 | 80554362 | 80556452 | 4.0401 | 3.62208 | 2.24668 |
| chr7 | 80402019 | 80403201 | 1000 | Promoter (<=1kb) | 7 | 80389194 | 80402762 | 13569 | 2 | ENSMUST00 | 0 | Furin | ENSMUSG0( | false | chr7 | 80401665 | 80403431 | 28.1615 | 93.53628 | 90.11485 | chr7 | 80403105 | 80403961 | 4.13488 | 3.64565 | 2.28953 |
| chr7 | 80232400 | 80232954 | 1000 | Promoter (<=1kb) | 7 | 80227156 | 80232813 | 5658 | 2 | ENSMUST00 | 0 | Cib1 | ENSMUSG0( | false | chr7 | 80231843 | 80233760 | 34.92026 | 124.22972 | 119.94103 | . | . | . | . | . | . |
| chr7 | 80192307 | 80192946 | 1000 | Intron (ENSMUST00000107383.2 | 7 | 80186841 | 80226394 | 39554 | 1 | ENSMUST00 | 5467 | Sema4b | ENSMUSG0( | false | chr7 | 80192373 | 80192939 | 10.4407 | 21.06721 | 18.71816 | chr7 | 80190524 | 80193223 | 4.86805 | 3.79798 | 2.93745 |
| chr7 | 80114337 | 80114837 | 1000 | Promoter (<=1kb) | 7 | 80094850 | 80115332 | 20483 | 2 | ENSMUST00 | 495 | Idh2 | ENSMUSG0( | false | chr7 | 80113906 | 80115785 | 33.40766 | 121.45458 | 117.2672 | chr7 | 80107135 | 80114500 | 4.80219 | 3.85138 | 2.88731 |
| chr7 | 70360317 | 70361129 | 1000 | Promoter (<=1kb) | 7 | 70351946 | 70360593 | 8648 | 2 | ENSMUST00 | 0 | Nr2f2 | ENSMUSG0( | false | chr7 | 70357618 | 70362933 | 29.75681 | 92.45776 | 89.05831 | . | . | . | . | . | . |
| chr7 | 68264469 | 68264985 | 1000 | Promoter (<=1kb) | 7 | 68236608 | 68264233 | 27626 | 2 | ENSMUST00 | -237 | Pgpep1l | ENSMUSG0( | false | chr7 | 68263831 | 68264961 | 11.56301 | 29.49975 | 27.03959 | chr7 | 68262613 | 68266799 | 4.09256 | 3.53896 | 2.3125 |
| chr7 | 29471060 | 29471695 | 1000 | Intron (ENSMUST00000085809.5 | 7 | 29384256 | 29505452 | 121197 | 2 | ENSMUST00 | 33757 | Sipa1l3 | ENSMUSG0( | false | . | . | . | . | . | . | chr7 | 29469527 | 29474693 | 6.76731 | 4.87889 | 4.50482 |
| chr7 | 28467383 | 28467766 | 1000 | 3' UTR | 7 | 28458658 | 28468242 | 9585 | 1 | ENSMUST00 | 8726 | Lrfn1 | ENSMUSG0( | false | chr7 | 28466889 | 28467949 | 25.22864 | 75.06109 | 71.99223 | chr7 | 28466521 | 28468188 | 5.43415 | 4.07545 | 3.38562 |
| chr7 | 79148246 | 79149311 | 1000 | Promoter (<=1kb) | 7 | 79133769 | 79149060 | 15292 | 2 | ENSMUST00 | 0 | Mfge8 | ENSMUSG0( | false | chr7 | 79148383 | 79149554 | 30.945 | 94.52721 | 91.08539 | chr7 | 79148251 | 79149435 | 4.03181 | 3.367 | 2.24958 |
| chr7 | 78690876 | 78691531 | 1000 | Distal Intergenic | 7 | 78775236 | 78783066 | 7831 | 2 | ENSMUST00 | 91535 | Mrpl46 | ENSMUSG0( | false | . | . | . | . | . | . | chr7 | 78689864 | 78693377 | 5.11876 | 4.05109 | 3.13481 |
| chr7 | 90475906 | 90476307 | 1000 | Promoter (<=1kb) | 7 | 90468194 | 90476001 | 7808 | 2 | ENSMUST00 | 0 | Tmem126b | ENSMUSG0( | false | chr7 | 90475283 | 90477161 | 22.06604 | 78.32127 | 75.1984 | . | . | . | . | . | . |
| chr7 | 89980146 | 89980967 | 1000 | Promoter (<=1kb) | 7 | 89976744 | 89980559 | 3816 | 2 | ENSMUST00 | 0 | Eed | ENSMUSG0( | false | chr7 | 89979712 | 89981930 | 23.85991 | 99.51073 | 95.95212 | . | . | . | . | . | . |
| chr7 | 89845766 | 89846019 | 1000 | Exon (ENSMUST00000032856.7/ | 7 | 89829282 | 89851962 | 22681 | 1 | ENSMUST00 | 16485 | Me3 | ENSMUSG0( | false | . | . | . | . | . | . | chr7 | 89845751 | 89846394 | 3.52395 | 3.27943 | 1.79672 |
| chr7 | 118541865 | 118542724 | 1000 | Exon (ENSMUST00000044195.4/ | 7 | 118525062 | 118533356 | 8295 | 2 | ENSMUST00 | -8510 | Coq7 | ENSMUSG0( | false | chr7 | 118539979 | 118542150 | 12.46556 | 27.94533 | 25.5059 | chr7 | 118536275 | 118546239 | 6.06229 | 4.47014 | 3.88658 |
| chr7 | 118242952 | 118243951 | 1000 | Promoter (<=1kb) | 7 | 118131308 | 118243637 | 112330 | 2 | ENSMUST00 | 0 | Smg1 | ENSMUSG0( | false | chr7 | 118242001 | 118243629 | 28.22734 | 101.53345 | 97.92473 | . | . | . | . | . | . |
| chr7 | 117380514 | 117381656 | 1000 | Promoter (<=1kb) | 7 | 117380979 | 117635185 | 254207 | 1 | ENSMUST00 | 0 | Xylt1 | ENSMUSG0( | false | chr7 | 117380196 | 117382139 | 19.74353 | 71.90538 | 68.88615 | chr7 | 117381543 | 117383388 | 5.64584 | 4.30275 | 3.55456 |
| chr14 | 120910753 | 120911468 | 1000 | Promoter (<=1kb) | 14 | 120911194 | 120948042 | 36849 | 1 | ENSMUST00 | 0 | Ipo5 | ENSMUSG0( | false | chr14 | 120910190 | 120912364 | 16.0743 | 67.86449 | 64.90525 | . | . | . | . | . | . |
| chr7 | 114414938 | 114415587 | 1000 | Promoter (<=1kb) | 7 | 114415281 | 114539251 | 123971 | 1 | ENSMUST00 | 0 | Pde3b | ENSMUSG0( | false | chr7 | 114414652 | 114417189 | 26.18681 | 91.82585 | 88.44535 | chr7 | 114412548 | 114415004 | 3.77464 | 3.45282 | 1.98656 |
| chr7 | 127025767 | 127027722 | 1000 | Promoter (<=1kb) | 7 | 127022143 | 127026479 | 4337 | 2 | ENSMUST00 | 0 | Maz | ENSMUSG0( | false | chr7 | 127025218 | 127028304 | 35.65887 | 112.85833 | 108.94193 | chr7 | 127020663 | 127025916 | 4.87603 | 3.99076 | 2.92065 |
| chr7 | 101323130 | 101323632 | 1000 | Promoter (1-2kb) | 7 | 101321971 | 101344332 | 22362 | 1 | ENSMUST00 | 1160 | Stard10 | ENSMUSG0( | false | . | . | . | . | . | . | chr7 | 101320561 | 101323782 | 4.03379 | 3.60603 | 2.23446 |
| chr7 | 101107894 | 101109722 | 1000 | Promoter (<=1kb) | 7 | 101108775 | 101284382 | 175608 | 1 | ENSMUST00 | 0 | Fchsd2 | ENSMUSG0( | true | chr7 | 101107784 | 101110527 | 43.94354 | 143.44185 | 138.423 | chr7 | 101107188 | 101108252 | 5.10392 | 4.17177 | 3.07786 |
| chr7 | 101119211 | 101119774 | 1000 | Intron (ENSMUST00000098250.4 | 7 | 101108882 | 101141129 | 32248 | 1 | ENSMUST00 | 10330 | Fchsd2 | ENSMUSG0( | true | . | . | . | . | . | . | chr7 | 101110387 | 101140869 | 4.70467 | 3.91558 | 2.76499 |
| chr7 | 126799085 | 126800488 | 1000 | Promoter (<=1kb) | 7 | 126795234 | 126799254 | 4021 | 2 | ENSMUST00 | 0 | Aldoa | ENSMUSG0( | false | chr7 | 126799476 | 126800481 | 23.69868 | 67.90152 | 64.94212 | chr7 | 126800202 | 126800945 | 3.90923 | 3.36395 | 2.12884 |
| chr7 | 100607107 | 100608675 | 1000 | Promoter (<=1kb) | 7 | 100607410 | 100641268 | 33859 | 1 | ENSMUST00 | 0 | Rab6a | ENSMUSG0( | false | chr7 | 100607615 | 100609354 | 38.42999 | 152.65175 | 147.26898 | . | . | . | . | . | . |
| chr7 | 100590732 | 100591319 | 1000 | Promoter (<=1kb) | 7 | 100550583 | 100591426 | 40844 | 2 | ENSMUST00 | 107 | Mrpl48 | ENSMUSG0( | false | chr7 | 100590651 | 100591651 | 19.14982 | 55.63055 | 52.84086 | . | . | . | . | . | . |
| chr7 | 126627955 | 126628616 | 1000 | Promoter (<=1kb) | 7 | 126623252 | 126627010 | 3759 | 2 | ENSMUST00 | -946 | Nupr1 | ENSMUSG0( | false | . | . | . | . | . | . | chr7 | 126626006 | 126631554 | 7.79469 | 5.453 | 5.28128 |
| chr7 | 126625378 | 126625833 | 1000 | Promoter (<=1kb) | 7 | 126623249 | 126625676 | 2428 | 2 | ENSMUST00 | 0 | Nupr1 | ENSMUSG0( | false | chr7 | 126622952 | 126625762 | 23.39993 | 73.62621 | 70.58024 | . | . | . | . | . | . |
| chr7 | 100159139 | 100159868 | 1000 | Promoter (<=1kb) | 7 | 100159277 | 100161365 | 2089 | 1 | ENSMUST00 | 0 | Lipt2 | ENSMUSG0( | false | chr7 | 100159002 | 100160115 | 36.25406 | 119.15681 | 115.04134 | . | . | . | . | . | . |
| chr7 | 100227239 | 100227801 | 1000 | Promoter (<=1kb) | 7 | 100227394 | 100275358 | 47965 | 1 | ENSMUST00 | 0 | Pgm2l1 | ENSMUSG0( | false | chr7 | 100226921 | 100228355 | 31.31394 | 100.15002 | 96.57744 | . | . | . | . | . | . |
| chr7 | 99171730 | 99172133 | 1000 | Intron (ENSMUST00000033001.4 | 7 | 99153663 | 99182713 | 29051 | 2 | ENSMUST00 | 10580 | Dgat2 | ENSMUSG0( | false | . | . | . | . | . | . | chr7 | 99170942 | 99172421 | 5.18767 | 4.12604 | 3.20751 |
| chr7 | 114322821 | 114323421 | 1000 | Intron (ENSMUST00000163996.1 | 7 | 114264606 | 114276118 | 11513 | 2 | ENSMUST00 | -46704 | Psma1 | ENSMUSG0( | false | . | . | . | . | . | . | chr7 | 114321160 | 114324452 | 4.23964 | 3.73609 | 2.38738 |
| chr7 | 125434719 | 125435159 | 1000 | Distal Intergenic | 7 | 125444676 | 125462265 | 17590 | 1 | ENSMUST00 | -9517 | Kdm8 | ENSMUSG0( | false | . | . | . | . | . | . | chr7 | 125434516 | 125435739 | 6.55131 | 4.64037 | 4.2955 |
| chr7 | 98702903 | 98703512 | 1000 | Promoter (<=1kb) | 7 | 98703103 | 98715367 | 12265 | 1 | ENSMUST00 | 0 | Prkrir | ENSMUSG0( | false | chr7 | 98702178 | 98704054 | 25.20037 | 82.41283 | 79.21442 | . | . | . | . | . | . |
| chr7 | 114203723 | 114204485 | 1000 | Distal Intergenic | 7 | 114215559 | 114254680 | 39122 | 2 | ENSMUST00 | 50195 | Copb1 | ENSMUSG0( | false | . | . | . | . | . | . | chr7 | 114199145 | 114205819 | 5.60331 | 4.36033 | 3.48713 |
| chr7 | 113513260 | 113514308 | 1000 | Promoter (<=1kb) | 7 | 113513834 | 113571511 | 57678 | 1 | ENSMUST00 | 0 | Far1 | ENSMUSG0( | false | chr7 | 113512797 | 113514889 | 33.04283 | 115.53907 | 111.53828 | chr7 | 113514261 | 113514954 | 3.34994 | 3.20224 | 1.65448 |
| chr7 | 123350212 | 123350692 | 1000 | Intron (ENSMUST00000098060.3 | 7 | 123279150 | 123369915 | 90766 | 2 | ENSMUST00 | 19223 | Arhgap17 | ENSMUSG0( | true | . | . | . | . | . | . | chr7 | 123327623 | 123369156 | 5.61251 | 4.27574 | 3.52744 |
| chr1 | 183077373 | 183077732 | 1000 | Distal Intergenic | 1 | 183086270 | 183147461 | 61192 | 2 | ENSMUST00 | 69729 | Disp1 | ENSMUSG0( | false | . | . | . | . | . | . | chr1 | 183075034 | 183077939 | 4.38135 | 3.67468 | 2.51838 |
| chr1 | 183116393 | 183116798 | 1000 | Intron (ENSMUST00000003035.6 | 1 | 183086270 | 183147461 | 61192 | 2 | ENSMUST00 | 30663 | Disp1 | ENSMUSG0( | false | . | . | . | . | . | . | chr1 | 183111643 | 183121343 | 6.66914 | 4.87138 | 4.37055 |
| chr1 | 183108675 | 183109012 | 1000 | Intron (ENSMUST00000003035.6 | 1 | 183086270 | 183147461 | 61192 | 2 | ENSMUST00 | 38449 | Disp1 | ENSMUSG0( | false | . | . | . | . | . | . | chr1 | 183107659 | 183110008 | 4.21369 | 3.61316 | 2.38184 |
| chr1 | 183151445 | 183151930 | 1000 | Intron (ENSMUST00000003035.6 | 1 | 183086270 | 183147461 | 61192 | 2 | ENSMUST00 | -3985 | Disp1 | ENSMUSG0( | false | . | . | . | . | . | . | chr1 | 183138001 | 183155887 | 4.39977 | 3.65871 | 2.54516 |
| chr1 | 183150485 | 183150915 | 1000 | Intron (ENSMUST00000003035.6 | 1 | 183086270 | 183147461 | 61192 | 2 | ENSMUST00 | -3025 | Disp1 | ENSMUSG0( | false | . | . | . | . | . | . | chr1 | 183138001 | 183155887 | 4.39977 | 3.65871 | 2.54516 |
| chr7 | 112462975 | 112463393 | 1000 | Intron (ENSMUST00000106643.2 | 7 | 112427905 | 112552929 | 125025 | 1 | ENSMUST00 | 35071 | Parva | ENSMUSG0( | true | . | . | . | . | . | . | chr7 | 112434747 | 112475955 | 6.44046 | 4.72904 | 4.19131 |
| chr7 | 112467276 | 112467955 | 1000 | Intron (ENSMUST00000106643.2 | 7 | 112501177 | 112547610 | 46434 | 1 | ENSMUST00 | -33222 | Parva | ENSMUSG0( | true | . | . | . | . | . | . | chr7 | 112434747 | 112475955 | 6.44046 | 4.72904 | 4.19131 |
| chr7 | 112427651 | 112428085 | 1000 | Promoter (<=1kb) | 7 | 112427706 | 112591692 | 163987 | 1 | ENSMUST00 | 0 | Parva | ENSMUSG0( | true | chr7 | 112427013 | 112429707 | 31.16539 | 106.97112 | 103.21663 | . | . | . | . | . | . |
| chr7 | 112572409 | 112572698 | 1000 | Intron (ENSMUST00000106643.2 | 7 | 112569250 | 112579659 | 10410 | 1 | ENSMUST00 | 3160 | Parva | ENSMUSG0( | true | . | . | . | . | . | . | chr7 | 112569473 | 112576741 | 4.79006 | 3.96332 | 2.83716 |
| chr7 | 112426827 | 112427139 | 1000 | Promoter (<=1kb) | 7 | 112427505 | 112589629 | 162125 | 1 | ENSMUST00 | -366 | Parva | ENSMUSG0( | true | chr7 | 112427013 | 112429707 | 31.16539 | 106.97112 | 103.21663 | chr7 | 112425680 | 112427351 | 5.76115 | 4.46539 | 3.62303 |
| chr7 | 97850317 | 97850954 | 1000 | Intron (ENSMUST00000033040.6 | 7 | 97842935 | 97912381 | 69447 | 1 | ENSMUST00 | 7383 | Pak1 | ENSMUSG0( | true | . | . | . | . | . | . | chr7 | 97847622 | 97867047 | 4.43625 | 3.78799 | 2.54474 |
| chr7 | 128311871 | 128312234 | 1000 | Distal Intergenic | 7 | 128271379 | 128298170 | 26792 | 2 | ENSMUST00 | -13702 | BC017158 | ENSMUSG0( | false | . | . | . | . | . | . | chr7 | 128310694 | 128312540 | 4.22843 | 3.68938 | 2.41458 |
| chr7 | 110630271 | 110630714 | 1000 | Promoter (2-3kb) | 7 | 110628028 | 110628551 | 524 | 1 | ENSMUST00 | 2244 | Adm | ENSMUSG0( | false | chr7 | 110629711 | 110631062 | 15.85026 | 49.49464 | 46.78799 | chr7 | 110630591 | 110631831 | 4.81083 | 3.95312 | 2.84064 |
| chr7 | 127745743 | 127746471 | 1000 | Promoter (<=1kb) | 7 | 127745770 | 127749717 | 3948 | 1 | ENSMUST00 | 0 | Fbxl19 | ENSMUSG0( | false | chr7 | 127744610 | 127747241 | 26.7916 | 76.85461 | 73.75612 | . | . | . | . | . | . |
| chr7 | 127708295 | 127708923 | 1000 | Promoter (<=1kb) | 7 | 127663353 | 127708785 | 45433 | 2 | ENSMUST00 | 0 | Bcl7c | ENSMUSG0( | false | chr7 | 127707789 | 127709220 | 34.47702 | 107.70252 | 103.93674 | . | . | . | . | . | . |
| chr7 | 45567828 | 45568662 | 1000 | Promoter (1-2kb) | 7 | 45570153 | 45589710 | 19558 | 1 | ENSMUST00 | -1491 | Bcat2 | ENSMUSG0( | false | . | . | . | . | . | . | chr7 | 45566728 | 45575570 | 5.46963 | 4.20393 | 3.40899 |
| chr7 | 130209541 | 130209878 | 1000 | Intron (ENSMUST00000124096.2 | 7 | 130180471 | 130196845 | 16375 | 2 | ENSMUST00 | -12697 | Fgfr2 | ENSMUSG0( | false | . | . | . | . | . | . | chr7 | 130199223 | 130211834 | 4.56673 | 3.87122 | 2.65732 |
| chr7 | 130167265 | 130167659 | 1000 | Promoter (<=1kb) | 7 | 130163608 | 130167884 | 4277 | 2 | ENSMUST00 | 225 | Fgfr2 | ENSMUSG0( | false | . | . | . | . | . | . | chr7 | 130164917 | 130170706 | 5.11872 | 4.14293 | 3.10716 |
| chr7 | 133121771 | 133124454 | 1000 | Promoter (<=1kb) | 7 | 130162451 | 133123350 | 2960900 | 2 | ENSMUST00 | 0 | Fgfr2 | ENSMUSG0( | false | chr7 | 133121202 | 133124729 | 36.31995 | 116.10173 | 112.08785 | . | . | . | . | . | . |
| chr7 | 130246512 | 130246833 | 1000 | Intron (ENSMUST00000124096.2 | 7 | 130163959 | 130261857 | 97899 | 2 | ENSMUST00 | 15024 | Fgfr2 | ENSMUSG0( | false | . | . | . | . | . | . | chr7 | 130228840 | 130264807 | 4.60924 | 3.83033 | 2.69727 |
| chr7 | 130258473 | 130258858 | 1000 | Promoter (2-3kb) | 7 | 130163959 | 130261857 | 97899 | 2 | ENSMUST00 | 2999 | Fgfr2 | ENSMUSG0( | false | . | . | . | . | . | . | chr7 | 130228840 | 130264807 | 4.60924 | 3.83033 | 2.69727 |
| chr7 | 130264987 | 130266566 | 1000 | Promoter (<=1kb) | 7 | 130162512 | 130266245 | 103734 | 2 | ENSMUST00 | 0 | Fgfr2 | ENSMUSG0( | false | chr7 | 130263579 | 130266780 | 19.35304 | 74.51352 | 71.45322 | . | . | . | . | . | . |
| chr7 | 130221003 | 130221458 | 1000 | Intron (ENSMUST00000124096.2 | 7 | 130199807 | 130228756 | 28950 | 2 | ENSMUST00 | 7298 | Fgfr2 | ENSMUSG0( | false | . | . | . | . | . | . | chr7 | 130219035 | 130227567 | 6.62101 | 4.76578 | 4.34548 |
| chr7 | 130121458 | 130122316 | 1000 | Distal Intergenic | 7 | 130163608 | 130167884 | 4277 | 2 | ENSMUST00 | 45568 | Fgfr2 | ENSMUSG0( | false | . | . | . | . | . | . | chr7 | 130121140 | 130123213 | 5.81139 | 4.41719 | 3.69887 |
| chr7 | 130654500 | 130655014 | 1000 | Intron (ENSMUST00000084513.5 | 7 | 130692556 | 130764784 | 72229 | 1 | ENSMUST00 | -37542 | Tacc2 | ENSMUSG0( | false | . | . | . | . | . | . | chr7 | 130652005 | 130660447 | 6.76004 | 4.77622 | 4.46982 |
| chr7 | 130631595 | 130631894 | 1000 | Intron (ENSMUST00000084513.5 | 7 | 130577484 | 130764784 | 187301 | 1 | ENSMUST00 | 54112 | Tacc2 | ENSMUSG0( | false | . | . | . | . | . | . | chr7 | 130631062 | 130632349 | 4.74689 | 4.04678 | 2.78436 |
| chr7 | 130645752 | 130646513 | 1000 | Intron (ENSMUST00000084513.5 | 7 | 130692556 | 130764784 | 72229 | 1 | ENSMUST00 | -46043 | Tacc2 | ENSMUSG0( | false | chr7 | 130645344 | 130646821 | 25.53309 | 82.08177 | 78.89293 | chr7 | 130643406 | 130645800 | 4.61393 | 3.92519 | 2.67826 |
| chr7 | 131436660 | 131437038 | 1000 | Promoter (2-3kb) | 7 | 131434246 | 131437519 | 3274 | 1 | ENSMUST00 | 2415 | Acadsb | ENSMUSG0( | false | . | . | . | . | . | . | chr7 | 131435991 | 131437744 | 4.27043 | 3.78341 | 2.39744 |
| chr7 | 131453126 | 131453484 | 1000 | Intron (ENSMUST00000124096.2 | 7 | 131434246 | 131437519 | 3274 | 1 | ENSMUST00 | 18881 | Acadsb | ENSMUSG0( | false | . | . | . | . | . | . | chr7 | 131452136 | 131454171 | 3.62031 | 3.42371 | 1.89106 |
| chr7 | 122066689 | 122067564 | 1000 | Promoter (<=1kb) | 7 | 122067175 | 122082194 | 15020 | 1 | ENSMUST00 | 0 | Ubfd1 | ENSMUSG0( | false | chr7 | 122066447 | 122068300 | 39.29051 | 134.21745 | 129.56854 | . | . | . | . | . | . |
| chr7 | 121034276 | 121035270 | 1000 | Promoter (<=1kb) | 7 | 121034445 | 121076834 | 42390 | 1 | ENSMUST00 | 0 | Mettl9 | ENSMUSG0( | false | chr7 | 121033856 | 121035911 | 34.54477 | 122.48211 | 118.25522 | . | . | . | . | . | . |
| chr7 | 120979283 | 120979980 | 1000 | Promoter (1-2kb) | 7 | 120957036 | 120981790 | 24755 | 2 | ENSMUST00 | 1810 | Cdr2 | ENSMUSG0( | false | . | . | . | . | . | . | chr7 | 120961858 | 120981649 | 5.33893 | 4.28516 | 3.2901 |
| chr7 | 133088792 | 133089384 | 1000 | Intron (ENSMUST00000124096.2 | 7 | 133025720 | 133111791 | 86072 | 2 | ENSMUST00 | 22407 | Ctbp2 | ENSMUSG0( | false | chr7 | 133088509 | 133089390 | 9.38717 | 20.36066 | 18.02005 | chr7 | 133085601 | 133091021 | 4.9669 | 4.01465 | 2.98399 |
| chr7 | 133750699 | 133751052 | 1000 | Intron (ENSMUST00000033290.6 | 7 | 133748564 | 133759472 | 10909 | 2 | ENSMUST00 | 8420 | Dhx32 | ENSMUSG0( | false | . | . | . | . | . | . | chr7 | 133743183 | 133753176 | 5.21662 | 4.08927 | 3.21065 |
| chrX | 13071266 | 13073042 | 1000 | Promoter (<=1kb) | 20 | 13071498 | 13123449 | 51952 | 1 | ENSMUST00 | 0 | Usp9x | ENSMUSG0( | false | chrX | 13070252 | 13073605 | 25.27427 | 69.73028 | 66.74294 | . | . | . | . | . | . |
| chrX | 13846196 | 13846965 | 1000 | Promoter (<=1kb) | 20 | 13517080 | 13846541 | 329462 | 2 | ENSMUST00 | 0 | Cask | ENSMUSG0( | false | chrX | 13845424 | 13847667 | 18.84082 | 47.13427 | 44.45482 | . | . | . | . | . | . |
| chr7 | 110121450 | 110123059 | 1000 | Promoter (<=1kb) | 7 | 110122350 | 110142112 | 19763 | 1 | ENSMUST00 | 0 | Wee1 | ENSMUSG0( | false | chr7 | 110121226 | 110123539 | 33.41831 | 117.26761 | 113.21259 | chr7 | 110121243 | 110121589 | 4.00576 | 3.64474 | 2.17118 |
| chr7 | 145302535 | 145303125 | 1000 | Promoter (1-2kb) | 7 | 145300899 | 145309349 | 8451 | 1 | ENSMUST00 | 1637 | Mrgprf | ENSMUSG0( | false | chr7 | 145302113 | 145302634 | 13.14203 | 32.98289 | 30.47944 | chr7 | 145303051 | 145309171 | 5.11867 | 4.06121 | 3.11909 |
| chr7 | 145300457 | 145300984 | 1000 | Promoter (<=1kb) | 7 | 145300828 | 145309557 | 8730 | 1 | ENSMUST00 | 0 | Mrgprf | ENSMUSG0( | false | chr7 | 145299716 | 145300691 | 11.33197 | 24.23275 | 21.84028 | . | . | . | . | . | . |
| chr7 | 144762842 | 144763218 | 1000 | Distal Intergenic | 7 | 144650503 | 144751974 | 101472 | 2 | ENSMUST00 | -10869 | Ano1 | ENSMUSG0( | false | . | . | . | . | . | . | . | . | . | . | . | . |
| chr7 | 144689291 | 144690407 | 1000 | Intron (ENSMUST00000121758.2 | 7 | 144655601 | 144678885 | 23285 | 2 | ENSMUST00 | -10407 | Ano1 | ENSMUSG0( | false | . | . | . | . | . | . | . | . | . | . | . | . |
| chrX | 48453607 | 48454607 | 1000 | Promoter (<=1kb) | 20 | 48411046 | 48454169 | 43124 | 2 | ENSMUST00 | 0 | Elf4 | ENSMUSG0( | true | chrX | 48453190 | 48454762 | 21.79372 | 61.12524 | 58.26245 | . | . | . | . | . | . |
| chrX | 48432897 | 48433195 | 1000 | Intron (ENSMUST00000114958.2 | 20 | 48411046 | 48454169 | 43124 | 2 | ENSMUST00 | 20974 | Elf4 | ENSMUSG0( | true | . | . | . | . | . | . | . | . | . | . | . | . |
| chrX | 52164744 | 52165658 | 1000 | Promoter (<=1kb) | 20 | 52053021 | 52165252 | 112232 | 2 | ENSMUST00 | 0 | Gpc4 | ENSMUSG0( | false | chrX | 52163494 | 52165503 | 17.00269 | 41.02908 | 38.42754 | . | . | . | . | . | . |
| chr9 | 95192366 | 95192736 | 1000 | Intron (ENSMUST00000033463.9 | 9 | 95121686 | 95124551 | 2866 | 1 | ENSMUST00 | 70681 | Slc9a9 | ENSMUSG0( | false | . | . | . | . | . | . | . | . | . | . | . | . |
| chr9 | 94681098 | 94681501 | 1000 | Intron (ENSMUST00000033463.9 | 9 | 94669932 | 94718846 | 48915 | 1 | ENSMUST00 | 11167 | Slc9a9 | ENSMUSG0( | false | . | . | . | . | . | . | . | . | . | . | . | . |
| chrX | 7658264 | 7658727 | 1000 | Promoter (<=1kb) | 20 | 7658533 | 7668186 | 9654 | 1 | ENSMUST00 | 0 | Prickle3 | ENSMUSG0( | false | chrX | 7657113 | 7658374 | 15.62409 | 36.57069 | 34.02363 | . | . | . | . | . | . |
| chrX | 8206312 | 8206675 | 1000 | Promoter (<=1kb) | 20 | 8193848 | 8206525 | 12678 | 2 | ENSMUST00 | 0 | Porcn | ENSMUSG0( | false | chrX | 8206074 | 8207019 | 17.92175 | 44.05972 | 41.41886 | . | . | . | . | . | . |
| chrX | 99218250 | 99218599 | 1000 | Distal Intergenic | 20 | 99138131 | 99147230 | 9100 | 1 | ENSMUST00 | 80120 | Efnb1 | ENSMUSG0( | false | . | . | . | . | . | . | . | . | . | . | . | . |
| chrX | 99172248 | 99172706 | 1000 | Distal Intergenic | 20 | 99138131 | 99147230 | 9100 | 1 | ENSMUST00 | 34118 | Efnb1 | ENSMUSG0( | false | . | . | . | . | . | . | . | . | . | . | . | . |
| chrX | 99149830 | 99150239 | 1000 | Distal Intergenic | 20 | 99138131 | 99147230 | 9100 | 1 | ENSMUST00 | 11700 | Efnb1 | ENSMUSG0( | false | . | . | . | . | . | . | . | . | . | . | . | . |
| chrX | 99141193 | 99141575 | 1000 | Intron (ENSMUST00000052839.6 | 20 | 99138131 | 99147230 | 9100 | 1 | ENSMUST00 | 3063 | Efnb1 | ENSMUSG0( | false | chrX | 99141344 | 99144749 | 10.10971 | 19.98882 | 17.65231 | . | . | . | . | . | . |
| chrX | 99221127 | 99221417 | 1000 | Distal Intergenic | 20 | 99138131 | 99147230 | 9100 | 1 | ENSMUST00 | 82997 | Efnb1 | ENSMUSG0( | false | . | . | . | . | . | . | . | . | . | . | . | . |
| chrX | 99239560 | 99239981 | 1000 | Distal Intergenic | 20 | 99138131 | 99147230 | 9100 | 1 | ENSMUST00 | 101430 | Efnb1 | ENSMUSG0( | false | . | . | . | . | . | . | . | . | . | . | . | . |
| chrX | 141475111 | 141476133 | 1000 | Promoter (<=1kb) | 20 | 141475385 | 141689234 | 213850 | 1 | ENSMUST00 | 0 | Col4a5 | ENSMUSG0( | true | chrX | 141475093 | 141476973 | 18.38128 | 45.59164 | 42.93153 | . | . | . | . | . | . |
| chrX | 142390373 | 142390920 | 1000 | Promoter (<=1kb) | 20 | 142317993 | 142390504 | 72512 | 2 | ENSMUST00 | 0 | Acsl4 | ENSMUSG0( | true | chrX | 142388849 | 142391115 | 22.97661 | 61.46103 | 58.5941 | . | . | . | . | . | . |
| chrX | 160502036 | 160502811 | 1000 | Promoter (<=1kb) | 20 | 160502166 | 160591141 | 88976 | 1 | ENSMUST00 | 0 | Phka2 | ENSMUSG0( | true | chrX | 160501750 | 160503740 | 24.3552 | 66.39809 | 63.45943 | . | . | . | . | . | . |
| chrX | 160598535 | 160599261 | 1000 | 3' UTR | 20 | 160557787 | 160562749 | 4963 | 1 | ENSMUST00 | 40749 | Phka2 | ENSMUSG0( | true | . | . | . | . | . | . | . | . | . | . | . | . |
| chrX | 160516216 | 160516739 | 1000 | Intron (ENSMUST00000112377.2 | 20 | 160523348 | 160541488 | 18141 | 1 | ENSMUST00 | -6609 | Phka2 | ENSMUSG0( | true | . | . | . | . | . | . | . | . | . | . | . | . |
| chrX | 159257069 | 159257517 | 1000 | Promoter (<=1kb) | 20 | 159256126 | 159328464 | 72339 | 1 | ENSMUST00 | 944 | Rps6ka3 | ENSMUSG0( | false | chrX | 159255753 | 159257406 | 18.84082 | 47.13427 | 44.45482 | . | . | . | . | . | . |
| chrX | 159254579 | 159256533 | 1000 | Promoter (<=1kb) | 20 | 159255782 | 159311525 | 55744 | 1 | ENSMUST00 | 0 | Rps6ka3 | ENSMUSG0( | false | chrX | 159254523 | 159255547 | 11.02877 | 22.59171 | 20.22099 | . | . | . | . | . | . |
| chrX | 159254114 | 159254436 | 1000 | Promoter (1-2kb) | 20 | 159255782 | 159311525 | 55744 | 1 | ENSMUST00 | -1346 | Rps6ka3 | ENSMUSG0( | false | chrX | 159254062 | 159254320 | 6.43345 | 10.42957 | 8.24651 | . | . | . | . | . | . |

| | | | | | | | | | | | | | | | | | | | | | | | | | | |
|---|---|---|---|---|---|---|---|---|---|---|---|---|---|---|---|---|---|---|---|---|---|---|---|---|---|---|
| chrX | 162759891 | 162760542 | 1000 | Promoter (<=1kb) | 20 | 162760402 | 162779092 | 18691 | 1 | ENSMUST00 | 0 | Rbbp7 | ENSMUSG0C | false | chrX | 162759606 | 162759971 | 4.13035 | 5.82909 | 3.75473 | . | . | . | . | . | . |
| chrX | 169123656 | 169124015 | 1000 | Intron (ENSMUST00000033721.7 | 20 | 169113047 | 169304435 | 191389 | 1 | ENSMUST00 | 10610 | Arhgap6 | ENSMUSG0C | false | . | . | . | . | . | . | . | . | . | . | . | . |
| chrX | 168673478 | 168674399 | 1000 | Promoter (<=1kb) | 20 | 168654117 | 168673898 | 19782 | 2 | ENSMUST00 | 0 | Msl3 | ENSMUSG0C | false | chrX | 168672052 | 168674729 | 21.59801 | 56.60135 | 53.80005 | . | . | . | . | . | . |
| chrX | 73940592 | 73940947 | 1000 | Downstream (1-2kb) | 20 | 73945321 | 73947432 | 2112 | 2 | ENSMUST00 | 6485 | Hcfc1 | ENSMUSG0C | false | . | . | . | . | . | . | . | . | . | . | . | . |
| chrX | 75095626 | 75096117 | 1000 | Promoter (<=1kb) | 20 | 75095854 | 75109777 | 13924 | 1 | ENSMUST00 | 0 | Dkc1 | ENSMUSG0C | false | chrX | 75095132 | 75096641 | 16.08362 | 38.04472 | 35.47989 | . | . | . | . | . | . |
| chrX | 136741476 | 136742174 | 1000 | Promoter (<=1kb) | 20 | 136732942 | 136741661 | 8720 | 2 | ENSMUST00 | 0 | Morf4l2 | ENSMUSG0C | false | chrX | 136741826 | 136742872 | 11.94783 | 25.26529 | 22.85895 | . | . | . | . | . | . |
| chrX | 140511162 | 140511653 | 1000 | Distal Intergenic | 20 | 140539528 | 140542668 | 3141 | 2 | ENSMUST00 | 31015 | Tsc22d3 | ENSMUSG0C | false | . | . | . | . | . | . | . | . | . | . | . | . |
| chrX | 140598617 | 140599762 | 1000 | Promoter (<=1kb) | 20 | 140540262 | 140600659 | 60398 | 2 | ENSMUST00 | 897 | Tsc22d3 | ENSMUSG0C | false | chrX | 140598132 | 140600658 | 17.46222 | 42.53877 | 39.91948 | . | . | . | . | . | . |
| chrX | 140520114 | 140520691 | 1000 | Distal Intergenic | 20 | 140539528 | 140542668 | 3141 | 2 | ENSMUST00 | 21977 | Tsc22d3 | ENSMUSG0C | false | . | . | . | . | . | . | . | . | . | . | . | . |
| chrX | 140529169 | 140529502 | 1000 | Distal Intergenic | 20 | 140539528 | 140542668 | 3141 | 2 | ENSMUST00 | 13166 | Tsc22d3 | ENSMUSG0C | false | . | . | . | . | . | . | . | . | . | . | . | . |
| chrX | 140516596 | 140516904 | 1000 | Distal Intergenic | 20 | 140539528 | 140542668 | 3141 | 2 | ENSMUST00 | 25764 | Tsc22d3 | ENSMUSG0C | false | . | . | . | . | . | . | . | . | . | . | . | . |
| chr8 | 12818918 | 12819205 | 1000 | Intron (ENSMUST00000033818.4 | 8 | 12822400 | 12828537 | 6138 | 1 | ENSMUST00 | -3195 | Atp11a | ENSMUSG0C | false | . | . | . | . | . | . | . | . | . | . | . | . |
| chr8 | 12756645 | 12757583 | 1000 | Promoter (<=1kb) | 8 | 12757014 | 12868728 | 111715 | 1 | ENSMUST00 | 0 | Atp11a | ENSMUSG0C | false | chr8 | 12756624 | 12758223 | 33.08631 | 99.21803 | 95.66874 | chr8 | 12756079 | 12756668 | 3.71578 | 3.48067 | 1.94102 |
| chr8 | 18728858 | 18729292 | 1000 | Intron (ENSMUST00000039412.9 | 8 | 18690263 | 18741562 | 51300 | 2 | ENSMUST00 | 12270 | Angpt2 | ENSMUSG0C | false | . | . | . | . | . | . | chr8 | 18728616 | 18729465 | 4.85365 | 3.849 | 2.91846 |
| chr8 | 22398207 | 22398845 | 1000 | Promoter (<=1kb) | 8 | 22375554 | 22398621 | 23068 | 2 | ENSMUST00 | 0 | Slc25a15 | ENSMUSG0C | false | chr8 | 22397785 | 22399225 | 23.61999 | 85.9385 | 82.67356 | . | . | . | . | . | . |
| chr8 | 27261128 | 27261543 | 1000 | Promoter (<=1kb) | 8 | 27260327 | 27275656 | 15330 | 1 | ENSMUST00 | 802 | Eif4ebp1 | ENSMUSG0C | false | chr8 | 27259721 | 27261776 | 30.46894 | 111.80851 | 107.92088 | chr8 | 27259685 | 27262143 | 4.81874 | 4.01756 | 2.86131 |
| chr8 | 11203244 | 11203569 | 1000 | Intron (ENSMUST00000151014.1 | 8 | 11212401 | 11216801 | 4401 | 2 | ENSMUST00 | 13232 | Col4a1 | ENSMUSG0C | false | . | . | . | . | . | . | chr8 | 11202860 | 11204404 | 4.40786 | 3.85948 | 2.50402 |
| chr8 | 11634926 | 11635470 | 1000 | Promoter (<=1kb) | 8 | 11611583 | 11635754 | 24172 | 2 | ENSMUST00 | 284 | Ankrd10 | ENSMUSG0C | false | chr8 | 11634291 | 11636731 | 43.41453 | 148.15424 | 142.93486 | . | . | . | . | . | . |
| chr8 | 34146560 | 34147242 | 1000 | Promoter (<=1kb) | 8 | 34135572 | 34147033 | 11462 | 2 | ENSMUST00 | 0 | Leprotl1 | ENSMUSG0C | false | chr8 | 34145755 | 34147509 | 35.67729 | 114.98298 | 111.00362 | chr8 | 34142564 | 34147869 | 4.04149 | 3.52066 | 2.22818 |
| chr8 | 34107958 | 34108593 | 1000 | Promoter (<=1kb) | 8 | 34090563 | 34108536 | 17974 | 2 | ENSMUST00 | 0 | Dctn6 | ENSMUSG0C | false | chr8 | 34107409 | 34109112 | 33.26678 | 112.81776 | 108.90401 | . | . | . | . | . | . |
| chr8 | 36577043 | 36577674 | 1000 | Promoter (<=1kb) | 8 | 36572145 | 36577269 | 5125 | 2 | ENSMUST00 | 0 | Dlc1 | ENSMUSG0C | true | . | . | . | . | . | . | chr8 | 36575944 | 36612020 | 5.64773 | 4.3412 | 3.54039 |
| chr8 | 34965332 | 34965861 | 1000 | Promoter (<=1kb) | 8 | 34829179 | 34965690 | 136512 | 2 | ENSMUST00 | 0 | Tnks | ENSMUSG0C | false | chr8 | 34964185 | 34966282 | 48.57222 | 166.35635 | 160.49539 | . | . | . | . | . | . |
| chr8 | 22758178 | 22758711 | 1000 | Promoter (<=1kb) | 8 | 22757744 | 22782847 | 25104 | 1 | ENSMUST00 | 435 | Plat | ENSMUSG0C | true | chr8 | 22756921 | 22764179 | 35.19295 | 114.13696 | 110.1796 | . | . | . | . | . | . |
| chr8 | 22859055 | 22860101 | 1000 | Promoter (<=1kb) | 8 | 22859535 | 22943259 | 83725 | 1 | ENSMUST00 | 0 | Kat6a | ENSMUSG0C | true | chr8 | 22858604 | 22859344 | 18.47733 | 48.83834 | 46.1367 | . | . | . | . | . | . |
| chr8 | 22847262 | 22847574 | 1000 | Distal Intergenic | 8 | 22859535 | 22943259 | 83725 | 1 | ENSMUST00 | -11961 | Kat6a | ENSMUSG0C | true | . | . | . | . | . | . | chr8 | 22847041 | 22847761 | 3.27833 | 3.15596 | 1.5829 |
| chr8 | 24531909 | 24532284 | 1000 | Exon (ENSMUST00000140417.2/ | 8 | 24533456 | 24540796 | 7341 | 2 | ENSMUST00 | 8512 | Ido2 | ENSMUSG0C | false | . | . | . | . | . | . | . | . | . | . | . | . |
| chr8 | 24787103 | 24787469 | 1000 | Intron (ENSMUST00000132180.2 | 8 | 24727093 | 24815473 | 88381 | 2 | ENSMUST00 | 28004 | Adam5 | ENSMUSG0C | false | . | . | . | . | . | . | . | . | . | . | . | . |
| chr8 | 25098127 | 25098716 | 1000 | Intron (ENSMUST00000064883.7 | 8 | 25042944 | 25101771 | 58828 | 2 | ENSMUST00 | 3055 | Plekha2 | ENSMUSG0C | true | chr8 | 25098560 | 25099109 | 8.34762 | 15.59896 | 13.32767 | chr8 | 25082618 | 25100838 | 5.3544 | 4.1622 | 3.30614 |
| chr8 | 48449562 | 48450011 | 1000 | Intron (ENSMUST00000190840.1 | 8 | 48227682 | 48367363 | 139682 | 2 | ENSMUST00 | -82200 | Tenm3 | ENSMUSG0C | true | . | . | . | . | . | . | chr8 | 48448782 | 48451764 | 5.22379 | 4.20564 | 3.22176 |
| chr8 | 48109877 | 48110596 | 1000 | Promoter (<=1kb) | 8 | 48109958 | 48141665 | 31708 | 1 | ENSMUST00 | 0 | Dctd | ENSMUSG0C | false | chr8 | 48109196 | 48110812 | 31.07761 | 98.73238 | 95.19199 | chr8 | 48108308 | 48110897 | 6.02469 | 4.57726 | 3.89256 |
| chr8 | 48126811 | 48127474 | 1000 | Intron (ENSMUST00000174278.2 | 8 | 48110211 | 48153233 | 43023 | 1 | ENSMUST00 | 16601 | Dctd | ENSMUSG0C | false | . | . | . | . | . | . | chr8 | 48121625 | 48130161 | 5.19553 | 4.10877 | 3.19299 |
| chr8 | 47990223 | 47991293 | 1000 | Promoter (<=1kb) | 8 | 47826083 | 47990543 | 164461 | 2 | ENSMUST00 | 0 | Wwc2 | ENSMUSG0C | false | chr8 | 47989082 | 47991874 | 41.76281 | 135.22986 | 130.53618 | . | . | . | . | . | . |
| chr8 | 25518582 | 25519766 | 1000 | Promoter (<=1kb) | 8 | 25518772 | 25575714 | 56943 | 1 | ENSMUST00 | 0 | Fgfr1 | ENSMUSG0C | true | chr8 | 25518487 | 25520589 | 36.25406 | 119.15681 | 115.04134 | . | . | . | . | . | . |
| chr8 | 25720040 | 25720542 | 1000 | Promoter (<=1kb) | 8 | 25720061 | 25724887 | 4827 | 1 | ENSMUST00 | 0 | Ppapdc1b | ENSMUSG0C | false | chr8 | 25719928 | 25720990 | 23.80619 | 80.75649 | 77.58801 | chr8 | 25719449 | 25721228 | 3.7613 | 3.38514 | 2.01257 |
| chr8 | 25796923 | 25797426 | 1000 | Intron (ENSMUST00000038421.6 | 8 | 25808474 | 25814723 | 6250 | 1 | ENSMUST00 | -11048 | Star | ENSMUSG0C | false | . | . | . | . | . | . | chr8 | 25793589 | 25799299 | 4.23404 | 3.50483 | 2.41744 |
| chr8 | 25806010 | 25806681 | 1000 | Promoter (1-2kb) | 8 | 25808474 | 25814723 | 6250 | 1 | ENSMUST00 | -1793 | Star | ENSMUSG0C | false | . | . | . | . | . | . | chr8 | 25804290 | 25809630 | 5.71806 | 4.3971 | 3.60241 |
| chr8 | 33653360 | 33654314 | 1000 | Promoter (<=1kb) | 8 | 33653238 | 33698163 | 44926 | 1 | ENSMUST00 | 123 | Gsr | ENSMUSG0C | false | chr8 | 33653035 | 33654537 | 30.17592 | 94.17345 | 90.73757 | chr8 | 33654267 | 33655792 | 3.22985 | 3.02093 | 1.60491 |
| chr8 | 33901340 | 33901670 | 1000 | Intron (ENSMUST00000053251.6 | 8 | 33863007 | 33891764 | 28758 | 2 | ENSMUST00 | -9577 | Rbpms | ENSMUSG0C | true | . | . | . | . | . | . | chr8 | 33863865 | 33927704 | 6.32022 | 4.67603 | 4.08842 |
| chr8 | 33902049 | 33903130 | 1000 | Intron (ENSMUST00000053251.6 | 8 | 33863007 | 33891764 | 28758 | 2 | ENSMUST00 | -10286 | Rbpms | ENSMUSG0C | true | . | . | . | . | . | . | chr8 | 33863865 | 33927704 | 6.32022 | 4.67603 | 4.08842 |
| chr8 | 33905810 | 33906496 | 1000 | Intron (ENSMUST00000053251.6 | 8 | 33863007 | 33891764 | 28758 | 2 | ENSMUST00 | -14047 | Rbpms | ENSMUSG0C | true | . | . | . | . | . | . | chr8 | 33863865 | 33927704 | 6.32022 | 4.67603 | 4.08842 |
| chr8 | 33842172 | 33842537 | 1000 | Promoter (1-2kb) | 8 | 33806742 | 33843671 | 36930 | 2 | ENSMUST00 | 1134 | Rbpms | ENSMUSG0C | true | . | . | . | . | . | . | chr8 | 33822415 | 33859327 | 6.05392 | 4.56707 | 3.87043 |
| chr8 | 33869128 | 33869678 | 1000 | Intron (ENSMUST00000053251.6 | 8 | 33784066 | 33853673 | 69608 | 2 | ENSMUST00 | -15456 | Rbpms | ENSMUSG0C | true | . | . | . | . | . | . | chr8 | 33863865 | 33927704 | 6.32022 | 4.67603 | 4.08842 |
| chr8 | 33929052 | 33930584 | 1000 | Promoter (<=1kb) | 8 | 33782643 | 33929863 | 147221 | 2 | ENSMUST00 | 0 | Rbpms | ENSMUSG0C | true | chr8 | 33927502 | 33930859 | 33.75878 | 107.48165 | 103.71877 | chr8 | 33930469 | 33931960 | 3.48954 | 3.33386 | 1.75051 |
| chr8 | 33916508 | 33916914 | 1000 | Intron (ENSMUST00000053251.6 | 8 | 33803939 | 33929805 | 125867 | 2 | ENSMUST00 | 12891 | Rbpms | ENSMUSG0C | true | . | . | . | . | . | . | chr8 | 33863865 | 33927704 | 6.32022 | 4.67603 | 4.08842 |
| chr8 | 64770823 | 64771194 | 1000 | Intron (ENSMUST00000034017.7 | 8 | 64718139 | 64733792 | 15654 | 2 | ENSMUST00 | -37032 | Msmo1 | ENSMUSG0C | false | . | . | . | . | . | . | chr8 | 64769610 | 64772118 | 4.45707 | 3.85413 | 2.56084 |
| chr8 | 64849448 | 64850402 | 1000 | Promoter (<=1kb) | 8 | 64739863 | 64849818 | 109956 | 2 | ENSMUST00 | 0 | Klhl2 | ENSMUSG0C | false | chr8 | 64848977 | 64850628 | 32.69058 | 104.59052 | 100.90593 | . | . | . | . | . | . |
| chr8 | 57530650 | 57531171 | 1000 | Intron (ENSMUST00000034021.6 | 8 | 57526024 | 57538453 | 12430 | 2 | ENSMUST00 | 7282 | Galnt7 | ENSMUSG0C | false | chr8 | 57530400 | 57531191 | 13.34131 | 30.05484 | 27.59022 | chr8 | 57530409 | 57531527 | 3.53204 | 3.3567 | 1.8627 |
| chr8 | 57409348 | 57409647 | 1000 | Distal Intergenic | 8 | 57455923 | 57477585 | 21663 | 1 | ENSMUST00 | -46276 | Scrg1 | ENSMUSG0C | false | . | . | . | . | . | . | chr8 | 57408138 | 57410679 | 4.33225 | 3.75843 | 2.48072 |
| chr8 | 76898886 | 76899700 | 1000 | Promoter (<=1kb) | 8 | 76899442 | 76908413 | 8972 | 1 | ENSMUST00 | 0 | Nr3c2 | ENSMUSG0C | false | chr8 | 76898535 | 76903248 | 21.17422 | 79.20749 | 76.06625 | . | . | . | . | . | . |
| chr8 | 46739372 | 46740293 | 1000 | Promoter (<=1kb) | 8 | 46739745 | 46847458 | 107714 | 1 | ENSMUST00 | 0 | Irf2 | ENSMUSG0C | false | chr8 | 46738444 | 46742094 | 33.45771 | 103.89173 | 100.22243 | . | . | . | . | . | . |
| chr8 | 46163395 | 46163940 | 1000 | Promoter (<=1kb) | 8 | 46163651 | 46195590 | 31940 | 1 | ENSMUST00 | 0 | 4933411K20I | ENSMUSG0C | false | chr8 | 46162924 | 46164395 | 33.94791 | 109.87469 | 106.04007 | chr8 | 46162665 | 46165162 | 5.39999 | 4.25765 | 3.3917 |
| chr8 | 61223750 | 61224661 | 1000 | Promoter (<=1kb) | 8 | 61224171 | 61396072 | 171902 | 1 | ENSMUST00 | 0 | Sh3rf1 | ENSMUSG0C | false | chr8 | 61223517 | 61225207 | 33.7938 | 119.00099 | 114.88904 | . | . | . | . | . | . |
| chr8 | 61008386 | 61009136 | 1000 | Intron (ENSMUST00000120689.2 | 8 | 60993241 | 61030600 | 37360 | 1 | ENSMUST00 | 15146 | Nek1 | ENSMUSG0C | true | . | . | . | . | . | . | chr8 | 61007494 | 61010706 | 4.67783 | 3.84507 | 2.76552 |
| chr8 | 89042238 | 89043913 | 1000 | Promoter (<=1kb) | 8 | 89027248 | 89044162 | 16915 | 2 | ENSMUST00 | 249 | Sall1 | ENSMUSG0C | false | chr8 | 89040662 | 89045860 | 28.17892 | 90.64638 | 87.2878 | . | . | . | . | . | . |
| chr8 | 102784943 | 102785320 | 1000 | Promoter (<=1kb) | 8 | 102632995 | 102785111 | 152117 | 2 | ENSMUST00 | 0 | Cdh11 | ENSMUSG0C | false | chr8 | 102783585 | 102786947 | 24.78212 | 78.86476 | 75.73294 | chr8 | 102785256 | 102786834 | 4.04549 | 3.54118 | 2.26587 |
| chr8 | 85379112 | 85379900 | 1000 | Promoter (1-2kb) | 8 | 85324447 | 85380978 | 56532 | 2 | ENSMUST00 | 1078 | Mylk3 | ENSMUSG0C | false | . | . | . | . | . | . | chr8 | 85373675 | 85384939 | 5.70069 | 4.34167 | 3.58874 |
| chr8 | 84066814 | 84067501 | 1000 | Promoter (<=1kb) | 8 | 84066836 | 84096992 | 30157 | 1 | ENSMUST00 | 0 | Rfx1 | ENSMUSG0C | false | chr8 | 84066764 | 84067958 | 24.40663 | 77.266 | 74.1616 | chr8 | 84067446 | 84075532 | 6.20464 | 4.49658 | 4.02581 |
| chr8 | 83079223 | 83079816 | 1000 | Intron (ENSMUST00000078525.5 | 8 | 83165352 | 83272422 | 107071 | 1 | ENSMUST00 | -85536 | Tbc1d9 | ENSMUSG0C | false | . | . | . | . | . | . | chr8 | 83076696 | 83081675 | 5.40594 | 4.28249 | 3.34276 |
| chr8 | 80823304 | 80823686 | 1000 | Intron (ENSMUST00000034150.8 | 8 | 80764434 | 80880479 | 116046 | 2 | ENSMUST00 | 56793 | Gab1 | ENSMUSG0C | false | . | . | . | . | . | . | chr8 | 80802749 | 80878363 | 5.64706 | 4.31183 | 3.55694 |
| chr8 | 80879650 | 80880658 | 1000 | Promoter (<=1kb) | 8 | 80764434 | 80880479 | 116046 | 2 | ENSMUST00 | 0 | Gab1 | ENSMUSG0C | false | chr8 | 80877775 | 80881139 | 42.61679 | 142.26089 | 137.289 | . | . | . | . | . | . |
| chr8 | 109565505 | 109565996 | 1000 | Promoter (<=1kb) | 8 | 109565892 | 109574051 | 8160 | 1 | ENSMUST00 | 0 | Txnl4b | ENSMUSG0C | false | chr8 | 109564705 | 109566641 | 33.65816 | 114.56059 | 110.59126 | . | . | . | . | . | . |
| chr8 | 91801832 | 91802285 | 1000 | Promoter (<=1kb) | 8 | 91798525 | 91801915 | 3391 | 2 | ENSMUST00 | 0 | Irx3 | ENSMUSG0C | false | chr8 | 91799641 | 91802556 | 28.29403 | 89.8816 | 86.54237 | chr8 | 91797521 | 91802659 | 4.92604 | 3.98943 | 2.9759 |
| chr8 | 93810161 | 93811006 | 1000 | Promoter (<=1kb) | 8 | 93810636 | 93811767 | 1132 | 1 | ENSMUST00 | 0 | Gnao1 | ENSMUSG0C | false | chr8 | 93809849 | 93811435 | 28.93225 | 84.27598 | 81.04385 | chr8 | 93810958 | 93813862 | 4.03905 | 3.53805 | 2.26254 |
| chr8 | 94178931 | 94179343 | 1000 | Promoter (<=1kb) | 8 | 94179089 | 94180325 | 1237 | 1 | ENSMUST00 | 0 | Mt1 | ENSMUSG0C | false | chr8 | 94178656 | 94180513 | 23.29623 | 62.85914 | 59.97368 | chr8 | 94178318 | 94180250 | 3.55431 | 3.32427 | 1.80613 |
| chr8 | 114098746 | 114099168 | 1000 | Distal Intergenic | 8 | 114133557 | 114154739 | 21183 | 1 | ENSMUST00 | -34389 | Nudt7 | ENSMUSG0C | false | . | . | . | . | . | . | chr8 | 114098281 | 114099825 | 3.91349 | 3.56594 | 2.11994 |
| chr8 | 94822749 | 94823283 | 1000 | Promoter (<=1kb) | 8 | 94822935 | 94824193 | 1259 | 2 | ENSMUST00 | 910 | Ciapin1 | ENSMUSG0C | false | . | . | . | . | . | . | chr8 | 94822288 | 94824586 | 6.5709 | 4.76387 | 4.32014 |
| chr8 | 71592078 | 71592727 | 1000 | Promoter (<=1kb) | 8 | 71592176 | 71596269 | 4094 | 1 | ENSMUST00 | 0 | Pgls | ENSMUSG0C | false | chr8 | 71591371 | 71593108 | 39.11606 | 129.53703 | 125.06399 | chr8 | 71591116 | 71594741 | 4.90266 | 3.94518 | 2.96352 |
| chr8 | 121526642 | 121527249 | 1000 | Intron (ENSMUST00000127664.1 | 8 | 121530828 | 121541869 | 11042 | 2 | ENSMUST00 | 14620 | 1700018B08I | ENSMUSG0C | false | . | . | . | . | . | . | chr8 | 121523588 | 121529488 | 6.52995 | 4.79127 | 4.2509 |
| chr8 | 120382371 | 120382890 | 1000 | Intron (ENSMUST00000127664.1 | 8 | 120488447 | 120581390 | 92944 | 1 | ENSMUST00 | -105557 | Gse1 | ENSMUSG0C | false | . | . | . | . | . | . | chr8 | 120360953 | 120384219 | 7.36608 | 4.94236 | 4.97938 |
| chr8 | 120368825 | 120369283 | 1000 | Intron (ENSMUST00000127664.1 | 8 | 120488447 | 120581390 | 92944 | 1 | ENSMUST00 | -119164 | Gse1 | ENSMUSG0C | false | . | . | . | . | . | . | chr8 | 120360953 | 120384219 | 7.36608 | 4.94236 | 4.97938 |
| chr8 | 120268465 | 120268831 | 1000 | Intron (ENSMUST00000127664.1 | 8 | 120230536 | 120553634 | 323099 | 1 | ENSMUST00 | 37930 | Gse1 | ENSMUSG0C | false | . | . | . | . | . | . | chr8 | 120263198 | 120270766 | 5.71876 | 4.35069 | 3.62232 |
| chr8 | 120376859 | 120377478 | 1000 | Intron (ENSMUST00000127664.1 | 8 | 120488447 | 120581390 | 92944 | 1 | ENSMUST00 | -110969 | Gse1 | ENSMUSG0C | false | . | . | . | . | . | . | chr8 | 120360953 | 120384219 | 7.36608 | 4.94236 | 4.97938 |
| chr8 | 120535780 | 120536561 | 1000 | Promoter (<=1kb) | 8 | 120535857 | 120579480 | 43624 | 1 | ENSMUST00 | 0 | Gse1 | ENSMUSG0C | false | chr8 | 120535456 | 120538941 | 22.15371 | 67.81625 | 64.85734 | chr8 | 120525068 | 120536956 | 4.78295 | 3.95 | 2.82791 |
| chr8 | 120536685 | 120538269 | 1000 | Promoter (<=1kb) | 8 | 120537429 | 120579326 | 41898 | 1 | ENSMUST00 | 0 | Gse1 | ENSMUSG0C | false | chr8 | 120535456 | 120538941 | 22.15371 | 67.81625 | 64.85734 | chr8 | 120537951 | 120565808 | 7.48404 | 5.10526 | 5.05119 |
| chr8 | 120487099 | 120488338 | 1000 | Promoter (<=1kb) | 8 | 120488447 | 120581390 | 92944 | 1 | ENSMUST00 | -109 | Gse1 | ENSMUSG0C | false | chr8 | 120486986 | 120490241 | 34.92444 | 106.42502 | 102.68874 | chr8 | 120486787 | 120488206 | 3.37475 | 3.21801 | 1.69604 |
| chr8 | 120265469 | 120265931 | 1000 | Intron (ENSMUST00000127664.1 | 8 | 120230536 | 120553634 | 323099 | 1 | ENSMUST00 | 34934 | Gse1 | ENSMUSG0C | false | . | . | . | . | . | . | chr8 | 120263198 | 120270766 | 5.71876 | 4.35069 | 3.62232 |
| chr8 | 120151312 | 120151717 | 1000 | Exon (ENSMUST00000108948.2/ | 8 | 120155864 | 120161386 | 5523 | 1 | ENSMUST00 | -4147 | 6430548M08 | ENSMUSG0C | false | . | . | . | . | . | . | . | . | . | . | . | . |
| chr8 | 120113455 | 120114211 | 1000 | Promoter (<=1kb) | 8 | 120114152 | 120154574 | 40423 | 1 | ENSMUST00 | 0 | 6430548M08 | ENSMUSG0C | false | chr8 | 120113193 | 120115008 | 38.2106 | 120.43382 | 116.28301 | chr8 | 120112499 | 120113529 | 5.39869 | 4.34015 | 3.31943 |
| chr8 | 120017269 | 120017721 | 1000 | Intron (ENSMUST00000127664.1 | 8 | 120013208 | 120015690 | 2483 | 1 | ENSMUST00 | 4062 | Crispld2 | ENSMUSG0C | false | . | . | . | . | . | . | chr8 | 120016592 | 120018240 | 4.42574 | 3.79934 | 2.54347 |
| chr8 | 119824868 | 119825320 | 1000 | Intron (ENSMUST00000168698.1 | 8 | 119809225 | 119840522 | 31298 | 2 | ENSMUST00 | 15202 | Cotl1 | ENSMUSG0C | false | . | . | . | . | . | . | chr8 | 119806261 | 119841342 | 7.1039 | 5.00847 | 4.72992 |
| chr8 | 70754489 | 70755062 | 1000 | Promoter (<=1kb) | 8 | 70754679 | 70758677 | 3999 | 1 | ENSMUST00 | 0 | Rab3a | ENSMUSG0C | false | chr8 | 70754657 | 70755419 | 13.82983 | 49.0377 | 46.33454 | chr8 | 70749419 | 70756647 | 5.88874 | 4.48997 | 3.73674 |
| chr8 | 70753126 | 70753567 | 1000 | Promoter (1-2kb) | 8 | 70754679 | 70758677 | 3999 | 1 | ENSMUST00 | -1112 | Rab3a | ENSMUSG0C | false | chr8 | 70753118 | 70753812 | 5.76737 | 9.70104 | 7.52928 | chr8 | 70749419 | 70756647 | 5.88874 | 4.48997 | 3.73674 |
| chr8 | 70742205 | 70742543 | 1000 | Promoter (<=1kb) | 8 | 70743079 | 70750473 | 7395 | 1 | ENSMUST00 | -536 | Pde4c | ENSMUSG0C | false | . | . | . | . | . | . | chr8 | 70740750 | 70748114 | 8.21466 | 5.47394 | 5.63599 |
| chr8 | 70673128 | 70673898 | 1000 | Promoter (<=1kb) | 8 | 70673248 | 70678752 | 5505 | 1 | ENSMUST00 | 0 | Lsm4 | ENSMUSG0C | false | chr8 | 70673266 | 70673970 | 26.6505 | 78.45867 | 75.33303 | chr8 | 70670054 | 70675670 | 8.834 | 5.78677 | 6.13804 |
| chr8 | 70682030 | 70682500 | 1000 | Distal Intergenic | 8 | 70677665 | 70678116 | 452 | 1 | ENSMUST00 | 4366 | Lsm4 | ENSMUSG0C | false | . | . | . | . | . | . | chr8 | 70678490 | 70683293 | 6.70585 | 4.76208 | 4.42699 |
| chr8 | 70671564 | 70672053 | 1000 | Promoter (1-2kb) | 8 | 70673248 | 70678752 | 5505 | 1 | ENSMUST00 | -1195 | Lsm4 | ENSMUSG0C | false | . | . | . | . | . | . | chr8 | 70670054 | 70675670 | 8.834 | 5.78677 | 6.13804 |
| chr9 | 8740188 | 8740504 | 1000 | Distal Intergenic | 9 | 8899833 | 8968611 | 68779 | 1 | ENSMUST00 | -159329 | Pgr | ENSMUSG0C | false | . | . | . | . | . | . | chr9 | 8739751 | 8741025 | 3.98213 | 3.50907 | 2.20055 |
| chr8 | 103717845 | 103718150 | 1000 | Distal Intergenic | 8 | 104101625 | 104144502 | 42878 | 1 | ENSMUST00 | -383475 | Cdh5 | ENSMUSG0C | false | . | . | . | . | . | . | chr8 | 103714124 | 103718890 | 4.39973 | 3.82564 | 2.50581 |
| chr8 | 104007261 | 104007594 | 1000 | Distal Intergenic | 8 | 104101625 | 104144502 | 42878 | 1 | ENSMUST00 | -94031 | Cdh5 | ENSMUSG0C | false | . | . | . | . | . | . | chr8 | 104004284 | 104009204 | 4.09691 | 3.51249 | 2.31302 |
| chr8 | 103880583 | 103881178 | 1000 | Distal Intergenic | 8 | 104101625 | 104144502 | 42878 | 1 | ENSMUST00 | -220447 | Cdh5 | ENSMUSG0C | false | . | . | . | . | . | . | chr8 | 103879704 | 103883395 | 4.18853 | 3.61747 | 2.3476 |
| chr8 | 103715866 | 103716400 | 1000 | Distal Intergenic | 8 | 104101625 | 104144502 | 42878 | 1 | ENSMUST00 | -385225 | Cdh5 | ENSMUSG0C | false | . | . | . | . | . | . | chr8 | 103714124 | 103718890 | 4.39973 | 3.82564 | 2.50581 |
| chr8 | 104340336 | 104340809 | 1000 | Promoter (<=1kb) | 8 | 104340594 | 104347672 | 7079 | 1 | ENSMUST00 | 0 | Cmtm3 | ENSMUSG0C | false | chr8 | 104340179 | 104341479 | 24.91517 | 69.11501 | 66.14118 | chr8 | 104339769 | 104341914 | 3.91904 | 3.5879 | 2.16318 |
| chr8 | 104886708 | 104887221 | 1000 | Distal Intergenic | 8 | 104867488 | 104874080 | 6593 | 1 | ENSMUST00 | 19221 | Ces2d-ps | ENSMUSG0C | false | . | . | . | . | . | . | chr8 | 104884771 | 104889614 | 5.64618 | 4.36566 | 3.53177 |
| chr8 | 105170347 | 105171691 | 1000 | Promoter (<=1kb) | 8 | 105170674 | 105217989 | 47316 | 1 | ENSMUST00 | 0 | Cbfb | ENSMUSG0C | false | chr8 | 105170151 | 105172219 | 39.11606 | 129.53703 | 125.06399 | . | . | . | . | . | . |
| chr8 | 105514675 | 105515034 | 1000 | Distal Intergenic | 8 | 105518746 | 105523984 | 5239 | 1 | ENSMUST00 | -3712 | Hsd11b2 | ENSMUSG0C | false | . | . | . | . | . | . | chr8 | 105513771 | 105519552 | 6.19344 | 4.58209 | 3.98829 |
| chr9 | 13749000 | 13749339 | 1000 | Promoter (<=1kb) | 9 | 13749181 | 13806481 | 57301 | 1 | ENSMUST00 | 0 | Mtmr2 | ENSMUSG0C | false | chr9 | 13748500 | 13750046 | 44.84737 | 157.52063 | 151.9514 | . | . | . | . | . | . |
| chr8 | 111729671 | 111730182 | 1000 | Intron (ENSMUST00000166232.2 | 8 | 111710485 | 111743849 | 33365 | 2 | ENSMUST00 | 13667 | Bcar1 | ENSMUSG0C | false | . | . | . | . | . | . | chr8 | 111710939 | 111747327 | 8.20913 | 5.4583 | 5.64085 |
| chr8 | 111769781 | 111770231 | 1000 | Intron (ENSMUST00000034432.5 | 8 | 111710485 | 111743849 | 33365 | 2 | ENSMUST00 | -25933 | Bcar1 | ENSMUSG0C | false | . | . | . | . | . | . | chr8 | 111768244 | 111772378 | 4.83429 | 3.89484 | 2.8868 |
| chr8 | 111716542 | 111717056 | 1000 | Intron (ENSMUST00000166232.2 | 8 | 111686510 | 111691010 | 4501 | 2 | ENSMUST00 | -25533 | Ctrb1 | ENSMUSG0C | false | . | . | . | . | . | . | chr8 | 111710939 | 111747327 | 8.20913 | 5.4583 | 5.64085 |
| chr8 | 123477744 | 123478241 | 1000 | Promoter (<=1kb) | 8 | 123477903 | 123503916 | 26014 | 1 | ENSMUST00 | 0 | Afg3l1 | ENSMUSG0C | false | chr8 | 123477139 | 123479106 | 18.02928 | 79.93274 | 76.78327 | . | . | . | . | . | . |
| chr8 | 124020778 | 124021646 | 1000 | Promoter (<=1kb) | 8 | 124021508 | 124048504 | 26997 | 1 | ENSMUST00 | 0 | Urb2 | ENSMUSG0C | false | chr8 | 124019985 | 124022062 | 29.03158 | 89.24218 | 85.91109 | . | . | . | . | . | . |
| chr8 | 124662707 | 124663513 | 1000 | Promoter (<=1kb) | 8 | 124635766 | 124663369 | 27604 | 2 | ENSMUST00 | 0 | 2310022B05I | ENSMUSG0C | false | chr8 | 124661452 | 124663780 | 36.46257 | 120.58523 | 116.42456 | chr8 | 124641308 | 124664635 | 6.64691 | 4.65387 | 4.38894 |
| chr8 | 124643917 | 124644866 | 1000 | Intron (ENSMUST00000034464.6 | 8 | 124635766 | 124663369 | 27604 | 2 | ENSMUST00 | 18503 | 2310022B05I | ENSMUSG0C | false | . | . | . | . | . | . | chr8 | 124641308 | 124664635 | 6.64691 | 4.65387 | 4.38894 |
| chr8 | 124949233 | 124949677 | 1000 | Promoter (<=1kb) | 8 | 124908596 | 124949254 | 40659 | 2 | ENSMUST00 | 0 | Egln1 | ENSMUSG0C | false | chr8 | 124946790 | 124950147 | 26.18681 | 91.82585 | 88.44535 | . | . | . | . | . | . |
| chr9 | 43982408 | 43982836 | 1000 | Distal Intergenic | 9 | 44043384 | 44048579 | 5196 | 1 | ENSMUST00 | -60548 | Thy1 | ENSMUSG0C | false | . | . | . | . | . | . | chr9 | 43981458 | 43985255 | 4.79246 | 3.99038 | 2.85353 |
| chr9 | 43239807 | 43240261 | 1000 | Promoter (<=1kb) | 9 | 43221278 | 43239816 | 18539 | 2 | ENSMUST00 | 0 | Oaf | ENSMUSG0C | false | chr9 | 43239020 | 43240233 | 31.38293 | 95.2858 | 91.82495 | . | . | . | . | . | . |
| chr9 | 48480402 | 48480919 | 1000 | Promoter (<=1kb) | 9 | 48468514 | 48480611 | 12098 | 2 | ENSMUST00 | 0 | Rexo2 | ENSMUSG0C | false | chr9 | 48479750 | 48481477 | 27.20707 | 96.65539 | 93.16586 | . | . | . | . | . | . |
| chr9 | 53667004 | 53667709 | 1000 | Promoter (<=1kb) | 9 | 53614582 | 53667507 | 52926 | 2 | ENSMUST00 | 0 | Cul5 | ENSMUSG0C | false | chr9 | 53666273 | 53667755 | 38.29964 | 140.16156 | 135.27679 | . | . | . | . | . | . |
| chr9 | 32646622 | 32647268 | 1000 | Intron (ENSMUST00000183879.1 | 9 | 32636239 | 32651476 | 15238 | 1 | ENSMUST00 | 10384 | Ets1 | ENSMUSG0C | true | . | . | . | . | . | . | chr9 | 32645640 | 32647917 | 6.06489 | 4.58743 | 3.88733 |
| chr9 | 32695572 | 32696464 | 1000 | Promoter (<=1kb) | 9 | 32696022 | 32757820 | 61799 | 1 | ENSMUST00 | 0 | Ets1 | ENSMUSG0C | true | chr9 | 32695428 | 32698295 | 24.03115 | 75.67385 | 72.59326 | chr9 | 32693762 | 32695670 | 4.34549 | 3.75055 | 2.50164 |
| chr9 | 32676023 | 32676403 | 1000 | Intron (ENSMUST00000183879.1 | 9 | 32696022 | 32757820 | 61799 | 1 | ENSMUST00 | -19619 | Ets1 | ENSMUSG0C | true | . | . | . | . | . | . | chr9 | 32666505 | 32679281 | 4.67115 | 3.90561 | 2.75084 |
| chr9 | 32696751 | 32697133 | 1000 | Promoter (<=1kb) | 9 | 32696384 | 32754428 | 58045 | 1 | ENSMUST00 | 368 | Ets1 | ENSMUSG0C | true | chr9 | 32695428 | 32698295 | 24.03115 | 75.67385 | 72.59326 | . | . | . | . | . | . |
| chr9 | 32671858 | 32672986 | 1000 | Intron (ENSMUST00000183879.1 | 9 | 32696022 | 32757820 | 61799 | 1 | ENSMUST00 | -23036 | Ets1 | ENSMUSG0C | true | chr9 | 32671954 | 32672786 | 8.53631 | 15.91019 | 13.6344 | chr9 | 32666505 | 32679281 | 4.67115 | 3.90561 | 2.75084 |
| chr9 | 34350116 | 34350432 | 1000 | Distal Intergenic | 9 | 34485894 | 34489158 | 3265 | 1 | ENSMUST00 | -135462 | Kirrel3 | ENSMUSG0C | false | . | . | . | . | . | . | chr9 | 34348404 | 34351921 | 4.76064 | 3.99049 | 2.80633 |
| chr9 | 35175661 | 35176240 | 1000 | Promoter (<=1kb) | 9 | 35124408 | 35176061 | 51654 | 2 | ENSMUST00 | 0 | Dcps | ENSMUSG0C | false | chr9 | 35175171 | 35176500 | 21.02725 | 63.18829 | 60.29588 | . | . | . | . | . | . |
| chr9 | 52048023 | 52049209 | 1000 | Promoter (<=1kb) | 9 | 52047941 | 52088737 | 40797 | 1 | ENSMUST00 | 83 | Rdx | ENSMUSG0C | false | chr9 | 52047271 | 52050025 | 33.7247 | 108.45113 | 104.65779 | . | . | . | . | . | . |
| chr9 | 44604425 | 44605440 | 1000 | Promoter (<=1kb) | 9 | 44604892 | 44640731 | 35840 | 1 | ENSMUST00 | 0 | Ddx6 | ENSMUSG0C | false | chr9 | 44603277 | 44606301 | 40.0545 | 133.36163 | 128.74257 | . | . | . | . | . | . |
| chr9 | 44406883 | 44407491 | 1000 | Promoter (<=1kb) | 9 | 44403761 | 44407548 | 3788 | 2 | ENSMUST00 | 57 | Trappc4 | ENSMUSG0C | false | chr9 | 44406531 | 44408619 | 29.2476 | 106.45832 | 102.71833 | . | . | . | . | . | . |
| chr9 | 44144046 | 44144313 | 1000 | Distal Intergenic | 9 | 44136861 | 44139682 | 2822 | 1 | ENSMUST00 | 7186 | Mcam | ENSMUSG0C | false | . | . | . | . | . | . | chr9 | 44143376 | 44145281 | 4.96875 | 4.01516 | 2.99211 |
| chr9 | 44129690 | 44130395 | 1000 | Intron (ENSMUST00000180670.1 | 9 | 44134469 | 44138934 | 4466 | 1 | ENSMUST00 | -4074 | Mcam | ENSMUSG0C | false | chr9 | 44129765 | 44130483 | 12.40737 | 26.6269 | 24.20249 | chr9 | 44124698 | 44131925 | 4.45799 | 3.82141 | 2.5616 |
| chr9 | 20651744 | 20652228 | 1000 | Promoter (<=1kb) | 9 | 20652130 | 20666584 | 14455 | 1 | ENSMUST00 | 0 | Pin1 | ENSMUSG0C | false | chr9 | 20651171 | 20651879 | 13.26345 | 35.90286 | 33.3624 | . | . | . | . | . | . |
| chr9 | 21354226 | 21354585 | 1000 | 3' UTR | 9 | 21368019 | 21401803 | 33785 | 1 | ENSMUST00 | -13434 | Ilf3 | ENSMUSG0C | false | . | . | . | . | . | . | chr9 | 21352092 | 21357162 | 6.16136 | 4.42154 | 3.97826 |
| chr9 | 21367553 | 21368501 | 1000 | Promoter (<=1kb) | 9 | 21368019 | 21401803 | 33785 | 1 | ENSMUST00 | 0 | Ilf3 | ENSMUSG0C | false | chr9 | 21367262 | 21369079 | 28.91247 | 96.86193 | 93.36482 | . | . | . | . | . | . |
| chr9 | 21354880 | 21355236 | 1000 | 3' UTR | 9 | 21368019 | 21401803 | 33785 | 1 | ENSMUST00 | -12783 | Ilf3 | ENSMUSG0C | false | . | . | . | . | . | . | chr9 | 21352092 | 21357162 | 6.16136 | 4.42154 | 3.97826 |
| chr9 | 75625433 | 75626231 | 1000 | Promoter (<=1kb) | 9 | 75625732 | 75637773 | 12042 | 1 | ENSMUST00 | 0 | Lysmd2 | ENSMUSG0C | false | chr9 | 75625127 | 75626647 | 32.27147 | 102.84008 | 99.19724 | chr9 | 75625196 | 75626761 | 4.35699 | 3.81448 | 2.50122 |
| chr9 | 75578633 | 75578951 | 1000 | Intron (ENSMUST00000098552.3 | 9 | 75573816 | 75597651 | 23836 | 2 | ENSMUST00 | 18700 | Tmod2 | ENSMUSG0C | false | . | . | . | . | . | . | chr9 | 75576366 | 75582593 | 4.78783 | 4.00601 | 2.82581 |
| chr9 | 71399643 | 71400041 | 1000 | Distal Intergenic | 9 | 71478471 | 71484958 | 6488 | 2 | ENSMUST00 | 84917 | Polr2m | ENSMUSG0C | false | . | . | . | . | . | . | chr9 | 71399565 | 71400400 | 3.58773 | 3.38549 | 1.87737 |
| chr9 | 70542413 | 70543316 | 1000 | Promoter (<=1kb) | 9 | 70542920 | 70592237 | 49318 | 1 | ENSMUST00 | 0 | Sltm | ENSMUSG0C | false | chr9 | 70541924 | 70543925 | 39.39634 | 133.30592 | 128.68834 | . | . | . | . | . | . |
| chr9 | 72661747 | 72662971 | 1000 | Promoter (<=1kb) | 9 | 72662346 | 72749852 | 87507 | 1 | ENSMUST00 | 0 | Nedd4 | ENSMUSG0C | true | chr9 | 72661484 | 72665268 | 29.58769 | 108.11006 | 104.32592 | . | . | . | . | . | . |
| chr9 | 70503141 | 70503956 | 1000 | Promoter (<=1kb) | 9 | 70425429 | 70503725 | 78297 | 2 | ENSMUST00 | 0 | Rnf111 | ENSMUSG0C | false | chr9 | 70502669 | 70504229 | 33.7938 | 119.00099 | 114.88904 | . | . | . | . | . | . |
| chr9 | 70221942 | 70222453 | 1000 | Intron (ENSMUST00000034745.7 | 9 | 70207350 | 70400067 | 192718 | 1 | ENSMUST00 | 14593 | Myo1e | ENSMUSG0C | true | . | . | . | . | . | . | chr9 | 70209471 | 70285922 | 6.22404 | 4.59358 | 4.02448 |
| chr9 | 70243210 | 70243897 | 1000 | Intron (ENSMUST00000034745.7 | 9 | 70207350 | 70400067 | 192718 | 1 | ENSMUST00 | 35861 | Myo1e | ENSMUSG0C | true | . | . | . | . | . | . | chr9 | 70209471 | 70285922 | 6.22404 | 4.59358 | 4.02448 |
| chr9 | 70111827 | 70112458 | 1000 | Intron (ENSMUST00000034749.9 | 9 | 70102550 | 70103076 | 527 | 2 | ENSMUST00 | -8752 | Fam81a | ENSMUSG0C | true | . | . | . | . | . | . | chr9 | 70107386 | 70116753 | 4.95028 | 3.89756 | 3.00574 |
| chr9 | 70132053 | 70132681 | 1000 | Intron (ENSMUST00000034749.9 | 9 | 70089310 | 70141578 | 52269 | 2 | ENSMUST00 | 8897 | Fam81a | ENSMUSG0C | true | . | . | . | . | . | . | chr9 | 70126965 | 70143919 | 5.28781 | 4.22747 | 3.24757 |
| chr9 | 72110951 | 72112202 | 1000 | Promoter (<=1kb) | 9 | 71842688 | 72111847 | 269160 | 2 | ENSMUST00 | 0 | Tcf12 | ENSMUSG0C | false | chr9 | 72110445 | 72112518 | 36.46257 | 120.58523 | 116.42456 | . | . | . | . | . | . |
| chr9 | 69469039 | 69469455 | 1000 | Intron (ENSMUST00000034756.9 | 9 | 69478465 | 69491795 | 13331 | 1 | ENSMUST00 | -9010 | Anxa2 | ENSMUSG0C | false | . | . | . | . | . | . | chr9 | 69455103 | 69480655 | 7.58546 | 5.20708 | 5.13718 |
| chr9 | 69453157 | 69453852 | 1000 | Promoter (<=1kb) | 9 | 69453620 | 69491795 | 38176 | 1 | ENSMUST00 | 0 | Anxa2 | ENSMUSG0C | false | chr9 | 69453593 | 69455744 | 36.10714 | 116.78466 | 112.7497 | chr9 | 69451818 | 69453430 | 3.75225 | 3.23583 | 2.01039 |
| chr9 | 71674434 | 71674867 | 1000 | Intron (ENSMUST00000072899.3 | 9 | 71632633 | 71648350 | 15718 | 2 | ENSMUST00 | -26085 | Cgnl1 | ENSMUSG0C | false | . | . | . | . | . | . | chr9 | 71672716 | 71681830 | 5.44209 | 4.19703 | 3.39991 |
| chr9 | 71625311 | 71625744 | 1000 | Downstream (<1kb) | 9 | 71628859 | 71631109 | 2251 | 2 | ENSMUST00 | 5365 | Cgnl1 | ENSMUSG0C | false | . | . | . | . | . | . | chr9 | 71621822 | 71628252 | 6.34849 | 4.65584 | 4.12076 |
| chr9 | 71753162 | 71753634 | 1000 | Intron (ENSMUST00000072899.3 | 9 | 71629211 | 71771560 | 142350 | 2 | ENSMUST00 | 17926 | Cgnl1 | ENSMUSG0C | false | . | . | . | . | . | . | chr9 | 71736274 | 71770833 | 5.2398 | 4.14793 | 3.22386 |
| chr9 | 69289584 | 69290163 | 1000 | Promoter (<=1kb) | 9 | 69289682 | 69379809 | 90128 | 1 | ENSMUST00 | 0 | Rora | ENSMUSG0C | false | chr9 | 69289292 | 69291949 | 26.72788 | 101.58999 | 97.97913 | . | . | . | . | . | . |
| chr9 | 62811337 | 62812177 | 1000 | Promoter (<=1kb) | 9 | 62791832 | 62811648 | 19817 | 2 | ENSMUST00 | 0 | Fem1b | ENSMUSG0C | false | chr9 | 62810356 | 62812424 | 33.41831 | 117.26761 | 113.21259 | . | . | . | . | . | . |
| chr9 | 62838411 | 62839057 | 1000 | Promoter (<=1kb) | 9 | 62838785 | 62852006 | 13222 | 1 | ENSMUST00 | 0 | Cln6 | ENSMUSG0C | false | chr9 | 62838121 | 62839384 | 38.14117 | 119.25307 | 115.13219 | chr9 | 62837714 | 62839802 | 4.49504 | 3.80765 | 2.63187 |
| chr9 | 82974723 | 82975949 | 1000 | Promoter (<=1kb) | 9 | 82866159 | 82975516 | 109358 | 2 | ENSMUST00 | 0 | Phip | ENSMUSG0C | false | chr9 | 82972816 | 82976631 | 23.3092 | 84.4034 | 81.16875 | . | . | . | . | . | . |
| chr9 | 83551167 | 83551585 | 1000 | Promoter (2-3kb) | 9 | 83548350 | 83638299 | 89950 | 1 | ENSMUST00 | 2818 | Sh3bgrl2 | ENSMUSG0C | true | . | . | . | . | . | . | chr9 | 83550325 | 83577249 | 5.75859 | 4.3064 | 3.65539 |
| chr9 | 83924973 | 83925363 | 1000 | Promoter (<=1kb) | 9 | 83925145 | 84124239 | 199095 | 1 | ENSMUST00 | 0 | Bckdhb | ENSMUSG0C | false | chr9 | 83924637 | 83925761 | 28.8526 | 98.7319 | 95.19158 | chr9 | 83924447 | 83926098 | 3.48175 | 3.16003 | 1.80136 |
| chr9 | 48984836 | 48985672 | 1000 | Promoter (<=1kb) | 9 | 48985385 | 49042517 | 57133 | 1 | ENSMUST00 | 0 | Usp28 | ENSMUSG0C | false | chr9 | 48984533 | 48986205 | 48.25087 | 161.14223 | 155.45619 | . | . | . | . | . | . |
| chr9 | 56994667 | 56995744 | 1000 | Promoter (<=1kb) | 9 | 56994968 | 57062805 | 67838 | 1 | ENSMUST00 | 0 | Ptpn9 | ENSMUSG0C | false | chr9 | 56994400 | 56996703 | 30.26787 | 111.42836 | 107.5545 | chr9 | 56994373 | 56994870 | 4.41206 | 3.88198 | 2.50908 |
| chr9 | 57262268 | 57262836 | 1000 | Promoter (<=1kb) | 9 | 57252322 | 57262599 | 10278 | 2 | ENSMUST00 | 0 | 1700017B05I | ENSMUSG0C | false | chr9 | 57261442 | 57262990 | 28.90751 | 104.81161 | 101.11721 | chr9 | 57260287 | 57263239 | 4.74791 | 3.76954 | 2.81045 |
| chr9 | 56041717 | 56042201 | 1000 | Promoter (<=1kb) | 9 | 56041845 | 56061882 | 20038 | 1 | ENSMUST00 | 0 | Rcn2 | ENSMUSG0C | false | chr9 | 56041506 | 56042817 | 34.36702 | 111.64683 | 107.76208 | . | . | . | . | . | . |
| chr9 | 58013059 | 58013444 | 1000 | Promoter (1-2kb) | 9 | 58014671 | 58027023 | 12353 | 1 | ENSMUST00 | -1227 | Cyp11a1 | ENSMUSG0C | false | . | . | . | . | . | . | chr9 | 58008267 | 58014975 | 6.15797 | 4.62219 | 3.96241 |
| chr9 | 58009100 | 58009528 | 1000 | Promoter (2-3kb) | 9 | 58006411 | 58020742 | 14332 | 1 | ENSMUST00 | 2690 | Cyp11a1 | ENSMUSG0C | false | . | . | . | . | . | . | chr9 | 58008267 | 58014975 | 6.15797 | 4.62219 | 3.96241 |
| chr9 | 79793173 | 79793709 | 1000 | Promoter (<=1kb) | 9 | 79768943 | 79793430 | 24488 | 2 | ENSMUST00 | 0 | Tmem30a | ENSMUSG0C | false | chr9 | 79792262 | 79793432 | 40.31151 | 144.91344 | 139.82919 | . | . | . | . | . | . |
| chr9 | 58295835 | 58296172 | 1000 | Intron (ENSMUST00000061799.8 | 9 | 58287723 | 58313212 | 25490 | 2 | ENSMUST00 | 17040 | Loxl1 | ENSMUSG0C | false | . | . | . | . | . | . | chr9 | 58294749 | 58300318 | 5.3165 | 4.1375 | 3.29392 |
| chr9 | 58581925 | 58582907 | 1000 | Promoter (<=1kb) | 9 | 58582240 | 58651318 | 69079 | 1 | ENSMUST00 | 0 | Nptn | ENSMUSG0C | false | chr9 | 58581598 | 58582102 | 10.01484 | 27.17025 | 24.74075 | . | . | . | . | . | . |
| chr9 | 59035533 | 59036912 | 1000 | Promoter (<=1kb) | 9 | 58874679 | 59036441 | 161763 | 2 | ENSMUST00 | 0 | Neo1 | ENSMUSG0C | false | chr9 | 59035272 | 59037558 | 38.48335 | 126.265 | 121.90878 | chr9 | 59032750 | 59035634 | 3.74082 | 3.50367 | 1.95985 |
| chr9 | 77910177 | 77911360 | 1000 | Distal Intergenic | 9 | 77917364 | 77984519 | 67156 | 1 | ENSMUST00 | -6004 | Elovl5 | ENSMUSG0C | false | chr9 | 77909795 | 77911783 | 25.89225 | 92.22567 | 88.8329 | chr9 | 77907575 | 77910279 | 4.68321 | 3.7783 | 2.77742 |
| chr9 | 77970891 | 77971227 | 1000 | Intron (ENSMUST00000034904.8 | 9 | 77941291 | 77961021 | 19731 | 1 | ENSMUST00 | 29601 | Elovl5 | ENSMUSG0C | false | . | . | . | . | . | . | chr9 | 77969136 | 77972087 | 5.11229 | 4.06506 | 3.14707 |
| chr9 | 77909274 | 77909895 | 1000 | Distal Intergenic | 9 | 77917364 | 77984519 | 67156 | 1 | ENSMUST00 | -7469 | Elovl5 | ENSMUSG0C | false | chr9 | 77909795 | 77911783 | 25.89225 | 92.22567 | 88.8329 | chr9 | 77907575 | 77910279 | 4.68321 | 3.7783 | 2.77742 |
| chr9 | 77461446 | 77462057 | 1000 | Intron (ENSMUST00000183873.2 | 9 | 77431074 | 77448686 | 17613 | 2 | ENSMUST00 | -12761 | Lrrc1 | ENSMUSG0C | true | . | . | . | . | . | . | chr9 | 77455835 | 77465492 | 5.32461 | 4.12133 | 3.29308 |
| chr9 | 77469811 | 77470168 | 1000 | Intron (ENSMUST00000183873.2 | 9 | 77431074 | 77448686 | 17613 | 2 | ENSMUST00 | -21126 | Lrrc1 | ENSMUSG0C | true | . | . | . | . | . | . | chr9 | 77468944 | 77487946 | 5.37817 | 4.20727 | 3.32875 |
| chr9 | 77544570 | 77545304 | 1000 | Promoter (<=1kb) | 9 | 77430823 | 77544870 | 114048 | 2 | ENSMUST00 | 0 | Lrrc1 | ENSMUSG0C | true | chr9 | 77542586 | 77546045 | 36.32541 | 120.3443 | 116.1959 | . | . | . | . | . | . |
| chr9 | 67049147 | 67049523 | 1000 | Promoter (<=1kb) | 9 | 67022590 | 67049267 | 26678 | 2 | ENSMUST00 | 0 | Tpm1 | ENSMUSG0C | true | chr9 | 67046899 | 67049946 | 34.78864 | 106.1911 | 102.46782 | . | . | . | . | . | . |
| chr9 | 66733154 | 66733751 | 1000 | Intron (ENSMUST00000071889.7 | 9 | 66747710 | 66764317 | 16608 | 1 | ENSMUST00 | -13959 | Car12 | ENSMUSG0C | false | . | . | . | . | . | . | chr9 | 66731592 | 66734285 | 6.22576 | 4.73567 | 4.00015 |
| chr9 | 66716471 | 66716917 | 1000 | Promoter (2-3kb) | 9 | 66713730 | 66766842 | 53113 | 1 | ENSMUST00 | 2742 | Car12 | ENSMUSG0C | false | . | . | . | . | . | . | chr9 | 66713595 | 66717842 | 3.81922 | 3.40543 | 2.0547 |
| chr9 | 92570175 | 92570679 | 1000 | Promoter (2-3kb) | 9 | 92573588 | 92591684 | 18097 | 1 | ENSMUST00 | -2909 | Plod2 | ENSMUSG0C | true | . | . | . | . | . | . | chr9 | 92543215 | 92573465 | 5.80463 | 4.42577 | 3.66882 |
| chr9 | 92572322 | 92572777 | 1000 | Promoter (<=1kb) | 9 | 92573588 | 92591684 | 18097 | 1 | ENSMUST00 | -811 | Plod2 | ENSMUSG0C | true | . | . | . | . | . | . | chr9 | 92543215 | 92573465 | 5.80463 | 4.42577 | 3.66882 |
| chr9 | 66124621 | 66125062 | 1000 | Promoter (<=1kb) | 9 | 66088133 | 66124886 | 36754 | 2 | ENSMUST00 | 0 | Snx1 | ENSMUSG0C | false | chr9 | 66123546 | 66125254 | 31.96831 | 119.80809 | 115.67081 | chr9 | 66122633 | 66127885 | 5.40174 | 4.13967 | 3.37486 |
| chr9 | 65908719 | 65909511 | 1000 | Promoter (<=1kb) | 9 | 65908974 | 66004168 | 95195 | 1 | ENSMUST00 | 0 | Csnk1g1 | ENSMUSG0C | false | chr9 | 65908312 | 65910257 | 22.44583 | 74.56482 | 71.50171 | . | . | . | . | . | . |
| chr9 | 65159030 | 65159464 | 1000 | Intron (ENSMUST00000034961.4 | 9 | 65141154 | 65185857 | 44704 | 1 | ENSMUST00 | 17877 | Igdcc3 | ENSMUSG0C | false | . | . | . | . | . | . | chr9 | 65158090 | 65159145 | 6.31281 | 4.6322 | 4.13138 |
| chr9 | 65162913 | 65163592 | 1000 | Intron (ENSMUST00000034961.4 | 9 | 65141154 | 65185857 | 44704 | 1 | ENSMUST00 | 21760 | Igdcc3 | ENSMUSG0C | false | chr9 | 65162638 | 65163554 | 8.26071 | 16.87976 | 14.58833 | chr9 | 65162295 | 65163612 | 5.35565 | 4.17933 | 3.33882 |
| chr9 | 65154583 | 65155327 | 1000 | Intron (ENSMUST00000034961.4 | 9 | 65141154 | 65185857 | 44704 | 1 | ENSMUST00 | 13430 | Igdcc3 | ENSMUSG0C | false | . | . | . | . | . | . | . | . | . | . | . | . |
| chr9 | 64116553 | 64117216 | 1000 | Promoter (<=1kb) | 9 | 64117147 | 64138118 | 20972 | 1 | ENSMUST00 | 0 | Lctl | ENSMUSG0C | true | . | . | . | . | . | . | chr9 | 64111545 | 64122374 | 5.31576 | 4.18523 | 3.29106 |
| chr9 | 63791196 | 63791700 | 1000 | Distal Intergenic | 9 | 63646767 | 63757994 | 111228 | 2 | ENSMUST00 | -33203 | Smad3 | ENSMUSG0C | false | . | . | . | . | . | . | . | . | . | . | . | . |
| chr9 | 63757609 | 63758733 | 1000 | Promoter (<=1kb) | 9 | 63646767 | 63757994 | 111228 | 2 | ENSMUST00 | 0 | Smad3 | ENSMUSG0C | false | chr9 | 63756312 | 63758698 | 29.14352 | 87.50663 | 84.2095 | chr9 | 63758376 | 63759683 | 3.93468 | 3.52104 | 2.14023 |
| chr9 | 63664958 | 63665334 | 1000 | Promoter (1-2kb) | 9 | 63654771 | 63666547 | 11777 | 2 | ENSMUST00 | 1213 | Smad3 | ENSMUSG0C | false | . | . | . | . | . | . | chr9 | 63662341 | 63669966 | 7.25169 | 5.00736 | 4.88788 |
| chr9 | 63716121 | 63716446 | 1000 | Intron (ENSMUST00000034973.4 | 9 | 63654799 | 63711969 | 57171 | 2 | ENSMUST00 | -4153 | Smad3 | ENSMUSG0C | false | . | . | . | . | . | . | chr9 | 63695180 | 63731540 | 5.72957 | 4.34734 | 3.61571 |
| chr9 | 63001591 | 63002005 | 1000 | Distal Intergenic | 9 | 62880077 | 62980879 | 100803 | 2 | ENSMUST00 | -20713 | Pias1 | ENSMUSG0C | false | . | . | . | . | . | . | chr9 | 63000023 | 63002831 | 5.48042 | 4.35112 | 3.41009 |
| chr9 | 96195875 | 96196710 | 1000 | Promoter (<=1kb) | 9 | 96196275 | 96318878 | 122604 | 1 | ENSMUST00 | 0 | Tfdp2 | ENSMUSG0C | false | chr9 | 96195685 | 96197584 | 36.04346 | 118.78738 | 114.68494 | . | . | . | . | . | . |
| chr9 | 96543692 | 96544360 | 1000 | Promoter (<=1kb) | 9 | 96542305 | 96545179 | 2875 | 2 | ENSMUST00 | 819 | Rasa2 | ENSMUSG0C | true | . | . | . | . | . | . | chr9 | 96542835 | 96544530 | 4.59718 | 3.68972 | 2.71486 |
| chr9 | 96630713 | 96631830 | 1000 | Promoter (<=1kb) | 9 | 96539300 | 96631617 | 92318 | 2 | ENSMUST00 | 0 | Rasa2 | ENSMUSG0C | true | chr9 | 96630303 | 96631963 | 24.60788 | 97.30984 | 93.80507 | . | . | . | . | . | . |
| chr9 | 86695598 | 86696056 | 1000 | Promoter (<=1kb) | 9 | 86581371 | 86695953 | 114583 | 2 | ENSMUST00 | 0 | Me1 | ENSMUSG0C | false | chr9 | 86695180 | 86696353 | 28.1615 | 93.53628 | 90.11485 | chr9 | 86695232 | 86696407 | 3.83197 | 3.43755 | 2.08631 |
| chr9 | 87730234 | 87730849 | 1000 | Promoter (<=1kb) | 9 | 87724397 | 87730497 | 6101 | 2 | ENSMUST00 | 0 | Tbx18 | ENSMUSG0C | false | chr9 | 87729621 | 87731071 | 10.56684 | 27.64477 | 25.20979 | chr9 | 87729191 | 87733968 | 4.0777 | 3.50571 | 2.26058 |
| chr9 | 88438843 | 88439219 | 1000 | Promoter (<=1kb) | 9 | 88376750 | 88438951 | 62202 | 2 | ENSMUST00 | 0 | Snx14 | ENSMUSG0C | false | chr9 | 88438145 | 88439793 | 37.78464 | 137.01973 | 132.25398 | . | . | . | . | . | . |
| chr9 | 88481541 | 88483288 | 1000 | Promoter (<=1kb) | 9 | 88449731 | 88482248 | 32518 | 2 | ENSMUST00 | 0 | Syncrip | ENSMUSG0C | false | chr9 | 88480664 | 88483536 | 40.19742 | 128.15663 | 123.73233 | . | . | . | . | . | . |
| chr9 | 114688595 | 114689001 | 1000 | Promoter (<=1kb) | 9 | 114688790 | 114724302 | 35513 | 1 | ENSMUST00 | 0 | Dync1li1 | ENSMUSG0C | false | chr9 | 114688118 | 114690077 | 33.10969 | 106.34647 | 102.61562 | chr9 | 114687972 | 114692983 | 4.36146 | 3.77838 | 2.51239 |
| chr9 | 114803037 | 114803373 | 1000 | Intron (ENSMUST00000047013.3 | 9 | 114756837 | 114781993 | 25157 | 2 | ENSMUST00 | -21045 | Cmtm7 | ENSMUSG0C | true | . | . | . | . | . | . | chr9 | 114801208 | 114806677 | 5.65143 | 4.12227 | 3.58214 |
| chr9 | 115309809 | 115310761 | 1000 | Promoter (<=1kb) | 9 | 115242607 | 115310421 | 67815 | 2 | ENSMUST00 | 0 | Stt3b | ENSMUSG0C | false | chr9 | 115307644 | 115311336 | 30.14656 | 119.38538 | 115.2599 | . | . | . | . | . | . |
| chr9 | 118478658 | 118478936 | 1000 | Promoter (<=1kb) | 9 | 118478861 | 118482757 | 3897 | 1 | ENSMUST00 | 0 | Eomes | ENSMUSG0C | false | chr9 | 118478380 | 118479448 | 17.28575 | 42.23933 | 39.6242 | chr9 | 118476506 | 118479487 | 5.23656 | 3.9469 | 3.22689 |
| chr9 | 100507633 | 100507953 | 1000 | Promoter (<=1kb) | 9 | 100495003 | 100506856 | 11854 | 2 | ENSMUST00 | -778 | Nck1 | ENSMUSG0C | false | . | . | . | . | . | . | chr9 | 100501215 | 100515854 | 3.93432 | 3.52119 | 2.13151 |
| chr9 | 110019211 | 110019593 | 1000 | Intron (ENSMUST00000035055.9 | 9 | 110036742 | 110072705 | 35964 | 1 | ENSMUST00 | -17149 | Map4 | ENSMUSG0C | false | . | . | . | . | . | . | chr9 | 110013879 | 110026147 | 5.4075 | 4.29297 | 3.32753 |
| chr9 | 110736955 | 110737338 | 1000 | Intron (ENSMUST00000006005.7 | 9 | 110722088 | 110742610 | 20523 | 2 | ENSMUST00 | 5272 | Pth1r | ENSMUSG0C | true | chr9 | 110735450 | 110737617 | 9.01168 | 19.18248 | 16.85847 | chr9 | 110716295 | 110740574 | 6.49295 | 4.67204 | 4.24267 |
| chr9 | 110736439 | 110736851 | 1000 | Intron (ENSMUST00000006005.7 | 9 | 110722088 | 110742610 | 20523 | 2 | ENSMUST00 | 5759 | Pth1r | ENSMUSG0C | true | chr9 | 110735450 | 110737617 | 9.01168 | 19.18248 | 16.85847 | chr9 | 110716295 | 110740574 | 6.49295 | 4.67204 | 4.24267 |
| chr9 | 110742248 | 110743910 | 1000 | Promoter (<=1kb) | 9 | 110722088 | 110742610 | 20523 | 2 | ENSMUST00 | 0 | Pth1r | ENSMUSG0C | true | chr9 | 110738126 | 110744546 | 29.35304 | 95.67908 | 92.21078 | . | . | . | . | . | . |
| chr9 | 111170430 | 111170896 | 1000 | Intron (ENSMUST00000035078.7 | 9 | 111118111 | 111225668 | 107558 | 1 | ENSMUST00 | 52320 | Lrrfip2 | ENSMUSG0C | false | . | . | . | . | . | . | chr9 | 111169001 | 111174442 | 5.19731 | 4.22965 | 3.16858 |

| | | | | | | | | | | | | | | | | | | | | | | | | | | |
|---|---|---|---|---|---|---|---|---|---|---|---|---|---|---|---|---|---|---|---|---|---|---|---|---|---|---|
| chr9 | 111205060 | 111205370 | 1000 | Intron (ENSMUST00000035078.7 | 9 | 111229870 | 111230948 | 1079 | 2 | ENSMUST00 | 25578 | Mlh1 | ENSMUSG0C | false | . | . | . | . | . | . | chr9 | 111203735 | 111205595 | 4.2828 | 3.61238 | 2.46694 |
| chr9 | 113707556 | 113708376 | 1000 | Promoter (<=1kb) | 9 | 113651744 | 113708259 | 56516 | 2 | ENSMUST00 | 0 | Pdcd6ip | ENSMUSG0C | false | chr9 | 113706732 | 113708611 | 37.92415 | 138.37384 | 133.55817 | . | . | . | . | . | . |
| chr9 | 119631844 | 119632440 | 1000 | Intron (ENSMUST00000084787.5 | 9 | 119484324 | 119579016 | 94693 | 2 | ENSMUST00 | -52829 | Scn5a | ENSMUSG0C | false | . | . | . | . | . | . | chr9 | 119630900 | 119634710 | 5.60192 | 4.23392 | 3.53631 |
| chr9 | 119984507 | 119985019 | 1000 | Intron (ENSMUST00000177637.1 | 9 | 119971166 | 119977250 | 6085 | 2 | ENSMUST00 | -7258 | Csrnp1 | ENSMUSG0C | false | chr9 | 119983901 | 119984798 | 18.39885 | 52.66533 | 49.91441 | chr9 | 119984393 | 119985457 | 4.44164 | 3.92098 | 2.53164 |
| chr9 | 119983561 | 119984181 | 1000 | Intron (ENSMUST00000177637.1 | 9 | 119971166 | 119977250 | 6085 | 2 | ENSMUST00 | -6312 | Csrnp1 | ENSMUSG0C | false | chr9 | 119982428 | 119983802 | 31.01414 | 97.6227 | 94.10706 | . | . | . | . | . | . |
| chr9 | 101036964 | 101037458 | 1000 | Promoter (2-3kb) | 9 | 100982032 | 101034898 | 52867 | 2 | ENSMUST00 | -2067 | Pccb | ENSMUSG0C | false | . | . | . | . | . | . | chr9 | 101033076 | 101041068 | 4.72591 | 3.79913 | 2.80177 |
| chr9 | 102726181 | 102726752 | 1000 | Promoter (1-2kb) | 9 | 102728331 | 102732341 | 4011 | 1 | ENSMUST00 | -1579 | Amotl2 | ENSMUSG0C | false | . | . | . | . | . | . | chr9 | 102720260 | 102734975 | 8.68688 | 5.64144 | 6.02662 |
| chr9 | 102721986 | 102722441 | 1000 | Promoter (<=1kb) | 9 | 102722316 | 102728326 | 6011 | 1 | ENSMUST00 | 0 | Amotl2 | ENSMUSG0C | false | chr9 | 102721728 | 102723352 | 11.64009 | 27.76484 | 25.32713 | chr9 | 102720260 | 102734975 | 8.68688 | 5.64144 | 6.02662 |
| chr9 | 102717110 | 102717964 | 1000 | Promoter (<=1kb) | 9 | 102717289 | 102720733 | 3445 | 1 | ENSMUST00 | 0 | Amotl2 | ENSMUSG0C | false | chr9 | 102716871 | 102721558 | 32.23346 | 129.76328 | 125.27943 | chr9 | 102714194 | 102717397 | 7.62791 | 5.22899 | 5.15878 |
| chr9 | 102354247 | 102354997 | 1000 | Promoter (<=1kb) | 9 | 101922128 | 102354693 | 432566 | 2 | ENSMUST00 | 0 | Ephb1 | ENSMUSG0C | true | chr9 | 102352850 | 102354848 | 31.54088 | 108.67455 | 104.87239 | . | . | . | . | . | . |
| chr9 | 107633203 | 107633806 | 1000 | Promoter (1-2kb) | 9 | 107614138 | 107635342 | 21205 | 2 | ENSMUST00 | 1536 | Gnai2 | ENSMUSG0C | false | chr9 | 107632708 | 107635411 | 27.71015 | 85.00858 | 81.76643 | chr9 | 107617961 | 107633388 | 7.13311 | 4.9094 | 4.77996 |
| chr9 | 105494732 | 105495650 | 1000 | Promoter (<=1kb) | 9 | 105410443 | 105495094 | 84652 | 2 | ENSMUST00 | 0 | Atp2c1 | ENSMUSG0C | false | chr9 | 105494013 | 105496076 | 36.809 | 123.33047 | 119.07505 | . | . | . | . | . | . |
| chr9 | 108796344 | 108797016 | 1000 | Promoter (<=1kb) | 9 | 108795994 | 108806333 | 10340 | 1 | ENSMUST00 | 351 | Ip6k2 | ENSMUSG0C | false | chr9 | 108795574 | 108798004 | 15.53947 | 48.13258 | 45.44172 | chr9 | 108784542 | 108804834 | 4.97749 | 3.96237 | 3.01028 |
| chr9 | 108693910 | 108694268 | 1000 | Promoter (1-2kb) | 9 | 108692143 | 108749512 | 57370 | 1 | ENSMUST00 | 1768 | Prkar2a | ENSMUSG0C | true | chr9 | 108691764 | 108693954 | 32.27147 | 102.84008 | 99.19724 | chr9 | 108693247 | 108705103 | 4.75631 | 3.90631 | 2.81912 |
| chr9 | 108347461 | 108348186 | 1000 | Promoter (<=1kb) | 9 | 108347831 | 108392529 | 44699 | 1 | ENSMUST00 | 0 | Usp4 | ENSMUSG0C | true | chr9 | 108346953 | 108348997 | 42.27695 | 136.11826 | 131.39047 | . | . | . | . | . | . |
| chr17 | 83350691 | 83351234 | 1000 | Promoter (<=1kb) | 17 | 83350931 | 83480361 | 129431 | 1 | ENSMUST00 | 0 | Eml4 | ENSMUSG0C | true | chr17 | 83350276 | 83350807 | 17.92175 | 44.05972 | 41.41886 | . | . | . | . | . | . |
| chr7 | 126503002 | 126503916 | 1000 | Promoter (<=1kb) | 7 | 126491708 | 126503437 | 11730 | 2 | ENSMUST00 | 0 | Atxn2l | ENSMUSG0C | false | chr7 | 126501677 | 126504111 | 42.82515 | 147.12178 | 141.95673 | . | . | . | . | . | . |
| chr6 | 134829969 | 134830599 | 1000 | Promoter (<=1kb) | 6 | 134830154 | 134858931 | 28778 | 1 | ENSMUST00 | 0 | Crebl2 | ENSMUSG0C | false | chr6 | 134829342 | 134829976 | 14.29631 | 42.77194 | 40.14703 | . | . | . | . | . | . |
| chr3 | 89182894 | 89183254 | 1000 | Promoter (<=1kb) | 3 | 89183143 | 89184679 | 1537 | 1 | ENSMUST00 | 0 | Fam189b | ENSMUSG0C | false | chr3 | 89182855 | 89183747 | 19.31422 | 56.74648 | 53.94233 | chr3 | 89182331 | 89183922 | 4.73757 | 3.89725 | 2.82964 |
| chr1 | 151976062 | 151976435 | 1000 | Intron (ENSMUST00000187603.1 | 1 | 151892903 | 151921227 | 28325 | 2 | ENSMUST00 | -54836 | 1700025G04 | ENSMUSG0C | true | . | . | . | . | . | . | chr1 | 151972781 | 151980078 | 5.57591 | 4.3299 | 3.48688 |
| chr1 | 152089060 | 152090603 | 1000 | Promoter (<=1kb) | 1 | 151852403 | 152089970 | 237568 | 2 | ENSMUST00 | 0 | 1700025G04 | ENSMUSG0C | true | chr1 | 152088689 | 152091255 | 23.5948 | 97.99914 | 94.47849 | chr1 | 152087132 | 152089110 | 4.62787 | 3.95637 | 2.69962 |
| chr2 | 104096439 | 104096866 | 1000 | Promoter (<=1kb) | 2 | 104095840 | 104114263 | 18424 | 1 | ENSMUST00 | 600 | Cd59a | ENSMUSG0C | true | chr2 | 104095745 | 104097982 | 31.16539 | 106.97112 | 103.21663 | . | . | . | . | . | . |
| chr18 | 65430365 | 65431433 | 1000 | Promoter (<=1kb) | 18 | 65430963 | 65478823 | 47861 | 1 | ENSMUST00 | 0 | Malt1 | ENSMUSG0C | true | chr18 | 65430226 | 65430779 | 14.26849 | 37.02594 | 34.4737 | . | . | . | . | . | . |
| chr19 | 25276619 | 25276991 | 1000 | Intron (ENSMUST00000049400.9 | 19 | 25236975 | 25434496 | 197522 | 1 | ENSMUST00 | 39645 | Kank1 | ENSMUSG0C | true | . | . | . | . | . | . | chr19 | 25261572 | 25293097 | 5.29409 | 4.21889 | 3.24888 |
| chr19 | 25307046 | 25307552 | 1000 | Intron (ENSMUST00000049400.9 | 19 | 25236975 | 25434496 | 197522 | 1 | ENSMUST00 | 70072 | Kank1 | ENSMUSG0C | true | . | . | . | . | . | . | chr19 | 25294157 | 25326429 | 5.37785 | 4.23317 | 3.31781 |
| chr19 | 25273348 | 25273646 | 1000 | Intron (ENSMUST00000049400.9 | 19 | 25236975 | 25434496 | 197522 | 1 | ENSMUST00 | 36374 | Kank1 | ENSMUSG0C | true | . | . | . | . | . | . | chr19 | 25261572 | 25293097 | 5.29409 | 4.21889 | 3.24888 |
| chr19 | 25238101 | 25238501 | 1000 | Promoter (1-2kb) | 19 | 25236975 | 25434496 | 197522 | 1 | ENSMUST00 | 1127 | Kank1 | ENSMUSG0C | true | chr19 | 25236314 | 25239245 | 23.12601 | 77.64799 | 74.53377 | . | . | . | . | . | . |
| chr13 | 111685789 | 111687074 | 1000 | Promoter (<=1kb) | 13 | 111686178 | 111718596 | 32419 | 1 | ENSMUST00 | 0 | Mier3 | ENSMUSG0C | false | chr13 | 111685596 | 111687859 | 31.95414 | 97.26555 | 93.76166 | . | . | . | . | . | . |
| chr9 | 56927910 | 56928510 | 1000 | Promoter (<=1kb) | 9 | 56917200 | 56928371 | 11172 | 2 | ENSMUST00 | 0 | Snx33 | ENSMUSG0C | false | chr9 | 56926890 | 56929065 | 19.22456 | 54.94707 | 52.16912 | . | . | . | . | . | . |
| chr9 | 56924615 | 56924985 | 1000 | Intron (ENSMUST00000050916.5 | 9 | 56917200 | 56928371 | 11172 | 2 | ENSMUST00 | 3386 | Snx33 | ENSMUSG0C | false | chr9 | 56924859 | 56925753 | 8.50221 | 18.80243 | 16.48241 | chr9 | 56919789 | 56927165 | 7.04198 | 4.86227 | 4.70521 |
| chr18 | 61888118 | 61888539 | 1000 | Intron (ENSMUST00000049378.9 | 18 | 61799395 | 61911824 | 112430 | 2 | ENSMUST00 | 23285 | Ablim3 | ENSMUSG0C | false | . | . | . | . | . | . | chr18 | 61877692 | 61898809 | 5.45644 | 4.15279 | 3.39554 |
| chr18 | 61880725 | 61881250 | 1000 | Intron (ENSMUST00000049378.9 | 18 | 61799395 | 61911824 | 112430 | 2 | ENSMUST00 | 30574 | Ablim3 | ENSMUSG0C | false | . | . | . | . | . | . | chr18 | 61877692 | 61898809 | 5.45644 | 4.15279 | 3.39554 |
| chr18 | 61894618 | 61894997 | 1000 | Intron (ENSMUST00000049378.9 | 18 | 61799395 | 61911824 | 112430 | 2 | ENSMUST00 | 16827 | Ablim3 | ENSMUSG0C | false | . | . | . | . | . | . | chr18 | 61877692 | 61898809 | 5.45644 | 4.15279 | 3.39554 |
| chr7 | 101837607 | 101838511 | 1000 | Promoter (<=1kb) | 7 | 101822632 | 101838076 | 15445 | 2 | ENSMUST00 | 0 | Inppl1 | ENSMUSG0C | false | chr7 | 101835409 | 101838921 | 30.94804 | 114.76607 | 110.79114 | chr7 | 101838506 | 101839917 | 5.06909 | 4.12598 | 3.10219 |
| chr11 | 29373322 | 29374191 | 1000 | Promoter (<=1kb) | 11 | 29373658 | 29510808 | 137151 | 1 | ENSMUST00 | 0 | Ccdc88a | ENSMUSG0C | false | chr11 | 29373114 | 29374752 | 34.36702 | 111.64683 | 107.76208 | . | . | . | . | . | . |
| chr5 | 120578567 | 120579811 | 1000 | Intron (ENSMUST00000046426.8 | 5 | 120534160 | 120588613 | 54454 | 2 | ENSMUST00 | 8802 | Tpcn1 | ENSMUSG0C | false | . | . | . | . | . | . | chr5 | 120558690 | 120580640 | 7.14372 | 4.95764 | 4.77953 |
| chr5 | 120563509 | 120563953 | 1000 | Intron (ENSMUST00000046426.8 | 5 | 120534160 | 120588613 | 54454 | 2 | ENSMUST00 | 24660 | Tpcn1 | ENSMUSG0C | false | . | . | . | . | . | . | chr5 | 120558690 | 120580640 | 7.14372 | 4.95764 | 4.77953 |
| chr5 | 120560614 | 120561297 | 1000 | Intron (ENSMUST00000046426.8 | 5 | 120534160 | 120588613 | 54454 | 2 | ENSMUST00 | 27316 | Tpcn1 | ENSMUSG0C | false | . | . | . | . | . | . | chr5 | 120558690 | 120580640 | 7.14372 | 4.95764 | 4.77953 |
| chr4 | 123239357 | 123240049 | 1000 | Promoter (<=1kb) | 4 | 123239757 | 123245989 | 6233 | 1 | ENSMUST00 | 0 | Heyl | ENSMUSG0C | false | . | . | . | . | . | . | chr4 | 123239083 | 123240238 | 4.9555 | 4.07409 | 2.98241 |
| chr4 | 123233523 | 123234174 | 1000 | Promoter (<=1kb) | 4 | 123233556 | 123249875 | 16320 | 1 | ENSMUST00 | 0 | Heyl | ENSMUSG0C | false | chr4 | 123233410 | 123234037 | 14.34627 | 35.7714 | 33.23358 | chr4 | 123233346 | 123234851 | 4.82833 | 4.06966 | 2.90534 |
| chr13 | 111489430 | 111490715 | 1000 | Promoter (<=1kb) | 13 | 111425680 | 111490111 | 64432 | 2 | ENSMUST00 | 0 | Gpbp1 | ENSMUSG0C | false | chr13 | 111488368 | 111489932 | 28.89995 | 120.94616 | 116.77392 | . | . | . | . | . | . |
| chr7 | 71960646 | 71961241 | 1000 | Distal Intergenic | 7 | 72077830 | 72306595 | 228766 | 2 | ENSMUST00 | 345354 | Mctp2 | ENSMUSG0C | false | . | . | . | . | . | . | chr7 | 71960252 | 71961607 | 3.40801 | 3.07749 | 1.7309 |
| chr7 | 72300290 | 72300604 | 1000 | Intron (ENSMUST00000079323.6 | 7 | 72077830 | 72306595 | 228766 | 2 | ENSMUST00 | 5991 | Mctp2 | ENSMUSG0C | false | . | . | . | . | . | . | . | . | . | . | . | . |
| chr11 | 69168186 | 69168595 | 1000 | Intron (ENSMUST00000036424.2 | 11 | 69156989 | 69169792 | 12804 | 1 | ENSMUST00 | 11198 | Alox12b | ENSMUSG0C | false | . | . | . | . | . | . | chr11 | 69167146 | 69170622 | 4.9796 | 4.04352 | 2.99715 |
| chr13 | 107888672 | 107889478 | 1000 | Promoter (<=1kb) | 13 | 107724618 | 107890064 | 165447 | 2 | ENSMUST00 | 586 | Zswim6 | ENSMUSG0C | false | chr13 | 107887775 | 107890104 | 33.94791 | 109.87469 | 106.04007 | . | . | . | . | . | . |
| chr13 | 107889688 | 107891515 | 1000 | Promoter (<=1kb) | 13 | 107724618 | 107890064 | 165447 | 2 | ENSMUST00 | 0 | Zswim6 | ENSMUSG0C | false | chr13 | 107887775 | 107890104 | 33.94791 | 109.87469 | 106.04007 | . | . | . | . | . | . |
| chr13 | 107887273 | 107887661 | 1000 | Promoter (2-3kb) | 13 | 107724618 | 107890064 | 165447 | 2 | ENSMUST00 | 2403 | Zswim6 | ENSMUSG0C | false | . | . | . | . | . | . | chr13 | 107859854 | 107887411 | 4.83407 | 3.98235 | 2.87425 |
| chr7 | 100911139 | 100911698 | 1000 | Intron (ENSMUST00000107032.1 | 7 | 100869746 | 100932161 | 62416 | 2 | ENSMUST00 | 20463 | Arhgef17 | ENSMUSG0C | false | chr7 | 100910881 | 100911312 | 4.88133 | 7.58987 | 5.46511 | chr7 | 100901870 | 100926579 | 7.14639 | 4.97053 | 4.78972 |
| chr7 | 100930713 | 100932961 | 1000 | Promoter (<=1kb) | 7 | 100869746 | 100932161 | 62416 | 2 | ENSMUST00 | 0 | Arhgef17 | ENSMUSG0C | false | chr7 | 100925652 | 100932907 | 25.18016 | 100.40004 | 96.81754 | chr7 | 100932304 | 100935855 | 4.57644 | 3.72921 | 2.69387 |
| chr7 | 100918117 | 100918576 | 1000 | Intron (ENSMUST00000107032.1 | 7 | 100869746 | 100932161 | 62416 | 2 | ENSMUST00 | 13585 | Arhgef17 | ENSMUSG0C | false | . | . | . | . | . | . | chr7 | 100901870 | 100926579 | 7.14639 | 4.97053 | 4.78972 |
| chr7 | 100926444 | 100927211 | 1000 | Intron (ENSMUST00000107032.1 | 7 | 100869746 | 100932161 | 62416 | 2 | ENSMUST00 | 4950 | Arhgef17 | ENSMUSG0C | false | chr7 | 100925652 | 100932907 | 25.18016 | 100.40004 | 96.81754 | chr7 | 100901870 | 100926579 | 7.14639 | 4.97053 | 4.78972 |
| chr16 | 75766743 | 75767126 | 1000 | Promoter (<=1kb) | 16 | 75755190 | 75766821 | 11632 | 2 | ENSMUST00 | 0 | Hspa13 | ENSMUSG0C | false | chr16 | 75766135 | 75767980 | 18.77433 | 54.14365 | 51.37372 | . | . | . | . | . | . |
| chr19 | 6401006 | 6402307 | 1000 | Promoter (<=1kb) | 19 | 6401695 | 6402446 | 752 | 1 | ENSMUST00 | 0 | Rasgrp2 | ENSMUSG0C | false | chr19 | 6400784 | 6402082 | 8.74864 | 23.49345 | 21.11253 | chr19 | 6398167 | 6401812 | 5.57956 | 4.20134 | 3.51519 |
| chr16 | 49698889 | 49699284 | 1000 | Promoter (<=1kb) | 16 | 49699233 | 49765126 | 65894 | 1 | ENSMUST00 | 0 | Ift57 | ENSMUSG0C | false | chr16 | 49698661 | 49699981 | 31.01882 | 98.6291 | 95.09013 | chr16 | 49698426 | 49700123 | 3.74087 | 3.52105 | 1.99242 |
| chr1 | 75478857 | 75479600 | 1000 | Promoter (<=1kb) | 1 | 75474569 | 75479271 | 4703 | 2 | ENSMUST00 | 0 | Chpf | ENSMUSG0C | false | chr1 | 75478421 | 75480081 | 27.88725 | 99.90214 | 96.33501 | chr1 | 75477205 | 75480428 | 7.75191 | 5.367 | 5.26863 |
| chr4 | 119539260 | 119540604 | 1000 | Promoter (<=1kb) | 4 | 119539661 | 119629119 | 89459 | 1 | ENSMUST00 | 0 | Foxj3 | ENSMUSG0C | true | chr4 | 119538635 | 119541529 | 41.61333 | 140.51047 | 135.61182 | . | . | . | . | . | . |
| chr14 | 103346357 | 103347073 | 1000 | Promoter (<=1kb) | 14 | 103113411 | 103346418 | 233008 | 2 | ENSMUST00 | 0 | Mycbp2 | ENSMUSG0C | false | chr14 | 103345093 | 103347425 | 30.50916 | 92.78792 | 89.38336 | . | . | . | . | . | . |
| chr3 | 103279285 | 103280330 | 1000 | Promoter (<=1kb) | 3 | 103279293 | 103358768 | 79476 | 1 | ENSMUST00 | 0 | Trim33 | ENSMUSG0C | false | chr3 | 103278682 | 103281247 | 32.64297 | 100.45817 | 96.87489 | . | . | . | . | . | . |
| chr18 | 80707993 | 80708608 | 1000 | Promoter (<=1kb) | 18 | 80606205 | 80708173 | 101969 | 2 | ENSMUST00 | 0 | Nfatc1 | ENSMUSG0C | false | chr18 | 80707078 | 80708822 | 33.04283 | 115.53907 | 111.53828 | chr18 | 80704695 | 80708710 | 3.39357 | 3.23629 | 1.72061 |
| chr18 | 80588991 | 80589415 | 1000 | Distal Intergenic | 18 | 80606205 | 80708173 | 101969 | 2 | ENSMUST00 | 118758 | Nfatc1 | ENSMUSG0C | false | . | . | . | . | . | . | chr18 | 80587168 | 80591017 | 5.94882 | 4.47979 | 3.82536 |
| chr18 | 61767769 | 61768484 | 1000 | Intron (ENSMUST00000154876.2 | 18 | 61730261 | 61786611 | 56351 | 2 | ENSMUST00 | 18127 | Afap1l1 | ENSMUSG0C | false | . | . | . | . | . | . | . | . | . | . | . | . |
| chr15 | 79108541 | 79109384 | 1000 | Promoter (<=1kb) | 15 | 79108972 | 79136897 | 27926 | 1 | ENSMUST00 | 0 | Micall1 | ENSMUSG0C | false | chr15 | 79108405 | 79110109 | 42.60553 | 141.12407 | 136.19626 | . | . | . | . | . | . |
| chr11 | 67649781 | 67650202 | 1000 | Promoter (2-3kb) | 11 | 67652577 | 67683688 | 31112 | 1 | ENSMUST00 | -2375 | Gas7 | ENSMUSG0C | true | . | . | . | . | . | . | chr11 | 67646563 | 67657817 | 7.67901 | 5.19007 | 5.23014 |
| chr11 | 67526630 | 67526946 | 1000 | Intron (ENSMUST00000108681.2 | 11 | 67547154 | 67684974 | 137821 | 1 | ENSMUST00 | -20208 | Gas7 | ENSMUSG0C | true | . | . | . | . | . | . | chr11 | 67521726 | 67547234 | 6.38934 | 4.66296 | 4.16347 |
| chr11 | 67454847 | 67456062 | 1000 | Promoter (<=1kb) | 11 | 67455437 | 67683460 | 228024 | 1 | ENSMUST00 | 0 | Gas7 | ENSMUSG0C | true | chr11 | 67454716 | 67457354 | 29.92778 | 109.76675 | 105.9399 | chr11 | 67454418 | 67455023 | 4.70932 | 4.03691 | 2.78841 |
| chr11 | 67628492 | 67628873 | 1000 | Intron (ENSMUST00000108681.2 | 11 | 67652577 | 67683688 | 31112 | 1 | ENSMUST00 | -23704 | Gas7 | ENSMUSG0C | true | . | . | . | . | . | . | chr11 | 67625478 | 67640419 | 5.80423 | 4.41975 | 3.66967 |
| chr11 | 67648681 | 67649124 | 1000 | Intron (ENSMUST00000108681.2 | 11 | 67652577 | 67683688 | 31112 | 1 | ENSMUST00 | -3453 | Gas7 | ENSMUSG0C | true | . | . | . | . | . | . | chr11 | 67646563 | 67657817 | 7.67901 | 5.19007 | 5.23014 |
| chr11 | 67515011 | 67515693 | 1000 | Intron (ENSMUST00000108681.2 | 11 | 67547154 | 67684974 | 137821 | 1 | ENSMUST00 | -31461 | Gas7 | ENSMUSG0C | true | . | . | . | . | . | . | chr11 | 67511965 | 67517600 | 7.10268 | 5.0078 | 4.75923 |
| chr11 | 67494056 | 67494838 | 1000 | Intron (ENSMUST00000108681.2 | 11 | 67455724 | 67665449 | 209726 | 1 | ENSMUST00 | 38333 | Gas7 | ENSMUSG0C | true | . | . | . | . | . | . | chr11 | 67489685 | 67495862 | 5.1759 | 4.0485 | 3.19077 |
| chr15 | 98093204 | 98093730 | 1000 | Promoter (<=1kb) | 15 | 98041393 | 98093569 | 52177 | 2 | ENSMUST00 | 0 | Senp1 | ENSMUSG0C | false | chr15 | 98092443 | 98094045 | 32.87838 | 107.99121 | 104.21945 | . | . | . | . | . | . |
| chr14 | 101608669 | 101609392 | 1000 | Promoter (<=1kb) | 14 | 101442360 | 101609033 | 166674 | 2 | ENSMUST00 | 0 | Tbc1d4 | ENSMUSG0C | false | chr14 | 101607475 | 101609516 | 33.06624 | 113.49754 | 109.55873 | . | . | . | . | . | . |
| chr14 | 101568742 | 101569179 | 1000 | Intron (ENSMUST00000161991.2 | 14 | 101443723 | 101608486 | 164764 | 2 | ENSMUST00 | 39307 | Tbc1d4 | ENSMUSG0C | false | . | . | . | . | . | . | chr14 | 101568012 | 101569991 | 4.11892 | 3.62781 | 2.29311 |
| chr14 | 101538034 | 101538383 | 1000 | Intron (ENSMUST00000161991.2 | 14 | 101442368 | 101507326 | 64959 | 2 | ENSMUST00 | -30709 | Tbc1d4 | ENSMUSG0C | false | . | . | . | . | . | . | chr14 | 101536669 | 101540662 | 4.8238 | 4.00444 | 2.86182 |
| chr15 | 78984057 | 78984410 | 1000 | Promoter (<=1kb) | 15 | 78983705 | 78991438 | 7734 | 1 | ENSMUST00 | 353 | Triobp | ENSMUSG0C | false | chr15 | 78982979 | 78984171 | 28.89701 | 129.08754 | 124.63428 | chr15 | 78981999 | 78988873 | 5.15157 | 4.09348 | 3.15503 |
| chr10 | 76501106 | 76501592 | 1000 | Exon (ENSMUST00000170795.1/ | 10 | 76531588 | 76535568 | 3981 | 1 | ENSMUST00 | -29996 | Lss | ENSMUSG0C | false | chr10 | 76500859 | 76501338 | 4.76124 | 7.69302 | 5.56588 | chr10 | 76499410 | 76507258 | 5.8758 | 4.41151 | 3.72672 |
| chr16 | 45844086 | 45844740 | 1000 | Promoter (<=1kb) | 16 | 45746243 | 45844378 | 98136 | 2 | ENSMUST00 | 0 | Phldb2 | ENSMUSG0C | false | chr16 | 45842591 | 45845351 | 29.36448 | 91.77025 | 88.3935 | . | . | . | . | . | . |
| chr16 | 45893633 | 45894153 | 1000 | Intron (ENSMUST00000134802.2 | 16 | 45901041 | 45901967 | 927 | 2 | ENSMUST00 | 7814 | Phldb2 | ENSMUSG0C | false | . | . | . | . | . | . | chr16 | 45888935 | 45896919 | 5.27673 | 4.2429 | 3.23254 |
| chr3 | 101603890 | 101604939 | 1000 | Promoter (<=1kb) | 3 | 101576219 | 101604603 | 28385 | 2 | ENSMUST00 | 0 | Atp1a1 | ENSMUSG0C | false | chr3 | 101602862 | 101605252 | 34.04953 | 116.30843 | 112.28568 | chr3 | 101604927 | 101613489 | 5.50887 | 4.25497 | 3.44478 |
| chr15 | 78796295 | 78797003 | 1000 | Intron (ENSMUST00000164826.2 | 15 | 78776493 | 78802669 | 26177 | 2 | ENSMUST00 | 5666 | Card10 | ENSMUSG0C | true | chr15 | 78794341 | 78796577 | 8.64794 | 16.56503 | 14.27926 | chr15 | 78770626 | 78800649 | 7.36271 | 5.05378 | 4.95398 |
| chr15 | 78803549 | 78804191 | 1000 | Promoter (<=1kb) | 15 | 78775138 | 78803042 | 27905 | 2 | ENSMUST00 | -508 | Card10 | ENSMUSG0C | true | chr15 | 78796787 | 78803936 | 36.38946 | 142.11884 | 137.14919 | chr15 | 78803127 | 78808136 | 6.88225 | 4.93932 | 4.56294 |
| chr15 | 78792163 | 78792871 | 1000 | Promoter (1-2kb) | 15 | 78787411 | 78790515 | 3105 | 2 | ENSMUST00 | -1649 | Card10 | ENSMUSG0C | true | . | . | . | . | . | . | chr15 | 78770626 | 78800649 | 7.36271 | 5.05378 | 4.95398 |
| chr15 | 78802583 | 78803343 | 1000 | Promoter (<=1kb) | 15 | 78775138 | 78803042 | 27905 | 2 | ENSMUST00 | 0 | Card10 | ENSMUSG0C | true | chr15 | 78796787 | 78803936 | 36.38946 | 142.11884 | 137.14919 | chr15 | 78803127 | 78808136 | 6.88225 | 4.93932 | 4.56294 |
| chr6 | 88589388 | 88589744 | 1000 | Promoter (<=1kb) | 6 | 88586528 | 88590659 | 4132 | 2 | ENSMUST00 | 915 | Kbtbd12 | ENSMUSG0C | false | . | . | . | . | . | . | chr6 | 88588887 | 88589903 | 4.56222 | 3.70147 | 2.67809 |
| chr18 | 46597324 | 46597922 | 1000 | Promoter (<=1kb) | 18 | 46585939 | 46597516 | 11578 | 2 | ENSMUST00 | 0 | Tmed7 | ENSMUSG0C | false | chr18 | 46596626 | 46598838 | 35.15467 | 116.16882 | 112.14974 | . | . | . | . | . | . |
| chr8 | 92846588 | 92847867 | 1000 | Intron (ENSMUST00000034187.7 | 8 | 92855350 | 92919279 | 63930 | 1 | ENSMUST00 | -7483 | Lpcat2 | ENSMUSG0C | false | . | . | . | . | . | . | chr8 | 92845551 | 92848245 | 7.2751 | 5.14753 | 4.87586 |
| chr15 | 78579586 | 78579891 | 1000 | Distal Intergenic | 15 | 78559169 | 78572783 | 13615 | 2 | ENSMUST00 | -6804 | Rac2 | ENSMUSG0C | false | . | . | . | . | . | . | chr15 | 78579344 | 78582264 | 4.73333 | 3.95093 | 2.78737 |
| chr15 | 96460012 | 96461575 | 1000 | Promoter (<=1kb) | 15 | 96411698 | 96460843 | 49146 | 2 | ENSMUST00 | 0 | Scaf11 | ENSMUSG0C | false | chr15 | 96459601 | 96460930 | 38.5588 | 123.16969 | 118.923 | . | . | . | . | . | . |
| chr15 | 96286507 | 96287927 | 1000 | Promoter (<=1kb) | 15 | 96287518 | 96352038 | 64521 | 1 | ENSMUST00 | 0 | Arid2 | ENSMUSG0C | false | chr15 | 96283762 | 96289319 | 34.73729 | 107.12603 | 103.36953 | . | . | . | . | . | . |
| chr8 | 105269557 | 105269959 | 1000 | Promoter (<=1kb) | 8 | 105269801 | 105275686 | 5886 | 1 | ENSMUST00 | 0 | Hsf4 | ENSMUSG0C | false | chr8 | 105268532 | 105270228 | 10.86952 | 32.54407 | 30.04659 | chr8 | 105268367 | 105271002 | 4.54883 | 3.84708 | 2.64998 |
| chr15 | 78436900 | 78437337 | 1000 | Promoter (2-3kb) | 15 | 78433982 | 78437579 | 3598 | 1 | ENSMUST00 | 2919 | Kctd17 | ENSMUSG0C | false | . | . | . | . | . | . | chr15 | 78435608 | 78438211 | 4.24809 | 3.76385 | 2.37764 |
| chr16 | 24393100 | 24394163 | 1000 | Promoter (<=1kb) | 16 | 24393350 | 24992576 | 599227 | 1 | ENSMUST00 | 0 | Lpp | ENSMUSG0C | false | chr16 | 24392713 | 24394688 | 34.00537 | 102.80987 | 99.17244 | . | . | . | . | . | . |
| chr16 | 24282373 | 24282785 | 1000 | Distal Intergenic | 16 | 24393350 | 24992576 | 599227 | 1 | ENSMUST00 | -110565 | Lpp | ENSMUSG0C | false | . | . | . | . | . | . | chr16 | 24282109 | 24283234 | 4.67663 | 3.85165 | 2.77965 |
| chr16 | 24506886 | 24507215 | 1000 | Intron (ENSMUST00000038053.7 | 16 | 24448091 | 24992576 | 544486 | 1 | ENSMUST00 | 58796 | Lpp | ENSMUSG0C | false | . | . | . | . | . | . | chr16 | 24505234 | 24508236 | 3.55819 | 3.32179 | 1.83089 |
| chr5 | 110598394 | 110599170 | 1000 | Intron (ENSMUST00000040001.8 | 5 | 110603505 | 110621382 | 17878 | 1 | ENSMUST00 | -4335 | Galnt9 | ENSMUSG0C | false | . | . | . | . | . | . | chr5 | 110596218 | 110601588 | 5.80345 | 4.3461 | 3.67267 |
| chr4 | 118179390 | 118179923 | 1000 | Promoter (<=1kb) | 4 | 118136959 | 118179654 | 42696 | 2 | ENSMUST00 | 0 | Kdm4a | ENSMUSG0C | false | chr4 | 118178793 | 118180839 | 33.94791 | 109.87469 | 106.04007 | . | . | . | . | . | . |
| chr11 | 65787583 | 65788577 | 1000 | Promoter (<=1kb) | 11 | 65688243 | 65788297 | 100055 | 2 | ENSMUST00 | 0 | Map2k4 | ENSMUSG0C | false | chr11 | 65787211 | 65788689 | 29.92778 | 109.76675 | 105.9399 | chr11 | 65781427 | 65787632 | 4.44144 | 3.77766 | 2.54904 |
| chr11 | 65723951 | 65724479 | 1000 | Intron (ENSMUST00000046963.4 | 11 | 65690333 | 65719603 | 29271 | 2 | ENSMUST00 | -4349 | Map2k4 | ENSMUSG0C | false | . | . | . | . | . | . | chr11 | 65723685 | 65726922 | 5.48321 | 4.25086 | 3.40856 |
| chr2 | 121955744 | 121956621 | 1000 | Promoter (<=1kb) | 2 | 121956001 | 122013642 | 57642 | 1 | ENSMUST00 | 0 | Ctdspl2 | ENSMUSG0C | false | chr2 | 121955227 | 121957775 | 29.8354 | 90.64201 | 87.2842 | . | . | . | . | . | . |
| chr19 | 60580359 | 60581209 | 1000 | Promoter (<=1kb) | 19 | 60524696 | 60581023 | 56328 | 2 | ENSMUST00 | 0 | Cacul1 | ENSMUSG0C | false | chr19 | 60579332 | 60581756 | 28.32768 | 117.73571 | 113.66808 | . | . | . | . | . | . |
| chr5 | 110107648 | 110108424 | 1000 | Promoter (<=1kb) | 5 | 110103975 | 110108197 | 4223 | 2 | ENSMUST00 | 0 | Gtpbp6 | ENSMUSG0C | false | chr5 | 110106963 | 110108282 | 34.83228 | 119.81895 | 115.68042 | . | . | . | . | . | . |
| chr12 | 76369169 | 76371482 | 1000 | Promoter (<=1kb) | 12 | 76370266 | 76396950 | 26685 | 1 | ENSMUST00 | 0 | Zbtb1 | ENSMUSG0C | false | chr12 | 76368766 | 76371973 | 35.02911 | 135.1772 | 130.48503 | chr12 | 76366511 | 76369425 | 5.78919 | 4.52618 | 3.64125 |
| chr19 | 57360650 | 57361015 | 1000 | Promoter (<=1kb) | 19 | 57361009 | 57389594 | 28586 | 1 | ENSMUST00 | 0 | Fam160b1 | ENSMUSG0C | false | chr19 | 57359910 | 57360885 | 14.3181 | 33.00736 | 30.50354 | . | . | . | . | . | . |
| chr19 | 57362822 | 57363317 | 1000 | Promoter (1-2kb) | 19 | 57361009 | 57389594 | 28586 | 1 | ENSMUST00 | 1814 | Fam160b1 | ENSMUSG0C | false | chr19 | 57361054 | 57363157 | 39.29051 | 134.21745 | 129.56854 | chr19 | 57361796 | 57370102 | 6.5788 | 4.76166 | 4.30497 |
| chr14 | 72638294 | 72638886 | 1000 | Intron (ENSMUST00000089017.6 | 14 | 72574566 | 72652076 | 77511 | 2 | ENSMUST00 | 13190 | Fndc3a | ENSMUSG0C | false | . | . | . | . | . | . | chr14 | 72637984 | 72638741 | 3.68032 | 3.1887 | 1.94975 |
| chr1 | 157472076 | 157472564 | 1000 | Exon (ENSMUST00000046743.5/ | 1 | 157458580 | 157489060 | 30481 | 1 | ENSMUST00 | 13497 | BC026585 | ENSMUSG0C | false | . | . | . | . | . | . | chr1 | 157470354 | 157473935 | 4.0188 | 3.64281 | 2.19531 |
| chr1 | 157450068 | 157450498 | 1000 | Distal Intergenic | 1 | 157458580 | 157489060 | 30481 | 1 | ENSMUST00 | -8082 | BC026585 | ENSMUSG0C | false | . | . | . | . | . | . | chr1 | 157448774 | 157450474 | 4.21147 | 3.6708 | 2.40544 |
| chr3 | 116345704 | 116346219 | 1000 | Intron (ENSMUST00000106491.2 | 3 | 116400186 | 116423024 | 22839 | 2 | ENSMUST00 | 76805 | Cdc14a | ENSMUSG0C | true | . | . | . | . | . | . | chr3 | 116319808 | 116393211 | 5.81811 | 4.44924 | 3.68159 |
| chr3 | 116338577 | 116339445 | 1000 | Intron (ENSMUST00000106491.2 | 3 | 116400186 | 116423024 | 22839 | 2 | ENSMUST00 | 83579 | Cdc14a | ENSMUSG0C | true | . | . | . | . | . | . | chr3 | 116319808 | 116393211 | 5.81811 | 4.44924 | 3.68159 |
| chr3 | 116423412 | 116424364 | 1000 | Promoter (<=1kb) | 3 | 116272553 | 116423986 | 151434 | 2 | ENSMUST00 | 0 | Cdc14a | ENSMUSG0C | true | chr3 | 116421128 | 116425134 | 42.70261 | 142.41019 | 137.43451 | . | . | . | . | . | . |
| chr3 | 116382626 | 116382908 | 1000 | Intron (ENSMUST00000106491.2 | 3 | 116400186 | 116423024 | 22839 | 2 | ENSMUST00 | 40116 | Cdc14a | ENSMUSG0C | true | . | . | . | . | . | . | chr3 | 116319808 | 116393211 | 5.81811 | 4.44924 | 3.68159 |
| chr8 | 111536269 | 111536966 | 1000 | Promoter (<=1kb) | 8 | 111536640 | 111626030 | 89391 | 1 | ENSMUST00 | 0 | Znrf1 | ENSMUSG0C | false | chr8 | 111536021 | 111538448 | 42.22279 | 143.81401 | 138.77701 | . | . | . | . | . | . |
| chr8 | 111555223 | 111555690 | 1000 | Intron (ENSMUST00000172856.2 | 8 | 111561840 | 111606796 | 44957 | 1 | ENSMUST00 | -6150 | Znrf1 | ENSMUSG0C | false | . | . | . | . | . | . | chr8 | 111553685 | 111560312 | 4.99215 | 3.92636 | 3.02966 |
| chr1 | 156718329 | 156719347 | 1000 | Promoter (<=1kb) | 1 | 156678532 | 156719072 | 40541 | 2 | ENSMUST00 | 0 | Fam20b | ENSMUSG0C | false | chr1 | 156717889 | 156719597 | 39.03915 | 124.0004 | 119.73064 | chr1 | 156719267 | 156719726 | 4.69023 | 3.91576 | 2.77191 |
| chr15 | 77306395 | 77307308 | 1000 | Promoter (<=1kb) | 15 | 77078990 | 77307053 | 228064 | 2 | ENSMUST00 | 0 | Rbfox2 | ENSMUSG0C | false | chr15 | 77305678 | 77307532 | 34.5013 | 111.88224 | 107.9941 | chr15 | 77307250 | 77307717 | 4.97565 | 4.144 | 2.96909 |
| chr15 | 77037634 | 77038245 | 1000 | Intron (ENSMUST00000166179.2 | 15 | 77044729 | 77056298 | 11570 | 1 | ENSMUST00 | -6484 | Apol6 | ENSMUSG0C | false | . | . | . | . | . | . | chr15 | 77032360 | 77040092 | 6.45197 | 4.68595 | 4.18859 |
| chr15 | 77036717 | 77037359 | 1000 | Intron (ENSMUST00000166179.2 | 15 | 77044729 | 77056298 | 11570 | 1 | ENSMUST00 | -7370 | Apol6 | ENSMUSG0C | false | . | . | . | . | . | . | chr15 | 77032360 | 77040092 | 6.45197 | 4.68595 | 4.18859 |
| chr9 | 80173492 | 80174086 | 1000 | Intron (ENSMUST00000035889.9 | 9 | 80165227 | 80308097 | 142871 | 1 | ENSMUST00 | 8266 | Myo6 | ENSMUSG0C | true | . | . | . | . | . | . | chr9 | 80167116 | 80198379 | 5.43108 | 4.25219 | 3.37051 |
| chr8 | 111360503 | 111361363 | 1000 | Intron (ENSMUST00000159336.2 | 8 | 111345135 | 111366477 | 21343 | 2 | ENSMUST00 | 5114 | Fa2h | ENSMUSG0C | true | . | . | . | . | . | . | chr8 | 111356598 | 111381726 | 5.44823 | 4.21871 | 3.39859 |
| chr8 | 111363506 | 111363936 | 1000 | Promoter (2-3kb) | 8 | 111345135 | 111366477 | 21343 | 2 | ENSMUST00 | 2541 | Fa2h | ENSMUSG0C | true | . | . | . | . | . | . | chr8 | 111356598 | 111381726 | 5.44823 | 4.21871 | 3.39859 |
| chr16 | 22162594 | 22163759 | 1000 | Promoter (<=1kb) | 16 | 22059009 | 22163299 | 104291 | 2 | ENSMUST00 | 0 | Igf2bp2 | ENSMUSG0C | false | chr16 | 22160989 | 22164041 | 33.94791 | 109.87469 | 106.04007 | . | . | . | . | . | . |
| chr19 | 34885796 | 34886247 | 1000 | Distal Intergenic | 19 | 34810894 | 34879455 | 68562 | 2 | ENSMUST00 | -6342 | Pank1 | ENSMUSG0C | false | . | . | . | . | . | . | chr19 | 34882848 | 34887489 | 4.53196 | 3.80863 | 2.62397 |
| chr19 | 34877938 | 34879688 | 1000 | Promoter (<=1kb) | 19 | 34810894 | 34879455 | 68562 | 2 | ENSMUST00 | 0 | Pank1 | ENSMUSG0C | false | chr19 | 34877357 | 34879837 | 39.29051 | 134.21745 | 129.56854 | chr19 | 34879564 | 34881765 | 4.92595 | 4.10418 | 2.95341 |
| chr9 | 65021528 | 65021936 | 1000 | Promoter (<=1kb) | 9 | 64986985 | 65021717 | 34733 | 2 | ENSMUST00 | 0 | Ptplad1 | ENSMUSG0C | false | chr9 | 65020724 | 65022246 | 36.6511 | 131.68239 | 127.12846 | chr9 | 65019477 | 65022575 | 5.10548 | 4.12581 | 3.15588 |
| chr14 | 70153221 | 70153863 | 1000 | Promoter (<=1kb) | 14 | 70138177 | 70153811 | 15635 | 2 | ENSMUST00 | 0 | Ccar2 | ENSMUSG0C | false | chr14 | 70152889 | 70154166 | 32.83827 | 106.89391 | 103.14682 | chr14 | 70152127 | 70154720 | 4.22016 | 3.67086 | 2.40058 |
| chr12 | 99391850 | 99392350 | 1000 | Promoter (<=1kb) | 12 | 99195094 | 99393072 | 197979 | 2 | ENSMUST00 | 722 | Foxn3 | ENSMUSG0C | false | chr12 | 99391404 | 99393876 | 34.17318 | 106.15081 | 102.42828 | . | . | . | . | . | . |
| chr12 | 99378361 | 99378786 | 1000 | Intron (ENSMUST00000046859.5 | 12 | 99195094 | 99393072 | 197979 | 2 | ENSMUST00 | 14286 | Foxn3 | ENSMUSG0C | false | . | . | . | . | . | . | chr12 | 99365961 | 99383588 | 5.3389 | 4.19815 | 3.29478 |
| chr12 | 99392559 | 99393496 | 1000 | Promoter (<=1kb) | 12 | 99195094 | 99393072 | 197979 | 2 | ENSMUST00 | 0 | Foxn3 | ENSMUSG0C | false | chr12 | 99391404 | 99393876 | 34.17318 | 106.15081 | 102.42828 | . | . | . | . | . | . |
| chr14 | 70074526 | 70075902 | 1000 | Promoter (1-2kb) | 14 | 70077445 | 70080157 | 2713 | 1 | ENSMUST00 | -1543 | Egr3 | ENSMUSG0C | false | chr14 | 70074380 | 70079100 | 25.84672 | 90.2271 | 86.88143 | chr14 | 70074142 | 70080178 | 6.65496 | 4.83303 | 4.35913 |
| chr14 | 69997517 | 69997917 | 1000 | Intron (ENSMUST00000022678.3 | 14 | 70077445 | 70080157 | 2713 | 1 | ENSMUST00 | -79528 | Egr3 | ENSMUSG0C | false | . | . | . | . | . | . | chr14 | 69996831 | 69999361 | 4.72732 | 3.82211 | 2.80906 |
| chr14 | 70001690 | 70002127 | 1000 | Intron (ENSMUST00000022678.3 | 14 | 70077445 | 70080157 | 2713 | 1 | ENSMUST00 | -75318 | Egr3 | ENSMUSG0C | false | . | . | . | . | . | . | chr14 | 70000686 | 70003126 | 5.81773 | 4.49428 | 3.71712 |
| chr14 | 69994756 | 69995423 | 1000 | Intron (ENSMUST00000022678.3 | 14 | 70077445 | 70080157 | 2713 | 1 | ENSMUST00 | -82022 | Egr3 | ENSMUSG0C | false | . | . | . | . | . | . | chr14 | 69994282 | 69995807 | 3.71747 | 3.42819 | 1.96605 |
| chr3 | 84135336 | 84135719 | 1000 | Intron (ENSMUST00000047368.6 | 3 | 84087934 | 84155786 | 67853 | 2 | ENSMUST00 | 20067 | Mnd1 | ENSMUSG0C | false | . | . | . | . | . | . | chr3 | 84135405 | 84135906 | 3.06074 | 3.05624 | 1.45685 |
| chr19 | 4756261 | 4756766 | 1000 | Promoter (<=1kb) | 19 | 4756543 | 4762876 | 6334 | 1 | ENSMUST00 | 0 | Rbm4b | ENSMUSG0C | false | chr19 | 4755824 | 4757634 | 30.90191 | 98.42355 | 94.89227 | . | . | . | . | . | . |
| chr3 | 84039501 | 84040482 | 1000 | Promoter (<=1kb) | 3 | 83898287 | 84040161 | 141875 | 2 | ENSMUST00 | 0 | D930015E06 | ENSMUSG0C | false | chr3 | 84038826 | 84040973 | 29.92778 | 109.76675 | 105.9399 | chr3 | 84038621 | 84040983 | 4.42526 | 3.71524 | 2.57528 |
| chr17 | 88627157 | 88627459 | 1000 | Promoter (<=1kb) | 17 | 88627364 | 88637634 | 10271 | 1 | ENSMUST00 | 0 | Ston1 | ENSMUSG0C | false | chr17 | 88626424 | 88629225 | 13.60354 | 37.21757 | 34.66072 | . | . | . | . | . | . |
| chr17 | 88597263 | 88597961 | 1000 | Promoter (<=1kb) | 17 | 88597684 | 88635829 | 38146 | 1 | ENSMUST00 | 0 | Ston1 | ENSMUSG0C | false | chr17 | 88597214 | 88598641 | 21.40274 | 64.72344 | 61.81032 | chr17 | 88595166 | 88604773 | 4.12794 | 3.59375 | 2.29889 |
| chr19 | 23140670 | 23141749 | 1000 | Promoter (<=1kb) | 19 | 23141226 | 23166911 | 25686 | 1 | ENSMUST00 | 0 | Klf9 | ENSMUSG0C | false | chr19 | 23140239 | 23143387 | 30.78991 | 105.27288 | 101.56436 | . | . | . | . | . | . |
| chr18 | 61400121 | 61400884 | 1000 | Promoter (<=1kb) | 18 | 61298136 | 61400431 | 102296 | 2 | ENSMUST00 | 0 | Ppargc1b | ENSMUSG0C | false | chr18 | 61398143 | 61401028 | 41.07278 | 140.67711 | 135.76277 | chr18 | 61399853 | 61401196 | 3.31531 | 3.19643 | 1.6024 |
| chr8 | 126961038 | 126961911 | 1000 | Intron (ENSMUST00000045994.6 | 8 | 126947175 | 126971079 | 23905 | 2 | ENSMUST00 | 9168 | Rbm34 | ENSMUSG0C | false | . | . | . | . | . | . | chr8 | 126957410 | 126968218 | 5.46408 | 4.25341 | 3.40496 |
| chr7 | 12977857 | 12978381 | 1000 | Promoter (<=1kb) | 7 | 12977893 | 12981463 | 3571 | 1 | ENSMUST00 | 0 | Zfp446 | ENSMUSG0C | false | chr7 | 12977339 | 12978767 | 32.46003 | 103.17023 | 99.51957 | . | . | . | . | . | . |
| chr17 | 80638081 | 80638460 | 1000 | Intron (ENSMUST00000025089.7 | 17 | 80523550 | 80563834 | 40285 | 2 | ENSMUST00 | -74248 | Cdkl4 | ENSMUSG0C | false | . | . | . | . | . | . | chr17 | 80637441 | 80638941 | 5.15947 | 4.11315 | 3.17036 |
| chr8 | 125971387 | 125971751 | 1000 | Distal Intergenic | 8 | 125995102 | 126030683 | 35582 | 1 | ENSMUST00 | -23351 | Kcnk1 | ENSMUSG0C | false | . | . | . | . | . | . | chr8 | 125971363 | 125972351 | 3.9152 | 3.57783 | 2.10373 |
| chr18 | 80285268 | 80285947 | 1000 | Intron (ENSMUST00000131780.2 | 18 | 80262840 | 80263485 | 646 | 1 | ENSMUST00 | 22429 | Pqlc1 | ENSMUSG0C | false | . | . | . | . | . | . | chr18 | 80280088 | 80289295 | 5.13437 | 4.0721 | 3.14211 |
| chr5 | 150673523 | 150674442 | 1000 | Promoter (<=1kb) | 5 | 150673739 | 150810690 | 136952 | 1 | ENSMUST00 | 0 | Pds5b | ENSMUSG0C | false | chr5 | 150673763 | 150674767 | 27.54716 | 98.2761 | 94.74957 | . | . | . | . | . | . |
| chr5 | 131241622 | 131241979 | 1000 | Intron (ENSMUST00000086023.6 | 5 | 131085272 | 131306915 | 221644 | 2 | ENSMUST00 | 64936 | Wbscr17 | ENSMUSG0C | false | . | . | . | . | . | . | chr5 | 131241296 | 131242890 | 4.15873 | 3.60701 | 2.31729 |
| chr5 | 131089840 | 131090216 | 1000 | Intron (ENSMUST00000086023.6 | 5 | 131085272 | 131306915 | 221644 | 2 | ENSMUST00 | 216699 | Wbscr17 | ENSMUSG0C | false | . | . | . | . | . | . | chr5 | 131088640 | 131090696 | 5.27695 | 3.95534 | 3.26865 |
| chrX | 102644009 | 102644505 | 1000 | Promoter (<=1kb) | 20 | 102513975 | 102644246 | 130272 | 2 | ENSMUST00 | 0 | Phka1 | ENSMUSG0C | false | chrX | 102643741 | 102645047 | 21.59801 | 56.60135 | 53.80005 | . | . | . | . | . | . |
| chr1 | 93511829 | 93512463 | 1000 | Promoter (<=1kb) | 1 | 93512079 | 93621976 | 109898 | 1 | ENSMUST00 | 0 | Farp2 | ENSMUSG0C | false | chr1 | 93511315 | 93512892 | 27.20707 | 96.65539 | 93.16586 | . | . | . | . | . | . |
| chr2 | 84635068 | 84636030 | 1000 | Promoter (2-3kb) | 2 | 84624378 | 84638105 | 13728 | 2 | ENSMUST00 | 2075 | Ctnnd1 | ENSMUSG0C | true | . | . | . | . | . | . | chr2 | 84621405 | 84648911 | 6.04389 | 4.54416 | 3.87093 |
| chr3 | 75930714 | 75931247 | 1000 | Intron (ENSMUST00000117242.2 | 3 | 75909856 | 75956650 | 46795 | 2 | ENSMUST00 | 25403 | Golim4 | ENSMUSG0C | true | . | . | . | . | . | . | chr3 | 75908736 | 75932827 | 4.90077 | 3.96047 | 2.93621 |
| chr3 | 75918161 | 75918539 | 1000 | Intron (ENSMUST00000117242.2 | 3 | 75894853 | 75902446 | 7594 | 2 | ENSMUST00 | -15716 | Golim4 | ENSMUSG0C | true | . | . | . | . | . | . | chr3 | 75908736 | 75932827 | 4.90077 | 3.96047 | 2.93621 |
| chr3 | 75860720 | 75861216 | 1000 | Distal Intergenic | 3 | 75876479 | 75879008 | 2530 | 2 | ENSMUST00 | 17792 | Golim4 | ENSMUSG0C | true | . | . | . | . | . | . | chr3 | 75860318 | 75861683 | 4.14398 | 3.30347 | 2.34904 |
| chr9 | 46110664 | 46111287 | 1000 | Intron (ENSMUST00000126865.2 | 9 | 46123237 | 46202885 | 79649 | 1 | ENSMUST00 | -11950 | Sik3 | ENSMUSG0C | false | . | . | . | . | . | . | chr9 | 46103496 | 46117248 | 5.7315 | 4.352 | 3.62195 |
| chr2 | 119476915 | 119478357 | 1000 | Promoter (<=1kb) | 2 | 119373042 | 119477687 | 104646 | 2 | ENSMUST00 | 0 | Ino80 | ENSMUSG0C | false | chr2 | 119476247 | 119478890 | 45.03415 | 147.57545 | 142.38377 | . | . | . | . | . | . |
| chr16 | 37869781 | 37870102 | 1000 | Promoter (1-2kb) | 16 | 37868389 | 37888858 | 20470 | 1 | ENSMUST00 | 1393 | Lrrc58 | ENSMUSG0C | false | chr16 | 37867771 | 37870669 | 34.00884 | 130.01486 | 125.52227 | chr16 | 37870054 | 37882292 | 5.93789 | 4.46579 | 3.79659 |
| chr1 | 93176476 | 93176957 | 1000 | Distal Intergenic | 1 | 93151349 | 93160948 | 9600 | 2 | ENSMUST00 | -15529 | 2310007B03 | ENSMUSG0C | false | chr1 | 93176368 | 93176997 | 12.3333 | 26.50278 | 24.08123 | chr1 | 93174946 | 93177186 | 4.65782 | 3.73478 | 2.76801 |
| chr15 | 76457181 | 76457795 | 1000 | Promoter (<=1kb) | 15 | 76457438 | 76459468 | 2031 | 1 | ENSMUST00 | 0 | Scx | ENSMUSG0C | false | chr15 | 76457167 | 76458476 | 25.04793 | 77.4957 | 74.38671 | chr15 | 76455205 | 76469027 | 6.45319 | 4.6614 | 4.20973 |
| chr15 | 76456207 | 76456746 | 1000 | Promoter (<=1kb) | 15 | 76457438 | 76459468 | 2031 | 1 | ENSMUST00 | -692 | Scx | ENSMUSG0C | false | . | . | . | . | . | . | chr15 | 76455205 | 76469027 | 6.45319 | 4.6614 | 4.20973 |
| chr17 | 47504900 | 47505238 | 1000 | Promoter (<=1kb) | 17 | 47505051 | 47598070 | 93020 | 1 | ENSMUST00 | 0 | Ccnd3 | ENSMUSG0C | false | chr17 | 47504679 | 47505280 | 11.4883 | 23.92001 | 21.53215 | chr17 | 47504225 | 47505666 | 6.10213 | 4.69576 | 3.89988 |
| chr12 | 86883783 | 86885222 | 1000 | Promoter (<=1kb) | 12 | 86880703 | 86884814 | 4112 | 2 | ENSMUST00 | 0 | Irf2bpl | ENSMUSG0C | false | chr12 | 86881379 | 86885421 | 34.90791 | 116.7856 | 112.7497 | chr12 | 86885110 | 86901194 | 5.68305 | 4.41688 | 3.56559 |
| chr6 | 91546206 | 91546798 | 1000 | Distal Intergenic | 6 | 91516047 | 91517481 | 1435 | 1 | ENSMUST00 | 30160 | Lsm3 | ENSMUSG0C | false | . | . | . | . | . | . | chr6 | 91545987 | 91549993 | 4.11226 | 3.49884 | 2.31598 |
| chr6 | 91566390 | 91566762 | 1000 | Distal Intergenic | 6 | 91516047 | 91517481 | 1435 | 1 | ENSMUST00 | 50344 | Lsm3 | ENSMUSG0C | false | . | . | . | . | . | . | chr6 | 91561703 | 91573814 | 5.57785 | 4.24835 | 3.49935 |
| chr1 | 92831227 | 92832006 | 1000 | Promoter (<=1kb) | 1 | 92831645 | 92860779 | 29135 | 1 | ENSMUST00 | 0 | Gpc1 | ENSMUSG0C | false | chr1 | 92831029 | 92833403 | 25.56571 | 75.64988 | 72.57267 | chr1 | 92830851 | 92835428 | 4.72324 | 3.90122 | 2.81783 |
| chr14 | 69284095 | 69285278 | 1000 | Promoter (<=1kb) | 14 | 69241848 | 69285112 | 43265 | 2 | ENSMUST00 | 0 | Slc25a37 | ENSMUSG0C | false | chr14 | 69283729 | 69285101 | 23.43614 | 63.09834 | 60.20888 | chr14 | 69283344 | 69284243 | 4.23744 | 3.7655 | 2.36907 |
| chr9 | 80066585 | 80067977 | 1000 | Promoter (<=1kb) | 9 | 80066903 | 80094484 | 27582 | 1 | ENSMUST00 | 0 | Senp6 | ENSMUSG0C | false | chr9 | 80066534 | 80068862 | 35.62435 | 116.99458 | 112.94973 | . | . | . | . | . | . |
| chr17 | 5593088 | 5593587 | 1000 | Intron (ENSMUST00000089185.4 | 17 | 5492600 | 5753891 | 261292 | 1 | ENSMUST00 | 100489 | Zdhhc14 | ENSMUSG0C | true | . | . | . | . | . | . | chr17 | 5587797 | 5599687 | 5.93548 | 4.48203 | 3.77591 |
| chr17 | 5492395 | 5492858 | 1000 | Promoter (<=1kb) | 17 | 5492600 | 5753891 | 261292 | 1 | ENSMUST00 | 0 | Zdhhc14 | ENSMUSG0C | true | chr17 | 5491522 | 5494381 | 31.29462 | 124.76181 | 120.45966 | . | . | . | . | . | . |
| chr17 | 5635275 | 5635629 | 1000 | Intron (ENSMUST00000089185.4 | 17 | 5492600 | 5753891 | 261292 | 1 | ENSMUST00 | 142676 | Zdhhc14 | ENSMUSG0C | true | . | . | . | . | . | . | chr17 | 5632857 | 5639534 | 4.55891 | 3.79533 | 2.67999 |
| chr6 | 113075987 | 113077645 | 1000 | Promoter (<=1kb) | 6 | 113077365 | 113107561 | 30197 | 1 | ENSMUST00 | 0 | Setd5 | ENSMUSG0C | false | chr6 | 113076843 | 113079361 | 35.66819 | 113.91924 | 109.96352 | . | . | . | . | . | . |
| chr12 | 85621696 | 85622039 | 1000 | Intron (ENSMUST00000177587.2 | 12 | 85600018 | 85639878 | 39861 | 1 | ENSMUST00 | 21679 | Jdp2 | ENSMUSG0C | true | . | . | . | . | . | . | chr12 | 85617726 | 85631513 | 6.15226 | 4.50635 | 3.95746 |
| chr12 | 85612597 | 85613065 | 1000 | Intron (ENSMUST00000177587.2 | 12 | 85600018 | 85639878 | 39861 | 1 | ENSMUST00 | 12580 | Jdp2 | ENSMUSG0C | true | . | . | . | . | . | . | chr12 | 85599922 | 85615323 | 5.04005 | 3.93431 | 3.0761 |
| chr12 | 85598670 | 85600041 | 1000 | Promoter (<=1kb) | 12 | 85599105 | 85639878 | 40774 | 1 | ENSMUST00 | 0 | Jdp2 | ENSMUSG0C | true | chr12 | 85598428 | 85600604 | 25.16654 | 87.04672 | 83.76057 | chr12 | 85599922 | 85615323 | 5.04005 | 3.93431 | 3.0761 |
| chr12 | 85608000 | 85608396 | 1000 | 5' UTR | 12 | 85600018 | 85639878 | 39861 | 1 | ENSMUST00 | 7983 | Jdp2 | ENSMUSG0C | true | . | . | . | . | . | . | chr12 | 85599922 | 85615323 | 5.04005 | 3.93431 | 3.0761 |
| chr11 | 116237973 | 116238420 | 1000 | Promoter (<=1kb) | 11 | 116220559 | 116238077 | 17519 | 2 | ENSMUST00 | 0 | Evpl | ENSMUSG0C | false | chr11 | 116237752 | 116238512 | 7.50973 | 14.65382 | 12.39649 | chr11 | 116237636 | 116238717 | 3.94596 | 3.56787 | 2.1394 |
| chr18 | 24709228 | 24709729 | 1000 | Promoter (<=1kb) | 18 | 24709445 | 25133500 | 424056 | 1 | ENSMUST00 | 0 | Fhod3 | ENSMUSG0C | false | chr18 | 24708503 | 24710423 | 18.36477 | 56.64417 | 53.84045 | chr18 | 24708241 | 24710605 | 4.62257 | 3.82887 | 2.73802 |
| chr9 | 37355699 | 37356023 | 1000 | Distal Intergenic | 9 | 37275835 | 37348392 | 72558 | 2 | ENSMUST00 | -7308 | Ccdc15 | ENSMUSG0C | false | . | . | . | . | . | . | chr9 | 37354560 | 37366659 | 5.20726 | 4.11038 | 3.19089 |
| chr6 | 90736078 | 90736940 | 1000 | Promoter (<=1kb) | 6 | 90690662 | 90736687 | 46026 | 2 | ENSMUST00 | 0 | Iqsec1 | ENSMUSG0C | false | chr6 | 90735442 | 90737552 | 35.51585 | 112.61008 | 108.7009 | chr6 | 90695436 | 90737892 | 6.93129 | 4.85273 | 4.61297 |
| chr9 | 44233593 | 44234348 | 1000 | Promoter (<=1kb) | 9 | 44149262 | 44234046 | 84785 | 2 | ENSMUST00 | 0 | Cbl | ENSMUSG0C | false | chr9 | 44232995 | 44234966 | 36.67561 | 120.95901 | 116.78313 | . | . | . | . | . | . |
| chr8 | 94532876 | 94533203 | 1000 | Promoter (<=1kb) | 8 | 94532990 | 94570529 | 37540 | 1 | ENSMUST00 | 0 | Cpne2 | ENSMUSG0C | false | chr8 | 94532544 | 94533595 | 23.60729 | 67.74274 | 64.78755 | chr8 | 94532359 | 94534704 | 4.65853 | 3.9104 | 2.72305 |
| chr19 | 10604164 | 10604628 | 1000 | Promoter (<=1kb) | 19 | 10592200 | 10604258 | 12059 | 2 | ENSMUST00 | 0 | Dak | ENSMUSG0C | false | chr19 | 10603045 | 10606602 | 41.22328 | 133.19821 | 128.59248 | chr19 | 10601470 | 10605052 | 4.9759 | 3.96533 | 3.02446 |
| chr19 | 10603569 | 10604045 | 1000 | Promoter (<=1kb) | 19 | 10592200 | 10604258 | 12059 | 2 | ENSMUST00 | 213 | Dak | ENSMUSG0C | false | chr19 | 10603045 | 10606602 | 41.22328 | 133.19821 | 128.59248 | chr19 | 10601470 | 10605052 | 4.9759 | 3.96533 | 3.02446 |
| chr17 | 6106383 | 6107537 | 1000 | Promoter (<=1kb) | 17 | 6106437 | 6138792 | 32356 | 1 | ENSMUST00 | 0 | Tulp4 | ENSMUSG0C | false | chr17 | 6106150 | 6108092 | 31.00994 | 95.62833 | 92.16279 | . | . | . | . | . | . |
| chr8 | 117256382 | 117257218 | 1000 | Promoter (<=1kb) | 8 | 117257019 | 117461503 | 204485 | 1 | ENSMUST00 | 0 | Cmip | ENSMUSG0C | false | chr8 | 117256201 | 117258587 | 38.6007 | 121.10697 | 116.92481 | . | . | . | . | . | . |
| chrX | 99470552 | 99470929 | 1000 | Promoter (<=1kb) | 20 | 99465738 | 99470687 | 4950 | 2 | ENSMUST00 | 0 | Pja1 | ENSMUSG0C | false | chrX | 99469695 | 99470948 | 24.3552 | 66.39809 | 63.45943 | . | . | . | . | . | . |
| chr11 | 4186567 | 4187045 | 1000 | Promoter (<=1kb) | 11 | 4186789 | 4214885 | 28097 | 1 | ENSMUST00 | 0 | Tbc1d10a | ENSMUSG0C | false | chr11 | 4186221 | 4188124 | 38.58521 | 122.14608 | 117.93867 | . | . | . | . | . | . |
| chr17 | 26397357 | 26397795 | 1000 | Distal Intergenic | 17 | 26414829 | 26441795 | 26967 | 1 | ENSMUST00 | -17034 | Neurl1b | ENSMUSG0C | false | . | . | . | . | . | . | chr17 | 26396231 | 26397904 | 4.48303 | 3.63606 | 2.61673 |
| chr6 | 90369326 | 90369814 | 1000 | Promoter (<=1kb) | 6 | 90369492 | 90385084 | 15593 | 1 | ENSMUST00 | 0 | Zxdc | ENSMUSG0C | false | chr6 | 90369051 | 90371189 | 40.8912 | 139.24503 | 134.39108 | . | . | . | . | . | . |
| chr1 | 90538259 | 90538627 | 1000 | Distal Intergenic | 1 | 90602981 | 90613341 | 10361 | 1 | ENSMUST00 | -64354 | Cops8 | ENSMUSG0C | false | . | . | . | . | . | . | chr1 | 90535451 | 90539952 | 4.47194 | 3.80284 | 2.61432 |
| chr5 | 105058898 | 105059273 | 1000 | Intron (ENSMUST00000031235.7 | 5 | 105014927 | 105051047 | 36121 | 2 | ENSMUST00 | -7852 | Gbp8 | ENSMUSG0C | false | . | . | . | . | . | . | chr5 | 105058800 | 105059357 | 2.51983 | 2.66008 | 1.02466 |
| chr14 | 65358270 | 65359055 | 1000 | Promoter (<=1kb) | 14 | 65358676 | 65398930 | 40255 | 1 | ENSMUST00 | 0 | Zfp395 | ENSMUSG0C | false | chr14 | 65357764 | 65359579 | 33.41831 | 117.26761 | 113.21259 | chr14 | 65358729 | 65364088 | 5.27047 | 4.22236 | 3.22306 |
| chr14 | 65343547 | 65344240 | 1000 | Distal Intergenic | 14 | 65358676 | 65398930 | 40255 | 1 | ENSMUST00 | -14436 | Zfp395 | ENSMUSG0C | false | chr14 | 65343139 | 65343787 | 20.26563 | 55.95697 | 53.16479 | chr14 | 65342449 | 65346186 | 4.72594 | 3.90525 | 2.78743 |
| chr10 | 83543754 | 83544370 | 1000 | Promoter (<=1kb) | 10 | 83543941 | 83596469 | 52529 | 1 | ENSMUST00 | 0 | A230046K03 | ENSMUSG0C | false | chr10 | 83543505 | 83544857 | 26.52689 | 93.43019 | 90.01563 | . | . | . | . | . | . |
| chr5 | 103469471 | 103469828 | 1000 | Intron (ENSMUST00000048957.9 | 5 | 103425192 | 103598359 | 173168 | 1 | ENSMUST00 | 44280 | Ptpn13 | ENSMUSG0C | false | . | . | . | . | . | . | chr5 | 103469028 | 103470676 | 3.66164 | 3.4441 | 1.89572 |
| chr5 | 103424610 | 103425408 | 1000 | Promoter (<=1kb) | 5 | 103425192 | 103598359 | 173168 | 1 | ENSMUST00 | 0 | Ptpn13 | ENSMUSG0C | false | chr5 | 103424449 | 103426532 | 35.36919 | 136.9064 | 132.15405 | chr5 | 103423670 | 103424731 | 6.06667 | 4.62214 | 3.89016 |
| chr5 | 103376770 | 103377408 | 1000 | Distal Intergenic | 5 | 103425192 | 103598359 | 173168 | 1 | ENSMUST00 | -47784 | Ptpn13 | ENSMUSG0C | false | chr5 | 103376727 | 103377469 | 5.59421 | 10.48996 | 8.30244 | chr5 | 103376057 | 103378714 | 5.86262 | 4.48493 | 3.69682 |

| | | | | | | | | | | | | | | | | | | | | | | | | | | |
|---|---|---|---|---|---|---|---|---|---|---|---|---|---|---|---|---|---|---|---|---|---|---|---|---|---|---|
| chr5 | 103365328 | 103365753 | 1000 | Distal Intergenic | 5 | 103425192 | 103598359 | 173168 | 1 | ENSMUST00 | -59439 | Ptpn13 | ENSMUSG0C | false | . | . | . | . | . | . | chr5 | 103364664 | 103365577 | 3.15401 | 3.08067 | 1.48598 |
| chr10 | 83301725 | 83302245 | 1000 | Promoter (2-3kb) | 10 | 83301016 | 83304375 | 3360 | 2 | ENSMUST00 | 2130 | Slc41a2 | ENSMUSG0C | true | . | . | . | . | . | . | chr10 | 83290999 | 83303347 | 4.92593 | 4.03489 | 2.94266 |
| chr10 | 83321826 | 83322140 | 1000 | Promoter (<=1kb) | 10 | 83316215 | 83322883 | 6669 | 2 | ENSMUST00 | 743 | Slc41a2 | ENSMUSG0C | true | . | . | . | . | . | . | chr10 | 83313612 | 83325459 | 3.72419 | 3.41803 | 1.99556 |
| chr10 | 83291755 | 83292234 | 1000 | Intron (ENSMUST00000039956.5 | 10 | 83301016 | 83304375 | 3360 | 2 | ENSMUST00 | 12141 | Slc41a2 | ENSMUSG0C | true | . | . | . | . | . | . | chr10 | 83290999 | 83303347 | 4.92593 | 4.03489 | 2.94266 |
| chr10 | 83337187 | 83338084 | 1000 | Promoter (<=1kb) | 10 | 83230848 | 83337882 | 107035 | 2 | ENSMUST00 | 0 | Slc41a2 | ENSMUSG0C | true | chr10 | 83336897 | 83338171 | 18.02578 | 59.25625 | 56.42029 | chr10 | 83335897 | 83338828 | 4.34943 | 3.82014 | 2.45676 |
| chr4 | 108459110 | 108460803 | 1000 | Promoter (<=1kb) | 4 | 108459426 | 108479153 | 19728 | 1 | ENSMUST00 | 0 | Zcchc11 | ENSMUSG0C | false | chr4 | 108458983 | 108461770 | 33.7938 | 119.00099 | 114.88904 | chr4 | 108458192 | 108459358 | 4.11475 | 3.66334 | 2.26967 |
| chr10 | 122770926 | 122771290 | 1000 | Intron (ENSMUST00000162853.2 | 10 | 122757198 | 122811471 | 54274 | 1 | ENSMUST00 | 13729 | Ppm1h | ENSMUSG0C | false | . | . | . | . | . | . | chr10 | 122767763 | 122772724 | 5.304 | 4.23982 | 3.25539 |
| chr11 | 3331964 | 3332672 | 1000 | Promoter (<=1kb) | 11 | 3332072 | 3333679 | 1608 | 1 | ENSMUST00 | 0 | Pik3ip1 | ENSMUSG0C | false | chr11 | 3331738 | 3332542 | 18.91326 | 49.58984 | 46.88035 | chr11 | 3330471 | 3335666 | 4.18582 | 3.64114 | 2.34797 |
| chr11 | 102555680 | 102556496 | 1000 | Promoter (<=1kb) | 11 | 102475915 | 102556199 | 80285 | 2 | ENSMUST00 | 0 | Gpatch8 | ENSMUSG0C | false | chr11 | 102554733 | 102556749 | 43.93484 | 147.93027 | 142.7272 | . | . | . | . | . | . |
| chr3 | 65527724 | 65528575 | 1000 | Promoter (<=1kb) | 3 | 65528410 | 65555145 | 26736 | 1 | ENSMUST00 | 0 | Tiparp | ENSMUSG0C | false | chr3 | 65527589 | 65530267 | 26.03857 | 105.07202 | 101.37238 | . | . | . | . | . | . |
| chr17 | 66076304 | 66077584 | 1000 | Promoter (<=1kb) | 17 | 65967501 | 66077046 | 109546 | 2 | ENSMUST00 | 0 | Ankrd12 | ENSMUSG0C | false | chr17 | 66075855 | 66077884 | 33.36056 | 104.74094 | 101.05196 | chr17 | 66077570 | 66077948 | 5.11581 | 4.12178 | 3.15794 |
| chr18 | 44828371 | 44829121 | 1000 | Promoter (<=1kb) | 18 | 44828665 | 44889720 | 61056 | 1 | ENSMUST00 | 0 | Ythdc2 | ENSMUSG0C | false | chr18 | 44827764 | 44829696 | 24.39013 | 102.54827 | 98.91666 | . | . | . | . | . | . |
| chr8 | 84388771 | 84389099 | 1000 | Promoter (<=1kb) | 8 | 84388440 | 84519039 | 130600 | 1 | ENSMUST00 | 332 | Cacna1a | ENSMUSG0C | false | . | . | . | . | . | . | chr8 | 84387638 | 84389484 | 4.51989 | 3.8849 | 2.60399 |
| chr8 | 84415152 | 84415641 | 1000 | Promoter (<=1kb) | 8 | 84415364 | 84640246 | 224883 | 1 | ENSMUST00 | 0 | Cacna1a | ENSMUSG0C | false | chr8 | 84414241 | 84415968 | 20.27628 | 60.14116 | 57.29097 | chr8 | 84414464 | 84416306 | 5.1587 | 3.95274 | 3.15369 |
| chr5 | 96997038 | 96998723 | 1000 | Promoter (<=1kb) | 5 | 96997689 | 97069649 | 71961 | 1 | ENSMUST00 | 0 | Bmp2k | ENSMUSG0C | false | chr5 | 96996701 | 96998913 | 21.26469 | 95.5017 | 92.03861 | chr5 | 96998713 | 97006670 | 4.23322 | 3.63518 | 2.41115 |
| chr11 | 102466983 | 102467514 | 1000 | Promoter (<=1kb) | 11 | 102456472 | 102467199 | 10728 | 2 | ENSMUST00 | 0 | Itga2b | ENSMUSG0C | false | chr11 | 102466964 | 102467901 | 14.99812 | 35.51057 | 32.97813 | chr11 | 102465676 | 102468730 | 5.13924 | 4.12194 | 3.12764 |
| chr17 | 34591408 | 34592727 | 1000 | Promoter (<=1kb) | 17 | 34592263 | 34597400 | 5138 | 1 | ENSMUST00 | 0 | Pbx2 | ENSMUSG0C | true | chr17 | 34592255 | 34596534 | 34.71595 | 106.06588 | 102.34386 | chr17 | 34586035 | 34591596 | 5.00173 | 4.02952 | 3.00733 |
| chr13 | 55487504 | 55488279 | 1000 | Promoter (<=1kb) | 13 | 55473429 | 55488111 | 14683 | 2 | ENSMUST00 | 0 | Dbn1 | ENSMUSG0C | false | chr13 | 55487090 | 55488351 | 18.33657 | 60.6815 | 57.82544 | chr13 | 55486940 | 55488662 | 4.42569 | 3.83808 | 2.53523 |
| chr11 | 114822865 | 114823258 | 1000 | Distal Intergenic | 11 | 114798924 | 114806745 | 7822 | 1 | ENSMUST00 | 23942 | Gpr142 | ENSMUSG0C | false | . | . | . | . | . | . | chr11 | 114818293 | 114824576 | 5.23544 | 4.14249 | 3.22448 |
| chr9 | 107707611 | 107708086 | 1000 | Promoter (2-3kb) | 9 | 107681502 | 107710475 | 28974 | 2 | ENSMUST00 | 2389 | Sema3f | ENSMUSG0C | false | chr9 | 107707503 | 107710482 | 34.42823 | 105.56979 | 101.85753 | chr9 | 107693726 | 107710535 | 7.184 | 5.04064 | 4.79532 |
| chr9 | 107699312 | 107699726 | 1000 | Intron (ENSMUST00000080560.3 | 9 | 107681502 | 107710475 | 28974 | 2 | ENSMUST00 | 10749 | Sema3f | ENSMUSG0C | false | chr9 | 107698856 | 107699487 | 6.93923 | 13.23942 | 11.00697 | chr9 | 107693726 | 107710535 | 7.184 | 5.04064 | 4.79532 |
| chr11 | 102430147 | 102430516 | 1000 | Promoter (<=1kb) | 11 | 102430315 | 102436809 | 6495 | 1 | ENSMUST00 | 0 | Grn | ENSMUSG0C | false | chr11 | 102429983 | 102431352 | 32.28179 | 97.83163 | 94.31397 | . | . | . | . | . | . |
| chr5 | 96160974 | 96162845 | 1000 | Promoter (<=1kb) | 5 | 96070333 | 96161990 | 91658 | 2 | ENSMUST00 | 0 | Cnot6l | ENSMUSG0C | false | chr5 | 96160231 | 96163458 | 32.11694 | 110.747 | 106.89115 | . | . | . | . | . | . |
| chr15 | 74516141 | 74516499 | 1000 | Promoter (<=1kb) | 15 | 74516195 | 74589464 | 73270 | 1 | ENSMUST00 | 0 | Bai1 | ENSMUSG0C | false | chr15 | 74516050 | 74517226 | 12.58327 | 33.30664 | 30.79935 | chr15 | 74515885 | 74520114 | 5.85295 | 4.38782 | 3.72551 |
| chr15 | 74449532 | 74449859 | 1000 | Distal Intergenic | 15 | 74516195 | 74589464 | 73270 | 1 | ENSMUST00 | -66336 | Bai1 | ENSMUSG0C | false | . | . | . | . | . | . | chr15 | 74448047 | 74450640 | 4.79221 | 3.91535 | 2.85034 |
| chr14 | 78731984 | 78733018 | 1000 | Distal Intergenic | 14 | 78569609 | 78725089 | 155481 | 2 | ENSMUST00 | -6896 | Dgkh | ENSMUSG0C | false | chr14 | 78731087 | 78733615 | 20.63592 | 96.67229 | 93.18274 | . | . | . | . | . | . |
| chr2 | 69104737 | 69105086 | 1000 | Exon (ENSMUST00000028426.3/ | 2 | 69135800 | 69189330 | 53531 | 1 | ENSMUST00 | -30714 | Nostrin | ENSMUSG0C | false | . | . | . | . | . | . | chr2 | 69095809 | 69110415 | 4.4714 | 3.76397 | 2.5933 |
| chr2 | 69084377 | 69084868 | 1000 | Intron (ENSMUST00000028426.3 | 2 | 69135800 | 69189330 | 53531 | 1 | ENSMUST00 | -50932 | Nostrin | ENSMUSG0C | false | . | . | . | . | . | . | chr2 | 69076857 | 69090376 | 5.44132 | 4.24372 | 3.37539 |
| chr2 | 69080533 | 69081236 | 1000 | Intron (ENSMUST00000028426.3 | 2 | 69135800 | 69189330 | 53531 | 1 | ENSMUST00 | -54564 | Nostrin | ENSMUSG0C | false | . | . | . | . | . | . | chr2 | 69076857 | 69090376 | 5.44132 | 4.24372 | 3.37539 |
| chr13 | 102864920 | 102865476 | 1000 | Intron (ENSMUST00000166726.2 | 13 | 102741969 | 102905786 | 163818 | 2 | ENSMUST00 | 40310 | Mast4 | ENSMUSG0C | true | . | . | . | . | . | . | chr13 | 102834339 | 102867396 | 5.64964 | 4.33081 | 3.54328 |
| chr10 | 81596666 | 81597558 | 1000 | Promoter (<=1kb) | 10 | 81590907 | 81597345 | 6439 | 2 | ENSMUST00 | 0 | Tle6 | ENSMUSG0C | false | chr10 | 81596760 | 81596971 | 7.81205 | 13.8349 | 11.58824 | chr10 | 81595405 | 81598950 | 6.4271 | 4.77786 | 4.17257 |
| chr4 | 107468205 | 107468567 | 1000 | Intron (ENSMUST00000135835.2 | 4 | 107438998 | 107635061 | 196064 | 1 | ENSMUST00 | 29208 | Glis1 | ENSMUSG0C | false | . | . | . | . | . | . | chr4 | 107467263 | 107470037 | 5.98773 | 4.52872 | 3.81005 |
| chr4 | 107434251 | 107435145 | 1000 | Promoter (<=1kb) | 4 | 107434591 | 107436036 | 1446 | 1 | ENSMUST00 | 0 | Glis1 | ENSMUSG0C | false | chr4 | 107434005 | 107435717 | 26.28407 | 85.32412 | 82.07336 | chr4 | 107433262 | 107435702 | 3.27817 | 3.10385 | 1.64505 |
| chr19 | 53507382 | 53507810 | 1000 | Distal Intergenic | 19 | 53529109 | 53542431 | 13323 | 1 | ENSMUST00 | -21299 | Dusp5 | ENSMUSG0C | false | . | . | . | . | . | . | chr19 | 53507394 | 53507864 | 3.15936 | 2.95985 | 1.53633 |
| chr10 | 81575092 | 81575495 | 1000 | Promoter (<=1kb) | 10 | 81575260 | 81590470 | 15211 | 1 | ENSMUST00 | 0 | Tle2 | ENSMUSG0C | false | chr10 | 81574805 | 81575907 | 27.51607 | 79.95555 | 76.80458 | chr10 | 81574695 | 81576594 | 5.0419 | 4.13264 | 3.0544 |
| chr11 | 102251494 | 102251930 | 1000 | Promoter (2-3kb) | 11 | 102248882 | 102265187 | 16306 | 1 | ENSMUST00 | 2613 | BC030867 | ENSMUSG0C | false | . | . | . | . | . | . | chr11 | 102250749 | 102252022 | 5.05333 | 4.08027 | 3.06065 |
| chr10 | 81545042 | 81545513 | 1000 | Promoter (<=1kb) | 10 | 81528724 | 81545190 | 16467 | 2 | ENSMUST00 | 0 | Gna11 | ENSMUSG0C | false | chr10 | 81543738 | 81545170 | 42.50686 | 139.83865 | 134.96973 | chr10 | 81540018 | 81545966 | 5.77146 | 4.42913 | 3.64778 |
| chr8 | 123122042 | 123122417 | 1000 | Intron (ENSMUST00000127664.1 | 8 | 123126517 | 123127373 | 857 | 1 | ENSMUST00 | -4100 | Cpne7 | ENSMUSG0C | false | . | . | . | . | . | . | chr8 | 123121872 | 123122913 | 4.11881 | 3.73548 | 2.26589 |
| chr12 | 69681306 | 69682377 | 1000 | Promoter (<=1kb) | 12 | 69583766 | 69681852 | 98087 | 2 | ENSMUST00 | 0 | Sos2 | ENSMUSG0C | false | chr12 | 69680563 | 69683199 | 37.7199 | 126.00829 | 121.65485 | . | . | . | . | . | . |
| chr6 | 83455892 | 83457251 | 1000 | Promoter (<=1kb) | 6 | 83368042 | 83456392 | 88351 | 2 | ENSMUST00 | 0 | Tet3 | ENSMUSG0C | true | chr6 | 83455308 | 83457331 | 30.69271 | 93.10602 | 89.69717 | chr6 | 83442085 | 83458124 | 5.8895 | 4.34304 | 3.76187 |
| chr6 | 83440864 | 83441177 | 1000 | Promoter (<=1kb) | 6 | 83362373 | 83441675 | 79303 | 2 | ENSMUST00 | 498 | Tet3 | ENSMUSG0C | true | chr6 | 83440905 | 83443443 | 28.36989 | 85.2065 | 81.96005 | . | . | . | . | . | . |
| chr6 | 83441256 | 83442027 | 1000 | Promoter (<=1kb) | 6 | 83362373 | 83441675 | 79303 | 2 | ENSMUST00 | 0 | Tet3 | ENSMUSG0C | true | chr6 | 83440905 | 83443443 | 28.36989 | 85.2065 | 81.96005 | . | . | . | . | . | . |
| chr9 | 60712813 | 60713383 | 1000 | Promoter (<=1kb) | 9 | 60712989 | 60738801 | 25813 | 1 | ENSMUST00 | 0 | Larp6 | ENSMUSG0C | false | chr9 | 60712746 | 60713581 | 25.56571 | 75.64988 | 72.57267 | chr9 | 60712521 | 60713957 | 4.10905 | 3.59383 | 2.29363 |
| chr9 | 74911746 | 74912300 | 1000 | Distal Intergenic | 9 | 74953053 | 75032468 | 79416 | 1 | ENSMUST00 | -40753 | Fam214a | ENSMUSG0C | false | . | . | . | . | . | . | chr9 | 74911404 | 74912756 | 4.56839 | 3.85187 | 2.68396 |
| chr10 | 81329461 | 81330017 | 1000 | Promoter (<=1kb) | 10 | 81328732 | 81335171 | 6440 | 1 | ENSMUST00 | 730 | Tbxa2r | ENSMUSG0C | false | chr10 | 81328327 | 81330303 | 6.89286 | 16.24995 | 13.96717 | chr10 | 81325851 | 81336202 | 6.94026 | 4.91316 | 4.59344 |
| chr10 | 81326885 | 81327186 | 1000 | Promoter (1-2kb) | 10 | 81328732 | 81335171 | 6440 | 1 | ENSMUST00 | -1546 | Tbxa2r | ENSMUSG0C | false | . | . | . | . | . | . | chr10 | 81325851 | 81336202 | 6.94026 | 4.91316 | 4.59344 |
| chr10 | 81317263 | 81317608 | 1000 | Promoter (<=1kb) | 10 | 81316842 | 81318013 | 1172 | 1 | ENSMUST00 | 422 | Pip5k1c | ENSMUSG0C | false | chr10 | 81316916 | 81317538 | 13.98552 | 41.45461 | 38.84487 | chr10 | 81316106 | 81319993 | 6.5738 | 4.78337 | 4.29633 |
| chr2 | 65238237 | 65239375 | 1000 | Promoter (<=1kb) | 2 | 65088339 | 65238626 | 150288 | 2 | ENSMUST00 | 0 | Cobll1 | ENSMUSG0C | false | chr2 | 65237620 | 65239624 | 36.84594 | 114.91216 | 110.93493 | chr2 | 65237612 | 65239672 | 3.66899 | 3.39458 | 1.95995 |
| chr9 | 72925263 | 72926006 | 1000 | Promoter (<=1kb) | 9 | 72925645 | 72952181 | 26537 | 1 | ENSMUST00 | 0 | Pygo1 | ENSMUSG0C | false | chr9 | 72925014 | 72926508 | 27.02239 | 79.1029 | 75.96719 | . | . | . | . | . | . |
| chr4 | 106633302 | 106633930 | 1000 | Promoter (2-3kb) | 4 | 106635977 | 106637628 | 1652 | 1 | ENSMUST00 | -2047 | Ttc22 | ENSMUSG0C | false | . | . | . | . | . | . | chr4 | 106630565 | 106635216 | 5.02664 | 4.13737 | 3.01085 |
| chr13 | 54693740 | 54694503 | 1000 | Promoter (<=1kb) | 13 | 54679410 | 54693847 | 14438 | 2 | ENSMUST00 | 0 | Rnf44 | ENSMUSG0C | false | chr13 | 54694268 | 54694933 | 18.31284 | 64.75944 | 61.84419 | . | . | . | . | . | . |
| chr13 | 54687110 | 54687972 | 1000 | Promoter (<=1kb) | 13 | 54679399 | 54687808 | 8410 | 2 | ENSMUST00 | 0 | Rnf44 | ENSMUSG0C | false | chr13 | 54686089 | 54688786 | 43.58744 | 151.86528 | 146.50356 | . | . | . | . | . | . |
| chr6 | 83136985 | 83137981 | 1000 | Promoter (<=1kb) | 6 | 83137089 | 83152579 | 15491 | 1 | ENSMUST00 | 0 | Rtkn | ENSMUSG0C | false | chr6 | 83136694 | 83138101 | 20.4053 | 65.4752 | 62.55133 | chr6 | 83135038 | 83138767 | 4.88184 | 4.03716 | 2.91481 |
| chr11 | 101720020 | 101720451 | 1000 | Distal Intergenic | 11 | 101732919 | 101735696 | 2778 | 1 | ENSMUST00 | -12468 | Dhx8 | ENSMUSG0C | false | . | . | . | . | . | . | chr11 | 101717646 | 101721997 | 5.52652 | 4.34966 | 3.43987 |
| chr11 | 101700278 | 101700813 | 1000 | Distal Intergenic | 11 | 101732919 | 101735696 | 2778 | 1 | ENSMUST00 | -32106 | Dhx8 | ENSMUSG0C | false | . | . | . | . | . | . | chr11 | 101699078 | 101705226 | 5.43127 | 4.24835 | 3.38069 |
| chr11 | 101686983 | 101687983 | 1000 | Distal Intergenic | 11 | 101665541 | 101667832 | 2292 | 1 | ENSMUST00 | 21443 | Arl4d | ENSMUSG0C | false | . | . | . | . | . | . | chr11 | 101685526 | 101693622 | 5.48414 | 4.26664 | 3.42353 |
| chr11 | 101675268 | 101675702 | 1000 | Distal Intergenic | 11 | 101665541 | 101667832 | 2292 | 1 | ENSMUST00 | 9728 | Arl4d | ENSMUSG0C | false | . | . | . | . | . | . | chr11 | 101669121 | 101683699 | 5.5434 | 4.31925 | 3.45858 |
| chr17 | 88440271 | 88441853 | 1000 | Promoter (<=1kb) | 17 | 88440711 | 88490533 | 49823 | 1 | ENSMUST00 | 0 | Foxn2 | ENSMUSG0C | false | chr17 | 88440015 | 88442495 | 37.2221 | 115.56093 | 111.55734 | . | . | . | . | . | . |
| chr2 | 62423097 | 62423748 | 1000 | Intron (ENSMUST00000136879.1 | 2 | 62330073 | 62412231 | 82159 | 2 | ENSMUST00 | -10867 | Dpp4 | ENSMUSG0C | true | . | . | . | . | . | . | chr2 | 62417275 | 62426614 | 4.96374 | 4.01867 | 2.98047 |
| chr2 | 62407284 | 62407602 | 1000 | Intron (ENSMUST00000136879.1 | 2 | 62407702 | 62411859 | 4158 | 2 | ENSMUST00 | 4257 | Dpp4 | ENSMUSG0C | true | . | . | . | . | . | . | chr2 | 62405593 | 62408853 | 4.86632 | 4.07627 | 2.88617 |
| chr2 | 62403768 | 62404201 | 1000 | Intron (ENSMUST00000136879.1 | 2 | 62407702 | 62411859 | 4158 | 2 | ENSMUST00 | 7658 | Dpp4 | ENSMUSG0C | true | . | . | . | . | . | . | chr2 | 62375333 | 62404604 | 4.73054 | 3.90376 | 2.80145 |
| chr10 | 81135990 | 81137154 | 1000 | Promoter (<=1kb) | 10 | 81136271 | 81152222 | 15952 | 1 | ENSMUST00 | 0 | Zbtb7a | ENSMUSG0C | false | chr10 | 81135087 | 81137270 | 45.56376 | 157.62607 | 152.05525 | chr10 | 81134083 | 81136069 | 5.94765 | 4.28225 | 3.82742 |
| chr10 | 81137274 | 81138794 | 1000 | Promoter (<=1kb) | 10 | 81137953 | 81148458 | 10506 | 1 | ENSMUST00 | 0 | Zbtb7a | ENSMUSG0C | false | chr10 | 81137634 | 81138894 | 27.66126 | 83.97217 | 80.74496 | chr10 | 81138291 | 81147299 | 5.78848 | 4.25289 | 3.67763 |
| chr12 | 54984681 | 54986493 | 1000 | Promoter (<=1kb) | 12 | 54892989 | 54986336 | 93348 | 2 | ENSMUST00 | 0 | Baz1a | ENSMUSG0C | false | chr12 | 54984607 | 54986836 | 26.72788 | 101.58999 | 97.97913 | . | . | . | . | . | . |
| chr2 | 125782248 | 125783118 | 1000 | Promoter (<=1kb) | 2 | 125736986 | 125782870 | 45885 | 2 | ENSMUST00 | 0 | Secisbp2l | ENSMUSG0C | false | chr2 | 125781288 | 125783313 | 27.03504 | 88.5911 | 85.27629 | . | . | . | . | . | . |
| chr16 | 58408016 | 58409075 | 1000 | Promoter (<=1kb) | 16 | 58408443 | 58469727 | 61285 | 1 | ENSMUST00 | 0 | Dcbld2 | ENSMUSG0C | false | chr16 | 58407933 | 58410191 | 35.02417 | 111.75519 | 107.8695 | . | . | . | . | . | . |
| chr16 | 58421959 | 58422445 | 1000 | Intron (ENSMUST00000046663.7 | 16 | 58408443 | 58469727 | 61285 | 1 | ENSMUST00 | 13517 | Dcbld2 | ENSMUSG0C | false | . | . | . | . | . | . | chr16 | 58419662 | 58426719 | 4.21043 | 3.69053 | 2.36294 |
| chr11 | 101260145 | 101260821 | 1000 | Promoter (<=1kb) | 11 | 101260567 | 101277409 | 16843 | 1 | ENSMUST00 | 0 | Wnk4 | ENSMUSG0C | false | chr11 | 101259782 | 101261361 | 21.13848 | 54.99947 | 52.22001 | chr11 | 101252869 | 101266377 | 5.45094 | 4.22522 | 3.38706 |
| chr11 | 101259039 | 101259951 | 1000 | Promoter (<=1kb) | 11 | 101260567 | 101277409 | 16843 | 1 | ENSMUST00 | -616 | Wnk4 | ENSMUSG0C | false | chr11 | 101259782 | 101261361 | 21.13848 | 54.99947 | 52.22001 | chr11 | 101252869 | 101266377 | 5.45094 | 4.22522 | 3.38706 |
| chr4 | 103050772 | 103051383 | 1000 | Intron (ENSMUST00000138960.2 | 4 | 103039272 | 103041617 | 2346 | 2 | ENSMUST00 | -9156 | Wdr78 | ENSMUSG0C | false | . | . | . | . | . | . | chr4 | 103048889 | 103051428 | 3.78775 | 3.44678 | 2.01269 |
| chr12 | 52503429 | 52505288 | 1000 | Distal Intergenic | 12 | 52516077 | 52567852 | 51776 | 1 | ENSMUST00 | -10789 | Arhgap5 | ENSMUSG0C | false | chr12 | 52502712 | 52505830 | 36.42221 | 131.26616 | 126.73016 | . | . | . | . | . | . |
| chrX | 94177875 | 94178249 | 1000 | Distal Intergenic | 20 | 94199641 | 94212651 | 13011 | 2 | ENSMUST00 | 34402 | Eif2s3x | ENSMUSG0C | false | chrX | 94177439 | 94178702 | 17.92175 | 44.05972 | 41.41886 | . | . | . | . | . | . |
| chr2 | 59608117 | 59608644 | 1000 | Distal Intergenic | 2 | 59612042 | 59845717 | 233676 | 1 | ENSMUST00 | -3398 | Tanc1 | ENSMUSG0C | true | . | . | . | . | . | . | chr2 | 59603840 | 59611869 | 4.71859 | 3.92118 | 2.78796 |
| chr2 | 59613366 | 59613837 | 1000 | Promoter (1-2kb) | 2 | 59612075 | 59649455 | 37381 | 1 | ENSMUST00 | 1292 | Tanc1 | ENSMUSG0C | true | chr2 | 59611415 | 59614138 | 45.5631 | 163.41409 | 157.64189 | chr2 | 59613497 | 59619964 | 4.85927 | 3.83189 | 2.93047 |
| chr2 | 59611593 | 59612631 | 1000 | Promoter (<=1kb) | 2 | 59612042 | 59845717 | 233676 | 1 | ENSMUST00 | 0 | Tanc1 | ENSMUSG0C | true | chr2 | 59611415 | 59614138 | 45.5631 | 163.41409 | 157.64189 | chr2 | 59603840 | 59611869 | 4.71859 | 3.92118 | 2.78796 |
| chr11 | 101167780 | 101168616 | 1000 | Promoter (1-2kb) | 11 | 101167474 | 101170354 | 2881 | 2 | ENSMUST00 | 1738 | Plekhh3 | ENSMUSG0C | false | . | . | . | . | . | . | chr11 | 101162175 | 101170220 | 6.22712 | 4.64855 | 4.01342 |
| chr12 | 51970752 | 51971523 | 1000 | Promoter (<=1kb) | 12 | 51875873 | 51971321 | 95449 | 2 | ENSMUST00 | 0 | Heatr5a | ENSMUSG0C | false | chr12 | 51970357 | 51971857 | 33.40766 | 121.45458 | 117.2672 | . | . | . | . | . | . |
| chr12 | 51952304 | 51952751 | 1000 | Exon (ENSMUST00000040583.5/ | 12 | 51875873 | 51971321 | 95449 | 2 | ENSMUST00 | 18570 | Heatr5a | ENSMUSG0C | false | . | . | . | . | . | . | chr12 | 51951674 | 51954426 | 4.96491 | 4.04128 | 2.98534 |
| chrX | 93831954 | 93832476 | 1000 | Promoter (<=1kb) | 20 | 93764607 | 93832201 | 67595 | 2 | ENSMUST00 | 0 | Pdk3 | ENSMUSG0C | false | chrX | 93831268 | 93832563 | 26.19333 | 73.09374 | 70.054 | . | . | . | . | . | . |
| chr13 | 59859415 | 59859881 | 1000 | Distal Intergenic | 13 | 59771879 | 59823147 | 51269 | 2 | ENSMUST00 | -36269 | Zcchc6 | ENSMUSG0C | false | . | . | . | . | . | . | chr13 | 59856293 | 59862398 | 4.54569 | 3.77915 | 2.64326 |
| chr13 | 59834663 | 59835840 | 1000 | Distal Intergenic | 13 | 59771879 | 59823147 | 51269 | 2 | ENSMUST00 | -11517 | Zcchc6 | ENSMUSG0C | false | . | . | . | . | . | . | chr13 | 59832928 | 59838791 | 4.70155 | 3.94381 | 2.76868 |
| chr2 | 163661426 | 163661914 | 1000 | Promoter (<=1kb) | 2 | 163661499 | 163725473 | 63975 | 1 | ENSMUST00 | 0 | Pkig | ENSMUSG0C | false | . | . | . | . | . | . | chr2 | 163658175 | 163666027 | 5.63736 | 4.28032 | 3.5511 |
| chr4 | 101068686 | 101069652 | 1000 | Promoter (<=1kb) | 4 | 101068983 | 101135356 | 66374 | 1 | ENSMUST00 | 0 | Raver2 | ENSMUSG0C | false | chr4 | 101068633 | 101070848 | 38.63803 | 123.30683 | 119.05459 | chr4 | 101064869 | 101068824 | 4.20578 | 3.73464 | 2.33867 |
| chr19 | 56660628 | 56661172 | 1000 | Distal Intergenic | 19 | 56722372 | 56724862 | 2491 | 1 | ENSMUST00 | -61200 | Adrb1 | ENSMUSG0C | false | . | . | . | . | . | . | . | . | . | . | . | . |
| chr12 | 51347882 | 51348350 | 1000 | Promoter (<=1kb) | 12 | 51348061 | 51376986 | 28926 | 1 | ENSMUST00 | 0 | G2e3 | ENSMUSG0C | false | chr12 | 51347377 | 51348896 | 31.48209 | 103.47416 | 99.8177 | . | . | . | . | . | . |
| chrX | 169882360 | 169882730 | 1000 | Promoter (2-3kb) | 20 | 169879664 | 169990571 | 110908 | 1 | ENSMUST00 | 2697 | Mid1 | ENSMUSG0C | false | . | . | . | . | . | . | . | . | . | . | . | . |
| chrX | 169881448 | 169881939 | 1000 | Promoter (1-2kb) | 20 | 169879664 | 169990571 | 110908 | 1 | ENSMUST00 | 1785 | Mid1 | ENSMUSG0C | false | chrX | 169879551 | 169881706 | 14.70503 | 33.66069 | 31.14909 | . | . | . | . | . | . |
| chr5 | 100491676 | 100492017 | 1000 | Intron (ENSMUST00000149714.2 | 5 | 100459586 | 100485411 | 25826 | 2 | ENSMUST00 | -6266 | Lin54 | ENSMUSG0C | false | . | . | . | . | . | . | chr5 | 100488222 | 100492384 | 4.58228 | 3.82928 | 2.68152 |
| chr5 | 100498456 | 100498840 | 1000 | Promoter (<=1kb) | 5 | 100441928 | 100498591 | 56664 | 2 | ENSMUST00 | 0 | Lin54 | ENSMUSG0C | false | chr5 | 100497647 | 100499112 | 37.68163 | 117.40437 | 113.3432 | chr5 | 100494362 | 100500216 | 4.84649 | 4.03773 | 2.87368 |
| chr5 | 100500524 | 100500902 | 1000 | Promoter (<=1kb) | 5 | 100441918 | 100500592 | 58675 | 2 | ENSMUST00 | 0 | Lin54 | ENSMUSG0C | false | chr5 | 100499667 | 100501061 | 18.85995 | 50.28701 | 47.56678 | . | . | . | . | . | . |
| chr9 | 64232545 | 64232918 | 1000 | Promoter (2-3kb) | 9 | 64235201 | 64236362 | 1162 | 1 | ENSMUST00 | -2283 | Uchl4 | ENSMUSG0C | false | . | . | . | . | . | . | chr9 | 64227997 | 64233257 | 6.97429 | 4.9005 | 4.65834 |
| chr19 | 45017833 | 45018288 | 1000 | Promoter (<=1kb) | 19 | 45018135 | 45027104 | 8970 | 1 | ENSMUST00 | 0 | Lzts2 | ENSMUSG0C | false | chr19 | 45017892 | 45019345 | 21.77823 | 66.26614 | 63.32996 | . | . | . | . | . | . |
| chr9 | 45906047 | 45906694 | 1000 | Promoter (<=1kb) | 9 | 45906569 | 45929692 | 23124 | 1 | ENSMUST00 | 0 | Pcsk7 | ENSMUSG0C | false | chr9 | 45905634 | 45907785 | 37.30079 | 124.19571 | 119.91465 | chr9 | 45904756 | 45906053 | 4.50578 | 3.81256 | 2.61236 |
| chr2 | 38286926 | 38287549 | 1000 | Promoter (<=1kb) | 2 | 37798991 | 38287375 | 488385 | 2 | ENSMUST00 | 0 | Dennd1a | ENSMUSG0C | false | chr2 | 38286358 | 38287858 | 41.48799 | 138.06813 | 133.26578 | chr2 | 38285628 | 38287930 | 2.98376 | 2.96948 | 1.41357 |
| chr10 | 80576477 | 80578180 | 1000 | Promoter (<=1kb) | 10 | 80567124 | 80577321 | 10198 | 2 | ENSMUST00 | 0 | Klf16 | ENSMUSG0C | false | chr10 | 80576172 | 80578703 | 38.46334 | 130.56975 | 126.05653 | chr10 | 80578116 | 80580104 | 4.76959 | 4.00221 | 2.81148 |
| chr2 | 37775954 | 37776383 | 1000 | Promoter (<=1kb) | 2 | 37776249 | 37799103 | 22855 | 1 | ENSMUST00 | 0 | Crb2 | ENSMUSG0C | false | chr2 | 37775966 | 37778479 | 22.18388 | 60.95028 | 58.09254 | chr2 | 37774517 | 37776214 | 5.40051 | 4.03254 | 3.36795 |
| chr2 | 37788203 | 37789013 | 1000 | Promoter (2-3kb) | 2 | 37791171 | 37793523 | 2353 | 1 | ENSMUST00 | -2158 | Crb2 | ENSMUSG0C | false | chr2 | 37788131 | 37791502 | 19.02488 | 51.36755 | 48.63442 | chr2 | 37777133 | 37803419 | 7.19536 | 4.98761 | 4.82001 |
| chr2 | 37784511 | 37784955 | 1000 | Intron (ENSMUST00000050372.7 | 2 | 37791171 | 37793523 | 2353 | 1 | ENSMUST00 | -6216 | Crb2 | ENSMUSG0C | false | . | . | . | . | . | . | chr2 | 37777133 | 37803419 | 7.19536 | 4.98761 | 4.82001 |
| chr2 | 37796074 | 37796598 | 1000 | Promoter (<=1kb) | 2 | 37795169 | 37796616 | 1448 | 1 | ENSMUST00 | 906 | Crb2 | ENSMUSG0C | false | . | . | . | . | . | . | chr2 | 37777133 | 37803419 | 7.19536 | 4.98761 | 4.82001 |
| chr2 | 37452031 | 37452573 | 1000 | Promoter (<=1kb) | 2 | 37452245 | 37475363 | 23119 | 1 | ENSMUST00 | 0 | Rabgap1 | ENSMUSG0C | false | chr2 | 37451563 | 37453152 | 43.50657 | 151.72289 | 146.37334 | . | . | . | . | . | . |
| chr11 | 80682526 | 80683157 | 1000 | Intron (ENSMUST00000041065.8 | 11 | 80673667 | 80780003 | 106337 | 2 | ENSMUST00 | 96846 | Myo1d | ENSMUSG0C | true | chr11 | 80681163 | 80682790 | 14.28371 | 39.87881 | 37.29034 | chr11 | 80677716 | 80688108 | 5.01363 | 4.04514 | 3.03102 |
| chr11 | 80679780 | 80680130 | 1000 | Exon (ENSMUST00000041065.8/ | 11 | 80673667 | 80780003 | 106337 | 2 | ENSMUST00 | 99873 | Myo1d | ENSMUSG0C | true | . | . | . | . | . | . | chr11 | 80677716 | 80688108 | 5.01363 | 4.04514 | 3.03102 |
| chr11 | 80700567 | 80700873 | 1000 | Intron (ENSMUST00000041065.8 | 11 | 80673667 | 80780003 | 106337 | 2 | ENSMUST00 | 79130 | Myo1d | ENSMUSG0C | true | . | . | . | . | . | . | chr11 | 80692188 | 80719976 | 5.39042 | 4.17842 | 3.33942 |
| chr14 | 59200866 | 59201870 | 1000 | Promoter (<=1kb) | 14 | 59201209 | 59223214 | 22006 | 1 | ENSMUST00 | 0 | Rcbtb1 | ENSMUSG0C | false | chr14 | 59200711 | 59202259 | 38.14666 | 123.52654 | 119.2661 | . | . | . | . | . | . |
| chrX | 85573608 | 85574435 | 1000 | Promoter (<=1kb) | 20 | 85574022 | 85634469 | 60448 | 1 | ENSMUST00 | 0 | Tab3 | ENSMUSG0C | false | chrX | 85573408 | 85573813 | 15.62409 | 36.57069 | 34.02363 | . | . | . | . | . | . |
| chr16 | 33580558 | 33581063 | 1000 | Intron (ENSMUST00000121925.2 | 16 | 33596562 | 33664135 | 67574 | 1 | ENSMUST00 | -15499 | Slc12a8 | ENSMUSG0C | false | . | . | . | . | . | . | chr16 | 33580365 | 33581663 | 2.84082 | 2.87605 | 1.27101 |
| chr16 | 33531276 | 33531722 | 1000 | Intron (ENSMUST00000121925.2 | 16 | 33518536 | 33663408 | 144873 | 1 | ENSMUST00 | 12741 | Slc12a8 | ENSMUSG0C | false | . | . | . | . | . | . | chr16 | 33529138 | 33532088 | 4.75546 | 3.95651 | 2.83762 |
| chr2 | 29245323 | 29245891 | 1000 | Promoter (2-3kb) | 2 | 29194821 | 29248040 | 53220 | 2 | ENSMUST00 | 2149 | Ntng2 | ENSMUSG0C | true | . | . | . | . | . | . | chr2 | 29230623 | 29250828 | 7.61258 | 5.25891 | 5.15208 |
| chr8 | 122917715 | 122918471 | 1000 | Promoter (1-2kb) | 8 | 122884151 | 122915987 | 31837 | 2 | ENSMUST00 | -1729 | Ankrd11 | ENSMUSG0C | false | . | . | . | . | . | . | chr8 | 122908962 | 122946959 | 5.2818 | 4.08553 | 3.26755 |
| chr7 | 3692671 | 3693121 | 1000 | Promoter (<=1kb) | 7 | 3683358 | 3692952 | 9595 | 2 | ENSMUST00 | 0 | Mboat7 | ENSMUSG0C | false | chr7 | 3692185 | 3695281 | 28.33848 | 93.85651 | 90.42933 | . | . | . | . | . | . |
| chr12 | 65036212 | 65036784 | 1000 | Promoter (<=1kb) | 12 | 65036333 | 65063386 | 27054 | 1 | ENSMUST00 | 0 | Prpf39 | ENSMUSG0C | false | chr12 | 65035953 | 65037435 | 38.6007 | 121.10697 | 116.92481 | . | . | . | . | . | . |
| chr10 | 84976697 | 84977164 | 1000 | Intron (ENSMUST00000038523.8 | 10 | 84917656 | 85016446 | 98791 | 1 | ENSMUST00 | 59042 | Ric8b | ENSMUSG0C | false | . | . | . | . | . | . | chr10 | 84974135 | 84978245 | 5.50264 | 4.22528 | 3.43815 |
| chr10 | 80150744 | 80151549 | 1000 | Promoter (<=1kb) | 10 | 80151154 | 80154472 | 3319 | 1 | ENSMUST00 | 0 | Midn | ENSMUSG0C | false | chr10 | 80147630 | 80154081 | 36.09911 | 114.6679 | 110.69713 | chr10 | 80150859 | 80167428 | 9.09735 | 5.82258 | 6.36001 |
| chr10 | 80151667 | 80153374 | 1000 | Promoter (<=1kb) | 10 | 80151238 | 80153696 | 2459 | 1 | ENSMUST00 | 430 | Midn | ENSMUSG0C | false | chr10 | 80147630 | 80154081 | 36.09911 | 114.6679 | 110.69713 | chr10 | 80150859 | 80167428 | 9.09735 | 5.82258 | 6.36001 |
| chr10 | 80147928 | 80150563 | 1000 | Promoter (<=1kb) | 10 | 80148272 | 80158367 | 10096 | 1 | ENSMUST00 | 0 | Midn | ENSMUSG0C | false | chr10 | 80147630 | 80154081 | 36.09911 | 114.6679 | 110.69713 | chr10 | 80140751 | 80148113 | 6.54123 | 4.83987 | 4.26386 |
| chr10 | 80158329 | 80158840 | 1000 | 3' UTR | 10 | 80151238 | 80153696 | 2459 | 1 | ENSMUST00 | 7092 | Midn | ENSMUSG0C | false | . | . | . | . | . | . | chr10 | 80150859 | 80167428 | 9.09735 | 5.82258 | 6.36001 |
| chr10 | 80138900 | 80139529 | 1000 | Promoter (<=1kb) | 10 | 80130551 | 80139365 | 8815 | 2 | ENSMUST00 | 0 | Dos | ENSMUSG0C | false | chr10 | 80138590 | 80139651 | 22.33355 | 63.77912 | 60.87984 | chr10 | 80136655 | 80139809 | 5.80968 | 4.55211 | 3.66506 |
| chr4 | 44776627 | 44776986 | 1000 | Intron (ENSMUST00000107824.3 | 4 | 44760444 | 44912059 | 151616 | 1 | ENSMUST00 | 16184 | Zcchc7 | ENSMUSG0C | false | . | . | . | . | . | . | chr4 | 44773552 | 44777329 | 4.25951 | 3.65718 | 2.40466 |
| chr8 | 84236953 | 84237559 | 1000 | Promoter (<=1kb) | 8 | 84210942 | 84237042 | 26101 | 2 | ENSMUST00 | 0 | Zswim4 | ENSMUSG0C | false | chr8 | 84235808 | 84237146 | 34.92444 | 106.42502 | 102.68874 | chr8 | 84235465 | 84238489 | 3.98234 | 3.4793 | 2.22273 |
| chr4 | 44167425 | 44168586 | 1000 | Promoter (<=1kb) | 4 | 44126210 | 44167608 | 41399 | 2 | ENSMUST00 | 0 | Rnf38 | ENSMUSG0C | false | chr4 | 44166984 | 44168121 | 29.2142 | 114.47596 | 110.50954 | . | . | . | . | . | . |
| chr4 | 44233665 | 44234414 | 1000 | Promoter (<=1kb) | 4 | 44152434 | 44233789 | 81356 | 2 | ENSMUST00 | 0 | Rnf38 | ENSMUSG0C | false | chr4 | 44232600 | 44234015 | 32.70897 | 100.57243 | 96.98774 | . | . | . | . | . | . |
| chr10 | 80018805 | 80019380 | 1000 | Promoter (<=1kb) | 10 | 80019653 | 80031468 | 11816 | 1 | ENSMUST00 | -273 | Hmha1 | ENSMUSG0C | false | . | . | . | . | . | . | chr10 | 80017834 | 80020155 | 4.67356 | 3.87903 | 2.75048 |
| chr7 | 97248082 | 97248490 | 1000 | Intron (ENSMUST00000004622.5 | 7 | 97312473 | 97315025 | 2553 | 2 | ENSMUST00 | 66535 | Usp35 | ENSMUSG0C | false | . | . | . | . | . | . | chr7 | 97245066 | 97250061 | 4.3972 | 3.71963 | 2.52358 |
| chr19 | 10286495 | 10287072 | 1000 | Intron (ENSMUST00000039327.5 | 19 | 10245268 | 10304857 | 59590 | 2 | ENSMUST00 | 17785 | Dagla | ENSMUSG0C | false | . | . | . | . | . | . | chr19 | 10270370 | 10305865 | 8.32241 | 5.52593 | 5.72899 |
| chr19 | 10266277 | 10267128 | 1000 | Intron (ENSMUST00000039327.5 | 19 | 10253330 | 10256871 | 3542 | 2 | ENSMUST00 | -9407 | Dagla | ENSMUSG0C | false | . | . | . | . | . | . | chr19 | 10264421 | 10268946 | 5.9519 | 4.50434 | 3.80579 |
| chr19 | 10296986 | 10297359 | 1000 | Intron (ENSMUST00000039327.5 | 19 | 10245268 | 10304857 | 59590 | 2 | ENSMUST00 | 7498 | Dagla | ENSMUSG0C | false | . | . | . | . | . | . | chr19 | 10270370 | 10305865 | 8.32241 | 5.52593 | 5.72899 |
| chr19 | 10289944 | 10290505 | 1000 | Intron (ENSMUST00000039327.5 | 19 | 10245268 | 10304857 | 59590 | 2 | ENSMUST00 | 14352 | Dagla | ENSMUSG0C | false | . | . | . | . | . | . | chr19 | 10270370 | 10305865 | 8.32241 | 5.52593 | 5.72899 |
| chr10 | 111376397 | 111377311 | 1000 | Distal Intergenic | 10 | 111298679 | 111301736 | 3058 | 1 | ENSMUST00 | 77719 | Bbs10 | ENSMUSG0C | false | . | . | . | . | . | . | chr10 | 111372873 | 111379701 | 4.78915 | 4.01987 | 2.82418 |
| chr10 | 111234446 | 111234977 | 1000 | Intron (ENSMUST00000095310.1 | 10 | 111298679 | 111301736 | 3058 | 1 | ENSMUST00 | -63702 | Bbs10 | ENSMUSG0C | false | . | . | . | . | . | . | chr10 | 111232705 | 111239147 | 4.92603 | 3.95685 | 2.97209 |
| chrX | 71492511 | 71492910 | 1000 | Promoter (<=1kb) | 20 | 71420060 | 71492645 | 72586 | 2 | ENSMUST00 | 0 | Cd99l2 | ENSMUSG0C | false | chrX | 71491872 | 71493555 | 21.13848 | 54.99947 | 52.22001 | . | . | . | . | . | . |
| chr10 | 111009533 | 111010662 | 1000 | Promoter (<=1kb) | 10 | 110941782 | 111010066 | 68285 | 2 | ENSMUST00 | 0 | Zdhhc17 | ENSMUSG0C | false | chr10 | 111008979 | 111011604 | 37.74925 | 129.3071 | 124.84747 | . | . | . | . | . | . |
| chr12 | 33957195 | 33958426 | 1000 | Promoter (<=1kb) | 12 | 33957671 | 33959817 | 2147 | 1 | ENSMUST00 | 0 | Twist1 | ENSMUSG0C | false | chr12 | 33957327 | 33958679 | 12.16433 | 45.21117 | 42.5593 | chr12 | 33956060 | 33960076 | 3.54435 | 3.27471 | 1.83935 |
| chr15 | 88861822 | 88862525 | 1000 | Promoter (<=1kb) | 15 | 88862186 | 88865718 | 3533 | 1 | ENSMUST00 | 0 | Pim3 | ENSMUSG0C | false | chr15 | 88861608 | 88864034 | 31.00681 | 94.63441 | 91.19181 | chr15 | 88861232 | 88861858 | 3.89064 | 3.49449 | 2.10157 |
| chr15 | 88863579 | 88864317 | 1000 | Promoter (1-2kb) | 15 | 88862186 | 88865718 | 3533 | 1 | ENSMUST00 | 1394 | Pim3 | ENSMUSG0C | false | chr15 | 88861608 | 88864034 | 31.00681 | 94.63441 | 91.19181 | chr15 | 88863272 | 88871136 | 9.58718 | 5.83938 | 6.75823 |
| chr13 | 81710826 | 81711652 | 1000 | Promoter (<=1kb) | 13 | 81673843 | 81711013 | 37171 | 2 | ENSMUST00 | 0 | Polr3g | ENSMUSG0C | false | chr13 | 81710251 | 81712428 | 26.89699 | 109.78668 | 105.95901 | . | . | . | . | . | . |
| chr13 | 81657486 | 81658233 | 1000 | Promoter (<=1kb) | 13 | 81657806 | 81671888 | 14083 | 1 | ENSMUST00 | 0 | Lysmd3 | ENSMUSG0C | false | chr13 | 81656629 | 81659028 | 28.59262 | 111.22587 | 107.35971 | . | . | . | . | . | . |
| chr17 | 66123383 | 66123982 | 1000 | Promoter (<=1kb) | 17 | 66123520 | 66152167 | 28648 | 1 | ENSMUST00 | 0 | Ddx11 | ENSMUSG0C | false | chr17 | 66123165 | 66124386 | 23.94845 | 71.00658 | 67.99948 | chr17 | 66122945 | 66125790 | 2.7261 | 2.82051 | 1.19178 |
| chr10 | 108336670 | 108337025 | 1000 | Intron (ENSMUST00000095313.3 | 10 | 108332189 | 108414391 | 82203 | 1 | ENSMUST00 | 4482 | Pawr | ENSMUSG0C | false | . | . | . | . | . | . | chr10 | 108334141 | 108343133 | 4.77514 | 3.97163 | 2.82359 |
| chr10 | 79766540 | 79767254 | 1000 | Promoter (<=1kb) | 10 | 79758515 | 79766952 | 8438 | 2 | ENSMUST00 | 0 | Rnf126 | ENSMUSG0C | false | chr10 | 79766064 | 79768018 | 26.2242 | 93.81972 | 90.39303 | chr10 | 79763178 | 79766687 | 4.04398 | 3.4522 | 2.26309 |
| chr15 | 86188864 | 86189431 | 1000 | Promoter (2-3kb) | 15 | 86139129 | 86186141 | 47013 | 2 | ENSMUST00 | -2724 | Cerk | ENSMUSG0C | false | . | . | . | . | . | . | chr15 | 86185990 | 86192464 | 8.70941 | 5.50823 | 6.04495 |
| chr15 | 86078801 | 86079288 | 1000 | Intron (ENSMUST00000088931.4 | 15 | 86074733 | 86091434 | 16702 | 1 | ENSMUST00 | 4069 | Gramd4 | ENSMUSG0C | false | . | . | . | . | . | . | chr15 | 86074674 | 86092519 | 7.22651 | 5.10744 | 4.82194 |
| chr15 | 86058493 | 86059473 | 1000 | Promoter (<=1kb) | 15 | 86058727 | 86137618 | 78892 | 1 | ENSMUST00 | 0 | Gramd4 | ENSMUSG0C | false | chr15 | 86058221 | 86059708 | 38.04976 | 128.75285 | 124.30675 | . | . | . | . | . | . |
| chr13 | 25069837 | 25070356 | 1000 | Intron (ENSMUST00000036932.9 | 13 | 25056253 | 25210706 | 154454 | 1 | ENSMUST00 | 13585 | Dcdc2a | ENSMUSG0C | true | . | . | . | . | . | . | chr13 | 25068613 | 25089235 | 4.80481 | 3.93018 | 2.85848 |
| chr13 | 25088555 | 25088875 | 1000 | Intron (ENSMUST00000036932.9 | 13 | 25096347 | 25119406 | 23060 | 1 | ENSMUST00 | -7472 | Dcdc2a | ENSMUSG0C | true | . | . | . | . | . | . | chr13 | 25068613 | 25089235 | 4.80481 | 3.93018 | 2.85848 |
| chr13 | 25098403 | 25099099 | 1000 | Promoter (2-3kb) | 13 | 25096347 | 25119406 | 23060 | 1 | ENSMUST00 | 2057 | Dcdc2a | ENSMUSG0C | true | . | . | . | . | . | . | chr13 | 25090221 | 25101703 | 5.1893 | 4.18623 | 3.1625 |
| chr9 | 37072214 | 37072903 | 1000 | Intron (ENSMUST00000177218.2 | 9 | 36892642 | 37083279 | 190638 | 2 | ENSMUST00 | 10376 | Pknox2 | ENSMUSG0C | true | . | . | . | . | . | . | chr9 | 37051691 | 37078073 | 5.33519 | 4.13112 | 3.31406 |
| chr9 | 36973222 | 36973781 | 1000 | Intron (ENSMUST00000177218.2 | 9 | 36892478 | 36968490 | 76013 | 2 | ENSMUST00 | -4733 | Pknox2 | ENSMUSG0C | true | . | . | . | . | . | . | chr9 | 36970192 | 36981240 | 5.71459 | 4.43768 | 3.58495 |
| chr9 | 37054599 | 37055130 | 1000 | Intron (ENSMUST00000177218.2 | 9 | 36892642 | 37083279 | 190638 | 2 | ENSMUST00 | 28149 | Pknox2 | ENSMUSG0C | true | chr9 | 37054394 | 37054779 | 4.42115 | 6.83029 | 4.72578 | chr9 | 37051691 | 37078073 | 5.33519 | 4.13112 | 3.31406 |
| chr9 | 37146726 | 37147307 | 1000 | Promoter (<=1kb) | 9 | 36890982 | 37147289 | 256308 | 2 | ENSMUST00 | 0 | Pknox2 | ENSMUSG0C | true | chr9 | 37144737 | 37147348 | 34.92444 | 106.42502 | 102.68874 | . | . | . | . | . | . |
| chr9 | 37144225 | 37144614 | 1000 | Promoter (1-2kb) | 9 | 36954772 | 37145660 | 190889 | 2 | ENSMUST00 | 1046 | Pknox2 | ENSMUSG0C | true | chr9 | 37144060 | 37144640 | 9.76265 | 21.55577 | 19.19873 | chr9 | 37137127 | 37144958 | 5.79298 | 4.47095 | 3.65934 |
| chr12 | 40445772 | 40447007 | 1000 | Promoter (<=1kb) | 12 | 40446053 | 40846486 | 400434 | 1 | ENSMUST00 | 0 | Dock4 | ENSMUSG0C | true | chr12 | 40444767 | 40448342 | 40.65367 | 138.82761 | 133.98773 | . | . | . | . | . | . |
| chr12 | 40472153 | 40472477 | 1000 | Intron (ENSMUST00000037488.6 | 12 | 40446053 | 40846486 | 400434 | 1 | ENSMUST00 | 26101 | Dock4 | ENSMUSG0C | true | . | . | . | . | . | . | chr12 | 40448062 | 40493881 | 5.07226 | 4.00134 | 3.09271 |
| chr4 | 43394475 | 43394955 | 1000 | Intron (ENSMUST00000035645.6 | 4 | 43401238 | 43427073 | 25836 | 1 | ENSMUST00 | -6283 | Rusc2 | ENSMUSG0C | false | . | . | . | . | . | . | chr4 | 43382882 | 43406168 | 6.80327 | 4.94439 | 4.48074 |
| chr10 | 105573787 | 105574793 | 1000 | Promoter (<=1kb) | 10 | 105187663 | 105574435 | 386773 | 2 | ENSMUST00 | 0 | Tmtc2 | ENSMUSG0C | false | chr10 | 105572740 | 105575603 | 30.6396 | 93.99698 | 90.56607 | chr10 | 105574626 | 105575006 | 3.19834 | 3.16308 | 1.55222 |
| chr14 | 50807320 | 50807989 | 1000 | Promoter (<=1kb) | 14 | 50807946 | 50821300 | 13355 | 1 | ENSMUST00 | 0 | Parp2 | ENSMUSG0C | false | chr14 | 50807147 | 50808551 | 20.26563 | 55.95697 | 53.16479 | chr14 | 50806822 | 50809702 | 3.01701 | 2.99972 | 1.43525 |
| chr9 | 51312890 | 51313335 | 1000 | Promoter (<=1kb) | 9 | 51290444 | 51312669 | 22226 | 2 | ENSMUST00 | -222 | 1810046K07I | ENSMUSG0C | false | . | . | . | . | . | . | chr9 | 51312405 | 51313768 | 4.48688 | 3.75816 | 2.62871 |
| chr9 | 51407970 | 51408404 | 1000 | Distal Intergenic | 9 | 51289686 | 51328917 | 39232 | 2 | ENSMUST00 | -79054 | 1810046K07I | ENSMUSG0C | false | . | . | . | . | . | . | chr9 | 51404996 | 51411058 | 4.97376 | 4.06092 | 2.98267 |
| chr4 | 42949648 | 42950568 | 1000 | Promoter (<=1kb) | 4 | 42949814 | 42959425 | 9612 | 1 | ENSMUST00 | 0 | Dnajb5 | ENSMUSG0C | false | chr4 | 42949218 | 42950597 | 27.18497 | 79.384 | 76.2402 | chr4 | 42949108 | 42951599 | 4.24148 | 3.73453 | 2.39126 |
| chr5 | 72913846 | 72914792 | 1000 | Promoter (<=1kb) | 5 | 72914304 | 72978829 | 64526 | 1 | ENSMUST00 | 0 | Slain2 | ENSMUSG0C | true | chr5 | 72913155 | 72914239 | 23.65566 | 74.08845 | 71.03342 | . | . | . | . | . | . |
| chr5 | 72933559 | 72934158 | 1000 | Intron (ENSMUST00000144843.2 | 5 | 72948499 | 72974702 | 26204 | 1 | ENSMUST00 | -14341 | Slain2 | ENSMUSG0C | true | . | . | . | . | . | . | chr5 | 72915763 | 72933978 | 5.15676 | 4.0985 | 3.1473 |
| chr2 | 52424550 | 52425152 | 1000 | Promoter (<=1kb) | 2 | 52397951 | 52424901 | 26951 | 2 | ENSMUST00 | 0 | Arl5a | ENSMUSG0C | false | chr2 | 52422513 | 52425752 | 30.33064 | 129.04645 | 124.59567 | . | . | . | . | . | . |
| chr19 | 44930612 | 44931648 | 1000 | Promoter (<=1kb) | 19 | 44931119 | 44983787 | 52669 | 1 | ENSMUST00 | 0 | Fam178a | ENSMUSG0C | false | chr19 | 44930334 | 44932028 | 46.79302 | 165.58527 | 159.74297 | chr19 | 44931583 | 44933997 | 3.95359 | 3.52585 | 2.16278 |
| chr18 | 9707362 | 9708024 | 1000 | Promoter (<=1kb) | 18 | 9707648 | 9877995 | 170348 | 1 | ENSMUST00 | 0 | Colec12 | ENSMUSG0C | true | chr18 | 9707115 | 9709019 | 34.59452 | 106.87945 | 103.13243 | . | . | . | . | . | . |
| chr15 | 84680768 | 84681956 | 1000 | Promoter (<=1kb) | 15 | 84681002 | 84703672 | 22671 | 1 | ENSMUST00 | 0 | Prr5 | ENSMUSG0C | false | chr15 | 84680404 | 84684554 | 34.34893 | 131.73137 | 127.17394 | chr15 | 84677966 | 84680818 | 4.88786 | 4.0225 | 2.91391 |
| chr15 | 84695997 | 84696622 | 1000 | Intron (ENSMUST00000171460.2 | 15 | 84681002 | 84703672 | 22671 | 1 | ENSMUST00 | 14996 | Prr5 | ENSMUSG0C | false | . | . | . | . | . | . | chr15 | 84682178 | 84700461 | 6.48581 | 4.70787 | 4.24565 |
| chr15 | 102975903 | 102976692 | 1000 | Promoter (<=1kb) | 15 | 102976439 | 102981669 | 5231 | 1 | ENSMUST00 | 0 | Hoxc9 | ENSMUSG0C | false | chr15 | 102975608 | 102978084 | 18.61129 | 49.85614 | 47.14222 | chr15 | 102975429 | 102981199 | 5.98646 | 4.4723 | 3.8277 |
| chr15 | 102976828 | 102977883 | 1000 | Promoter (<=1kb) | 15 | 102977032 | 102985099 | 8068 | 1 | ENSMUST00 | 0 | Hoxc9 | ENSMUSG0C | false | chr15 | 102975608 | 102978084 | 18.61129 | 49.85614 | 47.14222 | chr15 | 102975429 | 102981199 | 5.98646 | 4.4723 | 3.8277 |
| chr1 | 127213012 | 127213365 | 1000 | Intron (ENSMUST00000189427.1 | 1 | 127205028 | 127398792 | 193765 | 1 | ENSMUST00 | 7985 | Mgat5 | ENSMUSG0C | true | . | . | . | . | . | . | chr1 | 127206353 | 127222843 | 5.20875 | 4.0815 | 3.19162 |
| chr1 | 127241343 | 127241727 | 1000 | Intron (ENSMUST00000189427.1 | 1 | 127205028 | 127398792 | 193765 | 1 | ENSMUST00 | 36316 | Mgat5 | ENSMUSG0C | true | . | . | . | . | . | . | chr1 | 127237789 | 127252930 | 5.60995 | 4.26347 | 3.52359 |
| chr1 | 127307428 | 127307869 | 1000 | Promoter (<=1kb) | 1 | 127306762 | 127483704 | 176943 | 1 | ENSMUST00 | 667 | Mgat5 | ENSMUSG0C | true | chr1 | 127306652 | 127307946 | 25.3802 | 72.59858 | 69.5683 | . | . | . | . | . | . |
| chr1 | 127306400 | 127306861 | 1000 | Promoter (<=1kb) | 1 | 127306762 | 127483704 | 176943 | 1 | ENSMUST00 | 0 | Mgat5 | ENSMUSG0C | true | chr1 | 127306652 | 127307946 | 25.3802 | 72.59858 | 69.5683 | . | . | . | . | . | . |
| chr15 | 93595504 | 93596173 | 1000 | Promoter (<=1kb) | 15 | 93499114 | 93595891 | 96778 | 2 | ENSMUST00 | 0 | Prickle1 | ENSMUSG0C | false | chr15 | 93594446 | 93596861 | 38.6007 | 121.10697 | 116.92481 | chr15 | 93591141 | 93596895 | 3.96846 | 3.46499 | 2.17353 |
| chr2 | 26902809 | 26903277 | 1000 | Promoter (<=1kb) | 2 | 26890435 | 26902879 | 12445 | 2 | ENSMUST00 | 0 | Surf6 | ENSMUSG0C | false | chr2 | 26902239 | 26903850 | 28.69254 | 91.56027 | 88.18607 | . | . | . | . | . | . |
| chr2 | 52072564 | 52073373 | 1000 | Promoter (<=1kb) | 2 | 52072837 | 52122381 | 49545 | 1 | ENSMUST00 | 0 | Rif1 | ENSMUSG0C | false | chr2 | 52072429 | 52073853 | 41.911 | 144.38934 | 139.32462 | chr2 | 52072354 | 52074691 | 3.45561 | 3.36098 | 1.71715 |
| chr16 | 44724119 | 44724739 | 1000 | Promoter (<=1kb) | 16 | 44724301 | 44737284 | 12984 | 1 | ENSMUST00 | 0 | BC027231 | ENSMUSG0C | false | chr16 | 44723737 | 44725506 | 39.39634 | 133.30592 | 128.68834 | . | . | . | . | . | . |
| chr4 | 41135083 | 41136799 | 1000 | Promoter (<=1kb) | 4 | 41135743 | 41193380 | 57638 | 1 | ENSMUST00 | 0 | Ube2r2 | ENSMUSG0C | false | chr4 | 41134685 | 41137061 | 38.2674 | 121.59672 | 117.39927 | . | . | . | . | . | . |
| chr5 | 77407331 | 77408313 | 1000 | Promoter (<=1kb) | 5 | 77349240 | 77408045 | 58806 | 2 | ENSMUST00 | 0 | Igfbp7 | ENSMUSG0C | false | chr5 | 77405039 | 77408352 | 25.08176 | 91.61557 | 88.24099 | . | . | . | . | . | . |
| chr5 | 77384148 | 77384556 | 1000 | Intron (ENSMUST00000046746.6 | 5 | 77349240 | 77408045 | 58806 | 2 | ENSMUST00 | 23489 | Igfbp7 | ENSMUSG0C | false | . | . | . | . | . | . | chr5 | 77353137 | 77404361 | 6.51707 | 4.75542 | 4.25399 |
| chr5 | 77387032 | 77387513 | 1000 | Intron (ENSMUST00000046746.6 | 5 | 77349240 | 77408045 | 58806 | 2 | ENSMUST00 | 20532 | Igfbp7 | ENSMUSG0C | false | . | . | . | . | . | . | chr5 | 77353137 | 77404361 | 6.51707 | 4.75542 | 4.25399 |
| chr14 | 49066019 | 49066763 | 1000 | Promoter (<=1kb) | 14 | 49066495 | 49087723 | 21229 | 1 | ENSMUST00 | 0 | Ap5m1 | ENSMUSG0C | false | chr14 | 49065401 | 49067299 | 40.14342 | 141.26135 | 136.33099 | chr14 | 49062487 | 49066053 | 3.44546 | 3.23282 | 1.77409 |
| chr3 | 59228921 | 59229254 | 1000 | Intron (ENSMUST00000040325.8 | 3 | 59216360 | 59220259 | 3900 | 2 | ENSMUST00 | -8663 | P2ry12 | ENSMUSG0C | false | . | . | . | . | . | . | chr3 | 59228620 | 59232023 | 4.27046 | 3.66389 | 2.41675 |
| chr3 | 59243444 | 59243756 | 1000 | Intron (ENSMUST00000040325.8 | 3 | 59216271 | 59262831 | 46561 | 2 | ENSMUST00 | 19075 | P2ry12 | ENSMUSG0C | false | . | . | . | . | . | . | chr3 | 59242905 | 59244391 | 4.17663 | 3.73392 | 2.34566 |
| chr6 | 67075973 | 67076612 | 1000 | Intron (ENSMUST00000130657.2 | 6 | 67036897 | 67037457 | 561 | 2 | ENSMUST00 | -38517 | Gadd45a | ENSMUSG0C | false | . | . | . | . | . | . | chr6 | 67069252 | 67079338 | 6.07124 | 4.48472 | 3.90237 |
| chr6 | 66896100 | 66896945 | 1000 | Promoter (<=1kb) | 6 | 66896397 | 67021350 | 124954 | 1 | ENSMUST00 | 0 | Gng12 | ENSMUSG0C | false | chr6 | 66895983 | 66897619 | 41.73432 | 142.95721 | 137.9541 | chr6 | 66895817 | 66898778 | 4.66841 | 3.84133 | 2.75619 |
| chr5 | 76588675 | 76589214 | 1000 | Promoter (<=1kb) | 5 | 76588698 | 76644908 | 56211 | 1 | ENSMUST00 | 0 | Cep135 | ENSMUSG0C | false | chr5 | 76588487 | 76589753 | 17.56648 | 73.03062 | 69.99512 | chr5 | 76588285 | 76592340 | 3.67232 | 3.32386 | 1.94262 |
| chr17 | 46889452 | 46890956 | 1000 | Promoter (<=1kb) | 17 | 46890621 | 46892463 | 1843 | 1 | ENSMUST00 | 0 | Tbcc | ENSMUSG0C | false | chr17 | 46890621 | 46892494 | 28.56742 | 103.16996 | 99.51957 | chr17 | 46887820 | 46889476 | 4.12367 | 3.56122 | 2.31162 |
| chr3 | 58692044 | 58692757 | 1000 | Promoter (<=1kb) | 3 | 58674938 | 58692400 | 17463 | 2 | ENSMUST00 | 0 | Siah2 | ENSMUSG0C | false | chr3 | 58690452 | 58692965 | 31.43325 | 99.35611 | 95.79983 | chr3 | 58692727 | 58693486 | 5.56646 | 4.13223 | 3.50047 |
| chr5 | 76531943 | 76532214 | 1000 | Promoter (2-3kb) | 5 | 76529358 | 76570287 | 40930 | 1 | ENSMUST00 | 2586 | Exoc1 | ENSMUSG0C | false | . | . | . | . | . | . | chr5 | 76529877 | 76535256 | 5.4199 | 4.28416 | 3.35627 |
| chr17 | 87445933 | 87447311 | 1000 | Promoter (<=1kb) | 17 | 87433412 | 87446935 | 13524 | 2 | ENSMUST00 | 0 | Calm2 | ENSMUSG0C | true | chr17 | 87442999 | 87447868 | 33.21814 | 102.46268 | 98.83372 | . | . | . | . | . | . |
| chr18 | 38631499 | 38632156 | 1000 | Intron (ENSMUST00000145090.2 | 18 | 38627287 | 38642930 | 15644 | 1 | ENSMUST00 | 4213 | Arhgap26 | ENSMUSG0C | false | chr18 | 38631193 | 38632129 | 15.88037 | 38.4063 | 35.83701 | chr18 | 38631108 | 38632417 | 5.65459 | 4.47847 | 3.53779 |
| chr18 | 38611589 | 38611964 | 1000 | Intron (ENSMUST00000145090.2 | 18 | 38601603 | 38789045 | 187443 | 1 | ENSMUST00 | 9987 | Arhgap26 | ENSMUSG0C | false | . | . | . | . | . | . | chr18 | 38610902 | 38612576 | 4.55593 | 3.8773 | 2.63861 |
| chr7 | 34132976 | 34133597 | 1000 | Promoter (<=1kb) | 7 | 34109543 | 34133268 | 23726 | 2 | ENSMUST00 | 0 | Wtip | ENSMUSG0C | true | chr7 | 34131933 | 34133828 | 29.41006 | 85.09721 | 81.85307 | chr7 | 34133361 | 34134492 | 5.09532 | 4.17496 | 3.08034 |
| chr10 | 96616415 | 96617614 | 1000 | Promoter (<=1kb) | 10 | 96617001 | 96622813 | 5813 | 1 | ENSMUST00 | 0 | Btg1 | ENSMUSG0C | false | chr10 | 96616250 | 96616891 | 19.35304 | 74.51352 | 71.45322 | chr10 | 96614291 | 96616469 | 5.11022 | 4.16996 | 3.0839 |
| chr10 | 96690903 | 96691357 | 1000 | Distal Intergenic | 10 | 96617001 | 96622813 | 5813 | 1 | ENSMUST00 | 73903 | Btg1 | ENSMUSG0C | false | . | . | . | . | . | . | chr10 | 96689910 | 96692319 | 5.19629 | 4.13069 | 3.21553 |
| chr18 | 34506436 | 34507186 | 1000 | Promoter (<=1kb) | 18 | 34442352 | 34506823 | 64472 | 2 | ENSMUST00 | 0 | Fam13b | ENSMUSG0C | false | chr18 | 34505681 | 34507604 | 43.23485 | 145.58217 | 140.47385 | . | . | . | . | . | . |

| | | | | | | | | | | | | | | | | | | | | | | | | | | |
|---|---|---|---|---|---|---|---|---|---|---|---|---|---|---|---|---|---|---|---|---|---|---|---|---|---|---|
| chr17 | 79451540 | 79451866 | 1000 | Distal Intergenic | 17 | 79334025 | 79355091 | 21067 | 2 | ENSMUST00 | -96450 | Cdc42ep3 | ENSMUSG0( | false | . | . | . | . | . | . | chr17 | 79450922 | 79456272 | 5.18318 | 4.20836 | 3.14751 |
| chr11 | 50598541 | 50598861 | 1000 | Downstream (<1kb) | 11 | 50602084 | 50807573 | 205490 | 1 | ENSMUST00 | -3223 | Adamts2 | ENSMUSG0( | false | . | . | . | . | . | . | chr11 | 50597912 | 50598886 | 2.84094 | 2.69868 | 1.27561 |
| chr11 | 50601787 | 50602304 | 1000 | Promoter (<=1kb) | 11 | 50602084 | 50807573 | 205490 | 1 | ENSMUST00 | 0 | Adamts2 | ENSMUSG0( | false | chr11 | 50601623 | 50603606 | 25.98482 | 77.30067 | 74.19341 | chr11 | 50601226 | 50603484 | 5.54387 | 4.1633 | 3.50586 |
| chr17 | 46842795 | 46843469 | 1000 | Distal Intergenic | 17 | 46798116 | 46831413 | 33298 | 2 | ENSMUST00 | -11383 | Gltscr1l | ENSMUSG0( | false | . | . | . | . | . | . | chr17 | 46837300 | 46846637 | 4.69005 | 3.81936 | 2.79322 |
| chr17 | 46856832 | 46858755 | 1000 | Distal Intergenic | 17 | 46798116 | 46831413 | 33298 | 2 | ENSMUST00 | -25420 | Gltscr1l | ENSMUSG0( | false | chr17 | 46855436 | 46858855 | 40.61456 | 134.33821 | 129.68364 | . | . | . | . | . | . |
| chr1 | 86441196 | 86442895 | 1000 | Distal Intergenic | 1 | 86426329 | 86428052 | 1724 | 1 | ENSMUST00 | 14868 | 1700019O17 | ENSMUSG0( | false | chr1 | 86441607 | 86443281 | 19.04495 | 59.55748 | 56.71581 | chr1 | 86442357 | 86444207 | 4.38989 | 3.6795 | 2.50903 |
| chr18 | 38907344 | 38907848 | 1000 | Intron (ENSMUST00000148850.2 | 18 | 38838673 | 38918771 | 80099 | 2 | ENSMUST00 | 10923 | Fgf1 | ENSMUSG0( | false | . | . | . | . | . | . | chr18 | 38905811 | 38908304 | 3.57846 | 3.39889 | 1.82251 |
| chr2 | 20968024 | 20969832 | 1000 | Promoter (<=1kb) | 2 | 20847919 | 20968881 | 120963 | 2 | ENSMUST00 | 0 | Arhgap21 | ENSMUSG0( | false | chr2 | 20966651 | 20970031 | 34.52349 | 109.84589 | 106.01766 | . | . | . | . | . | . |
| chr15 | 89169489 | 89169835 | 1000 | Promoter (<=1kb) | 15 | 89155549 | 89170688 | 15140 | 2 | ENSMUST00 | 853 | Plxnb2 | ENSMUSG0( | false | chr15 | 89169012 | 89169636 | 11.73508 | 26.0954 | 23.67916 | chr15 | 89167407 | 89179959 | 7.78402 | 5.30321 | 5.27943 |
| chr9 | 25481381 | 25481790 | 1000 | Promoter (<=1kb) | 9 | 25481547 | 25604110 | 122564 | 1 | ENSMUST00 | 0 | Eepd1 | ENSMUSG0( | false | chr9 | 25481120 | 25482139 | 28.90617 | 85.18155 | 81.93615 | . | . | . | . | . | . |
| chr12 | 28874888 | 28875382 | 1000 | Distal Intergenic | 12 | 28751828 | 28867491 | 115664 | 1 | ENSMUST00 | 123061 | Tssc1 | ENSMUSG0( | false | . | . | . | . | . | . | chr12 | 28872140 | 28876378 | 5.8899 | 4.40225 | 3.76611 |
| chr2 | 20603390 | 20603768 | 1000 | Intron (ENSMUST00000114610.2 | 2 | 20559935 | 20709635 | 149701 | 1 | ENSMUST00 | 43456 | Etl4 | ENSMUSG0( | true | . | . | . | . | . | . | chr2 | 20596846 | 20605046 | 4.75299 | 3.86034 | 2.84874 |
| chr2 | 20551068 | 20551358 | 1000 | Intron (ENSMUST00000114610.2 | 2 | 20548022 | 20662316 | 114295 | 1 | ENSMUST00 | 3047 | Etl4 | ENSMUSG0( | true | . | . | . | . | . | . | chr2 | 20549869 | 20552702 | 5.15133 | 4.19742 | 3.11894 |
| chr2 | 20519461 | 20519865 | 1000 | Promoter (<=1kb) | 2 | 20519791 | 20809885 | 290095 | 1 | ENSMUST00 | 0 | Etl4 | ENSMUSG0( | true | chr2 | 20519278 | 20521509 | 21.87917 | 60.42461 | 57.57317 | chr2 | 20518420 | 20519470 | 4.37789 | 3.76655 | 2.51007 |
| chr11 | 50153643 | 50153970 | 1000 | Intron (ENSMUST00000093138.7 | 11 | 50148132 | 50149965 | 1834 | 1 | ENSMUST00 | 5512 | Tbc1d9b | ENSMUSG0( | false | . | . | . | . | . | . | chr11 | 50152887 | 50155780 | 6.3472 | 4.63814 | 4.14588 |
| chr11 | 50163279 | 50163565 | 1000 | Promoter (1-2kb) | 11 | 50161663 | 50164095 | 2433 | 1 | ENSMUST00 | 1617 | Tbc1d9b | ENSMUSG0( | false | . | . | . | . | . | . | chr11 | 50161791 | 50168280 | 6.80256 | 4.8557 | 4.49881 |
| chr15 | 73184433 | 73185436 | 1000 | Promoter (<=1kb) | 15 | 73095844 | 73184840 | 88997 | 2 | ENSMUST00 | 0 | Ago2 | ENSMUSG0( | false | chr15 | 73183476 | 73185498 | 26.61085 | 108.21047 | 104.42454 | . | . | . | . | . | . |
| chr15 | 73154562 | 73155083 | 1000 | Intron (ENSMUST00000044113.1 | 15 | 73095844 | 73184840 | 88997 | 2 | ENSMUST00 | 29757 | Ago2 | ENSMUSG0( | false | . | . | . | . | . | . | chr15 | 73144959 | 73166179 | 6.45968 | 4.61688 | 4.22831 |
| chr1 | 85822346 | 85822709 | 1000 | Intron (ENSMUST00000113360.2 | 1 | 85843080 | 85846070 | 2991 | 1 | ENSMUST00 | -20371 | Cab39 | ENSMUSG0( | false | . | . | . | . | . | . | chr1 | 85821124 | 85831850 | 5.01423 | 4.04862 | 3.04524 |
| chr1 | 84619198 | 84619639 | 1000 | Intron (ENSMUST00000049126.7 | 1 | 84583004 | 84623841 | 40838 | 2 | ENSMUST00 | 4202 | Dner | ENSMUSG0( | false | . | . | . | . | . | . | chr1 | 84617512 | 84619270 | 3.90271 | 3.42683 | 2.12302 |
| chr8 | 88118496 | 88119449 | 1000 | Promoter (<=1kb) | 8 | 88118769 | 88135106 | 16338 | 1 | ENSMUST00 | 0 | Cnep1r1 | ENSMUSG0( | false | chr8 | 88118680 | 88119684 | 34.54477 | 122.48211 | 118.25522 | . | . | . | . | . | . |
| chr5 | 139202176 | 139202453 | 1000 | Promoter (1-2kb) | 5 | 139200685 | 139226604 | 25920 | 1 | ENSMUST00 | 1492 | Sun1 | ENSMUSG0( | false | . | . | . | . | . | . | chr5 | 139199896 | 139210866 | 5.02005 | 4.00761 | 3.04428 |
| chr5 | 139200266 | 139201068 | 1000 | Promoter (<=1kb) | 5 | 139200637 | 139249840 | 49204 | 1 | ENSMUST00 | 0 | Sun1 | ENSMUSG0( | false | chr5 | 139199935 | 139201685 | 24.86315 | 92.1349 | 88.74321 | chr5 | 139199896 | 139210866 | 5.02005 | 4.00761 | 3.04428 |
| chr3 | 146404303 | 146405022 | 1000 | Promoter (<=1kb) | 3 | 146404642 | 146440139 | 35498 | 1 | ENSMUST00 | 0 | Ssx2ip | ENSMUSG0( | false | chr3 | 146404092 | 146405945 | 31.85236 | 101.09525 | 97.496 | . | . | . | . | . | . |
| chr3 | 145987585 | 145988877 | 1000 | Promoter (<=1kb) | 3 | 145987870 | 146021720 | 33851 | 1 | ENSMUST00 | 0 | Syde2 | ENSMUSG0( | false | chr3 | 145987290 | 145989994 | 29.3377 | 90.74815 | 87.38457 | chr3 | 145988714 | 145994049 | 4.9728 | 4.0991 | 2.96728 |
| chr9 | 64011883 | 64012261 | 1000 | Exon (ENSMUST00000041029.5/ | 9 | 63954009 | 64016987 | 62979 | 2 | ENSMUST00 | 4726 | Smad6 | ENSMUSG0( | true | . | . | . | . | . | . | chr9 | 63948352 | 64020083 | 7.27873 | 5.05859 | 4.874 |
| chr9 | 63960221 | 63960800 | 1000 | Intron (ENSMUST00000041029.5 | 9 | 63954009 | 64016987 | 62979 | 2 | ENSMUST00 | 56187 | Smad6 | ENSMUSG0( | true | . | . | . | . | . | . | chr9 | 63948352 | 64020083 | 7.27873 | 5.05859 | 4.874 |
| chr9 | 63965480 | 63966057 | 1000 | Intron (ENSMUST00000041029.5 | 9 | 63954009 | 64016987 | 62979 | 2 | ENSMUST00 | 50930 | Smad6 | ENSMUSG0( | true | . | . | . | . | . | . | chr9 | 63948352 | 64020083 | 7.27873 | 5.05859 | 4.874 |
| chr9 | 63959268 | 63959650 | 1000 | Intron (ENSMUST00000041029.5 | 9 | 63954009 | 64016987 | 62979 | 2 | ENSMUST00 | 57337 | Smad6 | ENSMUSG0( | true | . | . | . | . | . | . | chr9 | 63948352 | 64020083 | 7.27873 | 5.05859 | 4.874 |
| chr9 | 63958109 | 63958698 | 1000 | Intron (ENSMUST00000041029.5 | 9 | 63954009 | 64016987 | 62979 | 2 | ENSMUST00 | 58289 | Smad6 | ENSMUSG0( | true | . | . | . | . | . | . | chr9 | 63948352 | 64020083 | 7.27873 | 5.05859 | 4.874 |
| chr3 | 145947462 | 145947818 | 1000 | Distal Intergenic | 3 | 145939086 | 145944198 | 5113 | 1 | ENSMUST00 | 8377 | 2410004B18 | ENSMUSG0( | false | . | . | . | . | . | . | chr3 | 145937374 | 145952538 | 5.27117 | 4.192 | 3.23206 |
| chr3 | 62338718 | 62339929 | 1000 | Promoter (<=1kb) | 3 | 62338999 | 62379719 | 40721 | 1 | ENSMUST00 | 0 | Arhgef26 | ENSMUSG0( | true | chr3 | 62338524 | 62341788 | 33.83522 | 116.97303 | 112.93475 | chr3 | 62338506 | 62338776 | 3.44654 | 3.36317 | 1.70888 |
| chr2 | 44926695 | 44927524 | 1000 | Promoter (<=1kb) | 2 | 44575478 | 44927161 | 351684 | 2 | ENSMUST00 | 0 | Gtdc1 | ENSMUSG0( | false | chr2 | 44926157 | 44927009 | 20.4053 | 65.4752 | 62.55133 | . | . | . | . | . | . |
| chr7 | 30456970 | 30458061 | 1000 | Promoter (<=1kb) | 7 | 30447534 | 30457652 | 10119 | 2 | ENSMUST00 | 0 | Kirrel2 | ENSMUSG0( | false | chr7 | 30456714 | 30457803 | 11.4883 | 23.92001 | 21.53215 | chr7 | 30456551 | 30458962 | 4.06036 | 2.72883 | 2.26132 |
| chr17 | 87379898 | 87380400 | 1000 | Promoter (1-2kb) | 17 | 87378853 | 87381769 | 2917 | 1 | ENSMUST00 | 1046 | Ttc7 | ENSMUSG0( | false | . | . | . | . | . | . | chr17 | 87375030 | 87381879 | 8.86103 | 5.81864 | 6.15311 |
| chr17 | 87387100 | 87387434 | 1000 | Downstream (2-3kb) | 17 | 87378853 | 87381769 | 2917 | 1 | ENSMUST00 | 8248 | Ttc7 | ENSMUSG0( | false | . | . | . | . | . | . | chr17 | 87385214 | 87399910 | 5.00955 | 4.02526 | 3.02481 |
| chr4 | 136602174 | 136602896 | 1000 | Promoter (<=1kb) | 4 | 136550540 | 136602723 | 52184 | 2 | ENSMUST00 | 0 | Kdm1a | ENSMUSG0( | false | chr4 | 136601489 | 136602829 | 37.62274 | 127.9987 | 123.57949 | . | . | . | . | . | . |
| chrX | 48343363 | 48344002 | 1000 | Promoter (<=1kb) | 20 | 48343770 | 48408049 | 64280 | 1 | ENSMUST00 | 0 | Bcorl1 | ENSMUSG0( | false | chrX | 48343229 | 48343507 | 7.81205 | 13.8349 | 11.58824 | . | . | . | . | . | . |
| chrX | 48341214 | 48342116 | 1000 | Promoter (<=1kb) | 20 | 48341358 | 48408049 | 66692 | 1 | ENSMUST00 | 0 | Bcorl1 | ENSMUSG0( | false | chrX | 48340813 | 48342607 | 16.08362 | 38.04472 | 35.47989 | . | . | . | . | . | . |
| chr15 | 102135985 | 102136579 | 1000 | Promoter (<=1kb) | 15 | 102116528 | 102136192 | 19665 | 2 | ENSMUST00 | 0 | Spryd3 | ENSMUSG0( | false | chr15 | 102135500 | 102136850 | 33.04283 | 115.53907 | 111.53828 | chr15 | 102135156 | 102135995 | 8.61367 | 5.80951 | 5.93894 |
| chr17 | 3597914 | 3598332 | 1000 | Distal Intergenic | 17 | 3519263 | 3557713 | 38451 | 2 | ENSMUST00 | -40202 | Tfb1m | ENSMUSG0( | false | . | . | . | . | . | . | chr17 | 3596798 | 3599226 | 4.166 | 3.58578 | 2.36535 |
| chr7 | 105633229 | 105633834 | 1000 | Promoter (<=1kb) | 7 | 105604463 | 105633508 | 29046 | 2 | ENSMUST00 | 0 | Trim3 | ENSMUSG0( | false | chr7 | 105632662 | 105634117 | 33.94791 | 109.87469 | 106.04007 | . | . | . | . | . | . |
| chr15 | 102105409 | 102105842 | 1000 | Promoter (2-3kb) | 15 | 102102988 | 102116401 | 13414 | 1 | ENSMUST00 | 2422 | Tenc1 | ENSMUSG0( | false | chr15 | 102099897 | 102107616 | 31.25476 | 101.05123 | 97.45738 | chr15 | 102105799 | 102117464 | 7.8319 | 5.25795 | 5.32662 |
| chr15 | 102106274 | 102106836 | 1000 | Exon (ENSMUST00000046144.9/ | 15 | 102102988 | 102116401 | 13414 | 1 | ENSMUST00 | 3287 | Tenc1 | ENSMUSG0( | false | chr15 | 102099897 | 102107616 | 31.25476 | 101.05123 | 97.45738 | chr15 | 102105799 | 102117464 | 7.8319 | 5.25795 | 5.32662 |
| chr15 | 102102435 | 102103091 | 1000 | Promoter (<=1kb) | 15 | 102102988 | 102116391 | 13404 | 1 | ENSMUST00 | 0 | Tenc1 | ENSMUSG0( | false | chr15 | 102099897 | 102107616 | 31.25476 | 101.05123 | 97.45738 | chr15 | 102097473 | 102102796 | 4.79775 | 3.79288 | 2.84744 |
| chr15 | 102099992 | 102101362 | 1000 | Promoter (1-2kb) | 15 | 102102988 | 102116391 | 13404 | 1 | ENSMUST00 | -1626 | Tenc1 | ENSMUSG0( | false | chr15 | 102099897 | 102107616 | 31.25476 | 101.05123 | 97.45738 | chr15 | 102097473 | 102102796 | 4.79775 | 3.79288 | 2.84744 |
| chr15 | 102103342 | 102104511 | 1000 | Promoter (<=1kb) | 15 | 102102988 | 102116401 | 13414 | 1 | ENSMUST00 | 355 | Tenc1 | ENSMUSG0( | false | chr15 | 102099897 | 102107616 | 31.25476 | 101.05123 | 97.45738 | . | . | . | . | . | . |
| chr10 | 62339776 | 62340739 | 1000 | Promoter (<=1kb) | 10 | 62268855 | 62340542 | 71688 | 2 | ENSMUST00 | 0 | Hk1 | ENSMUSG0( | false | chr10 | 62338804 | 62341149 | 28.59476 | 85.59679 | 82.34128 | . | . | . | . | . | . |
| chr8 | 79294625 | 79295227 | 1000 | Promoter (<=1kb) | 8 | 79266424 | 79294956 | 28533 | 2 | ENSMUST00 | 0 | Mmaa | ENSMUSG0( | false | chr8 | 79293953 | 79295699 | 28.83278 | 87.92931 | 84.62811 | . | . | . | . | . | . |
| chr7 | 30169619 | 30170189 | 1000 | Promoter (<=1kb) | 7 | 30161274 | 30169727 | 8454 | 2 | ENSMUST00 | 0 | Zfp146 | ENSMUSG0( | false | chr7 | 30168946 | 30170898 | 24.99193 | 78.31828 | 75.19689 | . | . | . | . | . | . |
| chr10 | 62216544 | 62217199 | 1000 | Intron (ENSMUST00000047883.9 | 10 | 62185396 | 62231218 | 45823 | 2 | ENSMUST00 | 14019 | Tspan15 | ENSMUSG0( | false | . | . | . | . | . | . | chr10 | 62185685 | 62229768 | 8.28034 | 5.46771 | 5.70297 |
| chr1 | 119421513 | 119422500 | 1000 | Promoter (<=1kb) | 1 | 119415465 | 119422248 | 6784 | 2 | ENSMUST00 | 0 | Inhbb | ENSMUSG0( | false | chr1 | 119420266 | 119422956 | 32.43201 | 97.0959 | 93.5947 | chr1 | 119419981 | 119423043 | 4.45543 | 3.64568 | 2.59494 |
| chr7 | 105554073 | 105554511 | 1000 | Promoter (<=1kb) | 7 | 105554360 | 105558388 | 4029 | 1 | ENSMUST00 | 0 | Smpd1 | ENSMUSG0( | false | chr7 | 105553586 | 105555108 | 25.1466 | 74.00616 | 70.95453 | . | . | . | . | . | . |
| chr18 | 35598458 | 35599320 | 1000 | Promoter (<=1kb) | 18 | 35598617 | 35617186 | 18570 | 1 | ENSMUST00 | 0 | Paip2 | ENSMUSG0( | false | chr18 | 35598305 | 35599706 | 31.91637 | 110.38313 | 106.53261 | chr18 | 35597515 | 35598513 | 5.20445 | 4.25377 | 3.16315 |
| chr3 | 144719425 | 144720725 | 1000 | Promoter (<=1kb) | 3 | 144688844 | 144720323 | 31480 | 2 | ENSMUST00 | 0 | Sh3glb1 | ENSMUSG0( | false | chr3 | 144718529 | 144720795 | 29.95388 | 90.84766 | 87.48237 | chr3 | 144720604 | 144721592 | 4.82332 | 4.05267 | 2.85267 |
| chr3 | 144685099 | 144685782 | 1000 | Distal Intergenic | 3 | 144688844 | 144720323 | 31480 | 2 | ENSMUST00 | 34541 | Sh3glb1 | ENSMUSG0( | false | . | . | . | . | . | . | chr3 | 144681472 | 144689973 | 5.50423 | 4.31451 | 3.423 |
| chr15 | 58888415 | 58889640 | 1000 | Promoter (<=1kb) | 15 | 58889229 | 58902390 | 13162 | 1 | ENSMUST00 | 0 | Rnf139 | ENSMUSG0( | false | chr15 | 58888090 | 58890604 | 39.05373 | 137.10143 | 132.33189 | . | . | . | . | . | . |
| chr17 | 87107024 | 87107962 | 1000 | Promoter (<=1kb) | 17 | 87107679 | 87137583 | 29905 | 1 | ENSMUST00 | 0 | Socs5 | ENSMUSG0( | false | chr17 | 87106372 | 87108483 | 40.89836 | 130.45236 | 125.94646 | . | . | . | . | . | . |
| chr3 | 51560158 | 51561155 | 1000 | Promoter (<=1kb) | 3 | 51515319 | 51560878 | 45560 | 2 | ENSMUST00 | 0 | Setd7 | ENSMUSG0( | true | chr3 | 51558835 | 51561958 | 30.28657 | 98.33401 | 94.80609 | . | . | . | . | . | . |
| chr9 | 51031571 | 51031872 | 1000 | Distal Intergenic | 9 | 50892801 | 51009073 | 116273 | 2 | ENSMUST00 | -22499 | Sik2 | ENSMUSG0( | false | . | . | . | . | . | . | chr9 | 51030297 | 51033186 | 4.59576 | 3.8556 | 2.67702 |
| chr9 | 51008642 | 51009362 | 1000 | Promoter (<=1kb) | 9 | 50892801 | 51008936 | 116136 | 2 | ENSMUST00 | 0 | Sik2 | ENSMUSG0( | false | chr9 | 51009289 | 51009537 | 8.26071 | 16.87976 | 14.58833 | . | . | . | . | . | . |
| chr1 | 38627087 | 38627840 | 1000 | Promoter (<=1kb) | 1 | 38177326 | 38627185 | 449860 | 2 | ENSMUST00 | 0 | Aff3 | ENSMUSG0( | true | chr1 | 38626226 | 38628260 | 26.28407 | 85.32412 | 82.07336 | chr1 | 38625765 | 38629542 | 4.04709 | 3.59692 | 2.22598 |
| chr2 | 146047223 | 146047589 | 1000 | Promoter (<=1kb) | 2 | 146047251 | 146215035 | 167785 | 1 | ENSMUST00 | 0 | 4930529M08 | ENSMUSG0( | false | . | . | . | . | . | . | chr2 | 146044913 | 146047912 | 3.26816 | 3.08746 | 1.61884 |
| chr8 | 77517220 | 77518270 | 1000 | Promoter (<=1kb) | 8 | 77250366 | 77517907 | 267542 | 2 | ENSMUST00 | 0 | Arhgap10 | ENSMUSG0( | false | chr8 | 77517035 | 77518647 | 38.03245 | 121.19012 | 117.00504 | chr8 | 77515984 | 77518887 | 4.34071 | 3.79342 | 2.51663 |
| chr12 | 12966827 | 12967238 | 1000 | Distal Intergenic | 12 | 12936096 | 12941914 | 5819 | 2 | ENSMUST00 | -24914 | Mycn | ENSMUSG0( | false | . | . | . | . | . | . | chr12 | 12966075 | 12968824 | 4.48136 | 3.62934 | 2.62043 |
| chr12 | 13048359 | 13048690 | 1000 | Distal Intergenic | 12 | 12936096 | 12941914 | 5819 | 2 | ENSMUST00 | -106446 | Mycn | ENSMUSG0( | false | . | . | . | . | . | . | chr12 | 13047549 | 13049868 | 4.4634 | 3.81848 | 2.58992 |
| chr6 | 40429541 | 40430065 | 1000 | Intron (ENSMUST00000101492.4 | 6 | 40407500 | 40436133 | 28634 | 2 | ENSMUST00 | 6068 | E330009J07F | ENSMUSG0( | false | . | . | . | . | . | . | chr6 | 40428230 | 40432053 | 6.33133 | 4.73863 | 4.08389 |
| chr3 | 51275955 | 51277685 | 1000 | Promoter (<=1kb) | 3 | 51255665 | 51277514 | 21850 | 2 | ENSMUST00 | 0 | Elf2 | ENSMUSG0( | false | chr3 | 51275540 | 51278105 | 36.42221 | 131.26616 | 126.73016 | chr3 | 51274141 | 51276004 | 5.20196 | 4.197 | 3.18969 |
| chr3 | 51339934 | 51342130 | 1000 | Promoter (<=1kb) | 3 | 51255665 | 51340566 | 84902 | 2 | ENSMUST00 | 0 | Elf2 | ENSMUSG0( | false | chr3 | 51339237 | 51342406 | 37.30079 | 124.19571 | 119.91465 | chr3 | 51337503 | 51339957 | 3.65118 | 3.41338 | 1.89628 |
| chr17 | 10581836 | 10582268 | 1000 | Intron (ENSMUST00000041463.6 | 17 | 10576478 | 10631067 | 54590 | 2 | ENSMUST00 | 48799 | Pacrg | ENSMUSG0( | false | . | . | . | . | . | . | chr17 | 10578259 | 10587776 | 4.97694 | 4.03138 | 2.99727 |
| chr17 | 10677179 | 10677474 | 1000 | Intron (ENSMUST00000041463.6 | 17 | 10576478 | 10631067 | 54590 | 2 | ENSMUST00 | -46113 | Pacrg | ENSMUSG0( | false | . | . | . | . | . | . | chr17 | 10673669 | 10678435 | 5.44605 | 4.27759 | 3.37178 |
| chr2 | 11553509 | 11554214 | 1000 | Distal Intergenic | 2 | 11585439 | 11585990 | 552 | 2 | ENSMUST00 | 31776 | Rbm17 | ENSMUSG0( | false | chr2 | 11552855 | 11554333 | 29.3377 | 90.74815 | 87.38457 | chr2 | 11552415 | 11554325 | 5.11798 | 4.14934 | 3.09454 |
| chr5 | 34369639 | 34370130 | 1000 | Promoter (<=1kb) | 5 | 34369933 | 34486456 | 116524 | 1 | ENSMUST00 | 0 | Fam193a | ENSMUSG0( | false | chr5 | 34369227 | 34371027 | 33.69293 | 107.36673 | 103.60477 | . | . | . | . | . | . |
| chr3 | 37666748 | 37667383 | 1000 | Distal Intergenic | 3 | 37640080 | 37644598 | 4519 | 1 | ENSMUST00 | 26669 | Spry1 | ENSMUSG0( | false | . | . | . | . | . | . | chr3 | 37665617 | 37667922 | 6.36996 | 4.77213 | 4.16105 |
| chr8 | 26158369 | 26158971 | 1000 | Promoter (<=1kb) | 8 | 26158255 | 26159412 | 1158 | 1 | ENSMUST00 | 115 | Thap1 | ENSMUSG0( | false | chr8 | 26157472 | 26159346 | 22.72043 | 98.85921 | 95.31703 | . | . | . | . | . | . |
| chr5 | 34187007 | 34187891 | 1000 | Promoter (<=1kb) | 5 | 34173887 | 34187710 | 13824 | 2 | ENSMUST00 | 0 | Mxd4 | ENSMUSG0( | false | chr5 | 34186146 | 34187794 | 31.96831 | 119.80809 | 115.67081 | . | . | . | . | . | . |
| chr18 | 35561907 | 35563012 | 1000 | Promoter (<=1kb) | 18 | 35562146 | 35572490 | 10345 | 1 | ENSMUST00 | 0 | Matr3 | ENSMUSG0( | false | chr18 | 35561702 | 35564147 | 30.76814 | 122.68065 | 118.44827 | chr18 | 35561410 | 35561937 | 3.28211 | 3.07909 | 1.64521 |
| chr4 | 135235383 | 135235919 | 1000 | Intron (ENSMUST00000037099.8 | 4 | 135213969 | 135272814 | 58846 | 2 | ENSMUST00 | 36895 | Clic4 | ENSMUSG0( | false | . | . | . | . | . | . | chr4 | 135234407 | 135236534 | 5.05622 | 3.86999 | 3.07715 |
| chr4 | 135271933 | 135273336 | 1000 | Promoter (<=1kb) | 4 | 135213969 | 135272814 | 58846 | 2 | ENSMUST00 | 0 | Clic4 | ENSMUSG0( | false | chr4 | 135271193 | 135272820 | 27.66025 | 106.38611 | 102.65278 | chr4 | 135273275 | 135273904 | 6.7133 | 4.92367 | 4.38454 |
| chr18 | 73571890 | 73573757 | 1000 | Promoter (<=1kb) | 18 | 73573044 | 73592575 | 19532 | 1 | ENSMUST00 | 0 | Mex3c | ENSMUSG0( | false | chr18 | 73571379 | 73572465 | 18.02578 | 59.25625 | 56.42029 | . | . | . | . | . | . |
| chr9 | 63610824 | 63611500 | 1000 | Intron (ENSMUST00000041551.7 | 9 | 63602655 | 63641889 | 39235 | 1 | ENSMUST00 | 8170 | Aagab | ENSMUSG0( | false | chr9 | 63611389 | 63611663 | 4.77198 | 8.97201 | 6.81752 | chr9 | 63603357 | 63616669 | 6.44347 | 4.69808 | 4.19638 |
| chr8 | 25964350 | 25964726 | 1000 | Intron (ENSMUST00000037609.6 | 8 | 25944459 | 25976744 | 32286 | 2 | ENSMUST00 | 12018 | Hgsnat | ENSMUSG0( | false | . | . | . | . | . | . | chr8 | 25961491 | 25965675 | 6.4418 | 4.64993 | 4.22382 |
| chr9 | 100642794 | 100644427 | 1000 | Promoter (<=1kb) | 9 | 100643448 | 100796791 | 153344 | 1 | ENSMUST00 | 0 | Stag1 | ENSMUSG0( | false | chr9 | 100643537 | 100645131 | 36.08632 | 111.51659 | 107.63924 | . | . | . | . | . | . |
| chr9 | 42463708 | 42463997 | 1000 | Promoter (2-3kb) | 9 | 42413418 | 42461461 | 48044 | 2 | ENSMUST00 | -2248 | Tbcel | ENSMUSG0( | false | . | . | . | . | . | . | chr9 | 42458861 | 42464772 | 4.87425 | 3.90908 | 2.92868 |
| chr4 | 134767691 | 134768202 | 1000 | Promoter (<=1kb) | 4 | 134744735 | 134768024 | 23290 | 2 | ENSMUST00 | 0 | Ldlrap1 | ENSMUSG0( | false | chr4 | 134766368 | 134768617 | 40.29262 | 133.77719 | 129.14264 | . | . | . | . | . | . |
| chr4 | 134760730 | 134761377 | 1000 | Intron (ENSMUST00000037828.6 | 4 | 134744735 | 134768024 | 23290 | 2 | ENSMUST00 | 6647 | Ldlrap1 | ENSMUSG0( | false | . | . | . | . | . | . | chr4 | 134750748 | 134766877 | 7.637 | 5.15315 | 5.18044 |
| chr4 | 134683886 | 134684339 | 1000 | Intron (ENSMUST00000038628.3 | 4 | 134567889 | 134703767 | 135879 | 2 | ENSMUST00 | 19428 | Man1c1 | ENSMUSG0( | false | . | . | . | . | . | . | chr4 | 134682467 | 134687528 | 5.9319 | 4.46549 | 3.78246 |
| chr4 | 134685902 | 134686457 | 1000 | Intron (ENSMUST00000038628.3 | 4 | 134567889 | 134703767 | 135879 | 2 | ENSMUST00 | 17310 | Man1c1 | ENSMUSG0( | false | . | . | . | . | . | . | chr4 | 134682467 | 134687528 | 5.9319 | 4.46549 | 3.78246 |
| chrX | 20425325 | 20425900 | 1000 | Promoter (<=1kb) | 20 | 20425688 | 20519939 | 94252 | 1 | ENSMUST00 | 0 | Jade3 | ENSMUSG0( | false | chrX | 20424963 | 20426636 | 22.51707 | 59.83231 | 56.98677 | . | . | . | . | . | . |
| chr11 | 58009684 | 58010203 | 1000 | Promoter (<=1kb) | 11 | 58009064 | 58062034 | 52971 | 1 | ENSMUST00 | 621 | Larp1 | ENSMUSG0( | false | chr11 | 58008900 | 58010805 | 30.65557 | 96.00109 | 92.52367 | . | . | . | . | . | . |
| chr4 | 134496730 | 134497459 | 1000 | Promoter (<=1kb) | 4 | 134496975 | 134507215 | 10241 | 1 | ENSMUST00 | 0 | Paqr7 | ENSMUSG0( | false | chr4 | 134496595 | 134497573 | 25.16617 | 75.86667 | 72.78426 | chr4 | 134494929 | 134498560 | 5.35258 | 4.14792 | 3.3166 |
| chrX | 18164040 | 18164385 | 1000 | Promoter (1-2kb) | 20 | 18162648 | 18258968 | 96321 | 1 | ENSMUST00 | 1393 | Kdm6a | ENSMUSG0( | false | chrX | 18162564 | 18165181 | 23.43614 | 63.09834 | 60.20888 | . | . | . | . | . | . |
| chr10 | 24711851 | 24712222 | 1000 | Promoter (<=1kb) | 10 | 24637914 | 24712158 | 74245 | 2 | ENSMUST00 | 0 | Enpp1 | ENSMUSG0( | false | chr10 | 24711471 | 24712723 | 30.42596 | 90.69117 | 87.33178 | chr10 | 24711250 | 24712481 | 4.5993 | 3.86544 | 2.68248 |
| chr1 | 192735946 | 192736400 | 1000 | Promoter (<=1kb) | 1 | 192726197 | 192737279 | 11083 | 2 | ENSMUST00 | 879 | Hhat | ENSMUSG0( | false | . | . | . | . | . | . | chr1 | 192735527 | 192737336 | 6.3836 | 4.57699 | 4.17778 |
| chr1 | 192770525 | 192771361 | 1000 | Promoter (<=1kb) | 1 | 192512828 | 192771213 | 258386 | 2 | ENSMUST00 | 0 | Hhat | ENSMUSG0( | false | chr1 | 192769875 | 192771502 | 28.91247 | 96.86193 | 93.36482 | chr1 | 192771277 | 192771876 | 3.13421 | 3.09077 | 1.46216 |
| chr11 | 56023963 | 56024646 | 1000 | Intron (ENSMUST00000134285.1 | 11 | 56024987 | 56041010 | 16024 | 2 | ENSMUST00 | 16364 | Nmur2 | ENSMUSG0( | false | . | . | . | . | . | . | chr11 | 56023459 | 56026861 | 4.02889 | 3.48536 | 2.26023 |
| chr1 | 192136679 | 192138593 | 1000 | Promoter (<=1kb) | 1 | 192098656 | 192138010 | 39355 | 2 | ENSMUST00 | 0 | Rcor3 | ENSMUSG0( | false | chr1 | 192136182 | 192138954 | 25.18016 | 100.40004 | 96.81754 | . | . | . | . | . | . |
| chr1 | 36471218 | 36471829 | 1000 | Promoter (<=1kb) | 1 | 36471620 | 36508764 | 37145 | 1 | ENSMUST00 | 0 | Cnnm4 | ENSMUSG0( | false | chr1 | 36470838 | 36472511 | 30.19165 | 95.18787 | 91.73401 | chr1 | 36469968 | 36476122 | 3.74013 | 3.42249 | 2.01434 |
| chr9 | 90270510 | 90271047 | 1000 | Promoter (<=1kb) | 9 | 90202027 | 90270769 | 68743 | 2 | ENSMUST00 | 0 | Tbc1d2b | ENSMUSG0( | false | chr9 | 90269919 | 90271221 | 30.86817 | 91.45322 | 88.08056 | . | . | . | . | . | . |
| chr5 | 32991280 | 32991781 | 1000 | 3' UTR | 5 | 32967719 | 32974176 | 6458 | 1 | ENSMUST00 | 23562 | Depdc5 | ENSMUSG0( | false | . | . | . | . | . | . | chr5 | 32989461 | 32996528 | 5.20315 | 4.09317 | 3.21514 |
| chr5 | 137032462 | 137033062 | 1000 | Promoter (1-2kb) | 5 | 137030701 | 137031407 | 707 | 1 | ENSMUST00 | 1762 | Vgf | ENSMUSG0( | false | chr5 | 137031245 | 137033291 | 15.50707 | 38.47921 | 35.908 | chr5 | 137029894 | 137033165 | 7.20686 | 4.82218 | 4.83474 |
| chr1 | 191906144 | 191907000 | 1000 | Promoter (<=1kb) | 1 | 191906767 | 191913247 | 6481 | 1 | ENSMUST00 | 0 | Slc30a1 | ENSMUSG0( | false | chr1 | 191905836 | 191908156 | 38.67513 | 141.95352 | 136.99454 | . | . | . | . | . | . |
| chr4 | 134187014 | 134187392 | 1000 | Promoter (<=1kb) | 4 | 134129858 | 134187112 | 57255 | 2 | ENSMUST00 | 0 | Cep85 | ENSMUSG0( | false | chr4 | 134186247 | 134187154 | 25.15761 | 80.47005 | 77.31023 | . | . | . | . | . | . |
| chr1 | 36307559 | 36308268 | 1000 | Promoter (<=1kb) | 1 | 36307733 | 36319203 | 11471 | 1 | ENSMUST00 | 0 | Arid5a | ENSMUSG0( | false | chr1 | 36307691 | 36309096 | 38.6007 | 121.10697 | 116.92481 | chr1 | 36307219 | 36312041 | 4.43854 | 3.74332 | 2.57321 |
| chr15 | 38518214 | 38518887 | 1000 | Promoter (<=1kb) | 15 | 38500436 | 38518221 | 17786 | 2 | ENSMUST00 | 0 | Azin1 | ENSMUSG0( | false | chr15 | 38517684 | 38520107 | 40.21293 | 133.6382 | 129.0087 | . | . | . | . | . | . |
| chr15 | 38298701 | 38300161 | 1000 | Promoter (<=1kb) | 15 | 38294413 | 38300707 | 6295 | 2 | ENSMUST00 | 546 | Klf10 | ENSMUSG0( | false | chr15 | 38298493 | 38298933 | 9.32368 | 22.97803 | 20.60213 | chr15 | 38297535 | 38302848 | 5.70678 | 4.40106 | 3.58096 |
| chr15 | 38300649 | 38301545 | 1000 | Promoter (<=1kb) | 15 | 38294413 | 38300707 | 6295 | 2 | ENSMUST00 | 0 | Klf10 | ENSMUSG0( | false | chr15 | 38299118 | 38302130 | 30.08562 | 102.9821 | 99.33765 | chr15 | 38297535 | 38302848 | 5.70678 | 4.40106 | 3.58096 |
| chr15 | 38077829 | 38079134 | 1000 | Promoter (<=1kb) | 15 | 37967328 | 38078853 | 111526 | 2 | ENSMUST00 | 0 | Ubr5 | ENSMUSG0( | false | chr15 | 38077360 | 38079609 | 36.01109 | 111.38685 | 107.51837 | . | . | . | . | . | . |
| chr10 | 22674621 | 22674986 | 1000 | Intron (ENSMUST00000042261.4 | 10 | 22649008 | 22664690 | 15683 | 1 | ENSMUST00 | 25614 | Slc2a12 | ENSMUSG0( | false | . | . | . | . | . | . | chr10 | 22670879 | 22677260 | 4.29046 | 3.74548 | 2.42265 |
| chr10 | 22655631 | 22656221 | 1000 | Intron (ENSMUST00000042261.4 | 10 | 22649008 | 22664690 | 15683 | 1 | ENSMUST00 | 6624 | Slc2a12 | ENSMUSG0( | false | . | . | . | . | . | . | . | . | . | . | . | . |
| chr10 | 22644969 | 22645638 | 1000 | Promoter (<=1kb) | 10 | 22645011 | 22704285 | 59275 | 1 | ENSMUST00 | 0 | Slc2a12 | ENSMUSG0( | false | chr10 | 22644821 | 22645805 | 22.44027 | 62.24483 | 59.36732 | chr10 | 22643381 | 22650117 | 5.38368 | 4.25134 | 3.32798 |
| chr1 | 34842505 | 34843202 | 1000 | Promoter (<=1kb) | 1 | 34813219 | 34843050 | 29832 | 2 | ENSMUST00 | 0 | Fam168b | ENSMUSG0( | false | chr1 | 34841471 | 34843649 | 32.03847 | 101.42104 | 97.81718 | . | . | . | . | . | . |
| chr7 | 144531964 | 144532333 | 1000 | Intron (ENSMUST00000168134.4 | 7 | 144520283 | 144524370 | 4088 | 2 | ENSMUST00 | -7595 | Ppfia1 | ENSMUSG0( | false | . | . | . | . | . | . | chr7 | 144526111 | 144534909 | 5.1889 | 4.026 | 3.18747 |
| chr11 | 54522560 | 54523234 | 1000 | Promoter (<=1kb) | 11 | 54522847 | 54699285 | 176439 | 1 | ENSMUST00 | 0 | Rapgef6 | ENSMUSG0( | true | chr11 | 54521936 | 54523905 | 30.08562 | 102.9821 | 99.33765 | . | . | . | . | . | . |
| chr10 | 21399031 | 21399344 | 1000 | Distal Intergenic | 10 | 21419358 | 21426565 | 7208 | 1 | ENSMUST00 | -20014 | 1700021A07 | ENSMUSG0( | false | . | . | . | . | . | . | chr10 | 21397899 | 21400506 | 3.74931 | 3.3396 | 2.01461 |
| chr7 | 28375370 | 28376018 | 1000 | Promoter (2-3kb) | 7 | 28361092 | 28372599 | 11508 | 2 | ENSMUST00 | -2772 | Plekhg2 | ENSMUSG0( | true | chr7 | 28375275 | 28379742 | 28.38021 | 81.44218 | 78.26595 | chr7 | 28375103 | 28383860 | 6.23801 | 4.67039 | 4.01439 |
| chr7 | 28372277 | 28373080 | 1000 | Promoter (<=1kb) | 7 | 28359605 | 28372494 | 12890 | 2 | ENSMUST00 | 0 | Plekhg2 | ENSMUSG0( | true | chr7 | 28370828 | 28372772 | 30.78991 | 105.27288 | 101.56436 | . | . | . | . | . | . |
| chr11 | 94210317 | 94212400 | 1000 | Promoter (<=1kb) | 11 | 94211454 | 94215495 | 4042 | 1 | ENSMUST00 | 0 | Tob1 | ENSMUSG0( | false | chr11 | 94211421 | 94215068 | 50.8003 | 176.12823 | 170.0448 | chr11 | 94209745 | 94210697 | 3.98673 | 3.64849 | 2.15381 |
| chr5 | 81519330 | 81519942 | 1000 | Intron (ENSMUST00000132375.2 | 5 | 81645989 | 81688522 | 42534 | 1 | ENSMUST00 | -126047 | Lphn3 | ENSMUSG0( | false | chr5 | 81519672 | 81520026 | 5.15049 | 9.68248 | 7.51107 | chr5 | 81518468 | 81520895 | 3.83839 | 3.51106 | 2.04204 |
| chr5 | 81345360 | 81345620 | 1000 | Intron (ENSMUST00000147226.2 | 5 | 81310025 | 81796634 | 486610 | 1 | ENSMUST00 | 35336 | Lphn3 | ENSMUSG0( | false | . | . | . | . | . | . | chr5 | 81344436 | 81346072 | 3.81667 | 3.45707 | 2.04561 |
| chr7 | 143739930 | 143740628 | 1000 | Promoter (<=1kb) | 7 | 143688762 | 143740341 | 51580 | 2 | ENSMUST00 | 0 | Osbpl5 | ENSMUSG0( | false | chr7 | 143738218 | 143740816 | 29.58769 | 108.11006 | 104.32592 | chr7 | 143710495 | 143741311 | 6.41483 | 4.62224 | 4.19611 |
| chr7 | 143738490 | 143739050 | 1000 | Promoter (1-2kb) | 7 | 143713496 | 143740299 | 26804 | 2 | ENSMUST00 | 1249 | Osbpl5 | ENSMUSG0( | false | chr7 | 143738218 | 143740816 | 29.58769 | 108.11006 | 104.32592 | chr7 | 143710495 | 143741311 | 6.41483 | 4.62224 | 4.19611 |
| chr6 | 72216634 | 72217267 | 1000 | Intron (ENSMUST00000042646.7 | 6 | 72206177 | 72235577 | 29401 | 2 | ENSMUST00 | 18310 | Atoh8 | ENSMUSG0( | false | . | . | . | . | . | . | chr6 | 72211696 | 72224884 | 7.46431 | 5.12594 | 5.03432 |
| chr6 | 72235038 | 72235809 | 1000 | Promoter (<=1kb) | 6 | 72206177 | 72235577 | 29401 | 2 | ENSMUST00 | 0 | Atoh8 | ENSMUSG0( | false | chr6 | 72233454 | 72235881 | 33.76198 | 105.43819 | 101.727 | chr6 | 72235545 | 72237554 | 5.86328 | 4.31826 | 3.75114 |
| chr4 | 133338665 | 133339686 | 1000 | Promoter (<=1kb) | 4 | 133292459 | 133339324 | 46866 | 2 | ENSMUST00 | 0 | Wdtc1 | ENSMUSG0( | false | chr4 | 133338191 | 133339666 | 33.66875 | 128.30269 | 123.87101 | chr4 | 133337835 | 133339844 | 3.28447 | 3.11896 | 1.63239 |
| chr3 | 30995443 | 30996305 | 1000 | Promoter (<=1kb) | 3 | 30995747 | 31052741 | 56995 | 1 | ENSMUST00 | 0 | Prkci | ENSMUSG0( | false | chr3 | 30995072 | 30997340 | 37.7523 | 123.91259 | 119.6447 | . | . | . | . | . | . |
| chr15 | 35371101 | 35371919 | 1000 | Promoter (<=1kb) | 15 | 35371546 | 35931228 | 559683 | 1 | ENSMUST00 | 0 | Vps13b | ENSMUSG0( | false | chr15 | 35370761 | 35372510 | 33.94791 | 109.87469 | 106.04007 | . | . | . | . | . | . |
| chr7 | 143460819 | 143461334 | 1000 | Promoter (<=1kb) | 7 | 143458350 | 143461050 | 2701 | 2 | ENSMUST00 | 0 | Cdkn1c | ENSMUSG0( | true | chr7 | 143458479 | 143460972 | 21.40274 | 64.72344 | 61.81032 | . | . | . | . | . | . |
| chr7 | 143459436 | 143459859 | 1000 | Promoter (1-2kb) | 7 | 143458350 | 143461050 | 2701 | 2 | ENSMUST00 | 1191 | Cdkn1c | ENSMUSG0( | true | chr7 | 143458479 | 143460972 | 21.40274 | 64.72344 | 61.81032 | . | . | . | . | . | . |
| chr12 | 8154568 | 8155146 | 1000 | Distal Intergenic | 12 | 8208107 | 8285758 | 77652 | 1 | ENSMUST00 | -52961 | 1110057K04I | ENSMUSG0( | false | . | . | . | . | . | . | chr12 | 8153962 | 8156350 | 4.18808 | 3.6492 | 2.38285 |
| chr9 | 72531777 | 72532956 | 1000 | Promoter (<=1kb) | 9 | 72532240 | 72622829 | 90590 | 1 | ENSMUST00 | 0 | Rfx7 | ENSMUSG0( | false | chr9 | 72531033 | 72533138 | 41.81742 | 134.22485 | 129.57179 | chr9 | 72532839 | 72535864 | 3.47692 | 3.30893 | 1.75049 |
| chr12 | 112588286 | 112588981 | 1000 | Promoter (<=1kb) | 12 | 112588784 | 112615556 | 26773 | 1 | ENSMUST00 | 0 | Inf2 | ENSMUSG0( | false | chr12 | 112588066 | 112588495 | 7.01548 | 12.12394 | 9.90998 | chr12 | 112585329 | 112600582 | 8.19005 | 5.40781 | 5.63321 |
| chr12 | 112576295 | 112577092 | 1000 | Distal Intergenic | 12 | 112588784 | 112615556 | 26773 | 1 | ENSMUST00 | -11692 | Inf2 | ENSMUSG0( | false | . | . | . | . | . | . | chr12 | 112575007 | 112581917 | 7.31209 | 4.88694 | 4.93798 |
| chr5 | 67847120 | 67847627 | 1000 | Promoter (<=1kb) | 5 | 67618140 | 67847432 | 229293 | 2 | ENSMUST00 | 0 | Atp8a1 | ENSMUSG0( | true | chr5 | 67845830 | 67848432 | 32.29185 | 112.09676 | 108.19604 | . | . | . | . | . | . |
| chr12 | 112088468 | 112088983 | 1000 | Distal Intergenic | 12 | 112106683 | 112127573 | 20891 | 1 | ENSMUST00 | -17700 | Aspg | ENSMUSG0( | false | . | . | . | . | . | . | chr12 | 112085557 | 112091113 | 7.01668 | 5.01483 | 4.65162 |
| chr12 | 112096053 | 112096572 | 1000 | Distal Intergenic | 12 | 112106683 | 112127573 | 20891 | 1 | ENSMUST00 | -10111 | Aspg | ENSMUSG0( | false | . | . | . | . | . | . | chr12 | 112095270 | 112097183 | 5.87095 | 4.4284 | 3.73789 |
| chr12 | 112086714 | 112087379 | 1000 | Distal Intergenic | 12 | 112106683 | 112127573 | 20891 | 1 | ENSMUST00 | -19304 | Aspg | ENSMUSG0( | false | . | . | . | . | . | . | chr12 | 112085557 | 112091113 | 7.01668 | 5.01483 | 4.65162 |
| chr4 | 133068448 | 133068832 | 1000 | Promoter (2-3kb) | 4 | 133066249 | 133072063 | 5815 | 1 | ENSMUST00 | 2200 | Ahdc1 | ENSMUSG0( | false | . | . | . | . | . | . | chr4 | 133065767 | 133071849 | 5.14372 | 3.86248 | 3.15531 |
| chr4 | 133012342 | 133013719 | 1000 | Promoter (<=1kb) | 4 | 133012204 | 133077645 | 65442 | 1 | ENSMUST00 | 139 | Ahdc1 | ENSMUSG0( | false | chr4 | 133010598 | 133013973 | 43.65555 | 141.82639 | 136.87535 | chr4 | 133013189 | 133060125 | 8.03751 | 5.40542 | 5.50183 |
| chr4 | 133038513 | 133039677 | 1000 | Promoter (<=1kb) | 4 | 133039493 | 133077242 | 37750 | 1 | ENSMUST00 | 0 | Ahdc1 | ENSMUSG0( | false | chr4 | 133038350 | 133039857 | 24.03115 | 75.67385 | 72.59326 | chr4 | 133013189 | 133060125 | 8.03751 | 5.40542 | 5.50183 |
| chr4 | 133043326 | 133044099 | 1000 | Intron (ENSMUST00000154646.2 | 4 | 133039493 | 133077242 | 37750 | 1 | ENSMUST00 | 3834 | Ahdc1 | ENSMUSG0( | false | . | . | . | . | . | . | chr4 | 133013189 | 133060125 | 8.03751 | 5.40542 | 5.50183 |
| chr4 | 133046993 | 133047363 | 1000 | Intron (ENSMUST00000154646.2 | 4 | 133039493 | 133077242 | 37750 | 1 | ENSMUST00 | 7501 | Ahdc1 | ENSMUSG0( | false | . | . | . | . | . | . | chr4 | 133013189 | 133060125 | 8.03751 | 5.40542 | 5.50183 |
| chr4 | 133056055 | 133056735 | 1000 | Intron (ENSMUST00000154646.2 | 4 | 133066249 | 133072063 | 5815 | 1 | ENSMUST00 | -9514 | Ahdc1 | ENSMUSG0( | false | . | . | . | . | . | . | chr4 | 133013189 | 133060125 | 8.03751 | 5.40542 | 5.50183 |
| chr4 | 133020742 | 133021301 | 1000 | Intron (ENSMUST00000154646.2 | 4 | 133012204 | 133077645 | 65442 | 1 | ENSMUST00 | 8539 | Ahdc1 | ENSMUSG0( | false | . | . | . | . | . | . | chr4 | 133013189 | 133060125 | 8.03751 | 5.40542 | 5.50183 |
| chr14 | 45657790 | 45658443 | 1000 | Promoter (<=1kb) | 14 | 45593174 | 45658143 | 64970 | 2 | ENSMUST00 | 0 | Ddhd1 | ENSMUSG0( | false | chr14 | 45656729 | 45658743 | 27.88725 | 99.90214 | 96.33501 | chr14 | 45655751 | 45659281 | 5.10614 | 4.10597 | 3.14417 |
| chr14 | 45506948 | 45507454 | 1000 | Intron (ENSMUST00000045905.7 | 14 | 45504672 | 45529786 | 25115 | 2 | ENSMUST00 | 22332 | Fermt2 | ENSMUSG0( | true | . | . | . | . | . | . | chr14 | 45489964 | 45528839 | 6.12493 | 4.54024 | 3.93691 |
| chr14 | 45529369 | 45530579 | 1000 | Promoter (<=1kb) | 14 | 45458792 | 45530118 | 71327 | 2 | ENSMUST00 | 0 | Fermt2 | ENSMUSG0( | true | chr14 | 45528002 | 45530952 | 36.31995 | 116.10173 | 112.08785 | . | . | . | . | . | . |
| chr14 | 45388668 | 45389201 | 1000 | Promoter (<=1kb) | 14 | 45376334 | 45388802 | 12469 | 2 | ENSMUST00 | 0 | Gnpnat1 | ENSMUSG0( | false | chr14 | 45388277 | 45389470 | 34.60421 | 120.46815 | 116.3158 | chr14 | 45387660 | 45389486 | 4.1279 | 3.63421 | 2.29499 |
| chr11 | 77984310 | 77985184 | 1000 | Promoter (1-2kb) | 11 | 77982802 | 78030539 | 47738 | 1 | ENSMUST00 | 1509 | Phf12 | ENSMUSG0( | false | chr11 | 77982697 | 77985582 | 42.24747 | 139.38866 | 134.53271 | . | . | . | . | . | . |
| chr5 | 65763331 | 65765147 | 1000 | Promoter (<=1kb) | 5 | 65763521 | 65830108 | 66588 | 1 | ENSMUST00 | 0 | N4bp2 | ENSMUSG0( | false | chr5 | 65763081 | 65765578 | 32.49479 | 112.46503 | 108.55889 | . | . | . | . | . | . |
| chr9 | 63575900 | 63576287 | 1000 | Intron (ENSMUST00000042322.5 | 9 | 63421835 | 63602417 | 180583 | 2 | ENSMUST00 | 26130 | Iqch | ENSMUSG0( | false | . | . | . | . | . | . | chr9 | 63568228 | 63577350 | 4.15727 | 3.62156 | 2.32961 |
| chr18 | 35118479 | 35119529 | 1000 | Promoter (<=1kb) | 18 | 35118888 | 35254773 | 135886 | 1 | ENSMUST00 | 0 | Ctnna1 | ENSMUSG0( | false | chr18 | 35118136 | 35120892 | 33.07612 | 110.40385 | 106.55179 | . | . | . | . | . | . |
| chr18 | 35068887 | 35069303 | 1000 | Distal Intergenic | 18 | 35118888 | 35254773 | 135886 | 1 | ENSMUST00 | -49585 | Ctnna1 | ENSMUSG0( | false | . | . | . | . | . | . | chr18 | 35067816 | 35070475 | 5.20042 | 4.14643 | 3.16207 |
| chr5 | 65492579 | 65493696 | 1000 | Promoter (<=1kb) | 5 | 65446844 | 65493051 | 46208 | 2 | ENSMUST00 | 0 | Smim14 | ENSMUSG0( | false | chr5 | 65491649 | 65493977 | 36.42221 | 131.26616 | 126.73016 | chr5 | 65493414 | 65494645 | 4.94719 | 4.01524 | 2.97583 |
| chr2 | 157924515 | 157924835 | 1000 | Intron (ENSMUST00000109523.1 | 2 | 157914653 | 157944719 | 30067 | 1 | ENSMUST00 | 9863 | Vstm2l | ENSMUSG0( | false | . | . | . | . | . | . | . | . | . | . | . | . |
| chr10 | 68104042 | 68105026 | 1000 | Intron (ENSMUST00000020106.7 | 10 | 68043968 | 68051046 | 7079 | 1 | ENSMUST00 | 60075 | Rtkn2 | ENSMUSG0( | false | . | . | . | . | . | . | chr10 | 68101409 | 68108261 | 4.9792 | 4.05161 | 2.98005 |
| chr10 | 68136139 | 68136986 | 1000 | Intron (ENSMUST00000020106.7 | 10 | 68043968 | 68051046 | 7079 | 1 | ENSMUST00 | 92172 | Rtkn2 | ENSMUSG0( | false | chr10 | 68134465 | 68136910 | 16.165 | 45.52484 | 42.86639 | chr10 | 68136755 | 68141633 | 4.96273 | 4.05082 | 2.98182 |
| chr10 | 68060362 | 68060687 | 1000 | Distal Intergenic | 10 | 68043968 | 68051046 | 7079 | 1 | ENSMUST00 | 16395 | Rtkn2 | ENSMUSG0( | false | . | . | . | . | . | . | chr10 | 68060243 | 68061491 | 4.04804 | 3.60693 | 2.21567 |
| chr19 | 18641097 | 18641558 | 1000 | 3' UTR | 19 | 18631962 | 18637299 | 5338 | 1 | ENSMUST00 | 9136 | Nmrk1 | ENSMUSG0( | false | . | . | . | . | . | . | chr19 | 18639070 | 18643470 | 5.49478 | 4.36176 | 3.40026 |
| chr10 | 67126631 | 67127619 | 1000 | Promoter (<=1kb) | 10 | 67127258 | 67256318 | 129061 | 1 | ENSMUST00 | 0 | Jmjd1c | ENSMUSG0( | false | chr10 | 67126347 | 67127904 | 37.63272 | 121.56319 | 117.36774 | chr10 | 67123191 | 67126746 | 4.43338 | 3.83044 | 2.53845 |
| chr2 | 129832917 | 129833401 | 1000 | Distal Intergenic | 2 | 129800517 | 129832287 | 31771 | 1 | ENSMUST00 | 32401 | Stk35 | ENSMUSG0( | false | . | . | . | . | . | . | chr2 | 129832175 | 129835802 | 4.82808 | 3.99632 | 2.9026 |
| chr3 | 49908872 | 49909307 | 1000 | Distal Intergenic | 3 | 49743296 | 49757316 | 14021 | 2 | ENSMUST00 | -151557 | Pcdh18 | ENSMUSG0( | false | . | . | . | . | . | . | . | . | . | . | . | . |
| chr3 | 137863983 | 137864599 | 1000 | Promoter (<=1kb) | 3 | 137864487 | 137866922 | 2436 | 1 | ENSMUST00 | 0 | H2afz | ENSMUSG0( | false | chr3 | 137863702 | 137865762 | 33.76688 | 123.17645 | 118.92934 | . | . | . | . | . | . |
| chr12 | 111038760 | 111040144 | 1000 | Promoter (<=1kb) | 12 | 111039351 | 111115901 | 76551 | 1 | ENSMUST00 | 0 | Rcor1 | ENSMUSG0( | false | chr12 | 111038168 | 111040621 | 36.9999 | 115.1778 | 111.19392 | . | . | . | . | . | . |
| chr12 | 111089170 | 111089620 | 1000 | Intron (ENSMUST00000084968.8 | 12 | 111039798 | 111113384 | 73587 | 1 | ENSMUST00 | 49373 | Rcor1 | ENSMUSG0( | false | . | . | . | . | . | . | chr12 | 111088176 | 111092273 | 4.66613 | 3.75675 | 2.76726 |
| chr2 | 129592393 | 129593749 | 1000 | Promoter (<=1kb) | 2 | 129592835 | 129608992 | 16158 | 1 | ENSMUST00 | 0 | Sirpa | ENSMUSG0( | true | chr2 | 129592138 | 129595051 | 33.10969 | 106.34647 | 102.61562 | chr2 | 129588228 | 129592519 | 5.03543 | 4.10822 | 3.02491 |
| chr2 | 129624840 | 129625790 | 1000 | Intron (ENSMUST00000161620.2 | 2 | 129620685 | 129628044 | 7360 | 1 | ENSMUST00 | 4156 | Sirpa | ENSMUSG0( | true | . | . | . | . | . | . | chr2 | 129621436 | 129635375 | 5.74226 | 4.37432 | 3.64032 |
| chr11 | 99147227 | 99147759 | 1000 | Intron (ENSMUST00000103134.3 | 11 | 99144196 | 99155077 | 10882 | 2 | ENSMUST00 | 7318 | Ccr7 | ENSMUSG0( | false | . | . | . | . | . | . | . | . | . | . | . | . |
| chr13 | 49187220 | 49188166 | 1000 | Promoter (<=1kb) | 13 | 49187485 | 49196244 | 8760 | 1 | ENSMUST00 | 0 | Ninj1 | ENSMUSG0( | false | chr13 | 49187428 | 49188528 | 36.04937 | 140.37729 | 135.48044 | chr13 | 49185866 | 49191223 | 6.35279 | 4.64168 | 4.15734 |
| chr5 | 124861819 | 124863264 | 1000 | Promoter (<=1kb) | 5 | 124834418 | 124861885 | 27468 | 2 | ENSMUST00 | 0 | Ccdc92 | ENSMUSG0( | true | chr5 | 124860882 | 124863843 | 36.04672 | 129.50026 | 125.02953 | chr5 | 124839552 | 124861868 | 6.02936 | 4.57753 | 3.85106 |
| chr6 | 55648851 | 55649320 | 1000 | Distal Intergenic | 6 | 55677818 | 55681263 | 3446 | 2 | ENSMUST00 | 31943 | Neurod6 | ENSMUSG0( | false | . | . | . | . | . | . | chr6 | 55648400 | 55651115 | 4.67151 | 3.88099 | 2.76616 |
| chr11 | 98959438 | 98959943 | 1000 | Promoter (<=1kb) | 11 | 98960412 | 98974942 | 14531 | 1 | ENSMUST00 | -469 | Rara | ENSMUSG0( | false | chr11 | 98959620 | 98961343 | 26.65955 | 86.95457 | 83.67183 | chr11 | 98958785 | 98973618 | 4.55278 | 3.71427 | 2.6612 |
| chr11 | 98929702 | 98930037 | 1000 | Promoter (1-2kb) | 11 | 98927818 | 98974162 | 46345 | 1 | ENSMUST00 | 1885 | Rara | ENSMUSG0( | false | . | . | . | . | . | . | chr11 | 98928668 | 98957642 | 6.70032 | 4.70048 | 4.41905 |
| chr11 | 98963164 | 98964050 | 1000 | Promoter (2-3kb) | 11 | 98960412 | 98974942 | 14531 | 1 | ENSMUST00 | 2753 | Rara | ENSMUSG0( | false | chr11 | 98963188 | 98964126 | 8.50221 | 18.80243 | 16.48241 | chr11 | 98958785 | 98973618 | 4.55278 | 3.71427 | 2.6612 |
| chr11 | 98941904 | 98943154 | 1000 | Promoter (2-3kb) | 11 | 98939712 | 98950490 | 10779 | 1 | ENSMUST00 | 2193 | Rara | ENSMUSG0( | false | chr11 | 98941436 | 98942169 | 7.12487 | 12.70834 | 10.47977 | chr11 | 98928668 | 98957642 | 6.70032 | 4.70048 | 4.41905 |
| chr11 | 98966583 | 98966884 | 1000 | Intron (ENSMUST00000107474.2 | 11 | 98960412 | 98974942 | 14531 | 1 | ENSMUST00 | 6172 | Rara | ENSMUSG0( | false | . | . | . | . | . | . | chr11 | 98958785 | 98973618 | 4.55278 | 3.71427 | 2.6612 |
| chr6 | 127493751 | 127494100 | 1000 | 3' UTR | 6 | 127477655 | 127490047 | 12393 | 1 | ENSMUST00 | 16097 | Parp11 | ENSMUSG0( | false | . | . | . | . | . | . | chr6 | 127493345 | 127494806 | 6.66149 | 4.7064 | 4.43309 |
| chr4 | 86874178 | 86874654 | 1000 | Promoter (<=1kb) | 4 | 86874396 | 86920875 | 46480 | 1 | ENSMUST00 | 0 | Acer2 | ENSMUSG0( | false | chr4 | 86873870 | 86875346 | 29.2476 | 106.45832 | 102.71833 | chr4 | 86873840 | 86875167 | 4.68898 | 3.92095 | 2.80365 |
| chr15 | 99093425 | 99093969 | 1000 | Promoter (<=1kb) | 15 | 99093170 | 99100917 | 7748 | 1 | ENSMUST00 | 256 | Dnajc22 | ENSMUSG0( | false | chr15 | 99093529 | 99094203 | 17.93284 | 48.67312 | 45.97572 | chr15 | 99093546 | 99099596 | 5.88288 | 4.52988 | 3.73597 |
| chr13 | 48967922 | 48968593 | 1000 | Promoter (<=1kb) | 13 | 48879219 | 48968017 | 88799 | 2 | ENSMUST00 | 0 | Fam120a | ENSMUSG0( | false | chr13 | 48966128 | 48969320 | 37.30079 | 124.19571 | 119.91465 | . | . | . | . | . | . |
| chr4 | 86748260 | 86749159 | 1000 | Promoter (<=1kb) | 4 | 86748555 | 86830167 | 81613 | 1 | ENSMUST00 | 0 | Dennd4c | ENSMUSG0( | false | chr4 | 86747876 | 86749925 | 33.22105 | 122.16254 | 117.95486 | . | . | . | . | . | . |
| chr1 | 172312124 | 172312626 | 1000 | Promoter (<=1kb) | 1 | 172312367 | 172319841 | 7475 | 1 | ENSMUST00 | 0 | Igsf8 | ENSMUSG0( | false | chr1 | 172311315 | 172313237 | 31.01414 | 97.6227 | 94.10706 | . | . | . | . | . | . |
| chr13 | 48641333 | 48641639 | 1000 | Distal Intergenic | 13 | 48577873 | 48625573 | 47701 | 2 | ENSMUST00 | -15761 | Ptpdc1 | ENSMUSG0( | false | . | . | . | . | . | . | chr13 | 48639560 | 48641820 | 3.88183 | 3.32906 | 2.10531 |
| chr11 | 76294960 | 76295413 | 1000 | Intron (ENSMUST00000021204.3 | 11 | 76249212 | 76250508 | 1297 | 1 | ENSMUST00 | 45749 | Rnmtl1 | ENSMUSG0( | false | . | . | . | . | . | . | chr11 | 76283200 | 76297836 | 5.96193 | 4.42599 | 3.81873 |
| chr5 | 25498350 | 25499970 | 1000 | Promoter (<=1kb) | 5 | 25273609 | 25498745 | 225137 | 2 | ENSMUST00 | 0 | Kmt2c | ENSMUSG0( | false | chr5 | 25497212 | 25498824 | 47.35447 | 163.06985 | 157.31989 | . | . | . | . | . | . |
| chr6 | 54972256 | 54972755 | 1000 | Promoter (<=1kb) | 6 | 54923942 | 54972612 | 48671 | 2 | ENSMUST00 | 0 | Nod1 | ENSMUSG0( | false | chr6 | 54971060 | 54973296 | 35.78724 | 111.00061 | 107.13931 | . | . | . | . | . | . |
| chr6 | 54681540 | 54682026 | 1000 | Promoter (<=1kb) | 6 | 54681624 | 54703851 | 22228 | 1 | ENSMUST00 | 0 | Mturn | ENSMUSG0( | false | chr6 | 54680902 | 54682590 | 31.91637 | 110.38313 | 106.53261 | . | . | . | . | . | . |
| chr13 | 47178301 | 47178635 | 1000 | Intron (ENSMUST00000068891.5 | 13 | 47194003 | 47247991 | 53989 | 1 | ENSMUST00 | -15368 | Rnf144b | ENSMUSG0( | true | . | . | . | . | . | . | chr13 | 47176960 | 47179745 | 5.07277 | 4.14244 | 3.06516 |
| chr13 | 47122458 | 47122942 | 1000 | Promoter (<=1kb) | 13 | 47122720 | 47247991 | 125272 | 1 | ENSMUST00 | 0 | Rnf144b | ENSMUSG0( | true | chr13 | 47122064 | 47123515 | 27.24215 | 82.29425 | 79.10038 | chr13 | 47121911 | 47127413 | 4.95977 | 3.98421 | 3.01017 |
| chr5 | 25222731 | 25223347 | 1000 | Promoter (<=1kb) | 5 | 25222882 | 25265918 | 43037 | 1 | ENSMUST00 | 0 | Galnt11 | ENSMUSG0( | false | chr5 | 25222552 | 25223940 | 15.45146 | 50.98658 | 48.25883 | chr5 | 25222472 | 25225251 | 3.52889 | 3.34412 | 1.81157 |
| chr2 | 156196473 | 156197205 | 1000 | Promoter (<=1kb) | 2 | 156196642 | 156309952 | 113311 | 1 | ENSMUST00 | 0 | Phf20 | ENSMUSG0( | false | chr2 | 156196153 | 156197761 | 25.15761 | 80.47005 | 77.31023 | . | . | . | . | . | . |
| chr9 | 35421059 | 35422055 | 1000 | Promoter (<=1kb) | 9 | 35421128 | 35455365 | 34238 | 1 | ENSMUST00 | 0 | Cdon | ENSMUSG0( | false | chr9 | 35420777 | 35422586 | 47.36903 | 159.61537 | 153.98001 | . | . | . | . | . | . |
| chr8 | 47380256 | 47380568 | 1000 | Intron (ENSMUST00000181417.1 | 8 | 47180048 | 47352348 | 172301 | 2 | ENSMUST00 | -27909 | Stox2 | ENSMUSG0( | false | . | . | . | . | . | . | chr8 | 47379298 | 47381737 | 5.19532 | 4.21745 | 3.16144 |
| chr11 | 98588010 | 98588786 | 1000 | Promoter (<=1kb) | 11 | 98581256 | 98587368 | 6113 | 2 | ENSMUST00 | -643 | Ormdl3 | ENSMUSG0( | false | . | . | . | . | . | . | chr11 | 98580618 | 98591485 | 6.41628 | 4.67128 | 4.18007 |
| chr13 | 45964846 | 45965362 | 1000 | Promoter (<=1kb) | 13 | 45965253 | 45966508 | 1256 | 1 | ENSMUST00 | 0 | 5033430I15R | ENSMUSG0( | false | chr13 | 45962952 | 45966515 | 39.39634 | 133.30592 | 128.68834 | . | . | . | . | . | . |

| | | | | | | | | | | | | | | | | | | | | | | | | | | |
|---|---|---|---|---|---|---|---|---|---|---|---|---|---|---|---|---|---|---|---|---|---|---|---|---|---|---|
| chr16 | 26181692 | 26182072 | 1000 | Distal Intergenic | 16 | 25959288 | 26105784 | 146497 | 2 | ENSMUST00 | -75909 | Leprel1 | ENSMUSG0C | false | . | . | . | . | . | . | chr16 | 26180809 | 26183244 | 4.7381 | 3.91105 | 2.79506 |
| chr3 | 97749373 | 97749912 | 1000 | Intron (ENSMUST00000168438.3 | 3 | 97740817 | 97767931 | 27115 | 2 | ENSMUST00 | 18019 | Pde4dip | ENSMUSG0C | true | . | . | . | . | . | . | chr3 | 97748396 | 97750862 | 4.72134 | 3.82356 | 2.81338 |
| chr4 | 83251890 | 83252347 | 1000 | Intron (ENSMUST00000102823.4 | 4 | 83247613 | 83266039 | 18427 | 2 | ENSMUST00 | 13692 | Ttc39b | ENSMUSG0C | false | . | . | . | . | . | . | chr4 | 83250588 | 83252444 | 3.13021 | 3.07243 | 1.50959 |
| chr5 | 24550822 | 24551267 | 1000 | Intron (ENSMUST00000088302.5 | 5 | 24543434 | 24555469 | 12036 | 2 | ENSMUST00 | 4202 | 4931409K22I | ENSMUSG0C | false | chr5 | 24551250 | 24551521 | 4.82314 | 6.92697 | 4.82056 | chr5 | 24549786 | 24551853 | 5.25046 | 4.24181 | 3.20824 |
| chr10 | 43478516 | 43480116 | 1000 | Promoter (<=1kb) | 10 | 43479140 | 43515396 | 36257 | 1 | ENSMUST00 | 0 | Bend3 | ENSMUSG0C | false | chr10 | 43477903 | 43480632 | 36.90527 | 118.17374 | 114.08985 | . | . | . | . | . | . |
| chr11 | 75443582 | 75443939 | 1000 | Intron (ENSMUST00000173320.2 | 11 | 75435914 | 75439591 | 3678 | 2 | ENSMUST00 | -3992 | Serpinf2 | ENSMUSG0C | false | . | . | . | . | . | . | chr11 | 75443385 | 75444274 | 5.44376 | 4.03444 | 3.39935 |
| chr6 | 52230377 | 52230940 | 1000 | Promoter (<=1kb) | 6 | 52228247 | 52230532 | 2286 | 2 | ENSMUST00 | 0 | Hoxa9 | ENSMUSG0C | false | chr6 | 52227601 | 52235632 | 41.69514 | 140.65352 | 135.74945 | . | . | . | . | . | . |
| chr7 | 112217159 | 112217558 | 1000 | Distal Intergenic | 7 | 112225856 | 112355194 | 129339 | 1 | ENSMUST00 | -8298 | Mical2 | ENSMUSG0C | false | . | . | . | . | . | . | chr7 | 112216889 | 112217796 | 2.97956 | 3.0069 | 1.37034 |
| chr7 | 112316164 | 112316753 | 1000 | Promoter (<=1kb) | 7 | 112316029 | 112347491 | 31463 | 1 | ENSMUST00 | 136 | Mical2 | ENSMUSG0C | false | . | . | . | . | . | . | chr7 | 112315227 | 112322046 | 5.14867 | 4.13411 | 3.15663 |
| chr3 | 97227040 | 97227962 | 1000 | Promoter (<=1kb) | 3 | 97203662 | 97227364 | 23703 | 2 | ENSMUST00 | 0 | Bcl9 | ENSMUSG0C | false | chr3 | 97226662 | 97228545 | 27.03504 | 88.5911 | 85.27629 | . | . | . | . | . | . |
| chr7 | 45332644 | 45333080 | 1000 | Promoter (<=1kb) | 7 | 45303155 | 45333780 | 30626 | 2 | ENSMUST00 | 700 | Trpm4 | ENSMUSG0C | false | chr7 | 45332526 | 45333989 | 25.50663 | 88.63404 | 85.31516 | chr7 | 45332151 | 45334090 | 4.85422 | 3.72964 | 2.90984 |
| chr9 | 57967538 | 57967936 | 1000 | Distal Intergenic | 9 | 57940113 | 57962864 | 22752 | 1 | ENSMUST00 | 27426 | Sema7a | ENSMUSG0C | false | . | . | . | . | . | . | chr9 | 57965942 | 57968510 | 5.10102 | 4.07589 | 3.11971 |
| chr6 | 125148114 | 125148585 | 1000 | Promoter (<=1kb) | 6 | 125149007 | 125151487 | 2481 | 1 | ENSMUST00 | -422 | Iffo1 | ENSMUSG0C | false | . | . | . | . | . | . | chr6 | 125144609 | 125151122 | 6.70514 | 4.7951 | 4.43014 |
| chr7 | 111638371 | 111638835 | 1000 | Intron (ENSMUST00000049430.9 | 7 | 111471661 | 111779977 | 308317 | 2 | ENSMUST00 | 141142 | Galnt18 | ENSMUSG0C | false | . | . | . | . | . | . | chr7 | 111637634 | 111642613 | 4.46129 | 3.8031 | 2.60155 |
| chr1 | 57774672 | 57775315 | 1000 | Promoter (<=1kb) | 1 | 57774735 | 57946965 | 172231 | 1 | ENSMUST00 | 0 | Spats2l | ENSMUSG0C | false | chr1 | 57774051 | 57776163 | 31.70051 | 127.6562 | 123.25787 | . | . | . | . | . | . |
| chr5 | 24320428 | 24320865 | 1000 | Promoter (1-2kb) | 5 | 24320083 | 24322254 | 2172 | 2 | ENSMUST00 | 1389 | Kcnh2 | ENSMUSG0C | false | . | . | . | . | . | . | chr5 | 24320297 | 24322856 | 4.46105 | 3.73775 | 2.59271 |
| chr10 | 41866275 | 41866617 | 1000 | Intron (ENSMUST00000099931.4 | 10 | 41887439 | 41908424 | 20986 | 1 | ENSMUST00 | -20822 | Sesn1 | ENSMUSG0C | false | . | . | . | . | . | . | chr10 | 41864321 | 41868716 | 5.04981 | 4.09047 | 3.0579 |
| chr10 | 41809905 | 41810709 | 1000 | Promoter (<=1kb) | 10 | 41810574 | 41908424 | 97851 | 1 | ENSMUST00 | 0 | Sesn1 | ENSMUSG0C | false | chr10 | 41809166 | 41812188 | 38.97949 | 131.47847 | 126.9297 | . | . | . | . | . | . |
| chr1 | 55744159 | 55744613 | 1000 | Intron (ENSMUST00000042986.9 | 1 | 55702025 | 55735227 | 33203 | 1 | ENSMUST00 | 42135 | Plcl1 | ENSMUSG0C | true | . | . | . | . | . | . | chr1 | 55738786 | 55752458 | 4.85141 | 3.98095 | 2.88878 |
| chr1 | 55576137 | 55576635 | 1000 | Intron (ENSMUST00000042986.9 | 1 | 55702025 | 55735227 | 33203 | 1 | ENSMUST00 | -125390 | Plcl1 | ENSMUSG0C | true | . | . | . | . | . | . | chr1 | 55565857 | 55577601 | 5.51018 | 4.28641 | 3.43047 |
| chr1 | 55736760 | 55737301 | 1000 | Intron (ENSMUST00000042986.9 | 1 | 55702025 | 55735227 | 33203 | 1 | ENSMUST00 | 34736 | Plcl1 | ENSMUSG0C | true | chr1 | 55737106 | 55739417 | 8.08983 | 23.14769 | 20.77093 | . | . | . | . | . | . |
| chr11 | 97799050 | 97800653 | 1000 | Promoter (<=1kb) | 11 | 97799565 | 97838764 | 39200 | 1 | ENSMUST00 | 0 | Lasp1 | ENSMUSG0C | false | chr11 | 97798972 | 97802073 | 32.0113 | 129.32326 | 124.86235 | . | . | . | . | . | . |
| chr11 | 97813692 | 97814165 | 1000 | Promoter (1-2kb) | 11 | 97815248 | 97824949 | 9702 | 1 | ENSMUST00 | -1083 | Lasp1 | ENSMUSG0C | false | . | . | . | . | . | . | chr11 | 97801064 | 97833244 | 8.45203 | 5.49885 | 5.84074 |
| chr11 | 97807978 | 97808339 | 1000 | Promoter (1-2kb) | 11 | 97806502 | 97815782 | 9281 | 1 | ENSMUST00 | 1477 | Lasp1 | ENSMUSG0C | false | . | . | . | . | . | . | chr11 | 97801064 | 97833244 | 8.45203 | 5.49885 | 5.84074 |
| chr11 | 97816878 | 97817540 | 1000 | Promoter (1-2kb) | 11 | 97815248 | 97824949 | 9702 | 1 | ENSMUST00 | 1631 | Lasp1 | ENSMUSG0C | false | . | . | . | . | . | . | chr11 | 97801064 | 97833244 | 8.45203 | 5.49885 | 5.84074 |
| chr7 | 110613978 | 110615726 | 1000 | Promoter (<=1kb) | 7 | 110308013 | 110614922 | 306910 | 2 | ENSMUST00 | 0 | Sbf2 | ENSMUSG0C | true | chr7 | 110613377 | 110615752 | 31.19317 | 93.97161 | 90.54182 | . | . | . | . | . | . |
| chr7 | 110391982 | 110392434 | 1000 | Intron (ENSMUST00000164759.2 | 7 | 110341327 | 110364586 | 23260 | 2 | ENSMUST00 | -27397 | Sbf2 | ENSMUSG0C | true | . | . | . | . | . | . | chr7 | 110390902 | 110396335 | 4.31345 | 3.6685 | 2.45745 |
| chr13 | 32080366 | 32080911 | 1000 | Intron (ENSMUST00000041859.7 | 13 | 32157330 | 32227241 | 69912 | 2 | ENSMUST00 | 146330 | Gmds | ENSMUSG0C | true | . | . | . | . | . | . | chr13 | 32079788 | 32081720 | 4.1648 | 3.53747 | 2.342 |
| chr13 | 32298331 | 32298772 | 1000 | Intron (ENSMUST00000041859.7 | 13 | 32099227 | 32338585 | 239359 | 2 | ENSMUST00 | 39813 | Gmds | ENSMUSG0C | true | . | . | . | . | . | . | chr13 | 32293198 | 32299443 | 4.80818 | 3.9396 | 2.85944 |
| chr5 | 123141265 | 123142753 | 1000 | Promoter (<=1kb) | 5 | 123142193 | 123149981 | 7789 | 1 | ENSMUST00 | 0 | Setd1b | ENSMUSG0C | false | chr5 | 123136690 | 123143966 | 31.66066 | 102.77801 | 99.14294 | . | . | . | . | . | . |
| chr2 | 173226629 | 173227064 | 1000 | 3' UTR | 2 | 173228211 | 173234722 | 6512 | 2 | ENSMUST00 | 7658 | Pmepa1 | ENSMUSG0C | true | . | . | . | . | . | . | chr2 | 173223515 | 173275069 | 7.01359 | 4.89215 | 4.68115 |
| chr2 | 173275503 | 173276938 | 1000 | Promoter (<=1kb) | 2 | 173224458 | 173276533 | 52076 | 2 | ENSMUST00 | 0 | Pmepa1 | ENSMUSG0C | true | chr2 | 173274005 | 173277330 | 35.62435 | 116.99458 | 112.94973 | . | . | . | . | . | . |
| chr18 | 34887090 | 34887645 | 1000 | Distal Intergenic | 18 | 34861207 | 34864956 | 3750 | 1 | ENSMUST00 | 25884 | Egr1 | ENSMUSG0C | false | chr18 | 34887042 | 34887995 | 20.40337 | 54.55598 | 51.78317 | chr18 | 34884206 | 34889383 | 4.05361 | 3.48319 | 2.27183 |
| chr18 | 34884386 | 34884832 | 1000 | Distal Intergenic | 18 | 34861207 | 34864956 | 3750 | 1 | ENSMUST00 | 23180 | Egr1 | ENSMUSG0C | false | . | . | . | . | . | . | chr18 | 34884206 | 34889383 | 4.05361 | 3.48319 | 2.27183 |
| chr18 | 34860303 | 34861143 | 1000 | Promoter (<=1kb) | 18 | 34861207 | 34864956 | 3750 | 1 | ENSMUST00 | -64 | Egr1 | ENSMUSG0C | false | chr18 | 34859236 | 34862643 | 33.41831 | 117.26761 | 113.21259 | chr18 | 34858199 | 34863782 | 4.89469 | 4.00673 | 2.91424 |
| chr4 | 62496849 | 62497231 | 1000 | 3' UTR | 4 | 62499008 | 62502276 | 3269 | 2 | ENSMUST00 | 5045 | Hdhd3 | ENSMUSG0C | false | . | . | . | . | . | . | chr4 | 62496570 | 62497184 | 4.08737 | 3.31995 | 2.28521 |
| chr18 | 70529953 | 70530479 | 1000 | Promoter (<=1kb) | 18 | 70508680 | 70530138 | 21459 | 2 | ENSMUST00 | 0 | Poli | ENSMUSG0C | false | chr18 | 70529710 | 70531341 | 24.40663 | 77.266 | 74.1616 | chr18 | 70528961 | 70531927 | 2.79708 | 2.73168 | 1.24684 |
| chr11 | 97669029 | 97670042 | 1000 | Exon (ENSMUST00000044730.6/ | 11 | 97665001 | 97673548 | 8548 | 1 | ENSMUST00 | 4029 | Mllt6 | ENSMUSG0C | false | chr11 | 97669531 | 97669873 | 9.01168 | 19.18248 | 16.85847 | chr11 | 97664715 | 97673941 | 6.52303 | 4.69373 | 4.25389 |
| chr11 | 97662917 | 97664308 | 1000 | Promoter (<=1kb) | 11 | 97663414 | 97685463 | 22050 | 1 | ENSMUST00 | 0 | Mllt6 | ENSMUSG0C | false | chr11 | 97661507 | 97664564 | 31.01414 | 97.6227 | 94.10706 | chr11 | 97660731 | 97662935 | 5.22491 | 4.26722 | 3.18868 |
| chr6 | 124808661 | 124809040 | 1000 | Promoter (<=1kb) | 6 | 124808885 | 124810336 | 1452 | 1 | ENSMUST00 | 0 | Spsb2 | ENSMUSG0C | false | chr6 | 124808299 | 124810012 | 20.82289 | 72.3363 | 69.31286 | . | . | . | . | . | . |
| chr6 | 39543113 | 39543683 | 1000 | Exon (ENSMUST00000135363.1/ | 6 | 39462378 | 39557867 | 95490 | 2 | ENSMUST00 | 14184 | Dennd2a | ENSMUSG0C | false | . | . | . | . | . | . | chr6 | 39539485 | 39545229 | 6.37896 | 4.69531 | 4.14407 |
| chr7 | 84151569 | 84152332 | 1000 | Promoter (<=1kb) | 7 | 84109356 | 84151893 | 42538 | 2 | ENSMUST00 | 0 | Abhd17c | ENSMUSG0C | false | chr7 | 84150810 | 84152691 | 34.36702 | 111.64683 | 107.76208 | . | . | . | . | . | . |
| chr7 | 84135741 | 84136019 | 1000 | Intron (ENSMUST00000117085.1 | 7 | 84114251 | 84147815 | 33565 | 2 | ENSMUST00 | 11796 | Abhd17c | ENSMUSG0C | false | . | . | . | . | . | . | chr7 | 84125466 | 84151297 | 5.4347 | 4.25306 | 3.37492 |
| chr2 | 154656801 | 154657311 | 1000 | Promoter (<=1kb) | 2 | 154657018 | 154694785 | 37768 | 1 | ENSMUST00 | 0 | Chmp4b | ENSMUSG0C | false | chr2 | 154656983 | 154658473 | 38.95827 | 123.86062 | 119.59576 | . | . | . | . | . | . |
| chr1 | 170485243 | 170485777 | 1000 | Intron (ENSMUST00000160456.2 | 1 | 170338209 | 170454914 | 116706 | 2 | ENSMUST00 | -30330 | Nos1ap | ENSMUSG0C | false | . | . | . | . | . | . | chr1 | 170476687 | 170507715 | 5.67077 | 4.33326 | 3.56816 |
| chr1 | 170361190 | 170361596 | 1000 | Intron (ENSMUST00000160456.2 | 1 | 170302668 | 170327654 | 24987 | 2 | ENSMUST00 | -33537 | Nos1ap | ENSMUSG0C | false | . | . | . | . | . | . | chr1 | 170359992 | 170362162 | 6.63732 | 4.63045 | 4.37561 |
| chr1 | 170499874 | 170500234 | 1000 | Intron (ENSMUST00000160456.2 | 1 | 170338209 | 170454914 | 116706 | 2 | ENSMUST00 | -44961 | Nos1ap | ENSMUSG0C | false | . | . | . | . | . | . | chr1 | 170476687 | 170507715 | 5.67077 | 4.33326 | 3.56816 |
| chr1 | 170489419 | 170489748 | 1000 | Intron (ENSMUST00000160456.2 | 1 | 170338209 | 170454914 | 116706 | 2 | ENSMUST00 | -34506 | Nos1ap | ENSMUSG0C | false | . | . | . | . | . | . | chr1 | 170476687 | 170507715 | 5.67077 | 4.33326 | 3.56816 |
| chr10 | 40348934 | 40350067 | 1000 | Promoter (<=1kb) | 10 | 40349308 | 40483818 | 134511 | 1 | ENSMUST00 | 0 | Cdk19 | ENSMUSG0C | false | chr10 | 40348278 | 40350333 | 36.66269 | 126.29414 | 121.93508 | . | . | . | . | . | . |
| chr11 | 97362155 | 97363331 | 1000 | Promoter (<=1kb) | 11 | 97362551 | 97398542 | 35992 | 1 | ENSMUST00 | 0 | Socs7 | ENSMUSG0C | false | chr11 | 97361882 | 97364182 | 41.07278 | 140.67711 | 135.76277 | . | . | . | . | . | . |
| chr3 | 96181007 | 96181378 | 1000 | Promoter (<=1kb) | 3 | 96181151 | 96195521 | 14371 | 1 | ENSMUST00 | 0 | Sv2a | ENSMUSG0C | false | chr3 | 96180864 | 96181436 | 14.88741 | 34.64093 | 32.11682 | chr3 | 96179747 | 96182852 | 3.53565 | 3.29302 | 1.83412 |
| chr3 | 96104179 | 96104946 | 1000 | Promoter (<=1kb) | 3 | 96104527 | 96141518 | 36992 | 1 | ENSMUST00 | 0 | Otud7b | ENSMUSG0C | true | chr3 | 96103895 | 96107098 | 33.65816 | 114.56059 | 110.59126 | . | . | . | . | . | . |
| chr3 | 96128027 | 96128568 | 1000 | Intron (ENSMUST00000132980.2 | 3 | 96107087 | 96161129 | 54043 | 1 | ENSMUST00 | 20941 | Otud7b | ENSMUSG0C | true | . | . | . | . | . | . | chr3 | 96124412 | 96138707 | 5.31471 | 4.19209 | 3.2704 |
| chr7 | 44868787 | 44869979 | 1000 | Promoter (<=1kb) | 7 | 44863068 | 44869788 | 6721 | 2 | ENSMUST00 | 0 | Ptov1 | ENSMUSG0C | false | chr7 | 44868285 | 44870536 | 42.75948 | 139.16466 | 134.31596 | chr7 | 44860730 | 44868940 | 6.16567 | 4.38467 | 3.98867 |
| chr7 | 83883517 | 83884738 | 1000 | Promoter (<=1kb) | 7 | 83884466 | 83897940 | 13475 | 1 | ENSMUST00 | 0 | Mesdc2 | ENSMUSG0C | false | chr7 | 83882272 | 83884889 | 41.56779 | 133.79375 | 129.15814 | . | . | . | . | . | . |
| chr13 | 44623984 | 44624411 | 1000 | Distal Intergenic | 13 | 44729474 | 44731574 | 2101 | 1 | ENSMUST00 | -105063 | Jarid2 | ENSMUSG0C | false | . | . | . | . | . | . | chr13 | 44621314 | 44631701 | 5.57294 | 4.3396 | 3.4838 |
| chr13 | 44658009 | 44658678 | 1000 | Distal Intergenic | 13 | 44729474 | 44731574 | 2101 | 1 | ENSMUST00 | -70796 | Jarid2 | ENSMUSG0C | false | . | . | . | . | . | . | chr13 | 44656720 | 44658738 | 5.5017 | 4.23385 | 3.43196 |
| chr13 | 44839827 | 44840923 | 1000 | Promoter (<=1kb) | 13 | 44840691 | 44914319 | 73629 | 1 | ENSMUST00 | 0 | Jarid2 | ENSMUSG0C | false | chr13 | 44839940 | 44841021 | 14.03095 | 32.52118 | 30.02454 | chr13 | 44826187 | 44846053 | 5.23741 | 4.10613 | 3.22861 |
| chr13 | 44730194 | 44731475 | 1000 | Promoter (<=1kb) | 13 | 44730774 | 44920108 | 189335 | 1 | ENSMUST00 | 0 | Jarid2 | ENSMUSG0C | false | chr13 | 44730600 | 44734844 | 29.75681 | 92.45776 | 89.05831 | . | . | . | . | . | . |
| chr9 | 72274595 | 72275364 | 1000 | Promoter (<=1kb) | 9 | 72274860 | 72362891 | 88032 | 1 | ENSMUST00 | 0 | Zfp280d | ENSMUSG0C | false | chr9 | 72274530 | 72276395 | 13.98552 | 41.45461 | 38.84487 | . | . | . | . | . | . |
| chr6 | 38433221 | 38434878 | 1000 | Promoter (<=1kb) | 6 | 38433950 | 38512763 | 78814 | 1 | ENSMUST00 | 0 | Ubn2 | ENSMUSG0C | false | chr6 | 38432966 | 38435255 | 41.80831 | 147.58597 | 142.39091 | . | . | . | . | . | . |
| chr4 | 59761425 | 59761746 | 1000 | 3' UTR | 4 | 59771297 | 59783808 | 12512 | 2 | ENSMUST00 | 22062 | Inip | ENSMUSG0C | false | . | . | . | . | . | . | chr4 | 59756611 | 59762237 | 4.44224 | 3.62486 | 2.5942 |
| chr10 | 53422219 | 53422578 | 1000 | Distal Intergenic | 10 | 53278081 | 53379851 | 101771 | 2 | ENSMUST00 | -42369 | Cep85l | ENSMUSG0C | true | . | . | . | . | . | . | chr10 | 53407459 | 53424022 | 5.24828 | 4.19397 | 3.20837 |
| chr10 | 53410898 | 53411519 | 1000 | Distal Intergenic | 10 | 53278081 | 53379851 | 101771 | 2 | ENSMUST00 | -31048 | Cep85l | ENSMUSG0C | true | . | . | . | . | . | . | chr10 | 53407459 | 53424022 | 5.24828 | 4.19397 | 3.20837 |
| chr8 | 105594818 | 105595272 | 1000 | Distal Intergenic | 8 | 105605229 | 105622194 | 16966 | 1 | ENSMUST00 | -9957 | Fam65a | ENSMUSG0C | false | chr8 | 105593587 | 105594870 | 8.29801 | 20.47831 | 18.13586 | . | . | . | . | . | . |
| chr8 | 105606226 | 105606918 | 1000 | Promoter (<=1kb) | 8 | 105605229 | 105622194 | 16966 | 1 | ENSMUST00 | 998 | Fam65a | ENSMUSG0C | false | chr8 | 105600584 | 105607833 | 28.7777 | 83.06458 | 79.8585 | chr8 | 105598855 | 105624293 | 5.6449 | 4.36488 | 3.54309 |
| chr11 | 96824942 | 96825615 | 1000 | Promoter (<=1kb) | 11 | 96822223 | 96824552 | 2330 | 2 | ENSMUST00 | -391 | Nfe2l1 | ENSMUSG0C | false | chr11 | 96823537 | 96825054 | 12.76655 | 31.65917 | 29.17213 | chr11 | 96821899 | 96829042 | 7.7991 | 5.17723 | 5.30668 |
| chr11 | 96823928 | 96824626 | 1000 | Promoter (<=1kb) | 11 | 96817414 | 96824013 | 6600 | 2 | ENSMUST00 | 0 | Nfe2l1 | ENSMUSG0C | false | chr11 | 96823537 | 96825054 | 12.76655 | 31.65917 | 29.17213 | chr11 | 96821899 | 96829042 | 7.7991 | 5.17723 | 5.30668 |
| chr3 | 95627816 | 95628226 | 1000 | Promoter (2-3kb) | 3 | 95624980 | 95632102 | 7123 | 1 | ENSMUST00 | 2837 | Ensa | ENSMUSG0C | false | . | . | . | . | . | . | chr3 | 95625718 | 95639071 | 6.33259 | 4.60953 | 4.14581 |
| chr3 | 95635813 | 95636387 | 1000 | Distal Intergenic | 3 | 95624980 | 95632102 | 7123 | 1 | ENSMUST00 | 10834 | Ensa | ENSMUSG0C | false | . | . | . | . | . | . | chr3 | 95625718 | 95639071 | 6.33259 | 4.60953 | 4.14581 |
| chr1 | 182282200 | 182283044 | 1000 | Promoter (<=1kb) | 1 | 182275772 | 182282804 | 7033 | 2 | ENSMUST00 | 0 | Degs1 | ENSMUSG0C | false | chr1 | 182281371 | 182283460 | 26.1063 | 98.41757 | 94.88632 | . | . | . | . | . | . |
| chr6 | 37430540 | 37431785 | 1000 | Intron (ENSMUST00000041093.5 | 6 | 37327255 | 37442146 | 114892 | 2 | ENSMUST00 | 10361 | Creb3l2 | ENSMUSG0C | true | chr6 | 37431077 | 37431538 | 6.70576 | 11.61755 | 9.40775 | chr6 | 37373151 | 37440158 | 6.00246 | 4.49234 | 3.83883 |
| chr6 | 37484568 | 37485003 | 1000 | Distal Intergenic | 6 | 37327255 | 37442146 | 114892 | 2 | ENSMUST00 | -42423 | Creb3l2 | ENSMUSG0C | true | . | . | . | . | . | . | chr6 | 37483214 | 37487532 | 3.25926 | 3.12161 | 1.58017 |
| chr6 | 37419327 | 37419949 | 1000 | Intron (ENSMUST00000041093.5 | 6 | 37327255 | 37442146 | 114892 | 2 | ENSMUST00 | 22197 | Creb3l2 | ENSMUSG0C | true | . | . | . | . | . | . | chr6 | 37373151 | 37440158 | 6.00246 | 4.49234 | 3.83883 |
| chr11 | 96318834 | 96319686 | 1000 | Promoter (<=1kb) | 11 | 96318267 | 96321638 | 3372 | 1 | ENSMUST00 | 568 | Hoxb4 | ENSMUSG0C | false | chr11 | 96318131 | 96320429 | 33.10969 | 106.34647 | 102.61562 | chr11 | 96319587 | 96320962 | 3.97048 | 3.5261 | 2.17935 |
| chr2 | 34371560 | 34372997 | 1000 | Promoter (<=1kb) | 2 | 34171457 | 34372038 | 200582 | 2 | ENSMUST00 | 0 | Pbx3 | ENSMUSG0C | true | chr2 | 34370530 | 34373633 | 27.54716 | 98.2761 | 94.74957 | chr2 | 34372993 | 34373921 | 2.83506 | 2.8673 | 1.26113 |
| chr4 | 57809507 | 57809852 | 1000 | Intron (ENSMUST00000098066.3 | 4 | 57829245 | 57855049 | 25805 | 1 | ENSMUST00 | -19393 | Akap2 | ENSMUSG0C | false | . | . | . | . | . | . | chr4 | 57809325 | 57810841 | 4.19674 | 3.73449 | 2.35391 |
| chr4 | 57811731 | 57812450 | 1000 | Intron (ENSMUST00000098066.3 | 4 | 57829245 | 57855049 | 25805 | 1 | ENSMUST00 | -16795 | Akap2 | ENSMUSG0C | false | . | . | . | . | . | . | chr4 | 57811973 | 57814199 | 5.6232 | 4.34437 | 3.51202 |
| chr4 | 57851771 | 57852245 | 1000 | Promoter (1-2kb) | 4 | 57854159 | 57896982 | 42824 | 1 | ENSMUST00 | -1914 | Akap2 | ENSMUSG0C | false | . | . | . | . | . | . | chr4 | 57846708 | 57868207 | 6.85521 | 4.86004 | 4.55185 |
| chr4 | 57861173 | 57861536 | 1000 | Intron (ENSMUST00000098066.3 | 4 | 57854159 | 57896982 | 42824 | 1 | ENSMUST00 | 7015 | Akap2 | ENSMUSG0C | false | . | . | . | . | . | . | chr4 | 57846708 | 57868207 | 6.85521 | 4.86004 | 4.55185 |
| chr1 | 181211074 | 181212263 | 1000 | Promoter (<=1kb) | 1 | 181173228 | 181211552 | 38325 | 2 | ENSMUST00 | 0 | Wdr26 | ENSMUSG0C | false | chr1 | 181210182 | 181212891 | 39.20496 | 131.87447 | 127.31225 | . | . | . | . | . | . |
| chr4 | 57258259 | 57259053 | 1000 | Intron (ENSMUST00000075637.5 | 4 | 57218347 | 57222054 | 3708 | 2 | ENSMUST00 | -36206 | Ptpn3 | ENSMUSG0C | false | . | . | . | . | . | . | chr4 | 57256079 | 57260126 | 4.37335 | 3.7161 | 2.51884 |
| chr2 | 33631712 | 33632682 | 1000 | Promoter (<=1kb) | 2 | 33567156 | 33632283 | 65128 | 2 | ENSMUST00 | 0 | Lmx1b | ENSMUSG0C | true | chr2 | 33631177 | 33632414 | 18.44084 | 48.77526 | 46.0739 | chr2 | 33565260 | 33638297 | 6.78481 | 4.81457 | 4.48235 |
| chr2 | 33594259 | 33594629 | 1000 | Intron (ENSMUST00000041730.5 | 2 | 33567156 | 33632283 | 65128 | 2 | ENSMUST00 | 37654 | Lmx1b | ENSMUSG0C | true | . | . | . | . | . | . | chr2 | 33565260 | 33638297 | 6.78481 | 4.81457 | 4.48235 |
| chr2 | 33605081 | 33606316 | 1000 | Intron (ENSMUST00000041730.5 | 2 | 33567156 | 33632283 | 65128 | 2 | ENSMUST00 | 25967 | Lmx1b | ENSMUSG0C | true | . | . | . | . | . | . | chr2 | 33565260 | 33638297 | 6.78481 | 4.81457 | 4.48235 |
| chr3 | 95199710 | 95200081 | 1000 | Exon (ENSMUST00000155555.2/ | 3 | 95187589 | 95191898 | 4310 | 2 | ENSMUST00 | -7813 | Gabpb2 | ENSMUSG0C | true | . | . | . | . | . | . | chr3 | 95193311 | 95204022 | 6.01146 | 4.56637 | 3.8326 |
| chr10 | 50592388 | 50593064 | 1000 | Promoter (<=1kb) | 10 | 50592669 | 50851202 | 258534 | 1 | ENSMUST00 | 0 | Ascc3 | ENSMUSG0C | false | chr10 | 50591981 | 50593604 | 28.22734 | 101.53345 | 97.92473 | . | . | . | . | . | . |
| chr1 | 181016524 | 181017065 | 1000 | Promoter (<=1kb) | 1 | 180989554 | 181017569 | 28016 | 2 | ENSMUST00 | 504 | Ephx1 | ENSMUSG0C | false | chr1 | 181015848 | 181017585 | 28.9509 | 90.07051 | 86.72839 | chr1 | 181000567 | 181016642 | 7.46536 | 5.17908 | 5.02982 |
| chr1 | 181010542 | 181011208 | 1000 | Intron (ENSMUST00000036928.6 | 1 | 180989554 | 181017569 | 28016 | 2 | ENSMUST00 | 6361 | Ephx1 | ENSMUSG0C | false | . | . | . | . | . | . | chr1 | 181000567 | 181016642 | 7.46536 | 5.17908 | 5.02982 |
| chr3 | 95179927 | 95180462 | 1000 | Downstream (1-2kb) | 3 | 95170805 | 95171976 | 1172 | 1 | ENSMUST00 | 9123 | Sema6c | ENSMUSG0C | false | . | . | . | . | . | . | chr3 | 95169782 | 95186009 | 5.67371 | 4.33281 | 3.57492 |
| chr4 | 56828805 | 56829302 | 1000 | Promoter (<=1kb) | 4 | 56826266 | 56829853 | 3588 | 2 | ENSMUST00 | 551 | Ctnnal1 | ENSMUSG0C | true | . | . | . | . | . | . | chr4 | 56827266 | 56853148 | 5.48236 | 4.22607 | 3.42275 |
| chr4 | 56829560 | 56830114 | 1000 | Promoter (<=1kb) | 4 | 56826266 | 56829853 | 3588 | 2 | ENSMUST00 | 0 | Ctnnal1 | ENSMUSG0C | true | . | . | . | . | . | . | chr4 | 56827266 | 56853148 | 5.48236 | 4.22607 | 3.42275 |
| chr2 | 33165609 | 33166278 | 1000 | Promoter (1-2kb) | 2 | 33137636 | 33168222 | 30587 | 2 | ENSMUST00 | 1944 | Ralgps1 | ENSMUSG0C | false | . | . | . | . | . | . | chr2 | 33154669 | 33171007 | 7.40171 | 5.13174 | 4.97591 |
| chr2 | 33160296 | 33160800 | 1000 | Promoter (2-3kb) | 2 | 33143307 | 33158170 | 14864 | 2 | ENSMUST00 | -2127 | Ralgps1 | ENSMUSG0C | false | . | . | . | . | . | . | chr2 | 33154669 | 33171007 | 7.40171 | 5.13174 | 4.97591 |
| chr1 | 105990194 | 105990995 | 1000 | Promoter (<=1kb) | 1 | 105990406 | 106034074 | 43669 | 1 | ENSMUST00 | 0 | Zcchc2 | ENSMUSG0C | false | chr1 | 105989454 | 105992404 | 33.87313 | 113.90014 | 109.94483 | . | . | . | . | . | . |
| chr7 | 80370546 | 80371560 | 1000 | Promoter (<=1kb) | 7 | 80349098 | 80371375 | 22278 | 2 | ENSMUST00 | 0 | Man2a2 | ENSMUSG0C | false | chr7 | 80370176 | 80371869 | 31.16539 | 106.97112 | 103.21663 | chr7 | 80368144 | 80371674 | 5.59506 | 4.057 | 3.5213 |
| chr11 | 95369947 | 95370549 | 1000 | Intron (ENSMUST00000037502.6 | 11 | 95375467 | 95377585 | 2119 | 1 | ENSMUST00 | -4918 | Fam117a | ENSMUSG0C | false | . | . | . | . | . | . | chr11 | 95364434 | 95372547 | 6.52296 | 4.667 | 4.26701 |
| chr8 | 11008863 | 11009856 | 1000 | Promoter (<=1kb) | 8 | 10986980 | 11008458 | 21479 | 2 | ENSMUST00 | -406 | Irs2 | ENSMUSG0C | false | chr8 | 11006639 | 11010129 | 40.16868 | 134.66035 | 129.99733 | chr8 | 11009770 | 11013754 | 4.1229 | 3.64225 | 2.28814 |
| chr3 | 94837216 | 94838474 | 1000 | Promoter (<=1kb) | 3 | 94837567 | 94882326 | 44760 | 1 | ENSMUST00 | 0 | Pogz | ENSMUSG0C | false | chr3 | 94837352 | 94839024 | 26.28407 | 85.32412 | 82.07336 | chr3 | 94838440 | 94841661 | 3.88768 | 3.47393 | 2.10452 |
| chr11 | 95283031 | 95283362 | 1000 | Promoter (1-2kb) | 11 | 95275323 | 95281894 | 6572 | 2 | ENSMUST00 | -1138 | Kat7 | ENSMUSG0C | false | . | . | . | . | . | . | chr11 | 95282358 | 95285796 | 5.32177 | 4.25242 | 3.28749 |
| chr11 | 95309484 | 95309896 | 1000 | Promoter (<=1kb) | 11 | 95274259 | 95309602 | 35344 | 2 | ENSMUST00 | 0 | Kat7 | ENSMUSG0C | false | chr11 | 95309015 | 95309668 | 19.81326 | 53.53814 | 50.77763 | . | . | . | . | . | . |
| chr8 | 10945925 | 10946447 | 1000 | Distal Intergenic | 8 | 10924427 | 10928457 | 4031 | 2 | ENSMUST00 | -17469 | 3930402G23 | ENSMUSG0C | false | . | . | . | . | . | . | chr8 | 10944922 | 10946979 | 5.04945 | 4.15994 | 3.04116 |
| chr11 | 95037301 | 95037758 | 1000 | Promoter (2-3kb) | 11 | 95029654 | 95040558 | 10905 | 2 | ENSMUST00 | 2800 | Pdk2 | ENSMUSG0C | false | . | . | . | . | . | . | chr11 | 95031837 | 95049061 | 6.83336 | 4.76219 | 4.52264 |
| chr3 | 153912747 | 153913194 | 1000 | Promoter (<=1kb) | 3 | 153907289 | 153912966 | 5678 | 2 | ENSMUST00 | 0 | Rabggtb | ENSMUSG0C | false | chr3 | 153911873 | 153913720 | 26.01842 | 90.55051 | 87.195 | . | . | . | . | . | . |
| chr11 | 94992595 | 94993581 | 1000 | Promoter (<=1kb) | 11 | 94992880 | 95001326 | 8447 | 1 | ENSMUST00 | 0 | Ppp1r9b | ENSMUSG0C | false | chr11 | 94989305 | 94994756 | 33.94791 | 109.87469 | 106.04007 | chr11 | 94992312 | 95001916 | 5.70711 | 4.31596 | 3.59817 |
| chr11 | 94990678 | 94991356 | 1000 | Promoter (<=1kb) | 11 | 94991035 | 95006899 | 15865 | 1 | ENSMUST00 | 0 | Ppp1r9b | ENSMUSG0C | false | chr11 | 94989305 | 94994756 | 33.94791 | 109.87469 | 106.04007 | chr11 | 94988880 | 94990882 | 5.93865 | 4.53392 | 3.77484 |
| chr11 | 94989561 | 94990063 | 1000 | Promoter (<=1kb) | 11 | 94991035 | 95006899 | 15865 | 1 | ENSMUST00 | -972 | Ppp1r9b | ENSMUSG0C | false | chr11 | 94989305 | 94994756 | 33.94791 | 109.87469 | 106.04007 | chr11 | 94988880 | 94990882 | 5.93865 | 4.53392 | 3.77484 |
| chr13 | 38633273 | 38633588 | 1000 | Promoter (1-2kb) | 13 | 38602709 | 38635109 | 32401 | 2 | ENSMUST00 | 1521 | Bloc1s5 | ENSMUSG0C | false | . | . | . | . | . | . | chr13 | 38632294 | 38635780 | 4.70737 | 4.01539 | 2.74686 |
| chr13 | 38527786 | 38528666 | 1000 | Promoter (<=1kb) | 13 | 38500273 | 38527914 | 27642 | 2 | ENSMUST00 | 0 | Txndc5 | ENSMUSG0C | false | chr13 | 38527177 | 38528831 | 28.45235 | 93.08366 | 89.67602 | chr13 | 38528592 | 38528883 | 3.00925 | 2.84946 | 1.44689 |
| chr10 | 26776491 | 26776820 | 1000 | Intron (ENSMUST00000176060.2 | 10 | 26772477 | 26918648 | 146172 | 1 | ENSMUST00 | 4015 | Arhgap18 | ENSMUSG0C | false | . | . | . | . | . | . | chr10 | 26773957 | 26793135 | 5.02763 | 4.03577 | 3.04298 |
| chr10 | 26772317 | 26773480 | 1000 | Promoter (<=1kb) | 10 | 26772477 | 26918648 | 146172 | 1 | ENSMUST00 | 0 | Arhgap18 | ENSMUSG0C | false | chr10 | 26772092 | 26774866 | 26.85422 | 77.8849 | 74.7682 | chr10 | 26772137 | 26772380 | 4.72053 | 4.12055 | 2.75629 |
| chr2 | 6322451 | 6323496 | 1000 | Promoter (<=1kb) | 2 | 6322667 | 6446390 | 123724 | 1 | ENSMUST00 | 0 | Usp6nl | ENSMUSG0C | false | chr2 | 6322201 | 6324135 | 37.39668 | 122.22031 | 118.01075 | chr2 | 6323493 | 6324028 | 3.38179 | 3.177 | 1.7093 |
| chr2 | 6212936 | 6213582 | 1000 | Promoter (<=1kb) | 2 | 6188465 | 6213033 | 24569 | 2 | ENSMUST00 | 0 | Echdc3 | ENSMUSG0C | false | chr2 | 6212609 | 6213761 | 21.08548 | 68.47759 | 65.51048 | chr2 | 6212067 | 6213903 | 4.52741 | 3.79982 | 2.67748 |
| chr3 | 152478898 | 152479355 | 1000 | Intron (ENSMUST00000045262.6 | 3 | 152457417 | 152458535 | 1119 | 1 | ENSMUST00 | 21482 | Zzz3 | ENSMUSG0C | false | chr3 | 152477894 | 152479035 | 22.97661 | 61.46103 | 58.5941 | chr3 | 152477665 | 152479621 | 3.70285 | 3.39616 | 1.9771 |
| chr14 | 21987395 | 21988226 | 1000 | Promoter (1-2kb) | 14 | 21983962 | 21989601 | 5640 | 2 | ENSMUST00 | 1375 | Zfp503 | ENSMUSG0C | false | chr14 | 21984527 | 21987918 | 37.75571 | 118.58805 | 114.49699 | . | . | . | . | . | . |
| chr14 | 21985981 | 21986504 | 1000 | Exon (ENSMUST00000043409.7/ | 14 | 21983962 | 21989601 | 5640 | 2 | ENSMUST00 | 3097 | Zfp503 | ENSMUSG0C | false | chr14 | 21984527 | 21987918 | 37.75571 | 118.58805 | 114.49699 | chr14 | 21981011 | 21986083 | 4.65699 | 3.79582 | 2.74477 |
| chr13 | 37853937 | 37854348 | 1000 | Intron (ENSMUST00000122842.2 | 13 | 37857934 | 37893914 | 35981 | 1 | ENSMUST00 | -3586 | Rreb1 | ENSMUSG0C | false | chr13 | 37854017 | 37854797 | 8.80131 | 17.30502 | 15.00832 | chr13 | 37836337 | 37889224 | 5.54937 | 4.22662 | 3.48404 |
| chr13 | 37862612 | 37862953 | 1000 | Intron (ENSMUST00000122842.2 | 13 | 37857934 | 37893914 | 35981 | 1 | ENSMUST00 | 4679 | Rreb1 | ENSMUSG0C | false | . | . | . | . | . | . | chr13 | 37836337 | 37889224 | 5.54937 | 4.22662 | 3.48404 |
| chr13 | 37825738 | 37828038 | 1000 | Promoter (<=1kb) | 13 | 37825975 | 37893826 | 67852 | 1 | ENSMUST00 | 0 | Rreb1 | ENSMUSG0C | false | chr13 | 37825344 | 37828365 | 39.80159 | 147.3544 | 142.17488 | chr13 | 37825056 | 37825840 | 3.49771 | 3.34984 | 1.76698 |
| chr13 | 37778116 | 37779160 | 1000 | Promoter (<=1kb) | 13 | 37778400 | 37893849 | 115450 | 1 | ENSMUST00 | 0 | Rreb1 | ENSMUSG0C | false | chr13 | 37778006 | 37779602 | 24.87197 | 72.62122 | 69.58942 | chr13 | 37776941 | 37779645 | 3.57121 | 3.29393 | 1.88077 |
| chr10 | 26465338 | 26465754 | 1000 | Distal Intergenic | 10 | 26275452 | 26375185 | 99734 | 2 | ENSMUST00 | -90154 | L3mbtl3 | ENSMUSG0C | true | . | . | . | . | . | . | chr10 | 26464174 | 26466295 | 4.03283 | 3.45552 | 2.24048 |
| chr15 | 31540335 | 31540699 | 1000 | Distal Intergenic | 15 | 31455899 | 31531037 | 75139 | 2 | ENSMUST00 | -9299 | 38777 | ENSMUSG0C | false | . | . | . | . | . | . | chr15 | 31539046 | 31544069 | 5.25379 | 4.14739 | 3.21602 |
| chr4 | 63547821 | 63548190 | 1000 | Intron (ENSMUST00000035301.6 | 4 | 63544772 | 63550750 | 5979 | 1 | ENSMUST00 | 3050 | Atp6v1g1 | ENSMUSG0C | false | . | . | . | . | . | . | chr4 | 63544073 | 63548639 | 5.74686 | 4.38888 | 3.63689 |
| chr2 | 180024372 | 180025521 | 1000 | Promoter (<=1kb) | 2 | 180024987 | 180035461 | 10475 | 1 | ENSMUST00 | 0 | Lsm14b | ENSMUSG0C | false | chr2 | 180024135 | 180026031 | 36.04346 | 118.78738 | 114.68494 | chr2 | 180025440 | 180027711 | 5.60474 | 4.27644 | 3.5457 |
| chr11 | 94522892 | 94523343 | 1000 | Promoter (1-2kb) | 11 | 94501347 | 94521742 | 20396 | 2 | ENSMUST00 | -1151 | Mycbpap | ENSMUSG0C | false | . | . | . | . | . | . | chr11 | 94515793 | 94529022 | 5.67014 | 4.37838 | 3.56699 |
| chr2 | 179976392 | 179977317 | 1000 | Promoter (<=1kb) | 2 | 179912146 | 179976646 | 64501 | 2 | ENSMUST00 | 0 | Taf4a | ENSMUSG0C | false | chr2 | 179975983 | 179977691 | 42.9698 | 143.99599 | 138.94749 | . | . | . | . | . | . |
| chr17 | 44735688 | 44736789 | 1000 | Promoter (<=1kb) | 17 | 44495987 | 44736176 | 240190 | 2 | ENSMUST00 | 0 | Runx2 | ENSMUSG0C | false | chr17 | 44734011 | 44736771 | 24.48636 | 83.88964 | 80.66612 | chr17 | 44733614 | 44738430 | 5.14462 | 3.97095 | 3.15882 |
| chr5 | 53998128 | 53998932 | 1000 | Promoter (<=1kb) | 5 | 53998417 | 54121057 | 122641 | 1 | ENSMUST00 | 0 | Stim2 | ENSMUSG0C | false | chr5 | 53997850 | 54000404 | 26.41709 | 100.00125 | 96.43182 | . | . | . | . | . | . |
| chr6 | 30303722 | 30304574 | 1000 | Promoter (<=1kb) | 6 | 30211289 | 30304539 | 93251 | 2 | ENSMUST00 | 0 | Ube2h | ENSMUSG0C | false | chr6 | 30302352 | 30304619 | 30.76103 | 94.20794 | 90.76972 | . | . | . | . | . | . |
| chr10 | 25296665 | 25297381 | 1000 | Promoter (<=1kb) | 10 | 25283923 | 25297073 | 13151 | 2 | ENSMUST00 | 0 | Akap7 | ENSMUSG0C | false | chr10 | 25296481 | 25297459 | 26.1454 | 83.18384 | 79.97624 | . | . | . | . | . | . |
| chr7 | 79465747 | 79466492 | 1000 | Promoter (<=1kb) | 7 | 79448791 | 79466362 | 17572 | 2 | ENSMUST00 | 0 | Polg | ENSMUSG0C | false | chr7 | 79464733 | 79467421 | 37.51873 | 124.57824 | 120.27934 | . | . | . | . | . | . |
| chr14 | 21144297 | 21144730 | 1000 | Intron (ENSMUST00000045376.9 | 14 | 21052574 | 21448568 | 395995 | 1 | ENSMUST00 | 91724 | Adk | ENSMUSG0C | false | . | . | . | . | . | . | chr14 | 21143498 | 21146250 | 3.83323 | 3.50504 | 2.03426 |
| chr11 | 121702153 | 121702895 | 1000 | Promoter (<=1kb) | 11 | 121702411 | 121716306 | 13896 | 1 | ENSMUST00 | 0 | Metrnl | ENSMUSG0C | true | chr11 | 121701795 | 121706690 | 36.26418 | 124.50906 | 120.21295 | chr11 | 121701133 | 121702188 | 4.6375 | 3.96424 | 2.72918 |
| chr17 | 23802679 | 23803240 | 1000 | Promoter (<=1kb) | 17 | 23803187 | 23824736 | 21550 | 1 | ENSMUST00 | 0 | Srrm2 | ENSMUSG0C | false | chr17 | 23802526 | 23802961 | 10.89686 | 23.4988 | 21.11616 | . | . | . | . | . | . |
| chr13 | 13954353 | 13955121 | 1000 | Promoter (<=1kb) | 13 | 13954674 | 13999068 | 44395 | 1 | ENSMUST00 | 0 | B3galnt2 | ENSMUSG0C | false | chr13 | 13953612 | 13955626 | 33.41831 | 117.26761 | 113.21259 | . | . | . | . | . | . |
| chr2 | 32150758 | 32151689 | 1000 | Promoter (<=1kb) | 2 | 32151082 | 32234537 | 83456 | 1 | ENSMUST00 | 0 | Prrc2b | ENSMUSG0C | false | chr2 | 32150787 | 32152576 | 37.7199 | 126.00829 | 121.65485 | . | . | . | . | . | . |
| chr2 | 174112809 | 174113432 | 1000 | Promoter (1-2kb) | 2 | 174111573 | 174116103 | 4531 | 1 | ENSMUST00 | 1237 | Npepl1 | ENSMUSG0C | false | . | . | . | . | . | . | chr2 | 174109729 | 174116112 | 5.48827 | 4.29987 | 3.41377 |
| chr3 | 27736118 | 27736645 | 1000 | Distal Intergenic | 3 | 27416162 | 27710439 | 294278 | 2 | ENSMUST00 | -25680 | Fndc3b | ENSMUSG0C | true | . | . | . | . | . | . | chr3 | 27734964 | 27737253 | 4.76439 | 3.8955 | 2.86096 |
| chr3 | 27710003 | 27711419 | 1000 | Promoter (<=1kb) | 3 | 27416162 | 27710439 | 294278 | 2 | ENSMUST00 | 0 | Fndc3b | ENSMUSG0C | true | chr3 | 27708580 | 27711670 | 41.52505 | 152.7413 | 147.35226 | . | . | . | . | . | . |
| chr7 | 90129480 | 90130341 | 1000 | Promoter (<=1kb) | 7 | 90130232 | 90209447 | 79216 | 1 | ENSMUST00 | 0 | Picalm | ENSMUSG0C | false | chr7 | 90129152 | 90132776 | 34.36702 | 111.64683 | 107.76208 | chr7 | 90128685 | 90129796 | 3.92031 | 3.53691 | 2.10386 |
| chr7 | 89403433 | 89404484 | 1000 | Promoter (<=1kb) | 7 | 89156448 | 89404222 | 247775 | 2 | ENSMUST00 | 0 | Tmem135 | ENSMUSG0C | false | chr7 | 89403160 | 89405751 | 38.07755 | 144.18416 | 139.13362 | . | . | . | . | . | . |
| chr16 | 93832072 | 93832811 | 1000 | Promoter (<=1kb) | 16 | 93832121 | 93876072 | 43952 | 1 | ENSMUST00 | 0 | Morc3 | ENSMUSG0C | false | chr16 | 93832171 | 93833436 | 24.0356 | 94.23969 | 90.80066 | . | . | . | . | . | . |
| chr8 | 40423319 | 40424118 | 1000 | Promoter (<=1kb) | 8 | 40423815 | 40485725 | 61911 | 1 | ENSMUST00 | 0 | Zdhhc2 | ENSMUSG0C | false | chr8 | 40423157 | 40424706 | 25.84672 | 90.2271 | 86.88143 | chr8 | 40423047 | 40424974 | 4.39753 | 3.72101 | 2.52757 |
| chr5 | 142816994 | 142819030 | 1000 | Promoter (<=1kb) | 5 | 142754789 | 142817662 | 62874 | 2 | ENSMUST00 | 0 | Tnrc18 | ENSMUSG0C | false | chr5 | 142813148 | 142817670 | 35.38397 | 108.24114 | 104.45254 | . | . | . | . | . | . |
| chr5 | 142821448 | 142822042 | 1000 | Distal Intergenic | 5 | 142754789 | 142817662 | 62874 | 2 | ENSMUST00 | -3787 | Tnrc18 | ENSMUSG0C | false | chr5 | 142817908 | 142824082 | 29.69128 | 86.53342 | 83.26198 | chr5 | 142821951 | 142824303 | 4.24728 | 3.72552 | 2.37961 |
| chr2 | 30445275 | 30445839 | 1000 | Promoter (<=1kb) | 2 | 30445656 | 30447074 | 1419 | 1 | ENSMUST00 | 0 | Ppp2r4 | ENSMUSG0C | false | . | . | . | . | . | . | chr2 | 30439594 | 30459395 | 7.81508 | 5.29532 | 5.31547 |
| chr2 | 30441671 | 30442119 | 1000 | Promoter (<=1kb) | 2 | 30441853 | 30447131 | 5279 | 1 | ENSMUST00 | 0 | Ppp2r4 | ENSMUSG0C | false | chr2 | 30441461 | 30441830 | 9.39297 | 19.83805 | 17.50559 | chr2 | 30439594 | 30459395 | 7.81508 | 5.29532 | 5.31547 |
| chr17 | 35549391 | 35550127 | 1000 | Promoter (2-3kb) | 17 | 35552128 | 35557180 | 5053 | 1 | ENSMUST00 | -2001 | Cdsn | ENSMUSG0C | false | . | . | . | . | . | . | chr17 | 35546287 | 35552496 | 5.78427 | 4.44046 | 3.65751 |
| chr18 | 64643599 | 64644096 | 1000 | Intron (ENSMUST00000025482.8 | 18 | 64528979 | 64661000 | 132022 | 2 | ENSMUST00 | 16904 | Atp8b1 | ENSMUSG0C | true | chr18 | 64644031 | 64644507 | 5.82866 | 9.11851 | 6.96196 | chr18 | 64613324 | 64659141 | 5.21932 | 4.12513 | 3.19621 |
| chr18 | 64661588 | 64662046 | 1000 | Promoter (<=1kb) | 18 | 64528979 | 64661000 | 132022 | 2 | ENSMUST00 | -589 | Atp8b1 | ENSMUSG0C | true | chr18 | 64658246 | 64661826 | 35.98183 | 119.73972 | 115.60696 | chr18 | 64660486 | 64662366 | 3.41136 | 3.31646 | 1.67894 |
| chr18 | 64660356 | 64661443 | 1000 | Promoter (<=1kb) | 18 | 64528979 | 64661000 | 132022 | 2 | ENSMUST00 | 0 | Atp8b1 | ENSMUSG0C | true | chr18 | 64658246 | 64661826 | 35.98183 | 119.73972 | 115.60696 | chr18 | 64660486 | 64662366 | 3.41136 | 3.31646 | 1.67894 |
| chr18 | 64605935 | 64606429 | 1000 | Intron (ENSMUST00000025482.8 | 18 | 64528979 | 64661000 | 132022 | 2 | ENSMUST00 | 54571 | Atp8b1 | ENSMUSG0C | true | . | . | . | . | . | . | chr18 | 64600002 | 64611678 | 4.6748 | 3.82852 | 2.75176 |
| chr8 | 39005708 | 39006509 | 1000 | Promoter (<=1kb) | 8 | 39005867 | 39165112 | 159246 | 1 | ENSMUST00 | 0 | Tusc3 | ENSMUSG0C | false | chr8 | 39005471 | 39006885 | 38.67513 | 141.95352 | 136.99454 | chr8 | 39006413 | 39007211 | 3.40613 | 3.30984 | 1.67845 |
| chr16 | 4879363 | 4880774 | 1000 | Promoter (<=1kb) | 16 | 4874778 | 4880315 | 5538 | 2 | ENSMUST00 | 0 | Ubald1 | ENSMUSG0C | false | chr16 | 4880656 | 4881356 | 23.23877 | 64.48248 | 61.57465 | . | . | . | . | . | . |
| chr9 | 59750319 | 59751666 | 1000 | Promoter (<=1kb) | 9 | 59750896 | 59928866 | 177971 | 1 | ENSMUST00 | 0 | Myo9a | ENSMUSG0C | false | chr9 | 59749942 | 59750763 | 21.74415 | 57.67707 | 54.86403 | . | . | . | . | . | . |
| chr9 | 117192584 | 117192905 | 1000 | Intron (ENSMUST00000111773.4 | 9 | 116943123 | 117251597 | 308475 | 2 | ENSMUST00 | 58692 | Rbms3 | ENSMUSG0C | true | . | . | . | . | . | . | chr9 | 117191721 | 117196308 | 4.8312 | 3.96054 | 2.86504 |
| chr5 | 142440236 | 142440780 | 1000 | Intron (ENSMUST00000072837.5 | 5 | 142463931 | 142478715 | 14785 | 1 | ENSMUST00 | -23151 | Ap5z1 | ENSMUSG0C | false | chr5 | 142439877 | 142440543 | 7.13425 | 13.57188 | 11.33167 | chr5 | 142438011 | 142446243 | 6.37507 | 4.47997 | 4.17057 |
| chr5 | 142463687 | 142464113 | 1000 | Promoter (<=1kb) | 5 | 142463931 | 142478715 | 14785 | 1 | ENSMUST00 | 0 | Ap5z1 | ENSMUSG0C | false | chr5 | 142463474 | 142464783 | 35.38397 | 108.24114 | 104.45254 | chr5 | 142463287 | 142464993 | 3.99302 | 3.45645 | 2.21184 |
| chr17 | 24761140 | 24761599 | 1000 | Distal Intergenic | 17 | 24753003 | 24758683 | 5681 | 1 | ENSMUST00 | 8138 | Hs3st6 | ENSMUSG0C | false | . | . | . | . | . | . | chr17 | 24755141 | 24772276 | 6.3257 | 4.70291 | 4.09799 |
| chr16 | 4658618 | 4659342 | 1000 | Promoter (2-3kb) | 16 | 4634598 | 4661922 | 27325 | 2 | ENSMUST00 | 2580 | Coro7 | ENSMUSG0C | false | . | . | . | . | . | . | chr16 | 4642609 | 4661657 | 7.56188 | 5.16905 | 5.12366 |
| chr16 | 4656177 | 4656728 | 1000 | Intron (ENSMUST00000144815.1 | 16 | 4634598 | 4661922 | 27325 | 2 | ENSMUST00 | 5194 | Coro7 | ENSMUSG0C | false | . | . | . | . | . | . | chr16 | 4642609 | 4661657 | 7.56188 | 5.16905 | 5.12366 |
| chr19 | 37174056 | 37174986 | 1000 | Promoter (<=1kb) | 19 | 37021297 | 37174095 | 152799 | 2 | ENSMUST00 | 0 | Cpeb3 | ENSMUSG0C | false | chr19 | 37173708 | 37176086 | 24.14628 | 82.32003 | 79.12344 | chr19 | 37171746 | 37174122 | 3.60545 | 3.34591 | 1.88566 |
| chr19 | 37177881 | 37178317 | 1000 | Promoter (<=1kb) | 19 | 37174437 | 37178017 | 3581 | 2 | ENSMUST00 | 0 | Cpeb3 | ENSMUSG0C | false | chr19 | 37177484 | 37178568 | 11.02877 | 22.59171 | 20.22099 | chr19 | 37175901 | 37178526 | 3.48593 | 3.19866 | 1.78277 |
| chr2 | 165831558 | 165832805 | 1000 | Intron (ENSMUST00000109266.5 | 2 | 165845744 | 165852122 | 6379 | 2 | ENSMUST00 | 19317 | Zmynd8 | ENSMUSG0C | false | . | . | . | . | . | . | chr2 | 165823765 | 165834539 | 5.16576 | 4.03336 | 3.16898 |
| chr2 | 30133544 | 30133929 | 1000 | Promoter (<=1kb) | 2 | 30133746 | 30137296 | 3551 | 1 | ENSMUST00 | 0 | Tbc1d13 | ENSMUSG0C | false | chr2 | 30133098 | 30134976 | 35.34781 | 110.24136 | 106.39787 | chr2 | 30132392 | 30135577 | 5.08064 | 4.02768 | 3.09838 |
| chr16 | 92183731 | 92184238 | 1000 | Distal Intergenic | 16 | 92106261 | 92112227 | 5967 | 1 | ENSMUST00 | 77471 | Mrps6 | ENSMUSG0C | false | . | . | . | . | . | . | chr16 | 92182098 | 92185065 | 7.08408 | 5.14129 | 4.70455 |
| chr16 | 92157695 | 92158798 | 1000 | Distal Intergenic | 16 | 92106261 | 92112227 | 5967 | 1 | ENSMUST00 | 51435 | Mrps6 | ENSMUSG0C | false | chr16 | 92158227 | 92159042 | 13.18045 | 29.78312 | 27.32097 | chr16 | 92156045 | 92167743 | 5.46378 | 4.33054 | 3.39881 |
| chr5 | 141241259 | 141241792 | 1000 | Promoter (<=1kb) | 5 | 141241490 | 142215586 | 974097 | 1 | ENSMUST00 | 0 | Sdk1 | ENSMUSG0C | false | chr5 | 141240751 | 141242376 | 21.77823 | 66.26614 | 63.32996 | chr5 | 141240461 | 141243936 | 5.0214 | 4.01067 | 3.03991 |
| chr5 | 141946816 | 141947197 | 1000 | Intron (ENSMUST00000085774.5 | 5 | 141856787 | 142213194 | 356408 | 1 | ENSMUST00 | 90030 | Sdk1 | ENSMUSG0C | false | . | . | . | . | . | . | chr5 | 141946431 | 141947807 | 4.05217 | 3.57104 | 2.26932 |
| chr10 | 30802655 | 30803402 | 1000 | Promoter (<=1kb) | 10 | 30645584 | 30803107 | 157524 | 2 | ENSMUST00 | 0 | Ncoa7 | ENSMUSG0C | false | chr10 | 30802117 | 30803889 | 31.70129 | 105.8978 | 102.17969 | . | . | . | . | . | . |
| chr3 | 122983814 | 122984944 | 1000 | Promoter (<=1kb) | 3 | 122933601 | 122984447 | 50847 | 2 | ENSMUST00 | 0 | Usp53 | ENSMUSG0C | true | chr3 | 122983401 | 122985518 | 42.57743 | 146.68684 | 141.53157 | . | . | . | . | . | . |
| chr3 | 122986234 | 122986807 | 1000 | Promoter (1-2kb) | 3 | 122933601 | 122984447 | 50847 | 2 | ENSMUST00 | -1788 | Usp53 | ENSMUSG0C | true | . | . | . | . | . | . | chr3 | 122985135 | 122990580 | 4.5336 | 3.88052 | 2.61649 |
| chr15 | 9140324 | 9140720 | 1000 | Promoter (<=1kb) | 15 | 9140621 | 9198318 | 57698 | 1 | ENSMUST00 | 0 | Lmbrd2 | ENSMUSG0C | false | chr15 | 9139812 | 9141459 | 36.73951 | 123.20795 | 118.95689 | . | . | . | . | . | . |
| chr4 | 152110654 | 152111046 | 1000 | Promoter (2-3kb) | 4 | 152107729 | 152108209 | 481 | 1 | ENSMUST00 | 2926 | Plekhg5 | ENSMUSG0C | false | . | . | . | . | . | . | chr4 | 152106823 | 152117667 | 6.06834 | 4.4954 | 3.899 |
| chr4 | 152107427 | 152107861 | 1000 | Promoter (<=1kb) | 4 | 152107636 | 152115394 | 7759 | 1 | ENSMUST00 | 0 | Plekhg5 | ENSMUSG0C | false | chr4 | 152107604 | 152109436 | 25.32445 | 75.22865 | 72.15769 | chr4 | 152106823 | 152117667 | 6.06834 | 4.4954 | 3.899 |
| chr3 | 122619036 | 122620112 | 1000 | Promoter (<=1kb) | 3 | 122538719 | 122619667 | 80949 | 2 | ENSMUST00 | 0 | Fnbp1l | ENSMUSG0C | true | chr3 | 122617382 | 122620944 | 33.69019 | 111.49343 | 107.61708 | . | . | . | . | . | . |
| chr3 | 122567223 | 122567683 | 1000 | Intron (ENSMUST00000162409.2 | 3 | 122557910 | 122559244 | 1335 | 2 | ENSMUST00 | -7980 | Fnbp1l | ENSMUSG0C | true | . | . | . | . | . | . | chr3 | 122560136 | 122582388 | 4.30049 | 3.73293 | 2.42998 |
| chr11 | 120316896 | 120317392 | 1000 | Distal Intergenic | 11 | 120289793 | 120292167 | 2375 | 1 | ENSMUST00 | 27104 | Bahcc1 | ENSMUSG0C | false | chr11 | 120315894 | 120317217 | 16.34629 | 47.3758 | 44.69273 | chr11 | 120314427 | 120317826 | 3.86972 | 3.48335 | 2.10014 |
| chr11 | 120252620 | 120253303 | 1000 | Intron (ENSMUST00000122148.2 | 11 | 120236699 | 120292282 | 55584 | 1 | ENSMUST00 | 15922 | Bahcc1 | ENSMUSG0C | false | . | . | . | . | . | . | chr11 | 120250632 | 120257317 | 7.25676 | 4.82908 | 4.89578 |
| chr5 | 43781664 | 43782365 | 1000 | Promoter (<=1kb) | 5 | 43744616 | 43782118 | 37503 | 2 | ENSMUST00 | 0 | Fbxl5 | ENSMUSG0C | false | chr5 | 43780884 | 43782943 | 33.07612 | 110.40385 | 106.55179 | chr5 | 43774753 | 43783065 | 4.0027 | 3.52642 | 2.20871 |
| chr4 | 47091832 | 47092483 | 1000 | Promoter (<=1kb) | 4 | 47091909 | 47123070 | 31162 | 1 | ENSMUST00 | 0 | Galnt12 | ENSMUSG0C | false | chr4 | 47091669 | 47092843 | 13.73463 | 43.09688 | 40.46774 | chr4 | 47091909 | 47093171 | 6.16823 | 4.54669 | 3.9964 |
| chr5 | 43233489 | 43234913 | 1000 | Promoter (<=1kb) | 5 | 43233956 | 43287022 | 53067 | 1 | ENSMUST00 | 0 | Cpeb2 | ENSMUSG0C | false | chr5 | 43232302 | 43236237 | 40.20312 | 135.82269 | 131.11238 | . | . | . | . | . | . |
| chr3 | 121953016 | 121953434 | 1000 | Promoter (<=1kb) | 3 | 121953229 | 122016677 | 63449 | 1 | ENSMUST00 | 0 | Arhgap29 | ENSMUSG0C | false | chr3 | 121953085 | 121955451 | 20.109 | 88.33971 | 85.03291 | . | . | . | . | . | . |
| chr3 | 121887156 | 121887502 | 1000 | Distal Intergenic | 3 | 121953229 | 122016677 | 63449 | 1 | ENSMUST00 | -65727 | Arhgap29 | ENSMUSG0C | false | chr3 | 121887147 | 121887471 | 5.25681 | 8.51884 | 6.37244 | chr3 | 121885243 | 121894006 | 5.82719 | 4.55235 | 3.67654 |
| chr2 | 164910979 | 164911958 | 1000 | Promoter (<=1kb) | 2 | 164891882 | 164911757 | 19876 | 2 | ENSMUST00 | 0 | Zfp335 | ENSMUSG0C | false | chr2 | 164910073 | 164912468 | 39.24862 | 125.4371 | 121.11293 | . | . | . | . | . | . |
| chr10 | 18318361 | 18318835 | 1000 | Distal Intergenic | 10 | 18390322 | 18524289 | 133968 | 1 | ENSMUST00 | -71487 | Nhsl1 | ENSMUSG0C | false | chr10 | 18318175 | 18319701 | 26.03857 | 105.07202 | 101.37238 | chr10 | 18318031 | 18319686 | 3.44861 | 3.24031 | 1.76904 |
| chr6 | 28261727 | 28262460 | 1000 | Promoter (<=1kb) | 6 | 28239931 | 28261911 | 21981 | 2 | ENSMUST00 | 0 | Zfp800 | ENSMUSG0C | false | chr6 | 28260448 | 28262923 | 35.62435 | 116.99458 | 112.94973 | . | . | . | . | . | . |
| chr2 | 29696216 | 29696821 | 1000 | Intron (ENSMUST00000095087.5 | 2 | 29702482 | 29740363 | 37882 | 1 | ENSMUST00 | -5661 | Rapgef1 | ENSMUSG0C | false | . | . | . | . | . | . | chr2 | 29690471 | 29698739 | 5.92345 | 4.4721 | 3.78221 |
| chr2 | 164879384 | 164880323 | 1000 | Promoter (<=1kb) | 2 | 164880052 | 164883110 | 3059 | 1 | ENSMUST00 | 0 | Pcif1 | ENSMUSG0C | false | chr2 | 164880071 | 164881078 | 12.8669 | 28.00426 | 25.56298 | chr2 | 164880179 | 164886599 | 6.28839 | 4.71258 | 4.05871 |
| chr4 | 150297168 | 150297539 | 1000 | Intron (ENSMUST00000105682.3 | 4 | 150281812 | 150282466 | 655 | 1 | ENSMUST00 | 15357 | Rere | ENSMUSG0C | false | . | . | . | . | . | . | chr4 | 150292355 | 150301062 | 3.96838 | 3.54879 | 2.16729 |
| chr4 | 150457856 | 150458636 | 1000 | Intron (ENSMUST00000105682.3 | 4 | 150488393 | 150621962 | 133570 | 1 | ENSMUST00 | -29757 | Rere | ENSMUSG0C | false | . | . | . | . | . | . | chr4 | 150440410 | 150463295 | 5.62337 | 4.29719 | 3.53587 |
| chr4 | 150444078 | 150444380 | 1000 | Intron (ENSMUST00000105682.3 | 4 | 150406394 | 150470381 | 63988 | 1 | ENSMUST00 | 37685 | Rere | ENSMUSG0C | false | . | . | . | . | . | . | chr4 | 150440410 | 150463295 | 5.62337 | 4.29719 | 3.53587 |
| chr4 | 150371594 | 150372486 | 1000 | Intron (ENSMUST00000105682.3 | 4 | 150406394 | 150470381 | 63988 | 1 | ENSMUST00 | -33908 | Rere | ENSMUSG0C | false | chr4 | 150371430 | 150372679 | 27.50468 | 83.69844 | 80.48042 | chr4 | 150361010 | 150371752 | 4.90537 | 3.95042 | 2.93904 |
| chr4 | 150569148 | 150570181 | 1000 | Promoter (<=1kb) | 4 | 150569588 | 150570495 | 908 | 1 | ENSMUST00 | 0 | Rere | ENSMUSG0C | false | chr4 | 150568860 | 150571152 | 34.67015 | 117.41676 | 113.35385 | chr4 | 150555649 | 150569253 | 4.54721 | 3.83298 | 2.65135 |
| chr5 | 21529320 | 21529778 | 1000 | Intron (ENSMUST00000051358.5 | 5 | 21543527 | 21575900 | 32374 | 1 | ENSMUST00 | -13749 | Lrrc17 | ENSMUSG0C | false | . | . | . | . | . | . | . | . | . | . | . | . |

| | | | | | | | | | | | | | | | | | | | | | | | | | | |
|---|---|---|---|---|---|---|---|---|---|---|---|---|---|---|---|---|---|---|---|---|---|---|---|---|---|---|
| chr10 | 17511944 | 17512310 | 1000 | Distal Intergenic | 10 | 17723228 | 17725673 | 2446 | 1 | ENSMUST00 | -210918 | Cited2 | ENSMUSG0( | false | . | . | . | . | . | . | chr10 | 17511729 | 17513116 | 4.17781 | 3.51016 | 2.37139 |
| chr4 | 149968587 | 149968866 | 1000 | Distal Intergenic | 4 | 149907142 | 149955043 | 47902 | 2 | ENSMUST00 | -13545 | Spsb1 | ENSMUSG0( | false | . | . | . | . | . | . | . | . | . | . | . | . |
| chr4 | 149958172 | 149958523 | 1000 | Distal Intergenic | 4 | 149907142 | 149955043 | 47902 | 2 | ENSMUST00 | -3130 | Spsb1 | ENSMUSG0( | false | chr4 | 149958450 | 149958614 | 4.54943 | 6.49485 | 4.40153 | chr4 | 149957919 | 149958752 | 4.60919 | 3.95063 | 2.67584 |
| chr15 | 5343684 | 5344238 | 1000 | Distal Intergenic | 15 | 5233399 | 5244187 | 10789 | 2 | ENSMUST00 | -99498 | Ptger4 | ENSMUSG0( | false | . | . | . | . | . | . | chr15 | 5343403 | 5345027 | 4.35071 | 3.65208 | 2.48313 |
| chr15 | 5493456 | 5493880 | 1000 | Distal Intergenic | 15 | 5233399 | 5244187 | 10789 | 2 | ENSMUST00 | -249270 | Ptger4 | ENSMUSG0( | false | . | . | . | . | . | . | chr15 | 5493227 | 5494113 | 3.56363 | 3.38686 | 1.81453 |
| chr9 | 108275315 | 108275691 | 1000 | Distal Intergenic | 9 | 108205958 | 108263924 | 57967 | 2 | ENSMUST00 | -11392 | Dag1 | ENSMUSG0( | true | . | . | . | . | . | . | chr9 | 108264367 | 108278155 | 5.4782 | 4.31244 | 3.40132 |
| chr5 | 135488735 | 135489841 | 1000 | Intron (ENSMUST00000060311.8 | 5 | 135406523 | 135545122 | 138600 | 2 | ENSMUST00 | 55281 | Hip1 | ENSMUSG0( | true | chr5 | 135488346 | 135489702 | 31.70772 | 93.8754 | 90.44607 | chr5 | 135484962 | 135489884 | 4.50384 | 3.87639 | 2.59998 |
| chr11 | 119228263 | 119228739 | 1000 | Promoter (<=1kb) | 11 | 119228572 | 119265212 | 36641 | 1 | ENSMUST00 | 0 | Ccdc40 | ENSMUSG0( | false | chr11 | 119227871 | 119229195 | 23.73948 | 76.05544 | 72.96951 | chr11 | 119227688 | 119229383 | 4.51495 | 3.84638 | 2.60986 |
| chr4 | 34886551 | 34887131 | 1000 | Distal Intergenic | 4 | 34803113 | 34882960 | 79848 | 2 | ENSMUST00 | -3592 | Zfp292 | ENSMUSG0( | false | chr4 | 34886470 | 34887745 | 20.74539 | 66.97287 | 64.02668 | chr4 | 34886073 | 34886936 | 3.71606 | 3.26901 | 1.9892 |
| chr5 | 20950736 | 20952092 | 1000 | Promoter (<=1kb) | 5 | 20893028 | 20951822 | 58795 | 2 | ENSMUST00 | 0 | Rsbn1l | ENSMUSG0( | false | chr5 | 20950039 | 20952699 | 37.26972 | 119.86725 | 115.72688 | . | . | . | . | . | . |
| chr19 | 12500066 | 12500399 | 1000 | Promoter (1-2kb) | 19 | 12466341 | 12501996 | 35656 | 2 | ENSMUST00 | 1597 | Dtx4 | ENSMUSG0( | false | . | . | . | . | . | . | chr19 | 12499944 | 12502455 | 4.79267 | 4.01787 | 2.84167 |
| chr19 | 12486918 | 12487293 | 1000 | Intron (ENSMUST00000045521.7 | 19 | 12466341 | 12501996 | 35656 | 2 | ENSMUST00 | 14703 | Dtx4 | ENSMUSG0( | false | . | . | . | . | . | . | chr19 | 12486536 | 12489221 | 5.35438 | 4.11221 | 3.33279 |
| chr5 | 20881440 | 20882565 | 1000 | Promoter (<=1kb) | 5 | 20758663 | 20882124 | 123462 | 2 | ENSMUST00 | 0 | Phtf2 | ENSMUSG0( | false | chr5 | 20881099 | 20882123 | 28.45235 | 93.08366 | 89.67602 | . | . | . | . | . | . |
| chr11 | 119085347 | 119086713 | 1000 | Promoter (<=1kb) | 11 | 119077573 | 119086221 | 8649 | 2 | ENSMUST00 | 0 | Cbx4 | ENSMUSG0( | false | chr11 | 119084733 | 119086327 | 34.92444 | 106.42502 | 102.68874 | chr11 | 119086494 | 119091964 | 6.84315 | 4.95819 | 4.51269 |
| chr11 | 119090834 | 119091300 | 1000 | Distal Intergenic | 11 | 119081373 | 119086221 | 4849 | 2 | ENSMUST00 | -4614 | Cbx4 | ENSMUSG0( | false | chr11 | 119090174 | 119091704 | 27.01742 | 78.16648 | 75.04827 | chr11 | 119086494 | 119091964 | 6.84315 | 4.95819 | 4.51269 |
| chr10 | 128231874 | 128232366 | 1000 | Promoter (<=1kb) | 10 | 128232065 | 128252941 | 20877 | 1 | ENSMUST00 | 0 | Timeless | ENSMUSG0( | false | chr10 | 128231529 | 128232831 | 33.50033 | 109.08844 | 105.27855 | . | . | . | . | . | . |
| chr5 | 19226372 | 19227190 | 1000 | Promoter (<=1kb) | 5 | 19227046 | 20702903 | 1475858 | 1 | ENSMUST00 | 0 | Magi2 | ENSMUSG0( | true | chr5 | 19223861 | 19232405 | 34.78864 | 106.1911 | 102.46782 | . | . | . | . | . | . |
| chr2 | 118901091 | 118902966 | 1000 | Promoter (<=1kb) | 2 | 118901453 | 118924528 | 23076 | 1 | ENSMUST00 | 0 | Bahd1 | ENSMUSG0( | false | chr2 | 118900502 | 118903046 | 26.51101 | 114.88218 | 110.90597 | chr2 | 118902659 | 118904848 | 4.02427 | 3.62916 | 2.25107 |
| chr8 | 121875020 | 121875357 | 1000 | Intron (ENSMUST00000127664.1 | 8 | 121882802 | 121883914 | 1113 | 2 | ENSMUST00 | 8557 | Slc7a5 | ENSMUSG0( | false | . | . | . | . | . | . | chr8 | 121874189 | 121875792 | 5.83188 | 4.37677 | 3.73195 |
| chr19 | 59943160 | 59943915 | 1000 | Promoter (<=1kb) | 19 | 59902884 | 59943654 | 40771 | 2 | ENSMUST00 | 0 | Rab11fip2 | ENSMUSG0( | false | chr19 | 59942619 | 59944211 | 28.71633 | 95.52087 | 92.05702 | . | . | . | . | . | . |
| chr4 | 132211840 | 132212463 | 1000 | Promoter (<=1kb) | 4 | 132184912 | 132212303 | 27392 | 2 | ENSMUST00 | 0 | Ythdf2 | ENSMUSG0( | false | chr4 | 132210818 | 132212608 | 36.04346 | 118.78738 | 114.68494 | . | . | . | . | . | . |
| chr6 | 148831268 | 148831712 | 1000 | Promoter (<=1kb) | 6 | 148770683 | 148831467 | 60785 | 2 | ENSMUST00 | 0 | Ipo8 | ENSMUSG0( | false | chr6 | 148830615 | 148831891 | 27.41053 | 90.23362 | 86.88379 | chr6 | 148830046 | 148832310 | 3.59669 | 3.32202 | 1.8689 |
| chr10 | 128176355 | 128176733 | 1000 | Promoter (<=1kb) | 10 | 128129470 | 128176568 | 47099 | 2 | ENSMUST00 | 0 | Rbms2 | ENSMUSG0( | true | chr10 | 128174944 | 128177059 | 14.02077 | 44.3905 | 41.74443 | . | . | . | . | . | . |
| chr10 | 128092506 | 128093538 | 1000 | Promoter (<=1kb) | 10 | 128092783 | 128129303 | 36521 | 1 | ENSMUST00 | 0 | Baz2a | ENSMUSG0( | false | chr10 | 128091204 | 128094095 | 33.28429 | 109.73968 | 105.91746 | chr10 | 128093379 | 128103131 | 5.08043 | 4.07895 | 3.08556 |
| chr2 | 118728364 | 118728751 | 1000 | Promoter (<=1kb) | 2 | 118707517 | 118728438 | 20922 | 2 | ENSMUST00 | 0 | Plcb2 | ENSMUSG0( | false | . | . | . | . | . | . | chr2 | 118727772 | 118729075 | 4.09593 | 3.69881 | 2.27285 |
| chr2 | 118529476 | 118529953 | 1000 | 3' UTR | 2 | 118531974 | 118544587 | 12614 | 2 | ENSMUST00 | 14634 | Bmf | ENSMUSG0( | false | . | . | . | . | . | . | . | . | . | . | . | . |
| chr19 | 28963685 | 28964179 | 1000 | Promoter (<=1kb) | 19 | 28963953 | 28966811 | 2859 | 1 | ENSMUST00 | 0 | Ppapdc2 | ENSMUSG0( | false | chr19 | 28963503 | 28964799 | 37.68163 | 117.40437 | 113.3432 | . | . | . | . | . | . |
| chr9 | 40346027 | 40346376 | 1000 | Promoter (<=1kb) | 9 | 40293233 | 40346290 | 53058 | 2 | ENSMUST00 | 0 | Gramd1b | ENSMUSG0( | false | chr9 | 40344526 | 40346292 | 11.90309 | 30.75663 | 28.28125 | . | . | . | . | . | . |
| chr5 | 15934327 | 15935081 | 1000 | Promoter (<=1kb) | 5 | 15934691 | 16374511 | 439821 | 1 | ENSMUST00 | 0 | Cacna2d1 | ENSMUSG0( | false | chr5 | 15933989 | 15935703 | 23.60729 | 67.74274 | 64.78755 | chr5 | 15934055 | 15935799 | 3.88004 | 3.32264 | 2.11761 |
| chr14 | 56811075 | 56812512 | 1000 | Promoter (<=1kb) | 14 | 56790585 | 56811716 | 21132 | 2 | ENSMUST00 | 0 | Zmym5 | ENSMUSG0( | false | chr14 | 56810418 | 56811481 | 30.59503 | 95.89513 | 92.42037 | . | . | . | . | . | . |
| chr4 | 33247386 | 33248840 | 1000 | Promoter (<=1kb) | 4 | 33245423 | 33248510 | 3088 | 2 | ENSMUST00 | 0 | Pnrc1 | ENSMUSG0( | false | chr4 | 33246794 | 33249051 | 49.16994 | 165.05611 | 159.23541 | . | . | . | . | . | . |
| chr3 | 144479492 | 144479837 | 1000 | Intron (ENSMUST00000043325.7 | 3 | 144453963 | 144569881 | 115919 | 2 | ENSMUST00 | 90044 | Hs2st1 | ENSMUSG0( | false | . | . | . | . | . | . | chr3 | 144477657 | 144480938 | 5.02757 | 4.09812 | 3.04712 |
| chr3 | 144484581 | 144484879 | 1000 | Intron (ENSMUST00000043325.7 | 3 | 144453963 | 144569881 | 115919 | 2 | ENSMUST00 | 85002 | Hs2st1 | ENSMUSG0( | false | . | . | . | . | . | . | chr3 | 144483131 | 144485500 | 3.15359 | 3.06167 | 1.49846 |
| chr2 | 118111539 | 118111907 | 1000 | Promoter (<=1kb) | 2 | 118111876 | 118127133 | 15258 | 1 | ENSMUST00 | 0 | Thbs1 | ENSMUSG0( | false | chr2 | 118110556 | 118112456 | 21.4849 | 58.06458 | 55.24457 | chr2 | 118110316 | 118111567 | 3.84306 | 3.47438 | 2.09571 |
| chr11 | 73176817 | 73177849 | 1000 | Promoter (<=1kb) | 11 | 73177083 | 73182152 | 5070 | 1 | ENSMUST00 | 0 | Tax1bp3 | ENSMUSG0( | false | chr11 | 73175882 | 73178809 | 33.66875 | 128.30269 | 123.87101 | . | . | . | . | . | . |
| chr3 | 9609475 | 9610270 | 1000 | Promoter (<=1kb) | 3 | 9427011 | 9610085 | 183075 | 2 | ENSMUST00 | 0 | Zfp704 | ENSMUSG0( | false | chr3 | 9608405 | 9610425 | 28.1615 | 93.53628 | 90.11485 | chr3 | 9610106 | 9610551 | 3.56901 | 3.22046 | 1.87843 |
| chr3 | 18643647 | 18643989 | 1000 | Distal Intergenic | 3 | 18533861 | 18534289 | 429 | 1 | ENSMUST00 | 109787 | Gm6369 | ENSMUSG0( | false | . | . | . | . | . | . | chr3 | 18643088 | 18646122 | 4.75118 | 3.98987 | 2.805 |
| chr8 | 3674647 | 3675099 | 1000 | Promoter (1-2kb) | 8 | 3676299 | 3681255 | 4957 | 1 | ENSMUST00 | -1200 | Trappc5 | ENSMUSG0( | false | . | . | . | . | . | . | chr8 | 3674263 | 3676201 | 3.91835 | 3.5378 | 2.10898 |
| chr17 | 49543362 | 49543917 | 1000 | Intron (ENSMUST00000057610.6 | 17 | 49456022 | 49564337 | 108316 | 2 | ENSMUST00 | 20420 | Daam2 | ENSMUSG0( | true | . | . | . | . | . | . | chr17 | 49511234 | 49548441 | 5.06175 | 4.03075 | 3.07349 |
| chr17 | 49563329 | 49564495 | 1000 | Promoter (<=1kb) | 17 | 49456022 | 49564337 | 108316 | 2 | ENSMUST00 | 0 | Daam2 | ENSMUSG0( | true | chr17 | 49562125 | 49564376 | 19.02488 | 51.36755 | 48.63442 | chr17 | 49564293 | 49564687 | 4.37422 | 3.88114 | 2.48299 |
| chr17 | 49545447 | 49545861 | 1000 | Intron (ENSMUST00000057610.6 | 17 | 49456022 | 49564337 | 108316 | 2 | ENSMUST00 | 18476 | Daam2 | ENSMUSG0( | true | . | . | . | . | . | . | chr17 | 49511234 | 49548441 | 5.06175 | 4.03075 | 3.07349 |
| chr4 | 32238482 | 32239797 | 1000 | Promoter (<=1kb) | 4 | 32238804 | 32586104 | 347301 | 1 | ENSMUST00 | 0 | Bach2 | ENSMUSG0( | false | chr4 | 32238283 | 32240386 | 37.18615 | 120.78587 | 116.61589 | . | . | . | . | . | . |
| chrX | 12091051 | 12091467 | 1000 | Intron (ENSMUST00000043441.7 | 20 | 12036740 | 12080553 | 43814 | 2 | ENSMUST00 | -10499 | Bcor | ENSMUSG0( | false | chrX | 12090802 | 12091405 | 9.19064 | 17.46287 | 15.16242 | . | . | . | . | . | . |
| chrX | 12077069 | 12077624 | 1000 | Promoter (2-3kb) | 20 | 12036740 | 12080553 | 43814 | 2 | ENSMUST00 | 2929 | Bcor | ENSMUSG0( | false | chrX | 12076787 | 12081284 | 21.13848 | 54.99947 | 52.22001 | . | . | . | . | . | . |
| chrX | 12088096 | 12088926 | 1000 | Intron (ENSMUST00000043441.7 | 20 | 12036740 | 12080553 | 43814 | 2 | ENSMUST00 | -7544 | Bcor | ENSMUSG0( | false | chrX | 12087071 | 12088799 | 10.20265 | 24.60154 | 22.20526 | . | . | . | . | . | . |
| chrX | 12079338 | 12081391 | 1000 | Promoter (<=1kb) | 20 | 12036740 | 12080553 | 43814 | 2 | ENSMUST00 | 0 | Bcor | ENSMUSG0( | false | chrX | 12076787 | 12081284 | 21.13848 | 54.99947 | 52.22001 | . | . | . | . | . | . |
| chr1 | 160906163 | 160907196 | 1000 | Promoter (<=1kb) | 1 | 160906418 | 160974978 | 68561 | 1 | ENSMUST00 | 0 | Rc3h1 | ENSMUSG0( | false | chr1 | 160905985 | 160907996 | 41.77201 | 145.26822 | 140.17451 | . | . | . | . | . | . |
| chr11 | 107355386 | 107355904 | 1000 | Intron (ENSMUST00000103064.4 | 11 | 107226257 | 107348130 | 121874 | 2 | ENSMUST00 | -7257 | Pitpnc1 | ENSMUSG0( | false | . | . | . | . | . | . | chr11 | 107351556 | 107359305 | 6.08678 | 4.53131 | 3.89427 |
| chr11 | 107469630 | 107470916 | 1000 | Promoter (<=1kb) | 11 | 107207892 | 107470699 | 262808 | 2 | ENSMUST00 | 0 | Pitpnc1 | ENSMUSG0( | false | chr11 | 107468617 | 107471214 | 32.66734 | 113.81544 | 109.86345 | chr11 | 107470809 | 107473706 | 3.51775 | 3.33403 | 1.78864 |
| chr18 | 24529909 | 24530454 | 1000 | Promoter (<=1kb) | 18 | 24484962 | 24530204 | 45243 | 2 | ENSMUST00 | 0 | Rprd1a | ENSMUSG0( | false | chr18 | 24528898 | 24530722 | 33.1106 | 101.26704 | 97.66567 | chr18 | 24528175 | 24530863 | 3.31119 | 3.2171 | 1.6098 |
| chr11 | 72469068 | 72469548 | 1000 | Intron (ENSMUST00000045303.4 | 11 | 72459029 | 72475027 | 15999 | 2 | ENSMUST00 | 5479 | Spns2 | ENSMUSG0( | false | . | . | . | . | . | . | chr11 | 72459025 | 72472631 | 6.4619 | 4.65068 | 4.22104 |
| chr11 | 72467069 | 72467473 | 1000 | Intron (ENSMUST00000045303.4 | 11 | 72459029 | 72475027 | 15999 | 2 | ENSMUST00 | 7554 | Spns2 | ENSMUSG0( | false | . | . | . | . | . | . | chr11 | 72459025 | 72472631 | 6.4619 | 4.65068 | 4.22104 |
| chr11 | 72461369 | 72461861 | 1000 | Intron (ENSMUST00000045303.4 | 11 | 72452732 | 72456621 | 3890 | 2 | ENSMUST00 | -4749 | Spns2 | ENSMUSG0( | false | . | . | . | . | . | . | chr11 | 72459025 | 72472631 | 6.4619 | 4.65068 | 4.22104 |
| chr19 | 32346190 | 32346914 | 1000 | Promoter (2-3kb) | 19 | 32122730 | 32349316 | 226587 | 2 | ENSMUST00 | 2402 | Sgms1 | ENSMUSG0( | false | . | . | . | . | . | . | chr19 | 32332489 | 32359554 | 5.40081 | 4.18652 | 3.34993 |
| chr19 | 32349171 | 32349646 | 1000 | Promoter (<=1kb) | 19 | 32122730 | 32349316 | 226587 | 2 | ENSMUST00 | 0 | Sgms1 | ENSMUSG0( | false | . | . | . | . | . | . | chr19 | 32332489 | 32359554 | 5.40081 | 4.18652 | 3.34993 |
| chr19 | 32377318 | 32377680 | 1000 | Intron (ENSMUST00000099514.4 | 19 | 32289279 | 32388463 | 99185 | 2 | ENSMUST00 | 10783 | Sgms1 | ENSMUSG0( | false | . | . | . | . | . | . | chr19 | 32360669 | 32387358 | 5.88383 | 4.38649 | 3.75327 |
| chr19 | 32381260 | 32381730 | 1000 | Intron (ENSMUST00000099514.4 | 19 | 32289279 | 32388463 | 99185 | 2 | ENSMUST00 | 6733 | Sgms1 | ENSMUSG0( | false | . | . | . | . | . | . | chr19 | 32360669 | 32387358 | 5.88383 | 4.38649 | 3.75327 |
| chr19 | 32387707 | 32389641 | 1000 | Promoter (<=1kb) | 19 | 32122727 | 32388512 | 265786 | 2 | ENSMUST00 | 0 | Sgms1 | ENSMUSG0( | false | chr19 | 32386186 | 32390069 | 35.24744 | 113.18649 | 109.26083 | . | . | . | . | . | . |
| chr8 | 8994566 | 8995294 | 1000 | Distal Intergenic | 8 | 8666577 | 8690537 | 23961 | 2 | ENSMUST00 | -304030 | Arglu1 | ENSMUSG0( | false | chr8 | 8994672 | 8995271 | 9.78363 | 19.44713 | 17.12045 | chr8 | 8990485 | 8995162 | 5.68736 | 4.33322 | 3.56584 |
| chr11 | 107131069 | 107132422 | 1000 | Promoter (<=1kb) | 11 | 107033081 | 107131922 | 98842 | 2 | ENSMUST00 | 0 | Bptf | ENSMUSG0( | false | chr11 | 107130199 | 107132795 | 32.03048 | 96.40417 | 92.92239 | chr11 | 107132385 | 107133005 | 3.85936 | 3.47814 | 2.11204 |
| chr7 | 27332311 | 27332676 | 1000 | Promoter (<=1kb) | 7 | 27305998 | 27333613 | 27616 | 2 | ENSMUST00 | 937 | Ltbp4 | ENSMUSG0( | false | chr7 | 27332275 | 27333899 | 23.50745 | 64.07966 | 61.17685 | chr7 | 27328853 | 27337953 | 4.74964 | 3.95019 | 2.80772 |
| chr7 | 27330919 | 27331766 | 1000 | Promoter (1-2kb) | 7 | 27305998 | 27333613 | 27616 | 2 | ENSMUST00 | 1847 | Ltbp4 | ENSMUSG0( | false | chr7 | 27330451 | 27331846 | 14.6071 | 44.0986 | 41.45673 | chr7 | 27328853 | 27337953 | 4.74964 | 3.95019 | 2.80772 |
| chr11 | 72135383 | 72136145 | 1000 | Promoter (<=1kb) | 11 | 72047528 | 72135721 | 88194 | 2 | ENSMUST00 | 0 | Pitpnm3 | ENSMUSG0( | false | chr11 | 72134751 | 72136239 | 33.22886 | 104.51182 | 100.83223 | chr11 | 72123517 | 72139334 | 7.25529 | 5.05885 | 4.85135 |
| chr11 | 72127502 | 72127792 | 1000 | Intron (ENSMUST00000108508.2 | 11 | 72047528 | 72135721 | 88194 | 2 | ENSMUST00 | 7929 | Pitpnm3 | ENSMUSG0( | false | . | . | . | . | . | . | chr11 | 72123517 | 72139334 | 7.25529 | 5.05885 | 4.85135 |
| chr11 | 72129835 | 72130476 | 1000 | Intron (ENSMUST00000108508.2 | 11 | 72047528 | 72135721 | 88194 | 2 | ENSMUST00 | 5245 | Pitpnm3 | ENSMUSG0( | false | . | . | . | . | . | . | chr11 | 72123517 | 72139334 | 7.25529 | 5.05885 | 4.85135 |
| chr11 | 106612477 | 106613343 | 1000 | Promoter (<=1kb) | 11 | 106502147 | 106612930 | 110784 | 2 | ENSMUST00 | 0 | Tex2 | ENSMUSG0( | false | chr11 | 106611514 | 106613072 | 29.6869 | 105.26892 | 101.56436 | . | . | . | . | . | . |
| chr5 | 134932357 | 134932898 | 1000 | Promoter (<=1kb) | 5 | 134932368 | 134942637 | 10270 | 1 | ENSMUST00 | 0 | Wbscr27 | ENSMUSG0( | false | chr5 | 134932181 | 134932859 | 19.27906 | 51.80829 | 49.07 | chr5 | 134931501 | 134932953 | 4.24251 | 3.7301 | 2.42746 |
| chr19 | 36927186 | 36927560 | 1000 | Promoter (1-2kb) | 19 | 36926079 | 37012752 | 86674 | 1 | ENSMUST00 | 1108 | Btaf1 | ENSMUSG0( | false | chr19 | 36925411 | 36928022 | 31.66066 | 102.77801 | 99.14294 | chr19 | 36927194 | 36935272 | 4.66968 | 3.94009 | 2.73987 |
| chr19 | 37009102 | 37009397 | 1000 | Intron (ENSMUST00000099494.3 | 19 | 36926079 | 37012752 | 86674 | 1 | ENSMUST00 | 83024 | Btaf1 | ENSMUSG0( | false | . | . | . | . | . | . | chr19 | 37008385 | 37009571 | 3.32387 | 3.23788 | 1.68211 |
| chr4 | 14668366 | 14668759 | 1000 | Intron (ENSMUST00000023917.2 | 4 | 14502430 | 14621805 | 119376 | 2 | ENSMUST00 | -46562 | Slc26a7 | ENSMUSG0( | false | . | . | . | . | . | . | chr4 | 14667655 | 14670452 | 3.67807 | 3.34772 | 1.93722 |
| chr4 | 142221226 | 142221854 | 1000 | Intron (ENSMUST00000036476.4 | 4 | 142141541 | 142210206 | 68666 | 2 | ENSMUST00 | -11021 | Kazn | ENSMUSG0( | false | . | . | . | . | . | . | chr4 | 142218898 | 142232146 | 6.00295 | 4.43269 | 3.85028 |
| chr1 | 166253841 | 166254433 | 1000 | Promoter (<=1kb) | 1 | 166254139 | 166316823 | 62685 | 1 | ENSMUST00 | 0 | Ildr2 | ENSMUSG0( | true | chr1 | 166253860 | 166256724 | 27.62344 | 81.07606 | 77.90333 | . | . | . | . | . | . |
| chrX | 166170297 | 166170629 | 1000 | Promoter (<=1kb) | 20 | 166170454 | 166189804 | 19351 | 1 | ENSMUST00 | 0 | Gemin8 | ENSMUSG0( | false | chrX | 166170399 | 166170959 | 12.8669 | 28.00426 | 25.56298 | . | . | . | . | . | . |
| chr10 | 3755942 | 3756437 | 1000 | Intron (ENSMUST00000120274.2 | 10 | 3740665 | 3938577 | 197913 | 1 | ENSMUST00 | 15278 | Plekhg1 | ENSMUSG0( | true | . | . | . | . | . | . | chr10 | 3742850 | 3772389 | 5.62976 | 4.36437 | 3.52607 |
| chr10 | 3740097 | 3740532 | 1000 | Promoter (<=1kb) | 10 | 3740364 | 3967303 | 226940 | 1 | ENSMUST00 | 0 | Plekhg1 | ENSMUSG0( | true | chr10 | 3739900 | 3742294 | 41.69731 | 136.21887 | 131.48979 | . | . | . | . | . | . |
| chr10 | 3645070 | 3645384 | 1000 | Distal Intergenic | 10 | 3740364 | 3967303 | 226940 | 1 | ENSMUST00 | -94980 | Plekhg1 | ENSMUSG0( | true | . | . | . | . | . | . | chr10 | 3644396 | 3647183 | 4.34481 | 3.74487 | 2.51404 |
| chr14 | 27428666 | 27429768 | 1000 | Promoter (<=1kb) | 14 | 27428847 | 27483552 | 54706 | 1 | ENSMUST00 | 0 | Fam208a | ENSMUSG0( | false | chr14 | 27428244 | 27430149 | 37.30079 | 124.19571 | 119.91465 | . | . | . | . | . | . |
| chr9 | 65679503 | 65679922 | 1000 | Promoter (2-3kb) | 9 | 65676591 | 65688538 | 11948 | 1 | ENSMUST00 | 2913 | Oaz2 | ENSMUSG0( | false | . | . | . | . | . | . | chr9 | 65675551 | 65685323 | 4.69201 | 3.84119 | 2.79144 |
| chr4 | 141866961 | 141867733 | 1000 | Intron (ENSMUST00000036854.3 | 4 | 141858142 | 141874920 | 16779 | 2 | ENSMUST00 | 7187 | Efhd2 | ENSMUSG0( | false | . | . | . | . | . | . | chr4 | 141861816 | 141873890 | 7.33763 | 5.08181 | 4.9258 |
| chr19 | 4214071 | 4215578 | 1000 | Promoter (<=1kb) | 19 | 4214238 | 4223490 | 9253 | 1 | ENSMUST00 | 0 | Clcf1 | ENSMUSG0( | false | chr19 | 4213990 | 4215760 | 32.27147 | 102.84008 | 99.19724 | chr19 | 4213419 | 4218153 | 6.04705 | 4.60219 | 3.85122 |
| chr10 | 3972962 | 3973977 | 1000 | Promoter (<=1kb) | 10 | 3973118 | 4156111 | 182994 | 1 | ENSMUST00 | 0 | Mthfd1l | ENSMUSG0( | false | chr10 | 3972798 | 3974211 | 19.45739 | 70.46334 | 67.46697 | chr10 | 3972826 | 3974285 | 5.32101 | 4.24921 | 3.27241 |
| chr7 | 25802047 | 25803136 | 1000 | 3' UTR | 7 | 25809324 | 25815392 | 6069 | 2 | ENSMUST00 | 12256 | Cyp2s1 | ENSMUSG0( | false | . | . | . | . | . | . | chr7 | 25801478 | 25806139 | 7.18291 | 4.99145 | 4.80153 |
| chr7 | 25804713 | 25805294 | 1000 | Exon (ENSMUST00000152560.2/ | 7 | 25809324 | 25815392 | 6069 | 2 | ENSMUST00 | 10098 | Cyp2s1 | ENSMUSG0( | false | . | . | . | . | . | . | chr7 | 25801478 | 25806139 | 7.18291 | 4.99145 | 4.80153 |
| chr7 | 25815503 | 25815969 | 1000 | Promoter (<=1kb) | 7 | 25809324 | 25815392 | 6069 | 2 | ENSMUST00 | -112 | Cyp2s1 | ENSMUSG0( | false | chr7 | 25815840 | 25817023 | 18.23312 | 46.09947 | 43.43353 | chr7 | 25813779 | 25817911 | 6.75484 | 4.68651 | 4.47204 |
| chr11 | 32383770 | 32384358 | 1000 | Intron (ENSMUST00000038753.5 | 11 | 32347840 | 32390300 | 42461 | 1 | ENSMUST00 | 35931 | Sh3pxd2b | ENSMUSG0( | false | . | . | . | . | . | . | chr11 | 32383288 | 32386252 | 5.8032 | 4.27592 | 3.68113 |
| chr14 | 27044908 | 27045261 | 1000 | Intron (ENSMUST00000035336.3 | 14 | 27039001 | 27107286 | 68286 | 1 | ENSMUST00 | 5908 | Il17rd | ENSMUSG0( | false | . | . | . | . | . | . | chr14 | 27041643 | 27048842 | 5.16084 | 3.99802 | 3.16031 |
| chr1 | 165717706 | 165718700 | 1000 | Distal Intergenic | 1 | 165648945 | 165708094 | 59150 | 2 | ENSMUST00 | -9613 | Rcsd1 | ENSMUSG0( | true | chr1 | 165718458 | 165718951 | 7.88522 | 15.75676 | 13.48224 | chr1 | 165717334 | 165721559 | 5.59814 | 4.25202 | 3.52106 |
| chr1 | 165709769 | 165710302 | 1000 | Promoter (1-2kb) | 1 | 165648945 | 165708094 | 59150 | 2 | ENSMUST00 | -1676 | Rcsd1 | ENSMUSG0( | true | chr1 | 165709651 | 165709945 | 4.46497 | 6.3612 | 4.26978 | chr1 | 165708619 | 165712261 | 6.63203 | 4.73365 | 4.35395 |
| chr1 | 165708605 | 165709077 | 1000 | Promoter (<=1kb) | 1 | 165648945 | 165708094 | 59150 | 2 | ENSMUST00 | -512 | Rcsd1 | ENSMUSG0( | true | chr1 | 165706914 | 165708815 | 26.60951 | 79.31685 | 76.1741 | chr1 | 165708619 | 165712261 | 6.63203 | 4.73365 | 4.35395 |
| chr1 | 165707688 | 165708438 | 1000 | Promoter (<=1kb) | 1 | 165648945 | 165708094 | 59150 | 2 | ENSMUST00 | 0 | Rcsd1 | ENSMUSG0( | true | chr1 | 165706914 | 165708815 | 26.60951 | 79.31685 | 76.1741 | . | . | . | . | . | . |
| chr4 | 141538218 | 141539087 | 1000 | Promoter (<=1kb) | 4 | 141467890 | 141538449 | 70560 | 2 | ENSMUST00 | 0 | Spen | ENSMUSG0( | false | chr4 | 141535704 | 141538627 | 38.13901 | 127.82562 | 123.40861 | . | . | . | . | . | . |
| chr4 | 141535374 | 141535718 | 1000 | Promoter (2-3kb) | 4 | 141521803 | 141538253 | 16451 | 2 | ENSMUST00 | 2535 | Spen | ENSMUSG0( | false | chr4 | 141535704 | 141538627 | 38.13901 | 127.82562 | 123.40861 | chr4 | 141531641 | 141536451 | 4.1853 | 3.62885 | 2.34826 |
| chr6 | 121510049 | 121510700 | 1000 | Distal Intergenic | 6 | 121404801 | 121473678 | 68878 | 2 | ENSMUST00 | -36372 | Iqsec3 | ENSMUSG0( | false | . | . | . | . | . | . | chr6 | 121508713 | 121511567 | 4.89525 | 4.02035 | 2.91217 |
| chr7 | 19186568 | 19186934 | 1000 | Intron (ENSMUST00000117338.2 | 7 | 19181606 | 19186371 | 4766 | 1 | ENSMUST00 | 4963 | Eml2 | ENSMUSG0( | true | . | . | . | . | . | . | chr7 | 19178043 | 19190895 | 5.91671 | 4.49499 | 3.76502 |
| chr9 | 106685280 | 106686218 | 1000 | Promoter (<=1kb) | 9 | 106658746 | 106685682 | 26937 | 2 | ENSMUST00 | 0 | Tex264 | ENSMUSG0( | false | chr9 | 106684656 | 106686403 | 27.66025 | 106.38611 | 102.65278 | chr9 | 106686120 | 106687617 | 5.41497 | 4.26629 | 3.37616 |
| chr14 | 26579384 | 26580246 | 1000 | Promoter (<=1kb) | 14 | 26579578 | 26633394 | 53817 | 1 | ENSMUST00 | 0 | Dennd6a | ENSMUSG0( | false | chr14 | 26579063 | 26580603 | 44.00655 | 153.74478 | 148.311 | . | . | . | . | . | . |
| chr4 | 141139544 | 141139961 | 1000 | Promoter (<=1kb) | 4 | 141113001 | 141139727 | 26727 | 2 | ENSMUST00 | 0 | Szrd1 | ENSMUSG0( | false | chr4 | 141138228 | 141139871 | 29.94827 | 103.73744 | 100.07098 | . | . | . | . | . | . |
| chr17 | 84510407 | 84511213 | 1000 | Promoter (<=1kb) | 17 | 84511895 | 84622142 | 110248 | 1 | ENSMUST00 | -682 | Plekhh2 | ENSMUSG0( | true | . | . | . | . | . | . | chr17 | 84497894 | 84511794 | 4.6554 | 3.81889 | 2.73035 |
| chr17 | 84503762 | 84504550 | 1000 | Distal Intergenic | 17 | 84511895 | 84622142 | 110248 | 1 | ENSMUST00 | -7345 | Plekhh2 | ENSMUSG0( | true | chr17 | 84502135 | 84504110 | 11.32191 | 27.20368 | 24.77393 | chr17 | 84497894 | 84511794 | 4.6554 | 3.81889 | 2.73035 |
| chr7 | 25248000 | 25248454 | 1000 | Promoter (2-3kb) | 7 | 25245447 | 25250730 | 5284 | 2 | ENSMUST00 | 2276 | Erf | ENSMUSG0( | false | chr7 | 25247802 | 25251702 | 35.38397 | 108.24114 | 104.45254 | chr7 | 25243126 | 25249505 | 7.31348 | 5.08626 | 4.89724 |
| chr7 | 25249120 | 25251037 | 1000 | Promoter (<=1kb) | 7 | 25242561 | 25250761 | 8201 | 2 | ENSMUST00 | 0 | Erf | ENSMUSG0( | false | chr7 | 25247802 | 25251702 | 35.38397 | 108.24114 | 104.45254 | chr7 | 25251032 | 25252240 | 6.67549 | 4.86489 | 4.3681 |
| chr1 | 63113619 | 63114764 | 1000 | Promoter (<=1kb) | 1 | 63047801 | 63114267 | 66467 | 2 | ENSMUST00 | 0 | Ino80d | ENSMUSG0( | false | chr1 | 63112781 | 63116118 | 29.2476 | 106.45832 | 102.71833 | . | . | . | . | . | . |
| chr3 | 105452079 | 105453050 | 1000 | Promoter (<=1kb) | 3 | 105452330 | 105674002 | 221673 | 1 | ENSMUST00 | 0 | Kcnd3 | ENSMUSG0( | false | chr3 | 105450974 | 105453152 | 23.28017 | 72.50986 | 69.48009 | chr3 | 105450707 | 105453484 | 4.85717 | 3.92004 | 2.9323 |
| chr19 | 45696237 | 45696570 | 1000 | Distal Intergenic | 19 | 45578254 | 45660312 | 82059 | 2 | ENSMUST00 | -35926 | Fbxw4 | ENSMUSG0( | false | . | . | . | . | . | . | chr19 | 45695544 | 45696882 | 5.0982 | 3.90386 | 3.1256 |
| chr1 | 164253134 | 164253519 | 1000 | Intron (ENSMUST00000044021.6 | 1 | 164249253 | 164263518 | 14266 | 1 | ENSMUST00 | 3882 | Slc19a2 | ENSMUSG0( | false | . | . | . | . | . | . | chr1 | 164252466 | 164254381 | 5.17194 | 4.12364 | 3.20425 |
| chr19 | 28010609 | 28012004 | 1000 | Promoter (<=1kb) | 19 | 27766441 | 28011048 | 244608 | 2 | ENSMUST00 | 0 | Rfx3 | ENSMUSG0( | false | chr19 | 28009765 | 28012498 | 44.57462 | 145.65463 | 140.53799 | . | . | . | . | . | . |
| chr9 | 121313978 | 121314386 | 1000 | Distal Intergenic | 9 | 121102544 | 121277197 | 174654 | 2 | ENSMUST00 | -36782 | Ulk4 | ENSMUSG0( | false | chr9 | 121313886 | 121314840 | 13.75803 | 31.40302 | 28.92136 | chr9 | 121306967 | 121321326 | 5.57119 | 4.26333 | 3.49442 |
| chr3 | 133544186 | 133545105 | 1000 | Promoter (<=1kb) | 3 | 133463679 | 133544390 | 80712 | 2 | ENSMUST00 | 0 | Tet2 | ENSMUSG0( | false | chr3 | 133542885 | 133545353 | 27.9498 | 82.58102 | 79.37986 | . | . | . | . | . | . |
| chr4 | 140701864 | 140702922 | 1000 | Promoter (<=1kb) | 4 | 140701473 | 140723220 | 21748 | 1 | ENSMUST00 | 392 | Rcc2 | ENSMUSG0( | false | chr4 | 140700984 | 140703222 | 24.0356 | 94.23969 | 90.80066 | . | . | . | . | . | . |
| chr18 | 11839197 | 11839905 | 1000 | Promoter (<=1kb) | 18 | 11839220 | 11945621 | 106402 | 1 | ENSMUST00 | 0 | Cables1 | ENSMUSG0( | true | chr18 | 11838767 | 11842359 | 28.35087 | 106.74432 | 102.99886 | . | . | . | . | . | . |
| chr18 | 11810592 | 11811119 | 1000 | Distal Intergenic | 18 | 11839220 | 11945621 | 106402 | 1 | ENSMUST00 | -28101 | Cables1 | ENSMUSG0( | true | . | . | . | . | . | . | chr18 | 11809354 | 11813050 | 4.69669 | 3.97779 | 2.75808 |
| chr4 | 140610824 | 140611584 | 1000 | Exon (ENSMUST00000039204.4/ | 4 | 140593169 | 140617065 | 23897 | 2 | ENSMUST00 | 5481 | Arhgef10l | ENSMUSG0( | false | chr4 | 140610139 | 140611652 | 14.61261 | 36.2332 | 33.69079 | chr4 | 140597945 | 140613291 | 5.66962 | 4.2683 | 3.5931 |
| chr4 | 140622838 | 140623246 | 1000 | Intron (ENSMUST00000039204.4 | 4 | 140593169 | 140617065 | 23897 | 2 | ENSMUST00 | -5774 | Arhgef10l | ENSMUSG0( | false | . | . | . | . | . | . | chr4 | 140622957 | 140624266 | 2.99673 | 2.98674 | 1.39454 |
| chr4 | 140612020 | 140612478 | 1000 | Intron (ENSMUST00000039204.4 | 4 | 140593169 | 140617065 | 23897 | 2 | ENSMUST00 | 4587 | Arhgef10l | ENSMUSG0( | false | . | . | . | . | . | . | chr4 | 140597945 | 140613291 | 5.66962 | 4.2683 | 3.5931 |
| chr7 | 18844889 | 18845197 | 1000 | Intron (ENSMUST00000072415.6 | 7 | 18839966 | 18858654 | 18689 | 1 | ENSMUST00 | 4924 | Mill2 | ENSMUSG0( | true | . | . | . | . | . | . | chr7 | 18844491 | 18845223 | 4.5444 | 3.91957 | 2.66598 |
| chr3 | 132882730 | 132883394 | 1000 | 3' UTR | 3 | 132891176 | 132906359 | 15184 | 2 | ENSMUST00 | 22965 | Npnt | ENSMUSG0( | true | . | . | . | . | . | . | chr3 | 132880303 | 132885324 | 5.40591 | 4.23547 | 3.34377 |
| chr3 | 132911700 | 132912021 | 1000 | Intron (ENSMUST00000042744.1 | 3 | 132891176 | 132906359 | 15184 | 2 | ENSMUST00 | -5342 | Npnt | ENSMUSG0( | true | . | . | . | . | . | . | chr3 | 132908464 | 132914305 | 4.89816 | 4.00177 | 2.94071 |
| chr3 | 132893292 | 132893643 | 1000 | Intron (ENSMUST00000042744.1 | 3 | 132891176 | 132906359 | 15184 | 2 | ENSMUST00 | 12716 | Npnt | ENSMUSG0( | true | . | . | . | . | . | . | chr3 | 132890067 | 132898175 | 5.27144 | 4.147 | 3.22072 |
| chr4 | 128883333 | 128883813 | 1000 | Promoter (<=1kb) | 4 | 128883580 | 128903565 | 19986 | 1 | ENSMUST00 | 0 | Trim62 | ENSMUSG0( | false | chr4 | 128883298 | 128884769 | 28.48738 | 97.07422 | 93.57343 | chr4 | 128883147 | 128884887 | 3.62856 | 3.34723 | 1.89535 |
| chr4 | 139580774 | 139581306 | 1000 | Intron (ENSMUST00000123827.3 | 4 | 139574720 | 139620382 | 45663 | 1 | ENSMUST00 | 6055 | Iffo2 | ENSMUSG0( | false | chr4 | 139580247 | 139582371 | 12.5733 | 28.75278 | 26.30366 | chr4 | 139576299 | 139606726 | 8.92638 | 5.72021 | 6.23508 |
| chr4 | 139561651 | 139562187 | 1000 | Intron (ENSMUST00000123827.3 | 4 | 139574720 | 139620382 | 45663 | 1 | ENSMUST00 | -12533 | Iffo2 | ENSMUSG0( | false | . | . | . | . | . | . | chr4 | 139561788 | 139562567 | 4.17139 | 3.62108 | 2.36123 |
| chr1 | 59481881 | 59483138 | 1000 | Promoter (<=1kb) | 1 | 59482424 | 59486955 | 4532 | 1 | ENSMUST00 | 0 | Fzd7 | ENSMUSG0( | false | chr1 | 59482026 | 59483998 | 16.96705 | 61.94942 | 59.07605 | chr1 | 59482870 | 59486151 | 5.76271 | 4.44814 | 3.66227 |
| chrX | 152336493 | 152336888 | 1000 | 3' UTR | 20 | 152340265 | 152341801 | 1537 | 2 | ENSMUST00 | 4913 | Tspyl2 | ENSMUSG0( | false | . | . | . | . | . | . | . | . | . | . | . | . |
| chr13 | 20108950 | 20109346 | 1000 | Promoter (2-3kb) | 13 | 20111512 | 20342888 | 231377 | 1 | ENSMUST00 | -2166 | Elmo1 | ENSMUSG0( | true | . | . | . | . | . | . | chr13 | 20104132 | 20118020 | 3.95526 | 3.43632 | 2.18524 |
| chr13 | 20052437 | 20053144 | 1000 | Distal Intergenic | 13 | 20090596 | 20264007 | 173412 | 1 | ENSMUST00 | -37452 | Elmo1 | ENSMUSG0( | true | . | . | . | . | . | . | chr13 | 20044769 | 20055516 | 4.57843 | 3.87081 | 2.66499 |
| chr13 | 20050108 | 20050693 | 1000 | Distal Intergenic | 13 | 20090596 | 20264007 | 173412 | 1 | ENSMUST00 | -39903 | Elmo1 | ENSMUSG0( | true | . | . | . | . | . | . | chr13 | 20044769 | 20055516 | 4.57843 | 3.87081 | 2.66499 |
| chr13 | 20090276 | 20091420 | 1000 | Promoter (<=1kb) | 13 | 20090596 | 20264007 | 173412 | 1 | ENSMUST00 | 0 | Elmo1 | ENSMUSG0( | true | chr13 | 20089916 | 20091932 | 37.30079 | 124.19571 | 119.91465 | chr13 | 20089626 | 20090293 | 3.87824 | 3.51859 | 2.09414 |
| chr4 | 139092460 | 139093350 | 1000 | Promoter (<=1kb) | 4 | 139082284 | 139092993 | 10710 | 2 | ENSMUST00 | 0 | Nbl1 | ENSMUSG0( | false | chr4 | 139091610 | 139093306 | 40.89836 | 130.45236 | 125.94646 | chr4 | 139088999 | 139098811 | 5.83874 | 4.33468 | 3.72707 |
| chr4 | 16013681 | 16014331 | 1000 | Promoter (<=1kb) | 4 | 15997121 | 16013888 | 16768 | 2 | ENSMUST00 | 0 | Osgin2 | ENSMUSG0( | false | chr4 | 16013032 | 16014530 | 29.36448 | 91.77025 | 88.3935 | chr4 | 16014323 | 16014893 | 3.32821 | 3.24873 | 1.62651 |
| chrX | 151520170 | 151520723 | 1000 | Promoter (<=1kb) | 20 | 151520672 | 151570807 | 50136 | 1 | ENSMUST00 | 0 | Phf8 | ENSMUSG0( | false | chrX | 151520665 | 151522628 | 19.30035 | 48.68736 | 45.98855 | . | . | . | . | . | . |
| chr18 | 82691706 | 82692745 | 1000 | Promoter (<=1kb) | 18 | 82597155 | 82692734 | 95580 | 2 | ENSMUST00 | 0 | Zfp236 | ENSMUSG0( | false | chr18 | 82691477 | 82693572 | 23.88927 | 69.11851 | 66.14216 | . | . | . | . | . | . |
| chr9 | 54500976 | 54502152 | 1000 | Promoter (<=1kb) | 9 | 54365158 | 54501626 | 136469 | 2 | ENSMUST00 | 0 | Dmxl2 | ENSMUSG0( | false | chr9 | 54500833 | 54502265 | 29.18609 | 122.55783 | 118.32927 | chr9 | 54500321 | 54501021 | 3.00686 | 3.0304 | 1.35975 |
| chr13 | 17803918 | 17805768 | 1000 | Promoter (<=1kb) | 13 | 17715962 | 17805097 | 89136 | 2 | ENSMUST00 | 0 | Cdk13 | ENSMUSG0( | false | chr13 | 17802516 | 17806006 | 35.02911 | 135.1772 | 130.48503 | . | . | . | . | . | . |
| chr8 | 106935498 | 106936461 | 1000 | Promoter (<=1kb) | 8 | 106935750 | 107014192 | 78443 | 1 | ENSMUST00 | 0 | Sntb2 | ENSMUSG0( | false | chr8 | 106935129 | 106937196 | 31.1999 | 108.05416 | 104.27962 | . | . | . | . | . | . |
| chr5 | 148342416 | 148342813 | 1000 | Exon (ENSMUST00000048116.9/ | 5 | 148352153 | 148371546 | 19394 | 2 | ENSMUST00 | 28733 | Slc7a1 | ENSMUSG0( | false | chr5 | 148341821 | 148342904 | 13.83063 | 32.8443 | 30.34371 | chr5 | 148341752 | 148343231 | 4.85391 | 3.92861 | 2.89924 |
| chr17 | 23675173 | 23675550 | 1000 | Promoter (1-2kb) | 17 | 23668618 | 23673882 | 5265 | 2 | ENSMUST00 | -1292 | Thoc6 | ENSMUSG0( | false | . | . | . | . | . | . | chr17 | 23673161 | 23678044 | 6.00185 | 4.61543 | 3.82218 |
| chr12 | 105685079 | 105685659 | 1000 | Promoter (<=1kb) | 12 | 105613539 | 105685241 | 71703 | 2 | ENSMUST00 | 0 | Atg2b | ENSMUSG0( | false | chr12 | 105683809 | 105686329 | 38.8867 | 129.13618 | 124.68224 | . | . | . | . | . | . |
| chr9 | 71578566 | 71579532 | 1000 | Promoter (<=1kb) | 9 | 71561038 | 71579885 | 18848 | 2 | ENSMUST00 | 353 | Myzap | ENSMUSG0( | false | . | . | . | . | . | . | chr9 | 71561341 | 71584420 | 5.41782 | 4.13897 | 3.38306 |
| chr9 | 71592082 | 71592564 | 1000 | Promoter (<=1kb) | 9 | 71504347 | 71592360 | 88014 | 2 | ENSMUST00 | 0 | Myzap | ENSMUSG0( | false | chr9 | 71590867 | 71592639 | 33.30314 | 106.68535 | 102.93993 | chr9 | 71586004 | 71596232 | 6.06407 | 4.60974 | 3.87823 |
| chr9 | 71567453 | 71567788 | 1000 | Intron (ENSMUST00000163998.1 | 9 | 71561038 | 71579885 | 18848 | 2 | ENSMUST00 | 12097 | Myzap | ENSMUSG0( | false | . | . | . | . | . | . | chr9 | 71561341 | 71584420 | 5.41782 | 4.13897 | 3.38306 |
| chr7 | 16286070 | 16286907 | 1000 | Promoter (<=1kb) | 7 | 16274045 | 16286790 | 12746 | 2 | ENSMUST00 | 0 | Ccdc9 | ENSMUSG0( | false | chr7 | 16286309 | 16287057 | 30.78865 | 90.34473 | 86.99165 | . | . | . | . | . | . |
| chr7 | 16296683 | 16297252 | 1000 | Distal Intergenic | 7 | 16282755 | 16286795 | 4041 | 2 | ENSMUST00 | -9889 | Ccdc9 | ENSMUSG0( | false | chr7 | 16297079 | 16297339 | 7.55891 | 13.84842 | 11.60173 | chr7 | 16295229 | 16298235 | 6.20923 | 4.6252 | 3.99375 |
| chr6 | 120188263 | 120188651 | 1000 | Intron (ENSMUST00000112711.3 | 6 | 120193823 | 120200334 | 6512 | 1 | ENSMUST00 | -5172 | Ninj2 | ENSMUSG0( | false | . | . | . | . | . | . | chr6 | 120187663 | 120189185 | 5.31131 | 4.30317 | 3.25488 |
| chr6 | 120109198 | 120109672 | 1000 | Intron (ENSMUST00000112711.3 | 6 | 120093350 | 120200339 | 106990 | 1 | ENSMUST00 | 15849 | Ninj2 | ENSMUSG0( | false | . | . | . | . | . | . | chr6 | 120107892 | 120116189 | 5.35272 | 4.21388 | 3.29824 |
| chr16 | 18781246 | 18782304 | 1000 | 3' UTR | 16 | 18776847 | 18778260 | 1414 | 1 | ENSMUST00 | 4400 | Cldn5 | ENSMUSG0( | true | . | . | . | . | . | . | chr16 | 18779014 | 18787308 | 5.37592 | 4.15125 | 3.35319 |
| chr16 | 18776623 | 18777031 | 1000 | Promoter (<=1kb) | 16 | 18776847 | 18778260 | 1414 | 1 | ENSMUST00 | 0 | Cldn5 | ENSMUSG0( | true | chr16 | 18776124 | 18780444 | 33.04283 | 115.53907 | 111.53828 | . | . | . | . | . | . |
| chr1 | 163991998 | 163992807 | 1000 | Promoter (1-2kb) | 1 | 163989738 | 163994541 | 4804 | 2 | ENSMUST00 | 1734 | BC055324 | ENSMUSG0( | false | . | . | . | . | . | . | chr1 | 163991574 | 163996008 | 5.65475 | 4.33948 | 3.57649 |
| chr12 | 104751536 | 104752349 | 1000 | Promoter (<=1kb) | 12 | 104687742 | 104751952 | 64211 | 2 | ENSMUST00 | 0 | Dicer1 | ENSMUSG0( | false | chr12 | 104750854 | 104752922 | 38.55812 | 129.64772 | 125.16568 | . | . | . | . | . | . |
| chr13 | 101739825 | 101740300 | 1000 | Intron (ENSMUST00000055518.7 | 13 | 101755571 | 101768213 | 12643 | 2 | ENSMUST00 | 27913 | Pik3r1 | ENSMUSG0( | false | . | . | . | . | . | . | chr13 | 101735550 | 101742952 | 3.81937 | 3.46776 | 2.06025 |
| chr13 | 101694094 | 101695255 | 1000 | Promoter (<=1kb) | 13 | 101689040 | 101694600 | 5561 | 2 | ENSMUST00 | 0 | Pik3r1 | ENSMUSG0( | false | chr13 | 101693541 | 101695297 | 31.70772 | 93.8754 | 90.44607 | chr13 | 101690067 | 101698367 | 4.61928 | 3.86189 | 2.69344 |
| chr17 | 24636721 | 24637137 | 1000 | Intron (ENSMUST00000047611.2 | 17 | 24632682 | 24638838 | 6157 | 1 | ENSMUST00 | 4040 | Nthl1 | ENSMUSG0( | false | . | . | . | . | . | . | chr17 | 24630730 | 24649025 | 8.70874 | 5.59711 | 6.05272 |
| chr8 | 106893006 | 106893755 | 1000 | Promoter (<=1kb) | 8 | 106893640 | 106923093 | 29454 | 1 | ENSMUST00 | 0 | Cirh1a | ENSMUSG0( | false | chr8 | 106892093 | 106894289 | 38.08716 | 126.65268 | 122.28292 | . | . | . | . | . | . |
| chr9 | 32243446 | 32244153 | 1000 | Promoter (1-2kb) | 9 | 32245164 | 32249464 | 4301 | 1 | ENSMUST00 | -1011 | Arhgap32 | ENSMUSG0( | false | . | . | . | . | . | . | chr9 | 32232319 | 32247487 | 4.87198 | 3.98331 | 2.90407 |
| chr6 | 119175139 | 119175884 | 1000 | Promoter (<=1kb) | 6 | 119175253 | 119221616 | 46364 | 1 | ENSMUST00 | 0 | Dcp1b | ENSMUSG0( | false | chr6 | 119174737 | 119176198 | 36.04346 | 118.78738 | 114.68494 | . | . | . | . | . | . |
| chr12 | 104114915 | 104115229 | 1000 | Promoter (<=1kb) | 12 | 104115970 | 104121518 | 5549 | 1 | ENSMUST00 | -741 | Serpina3a | ENSMUSG0( | false | . | . | . | . | . | . | . | . | . | . | . | . |
| chr6 | 144781392 | 144782490 | 1000 | Promoter (<=1kb) | 6 | 143843739 | 144781977 | 938239 | 2 | ENSMUST00 | 0 | Sox5 | ENSMUSG0( | true | chr6 | 144780525 | 144782962 | 19.57973 | 66.4542 | 63.51413 | chr6 | 144782320 | 144784334 | 3.50482 | 3.28751 | 1.7921 |
| chr5 | 116422775 | 116423322 | 1000 | Promoter (<=1kb) | 5 | 116408491 | 116422864 | 14374 | 2 | ENSMUST00 | 0 | Hspb8 | ENSMUSG0( | false | chr5 | 116421455 | 116423156 | 17.74057 | 61.94745 | 59.07436 | chr5 | 116422275 | 116423843 | 4.22688 | 3.7331 | 2.38607 |
| chr4 | 148163982 | 148164343 | 1000 | Exon (ENSMUST00000047951.8/ | 4 | 148160621 | 148166424 | 5804 | 1 | ENSMUST00 | 3362 | Fbxo2 | ENSMUSG0( | false | chr4 | 148163958 | 148164618 | 6.75876 | 12.51203 | 10.2897 | chr4 | 148163636 | 148165113 | 4.295 | 3.75673 | 2.44477 |
| chr11 | 114043491 | 114044061 | 1000 | Intron (ENSMUST00000041627.8 | 11 | 113848562 | 114065975 | 217414 | 2 | ENSMUST00 | 21914 | Sdk2 | ENSMUSG0( | false | . | . | . | . | . | . | chr11 | 114042166 | 114045202 | 6.07085 | 4.58888 | 3.88909 |
| chr11 | 114066749 | 114067278 | 1000 | Promoter (<=1kb) | 11 | 113865600 | 114067046 | 201447 | 2 | ENSMUST00 | 0 | Sdk2 | ENSMUSG0( | false | chr11 | 114065630 | 114067369 | 21.76566 | 71.50774 | 68.49327 | chr11 | 114064810 | 114067260 | 5.50609 | 4.19583 | 3.44854 |
| chr11 | 113806963 | 113807452 | 1000 | Promoter (2-3kb) | 11 | 113793839 | 113810423 | 16585 | 2 | ENSMUST00 | 2971 | Sdk2 | ENSMUSG0( | false | . | . | . | . | . | . | . | . | . | . | . | . |
| chr14 | 122982785 | 122983589 | 1000 | Promoter (<=1kb) | 14 | 122918971 | 122983283 | 64313 | 2 | ENSMUST00 | 0 | Tmtc4 | ENSMUSG0( | false | chr14 | 122982323 | 122984015 | 28.53699 | 95.19626 | 91.7378 | chr14 | 122982505 | 122983871 | 3.45229 | 3.28532 | 1.78617 |
| chr11 | 113683462 | 113684776 | 1000 | Promoter (<=1kb) | 11 | 113684412 | 113694647 | 10236 | 1 | ENSMUST00 | 0 | D11Wsu47e | ENSMUSG0( | false | chr11 | 113682857 | 113685142 | 26.86698 | 95.04005 | 91.5885 | . | . | . | . | . | . |
| chr1 | 136140168 | 136140677 | 1000 | Intron (ENSMUST00000075164.5 | 1 | 136131454 | 136149993 | 18540 | 1 | ENSMUST00 | 8715 | Kif21b | ENSMUSG0( | false | chr1 | 136139975 | 136140710 | 9.32368 | 22.97803 | 20.60213 | chr1 | 136136450 | 136141742 | 3.35814 | 3.22403 | 1.65544 |
| chr14 | 122564363 | 122564697 | 1000 | Intron (ENSMUST00000148172.1 | 14 | 122582684 | 122889828 | 307145 | 1 | ENSMUST00 | -17987 | Pcca | ENSMUSG0( | false | . | . | . | . | . | . | chr14 | 122563777 | 122566340 | 3.68769 | 3.36492 | 1.96488 |
| chr14 | 32950624 | 32951404 | 1000 | Distal Intergenic | 14 | 32991382 | 33015292 | 23911 | 1 | ENSMUST00 | -39978 | Lrrc18 | ENSMUSG0( | false | . | . | . | . | . | . | chr14 | 32945937 | 32960190 | 5.28697 | 4.21543 | 3.24118 |
| chr11 | 112772914 | 112773240 | 1000 | Intron (ENSMUST00000137081.2 | 11 | 112687645 | 112781976 | 94332 | 2 | ENSMUST00 | 8736 | BC006965 | ENSMUSG0( | false | . | . | . | . | . | . | chr11 | 112772876 | 112773912 | 4.16942 | 3.35151 | 2.35911 |
| chr13 | 98814532 | 98815745 | 1000 | Promoter (<=1kb) | 13 | 98723407 | 98815449 | 92043 | 2 | ENSMUST00 | 0 | Fcho2 | ENSMUSG0( | false | chr13 | 98813742 | 98815937 | 32.69058 | 104.59052 | 100.90593 | chr13 | 98815688 | 98816790 | 2.85783 | 2.88839 | 1.32679 |
| chr12 | 102876034 | 102876462 | 1000 | Promoter (1-2kb) | 12 | 102784656 | 102878406 | 93751 | 2 | ENSMUST00 | 1944 | Btbd7 | ENSMUSG0( | false | chr12 | 102876356 | 102879146 | 34.81759 | 111.39526 | 107.52535 | chr12 | 102869011 | 102877614 | 5.31821 | 4.25336 | 3.26871 |
| chr12 | 102877476 | 102878813 | 1000 | Promoter (<=1kb) | 12 | 102784656 | 102878406 | 93751 | 2 | ENSMUST00 | 0 | Btbd7 | ENSMUSG0( | false | chr12 | 102876356 | 102879146 | 34.81759 | 111.39526 | 107.52535 | chr12 | 102869011 | 102877614 | 5.31821 | 4.25336 | 3.26871 |
| chr9 | 62435708 | 62436289 | 1000 | Intron (ENSMUST00000048043.6 | 9 | 62428263 | 62429250 | 988 | 2 | ENSMUST00 | -6459 | Coro2b | ENSMUSG0( | true | . | . | . | . | . | . | chr9 | 62432367 | 62439369 | 7.25333 | 4.95646 | 4.88893 |
| chr9 | 62488715 | 62488989 | 1000 | Intron (ENSMUST00000048043.6 | 9 | 62427936 | 62510498 | 82563 | 2 | ENSMUST00 | 21509 | Coro2b | ENSMUSG0( | true | . | . | . | . | . | . | chr9 | 62449515 | 62535966 | 6.98492 | 4.90408 | 4.64549 |
| chr9 | 62531498 | 62532110 | 1000 | Intron (ENSMUST00000048043.6 | 9 | 62453283 | 62536960 | 83678 | 2 | ENSMUST00 | 4850 | Coro2b | ENSMUSG0( | true | . | . | . | . | . | . | chr9 | 62449515 | 62535966 | 6.98492 | 4.90408 | 4.64549 |
| chr9 | 62536649 | 62537183 | 1000 | Promoter (<=1kb) | 9 | 62419492 | 62537044 | 117553 | 2 | ENSMUST00 | 0 | Coro2b | ENSMUSG0( | true | chr9 | 62534315 | 62537345 | 30.01348 | 90.95103 | 87.5851 | chr9 | 62536973 | 62542648 | 4.83986 | 4.06256 | 2.86567 |
| chr9 | 62456013 | 62456322 | 1000 | Intron (ENSMUST00000048043.6 | 9 | 62429334 | 62449148 | 19815 | 2 | ENSMUST00 | -6866 | Coro2b | ENSMUSG0( | true | . | . | . | . | . | . | chr9 | 62449515 | 62535966 | 6.98492 | 4.90408 | 4.64549 |
| chr3 | 87174374 | 87174964 | 1000 | Promoter (<=1kb) | 3 | 87078593 | 87174700 | 96108 | 2 | ENSMUST00 | 0 | Kirrel | ENSMUSG0( | true | chr3 | 87171121 | 87175434 | 30.17592 | 94.17345 | 90.73757 | chr3 | 87174920 | 87175717 | 5.21527 | 3.95515 | 3.2152 |
| chr3 | 87171837 | 87172677 | 1000 | Promoter (1-2kb) | 3 | 87084628 | 87174550 | 89923 | 2 | ENSMUST00 | 1873 | Kirrel | ENSMUSG0( | true | chr3 | 87171121 | 87175434 | 30.17592 | 94.17345 | 90.73757 | . | . | . | . | . | . |
| chr3 | 87160789 | 87161168 | 1000 | Intron (ENSMUST00000159976.2 | 3 | 87084628 | 87174550 | 89923 | 2 | ENSMUST00 | 13382 | Kirrel | ENSMUSG0( | true | . | . | . | . | . | . | chr3 | 87139743 | 87171483 | 6.81014 | 4.88372 | 4.49361 |
| chr3 | 87165159 | 87166022 | 1000 | Intron (ENSMUST00000159976.2 | 3 | 87084628 | 87174550 | 89923 | 2 | ENSMUST00 | 8528 | Kirrel | ENSMUSG0( | true | chr3 | 87165808 | 87166737 | 7.13425 | 13.57188 | 11.33167 | chr3 | 87139743 | 87171483 | 6.81014 | 4.88372 | 4.49361 |
| chr1 | 133222505 | 133223148 | 1000 | Intron (ENSMUST00000187285.1 | 1 | 133246097 | 133303435 | 57339 | 1 | ENSMUST00 | -22949 | Plekha6 | ENSMUSG0( | false | . | . | . | . | . | . | chr1 | 133221862 | 133228348 | 6.32582 | 4.78563 | 4.08006 |
| chr7 | 138788017 | 138788354 | 1000 | Distal Intergenic | 7 | 138846079 | 138871702 | 25624 | 1 | ENSMUST00 | -57725 | Ppp2r2d | ENSMUSG0( | false | . | . | . | . | . | . | chr7 | 138787248 | 138790593 | 5.14773 | 4.02367 | 3.15774 |
| chr7 | 138846154 | 138846674 | 1000 | Promoter (<=1kb) | 7 | 138846359 | 138883057 | 36699 | 1 | ENSMUST00 | 0 | Ppp2r2d | ENSMUSG0( | false | chr7 | 138845531 | 138847675 | 37.04683 | 119.47973 | 115.35273 | . | . | . | . | . | . |
| chr13 | 97240978 | 97241625 | 1000 | Promoter (<=1kb) | 13 | 97241105 | 97253039 | 11935 | 1 | ENSMUST00 | 0 | Enc1 | ENSMUSG0( | false | chr13 | 97240304 | 97242323 | 24.48636 | 83.88964 | 80.66612 | chr13 | 97239205 | 97242784 | 4.46023 | 3.744 | 2.60558 |
| chr1 | 21034382 | 21034685 | 1000 | Intron (ENSMUST00000037998.5 | 1 | 21004313 | 21043656 | 39344 | 2 | ENSMUST00 | 8971 | Tram2 | ENSMUSG0( | false | . | . | . | . | . | . | chr1 | 21033430 | 21035825 | 4.21464 | 3.5031 | 2.39631 |
| chr1 | 21052652 | 21053106 | 1000 | Intron (ENSMUST00000037998.5 | 1 | 21004313 | 21043656 | 39344 | 2 | ENSMUST00 | -8997 | Tram2 | ENSMUSG0( | false | . | . | . | . | . | . | chr1 | 21036770 | 21057423 | 5.9841 | 4.45655 | 3.82459 |
| chr1 | 21079053 | 21079666 | 1000 | Promoter (<=1kb) | 1 | 20996299 | 21079229 | 82931 | 2 | ENSMUST00 | 0 | Tram2 | ENSMUSG0( | false | chr1 | 21077190 | 21079961 | 26.86698 | 95.04005 | 91.5885 | . | . | . | . | . | . |
| chr1 | 21047636 | 21048157 | 1000 | Intron (ENSMUST00000037998.5 | 1 | 21004313 | 21043656 | 39344 | 2 | ENSMUST00 | -3981 | Tram2 | ENSMUSG0( | false | . | . | . | . | . | . | chr1 | 21036770 | 21057423 | 5.9841 | 4.45655 | 3.82459 |
| chr1 | 20951347 | 20951756 | 1000 | Promoter (<=1kb) | 1 | 20951626 | 20955527 | 3902 | 1 | ENSMUST00 | 0 | Efhc1 | ENSMUSG0( | false | chr1 | 20951200 | 20952243 | 15.9403 | 41.39913 | 38.79041 | chr1 | 20951169 | 20952171 | 3.70481 | 3.23135 | 1.96562 |
| chr12 | 101083166 | 101084015 | 1000 | Promoter (<=1kb) | 12 | 101039409 | 101083702 | 44294 | 2 | ENSMUST00 | 0 | Smek1 | ENSMUSG0( | false | chr12 | 101081809 | 101084598 | 31.07893 | 124.33488 | 120.0426 | . | . | . | . | . | . |
| chr15 | 82932888 | 82933326 | 1000 | Distal Intergenic | 15 | 82808629 | 82912134 | 103506 | 2 | ENSMUST00 | -20755 | Tcf20 | ENSMUSG0( | false | . | . | . | . | . | . | chr15 | 82933085 | 82933950 | 9.08237 | 5.98297 | 6.32509 |
| chr5 | 114588424 | 114588996 | 1000 | Intron (ENSMUST00000043650.7 | 5 | 114568017 | 114594678 | 26662 | 1 | ENSMUST00 | 20408 | Fam222a | ENSMUSG0( | true | chr5 | 114588116 | 114588585 | 11.51562 | 24.54127 | 22.1466 | chr5 | 114571033 | 114614394 | 7.1045 | 4.9325 | 4.74958 |
| chr5 | 114593089 | 114593630 | 1000 | Intron (ENSMUST00000043650.7 | 5 | 114568017 | 114594678 | 26662 | 1 | ENSMUST00 | 25073 | Fam222a | ENSMUSG0( | true | . | . | . | . | . | . | chr5 | 114571033 | 114614394 | 7.1045 | 4.9325 | 4.74958 |
| chr5 | 114580937 | 114581899 | 1000 | Intron (ENSMUST00000043650.7 | 5 | 114568017 | 114594678 | 26662 | 1 | ENSMUST00 | 12921 | Fam222a | ENSMUSG0( | true | . | . | . | . | . | . | chr5 | 114571033 | 114614394 | 7.1045 | 4.9325 | 4.74958 |
| chr5 | 114570066 | 114570505 | 1000 | Promoter (2-3kb) | 5 | 114568017 | 114594678 | 26662 | 1 | ENSMUST00 | 2050 | Fam222a | ENSMUSG0( | true | chr5 | 114570268 | 114572884 | 12.92336 | 34.59913 | 32.07519 | . | . | . | . | . | . |
| chr5 | 114571341 | 114571956 | 1000 | Intron (ENSMUST00000043650.7 | 5 | 114568017 | 114594678 | 26662 | 1 | ENSMUST00 | 3325 | Fam222a | ENSMUSG0( | true | chr5 | 114570268 | 114572884 | 12.92336 | 34.59913 | 32.07519 | chr5 | 114571033 | 114614394 | 7.1045 | 4.9325 | 4.74958 |

| | | | | | | | | | | | | | | | | | | | | | | | | | | |
|---|---|---|---|---|---|---|---|---|---|---|---|---|---|---|---|---|---|---|---|---|---|---|---|---|---|---|
| chr5 | 114587866 | 114588329 | 1000 | Intron (ENSMUST00000043650.7 | 5 | 114568017 | 114594678 | 26662 | 1 | ENSMUST00 | 19850 | Fam222a | ENSMUSG0( | true | chr5 | 114588116 | 114588585 | 11.51562 | 24.54127 | 22.1466 | chr5 | 114571033 | 114614394 | 7.1045 | 4.9325 | 4.74958 |
| chr4 | 156197220 | 156198180 | 1000 | Promoter (<=1kb) | 4 | 156165778 | 156197488 | 31711 | 2 | ENSMUST00 | 0 | Agrn | ENSMUSG0( | false | chr4 | 156196002 | 156198318 | 31.28813 | 116.44205 | 112.41405 | . | . | . | . | . | . |
| chr4 | 156192140 | 156192941 | 1000 | Intron (ENSMUST00000180572.1 | 4 | 156165778 | 156197488 | 31711 | 2 | ENSMUST00 | 4547 | Agrn | ENSMUSG0( | false | chr4 | 156191798 | 156192581 | 9.73822 | 22.62381 | 20.25297 | chr4 | 156183439 | 156196896 | 6.66871 | 4.82292 | 4.38957 |
| chr16 | 16244690 | 16245117 | 1000 | Intron (ENSMUST00000161342.2 | 16 | 16213318 | 16272712 | 59395 | 1 | ENSMUST00 | 31373 | Pkp2 | ENSMUSG0( | false | . | . | . | . | . | . | chr16 | 16243056 | 16246974 | 4.27978 | 3.64678 | 2.4386 |
| chr3 | 93554980 | 93555370 | 1000 | Promoter (<=1kb) | 3 | 93555080 | 93564643 | 9564 | 1 | ENSMUST00 | 0 | S100a10 | ENSMUSG0( | false | chr3 | 93554542 | 93557324 | 37.4081 | 121.17245 | 116.98877 | . | . | . | . | . | . |
| chr11 | 5444044 | 5445197 | 1000 | Promoter (<=1kb) | 11 | 5276324 | 5444847 | 168524 | 2 | ENSMUST00 | 0 | Znrf3 | ENSMUSG0( | false | chr11 | 5443230 | 5445570 | 28.08037 | 85.65662 | 82.39679 | . | . | . | . | . | . |
| chr3 | 87617179 | 87617771 | 1000 | Promoter (<=1kb) | 3 | 87617559 | 87699145 | 81587 | 1 | ENSMUST00 | 0 | Arhgef11 | ENSMUSG0( | false | chr3 | 87617020 | 87619659 | 33.41831 | 117.26761 | 113.21259 | . | . | . | . | . | . |
| chr2 | 70824995 | 70826144 | 1000 | Promoter (<=1kb) | 2 | 70712407 | 70825397 | 112991 | 2 | ENSMUST00 | 0 | Tlk1 | ENSMUSG0( | false | chr2 | 70823887 | 70826522 | 27.46926 | 112.95284 | 109.03484 | . | . | . | . | . | . |
| chr13 | 94976195 | 94976743 | 1000 | Promoter (<=1kb) | 13 | 94976344 | 95014531 | 38188 | 1 | ENSMUST00 | 0 | Wdr41 | ENSMUSG0( | false | chr13 | 94975953 | 94977107 | 21.17422 | 79.20749 | 76.06625 | . | . | . | . | . | . |
| chr11 | 40692144 | 40692814 | 1000 | Promoter (<=1kb) | 11 | 40679314 | 40692774 | 13461 | 2 | ENSMUST00 | 0 | Mat2b | ENSMUSG0( | false | chr11 | 40691371 | 40693824 | 35.56825 | 117.95176 | 113.87694 | chr11 | 40688035 | 40693673 | 4.65165 | 3.82616 | 2.7271 |
| chr1 | 132417244 | 132418205 | 1000 | Promoter (<=1kb) | 1 | 132417555 | 132466958 | 49404 | 1 | ENSMUST00 | 0 | Dstyk | ENSMUSG0( | false | chr1 | 132416868 | 132418544 | 37.38551 | 123.27131 | 119.01954 | chr1 | 132416647 | 132417288 | 3.48045 | 3.33825 | 1.80575 |
| chr7 | 129509639 | 129510055 | 1000 | Distal Intergenic | 7 | 129591863 | 129635738 | 43876 | 1 | ENSMUST00 | -81808 | Wdr11 | ENSMUSG0( | false | . | . | . | . | . | . | . | . | . | . | . | . |
| chr1 | 132353387 | 132353781 | 1000 | Downstream (2-3kb) | 1 | 132361026 | 132364948 | 3923 | 2 | ENSMUST00 | 11167 | Tmcc2 | ENSMUSG0( | false | . | . | . | . | . | . | chr1 | 132352707 | 132356435 | 8.61007 | 5.59858 | 5.98871 |
| chr6 | 117916803 | 117917462 | 1000 | Promoter (<=1kb) | 6 | 117916981 | 117924382 | 7402 | 1 | ENSMUST00 | 0 | Hnrnpf | ENSMUSG0( | false | chr6 | 117916314 | 117918503 | 25.14555 | 83.24793 | 80.0387 | . | . | . | . | . | . |
| chr6 | 117907425 | 117908008 | 1000 | Promoter (<=1kb) | 6 | 117907110 | 117907760 | 651 | 2 | ENSMUST00 | 0 | 4933440N22 | ENSMUSG0( | false | chr6 | 117906713 | 117908804 | 44.72366 | 151.575 | 146.22792 | . | . | . | . | . | . |
| chr7 | 128610849 | 128611897 | 1000 | Promoter (<=1kb) | 7 | 128611328 | 128671622 | 60295 | 1 | ENSMUST00 | 0 | Inpp5f | ENSMUSG0( | false | chr7 | 128611271 | 128612373 | 35.37319 | 123.98246 | 119.71299 | . | . | . | . | . | . |
| chr5 | 113993363 | 113994136 | 1000 | Promoter (<=1kb) | 5 | 113937094 | 113993894 | 56801 | 2 | ENSMUST00 | 0 | Ssh1 | ENSMUSG0( | false | chr5 | 113993025 | 113994545 | 34.26917 | 107.34218 | 103.58047 | chr5 | 113983235 | 113993428 | 4.81586 | 3.83682 | 2.88443 |
| chr5 | 113986562 | 113986989 | 1000 | Promoter (2-3kb) | 5 | 113942192 | 113989743 | 47552 | 2 | ENSMUST00 | 2754 | Ssh1 | ENSMUSG0( | false | . | . | . | . | . | . | chr5 | 113983235 | 113993428 | 4.81586 | 3.83682 | 2.88443 |
| chrX | 142966272 | 142967029 | 1000 | Promoter (<=1kb) | 20 | 142851146 | 142966728 | 115583 | 2 | ENSMUST00 | 0 | Ammecr1 | ENSMUSG0( | false | chrX | 142965604 | 142967368 | 20.21941 | 51.82391 | 49.08421 | . | . | . | . | . | . |
| chr4 | 3678459 | 3679158 | 1000 | Promoter (<=1kb) | 4 | 3678130 | 3739176 | 61047 | 1 | ENSMUST00 | 330 | Lyn | ENSMUSG0( | false | chr4 | 3677777 | 3679180 | 30.26787 | 111.42836 | 107.5545 | chr4 | 3678882 | 3683092 | 4.05759 | 3.60604 | 2.27807 |
| chr17 | 42358441 | 42358917 | 1000 | Intron (ENSMUST00000048691.4 | 17 | 42315947 | 42507741 | 191795 | 1 | ENSMUST00 | 42495 | Ptchd4 | ENSMUSG0( | true | . | . | . | . | . | . | chr17 | 42356956 | 42366187 | 4.47326 | 3.86148 | 2.56747 |
| chr17 | 42113713 | 42114116 | 1000 | Distal Intergenic | 17 | 42315947 | 42507741 | 191795 | 1 | ENSMUST00 | -201831 | Ptchd4 | ENSMUSG0( | true | . | . | . | . | . | . | chr17 | 42112614 | 42115647 | 3.70808 | 3.40834 | 1.98723 |
| chr17 | 42316178 | 42316527 | 1000 | Promoter (<=1kb) | 17 | 42315947 | 42507741 | 191795 | 1 | ENSMUST00 | 232 | Ptchd4 | ENSMUSG0( | true | chr17 | 42315825 | 42317062 | 24.93112 | 71.82204 | 68.80418 | chr17 | 42315188 | 42317391 | 4.25142 | 3.70747 | 2.39228 |
| chr15 | 81190529 | 81191097 | 1000 | Promoter (<=1kb) | 15 | 81012281 | 81190733 | 178453 | 2 | ENSMUST00 | 0 | Mkl1 | ENSMUSG0( | false | chr15 | 81190019 | 81190846 | 31.62822 | 118.12274 | 114.03909 | . | . | . | . | . | . |
| chr11 | 22286399 | 22287369 | 1000 | Promoter (<=1kb) | 11 | 22005828 | 22286795 | 280968 | 2 | ENSMUST00 | 0 | Ehbp1 | ENSMUSG0( | false | chr11 | 22286225 | 22287137 | 38.37755 | 123.92766 | 119.65938 | chr11 | 22285600 | 22286512 | 3.6345 | 3.46018 | 1.86885 |
| chr7 | 127776651 | 127777885 | 1000 | Promoter (<=1kb) | 7 | 127776670 | 127800122 | 23453 | 1 | ENSMUST00 | 0 | Setd1a | ENSMUSG0( | false | chr7 | 127776169 | 127778338 | 22.99841 | 82.87405 | 79.66897 | . | . | . | . | . | . |
| chr4 | 154941978 | 154942299 | 1000 | Distal Intergenic | 4 | 154926628 | 154928563 | 1936 | 2 | ENSMUST00 | -13416 | Tnfrsf14 | ENSMUSG0( | false | . | . | . | . | . | . | . | . | . | . | . | . |
| chr4 | 154942493 | 154942777 | 1000 | Distal Intergenic | 4 | 154926628 | 154928563 | 1936 | 2 | ENSMUST00 | -13931 | Tnfrsf14 | ENSMUSG0( | false | . | . | . | . | . | . | . | . | . | . | . | . |
| chr13 | 113621977 | 113622779 | 1000 | Distal Intergenic | 13 | 113592180 | 113618564 | 26385 | 2 | ENSMUST00 | -3414 | Snx18 | ENSMUSG0( | false | . | . | . | . | . | . | chr13 | 113620854 | 113624004 | 4.25547 | 3.67701 | 2.40082 |
| chr13 | 113618341 | 113618776 | 1000 | Promoter (<=1kb) | 13 | 113592180 | 113618564 | 26385 | 2 | ENSMUST00 | 0 | Snx18 | ENSMUSG0( | false | chr13 | 113614178 | 113618647 | 32.67316 | 105.57997 | 101.8659 | . | . | . | . | . | . |
| chr3 | 90265060 | 90266073 | 1000 | Promoter (<=1kb) | 3 | 90265185 | 90280669 | 15485 | 1 | ENSMUST00 | 0 | Dennd4b | ENSMUSG0( | false | chr3 | 90264901 | 90266211 | 38.61126 | 124.33321 | 120.0426 | chr3 | 90263785 | 90266809 | 4.83883 | 4.11015 | 2.86112 |
| chr3 | 90292752 | 90294013 | 1000 | Distal Intergenic | 3 | 90278008 | 90278670 | 663 | 1 | ENSMUST00 | 14745 | Dennd4b | ENSMUSG0( | false | chr3 | 90293103 | 90294459 | 33.19039 | 103.42804 | 99.77289 | . | . | . | . | . | . |
| chr13 | 112987451 | 112988132 | 1000 | Promoter (<=1kb) | 13 | 112987802 | 112990778 | 2977 | 1 | ENSMUST00 | 0 | Ccno | ENSMUSG0( | false | chr13 | 112987057 | 112988572 | 27.11239 | 76.48743 | 73.39479 | chr13 | 112986424 | 112990826 | 4.22206 | 3.70622 | 2.3781 |
| chr9 | 70657056 | 70657467 | 1000 | Promoter (<=1kb) | 9 | 70599014 | 70657174 | 58161 | 2 | ENSMUST00 | 0 | Fam63b | ENSMUSG0( | false | chr9 | 70655957 | 70657918 | 34.04953 | 116.30843 | 112.28568 | . | . | . | . | . | . |
| chr6 | 8209104 | 8209840 | 1000 | Promoter (<=1kb) | 6 | 8209222 | 8236274 | 27053 | 1 | ENSMUST00 | 0 | Mios | ENSMUSG0( | false | chr6 | 8208860 | 8210167 | 35.79183 | 131.19528 | 126.66647 | . | . | . | . | . | . |
| chr7 | 127266230 | 127266759 | 1000 | Distal Intergenic | 7 | 127256959 | 127260709 | 3751 | 2 | ENSMUST00 | -5522 | Dctpp1 | ENSMUSG0( | false | . | . | . | . | . | . | chr7 | 127265106 | 127268380 | 4.66021 | 3.86831 | 2.76589 |
| chrX | 139684768 | 139685252 | 1000 | Promoter (<=1kb) | 20 | 139684996 | 139753405 | 68410 | 1 | ENSMUST00 | 0 | Tbc1d8b | ENSMUSG0( | true | chrX | 139684244 | 139686102 | 17.46222 | 42.53877 | 39.91948 | . | . | . | . | . | . |
| chr12 | 84192553 | 84194131 | 1000 | Promoter (<=1kb) | 12 | 84149278 | 84194015 | 44738 | 2 | ENSMUST00 | 0 | Elmsan1 | ENSMUSG0( | false | chr12 | 84192322 | 84194307 | 26.65955 | 86.95457 | 83.67183 | chr12 | 84184648 | 84196462 | 7.46806 | 5.03703 | 5.02687 |
| chr15 | 79741826 | 79742912 | 1000 | Promoter (<=1kb) | 15 | 79724070 | 79742456 | 18387 | 2 | ENSMUST00 | 0 | Sun2 | ENSMUSG0( | false | chr15 | 79741496 | 79743083 | 26.72788 | 101.58999 | 97.97913 | . | . | . | . | . | . |
| chr15 | 79735412 | 79735782 | 1000 | Promoter (<=1kb) | 15 | 79734108 | 79735951 | 1844 | 2 | ENSMUST00 | 169 | Sun2 | ENSMUSG0( | false | . | . | . | . | . | . | chr15 | 79734747 | 79736203 | 5.00055 | 4.06353 | 3.02064 |
| chr2 | 153345431 | 153346727 | 1000 | Promoter (<=1kb) | 2 | 153345845 | 153404007 | 58163 | 1 | ENSMUST00 | 0 | Asxl1 | ENSMUSG0( | false | chr2 | 153345733 | 153347755 | 30.91854 | 102.46954 | 98.8397 | . | . | . | . | . | . |
| chr5 | 121833785 | 121834292 | 1000 | Promoter (1-2kb) | 5 | 121817010 | 121836104 | 19095 | 2 | ENSMUST00 | 1812 | Sh2b3 | ENSMUSG0( | false | . | . | . | . | . | . | chr5 | 121827805 | 121836228 | 4.97092 | 3.99548 | 3.00296 |
| chr6 | 39205129 | 39207296 | 1000 | Promoter (<=1kb) | 6 | 39136623 | 39206789 | 70167 | 2 | ENSMUST00 | 0 | Kdm7a | ENSMUSG0( | false | chr6 | 39204806 | 39207527 | 39.05061 | 143.74966 | 138.71384 | . | . | . | . | . | . |
| chr5 | 121711351 | 121712662 | 1000 | Promoter (<=1kb) | 5 | 121711609 | 121814949 | 103341 | 1 | ENSMUST00 | 0 | Atxn2 | ENSMUSG0( | false | chr5 | 121710758 | 121713663 | 37.78396 | 121.82604 | 117.62474 | . | . | . | . | . | . |
| chr5 | 121712684 | 121713514 | 1000 | Promoter (<=1kb) | 5 | 121712969 | 121772787 | 59819 | 1 | ENSMUST00 | 0 | Atxn2 | ENSMUSG0( | false | chr5 | 121710758 | 121713663 | 37.78396 | 121.82604 | 117.62474 | chr5 | 121713372 | 121725102 | 4.18753 | 3.69327 | 2.34616 |
| chr4 | 126110736 | 126111533 | 1000 | Intron (ENSMUST00000116286.3 | 4 | 126104035 | 126117698 | 13664 | 1 | ENSMUST00 | 6702 | Stk40 | ENSMUSG0( | false | chr4 | 126110399 | 126111581 | 8.3822 | 16.12319 | 13.84377 | chr4 | 126104618 | 126117854 | 6.47728 | 4.72001 | 4.23059 |
| chr4 | 126133223 | 126133658 | 1000 | Intron (ENSMUST00000116286.3 | 4 | 126128819 | 126132798 | 3980 | 1 | ENSMUST00 | 4405 | Stk40 | ENSMUSG0( | false | . | . | . | . | . | . | chr4 | 126129881 | 126135597 | 7.91466 | 5.29987 | 5.45182 |
| chr4 | 126130484 | 126130858 | 1000 | Promoter (1-2kb) | 4 | 126128819 | 126132798 | 3980 | 1 | ENSMUST00 | 1666 | Stk40 | ENSMUSG0( | false | . | . | . | . | . | . | chr4 | 126129881 | 126135597 | 7.91466 | 5.29987 | 5.45182 |
| chr12 | 83596917 | 83597415 | 1000 | Promoter (<=1kb) | 12 | 83546941 | 83597147 | 50207 | 2 | ENSMUST00 | 0 | Zfyve1 | ENSMUSG0( | false | chr12 | 83597366 | 83597737 | 8.63619 | 18.02194 | 15.71366 | . | . | . | . | . | . |
| chr17 | 27048534 | 27048966 | 1000 | Distal Intergenic | 17 | 27057304 | 27122223 | 64920 | 1 | ENSMUST00 | -8338 | Itpr3 | ENSMUSG0( | false | . | . | . | . | . | . | chr17 | 27047791 | 27052228 | 4.49279 | 3.68146 | 2.62823 |
| chr17 | 27056195 | 27056615 | 1000 | Promoter (<=1kb) | 17 | 27057304 | 27122223 | 64920 | 1 | ENSMUST00 | -689 | Itpr3 | ENSMUSG0( | false | chr17 | 27056273 | 27059567 | 32.33232 | 98.9174 | 95.3739 | . | . | . | . | . | . |
| chr17 | 27056924 | 27057540 | 1000 | Promoter (<=1kb) | 17 | 27057304 | 27122223 | 64920 | 1 | ENSMUST00 | 0 | Itpr3 | ENSMUSG0( | false | chr17 | 27056273 | 27059567 | 32.33232 | 98.9174 | 95.3739 | chr17 | 27056774 | 27076697 | 5.08038 | 3.96808 | 3.08893 |
| chr17 | 27058498 | 27059191 | 1000 | Promoter (1-2kb) | 17 | 27057304 | 27122223 | 64920 | 1 | ENSMUST00 | 1195 | Itpr3 | ENSMUSG0( | false | chr17 | 27056273 | 27059567 | 32.33232 | 98.9174 | 95.3739 | chr17 | 27056774 | 27076697 | 5.08038 | 3.96808 | 3.08893 |
| chr2 | 152951456 | 152952033 | 1000 | Promoter (<=1kb) | 2 | 152944933 | 152951582 | 6650 | 2 | ENSMUST00 | 0 | Dusp15 | ENSMUSG0( | false | chr2 | 152949781 | 152952321 | 21.13367 | 73.82313 | 70.77491 | chr2 | 152946044 | 152955187 | 5.87363 | 4.46622 | 3.72171 |
| chr2 | 152950309 | 152950651 | 1000 | Promoter (<=1kb) | 2 | 152940997 | 152951456 | 10460 | 2 | ENSMUST00 | 805 | Dusp15 | ENSMUSG0( | false | chr2 | 152949781 | 152952321 | 21.13367 | 73.82313 | 70.77491 | chr2 | 152946044 | 152955187 | 5.87363 | 4.46622 | 3.72171 |
| chr2 | 152952260 | 152952753 | 1000 | Promoter (<=1kb) | 2 | 152944977 | 152951698 | 6722 | 2 | ENSMUST00 | -563 | Dusp15 | ENSMUSG0( | false | chr2 | 152949781 | 152952321 | 21.13367 | 73.82313 | 70.77491 | chr2 | 152946044 | 152955187 | 5.87363 | 4.46622 | 3.72171 |
| chr4 | 125122523 | 125123585 | 1000 | Promoter (<=1kb) | 4 | 125120094 | 125122520 | 2427 | 2 | ENSMUST00 | -4 | Zc3h12a | ENSMUSG0( | false | . | . | . | . | . | . | chr4 | 125106469 | 125127334 | 7.64816 | 5.22391 | 5.18478 |
| chr1 | 153535582 | 153535976 | 1000 | Exon (ENSMUST00000041874.8/ | 1 | 153548422 | 153549711 | 1290 | 2 | ENSMUST00 | 13735 | Npl | ENSMUSG0( | false | . | . | . | . | . | . | chr1 | 153531685 | 153537651 | 4.35455 | 3.71394 | 2.519 |
| chr1 | 17097507 | 17098183 | 1000 | Promoter (<=1kb) | 1 | 16994938 | 17097889 | 102952 | 2 | ENSMUST00 | 0 | Jph1 | ENSMUSG0( | false | chr1 | 17096332 | 17098208 | 29.28716 | 88.72003 | 85.40064 | . | . | . | . | . | . |
| chr9 | 75409299 | 75410390 | 1000 | Promoter (<=1kb) | 9 | 75386902 | 75409359 | 22458 | 2 | ENSMUST00 | 0 | Mapk6 | ENSMUSG0( | false | chr9 | 75408649 | 75410600 | 27.34946 | 104.78247 | 101.09169 | . | . | . | . | . | . |
| chr19 | 47536930 | 47537317 | 1000 | Promoter (<=1kb) | 19 | 47501033 | 47537119 | 36087 | 2 | ENSMUST00 | 0 | Obfc1 | ENSMUSG0( | false | chr19 | 47536467 | 47537802 | 17.60932 | 44.27721 | 41.6341 | chr19 | 47536115 | 47539172 | 6.59782 | 4.88346 | 4.29922 |
| chr12 | 82169225 | 82170275 | 1000 | Promoter (<=1kb) | 12 | 82170016 | 82451782 | 281767 | 1 | ENSMUST00 | 0 | Sipa1l1 | ENSMUSG0( | false | chr12 | 82168802 | 82171414 | 42.12499 | 142.52214 | 137.54099 | . | . | . | . | . | . |
| chr5 | 121385184 | 121385776 | 1000 | Promoter (<=1kb) | 5 | 121371725 | 121385632 | 13908 | 2 | ENSMUST00 | 0 | Trafd1 | ENSMUSG0( | false | chr5 | 121384670 | 121386247 | 22.35204 | 62.09245 | 59.21756 | . | . | . | . | . | . |
| chr6 | 13677444 | 13678399 | 1000 | Promoter (<=1kb) | 6 | 13625677 | 13677966 | 52290 | 2 | ENSMUST00 | 0 | B630005N14 | ENSMUSG0( | false | chr6 | 13676723 | 13678457 | 29.35304 | 95.67908 | 92.21078 | chr6 | 13678191 | 13678646 | 5.53238 | 4.4088 | 3.50718 |
| chr9 | 103708921 | 103709257 | 1000 | Intron (ENSMUST00000189588.1 | 9 | 103482947 | 103692807 | 209861 | 2 | ENSMUST00 | -16115 | Tmem108 | ENSMUSG0( | true | . | . | . | . | . | . | chr9 | 103706426 | 103718720 | 5.35544 | 4.27413 | 3.29799 |
| chr9 | 103709657 | 103710091 | 1000 | Intron (ENSMUST00000189588.1 | 9 | 103482947 | 103692807 | 209861 | 2 | ENSMUST00 | -16851 | Tmem108 | ENSMUSG0( | true | . | . | . | . | . | . | chr9 | 103706426 | 103718720 | 5.35544 | 4.27413 | 3.29799 |
| chr9 | 103712523 | 103712930 | 1000 | Intron (ENSMUST00000189588.1 | 9 | 103482947 | 103692807 | 209861 | 2 | ENSMUST00 | -19717 | Tmem108 | ENSMUSG0( | true | . | . | . | . | . | . | chr9 | 103706426 | 103718720 | 5.35544 | 4.27413 | 3.29799 |
| chr9 | 103679212 | 103679625 | 1000 | Intron (ENSMUST00000049452.9 | 9 | 103482947 | 103692807 | 209861 | 2 | ENSMUST00 | 13182 | Tmem108 | ENSMUSG0( | true | . | . | . | . | . | . | chr9 | 103675009 | 103695608 | 5.07737 | 4.10994 | 3.07001 |
| chr1 | 152902140 | 152903618 | 1000 | Promoter (<=1kb) | 1 | 152836995 | 152902622 | 65628 | 2 | ENSMUST00 | 0 | Smg7 | ENSMUSG0( | false | chr1 | 152901741 | 152904742 | 36.83603 | 116.99731 | 112.9521 | . | . | . | . | . | . |
| chr7 | 97737866 | 97738365 | 1000 | Promoter (<=1kb) | 7 | 97726379 | 97738247 | 11869 | 2 | ENSMUST00 | 0 | Aqp11 | ENSMUSG0( | false | chr7 | 97737410 | 97738709 | 20.20131 | 69.382 | 66.39919 | chr7 | 97736160 | 97738686 | 4.28121 | 3.77143 | 2.41748 |
| chr8 | 121083617 | 121084488 | 1000 | Promoter (<=1kb) | 8 | 121084386 | 121088144 | 3759 | 1 | ENSMUST00 | 0 | Foxf1 | ENSMUSG0( | false | chr8 | 121082669 | 121085168 | 40.23391 | 129.30441 | 124.84723 | chr8 | 121080807 | 121085331 | 4.65031 | 3.72632 | 2.75058 |
| chr2 | 167542979 | 167543681 | 1000 | Distal Intergenic | 2 | 167538195 | 167542814 | 4620 | 1 | ENSMUST00 | 4785 | Snai1 | ENSMUSG0( | false | . | . | . | . | . | . | chr2 | 167541955 | 167544771 | 7.69518 | 5.21237 | 5.24384 |
| chr18 | 33463727 | 33464497 | 1000 | Promoter (<=1kb) | 18 | 33437019 | 33463747 | 26729 | 2 | ENSMUST00 | 0 | Nrep | ENSMUSG0( | false | chr18 | 33462104 | 33464772 | 29.3377 | 90.74815 | 87.38457 | . | . | . | . | . | . |
| chr13 | 32964958 | 32965655 | 1000 | Promoter (<=1kb) | 13 | 32965209 | 32979067 | 13859 | 1 | ENSMUST00 | 0 | Serpinb6b | ENSMUSG0( | false | chr13 | 32965180 | 32966767 | 16.23624 | 41.91433 | 39.30089 | chr13 | 32957470 | 32971328 | 4.32621 | 3.59198 | 2.48777 |
| chr2 | 128967078 | 128968336 | 1000 | Promoter (<=1kb) | 2 | 128967402 | 129018563 | 51162 | 1 | ENSMUST00 | 0 | Zc3h6 | ENSMUSG0( | false | chr2 | 128966468 | 128968509 | 41.911 | 144.38934 | 139.32462 | chr2 | 128968232 | 128969041 | 3.00098 | 2.92419 | 1.35513 |
| chr8 | 75063015 | 75063586 | 1000 | Intron (ENSMUST00000078847.6 | 8 | 75033686 | 75070121 | 36436 | 1 | ENSMUST00 | 29330 | Tom1 | ENSMUSG0( | false | . | . | . | . | . | . | chr8 | 75061373 | 75065112 | 5.02936 | 4.08771 | 3.02529 |
| chrX | 101254005 | 101254671 | 1000 | Promoter (<=1kb) | 20 | 101254528 | 101260873 | 6346 | 1 | ENSMUST00 | 0 | Foxo4 | ENSMUSG0( | false | chrX | 101253735 | 101255612 | 16.14593 | 43.98602 | 41.34693 | . | . | . | . | . | . |
| chr15 | 7203891 | 7204307 | 1000 | Downstream (1-2kb) | 15 | 7212191 | 7223069 | 10879 | 2 | ENSMUST00 | 18762 | Egflam | ENSMUSG0( | false | . | . | . | . | . | . | chr15 | 7200566 | 7204608 | 3.39469 | 3.13123 | 1.72761 |
| chr4 | 35138036 | 35138365 | 1000 | Intron (ENSMUST00000102975.4 | 4 | 35152056 | 35154005 | 1950 | 1 | ENSMUST00 | -13691 | Ifnk | ENSMUSG0( | false | . | . | . | . | . | . | . | . | . | . | . | . |
| chr7 | 16905379 | 16905914 | 1000 | Promoter (<=1kb) | 7 | 16906490 | 16910905 | 4416 | 1 | ENSMUST00 | -576 | Ptgir | ENSMUSG0( | false | . | . | . | . | . | . | chr7 | 16900106 | 16910944 | 5.45499 | 4.23801 | 3.39476 |
| chr6 | 108531881 | 108532408 | 1000 | Intron (ENSMUST00000032192.6 | 6 | 108563445 | 108565693 | 2249 | 2 | ENSMUST00 | 33285 | Gm16433 | ENSMUSG0( | false | . | . | . | . | . | . | chr6 | 108530788 | 108538190 | 6.66183 | 4.72814 | 4.3796 |
| chr6 | 108570921 | 108571663 | 1000 | Distal Intergenic | 6 | 108563445 | 108565693 | 2249 | 2 | ENSMUST00 | -5229 | Gm16433 | ENSMUSG0( | false | chr6 | 108570723 | 108571632 | 14.00287 | 32.47356 | 29.97816 | chr6 | 108569788 | 108574684 | 5.11474 | 4.14126 | 3.10691 |
| chr5 | 122145789 | 122146296 | 1000 | Distal Intergenic | 5 | 122134437 | 122138957 | 4521 | 2 | ENSMUST00 | -6833 | Ccdc63 | ENSMUSG0( | false | . | . | . | . | . | . | chr5 | 122145419 | 122146901 | 3.84848 | 3.34299 | 2.088 |
| chr18 | 60609754 | 60610410 | 1000 | Promoter (<=1kb) | 18 | 60600207 | 60610105 | 9899 | 2 | ENSMUST00 | 0 | Synpo | ENSMUSG0( | true | chr18 | 60603340 | 60613557 | 27.39171 | 92.13462 | 88.74321 | . | . | . | . | . | . |
| chr18 | 60606047 | 60606850 | 1000 | Exon (ENSMUST00000137894.1/ | 18 | 60600207 | 60610105 | 9899 | 2 | ENSMUST00 | 3255 | Synpo | ENSMUSG0( | true | chr18 | 60603340 | 60613557 | 27.39171 | 92.13462 | 88.74321 | chr18 | 60593903 | 60608752 | 6.45808 | 4.76452 | 4.19811 |
| chr18 | 60619051 | 60619903 | 1000 | Intron (ENSMUST00000097566.4 | 18 | 60604109 | 60623325 | 19217 | 2 | ENSMUST00 | 3422 | Synpo | ENSMUSG0( | true | . | . | . | . | . | . | chr18 | 60613738 | 60622693 | 7.427 | 5.19702 | 4.99228 |
| chr11 | 75170144 | 75170675 | 1000 | Promoter (<=1kb) | 11 | 75164565 | 75169519 | 4955 | 2 | ENSMUST00 | -626 | Hic1 | ENSMUSG0( | false | chr11 | 75170188 | 75170675 | 19.14982 | 55.63055 | 52.84086 | chr11 | 75165128 | 75174332 | 5.71251 | 4.38871 | 3.59566 |
| chr12 | 112490528 | 112490882 | 1000 | Promoter (1-2kb) | 12 | 112489448 | 112499927 | 10480 | 1 | ENSMUST00 | 1081 | A530016L24 | ENSMUSG0( | false | . | . | . | . | . | . | chr12 | 112489823 | 112492253 | 6.85166 | 4.73114 | 4.5344 |
| chr10 | 61495488 | 61495821 | 1000 | Intron (ENSMUST00000049242.7 | 10 | 61475245 | 61476937 | 1693 | 2 | ENSMUST00 | -18552 | D830039M14 | ENSMUSG0( | false | . | . | . | . | . | . | chr10 | 61494269 | 61496302 | 4.65675 | 3.94834 | 2.72799 |
| chr9 | 101251283 | 101252058 | 1000 | Promoter (<=1kb) | 9 | 101105084 | 101251795 | 146712 | 2 | ENSMUST00 | 0 | Ppp2r3a | ENSMUSG0( | false | chr9 | 101250702 | 101251977 | 26.52689 | 93.43019 | 90.01563 | . | . | . | . | . | . |
| chr3 | 92080443 | 92081995 | 1000 | Promoter (1-2kb) | 3 | 92080271 | 92083142 | 2872 | 2 | ENSMUST00 | 1147 | Lor | ENSMUSG0( | false | chr3 | 92080275 | 92082186 | 21.02725 | 63.18829 | 60.29588 | chr3 | 92081855 | 92082684 | 5.21425 | 4.22495 | 3.24704 |
| chr2 | 5950926 | 5951936 | 1000 | Promoter (<=1kb) | 2 | 5951469 | 6056703 | 105235 | 1 | ENSMUST00 | 0 | Upf2 | ENSMUSG0( | false | chr2 | 5950319 | 5951076 | 20.29158 | 53.552 | 50.79138 | . | . | . | . | . | . |
| chr4 | 133690249 | 133690621 | 1000 | Exon (ENSMUST00000138473.2/ | 4 | 133668757 | 133672647 | 3891 | 2 | ENSMUST00 | -17603 | Pigv | ENSMUSG0( | false | . | . | . | . | . | . | chr4 | 133686744 | 133692777 | 5.78466 | 4.23388 | 3.6953 |
| chr10 | 70440727 | 70441525 | 1000 | Promoter (<=1kb) | 10 | 70440880 | 70558729 | 117850 | 1 | ENSMUST00 | 0 | Fam13c | ENSMUSG0( | false | chr10 | 70440157 | 70441617 | 25.90858 | 83.69981 | 80.48042 | chr10 | 70441443 | 70443202 | 3.55181 | 3.32031 | 1.80507 |
| chr5 | 110448378 | 110448763 | 1000 | Promoter (<=1kb) | 5 | 110361754 | 110448503 | 86750 | 2 | ENSMUST00 | 0 | Fbrsl1 | ENSMUSG0( | false | chr5 | 110447113 | 110449296 | 37.30079 | 124.19571 | 119.91465 | . | . | . | . | . | . |
| chr2 | 74697727 | 74698810 | 1000 | Promoter (<=1kb) | 2 | 74697727 | 74700208 | 2482 | 1 | ENSMUST00 | 1 | Hoxd9 | ENSMUSG0( | false | chr2 | 74697475 | 74699399 | 28.56742 | 103.16996 | 99.51957 | . | . | . | . | . | . |
| chr11 | 103142400 | 103142965 | 1000 | Distal Intergenic | 11 | 103133865 | 103138279 | 4415 | 1 | ENSMUST00 | 8536 | Hexim2 | ENSMUSG0( | false | chr11 | 103142280 | 103143074 | 7.82203 | 16.59716 | 14.30959 | chr11 | 103138329 | 103150034 | 6.87245 | 4.95791 | 4.53389 |
| chr11 | 103147814 | 103148208 | 1000 | Distal Intergenic | 11 | 103133865 | 103138279 | 4415 | 1 | ENSMUST00 | 13950 | Hexim2 | ENSMUSG0( | false | . | . | . | . | . | . | chr11 | 103138329 | 103150034 | 6.87245 | 4.95791 | 4.53389 |
| chr19 | 5845418 | 5847360 | 1000 | Promoter (<=1kb) | 19 | 5845623 | 5851388 | 5766 | 1 | ENSMUST00 | 0 | Frmd8os | ENSMUSG0( | false | chr19 | 5846110 | 5846457 | 10.55355 | 22.91593 | 20.54101 | chr19 | 5846278 | 5852772 | 9.4084 | 5.8562 | 6.60958 |
| chr5 | 110839528 | 110839931 | 1000 | Promoter (<=1kb) | 5 | 110829070 | 110839777 | 10708 | 2 | ENSMUST00 | 0 | Hscb | ENSMUSG0( | false | chr5 | 110839100 | 110840690 | 24.78212 | 78.86476 | 75.73294 | . | . | . | . | . | . |
| chr2 | 29124894 | 29125653 | 1000 | Promoter (<=1kb) | 2 | 29124992 | 29182471 | 57480 | 1 | ENSMUST00 | 0 | Setx | ENSMUSG0( | false | chr2 | 29124443 | 29126022 | 29.3377 | 90.74815 | 87.38457 | . | . | . | . | . | . |
| chr3 | 7503263 | 7503826 | 1000 | Promoter (<=1kb) | 3 | 7503426 | 7553848 | 50423 | 1 | ENSMUST00 | 0 | Zc2hc1a | ENSMUSG0( | false | chr3 | 7503032 | 7504295 | 30.94804 | 114.76607 | 110.79114 | chr3 | 7502261 | 7508301 | 3.955 | 3.46223 | 2.19023 |
| chr11 | 70763938 | 70764690 | 1000 | Promoter (<=1kb) | 11 | 70764209 | 70772928 | 8720 | 1 | ENSMUST00 | 0 | Zfp3 | ENSMUSG0( | false | chr11 | 70763721 | 70765043 | 30.17466 | 106.17292 | 102.44986 | chr11 | 70763427 | 70763972 | 5.62231 | 4.40221 | 3.51544 |
| chr5 | 89882034 | 89882746 | 1000 | Promoter (<=1kb) | 5 | 89673841 | 89883334 | 209494 | 2 | ENSMUST00 | 588 | Adamts3 | ENSMUSG0( | false | chr5 | 89882037 | 89883791 | 27.08035 | 80.13548 | 76.98148 | chr5 | 89881940 | 89883731 | 5.49256 | 4.33474 | 3.40025 |
| chr5 | 89883020 | 89883713 | 1000 | Promoter (<=1kb) | 5 | 89673841 | 89883334 | 209494 | 2 | ENSMUST00 | 0 | Adamts3 | ENSMUSG0( | false | chr5 | 89882037 | 89883791 | 27.08035 | 80.13548 | 76.98148 | chr5 | 89881940 | 89883731 | 5.49256 | 4.33474 | 3.40025 |
| chr11 | 120348149 | 120349022 | 1000 | Promoter (<=1kb) | 11 | 120348678 | 120351190 | 2513 | 1 | ENSMUST00 | 0 | 0610009L18 | ENSMUSG0( | false | chr11 | 120347215 | 120350938 | 34.00618 | 119.38669 | 115.26049 | . | . | . | . | . | . |
| chr11 | 120349907 | 120350655 | 1000 | Promoter (1-2kb) | 11 | 120348678 | 120351190 | 2513 | 1 | ENSMUST00 | 1230 | 0610009L18 | ENSMUSG0( | false | chr11 | 120347215 | 120350938 | 34.00618 | 119.38669 | 115.26049 | chr11 | 120349642 | 120356034 | 5.56478 | 4.37057 | 3.45374 |
| chr11 | 59787237 | 59787814 | 1000 | Promoter (<=1kb) | 11 | 59783897 | 59787645 | 3749 | 2 | ENSMUST00 | 0 | Pld6 | ENSMUSG0( | false | chr11 | 59786975 | 59787807 | 17.37054 | 45.37996 | 42.72523 | chr11 | 59785037 | 59788001 | 4.856 | 3.9381 | 2.91233 |
| chr11 | 59786417 | 59786881 | 1000 | Promoter (<=1kb) | 11 | 59784703 | 59787642 | 2940 | 2 | ENSMUST00 | 761 | Pld6 | ENSMUSG0( | false | . | . | . | . | . | . | chr11 | 59785037 | 59788001 | 4.856 | 3.9381 | 2.91233 |
| chr8 | 120634253 | 120634720 | 1000 | Promoter (<=1kb) | 8 | 120608602 | 120634492 | 25891 | 2 | ENSMUST00 | 0 | 1190005I06R | ENSMUSG0( | false | chr8 | 120633713 | 120634858 | 15.77044 | 42.5731 | 39.95169 | chr8 | 120633518 | 120634951 | 3.88209 | 3.43348 | 2.0992 |
| chr11 | 116917403 | 116917716 | 1000 | Promoter (1-2kb) | 11 | 116918863 | 116986948 | 68086 | 1 | ENSMUST00 | -1147 | Mgat5b | ENSMUSG0( | false | . | . | . | . | . | . | chr11 | 116917199 | 116923338 | 4.7567 | 3.98532 | 2.79837 |
| chr11 | 116967513 | 116967853 | 1000 | Promoter (<=1kb) | 11 | 116966796 | 116978393 | 11598 | 1 | ENSMUST00 | 718 | Mgat5b | ENSMUSG0( | false | . | . | . | . | . | . | chr11 | 116958378 | 116973301 | 7.07011 | 4.94661 | 4.71174 |
| chr9 | 15771582 | 15772214 | 1000 | Distal Intergenic | 9 | 15709739 | 15742384 | 32646 | 1 | ENSMUST00 | 61844 | Slc36a4 | ENSMUSG0( | false | . | . | . | . | . | . | chr9 | 15770402 | 15776723 | 6.03027 | 4.5753 | 3.83562 |
| chr9 | 15782124 | 15782476 | 1000 | Distal Intergenic | 9 | 15709739 | 15742384 | 32646 | 1 | ENSMUST00 | 72386 | Slc36a4 | ENSMUSG0( | false | . | . | . | . | . | . | chr9 | 15781534 | 15783890 | 4.44284 | 3.86424 | 2.54275 |
| chr9 | 20976488 | 20977017 | 1000 | Promoter (<=1kb) | 9 | 20965952 | 20976793 | 10842 | 2 | ENSMUST00 | 0 | S1pr2 | ENSMUSG0( | false | chr9 | 20975370 | 20977071 | 29.28796 | 98.53321 | 94.99628 | . | . | . | . | . | . |
| chr9 | 44417951 | 44418433 | 1000 | Promoter (<=1kb) | 9 | 44410160 | 44418003 | 7844 | 2 | ENSMUST00 | 0 | Ccdc84 | ENSMUSG0( | false | chr9 | 44417144 | 44418636 | 23.17719 | 89.67316 | 86.33752 | chr9 | 44416177 | 44418741 | 4.29055 | 3.60635 | 2.46344 |
| chr5 | 101968501 | 101968964 | 1000 | Exon (ENSMUST00000053177.8/ | 5 | 101836158 | 101981558 | 145401 | 2 | ENSMUST00 | 12594 | Wdfy3 | ENSMUSG0( | false | . | . | . | . | . | . | chr5 | 101962254 | 101974242 | 5.86009 | 4.43256 | 3.74825 |
| chr5 | 102069115 | 102070284 | 1000 | Promoter (<=1kb) | 5 | 101832956 | 102069921 | 236966 | 2 | ENSMUST00 | 0 | Wdfy3 | ENSMUSG0( | false | chr5 | 102068756 | 102070662 | 28.8526 | 98.7319 | 95.19158 | chr5 | 102070197 | 102071128 | 4.08112 | 3.70737 | 2.26712 |
| chr5 | 101992805 | 101993322 | 1000 | Intron (ENSMUST00000053177.8 | 5 | 101836158 | 101981558 | 145401 | 2 | ENSMUST00 | -11248 | Wdfy3 | ENSMUSG0( | false | . | . | . | . | . | . | chr5 | 101981137 | 102004930 | 5.09217 | 4.0974 | 3.08644 |
| chr5 | 101993892 | 101994478 | 1000 | Intron (ENSMUST00000053177.8 | 5 | 101836158 | 101981558 | 145401 | 2 | ENSMUST00 | -12335 | Wdfy3 | ENSMUSG0( | false | . | . | . | . | . | . | chr5 | 101981137 | 102004930 | 5.09217 | 4.0974 | 3.08644 |
| chr4 | 126202281 | 126203087 | 1000 | Promoter (<=1kb) | 4 | 126164082 | 126202760 | 38679 | 2 | ENSMUST00 | 0 | Thrap3 | ENSMUSG0( | false | chr4 | 126201231 | 126202738 | 28.1615 | 93.53628 | 90.11485 | . | . | . | . | . | . |
| chr18 | 36280407 | 36281635 | 1000 | Promoter (<=1kb) | 18 | 36281097 | 36301521 | 20425 | 1 | ENSMUST00 | 0 | Pura | ENSMUSG0( | false | chr18 | 36280325 | 36282308 | 37.22259 | 125.13314 | 120.81451 | . | . | . | . | . | . |
| chr18 | 36286237 | 36287428 | 1000 | 5' UTR | 18 | 36281097 | 36301521 | 20425 | 1 | ENSMUST00 | 5141 | Pura | ENSMUSG0( | false | chr18 | 36285948 | 36288201 | 33.41831 | 117.26761 | 113.21259 | chr18 | 36284770 | 36286535 | 4.04344 | 3.66091 | 2.2014 |
| chr11 | 30885136 | 30885607 | 1000 | Promoter (<=1kb) | 11 | 30885358 | 30893729 | 8372 | 1 | ENSMUST00 | 0 | Gpr75 | ENSMUSG0( | false | chr11 | 30884899 | 30885770 | 17.55798 | 43.44164 | 40.81004 | chr11 | 30883429 | 30885501 | 2.912 | 2.86996 | 1.32398 |
| chr11 | 101173229 | 101173992 | 1000 | Promoter (1-2kb) | 11 | 101172997 | 101175443 | 2447 | 2 | ENSMUST00 | 1451 | Ccr10 | ENSMUSG0( | false | chr11 | 101173012 | 101174627 | 12.76655 | 31.65917 | 29.17213 | chr11 | 101171694 | 101177404 | 5.70547 | 4.40178 | 3.59444 |
| chr3 | 103914042 | 103915140 | 1000 | Promoter (<=1kb) | 3 | 103914120 | 103966613 | 52494 | 1 | ENSMUST00 | 0 | Rsbn1 | ENSMUSG0( | false | chr3 | 103913684 | 103916036 | 34.00884 | 130.01486 | 125.52227 | . | . | . | . | . | . |
| chr12 | 69371454 | 69372669 | 1000 | Promoter (<=1kb) | 12 | 69372150 | 69375979 | 3830 | 1 | ENSMUST00 | 0 | Arf6 | ENSMUSG0( | false | chr12 | 69371247 | 69375626 | 29.75837 | 125.79383 | 121.46131 | . | . | . | . | . | . |
| chrX | 12673428 | 12673798 | 1000 | Promoter (<=1kb) | 20 | 12654879 | 12673668 | 18790 | 2 | ENSMUST00 | 0 | 1810030O07 | ENSMUSG0( | false | chrX | 12672910 | 12674275 | 26.19333 | 73.09374 | 70.054 | . | . | . | . | . | . |
| chrX | 36904216 | 36904610 | 1000 | Promoter (1-2kb) | 20 | 36887540 | 36902899 | 15360 | 2 | ENSMUST00 | -1318 | Nkrf | ENSMUSG0( | false | . | . | . | . | . | . | . | . | . | . | . | . |
| chr6 | 65516804 | 65517238 | 1000 | Distal Intergenic | 6 | 65590398 | 65634040 | 43643 | 1 | ENSMUST00 | -73160 | Tnip3 | ENSMUSG0( | false | . | . | . | . | . | . | . | . | . | . | . | . |
| chr18 | 62652606 | 62652922 | 1000 | Promoter (2-3kb) | 18 | 62650504 | 62653475 | 2972 | 1 | ENSMUST00 | 2103 | Spink10 | ENSMUSG0( | false | . | . | . | . | . | . | chr18 | 62652593 | 62652877 | 3.07723 | 2.94952 | 1.43298 |
| chr11 | 94231657 | 94232489 | 1000 | Distal Intergenic | 11 | 94235956 | 94242707 | 6752 | 2 | ENSMUST00 | 10218 | Wfikkn2 | ENSMUSG0( | false | . | . | . | . | . | . | chr11 | 94224699 | 94234379 | 6.03156 | 4.54506 | 3.85595 |
| chr5 | 139389607 | 139390054 | 1000 | Promoter (<=1kb) | 5 | 139389785 | 139396414 | 6630 | 1 | ENSMUST00 | 0 | Gpr146 | ENSMUSG0( | false | . | . | . | . | . | . | chr5 | 139381256 | 139395525 | 8.99937 | 5.68291 | 6.29852 |
| chr5 | 139380244 | 139380979 | 1000 | Promoter (<=1kb) | 5 | 139380581 | 139396415 | 15835 | 1 | ENSMUST00 | 0 | Gpr146 | ENSMUSG0( | false | chr5 | 139380120 | 139383385 | 24.55236 | 90.57755 | 87.22021 | chr5 | 139375727 | 139380310 | 6.52375 | 4.83766 | 4.24296 |
| chr5 | 139378968 | 139379470 | 1000 | Promoter (<=1kb) | 5 | 139378220 | 139379259 | 1040 | 1 | ENSMUST00 | 749 | Gpr146 | ENSMUSG0( | false | . | . | . | . | . | . | chr5 | 139375727 | 139380310 | 6.52375 | 4.83766 | 4.24296 |
| chr5 | 139391540 | 139391981 | 1000 | Promoter (1-2kb) | 5 | 139389785 | 139396414 | 6630 | 1 | ENSMUST00 | 1756 | Gpr146 | ENSMUSG0( | false | . | . | . | . | . | . | chr5 | 139381256 | 139395525 | 8.99937 | 5.68291 | 6.29852 |
| chr15 | 10470102 | 10470813 | 1000 | Promoter (<=1kb) | 15 | 10446756 | 10470516 | 23761 | 2 | ENSMUST00 | 0 | Dnajc21 | ENSMUSG0( | false | chr15 | 10469446 | 10471180 | 33.62733 | 107.25218 | 103.49312 | . | . | . | . | . | . |
| chr3 | 85924170 | 85924766 | 1000 | Distal Intergenic | 3 | 85915783 | 85916070 | 288 | 1 | ENSMUST00 | 8388 | Gm9790 | ENSMUSG0( | false | . | . | . | . | . | . | chr3 | 85922138 | 85925692 | 5.66962 | 4.38034 | 3.6019 |
| chr19 | 42147124 | 42147898 | 1000 | Promoter (<=1kb) | 19 | 42147400 | 42151703 | 4304 | 1 | ENSMUST00 | 0 | Marveld1 | ENSMUSG0( | false | chr19 | 42147079 | 42151271 | 36.24507 | 114.92111 | 110.94343 | chr19 | 42144080 | 42147385 | 6.5634 | 4.79797 | 4.29474 |
| chr19 | 42148234 | 42148739 | 1000 | Promoter (<=1kb) | 19 | 42147708 | 42151462 | 3755 | 1 | ENSMUST00 | 527 | Marveld1 | ENSMUSG0( | false | chr19 | 42147079 | 42151271 | 36.24507 | 114.92111 | 110.94343 | . | . | . | . | . | . |
| chr14 | 76927465 | 76928149 | 1000 | Distal Intergenic | 14 | 77034950 | 77036081 | 1132 | 2 | ENSMUST00 | 107932 | Lacc1 | ENSMUSG0( | false | . | . | . | . | . | . | chr14 | 76926280 | 76931806 | 5.3758 | 4.23387 | 3.33603 |
| chr14 | 76912446 | 76913064 | 1000 | Distal Intergenic | 14 | 77034950 | 77036081 | 1132 | 2 | ENSMUST00 | 123017 | Lacc1 | ENSMUSG0( | false | . | . | . | . | . | . | chr14 | 76911062 | 76914215 | 5.4674 | 4.18211 | 3.419 |
| chr19 | 23134056 | 23136303 | 1000 | Promoter (<=1kb) | 19 | 23134900 | 23135717 | 818 | 1 | ENSMUST00 | 0 | 2410080I02R | ENSMUSG0( | false | chr19 | 23134219 | 23136818 | 17.93284 | 48.67312 | 45.97572 | chr19 | 23132889 | 23137841 | 4.72354 | 3.90939 | 2.78444 |
| chr17 | 10319175 | 10320507 | 1000 | Promoter (<=1kb) | 17 | 10319945 | 10320229 | 285 | 2 | ENSMUST00 | 0 | Qk | ENSMUSG0( | true | chr17 | 10319435 | 10321332 | 33.41831 | 117.26761 | 113.21259 | . | . | . | . | . | . |
| chr14 | 67924112 | 67924449 | 1000 | Intron (ENSMUST00000039135.4 | 14 | 67751928 | 67933572 | 181645 | 2 | ENSMUST00 | 9123 | Dock5 | ENSMUSG0( | true | . | . | . | . | . | . | chr14 | 67901014 | 67931267 | 5.24297 | 4.14115 | 3.21827 |
| chr14 | 67933175 | 67934003 | 1000 | Promoter (<=1kb) | 14 | 67751928 | 67933572 | 181645 | 2 | ENSMUST00 | 0 | Dock5 | ENSMUSG0( | true | chr14 | 67930568 | 67934296 | 26.72788 | 101.58999 | 97.97913 | chr14 | 67933844 | 67935659 | 3.93863 | 3.63998 | 2.11574 |
| chr3 | 100488593 | 100489683 | 1000 | Promoter (<=1kb) | 3 | 100451628 | 100489198 | 37571 | 2 | ENSMUST00 | 0 | Fam46c | ENSMUSG0( | false | chr3 | 100488212 | 100489978 | 34.42823 | 105.56979 | 101.85753 | . | . | . | . | . | . |
| chr6 | 31216243 | 31217116 | 1000 | Promoter (1-2kb) | 6 | 31166524 | 31218346 | 51823 | 2 | ENSMUST00 | 1230 | Lincpint | ENSMUSG0( | false | chr6 | 31215613 | 31219210 | 29.2142 | 114.47596 | 110.50954 | chr6 | 31203970 | 31216348 | 5.02803 | 4.03958 | 3.05299 |
| chr17 | 30156490 | 30156860 | 1000 | Intron (ENSMUST00000057897.7 | 17 | 30005087 | 30210019 | 204933 | 1 | ENSMUST00 | 151404 | Zfand3 | ENSMUSG0( | false | . | . | . | . | . | . | chr17 | 30154012 | 30159141 | 6.6536 | 4.8367 | 4.38922 |
| chr17 | 30004309 | 30005926 | 1000 | Promoter (<=1kb) | 17 | 30005087 | 30210019 | 204933 | 1 | ENSMUST00 | 0 | Zfand3 | ENSMUSG0( | false | chr17 | 30003745 | 30005950 | 37.83443 | 118.72395 | 114.62898 | . | . | . | . | . | . |
| chr4 | 147940780 | 147941035 | 1000 | Promoter (<=1kb) | 4 | 147940895 | 147947314 | 6420 | 1 | ENSMUST00 | 0 | 2510039O18 | ENSMUSG0( | false | chr4 | 147940819 | 147942100 | 42.27695 | 136.11826 | 131.39047 | chr4 | 147937366 | 147944112 | 5.71174 | 4.27591 | 3.6236 |
| chr11 | 31541135 | 31541416 | 1000 | Distal Intergenic | 11 | 31666487 | 31671727 | 5241 | 2 | ENSMUST00 | 130311 | Bod1 | ENSMUSG0( | false | . | . | . | . | . | . | chr11 | 31540815 | 31541446 | 4.04748 | 3.34406 | 2.26308 |
| chr17 | 56509777 | 56510130 | 1000 | Promoter (2-3kb) | 17 | 56511248 | 56512483 | 1236 | 2 | ENSMUST00 | 2353 | Znrf4 | ENSMUSG0( | false | . | . | . | . | . | . | chr17 | 56506710 | 56511612 | 4.49716 | 3.82906 | 2.61882 |
| chr14 | 70160632 | 70161121 | 1000 | Promoter (1-2kb) | 14 | 70154405 | 70159502 | 5098 | 2 | ENSMUST00 | -1131 | 9930012K11 | ENSMUSG0( | false | . | . | . | . | . | . | chr14 | 70156516 | 70175803 | 7.12683 | 5 | 4.75261 |
| chr6 | 112292971 | 112293754 | 1000 | Intron (ENSMUST00000032376.5 | 6 | 112285857 | 112288837 | 2981 | 2 | ENSMUST00 | -4135 | 5031434C07 | ENSMUSG0( | false | . | . | . | . | . | . | chr6 | 112291236 | 112293802 | 2.95763 | 2.91514 | 1.37364 |
| chr10 | 82319814 | 82320266 | 1000 | Distal Intergenic | 10 | 82282116 | 82285278 | 3163 | 2 | ENSMUST00 | -34537 | 4932415D10 | ENSMUSG0( | false | . | . | . | . | . | . | chr10 | 82319562 | 82320570 | 3.1736 | 3.11195 | 1.52132 |
| chr16 | 21984240 | 21984555 | 1000 | Exon (ENSMUST00000060673.6/ | 16 | 21953826 | 21995542 | 41717 | 2 | ENSMUST00 | 10987 | Liph | ENSMUSG0( | false | . | . | . | . | . | . | . | . | . | . | . | . |
| chr2 | 168096699 | 168097220 | 1000 | Intron (ENSMUST00000052125.6 | 2 | 168081004 | 168101203 | 20200 | 1 | ENSMUST00 | 15696 | Pard6b | ENSMUSG0( | false | . | . | . | . | . | . | chr2 | 168084045 | 168104695 | 5.60663 | 4.26449 | 3.52969 |
| chr2 | 168080499 | 168081390 | 1000 | Promoter (<=1kb) | 2 | 168081004 | 168101203 | 20200 | 1 | ENSMUST00 | 0 | Pard6b | ENSMUSG0( | false | chr2 | 168080352 | 168080877 | 13.37139 | 34.05986 | 31.54458 | chr2 | 168063587 | 168080595 | 6.53927 | 4.82888 | 4.26326 |
| chr18 | 76063025 | 76063523 | 1000 | Intron (ENSMUST00000058997.8 | 18 | 76059458 | 76146443 | 86986 | 1 | ENSMUST00 | 3568 | Zbtb7c | ENSMUSG0( | true | chr18 | 76063435 | 76064324 | 7.88522 | 15.75676 | 13.48224 | chr18 | 76061445 | 76132293 | 6.70015 | 4.79833 | 4.4158 |
| chr18 | 76022132 | 76022588 | 1000 | Intron (ENSMUST00000058997.8 | 18 | 76059458 | 76146443 | 86986 | 1 | ENSMUST00 | -36870 | Zbtb7c | ENSMUSG0( | true | . | . | . | . | . | . | chr18 | 76020071 | 76023231 | 4.04971 | 3.64011 | 2.21804 |
| chr18 | 76087465 | 76088530 | 1000 | Intron (ENSMUST00000058997.8 | 18 | 76059458 | 76146443 | 86986 | 1 | ENSMUST00 | 28008 | Zbtb7c | ENSMUSG0( | true | . | . | . | . | . | . | chr18 | 76061445 | 76132293 | 6.70015 | 4.79833 | 4.4158 |
| chr18 | 76083987 | 76085233 | 1000 | Intron (ENSMUST00000058997.8 | 18 | 76059458 | 76146443 | 86986 | 1 | ENSMUST00 | 24530 | Zbtb7c | ENSMUSG0( | true | . | . | . | . | . | . | chr18 | 76061445 | 76132293 | 6.70015 | 4.79833 | 4.4158 |
| chr18 | 76056990 | 76057887 | 1000 | Promoter (1-2kb) | 18 | 76059458 | 76146443 | 86986 | 1 | ENSMUST00 | -1571 | Zbtb7c | ENSMUSG0( | true | . | . | . | . | . | . | chr18 | 76050774 | 76058938 | 6.53532 | 4.72087 | 4.27307 |
| chr18 | 76096163 | 76096544 | 1000 | Intron (ENSMUST00000058997.8 | 18 | 76059458 | 76146443 | 86986 | 1 | ENSMUST00 | 36706 | Zbtb7c | ENSMUSG0( | true | . | . | . | . | . | . | chr18 | 76061445 | 76132293 | 6.70015 | 4.79833 | 4.4158 |
| chr18 | 75984277 | 75984658 | 1000 | Intron (ENSMUST00000058997.8 | 18 | 76059458 | 76146443 | 86986 | 1 | ENSMUST00 | -74800 | Zbtb7c | ENSMUSG0( | true | . | . | . | . | . | . | chr18 | 75979932 | 75986273 | 5.94014 | 4.47149 | 3.77873 |
| chr18 | 76010940 | 76011641 | 1000 | 5' UTR | 18 | 76059458 | 76146443 | 86986 | 1 | ENSMUST00 | -47817 | Zbtb7c | ENSMUSG0( | true | . | . | . | . | . | . | chr18 | 75999046 | 76019015 | 7.41965 | 5.13063 | 4.98856 |
| chr18 | 75819651 | 75820846 | 1000 | Promoter (<=1kb) | 18 | 75820178 | 76148561 | 328384 | 1 | ENSMUST00 | 0 | Zbtb7c | ENSMUSG0( | true | chr18 | 75818942 | 75821711 | 21.73903 | 87.55687 | 84.25908 | chr18 | 75820689 | 75827583 | 4.6164 | 3.88748 | 2.69497 |
| chr18 | 75983729 | 75984145 | 1000 | Intron (ENSMUST00000058997.8 | 18 | 76059458 | 76146443 | 86986 | 1 | ENSMUST00 | -75313 | Zbtb7c | ENSMUSG0( | true | . | . | . | . | . | . | chr18 | 75979932 | 75986273 | 5.94014 | 4.47149 | 3.77873 |
| chr1 | 63978768 | 63979174 | 1000 | Distal Intergenic | 1 | 63964727 | 63968999 | 4273 | 1 | ENSMUST00 | 14042 | Gm13749 | ENSMUSG0( | false | . | . | . | . | . | . | chr1 | 63977049 | 63981659 | 4.15039 | 3.64414 | 2.3046 |
| chr11 | 43549622 | 43550111 | 1000 | Intron (ENSMUST00000050574.6 | 11 | 43529246 | 43556689 | 27444 | 1 | ENSMUST00 | 20377 | Ccnjl | ENSMUSG0( | false | . | . | . | . | . | . | chr11 | 43548137 | 43550926 | 4.66358 | 3.65089 | 2.76867 |
| chr1 | 97660942 | 97662344 | 1000 | Promoter (<=1kb) | 1 | 97606318 | 97662074 | 55757 | 2 | ENSMUST00 | 0 | D1Ertd622e | ENSMUSG0( | false | chr1 | 97660633 | 97662645 | 37.09717 | 122.76616 | 118.52905 | . | . | . | . | . | . |
| chr10 | 42855245 | 42855657 | 1000 | Distal Intergenic | 10 | 42860370 | 42960780 | 100411 | 1 | ENSMUST00 | -4713 | Scml4 | ENSMUSG0( | false | . | . | . | . | . | . | chr10 | 42854998 | 42855784 | 3.97861 | 3.50352 | 2.20689 |
| chr7 | 28376305 | 28376687 | 1000 | Promoter (2-3kb) | 7 | 28376784 | 28379255 | 2472 | 2 | ENSMUST00 | 2568 | Zfp36 | ENSMUSG0( | false | chr7 | 28375275 | 28379742 | 28.38021 | 81.44218 | 78.26595 | chr7 | 28375103 | 28383860 | 6.23801 | 4.67039 | 4.01439 |
| chr7 | 28379180 | 28379753 | 1000 | Promoter (<=1kb) | 7 | 28376784 | 28379255 | 2472 | 2 | ENSMUST00 | 0 | Zfp36 | ENSMUSG0( | false | chr7 | 28375275 | 28379742 | 28.38021 | 81.44218 | 78.26595 | chr7 | 28375103 | 28383860 | 6.23801 | 4.67039 | 4.01439 |
| chr7 | 28381838 | 28382372 | 1000 | Promoter (2-3kb) | 7 | 28376784 | 28379255 | 2472 | 2 | ENSMUST00 | -2584 | Zfp36 | ENSMUSG0( | false | . | . | . | . | . | . | chr7 | 28375103 | 28383860 | 6.23801 | 4.67039 | 4.01439 |
| chr9 | 110531934 | 110533177 | 1000 | Promoter (<=1kb) | 9 | 110532597 | 110618633 | 86037 | 1 | ENSMUST00 | 0 | Setd2 | ENSMUSG0( | false | chr9 | 110531641 | 110533828 | 33.41831 | 117.26761 | 113.21259 | . | . | . | . | . | . |
| chr4 | 45473105 | 45473549 | 1000 | Intron (ENSMUST00000147926.2 | 4 | 45423278 | 45482802 | 59525 | 2 | ENSMUST00 | 9253 | Shb | ENSMUSG0( | false | . | . | . | . | . | . | chr4 | 45467198 | 45475428 | 4.97897 | 3.96606 | 3.0261 |
| chr5 | 64876111 | 64876414 | 1000 | Distal Intergenic | 5 | 64924811 | 64932761 | 7951 | 2 | ENSMUST00 | 56347 | Tlr1 | ENSMUSG0( | false | . | . | . | . | . | . | chr5 | 64875130 | 64879652 | 5.60891 | 4.1449 | 3.54722 |
| chr5 | 64884295 | 64884693 | 1000 | Distal Intergenic | 5 | 64924811 | 64932761 | 7951 | 2 | ENSMUST00 | 48068 | Tlr1 | ENSMUSG0( | false | . | . | . | . | . | . | chr5 | 64883738 | 64885565 | 4.50228 | 3.82342 | 2.59184 |
| chr17 | 27203983 | 27204638 | 1000 | Promoter (<=1kb) | 17 | 27189601 | 27204438 | 14838 | 2 | ENSMUST00 | 0 | Lemd2 | ENSMUSG0( | false | chr17 | 27202960 | 27204531 | 33.7938 | 119.00099 | 114.88904 | . | . | . | . | . | . |
| chr19 | 47918839 | 47919428 | 1000 | Promoter (<=1kb) | 19 | 47786580 | 47919287 | 132708 | 2 | ENSMUST00 | 0 | Wdr96 | ENSMUSG0( | false | chr19 | 47917779 | 47920263 | 28.22734 | 101.53345 | 97.92473 | . | . | . | . | . | . |
| chr11 | 32454975 | 32455810 | 1000 | Promoter (<=1kb) | 11 | 32455370 | 32516687 | 61318 | 1 | ENSMUST00 | 0 | Ubtd2 | ENSMUSG0( | false | chr11 | 32454239 | 32456649 | 46.03907 | 157.30389 | 151.74776 | . | . | . | . | . | . |
| chr11 | 32435656 | 32436031 | 1000 | Distal Intergenic | 11 | 32455370 | 32516687 | 61318 | 1 | ENSMUST00 | -19339 | Ubtd2 | ENSMUSG0( | false | . | . | . | . | . | . | chr11 | 32434576 | 32437630 | 3.79569 | 3.44362 | 2.05309 |
| chr1 | 64737337 | 64738067 | 1000 | Promoter (<=1kb) | 1 | 64730558 | 64737751 | 7194 | 2 | ENSMUST00 | 0 | Fzd5 | ENSMUSG0( | false | chr1 | 64737999 | 64739348 | 18.1972 | 47.5761 | 44.89159 | chr1 | 64738066 | 64739517 | 4.37992 | 3.84802 | 2.49167 |
| chr17 | 86167207 | 86168383 | 1000 | Promoter (<=1kb) | 17 | 86167785 | 86657919 | 490135 | 1 | ENSMUST00 | 0 | Prkce | ENSMUSG0( | true | chr17 | 86166332 | 86169593 | 32.75195 | 98.6424 | 95.10313 | . | . | . | . | . | . |
| chr17 | 86203694 | 86204534 | 1000 | Intron (ENSMUST00000097275.3 | 17 | 86168174 | 86655415 | 487242 | 1 | ENSMUST00 | 35521 | Prkce | ENSMUSG0( | true | . | . | . | . | . | . | chr17 | 86200784 | 86212339 | 6.33511 | 4.67161 | 4.1119 |
| chr7 | 25319124 | 25319454 | 1000 | Promoter (1-2kb) | 7 | 25317164 | 25365917 | 48754 | 1 | ENSMUST00 | 1961 | Megf8 | ENSMUSG0( | false | . | . | . | . | . | . | chr7 | 25314793 | 25322508 | 5.81353 | 4.33708 | 3.6977 |
| chr7 | 25316637 | 25317146 | 1000 | Promoter (<=1kb) | 7 | 25317164 | 25365917 | 48754 | 1 | ENSMUST00 | -18 | Megf8 | ENSMUSG0( | false | chr7 | 25316780 | 25318001 | 32.44658 | 100.11784 | 96.54536 | chr7 | 25314793 | 25322508 | 5.81353 | 4.33708 | 3.6977 |
| chr7 | 25322011 | 25322767 | 1000 | Intron (ENSMUST00000128119.1 | 7 | 25317164 | 25365917 | 48754 | 1 | ENSMUST00 | 4848 | Megf8 | ENSMUSG0( | false | . | . | . | . | . | . | chr7 | 25314793 | 25322508 | 5.81353 | 4.33708 | 3.6977 |
| chr12 | 85289205 | 85290065 | 1000 | Promoter (<=1kb) | 12 | 85288591 | 85299358 | 10768 | 1 | ENSMUST00 | 615 | Zc2hc1c | ENSMUSG0( | false | chr12 | 85287825 | 85289333 | 19.06563 | 48.28789 | 45.59428 | chr12 | 85287483 | 85292663 | 4.73802 | 4.00512 | 2.78959 |
| chr4 | 59625954 | 59627003 | 1000 | Promoter (<=1kb) | 4 | 59626211 | 59761439 | 135229 | 1 | ENSMUST00 | 0 | E130308A19 | ENSMUSG0( | true | chr4 | 59625627 | 59627522 | 46.54264 | 160.49254 | 154.82993 | chr4 | 59626815 | 59631370 | 3.34043 | 3.24063 | 1.64431 |
| chr5 | 143053633 | 143053995 | 1000 | Intron (ENSMUST00000053498.8 | 5 | 142990893 | 143113020 | 122128 | 2 | ENSMUST00 | 59025 | Rnf216 | ENSMUSG0( | false | . | . | . | . | . | . | chr5 | 143046832 | 143061805 | 7.05595 | 4.92101 | 4.70914 |
| chr4 | 36907825 | 36908172 | 1000 | Intron (ENSMUST00000108122.2 | 4 | 36718557 | 36950467 | 231911 | 2 | ENSMUST00 | 42295 | Lingo2 | ENSMUSG0( | true | . | . | . | . | . | . | chr4 | 36906724 | 36909523 | 4.2908 | 3.72166 | 2.41673 |
| chr6 | 94283587 | 94284131 | 1000 | Promoter (<=1kb) | 6 | 93675455 | 94283322 | 607868 | 2 | ENSMUST00 | -266 | Magi1 | ENSMUSG0( | false | chr6 | 94282364 | 94284879 | 24.86315 | 92.1349 | 88.74321 | . | . | . | . | . | . |

| | | | | | | | | | | | | | | | | | | | | | | | | | | |
|---|---|---|---|---|---|---|---|---|---|---|---|---|---|---|---|---|---|---|---|---|---|---|---|---|---|---|
| chr5 | 34169261 | 34169878 | 1000 | Promoter (<=1kb) | 5 | 34007198 | 34169448 | 162251 | 2 | ENSMUST00 | 0 | Poln | ENSMUSG0C | false | chr5 | 34169672 | 34169994 | 10.75319 | 23.25534 | 20.87728 | chr5 | 34168693 | 34173571 | 5.15082 | 4.13032 | 3.13292 |
| chr5 | 34170782 | 34171111 | 1000 | Promoter (1-2kb) | 5 | 34007198 | 34169448 | 162251 | 2 | ENSMUST00 | -1335 | Poln | ENSMUSG0C | false | . | . | . | . | . | . | chr5 | 34168693 | 34173571 | 5.15082 | 4.13032 | 3.13292 |
| chrX | 85048121 | 85048496 | 1000 | Promoter (<=1kb) | 20 | 85048295 | 85055597 | 7303 | 1 | ENSMUST00 | 0 | Dmd | ENSMUSG0C | false | chrX | 85047907 | 85049126 | 17.46222 | 42.53877 | 39.91948 | . | . | . | . | . | . |
| chr13 | 34118318 | 34118678 | 1000 | Distal Intergenic | 13 | 34127008 | 34130354 | 3347 | 2 | ENSMUST00 | 11676 | Tubb2b | ENSMUSG0C | false | . | . | . | . | . | . | chr13 | 34109058 | 34120340 | 5.26535 | 4.17937 | 3.22481 |
| chr10 | 80159666 | 80160412 | 1000 | Distal Intergenic | 10 | 80165985 | 80170316 | 4332 | 1 | ENSMUST00 | -5573 | Cirbp | ENSMUSG0C | false | chr10 | 80159097 | 80159881 | 9.74399 | 19.38118 | 17.0552 | chr10 | 80150859 | 80167428 | 9.09735 | 5.82258 | 6.36001 |
| chr10 | 80160810 | 80162100 | 1000 | Distal Intergenic | 10 | 80165985 | 80170316 | 4332 | 1 | ENSMUST00 | -3885 | Cirbp | ENSMUSG0C | false | chr10 | 80161186 | 80161536 | 10.48118 | 21.67722 | 19.31875 | chr10 | 80150859 | 80167428 | 9.09735 | 5.82258 | 6.36001 |
| chr14 | 70583705 | 70584042 | 1000 | 3' UTR | 14 | 70577847 | 70582571 | 4725 | 1 | ENSMUST00 | 5859 | Nudt18 | ENSMUSG0C | false | . | . | . | . | . | . | chr14 | 70583047 | 70585331 | 6.34664 | 4.72941 | 4.11177 |
| chr1 | 36068118 | 36068651 | 1000 | Promoter (<=1kb) | 1 | 36068400 | 36106446 | 38047 | 1 | ENSMUST00 | 0 | Hs6st1 | ENSMUSG0C | false | chr1 | 36067902 | 36069968 | 29.20317 | 85.69358 | 82.43262 | . | . | . | . | . | . |
| chr15 | 32042226 | 32042897 | 1000 | Distal Intergenic | 15 | 32177289 | 32178296 | 1008 | 1 | ENSMUST00 | -134392 | Tas2r119 | ENSMUSG0C | false | . | . | . | . | . | . | chr15 | 32033065 | 32044868 | 4.832 | 3.92577 | 2.87384 |
| chr11 | 75193494 | 75193999 | 1000 | Promoter (<=1kb) | 11 | 75193783 | 75267769 | 73987 | 1 | ENSMUST00 | 0 | Rtn4rl1 | ENSMUSG0C | false | chr11 | 75193308 | 75195378 | 19.27906 | 51.80829 | 49.07 | chr11 | 75192567 | 75195444 | 6.01616 | 4.51859 | 3.86626 |
| chr11 | 75255312 | 75255691 | 1000 | Intron (ENSMUST00000102514.3 | 11 | 75193783 | 75267769 | 73987 | 1 | ENSMUST00 | 61530 | Rtn4rl1 | ENSMUSG0C | false | . | . | . | . | . | . | . | . | . | . | . | . |
| chr5 | 35233227 | 35233598 | 1000 | Distal Intergenic | 5 | 35278566 | 35281761 | 3196 | 1 | ENSMUST00 | -44968 | Adra2c | ENSMUSG0C | false | . | . | . | . | . | . | chr5 | 35229105 | 35239186 | 4.86991 | 4.0349 | 2.9021 |
| chr5 | 35221202 | 35221638 | 1000 | Distal Intergenic | 5 | 35278566 | 35281761 | 3196 | 1 | ENSMUST00 | -56928 | Adra2c | ENSMUSG0C | false | . | . | . | . | . | . | chr5 | 35220655 | 35222602 | 4.29049 | 3.74688 | 2.41995 |
| chr3 | 96618184 | 96618663 | 1000 | Exon (ENSMUST00000062058.3/ | 3 | 96607624 | 96608725 | 1102 | 2 | ENSMUST00 | -9460 | 6330549D23 | ENSMUSG0C | false | . | . | . | . | . | . | chr3 | 96616884 | 96618370 | 3.67272 | 3.40311 | 1.94875 |
| chr8 | 87741930 | 87742420 | 1000 | Intron (ENSMUST00000165770.3 | 8 | 87783147 | 87803823 | 20677 | 2 | ENSMUST00 | 61403 | Zfp423 | ENSMUSG0C | true | . | . | . | . | . | . | chr8 | 87729485 | 87749127 | 6.31805 | 4.63416 | 4.1004 |
| chr8 | 87938352 | 87939453 | 1000 | Intron (ENSMUST00000109655.3 | 8 | 87783316 | 87944052 | 160737 | 2 | ENSMUST00 | 4599 | Zfp423 | ENSMUSG0C | true | chr8 | 87939025 | 87939348 | 6.38171 | 10.34615 | 8.16704 | chr8 | 87909819 | 87941142 | 6.06882 | 4.48607 | 3.90276 |
| chr8 | 87958461 | 87959867 | 1000 | Promoter (<=1kb) | 8 | 87661810 | 87959595 | 297786 | 2 | ENSMUST00 | 0 | Zfp423 | ENSMUSG0C | true | chr8 | 87957648 | 87961172 | 32.66734 | 113.81544 | 109.86345 | . | . | . | . | . | . |
| chr8 | 87933550 | 87934511 | 1000 | Intron (ENSMUST00000109655.3 | 8 | 87783316 | 87944052 | 160737 | 2 | ENSMUST00 | 9541 | Zfp423 | ENSMUSG0C | true | . | . | . | . | . | . | chr8 | 87909819 | 87941142 | 6.06882 | 4.48607 | 3.90276 |
| chr8 | 87960019 | 87960830 | 1000 | Promoter (<=1kb) | 8 | 87661810 | 87959595 | 297786 | 2 | ENSMUST00 | -425 | Zfp423 | ENSMUSG0C | true | chr8 | 87957648 | 87961172 | 32.66734 | 113.81544 | 109.86345 | chr8 | 87960719 | 87962469 | 6.18384 | 4.52485 | 4.00339 |
| chr8 | 87740926 | 87741289 | 1000 | Intron (ENSMUST00000165770.3 | 8 | 87783147 | 87803823 | 20677 | 2 | ENSMUST00 | 62534 | Zfp423 | ENSMUSG0C | true | . | . | . | . | . | . | chr8 | 87729485 | 87749127 | 6.31805 | 4.63416 | 4.1004 |
| chr8 | 87914200 | 87914791 | 1000 | Intron (ENSMUST00000109655.3 | 8 | 87661812 | 87895735 | 233924 | 2 | ENSMUST00 | -18466 | Zfp423 | ENSMUSG0C | true | . | . | . | . | . | . | chr8 | 87909819 | 87941142 | 6.06882 | 4.48607 | 3.90276 |
| chr12 | 99940771 | 99941124 | 1000 | Intron (ENSMUST00000153627.2 | 12 | 99964499 | 100062682 | 98184 | 1 | ENSMUST00 | -23375 | Kcnk13 | ENSMUSG0C | false | . | . | . | . | . | . | chr12 | 99937595 | 99942444 | 4.80356 | 3.84613 | 2.87857 |
| chr9 | 94537581 | 94538840 | 1000 | Promoter (<=1kb) | 9 | 94517864 | 94538081 | 20218 | 2 | ENSMUST00 | 0 | 1190002N15 | ENSMUSG0C | false | chr9 | 94536192 | 94539228 | 32.67316 | 105.57997 | 101.8659 | . | . | . | . | . | . |
| chr2 | 164454836 | 164456239 | 1000 | Promoter (<=1kb) | 2 | 164456964 | 164460936 | 3973 | 1 | ENSMUST00 | -725 | Sys1 | ENSMUSG0C | false | . | . | . | . | . | . | chr2 | 164447103 | 164460527 | 5.1383 | 4.18055 | 3.12116 |
| chr17 | 63863292 | 63864441 | 1000 | Promoter (<=1kb) | 17 | 63863300 | 63863791 | 492 | 2 | ENSMUST00 | 0 | A930002H24 | ENSMUSG0C | false | chr17 | 63862964 | 63864912 | 32.04747 | 138.90092 | 134.05882 | . | . | . | . | . | . |
| chr11 | 102946595 | 102946927 | 1000 | Promoter (<=1kb) | 11 | 102939264 | 102946688 | 7425 | 2 | ENSMUST00 | 0 | C1ql1 | ENSMUSG0C | false | chr11 | 102946065 | 102947328 | 11.31597 | 24.78923 | 22.38983 | chr11 | 102945019 | 102949394 | 5.6668 | 4.35893 | 3.56122 |
| chr18 | 61952543 | 61953511 | 1000 | Promoter (<=1kb) | 18 | 61953075 | 62015715 | 62641 | 1 | ENSMUST00 | 0 | Sh3tc2 | ENSMUSG0C | false | chr18 | 61952994 | 61954105 | 11.64009 | 27.76484 | 25.32713 | chr18 | 61946744 | 61959660 | 5.23742 | 4.1986 | 3.20658 |
| chr8 | 40963048 | 40963704 | 1000 | Intron (ENSMUST00000034004.9 | 8 | 40998770 | 41000205 | 1436 | 2 | ENSMUST00 | 36501 | Mtus1 | ENSMUSG0C | false | . | . | . | . | . | . | chr8 | 40961973 | 40966247 | 5.49667 | 4.32494 | 3.42147 |
| chr8 | 41039479 | 41039876 | 1000 | Promoter (<=1kb) | 8 | 41014376 | 41039587 | 25212 | 2 | ENSMUST00 | 0 | Mtus1 | ENSMUSG0C | false | chr8 | 41038344 | 41040438 | 17.22774 | 43.62438 | 40.99156 | chr8 | 41020996 | 41040121 | 5.92676 | 4.49745 | 3.7648 |
| chr8 | 41016456 | 41016905 | 1000 | Promoter (<=1kb) | 8 | 41000675 | 41016755 | 16081 | 2 | ENSMUST00 | 0 | Mtus1 | ENSMUSG0C | false | chr8 | 41016234 | 41017093 | 7.48194 | 15.52168 | 13.25109 | chr8 | 41003261 | 41019903 | 5.33155 | 4.20563 | 3.29685 |
| chr11 | 19923827 | 19924793 | 1000 | Promoter (<=1kb) | 11 | 19924375 | 20024026 | 99652 | 1 | ENSMUST00 | 0 | Spred2 | ENSMUSG0C | true | chr11 | 19923664 | 19926262 | 36.39542 | 121.53264 | 117.33772 | chr11 | 19923171 | 19923847 | 3.31596 | 3.15195 | 1.62218 |
| chr11 | 19919852 | 19920708 | 1000 | Distal Intergenic | 11 | 19924375 | 20024026 | 99652 | 1 | ENSMUST00 | -3667 | Spred2 | ENSMUSG0C | true | chr11 | 19919639 | 19920788 | 26.61343 | 84.02032 | 80.79139 | chr11 | 19919370 | 19921926 | 5.01188 | 4.15253 | 3.01706 |
| chr10 | 22875134 | 22875635 | 1000 | Distal Intergenic | 10 | 22817279 | 22820128 | 2850 | 2 | ENSMUST00 | -55007 | Tcf21 | ENSMUSG0C | true | . | . | . | . | . | . | chr10 | 22875585 | 22880104 | 5.38004 | 4.32991 | 3.30597 |
| chr10 | 22877795 | 22878235 | 1000 | Distal Intergenic | 10 | 22817279 | 22820128 | 2850 | 2 | ENSMUST00 | -57668 | Tcf21 | ENSMUSG0C | true | . | . | . | . | . | . | chr10 | 22875585 | 22880104 | 5.38004 | 4.32991 | 3.30597 |
| chr10 | 22813611 | 22813935 | 1000 | Distal Intergenic | 10 | 22817279 | 22820128 | 2850 | 2 | ENSMUST00 | 6193 | Tcf21 | ENSMUSG0C | true | . | . | . | . | . | . | chr10 | 22811405 | 22814144 | 3.60771 | 3.44295 | 1.84123 |
| chr10 | 22868297 | 22868932 | 1000 | Distal Intergenic | 10 | 22817279 | 22820128 | 2850 | 2 | ENSMUST00 | -48170 | Tcf21 | ENSMUSG0C | true | . | . | . | . | . | . | chr10 | 22866220 | 22874539 | 4.6225 | 3.87067 | 2.69769 |
| chr10 | 22843941 | 22844340 | 1000 | Distal Intergenic | 10 | 22817279 | 22820128 | 2850 | 2 | ENSMUST00 | -23814 | Tcf21 | ENSMUSG0C | true | chr10 | 22842940 | 22844360 | 10.44916 | 25.63548 | 23.2261 | chr10 | 22841048 | 22847295 | 4.04369 | 3.59873 | 2.2163 |
| chr7 | 142330156 | 142330726 | 1000 | Intron (ENSMUST00000131791.1 | 7 | 142325837 | 142338022 | 12186 | 2 | ENSMUST00 | 7296 | Ifitm10 | ENSMUSG0C | false | . | . | . | . | . | . | chr7 | 142329940 | 142330848 | 2.97938 | 2.87125 | 1.38149 |
| chr17 | 84187296 | 84188877 | 1000 | Promoter (<=1kb) | 17 | 84183931 | 84187947 | 4017 | 2 | ENSMUST00 | 0 | Zfp36l2 | ENSMUSG0C | false | chr17 | 84184922 | 84188829 | 24.14628 | 82.32003 | 79.12344 | chr17 | 84188257 | 84189216 | 4.57096 | 3.7402 | 2.66061 |
| chr10 | 95236164 | 95236526 | 1000 | Intron (ENSMUST00000053594.5 | 10 | 95174746 | 95324097 | 149352 | 2 | ENSMUST00 | 87571 | Cradd | ENSMUSG0C | false | . | . | . | . | . | . | chr10 | 95228500 | 95239876 | 5.61536 | 4.35143 | 3.5079 |
| chr10 | 95262470 | 95263244 | 1000 | Intron (ENSMUST00000053594.5 | 10 | 95174746 | 95324097 | 149352 | 2 | ENSMUST00 | 60853 | Cradd | ENSMUSG0C | false | . | . | . | . | . | . | chr10 | 95253598 | 95264609 | 4.86524 | 3.97711 | 2.9078 |
| chr6 | 120037683 | 120039433 | 1000 | Promoter (<=1kb) | 6 | 119923969 | 120038672 | 114704 | 2 | ENSMUST00 | 0 | Wnk1 | ENSMUSG0C | false | chr6 | 120035822 | 120038688 | 36.53154 | 123.91184 | 119.64436 | . | . | . | . | . | . |
| chr6 | 119981854 | 119982193 | 1000 | Promoter (<=1kb) | 6 | 119969565 | 119981448 | 11884 | 2 | ENSMUST00 | -407 | Wnk1 | ENSMUSG0C | false | . | . | . | . | . | . | chr6 | 119981378 | 119983281 | 4.27754 | 3.73987 | 2.42868 |
| chr2 | 153241259 | 153241725 | 1000 | Promoter (<=1kb) | 2 | 153241533 | 153270247 | 28715 | 1 | ENSMUST00 | 0 | Pofut1 | ENSMUSG0C | true | chr2 | 153239613 | 153242107 | 24.32174 | 95.77222 | 92.30251 | . | . | . | . | . | . |
| chr1 | 133130983 | 133131577 | 1000 | Promoter (<=1kb) | 1 | 133131143 | 133139783 | 8641 | 1 | ENSMUST00 | 0 | Ppp1r15b | ENSMUSG0C | false | chr1 | 133130586 | 133135428 | 34.54275 | 123.54154 | 119.28033 | . | . | . | . | . | . |
| chr13 | 98608059 | 98608518 | 1000 | Distal Intergenic | 13 | 98634978 | 98637410 | 2433 | 2 | ENSMUST00 | 28892 | Tmem174 | ENSMUSG0C | false | . | . | . | . | . | . | chr13 | 98607528 | 98608731 | 5.17215 | 4.22411 | 3.14768 |
| chr12 | 78983992 | 78984441 | 1000 | Promoter (<=1kb) | 12 | 78961795 | 78983478 | 21684 | 2 | ENSMUST00 | -515 | Tmem229b | ENSMUSG0C | false | . | . | . | . | . | . | chr12 | 78980486 | 78985561 | 5.38501 | 4.26618 | 3.32823 |
| chr17 | 9895172 | 9895643 | 1000 | Distal Intergenic | 17 | 9666497 | 9669704 | 3208 | 2 | ENSMUST00 | -225469 | Pabpc6 | ENSMUSG0C | false | . | . | . | . | . | . | chr17 | 9893324 | 9898140 | 4.38128 | 3.74587 | 2.51066 |
| chr17 | 3114515 | 3115758 | 1000 | Promoter (<=1kb) | 17 | 3114972 | 3198855 | 83884 | 1 | ENSMUST00 | 0 | Scaf8 | ENSMUSG0C | false | chr17 | 3113719 | 3116459 | 36.98618 | 117.25745 | 113.20857 | chr17 | 3115708 | 3120462 | 4.42595 | 3.77349 | 2.5382 |
| chr19 | 16780746 | 16781256 | 1000 | Promoter (<=1kb) | 19 | 16616763 | 16780822 | 164060 | 2 | ENSMUST00 | 0 | Vps13a | ENSMUSG0C | false | chr19 | 16779373 | 16781820 | 31.70051 | 127.6562 | 123.25787 | . | . | . | . | . | . |
| chrX | 7375701 | 7376165 | 1000 | Promoter (<=1kb) | 20 | 7371276 | 7375826 | 4551 | 2 | ENSMUST00 | 0 | Usp27x | ENSMUSG0C | false | chrX | 7374969 | 7376714 | 19.30035 | 48.68736 | 45.98855 | . | . | . | . | . | . |
| chr3 | 89865422 | 89865815 | 1000 | Distal Intergenic | 3 | 89831370 | 89858834 | 27465 | 1 | ENSMUST00 | 34053 | She | ENSMUSG0C | false | . | . | . | . | . | . | chr3 | 89858315 | 89886122 | 6.15651 | 4.57192 | 3.96514 |
| chr3 | 89858934 | 89859319 | 1000 | Distal Intergenic | 3 | 89831370 | 89858834 | 27465 | 1 | ENSMUST00 | 27565 | She | ENSMUSG0C | false | . | . | . | . | . | . | chr3 | 89858315 | 89886122 | 6.15651 | 4.57192 | 3.96514 |
| chr11 | 49203211 | 49203946 | 1000 | Promoter (<=1kb) | 11 | 49203292 | 49218553 | 15262 | 1 | ENSMUST00 | 0 | Zfp62 | ENSMUSG0C | false | chr11 | 49202015 | 49204776 | 21.75525 | 76.8157 | 73.7174 | . | . | . | . | . | . |
| chr11 | 62648082 | 62648900 | 1000 | Promoter (<=1kb) | 11 | 62604884 | 62648523 | 43640 | 2 | ENSMUST00 | 0 | Lrrc75a | ENSMUSG0C | false | chr11 | 62647832 | 62649284 | 26.84644 | 76.94898 | 73.84959 | chr11 | 62645300 | 62649776 | 3.45477 | 3.20498 | 1.78642 |
| chr11 | 87358399 | 87359255 | 1000 | Promoter (<=1kb) | 11 | 87226906 | 87359023 | 132118 | 2 | ENSMUST00 | 0 | Ppm1e | ENSMUSG0C | false | chr11 | 87357598 | 87359414 | 35.34781 | 110.24136 | 106.39787 | chr11 | 87357396 | 87359628 | 4.36603 | 3.77578 | 2.49651 |
| chr4 | 138453711 | 138455318 | 1000 | Promoter (<=1kb) | 4 | 138454314 | 138460123 | 5810 | 1 | ENSMUST00 | 0 | Camk2n1 | ENSMUSG0C | false | chr4 | 138453728 | 138456452 | 24.86315 | 92.1349 | 88.74321 | chr4 | 138453459 | 138454071 | 5.95408 | 4.54267 | 3.77407 |
| chr4 | 138463579 | 138464238 | 1000 | Distal Intergenic | 4 | 138454314 | 138460123 | 5810 | 1 | ENSMUST00 | 9266 | Camk2n1 | ENSMUSG0C | false | . | . | . | . | . | . | chr4 | 138463360 | 138465484 | 5.99771 | 4.45783 | 3.84992 |
| chr12 | 33717595 | 33717978 | 1000 | Distal Intergenic | 12 | 33928425 | 33929310 | 886 | 1 | ENSMUST00 | -210447 | Ferd3l | ENSMUSG0C | false | chr12 | 33717637 | 33718380 | 4.04026 | 6.18694 | 4.10202 | chr12 | 33714598 | 33719432 | 3.49213 | 3.2133 | 1.80507 |
| chr16 | 30654267 | 30654588 | 1000 | Distal Intergenic | 16 | 30599723 | 30602797 | 3075 | 1 | ENSMUST00 | 54545 | Fam43a | ENSMUSG0C | false | . | . | . | . | . | . | . | . | . | . | . | . |
| chr16 | 30691208 | 30691530 | 1000 | Distal Intergenic | 16 | 30599723 | 30602797 | 3075 | 1 | ENSMUST00 | 91486 | Fam43a | ENSMUSG0C | false | . | . | . | . | . | . | chr16 | 30691145 | 30692703 | 4.72215 | 4.00507 | 2.76653 |
| chr16 | 30599460 | 30599915 | 1000 | Promoter (<=1kb) | 16 | 30599723 | 30602797 | 3075 | 1 | ENSMUST00 | 0 | Fam43a | ENSMUSG0C | false | chr16 | 30599184 | 30602189 | 29.75681 | 92.45776 | 89.05831 | . | . | . | . | . | . |
| chr7 | 45052670 | 45053630 | 1000 | Promoter (<=1kb) | 7 | 45027707 | 45052881 | 25175 | 2 | ENSMUST00 | 0 | Prr12 | ENSMUSG0C | false | chr7 | 45052080 | 45054223 | 37.68163 | 117.40437 | 113.3432 | chr7 | 45051084 | 45052773 | 4.93289 | 4.12509 | 2.96638 |
| chr2 | 38933838 | 38934511 | 1000 | Promoter (1-2kb) | 2 | 38931978 | 38963753 | 31776 | 1 | ENSMUST00 | 1861 | Olfml2a | ENSMUSG0C | false | . | . | . | . | . | . | chr2 | 38931748 | 38936400 | 4.94393 | 3.83872 | 2.99043 |
| chr18 | 27292099 | 27292493 | 1000 | Distal Intergenic | 18 | 27598644 | 27599756 | 1113 | 1 | ENSMUST00 | -306151 | Gm7729 | ENSMUSG0C | false | . | . | . | . | . | . | chr18 | 27291415 | 27293201 | 3.2296 | 3.12301 | 1.58181 |
| chr18 | 35815452 | 35815931 | 1000 | Distal Intergenic | 18 | 35829818 | 35861687 | 31870 | 1 | ENSMUST00 | -13887 | Cxxc5 | ENSMUSG0C | false | chr18 | 35814542 | 35815564 | 10.17627 | 21.16558 | 18.81479 | chr18 | 35812838 | 35818671 | 7.77207 | 5.31683 | 5.31473 |
| chr18 | 35829477 | 35831321 | 1000 | Promoter (<=1kb) | 18 | 35829818 | 35861687 | 31870 | 1 | ENSMUST00 | 0 | Cxxc5 | ENSMUSG0C | false | chr18 | 35829308 | 35833126 | 34.46491 | 104.61457 | 100.92722 | chr18 | 35827671 | 35829647 | 7.07364 | 5.10378 | 4.71506 |
| chr4 | 98321324 | 98321637 | 1000 | Promoter (<=1kb) | 4 | 98321398 | 98322850 | 1453 | 1 | ENSMUST00 | 0 | 0610025J13 | ENSMUSG0C | false | chr4 | 98321436 | 98322450 | 23.23877 | 64.48248 | 61.57465 | chr4 | 98320771 | 98321500 | 3.44331 | 3.29961 | 1.7864 |
| chr3 | 127789762 | 127790810 | 1000 | Promoter (<=1kb) | 3 | 127789872 | 127798392 | 8521 | 1 | ENSMUST00 | 0 | Tifa | ENSMUSG0C | false | chr3 | 127789334 | 127791477 | 22.78592 | 76.10319 | 73.01487 | . | . | . | . | . | . |
| chr8 | 106892338 | 106892670 | 1000 | Promoter (<=1kb) | 8 | 106883894 | 106893215 | 9322 | 2 | ENSMUST00 | 545 | Chtf8 | ENSMUSG0C | false | chr8 | 106892093 | 106894289 | 38.08716 | 126.65268 | 122.28292 | chr8 | 106891657 | 106892371 | 6.24018 | 4.56004 | 4.07445 |
| chr8 | 95488533 | 95489059 | 1000 | Promoter (<=1kb) | 8 | 95446096 | 95488820 | 42725 | 2 | ENSMUST00 | 0 | Csnk2a2 | ENSMUSG0C | false | chr8 | 95487773 | 95489217 | 34.10395 | 106.03094 | 102.3112 | chr8 | 95489042 | 95489472 | 4.79859 | 3.81089 | 2.87016 |
| chr11 | 95508228 | 95508723 | 1000 | Downstream (1-2kb) | 11 | 95509845 | 95514570 | 4726 | 2 | ENSMUST00 | 5847 | Nxph3 | ENSMUSG0C | false | . | . | . | . | . | . | chr11 | 95489084 | 95513208 | 6.11718 | 4.48759 | 3.93601 |
| chr11 | 95504247 | 95505028 | 1000 | Distal Intergenic | 11 | 95509845 | 95514570 | 4726 | 2 | ENSMUST00 | 9542 | Nxph3 | ENSMUSG0C | false | . | . | . | . | . | . | chr11 | 95489084 | 95513208 | 6.11718 | 4.48759 | 3.93601 |
| chr11 | 95513479 | 95514892 | 1000 | Promoter (<=1kb) | 11 | 95509845 | 95514570 | 4726 | 2 | ENSMUST00 | 0 | Nxph3 | ENSMUSG0C | false | chr11 | 95512463 | 95515642 | 20.31581 | 74.8081 | 71.74177 | chr11 | 95514812 | 95517733 | 7.19932 | 5.16539 | 4.79467 |
| chr7 | 45313126 | 45313598 | 1000 | Intron (ENSMUST00000042194.8 | 7 | 45324941 | 45325651 | 711 | 1 | ENSMUST00 | -11343 | Rpl14-ps1 | ENSMUSG0C | false | . | . | . | . | . | . | chr7 | 45311729 | 45314737 | 5.49406 | 4.15388 | 3.45285 |
| chr18 | 36353691 | 36354153 | 1000 | Intron (ENSMUST00000139727.1 | 18 | 36348774 | 36393370 | 44597 | 1 | ENSMUST00 | 4918 | Cystm1 | ENSMUSG0C | false | . | . | . | . | . | . | chr18 | 36350295 | 36357415 | 4.45104 | 3.63325 | 2.59464 |
| chr11 | 69880837 | 69881570 | 1000 | Promoter (<=1kb) | 11 | 69878261 | 69881406 | 3146 | 2 | ENSMUST00 | 0 | Kctd11 | ENSMUSG0C | false | chr11 | 69879287 | 69881992 | 16.78262 | 53.63071 | 50.86883 | . | . | . | . | . | . |
| chr6 | 135048872 | 135049283 | 1000 | Distal Intergenic | 6 | 135065651 | 135084709 | 19059 | 1 | ENSMUST00 | -16368 | Gprc5a | ENSMUSG0C | false | . | . | . | . | . | . | chr6 | 135048398 | 135049564 | 4.94521 | 4.06897 | 2.95373 |
| chr3 | 38723445 | 38723932 | 1000 | Distal Intergenic | 3 | 38886940 | 39011985 | 125046 | 1 | ENSMUST00 | -163008 | Fat4 | ENSMUSG0C | false | . | . | . | . | . | . | . | . | . | . | . | . |
| chr12 | 83950439 | 83951126 | 1000 | Promoter (<=1kb) | 12 | 83950608 | 83952953 | 2346 | 1 | ENSMUST00 | 0 | 2410016O06 | ENSMUSG0C | false | chr12 | 83949954 | 83951946 | 28.56742 | 103.16996 | 99.51957 | . | . | . | . | . | . |
| chr5 | 5514288 | 5514937 | 1000 | Promoter (<=1kb) | 5 | 5505015 | 5514789 | 9775 | 2 | ENSMUST00 | 0 | Cldn12 | ENSMUSG0C | false | chr5 | 5513956 | 5515239 | 28.22734 | 101.53345 | 97.92473 | chr5 | 5510287 | 5515254 | 3.98273 | 3.53052 | 2.15744 |
| chr6 | 34874265 | 34874817 | 1000 | Promoter (2-3kb) | 6 | 34875346 | 34876980 | 1635 | 2 | ENSMUST00 | 2163 | 3110062M04 | ENSMUSG0C | false | . | . | . | . | . | . | chr6 | 34863195 | 34879034 | 5.53817 | 4.34013 | 3.44642 |
| chr6 | 34889181 | 34889561 | 1000 | Exon (ENSMUST00000081214.6/ | 6 | 34871840 | 34878060 | 6221 | 2 | ENSMUST00 | -11122 | 3110062M04 | ENSMUSG0C | false | . | . | . | . | . | . | chr6 | 34887903 | 34891132 | 3.62154 | 3.29537 | 1.91912 |
| chr5 | 72282943 | 72283348 | 1000 | Promoter (2-3kb) | 5 | 72280047 | 72298762 | 18716 | 1 | ENSMUST00 | 2897 | Atp10d | ENSMUSG0C | false | . | . | . | . | . | . | chr5 | 72280129 | 72285791 | 4.67903 | 3.84672 | 2.78486 |
| chr13 | 45963390 | 45964507 | 1000 | Promoter (<=1kb) | 13 | 45549758 | 45964991 | 415234 | 2 | ENSMUST00 | 484 | Atxn1 | ENSMUSG0C | false | chr13 | 45962952 | 45966515 | 39.39634 | 133.30592 | 128.68834 | . | . | . | . | . | . |
| chr18 | 83936540 | 83936867 | 1000 | Distal Intergenic | 18 | 84012082 | 84086390 | 74309 | 2 | ENSMUST00 | 149523 | Tshz1 | ENSMUSG0C | false | . | . | . | . | . | . | chr18 | 83936399 | 83937397 | 4.00391 | 3.54865 | 2.24524 |
| chr18 | 83840879 | 83841322 | 1000 | Distal Intergenic | 18 | 84012082 | 84086390 | 74309 | 2 | ENSMUST00 | 245068 | Tshz1 | ENSMUSG0C | false | . | . | . | . | . | . | chr18 | 83839199 | 83842091 | 5.41539 | 4.11829 | 3.38994 |
| chr5 | 44226351 | 44227202 | 1000 | Promoter (<=1kb) | 5 | 44175164 | 44226606 | 51443 | 2 | ENSMUST00 | 0 | Tapt1 | ENSMUSG0C | false | chr5 | 44225630 | 44227675 | 37.24078 | 123.01789 | 118.77389 | . | . | . | . | . | . |
| chr1 | 55236912 | 55237345 | 1000 | Promoter (<=1kb) | 1 | 55237177 | 55248470 | 11294 | 1 | ENSMUST00 | 0 | Mar-02 | ENSMUSG0C | false | chr1 | 55236585 | 55238734 | 35.37319 | 123.98246 | 119.71299 | . | . | . | . | . | . |
| chr9 | 96977888 | 96978512 | 1000 | Intron (ENSMUST00000055433.4 | 9 | 96943485 | 96984195 | 40711 | 2 | ENSMUST00 | 5683 | Spsb4 | ENSMUSG0C | false | . | . | . | . | . | . | chr9 | 96975779 | 96994993 | 5.8258 | 4.40484 | 3.6945 |
| chr9 | 97018102 | 97019128 | 1000 | Promoter (<=1kb) | 9 | 96943482 | 97018843 | 75362 | 2 | ENSMUST00 | 0 | Spsb4 | ENSMUSG0C | false | chr9 | 97017209 | 97019396 | 27.6016 | 85.77155 | 82.50918 | chr9 | 97016784 | 97019499 | 3.80073 | 3.30715 | 2.05208 |
| chr2 | 167476352 | 167476704 | 1000 | 3' UTR | 2 | 167484520 | 167485830 | 1311 | 2 | ENSMUST00 | 9126 | Spata2 | ENSMUSG0C | false | . | . | . | . | . | . | chr2 | 167475480 | 167478411 | 6.58875 | 4.60214 | 4.34695 |
| chrX | 142680689 | 142682068 | 1000 | Promoter (<=1kb) | 20 | 142681406 | 142843494 | 162089 | 1 | ENSMUST00 | 0 | Tmem164 | ENSMUSG0C | true | chrX | 142680507 | 142681054 | 9.65017 | 18.71579 | 16.39739 | . | . | . | . | . | . |
| chrX | 142834927 | 142835196 | 1000 | 3' UTR | 20 | 142825741 | 142838238 | 12498 | 1 | ENSMUST00 | 9187 | Tmem164 | ENSMUSG0C | true | . | . | . | . | . | . | . | . | . | . | . | . |
| chr17 | 56276605 | 56276996 | 1000 | Promoter (<=1kb) | 17 | 56269319 | 56276786 | 7468 | 2 | ENSMUST00 | 0 | Ticam1 | ENSMUSG0C | false | chr17 | 56276023 | 56277505 | 18.77877 | 48.57565 | 45.87926 | chr17 | 56272181 | 56277752 | 4.07905 | 3.60522 | 2.27287 |
| chr11 | 86757220 | 86758369 | 1000 | Promoter (<=1kb) | 11 | 86694351 | 86757565 | 63215 | 2 | ENSMUST00 | 0 | Cltc | ENSMUSG0C | false | chr11 | 86755991 | 86758872 | 42.56624 | 145.53995 | 140.43991 | . | . | . | . | . | . |
| chr14 | 55884708 | 55885267 | 1000 | Promoter (<=1kb) | 14 | 55884969 | 55896781 | 11813 | 1 | ENSMUST00 | 0 | Khnyn | ENSMUSG0C | false | chr14 | 55884665 | 55886221 | 35.71972 | 120.33891 | 116.19067 | chr14 | 55883960 | 55886527 | 3.22342 | 3.09535 | 1.59174 |
| chr3 | 16182938 | 16183475 | 1000 | Promoter (<=1kb) | 3 | 16183183 | 16217035 | 33853 | 1 | ENSMUST00 | 0 | Ythdf3 | ENSMUSG0C | false | chr3 | 16182684 | 16184590 | 38.11044 | 121.32513 | 117.13898 | . | . | . | . | . | . |
| chr12 | 64917712 | 64918316 | 1000 | Promoter (<=1kb) | 12 | 64917911 | 64924591 | 6681 | 1 | ENSMUST00 | 0 | Gm527 | ENSMUSG0C | false | chr12 | 64917326 | 64918820 | 32.41345 | 102.0761 | 98.45256 | . | . | . | . | . | . |
| chr2 | 36230289 | 36230710 | 1000 | Promoter (<=1kb) | 2 | 36230426 | 36252272 | 21847 | 1 | ENSMUST00 | 0 | Ptgs1 | ENSMUSG0C | false | chr2 | 36230166 | 36231210 | 12.74236 | 36.28267 | 33.73857 | chr2 | 36229934 | 36231701 | 4.98851 | 3.92601 | 3.03277 |
| chr11 | 69901473 | 69902022 | 1000 | Promoter (<=1kb) | 11 | 69901816 | 69913515 | 11700 | 1 | ENSMUST00 | 0 | Neurl4 | ENSMUSG0C | false | chr11 | 69900342 | 69902635 | 23.4661 | 79.19912 | 76.05865 | . | . | . | . | . | . |
| chr13 | 48027748 | 48028124 | 1000 | Distal Intergenic | 13 | 47960330 | 48018608 | 58279 | 2 | ENSMUST00 | -9141 | 4931429P17 | ENSMUSG0C | false | . | . | . | . | . | . | chr13 | 48026731 | 48028918 | 4.04056 | 3.64919 | 2.22115 |
| chr1 | 59602841 | 59603183 | 1000 | Intron (ENSMUST00000114243.2 | 1 | 59606782 | 59628261 | 21480 | 1 | ENSMUST00 | -3599 | Gm973 | ENSMUSG0C | false | . | . | . | . | . | . | chr1 | 59601911 | 59604957 | 5.32432 | 4.24379 | 3.27412 |
| chr1 | 59590678 | 59591023 | 1000 | Intron (ENSMUST00000114243.2 | 1 | 59606782 | 59628261 | 21480 | 1 | ENSMUST00 | -15759 | Gm973 | ENSMUSG0C | false | . | . | . | . | . | . | chr1 | 59588365 | 59593782 | 4.54249 | 3.78158 | 2.65371 |
| chr9 | 118926133 | 118927285 | 1000 | Promoter (<=1kb) | 9 | 118926453 | 119043998 | 117546 | 1 | ENSMUST00 | 0 | Ctdspl | ENSMUSG0C | true | chr9 | 118925107 | 118929928 | 34.45293 | 106.63475 | 102.89172 | . | . | . | . | . | . |
| chr9 | 118960288 | 118960814 | 1000 | Intron (ENSMUST00000073109.6 | 9 | 118929867 | 119033628 | 103762 | 1 | ENSMUST00 | 30422 | Ctdspl | ENSMUSG0C | true | . | . | . | . | . | . | chr9 | 118927678 | 119010843 | 6.89652 | 4.89961 | 4.57102 |
| chr9 | 118964479 | 118964823 | 1000 | Intron (ENSMUST00000073109.6 | 9 | 118929867 | 119033628 | 103762 | 1 | ENSMUST00 | 34613 | Ctdspl | ENSMUSG0C | true | . | . | . | . | . | . | chr9 | 118927678 | 119010843 | 6.89652 | 4.89961 | 4.57102 |
| chr9 | 118975065 | 118975810 | 1000 | Intron (ENSMUST00000073109.6 | 9 | 118929867 | 119033628 | 103762 | 1 | ENSMUST00 | 45199 | Ctdspl | ENSMUSG0C | true | . | . | . | . | . | . | chr9 | 118927678 | 119010843 | 6.89652 | 4.89961 | 4.57102 |
| chr9 | 118937477 | 118937891 | 1000 | Intron (ENSMUST00000073109.6 | 9 | 118929867 | 119033628 | 103762 | 1 | ENSMUST00 | 7611 | Ctdspl | ENSMUSG0C | true | . | . | . | . | . | . | chr9 | 118927678 | 119010843 | 6.89652 | 4.89961 | 4.57102 |
| chr12 | 95694940 | 95696159 | 1000 | Promoter (<=1kb) | 12 | 95695357 | 95781107 | 85751 | 1 | ENSMUST00 | 0 | Flrt2 | ENSMUSG0C | true | chr12 | 95691821 | 95698644 | 22.60491 | 86.65532 | 83.38163 | . | . | . | . | . | . |
| chr12 | 95746882 | 95747647 | 1000 | Intron (ENSMUST00000057324.3 | 12 | 95695357 | 95781107 | 85751 | 1 | ENSMUST00 | 51526 | Flrt2 | ENSMUSG0C | true | . | . | . | . | . | . | chr12 | 95729094 | 95751550 | 5.29594 | 4.20584 | 3.24965 |
| chr13 | 93138065 | 93138484 | 1000 | Intron (ENSMUST00000062122.3 | 13 | 93040713 | 93144724 | 104012 | 2 | ENSMUST00 | 6240 | Cmya5 | ENSMUSG0C | false | . | . | . | . | . | . | chr13 | 93136866 | 93139162 | 4.8944 | 3.94341 | 2.9335 |
| chr12 | 40037370 | 40038484 | 1000 | Promoter (<=1kb) | 12 | 40005447 | 40037376 | 31930 | 2 | ENSMUST00 | 0 | Arl4a | ENSMUSG0C | false | chr12 | 40036276 | 40038537 | 36.73951 | 123.20795 | 118.95689 | . | . | . | . | . | . |
| chr12 | 78225949 | 78226836 | 1000 | Promoter (<=1kb) | 12 | 78226655 | 78684767 | 458113 | 1 | ENSMUST00 | 0 | Gphn | ENSMUSG0C | false | chr12 | 78225441 | 78228015 | 33.69441 | 105.32094 | 101.61211 | . | . | . | . | . | . |
| chr15 | 11165867 | 11166167 | 1000 | Intron (ENSMUST00000061318.7 | 15 | 11064790 | 11346867 | 282078 | 1 | ENSMUST00 | 101078 | Adamts12 | ENSMUSG0C | false | . | . | . | . | . | . | chr15 | 11165308 | 11167383 | 5.46544 | 4.34769 | 3.38022 |
| chr19 | 41847456 | 41848549 | 1000 | Promoter (<=1kb) | 19 | 41845978 | 41848132 | 2155 | 2 | ENSMUST00 | 0 | Frat2 | ENSMUSG0C | false | chr19 | 41846750 | 41848862 | 31.91637 | 110.38313 | 106.53261 | . | . | . | . | . | . |
| chr10 | 8518329 | 8519183 | 1000 | Promoter (<=1kb) | 10 | 8204753 | 8518825 | 314073 | 2 | ENSMUST00 | 0 | Ust | ENSMUSG0C | true | chr10 | 8517801 | 8519623 | 39.51976 | 124.83012 | 120.52397 | . | . | . | . | . | . |
| chr7 | 126695234 | 126695535 | 1000 | Promoter (<=1kb) | 7 | 126695401 | 126696687 | 1287 | 1 | ENSMUST00 | 0 | Bola2 | ENSMUSG0C | false | chr7 | 126694880 | 126696774 | 34.0961 | 113.25239 | 109.32404 | chr7 | 126693450 | 126697330 | 4.90769 | 3.95971 | 2.94743 |
| chr19 | 46623109 | 46623529 | 1000 | Promoter (<=1kb) | 19 | 46623401 | 46652522 | 29122 | 1 | ENSMUST00 | 0 | Wbp1l | ENSMUSG0C | false | chr19 | 46623471 | 46624159 | 6.52658 | 13.36193 | 11.1274 | chr19 | 46599683 | 46650166 | 6.55215 | 4.7277 | 4.28832 |
| chr8 | 82939856 | 82940389 | 1000 | Intron (ENSMUST00000078525.5 | 8 | 82863356 | 83091271 | 227916 | 1 | ENSMUST00 | 76501 | Rnf150 | ENSMUSG0C | true | . | . | . | . | . | . | chr8 | 82936463 | 82941511 | 3.87884 | 3.42569 | 2.13119 |
| chr11 | 64434867 | 64435397 | 1000 | Promoter (<=1kb) | 11 | 64435332 | 64522841 | 87510 | 1 | ENSMUST00 | 0 | Hs3st3a1 | ENSMUSG0C | true | chr11 | 64434865 | 64438804 | 22.72043 | 98.85921 | 95.31703 | chr11 | 64433837 | 64435000 | 2.85084 | 2.77615 | 1.25095 |
| chr9 | 60687850 | 60688337 | 1000 | Promoter (<=1kb) | 9 | 60568859 | 60688124 | 119266 | 2 | ENSMUST00 | 0 | Lrrc49 | ENSMUSG0C | true | chr9 | 60687282 | 60688502 | 15.39584 | 39.00168 | 36.42487 | chr9 | 60687221 | 60688658 | 4.7687 | 4.02607 | 2.8171 |
| chr9 | 60649530 | 60649912 | 1000 | Promoter (<=1kb) | 9 | 60569172 | 60649801 | 80630 | 2 | ENSMUST00 | 0 | Lrrc49 | ENSMUSG0C | true | chr9 | 60648251 | 60649883 | 12.58327 | 33.30664 | 30.79935 | chr9 | 60643773 | 60650965 | 3.96179 | 3.59314 | 2.14865 |
| chr9 | 60637042 | 60637459 | 1000 | Intron (ENSMUST00000150060.2 | 9 | 60610270 | 60649710 | 39441 | 2 | ENSMUST00 | 12251 | Lrrc49 | ENSMUSG0C | true | . | . | . | . | . | . | chr9 | 60623926 | 60640440 | 4.81608 | 3.88284 | 2.88797 |
| chr4 | 151996043 | 151996957 | 1000 | Promoter (<=1kb) | 4 | 151989633 | 151996258 | 6626 | 2 | ENSMUST00 | 0 | Phf13 | ENSMUSG0C | false | chr4 | 151994455 | 151997533 | 32.98857 | 124.89146 | 120.57959 | . | . | . | . | . | . |
| chr19 | 7122971 | 7123374 | 1000 | Intron (ENSMUST00000040261.5 | 19 | 7092014 | 7105729 | 13716 | 2 | ENSMUST00 | -17243 | Flrt1 | ENSMUSG0C | false | . | . | . | . | . | . | chr19 | 7120136 | 7123619 | 8.00868 | 5.27125 | 5.51846 |
| chr19 | 7148242 | 7148584 | 1000 | Intron (ENSMUST00000040261.5 | 19 | 7092014 | 7105729 | 13716 | 2 | ENSMUST00 | -42514 | Flrt1 | ENSMUSG0C | false | . | . | . | . | . | . | chr19 | 7127706 | 7162730 | 7.6862 | 5.1249 | 5.24023 |
| chr19 | 7121248 | 7121707 | 1000 | Intron (ENSMUST00000040261.5 | 19 | 7092014 | 7105729 | 13716 | 2 | ENSMUST00 | -15520 | Flrt1 | ENSMUSG0C | false | . | . | . | . | . | . | chr19 | 7120136 | 7123619 | 8.00868 | 5.27125 | 5.51846 |
| chr19 | 7135415 | 7136574 | 1000 | Intron (ENSMUST00000040261.5 | 19 | 7092014 | 7105729 | 13716 | 2 | ENSMUST00 | -29687 | Flrt1 | ENSMUSG0C | false | . | . | . | . | . | . | chr19 | 7127706 | 7162730 | 7.6862 | 5.1249 | 5.24023 |
| chr15 | 57793289 | 57794108 | 1000 | Intron (ENSMUST00000096430.5 | 15 | 57736563 | 57738666 | 2104 | 2 | ENSMUST00 | -54624 | 9330154K18 | ENSMUSG0C | false | . | . | . | . | . | . | chr15 | 57790092 | 57797131 | 5.18328 | 3.90888 | 3.19255 |
| chr5 | 144254236 | 144256428 | 1000 | Promoter (<=1kb) | 5 | 144255225 | 144446757 | 191533 | 1 | ENSMUST00 | 0 | Bri3 | ENSMUSG0C | false | chr5 | 144254325 | 144259060 | 27.88725 | 99.90214 | 96.33501 | chr5 | 144253638 | 144254273 | 4.03563 | 3.594 | 2.25125 |
| chr18 | 10029677 | 10030269 | 1000 | Promoter (<=1kb) | 18 | 9995432 | 10030141 | 34710 | 2 | ENSMUST00 | 0 | Usp14 | ENSMUSG0C | true | chr18 | 10028791 | 10030110 | 26.86698 | 95.04005 | 91.5885 | . | . | . | . | . | . |
| chr5 | 63936661 | 63937138 | 1000 | Intron (ENSMUST00000154169.1 | 5 | 63924735 | 63930912 | 6178 | 2 | ENSMUST00 | -5750 | Rell1 | ENSMUSG0C | false | . | . | . | . | . | . | chr5 | 63936316 | 63938627 | 4.50457 | 3.60183 | 2.64051 |
| chr5 | 63968634 | 63969289 | 1000 | Promoter (<=1kb) | 5 | 63908896 | 63968905 | 60010 | 2 | ENSMUST00 | 0 | Rell1 | ENSMUSG0C | false | chr5 | 63967664 | 63969562 | 32.3084 | 121.498 | 117.30614 | . | . | . | . | . | . |
| chr15 | 80780930 | 80781228 | 1000 | Intron (ENSMUST00000067689.7 | 15 | 80711319 | 80941086 | 229768 | 1 | ENSMUST00 | 69612 | Tnrc6b | ENSMUSG0C | false | . | . | . | . | . | . | chr15 | 80779168 | 80781970 | 6.0326 | 4.52152 | 3.87951 |
| chr2 | 169632484 | 169632776 | 1000 | Promoter (<=1kb) | 2 | 169633013 | 169913093 | 280081 | 1 | ENSMUST00 | -237 | Tshz2 | ENSMUSG0C | true | . | . | . | . | . | . | chr2 | 169631871 | 169632608 | 4.2515 | 3.66469 | 2.39639 |
| chr2 | 169728595 | 169729098 | 1000 | Intron (ENSMUST00000109159.2 | 2 | 169633676 | 169888504 | 254829 | 1 | ENSMUST00 | 94920 | Tshz2 | ENSMUSG0C | true | . | . | . | . | . | . | chr2 | 169727082 | 169731791 | 5.74299 | 4.41842 | 3.62669 |
| chr15 | 73060931 | 73061381 | 1000 | Promoter (<=1kb) | 15 | 72589621 | 73061204 | 471584 | 2 | ENSMUST00 | 0 | Trappc9 | ENSMUSG0C | false | chr15 | 73060364 | 73061115 | 30.33627 | 112.57923 | 108.67202 | chr15 | 73059908 | 73062963 | 4.77387 | 3.96831 | 2.82032 |
| chr3 | 107035939 | 107036511 | 1000 | Promoter (<=1kb) | 3 | 107036162 | 107038129 | 1968 | 1 | ENSMUST00 | 0 | Kcna3 | ENSMUSG0C | false | chr3 | 107035703 | 107037207 | 20.65177 | 61.66082 | 58.78991 | chr3 | 107035744 | 107037240 | 4.3579 | 3.47714 | 2.51732 |
| chr7 | 107708636 | 107709076 | 1000 | Exon (ENSMUST00000040056.8/ | 7 | 107748455 | 107758032 | 9578 | 2 | ENSMUST00 | 48956 | Cyb5r2 | ENSMUSG0C | false | . | . | . | . | . | . | chr7 | 107704956 | 107711453 | 4.50562 | 3.70308 | 2.63801 |
| chr7 | 107752030 | 107752423 | 1000 | Exon (ENSMUST00000052438.7/ | 7 | 107748455 | 107758032 | 9578 | 2 | ENSMUST00 | 5609 | Cyb5r2 | ENSMUSG0C | false | . | . | . | . | . | . | chr7 | 107748148 | 107752802 | 5.36677 | 4.0977 | 3.32424 |
| chr7 | 107736343 | 107736910 | 1000 | Intron (ENSMUST00000040056.8 | 7 | 107748455 | 107758032 | 9578 | 2 | ENSMUST00 | 21122 | Cyb5r2 | ENSMUSG0C | false | . | . | . | . | . | . | chr7 | 107734046 | 107737802 | 4.69069 | 3.88568 | 2.75987 |
| chr7 | 96745955 | 96746372 | 1000 | Intron (ENSMUST00000107162.2 | 7 | 96777934 | 96797733 | 19800 | 1 | ENSMUST00 | -31562 | Tenm4 | ENSMUSG0C | false | . | . | . | . | . | . | chr7 | 96745559 | 96749957 | 5.32918 | 4.06898 | 3.31309 |
| chr7 | 96796438 | 96796740 | 1000 | Promoter (<=1kb) | 7 | 96796641 | 96798576 | 1936 | 1 | ENSMUST00 | 0 | Tenm4 | ENSMUSG0C | false | . | . | . | . | . | . | chr7 | 96795768 | 96798344 | 4.1797 | 3.60757 | 2.37214 |
| chr7 | 96747360 | 96747804 | 1000 | Intron (ENSMUST00000107162.2 | 7 | 96777934 | 96797733 | 19800 | 1 | ENSMUST00 | -30130 | Tenm4 | ENSMUSG0C | false | . | . | . | . | . | . | chr7 | 96745559 | 96749957 | 5.32918 | 4.06898 | 3.31309 |
| chr12 | 71016043 | 71017075 | 1000 | Promoter (<=1kb) | 12 | 71016708 | 71047339 | 30632 | 1 | ENSMUST00 | 0 | Arid4a | ENSMUSG0C | false | chr12 | 71015870 | 71017300 | 46.37146 | 163.67729 | 157.89816 | . | . | . | . | . | . |
| chr7 | 128739994 | 128740680 | 1000 | Promoter (<=1kb) | 7 | 128696441 | 128740495 | 44055 | 2 | ENSMUST00 | 0 | Mcmbp | ENSMUSG0C | false | chr7 | 128739046 | 128740883 | 37.76794 | 122.86759 | 118.62959 | . | . | . | . | . | . |
| chr2 | 4717440 | 4718706 | 1000 | Promoter (<=1kb) | 2 | 4717831 | 4800563 | 82733 | 1 | ENSMUST00 | 0 | Bend7 | ENSMUSG0C | false | chr2 | 4717210 | 4718922 | 37.60472 | 123.65472 | 119.39239 | chr2 | 4718549 | 4719495 | 3.5749 | 3.40003 | 1.86691 |
| chr7 | 141463780 | 141464178 | 1000 | Promoter (<=1kb) | 7 | 141464044 | 141464832 | 789 | 1 | ENSMUST00 | 0 | Efcab4a | ENSMUSG0C | false | . | . | . | . | . | . | chr7 | 141456108 | 141486560 | 6.62038 | 4.76228 | 4.33989 |
| chr15 | 97376699 | 97377271 | 1000 | Intron (ENSMUST00000059433.6 | 15 | 97244074 | 97247287 | 3214 | 2 | ENSMUST00 | -129413 | Amigo2 | ENSMUSG0C | false | chr15 | 97376804 | 97377279 | 8.63619 | 18.02194 | 15.71366 | chr15 | 97376312 | 97377838 | 5.0596 | 4.10055 | 3.11143 |
| chr11 | 85233597 | 85234196 | 1000 | Promoter (<=1kb) | 11 | 85234775 | 85235223 | 449 | 1 | ENSMUST00 | -579 | Appbp2os | ENSMUSG0C | false | chr11 | 85233236 | 85235725 | 29.28796 | 98.53321 | 94.99628 | chr11 | 85226558 | 85234087 | 4.20347 | 3.62744 | 2.38549 |
| chr11 | 85234377 | 85235093 | 1000 | Promoter (<=1kb) | 11 | 85234775 | 85235223 | 449 | 1 | ENSMUST00 | 0 | Appbp2os | ENSMUSG0C | false | chr11 | 85233236 | 85235725 | 29.28796 | 98.53321 | 94.99628 | . | . | . | . | . | . |
| chr17 | 26715270 | 26716774 | 1000 | Promoter (<=1kb) | 17 | 26715650 | 26776635 | 60986 | 1 | ENSMUST00 | 0 | Crebrf | ENSMUSG0C | false | chr17 | 26714994 | 26715492 | 16.04633 | 42.32132 | 39.70501 | . | . | . | . | . | . |
| chr12 | 70824719 | 70825797 | 1000 | Promoter (<=1kb) | 12 | 70825514 | 70902232 | 76719 | 1 | ENSMUST00 | 0 | Frmd6 | ENSMUSG0C | true | chr12 | 70824542 | 70825277 | 15.01947 | 39.77812 | 37.19162 | chr12 | 70823862 | 70824904 | 4.95523 | 4.05833 | 2.9524 |
| chr12 | 70799503 | 70799860 | 1000 | Distal Intergenic | 12 | 70825514 | 70902232 | 76719 | 1 | ENSMUST00 | -25654 | Frmd6 | ENSMUSG0C | true | . | . | . | . | . | . | chr12 | 70798228 | 70801529 | 6.77722 | 4.82948 | 4.46758 |
| chr15 | 36500420 | 36501100 | 1000 | Downstream (2-3kb) | 15 | 36477668 | 36496791 | 19124 | 2 | ENSMUST00 | -3630 | Ankrd46 | ENSMUSG0C | false | . | . | . | . | . | . | chr15 | 36487947 | 36503116 | 6.4169 | 4.70464 | 4.17261 |
| chr14 | 104336844 | 104337145 | 1000 | Distal Intergenic | 14 | 104463973 | 104465365 | 1393 | 2 | ENSMUST00 | 128220 | Pou4f1 | ENSMUSG0C | false | chr14 | 104336851 | 104337137 | 22.5292 | 69.37365 | 66.39268 | . | . | . | . | . | . |
| chr4 | 108327917 | 108328443 | 1000 | Promoter (<=1kb) | 4 | 108328140 | 108341544 | 13405 | 1 | ENSMUST00 | 0 | Coa7 | ENSMUSG0C | false | chr4 | 108327397 | 108328719 | 34.49972 | 105.69308 | 101.97717 | . | . | . | . | . | . |
| chr13 | 95638892 | 95639351 | 1000 | Intron (ENSMUST00000068603.6 | 13 | 95601804 | 95618459 | 16656 | 2 | ENSMUST00 | -20434 | F2r | ENSMUSG0C | false | . | . | . | . | . | . | chr13 | 95637625 | 95641811 | 6.80292 | 4.91506 | 4.48347 |
| chr16 | 25417942 | 25418633 | 1000 | Intron (ENSMUST00000056087.3 | 16 | 25286821 | 25318235 | 31415 | 1 | ENSMUST00 | 131122 | Tprg | ENSMUSG0C | false | . | . | . | . | . | . | chr16 | 25417874 | 25419475 | 4.33733 | 3.46919 | 2.49501 |
| chr4 | 141545852 | 141547184 | 1000 | Promoter (<=1kb) | 4 | 141546189 | 141547892 | 1704 | 1 | ENSMUST00 | 0 | B330016D10 | ENSMUSG0C | false | chr4 | 141544698 | 141545900 | 24.99264 | 71.03027 | 68.02251 | . | . | . | . | . | . |
| chr3 | 105704464 | 105705662 | 1000 | Promoter (<=1kb) | 3 | 105704599 | 105720842 | 16244 | 1 | ENSMUST00 | 0 | Fam212b | ENSMUSG0C | false | chr3 | 105704253 | 105705864 | 19.22456 | 54.94707 | 52.16912 | chr3 | 105704128 | 105707922 | 3.38042 | 3.08641 | 1.72251 |
| chr12 | 86974195 | 86974751 | 1000 | Distal Intergenic | 12 | 86983381 | 86988676 | 5296 | 2 | ENSMUST00 | 13925 | Zdhhc22 | ENSMUSG0C | false | chr12 | 86974173 | 86974966 | 13.56528 | 31.07685 | 28.5984 | chr12 | 86971639 | 86975722 | 5.78025 | 4.45948 | 3.65006 |
| chr4 | 129445166 | 129445740 | 1000 | Distal Intergenic | 4 | 129425765 | 129440853 | 15089 | 2 | ENSMUST00 | -4314 | Zbtb8b | ENSMUSG0C | false | chr4 | 129445146 | 129445784 | 8.69484 | 19.14606 | 16.82409 | chr4 | 129443250 | 129448315 | 4.28841 | 3.66336 | 2.43158 |
| chr16 | 76369939 | 76370332 | 1000 | Promoter (2-3kb) | 16 | 76328283 | 76373037 | 44755 | 2 | ENSMUST00 | 2705 | Nrip1 | ENSMUSG0C | false | . | . | . | . | . | . | chr16 | 76347307 | 76371866 | 5.01893 | 4.04265 | 3.02548 |
| chr16 | 76369369 | 76369657 | 1000 | Intron (ENSMUST00000121927.2 | 16 | 76328283 | 76373037 | 44755 | 2 | ENSMUST00 | 3380 | Nrip1 | ENSMUSG0C | false | . | . | . | . | . | . | chr16 | 76347307 | 76371866 | 5.01893 | 4.04265 | 3.02548 |
| chr11 | 115071839 | 115072433 | 1000 | Distal Intergenic | 11 | 115051917 | 115062177 | 10261 | 2 | ENSMUST00 | -9663 | Cd300e | ENSMUSG0C | false | . | . | . | . | . | . | chr11 | 115072046 | 115072990 | 4.5308 | 3.49172 | 2.65366 |
| chr9 | 44721213 | 44721643 | 1000 | Promoter (<=1kb) | 9 | 44716124 | 44721424 | 5301 | 2 | ENSMUST00 | 0 | Phldb1 | ENSMUSG0C | false | chr9 | 44720798 | 44721735 | 23.98789 | 70.18118 | 67.18839 | chr9 | 44719663 | 44729806 | 5.68645 | 4.29249 | 3.57992 |
| chr12 | 118836587 | 118837143 | 1000 | Distal Intergenic | 12 | 118846329 | 118852578 | 6250 | 1 | ENSMUST00 | -9186 | Sp8 | ENSMUSG0C | false | chr12 | 118836477 | 118837548 | 12.87622 | 39.27038 | 36.69063 | chr12 | 118836450 | 118837708 | 7.03546 | 4.96712 | 4.67671 |
| chr5 | 115157941 | 115158651 | 1000 | Promoter (<=1kb) | 5 | 115142981 | 115158179 | 15199 | 2 | ENSMUST00 | 0 | Mlec | ENSMUSG0C | false | chr5 | 115156953 | 115159326 | 36.42221 | 131.26616 | 126.73016 | . | . | . | . | . | . |
| chr14 | 57057486 | 57058272 | 1000 | Promoter (<=1kb) | 14 | 57035608 | 57058030 | 22423 | 2 | ENSMUST00 | 0 | Gja3 | ENSMUSG0C | false | chr14 | 57056396 | 57058454 | 32.01914 | 97.37792 | 93.87154 | chr14 | 57057793 | 57058946 | 5.34842 | 4.23338 | 3.29185 |
| chrX | 142656533 | 142656804 | 1000 | Distal Intergenic | 20 | 142631483 | 142634015 | 2533 | 2 | ENSMUST00 | -22519 | Gm5763 | ENSMUSG0C | false | . | . | . | . | . | . | . | . | . | . | . | . |
| chr19 | 38038190 | 38038556 | 1000 | Intron (ENSMUST00000041475.9 | 19 | 37899036 | 38043577 | 144542 | 2 | ENSMUST00 | 5021 | Myof | ENSMUSG0C | true | . | . | . | . | . | . | chr19 | 38020047 | 38042075 | 6.29391 | 4.65304 | 4.06409 |
| chr19 | 38025088 | 38025689 | 1000 | Intron (ENSMUST00000041475.9 | 19 | 37899036 | 38043577 | 144542 | 2 | ENSMUST00 | 17888 | Myof | ENSMUSG0C | true | . | . | . | . | . | . | chr19 | 38020047 | 38042075 | 6.29391 | 4.65304 | 4.06409 |
| chr19 | 38030671 | 38031177 | 1000 | Intron (ENSMUST00000041475.9 | 19 | 37899036 | 38043577 | 144542 | 2 | ENSMUST00 | 12400 | Myof | ENSMUSG0C | true | . | . | . | . | . | . | chr19 | 38020047 | 38042075 | 6.29391 | 4.65304 | 4.06409 |
| chr19 | 37881356 | 37881998 | 1000 | Distal Intergenic | 19 | 37899036 | 38043577 | 144542 | 2 | ENSMUST00 | 161579 | Myof | ENSMUSG0C | true | . | . | . | . | . | . | chr19 | 37875685 | 37888069 | 5.45448 | 4.31969 | 3.38198 |
| chr19 | 38041980 | 38042606 | 1000 | Promoter (<=1kb) | 19 | 37899036 | 38043577 | 144542 | 2 | ENSMUST00 | 971 | Myof | ENSMUSG0C | true | chr19 | 38041811 | 38043957 | 14.31595 | 48.63377 | 45.93701 | chr19 | 38020047 | 38042075 | 6.29391 | 4.65304 | 4.06409 |
| chr19 | 38043200 | 38043641 | 1000 | Promoter (<=1kb) | 19 | 37899036 | 38043577 | 144542 | 2 | ENSMUST00 | 0 | Myof | ENSMUSG0C | true | chr19 | 38041811 | 38043957 | 14.31595 | 48.63377 | 45.93701 | . | . | . | . | . | . |
| chr3 | 55132160 | 55132465 | 1000 | Intron (ENSMUST00000044116.8 | 3 | 55137339 | 55172935 | 35597 | 1 | ENSMUST00 | -4874 | Ccdc169 | ENSMUSG0C | false | . | . | . | . | . | . | chr3 | 55129111 | 55132546 | 4.68434 | 3.81851 | 2.79247 |
| chr6 | 67142133 | 67142544 | 1000 | Intron (ENSMUST00000181304.1 | 6 | 67165203 | 67165616 | 414 | 1 | ENSMUST00 | -22659 | Gm27015 | ENSMUSG0C | false | . | . | . | . | . | . | chr6 | 67123607 | 67146074 | 5.6205 | 4.38535 | 3.51128 |
| chr7 | 145219781 | 145220039 | 1000 | Distal Intergenic | 7 | 145253923 | 145283927 | 30005 | 2 | ENSMUST00 | 63888 | Tpcn2 | ENSMUSG0C | false | . | . | . | . | . | . | chr7 | 145219038 | 145220138 | 4.26137 | 3.6235 | 2.41345 |
| chr10 | 80387611 | 80388042 | 1000 | Promoter (<=1kb) | 10 | 80380355 | 80387659 | 7305 | 2 | ENSMUST00 | 0 | Mex3d | ENSMUSG0C | false | chr10 | 80385919 | 80387636 | 24.61517 | 67.71854 | 64.76461 | chr10 | 80383268 | 80388494 | 5.36762 | 4.25827 | 3.32111 |
| chr10 | 70149169 | 70149558 | 1000 | Intron (ENSMUST00000147545.2 | 10 | 70174509 | 70182206 | 7698 | 1 | ENSMUST00 | -24951 | Ccdc6 | ENSMUSG0C | false | . | . | . | . | . | . | chr10 | 70148720 | 70154033 | 6.46015 | 4.65202 | 4.2548 |
| chr10 | 70096480 | 70097594 | 1000 | Promoter (<=1kb) | 10 | 70097121 | 70193200 | 96080 | 1 | ENSMUST00 | 0 | Ccdc6 | ENSMUSG0C | false | chr10 | 70095817 | 70098606 | 30.78991 | 105.27288 | 101.56436 | chr10 | 70094512 | 70096518 | 5.05273 | 4.16827 | 3.0413 |
| chr10 | 42273888 | 42274726 | 1000 | Promoter (1-2kb) | 10 | 42196268 | 42276696 | 80429 | 2 | ENSMUST00 | 1970 | Foxo3 | ENSMUSG0C | false | chr10 | 42273937 | 42278830 | 36.72955 | 143.86444 | 138.82292 | chr10 | 42243354 | 42275609 | 6.05342 | 4.50564 | 3.88797 |
| chr10 | 42275212 | 42277218 | 1000 | Promoter (<=1kb) | 10 | 42181841 | 42276752 | 94912 | 2 | ENSMUST00 | 0 | Foxo3 | ENSMUSG0C | false | chr10 | 42273937 | 42278830 | 36.72955 | 143.86444 | 138.82292 | chr10 | 42243354 | 42275609 | 6.05342 | 4.50564 | 3.88797 |
| chr10 | 42204993 | 42205501 | 1000 | Intron (ENSMUST00000105502.2 | 10 | 42197554 | 42258366 | 60813 | 2 | ENSMUST00 | 52865 | Foxo3 | ENSMUSG0C | false | . | . | . | . | . | . | chr10 | 42198346 | 42216285 | 5.12772 | 4.0207 | 3.1378 |
| chr9 | 106418092 | 106418642 | 1000 | Distal Intergenic | 9 | 106429454 | 106431568 | 2115 | 1 | ENSMUST00 | -10812 | Rpl29 | ENSMUSG0C | false | chr9 | 106418104 | 106418716 | 8.85273 | 17.39095 | 15.09274 | chr9 | 106417277 | 106419225 | 6.67888 | 4.85392 | 4.41112 |
| chr4 | 117251718 | 117252294 | 1000 | Promoter (<=1kb) | 4 | 117251951 | 117268580 | 16630 | 1 | ENSMUST00 | 0 | Tmem53 | ENSMUSG0C | false | chr4 | 117251477 | 117252620 | 29.66345 | 100.21 | 96.63225 | chr4 | 117250852 | 117254641 | 4.1612 | 3.54286 | 2.35769 |
| chr3 | 108197033 | 108197603 | 1000 | Promoter (2-3kb) | 3 | 108198273 | 108200595 | 2323 | 2 | ENSMUST00 | 2992 | Cyb561d1 | ENSMUSG0C | false | . | . | . | . | . | . | chr3 | 108193497 | 108198144 | 7.42068 | 5.10456 | 4.99118 |
| chr18 | 53781364 | 53781742 | 1000 | Distal Intergenic | 18 | 53681724 | 53744547 | 62824 | 2 | ENSMUST00 | -36818 | Cep120 | ENSMUSG0C | false | . | . | . | . | . | . | chr18 | 53780123 | 53781420 | 3.44756 | 3.24127 | 1.74303 |
| chr18 | 53744340 | 53745098 | 1000 | Promoter (<=1kb) | 18 | 53681724 | 53744547 | 62824 | 2 | ENSMUST00 | 0 | Cep120 | ENSMUSG0C | false | chr18 | 53743447 | 53745668 | 31.14379 | 104.89771 | 101.20105 | . | . | . | . | . | . |
| chr11 | 103115948 | 103116639 | 1000 | Promoter (<=1kb) | 11 | 103116231 | 103119725 | 3495 | 1 | ENSMUST00 | 0 | Hexim1 | ENSMUSG0C | false | chr11 | 103114825 | 103116009 | 22.63194 | 64.29881 | 61.39256 | . | . | . | . | . | . |
| chr11 | 80476466 | 80477204 | 1000 | Promoter (<=1kb) | 11 | 80477023 | 80481184 | 4162 | 1 | ENSMUST00 | 0 | Cdk5r1 | ENSMUSG0C | false | chr11 | 80476339 | 80478051 | 22.78592 | 76.10319 | 73.01487 | chr11 | 80476244 | 80480069 | 5.82369 | 4.25853 | 3.71416 |
| chr7 | 80024653 | 80025492 | 1000 | Promoter (<=1kb) | 7 | 80024814 | 80092751 | 67938 | 1 | ENSMUST00 | 0 | Zfp710 | ENSMUSG0C | false | chr7 | 80024655 | 80028372 | 33.9579 | 107.82887 | 104.06033 | chr7 | 80023473 | 80025921 | 3.47992 | 3.28125 | 1.77065 |

| | | | | | | | | | | | | | | | | | | | | | | | | | | |
|---|---|---|---|---|---|---|---|---|---|---|---|---|---|---|---|---|---|---|---|---|---|---|---|---|---|---|
| chr7 | 80026107 | 80027673 | 1000 | Promoter (<=1kb) | 7 | 80026228 | 80092751 | 66524 | 1 | ENSMUST00 | 0 | Zfp710 | ENSMUSG0C | false | chr7 | 80024655 | 80028372 | 33.9579 | 107.82887 | 104.06033 | chr7 | 80027509 | 80037681 | 5.83309 | 4.41255 | 3.696 |
| chr4 | 119492386 | 119492838 | 1000 | Promoter (<=1kb) | 4 | 119465283 | 119492628 | 27346 | 2 | ENSMUST00 | 0 | Rimkla | ENSMUSG0C | false | chr4 | 119492004 | 119493020 | 23.14593 | 64.32256 | 61.41582 | chr4 | 119491899 | 119493046 | 3.39513 | 3.25875 | 1.72453 |
| chr2 | 131352336 | 131353121 | 1000 | Promoter (<=1kb) | 2 | 131298064 | 131352881 | 54818 | 2 | ENSMUST00 | 0 | Rnf24 | ENSMUSG0C | false | chr2 | 131351759 | 131353433 | 45.08069 | 160.24072 | 154.5827 | . | . | . | . | . | . |
| chr17 | 62882388 | 62883340 | 1000 | Promoter (1-2kb) | 17 | 62604184 | 62881317 | 277134 | 2 | ENSMUST00 | -1072 | Efna5 | ENSMUSG0C | false | chr17 | 62879599 | 62884335 | 19.57973 | 66.4542 | 63.51413 | . | . | . | . | . | . |
| chr7 | 143783879 | 143784415 | 1000 | Promoter (<=1kb) | 7 | 143778365 | 143784500 | 6136 | 2 | ENSMUST00 | 85 | Mrgpre | ENSMUSG0C | false | chr7 | 143783582 | 143784666 | 8.43118 | 16.20477 | 13.92443 | chr7 | 143782844 | 143786651 | 6.54139 | 4.84039 | 4.2661 |
| chr8 | 69935377 | 69935810 | 1000 | Exon (ENSMUST00000116463.2/ | 8 | 69887788 | 69902712 | 14925 | 2 | ENSMUST00 | -32666 | Yjefn3 | ENSMUSG0C | false | . | . | . | . | . | . | chr8 | 69918767 | 69945895 | 6.06816 | 4.46404 | 3.90949 |
| chr8 | 15046708 | 15047215 | 1000 | Promoter (<=1kb) | 8 | 15042502 | 15046078 | 3577 | 2 | ENSMUST00 | -631 | BB014433 | ENSMUSG0C | false | . | . | . | . | . | . | chr8 | 15043191 | 15051415 | 5.67584 | 4.41005 | 3.56153 |
| chr2 | 72238972 | 72239377 | 1000 | Promoter (2-3kb) | 2 | 72242294 | 72256655 | 14362 | 1 | ENSMUST00 | -2917 | Rapgef4 | ENSMUSG0C | false | . | . | . | . | . | . | chr2 | 72238558 | 72240566 | 5.53304 | 4.29348 | 3.4567 |
| chr18 | 84087327 | 84088732 | 1000 | Promoter (<=1kb) | 18 | 84088158 | 84097514 | 9357 | 1 | ENSMUST00 | 0 | Zadh2 | ENSMUSG0C | false | chr18 | 84084710 | 84090178 | 35.29575 | 125.98199 | 121.63858 | . | . | . | . | . | . |
| chr18 | 84089351 | 84089654 | 1000 | Promoter (1-2kb) | 18 | 84088158 | 84097514 | 9357 | 1 | ENSMUST00 | 1194 | Zadh2 | ENSMUSG0C | false | chr18 | 84084710 | 84090178 | 35.29575 | 125.98199 | 121.63858 | chr18 | 84089499 | 84093736 | 4.61079 | 3.91659 | 2.69639 |
| chr6 | 67480581 | 67480867 | 1000 | Intron (ENSMUST00000118364.1 | 6 | 67422932 | 67491855 | 68924 | 2 | ENSMUST00 | 10988 | Il23r | ENSMUSG0C | false | . | . | . | . | . | . | chr6 | 67479931 | 67481848 | 4.07624 | 3.65751 | 2.24293 |
| chr6 | 126034818 | 126035140 | 1000 | Intron (ENSMUST00000160496.2 | 6 | 126101413 | 126166744 | 65332 | 2 | ENSMUST00 | 131604 | Ntf3 | ENSMUSG0C | false | . | . | . | . | . | . | chr6 | 126033315 | 126036007 | 4.26705 | 3.72837 | 2.4162 |
| chr4 | 5643962 | 5644873 | 1000 | Promoter (<=1kb) | 4 | 5644090 | 5798382 | 154293 | 1 | ENSMUST00 | 0 | Fam110b | ENSMUSG0C | false | chr4 | 5643981 | 5645206 | 30.17592 | 94.17345 | 90.73757 | chr4 | 5644280 | 5645949 | 3.44295 | 3.28432 | 1.73561 |
| chr17 | 26101425 | 26101868 | 1000 | Promoter (2-3kb) | 17 | 26098872 | 26099257 | 386 | 2 | ENSMUST00 | -2169 | Gm8186 | ENSMUSG0C | false | . | . | . | . | . | . | chr17 | 26099235 | 26102429 | 4.01625 | 3.54732 | 2.21402 |
| chr18 | 60581435 | 60581760 | 1000 | Intron (ENSMUST00000056533.7 | 18 | 60576316 | 60591788 | 15473 | 2 | ENSMUST00 | 10028 | Myoz3 | ENSMUSG0C | false | . | . | . | . | . | . | chr18 | 60581472 | 60581702 | 3.7446 | 3.48658 | 1.9863 |
| chr4 | 11965651 | 11966814 | 1000 | Promoter (<=1kb) | 4 | 11958184 | 11966452 | 8269 | 2 | ENSMUST00 | 0 | Pdp1 | ENSMUSG0C | false | chr4 | 11965470 | 11967183 | 32.3084 | 121.498 | 117.30614 | . | . | . | . | . | . |
| chr6 | 59208687 | 59209151 | 1000 | Promoter (<=1kb) | 6 | 59208870 | 59212033 | 3164 | 1 | ENSMUST00 | 0 | Tigd2 | ENSMUSG0C | false | chr6 | 59208815 | 59209796 | 27.03867 | 103.18376 | 99.53125 | chr6 | 59208002 | 59210677 | 3.7582 | 3.44506 | 2.00064 |
| chr5 | 123879894 | 123880369 | 1000 | Promoter (<=1kb) | 5 | 123876736 | 123880020 | 3285 | 2 | ENSMUST00 | 0 | Hcar1 | ENSMUSG0C | false | chr5 | 123879475 | 123880321 | 7.50973 | 14.65382 | 12.39649 | chr5 | 123877917 | 123880369 | 5.53335 | 4.36017 | 3.4448 |
| chr5 | 30587892 | 30588324 | 1000 | Promoter (<=1kb) | 5 | 30588170 | 30625270 | 37101 | 1 | ENSMUST00 | 0 | Kcnk3 | ENSMUSG0C | false | chr5 | 30588173 | 30589030 | 14.66885 | 35.63733 | 33.10225 | chr5 | 30587932 | 30589804 | 8.30367 | 5.27746 | 5.73073 |
| chr9 | 40241561 | 40241862 | 1000 | Distal Intergenic | 9 | 40269216 | 40291617 | 22402 | 1 | ENSMUST00 | -27354 | Scn3b | ENSMUSG0C | false | chr9 | 40241510 | 40242244 | 13.14203 | 32.98289 | 30.47944 | chr9 | 40241043 | 40242388 | 4.07735 | 3.58083 | 2.30692 |
| chr11 | 59163302 | 59163973 | 1000 | Promoter (<=1kb) | 11 | 59155369 | 59163739 | 8371 | 2 | ENSMUST00 | 0 | Iba57 | ENSMUSG0C | false | chr11 | 59162784 | 59164239 | 34.32027 | 109.49174 | 105.67705 | chr11 | 59161523 | 59165816 | 3.79328 | 3.4717 | 2.00229 |
| chr5 | 124453691 | 124454073 | 1000 | Promoter (2-3kb) | 5 | 124451171 | 124460837 | 9667 | 1 | ENSMUST00 | 2521 | Setd8 | ENSMUSG0C | false | . | . | . | . | . | . | chr5 | 124446227 | 124457002 | 6.32983 | 4.7089 | 4.09735 |
| chr11 | 36495133 | 36495715 | 1000 | Promoter (<=1kb) | 11 | 36163719 | 36495139 | 331421 | 2 | ENSMUST00 | 0 | Tenm2 | ENSMUSG0C | false | . | . | . | . | . | . | chr11 | 36494200 | 36495977 | 3.86283 | 3.55946 | 2.07276 |
| chr11 | 37047676 | 37048033 | 1000 | Intron (ENSMUST00000102801.2 | 11 | 36139480 | 36944166 | 804687 | 2 | ENSMUST00 | -103511 | Tenm2 | ENSMUSG0C | false | . | . | . | . | . | . | chr11 | 37047178 | 37048618 | 3.17486 | 3.0821 | 1.55553 |
| chr2 | 180589246 | 180589648 | 1000 | Promoter (<=1kb) | 2 | 180589462 | 180593849 | 4388 | 1 | ENSMUST00 | 0 | Ogfr | ENSMUSG0C | false | chr2 | 180589025 | 180590552 | 43.41452 | 148.15424 | 142.93486 | . | . | . | . | . | . |
| chr18 | 12074009 | 12074419 | 1000 | Intron (ENSMUST00000050228.8 | 18 | 11981303 | 12121480 | 140178 | 2 | ENSMUST00 | 47061 | Tmem241 | ENSMUSG0C | false | . | . | . | . | . | . | chr18 | 12073733 | 12076817 | 3.4813 | 3.31016 | 1.80976 |
| chr18 | 11995917 | 11996463 | 1000 | Intron (ENSMUST00000050228.8 | 18 | 11981303 | 12121480 | 140178 | 2 | ENSMUST00 | 125017 | Tmem241 | ENSMUSG0C | false | . | . | . | . | . | . | chr18 | 11988526 | 11997770 | 5.05489 | 4.03677 | 3.06287 |
| chr11 | 53350238 | 53351235 | 1000 | Promoter (<=1kb) | 11 | 53350833 | 53421830 | 70998 | 1 | ENSMUST00 | 0 | Aff4 | ENSMUSG0C | false | chr11 | 53350131 | 53352443 | 33.07612 | 110.40385 | 106.55179 | . | . | . | . | . | . |
| chr7 | 47008090 | 47008555 | 1000 | Promoter (<=1kb) | 7 | 46990396 | 47008411 | 18016 | 2 | ENSMUST00 | 0 | Spty2d1 | ENSMUSG0C | false | chr7 | 47007232 | 47009188 | 33.08674 | 107.33151 | 103.57004 | . | . | . | . | . | . |
| chr15 | 78842257 | 78843165 | 1000 | Promoter (<=1kb) | 15 | 78842647 | 78850902 | 8256 | 1 | ENSMUST00 | 0 | Cdc42ep1 | ENSMUSG0C | false | chr15 | 78842114 | 78845803 | 33.10969 | 106.34647 | 102.61562 | chr15 | 78830112 | 78842533 | 8.19317 | 5.56805 | 5.60698 |
| chr15 | 78834211 | 78835064 | 1000 | Distal Intergenic | 15 | 78842647 | 78850902 | 8256 | 1 | ENSMUST00 | -7583 | Cdc42ep1 | ENSMUSG0C | false | . | . | . | . | . | . | chr15 | 78830112 | 78842533 | 8.19317 | 5.56805 | 5.60698 |
| chr14 | 52315916 | 52316515 | 1000 | Promoter (<=1kb) | 14 | 52311172 | 52316323 | 5152 | 2 | ENSMUST00 | 0 | Sall2 | ENSMUSG0C | false | chr14 | 52315306 | 52316762 | 28.95052 | 83.36136 | 80.14891 | chr14 | 52314721 | 52317160 | 4.64942 | 3.9716 | 2.71402 |
| chr13 | 70789108 | 70789442 | 1000 | Intron (ENSMUST00000080145.7 | 13 | 70727810 | 70841775 | 113966 | 2 | ENSMUST00 | 52333 | Adamts16 | ENSMUSG0C | false | . | . | . | . | . | . | chr13 | 70786727 | 70790846 | 5.46026 | 4.25496 | 3.39558 |
| chr6 | 71934229 | 71934557 | 1000 | Intron (ENSMUST00000055296.8 | 6 | 71909053 | 71979356 | 70304 | 1 | ENSMUST00 | 25177 | Polr1a | ENSMUSG0C | false | . | . | . | . | . | . | chr6 | 71933375 | 71935580 | 6.31518 | 4.52591 | 4.12079 |
| chr19 | 5567566 | 5568272 | 1000 | Promoter (<=1kb) | 19 | 5568074 | 5571261 | 3188 | 1 | ENSMUST00 | 0 | Ap5b1 | ENSMUSG0C | false | chr19 | 5566994 | 5568966 | 31.16539 | 106.97112 | 103.21663 | . | . | . | . | . | . |
| chr3 | 108770783 | 108771349 | 1000 | Intron (ENSMUST00000133931.3 | 3 | 108760214 | 108775002 | 14789 | 1 | ENSMUST00 | 10570 | Aknad1 | ENSMUSG0C | false | . | . | . | . | . | . | chr3 | 108769481 | 108771524 | 3.75878 | 3.41316 | 1.97538 |
| chr11 | 55077265 | 55077907 | 1000 | Promoter (<=1kb) | 11 | 55049731 | 55078107 | 28377 | 2 | ENSMUST00 | 200 | Ccdc69 | ENSMUSG0C | false | chr11 | 55077234 | 55078353 | 15.98415 | 46.70646 | 44.03467 | chr11 | 55075570 | 55078747 | 6.25261 | 4.74604 | 4.02264 |
| chr4 | 45012080 | 45012634 | 1000 | Promoter (<=1kb) | 4 | 44991242 | 45012394 | 21153 | 2 | ENSMUST00 | 0 | Zbtb5 | ENSMUSG0C | false | chr4 | 45010792 | 45012460 | 41.07773 | 137.35284 | 132.57864 | . | . | . | . | . | . |
| chr13 | 100103789 | 100104188 | 1000 | Promoter (<=1kb) | 13 | 100017994 | 100104070 | 86077 | 2 | ENSMUST00 | 0 | Bdp1 | ENSMUSG0C | false | chr13 | 100102898 | 100104640 | 31.16539 | 106.97112 | 103.21663 | chr13 | 100101947 | 100104857 | 4.3775 | 3.58797 | 2.52868 |
| chr11 | 20740933 | 20741993 | 1000 | Promoter (<=1kb) | 11 | 20685084 | 20741589 | 56506 | 2 | ENSMUST00 | 0 | Aftph | ENSMUSG0C | false | chr11 | 20740526 | 20742745 | 41.35789 | 132.3362 | 127.75856 | . | . | . | . | . | . |
| chr17 | 69382540 | 69384149 | 1000 | Promoter (<=1kb) | 17 | 69383050 | 69390750 | 7701 | 1 | ENSMUST00 | 0 | Zbtb14 | ENSMUSG0C | false | chr17 | 69382406 | 69384678 | 32.50483 | 115.59841 | 111.59217 | chr17 | 69382295 | 69382614 | 4.38243 | 3.68718 | 2.53186 |
| chr7 | 107664066 | 107665016 | 1000 | Intron (ENSMUST00000040056.8 | 7 | 107623971 | 107625161 | 1191 | 2 | ENSMUST00 | -38906 | 5330417H12I | ENSMUSG0C | false | chr7 | 107664446 | 107667691 | 20.9555 | 57.98476 | 55.16582 | chr7 | 107661425 | 107665686 | 5.62473 | 4.32581 | 3.51662 |
| chr7 | 107683197 | 107683525 | 1000 | Intron (ENSMUST00000040056.8 | 7 | 107623971 | 107625161 | 1191 | 2 | ENSMUST00 | -58037 | 5330417H12I | ENSMUSG0C | false | . | . | . | . | . | . | chr7 | 107666979 | 107703704 | 5.18106 | 4.08532 | 3.17145 |
| chrX | 167207645 | 167208059 | 1000 | Promoter (<=1kb) | 20 | 167207292 | 167208827 | 1536 | 2 | ENSMUST00 | 768 | Tmsb4x | ENSMUSG0C | false | chrX | 167206266 | 167210665 | 18.38128 | 45.59164 | 42.93153 | . | . | . | . | . | . |
| chr7 | 89434154 | 89434520 | 1000 | Distal Intergenic | 7 | 89404366 | 89410110 | 5745 | 1 | ENSMUST00 | 29789 | Fzd4 | ENSMUSG0C | false | . | . | . | . | . | . | chr7 | 89432530 | 89434699 | 4.06012 | 3.59748 | 2.2348 |
| chr11 | 20542534 | 20543427 | 1000 | Promoter (<=1kb) | 11 | 20543253 | 20597020 | 53768 | 1 | ENSMUST00 | 0 | Sertad2 | ENSMUSG0C | true | chr11 | 20543047 | 20544428 | 26.61085 | 108.21047 | 104.42454 | . | . | . | . | . | . |
| chr11 | 97441570 | 97442185 | 1000 | Intron (ENSMUST00000121799.2 | 11 | 97436285 | 97444228 | 7944 | 1 | ENSMUST00 | 5286 | Arhgap23 | ENSMUSG0C | true | chr11 | 97439863 | 97443213 | 40.55256 | 150.97545 | 145.64915 | chr11 | 97441626 | 97469759 | 6.61137 | 4.61316 | 4.36299 |
| chr11 | 97437657 | 97438334 | 1000 | Promoter (1-2kb) | 11 | 97436285 | 97444228 | 7944 | 1 | ENSMUST00 | 1373 | Arhgap23 | ENSMUSG0C | true | . | . | . | . | . | . | chr11 | 97431804 | 97440135 | 9.81823 | 6.18842 | 6.94298 |
| chr11 | 97404174 | 97404652 | 1000 | Distal Intergenic | 11 | 97415533 | 97502402 | 86870 | 1 | ENSMUST00 | -10881 | Arhgap23 | ENSMUSG0C | true | chr11 | 97404289 | 97405289 | 23.82055 | 66.35441 | 63.41718 | chr11 | 97403912 | 97406808 | 7.80875 | 5.43335 | 5.28241 |
| chr11 | 97449632 | 97450819 | 1000 | Promoter (<=1kb) | 11 | 97450160 | 97502400 | 52241 | 1 | ENSMUST00 | 0 | Arhgap23 | ENSMUSG0C | true | chr11 | 97450104 | 97452307 | 17.16547 | 48.86851 | 46.16669 | chr11 | 97441626 | 97469759 | 6.61137 | 4.61316 | 4.36299 |
| chr11 | 97439942 | 97440647 | 1000 | Intron (ENSMUST00000121799.2 | 11 | 97436285 | 97444228 | 7944 | 1 | ENSMUST00 | 3658 | Arhgap23 | ENSMUSG0C | true | chr11 | 97439863 | 97443213 | 40.55256 | 150.97545 | 145.64915 | chr11 | 97431804 | 97440135 | 9.81823 | 6.18842 | 6.94298 |
| chr17 | 34894039 | 34895224 | 1000 | Promoter (<=1kb) | 17 | 34894559 | 34896867 | 2309 | 1 | ENSMUST00 | 0 | Zbtb12 | ENSMUSG0C | false | chr17 | 34893816 | 34895421 | 24.3552 | 66.39809 | 63.45943 | chr17 | 34894965 | 34898038 | 4.84838 | 3.93846 | 2.88461 |
| chr1 | 157411767 | 157412962 | 1000 | Promoter (<=1kb) | 1 | 157412352 | 157420236 | 7885 | 1 | ENSMUST00 | 0 | 2810025M15 | ENSMUSG0C | false | chr1 | 157411007 | 157413312 | 36.53154 | 123.91184 | 119.64436 | . | . | . | . | . | . |
| chr12 | 69582648 | 69583293 | 1000 | Promoter (<=1kb) | 12 | 69577628 | 69583028 | 5401 | 2 | ENSMUST00 | 0 | Vcpkmt | ENSMUSG0C | false | chr12 | 69580317 | 69583372 | 28.69329 | 112.43184 | 108.52676 | chr12 | 69583141 | 69583630 | 3.06982 | 3.08617 | 1.41005 |
| chr12 | 69580851 | 69581178 | 1000 | Promoter (1-2kb) | 12 | 69577628 | 69583028 | 5401 | 2 | ENSMUST00 | 1850 | Vcpkmt | ENSMUSG0C | false | chr12 | 69580317 | 69583372 | 28.69329 | 112.43184 | 108.52676 | chr12 | 69573229 | 69581639 | 5.27043 | 4.12296 | 3.23088 |
| chr11 | 59964806 | 59965225 | 1000 | Promoter (<=1kb) | 11 | 59963181 | 59964944 | 1764 | 2 | ENSMUST00 | 0 | Rasd1 | ENSMUSG0C | false | chr11 | 59963913 | 59965603 | 25.81463 | 76.08353 | 72.99751 | chr11 | 59963508 | 59965831 | 4.33867 | 3.64315 | 2.47415 |
| chr6 | 28832112 | 28832912 | 1000 | Promoter (<=1kb) | 6 | 28828133 | 28831747 | 3615 | 2 | ENSMUST00 | -366 | Lrrc4 | ENSMUSG0C | false | chr6 | 28831812 | 28833824 | 33.66105 | 104.24397 | 100.57021 | chr6 | 28830957 | 28833786 | 3.57503 | 3.3004 | 1.88157 |
| chr14 | 20393300 | 20393902 | 1000 | Promoter (<=1kb) | 14 | 20391231 | 20393555 | 2325 | 2 | ENSMUST00 | 0 | Mrps16 | ENSMUSG0C | false | chr14 | 20392850 | 20394150 | 28.93225 | 84.27598 | 81.04385 | chr14 | 20392476 | 20394473 | 3.49408 | 3.35996 | 1.77558 |
| chr4 | 11021769 | 11022160 | 1000 | Distal Intergenic | 4 | 10990749 | 11007927 | 17179 | 2 | ENSMUST00 | -13843 | Plekhf2 | ENSMUSG0C | false | . | . | . | . | . | . | chr4 | 11020775 | 11023017 | 4.15624 | 3.59718 | 2.34125 |
| chr13 | 58157455 | 58157754 | 1000 | Promoter (<=1kb) | 13 | 58157649 | 58164693 | 7045 | 1 | ENSMUST00 | 0 | Idnk | ENSMUSG0C | false | chr13 | 58157315 | 58158795 | 29.66345 | 100.21 | 96.63225 | chr13 | 58156945 | 58158664 | 4.6555 | 3.73413 | 2.75906 |
| chr5 | 67607616 | 67608512 | 1000 | Promoter (<=1kb) | 5 | 67607883 | 67613992 | 6110 | 1 | ENSMUST00 | 0 | Shisa3 | ENSMUSG0C | false | chr5 | 67606389 | 67611595 | 34.3275 | 112.61823 | 108.70669 | . | . | . | . | . | . |
| chr5 | 67605911 | 67606282 | 1000 | Promoter (1-2kb) | 5 | 67607883 | 67613992 | 6110 | 1 | ENSMUST00 | -1601 | Shisa3 | ENSMUSG0C | false | . | . | . | . | . | . | . | . | . | . | . | . |
| chr5 | 67583526 | 67583858 | 1000 | Distal Intergenic | 5 | 67607883 | 67613992 | 6110 | 1 | ENSMUST00 | -24025 | Shisa3 | ENSMUSG0C | false | . | . | . | . | . | . | chr5 | 67578715 | 67584576 | 4.51681 | 3.77744 | 2.62069 |
| chr5 | 67581269 | 67581735 | 1000 | Distal Intergenic | 5 | 67607883 | 67613992 | 6110 | 1 | ENSMUST00 | -26148 | Shisa3 | ENSMUSG0C | false | . | . | . | . | . | . | chr5 | 67578715 | 67584576 | 4.51681 | 3.77744 | 2.62069 |
| chr5 | 111330395 | 111330864 | 1000 | Promoter (<=1kb) | 5 | 111330763 | 111388359 | 57597 | 1 | ENSMUST00 | 0 | Pitpnb | ENSMUSG0C | false | chr5 | 111330131 | 111331640 | 35.49035 | 116.75928 | 112.72716 | . | . | . | . | . | . |
| chr5 | 140777048 | 140777488 | 1000 | Intron (ENSMUST00000000153.4 | 5 | 140749283 | 140761439 | 12157 | 1 | ENSMUST00 | 27766 | Amz1 | ENSMUSG0C | false | . | . | . | . | . | . | chr5 | 140761728 | 140779292 | 6.36667 | 4.61767 | 4.14055 |
| chr10 | 85102265 | 85103574 | 1000 | Promoter (<=1kb) | 10 | 85097019 | 85102495 | 5477 | 2 | ENSMUST00 | 0 | Fhl4 | ENSMUSG0C | false | chr10 | 85101964 | 85103897 | 40.3785 | 139.45323 | 134.59462 | chr10 | 85103511 | 85105206 | 3.87055 | 3.54556 | 2.08109 |
| chr8 | 13974368 | 13975169 | 1000 | Promoter (<=1kb) | 8 | 13952009 | 13974777 | 22769 | 2 | ENSMUST00 | 0 | Tdrp | ENSMUSG0C | false | chr8 | 13972915 | 13975381 | 32.98857 | 124.89146 | 120.57959 | . | . | . | . | . | . |
| chr8 | 13957642 | 13958012 | 1000 | Intron (ENSMUST00000062613.5 | 8 | 13972616 | 13974743 | 2128 | 2 | ENSMUST00 | 16731 | Tdrp | ENSMUSG0C | false | . | . | . | . | . | . | chr8 | 13954846 | 13966705 | 5.11039 | 3.95144 | 3.12986 |
| chrX | 153497858 | 153498401 | 1000 | Promoter (<=1kb) | 20 | 153498227 | 153501570 | 3344 | 1 | ENSMUST00 | 0 | Ubqln2 | ENSMUSG0C | false | chrX | 153497465 | 153499227 | 17.46222 | 42.53877 | 39.91948 | . | . | . | . | . | . |
| chr4 | 126147467 | 126148954 | 1000 | Promoter (<=1kb) | 4 | 126147744 | 126149874 | 2131 | 1 | ENSMUST00 | 0 | Eva1b | ENSMUSG0C | false | chr4 | 126147596 | 126150575 | 31.96693 | 102.30585 | 98.67843 | chr4 | 126145657 | 126153251 | 4.91678 | 3.70661 | 2.96666 |
| chr8 | 36094598 | 36095025 | 1000 | Promoter (<=1kb) | 8 | 36094828 | 36147787 | 52960 | 1 | ENSMUST00 | 0 | D8Ertd82e | ENSMUSG0C | false | chr8 | 36094331 | 36095970 | 27.29656 | 83.33401 | 80.12321 | chr8 | 36093863 | 36096527 | 4.54244 | 3.90883 | 2.62567 |
| chr11 | 102606624 | 102607305 | 1000 | Promoter (2-3kb) | 11 | 102604396 | 102608058 | 3663 | 1 | ENSMUST00 | 2229 | Fzd2 | ENSMUSG0C | false | chr11 | 102603096 | 102607889 | 33.76198 | 105.43819 | 101.727 | . | . | . | . | . | . |
| chr13 | 31806286 | 31807373 | 1000 | Promoter (<=1kb) | 13 | 31806691 | 31810643 | 3953 | 1 | ENSMUST00 | 0 | Foxc1 | ENSMUSG0C | false | chr13 | 31805773 | 31812441 | 35.46717 | 120.9532 | 116.77927 | . | . | . | . | . | . |
| chr13 | 31811035 | 31811603 | 1000 | Distal Intergenic | 13 | 31806691 | 31810643 | 3953 | 1 | ENSMUST00 | 4345 | Foxc1 | ENSMUSG0C | false | chr13 | 31805773 | 31812441 | 35.46717 | 120.9532 | 116.77927 | . | . | . | . | . | . |
| chr13 | 31818484 | 31818955 | 1000 | Downstream (<1kb) | 13 | 31806691 | 31810643 | 3953 | 1 | ENSMUST00 | 11794 | Foxc1 | ENSMUSG0C | false | . | . | . | . | . | . | chr13 | 31812565 | 31835928 | 5.28848 | 4.1855 | 3.24535 |
| chr15 | 6707539 | 6708523 | 1000 | Promoter (<=1kb) | 15 | 6708381 | 6800398 | 92018 | 1 | ENSMUST00 | 0 | Rictor | ENSMUSG0C | false | chr15 | 6707265 | 6709859 | 31.49545 | 99.46503 | 95.90771 | chr15 | 6706764 | 6707709 | 3.00098 | 2.87493 | 1.35513 |
| chrX | 95444134 | 95445227 | 1000 | Promoter (<=1kb) | 20 | 95420318 | 95444872 | 24555 | 2 | ENSMUST00 | 0 | Amer1 | ENSMUSG0C | false | chrX | 95443457 | 95445327 | 17.46222 | 42.53877 | 39.91948 | . | . | . | . | . | . |
| chr2 | 164785532 | 164786307 | 1000 | Promoter (<=1kb) | 2 | 164785823 | 164792487 | 6665 | 1 | ENSMUST00 | 0 | Snx21 | ENSMUSG0C | false | chr2 | 164784984 | 164786753 | 24.24157 | 89.02564 | 85.70093 | . | . | . | . | . | . |
| chr4 | 129248141 | 129248708 | 1000 | Promoter (<=1kb) | 4 | 129219578 | 129248443 | 28866 | 2 | ENSMUST00 | 0 | C77080 | ENSMUSG0C | false | chr4 | 129247693 | 129249005 | 28.6362 | 83.76622 | 80.54601 | chr4 | 129245280 | 129249936 | 4.20173 | 3.67127 | 2.35538 |
| chr7 | 19131312 | 19131810 | 1000 | Promoter (<=1kb) | 7 | 19131693 | 19137812 | 6120 | 1 | ENSMUST00 | 0 | Fbxo46 | ENSMUSG0C | false | chr7 | 19131078 | 19131373 | 6.83737 | 14.36054 | 12.10883 | chr7 | 19128141 | 19133343 | 4.70851 | 3.90871 | 2.78695 |
| chr2 | 50066202 | 50066745 | 1000 | Promoter (<=1kb) | 2 | 50066429 | 50193569 | 127141 | 1 | ENSMUST00 | 0 | Lypd6 | ENSMUSG0C | false | chr2 | 50065979 | 50067249 | 21.51301 | 61.48686 | 58.61925 | chr2 | 50065907 | 50067245 | 4.47634 | 3.72009 | 2.61746 |
| chr1 | 121540495 | 121540831 | 1000 | Distal Intergenic | 1 | 121509842 | 121528465 | 18624 | 2 | ENSMUST00 | -12031 | Htr5b | ENSMUSG0C | false | . | . | . | . | . | . | chr1 | 121540319 | 121543510 | 4.07973 | 3.50017 | 2.29143 |
| chr3 | 127895737 | 127896543 | 1000 | Promoter (<=1kb) | 3 | 127869688 | 127896323 | 26636 | 2 | ENSMUST00 | 0 | 5730508B09I | ENSMUSG0C | false | chr3 | 127895360 | 127896376 | 24.14628 | 82.32003 | 79.12344 | . | . | . | . | . | . |
| chr11 | 116433763 | 116434809 | 1000 | Promoter (<=1kb) | 11 | 116434094 | 116439077 | 4984 | 1 | ENSMUST00 | 0 | Ubald2 | ENSMUSG0C | false | chr11 | 116434028 | 116435508 | 17.23216 | 63.31984 | 60.4264 | . | . | . | . | . | . |
| chr5 | 100158645 | 100159669 | 1000 | Promoter (<=1kb) | 5 | 100083946 | 100159432 | 75487 | 2 | ENSMUST00 | 0 | Tmem150c | ENSMUSG0C | true | chr5 | 100156800 | 100160103 | 34.80912 | 107.25002 | 103.49115 | chr5 | 100159061 | 100166883 | 5.23434 | 4.17007 | 3.22231 |
| chr15 | 5143488 | 5144411 | 1000 | Promoter (<=1kb) | 15 | 5143861 | 5181899 | 38039 | 1 | ENSMUST00 | 0 | Prkaa1 | ENSMUSG0C | false | chr15 | 5143151 | 5144825 | 35.94841 | 113.36032 | 109.43 | chr15 | 5142410 | 5146323 | 4.09737 | 3.45193 | 2.27341 |
| chr9 | 65579933 | 65580598 | 1000 | Promoter (<=1kb) | 9 | 65554386 | 65580040 | 25655 | 2 | ENSMUST00 | 0 | Plekho2 | ENSMUSG0C | false | chr9 | 65579586 | 65581133 | 21.92336 | 59.65632 | 56.81357 | chr9 | 65578267 | 65580867 | 2.97788 | 2.92823 | 1.36512 |
| chr6 | 72400150 | 72400495 | 1000 | Distal Intergenic | 6 | 72385223 | 72390703 | 5481 | 2 | ENSMUST00 | -9448 | Vamp8 | ENSMUSG0C | false | . | . | . | . | . | . | chr6 | 72395743 | 72402273 | 5.15999 | 4.09221 | 3.15674 |
| chr6 | 72389891 | 72390756 | 1000 | Promoter (<=1kb) | 6 | 72385223 | 72390703 | 5481 | 2 | ENSMUST00 | 0 | Vamp8 | ENSMUSG0C | false | chr6 | 72389183 | 72391497 | 20.5555 | 53.19651 | 50.4419 | chr6 | 72385690 | 72394714 | 4.39484 | 3.72006 | 2.54009 |
| chr4 | 58911616 | 58913016 | 1000 | Promoter (<=1kb) | 4 | 58798911 | 58912749 | 113839 | 2 | ENSMUST00 | 0 | AI314180 | ENSMUSG0C | false | chr4 | 58910802 | 58913238 | 34.54477 | 122.48211 | 118.25522 | . | . | . | . | . | . |
| chr17 | 25518261 | 25518559 | 1000 | Distal Intergenic | 17 | 25489875 | 25497288 | 7414 | 2 | ENSMUST00 | -20974 | Sstr5 | ENSMUSG0C | false | . | . | . | . | . | . | chr17 | 25516170 | 25521099 | 4.97676 | 3.80972 | 3.0233 |
| chr4 | 118553070 | 118553662 | 1000 | Downstream (<1kb) | 4 | 118541859 | 118544044 | 2186 | 2 | ENSMUST00 | -9027 | Tmem125 | ENSMUSG0C | false | . | . | . | . | . | . | chr4 | 118552510 | 118555443 | 4.99368 | 3.92604 | 3.02233 |
| chr4 | 134287159 | 134287567 | 1000 | Promoter (<=1kb) | 4 | 134275002 | 134287239 | 12238 | 2 | ENSMUST00 | 0 | Pdik1l | ENSMUSG0C | false | chr4 | 134286315 | 134288340 | 26.77612 | 116.44383 | 112.41551 | . | . | . | . | . | . |
| chr18 | 5390171 | 5390655 | 1000 | Distal Intergenic | 18 | 5210031 | 5334439 | 124409 | 2 | ENSMUST00 | -55733 | Zfp438 | ENSMUSG0C | false | . | . | . | . | . | . | chr18 | 5389082 | 5390893 | 3.12696 | 2.99917 | 1.50556 |
| chr10 | 56474981 | 56475349 | 1000 | Distal Intergenic | 10 | 56377300 | 56390419 | 13120 | 1 | ENSMUST00 | 97682 | Gja1 | ENSMUSG0C | false | . | . | . | . | . | . | chr10 | 56474298 | 56475126 | 6.3993 | 4.85826 | 4.12913 |
| chr11 | 108343671 | 108344409 | 1000 | Promoter (<=1kb) | 11 | 107933387 | 108343928 | 410542 | 2 | ENSMUST00 | 0 | Prkca | ENSMUSG0C | false | chr11 | 108342513 | 108344600 | 38.27806 | 124.82941 | 120.52397 | . | . | . | . | . | . |
| chr7 | 141413978 | 141414531 | 1000 | Promoter (<=1kb) | 7 | 141408135 | 141414090 | 5956 | 2 | ENSMUST00 | 0 | Pddc1 | ENSMUSG0C | false | chr7 | 141413270 | 141414145 | 24.55236 | 90.57755 | 87.22021 | chr7 | 141412445 | 141415130 | 4.2922 | 3.59313 | 2.46905 |
| chr11 | 60115022 | 60115557 | 1000 | Promoter (<=1kb) | 11 | 60112219 | 60115230 | 3012 | 2 | ENSMUST00 | 0 | 4930412M03I | ENSMUSG0C | false | chr11 | 60114432 | 60115777 | 14.00966 | 37.26205 | 34.70494 | chr11 | 60106353 | 60117543 | 5.28818 | 4.15483 | 3.25623 |
| chr11 | 114846409 | 114846739 | 1000 | Distal Intergenic | 11 | 114851152 | 114863733 | 12582 | 1 | ENSMUST00 | -4413 | Gprc5c | ENSMUSG0C | false | . | . | . | . | . | . | . | . | . | . | . | . |
| chr11 | 114866921 | 114867532 | 1000 | Promoter (2-3kb) | 11 | 114864492 | 114870607 | 6116 | 1 | ENSMUST00 | 2430 | Gprc5c | ENSMUSG0C | false | . | . | . | . | . | . | chr11 | 114865142 | 114868789 | 5.38654 | 4.23201 | 3.33703 |
| chr16 | 28929488 | 28930119 | 1000 | Promoter (<=1kb) | 16 | 28826176 | 28929698 | 103523 | 2 | ENSMUST00 | 0 | Mb21d2 | ENSMUSG0C | false | chr16 | 28928458 | 28930569 | 36.89429 | 120.27677 | 116.13108 | . | . | . | . | . | . |
| chr3 | 100796578 | 100797024 | 1000 | Distal Intergenic | 3 | 100825459 | 100895679 | 70221 | 1 | ENSMUST00 | -28435 | Vtcn1 | ENSMUSG0C | false | . | . | . | . | . | . | chr3 | 100794605 | 100798870 | 5.16708 | 4.05467 | 3.17162 |
| chr16 | 20621109 | 20621891 | 1000 | Promoter (<=1kb) | 16 | 20619215 | 20621287 | 2073 | 2 | ENSMUST00 | 0 | Camk2n2 | ENSMUSG0C | false | chr16 | 20620717 | 20622730 | 29.69128 | 86.53342 | 83.26198 | chr16 | 20619621 | 20622569 | 4.7334 | 3.74519 | 2.81748 |
| chr7 | 5015082 | 5015701 | 1000 | Promoter (<=1kb) | 7 | 5015501 | 5018486 | 2986 | 1 | ENSMUST00 | 0 | Zfp524 | ENSMUSG0C | false | chr7 | 5014072 | 5016177 | 33.5288 | 108.10788 | 104.32592 | . | . | . | . | . | . |
| chr1 | 58392630 | 58393808 | 1000 | Promoter (<=1kb) | 1 | 58392898 | 58407353 | 14456 | 1 | ENSMUST00 | 0 | Bzw1 | ENSMUSG0C | false | chr1 | 58391838 | 58394196 | 30.41442 | 103.5799 | 99.9162 | . | . | . | . | . | . |
| chr10 | 120898470 | 120899317 | 1000 | Promoter (<=1kb) | 10 | 120781096 | 120899101 | 118006 | 2 | ENSMUST00 | 0 | Msrb3 | ENSMUSG0C | true | chr10 | 120897818 | 120899446 | 44.57462 | 145.65463 | 140.53799 | . | . | . | . | . | . |
| chr1 | 7088611 | 7089451 | 1000 | Promoter (<=1kb) | 1 | 7088920 | 7173628 | 84709 | 1 | ENSMUST00 | 0 | Pcmtd1 | ENSMUSG0C | true | chr1 | 7089010 | 7090932 | 17.64787 | 49.73517 | 47.02245 | . | . | . | . | . | . |
| chr13 | 43303905 | 43304649 | 1000 | Promoter (<=1kb) | 13 | 43195519 | 43304172 | 108654 | 2 | ENSMUST00 | 0 | Gfod1 | ENSMUSG0C | false | chr13 | 43302084 | 43304284 | 34.3275 | 112.61823 | 108.70669 | chr13 | 43304469 | 43305753 | 6.07098 | 4.63333 | 3.88705 |
| chr1 | 64903561 | 64903869 | 1000 | Intron (ENSMUST00000139649.2 | 1 | 64860298 | 64928097 | 67800 | 2 | ENSMUST00 | 24228 | Plekhm3 | ENSMUSG0C | false | . | . | . | . | . | . | chr1 | 64902486 | 64906683 | 3.76074 | 3.42786 | 1.99454 |
| chr1 | 64852207 | 64852569 | 1000 | Intron (ENSMUST00000139649.2 | 1 | 64793809 | 64836764 | 42956 | 2 | ENSMUST00 | -15444 | Plekhm3 | ENSMUSG0C | false | . | . | . | . | . | . | chr1 | 64839724 | 64855177 | 5.20733 | 4.17383 | 3.17425 |
| chr4 | 136285404 | 136286374 | 1000 | Promoter (<=1kb) | 4 | 136286067 | 136293942 | 7876 | 1 | ENSMUST00 | 0 | Zfp46 | ENSMUSG0C | false | chr4 | 136284651 | 136286770 | 23.12601 | 77.64799 | 74.53377 | chr4 | 136286340 | 136287316 | 6.10365 | 4.57119 | 3.91068 |
| chr15 | 37803278 | 37804305 | 1000 | Distal Intergenic | 15 | 37397171 | 37792570 | 395400 | 2 | ENSMUST00 | -10709 | Ncald | ENSMUSG0C | true | chr15 | 37803434 | 37804046 | 7.54398 | 13.82373 | 11.57928 | chr15 | 37801944 | 37809187 | 4.93525 | 4.03004 | 2.96091 |
| chr2 | 32120403 | 32120737 | 1000 | Distal Intergenic | 2 | 32096155 | 32110808 | 14654 | 1 | ENSMUST00 | 24249 | Ppapdc3 | ENSMUSG0C | false | . | . | . | . | . | . | chr2 | 32120028 | 32122631 | 3.97307 | 3.45805 | 2.20756 |
| chr18 | 38215124 | 38215461 | 1000 | Intron (ENSMUST00000160935.1 | 18 | 38185915 | 38212053 | 26139 | 2 | ENSMUST00 | -3072 | Pcdh1 | ENSMUSG0C | false | . | . | . | . | . | . | . | . | . | . | . | . |
| chr18 | 38185100 | 38185608 | 1000 | Downstream (<1kb) | 18 | 38188888 | 38203163 | 14276 | 2 | ENSMUST00 | 17555 | Pcdh1 | ENSMUSG0C | false | chr18 | 38185112 | 38186528 | 20.38514 | 52.9059 | 50.15308 | . | . | . | . | . | . |
| chr18 | 38172030 | 38172730 | 1000 | Distal Intergenic | 18 | 38188888 | 38203163 | 14276 | 2 | ENSMUST00 | 30433 | Pcdh1 | ENSMUSG0C | false | . | . | . | . | . | . | chr18 | 38170740 | 38172822 | 4.03414 | 3.38269 | 2.24983 |
| chr6 | 67533044 | 67533455 | 1000 | Promoter (2-3kb) | 6 | 67534059 | 67535822 | 1764 | 2 | ENSMUST00 | 2367 | Tacstd2 | ENSMUSG0C | false | chr6 | 67532797 | 67533769 | 8.16212 | 17.69099 | 15.38708 | chr6 | 67531372 | 67534276 | 4.69776 | 3.92918 | 2.75749 |
| chr7 | 90442591 | 90443477 | 1000 | Promoter (<=1kb) | 7 | 90442729 | 90446104 | 3376 | 1 | ENSMUST00 | 0 | Crebzf | ENSMUSG0C | false | chr7 | 90442171 | 90445003 | 37.34335 | 127.50397 | 123.10989 | . | . | . | . | . | . |
| chr18 | 24052869 | 24053245 | 1000 | Distal Intergenic | 18 | 24009702 | 24020810 | 11109 | 2 | ENSMUST00 | -32060 | Zfp191 | ENSMUSG0C | false | . | . | . | . | . | . | chr18 | 24052236 | 24054659 | 5.06147 | 4.15061 | 3.06045 |
| chr18 | 24020293 | 24021085 | 1000 | Promoter (<=1kb) | 18 | 24009702 | 24020810 | 11109 | 2 | ENSMUST00 | 0 | Zfp191 | ENSMUSG0C | false | chr18 | 24019506 | 24021414 | 31.98759 | 106.40915 | 102.67528 | . | . | . | . | . | . |
| chr8 | 126666043 | 126666845 | 1000 | Distal Intergenic | 8 | 126588296 | 126593986 | 5691 | 2 | ENSMUST00 | -72058 | Irf2bp2 | ENSMUSG0C | false | . | . | . | . | . | . | chr8 | 126662916 | 126668496 | 7.0037 | 4.98697 | 4.64274 |
| chr8 | 126595960 | 126596503 | 1000 | Promoter (1-2kb) | 8 | 126588296 | 126593986 | 5691 | 2 | ENSMUST00 | -1975 | Irf2bp2 | ENSMUSG0C | false | . | . | . | . | . | . | chr8 | 126594816 | 126596983 | 8.16677 | 5.62022 | 5.58389 |
| chr8 | 126593749 | 126594301 | 1000 | Promoter (<=1kb) | 8 | 126588296 | 126593986 | 5691 | 2 | ENSMUST00 | 0 | Irf2bp2 | ENSMUSG0C | false | chr8 | 126589312 | 126594549 | 35.95432 | 114.41653 | 110.45183 | . | . | . | . | . | . |
| chr8 | 126735706 | 126736193 | 1000 | Distal Intergenic | 8 | 126588296 | 126593986 | 5691 | 2 | ENSMUST00 | -141721 | Irf2bp2 | ENSMUSG0C | false | . | . | . | . | . | . | chr8 | 126733289 | 126737272 | 7.19033 | 5.00384 | 4.80902 |
| chr6 | 120999901 | 121000577 | 1000 | Promoter (2-3kb) | 6 | 120931707 | 121003152 | 71446 | 2 | ENSMUST00 | 2575 | Mical3 | ENSMUSG0C | false | . | . | . | . | . | . | chr6 | 120994546 | 121005852 | 6.94439 | 4.93247 | 4.60074 |
| chr6 | 120987355 | 120988021 | 1000 | Intron (ENSMUST00000098457.3 | 6 | 120931707 | 121003152 | 71446 | 2 | ENSMUST00 | 15131 | Mical3 | ENSMUSG0C | false | . | . | . | . | . | . | chr6 | 120985290 | 120992730 | 7.03957 | 5.08701 | 4.66881 |
| chr14 | 120477439 | 120478699 | 1000 | Promoter (<=1kb) | 14 | 120478461 | 120507194 | 28734 | 1 | ENSMUST00 | 0 | Rap2a | ENSMUSG0C | true | chr14 | 120476887 | 120479540 | 20.41348 | 80.25478 | 77.09972 | . | . | . | . | . | . |
| chr11 | 22859376 | 22860197 | 1000 | Promoter (<=1kb) | 11 | 22834747 | 22860101 | 25355 | 2 | ENSMUST00 | 0 | B3gnt2 | ENSMUSG0C | false | chr11 | 22858157 | 22860897 | 30.41442 | 103.5799 | 99.9162 | . | . | . | . | . | . |
| chr4 | 65496369 | 65496694 | 1000 | Intron (ENSMUST00000084496.2 | 4 | 65604986 | 65616238 | 11253 | 1 | ENSMUST00 | -108292 | Trim32 | ENSMUSG0C | false | . | . | . | . | . | . | chr4 | 65494334 | 65496964 | 5.24895 | 4.09941 | 3.20772 |
| chr9 | 59749018 | 59749335 | 1000 | Promoter (1-2kb) | 9 | 59734259 | 59750649 | 16391 | 2 | ENSMUST00 | 1314 | Senp8 | ENSMUSG0C | false | . | . | . | . | . | . | chr9 | 59748886 | 59750132 | 4.96505 | 4.08269 | 3.00593 |
| chr3 | 67975087 | 67975437 | 1000 | Intron (ENSMUST00000063263.4 | 3 | 68040729 | 68056593 | 15865 | 1 | ENSMUST00 | -65292 | Iqcj | ENSMUSG0C | false | . | . | . | . | . | . | chr3 | 67974606 | 67977604 | 3.9094 | 3.41504 | 2.13469 |
| chr11 | 69837534 | 69838141 | 1000 | Promoter (<=1kb) | 11 | 69823122 | 69837784 | 14663 | 2 | ENSMUST00 | 0 | Nlgn2 | ENSMUSG0C | false | chr11 | 69836901 | 69838304 | 26.80025 | 77.79173 | 74.67541 | chr11 | 69833818 | 69837639 | 4.21706 | 3.65774 | 2.36499 |
| chr13 | 69541739 | 69542204 | 1000 | Promoter (1-2kb) | 13 | 69543046 | 69544023 | 978 | 2 | ENSMUST00 | 1819 | A530095I07F | ENSMUSG0C | false | . | . | . | . | . | . | chr13 | 69535499 | 69543951 | 7.03513 | 4.89229 | 4.72068 |
| chr2 | 152396123 | 152398275 | 1000 | Promoter (<=1kb) | 2 | 152393611 | 152398063 | 4453 | 2 | ENSMUST00 | 0 | Sox12 | ENSMUSG0C | false | chr2 | 152395767 | 152399538 | 37.89522 | 119.88892 | 115.74806 | chr2 | 152393058 | 152396892 | 3.78225 | 3.17389 | 2.04248 |
| chr2 | 152406166 | 152406541 | 1000 | Distal Intergenic | 2 | 152393611 | 152398063 | 4453 | 2 | ENSMUST00 | -8104 | Sox12 | ENSMUSG0C | false | chr2 | 152406280 | 152406745 | 10.10971 | 19.98882 | 17.65231 | chr2 | 152405565 | 152407041 | 4.35529 | 3.6262 | 2.50668 |
| chr6 | 83443246 | 83444155 | 1000 | Promoter (1-2kb) | 6 | 83441812 | 83442640 | 829 | 1 | ENSMUST00 | 1435 | B230319C09 | ENSMUSG0C | false | chr6 | 83440905 | 83443443 | 28.36989 | 85.2065 | 81.96005 | chr6 | 83442085 | 83458124 | 5.8895 | 4.34304 | 3.76187 |
| chr15 | 98762117 | 98763422 | 1000 | Promoter (<=1kb) | 15 | 98737621 | 98763211 | 25591 | 2 | ENSMUST00 | 0 | Arf3 | ENSMUSG0C | false | chr15 | 98762094 | 98763689 | 41.49189 | 142.53105 | 137.54099 | . | . | . | . | . | . |
| chr15 | 86219362 | 86219927 | 1000 | Intron (ENSMUST00000063414.8 | 15 | 86214459 | 86498503 | 284045 | 1 | ENSMUST00 | 4904 | Tbc1d22a | ENSMUSG0C | false | . | . | . | . | . | . | chr15 | 86214821 | 86223016 | 7.24796 | 4.98603 | 4.84278 |
| chr15 | 99126430 | 99126926 | 1000 | Promoter (<=1kb) | 15 | 99126578 | 99213206 | 86629 | 1 | ENSMUST00 | 0 | Spats2 | ENSMUSG0C | false | chr15 | 99125784 | 99127374 | 33.10969 | 106.34647 | 102.61562 | . | . | . | . | . | . |
| chr5 | 149678112 | 149678643 | 1000 | Promoter (<=1kb) | 5 | 149678230 | 149762599 | 84370 | 1 | ENSMUST00 | 0 | B3glct | ENSMUSG0C | false | chr5 | 149677956 | 149679210 | 36.76257 | 113.72284 | 109.77938 | chr5 | 149678011 | 149679562 | 4.09977 | 3.52139 | 2.32066 |
| chr5 | 149868285 | 149868806 | 1000 | Distal Intergenic | 5 | 149753942 | 149760607 | 6666 | 1 | ENSMUST00 | 114344 | B3glct | ENSMUSG0C | false | . | . | . | . | . | . | chr5 | 149864198 | 149869407 | 4.14149 | 3.61486 | 2.32443 |
| chr19 | 44514457 | 44515233 | 1000 | Downstream (1-2kb) | 19 | 44517644 | 44520593 | 2950 | 2 | ENSMUST00 | 5360 | Sec31b | ENSMUSG0C | false | . | . | . | . | . | . | chr19 | 44509183 | 44520604 | 5.97554 | 4.52339 | 3.81807 |
| chr1 | 133692232 | 133692593 | 1000 | Promoter (2-3kb) | 1 | 133679091 | 133690108 | 11018 | 2 | ENSMUST00 | -2125 | Lax1 | ENSMUSG0C | false | . | . | . | . | . | . | chr1 | 133689861 | 133697286 | 7.10741 | 4.96538 | 4.74647 |
| chr2 | 107519295 | 107519835 | 1000 | Distal Intergenic | 2 | 107595630 | 107600539 | 4910 | 2 | ENSMUST00 | 80704 | Gm9864 | ENSMUSG0C | false | . | . | . | . | . | . | chr2 | 107517224 | 107520985 | 4.43495 | 3.75429 | 2.55123 |
| chr7 | 63924755 | 63926347 | 1000 | Promoter (<=1kb) | 7 | 63891400 | 63924870 | 33471 | 2 | ENSMUST00 | 0 | Klf13 | ENSMUSG0C | false | . | . | . | . | . | . | chr7 | 63885397 | 63936279 | 7.51388 | 5.18869 | 5.06896 |
| chr7 | 63937137 | 63939403 | 1000 | Promoter (<=1kb) | 7 | 63886347 | 63938915 | 52569 | 2 | ENSMUST00 | 0 | Klf13 | ENSMUSG0C | false | chr7 | 63935484 | 63939001 | 41.67902 | 156.4369 | 150.90947 | . | . | . | . | . | . |
| chr1 | 61641876 | 61642385 | 1000 | Promoter (2-3kb) | 1 | 61639243 | 62642284 | 1003042 | 1 | ENSMUST00 | 2634 | Pard3b | ENSMUSG0C | true | . | . | . | . | . | . | chr1 | 61642258 | 61660533 | 4.74548 | 3.9194 | 2.80283 |
| chr1 | 61638632 | 61639295 | 1000 | Promoter (<=1kb) | 1 | 61638824 | 62078259 | 439436 | 1 | ENSMUST00 | 0 | Pard3b | ENSMUSG0C | true | chr1 | 61638253 | 61641326 | 26.89699 | 109.78668 | 105.95901 | . | . | . | . | . | . |
| chr17 | 66352813 | 66353284 | 1000 | Intron (ENSMUST00000086693.6 | 17 | 66343946 | 66346545 | 2600 | 2 | ENSMUST00 | -6269 | Mtcl1 | ENSMUSG0C | true | . | . | . | . | . | . | chr17 | 66349054 | 66377196 | 6.56105 | 4.76301 | 4.29736 |
| chr17 | 66449105 | 66450240 | 1000 | Promoter (<=1kb) | 17 | 66336982 | 66449750 | 112769 | 2 | ENSMUST00 | 0 | Mtcl1 | ENSMUSG0C | true | chr17 | 66446496 | 66450714 | 31.54088 | 108.67455 | 104.87239 | chr17 | 66450004 | 66451795 | 6.85649 | 4.88815 | 4.54732 |
| chr17 | 66357197 | 66357569 | 1000 | Intron (ENSMUST00000086693.6 | 17 | 66354205 | 66362726 | 8522 | 2 | ENSMUST00 | 5157 | Mtcl1 | ENSMUSG0C | true | . | . | . | . | . | . | chr17 | 66349054 | 66377196 | 6.56105 | 4.76301 | 4.29736 |
| chr17 | 66469408 | 66470457 | 1000 | Distal Intergenic | 17 | 66336986 | 66449750 | 112765 | 2 | ENSMUST00 | -19659 | Mtcl1 | ENSMUSG0C | true | . | . | . | . | . | . | chr17 | 66469072 | 66472170 | 3.92699 | 3.56983 | 2.10881 |
| chr4 | 120282673 | 120283513 | 1000 | Intron (ENSMUST00000102656.3 | 4 | 120267079 | 120287349 | 20271 | 2 | ENSMUST00 | 3836 | Foxo6 | ENSMUSG0C | false | chr4 | 120282409 | 120283440 | 29.86959 | 86.83979 | 83.56236 | chr4 | 120281377 | 120283888 | 3.9801 | 3.56023 | 2.1925 |
| chr5 | 32068554 | 32069226 | 1000 | Intron (ENSMUST00000080598.8 | 5 | 32079697 | 32084746 | 5050 | 1 | ENSMUST00 | -10471 | Bre | ENSMUSG0C | false | . | . | . | . | . | . | chr5 | 32061891 | 32071385 | 4.67719 | 3.8544 | 2.75917 |
| chr9 | 60838303 | 60838810 | 1000 | Promoter (<=1kb) | 9 | 60821855 | 60838200 | 16346 | 2 | ENSMUST00 | -104 | Gm9869 | ENSMUSG0C | false | chr9 | 60836859 | 60842659 | 24.93112 | 71.82204 | 68.80418 | . | . | . | . | . | . |
| chr9 | 60848748 | 60849141 | 1000 | Intron (ENSMUST00000050183.6 | 9 | 60821855 | 60838200 | 16346 | 2 | ENSMUST00 | -10549 | Gm9869 | ENSMUSG0C | false | . | . | . | . | . | . | chr9 | 60839714 | 60851928 | 5.61355 | 4.39804 | 3.5066 |
| chr6 | 100833493 | 100834200 | 1000 | Promoter (<=1kb) | 6 | 100833638 | 100868717 | 35080 | 1 | ENSMUST00 | 0 | Ppp4r2 | ENSMUSG0C | false | chr6 | 100833184 | 100834843 | 24.03115 | 75.67385 | 72.59326 | . | . | . | . | . | . |
| chr2 | 48882618 | 48882983 | 1000 | Intron (ENSMUST00000063886.3 | 2 | 48890389 | 48900248 | 9860 | 1 | ENSMUST00 | -7406 | Acvr2a | ENSMUSG0C | false | . | . | . | . | . | . | chr2 | 48882072 | 48884114 | 4.57086 | 3.88837 | 2.63984 |
| chr19 | 6528234 | 6528571 | 1000 | Intron (ENSMUST00000113462.2 | 19 | 6530789 | 6531788 | 1000 | 2 | ENSMUST00 | 3217 | Gm14964 | ENSMUSG0C | false | . | . | . | . | . | . | chr19 | 6527406 | 6533037 | 4.81432 | 3.87913 | 2.86713 |
| chr5 | 144182592 | 144183040 | 1000 | Exon (ENSMUST00000041804.7/ | 5 | 144190286 | 144194441 | 4156 | 1 | ENSMUST00 | -7246 | Bhlha15 | ENSMUSG0C | false | . | . | . | . | . | . | chr5 | 144181835 | 144185686 | 5.69891 | 4.32402 | 3.61705 |
| chr1 | 54897294 | 54897668 | 1000 | Intron (ENSMUST00000179030.2 | 1 | 54766792 | 54884711 | 117920 | 2 | ENSMUST00 | -12584 | Ankrd44 | ENSMUSG0C | true | . | . | . | . | . | . | chr1 | 54896196 | 54904292 | 5.4736 | 4.28869 | 3.39741 |
| chr1 | 54867244 | 54867642 | 1000 | Intron (ENSMUST00000179030.2 | 1 | 54766792 | 54884711 | 117920 | 2 | ENSMUST00 | 17069 | Ankrd44 | ENSMUSG0C | true | . | . | . | . | . | . | chr1 | 54864456 | 54889599 | 5.98614 | 4.56539 | 3.81071 |
| chr1 | 54922254 | 54922597 | 1000 | Intron (ENSMUST00000179030.2 | 1 | 54735302 | 54926354 | 191053 | 2 | ENSMUST00 | 3757 | Ankrd44 | ENSMUSG0C | true | . | . | . | . | . | . | chr1 | 54909245 | 54924829 | 5.57394 | 4.32237 | 3.47681 |
| chr7 | 83921561 | 83922040 | 1000 | Distal Intergenic | 7 | 83934783 | 83944340 | 9558 | 2 | ENSMUST00 | 22300 | Cemip | ENSMUSG0C | false | . | . | . | . | . | . | chr7 | 83920330 | 83922379 | 4.37735 | 3.73784 | 2.50175 |
| chrX | 8892816 | 8893180 | 1000 | Promoter (<=1kb) | 20 | 8892382 | 8894964 | 2583 | 1 | ENSMUST00 | 435 | B630019K06 | ENSMUSG0C | false | chrX | 8892266 | 8893915 | 18.38128 | 45.59164 | 42.93153 | . | . | . | . | . | . |
| chr15 | 97950310 | 97950650 | 1000 | Distal Intergenic | 15 | 97964229 | 97970286 | 6058 | 1 | ENSMUST00 | -13579 | Tmem106c | ENSMUSG0C | false | . | . | . | . | . | . | chr15 | 97948754 | 97950917 | 4.80262 | 3.82762 | 2.88558 |
| chr11 | 102026564 | 102027606 | 1000 | Promoter (<=1kb) | 11 | 101999652 | 102026934 | 27283 | 2 | ENSMUST00 | 0 | Mpp3 | ENSMUSG0C | false | chr11 | 102025982 | 102027728 | 33.54585 | 101.01101 | 97.41918 | chr11 | 102023623 | 102028163 | 3.81611 | 3.52131 | 2.02106 |
| chr2 | 13554033 | 13554585 | 1000 | Distal Intergenic | 2 | 13557848 | 13560324 | 2477 | 1 | ENSMUST00 | -3263 | Gm9875 | ENSMUSG0C | false | . | . | . | . | . | . | chr2 | 13552881 | 13559080 | 4.50577 | 3.85199 | 2.58778 |
| chr17 | 6736866 | 6737362 | 1000 | Intron (ENSMUST00000097430.3 | 17 | 6740837 | 6742821 | 1985 | 2 | ENSMUST00 | 5459 | Ezr | ENSMUSG0C | false | . | . | . | . | . | . | chr17 | 6734583 | 6749633 | 6.00896 | 4.44798 | 3.8635 |
| chr17 | 6742151 | 6742540 | 1000 | Promoter (<=1kb) | 17 | 6740837 | 6742821 | 1985 | 2 | ENSMUST00 | 281 | Ezr | ENSMUSG0C | false | . | . | . | . | . | . | chr17 | 6734583 | 6749633 | 6.00896 | 4.44798 | 3.8635 |
| chr15 | 10615264 | 10615607 | 1000 | Intron (ENSMUST00000090339.4 | 15 | 10560524 | 10561524 | 1001 | 2 | ENSMUST00 | -53741 | Gm8174 | ENSMUSG0C | false | . | . | . | . | . | . | chr15 | 10614288 | 10618293 | 4.31146 | 3.74307 | 2.44769 |
| chr7 | 49262355 | 49262743 | 1000 | Intron (ENSMUST00000183659.2 | 7 | 49246845 | 49494000 | 247156 | 1 | ENSMUST00 | 15511 | Nav2 | ENSMUSG0C | false | . | . | . | . | . | . | chr7 | 49261971 | 49264613 | 6.7792 | 4.83613 | 4.4635 |
| chr7 | 49444178 | 49444599 | 1000 | Intron (ENSMUST00000183659.2 | 7 | 49450760 | 49464580 | 13821 | 1 | ENSMUST00 | -6161 | Nav2 | ENSMUSG0C | false | . | . | . | . | . | . | chr7 | 49442742 | 49445288 | 5.83031 | 4.19531 | 3.72205 |
| chr7 | 49247946 | 49248803 | 1000 | Promoter (1-2kb) | 7 | 49246845 | 49494000 | 247156 | 1 | ENSMUST00 | 1102 | Nav2 | ENSMUSG0C | false | chr7 | 49245452 | 49248461 | 23.73948 | 76.05544 | 72.96951 | chr7 | 49245394 | 49251528 | 6.01396 | 4.39599 | 3.85646 |
| chr7 | 49448535 | 49448976 | 1000 | Promoter (1-2kb) | 7 | 49450760 | 49464580 | 13821 | 1 | ENSMUST00 | -1784 | Nav2 | ENSMUSG0C | false | . | . | . | . | . | . | chr7 | 49446529 | 49453829 | 5.00391 | 3.99232 | 3.02197 |
| chr7 | 49437783 | 49438493 | 1000 | Exon (ENSMUST00000184955.2/ | 7 | 49450760 | 49464580 | 13821 | 1 | ENSMUST00 | -12267 | Nav2 | ENSMUSG0C | false | chr7 | 49437771 | 49438084 | 7.76779 | 14.19323 | 11.94406 | chr7 | 49438252 | 49439675 | 8.12968 | 5.19317 | 5.60514 |
| chr7 | 49245844 | 49246371 | 1000 | Promoter (<=1kb) | 7 | 49246189 | 49409643 | 163455 | 1 | ENSMUST00 | 0 | Nav2 | ENSMUSG0C | false | chr7 | 49245452 | 49248461 | 23.73948 | 76.05544 | 72.96951 | chr7 | 49245394 | 49251528 | 6.01396 | 4.39599 | 3.85646 |
| chr16 | 74411311 | 74412224 | 1000 | Promoter (<=1kb) | 16 | 73920206 | 74411785 | 491580 | 2 | ENSMUST00 | 0 | Robo2 | ENSMUSG0C | true | chr16 | 74408228 | 74412370 | 19.04495 | 59.55748 | 56.71581 | . | . | . | . | . | . |
| chr1 | 168431528 | 168432590 | 1000 | Promoter (<=1kb) | 1 | 168119364 | 168432270 | 312907 | 2 | ENSMUST00 | 0 | Pbx1 | ENSMUSG0C | false | chr1 | 168432526 | 168432982 | 17.37054 | 45.37996 | 42.72523 | . | . | . | . | . | . |
| chr3 | 104220111 | 104220900 | 1000 | Promoter (<=1kb) | 3 | 104013259 | 104220374 | 207116 | 2 | ENSMUST00 | 0 | Magi3 | ENSMUSG0C | false | chr3 | 104218748 | 104221302 | 31.16539 | 106.97112 | 103.21663 | . | . | . | . | . | . |
| chr8 | 117157607 | 117158139 | 1000 | Promoter (<=1kb) | 8 | 117158135 | 117205186 | 47052 | 1 | ENSMUST00 | 0 | Gan | ENSMUSG0C | false | chr8 | 117157074 | 117158922 | 44.57026 | 147.90271 | 142.70154 | . | . | . | . | . | . |
| chr14 | 76569401 | 76569927 | 1000 | Distal Intergenic | 14 | 76532814 | 76556687 | 23874 | 2 | ENSMUST00 | -12715 | Serp2 | ENSMUSG0C | false | . | . | . | . | . | . | chr14 | 76566818 | 76570527 | 5.20463 | 3.99775 | 3.20905 |
| chr7 | 78614648 | 78614940 | 1000 | Distal Intergenic | 7 | 78578830 | 78581038 | 2209 | 1 | ENSMUST00 | 35819 | Gm9885 | ENSMUSG0C | false | . | . | . | . | . | . | chr7 | 78613393 | 78625373 | 6.15156 | 4.58262 | 3.94687 |
| chr7 | 78651172 | 78651929 | 1000 | Distal Intergenic | 7 | 78578830 | 78581038 | 2209 | 1 | ENSMUST00 | 72343 | Gm9885 | ENSMUSG0C | false | . | . | . | . | . | . | chr7 | 78647720 | 78662742 | 5.70256 | 4.4354 | 3.58296 |
| chr7 | 78660696 | 78661345 | 1000 | Distal Intergenic | 7 | 78578830 | 78581038 | 2209 | 1 | ENSMUST00 | 81867 | Gm9885 | ENSMUSG0C | false | . | . | . | . | . | . | chr7 | 78647720 | 78662742 | 5.70256 | 4.4354 | 3.58296 |
| chr6 | 72487333 | 72487610 | 1000 | Distal Intergenic | 6 | 72513644 | 72515290 | 1647 | 2 | ENSMUST00 | 27680 | Sh2d6 | ENSMUSG0C | false | . | . | . | . | . | . | chr6 | 72480228 | 72490717 | 5.30413 | 4.25075 | 3.25547 |
| chr12 | 21134068 | 21134610 | 1000 | Intron (ENSMUST00000174530.1 | 12 | 21111954 | 21268179 | 156226 | 1 | ENSMUST00 | 22115 | Asap2 | ENSMUSG0C | false | . | . | . | . | . | . | chr12 | 21122843 | 21147439 | 6.77508 | 4.82373 | 4.46417 |
| chr12 | 21135886 | 21136420 | 1000 | Intron (ENSMUST00000174530.1 | 12 | 21111954 | 21268179 | 156226 | 1 | ENSMUST00 | 23933 | Asap2 | ENSMUSG0C | false | . | . | . | . | . | . | chr12 | 21122843 | 21147439 | 6.77508 | 4.82373 | 4.46417 |
| chr12 | 69458146 | 69458844 | 1000 | Distal Intergenic | 12 | 69371835 | 69372458 | 624 | 2 | ENSMUST00 | -85689 | Gm9887 | ENSMUSG0C | false | . | . | . | . | . | . | chr12 | 69456245 | 69463797 | 5.86899 | 4.48095 | 3.71706 |
| chr12 | 69404015 | 69404563 | 1000 | Distal Intergenic | 12 | 69371835 | 69372458 | 624 | 2 | ENSMUST00 | -31558 | Gm9887 | ENSMUSG0C | false | chr12 | 69404298 | 69406539 | 18.70486 | 58.09692 | 55.27494 | chr12 | 69400543 | 69404256 | 6.14189 | 4.63503 | 3.94789 |
| chr12 | 69434288 | 69434685 | 1000 | Distal Intergenic | 12 | 69371835 | 69372458 | 624 | 2 | ENSMUST00 | -61831 | Gm9887 | ENSMUSG0C | false | . | . | . | . | . | . | chr12 | 69433996 | 69434868 | 5.64876 | 4.48093 | 3.52441 |
| chr12 | 69442215 | 69442846 | 1000 | Distal Intergenic | 12 | 69371835 | 69372458 | 624 | 2 | ENSMUST00 | -69758 | Gm9887 | ENSMUSG0C | false | . | . | . | . | . | . | chr12 | 69436684 | 69444561 | 5.15172 | 4.15476 | 3.12711 |
| chr12 | 69438162 | 69438474 | 1000 | Distal Intergenic | 12 | 69371835 | 69372458 | 624 | 2 | ENSMUST00 | -65705 | Gm9887 | ENSMUSG0C | false | . | . | . | . | . | . | chr12 | 69436684 | 69444561 | 5.15172 | 4.15476 | 3.12711 |
| chr10 | 117940985 | 117941315 | 1000 | Distal Intergenic | 10 | 117814471 | 117845935 | 31465 | 2 | ENSMUST00 | -95051 | Rap1b | ENSMUSG0C | false | . | . | . | . | . | . | chr10 | 117940267 | 117941741 | 4.32151 | 3.81416 | 2.43242 |

| | | | | | | | | | | | | | | | | | | | | | | | | | | |
|---|---|---|---|---|---|---|---|---|---|---|---|---|---|---|---|---|---|---|---|---|---|---|---|---|---|---|
| chr10 | 117905705 | 117906822 | 1000 | Distal Intergenic | 10 | 117814471 | 117845935 | 31465 | 2 | ENSMUST00 | -59771 | Rap1b | ENSMUSG0C | false | chr10 | 117905555 | 117906960 | 14.48567 | 38.11502 | 35.5495 | chr10 | 117905498 | 117907126 | 3.4126 | 3.29812 | 1.75299 |
| chr7 | 123097932 | 123098386 | 1000 | Distal Intergenic | 7 | 123123885 | 123195288 | 71404 | 1 | ENSMUST00 | -25499 | Tnrc6a | ENSMUSG0C | false | . | . | . | . | . | . | chr7 | 123097695 | 123099674 | 7.86312 | 5.3651 | 5.35389 |
| chr17 | 29297093 | 29297636 | 1000 | Promoter (<=1kb) | 17 | 29298284 | 29301525 | 3242 | 1 | ENSMUST00 | -648 | BC004004 | ENSMUSG0C | false | . | . | . | . | . | . | chr17 | 29292484 | 29307926 | 7.62734 | 5.21808 | 5.16527 |
| chr17 | 29302096 | 29302616 | 1000 | 3' UTR | 17 | 29298284 | 29301525 | 3242 | 1 | ENSMUST00 | 3813 | BC004004 | ENSMUSG0C | false | . | . | . | . | . | . | chr17 | 29292484 | 29307926 | 7.62734 | 5.21808 | 5.16527 |
| chr9 | 114767530 | 114768281 | 1000 | Intron (ENSMUST00000035009.9 | 9 | 114780100 | 114782257 | 2158 | 1 | ENSMUST00 | -11819 | Gm9888 | ENSMUSG0C | false | . | . | . | . | . | . | chr9 | 114758262 | 114780983 | 6.81555 | 4.89533 | 4.50006 |
| chr13 | 99514832 | 99515607 | 1000 | Promoter (<=1kb) | 13 | 99421464 | 99516602 | 95139 | 2 | ENSMUST00 | 995 | Map1b | ENSMUSG0C | false | chr13 | 99514671 | 99517115 | 21.46036 | 80.68572 | 77.52129 | chr13 | 99513083 | 99514842 | 3.68618 | 3.4299 | 1.92073 |
| chr13 | 99516131 | 99516861 | 1000 | Promoter (<=1kb) | 13 | 99421464 | 99516602 | 95139 | 2 | ENSMUST00 | 0 | Map1b | ENSMUSG0C | false | chr13 | 99514671 | 99517115 | 21.46036 | 80.68572 | 77.52129 | chr13 | 99515980 | 99516907 | 3.01922 | 3.01727 | 1.3782 |
| chr6 | 48593688 | 48594442 | 1000 | Promoter (<=1kb) | 6 | 48593883 | 48597387 | 3505 | 1 | ENSMUST00 | 0 | Repin1 | ENSMUSG0C | false | chr6 | 48593384 | 48594992 | 39.06023 | 122.966 | 118.72274 | chr6 | 48593220 | 48595793 | 3.6894 | 3.26742 | 1.96454 |
| chr3 | 115789000 | 115789598 | 1000 | Distal Intergenic | 3 | 115715150 | 115716727 | 1578 | 1 | ENSMUST00 | 73851 | Gm9889 | ENSMUSG0C | false | . | . | . | . | . | . | . | . | . | . | . | . |
| chr7 | 16361415 | 16361721 | 1000 | Intron (ENSMUST00000094815.3 | 7 | 16327053 | 16387797 | 60745 | 2 | ENSMUST00 | 26076 | Sae1 | ENSMUSG0C | false | . | . | . | . | . | . | chr7 | 16360337 | 16366203 | 4.80253 | 3.95138 | 2.85963 |
| chr9 | 108478666 | 108479191 | 1000 | Promoter (<=1kb) | 9 | 108479862 | 108490530 | 10669 | 1 | ENSMUST00 | -671 | Lamb2 | ENSMUSG0C | false | chr9 | 108478517 | 108480814 | 19.47644 | 60.36021 | 57.50917 | chr9 | 108477974 | 108483462 | 4.94861 | 3.89359 | 3.00507 |
| chr11 | 98788887 | 98789722 | 1000 | Distal Intergenic | 11 | 98795516 | 98807859 | 12344 | 1 | ENSMUST00 | -5794 | Msl1 | ENSMUSG0C | false | chr11 | 98789230 | 98789849 | 11.4883 | 23.92001 | 21.53215 | chr11 | 98780022 | 98794752 | 7.02549 | 5.01464 | 4.66092 |
| chr11 | 98794727 | 98796242 | 1000 | Promoter (<=1kb) | 11 | 98795516 | 98807859 | 12344 | 1 | ENSMUST00 | 0 | Msl1 | ENSMUSG0C | false | chr11 | 98794305 | 98797777 | 41.07278 | 140.67711 | 135.76277 | chr11 | 98780022 | 98794752 | 7.02549 | 5.01464 | 4.66092 |
| chr19 | 30623651 | 30624121 | 1000 | Promoter (1-2kb) | 19 | 30619371 | 30625491 | 6121 | 2 | ENSMUST00 | 1370 | Prkg1 | ENSMUSG0C | true | . | . | . | . | . | . | chr19 | 30623051 | 30626449 | 3.76341 | 3.39912 | 2.0139 |
| chr19 | 31444109 | 31444441 | 1000 | Intron (ENSMUST00000065067.7 | 19 | 30568547 | 31664370 | 1095824 | 2 | ENSMUST00 | 219929 | Prkg1 | ENSMUSG0C | true | . | . | . | . | . | . | chr19 | 31443357 | 31447049 | 4.4722 | 3.68324 | 2.60379 |
| chr2 | 33640621 | 33641406 | 1000 | Promoter (<=1kb) | 2 | 33641193 | 33642593 | 1401 | 1 | ENSMUST00 | 0 | C130021I20R | ENSMUSG0C | false | chr2 | 33637941 | 33643219 | 26.18681 | 91.82585 | 88.44535 | chr2 | 33640996 | 33643597 | 3.43705 | 3.26878 | 1.72381 |
| chr13 | 60177049 | 60178017 | 1000 | Promoter (<=1kb) | 13 | 60174405 | 60177365 | 2961 | 2 | ENSMUST00 | 0 | Gas1 | ENSMUSG0C | false | chr13 | 60177660 | 60178651 | 13.87066 | 31.59332 | 29.10884 | . | . | . | . | . | . |
| chr13 | 60047809 | 60048252 | 1000 | Distal Intergenic | 13 | 60174405 | 60177365 | 2961 | 2 | ENSMUST00 | 129113 | Gas1 | ENSMUSG0C | false | . | . | . | . | . | . | chr13 | 60046981 | 60050078 | 6.40268 | 4.69743 | 4.16328 |
| chr13 | 69739662 | 69740203 | 1000 | Promoter (<=1kb) | 13 | 69702835 | 69739897 | 37063 | 2 | ENSMUST00 | 0 | Ube2ql1 | ENSMUSG0C | false | chr13 | 69738105 | 69740890 | 33.0268 | 99.11563 | 95.56912 | chr13 | 69739580 | 69740700 | 4.92268 | 3.98243 | 2.94987 |
| chr13 | 69692158 | 69692671 | 1000 | Distal Intergenic | 13 | 69702835 | 69739897 | 37063 | 2 | ENSMUST00 | 47226 | Ube2ql1 | ENSMUSG0C | false | . | . | . | . | . | . | . | . | . | . | . | . |
| chr6 | 114397666 | 114398195 | 1000 | Promoter (<=1kb) | 6 | 114397936 | 114483296 | 85361 | 1 | ENSMUST00 | 0 | Hrh1 | ENSMUSG0C | false | chr6 | 114397453 | 114398494 | 17.85986 | 47.76698 | 45.0813 | chr6 | 114397098 | 114398772 | 4.82554 | 4.03226 | 2.8452 |
| chr6 | 114450714 | 114451113 | 1000 | Intron (ENSMUST00000088987.2 | 6 | 114458519 | 114482468 | 23950 | 1 | ENSMUST00 | -7406 | Hrh1 | ENSMUSG0C | false | . | . | . | . | . | . | chr6 | 114450532 | 114454426 | 5.99252 | 4.5118 | 3.83078 |
| chr7 | 142021709 | 142022091 | 1000 | Intron (ENSMUST00000084418.2 | 7 | 141998661 | 142001441 | 2781 | 1 | ENSMUST00 | 23049 | Brsk2 | ENSMUSG0C | false | chr7 | 142021191 | 142026097 | 20.8177 | 63.66574 | 60.7686 | chr7 | 142014396 | 142028430 | 5.88599 | 4.36974 | 3.76503 |
| chr19 | 7417457 | 7418236 | 1000 | Promoter (<=1kb) | 19 | 7417625 | 7425904 | 8280 | 1 | ENSMUST00 | 0 | 2700081O15 | ENSMUSG0C | false | chr19 | 7416554 | 7418702 | 27.66286 | 83.97498 | 80.7477 | . | . | . | . | . | . |
| chr9 | 8004384 | 8004925 | 1000 | Promoter (<=1kb) | 9 | 7931999 | 8004588 | 72590 | 2 | ENSMUST00 | 0 | Yap1 | ENSMUSG0C | false | chr9 | 8002688 | 8005179 | 35.84351 | 110.06287 | 106.22365 | . | . | . | . | . | . |
| chr11 | 117962196 | 117962632 | 1000 | Distal Intergenic | 11 | 117966079 | 117969186 | 3108 | 2 | ENSMUST00 | 6554 | Socs3 | ENSMUSG0C | false | . | . | . | . | . | . | chr11 | 117954133 | 117968344 | 6.92433 | 4.93094 | 4.58678 |
| chr11 | 117969128 | 117969846 | 1000 | Promoter (<=1kb) | 11 | 117966079 | 117969186 | 3108 | 2 | ENSMUST00 | 0 | Socs3 | ENSMUSG0C | false | chr11 | 117967602 | 117970051 | 29.83578 | 117.74437 | 113.67492 | chr11 | 117969624 | 117971164 | 4.27282 | 3.78764 | 2.40711 |
| chr11 | 117956920 | 117957263 | 1000 | Distal Intergenic | 11 | 117966079 | 117969186 | 3108 | 2 | ENSMUST00 | 11923 | Socs3 | ENSMUSG0C | false | . | . | . | . | . | . | chr11 | 117954133 | 117968344 | 6.92433 | 4.93094 | 4.58678 |
| chr11 | 117963034 | 117963903 | 1000 | Downstream (2-3kb) | 11 | 117966079 | 117969186 | 3108 | 2 | ENSMUST00 | 5283 | Socs3 | ENSMUSG0C | false | chr11 | 117963726 | 117964052 | 9.63447 | 24.12454 | 21.73289 | chr11 | 117954133 | 117968344 | 6.92433 | 4.93094 | 4.58678 |
| chr7 | 27507045 | 27507436 | 1000 | Promoter (<=1kb) | 7 | 27507801 | 27520034 | 12234 | 1 | ENSMUST00 | -365 | Prx | ENSMUSG0C | false | chr7 | 27507246 | 27508158 | 22.15371 | 67.81625 | 64.85734 | chr7 | 27488220 | 27512085 | 6.36425 | 4.68802 | 4.12603 |
| chr5 | 135465739 | 135466298 | 1000 | Intron (ENSMUST00000060311.8 | 5 | 135376141 | 135394546 | 18406 | 2 | ENSMUST00 | -71194 | Pom121 | ENSMUSG0C | false | chr5 | 135465527 | 135465869 | 7.82748 | 15.19303 | 12.92724 | chr5 | 135458659 | 135480128 | 5.22779 | 4.14143 | 3.19665 |
| chr9 | 37567980 | 37568867 | 1000 | Distal Intergenic | 9 | 37544492 | 37552904 | 8413 | 2 | ENSMUST00 | -15077 | Nrgn | ENSMUSG0C | false | . | . | . | . | . | . | chr9 | 37567805 | 37569121 | 4.61613 | 3.89554 | 2.67639 |
| chr5 | 150152047 | 150152565 | 1000 | Distal Intergenic | 5 | 150080622 | 150082184 | 1563 | 1 | ENSMUST00 | 71426 | Rxfp2 | ENSMUSG0C | false | . | . | . | . | . | . | chr5 | 150148830 | 150161159 | 5.72506 | 4.43285 | 3.60287 |
| chr5 | 150051523 | 150051812 | 1000 | Exon (ENSMUST00000143989.1/ | 5 | 150065870 | 150069355 | 3486 | 1 | ENSMUST00 | -14058 | Rxfp2 | ENSMUSG0C | false | . | . | . | . | . | . | chr5 | 150050277 | 150052011 | 3.39984 | 3.25941 | 1.70478 |
| chr5 | 150130005 | 150130324 | 1000 | Distal Intergenic | 5 | 150080622 | 150082184 | 1563 | 1 | ENSMUST00 | 49384 | Rxfp2 | ENSMUSG0C | false | . | . | . | . | . | . | chr5 | 150127781 | 150137123 | 4.81874 | 3.97228 | 2.87137 |
| chr5 | 150118451 | 150119275 | 1000 | Distal Intergenic | 5 | 150080622 | 150082184 | 1563 | 1 | ENSMUST00 | 37830 | Rxfp2 | ENSMUSG0C | false | chr5 | 150118265 | 150119526 | 30.94091 | 93.53562 | 90.11485 | chr5 | 150117949 | 150123701 | 5.27343 | 4.24721 | 3.2367 |
| chr6 | 71632104 | 71633681 | 1000 | Promoter (<=1kb) | 6 | 71588974 | 71632686 | 43713 | 2 | ENSMUST00 | 0 | Kdm3a | ENSMUSG0C | false | chr6 | 71631634 | 71633659 | 36.04346 | 118.78738 | 114.68494 | chr6 | 71628187 | 71632106 | 3.68186 | 3.40338 | 1.94115 |
| chr4 | 109000308 | 109000927 | 1000 | Promoter (<=1kb) | 4 | 109000655 | 109061777 | 61123 | 1 | ENSMUST00 | 0 | Nrd1 | ENSMUSG0C | false | chr4 | 108999883 | 109000456 | 15.64407 | 45.32187 | 42.668 | . | . | . | . | . | . |
| chr11 | 33791347 | 33791857 | 1000 | Intron (ENSMUST00000101368.3 | 11 | 33629339 | 33843461 | 214123 | 2 | ENSMUST00 | 51604 | Kcnip1 | ENSMUSG0C | false | . | . | . | . | . | . | chr11 | 33790884 | 33792357 | 3.28088 | 3.11473 | 1.61865 |
| chr8 | 84661167 | 84661899 | 1000 | Promoter (<=1kb) | 8 | 84661331 | 84662852 | 1522 | 2 | ENSMUST00 | 953 | Ier2 | ENSMUSG0C | false | chr8 | 84659950 | 84663752 | 33.52805 | 106.05383 | 102.33234 | . | . | . | . | . | . |
| chr11 | 105589399 | 105590427 | 1000 | Promoter (<=1kb) | 11 | 105589986 | 105776834 | 186849 | 1 | ENSMUST00 | 0 | Tanc2 | ENSMUSG0C | false | chr11 | 105589174 | 105590737 | 36.04346 | 118.78738 | 114.68494 | chr11 | 105590409 | 105591309 | 3.50216 | 3.1406 | 1.81876 |
| chr6 | 70749954 | 70750431 | 1000 | Distal Intergenic | 6 | 70766273 | 70773692 | 7420 | 2 | ENSMUST00 | 23261 | Rpia | ENSMUSG0C | false | . | . | . | . | . | . | chr6 | 70749063 | 70752667 | 4.61576 | 3.87545 | 2.69373 |
| chr6 | 70791844 | 70792307 | 1000 | Promoter (<=1kb) | 6 | 70765720 | 70792232 | 26513 | 2 | ENSMUST00 | 0 | Rpia | ENSMUSG0C | false | chr6 | 70790221 | 70792994 | 33.84988 | 110.73796 | 106.88306 | . | . | . | . | . | . |
| chr9 | 109095029 | 109096085 | 1000 | Promoter (<=1kb) | 9 | 109095436 | 109119911 | 24476 | 1 | ENSMUST00 | 0 | Plxnb1 | ENSMUSG0C | false | chr9 | 109094798 | 109099441 | 38.03245 | 121.19012 | 117.00504 | chr9 | 109092590 | 109095286 | 5.26963 | 4.23227 | 3.23275 |
| chr9 | 109096403 | 109097618 | 1000 | Promoter (<=1kb) | 9 | 109096710 | 109100498 | 3789 | 1 | ENSMUST00 | 0 | Plxnb1 | ENSMUSG0C | false | chr9 | 109094798 | 109099441 | 38.03245 | 121.19012 | 117.00504 | chr9 | 109096591 | 109107997 | 6.47539 | 4.76075 | 4.21763 |
| chr1 | 167182282 | 167182699 | 1000 | Distal Intergenic | 1 | 167223827 | 167223919 | 93 | 2 | ENSMUST00 | 41220 | Uck2 | ENSMUSG0C | true | . | . | . | . | . | . | chr1 | 167181488 | 167186065 | 5.56647 | 4.31802 | 3.46843 |
| chr1 | 167249247 | 167249782 | 1000 | Intron (ENSMUST00000027839.8 | 1 | 167223827 | 167223919 | 93 | 2 | ENSMUST00 | -25329 | Uck2 | ENSMUSG0C | true | . | . | . | . | . | . | chr1 | 167235376 | 167257440 | 5.06825 | 4.01124 | 3.08857 |
| chr1 | 167159564 | 167160279 | 1000 | Distal Intergenic | 1 | 167223827 | 167223919 | 93 | 2 | ENSMUST00 | 63640 | Uck2 | ENSMUSG0C | true | . | . | . | . | . | . | chr1 | 167158891 | 167160202 | 5.37935 | 4.2586 | 3.35897 |
| chr2 | 17714677 | 17715722 | 1000 | Intron (ENSMUST00000028080.6 | 2 | 17343909 | 17731043 | 387135 | 2 | ENSMUST00 | 15321 | Nebl | ENSMUSG0C | true | . | . | . | . | . | . | chr2 | 17675160 | 17718684 | 4.75621 | 3.88836 | 2.82315 |
| chr2 | 17658208 | 17658596 | 1000 | Intron (ENSMUST00000028080.6 | 2 | 17343909 | 17731043 | 387135 | 2 | ENSMUST00 | 72447 | Nebl | ENSMUSG0C | true | . | . | . | . | . | . | chr2 | 17655501 | 17662021 | 4.58993 | 3.73654 | 2.70414 |
| chr9 | 106371728 | 106372228 | 1000 | Promoter (<=1kb) | 9 | 106370744 | 106373856 | 3113 | 1 | ENSMUST00 | 985 | Dusp7 | ENSMUSG0C | false | chr9 | 106371249 | 106372180 | 7.96309 | 14.9624 | 12.70017 | chr9 | 106369383 | 106374003 | 5.91164 | 4.40656 | 3.78976 |
| chr9 | 106368240 | 106369102 | 1000 | Promoter (<=1kb) | 9 | 106368632 | 106375724 | 7093 | 1 | ENSMUST00 | 0 | Dusp7 | ENSMUSG0C | false | chr9 | 106367929 | 106370134 | 27.97104 | 107.99458 | 104.2211 | . | . | . | . | . | . |
| chr4 | 129696431 | 129696928 | 1000 | Promoter (<=1kb) | 4 | 129676355 | 129696796 | 20442 | 2 | ENSMUST00 | 0 | Tmem39b | ENSMUSG0C | false | chr4 | 129695887 | 129697149 | 24.55236 | 90.57755 | 87.22021 | chr4 | 129695512 | 129697155 | 4.00503 | 3.62079 | 2.17362 |
| chr19 | 37550295 | 37551170 | 1000 | Promoter (<=1kb) | 19 | 37550418 | 37683245 | 132828 | 1 | ENSMUST00 | 0 | Exoc6 | ENSMUSG0C | false | chr19 | 37550051 | 37551353 | 33.08674 | 107.33151 | 103.57004 | chr19 | 37549871 | 37552859 | 4.11065 | 3.70153 | 2.25912 |
| chr3 | 126596612 | 126597502 | 1000 | Promoter (<=1kb) | 3 | 126596951 | 126844913 | 247963 | 1 | ENSMUST00 | 0 | Camk2d | ENSMUSG0C | false | chr3 | 126595914 | 126598682 | 36.46498 | 119.52629 | 115.39857 | . | . | . | . | . | . |
| chr17 | 36023596 | 36024405 | 1000 | Intron (ENSMUST00000173900.1 | 17 | 36005695 | 36020560 | 14866 | 2 | ENSMUST00 | -3037 | H2-T24 | ENSMUSG0C | false | . | . | . | . | . | . | chr17 | 36019995 | 36026641 | 6.63467 | 4.74288 | 4.36704 |
| chr7 | 127511936 | 127513084 | 1000 | Promoter (<=1kb) | 7 | 127511983 | 127527040 | 15058 | 1 | ENSMUST00 | 0 | Srcap | ENSMUSG0C | false | chr7 | 127511504 | 127513494 | 25.79552 | 96.83904 | 93.34669 | . | . | . | . | . | . |
| chr8 | 68279440 | 68280081 | 1000 | Promoter (2-3kb) | 8 | 68276567 | 68347694 | 71128 | 1 | ENSMUST00 | 2874 | Sh2d4a | ENSMUSG0C | true | chr8 | 68276198 | 68280543 | 31.1999 | 108.05416 | 104.27962 | chr8 | 68277725 | 68299894 | 4.63858 | 3.82888 | 2.72658 |
| chr8 | 68307526 | 68307979 | 1000 | Intron (ENSMUST00000066594.3 | 8 | 68276567 | 68347694 | 71128 | 1 | ENSMUST00 | 30960 | Sh2d4a | ENSMUSG0C | true | . | . | . | . | . | . | chr8 | 68303974 | 68315122 | 5.09606 | 4.07709 | 3.08935 |
| chr8 | 68260317 | 68260716 | 1000 | Distal Intergenic | 8 | 68276567 | 68347694 | 71128 | 1 | ENSMUST00 | -15851 | Sh2d4a | ENSMUSG0C | true | . | . | . | . | . | . | chr8 | 68259255 | 68260869 | 4.20874 | 3.57837 | 2.35849 |
| chr8 | 68276367 | 68277684 | 1000 | Promoter (<=1kb) | 8 | 68276567 | 68347694 | 71128 | 1 | ENSMUST00 | 0 | Sh2d4a | ENSMUSG0C | true | chr8 | 68276198 | 68280543 | 31.1999 | 108.05416 | 104.27962 | chr8 | 68271010 | 68276729 | 5.1542 | 4.08626 | 3.14495 |
| chr7 | 28827396 | 28827687 | 1000 | Promoter (1-2kb) | 7 | 28826204 | 28832247 | 6044 | 1 | ENSMUST00 | 1193 | Ech1 | ENSMUSG0C | false | . | . | . | . | . | . | chr7 | 28811756 | 28829379 | 5.36936 | 4.22722 | 3.31467 |
| chr6 | 72452261 | 72452617 | 1000 | Distal Intergenic | 6 | 72432799 | 72439558 | 6760 | 2 | ENSMUST00 | -12704 | Mat2a | ENSMUSG0C | false | . | . | . | . | . | . | chr6 | 72451261 | 72456501 | 5.99396 | 4.56055 | 3.8382 |
| chr3 | 121426117 | 121426661 | 1000 | Promoter (<=1kb) | 3 | 121426541 | 121458205 | 31665 | 1 | ENSMUST00 | 0 | Cnn3 | ENSMUSG0C | false | chr3 | 121425851 | 121428534 | 43.84918 | 145.52415 | 140.42462 | chr3 | 121425309 | 121426133 | 4.07288 | 3.68561 | 2.22889 |
| chr3 | 121426873 | 121427511 | 1000 | Promoter (<=1kb) | 3 | 121426541 | 121458205 | 31665 | 1 | ENSMUST00 | 333 | Cnn3 | ENSMUSG0C | false | chr3 | 121425851 | 121428534 | 43.84918 | 145.52415 | 140.42462 | . | . | . | . | . | . |
| chr1 | 180417078 | 180417490 | 1000 | Intron (ENSMUST00000070181.6 | 1 | 180432387 | 180483467 | 51081 | 2 | ENSMUST00 | 65977 | 6330403A02I | ENSMUSG0C | false | . | . | . | . | . | . | . | . | . | . | . | . |
| chr4 | 93381735 | 93382052 | 1000 | Distal Intergenic | 4 | 93334138 | 93335511 | 1374 | 2 | ENSMUST00 | -46225 | Tusc1 | ENSMUSG0C | false | . | . | . | . | . | . | chr4 | 93381595 | 93382675 | 3.36709 | 3.16382 | 1.70221 |
| chr12 | 112511056 | 112511359 | 1000 | Promoter (<=1kb) | 12 | 112500184 | 112511160 | 10977 | 2 | ENSMUST00 | 0 | Tmem179 | ENSMUSG0C | false | . | . | . | . | . | . | chr12 | 112510653 | 112511584 | 4.21475 | 3.68696 | 2.35086 |
| chr11 | 93885547 | 93886292 | 1000 | Promoter (<=1kb) | 11 | 93859243 | 93885766 | 26524 | 2 | ENSMUST00 | 0 | Utp18 | ENSMUSG0C | false | chr11 | 93884540 | 93887391 | 41.19049 | 135.33968 | 130.64169 | . | . | . | . | . | . |
| chr3 | 122924025 | 122924497 | 1000 | Promoter (<=1kb) | 3 | 122924242 | 122926194 | 1953 | 1 | ENSMUST00 | 0 | 1810037I17R | ENSMUSG0C | false | chr3 | 122923822 | 122925193 | 27.08035 | 80.13548 | 76.98148 | . | . | . | . | . | . |
| chr12 | 105009587 | 105010272 | 1000 | Promoter (<=1kb) | 12 | 104929949 | 105009809 | 79861 | 2 | ENSMUST00 | 0 | Syne3 | ENSMUSG0C | false | chr12 | 105008329 | 105010500 | 19.08793 | 73.09275 | 70.054 | chr12 | 105000163 | 105017783 | 5.53931 | 4.22324 | 3.47881 |
| chr5 | 33721567 | 33722460 | 1000 | Promoter (<=1kb) | 5 | 33721724 | 33737067 | 15344 | 1 | ENSMUST00 | 0 | Fgfr3 | ENSMUSG0C | false | chr5 | 33721432 | 33723845 | 28.24444 | 85.94312 | 82.67722 | chr5 | 33720971 | 33724013 | 4.45267 | 3.77858 | 2.6086 |
| chr13 | 41689428 | 41690151 | 1000 | Distal Intergenic | 13 | 41655697 | 41656524 | 828 | 1 | ENSMUST00 | 33732 | Gm5082 | ENSMUSG0C | false | . | . | . | . | . | . | chr13 | 41681725 | 41693613 | 4.99308 | 4.08684 | 2.99341 |
| chr13 | 41705556 | 41705960 | 1000 | Distal Intergenic | 13 | 41655697 | 41656524 | 828 | 1 | ENSMUST00 | 49860 | Gm5082 | ENSMUSG0C | false | . | . | . | . | . | . | chr13 | 41700288 | 41710100 | 5.7931 | 4.46811 | 3.65893 |
| chr15 | 7128908 | 7130165 | 1000 | Promoter (<=1kb) | 15 | 7129559 | 7191743 | 62185 | 1 | ENSMUST00 | 0 | Lifr | ENSMUSG0C | false | chr15 | 7128546 | 7131106 | 37.17318 | 134.81126 | 130.13722 | . | . | . | . | . | . |
| chr5 | 32854090 | 32854514 | 1000 | Promoter (<=1kb) | 5 | 32789218 | 32854256 | 65039 | 2 | ENSMUST00 | 0 | Prr14l | ENSMUSG0C | false | chr5 | 32853439 | 32854321 | 43.14866 | 145.43195 | 140.33446 | chr5 | 32851198 | 32855103 | 5.56523 | 4.26263 | 3.47964 |
| chr8 | 105445882 | 105446515 | 1000 | Exon (ENSMUST00000109355.3/ | 8 | 105427640 | 105464084 | 36445 | 1 | ENSMUST00 | 18243 | Lrrc36 | ENSMUSG0C | false | . | . | . | . | . | . | chr8 | 105445249 | 105446373 | 3.93552 | 3.43196 | 2.17098 |
| chr12 | 8428606 | 8429000 | 1000 | Distal Intergenic | 12 | 8497763 | 8499985 | 2223 | 2 | ENSMUST00 | 70985 | Rhob | ENSMUSG0C | false | . | . | . | . | . | . | chr12 | 8427817 | 8429728 | 4.09208 | 3.59554 | 2.30498 |
| chr8 | 54529357 | 54530352 | 1000 | Promoter (<=1kb) | 8 | 54520433 | 54529998 | 9566 | 2 | ENSMUST00 | 0 | Spcs3 | ENSMUSG0C | false | chr8 | 54529007 | 54530608 | 35.33424 | 109.18479 | 105.37405 | . | . | . | . | . | . |
| chr10 | 81560875 | 81561392 | 1000 | Promoter (1-2kb) | 10 | 81559494 | 81566362 | 6869 | 1 | ENSMUST00 | 1382 | Aes | ENSMUSG0C | false | chr10 | 81559124 | 81561616 | 23.12601 | 77.64799 | 74.53377 | chr10 | 81560928 | 81565156 | 7.15643 | 4.80865 | 4.80181 |
| chr17 | 73107715 | 73108156 | 1000 | Promoter (<=1kb) | 17 | 73107985 | 73161991 | 54007 | 1 | ENSMUST00 | 0 | Lclat1 | ENSMUSG0C | false | chr17 | 73107245 | 73108937 | 43.83018 | 166.11598 | 160.25778 | . | . | . | . | . | . |
| chr6 | 117437647 | 117438024 | 1000 | Distal Intergenic | 6 | 117559697 | 117568193 | 8497 | 1 | ENSMUST00 | -121673 | Gm9946 | ENSMUSG0C | false | . | . | . | . | . | . | chr6 | 117434327 | 117438832 | 4.13968 | 3.59187 | 2.31244 |
| chr7 | 134026259 | 134026556 | 1000 | Intron (ENSMUST00000067680.5 | 7 | 133886196 | 133961265 | 75070 | 2 | ENSMUST00 | -64995 | Adam12 | ENSMUSG0C | false | . | . | . | . | . | . | chr7 | 134026117 | 134028829 | 3.75726 | 3.46209 | 2.00995 |
| chr19 | 4396377 | 4397833 | 1000 | Promoter (<=1kb) | 19 | 4316170 | 4397077 | 80908 | 2 | ENSMUST00 | 0 | Kdm2a | ENSMUSG0C | false | chr19 | 4395327 | 4397114 | 41.97713 | 143.38342 | 138.36984 | . | . | . | . | . | . |
| chr7 | 137048117 | 137048457 | 1000 | Intron (ENSMUST00000081510.2 | 7 | 136894611 | 137128187 | 233577 | 1 | ENSMUST00 | 153507 | Mgmt | ENSMUSG0C | false | . | . | . | . | . | . | chr7 | 137047111 | 137049679 | 4.64201 | 3.9467 | 2.69823 |
| chr7 | 137048553 | 137049013 | 1000 | Intron (ENSMUST00000081510.2 | 7 | 136894611 | 137128187 | 233577 | 1 | ENSMUST00 | 153943 | Mgmt | ENSMUSG0C | false | . | . | . | . | . | . | chr7 | 137047111 | 137049679 | 4.64201 | 3.9467 | 2.69823 |
| chr10 | 81643362 | 81643909 | 1000 | Promoter (<=1kb) | 10 | 81643803 | 81647474 | 3672 | 1 | ENSMUST00 | 0 | Ankrd24 | ENSMUSG0C | false | chr10 | 81642746 | 81644258 | 32.06079 | 99.44822 | 95.89121 | chr10 | 81642077 | 81644665 | 3.40147 | 3.09571 | 1.71882 |
| chr5 | 105519086 | 105519671 | 1000 | Promoter (<=1kb) | 5 | 105519388 | 105613018 | 93631 | 1 | ENSMUST00 | 0 | Lrrc8c | ENSMUSG0C | false | chr5 | 105518730 | 105520178 | 15.94852 | 63.64039 | 60.74356 | chr5 | 105517662 | 105520105 | 4.37462 | 3.74225 | 2.50101 |
| chr2 | 30065981 | 30067459 | 1000 | Promoter (<=1kb) | 2 | 30066456 | 30072577 | 6122 | 1 | ENSMUST00 | 0 | Set | ENSMUSG0C | false | chr2 | 30065820 | 30067950 | 34.78613 | 113.42423 | 109.48744 | chr2 | 30067436 | 30068329 | 4.26347 | 3.63475 | 2.42072 |
| chr7 | 28939527 | 28940169 | 1000 | Intron (ENSMUST00000068045.8 | 7 | 28913378 | 28949759 | 36382 | 2 | ENSMUST00 | 9590 | Actn4 | ENSMUSG0C | true | . | . | . | . | . | . | chr7 | 28910727 | 28959574 | 8.49369 | 5.62595 | 5.86532 |
| chr7 | 28923885 | 28924304 | 1000 | Intron (ENSMUST00000068045.8 | 7 | 28907014 | 28919221 | 12208 | 2 | ENSMUST00 | -4665 | Actn4 | ENSMUSG0C | true | . | . | . | . | . | . | chr7 | 28910727 | 28959574 | 8.49369 | 5.62595 | 5.86532 |
| chr7 | 28930737 | 28931074 | 1000 | Intron (ENSMUST00000068045.8 | 7 | 28907014 | 28919221 | 12208 | 2 | ENSMUST00 | -11517 | Actn4 | ENSMUSG0C | true | . | . | . | . | . | . | chr7 | 28910727 | 28959574 | 8.49369 | 5.62595 | 5.86532 |
| chr7 | 28962200 | 28962612 | 1000 | Promoter (<=1kb) | 7 | 28893248 | 28962340 | 69093 | 2 | ENSMUST00 | 0 | Actn4 | ENSMUSG0C | true | chr7 | 28958851 | 28962304 | 37.17318 | 134.81126 | 130.13722 | . | . | . | . | . | . |
| chr7 | 28925287 | 28925724 | 1000 | Intron (ENSMUST00000068045.8 | 7 | 28907014 | 28919221 | 12208 | 2 | ENSMUST00 | -6067 | Actn4 | ENSMUSG0C | true | . | . | . | . | . | . | chr7 | 28910727 | 28959574 | 8.49369 | 5.62595 | 5.86532 |
| chr7 | 28964387 | 28964881 | 1000 | Promoter (2-3kb) | 7 | 28893248 | 28962340 | 69093 | 2 | ENSMUST00 | -2048 | Actn4 | ENSMUSG0C | true | . | . | . | . | . | . | chr7 | 28962675 | 28973922 | 7.08725 | 4.96018 | 4.72253 |
| chr7 | 28945529 | 28946212 | 1000 | Intron (ENSMUST00000068045.8 | 7 | 28913378 | 28949759 | 36382 | 2 | ENSMUST00 | 3547 | Actn4 | ENSMUSG0C | true | . | . | . | . | . | . | chr7 | 28910727 | 28959574 | 8.49369 | 5.62595 | 5.86532 |
| chr7 | 28894868 | 28895277 | 1000 | Promoter (<=1kb) | 7 | 28894232 | 28894852 | 621 | 2 | ENSMUST00 | -17 | Actn4 | ENSMUSG0C | true | . | . | . | . | . | . | chr7 | 28892787 | 28898188 | 5.46695 | 4.19683 | 3.43075 |
| chr19 | 57610368 | 57611202 | 1000 | Promoter (<=1kb) | 19 | 57611034 | 58133338 | 522305 | 1 | ENSMUST00 | 0 | Atrnl1 | ENSMUSG0C | false | chr19 | 57609779 | 57612310 | 33.92788 | 125.59804 | 121.27206 | chr19 | 57609822 | 57610372 | 3.46872 | 3.15578 | 1.78369 |
| chr17 | 80760894 | 80761188 | 1000 | Distal Intergenic | 17 | 80728131 | 80728522 | 392 | 1 | ENSMUST00 | 32764 | Gm9959 | ENSMUSG0C | false | . | . | . | . | . | . | . | . | . | . | . | . |
| chr9 | 65394313 | 65394738 | 1000 | Promoter (2-3kb) | 9 | 65388684 | 65391652 | 2969 | 2 | ENSMUST00 | -2662 | Kbtbd13 | ENSMUSG0C | false | . | . | . | . | . | . | . | . | . | . | . | . |
| chr1 | 89454290 | 89455256 | 1000 | Promoter (<=1kb) | 1 | 89454806 | 89897617 | 442812 | 1 | ENSMUST00 | 0 | Agap1 | ENSMUSG0C | false | chr1 | 89454009 | 89456011 | 24.78212 | 78.86476 | 75.73294 | . | . | . | . | . | . |
| chr15 | 81584994 | 81586476 | 1000 | Promoter (<=1kb) | 15 | 81585351 | 81588344 | 2994 | 1 | ENSMUST00 | 0 | Ep300 | ENSMUSG0C | false | chr15 | 81584642 | 81585068 | 14.66885 | 35.63733 | 33.10225 | . | . | . | . | . | . |
| chr10 | 81401790 | 81402539 | 1000 | Promoter (1-2kb) | 10 | 81399196 | 81400554 | 1359 | 2 | ENSMUST00 | -1237 | Nfic | ENSMUSG0C | false | chr10 | 81401906 | 81402504 | 5.97392 | 9.35129 | 7.18948 | chr10 | 81392649 | 81426205 | 8.72751 | 5.63067 | 6.0679 |
| chr10 | 81426041 | 81427408 | 1000 | Promoter (<=1kb) | 10 | 81396186 | 81427187 | 31002 | 2 | ENSMUST00 | 0 | Nfic | ENSMUSG0C | false | chr10 | 81425223 | 81427495 | 34.46491 | 104.61457 | 100.92722 | chr10 | 81427223 | 81436730 | 7.9801 | 5.39294 | 5.46678 |
| chr10 | 81435392 | 81435753 | 1000 | Distal Intergenic | 10 | 81400953 | 81431005 | 30053 | 2 | ENSMUST00 | -4388 | Nfic | ENSMUSG0C | false | . | . | . | . | . | . | chr10 | 81427223 | 81436730 | 7.9801 | 5.39294 | 5.46678 |
| chr10 | 81429769 | 81431234 | 1000 | Promoter (<=1kb) | 10 | 81400953 | 81431005 | 30053 | 2 | ENSMUST00 | 0 | Nfic | ENSMUSG0C | false | chr10 | 81429715 | 81431421 | 37.37859 | 116.88169 | 112.84478 | chr10 | 81427223 | 81436730 | 7.9801 | 5.39294 | 5.46678 |
| chr7 | 95789482 | 95789886 | 1000 | Distal Intergenic | 7 | 95958215 | 95961613 | 3399 | 1 | ENSMUST00 | -168329 | Gm9966 | ENSMUSG0C | false | . | . | . | . | . | . | chr7 | 95784545 | 95792909 | 5.1339 | 4.17838 | 3.10723 |
| chr13 | 110845695 | 110845997 | 1000 | Distal Intergenic | 13 | 111255013 | 111257749 | 2737 | 1 | ENSMUST00 | -409016 | Actbl2 | ENSMUSG0C | false | . | . | . | . | . | . | chr13 | 110844252 | 110846792 | 3.88323 | 3.49356 | 2.09543 |
| chr13 | 110913376 | 110913753 | 1000 | Distal Intergenic | 13 | 111255013 | 111257749 | 2737 | 1 | ENSMUST00 | -341260 | Actbl2 | ENSMUSG0C | false | . | . | . | . | . | . | chr13 | 110913081 | 110914475 | 4.20555 | 3.58008 | 2.37476 |
| chr4 | 114821569 | 114822397 | 1000 | Promoter (<=1kb) | 4 | 114821722 | 114856166 | 34445 | 1 | ENSMUST00 | 0 | Gm12830 | ENSMUSG0C | false | chr4 | 114821453 | 114822346 | 22.05606 | 59.8844 | 57.03867 | chr4 | 114821020 | 114823896 | 3.44732 | 3.17501 | 1.76058 |
| chr4 | 114779795 | 114780366 | 1000 | Distal Intergenic | 4 | 114821722 | 114856166 | 34445 | 1 | ENSMUST00 | -41356 | Gm12830 | ENSMUSG0C | false | . | . | . | . | . | . | chr4 | 114777307 | 114780683 | 6.16318 | 4.61474 | 3.98775 |
| chr5 | 90365253 | 90367027 | 1000 | Promoter (<=1kb) | 5 | 90227190 | 90366185 | 138996 | 2 | ENSMUST00 | 0 | Ankrd17 | ENSMUSG0C | false | chr5 | 90364700 | 90367410 | 33.04283 | 115.53907 | 111.53828 | . | . | . | . | . | . |
| chr6 | 72898810 | 72900340 | 1000 | Promoter (<=1kb) | 6 | 72841114 | 72899979 | 58866 | 2 | ENSMUST00 | 0 | Kcmf1 | ENSMUSG0C | false | chr6 | 72898418 | 72900908 | 37.2221 | 115.56093 | 111.55734 | . | . | . | . | . | . |
| chr13 | 59009501 | 59010110 | 1000 | Intron (ENSMUST00000079828.5 | 13 | 58807900 | 59133970 | 326071 | 1 | ENSMUST00 | 201602 | Ntrk2 | ENSMUSG0C | false | . | . | . | . | . | . | . | . | . | . | . | . |
| chr4 | 56946685 | 56947694 | 1000 | Promoter (<=1kb) | 4 | 56866923 | 56947437 | 80515 | 2 | ENSMUST00 | 0 | Tmem245 | ENSMUSG0C | false | chr4 | 56945931 | 56947685 | 25.17393 | 93.69764 | 90.27328 | . | . | . | . | . | . |
| chr7 | 112678900 | 112680254 | 1000 | Promoter (<=1kb) | 7 | 112679318 | 112899459 | 220142 | 1 | ENSMUST00 | 0 | Tead1 | ENSMUSG0C | false | chr7 | 112678205 | 112681471 | 36.76257 | 113.72284 | 109.77938 | chr7 | 112677714 | 112678974 | 5.79943 | 4.50423 | 3.65278 |
| chr7 | 112697158 | 112697845 | 1000 | Intron (ENSMUST00000084705.6 | 7 | 112691698 | 112759529 | 67832 | 1 | ENSMUST00 | 5461 | Tead1 | ENSMUSG0C | false | . | . | . | . | . | . | chr7 | 112684336 | 112703707 | 6.1881 | 4.54631 | 4.00467 |
| chr7 | 112728462 | 112728806 | 1000 | Intron (ENSMUST00000084705.6 | 7 | 112742045 | 112898660 | 156616 | 1 | ENSMUST00 | -13239 | Tead1 | ENSMUSG0C | false | . | . | . | . | . | . | chr7 | 112726136 | 112737477 | 6.09225 | 4.56315 | 3.92226 |
| chr7 | 112731547 | 112732035 | 1000 | Intron (ENSMUST00000084705.6 | 7 | 112742045 | 112898660 | 156616 | 1 | ENSMUST00 | -10010 | Tead1 | ENSMUSG0C | false | . | . | . | . | . | . | chr7 | 112726136 | 112737477 | 6.09225 | 4.56315 | 3.92226 |
| chr1 | 73930747 | 73931118 | 1000 | Promoter (<=1kb) | 1 | 73925417 | 73930811 | 5395 | 2 | ENSMUST00 | 0 | Tns1 | ENSMUSG0C | false | . | . | . | . | . | . | chr1 | 73927832 | 73933110 | 4.94644 | 3.95251 | 2.98269 |
| chr1 | 74124369 | 74124783 | 1000 | Promoter (<=1kb) | 1 | 73910231 | 74124447 | 214217 | 2 | ENSMUST00 | 0 | Tns1 | ENSMUSG0C | false | chr1 | 74124171 | 74125317 | 22.10575 | 73.03297 | 69.99519 | chr1 | 74124128 | 74125590 | 4.70474 | 4.01886 | 2.75832 |
| chr1 | 74081634 | 74082061 | 1000 | Intron (ENSMUST00000169786.2 | 1 | 73910232 | 74091351 | 181120 | 2 | ENSMUST00 | 9290 | Tns1 | ENSMUSG0C | false | . | . | . | . | . | . | chr1 | 74081332 | 74083308 | 5.0869 | 3.92899 | 3.09986 |
| chr4 | 148151106 | 148151525 | 1000 | Promoter (<=1kb) | 4 | 148149444 | 148151646 | 2203 | 2 | ENSMUST00 | 121 | Fbxo6 | ENSMUSG0C | false | chr4 | 148150673 | 148152220 | 31.96453 | 96.29042 | 92.80983 | chr4 | 148148886 | 148152963 | 5.22456 | 4.18047 | 3.19373 |
| chr6 | 89050894 | 89051318 | 1000 | Intron (ENSMUST00000168837.3 | 6 | 89096110 | 89107834 | 11725 | 1 | ENSMUST00 | -44792 | 4933427D06I | ENSMUSG0C | false | . | . | . | . | . | . | . | . | . | . | . | . |
| chr14 | 93734248 | 93734885 | 1000 | Intron (ENSMUST00000068992.2 | 14 | 93015512 | 93888732 | 873221 | 2 | ENSMUST00 | 153847 | Pcdh9 | ENSMUSG0C | false | . | . | . | . | . | . | chr14 | 93733460 | 93735794 | 4.11716 | 3.6607 | 2.3151 |
| chr2 | 157078752 | 157079586 | 1000 | Promoter (<=1kb) | 2 | 157015799 | 157079254 | 63456 | 2 | ENSMUST00 | 0 | Soga1 | ENSMUSG0C | false | chr2 | 157077375 | 157079761 | 27.41053 | 90.23362 | 86.88379 | chr2 | 157074030 | 157079796 | 4.31246 | 3.68722 | 2.46593 |
| chr19 | 55244768 | 55245417 | 1000 | Distal Intergenic | 19 | 55198297 | 55241236 | 42940 | 2 | ENSMUST00 | -3533 | Gucy2g | ENSMUSG0C | false | . | . | . | . | . | . | chr19 | 55244366 | 55246544 | 5.44545 | 4.08015 | 3.41307 |
| chr19 | 55217497 | 55217841 | 1000 | Intron (ENSMUST00000069183.6 | 19 | 55198297 | 55241236 | 42940 | 2 | ENSMUST00 | 23395 | Gucy2g | ENSMUSG0C | false | . | . | . | . | . | . | chr19 | 55216941 | 55218667 | 5.89084 | 4.38225 | 3.79487 |
| chr14 | 25725074 | 25725616 | 1000 | Intron (ENSMUST00000069180.7 | 14 | 25711644 | 25719792 | 8149 | 2 | ENSMUST00 | -5283 | Zcchc24 | ENSMUSG0C | false | . | . | . | . | . | . | chr14 | 25724370 | 25726567 | 5.80514 | 4.21385 | 3.69433 |
| chr14 | 25718419 | 25718908 | 1000 | Promoter (<=1kb) | 14 | 25711644 | 25719792 | 8149 | 2 | ENSMUST00 | 884 | Zcchc24 | ENSMUSG0C | false | . | . | . | . | . | . | chr14 | 25717907 | 25719376 | 4.94144 | 3.92249 | 2.99753 |
| chr7 | 141047740 | 141048299 | 1000 | Distal Intergenic | 7 | 141061274 | 141072119 | 10846 | 1 | ENSMUST00 | -12975 | B4galnt4 | ENSMUSG0C | false | chr7 | 141047937 | 141048901 | 11.87651 | 25.14597 | 22.74365 | chr7 | 141046795 | 141048933 | 3.50285 | 3.33276 | 1.77277 |
| chr1 | 180343814 | 180344244 | 1000 | Intron (ENSMUST00000070181.6 | 1 | 180326970 | 180330550 | 3581 | 2 | ENSMUST00 | -13265 | Gm5069 | ENSMUSG0C | false | . | . | . | . | . | . | chr1 | 180343468 | 180344577 | 2.95156 | 2.97523 | 1.34344 |
| chr1 | 180365094 | 180365826 | 1000 | Intron (ENSMUST00000070181.6 | 1 | 180326970 | 180330550 | 3581 | 2 | ENSMUST00 | -34545 | Gm5069 | ENSMUSG0C | false | chr1 | 180365023 | 180365940 | 12.34421 | 27.73961 | 25.30389 | chr1 | 180364576 | 180367327 | 4.22095 | 3.66911 | 2.38032 |
| chr7 | 114086341 | 114086921 | 1000 | Intron (ENSMUST00000069449.5 | 7 | 114046785 | 114117781 | 70997 | 2 | ENSMUST00 | 30860 | Rras2 | ENSMUSG0C | true | . | . | . | . | . | . | chr7 | 114058507 | 114116129 | 5.64505 | 4.33683 | 3.54844 |
| chr7 | 114087308 | 114087705 | 1000 | Intron (ENSMUST00000069449.5 | 7 | 114046785 | 114117781 | 70997 | 2 | ENSMUST00 | 30076 | Rras2 | ENSMUSG0C | true | . | . | . | . | . | . | chr7 | 114058507 | 114116129 | 5.64505 | 4.33683 | 3.54844 |
| chr7 | 114116839 | 114117396 | 1000 | Promoter (<=1kb) | 7 | 114046785 | 114117781 | 70997 | 2 | ENSMUST00 | 385 | Rras2 | ENSMUSG0C | true | chr7 | 114115691 | 114118263 | 32.67761 | 108.66347 | 104.86716 | . | . | . | . | . | . |
| chr15 | 3433166 | 3433622 | 1000 | Intron (ENSMUST00000069451.5 | 15 | 3327539 | 3471644 | 144106 | 2 | ENSMUST00 | 38022 | Ghr | ENSMUSG0C | false | . | . | . | . | . | . | chr15 | 3424973 | 3437760 | 4.97978 | 4.03653 | 3.00524 |
| chr15 | 3582981 | 3583639 | 1000 | Promoter (<=1kb) | 15 | 3317760 | 3583230 | 265471 | 2 | ENSMUST00 | 0 | Ghr | ENSMUSG0C | false | chr15 | 3582852 | 3583587 | 33.01748 | 102.11515 | 98.48943 | . | . | . | . | . | . |
| chr18 | 30487175 | 30487491 | 1000 | Distal Intergenic | 18 | 30495001 | 30495375 | 375 | 2 | ENSMUST00 | 7884 | Gm7936 | ENSMUSG0C | false | . | . | . | . | . | . | chr18 | 30486688 | 30488192 | 4.47405 | 3.84126 | 2.62731 |
| chr6 | 72788718 | 72789679 | 1000 | Promoter (<=1kb) | 6 | 72626378 | 72789254 | 162877 | 2 | ENSMUST00 | 0 | Tcf7l1 | ENSMUSG0C | false | chr6 | 72787705 | 72789802 | 36.88168 | 122.38801 | 118.16834 | chr6 | 72789635 | 72790122 | 8.14537 | 5.60902 | 5.56039 |
| chr6 | 72743683 | 72744402 | 1000 | Intron (ENSMUST00000069536.6 | 6 | 72632976 | 72788598 | 155623 | 2 | ENSMUST00 | 44196 | Tcf7l1 | ENSMUSG0C | false | . | . | . | . | . | . | chr6 | 72739919 | 72749221 | 7.65916 | 5.17359 | 5.21352 |
| chr11 | 103158600 | 103159193 | 1000 | Distal Intergenic | 11 | 103171107 | 103198901 | 27795 | 1 | ENSMUST00 | -11914 | Fmnl1 | ENSMUSG0C | false | chr11 | 103158897 | 103159545 | 11.18601 | 23.41478 | 21.03445 | chr11 | 103152989 | 103159932 | 4.37918 | 3.77334 | 2.50587 |
| chr17 | 23828944 | 23829411 | 1000 | Promoter (<=1kb) | 17 | 23824740 | 23829109 | 4370 | 2 | ENSMUST00 | 0 | Tceb2 | ENSMUSG0C | false | chr17 | 23828084 | 23829590 | 33.94791 | 109.87469 | 106.04007 | chr17 | 23826449 | 23829983 | 3.36669 | 3.12595 | 1.68099 |
| chr1 | 91456524 | 91456905 | 1000 | Promoter (2-3kb) | 1 | 91415982 | 91459324 | 43343 | 2 | ENSMUST00 | 2419 | Per2 | ENSMUSG0C | false | . | . | . | . | . | . | chr1 | 91451145 | 91461043 | 6.24965 | 4.62037 | 4.05296 |
| chr2 | 181483629 | 181483908 | 1000 | Distal Intergenic | 2 | 181493206 | 181494980 | 1775 | 1 | ENSMUST00 | -9298 | Abhd16b | ENSMUSG0C | false | . | . | . | . | . | . | chr2 | 181482359 | 181484246 | 4.57936 | 3.84452 | 2.72151 |
| chr12 | 40387690 | 40388161 | 1000 | Intron (ENSMUST00000069637.8 | 12 | 40315558 | 40445769 | 130212 | 2 | ENSMUST00 | 57608 | Zfp277 | ENSMUSG0C | false | . | . | . | . | . | . | chr12 | 40378830 | 40393499 | 4.81877 | 3.95373 | 2.86926 |
| chr12 | 40438043 | 40438676 | 1000 | Intron (ENSMUST00000069637.8 | 12 | 40315558 | 40445769 | 130212 | 2 | ENSMUST00 | 7093 | Zfp277 | ENSMUSG0C | false | . | . | . | . | . | . | chr12 | 40422298 | 40443652 | 4.88409 | 4.00258 | 2.915 |
| chr8 | 91376662 | 91377206 | 1000 | Intron (ENSMUST00000069718.8 | 8 | 91387736 | 91409840 | 22105 | 1 | ENSMUST00 | -10530 | Fto | ENSMUSG0C | false | . | . | . | . | . | . | chr8 | 91370620 | 91386905 | 5.91421 | 4.41152 | 3.77852 |
| chr8 | 91522206 | 91522591 | 1000 | Intron (ENSMUST00000069718.8 | 8 | 91467685 | 91666946 | 199262 | 1 | ENSMUST00 | 54522 | Fto | ENSMUSG0C | false | . | . | . | . | . | . | chr8 | 91521033 | 91528017 | 5.83853 | 4.42897 | 3.70928 |
| chr8 | 91338748 | 91339148 | 1000 | Intron (ENSMUST00000069718.8 | 8 | 91313568 | 91487791 | 174224 | 1 | ENSMUST00 | 25181 | Fto | ENSMUSG0C | false | . | . | . | . | . | . | chr8 | 91337651 | 91367410 | 5.62588 | 4.32828 | 3.53466 |
| chr1 | 82290683 | 82292030 | 1000 | Promoter (<=1kb) | 1 | 82233101 | 82291416 | 58316 | 2 | ENSMUST00 | 0 | Irs1 | ENSMUSG0C | false | chr1 | 82289450 | 82292285 | 25.50663 | 88.63404 | 85.31516 | chr1 | 82291780 | 82294158 | 5.60696 | 4.25715 | 3.53276 |
| chr1 | 82274145 | 82274706 | 1000 | Intron (ENSMUST00000069799.2 | 1 | 82233101 | 82291416 | 58316 | 2 | ENSMUST00 | 16710 | Irs1 | ENSMUSG0C | false | . | . | . | . | . | . | chr1 | 82270621 | 82285607 | 3.93745 | 3.56507 | 2.13398 |
| chr1 | 82283089 | 82283716 | 1000 | Intron (ENSMUST00000069799.2 | 1 | 82233101 | 82291416 | 58316 | 2 | ENSMUST00 | 7700 | Irs1 | ENSMUSG0C | false | chr1 | 82282881 | 82284308 | 16.8969 | 46.84153 | 44.16631 | chr1 | 82270621 | 82285607 | 3.93745 | 3.56507 | 2.13398 |
| chr8 | 88666423 | 88666780 | 1000 | Intron (ENSMUST00000054324.9 | 8 | 88651341 | 88688474 | 37134 | 1 | ENSMUST00 | 15083 | Nod2 | ENSMUSG0C | false | . | . | . | . | . | . | . | . | . | . | . | . |
| chr5 | 9265909 | 9266244 | 1000 | Promoter (<=1kb) | 5 | 9266118 | 9479736 | 213619 | 1 | ENSMUST00 | 0 | 9330182L06I | ENSMUSG0C | false | chr5 | 9265761 | 9266601 | 21.5194 | 62.35189 | 59.47348 | chr5 | 9265735 | 9266708 | 3.00757 | 2.86347 | 1.41548 |
| chr11 | 94716893 | 94717218 | 1000 | Distal Intergenic | 11 | 94694498 | 94704499 | 10002 | 2 | ENSMUST00 | -12395 | Gm11541 | ENSMUSG0C | false | . | . | . | . | . | . | chr11 | 94715033 | 94719556 | 5.32725 | 4.25479 | 3.2741 |
| chr1 | 183369441 | 183369803 | 1000 | Promoter (<=1kb) | 1 | 183326725 | 183369529 | 42805 | 2 | ENSMUST00 | 0 | Mia3 | ENSMUSG0C | false | chr1 | 183368584 | 183370028 | 38.04976 | 128.75285 | 124.30675 | chr1 | 183367338 | 183370220 | 4.85798 | 3.90169 | 2.92728 |
| chr1 | 183345229 | 183345755 | 1000 | Promoter (<=1kb) | 1 | 183326725 | 183345296 | 18572 | 2 | ENSMUST00 | 0 | Mia3 | ENSMUSG0C | false | chr1 | 183344773 | 183346260 | 37.7199 | 126.00829 | 121.65485 | chr1 | 183344788 | 183346266 | 4.10935 | 3.70331 | 2.307 |
| chr15 | 27782674 | 27783294 | 1000 | Intron (ENSMUST00000090247.5 | 15 | 27655071 | 27681542 | 26472 | 2 | ENSMUST00 | -101133 | Fam105a | ENSMUSG0C | false | . | . | . | . | . | . | chr15 | 27777264 | 27789305 | 7.24061 | 5.11987 | 4.8476 |
| chr15 | 25510049 | 25510526 | 1000 | Distal Intergenic | 15 | 25414192 | 25414844 | 653 | 1 | ENSMUST00 | 95858 | Gm5468 | ENSMUSG0C | false | . | . | . | . | . | . | chr15 | 25508591 | 25510817 | 3.653 | 3.09453 | 1.9373 |
| chr6 | 72097301 | 72098035 | 1000 | Promoter (<=1kb) | 6 | 72097592 | 72154571 | 56980 | 1 | ENSMUST00 | 0 | St3gal5 | ENSMUSG0C | false | chr6 | 72096911 | 72098643 | 37.13757 | 117.51956 | 113.45509 | chr6 | 72095758 | 72100529 | 3.80657 | 3.46166 | 2.02715 |
| chr6 | 72092757 | 72093081 | 1000 | Distal Intergenic | 6 | 72097592 | 72154571 | 56980 | 1 | ENSMUST00 | -4511 | St3gal5 | ENSMUSG0C | false | . | . | . | . | . | . | chr6 | 72091246 | 72093533 | 4.49212 | 3.74748 | 2.60796 |
| chr10 | 53434425 | 53434949 | 1000 | Distal Intergenic | 10 | 53474964 | 53475467 | 504 | 1 | ENSMUST00 | -40015 | Gm5423 | ENSMUSG0C | false | . | . | . | . | . | . | chr10 | 53434894 | 53439749 | 5.3576 | 4.21111 | 3.30184 |
| chr1 | 138930702 | 138931159 | 1000 | Distal Intergenic | 1 | 138963436 | 139063719 | 100284 | 1 | ENSMUST00 | -32277 | Dennd1b | ENSMUSG0C | false | . | . | . | . | . | . | chr1 | 138929024 | 138931119 | 3.34952 | 3.14719 | 1.69616 |
| chr1 | 138937891 | 138938428 | 1000 | Distal Intergenic | 1 | 138963436 | 139063719 | 100284 | 1 | ENSMUST00 | -25008 | Dennd1b | ENSMUSG0C | false | . | . | . | . | . | . | chr1 | 138936117 | 138938994 | 4.42918 | 3.8072 | 2.53869 |
| chr11 | 23306024 | 23307260 | 1000 | Promoter (<=1kb) | 11 | 23306895 | 23490560 | 183666 | 1 | ENSMUST00 | 0 | Usp34 | ENSMUSG0C | false | chr11 | 23305515 | 23307804 | 36.53698 | 118.59147 | 114.49885 | . | . | . | . | . | . |
| chr11 | 35142743 | 35143195 | 1000 | Intron (ENSMUST00000069837.3 | 11 | 35121224 | 35708507 | 587284 | 1 | ENSMUST00 | 21520 | Slit3 | ENSMUSG0C | false | . | . | . | . | . | . | chr11 | 35141575 | 35143716 | 5.27437 | 4.14805 | 3.26167 |
| chr2 | 167683537 | 167684013 | 1000 | Intron (ENSMUST00000148262.1 | 2 | 167688915 | 167690418 | 1504 | 1 | ENSMUST00 | -4902 | Cebpb | ENSMUSG0C | false | . | . | . | . | . | . | chr2 | 167682328 | 167686587 | 4.12823 | 3.6474 | 2.30102 |
| chr17 | 27820421 | 27821083 | 1000 | Promoter (<=1kb) | 17 | 27751232 | 27820558 | 69327 | 2 | ENSMUST00 | 0 | D17Wsu92e | ENSMUSG0C | false | chr17 | 27819348 | 27820669 | 31.14379 | 104.89771 | 101.20105 | chr17 | 27821022 | 27821718 | 6.14206 | 4.59952 | 3.9878 |
| chr9 | 110653844 | 110654340 | 1000 | Promoter (<=1kb) | 9 | 110624789 | 110654161 | 29373 | 2 | ENSMUST00 | 0 | Nbeal2 | ENSMUSG0C | false | chr9 | 110653365 | 110654625 | 23.93078 | 87.47927 | 84.18233 | chr9 | 110651086 | 110659022 | 5.37784 | 4.15081 | 3.31581 |
| chr6 | 72552673 | 72553239 | 1000 | Promoter (<=1kb) | 6 | 72552938 | 72558257 | 5320 | 1 | ENSMUST00 | 0 | Capg | ENSMUSG0C | false | chr6 | 72552561 | 72553439 | 19.75988 | 50.25065 | 47.53236 | chr6 | 72552300 | 72553486 | 3.53422 | 3.37582 | 1.79671 |
| chr11 | 66067560 | 66068125 | 1000 | Intron (ENSMUST00000080665.4 | 11 | 65831403 | 66005195 | 173793 | 2 | ENSMUST00 | -62366 | Dnah9 | ENSMUSG0C | false | . | . | . | . | . | . | . | . | . | . | . | . |
| chr11 | 66046677 | 66047060 | 1000 | Intron (ENSMUST00000080665.4 | 11 | 65831403 | 66005195 | 173793 | 2 | ENSMUST00 | -41483 | Dnah9 | ENSMUSG0C | false | . | . | . | . | . | . | chr11 | 66044814 | 66048224 | 4.09724 | 3.67211 | 2.25748 |
| chr8 | 84071799 | 84072206 | 1000 | Intron (ENSMUST00000005600.5 | 8 | 84065236 | 84067287 | 2052 | 2 | ENSMUST00 | -4513 | C330011M18 | ENSMUSG0C | false | . | . | . | . | . | . | chr8 | 84067446 | 84075532 | 6.20464 | 4.49658 | 4.02581 |
| chr7 | 40192358 | 40192750 | 1000 | Promoter (1-2kb) | 7 | 40190935 | 40191324 | 390 | 1 | ENSMUST00 | 1424 | Gm6851 | ENSMUSG0C | false | . | . | . | . | . | . | . | . | . | . | . | . |
| chr3 | 32822658 | 32823145 | 1000 | Intron (ENSMUST00000072312.6 | 3 | 32817711 | 32915033 | 97323 | 1 | ENSMUST00 | 4948 | Usp13 | ENSMUSG0C | true | . | . | . | . | . | . | chr3 | 32819248 | 32827108 | 6.69053 | 4.68007 | 4.42218 |
| chr6 | 113696902 | 113697493 | 1000 | Promoter (<=1kb) | 6 | 113697050 | 113711069 | 14020 | 1 | ENSMUST00 | 0 | Tatdn2 | ENSMUSG0C | false | chr6 | 113696360 | 113698200 | 36.76257 | 113.72284 | 109.77938 | . | . | . | . | . | . |
| chr2 | 35109318 | 35109979 | 1000 | Promoter (<=1kb) | 2 | 35109492 | 35130713 | 21222 | 1 | ENSMUST00 | 0 | Cntrl | ENSMUSG0C | false | chr2 | 35108876 | 35110289 | 25.50663 | 88.63404 | 85.31516 | chr2 | 35108791 | 35110833 | 4.17046 | 3.76631 | 2.3065 |
| chr2 | 161108319 | 161108928 | 1000 | Promoter (<=1kb) | 2 | 161035031 | 161109004 | 73974 | 2 | ENSMUST00 | 76 | Chd6 | ENSMUSG0C | false | chr2 | 161107862 | 161109129 | 31.28813 | 116.44205 | 112.41405 | . | . | . | . | . | . |
| chr2 | 161102181 | 161102610 | 1000 | Intron (ENSMUST00000039782.8 | 2 | 161035031 | 161109004 | 73974 | 2 | ENSMUST00 | 6394 | Chd6 | ENSMUSG0C | false | . | . | . | . | . | . | chr2 | 161099721 | 161107860 | 4.98733 | 3.91915 | 3.02701 |
| chr7 | 104352950 | 104353562 | 1000 | Promoter (<=1kb) | 7 | 104338754 | 104353362 | 14609 | 2 | ENSMUST00 | 0 | Trim12c | ENSMUSG0C | false | chr7 | 104352353 | 104353745 | 20.52563 | 54.76614 | 51.99178 | chr7 | 104351295 | 104354082 | 3.53522 | 3.34867 | 1.80738 |
| chr2 | 114654489 | 114655030 | 1000 | Promoter (<=1kb) | 2 | 114516416 | 114654953 | 138538 | 2 | ENSMUST00 | 0 | Dph6 | ENSMUSG0C | false | chr2 | 114654182 | 114655559 | 23.09106 | 69.49222 | 66.50756 | chr2 | 114653066 | 114655589 | 4.32539 | 3.75726 | 2.439 |
| chr1 | 39713848 | 39714592 | 1000 | Intron (ENSMUST00000151913.1 | 1 | 39665306 | 39720997 | 55692 | 2 | ENSMUST00 | 6405 | Rfx8 | ENSMUSG0C | false | . | . | . | . | . | . | chr1 | 39713045 | 39715573 | 4.67341 | 3.81045 | 2.74942 |
| chr4 | 129335033 | 129335711 | 1000 | Promoter (<=1kb) | 4 | 129307100 | 129335370 | 28271 | 2 | ENSMUST00 | 0 | Rbbp4 | ENSMUSG0C | false | chr4 | 129335566 | 129336753 | 21.91892 | 63.05404 | 60.16603 | . | . | . | . | . | . |
| chr4 | 58304540 | 58304903 | 1000 | Intron (ENSMUST00000081919.6 | 4 | 58286175 | 58304324 | 18150 | 1 | ENSMUST00 | 18366 | Musk | ENSMUSG0C | false | . | . | . | . | . | . | chr4 | 58304393 | 58305041 | 4.2556 | 3.71526 | 2.45473 |
| chr4 | 58254520 | 58254900 | 1000 | Distal Intergenic | 4 | 58285960 | 58374302 | 88343 | 1 | ENSMUST00 | -31060 | Musk | ENSMUSG0C | false | . | . | . | . | . | . | chr4 | 58253294 | 58254777 | 4.33793 | 3.51135 | 2.48095 |
| chr5 | 102829730 | 102830160 | 1000 | Intron (ENSMUST00000094559.3 | 5 | 102845007 | 102897829 | 52823 | 1 | ENSMUST00 | -14847 | Arhgap24 | ENSMUSG0C | true | . | . | . | . | . | . | chr5 | 102806003 | 102845305 | 4.78306 | 3.97005 | 2.82906 |
| chr5 | 102739775 | 102740172 | 1000 | Intron (ENSMUST00000094559.3 | 5 | 102724973 | 102897829 | 172857 | 1 | ENSMUST00 | 14803 | Arhgap24 | ENSMUSG0C | true | . | . | . | . | . | . | chr5 | 102734086 | 102753392 | 4.81032 | 3.97837 | 2.85457 |
| chr5 | 102833114 | 102833400 | 1000 | Intron (ENSMUST00000094559.3 | 5 | 102845007 | 102897829 | 52823 | 1 | ENSMUST00 | -11607 | Arhgap24 | ENSMUSG0C | true | . | . | . | . | . | . | chr5 | 102806003 | 102845305 | 4.78306 | 3.97005 | 2.82906 |
| chr5 | 102751780 | 102752662 | 1000 | Intron (ENSMUST00000094559.3 | 5 | 102768771 | 102897829 | 129059 | 1 | ENSMUST00 | -16109 | Arhgap24 | ENSMUSG0C | true | . | . | . | . | . | . | chr5 | 102734086 | 102753392 | 4.81032 | 3.97837 | 2.85457 |
| chr5 | 102867984 | 102868502 | 1000 | Intron (ENSMUST00000094559.3 | 5 | 102845007 | 102897829 | 52823 | 1 | ENSMUST00 | 22978 | Arhgap24 | ENSMUSG0C | true | . | . | . | . | . | . | chr5 | 102861645 | 102881373 | 4.89607 | 4.00302 | 2.92218 |
| chr5 | 102845021 | 102846115 | 1000 | Promoter (<=1kb) | 5 | 102845007 | 102897829 | 52823 | 1 | ENSMUST00 | 15 | Arhgap24 | ENSMUSG0C | true | chr5 | 102845002 | 102848980 | 15.98415 | 46.70646 | 44.03467 | chr5 | 102806003 | 102845305 | 4.78306 | 3.97005 | 2.82906 |
| chr1 | 106719156 | 106719559 | 1000 | Downstream (<1kb) | 1 | 106538178 | 106714274 | 176097 | 2 | ENSMUST00 | -4883 | Bcl2 | ENSMUSG0C | false | . | . | . | . | . | . | chr1 | 106714557 | 106720254 | 4.66173 | 3.83764 | 2.74504 |
| chr1 | 106713846 | 106714739 | 1000 | Promoter (<=1kb) | 1 | 106538178 | 106714274 | 176097 | 2 | ENSMUST00 | 0 | Bcl2 | ENSMUSG0C | false | chr1 | 106712428 | 106714820 | 29.05819 | 88.32191 | 85.01541 | chr1 | 106714557 | 106720254 | 4.66173 | 3.83764 | 2.74504 |
| chr4 | 107318731 | 107319495 | 1000 | Promoter (<=1kb) | 4 | 107318971 | 107359823 | 40853 | 1 | ENSMUST00 | 0 | Yipf1 | ENSMUSG0C | false | . | . | . | . | . | . | chr4 | 107313492 | 107323582 | 6.11384 | 4.58418 | 3.92071 |

| | | | | | | | | | | | | | | | | | | | | | | | | | | |
|---|---|---|---|---|---|---|---|---|---|---|---|---|---|---|---|---|---|---|---|---|---|---|---|---|---|---|
| chr11 | 95402771 | 95403284 | 1000 | Distal Intergenic | 11 | 95414080 | 95493406 | 79327 | 1 | ENSMUST00 | -10796 | Spop | ENSMUSG0( | false | . | . | . | . | . | . | chr11 | 95401550 | 95404301 | 5.59151 | 4.36791 | 3.49423 |
| chr11 | 95494024 | 95494791 | 1000 | Distal Intergenic | 11 | 95474319 | 95490315 | 15997 | 1 | ENSMUST00 | 19706 | Spop | ENSMUSG0( | false | . | . | . | . | . | . | chr11 | 95489084 | 95513208 | 6.11718 | 4.48759 | 3.93601 |
| chr13 | 44939408 | 44939977 | 1000 | Intron (ENSMUST00000072329.8 | 13 | 44922079 | 45002096 | 80018 | 2 | ENSMUST00 | 62119 | Dtnbp1 | ENSMUSG0( | false | . | . | . | . | . | . | chr13 | 44937128 | 44941690 | 5.21885 | 4.08717 | 3.19078 |
| chr5 | 18436377 | 18436798 | 1000 | Distal Intergenic | 5 | 18265135 | 18360413 | 95279 | 2 | ENSMUST00 | -75965 | Gnai1 | ENSMUSG0( | false | . | . | . | . | . | . | chr5 | 18435847 | 18436767 | 4.3346 | 3.71507 | 2.49874 |
| chr8 | 83698385 | 83699492 | 1000 | Promoter (<=1kb) | 8 | 83669762 | 83699179 | 29418 | 2 | ENSMUST00 | 0 | Pkn1 | ENSMUSG0( | false | chr8 | 83697311 | 83699766 | 46.69001 | 154.98724 | 149.51399 | chr8 | 83692808 | 83698419 | 7.74578 | 5.12799 | 5.29944 |
| chr7 | 82866004 | 82868194 | 1000 | Promoter (<=1kb) | 7 | 82867333 | 82871563 | 4231 | 1 | ENSMUST00 | 0 | Mex3b | ENSMUSG0( | false | chr7 | 82865501 | 82869284 | 27.54716 | 98.2761 | 94.74957 | . | . | . | . | . | . |
| chr2 | 29965017 | 29966062 | 1000 | Promoter (<=1kb) | 2 | 29965560 | 30031448 | 65889 | 1 | ENSMUST00 | 0 | Sptan1 | ENSMUSG0( | false | chr2 | 29964945 | 29967047 | 38.08445 | 130.98944 | 126.4684 | . | . | . | . | . | . |
| chr16 | 8579633 | 8579957 | 1000 | Exon (ENSMUST00000140795.1/ | 16 | 8583627 | 8603047 | 19421 | 1 | ENSMUST00 | -3670 | Abat | ENSMUSG0( | false | . | . | . | . | . | . | chr16 | 8579226 | 8580391 | 3.86449 | 3.51369 | 2.10372 |
| chr16 | 8513175 | 8513611 | 1000 | Promoter (<=1kb) | 16 | 8513429 | 8621568 | 108140 | 1 | ENSMUST00 | 0 | Abat | ENSMUSG0( | false | chr16 | 8513058 | 8514438 | 27.40733 | 77.91289 | 74.79585 | chr16 | 8513054 | 8516114 | 4.31331 | 3.7593 | 2.49116 |
| chr16 | 8555548 | 8555852 | 1000 | Intron (ENSMUST00000065987.8 | 16 | 8572902 | 8605633 | 32732 | 1 | ENSMUST00 | -17050 | Abat | ENSMUSG0( | false | . | . | . | . | . | . | chr16 | 8554063 | 8556165 | 3.94313 | 3.48179 | 2.16143 |
| chr14 | 66826190 | 66826626 | 1000 | Intron (ENSMUST00000022629.8 | 14 | 66784529 | 66793758 | 9230 | 1 | ENSMUST00 | 41662 | Gm10032 | ENSMUSG0( | false | . | . | . | . | . | . | chr14 | 66815115 | 66830778 | 7.77502 | 5.24019 | 5.30996 |
| chr14 | 66816762 | 66817348 | 1000 | Intron (ENSMUST00000022629.8 | 14 | 66784529 | 66793758 | 9230 | 1 | ENSMUST00 | 32234 | Gm10032 | ENSMUSG0( | false | . | . | . | . | . | . | chr14 | 66815115 | 66830778 | 7.77502 | 5.24019 | 5.30996 |
| chr11 | 116077915 | 116078359 | 1000 | Promoter (<=1kb) | 11 | 116062096 | 116077961 | 15866 | 2 | ENSMUST00 | 0 | Unc13d | ENSMUSG0( | false | . | . | . | . | . | . | chr11 | 116075681 | 116079564 | 5.41266 | 4.23435 | 3.3688 |
| chr5 | 93123927 | 93124294 | 1000 | Intron (ENSMUST00000074733.7 | 5 | 93093457 | 93174958 | 81502 | 1 | ENSMUST00 | 30471 | Sep-11 | ENSMUSG0( | false | . | . | . | . | . | . | chr5 | 93095018 | 93154711 | 6.59326 | 4.78158 | 4.31782 |
| chr5 | 93115157 | 93115511 | 1000 | Intron (ENSMUST00000074733.7 | 5 | 93093457 | 93174958 | 81502 | 1 | ENSMUST00 | 21701 | Sep-11 | ENSMUSG0( | false | . | . | . | . | . | . | chr5 | 93095018 | 93154711 | 6.59326 | 4.78158 | 4.31782 |
| chr5 | 93124406 | 93125068 | 1000 | Intron (ENSMUST00000074733.7 | 5 | 93093457 | 93174958 | 81502 | 1 | ENSMUST00 | 30950 | Sep-11 | ENSMUSG0( | false | . | . | . | . | . | . | chr5 | 93095018 | 93154711 | 6.59326 | 4.78158 | 4.31782 |
| chr5 | 93127745 | 93128127 | 1000 | Intron (ENSMUST00000074733.7 | 5 | 93093457 | 93174958 | 81502 | 1 | ENSMUST00 | 34289 | Sep-11 | ENSMUSG0( | false | . | . | . | . | . | . | chr5 | 93095018 | 93154711 | 6.59326 | 4.78158 | 4.31782 |
| chr5 | 93092934 | 93094360 | 1000 | Promoter (<=1kb) | 5 | 93093457 | 93174958 | 81502 | 1 | ENSMUST00 | 0 | Sep-11 | ENSMUSG0( | false | chr5 | 93091910 | 93098273 | 29.81577 | 89.63673 | 86.30187 | . | . | . | . | . | . |
| chr8 | 61877582 | 61877991 | 1000 | 5' UTR | 8 | 61513246 | 61902669 | 389424 | 2 | ENSMUST00 | 24678 | Palld | ENSMUSG0( | true | . | . | . | . | . | . | chr8 | 61877408 | 61878237 | 3.32957 | 2.97816 | 1.66707 |
| chr8 | 61642309 | 61642891 | 1000 | Intron (ENSMUST00000121785.2 | 8 | 61513248 | 61591142 | 77895 | 2 | ENSMUST00 | -51168 | Palld | ENSMUSG0( | true | . | . | . | . | . | . | chr8 | 61639963 | 61648596 | 5.13767 | 4.13724 | 3.11858 |
| chr8 | 61679597 | 61679908 | 1000 | Intron (ENSMUST00000121785.2 | 8 | 61513252 | 61760089 | 246838 | 2 | ENSMUST00 | 80181 | Palld | ENSMUSG0( | true | . | . | . | . | . | . | chr8 | 61678832 | 61684610 | 4.89827 | 4.02741 | 2.92515 |
| chr8 | 61555463 | 61555794 | 1000 | Intron (ENSMUST00000121785.2 | 8 | 61525121 | 61533841 | 8721 | 2 | ENSMUST00 | -21623 | Palld | ENSMUSG0( | true | . | . | . | . | . | . | chr8 | 61550887 | 61568090 | 5.03432 | 4.07287 | 3.05874 |
| chr8 | 61681276 | 61681660 | 1000 | Intron (ENSMUST00000121785.2 | 8 | 61513252 | 61760089 | 246838 | 2 | ENSMUST00 | 78429 | Palld | ENSMUSG0( | true | . | . | . | . | . | . | chr8 | 61678832 | 61684610 | 4.89827 | 4.02741 | 2.92515 |
| chr8 | 61645474 | 61645852 | 1000 | Intron (ENSMUST00000121785.2 | 8 | 61513248 | 61591142 | 77895 | 2 | ENSMUST00 | -54333 | Palld | ENSMUSG0( | true | . | . | . | . | . | . | chr8 | 61639963 | 61648596 | 5.13767 | 4.13724 | 3.11858 |
| chr12 | 108493972 | 108494489 | 1000 | Intron (ENSMUST00000138456.2 | 12 | 108471853 | 108539564 | 67712 | 1 | ENSMUST00 | 22120 | Eml1 | ENSMUSG0( | false | . | . | . | . | . | . | chr12 | 108488039 | 108499693 | 6.71751 | 4.82027 | 4.43554 |
| chr12 | 108495428 | 108495948 | 1000 | Intron (ENSMUST00000138456.2 | 12 | 108471853 | 108539564 | 67712 | 1 | ENSMUST00 | 23576 | Eml1 | ENSMUSG0( | false | . | . | . | . | . | . | chr12 | 108488039 | 108499693 | 6.71751 | 4.82027 | 4.43554 |
| chr7 | 66960197 | 66960942 | 1000 | Intron (ENSMUST00000156396.2 | 7 | 66909468 | 66969555 | 60088 | 1 | ENSMUST00 | 50730 | Adamts17 | ENSMUSG0( | false | . | . | . | . | . | . | . | . | . | . | . | . |
| chr18 | 59174997 | 59175613 | 1000 | Promoter (<=1kb) | 18 | 59175401 | 59410446 | 235046 | 1 | ENSMUST00 | 0 | Chsy3 | ENSMUSG0( | false | chr18 | 59174453 | 59176473 | 38.30962 | 127.04226 | 122.65842 | chr18 | 59174406 | 59176933 | 4.95916 | 4.01356 | 3.01828 |
| chr7 | 16614122 | 16615542 | 1000 | Promoter (<=1kb) | 7 | 16493719 | 16614993 | 121275 | 2 | ENSMUST00 | 0 | Arhgap35 | ENSMUSG0( | false | chr7 | 16613597 | 16615423 | 36.32796 | 122.48113 | 118.25522 | . | . | . | . | . | . |
| chr7 | 30955870 | 30957176 | 1000 | Promoter (<=1kb) | 7 | 30945252 | 30956176 | 10925 | 2 | ENSMUST00 | 0 | Usf2 | ENSMUSG0( | false | chr7 | 30955688 | 30958528 | 36.31549 | 112.95179 | 109.03448 | chr7 | 30954839 | 30956027 | 4.09835 | 3.58019 | 2.29053 |
| chr2 | 103113032 | 103113270 | 1000 | Distal Intergenic | 2 | 103099155 | 103099601 | 447 | 1 | ENSMUST00 | 13878 | Gm10038 | ENSMUSG0( | false | . | . | . | . | . | . | . | . | . | . | . | . |
| chr5 | 138620903 | 138621237 | 1000 | Promoter (1-2kb) | 5 | 138604616 | 138619761 | 15146 | 2 | ENSMUST00 | -1143 | Zfp68 | ENSMUSG0( | false | . | . | . | . | . | . | chr5 | 138620065 | 138624832 | 4.56009 | 3.89358 | 2.6416 |
| chr2 | 92183611 | 92184912 | 1000 | Promoter (<=1kb) | 2 | 92184106 | 92361829 | 177724 | 1 | ENSMUST00 | 0 | Phf21a | ENSMUSG0( | false | chr2 | 92182752 | 92186025 | 28.03146 | 79.91027 | 76.76085 | . | . | . | . | . | . |
| chr2 | 92208394 | 92208749 | 1000 | Intron (ENSMUST00000044036.9 | 2 | 92221562 | 92364666 | 143105 | 1 | ENSMUST00 | -12813 | Phf21a | ENSMUSG0( | false | . | . | . | . | . | . | chr2 | 92198461 | 92211374 | 4.94471 | 3.98252 | 2.99295 |
| chr10 | 127748964 | 127749572 | 1000 | Promoter (2-3kb) | 10 | 127747276 | 127751732 | 4457 | 2 | ENSMUST00 | 2160 | Gpr182 | ENSMUSG0( | false | . | . | . | . | . | . | chr10 | 127743127 | 127759693 | 6.17884 | 4.54629 | 3.98922 |
| chr7 | 118447372 | 118448612 | 1000 | Distal Intergenic | 7 | 118381856 | 118443552 | 61697 | 2 | ENSMUST00 | -3821 | Syt17 | ENSMUSG0( | false | chr7 | 118447355 | 118448957 | 30.78991 | 105.27288 | 101.56436 | chr7 | 118441543 | 118449329 | 4.02821 | 3.5716 | 2.21247 |
| chr6 | 30047626 | 30048776 | 1000 | Promoter (<=1kb) | 6 | 30047988 | 30153458 | 105471 | 1 | ENSMUST00 | 0 | Nrf1 | ENSMUSG0( | false | chr6 | 30047101 | 30049195 | 33.04283 | 115.53907 | 111.53828 | . | . | . | . | . | . |
| chr6 | 54816384 | 54817094 | 1000 | Promoter (<=1kb) | 6 | 54816916 | 54893500 | 76585 | 1 | ENSMUST00 | 0 | Znrf2 | ENSMUSG0( | false | chr6 | 54816612 | 54818738 | 32.83994 | 103.83406 | 100.16508 | chr6 | 54815506 | 54816492 | 4.39055 | 3.82362 | 2.49081 |
| chr7 | 143832279 | 143832636 | 1000 | Promoter (2-3kb) | 7 | 143830258 | 143837888 | 7631 | 1 | ENSMUST00 | 2022 | Dhcr7 | ENSMUSG0( | false | . | . | . | . | . | . | chr7 | 143830649 | 143839948 | 5.20091 | 4.06122 | 3.19483 |
| chr9 | 75561268 | 75561629 | 1000 | Promoter (1-2kb) | 9 | 75497796 | 75559657 | 61862 | 2 | ENSMUST00 | -1612 | Tmod3 | ENSMUSG0( | true | . | . | . | . | . | . | chr9 | 75559804 | 75567102 | 4.3979 | 3.82685 | 2.50388 |
| chr9 | 75548932 | 75549352 | 1000 | Intron (ENSMUST00000072232.7 | 9 | 75497796 | 75559657 | 61862 | 2 | ENSMUST00 | 10305 | Tmod3 | ENSMUSG0( | true | chr9 | 75549201 | 75549725 | 7.13425 | 13.57188 | 11.33167 | chr9 | 75524164 | 75555861 | 5.5746 | 4.35546 | 3.47394 |
| chr9 | 75527362 | 75527752 | 1000 | Intron (ENSMUST00000072232.7 | 9 | 75497796 | 75559657 | 61862 | 2 | ENSMUST00 | 31905 | Tmod3 | ENSMUSG0( | true | . | . | . | . | . | . | chr9 | 75524164 | 75555861 | 5.5746 | 4.35546 | 3.47394 |
| chr9 | 75523558 | 75524029 | 1000 | Intron (ENSMUST00000072232.7 | 9 | 75497796 | 75559657 | 61862 | 2 | ENSMUST00 | 35628 | Tmod3 | ENSMUSG0( | true | . | . | . | . | . | . | . | . | . | . | . | . |
| chr10 | 90576373 | 90576845 | 1000 | Promoter (<=1kb) | 10 | 90576377 | 90970258 | 393882 | 1 | ENSMUST00 | 0 | Anks1b | ENSMUSG0( | true | chr10 | 90575990 | 90578186 | 10.01484 | 27.17025 | 24.74075 | . | . | . | . | . | . |
| chr10 | 90637423 | 90637820 | 1000 | Intron (ENSMUST00000183156.2 | 10 | 90576992 | 90969820 | 392829 | 1 | ENSMUST00 | 60432 | Anks1b | ENSMUSG0( | true | . | . | . | . | . | . | chr10 | 90636337 | 90638952 | 3.93366 | 3.54635 | 2.17633 |
| chr15 | 53896823 | 53897273 | 1000 | Intron (ENSMUST00000154119.2 | 15 | 53462194 | 53902381 | 440188 | 2 | ENSMUST00 | 5108 | Samd12 | ENSMUSG0( | true | . | . | . | . | . | . | chr15 | 53890832 | 53902905 | 4.62295 | 3.89484 | 2.70376 |
| chr14 | 36892965 | 36893431 | 1000 | Intron (ENSMUST00000067700.7 | 14 | 36879570 | 36886360 | 6791 | 2 | ENSMUST00 | -6606 | Ccser2 | ENSMUSG0( | false | . | . | . | . | . | . | chr14 | 36889985 | 36897083 | 4.72464 | 3.95482 | 2.77604 |
| chr14 | 36942663 | 36943043 | 1000 | Promoter (2-3kb) | 14 | 36900790 | 36939945 | 39156 | 2 | ENSMUST00 | -2719 | Ccser2 | ENSMUSG0( | false | . | . | . | . | . | . | chr14 | 36941939 | 36944754 | 4.10116 | 3.43305 | 2.29273 |
| chr17 | 74211722 | 74212014 | 1000 | Intron (ENSMUST00000078459.6 | 17 | 74200706 | 74294883 | 94178 | 2 | ENSMUST00 | 82869 | Memo1 | ENSMUSG0( | false | . | . | . | . | . | . | chr17 | 74211255 | 74212877 | 5.44764 | 4.38547 | 3.35871 |
| chr17 | 74294373 | 74295859 | 1000 | Promoter (<=1kb) | 17 | 74200706 | 74294883 | 94178 | 2 | ENSMUST00 | 0 | Memo1 | ENSMUSG0( | false | chr17 | 74293736 | 74296377 | 31.60199 | 102.67427 | 99.04176 | . | . | . | . | . | . |
| chr1 | 180640796 | 180641534 | 1000 | Promoter (<=1kb) | 1 | 180641150 | 180690518 | 49369 | 1 | ENSMUST00 | 0 | Lin9 | ENSMUSG0( | false | chr1 | 180640589 | 180642144 | 35.11919 | 146.21799 | 141.08704 | . | . | . | . | . | . |
| chr11 | 98746811 | 98748059 | 1000 | Promoter (2-3kb) | 11 | 98744233 | 98753777 | 9545 | 1 | ENSMUST00 | 2579 | Thra | ENSMUSG0( | false | chr11 | 98746952 | 98747514 | 7.45894 | 16.41436 | 14.12962 | chr11 | 98739896 | 98756634 | 7.38853 | 5.10697 | 4.96875 |
| chr11 | 98750427 | 98751571 | 1000 | Promoter (2-3kb) | 11 | 98753586 | 98768403 | 14818 | 1 | ENSMUST00 | -2015 | Thra | ENSMUSG0( | false | chr11 | 98750726 | 98751443 | 8.98559 | 17.12414 | 14.83055 | chr11 | 98739896 | 98756634 | 7.38853 | 5.10697 | 4.96875 |
| chr11 | 98741587 | 98742217 | 1000 | Promoter (<=1kb) | 11 | 98741811 | 98757695 | 15885 | 1 | ENSMUST00 | 0 | Thra | ENSMUSG0( | false | chr11 | 98740244 | 98743094 | 32.66734 | 113.81544 | 109.86345 | chr11 | 98739896 | 98756634 | 7.38853 | 5.10697 | 4.96875 |
| chr10 | 111419658 | 111420110 | 1000 | Distal Intergenic | 10 | 111473192 | 111496886 | 23695 | 1 | ENSMUST00 | -53082 | Nap1l1 | ENSMUSG0( | false | chr10 | 111419043 | 111420465 | 9.65017 | 18.71579 | 16.39739 | chr10 | 111416547 | 111423445 | 5.41255 | 4.32457 | 3.33421 |
| chr10 | 111473015 | 111473799 | 1000 | Promoter (<=1kb) | 10 | 111473192 | 111496886 | 23695 | 1 | ENSMUST00 | 0 | Nap1l1 | ENSMUSG0( | false | chr10 | 111472714 | 111474509 | 29.28796 | 98.53321 | 94.99628 | . | . | . | . | . | . |
| chr2 | 23039647 | 23040395 | 1000 | Promoter (<=1kb) | 2 | 22940073 | 23040195 | 100123 | 2 | ENSMUST00 | 0 | Abi1 | ENSMUSG0( | false | chr2 | 23038901 | 23040223 | 39.05373 | 137.10143 | 132.33189 | . | . | . | . | . | . |
| chr18 | 82914079 | 82915042 | 1000 | Promoter (<=1kb) | 18 | 82914632 | 83005314 | 90683 | 1 | ENSMUST00 | 0 | Zfp516 | ENSMUSG0( | false | chr18 | 82909434 | 82915656 | 29.53108 | 87.21563 | 83.92796 | . | . | . | . | . | . |
| chr18 | 57570451 | 57570757 | 1000 | Intron (ENSMUST00000079738.4 | 18 | 57563176 | 57731006 | 167831 | 1 | ENSMUST00 | 7276 | 1700011I03R | ENSMUSG0( | false | . | . | . | . | . | . | chr18 | 57570470 | 57572004 | 4.69314 | 3.95343 | 2.80601 |
| chr6 | 120531108 | 120531580 | 1000 | Promoter (<=1kb) | 6 | 120509494 | 120531319 | 21826 | 2 | ENSMUST00 | 0 | Cecr5 | ENSMUSG0( | false | chr6 | 120530741 | 120532312 | 30.14656 | 119.38538 | 115.2599 | . | . | . | . | . | . |
| chr14 | 78848434 | 78848777 | 1000 | Promoter (<=1kb) | 14 | 78849178 | 79202310 | 353133 | 1 | ENSMUST00 | -401 | Vwa8 | ENSMUSG0( | false | chr14 | 78848611 | 78849970 | 28.71633 | 95.52087 | 92.05702 | chr14 | 78846578 | 78848524 | 4.81829 | 4.09986 | 2.84496 |
| chr14 | 78848912 | 78849502 | 1000 | Promoter (<=1kb) | 14 | 78849178 | 79202310 | 353133 | 1 | ENSMUST00 | 0 | Vwa8 | ENSMUSG0( | false | chr14 | 78848611 | 78849970 | 28.71633 | 95.52087 | 92.05702 | . | . | . | . | . | . |
| chr16 | 9670643 | 9671005 | 1000 | Intron (ENSMUST00000115835.2 | 16 | 9577827 | 9992533 | 414707 | 2 | ENSMUST00 | 321528 | Grin2a | ENSMUSG0( | false | . | . | . | . | . | . | chr16 | 9670194 | 9672253 | 3.6435 | 3.40332 | 1.91444 |
| chr2 | 75658776 | 75659927 | 1000 | Promoter (<=1kb) | 2 | 75659261 | 75665839 | 6579 | 1 | ENSMUST00 | 0 | Hnrnpa3 | ENSMUSG0( | false | chr2 | 75659232 | 75661067 | 27.88725 | 99.90214 | 96.33501 | . | . | . | . | . | . |
| chr2 | 32727411 | 32727800 | 1000 | Promoter (<=1kb) | 2 | 32727706 | 32744844 | 17139 | 1 | ENSMUST00 | 0 | Sh2d3c | ENSMUSG0( | false | chr2 | 32727689 | 32728790 | 6.38327 | 11.47542 | 9.27082 | chr2 | 32726994 | 32728582 | 4.95074 | 4.15085 | 2.98551 |
| chr6 | 29066511 | 29066835 | 1000 | Promoter (1-2kb) | 6 | 29065462 | 29070972 | 5511 | 1 | ENSMUST00 | 1050 | Lep | ENSMUSG0( | false | . | . | . | . | . | . | chr6 | 29065125 | 29067495 | 5.28294 | 4.27116 | 3.22981 |
| chr15 | 98812724 | 98813404 | 1000 | 3' UTR | 15 | 98803782 | 98807925 | 4144 | 2 | ENSMUST00 | -4800 | Ddn | ENSMUSG0( | true | chr15 | 98812788 | 98813788 | 7.14186 | 14.46535 | 12.21151 | chr15 | 98809760 | 98818473 | 4.97708 | 4.01159 | 3.00452 |
| chr15 | 98809287 | 98809811 | 1000 | Promoter (1-2kb) | 15 | 98803782 | 98807925 | 4144 | 2 | ENSMUST00 | -1363 | Ddn | ENSMUSG0( | true | chr15 | 98803793 | 98809569 | 29.66345 | 100.21 | 96.63225 | chr15 | 98809760 | 98818473 | 4.97708 | 4.01159 | 3.00452 |
| chr11 | 117265948 | 117266481 | 1000 | Promoter (<=1kb) | 11 | 117266246 | 117362313 | 96068 | 1 | ENSMUST00 | 0 | Sep-09 | ENSMUSG0( | false | chr11 | 117265778 | 117267655 | 22.10065 | 59.96099 | 57.11441 | chr11 | 117264529 | 117268124 | 5.54315 | 4.37602 | 3.43357 |
| chr7 | 16400641 | 16401567 | 1000 | Promoter (<=1kb) | 7 | 16401196 | 16437694 | 36499 | 1 | ENSMUST00 | 0 | Zc3h4 | ENSMUSG0( | false | chr7 | 16399095 | 16403914 | 21.20881 | 84.61981 | 81.38317 | . | . | . | . | . | . |
| chr7 | 16399850 | 16400417 | 1000 | Promoter (<=1kb) | 7 | 16401196 | 16437694 | 36499 | 1 | ENSMUST00 | -779 | Zc3h4 | ENSMUSG0( | false | chr7 | 16399095 | 16403914 | 21.20881 | 84.61981 | 81.38317 | . | . | . | . | . | . |
| chr13 | 35740638 | 35741911 | 1000 | Promoter (<=1kb) | 13 | 35741313 | 35874058 | 132746 | 1 | ENSMUST00 | 0 | Cdyl | ENSMUSG0( | true | chr13 | 35740437 | 35743017 | 37.4318 | 119.08698 | 114.97395 | . | . | . | . | . | . |
| chr13 | 35486093 | 35486471 | 1000 | Distal Intergenic | 13 | 35659862 | 35874064 | 214203 | 1 | ENSMUST00 | -173391 | Cdyl | ENSMUSG0( | true | . | . | . | . | . | . | chr13 | 35485326 | 35487625 | 3.4637 | 3.32573 | 1.73258 |
| chr2 | 29802504 | 29802782 | 1000 | Promoter (<=1kb) | 2 | 29802634 | 29817522 | 14889 | 1 | ENSMUST00 | 0 | Slc27a4 | ENSMUSG0( | false | chr2 | 29802524 | 29803400 | 30.17592 | 94.17345 | 90.73757 | chr2 | 29802430 | 29804169 | 3.68974 | 3.37797 | 1.94784 |
| chr10 | 80887270 | 80887669 | 1000 | Exon (ENSMUST00000105333.2/ | 10 | 80879720 | 80899441 | 19722 | 1 | ENSMUST00 | 7551 | Tmprss9 | ENSMUSG0( | false | . | . | . | . | . | . | chr10 | 80886665 | 80888408 | 5.2216 | 4.16407 | 3.19372 |
| chr11 | 85640625 | 85641065 | 1000 | Promoter (<=1kb) | 11 | 85640704 | 85826055 | 185352 | 1 | ENSMUST00 | 0 | Bcas3 | ENSMUSG0( | false | . | . | . | . | . | . | chr11 | 85639815 | 85643227 | 4.23086 | 3.69506 | 2.37239 |
| chr11 | 85523752 | 85524135 | 1000 | Intron (ENSMUST00000108061.2 | 11 | 85509638 | 85826058 | 316421 | 1 | ENSMUST00 | 14115 | Bcas3 | ENSMUSG0( | false | . | . | . | . | . | . | chr11 | 85523184 | 85532294 | 6.02645 | 4.52951 | 3.85324 |
| chr11 | 85541413 | 85541755 | 1000 | Intron (ENSMUST00000108061.2 | 11 | 85509638 | 85826058 | 316421 | 1 | ENSMUST00 | 31776 | Bcas3 | ENSMUSG0( | false | . | . | . | . | . | . | chr11 | 85540288 | 85543551 | 4.77125 | 3.94401 | 2.85621 |
| chr7 | 25627371 | 25627788 | 1000 | Promoter (<=1kb) | 7 | 25627624 | 25629490 | 1867 | 1 | ENSMUST00 | 0 | B3gnt8 | ENSMUSG0( | false | chr7 | 25626852 | 25628244 | 15.50707 | 38.47921 | 35.908 | chr7 | 25626394 | 25629065 | 4.72369 | 3.86396 | 2.78004 |
| chr6 | 56797204 | 56797894 | 1000 | Promoter (<=1kb) | 6 | 56777524 | 56797736 | 20213 | 2 | ENSMUST00 | 0 | Kbtbd2 | ENSMUSG0( | false | chr6 | 56796286 | 56798543 | 30.35223 | 94.4818 | 91.04041 | . | . | . | . | . | . |
| chrX | 162159906 | 162160239 | 1000 | Promoter (<=1kb) | 20 | 161833296 | 162159730 | 326435 | 2 | ENSMUST00 | -177 | Nhs | ENSMUSG0( | false | chrX | 162158562 | 162160279 | 13.78596 | 30.80405 | 28.32676 | . | . | . | . | . | . |
| chr9 | 43076138 | 43076577 | 1000 | Intron (ENSMUST00000165665.2 | 9 | 42963843 | 43105486 | 141644 | 2 | ENSMUST00 | 28909 | Arhgef12 | ENSMUSG0( | true | . | . | . | . | . | . | chr9 | 43060024 | 43103547 | 5.90513 | 4.47373 | 3.75735 |
| chr9 | 43105488 | 43106409 | 1000 | Promoter (<=1kb) | 9 | 42963842 | 43105718 | 141877 | 2 | ENSMUST00 | 0 | Arhgef12 | ENSMUSG0( | true | chr9 | 43102050 | 43107425 | 27.39171 | 92.13462 | 88.74321 | . | . | . | . | . | . |
| chr9 | 43071563 | 43072077 | 1000 | Intron (ENSMUST00000165665.2 | 9 | 42963843 | 43105486 | 141644 | 2 | ENSMUST00 | 33409 | Arhgef12 | ENSMUSG0( | true | . | . | . | . | . | . | chr9 | 43060024 | 43103547 | 5.90513 | 4.47373 | 3.75735 |
| chr9 | 43036554 | 43036946 | 1000 | Intron (ENSMUST00000165665.2 | 9 | 42963843 | 43105486 | 141644 | 2 | ENSMUST00 | 68540 | Arhgef12 | ENSMUSG0( | true | . | . | . | . | . | . | chr9 | 43027603 | 43053830 | 5.31283 | 4.19603 | 3.26716 |
| chr9 | 43034702 | 43035018 | 1000 | Intron (ENSMUST00000165665.2 | 9 | 42963843 | 43105486 | 141644 | 2 | ENSMUST00 | 70468 | Arhgef12 | ENSMUSG0( | true | . | . | . | . | . | . | chr9 | 43027603 | 43053830 | 5.31283 | 4.19603 | 3.26716 |
| chr12 | 24498471 | 24498967 | 1000 | Promoter (<=1kb) | 12 | 24498581 | 24558571 | 59991 | 1 | ENSMUST00 | 0 | Taf1b | ENSMUSG0( | false | chr12 | 24498062 | 24499146 | 29.38432 | 86.00549 | 82.73921 | chr12 | 24497705 | 24499371 | 3.63773 | 3.38377 | 1.87819 |
| chr14 | 54641157 | 54641647 | 1000 | Promoter (<=1kb) | 14 | 54631992 | 54641364 | 9373 | 2 | ENSMUST00 | 0 | Cdh24 | ENSMUSG0( | false | chr14 | 54640971 | 54641938 | 22.68762 | 81.35053 | 78.1749 | chr14 | 54640784 | 54642016 | 3.00098 | 2.90284 | 1.35513 |
| chr9 | 110778861 | 110779506 | 1000 | Distal Intergenic | 9 | 110769001 | 110769794 | 794 | 1 | ENSMUST00 | 9861 | Myl3 | ENSMUSG0( | false | . | . | . | . | . | . | chr9 | 110772879 | 110782615 | 6.7891 | 4.9169 | 4.4717 |
| chr11 | 69671155 | 69671763 | 1000 | Promoter (<=1kb) | 11 | 69670180 | 69671730 | 1551 | 2 | ENSMUST00 | 0 | Eif4a1 | ENSMUSG0( | false | chr11 | 69670578 | 69672426 | 37.30296 | 124.19952 | 119.9176 | . | . | . | . | . | . |
| chr7 | 45704815 | 45705400 | 1000 | Promoter (<=1kb) | 7 | 45705088 | 45710155 | 5068 | 1 | ENSMUST00 | 0 | Dbp | ENSMUSG0( | false | chr7 | 45703819 | 45709044 | 27.54716 | 98.2761 | 94.74957 | . | . | . | . | . | . |
| chr7 | 45706597 | 45707833 | 1000 | Promoter (<=1kb) | 7 | 45707104 | 45710028 | 2925 | 1 | ENSMUST00 | 0 | Dbp | ENSMUSG0( | false | chr7 | 45703819 | 45709044 | 27.54716 | 98.2761 | 94.74957 | chr7 | 45705481 | 45717582 | 6.81134 | 4.85162 | 4.49354 |
| chr3 | 41602637 | 41603089 | 1000 | Intron (ENSMUST00000026865.9 | 3 | 41627117 | 41630123 | 3007 | 2 | ENSMUST00 | 27034 | Sclt1 | ENSMUSG0( | false | . | . | . | . | . | . | chr3 | 41598517 | 41607869 | 6.56571 | 4.58882 | 4.32841 |
| chr18 | 80349178 | 80349658 | 1000 | Intron (ENSMUST00000077962.7 | 18 | 80322679 | 80363336 | 40658 | 2 | ENSMUST00 | 13678 | Kcng2 | ENSMUSG0( | false | . | . | . | . | . | . | chr18 | 80344101 | 80352099 | 5.09327 | 3.95276 | 3.12772 |
| chr3 | 141464695 | 141465689 | 1000 | Promoter (<=1kb) | 3 | 141465564 | 141834924 | 369361 | 1 | ENSMUST00 | 0 | Unc5c | ENSMUSG0( | true | chr3 | 141464561 | 141466743 | 39.20496 | 131.87447 | 127.31225 | . | . | . | . | . | . |
| chr11 | 115708388 | 115708817 | 1000 | Promoter (<=1kb) | 11 | 115644045 | 115708597 | 64553 | 2 | ENSMUST00 | 0 | Grb2 | ENSMUSG0( | true | chr11 | 115707334 | 115709306 | 23.46131 | 103.32642 | 99.67202 | . | . | . | . | . | . |
| chr9 | 29459008 | 29459323 | 1000 | Intron (ENSMUST00000075069.5 | 9 | 28996012 | 29412204 | 416193 | 2 | ENSMUST00 | -46805 | Ntm | ENSMUSG0( | false | . | . | . | . | . | . | . | . | . | . | . | . |
| chr11 | 102296185 | 102297466 | 1000 | Promoter (<=1kb) | 11 | 102289300 | 102296631 | 7332 | 2 | ENSMUST00 | 0 | Atxn7l3 | ENSMUSG0( | false | chr11 | 102295679 | 102297953 | 33.1785 | 101.38432 | 97.78168 | . | . | . | . | . | . |
| chr14 | 75844870 | 75845936 | 1000 | Promoter (<=1kb) | 14 | 75845093 | 75848415 | 3323 | 1 | ENSMUST00 | 0 | Tpt1 | ENSMUSG0( | false | chr14 | 75845256 | 75846538 | 27.47569 | 96.18372 | 92.70509 | . | . | . | . | . | . |
| chr14 | 105732192 | 105732896 | 1000 | Distal Intergenic | 14 | 105681828 | 105682211 | 384 | 1 | ENSMUST00 | 50365 | Gm10076 | ENSMUSG0( | false | chr14 | 105731973 | 105733249 | 12.81979 | 29.17209 | 26.71728 | chr14 | 105730765 | 105733801 | 5.29484 | 4.2319 | 3.23643 |
| chr16 | 18234926 | 18235450 | 1000 | Promoter (<=1kb) | 16 | 18220754 | 18235136 | 14383 | 2 | ENSMUST00 | 0 | Zdhhc8 | ENSMUSG0( | false | chr16 | 18234256 | 18236098 | 20.74539 | 66.97287 | 64.02668 | . | . | . | . | . | . |
| chr13 | 58358804 | 58359339 | 1000 | Distal Intergenic | 13 | 58287516 | 58354862 | 67347 | 2 | ENSMUST00 | -3943 | Kif27 | ENSMUSG0( | false | chr13 | 58358554 | 58359564 | 18.44084 | 48.77526 | 46.0739 | chr13 | 58358557 | 58359535 | 3.81371 | 3.49746 | 2.01179 |
| chr10 | 117049614 | 117050044 | 1000 | Downstream (<1kb) | 10 | 117045341 | 117046768 | 1428 | 1 | ENSMUST00 | 4274 | Lrrc10 | ENSMUSG0( | false | . | . | . | . | . | . | chr10 | 117045739 | 117050616 | 4.20138 | 3.5215 | 2.382 |
| chr4 | 54944915 | 54946341 | 1000 | Promoter (<=1kb) | 4 | 54945048 | 55008314 | 63267 | 1 | ENSMUST00 | 0 | Zfp462 | ENSMUSG0( | true | chr4 | 54944740 | 54946604 | 25.1466 | 74.00616 | 70.95453 | chr4 | 54944398 | 54945454 | 3.20552 | 3.13423 | 1.52483 |
| chr4 | 54947182 | 54948752 | 1000 | Promoter (<=1kb) | 4 | 54947976 | 55083563 | 135588 | 1 | ENSMUST00 | 0 | Zfp462 | ENSMUSG0( | true | chr4 | 54946727 | 54950127 | 24.72749 | 72.3696 | 69.3433 | . | . | . | . | . | . |
| chr11 | 70432146 | 70432782 | 1000 | Promoter (<=1kb) | 11 | 70432635 | 70435996 | 3362 | 1 | ENSMUST00 | 0 | Arrb2 | ENSMUSG0( | false | chr11 | 70432049 | 70433420 | 20.14837 | 78.81087 | 75.68276 | . | . | . | . | . | . |
| chr7 | 139251869 | 139252315 | 1000 | Intron (ENSMUST00000093993.3 | 7 | 139248482 | 139267253 | 18772 | 1 | ENSMUST00 | 3388 | Pwwp2b | ENSMUSG0( | false | . | . | . | . | . | . | chr7 | 139246761 | 139254099 | 8.53008 | 5.68016 | 5.90094 |
| chr7 | 139247969 | 139249269 | 1000 | Promoter (<=1kb) | 7 | 139248482 | 139267253 | 18772 | 1 | ENSMUST00 | 0 | Pwwp2b | ENSMUSG0( | false | chr7 | 139247202 | 139249854 | 22.89105 | 88.16154 | 84.85722 | chr7 | 139246761 | 139254099 | 8.53008 | 5.68016 | 5.90094 |
| chr7 | 139273321 | 139273768 | 1000 | Distal Intergenic | 7 | 139248482 | 139267253 | 18772 | 1 | ENSMUST00 | 24840 | Pwwp2b | ENSMUSG0( | false | . | . | . | . | . | . | chr7 | 139270937 | 139275472 | 8.52846 | 5.56821 | 5.90402 |
| chr7 | 139285583 | 139285965 | 1000 | Distal Intergenic | 7 | 139248482 | 139267253 | 18772 | 1 | ENSMUST00 | 37102 | Pwwp2b | ENSMUSG0( | false | . | . | . | . | . | . | chr7 | 139284286 | 139286649 | 4.76593 | 3.85372 | 2.86239 |
| chr7 | 44929144 | 44930119 | 1000 | Promoter (<=1kb) | 7 | 44900373 | 44929496 | 29124 | 2 | ENSMUST00 | 0 | Ap2a1 | ENSMUSG0( | false | chr7 | 44929975 | 44930461 | 17.78412 | 46.86213 | 44.18626 | . | . | . | . | . | . |
| chr9 | 44483732 | 44484080 | 1000 | Distal Intergenic | 9 | 44478778 | 44482180 | 3403 | 1 | ENSMUST00 | 4955 | C030014I23R | ENSMUSG0( | false | chr9 | 44482529 | 44484376 | 36.04672 | 129.50026 | 125.02953 | chr9 | 44483700 | 44502510 | 6.77697 | 4.85012 | 4.46719 |
| chr5 | 129163892 | 129164506 | 1000 | Intron (ENSMUST00000056617.8 | 5 | 129128077 | 129128517 | 441 | 2 | ENSMUST00 | -35376 | Rps16-ps2 | ENSMUSG0( | false | . | . | . | . | . | . | chr5 | 129163439 | 129166604 | 6.24778 | 4.67812 | 4.02947 |
| chr15 | 55810702 | 55811068 | 1000 | Intron (ENSMUST00000039769.6 | 15 | 55791938 | 55906310 | 114373 | 2 | ENSMUST00 | 95242 | Sntb1 | ENSMUSG0( | false | . | . | . | . | . | . | . | . | . | . | . | . |
| chr15 | 55791327 | 55791642 | 1000 | Intron (ENSMUST00000039769.6 | 15 | 55791938 | 55906310 | 114373 | 2 | ENSMUST00 | 114668 | Sntb1 | ENSMUSG0( | false | . | . | . | . | . | . | . | . | . | . | . | . |
| chr10 | 9647657 | 9648251 | 1000 | Intron (ENSMUST00000100070.2 | 10 | 9627259 | 9675208 | 47950 | 2 | ENSMUST00 | 26957 | Samd5 | ENSMUSG0( | false | . | . | . | . | . | . | chr10 | 9646052 | 9652313 | 4.83709 | 3.95568 | 2.88396 |
| chr9 | 20521064 | 20521764 | 1000 | Promoter (<=1kb) | 9 | 20495068 | 20521417 | 26350 | 2 | ENSMUST00 | 0 | Zfp266 | ENSMUSG0( | false | chr9 | 20520577 | 20522228 | 26.52689 | 93.43019 | 90.01563 | . | . | . | . | . | . |
| chr14 | 60913064 | 60913502 | 1000 | Distal Intergenic | 14 | 60963834 | 61037987 | 74154 | 2 | ENSMUST00 | 124485 | Tnfrsf19 | ENSMUSG0( | false | . | . | . | . | . | . | chr14 | 60909576 | 60917829 | 5.09447 | 4.13941 | 3.083 |
| chr1 | 167052971 | 167053391 | 1000 | Intron (ENSMUST00000165874.2 | 1 | 167001589 | 167091009 | 89421 | 1 | ENSMUST00 | 51383 | Fam78b | ENSMUSG0( | true | . | . | . | . | . | . | chr1 | 167042661 | 167060137 | 5.59987 | 4.30177 | 3.51451 |
| chr1 | 167000544 | 167001598 | 1000 | Promoter (<=1kb) | 1 | 167001417 | 167091302 | 89886 | 1 | ENSMUST00 | 0 | Fam78b | ENSMUSG0( | true | chr1 | 167000262 | 167003097 | 33.34559 | 100.66625 | 97.08096 | . | . | . | . | . | . |
| chr7 | 140954133 | 140954485 | 1000 | Promoter (1-2kb) | 7 | 140954839 | 140955961 | 1123 | 2 | ENSMUST00 | 1476 | Ifitm2 | ENSMUSG0( | false | chr7 | 140953561 | 140956714 | 27.66126 | 83.97217 | 80.74496 | chr7 | 140948249 | 140954254 | 8.1757 | 5.53651 | 5.60556 |
| chr7 | 44560253 | 44560743 | 1000 | Distal Intergenic | 7 | 44551057 | 44553951 | 2895 | 2 | ENSMUST00 | -6303 | Nr1h2 | ENSMUSG0( | false | . | . | . | . | . | . | chr7 | 44558997 | 44561380 | 6.67567 | 4.90716 | 4.37039 |
| chr3 | 89951803 | 89952309 | 1000 | Promoter (1-2kb) | 3 | 89939489 | 89953365 | 13877 | 2 | ENSMUST00 | 1056 | Atp8b2 | ENSMUSG0( | false | . | . | . | . | . | . | chr3 | 89950392 | 89954155 | 5.25526 | 4.19933 | 3.22585 |
| chr2 | 60283723 | 60284448 | 1000 | Promoter (<=1kb) | 2 | 60251993 | 60284418 | 32426 | 2 | ENSMUST00 | 0 | Cd302 | ENSMUSG0( | false | chr2 | 60283735 | 60284626 | 16.8969 | 46.84153 | 44.16631 | chr2 | 60282979 | 60284362 | 4.04953 | 3.64112 | 2.21417 |
| chr1 | 180813386 | 180814330 | 1000 | Promoter (<=1kb) | 1 | 180800832 | 180813632 | 12801 | 2 | ENSMUST00 | 0 | H3f3a | ENSMUSG0( | false | chr1 | 180811048 | 180813568 | 35.54802 | 115.80763 | 111.79785 | . | . | . | . | . | . |
| chr10 | 63541705 | 63542142 | 1000 | Intron (ENSMUST00000105441.2 | 10 | 63574506 | 63585641 | 11136 | 1 | ENSMUST00 | -32364 | Ctnna3 | ENSMUSG0( | false | . | . | . | . | . | . | chr10 | 63541072 | 63542936 | 4.60674 | 3.85482 | 2.71754 |
| chr10 | 88730862 | 88731587 | 1000 | Promoter (<=1kb) | 10 | 88731068 | 88744094 | 13027 | 1 | ENSMUST00 | 0 | Arl1 | ENSMUSG0( | false | chr10 | 88730430 | 88732564 | 32.98857 | 124.89146 | 120.57959 | . | . | . | . | . | . |
| chr1 | 156306309 | 156306952 | 1000 | Promoter (2-3kb) | 1 | 156255296 | 156303664 | 48369 | 2 | ENSMUST00 | -2646 | Tdrd5 | ENSMUSG0( | true | . | . | . | . | . | . | . | . | . | . | . | . |
| chr7 | 110061084 | 110062322 | 1000 | Promoter (<=1kb) | 7 | 110061702 | 110095392 | 33691 | 1 | ENSMUST00 | 0 | Zfp143 | ENSMUSG0( | false | chr7 | 110061710 | 110062924 | 20.02967 | 73.35366 | 70.31202 | . | . | . | . | . | . |
| chrX | 53131470 | 53131939 | 1000 | Intron (ENSMUST00000074861.3 | 20 | 53069996 | 53114530 | 44535 | 2 | ENSMUST00 | -16941 | Plac1 | ENSMUSG0( | false | . | . | . | . | . | . | . | . | . | . | . | . |
| chrX | 53212103 | 53212392 | 1000 | Promoter (2-3kb) | 20 | 53070172 | 53209724 | 139553 | 2 | ENSMUST00 | -2380 | Plac1 | ENSMUSG0( | false | . | . | . | . | . | . | . | . | . | . | . | . |
| chr17 | 84957823 | 84958648 | 1000 | Promoter (<=1kb) | 17 | 84958001 | 85014776 | 56776 | 1 | ENSMUST00 | 0 | Ppm1b | ENSMUSG0( | false | chr17 | 84957997 | 84959482 | 32.55857 | 101.31886 | 97.7165 | chr17 | 84958632 | 84963957 | 4.09512 | 3.59539 | 2.27727 |
| chr3 | 52104531 | 52105229 | 1000 | Promoter (<=1kb) | 3 | 51687611 | 52104891 | 417281 | 2 | ENSMUST00 | 0 | Maml3 | ENSMUSG0( | true | chr3 | 52102288 | 52105222 | 37.14626 | 116.48058 | 112.45192 | . | . | . | . | . | . |
| chr3 | 51771773 | 51772353 | 1000 | Intron (ENSMUST00000121440.2 | 3 | 51689437 | 51796535 | 107099 | 2 | ENSMUST00 | 24182 | Maml3 | ENSMUSG0( | true | . | . | . | . | . | . | chr3 | 51767223 | 51782456 | 5.75718 | 4.44357 | 3.63488 |
| chr3 | 52034895 | 52035846 | 1000 | Intron (ENSMUST00000121440.2 | 3 | 51687611 | 52104891 | 417281 | 2 | ENSMUST00 | 69045 | Maml3 | ENSMUSG0( | true | . | . | . | . | . | . | chr3 | 52009386 | 52064052 | 5.58271 | 4.32835 | 3.48679 |
| chr3 | 52101798 | 52103185 | 1000 | Promoter (1-2kb) | 3 | 51687611 | 52104891 | 417281 | 2 | ENSMUST00 | 1706 | Maml3 | ENSMUSG0( | true | chr3 | 52101179 | 52102153 | 17.46222 | 42.53877 | 39.91948 | chr3 | 52077242 | 52101863 | 5.27763 | 4.15049 | 3.25061 |
| chr3 | 52033771 | 52034184 | 1000 | Intron (ENSMUST00000121440.2 | 3 | 51687611 | 52104891 | 417281 | 2 | ENSMUST00 | 70707 | Maml3 | ENSMUSG0( | true | . | . | . | . | . | . | chr3 | 52009386 | 52064052 | 5.58271 | 4.32835 | 3.48679 |
| chr3 | 52029668 | 52030027 | 1000 | Intron (ENSMUST00000121440.2 | 3 | 51687611 | 52104891 | 417281 | 2 | ENSMUST00 | 74864 | Maml3 | ENSMUSG0( | true | . | . | . | . | . | . | chr3 | 52009386 | 52064052 | 5.58271 | 4.32835 | 3.48679 |
| chr3 | 79567428 | 79568349 | 1000 | Promoter (<=1kb) | 3 | 79455971 | 79567679 | 111709 | 2 | ENSMUST00 | 0 | Fnip2 | ENSMUSG0( | false | chr3 | 79566758 | 79568522 | 34.78613 | 113.42423 | 109.48744 | . | . | . | . | . | . |
| chr4 | 111517118 | 111517430 | 1000 | Intron (ENSMUST00000106591.2 | 4 | 111566695 | 111656569 | 89875 | 1 | ENSMUST00 | -49265 | Agbl4 | ENSMUSG0( | false | . | . | . | . | . | . | chr4 | 111513434 | 111520497 | 5.48354 | 4.34928 | 3.40316 |
| chr4 | 111515173 | 111515726 | 1000 | Intron (ENSMUST00000106591.2 | 4 | 111566695 | 111656569 | 89875 | 1 | ENSMUST00 | -50969 | Agbl4 | ENSMUSG0( | false | . | . | . | . | . | . | chr4 | 111513434 | 111520497 | 5.48354 | 4.34928 | 3.40316 |
| chr11 | 120144079 | 120144501 | 1000 | Promoter (1-2kb) | 11 | 120139109 | 120142784 | 3676 | 2 | ENSMUST00 | -1296 | Slc38a10 | ENSMUSG0( | false | . | . | . | . | . | . | chr11 | 120142061 | 120146294 | 8.6033 | 5.5533 | 5.96569 |
| chr8 | 25753943 | 25754663 | 1000 | Promoter (<=1kb) | 8 | 25725324 | 25754280 | 28957 | 2 | ENSMUST00 | 0 | Ddhd2 | ENSMUSG0( | false | chr8 | 25753663 | 25755164 | 23.80619 | 80.75649 | 77.58801 | . | . | . | . | . | . |
| chr14 | 65615045 | 65615368 | 1000 | Intron (ENSMUST00000150897.2 | 14 | 65605267 | 65648444 | 43178 | 1 | ENSMUST00 | 9779 | Nuggc | ENSMUSG0( | false | . | . | . | . | . | . | chr14 | 65614629 | 65615359 | 3.44675 | 3.23247 | 1.77815 |
| chr9 | 119401904 | 119402407 | 1000 | Promoter (<=1kb) | 9 | 119402444 | 119433501 | 31058 | 1 | ENSMUST00 | -37 | Acvr2b | ENSMUSG0( | false | chr9 | 119401529 | 119403875 | 32.48404 | 109.34737 | 105.53431 | . | . | . | . | . | . |
| chr2 | 153529463 | 153530439 | 1000 | Promoter (<=1kb) | 2 | 153407462 | 153529971 | 122510 | 2 | ENSMUST00 | 0 | Nol4l | ENSMUSG0( | false | chr2 | 153527885 | 153530666 | 30.59503 | 95.89513 | 92.42037 | chr2 | 153527739 | 153530465 | 5.1686 | 4.23542 | 3.13389 |
| chr2 | 153443733 | 153445304 | 1000 | Promoter (<=1kb) | 2 | 153407466 | 153444479 | 37014 | 2 | ENSMUST00 | 0 | Nol4l | ENSMUSG0( | false | chr2 | 153444761 | 153446242 | 25.50448 | 78.30592 | 75.18496 | . | . | . | . | . | . |
| chr2 | 153469816 | 153470427 | 1000 | Intron (ENSMUST00000035346.8 | 2 | 153463545 | 153479726 | 16182 | 2 | ENSMUST00 | 9299 | Nol4l | ENSMUSG0( | false | . | . | . | . | . | . | chr2 | 153467463 | 153471752 | 6.49225 | 4.66409 | 4.27861 |
| chr2 | 153527956 | 153528628 | 1000 | Promoter (1-2kb) | 2 | 153407462 | 153529971 | 122510 | 2 | ENSMUST00 | 1343 | Nol4l | ENSMUSG0( | false | chr2 | 153527885 | 153530666 | 30.59503 | 95.89513 | 92.42037 | chr2 | 153527739 | 153530465 | 5.1686 | 4.23542 | 3.13389 |
| chr2 | 153561095 | 153561457 | 1000 | Distal Intergenic | 2 | 153407462 | 153529971 | 122510 | 2 | ENSMUST00 | -31125 | Nol4l | ENSMUSG0( | false | . | . | . | . | . | . | chr2 | 153560401 | 153562671 | 4.60039 | 3.57759 | 2.71448 |
| chr2 | 153436647 | 153437010 | 1000 | Intron (ENSMUST00000035346.8 | 2 | 153407466 | 153444479 | 37014 | 2 | ENSMUST00 | 7469 | Nol4l | ENSMUSG0( | false | . | . | . | . | . | . | chr2 | 153415054 | 153443585 | 6.91758 | 4.79199 | 4.58849 |
| chr6 | 38821759 | 38822148 | 1000 | Intron (ENSMUST00000161779.2 | 6 | 38745457 | 38818346 | 72890 | 2 | ENSMUST00 | -3414 | Hipk2 | ENSMUSG0( | false | . | . | . | . | . | . | chr6 | 38820350 | 38825277 | 4.48636 | 3.79375 | 2.59668 |
| chr19 | 5071897 | 5072621 | 1000 | Promoter (<=1kb) | 19 | 5070859 | 5072132 | 1274 | 2 | ENSMUST00 | 0 | Tmem151a | ENSMUSG0( | false | chr19 | 5071646 | 5072829 | 19.14982 | 55.63055 | 52.84086 | chr19 | 5072545 | 5073228 | 4.73656 | 3.86042 | 2.81432 |
| chr19 | 5085326 | 5085770 | 1000 | Promoter (<=1kb) | 19 | 5071035 | 5085487 | 14453 | 2 | ENSMUST00 | 0 | Tmem151a | ENSMUSG0( | false | chr19 | 5084979 | 5085826 | 17.16829 | 59.16322 | 56.32936 | chr19 | 5084840 | 5086316 | 4.87534 | 3.91084 | 2.92156 |
| chr9 | 101592629 | 101593031 | 1000 | Distal Intergenic | 9 | 101448501 | 101449353 | 853 | 2 | ENSMUST00 | -143277 | Gm5161 | ENSMUSG0( | false | . | . | . | . | . | . | chr9 | 101592829 | 101593988 | 2.95835 | 2.94 | 1.39281 |
| chr5 | 43441164 | 43441837 | 1000 | Distal Intergenic | 5 | 43515569 | 43616585 | 101017 | 1 | ENSMUST00 | -73732 | C1qtnf7 | ENSMUSG0( | true | . | . | . | . | . | . | . | . | . | . | . | . |
| chr9 | 121704659 | 121705508 | 1000 | Promoter (<=1kb) | 9 | 121680047 | 121705029 | 24983 | 2 | ENSMUST00 | 0 | Sec22c | ENSMUSG0( | false | chr9 | 121704441 | 121705820 | 31.61691 | 99.67752 | 96.11594 | chr9 | 121703875 | 121712716 | 6.06469 | 4.507 | 3.89251 |
| chr5 | 117551053 | 117551361 | 1000 | Intron (ENSMUST00000073347.4 | 5 | 117414000 | 117775003 | 361004 | 1 | ENSMUST00 | 137054 | Ksr2 | ENSMUSG0( | false | . | . | . | . | . | . | chr5 | 117540999 | 117552608 | 4.94641 | 4.02258 | 2.97857 |
| chr1 | 37918130 | 37918496 | 1000 | Promoter (1-2kb) | 1 | 37905921 | 37916493 | 10573 | 2 | ENSMUST00 | -1638 | Lyg2 | ENSMUSG0( | false | . | . | . | . | . | . | chr1 | 37916050 | 37923921 | 5.19706 | 4.03622 | 3.21124 |
| chr10 | 80754852 | 80755536 | 1000 | Promoter (<=1kb) | 10 | 80755206 | 80795461 | 40256 | 1 | ENSMUST00 | 0 | Dot1l | ENSMUSG0( | false | chr10 | 80754650 | 80756360 | 41.80276 | 137.50731 | 132.72684 | . | . | . | . | . | . |
| chr5 | 14514793 | 14515339 | 1000 | Promoter (<=1kb) | 5 | 14514918 | 14795993 | 281076 | 1 | ENSMUST00 | 0 | Pclo | ENSMUSG0( | false | chr5 | 14514361 | 14515748 | 31.91637 | 110.38313 | 106.53261 | chr5 | 14515316 | 14516085 | 4.49688 | 3.92898 | 2.57647 |
| chr17 | 42943115 | 42943452 | 1000 | Distal Intergenic | 17 | 42792951 | 42876424 | 83474 | 2 | ENSMUST00 | -66692 | Cd2ap | ENSMUSG0( | true | . | . | . | . | . | . | chr17 | 42938699 | 42944992 | 4.03776 | 3.57328 | 2.22013 |
| chr17 | 42916669 | 42917282 | 1000 | Distal Intergenic | 17 | 42792951 | 42876424 | 83474 | 2 | ENSMUST00 | -40246 | Cd2ap | ENSMUSG0( | true | . | . | . | . | . | . | chr17 | 42914306 | 42920043 | 5.49156 | 4.20042 | 3.41297 |
| chr17 | 42876081 | 42877126 | 1000 | Promoter (<=1kb) | 17 | 42792951 | 42876424 | 83474 | 2 | ENSMUST00 | 0 | Cd2ap | ENSMUSG0( | true | chr17 | 42874504 | 42877282 | 43.42055 | 150.43358 | 145.13264 | chr17 | 42876954 | 42878166 | 3.99546 | 3.63629 | 2.17815 |
| chr15 | 53322792 | 53323242 | 1000 | Intron (ENSMUST00000077273.3 | 15 | 53068664 | 53345659 | 276996 | 2 | ENSMUST00 | 22417 | Ext1 | ENSMUSG0( | false | . | . | . | . | . | . | chr15 | 53321519 | 53328428 | 3.95744 | 3.46118 | 2.18413 |
| chr15 | 53347120 | 53347530 | 1000 | Promoter (<=1kb) | 15 | 53064038 | 53346159 | 282122 | 2 | ENSMUST00 | -962 | Ext1 | ENSMUSG0( | false | . | . | . | . | . | . | chr15 | 53346743 | 53348076 | 4.0472 | 3.52755 | 2.25734 |
| chr15 | 82379877 | 82380706 | 1000 | Promoter (<=1kb) | 15 | 82370528 | 82380260 | 9733 | 2 | ENSMUST00 | 0 | Cyp2d22 | ENSMUSG0( | false | chr15 | 82379548 | 82380266 | 16.08362 | 38.04472 | 35.47989 | chr15 | 82376483 | 82381308 | 5.16064 | 4.22652 | 3.12826 |
| chr16 | 34469459 | 34469929 | 1000 | Intron (ENSMUST00000114961.2 | 16 | 33975495 | 34456917 | 481423 | 2 | ENSMUST00 | -12543 | Kalrn | ENSMUSG0( | false | . | . | . | . | . | . | chr16 | 34469036 | 34470604 | 3.67282 | 3.3187 | 1.91331 |
| chr16 | 34573406 | 34574093 | 1000 | Promoter (<=1kb) | 16 | 34049689 | 34573532 | 523844 | 2 | ENSMUST00 | 0 | Kalrn | ENSMUSG0( | false | chr16 | 34572257 | 34574303 | 30.17592 | 94.17345 | 90.73757 | chr16 | 34572088 | 34574259 | 4.73137 | 3.76861 | 2.81847 |
| chr6 | 95718072 | 95718990 | 1000 | Promoter (<=1kb) | 6 | 95474134 | 95718837 | 244704 | 2 | ENSMUST00 | 0 | Suclg2 | ENSMUSG0( | false | chr6 | 95718032 | 95719391 | 28.91247 | 96.86193 | 93.36482 | chr6 | 95718925 | 95721895 | 5.74506 | 4.21429 | 3.63104 |
| chr6 | 95675811 | 95676087 | 1000 | Intron (ENSMUST00000079847.5 | 6 | 95474134 | 95718837 | 244704 | 2 | ENSMUST00 | 42750 | Suclg2 | ENSMUSG0( | false | . | . | . | . | . | . | chr6 | 95674582 | 95680486 | 5.66179 | 4.26895 | 3.59002 |
| chr5 | 123946579 | 123947410 | 1000 | Promoter (<=1kb) | 5 | 123946386 | 123954515 | 8130 | 1 | ENSMUST00 | 194 | Ccdc62 | ENSMUSG0( | false | . | . | . | . | . | . | chr5 | 123945336 | 123948582 | 5.63933 | 4.3243 | 3.52965 |
| chr4 | 106911022 | 106911875 | 1000 | Promoter (<=1kb) | 4 | 106911470 | 107049694 | 138225 | 1 | ENSMUST00 | 0 | Ssbp3 | ENSMUSG0( | false | chr4 | 106909808 | 106912767 | 41.54923 | 137.06703 | 132.30061 | . | . | . | . | . | . |
| chr4 | 106978336 | 106978784 | 1000 | Intron (ENSMUST00000150915.2 | 4 | 106966426 | 107031067 | 64642 | 1 | ENSMUST00 | 11911 | Ssbp3 | ENSMUSG0( | false | . | . | . | . | . | . | chr4 | 106975874 | 106982715 | 4.16048 | 3.52562 | 2.34565 |
| chr4 | 106909997 | 106910474 | 1000 | Promoter (<=1kb) | 4 | 106910701 | 107031069 | 120369 | 1 | ENSMUST00 | -227 | Ssbp3 | ENSMUSG0( | false | chr4 | 106909808 | 106912767 | 41.54923 | 137.06703 | 132.30061 | chr4 | 106907586 | 106910135 | 7.86904 | 5.27731 | 5.36812 |
| chr15 | 38198717 | 38199121 | 1000 | Distal Intergenic | 15 | 38219203 | 38226735 | 7533 | 1 | ENSMUST00 | -20082 | Odf1 | ENSMUSG0( | false | chr15 | 38198637 | 38199902 | 7.88522 | 15.75676 | 13.48224 | chr15 | 38198092 | 38200961 | 4.67326 | 3.96131 | 2.73207 |
| chr14 | 46821899 | 46822465 | 1000 | Promoter (<=1kb) | 14 | 46808149 | 46822242 | 14094 | 2 | ENSMUST00 | 0 | Gmfb | ENSMUSG0( | false | chr14 | 46820959 | 46822650 | 40.9965 | 143.89526 | 138.84959 | . | . | . | . | . | . |
| chr12 | 21316205 | 21316807 | 1000 | Promoter (<=1kb) | 12 | 21316392 | 21323605 | 7214 | 1 | ENSMUST00 | 0 | Iah1 | ENSMUSG0( | false | chr12 | 21316058 | 21317124 | 35.15467 | 116.16882 | 112.14974 | chr12 | 21316035 | 21317641 | 4.09002 | 3.72194 | 2.24908 |
| chr10 | 39185212 | 39185680 | 1000 | Distal Intergenic | 10 | 39150971 | 39163794 | 12824 | 2 | ENSMUST00 | -21419 | Wisp3 | ENSMUSG0( | false | . | . | . | . | . | . | chr10 | 39182951 | 39186097 | 4.62693 | 3.88235 | 2.70051 |
| chr17 | 10093886 | 10094415 | 1000 | Distal Intergenic | 17 | 10206471 | 10319361 | 112891 | 2 | ENSMUST00 | 224946 | Qk | ENSMUSG0( | true | . | . | . | . | . | . | chr17 | 10092887 | 10106229 | 5.44202 | 4.21288 | 3.38953 |
| chr17 | 10085297 | 10085835 | 1000 | Distal Intergenic | 17 | 10206471 | 10319361 | 112891 | 2 | ENSMUST00 | 233526 | Qk | ENSMUSG0( | true | . | . | . | . | . | . | chr17 | 10078916 | 10088169 | 4.84297 | 3.88168 | 2.89341 |
| chr17 | 10105315 | 10105617 | 1000 | Distal Intergenic | 17 | 10206471 | 10319361 | 112891 | 2 | ENSMUST00 | 213744 | Qk | ENSMUSG0( | true | . | . | . | . | . | . | chr17 | 10092887 | 10106229 | 5.44202 | 4.21288 | 3.38953 |
| chr5 | 74256779 | 74257182 | 1000 | Intron (ENSMUST00000072857.7 | 5 | 74206089 | 74225484 | 19396 | 2 | ENSMUST00 | -31296 | Scfd2 | ENSMUSG0( | true | . | . | . | . | . | . | chr5 | 74256604 | 74257588 | 4.01602 | 3.42041 | 2.2128 |
| chr11 | 60175279 | 60177315 | 1000 | Promoter (<=1kb) | 11 | 60175879 | 60199197 | 23319 | 1 | ENSMUST00 | 0 | Rai1 | ENSMUSG0( | false | chr11 | 60175495 | 60177680 | 20.5121 | 70.85588 | 67.85384 | chr11 | 60176890 | 60178980 | 8.09767 | 5.33041 | 5.55498 |
| chr2 | 156840976 | 156841574 | 1000 | Promoter (<=1kb) | 2 | 156841022 | 156847674 | 6653 | 1 | ENSMUST00 | 0 | Tgif2 | ENSMUSG0( | false | chr2 | 156839609 | 156842066 | 20.14837 | 78.81087 | 75.68276 | chr2 | 156841021 | 156842683 | 5.13196 | 3.89585 | 3.14963 |
| chr2 | 156839743 | 156840522 | 1000 | Promoter (<=1kb) | 2 | 156840007 | 156855570 | 15564 | 1 | ENSMUST00 | 0 | Tgif2 | ENSMUSG0( | false | chr2 | 156839609 | 156842066 | 20.14837 | 78.81087 | 75.68276 | chr2 | 156839584 | 156840094 | 3.38562 | 3.13767 | 1.71549 |
| chr17 | 30471235 | 30471856 | 1000 | Intron (ENSMUST00000079924.5 | 17 | 30215524 | 30576287 | 360764 | 2 | ENSMUST00 | 104431 | Btbd9 | ENSMUSG0( | false | . | . | . | . | . | . | chr17 | 30466527 | 30472388 | 5.79007 | 4.41639 | 3.69487 |
| chr18 | 50053189 | 50053773 | 1000 | Promoter (<=1kb) | 18 | 50053282 | 50091782 | 38501 | 1 | ENSMUST00 | 0 | Tnfaip8 | ENSMUSG0( | false | chr18 | 50052968 | 50055720 | 27.88725 | 99.90214 | 96.33501 | . | . | . | . | . | . |
| chr7 | 19746095 | 19746826 | 1000 | Promoter (2-3kb) | 7 | 19716644 | 19749573 | 32930 | 2 | ENSMUST00 | 2747 | Pvrl2 | ENSMUSG0( | false | . | . | . | . | . | . | chr7 | 19740552 | 19751186 | 6.7145 | 4.82874 | 4.41566 |
| chr11 | 98412205 | 98412937 | 1000 | Promoter (<=1kb) | 11 | 98412470 | 98437716 | 25247 | 1 | ENSMUST00 | 0 | Erbb2 | ENSMUSG0( | false | chr11 | 98412075 | 98414297 | 36.26418 | 124.50906 | 120.21295 | chr11 | 98411669 | 98412379 | 3.75414 | 3.44188 | 1.97615 |
| chr18 | 67252300 | 67252753 | 1000 | Distal Intergenic | 18 | 67221530 | 67245830 | 20301 | 2 | ENSMUST00 | -6471 | Mppe1 | ENSMUSG0( | false | . | . | . | . | . | . | chr18 | 67248989 | 67256141 | 4.10353 | 3.53376 | 2.30208 |
| chr7 | 107521405 | 107521850 | 1000 | Intron (ENSMUST00000130414.2 | 7 | 107504096 | 107548653 | 44558 | 1 | ENSMUST00 | 17310 | Syt9 | ENSMUSG0( | false | . | . | . | . | . | . | chr7 | 107515930 | 107522914 | 4.4782 | 3.78829 | 2.60858 |
| chr7 | 107455100 | 107455516 | 1000 | Intron (ENSMUST00000130414.2 | 7 | 107504062 | 107548653 | 44592 | 1 | ENSMUST00 | -48546 | Syt9 | ENSMUSG0( | false | . | . | . | . | . | . | chr7 | 107454058 | 107456447 | 4.89607 | 4.01244 | 2.95389 |
| chr12 | 99116423 | 99116882 | 1000 | Distal Intergenic | 12 | 99047230 | 99047577 | 348 | 1 | ENSMUST00 | 69194 | Rpl30-ps8 | ENSMUSG0( | false | . | . | . | . | . | . | chr12 | 99113145 | 99121617 | 5.3804 | 4.24636 | 3.34008 |
| chr14 | 87799755 | 87800179 | 1000 | Distal Intergenic | 14 | 88123017 | 88123808 | 792 | 2 | ENSMUST00 | 323629 | Rps3a2 | ENSMUSG0( | false | . | . | . | . | . | . | chr14 | 87799559 | 87800938 | 3.80281 | 3.30993 | 2.05281 |

| | | | | | | | | | | | | | | | | | | | | | | | | | | |
|---|---|---|---|---|---|---|---|---|---|---|---|---|---|---|---|---|---|---|---|---|---|---|---|---|---|---|
| chr2 | 31263732 | 31264216 | 1000 | Intron (ENSMUST00000000199.7 | 2 | 31246166 | 31292319 | 46154 | 1 | ENSMUST00 | 17567 | Ncs1 | ENSMUSG0C | false | . | . | . | . | . | . | chr2 | 31263119 | 31269366 | 6.14924 | 4.58843 | 3.93853 |
| chr15 | 84450436 | 84450977 | 1000 | Distal Intergenic | 15 | 84379203 | 84447097 | 67895 | 2 | ENSMUST00 | -3340 | 1810041L15F | ENSMUSG0C | false | . | . | . | . | . | . | chr15 | 84450267 | 84451391 | 6.99467 | 5.06682 | 4.62736 |
| chr5 | 31452176 | 31452616 | 1000 | Promoter (<=1kb) | 5 | 31452436 | 31481749 | 29314 | 1 | ENSMUST00 | 0 | Zfp512 | ENSMUSG0C | false | chr5 | 31451928 | 31453552 | 19.38504 | 61.02572 | 58.1656 | chr5 | 31451742 | 31455192 | 4.83541 | 3.98435 | 2.88541 |
| chr9 | 108560015 | 108560436 | 1000 | Promoter (<=1kb) | 9 | 108560534 | 108565519 | 4986 | 1 | ENSMUST00 | -98 | Impdh2 | ENSMUSG0C | false | chr9 | 108560313 | 108561124 | 38.89715 | 125.90571 | 121.56978 | chr9 | 108558780 | 108562207 | 4.29051 | 3.64048 | 2.46849 |
| chr16 | 20097426 | 20098374 | 1000 | Promoter (<=1kb) | 16 | 20097554 | 20127744 | 30191 | 1 | ENSMUST00 | 0 | Klhl24 | ENSMUSG0C | false | chr16 | 20097199 | 20098659 | 32.34344 | 150.1674 | 144.87335 | . | . | . | . | . | . |
| chrX | 60403617 | 60404584 | 1000 | Promoter (<=1kb) | 20 | 60223293 | 60403981 | 180689 | 2 | ENSMUST00 | 0 | Atp11c | ENSMUSG0C | false | chrX | 60403062 | 60404044 | 25.27427 | 69.73028 | 66.74294 | . | . | . | . | . | . |
| chrX | 7416451 | 7416886 | 1000 | Intron (ENSMUST00000154346.1 | 20 | 7412087 | 7412825 | 739 | 2 | ENSMUST00 | -3627 | 2010204K13I | ENSMUSG0C | false | . | . | . | . | . | . | . | . | . | . | . | . |
| chr8 | 47674622 | 47675920 | 1000 | Promoter (<=1kb) | 8 | 47667178 | 47675158 | 7981 | 2 | ENSMUST00 | 0 | Ing2 | ENSMUSG0C | false | chr8 | 47673181 | 47676469 | 40.65367 | 138.82761 | 133.98773 | . | . | . | . | . | . |
| chr7 | 126759298 | 126759794 | 1000 | Promoter (<=1kb) | 7 | 126759601 | 126764845 | 5245 | 1 | ENSMUST00 | 0 | Mapk3 | ENSMUSG0C | false | chr7 | 126759577 | 126760602 | 28.90617 | 85.18155 | 81.93615 | chr7 | 126759098 | 126762191 | 6.21078 | 4.63256 | 4.02152 |
| chr4 | 149196163 | 149196525 | 1000 | Promoter (1-2kb) | 4 | 149178863 | 149198494 | 19632 | 2 | ENSMUST00 | 1969 | Kif1b | ENSMUSG0C | false | . | . | . | . | . | . | chr4 | 149196186 | 149196445 | 3.25478 | 3.18482 | 1.59265 |
| chr18 | 5441156 | 5441554 | 1000 | Distal Intergenic | 18 | 5518415 | 5521109 | 2695 | 2 | ENSMUST00 | 79555 | Gm10125 | ENSMUSG0C | false | . | . | . | . | . | . | . | . | . | . | . | . |
| chr5 | 134552043 | 134552651 | 1000 | Promoter (<=1kb) | 5 | 134489386 | 134552434 | 63049 | 2 | ENSMUST00 | 0 | Clip2 | ENSMUSG0C | false | chr5 | 134550699 | 134552810 | 36.27325 | 129.91212 | 125.42401 | . | . | . | . | . | . |
| chr5 | 134548309 | 134548645 | 1000 | Intron (ENSMUST00000036999.6 | 5 | 134489386 | 134552434 | 63049 | 2 | ENSMUST00 | 3789 | Clip2 | ENSMUSG0C | false | . | . | . | . | . | . | chr5 | 134538830 | 134551710 | 6.68849 | 4.77556 | 4.40881 |
| chr7 | 27258326 | 27259078 | 1000 | Promoter (<=1kb) | 7 | 27258433 | 27266801 | 8369 | 1 | ENSMUST00 | 0 | Numbl | ENSMUSG0C | false | chr7 | 27258187 | 27259515 | 25.16654 | 87.04672 | 83.76057 | chr7 | 27256723 | 27259803 | 4.43709 | 3.67834 | 2.58182 |
| chr6 | 148315344 | 148315767 | 1000 | Intron (ENSMUST00000060095.9 | 6 | 148354656 | 148355598 | 943 | 1 | ENSMUST00 | -38889 | Rps4l | ENSMUSG0C | false | . | . | . | . | . | . | chr6 | 148308870 | 148320731 | 5.77782 | 4.49565 | 3.63716 |
| chr15 | 103304960 | 103305618 | 1000 | Downstream (2-3kb) | 15 | 103308236 | 103310438 | 2203 | 2 | ENSMUST00 | 4820 | Gpr84 | ENSMUSG0C | false | chr15 | 103304505 | 103305880 | 24.3552 | 66.39809 | 63.45943 | chr15 | 103298885 | 103310553 | 5.16019 | 4.10407 | 3.16269 |
| chr16 | 16983011 | 16983690 | 1000 | Promoter (<=1kb) | 16 | 16983382 | 17044749 | 61368 | 1 | ENSMUST00 | 0 | Mapk1 | ENSMUSG0C | false | chr16 | 16982800 | 16984633 | 29.2476 | 106.45832 | 102.71833 | chr16 | 16982743 | 16983110 | 4.75672 | 4.08911 | 2.82647 |
| chr9 | 44488846 | 44489439 | 1000 | Distal Intergenic | 9 | 44499136 | 44510388 | 11253 | 1 | ENSMUST00 | -9697 | Bcl9l | ENSMUSG0C | false | . | . | . | . | . | . | chr9 | 44483700 | 44502510 | 6.77697 | 4.85012 | 4.46719 |
| chr14 | 121339186 | 121339612 | 1000 | Intron (ENSMUST00000079817.7 | 14 | 121286343 | 121379334 | 92992 | 2 | ENSMUST00 | 39722 | Stk24 | ENSMUSG0C | false | . | . | . | . | . | . | chr14 | 121336845 | 121343023 | 6.56058 | 4.61176 | 4.28643 |
| chr14 | 121378768 | 121379732 | 1000 | Promoter (<=1kb) | 14 | 121286343 | 121379334 | 92992 | 2 | ENSMUST00 | 0 | Stk24 | ENSMUSG0C | false | chr14 | 121377820 | 121380208 | 31.1999 | 108.05416 | 104.27962 | . | . | . | . | . | . |
| chr10 | 40638332 | 40638667 | 1000 | Intron (ENSMUST00000019977.7 | 10 | 40630011 | 40649931 | 19921 | 1 | ENSMUST00 | 8322 | Ddo | ENSMUSG0C | false | . | . | . | . | . | . | chr10 | 40637480 | 40639273 | 4.14029 | 3.52216 | 2.34648 |
| chr12 | 75818032 | 75818747 | 1000 | Promoter (<=1kb) | 12 | 75818134 | 75880506 | 62373 | 1 | ENSMUST00 | 0 | Syne2 | ENSMUSG0C | false | chr12 | 75817574 | 75819634 | 34.14626 | 110.2223 | 106.38018 | . | . | . | . | . | . |
| chr6 | 38253893 | 38254535 | 1000 | Promoter (<=1kb) | 6 | 38123176 | 38254009 | 130834 | 2 | ENSMUST00 | 0 | D630045J12I | ENSMUSG0C | false | chr6 | 38253205 | 38254651 | 22.30831 | 66.34106 | 63.40397 | chr6 | 38252875 | 38254770 | 3.68155 | 3.43576 | 1.93055 |
| chr14 | 22329237 | 22329555 | 1000 | Intron (ENSMUST00000075639.5 | 14 | 22310165 | 22584439 | 274275 | 1 | ENSMUST00 | 19073 | 1700112E06I | ENSMUSG0C | false | . | . | . | . | . | . | chr14 | 22328921 | 22330403 | 3.38196 | 3.17319 | 1.72714 |
| chr14 | 22745860 | 22746490 | 1000 | Intron (ENSMUST00000075639.5 | 14 | 22577727 | 22599268 | 21542 | 1 | ENSMUST00 | 168134 | 1700112E06I | ENSMUSG0C | false | . | . | . | . | . | . | chr14 | 22744456 | 22748117 | 4.97368 | 4.08352 | 3.00438 |
| chr14 | 22757674 | 22758181 | 1000 | Intron (ENSMUST00000075639.5 | 14 | 22577727 | 22599268 | 21542 | 1 | ENSMUST00 | 179948 | 1700112E06I | ENSMUSG0C | false | . | . | . | . | . | . | chr14 | 22753942 | 22761145 | 5.35278 | 4.22408 | 3.30424 |
| chr5 | 14025144 | 14025782 | 1000 | Promoter (<=1kb) | 5 | 14025276 | 14254314 | 229039 | 1 | ENSMUST00 | 0 | Sema3e | ENSMUSG0C | true | chr5 | 14025087 | 14028582 | 22.32483 | 64.62918 | 61.71925 | . | . | . | . | . | . |
| chr5 | 14039199 | 14039644 | 1000 | Intron (ENSMUST00000130116.1 | 5 | 14025317 | 14256689 | 231373 | 1 | ENSMUST00 | 13883 | Sema3e | ENSMUSG0C | true | . | . | . | . | . | . | chr5 | 14038084 | 14051077 | 3.98378 | 3.51352 | 2.17931 |
| chr6 | 53067821 | 53068880 | 1000 | Promoter (<=1kb) | 6 | 52768797 | 53068631 | 299835 | 2 | ENSMUST00 | 0 | Jazf1 | ENSMUSG0C | false | chr6 | 53067479 | 53069129 | 33.1785 | 101.38432 | 97.78168 | chr6 | 53068777 | 53070301 | 3.29046 | 3.05985 | 1.6409 |
| chr8 | 103290127 | 103290480 | 1000 | Distal Intergenic | 8 | 102864779 | 102865853 | 1075 | 2 | ENSMUST00 | -424275 | Gm8730 | ENSMUSG0C | false | . | . | . | . | . | . | chr8 | 103288175 | 103293261 | 5.7653 | 4.41075 | 3.63503 |
| chr8 | 103408966 | 103409649 | 1000 | Distal Intergenic | 8 | 102864779 | 102865853 | 1075 | 2 | ENSMUST00 | -543114 | Gm8730 | ENSMUSG0C | false | . | . | . | . | . | . | chr8 | 103405303 | 103411536 | 4.39795 | 3.72409 | 2.54416 |
| chr2 | 73038288 | 73038714 | 1000 | Distal Intergenic | 2 | 72987241 | 72988607 | 1367 | 1 | ENSMUST00 | 51048 | Sp3os | ENSMUSG0C | false | chr2 | 73037974 | 73038856 | 7.14186 | 14.46535 | 12.21151 | chr2 | 73037720 | 73038477 | 4.5461 | 3.66585 | 2.66698 |
| chr9 | 57433105 | 57433849 | 1000 | Intron (ENSMUST00000085709.4 | 9 | 57412668 | 57440114 | 27447 | 2 | ENSMUST00 | 6265 | Ppcdc | ENSMUSG0C | false | . | . | . | . | . | . | chr9 | 57431460 | 57434338 | 5.17477 | 3.93273 | 3.17222 |
| chr6 | 125088677 | 125090275 | 1000 | Distal Intergenic | 6 | 125095981 | 125130585 | 34605 | 1 | ENSMUST00 | -5706 | Chd4 | ENSMUSG0C | false | . | . | . | . | . | . | chr6 | 125088715 | 125092715 | 6.09443 | 4.53423 | 3.91246 |
| chr6 | 125095410 | 125097423 | 1000 | Promoter (<=1kb) | 6 | 125095981 | 125130585 | 34605 | 1 | ENSMUST00 | 0 | Chd4 | ENSMUSG0C | false | chr6 | 125096062 | 125099104 | 36.7977 | 133.03651 | 128.43381 | . | . | . | . | . | . |
| chr2 | 145167449 | 145168131 | 1000 | Promoter (<=1kb) | 2 | 145167754 | 145642166 | 474413 | 1 | ENSMUST00 | 0 | Slc24a3 | ENSMUSG0C | false | chr2 | 145167300 | 145168685 | 18.85995 | 50.28701 | 47.56678 | chr2 | 145167216 | 145169104 | 6.65365 | 4.6636 | 4.41446 |
| chr17 | 12199787 | 12200120 | 1000 | Intron (ENSMUST00000164143.2 | 17 | 12189921 | 12190456 | 536 | 1 | ENSMUST00 | 9867 | Tdgf1-ps2 | ENSMUSG0C | false | . | . | . | . | . | . | chr17 | 12189838 | 12208021 | 6.13066 | 4.60409 | 3.94082 |
| chr5 | 116446661 | 116447086 | 1000 | Promoter (<=1kb) | 5 | 116442432 | 116446844 | 4413 | 2 | ENSMUST00 | 0 | Srrm4 | ENSMUSG0C | false | chr5 | 116445494 | 116447174 | 17.46222 | 42.53877 | 39.91948 | . | . | . | . | . | . |
| chr17 | 28800631 | 28801815 | 1000 | Promoter (<=1kb) | 17 | 28801090 | 28838546 | 37457 | 1 | ENSMUST00 | 0 | Brpf3 | ENSMUSG0C | false | chr17 | 28800335 | 28802507 | 32.16725 | 95.65015 | 92.18271 | chr17 | 28799726 | 28800792 | 5.89616 | 4.08395 | 3.78567 |
| chr18 | 73899869 | 73900346 | 1000 | Distal Intergenic | 18 | 73863672 | 73871980 | 8309 | 1 | ENSMUST00 | 36198 | Mro | ENSMUSG0C | false | . | . | . | . | . | . | chr18 | 73898686 | 73902317 | 4.23185 | 3.58169 | 2.41095 |
| chr13 | 67864944 | 67865754 | 1000 | Promoter (1-2kb) | 13 | 67863326 | 67863923 | 598 | 1 | ENSMUST00 | 1619 | BC048507 | ENSMUSG0C | false | . | . | . | . | . | . | chr13 | 67865615 | 67870262 | 5.53014 | 4.37784 | 3.42705 |
| chr17 | 49472327 | 49472973 | 1000 | Intron (ENSMUST00000057610.6 | 17 | 49452284 | 49455435 | 3152 | 1 | ENSMUST00 | 20044 | Mocs1 | ENSMUSG0C | false | . | . | . | . | . | . | chr17 | 49468167 | 49480126 | 6.43614 | 4.71108 | 4.19031 |
| chr17 | 49485562 | 49486056 | 1000 | Intron (ENSMUST00000057610.6 | 17 | 49452284 | 49455435 | 3152 | 1 | ENSMUST00 | 33279 | Mocs1 | ENSMUSG0C | false | chr17 | 49485083 | 49485590 | 6.85183 | 11.47078 | 9.26929 | chr17 | 49481205 | 49490260 | 5.66436 | 4.31035 | 3.55927 |
| chr17 | 49477271 | 49477791 | 1000 | Intron (ENSMUST00000057610.6 | 17 | 49452284 | 49455435 | 3152 | 1 | ENSMUST00 | 24988 | Mocs1 | ENSMUSG0C | false | . | . | . | . | . | . | chr17 | 49468167 | 49480126 | 6.43614 | 4.71108 | 4.19031 |
| chr17 | 49432332 | 49432720 | 1000 | Promoter (<=1kb) | 17 | 49433196 | 49435532 | 2337 | 1 | ENSMUST00 | -476 | Mocs1 | ENSMUSG0C | false | . | . | . | . | . | . | chr17 | 49427791 | 49436927 | 6.84895 | 4.83754 | 4.51735 |
| chrX | 12761546 | 12762541 | 1000 | Promoter (<=1kb) | 20 | 12675369 | 12762073 | 86705 | 2 | ENSMUST00 | 0 | Med14 | ENSMUSG0C | false | chrX | 12760788 | 12762958 | 20.21941 | 51.82391 | 49.08421 | . | . | . | . | . | . |
| chr10 | 116949812 | 116950635 | 1000 | Promoter (<=1kb) | 10 | 116905784 | 116950380 | 44597 | 2 | ENSMUST00 | 0 | Rab3ip | ENSMUSG0C | false | chr10 | 116949413 | 116951139 | 34.56959 | 108.89287 | 105.0881 | . | . | . | . | . | . |
| chr15 | 101827368 | 101827686 | 1000 | Distal Intergenic | 15 | 101810689 | 101818169 | 7481 | 2 | ENSMUST00 | -9200 | Krt2 | ENSMUSG0C | false | . | . | . | . | . | . | chr15 | 101826714 | 101827825 | 5.76195 | 4.47445 | 3.67516 |
| chr15 | 95790488 | 95791138 | 1000 | Promoter (<=1kb) | 15 | 95790843 | 95975464 | 184622 | 1 | ENSMUST00 | 0 | Ano6 | ENSMUSG0C | false | chr15 | 95790185 | 95792142 | 36.88168 | 122.38801 | 118.16834 | . | . | . | . | . | . |
| chr5 | 21271760 | 21272447 | 1000 | Intron (ENSMUST00000036031.8 | 5 | 21319023 | 21322746 | 3724 | 2 | ENSMUST00 | 50299 | Ccdc146 | ENSMUSG0C | false | . | . | . | . | . | . | chr5 | 21270486 | 21275517 | 5.32014 | 4.12064 | 3.30837 |
| chr2 | 61656438 | 61657016 | 1000 | Distal Intergenic | 2 | 61643831 | 61649977 | 6147 | 1 | ENSMUST00 | 12608 | Tank | ENSMUSG0C | false | chr2 | 61655178 | 61657128 | 24.56396 | 67.6308 | 64.67719 | chr2 | 61655080 | 61657186 | 4.58965 | 3.8314 | 2.66306 |
| chr12 | 112644639 | 112645723 | 1000 | Promoter (<=1kb) | 12 | 112644828 | 112649152 | 4325 | 1 | ENSMUST00 | 0 | Siva1 | ENSMUSG0C | false | chr12 | 112644742 | 112646905 | 35.13211 | 109.86813 | 106.03947 | chr12 | 112645492 | 112648303 | 5.55915 | 4.25614 | 3.49523 |
| chr11 | 98950988 | 98951993 | 1000 | Intron (ENSMUST00000107474.2 | 11 | 98946444 | 98946550 | 107 | 1 | ENSMUST00 | 4545 | Gm22061 | ENSMUSG0C | false | chr11 | 98951833 | 98952000 | 5.6323 | 9.47728 | 7.31216 | chr11 | 98928668 | 98957642 | 6.70032 | 4.70048 | 4.41905 |
| chr8 | 47764126 | 47764503 | 1000 | Distal Intergenic | 8 | 47719246 | 47719369 | 124 | 1 | ENSMUST00 | 44881 | n-R5s96 | ENSMUSG0C | false | chr8 | 47763766 | 47764916 | 6.89298 | 11.53747 | 9.33025 | chr8 | 47762950 | 47764955 | 3.9038 | 3.48015 | 2.15657 |
| chr5 | 96912367 | 96912675 | 1000 | Distal Intergenic | 5 | 96907632 | 96907825 | 194 | 1 | ENSMUST00 | 4736 | Gm24623 | ENSMUSG0C | false | . | . | . | . | . | . | chr5 | 96911502 | 96913313 | 3.21852 | 3.09399 | 1.53272 |
| chr7 | 138121491 | 138121826 | 1000 | Distal Intergenic | 7 | 138101433 | 138101539 | 107 | 2 | ENSMUST00 | -19953 | Gm25798 | ENSMUSG0C | false | . | . | . | . | . | . | chr7 | 138119026 | 138122562 | 5.3525 | 4.22418 | 3.34817 |
| chr3 | 79244308 | 79244686 | 1000 | Distal Intergenic | 3 | 79196374 | 79196477 | 104 | 1 | ENSMUST00 | 47935 | Gm24150 | ENSMUSG0C | false | . | . | . | . | . | . | chr3 | 79238335 | 79244955 | 4.79647 | 3.87169 | 2.87846 |
| chr10 | 113161469 | 113161885 | 1000 | Distal Intergenic | 10 | 112931424 | 112931583 | 160 | 1 | ENSMUST00 | 230046 | Gm22507 | ENSMUSG0C | false | . | . | . | . | . | . | . | . | . | . | . | . |
| chrX | 58368212 | 58368612 | 1000 | Distal Intergenic | 20 | 58357952 | 58358058 | 107 | 2 | ENSMUST00 | -10155 | Gm25851 | ENSMUSG0C | false | . | . | . | . | . | . | . | . | . | . | . | . |
| chr9 | 72640269 | 72640829 | 1000 | Distal Intergenic | 9 | 72633447 | 72633553 | 107 | 1 | ENSMUST00 | 6823 | Gm23057 | ENSMUSG0C | false | . | . | . | . | . | . | chr9 | 72635407 | 72647016 | 5.05667 | 4.13544 | 3.04666 |
| chr14 | 54592187 | 54592830 | 1000 | Intron (ENSMUST00000022786.4 | 14 | 54581177 | 54581283 | 107 | 1 | ENSMUST00 | 11011 | Gm24705 | ENSMUSG0C | false | . | . | . | . | . | . | chr14 | 54592230 | 54595906 | 3.81373 | 3.30769 | 2.043 |
| chr15 | 85356180 | 85356498 | 1000 | Intron (ENSMUST00000163242.1 | 15 | 85351063 | 85351254 | 192 | 2 | ENSMUST00 | -4927 | Gm23517 | ENSMUSG0C | false | . | . | . | . | . | . | chr15 | 85353511 | 85360452 | 4.69954 | 3.8588 | 2.77487 |
| chr4 | 49253791 | 49254074 | 1000 | Intron (ENSMUST00000076670.2 | 4 | 49289298 | 49289403 | 106 | 1 | ENSMUST00 | -35224 | Gm22685 | ENSMUSG0C | false | . | . | . | . | . | . | chr4 | 49250271 | 49254046 | 4.84306 | 3.98037 | 2.88173 |
| chr4 | 149860304 | 149860665 | 1000 | Distal Intergenic | 4 | 149817732 | 149817840 | 109 | 1 | ENSMUST00 | 42573 | n-R5s193 | ENSMUSG0C | false | . | . | . | . | . | . | chr4 | 149859847 | 149861376 | 5.80362 | 4.28726 | 3.6996 |
| chr16 | 73647189 | 73647686 | 1000 | Distal Intergenic | 16 | 73422793 | 73422959 | 167 | 2 | ENSMUST00 | -224231 | Gm25699 | ENSMUSG0C | false | chr16 | 73647566 | 73648413 | 8.43118 | 16.20477 | 13.92443 | chr16 | 73645928 | 73649591 | 4.13714 | 3.67768 | 2.31893 |
| chr9 | 63043291 | 63043759 | 1000 | Distal Intergenic | 9 | 63027560 | 63027717 | 158 | 1 | ENSMUST00 | 15732 | Gm24526 | ENSMUSG0C | false | . | . | . | . | . | . | chr9 | 63042952 | 63044209 | 4.38539 | 3.73421 | 2.53764 |
| chr8 | 66049935 | 66050219 | 1000 | Intron (ENSMUST00000152320.2 | 8 | 66002346 | 66002452 | 107 | 2 | ENSMUST00 | -47484 | Gm24205 | ENSMUSG0C | false | . | . | . | . | . | . | chr8 | 66048589 | 66053641 | 3.94247 | 3.53062 | 2.16511 |
| chr7 | 77338333 | 77338987 | 1000 | Distal Intergenic | 7 | 77910140 | 77910320 | 181 | 2 | ENSMUST00 | 571333 | Gm23239 | ENSMUSG0C | false | . | . | . | . | . | . | chr7 | 77337779 | 77339641 | 3.05783 | 3.03347 | 1.40215 |
| chr6 | 97568500 | 97568950 | 1000 | Intron (ENSMUST00000113355.3 | 6 | 97572076 | 97572238 | 163 | 2 | ENSMUST00 | 3288 | Gm23510 | ENSMUSG0C | false | . | . | . | . | . | . | chr6 | 97563267 | 97570348 | 5.20999 | 4.19537 | 3.17679 |
| chr3 | 51916079 | 51916782 | 1000 | Intron (ENSMUST00000121440.2 | 3 | 51828912 | 51829028 | 117 | 1 | ENSMUST00 | 87168 | n-R5s196 | ENSMUSG0C | false | . | . | . | . | . | . | chr3 | 51904229 | 51919287 | 6.09838 | 4.57081 | 3.90639 |
| chr3 | 51965730 | 51966279 | 1000 | Intron (ENSMUST00000121440.2 | 3 | 51828912 | 51829028 | 117 | 1 | ENSMUST00 | 136819 | n-R5s196 | ENSMUSG0C | false | . | . | . | . | . | . | chr3 | 51962114 | 51970468 | 5.40373 | 4.26175 | 3.34479 |
| chr3 | 51879766 | 51880430 | 1000 | Intron (ENSMUST00000121440.2 | 3 | 51828912 | 51829028 | 117 | 1 | ENSMUST00 | 50855 | n-R5s196 | ENSMUSG0C | false | . | . | . | . | . | . | chr3 | 51877616 | 51896424 | 5.58413 | 4.30162 | 3.49753 |
| chr8 | 122526543 | 122526867 | 1000 | Intron (ENSMUST00000127664.1 | 8 | 122520568 | 122520674 | 107 | 2 | ENSMUST00 | -5870 | Gm26497 | ENSMUSG0C | false | . | . | . | . | . | . | chr8 | 122522260 | 122534895 | 7.72566 | 5.23855 | 5.24489 |
| chr8 | 115529731 | 115530199 | 1000 | Distal Intergenic | 8 | 115591170 | 115591273 | 104 | 2 | ENSMUST00 | 61074 | Gm23677 | ENSMUSG0C | false | . | . | . | . | . | . | chr8 | 115529587 | 115530474 | 3.18627 | 3.14017 | 1.56013 |
| chr3 | 151644398 | 151644869 | 1000 | Distal Intergenic | 3 | 151608343 | 151608449 | 107 | 1 | ENSMUST00 | 36056 | Gm24458 | ENSMUSG0C | false | . | . | . | . | . | . | chr3 | 151643550 | 151645981 | 3.67625 | 3.42856 | 1.94544 |
| chr7 | 84358236 | 84358692 | 1000 | Intron (ENSMUST00000085077.3 | 7 | 84323309 | 84323415 | 107 | 2 | ENSMUST00 | -34822 | Gm25790 | ENSMUSG0C | false | . | . | . | . | . | . | chr7 | 84355508 | 84365417 | 5.93716 | 4.48435 | 3.77591 |
| chr4 | 108498799 | 108499122 | 1000 | Promoter (<=1kb) | 4 | 108498044 | 108498146 | 103 | 1 | ENSMUST00 | 756 | Gm23138 | ENSMUSG0C | false | . | . | . | . | . | . | chr4 | 108495296 | 108501329 | 3.61914 | 3.39824 | 1.85771 |
| chr11 | 18745355 | 18746386 | 1000 | Distal Intergenic | 11 | 18738695 | 18738796 | 102 | 1 | ENSMUST00 | 6661 | Gm23093 | ENSMUSG0C | false | . | . | . | . | . | . | chr11 | 18745255 | 18746574 | 3.29718 | 3.07195 | 1.64097 |
| chr17 | 17825949 | 17827422 | 1000 | Promoter (2-3kb) | 17 | 17830188 | 17830257 | 70 | 1 | ENSMUST00 | -2766 | Mir99b | ENSMUSG0C | false | chr17 | 17825989 | 17829531 | 29.75681 | 92.45776 | 89.05831 | chr17 | 17825658 | 17826088 | 4.98022 | 4.18295 | 2.98106 |
| chr1 | 23263863 | 23264740 | 1000 | Distal Intergenic | 1 | 23272264 | 23272345 | 82 | 1 | ENSMUST00 | -7524 | Mir30a | ENSMUSG0C | false | . | . | . | . | . | . | chr1 | 23258890 | 23271405 | 4.63528 | 3.84316 | 2.71286 |
| chr10 | 122985417 | 122986604 | 1000 | Promoter (<=1kb) | 10 | 122985634 | 122985723 | 90 | 2 | ENSMUST00 | 0 | Mirlet7i | ENSMUSG0C | false | chr10 | 122984594 | 122987014 | 36.09495 | 113.61411 | 109.67428 | . | . | . | . | . | . |
| chr13 | 48552216 | 48552906 | 1000 | Distal Intergenic | 13 | 48538171 | 48538270 | 100 | 2 | ENSMUST00 | -13947 | Mirlet7a-1 | ENSMUSG0C | false | chr13 | 48552169 | 48552951 | 6.89286 | 16.24995 | 13.96717 | chr13 | 48550225 | 48557015 | 4.02425 | 3.56278 | 2.21826 |
| chr13 | 48545452 | 48546870 | 1000 | Distal Intergenic | 13 | 48538171 | 48538270 | 100 | 2 | ENSMUST00 | -7183 | Mirlet7a-1 | ENSMUSG0C | false | chr13 | 48545980 | 48546574 | 36.38946 | 142.11884 | 137.14919 | . | . | . | . | . | . |
| chr18 | 61653200 | 61653822 | 1000 | Intron (ENSMUST00000182244.2 | 18 | 61649190 | 61649269 | 80 | 2 | ENSMUST00 | -3932 | Mir143 | ENSMUSG0C | false | . | . | . | . | . | . | chr18 | 61652151 | 61656674 | 6.13635 | 4.45152 | 3.96911 |
| chr6 | 29022128 | 29022551 | 1000 | Promoter (<=1kb) | 6 | 29022619 | 29022691 | 73 | 1 | ENSMUST00 | -68 | Mir129-1 | ENSMUSG0C | false | . | . | . | . | . | . | chr6 | 29020988 | 29024361 | 6.3597 | 4.65726 | 4.12807 |
| chr4 | 150070059 | 150070608 | 1000 | Promoter (1-2kb) | 4 | 150068454 | 150068555 | 102 | 1 | ENSMUST00 | 1606 | Mir34a | ENSMUSG0C | false | . | . | . | . | . | . | chr4 | 150065938 | 150072940 | 5.00292 | 4.0869 | 3.01476 |
| chr9 | 119002259 | 119002661 | 1000 | Intron (ENSMUST00000073109.6 | 9 | 119031796 | 119031885 | 90 | 1 | ENSMUST00 | -29135 | Mir26a-1 | ENSMUSG0C | false | . | . | . | . | . | . | chr9 | 118927678 | 119010843 | 6.89652 | 4.89961 | 4.57102 |
| chr11 | 75462947 | 75463459 | 1000 | Promoter (<=1kb) | 11 | 75463716 | 75463810 | 95 | 1 | ENSMUST00 | -257 | Mir22 | ENSMUSG0C | false | chr11 | 75461573 | 75463115 | 48.67142 | 166.52875 | 160.66557 | chr11 | 75462625 | 75465901 | 6.68681 | 4.87848 | 4.38162 |
| chr7 | 141221719 | 141222112 | 1000 | Promoter (<=1kb) | 7 | 141221384 | 141221493 | 110 | 2 | ENSMUST00 | -227 | Mir210 | ENSMUSG0C | false | chr7 | 141221948 | 141222415 | 7.76973 | 17.46802 | 15.16681 | chr7 | 141221021 | 141222870 | 5.44223 | 4.09447 | 3.39645 |
| chr19 | 29147260 | 29147602 | 1000 | Distal Intergenic | 19 | 29135279 | 29135375 | 97 | 1 | ENSMUST00 | 11982 | Mir101b | ENSMUSG0C | false | . | . | . | . | . | . | . | . | . | . | . | . |
| chr7 | 99547984 | 99548490 | 1000 | Intron (ENSMUST00000098266.3 | 7 | 99552269 | 99552363 | 95 | 1 | ENSMUST00 | -3779 | Mir326 | ENSMUSG0C | false | . | . | . | . | . | . | chr7 | 99536482 | 99576026 | 7.32182 | 4.95837 | 4.9327 |
| chr7 | 99562374 | 99562875 | 1000 | Intron (ENSMUST00000098266.3 | 7 | 99552269 | 99552363 | 95 | 1 | ENSMUST00 | 10106 | Mir326 | ENSMUSG0C | false | . | . | . | . | . | . | chr7 | 99536482 | 99576026 | 7.32182 | 4.95837 | 4.9327 |
| chr7 | 99546093 | 99546763 | 1000 | Intron (ENSMUST00000098266.3 | 7 | 99552269 | 99552363 | 95 | 1 | ENSMUST00 | -5506 | Mir326 | ENSMUSG0C | false | . | . | . | . | . | . | chr7 | 99536482 | 99576026 | 7.32182 | 4.95837 | 4.9327 |
| chr12 | 112130239 | 112130886 | 1000 | Promoter (<=1kb) | 12 | 112130880 | 112130955 | 76 | 1 | ENSMUST00 | 0 | Mir203 | ENSMUSG0C | false | chr12 | 112130010 | 112131148 | 31.85236 | 101.09525 | 97.496 | chr12 | 112129304 | 112131444 | 4.13304 | 3.71722 | 2.27819 |
| chr18 | 61647595 | 61648064 | 1000 | Promoter (<=1kb) | 18 | 61647816 | 61647903 | 88 | 2 | ENSMUST00 | 0 | Mir145a | ENSMUSG0C | false | chr18 | 61647373 | 61648232 | 13.24688 | 29.26311 | 26.80765 | chr18 | 61645922 | 61648647 | 4.10949 | 3.54343 | 2.31315 |
| chr15 | 85675911 | 85676305 | 1000 | Distal Intergenic | 15 | 85706603 | 85706697 | 95 | 1 | ENSMUST00 | -30298 | Mirlet7c-2 | ENSMUSG0C | false | . | . | . | . | . | . | chr15 | 85673573 | 85693849 | 7.40599 | 5.12361 | 4.97896 |
| chr15 | 85705719 | 85706318 | 1000 | Promoter (<=1kb) | 15 | 85706603 | 85706697 | 95 | 1 | ENSMUST00 | -285 | Mirlet7c-2 | ENSMUSG0C | false | . | . | . | . | . | . | chr15 | 85698120 | 85707482 | 9.04895 | 5.54233 | 6.37394 |
| chr8 | 84203468 | 84203951 | 1000 | Distal Intergenic | 8 | 84208518 | 84208592 | 75 | 1 | ENSMUST00 | -4567 | Mir23a | ENSMUSG0C | false | chr8 | 84200501 | 84204316 | 27.20707 | 96.65539 | 93.16586 | chr8 | 84197000 | 84210598 | 6.06328 | 4.55601 | 3.88169 |
| chr11 | 4339913 | 4340415 | 1000 | Distal Intergenic | 11 | 4324149 | 4324255 | 107 | 2 | ENSMUST00 | -15659 | Gm24803 | ENSMUSG0C | false | . | . | . | . | . | . | chr11 | 4339487 | 4342614 | 6.34427 | 4.70465 | 4.10388 |
| chr19 | 25174638 | 25175139 | 1000 | Intron (ENSMUST00000025831.6 | 19 | 25166085 | 25166216 | 132 | 2 | ENSMUST00 | -8423 | Gm24252 | ENSMUSG0C | false | . | . | . | . | . | . | chr19 | 25162630 | 25181812 | 5.3445 | 4.22539 | 3.29608 |
| chr19 | 25169417 | 25169823 | 1000 | Exon (ENSMUST00000025831.6/ | 19 | 25166085 | 25166216 | 132 | 2 | ENSMUST00 | -3202 | Gm24252 | ENSMUSG0C | false | . | . | . | . | . | . | chr19 | 25162630 | 25181812 | 5.3445 | 4.22539 | 3.29608 |
| chr19 | 25187625 | 25188251 | 1000 | Intron (ENSMUST00000025831.6 | 19 | 25166085 | 25166216 | 132 | 2 | ENSMUST00 | -21410 | Gm24252 | ENSMUSG0C | false | . | . | . | . | . | . | chr19 | 25182964 | 25191080 | 6.43266 | 4.75895 | 4.19285 |
| chr8 | 126832888 | 126833251 | 1000 | Distal Intergenic | 8 | 126944847 | 126944980 | 134 | 2 | ENSMUST00 | 111729 | Gm26397 | ENSMUSG0C | false | . | . | . | . | . | . | chr8 | 126824457 | 126835290 | 5.5694 | 4.27648 | 3.48091 |
| chr8 | 126944720 | 126945087 | 1000 | Promoter (<=1kb) | 8 | 126944847 | 126944980 | 134 | 2 | ENSMUST00 | 0 | Gm26397 | ENSMUSG0C | false | chr8 | 126943948 | 126946384 | 23.3092 | 84.4034 | 81.16875 | chr8 | 126939915 | 126945088 | 4.81672 | 3.88889 | 2.89487 |
| chr10 | 108385774 | 108386109 | 1000 | Intron (ENSMUST00000095313.3 | 10 | 108363665 | 108363798 | 134 | 1 | ENSMUST00 | 22110 | Gm23105 | ENSMUSG0C | false | . | . | . | . | . | . | chr10 | 108383715 | 108391831 | 4.99674 | 4.04583 | 3.02244 |
| chr10 | 108350413 | 108351067 | 1000 | Intron (ENSMUST00000095313.3 | 10 | 108363665 | 108363798 | 134 | 1 | ENSMUST00 | -12598 | Gm23105 | ENSMUSG0C | false | . | . | . | . | . | . | chr10 | 108347480 | 108358834 | 5.11833 | 4.07651 | 3.11993 |
| chr10 | 8752543 | 8752945 | 1000 | Intron (ENSMUST00000015449.5 | 10 | 8824011 | 8824147 | 137 | 2 | ENSMUST00 | 71202 | Gm25410 | ENSMUSG0C | false | . | . | . | . | . | . | chr10 | 8751277 | 8755790 | 6.42059 | 4.71149 | 4.18065 |
| chr7 | 73617844 | 73618928 | 1000 | Distal Intergenic | 7 | 73645985 | 73646089 | 105 | 2 | ENSMUST00 | 27161 | Gm26176 | ENSMUSG0C | false | chr7 | 73616872 | 73618375 | 27.78601 | 91.88203 | 88.49705 | . | . | . | . | . | . |
| chr7 | 73606770 | 73607271 | 1000 | Distal Intergenic | 7 | 73645985 | 73646089 | 105 | 2 | ENSMUST00 | 38818 | Gm26176 | ENSMUSG0C | false | chr7 | 73606918 | 73608349 | 12.0809 | 26.07937 | 23.66483 | chr7 | 73605792 | 73607723 | 3.97831 | 3.59295 | 2.18304 |
| chr17 | 45619363 | 45619723 | 1000 | Promoter (<=1kb) | 17 | 45620679 | 45620759 | 81 | 1 | ENSMUST00 | -956 | Gm25008 | ENSMUSG0C | false | . | . | . | . | . | . | chr17 | 45618799 | 45620939 | 5.71313 | 4.51821 | 3.5875 |
| chr15 | 49535064 | 49535403 | 1000 | Promoter (2-3kb) | 15 | 49538195 | 49538300 | 106 | 1 | ENSMUST00 | -2792 | Gm23848 | ENSMUSG0C | false | . | . | . | . | . | . | chr15 | 49534226 | 49538322 | 3.86398 | 3.4808 | 2.0709 |
| chr11 | 101604456 | 101605002 | 1000 | Promoter (<=1kb) | 11 | 101604850 | 101605040 | 191 | 1 | ENSMUST00 | 0 | Gm26316 | ENSMUSG0C | false | chr11 | 101604364 | 101605060 | 10.74617 | 22.67706 | 20.30526 | chr11 | 101604296 | 101604837 | 2.98405 | 2.98543 | 1.39819 |
| chrX | 11807053 | 11807877 | 1000 | Distal Intergenic | 20 | 11850484 | 11850590 | 107 | 1 | ENSMUST00 | -42607 | Gm26314 | ENSMUSG0C | false | chrX | 11807126 | 11808092 | 12.40737 | 26.6269 | 24.20249 | . | . | . | . | . | . |
| chr2 | 63702447 | 63703137 | 1000 | Distal Intergenic | 2 | 63432088 | 63432194 | 107 | 2 | ENSMUST00 | -270254 | Gm23503 | ENSMUSG0C | false | chr2 | 63701903 | 63703074 | 8.50137 | 15.8526 | 13.57707 | chr2 | 63701955 | 63703908 | 4.28156 | 3.79438 | 2.40494 |
| chr6 | 140337295 | 140337964 | 1000 | Distal Intergenic | 6 | 140319630 | 140319733 | 104 | 1 | ENSMUST00 | 17666 | n-R5s168 | ENSMUSG0C | false | chr6 | 140337804 | 140338672 | 10.47775 | 22.22469 | 19.86031 | chr6 | 140336296 | 140338685 | 2.89794 | 2.91378 | 1.33499 |
| chr6 | 140300481 | 140300812 | 1000 | Distal Intergenic | 6 | 140319630 | 140319733 | 104 | 1 | ENSMUST00 | -18818 | n-R5s168 | ENSMUSG0C | false | . | . | . | . | . | . | chr6 | 140299913 | 140301098 | 3.06391 | 3.00392 | 1.4747 |
| chr9 | 35488533 | 35488940 | 1000 | Intron (ENSMUST00000119129.3 | 9 | 35474885 | 35475046 | 162 | 1 | ENSMUST00 | 13649 | Gm25630 | ENSMUSG0C | false | . | . | . | . | . | . | chr9 | 35486585 | 35489810 | 4.40428 | 3.73032 | 2.53451 |
| chr9 | 35467388 | 35467885 | 1000 | Intron (ENSMUST00000119129.3 | 9 | 35474885 | 35475046 | 162 | 1 | ENSMUST00 | -7000 | Gm25630 | ENSMUSG0C | false | . | . | . | . | . | . | chr9 | 35466445 | 35468786 | 4.8709 | 4.08684 | 2.90897 |
| chr8 | 25240467 | 25241113 | 1000 | Distal Intergenic | 8 | 25154552 | 25201449 | 46898 | 2 | ENSMUST00 | -39019 | Tacc1 | ENSMUSG0C | false | chr8 | 25240215 | 25241431 | 33.04283 | 115.53907 | 111.53828 | chr8 | 25239982 | 25242224 | 3.82277 | 3.52298 | 2.02578 |
| chr8 | 25323664 | 25323924 | 1000 | Distal Intergenic | 8 | 25154552 | 25201449 | 46898 | 2 | ENSMUST00 | -122216 | Tacc1 | ENSMUSG0C | false | . | . | . | . | . | . | chr8 | 25323685 | 25324031 | 3.50232 | 3.42417 | 1.80071 |
| chr8 | 25311856 | 25312207 | 1000 | Distal Intergenic | 8 | 25154552 | 25201449 | 46898 | 2 | ENSMUST00 | -110408 | Tacc1 | ENSMUSG0C | false | . | . | . | . | . | . | chr8 | 25304305 | 25313043 | 5.09789 | 4.06048 | 3.12494 |
| chr8 | 25201179 | 25201832 | 1000 | Promoter (<=1kb) | 8 | 25154552 | 25201449 | 46898 | 2 | ENSMUST00 | 0 | Tacc1 | ENSMUSG0C | false | chr8 | 25200274 | 25202264 | 27.20707 | 96.65539 | 93.16586 | . | . | . | . | . | . |
| chr4 | 132387771 | 132388275 | 1000 | Intron (ENSMUST00000102568.4 | 4 | 132377461 | 132398199 | 20739 | 2 | ENSMUST00 | 9924 | Phactr4 | ENSMUSG0C | true | . | . | . | . | . | . | chr4 | 132379835 | 132397156 | 5.13856 | 4.06122 | 3.15072 |
| chr4 | 132389345 | 132389693 | 1000 | Intron (ENSMUST00000102568.4 | 4 | 132377461 | 132398199 | 20739 | 2 | ENSMUST00 | 8506 | Phactr4 | ENSMUSG0C | true | . | . | . | . | . | . | chr4 | 132379835 | 132397156 | 5.13856 | 4.06122 | 3.15072 |
| chr4 | 62287429 | 62288179 | 1000 | Promoter (<=1kb) | 4 | 62286449 | 62298411 | 11963 | 1 | ENSMUST00 | 981 | Slc31a2 | ENSMUSG0C | false | chr4 | 62288080 | 62288869 | 8.26071 | 16.87976 | 14.58833 | chr4 | 62287890 | 62304836 | 6.16439 | 4.4648 | 3.98538 |
| chr17 | 28522959 | 28523537 | 1000 | Promoter (<=1kb) | 17 | 28523257 | 28525347 | 2091 | 1 | ENSMUST00 | 0 | E230001N04 | ENSMUSG0C | false | chr17 | 28522419 | 28523541 | 24.0633 | 66.77184 | 63.83004 | chr17 | 28522194 | 28523718 | 5.2971 | 4.25669 | 3.25596 |
| chr9 | 123021050 | 123021831 | 1000 | Promoter (<=1kb) | 9 | 123021326 | 123023490 | 2165 | 1 | ENSMUST00 | 0 | Tmem42 | ENSMUSG0C | false | chr9 | 123020357 | 123022072 | 28.71633 | 95.52087 | 92.05702 | chr9 | 123020080 | 123022094 | 3.93812 | 3.59546 | 2.12604 |
| chr7 | 101969741 | 101971294 | 1000 | Promoter (<=1kb) | 7 | 101969843 | 102014959 | 45117 | 1 | ENSMUST00 | 0 | Numa1 | ENSMUSG0C | false | chr7 | 101969394 | 101971896 | 25.16654 | 87.04672 | 83.76057 | chr7 | 101971197 | 101976518 | 4.93622 | 4.03726 | 2.95537 |
| chr7 | 75642250 | 75642833 | 1000 | Promoter (<=1kb) | 7 | 75643319 | 75704422 | 61104 | 1 | ENSMUST00 | -486 | Akap13 | ENSMUSG0C | false | chr7 | 75642319 | 75642769 | 7.13425 | 13.57188 | 11.33167 | chr7 | 75639109 | 75646053 | 5.29783 | 4.15948 | 3.25907 |
| chr9 | 101073400 | 101075102 | 1000 | Promoter (<=1kb) | 9 | 101074101 | 101101785 | 27685 | 1 | ENSMUST00 | 0 | Msl2 | ENSMUSG0C | true | chr9 | 101073983 | 101079109 | 33.41831 | 117.26761 | 113.21259 | . | . | . | . | . | . |
| chr5 | 149051230 | 149052400 | 1000 | Promoter (<=1kb) | 5 | 149048228 | 149051300 | 3073 | 2 | ENSMUST00 | 0 | Hmgb1 | ENSMUSG0C | false | chr5 | 149050983 | 149053595 | 30.86329 | 102.37074 | 98.74274 | . | . | . | . | . | . |
| chr5 | 129941464 | 129942212 | 1000 | Promoter (<=1kb) | 5 | 129942109 | 129984745 | 42637 | 1 | ENSMUST00 | 0 | Vkorc1l1 | ENSMUSG0C | false | chr5 | 129941943 | 129942136 | 5.50447 | 8.289 | 6.15165 | . | . | . | . | . | . |
| chr1 | 43445075 | 43446579 | 1000 | Promoter (<=1kb) | 1 | 43445751 | 43462194 | 16444 | 1 | ENSMUST00 | 0 | Nck2 | ENSMUSG0C | true | chr1 | 43444845 | 43448373 | 43.42055 | 150.43358 | 145.13264 | chr1 | 43444516 | 43445300 | 4.35943 | 3.84373 | 2.47308 |
| chr1 | 43474495 | 43474944 | 1000 | Intron (ENSMUST00000187435.1 | 1 | 43457273 | 43462222 | 4950 | 1 | ENSMUST00 | 17223 | Nck2 | ENSMUSG0C | true | . | . | . | . | . | . | chr1 | 43446894 | 43509082 | 5.99677 | 4.48008 | 3.83622 |
| chr1 | 43469017 | 43469358 | 1000 | Intron (ENSMUST00000187435.1 | 1 | 43457273 | 43462222 | 4950 | 1 | ENSMUST00 | 11745 | Nck2 | ENSMUSG0C | true | . | . | . | . | . | . | chr1 | 43446894 | 43509082 | 5.99677 | 4.48008 | 3.83622 |
| chr8 | 71922483 | 71922866 | 1000 | Promoter (<=1kb) | 8 | 71922852 | 71934629 | 11778 | 1 | ENSMUST00 | 0 | Zfp617 | ENSMUSG0C | false | chr8 | 71922378 | 71923838 | 29.28796 | 98.53321 | 94.99628 | . | . | . | . | . | . |
| chr5 | 117115675 | 117116136 | 1000 | Promoter (<=1kb) | 5 | 117091682 | 117115993 | 24312 | 2 | ENSMUST00 | 0 | Suds3 | ENSMUSG0C | false | chr5 | 117114778 | 117116319 | 35.62435 | 116.99458 | 112.94973 | . | . | . | . | . | . |
| chr1 | 82586631 | 82587258 | 1000 | Promoter (<=1kb) | 1 | 82448423 | 82586849 | 138427 | 2 | ENSMUST00 | 0 | Col4a4 | ENSMUSG0C | true | chr1 | 82583771 | 82590897 | 23.80619 | 80.75649 | 77.58801 | . | . | . | . | . | . |
| chr19 | 41829554 | 41830558 | 1000 | Promoter (<=1kb) | 19 | 41830168 | 41830809 | 642 | 1 | ENSMUST00 | 0 | Frat1 | ENSMUSG0C | false | chr19 | 41829166 | 41831508 | 30.60556 | 103.92632 | 100.25623 | chr19 | 41829110 | 41829606 | 3.58793 | 3.11354 | 1.89254 |
| chrX | 153007444 | 153007792 | 1000 | Promoter (<=1kb) | 20 | 153006469 | 153009416 | 2948 | 1 | ENSMUST00 | 976 | Usp51 | ENSMUSG0C | false | chrX | 153006655 | 153007849 | 17.92175 | 44.05972 | 41.41886 | . | . | . | . | . | . |
| chrX | 152974177 | 152974649 | 1000 | Distal Intergenic | 20 | 153006469 | 153009416 | 2948 | 1 | ENSMUST00 | -31820 | Usp51 | ENSMUSG0C | false | chrX | 152973883 | 152974272 | 8.73111 | 16.23101 | 13.94854 | . | . | . | . | . | . |
| chr5 | 45040796 | 45041112 | 1000 | Distal Intergenic | 5 | 44983861 | 44984242 | 382 | 2 | ENSMUST00 | -56555 | Gm10205 | ENSMUSG0C | false | . | . | . | . | . | . | chr5 | 45040088 | 45042578 | 4.07514 | 3.62745 | 2.24402 |
| chr1 | 59833725 | 59834099 | 1000 | Intron (ENSMUST00000087435.6 | 1 | 59850665 | 59852351 | 1687 | 1 | ENSMUST00 | -16566 | Bmpr2 | ENSMUSG0C | true | . | . | . | . | . | . | chr1 | 59829971 | 59834424 | 4.26072 | 3.73225 | 2.38707 |
| chr1 | 59763128 | 59764152 | 1000 | Promoter (<=1kb) | 1 | 59763400 | 59879014 | 115615 | 1 | ENSMUST00 | 0 | Bmpr2 | ENSMUSG0C | true | chr1 | 59762382 | 59765962 | 25.48472 | 95.2657 | 91.8051 | . | . | . | . | . | . |
| chr1 | 59762138 | 59762747 | 1000 | Promoter (<=1kb) | 1 | 59763400 | 59879014 | 115615 | 1 | ENSMUST00 | -653 | Bmpr2 | ENSMUSG0C | true | chr1 | 59762382 | 59765962 | 25.48472 | 95.2657 | 91.8051 | chr1 | 59760312 | 59762827 | 3.33274 | 3.24585 | 1.64988 |
| chr17 | 45584458 | 45584843 | 1000 | Promoter (1-2kb) | 17 | 45586120 | 45588377 | 2258 | 1 | ENSMUST00 | -1277 | Gm17080 | ENSMUSG0C | false | . | . | . | . | . | . | chr17 | 45580554 | 45586436 | 6.35903 | 4.63318 | 4.14002 |
| chr13 | 51328251 | 51328705 | 1000 | Distal Intergenic | 13 | 51408618 | 51422797 | 14180 | 1 | ENSMUST00 | -79913 | S1pr3 | ENSMUSG0C | false | . | . | . | . | . | . | chr13 | 51327944 | 51329287 | 3.01346 | 2.94697 | 1.40327 |
| chr15 | 98832231 | 98832596 | 1000 | Promoter (<=1kb) | 15 | 98812798 | 98831508 | 18711 | 2 | ENSMUST00 | -724 | Prkag1 | ENSMUSG0C | false | . | . | . | . | . | . | chr15 | 98831563 | 98833615 | 5.51648 | 4.29781 | 3.45636 |
| chr5 | 24526511 | 24527318 | 1000 | Promoter (<=1kb) | 5 | 24502757 | 24527202 | 24446 | 2 | ENSMUST00 | 0 | Gbx1 | ENSMUSG0C | false | chr5 | 24525602 | 24527538 | 25.56571 | 75.64988 | 72.57267 | chr5 | 24527175 | 24528054 | 3.52 | 3.36018 | 1.79647 |
| chr5 | 24497596 | 24498242 | 1000 | Exon (ENSMUST00000024123.4/ | 5 | 24502757 | 24527202 | 24446 | 2 | ENSMUST00 | 28960 | Gbx1 | ENSMUSG0C | false | chr5 | 24497977 | 24499046 | 8.90104 | 20.03618 | 17.69957 | chr5 | 24495765 | 24506361 | 5.85559 | 4.41542 | 3.73746 |
| chr5 | 24512314 | 24512832 | 1000 | Intron (ENSMUST00000088311.5 | 5 | 24502757 | 24527202 | 24446 | 2 | ENSMUST00 | 14370 | Gbx1 | ENSMUSG0C | false | . | . | . | . | . | . | chr5 | 24511686 | 24513888 | 6.32215 | 4.56472 | 4.12943 |
| chr5 | 24505388 | 24505758 | 1000 | Intron (ENSMUST00000088311.5 | 5 | 24502757 | 24527202 | 24446 | 2 | ENSMUST00 | 21444 | Gbx1 | ENSMUSG0C | false | . | . | . | . | . | . | chr5 | 24495765 | 24506361 | 5.85559 | 4.41542 | 3.73746 |
| chr6 | 121160215 | 121160621 | 1000 | Distal Intergenic | 6 | 121183667 | 121196185 | 12519 | 1 | ENSMUST00 | -23046 | Pex26 | ENSMUSG0C | false | . | . | . | . | . | . | chr6 | 121159156 | 121161151 | 4.66549 | 3.96415 | 2.78617 |
| chr1 | 10232089 | 10233326 | 1000 | Promoter (<=1kb) | 1 | 10137571 | 10232670 | 95100 | 2 | ENSMUST00 | 0 | Arfgef1 | ENSMUSG0C | false | chr1 | 10231713 | 10233939 | 43.16833 | 149.98996 | 144.69951 | chr1 | 10228067 | 10232125 | 4.04298 | 3.56329 | 2.26326 |
| chr17 | 21566745 | 21567330 | 1000 | Promoter (<=1kb) | 17 | 21567046 | 21588697 | 21652 | 1 | ENSMUST00 | 0 | Zfp948 | ENSMUSG0C | false | chr17 | 21566408 | 21567855 | 16.96705 | 61.94942 | 59.07605 | . | . | . | . | . | . |
| chr14 | 74947601 | 74948131 | 1000 | Promoter (<=1kb) | 14 | 74754674 | 74947877 | 193204 | 2 | ENSMUST00 | 0 | Lrch1 | ENSMUSG0C | true | chr14 | 74944710 | 74948380 | 36.93744 | 125.70555 | 121.37619 | . | . | . | . | . | . |
| chr14 | 74925979 | 74926379 | 1000 | Intron (ENSMUST00000088970.5 | 14 | 74754674 | 74947877 | 193204 | 2 | ENSMUST00 | 21498 | Lrch1 | ENSMUSG0C | true | . | . | . | . | . | . | chr14 | 74857233 | 74945366 | 6.29465 | 4.64624 | 4.07764 |
| chr14 | 74910208 | 74910753 | 1000 | Intron (ENSMUST00000088970.5 | 14 | 74754674 | 74947877 | 193204 | 2 | ENSMUST00 | 37124 | Lrch1 | ENSMUSG0C | true | . | . | . | . | . | . | chr14 | 74857233 | 74945366 | 6.29465 | 4.64624 | 4.07764 |
| chr14 | 74863020 | 74863370 | 1000 | Intron (ENSMUST00000088970.5 | 14 | 74754674 | 74947877 | 193204 | 2 | ENSMUST00 | 84507 | Lrch1 | ENSMUSG0C | true | . | . | . | . | . | . | chr14 | 74857233 | 74945366 | 6.29465 | 4.64624 | 4.07764 |
| chr14 | 74946453 | 74947077 | 1000 | Promoter (<=1kb) | 14 | 74754674 | 74947877 | 193204 | 2 | ENSMUST00 | 800 | Lrch1 | ENSMUSG0C | true | chr14 | 74944710 | 74948380 | 36.93744 | 125.70555 | 121.37619 | . | . | . | . | . | . |
| chr14 | 74918454 | 74918882 | 1000 | Intron (ENSMUST00000088970.5 | 14 | 74754674 | 74947877 | 193204 | 2 | ENSMUST00 | 28995 | Lrch1 | ENSMUSG0C | true | . | . | . | . | . | . | chr14 | 74857233 | 74945366 | 6.29465 | 4.64624 | 4.07764 |
| chr14 | 74882765 | 74883482 | 1000 | Intron (ENSMUST00000088970.5 | 14 | 74754674 | 74947877 | 193204 | 2 | ENSMUST00 | 64395 | Lrch1 | ENSMUSG0C | true | . | . | . | . | . | . | chr14 | 74857233 | 74945366 | 6.29465 | 4.64624 | 4.07764 |
| chr14 | 74945082 | 74945726 | 1000 | Promoter (2-3kb) | 14 | 74754674 | 74947877 | 193204 | 2 | ENSMUST00 | 2151 | Lrch1 | ENSMUSG0C | true | chr14 | 74944710 | 74948380 | 36.93744 | 125.70555 | 121.37619 | chr14 | 74857233 | 74945366 | 6.29465 | 4.64624 | 4.07764 |
| chr17 | 13759899 | 13760574 | 1000 | Promoter (<=1kb) | 17 | 13760539 | 13810518 | 49980 | 1 | ENSMUST00 | 0 | Mllt4 | ENSMUSG0C | false | chr17 | 13759895 | 13761892 | 20.4053 | 65.4752 | 62.55133 | chr17 | 13758839 | 13761757 | 4.04384 | 3.35224 | 2.26336 |
| chr2 | 153186217 | 153186561 | 1000 | Intron (ENSMUST00000137446.2 | 2 | 153190399 | 153192187 | 1789 | 1 | ENSMUST00 | -3838 | Tm9sf4 | ENSMUSG0C | false | . | . | . | . | . | . | chr2 | 153184117 | 153189002 | 5.31615 | 4.17455 | 3.27624 |
| chr2 | 153191723 | 153192515 | 1000 | Promoter (<=1kb) | 2 | 153190926 | 153200761 | 9836 | 1 | ENSMUST00 | 798 | Tm9sf4 | ENSMUSG0C | false | . | . | . | . | . | . | chr2 | 153190018 | 153193556 | 4.20478 | 3.4605 | 2.39267 |
| chr15 | 78928085 | 78928560 | 1000 | Promoter (1-2kb) | 15 | 78926725 | 78930465 | 3741 | 1 | ENSMUST00 | 1361 | Lgals1 | ENSMUSG0C | false | chr15 | 78927804 | 78928429 | 9.01168 | 19.18248 | 16.85847 | chr15 | 78925753 | 78928195 | 5.28148 | 4.19836 | 3.23857 |
| chrX | 6399894 | 6400592 | 1000 | Promoter (<=1kb) | 20 | 6399854 | 6633837 | 233984 | 1 | ENSMUST00 | 41 | Shroom4 | ENSMUSG0C | false | chrX | 6399688 | 6401191 | 25.7338 | 71.40817 | 68.39619 | . | . | . | . | . | . |
| chr16 | 44172931 | 44173844 | 1000 | Promoter (<=1kb) | 16 | 44173246 | 44227465 | 54220 | 1 | ENSMUST00 | 0 | Gm608 | ENSMUSG0C | false | chr16 | 44173160 | 44174315 | 31.91637 | 110.38313 | 106.53261 | . | . | . | . | . | . |
| chr6 | 83268155 | 83268499 | 1000 | Exon (ENSMUST00000113900.2/ | 6 | 83277393 | 83288451 | 11059 | 1 | ENSMUST00 | -8894 | Slc4a5 | ENSMUSG0C | false | chr6 | 83268078 | 83268644 | 15.04464 | 36.28028 | 33.73701 | . | . | . | . | . | . |
| chr2 | 104267020 | 104267287 | 1000 | Intron (ENSMUST00000150450.1 | 2 | 104207666 | 104257984 | 50319 | 2 | ENSMUST00 | -9037 | D430041D05 | ENSMUSG0C | false | . | . | . | . | . | . | chr2 | 104266882 | 104267817 | 3.22699 | 3.14713 | 1.58996 |
| chr15 | 73090249 | 73090957 | 1000 | Promoter (<=1kb) | 15 | 73090392 | 73094075 | 3684 | 1 | ENSMUST00 | 0 | Chrac1 | ENSMUSG0C | false | chr15 | 73089864 | 73091428 | 36.76257 | 113.72284 | 109.77938 | . | . | . | . | . | . |
| chr6 | 48444933 | 48446572 | 1000 | Promoter (<=1kb) | 6 | 48427697 | 48445464 | 17768 | 2 | ENSMUST00 | 0 | Zfp467 | ENSMUSG0C | false | chr6 | 48444505 | 48446173 | 25.79552 | 96.83904 | 93.34669 | . | . | . | . | . | . |
| chr6 | 48443931 | 48444874 | 1000 | Promoter (<=1kb) | 6 | 48427697 | 48444036 | 16340 | 2 | ENSMUST00 | 0 | Zfp467 | ENSMUSG0C | false | chr6 | 48443941 | 48444382 | 16.22329 | 39.70485 | 37.12044 | chr6 | 48442665 | 48444265 | 3.71782 | 3.50123 | 1.94259 |
| chr7 | 3292784 | 3293261 | 1000 | Promoter (2-3kb) | 7 | 3290556 | 3299345 | 8790 | 1 | ENSMUST00 | 2229 | Myadm | ENSMUSG0C | false | chr7 | 3293062 | 3293649 | 9.68513 | 19.2832 | 16.95758 | chr7 | 3290873 | 3298443 | 4.7394 | 4.01231 | 2.78151 |
| chr6 | 40728504 | 40729064 | 1000 | Intron (ENSMUST00000071535.6 | 6 | 40628831 | 40769123 | 140293 | 1 | ENSMUST00 | 99674 | Mgam | ENSMUSG0C | false | . | . | . | . | . | . | chr6 | 40727806 | 40731788 | 3.75798 | 3.42302 | 1.98993 |
| chr6 | 40691920 | 40692464 | 1000 | Intron (ENSMUST00000071535.6 | 6 | 40628831 | 40769123 | 140293 | 1 | ENSMUST00 | 63090 | Mgam | ENSMUSG0C | false | . | . | . | . | . | . | chr6 | 40691104 | 40695012 | 3.79191 | 3.50535 | 2.00539 |
| chr6 | 40757349 | 40757805 | 1000 | Exon (ENSMUST00000071535.6/ | 6 | 40628831 | 40769123 | 140293 | 1 | ENSMUST00 | 128519 | Mgam | ENSMUSG0C | false | . | . | . | . | . | . | chr6 | 40756374 | 40758065 | 4.58975 | 3.84803 | 2.67922 |
| chr16 | 15052826 | 15053235 | 1000 | Distal Intergenic | 16 | 14906637 | 14925879 | 19243 | 1 | ENSMUST00 | 146190 | Efcab1 | ENSMUSG0C | false | . | . | . | . | . | . | chr16 | 15051907 | 15054883 | 3.97672 | 3.49044 | 2.17804 |
| chr2 | 93187285 | 93187908 | 1000 | Promoter (<=1kb) | 2 | 93187548 | 93198919 | 11372 | 1 | ENSMUST00 | 0 | Trp53i11 | ENSMUSG0C | false | chr2 | 93187257 | 93189185 | 33.10969 | 106.34647 | 102.61562 | chr2 | 93186945 | 93189255 | 3.52931 | 3.32109 | 1.81867 |
| chr2 | 92434041 | 92434624 | 1000 | Promoter (<=1kb) | 2 | 92403646 | 92434043 | 30398 | 2 | ENSMUST00 | 0 | Cry2 | ENSMUSG0C | false | chr2 | 92431256 | 92434116 | 30.13579 | 99.06311 | 95.51676 | chr2 | 92434470 | 92435903 | 6.09838 | 4.41148 | 3.93863 |
| chr2 | 92423634 | 92424116 | 1000 | Intron (ENSMUST00000126002.1 | 2 | 92426904 | 92432392 | 5489 | 2 | ENSMUST00 | 8276 | Cry2 | ENSMUSG0C | false | . | . | . | . | . | . | chr2 | 92421995 | 92433198 | 7.705 | 5.3487 | 5.2209 |
| chr14 | 14345571 | 14345849 | 1000 | Promoter (<=1kb) | 14 | 14346495 | 14355490 | 8996 | 1 | ENSMUST00 | -646 | Il3ra | ENSMUSG0C | false | chr14 | 14345517 | 14346893 | 32.16725 | 95.65015 | 92.18271 | chr14 | 14345200 | 14346551 | 3.37383 | 3.28911 | 1.67052 |
| chr2 | 69861337 | 69862060 | 1000 | Promoter (<=1kb) | 2 | 69861562 | 69871786 | 10225 | 1 | ENSMUST00 | 0 | Ssb | ENSMUSG0C | false | chr2 | 69860961 | 69862760 | 27.88725 | 99.90214 | 96.33501 | . | . | . | . | . | . |
| chr15 | 31430174 | 31430653 | 1000 | Distal Intergenic | 15 | 31446380 | 31446817 | 438 | 2 | ENSMUST00 | 16164 | Gm6361 | ENSMUSG0C | false | . | . | . | . | . | . | chr15 | 31428473 | 31434591 | 4.90088 | 3.9058 | 2.96801 |
| chrY | 809039 | 809632 | 1000 | Distal Intergenic | 21 | 818649 | 843684 | 25036 | 1 | ENSMUST00 | -9017 | Uba1y | ENSMUSG0C | false | . | . | . | . | . | . | . | . | . | . | . | . |
| chr13 | 100724300 | 100724747 | 1000 | Intron (ENSMUST00000091299.6 | 13 | 100697027 | 100730942 | 33916 | 2 | ENSMUST00 | 6195 | Cdk7 | ENSMUSG0C | false | . | . | . | . | . | . | chr13 | 100723447 | 100730243 | 5.44102 | 4.21894 | 3.38477 |
| chr3 | 9249695 | 9251896 | 1000 | Promoter (<=1kb) | 3 | 9250602 | 9285333 | 34732 | 1 | ENSMUST00 | 0 | Zbtb10 | ENSMUSG0C | false | chr3 | 9249532 | 9253181 | 39.24862 | 125.4371 | 121.11293 | . | . | . | . | . | . |
| chr13 | 97912370 | 97912771 | 1000 | Intron (ENSMUST00000109426.1 | 13 | 97760130 | 97760588 | 459 | 2 | ENSMUST00 | -151783 | Gm10260 | ENSMUSG0C | false | . | . | . | . | . | . | chr13 | 97909477 | 97929349 | 4.81516 | 3.96306 | 2.86901 |
| chr13 | 73971030 | 73971583 | 1000 | Intron (ENSMUST00000091493.4 | 13 | 73963851 | 73992989 | 29139 | 1 | ENSMUST00 | 7180 | Zdhhc11 | ENSMUSG0C | false | . | . | . | . | . | . | chr13 | 73970427 | 73971732 | 4.89859 | 4.02044 | 2.96132 |
| chr13 | 30659782 | 30660385 | 1000 | Promoter (<=1kb) | 13 | 30659999 | 30711231 | 51233 | 1 | ENSMUST00 | 0 | Dusp22 | ENSMUSG0C | false | chr13 | 30659404 | 30660874 | 31.01414 | 97.6227 | 94.10706 | . | . | . | . | . | . |
| chr16 | 15195295 | 15195736 | 1000 | Distal Intergenic | 16 | 15216859 | 15217328 | 470 | 2 | ENSMUST00 | 21592 | Gm7731 | ENSMUSG0C | false | . | . | . | . | . | . | chr16 | 15195049 | 15195982 | 5.65738 | 4.41928 | 3.54094 |
| chr16 | 15252939 | 15253346 | 1000 | Distal Intergenic | 16 | 15216859 | 15217328 | 470 | 2 | ENSMUST00 | -35612 | Gm7731 | ENSMUSG0C | false | . | . | . | . | . | . | chr16 | 15251497 | 15254440 | 5.11316 | 4.13125 | 3.10432 |
| chr18 | 53993739 | 53994492 | 1000 | Distal Intergenic | 18 | 53984321 | 53985328 | 1008 | 1 | ENSMUST00 | 9419 | Gm5507 | ENSMUSG0C | false | . | . | . | . | . | . | chr18 | 53992848 | 53996380 | 3.84914 | 3.47533 | 2.06311 |
| chr18 | 54118953 | 54120131 | 1000 | Distal Intergenic | 18 | 53984321 | 53985328 | 1008 | 1 | ENSMUST00 | 134633 | Gm5507 | ENSMUSG0C | false | . | . | . | . | . | . | chr18 | 54117000 | 54127216 | 4.98589 | 4.05256 | 3.0024 |

| | | | | | | | | | | | | | | | | | | | | | | | | | | |
|---|---|---|---|---|---|---|---|---|---|---|---|---|---|---|---|---|---|---|---|---|---|---|---|---|---|---|
| chr15 | 81836373 | 81836909 | 1000 | Distal Intergenic | 15 | 81843343 | 81843699 | 357 | 2 | ENSMUST00 | 6790 | Gm8444 | ENSMUSG0( | false | chr15 | 81836244 | 81836833 | 8.8423 | 19.93079 | 17.59637 | chr15 | 81835467 | 81837874 | 5.11459 | 4.10976 | 3.1026 |
| chr2 | 49450952 | 49452249 | 1000 | Promoter (<=1kb) | 2 | 49451486 | 49551948 | 100463 | 1 | ENSMUST00 | 0 | Epc2 | ENSMUSG0( | false | chr2 | 49450752 | 49453348 | 29.58769 | 108.11006 | 104.32592 | . | . | . | . | . | . |
| chr10 | 115361941 | 115362687 | 1000 | Promoter (<=1kb) | 10 | 115340739 | 115362262 | 21524 | 2 | ENSMUST00 | 0 | Tmem19 | ENSMUSG0( | false | chr10 | 115361706 | 115362980 | 16.8969 | 46.84153 | 44.16631 | chr10 | 115360380 | 115370857 | 5.25252 | 4.21307 | 3.20827 |
| chr11 | 117482282 | 117483470 | 1000 | Promoter (<=1kb) | 11 | 117484368 | 117489014 | 4647 | 1 | ENSMUST00 | -898 | Gm11733 | ENSMUSG0( | false | chr11 | 117482135 | 117483059 | 13.34131 | 30.05484 | 27.59022 | chr11 | 117480350 | 117485365 | 5.8868 | 4.46037 | 3.74474 |
| chr10 | 69812762 | 69813240 | 1000 | Intron (ENSMUST00000182194.2 | 10 | 69787456 | 69850148 | 62693 | 1 | ENSMUST00 | 25307 | Ank3 | ENSMUSG0( | false | . | . | . | . | . | . | chr10 | 69810964 | 69814715 | 3.99588 | 3.61973 | 2.19058 |
| chr10 | 37138788 | 37139134 | 1000 | Promoter (<=1kb) | 10 | 37133375 | 37138920 | 5546 | 2 | ENSMUST00 | 0 | Marcks | ENSMUSG0( | false | chr10 | 37138147 | 37141657 | 19.27906 | 51.80829 | 49.07 | chr10 | 37136051 | 37141791 | 4.47232 | 3.83588 | 2.60562 |
| chr10 | 37135992 | 37137358 | 1000 | Promoter (1-2kb) | 10 | 37133375 | 37138920 | 5546 | 2 | ENSMUST00 | 1562 | Marcks | ENSMUSG0( | false | chr10 | 37135907 | 37137976 | 16.70194 | 60.58573 | 57.73199 | chr10 | 37136051 | 37141791 | 4.47232 | 3.83588 | 2.60562 |
| chr17 | 4907782 | 4908578 | 1000 | Distal Intergenic | 17 | 4994332 | 5097733 | 103402 | 1 | ENSMUST00 | -85754 | Arid1b | ENSMUSG0( | false | chr17 | 4907803 | 4909035 | 19.43846 | 52.88828 | 50.13554 | . | . | . | . | . | . |
| chr17 | 4995322 | 4996865 | 1000 | Promoter (<=1kb) | 17 | 4996421 | 5347656 | 351236 | 1 | ENSMUST00 | 0 | Arid1b | ENSMUSG0( | false | chr17 | 4993664 | 4997887 | 36.69013 | 115.69205 | 111.68276 | . | . | . | . | . | . |
| chr11 | 88717775 | 88718944 | 1000 | Promoter (<=1kb) | 11 | 88339382 | 88718229 | 378848 | 2 | ENSMUST00 | 0 | Msi2 | ENSMUSG0( | false | chr11 | 88715830 | 88719254 | 33.54585 | 101.01101 | 97.41918 | chr11 | 88718897 | 88719269 | 4.42611 | 3.77716 | 2.54237 |
| chr11 | 88620000 | 88620844 | 1000 | Intron (ENSMUST00000107909.2 | 11 | 88365515 | 88590147 | 224633 | 2 | ENSMUST00 | -29854 | Msi2 | ENSMUSG0( | false | . | . | . | . | . | . | chr11 | 88603656 | 88625501 | 5.32676 | 4.10238 | 3.30904 |
| chr11 | 76201727 | 76202378 | 1000 | Promoter (<=1kb) | 11 | 76202015 | 76209416 | 7402 | 1 | ENSMUST00 | 0 | Fam57a | ENSMUSG0( | false | chr11 | 76201511 | 76202751 | 27.34946 | 104.78247 | 101.09169 | chr11 | 76201329 | 76203072 | 2.96184 | 2.96238 | 1.39188 |
| chr19 | 8989871 | 8990615 | 1000 | Promoter (<=1kb) | 19 | 8989284 | 9076919 | 87636 | 1 | ENSMUST00 | 588 | Ahnak | ENSMUSG0( | false | chr19 | 8988723 | 8990788 | 33.41831 | 117.26761 | 113.21259 | chr19 | 8990337 | 9001009 | 6.29078 | 4.5762 | 4.06914 |
| chr8 | 70609379 | 70611412 | 1000 | Promoter (1-2kb) | 8 | 70597491 | 70608314 | 10824 | 2 | ENSMUST00 | -1066 | Ssbp4 | ENSMUSG0( | false | chr8 | 70610927 | 70612123 | 16.91761 | 46.11281 | 43.44656 | chr8 | 70604127 | 70612475 | 6.23684 | 4.68532 | 4.01553 |
| chr8 | 44937884 | 44938722 | 1000 | Promoter (2-3kb) | 8 | 44935447 | 45052257 | 116811 | 1 | ENSMUST00 | 2438 | Fat1 | ENSMUSG0( | true | chr8 | 44936734 | 44939539 | 30.08562 | 102.9821 | 99.33765 | . | . | . | . | . | . |
| chr8 | 44959456 | 44960137 | 1000 | Intron (ENSMUST00000191428.1 | 8 | 44950208 | 45052257 | 102050 | 1 | ENSMUST00 | 9249 | Fat1 | ENSMUSG0( | true | . | . | . | . | . | . | chr8 | 44953863 | 44987890 | 5.49999 | 4.27545 | 3.43184 |
| chr8 | 44960524 | 44960861 | 1000 | Intron (ENSMUST00000191428.1 | 8 | 44950208 | 45052257 | 102050 | 1 | ENSMUST00 | 10317 | Fat1 | ENSMUSG0( | true | . | . | . | . | . | . | chr8 | 44953863 | 44987890 | 5.49999 | 4.27545 | 3.43184 |
| chr1 | 162218726 | 162219141 | 1000 | Promoter (<=1kb) | 1 | 162217814 | 162217923 | 110 | 1 | ENSMUST00 | 913 | Mir199a-2 | ENSMUSG0( | false | . | . | . | . | . | . | chr1 | 162216544 | 162219357 | 4.37103 | 3.64578 | 2.51702 |
| chr10 | 58341752 | 58342103 | 1000 | Intron (ENSMUST00000020078.8 | 10 | 58349498 | 58349604 | 107 | 2 | ENSMUST00 | 7501 | Gm23058 | ENSMUSG0( | false | . | . | . | . | . | . | chr10 | 58340746 | 58343999 | 6.03255 | 4.57621 | 3.84704 |
| chr11 | 97883504 | 97884534 | 1000 | Promoter (<=1kb) | 11 | 97853826 | 97884154 | 30329 | 2 | ENSMUST00 | 0 | Fbxo47 | ENSMUSG0( | false | chr11 | 97884236 | 97884966 | 15.84839 | 38.35189 | 35.78466 | . | . | . | . | . | . |
| chr11 | 97350156 | 97350456 | 1000 | Promoter (1-2kb) | 11 | 97332109 | 97352016 | 19908 | 2 | ENSMUST00 | 1560 | Gpr179 | ENSMUSG0( | false | . | . | . | . | . | . | chr11 | 97350260 | 97351667 | 4.10304 | 3.41554 | 2.30464 |
| chr7 | 144941452 | 144941941 | 1000 | Promoter (1-2kb) | 7 | 144929931 | 144939869 | 9939 | 2 | ENSMUST00 | -1584 | Ccnd1 | ENSMUSG0( | false | chr7 | 144941140 | 144943434 | 27.88725 | 99.90214 | 96.33501 | chr7 | 144938154 | 144943823 | 5.32689 | 4.2503 | 3.27022 |
| chr3 | 96246382 | 96246772 | 1000 | Promoter (<=1kb) | 3 | 96246685 | 96248509 | 1825 | 1 | ENSMUST00 | 0 | Gm20634 | ENSMUSG0( | false | chr3 | 96246039 | 96246579 | 28.61382 | 119.33878 | 115.2169 | . | . | . | . | . | . |
| chr11 | 63915681 | 63916147 | 1000 | Intron (ENSMUST00000094103.3 | 11 | 63885792 | 63922290 | 36499 | 2 | ENSMUST00 | 6143 | Hs3st3b1 | ENSMUSG0( | false | . | . | . | . | . | . | chr11 | 63904532 | 63920020 | 5.40463 | 4.1524 | 3.35604 |
| chr3 | 58636503 | 58637224 | 1000 | Promoter (<=1kb) | 3 | 58616300 | 58637207 | 20908 | 2 | ENSMUST00 | 0 | Erich6 | ENSMUSG0( | false | chr3 | 58636252 | 58637357 | 17.00269 | 41.02908 | 38.42754 | chr3 | 58636129 | 58638685 | 3.21468 | 3.0814 | 1.58029 |
| chr7 | 73372361 | 73372698 | 1000 | Promoter (2-3kb) | 7 | 73375520 | 73419899 | 44380 | 1 | ENSMUST00 | -2822 | Rgma | ENSMUSG0( | false | chr7 | 73372401 | 73372945 | 10.91731 | 22.4059 | 20.03854 | . | . | . | . | . | . |
| chr7 | 73374767 | 73375770 | 1000 | Promoter (<=1kb) | 7 | 73375520 | 73419899 | 44380 | 1 | ENSMUST00 | 0 | Rgma | ENSMUSG0( | false | chr7 | 73374798 | 73376759 | 25.15761 | 80.47005 | 77.31023 | chr7 | 73372782 | 73376627 | 5.46181 | 4.19274 | 3.44118 |
| chr2 | 160645961 | 160646928 | 1000 | Promoter (<=1kb) | 2 | 160645980 | 160650107 | 4128 | 1 | ENSMUST00 | 0 | Top1 | ENSMUSG0( | false | chr2 | 160645955 | 160647510 | 32.53129 | 125.07367 | 120.7602 | . | . | . | . | . | . |
| chr1 | 157143251 | 157143759 | 1000 | Intron (ENSMUST00000078308.7 | 1 | 157157651 | 157163933 | 6283 | 2 | ENSMUST00 | 20174 | Rasal2 | ENSMUSG0( | false | . | . | . | . | . | . | chr1 | 157139568 | 157146134 | 4.7708 | 3.94145 | 2.83546 |
| chr5 | 107437752 | 107438314 | 1000 | Promoter (<=1kb) | 5 | 107437997 | 107491628 | 53632 | 1 | ENSMUST00 | 0 | Btbd8 | ENSMUSG0( | false | chr5 | 107437510 | 107438662 | 39.39634 | 133.30592 | 128.68834 | . | . | . | . | . | . |
| chr5 | 105415329 | 105416077 | 1000 | Promoter (<=1kb) | 5 | 105415775 | 105486189 | 70415 | 1 | ENSMUST00 | 0 | Lrrc8b | ENSMUSG0( | false | chr5 | 105414998 | 105416445 | 34.54477 | 122.48211 | 118.25522 | chr5 | 105414802 | 105417367 | 3.75656 | 3.39015 | 1.99796 |
| chr1 | 133468897 | 133469559 | 1000 | Distal Intergenic | 1 | 133382303 | 133424377 | 42075 | 2 | ENSMUST00 | -44521 | Sox13 | ENSMUSG0( | false | . | . | . | . | . | . | chr1 | 133468626 | 133471492 | 4.0268 | 3.49346 | 2.23237 |
| chr1 | 133423751 | 133424826 | 1000 | Promoter (<=1kb) | 1 | 133382303 | 133424377 | 42075 | 2 | ENSMUST00 | 0 | Sox13 | ENSMUSG0( | false | chr1 | 133421916 | 133424490 | 24.0356 | 94.23969 | 90.80066 | . | . | . | . | . | . |
| chr8 | 72303638 | 72304034 | 1000 | Promoter (1-2kb) | 8 | 72304118 | 72305276 | 1159 | 2 | ENSMUST00 | 1242 | Gm10282 | ENSMUSG0( | false | . | . | . | . | . | . | chr8 | 72302951 | 72304772 | 3.41569 | 3.20808 | 1.75616 |
| chr4 | 131921184 | 131922061 | 1000 | Promoter (<=1kb) | 4 | 131921771 | 131923140 | 1370 | 1 | ENSMUST00 | 0 | Tmem200b | ENSMUSG0( | false | chr4 | 131920283 | 131922489 | 21.79372 | 61.12524 | 58.26245 | chr4 | 131920243 | 131922772 | 5.42572 | 3.9801 | 3.38953 |
| chr7 | 16940343 | 16940678 | 1000 | Distal Intergenic | 7 | 16944682 | 16948828 | 4147 | 1 | ENSMUST00 | -4004 | Pnmal2 | ENSMUSG0( | false | . | . | . | . | . | . | chr7 | 16937974 | 16950184 | 4.88741 | 4.01233 | 2.92708 |
| chr4 | 119462837 | 119463202 | 1000 | Intron (ENSMUST00000129150.1 | 4 | 119444720 | 119467444 | 22725 | 1 | ENSMUST00 | 18118 | Zmynd12 | ENSMUSG0( | false | . | . | . | . | . | . | . | . | . | . | . | . |
| chr4 | 114405740 | 114406469 | 1000 | Promoter (<=1kb) | 4 | 114406724 | 114615098 | 208375 | 1 | ENSMUST00 | -255 | Trabd2b | ENSMUSG0( | false | chr4 | 114405755 | 114407430 | 42.51921 | 140.97417 | 136.05247 | chr4 | 114405103 | 114411487 | 5.01818 | 3.99702 | 3.06193 |
| chr4 | 114504583 | 114504955 | 1000 | Intron (ENSMUST00000094894.3 | 4 | 114406724 | 114615098 | 208375 | 1 | ENSMUST00 | 97860 | Trabd2b | ENSMUSG0( | false | . | . | . | . | . | . | chr4 | 114495475 | 114506146 | 5.17812 | 4.15053 | 3.15948 |
| chr1 | 43067413 | 43067876 | 1000 | Exon (ENSMUST00000095014.2/ | 1 | 43054708 | 43060933 | 6226 | 2 | ENSMUST00 | -6481 | Tgfbrap1 | ENSMUSG0( | false | . | . | . | . | . | . | chr1 | 43065837 | 43068517 | 6.89831 | 5.00604 | 4.56403 |
| chr4 | 43726893 | 43727371 | 1000 | Promoter (<=1kb) | 4 | 43727188 | 43728639 | 1452 | 1 | ENSMUST00 | 0 | Hrct1 | ENSMUSG0( | false | . | . | . | . | . | . | chr4 | 43725362 | 43730847 | 5.67024 | 4.30531 | 3.60642 |
| chr8 | 79028724 | 79031083 | 1000 | Promoter (<=1kb) | 8 | 79028587 | 79190856 | 162270 | 1 | ENSMUST00 | 138 | Zfp827 | ENSMUSG0( | false | chr8 | 79028069 | 79031248 | 27.54716 | 98.2761 | 94.74957 | chr8 | 79030999 | 79031964 | 4.09899 | 3.6431 | 2.26094 |
| chr8 | 79111967 | 79112864 | 1000 | Intron (ENSMUST00000098614.3 | 8 | 79118327 | 79193766 | 75440 | 1 | ENSMUST00 | -5463 | Zfp827 | ENSMUSG0( | false | . | . | . | . | . | . | chr8 | 79111391 | 79116090 | 5.16683 | 3.99061 | 3.14481 |
| chr8 | 70698219 | 70699367 | 1000 | Promoter (<=1kb) | 8 | 70697739 | 70700616 | 2878 | 1 | ENSMUST00 | 481 | Jund | ENSMUSG0( | false | chr8 | 70697986 | 70698758 | 15.08796 | 37.05233 | 34.49837 | chr8 | 70693345 | 70698398 | 5.3592 | 4.1184 | 3.30975 |
| chr6 | 120665892 | 120666913 | 1000 | Promoter (<=1kb) | 6 | 120666369 | 120724920 | 58552 | 1 | ENSMUST00 | 0 | Cecr2 | ENSMUSG0( | false | chr6 | 120665084 | 120668136 | 38.87692 | 132.39133 | 127.80991 | . | . | . | . | . | . |
| chr17 | 23564002 | 23564375 | 1000 | Promoter (<=1kb) | 17 | 23556769 | 23564226 | 7458 | 2 | ENSMUST00 | 0 | Zfp213 | ENSMUSG0( | false | chr17 | 23563495 | 23564961 | 40.29262 | 133.77719 | 129.14264 | chr17 | 23562779 | 23565245 | 5.41453 | 4.19981 | 3.38218 |
| chr10 | 59547354 | 59547870 | 1000 | Intron (ENSMUST00000020312.7 | 10 | 59616063 | 59616398 | 336 | 1 | ENSMUST00 | -68193 | Gm10322 | ENSMUSG0( | false | . | . | . | . | . | . | chr10 | 59547090 | 59550035 | 4.00881 | 3.59668 | 2.23802 |
| chr10 | 59588196 | 59588536 | 1000 | Intron (ENSMUST00000020312.7 | 10 | 59616063 | 59616398 | 336 | 1 | ENSMUST00 | -27527 | Gm10322 | ENSMUSG0( | false | . | . | . | . | . | . | chr10 | 59584028 | 59593191 | 4.02155 | 3.528 | 2.20777 |
| chr10 | 59537408 | 59538014 | 1000 | Intron (ENSMUST00000020312.7 | 10 | 59616063 | 59616398 | 336 | 1 | ENSMUST00 | -78049 | Gm10322 | ENSMUSG0( | false | . | . | . | . | . | . | chr10 | 59536235 | 59540855 | 6.98872 | 4.97826 | 4.65808 |
| chr10 | 41934512 | 41935554 | 1000 | Intron (ENSMUST00000160262.3 | 10 | 41916051 | 41976122 | 60072 | 2 | ENSMUST00 | 40568 | Armc2 | ENSMUSG0( | false | chr10 | 41934492 | 41935362 | 5.43663 | 10.5674 | 8.37697 | chr10 | 41932117 | 41937048 | 4.69283 | 3.92408 | 2.78135 |
| chr10 | 22696269 | 22696604 | 1000 | Intron (ENSMUST00000042261.4 | 10 | 22705523 | 22706385 | 863 | 2 | ENSMUST00 | 9781 | Tbpl1 | ENSMUSG0( | false | . | . | . | . | . | . | chr10 | 22695577 | 22702618 | 5.6952 | 4.24719 | 3.6107 |
| chr10 | 22710307 | 22710732 | 1000 | Intron (ENSMUST00000127698.2 | 10 | 22705523 | 22706385 | 863 | 2 | ENSMUST00 | -3923 | Tbpl1 | ENSMUSG0( | false | . | . | . | . | . | . | chr10 | 22710291 | 22711125 | 4.67629 | 3.70268 | 2.77869 |
| chr10 | 19987797 | 19988182 | 1000 | Intron (ENSMUST00000095806.4 | 10 | 19951055 | 20023699 | 72645 | 1 | ENSMUST00 | 36743 | Map3k5 | ENSMUSG0( | true | . | . | . | . | . | . | chr10 | 19985983 | 19992169 | 3.73319 | 3.3726 | 1.98609 |
| chr10 | 20040114 | 20040467 | 1000 | Intron (ENSMUST00000095806.4 | 10 | 20099525 | 20105437 | 5913 | 1 | ENSMUST00 | -59058 | Map3k5 | ENSMUSG0( | true | . | . | . | . | . | . | chr10 | 20036920 | 20042145 | 4.52371 | 3.81961 | 2.62584 |
| chr10 | 20031946 | 20032341 | 1000 | Intron (ENSMUST00000095806.4 | 10 | 20099525 | 20105437 | 5913 | 1 | ENSMUST00 | -67184 | Map3k5 | ENSMUSG0( | true | . | . | . | . | . | . | chr10 | 20031592 | 20033293 | 5.66136 | 4.16022 | 3.57934 |
| chr12 | 17733253 | 17733768 | 1000 | Intron (ENSMUST00000071858.3 | 12 | 17690814 | 17791926 | 101113 | 1 | ENSMUST00 | 42440 | Hpcal1 | ENSMUSG0( | false | . | . | . | . | . | . | chr12 | 17731846 | 17735069 | 5.54101 | 4.2777 | 3.50005 |
| chr6 | 64238776 | 64239101 | 1000 | Intron (ENSMUST00000095852.3 | 6 | 64076792 | 64094748 | 17957 | 1 | ENSMUST00 | 161985 | Grid2 | ENSMUSG0( | false | . | . | . | . | . | . | . | . | . | . | . | . |
| chr12 | 3572084 | 3572917 | 1000 | Promoter (<=1kb) | 12 | 3572381 | 3781796 | 209416 | 1 | ENSMUST00 | 0 | Dtnb | ENSMUSG0( | false | chr12 | 3571809 | 3573315 | 41.59238 | 139.36057 | 134.50562 | chr12 | 3571974 | 3573779 | 3.14417 | 3.08594 | 1.48793 |
| chr10 | 14481000 | 14481384 | 1000 | Intron (ENSMUST00000041168.4 | 10 | 14523078 | 14523545 | 468 | 1 | ENSMUST00 | -41694 | Gm10335 | ENSMUSG0( | false | . | . | . | . | . | . | chr10 | 14480601 | 14481567 | 4.22322 | 3.61654 | 2.39472 |
| chr19 | 23959796 | 23960593 | 1000 | Distal Intergenic | 19 | 23972751 | 24031019 | 58269 | 2 | ENSMUST00 | 70426 | Fam189a2 | ENSMUSG0( | false | . | . | . | . | . | . | chr19 | 23958921 | 23962405 | 6.62285 | 4.82299 | 4.33144 |
| chr19 | 23925478 | 23925927 | 1000 | Intron (ENSMUST00000025830.7 | 19 | 23972751 | 24031019 | 58269 | 2 | ENSMUST00 | 105092 | Fam189a2 | ENSMUSG0( | false | . | . | . | . | . | . | . | . | . | . | . | . |
| chr19 | 8966827 | 8967315 | 1000 | Promoter (<=1kb) | 19 | 8967041 | 8978479 | 11439 | 1 | ENSMUST00 | 0 | Eef1g | ENSMUSG0( | false | chr19 | 8965979 | 8966930 | 18.95815 | 63.55374 | 60.65727 | . | . | . | . | . | . |
| chr19 | 8871342 | 8871868 | 1000 | Promoter (<=1kb) | 19 | 8871559 | 8875655 | 4097 | 1 | ENSMUST00 | 0 | Ubxn1 | ENSMUSG0( | false | chr19 | 8870791 | 8872513 | 31.62822 | 118.12274 | 114.03909 | chr19 | 8870632 | 8874980 | 4.20911 | 3.69101 | 2.37602 |
| chr19 | 8819715 | 8820725 | 1000 | Promoter (<=1kb) | 19 | 8819401 | 8834142 | 14742 | 1 | ENSMUST00 | 315 | Hnrnpul2 | ENSMUSG0( | false | chr19 | 8819501 | 8822631 | 40.17707 | 149.16289 | 143.90889 | . | . | . | . | . | . |
| chr15 | 78416899 | 78417515 | 1000 | Distal Intergenic | 15 | 78407104 | 78414002 | 6899 | 1 | ENSMUST00 | 9796 | Mpst | ENSMUSG0( | false | . | . | . | . | . | . | chr15 | 78415224 | 78417690 | 3.81229 | 3.28501 | 2.07468 |
| chr15 | 57695344 | 57696278 | 1000 | Promoter (<=1kb) | 15 | 57695729 | 57821110 | 125382 | 1 | ENSMUST00 | 0 | Zhx2 | ENSMUSG0( | true | chr15 | 57692968 | 57696625 | 40.46864 | 131.89105 | 127.3278 | chr15 | 57695806 | 57714739 | 5.09262 | 4.12618 | 3.0824 |
| chr15 | 57659663 | 57660381 | 1000 | Distal Intergenic | 15 | 57694665 | 57839832 | 145168 | 1 | ENSMUST00 | -34284 | Zhx2 | ENSMUSG0( | true | . | . | . | . | . | . | chr15 | 57659154 | 57662270 | 5.11923 | 4.14962 | 3.10605 |
| chr15 | 57693873 | 57694610 | 1000 | Promoter (<=1kb) | 15 | 57694665 | 57839832 | 145168 | 1 | ENSMUST00 | -55 | Zhx2 | ENSMUSG0( | true | chr15 | 57692968 | 57696625 | 40.46864 | 131.89105 | 127.3278 | . | . | . | . | . | . |
| chr18 | 44686639 | 44687052 | 1000 | Intron (ENSMUST00000089874.5 | 18 | 44425061 | 44662266 | 237206 | 2 | ENSMUST00 | -24374 | Mcc | ENSMUSG0( | true | . | . | . | . | . | . | chr18 | 44685683 | 44687601 | 4.71787 | 3.95657 | 2.82787 |
| chr18 | 43809659 | 43809943 | 1000 | Distal Intergenic | 18 | 43777196 | 43792878 | 15683 | 2 | ENSMUST00 | -16782 | Gm94 | ENSMUSG0( | false | . | . | . | . | . | . | chr18 | 43809007 | 43810688 | 3.47052 | 3.30399 | 1.73543 |
| chr11 | 6415891 | 6416904 | 1000 | Promoter (<=1kb) | 11 | 6415876 | 6419812 | 3937 | 1 | ENSMUST00 | 16 | Ppia | ENSMUSG0( | false | chr11 | 6415138 | 6417437 | 36.39639 | 123.67226 | 119.40948 | chr11 | 6416820 | 6419020 | 4.13032 | 3.51607 | 2.32578 |
| chr17 | 24267056 | 24267555 | 1000 | Exon (ENSMUST00000121226.2/ | 17 | 24223232 | 24251409 | 28178 | 2 | ENSMUST00 | -15648 | Ccnf | ENSMUSG0( | false | . | . | . | . | . | . | chr17 | 24266534 | 24269452 | 3.66235 | 3.43735 | 1.89171 |
| chr16 | 18686898 | 18687342 | 1000 | Distal Intergenic | 16 | 18621811 | 18629938 | 8128 | 2 | ENSMUST00 | -56961 | Sep-05 | ENSMUSG0( | false | . | . | . | . | . | . | chr16 | 18686053 | 18687913 | 5.89956 | 4.48206 | 3.75077 |
| chr16 | 48304131 | 48304464 | 1000 | Distal Intergenic | 16 | 48310477 | 48319723 | 9247 | 1 | ENSMUST00 | -6013 | Dppa2 | ENSMUSG0( | false | . | . | . | . | . | . | chr16 | 48303053 | 48304568 | 3.58167 | 3.34761 | 1.8427 |
| chr5 | 134676003 | 134676797 | 1000 | Promoter (<=1kb) | 5 | 134676490 | 134680145 | 3656 | 1 | ENSMUST00 | 0 | Gm10369 | ENSMUSG0( | false | chr5 | 134675807 | 134676758 | 9.52248 | 22.23582 | 19.8701 | chr5 | 134675072 | 134677208 | 6.30627 | 4.63071 | 4.08252 |
| chr15 | 53826821 | 53827105 | 1000 | Intron (ENSMUST00000154119.2 | 15 | 53884906 | 53886597 | 1692 | 1 | ENSMUST00 | -57801 | Gm7489 | ENSMUSG0( | false | . | . | . | . | . | . | . | . | . | . | . | . |
| chr15 | 36938417 | 36939064 | 1000 | Distal Intergenic | 15 | 36870501 | 36879816 | 9316 | 2 | ENSMUST00 | -58602 | Gm10384 | ENSMUSG0( | false | chr15 | 36938198 | 36939286 | 22.90469 | 70.93823 | 67.93287 | chr15 | 36935585 | 36938644 | 6.1812 | 4.6712 | 4.00794 |
| chr11 | 78827543 | 78827999 | 1000 | Promoter (<=1kb) | 11 | 78826629 | 78839275 | 12647 | 1 | ENSMUST00 | 915 | Lyrm9 | ENSMUSG0( | false | . | . | . | . | . | . | chr11 | 78823496 | 78836454 | 6.00182 | 4.51146 | 3.83406 |
| chr11 | 78826417 | 78827124 | 1000 | Promoter (<=1kb) | 11 | 78826584 | 78846870 | 20287 | 1 | ENSMUST00 | 0 | Lyrm9 | ENSMUSG0( | false | chr11 | 78826527 | 78827336 | 21.50638 | 60.6253 | 57.77077 | chr11 | 78823496 | 78836454 | 6.00182 | 4.51146 | 3.83406 |
| chr11 | 78829319 | 78829995 | 1000 | Promoter (2-3kb) | 11 | 78826629 | 78839275 | 12647 | 1 | ENSMUST00 | 2691 | Lyrm9 | ENSMUSG0( | false | . | . | . | . | . | . | chr11 | 78823496 | 78836454 | 6.00182 | 4.51146 | 3.83406 |
| chr6 | 144647941 | 144648711 | 1000 | Intron (ENSMUST00000141102.2 | 6 | 144670427 | 144672720 | 2294 | 2 | ENSMUST00 | 24009 | Sox5it | ENSMUSG0( | false | . | . | . | . | . | . | chr6 | 144638785 | 144651275 | 5.59752 | 4.38604 | 3.49804 |
| chr14 | 24823039 | 24823682 | 1000 | Distal Intergenic | 14 | 24909414 | 24913440 | 4027 | 1 | ENSMUST00 | -85732 | Gm10398 | ENSMUSG0( | false | . | . | . | . | . | . | chr14 | 24822563 | 24829369 | 4.4396 | 3.69842 | 2.58436 |
| chr6 | 100109755 | 100110355 | 1000 | Distal Intergenic | 6 | 100228566 | 100287441 | 58876 | 2 | ENSMUST00 | 177086 | Rybp | ENSMUSG0( | false | . | . | . | . | . | . | chr6 | 100109412 | 100111163 | 4.48774 | 3.58181 | 2.62125 |
| chr6 | 100286603 | 100288009 | 1000 | Promoter (<=1kb) | 6 | 100228566 | 100287441 | 58876 | 2 | ENSMUST00 | 0 | Rybp | ENSMUSG0( | false | chr6 | 100286055 | 100287492 | 25.0055 | 70.15867 | 67.16741 | chr6 | 100287772 | 100289184 | 5.57913 | 4.23799 | 3.50814 |
| chr5 | 52363509 | 52364114 | 1000 | Promoter (<=1kb) | 5 | 52363946 | 52371405 | 7460 | 1 | ENSMUST00 | 0 | Sod3 | ENSMUSG0( | false | . | . | . | . | . | . | chr5 | 52363284 | 52365001 | 4.45803 | 3.78203 | 2.60217 |
| chr12 | 83998587 | 83998926 | 1000 | Distal Intergenic | 12 | 84009502 | 84017670 | 8169 | 1 | ENSMUST00 | -10576 | Acot1 | ENSMUSG0( | false | . | . | . | . | . | . | chr12 | 83998034 | 84000817 | 8.05826 | 5.53182 | 5.48453 |
| chr17 | 9407703 | 9408061 | 1000 | Distal Intergenic | 17 | 9422060 | 9422458 | 399 | 1 | ENSMUST00 | -13999 | Gm17728 | ENSMUSG0( | false | . | . | . | . | . | . | chr17 | 9407019 | 9408303 | 3.711 | 3.46758 | 1.94536 |
| chr6 | 72636137 | 72636415 | 1000 | Promoter (<=1kb) | 6 | 72636253 | 72637717 | 1465 | 1 | ENSMUST00 | 0 | Gm15401 | ENSMUSG0( | false | . | . | . | . | . | . | chr6 | 72635812 | 72637074 | 6.34356 | 4.68544 | 4.14654 |
| chr6 | 54393795 | 54394288 | 1000 | Intron (ENSMUST00000145111.2 | 6 | 54424564 | 54429537 | 4974 | 2 | ENSMUST00 | 35249 | 9130019P16I | ENSMUSG0( | false | . | . | . | . | . | . | chr6 | 54391673 | 54403137 | 6.55024 | 4.83886 | 4.27494 |
| chr6 | 54328974 | 54329395 | 1000 | Promoter (1-2kb) | 6 | 54302652 | 54327341 | 24690 | 2 | ENSMUST00 | -1634 | 9130019P16I | ENSMUSG0( | false | . | . | . | . | . | . | chr6 | 54328542 | 54331278 | 6.93202 | 5.0487 | 4.56667 |
| chr6 | 54398474 | 54398865 | 1000 | Intron (ENSMUST00000145111.2 | 6 | 54424564 | 54429537 | 4974 | 2 | ENSMUST00 | 30672 | 9130019P16I | ENSMUSG0( | false | . | . | . | . | . | . | chr6 | 54391673 | 54403137 | 6.55024 | 4.83886 | 4.27494 |
| chr6 | 54351792 | 54352412 | 1000 | Intron (ENSMUST00000145111.2 | 6 | 54302652 | 54327341 | 24690 | 2 | ENSMUST00 | -24452 | 9130019P16I | ENSMUSG0( | false | . | . | . | . | . | . | chr6 | 54350138 | 54356215 | 5.61723 | 4.41849 | 3.50454 |
| chr6 | 54429335 | 54429836 | 1000 | Promoter (<=1kb) | 6 | 54424564 | 54429537 | 4974 | 2 | ENSMUST00 | 0 | 9130019P16I | ENSMUSG0( | false | chr6 | 54429012 | 54432013 | 22.68762 | 81.35053 | 78.1749 | chr6 | 54428707 | 54430076 | 3.49164 | 3.21009 | 1.77445 |
| chr6 | 54327591 | 54328198 | 1000 | Promoter (<=1kb) | 6 | 54302652 | 54327341 | 24690 | 2 | ENSMUST00 | -251 | 9130019P16I | ENSMUSG0( | false | chr6 | 54326642 | 54328336 | 20.06521 | 63.98471 | 61.08233 | . | . | . | . | . | . |
| chr5 | 23432581 | 23433202 | 1000 | Promoter (<=1kb) | 5 | 23422404 | 23433309 | 10906 | 2 | ENSMUST00 | 107 | 5031425E22I | ENSMUSG0( | false | chr5 | 23431644 | 23437822 | 48.25087 | 161.14223 | 155.45619 | . | . | . | . | . | . |
| chr5 | 23413644 | 23413999 | 1000 | Intron (ENSMUST00000185943.1 | 5 | 23422404 | 23433309 | 10906 | 2 | ENSMUST00 | 19310 | 5031425E22I | ENSMUSG0( | false | . | . | . | . | . | . | chr5 | 23407933 | 23417945 | 6.48738 | 4.78205 | 4.22896 |
| chr5 | 20701797 | 20702659 | 1000 | Promoter (<=1kb) | 5 | 20702277 | 20706049 | 3773 | 1 | ENSMUST00 | 0 | Magi2 | ENSMUSG0( | true | chr5 | 20701623 | 20702915 | 15.07427 | 36.33085 | 33.78652 | chr5 | 20701192 | 20702866 | 4.83648 | 3.83177 | 2.92769 |
| chr17 | 46864535 | 46864972 | 1000 | Distal Intergenic | 17 | 46889216 | 46890405 | 1190 | 2 | ENSMUST00 | 25433 | A330017A19 | ENSMUSG0( | false | chr17 | 46864321 | 46864861 | 7.54398 | 13.82373 | 11.57928 | chr17 | 46863713 | 46867626 | 5.95403 | 4.41683 | 3.82696 |
| chr17 | 35417999 | 35418851 | 1000 | Distal Intergenic | 17 | 35424850 | 35430055 | 5206 | 1 | ENSMUST00 | -5999 | H2-Q6 | ENSMUSG0( | true | . | . | . | . | . | . | chr17 | 35412151 | 35429685 | 5.1178 | 3.62349 | 3.13713 |
| chr17 | 27389116 | 27389532 | 1000 | Distal Intergenic | 17 | 27342460 | 27344825 | 2366 | 1 | ENSMUST00 | 46657 | Gm10505 | ENSMUSG0( | false | . | . | . | . | . | . | chr17 | 27386011 | 27390573 | 5.14515 | 4.03563 | 3.14987 |
| chr17 | 7945406 | 7946076 | 1000 | Promoter (<=1kb) | 17 | 7945653 | 7979824 | 34172 | 1 | ENSMUST00 | 0 | Rsph3a | ENSMUSG0( | false | chr17 | 7945193 | 7946321 | 22.90469 | 70.93823 | 67.93287 | chr17 | 7944889 | 7945456 | 4.39235 | 3.79359 | 2.537 |
| chr1 | 184851198 | 184851639 | 1000 | Distal Intergenic | 1 | 184822375 | 184846451 | 24077 | 2 | ENSMUST00 | -4748 | 37316 | ENSMUSG0( | false | . | . | . | . | . | . | chr1 | 184846562 | 184855069 | 5.5375 | 4.29731 | 3.4563 |
| chr18 | 53861763 | 53863120 | 1000 | Promoter (<=1kb) | 18 | 53862113 | 53955684 | 93572 | 1 | ENSMUST00 | 0 | Csnk1g3 | ENSMUSG0( | false | chr18 | 53861654 | 53863702 | 28.22848 | 94.63612 | 91.19299 | . | . | . | . | . | . |
| chr18 | 53917029 | 53917418 | 1000 | Exon (ENSMUST00000069597.6/ | 18 | 53862113 | 53955684 | 93572 | 1 | ENSMUST00 | 54917 | Csnk1g3 | ENSMUSG0( | false | . | . | . | . | . | . | chr18 | 53913134 | 53917201 | 5.08097 | 4.12194 | 3.07956 |
| chr18 | 33470996 | 33471449 | 1000 | Intron (ENSMUST00000097634.3 | 18 | 33464163 | 33472448 | 8286 | 1 | ENSMUST00 | 6834 | Gm10549 | ENSMUSG0( | false | . | . | . | . | . | . | chr18 | 33468172 | 33471896 | 4.0663 | 3.65807 | 2.24112 |
| chr1 | 79761331 | 79761894 | 1000 | Promoter (<=1kb) | 1 | 79702262 | 79761769 | 59508 | 2 | ENSMUST00 | 0 | Wdfy1 | ENSMUSG0( | false | chr1 | 79760642 | 79762273 | 23.17719 | 89.67316 | 86.33752 | chr1 | 79751745 | 79763284 | 5.0777 | 4.06964 | 3.07453 |
| chr1 | 60180217 | 60181255 | 1000 | Promoter (<=1kb) | 1 | 60180599 | 60338328 | 157730 | 1 | ENSMUST00 | 0 | Nbeal1 | ENSMUSG0( | true | chr1 | 60179757 | 60180364 | 23.94845 | 71.00658 | 67.99948 | . | . | . | . | . | . |
| chr4 | 155249669 | 155250334 | 1000 | Promoter (<=1kb) | 4 | 155249802 | 155256419 | 6618 | 1 | ENSMUST00 | 0 | 2610002J02F | ENSMUSG0( | false | chr4 | 155249526 | 155251162 | 24.86209 | 73.50948 | 70.465 | chr4 | 155248891 | 155252103 | 5.08587 | 4.11596 | 3.05981 |
| chr1 | 24678372 | 24678974 | 1000 | Promoter (<=1kb) | 1 | 24678630 | 24765448 | 86819 | 1 | ENSMUST00 | 0 | Lmbrd1 | ENSMUSG0( | false | chr1 | 24677760 | 24679595 | 43.58744 | 151.86528 | 146.50356 | . | . | . | . | . | . |
| chr4 | 124575287 | 124576359 | 1000 | Intron (ENSMUST00000184062.1 | 4 | 124569190 | 124574645 | 5456 | 1 | ENSMUST00 | 6098 | 4933407E24I | ENSMUSG0( | false | chr4 | 124575138 | 124576197 | 15.16456 | 35.10921 | 32.57929 | chr4 | 124568013 | 124576752 | 4.65263 | 3.69297 | 2.75743 |
| chr7 | 142229512 | 142229956 | 1000 | Promoter (<=1kb) | 7 | 142228795 | 142229971 | 1177 | 2 | ENSMUST00 | 15 | Krtap5-5 | ENSMUSG0( | false | . | . | . | . | . | . | chr7 | 142228878 | 142230918 | 3.68476 | 3.37935 | 1.9741 |
| chr4 | 99773096 | 99773390 | 1000 | Intron (ENSMUST00000146258.1 | 4 | 99719152 | 99739473 | 20322 | 1 | ENSMUST00 | 53945 | Alg6 | ENSMUSG0( | false | . | . | . | . | . | . | chr4 | 99772461 | 99775124 | 3.84788 | 3.41573 | 2.09538 |
| chr7 | 134944781 | 134945064 | 1000 | Intron (ENSMUST00000084488.4 | 7 | 134881926 | 134938430 | 56505 | 2 | ENSMUST00 | -6352 | Fam196a | ENSMUSG0( | false | . | . | . | . | . | . | chr7 | 134943649 | 134945007 | 7.07136 | 4.94106 | 4.75321 |
| chr4 | 41773226 | 41773902 | 1000 | Promoter (<=1kb) | 4 | 41769467 | 41773537 | 4071 | 2 | ENSMUST00 | 0 | Ccl27a | ENSMUSG0( | false | chr4 | 41773381 | 41773915 | 13.20699 | 29.82797 | 27.36516 | . | . | . | . | . | . |
| chr7 | 102249750 | 102250936 | 1000 | Promoter (<=1kb) | 7 | 102239123 | 102250088 | 10966 | 2 | ENSMUST00 | 0 | Rhog | ENSMUSG0( | false | chr7 | 102249024 | 102251102 | 40.75532 | 136.78966 | 132.04059 | . | . | . | . | . | . |
| chr8 | 126497603 | 126497917 | 1000 | Distal Intergenic | 8 | 126476248 | 126476412 | 165 | 2 | ENSMUST00 | -21192 | Gm6091 | ENSMUSG0( | false | chr8 | 126497748 | 126498330 | 8.26071 | 16.87976 | 14.58833 | chr8 | 126497216 | 126499328 | 7.04565 | 4.97782 | 4.68136 |
| chr8 | 119416541 | 119417439 | 1000 | Distal Intergenic | 8 | 119434124 | 119443077 | 8954 | 1 | ENSMUST00 | -16685 | Osgin1 | ENSMUSG0( | false | . | . | . | . | . | . | chr8 | 119415900 | 119416559 | 3.57196 | 3.21228 | 1.87331 |
| chr7 | 38227765 | 38228347 | 1000 | Promoter (<=1kb) | 7 | 38220654 | 38227994 | 7341 | 2 | ENSMUST00 | 0 | Plekhf1 | ENSMUSG0( | false | chr7 | 38227135 | 38228402 | 28.666 | 87.63837 | 84.3392 | chr7 | 38223264 | 38228403 | 4.20626 | 3.5591 | 2.38787 |
| chr8 | 90907829 | 90909475 | 1000 | Promoter (<=1kb) | 8 | 90907939 | 90908415 | 477 | 2 | ENSMUST00 | 0 | Gm6658 | ENSMUSG0( | false | chr8 | 90907503 | 90909657 | 36.76141 | 116.86796 | 112.83147 | . | . | . | . | . | . |
| chr8 | 90869133 | 90869778 | 1000 | Intron (ENSMUST00000048665.6 | 8 | 90907939 | 90908415 | 477 | 2 | ENSMUST00 | 38637 | Gm6658 | ENSMUSG0( | false | . | . | . | . | . | . | chr8 | 90867174 | 90871561 | 5.45736 | 4.12147 | 3.3984 |
| chr7 | 30121657 | 30122192 | 1000 | Promoter (<=1kb) | 7 | 30121942 | 30134665 | 12724 | 1 | ENSMUST00 | 0 | Zfp382 | ENSMUSG0( | false | chr7 | 30121250 | 30122569 | 29.35304 | 95.67908 | 92.21078 | . | . | . | . | . | . |
| chr3 | 128145224 | 128145602 | 1000 | Distal Intergenic | 3 | 128038694 | 128040661 | 1968 | 1 | ENSMUST00 | 106531 | Gm10650 | ENSMUSG0( | false | . | . | . | . | . | . | chr3 | 128144690 | 128146097 | 5.51962 | 4.27521 | 3.44708 |
| chr3 | 121663176 | 121663664 | 1000 | Distal Intergenic | 3 | 121671317 | 121682835 | 11519 | 1 | ENSMUST00 | -7653 | 4930432M17 | ENSMUSG0( | false | . | . | . | . | . | . | chr3 | 121661916 | 121664136 | 4.80771 | 3.87568 | 2.86593 |
| chr9 | 61361075 | 61361643 | 1000 | Distal Intergenic | 9 | 61370878 | 61371627 | 750 | 2 | ENSMUST00 | 9984 | Gm10655 | ENSMUSG0( | false | . | . | . | . | . | . | chr9 | 61360198 | 61362657 | 5.64751 | 4.37108 | 3.54161 |
| chr7 | 25461281 | 25461989 | 1000 | 3' UTR | 7 | 25461702 | 25477625 | 15924 | 2 | ENSMUST00 | 15636 | Ceacam1 | ENSMUSG0( | false | . | . | . | . | . | . | chr7 | 25459583 | 25465057 | 6.73544 | 4.89232 | 4.43593 |
| chr7 | 15962306 | 15963176 | 1000 | Intron (ENSMUST00000098799.3 | 7 | 15957683 | 15967475 | 9793 | 2 | ENSMUST00 | 4299 | Ehd2 | ENSMUSG0( | false | . | . | . | . | . | . | chr7 | 15953828 | 15968448 | 7.04869 | 5.00331 | 4.66896 |
| chr7 | 15960676 | 15961234 | 1000 | Intron (ENSMUST00000098799.3 | 7 | 15957683 | 15967475 | 9793 | 2 | ENSMUST00 | 6241 | Ehd2 | ENSMUSG0( | false | . | . | . | . | . | . | chr7 | 15953828 | 15968448 | 7.04869 | 5.00331 | 4.66896 |
| chr8 | 25388921 | 25389297 | 1000 | Distal Intergenic | 8 | 25454273 | 25454704 | 432 | 1 | ENSMUST00 | -64976 | Gm10689 | ENSMUSG0( | false | . | . | . | . | . | . | chr8 | 25388495 | 25389912 | 4.75797 | 3.98662 | 2.8439 |
| chr8 | 25449609 | 25449951 | 1000 | Distal Intergenic | 8 | 25454273 | 25454704 | 432 | 1 | ENSMUST00 | -4322 | Gm10689 | ENSMUSG0( | false | . | . | . | . | . | . | chr8 | 25447523 | 25451206 | 5.02392 | 4.0535 | 3.03891 |
| chr8 | 25437131 | 25437571 | 1000 | Distal Intergenic | 8 | 25454273 | 25454704 | 432 | 1 | ENSMUST00 | -16702 | Gm10689 | ENSMUSG0( | false | . | . | . | . | . | . | chr8 | 25435682 | 25441738 | 6.54241 | 4.78324 | 4.28383 |
| chr8 | 25440561 | 25441133 | 1000 | Distal Intergenic | 8 | 25454273 | 25454704 | 432 | 1 | ENSMUST00 | -13140 | Gm10689 | ENSMUSG0( | false | chr8 | 25439977 | 25440949 | 7.35251 | 12.67314 | 10.44663 | chr8 | 25435682 | 25441738 | 6.54241 | 4.78324 | 4.28383 |
| chr9 | 41611903 | 41612324 | 1000 | Promoter (1-2kb) | 9 | 41614237 | 41617772 | 3536 | 1 | ENSMUST00 | -1913 | 2610203C20I | ENSMUSG0( | false | chr9 | 41611913 | 41612849 | 8.24817 | 15.43468 | 13.16567 | . | . | . | . | . | . |
| chr9 | 21767849 | 21768214 | 1000 | 3' UTR | 9 | 21755442 | 21760286 | 4845 | 2 | ENSMUST00 | -7564 | Spc24 | ENSMUSG0( | false | . | . | . | . | . | . | chr9 | 21767075 | 21769846 | 6.65311 | 4.68506 | 4.41065 |
| chr9 | 21752304 | 21752863 | 1000 | Downstream (2-3kb) | 9 | 21755442 | 21760286 | 4845 | 2 | ENSMUST00 | 7423 | Spc24 | ENSMUSG0( | false | . | . | . | . | . | . | chr9 | 21751246 | 21755376 | 6.77488 | 4.93372 | 4.46002 |
| chr3 | 84581998 | 84582922 | 1000 | Promoter (<=1kb) | 3 | 84496093 | 84582534 | 86442 | 2 | ENSMUST00 | 0 | Arfip1 | ENSMUSG0( | false | chr3 | 84580512 | 84583491 | 33.1106 | 101.26704 | 97.66567 | . | . | . | . | . | . |
| chr2 | 160366376 | 160367575 | 1000 | Promoter (<=1kb) | 2 | 160363703 | 160367065 | 3363 | 2 | ENSMUST00 | 0 | Mafb | ENSMUSG0( | true | chr2 | 160367241 | 160369416 | 27.62823 | 80.1489 | 76.99468 | chr2 | 160367527 | 160369510 | 4.90013 | 3.92828 | 2.92855 |
| chr2 | 160201971 | 160202397 | 1000 | Distal Intergenic | 2 | 160311393 | 160313616 | 2224 | 2 | ENSMUST00 | 111219 | Gm826 | ENSMUSG0( | false | . | . | . | . | . | . | chr2 | 160199744 | 160203956 | 5.01478 | 4.0483 | 3.01431 |
| chr2 | 160288579 | 160289202 | 1000 | Distal Intergenic | 2 | 160311393 | 160313616 | 2224 | 2 | ENSMUST00 | 24414 | Gm826 | ENSMUSG0( | false | . | . | . | . | . | . | chr2 | 160286587 | 160290821 | 5.12701 | 4.23469 | 3.10338 |
| chr2 | 152928181 | 152928705 | 1000 | Distal Intergenic | 2 | 152931898 | 152933208 | 1311 | 1 | ENSMUST00 | -4503 | Foxs1 | ENSMUSG0( | false | . | . | . | . | . | . | chr2 | 152925247 | 152934463 | 6.47975 | 4.78702 | 4.2189 |
| chr2 | 152410829 | 152411546 | 1000 | Downstream (<1kb) | 2 | 152411955 | 152415207 | 3073 | 2 | ENSMUST00 | 3481 | Zcchc3 | ENSMUSG0( | false | . | . | . | . | . | . | chr2 | 152408919 | 152413746 | 5.68724 | 4.10309 | 3.60771 |
| chr2 | 152414407 | 152415184 | 1000 | Promoter (<=1kb) | 2 | 152411955 | 152415207 | 3073 | 2 | ENSMUST00 | 0 | Zcchc3 | ENSMUSG0( | false | chr2 | 152413063 | 152415644 | 35.70928 | 138.63979 | 133.81488 | . | . | . | . | . | . |
| chr2 | 152226448 | 152227446 | 1000 | Promoter (<=1kb) | 2 | 152226175 | 152281852 | 55014 | 1 | ENSMUST00 | 0 | Csnk2a1 | ENSMUSG0( | false | chr2 | 152226184 | 152227818 | 35.2856 | 127.03558 | 122.65217 | . | . | . | . | . | . |
| chr19 | 59345252 | 59346333 | 1000 | Promoter (<=1kb) | 19 | 59296084 | 59345780 | 49697 | 2 | ENSMUST00 | 0 | Pdzd8 | ENSMUSG0( | false | chr19 | 59343941 | 59346591 | 37.72364 | 120.6551 | 116.49378 | . | . | . | . | . | . |
| chr2 | 146888700 | 146889142 | 1000 | Exon (ENSMUST00000099278.3/ | 2 | 146855909 | 146889295 | 33387 | 2 | ENSMUST00 | 32792 | Kiz | ENSMUSG0( | false | . | . | . | . | . | . | chr2 | 146885370 | 146890848 | 4.9551 | 3.86874 | 2.9915 |
| chr10 | 95514608 | 95515270 | 1000 | Promoter (<=1kb) | 10 | 95515162 | 95545658 | 30497 | 1 | ENSMUST00 | 0 | Ube2n | ENSMUSG0( | false | chr10 | 95514318 | 95516303 | 45.54573 | 153.00453 | 147.60376 | . | . | . | . | . | . |
| chr13 | 80882942 | 80883457 | 1000 | Promoter (<=1kb) | 13 | 80883384 | 80889643 | 6260 | 1 | ENSMUST00 | 0 | Arrdc3 | ENSMUSG0( | false | chr13 | 80882761 | 80883175 | 33.47463 | 112.14942 | 108.24687 | . | . | . | . | . | . |
| chr19 | 46003270 | 46003836 | 1000 | Promoter (<=1kb) | 19 | 46003478 | 46006173 | 2696 | 1 | ENSMUST00 | 0 | Hps6 | ENSMUSG0( | false | chr19 | 46003066 | 46004412 | 37.68163 | 117.40437 | 113.3432 | chr19 | 46002951 | 46006112 | 4.60993 | 3.76933 | 2.73023 |
| chr2 | 122625876 | 122626225 | 1000 | Distal Intergenic | 2 | 122630625 | 122632704 | 2080 | 1 | ENSMUST00 | -4297 | Spata5l1 | ENSMUSG0( | false | . | . | . | . | . | . | chr2 | 122625654 | 122627162 | 4.21677 | 3.68897 | 2.37116 |
| chr2 | 121448846 | 121450046 | 1000 | Promoter (<=1kb) | 2 | 121449195 | 121453420 | 4226 | 1 | ENSMUST00 | 0 | Serf2 | ENSMUSG0( | false | chr2 | 121448585 | 121449040 | 11.60718 | 37.9516 | 35.3891 | . | . | . | . | . | . |
| chr13 | 55444402 | 55445484 | 1000 | Promoter (<=1kb) | 13 | 55445334 | 55459577 | 14244 | 1 | ENSMUST00 | 0 | Grk6 | ENSMUSG0( | false | chr13 | 55444309 | 55446120 | 36.66269 | 126.29414 | 121.93508 | chr13 | 55444302 | 55444618 | 3.00912 | 2.93524 | 1.36152 |
| chr2 | 120086029 | 120086412 | 1000 | Promoter (<=1kb) | 2 | 120076214 | 120085772 | 9559 | 2 | ENSMUST00 | -258 | Sptbn5 | ENSMUSG0( | false | . | . | . | . | . | . | chr2 | 120084728 | 120087163 | 5.9071 | 4.31805 | 3.77927 |
| chr19 | 26942122 | 26942535 | 1000 | Distal Intergenic | 19 | 26885925 | 26888639 | 2715 | 1 | ENSMUST00 | 56198 | Gm815 | ENSMUSG0( | false | . | . | . | . | . | . | . | . | . | . | . | . |
| chr2 | 118745443 | 118746077 | 1000 | Promoter (<=1kb) | 2 | 118745758 | 118748810 | 3053 | 1 | ENSMUST00 | 0 | 5430417L2I | ENSMUSG0( | false | chr2 | 118745142 | 118747076 | 33.7247 | 108.45113 | 104.65779 | chr2 | 118743438 | 118745488 | 4.9534 | 4.13927 | 2.96732 |
| chr19 | 24469167 | 24469623 | 1000 | Intron (ENSMUST00000112673.3 | 19 | 24476502 | 24477356 | 855 | 2 | ENSMUST00 | 7733 | Fam122a | ENSMUSG0( | false | . | . | . | . | . | . | chr19 | 24467482 | 24471892 | 4.06001 | 3.5939 | 2.26098 |
| chr13 | 32924549 | 32925304 | 1000 | Promoter (<=1kb) | 13 | 32938524 | 32938691 | 168 | 2 | ENSMUST00 | 13387 | Gm11382 | ENSMUSG0( | false | . | . | . | . | . | . | chr13 | 32922423 | 32930295 | 4.39099 | 3.75239 | 2.52126 |
| chr2 | 105058676 | 105059145 | 1000 | Distal Intergenic | 2 | 105077895 | 105078107 | 213 | 2 | ENSMUST00 | 18962 | Wt1os | ENSMUSG0( | true | . | . | . | . | . | . | chr2 | 105057120 | 105080197 | 5.15057 | 4.16934 | 3.12788 |
| chr2 | 104816468 | 104817307 | 1000 | Promoter (<=1kb) | 2 | 104754795 | 104816696 | 61902 | 2 | ENSMUST00 | 0 | Qser1 | ENSMUSG0( | false | chr2 | 104815665 | 104818377 | 33.32351 | 103.65904 | 99.99326 | chr2 | 104814358 | 104818966 | 3.92436 | 3.53073 | 2.12871 |
| chr2 | 94273145 | 94273986 | 1000 | Promoter (<=1kb) | 2 | 94260966 | 94264758 | 3793 | 2 | ENSMUST00 | -8388 | E530001K10I | ENSMUSG0( | false | chr2 | 94273044 | 94273945 | 21.02725 | 63.18829 | 60.29588 | chr2 | 94273027 | 94274135 | 4.12728 | 3.68798 | 2.27447 |
| chr2 | 93045497 | 93046457 | 1000 | Promoter (<=1kb) | 2 | 92965151 | 93045571 | 80421 | 2 | ENSMUST00 | 0 | Prdm11 | ENSMUSG0( | false | chr2 | 93044878 | 93046685 | 36.76257 | 113.72284 | 109.77938 | . | . | . | . | . | . |
| chr10 | 30327914 | 30328268 | 1000 | Distal Intergenic | 10 | 30196011 | 30200540 | 4530 | 2 | ENSMUST00 | -127375 | Cenpw | ENSMUSG0( | false | . | . | . | . | . | . | chr10 | 30326259 | 30329046 | 4.29691 | 3.70523 | 2.43187 |
| chr2 | 74748551 | 74748912 | 1000 | Promoter (1-2kb) | 2 | 74748422 | 74750506 | 2085 | 1 | ENSMUST00 | 1594 | Haglr | ENSMUSG0( | false | . | . | . | . | . | . | chr2 | 74744931 | 74751115 | 5.56017 | 4.28612 | 3.49137 |
| chr2 | 74749276 | 74750307 | 1000 | Promoter (<=1kb) | 2 | 74748422 | 74750506 | 2085 | 1 | ENSMUST00 | 199 | Haglr | ENSMUSG0( | false | chr2 | 74749104 | 74750334 | 12.46893 | 30.48504 | 28.01359 | chr2 | 74744931 | 74751115 | 5.56017 | 4.28612 | 3.49137 |
| chr19 | 4175266 | 4175938 | 1000 | Promoter (<=1kb) | 19 | 4164324 | 4175479 | 11156 | 2 | ENSMUST00 | 0 | Carns1 | ENSMUSG0( | false | . | . | . | . | . | . | chr19 | 4166946 | 4176432 | 4.77668 | 3.94064 | 2.82677 |
| chr10 | 4386629 | 4388388 | 1000 | Promoter (<=1kb) | 10 | 4367074 | 4388108 | 21035 | 2 | ENSMUST00 | 0 | Zbtb2 | ENSMUSG0( | false | chr10 | 4386149 | 4389234 | 34.90791 | 116.7856 | 112.7497 | chr10 | 4385764 | 4386648 | 5.65304 | 4.39657 | 3.53013 |
| chr2 | 37422519 | 37423179 | 1000 | Promoter (<=1kb) | 2 | 37370069 | 37422943 | 52835 | 2 | ENSMUST00 | 0 | Rc3h2 | ENSMUSG0( | false | chr2 | 37422026 | 37423429 | 32.87541 | 111.07999 | 107.21675 | . | . | . | . | . | . |
| chr11 | 120189395 | 120190233 | 1000 | Promoter (<=1kb) | 11 | 120187707 | 120189852 | 2146 | 2 | ENSMUST00 | 0 | 2810410L24F | ENSMUSG0( | false | chr11 | 120186648 | 120190389 | 30.60583 | 111.04385 | 107.18215 | . | . | . | . | . | . |
| chr15 | 103060170 | 103061252 | 1000 | Promoter (<=1kb) | 15 | 103034395 | 103036852 | 2458 | 1 | ENSMUST00 | 25776 | Hoxc4 | ENSMUSG0( | false | chr15 | 103060301 | 103061298 | 13.64722 | 31.21559 | 28.73584 | chr15 | 103061190 | 103063160 | 3.25054 | 3.0956 | 1.59346 |
| chr15 | 103061458 | 103061775 | 1000 | Promoter (<=1kb) | 15 | 103034395 | 103036852 | 2458 | 1 | ENSMUST00 | 27064 | Hoxc4 | ENSMUSG0( | false | chr15 | 103061395 | 103061996 | 6.94583 | 12.01026 | 9.79738 | chr15 | 103061190 | 103063160 | 3.25054 | 3.0956 | 1.59346 |
| chr15 | 101293994 | 101294407 | 1000 | Promoter (<=1kb) | 15 | 101293232 | 101297426 | 4195 | 1 | ENSMUST00 | 763 | 6030408B16I | ENSMUSG0( | false | . | . | . | . | . | . | chr15 | 101291483 | 101296758 | 5.6438 | 4.08858 | 3.5627 |
| chr2 | 31135536 | 31135985 | 1000 | Intron (ENSMUST00000073879.6 | 2 | 31026206 | 31141802 | 115597 | 2 | ENSMUST00 | 5817 | Fnbp1 | ENSMUSG0( | false | . | . | . | . | . | . | chr2 | 31124399 | 31141165 | 6.19999 | 4.65578 | 3.97992 |
| chr2 | 31087725 | 31088606 | 1000 | Intron (ENSMUST00000073879.6 | 2 | 31054200 | 31096498 | 42299 | 2 | ENSMUST00 | 7892 | Fnbp1 | ENSMUSG0( | false | . | . | . | . | . | . | chr2 | 31086368 | 31093289 | 6.36762 | 4.66528 | 4.15642 |
| chr2 | 31132679 | 31133071 | 1000 | Intron (ENSMUST00000073879.6 | 2 | 31026206 | 31141802 | 115597 | 2 | ENSMUST00 | 8731 | Fnbp1 | ENSMUSG0( | false | . | . | . | . | . | . | chr2 | 31124399 | 31141165 | 6.19999 | 4.65578 | 3.97992 |
| chr2 | 30711681 | 30712411 | 1000 | Downstream (<1kb) | 2 | 30713150 | 30717043 | 3894 | 2 | ENSMUST00 | 4632 | Gm14488 | ENSMUSG0( | false | chr2 | 30711573 | 30712506 | 17.45395 | 46.28863 | 43.62135 | chr2 | 30711079 | 30713834 | 5.05675 | 4.11298 | 3.0463 |
| chr2 | 30707408 | 30708602 | 1000 | Distal Intergenic | 2 | 30713150 | 30717043 | 3894 | 2 | ENSMUST00 | 8441 | Gm14488 | ENSMUSG0( | false | chr2 | 30707595 | 30708517 | 22.63194 | 64.29881 | 61.39256 | chr2 | 30707175 | 30709614 | 4.78885 | 3.80828 | 2.85067 |
| chr9 | 107063366 | 107063635 | 1000 | Intron (ENSMUST00000044532.5 | 9 | 107047803 | 107049009 | 1207 | 1 | ENSMUST00 | 15564 | Gm17205 | ENSMUSG0( | false | . | . | . | . | . | . | . | . | . | . | . | . |
| chr2 | 16148096 | 16148405 | 1000 | Intron (ENSMUST00000146205.1 | 2 | 16042126 | 16150806 | 108681 | 1 | ENSMUST00 | 105971 | Malrd1 | ENSMUSG0( | false | . | . | . | . | . | . | chr2 | 16147619 | 16149085 | 4.09585 | 3.59207 | 2.2868 |
| chr15 | 76091375 | 76091914 | 1000 | Promoter (1-2kb) | 15 | 76085595 | 76090013 | 4419 | 2 | ENSMUST00 | -1363 | Nrbp2 | ENSMUSG0( | false | chr15 | 76090740 | 76092009 | 30.05689 | 90.05383 | 86.71302 | chr15 | 76090235 | 76092300 | 5.26433 | 4.19625 | 3.22646 |
| chr16 | 18448079 | 18448468 | 1000 | Intron (ENSMUST00000115605.2 | 16 | 18452039 | 18472249 | 20211 | 1 | ENSMUST00 | -3571 | Txnrd2 | ENSMUSG0( | false | . | . | . | . | . | . | chr16 | 18446552 | 18452802 | 5.74275 | 4.36487 | 3.63646 |
| chr10 | 80048161 | 80048676 | 1000 | Promoter (<=1kb) | 10 | 80047166 | 80056439 | 9274 | 1 | ENSMUST00 | 996 | Gpx4 | ENSMUSG0( | false | . | . | . | . | . | . | chr10 | 80046083 | 80053338 | 6.68412 | 4.69614 | 4.42765 |
| chr8 | 128608929 | 128609336 | 1000 | Distal Intergenic | 8 | 128650997 | 128651103 | 107 | 1 | ENSMUST00 | -41661 | Gm26002 | ENSMUSG0( | false | . | . | . | . | . | . | chr8 | 128604938 | 128611438 | 5.48899 | 4.2322 | 3.41813 |
| chr4 | 9220684 | 9221247 | 1000 | Distal Intergenic | 4 | 9210003 | 9210108 | 106 | 1 | ENSMUST00 | 10682 | Gm23423 | ENSMUSG0( | false | . | . | . | . | . | . | chr4 | 9220261 | 9221710 | 4.14072 | 3.62531 | 2.3481 |
| chr11 | 102655740 | 102656320 | 1000 | Downstream (2-3kb) | 11 | 102659572 | 102659680 | 109 | 2 | ENSMUST00 | 3360 | Gm22920 | ENSMUSG0( | false | . | . | . | . | . | . | chr11 | 102653489 | 102657768 | 4.98684 | 4.0937 | 3.00781 |
| chr18 | 66187864 | 66188204 | 1000 | Intron (ENSMUST00000146610.1 | 18 | 66219264 | 66219333 | 70 | 2 | ENSMUST00 | 31129 | Mir694 | ENSMUSG0( | false | . | . | . | . | . | . | chr18 | 66187099 | 66189734 | 3.75494 | 3.33367 | 2.01261 |
| chr2 | 156324046 | 156324488 | 1000 | Promoter (1-2kb) | 2 | 156322552 | 156322647 | 96 | 2 | ENSMUST00 | 1495 | Gm22759 | ENSMUSG0( | false | . | . | . | . | . | . | chr2 | 156323497 | 156325472 | 5.42305 | 4.2172 | 3.37645 |
| chr1 | 131486431 | 131487432 | 1000 | Intron (ENSMUST00000097588.3 | 1 | 131498639 | 131498802 | 164 | 2 | ENSMUST00 | 11370 | Gm24273 | ENSMUSG0( | false | . | . | . | . | . | . | chr1 | 131451954 | 131498502 | 5.16635 | 4.13104 | 3.14877 |
| chr15 | 35618936 | 35619247 | 1000 | Intron (ENSMUST00000048646.7 | 15 | 35660831 | 35660918 | 88 | 2 | ENSMUST00 | 41671 | Mir599 | ENSMUSG0( | false | . | . | . | . | . | . | chr15 | 35617582 | 35620501 | 3.61383 | 3.24791 | 1.88439 |
| chr15 | 35649758 | 35650130 | 1000 | Intron (ENSMUST00000048646.7 | 15 | 35660831 | 35660918 | 88 | 2 | ENSMUST00 | 10788 | Mir599 | ENSMUSG0( | false | . | . | . | . | . | . | chr15 | 35649071 | 35651664 | 3.55525 | 3.33258 | 1.81914 |
| chr9 | 42897261 | 42897979 | 1000 | Intron (ENSMUST00000114865.2 | 9 | 42862051 | 42862129 | 79 | 2 | ENSMUST00 | -35133 | Gm27509 | ENSMUSG0( | false | chr9 | 42896787 | 42898140 | 20.9555 | 57.98476 | 55.16582 | chr9 | 42896902 | 42898536 | 5.59432 | 4.41031 | 3.4971 |
| chr3 | 38697963 | 38698477 | 1000 | Distal Intergenic | 3 | 38558109 | 38558214 | 106 | 2 | ENSMUST00 | -139750 | Gm24462 | ENSMUSG0( | false | . | . | . | . | . | . | . | . | . | . | . | . |
| chr11 | 65081052 | 65081286 | 1000 | Intron (ENSMUST00000047463.9 | 11 | 65067709 | 65067807 | 99 | 2 | ENSMUST00 | -13246 | Gm25991 | ENSMUSG0( | false | . | . | . | . | . | . | chr11 | 65079421 | 65084181 | 3.80681 | 3.44976 | 2.01799 |
| chr12 | 21417110 | 21418028 | 1000 | Promoter (<=1kb) | 12 | 21390071 | 21417637 | 27567 | 2 | ENSMUST00 | 0 | Ywhaq | ENSMUSG0( | false | chr12 | 21416080 | 21418715 | 26.74277 | 88.06109 | 84.75893 | . | . | . | . | . | . |
| chr11 | 118326962 | 118328028 | 1000 | Promoter (<=1kb) | 11 | 118332415 | 118334016 | 1602 | 1 | ENSMUST00 | 5988 | BC100451 | ENSMUSG0( | false | chr11 | 118327240 | 118327487 | 4.72831 | 6.77738 | 4.67505 | chr11 | 118319919 | 118354581 | 6.53665 | 4.67889 | 4.28866 |
| chr6 | 70073781 | 70074409 | 1000 | Promoter (<=1kb) | 6 | 70074024 | 70074584 | 561 | 2 | ENSMUST00 | 175 | Igkv6-32 | ENSMUSG0( | false | . | . | . | . | . | . | chr6 | 70072263 | 70075068 | 3.76337 | 3.46811 | 1.98668 |
| chr6 | 70116852 | 70117284 | 1000 | Promoter (<=1kb) | 6 | 70117061 | 70117617 | 557 | 2 | ENSMUST00 | 333 | Igkv8-30 | ENSMUSG0( | false | . | . | . | . | . | . | chr6 | 70115225 | 70118243 | 3.61941 | 3.39411 | 1.85895 |
| chr6 | 70105671 | 70105981 | 1000 | Distal Intergenic | 6 | 70117061 | 70117617 | 557 | 2 | ENSMUST00 | 11636 | Igkv8-30 | ENSMUSG0( | false | chr6 | . | . | . | . | . | chr6 | 70102385 | 70108287 | 4.05658 | 3.56049 | 2.23003 |

| | | | | | | | | | | | | | | | | | | | | | | | | | |
|---|---|---|---|---|---|---|---|---|---|---|---|---|---|---|---|---|---|---|---|---|---|---|---|---|---|
| chr6 | 70193652 | 70194020 | 1000 | Promoter (<=1kb) | 6 | 70193228 | 70193797 | 570 | 1 | ENSMUST00 | 425 | Igkv8-26 | ENSMUSG0C | false | . | . | . | . | . | . | chr6 | 70193532 | 70195164 | 3.62092 | 3.25752 | 1.91305 |
| chr6 | 70216658 | 70217036 | 1000 | Promoter (<=1kb) | 6 | 70216858 | 70217421 | 564 | 2 | ENSMUST00 | 385 | Igkv8-24 | ENSMUSG0C | false | . | . | . | . | . | . | chr6 | 70213474 | 70219351 | 5.14256 | 4.19463 | 3.11118 |
| chr6 | 70356240 | 70356616 | 1000 | Promoter (<=1kb) | 6 | 70355877 | 70356445 | 569 | 1 | ENSMUST00 | 364 | Igkv8-18 | ENSMUSG0C | false | . | . | . | . | . | . | chr6 | 70355060 | 70358231 | 3.42674 | 3.18127 | 1.76216 |
| chr6 | 70386409 | 70386889 | 1000 | Promoter (<=1kb) | 6 | 70386672 | 70387238 | 567 | 2 | ENSMUST00 | 349 | Igkv8-16 | ENSMUSG0C | false | . | . | . | . | . | . | chr6 | 70385956 | 70388311 | 6.31575 | 4.65551 | 4.07275 |
| chr11 | 57751542 | 57751961 | 1000 | Intron (ENSMUST00000066987.8 | 11 | 57741300 | 57741402 | 103 | 1 | ENSMUST00 | 10243 | Gm24582 | ENSMUSG0C | false | . | . | . | . | . | . | chr11 | 57744780 | 57769452 | 6.61093 | 4.7685 | 4.34185 |
| chr11 | 57732528 | 57733049 | 1000 | Intron (ENSMUST00000066987.8 | 11 | 57741300 | 57741402 | 103 | 1 | ENSMUST00 | -8251 | Gm24582 | ENSMUSG0C | false | . | . | . | . | . | . | chr11 | 57724712 | 57734543 | 6.29874 | 4.67914 | 4.07864 |
| chr4 | 62862974 | 62863640 | 1000 | Distal Intergenic | 4 | 62812428 | 62812560 | 133 | 1 | ENSMUST00 | 50547 | Gm24117 | ENSMUSG0C | false | . | . | . | . | . | . | chr4 | 62862856 | 62863824 | 5.20256 | 4.19938 | 3.15047 |
| chr7 | 70078546 | 70078865 | 1000 | Distal Intergenic | 7 | 69957776 | 69957904 | 129 | 2 | ENSMUST00 | -120643 | Gm24120 | ENSMUSG0C | false | . | . | . | . | . | . | chr7 | 70078245 | 70079244 | 4.2612 | 3.7079 | 2.43178 |
| chr2 | 159270194 | 159271091 | 1000 | Distal Intergenic | 2 | 159139361 | 159139492 | 132 | 1 | ENSMUST00 | 130834 | Gm25090 | ENSMUSG0C | false | . | . | . | . | . | . | chr2 | 159269480 | 159284862 | 4.11414 | 3.61003 | 2.27932 |
| chr11 | 45693186 | 45693636 | 1000 | Distal Intergenic | 11 | 45639383 | 45639514 | 132 | 1 | ENSMUST00 | 53804 | Gm22284 | ENSMUSG0C | false | . | . | . | . | . | . | . | . | . | . | . | . |
| chr9 | 37054179 | 37054531 | 1000 | Intron (ENSMUST00000177218.2 | 9 | 37026399 | 37026507 | 109 | 1 | ENSMUST00 | 27781 | n-R5s82 | ENSMUSG0C | false | chr9 | 37054394 | 37054779 | 4.42115 | 6.83029 | 4.72578 | chr9 | 37051691 | 37078073 | 5.33519 | 4.13112 | 3.31406 |
| chr9 | 37024026 | 37024412 | 1000 | Promoter (1-2kb) | 9 | 37026399 | 37026507 | 109 | 1 | ENSMUST00 | -1987 | n-R5s82 | ENSMUSG0C | false | . | . | . | . | . | . | chr9 | 37021034 | 37031907 | 6.40408 | 4.65381 | 4.17785 |
| chr15 | 75778178 | 75778453 | 1000 | Intron (ENSMUST00000100538.2 | 15 | 75759859 | 75759989 | 131 | 1 | ENSMUST00 | 18320 | Gm23747 | ENSMUSG0C | false | . | . | . | . | . | . | chr15 | 75777173 | 75779080 | 5.2118 | 3.99176 | 3.21498 |
| chr16 | 78213799 | 78214289 | 1000 | Distal Intergenic | 16 | 78240258 | 78240388 | 131 | 1 | ENSMUST00 | -25969 | Gm25916 | ENSMUSG0C | false | . | . | . | . | . | . | . | . | . | . | . | . |
| chr3 | 67557830 | 67558310 | 1000 | Distal Intergenic | 3 | 67573160 | 67573291 | 132 | 2 | ENSMUST00 | 14981 | Gm25919 | ENSMUSG0C | false | . | . | . | . | . | . | chr3 | 67555913 | 67559354 | 5.28251 | 4.16844 | 3.24229 |
| chr17 | 12292922 | 12293264 | 1000 | Intron (ENSMUST00000089058.5 | 17 | 12301036 | 12301175 | 140 | 1 | ENSMUST00 | -7772 | Gm24794 | ENSMUSG0C | false | . | . | . | . | . | . | chr17 | 12290254 | 12298600 | 5.97528 | 4.50765 | 3.81519 |
| chr13 | 70277699 | 70278777 | 1000 | Distal Intergenic | 13 | 70168832 | 70168959 | 128 | 1 | ENSMUST00 | 108868 | Gm26018 | ENSMUSG0C | false | . | . | . | . | . | . | chr13 | 70276634 | 70283449 | 5.13818 | 4.18533 | 3.1123 |
| chr4 | 86498737 | 86499126 | 1000 | Intron (ENSMUST00000151481.2 | 4 | 86509152 | 86509225 | 74 | 1 | ENSMUST00 | -10026 | Gm26017 | ENSMUSG0C | false | . | . | . | . | . | . | chr4 | 86498540 | 86499041 | 2.54089 | 2.56713 | 1.03618 |
| chr7 | 28356931 | 28357392 | 1000 | Downstream (2-3kb) | 7 | 28351508 | 28351632 | 125 | 1 | ENSMUST00 | 5424 | AF357399 | ENSMUSG0C | false | . | . | . | . | . | . | chr7 | 28351351 | 28371775 | 6.79815 | 4.79627 | 4.48731 |
| chr3 | 37999677 | 38000242 | 1000 | Distal Intergenic | 3 | 37995105 | 37995235 | 131 | 2 | ENSMUST00 | -4443 | Gm22899 | ENSMUSG0C | false | . | . | . | . | . | . | chr3 | 37998031 | 38001135 | 4.97791 | 4.02404 | 3.02162 |
| chr12 | 33626059 | 33626430 | 1000 | Distal Intergenic | 12 | 33452712 | 33452843 | 132 | 2 | ENSMUST00 | -173217 | Gm23529 | ENSMUSG0C | false | chr12 | 33625974 | 33626252 | 5.7815 | 10.45037 | 8.26584 | . | . | . | . | . | . |
| chr1 | 19899315 | 19899825 | 1000 | Distal Intergenic | 1 | 19921310 | 19921399 | 90 | 2 | ENSMUST00 | 21574 | Gm25568 | ENSMUSG0C | false | . | . | . | . | . | . | chr1 | 19898820 | 19902584 | 4.3215 | 3.77516 | 2.44276 |
| chr13 | 58029973 | 58030398 | 1000 | Intron (ENSMUST00000160860.2 | 13 | 58023125 | 58023200 | 76 | 2 | ENSMUST00 | -6774 | Mir874 | ENSMUSG0C | false | . | . | . | . | . | . | chr13 | 58029180 | 58032225 | 5.11035 | 4.12179 | 3.13401 |
| chr13 | 58035422 | 58036097 | 1000 | Intron (ENSMUST00000160860.2 | 13 | 58023125 | 58023200 | 76 | 2 | ENSMUST00 | -12223 | Mir874 | ENSMUSG0C | false | . | . | . | . | . | . | chr13 | 58034253 | 58039192 | 5.68384 | 4.1441 | 3.60254 |
| chr13 | 12702907 | 12703863 | 1000 | Promoter (<=1kb) | 13 | 12702362 | 12702589 | 228 | 1 | ENSMUST00 | 546 | Gm2399 | ENSMUSG0C | false | chr13 | 12702430 | 12703024 | 18.33657 | 60.6815 | 57.82544 | chr13 | 12700774 | 12706233 | 5.06138 | 3.93361 | 3.0952 |
| chr1 | 176092040 | 176092481 | 1000 | Intron (ENSMUST00000065967.8 | 1 | 175977777 | 175979114 | 1338 | 2 | ENSMUST00 | -112927 | B020018G12 | ENSMUSG0C | false | . | . | . | . | . | . | chr1 | 176089383 | 176099329 | 4.1765 | 3.68436 | 2.32939 |
| chr1 | 176064657 | 176065011 | 1000 | Intron (ENSMUST00000065967.8 | 1 | 175977777 | 175979114 | 1338 | 2 | ENSMUST00 | -85544 | B020018G12 | ENSMUSG0C | false | . | . | . | . | . | . | chr1 | 176063189 | 176065619 | 5.54365 | 4.28504 | 3.45838 |
| chr13 | 98354238 | 98355168 | 1000 | Promoter (<=1kb) | 13 | 98354242 | 98356703 | 2462 | 1 | ENSMUST00 | 0 | Foxd1 | ENSMUSG0C | false | chr13 | 98353706 | 98359675 | 27.78601 | 91.88203 | 88.49705 | . | . | . | . | . | . |
| chr13 | 98365671 | 98366626 | 1000 | Distal Intergenic | 13 | 98354242 | 98356703 | 2462 | 1 | ENSMUST00 | 11430 | Foxd1 | ENSMUSG0C | false | chr13 | 98365666 | 98367402 | 22.21283 | 62.70797 | 59.8237 | . | . | . | . | . | . |
| chr4 | 154024099 | 154024558 | 1000 | Promoter (<=1kb) | 4 | 154023311 | 154024355 | 1045 | 2 | ENSMUST00 | 0 | Smim1 | ENSMUSG0C | false | chr4 | 154023440 | 154024806 | 7.14186 | 14.46535 | 12.21151 | chr4 | 154021005 | 154028094 | 7.34568 | 5.07579 | 4.95328 |
| chr4 | 154022447 | 154022926 | 1000 | Promoter (1-2kb) | 4 | 154023724 | 154024323 | 600 | 2 | ENSMUST00 | 1397 | Smim1 | ENSMUSG0C | false | . | . | . | . | . | . | chr4 | 154021005 | 154028094 | 7.34568 | 5.07579 | 4.95328 |
| chr10 | 126971778 | 126972472 | 1000 | Distal Intergenic | 10 | 126978717 | 126999975 | 21259 | 1 | ENSMUST00 | -6245 | Ctdsp2 | ENSMUSG0C | false | . | . | . | . | . | . | chr10 | 126967116 | 126978243 | 5.23135 | 4.17886 | 3.19381 |
| chr10 | 126978288 | 126980237 | 1000 | Promoter (<=1kb) | 10 | 126978717 | 126999975 | 21259 | 1 | ENSMUST00 | 0 | Ctdsp2 | ENSMUSG0C | false | chr10 | 126977989 | 126978495 | 9.20781 | 17.98236 | 15.67656 | . | . | . | . | . | . |
| chr4 | 155793733 | 155794084 | 1000 | Promoter (2-3kb) | 4 | 155791172 | 155799503 | 8332 | 1 | ENSMUST00 | 2562 | Ankrd65 | ENSMUSG0C | false | . | . | . | . | . | . | . | . | . | . | . | . |
| chr4 | 141722946 | 141723653 | 1000 | Promoter (<=1kb) | 4 | 141677549 | 141723419 | 45871 | 2 | ENSMUST00 | 0 | Ddi2 | ENSMUSG0C | false | chr4 | 141722638 | 141723984 | 29.2476 | 106.45832 | 102.71833 | chr4 | 141717923 | 141725687 | 4.52579 | 3.82801 | 2.61933 |
| chr4 | 131405346 | 131405816 | 1000 | Distal Intergenic | 4 | 131487197 | 131505527 | 18331 | 1 | ENSMUST00 | -81381 | Gm831 | ENSMUSG0C | false | . | . | . | . | . | . | chr4 | 131403590 | 131411016 | 4.73067 | 3.88614 | 2.78676 |
| chr4 | 131381082 | 131381450 | 1000 | Distal Intergenic | 4 | 131487197 | 131505527 | 18331 | 1 | ENSMUST00 | -105747 | Gm831 | ENSMUSG0C | false | . | . | . | . | . | . | chr4 | 131378929 | 131382281 | 6.34379 | 4.71406 | 4.11296 |
| chr4 | 131489078 | 131489466 | 1000 | Promoter (1-2kb) | 4 | 131487197 | 131505527 | 18331 | 1 | ENSMUST00 | 1882 | Gm831 | ENSMUSG0C | false | . | . | . | . | . | . | chr4 | 131488351 | 131489644 | 4.66453 | 3.88473 | 2.76991 |
| chr7 | 138909257 | 138909981 | 1000 | Promoter (<=1kb) | 7 | 138890836 | 138909506 | 18671 | 2 | ENSMUST00 | 0 | Bnip3 | ENSMUSG0C | false | chr7 | 138908947 | 138910265 | 34.90036 | 108.43528 | 104.64344 | chr7 | 138907509 | 138910513 | 3.84455 | 3.45805 | 2.09793 |
| chr3 | 135437858 | 135439584 | 1000 | Promoter (<=1kb) | 3 | 135438297 | 135466058 | 27762 | 1 | ENSMUST00 | 0 | Ube2d3 | ENSMUSG0C | false | chr3 | 135438324 | 135440830 | 34.16929 | 120.7392 | 116.5701 | . | . | . | . | . | . |
| chr11 | 114198370 | 114198772 | 1000 | Promoter (<=1kb) | 11 | 114198258 | 114199199 | 942 | 1 | ENSMUST00 | 113 | 1700092K14I | ENSMUSG0C | false | . | . | . | . | . | . | chr11 | 114198195 | 114199939 | 6.75303 | 4.88829 | 4.48258 |
| chr7 | 73513940 | 73514418 | 1000 | Exon (ENSMUST00000169922.3/ | 7 | 73501920 | 73519588 | 17669 | 2 | ENSMUST00 | 5170 | Chd2 | ENSMUSG0C | false | . | . | . | . | . | . | chr7 | 73507322 | 73518557 | 5.44869 | 4.30439 | 3.36653 |
| chr11 | 97685461 | 97686057 | 1000 | Promoter (<=1kb) | 11 | 97685826 | 97688621 | 2796 | 1 | ENSMUST00 | 0 | Cisd3 | ENSMUSG0C | false | chr11 | 97685345 | 97686413 | 9.65017 | 18.71579 | 16.39739 | chr11 | 97683689 | 97687456 | 6.46404 | 4.74717 | 4.22185 |
| chr11 | 96953723 | 96954138 | 1000 | 3' UTR | 11 | 96944312 | 96963895 | 19584 | 1 | ENSMUST00 | 9412 | D030028A08 | ENSMUSG0C | false | . | . | . | . | . | . | chr11 | 96951250 | 96960443 | 6.86299 | 4.78743 | 4.54283 |
| chr7 | 27838362 | 27838704 | 1000 | Distal Intergenic | 7 | 27842855 | 27855086 | 12232 | 1 | ENSMUST00 | -4151 | Zfp59 | ENSMUSG0C | false | chr7 | 27837846 | 27839317 | 21.92587 | 57.98755 | 55.16829 | . | . | . | . | . | . |
| chr11 | 69921216 | 69921863 | 1000 | Promoter (<=1kb) | 11 | 69916720 | 69921386 | 4667 | 2 | ENSMUST00 | 0 | Eif5a | ENSMUSG0C | false | chr11 | 69921657 | 69922644 | 19.93088 | 52.12947 | 49.38807 | . | . | . | . | . | . |
| chr2 | 167631492 | 167632429 | 1000 | Promoter (<=1kb) | 2 | 167607638 | 167632095 | 24458 | 2 | ENSMUST00 | 0 | Ube2v1 | ENSMUSG0C | false | chr2 | 167630879 | 167632622 | 37.7199 | 126.00829 | 121.65485 | chr2 | 167632351 | 167635256 | 7.37954 | 5.22271 | 4.9537 |
| chr15 | 96056059 | 96056436 | 1000 | Intron (ENSMUST00000100271.3 | 15 | 96051616 | 96078134 | 26519 | 2 | ENSMUST00 | 21698 | D030018L15I | ENSMUSG0C | false | . | . | . | . | . | . | chr15 | 96054907 | 96057007 | 4.66694 | 3.96012 | 2.72621 |
| chr13 | 97548451 | 97548992 | 1000 | Distal Intergenic | 13 | 97482955 | 97489700 | 6746 | 1 | ENSMUST00 | 65497 | Lincenc1 | ENSMUSG0C | false | . | . | . | . | . | . | chr13 | 97545926 | 97548907 | 3.1867 | 3.12648 | 1.50253 |
| chr13 | 97435303 | 97435576 | 1000 | Promoter (<=1kb) | 13 | 97434970 | 97497007 | 62038 | 1 | ENSMUST00 | 334 | Lincenc1 | ENSMUSG0C | false | chr13 | 97435351 | 97436281 | 9.92602 | 20.74379 | 18.3991 | chr13 | 97434721 | 97436585 | 4.12724 | 3.60969 | 2.32507 |
| chr13 | 97607469 | 97607849 | 1000 | Distal Intergenic | 13 | 97482955 | 97489700 | 6746 | 1 | ENSMUST00 | 124515 | Lincenc1 | ENSMUSG0C | false | . | . | . | . | . | . | chr13 | 97607207 | 97607783 | 3.53264 | 3.22615 | 1.84292 |
| chr15 | 84714707 | 84715130 | 1000 | Distal Intergenic | 15 | 84720052 | 84772207 | 52156 | 1 | ENSMUST00 | -4922 | Arhgap8 | ENSMUSG0C | false | . | . | . | . | . | . | chr15 | 84711388 | 84717566 | 4.51941 | 3.88236 | 2.6032 |
| chr2 | 160507740 | 160508294 | 1000 | Distal Intergenic | 2 | 160583357 | 160585191 | 1835 | 1 | ENSMUST00 | -75063 | Gm14222 | ENSMUSG0C | false | . | . | . | . | . | . | chr2 | 160507144 | 160511138 | 4.2908 | 3.67861 | 2.46178 |
| chr8 | 83996683 | 83998043 | 1000 | Promoter (<=1kb) | 8 | 83997672 | 84000386 | 2715 | 1 | ENSMUST00 | 0 | Samd1 | ENSMUSG0C | false | chr8 | 83996357 | 83998054 | 31.32211 | 93.21193 | 89.80268 | . | . | . | . | . | . |
| chr2 | 131909645 | 131910293 | 1000 | Promoter (<=1kb) | 2 | 131909928 | 131938429 | 28502 | 1 | ENSMUST00 | 0 | Prnp | ENSMUSG0C | false | chr2 | 131909450 | 131910584 | 24.78212 | 78.86476 | 75.73294 | . | . | . | . | . | . |
| chr1 | 193173102 | 193173785 | 1000 | Promoter (<=1kb) | 1 | 193173469 | 193176298 | 2830 | 1 | ENSMUST00 | 0 | A130010J15I | ENSMUSG0C | false | chr1 | 193172724 | 193174122 | 22.64381 | 71.35664 | 68.34631 | . | . | . | . | . | . |
| chr1 | 182986732 | 182987164 | 1000 | Distal Intergenic | 1 | 182954788 | 182976044 | 21257 | 1 | ENSMUST00 | 31945 | Tlr5 | ENSMUSG0C | false | . | . | . | . | . | . | chr1 | 182985930 | 182987446 | 5.4881 | 4.3966 | 3.39416 |
| chr2 | 105097024 | 105097784 | 1000 | Intron (ENSMUST00000172701.2 | 2 | 105092018 | 105093760 | 1743 | 1 | ENSMUST00 | 5007 | Gm11060 | ENSMUSG0C | false | . | . | . | . | . | . | chr2 | 105095065 | 105107198 | 5.21123 | 4.18238 | 3.18288 |
| chr14 | 56667977 | 56668890 | 1000 | Promoter (<=1kb) | 14 | 56668248 | 56697430 | 29183 | 1 | ENSMUST00 | 0 | Mphosph8 | ENSMUSG0C | false | chr14 | 56667534 | 56669410 | 29.31851 | 87.80975 | 84.5099 | . | . | . | . | . | . |
| chr9 | 120010964 | 120011378 | 1000 | Intron (ENSMUST00000177637.1 | 9 | 120013755 | 120023598 | 9844 | 2 | ENSMUST00 | 12220 | Xirp1 | ENSMUSG0C | false | chr9 | 120010981 | 120011832 | 18.77877 | 48.57565 | 45.87926 | chr9 | 120010413 | 120012164 | 5.63569 | 4.41725 | 3.58143 |
| chr9 | 114563935 | 114564758 | 1000 | Promoter (<=1kb) | 9 | 114507133 | 114564369 | 57237 | 2 | ENSMUST00 | 0 | Trim71 | ENSMUSG0C | false | chr9 | 114562275 | 114564649 | 21.59801 | 56.60135 | 53.80005 | chr9 | 114560214 | 114565323 | 5.70594 | 4.35756 | 3.60901 |
| chr2 | 74746446 | 74747419 | 1000 | Promoter (<=1kb) | 2 | 74745722 | 74748442 | 2721 | 1 | ENSMUST00 | 725 | Hoxd3 | ENSMUSG0C | false | chr2 | 74746552 | 74747523 | 15.14328 | 38.56274 | 35.98906 | chr2 | 74744931 | 74751115 | 5.56017 | 4.28612 | 3.49137 |
| chr2 | 74740294 | 74740907 | 1000 | Promoter (<=1kb) | 2 | 74739619 | 74747900 | 8282 | 1 | ENSMUST00 | 676 | Hoxd3 | ENSMUSG0C | false | chr2 | 74738622 | 74741151 | 20.65177 | 61.66082 | 58.78991 | chr2 | 74737574 | 74742233 | 4.86524 | 4.00461 | 2.89696 |
| chr9 | 104101407 | 104101905 | 1000 | Promoter (<=1kb) | 9 | 104084157 | 104102560 | 18404 | 2 | ENSMUST00 | 655 | Ackr4 | ENSMUSG0C | false | chr9 | 104100927 | 104101613 | 6.38327 | 11.47542 | 9.27082 | chr9 | 104100336 | 104102534 | 4.12157 | 3.52496 | 2.34956 |
| chr1 | 82603277 | 82603815 | 1000 | Intron (ENSMUST00000141994.1 | 1 | 82586976 | 82722059 | 135084 | 1 | ENSMUST00 | 16302 | Col4a3 | ENSMUSG0C | true | . | . | . | . | . | . | chr1 | 82589332 | 82610610 | 5.25431 | 4.10973 | 3.23867 |
| chr6 | 88027410 | 88027754 | 1000 | Intron (ENSMUST00000113600.4 | 6 | 87999940 | 88019783 | 19844 | 2 | ENSMUST00 | -7628 | Rab7 | ENSMUSG0C | false | . | . | . | . | . | . | chr6 | 88024679 | 88044328 | 5.33784 | 4.18144 | 3.30124 |
| chrX | 101954147 | 101954577 | 1000 | Intron (ENSMUST00000124279.1 | 20 | 101961797 | 102077881 | 116085 | 1 | ENSMUST00 | -7220 | Nhsl2 | ENSMUSG0C | false | . | . | . | . | . | . | . | . | . | . | . | . |
| chr2 | 29251048 | 29251988 | 1000 | Promoter (1-2kb) | 2 | 29245107 | 29252993 | 7887 | 2 | ENSMUST00 | 1005 | 6530402F18I | ENSMUSG0C | false | chr2 | 29249202 | 29253779 | 38.46334 | 130.56975 | 126.05653 | . | . | . | . | . | . |
| chr2 | 29253547 | 29253861 | 1000 | Promoter (<=1kb) | 2 | 29250910 | 29253006 | 2097 | 2 | ENSMUST00 | -542 | 6530402F18I | ENSMUSG0C | false | chr2 | 29249202 | 29253779 | 38.46334 | 130.56975 | 126.05653 | chr2 | 29253440 | 29256224 | 4.76042 | 3.89351 | 2.82399 |
| chr6 | 73013781 | 73014118 | 1000 | Distal Intergenic | 6 | 72957361 | 72958748 | 1388 | 2 | ENSMUST00 | -55034 | Tmsb10 | ENSMUSG0C | false | . | . | . | . | . | . | chr6 | 73012974 | 73015121 | 4.39161 | 3.74498 | 2.52283 |
| chr6 | 72981182 | 72981775 | 1000 | Distal Intergenic | 6 | 72957361 | 72958748 | 1388 | 2 | ENSMUST00 | -22435 | Tmsb10 | ENSMUSG0C | false | . | . | . | . | . | . | chr6 | 72980327 | 72983165 | 5.84177 | 4.43182 | 3.74919 |
| chr6 | 72957144 | 72957451 | 1000 | Promoter (<=1kb) | 6 | 72957353 | 72958302 | 950 | 2 | ENSMUST00 | 851 | Tmsb10 | ENSMUSG0C | false | chr6 | 72956611 | 72959041 | 38.01312 | 122.22395 | 118.01301 | . | . | . | . | . | . |
| chr17 | 33890655 | 33891098 | 1000 | Promoter (<=1kb) | 17 | 33875666 | 33890661 | 14996 | 2 | ENSMUST00 | 0 | Kifc1 | ENSMUSG0C | false | chr17 | 33890057 | 33890723 | 37.64722 | 120.52259 | 116.3698 | chr17 | 33889700 | 33891136 | 4.70022 | 3.7574 | 2.79528 |
| chr6 | 38662735 | 38663414 | 1000 | Promoter (<=1kb) | 6 | 38663069 | 38680864 | 17796 | 1 | ENSMUST00 | 0 | Clec2l | ENSMUSG0C | false | chr6 | 38662591 | 38663553 | 20.83814 | 54.48703 | 51.7144 | chr6 | 38662835 | 38663614 | 3.38048 | 3.15445 | 1.72072 |
| chr18 | 67774668 | 67775548 | 1000 | Promoter (<=1kb) | 18 | 67774876 | 67795461 | 20586 | 1 | ENSMUST00 | 0 | Seh1l | ENSMUSG0C | false | chr18 | 67774259 | 67776254 | 34.16929 | 120.7392 | 116.5701 | . | . | . | . | . | . |
| chr9 | 117253360 | 117253770 | 1000 | Promoter (<=1kb) | 9 | 117252536 | 117254696 | 2161 | 1 | ENSMUST00 | 825 | Gm17396 | ENSMUSG0C | false | chr9 | 117249955 | 117253836 | 26.19333 | 73.09374 | 70.054 | . | . | . | . | . | . |
| chr17 | 12266301 | 12267097 | 1000 | Intron (ENSMUST00000115691.2 | 17 | 12232481 | 12271184 | 38704 | 1 | ENSMUST00 | 33821 | Map3k4 | ENSMUSG0C | false | . | . | . | . | . | . | chr17 | 12265068 | 12269730 | 6.88449 | 4.92433 | 4.57054 |
| chr8 | 70700480 | 70700896 | 1000 | Promoter (<=1kb) | 8 | 70699538 | 70700080 | 543 | 2 | ENSMUST00 | -401 | Gm11175 | ENSMUSG0C | false | chr8 | 70698975 | 70701534 | 23.70827 | 104.82465 | 101.13002 | chr8 | 70700836 | 70705482 | 4.16546 | 3.64006 | 2.32667 |
| chr8 | 70707200 | 70707697 | 1000 | Distal Intergenic | 8 | 70699538 | 70700080 | 543 | 2 | ENSMUST00 | -7121 | Gm11175 | ENSMUSG0C | false | . | . | . | . | . | . | chr8 | 70706856 | 70708594 | 5.04803 | 4.07593 | 3.06811 |
| chr17 | 17838119 | 17838488 | 1000 | Promoter (<=1kb) | 17 | 17838281 | 17839071 | 791 | 1 | ENSMUST00 | 0 | Spaca6 | ENSMUSG0C | true | chr17 | 17837498 | 17839072 | 14.45496 | 36.65565 | 34.10837 | chr17 | 17828149 | 17841746 | 4.51187 | 3.80845 | 2.61198 |
| chr10 | 96751501 | 96751989 | 1000 | Distal Intergenic | 10 | 96855582 | 96855670 | 89 | 2 | ENSMUST00 | 103681 | Gm24587 | ENSMUSG0C | false | chr10 | 96751517 | 96752077 | 11.94783 | 25.26529 | 22.85895 | chr10 | 96750862 | 96753805 | 4.41434 | 3.79422 | 2.55352 |
| chr9 | 115355903 | 115356312 | 1000 | Distal Intergenic | 9 | 115381819 | 115381939 | 121 | 1 | ENSMUST00 | -25507 | Mir467h | ENSMUSG0C | false | . | . | . | . | . | . | chr9 | 115354829 | 115357022 | 4.77352 | 3.89621 | 2.86366 |
| chr3 | 85482550 | 85483043 | 1000 | Intron (ENSMUST00000154148.2 | 3 | 85467377 | 85467498 | 122 | 1 | ENSMUST00 | 15174 | Mir466k | ENSMUSG0C | false | . | . | . | . | . | . | chr3 | 85482189 | 85483445 | 3.66723 | 3.34411 | 1.93732 |
| chr15 | 37568714 | 37569437 | 1000 | Intron (ENSMUST00000168992.2 | 15 | 37603415 | 37603534 | 120 | 2 | ENSMUST00 | 34097 | Gm24833 | ENSMUSG0C | false | . | . | . | . | . | . | chr15 | 37567895 | 37570984 | 4.89045 | 3.96584 | 2.91914 |
| chr11 | 64544268 | 64544644 | 1000 | Distal Intergenic | 11 | 64590495 | 64590640 | 146 | 2 | ENSMUST00 | 45996 | Gm24275 | ENSMUSG0C | false | . | . | . | . | . | . | chr11 | 64539605 | 64548358 | 5.34347 | 4.23728 | 3.29791 |
| chr18 | 35921409 | 35921827 | 1000 | Distal Intergenic | 18 | 35952824 | 35952907 | 84 | 1 | ENSMUST00 | -30997 | Gm23882 | ENSMUSG0C | false | . | . | . | . | . | . | chr18 | 35918346 | 35921757 | 4.57948 | 3.8339 | 2.68759 |
| chr18 | 35927170 | 35927570 | 1000 | Distal Intergenic | 18 | 35952824 | 35952907 | 84 | 1 | ENSMUST00 | -25254 | Gm23882 | ENSMUSG0C | false | . | . | . | . | . | . | chr18 | 35926400 | 35928452 | 5.67483 | 4.24485 | 3.58221 |
| chr18 | 22393817 | 22394338 | 1000 | Promoter (1-2kb) | 18 | 22395371 | 22395458 | 88 | 1 | ENSMUST00 | -1033 | Gm24483 | ENSMUSG0C | false | . | . | . | . | . | . | chr18 | 22392200 | 22396748 | 4.04234 | 3.56032 | 2.23926 |
| chr13 | 60455096 | 60455503 | 1000 | Distal Intergenic | 13 | 60529296 | 60529409 | 114 | 2 | ENSMUST00 | 73906 | Gm25000 | ENSMUSG0C | false | . | . | . | . | . | . | chr13 | 60450423 | 60457848 | 4.93919 | 4.02175 | 2.96511 |
| chr13 | 60488591 | 60488970 | 1000 | Distal Intergenic | 13 | 60529296 | 60529409 | 114 | 2 | ENSMUST00 | 40439 | Gm25000 | ENSMUSG0C | false | . | . | . | . | . | . | chr13 | 60487671 | 60489300 | 4.26373 | 3.37698 | 2.43276 |
| chr13 | 60343498 | 60343828 | 1000 | Distal Intergenic | 13 | 60293672 | 60293772 | 101 | 1 | ENSMUST00 | 49827 | Gm24999 | ENSMUSG0C | false | . | . | . | . | . | . | chr13 | 60338549 | 60344628 | 4.38308 | 3.79503 | 2.48565 |
| chr13 | 60267524 | 60268065 | 1000 | Distal Intergenic | 13 | 60293672 | 60293772 | 101 | 1 | ENSMUST00 | -25607 | Gm24999 | ENSMUSG0C | false | . | . | . | . | . | . | chr13 | 60265916 | 60273445 | 5.63968 | 4.31967 | 3.54273 |
| chr13 | 60332568 | 60332959 | 1000 | Distal Intergenic | 13 | 60293672 | 60293772 | 101 | 1 | ENSMUST00 | 38897 | Gm24999 | ENSMUSG0C | false | . | . | . | . | . | . | chr13 | 60331876 | 60337138 | 4.59063 | 3.63495 | 2.69691 |
| chr5 | 74315771 | 74316243 | 1000 | Promoter (<=1kb) | 5 | 74316566 | 74317029 | 464 | 2 | ENSMUST00 | 786 | Gm15981 | ENSMUSG0C | false | . | . | . | . | . | . | chr5 | 74314743 | 74317419 | 5.46294 | 4.35609 | 3.38771 |
| chr4 | 76386972 | 76387462 | 1000 | Intron (ENSMUST00000107289.3 | 4 | 76393154 | 76394092 | 939 | 1 | ENSMUST00 | -5692 | Gm11252 | ENSMUSG0C | false | . | . | . | . | . | . | chr4 | 76377583 | 76403972 | 4.17482 | 3.66873 | 2.32976 |
| chr1 | 60960481 | 60960903 | 1000 | Distal Intergenic | 1 | 60948230 | 60948465 | 236 | 2 | ENSMUST00 | -12017 | Gm11582 | ENSMUSG0C | false | . | . | . | . | . | . | . | . | . | . | . | . |
| chrX | 11664341 | 11665105 | 1000 | Distal Intergenic | 20 | 11684002 | 11684124 | 123 | 2 | ENSMUST00 | 19019 | Gm14513 | ENSMUSG0C | false | chrX | 11664285 | 11665249 | 14.2455 | 32.22549 | 29.73087 | . | . | . | . | . | . |
| chrX | 11703650 | 11703978 | 1000 | Distal Intergenic | 20 | 11684002 | 11684124 | 123 | 2 | ENSMUST00 | -19527 | Gm14513 | ENSMUSG0C | false | . | . | . | . | . | . | . | . | . | . | . | . |
| chr4 | 59528321 | 59528811 | 1000 | Intron (ENSMUST00000030076.6 | 4 | 59532747 | 59533014 | 268 | 1 | ENSMUST00 | -3936 | Gm12529 | ENSMUSG0C | false | . | . | . | . | . | . | chr4 | 59518606 | 59536295 | 5.04232 | 4.08706 | 3.03874 |
| chr5 | 123057824 | 123058351 | 1000 | Intron (ENSMUST00000181022.1 | 5 | 123066242 | 123066699 | 458 | 1 | ENSMUST00 | -7891 | Gm6444 | ENSMUSG0C | false | chr5 | 123058082 | 123058657 | 12.24318 | 32.0257 | 29.53485 | chr5 | 123057396 | 123058755 | 3.98558 | 3.63204 | 2.16269 |
| chr9 | 116091179 | 116091596 | 1000 | Intron (ENSMUST00000035014.6 | 9 | 116040593 | 116041064 | 472 | 2 | ENSMUST00 | -50116 | Gm9385 | ENSMUSG0C | false | . | . | . | . | . | . | chr9 | 116087970 | 116094271 | 5.91845 | 4.43298 | 3.76556 |
| chr1 | 152454585 | 152454995 | 1000 | Intron (ENSMUST00000127586.2 | 1 | 152407074 | 152407525 | 452 | 2 | ENSMUST00 | -47061 | Gm15618 | ENSMUSG0C | false | . | . | . | . | . | . | chr1 | 152452441 | 152462754 | 5.57071 | 4.29497 | 3.48344 |
| chrX | 94562338 | 94562695 | 1000 | Distal Intergenic | 20 | 94557341 | 94557749 | 409 | 1 | ENSMUST00 | 4998 | Gm14831 | ENSMUSG0C | false | . | . | . | . | . | . | . | . | . | . | . | . |
| chr5 | 130194228 | 130194600 | 1000 | Intron (ENSMUST00000119797.2 | 5 | 130188395 | 130188796 | 402 | 1 | ENSMUST00 | 5834 | Gm15920 | ENSMUSG0C | false | . | . | . | . | . | . | chr5 | 130172335 | 130207681 | 5.43982 | 4.22227 | 3.37739 |
| chrX | 11254166 | 11254935 | 1000 | Distal Intergenic | 20 | 11220607 | 11221235 | 629 | 1 | ENSMUST00 | 33560 | Gm14481 | ENSMUSG0C | false | . | . | . | . | . | . | . | . | . | . | . | . |
| chr4 | 100307324 | 100307785 | 1000 | Intron (ENSMUST00000039630.5 | 4 | 100361362 | 100362059 | 698 | 2 | ENSMUST00 | 54274 | Gm12706 | ENSMUSG0C | false | . | . | . | . | . | . | chr4 | 100306284 | 100309516 | 5.31272 | 4.02354 | 3.31768 |
| chr4 | 100350017 | 100350589 | 1000 | Intron (ENSMUST00000039630.5 | 4 | 100361362 | 100362059 | 698 | 2 | ENSMUST00 | 11470 | Gm12706 | ENSMUSG0C | false | . | . | . | . | . | . | chr4 | 100347301 | 100354535 | 6.63157 | 4.80816 | 4.33798 |
| chr2 | 152180315 | 152180824 | 1000 | Distal Intergenic | 2 | 152185291 | 152185682 | 392 | 2 | ENSMUST00 | 4858 | Rps15a-ps7 | ENSMUSG0C | false | . | . | . | . | . | . | chr2 | 152179241 | 152181916 | 4.23696 | 3.65342 | 2.4024 |
| chr14 | 13399475 | 13399970 | 1000 | Intron (ENSMUST00000070323.6 | 14 | 13353876 | 13354007 | 132 | 2 | ENSMUST00 | -45469 | Gm15913 | ENSMUSG0C | false | . | . | . | . | . | . | chr14 | 13398464 | 13400493 | 4.67426 | 3.92704 | 2.77797 |
| chr4 | 135404024 | 135404388 | 1000 | Promoter (<=1kb) | 4 | 135403189 | 135403656 | 468 | 2 | ENSMUST00 | -369 | Gm12990 | ENSMUSG0C | false | . | . | . | . | . | . | chr4 | 135402165 | 135406081 | 4.50505 | 3.80454 | 2.62652 |
| chr2 | 59534391 | 59534757 | 1000 | Distal Intergenic | 2 | 59564268 | 59564684 | 417 | 2 | ENSMUST00 | 29927 | Gm13552 | ENSMUSG0C | false | . | . | . | . | . | . | chr2 | 59533698 | 59534682 | 5.30634 | 3.92121 | 3.29777 |
| chr2 | 59526893 | 59527345 | 1000 | Distal Intergenic | 2 | 59564268 | 59564684 | 417 | 2 | ENSMUST00 | 37339 | Gm13552 | ENSMUSG0C | false | . | . | . | . | . | . | chr2 | 59526252 | 59528488 | 4.45107 | 3.73925 | 2.59228 |
| chr5 | 115497059 | 115497375 | 1000 | Distal Intergenic | 5 | 115491891 | 115492455 | 565 | 2 | ENSMUST00 | -4605 | Gm13836 | ENSMUSG0C | false | . | . | . | . | . | . | chr5 | 115495856 | 115500926 | 5.04255 | 4.05763 | 3.05495 |
| chr4 | 40884692 | 40885221 | 1000 | Distal Intergenic | 4 | 40912741 | 40913355 | 615 | 2 | ENSMUST00 | 28134 | Gm12397 | ENSMUSG0C | false | . | . | . | . | . | . | chr4 | 40882325 | 40890611 | 4.88004 | 4.00478 | 2.90476 |
| chr2 | 93759136 | 93759528 | 1000 | Intron (ENSMUST00000184931.2 | 2 | 93794570 | 93795080 | 511 | 1 | ENSMUST00 | -35042 | Gm13818 | ENSMUSG0C | false | . | . | . | . | . | . | chr2 | 93757685 | 93760236 | 4.90066 | 3.87491 | 2.97166 |
| chrX | 77720884 | 77721201 | 1000 | Promoter (2-3kb) | 20 | 77717052 | 77717983 | 932 | 2 | ENSMUST00 | -2902 | Gm14746 | ENSMUSG0C | false | . | . | . | . | . | . | . | . | . | . | . | . |
| chr7 | 133747343 | 133747718 | 1000 | Intron (ENSMUST00000033290.6 | 7 | 133738479 | 133738934 | 456 | 2 | ENSMUST00 | -8410 | Gm15483 | ENSMUSG0C | false | . | . | . | . | . | . | chr7 | 133743183 | 133753176 | 5.21662 | 4.08927 | 3.21065 |
| chr2 | 167594395 | 167594884 | 1000 | Intron (ENSMUST00000127006.1 | 2 | 167591395 | 167591870 | 476 | 1 | ENSMUST00 | 3001 | Gm11475 | ENSMUSG0C | false | . | . | . | . | . | . | chr2 | 167594068 | 167598191 | 6.05431 | 4.56249 | 3.89098 |
| chr15 | 97781923 | 97782981 | 1000 | Promoter (1-2kb) | 15 | 97784355 | 97792692 | 8338 | 1 | ENSMUST00 | -1374 | Slc48a1 | ENSMUSG0C | false | chr15 | 97782222 | 97782979 | 8.94881 | 17.06333 | 14.77052 | chr15 | 97771183 | 97790304 | 7.43452 | 5.14328 | 5.01036 |
| chr15 | 97778571 | 97779129 | 1000 | Distal Intergenic | 15 | 97784355 | 97792692 | 8338 | 1 | ENSMUST00 | -5226 | Slc48a1 | ENSMUSG0C | false | . | . | . | . | . | . | chr15 | 97771183 | 97790304 | 7.43452 | 5.14328 | 5.01036 |
| chr6 | 29744046 | 29744405 | 1000 | Intron (ENSMUST00000001812.4 | 6 | 29750761 | 29751125 | 365 | 1 | ENSMUST00 | -6356 | Gm13717 | ENSMUSG0C | false | . | . | . | . | . | . | chr6 | 29736983 | 29747483 | 4.58825 | 3.76848 | 2.68875 |
| chr11 | 63287556 | 63288188 | 1000 | Distal Intergenic | 11 | 63312310 | 63313311 | 1002 | 2 | ENSMUST00 | 25123 | Gm12286 | ENSMUSG0C | false | . | . | . | . | . | . | chr11 | 63285912 | 63290836 | 3.92834 | 3.47453 | 2.13204 |
| chr11 | 63293515 | 63293994 | 1000 | Distal Intergenic | 11 | 63312310 | 63313311 | 1002 | 2 | ENSMUST00 | 19317 | Gm12286 | ENSMUSG0C | false | . | . | . | . | . | . | chr11 | 63292091 | 63294920 | 5.05367 | 4.14002 | 3.04222 |
| chr4 | 149798710 | 149799021 | 1000 | Distal Intergenic | 4 | 149793279 | 149793419 | 141 | 2 | ENSMUST00 | -5292 | Gm13065 | ENSMUSG0C | false | . | . | . | . | . | . | chr4 | 149797276 | 149802576 | 5.06815 | 4.07327 | 3.06703 |
| chrX | 77409417 | 77409842 | 1000 | Distal Intergenic | 20 | 77456586 | 77457228 | 643 | 1 | ENSMUST00 | -46744 | Gm5937 | ENSMUSG0C | false | . | . | . | . | . | . | . | . | . | . | . | . |
| chrX | 77433251 | 77433789 | 1000 | Distal Intergenic | 20 | 77456586 | 77457228 | 643 | 1 | ENSMUST00 | -22797 | Gm5937 | ENSMUSG0C | false | . | . | . | . | . | . | . | . | . | . | . | . |
| chr2 | 132676838 | 132677181 | 1000 | Promoter (1-2kb) | 2 | 132678269 | 132679239 | 971 | 1 | ENSMUST00 | -1088 | Gm14098 | ENSMUSG0C | false | . | . | . | . | . | . | chr2 | 132676150 | 132677332 | 4.67439 | 3.93697 | 2.77131 |
| chr11 | 80817933 | 80818485 | 1000 | Intron (ENSMUST00000040865.8 | 11 | 80822780 | 80823193 | 414 | 2 | ENSMUST00 | 4708 | Gm11207 | ENSMUSG0C | false | . | . | . | . | . | . | chr11 | 80813757 | 80819265 | 5.24947 | 4.17377 | 3.2186 |
| chr2 | 103554022 | 103554551 | 1000 | Distal Intergenic | 2 | 103547259 | 103547688 | 430 | 1 | ENSMUST00 | 6764 | Gm13996 | ENSMUSG0C | false | . | . | . | . | . | . | chr2 | 103552142 | 103556696 | 5.10403 | 4.16116 | 3.08654 |
| chrX | 77050896 | 77051401 | 1000 | Distal Intergenic | 20 | 76980706 | 76981144 | 439 | 1 | ENSMUST00 | 70191 | Gm14714 | ENSMUSG0C | false | . | . | . | . | . | . | . | . | . | . | . | . |
| chrX | 142516312 | 142516737 | 1000 | Distal Intergenic | 20 | 142533439 | 142533870 | 432 | 2 | ENSMUST00 | 17133 | Gm4995 | ENSMUSG0C | false | . | . | . | . | . | . | . | . | . | . | . | . |
| chr11 | 64711700 | 64712121 | 1000 | Distal Intergenic | 11 | 64759717 | 64760027 | 311 | 2 | ENSMUST00 | 47906 | Gm12291 | ENSMUSG0C | false | . | . | . | . | . | . | chr11 | 64706948 | 64713410 | 4.30838 | 3.74761 | 2.44128 |
| chr17 | 5189621 | 5190365 | 1000 | Intron (ENSMUST00000092723.5 | 17 | 5110781 | 5112119 | 1339 | 2 | ENSMUST00 | -77503 | Gm15599 | ENSMUSG0C | false | chr17 | 5189499 | 5190676 | 15.70458 | 43.19576 | 40.56621 | chr17 | 5188721 | 5199127 | 4.36014 | 3.6534 | 2.51291 |
| chr2 | 83742845 | 83743161 | 1000 | Promoter (2-3kb) | 2 | 83744484 | 83745736 | 1253 | 2 | ENSMUST00 | 2575 | Dnmt3l-ps1 | ENSMUSG0C | false | . | . | . | . | . | . | chr2 | 83738044 | 83745203 | 5.80522 | 4.36226 | 3.6965 |
| chr2 | 83739716 | 83740200 | 1000 | Exon (ENSMUST00000155521.1/ | 2 | 83744484 | 83745736 | 1253 | 2 | ENSMUST00 | 5536 | Dnmt3l-ps1 | ENSMUSG0C | false | . | . | . | . | . | . | chr2 | 83738044 | 83745203 | 5.80522 | 4.36226 | 3.6965 |
| chr11 | 19851196 | 19851672 | 1000 | Distal Intergenic | 11 | 19783652 | 19784483 | 832 | 1 | ENSMUST00 | 67545 | Gm12029 | ENSMUSG0C | false | . | . | . | . | . | . | chr11 | 19850150 | 19851311 | 6.16042 | 4.47517 | 4.00729 |
| chr3 | 137894496 | 137894800 | 1000 | Exon (ENSMUST00000125226.1/ | 3 | 137905009 | 137905399 | 391 | 1 | ENSMUST00 | -10209 | Gm15460 | ENSMUSG0C | false | . | . | . | . | . | . | chr3 | 137894420 | 137896622 | 3.56215 | 3.36562 | 1.87492 |
| chr2 | 117240645 | 117240990 | 1000 | Promoter (<=1kb) | 2 | 117240883 | 117241441 | 559 | 1 | ENSMUST00 | 0 | Rap1a-ps2 | ENSMUSG0C | false | . | . | . | . | . | . | chr2 | 117240633 | 117241558 | 2.82635 | 2.79289 | 1.27109 |
| chr7 | 34485330 | 34486036 | 1000 | Intron (ENSMUST00000140298.1 | 7 | 34495421 | 34496065 | 645 | 2 | ENSMUST00 | 10029 | Gm12780 | ENSMUSG0C | false | . | . | . | . | . | . | . | . | . | . | . | . |
| chrX | 11998935 | 11999455 | 1000 | Distal Intergenic | 20 | 11984475 | 11984650 | 176 | 1 | ENSMUST00 | 14461 | Gm14512 | ENSMUSG0C | false | chrX | 11998979 | 11999498 | 5.44843 | 8.50832 | 6.36429 | . | . | . | . | . | . |
| chr11 | 48176519 | 48176857 | 1000 | Distal Intergenic | 11 | 48289984 | 48290313 | 330 | 2 | ENSMUST00 | 113456 | Gm12178 | ENSMUSG0C | false | . | . | . | . | . | . | chr11 | 48175810 | 48177199 | 3.40717 | 3.22297 | 1.72733 |
| chr11 | 107292028 | 107292483 | 1000 | Intron (ENSMUST00000040380.7 | 11 | 107330708 | 107331058 | 351 | 1 | ENSMUST00 | -38225 | Gm11719 | ENSMUSG0C | false | . | . | . | . | . | . | chr11 | 107290753 | 107293209 | 4.48871 | 3.71724 | 2.62207 |
| chr11 | 78600176 | 78600628 | 1000 | Intron (ENSMUST00000142739.2 | 11 | 78585161 | 78585555 | 395 | 2 | ENSMUST00 | -14622 | Gm11196 | ENSMUSG0C | false | . | . | . | . | . | . | chr11 | 78596906 | 78604203 | 5.14453 | 4.16406 | 3.12824 |
| chr13 | 25639949 | 25640325 | 1000 | Distal Intergenic | 13 | 25680274 | 25681143 | 870 | 2 | ENSMUST00 | 40818 | Gm11350 | ENSMUSG0C | false | . | . | . | . | . | . | chr13 | 25639402 | 25641908 | 3.88223 | 3.53077 | 2.06878 |
| chrX | 60547417 | 60548139 | 1000 | Promoter (<=1kb) | 20 | 60548019 | 60548849 | 831 | 1 | ENSMUST00 | 0 | Gm715 | ENSMUSG0C | false | chrX | 60547102 | 60548340 | 16.54316 | 39.53096 | 36.94807 | . | . | . | . | . | . |
| chr2 | 153418623 | 153418935 | 1000 | Intron (ENSMUST00000035346.8 | 2 | 153413849 | 153414130 | 282 | 1 | ENSMUST00 | 4775 | Gm14472 | ENSMUSG0C | false | . | . | . | . | . | . | chr2 | 153415054 | 153443585 | 6.91758 | 4.79199 | 4.58849 |
| chr2 | 11429596 | 11430002 | 1000 | Distal Intergenic | 2 | 11420222 | 11421236 | 1015 | 1 | ENSMUST00 | 9375 | Gm13296 | ENSMUSG0C | false | . | . | . | . | . | . | chr2 | 11423400 | 11433608 | 4.58291 | 3.77807 | 2.6805 |
| chr2 | 11425765 | 11426324 | 1000 | Distal Intergenic | 2 | 11420222 | 11421236 | 1015 | 1 | ENSMUST00 | 5544 | Gm13296 | ENSMUSG0C | false | . | . | . | . | . | . | chr2 | 11423400 | 11433608 | 4.58291 | 3.77807 | 2.6805 |
| chr13 | 111894866 | 111895191 | 1000 | Distal Intergenic | 13 | 111882393 | 111884177 | 1785 | 2 | ENSMUST00 | -10690 | Gm9025 | ENSMUSG0C | false | . | . | . | . | . | . | . | . | . | . | . | . |
| chr4 | 129384335 | 129384803 | 1000 | Distal Intergenic | 4 | 129379844 | 129380179 | 336 | 1 | ENSMUST00 | 4492 | Gm12980 | ENSMUSG0C | false | . | . | . | . | . | . | chr4 | 129378069 | 129387044 | 4.79047 | 3.93835 | 2.85504 |
| chr4 | 129382423 | 129382915 | 1000 | Promoter (2-3kb) | 4 | 129379844 | 129380179 | 336 | 1 | ENSMUST00 | 2580 | Gm12980 | ENSMUSG0C | false | chr4 | 129382531 | 129383931 | 10.65138 | 21.96196 | 19.60116 | chr4 | 129378069 | 129387044 | 4.79047 | 3.93835 | 2.85504 |
| chr11 | 108419958 | 108420655 | 1000 | Promoter (2-3kb) | 11 | 108423273 | 108423758 | 486 | 1 | ENSMUST00 | -2618 | Gm11667 | ENSMUSG0C | false | . | . | . | . | . | . | . | . | . | . | . | . |
| chr13 | 30213315 | 30213660 | 1000 | Intron (ENSMUST00000047311.9 | 13 | 30181716 | 30182144 | 429 | 1 | ENSMUST00 | 31600 | Gm11366 | ENSMUSG0C | false | . | . | . | . | . | . | chr13 | 30212223 | 30214525 | 3.52086 | 3.26537 | 1.82369 |
| chr16 | 11526956 | 11527401 | 1000 | Intron (ENSMUST00000180792.2 | 16 | 11494821 | 11496306 | 1486 | 2 | ENSMUST00 | -30651 | Gm15897 | ENSMUSG0C | false | . | . | . | . | . | . | chr16 | 11526093 | 11533919 | 5.11794 | 4.15474 | 3.11474 |
| chr2 | 128512786 | 128513137 | 1000 | Distal Intergenic | 2 | 128557254 | 128558234 | 981 | 2 | ENSMUST00 | 45097 | Gm14006 | ENSMUSG0C | false | . | . | . | . | . | . | chr2 | 128511225 | 128516189 | 4.79079 | 3.90535 | 2.87325 |
| chr5 | 35921620 | 35922800 | 1000 | Intron (ENSMUST00000064571.5 | 5 | 35917897 | 35918201 | 305 | 1 | ENSMUST00 | 3724 | H2af-ps | ENSMUSG0C | false | . | . | . | . | . | . | chr5 | 35902761 | 35945832 | 6.32118 | 4.58806 | 4.1083 |
| chr5 | 35973704 | 35974260 | 1000 | Intron (ENSMUST00000064571.5 | 5 | 35917897 | 35918201 | 305 | 1 | ENSMUST00 | 55808 | H2af-ps | ENSMUSG0C | false | . | . | . | . | . | . | chr5 | 35969643 | 35976391 | 7.51744 | 5.22127 | 5.05303 |
| chr5 | 35924020 | 35924417 | 1000 | Intron (ENSMUST00000064571.5 | 5 | 35917897 | 35918201 | 305 | 1 | ENSMUST00 | 6124 | H2af-ps | ENSMUSG0C | false | . | . | . | . | . | . | chr5 | 35902761 | 35945832 | 6.32118 | 4.58806 | 4.1083 |
| chr4 | 106092307 | 106092776 | 1000 | Intron (ENSMUST00000130088.1 | 4 | 106098924 | 106099282 | 359 | 1 | ENSMUST00 | -6148 | Gm12727 | ENSMUSG0C | false | . | . | . | . | . | . | chr4 | 106089154 | 106096006 | 4.26363 | 3.67795 | 2.43447 |
| chr7 | 96408468 | 96408837 | 1000 | Intron (ENSMUST00000107162.2 | 7 | 96435546 | 96435999 | 454 | 2 | ENSMUST00 | 27162 | Rps11-ps5 | ENSMUSG0C | false | . | . | . | . | . | . | . | . | . | . | . | . |
| chr11 | 44349849 | 44350262 | 1000 | Distal Intergenic | 11 | 44387115 | 44387274 | 160 | 2 | ENSMUST00 | 37012 | Gm12155 | ENSMUSG0C | false | . | . | . | . | . | . | chr11 | 44347620 | 44351377 | 4.90686 | 4.08367 | 2.93913 |
| chr2 | 68805771 | 68806227 | 1000 | Distal Intergenic | 2 | 68831017 | 68831406 | 390 | 1 | ENSMUST00 | -24790 | Gm13612 | ENSMUSG0C | false | . | . | . | . | . | . | chr2 | 68805653 | 68807102 | 4.99964 | 4.10695 | 3.02683 |
| chr2 | 68815537 | 68815847 | 1000 | Distal Intergenic | 2 | 68831017 | 68831406 | 390 | 1 | ENSMUST00 | -15170 | Gm13612 | ENSMUSG0C | false | . | . | . | . | . | . | chr2 | 68814340 | 68815754 | 5.82294 | 4.41485 | 3.70889 |
| chrX | 77734099 | 77734637 | 1000 | Promoter (<=1kb) | 20 | 77733668 | 77737745 | 4078 | 1 | ENSMUST00 | 432 | Gm14748 | ENSMUSG0C | false | . | . | . | . | . | . | . | . | . | . | . | . |
| chr10 | 120920345 | 120920832 | 1000 | Downstream (2-3kb) | 10 | 120928186 | 120928972 | 787 | 1 | ENSMUST00 | -7354 | Gm16166 | ENSMUSG0C | false | . | . | . | . | . | . | chr10 | 120919810 | 120921186 | 3.61558 | 3.30238 | 1.90575 |
| chr2 | 8877114 | 8877427 | 1000 | Distal Intergenic | 2 | 9041124 | 9042484 | 1361 | 2 | ENSMUST00 | 165057 | Gm13217 | ENSMUSG0C | false | . | . | . | . | . | . | . | . | . | . | . | . |
| chr2 | 145888322 | 145888861 | 1000 | Distal Intergenic | 2 | 145892082 | 145892704 | 623 | 1 | ENSMUST00 | -3221 | Gm11764 | ENSMUSG0C | false | . | . | . | . | . | . | chr2 | 145866961 | 145893931 | 5.63035 | 4.25092 | 3.54043 |
| chr4 | 66202146 | 66202517 | 1000 | Intron (ENSMUST00000084496.2 | 4 | 66353849 | 66354186 | 338 | 1 | ENSMUST00 | -151332 | Gm11484 | ENSMUSG0C | false | . | . | . | . | . | . | chr4 | 66199366 | 66203835 | 4.35757 | 3.74325 | 2.47897 |
| chr11 | 63785315 | 63785800 | 1000 | Distal Intergenic | 11 | 63817019 | 63818300 | 1282 | 2 | ENSMUST00 | 32500 | Gm12288 | ENSMUSG0C | false | . | . | . | . | . | . | chr11 | 63784346 | 63787307 | 5.19423 | 4.00117 | 3.18789 |
| chr11 | 63775355 | 63775827 | 1000 | Distal Intergenic | 11 | 63817019 | 63818300 | 1282 | 2 | ENSMUST00 | 42473 | Gm12288 | ENSMUSG0C | false | . | . | . | . | . | . | chr11 | 63774004 | 63780566 | 4.55324 | 3.87984 | 2.63947 |
| chr2 | 61138747 | 61139563 | 1000 | Distal Intergenic | 2 | 61224792 | 61225082 | 291 | 1 | ENSMUST00 | -85229 | Gm13581 | ENSMUSG0C | false | . | . | . | . | . | . | chr2 | 61136622 | 61140725 | 5.9439 | 4.56787 | 3.77059 |
| chrX | 109137359 | 109137671 | 1000 | Intron (ENSMUST00000033598.8 | 20 | 109120801 | 109121684 | 884 | 2 | ENSMUST00 | -15676 | Gm14869 | ENSMUSG0C | false | . | . | . | . | . | . | . | . | . | . | . | . |
| chr16 | 90891706 | 90892079 | 1000 | Intron (ENSMUST00000099548.3 | 16 | 90915180 | 90915731 | 552 | 1 | ENSMUST00 | -23101 | Gm7831 | ENSMUSG0C | false | . | . | . | . | . | . | . | . | . | . | . | . |
| chr4 | 133292903 | 133293971 | 1000 | 3' UTR | 4 | 133280857 | 133281312 | 456 | 2 | ENSMUST00 | -11592 | Gm13259 | ENSMUSG0C | false | chr4 | 133293289 | 133294010 | 13.51752 | 34.31881 | 31.79871 | chr4 | 133292100 | 133300204 | 5.11987 | 4.07043 | 3.13309 |
| chr4 | 14775964 | 14776542 | 1000 | Intron (ENSMUST00000023917.2 | 4 | 14766335 | 14766565 | 231 | 1 | ENSMUST00 | 9630 | Gm11838 | ENSMUSG0C | false | . | . | . | . | . | . | chr4 | 14772277 | 14777374 | 3.67793 | 3.23823 | 1.95792 |
| chr2 | 61100853 | 61101290 | 1000 | Distal Intergenic | 2 | 61030351 | 61030970 | 620 | 2 | ENSMUST00 | -69884 | Gm13579 | ENSMUSG0C | false | . | . | . | . | . | . | chr2 | 61099341 | 61105801 | 4.0096 | 3.59734 | 2.20741 |
| chr2 | 61110616 | 61110978 | 1000 | Distal Intergenic | 2 | 61030351 | 61030970 | 620 | 2 | ENSMUST00 | -79647 | Gm13579 | ENSMUSG0C | false | . | . | . | . | . | . | chr2 | 61107692 | 61115144 | 4.44587 | 3.7065 | 2.56286 |
| chr2 | 61058565 | 61059273 | 1000 | Distal Intergenic | 2 | 61030351 | 61030970 | 620 | 2 | ENSMUST00 | -27596 | Gm13579 | ENSMUSG0C | false | . | . | . | . | . | . | chr2 | 61057908 | 61060199 | 4.90915 | 3.96028 | 2.94481 |
| chr16 | 37881139 | 37881503 | 1000 | Promoter (1-2kb) | 16 | 37882542 | 37883631 | 1090 | 1 | ENSMUST00 | -1039 | Gm15725 | ENSMUSG0C | false | . | . | . | . | . | . | chr16 | 37870054 | 37882292 | 5.93789 | 4.46579 | 3.79659 |
| chr2 | 159646568 | 159646945 | 1000 | Distal Intergenic | 2 | 159777676 | 159778207 | 532 | 2 | ENSMUST00 | 131262 | Gm11445 | ENSMUSG0C | false | . | . | . | . | . | . | chr2 | 159645432 | 159647932 | 4.5832 | 3.86829 | 2.65457 |
| chr2 | 105026985 | 105027830 | 1000 | Distal Intergenic | 2 | 105031866 | 105032287 | 422 | 2 | ENSMUST00 | 4457 | Gm13888 | ENSMUSG0C | false | . | . | . | . | . | . | chr2 | 105020279 | 105029356 | 4.59547 | 3.90746 | 2.6737 |
| chr2 | 45514817 | 45515200 | 1000 | Distal Intergenic | 2 | 45647191 | 45647603 | 413 | 1 | ENSMUST00 | -131991 | Gm13478 | ENSMUSG0C | false | . | . | . | . | . | . | chr2 | 45514704 | 45515266 | 3.83094 | 3.51925 | 2.05506 |
| chr3 | 57496598 | 57496887 | 1000 | Intron (ENSMUST00000029380.8 | 3 | 57528232 | 57528781 | 550 | 1 | ENSMUST00 | -31345 | Gm16016 | ENSMUSG0C | false | . | . | . | . | . | . | chr3 | 57491647 | 57498697 | 4.51463 | 3.80006 | 2.60787 |
| chr6 | 12892966 | 12893636 | 1000 | Distal Intergenic | 6 | 12857397 | 12858390 | 994 | 1 | ENSMUST00 | 35570 | Gm15529 | ENSMUSG0C | false | . | . | . | . | . | . | chr6 | 12890342 | 12893131 | 3.6939 | 3.43811 | 1.91827 |
| chr4 | 123250099 | 123250610 | 1000 | Distal Intergenic | 4 | 123258046 | 123258904 | 859 | 1 | ENSMUST00 | -7436 | Gm12901 | ENSMUSG0C | false | . | . | . | . | . | . | chr4 | 123249475 | 123251134 | 4.27297 | 3.70879 | 2.42369 |
| chrX | 12950918 | 12951608 | 1000 | Distal Intergenic | 20 | 12944610 | 12944966 | 357 | 2 | ENSMUST00 | -5953 | Gm14534 | ENSMUSG0C | false | . | . | . | . | . | . | . | . | . | . | . | . |
| chr2 | 143310309 | 143310659 | 1000 | Distal Intergenic | 2 | 143349512 | 143349661 | 150 | 1 | ENSMUST00 | -38853 | Gm11644 | ENSMUSG0C | false | . | . | . | . | . | . | chr2 | 143310448 | 143311251 | 2.79364 | 2.79821 | 1.23543 |
| chr10 | 80961014 | 80961689 | 1000 | Intron (ENSMUST00000099462.2 | 10 | 80954454 | 80954710 | 257 | 2 | ENSMUST00 | -6305 | Gm3828 | ENSMUSG0C | false | . | . | . | . | . | . | chr10 | 80960060 | 80966904 | 7.38858 | 5.12986 | 4.98152 |
| chr4 | 143161670 | 143162418 | 1000 | Exon (ENSMUST00000105778.2/ | 4 | 143166603 | 143167396 | 794 | 1 | ENSMUST00 | -4185 | Gm13039 | ENSMUSG0C | false | chr4 | 143160943 | 143162268 | 24.82645 | 85.46523 | 82.21117 | chr4 | 143150104 | 143163160 | 5.00338 | 4.03286 | 3.02545 |
| chr2 | 159356564 | 159357068 | 1000 | Distal Intergenic | 2 | 159456170 | 159456457 | 288 | 2 | ENSMUST00 | 99389 | Gm14219 | ENSMUSG0C | false | . | . | . | . | . | . | chr2 | 159353426 | 159358106 | 4.96229 | 3.96932 | 2.99075 |
| chr2 | 159607101 | 159607685 | 1000 | Distal Intergenic | 2 | 159456170 | 159456457 | 288 | 2 | ENSMUST00 | -150645 | Gm14219 | ENSMUSG0C | false | . | . | . | . | . | . | chr2 | 159604547 | 159611310 | 5.03599 | 4.15595 | 3.02688 |
| chr11 | 117135500 | 117136087 | 1000 | Promoter (<=1kb) | 11 | 117134598 | 117135045 | 448 | 1 | ENSMUST00 | 903 | Gm11745 | ENSMUSG0C | false | chr11 | 117135886 | 117136235 | 4.76124 | 7.69302 | 5.56588 | chr11 | 117116362 | 117143393 | 7.76107 | 5.12248 | 5.29175 |
| chr5 | 103617128 | 103617586 | 1000 | Intron (ENSMUST00000031263.1 | 5 | 103624471 | 103625046 | 576 | 1 | ENSMUST00 | -6885 | Gm15844 | ENSMUSG0C | false | . | . | . | . | . | . | chr5 | 103617128 | 103621618 | 4.46422 | 3.80493 | 2.5717 |
| chr2 | 72390434 | 72391083 | 1000 | Intron (ENSMUST00000090824.6 | 2 | 72360247 | 72360542 | 296 | 1 | ENSMUST00 | 30188 | Rpl36a-ps4 | ENSMUSG0C | false | . | . | . | . | . | . | chr2 | 72388582 | 72393929 | 4.9635 | 4.09297 | 2.98642 |
| chr15 | 37405934 | 37406291 | 1000 | Intron (ENSMUST00000168992.2 | 15 | 37421248 | 37422694 | 1447 | 1 | ENSMUST00 | -14957 | Gm8668 | ENSMUSG0C | false | . | . | . | . | . | . | chr15 | 37404787 | 37408305 | 4.85723 | 3.87348 | 2.91731 |

| | | | | | | | | | | | | | | | | | | | | | | | | | | |
|---|---|---|---|---|---|---|---|---|---|---|---|---|---|---|---|---|---|---|---|---|---|---|---|---|---|---|
| chr6 | 99833495 | 99833776 | 1000 | Distal Intergenic | 6 | 99877816 | 99878334 | 519 | 2 | ENSMUST00 | 44558 | Tpt1-ps3 | ENSMUSG0( | false | . | . | . | . | . | . | chr6 | 99832689 | 99833660 | 3.46777 | 3.36237 | 1.72772 |
| chr2 | 57356214 | 57356539 | 1000 | Exon (ENSMUST00000169687.2/ | 2 | 57425167 | 57425709 | 543 | 2 | ENSMUST00 | 69170 | Gm13531 | ENSMUSG0( | false | . | . | . | . | . | . | chr2 | 57355659 | 57357243 | 4.34325 | 3.7965 | 2.45481 |
| chr11 | 88404814 | 88405555 | 1000 | Intron (ENSMUST00000107909.2 | 11 | 88432986 | 88433514 | 529 | 2 | ENSMUST00 | 27959 | Gm11510 | ENSMUSG0( | false | . | . | . | . | . | . | chr11 | 88403468 | 88407670 | 5.70199 | 4.27467 | 3.61961 |
| chr11 | 88463958 | 88464254 | 1000 | Intron (ENSMUST00000107909.2 | 11 | 88432986 | 88433514 | 529 | 2 | ENSMUST00 | -30445 | Gm11510 | ENSMUSG0( | false | . | . | . | . | . | . | chr11 | 88445933 | 88476041 | 6.2627 | 4.6354 | 4.05065 |
| chr11 | 88470349 | 88471116 | 1000 | Intron (ENSMUST00000107909.2 | 11 | 88432986 | 88433514 | 529 | 2 | ENSMUST00 | -36836 | Gm11510 | ENSMUSG0( | false | . | . | . | . | . | . | chr11 | 88445933 | 88476041 | 6.2627 | 4.6354 | 4.05065 |
| chr11 | 88467707 | 88468242 | 1000 | Intron (ENSMUST00000107909.2 | 11 | 88432986 | 88433514 | 529 | 2 | ENSMUST00 | -34194 | Gm11510 | ENSMUSG0( | false | . | . | . | . | . | . | chr11 | 88445933 | 88476041 | 6.2627 | 4.6354 | 4.05065 |
| chr9 | 73854234 | 73854580 | 1000 | Intron (ENSMUST00000184666.2 | 9 | 73795128 | 73795257 | 130 | 2 | ENSMUST00 | -58978 | Gm25105 | ENSMUSG0( | false | . | . | . | . | . | . | . | . | . | . | . | . |
| chr16 | 23401284 | 23401870 | 1000 | Distal Intergenic | 16 | 23423422 | 23423735 | 314 | 2 | ENSMUST00 | 21865 | Gm23430 | ENSMUSG0( | false | . | . | . | . | . | . | chr16 | 23400498 | 23403754 | 4.56425 | 3.90076 | 2.6624 |
| chr14 | 78662659 | 78663037 | 1000 | Intron (ENSMUST00000074729.5 | 14 | 78603710 | 78603831 | 122 | 1 | ENSMUST00 | 58950 | Gm26251 | ENSMUSG0( | false | . | . | . | . | . | . | chr14 | 78659799 | 78663968 | 3.74336 | 3.40827 | 2.00197 |
| chr10 | 51175037 | 51175376 | 1000 | Distal Intergenic | 10 | 51182299 | 51182405 | 107 | 1 | ENSMUST00 | -6923 | Gm26257 | ENSMUSG0( | false | . | . | . | . | . | . | chr10 | 51174675 | 51175916 | 3.32303 | 3.16968 | 1.65822 |
| chr6 | 140576177 | 140576509 | 1000 | Intron (ENSMUST00000087622.3 | 6 | 140530963 | 140531215 | 253 | 1 | ENSMUST00 | 45215 | Gm25780 | ENSMUSG0( | false | . | . | . | . | . | . | chr6 | 140574226 | 140577474 | 4.67928 | 3.77086 | 2.78431 |
| chr8 | 128387209 | 128387608 | 1000 | Intron (ENSMUST00000026917.8 | 8 | 128359241 | 128359320 | 80 | 1 | ENSMUST00 | 27969 | Mir1903 | ENSMUSG0( | false | . | . | . | . | . | . | chr8 | 128386761 | 128392770 | 5.18649 | 3.90151 | 3.1901 |
| chr9 | 62517286 | 62517810 | 1000 | Promoter (2-3kb) | 9 | 62515029 | 62515137 | 109 | 1 | ENSMUST00 | 2258 | n-R5s84 | ENSMUSG0( | false | . | . | . | . | . | . | chr9 | 62449515 | 62535966 | 6.98492 | 4.90408 | 4.64549 |
| chr5 | 66323637 | 66323959 | 1000 | Promoter (2-3kb) | 5 | 66325740 | 66326062 | 323 | 2 | ENSMUST00 | 2103 | Gm23183 | ENSMUSG0( | false | . | . | . | . | . | . | chr5 | 66319472 | 66333636 | 6.11352 | 4.4789 | 3.95234 |
| chr5 | 119325757 | 119326206 | 1000 | Distal Intergenic | 5 | 119297245 | 119297361 | 117 | 1 | ENSMUST00 | 28513 | n-R5s175 | ENSMUSG0( | false | . | . | . | . | . | . | . | . | . | . | . | . |
| chr3 | 134239645 | 134240931 | 1000 | Promoter (<=1kb) | 3 | 134240505 | 134240583 | 79 | 2 | ENSMUST00 | 0 | Mir1895 | ENSMUSG0( | false | chr3 | 134239268 | 134240836 | 28.18876 | 82.05154 | 78.86629 | chr3 | 134239254 | 134240687 | 4.41286 | 3.61362 | 2.54783 |
| chr18 | 11973679 | 11974147 | 1000 | Distal Intergenic | 18 | 11840361 | 11840438 | 78 | 2 | ENSMUST00 | -133242 | Mir1901 | ENSMUSG0( | false | . | . | . | . | . | . | chr18 | 11971839 | 11979509 | 5.50442 | 4.29031 | 3.42052 |
| chr18 | 11939752 | 11940270 | 1000 | Exon (ENSMUST00000171109.2/ | 18 | 11840361 | 11840438 | 78 | 2 | ENSMUST00 | -99315 | Mir1901 | ENSMUSG0( | false | . | . | . | . | . | . | chr18 | 11936954 | 11942682 | 6.83828 | 4.75724 | 4.53728 |
| chr18 | 11877749 | 11878048 | 1000 | Intron (ENSMUST00000171109.2 | 18 | 11840361 | 11840438 | 78 | 2 | ENSMUST00 | -37312 | Mir1901 | ENSMUSG0( | false | . | . | . | . | . | . | chr18 | 11869945 | 11889578 | 6.23298 | 4.56582 | 4.04247 |
| chr18 | 11915508 | 11915844 | 1000 | Intron (ENSMUST00000171109.2 | 18 | 11840361 | 11840438 | 78 | 2 | ENSMUST00 | -75071 | Mir1901 | ENSMUSG0( | false | . | . | . | . | . | . | chr18 | 11896077 | 11922616 | 5.15405 | 3.99993 | 3.17378 |
| chr18 | 11967159 | 11967674 | 1000 | Distal Intergenic | 18 | 11840361 | 11840438 | 78 | 2 | ENSMUST00 | -126722 | Mir1901 | ENSMUSG0( | false | . | . | . | . | . | . | chr18 | 11964714 | 11968820 | 4.82783 | 3.83752 | 2.88025 |
| chr16 | 35199684 | 35200096 | 1000 | Intron (ENSMUST00000114913.1 | 16 | 35234796 | 35234923 | 128 | 2 | ENSMUST00 | 34827 | Gm25179 | ENSMUSG0( | false | . | . | . | . | . | . | chr16 | 35198472 | 35207433 | 6.25906 | 4.64811 | 4.02553 |
| chr6 | 88321977 | 88322297 | 1000 | Intron (ENSMUST00000165242.1 | 6 | 88405977 | 88406301 | 325 | 1 | ENSMUST00 | -83680 | Gm25178 | ENSMUSG0( | false | . | . | . | . | . | . | chr6 | 88319571 | 88323455 | 6.45451 | 4.73731 | 4.20763 |
| chr19 | 26351678 | 26352011 | 1000 | Distal Intergenic | 19 | 26260369 | 26260493 | 125 | 2 | ENSMUST00 | -91186 | Gm25176 | ENSMUSG0( | false | . | . | . | . | . | . | chr19 | 26351132 | 26356191 | 4.5733 | 3.82867 | 2.65437 |
| chr4 | 82442955 | 82443365 | 1000 | Intron (ENSMUST00000050872.9 | 4 | 82439410 | 82439514 | 105 | 1 | ENSMUST00 | 3546 | n-R5s188 | ENSMUSG0( | false | . | . | . | . | . | . | chr4 | 82442500 | 82444214 | 4.27587 | 3.66937 | 2.44344 |
| chr15 | 34583145 | 34583468 | 1000 | Intron (ENSMUST00000040791.7 | 15 | 34531714 | 34531823 | 110 | 1 | ENSMUST00 | 51432 | Gm22208 | ENSMUSG0( | false | . | . | . | . | . | . | chr15 | 34583044 | 34583578 | 3.65477 | 3.25534 | 1.9355 |
| chr3 | 60120953 | 60121242 | 1000 | Distal Intergenic | 3 | 60127641 | 60127747 | 107 | 2 | ENSMUST00 | 6505 | Gm24382 | ENSMUSG0( | false | . | . | . | . | . | . | chr3 | 60120518 | 60128464 | 4.1733 | 3.66476 | 2.33216 |
| chr13 | 102434361 | 102434719 | 1000 | Distal Intergenic | 13 | 102228669 | 102228952 | 284 | 1 | ENSMUST00 | 205693 | Gm25808 | ENSMUSG0( | false | . | . | . | . | . | . | . | . | . | . | . | . |
| chr4 | 124388862 | 124389253 | 1000 | Distal Intergenic | 4 | 124414416 | 124418858 | 4443 | 2 | ENSMUST00 | 29605 | 1700057H15 | ENSMUSG0( | false | . | . | . | . | . | . | chr4 | 124384269 | 124390248 | 5.94125 | 4.46125 | 3.82162 |
| chr4 | 124380862 | 124381226 | 1000 | Distal Intergenic | 4 | 124414416 | 124418858 | 4443 | 2 | ENSMUST00 | 37632 | 1700057H15 | ENSMUSG0( | false | . | . | . | . | . | . | chr4 | 124379796 | 124383107 | 4.52012 | 3.82789 | 2.66381 |
| chr4 | 124385022 | 124385548 | 1000 | Distal Intergenic | 4 | 124414416 | 124418858 | 4443 | 2 | ENSMUST00 | 33310 | 1700057H15 | ENSMUSG0( | false | . | . | . | . | . | . | chr4 | 124384269 | 124390248 | 5.94125 | 4.46125 | 3.82162 |
| chr2 | 35245072 | 35245417 | 1000 | Distal Intergenic | 2 | 35224535 | 35226254 | 1720 | 1 | ENSMUST00 | 20538 | Gm13605 | ENSMUSG0( | false | . | . | . | . | . | . | chr2 | 35244330 | 35250667 | 6.28493 | 4.62322 | 4.08155 |
| chr4 | 133813054 | 133814790 | 1000 | Promoter (1-2kb) | 4 | 133816420 | 133817614 | 1195 | 1 | ENSMUST00 | -1630 | Gm12974 | ENSMUSG0( | false | chr4 | 133812841 | 133815309 | 30.78865 | 90.34473 | 86.99165 | chr4 | 133812161 | 133813069 | 3.00098 | 2.91197 | 1.35513 |
| chr2 | 167777233 | 167778139 | 1000 | Promoter (<=1kb) | 2 | 167777467 | 167782569 | 5103 | 1 | ENSMUST00 | 0 | Gm14321 | ENSMUSG0( | false | chr2 | 167777365 | 167778095 | 19.85205 | 54.41815 | 51.64575 | chr2 | 167773374 | 167783003 | 5.39085 | 4.1895 | 3.35037 |
| chr4 | 138131214 | 138131596 | 1000 | Intron (ENSMUST00000084215.6 | 4 | 138137595 | 138138518 | 924 | 2 | ENSMUST00 | 6922 | 1700095J12F | ENSMUSG0( | false | . | . | . | . | . | . | chr4 | 138129605 | 138131871 | 3.58358 | 3.17887 | 1.87757 |
| chrX | 167099437 | 167099863 | 1000 | Promoter (2-3kb) | 20 | 167088093 | 167102090 | 13998 | 2 | ENSMUST00 | 2227 | Gm15232 | ENSMUSG0( | false | . | . | . | . | . | . | . | . | . | . | . | . |
| chr4 | 149780792 | 149781347 | 1000 | Promoter (<=1kb) | 4 | 149780024 | 149780714 | 691 | 2 | ENSMUST00 | -79 | Gm13073 | ENSMUSG0( | false | . | . | . | . | . | . | chr4 | 149780418 | 149783610 | 5.7226 | 4.49571 | 3.59199 |
| chr8 | 111155720 | 111156210 | 1000 | 3' UTR | 8 | 111129076 | 111145480 | 16405 | 2 | ENSMUST00 | -10241 | 9430091E24 | ENSMUSG0( | false | . | . | . | . | . | . | chr8 | 111152542 | 111158131 | 5.98348 | 4.29022 | 3.84979 |
| chrX | 76077853 | 76078226 | 1000 | Distal Intergenic | 20 | 76153331 | 76162605 | 9275 | 2 | ENSMUST00 | 84379 | Gm14710 | ENSMUSG0( | false | . | . | . | . | . | . | . | . | . | . | . | . |
| chr4 | 155734465 | 155735225 | 1000 | Promoter (<=1kb) | 4 | 155734800 | 155740142 | 5343 | 1 | ENSMUST00 | 0 | Tmem240 | ENSMUSG0( | false | chr4 | 155734052 | 155735926 | 25.90858 | 83.69981 | 80.48042 | chr4 | 155734075 | 155736117 | 5.03231 | 3.98697 | 3.07912 |
| chr13 | 104540634 | 104541219 | 1000 | Distal Intergenic | 13 | 104684931 | 104694994 | 10064 | 1 | ENSMUST00 | -143712 | 2610204G07 | ENSMUSG0( | false | . | . | . | . | . | . | chr13 | 104540160 | 104543750 | 4.92729 | 4.08171 | 2.945 |
| chr16 | 91459060 | 91459408 | 1000 | Distal Intergenic | 16 | 91465104 | 91470121 | 5018 | 1 | ENSMUST00 | -5696 | A930006K02 | ENSMUSG0( | false | . | . | . | . | . | . | chr16 | 91458549 | 91460163 | 3.40479 | 3.30555 | 1.70173 |
| chr15 | 18818473 | 18818776 | 1000 | Promoter (<=1kb) | 15 | 18698194 | 18818772 | 120579 | 2 | ENSMUST00 | 0 | C030047K22 | ENSMUSG0( | false | . | . | . | . | . | . | . | . | . | . | . | . |
| chr12 | 108244886 | 108245232 | 1000 | Intron (ENSMUST00000136175.1 | 12 | 108206345 | 108275417 | 69073 | 2 | ENSMUST00 | 30185 | Ccdc85c | ENSMUSG0( | false | . | . | . | . | . | . | chr12 | 108233398 | 108274893 | 6.76084 | 4.78759 | 4.47033 |
| chr12 | 108239597 | 108240150 | 1000 | Intron (ENSMUST00000136175.1 | 12 | 108206345 | 108275417 | 69073 | 2 | ENSMUST00 | 35267 | Ccdc85c | ENSMUSG0( | false | . | . | . | . | . | . | chr12 | 108233398 | 108274893 | 6.76084 | 4.78759 | 4.47033 |
| chr12 | 108275032 | 108275977 | 1000 | Promoter (<=1kb) | 12 | 108206345 | 108275417 | 69073 | 2 | ENSMUST00 | 0 | Ccdc85c | ENSMUSG0( | false | chr12 | 108272978 | 108276795 | 32.66734 | 113.81544 | 109.86345 | . | . | . | . | . | . |
| chrX | 152368901 | 152369232 | 1000 | Promoter (<=1kb) | 20 | 152369028 | 152403045 | 34018 | 1 | ENSMUST00 | 0 | 3010001F23 | ENSMUSG0( | false | chrX | 152368836 | 152369430 | 8.27158 | 15.0213 | 12.75664 | . | . | . | . | . | . |
| chr11 | 3535406 | 3536010 | 1000 | Promoter (1-2kb) | 11 | 3534238 | 3534907 | 670 | 1 | ENSMUST00 | 1169 | Gm11947 | ENSMUSG0( | false | chr11 | 3535468 | 3536634 | 21.91997 | 62.19815 | 59.32085 | chr11 | 3532889 | 3538526 | 7.62005 | 5.18049 | 5.17074 |
| chr19 | 12714481 | 12714914 | 1000 | Exon (ENSMUST00000025598.4/ | 19 | 12706055 | 12721203 | 15149 | 2 | ENSMUST00 | 6289 | Gm15962 | ENSMUSG0( | false | . | . | . | . | . | . | chr19 | 12711318 | 12720714 | 4.33765 | 3.73227 | 2.4689 |
| chr11 | 120518884 | 120519198 | 1000 | Distal Intergenic | 11 | 120526855 | 120530554 | 3700 | 2 | ENSMUST00 | 11356 | Gm11789 | ENSMUSG0( | false | . | . | . | . | . | . | chr11 | 120517836 | 120520273 | 5.14815 | 4.17924 | 3.12897 |
| chr1 | 93211355 | 93211932 | 1000 | Distal Intergenic | 1 | 93215899 | 93231072 | 15174 | 1 | ENSMUST00 | -3967 | E030010N08 | ENSMUSG0( | false | . | . | . | . | . | . | chr1 | 93210088 | 93211973 | 5.50157 | 4.34556 | 3.43814 |
| chr2 | 128753134 | 128753820 | 1000 | Promoter (<=1kb) | 2 | 128751271 | 128753705 | 2435 | 2 | ENSMUST00 | 0 | Gm14011 | ENSMUSG0( | false | . | . | . | . | . | . | chr2 | 128721655 | 128769691 | 5.949 | 4.48828 | 3.78473 |
| chr7 | 112533528 | 112534079 | 1000 | Promoter (1-2kb) | 7 | 112526972 | 112535683 | 8712 | 2 | ENSMUST00 | 1604 | Parvaos | ENSMUSG0( | false | . | . | . | . | . | . | chr7 | 112487030 | 112537122 | 5.64568 | 4.33316 | 3.55068 |
| chr7 | 112534586 | 112534916 | 1000 | Promoter (<=1kb) | 7 | 112526972 | 112535683 | 8712 | 2 | ENSMUST00 | 767 | Parvaos | ENSMUSG0( | false | . | . | . | . | . | . | chr7 | 112487030 | 112537122 | 5.64568 | 4.33316 | 3.55068 |
| chrX | 99316624 | 99317049 | 1000 | Distal Intergenic | 20 | 99409571 | 99418895 | 9325 | 2 | ENSMUST00 | 101846 | Gm14809 | ENSMUSG0( | false | . | . | . | . | . | . | . | . | . | . | . | . |
| chrX | 99388584 | 99389025 | 1000 | Distal Intergenic | 20 | 99409571 | 99418895 | 9325 | 2 | ENSMUST00 | 29870 | Gm14809 | ENSMUSG0( | false | . | . | . | . | . | . | . | . | . | . | . | . |
| chrX | 99391628 | 99391947 | 1000 | Distal Intergenic | 20 | 99409571 | 99418895 | 9325 | 2 | ENSMUST00 | 26948 | Gm14809 | ENSMUSG0( | false | . | . | . | . | . | . | . | . | . | . | . | . |
| chrX | 99294633 | 99295232 | 1000 | Distal Intergenic | 20 | 99409571 | 99418895 | 9325 | 2 | ENSMUST00 | 123663 | Gm14809 | ENSMUSG0( | false | . | . | . | . | . | . | . | . | . | . | . | . |
| chrX | 99389950 | 99390397 | 1000 | Distal Intergenic | 20 | 99409571 | 99418895 | 9325 | 2 | ENSMUST00 | 28498 | Gm14809 | ENSMUSG0( | false | . | . | . | . | . | . | . | . | . | . | . | . |
| chrX | 99292712 | 99293074 | 1000 | Distal Intergenic | 20 | 99409571 | 99418895 | 9325 | 2 | ENSMUST00 | 125821 | Gm14809 | ENSMUSG0( | false | . | . | . | . | . | . | . | . | . | . | . | . |
| chrX | 99291343 | 99292019 | 1000 | Distal Intergenic | 20 | 99409571 | 99418895 | 9325 | 2 | ENSMUST00 | 126876 | Gm14809 | ENSMUSG0( | false | . | . | . | . | . | . | . | . | . | . | . | . |
| chrX | 99319195 | 99319659 | 1000 | Distal Intergenic | 20 | 99409571 | 99418895 | 9325 | 2 | ENSMUST00 | 99236 | Gm14809 | ENSMUSG0( | false | . | . | . | . | . | . | . | . | . | . | . | . |
| chrX | 99326169 | 99326557 | 1000 | Distal Intergenic | 20 | 99409571 | 99418895 | 9325 | 2 | ENSMUST00 | 92338 | Gm14809 | ENSMUSG0( | false | . | . | . | . | . | . | . | . | . | . | . | . |
| chrX | 99392548 | 99392856 | 1000 | Distal Intergenic | 20 | 99409571 | 99418895 | 9325 | 2 | ENSMUST00 | 26039 | Gm14809 | ENSMUSG0( | false | . | . | . | . | . | . | . | . | . | . | . | . |
| chr6 | 39539551 | 39540198 | 1000 | Intron (ENSMUST00000128919.1 | 6 | 39527522 | 39544486 | 16965 | 1 | ENSMUST00 | 12030 | 8030453O22 | ENSMUSG0( | false | . | . | . | . | . | . | chr6 | 39539485 | 39545229 | 6.37896 | 4.69531 | 4.14407 |
| chr11 | 80532893 | 80533316 | 1000 | Intron (ENSMUST00000041065.8 | 11 | 80507410 | 80509106 | 1697 | 2 | ENSMUST00 | -23788 | C030013C21 | ENSMUSG0( | false | . | . | . | . | . | . | chr11 | 80530643 | 80535922 | 6.03308 | 4.31286 | 3.89746 |
| chr11 | 80506899 | 80507262 | 1000 | Promoter (1-2kb) | 11 | 80507410 | 80509106 | 1697 | 2 | ENSMUST00 | 1844 | C030013C21 | ENSMUSG0( | false | . | . | . | . | . | . | chr11 | 80504064 | 80520376 | 4.41202 | 3.72882 | 2.53764 |
| chr11 | 119834473 | 119834892 | 1000 | Promoter (<=1kb) | 11 | 119820812 | 119834365 | 13554 | 2 | ENSMUST00 | -109 | Rptoros | ENSMUSG0( | false | . | . | . | . | . | . | chr11 | 119831481 | 119836988 | 5.44333 | 4.10671 | 3.40297 |
| chr4 | 129048679 | 129049420 | 1000 | Downstream (<1kb) | 4 | 129050097 | 129057983 | 7887 | 2 | ENSMUST00 | 8563 | 1700086P04 | ENSMUSG0( | false | chr4 | 129048436 | 129049847 | 25.84577 | 73.40098 | 70.35904 | chr4 | 129048159 | 129050161 | 3.99752 | 3.42649 | 2.21515 |
| chr4 | 129035205 | 129035470 | 1000 | Distal Intergenic | 4 | 129050097 | 129057983 | 7887 | 2 | ENSMUST00 | 22513 | 1700086P04 | ENSMUSG0( | false | . | . | . | . | . | . | chr4 | 129032304 | 129035991 | 5.85449 | 4.44408 | 3.71528 |
| chr2 | 33179873 | 33180680 | 1000 | Promoter (1-2kb) | 2 | 33177902 | 33181881 | 3980 | 1 | ENSMUST00 | 1972 | Gm13528 | ENSMUSG0( | false | . | . | . | . | . | . | chr2 | 33175823 | 33189808 | 7.09799 | 4.90853 | 4.74291 |
| chr3 | 81418759 | 81419321 | 1000 | Distal Intergenic | 3 | 81037408 | 81040437 | 3030 | 2 | ENSMUST00 | -378323 | Gm16000 | ENSMUSG0( | false | . | . | . | . | . | . | . | . | . | . | . | . |
| chr11 | 121763494 | 121763900 | 1000 | Distal Intergenic | 11 | 121803107 | 121810319 | 7213 | 1 | ENSMUST00 | -39207 | Gm12589 | ENSMUSG0( | false | . | . | . | . | . | . | chr11 | 121762232 | 121766311 | 4.44206 | 3.79793 | 2.56192 |
| chr11 | 121794576 | 121795090 | 1000 | Distal Intergenic | 11 | 121803107 | 121810319 | 7213 | 1 | ENSMUST00 | -8017 | Gm12589 | ENSMUSG0( | false | . | . | . | . | . | . | chr11 | 121787526 | 121808717 | 6.67832 | 4.85129 | 4.38811 |
| chr2 | 144224111 | 144224816 | 1000 | Distal Intergenic | 2 | 144243241 | 144244849 | 1609 | 1 | ENSMUST00 | -18425 | 4930444E06 | ENSMUSG0( | false | chr2 | 144224128 | 144224938 | 7.82203 | 16.59716 | 14.30959 | chr2 | 144223213 | 144227135 | 3.77986 | 3.52638 | 2.00101 |
| chr11 | 81096003 | 81096411 | 1000 | Intron (ENSMUST00000021045.7 | 11 | 81072403 | 81083358 | 10956 | 1 | ENSMUST00 | 23601 | Gm11417 | ENSMUSG0( | false | . | . | . | . | . | . | chr11 | 81095149 | 81098636 | 4.57174 | 3.86096 | 2.67966 |
| chr11 | 65386757 | 65387153 | 1000 | Distal Intergenic | 11 | 65285936 | 65365798 | 79863 | 2 | ENSMUST00 | -20960 | Gm12295 | ENSMUSG0( | false | . | . | . | . | . | . | chr11 | 65386261 | 65387823 | 5.06573 | 4.08458 | 3.04852 |
| chr4 | 62711170 | 62711931 | 1000 | Intron (ENSMUST00000150203.1 | 4 | 62705981 | 62725737 | 19757 | 1 | ENSMUST00 | 5190 | Gm11210 | ENSMUSG0( | false | . | . | . | . | . | . | chr4 | 62700580 | 62712268 | 6.69274 | 4.86565 | 4.38997 |
| chr1 | 61321675 | 61322157 | 1000 | Distal Intergenic | 1 | 61301909 | 61307746 | 5838 | 1 | ENSMUST00 | 19767 | Gm11587 | ENSMUSG0( | false | . | . | . | . | . | . | chr1 | 61321473 | 61322445 | 4.89327 | 3.8512 | 2.94361 |
| chr2 | 72674233 | 72674666 | 1000 | Distal Intergenic | 2 | 72703510 | 72729468 | 25959 | 1 | ENSMUST00 | -28844 | 8430437L04F | ENSMUSG0( | false | . | . | . | . | . | . | chr2 | 72672965 | 72675132 | 3.38344 | 3.15544 | 1.728 |
| chr2 | 92491864 | 92492511 | 1000 | Intron (ENSMUST00000141341.1 | 2 | 92483926 | 92529809 | 45884 | 1 | ENSMUST00 | 7939 | Gm13814 | ENSMUSG0( | false | . | . | . | . | . | . | chr2 | 92484342 | 92499595 | 6.72219 | 4.87617 | 4.42323 |
| chr2 | 92490843 | 92491372 | 1000 | Intron (ENSMUST00000141341.1 | 2 | 92483926 | 92529809 | 45884 | 1 | ENSMUST00 | 6918 | Gm13814 | ENSMUSG0( | false | . | . | . | . | . | . | chr2 | 92484342 | 92499595 | 6.72219 | 4.87617 | 4.42323 |
| chr2 | 92477638 | 92478004 | 1000 | Distal Intergenic | 2 | 92483926 | 92529809 | 45884 | 1 | ENSMUST00 | -5922 | Gm13814 | ENSMUSG0( | false | . | . | . | . | . | . | chr2 | 92477505 | 92478556 | 4.30308 | 3.63915 | 2.45629 |
| chr2 | 92512171 | 92512683 | 1000 | Intron (ENSMUST00000141341.1 | 2 | 92483926 | 92529809 | 45884 | 1 | ENSMUST00 | 28246 | Gm13814 | ENSMUSG0( | false | . | . | . | . | . | . | chr2 | 92506711 | 92530936 | 5.29916 | 4.18157 | 3.26309 |
| chr2 | 92514976 | 92515411 | 1000 | Intron (ENSMUST00000141341.1 | 2 | 92529694 | 92540980 | 11287 | 2 | ENSMUST00 | 25569 | Gm13817 | ENSMUSG0( | false | . | . | . | . | . | . | chr2 | 92506711 | 92530936 | 5.29916 | 4.18157 | 3.26309 |
| chr4 | 114886870 | 114887302 | 1000 | Intron (ENSMUST00000155499.1 | 4 | 114817161 | 114902859 | 85699 | 2 | ENSMUST00 | 15557 | 9130410C08 | ENSMUSG0( | false | . | . | . | . | . | . | chr4 | 114884610 | 114889294 | 6.06084 | 4.59598 | 3.87048 |
| chr10 | 81410851 | 81411387 | 1000 | Promoter (1-2kb) | 10 | 81409575 | 81411579 | 2005 | 1 | ENSMUST00 | 1277 | Gm16104 | ENSMUSG0( | false | . | . | . | . | . | . | chr10 | 81392649 | 81426205 | 8.72751 | 5.63067 | 6.0679 |
| chr10 | 81416878 | 81417398 | 1000 | Intron (ENSMUST00000105321.4 | 10 | 81409575 | 81411579 | 2005 | 1 | ENSMUST00 | 7304 | Gm16104 | ENSMUSG0( | false | . | . | . | . | . | . | chr10 | 81392649 | 81426205 | 8.72751 | 5.63067 | 6.0679 |
| chr5 | 127910363 | 127910755 | 1000 | Intron (ENSMUST00000044441.6 | 5 | 127878531 | 127882376 | 3846 | 1 | ENSMUST00 | 31833 | Gm17131 | ENSMUSG0( | false | . | . | . | . | . | . | chr5 | 127909852 | 127912806 | 5.33806 | 4.24441 | 3.29039 |
| chr2 | 49804963 | 49805437 | 1000 | Intron (ENSMUST00000028103.7 | 2 | 49805976 | 49812948 | 6973 | 2 | ENSMUST00 | 7511 | Gm13480 | ENSMUSG0( | false | . | . | . | . | . | . | . | . | . | . | . | . |
| chr2 | 49813923 | 49814320 | 1000 | Promoter (<=1kb) | 2 | 49805976 | 49812948 | 6973 | 2 | ENSMUST00 | -976 | Gm13480 | ENSMUSG0( | false | . | . | . | . | . | . | chr2 | 49811428 | 49816755 | 3.89955 | 3.44869 | 2.15154 |
| chr6 | 34637616 | 34638003 | 1000 | Intron (ENSMUST00000079391.4 | 6 | 34651354 | 34653894 | 2541 | 2 | ENSMUST00 | 15891 | Gm13861 | ENSMUSG0( | false | . | . | . | . | . | . | chr6 | 34636828 | 34643532 | 4.67449 | 3.94616 | 2.73484 |
| chr1 | 38268902 | 38269555 | 1000 | Intron (ENSMUST00000095027.3 | 1 | 38239700 | 38246666 | 6967 | 1 | ENSMUST00 | 29203 | Gm16150 | ENSMUSG0( | false | . | . | . | . | . | . | chr1 | 38267333 | 38268903 | 5.03434 | 4.11158 | 3.08535 |
| chr4 | 141281818 | 141282149 | 1000 | Distal Intergenic | 4 | 141278439 | 141279794 | 1356 | 1 | ENSMUST00 | 3380 | Gm13056 | ENSMUSG0( | false | . | . | . | . | . | . | chr4 | 141277124 | 141282934 | 7.25702 | 5.14963 | 4.88042 |
| chr4 | 114910961 | 114911604 | 1000 | Promoter (1-2kb) | 4 | 114909257 | 114921967 | 12711 | 1 | ENSMUST00 | 1705 | Foxd2os | ENSMUSG0( | false | chr4 | 114904677 | 114911326 | 45.47067 | 151.73354 | 146.38181 | chr4 | 114911555 | 114917656 | 4.83581 | 4.00866 | 2.87673 |
| chr10 | 94409637 | 94410410 | 1000 | Promoter (<=1kb) | 10 | 94408891 | 94410732 | 1842 | 1 | ENSMUST00 | 747 | Gm16155 | ENSMUSG0( | false | . | . | . | . | . | . | chr10 | 94408503 | 94417576 | 5.71193 | 4.27942 | 3.61952 |
| chr2 | 167864105 | 167864706 | 1000 | Distal Intergenic | 2 | 167858585 | 167862542 | 3958 | 1 | ENSMUST00 | 5521 | Gm14319 | ENSMUSG0( | false | . | . | . | . | . | . | chr2 | 167855418 | 167869855 | 6.99032 | 4.95783 | 4.63791 |
| chr2 | 167866719 | 167867212 | 1000 | Distal Intergenic | 2 | 167858585 | 167862542 | 3958 | 1 | ENSMUST00 | 8135 | Gm14319 | ENSMUSG0( | false | . | . | . | . | . | . | chr2 | 167855418 | 167869855 | 6.99032 | 4.95783 | 4.63791 |
| chr2 | 167871487 | 167871956 | 1000 | Distal Intergenic | 2 | 167858585 | 167862542 | 3958 | 1 | ENSMUST00 | 12903 | Gm14319 | ENSMUSG0( | false | . | . | . | . | . | . | chr2 | 167871188 | 167873275 | 6.24944 | 4.65092 | 4.04596 |
| chr2 | 110192295 | 110193004 | 1000 | Distal Intergenic | 2 | 110196889 | 110256131 | 59243 | 1 | ENSMUST00 | -3885 | Gm13936 | ENSMUSG0( | false | . | . | . | . | . | . | chr2 | 110172226 | 110205814 | 5.39899 | 4.25006 | 3.33729 |
| chr2 | 110179520 | 110180361 | 1000 | Distal Intergenic | 2 | 110196889 | 110256131 | 59243 | 1 | ENSMUST00 | -16528 | Gm13936 | ENSMUSG0( | false | . | . | . | . | . | . | chr2 | 110172226 | 110205814 | 5.39899 | 4.25006 | 3.33729 |
| chr2 | 110196694 | 110197040 | 1000 | Promoter (<=1kb) | 2 | 110196889 | 110256131 | 59243 | 1 | ENSMUST00 | 0 | Gm13936 | ENSMUSG0( | false | . | . | . | . | . | . | chr2 | 110172226 | 110205814 | 5.39899 | 4.25006 | 3.33729 |
| chr1 | 61407165 | 61407531 | 1000 | Distal Intergenic | 1 | 61378432 | 61395904 | 17473 | 1 | ENSMUST00 | 28734 | 9530026F06 | ENSMUSG0( | false | . | . | . | . | . | . | chr1 | 61402840 | 61409167 | 4.3385 | 3.6894 | 2.48712 |
| chr12 | 30069195 | 30069786 | 1000 | Intron (ENSMUST00000021005.9 | 12 | 30059657 | 30062113 | 2457 | 1 | ENSMUST00 | 9539 | Gm15691 | ENSMUSG0( | false | . | . | . | . | . | . | chr12 | 30069018 | 30071689 | 3.85844 | 3.45894 | 2.09223 |
| chr12 | 30054757 | 30055073 | 1000 | 3' UTR | 12 | 30059657 | 30062113 | 2457 | 1 | ENSMUST00 | -4584 | Gm15691 | ENSMUSG0( | false | . | . | . | . | . | . | chr12 | 30053642 | 30055101 | 3.05868 | 2.97321 | 1.44259 |
| chr11 | 4294771 | 4295399 | 1000 | Distal Intergenic | 11 | 4285460 | 4286939 | 1480 | 1 | ENSMUST00 | 9312 | Gm11958 | ENSMUSG0( | false | . | . | . | . | . | . | chr11 | 4291014 | 4306323 | 6.81252 | 4.86662 | 4.50325 |
| chr5 | 43561620 | 43562246 | 1000 | Intron (ENSMUST00000167522.1 | 5 | 43549930 | 43557769 | 7840 | 2 | ENSMUST00 | -3852 | Gm15865 | ENSMUSG0( | false | . | . | . | . | . | . | chr5 | 43560201 | 43596005 | 4.90131 | 4.0072 | 2.93052 |
| chr9 | 97013120 | 97013527 | 1000 | Promoter (2-3kb) | 9 | 97015966 | 97022043 | 6078 | 1 | ENSMUST00 | -2439 | Gm16010 | ENSMUSG0( | false | . | . | . | . | . | . | chr9 | 97009875 | 97014254 | 4.87082 | 3.81409 | 2.93633 |
| chrX | 99463890 | 99464270 | 1000 | Downstream (1-2kb) | 20 | 99459025 | 99460753 | 1729 | 1 | ENSMUST00 | 4866 | Gm14808 | ENSMUSG0( | false | . | . | . | . | . | . | . | . | . | . | . | . |
| chr7 | 96967155 | 96967649 | 1000 | Intron (ENSMUST00000044466.6 | 7 | 96947910 | 96951886 | 3977 | 2 | ENSMUST00 | -15270 | C230038L03F | ENSMUSG0( | false | . | . | . | . | . | . | chr7 | 96952896 | 96991589 | 4.87493 | 3.9704 | 2.91243 |
| chr8 | 11155558 | 11156011 | 1000 | Distal Intergenic | 8 | 11187755 | 11204503 | 16749 | 1 | ENSMUST00 | -31744 | Gm15418 | ENSMUSG0( | false | . | . | . | . | . | . | chr8 | 11154514 | 11156580 | 5.64845 | 4.21781 | 3.56775 |
| chr4 | 107011812 | 107012157 | 1000 | Intron (ENSMUST00000150915.2 | 4 | 107017935 | 107022499 | 4565 | 2 | ENSMUST00 | 10342 | Gm12786 | ENSMUSG0( | false | . | . | . | . | . | . | chr4 | 107009465 | 107020061 | 6.67324 | 4.69024 | 4.42102 |
| chr11 | 97659822 | 97660805 | 1000 | Promoter (<=1kb) | 11 | 97644184 | 97660367 | 16184 | 2 | ENSMUST00 | 0 | Gm11613 | ENSMUSG0( | false | chr11 | 97659512 | 97661428 | 28.83795 | 83.16807 | 79.96098 | chr11 | 97660731 | 97662935 | 5.22491 | 4.26722 | 3.18868 |
| chr6 | 86472260 | 86472946 | 1000 | 3' UTR | 6 | 86470805 | 86480770 | 9966 | 2 | ENSMUST00 | 7824 | A430078I02F | ENSMUSG0( | false | chr6 | 86471752 | 86472261 | 13.16437 | 29.12438 | 26.66979 | chr6 | 86472845 | 86482455 | 6.5006 | 4.7144 | 4.25243 |
| chr4 | 151339347 | 151339640 | 1000 | Intron (ENSMUST00000169423.3 | 4 | 151376470 | 151379582 | 3113 | 1 | ENSMUST00 | -36830 | 4930589P08 | ENSMUSG0( | false | . | . | . | . | . | . | . | . | . | . | . | . |
| chr4 | 151385995 | 151386924 | 1000 | Intron (ENSMUST00000169423.3 | 4 | 151376470 | 151379582 | 3113 | 1 | ENSMUST00 | 9526 | 4930589P08 | ENSMUSG0( | false | chr4 | 151386347 | 151387508 | 14.26849 | 37.02594 | 34.4737 | chr4 | 151384634 | 151392014 | 4.60253 | 3.82226 | 2.70339 |
| chr2 | 135578095 | 135578514 | 1000 | Intron (ENSMUST00000153402.1 | 2 | 135580530 | 135583296 | 2767 | 2 | ENSMUST00 | 4782 | 9630028H03 | ENSMUSG0( | false | . | . | . | . | . | . | . | . | . | . | . | . |
| chr4 | 56070644 | 56070938 | 1000 | Distal Intergenic | 4 | 55969023 | 55993739 | 24717 | 2 | ENSMUST00 | -76906 | Gm12519 | ENSMUSG0( | false | . | . | . | . | . | . | . | . | . | . | . | . |
| chr4 | 153175723 | 153176189 | 1000 | Distal Intergenic | 4 | 153145590 | 153146943 | 1354 | 2 | ENSMUST00 | -28781 | Gm13173 | ENSMUSG0( | false | . | . | . | . | . | . | chr4 | 153175502 | 153176427 | 5.08604 | 3.89428 | 3.12486 |
| chr4 | 153110324 | 153110934 | 1000 | Distal Intergenic | 4 | 153145590 | 153146943 | 1354 | 2 | ENSMUST00 | 36009 | Gm13173 | ENSMUSG0( | false | . | . | . | . | . | . | chr4 | 153109725 | 153115297 | 5.19331 | 4.20298 | 3.16863 |
| chr11 | 23067718 | 23068219 | 1000 | Promoter (<=1kb) | 11 | 23067998 | 23075574 | 7577 | 1 | ENSMUST00 | 0 | Gm12059 | ENSMUSG0( | false | chr11 | 23067253 | 23068481 | 27.34985 | 78.73917 | 75.61149 | chr11 | 23067141 | 23068590 | 4.89236 | 3.95874 | 2.96174 |
| chr10 | 42038528 | 42039023 | 1000 | Distal Intergenic | 10 | 42054113 | 42055306 | 1194 | 2 | ENSMUST00 | 16283 | Gm9855 | ENSMUSG0( | false | . | . | . | . | . | . | chr10 | 42036723 | 42039074 | 4.18639 | 3.41457 | 2.36286 |
| chr2 | 117381237 | 117381708 | 1000 | Distal Intergenic | 2 | 117414392 | 117417697 | 3306 | 2 | ENSMUST00 | 35989 | Gm13982 | ENSMUSG0( | false | . | . | . | . | . | . | chr2 | 117379234 | 117382776 | 4.305 | 3.52039 | 2.47014 |
| chr6 | 65793010 | 65793326 | 1000 | Intron (ENSMUST00000031976.8 | 6 | 65768544 | 65779249 | 10706 | 2 | ENSMUST00 | -13762 | Gm16838 | ENSMUSG0( | false | . | . | . | . | . | . | chr6 | 65792044 | 65793495 | 3.41819 | 3.29429 | 1.72069 |
| chr6 | 52175607 | 52176277 | 1000 | Promoter (<=1kb) | 6 | 52175235 | 52215076 | 39842 | 1 | ENSMUST00 | 373 | Hoxaas3 | ENSMUSG0( | false | chr6 | 52175412 | 52176523 | 20.67895 | 53.40692 | 50.64761 | chr6 | 52174986 | 52186369 | 4.8401 | 4.02656 | 2.87141 |
| chr2 | 135711858 | 135712243 | 1000 | Intron (ENSMUST00000144931.1 | 2 | 135686174 | 135693154 | 6981 | 2 | ENSMUST00 | -18705 | Gm14211 | ENSMUSG0( | false | chr2 | 135711966 | 135712232 | 3.37938 | 4.21603 | 2.20503 | chr2 | 135708887 | 135713201 | 4.38898 | 3.77563 | 2.5066 |
| chr4 | 101317475 | 101317817 | 1000 | Intron (ENSMUST00000131864.1 | 4 | 101292414 | 101321745 | 29332 | 2 | ENSMUST00 | 3928 | Gm12796 | ENSMUSG0( | false | . | . | . | . | . | . | chr4 | 101316720 | 101318846 | 4.17514 | 3.70301 | 2.3191 |
| chr10 | 127661833 | 127662724 | 1000 | Promoter (1-2kb) | 10 | 127663839 | 127665484 | 1646 | 1 | ENSMUST00 | -1115 | Gm16229 | ENSMUSG0( | false | chr10 | 127662238 | 127666785 | 29.66345 | 100.21 | 96.63225 | chr10 | 127658015 | 127665178 | 5.91846 | 4.4802 | 3.77282 |
| chr5 | 148789204 | 148789521 | 1000 | Distal Intergenic | 5 | 148777963 | 148783080 | 5118 | 1 | ENSMUST00 | 11242 | 4930505K14 | ENSMUSG0( | false | . | . | . | . | . | . | chr5 | 148787312 | 148790648 | 3.52762 | 3.22001 | 1.82091 |
| chr5 | 148768390 | 148768803 | 1000 | Distal Intergenic | 5 | 148777963 | 148783080 | 5118 | 1 | ENSMUST00 | -9160 | 4930505K14 | ENSMUSG0( | false | . | . | . | . | . | . | chr5 | 148768023 | 148769150 | 5.32725 | 4.16001 | 3.28485 |
| chr11 | 75051428 | 75052033 | 1000 | Intron (ENSMUST00000045281.7 | 11 | 75036476 | 75039348 | 2873 | 2 | ENSMUST00 | -12081 | Gm12333 | ENSMUSG0( | false | . | . | . | . | . | . | chr11 | 75046409 | 75055521 | 5.55534 | 4.32005 | 3.48408 |
| chr13 | 32518727 | 32519364 | 1000 | Distal Intergenic | 13 | 32379907 | 32383688 | 3782 | 2 | ENSMUST00 | -135040 | Gm11381 | ENSMUSG0( | false | . | . | . | . | . | . | chr13 | 32517308 | 32520163 | 5.41626 | 4.29262 | 3.3411 |
| chr11 | 31824375 | 31824673 | 1000 | Promoter (<=1kb) | 11 | 31790113 | 31824473 | 34361 | 2 | ENSMUST00 | 0 | D630024D03 | ENSMUSG0( | false | . | . | . | . | . | . | chr11 | 31821641 | 31832598 | 5.80216 | 4.3489 | 3.6737 |
| chr11 | 31827994 | 31828381 | 1000 | Distal Intergenic | 11 | 31800688 | 31824524 | 23837 | 2 | ENSMUST00 | -3471 | D630024D03 | ENSMUSG0( | false | . | . | . | . | . | . | chr11 | 31821641 | 31832598 | 5.80216 | 4.3489 | 3.6737 |
| chr11 | 31787868 | 31788228 | 1000 | Downstream (1-2kb) | 11 | 31790113 | 31824473 | 34361 | 2 | ENSMUST00 | 36245 | D630024D03 | ENSMUSG0( | false | . | . | . | . | . | . | chr11 | 31787695 | 31788564 | 4.67866 | 3.93455 | 2.73407 |
| chr8 | 26978389 | 26979286 | 1000 | Promoter (1-2kb) | 8 | 26977336 | 26981461 | 4126 | 1 | ENSMUST00 | 1054 | Zfp703 | ENSMUSG0( | false | chr8 | 26976009 | 26979489 | 27.41053 | 90.23362 | 86.88379 | chr8 | 26978900 | 26985294 | 4.32877 | 3.61461 | 2.48483 |
| chr8 | 26976635 | 26977477 | 1000 | Promoter (<=1kb) | 8 | 26977336 | 26981461 | 4126 | 1 | ENSMUST00 | 0 | Zfp703 | ENSMUSG0( | false | chr8 | 26976009 | 26979489 | 27.41053 | 90.23362 | 86.88379 | . | . | . | . | . | . |
| chr6 | 127917039 | 127917509 | 1000 | Intron (ENSMUST00000032501.4 | 6 | 127934689 | 127944839 | 10151 | 2 | ENSMUST00 | 27330 | Gm15870 | ENSMUSG0( | false | . | . | . | . | . | . | chr6 | 127916854 | 127918694 | 4.47232 | 3.76212 | 2.59248 |
| chr6 | 127932813 | 127933224 | 1000 | Intron (ENSMUST00000032501.4 | 6 | 127934689 | 127944839 | 10151 | 2 | ENSMUST00 | 11615 | Gm15870 | ENSMUSG0( | false | . | . | . | . | . | . | chr6 | 127931822 | 127933603 | 6.61563 | 4.51221 | 4.38732 |
| chr4 | 137897344 | 137897686 | 1000 | Intron (ENSMUST00000102518.4 | 4 | 137882370 | 137883937 | 1568 | 2 | ENSMUST00 | -13408 | Gm13012 | ENSMUSG0( | false | . | . | . | . | . | . | chr4 | 137896034 | 137898992 | 5.021 | 3.92895 | 3.0585 |
| chr2 | 25292117 | 25292493 | 1000 | Promoter (<=1kb) | 2 | 25291220 | 25298925 | 7706 | 1 | ENSMUST00 | 898 | Grin1os | ENSMUSG0( | false | chr2 | 25291872 | 25292980 | 16.95696 | 43.908 | 41.27272 | chr2 | 25290088 | 25293205 | 5.62432 | 4.39082 | 3.51618 |
| chr17 | 27945966 | 27946321 | 1000 | Intron (ENSMUST00000114842.2 | 17 | 27959615 | 27965316 | 5702 | 2 | ENSMUST00 | 18995 | Gm15597 | ENSMUSG0( | false | . | . | . | . | . | . | chr17 | 27945027 | 27950123 | 5.05743 | 3.9964 | 3.06404 |
| chr17 | 27938355 | 27938701 | 1000 | Intron (ENSMUST00000114842.2 | 17 | 27959615 | 27965316 | 5702 | 2 | ENSMUST00 | 26615 | Gm15597 | ENSMUSG0( | false | . | . | . | . | . | . | chr17 | 27930351 | 27942547 | 4.95598 | 3.97496 | 2.99042 |
| chr4 | 101291934 | 101292443 | 1000 | Exon (ENSMUST00000131864.1/ | 4 | 101284221 | 101285203 | 983 | 1 | ENSMUST00 | 7714 | Gm12801 | ENSMUSG0( | false | chr4 | 101292040 | 101293747 | 23.60507 | 64.24684 | 61.34294 | chr4 | 101291052 | 101294245 | 4.50505 | 3.85025 | 2.60632 |
| chr4 | 114688329 | 114689011 | 1000 | Distal Intergenic | 4 | 114680804 | 114687595 | 6792 | 1 | ENSMUST00 | 7526 | Gm12829 | ENSMUSG0( | false | chr4 | 114688177 | 114688804 | 8.63619 | 18.02194 | 15.71366 | chr4 | 114677087 | 114691330 | 5.31309 | 4.19527 | 3.28131 |
| chr4 | 12490764 | 12491165 | 1000 | Distal Intergenic | 4 | 12271399 | 12314583 | 43185 | 1 | ENSMUST00 | 219366 | Gm11846 | ENSMUSG0( | false | . | . | . | . | . | . | chr4 | 12489611 | 12492934 | 3.37407 | 3.23783 | 1.69022 |
| chr4 | 141881268 | 141882000 | 1000 | Distal Intergenic | 4 | 141885654 | 141897282 | 11629 | 1 | ENSMUST00 | -3654 | Fhad1os2 | ENSMUSG0( | false | . | . | . | . | . | . | chr4 | 141877986 | 141887420 | 6.94416 | 4.90069 | 4.61404 |
| chr2 | 27540078 | 27540548 | 1000 | Promoter (<=1kb) | 2 | 27540426 | 27543362 | 2937 | 1 | ENSMUST00 | 0 | Gm13421 | ENSMUSG0( | false | chr2 | 27540322 | 27540901 | 15.01131 | 34.85038 | 32.32499 | chr2 | 27538977 | 27541379 | 3.82422 | 3.52624 | 2.04577 |
| chr13 | 28417760 | 28418458 | 1000 | Distal Intergenic | 13 | 28460778 | 28480841 | 20064 | 2 | ENSMUST00 | 62383 | 2610307P16 | ENSMUSG0( | false | chr13 | 28417689 | 28417950 | 9.06242 | 17.74057 | 15.43651 | chr13 | 28413931 | 28420800 | 5.81821 | 4.35832 | 3.69744 |
| chr2 | 173858654 | 173858920 | 1000 | Distal Intergenic | 2 | 173793437 | 173847670 | 54234 | 2 | ENSMUST00 | -10985 | Gm14371 | ENSMUSG0( | false | . | . | . | . | . | . | chr2 | 173847661 | 173860588 | 5.03291 | 4.0817 | 3.03782 |
| chr2 | 92751205 | 92751753 | 1000 | Intron (ENSMUST00000152230.1 | 2 | 92742230 | 92759208 | 16979 | 1 | ENSMUST00 | 8976 | Gm13816 | ENSMUSG0( | false | . | . | . | . | . | . | chr2 | 92749887 | 92754068 | 7.8726 | 5.31163 | 5.37358 |
| chr11 | 94133952 | 94134753 | 1000 | Promoter (<=1kb) | 11 | 94133502 | 94155776 | 22275 | 1 | ENSMUST00 | 451 | B230206L02 | ENSMUSG0( | false | . | . | . | . | . | . | chr11 | 94129611 | 94138962 | 5.93933 | 4.52942 | 3.772 |
| chr11 | 54735127 | 54735458 | 1000 | Intron (ENSMUST00000064104.7 | 11 | 54728677 | 54734323 | 5647 | 1 | ENSMUST00 | 6451 | Gm12227 | ENSMUSG0( | false | . | . | . | . | . | . | chr11 | 54722256 | 54750513 | 5.8032 | 4.39 | 3.67574 |
| chr6 | 54450452 | 54450965 | 1000 | Promoter (1-2kb) | 6 | 54452883 | 54503768 | 50886 | 1 | ENSMUST00 | -1918 | Wipf3 | ENSMUSG0( | true | . | . | . | . | . | . | chr6 | 54431186 | 54455170 | 5.36229 | 4.20786 | 3.31695 |
| chr1 | 75392933 | 75393376 | 1000 | Promoter (2-3kb) | 1 | 75392131 | 75396363 | 4233 | 2 | ENSMUST00 | 2987 | Gm15178 | ENSMUSG0( | false | . | . | . | . | . | . | chr1 | 75386249 | 75400648 | 7.0728 | 5.02664 | 4.70473 |
| chr1 | 75395214 | 75395597 | 1000 | Promoter (<=1kb) | 1 | 75392131 | 75396363 | 4233 | 2 | ENSMUST00 | 766 | Gm15178 | ENSMUSG0( | false | . | . | . | . | . | . | chr1 | 75386249 | 75400648 | 7.0728 | 5.02664 | 4.70473 |
| chr14 | 98105579 | 98105903 | 1000 | Intron (ENSMUST00000069334.7 | 14 | 98118509 | 98119006 | 498 | 1 | ENSMUST00 | -12606 | Gm16331 | ENSMUSG0( | false | . | . | . | . | . | . | chr14 | 98103378 | 98106625 | 5.02834 | 3.92695 | 3.04488 |
| chr8 | 25546900 | 25547328 | 1000 | Intron (ENSMUST00000155564.2 | 8 | 25552678 | 25553560 | 883 | 2 | ENSMUST00 | 6232 | Gm16159 | ENSMUSG0( | false | . | . | . | . | . | . | chr8 | 25521452 | 25558014 | 5.35885 | 4.21426 | 3.3104 |
| chr4 | 62727566 | 62728046 | 1000 | Promoter (<=1kb) | 4 | 62723516 | 62727537 | 4022 | 2 | ENSMUST00 | -30 | Gm11211 | ENSMUSG0( | false | . | . | . | . | . | . | chr4 | 62726795 | 62739050 | 7.64763 | 5.19923 | 5.18438 |
| chr10 | 24379406 | 24379770 | 1000 | Distal Intergenic | 10 | 24447490 | 24452657 | 5168 | 1 | ENSMUST00 | -67720 | Gm15271 | ENSMUSG0( | false | . | . | . | . | . | . | chr10 | 24378933 | 24381355 | 5.20558 | 3.92704 | 3.21313 |
| chr17 | 28018057 | 28018788 | 1000 | Intron (ENSMUST00000114842.2 | 17 | 27988028 | 28008432 | 20405 | 2 | ENSMUST00 | -9626 | Gm15598 | ENSMUSG0( | false | . | . | . | . | . | . | chr17 | 28005688 | 28028167 | 8.71897 | 5.48322 | 6.09085 |
| chr17 | 28013373 | 28013763 | 1000 | Intron (ENSMUST00000114842.2 | 17 | 27988028 | 28008432 | 20405 | 2 | ENSMUST00 | -4942 | Gm15598 | ENSMUSG0( | false | . | . | . | . | . | . | chr17 | 28005688 | 28028167 | 8.71897 | 5.48322 | 6.09085 |
| chr17 | 28025406 | 28025894 | 1000 | Intron (ENSMUST00000114842.2 | 17 | 27988028 | 28008432 | 20405 | 2 | ENSMUST00 | -16975 | Gm15598 | ENSMUSG0( | false | . | . | . | . | . | . | chr17 | 28005688 | 28028167 | 8.71897 | 5.48322 | 6.09085 |
| chr4 | 32560282 | 32560606 | 1000 | Exon (ENSMUST00000126225.1/ | 4 | 32563097 | 32571617 | 8521 | 2 | ENSMUST00 | 11011 | Bach2os | ENSMUSG0( | false | . | . | . | . | . | . | chr4 | 32560057 | 32561997 | 3.65302 | 3.46599 | 1.89636 |
| chr2 | 72253999 | 72254302 | 1000 | Intron (ENSMUST00000090826.6 | 2 | 72244965 | 72248975 | 4011 | 2 | ENSMUST00 | -5025 | Rapgef4os3 | ENSMUSG0( | false | . | . | . | . | . | . | chr2 | 72253306 | 72255856 | 5.20614 | 4.2309 | 3.20005 |
| chr4 | 106232598 | 106233013 | 1000 | Distal Intergenic | 4 | 106241216 | 106290026 | 48811 | 1 | ENSMUST00 | -8203 | Gm12724 | ENSMUSG0( | false | . | . | . | . | . | . | chr4 | 106232380 | 106233132 | 4.25666 | 3.74435 | 2.44135 |
| chr7 | 124707122 | 124707520 | 1000 | Promoter (1-2kb) | 7 | 124625670 | 124708935 | 83266 | 2 | ENSMUST00 | 1415 | 3100003L05F | ENSMUSG0( | false | . | . | . | . | . | . | . | . | . | . | . | . |
| chr11 | 102624225 | 102624661 | 1000 | Promoter (<=1kb) | 11 | 102619796 | 102624416 | 4621 | 2 | ENSMUST00 | 0 | 2810433D01 | ENSMUSG0( | false | chr11 | 102623539 | 102624414 | 17.19387 | 43.56631 | 40.93357 | chr11 | 102621435 | 102629837 | 4.81998 | 3.90168 | 2.88673 |
| chr18 | 74498695 | 74499101 | 1000 | Intron (ENSMUST00000125882.2 | 18 | 74525253 | 74531106 | 5854 | 2 | ENSMUST00 | 32005 | 1700120E14 | ENSMUSG0( | false | . | . | . | . | . | . | chr18 | 74494956 | 74502808 | 5.10387 | 4.10112 | 3.10085 |
| chr4 | 129500767 | 129501320 | 1000 | Intron (ENSMUST00000145488.1 | 4 | 129491969 | 129509746 | 17778 | 2 | ENSMUST00 | 8426 | Gm12979 | ENSMUSG0( | false | chr4 | 129500396 | 129501680 | 25.98482 | 77.30067 | 74.19341 | chr4 | 129501210 | 129502253 | 4.52939 | 3.92768 | 2.61762 |
| chr2 | 18056131 | 18057333 | 1000 | Promoter (<=1kb) | 2 | 18054193 | 18056613 | 2421 | 2 | ENSMUST00 | 0 | 1810059C17 | ENSMUSG0( | false | chr2 | 18055597 | 18058275 | 30.32912 | 88.589 | 85.27629 | . | . | . | . | . | . |
| chr2 | 48441256 | 48441637 | 1000 | Intron (ENSMUST00000127672.1 | 2 | 48062548 | 48457245 | 394698 | 2 | ENSMUST00 | 15608 | Gm13481 | ENSMUSG0( | false | . | . | . | . | . | . | chr2 | 48441121 | 48442154 | 4.25067 | 3.59399 | 2.4484 |
| chr4 | 100214807 | 100215280 | 1000 | Intron (ENSMUST00000039630.5 | 4 | 100119621 | 100120841 | 1221 | 1 | ENSMUST00 | 95187 | Gm12701 | ENSMUSG0( | false | . | . | . | . | . | . | chr4 | 100214112 | 100219382 | 5.3099 | 4.17199 | 3.28584 |
| chr2 | 30648507 | 30649269 | 1000 | Intron (ENSMUST00000146702.1 | 2 | 30629655 | 30689210 | 59556 | 1 | ENSMUST00 | 18853 | 9330198N18 | ENSMUSG0( | false | . | . | . | . | . | . | chr2 | 30647143 | 30648739 | 3.19124 | 3.1382 | 1.54372 |
| chr2 | 5045305 | 5045896 | 1000 | Intron (ENSMUST00000114996.2 | 2 | 5037280 | 5041133 | 3854 | 1 | ENSMUST00 | 8026 | 4930551O13 | ENSMUSG0( | false | . | . | . | . | . | . | chr2 | 5041736 | 5062961 | 5.08721 | 4.02119 | 3.09837 |
| chr10 | 127600073 | 127600761 | 1000 | 3' UTR | 10 | 127596548 | 127597870 | 1323 | 1 | ENSMUST00 | 3526 | Gm16217 | ENSMUSG0( | false | . | . | . | . | . | . | chr10 | 127599312 | 127601246 | 6.5695 | 4.7699 | 4.33981 |
| chr6 | 121224964 | 121225427 | 1000 | Promoter (<=1kb) | 6 | 121223692 | 121226153 | 2462 | 2 | ENSMUST00 | 726 | Gm15856 | ENSMUSG0( | false | . | . | . | . | . | . | chr6 | 121224991 | 121225253 | 4.7038 | 4.01322 | 2.74337 |
| chr2 | 73592801 | 73593149 | 1000 | Distal Intergenic | 2 | 73596526 | 73616012 | 19487 | 1 | ENSMUST00 | -3377 | Chn1os3 | ENSMUSG0( | false | . | . | . | . | . | . | chr2 | 73592046 | 73597733 | 5.60047 | 4.20841 | 3.53877 |
| chr2 | 73596747 | 73597209 | 1000 | Promoter (<=1kb) | 2 | 73596824 | 73599200 | 2377 | 1 | ENSMUST00 | 0 | Chn1os3 | ENSMUSG0( | false | chr2 | 73596734 | 73596915 | 5.6323 | 9.47728 | 7.31216 | chr2 | 73592046 | 73597733 | 5.60047 | 4.20841 | 3.53877 |
| chr4 | 117982682 | 117983107 | 1000 | Intron (ENSMUST00000146982.1 | 4 | 117975036 | 117993972 | 18937 | 1 | ENSMUST00 | 7647 | 9530034E10 | ENSMUSG0( | false | . | . | . | . | . | . | chr4 | 117972459 | 117988550 | 5.57621 | 4.24526 | 3.52237 |
| chr18 | 50064299 | 50064637 | 1000 | Intron (ENSMUST00000148989.2 | 18 | 50052299 | 50053844 | 1546 | 2 | ENSMUST00 | -10456 | C030005K06 | ENSMUSG0( | false | chr18 | 50064550 | 50064868 | 7.81406 | 14.26947 | 12.02001 | chr18 | 50054967 | 50071969 | 5.75206 | 4.39775 | 3.62765 |
| chr18 | 50061844 | 50062226 | 1000 | Intron (ENSMUST00000148989.2 | 18 | 50052299 | 50053844 | 1546 | 2 | ENSMUST00 | -8001 | C030005K06 | ENSMUSG0( | false | . | . | . | . | . | . | chr18 | 50054967 | 50071969 | 5.75206 | 4.39775 | 3.62765 |
| chr2 | 160017098 | 160017415 | 1000 | Promoter (<=1kb) | 2 | 160016653 | 160040619 | 23967 | 1 | ENSMUST00 | 446 | 9130015L21F | ENSMUSG0( | false | . | . | . | . | . | . | chr2 | 160016048 | 160022298 | 5.11924 | 4.10769 | 3.10369 |
| chr2 | 160028037 | 160028428 | 1000 | Intron (ENSMUST00000140319.1 | 2 | 160016653 | 160040619 | 23967 | 1 | ENSMUST00 | 11385 | 9130015L21F | ENSMUSG0( | false | . | . | . | . | . | . | chr2 | 160025783 | 160029634 | 4.45414 | 3.76448 | 2.58441 |
| chr7 | 65500732 | 65501267 | 1000 | Distal Intergenic | 7 | 65582031 | 65618869 | 36839 | 1 | ENSMUST00 | -80764 | BC046251 | ENSMUSG0( | false | . | . | . | . | . | . | chr7 | 65500002 | 65503380 | 6.61736 | 4.8051 | 4.33052 |
| chr2 | 27327052 | 27327519 | 1000 | Promoter (<=1kb) | 2 | 27326704 | 27331297 | 4594 | 1 | ENSMUST00 | 349 | AA645442 | ENSMUSG0( | false | . | . | . | . | . | . | chr2 | 27326070 | 27327407 | 3.20725 | 3.13441 | 1.57971 |
| chr6 | 32388413 | 32388823 | 1000 | Intron (ENSMUST00000115096.3 | 6 | 32451170 | 32454750 | 3581 | 1 | ENSMUST00 | -62347 | Plxna4os2 | ENSMUSG0( | false | . | . | . | . | . | . | chr6 | 32386110 | 32389324 | 4.94933 | 3.90023 | 2.9937 |
| chr6 | 32397021 | 32397584 | 1000 | Intron (ENSMUST00000115096.3 | 6 | 32451170 | 32454750 | 3581 | 1 | ENSMUST00 | -53586 | Plxna4os2 | ENSMUSG0( | false | . | . | . | . | . | . | chr6 | 32395935 | 32398151 | 4.98785 | 4.02539 | 3.03457 |
| chr11 | 62920758 | 62921877 | 1000 | Promoter (<=1kb) | 11 | 62919079 | 62921592 | 2514 | 2 | ENSMUST00 | 0 | Tvp23bos | ENSMUSG0( | false | chr11 | 62921017 | 62921848 | 5.86754 | 9.51579 | 7.34688 | chr11 | 62920175 | 62922639 | 4.05062 | 3.66091 | 2.21182 |
| chr8 | 125342588 | 125343351 | 1000 | Distal Intergenic | 8 | 125224400 | 125227323 | 2924 | 2 | ENSMUST00 | -115266 | Gm16237 | ENSMUSG0( | false | . | . | . | . | . | . | chr8 | 125335216 | 125352727 | 6.82037 | 4.91647 | 4.50391 |
| chr11 | 44232673 | 44233100 | 1000 | Distal Intergenic | 11 | 44265563 | 44267209 | 1647 | 2 | ENSMUST00 | 34109 | Gm12154 | ENSMUSG0( | false | chr11 | 44232741 | 44233137 | 9.62697 | 19.18633 | 16.86223 | chr11 | 44232041 | 44233438 | 4.37741 | 3.57635 | 2.52369 |
| chr4 | 148967656 | 148968128 | 1000 | Exon (ENSMUST00000145143.1/ | 4 | 148960360 | 148966998 | 6639 | 1 | ENSMUST00 | 7297 | Gm17029 | ENSMUSG0( | false | chr4 | 148966100 | 148968885 | 16.91761 | 46.11281 | 43.44656 | . | . | . | . | . | . |
| chr4 | 148969421 | 148969977 | 1000 | Exon (ENSMUST00000128067.2/ | 4 | 148960360 | 148966998 | 6639 | 1 | ENSMUST00 | 9062 | Gm17029 | ENSMUSG0( | false | . | . | . | . | . | . | chr4 | 148968291 | 148974289 | 7.92118 | 5.40621 | 5.39072 |
| chr3 | 102747134 | 102747881 | 1000 | Intron (ENSMUST00000029451.6 | 3 | 102720231 | 102735417 | 15187 | 2 | ENSMUST00 | -11718 | Tspan2os | ENSMUSG0( | false | . | . | . | . | . | . | chr3 | 102735904 | 102757857 | 6.42245 | 4.63759 | 4.19001 |
| chr3 | 102736490 | 102736866 | 1000 | Promoter (1-2kb) | 3 | 102720231 | 102735417 | 15187 | 2 | ENSMUST00 | -1074 | Tspan2os | ENSMUSG0( | false | . | . | . | . | . | . | chr3 | 102735904 | 102757857 | 6.42245 | 4.63759 | 4.19001 |

| | | | | | | | | | | | | | | | | | | | | | | | | | | |
|---|---|---|---|---|---|---|---|---|---|---|---|---|---|---|---|---|---|---|---|---|---|---|---|---|---|---|
| chr3 | 102748077 | 102748462 | 1000 | Intron (ENSMUST00000029451.6 | 3 | 102720231 | 102735417 | 15187 | 2 | ENSMUST00 | -12661 | Tspan2os | ENSMUSG0C | false | . | . | . | . | . | . | chr3 | 102735904 | 102757857 | 6.42245 | 4.63759 | 4.19001 |
| chr14 | 46387271 | 46387647 | 1000 | Promoter (<=1kb) | 14 | 46387519 | 46389282 | 1764 | 1 | ENSMUST00 | 0 | Gm15222 | ENSMUSG0C | false | chr14 | 46386588 | 46388392 | 26.19333 | 73.09374 | 70.054 | chr14 | 46385786 | 46390843 | 3.99631 | 3.58417 | 2.19112 |
| chr11 | 20833226 | 20833621 | 1000 | Promoter (1-2kb) | 11 | 20831481 | 20831919 | 439 | 2 | ENSMUST00 | -1308 | Gm12037 | ENSMUSG0C | false | . | . | . | . | . | . | chr11 | 20831700 | 20835450 | 5.08603 | 4.1192 | 3.07417 |
| chr11 | 20832671 | 20832975 | 1000 | Promoter (<=1kb) | 11 | 20831481 | 20831919 | 439 | 2 | ENSMUST00 | -753 | Gm12037 | ENSMUSG0C | false | . | . | . | . | . | . | chr11 | 20831700 | 20835450 | 5.08603 | 4.1192 | 3.07417 |
| chr4 | 148918988 | 148919540 | 1000 | Intron (ENSMUST00000094464.4 | 4 | 148913511 | 148914324 | 814 | 2 | ENSMUST00 | -4665 | Gm15969 | ENSMUSG0C | false | . | . | . | . | . | . | chr4 | 148901819 | 148933146 | 7.00771 | 4.91323 | 4.66351 |
| chr6 | 31629928 | 31630444 | 1000 | Intron (ENSMUST00000124919.2 | 6 | 31651041 | 31662021 | 10981 | 1 | ENSMUST00 | -20597 | Gm13846 | ENSMUSG0C | false | . | . | . | . | . | . | chr6 | 31612785 | 31643065 | 5.56305 | 4.31748 | 3.47573 |
| chr6 | 31627854 | 31628506 | 1000 | Intron (ENSMUST00000124919.2 | 6 | 31651041 | 31662021 | 10981 | 1 | ENSMUST00 | -22535 | Gm13846 | ENSMUSG0C | false | . | . | . | . | . | . | chr6 | 31612785 | 31643065 | 5.56305 | 4.31748 | 3.47573 |
| chr13 | 29954725 | 29955074 | 1000 | Intron (ENSMUST00000102948.4 | 13 | 29940283 | 29942261 | 1979 | 1 | ENSMUST00 | 14443 | Gm11365 | ENSMUSG0C | false | . | . | . | . | . | . | chr13 | 29953873 | 29962790 | 5.9859 | 4.53593 | 3.82195 |
| chr13 | 29957663 | 29958295 | 1000 | Intron (ENSMUST00000102948.4 | 13 | 29940283 | 29942261 | 1979 | 1 | ENSMUST00 | 17381 | Gm11365 | ENSMUSG0C | false | . | . | . | . | . | . | chr13 | 29953873 | 29962790 | 5.9859 | 4.53593 | 3.82195 |
| chr2 | 67242098 | 67242474 | 1000 | Distal Intergenic | 2 | 67157646 | 67194735 | 37090 | 2 | ENSMUST00 | -47364 | Gm13598 | ENSMUSG0C | false | . | . | . | . | . | . | . | . | . | . | . | . |
| chr6 | 31148305 | 31148748 | 1000 | Intron (ENSMUST00000187876.1 | 6 | 31141979 | 31142663 | 685 | 2 | ENSMUST00 | -5643 | Gm13835 | ENSMUSG0C | false | . | . | . | . | . | . | chr6 | 31141046 | 31151719 | 5.50019 | 4.22494 | 3.43224 |
| chr11 | 119938167 | 119938495 | 1000 | Promoter (<=1kb) | 11 | 119938511 | 119942074 | 3564 | 1 | ENSMUST00 | -16 | Gm11766 | ENSMUSG0C | false | . | . | . | . | . | . | chr11 | 119937453 | 119942377 | 6.8796 | 4.95571 | 4.54178 |
| chr11 | 57682978 | 57683508 | 1000 | Intron (ENSMUST00000066987.8 | 11 | 57655415 | 57657826 | 2412 | 2 | ENSMUST00 | -25153 | 4933426K07I | ENSMUSG0C | false | . | . | . | . | . | . | chr11 | 57646153 | 57704104 | 7.42247 | 5.08195 | 5.00645 |
| chr11 | 57672957 | 57673284 | 1000 | Intron (ENSMUST00000066987.8 | 11 | 57655415 | 57657826 | 2412 | 2 | ENSMUST00 | -15132 | 4933426K07I | ENSMUSG0C | false | . | . | . | . | . | . | chr11 | 57646153 | 57704104 | 7.42247 | 5.08195 | 5.00645 |
| chr4 | 106839798 | 106840529 | 1000 | Downstream (1-2kb) | 4 | 106841761 | 106847920 | 6160 | 2 | ENSMUST00 | 7391 | Gm12746 | ENSMUSG0C | false | chr4 | 106839087 | 106840723 | 27.09851 | 84.88239 | 81.64215 | chr4 | 106837700 | 106843971 | 4.90694 | 4.01968 | 2.94365 |
| chr4 | 106827725 | 106828409 | 1000 | Distal Intergenic | 4 | 106841761 | 106847920 | 6160 | 2 | ENSMUST00 | 19511 | Gm12746 | ENSMUSG0C | false | chr4 | 106827796 | 106828358 | 9.22042 | 18.50693 | 16.19293 | chr4 | 106827481 | 106829093 | 5.48912 | 4.25467 | 3.41499 |
| chr4 | 106858356 | 106858730 | 1000 | Distal Intergenic | 4 | 106841761 | 106847920 | 6160 | 2 | ENSMUST00 | -10437 | Gm12746 | ENSMUSG0C | false | chr4 | 106858151 | 106858528 | 4.88133 | 7.58987 | 5.46511 | chr4 | 106853465 | 106864205 | 5.81032 | 4.40131 | 3.67789 |
| chr10 | 39557798 | 39558346 | 1000 | Promoter (<=1kb) | 10 | 39556071 | 39558062 | 1992 | 2 | ENSMUST00 | 0 | Gm16364 | ENSMUSG0C | false | . | . | . | . | . | . | chr10 | 39550959 | 39564232 | 7.35607 | 5.07236 | 4.95132 |
| chr10 | 39561652 | 39562583 | 1000 | Exon (ENSMUST00000122987.1/ | 10 | 39556071 | 39558062 | 1992 | 2 | ENSMUST00 | -3591 | Gm16364 | ENSMUSG0C | false | . | . | . | . | . | . | chr10 | 39550959 | 39564232 | 7.35607 | 5.07236 | 4.95132 |
| chr4 | 148327223 | 148327858 | 1000 | Distal Intergenic | 4 | 148306853 | 148316569 | 9717 | 1 | ENSMUST00 | 20371 | Gm13206 | ENSMUSG0C | false | . | . | . | . | . | . | chr4 | 148324659 | 148328387 | 5.58334 | 4.25719 | 3.51337 |
| chr11 | 117103605 | 117103957 | 1000 | Distal Intergenic | 11 | 117094948 | 117097067 | 2120 | 2 | ENSMUST00 | -6539 | Gm11730 | ENSMUSG0C | false | . | . | . | . | . | . | chr11 | 117097696 | 117104768 | 5.85805 | 4.31896 | 3.74714 |
| chr11 | 97025702 | 97026182 | 1000 | Promoter (<=1kb) | 11 | 97020508 | 97026248 | 5741 | 2 | ENSMUST00 | 66 | Gm11532 | ENSMUSG0C | false | chr11 | 97025529 | 97026251 | 14.11588 | 32.66511 | 30.16695 | chr11 | 97020867 | 97026684 | 5.07759 | 4.06029 | 3.06843 |
| chr12 | 111406638 | 111407047 | 1000 | Promoter (<=1kb) | 12 | 111406809 | 111412077 | 5269 | 1 | ENSMUST00 | 0 | A230065H16 | ENSMUSG0C | false | . | . | . | . | . | . | chr12 | 111406302 | 111407412 | 6.60325 | 4.90675 | 4.30432 |
| chr2 | 143682145 | 143682474 | 1000 | Intron (ENSMUST00000028905.9 | 2 | 143747583 | 143789186 | 41604 | 2 | ENSMUST00 | 106712 | Pcsk2os2 | ENSMUSG0C | false | . | . | . | . | . | . | chr2 | 143678793 | 143683818 | 4.17322 | 3.54281 | 2.35985 |
| chr2 | 6164704 | 6165126 | 1000 | Distal Intergenic | 2 | 6193251 | 6321611 | 128361 | 1 | ENSMUST00 | -28125 | A230108P19 | ENSMUSG0C | false | . | . | . | . | . | . | chr2 | 6162576 | 6166166 | 4.08213 | 3.61767 | 2.25524 |
| chr2 | 163691944 | 163692437 | 1000 | Promoter (<=1kb) | 2 | 163685004 | 163692279 | 7276 | 2 | ENSMUST00 | 0 | Gm16316 | ENSMUSG0C | false | . | . | . | . | . | . | chr2 | 163667073 | 163719465 | 6.42152 | 4.6796 | 4.1796 |
| chr2 | 163685457 | 163686062 | 1000 | Exon (ENSMUST00000154065.1/ | 2 | 163685004 | 163692279 | 7276 | 2 | ENSMUST00 | 6217 | Gm16316 | ENSMUSG0C | false | . | . | . | . | . | . | chr2 | 163667073 | 163719465 | 6.42152 | 4.6796 | 4.1796 |
| chr1 | 38368533 | 38368922 | 1000 | Intron (ENSMUST00000095027.3 | 1 | 38384949 | 38393974 | 9026 | 1 | ENSMUST00 | -16027 | Gm16152 | ENSMUSG0C | false | . | . | . | . | . | . | chr1 | 38366741 | 38371656 | 5.95599 | 4.51574 | 3.80421 |
| chr11 | 57786370 | 57786998 | 1000 | 3' UTR | 11 | 57795606 | 57801398 | 5793 | 2 | ENSMUST00 | 14400 | 2010001A14I | ENSMUSG0C | false | . | . | . | . | . | . | chr11 | 57782717 | 57788725 | 7.44149 | 5.10953 | 5.01045 |
| chr2 | 18693917 | 18694727 | 1000 | Promoter (<=1kb) | 2 | 18694032 | 18698884 | 4853 | 1 | ENSMUST00 | 0 | Gm13334 | ENSMUSG0C | false | chr2 | 18693722 | 18694863 | 39.38308 | 128.91518 | 124.46609 | chr2 | 18693178 | 18695519 | 3.34231 | 3.21011 | 1.67202 |
| chr2 | 36224686 | 36225098 | 1000 | Distal Intergenic | 2 | 36220490 | 36222865 | 2376 | 1 | ENSMUST00 | 4197 | Gm13431 | ENSMUSG0C | false | . | . | . | . | . | . | . | . | . | . | . | . |
| chr2 | 92139422 | 92139777 | 1000 | Intron (ENSMUST00000044036.9 | 2 | 92043373 | 92096042 | 52670 | 2 | ENSMUST00 | -43381 | Gm13780 | ENSMUSG0C | false | . | . | . | . | . | . | chr2 | 92138729 | 92141360 | 6.1788 | 4.69447 | 3.95993 |
| chr15 | 4027024 | 4027736 | 1000 | Promoter (<=1kb) | 15 | 4021686 | 4027393 | 5708 | 2 | ENSMUST00 | 0 | BC037032 | ENSMUSG0C | false | chr15 | 4025971 | 4028201 | 33.65816 | 114.56059 | 110.59126 | . | . | . | . | . | . |
| chr5 | 74003930 | 74004399 | 1000 | Intron (ENSMUST00000119154.2 | 5 | 74014216 | 74029680 | 15465 | 1 | ENSMUST00 | -9817 | Usp46os1 | ENSMUSG0C | false | . | . | . | . | . | . | chr5 | 74001579 | 74008551 | 5.27799 | 4.06988 | 3.27177 |
| chr10 | 86164834 | 86165365 | 1000 | Intron (ENSMUST00000120638.2 | 10 | 86125827 | 86129854 | 4028 | 1 | ENSMUST00 | 39008 | Gm15990 | ENSMUSG0C | false | . | . | . | . | . | . | chr10 | 86157796 | 86167524 | 6.43176 | 4.7099 | 4.1851 |
| chr10 | 86123949 | 86124435 | 1000 | Promoter (1-2kb) | 10 | 86125827 | 86129854 | 4028 | 1 | ENSMUST00 | -1392 | Gm15990 | ENSMUSG0C | false | . | . | . | . | . | . | chr10 | 86122373 | 86133417 | 5.202 | 4.08244 | 3.19933 |
| chr10 | 86114385 | 86115009 | 1000 | Intron (ENSMUST00000120638.2 | 10 | 86125827 | 86129854 | 4028 | 1 | ENSMUST00 | -10818 | Gm15990 | ENSMUSG0C | false | . | . | . | . | . | . | chr10 | 86112561 | 86116672 | 5.219 | 4.14441 | 3.19963 |
| chr11 | 113165591 | 113165935 | 1000 | Intron (ENSMUST00000136438.2 | 11 | 113043895 | 113173077 | 129183 | 2 | ENSMUST00 | 7142 | 2610035D17I | ENSMUSG0C | false | . | . | . | . | . | . | chr11 | 113163715 | 113168269 | 5.39105 | 4.28786 | 3.31527 |
| chr2 | 30658191 | 30659443 | 1000 | Exon (ENSMUST00000147959.1/ | 2 | 30658468 | 30669908 | 11441 | 2 | ENSMUST00 | 10465 | Gm14486 | ENSMUSG0C | false | chr2 | 30658018 | 30659016 | 18.88512 | 67.59834 | 64.64505 | chr2 | 30656689 | 30660996 | 5.17631 | 4.15173 | 3.18095 |
| chr4 | 148710582 | 148710881 | 1000 | Promoter (2-3kb) | 4 | 148713703 | 148716581 | 2879 | 1 | ENSMUST00 | -2822 | Gm13203 | ENSMUSG0C | false | . | . | . | . | . | . | chr4 | 148709604 | 148711445 | 4.91207 | 4.11321 | 2.94661 |
| chr5 | 43586934 | 43587252 | 1000 | Intron (ENSMUST00000167522.1 | 5 | 43557285 | 43601730 | 44446 | 2 | ENSMUST00 | 14478 | Gm15866 | ENSMUSG0C | false | . | . | . | . | . | . | chr5 | 43560201 | 43596005 | 4.90131 | 4.0072 | 2.93052 |
| chr6 | 144105182 | 144105742 | 1000 | Exon (ENSMUST00000149451.2/ | 6 | 144021908 | 144023778 | 1871 | 1 | ENSMUST00 | 83275 | Sox5os1 | ENSMUSG0C | false | . | . | . | . | . | . | chr6 | 144103132 | 144107582 | 5.73038 | 4.43689 | 3.60871 |
| chr6 | 8473502 | 8474644 | 1000 | Intron (ENSMUST00000162564.2 | 6 | 8489933 | 8497979 | 8047 | 2 | ENSMUST00 | 23335 | A430035B10 | ENSMUSG0C | false | . | . | . | . | . | . | chr6 | 8462693 | 8475616 | 4.97986 | 4.07957 | 2.98249 |
| chr4 | 134446005 | 134446411 | 1000 | Downstream (2-3kb) | 4 | 134448766 | 134450171 | 1406 | 2 | ENSMUST00 | 3760 | 1700021N21 | ENSMUSG0C | false | . | . | . | . | . | . | chr4 | 134444756 | 134452233 | 5.18386 | 4.23712 | 3.15068 |
| chr4 | 134450835 | 134451310 | 1000 | Promoter (<=1kb) | 4 | 134448766 | 134450171 | 1406 | 2 | ENSMUST00 | -665 | 1700021N21 | ENSMUSG0C | false | . | . | . | . | . | . | chr4 | 134444756 | 134452233 | 5.18386 | 4.23712 | 3.15068 |
| chr7 | 132714307 | 132714742 | 1000 | 3' UTR | 7 | 132657378 | 132659884 | 2507 | 2 | ENSMUST00 | -54424 | Gm15582 | ENSMUSG0C | false | . | . | . | . | . | . | chr7 | 132713756 | 132715475 | 5.40793 | 4.13039 | 3.3711 |
| chr6 | 34145973 | 34146444 | 1000 | Distal Intergenic | 6 | 34151929 | 34174235 | 22307 | 1 | ENSMUST00 | -5485 | Gm13856 | ENSMUSG0C | false | . | . | . | . | . | . | chr6 | 34145432 | 34157774 | 5.20587 | 4.07382 | 3.18968 |
| chr6 | 34101931 | 34102330 | 1000 | Intron (ENSMUST00000070189.4 | 6 | 34151929 | 34174235 | 22307 | 1 | ENSMUST00 | -49599 | Gm13856 | ENSMUSG0C | false | . | . | . | . | . | . | chr6 | 34101369 | 34102259 | 4.24429 | 3.72649 | 2.37464 |
| chr4 | 95052246 | 95053421 | 1000 | Promoter (<=1kb) | 4 | 95052951 | 95060658 | 7708 | 1 | ENSMUST00 | 0 | Junos | ENSMUSG0C | false | chr4 | 95052498 | 95053357 | 25.86957 | 74.34999 | 71.29122 | . | . | . | . | . | . |
| chr13 | 41605508 | 41606174 | 1000 | Promoter (<=1kb) | 13 | 41606216 | 41641347 | 35132 | 1 | ENSMUST00 | -42 | Tmem170b | ENSMUSG0C | false | chr13 | 41605028 | 41605689 | 25.1466 | 74.00616 | 70.95453 | . | . | . | . | . | . |
| chr6 | 31288338 | 31288723 | 1000 | Intron (ENSMUST00000144612.2 | 6 | 31285004 | 31337403 | 52400 | 1 | ENSMUST00 | 3335 | 2210408F21I | ENSMUSG0C | false | . | . | . | . | . | . | chr6 | 31287055 | 31289853 | 5.83326 | 4.44407 | 3.71842 |
| chr6 | 31220218 | 31220637 | 1000 | Promoter (<=1kb) | 6 | 31220351 | 31337247 | 116897 | 1 | ENSMUST00 | 0 | 2210408F21I | ENSMUSG0C | false | chr6 | 31219869 | 31221288 | 31.84599 | 102.09332 | 98.46812 | chr6 | 31218871 | 31221916 | 4.82639 | 3.95698 | 2.89638 |
| chr10 | 24613537 | 24613891 | 1000 | Distal Intergenic | 10 | 24595249 | 24597108 | 1860 | 2 | ENSMUST00 | -16430 | Gm15270 | ENSMUSG0C | false | . | . | . | . | . | . | . | . | . | . | . | . |
| chr10 | 24591236 | 24591693 | 1000 | Promoter (2-3kb) | 10 | 24580048 | 24593758 | 13711 | 2 | ENSMUST00 | 2065 | Gm15270 | ENSMUSG0C | false | . | . | . | . | . | . | chr10 | 24589538 | 24595113 | 3.89644 | 3.5432 | 2.09048 |
| chr3 | 90446170 | 90446932 | 1000 | Distal Intergenic | 3 | 90451583 | 90457095 | 5513 | 1 | ENSMUST00 | -4651 | Gm16048 | ENSMUSG0C | false | . | . | . | . | . | . | chr3 | 90444443 | 90452742 | 6.22784 | 4.66702 | 4.02057 |
| chr3 | 90447167 | 90447613 | 1000 | Downstream (2-3kb) | 3 | 90451583 | 90457095 | 5513 | 1 | ENSMUST00 | -3970 | Gm16048 | ENSMUSG0C | false | . | . | . | . | . | . | chr3 | 90444443 | 90452742 | 6.22784 | 4.66702 | 4.02057 |
| chr4 | 129129362 | 129129700 | 1000 | Distal Intergenic | 4 | 129135091 | 129136910 | 1820 | 1 | ENSMUST00 | -5391 | Gm15906 | ENSMUSG0C | false | . | . | . | . | . | . | chr4 | 129128207 | 129131816 | 5.97344 | 4.48503 | 3.80935 |
| chr4 | 129130612 | 129131037 | 1000 | Distal Intergenic | 4 | 129135091 | 129136910 | 1820 | 1 | ENSMUST00 | -4054 | Gm15906 | ENSMUSG0C | false | . | . | . | . | . | . | chr4 | 129128207 | 129131816 | 5.97344 | 4.48503 | 3.80935 |
| chr4 | 137133795 | 137134155 | 1000 | Distal Intergenic | 4 | 137117143 | 137118143 | 1001 | 2 | ENSMUST00 | -15653 | Gm13001 | ENSMUSG0C | false | . | . | . | . | . | . | chr4 | 137132391 | 137134931 | 5.71815 | 4.2776 | 3.65042 |
| chr4 | 153263340 | 153263737 | 1000 | Distal Intergenic | 4 | 153218176 | 153218595 | 420 | 1 | ENSMUST00 | 45165 | Gm13174 | ENSMUSG0C | false | . | . | . | . | . | . | chr4 | 153261073 | 153271626 | 5.04729 | 4.02401 | 3.0775 |
| chr4 | 153230821 | 153231154 | 1000 | Distal Intergenic | 4 | 153218176 | 153218595 | 420 | 1 | ENSMUST00 | 12646 | Gm13174 | ENSMUSG0C | false | . | . | . | . | . | . | chr4 | 153219753 | 153237639 | 5.81029 | 4.42202 | 3.67688 |
| chr4 | 153236172 | 153236651 | 1000 | Distal Intergenic | 4 | 153218176 | 153218595 | 420 | 1 | ENSMUST00 | 17997 | Gm13174 | ENSMUSG0C | false | . | . | . | . | . | . | chr4 | 153219753 | 153237639 | 5.81029 | 4.42202 | 3.67688 |
| chr4 | 153268968 | 153269329 | 1000 | Distal Intergenic | 4 | 153218176 | 153218595 | 420 | 1 | ENSMUST00 | 50793 | Gm13174 | ENSMUSG0C | false | . | . | . | . | . | . | chr4 | 153261073 | 153271626 | 5.04729 | 4.02401 | 3.0775 |
| chr4 | 153224619 | 153225105 | 1000 | Distal Intergenic | 4 | 153218176 | 153218595 | 420 | 1 | ENSMUST00 | 6444 | Gm13174 | ENSMUSG0C | false | . | . | . | . | . | . | chr4 | 153219753 | 153237639 | 5.81029 | 4.42202 | 3.67688 |
| chr4 | 153207213 | 153208043 | 1000 | Distal Intergenic | 4 | 153218176 | 153218595 | 420 | 1 | ENSMUST00 | -10133 | Gm13174 | ENSMUSG0C | false | . | . | . | . | . | . | chr4 | 153205812 | 153211213 | 5.02003 | 4.09646 | 3.02461 |
| chr3 | 37976356 | 37977041 | 1000 | Distal Intergenic | 3 | 37959612 | 37960087 | 476 | 2 | ENSMUST00 | -16270 | Gm7799 | ENSMUSG0C | false | . | . | . | . | . | . | chr3 | 37970670 | 37979109 | 4.79981 | 3.95584 | 2.8516 |
| chr14 | 25460176 | 25461770 | 1000 | Promoter (<=1kb) | 14 | 25455737 | 25461128 | 5392 | 2 | ENSMUST00 | 0 | Zmiz1os1 | ENSMUSG0C | false | chr14 | 25457181 | 25461653 | 31.54088 | 108.67455 | 104.87239 | chr14 | 25460928 | 25485304 | 5.40667 | 4.23635 | 3.3428 |
| chr4 | 129263438 | 129263874 | 1000 | Promoter (<=1kb) | 4 | 129263730 | 129265010 | 1281 | 1 | ENSMUST00 | 0 | Gm12976 | ENSMUSG0C | false | . | . | . | . | . | . | chr4 | 129260820 | 129264597 | 5.77718 | 4.46944 | 3.66067 |
| chr19 | 36688113 | 36688757 | 1000 | Promoter (<=1kb) | 19 | 36645683 | 36689273 | 43591 | 2 | ENSMUST00 | 516 | 1500017E21I | ENSMUSG0C | false | . | . | . | . | . | . | chr19 | 36687857 | 36690681 | 7.37966 | 5.24694 | 4.94518 |
| chr19 | 36680175 | 36680594 | 1000 | Intron (ENSMUST00000149601.2 | 19 | 36645683 | 36689273 | 43591 | 2 | ENSMUST00 | 8679 | 1500017E21I | ENSMUSG0C | false | . | . | . | . | . | . | chr19 | 36677521 | 36681530 | 4.66501 | 3.92586 | 2.73896 |
| chr18 | 33757729 | 33758088 | 1000 | Distal Intergenic | 18 | 33794892 | 33795986 | 1095 | 1 | ENSMUST00 | -36804 | 2410004N09 | ENSMUSG0C | false | . | . | . | . | . | . | chr18 | 33756350 | 33758760 | 3.69256 | 3.38301 | 1.95191 |
| chr7 | 19816246 | 19816721 | 1000 | Promoter (<=1kb) | 7 | 19816553 | 19818625 | 2073 | 1 | ENSMUST00 | 0 | Gm16174 | ENSMUSG0C | false | chr7 | 19816114 | 19817765 | 13.64722 | 31.21559 | 28.73584 | chr7 | 19815747 | 19818650 | 6.26859 | 4.57703 | 4.06494 |
| chr7 | 19817349 | 19817849 | 1000 | Promoter (<=1kb) | 7 | 19816553 | 19818625 | 2073 | 1 | ENSMUST00 | 797 | Gm16174 | ENSMUSG0C | false | chr7 | 19816114 | 19817765 | 13.64722 | 31.21559 | 28.73584 | chr7 | 19815747 | 19818650 | 6.26859 | 4.57703 | 4.06494 |
| chr2 | 145682834 | 145683339 | 1000 | Intron (ENSMUST00000144661.1 | 2 | 145675367 | 145714989 | 39623 | 1 | ENSMUST00 | 7468 | BC039771 | ENSMUSG0C | false | . | . | . | . | . | . | chr2 | 145676010 | 145686155 | 5.31951 | 4.23417 | 3.27212 |
| chr18 | 56756106 | 56756473 | 1000 | Distal Intergenic | 18 | 56769717 | 56801688 | 31972 | 1 | ENSMUST00 | -13244 | Gm15345 | ENSMUSG0C | false | chr18 | 56755851 | 56756882 | 16.14593 | 43.98602 | 41.34693 | chr18 | 56753533 | 56757612 | 3.84422 | 3.46396 | 2.09571 |
| chr2 | 173277105 | 173277517 | 1000 | Promoter (<=1kb) | 2 | 173276698 | 173278428 | 1731 | 1 | ENSMUST00 | 408 | Pmepa1os | ENSMUSG0C | false | chr2 | 173274005 | 173277330 | 35.62435 | 116.99458 | 112.94973 | chr2 | 173276948 | 173281914 | 5.70628 | 4.41535 | 3.58938 |
| chr2 | 173307080 | 173307943 | 1000 | Distal Intergenic | 2 | 173276698 | 173278428 | 1731 | 1 | ENSMUST00 | 30383 | Pmepa1os | ENSMUSG0C | false | . | . | . | . | . | . | chr2 | 173304893 | 173308639 | 4.66531 | 3.76203 | 2.75195 |
| chr16 | 30964601 | 30965292 | 1000 | Intron (ENSMUST00000055389.7 | 16 | 30969393 | 30973518 | 4126 | 2 | ENSMUST00 | 8226 | Gm15742 | ENSMUSG0C | false | . | . | . | . | . | . | chr16 | 30959851 | 30970783 | 4.98026 | 4.05336 | 2.99614 |
| chr3 | 96576689 | 96577338 | 1000 | Promoter (<=1kb) | 3 | 96576984 | 96580663 | 3680 | 1 | ENSMUST00 | 0 | Gm16253 | ENSMUSG0C | false | chr3 | 96576347 | 96576879 | 8.00962 | 15.03963 | 12.77451 | chr3 | 96568864 | 96577867 | 5.10498 | 4.12212 | 3.09255 |
| chr11 | 95034862 | 95035369 | 1000 | Promoter (2-3kb) | 11 | 95032759 | 95038283 | 5525 | 1 | ENSMUST00 | 2104 | Gm11513 | ENSMUSG0C | false | . | . | . | . | . | . | chr11 | 95031837 | 95049061 | 6.83336 | 4.76219 | 4.52264 |
| chr2 | 168154456 | 168154822 | 1000 | Promoter (<=1kb) | 2 | 168152603 | 168154513 | 1911 | 2 | ENSMUST00 | 0 | E130018N17 | ENSMUSG0C | false | . | . | . | . | . | . | chr2 | 168154208 | 168155668 | 3.10814 | 3.0829 | 1.47337 |
| chr2 | 26139938 | 26140819 | 1000 | Promoter (<=1kb) | 2 | 26136814 | 26140521 | 3708 | 2 | ENSMUST00 | 0 | C330006A16I | ENSMUSG0C | false | chr2 | 26139348 | 26141046 | 31.94275 | 111.4613 | 107.58682 | chr2 | 26140802 | 26142075 | 5.21147 | 4.12897 | 3.17997 |
| chr2 | 62448408 | 62448847 | 1000 | Intron (ENSMUST00000136879.1 | 2 | 62436880 | 62441142 | 4263 | 2 | ENSMUST00 | -7267 | Gm13565 | ENSMUSG0C | false | . | . | . | . | . | . | chr2 | 62446586 | 62450644 | 3.81602 | 3.45995 | 2.04157 |
| chr3 | 94917653 | 94918111 | 1000 | Promoter (2-3kb) | 3 | 94920322 | 94920423 | 102 | 2 | ENSMUST00 | 2312 | Gm26279 | ENSMUSG0C | false | . | . | . | . | . | . | chr3 | 94916017 | 94919084 | 4.21207 | 3.71027 | 2.36291 |
| chr3 | 122509747 | 122510185 | 1000 | Intron (ENSMUST00000029766.4 | 3 | 122464727 | 122464882 | 156 | 1 | ENSMUST00 | 45021 | Gm25153 | ENSMUSG0C | false | . | . | . | . | . | . | chr3 | 122468157 | 122521853 | 6.29611 | 4.62489 | 4.08524 |
| chr3 | 122482273 | 122483421 | 1000 | Intron (ENSMUST00000029766.4 | 3 | 122464727 | 122464882 | 156 | 1 | ENSMUST00 | 17547 | Gm25153 | ENSMUSG0C | false | chr3 | 122483254 | 122485690 | 12.74236 | 36.28267 | 33.73857 | chr3 | 122468157 | 122521853 | 6.29611 | 4.62489 | 4.08524 |
| chr8 | 77842344 | 77842757 | 1000 | Distal Intergenic | 8 | 77807535 | 77807640 | 106 | 1 | ENSMUST00 | 34810 | Gm23981 | ENSMUSG0C | false | . | . | . | . | . | . | chr8 | 77841927 | 77843389 | 4.2209 | 3.68629 | 2.36683 |
| chr5 | 20256469 | 20256800 | 1000 | Intron (ENSMUST00000088516.4 | 5 | 20264341 | 20264440 | 100 | 1 | ENSMUST00 | -7541 | Gm22755 | ENSMUSG0C | false | . | . | . | . | . | . | chr5 | 20247256 | 20259329 | 3.85493 | 3.37072 | 2.09981 |
| chr10 | 70494505 | 70494856 | 1000 | Intron (ENSMUST00000105436.3 | 10 | 70501192 | 70501503 | 312 | 2 | ENSMUST00 | 6647 | Gm22320 | ENSMUSG0C | false | . | . | . | . | . | . | chr10 | 70492552 | 70498239 | 4.70976 | 3.88142 | 2.80749 |
| chr6 | 89239477 | 89240380 | 1000 | Distal Intergenic | 6 | 89249468 | 89249602 | 135 | 2 | ENSMUST00 | 9222 | Gm25961 | ENSMUSG0C | false | . | . | . | . | . | . | chr6 | 89239017 | 89240690 | 5.25248 | 3.97781 | 3.24423 |
| chr9 | 113340798 | 113341237 | 1000 | Distal Intergenic | 9 | 113403475 | 113403771 | 297 | 1 | ENSMUST00 | -62238 | Gm22492 | ENSMUSG0C | false | . | . | . | . | . | . | chr9 | 113339166 | 113343758 | 4.62012 | 3.83176 | 2.72203 |
| chr3 | 65482687 | 65483056 | 1000 | Distal Intergenic | 3 | 65525701 | 65525793 | 93 | 1 | ENSMUST00 | -42645 | Gm22279 | ENSMUSG0C | false | . | . | . | . | . | . | chr3 | 65481033 | 65483719 | 6.2348 | 4.57775 | 4.05042 |
| chr4 | 114842927 | 114843824 | 1000 | Promoter (1-2kb) | 4 | 114844985 | 114845117 | 133 | 2 | ENSMUST00 | 1293 | Gm23230 | ENSMUSG0C | false | chr4 | 114842462 | 114842938 | 10.56924 | 21.28106 | 18.92771 | chr4 | 114841210 | 114845011 | 4.22202 | 3.73999 | 2.3678 |
| chr4 | 114869859 | 114870519 | 1000 | Intron (ENSMUST00000155499.1 | 4 | 114844985 | 114845117 | 133 | 2 | ENSMUST00 | -24743 | Gm23230 | ENSMUSG0C | false | . | . | . | . | . | . | chr4 | 114865811 | 114874373 | 4.93378 | 4.02281 | 2.96897 |
| chr4 | 62878933 | 62879311 | 1000 | Distal Intergenic | 4 | 62939341 | 62939474 | 134 | 1 | ENSMUST00 | -60030 | Gm25391 | ENSMUSG0C | false | . | . | . | . | . | . | chr4 | 62878464 | 62879969 | 3.51867 | 3.22245 | 1.81692 |
| chr3 | 151894760 | 151895214 | 1000 | Distal Intergenic | 3 | 151926348 | 151926409 | 62 | 2 | ENSMUST00 | 31195 | Gm25056 | ENSMUSG0C | false | . | . | . | . | . | . | chr3 | 151893340 | 151895481 | 4.36795 | 3.73034 | 2.53421 |
| chr4 | 131133664 | 131133938 | 1000 | Distal Intergenic | 4 | 131239277 | 131239410 | 134 | 1 | ENSMUST00 | -105339 | Gm25261 | ENSMUSG0C | false | . | . | . | . | . | . | chr4 | 131132400 | 131133974 | 4.45101 | 3.85614 | 2.55224 |
| chr4 | 131270499 | 131270964 | 1000 | Distal Intergenic | 4 | 131239277 | 131239410 | 134 | 1 | ENSMUST00 | 31223 | Gm25261 | ENSMUSG0C | false | . | . | . | . | . | . | chr4 | 131270622 | 131271027 | 5.55617 | 4.28572 | 3.50853 |
| chr15 | 33499528 | 33499942 | 1000 | Intron (ENSMUST00000042167.9 | 15 | 33589606 | 33589736 | 131 | 2 | ENSMUST00 | 89794 | Gm24476 | ENSMUSG0C | false | . | . | . | . | . | . | chr15 | 33493452 | 33502805 | 4.41313 | 3.78751 | 2.51715 |
| chr5 | 132673652 | 132674112 | 1000 | Distal Intergenic | 5 | 132801678 | 132801956 | 279 | 1 | ENSMUST00 | -127566 | Gm25722 | ENSMUSG0C | false | . | . | . | . | . | . | chr5 | 132672945 | 132674667 | 3.56525 | 3.24953 | 1.85486 |
| chr5 | 132923347 | 132923757 | 1000 | Distal Intergenic | 5 | 132801678 | 132801956 | 279 | 1 | ENSMUST00 | 121670 | Gm25722 | ENSMUSG0C | false | . | . | . | . | . | . | chr5 | 132922177 | 132925215 | 4.40417 | 3.7649 | 2.51081 |
| chr5 | 132933451 | 132933840 | 1000 | Distal Intergenic | 5 | 132801678 | 132801956 | 279 | 1 | ENSMUST00 | 131774 | Gm25722 | ENSMUSG0C | false | . | . | . | . | . | . | chr5 | 132932438 | 132935292 | 3.41124 | 3.27103 | 1.75438 |
| chr17 | 83983027 | 83983688 | 1000 | Distal Intergenic | 17 | 83947588 | 83947696 | 109 | 1 | ENSMUST00 | 35440 | Gm23905 | ENSMUSG0C | false | . | . | . | . | . | . | chr17 | 83980852 | 83984203 | 3.79057 | 3.46339 | 2.01806 |
| chr17 | 83938696 | 83939197 | 1000 | Distal Intergenic | 17 | 83947588 | 83947696 | 109 | 1 | ENSMUST00 | -8391 | Gm23905 | ENSMUSG0C | false | . | . | . | . | . | . | chr17 | 83935321 | 83943115 | 7.64209 | 5.27918 | 5.16816 |
| chr17 | 83979046 | 83979493 | 1000 | Distal Intergenic | 17 | 83947588 | 83947696 | 109 | 1 | ENSMUST00 | 31459 | Gm23905 | ENSMUSG0C | false | . | . | . | . | . | . | chr17 | 83978817 | 83979462 | 3.85493 | 3.59904 | 2.03923 |
| chr17 | 83950138 | 83950435 | 1000 | Promoter (2-3kb) | 17 | 83947588 | 83947696 | 109 | 1 | ENSMUST00 | 2551 | Gm23905 | ENSMUSG0C | false | . | . | . | . | . | . | chr17 | 83949461 | 83951783 | 5.37712 | 4.24569 | 3.31776 |
| chr12 | 108865022 | 108865416 | 1000 | Promoter (2-3kb) | 12 | 108862564 | 108862669 | 106 | 2 | ENSMUST00 | -2354 | Gm22079 | ENSMUSG0C | false | . | . | . | . | . | . | chr12 | 108864469 | 108866741 | 5.85117 | 4.17531 | 3.74867 |
| chr18 | 55209870 | 55210220 | 1000 | Distal Intergenic | 18 | 55179840 | 55179919 | 80 | 1 | ENSMUST00 | 30031 | Gm22597 | ENSMUSG0C | false | . | . | . | . | . | . | chr18 | 55206940 | 55211167 | 4.47449 | 3.8428 | 2.58865 |
| chr18 | 55256823 | 55257322 | 1000 | Distal Intergenic | 18 | 55179840 | 55179919 | 80 | 1 | ENSMUST00 | 76984 | Gm22597 | ENSMUSG0C | false | . | . | . | . | . | . | chr18 | 55255955 | 55261639 | 5.0948 | 4.14456 | 3.07328 |
| chr18 | 55258484 | 55258969 | 1000 | Distal Intergenic | 18 | 55179840 | 55179919 | 80 | 1 | ENSMUST00 | 78645 | Gm22597 | ENSMUSG0C | false | . | . | . | . | . | . | chr18 | 55255955 | 55261639 | 5.0948 | 4.14456 | 3.07328 |
| chr15 | 102377751 | 102378136 | 1000 | Distal Intergenic | 15 | 102373880 | 102373983 | 104 | 1 | ENSMUST00 | 3872 | Gm23590 | ENSMUSG0C | false | . | . | . | . | . | . | chr15 | 102376196 | 102377784 | 4.42101 | 3.66961 | 2.56574 |
| chr2 | 61402128 | 61402465 | 1000 | Distal Intergenic | 2 | 61492766 | 61492894 | 129 | 1 | ENSMUST00 | -90301 | Gm22338 | ENSMUSG0C | false | chr2 | 61401859 | 61402201 | 9.63953 | 19.72805 | 17.39678 | chr2 | 61398332 | 61406140 | 5.10401 | 4.1067 | 3.09609 |
| chr15 | 72687175 | 72687707 | 1000 | Intron (ENSMUST00000023276.9 | 15 | 72721321 | 72721387 | 67 | 2 | ENSMUST00 | 33680 | Gm28020 | ENSMUSG0C | false | . | . | . | . | . | . | chr15 | 72686038 | 72689611 | 5.42595 | 4.11861 | 3.39229 |
| chr1 | 89555249 | 89555797 | 1000 | Intron (ENSMUST00000027521.9 | 1 | 89563360 | 89563474 | 115 | 1 | ENSMUST00 | -7563 | Gm25180 | ENSMUSG0C | false | . | . | . | . | . | . | chr1 | 89553718 | 89560143 | 7.51776 | 5.25618 | 5.0628 |
| chr18 | 38353614 | 38353966 | 1000 | Promoter (2-3kb) | 18 | 38356578 | 38356674 | 97 | 1 | ENSMUST00 | -2612 | Gm24690 | ENSMUSG0C | false | . | . | . | . | . | . | chr18 | 38353319 | 38362122 | 5.91487 | 4.46248 | 3.75704 |
| chr18 | 38354307 | 38354702 | 1000 | Promoter (1-2kb) | 18 | 38356578 | 38356674 | 97 | 1 | ENSMUST00 | -1876 | Gm24690 | ENSMUSG0C | false | . | . | . | . | . | . | chr18 | 38353319 | 38362122 | 5.91487 | 4.46248 | 3.75704 |
| chr12 | 54265380 | 54265899 | 1000 | Distal Intergenic | 12 | 54240755 | 54240900 | 146 | 2 | ENSMUST00 | -24481 | Gm24692 | ENSMUSG0C | false | . | . | . | . | . | . | chr12 | 54259658 | 54266087 | 5.59493 | 4.35111 | 3.49425 |
| chr12 | 86800753 | 86801881 | 1000 | Distal Intergenic | 12 | 86809957 | 86810258 | 302 | 1 | ENSMUST00 | -8076 | Gm25888 | ENSMUSG0C | false | chr12 | 86801398 | 86802297 | 8.73111 | 16.23101 | 13.94854 | chr12 | 86797890 | 86804646 | 7.89277 | 5.3352 | 5.36976 |
| chr12 | 86824877 | 86825381 | 1000 | Distal Intergenic | 12 | 86809957 | 86810258 | 302 | 1 | ENSMUST00 | 14921 | Gm25888 | ENSMUSG0C | false | . | . | . | . | . | . | chr12 | 86820408 | 86827213 | 6.55613 | 4.76961 | 4.27793 |
| chr17 | 83275926 | 83276735 | 1000 | Distal Intergenic | 17 | 83269971 | 83270105 | 135 | 1 | ENSMUST00 | 5956 | Gm24240 | ENSMUSG0C | false | chr17 | 83276028 | 83277166 | 16.77503 | 49.70845 | 46.99806 | chr17 | 83272007 | 83278538 | 5.76726 | 4.43266 | 3.64322 |
| chr10 | 17299531 | 17300074 | 1000 | Distal Intergenic | 10 | 17116804 | 17116900 | 97 | 1 | ENSMUST00 | 182728 | Gm25382 | ENSMUSG0C | false | . | . | . | . | . | . | chr10 | 17298296 | 17301518 | 3.72315 | 3.35455 | 1.99403 |
| chr10 | 110855899 | 110856280 | 1000 | Intron (ENSMUST00000188776.1 | 10 | 110892298 | 110892419 | 122 | 2 | ENSMUST00 | 36139 | Gm22186 | ENSMUSG0C | false | . | . | . | . | . | . | chr10 | 110854868 | 110856782 | 4.16304 | 3.58455 | 2.36137 |
| chr12 | 97612410 | 97613025 | 1000 | Distal Intergenic | 12 | 97371034 | 97371113 | 80 | 2 | ENSMUST00 | -241298 | Gm23558 | ENSMUSG0C | false | . | . | . | . | . | . | chr12 | 97610637 | 97613967 | 3.77716 | 3.48837 | 1.99328 |
| chr5 | 103606238 | 103606750 | 1000 | 3' UTR | 5 | 103602587 | 103602879 | 293 | 1 | ENSMUST00 | 3652 | Gm23085 | ENSMUSG0C | false | . | . | . | . | . | . | chr5 | 103602894 | 103607755 | 4.47885 | 3.74726 | 2.61032 |
| chr13 | 3695075 | 3695447 | 1000 | Distal Intergenic | 13 | 3648537 | 3648650 | 114 | 1 | ENSMUST00 | 46539 | Gm23084 | ENSMUSG0C | false | . | . | . | . | . | . | chr13 | 3693960 | 3695755 | 2.85427 | 2.87825 | 1.29796 |
| chr8 | 127719723 | 127720263 | 1000 | Distal Intergenic | 8 | 127597706 | 127597823 | 118 | 1 | ENSMUST00 | 122018 | Gm22968 | ENSMUSG0C | false | . | . | . | . | . | . | chr8 | 127719334 | 127721741 | 4.98893 | 4.08553 | 2.9845 |
| chr8 | 127638738 | 127639122 | 1000 | Distal Intergenic | 8 | 127597706 | 127597823 | 118 | 1 | ENSMUST00 | 41033 | Gm22968 | ENSMUSG0C | false | . | . | . | . | . | . | chr8 | 127638014 | 127642276 | 4.29978 | 3.69257 | 2.4376 |
| chr8 | 127915787 | 127916148 | 1000 | Distal Intergenic | 8 | 127597706 | 127597823 | 118 | 1 | ENSMUST00 | 318082 | Gm22968 | ENSMUSG0C | false | . | . | . | . | . | . | chr8 | 127913940 | 127916729 | 5.22882 | 4.19723 | 3.18969 |
| chr8 | 127768368 | 127768747 | 1000 | Distal Intergenic | 8 | 127597706 | 127597823 | 118 | 1 | ENSMUST00 | 170663 | Gm22968 | ENSMUSG0C | false | . | . | . | . | . | . | chr8 | 127767175 | 127769301 | 4.25431 | 3.76911 | 2.38873 |
| chr3 | 75696292 | 75697027 | 1000 | Distal Intergenic | 3 | 75714790 | 75714924 | 135 | 1 | ENSMUST00 | -17763 | Gm25846 | ENSMUSG0C | false | . | . | . | . | . | . | . | . | . | . | . | . |
| chr6 | 99485131 | 99485642 | 1000 | Promoter (<=1kb) | 6 | 99484591 | 99484702 | 112 | 2 | ENSMUST00 | -430 | Gm22328 | ENSMUSG0C | false | chr6 | 99484863 | 99485599 | 8.36866 | 16.57842 | 14.29226 | chr6 | 99483768 | 99487728 | 5.64568 | 4.35342 | 3.5421 |
| chr14 | 40349900 | 40350405 | 1000 | Distal Intergenic | 14 | 40318072 | 40318250 | 179 | 2 | ENSMUST00 | -31651 | Gm22726 | ENSMUSG0C | false | . | . | . | . | . | . | . | . | . | . | . | . |
| chr10 | 59194915 | 59195323 | 1000 | Promoter (<=1kb) | 10 | 59195699 | 59195825 | 127 | 1 | ENSMUST00 | -376 | Gm23223 | ENSMUSG0C | false | . | . | . | . | . | . | chr10 | 59193222 | 59198884 | 4.05369 | 3.56936 | 2.23565 |
| chr18 | 67460719 | 67461278 | 1000 | Promoter (1-2kb) | 18 | 67462342 | 67462475 | 134 | 1 | ENSMUST00 | -1064 | Gm24900 | ENSMUSG0C | false | . | . | . | . | . | . | chr18 | 67457849 | 67462154 | 5.76781 | 4.35389 | 3.65015 |
| chr14 | 45456455 | 45456806 | 1000 | Downstream (1-2kb) | 14 | 45461363 | 45461645 | 283 | 2 | ENSMUST00 | 4839 | Gm22637 | ENSMUSG0C | false | . | . | . | . | . | . | chr14 | 45452174 | 45458680 | 5.35592 | 3.97788 | 3.3317 |
| chr14 | 45463591 | 45464068 | 1000 | Promoter (1-2kb) | 14 | 45461363 | 45461645 | 283 | 2 | ENSMUST00 | -1947 | Gm22637 | ENSMUSG0C | false | . | . | . | . | . | . | chr14 | 45462387 | 45466113 | 6.6239 | 4.57735 | 4.37616 |
| chr14 | 45452697 | 45453142 | 1000 | Distal Intergenic | 14 | 45461363 | 45461645 | 283 | 2 | ENSMUST00 | 8503 | Gm22637 | ENSMUSG0C | false | . | . | . | . | . | . | chr14 | 45452174 | 45458680 | 5.35592 | 3.97788 | 3.3317 |
| chr1 | 89750801 | 89751225 | 1000 | Intron (ENSMUST00000027521.9 | 1 | 89684802 | 89685112 | 311 | 1 | ENSMUST00 | 66000 | Gm26037 | ENSMUSG0C | false | chr1 | 89750882 | 89751404 | 6.08969 | 9.87507 | 7.70114 | chr1 | 89750412 | 89758303 | 5.13716 | 4.06556 | 3.12402 |
| chr1 | 89691043 | 89691426 | 1000 | Intron (ENSMUST00000027521.9 | 1 | 89684802 | 89685112 | 311 | 1 | ENSMUST00 | 6242 | Gm26037 | ENSMUSG0C | false | . | . | . | . | . | . | chr1 | 89688712 | 89693004 | 6.10578 | 4.57559 | 3.94844 |
| chr6 | 145482785 | 145483293 | 1000 | Intron (ENSMUST00000148739.2 | 6 | 145464189 | 145464303 | 115 | 1 | ENSMUST00 | 18597 | Gm25373 | ENSMUSG0C | false | . | . | . | . | . | . | chr6 | 145480832 | 145485642 | 4.34444 | 3.64479 | 2.48815 |
| chr17 | 88258998 | 88259474 | 1000 | Distal Intergenic | 17 | 88342388 | 88342482 | 95 | 2 | ENSMUST00 | 83008 | Gm22542 | ENSMUSG0C | false | . | . | . | . | . | . | chr17 | 88255594 | 88263362 | 4.52852 | 3.7778 | 2.62681 |
| chr17 | 88326266 | 88326938 | 1000 | Distal Intergenic | 17 | 88342388 | 88342482 | 95 | 2 | ENSMUST00 | 15544 | Gm22542 | ENSMUSG0C | false | . | . | . | . | . | . | chr17 | 88323758 | 88330195 | 6.07828 | 4.50218 | 3.92039 |
| chr17 | 88317299 | 88317761 | 1000 | Distal Intergenic | 17 | 88342388 | 88342482 | 95 | 2 | ENSMUST00 | 24721 | Gm22542 | ENSMUSG0C | false | . | . | . | . | . | . | chr17 | 88316889 | 88319209 | 5.25399 | 4.3093 | 3.20008 |
| chr1 | 15570354 | 15570647 | 1000 | Intron (ENSMUST00000175681.2 | 1 | 15685935 | 15686046 | 112 | 2 | ENSMUST00 | 115399 | Gm25227 | ENSMUSG0C | false | . | . | . | . | . | . | . | . | . | . | . | . |
| chr4 | 45513798 | 45514282 | 1000 | Intron (ENSMUST00000061986.6 | 4 | 45500485 | 45500552 | 68 | 2 | ENSMUST00 | -13247 | Gm22518 | ENSMUSG0C | false | . | . | . | . | . | . | chr4 | 45486900 | 45516085 | 7.16373 | 5.03069 | 4.78278 |
| chr4 | 45502444 | 45502857 | 1000 | Promoter (1-2kb) | 4 | 45500485 | 45500552 | 68 | 2 | ENSMUST00 | -1893 | Gm22518 | ENSMUSG0C | false | . | . | . | . | . | . | chr4 | 45486900 | 45516085 | 7.16373 | 5.03069 | 4.78278 |
| chr19 | 61022205 | 61022693 | 1000 | Intron (ENSMUST00000003313.8 | 19 | 61013417 | 61013545 | 129 | 2 | ENSMUST00 | -8661 | Gm22520 | ENSMUSG0C | false | . | . | . | . | . | . | chr19 | 61016804 | 61032271 | 5.33215 | 4.18451 | 3.28069 |
| chr9 | 80012772 | 80013234 | 1000 | Distal Intergenic | 9 | 80037429 | 80037714 | 286 | 1 | ENSMUST00 | -24195 | Gm25433 | ENSMUSG0C | false | . | . | . | . | . | . | chr9 | 80010983 | 80013740 | 4.24228 | 3.76128 | 2.39673 |
| chr4 | 124349640 | 124350005 | 1000 | Distal Intergenic | 4 | 124335578 | 124335681 | 104 | 2 | ENSMUST00 | -13960 | Gm22983 | ENSMUSG0C | false | . | . | . | . | . | . | chr4 | 124349684 | 124350668 | 3.4652 | 3.15035 | 1.76615 |
| chr3 | 63413340 | 63413695 | 1000 | Distal Intergenic | 3 | 63488993 | 63489098 | 106 | 1 | ENSMUST00 | -75298 | Gm24946 | ENSMUSG0C | false | . | . | . | . | . | . | chr3 | 63411954 | 63414537 | 4.22957 | 3.73827 | 2.36397 |
| chr1 | 60652638 | 60653054 | 1000 | Distal Intergenic | 1 | 60613698 | 60613803 | 106 | 2 | ENSMUST00 | -38836 | Gm23762 | ENSMUSG0C | false | . | . | . | . | . | . | . | . | . | . | . | . |
| chr5 | 92997726 | 92998171 | 1000 | Distal Intergenic | 5 | 93013032 | 93013139 | 108 | 2 | ENSMUST00 | 14968 | Gm25521 | ENSMUSG0C | false | . | . | . | . | . | . | . | . | . | . | . | . |
| chr16 | 74068961 | 74069293 | 1000 | Intron (ENSMUST00000117785.2 | 16 | 74058899 | 74059033 | 135 | 2 | ENSMUST00 | -9929 | Gm22163 | ENSMUSG0C | false | . | . | . | . | . | . | chr16 | 74067800 | 74073254 | 3.87731 | 3.51735 | 2.07937 |
| chr3 | 64112486 | 64113100 | 1000 | Promoter (2-3kb) | 3 | 64109635 | 64109737 | 103 | 1 | ENSMUST00 | 2852 | Gm22165 | ENSMUSG0C | false | . | . | . | . | . | . | chr3 | 64110074 | 64115529 | 4.6808 | 3.86905 | 2.73646 |
| chr1 | 186444739 | 186445152 | 1000 | Distal Intergenic | 1 | 186676742 | 186676876 | 135 | 1 | ENSMUST00 | -231590 | Gm26169 | ENSMUSG0C | false | . | . | . | . | . | . | . | . | . | . | . | . |
| chr3 | 37859705 | 37860479 | 1000 | Distal Intergenic | 3 | 37802544 | 37802652 | 109 | 2 | ENSMUST00 | -57054 | Gm26404 | ENSMUSG0C | false | . | . | . | . | . | . | chr3 | 37856296 | 37862135 | 4.25522 | 3.61324 | 2.4186 |
| chr3 | 37846638 | 37847071 | 1000 | Distal Intergenic | 3 | 37802544 | 37802652 | 109 | 2 | ENSMUST00 | -43987 | Gm26404 | ENSMUSG0C | false | . | . | . | . | . | . | chr3 | 37844205 | 37849062 | 4.77419 | 3.94961 | 2.82955 |
| chr6 | 115085592 | 115085928 | 1000 | Distal Intergenic | 6 | 115099554 | 115099648 | 95 | 1 | ENSMUST00 | -13626 | Gm24767 | ENSMUSG0C | false | . | . | . | . | . | . | chr6 | 115085205 | 115086049 | 3.1531 | 2.73923 | 1.52127 |
| chr2 | 46669400 | 46669810 | 1000 | Distal Intergenic | 2 | 46606967 | 46607083 | 117 | 2 | ENSMUST00 | -62318 | Gm25264 | ENSMUSG0C | false | . | . | . | . | . | . | chr2 | 46664787 | 46671476 | 3.53483 | 3.36759 | 1.80184 |
| chr3 | 63128481 | 63128882 | 1000 | Distal Intergenic | 3 | 63152217 | 63152358 | 142 | 1 | ENSMUST00 | -23335 | Gm22433 | ENSMUSG0C | false | . | . | . | . | . | . | . | . | . | . | . | . |
| chr17 | 30269467 | 30269775 | 1000 | Intron (ENSMUST00000079924.5 | 17 | 30311079 | 30311151 | 73 | 1 | ENSMUST00 | -41304 | Gm25049 | ENSMUSG0C | false | . | . | . | . | . | . | chr17 | 30266579 | 30270803 | 4.56075 | 3.64149 | 2.68041 |
| chr3 | 153692044 | 153692653 | 1000 | Intron (ENSMUST00000064460.5 | 3 | 153708911 | 153709026 | 116 | 1 | ENSMUST00 | -16258 | Gm22206 | ENSMUSG0C | false | . | . | . | . | . | . | chr3 | 153686316 | 153712771 | 6.43925 | 4.66349 | 4.19773 |
| chr15 | 77212404 | 77212767 | 1000 | Intron (ENSMUST00000048145.6 | 15 | 77242406 | 77242551 | 146 | 2 | ENSMUST00 | 29784 | Gm24056 | ENSMUSG0C | false | . | . | . | . | . | . | chr15 | 77208986 | 77216071 | 4.99519 | 4.05687 | 3.00547 |
| chr15 | 84300941 | 84301388 | 1000 | Intron (ENSMUST00000023072.6 | 15 | 84323209 | 84323352 | 144 | 2 | ENSMUST00 | 21964 | Gm22890 | ENSMUSG0C | false | . | . | . | . | . | . | chr15 | 84297650 | 84306963 | 6.75434 | 4.89978 | 4.45766 |
| chr15 | 84313624 | 84314808 | 1000 | 3' UTR | 15 | 84323209 | 84323352 | 144 | 2 | ENSMUST00 | 8544 | Gm22890 | ENSMUSG0C | false | . | . | . | . | . | . | chr15 | 84310303 | 84316013 | 7.01159 | 5.06165 | 4.64911 |
| chr15 | 84298878 | 84299358 | 1000 | Intron (ENSMUST00000023072.6 | 15 | 84323209 | 84323352 | 144 | 2 | ENSMUST00 | 23994 | Gm22890 | ENSMUSG0C | false | . | . | . | . | . | . | chr15 | 84297650 | 84306963 | 6.75434 | 4.89978 | 4.45766 |
| chr5 | 92763491 | 92763919 | 1000 | Intron (ENSMUST00000113055.3 | 5 | 92788451 | 92788562 | 112 | 2 | ENSMUST00 | 24643 | Mir1961 | ENSMUSG0C | false | . | . | . | . | . | . | chr5 | 92759258 | 92768252 | 4.69185 | 3.93143 | 2.74833 |
| chr4 | 57761386 | 57761708 | 1000 | Promoter (<=1kb) | 4 | 57762607 | 57762710 | 104 | 1 | ENSMUST00 | -899 | Gm24277 | ENSMUSG0C | false | . | . | . | . | . | . | chr4 | 57761174 | 57763341 | 3.79208 | 3.37446 | 2.02964 |
| chr4 | 133166646 | 133167282 | 1000 | Intron (ENSMUST00000084241.6 | 4 | 133171427 | 133171714 | 288 | 2 | ENSMUST00 | 4432 | Gm24636 | ENSMUSG0C | false | . | . | . | . | . | . | chr4 | 133148299 | 133179985 | 6.32767 | 4.58392 | 4.12241 |
| chr10 | 3920461 | 3920803 | 1000 | Intron (ENSMUST00000120274.2 | 10 | 3934769 | 3934877 | 109 | 2 | ENSMUST00 | 14074 | Gm23023 | ENSMUSG0C | false | . | . | . | . | . | . | chr10 | 3917748 | 3924115 | 5.49509 | 4.26832 | 3.42809 |
| chr7 | 70690939 | 70691276 | 1000 | Distal Intergenic | 7 | 70752416 | 70752500 | 85 | 1 | ENSMUST00 | -61140 | Gm24880 | ENSMUSG0C | false | . | . | . | . | . | . | . | . | . | . | . | . |
| chr7 | 70813094 | 70813569 | 1000 | Distal Intergenic | 7 | 70752416 | 70752500 | 85 | 1 | ENSMUST00 | 60679 | Gm24880 | ENSMUSG0C | false | . | . | . | . | . | . | chr7 | 70811734 | 70814902 | 3.71397 | 3.26234 | 1.99392 |
| chr17 | 87459268 | 87459798 | 1000 | Distal Intergenic | 17 | 87453223 | 87453356 | 134 | 2 | ENSMUST00 | -5913 | Gm26005 | ENSMUSG0C | false | chr17 | 87459188 | 87459424 | 6.75876 | 12.51203 | 10.2897 | chr17 | 87457781 | 87461935 | 5.96389 | 4.48039 | 3.80987 |
| chr8 | 105613730 | 105614400 | 1000 | Promoter (1-2kb) | 8 | 105615466 | 105615573 | 108 | 1 | ENSMUST00 | -1066 | Mir1966 | ENSMUSG0C | false | chr8 | 105612839 | 105614792 | 22.51707 | 59.83231 | 56.98677 | chr8 | 105598855 | 105624293 | 5.6449 | 4.36488 | 3.54309 |
| chr8 | 73198295 | 73198644 | 1000 | Intron (ENSMUST00000004497.9 | 8 | 73175119 | 73175258 | 140 | 2 | ENSMUST00 | -23038 | Gm22532 | ENSMUSG0C | false | . | . | . | . | . | . | chr8 | 73193455 | 73204706 | 4.46876 | 3.78541 | 2.57026 |
| chr8 | 73178277 | 73178814 | 1000 | Intron (ENSMUST00000004497.9 | 8 | 73175119 | 73175258 | 140 | 2 | ENSMUST00 | -3020 | Gm22532 | ENSMUSG0C | false | . | . | . | . | . | . | chr8 | 73174904 | 73181947 | 5.02217 | 4.11881 | 3.0202 |
| chr9 | 66988937 | 66989396 | 1000 | Distal Intergenic | 9 | 66981088 | 66981185 | 98 | 2 | ENSMUST00 | -7753 | Gm24225 | ENSMUSG0C | false | . | . | . | . | . | . | chr9 | 66988525 | 66989703 | 5.41573 | 4.2984 | 3.33019 |
| chr17 | 72503573 | 72503949 | 1000 | Intron (ENSMUST00000086639.4 | 17 | 72499289 | 72499418 | 130 | 2 | ENSMUST00 | -4156 | Gm24736 | ENSMUSG0C | false | . | . | . | . | . | . | chr17 | 72501367 | 72505927 | 4.17628 | 3.68877 | 2.33579 |
| chr14 | 97791732 | 97792130 | 1000 | Intron (ENSMUST00000069334.7 | 14 | 97967829 | 97967991 | 163 | 2 | ENSMUST00 | 175861 | Gm25831 | ENSMUSG0C | false | . | . | . | . | . | . | chr14 | 97791486 | 97803403 | 3.98924 | 3.5968 | 2.16581 |
| chr1 | 23256631 | 23257094 | 1000 | Distal Intergenic | 1 | 23245211 | 23245331 | 121 | 1 | ENSMUST00 | 11421 | Gm24176 | ENSMUSG0C | false | chr1 | 23256193 | 23259443 | 33.26644 | 105.59763 | 101.88331 | . | . | . | . | . | . |
| chr12 | 29096073 | 29096471 | 1000 | Distal Intergenic | 12 | 29035180 | 29035294 | 115 | 2 | ENSMUST00 | -60780 | Gm23342 | ENSMUSG0C | false | . | . | . | . | . | . | chr12 | 29094881 | 29096934 | 4.64537 | 3.84481 | 2.70715 |
| chr12 | 29015387 | 29015996 | 1000 | Distal Intergenic | 12 | 29035180 | 29035294 | 115 | 2 | ENSMUST00 | 19298 | Gm23342 | ENSMUSG0C | false | . | . | . | . | . | . | chr12 | 29013966 | 29017219 | 3.81289 | 3.48874 | 2.07878 |
| chr10 | 124288594 | 124289112 | 1000 | Distal Intergenic | 10 | 124455059 | 124455317 | 259 | 1 | ENSMUST00 | -165947 | Gm23777 | ENSMUSG0C | false | chr10 | 124288068 | 124289218 | 16.14593 | 43.98602 | 41.34693 | chr10 | 124287939 | 124289356 | 3.71093 | 3.50249 | 1.94662 |
| chr15 | 86089650 | 86090542 | 1000 | Promoter (<=1kb) | 15 | 86087640 | 86090060 | 2421 | 2 | ENSMUST00 | 0 | Gm15722 | ENSMUSG0C | false | . | . | . | . | . | . | chr15 | 86074674 | 86092519 | 7.22651 | 5.10744 | 4.82194 |
| chr14 | 54883163 | 54883798 | 1000 | Promoter (<=1kb) | 14 | 54883377 | 54888234 | 4858 | 1 | ENSMUST00 | 0 | Bcl2l2 | ENSMUSG0C | false | chr14 | 54882829 | 54884007 | 20.74539 | 66.97287 | 64.02668 | chr14 | 54882139 | 54887241 | 4.16128 | 3.4964 | 2.33704 |
| chr15 | 79834010 | 79834641 | 1000 | Promoter (<=1kb) | 15 | 79826141 | 79834237 | 8097 | 2 | ENSMUST00 | 0 | Cbx6 | ENSMUSG0C | false | chr15 | 79833220 | 79834719 | 24.1157 | 67.74074 | 64.78616 | chr15 | 79832125 | 79835039 | 5.32077 | 4.19974 | 3.31063 |
| chr14 | 115040447 | 115042337 | 1000 | Promoter (<=1kb) | 14 | 115042879 | 115046727 | 3849 | 1 | ENSMUST00 | -542 | Mir17hg | ENSMUSG0C | false | chr14 | 115040106 | 115042806 | 33.08631 | 99.21803 | 95.66874 | chr14 | 115040204 | 115040670 | 5.64431 | 4.45292 | 3.52254 |
| chr2 | 167658756 | 167659129 | 1000 | Promoter (2-3kb) | 2 | 167607850 | 167661496 | 53647 | 2 | ENSMUST00 | 2367 | Gm20431 | ENSMUSG0C | false | chr2 | 167658261 | 167658813 | 8.66089 | 16.11541 | 13.83781 | chr2 | 167651959 | 167660835 | 7.77666 | 5.30918 | 5.27423 |
| chr1 | 37600734 | 37601416 | 1000 | Distal Intergenic | 1 | 37614259 | 37615372 | 1114 | 1 | ENSMUST00 | -12843 | 4930556I23R | ENSMUSG0C | false | . | . | . | . | . | . | chr1 | 37596513 | 37602675 | 6.10872 | 4.60071 | 3.91719 |
| chr3 | 9785266 | 9785662 | 1000 | Intron (ENSMUST00000160368.1 | 3 | 9757607 | 9786142 | 28536 | 1 | ENSMUST00 | 27660 | Gm16337 | ENSMUSG0C | false | . | . | . | . | . | . | chr3 | 9778824 | 9802371 | 5.7635 | 4.36373 | 3.65538 |
| chr3 | 9739289 | 9739702 | 1000 | Intron (ENSMUST00000161949.2 | 3 | 9757607 | 9786142 | 28536 | 1 | ENSMUST00 | -17905 | Gm16337 | ENSMUSG0C | false | . | . | . | . | . | . | chr3 | 9738186 | 9742454 | 5.59795 | 4.34083 | 3.54839 |
| chr13 | 44232235 | 44232822 | 1000 | Distal Intergenic | 13 | 44121294 | 44125179 | 3886 | 1 | ENSMUST00 | 110942 | Gm5083 | ENSMUSG0C | false | . | . | . | . | . | . | chr13 | 44226456 | 44236587 | 5.70374 | 4.41191 | 3.58124 |
| chr3 | 116662094 | 116662881 | 1000 | Promoter (<=1kb) | 3 | 116631164 | 116662677 | 31514 | 2 | ENSMUST00 | 0 | Hiat1 | ENSMUSG0C | false | chr3 | 116661343 | 116663117 | 37.34335 | 127.50397 | 123.10989 | . | . | . | . | . | . |
| chr3 | 141660622 | 141661103 | 1000 | Intron (ENSMUST00000106236.3 | 3 | 141662889 | 141682900 | 20012 | 2 | ENSMUST00 | 21797 | Gm15689 | ENSMUSG0C | false | . | . | . | . | . | . | chr3 | 141658950 | 141661342 | 4.63927 | 3.99012 | 2.69645 |
| chr17 | 10501475 | 10501803 | 1000 | Intron (ENSMUST00000161274.1 | 17 | 10496453 | 10553169 | 56717 | 1 | ENSMUST00 | 5023 | Gm16168 | ENSMUSG0C | false | . | . | . | . | . | . | chr17 | 10500241 | 10502548 | 5.12183 | 4.19327 | 3.09242 |

| | | | | | | | | | | | | | | | | | | | | | | | | | | |
|---|---|---|---|---|---|---|---|---|---|---|---|---|---|---|---|---|---|---|---|---|---|---|---|---|---|---|
| chr17 | 10529704 | 10530021 | 1000 | Intron (ENSMUST00000161274.1 | 17 | 10496453 | 10553169 | 56717 | 1 | ENSMUST00 | 33252 | Gm16168 | ENSMUSG0C | false | . | . | . | . | . | . | chr17 | 10526632 | 10533668 | 4.66575 | 3.85507 | 2.75675 |
| chr5 | 66359606 | 66360099 | 1000 | Promoter (2-3kb) | 5 | 66356918 | 66357185 | 268 | 1 | ENSMUST00 | 2689 | Gm16273 | ENSMUSG0C | false | . | . | . | . | . | . | chr5 | 66352002 | 66369698 | 7.18935 | 5.00706 | 4.81216 |
| chr5 | 99444703 | 99445289 | 1000 | Intron (ENSMUST00000146396.2 | 5 | 99509944 | 99510682 | 739 | 2 | ENSMUST00 | 65393 | Gm16227 | ENSMUSG0C | false | . | . | . | . | . | . | chr5 | 99442196 | 99449542 | 4.74151 | 3.92904 | 2.79783 |
| chr13 | 54211656 | 54212291 | 1000 | Intron (ENSMUST00000038101.3 | 13 | 54219453 | 54223023 | 3571 | 2 | ENSMUST00 | 10732 | Gm16578 | ENSMUSG0C | false | . | . | . | . | . | . | chr13 | 54208772 | 54213387 | 4.89932 | 4.10866 | 2.91146 |
| chr13 | 54231846 | 54232240 | 1000 | Distal Intergenic | 13 | 54219453 | 54223023 | 3571 | 2 | ENSMUST00 | -8824 | Gm16578 | ENSMUSG0C | false | . | . | . | . | . | . | chr13 | 54227203 | 54232943 | 5.05542 | 4.16332 | 3.04801 |
| chr8 | 110761172 | 110761603 | 1000 | Intron (ENSMUST00000150680.1 | 8 | 110754992 | 110758970 | 3979 | 1 | ENSMUST00 | 6181 | Gm15894 | ENSMUSG0C | false | . | . | . | . | . | . | chr8 | 110759571 | 110762731 | 4.77217 | 3.91612 | 2.8218 |
| chr4 | 57557138 | 57557511 | 1000 | Intron (ENSMUST00000102905.2 | 4 | 57568159 | 57711802 | 143644 | 1 | ENSMUST00 | -10648 | Palm2 | ENSMUSG0C | false | . | . | . | . | . | . | chr4 | 57554631 | 57561589 | 3.94037 | 3.54927 | 2.15417 |
| chr4 | 57559992 | 57560392 | 1000 | Intron (ENSMUST00000102905.2 | 4 | 57568159 | 57711802 | 143644 | 1 | ENSMUST00 | -7767 | Palm2 | ENSMUSG0C | false | . | . | . | . | . | . | chr4 | 57554631 | 57561589 | 3.94037 | 3.54927 | 2.15417 |
| chr8 | 127533746 | 127534060 | 1000 | Intron (ENSMUST00000162309.2 | 8 | 127561435 | 127561869 | 435 | 1 | ENSMUST00 | -27375 | Gm6921 | ENSMUSG0C | false | . | . | . | . | . | . | chr8 | 127531330 | 127535604 | 5.38019 | 4.18427 | 3.32877 |
| chr5 | 123136441 | 123136785 | 1000 | Promoter (1-2kb) | 5 | 123133723 | 123138109 | 4387 | 2 | ENSMUST00 | 1324 | AI480526 | ENSMUSG0C | false | chr5 | 123135355 | 123136555 | 12.76676 | 28.45456 | 26.00909 | chr5 | 123131739 | 123139661 | 6.68632 | 4.77641 | 4.39554 |
| chr16 | 26049166 | 26050069 | 1000 | Intron (ENSMUST00000039990.5 | 16 | 26031810 | 26040870 | 9061 | 2 | ENSMUST00 | -8297 | AU015336 | ENSMUSG0C | false | . | . | . | . | . | . | chr16 | 26035252 | 26056185 | 6.03158 | 4.46161 | 3.87334 |
| chr16 | 96757588 | 96758117 | 1000 | Intron (ENSMUST00000056102.7 | 16 | 96467606 | 96525793 | 58188 | 1 | ENSMUST00 | 289983 | Pcp4 | ENSMUSG0C | false | . | . | . | . | . | . | chr16 | 96757159 | 96758588 | 3.78436 | 3.30537 | 2.04389 |
| chr16 | 17863188 | 17863607 | 1000 | Intron (ENSMUST00000150068.1 | 16 | 17835276 | 17838364 | 3089 | 2 | ENSMUST00 | -24825 | Car15 | ENSMUSG0C | false | . | . | . | . | . | . | chr16 | 17860699 | 17864520 | 4.78184 | 3.9104 | 2.83477 |
| chr19 | 10457248 | 10457712 | 1000 | Promoter (<=1kb) | 19 | 10455371 | 10457447 | 2077 | 2 | ENSMUST00 | 0 | Lrrc10b | ENSMUSG0C | false | chr19 | 10456412 | 10458156 | 19.69817 | 53.33889 | 50.58166 | chr19 | 10454448 | 10459480 | 6.0164 | 4.43395 | 3.89018 |
| chr19 | 10469440 | 10469734 | 1000 | Distal Intergenic | 19 | 10455371 | 10457447 | 2077 | 2 | ENSMUST00 | -11994 | Lrrc10b | ENSMUSG0C | false | . | . | . | . | . | . | chr19 | 10468814 | 10470528 | 5.84089 | 4.55984 | 3.68423 |
| chr13 | 102764324 | 102764808 | 1000 | Promoter (1-2kb) | 13 | 102762435 | 102764785 | 2351 | 1 | ENSMUST00 | 1890 | Gm17160 | ENSMUSG0C | false | . | . | . | . | . | . | chr13 | 102747137 | 102771216 | 6.28304 | 4.64045 | 4.06151 |
| chr1 | 160036849 | 160037268 | 1000 | Distal Intergenic | 1 | 160044384 | 160071536 | 27153 | 1 | ENSMUST00 | -7116 | 4930523C07I | ENSMUSG0C | false | . | . | . | . | . | . | chr1 | 160035622 | 160038099 | 6.30681 | 4.66224 | 4.09676 |
| chr18 | 78312906 | 78313327 | 1000 | Distal Intergenic | 18 | 78349754 | 78350701 | 948 | 1 | ENSMUST00 | -36427 | Gm6133 | ENSMUSG0C | false | . | . | . | . | . | . | chr18 | 78311113 | 78313785 | 4.98111 | 4.09217 | 3.04672 |
| chr6 | 117332538 | 117332982 | 1000 | Distal Intergenic | 6 | 117213656 | 117214048 | 393 | 2 | ENSMUST00 | -118491 | Rpl28-ps4 | ENSMUSG0C | false | . | . | . | . | . | . | chr6 | 117332152 | 117334036 | 4.9276 | 4.07359 | 2.95832 |
| chr5 | 73209114 | 73209644 | 1000 | Promoter (2-3kb) | 5 | 73210109 | 73212634 | 2526 | 2 | ENSMUST00 | 2990 | Gm17207 | ENSMUSG0C | false | . | . | . | . | . | . | chr5 | 73196545 | 73254445 | 6.29302 | 4.62166 | 4.08669 |
| chr5 | 73227491 | 73227876 | 1000 | Intron (ENSMUST00000101127.6 | 5 | 73210109 | 73212634 | 2526 | 2 | ENSMUST00 | -14858 | Gm17207 | ENSMUSG0C | false | . | . | . | . | . | . | chr5 | 73196545 | 73254445 | 6.29302 | 4.62166 | 4.08669 |
| chr9 | 107307800 | 107308190 | 1000 | Distal Intergenic | 9 | 107300380 | 107301838 | 1459 | 2 | ENSMUST00 | -5963 | Gm17041 | ENSMUSG0C | false | . | . | . | . | . | . | chr9 | 107305847 | 107308682 | 5.22193 | 4.10815 | 3.23628 |
| chr2 | 174355596 | 174355904 | 1000 | Distal Intergenic | 2 | 174330319 | 174346712 | 16394 | 2 | ENSMUST00 | -8885 | Gm20721 | ENSMUSG0C | false | . | . | . | . | . | . | chr2 | 174354854 | 174356346 | 4.63733 | 3.67831 | 2.74722 |
| chr8 | 91558575 | 91558947 | 1000 | Intron (ENSMUST00000069718.8 | 8 | 91621448 | 91629729 | 8282 | 2 | ENSMUST00 | 70782 | Gm3235 | ENSMUSG0C | false | . | . | . | . | . | . | chr8 | 91549338 | 91568172 | 5.03197 | 4.01835 | 3.04808 |
| chr3 | 98427495 | 98427865 | 1000 | Exon (ENSMUST00000056096.9/ | 3 | 98447198 | 98448976 | 1779 | 2 | ENSMUST00 | 21111 | Gm4450 | ENSMUSG0C | false | chr3 | 98427166 | 98428732 | 29.28796 | 98.53321 | 94.99628 | chr3 | 98426930 | 98429301 | 4.55821 | 3.71485 | 2.66183 |
| chr13 | 41267856 | 41268232 | 1000 | Intron (ENSMUST00000165561.2 | 13 | 41273803 | 41273977 | 175 | 2 | ENSMUST00 | 5745 | Gm17364 | ENSMUSG0C | false | . | . | . | . | . | . | chr13 | 41266733 | 41269608 | 4.99376 | 4.07084 | 3.051 |
| chr12 | 32486740 | 32487093 | 1000 | Distal Intergenic | 12 | 32378704 | 32382943 | 4240 | 1 | ENSMUST00 | 108037 | Ccdc71l | ENSMUSG0C | false | . | . | . | . | . | . | chr12 | 32486961 | 32489937 | 5.41988 | 4.30762 | 3.33576 |
| chr1 | 182191696 | 182192511 | 1000 | Distal Intergenic | 1 | 182157144 | 182158319 | 1176 | 2 | ENSMUST00 | -33378 | Vmn1r1 | ENSMUSG0C | false | . | . | . | . | . | . | chr1 | 182189674 | 182193542 | 5.45512 | 4.3571 | 3.36845 |
| chr14 | 24222254 | 24222734 | 1000 | Intron (ENSMUST00000073687.7 | 14 | 24238172 | 24243391 | 5220 | 2 | ENSMUST00 | 20657 | Gm17105 | ENSMUSG0C | false | . | . | . | . | . | . | chr14 | 24191579 | 24244546 | 6.62149 | 4.76248 | 4.34728 |
| chr15 | 81514194 | 81514573 | 1000 | Distal Intergenic | 15 | 81499084 | 81499548 | 465 | 2 | ENSMUST00 | -14647 | Gm5218 | ENSMUSG0C | false | chr15 | 81514051 | 81514658 | 10.09152 | 19.95864 | 17.62411 | chr15 | 81514015 | 81515265 | 4.59133 | 3.84925 | 2.69069 |
| chr13 | 41249517 | 41250387 | 1000 | Promoter (<=1kb) | 13 | 41249860 | 41276574 | 26715 | 1 | ENSMUST00 | 0 | Smim13 | ENSMUSG0C | false | chr13 | 41249177 | 41250862 | 33.52805 | 106.05383 | 102.33234 | . | . | . | . | . | . |
| chr13 | 51221751 | 51222142 | 1000 | Distal Intergenic | 13 | 51202688 | 51203065 | 378 | 2 | ENSMUST00 | -18687 | Hist1h2al | ENSMUSG0C | false | . | . | . | . | . | . | chr13 | 51220218 | 51223470 | 6.28605 | 4.52708 | 4.11463 |
| chr13 | 51222710 | 51223260 | 1000 | Distal Intergenic | 13 | 51202688 | 51203065 | 378 | 2 | ENSMUST00 | -19646 | Hist1h2al | ENSMUSG0C | false | . | . | . | . | . | . | chr13 | 51220218 | 51223470 | 6.28605 | 4.52708 | 4.11463 |
| chr13 | 51250715 | 51251288 | 1000 | Distal Intergenic | 13 | 51202688 | 51203065 | 378 | 2 | ENSMUST00 | -47651 | Hist1h2al | ENSMUSG0C | false | . | . | . | . | . | . | chr13 | 51249447 | 51251303 | 3.77395 | 3.45027 | 2.04755 |
| chr13 | 51212191 | 51212776 | 1000 | Distal Intergenic | 13 | 51202688 | 51203065 | 378 | 2 | ENSMUST00 | -9127 | Hist1h2al | ENSMUSG0C | false | . | . | . | . | . | . | chr13 | 51210775 | 51213537 | 4.39382 | 3.78217 | 2.50839 |
| chr1 | 166304629 | 166305018 | 1000 | Intron (ENSMUST00000111416.1 | 1 | 166306741 | 166309585 | 2845 | 2 | ENSMUST00 | 4567 | 5330438I03R | ENSMUSG0C | false | . | . | . | . | . | . | chr1 | 166301548 | 166311479 | 5.01639 | 4.00677 | 3.03101 |
| chr1 | 166282566 | 166283042 | 1000 | Intron (ENSMUST00000111416.1 | 1 | 166306741 | 166309585 | 2845 | 2 | ENSMUST00 | 26543 | 5330438I03R | ENSMUSG0C | false | . | . | . | . | . | . | chr1 | 166277550 | 166299518 | 5.01596 | 4.06002 | 3.04182 |
| chr9 | 107456555 | 107457130 | 1000 | Intron (ENSMUST00000010210.7 | 9 | 107443950 | 107445031 | 1082 | 2 | ENSMUST00 | -11525 | Gm17118 | ENSMUSG0C | false | . | . | . | . | . | . | chr9 | 107452389 | 107458103 | 5.78644 | 4.23585 | 3.68997 |
| chr11 | 100364262 | 100364632 | 1000 | Promoter (2-3kb) | 11 | 100361716 | 100362126 | 411 | 2 | ENSMUST00 | -2137 | Gm10039 | ENSMUSG0C | false | . | . | . | . | . | . | chr11 | 100362708 | 100370334 | 5.36303 | 4.02698 | 3.33266 |
| chr17 | 3529302 | 3529765 | 1000 | Promoter (2-3kb) | 17 | 3532554 | 3533213 | 660 | 1 | ENSMUST00 | -2789 | Cldn20 | ENSMUSG0C | false | . | . | . | . | . | . | chr17 | 3527625 | 3530730 | 4.36386 | 3.63864 | 2.52559 |
| chr19 | 7306938 | 7307341 | 1000 | Intron (ENSMUST00000032557.9 | 19 | 7297734 | 7304350 | 6617 | 1 | ENSMUST00 | 9205 | Gm17227 | ENSMUSG0C | false | . | . | . | . | . | . | chr19 | 7304919 | 7311624 | 7.73326 | 5.08563 | 5.28086 |
| chr18 | 67597867 | 67598148 | 1000 | Distal Intergenic | 18 | 67562387 | 67562851 | 465 | 1 | ENSMUST00 | 35481 | Gm17669 | ENSMUSG0C | false | . | . | . | . | . | . | chr18 | 67596706 | 67598451 | 3.49959 | 3.27697 | 1.82031 |
| chr8 | 105010086 | 105010553 | 1000 | Intron (ENSMUST00000172032.2 | 8 | 105000853 | 105020410 | 19558 | 1 | ENSMUST00 | 9234 | Ces2h | ENSMUSG0C | false | . | . | . | . | . | . | chr8 | 105008733 | 105012084 | 4.01739 | 3.50668 | 2.22618 |
| chr9 | 107421565 | 107421862 | 1000 | Intron (ENSMUST00000010210.7 | 9 | 107423857 | 107430567 | 6711 | 2 | ENSMUST00 | 8705 | Gm17141 | ENSMUSG0C | false | . | . | . | . | . | . | chr9 | 107421188 | 107422096 | 3.87858 | 3.49103 | 2.14096 |
| chr18 | 35771295 | 35772193 | 1000 | Promoter (<=1kb) | 18 | 35771559 | 35807172 | 35614 | 1 | ENSMUST00 | 0 | Ube2d2a | ENSMUSG0C | false | chr18 | 35770990 | 35772691 | 32.0113 | 129.32326 | 124.86235 | . | . | . | . | . | . |
| chr13 | 102749156 | 102749611 | 1000 | Intron (ENSMUST00000166726.2 | 13 | 102760255 | 102761896 | 1642 | 1 | ENSMUST00 | -10644 | 1700099I09R | ENSMUSG0C | false | . | . | . | . | . | . | chr13 | 102747137 | 102771216 | 6.28304 | 4.64045 | 4.06151 |
| chr13 | 102756447 | 102757100 | 1000 | Intron (ENSMUST00000166726.2 | 13 | 102760255 | 102761896 | 1642 | 1 | ENSMUST00 | -3155 | 1700099I09R | ENSMUSG0C | false | . | . | . | . | . | . | chr13 | 102747137 | 102771216 | 6.28304 | 4.64045 | 4.06151 |
| chr13 | 102753543 | 102754094 | 1000 | Exon (ENSMUST00000167058.2/ | 13 | 102760255 | 102761896 | 1642 | 1 | ENSMUST00 | -6161 | 1700099I09R | ENSMUSG0C | false | . | . | . | . | . | . | chr13 | 102747137 | 102771216 | 6.28304 | 4.64045 | 4.06151 |
| chr7 | 24850927 | 24851268 | 1000 | Distal Intergenic | 7 | 24862213 | 24862697 | 485 | 1 | ENSMUST00 | -10945 | Gm9844 | ENSMUSG0C | false | . | . | . | . | . | . | chr7 | 24846056 | 24853926 | 4.30179 | 3.69763 | 2.44036 |
| chr7 | 24850080 | 24850514 | 1000 | Distal Intergenic | 7 | 24862213 | 24862697 | 485 | 1 | ENSMUST00 | -11699 | Gm9844 | ENSMUSG0C | false | . | . | . | . | . | . | chr7 | 24846056 | 24853926 | 4.30179 | 3.69763 | 2.44036 |
| chr5 | 67427107 | 67427940 | 1000 | Promoter (<=1kb) | 5 | 67392147 | 67427799 | 35653 | 2 | ENSMUST00 | 0 | Bend4 | ENSMUSG0C | false | chr5 | 67426148 | 67428221 | 24.40663 | 77.266 | 74.1616 | chr5 | 67425840 | 67429332 | 3.45759 | 3.25718 | 1.74931 |
| chr5 | 67440492 | 67440832 | 1000 | Distal Intergenic | 5 | 67392147 | 67427799 | 35653 | 2 | ENSMUST00 | -12694 | Bend4 | ENSMUSG0C | false | . | . | . | . | . | . | chr5 | 67440196 | 67441289 | 3.15532 | 3.10153 | 1.54269 |
| chr5 | 92742629 | 92742908 | 1000 | Intron (ENSMUST00000113055.3 | 5 | 92736205 | 92737442 | 1238 | 1 | ENSMUST00 | 6425 | Gm20500 | ENSMUSG0C | false | . | . | . | . | . | . | chr5 | 92741029 | 92749681 | 4.65041 | 3.88261 | 2.72811 |
| chr5 | 92761918 | 92762279 | 1000 | Intron (ENSMUST00000113055.3 | 5 | 92736205 | 92737442 | 1238 | 1 | ENSMUST00 | 25714 | Gm20500 | ENSMUSG0C | false | . | . | . | . | . | . | chr5 | 92759258 | 92768252 | 4.69185 | 3.93143 | 2.74833 |
| chr16 | 50786043 | 50786723 | 1000 | Promoter (2-3kb) | 16 | 50783478 | 50784065 | 588 | 1 | ENSMUST00 | 2566 | 1700116B05I | ENSMUSG0C | false | . | . | . | . | . | . | chr16 | 50781927 | 50788895 | 5.4458 | 4.28026 | 3.36614 |
| chr16 | 50783474 | 50784181 | 1000 | Promoter (<=1kb) | 16 | 50783478 | 50784065 | 588 | 1 | ENSMUST00 | 0 | 1700116B05I | ENSMUSG0C | false | . | . | . | . | . | . | chr16 | 50781927 | 50788895 | 5.4458 | 4.28026 | 3.36614 |
| chr16 | 50792937 | 50793391 | 1000 | Distal Intergenic | 16 | 50783478 | 50784065 | 588 | 1 | ENSMUST00 | 9460 | 1700116B05I | ENSMUSG0C | false | . | . | . | . | . | . | chr16 | 50790830 | 50799832 | 4.14172 | 3.59695 | 2.3089 |
| chr19 | 5803033 | 5803957 | 1000 | Promoter (<=1kb) | 19 | 5795690 | 5802672 | 6983 | 2 | ENSMUST00 | -362 | Malat1 | ENSMUSG0C | false | chr19 | 5796007 | 5804263 | 15.48624 | 61.01235 | 58.15405 | chr19 | 5803906 | 5840432 | 6.03605 | 4.50406 | 3.86334 |
| chr10 | 127290190 | 127290766 | 1000 | Promoter (<=1kb) | 10 | 127289696 | 127290724 | 1029 | 2 | ENSMUST00 | 0 | Gm4189 | ENSMUSG0C | false | chr10 | 127290751 | 127292540 | 30.19165 | 95.18787 | 91.73401 | chr10 | 127289605 | 127290323 | 5.06022 | 4.15261 | 3.03912 |
| chr9 | 117461901 | 117462229 | 1000 | Intron (ENSMUST00000164018.3 | 9 | 117402979 | 117404342 | 1364 | 1 | ENSMUST00 | 58923 | Gm20396 | ENSMUSG0C | false | . | . | . | . | . | . | chr9 | 117461808 | 117463880 | 4.06004 | 3.66618 | 2.22197 |
| chr5 | 25472063 | 25472402 | 1000 | Intron (ENSMUST00000045291.8 | 5 | 25450305 | 25450908 | 604 | 1 | ENSMUST00 | 21759 | Gm18343 | ENSMUSG0C | false | . | . | . | . | . | . | chr5 | 25467432 | 25476063 | 4.23244 | 3.6851 | 2.39341 |
| chr9 | 117117149 | 117117514 | 1000 | Intron (ENSMUST00000111773.4 | 9 | 117065527 | 117065888 | 362 | 1 | ENSMUST00 | 51623 | Gm20397 | ENSMUSG0C | false | . | . | . | . | . | . | chr9 | 117113138 | 117121657 | 4.97914 | 3.95422 | 3.00047 |
| chr9 | 79918385 | 79918715 | 1000 | Intron (ENSMUST00000093811.5 | 9 | 79913200 | 79913631 | 432 | 1 | ENSMUST00 | 5186 | Gm3211 | ENSMUSG0C | false | . | . | . | . | . | . | chr9 | 79911389 | 79923121 | 5.13391 | 4.11345 | 3.12085 |
| chr9 | 79915315 | 79915627 | 1000 | Promoter (2-3kb) | 9 | 79913200 | 79913631 | 432 | 1 | ENSMUST00 | 2116 | Gm3211 | ENSMUSG0C | false | . | . | . | . | . | . | chr9 | 79911389 | 79923121 | 5.13391 | 4.11345 | 3.12085 |
| chr9 | 79913355 | 79913669 | 1000 | Promoter (<=1kb) | 9 | 79913200 | 79913631 | 432 | 1 | ENSMUST00 | 156 | Gm3211 | ENSMUSG0C | false | . | . | . | . | . | . | chr9 | 79911389 | 79923121 | 5.13391 | 4.11345 | 3.12085 |
| chr13 | 81131344 | 81131669 | 1000 | Intron (ENSMUST00000128120.1 | 13 | 81065477 | 81065560 | 84 | 1 | ENSMUST00 | 65868 | Gm25241 | ENSMUSG0C | false | . | . | . | . | . | . | chr13 | 81128690 | 81133234 | 5.13344 | 4.16391 | 3.11287 |
| chr2 | 44999890 | 45000357 | 1000 | Intron (ENSMUST00000068415.5 | 2 | 45023100 | 45023177 | 78 | 2 | ENSMUST00 | 22820 | Mir5129 | ENSMUSG0C | false | . | . | . | . | . | . | chr2 | 44999483 | 45000933 | 4.43428 | 3.78197 | 2.59152 |
| chr3 | 129698852 | 129699271 | 1000 | Intron (ENSMUST00000029653.2 | 3 | 129684133 | 129684241 | 109 | 2 | ENSMUST00 | -14612 | Gm24515 | ENSMUSG0C | false | . | . | . | . | . | . | chr3 | 129694610 | 129701042 | 5.50271 | 4.23715 | 3.46885 |
| chr18 | 55548801 | 55549597 | 1000 | Distal Intergenic | 18 | 55622636 | 55622920 | 285 | 2 | ENSMUST00 | 73323 | Gm24514 | ENSMUSG0C | false | chr18 | 55548461 | 55549338 | 11.49921 | 31.27799 | 28.79692 | chr18 | 55546154 | 55550197 | 4.63718 | 3.88951 | 2.70527 |
| chr2 | 145878509 | 145878906 | 1000 | Promoter (<=1kb) | 2 | 145878286 | 145878473 | 188 | 1 | ENSMUST00 | 224 | Gm25026 | ENSMUSG0C | false | . | . | . | . | . | . | chr2 | 145866961 | 145893931 | 5.63035 | 4.25092 | 3.54043 |
| chr2 | 32317945 | 32318460 | 1000 | Promoter (<=1kb) | 2 | 32318249 | 32318356 | 108 | 1 | ENSMUST00 | 0 | Gm23363 | ENSMUSG0C | false | chr2 | 32318161 | 32318504 | 8.73111 | 16.23101 | 13.94854 | chr2 | 32315734 | 32320167 | 5.03703 | 3.88571 | 3.06941 |
| chr18 | 33863537 | 33863891 | 1000 | Intron (ENSMUST00000025234.5 | 18 | 33795164 | 33795295 | 132 | 1 | ENSMUST00 | 68374 | Gm23639 | ENSMUSG0C | false | . | . | . | . | . | . | chr18 | 33861621 | 33864746 | 5.36645 | 4.25909 | 3.30026 |
| chr2 | 168072483 | 168073295 | 1000 | Distal Intergenic | 2 | 168064820 | 168064897 | 78 | 2 | ENSMUST00 | -7587 | Gm24327 | ENSMUSG0C | false | . | . | . | . | . | . | chr2 | 168063587 | 168080595 | 6.53927 | 4.82888 | 4.26326 |
| chr1 | 34434900 | 34435205 | 1000 | Promoter (1-2kb) | 1 | 34433121 | 34433199 | 79 | 2 | ENSMUST00 | -1702 | Mir5103 | ENSMUSG0C | false | chr1 | 34434002 | 34435403 | 17.60262 | 45.78123 | 43.11721 | chr1 | 34433765 | 34435611 | 4.29264 | 3.76707 | 2.4137 |
| chr6 | 48227215 | 48227778 | 1000 | Intron (ENSMUST00000180741.1 | 6 | 48289228 | 48289427 | 200 | 1 | ENSMUST00 | -61450 | Gm22122 | ENSMUSG0C | false | . | . | . | . | . | . | chr6 | 48226682 | 48227901 | 4.90119 | 3.97673 | 2.9445 |
| chr14 | 102982220 | 102983186 | 1000 | Promoter (<=1kb) | 14 | 102982549 | 102982632 | 84 | 2 | ENSMUST00 | 0 | Mir5130 | ENSMUSG0C | false | chr14 | 102980932 | 102983668 | 31.01414 | 97.6227 | 94.10706 | chr14 | 102980620 | 102982291 | 4.03906 | 3.44113 | 2.24082 |
| chr7 | 112615248 | 112615856 | 1000 | Promoter (2-3kb) | 7 | 112618096 | 112618225 | 130 | 2 | ENSMUST00 | 2369 | Gm24154 | ENSMUSG0C | false | . | . | . | . | . | . | chr7 | 112613970 | 112618130 | 4.20584 | 3.67661 | 2.35462 |
| chr5 | 64574448 | 64574858 | 1000 | Distal Intergenic | 5 | 64647633 | 64647749 | 117 | 2 | ENSMUST00 | 72891 | Gm25306 | ENSMUSG0C | false | . | . | . | . | . | . | chr5 | 64574395 | 64576348 | 4.70992 | 3.82406 | 2.81128 |
| chr4 | 133659931 | 133660908 | 1000 | 3' UTR | 4 | 133654002 | 133654257 | 256 | 1 | ENSMUST00 | 5930 | Gm22662 | ENSMUSG0C | false | chr4 | 133659625 | 133660921 | 15.98415 | 46.70646 | 44.03467 | chr4 | 133658272 | 133663618 | 5.53059 | 4.29226 | 3.48895 |
| chr9 | 69458940 | 69459284 | 1000 | Promoter (1-2kb) | 9 | 69456944 | 69457031 | 88 | 1 | ENSMUST00 | 1997 | Mir3109 | ENSMUSG0C | false | . | . | . | . | . | . | chr9 | 69455103 | 69480655 | 7.58546 | 5.20708 | 5.13718 |
| chr1 | 190513893 | 190514314 | 1000 | Distal Intergenic | 1 | 190482845 | 190483157 | 313 | 2 | ENSMUST00 | -30737 | Gm23153 | ENSMUSG0C | false | . | . | . | . | . | . | . | . | . | . | . | . |
| chr2 | 30743900 | 30744638 | 1000 | Distal Intergenic | 2 | 30721210 | 30721293 | 84 | 1 | ENSMUST00 | 22691 | Mir3089 | ENSMUSG0C | false | . | . | . | . | . | . | chr2 | 30743149 | 30744848 | 8.39244 | 5.64319 | 5.7678 |
| chr13 | 98534105 | 98534861 | 1000 | Distal Intergenic | 13 | 98560354 | 98560461 | 108 | 1 | ENSMUST00 | -25493 | Gm22698 | ENSMUSG0C | false | . | . | . | . | . | . | chr13 | 98532862 | 98535918 | 4.11254 | 3.58103 | 2.30807 |
| chr13 | 98594735 | 98595112 | 1000 | Distal Intergenic | 13 | 98560354 | 98560461 | 108 | 1 | ENSMUST00 | 34382 | Gm22698 | ENSMUSG0C | false | . | . | . | . | . | . | chr13 | 98594594 | 98595590 | 5.25136 | 4.27411 | 3.20699 |
| chr13 | 98533417 | 98533842 | 1000 | Distal Intergenic | 13 | 98560354 | 98560461 | 108 | 1 | ENSMUST00 | -26512 | Gm22698 | ENSMUSG0C | false | . | . | . | . | . | . | chr13 | 98532862 | 98535918 | 4.11254 | 3.58103 | 2.30807 |
| chr17 | 26946569 | 26947087 | 1000 | Promoter (1-2kb) | 17 | 26948056 | 26948119 | 64 | 2 | ENSMUST00 | 1032 | Mir3083 | ENSMUSG0C | false | chr17 | 26946217 | 26947265 | 16.14593 | 43.98602 | 41.34693 | chr17 | 26946231 | 26949664 | 3.48984 | 3.31799 | 1.75783 |
| chr15 | 99262709 | 99263025 | 1000 | Distal Intergenic | 15 | 99254882 | 99255014 | 133 | 2 | ENSMUST00 | -7696 | Gm25999 | ENSMUSG0C | false | . | . | . | . | . | . | chr15 | 99262689 | 99263130 | 3.63657 | 3.45804 | 1.85995 |
| chr1 | 127322061 | 127322444 | 1000 | Intron (ENSMUST00000189427.1 | 1 | 127331813 | 127331941 | 129 | 1 | ENSMUST00 | -9369 | Gm23370 | ENSMUSG0C | false | . | . | . | . | . | . | chr1 | 127310467 | 127366303 | 5.56318 | 4.29763 | 3.47737 |
| chr1 | 127348692 | 127349206 | 1000 | Intron (ENSMUST00000189427.1 | 1 | 127331813 | 127331941 | 129 | 1 | ENSMUST00 | 16880 | Gm23370 | ENSMUSG0C | false | . | . | . | . | . | . | chr1 | 127310467 | 127366303 | 5.56318 | 4.29763 | 3.47737 |
| chr2 | 172862913 | 172863904 | 1000 | Promoter (<=1kb) | 2 | 172864264 | 172864341 | 78 | 2 | ENSMUST00 | 437 | Gm22773 | ENSMUSG0C | false | chr2 | 172862620 | 172863294 | 11.94783 | 25.26529 | 22.85895 | chr2 | 172861503 | 172865471 | 5.03829 | 4.03624 | 3.05316 |
| chr6 | 86585019 | 86585504 | 1000 | Distal Intergenic | 6 | 86574100 | 86574161 | 62 | 2 | ENSMUST00 | -10859 | Gm22184 | ENSMUSG0C | false | chr6 | 86584683 | 86585388 | 13.41152 | 31.46699 | 28.98322 | chr6 | 86584317 | 86586140 | 4.33258 | 3.68883 | 2.50122 |
| chr7 | 15948283 | 15948892 | 1000 | Promoter (2-3kb) | 7 | 15946248 | 15946495 | 248 | 1 | ENSMUST00 | 2036 | Gm24576 | ENSMUSG0C | false | . | . | . | . | . | . | chr7 | 15944204 | 15952855 | 5.15429 | 4.09924 | 3.13011 |
| chr11 | 85760674 | 85761119 | 1000 | Promoter (<=1kb) | 11 | 85760616 | 85760702 | 87 | 2 | ENSMUST00 | 0 | Mir5110 | ENSMUSG0C | false | . | . | . | . | . | . | chr11 | 85760058 | 85761337 | 6.71488 | 4.27359 | 4.44986 |
| chr7 | 100893082 | 100893588 | 1000 | Intron (ENSMUST00000107032.1 | 7 | 100882306 | 100882409 | 104 | 2 | ENSMUST00 | -10674 | Mir3102 | ENSMUSG0C | false | chr7 | 100893076 | 100894901 | 19.90079 | 58.62946 | 55.80003 | chr7 | 100889408 | 100898919 | 7.15353 | 5.01893 | 4.77985 |
| chr7 | 100894324 | 100894925 | 1000 | Intron (ENSMUST00000107032.1 | 7 | 100882306 | 100882409 | 104 | 2 | ENSMUST00 | -11916 | Mir3102 | ENSMUSG0C | false | chr7 | 100893076 | 100894901 | 19.90079 | 58.62946 | 55.80003 | chr7 | 100889408 | 100898919 | 7.15353 | 5.01893 | 4.77985 |
| chr19 | 44311840 | 44312342 | 1000 | Distal Intergenic | 19 | 44303171 | 44303231 | 61 | 1 | ENSMUST00 | 8670 | Mir5114 | ENSMUSG0C | false | . | . | . | . | . | . | chr19 | 44310172 | 44316564 | 5.73307 | 4.39799 | 3.62835 |
| chr13 | 107848907 | 107849237 | 1000 | Intron (ENSMUST00000105097.2 | 13 | 107829110 | 107829280 | 171 | 2 | ENSMUST00 | -19628 | Gm22118 | ENSMUSG0C | false | . | . | . | . | . | . | chr13 | 107830048 | 107858603 | 6.21396 | 4.54734 | 4.02799 |
| chr13 | 107838105 | 107838459 | 1000 | Intron (ENSMUST00000105097.2 | 13 | 107829110 | 107829280 | 171 | 2 | ENSMUST00 | -8826 | Gm22118 | ENSMUSG0C | false | chr13 | 107838288 | 107838613 | 6.34786 | 10.65255 | 8.46139 | chr13 | 107830048 | 107858603 | 6.21396 | 4.54734 | 4.02799 |
| chr2 | 32654525 | 32655275 | 1000 | Promoter (2-3kb) | 2 | 32652329 | 32652414 | 86 | 1 | ENSMUST00 | 2197 | Mir1954 | ENSMUSG0C | false | . | . | . | . | . | . | chr2 | 32652200 | 32659270 | 4.83732 | 3.88853 | 2.88316 |
| chr2 | 32655701 | 32656324 | 1000 | Intron (ENSMUST00000009705.8 | 2 | 32652329 | 32652414 | 86 | 1 | ENSMUST00 | 3373 | Mir1954 | ENSMUSG0C | false | . | . | . | . | . | . | chr2 | 32652200 | 32659270 | 4.83732 | 3.88853 | 2.88316 |
| chr15 | 68363196 | 68363633 | 1000 | Promoter (<=1kb) | 15 | 68340983 | 68363176 | 22194 | 2 | ENSMUST00 | -21 | Gm20732 | ENSMUSG0C | false | chr15 | 68360668 | 68363243 | 41.31276 | 141.09892 | 136.17311 | . | . | . | . | . | . |
| chr17 | 66860591 | 66860961 | 1000 | Intron (ENSMUST00000037974.8 | 17 | 67026147 | 67026235 | 89 | 2 | ENSMUST00 | 165274 | Mir5709 | ENSMUSG0C | false | . | . | . | . | . | . | . | . | . | . | . | . |
| chr7 | 64019885 | 64020255 | 1000 | Distal Intergenic | 7 | 64057602 | 64062958 | 5357 | 1 | ENSMUST00 | -37347 | Gm20670 | ENSMUSG0C | false | . | . | . | . | . | . | chr7 | 64015632 | 64024193 | 6.12605 | 4.59747 | 3.93071 |
| chr7 | 64021590 | 64022129 | 1000 | Distal Intergenic | 7 | 64057602 | 64062958 | 5357 | 1 | ENSMUST00 | -35473 | Gm20670 | ENSMUSG0C | false | . | . | . | . | . | . | chr7 | 64015632 | 64024193 | 6.12605 | 4.59747 | 3.93071 |
| chr14 | 79554320 | 79554628 | 1000 | Intron (ENSMUST00000040131.6 | 14 | 79562167 | 79562292 | 126 | 2 | ENSMUST00 | 7664 | Gm27857 | ENSMUSG0C | false | . | . | . | . | . | . | chr14 | 79551216 | 79555059 | 3.54755 | 3.32695 | 1.83305 |
| chr17 | 66419458 | 66419852 | 1000 | Intron (ENSMUST00000086693.6 | 17 | 66413631 | 66413984 | 354 | 2 | ENSMUST00 | -5475 | Gm4705 | ENSMUSG0C | false | . | . | . | . | . | . | chr17 | 66401889 | 66429482 | 6.16935 | 4.5489 | 3.97689 |
| chr8 | 108859957 | 108860377 | 1000 | Intron (ENSMUST00000043896.9 | 8 | 108918264 | 108921874 | 3611 | 2 | ENSMUST00 | 61497 | Gm20686 | ENSMUSG0C | false | . | . | . | . | . | . | chr8 | 108859356 | 108861181 | 4.81468 | 4.03975 | 2.85738 |
| chr9 | 64084634 | 64085484 | 1000 | Intron (ENSMUST00000176950.1 | 9 | 64080761 | 64086961 | 6201 | 1 | ENSMUST00 | 3874 | Scarletltr | ENSMUSG0C | false | chr9 | 64084430 | 64085342 | 18.16205 | 47.51548 | 44.83143 | chr9 | 64084037 | 64087913 | 5.30986 | 4.14497 | 3.30306 |
| chr10 | 108444528 | 108445014 | 1000 | Distal Intergenic | 10 | 108428067 | 108428160 | 94 | 1 | ENSMUST00 | 16462 | Gm25591 | ENSMUSG0C | false | chr10 | 108443290 | 108444613 | 15.42443 | 39.05124 | 36.47339 | chr10 | 108444796 | 108448791 | 5.7008 | 4.31379 | 3.59208 |
| chr9 | 70325915 | 70326539 | 1000 | Intron (ENSMUST00000034745.7 | 9 | 70405694 | 70405753 | 60 | 2 | ENSMUST00 | 79214 | Mir5626 | ENSMUSG0C | false | . | . | . | . | . | . | chr9 | 70312793 | 70333015 | 5.95088 | 4.44065 | 3.80175 |
| chr9 | 70322695 | 70323194 | 1000 | Intron (ENSMUST00000034745.7 | 9 | 70405694 | 70405753 | 60 | 2 | ENSMUST00 | 82559 | Mir5626 | ENSMUSG0C | false | . | . | . | . | . | . | chr9 | 70312793 | 70333015 | 5.95088 | 4.44065 | 3.80175 |
| chr13 | 119689871 | 119690599 | 1000 | Promoter (<=1kb) | 13 | 119690462 | 119708086 | 17625 | 1 | ENSMUST00 | 0 | Hmgcs1 | ENSMUSG0C | false | chr13 | 119689658 | 119690178 | 16.2384 | 40.45376 | 37.85896 | chr13 | 119689324 | 119689951 | 3.24855 | 3.15324 | 1.54928 |
| chr14 | 69502364 | 69503566 | 1000 | Promoter (<=1kb) | 14 | 69363504 | 69503320 | 139817 | 2 | ENSMUST00 | 0 | Gm28049 | ENSMUSG0C | false | chr14 | 69503539 | 69503826 | 7.81205 | 13.8349 | 11.58824 | . | . | . | . | . | . |
| chr16 | 78737232 | 78737520 | 1000 | Distal Intergenic | 16 | 78624747 | 78624853 | 107 | 1 | ENSMUST00 | 112486 | Gm24861 | ENSMUSG0C | false | . | . | . | . | . | . | chr16 | 78736075 | 78738558 | 3.66193 | 3.40149 | 1.89956 |
| chr17 | 8103037 | 8103366 | 1000 | Promoter (1-2kb) | 17 | 8038510 | 8101228 | 62719 | 2 | ENSMUST00 | -1810 | Gm1604A | ENSMUSG0C | false | . | . | . | . | . | . | . | . | . | . | . | . |
| chr9 | 63091887 | 63092639 | 1000 | Distal Intergenic | 9 | 63133283 | 63133389 | 107 | 1 | ENSMUST00 | -40644 | Gm25064 | ENSMUSG0C | false | chr9 | 63091662 | 63092777 | 6.52658 | 13.36193 | 11.1274 | chr9 | 63090225 | 63093748 | 4.55943 | 3.89487 | 2.64668 |
| chr4 | 142845401 | 142845862 | 1000 | Distal Intergenic | 4 | 142557390 | 142557836 | 447 | 2 | ENSMUST00 | -287566 | Gm13018 | ENSMUSG0C | false | . | . | . | . | . | . | . | . | . | . | . | . |
| chr9 | 49922134 | 49922501 | 1000 | Distal Intergenic | 9 | 49825869 | 49825975 | 107 | 2 | ENSMUST00 | -96160 | Gm25633 | ENSMUSG0C | false | . | . | . | . | . | . | chr9 | 49921974 | 49923173 | 4.10969 | 3.41881 | 2.30559 |
| chr9 | 49989908 | 49990333 | 1000 | Distal Intergenic | 9 | 49825869 | 49825975 | 107 | 2 | ENSMUST00 | -163934 | Gm25633 | ENSMUSG0C | false | . | . | . | . | . | . | chr9 | 49987558 | 49990811 | 5.24183 | 4.05424 | 3.24295 |
| chr16 | 51086649 | 51087171 | 1000 | Distal Intergenic | 16 | 50920542 | 50920648 | 107 | 2 | ENSMUST00 | -166002 | Gm24925 | ENSMUSG0C | false | . | . | . | . | . | . | chr16 | 51086334 | 51086791 | 3.04844 | 3.10198 | 1.39236 |
| chr16 | 51094898 | 51095192 | 1000 | Distal Intergenic | 16 | 50920542 | 50920648 | 107 | 2 | ENSMUST00 | -174251 | Gm24925 | ENSMUSG0C | false | . | . | . | . | . | . | . | . | . | . | . | . |
| chr6 | 70143377 | 70143788 | 1000 | Promoter (<=1kb) | 6 | 70143593 | 70143897 | 305 | 2 | ENSMUST00 | 109 | Igkv8-28 | ENSMUSG0C | false | . | . | . | . | . | . | chr6 | 70143123 | 70145035 | 3.54019 | 3.33533 | 1.80146 |
| chr4 | 139587349 | 139587956 | 1000 | Promoter (1-2kb) | 4 | 139588876 | 139588977 | 102 | 2 | ENSMUST00 | 1021 | Gm25571 | ENSMUSG0C | false | . | . | . | . | . | . | chr4 | 139576299 | 139606726 | 8.92638 | 5.72021 | 6.23508 |
| chr4 | 139582154 | 139582509 | 1000 | Intron (ENSMUST00000123827.3 | 4 | 139588876 | 139588977 | 102 | 2 | ENSMUST00 | 6468 | Gm25571 | ENSMUSG0C | false | chr4 | 139580247 | 139582371 | 12.5733 | 28.75278 | 26.30366 | chr4 | 139576299 | 139606726 | 8.92638 | 5.72021 | 6.23508 |
| chr15 | 6122916 | 6123468 | 1000 | Distal Intergenic | 15 | 6151217 | 6151348 | 132 | 2 | ENSMUST00 | 27880 | Gm23139 | ENSMUSG0C | false | . | . | . | . | . | . | chr15 | 6123043 | 6123632 | 3.02472 | 2.97787 | 1.37375 |
| chr4 | 139598984 | 139599634 | 1000 | Intron (ENSMUST00000123827.3 | 4 | 139603186 | 139662691 | 59506 | 1 | ENSMUST00 | -3552 | Gm21969 | ENSMUSG0C | false | chr4 | 139598866 | 139599073 | 6.70576 | 11.61755 | 9.40775 | chr4 | 139576299 | 139606726 | 8.92638 | 5.72021 | 6.23508 |
| chr6 | 51811833 | 51812393 | 1000 | Distal Intergenic | 6 | 51621632 | 51621708 | 77 | 1 | ENSMUST00 | 190202 | Gm22914 | ENSMUSG0C | false | . | . | . | . | . | . | chr6 | 51808893 | 51812949 | 5.35495 | 4.18951 | 3.30309 |
| chr6 | 120493206 | 120494201 | 1000 | Promoter (<=1kb) | 6 | 120488939 | 120493807 | 4869 | 2 | ENSMUST00 | 0 | Cecr6 | ENSMUSG0C | false | chr6 | 120492232 | 120494265 | 28.03146 | 79.91027 | 76.76085 | chr6 | 120492268 | 120494664 | 7.69762 | 5.16645 | 5.22414 |
| chr17 | 45733730 | 45734220 | 1000 | Promoter (<=1kb) | 17 | 45732864 | 45733844 | 981 | 2 | ENSMUST00 | 0 | 1600014C23I | ENSMUSG0C | false | chr17 | 45733451 | 45734265 | 11.40015 | 24.34736 | 21.95423 | chr17 | 45731816 | 45737308 | 6.56758 | 4.65756 | 4.31435 |
| chr8 | 108045228 | 108045588 | 1000 | Distal Intergenic | 8 | 107842451 | 107842551 | 101 | 1 | ENSMUST00 | 202778 | Gm22825 | ENSMUSG0C | false | . | . | . | . | . | . | chr8 | 108044290 | 108046363 | 5.29298 | 4.19586 | 3.28019 |
| chr5 | 22813445 | 22813770 | 1000 | Distal Intergenic | 5 | 22775630 | 22778702 | 3073 | 1 | ENSMUST00 | 37816 | Gm21846 | ENSMUSG0C | false | . | . | . | . | . | . | chr5 | 22813304 | 22813917 | 2.88102 | 2.93569 | 1.30462 |
| chr1 | 85758285 | 85758859 | 1000 | Intron (ENSMUST00000159122.2 | 1 | 85763794 | 85763871 | 78 | 1 | ENSMUST00 | -4935 | Gm25472 | ENSMUSG0C | false | . | . | . | . | . | . | chr1 | 85757569 | 85759528 | 5.46235 | 4.33237 | 3.394 |
| chr10 | 43765307 | 43765676 | 1000 | Distal Intergenic | 10 | 43821231 | 43822235 | 1005 | 1 | ENSMUST00 | -55555 | Gm9034 | ENSMUSG0C | false | . | . | . | . | . | . | . | . | . | . | . | . |
| chr17 | 48039469 | 48039937 | 1000 | Promoter (1-2kb) | 17 | 48036745 | 48038293 | 1549 | 2 | ENSMUST00 | -1177 | 1700122O11 | ENSMUSG0C | false | . | . | . | . | . | . | chr17 | 48039013 | 48040234 | 4.62489 | 3.96903 | 2.69237 |
| chr6 | 70215734 | 70216150 | 1000 | Promoter (<=1kb) | 6 | 70215433 | 70215957 | 525 | 1 | ENSMUST00 | 302 | Igkv6-25 | ENSMUSG0C | false | . | . | . | . | . | . | chr6 | 70213474 | 70219351 | 5.14256 | 4.19463 | 3.11118 |
| chr12 | 94257616 | 94258051 | 1000 | Distal Intergenic | 12 | 93927004 | 93929102 | 2099 | 2 | ENSMUST00 | -328515 | Gm9726 | ENSMUSG0C | false | . | . | . | . | . | . | . | . | . | . | . | . |
| chr12 | 94282894 | 94283422 | 1000 | Distal Intergenic | 12 | 93927004 | 93929102 | 2099 | 2 | ENSMUST00 | -353793 | Gm9726 | ENSMUSG0C | false | . | . | . | . | . | . | chr12 | 94279611 | 94284656 | 3.79808 | 3.5006 | 2.0053 |
| chr6 | 69858254 | 69858630 | 1000 | Promoter (<=1kb) | 6 | 69858420 | 69858884 | 465 | 2 | ENSMUST00 | 254 | Igkv12-41 | ENSMUSG0C | false | . | . | . | . | . | . | chr6 | 69857739 | 69859214 | 6.42763 | 4.81646 | 4.16397 |
| chr17 | 87185711 | 87186051 | 1000 | Promoter (1-2kb) | 17 | 87184105 | 87184225 | 121 | 2 | ENSMUST00 | -1487 | Gm22346 | ENSMUSG0C | false | . | . | . | . | . | . | chr17 | 87185193 | 87187241 | 5.36753 | 4.27934 | 3.30999 |
| chr7 | 63920432 | 63920718 | 1000 | 3' UTR | 7 | 63916857 | 63920539 | 3683 | 1 | ENSMUST00 | 3576 | E030018B13I | ENSMUSG0C | false | . | . | . | . | . | . | chr7 | 63885397 | 63936279 | 7.51388 | 5.18869 | 5.06896 |
| chr11 | 64511345 | 64511893 | 1000 | Intron (ENSMUST00000058652.5 | 11 | 64435156 | 64436653 | 1498 | 2 | ENSMUST00 | -74693 | F930015N05 | ENSMUSG0C | false | . | . | . | . | . | . | chr11 | 64509630 | 64515155 | 5.43544 | 4.26981 | 3.39776 |
| chr15 | 77068745 | 77069257 | 1000 | Promoter (<=1kb) | 15 | 77067791 | 77067897 | 107 | 1 | ENSMUST00 | 955 | Gm24753 | ENSMUSG0C | false | . | . | . | . | . | . | chr15 | 77068375 | 77069486 | 5.49084 | 4.13979 | 3.438 |
| chr6 | 128285738 | 128286168 | 1000 | Intron (ENSMUST00000006311.7 | 6 | 128294096 | 128294506 | 411 | 1 | ENSMUST00 | -7928 | Gm21893 | ENSMUSG0C | false | chr6 | 128285871 | 128286241 | 12.97578 | 28.80697 | 26.35731 | chr6 | 128284568 | 128292111 | 4.60797 | 3.90404 | 2.68631 |
| chr13 | 69988074 | 69988708 | 1000 | Distal Intergenic | 13 | 70007204 | 70007614 | 411 | 2 | ENSMUST00 | 18906 | Gm6132 | ENSMUSG0C | false | . | . | . | . | . | . | chr13 | 69981325 | 69991868 | 4.48541 | 3.78231 | 2.60299 |
| chr15 | 101046451 | 101047119 | 1000 | Distal Intergenic | 15 | 101050194 | 101054399 | 4206 | 2 | ENSMUST00 | 7280 | Fignl2 | ENSMUSG0C | false | chr15 | 101046148 | 101047343 | 17.00309 | 47.03089 | 44.35329 | chr15 | 101046007 | 101047332 | 6.75392 | 4.63557 | 4.47022 |
| chr5 | 51072765 | 51073111 | 1000 | Distal Intergenic | 5 | 51223587 | 51223680 | 94 | 1 | ENSMUST00 | -150476 | Gm22976 | ENSMUSG0C | false | . | . | . | . | . | . | chr5 | 51070089 | 51073348 | 4.24262 | 3.68278 | 2.39081 |
| chr6 | 70240204 | 70240610 | 1000 | Distal Intergenic | 6 | 70260409 | 70260933 | 525 | 2 | ENSMUST00 | 20323 | Igkv6-23 | ENSMUSG0C | false | . | . | . | . | . | . | chr6 | 70239615 | 70241403 | 4.75318 | 3.82127 | 2.83041 |
| chr14 | 53024832 | 53025587 | 1000 | Distal Intergenic | 14 | 53032902 | 53033418 | 517 | 1 | ENSMUST00 | -7315 | Trav13d-3 | ENSMUSG0C | false | . | . | . | . | . | . | . | . | . | . | . | . |
| chr17 | 6655261 | 6656080 | 1000 | Promoter (<=1kb) | 17 | 6648941 | 6655939 | 6999 | 2 | ENSMUST00 | 0 | Dynlt1f | ENSMUSG0C | false | chr17 | 6654460 | 6656315 | 20.93824 | 109.0712 | 105.26324 | chr17 | 6654090 | 6656424 | 4.66774 | 3.6516 | 2.76541 |
| chr12 | 69494452 | 69495014 | 1000 | Distal Intergenic | 12 | 69561229 | 69561337 | 109 | 2 | ENSMUST00 | 66323 | Gm23018 | ENSMUSG0C | false | . | . | . | . | . | . | chr12 | 69491433 | 69504349 | 6.88286 | 4.93684 | 4.54772 |
| chr12 | 69501753 | 69502014 | 1000 | Distal Intergenic | 12 | 69561229 | 69561337 | 109 | 2 | ENSMUST00 | 59323 | Gm23018 | ENSMUSG0C | false | . | . | . | . | . | . | chr12 | 69491433 | 69504349 | 6.88286 | 4.93684 | 4.54772 |
| chr12 | 69502192 | 69502734 | 1000 | Distal Intergenic | 12 | 69561229 | 69561337 | 109 | 2 | ENSMUST00 | 58603 | Gm23018 | ENSMUSG0C | false | chr12 | 69502212 | 69503540 | 17.6846 | 53.76268 | 50.99834 | chr12 | 69491433 | 69504349 | 6.88286 | 4.93684 | 4.54772 |
| chr16 | 30821192 | 30822203 | 1000 | Distal Intergenic | 16 | 30814162 | 30814268 | 107 | 1 | ENSMUST00 | 7031 | Gm26295 | ENSMUSG0C | false | . | . | . | . | . | . | chr16 | 30820618 | 30823387 | 4.71605 | 3.8569 | 2.80454 |
| chr6 | 70371814 | 70372273 | 1000 | Promoter (<=1kb) | 6 | 70371469 | 70371993 | 525 | 1 | ENSMUST00 | 346 | Igkv6-17 | ENSMUSG0C | false | . | . | . | . | . | . | chr6 | 70370347 | 70374796 | 4.78873 | 3.97689 | 2.85976 |
| chr5 | 50457701 | 50458119 | 1000 | Distal Intergenic | 5 | 50290380 | 50290486 | 107 | 2 | ENSMUST00 | -167216 | Gm22618 | ENSMUSG0C | false | . | . | . | . | . | . | . | . | . | . | . | . |
| chr16 | 58074915 | 58075316 | 1000 | Distal Intergenic | 16 | 57884008 | 57884139 | 132 | 2 | ENSMUST00 | -190777 | Gm24039 | ENSMUSG0C | false | . | . | . | . | . | . | chr16 | 58074687 | 58076552 | 2.73573 | 2.7665 | 1.18641 |
| chr9 | 60743040 | 60743731 | 1000 | Promoter (<=1kb) | 9 | 60743120 | 60770276 | 27157 | 1 | ENSMUST00 | 0 | 1700036A12I | ENSMUSG0C | false | chr9 | 60742982 | 60743346 | 7.88522 | 15.75676 | 13.48224 | chr9 | 60741220 | 60745216 | 5.18588 | 4.09349 | 3.20057 |
| chr17 | 43554448 | 43555011 | 1000 | Distal Intergenic | 17 | 43534174 | 43543639 | 9466 | 2 | ENSMUST00 | -10810 | Ankrd66 | ENSMUSG0C | false | . | . | . | . | . | . | chr17 | 43554410 | 43555332 | 4.40255 | 3.73011 | 2.56571 |
| chr17 | 43547447 | 43547842 | 1000 | Distal Intergenic | 17 | 43534174 | 43543639 | 9466 | 2 | ENSMUST00 | -3809 | Ankrd66 | ENSMUSG0C | false | . | . | . | . | . | . | chr17 | 43546947 | 43548208 | 3.729 | 3.39384 | 1.98866 |
| chr14 | 57034161 | 57034566 | 1000 | Downstream (1-2kb) | 14 | 57015134 | 57018982 | 3849 | 1 | ENSMUST00 | 19028 | B020004C17I | ENSMUSG0C | false | . | . | . | . | . | . | chr14 | 57032814 | 57035802 | 5.37941 | 4.25482 | 3.33652 |
| chr9 | 70862989 | 70863646 | 1000 | Intron (ENSMUST00000034731.8 | 9 | 70852576 | 70853418 | 843 | 1 | ENSMUST00 | 10414 | Gm3436 | ENSMUSG0C | false | . | . | . | . | . | . | chr9 | 70861061 | 70864431 | 5.58591 | 4.34029 | 3.49025 |
| chr15 | 72951177 | 72951634 | 1000 | Intron (ENSMUST00000023276.9 | 15 | 72913357 | 72913560 | 204 | 1 | ENSMUST00 | 37821 | Gm3150 | ENSMUSG0C | false | . | . | . | . | . | . | chr15 | 72949680 | 72953379 | 5.04063 | 3.98924 | 3.08344 |
| chr4 | 57867025 | 57867386 | 1000 | Intron (ENSMUST00000098066.3 | 4 | 57877389 | 57877495 | 107 | 2 | ENSMUST00 | 10109 | Gm26138 | ENSMUSG0C | false | . | . | . | . | . | . | chr4 | 57846708 | 57868207 | 6.85521 | 4.86004 | 4.55185 |
| chr3 | 119949006 | 119949396 | 1000 | Distal Intergenic | 3 | 119837904 | 119838024 | 121 | 1 | ENSMUST00 | 111103 | Gm26137 | ENSMUSG0C | false | . | . | . | . | . | . | chr3 | 119948466 | 119950445 | 3.75415 | 3.34156 | 2.01361 |
| chr17 | 4626079 | 4626671 | 1000 | Distal Intergenic | 17 | 4678020 | 4678126 | 107 | 1 | ENSMUST00 | -51349 | Gm23186 | ENSMUSG0C | false | chr17 | 4626080 | 4626682 | 79.22768 | 354.44785 | 346.18814 | . | . | . | . | . | . |
| chr15 | 79028179 | 79028716 | 1000 | Promoter (<=1kb) | 15 | 79028212 | 79030498 | 2287 | 1 | ENSMUST00 | 0 | H1f0 | ENSMUSG0C | false | chr15 | 79027416 | 79030756 | 27.03504 | 88.5911 | 85.27629 | . | . | . | . | . | . |
| chr8 | 10664882 | 10665249 | 1000 | Distal Intergenic | 8 | 10808730 | 10809170 | 441 | 2 | ENSMUST00 | 143921 | Rps16-ps3 | ENSMUSG0C | false | . | . | . | . | . | . | chr8 | 10663553 | 10665989 | 4.68753 | 3.83769 | 2.79655 |
| chr18 | 13221851 | 13222357 | 1000 | Distal Intergenic | 18 | 13209702 | 13209808 | 107 | 2 | ENSMUST00 | -12044 | Gm22251 | ENSMUSG0C | false | . | . | . | . | . | . | chr18 | 13220295 | 13222808 | 4.17372 | 3.48837 | 2.35995 |
| chr9 | 43858791 | 43859111 | 1000 | Distal Intergenic | 9 | 43829963 | 43832516 | 2554 | 1 | ENSMUST00 | 28829 | Gm3898 | ENSMUSG0C | false | . | . | . | . | . | . | . | . | . | . | . | . |
| chr9 | 43863773 | 43864424 | 1000 | Distal Intergenic | 9 | 43829963 | 43832516 | 2554 | 1 | ENSMUST00 | 33811 | Gm3898 | ENSMUSG0C | false | . | . | . | . | . | . | chr9 | 43863659 | 43865503 | 5.22983 | 4.13814 | 3.21075 |
| chr4 | 132409209 | 132409609 | 1000 | Promoter (<=1kb) | 4 | 132408309 | 132408404 | 96 | 1 | ENSMUST00 | 901 | Gm25707 | ENSMUSG0C | false | . | . | . | . | . | . | chr4 | 132398431 | 132412921 | 4.68778 | 3.92949 | 2.75307 |
| chr4 | 40818427 | 40818824 | 1000 | Intron (ENSMUST00000030121.7 | 4 | 40850056 | 40850138 | 83 | 2 | ENSMUST00 | 31314 | Mir5123 | ENSMUSG0C | false | . | . | . | . | . | . | chr4 | 40814142 | 40820715 | 6.71689 | 4.72651 | 4.42369 |
| chr10 | 5179807 | 5180557 | 1000 | Distal Intergenic | 10 | 5069192 | 5071975 | 2784 | 1 | ENSMUST00 | 110616 | Gm10097 | ENSMUSG0C | false | . | . | . | . | . | . | chr10 | 5174596 | 5182624 | 4.92543 | 4.00808 | 2.95185 |
| chr10 | 5139287 | 5140274 | 1000 | Distal Intergenic | 10 | 5069192 | 5071975 | 2784 | 1 | ENSMUST00 | 70096 | Gm10097 | ENSMUSG0C | false | . | . | . | . | . | . | chr10 | 5137874 | 5150928 | 4.52065 | 3.84085 | 2.61241 |
| chr6 | 70340649 | 70341069 | 1000 | Promoter (<=1kb) | 6 | 70340876 | 70341180 | 305 | 2 | ENSMUST00 | 111 | Igkv8-19 | ENSMUSG0C | false | . | . | . | . | . | . | chr6 | 70340487 | 70343444 | 4.70979 | 3.93738 | 2.77295 |
| chr1 | 151062397 | 151062744 | 1000 | Promoter (<=1kb) | 1 | 151062004 | 151062110 | 107 | 2 | ENSMUST00 | -288 | Gm22756 | ENSMUSG0C | false | . | . | . | . | . | . | chr1 | 151062011 | 151063610 | 4.13649 | 3.70907 | 2.28344 |
| chr16 | 21333093 | 21333529 | 1000 | Distal Intergenic | 16 | 21247518 | 21296872 | 49355 | 2 | ENSMUST00 | -36222 | Gm16863 | ENSMUSG0C | false | chr16 | 21332479 | 21334089 | 33.10969 | 106.34647 | 102.61562 | . | . | . | . | . | . |
| chr8 | 11044551 | 11045150 | 1000 | Exon (ENSMUST00000180750.2/ | 8 | 11049578 | 11054541 | 4964 | 1 | ENSMUST00 | -4428 | 9530052E02I | ENSMUSG0C | false | . | . | . | . | . | . | chr8 | 11043855 | 11047520 | 4.42731 | 3.73242 | 2.54378 |
| chr8 | 11000962 | 11001427 | 1000 | Intron (ENSMUST00000040514.6 | 8 | 11007850 | 11054536 | 46687 | 1 | ENSMUST00 | -6423 | 9530052E02I | ENSMUSG0C | false | . | . | . | . | . | . | chr8 | 10998695 | 11007069 | 6.48531 | 4.76202 | 4.22275 |
| chr1 | 155083059 | 155083961 | 1000 | Promoter (<=1kb) | 1 | 155063447 | 155083266 | 19820 | 2 | ENSMUST00 | 0 | Gm9530 | ENSMUSG0C | false | chr1 | 155082926 | 155084017 | 16.7644 | 42.82802 | 40.20103 | . | . | . | . | . | . |
| chr15 | 10720720 | 10721270 | 1000 | Exon (ENSMUST00000181739.1/ | 15 | 10717144 | 10722471 | 5328 | 1 | ENSMUST00 | 3577 | 4930556M19 | ENSMUSG0C | false | . | . | . | . | . | . | chr15 | 10719289 | 10724261 | 4.16849 | 3.55198 | 2.35744 |
| chr11 | 88040831 | 88041683 | 1000 | Intron (ENSMUST00000129337.1 | 11 | 88039578 | 88047360 | 7783 | 2 | ENSMUST00 | 5677 | 2010015M23 | ENSMUSG0C | false | . | . | . | . | . | . | chr11 | 88041018 | 88041765 | 3.12966 | 3.03052 | 1.48234 |
| chr13 | 69907072 | 69907442 | 1000 | Distal Intergenic | 13 | 69809914 | 69816094 | 6181 | 2 | ENSMUST00 | -90979 | Gm26819 | ENSMUSG0C | false | . | . | . | . | . | . | . | . | . | . | . | . |
| chr7 | 18957873 | 18958394 | 1000 | Exon (ENSMUST00000172835.2/ | 7 | 18962415 | 18965173 | 2759 | 2 | ENSMUST00 | 6779 | Gm26821 | ENSMUSG0C | false | chr7 | 18957376 | 18958691 | 21.42557 | 69.98924 | 66.99868 | chr7 | 18956721 | 18958744 | 4.24108 | 3.74386 | 2.38048 |
| chr8 | 30733395 | 30733700 | 1000 | Distal Intergenic | 8 | 30568393 | 30575573 | 7181 | 1 | ENSMUST00 | 165003 | 4933433F19I | ENSMUSG0C | false | . | . | . | . | . | . | chr8 | 30733014 | 30733993 | 3.50567 | 3.31441 | 1.77522 |
| chr8 | 126566278 | 126567119 | 1000 | Promoter (<=1kb) | 8 | 126566970 | 126590988 | 24019 | 1 | ENSMUST00 | 0 | Gm26759 | ENSMUSG0C | false | . | . | . | . | . | . | chr8 | 126562406 | 126572815 | 5.75135 | 4.37077 | 3.62239 |
| chr15 | 92276836 | 92277325 | 1000 | Intron (ENSMUST00000169825.2 | 15 | 92344359 | 92378088 | 33730 | 1 | ENSMUST00 | -67034 | Gm26760 | ENSMUSG0C | false | . | . | . | . | . | . | . | . | . | . | . | . |
| chr12 | 111430364 | 111431367 | 1000 | Promoter (<=1kb) | 12 | 111426874 | 111430946 | 4073 | 2 | ENSMUST00 | 0 | Gm10425 | ENSMUSG0C | false | . | . | . | . | . | . | chr12 | 111425142 | 111437454 | 7.80275 | 5.329 | 5.29325 |
| chr8 | 121124607 | 121125847 | 1000 | Promoter (2-3kb) | 8 | 121127940 | 121130644 | 2705 | 1 | ENSMUST00 | -2093 | Foxl1 | ENSMUSG0C | false | chr8 | 121125330 | 121126765 | 15.30398 | 43.94659 | 41.30965 | chr8 | 121121895 | 121136533 | 5.26018 | 4.19829 | 3.22208 |
| chr8 | 121134585 | 121134931 | 1000 | Intron (ENSMUST00000127664.1 | 8 | 121127940 | 121130644 | 2705 | 1 | ENSMUST00 | 6646 | Foxl1 | ENSMUSG0C | false | . | . | . | . | . | . | chr8 | 121121895 | 121136533 | 5.26018 | 4.19829 | 3.22208 |
| chr15 | 78863717 | 78864093 | 1000 | Promoter (2-3kb) | 15 | 78862956 | 78866889 | 3934 | 2 | ENSMUST00 | 2796 | Gm26634 | ENSMUSG0C | false | . | . | . | . | . | . | chr15 | 78863592 | 78864667 | 5.95596 | 4.38821 | 3.84349 |
| chr4 | 133095003 | 133095670 | 1000 | Distal Intergenic | 4 | 133085474 | 133089146 | 3673 | 1 | ENSMUST00 | 9530 | Gm26615 | ENSMUSG0C | false | chr4 | 133095363 | 133097697 | 19.17126 | 69.02763 | 66.05415 | chr4 | 133082180 | 133096297 | 7.96206 | 5.47868 | 5.42168 |
| chr4 | 133107599 | 133108131 | 1000 | Distal Intergenic | 4 | 133085474 | 133089146 | 3673 | 1 | ENSMUST00 | 22126 | Gm26615 | ENSMUSG0C | false | . | . | . | . | . | . | chr4 | 133105817 | 133110012 | 5.55758 | 4.31138 | 3.4668 |
| chr4 | 133093944 | 133094459 | 1000 | Distal Intergenic | 4 | 133085474 | 133089146 | 3673 | 1 | ENSMUST00 | 8471 | Gm26615 | ENSMUSG0C | false | . | . | . | . | . | . | chr4 | 133082180 | 133096297 | 7.96206 | 5.47868 | 5.42168 |
| chr4 | 133087785 | 133088243 | 1000 | Promoter (2-3kb) | 4 | 133085474 | 133089146 | 3673 | 1 | ENSMUST00 | 2312 | Gm26615 | ENSMUSG0C | false | . | . | . | . | . | . | chr4 | 133082180 | 133096297 | 7.96206 | 5.47868 | 5.42168 |

| | | | | | | | | | | | | | | | | | | | | | | | | | | |
|---|---|---|---|---|---|---|---|---|---|---|---|---|---|---|---|---|---|---|---|---|---|---|---|---|---|---|
| chr4 | 133085058 | 133085720 | 1000 | Promoter (<=1kb) | 4 | 133085474 | 133089146 | 3673 | 1 | ENSMUST00 | 0 | Gm26615 | ENSMUSG0C | false | chr4 | 133084656 | 133086042 | 12.01557 | 29.04976 | 26.5957 | chr4 | 133082180 | 133096297 | 7.96206 | 5.47868 | 5.42168 |
| chr4 | 133084155 | 133084837 | 1000 | Promoter (<=1kb) | 4 | 133085474 | 133089146 | 3673 | 1 | ENSMUST00 | -637 | Gm26615 | ENSMUSG0C | false | chr4 | 133084656 | 133086042 | 12.01557 | 29.04976 | 26.5957 | chr4 | 133082180 | 133096297 | 7.96206 | 5.47868 | 5.42168 |
| chr17 | 27194304 | 27195002 | 1000 | Intron (ENSMUST00000055117.7 | 17 | 27201032 | 27203071 | 2040 | 1 | ENSMUST00 | -6030 | Gm26724 | ENSMUSG0C | false | . | . | . | . | . | . | chr17 | 27191319 | 27203697 | 8.84294 | 5.47024 | 6.1733 |
| chr17 | 27199833 | 27200322 | 1000 | Promoter (<=1kb) | 17 | 27201032 | 27203071 | 2040 | 1 | ENSMUST00 | -710 | Gm26724 | ENSMUSG0C | false | chr17 | 27200284 | 27200479 | 5.34626 | 8.34382 | 6.2035 | chr17 | 27191319 | 27203697 | 8.84294 | 5.47024 | 6.1733 |
| chr17 | 27200780 | 27201317 | 1000 | Promoter (<=1kb) | 17 | 27201032 | 27203071 | 2040 | 1 | ENSMUST00 | 0 | Gm26724 | ENSMUSG0C | false | . | . | . | . | . | . | chr17 | 27191319 | 27203697 | 8.84294 | 5.47024 | 6.1733 |
| chr8 | 121264484 | 121264960 | 1000 | Intron (ENSMUST00000127664.1 | 8 | 121198714 | 121201835 | 3122 | 2 | ENSMUST00 | -62650 | 5033426O07 | ENSMUSG0C | false | . | . | . | . | . | . | chr8 | 121255503 | 121265207 | 5.96407 | 4.44635 | 3.80299 |
| chr8 | 121239215 | 121239544 | 1000 | Intron (ENSMUST00000127664.1 | 8 | 121198714 | 121201835 | 3122 | 2 | ENSMUST00 | -37381 | 5033426O07 | ENSMUSG0C | false | . | . | . | . | . | . | chr8 | 121236124 | 121247254 | 7.60766 | 5.2125 | 5.14118 |
| chr8 | 121241670 | 121242435 | 1000 | Intron (ENSMUST00000127664.1 | 8 | 121198714 | 121201835 | 3122 | 2 | ENSMUST00 | -39836 | 5033426O07 | ENSMUSG0C | false | . | . | . | . | . | . | chr8 | 121236124 | 121247254 | 7.60766 | 5.2125 | 5.14118 |
| chr8 | 121257810 | 121258974 | 1000 | Exon (ENSMUST00000181200.1/ | 8 | 121198714 | 121201835 | 3122 | 2 | ENSMUST00 | -55976 | 5033426O07 | ENSMUSG0C | false | . | . | . | . | . | . | chr8 | 121255503 | 121265207 | 5.96407 | 4.44635 | 3.80299 |
| chr17 | 6852507 | 6852972 | 1000 | Distal Intergenic | 17 | 6798390 | 6827990 | 29601 | 2 | ENSMUST00 | -24518 | Gm2885 | ENSMUSG0C | false | . | . | . | . | . | . | chr17 | 6850662 | 6853671 | 4.16631 | 3.40223 | 2.34843 |
| chr8 | 84200627 | 84201117 | 1000 | Promoter (2-3kb) | 8 | 84192774 | 84197667 | 4894 | 2 | ENSMUST00 | -2961 | Gm26887 | ENSMUSG0C | false | chr8 | 84200501 | 84204316 | 27.20707 | 96.65539 | 93.16586 | chr8 | 84197000 | 84210598 | 6.06328 | 4.55601 | 3.88169 |
| chr8 | 84192543 | 84192872 | 1000 | Exon (ENSMUST00000181282.1/ | 8 | 84192774 | 84197667 | 4894 | 2 | ENSMUST00 | 4795 | Gm26887 | ENSMUSG0C | false | chr8 | 84192248 | 84192996 | 8.63619 | 18.02194 | 15.71366 | chr8 | 84190694 | 84193192 | 4.40739 | 3.61139 | 2.5518 |
| chr8 | 84197300 | 84198821 | 1000 | Promoter (<=1kb) | 8 | 84192774 | 84197667 | 4894 | 2 | ENSMUST00 | 0 | Gm26887 | ENSMUSG0C | false | chr8 | 84197247 | 84198456 | 13.60354 | 37.21757 | 34.66072 | chr8 | 84197000 | 84210598 | 6.06328 | 4.55601 | 3.88169 |
| chr5 | 92834486 | 92834902 | 1000 | Intron (ENSMUST00000113055.3 | 5 | 92804220 | 92809488 | 5269 | 2 | ENSMUST00 | -24999 | 4932430I15R | ENSMUSG0C | false | . | . | . | . | . | . | chr5 | 92813516 | 92845987 | 5.7053 | 4.3919 | 3.59815 |
| chr5 | 92841525 | 92841863 | 1000 | Intron (ENSMUST00000113055.3 | 5 | 92804220 | 92809488 | 5269 | 2 | ENSMUST00 | -32038 | 4932430I15R | ENSMUSG0C | false | . | . | . | . | . | . | chr5 | 92813516 | 92845987 | 5.7053 | 4.3919 | 3.59815 |
| chr16 | 92632103 | 92632665 | 1000 | Intron (ENSMUST00000023673.8 | 16 | 92612824 | 92620032 | 7209 | 1 | ENSMUST00 | 19280 | Gm26626 | ENSMUSG0C | false | . | . | . | . | . | . | chr16 | 92630983 | 92633222 | 3.39298 | 3.22802 | 1.71797 |
| chr3 | 97297932 | 97298604 | 1000 | Promoter (<=1kb) | 3 | 97284908 | 97297778 | 12871 | 2 | ENSMUST00 | -155 | Gm17608 | ENSMUSG0C | false | chr3 | 97296446 | 97298749 | 21.79372 | 61.12524 | 58.26245 | chr3 | 97298263 | 97299440 | 3.6202 | 3.46859 | 1.85236 |
| chr5 | 52256687 | 52256976 | 1000 | Distal Intergenic | 5 | 52190681 | 52197984 | 7304 | 1 | ENSMUST00 | 66007 | 9230114K14I | ENSMUSG0C | false | . | . | . | . | . | . | chr5 | 52256324 | 52257444 | 4.02692 | 3.55467 | 2.23597 |
| chr12 | 51909206 | 51909512 | 1000 | Intron (ENSMUST00000040583.5 | 12 | 51888015 | 51889459 | 1445 | 1 | ENSMUST00 | 21192 | Gm26517 | ENSMUSG0C | false | . | . | . | . | . | . | chr12 | 51908906 | 51909426 | 3.72189 | 3.37926 | 1.98621 |
| chr16 | 30034044 | 30034586 | 1000 | Distal Intergenic | 16 | 30008667 | 30021430 | 12764 | 1 | ENSMUST00 | 25378 | 4632428C04I | ENSMUSG0C | false | . | . | . | . | . | . | chr16 | 30032619 | 30035262 | 3.83575 | 3.33687 | 2.10367 |
| chr16 | 30013107 | 30013599 | 1000 | Intron (ENSMUST00000181485.1 | 16 | 30008667 | 30021430 | 12764 | 1 | ENSMUST00 | 4441 | 4632428C04I | ENSMUSG0C | false | . | . | . | . | . | . | chr16 | 30013114 | 30013950 | 3.49308 | 3.31718 | 1.77294 |
| chr16 | 30008506 | 30009322 | 1000 | Promoter (<=1kb) | 16 | 30008667 | 30021430 | 12764 | 1 | ENSMUST00 | 0 | 4632428C04I | ENSMUSG0C | false | chr16 | 30008531 | 30009799 | 13.32643 | 29.39681 | 26.9387 | chr16 | 30008297 | 30010007 | 5.81527 | 4.54129 | 3.66158 |
| chr9 | 88522891 | 88523417 | 1000 | Promoter (<=1kb) | 9 | 88521050 | 88522929 | 1880 | 2 | ENSMUST00 | 0 | Snhg5 | ENSMUSG0C | false | chr9 | 88522164 | 88523627 | 26.86175 | 78.82483 | 75.69657 | . | . | . | . | . | . |
| chrX | 48284632 | 48285919 | 1000 | Promoter (<=1kb) | 20 | 48276792 | 48285426 | 8635 | 2 | ENSMUST00 | 0 | 9530027J09F | ENSMUSG0C | false | chrX | 48284800 | 48285343 | 16.08362 | 38.04472 | 35.47989 | . | . | . | . | . | . |
| chr9 | 40522094 | 40522558 | 1000 | Intron (ENSMUST00000119373.2 | 9 | 40531155 | 40533620 | 2466 | 1 | ENSMUST00 | -8597 | Gm17540 | ENSMUSG0C | false | . | . | . | . | . | . | chr9 | 40521881 | 40524616 | 4.13519 | 3.54361 | 2.33402 |
| chr18 | 11239676 | 11240024 | 1000 | Distal Intergenic | 18 | 11049931 | 11052567 | 2637 | 2 | ENSMUST00 | -187110 | 1010001N08 | ENSMUSG0C | false | chr18 | 11239657 | 11240387 | 7.04473 | 12.99314 | 10.76275 | chr18 | 11239266 | 11241078 | 4.99867 | 3.79646 | 3.04748 |
| chr19 | 4169391 | 4169743 | 1000 | Promoter (<=1kb) | 19 | 4169566 | 4174284 | 4719 | 1 | ENSMUST00 | 0 | Gm17552 | ENSMUSG0C | false | chr19 | 4169619 | 4170188 | 16.52141 | 45.4089 | 42.75185 | chr19 | 4166946 | 4176432 | 4.77668 | 3.94064 | 2.82677 |
| chr14 | 70203319 | 70203740 | 1000 | Promoter (1-2kb) | 14 | 70205324 | 70217659 | 12336 | 1 | ENSMUST00 | -1584 | Gm26908 | ENSMUSG0C | false | . | . | . | . | . | . | chr14 | 70202314 | 70210660 | 7.58237 | 5.24911 | 5.1119 |
| chr1 | 182385693 | 182386028 | 1000 | Distal Intergenic | 1 | 182392543 | 182393535 | 993 | 2 | ENSMUST00 | 7507 | Gm5706 | ENSMUSG0C | false | . | . | . | . | . | . | chr1 | 182384457 | 182389495 | 5.25206 | 4.10192 | 3.26079 |
| chr1 | 164543413 | 164543814 | 1000 | Distal Intergenic | 1 | 164503307 | 164514882 | 11576 | 1 | ENSMUST00 | 40107 | Gm26685 | ENSMUSG0C | false | . | . | . | . | . | . | chr1 | 164543411 | 164543930 | 3.87697 | 3.54337 | 2.14283 |
| chr10 | 14017899 | 14018454 | 1000 | Intron (ENSMUST00000187083.1 | 10 | 13967024 | 14007405 | 40382 | 2 | ENSMUST00 | -10495 | Gm26835 | ENSMUSG0C | false | chr10 | 14017825 | 14018758 | 10.88283 | 27.02401 | 24.59589 | chr10 | 14017579 | 14020834 | 5.99895 | 4.59741 | 3.82797 |
| chr3 | 37747641 | 37748114 | 1000 | Distal Intergenic | 3 | 37734730 | 37735104 | 375 | 1 | ENSMUST00 | 12912 | Rpl31-ps15 | ENSMUSG0C | false | . | . | . | . | . | . | chr3 | 37747239 | 37748058 | 3.63582 | 3.44551 | 1.87247 |
| chr16 | 29813899 | 29814379 | 1000 | Promoter (2-3kb) | 16 | 29811058 | 29837814 | 26757 | 1 | ENSMUST00 | 2842 | Gm26834 | ENSMUSG0C | false | chr16 | 29813613 | 29814424 | 12.0178 | 35.52963 | 32.99667 | chr16 | 29813284 | 29814316 | 4.55703 | 3.44605 | 2.69217 |
| chr16 | 29945766 | 29946552 | 1000 | Promoter (<=1kb) | 16 | 29907519 | 29946517 | 38999 | 2 | ENSMUST00 | 0 | Gm26569 | ENSMUSG0C | false | chr16 | 29945587 | 29946602 | 12.99241 | 30.10301 | 27.6364 | chr16 | 29943407 | 29947564 | 5.10658 | 4.06635 | 3.12869 |
| chr1 | 180768179 | 180768686 | 1000 | Distal Intergenic | 1 | 180802568 | 180809483 | 6916 | 1 | ENSMUST00 | -33882 | Gm17275 | ENSMUSG0C | false | chr1 | 180768151 | 180769015 | 15.76414 | 41.82454 | 39.21188 | chr1 | 180767095 | 180770353 | 4.8676 | 4.03706 | 2.90659 |
| chr17 | 12327785 | 12328448 | 1000 | Distal Intergenic | 17 | 12318885 | 12327250 | 8366 | 1 | ENSMUST00 | 8901 | 4732491K20I | ENSMUSG0C | false | . | . | . | . | . | . | chr17 | 12324606 | 12330601 | 6.56988 | 4.78246 | 4.31043 |
| chr5 | 120194651 | 120194937 | 1000 | Intron (ENSMUST00000031590.8 | 5 | 120131485 | 120139263 | 7779 | 2 | ENSMUST00 | -55389 | Gm10390 | ENSMUSG0C | false | . | . | . | . | . | . | chr5 | 120193651 | 120195560 | 5.25406 | 4.24789 | 3.21269 |
| chr8 | 121342883 | 121343393 | 1000 | Intron (ENSMUST00000127664.1 | 8 | 121256747 | 121356836 | 100090 | 2 | ENSMUST00 | 13443 | Gm26815 | ENSMUSG0C | false | . | . | . | . | . | . | chr8 | 121340922 | 121349877 | 6.81979 | 4.84985 | 4.53151 |
| chr8 | 121280463 | 121281037 | 1000 | Intron (ENSMUST00000127664.1 | 8 | 121256747 | 121356836 | 100090 | 2 | ENSMUST00 | 75799 | Gm26815 | ENSMUSG0C | false | . | . | . | . | . | . | chr8 | 121277448 | 121284034 | 5.99549 | 4.54477 | 3.8074 |
| chr8 | 121301359 | 121301777 | 1000 | Intron (ENSMUST00000127664.1 | 8 | 121256747 | 121356836 | 100090 | 2 | ENSMUST00 | 55059 | Gm26815 | ENSMUSG0C | false | . | . | . | . | . | . | chr8 | 121299982 | 121302568 | 7.20896 | 5.12344 | 4.81006 |
| chr8 | 121332510 | 121332890 | 1000 | Intron (ENSMUST00000127664.1 | 8 | 121256747 | 121356836 | 100090 | 2 | ENSMUST00 | 23946 | Gm26815 | ENSMUSG0C | false | . | . | . | . | . | . | chr8 | 121331338 | 121336444 | 7.7568 | 5.32028 | 5.25757 |
| chr8 | 121364643 | 121364991 | 1000 | Intron (ENSMUST00000127664.1 | 8 | 121256747 | 121356836 | 100090 | 2 | ENSMUST00 | -7808 | Gm26815 | ENSMUSG0C | false | . | . | . | . | . | . | chr8 | 121362805 | 121370856 | 6.80763 | 4.93598 | 4.48579 |
| chr8 | 121318714 | 121319036 | 1000 | Intron (ENSMUST00000127664.1 | 8 | 121256747 | 121356836 | 100090 | 2 | ENSMUST00 | 37800 | Gm26815 | ENSMUSG0C | false | . | . | . | . | . | . | chr8 | 121315121 | 121320000 | 4.45887 | 3.7818 | 2.55833 |
| chr17 | 46266042 | 46266807 | 1000 | Intron (ENSMUST00000012440.7 | 17 | 46273479 | 46280060 | 6582 | 1 | ENSMUST00 | -6672 | Gm26785 | ENSMUSG0C | false | . | . | . | . | . | . | chr17 | 46263490 | 46282075 | 6.71144 | 4.73752 | 4.44242 |
| chr8 | 109105613 | 109106135 | 1000 | Intron (ENSMUST00000180524.2 | 8 | 109075223 | 109231047 | 155825 | 1 | ENSMUST00 | 30391 | D030068K23 | ENSMUSG0C | false | chr8 | 109105405 | 109105849 | 5.44141 | 9.50478 | 7.33812 | chr8 | 109102953 | 109108862 | 5.75078 | 4.39185 | 3.61981 |
| chr1 | 88738904 | 88739373 | 1000 | Distal Intergenic | 1 | 88754890 | 88755736 | 847 | 1 | ENSMUST00 | -15517 | 2410088K16I | ENSMUSG0C | false | . | . | . | . | . | . | chr1 | 88738424 | 88740544 | 5.0809 | 4.02794 | 3.1253 |
| chr12 | 80083910 | 80084341 | 1000 | Distal Intergenic | 12 | 80108034 | 80113000 | 4967 | 1 | ENSMUST00 | -23693 | Gm26669 | ENSMUSG0C | false | . | . | . | . | . | . | chr12 | 80079992 | 80084942 | 5.65 | 4.35804 | 3.54363 |
| chr10 | 22491912 | 22492223 | 1000 | Distal Intergenic | 10 | 22452891 | 22453520 | 630 | 2 | ENSMUST00 | -38393 | Gm26585 | ENSMUSG0C | false | . | . | . | . | . | . | chr10 | 22491709 | 22492320 | 3.58378 | 3.26389 | 1.88582 |
| chr12 | 112950207 | 112950824 | 1000 | Promoter (<=1kb) | 12 | 112946363 | 112950512 | 4150 | 2 | ENSMUST00 | 0 | Gm26583 | ENSMUSG0C | false | chr12 | 112949963 | 112951035 | 22.21283 | 62.70797 | 59.8237 | chr12 | 112948716 | 112953474 | 5.69251 | 4.20255 | 3.60267 |
| chr8 | 46691348 | 46691971 | 1000 | Distal Intergenic | 8 | 46730972 | 46739515 | 8544 | 2 | ENSMUST00 | 47544 | Gm16675 | ENSMUSG0C | false | chr8 | 46691543 | 46691873 | 7.12487 | 12.70834 | 10.47977 | chr8 | 46690388 | 46692872 | 4.20632 | 3.61582 | 2.36788 |
| chr9 | 78176930 | 78177463 | 1000 | Promoter (1-2kb) | 9 | 78175914 | 78178879 | 2966 | 1 | ENSMUST00 | 1017 | C920006O11 | ENSMUSG0C | false | chr9 | 78175862 | 78177230 | 28.22848 | 94.63612 | 91.19299 | chr9 | 78176819 | 78178618 | 3.93596 | 3.55731 | 2.11808 |
| chr14 | 61769886 | 61770330 | 1000 | Distal Intergenic | 14 | 61602839 | 61682373 | 79535 | 2 | ENSMUST00 | -87514 | Dleu2 | ENSMUSG0C | false | . | . | . | . | . | . | chr14 | 61768459 | 61770808 | 3.99873 | 3.45162 | 2.22776 |
| chr14 | 61680948 | 61683298 | 1000 | Promoter (<=1kb) | 14 | 61602839 | 61682373 | 79535 | 2 | ENSMUST00 | 0 | Dleu2 | ENSMUSG0C | false | chr14 | 61680629 | 61683530 | 27.78601 | 91.88203 | 88.49705 | chr14 | 61683024 | 61685193 | 5.50185 | 4.36517 | 3.40928 |
| chr14 | 61648686 | 61649096 | 1000 | Promoter (<=1kb) | 14 | 61603589 | 61648822 | 45234 | 2 | ENSMUST00 | 0 | Dleu2 | ENSMUSG0C | false | chr14 | 61646907 | 61648830 | 35.15467 | 116.16882 | 112.14974 | . | . | . | . | . | . |
| chr14 | 25377905 | 25378708 | 1000 | Promoter (<=1kb) | 14 | 25355801 | 25378635 | 22835 | 2 | ENSMUST00 | 0 | Gm26660 | ENSMUSG0C | false | chr14 | 25377781 | 25378838 | 14.30691 | 45.69287 | 43.03132 | chr14 | 25377226 | 25379461 | 3.95555 | 3.55433 | 2.1307 |
| chr8 | 68205965 | 68206631 | 1000 | Distal Intergenic | 8 | 68218173 | 68228413 | 10241 | 2 | ENSMUST00 | 21782 | Gm26659 | ENSMUSG0C | false | . | . | . | . | . | . | chr8 | 68205617 | 68206761 | 3.7207 | 3.52798 | 1.98891 |
| chr18 | 44535179 | 44535643 | 1000 | Intron (ENSMUST00000164666.2 | 18 | 44661665 | 44676271 | 14607 | 1 | ENSMUST00 | -126022 | A930012L18I | ENSMUSG0C | false | . | . | . | . | . | . | chr18 | 44508889 | 44549417 | 5.66005 | 4.35225 | 3.55985 |
| chr18 | 44542537 | 44543144 | 1000 | Intron (ENSMUST00000164666.2 | 18 | 44661665 | 44676271 | 14607 | 1 | ENSMUST00 | -118521 | A930012L18I | ENSMUSG0C | false | chr18 | 44543051 | 44544910 | 8.40012 | 17.61332 | 15.31135 | chr18 | 44508889 | 44549417 | 5.66005 | 4.35225 | 3.55985 |
| chr18 | 44550826 | 44551217 | 1000 | Intron (ENSMUST00000164666.2 | 18 | 44661665 | 44676271 | 14607 | 1 | ENSMUST00 | -110448 | A930012L18I | ENSMUSG0C | false | . | . | . | . | . | . | chr18 | 44551200 | 44551435 | 3.3548 | 3.28289 | 1.69278 |
| chr1 | 185481807 | 185482369 | 1000 | Distal Intergenic | 1 | 185447116 | 185455568 | 8453 | 2 | ENSMUST00 | -26240 | Gm2061 | ENSMUSG0C | false | chr1 | 185481802 | 185482467 | 8.60706 | 16.02675 | 13.74988 | chr1 | 185480719 | 185483053 | 5.4903 | 4.40262 | 3.40878 |
| chr15 | 96233242 | 96233565 | 1000 | Intron (ENSMUST00000180852.1 | 15 | 96231870 | 96240588 | 8719 | 2 | ENSMUST00 | 7023 | 4833422M21 | ENSMUSG0C | false | . | . | . | . | . | . | . | . | . | . | . | . |
| chr5 | 19180792 | 19181379 | 1000 | Distal Intergenic | 5 | 19202368 | 19226555 | 24188 | 2 | ENSMUST00 | 45176 | 4921504A21 | ENSMUSG0C | true | . | . | . | . | . | . | chr5 | 19177836 | 19184145 | 4.31014 | 3.73966 | 2.43836 |
| chr5 | 19171258 | 19171684 | 1000 | Distal Intergenic | 5 | 19202368 | 19226555 | 24188 | 2 | ENSMUST00 | 54871 | 4921504A21 | ENSMUSG0C | true | . | . | . | . | . | . | chr5 | 19170700 | 19171959 | 4.21307 | 3.73549 | 2.3726 |
| chr5 | 19225320 | 19225771 | 1000 | Promoter (<=1kb) | 5 | 19202368 | 19226555 | 24188 | 2 | ENSMUST00 | 784 | 4921504A21 | ENSMUSG0C | true | chr5 | 19223861 | 19232405 | 34.78864 | 106.1911 | 102.46782 | . | . | . | . | . | . |
| chr19 | 55622132 | 55622491 | 1000 | Intron (ENSMUST00000095950.2 | 19 | 55640589 | 55702204 | 61616 | 2 | ENSMUST00 | 79713 | 4930552P12I | ENSMUSG0C | false | . | . | . | . | . | . | chr19 | 55621588 | 55623432 | 5.77033 | 4.05561 | 3.67413 |
| chr6 | 122826080 | 122826746 | 1000 | Intron (ENSMUST00000003238.8 | 6 | 122813626 | 122820614 | 6989 | 2 | ENSMUST00 | -5467 | Gm26826 | ENSMUSG0C | false | chr6 | 122826058 | 122826979 | 7.54398 | 13.82373 | 11.57928 | chr6 | 122818679 | 122828548 | 6.07039 | 4.57354 | 3.88889 |
| chr19 | 53420856 | 53421228 | 1000 | Distal Intergenic | 19 | 53444014 | 53447314 | 3301 | 2 | ENSMUST00 | 26086 | 5830416P10I | ENSMUSG0C | false | chr19 | 53420993 | 53421471 | 13.32643 | 29.39681 | 26.9387 | chr19 | 53420386 | 53425725 | 3.65118 | 3.39611 | 1.90064 |
| chr19 | 53437294 | 53437938 | 1000 | Distal Intergenic | 19 | 53444014 | 53447314 | 3301 | 2 | ENSMUST00 | 9376 | 5830416P10I | ENSMUSG0C | false | . | . | . | . | . | . | chr19 | 53437010 | 53438362 | 5.63677 | 4.32971 | 3.57787 |
| chr19 | 53482578 | 53482965 | 1000 | Distal Intergenic | 19 | 53441212 | 53464796 | 23585 | 2 | ENSMUST00 | -17783 | 5830416P10I | ENSMUSG0C | false | . | . | . | . | . | . | chr19 | 53482021 | 53482873 | 3.65198 | 3.33829 | 1.93267 |
| chr19 | 53452427 | 53452766 | 1000 | Promoter (1-2kb) | 19 | 53451641 | 53454094 | 2454 | 2 | ENSMUST00 | 1328 | 5830416P10I | ENSMUSG0C | false | . | . | . | . | . | . | chr19 | 53450950 | 53453385 | 3.92468 | 3.62056 | 2.11855 |
| chr15 | 38384179 | 38385122 | 1000 | Promoter (<=1kb) | 15 | 38384981 | 38386300 | 1320 | 1 | ENSMUST00 | 0 | Gm26863 | ENSMUSG0C | false | chr15 | 38384184 | 38385474 | 13.14203 | 32.98289 | 30.47944 | chr15 | 38383689 | 38386004 | 4.35056 | 3.74144 | 2.48199 |
| chr13 | 15426907 | 15427365 | 1000 | Distal Intergenic | 13 | 15459980 | 15461295 | 1316 | 2 | ENSMUST00 | 33930 | Gm26645 | ENSMUSG0C | false | chr13 | 15426617 | 15427491 | 4.4453 | 8.34 | 6.1997 | chr13 | 15426570 | 15427644 | 4.22526 | 3.63145 | 2.41306 |
| chr18 | 75502742 | 75503373 | 1000 | Intron (ENSMUST00000165559.1 | 18 | 75514645 | 75522806 | 8162 | 1 | ENSMUST00 | -11272 | Gm10532 | ENSMUSG0C | false | . | . | . | . | . | . | chr18 | 75500942 | 75504289 | 3.90055 | 3.49915 | 2.15306 |
| chr13 | 12650102 | 12651154 | 1000 | Promoter (<=1kb) | 13 | 12619084 | 12650297 | 31214 | 2 | ENSMUST00 | 0 | Gpr137b-ps | ENSMUSG0C | false | chr13 | 12649806 | 12651375 | 27.78601 | 91.88203 | 88.49705 | chr13 | 12651065 | 12651470 | 5.49429 | 4.45744 | 3.40649 |
| chr17 | 45886510 | 45887102 | 1000 | Distal Intergenic | 17 | 45756710 | 45788594 | 31885 | 1 | ENSMUST00 | 129801 | F630040K05I | ENSMUSG0C | false | . | . | . | . | . | . | chr17 | 45885661 | 45888081 | 7.57576 | 5.2058 | 5.16731 |
| chr17 | 45805670 | 45806160 | 1000 | Distal Intergenic | 17 | 45756710 | 45788594 | 31885 | 1 | ENSMUST00 | 48961 | F630040K05I | ENSMUSG0C | false | . | . | . | . | . | . | chr17 | 45804581 | 45807832 | 4.77899 | 3.85908 | 2.86622 |
| chr17 | 45736510 | 45737046 | 1000 | Promoter (2-3kb) | 17 | 45734506 | 45781251 | 46746 | 1 | ENSMUST00 | 2005 | F630040K05I | ENSMUSG0C | false | chr17 | 45736363 | 45737272 | 16.52141 | 45.4089 | 42.75185 | chr17 | 45731816 | 45737308 | 6.56758 | 4.65756 | 4.31435 |
| chr17 | 45787466 | 45787779 | 1000 | Exon (ENSMUST00000180519.1/ | 17 | 45756710 | 45788594 | 31885 | 1 | ENSMUST00 | 30757 | F630040K05I | ENSMUSG0C | false | . | . | . | . | . | . | chr17 | 45784588 | 45788810 | 6.07111 | 4.63598 | 3.87687 |
| chr12 | 80168087 | 80168590 | 1000 | 3' UTR | 12 | 80120548 | 80132844 | 12297 | 2 | ENSMUST00 | -35244 | 2310015A10I | ENSMUSG0C | false | chr12 | 80168115 | 80168828 | 16.32424 | 48.10018 | 45.4107 | chr12 | 80166901 | 80171473 | 3.22452 | 3.14317 | 1.53182 |
| chr12 | 80130608 | 80131061 | 1000 | Promoter (1-2kb) | 12 | 80120548 | 80132844 | 12297 | 2 | ENSMUST00 | 1783 | 2310015A10I | ENSMUSG0C | false | . | . | . | . | . | . | chr12 | 80116083 | 80133218 | 5.88241 | 4.4251 | 3.73792 |
| chr3 | 152585583 | 152585930 | 1000 | Intron (ENSMUST00000045262.6 | 3 | 152666913 | 152681432 | 14520 | 1 | ENSMUST00 | -80983 | AI115009 | ENSMUSG0C | false | . | . | . | . | . | . | chr3 | 152585649 | 152586499 | 2.79984 | 2.85668 | 1.25196 |
| chr2 | 5200790 | 5201384 | 1000 | Intron (ENSMUST00000027988.7 | 2 | 5137749 | 5230864 | 93116 | 2 | ENSMUST00 | 29480 | Gm26776 | ENSMUSG0C | false | . | . | . | . | . | . | chr2 | 5196720 | 5202535 | 4.35636 | 3.76045 | 2.46313 |
| chr8 | 121391311 | 121392105 | 1000 | Intron (ENSMUST00000127664.1 | 8 | 121421345 | 121427188 | 5844 | 1 | ENSMUST00 | -29240 | Gm26747 | ENSMUSG0C | false | . | . | . | . | . | . | chr8 | 121389629 | 121400811 | 5.77409 | 4.45821 | 3.63728 |
| chr8 | 121397108 | 121397588 | 1000 | Intron (ENSMUST00000127664.1 | 8 | 121421345 | 121427188 | 5844 | 1 | ENSMUST00 | -23757 | Gm26747 | ENSMUSG0C | false | . | . | . | . | . | . | chr8 | 121389629 | 121400811 | 5.77409 | 4.45821 | 3.63728 |
| chr13 | 99124770 | 99125289 | 1000 | Distal Intergenic | 13 | 99100706 | 99107580 | 6875 | 1 | ENSMUST00 | 24065 | Gm807 | ENSMUSG0C | false | . | . | . | . | . | . | chr13 | 99123445 | 99129979 | 5.58129 | 4.22085 | 3.50954 |
| chr13 | 99123883 | 99124400 | 1000 | Distal Intergenic | 13 | 99100706 | 99107580 | 6875 | 1 | ENSMUST00 | 23178 | Gm807 | ENSMUSG0C | false | . | . | . | . | . | . | chr13 | 99123445 | 99129979 | 5.58129 | 4.22085 | 3.50954 |
| chr2 | 52935859 | 52936366 | 1000 | Intron (ENSMUST00000090952.5 | 2 | 52859723 | 52863516 | 3794 | 2 | ENSMUST00 | -72344 | Gm26603 | ENSMUSG0C | false | . | . | . | . | . | . | chr2 | 52917882 | 52946369 | 5.21081 | 4.16706 | 3.18635 |
| chr4 | 3703468 | 3703952 | 1000 | Intron (ENSMUST00000041377.7 | 4 | 3678117 | 3689548 | 11432 | 2 | ENSMUST00 | -13921 | Gm26857 | ENSMUSG0C | false | . | . | . | . | . | . | chr4 | 3702725 | 3705314 | 5.83453 | 4.40274 | 3.70748 |
| chr9 | 56572602 | 56573359 | 1000 | Distal Intergenic | 9 | 56606693 | 56614471 | 7779 | 1 | ENSMUST00 | -33334 | Gm26868 | ENSMUSG0C | false | . | . | . | . | . | . | chr9 | 56572274 | 56577148 | 4.79706 | 3.95319 | 2.87713 |
| chr9 | 56604757 | 56605094 | 1000 | Promoter (1-2kb) | 9 | 56606693 | 56614471 | 7779 | 1 | ENSMUST00 | -1599 | Gm26868 | ENSMUSG0C | false | . | . | . | . | . | . | chr9 | 56604278 | 56606624 | 5.59341 | 4.3412 | 3.53891 |
| chr3 | 51559525 | 51560065 | 1000 | Promoter (<=1kb) | 3 | 51559757 | 51567117 | 7361 | 1 | ENSMUST00 | 0 | 5031434O11 | ENSMUSG0C | false | chr3 | 51558835 | 51561958 | 30.28657 | 98.33401 | 94.80609 | . | . | . | . | . | . |
| chr12 | 86001347 | 86001741 | 1000 | Intron (ENSMUST00000040179.8 | 12 | 85962598 | 85979014 | 16417 | 2 | ENSMUST00 | -22334 | Gm26531 | ENSMUSG0C | false | . | . | . | . | . | . | chr12 | 85999690 | 86002429 | 3.93686 | 3.37853 | 2.16761 |
| chr14 | 120336476 | 120336882 | 1000 | Intron (ENSMUST00000088419.7 | 14 | 120384263 | 120388985 | 4723 | 2 | ENSMUST00 | 52103 | Gm26679 | ENSMUSG0C | false | . | . | . | . | . | . | chr14 | 120329884 | 120339572 | 6.16737 | 4.46838 | 4.00867 |
| chr1 | 23443986 | 23444278 | 1000 | Distal Intergenic | 1 | 23479539 | 23480527 | 989 | 1 | ENSMUST00 | -35261 | Gm7784 | ENSMUSG0C | false | . | . | . | . | . | . | chr1 | 23443568 | 23445145 | 3.33211 | 3.18103 | 1.68401 |
| chr1 | 189820381 | 189820766 | 1000 | Intron (ENSMUST00000097442.3 | 1 | 189833141 | 189836620 | 3480 | 2 | ENSMUST00 | 15854 | Gm26574 | ENSMUSG0C | false | . | . | . | . | . | . | chr1 | 189811241 | 189822755 | 6.05631 | 4.50959 | 3.8969 |
| chr8 | 120228122 | 120228717 | 1000 | Promoter (<=1kb) | 8 | 120204434 | 120228230 | 23797 | 2 | ENSMUST00 | 0 | A330074K22 | ENSMUSG0C | false | chr8 | 120228411 | 120230255 | 25.66166 | 72.18176 | 69.16018 | chr8 | 120224195 | 120235757 | 5.14185 | 4.11684 | 3.13065 |
| chr8 | 121922055 | 121922410 | 1000 | Promoter (2-3kb) | 8 | 121919024 | 121919690 | 667 | 2 | ENSMUST00 | -2366 | Gm26999 | ENSMUSG0C | false | . | . | . | . | . | . | chr8 | 121918157 | 121923089 | 5.61939 | 4.40193 | 3.52271 |
| chr8 | 79502129 | 79502555 | 1000 | Distal Intergenic | 8 | 79549774 | 79550748 | 975 | 2 | ENSMUST00 | 48193 | Gm8079 | ENSMUSG0C | false | . | . | . | . | . | . | chr8 | 79500857 | 79503094 | 4.38181 | 3.72475 | 2.54039 |
| chr7 | 70505822 | 70506204 | 1000 | Distal Intergenic | 7 | 70490848 | 70491851 | 1004 | 1 | ENSMUST00 | 14975 | Gm7656 | ENSMUSG0C | false | . | . | . | . | . | . | chr7 | 70505230 | 70506733 | 4.72548 | 3.87385 | 2.82262 |
| chr1 | 135129068 | 135129507 | 1000 | Promoter (<=1kb) | 1 | 135126376 | 135128262 | 1887 | 2 | ENSMUST00 | -807 | Ptprv | ENSMUSG0C | false | . | . | . | . | . | . | chr1 | 135126075 | 135132880 | 5.21831 | 4.07933 | 3.22044 |
| chr18 | 55815741 | 55816281 | 1000 | Distal Intergenic | 18 | 55837972 | 55838949 | 978 | 2 | ENSMUST00 | 22668 | Gm8614 | ENSMUSG0C | false | . | . | . | . | . | . | chr18 | 55815404 | 55817596 | 4.4866 | 3.74871 | 2.62571 |
| chr9 | 50191125 | 50191559 | 1000 | Distal Intergenic | 9 | 50186043 | 50187045 | 1003 | 2 | ENSMUST00 | -4081 | Gm8907 | ENSMUSG0C | false | . | . | . | . | . | . | chr9 | 50187925 | 50193346 | 4.39357 | 3.76919 | 2.5395 |
| chr14 | 62021623 | 62022194 | 1000 | Distal Intergenic | 14 | 62201915 | 62204352 | 2438 | 2 | ENSMUST00 | 182158 | Gm26969 | ENSMUSG0C | false | . | . | . | . | . | . | chr14 | 62020569 | 62025093 | 6.58122 | 4.80655 | 4.30387 |
| chr18 | 61639511 | 61640159 | 1000 | Promoter (<=1kb) | 18 | 61639542 | 61647292 | 7751 | 1 | ENSMUST00 | 0 | Gm20748 | ENSMUSG0C | false | chr18 | 61639449 | 61639795 | 4.88133 | 7.58987 | 5.46511 | chr18 | 61630875 | 61644532 | 7.58082 | 5.1921 | 5.12657 |
| chr18 | 61634701 | 61635172 | 1000 | Distal Intergenic | 18 | 61639542 | 61647292 | 7751 | 1 | ENSMUST00 | -4370 | Gm20748 | ENSMUSG0C | false | . | . | . | . | . | . | chr18 | 61630875 | 61644532 | 7.58082 | 5.1921 | 5.12657 |
| chr18 | 61637765 | 61638466 | 1000 | Promoter (1-2kb) | 18 | 61639542 | 61647292 | 7751 | 1 | ENSMUST00 | -1076 | Gm20748 | ENSMUSG0C | false | . | . | . | . | . | . | chr18 | 61630875 | 61644532 | 7.58082 | 5.1921 | 5.12657 |
| chr13 | 44506965 | 44507395 | 1000 | Distal Intergenic | 13 | 44400436 | 44455303 | 54868 | 2 | ENSMUST00 | -51663 | Gm27007 | ENSMUSG0C | false | . | . | . | . | . | . | chr13 | 44501171 | 44510788 | 6.13728 | 4.58773 | 3.94959 |
| chr13 | 98390932 | 98391846 | 1000 | Distal Intergenic | 13 | 98402095 | 98403083 | 989 | 2 | ENSMUST00 | 11237 | Gm2445 | ENSMUSG0C | false | chr13 | 98390311 | 98391925 | 25.17599 | 69.56199 | 66.57701 | . | . | . | . | . | . |
| chr13 | 98393164 | 98393924 | 1000 | Distal Intergenic | 13 | 98402095 | 98403083 | 989 | 2 | ENSMUST00 | 9159 | Gm2445 | ENSMUSG0C | false | . | . | . | . | . | . | chr13 | 98391922 | 98395487 | 4.17542 | 3.68065 | 2.34863 |
| chr13 | 98384110 | 98384490 | 1000 | Distal Intergenic | 13 | 98402095 | 98403083 | 989 | 2 | ENSMUST00 | 18593 | Gm2445 | ENSMUSG0C | false | . | . | . | . | . | . | chr13 | 98382088 | 98385324 | 4.83837 | 3.87926 | 2.9137 |
| chr7 | 112957463 | 112958094 | 1000 | Distal Intergenic | 7 | 112953962 | 112957457 | 3496 | 1 | ENSMUST00 | 3502 | Rassf10 | ENSMUSG0C | false | chr7 | 112953937 | 112957582 | 22.03264 | 83.65932 | 80.44393 | chr7 | 112958081 | 112961136 | 4.90276 | 4.06988 | 2.91926 |
| chr8 | 36354913 | 36355213 | 1000 | Distal Intergenic | 8 | 36372599 | 36373642 | 1044 | 2 | ENSMUST00 | 18429 | Gm5787 | ENSMUSG0C | false | . | . | . | . | . | . | . | . | . | . | . | . |
| chr12 | 15681732 | 15682062 | 1000 | Distal Intergenic | 12 | 15728097 | 15729095 | 999 | 1 | ENSMUST00 | -46035 | Gm4804 | ENSMUSG0C | false | . | . | . | . | . | . | chr12 | 15678204 | 15686497 | 4.63142 | 3.94161 | 2.69502 |
| chr12 | 15721597 | 15722293 | 1000 | Distal Intergenic | 12 | 15728097 | 15729095 | 999 | 1 | ENSMUST00 | -5804 | Gm4804 | ENSMUSG0C | false | . | . | . | . | . | . | chr12 | 15715872 | 15727235 | 4.77348 | 3.97042 | 2.82697 |
| chr12 | 15704259 | 15704918 | 1000 | Distal Intergenic | 12 | 15728097 | 15729095 | 999 | 1 | ENSMUST00 | -23179 | Gm4804 | ENSMUSG0C | false | . | . | . | . | . | . | chr12 | 15695149 | 15708519 | 5.84812 | 4.504 | 3.69923 |
| chr12 | 15698518 | 15699412 | 1000 | Distal Intergenic | 12 | 15728097 | 15729095 | 999 | 1 | ENSMUST00 | -28685 | Gm4804 | ENSMUSG0C | false | . | . | . | . | . | . | chr12 | 15695149 | 15708519 | 5.84812 | 4.504 | 3.69923 |
| chr8 | 33875831 | 33876453 | 1000 | Intron (ENSMUST00000053251.6 | 8 | 33884774 | 33885747 | 974 | 2 | ENSMUST00 | 9294 | Gm26978 | ENSMUSG0C | false | . | . | . | . | . | . | chr8 | 33863865 | 33927704 | 6.32022 | 4.67603 | 4.08842 |
| chr6 | 86247761 | 86248105 | 1000 | Intron (ENSMUST00000032066.9 | 6 | 86284887 | 86285894 | 1008 | 1 | ENSMUST00 | -36782 | Gm7545 | ENSMUSG0C | false | . | . | . | . | . | . | chr6 | 86240704 | 86257707 | 6.40824 | 4.67669 | 4.16105 |
| chr6 | 86252186 | 86253000 | 1000 | Intron (ENSMUST00000032066.9 | 6 | 86284887 | 86285894 | 1008 | 1 | ENSMUST00 | -31887 | Gm7545 | ENSMUSG0C | false | . | . | . | . | . | . | chr6 | 86240704 | 86257707 | 6.40824 | 4.67669 | 4.16105 |
| chr15 | 75856830 | 75857256 | 1000 | Promoter (<=1kb) | 15 | 75854819 | 75856228 | 1410 | 2 | ENSMUST00 | -603 | Gm9568 | ENSMUSG0C | false | . | . | . | . | . | . | chr15 | 75856195 | 75858200 | 5.92258 | 4.44662 | 3.78102 |
| chr13 | 84347971 | 84348609 | 1000 | Distal Intergenic | 13 | 84339149 | 84340113 | 965 | 2 | ENSMUST00 | -7859 | Gm26927 | ENSMUSG0C | false | chr13 | 84347485 | 84349633 | 17.27343 | 42.95708 | 40.32916 | chr13 | 84345332 | 84349559 | 3.53018 | 3.21309 | 1.83238 |
| chr14 | 69251929 | 69252387 | 1000 | Intron (ENSMUST00000037064.4 | 14 | 69259146 | 69262037 | 2892 | 1 | ENSMUST00 | -6759 | Gm27222 | ENSMUSG0C | false | . | . | . | . | . | . | chr14 | 69247679 | 69257987 | 5.54859 | 4.38732 | 3.45181 |
| chr14 | 69255875 | 69256347 | 1000 | Promoter (2-3kb) | 14 | 69259146 | 69262037 | 2892 | 1 | ENSMUST00 | -2799 | Gm27222 | ENSMUSG0C | false | . | . | . | . | . | . | chr14 | 69247679 | 69257987 | 5.54859 | 4.38732 | 3.45181 |
| chr18 | 76153836 | 76154269 | 1000 | Distal Intergenic | 18 | 76170552 | 76170652 | 101 | 1 | ENSMUST00 | -16283 | Mir6358 | ENSMUSG0C | false | . | . | . | . | . | . | chr18 | 76133317 | 76155019 | 6.7969 | 4.75664 | 4.51499 |
| chr18 | 76204537 | 76204910 | 1000 | Distal Intergenic | 18 | 76170552 | 76170652 | 101 | 1 | ENSMUST00 | 33986 | Mir6358 | ENSMUSG0C | false | . | . | . | . | . | . | chr18 | 76203827 | 76205710 | 5.71624 | 4.26135 | 3.61594 |
| chr18 | 76140234 | 76141359 | 1000 | Intron (ENSMUST00000058997.8 | 18 | 76170552 | 76170652 | 101 | 1 | ENSMUST00 | -29193 | Mir6358 | ENSMUSG0C | false | . | . | . | . | . | . | chr18 | 76133317 | 76155019 | 6.7969 | 4.75664 | 4.51499 |
| chr9 | 72791118 | 72791495 | 1000 | Distal Intergenic | 9 | 72774063 | 72776225 | 2163 | 2 | ENSMUST00 | -14894 | Gm27204 | ENSMUSG0C | false | . | . | . | . | . | . | . | . | . | . | . | . |
| chr18 | 75040329 | 75040610 | 1000 | Intron (ENSMUST00000039608.7 | 18 | 75023856 | 75023947 | 92 | 1 | ENSMUST00 | 16474 | Gm27781 | ENSMUSG0C | false | . | . | . | . | . | . | chr18 | 75035531 | 75042495 | 4.4278 | 3.7892 | 2.54156 |
| chr5 | 100423682 | 100424104 | 1000 | Promoter (1-2kb) | 5 | 100421968 | 100422041 | 74 | 1 | ENSMUST00 | 1715 | Gm27553 | ENSMUSG0C | false | . | . | . | . | . | . | chr5 | 100419442 | 100425207 | 4.90144 | 3.80197 | 2.9662 |
| chr3 | 97712377 | 97712767 | 1000 | Intron (ENSMUST00000168438.3 | 3 | 97690509 | 97690562 | 54 | 2 | ENSMUST00 | -21816 | Mir7225 | ENSMUSG0C | false | . | . | . | . | . | . | chr3 | 97710479 | 97715210 | 4.70972 | 3.95774 | 2.77291 |
| chr3 | 97697530 | 97698037 | 1000 | Intron (ENSMUST00000168438.3 | 3 | 97690509 | 97690562 | 54 | 2 | ENSMUST00 | -6969 | Mir7225 | ENSMUSG0C | false | . | . | . | . | . | . | chr3 | 97696109 | 97700653 | 6.52386 | 4.79323 | 4.25712 |
| chr3 | 97683374 | 97684328 | 1000 | Distal Intergenic | 3 | 97690509 | 97690562 | 54 | 2 | ENSMUST00 | 6234 | Mir7225 | ENSMUSG0C | false | . | . | . | . | . | . | chr3 | 97680762 | 97692084 | 4.4508 | 3.75815 | 2.57136 |
| chr11 | 101028272 | 101028671 | 1000 | Promoter (1-2kb) | 11 | 101030340 | 101030409 | 70 | 1 | ENSMUST00 | -1669 | Mir6928 | ENSMUSG0C | false | . | . | . | . | . | . | chr11 | 101025610 | 101032059 | 5.06311 | 4.12357 | 3.05904 |
| chr4 | 116732309 | 116732745 | 1000 | Intron (ENSMUST00000124628.2 | 4 | 116725899 | 116725989 | 91 | 1 | ENSMUST00 | 6411 | Gm27688 | ENSMUSG0C | false | . | . | . | . | . | . | chr4 | 116731176 | 116741002 | 4.98596 | 4.04488 | 2.98621 |
| chr7 | 73558053 | 73558786 | 1000 | Promoter (<=1kb) | 7 | 73539801 | 73558395 | 18595 | 2 | ENSMUST00 | 0 | 1810026B05I | ENSMUSG0C | false | chr7 | 73555379 | 73558486 | 30.78991 | 105.27288 | 101.56436 | . | . | . | . | . | . |
| chr11 | 107651446 | 107651974 | 1000 | Intron (ENSMUST00000100305.2 | 11 | 107637722 | 107637779 | 58 | 1 | ENSMUST00 | 13725 | Mir6932 | ENSMUSG0C | false | . | . | . | . | . | . | chr11 | 107647908 | 107653872 | 5.72702 | 4.35515 | 3.61075 |
| chr11 | 107652505 | 107652890 | 1000 | Intron (ENSMUST00000100305.2 | 11 | 107637722 | 107637779 | 58 | 1 | ENSMUST00 | 14784 | Mir6932 | ENSMUSG0C | false | . | . | . | . | . | . | chr11 | 107647908 | 107653872 | 5.72702 | 4.35515 | 3.61075 |
| chr9 | 102299864 | 102300147 | 1000 | Intron (ENSMUST00000035129.8 | 9 | 102304924 | 102304975 | 52 | 2 | ENSMUST00 | 4828 | Mir7243 | ENSMUSG0C | false | . | . | . | . | . | . | chr9 | 102297810 | 102301329 | 4.27532 | 3.50194 | 2.45277 |
| chr14 | 22161320 | 22161825 | 1000 | Intron (ENSMUST00000075639.5 | 14 | 22021958 | 22022070 | 113 | 2 | ENSMUST00 | -139251 | Gm27532 | ENSMUSG0C | false | . | . | . | . | . | . | chr14 | 22158919 | 22162889 | 4.15949 | 3.66084 | 2.34715 |
| chr14 | 22059449 | 22059854 | 1000 | Intron (ENSMUST00000075639.5 | 14 | 22021958 | 22022070 | 113 | 2 | ENSMUST00 | -37380 | Gm27532 | ENSMUSG0C | false | . | . | . | . | . | . | chr14 | 22054966 | 22061026 | 5.0167 | 3.80015 | 3.05262 |
| chr11 | 60189425 | 60190187 | 1000 | Exon (ENSMUST00000064190.7/ | 11 | 60200538 | 60200617 | 80 | 2 | ENSMUST00 | 10430 | Mir6921 | ENSMUSG0C | false | . | . | . | . | . | . | chr11 | 60181134 | 60192821 | 7.23138 | 4.91369 | 4.86391 |
| chr15 | 5617705 | 5618239 | 1000 | Distal Intergenic | 15 | 5864306 | 5864419 | 114 | 1 | ENSMUST00 | -246067 | Gm27760 | ENSMUSG0C | false | . | . | . | . | . | . | chr15 | 5617091 | 5619625 | 4.3034 | 3.71772 | 2.48127 |
| chr10 | 128583724 | 128584025 | 1000 | Promoter (<=1kb) | 10 | 128584437 | 128584497 | 61 | 2 | ENSMUST00 | 472 | Mir6917 | ENSMUSG0C | false | . | . | . | . | . | . | chr10 | 128583168 | 128584845 | 4.53542 | 3.58143 | 2.65378 |
| chr16 | 85160153 | 85160521 | 1000 | Intron (ENSMUST00000005406.1 | 16 | 85168849 | 85169001 | 153 | 1 | ENSMUST00 | -8328 | Gm27295 | ENSMUSG0C | false | . | . | . | . | . | . | chr16 | 85126656 | 85166763 | 5.48022 | 4.2963 | 3.40197 |
| chr6 | 71268463 | 71268951 | 1000 | Promoter (2-3kb) | 6 | 71271671 | 71271801 | 131 | 1 | ENSMUST00 | -2720 | Mir8112 | ENSMUSG0C | false | . | . | . | . | . | . | chr6 | 71262133 | 71270664 | 5.23015 | 4.16906 | 3.19645 |
| chr15 | 89180353 | 89181210 | 1000 | Downstream (1-2kb) | 15 | 89185465 | 89185537 | 73 | 2 | ENSMUST00 | 4327 | Mir6958 | ENSMUSG0C | false | chr15 | 89178489 | 89181434 | 28.29403 | 89.8816 | 86.54237 | chr15 | 89181073 | 89182474 | 5.37297 | 4.33688 | 3.30039 |
| chr6 | 108448272 | 108448666 | 1000 | Intron (ENSMUST00000032192.6 | 6 | 108456183 | 108456536 | 354 | 1 | ENSMUST00 | -7517 | Gm27813 | ENSMUSG0C | false | . | . | . | . | . | . | chr6 | 108447009 | 108453900 | 4.80361 | 3.89721 | 2.88341 |
| chr1 | 36352355 | 36352761 | 1000 | Intron (ENSMUST00000185912.1 | 1 | 36348682 | 36348758 | 77 | 2 | ENSMUST00 | -3598 | Mir6898 | ENSMUSG0C | false | . | . | . | . | . | . | chr1 | 36350640 | 36368234 | 5.46208 | 4.18784 | 3.39932 |
| chr1 | 36355284 | 36355643 | 1000 | Intron (ENSMUST00000185912.1 | 1 | 36348682 | 36348758 | 77 | 2 | ENSMUST00 | -6527 | Mir6898 | ENSMUSG0C | false | . | . | . | . | . | . | chr1 | 36350640 | 36368234 | 5.46208 | 4.18784 | 3.39932 |
| chr7 | 63887033 | 63887444 | 1000 | 3' UTR | 7 | 63896533 | 63899698 | 3166 | 1 | ENSMUST00 | -9089 | Gm27252 | ENSMUSG0C | false | . | . | . | . | . | . | chr7 | 63885397 | 63936279 | 7.51388 | 5.18869 | 5.06896 |
| chr19 | 16366251 | 16366566 | 1000 | Intron (ENSMUST00000025541.5 | 19 | 16314892 | 16315004 | 113 | 2 | ENSMUST00 | -51248 | Mir496b | ENSMUSG0C | false | . | . | . | . | . | . | chr19 | 16365264 | 16373770 | 4.4736 | 3.84229 | 2.57444 |
| chr6 | 125099359 | 125099887 | 1000 | Promoter (2-3kb) | 6 | 125097024 | 125097086 | 63 | 1 | ENSMUST00 | 2336 | Mir7045 | ENSMUSG0C | false | chr6 | 125099615 | 125100404 | 4.63715 | 6.63346 | 4.53566 | chr6 | 125097452 | 125101880 | 7.168 | 4.97791 | 4.79006 |
| chr12 | 26356208 | 26356536 | 1000 | Intron (ENSMUST00000020971.7 | 12 | 26414901 | 26415010 | 110 | 2 | ENSMUST00 | 58474 | Mir6538 | ENSMUSG0C | false | . | . | . | . | . | . | . | . | . | . | . | . |
| chr19 | 47291290 | 47291672 | 1000 | Intron (ENSMUST00000081619.3 | 19 | 47273519 | 47273589 | 71 | 2 | ENSMUST00 | -17702 | Mir6995 | ENSMUSG0C | false | . | . | . | . | . | . | chr19 | 47286575 | 47295351 | 6.82895 | 4.98258 | 4.49361 |
| chr19 | 47320128 | 47320736 | 1000 | Exon (ENSMUST00000081619.3/ | 19 | 47273519 | 47273589 | 71 | 2 | ENSMUST00 | -46540 | Mir6995 | ENSMUSG0C | false | chr19 | 47319924 | 47320673 | 5.43663 | 10.5674 | 8.37697 | chr19 | 47311839 | 47321707 | 6.07643 | 4.48035 | 3.91145 |
| chr19 | 47276740 | 47277167 | 1000 | Intron (ENSMUST00000081619.3 | 19 | 47273519 | 47273589 | 71 | 2 | ENSMUST00 | -3152 | Mir6995 | ENSMUSG0C | false | . | . | . | . | . | . | chr19 | 47274861 | 47283196 | 8.38768 | 5.49381 | 5.82214 |
| chr19 | 47280763 | 47281346 | 1000 | Intron (ENSMUST00000081619.3 | 19 | 47273519 | 47273589 | 71 | 2 | ENSMUST00 | -7175 | Mir6995 | ENSMUSG0C | false | . | . | . | . | . | . | chr19 | 47274861 | 47283196 | 8.38768 | 5.49381 | 5.82214 |
| chr13 | 32752194 | 32752651 | 1000 | Intron (ENSMUST00000057428.5 | 13 | 32758202 | 32758310 | 109 | 1 | ENSMUST00 | -5551 | Gm27342 | ENSMUSG0C | false | . | . | . | . | . | . | chr13 | 32751192 | 32752903 | 5.72996 | 4.45819 | 3.64347 |
| chr2 | 27441927 | 27442585 | 1000 | Downstream (2-3kb) | 2 | 27450549 | 27450630 | 82 | 2 | ENSMUST00 | 8045 | Mir7578 | ENSMUSG0C | false | chr2 | 27441884 | 27442297 | 13.32643 | 29.39681 | 26.9387 | chr2 | 27441204 | 27443305 | 3.82158 | 3.52419 | 2.0789 |
| chr9 | 32717922 | 32718484 | 1000 | Intron (ENSMUST00000183879.1 | 9 | 32709652 | 32711171 | 1520 | 2 | ENSMUST00 | -6752 | Gm27240 | ENSMUSG0C | false | . | . | . | . | . | . | chr9 | 32697729 | 32732744 | 6.76714 | 4.77458 | 4.47728 |
| chr9 | 32709270 | 32709722 | 1000 | Promoter (1-2kb) | 9 | 32709652 | 32711171 | 1520 | 2 | ENSMUST00 | 1449 | Gm27240 | ENSMUSG0C | false | . | . | . | . | . | . | chr9 | 32697729 | 32732744 | 6.76714 | 4.77458 | 4.47728 |
| chr6 | 11355489 | 11356031 | 1000 | Distal Intergenic | 6 | 10858427 | 10858518 | 92 | 1 | ENSMUST00 | 497063 | Gm27894 | ENSMUSG0C | false | . | . | . | . | . | . | . | . | . | . | . | . |
| chr14 | 67877971 | 67878505 | 1000 | Intron (ENSMUST00000039135.4 | 14 | 67859636 | 67859745 | 110 | 2 | ENSMUST00 | -18227 | Mir6539 | ENSMUSG0C | false | . | . | . | . | . | . | chr14 | 67862320 | 67900048 | 6.11049 | 4.53842 | 3.92537 |
| chr3 | 31056920 | 31057496 | 1000 | Distal Intergenic | 3 | 31038966 | 31039028 | 63 | 1 | ENSMUST00 | 17955 | Mir7008 | ENSMUSG0C | false | chr3 | 31056300 | 31057454 | 20.63265 | 55.76951 | 52.97896 | . | . | . | . | . | . |
| chr3 | 65635947 | 65636314 | 1000 | Distal Intergenic | 3 | 65659288 | 65659426 | 139 | 1 | ENSMUST00 | -22974 | Mir8120 | ENSMUSG0C | false | . | . | . | . | . | . | chr3 | 65633452 | 65637634 | 5.73459 | 4.38136 | 3.62192 |
| chr16 | 93079463 | 93079935 | 1000 | Distal Intergenic | 16 | 93072755 | 93072822 | 68 | 1 | ENSMUST00 | 6709 | Gm28003 | ENSMUSG0C | false | chr16 | 93079347 | 93080307 | 6.80177 | 13.42905 | 11.19262 | chr16 | 93077773 | 93080926 | 5.41211 | 4.35728 | 3.32701 |
| chr10 | 127766507 | 127767128 | 1000 | 3' UTR | 10 | 127759787 | 127792694 | 32908 | 1 | ENSMUST00 | 6721 | Gm28046 | ENSMUSG0C | false | . | . | . | . | . | . | chr10 | 127761024 | 127770039 | 5.65995 | 4.35124 | 3.57259 |
| chr8 | 70801219 | 70801544 | 1000 | Distal Intergenic | 8 | 70798135 | 70798192 | 58 | 1 | ENSMUST00 | 3085 | Mir7240 | ENSMUSG0C | false | . | . | . | . | . | . | chr8 | 70800539 | 70805424 | 7.12963 | 5.06093 | 4.74105 |
| chr2 | 92389006 | 92389729 | 1000 | Promoter (1-2kb) | 2 | 92387384 | 92387446 | 63 | 2 | ENSMUST00 | -1561 | Mir7000 | ENSMUSG0C | false | chr2 | 92388689 | 92389914 | 27.18019 | 77.52287 | 74.41271 | chr2 | 92388364 | 92393091 | 4.25911 | 3.70679 | 2.40514 |
| chr9 | 77546004 | 77546470 | 1000 | Promoter (<=1kb) | 9 | 77545202 | 77546467 | 1266 | 1 | ENSMUST00 | 803 | Gm19569 | ENSMUSG0C | false | chr9 | 77542586 | 77546045 | 36.32541 | 120.3443 | 116.1959 | chr9 | 77545346 | 77549514 | 6.11128 | 4.58585 | 3.90858 |
| chr7 | 143389255 | 143389837 | 1000 | Intron (ENSMUST00000009689.5 | 7 | 143447605 | 143447665 | 61 | 1 | ENSMUST00 | -57768 | Gm27901 | ENSMUSG0C | false | . | . | . | . | . | . | chr7 | 143388402 | 143390632 | 5.79172 | 4.33762 | 3.70011 |
| chr15 | 38470131 | 38471083 | 1000 | Distal Intergenic | 15 | 38490473 | 38490547 | 75 | 2 | ENSMUST00 | 19464 | Mir6951 | ENSMUSG0C | false | chr15 | 38470059 | 38471389 | 13.87847 | 36.33116 | 33.78668 | chr15 | 38468227 | 38474123 | 4.55881 | 3.82285 | 2.65383 |
| chr3 | 153261615 | 153261987 | 1000 | Intron (ENSMUST00000064460.5 | 3 | 153313713 | 153313824 | 112 | 1 | ENSMUST00 | -51726 | Gm27405 | ENSMUSG0C | false | . | . | . | . | . | . | chr3 | 153258909 | 153264662 | 5.67433 | 4.35122 | 3.59934 |
| chr7 | 100474670 | 100474979 | 1000 | Promoter (<=1kb) | 7 | 100446737 | 100474841 | 28105 | 2 | ENSMUST00 | 0 | Gm10603 | ENSMUSG0C | false | . | . | . | . | . | . | chr7 | 100473183 | 100476622 | 5.42622 | 4.14194 | 3.38677 |
| chr11 | 53762881 | 53764332 | 1000 | Promoter (<=1kb) | 11 | 53763597 | 53763654 | 58 | 2 | ENSMUST00 | 0 | Mir7671 | ENSMUSG0C | false | chr11 | 53762730 | 53763974 | 7.72573 | 18.36004 | 16.04866 | chr11 | 53758641 | 53765102 | 7.01527 | 4.95888 | 4.6564 |
| chr11 | 53761428 | 53762190 | 1000 | Promoter (1-2kb) | 11 | 53763597 | 53763654 | 58 | 2 | ENSMUST00 | 1464 | Mir7671 | ENSMUSG0C | false | . | . | . | . | . | . | chr11 | 53758641 | 53765102 | 7.01527 | 4.95888 | 4.6564 |
| chr13 | 28757716 | 28758552 | 1000 | Intron (ENSMUST00000148885.2 | 13 | 28710781 | 28710873 | 93 | 2 | ENSMUST00 | -46844 | Mir6368 | ENSMUSG0C | false | chr13 | 28757850 | 28758518 | 5.44141 | 9.50478 | 7.33812 | chr13 | 28755695 | 28761334 | 5.21524 | 4.1013 | 3.18882 |
| chr10 | 62210600 | 62211309 | 1000 | Intron (ENSMUST00000047883.9 | 10 | 62194167 | 62194227 | 61 | 2 | ENSMUST00 | -16374 | Mir7662 | ENSMUSG0C | false | . | . | . | . | . | . | chr10 | 62185685 | 62229768 | 8.28034 | 5.46771 | 5.70297 |
| chr10 | 62192776 | 62193475 | 1000 | Promoter (<=1kb) | 10 | 62194167 | 62194227 | 61 | 2 | ENSMUST00 | 752 | Mir7662 | ENSMUSG0C | false | . | . | . | . | . | . | chr10 | 62185685 | 62229768 | 8.28034 | 5.46771 | 5.70297 |
| chr8 | 87198163 | 87198575 | 1000 | Promoter (1-2kb) | 8 | 87169747 | 87199839 | 30093 | 2 | ENSMUST00 | 1264 | Gm10637 | ENSMUSG0C | false | . | . | . | . | . | . | chr8 | 87197407 | 87200154 | 4.75611 | 3.82338 | 2.8338 |
| chr9 | 72132736 | 72133508 | 1000 | Distal Intergenic | 9 | 72128587 | 72129391 | 805 | 1 | ENSMUST00 | 4150 | Gm7863 | ENSMUSG0C | false | chr9 | 72132652 | 72133982 | 28.5703 | 92.31807 | 88.92419 | chr9 | 72132726 | 72134236 | 2.96242 | 2.95584 | 1.39473 |
| chr14 | 69474032 | 69474667 | 1000 | Intron (ENSMUST00000111062.3 | 14 | 69502856 | 69523605 | 20750 | 1 | ENSMUST00 | -28189 | Gm27175 | ENSMUSG0C | false | . | . | . | . | . | . | . | . | . | . | . | . |
| chr7 | 143905632 | 143905998 | 1000 | Distal Intergenic | 7 | 143933115 | 143933208 | 94 | 1 | ENSMUST00 | -27117 | Mir6404 | ENSMUSG0C | false | . | . | . | . | . | . | chr7 | 143905004 | 143907923 | 5.84981 | 4.39457 | 3.74354 |
| chr8 | 128290443 | 128290839 | 1000 | Distal Intergenic | 8 | 128278148 | 128278225 | 78 | 2 | ENSMUST00 | -12219 | Mir21c | ENSMUSG0C | false | . | . | . | . | . | . | chr8 | 128289162 | 128291220 | 5.15281 | 3.95163 | 3.16379 |
| chr8 | 13208973 | 13209362 | 1000 | Distal Intergenic | 8 | 13200820 | 13207707 | 6888 | 1 | ENSMUST00 | 8154 | RP23-185G9. | ENSMUSG0C | false | . | . | . | . | . | . | chr8 | 13208705 | 13211559 | 4.95412 | 3.93735 | 2.99787 |
| chr13 | 91315104 | 91315402 | 1000 | Distal Intergenic | 13 | 91373842 | 91374294 | 453 | 2 | ENSMUST00 | 58892 | RP23-241P22 | ENSMUSG0C | false | . | . | . | . | . | . | chr13 | 91313100 | 91315883 | 5.06399 | 4.08631 | 3.09892 |
| chr13 | 91370890 | 91371539 | 1000 | Promoter (2-3kb) | 13 | 91373842 | 91374294 | 453 | 2 | ENSMUST00 | 2755 | RP23-241P22 | ENSMUSG0C | false | chr13 | 91370313 | 91371539 | 17.7171 | 46.74591 | 44.0739 | chr13 | 91368047 | 91375238 | 5.87525 | 4.49782 | 3.72754 |
| chr1 | 88976074 | 88976522 | 1000 | Distal Intergenic | 1 | 89016113 | 89022210 | 6098 | 1 | ENSMUST00 | -39591 | RP24-423I20 | ENSMUSG0C | false | . | . | . | . | . | . | chr1 | 88975800 | 88977045 | 3.97902 | 3.62673 | 2.14992 |
| chr1 | 89012315 | 89012602 | 1000 | Distal Intergenic | 1 | 89016113 | 89022210 | 6098 | 1 | ENSMUST00 | -3511 | RP24-423I20 | ENSMUSG0C | false | . | . | . | . | . | . | chr1 | 89009654 | 89017093 | 4.35608 | 3.6675 | 2.50137 |
| chr1 | 89015410 | 89015781 | 1000 | Promoter (<=1kb) | 1 | 89016113 | 89022210 | 6098 | 1 | ENSMUST00 | -332 | RP24-423I20 | ENSMUSG0C | false | . | . | . | . | . | . | chr1 | 89009654 | 89017093 | 4.35608 | 3.6675 | 2.50137 |
| chr1 | 89011571 | 89012078 | 1000 | Distal Intergenic | 1 | 89016113 | 89022210 | 6098 | 1 | ENSMUST00 | -4035 | RP24-423I20 | ENSMUSG0C | false | . | . | . | . | . | . | chr1 | 89009654 | 89017093 | 4.35608 | 3.6675 | 2.50137 |
| chr17 | 10338035 | 10338591 | 1000 | Distal Intergenic | 17 | 10324602 | 10329307 | 4706 | 1 | ENSMUST00 | 13434 | RP24-162O2. | ENSMUSG0C | false | . | . | . | . | . | . | chr17 | 10330143 | 10344083 | 5.0812 | 4.05752 | 3.08199 |
| chr17 | 10335118 | 10336336 | 1000 | Distal Intergenic | 17 | 10324602 | 10329307 | 4706 | 1 | ENSMUST00 | 10517 | RP24-162O2. | ENSMUSG0C | false | chr17 | 10334584 | 10336344 | 13.14203 | 32.98289 | 30.47944 | chr17 | 10330143 | 10344083 | 5.0812 | 4.05752 | 3.08199 |
| chr8 | 61258884 | 61259180 | 1000 | Intron (ENSMUST00000034060.5 | 8 | 61281522 | 61288872 | 7351 | 2 | ENSMUST00 | 29692 | RP23-5F21.1 | ENSMUSG0C | false | . | . | . | . | . | . | . | . | . | . | . | . |
| chr4 | 54615852 | 54616474 | 1000 | Distal Intergenic | 4 | 54645811 | 54646280 | 470 | 2 | ENSMUST00 | 29806 | RP23-390I8.4 | ENSMUSG0C | false | . | . | . | . | . | . | chr4 | 54612332 | 54617800 | 4.99456 | 4.06573 | 3.03769 |

| | | | | | | | | | | | | | | | | | | | | | | | | | | |
|---|---|---|---|---|---|---|---|---|---|---|---|---|---|---|---|---|---|---|---|---|---|---|---|---|---|---|
| chr1 | 89041256 | 89041606 | 1000 | Distal Intergenic | 1 | 89048974 | 89049216 | 243 | 1 | ENSMUST00 | -7368 | RP24-423I20 | ENSMUSG0C | false | . | . | . | . | . | . | chr1 | 89041020 | 89043070 | 4.42448 | 3.72778 | 2.54235 |
| chr1 | 43310931 | 43311428 | 1000 | Distal Intergenic | 1 | 43250156 | 43302925 | 52770 | 2 | ENSMUST00 | -8007 | RP23-305F24 | ENSMUSG0C | false | . | . | . | . | . | . | chr1 | 43299283 | 43315017 | 5.27004 | 4.116 | 3.25173 |
| chr1 | 43321062 | 43321536 | 1000 | Distal Intergenic | 1 | 43250156 | 43302925 | 52770 | 2 | ENSMUST00 | -18138 | RP23-305F24 | ENSMUSG0C | false | . | . | . | . | . | . | chr1 | 43316042 | 43324876 | 5.43437 | 4.3107 | 3.35534 |
| chr1 | 43305309 | 43305622 | 1000 | Promoter (2-3kb) | 1 | 43250156 | 43302925 | 52770 | 2 | ENSMUST00 | -2385 | RP23-305F24 | ENSMUSG0C | false | . | . | . | . | . | . | chr1 | 43299283 | 43315017 | 5.27004 | 4.116 | 3.25173 |
| chr12 | 99123687 | 99124229 | 1000 | Distal Intergenic | 12 | 99190078 | 99191484 | 1407 | 1 | ENSMUST00 | -65849 | RP24-69I6.1 | ENSMUSG0C | false | . | . | . | . | . | . | chr12 | 99123319 | 99126013 | 5.77385 | 4.53587 | 3.63052 |
| chr17 | 11514001 | 11514471 | 1000 | Intron (ENSMUST00000066658.7 | 17 | 11545505 | 11545798 | 294 | 2 | ENSMUST00 | 31327 | RP23-297J19 | ENSMUSG0C | false | . | . | . | . | . | . | chr17 | 11511421 | 11515991 | 5.45303 | 4.22483 | 3.40938 |
| chr19 | 46348785 | 46349246 | 1000 | Promoter (1-2kb) | 19 | 46348201 | 46350302 | 2102 | 2 | ENSMUST00 | 1056 | RP23-62I23.3 | ENSMUSG0C | false | chr19 | 46348294 | 46349707 | 28.32091 | 81.34037 | 78.16626 | . | . | . | . | . | . |
| chr1 | 69437028 | 69437376 | 1000 | Distal Intergenic | 1 | 69441056 | 69444084 | 3029 | 1 | ENSMUST00 | -3680 | RP24-280N1 | ENSMUSG0C | false | . | . | . | . | . | . | chr1 | 69436771 | 69439099 | 4.84269 | 3.97486 | 2.93068 |
| chr7 | 71383261 | 71383983 | 1000 | Intron (ENSMUST00000190004.1 | 7 | 71370333 | 71379813 | 9481 | 1 | ENSMUST00 | 12929 | RP24-173C1 | ENSMUSG0C | false | . | . | . | . | . | . | chr7 | 71378101 | 71386659 | 5.91662 | 4.56543 | 3.76231 |
| chr7 | 71409025 | 71409431 | 1000 | Distal Intergenic | 7 | 71370333 | 71379813 | 9481 | 1 | ENSMUST00 | 38693 | RP24-173C1 | ENSMUSG0C | false | . | . | . | . | . | . | chr7 | 71408173 | 71411949 | 5.13174 | 4.08589 | 3.11505 |
| chr1 | 73425012 | 73425367 | 1000 | Intron (ENSMUST00000186582.1 | 1 | 73395334 | 73407569 | 12236 | 2 | ENSMUST00 | -17444 | Pinc | ENSMUSG0C | false | . | . | . | . | . | . | chr1 | 73424625 | 73425733 | 3.02677 | 2.90971 | 1.4046 |
| chr8 | 123041703 | 123042688 | 1000 | Promoter (<=1kb) | 8 | 123042466 | 123044602 | 2137 | 1 | ENSMUST00 | 0 | RP23-460F21 | ENSMUSG0C | false | chr8 | 123040842 | 123042311 | 30.41442 | 103.5799 | 99.9162 | . | . | . | . | . | . |
| chr3 | 85338429 | 85338796 | 1000 | Intron (ENSMUST00000154148.2 | 3 | 85317519 | 85332030 | 14512 | 1 | ENSMUST00 | 20911 | RP23-94M23 | ENSMUSG0C | false | . | . | . | . | . | . | chr3 | 85337362 | 85338947 | 6.23894 | 4.63263 | 4.06894 |
| chr1 | 52597173 | 52598257 | 1000 | Distal Intergenic | 1 | 52584518 | 52585159 | 642 | 1 | ENSMUST00 | 12656 | RP24-388H6. | ENSMUSG0C | false | . | . | . | . | . | . | chr1 | 52595947 | 52600230 | 5.92297 | 4.53953 | 3.76138 |
| chr13 | 37492199 | 37492627 | 1000 | Distal Intergenic | 13 | 37508940 | 37512074 | 3135 | 1 | ENSMUST00 | -16313 | RP23-21N15 | ENSMUSG0C | false | . | . | . | . | . | . | chr13 | 37489375 | 37494220 | 4.57256 | 3.81861 | 2.66162 |
| chr13 | 37531673 | 37532391 | 1000 | Distal Intergenic | 13 | 37508940 | 37512074 | 3135 | 1 | ENSMUST00 | 22734 | RP23-21N15 | ENSMUSG0C | false | . | . | . | . | . | . | chr13 | 37526947 | 37535595 | 4.83754 | 3.94 | 2.87988 |
| chr13 | 42049103 | 42049911 | 1000 | Promoter (<=1kb) | 13 | 42049334 | 42050287 | 954 | 1 | ENSMUST00 | 0 | RP24-87A4.1 | ENSMUSG0C | false | chr13 | 42049007 | 42050458 | 23.68814 | 67.0026 | 64.0563 | chr13 | 42048704 | 42051003 | 4.30453 | 3.77001 | 2.42817 |
| chr1 | 46933897 | 46934210 | 1000 | Distal Intergenic | 1 | 46896583 | 46896879 | 297 | 2 | ENSMUST00 | -37019 | RP23-461J11 | ENSMUSG0C | false | . | . | . | . | . | . | chr1 | 46931976 | 46937290 | 4.53037 | 3.74579 | 2.63431 |
| chr6 | 86822271 | 86822884 | 1000 | Distal Intergenic | 6 | 86848399 | 86849440 | 1042 | 2 | ENSMUST00 | 26556 | RP24-467O4 | ENSMUSG0C | false | chr6 | 86822508 | 86823042 | 7.60649 | 13.49871 | 11.2608 | chr6 | 86819907 | 86824438 | 5.22703 | 4.2192 | 3.18482 |
| chr5 | 73279704 | 73280194 | 1000 | Intron (ENSMUST00000188758.1 | 5 | 73268581 | 73284174 | 15594 | 2 | ENSMUST00 | 3980 | RP23-153H1 | ENSMUSG0C | false | . | . | . | . | . | . | chr5 | 73277459 | 73282289 | 4.62689 | 3.9179 | 2.69818 |
| chr4 | 95161056 | 95161872 | 1000 | Exon (ENSMUST00000130684.2/ | 4 | 95151740 | 95157503 | 5764 | 1 | ENSMUST00 | 9317 | RP23-77I17.4 | ENSMUSG0C | false | chr4 | 95160997 | 95162146 | 13.89301 | 35.6666 | 33.12951 | chr4 | 95159243 | 95162809 | 4.78211 | 3.95901 | 2.84821 |
| chr18 | 35849582 | 35850023 | 1000 | Intron (ENSMUST00000060722.6 | 18 | 35830389 | 35831093 | 705 | 2 | ENSMUST00 | -18490 | RP24-326K1 | ENSMUSG0C | false | . | . | . | . | . | . | chr18 | 35831800 | 35857612 | 7.70883 | 5.15861 | 5.24645 |
| chr18 | 35836976 | 35837482 | 1000 | Intron (ENSMUST00000060722.6 | 18 | 35830389 | 35831093 | 705 | 2 | ENSMUST00 | -5884 | RP24-326K1 | ENSMUSG0C | false | . | . | . | . | . | . | chr18 | 35831800 | 35857612 | 7.70883 | 5.15861 | 5.24645 |
| chr18 | 35887473 | 35888630 | 1000 | Distal Intergenic | 18 | 35830389 | 35831093 | 705 | 2 | ENSMUST00 | -56381 | RP24-326K1 | ENSMUSG0C | false | chr18 | 35887544 | 35890163 | 24.02885 | 69.3614 | 66.38335 | chr18 | 35887056 | 35890338 | 5.5032 | 4.25211 | 3.4607 |
| chr18 | 35831645 | 35832791 | 1000 | Promoter (<=1kb) | 18 | 35830389 | 35831093 | 705 | 2 | ENSMUST00 | -553 | RP24-326K1 | ENSMUSG0C | false | chr18 | 35829308 | 35833126 | 34.46491 | 104.61457 | 100.92722 | chr18 | 35831800 | 35857612 | 7.70883 | 5.15861 | 5.24645 |
| chr18 | 35888947 | 35890001 | 1000 | Distal Intergenic | 18 | 35830389 | 35831093 | 705 | 2 | ENSMUST00 | -57855 | RP24-326K1 | ENSMUSG0C | false | chr18 | 35887544 | 35890163 | 24.02885 | 69.3614 | 66.38335 | chr18 | 35887056 | 35890338 | 5.5032 | 4.25211 | 3.4607 |
| chr8 | 26882644 | 26882960 | 1000 | Distal Intergenic | 8 | 26814593 | 26824423 | 9831 | 2 | ENSMUST00 | -58222 | RP24-502D8. | ENSMUSG0C | false | . | . | . | . | . | . | chr8 | 26882006 | 26883091 | 3.04995 | 3.02384 | 1.3955 |
| chr1 | 56168693 | 56169079 | 1000 | Distal Intergenic | 1 | 56019023 | 56020137 | 1115 | 1 | ENSMUST00 | 149671 | RP23-258N1 | ENSMUSG0C | false | . | . | . | . | . | . | chr1 | 56168596 | 56170407 | 3.49176 | 3.34334 | 1.76962 |
| chr1 | 61546022 | 61546338 | 1000 | Promoter (1-2kb) | 1 | 61548122 | 61548511 | 390 | 1 | ENSMUST00 | -1784 | RP23-280M7 | ENSMUSG0C | false | . | . | . | . | . | . | . | . | . | . | . | . |
| chr1 | 61529564 | 61530199 | 1000 | Distal Intergenic | 1 | 61548122 | 61548511 | 390 | 1 | ENSMUST00 | -17923 | RP23-280M7 | ENSMUSG0C | false | . | . | . | . | . | . | chr1 | 61526896 | 61533551 | 4.85222 | 3.97276 | 2.89832 |
| chr4 | 141879211 | 141879574 | 1000 | Distal Intergenic | 4 | 141875707 | 141877033 | 1327 | 1 | ENSMUST00 | 3505 | RP23-423G1 | ENSMUSG0C | false | . | . | . | . | . | . | chr4 | 141877986 | 141887420 | 6.94416 | 4.90069 | 4.61404 |
| chr13 | 98476294 | 98476759 | 1000 | Promoter (<=1kb) | 13 | 98476598 | 98477459 | 862 | 2 | ENSMUST00 | 700 | RP23-369M1 | ENSMUSG0C | false | . | . | . | . | . | . | chr13 | 98474863 | 98480463 | 5.33341 | 4.04738 | 3.29871 |
| chr5 | 132667100 | 132667449 | 1000 | Distal Intergenic | 5 | 132544435 | 132545593 | 1159 | 1 | ENSMUST00 | 122666 | RP24-399A1 | ENSMUSG0C | false | . | . | . | . | . | . | chr5 | 132666225 | 132668977 | 4.15391 | 3.60284 | 2.34769 |
| chr1 | 62834800 | 62835192 | 1000 | Promoter (2-3kb) | 1 | 62821355 | 62832370 | 11016 | 2 | ENSMUST00 | -2431 | RP23-149A5. | ENSMUSG0C | false | . | . | . | . | . | . | chr1 | 62830064 | 62836891 | 4.65649 | 3.87137 | 2.75574 |
| chr1 | 62860719 | 62861258 | 1000 | Distal Intergenic | 1 | 62821355 | 62832370 | 11016 | 2 | ENSMUST00 | -28350 | RP23-149A5. | ENSMUSG0C | false | . | . | . | . | . | . | chr1 | 62857305 | 62862928 | 5.80429 | 4.51093 | 3.66956 |
| chr1 | 62887318 | 62887805 | 1000 | Distal Intergenic | 1 | 62821355 | 62832370 | 11016 | 2 | ENSMUST00 | -54949 | RP23-149A5. | ENSMUSG0C | false | . | . | . | . | . | . | chr1 | 62885282 | 62890313 | 4.6486 | 3.92393 | 2.70924 |
| chr18 | 83134878 | 83135474 | 1000 | Distal Intergenic | 18 | 83106528 | 83107860 | 1333 | 2 | ENSMUST00 | -27019 | RP23-450K16 | ENSMUSG0C | false | . | . | . | . | . | . | chr18 | 83134824 | 83135617 | 5.84373 | 4.45588 | 3.69864 |
| chr1 | 92686918 | 92687534 | 1000 | Distal Intergenic | 1 | 92714234 | 92717202 | 2969 | 2 | ENSMUST00 | 29668 | RP23-145J9. | ENSMUSG0C | false | . | . | . | . | . | . | chr1 | 92685766 | 92687883 | 7.08707 | 5.07287 | 4.70243 |
| chr1 | 92698980 | 92699551 | 1000 | Distal Intergenic | 1 | 92714234 | 92717202 | 2969 | 2 | ENSMUST00 | 17651 | RP23-145J9. | ENSMUSG0C | false | . | . | . | . | . | . | chr1 | 92697284 | 92701611 | 6.43184 | 4.48877 | 4.22381 |
| chr5 | 118967799 | 118968181 | 1000 | Distal Intergenic | 5 | 119108234 | 119114543 | 6310 | 1 | ENSMUST00 | -140053 | RP24-389L10 | ENSMUSG0C | false | . | . | . | . | . | . | . | . | . | . | . | . |
| chr7 | 16581329 | 16582026 | 1000 | Intron (ENSMUST00000075845.5 | 7 | 16613824 | 16632544 | 18721 | 1 | ENSMUST00 | -31798 | RP24-490A2 | ENSMUSG0C | false | . | . | . | . | . | . | chr7 | 16580388 | 16590016 | 6.49854 | 4.67421 | 4.27706 |
| chr16 | 34942906 | 34943209 | 1000 | Intron (ENSMUST00000023538.8 | 16 | 34935854 | 34939329 | 3476 | 2 | ENSMUST00 | -3578 | RP23-80O10 | ENSMUSG0C | false | . | . | . | . | . | . | chr16 | 34923663 | 34969430 | 6.94598 | 4.91232 | 4.61654 |
| chr16 | 34937669 | 34938127 | 1000 | Promoter (1-2kb) | 16 | 34935854 | 34939329 | 3476 | 2 | ENSMUST00 | 1202 | RP23-80O10 | ENSMUSG0C | false | . | . | . | . | . | . | chr16 | 34923663 | 34969430 | 6.94598 | 4.91232 | 4.61654 |
| chr16 | 34958616 | 34959125 | 1000 | Intron (ENSMUST00000023538.8 | 16 | 34935854 | 34939329 | 3476 | 2 | ENSMUST00 | -19288 | RP23-80O10 | ENSMUSG0C | false | . | . | . | . | . | . | chr16 | 34923663 | 34969430 | 6.94598 | 4.91232 | 4.61654 |
| chr16 | 34941898 | 34942405 | 1000 | Promoter (2-3kb) | 16 | 34935854 | 34939329 | 3476 | 2 | ENSMUST00 | -2570 | RP23-80O10 | ENSMUSG0C | false | . | . | . | . | . | . | chr16 | 34923663 | 34969430 | 6.94598 | 4.91232 | 4.61654 |
| chr16 | 34959358 | 34959704 | 1000 | Intron (ENSMUST00000023538.8 | 16 | 34935854 | 34939329 | 3476 | 2 | ENSMUST00 | -20030 | RP23-80O10 | ENSMUSG0C | false | . | . | . | . | . | . | chr16 | 34923663 | 34969430 | 6.94598 | 4.91232 | 4.61654 |
| chr16 | 34921213 | 34921713 | 1000 | Exon (ENSMUST00000023538.8/ | 16 | 34935854 | 34939329 | 3476 | 2 | ENSMUST00 | 17616 | RP23-80O10 | ENSMUSG0C | false | chr16 | 34920992 | 34925857 | 23.32542 | 76.20397 | 73.11496 | chr16 | 34919037 | 34921264 | 6.33072 | 4.5827 | 4.11867 |
| chr15 | 95209143 | 95209637 | 1000 | Distal Intergenic | 15 | 95097542 | 95162394 | 64853 | 2 | ENSMUST00 | -46750 | RP24-303N1 | ENSMUSG0C | false | . | . | . | . | . | . | chr15 | 95207375 | 95211171 | 4.74707 | 3.87834 | 2.83808 |
| chr15 | 95214728 | 95215049 | 1000 | Distal Intergenic | 15 | 95097542 | 95162394 | 64853 | 2 | ENSMUST00 | -52335 | RP24-303N1 | ENSMUSG0C | false | . | . | . | . | . | . | chr15 | 95213810 | 95221642 | 5.84437 | 4.48428 | 3.6948 |
| chr15 | 95217186 | 95217714 | 1000 | Downstream (1-2kb) | 15 | 95097542 | 95162394 | 64853 | 2 | ENSMUST00 | -54793 | RP24-303N1 | ENSMUSG0C | false | . | . | . | . | . | . | chr15 | 95213810 | 95221642 | 5.84437 | 4.48428 | 3.6948 |
| chr15 | 95280872 | 95281552 | 1000 | Intron (ENSMUST00000166170.2 | 15 | 95097542 | 95162394 | 64853 | 2 | ENSMUST00 | -118479 | RP24-303N1 | ENSMUSG0C | false | . | . | . | . | . | . | chr15 | 95280101 | 95284037 | 5.82739 | 4.36965 | 3.73075 |
| chr19 | 45304420 | 45305040 | 1000 | Promoter (1-2kb) | 19 | 45300463 | 45303034 | 2572 | 2 | ENSMUST00 | -1387 | RP23-188J8. | ENSMUSG0C | false | . | . | . | . | . | . | chr19 | 45300529 | 45315943 | 6.02178 | 4.57633 | 3.84298 |
| chr19 | 45310849 | 45311291 | 1000 | Promoter (<=1kb) | 19 | 45300454 | 45311011 | 10558 | 2 | ENSMUST00 | 0 | RP23-188J8. | ENSMUSG0C | false | . | . | . | . | . | . | chr19 | 45300529 | 45315943 | 6.02178 | 4.57633 | 3.84298 |
| chr19 | 45294669 | 45295510 | 1000 | Distal Intergenic | 19 | 45300463 | 45303034 | 2572 | 2 | ENSMUST00 | 7524 | RP23-188J8. | ENSMUSG0C | false | . | . | . | . | . | . | chr19 | 45292204 | 45298207 | 6.27899 | 4.63491 | 4.0501 |
| chr1 | 154150105 | 154150503 | 1000 | Distal Intergenic | 1 | 154048109 | 154117951 | 69843 | 2 | ENSMUST00 | -32155 | RP24-255G2 | ENSMUSG0C | false | . | . | . | . | . | . | chr1 | 154149398 | 154151121 | 5.2312 | 4.14349 | 3.2251 |
| chr7 | 71121577 | 71121918 | 1000 | Distal Intergenic | 7 | 71217049 | 71224409 | 7361 | 1 | ENSMUST00 | -95131 | RP23-257M2 | ENSMUSG0C | false | . | . | . | . | . | . | chr7 | 71120039 | 71124347 | 3.49057 | 3.31038 | 1.77707 |
| chr10 | 119065414 | 119065910 | 1000 | Distal Intergenic | 10 | 119092905 | 119103107 | 10203 | 1 | ENSMUST00 | -26995 | RP23-360L4. | ENSMUSG0C | false | . | . | . | . | . | . | chr10 | 119061401 | 119066309 | 4.92666 | 3.97999 | 2.96336 |
| chr8 | 48842990 | 48843680 | 1000 | Promoter (<=1kb) | 8 | 48843665 | 48869016 | 25352 | 1 | ENSMUST00 | 0 | RP23-61B8.2 | ENSMUSG0C | false | chr8 | 48842559 | 48844996 | 32.27147 | 102.84008 | 99.19724 | chr8 | 48842176 | 48845151 | 3.61777 | 3.28578 | 1.90608 |
| chr6 | 141249434 | 141250419 | 1000 | Promoter (<=1kb) | 6 | 141246920 | 141250009 | 3090 | 2 | ENSMUST00 | 0 | RP23-97L11. | ENSMUSG0C | false | chr6 | 141249145 | 141251922 | 34.07522 | 104.96031 | 101.26198 | . | . | . | . | . | . |
| chr17 | 86236903 | 86237353 | 1000 | Intron (ENSMUST00000097275.3 | 17 | 86285164 | 86287178 | 2015 | 2 | ENSMUST00 | 49825 | RP23-458O2 | ENSMUSG0C | false | . | . | . | . | . | . | chr17 | 86233898 | 86244079 | 5.46025 | 4.17001 | 3.40504 |
| chr17 | 86243025 | 86243362 | 1000 | Intron (ENSMUST00000097275.3 | 17 | 86285164 | 86287178 | 2015 | 2 | ENSMUST00 | 43816 | RP23-458O2 | ENSMUSG0C | false | . | . | . | . | . | . | chr17 | 86233898 | 86244079 | 5.46025 | 4.17001 | 3.40504 |
| chr17 | 86379379 | 86379973 | 1000 | Intron (ENSMUST00000097275.3 | 17 | 86285164 | 86287178 | 2015 | 2 | ENSMUST00 | -92202 | RP23-458O2 | ENSMUSG0C | false | . | . | . | . | . | . | chr17 | 86376181 | 86386191 | 6.60767 | 4.81689 | 4.34402 |
| chr13 | 37441912 | 37442470 | 1000 | Distal Intergenic | 13 | 37421945 | 37435149 | 13205 | 2 | ENSMUST00 | -6764 | RP23-21N15 | ENSMUSG0C | false | . | . | . | . | . | . | chr13 | 37438115 | 37444000 | 5.21214 | 4.12116 | 3.17794 |
| chr13 | 37440551 | 37441413 | 1000 | Distal Intergenic | 13 | 37421945 | 37435149 | 13205 | 2 | ENSMUST00 | -5403 | RP23-21N15 | ENSMUSG0C | false | . | . | . | . | . | . | chr13 | 37438115 | 37444000 | 5.21214 | 4.12116 | 3.17794 |
| chr5 | 149229011 | 149229583 | 1000 | Promoter (1-2kb) | 5 | 149231332 | 149237984 | 6653 | 1 | ENSMUST00 | -1749 | RP23-408A1. | ENSMUSG0C | false | chr5 | 149228600 | 149229612 | 15.98415 | 46.70646 | 44.03467 | . | . | . | . | . | . |
| chr1 | 62751202 | 62751677 | 1000 | Intron (ENSMUST00000075144.6 | 1 | 62714723 | 62718073 | 3351 | 2 | ENSMUST00 | -33130 | RP23-20G7.7 | ENSMUSG0C | false | . | . | . | . | . | . | chr1 | 62750416 | 62751822 | 3.71852 | 3.36903 | 1.97792 |
| chr3 | 85373034 | 85373477 | 1000 | Intron (ENSMUST00000154148.2 | 3 | 85360228 | 85361174 | 947 | 2 | ENSMUST00 | -11861 | RP23-94M23 | ENSMUSG0C | false | . | . | . | . | . | . | chr3 | 85370974 | 85374501 | 6.48463 | 4.69952 | 4.27954 |
| chr17 | 87473585 | 87474247 | 1000 | Distal Intergenic | 17 | 87490748 | 87502616 | 11869 | 1 | ENSMUST00 | -16501 | RP23-243O1 | ENSMUSG0C | false | chr17 | 87473025 | 87474630 | 9.76265 | 21.55577 | 19.19873 | chr17 | 87471014 | 87475647 | 6.19739 | 4.65842 | 3.98524 |
| chr5 | 64505188 | 64505943 | 1000 | Distal Intergenic | 5 | 64456328 | 64467946 | 11619 | 2 | ENSMUST00 | -37243 | RP23-195B3. | ENSMUSG0C | false | chr5 | 64505373 | 64506387 | 13.26345 | 35.90286 | 33.3624 | . | . | . | . | . | . |
| chr4 | 114550264 | 114550762 | 1000 | Intron (ENSMUST00000094894.3 | 4 | 114672697 | 114673340 | 644 | 2 | ENSMUST00 | 122578 | RP23-454F1 | ENSMUSG0C | false | . | . | . | . | . | . | chr4 | 114538576 | 114564774 | 5.66596 | 4.26794 | 3.56957 |
| chr4 | 114672427 | 114672888 | 1000 | Promoter (<=1kb) | 4 | 114672697 | 114673340 | 644 | 2 | ENSMUST00 | 452 | RP23-454F1 | ENSMUSG0C | false | . | . | . | . | . | . | chr4 | 114665906 | 114674187 | 5.6616 | 4.39937 | 3.54713 |
| chr4 | 114584659 | 114585356 | 1000 | Intron (ENSMUST00000094894.3 | 4 | 114672697 | 114673340 | 644 | 2 | ENSMUST00 | 87984 | RP23-454F1 | ENSMUSG0C | false | . | . | . | . | . | . | chr4 | 114581118 | 114602451 | 5.21128 | 4.07567 | 3.19659 |
| chr4 | 114593855 | 114594218 | 1000 | Intron (ENSMUST00000094894.3 | 4 | 114672697 | 114673340 | 644 | 2 | ENSMUST00 | 79122 | RP23-454F1 | ENSMUSG0C | false | . | . | . | . | . | . | chr4 | 114581118 | 114602451 | 5.21128 | 4.07567 | 3.19659 |
| chr1 | 124730416 | 124730765 | 1000 | Intron (ENSMUST00000187286.1 | 1 | 124615191 | 124615596 | 406 | 2 | ENSMUST00 | -114821 | RP24-63O23 | ENSMUSG0C | false | . | . | . | . | . | . | chr1 | 124729926 | 124732275 | 4.72892 | 3.92956 | 2.78946 |
| chr7 | 16618438 | 16618982 | 1000 | Promoter (<=1kb) | 7 | 16618627 | 16619832 | 1206 | 2 | ENSMUST00 | 850 | RP24-490A2 | ENSMUSG0C | false | . | . | . | . | . | . | chr7 | 16615607 | 16621870 | 5.3619 | 4.10589 | 3.33445 |
| chr14 | 76764223 | 76764866 | 1000 | Distal Intergenic | 14 | 76681754 | 76686887 | 5134 | 1 | ENSMUST00 | 82470 | RP23-421C2 | ENSMUSG0C | false | . | . | . | . | . | . | chr14 | 76762591 | 76768747 | 5.81188 | 4.37563 | 3.678 |
| chr14 | 76714693 | 76715138 | 1000 | Distal Intergenic | 14 | 76681754 | 76686887 | 5134 | 1 | ENSMUST00 | 32940 | RP23-421C2 | ENSMUSG0C | false | . | . | . | . | . | . | chr14 | 76713799 | 76716136 | 4.11408 | 3.68286 | 2.27647 |
| chr1 | 73200134 | 73200433 | 1000 | Distal Intergenic | 1 | 73269064 | 73306360 | 37297 | 1 | ENSMUST00 | -68631 | RP24-141F1 | ENSMUSG0C | false | . | . | . | . | . | . | chr1 | 73200086 | 73200792 | 2.64537 | 2.73869 | 1.13454 |
| chr1 | 73232423 | 73232800 | 1000 | Distal Intergenic | 1 | 73269064 | 73306360 | 37297 | 1 | ENSMUST00 | -36264 | RP24-141F1 | ENSMUSG0C | false | . | . | . | . | . | . | . | . | . | . | . | . |
| chr2 | 74727736 | 74728297 | 1000 | Promoter (<=1kb) | 2 | 74727074 | 74729123 | 2050 | 1 | ENSMUST00 | 663 | Hoxd4 | ENSMUSG0C | false | chr2 | 74727178 | 74727798 | 18.39885 | 52.66533 | 49.91441 | chr2 | 74726816 | 74729597 | 5.16616 | 4.19691 | 3.16396 |
| chr1 | 159393338 | 159393680 | 1000 | Distal Intergenic | 1 | 159358509 | 159358903 | 395 | 1 | ENSMUST00 | 34830 | RP23-145H3. | ENSMUSG0C | false | . | . | . | . | . | . | chr1 | 159392770 | 159394484 | 5.12048 | 3.99253 | 3.14242 |
| chr1 | 29182857 | 29183207 | 1000 | Distal Intergenic | 1 | 29219700 | 29220249 | 550 | 1 | ENSMUST00 | -36493 | RP24-430C1. | ENSMUSG0C | false | . | . | . | . | . | . | chr1 | 29181843 | 29183813 | 4.13141 | 3.51352 | 2.31698 |
| chr1 | 86472058 | 86472565 | 1000 | Promoter (1-2kb) | 1 | 86473659 | 86474006 | 348 | 1 | ENSMUST00 | -1094 | RP24-403I3.6 | ENSMUSG0C | false | . | . | . | . | . | . | chr1 | 86469522 | 86474676 | 6.26934 | 4.72595 | 4.04395 |
| chr1 | 88956983 | 88957516 | 1000 | Distal Intergenic | 1 | 88932319 | 88933573 | 1255 | 1 | ENSMUST00 | 24665 | RP23-358L5. | ENSMUSG0C | false | . | . | . | . | . | . | chr1 | 88954640 | 88960619 | 4.16838 | 3.61348 | 2.33082 |
| chr16 | 5661888 | 5662154 | 1000 | Distal Intergenic | 16 | 5975585 | 5998496 | 22912 | 1 | ENSMUST00 | -313431 | RP23-356A2 | ENSMUSG0C | false | . | . | . | . | . | . | chr16 | 5661693 | 5662817 | 3.86553 | 3.48483 | 2.1049 |
| chr1 | 184447205 | 184447657 | 1000 | Distal Intergenic | 1 | 184527841 | 184557691 | 29851 | 2 | ENSMUST00 | 110034 | RP23-172L16 | ENSMUSG0C | false | . | . | . | . | . | . | chr1 | 184445652 | 184449151 | 5.44014 | 4.28091 | 3.36067 |
| chr15 | 103115134 | 103115687 | 1000 | Distal Intergenic | 15 | 103102284 | 103104602 | 2319 | 2 | ENSMUST00 | -10533 | Gm10830 | ENSMUSG0C | false | . | . | . | . | . | . | chr15 | 103113119 | 103118268 | 4.97916 | 3.82566 | 3.02992 |
| chr9 | 96536777 | 96537373 | 1000 | Promoter (<=1kb) | 9 | 96537161 | 96537854 | 694 | 1 | ENSMUST00 | 0 | RP24-91A16. | ENSMUSG0C | false | chr9 | 96536896 | 96537429 | 14.96389 | 42.58086 | 39.95799 | chr9 | 96530728 | 96539290 | 4.63815 | 3.80816 | 2.74726 |
| chr5 | 112102134 | 112102651 | 1000 | Distal Intergenic | 5 | 112206763 | 112210002 | 3240 | 1 | ENSMUST00 | -104112 | RP24-317K1 | ENSMUSG0C | false | . | . | . | . | . | . | . | . | . | . | . | . |
| chr12 | 79949102 | 79949466 | 1000 | Distal Intergenic | 12 | 79963568 | 79963872 | 305 | 1 | ENSMUST00 | -14102 | RP23-179D3. | ENSMUSG0C | false | . | . | . | . | . | . | chr12 | 79942657 | 79955332 | 5.38416 | 4.22706 | 3.33447 |
| chr12 | 79856292 | 79856877 | 1000 | Distal Intergenic | 12 | 79963568 | 79963872 | 305 | 1 | ENSMUST00 | -106691 | RP23-179D3. | ENSMUSG0C | false | . | . | . | . | . | . | chr12 | 79854046 | 79861014 | 5.11326 | 4.07625 | 3.10953 |
| chr12 | 79661287 | 79661897 | 1000 | Distal Intergenic | 12 | 79963568 | 79963872 | 305 | 1 | ENSMUST00 | -301671 | RP23-179D3. | ENSMUSG0C | false | . | . | . | . | . | . | chr12 | 79660835 | 79664557 | 4.52908 | 3.69215 | 2.65996 |
| chr12 | 79975807 | 79976319 | 1000 | Distal Intergenic | 12 | 79963568 | 79963872 | 305 | 1 | ENSMUST00 | 12240 | RP23-179D3. | ENSMUSG0C | false | . | . | . | . | . | . | chr12 | 79973528 | 79977383 | 6.12958 | 4.63276 | 3.93422 |
| chr12 | 79993814 | 79994450 | 1000 | Distal Intergenic | 12 | 79963568 | 79963872 | 305 | 1 | ENSMUST00 | 30247 | RP23-179D3. | ENSMUSG0C | false | . | . | . | . | . | . | chr12 | 79993084 | 80001591 | 4.83553 | 3.97765 | 2.87564 |
| chr12 | 79905728 | 79906218 | 1000 | Distal Intergenic | 12 | 79963568 | 79963872 | 305 | 1 | ENSMUST00 | -57350 | RP23-179D3. | ENSMUSG0C | false | . | . | . | . | . | . | chr12 | 79902506 | 79908120 | 7.24129 | 5.11539 | 4.82473 |
| chr12 | 79755611 | 79756715 | 1000 | Distal Intergenic | 12 | 79963568 | 79963872 | 305 | 1 | ENSMUST00 | -206853 | RP23-179D3. | ENSMUSG0C | false | . | . | . | . | . | . | chr12 | 79753608 | 79761465 | 5.50685 | 4.27239 | 3.4259 |
| chr14 | 60558562 | 60558929 | 1000 | Intron (ENSMUST00000190392.1 | 14 | 60485208 | 60599031 | 113824 | 1 | ENSMUST00 | 73355 | RP24-127N1 | ENSMUSG0C | false | . | . | . | . | . | . | chr14 | 60552371 | 60562219 | 6.22671 | 4.66086 | 4.01671 |
| chr1 | 57528838 | 57529136 | 1000 | Distal Intergenic | 1 | 57518523 | 57519412 | 890 | 2 | ENSMUST00 | -9427 | RP23-433B1 | ENSMUSG0C | false | . | . | . | . | . | . | chr1 | 57523871 | 57531673 | 4.91427 | 4.0154 | 2.94536 |
| chr15 | 85610749 | 85611134 | 1000 | Distal Intergenic | 15 | 85639747 | 85645934 | 6188 | 1 | ENSMUST00 | -28613 | RP23-64J7.2 | ENSMUSG0C | false | . | . | . | . | . | . | chr15 | 85609381 | 85614104 | 7.03969 | 4.78593 | 4.70306 |
| chr15 | 85670578 | 85671517 | 1000 | Distal Intergenic | 15 | 85639747 | 85645934 | 6188 | 1 | ENSMUST00 | 30832 | RP23-64J7.2 | ENSMUSG0C | false | chr15 | 85668507 | 85673353 | 33.09821 | 104.28432 | 100.60858 | . | . | . | . | . | . |
| chr7 | 73250526 | 73250933 | 1000 | Distal Intergenic | 7 | 73185014 | 73185393 | 380 | 2 | ENSMUST00 | -65134 | RP24-136L4. | ENSMUSG0C | false | . | . | . | . | . | . | chr7 | 73250257 | 73251444 | 3.9181 | 3.44262 | 2.15166 |
| chr5 | 25529499 | 25530263 | 1000 | Promoter (<=1kb) | 5 | 25530013 | 25531466 | 1454 | 1 | ENSMUST00 | 0 | RP23-405O1 | ENSMUSG0C | false | chr5 | 25529018 | 25529675 | 23.14593 | 64.32256 | 61.41582 | . | . | . | . | . | . |
| chr2 | 118125346 | 118125743 | 1000 | Promoter (1-2kb) | 2 | 118123451 | 118124047 | 597 | 2 | ENSMUST00 | -1300 | RP23-75E19. | ENSMUSG0C | false | . | . | . | . | . | . | chr2 | 118124392 | 118126474 | 5.69378 | 4.33711 | 3.6005 |
| chr16 | 30187804 | 30188412 | 1000 | Distal Intergenic | 16 | 30195071 | 30200299 | 5229 | 1 | ENSMUST00 | -6659 | RP23-11P15. | ENSMUSG0C | false | chr16 | 30187625 | 30188119 | 11.2646 | 26.49366 | 24.07236 | chr16 | 30184555 | 30189184 | 7.55656 | 5.24836 | 5.09841 |
| chr3 | 21546453 | 21546789 | 1000 | Distal Intergenic | 3 | 21268372 | 21269090 | 719 | 2 | ENSMUST00 | -277364 | RP24-399D1 | ENSMUSG0C | false | . | . | . | . | . | . | chr3 | 21546172 | 21548054 | 3.74455 | 3.41227 | 1.99982 |
| chr3 | 21478696 | 21479525 | 1000 | Distal Intergenic | 3 | 21268372 | 21269090 | 719 | 2 | ENSMUST00 | -209607 | RP24-399D1 | ENSMUSG0C | false | . | . | . | . | . | . | chr3 | 21478289 | 21480106 | 3.80042 | 3.28673 | 2.05636 |
| chr1 | 43363074 | 43363573 | 1000 | Distal Intergenic | 1 | 43409584 | 43409963 | 380 | 1 | ENSMUST00 | -46011 | RP23-389C9. | ENSMUSG0C | false | . | . | . | . | . | . | chr1 | 43359500 | 43364100 | 5.81258 | 4.44548 | 3.69389 |
| chr9 | 94408628 | 94408968 | 1000 | Promoter (<=1kb) | 9 | 94407553 | 94408126 | 574 | 2 | ENSMUST00 | -503 | RP23-19M11 | ENSMUSG0C | false | . | . | . | . | . | . | chr9 | 94406623 | 94410448 | 4.34645 | 3.69301 | 2.47708 |
| chr10 | 17102055 | 17102426 | 1000 | Distal Intergenic | 10 | 16951294 | 17095782 | 144489 | 2 | ENSMUST00 | -6274 | Gm20125 | ENSMUSG0C | false | . | . | . | . | . | . | chr10 | 17100485 | 17103034 | 4.096 | 3.67824 | 2.25337 |
| chr10 | 17012433 | 17012735 | 1000 | Intron (ENSMUST00000185722.1 | 10 | 16951294 | 17095782 | 144489 | 2 | ENSMUST00 | 83047 | Gm20125 | ENSMUSG0C | false | . | . | . | . | . | . | chr10 | 17011739 | 17013273 | 3.67244 | 3.40132 | 1.89931 |
| chr10 | 16877568 | 16877911 | 1000 | Distal Intergenic | 10 | 16951294 | 17095782 | 144489 | 2 | ENSMUST00 | 217871 | Gm20125 | ENSMUSG0C | false | . | . | . | . | . | . | chr10 | 16876972 | 16880105 | 5.07643 | 4.14205 | 3.05115 |
| chr1 | 90506487 | 90506973 | 1000 | Distal Intergenic | 1 | 90453394 | 90454501 | 1108 | 2 | ENSMUST00 | -51987 | RP24-154L1 | ENSMUSG0C | false | chr1 | 90506776 | 90507928 | 12.12078 | 33.7584 | 31.24543 | chr1 | 90502443 | 90511262 | 5.2824 | 4.18363 | 3.25082 |
| chr1 | 90440719 | 90441636 | 1000 | Distal Intergenic | 1 | 90453394 | 90454501 | 1108 | 2 | ENSMUST00 | 12865 | RP24-154L1 | ENSMUSG0C | false | chr1 | 90441549 | 90441950 | 5.02932 | 7.53255 | 5.41116 | chr1 | 90438990 | 90445431 | 5.52456 | 4.20032 | 3.45273 |
| chr1 | 90449664 | 90450218 | 1000 | Distal Intergenic | 1 | 90453394 | 90454501 | 1108 | 2 | ENSMUST00 | 4283 | RP24-154L1 | ENSMUSG0C | false | . | . | . | . | . | . | chr1 | 90448827 | 90457765 | 4.51771 | 3.82793 | 2.63689 |
| chr1 | 90442551 | 90443219 | 1000 | Distal Intergenic | 1 | 90453394 | 90454501 | 1108 | 2 | ENSMUST00 | 11282 | RP24-154L1 | ENSMUSG0C | false | . | . | . | . | . | . | chr1 | 90438990 | 90445431 | 5.52456 | 4.20032 | 3.45273 |
| chr1 | 90470900 | 90471510 | 1000 | Distal Intergenic | 1 | 90453394 | 90454501 | 1108 | 2 | ENSMUST00 | -16400 | RP24-154L1 | ENSMUSG0C | false | . | . | . | . | . | . | chr1 | 90461309 | 90477894 | 5.37806 | 4.21863 | 3.31987 |
| chr13 | 107700370 | 107700643 | 1000 | Distal Intergenic | 13 | 107687397 | 107692173 | 4777 | 1 | ENSMUST00 | 12974 | RP23-125L8. | ENSMUSG0C | false | . | . | . | . | . | . | chr13 | 107698891 | 107701370 | 6.52131 | 4.62778 | 4.28903 |
| chr13 | 107676322 | 107677103 | 1000 | Distal Intergenic | 13 | 107687397 | 107692162 | 4766 | 1 | ENSMUST00 | -10294 | RP23-125L8. | ENSMUSG0C | false | . | . | . | . | . | . | chr13 | 107675689 | 107677287 | 5.25939 | 4.03293 | 3.25311 |
| chr1 | 125458515 | 125458890 | 1000 | Distal Intergenic | 1 | 125444567 | 125447398 | 2832 | 1 | ENSMUST00 | 13949 | RP23-411B1 | ENSMUSG0C | false | . | . | . | . | . | . | chr1 | 125457959 | 125460133 | 5.30427 | 4.25663 | 3.24883 |
| chr6 | 141268243 | 141268560 | 1000 | Intron (ENSMUST00000043259.8 | 6 | 141238278 | 141250561 | 12284 | 2 | ENSMUST00 | -17683 | RP23-97L11. | ENSMUSG0C | false | . | . | . | . | . | . | chr6 | 141263575 | 141273023 | 4.35401 | 3.66878 | 2.50488 |
| chr15 | 67206431 | 67206873 | 1000 | Distal Intergenic | 15 | 67226769 | 67377094 | 150326 | 1 | ENSMUST00 | -19896 | RP23-163N4 | ENSMUSG0C | false | . | . | . | . | . | . | chr15 | 67205228 | 67211603 | 4.50274 | 3.77383 | 2.6424 |
| chr15 | 67361114 | 67361678 | 1000 | Intron (ENSMUST00000187233.1 | 15 | 67226769 | 67377094 | 150326 | 1 | ENSMUST00 | 134346 | RP23-163N4 | ENSMUSG0C | false | . | . | . | . | . | . | chr15 | 67357562 | 67366150 | 5.52464 | 4.33965 | 3.4393 |
| chr15 | 67176520 | 67177183 | 1000 | Distal Intergenic | 15 | 67226769 | 67377094 | 150326 | 1 | ENSMUST00 | -49586 | RP23-163N4 | ENSMUSG0C | false | chr15 | 67173622 | 67176826 | 30.67871 | 107.10089 | 103.34502 | . | . | . | . | . | . |
| chr15 | 67303844 | 67304365 | 1000 | Intron (ENSMUST00000187233.1 | 15 | 67226769 | 67377094 | 150326 | 1 | ENSMUST00 | 77076 | RP23-163N4 | ENSMUSG0C | false | . | . | . | . | . | . | chr15 | 67300501 | 67306545 | 4.04245 | 3.58414 | 2.22633 |
| chr15 | 31563735 | 31564187 | 999 | Distal Intergenic | 15 | 31568791 | 31588259 | 19469 | 1 | ENSMUST00 | -4604 | Cmbl | ENSMUSG0C | false | chr15 | 31563739 | 31564155 | 6.20376 | 10.41759 | 8.23702 | chr15 | 31562637 | 31570270 | 5.23895 | 4.15286 | 3.24218 |
| chr15 | 41447096 | 41447663 | 999 | Promoter (<=1kb) | 15 | 41447482 | 41861045 | 413564 | 1 | ENSMUST00 | 0 | Oxr1 | ENSMUSG0C | false | chr15 | 41446789 | 41448543 | 26.22065 | 76.7888 | 73.6919 | chr15 | 41446681 | 41448800 | 4.05251 | 3.54005 | 2.23134 |
| chr4 | 150281113 | 150282158 | 999 | Promoter (<=1kb) | 4 | 150281646 | 150621966 | 340321 | 1 | ENSMUST00 | 0 | Rere | ENSMUSG0C | false | chr4 | 150280873 | 150282960 | 34.16929 | 120.7392 | 116.5701 | . | . | . | . | . | . |
| chr6 | 15720467 | 15721412 | 999 | Promoter (<=1kb) | 6 | 15720661 | 15802165 | 81505 | 1 | ENSMUST00 | 0 | Mdfic | ENSMUSG0C | true | chr6 | 15720519 | 15722168 | 31.7393 | 99.89149 | 96.32799 | . | . | . | . | . | . |
| chr15 | 38473429 | 38474016 | 999 | Distal Intergenic | 15 | 38490473 | 38490547 | 75 | 2 | ENSMUST00 | 16531 | Mir6951 | ENSMUSG0C | false | chr15 | 38473271 | 38473881 | 26.82304 | 80.62296 | 77.45952 | chr15 | 38468227 | 38474123 | 4.55881 | 3.82285 | 2.65383 |
| chr11 | 98021946 | 98022342 | 998 | Promoter (<=1kb) | 11 | 98002901 | 98022627 | 19727 | 2 | ENSMUST00 | 285 | Cacnb1 | ENSMUSG0C | false | chr11 | 98021697 | 98023289 | 34.78613 | 113.42423 | 109.48744 | chr11 | 98018411 | 98024278 | 4.45887 | 3.76503 | 2.58806 |
| chrX | 142637545 | 142637828 | 998 | Distal Intergenic | 20 | 142631483 | 142634015 | 2533 | 2 | ENSMUST00 | -3531 | Gm5763 | ENSMUSG0C | false | . | . | . | . | . | . | . | . | . | . | . | . |
| chr7 | 54892067 | 54892652 | 998 | Intron (ENSMUST00000082373.6 | 7 | 54835959 | 55265366 | 429408 | 1 | ENSMUST00 | 56109 | Luzp2 | ENSMUSG0C | false | . | . | . | . | . | . | chr7 | 54890842 | 54894291 | 3.28959 | 3.19476 | 1.58303 |
| chr2 | 104052092 | 104052416 | 998 | Intron (ENSMUST00000028603.4 | 2 | 104040669 | 104040807 | 139 | 1 | ENSMUST00 | 11424 | Gm24644 | ENSMUSG0C | false | . | . | . | . | . | . | chr2 | 104051883 | 104053438 | 4.02056 | 3.57317 | 2.19739 |
| chr15 | 72927454 | 72927824 | 998 | Intron (ENSMUST00000023276.9 | 15 | 72913357 | 72913560 | 204 | 1 | ENSMUST00 | 14098 | Gm3150 | ENSMUSG0C | false | . | . | . | . | . | . | chr15 | 72926813 | 72930315 | 6.49107 | 4.45488 | 4.26015 |
| chr3 | 50272275 | 50272533 | 998 | Intron (ENSMUST00000190272.1 | 3 | 50259932 | 50275128 | 15197 | 1 | ENSMUST00 | 12344 | RP24-377J3.1 | ENSMUSG0C | false | . | . | . | . | . | . | chr3 | 50271221 | 50273186 | 3.75318 | 3.43594 | 1.97235 |
| chr3 | 31088867 | 31089158 | 997 | Distal Intergenic | 3 | 31095058 | 31122577 | 27520 | 1 | ENSMUST00 | -5900 | Skil | ENSMUSG0C | false | . | . | . | . | . | . | chr3 | 31088958 | 31090566 | 4.59479 | 3.867 | 2.68827 |
| chr5 | 124480005 | 124480301 | 997 | Promoter (1-2kb) | 5 | 124463265 | 124478366 | 15102 | 2 | ENSMUST00 | -1640 | Rilpl2 | ENSMUSG0C | false | . | . | . | . | . | . | chr5 | 124470465 | 124482757 | 4.75304 | 3.83886 | 2.83352 |
| chr16 | 90402579 | 90402821 | 997 | Intron (ENSMUST00000065856.6 | 16 | 90386397 | 90499553 | 113157 | 1 | ENSMUST00 | 16183 | Hunk | ENSMUSG0C | false | . | . | . | . | . | . | chr16 | 90400784 | 90404150 | 4.52588 | 3.79392 | 2.63265 |
| chr5 | 74068173 | 74068714 | 997 | Promoter (<=1kb) | 5 | 73998463 | 74068420 | 69958 | 2 | ENSMUST00 | 0 | Usp46 | ENSMUSG0C | false | chr5 | 74067293 | 74069360 | 27.03867 | 103.18376 | 99.53125 | . | . | . | . | . | . |
| chr9 | 48737453 | 48737949 | 997 | Intron (ENSMUST00000093852.3 | 9 | 48654311 | 48835945 | 181635 | 2 | ENSMUST00 | 97996 | Zbtb16 | ENSMUSG0C | false | . | . | . | . | . | . | chr9 | 48731594 | 48741569 | 5.57909 | 4.32779 | 3.48116 |
| chr13 | 99202197 | 99202463 | 996 | Intron (ENSMUST00000056558.8 | 13 | 99184823 | 99247037 | 62215 | 1 | ENSMUST00 | 17375 | Zfp366 | ENSMUSG0C | false | . | . | . | . | . | . | chr13 | 99201930 | 99202746 | 3.43007 | 3.26558 | 1.73388 |
| chr7 | 107370570 | 107371116 | 996 | Promoter (<=1kb) | 7 | 107370728 | 107436336 | 65609 | 1 | ENSMUST00 | 0 | Syt9 | ENSMUSG0C | false | chr7 | 107370116 | 107371837 | 26.18681 | 91.82585 | 88.44535 | chr7 | 107369153 | 107372560 | 3.16739 | 3.12249 | 1.56646 |
| chr10 | 8755131 | 8755486 | 996 | Intron (ENSMUST00000015449.5 | 10 | 8824011 | 8824147 | 137 | 2 | ENSMUST00 | 68661 | Gm25410 | ENSMUSG0C | false | . | . | . | . | . | . | chr10 | 8751277 | 8755790 | 6.42059 | 4.71149 | 4.18065 |
| chr4 | 62294196 | 62294517 | 996 | Promoter (2-3kb) | 4 | 62291547 | 62297477 | 5931 | 1 | ENSMUST00 | 2650 | Slc31a2 | ENSMUSG0C | false | . | . | . | . | . | . | chr4 | 62287890 | 62304836 | 6.16439 | 4.4648 | 3.98538 |
| chr14 | 20572150 | 20572432 | 996 | Intron (ENSMUST00000022356.5 | 14 | 20565011 | 20570687 | 5677 | 1 | ENSMUST00 | 7140 | 1810062O18 | ENSMUSG0C | false | . | . | . | . | . | . | chr14 | 20571546 | 20573586 | 5.1237 | 4.06837 | 3.10761 |
| chr7 | 39544239 | 39544540 | 996 | Distal Intergenic | 7 | 39552496 | 39556305 | 3810 | 1 | ENSMUST00 | -7956 | RP23-402K2 | ENSMUSG0C | false | chr7 | 39543998 | 39545210 | 30.03704 | 95.90482 | 92.42858 | . | . | . | . | . | . |
| chr14 | 70176627 | 70177085 | 995 | Promoter (<=1kb) | 14 | 70164231 | 70176819 | 12589 | 2 | ENSMUST00 | 0 | Pdlim2 | ENSMUSG0C | true | chr14 | 70173369 | 70177377 | 26.1063 | 98.41757 | 94.88632 | chr14 | 70177011 | 70181867 | 6.30878 | 4.59442 | 4.09527 |
| chr5 | 146846230 | 146846776 | 995 | Promoter (1-2kb) | 5 | 146845071 | 146847726 | 2656 | 1 | ENSMUST00 | 1160 | Rasl11a | ENSMUSG0C | true | chr5 | 146844279 | 146848368 | 25.50663 | 88.63404 | 85.31516 | . | . | . | . | . | . |
| chr14 | 21062844 | 21063149 | 995 | Intron (ENSMUST00000045376.9 | 14 | 21052574 | 21448568 | 395995 | 1 | ENSMUST00 | 10271 | Adk | ENSMUSG0C | false | . | . | . | . | . | . | chr14 | 21061598 | 21064282 | 3.67159 | 3.37664 | 1.96132 |
| chr6 | 86360725 | 86361096 | 995 | Distal Intergenic | 6 | 86365683 | 86370057 | 4375 | 1 | ENSMUST00 | -4587 | Fam136a | ENSMUSG0C | false | . | . | . | . | . | . | chr6 | 86356929 | 86365074 | 4.66248 | 3.70767 | 2.75361 |
| chr4 | 82148368 | 82148734 | 995 | Distal Intergenic | 4 | 82065957 | 82102673 | 36717 | 1 | ENSMUST00 | 82412 | Gm5860 | ENSMUSG0C | false | . | . | . | . | . | . | chr4 | 82148082 | 82148595 | 3.19005 | 2.98955 | 1.55713 |
| chr17 | 34850115 | 34850486 | 994 | Promoter (<=1kb) | 17 | 34850391 | 34856369 | 5979 | 1 | ENSMUST00 | 0 | Nelfe | ENSMUSG0C | false | chr17 | 34849538 | 34851078 | 38.04976 | 128.75285 | 124.30675 | chr17 | 34849116 | 34851626 | 5.69036 | 4.11048 | 3.60335 |
| chr1 | 16535443 | 16535730 | 994 | Distal Intergenic | 1 | 16228816 | 16520112 | 291297 | 2 | ENSMUST00 | -15332 | Stau2 | ENSMUSG0C | false | . | . | . | . | . | . | chr1 | 16535401 | 16536238 | 3.5025 | 3.28407 | 1.7996 |
| chr2 | 58532176 | 58532715 | 994 | Intron (ENSMUST00000090935.3 | 2 | 58477758 | 58538438 | 60681 | 2 | ENSMUST00 | 5723 | Acvr1 | ENSMUSG0C | false | . | . | . | . | . | . | chr2 | 58529256 | 58537023 | 5.13878 | 4.14505 | 3.13264 |
| chr3 | 87768862 | 87769194 | 994 | Promoter (<=1kb) | 3 | 87747890 | 87768948 | 21059 | 2 | ENSMUST00 | 0 | Pear1 | ENSMUSG0C | false | chr3 | 87768580 | 87769243 | 11.73166 | 34.30278 | 31.78458 | chr3 | 87768535 | 87769483 | 3.42333 | 3.17446 | 1.7138 |
| chr11 | 30025819 | 30026356 | 994 | Promoter (<=1kb) | 11 | 29743048 | 30026033 | 282986 | 2 | ENSMUST00 | 0 | Eml6 | ENSMUSG0C | false | chr11 | 30024227 | 30026706 | 29.34276 | 99.62727 | 96.06711 | . | . | . | . | . | . |
| chr2 | 72663460 | 72663864 | 994 | Distal Intergenic | 2 | 72633936 | 72634607 | 672 | 1 | ENSMUST00 | 29525 | Ak3l2-ps | ENSMUSG0C | false | . | . | . | . | . | . | chr2 | 72662753 | 72664416 | 3.49571 | 3.32567 | 1.78088 |
| chr17 | 12205685 | 12206063 | 993 | Intron (ENSMUST00000164143.2 | 17 | 12215125 | 12219253 | 4129 | 1 | ENSMUST00 | -9062 | Agpat4 | ENSMUSG0C | true | . | . | . | . | . | . | chr17 | 12189838 | 12208021 | 6.13066 | 4.60409 | 3.94082 |
| chr19 | 46042029 | 46042286 | 993 | Promoter (2-3kb) | 19 | 46032594 | 46039621 | 7028 | 2 | ENSMUST00 | -2409 | Ldb1 | ENSMUSG0C | false | chr19 | 46041290 | 46043334 | 8.08703 | 14.71837 | 12.46039 | chr19 | 46036064 | 46044344 | 7.61565 | 5.16237 | 5.15378 |
| chr8 | 87731108 | 87731493 | 993 | Intron (ENSMUST00000165770.3 | 8 | 87783147 | 87803823 | 20677 | 2 | ENSMUST00 | 72330 | Zfp423 | ENSMUSG0C | true | . | . | . | . | . | . | chr8 | 87729485 | 87749127 | 6.31805 | 4.63416 | 4.1004 |
| chr1 | 39576940 | 39577575 | 993 | Promoter (<=1kb) | 1 | 39551296 | 39577405 | 26110 | 2 | ENSMUST00 | 0 | Rnf149 | ENSMUSG0C | false | chr1 | 39576052 | 39577945 | 29.47202 | 98.86652 | 95.32386 | chr1 | 39575536 | 39578032 | 4.12878 | 3.66056 | 2.30191 |
| chr13 | 28629610 | 28629991 | 993 | Intron (ENSMUST00000148885.2 | 13 | 28710781 | 28710873 | 93 | 2 | ENSMUST00 | 80882 | Mir6368 | ENSMUSG0C | false | . | . | . | . | . | . | chr13 | 28628983 | 28631243 | 5.14095 | 4.11234 | 3.16632 |
| chr8 | 73353098 | 73353898 | 992 | Promoter (<=1kb) | 8 | 72814599 | 73353540 | 538942 | 2 | ENSMUST00 | 0 | Large | ENSMUSG0C | false | chr8 | 73351993 | 73354929 | 30.60796 | 113.09483 | 109.17188 | . | . | . | . | . | . |
| chr10 | 7183132 | 7183591 | 992 | Intron (ENSMUST00000015346.6 | 10 | 7119063 | 7212237 | 93175 | 2 | ENSMUST00 | 28646 | Cnksr3 | ENSMUSG0C | false | chr10 | 7181972 | 7183538 | 15.39495 | 41.17039 | 38.56658 | chr10 | 7176560 | 7190519 | 5.92831 | 4.42258 | 3.77546 |
| chr2 | 174074559 | 174074986 | 992 | Promoter (1-2kb) | 2 | 174076308 | 174092433 | 16126 | 1 | ENSMUST00 | -1322 | Stx16 | ENSMUSG0C | false | . | . | . | . | . | . | chr2 | 174073295 | 174092361 | 5.14014 | 4.03613 | 3.13424 |
| chr4 | 148130535 | 148130969 | 992 | Promoter (<=1kb) | 4 | 148130915 | 148145699 | 14785 | 1 | ENSMUST00 | 0 | Mad2l2 | ENSMUSG0C | false | chr4 | 148130346 | 148131392 | 22.51707 | 59.83231 | 56.98677 | chr4 | 148130095 | 148131788 | 5.91656 | 4.47238 | 3.76007 |
| chr2 | 154436140 | 154436753 | 992 | Promoter (<=1kb) | 2 | 154436481 | 154539356 | 102876 | 1 | ENSMUST00 | 0 | Cbfa2t2 | ENSMUSG0C | false | chr2 | 154436395 | 154437605 | 20.80241 | 92.62246 | 89.22079 | . | . | . | . | . | . |
| chr6 | 95596993 | 95597460 | 992 | Intron (ENSMUST00000079847.5 | 6 | 95474134 | 95718837 | 244704 | 2 | ENSMUST00 | 121377 | Suclg2 | ENSMUSG0C | false | . | . | . | . | . | . | chr6 | 95596673 | 95598224 | 5.29044 | 4.1696 | 3.25456 |
| chr2 | 60963778 | 60964036 | 991 | Promoter (<=1kb) | 2 | 60750193 | 60963192 | 213000 | 2 | ENSMUST00 | -587 | Rbms1 | ENSMUSG0C | true | chr2 | 60963877 | 60964298 | 5.43663 | 10.5674 | 8.37697 | . | . | . | . | . | . |
| chr4 | 62965271 | 62965985 | 991 | Promoter (<=1kb) | 4 | 62965573 | 63115748 | 150176 | 1 | ENSMUST00 | 0 | Zfp618 | ENSMUSG0C | false | chr4 | 62965204 | 62966197 | 27.57193 | 78.19525 | 75.07539 | . | . | . | . | . | . |
| chr8 | 94098066 | 94098631 | 991 | Promoter (<=1kb) | 8 | 94067954 | 94098811 | 30858 | 2 | ENSMUST00 | 180 | Bbs2 | ENSMUSG0C | false | chr8 | 94097748 | 94099004 | 19.38504 | 61.02572 | 58.1656 | chr8 | 94097480 | 94099379 | 4.00493 | 3.40566 | 2.22407 |
| chr19 | 38836383 | 38837069 | 991 | Promoter (<=1kb) | 19 | 38836579 | 38919923 | 83345 | 1 | ENSMUST00 | 0 | Tbc1d12 | ENSMUSG0C | false | chr19 | 38836126 | 38837723 | 33.26678 | 112.81776 | 108.90401 | . | . | . | . | . | . |
| chr9 | 118982874 | 118983162 | 991 | Intron (ENSMUST00000073109.6 | 9 | 119031796 | 119031885 | 90 | 1 | ENSMUST00 | -48634 | Mir26a-1 | ENSMUSG0C | false | . | . | . | . | . | . | chr9 | 118927678 | 119010843 | 6.89652 | 4.89961 | 4.57102 |
| chr9 | 108517092 | 108517685 | 990 | Promoter (<=1kb) | 9 | 108517087 | 108560167 | 43081 | 1 | ENSMUST00 | 6 | Qrich1 | ENSMUSG0C | false | chr9 | 108516286 | 108518500 | 45.67889 | 150.95599 | 145.63583 | . | . | . | . | . | . |
| chr3 | 113629693 | 113630448 | 990 | Promoter (<=1kb) | 3 | 113605067 | 113629900 | 24834 | 2 | ENSMUST00 | 0 | Rnpc3 | ENSMUSG0C | false | chr3 | 113629253 | 113630972 | 32.03847 | 101.42104 | 97.81718 | . | . | . | . | . | . |
| chr1 | 74007542 | 74008103 | 990 | Intron (ENSMUST00000169786.2 | 1 | 73992205 | 74001567 | 9363 | 2 | ENSMUST00 | -5976 | Tns1 | ENSMUSG0C | false | . | . | . | . | . | . | chr1 | 74006078 | 74010176 | 3.31651 | 3.21932 | 1.61985 |
| chr7 | 44908741 | 44909262 | 990 | Promoter (<=1kb) | 7 | 44906114 | 44909829 | 3716 | 2 | ENSMUST00 | 567 | Ap2a1 | ENSMUSG0C | false | . | . | . | . | . | . | chr7 | 44908031 | 44909943 | 6.86677 | 4.89193 | 4.56239 |
| chr4 | 143125593 | 143125903 | 990 | Intron (ENSMUST00000105778.2 | 4 | 143152572 | 143153030 | 459 | 1 | ENSMUST00 | -26669 | Gm13038 | ENSMUSG0C | false | . | . | . | . | . | . | chr4 | 143124995 | 143129649 | 4.54626 | 3.63885 | 2.66401 |
| chr15 | 99818336 | 99818836 | 990 | Promoter (<=1kb) | 15 | 99819020 | 99834570 | 15551 | 1 | ENSMUST00 | -184 | Gm17057 | ENSMUSG0C | false | chr15 | 99817939 | 99820181 | 16.08362 | 38.04472 | 35.47989 | chr15 | 99811513 | 99819053 | 3.94956 | 3.48882 | 2.16316 |
| chr7 | 70361227 | 70362097 | 989 | Promoter (<=1kb) | 7 | 70351946 | 70360593 | 8648 | 2 | ENSMUST00 | -635 | Nr2f2 | ENSMUSG0C | false | chr7 | 70357618 | 70362933 | 29.75681 | 92.45776 | 89.05831 | . | . | . | . | . | . |
| chr9 | 75581177 | 75581484 | 989 | Intron (ENSMUST00000098552.3 | 9 | 75573816 | 75597651 | 23836 | 2 | ENSMUST00 | 16167 | Tmod2 | ENSMUSG0C | false | . | . | . | . | . | . | chr9 | 75576366 | 75582593 | 4.78783 | 4.00601 | 2.82581 |
| chr11 | 6443985 | 6444852 | 989 | Promoter (<=1kb) | 11 | 6427229 | 6444443 | 17215 | 2 | ENSMUST00 | 0 | H2afv | ENSMUSG0C | false | chr11 | 6443550 | 6444540 | 31.25687 | 94.08162 | 90.65025 | . | . | . | . | . | . |
| chr17 | 84574774 | 84575424 | 989 | Exon (ENSMUST00000047206.5/ | 17 | 84624402 | 84624510 | 109 | 1 | ENSMUST00 | -48978 | Gm26066 | ENSMUSG0C | false | . | . | . | . | . | . | chr17 | 84573511 | 84577398 | 4.00295 | 3.44887 | 2.2295 |
| chr1 | 92683161 | 92683853 | 989 | Distal Intergenic | 1 | 92714234 | 92717202 | 2969 | 2 | ENSMUST00 | 33349 | RP23-145J9. | ENSMUSG0C | false | . | . | . | . | . | . | chr1 | 92680001 | 92684688 | 6.40462 | 4.67491 | 4.16393 |
| chr16 | 13256183 | 13257057 | 988 | Promoter (<=1kb) | 16 | 13256481 | 13415063 | 158583 | 1 | ENSMUST00 | 0 | Mkl2 | ENSMUSG0C | false | chr16 | 13255775 | 13257573 | 33.41831 | 117.26761 | 113.21259 | chr16 | 13255800 | 13258637 | 3.73308 | 3.27952 | 2.01502 |
| chr10 | 75060345 | 75060846 | 988 | Promoter (<=1kb) | 10 | 75060896 | 75184923 | 124028 | 1 | ENSMUST00 | -50 | Bcr | ENSMUSG0C | false | chr10 | 75059656 | 75062040 | 36.01637 | 112.4352 | 108.52973 | chr10 | 75059548 | 75063902 | 3.94451 | 3.3221 | 2.17899 |
| chr4 | 126103766 | 126104348 | 988 | Promoter (<=1kb) | 4 | 126103957 | 126141028 | 37072 | 1 | ENSMUST00 | 0 | Stk40 | ENSMUSG0C | false | chr4 | 126102974 | 126103823 | 12.12078 | 33.7584 | 31.24543 | . | . | . | . | . | . |
| chr4 | 129702462 | 129702792 | 988 | Downstream (<1kb) | 4 | 129676355 | 129696838 | 20484 | 2 | ENSMUST00 | -5625 | Tmem39b | ENSMUSG0C | false | . | . | . | . | . | . | chr4 | 129701828 | 129702852 | 5.69617 | 4.30477 | 3.61801 |
| chr11 | 69715887 | 69716206 | 988 | Distal Intergenic | 11 | 69729995 | 69730584 | 590 | 2 | ENSMUST00 | 14378 | Gm12307 | ENSMUSG0C | false | . | . | . | . | . | . | chr11 | 69708924 | 69717022 | 4.65082 | 3.90064 | 2.7202 |
| chr19 | 61044528 | 61044797 | 988 | Intron (ENSMUST00000003313.8 | 19 | 61013417 | 61013545 | 129 | 2 | ENSMUST00 | -30984 | Gm22520 | ENSMUSG0C | false | . | . | . | . | . | . | chr19 | 61033925 | 61053045 | 5.06464 | 4.02759 | 3.06894 |
| chr2 | 170345075 | 170345673 | 987 | Downstream (1-2kb) | 2 | 170347931 | 170349740 | 1810 | 2 | ENSMUST00 | 4067 | Bcas1 | ENSMUSG0C | false | . | . | . | . | . | . | chr2 | 170343669 | 170347174 | 5.91388 | 4.45747 | 3.81613 |
| chr11 | 111065750 | 111066199 | 987 | Promoter (<=1kb) | 11 | 111066164 | 111076821 | 10658 | 1 | ENSMUST00 | 0 | Kcnj2 | ENSMUSG0C | false | chr11 | 111064881 | 111066677 | 21.40274 | 64.72344 | 61.81032 | chr11 | 111064811 | 111066979 | 5.5543 | 4.31218 | 3.4442 |
| chr15 | 80173220 | 80173832 | 987 | Promoter (<=1kb) | 15 | 80173721 | 80215519 | 41799 | 1 | ENSMUST00 | 0 | Mgat3 | ENSMUSG0C | false | chr15 | 80173046 | 80174497 | 21.02725 | 63.18829 | 60.29588 | chr15 | 80173132 | 80174847 | 6.20365 | 4.50592 | 4.03057 |
| chr3 | 93444372 | 93444964 | 987 | Promoter (2-3kb) | 3 | 93442330 | 93449077 | 6748 | 1 | ENSMUST00 | 2043 | Tchh | ENSMUSG0C | false | chr3 | 93443512 | 93448142 | 14.07679 | 50.30981 | 47.58934 | chr3 | 93443434 | 93448313 | 4.58012 | 3.85381 | 2.66881 |
| chr18 | 55660555 | 55660835 | 987 | Distal Intergenic | 18 | 55622636 | 55622920 | 285 | 2 | ENSMUST00 | -37636 | Gm24514 | ENSMUSG0C | false | . | . | . | . | . | . | chr18 | 55660285 | 55661073 | 4.5453 | 3.7652 | 2.64663 |

| | | | | | | | | | | | | | | | | | | | | | | | | | | |
|---|---|---|---|---|---|---|---|---|---|---|---|---|---|---|---|---|---|---|---|---|---|---|---|---|---|---|
| chr14 | 30735858 | 30736159 | 986 | Intron (ENSMUST00000112184.2 | 14 | 30716491 | 30818346 | 101856 | 1 | ENSMUST00 | 19368 | Sfmbt1 | ENSMUSG0( | false | . | . | . | . | . | . | chr14 | 30731435 | 30737566 | 4.24682 | 3.58331 | 2.43823 |
| chr16 | 30596798 | 30597086 | 986 | Promoter (2-3kb) | 16 | 30599723 | 30602797 | 3075 | 1 | ENSMUST00 | -2637 | Fam43a | ENSMUSG0( | false | chr16 | 30596816 | 30597074 | 6.57701 | 11.02508 | 8.83021 | chr16 | 30596541 | 30598911 | 5.21438 | 4.22344 | 3.18444 |
| chr15 | 3269431 | 3269804 | 986 | Promoter (<=1kb) | 15 | 3268547 | 3275782 | 7236 | 1 | ENSMUST00 | 885 | Sepp1 | ENSMUSG0( | false | . | . | . | . | . | . | chr15 | 3269193 | 3270212 | 4.6433 | 3.99837 | 2.71152 |
| chr12 | 74458584 | 74458929 | 986 | Distal Intergenic | 12 | 74315110 | 74316394 | 1285 | 2 | ENSMUST00 | -142191 | Gm11042 | ENSMUSG0( | false | . | . | . | . | . | . | chr12 | 74456614 | 74460058 | 5.4766 | 4.31612 | 3.39186 |
| chr14 | 69443104 | 69443422 | 986 | Intron (ENSMUST00000111062.3 | 14 | 69419394 | 69426477 | 7084 | 2 | ENSMUST00 | -16628 | Gm28049 | ENSMUSG0( | false | . | . | . | . | . | . | chr14 | 69440416 | 69448720 | 4.10395 | 3.65347 | 2.25467 |
| chr8 | 26756515 | 26756862 | 986 | Distal Intergenic | 8 | 26814593 | 26824423 | 9831 | 2 | ENSMUST00 | 67561 | RP24-502D8. | ENSMUSG0( | false | . | . | . | . | . | . | chr8 | 26755853 | 26758237 | 3.83496 | 3.34765 | 2.08448 |
| chr16 | 37776739 | 37777298 | 985 | Promoter (<=1kb) | 16 | 37776873 | 37836514 | 59642 | 1 | ENSMUST00 | 0 | Fstl1 | ENSMUSG0( | false | chr16 | 37776641 | 37777898 | 26.86698 | 95.04005 | 91.5885 | . | . | . | . | . | . |
| chr15 | 99865951 | 99866276 | 985 | Intron (ENSMUST00000073691.3 | 15 | 99841228 | 99875416 | 34189 | 2 | ENSMUST00 | 9140 | Lima1 | ENSMUSG0( | false | . | . | . | . | . | . | chr15 | 99864908 | 99867960 | 4.52042 | 3.84277 | 2.6189 |
| chr2 | 33629529 | 33629855 | 985 | Promoter (2-3kb) | 2 | 33567156 | 33632283 | 65128 | 2 | ENSMUST00 | 2428 | Lmx1b | ENSMUSG0( | true | . | . | . | . | . | . | chr2 | 33565260 | 33638297 | 6.78481 | 4.81457 | 4.48235 |
| chr15 | 68301138 | 68301490 | 985 | Distal Intergenic | 15 | 68309819 | 68310301 | 483 | 1 | ENSMUST00 | -8329 | Gm10240 | ENSMUSG0( | false | . | . | . | . | . | . | chr15 | 68298467 | 68303912 | 4.64547 | 3.96275 | 2.70162 |
| chr6 | 31079582 | 31079907 | 985 | Distal Intergenic | 6 | 31087609 | 31087912 | 304 | 2 | ENSMUST00 | 8005 | Gm13834 | ENSMUSG0( | false | . | . | . | . | . | . | . | . | . | . | . | . |
| chr16 | 52011573 | 52011996 | 984 | Distal Intergenic | 16 | 52031549 | 52208047 | 176499 | 1 | ENSMUST00 | -19553 | Cblb | ENSMUSG0( | true | . | . | . | . | . | . | chr16 | 52009607 | 52016621 | 4.28782 | 3.76463 | 2.41076 |
| chr19 | 55754077 | 55754403 | 984 | Intron (ENSMUST00000061496.1 | 19 | 55742845 | 55910681 | 167837 | 1 | ENSMUST00 | 11233 | Tcf7l2 | ENSMUSG0( | false | . | . | . | . | . | . | chr19 | 55744268 | 55760177 | 6.20416 | 4.6193 | 3.99773 |
| chr4 | 100776471 | 100777320 | 984 | Promoter (<=1kb) | 4 | 100776675 | 101004544 | 227870 | 1 | ENSMUST00 | 0 | Cachd1 | ENSMUSG0( | false | chr4 | 100776119 | 100778615 | 28.83684 | 94.75451 | 91.30991 | . | . | . | . | . | . |
| chr12 | 102291725 | 102292139 | 984 | Intron (ENSMUST00000056950.8 | 12 | 102283641 | 102390855 | 107215 | 1 | ENSMUST00 | 8085 | Rin3 | ENSMUSG0( | false | . | . | . | . | . | . | chr12 | 102282553 | 102299505 | 4.61003 | 3.86052 | 2.6933 |
| chr12 | 105012931 | 105013569 | 984 | Distal Intergenic | 12 | 104929949 | 105009809 | 79861 | 2 | ENSMUST00 | -3123 | Syne3 | ENSMUSG0( | false | . | . | . | . | . | . | chr12 | 105000163 | 105017783 | 5.53931 | 4.22324 | 3.47881 |
| chr19 | 8911830 | 8912118 | 984 | Exon (ENSMUST00000096246.3/ | 19 | 8920374 | 8927234 | 6861 | 1 | ENSMUST00 | -8256 | B3gat3 | ENSMUSG0( | false | . | . | . | . | . | . | chr19 | 8911015 | 8915692 | 5.21461 | 4.08859 | 3.21453 |
| chr11 | 5061789 | 5062205 | 983 | Promoter (2-3kb) | 11 | 5060470 | 5064851 | 4382 | 2 | ENSMUST00 | 2646 | Gas2l1 | ENSMUSG0( | false | chr11 | 5061516 | 5062377 | 11.38209 | 24.90129 | 22.50064 | chr11 | 5060962 | 5066029 | 6.062 | 4.51634 | 3.91645 |
| chr5 | 24750366 | 24751248 | 983 | Promoter (<=1kb) | 5 | 24751003 | 24751324 | 322 | 2 | ENSMUST00 | 76 | Crygn | ENSMUSG0( | false | chr5 | 24750232 | 24752112 | 18.39885 | 52.66533 | 49.91441 | chr5 | 24749817 | 24752596 | 3.96722 | 3.59791 | 2.16982 |
| chr3 | 122586985 | 122587352 | 983 | Intron (ENSMUST00000162409.2 | 3 | 122557910 | 122559244 | 1335 | 2 | ENSMUST00 | -27742 | Fnbp1l | ENSMUSG0( | true | . | . | . | . | . | . | chr3 | 122585238 | 122617925 | 4.82785 | 4.00989 | 2.86632 |
| chr8 | 22807848 | 22808471 | 983 | Promoter (<=1kb) | 8 | 22808275 | 22808732 | 458 | 1 | ENSMUST00 | 0 | 1700041G16 | ENSMUSG0( | false | chr8 | 22805201 | 22809095 | 30.03893 | 101.89226 | 98.2715 | chr8 | 22805049 | 22808872 | 4.2967 | 3.64033 | 2.46581 |
| chr6 | 86497581 | 86498308 | 983 | Promoter (<=1kb) | 6 | 86498781 | 86500537 | 1757 | 1 | ENSMUST00 | -473 | RP23-281H4. | ENSMUSG0( | false | chr6 | 86497025 | 86498372 | 8.56326 | 16.42445 | 14.13964 | chr6 | 86495850 | 86499944 | 4.87798 | 3.96129 | 2.95487 |
| chr11 | 74838211 | 74838766 | 982 | Promoter (<=1kb) | 11 | 74837468 | 74843775 | 6308 | 1 | ENSMUST00 | 744 | Mnt | ENSMUSG0( | false | chr11 | 74837494 | 74839155 | 33.08631 | 99.21803 | 95.66874 | . | . | . | . | . | . |
| chr8 | 72571916 | 72572480 | 982 | Promoter (<=1kb) | 8 | 72572103 | 72587282 | 15180 | 1 | ENSMUST00 | 0 | Tmem38a | ENSMUSG0( | false | chr8 | 72570135 | 72572933 | 22.15371 | 67.81625 | 64.85734 | . | . | . | . | . | . |
| chr9 | 75070954 | 75071657 | 982 | Promoter (<=1kb) | 9 | 75071015 | 75117308 | 46294 | 1 | ENSMUST00 | 0 | Myo5a | ENSMUSG0( | false | chr9 | 75070770 | 75072219 | 32.69058 | 104.59052 | 100.90593 | chr9 | 75070559 | 75072138 | 4.00295 | 3.67256 | 2.17007 |
| chr11 | 98937504 | 98937983 | 982 | Promoter (<=1kb) | 11 | 98937698 | 98974162 | 36465 | 1 | ENSMUST00 | 0 | Rara | ENSMUSG0( | false | chr11 | 98936314 | 98938742 | 41.35789 | 132.3362 | 127.75856 | chr11 | 98928668 | 98957642 | 6.70032 | 4.70048 | 4.41905 |
| chr2 | 157048462 | 157048881 | 982 | Intron (ENSMUST00000069098.6 | 2 | 157037050 | 157040239 | 3190 | 2 | ENSMUST00 | -8224 | Soga1 | ENSMUSG0( | false | . | . | . | . | . | . | chr2 | 157047505 | 157049845 | 3.42377 | 3.07497 | 1.74981 |
| chr5 | 93000790 | 93001565 | 982 | Distal Intergenic | 5 | 93013032 | 93013139 | 108 | 2 | ENSMUST00 | 11574 | Gm25521 | ENSMUSG0( | false | . | . | . | . | . | . | chr5 | 93000185 | 93002316 | 3.88932 | 3.45352 | 2.1187 |
| chr3 | 88022945 | 88023270 | 981 | Promoter (<=1kb) | 3 | 88022638 | 88023673 | 1036 | 2 | ENSMUST00 | 403 | Hapln2 | ENSMUSG0( | false | chr3 | 88022473 | 88024031 | 24.48636 | 83.88964 | 80.66612 | chr3 | 88022419 | 88024199 | 6.05367 | 4.5232 | 3.91103 |
| chr18 | 56572737 | 56573239 | 981 | Promoter (<=1kb) | 18 | 56572822 | 56587712 | 14891 | 1 | ENSMUST00 | 0 | Phax | ENSMUSG0( | false | chr18 | 56572380 | 56574181 | 24.14628 | 82.32003 | 79.12344 | . | . | . | . | . | . |
| chr15 | 84975776 | 84976103 | 981 | Intron (ENSMUST00000047144.7 | 15 | 84945720 | 84987971 | 42252 | 2 | ENSMUST00 | 11868 | 5031439G07 | ENSMUSG0( | false | . | . | . | . | . | . | chr15 | 84974957 | 84979237 | 5.23029 | 4.20331 | 3.19729 |
| chr10 | 127060120 | 127060504 | 981 | Promoter (<=1kb) | 10 | 127055011 | 127060184 | 5174 | 2 | ENSMUST00 | 0 | 39873 | ENSMUSG0( | false | chr10 | 127059084 | 127060802 | 32.66316 | 99.48935 | 95.93111 | chr10 | 127058568 | 127061029 | 4.59831 | 3.87696 | 2.68248 |
| chr2 | 113848847 | 113849171 | 981 | Promoter (<=1kb) | 2 | 113831492 | 113848661 | 17170 | 2 | ENSMUST00 | -187 | Arhgap11a | ENSMUSG0( | false | chr2 | 113847972 | 113849522 | 29.75681 | 92.45776 | 89.05831 | chr2 | 113847671 | 113849346 | 5.54748 | 4.19324 | 3.48314 |
| chr5 | 114568892 | 114569229 | 981 | Promoter (<=1kb) | 5 | 114568017 | 114594678 | 26662 | 1 | ENSMUST00 | 876 | Fam222a | ENSMUSG0( | true | chr5 | 114567128 | 114570164 | 43.41453 | 148.15424 | 142.93486 | . | . | . | . | . | . |
| chr17 | 44235089 | 44235531 | 980 | Intron (ENSMUST00000024755.5 | 17 | 44188572 | 44280168 | 91597 | 1 | ENSMUST00 | 46518 | Clic5 | ENSMUSG0( | true | . | . | . | . | . | . | chr17 | 44197784 | 44272727 | 5.95006 | 4.47222 | 3.79255 |
| chr5 | 3343626 | 3344353 | 980 | Promoter (<=1kb) | 5 | 3343893 | 3523218 | 179326 | 1 | ENSMUST00 | 0 | Cdk6 | ENSMUSG0( | false | chr5 | 3341034 | 3344969 | 32.55001 | 105.36325 | 101.65399 | chr5 | 3341133 | 3344421 | 4.3677 | 3.73533 | 2.50877 |
| chr19 | 7134538 | 7135048 | 980 | Intron (ENSMUST00000040261.5 | 19 | 7092014 | 7105729 | 13716 | 2 | ENSMUST00 | -28810 | Flrt1 | ENSMUSG0( | false | . | . | . | . | . | . | chr19 | 7127706 | 7162730 | 7.6862 | 5.1249 | 5.24023 |
| chr15 | 102470860 | 102471442 | 980 | Promoter (<=1kb) | 15 | 102470632 | 102500059 | 29428 | 1 | ENSMUST00 | 229 | Pcbp2 | ENSMUSG0( | false | chr15 | 102470489 | 102471870 | 22.78592 | 76.10319 | 73.01487 | . | . | . | . | . | . |
| chr11 | 22639153 | 22639687 | 980 | Distal Intergenic | 11 | 22626375 | 22626664 | 290 | 2 | ENSMUST00 | -12490 | Gm12053 | ENSMUSG0( | false | . | . | . | . | . | . | chr11 | 22635852 | 22640892 | 6.051 | 4.54017 | 3.86945 |
| chr10 | 95414301 | 95414647 | 979 | Promoter (<=1kb) | 10 | 95412420 | 95414827 | 2408 | 2 | ENSMUST00 | 180 | Socs2 | ENSMUSG0( | false | chr10 | 95414179 | 95418016 | 35.49308 | 110.49252 | 106.63927 | chr10 | 95413852 | 95418311 | 4.09983 | 3.68592 | 2.25149 |
| chr11 | 98022429 | 98022989 | 979 | Promoter (<=1kb) | 11 | 98001508 | 98022634 | 21127 | 2 | ENSMUST00 | 0 | Cacnb1 | ENSMUSG0( | false | chr11 | 98021697 | 98023289 | 34.78613 | 113.42423 | 109.48744 | chr11 | 98018411 | 98024278 | 4.45887 | 3.76503 | 2.58806 |
| chr12 | 108334066 | 108334473 | 979 | Promoter (<=1kb) | 12 | 108334377 | 108362231 | 27855 | 1 | ENSMUST00 | 0 | Cyp46a1 | ENSMUSG0( | false | chr12 | 108334256 | 108334858 | 23.61999 | 85.9385 | 82.67356 | chr12 | 108332878 | 108334952 | 6.75437 | 5.01921 | 4.43145 |
| chr9 | 37238937 | 37239238 | 979 | Exon (ENSMUST00000115068.3/ | 9 | 37230512 | 37235154 | 4643 | 2 | ENSMUST00 | -3784 | Slc37a2 | ENSMUSG0( | false | . | . | . | . | . | . | chr9 | 37237899 | 37240331 | 5.29119 | 4.18586 | 3.28128 |
| chr7 | 34812612 | 34813055 | 979 | Promoter (<=1kb) | 7 | 34674468 | 34812686 | 138219 | 2 | ENSMUST00 | 0 | Chst8 | ENSMUSG0( | false | chr7 | 34811244 | 34813377 | 19.04495 | 59.55748 | 56.71581 | chr7 | 34810803 | 34813269 | 4.14049 | 3.64326 | 2.30862 |
| chr3 | 51721587 | 51721886 | 979 | Intron (ENSMUST00000121440.2 | 3 | 51693771 | 51695090 | 1320 | 1 | ENSMUST00 | 27817 | Gm10729 | ENSMUSG0( | false | . | . | . | . | . | . | chr3 | 51705501 | 51729803 | 5.39003 | 4.2072 | 3.33836 |
| chr17 | 34121969 | 34122917 | 978 | Promoter (<=1kb) | 17 | 34112023 | 34122317 | 10295 | 2 | ENSMUST00 | 0 | Brd2 | ENSMUSG0( | false | chr17 | 34119478 | 34123860 | 39.82221 | 128.59171 | 124.15125 | . | . | . | . | . | . |
| chr11 | 76285468 | 76285804 | 978 | Intron (ENSMUST00000021204.3 | 11 | 76249212 | 76250508 | 1297 | 1 | ENSMUST00 | 36257 | Rnmtl1 | ENSMUSG0( | false | . | . | . | . | . | . | chr11 | 76283200 | 76297836 | 5.96193 | 4.42599 | 3.81873 |
| chr4 | 152253435 | 152253790 | 978 | Intron (ENSMUST00000075363.4 | 4 | 152274232 | 152285337 | 11106 | 1 | ENSMUST00 | -20442 | Gpr153 | ENSMUSG0( | false | . | . | . | . | . | . | chr4 | 152251696 | 152254534 | 5.57038 | 4.28494 | 3.47049 |
| chr7 | 99559221 | 99559574 | 978 | Intron (ENSMUST00000098266.3 | 7 | 99552269 | 99552363 | 95 | 1 | ENSMUST00 | 6953 | Mir326 | ENSMUSG0( | false | . | . | . | . | . | . | chr7 | 99536482 | 99576026 | 7.32182 | 4.95837 | 4.9327 |
| chr4 | 107130598 | 107130912 | 978 | Intron (ENSMUST00000145510.1 | 4 | 107126613 | 107136252 | 9640 | 1 | ENSMUST00 | 3986 | Gm12802 | ENSMUSG0( | false | . | . | . | . | . | . | chr4 | 107130414 | 107130874 | 4.65336 | 3.98193 | 2.75281 |
| chr10 | 121069871 | 121070480 | 977 | Intron (ENSMUST00000020439.5 | 10 | 121083755 | 121100637 | 16883 | 1 | ENSMUST00 | -13275 | Wif1 | ENSMUSG0( | false | . | . | . | . | . | . | . | . | . | . | . | . |
| chr2 | 166031161 | 166031506 | 977 | Intron (ENSMUST00000109252.2 | 2 | 166047918 | 166052398 | 4481 | 1 | ENSMUST00 | -16412 | Ncoa3 | ENSMUSG0( | false | chr2 | 166030178 | 166031780 | 12.3333 | 26.50278 | 24.08123 | chr2 | 166014974 | 166032775 | 4.52076 | 3.84017 | 2.61642 |
| chr5 | 121586664 | 121587068 | 977 | Intron (ENSMUST00000031411.9 | 5 | 121570312 | 121593580 | 23269 | 2 | ENSMUST00 | 6512 | Aldh2 | ENSMUSG0( | false | . | . | . | . | . | . | chr5 | 121586570 | 121587447 | 4.96793 | 4.02974 | 3.01644 |
| chr3 | 152395666 | 152396764 | 977 | Promoter (<=1kb) | 3 | 152396003 | 152462826 | 66824 | 1 | ENSMUST00 | 0 | Zzz3 | ENSMUSG0( | false | chr3 | 152395164 | 152397484 | 30.59503 | 95.89513 | 92.42037 | . | . | . | . | . | . |
| chr3 | 107332355 | 107332944 | 977 | Promoter (<=1kb) | 3 | 107326110 | 107333289 | 7180 | 2 | ENSMUST00 | 345 | Rbm15 | ENSMUSG0( | false | chr3 | 107331120 | 107333142 | 38.99987 | 144.77138 | 139.69241 | . | . | . | . | . | . |
| chr15 | 103085843 | 103086800 | 977 | Distal Intergenic | 15 | 103096151 | 103102111 | 5961 | 2 | ENSMUST00 | 15311 | RP24-89G21 | ENSMUSG0( | false | chr15 | 103085884 | 103086945 | 16.53312 | 44.67334 | 42.02541 | chr15 | 103085467 | 103091146 | 6.12281 | 4.30788 | 3.96408 |
| chr3 | 117868697 | 117869196 | 976 | Promoter (<=1kb) | 3 | 117781497 | 117868936 | 87440 | 2 | ENSMUST00 | 0 | Snx7 | ENSMUSG0( | false | chr3 | 117868381 | 117869632 | 24.03115 | 75.67385 | 72.59326 | . | . | . | . | . | . |
| chr8 | 92355696 | 92356422 | 976 | Promoter (<=1kb) | 8 | 92326038 | 92355831 | 29794 | 2 | ENSMUST00 | 0 | Crnde | ENSMUSG0( | false | chr8 | 92355053 | 92358743 | 27.29887 | 88.10642 | 84.80337 | chr8 | 92354184 | 92360798 | 6.77952 | 4.77718 | 4.47075 |
| chr9 | 116910018 | 116910363 | 976 | Intron (ENSMUST00000111773.4 | 9 | 116582822 | 116822819 | 239998 | 2 | ENSMUST00 | -87200 | Rbms3 | ENSMUSG0( | true | . | . | . | . | . | . | chr9 | 116903777 | 116913504 | 4.73906 | 3.97114 | 2.79562 |
| chr7 | 80044189 | 80044962 | 976 | Intron (ENSMUST00000164056.3 | 7 | 80026418 | 80026513 | 96 | 2 | ENSMUST00 | -17677 | Gm27983 | ENSMUSG0( | false | . | . | . | . | . | . | chr7 | 80043846 | 80073175 | 7.55169 | 5.20662 | 5.10629 |
| chr14 | 76648468 | 76648765 | 976 | Distal Intergenic | 14 | 76672963 | 76687169 | 14207 | 1 | ENSMUST00 | -24198 | RP23-421C2( | ENSMUSG0( | false | . | . | . | . | . | . | . | . | . | . | . | . |
| chr17 | 31277007 | 31277566 | 975 | Promoter (<=1kb) | 17 | 31255019 | 31277356 | 22338 | 2 | ENSMUST00 | 0 | Rsph1 | ENSMUSG0( | false | chr17 | 31276940 | 31277678 | 10.07419 | 29.06419 | 26.60987 | . | . | . | . | . | . |
| chr4 | 43267000 | 43267580 | 975 | Promoter (<=1kb) | 4 | 43267159 | 43344582 | 77424 | 1 | ENSMUST00 | 0 | Atp8b5 | ENSMUSG0( | false | chr4 | 43266935 | 43267554 | 9.06831 | 19.28032 | 16.9548 | chr4 | 43266860 | 43268125 | 4.28094 | 3.73493 | 2.45663 |
| chr18 | 80046689 | 80047189 | 975 | Promoter (<=1kb) | 18 | 80046895 | 80119640 | 72746 | 1 | ENSMUST00 | 0 | Pard6g | ENSMUSG0( | false | chr18 | 80046481 | 80048014 | 25.15761 | 80.47005 | 77.31023 | chr18 | 80046567 | 80049731 | 4.11742 | 3.63365 | 2.26569 |
| chr11 | 88488585 | 88488870 | 975 | Intron (ENSMUST00000107909.2 | 11 | 88432986 | 88433514 | 529 | 2 | ENSMUST00 | -55072 | Gm11510 | ENSMUSG0( | false | . | . | . | . | . | . | chr11 | 88480578 | 88492236 | 6.52217 | 4.75954 | 4.2595 |
| chr16 | 35215213 | 35215711 | 975 | Intron (ENSMUST00000114913.1 | 16 | 35234796 | 35234923 | 128 | 2 | ENSMUST00 | 19212 | Gm25179 | ENSMUSG0( | false | . | . | . | . | . | . | chr16 | 35214600 | 35217040 | 5.77354 | 4.38929 | 3.68192 |
| chr13 | 44228544 | 44228873 | 975 | Distal Intergenic | 13 | 44121294 | 44125179 | 3886 | 1 | ENSMUST00 | 107251 | Gm5083 | ENSMUSG0( | false | chr13 | 44228751 | 44229291 | 6.37612 | 11.07527 | 8.87976 | chr13 | 44226456 | 44236587 | 5.70374 | 4.41191 | 3.58124 |
| chr12 | 55491758 | 55492350 | 974 | Promoter (<=1kb) | 12 | 55489411 | 55492647 | 3237 | 2 | ENSMUST00 | 297 | Nfkbia | ENSMUSG0( | false | chr12 | 55490207 | 55493180 | 37.2221 | 115.56093 | 111.55734 | . | . | . | . | . | . |
| chr18 | 74688463 | 74689132 | 974 | Intron (ENSMUST00000074157.7 | 18 | 74707506 | 74717075 | 9570 | 1 | ENSMUST00 | -18374 | Myo5b | ENSMUSG0( | false | . | . | . | . | . | . | chr18 | 74687877 | 74689308 | 4.02817 | 3.62078 | 2.20685 |
| chr9 | 108795767 | 108796173 | 974 | Promoter (<=1kb) | 9 | 108795994 | 108806333 | 10340 | 1 | ENSMUST00 | 0 | Ip6k2 | ENSMUSG0( | false | chr9 | 108795574 | 108798004 | 15.53947 | 48.13258 | 45.44172 | chr9 | 108784542 | 108804834 | 4.97749 | 3.96237 | 3.01028 |
| chr11 | 5184224 | 5184521 | 974 | Distal Intergenic | 11 | 5106265 | 5152257 | 45993 | 2 | ENSMUST00 | -31968 | Emid1 | ENSMUSG0( | false | . | . | . | . | . | . | chr11 | 5181494 | 5198688 | 5.21662 | 4.12982 | 3.19453 |
| chr1 | 161034380 | 161035155 | 974 | Promoter (<=1kb) | 1 | 161008794 | 161034849 | 26056 | 2 | ENSMUST00 | 0 | Zbtb37 | ENSMUSG0( | false | chr1 | 161033938 | 161036613 | 33.32351 | 103.65904 | 99.99326 | . | . | . | . | . | . |
| chr2 | 173737392 | 173737825 | 974 | Promoter (<=1kb) | 2 | 173737511 | 173784336 | 46826 | 1 | ENSMUST00 | 0 | Vapb | ENSMUSG0( | false | chr2 | 173737113 | 173738489 | 36.30304 | 111.89011 | 107.99921 | chr2 | 173736858 | 173745093 | 5.38519 | 4.22381 | 3.32635 |
| chr6 | 124790187 | 124790515 | 973 | Distal Intergenic | 6 | 124769869 | 124779727 | 9859 | 2 | ENSMUST00 | -10461 | Lrrc23 | ENSMUSG0( | false | . | . | . | . | . | . | chr6 | 124788459 | 124792292 | 3.65803 | 3.448 | 1.93739 |
| chr5 | 103423523 | 103423886 | 973 | Promoter (1-2kb) | 5 | 103425192 | 103598359 | 173168 | 1 | ENSMUST00 | -1306 | Ptpn13 | ENSMUSG0( | false | . | . | . | . | . | . | chr5 | 103423670 | 103424731 | 6.06667 | 4.62214 | 3.89016 |
| chr11 | 104246312 | 104246772 | 973 | Intron (ENSMUST00000106992.4 | 11 | 104251012 | 104251144 | 133 | 2 | ENSMUST00 | 4372 | Gm22743 | ENSMUSG0( | false | . | . | . | . | . | . | chr11 | 104242646 | 104282246 | 6.16611 | 4.45267 | 3.98703 |
| chr18 | 66210579 | 66211003 | 973 | Intron (ENSMUST00000146610.1 | 18 | 66219264 | 66219333 | 70 | 2 | ENSMUST00 | 8330 | Mir694 | ENSMUSG0( | false | . | . | . | . | . | . | chr18 | 66208792 | 66211535 | 4.66329 | 3.94931 | 2.73468 |
| chr13 | 32601326 | 32601685 | 973 | Distal Intergenic | 13 | 32758202 | 32758310 | 109 | 1 | ENSMUST00 | -156517 | Gm27342 | ENSMUSG0( | false | . | . | . | . | . | . | chr13 | 32600225 | 32603919 | 5.20923 | 4.26392 | 3.17852 |
| chr11 | 7114381 | 7114730 | 972 | Intron (ENSMUST00000020706.4 | 11 | 7159882 | 7172879 | 12998 | 1 | ENSMUST00 | -45152 | Adcy1 | ENSMUSG0( | true | . | . | . | . | . | . | chr11 | 7111251 | 7145535 | 5.23684 | 4.17059 | 3.20426 |
| chr8 | 71485844 | 71486180 | 972 | Promoter (<=1kb) | 8 | 71476023 | 71486067 | 10045 | 2 | ENSMUST00 | 0 | Ano8 | ENSMUSG0( | false | chr8 | 71485346 | 71486066 | 22.44583 | 74.56482 | 71.50171 | chr8 | 71485439 | 71487703 | 4.21578 | 3.55568 | 2.39503 |
| chr18 | 65565311 | 65565983 | 972 | Distal Intergenic | 18 | 65580230 | 65689443 | 109214 | 1 | ENSMUST00 | -14247 | Zfp532 | ENSMUSG0( | false | . | . | . | . | . | . | chr18 | 65565238 | 65565917 | 4.23228 | 3.42875 | 2.40705 |
| chr7 | 35753968 | 35754593 | 972 | Promoter (<=1kb) | 7 | 35685165 | 35754454 | 69290 | 2 | ENSMUST00 | 0 | Dpy19l3 | ENSMUSG0( | false | chr7 | 35753775 | 35754912 | 27.88725 | 99.90214 | 96.33501 | chr7 | 35752993 | 35755046 | 4.29668 | 3.78596 | 2.41652 |
| chr14 | 56575394 | 56575895 | 972 | Promoter (<=1kb) | 14 | 56575619 | 56659794 | 84176 | 1 | ENSMUST00 | 0 | Parp4 | ENSMUSG0( | true | chr14 | 56575064 | 56576957 | 36.25406 | 119.15681 | 115.04134 | . | . | . | . | . | . |
| chr5 | 124875584 | 124875956 | 972 | Promoter (<=1kb) | 5 | 124875970 | 124885163 | 9194 | 1 | ENSMUST00 | -14 | Zfp664 | ENSMUSG0( | false | . | . | . | . | . | . | chr5 | 124870149 | 124879957 | 4.56966 | 3.79154 | 2.67149 |
| chr6 | 87653978 | 87654385 | 971 | Intron (ENSMUST00000032130.3 | 6 | 87628429 | 87672168 | 43740 | 2 | ENSMUST00 | 17783 | Aplf | ENSMUSG0( | false | . | . | . | . | . | . | chr6 | 87652581 | 87655651 | 5.36724 | 4.29294 | 3.33897 |
| chr1 | 74083098 | 74083547 | 971 | Intron (ENSMUST00000169786.2 | 1 | 73910232 | 74091351 | 181120 | 2 | ENSMUST00 | 7804 | Tns1 | ENSMUSG0( | false | . | . | . | . | . | . | chr1 | 74081332 | 74083308 | 5.0869 | 3.92899 | 3.09986 |
| chr5 | 150215994 | 150216320 | 971 | Distal Intergenic | 5 | 150259930 | 150497752 | 237823 | 1 | ENSMUST00 | -43610 | Fry | ENSMUSG0( | true | . | . | . | . | . | . | chr5 | 150213346 | 150216310 | 4.23389 | 3.66216 | 2.42818 |
| chr15 | 82212708 | 82213023 | 971 | Promoter (<=1kb) | 15 | 82206952 | 82212815 | 5864 | 2 | ENSMUST00 | 0 | AI848285 | ENSMUSG0( | false | chr15 | 82211767 | 82213072 | 18.38128 | 45.59164 | 42.93153 | chr15 | 82211519 | 82213435 | 5.33302 | 4.22274 | 3.2939 |
| chr7 | 143445658 | 143446207 | 971 | Promoter (1-2kb) | 7 | 143447605 | 143447665 | 61 | 1 | ENSMUST00 | -1398 | Gm27901 | ENSMUSG0( | false | chr7 | 143444764 | 143446666 | 17.60262 | 45.78123 | 43.11721 | chr7 | 143437483 | 143448788 | 5.47047 | 4.33141 | 3.38873 |
| chr3 | 85346766 | 85347138 | 971 | Intron (ENSMUST00000154148.2 | 3 | 85360228 | 85361174 | 947 | 2 | ENSMUST00 | 14036 | RP23-94M23 | ENSMUSG0( | false | . | . | . | . | . | . | chr3 | 85346536 | 85348549 | 2.99389 | 2.92302 | 1.36838 |
| chr4 | 94560005 | 94560296 | 970 | Distal Intergenic | 4 | 94500081 | 94556796 | 56716 | 2 | ENSMUST00 | -3210 | Caap1 | ENSMUSG0( | false | . | . | . | . | . | . | chr4 | 94558911 | 94563213 | 4.41548 | 3.77841 | 2.54747 |
| chr13 | 70794012 | 70794617 | 970 | Intron (ENSMUST00000080145.7 | 13 | 70727810 | 70841775 | 113966 | 2 | ENSMUST00 | 47158 | Adamts16 | ENSMUSG0( | false | . | . | . | . | . | . | chr13 | 70792267 | 70795677 | 4.55535 | 3.8229 | 2.687 |
| chr12 | 40798534 | 40798889 | 970 | Intron (ENSMUST00000037488.6 | 12 | 41024090 | 41955588 | 931499 | 1 | ENSMUST00 | -225201 | Immp2l | ENSMUSG0( | false | . | . | . | . | . | . | chr12 | 40796973 | 40799899 | 5.35922 | 4.23795 | 3.33367 |
| chr3 | 9800253 | 9800566 | 970 | Intron (ENSMUST00000161949.2 | 3 | 9829620 | 9829728 | 109 | 2 | ENSMUST00 | 29162 | Gm24786 | ENSMUSG0( | false | . | . | . | . | . | . | chr3 | 9778824 | 9802371 | 5.7635 | 4.36373 | 3.65538 |
| chr15 | 88837385 | 88837766 | 970 | Promoter (2-3kb) | 15 | 88835300 | 88835406 | 107 | 1 | ENSMUST00 | 2086 | Gm23144 | ENSMUSG0( | false | chr15 | 88837190 | 88838108 | 17.92175 | 44.05972 | 41.41886 | chr15 | 88836763 | 88838109 | 3.56248 | 3.22659 | 1.84462 |
| chr15 | 38661651 | 38662130 | 969 | Promoter (<=1kb) | 15 | 38661904 | 38692443 | 30540 | 1 | ENSMUST00 | 0 | Atp6v1c1 | ENSMUSG0( | false | chr15 | 38661292 | 38662773 | 42.74922 | 148.11885 | 142.90198 | chr15 | 38661162 | 38664135 | 4.10141 | 3.53423 | 2.29322 |
| chr19 | 56997260 | 56997641 | 969 | Intron (ENSMUST00000118800.2 | 19 | 56930098 | 57001046 | 70949 | 2 | ENSMUST00 | 3405 | Afap1l2 | ENSMUSG0( | false | . | . | . | . | . | . | chr19 | 56996211 | 56998015 | 4.21007 | 3.66743 | 2.35357 |
| chr6 | 143153685 | 143154055 | 969 | Distal Intergenic | 6 | 143167230 | 143208542 | 41313 | 1 | ENSMUST00 | -13175 | Etnk1 | ENSMUSG0( | false | . | . | . | . | . | . | chr6 | 143153717 | 143154213 | 2.96539 | 2.81999 | 1.38119 |
| chr17 | 33757175 | 33757538 | 969 | Promoter (2-3kb) | 17 | 33759992 | 33761870 | 1879 | 1 | ENSMUST00 | -2454 | Gm17251 | ENSMUSG0( | false | . | . | . | . | . | . | chr17 | 33750126 | 33759845 | 7.3113 | 4.95775 | 4.92322 |
| chr15 | 101078429 | 101079011 | 969 | Distal Intergenic | 15 | 101050194 | 101054399 | 4206 | 2 | ENSMUST00 | -24031 | Fignl2 | ENSMUSG0( | false | chr15 | 101077078 | 101079386 | 32.23851 | 100.76218 | 97.17527 | . | . | . | . | . | . |
| chr1 | 75250587 | 75250893 | 969 | Promoter (1-2kb) | 1 | 75252322 | 75257427 | 5106 | 1 | ENSMUST00 | -1429 | RP23-321N2. | ENSMUSG0( | false | . | . | . | . | . | . | chr1 | 75247333 | 75252393 | 6.77466 | 4.89628 | 4.45166 |
| chr11 | 95000498 | 95000865 | 968 | Intron (ENSMUST00000038696.6 | 11 | 94996064 | 95006899 | 10836 | 1 | ENSMUST00 | 4435 | Ppp1r9b | ENSMUSG0( | false | . | . | . | . | . | . | chr11 | 94992312 | 95001916 | 5.70711 | 4.31596 | 3.59817 |
| chr19 | 29805398 | 29806752 | 968 | Promoter (<=1kb) | 19 | 29714402 | 29805507 | 91106 | 2 | ENSMUST00 | 0 | 9930021J03F | ENSMUSG0( | false | chr19 | 29804622 | 29807012 | 40.11768 | 137.88335 | 133.09009 | . | . | . | . | . | . |
| chr19 | 34845204 | 34845481 | 968 | Intron (ENSMUST00000036584.6 | 19 | 34820687 | 34820773 | 87 | 2 | ENSMUST00 | -24432 | Mir107 | ENSMUSG0( | false | . | . | . | . | . | . | chr19 | 34841223 | 34848354 | 5.17461 | 4.07032 | 3.17219 |
| chr4 | 43651485 | 43651825 | 968 | Promoter (<=1kb) | 4 | 43651335 | 43652159 | 825 | 2 | ENSMUST00 | 334 | Spag8 | ENSMUSG0( | false | . | . | . | . | . | . | chr4 | 43651237 | 43652136 | 4.28341 | 3.76126 | 2.42291 |
| chr11 | 95765399 | 95765765 | 968 | 3' UTR | 11 | 95761491 | 95761665 | 175 | 1 | ENSMUST00 | 3909 | Polr2k-ps | ENSMUSG0( | false | . | . | . | . | . | . | chr11 | 95762899 | 95766177 | 4.89411 | 3.98606 | 2.93979 |
| chr13 | 37451664 | 37452729 | 968 | Promoter (<=1kb) | 13 | 37452212 | 37453208 | 997 | 2 | ENSMUST00 | 479 | RP23-21N15 | ENSMUSG0( | false | chr13 | 37452098 | 37452559 | 8.50221 | 18.80243 | 16.48241 | chr13 | 37449991 | 37457701 | 5.94961 | 4.52437 | 3.79226 |
| chr3 | 138767174 | 138767609 | 967 | Intron (ENSMUST00000029800.3 | 3 | 138778206 | 138902494 | 124289 | 1 | ENSMUST00 | -10597 | Tspan5 | ENSMUSG0( | true | . | . | . | . | . | . | chr3 | 138763693 | 138776582 | 5.61138 | 4.35746 | 3.51014 |
| chr4 | 84674719 | 84675302 | 967 | Promoter (<=1kb) | 4 | 84275095 | 84675034 | 399940 | 2 | ENSMUST00 | 0 | Bnc2 | ENSMUSG0( | false | chr4 | 84674350 | 84676199 | 18.33657 | 60.6815 | 57.82544 | chr4 | 84674065 | 84676172 | 3.68811 | 3.35717 | 1.9605 |
| chr9 | 56123045 | 56123333 | 967 | Promoter (<=1kb) | 9 | 56123055 | 56124047 | 993 | 1 | ENSMUST00 | 0 | Pstpip1 | ENSMUSG0( | false | . | . | . | . | . | . | chr9 | 56122553 | 56125978 | 4.82223 | 3.98475 | 2.86909 |
| chr15 | 32171741 | 32172101 | 967 | Distal Intergenic | 15 | 32177289 | 32178296 | 1008 | 1 | ENSMUST00 | -5188 | Tas2r119 | ENSMUSG0( | false | . | . | . | . | . | . | chr15 | 32169672 | 32172579 | 3.91046 | 3.50655 | 2.15888 |
| chr7 | 77303959 | 77304576 | 967 | Distal Intergenic | 7 | 77910140 | 77910320 | 181 | 2 | ENSMUST00 | 605744 | Gm23239 | ENSMUSG0( | false | . | . | . | . | . | . | chr7 | 77301863 | 77306899 | 5.60865 | 4.416 | 3.50558 |
| chr1 | 41431118 | 41431413 | 967 | Distal Intergenic | 1 | 41526931 | 41529543 | 2613 | 2 | ENSMUST00 | 98130 | RP23-59E7.1 | ENSMUSG0( | false | . | . | . | . | . | . | chr1 | 41430728 | 41431785 | 3.41351 | 3.26978 | 1.71744 |
| chr9 | 44320123 | 44320636 | 966 | Promoter (<=1kb) | 9 | 44309237 | 44320282 | 11046 | 2 | ENSMUST00 | 0 | C2cd2l | ENSMUSG0( | false | chr9 | 44318922 | 44320981 | 39.57526 | 126.00172 | 121.65485 | chr9 | 44317736 | 44321169 | 6.42376 | 4.74806 | 4.18868 |
| chr10 | 42260565 | 42260921 | 966 | Promoter (2-3kb) | 10 | 42197554 | 42258366 | 60813 | 2 | ENSMUST00 | -2200 | Foxo3 | ENSMUSG0( | false | . | . | . | . | . | . | chr10 | 42243354 | 42275609 | 6.05342 | 4.50564 | 3.88797 |
| chr9 | 56635592 | 56636056 | 966 | Promoter (<=1kb) | 9 | 56619088 | 56635628 | 16541 | 2 | ENSMUST00 | 0 | Lingo1 | ENSMUSG0( | false | chr9 | 56634833 | 56636173 | 23.4661 | 79.19912 | 76.05865 | chr9 | 56634857 | 56638086 | 4.71943 | 3.97843 | 2.76558 |
| chr2 | 160433223 | 160433669 | 966 | Distal Intergenic | 2 | 160363703 | 160367065 | 3363 | 2 | ENSMUST00 | -66159 | Mafb | ENSMUSG0( | true | . | . | . | . | . | . | chr2 | 160432610 | 160434900 | 5.31291 | 3.98525 | 3.30134 |
| chr15 | 37577106 | 37577338 | 966 | Intron (ENSMUST00000168992.2 | 15 | 37603415 | 37603534 | 120 | 2 | ENSMUST00 | 26196 | Gm24833 | ENSMUSG0( | false | . | . | . | . | . | . | chr15 | 37577005 | 37578071 | 4.69338 | 3.94528 | 2.77292 |
| chr7 | 19308526 | 19308969 | 965 | Promoter (1-2kb) | 7 | 19302721 | 19310045 | 7325 | 2 | ENSMUST00 | 1076 | Fosb | ENSMUSG0( | false | chr7 | 19307857 | 19310719 | 23.80619 | 80.75649 | 77.58801 | chr7 | 19307852 | 19310879 | 4.30679 | 3.70681 | 2.44358 |
| chr18 | 5063252 | 5063636 | 965 | Promoter (<=1kb) | 18 | 5064472 | 5119291 | 54820 | 1 | ENSMUST00 | -836 | Svil | ENSMUSG0( | false | . | . | . | . | . | . | chr18 | 5063023 | 5064824 | 5.13156 | 4.06977 | 3.17116 |
| chr1 | 162590460 | 162590775 | 965 | Intron (ENSMUST00000132158.2 | 1 | 162574821 | 162594753 | 19933 | 1 | ENSMUST00 | 15640 | Vamp4 | ENSMUSG0( | true | . | . | . | . | . | . | chr1 | 162587733 | 162591044 | 4.60166 | 3.90294 | 2.67356 |
| chr6 | 92446839 | 92447139 | 965 | Intron (ENSMUST00000032093.6 | 6 | 92375384 | 92481392 | 106009 | 2 | ENSMUST00 | 34253 | Prickle2 | ENSMUSG0( | false | . | . | . | . | . | . | chr6 | 92445231 | 92448610 | 4.30401 | 3.72548 | 2.44321 |
| chr14 | 61012365 | 61012703 | 965 | Intron (ENSMUST00000111236.2 | 14 | 60963834 | 61037987 | 74154 | 2 | ENSMUST00 | 25284 | Tnfrsf19 | ENSMUSG0( | false | chr14 | 61012395 | 61012987 | 10.7597 | 23.84104 | 21.45527 | chr14 | 61012247 | 61013761 | 3.36381 | 3.24929 | 1.6798 |
| chr2 | 159575346 | 159575722 | 965 | Distal Intergenic | 2 | 159456170 | 159456457 | 288 | 2 | ENSMUST00 | -118890 | Gm14219 | ENSMUSG0( | false | . | . | . | . | . | . | chr2 | 159574796 | 159576383 | 3.72618 | 3.37928 | 1.97394 |
| chr10 | 54075406 | 54076119 | 964 | Promoter (<=1kb) | 10 | 53904788 | 54075796 | 171009 | 2 | ENSMUST00 | 0 | Man1a | ENSMUSG0( | false | chr10 | 54074002 | 54076827 | 34.34893 | 131.73137 | 127.17394 | . | . | . | . | . | . |
| chr14 | 31494772 | 31495234 | 964 | Promoter (<=1kb) | 14 | 31495079 | 31509858 | 14780 | 1 | ENSMUST00 | 0 | Eaf1 | ENSMUSG0( | false | chr14 | 31493999 | 31495974 | 21.7465 | 82.16968 | 78.97962 | . | . | . | . | . | . |
| chr9 | 63856454 | 63856761 | 964 | Distal Intergenic | 9 | 63646767 | 63757994 | 111228 | 2 | ENSMUST00 | -98461 | Smad3 | ENSMUSG0( | false | . | . | . | . | . | . | chr9 | 63856070 | 63857723 | 4.65141 | 3.90351 | 2.71328 |
| chr5 | 33224526 | 33224862 | 964 | Distal Intergenic | 5 | 33213518 | 33218238 | 4721 | 2 | ENSMUST00 | -6289 | Spon2 | ENSMUSG0( | false | . | . | . | . | . | . | chr5 | 33221817 | 33232043 | 5.32048 | 4.18629 | 3.28315 |
| chr9 | 106788912 | 106789364 | 964 | Promoter (<=1kb) | 9 | 106688082 | 106789194 | 101113 | 2 | ENSMUST00 | 0 | Rad54l2 | ENSMUSG0( | false | chr9 | 106788240 | 106789925 | 26.2459 | 113.32486 | 109.39578 | chr9 | 106783504 | 106790006 | 3.25538 | 3.14663 | 1.56876 |
| chr18 | 84234687 | 84234992 | 964 | Intron (ENSMUST00000125450.3 | 18 | 84088158 | 84097514 | 9357 | 1 | ENSMUST00 | 146530 | Zadh2 | ENSMUSG0( | false | . | . | . | . | . | . | chr18 | 84234233 | 84237935 | 4.62628 | 3.9182 | 2.7086 |
| chr5 | 135040862 | 135041179 | 963 | Promoter (<=1kb) | 5 | 135041060 | 135051099 | 10040 | 1 | ENSMUST00 | 0 | Stx1a | ENSMUSG0( | false | . | . | . | . | . | . | chr5 | 135038993 | 135042779 | 5.94879 | 4.5474 | 3.78926 |
| chr12 | 80114098 | 80114698 | 963 | Promoter (1-2kb) | 12 | 80107760 | 80113013 | 5254 | 2 | ENSMUST00 | -1086 | Zfp36l1 | ENSMUSG0( | false | chr12 | 80113160 | 80116456 | 21.91337 | 68.25864 | 65.29428 | chr12 | 80113722 | 80114749 | 4.8306 | 4.07928 | 2.85837 |
| chr16 | 58727779 | 58728311 | 963 | Promoter (<=1kb) | 16 | 58727910 | 58734251 | 6342 | 1 | ENSMUST00 | 0 | Cldn25 | ENSMUSG0( | false | chr16 | 58727491 | 58729008 | 25.84672 | 90.2271 | 86.88143 | . | . | . | . | . | . |
| chr19 | 37207438 | 37207925 | 963 | Promoter (<=1kb) | 19 | 37207543 | 37222139 | 14597 | 1 | ENSMUST00 | 0 | 38412 | ENSMUSG0( | false | chr19 | 37206366 | 37209314 | 29.18609 | 122.55783 | 118.32927 | . | . | . | . | . | . |
| chr11 | 121510574 | 121510910 | 963 | Intron (ENSMUST00000103013.4 | 11 | 121510978 | 121519333 | 8356 | 2 | ENSMUST00 | 8423 | Zfp750 | ENSMUSG0( | false | . | . | . | . | . | . | chr11 | 121507341 | 121511263 | 4.40132 | 3.80131 | 2.55836 |
| chr10 | 86128180 | 86128454 | 963 | Promoter (2-3kb) | 10 | 86125827 | 86129854 | 4028 | 1 | ENSMUST00 | 2354 | Gm15990 | ENSMUSG0( | false | . | . | . | . | . | . | chr10 | 86122373 | 86133417 | 5.202 | 4.08244 | 3.19933 |
| chr11 | 88114771 | 88115182 | 962 | Intron (ENSMUST00000018522.7 | 11 | 88099290 | 88185365 | 86076 | 1 | ENSMUST00 | 15482 | Cuedc1 | ENSMUSG0( | true | . | . | . | . | . | . | chr11 | 88103459 | 88120719 | 6.48176 | 4.71748 | 4.23259 |
| chr11 | 105311398 | 105311889 | 962 | Intron (ENSMUST00000126931.1 | 11 | 105329052 | 105332299 | 3248 | 1 | ENSMUST00 | -17163 | Mrc2 | ENSMUSG0( | true | . | . | . | . | . | . | chr11 | 105306066 | 105319029 | 6.36006 | 4.57632 | 4.15147 |
| chr3 | 41082935 | 41083538 | 962 | Promoter (<=1kb) | 3 | 41066326 | 41083046 | 16721 | 2 | ENSMUST00 | 0 | Pgrmc2 | ENSMUSG0( | false | chr3 | 41081402 | 41083110 | 39.37477 | 126.73389 | 122.35824 | . | . | . | . | . | . |
| chr9 | 70678701 | 70679258 | 962 | Promoter (<=1kb) | 9 | 70678997 | 70778382 | 99386 | 1 | ENSMUST00 | 0 | Adam10 | ENSMUSG0( | false | chr9 | 70678383 | 70678768 | 21.75113 | 60.20329 | 57.35284 | . | . | . | . | . | . |
| chr2 | 152933116 | 152933409 | 962 | Promoter (<=1kb) | 2 | 152931898 | 152933208 | 1311 | 2 | ENSMUST00 | 0 | Foxs1 | ENSMUSG0( | false | chr2 | 152931092 | 152933192 | 16.52141 | 45.4089 | 42.75185 | chr2 | 152925247 | 152934463 | 6.47975 | 4.78702 | 4.2189 |
| chr11 | 103097181 | 103097798 | 961 | Intron (ENSMUST00000103077.1 | 11 | 103081730 | 103101591 | 19862 | 2 | ENSMUST00 | 3793 | Plcd3 | ENSMUSG0( | false | . | . | . | . | . | . | chr11 | 103094623 | 103106775 | 6.90747 | 4.97013 | 4.56297 |
| chr3 | 97609769 | 97610327 | 961 | Promoter (<=1kb) | 3 | 97560742 | 97610203 | 49462 | 2 | ENSMUST00 | 0 | Chd1l | ENSMUSG0( | false | chr3 | 97609302 | 97610592 | 32.29185 | 112.09676 | 108.19604 | . | . | . | . | . | . |
| chr7 | 97852228 | 97852567 | 961 | Intron (ENSMUST00000033040.6 | 7 | 97842935 | 97912381 | 69447 | 1 | ENSMUST00 | 9294 | Pak1 | ENSMUSG0( | true | . | . | . | . | . | . | chr7 | 97847622 | 97867047 | 4.43625 | 3.78799 | 2.54474 |
| chr8 | 27160679 | 27160991 | 961 | Intron (ENSMUST00000033878.7 | 8 | 27138773 | 27174646 | 35874 | 2 | ENSMUST00 | 13655 | Rab11fip1 | ENSMUSG0( | false | . | . | . | . | . | . | chr8 | 27157652 | 27167682 | 5.55328 | 4.34877 | 3.47001 |
| chr8 | 70658968 | 70659291 | 961 | Promoter (<=1kb) | 8 | 70646436 | 70659738 | 13303 | 2 | ENSMUST00 | 447 | Pgpep1 | ENSMUSG0( | false | chr8 | 70658804 | 70660437 | 32.16322 | 96.63297 | 93.14677 | chr8 | 70656983 | 70658981 | 3.8841 | 3.31462 | 2.13645 |
| chr16 | 85094373 | 85094677 | 961 | Intron (ENSMUST00000005406.1 | 16 | 85168849 | 85169001 | 153 | 1 | ENSMUST00 | -74172 | Gm27295 | ENSMUSG0( | false | . | . | . | . | . | . | chr16 | 85091293 | 85097157 | 4.84003 | 4.0199 | 2.87443 |
| chr19 | 26758657 | 26759025 | 960 | Intron (ENSMUST00000025862.8 | 19 | 26753600 | 26777651 | 24052 | 1 | ENSMUST00 | 5058 | Smarca2 | ENSMUSG0( | false | . | . | . | . | . | . | chr19 | 26758241 | 26758977 | 2.92441 | 2.93094 | 1.33955 |
| chr9 | 121414331 | 121414626 | 960 | Intron (ENSMUST00000045903.6 | 9 | 121366958 | 121474916 | 107959 | 1 | ENSMUST00 | 47374 | Trak1 | ENSMUSG0( | false | . | . | . | . | . | . | chr9 | 121414032 | 121417802 | 4.97692 | 3.93778 | 3.02735 |
| chr9 | 58541860 | 58542160 | 960 | Promoter (<=1kb) | 9 | 58524300 | 58540941 | 16642 | 2 | ENSMUST00 | -920 | Cd276 | ENSMUSG0( | false | . | . | . | . | . | . | chr9 | 58541298 | 58542496 | 5.43317 | 4.11368 | 3.40583 |
| chr1 | 38358732 | 38359131 | 960 | Intron (ENSMUST00000095027.3 | 1 | 38384949 | 38393974 | 9026 | 1 | ENSMUST00 | -25818 | Gm16152 | ENSMUSG0( | false | . | . | . | . | . | . | chr1 | 38357218 | 38359292 | 3.81428 | 3.31126 | 2.0622 |
| chr5 | 140194275 | 140194609 | 960 | Intron (ENSMUST00000110829.2 | 5 | 140207385 | 140210751 | 3367 | 1 | ENSMUST00 | -12776 | Gm16120 | ENSMUSG0( | false | . | . | . | . | . | . | chr5 | 140193717 | 140194785 | 2.85904 | 2.84265 | 1.30329 |
| chr1 | 73223613 | 73224030 | 960 | Distal Intergenic | 1 | 73269064 | 73306360 | 37297 | 1 | ENSMUST00 | -45034 | RP24-141F1( | ENSMUSG0( | false | . | . | . | . | . | . | . | . | . | . | . | . |
| chr2 | 32618097 | 32618506 | 959 | Exon (ENSMUST00000140983.2/ | 2 | 32621758 | 32635058 | 13301 | 1 | ENSMUST00 | -3252 | Ak1 | ENSMUSG0( | false | chr2 | 32618371 | 32618968 | 7.97015 | 15.43348 | 13.16457 | chr2 | 32605009 | 32639275 | 6.86759 | 4.92928 | 4.53618 |
| chr9 | 64022800 | 64023311 | 959 | Promoter (<=1kb) | 9 | 63953076 | 64022059 | 68984 | 2 | ENSMUST00 | -742 | Smad6 | ENSMUSG0( | true | chr9 | 64016675 | 64022981 | 24.82645 | 85.46523 | 82.21117 | chr9 | 64022727 | 64025433 | 4.45683 | 3.81073 | 2.55791 |
| chr18 | 35103214 | 35103658 | 959 | Distal Intergenic | 18 | 35118888 | 35254773 | 135886 | 1 | ENSMUST00 | -15230 | Ctnna1 | ENSMUSG0( | false | . | . | . | . | . | . | chr18 | 35102797 | 35103903 | 3.30665 | 3.18838 | 1.66623 |
| chr17 | 66374353 | 66375004 | 959 | Promoter (<=1kb) | 17 | 66366344 | 66374664 | 8321 | 2 | ENSMUST00 | 0 | Mtcl1 | ENSMUSG0( | true | . | . | . | . | . | . | chr17 | 66349054 | 66377196 | 6.56105 | 4.76301 | 4.29736 |
| chr6 | 140764873 | 140765159 | 959 | Distal Intergenic | 6 | 140746960 | 140747172 | 213 | 2 | ENSMUST00 | -17702 | Rpl38-ps2 | ENSMUSG0( | false | . | . | . | . | . | . | . | . | . | . | . | . |
| chr13 | 29769641 | 29770045 | 959 | Intron (ENSMUST00000140278.2 | 13 | 29732942 | 29733036 | 95 | 1 | ENSMUST00 | 36700 | Gm23729 | ENSMUSG0( | false | . | . | . | . | . | . | chr13 | 29767558 | 29771253 | 5.8376 | 4.50667 | 3.69547 |
| chr18 | 84885982 | 84886245 | 958 | Distal Intergenic | 18 | 84877601 | 84879864 | 2264 | 1 | ENSMUST00 | 8382 | Cyb5a | ENSMUSG0( | false | . | . | . | . | . | . | chr18 | 84884593 | 84887369 | 4.7517 | 4.01238 | 2.80867 |
| chr6 | 35133575 | 35133987 | 958 | Promoter (<=1kb) | 6 | 35022066 | 35133737 | 111672 | 2 | ENSMUST00 | 0 | Cnot4 | ENSMUSG0( | false | chr6 | 35132576 | 35134663 | 32.88158 | 104.92503 | 101.22703 | . | . | . | . | . | . |
| chr9 | 31030209 | 31031109 | 958 | Promoter (<=1kb) | 9 | 31030644 | 31075879 | 45236 | 1 | ENSMUST00 | 0 | Zbtb44 | ENSMUSG0( | false | chr9 | 31029400 | 31031742 | 28.04154 | 116.13699 | 112.1206 | . | . | . | . | . | . |
| chr8 | 91403640 | 91404005 | 958 | Intron (ENSMUST00000069718.8 | 8 | 91390555 | 91409464 | 18910 | 1 | ENSMUST00 | 13086 | Fto | ENSMUSG0( | false | . | . | . | . | . | . | chr8 | 91402710 | 91405681 | 6.12993 | 4.61083 | 3.95543 |
| chr5 | 64274718 | 64275048 | 958 | Intron (ENSMUST00000043893.7 | 5 | 64252578 | 64253206 | 629 | 2 | ENSMUST00 | -21513 | Gm15823 | ENSMUSG0( | false | . | . | . | . | . | . | chr5 | 64273894 | 64301350 | 6.13726 | 4.51007 | 3.95856 |
| chr14 | 78645659 | 78645992 | 958 | Intron (ENSMUST00000074729.5 | 14 | 78603710 | 78603831 | 122 | 1 | ENSMUST00 | 41950 | Gm26251 | ENSMUSG0( | false | . | . | . | . | . | . | chr14 | 78639208 | 78651275 | 4.93549 | 3.97381 | 2.99528 |
| chr7 | 111082040 | 111082843 | 957 | Promoter (<=1kb) | 7 | 111079338 | 111082195 | 2858 | 2 | ENSMUST00 | 0 | Eif4g2 | ENSMUSG0( | false | chr7 | 111081145 | 111083078 | 31.56964 | 96.59996 | 93.11521 | . | . | . | . | . | . |
| chr2 | 3730782 | 3731101 | 957 | Intron (ENSMUST00000115052.3 | 2 | 3713478 | 3779921 | 66444 | 1 | ENSMUST00 | 17305 | Fam107b | ENSMUSG0( | false | . | . | . | . | . | . | . | . | . | . | . | . |
| chr2 | 154565600 | 154565969 | 957 | Intron (ENSMUST00000103145.5 | 2 | 154559634 | 154569720 | 10087 | 2 | ENSMUST00 | 3751 | E2f1 | ENSMUSG0( | false | . | . | . | . | . | . | chr2 | 154555502 | 154571286 | 6.03478 | 4.51051 | 3.86945 |
| chr8 | 110774195 | 110774509 | 957 | Intron (ENSMUST00000150680.1 | 8 | 110755236 | 110790888 | 35653 | 2 | ENSMUST00 | 16379 | Il34 | ENSMUSG0( | false | . | . | . | . | . | . | chr8 | 110771936 | 110776315 | 5.81817 | 4.50309 | 3.66655 |
| chr9 | 51076967 | 51077360 | 957 | Promoter (<=1kb) | 9 | 51056780 | 51077094 | 20315 | 2 | ENSMUST00 | 0 | Layn | ENSMUSG0( | false | chr9 | 51076314 | 51077616 | 28.49948 | 87.34749 | 84.05397 | chr9 | 51075652 | 51079387 | 3.4741 | 3.29042 | 1.7771 |
| chr11 | 98705440 | 98705780 | 956 | Promoter (<=1kb) | 11 | 98704843 | 98706480 | 1638 | 2 | ENSMUST00 | 700 | Med24 | ENSMUSG0( | false | chr11 | 98703830 | 98705879 | 15.26358 | 39.48916 | 36.90792 | chr11 | 98700043 | 98711662 | 7.93936 | 5.23478 | 5.43738 |
| chr18 | 80715187 | 80715603 | 956 | Promoter (2-3kb) | 18 | 80606205 | 80713071 | 106867 | 2 | ENSMUST00 | -2117 | Nfatc1 | ENSMUSG0( | false | chr18 | 80714491 | 80715784 | 14.82622 | 34.53743 | 32.01512 | chr18 | 80711836 | 80715689 | 5.07579 | 4.0937 | 3.06953 |
| chr5 | 36457741 | 36458043 | 956 | Intron (ENSMUST00000149687.1 | 5 | 36455768 | 36464811 | 9044 | 2 | ENSMUST00 | 6768 | 2210406O10 | ENSMUSG0( | false | . | . | . | . | . | . | chr5 | 36457405 | 36458676 | 4.25166 | 3.73074 | 2.43449 |
| chr6 | 122339217 | 122340227 | 956 | Promoter (<=1kb) | 6 | 122317738 | 122339658 | 21921 | 2 | ENSMUST00 | 0 | Phc1 | ENSMUSG0( | false | chr6 | 122338950 | 122340676 | 34.73729 | 107.12603 | 103.36953 | chr6 | 122337348 | 122339282 | 3.80561 | 3.53539 | 2.02456 |
| chr18 | 84085013 | 84085484 | 956 | Promoter (<=1kb) | 18 | 84012082 | 84086390 | 74309 | 2 | ENSMUST00 | 906 | Tshz1 | ENSMUSG0( | false | chr18 | 84084710 | 84090178 | 35.29575 | 125.98199 | 121.63858 | . | . | . | . | . | . |
| chr4 | 106278813 | 106279147 | 956 | Intron (ENSMUST00000140196.1 | 4 | 106291269 | 106292255 | 987 | 1 | ENSMUST00 | -12122 | Gm12730 | ENSMUSG0( | false | . | . | . | . | . | . | chr4 | 106278204 | 106279495 | 3.9755 | 3.57803 | 2.16206 |
| chr10 | 83353899 | 83354346 | 955 | Distal Intergenic | 10 | 83360221 | 83368835 | 8615 | 1 | ENSMUST00 | -5875 | D10Wsu102e | ENSMUSG0( | false | . | . | . | . | . | . | chr10 | 83353415 | 83355440 | 6.22065 | 4.60643 | 4.02852 |
| chr5 | 146847701 | 146848036 | 955 | Promoter (2-3kb) | 5 | 146845071 | 146847726 | 2656 | 1 | ENSMUST00 | 2631 | Rasl11a | ENSMUSG0( | true | chr5 | 146844279 | 146848368 | 25.50663 | 88.63404 | 85.31516 | . | . | . | . | . | . |
| chr9 | 69243700 | 69244162 | 955 | Intron (ENSMUST00000034766.8 | 9 | 69289682 | 69379809 | 90128 | 1 | ENSMUST00 | -45520 | Rora | ENSMUSG0( | false | . | . | . | . | . | . | chr9 | 69242329 | 69246961 | 4.17138 | 3.70907 | 2.32261 |
| chr3 | 95216910 | 95217872 | 955 | Promoter (<=1kb) | 3 | 95181766 | 95217709 | 35944 | 2 | ENSMUST00 | 0 | Gabpb2 | ENSMUSG0( | true | chr3 | 95216676 | 95218334 | 28.50113 | 87.35039 | 84.05681 | . | . | . | . | . | . |
| chr7 | 134670495 | 134671068 | 955 | Promoter (<=1kb) | 7 | 134670687 | 135173639 | 502953 | 1 | ENSMUST00 | 0 | Dock1 | ENSMUSG0( | false | chr7 | 134670248 | 134672039 | 33.89795 | 105.67399 | 101.95845 | chr7 | 134669829 | 134680714 | 3.36907 | 3.13469 | 1.68815 |
| chr3 | 97337396 | 97337706 | 955 | Distal Intergenic | 3 | 97284908 | 97297778 | 12871 | 2 | ENSMUST00 | -39619 | Gm17608 | ENSMUSG0( | false | . | . | . | . | . | . | chr3 | 97336194 | 97339033 | 4.89652 | 3.82992 | 2.95388 |
| chr10 | 60831373 | 60831871 | 954 | Promoter (<=1kb) | 10 | 60762610 | 60831581 | 68972 | 2 | ENSMUST00 | 0 | Unc5b | ENSMUSG0( | false | chr10 | 60831673 | 60832323 | 14.20671 | 34.84183 | 32.31654 | . | . | . | . | . | . |
| chr15 | 6874214 | 6874555 | 954 | Promoter (<=1kb) | 15 | 6836758 | 6874268 | 37511 | 2 | ENSMUST00 | 0 | Osmr | ENSMUSG0( | false | chr15 | 6873356 | 6875149 | 21.91892 | 63.05404 | 60.16603 | chr15 | 6873438 | 6874974 | 4.64132 | 3.86951 | 2.7388 |
| chr13 | 48633643 | 48633991 | 954 | Distal Intergenic | 13 | 48577873 | 48625573 | 47701 | 2 | ENSMUST00 | -8071 | Ptpdc1 | ENSMUSG0( | false | . | . | . | . | . | . | chr13 | 48632270 | 48635009 | 4.89934 | 4.05855 | 2.92415 |
| chr3 | 8666930 | 8667765 | 954 | Promoter (<=1kb) | 3 | 8663362 | 8667039 | 3678 | 2 | ENSMUST00 | 0 | Hey1 | ENSMUSG0( | false | chr3 | 8666061 | 8668353 | 25.48472 | 95.2657 | 91.8051 | . | . | . | . | . | . |
| chr4 | 62699276 | 62699759 | 954 | Promoter (<=1kb) | 4 | 62699466 | 62703018 | 3553 | 1 | ENSMUST00 | 0 | Rgs3 | ENSMUSG0( | false | chr4 | 62699665 | 62702487 | 24.69334 | 83.34509 | 80.13418 | chr4 | 62694793 | 62699426 | 7.42731 | 5.01341 | 4.9919 |
| chr8 | 33883629 | 33884017 | 954 | Promoter (1-2kb) | 8 | 33884774 | 33885747 | 974 | 2 | ENSMUST00 | 1730 | Gm26978 | ENSMUSG0( | false | . | . | . | . | . | . | chr8 | 33863865 | 33927704 | 6.32022 | 4.67603 | 4.08842 |
| chr11 | 104232136 | 104233060 | 953 | Promoter (<=1kb) | 11 | 104231584 | 104328219 | 96636 | 1 | ENSMUST00 | 553 | Mapt | ENSMUSG0( | true | chr11 | 104231437 | 104234756 | 38.46334 | 130.56975 | 126.05653 | . | . | . | . | . | . |
| chr5 | 136568721 | 136569060 | 953 | Promoter (1-2kb) | 5 | 136482875 | 136567490 | 84616 | 2 | ENSMUST00 | -1232 | Cux1 | ENSMUSG0( | false | chr5 | 136565019 | 136569262 | 36.46257 | 120.58523 | 116.42456 | . | . | . | . | . | . |
| chr13 | 9093563 | 9094068 | 953 | Promoter (<=1kb) | 13 | 9093881 | 9171104 | 77224 | 1 | ENSMUST00 | 0 | Larp4b | ENSMUSG0( | false | chr13 | 9093707 | 9095228 | 44.57462 | 145.65463 | 140.53799 | . | . | . | . | . | . |
| chr4 | 22357384 | 22357989 | 953 | Promoter (<=1kb) | 4 | 22357543 | 22376725 | 19183 | 1 | ENSMUST00 | 0 | Fbxl4 | ENSMUSG0( | false | chr4 | 22357091 | 22358524 | 24.55236 | 90.57755 | 87.22021 | chr4 | 22356594 | 22360304 | 4.19656 | 3.73171 | 2.34258 |
| chr9 | 7763869 | 7764495 | 953 | Promoter (<=1kb) | 9 | 7764041 | 7794333 | 30293 | 1 | ENSMUST00 | 0 | Tmem123 | ENSMUSG0( | false | chr9 | 7763992 | 7764964 | 32.06657 | 104.51031 | 100.83098 | . | . | . | . | . | . |
| chr19 | 55841802 | 55842448 | 953 | Promoter (<=1kb) | 19 | 55841672 | 55845897 | 4226 | 1 | ENSMUST00 | 131 | RP24-210C4. | ENSMUSG0( | false | . | . | . | . | . | . | chr19 | 55838098 | 55844140 | 5.7222 | 4.3683 | 3.6165 |
| chr9 | 57644989 | 57645863 | 952 | Promoter (<=1kb) | 9 | 57626647 | 57645653 | 19007 | 2 | ENSMUST00 | 0 | Csk | ENSMUSG0( | false | chr9 | 57644457 | 57645948 | 38.97723 | 131.47449 | 126.92584 | chr9 | 57632762 | 57645064 | 6.57255 | 4.74185 | 4.30478 |

| | | | | | | | | | | | | | | | | | | | | | | | | | | |
|---|---|---|---|---|---|---|---|---|---|---|---|---|---|---|---|---|---|---|---|---|---|---|---|---|---|---|
| chr18 | 76071939 | 76072527 | 952 | Intron (ENSMUST00000058997.8 | 18 | 76059458 | 76146443 | 86986 | 1 | ENSMUST00 | 12482 | Zbtb7c | ENSMUSG0C | true | . | . | . | . | . | . | chr18 | 76061445 | 76132293 | 6.70015 | 4.79833 | 4.4158 |
| chr8 | 82914626 | 82914970 | 952 | Intron (ENSMUST00000078525.5 | 8 | 82863356 | 83091271 | 227916 | 1 | ENSMUST00 | 51271 | Rnf150 | ENSMUSG0C | true | . | . | . | . | . | . | chr8 | 82912309 | 82917956 | 3.76062 | 3.41205 | 1.99098 |
| chr1 | 43487156 | 43487523 | 952 | Intron (ENSMUST00000187435.1 | 1 | 43457273 | 43462222 | 4950 | 1 | ENSMUST00 | 29884 | Nck2 | ENSMUSG0C | true | . | . | . | . | . | . | chr1 | 43446894 | 43509082 | 5.99677 | 4.48008 | 3.83622 |
| chr11 | 53345418 | 53345693 | 952 | Promoter (<=1kb) | 11 | 53345290 | 53345677 | 388 | 2 | ENSMUST00 | 0 | Gm11188 | ENSMUSG0C | false | . | . | . | . | . | . | chr11 | 53341291 | 53346716 | 4.65096 | 3.97609 | 2.7137 |
| chr9 | 108263044 | 108264166 | 951 | Promoter (<=1kb) | 9 | 108205955 | 108263918 | 57964 | 2 | ENSMUST00 | 0 | Dag1 | ENSMUSG0C | true | chr9 | 108261324 | 108263997 | 35.98183 | 119.73972 | 115.60696 | . | . | . | . | . | . |
| chr9 | 15746368 | 15746808 | 951 | Distal Intergenic | 9 | 15709739 | 15742384 | 32646 | 1 | ENSMUST00 | 36630 | Slc36a4 | ENSMUSG0C | false | . | . | . | . | . | . | chr9 | 15743257 | 15748686 | 4.79895 | 3.89356 | 2.89308 |
| chr15 | 78405762 | 78406223 | 951 | Promoter (<=1kb) | 15 | 78399556 | 78405859 | 6304 | 2 | ENSMUST00 | 0 | Tst | ENSMUSG0C | false | chr15 | 78405455 | 78407074 | 31.91637 | 110.38313 | 106.53261 | chr15 | 78405406 | 78405876 | 5.06869 | 4.06825 | 3.11388 |
| chr9 | 99447265 | 99447575 | 951 | Distal Intergenic | 9 | 99456207 | 99470932 | 14726 | 1 | ENSMUST00 | -8632 | 1600029I14R | ENSMUSG0C | false | . | . | . | . | . | . | chr9 | 99447222 | 99450298 | 4.93571 | 3.99558 | 2.99805 |
| chr11 | 115899889 | 115900226 | 951 | Promoter (<=1kb) | 11 | 115899966 | 115906269 | 6304 | 1 | ENSMUST00 | 0 | Smim5 | ENSMUSG0C | false | . | . | . | . | . | . | chr11 | 115899613 | 115906730 | 7.08174 | 4.8342 | 4.74864 |
| chr13 | 9578152 | 9578682 | 951 | Intron (ENSMUST00000174552.2 | 13 | 9623380 | 9623627 | 248 | 2 | ENSMUST00 | 44945 | Gm23496 | ENSMUSG0C | false | . | . | . | . | . | . | chr13 | 9577500 | 9580497 | 3.94488 | 3.56059 | 2.13635 |
| chr2 | 119208529 | 119209142 | 950 | Promoter (<=1kb) | 2 | 119172500 | 119208795 | 36296 | 2 | ENSMUST00 | 0 | Dnajc17 | ENSMUSG0C | false | chr2 | 119207787 | 119209563 | 36.76537 | 115.82222 | 111.81207 | . | . | . | . | . | . |
| chr9 | 114978353 | 114979121 | 950 | Promoter (<=1kb) | 9 | 114978569 | 115232223 | 253655 | 1 | ENSMUST00 | 0 | Osbpl10 | ENSMUSG0C | false | chr9 | 114977880 | 114979450 | 31.70772 | 93.8754 | 90.44607 | chr9 | 114977805 | 114979514 | 4.83351 | 4.00041 | 2.86807 |
| chr1 | 106171434 | 106172177 | 950 | Promoter (<=1kb) | 1 | 106171752 | 106394250 | 222499 | 1 | ENSMUST00 | 0 | Phlpp1 | ENSMUSG0C | false | chr1 | 106171618 | 106174296 | 38.55812 | 129.64772 | 125.16568 | . | . | . | . | . | . |
| chr4 | 107336435 | 107336694 | 950 | Exon (ENSMUST00000075693.6/ | 4 | 107343111 | 107351589 | 8479 | 1 | ENSMUST00 | -6417 | Yipf1 | ENSMUSG0C | false | . | . | . | . | . | . | chr4 | 107334373 | 107338462 | 4.40841 | 3.53981 | 2.56287 |
| chr2 | 133467450 | 133467851 | 950 | Distal Intergenic | 2 | 133463118 | 133465784 | 2667 | 1 | ENSMUST00 | 4333 | Gm14100 | ENSMUSG0C | false | . | . | . | . | . | . | . | . | . | . | . | . |
| chr18 | 61640628 | 61641801 | 950 | Promoter (<=1kb) | 18 | 61639653 | 61647472 | 7820 | 1 | ENSMUST00 | 976 | Gm20748 | ENSMUSG0C | false | . | . | . | . | . | . | chr18 | 61630875 | 61644532 | 7.58082 | 5.1921 | 5.12657 |
| chr15 | 28024190 | 28026246 | 949 | Promoter (<=1kb) | 15 | 27730652 | 28025848 | 295197 | 2 | ENSMUST00 | 0 | Trio | ENSMUSG0C | false | chr15 | 28026111 | 28026553 | 9.1888 | 20.54895 | 18.20621 | chr15 | 28020404 | 28024202 | 3.60843 | 3.31574 | 1.8938 |
| chr8 | 15047421 | 15047812 | 949 | Promoter (1-2kb) | 8 | 15042502 | 15046078 | 3577 | 2 | ENSMUST00 | -1344 | BB014433 | ENSMUSG0C | false | . | . | . | . | . | . | chr8 | 15043191 | 15051415 | 5.67584 | 4.41005 | 3.56153 |
| chr7 | 64048358 | 64048724 | 949 | Distal Intergenic | 7 | 64057602 | 64062958 | 5357 | 1 | ENSMUST00 | -8878 | Gm20670 | ENSMUSG0C | false | chr7 | 64048343 | 64049162 | 8.27169 | 15.93882 | 13.66243 | chr7 | 64047583 | 64051824 | 7.77647 | 5.37371 | 5.28037 |
| chr15 | 89211305 | 89211671 | 949 | Exon (ENSMUST00000181384.1/ | 15 | 89211559 | 89215012 | 3454 | 2 | ENSMUST00 | 3341 | Gm26798 | ENSMUSG0C | false | chr15 | 89210792 | 89212473 | 39.44868 | 124.7075 | 120.4059 | . | . | . | . | . | . |
| chr19 | 38778770 | 38779146 | 949 | Intron (ENSMUST00000182481.2 | 19 | 38748760 | 38752645 | 3886 | 2 | ENSMUST00 | -26126 | RP23-221A9. | ENSMUSG0C | false | . | . | . | . | . | . | chr19 | 38774674 | 38780110 | 4.20404 | 3.6078 | 2.37963 |
| chr13 | 107692375 | 107692723 | 949 | Distal Intergenic | 13 | 107687397 | 107692173 | 4777 | 1 | ENSMUST00 | 4979 | RP23-125L8. | ENSMUSG0C | false | . | . | . | . | . | . | chr13 | 107688659 | 107693312 | 4.03041 | 3.39672 | 2.24558 |
| chr7 | 27596863 | 27597168 | 948 | Promoter (<=1kb) | 7 | 27596830 | 27618233 | 21404 | 1 | ENSMUST00 | 34 | Akt2 | ENSMUSG0C | false | . | . | . | . | . | . | chr7 | 27592472 | 27610994 | 7.14155 | 5.01543 | 4.77185 |
| chr8 | 3429994 | 3430289 | 948 | Intron (ENSMUST00000131773.2 | 8 | 3437011 | 3456600 | 19590 | 1 | ENSMUST00 | -6722 | Arhgef18 | ENSMUSG0C | true | . | . | . | . | . | . | chr8 | 3394895 | 3435896 | 6.08039 | 4.58876 | 3.89109 |
| chr16 | 14163019 | 14163393 | 948 | Promoter (<=1kb) | 16 | 14163275 | 14192928 | 29654 | 1 | ENSMUST00 | 0 | Nde1 | ENSMUSG0C | false | chr16 | 14162717 | 14164701 | 20.60195 | 76.26862 | 73.17842 | . | . | . | . | . | . |
| chr8 | 41239456 | 41240066 | 948 | Promoter (<=1kb) | 8 | 41239759 | 41334086 | 94328 | 1 | ENSMUST00 | 0 | Pcm1 | ENSMUSG0C | false | chr8 | 41239069 | 41241158 | 48.76767 | 169.04231 | 163.0952 | . | . | . | . | . | . |
| chr12 | 40449564 | 40449895 | 948 | Intron (ENSMUST00000037488.6 | 12 | 40446053 | 40846486 | 400434 | 1 | ENSMUST00 | 3512 | Dock4 | ENSMUSG0C | true | . | . | . | . | . | . | chr12 | 40448062 | 40493881 | 5.07226 | 4.00134 | 3.09271 |
| chr4 | 56837713 | 56838130 | 948 | Promoter (<=1kb) | 4 | 56829525 | 56838584 | 9060 | 2 | ENSMUST00 | 454 | Ctnnal1 | ENSMUSG0C | true | . | . | . | . | . | . | chr4 | 56827266 | 56853148 | 5.48236 | 4.22607 | 3.42275 |
| chr2 | 29239314 | 29239668 | 947 | Intron (ENSMUST00000183583.2 | 2 | 29194821 | 29248040 | 53220 | 2 | ENSMUST00 | 8372 | Ntng2 | ENSMUSG0C | true | . | . | . | . | . | . | chr2 | 29230623 | 29250828 | 7.61258 | 5.25891 | 5.15208 |
| chr15 | 102101622 | 102102324 | 947 | Promoter (<=1kb) | 15 | 102102988 | 102116391 | 13404 | 1 | ENSMUST00 | -664 | Tenc1 | ENSMUSG0C | false | chr15 | 102099897 | 102107616 | 31.25476 | 101.05123 | 97.45738 | chr15 | 102097473 | 102102796 | 4.79775 | 3.79288 | 2.84744 |
| chr6 | 34493708 | 34494028 | 947 | Intron (ENSMUST00000045372.5 | 6 | 34477142 | 34487848 | 10707 | 1 | ENSMUST00 | 16567 | Bpgm | ENSMUSG0C | false | . | . | . | . | . | . | chr6 | 34492938 | 34496247 | 4.168 | 3.67796 | 2.34698 |
| chr5 | 119378264 | 119378670 | 947 | Distal Intergenic | 5 | 119297245 | 119297361 | 117 | 1 | ENSMUST00 | 81020 | n-R5s175 | ENSMUSG0C | false | . | . | . | . | . | . | . | . | . | . | . | . |
| chr9 | 117393950 | 117394347 | 947 | Intron (ENSMUST00000164018.3 | 9 | 117402979 | 117404342 | 1364 | 1 | ENSMUST00 | -8632 | Gm20396 | ENSMUSG0C | false | . | . | . | . | . | . | . | . | . | . | . | . |
| chr15 | 103502664 | 103503324 | 946 | Promoter (<=1kb) | 15 | 103503298 | 103530052 | 26755 | 1 | ENSMUST00 | 0 | Pde1b | ENSMUSG0C | false | chr15 | 103502456 | 103503556 | 12.59008 | 38.01365 | 35.45054 | chr15 | 103502614 | 103503914 | 4.09216 | 3.69955 | 2.25073 |
| chr4 | 131743945 | 131744220 | 946 | Distal Intergenic | 4 | 131768458 | 131768921 | 464 | 2 | ENSMUST00 | 24701 | Ptpru | ENSMUSG0C | false | . | . | . | . | . | . | chr4 | 131743070 | 131745154 | 4.49998 | 3.85941 | 2.63763 |
| chr6 | 13607705 | 13608234 | 946 | Promoter (<=1kb) | 6 | 13580687 | 13608100 | 27414 | 2 | ENSMUST00 | 0 | Tmem168 | ENSMUSG0C | false | chr6 | 13607283 | 13608435 | 36.7977 | 133.03651 | 128.43381 | . | . | . | . | . | . |
| chr7 | 118543652 | 118543930 | 946 | Intron (ENSMUST00000044195.4 | 7 | 118525062 | 118533356 | 8295 | 2 | ENSMUST00 | -10297 | Coq7 | ENSMUSG0C | false | . | . | . | . | . | . | chr7 | 118536275 | 118546239 | 6.06229 | 4.47014 | 3.88658 |
| chr8 | 33803256 | 33803633 | 946 | Intron (ENSMUST00000053251.6 | 8 | 33784064 | 33806799 | 22736 | 2 | ENSMUST00 | 3166 | Rbpms | ENSMUSG0C | true | . | . | . | . | . | . | chr8 | 33797013 | 33821269 | 7.29084 | 5.02722 | 4.89898 |
| chr4 | 11321904 | 11322366 | 946 | Promoter (<=1kb) | 4 | 11261315 | 11322137 | 60823 | 2 | ENSMUST00 | 0 | Dpy19l4 | ENSMUSG0C | false | chr4 | 11321290 | 11322745 | 41.911 | 144.38934 | 139.32462 | . | . | . | . | . | . |
| chr13 | 63486721 | 63487191 | 945 | Intron (ENSMUST00000159024.1 | 13 | 63347512 | 63497278 | 149767 | 2 | ENSMUST00 | 10087 | Fancc | ENSMUSG0C | false | . | . | . | . | . | . | chr13 | 63484955 | 63493657 | 5.66759 | 4.30416 | 3.56899 |
| chr13 | 91223711 | 91224254 | 945 | Promoter (<=1kb) | 13 | 90935349 | 91223987 | 288639 | 2 | ENSMUST00 | 0 | Atg10 | ENSMUSG0C | false | chr13 | 91223518 | 91224518 | 29.2142 | 114.47596 | 110.50954 | . | . | . | . | . | . |
| chr19 | 46044862 | 46045714 | 945 | Promoter (<=1kb) | 19 | 46032593 | 46045214 | 12622 | 2 | ENSMUST00 | 0 | Ldb1 | ENSMUSG0C | false | chr19 | 46043949 | 46046177 | 34.10395 | 106.03094 | 102.3112 | chr19 | 46045644 | 46047114 | 3.8061 | 3.50605 | 2.01095 |
| chr4 | 130163373 | 130163823 | 945 | Promoter (2-3kb) | 4 | 130165604 | 130166611 | 1008 | 2 | ENSMUST00 | 2788 | Tinagl1 | ENSMUSG0C | false | . | . | . | . | . | . | chr4 | 130162993 | 130163722 | 2.86152 | 2.89489 | 1.32681 |
| chr4 | 155074512 | 155074796 | 945 | 3' UTR | 4 | 155067059 | 155069116 | 2058 | 1 | ENSMUST00 | 7454 | Pex10 | ENSMUSG0C | false | . | . | . | . | . | . | chr4 | 155065929 | 155080644 | 7.3659 | 5.06273 | 4.94236 |
| chr11 | 117414839 | 117415223 | 945 | Promoter (<=1kb) | 11 | 117414389 | 117415368 | 980 | 1 | ENSMUST00 | 451 | Gm11729 | ENSMUSG0C | false | chr11 | 117414833 | 117415381 | 11.02877 | 22.59171 | 20.22099 | chr11 | 117406406 | 117416739 | 6.0329 | 4.55882 | 3.85214 |
| chr1 | 171329096 | 171329890 | 944 | Promoter (<=1kb) | 1 | 171329145 | 171342331 | 13187 | 1 | ENSMUST00 | 0 | Dedd | ENSMUSG0C | false | chr1 | 171328255 | 171330422 | 28.50037 | 84.48055 | 81.24493 | . | . | . | . | . | . |
| chr11 | 80069653 | 80070068 | 944 | Intron (ENSMUST00000103233.4 | 11 | 80059312 | 80080861 | 21550 | 2 | ENSMUST00 | 10793 | Crlf3 | ENSMUSG0C | true | . | . | . | . | . | . | chr11 | 80064592 | 80080109 | 4.7628 | 3.97341 | 2.8246 |
| chr11 | 5261231 | 5262006 | 944 | Promoter (<=1kb) | 11 | 5191552 | 5261558 | 70007 | 2 | ENSMUST00 | 0 | Kremen1 | ENSMUSG0C | true | chr11 | 5260004 | 5262317 | 38.30962 | 127.04226 | 122.65842 | . | . | . | . | . | . |
| chr10 | 121364917 | 121365575 | 944 | Promoter (<=1kb) | 10 | 121365090 | 121397245 | 32156 | 1 | ENSMUST00 | 0 | Gns | ENSMUSG0C | false | chr10 | 121364588 | 121366171 | 25.98079 | 111.7719 | 107.88497 | . | . | . | . | . | . |
| chr18 | 60600304 | 60600846 | 944 | Promoter (<=1kb) | 18 | 60593991 | 60600497 | 6507 | 2 | ENSMUST00 | 0 | Synpo | ENSMUSG0C | true | . | . | . | . | . | . | chr18 | 60593903 | 60608752 | 6.45808 | 4.76452 | 4.19811 |
| chr2 | 156720611 | 156721456 | 944 | Promoter (<=1kb) | 2 | 156721031 | 156762725 | 41695 | 1 | ENSMUST00 | 0 | Dlgap4 | ENSMUSG0C | true | chr2 | 156720047 | 156723142 | 30.41442 | 103.5799 | 99.9162 | . | . | . | . | . | . |
| chr3 | 57846964 | 57848121 | 943 | Promoter (<=1kb) | 3 | 57841895 | 57847564 | 5670 | 2 | ENSMUST00 | 0 | Pfn2 | ENSMUSG0C | false | chr3 | 57846610 | 57848374 | 36.10713 | 116.78466 | 112.7497 | chr3 | 57846391 | 57848532 | 3.63303 | 3.36688 | 1.93638 |
| chr9 | 108221829 | 108222135 | 943 | Intron (ENSMUST00000166905.2 | 9 | 108096022 | 108190384 | 94363 | 2 | ENSMUST00 | -31446 | Bsn | ENSMUSG0C | false | . | . | . | . | . | . | chr9 | 108210099 | 108251502 | 5.76916 | 4.35802 | 3.65688 |
| chr8 | 104442411 | 104443115 | 943 | Promoter (<=1kb) | 8 | 104417674 | 104443047 | 25374 | 2 | ENSMUST00 | 0 | Dync1li2 | ENSMUSG0C | false | chr8 | 104441803 | 104443201 | 39.66383 | 132.67865 | 128.09186 | . | . | . | . | . | . |
| chr9 | 122805392 | 122805847 | 943 | Promoter (<=1kb) | 9 | 122805539 | 122836334 | 30796 | 1 | ENSMUST00 | 0 | Tcaim | ENSMUSG0C | false | chr9 | 122805110 | 122806474 | 27.39917 | 84.46252 | 81.22775 | chr9 | 122804781 | 122807034 | 3.94229 | 3.60782 | 2.12692 |
| chr10 | 81559274 | 81559720 | 943 | Promoter (<=1kb) | 10 | 81559494 | 81566362 | 6869 | 1 | ENSMUST00 | 0 | Aes | ENSMUSG0C | false | chr10 | 81559124 | 81561616 | 23.12601 | 77.64799 | 74.53377 | . | . | . | . | . | . |
| chr19 | 23964598 | 23964975 | 943 | Distal Intergenic | 19 | 23972751 | 24031019 | 58269 | 2 | ENSMUST00 | 66044 | Fam189a2 | ENSMUSG0C | false | . | . | . | . | . | . | chr19 | 23963776 | 23967797 | 5.4084 | 4.28789 | 3.37212 |
| chr19 | 47854794 | 47855150 | 942 | Promoter (<=1kb) | 19 | 47854970 | 47864790 | 9821 | 1 | ENSMUST00 | 0 | Gsto1 | ENSMUSG0C | false | chr19 | 47854432 | 47855663 | 17.28728 | 67.4904 | 64.54075 | chr19 | 47854117 | 47855634 | 5.87494 | 4.19464 | 3.75878 |
| chr8 | 33961601 | 33961850 | 942 | Distal Intergenic | 8 | 33782643 | 33929863 | 147221 | 2 | ENSMUST00 | -31739 | Rbpms | ENSMUSG0C | true | . | . | . | . | . | . | chr8 | 33960821 | 33962454 | 3.6485 | 3.42364 | 1.95023 |
| chr11 | 5740804 | 5741420 | 942 | Promoter (<=1kb) | 11 | 5713417 | 5741150 | 27734 | 2 | ENSMUST00 | 0 | Urgcp | ENSMUSG0C | false | chr11 | 5740331 | 5741629 | 30.14656 | 119.38538 | 115.2599 | . | . | . | . | . | . |
| chr5 | 97079564 | 97079873 | 942 | Intron (ENSMUST00000035635.8 | 5 | 97095932 | 97097771 | 1840 | 2 | ENSMUST00 | 17898 | Paqr3 | ENSMUSG0C | false | . | . | . | . | . | . | chr5 | 97078644 | 97082317 | 4.22791 | 3.67168 | 2.38959 |
| chr11 | 69692438 | 69692772 | 942 | Promoter (2-3kb) | 11 | 69687340 | 69695521 | 8182 | 2 | ENSMUST00 | 2749 | Tnfsf12 | ENSMUSG0C | false | . | . | . | . | . | . | chr11 | 69688439 | 69697396 | 5.01103 | 4.04632 | 3.01977 |
| chr2 | 152707531 | 152707981 | 942 | 3' UTR | 2 | 152694733 | 152696692 | 1960 | 2 | ENSMUST00 | -10840 | Gm26841 | ENSMUSG0C | false | chr2 | 152707371 | 152708201 | 9.56722 | 36.15532 | 33.61367 | chr2 | 152705988 | 152708196 | 4.64341 | 3.67836 | 2.74675 |
| chr3 | 95314900 | 95315680 | 941 | Promoter (<=1kb) | 3 | 95315084 | 95323599 | 8516 | 1 | ENSMUST00 | 0 | Cers2 | ENSMUSG0C | false | chr3 | 95314707 | 95316369 | 35.46717 | 120.9532 | 116.77927 | chr3 | 95314716 | 95315039 | 3.23208 | 3.13666 | 1.53636 |
| chr7 | 139364026 | 139364440 | 941 | Distal Intergenic | 7 | 139389109 | 139579652 | 190544 | 1 | ENSMUST00 | -24669 | Inpp5a | ENSMUSG0C | false | . | . | . | . | . | . | chr7 | 139363267 | 139368578 | 5.11722 | 4.13054 | 3.10207 |
| chr6 | 32588091 | 32588466 | 941 | Promoter (<=1kb) | 6 | 32144268 | 32588192 | 443925 | 2 | ENSMUST00 | 0 | Plxna4 | ENSMUSG0C | false | chr6 | 32586391 | 32588544 | 29.22959 | 88.61999 | 85.30408 | chr6 | 32586502 | 32588661 | 6.30071 | 4.65622 | 4.13929 |
| chr14 | 72709431 | 72710220 | 941 | Promoter (<=1kb) | 14 | 72537946 | 72710003 | 172058 | 2 | ENSMUST00 | 0 | Fndc3a | ENSMUSG0C | false | chr14 | 72709140 | 72711049 | 31.70051 | 127.6562 | 123.25787 | . | . | . | . | . | . |
| chr14 | 55853820 | 55854251 | 941 | Promoter (<=1kb) | 14 | 55854036 | 55874291 | 20256 | 1 | ENSMUST00 | 0 | Nynrin | ENSMUSG0C | false | chr14 | 55853819 | 55855746 | 21.02725 | 63.18829 | 60.29588 | chr14 | 55853457 | 55858185 | 3.82651 | 3.51688 | 2.06868 |
| chr6 | 100011620 | 100012206 | 941 | Distal Intergenic | 6 | 99893226 | 99893547 | 322 | 2 | ENSMUST00 | -118074 | Gm24248 | ENSMUSG0C | false | . | . | . | . | . | . | chr6 | 100010729 | 100012240 | 3.70152 | 3.26035 | 1.97137 |
| chr2 | 166073112 | 166073428 | 940 | Promoter (2-3kb) | 2 | 166074670 | 166076018 | 1349 | 2 | ENSMUST00 | 2590 | Sulf2 | ENSMUSG0C | false | . | . | . | . | . | . | chr2 | 166072077 | 166074593 | 4.99505 | 3.9057 | 3.0229 |
| chrX | 151017246 | 151017495 | 940 | Promoter (<=1kb) | 20 | 150983141 | 151017322 | 34182 | 2 | ENSMUST00 | 0 | Gnl3l | ENSMUSG0C | false | chrX | 151016646 | 151018076 | 17.92175 | 44.05972 | 41.41886 | . | . | . | . | . | . |
| chrX | 42148746 | 42149280 | 940 | Promoter (<=1kb) | 20 | 42149317 | 42277185 | 127869 | 1 | ENSMUST00 | -37 | Stag2 | ENSMUSG0C | false | chrX | 42148652 | 42149104 | 12.40737 | 26.6269 | 24.20249 | . | . | . | . | . | . |
| chr11 | 52396174 | 52396567 | 940 | Promoter (<=1kb) | 11 | 52396428 | 52404242 | 7815 | 1 | ENSMUST00 | 0 | 9530068E07I | ENSMUSG0C | false | chr11 | 52396013 | 52397309 | 37.4597 | 129.87831 | 125.39164 | chr11 | 52395783 | 52402328 | 4.20317 | 3.55173 | 2.37894 |
| chr7 | 7298380 | 7298807 | 940 | Promoter (<=1kb) | 7 | 7298891 | 7298946 | 56 | 1 | ENSMUST00 | -84 | Mir5620 | ENSMUSG0C | false | chr7 | 7297989 | 7299897 | 25.04113 | 84.92952 | 81.6889 | . | . | . | . | . | . |
| chr10 | 23349829 | 23350615 | 939 | Promoter (<=1kb) | 10 | 23104168 | 23349895 | 245728 | 2 | ENSMUST00 | 0 | Eya4 | ENSMUSG0C | true | chr10 | 23348112 | 23350902 | 24.07142 | 68.54774 | 65.57966 | chr10 | 23350408 | 23350966 | 3.79943 | 3.53113 | 2.05868 |
| chr5 | 119670441 | 119671206 | 939 | Promoter (<=1kb) | 5 | 119670669 | 119684561 | 13893 | 1 | ENSMUST00 | 0 | Tbx3 | ENSMUSG0C | false | chr5 | 119669890 | 119672818 | 8.7021 | 20.72798 | 18.38387 | chr5 | 119667260 | 119672434 | 3.90359 | 3.53401 | 2.11341 |
| chr19 | 47891588 | 47892319 | 939 | Intron (ENSMUST00000026048.4 | 19 | 47870262 | 47884704 | 14443 | 1 | ENSMUST00 | 21327 | Gsto2 | ENSMUSG0C | false | . | . | . | . | . | . | chr19 | 47889899 | 47894231 | 5.40065 | 4.08482 | 3.36637 |
| chr9 | 103364727 | 103366028 | 939 | Promoter (<=1kb) | 9 | 103353094 | 103365840 | 12747 | 2 | ENSMUST00 | 0 | Cdv3 | ENSMUSG0C | false | chr9 | 103363917 | 103365715 | 20.109 | 88.33971 | 85.03291 | . | . | . | . | . | . |
| chr4 | 49597137 | 49597903 | 939 | Promoter (<=1kb) | 4 | 49584506 | 49597876 | 13371 | 2 | ENSMUST00 | 0 | Tmem246 | ENSMUSG0C | false | chr4 | 49596702 | 49598204 | 35.46717 | 120.9532 | 116.77927 | . | . | . | . | . | . |
| chr7 | 70822523 | 70823235 | 939 | Distal Intergenic | 7 | 70752416 | 70752500 | 85 | 1 | ENSMUST00 | 70108 | Gm24880 | ENSMUSG0C | false | chr7 | 70822562 | 70822998 | 4.35105 | 6.99241 | 4.88445 | chr7 | 70822270 | 70823715 | 2.90356 | 2.91766 | 1.32364 |
| chr1 | 54866524 | 54866841 | 938 | Intron (ENSMUST00000179030.2 | 1 | 54766792 | 54884711 | 117920 | 2 | ENSMUST00 | 17870 | Ankrd44 | ENSMUSG0C | true | . | . | . | . | . | . | chr1 | 54864456 | 54889599 | 5.98614 | 4.56539 | 3.81071 |
| chr6 | 72380177 | 72380734 | 938 | Promoter (<=1kb) | 6 | 72368049 | 72380468 | 12420 | 2 | ENSMUST00 | 0 | Vamp5 | ENSMUSG0C | true | chr6 | 72379564 | 72380905 | 23.42648 | 72.77338 | 69.73964 | chr6 | 72369039 | 72384082 | 5.14581 | 4.03784 | 3.16124 |
| chr6 | 37617393 | 37618071 | 938 | Distal Intergenic | 6 | 37580335 | 37580427 | 93 | 1 | ENSMUST00 | 37059 | Gm25062 | ENSMUSG0C | false | . | . | . | . | . | . | chr6 | 37613519 | 37619590 | 4.74098 | 3.99613 | 2.78613 |
| chr2 | 169213427 | 169213733 | 938 | Distal Intergenic | 2 | 169234991 | 169237770 | 2780 | 1 | ENSMUST00 | -21258 | 9430093N23 | ENSMUSG0C | false | . | . | . | . | . | . | chr2 | 169212830 | 169214040 | 4.47564 | 3.85507 | 2.58577 |
| chr1 | 180777649 | 180778425 | 938 | Distal Intergenic | 1 | 180802568 | 180809483 | 6916 | 1 | ENSMUST00 | -24143 | Gm17275 | ENSMUSG0C | false | chr1 | 180777278 | 180778774 | 24.82645 | 85.46523 | 82.21117 | chr1 | 180775608 | 180777796 | 5.47738 | 4.21265 | 3.44239 |
| chr17 | 21488738 | 21489106 | 938 | Promoter (<=1kb) | 17 | 21485177 | 21488875 | 3699 | 2 | ENSMUST00 | 0 | Gm26753 | ENSMUSG0C | false | chr17 | 21488599 | 21489847 | 23.96561 | 66.60394 | 63.66297 | chr17 | 21488696 | 21490771 | 4.87252 | 4.05028 | 2.88651 |
| chr2 | 76693265 | 76693600 | 937 | Intron (ENSMUST00000111920.1 | 2 | 76689853 | 76692764 | 2912 | 1 | ENSMUST00 | 3413 | Plekha3 | ENSMUSG0C | false | . | . | . | . | . | . | chr2 | 76691542 | 76694460 | 4.19417 | 3.58683 | 2.35442 |
| chr2 | 90553846 | 90554196 | 937 | Intron (ENSMUST00000168621.2 | 2 | 90432889 | 90580647 | 147759 | 2 | ENSMUST00 | 26451 | Ptprj | ENSMUSG0C | false | . | . | . | . | . | . | chr2 | 90529589 | 90563488 | 5.67436 | 4.3718 | 3.56584 |
| chr10 | 61475562 | 61476084 | 937 | Promoter (<=1kb) | 10 | 61475833 | 61582226 | 106394 | 1 | ENSMUST00 | 0 | Lrrc20 | ENSMUSG0C | false | chr10 | 61475289 | 61476610 | 27.38781 | 96.99624 | 93.49658 | chr10 | 61474630 | 61477427 | 5.80316 | 4.50266 | 3.66366 |
| chr6 | 85374372 | 85374925 | 937 | Promoter (<=1kb) | 6 | 85334963 | 85374634 | 39672 | 2 | ENSMUST00 | 0 | Rab11fip5 | ENSMUSG0C | false | chr6 | 85373844 | 85375332 | 33.32866 | 126.59486 | 122.2257 | . | . | . | . | . | . |
| chr7 | 46838086 | 46838459 | 937 | Distal Intergenic | 7 | 46841475 | 46851994 | 10520 | 1 | ENSMUST00 | -3016 | Ldha | ENSMUSG0C | false | . | . | . | . | . | . | chr7 | 46837117 | 46841159 | 5.05688 | 4.03651 | 3.08308 |
| chr12 | 85252188 | 85252473 | 937 | Intron (ENSMUST00000019378.6 | 12 | 85245923 | 85245985 | 63 | 2 | ENSMUST00 | -6204 | Mir6938 | ENSMUSG0C | false | . | . | . | . | . | . | chr12 | 85251024 | 85257823 | 3.64793 | 3.37643 | 1.89211 |
| chr5 | 131600519 | 131600790 | 936 | Intron (ENSMUST00000161226.3 | 5 | 131437788 | 131616605 | 178818 | 2 | ENSMUST00 | 15815 | Auts2 | ENSMUSG0C | true | . | . | . | . | . | . | chr5 | 131599383 | 131601455 | 3.54569 | 3.18116 | 1.85269 |
| chr11 | 98884949 | 98885211 | 936 | 5' UTR | 11 | 98891173 | 98892097 | 925 | 1 | ENSMUST00 | -5962 | Wipf2 | ENSMUSG0C | false | . | . | . | . | . | . | chr11 | 98883993 | 98886819 | 4.53284 | 3.86155 | 2.6706 |
| chr3 | 51881943 | 51882251 | 936 | Intron (ENSMUST00000121440.2 | 3 | 51828912 | 51829028 | 117 | 1 | ENSMUST00 | 53032 | n-R5s196 | ENSMUSG0C | false | . | . | . | . | . | . | chr3 | 51877616 | 51896424 | 5.58413 | 4.30162 | 3.49753 |
| chr17 | 12734209 | 12734481 | 936 | Intron (ENSMUST00000024599.8 | 17 | 12741311 | 12860122 | 118812 | 1 | ENSMUST00 | -6830 | Airn | ENSMUSG0C | false | . | . | . | . | . | . | chr17 | 12733766 | 12734884 | 4.3457 | 3.82065 | 2.45428 |
| chr4 | 105338914 | 105339297 | 936 | Distal Intergenic | 4 | 105374454 | 105374946 | 493 | 2 | ENSMUST00 | 35649 | Gm12722 | ENSMUSG0C | false | . | . | . | . | . | . | chr4 | 105337701 | 105339941 | 3.15201 | 3.10188 | 1.52761 |
| chr3 | 94404398 | 94404691 | 936 | Promoter (1-2kb) | 3 | 94392971 | 94402848 | 9878 | 2 | ENSMUST00 | -1551 | Gm17179 | ENSMUSG0C | false | . | . | . | . | . | . | chr3 | 94403221 | 94406496 | 6.08922 | 4.53013 | 3.93833 |
| chr1 | 69685776 | 69686364 | 935 | Promoter (<=1kb) | 1 | 69531214 | 69685821 | 154608 | 2 | ENSMUST00 | 0 | Ikzf2 | ENSMUSG0C | false | chr1 | 69685182 | 69688560 | 36.31549 | 112.95179 | 109.03448 | . | . | . | . | . | . |
| chr4 | 119541289 | 119541542 | 935 | Promoter (<=1kb) | 4 | 119540522 | 119573745 | 33224 | 1 | ENSMUST00 | 768 | Foxj3 | ENSMUSG0C | true | chr4 | 119538635 | 119541529 | 41.61333 | 140.51047 | 135.61182 | . | . | . | . | . | . |
| chr3 | 144569973 | 144570480 | 935 | Promoter (<=1kb) | 3 | 144570304 | 144597257 | 26954 | 1 | ENSMUST00 | 0 | Sep-15 | ENSMUSG0C | false | chr3 | 144568873 | 144570248 | 35.02417 | 111.75519 | 107.8695 | . | . | . | . | . | . |
| chr19 | 28679540 | 28680434 | 935 | Promoter (<=1kb) | 19 | 28258851 | 28680077 | 421227 | 2 | ENSMUST00 | 0 | Glis3 | ENSMUSG0C | false | chr19 | 28677815 | 28680438 | 26.39899 | 75.26234 | 72.19086 | chr19 | 28677731 | 28680307 | 3.41802 | 3.25677 | 1.70356 |
| chr2 | 123397618 | 123397907 | 935 | Distal Intergenic | 2 | 123273924 | 123274211 | 288 | 1 | ENSMUST00 | 123695 | Gm13988 | ENSMUSG0C | false | . | . | . | . | . | . | chr2 | 123397317 | 123401334 | 4.1122 | 3.5192 | 2.30929 |
| chr14 | 45445570 | 45446035 | 935 | Distal Intergenic | 14 | 45461363 | 45461645 | 283 | 2 | ENSMUST00 | 15610 | Gm22637 | ENSMUSG0C | false | . | . | . | . | . | . | chr14 | 45444716 | 45447212 | 4.19073 | 3.50624 | 2.37003 |
| chr3 | 95356947 | 95357413 | 934 | Promoter (<=1kb) | 3 | 95323525 | 95357155 | 33631 | 2 | ENSMUST00 | 0 | Setdb1 | ENSMUSG0C | false | chr3 | 95356009 | 95357528 | 16.32424 | 48.10018 | 45.4107 | . | . | . | . | . | . |
| chr8 | 47713808 | 47714482 | 934 | Promoter (<=1kb) | 8 | 47709344 | 47713931 | 4588 | 2 | ENSMUST00 | 0 | Cdkn2aip | ENSMUSG0C | false | chr8 | 47712794 | 47714015 | 31.76134 | 137.24866 | 132.47624 | . | . | . | . | . | . |
| chr1 | 134560018 | 134561508 | 934 | Promoter (<=1kb) | 1 | 134560171 | 134635285 | 75115 | 1 | ENSMUST00 | 0 | Kdm5b | ENSMUSG0C | false | chr1 | 134559887 | 134561761 | 32.2491 | 132.9626 | 128.36652 | chr1 | 134559712 | 134560169 | 3.00098 | 3.05606 | 1.35513 |
| chr5 | 150465433 | 150465756 | 934 | Intron (ENSMUST00000087204.5 | 5 | 150509694 | 150511338 | 1645 | 2 | ENSMUST00 | 45582 | Gm15933 | ENSMUSG0C | false | . | . | . | . | . | . | chr5 | 150464615 | 150467435 | 7.08694 | 5.02005 | 4.70994 |
| chr2 | 30471855 | 30472552 | 934 | Promoter (1-2kb) | 2 | 30471537 | 30474219 | 2683 | 2 | ENSMUST00 | 1667 | Ier5l | ENSMUSG0C | false | chr2 | 30471582 | 30474710 | 31.56964 | 96.59996 | 93.11521 | chr2 | 30469252 | 30473405 | 4.8209 | 4.06708 | 2.84851 |
| chr4 | 130829182 | 130829657 | 934 | Promoter (2-3kb) | 4 | 130824394 | 130826313 | 1920 | 2 | ENSMUST00 | -2870 | Gm26716 | ENSMUSG0C | false | . | . | . | . | . | . | chr4 | 130828492 | 130830002 | 5.43886 | 4.24999 | 3.41315 |
| chr15 | 32539111 | 32539406 | 933 | Intron (ENSMUST00000067458.6 | 15 | 32244813 | 32696341 | 451529 | 1 | ENSMUST00 | 294299 | Sema5a | ENSMUSG0C | true | . | . | . | . | . | . | chr15 | 32499761 | 32540421 | 4.87542 | 4.01677 | 2.90332 |
| chr4 | 134914584 | 134915091 | 933 | Promoter (<=1kb) | 4 | 134897849 | 134915024 | 17176 | 2 | ENSMUST00 | 0 | Tmem50a | ENSMUSG0C | false | chr4 | 134913993 | 134915235 | 32.11694 | 110.747 | 106.89115 | chr4 | 134913137 | 134915909 | 3.91408 | 3.51827 | 2.1678 |
| chr9 | 57708396 | 57708753 | 933 | Promoter (<=1kb) | 9 | 57708540 | 57752499 | 43960 | 1 | ENSMUST00 | 0 | Edc3 | ENSMUSG0C | false | chr9 | 57708441 | 57709366 | 30.03893 | 101.89226 | 98.2715 | chr9 | 57708240 | 57710570 | 3.95507 | 3.62022 | 2.13813 |
| chr8 | 91444772 | 91445190 | 933 | Intron (ENSMUST00000069718.8 | 8 | 91467685 | 91666946 | 199262 | 1 | ENSMUST00 | -22495 | Fto | ENSMUSG0C | false | . | . | . | . | . | . | chr8 | 91437454 | 91445828 | 3.89467 | 3.50581 | 2.10835 |
| chr11 | 60104875 | 60106198 | 933 | Promoter (<=1kb) | 11 | 60105013 | 60199195 | 94183 | 1 | ENSMUST00 | 0 | Rai1 | ENSMUSG0C | false | chr11 | 60104773 | 60106757 | 36.76944 | 114.78011 | 110.80492 | chr11 | 60104188 | 60104909 | 4.31724 | 3.63497 | 2.48429 |
| chrX | 104464368 | 104464676 | 933 | Distal Intergenic | 20 | 104482818 | 104507490 | 24673 | 1 | ENSMUST00 | -18142 | Uprt | ENSMUSG0C | false | . | . | . | . | . | . | . | . | . | . | . | . |
| chr4 | 118131185 | 118131610 | 932 | Intron (ENSMUST00000030263.3 | 4 | 117932160 | 118134882 | 202723 | 2 | ENSMUST00 | 3272 | St3gal3 | ENSMUSG0C | true | . | . | . | . | . | . | chr4 | 118119621 | 118134187 | 5.32651 | 4.16451 | 3.29068 |
| chr9 | 103007700 | 103008084 | 932 | Promoter (<=1kb) | 9 | 103008002 | 103090451 | 82450 | 1 | ENSMUST00 | 0 | Slco2a1 | ENSMUSG0C | true | chr9 | 103007566 | 103008928 | 30.2403 | 90.37072 | 87.01736 | chr9 | 102991889 | 103008204 | 5.47516 | 4.26994 | 3.39839 |
| chr3 | 116807971 | 116808414 | 932 | Promoter (<=1kb) | 3 | 116739999 | 116808166 | 68168 | 2 | ENSMUST00 | 0 | Agl | ENSMUSG0C | false | chr3 | 116807428 | 116808817 | 21.76566 | 71.50774 | 68.49327 | . | . | . | . | . | . |
| chr15 | 58166176 | 58166501 | 932 | Distal Intergenic | 15 | 58146616 | 58146742 | 127 | 1 | ENSMUST00 | 19561 | Gm22488 | ENSMUSG0C | false | . | . | . | . | . | . | . | . | . | . | . | . |
| chr9 | 97813236 | 97813574 | 932 | Intron (ENSMUST00000035027.7 | 9 | 98003420 | 98005769 | 2350 | 1 | ENSMUST00 | -189846 | 4921534H16I | ENSMUSG0C | false | . | . | . | . | . | . | chr9 | 97812745 | 97813782 | 4.50717 | 3.91967 | 2.64549 |
| chr5 | 73231845 | 73232329 | 932 | Intron (ENSMUST00000101127.6 | 5 | 73210109 | 73212634 | 2526 | 2 | ENSMUST00 | -19212 | Gm17207 | ENSMUSG0C | false | . | . | . | . | . | . | chr5 | 73196545 | 73254445 | 6.29302 | 4.62166 | 4.08669 |
| chr6 | 92706102 | 92706489 | 931 | Promoter (<=1kb) | 6 | 92370892 | 92706145 | 335254 | 2 | ENSMUST00 | 0 | Prickle2 | ENSMUSG0C | false | chr6 | 92705404 | 92706488 | 21.51301 | 61.48686 | 58.61925 | chr6 | 92704892 | 92706483 | 4.94594 | 4.09281 | 3.00801 |
| chr13 | 103773965 | 103774817 | 931 | Distal Intergenic | 13 | 103741615 | 103764504 | 22890 | 2 | ENSMUST00 | -9462 | Srek1 | ENSMUSG0C | false | chr13 | 103773260 | 103775065 | 19.74353 | 71.90538 | 68.88615 | chr13 | 103774777 | 103775223 | 4.88796 | 4.08013 | 2.90636 |
| chr13 | 64153385 | 64153767 | 931 | Promoter (<=1kb) | 13 | 64140468 | 64153202 | 12735 | 2 | ENSMUST00 | -184 | Zfp367 | ENSMUSG0C | false | chr13 | 64153467 | 64154294 | 12.8449 | 29.2147 | 26.75964 | . | . | . | . | . | . |
| chr17 | 17827711 | 17828013 | 931 | Promoter (2-3kb) | 17 | 17830188 | 17830257 | 70 | 1 | ENSMUST00 | -2175 | Mir99b | ENSMUSG0C | false | chr17 | 17825989 | 17829531 | 29.75681 | 92.45776 | 89.05831 | . | . | . | . | . | . |
| chr2 | 30692966 | 30693573 | 931 | Distal Intergenic | 2 | 30658613 | 30678011 | 19399 | 2 | ENSMUST00 | -14956 | Gm14486 | ENSMUSG0C | false | chr2 | 30693330 | 30693904 | 8.26071 | 16.87976 | 14.58833 | chr2 | 30691850 | 30694277 | 4.53114 | 3.84302 | 2.64072 |
| chr11 | 69995346 | 69995975 | 930 | Promoter (<=1kb) | 11 | 69995777 | 70000009 | 4233 | 1 | ENSMUST00 | 0 | Phf23 | ENSMUSG0C | false | chr11 | 69995731 | 69997499 | 38.91574 | 128.10115 | 123.6785 | . | . | . | . | . | . |
| chr10 | 79934307 | 79934707 | 930 | Intron (ENSMUST00000019708.6 | 10 | 79930424 | 79931218 | 795 | 1 | ENSMUST00 | 3884 | Arid3a | ENSMUSG0C | false | . | . | . | . | . | . | chr10 | 79931436 | 79938436 | 8.97089 | 5.80763 | 6.24553 |
| chr11 | 69099806 | 69100243 | 930 | Promoter (<=1kb) | 11 | 69100152 | 69101048 | 897 | 1 | ENSMUST00 | 0 | Per1 | ENSMUSG0C | true | chr11 | 69098805 | 69100459 | 19.38504 | 61.02572 | 58.1656 | chr11 | 69097810 | 69105920 | 6.46779 | 4.59054 | 4.2103 |
| chr3 | 41605622 | 41606201 | 930 | Intron (ENSMUST00000026865.9 | 3 | 41627117 | 41630123 | 3007 | 2 | ENSMUST00 | 23922 | Sclt1 | ENSMUSG0C | false | . | . | . | . | . | . | chr3 | 41598517 | 41607869 | 6.56571 | 4.58882 | 4.32841 |
| chr8 | 116814521 | 116814915 | 930 | Distal Intergenic | 8 | 116764144 | 116765479 | 1336 | 2 | ENSMUST00 | -49043 | Gm10620 | ENSMUSG0C | false | . | . | . | . | . | . | chr8 | 116814302 | 116815183 | 4.48722 | 3.69215 | 2.58374 |
| chr4 | 97751644 | 97752322 | 930 | Intron (ENSMUST00000146447.2 | 4 | 97754168 | 97757389 | 3222 | 2 | ENSMUST00 | 5067 | Gm12676 | ENSMUSG0C | false | chr4 | 97751131 | 97752475 | 27.46033 | 79.85941 | 76.71169 | . | . | . | . | . | . |
| chr11 | 55419630 | 55420136 | 929 | Promoter (<=1kb) | 11 | 55394500 | 55419898 | 25399 | 2 | ENSMUST00 | 0 | Sparc | ENSMUSG0C | true | chr11 | 55417325 | 55420160 | 29.3377 | 90.74815 | 87.38457 | chr11 | 55419979 | 55420777 | 5.85316 | 4.56839 | 3.70909 |
| chr10 | 121962303 | 121962595 | 929 | Intron (ENSMUST00000020322.6 | 10 | 121780991 | 122047308 | 266318 | 2 | ENSMUST00 | 84713 | Srgap1 | ENSMUSG0C | true | . | . | . | . | . | . | chr10 | 121945243 | 122036389 | 5.74397 | 4.3951 | 3.62974 |
| chr19 | 42090137 | 42090688 | 929 | Promoter (<=1kb) | 19 | 42090435 | 42122218 | 31784 | 1 | ENSMUST00 | 0 | Pi4k2a | ENSMUSG0C | false | chr19 | 42090349 | 42091418 | 30.17592 | 94.17345 | 90.73757 | . | . | . | . | . | . |
| chr5 | 130200647 | 130200982 | 929 | Intron (ENSMUST00000119797.2 | 5 | 130207053 | 130210991 | 3939 | 1 | ENSMUST00 | -6071 | Rabgef1 | ENSMUSG0C | false | . | . | . | . | . | . | chr5 | 130172335 | 130207681 | 5.43982 | 4.22227 | 3.37739 |
| chr4 | 150778795 | 150779138 | 929 | Distal Intergenic | 4 | 150825733 | 150839575 | 13843 | 1 | ENSMUST00 | -46595 | Gm13049 | ENSMUSG0C | false | . | . | . | . | . | . | . | . | . | . | . | . |
| chr1 | 72944270 | 72944624 | 929 | Distal Intergenic | 1 | 72984844 | 73024724 | 39881 | 1 | ENSMUST00 | -40220 | RP23-445K8. | ENSMUSG0C | false | . | . | . | . | . | . | chr1 | 72944482 | 72946132 | 4.61488 | 3.67114 | 2.71941 |
| chr19 | 10164640 | 10164931 | 928 | Distal Intergenic | 19 | 10182888 | 10196870 | 13983 | 1 | ENSMUST00 | -17957 | Fads1 | ENSMUSG0C | false | . | . | . | . | . | . | . | . | . | . | . | . |
| chr1 | 43698445 | 43698735 | 928 | Distal Intergenic | 1 | 43730602 | 43742578 | 11977 | 1 | ENSMUST00 | -31867 | 1500015O10 | ENSMUSG0C | false | . | . | . | . | . | . | chr1 | 43698101 | 43699817 | 5.44098 | 4.223 | 3.39102 |
| chr19 | 32304077 | 32304408 | 928 | Intron (ENSMUST00000099514.4 | 19 | 32220587 | 32289579 | 68993 | 2 | ENSMUST00 | -14499 | Sgms1 | ENSMUSG0C | false | . | . | . | . | . | . | chr19 | 32301122 | 32307156 | 5.60323 | 4.40652 | 3.49303 |
| chr11 | 5421924 | 5422340 | 928 | Intron (ENSMUST00000109867.2 | 11 | 5276324 | 5444847 | 168524 | 2 | ENSMUST00 | 22507 | Znrf3 | ENSMUSG0C | false | . | . | . | . | . | . | chr11 | 5410686 | 5427806 | 5.74514 | 4.38198 | 3.63413 |
| chr3 | 121650275 | 121650734 | 928 | Distal Intergenic | 3 | 121637145 | 121643485 | 6341 | 2 | ENSMUST00 | -6791 | A730020M07 | ENSMUSG0C | false | . | . | . | . | . | . | chr3 | 121649086 | 121651504 | 5.48653 | 4.35187 | 3.40071 |
| chr4 | 148979319 | 148979590 | 928 | Intron (ENSMUST00000146124.1 | 4 | 148960360 | 148966998 | 6639 | 1 | ENSMUST00 | 18960 | Gm17029 | ENSMUSG0C | false | . | . | . | . | . | . | chr4 | 148978452 | 148981491 | 3.12155 | 3.05113 | 1.51451 |
| chr2 | 90940289 | 90941110 | 927 | Promoter (<=1kb) | 2 | 90940382 | 90987754 | 47373 | 1 | ENSMUST00 | 0 | Celf1 | ENSMUSG0C | false | chr2 | 90940019 | 90941495 | 34.63471 | 129.05151 | 124.59962 | . | . | . | . | . | . |
| chr14 | 54653552 | 54653847 | 927 | Promoter (<=1kb) | 14 | 54642161 | 54653616 | 11456 | 2 | ENSMUST00 | 0 | Acin1 | ENSMUSG0C | false | chr14 | 54652953 | 54653965 | 20.67921 | 57.50454 | 54.69232 | chr14 | 54650135 | 54654884 | 3.9357 | 3.27722 | 2.1717 |
| chr18 | 22343812 | 22344757 | 927 | Promoter (<=1kb) | 18 | 22344883 | 22378950 | 34068 | 1 | ENSMUST00 | -126 | Asxl3 | ENSMUSG0C | false | chr18 | 22342934 | 22345013 | 37.30079 | 124.19571 | 119.91465 | . | . | . | . | . | . |
| chr2 | 167188707 | 167189221 | 927 | Promoter (<=1kb) | 2 | 167103378 | 167188799 | 85422 | 2 | ENSMUST00 | 0 | Kcnb1 | ENSMUSG0C | false | chr2 | 167188384 | 167189902 | 22.97661 | 61.46103 | 58.5941 | chr2 | 167188136 | 167190629 | 4.63484 | 3.74978 | 2.73873 |
| chr8 | 92064794 | 92065149 | 927 | Distal Intergenic | 8 | 92039917 | 92040020 | 104 | 1 | ENSMUST00 | 24878 | Gm24254 | ENSMUSG0C | false | . | . | . | . | . | . | chr8 | 92064727 | 92067058 | 4.23558 | 3.63614 | 2.39963 |
| chr1 | 137354972 | 137355293 | 927 | Distal Intergenic | 1 | 137367151 | 137367247 | 97 | 2 | ENSMUST00 | 11954 | Gm23534 | ENSMUSG0C | false | . | . | . | . | . | . | chr1 | 137354458 | 137356867 | 4.4293 | 3.83358 | 2.54293 |
| chr15 | 88733136 | 88733888 | 926 | Promoter (<=1kb) | 15 | 88698851 | 88734230 | 35380 | 2 | ENSMUST00 | 342 | Brd1 | ENSMUSG0C | false | chr15 | 88732641 | 88734627 | 19.45739 | 70.46334 | 67.46697 | . | . | . | . | . | . |
| chrX | 38564473 | 38565138 | 926 | Promoter (<=1kb) | 20 | 38533274 | 38564525 | 31252 | 2 | ENSMUST00 | 0 | Cul4b | ENSMUSG0C | false | chrX | 38563569 | 38565117 | 19.30035 | 48.68736 | 45.98855 | . | . | . | . | . | . |
| chr18 | 73754275 | 73754667 | 926 | Promoter (<=1kb) | 18 | 73735038 | 73754479 | 19442 | 2 | ENSMUST00 | 0 | Elac1 | ENSMUSG0C | false | chr18 | 73753844 | 73755108 | 28.1615 | 93.53628 | 90.11485 | chr18 | 73753329 | 73755465 | 4.00046 | 3.59996 | 2.23539 |
| chr17 | 70851105 | 70852042 | 926 | Promoter (<=1kb) | 17 | 70844205 | 70851207 | 7003 | 2 | ENSMUST00 | 0 | Tgif1 | ENSMUSG0C | false | chr17 | 70848566 | 70853866 | 23.45396 | 71.92415 | 68.90385 | . | . | . | . | . | . |
| chr6 | 87294866 | 87295144 | 926 | Intron (ENSMUST00000042025.9 | 6 | 87281149 | 87281260 | 112 | 1 | ENSMUST00 | 13718 | n-R5s164 | ENSMUSG0C | false | . | . | . | . | . | . | chr6 | 87294617 | 87295618 | 4.86309 | 4.03378 | 2.90815 |
| chr1 | 132335457 | 132335909 | 926 | Distal Intergenic | 1 | 132342023 | 132349626 | 7604 | 1 | ENSMUST00 | -6114 | RP23-217K18 | ENSMUSG0C | false | . | . | . | . | . | . | chr1 | 132317064 | 132339308 | 6.74463 | 4.86917 | 4.43585 |
| chr7 | 31111880 | 31112376 | 925 | Promoter (<=1kb) | 7 | 31109140 | 31111958 | 2819 | 2 | ENSMUST00 | 0 | Hpn | ENSMUSG0C | false | . | . | . | . | . | . | chr7 | 31108643 | 31112914 | 3.88171 | 3.47007 | 2.1192 |
| chr4 | 136144558 | 136144973 | 925 | Promoter (<=1kb) | 4 | 136144974 | 136145755 | 782 | 1 | ENSMUST00 | -1 | Id3 | ENSMUSG0C | false | chr4 | 136142962 | 136145178 | 19.27017 | 55.84607 | 53.05495 | chr4 | 136142629 | 136145211 | 5.36542 | 4.15748 | 3.32061 |
| chrX | 35888508 | 35888930 | 925 | Promoter (<=1kb) | 20 | 35888832 | 36076562 | 187731 | 1 | ENSMUST00 | 0 | Dock11 | ENSMUSG0C | true | chrX | 35888509 | 35889983 | 23.43614 | 63.09834 | 60.20888 | . | . | . | . | . | . |
| chrX | 145505086 | 145505694 | 925 | Promoter (<=1kb) | 20 | 145446425 | 145505144 | 58720 | 2 | ENSMUST00 | 0 | Amot | ENSMUSG0C | false | chrX | 145504092 | 145506026 | 22.51707 | 59.83231 | 56.98677 | . | . | . | . | . | . |
| chr14 | 21164176 | 21164503 | 925 | Intron (ENSMUST00000045376.9 | 14 | 21263624 | 21263859 | 236 | 1 | ENSMUST00 | -99121 | Gm26372 | ENSMUSG0C | false | chr14 | 21164096 | 21164544 | 4.42115 | 6.83029 | 4.72578 | chr14 | 21163555 | 21165155 | 3.29544 | 3.25744 | 1.6005 |
| chr14 | 25371592 | 25372047 | 925 | Intron (ENSMUST00000181582.1 | 14 | 25355801 | 25378635 | 22835 | 2 | ENSMUST00 | 6588 | Gm26660 | ENSMUSG0C | false | . | . | . | . | . | . | chr14 | 25370182 | 25373552 | 4.50744 | 3.55192 | 2.63962 |
| chr7 | 25275767 | 25276250 | 924 | Promoter (2-3kb) | 7 | 25278422 | 25284969 | 6548 | 1 | ENSMUST00 | -2172 | Cic | ENSMUSG0C | false | . | . | . | . | . | . | chr7 | 25269284 | 25281823 | 5.87867 | 4.36785 | 3.75349 |
| chr17 | 74527738 | 74528849 | 924 | Promoter (<=1kb) | 17 | 74528295 | 74703356 | 175062 | 1 | ENSMUST00 | 0 | Birc6 | ENSMUSG0C | false | chr17 | 74527497 | 74528154 | 17.60262 | 45.78123 | 43.11721 | . | . | . | . | . | . |
| chr3 | 90079430 | 90079745 | 924 | Promoter (<=1kb) | 3 | 90079540 | 90100901 | 21362 | 1 | ENSMUST00 | 0 | Tpm3 | ENSMUSG0C | false | chr3 | 90079128 | 90080759 | 41.91343 | 144.39362 | 139.32506 | . | . | . | . | . | . |
| chr7 | 120102167 | 120102854 | 924 | Promoter (<=1kb) | 7 | 120102376 | 120120986 | 18611 | 1 | ENSMUST00 | 0 | Tmem159 | ENSMUSG0C | false | chr7 | 120101899 | 120103846 | 34.54477 | 122.48211 | 118.25522 | chr7 | 120100216 | 120102228 | 4.92806 | 3.94137 | 2.9664 |
| chrX | 56455744 | 56456194 | 924 | Promoter (<=1kb) | 20 | 56454885 | 56507843 | 52959 | 1 | ENSMUST00 | 860 | Ddx26b | ENSMUSG0C | false | chrX | 56454847 | 56456286 | 25.27427 | 69.73028 | 66.74294 | . | . | . | . | . | . |
| chr3 | 132949943 | 132950542 | 924 | Promoter (<=1kb) | 3 | 132881745 | 132950291 | 68547 | 2 | ENSMUST00 | 0 | Npnt | ENSMUSG0C | true | chr3 | 132948145 | 132950666 | 16.50874 | 70.50747 | 67.5107 | . | . | . | . | . | . |
| chr17 | 44105526 | 44106046 | 923 | Promoter (<=1kb) | 17 | 44096308 | 44105809 | 9502 | 2 | ENSMUST00 | 0 | Enpp4 | ENSMUSG0C | false | chr17 | 44105080 | 44106469 | 34.36702 | 111.64683 | 107.76208 | . | . | . | . | . | . |
| chr19 | 25267299 | 25268173 | 923 | Intron (ENSMUST00000049400.9 | 19 | 25236975 | 25434496 | 197522 | 1 | ENSMUST00 | 30325 | Kank1 | ENSMUSG0C | true | . | . | . | . | . | . | chr19 | 25261572 | 25293097 | 5.29409 | 4.21889 | 3.24888 |
| chr18 | 38634790 | 38635214 | 923 | Intron (ENSMUST00000145090.2 | 18 | 38627287 | 38642930 | 15644 | 1 | ENSMUST00 | 7504 | Arhgap26 | ENSMUSG0C | false | . | . | . | . | . | . | chr18 | 38633366 | 38637673 | 5.18756 | 4.19008 | 3.17393 |
| chr11 | 58008575 | 58008946 | 923 | Promoter (<=1kb) | 11 | 58009064 | 58062031 | 52968 | 1 | ENSMUST00 | -118 | Larp1 | ENSMUSG0C | false | chr11 | 58008900 | 58010805 | 30.65557 | 96.00109 | 92.52367 | chr11 | 58007897 | 58008657 | 4.4095 | 3.82137 | 2.51157 |
| chr5 | 144357687 | 144358630 | 923 | Promoter (<=1kb) | 5 | 144264526 | 144358112 | 93587 | 2 | ENSMUST00 | 0 | Baiap2l1 | ENSMUSG0C | false | chr5 | 144357066 | 144358946 | 33.41831 | 117.26761 | 113.21259 | . | . | . | . | . | . |
| chr2 | 32534938 | 32535818 | 923 | Promoter (<=1kb) | 2 | 32535332 | 32569750 | 34419 | 1 | ENSMUST00 | 0 | Fam102a | ENSMUSG0C | false | chr2 | 32534502 | 32536789 | 36.10317 | 115.72481 | 111.71513 | chr2 | 32535767 | 32537408 | 4.06627 | 3.70387 | 2.22048 |
| chr9 | 37613195 | 37613998 | 922 | Promoter (<=1kb) | 9 | 37613847 | 37648318 | 34472 | 1 | ENSMUST00 | 0 | Siae | ENSMUSG0C | false | chr9 | 37612835 | 37614176 | 26.65286 | 74.78688 | 71.72144 | chr9 | 37612768 | 37614282 | 4.47919 | 3.85366 | 2.57876 |
| chr11 | 78566695 | 78567188 | 922 | Promoter (1-2kb) | 11 | 78567402 | 78568916 | 1515 | 2 | ENSMUST00 | 1728 | Nlk | ENSMUSG0C | true | . | . | . | . | . | . | chr11 | 78565466 | 78571591 | 4.72666 | 3.87884 | 2.82952 |
| chr4 | 149990218 | 149990706 | 922 | Intron (ENSMUST00000084117.7 | 4 | 149995848 | 150001265 | 5418 | 2 | ENSMUST00 | 10559 | H6pd | ENSMUSG0C | false | . | . | . | . | . | . | chr4 | 149985077 | 150009610 | 6.9066 | 4.88029 | 4.57705 |
| chr9 | 63980374 | 63980812 | 922 | Intron (ENSMUST00000041029.5 | 9 | 63954009 | 64016987 | 62979 | 2 | ENSMUST00 | 36175 | Smad6 | ENSMUSG0C | true | . | . | . | . | . | . | chr9 | 63948352 | 64020083 | 7.27873 | 5.05859 | 4.874 |
| chr13 | 20077220 | 20077542 | 922 | Distal Intergenic | 13 | 20090596 | 20264007 | 173412 | 1 | ENSMUST00 | -13054 | Elmo1 | ENSMUSG0C | true | . | . | . | . | . | . | chr13 | 20076604 | 20077767 | 4.75772 | 3.98275 | 2.80149 |
| chr8 | 109415410 | 109415750 | 922 | Distal Intergenic | 8 | 109425644 | 109425744 | 101 | 2 | ENSMUST00 | 9994 | Gm23163 | ENSMUSG0C | false | . | . | . | . | . | . | chr8 | 109414917 | 109416101 | 3.14658 | 3.11285 | 1.48062 |
| chr5 | 148511261 | 148511652 | 921 | Intron (ENSMUST00000079324.8 | 5 | 148504631 | 148552788 | 48158 | 2 | ENSMUST00 | 41136 | Ubl3 | ENSMUSG0C | false | . | . | . | . | . | . | chr5 | 148509345 | 148513472 | 4.55819 | 3.69621 | 2.67521 |
| chr6 | 137440313 | 137440662 | 921 | Intron (ENSMUST00000077115.7 | 6 | 137410721 | 137464629 | 53909 | 1 | ENSMUST00 | 29593 | Ptpro | ENSMUSG0C | true | . | . | . | . | . | . | chr6 | 137437087 | 137441627 | 4.75901 | 4.0003 | 2.8115 |
| chr9 | 71660514 | 71660830 | 921 | Intron (ENSMUST00000072899.3 | 9 | 71632633 | 71648350 | 15718 | 2 | ENSMUST00 | -12165 | Cgnl1 | ENSMUSG0C | false | . | . | . | . | . | . | chr9 | 71657960 | 71671511 | 6.52791 | 4.76881 | 4.25927 |
| chr19 | 32322230 | 32322695 | 921 | Promoter (<=1kb) | 19 | 32160170 | 32323229 | 163060 | 2 | ENSMUST00 | 534 | Sgms1 | ENSMUSG0C | false | . | . | . | . | . | . | chr19 | 32318143 | 32331326 | 6.10815 | 4.62528 | 3.91491 |
| chr14 | 33015262 | 33015549 | 921 | 3' UTR | 14 | 33018284 | 33023337 | 5054 | 2 | ENSMUST00 | 7788 | Wdfy4 | ENSMUSG0C | false | . | . | . | . | . | . | chr14 | 33013795 | 33017224 | 6.58401 | 4.77985 | 4.30665 |
| chr13 | 51249088 | 51249473 | 921 | Distal Intergenic | 13 | 51202688 | 51203065 | 378 | 2 | ENSMUST00 | -46024 | Hist1h2al | ENSMUSG0C | false | . | . | . | . | . | . | chr13 | 51249447 | 51251303 | 3.77395 | 3.45027 | 2.04755 |
| chr10 | 13090494 | 13090868 | 920 | Promoter (<=1kb) | 10 | 13090691 | 13124860 | 34170 | 1 | ENSMUST00 | 0 | Plagl1 | ENSMUSG0C | true | chr10 | 13090292 | 13091986 | 23.26691 | 63.66726 | 60.76967 | . | . | . | . | . | . |
| chr13 | 13394004 | 13394364 | 920 | Promoter (<=1kb) | 13 | 13357623 | 13393624 | 36002 | 2 | ENSMUST00 | -381 | Gpr137b | ENSMUSG0C | false | chr13 | 13392538 | 13394560 | 25.29353 | 74.26213 | 71.20484 | chr13 | 13391893 | 13394867 | 5.33543 | 4.08284 | 3.32348 |
| chr5 | 76183780 | 76184560 | 920 | Promoter (<=1kb) | 5 | 76183880 | 76209245 | 25366 | 1 | ENSMUST00 | 0 | Tmem165 | ENSMUSG0C | false | chr5 | 76183382 | 76185050 | 40.73933 | 130.17783 | 125.67887 | chr5 | 76182437 | 76191091 | 3.91563 | 3.48666 | 2.14013 |
| chr14 | 70044059 | 70044455 | 920 | Intron (ENSMUST00000022678.3 | 14 | 70077445 | 70080157 | 2713 | 1 | ENSMUST00 | -32990 | Egr3 | ENSMUSG0C | false | . | . | . | . | . | . | chr14 | 70043319 | 70045778 | 8.71215 | 5.73982 | 6.03936 |
| chr18 | 88893937 | 88894485 | 920 | Promoter (<=1kb) | 18 | 88668695 | 88894339 | 225645 | 2 | ENSMUST00 | 0 | Socs6 | ENSMUSG0C | false | chr18 | 88893282 | 88894599 | 26.35494 | 78.87288 | 75.741 | chr18 | 88892303 | 88894478 | 4.43387 | 3.87033 | 2.52536 |
| chr15 | 73864881 | 73865191 | 920 | Distal Intergenic | 15 | 73796736 | 73839699 | 42964 | 2 | ENSMUST00 | -25183 | Mroh5 | ENSMUSG0C | false | . | . | . | . | . | . | . | . | . | . | . | . |
| chr17 | 23803409 | 23803771 | 919 | Promoter (<=1kb) | 17 | 23803221 | 23824741 | 21521 | 1 | ENSMUST00 | 189 | Srrm2 | ENSMUSG0C | false | chr17 | 23803067 | 23805197 | 35.30345 | 112.24108 | 108.33721 | . | . | . | . | . | . |
| chr15 | 78832378 | 78833155 | 919 | Distal Intergenic | 15 | 78842647 | 78850902 | 8256 | 1 | ENSMUST00 | -9492 | Cdc42ep1 | ENSMUSG0C | false | . | . | . | . | . | . | chr15 | 78830112 | 78842533 | 8.19317 | 5.56805 | 5.60698 |

| | | | | | | | | | | | | | | | | | | | | | | | | | | |
|---|---|---|---|---|---|---|---|---|---|---|---|---|---|---|---|---|---|---|---|---|---|---|---|---|---|---|
| chr5 | 88564716 | 88565156 | 919 | Distal Intergenic | 5 | 88554483 | 88556083 | 1601 | 1 | ENSMUST00 | 10234 | Utp3 | ENSMUSG0C | false | chr5 | 88564232 | 88566691 | 28.91247 | 96.86193 | 93.36482 | . | . | . | . | . | . |
| chr2 | 168123666 | 168124055 | 919 | Distal Intergenic | 2 | 168148017 | 168153533 | 5517 | 2 | ENSMUST00 | 29478 | Gm14235 | ENSMUSG0C | false | . | . | . | . | . | . | chr2 | 168121758 | 168126187 | 6.18138 | 4.66453 | 3.96064 |
| chr8 | 123041171 | 123041591 | 919 | Promoter (<=1kb) | 8 | 123042466 | 123044602 | 2137 | 1 | ENSMUST00 | -875 | RP23-460F2 | ENSMUSG0C | false | chr8 | 123040842 | 123042311 | 30.41442 | 103.5799 | 99.9162 | . | . | . | . | . | . |
| chr7 | 143367626 | 143368049 | 919 | Intron (ENSMUST00000009689.5 | 7 | 143322775 | 143330650 | 7876 | 2 | ENSMUST00 | -36977 | RP23-124B2. | ENSMUSG0C | false | chr7 | 143367648 | 143368462 | 13.02911 | 28.89677 | 26.44547 | chr7 | 143367227 | 143367737 | 3.13348 | 2.99869 | 1.5108 |
| chr17 | 28232534 | 28232887 | 918 | Promoter (<=1kb) | 17 | 28232754 | 28301469 | 68716 | 1 | ENSMUST00 | 0 | Ppard | ENSMUSG0C | false | chr17 | 28232691 | 28233753 | 43.4858 | 141.53311 | 136.59532 | chr17 | 28232288 | 28243092 | 6.25267 | 4.64244 | 4.03578 |
| chr11 | 65209522 | 65209836 | 918 | Intron (ENSMUST00000102635.4 | 11 | 65196102 | 65205467 | 9366 | 2 | ENSMUST00 | -4056 | Myocd | ENSMUSG0C | false | . | . | . | . | . | . | chr11 | 65209265 | 65211188 | 5.10319 | 4.08676 | 3.10012 |
| chr12 | 108835630 | 108836211 | 918 | Promoter (<=1kb) | 12 | 108825873 | 108835883 | 10011 | 2 | ENSMUST00 | 0 | Slc25a29 | ENSMUSG0C | false | chr12 | 108834590 | 108837062 | 32.32999 | 116.31703 | 112.294 | . | . | . | . | . | . |
| chr2 | 179947312 | 179947954 | 918 | Intron (ENSMUST00000041618.8 | 2 | 179922004 | 179929355 | 7352 | 2 | ENSMUST00 | -17958 | Taf4a | ENSMUSG0C | false | . | . | . | . | . | . | chr2 | 179942669 | 179955942 | 5.64155 | 4.18843 | 3.56812 |
| chr3 | 41172388 | 41172732 | 918 | Distal Intergenic | 3 | 41066326 | 41083046 | 16721 | 2 | ENSMUST00 | -89343 | Pgrmc2 | ENSMUSG0C | false | . | . | . | . | . | . | chr3 | 41171088 | 41174241 | 3.49791 | 3.29083 | 1.78296 |
| chr11 | 87471273 | 87471665 | 918 | Promoter (<=1kb) | 11 | 87471368 | 87471582 | 215 | 1 | ENSMUST00 | 0 | Rnu3b2 | ENSMUSG0C | false | . | . | . | . | . | . | . | . | . | . | . | . |
| chr11 | 94491057 | 94491358 | 917 | 3' UTR | 11 | 94484608 | 94486179 | 1572 | 2 | ENSMUST00 | -4879 | Spata20 | ENSMUSG0C | false | . | . | . | . | . | . | chr11 | 94489158 | 94497346 | 7.88949 | 5.284 | 5.39442 |
| chr12 | 111945099 | 111945419 | 917 | Distal Intergenic | 12 | 111961376 | 111966977 | 5602 | 2 | ENSMUST00 | 21558 | 2010107E04I | ENSMUSG0C | false | . | . | . | . | . | . | chr12 | 111942224 | 111946871 | 4.57899 | 3.59176 | 2.69776 |
| chr15 | 77793498 | 77793871 | 917 | Intron (ENSMUST00000016771.7 | 15 | 77794652 | 77813231 | 18580 | 2 | ENSMUST00 | 19360 | Myh9 | ENSMUSG0C | true | . | . | . | . | . | . | chr15 | 77781877 | 77840076 | 8.49833 | 5.61339 | 5.87304 |
| chr5 | 131537183 | 131537687 | 917 | Promoter (<=1kb) | 5 | 131475328 | 131537970 | 62643 | 2 | ENSMUST00 | 283 | Auts2 | ENSMUSG0C | true | chr5 | 131536784 | 131539135 | 30.78865 | 90.34473 | 86.99165 | . | . | . | . | . | . |
| chr6 | 82429898 | 82430236 | 917 | Intron (ENSMUST00000032122.8 | 6 | 82402475 | 82560103 | 157629 | 1 | ENSMUST00 | 27424 | Tacr1 | ENSMUSG0C | false | . | . | . | . | . | . | chr6 | 82429348 | 82431608 | 3.61359 | 3.30339 | 1.9042 |
| chr2 | 160225469 | 160225729 | 917 | Distal Intergenic | 2 | 160311393 | 160313616 | 2224 | 2 | ENSMUST00 | 87887 | Gm826 | ENSMUSG0C | false | . | . | . | . | . | . | chr2 | 160225122 | 160226692 | 5.05897 | 4.08108 | 3.10305 |
| chr17 | 12031268 | 12031547 | 916 | Promoter (1-2kb) | 17 | 12029916 | 12063358 | 33443 | 1 | ENSMUST00 | 1353 | Park2 | ENSMUSG0C | false | . | . | . | . | . | . | chr17 | 12029969 | 12032737 | 3.64787 | 3.39388 | 1.90076 |
| chr4 | 63528831 | 63529079 | 916 | Distal Intergenic | 4 | 63544772 | 63550750 | 5979 | 1 | ENSMUST00 | -15693 | Atp6v1g1 | ENSMUSG0C | false | . | . | . | . | . | . | chr4 | 63528040 | 63529535 | 4.22465 | 3.67006 | 2.4167 |
| chr13 | 93191805 | 93192658 | 916 | Promoter (<=1kb) | 13 | 93147400 | 93192283 | 44884 | 2 | ENSMUST00 | 0 | Papd4 | ENSMUSG0C | false | chr13 | 93191112 | 93192929 | 27.46926 | 112.95284 | 109.03484 | . | . | . | . | . | . |
| chr2 | 119740625 | 119740926 | 916 | Promoter (<=1kb) | 2 | 119740828 | 119740956 | 129 | 2 | ENSMUST00 | 30 | Gm25857 | ENSMUSG0C | false | . | . | . | . | . | . | chr2 | 119740863 | 119745199 | 5.39081 | 4.15988 | 3.37725 |
| chr2 | 60116009 | 60116358 | 916 | Intron (ENSMUST00000112550.2 | 2 | 60125096 | 60125819 | 724 | 1 | ENSMUST00 | -8738 | Gm13620 | ENSMUSG0C | false | . | . | . | . | . | . | chr2 | 60113393 | 60124063 | 4.90918 | 3.93506 | 2.95446 |
| chr11 | 117098188 | 117098668 | 916 | Promoter (1-2kb) | 11 | 117094948 | 117097067 | 2120 | 2 | ENSMUST00 | -1122 | Gm11730 | ENSMUSG0C | false | . | . | . | . | . | . | chr11 | 117097696 | 117104768 | 5.85805 | 4.31896 | 3.74714 |
| chr7 | 128591357 | 128591736 | 916 | Distal Intergenic | 7 | 128595979 | 128596278 | 300 | 2 | ENSMUST00 | 4542 | Gm7258 | ENSMUSG0C | false | . | . | . | . | . | . | chr7 | 128590684 | 128593255 | 4.75645 | 3.79214 | 2.84874 |
| chr1 | 125560270 | 125560650 | 915 | Promoter (<=1kb) | 1 | 125560595 | 125568573 | 7979 | 1 | ENSMUST00 | 0 | Slc35f5 | ENSMUSG0C | false | chr1 | 125559921 | 125561892 | 40.46864 | 131.89105 | 127.3278 | . | . | . | . | . | . |
| chr5 | 104046666 | 104047007 | 915 | Promoter (<=1kb) | 5 | 104046869 | 104065379 | 18511 | 1 | ENSMUST00 | 0 | Nudt9 | ENSMUSG0C | false | chr5 | 104046126 | 104048099 | 36.66269 | 126.29414 | 121.93508 | . | . | . | . | . | . |
| chr9 | 65646003 | 65646285 | 915 | Intron (ENSMUST00000169003.2 | 9 | 65630582 | 65660505 | 29924 | 1 | ENSMUST00 | 15422 | Rbpms2 | ENSMUSG0C | false | . | . | . | . | . | . | chr9 | 65645908 | 65646467 | 3.14835 | 3.06899 | 1.5213 |
| chr17 | 63829032 | 63829332 | 915 | Distal Intergenic | 17 | 63863300 | 63863791 | 492 | 2 | ENSMUST00 | 34459 | A930002H24 | ENSMUSG0C | false | . | . | . | . | . | . | . | . | . | . | . | . |
| chr15 | 72768816 | 72769176 | 915 | Intron (ENSMUST00000023276.9 | 15 | 72721321 | 72721387 | 67 | 2 | ENSMUST00 | -47430 | Gm28020 | ENSMUSG0C | false | . | . | . | . | . | . | chr15 | 72767818 | 72769639 | 4.18812 | 3.65634 | 2.39168 |
| chr9 | 83721055 | 83721372 | 915 | Distal Intergenic | 9 | 83691475 | 83693223 | 1749 | 1 | ENSMUST00 | 29581 | RP23-101D1 | ENSMUSG0C | false | . | . | . | . | . | . | chr9 | 83720114 | 83721677 | 4.51252 | 3.71186 | 2.62526 |
| chr10 | 105840994 | 105841577 | 914 | Promoter (<=1kb) | 10 | 105841067 | 105847833 | 6767 | 1 | ENSMUST00 | 0 | Ccdc59 | ENSMUSG0C | false | chr10 | 105840876 | 105842352 | 34.36702 | 111.64683 | 107.76208 | . | . | . | . | . | . |
| chr3 | 28262713 | 28263643 | 914 | Promoter (<=1kb) | 3 | 28263214 | 28670585 | 407372 | 1 | ENSMUST00 | 0 | Tnik | ENSMUSG0C | false | chr3 | 28262261 | 28263926 | 24.30838 | 70.74035 | 67.73997 | . | . | . | . | . | . |
| chr8 | 36608512 | 36608832 | 914 | Intron (ENSMUST00000033923.8 | 8 | 36593231 | 36613874 | 20644 | 2 | ENSMUST00 | 5042 | Dlc1 | ENSMUSG0C | true | . | . | . | . | . | . | chr8 | 36575944 | 36612020 | 5.64773 | 4.3412 | 3.54039 |
| chr7 | 80657943 | 80658247 | 914 | Intron (ENSMUST00000122255.2 | 7 | 80636017 | 80644384 | 8368 | 1 | ENSMUST00 | 21927 | Gm15880 | ENSMUSG0C | false | . | . | . | . | . | . | chr7 | 80644966 | 80674155 | 4.92417 | 4.02487 | 2.94906 |
| chr17 | 56425127 | 56425949 | 914 | Exon (ENSMUST00000067538.5/ | 17 | 56418846 | 56418909 | 64 | 2 | ENSMUST00 | -6219 | Mir6977 | ENSMUSG0C | false | chr17 | 56425075 | 56426001 | 27.57193 | 78.19525 | 75.07539 | chr17 | 56424857 | 56425882 | 5.00873 | 4.15198 | 3.00683 |
| chr18 | 35874380 | 35874691 | 914 | Distal Intergenic | 18 | 35830389 | 35831093 | 705 | 2 | ENSMUST00 | -43288 | RP24-326K1 | ENSMUSG0C | false | . | . | . | . | . | . | chr18 | 35870146 | 35877349 | 5.65833 | 4.25717 | 3.58139 |
| chr17 | 34127541 | 34127826 | 913 | Distal Intergenic | 17 | 34119867 | 34122634 | 2768 | 2 | ENSMUST00 | -4908 | Brd2 | ENSMUSG0C | false | . | . | . | . | . | . | chr17 | 34123233 | 34131489 | 5.36434 | 4.14852 | 3.33089 |
| chr18 | 44452788 | 44453145 | 913 | Intron (ENSMUST00000164666.2 | 18 | 44380500 | 44424969 | 44470 | 1 | ENSMUST00 | 72289 | Dcp2 | ENSMUSG0C | false | . | . | . | . | . | . | chr18 | 44451929 | 44456355 | 6.16447 | 4.40422 | 4.00607 |
| chrX | 99199156 | 99199438 | 913 | Distal Intergenic | 20 | 99138131 | 99147230 | 9100 | 1 | ENSMUST00 | 61026 | Efnb1 | ENSMUSG0C | false | chrX | 99198925 | 99199175 | 7.81205 | 13.8349 | 11.58824 | . | . | . | . | . | . |
| chr7 | 120327417 | 120327716 | 913 | Promoter (<=1kb) | 7 | 120328684 | 120361690 | 33007 | 1 | ENSMUST00 | -968 | Abca15 | ENSMUSG0C | false | . | . | . | . | . | . | chr7 | 120326969 | 120328673 | 4.74908 | 3.95864 | 2.855 |
| chr14 | 55386835 | 55387215 | 913 | Distal Intergenic | 14 | 55420269 | 55420724 | 456 | 2 | ENSMUST00 | 33509 | Gm8894 | ENSMUSG0C | false | . | . | . | . | . | . | chr14 | 55386972 | 55387630 | 3.61316 | 3.16541 | 1.90173 |
| chrX | 104499601 | 104499821 | 913 | Intron (ENSMUST00000087867.5 | 20 | 104482818 | 104507490 | 24673 | 1 | ENSMUST00 | 16784 | Uprt | ENSMUSG0C | false | . | . | . | . | . | . | . | . | . | . | . | . |
| chr10 | 84418042 | 84418338 | 912 | Intron (ENSMUST00000020220.9 | 10 | 84370905 | 84440597 | 69693 | 2 | ENSMUST00 | 22259 | Nuak1 | ENSMUSG0C | false | . | . | . | . | . | . | chr10 | 84414812 | 84419331 | 6.36294 | 4.69905 | 4.14802 |
| chr15 | 97830886 | 97831908 | 912 | Promoter (<=1kb) | 15 | 97792664 | 97831494 | 38831 | 2 | ENSMUST00 | 0 | Hdac7 | ENSMUSG0C | false | chr15 | 97830319 | 97832239 | 27.11239 | 76.48743 | 73.39479 | chr15 | 97819806 | 97833530 | 5.44469 | 4.22025 | 3.37668 |
| chr6 | 134450689 | 134451006 | 912 | 3' UTR | 6 | 134450483 | 134471293 | 20811 | 2 | ENSMUST00 | 20287 | Lrp6 | ENSMUSG0C | false | . | . | . | . | . | . | chr6 | 134449480 | 134451717 | 4.1718 | 3.70807 | 2.32071 |
| chr6 | 114915906 | 114916252 | 912 | Intron (ENSMUST00000032459.8 | 6 | 114861424 | 114921821 | 60398 | 2 | ENSMUST00 | 5569 | Vgll4 | ENSMUSG0C | true | . | . | . | . | . | . | chr6 | 114853844 | 114919520 | 8.05355 | 5.36011 | 5.51688 |
| chr9 | 63780159 | 63780752 | 912 | Distal Intergenic | 9 | 63646767 | 63757994 | 111228 | 2 | ENSMUST00 | -22166 | Smad3 | ENSMUSG0C | false | . | . | . | . | . | . | chr9 | 63779840 | 63781622 | 6.5781 | 4.83582 | 4.30576 |
| chr2 | 132646900 | 132647167 | 912 | Promoter (1-2kb) | 2 | 132645881 | 132652999 | 7119 | 1 | ENSMUST00 | 1020 | AU019990 | ENSMUSG0C | false | . | . | . | . | . | . | chr2 | 132645606 | 132648470 | 4.60592 | 3.97308 | 2.67712 |
| chr4 | 117288880 | 117289719 | 911 | Promoter (<=1kb) | 4 | 117285092 | 117289442 | 4351 | 2 | ENSMUST00 | 0 | Rnf220 | ENSMUSG0C | false | chr4 | 117288453 | 117289927 | 20.69629 | 56.7044 | 53.90051 | chr4 | 117288080 | 117289996 | 3.65348 | 3.28285 | 1.92036 |
| chrX | 101419731 | 101420053 | 911 | Promoter (<=1kb) | 20 | 101404384 | 101419798 | 15415 | 2 | ENSMUST00 | 0 | Zmym3 | ENSMUSG0C | false | chrX | 101419086 | 101420481 | 20.21941 | 51.82391 | 49.08421 | . | . | . | . | . | . |
| chr4 | 8689974 | 8690372 | 911 | Promoter (<=1kb) | 4 | 8690406 | 8755906 | 65501 | 1 | ENSMUST00 | -34 | Chd7 | ENSMUSG0C | false | chr4 | 8689673 | 8692342 | 39.81545 | 135.14195 | 130.45284 | . | . | . | . | . | . |
| chr12 | 29175818 | 29176118 | 911 | Distal Intergenic | 12 | 29035180 | 29035294 | 115 | 2 | ENSMUST00 | -140525 | Gm23342 | ENSMUSG0C | false | . | . | . | . | . | . | chr12 | 29172063 | 29176872 | 5.05824 | 4.08857 | 3.09154 |
| chr11 | 87448476 | 87448938 | 911 | Promoter (<=1kb) | 11 | 87448627 | 87448842 | 216 | 1 | ENSMUST00 | 0 | Rnu3b3 | ENSMUSG0C | false | chr11 | 87447966 | 87448754 | 21.90749 | 75.32819 | 72.25517 | chr11 | 87447777 | 87449281 | 3.9216 | 3.54316 | 2.11185 |
| chr15 | 85669591 | 85670246 | 911 | Distal Intergenic | 15 | 85639747 | 85645934 | 6188 | 1 | ENSMUST00 | 29845 | RP23-64J7.2 | ENSMUSG0C | false | chr15 | 85668507 | 85673353 | 33.09821 | 104.28432 | 100.60858 | chr15 | 85667739 | 85670381 | 4.42261 | 3.85806 | 2.51828 |
| chr11 | 96956147 | 96957085 | 910 | Exon (ENSMUST00000107626.2/ | 11 | 96953473 | 96968665 | 15193 | 2 | ENSMUST00 | 11580 | Sp2 | ENSMUSG0C | false | . | . | . | . | . | . | chr11 | 96951250 | 96960443 | 6.86299 | 4.78743 | 4.54283 |
| chr13 | 41352284 | 41352604 | 910 | Intron (ENSMUST00000163623.1 | 13 | 41309916 | 41359001 | 49086 | 2 | ENSMUST00 | 6397 | Nedd9 | ENSMUSG0C | false | . | . | . | . | . | . | chr13 | 41318388 | 41357685 | 5.92416 | 4.40318 | 3.78151 |
| chr16 | 26116351 | 26116689 | 910 | Distal Intergenic | 16 | 25959288 | 26105784 | 146497 | 2 | ENSMUST00 | -10568 | Leprel1 | ENSMUSG0C | false | . | . | . | . | . | . | chr16 | 26115051 | 26118846 | 4.32526 | 3.78072 | 2.43692 |
| chr18 | 60632706 | 60633090 | 910 | Promoter (2-3kb) | 18 | 60593990 | 60629853 | 35864 | 2 | ENSMUST00 | -2854 | Synpo | ENSMUSG0C | true | . | . | . | . | . | . | chr18 | 60631929 | 60634352 | 6.69729 | 4.868 | 4.40499 |
| chr9 | 43086607 | 43086915 | 910 | Intron (ENSMUST00000165665.2 | 9 | 42963843 | 43105486 | 141644 | 2 | ENSMUST00 | 18571 | Arhgef12 | ENSMUSG0C | true | . | . | . | . | . | . | chr9 | 43060024 | 43103547 | 5.90513 | 4.47373 | 3.75735 |
| chr9 | 48740136 | 48740500 | 910 | Intron (ENSMUST00000093852.3 | 9 | 48654311 | 48835945 | 181635 | 2 | ENSMUST00 | 95445 | Zbtb16 | ENSMUSG0C | false | . | . | . | . | . | . | chr9 | 48731594 | 48741569 | 5.57909 | 4.32779 | 3.48116 |
| chr12 | 106010052 | 106010404 | 909 | Promoter (<=1kb) | 12 | 106010263 | 106077410 | 67148 | 1 | ENSMUST00 | 0 | Vrk1 | ENSMUSG0C | false | chr12 | 106009676 | 106011099 | 39.11606 | 129.53703 | 125.06399 | chr12 | 106009773 | 106011744 | 4.34987 | 3.7643 | 2.49577 |
| chr13 | 63245288 | 63245989 | 909 | Intron (ENSMUST00000091560.5 | 13 | 63282142 | 63326096 | 43955 | 1 | ENSMUST00 | -36153 | 2010111I01R | ENSMUSG0C | false | chr13 | 63244274 | 63245622 | 16.14593 | 43.98602 | 41.34693 | chr13 | 63244614 | 63251804 | 3.9909 | 3.58662 | 2.16637 |
| chr11 | 120948876 | 120949201 | 909 | Promoter (<=1kb) | 11 | 120949053 | 120956747 | 7695 | 1 | ENSMUST00 | 0 | Slc16a3 | ENSMUSG0C | false | chr11 | 120948429 | 120949719 | 15.92618 | 39.91766 | 37.3274 | chr11 | 120948533 | 120951419 | 5.55747 | 4.1386 | 3.50262 |
| chr2 | 155473325 | 155474034 | 909 | Promoter (<=1kb) | 2 | 155390665 | 155473806 | 83142 | 2 | ENSMUST00 | 0 | Ncoa6 | ENSMUSG0C | false | chr2 | 155472648 | 155474429 | 27.57193 | 78.19525 | 75.07539 | . | . | . | . | . | . |
| chr14 | 58972051 | 58972518 | 909 | Distal Intergenic | 14 | 58893506 | 58894129 | 624 | 1 | ENSMUST00 | 78546 | Rpl13-ps3 | ENSMUSG0C | false | . | . | . | . | . | . | chr14 | 58971205 | 58973620 | 4.26018 | 3.75828 | 2.38711 |
| chr14 | 60506279 | 60506621 | 909 | Exon (ENSMUST00000190392.1/ | 14 | 60485208 | 60599031 | 113824 | 1 | ENSMUST00 | 21072 | RP24-127N1 | ENSMUSG0C | false | chr14 | 60506281 | 60507719 | 21.59801 | 56.60135 | 53.80005 | . | . | . | . | . | . |
| chr8 | 125507328 | 125507619 | 908 | Distal Intergenic | 8 | 125418063 | 125492710 | 74648 | 2 | ENSMUST00 | -14619 | Sipa1l2 | ENSMUSG0C | false | . | . | . | . | . | . | chr8 | 125502637 | 125510244 | 6.21729 | 4.41596 | 4.04306 |
| chr10 | 84285951 | 84286376 | 908 | Distal Intergenic | 10 | 84375296 | 84392388 | 17093 | 2 | ENSMUST00 | 106012 | Nuak1 | ENSMUSG0C | false | . | . | . | . | . | . | chr10 | 84285136 | 84287067 | 5.23493 | 4.23915 | 3.20785 |
| chr5 | 125506666 | 125506955 | 908 | Promoter (<=1kb) | 5 | 125506149 | 125512737 | 6589 | 1 | ENSMUST00 | 518 | Aacs | ENSMUSG0C | false | . | . | . | . | . | . | chr5 | 125504189 | 125507809 | 4.00135 | 3.4541 | 2.20743 |
| chr7 | 126272230 | 126272688 | 908 | Promoter (<=1kb) | 7 | 126272619 | 126294999 | 22381 | 1 | ENSMUST00 | 0 | Sbk1 | ENSMUSG0C | false | chr7 | 126271603 | 126273389 | 28.91247 | 96.86193 | 93.36482 | . | . | . | . | . | . |
| chr8 | 122281339 | 122282151 | 908 | Promoter (<=1kb) | 8 | 122282141 | 122337251 | 55111 | 1 | ENSMUST00 | 0 | Zfpm1 | ENSMUSG0C | false | chr8 | 122281032 | 122281852 | 11.22292 | 28.25542 | 25.81138 | . | . | . | . | . | . |
| chr3 | 58631719 | 58632068 | 908 | Intron (ENSMUST00000041115.6 | 3 | 58616300 | 58637207 | 20908 | 2 | ENSMUST00 | 5139 | Erich6 | ENSMUSG0C | false | . | . | . | . | . | . | chr3 | 58630230 | 58632046 | 2.8749 | 2.82064 | 1.30493 |
| chr5 | 142340250 | 142340726 | 908 | Distal Intergenic | 5 | 142283106 | 142284069 | 964 | 1 | ENSMUST00 | 57145 | Gm26970 | ENSMUSG0C | false | . | . | . | . | . | . | chr5 | 142337724 | 142341069 | 4.19262 | 3.66897 | 2.3405 |
| chr6 | 119452102 | 119452558 | 907 | Promoter (2-3kb) | 6 | 119432531 | 119449386 | 16856 | 2 | ENSMUST00 | -2717 | Wnt5b | ENSMUSG0C | false | chr6 | 119451936 | 119452726 | 14.24974 | 34.23454 | 31.71693 | chr6 | 119448074 | 119453219 | 5.22359 | 4.16949 | 3.21432 |
| chr14 | 49171862 | 49173168 | 907 | Promoter (<=1kb) | 14 | 49172226 | 49190866 | 18641 | 1 | ENSMUST00 | 0 | Naa30 | ENSMUSG0C | false | chr14 | 49171516 | 49174414 | 37.7199 | 126.00829 | 121.65485 | chr14 | 49173007 | 49179961 | 4.49705 | 3.84277 | 2.59358 |
| chr18 | 76061966 | 76062258 | 907 | Promoter (2-3kb) | 18 | 76059458 | 76146443 | 86986 | 1 | ENSMUST00 | 2509 | Zbtb7c | ENSMUSG0C | true | chr18 | 76059035 | 76063179 | 44.34918 | 154.34741 | 148.89743 | chr18 | 76061445 | 76132293 | 6.70015 | 4.79833 | 4.4158 |
| chr2 | 153179142 | 153179462 | 907 | Intron (ENSMUST00000137446.2 | 2 | 153190399 | 153192187 | 1789 | 1 | ENSMUST00 | -10937 | Tm9sf4 | ENSMUSG0C | false | . | . | . | . | . | . | chr2 | 153173775 | 153181365 | 6.89193 | 4.98213 | 4.56161 |
| chr13 | 60470932 | 60471263 | 907 | Distal Intergenic | 13 | 60529296 | 60529409 | 114 | 2 | ENSMUST00 | 58146 | Gm25000 | ENSMUSG0C | false | . | . | . | . | . | . | chr13 | 60470120 | 60471593 | 5.63808 | 4.38876 | 3.56202 |
| chr2 | 158113818 | 158114143 | 907 | Intron (ENSMUST00000152452.2 | 2 | 158117154 | 158117905 | 752 | 1 | ENSMUST00 | -3011 | Gm20412 | ENSMUSG0C | false | . | . | . | . | . | . | chr2 | 158113221 | 158117875 | 5.53387 | 4.29787 | 3.45991 |
| chr18 | 34220794 | 34221499 | 906 | Promoter (<=1kb) | 18 | 34220924 | 34313506 | 92583 | 1 | ENSMUST00 | 0 | Apc | ENSMUSG0C | false | chr18 | 34220262 | 34222439 | 25.48472 | 95.2657 | 91.8051 | . | . | . | . | . | . |
| chr2 | 93810731 | 93811256 | 906 | Intron (ENSMUST00000184931.2 | 2 | 93696127 | 93822545 | 126419 | 2 | ENSMUST00 | 11289 | Ext2 | ENSMUSG0C | true | . | . | . | . | . | . | chr2 | 93805378 | 93812725 | 6.22701 | 4.60265 | 4.02277 |
| chr6 | 137252148 | 137252517 | 906 | Promoter (<=1kb) | 6 | 137252299 | 137464629 | 212331 | 1 | ENSMUST00 | 0 | Ptpro | ENSMUSG0C | true | chr6 | 137251894 | 137256757 | 32.97563 | 101.03377 | 97.44133 | . | . | . | . | . | . |
| chr7 | 29271028 | 29271308 | 906 | Intron (ENSMUST00000098604.5 | 7 | 29256331 | 29281977 | 25647 | 2 | ENSMUST00 | 10669 | Spint2 | ENSMUSG0C | false | . | . | . | . | . | . | chr7 | 29270378 | 29278106 | 5.53936 | 4.34059 | 3.44839 |
| chr17 | 8621661 | 8622074 | 906 | Intron (ENSMUST00000115715.2 | 17 | 8564258 | 8568860 | 4603 | 2 | ENSMUST00 | -52802 | Gm15425 | ENSMUSG0C | false | . | . | . | . | . | . | chr17 | 8621094 | 8623257 | 5.13034 | 4.0407 | 3.14284 |
| chr9 | 70332051 | 70332392 | 906 | Intron (ENSMUST00000034745.7 | 9 | 70405694 | 70405753 | 60 | 2 | ENSMUST00 | 73361 | Mir5626 | ENSMUSG0C | false | . | . | . | . | . | . | chr9 | 70312793 | 70333015 | 5.95088 | 4.44065 | 3.80175 |
| chr11 | 63132474 | 63133191 | 905 | Promoter (<=1kb) | 11 | 63132569 | 63158484 | 25916 | 1 | ENSMUST00 | 0 | Pmp22 | ENSMUSG0C | false | chr11 | 63130660 | 63133452 | 20.74539 | 66.97287 | 64.02668 | chr11 | 63129453 | 63136936 | 5.01478 | 3.91056 | 3.06639 |
| chr7 | 78725303 | 78725558 | 905 | Distal Intergenic | 7 | 78775236 | 78783066 | 7831 | 2 | ENSMUST00 | 57508 | Mrpl46 | ENSMUSG0C | false | . | . | . | . | . | . | chr7 | 78725148 | 78726157 | 3.8772 | 3.56568 | 2.06416 |
| chr9 | 78489043 | 78489369 | 905 | Promoter (<=1kb) | 9 | 78479649 | 78489151 | 9503 | 2 | ENSMUST00 | 0 | Eef1a1 | ENSMUSG0C | false | chr9 | 78488672 | 78489472 | 15.01947 | 39.77812 | 37.19162 | chr9 | 78488258 | 78489656 | 3.64735 | 3.4138 | 1.88179 |
| chr6 | 135559469 | 135559932 | 905 | Distal Intergenic | 6 | 135584192 | 135584306 | 115 | 1 | ENSMUST00 | -24260 | Gm25136 | ENSMUSG0C | false | . | . | . | . | . | . | chr6 | 135558601 | 135561802 | 4.89403 | 3.99978 | 2.92782 |
| chr5 | 21144575 | 21145016 | 905 | Distal Intergenic | 5 | 21119149 | 21119522 | 374 | 2 | ENSMUST00 | -25054 | Rpl31-ps21 | ENSMUSG0C | false | . | . | . | . | . | . | chr5 | 21143537 | 21146177 | 4.07237 | 3.46635 | 2.2541 |
| chr17 | 6812566 | 6812857 | 905 | Intron (ENSMUST00000181895.1 | 17 | 6798390 | 6827990 | 29601 | 2 | ENSMUST00 | 15133 | Gm2885 | ENSMUSG0C | false | . | . | . | . | . | . | chr17 | 6811040 | 6816818 | 5.75765 | 4.4253 | 3.63624 |
| chr4 | 133910657 | 133911198 | 904 | Distal Intergenic | 4 | 133847290 | 133887797 | 40508 | 2 | ENSMUST00 | -22861 | Rps6ka1 | ENSMUSG0C | false | chr4 | 133910275 | 133911377 | 18.77433 | 54.14365 | 51.37372 | chr4 | 133910033 | 133911749 | 4.25688 | 3.62588 | 2.43743 |
| chr11 | 76398131 | 76399309 | 904 | Promoter (<=1kb) | 11 | 76257198 | 76399140 | 141943 | 2 | ENSMUST00 | 0 | Nxn | ENSMUSG0C | true | chr11 | 76397725 | 76399811 | 28.12215 | 95.42207 | 91.9593 | . | . | . | . | . | . |
| chr18 | 76241169 | 76242466 | 904 | Promoter (<=1kb) | 18 | 76241580 | 76305731 | 64152 | 1 | ENSMUST00 | 0 | Smad2 | ENSMUSG0C | false | chr18 | 76240774 | 76242782 | 27.24373 | 82.29701 | 79.10307 | . | . | . | . | . | . |
| chr8 | 25101475 | 25102498 | 904 | Promoter (<=1kb) | 8 | 25039144 | 25102194 | 63051 | 2 | ENSMUST00 | 0 | Plekha2 | ENSMUSG0C | true | chr8 | 25099212 | 25103419 | 35.56825 | 117.95176 | 113.87694 | . | . | . | . | . | . |
| chr9 | 44482503 | 44483133 | 904 | Distal Intergenic | 9 | 44478778 | 44482180 | 3403 | 1 | ENSMUST00 | 3726 | C030014I23R | ENSMUSG0C | false | chr9 | 44482529 | 44484376 | 36.04672 | 129.50026 | 125.02953 | chr9 | 44481741 | 44482719 | 3.92779 | 3.59782 | 2.11494 |
| chr16 | 33321234 | 33321795 | 904 | Distal Intergenic | 16 | 33379855 | 33380727 | 873 | 2 | ENSMUST00 | 58932 | 1700007L15F | ENSMUSG0C | false | . | . | . | . | . | . | . | . | . | . | . | . |
| chr12 | 70227544 | 70227889 | 903 | Promoter (<=1kb) | 12 | 70190811 | 70227683 | 36873 | 2 | ENSMUST00 | 0 | Pygl | ENSMUSG0C | false | chr12 | 70226918 | 70228127 | 18.44084 | 48.77526 | 46.0739 | chr12 | 70226936 | 70227738 | 3.26168 | 3.06696 | 1.62014 |
| chr16 | 10976351 | 10976680 | 903 | Promoter (<=1kb) | 16 | 10960824 | 10975579 | 14756 | 2 | ENSMUST00 | -773 | Litaf | ENSMUSG0C | false | . | . | . | . | . | . | chr16 | 10975662 | 10992001 | 5.99475 | 4.33209 | 3.84709 |
| chr7 | 81828811 | 81829831 | 903 | Promoter (<=1kb) | 7 | 81792075 | 81829431 | 37357 | 2 | ENSMUST00 | 0 | Btbd1 | ENSMUSG0C | false | chr7 | 81827970 | 81829981 | 27.34946 | 104.78247 | 101.09169 | chr7 | 81829801 | 81830443 | 3.47713 | 3.24702 | 1.74356 |
| chr2 | 73721847 | 73722212 | 903 | Intron (ENSMUST00000112024.4 | 2 | 73613084 | 73716409 | 103326 | 2 | ENSMUST00 | -5439 | Chn1 | ENSMUSG0C | false | . | . | . | . | . | . | chr2 | 73721720 | 73722896 | 3.71849 | 3.2032 | 1.98266 |
| chr12 | 59411482 | 59411793 | 903 | Distal Intergenic | 12 | 59291964 | 59292938 | 975 | 1 | ENSMUST00 | 119519 | Gm7719 | ENSMUSG0C | false | . | . | . | . | . | . | chr12 | 59410587 | 59412183 | 3.53871 | 3.19864 | 1.84654 |
| chr1 | 80091486 | 80091951 | 903 | Distal Intergenic | 1 | 80098622 | 80098885 | 264 | 1 | ENSMUST00 | -6671 | RP23-470G1 | ENSMUSG0C | false | . | . | . | . | . | . | . | . | . | . | . | . |
| chr5 | 21054433 | 21054927 | 902 | Promoter (<=1kb) | 5 | 21009444 | 21055725 | 46282 | 2 | ENSMUST00 | 798 | Ptpn12 | ENSMUSG0C | true | chr5 | 21053536 | 21057043 | 37.92415 | 138.37384 | 133.55817 | . | . | . | . | . | . |
| chr4 | 129188768 | 129190059 | 902 | Promoter (<=1kb) | 4 | 129189760 | 129219607 | 29848 | 1 | ENSMUST00 | 0 | Yars | ENSMUSG0C | false | chr4 | 129188679 | 129191045 | 43.41452 | 148.15424 | 142.93486 | chr4 | 129183108 | 129188971 | 3.74369 | 3.42065 | 1.97895 |
| chr14 | 62761210 | 62761861 | 902 | Promoter (<=1kb) | 14 | 62676330 | 62761112 | 84783 | 2 | ENSMUST00 | -99 | Ints6 | ENSMUSG0C | false | chr14 | 62761387 | 62762328 | 23.6529 | 67.82199 | 64.86302 | . | . | . | . | . | . |
| chr4 | 16163413 | 16164219 | 902 | Promoter (<=1kb) | 4 | 16122733 | 16163647 | 40915 | 2 | ENSMUST00 | 0 | Ripk2 | ENSMUSG0C | false | chr4 | 16164084 | 16164594 | 13.98552 | 41.45461 | 38.84487 | chr4 | 16164197 | 16167792 | 3.77251 | 3.43128 | 2.00264 |
| chr9 | 108072826 | 108073336 | 902 | Promoter (<=1kb) | 9 | 108071382 | 108073121 | 1740 | 2 | ENSMUST00 | 0 | Rnf123 | ENSMUSG0C | false | . | . | . | . | . | . | chr9 | 108072513 | 108075801 | 7.10708 | 5.05191 | 4.70896 |
| chr13 | 52978323 | 52978764 | 902 | Promoter (2-3kb) | 13 | 52967209 | 52981073 | 13865 | 2 | ENSMUST00 | 2309 | Nfil3 | ENSMUSG0C | false | chr13 | 52977991 | 52979106 | 20.60796 | 54.90754 | 52.13 | chr13 | 52975691 | 52979722 | 3.76935 | 3.36446 | 2.03766 |
| chr8 | 26865001 | 26865365 | 902 | Distal Intergenic | 8 | 26814593 | 26824423 | 9831 | 2 | ENSMUST00 | -40579 | RP24-502D8. | ENSMUSG0C | false | . | . | . | . | . | . | chr8 | 26864265 | 26865581 | 4.3272 | 3.72826 | 2.51234 |
| chr13 | 45709575 | 45709907 | 901 | Intron (ENSMUST00000091628.4 | 13 | 45538809 | 45542550 | 3742 | 1 | ENSMUST00 | 170767 | Gmpr | ENSMUSG0C | false | chr13 | 45709333 | 45709733 | 7.88522 | 15.75676 | 13.48224 | chr13 | 45707075 | 45710178 | 3.78216 | 3.45546 | 2.01498 |
| chr6 | 132463605 | 132463906 | 901 | Distal Intergenic | 6 | 132361041 | 132364134 | 3094 | 2 | ENSMUST00 | -99472 | Gm8882 | ENSMUSG0C | false | chr6 | 132463619 | 132463896 | 29.41006 | 85.09721 | 81.85307 | . | . | . | . | . | . |
| chr6 | 72477630 | 72478106 | 901 | Distal Intergenic | 6 | 72513644 | 72515290 | 1647 | 2 | ENSMUST00 | 37184 | Sh2d6 | ENSMUSG0C | false | . | . | . | . | . | . | chr6 | 72472745 | 72478497 | 5.05532 | 4.02854 | 3.06461 |
| chr12 | 76954885 | 76955532 | 901 | Intron (ENSMUST00000082136.5 | 12 | 76937269 | 76962248 | 24980 | 2 | ENSMUST00 | 6716 | Max | ENSMUSG0C | false | . | . | . | . | . | . | chr12 | 76953207 | 76961808 | 5.68558 | 4.38218 | 3.58599 |
| chr2 | 129186579 | 129186945 | 901 | Promoter (2-3kb) | 2 | 129182482 | 129183947 | 1466 | 2 | ENSMUST00 | -2633 | Gm14029 | ENSMUSG0C | false | . | . | . | . | . | . | chr2 | 129185462 | 129188434 | 4.25415 | 3.59661 | 2.42798 |
| chr3 | 8767191 | 8767859 | 901 | Distal Intergenic | 3 | 8679359 | 8679477 | 119 | 2 | ENSMUST00 | -87715 | Gm23670 | ENSMUSG0C | false | chr3 | 8766691 | 8767982 | 19.43846 | 52.88828 | 50.13554 | chr3 | 8764904 | 8768381 | 3.97957 | 3.54738 | 2.17292 |
| chr19 | 15924413 | 15924859 | 900 | Promoter (<=1kb) | 19 | 15905127 | 15924622 | 19496 | 2 | ENSMUST00 | 0 | Psat1 | ENSMUSG0C | false | chr19 | 15924003 | 15925843 | 19.5687 | 55.55706 | 52.77042 | chr19 | 15923492 | 15926257 | 3.88024 | 3.5774 | 2.07456 |
| chr2 | 158771355 | 158771786 | 900 | Intron (ENSMUST00000029183.2 | 2 | 158768093 | 158786637 | 18545 | 1 | ENSMUST00 | 3263 | Fam83d | ENSMUSG0C | false | . | . | . | . | . | . | chr2 | 158767460 | 158773109 | 4.66591 | 3.76493 | 2.78491 |
| chr9 | 63969998 | 63970658 | 900 | Intron (ENSMUST00000041029.5 | 9 | 63954009 | 64016987 | 62979 | 2 | ENSMUST00 | 46329 | Smad6 | ENSMUSG0C | true | . | . | . | . | . | . | chr9 | 63948352 | 64020083 | 7.27873 | 5.05859 | 4.874 |
| chr8 | 87836714 | 87837219 | 900 | Intron (ENSMUST00000109655.3 | 8 | 87661810 | 87804439 | 142630 | 2 | ENSMUST00 | -32276 | Zfp423 | ENSMUSG0C | true | . | . | . | . | . | . | chr8 | 87835485 | 87838634 | 6.13963 | 4.58154 | 3.93484 |
| chr3 | 133601010 | 133601261 | 900 | Distal Intergenic | 3 | 133588406 | 133588537 | 132 | 2 | ENSMUST00 | -12474 | Gm23011 | ENSMUSG0C | false | . | . | . | . | . | . | chr3 | 133599912 | 133602007 | 3.82672 | 3.27279 | 2.08009 |
| chr3 | 123503723 | 123504249 | 900 | Intron (ENSMUST00000029603.8 | 3 | 123507937 | 123508066 | 130 | 2 | ENSMUST00 | 3817 | Snora24 | ENSMUSG0C | false | . | . | . | . | . | . | chr3 | 123503484 | 123504716 | 4.60123 | 3.85325 | 2.68142 |
| chr8 | 105759291 | 105759637 | 899 | Promoter (<=1kb) | 8 | 105713854 | 105758664 | 44811 | 2 | ENSMUST00 | -628 | Gfod2 | ENSMUSG0C | false | chr8 | 105757781 | 105759401 | 28.22734 | 101.53345 | 97.92473 | chr8 | 105757102 | 105760913 | 4.29376 | 3.70013 | 2.47708 |
| chr17 | 67917039 | 67917307 | 899 | Intron (ENSMUST00000024840.6 | 17 | 67845735 | 67901471 | 55737 | 2 | ENSMUST00 | -15569 | Arhgap28 | ENSMUSG0C | true | . | . | . | . | . | . | chr17 | 67915917 | 67949162 | 5.38115 | 4.23981 | 3.32205 |
| chr10 | 70301161 | 70301914 | 899 | Distal Intergenic | 10 | 70285549 | 70288176 | 2628 | 2 | ENSMUST00 | -12986 | Gm9877 | ENSMUSG0C | false | . | . | . | . | . | . | chr10 | 70295718 | 70302457 | 4.49539 | 3.8568 | 2.58998 |
| chr6 | 113668467 | 113668888 | 899 | Promoter (<=1kb) | 6 | 113668850 | 113672939 | 4090 | 1 | ENSMUST00 | 0 | Irak2 | ENSMUSG0C | false | . | . | . | . | . | . | chr6 | 113667888 | 113670307 | 4.73112 | 3.72713 | 2.82465 |
| chr8 | 25635350 | 25635677 | 899 | Intron (ENSMUST00000155861.2 | 8 | 25631950 | 25632079 | 130 | 1 | ENSMUST00 | 3401 | Gm24992 | ENSMUSG0C | false | . | . | . | . | . | . | chr8 | 25632001 | 25639825 | 4.83789 | 3.86947 | 2.90597 |
| chr5 | 118950894 | 118951405 | 899 | Distal Intergenic | 5 | 119108234 | 119114543 | 6310 | 1 | ENSMUST00 | -156829 | RP24-389L1C | ENSMUSG0C | false | . | . | . | . | . | . | . | . | . | . | . | . |
| chr11 | 69846190 | 69846687 | 898 | Promoter (<=1kb) | 11 | 69846376 | 69851036 | 4661 | 1 | ENSMUST00 | 0 | Plscr3 | ENSMUSG0C | false | chr11 | 69845891 | 69847275 | 21.17422 | 79.20749 | 76.06625 | chr11 | 69845686 | 69847750 | 5.65406 | 4.48347 | 3.53564 |
| chr9 | 66059934 | 66060461 | 898 | Promoter (<=1kb) | 9 | 66060169 | 66066629 | 6461 | 1 | ENSMUST00 | 0 | Ppib | ENSMUSG0C | false | chr9 | 66059514 | 66061149 | 23.46333 | 91.19009 | 87.82124 | chr9 | 66059409 | 66061736 | 4.61649 | 3.61356 | 2.72081 |
| chr17 | 42918245 | 42918532 | 898 | Distal Intergenic | 17 | 42792951 | 42876424 | 83474 | 2 | ENSMUST00 | -41822 | Cd2ap | ENSMUSG0C | true | . | . | . | . | . | . | chr17 | 42914306 | 42920043 | 5.49156 | 4.20042 | 3.41297 |
| chr8 | 14911288 | 14911855 | 898 | Promoter (<=1kb) | 8 | 14911663 | 15001085 | 89423 | 1 | ENSMUST00 | 0 | Arhgef10 | ENSMUSG0C | false | chr8 | 14910723 | 14912537 | 24.40663 | 77.266 | 74.1616 | chr8 | 14910525 | 14913343 | 5.38264 | 4.17488 | 3.32821 |
| chr17 | 28033665 | 28033994 | 898 | Intron (ENSMUST00000114842.2 | 17 | 27988028 | 28008432 | 20405 | 2 | ENSMUST00 | -25234 | Gm15598 | ENSMUSG0C | false | . | . | . | . | . | . | chr17 | 28029226 | 28043441 | 7.08706 | 4.83268 | 4.74553 |
| chr2 | 170222244 | 170222995 | 898 | Distal Intergenic | 2 | 170192449 | 170194053 | 1605 | 2 | ENSMUST00 | -28192 | Gm17619 | ENSMUSG0C | false | . | . | . | . | . | . | chr2 | 170218554 | 170223575 | 5.12078 | 4.17757 | 3.10203 |
| chr15 | 27941359 | 27941644 | 897 | Intron (ENSMUST00000090247.5 | 15 | 27730652 | 28025848 | 295197 | 2 | ENSMUST00 | 84204 | Trio | ENSMUSG0C | false | . | . | . | . | . | . | chr15 | 27936585 | 27946559 | 5.44547 | 4.23139 | 3.36928 |
| chr9 | 110131599 | 110132217 | 897 | Promoter (<=1kb) | 9 | 110132037 | 110184303 | 52267 | 1 | ENSMUST00 | 0 | Smarcc1 | ENSMUSG0C | false | chr9 | 110131232 | 110132839 | 38.65522 | 135.28891 | 130.59163 | . | . | . | . | . | . |
| chr12 | 44304967 | 44305292 | 897 | Exon (ENSMUST00000122902.2/ | 12 | 44299277 | 44311844 | 12568 | 1 | ENSMUST00 | 5691 | Pnpla8 | ENSMUSG0C | false | . | . | . | . | . | . | chr12 | 44301384 | 44306896 | 4.3662 | 3.77333 | 2.49403 |
| chr3 | 87162436 | 87162756 | 897 | Intron (ENSMUST00000159976.2 | 3 | 87084628 | 87174550 | 89923 | 2 | ENSMUST00 | 11794 | Kirrel | ENSMUSG0C | true | . | . | . | . | . | . | chr3 | 87139743 | 87171483 | 6.81014 | 4.88372 | 4.49361 |
| chr7 | 105399867 | 105400470 | 897 | Promoter (<=1kb) | 7 | 105371211 | 105400054 | 28844 | 2 | ENSMUST00 | 0 | Fam160a2 | ENSMUSG0C | false | chr7 | 105399362 | 105400078 | 33.94791 | 109.87469 | 106.04007 | chr7 | 105398872 | 105401219 | 4.65327 | 3.78452 | 2.76388 |
| chr9 | 43048852 | 43049310 | 897 | Intron (ENSMUST00000165665.2 | 9 | 42963843 | 43105486 | 141644 | 2 | ENSMUST00 | 56176 | Arhgef12 | ENSMUSG0C | true | . | . | . | . | . | . | chr9 | 43027603 | 43053830 | 5.31283 | 4.19603 | 3.26716 |
| chr11 | 94265396 | 94265865 | 897 | Distal Intergenic | 11 | 94284652 | 94285985 | 1334 | 1 | ENSMUST00 | -18787 | Gm21885 | ENSMUSG0C | false | chr11 | 94265397 | 94265834 | 8.73111 | 16.23101 | 13.94854 | chr11 | 94265191 | 94266926 | 3.7219 | 3.42718 | 1.95106 |
| chr16 | 4603078 | 4603392 | 896 | Promoter (1-2kb) | 16 | 4601163 | 4610413 | 9251 | 1 | ENSMUST00 | 1916 | Glis2 | ENSMUSG0C | false | . | . | . | . | . | . | chr16 | 4591906 | 4609359 | 7.50331 | 5.19962 | 5.05636 |
| chr13 | 95775053 | 95775330 | 896 | Intron (ENSMUST00000068603.6 | 13 | 95696853 | 95702739 | 5887 | 1 | ENSMUST00 | 78201 | F2rl2 | ENSMUSG0C | false | . | . | . | . | . | . | chr13 | 95772302 | 95783884 | 6.26953 | 4.56663 | 4.0844 |
| chrX | 77522959 | 77523342 | 896 | Intron (ENSMUST00000088217.6 | 20 | 77511214 | 77659743 | 148530 | 1 | ENSMUST00 | 11746 | Tbl1x | ENSMUSG0C | false | . | . | . | . | . | . | . | . | . | . | . | . |
| chr2 | 3283730 | 3284571 | 896 | Promoter (<=1kb) | 2 | 3284212 | 3328877 | 44666 | 1 | ENSMUST00 | 0 | Nmt2 | ENSMUSG0C | false | chr2 | 3284114 | 3285676 | 43.40503 | 144.75304 | 139.67491 | . | . | . | . | . | . |
| chr5 | 147860955 | 147861338 | 896 | Promoter (<=1kb) | 5 | 147860628 | 147875778 | 15151 | 1 | ENSMUST00 | 328 | Pomp | ENSMUSG0C | false | chr5 | 147859955 | 147861658 | 38.67513 | 141.95352 | 136.99454 | chr5 | 147861324 | 147862598 | 4.21697 | 3.38749 | 2.39162 |
| chr17 | 47139720 | 47140172 | 896 | Promoter (<=1kb) | 17 | 47140875 | 47329536 | 188662 | 1 | ENSMUST00 | -703 | Trerf1 | ENSMUSG0C | false | chr17 | 47139299 | 47142406 | 27.6016 | 85.77155 | 82.50918 | chr17 | 47139017 | 47143146 | 4.41006 | 3.81485 | 2.5073 |
| chr4 | 133756637 | 133757748 | 895 | Promoter (<=1kb) | 4 | 133679941 | 133756769 | 76829 | 2 | ENSMUST00 | 0 | Arid1a | ENSMUSG0C | false | chr4 | 133756056 | 133757903 | 24.51421 | 78.37994 | 75.25539 | chr4 | 133754644 | 133758088 | 8.04793 | 5.46477 | 5.4798 |
| chr7 | 99534839 | 99535245 | 895 | Promoter (<=1kb) | 7 | 99535466 | 99606771 | 71306 | 1 | ENSMUST00 | -221 | Arrb1 | ENSMUSG0C | false | chr7 | 99535128 | 99537920 | 25.99992 | 84.80917 | 81.57088 | chr7 | 99534502 | 99535527 | 5.81443 | 4.51927 | 3.65705 |
| chr12 | 51829005 | 51830042 | 895 | Promoter (<=1kb) | 12 | 51743722 | 51829536 | 85815 | 2 | ENSMUST00 | 0 | Hectd1 | ENSMUSG0C | false | chr12 | 51828430 | 51830566 | 23.17719 | 89.67316 | 86.33752 | . | . | . | . | . | . |
| chr1 | 184827127 | 184827514 | 895 | Promoter (<=1kb) | 1 | 184813678 | 184827482 | 13805 | 2 | ENSMUST00 | 0 | 37316 | ENSMUSG0C | false | chr1 | 184826893 | 184827200 | 7.12487 | 12.70834 | 10.47977 | chr1 | 184822720 | 184831886 | 4.91956 | 3.91689 | 2.97507 |
| chr9 | 28196596 | 28197031 | 895 | Promoter (<=1kb) | 9 | 28197887 | 28199244 | 1358 | 1 | ENSMUST00 | -856 | Gm15606 | ENSMUSG0C | false | . | . | . | . | . | . | chr9 | 28193892 | 28197673 | 3.52639 | 3.299 | 1.83437 |
| chr1 | 118878321 | 118878650 | 895 | Intron (ENSMUST00000062483.9 | 1 | 118810136 | 118810453 | 318 | 1 | ENSMUST00 | 68186 | RP23-353I19 | ENSMUSG0C | false | . | . | . | . | . | . | . | . | . | . | . | . |
| chr10 | 80676342 | 80676762 | 894 | Distal Intergenic | 10 | 80665332 | 80671976 | 6645 | 2 | ENSMUST00 | -4367 | Mknk2 | ENSMUSG0C | false | chr10 | 80676724 | 80677626 | 15.86955 | 42.74938 | 40.12535 | chr10 | 80669195 | 80680494 | 7.16995 | 4.88679 | 4.8213 |
| chr12 | 85486787 | 85487131 | 894 | Distal Intergenic | 12 | 85474911 | 85476063 | 1153 | 1 | ENSMUST00 | 11877 | Fos | ENSMUSG0C | false | chr12 | 85485608 | 85487883 | 27.24215 | 82.29425 | 79.10038 | chr12 | 85485423 | 85488221 | 5.47077 | 4.36176 | 3.37833 |
| chr13 | 24949403 | 24949777 | 894 | Intron (ENSMUST00000021773.6 | 13 | 24955854 | 24962818 | 6965 | 1 | ENSMUST00 | -6077 | Gpld1 | ENSMUSG0C | false | . | . | . | . | . | . | chr13 | 24947308 | 24961124 | 4.97023 | 4.07319 | 2.98257 |
| chr13 | 30396876 | 30397305 | 894 | Promoter (2-3kb) | 13 | 30399617 | 30399934 | 318 | 2 | ENSMUST00 | 2629 | Gm11369 | ENSMUSG0C | false | . | . | . | . | . | . | . | . | . | . | . | . |
| chr1 | 131084345 | 131084711 | 894 | Intron (ENSMUST00000016672.5 | 1 | 131073658 | 131073751 | 94 | 2 | ENSMUST00 | -10595 | Gm25549 | ENSMUSG0C | false | . | . | . | . | . | . | chr1 | 131068512 | 131096874 | 6.05258 | 4.49318 | 3.88351 |
| chr18 | 75532751 | 75533134 | 894 | Intron (ENSMUST00000165559.1 | 18 | 75514645 | 75522806 | 8162 | 1 | ENSMUST00 | 18107 | Gm10532 | ENSMUSG0C | false | . | . | . | . | . | . | chr18 | 75533002 | 75533884 | 4.41617 | 3.66707 | 2.55778 |
| chr4 | 72199945 | 72201284 | 893 | Promoter (<=1kb) | 4 | 72117142 | 72200919 | 83778 | 2 | ENSMUST00 | 0 | Tle1 | ENSMUSG0C | false | chr4 | 72198600 | 72202177 | 26.52689 | 93.43019 | 90.01563 | . | . | . | . | . | . |
| chr16 | 23272566 | 23272893 | 893 | Promoter (2-3kb) | 16 | 23269629 | 23321179 | 51551 | 1 | ENSMUST00 | 2938 | St6gal1 | ENSMUSG0C | true | . | . | . | . | . | . | chr16 | 23227288 | 23275612 | 5.51887 | 4.26961 | 3.44483 |
| chr3 | 68493319 | 68494512 | 893 | Promoter (<=1kb) | 3 | 68493491 | 68626231 | 132741 | 1 | ENSMUST00 | 0 | Schip1 | ENSMUSG0C | false | chr3 | 68492555 | 68495979 | 22.26925 | 90.51499 | 87.16024 | . | . | . | . | . | . |
| chr18 | 61398721 | 61399932 | 893 | Promoter (<=1kb) | 18 | 61298510 | 61400400 | 101891 | 2 | ENSMUST00 | 468 | Ppargc1b | ENSMUSG0C | false | chr18 | 61398143 | 61401028 | 41.07278 | 140.67711 | 135.76277 | chr18 | 61399853 | 61401196 | 3.31531 | 3.19643 | 1.6024 |
| chr5 | 31750632 | 31751082 | 893 | Intron (ENSMUST00000080598.8 | 5 | 31785383 | 31820643 | 35261 | 1 | ENSMUST00 | -34301 | Bre | ENSMUSG0C | false | . | . | . | . | . | . | chr5 | 31750491 | 31751443 | 3.69115 | 3.18611 | 1.96445 |
| chr14 | 22751201 | 22751644 | 893 | Intron (ENSMUST00000075639.5 | 14 | 22577727 | 22599268 | 21542 | 1 | ENSMUST00 | 173475 | 1700112E06I | ENSMUSG0C | false | . | . | . | . | . | . | chr14 | 22749519 | 22752555 | 6.00692 | 4.62149 | 3.83379 |
| chr12 | 108309486 | 108309825 | 892 | Intron (ENSMUST00000021685.6 | 12 | 108306270 | 108328300 | 22031 | 1 | ENSMUST00 | 3217 | Hhipl1 | ENSMUSG0C | false | . | . | . | . | . | . | chr12 | 108309061 | 108310754 | 4.25748 | 3.81765 | 2.41559 |
| chr3 | 131564398 | 131565158 | 892 | Promoter (<=1kb) | 3 | 131564768 | 131643670 | 78903 | 1 | ENSMUST00 | 0 | Papss1 | ENSMUSG0C | false | chr3 | 131564253 | 131564610 | 7.14186 | 14.46535 | 12.21151 | . | . | . | . | . | . |
| chr1 | 54896434 | 54896796 | 892 | Intron (ENSMUST00000179030.2 | 1 | 54766792 | 54884711 | 117920 | 2 | ENSMUST00 | -11724 | Ankrd44 | ENSMUSG0C | true | . | . | . | . | . | . | chr1 | 54896196 | 54904292 | 5.4736 | 4.28869 | 3.39741 |
| chr18 | 80390900 | 80391147 | 892 | Distal Intergenic | 18 | 80294546 | 80364254 | 69709 | 2 | ENSMUST00 | -26647 | Kcng2 | ENSMUSG0C | false | . | . | . | . | . | . | chr18 | 80390740 | 80391597 | 3.29248 | 3.11927 | 1.61719 |
| chr11 | 62636882 | 62637214 | 892 | Promoter (2-3kb) | 11 | 62634704 | 62638788 | 4085 | 1 | ENSMUST00 | 2179 | Lrrc75aos1 | ENSMUSG0C | false | . | . | . | . | . | . | chr11 | 62634966 | 62638705 | 3.87081 | 3.49347 | 2.10132 |
| chr3 | 41321854 | 41322121 | 892 | Distal Intergenic | 3 | 41441343 | 41441467 | 125 | 2 | ENSMUST00 | 119346 | Gm25487 | ENSMUSG0C | false | . | . | . | . | . | . | chr3 | 41321397 | 41322623 | 3.45475 | 3.25926 | 1.76385 |
| chr2 | 31164948 | 31165370 | 891 | Intron (ENSMUST00000056433.6 | 2 | 31172267 | 31184851 | 12585 | 1 | ENSMUST00 | -6897 | Gpr107 | ENSMUSG0C | false | . | . | . | . | . | . | chr2 | 31158838 | 31168082 | 5.9417 | 4.14532 | 3.82034 |
| chr10 | 121586555 | 121587053 | 891 | Promoter (<=1kb) | 10 | 121546455 | 121586794 | 40340 | 2 | ENSMUST00 | 0 | Tbk1 | ENSMUSG0C | false | chr10 | 121585897 | 121587605 | 27.03504 | 88.5911 | 85.27629 | . | . | . | . | . | . |
| chr13 | 112751900 | 112752286 | 891 | Distal Intergenic | 13 | 112800894 | 112867881 | 66988 | 1 | ENSMUST00 | -48608 | Ppap2a | ENSMUSG0C | false | . | . | . | . | . | . | chr13 | 112751229 | 112752032 | 5.72966 | 4.51241 | 3.64385 |
| chr6 | 148373524 | 148373878 | 891 | Intron (ENSMUST00000060095.9 | 6 | 148354656 | 148355598 | 943 | 1 | ENSMUST00 | 18869 | Rps4l | ENSMUSG0C | false | . | . | . | . | . | . | chr6 | 148370427 | 148377290 | 5.88115 | 4.51351 | 3.72575 |
| chr8 | 83442231 | 83442716 | 891 | Promoter (<=1kb) | 8 | 83434493 | 83442648 | 8156 | 2 | ENSMUST00 | 0 | Scoc | ENSMUSG0C | false | chr8 | 83441742 | 83443146 | 26.61343 | 84.02032 | 80.79139 | . | . | . | . | . | . |
| chr6 | 67131616 | 67131940 | 891 | Intron (ENSMUST00000181304.1 | 6 | 67115673 | 67168426 | 52754 | 1 | ENSMUST00 | 15944 | A430010J10F | ENSMUSG0C | false | . | . | . | . | . | . | chr6 | 67123607 | 67146074 | 5.6205 | 4.38535 | 3.51128 |
| chr13 | 99412732 | 99413132 | 891 | Promoter (<=1kb) | 13 | 99408610 | 99412864 | 4255 | 2 | ENSMUST00 | 0 | 6430562O15 | ENSMUSG0C | false | chr13 | 99412495 | 99412862 | 4.97263 | 8.68936 | 6.53912 | chr13 | 99411965 | 99413323 | 4.47516 | 3.81338 | 2.61903 |
| chr17 | 69973235 | 69973554 | 890 | Intron (ENSMUST00000133983.2 | 17 | 69969421 | 70594408 | 624988 | 1 | ENSMUST00 | 3815 | Dlgap1 | ENSMUSG0C | false | . | . | . | . | . | . | . | . | . | . | . | . |
| chr2 | 32711979 | 32712455 | 890 | Promoter (<=1kb) | 2 | 32707444 | 32712267 | 4824 | 2 | ENSMUST00 | 0 | Cdk9 | ENSMUSG0C | false | chr2 | 32711568 | 32713700 | 30.94804 | 114.76607 | 110.79114 | chr2 | 32710967 | 32712041 | 8.09371 | 5.63132 | 5.52562 |
| chr3 | 116406452 | 116406769 | 890 | Intron (ENSMUST00000106491.2 | 3 | 116400186 | 116423024 | 22839 | 2 | ENSMUST00 | 16255 | Cdc14a | ENSMUSG0C | true | . | . | . | . | . | . | chr3 | 116394159 | 116421460 | 5.70692 | 4.42755 | 3.58909 |
| chr11 | 72478282 | 72478547 | 890 | Intron (ENSMUST00000045303.4 | 11 | 72459029 | 72475027 | 15999 | 2 | ENSMUST00 | -3256 | Spns2 | ENSMUSG0C | false | . | . | . | . | . | . | chr11 | 72473568 | 72481707 | 6.44299 | 4.72774 | 4.18703 |
| chr13 | 32493842 | 32494142 | 890 | Distal Intergenic | 13 | 32379907 | 32383688 | 3782 | 2 | ENSMUST00 | -110155 | Gm11381 | ENSMUSG0C | false | . | . | . | . | . | . | chr13 | 32484237 | 32502844 | 4.66222 | 3.8395 | 2.75657 |
| chr9 | 75496087 | 75496444 | 890 | Downstream (1-2kb) | 9 | 75441122 | 75441652 | 531 | 2 | ENSMUST00 | -54436 | A130057D12 | ENSMUSG0C | false | . | . | . | . | . | . | chr9 | 75493281 | 75497630 | 5.72539 | 4.39347 | 3.61269 |
| chr11 | 77806845 | 77807131 | 889 | Intron (ENSMUST00000108375.3 | 11 | 77811515 | 77865980 | 54466 | 1 | ENSMUST00 | -4384 | Myo18a | ENSMUSG0C | false | . | . | . | . | . | . | chr11 | 77799596 | 77817107 | 5.11096 | 4.06241 | 3.11534 |
| chr19 | 58817497 | 58817786 | 889 | Intron (ENSMUST00000066285.4 | 19 | 58785803 | 58794461 | 8659 | 2 | ENSMUST00 | -23037 | 1700019N19 | ENSMUSG0C | false | . | . | . | . | . | . | chr19 | 58814047 | 58859817 | 7.46659 | 5.12677 | 5.03638 |
| chr3 | 9727043 | 9727418 | 889 | Intron (ENSMUST00000161949.2 | 3 | 9692767 | 9705002 | 12236 | 2 | ENSMUST00 | -22042 | Pag1 | ENSMUSG0C | true | . | . | . | . | . | . | chr3 | 9723127 | 9731243 | 5.21145 | 4.10329 | 3.21588 |
| chr7 | 84669858 | 84670136 | 889 | Intron (ENSMUST00000069537.2 | 7 | 84615054 | 84679351 | 64298 | 2 | ENSMUST00 | 9215 | Zfand6 | ENSMUSG0C | false | . | . | . | . | . | . | chr7 | 84662142 | 84686408 | 4.53031 | 3.83251 | 2.62083 |
| chr19 | 29325243 | 29325614 | 889 | Promoter (<=1kb) | 19 | 29321344 | 29325356 | 4013 | 2 | ENSMUST00 | 0 | Insl6 | ENSMUSG0C | false | chr19 | 29325013 | 29325338 | 10.96168 | 23.03899 | 20.66252 | chr19 | 29323469 | 29326100 | 4.62638 | 3.94921 | 2.70535 |
| chr1 | 134099881 | 134100216 | 889 | Distal Intergenic | 1 | 134109368 | 134109799 | 432 | 1 | ENSMUST00 | -9152 | Gm8618 | ENSMUSG0C | false | chr1 | 134099435 | 134099888 | 10.36119 | 22.58756 | 20.21866 | chr1 | 134095503 | 134101645 | 7.68348 | 5.30363 | 5.20617 |
| chr11 | 88172992 | 88173362 | 888 | Intron (ENSMUST00000018522.7 | 11 | 88182303 | 88183280 | 978 | 1 | ENSMUST00 | -8941 | Cuedc1 | ENSMUSG0C | true | . | . | . | . | . | . | chr11 | 88146778 | 88183132 | 6.97604 | 4.89368 | 4.62779 |
| chr17 | 29660400 | 29660809 | 888 | Promoter (<=1kb) | 17 | 29660595 | 29705979 | 45385 | 1 | ENSMUST00 | 0 | Cmtr1 | ENSMUSG0C | false | chr17 | 29660044 | 29661644 | 26.72788 | 101.58999 | 97.97913 | . | . | . | . | . | . |
| chr7 | 70350866 | 70351578 | 888 | Downstream (<1kb) | 7 | 70351946 | 70360593 | 8648 | 2 | ENSMUST00 | 9015 | Nr2f2 | ENSMUSG0C | false | chr7 | 70350687 | 70352400 | 22.97661 | 61.46103 | 58.5941 | . | . | . | . | . | . |
| chr1 | 91211794 | 91212074 | 888 | Intron (ENSMUST00000097648.5 | 1 | 91180046 | 91225196 | 45151 | 1 | ENSMUST00 | 31749 | Ramp1 | ENSMUSG0C | false | . | . | . | . | . | . | chr1 | 91210661 | 91212446 | 4.71495 | 3.72845 | 2.80384 |
| chr13 | 44907692 | 44908164 | 888 | Intron (ENSMUST00000173704.2 | 13 | 44840783 | 44896481 | 55699 | 1 | ENSMUST00 | 66910 | Jarid2 | ENSMUSG0C | false | . | . | . | . | . | . | chr13 | 44903494 | 44909610 | 5.92541 | 4.18183 | 3.80911 |
| chr2 | 102451561 | 102452159 | 888 | Promoter (<=1kb) | 2 | 102449366 | 102452499 | 3134 | 2 | ENSMUST00 | 340 | Fjx1 | ENSMUSG0C | false | chr2 | 102450101 | 102452417 | 25.88708 | 77.13007 | 74.02996 | chr2 | 102449784 | 102452396 | 5.13159 | 3.96451 | 3.13453 |

| | | | | | | | | | | | | | | | | | | | | | | | | | | |
|---|---|---|---|---|---|---|---|---|---|---|---|---|---|---|---|---|---|---|---|---|---|---|---|---|---|---|
| chr2 | 180225596 | 180226083 | 887 | Promoter (<=1kb) | 2 | 180176373 | 180225859 | 49487 | 2 | ENSMUST00 | 0 | Lama5 | ENSMUSG0C | true | chr2 | 180223494 | 180226303 | 32.29185 | 112.09676 | 108.19604 | chr2 | 180225908 | 180236447 | 7.12665 | 5.06061 | 4.74583 |
| chr9 | 51239723 | 51240016 | 887 | 3' UTR | 9 | 51213690 | 51240079 | 26390 | 1 | ENSMUST00 | 26034 | Pou2af1 | ENSMUSG0C | false | . | . | . | . | . | . | chr9 | 51238831 | 51242460 | 5.56756 | 4.32807 | 3.47861 |
| chr17 | 86458418 | 86458756 | 887 | Intron (ENSMUST00000097275.3 | 17 | 86490562 | 86496112 | 5551 | 1 | ENSMUST00 | -31806 | Prkce | ENSMUSG0C | true | . | . | . | . | . | . | chr17 | 86455682 | 86461399 | 4.95475 | 3.9467 | 3.00207 |
| chr9 | 44735095 | 44735378 | 887 | Promoter (<=1kb) | 9 | 44686308 | 44735198 | 48891 | 2 | ENSMUST00 | 0 | Phldb1 | ENSMUSG0C | false | chr9 | 44733443 | 44735510 | 29.66345 | 100.21 | 96.63225 | chr9 | 44732060 | 44735669 | 3.74755 | 3.35522 | 1.98328 |
| chr10 | 82814143 | 82814691 | 887 | Promoter (2-3kb) | 10 | 82812123 | 82824242 | 12120 | 1 | ENSMUST00 | 2021 | 1700028I16R | ENSMUSG0C | false | . | . | . | . | . | . | chr10 | 82812050 | 82815775 | 5.74107 | 4.43512 | 3.61361 |
| chr5 | 93099436 | 93099726 | 887 | Intron (ENSMUST00000074733.7 | 5 | 93093457 | 93174958 | 81502 | 1 | ENSMUST00 | 5980 | Sep-11 | ENSMUSG0C | false | . | . | . | . | . | . | chr5 | 93095018 | 93154711 | 6.59326 | 4.78158 | 4.31782 |
| chr15 | 98776324 | 98776692 | 886 | Promoter (<=1kb) | 15 | 98772200 | 98777261 | 5062 | 2 | ENSMUST00 | 569 | Wnt10b | ENSMUSG0C | false | chr15 | 98776332 | 98776744 | 8.78697 | 16.7955 | 14.50663 | chr15 | 98775676 | 98778436 | 4.71083 | 3.81658 | 2.80395 |
| chr6 | 108537044 | 108537375 | 886 | Intron (ENSMUST00000032192.6 | 6 | 108563445 | 108565693 | 2249 | 2 | ENSMUST00 | 28318 | Gm16433 | ENSMUSG0C | false | . | . | . | . | . | . | chr6 | 108530788 | 108538190 | 6.66183 | 4.72814 | 4.3796 |
| chr4 | 144903899 | 144904266 | 886 | Intron (ENSMUST00000105744.2 | 4 | 144893219 | 144927639 | 34421 | 1 | ENSMUST00 | 10681 | Dhrs3 | ENSMUSG0C | false | . | . | . | . | . | . | chr4 | 144899456 | 144904697 | 3.84043 | 3.49289 | 2.05554 |
| chr4 | 109017429 | 109017777 | 886 | Promoter (<=1kb) | 4 | 109017655 | 109017730 | 76 | 1 | ENSMUST00 | 0 | Mir761 | ENSMUSG0C | false | . | . | . | . | . | . | chr4 | 109016641 | 109018428 | 4.7047 | 4.01243 | 2.75163 |
| chr10 | 105802143 | 105802440 | 886 | Promoter (1-2kb) | 10 | 105804087 | 105804197 | 111 | 2 | ENSMUST00 | 1757 | n-R5s80 | ENSMUSG0C | false | . | . | . | . | . | . | chr10 | 105801283 | 105802753 | 3.822 | 3.57102 | 2.02063 |
| chr5 | 21136373 | 21136788 | 886 | Distal Intergenic | 5 | 21119149 | 21119522 | 374 | 2 | ENSMUST00 | -16852 | Rpl31-ps21 | ENSMUSG0C | false | . | . | . | . | . | . | chr5 | 21135523 | 21137102 | 5.009 | 4.08637 | 3.02466 |
| chr1 | 62884983 | 62885519 | 886 | Distal Intergenic | 1 | 62821355 | 62832370 | 11016 | 2 | ENSMUST00 | -52614 | RP23-149A5. | ENSMUSG0C | false | chr1 | 62883755 | 62885574 | 13.51752 | 34.31881 | 31.79871 | chr1 | 62885282 | 62890313 | 4.6486 | 3.92393 | 2.70924 |
| chr17 | 56462091 | 56462694 | 885 | Promoter (<=1kb) | 17 | 56412431 | 56462216 | 49786 | 2 | ENSMUST00 | 0 | Ptprs | ENSMUSG0C | false | chr17 | 56461389 | 56463239 | 29.93067 | 92.76174 | 89.35763 | chr17 | 56460929 | 56463551 | 7.72585 | 5.25113 | 5.23609 |
| chr2 | 27515011 | 27515342 | 885 | Promoter (<=1kb) | 2 | 27515157 | 27520653 | 5497 | 1 | ENSMUST00 | 0 | Wdr5 | ENSMUSG0C | false | chr2 | 27514311 | 27516190 | 31.61691 | 99.67752 | 96.11594 | . | . | . | . | . | . |
| chr5 | 115259291 | 115259611 | 885 | Promoter (<=1kb) | 5 | 115251314 | 115260226 | 8913 | 2 | ENSMUST00 | 615 | Rnf10 | ENSMUSG0C | false | . | . | . | . | . | . | chr5 | 115257865 | 115271333 | 6.24384 | 4.53084 | 4.06041 |
| chr1 | 89580008 | 89580695 | 885 | Promoter (<=1kb) | 1 | 89580236 | 89895277 | 315042 | 1 | ENSMUST00 | 0 | Agap1 | ENSMUSG0C | false | chr1 | 89579858 | 89582737 | 22.99841 | 82.87405 | 79.66897 | chr1 | 89564740 | 89589718 | 5.63127 | 4.22514 | 3.56367 |
| chr2 | 92742033 | 92742443 | 885 | Promoter (<=1kb) | 2 | 92742230 | 92759208 | 16979 | 1 | ENSMUST00 | 0 | Gm13816 | ENSMUSG0C | false | . | . | . | . | . | . | chr2 | 92741714 | 92744170 | 6.09939 | 4.6297 | 3.92188 |
| chr2 | 44979046 | 44979564 | 885 | Distal Intergenic | 2 | 45023100 | 45023177 | 78 | 2 | ENSMUST00 | 43613 | Mir5129 | ENSMUSG0C | false | . | . | . | . | . | . | chr2 | 44975552 | 44981626 | 5.83223 | 4.48649 | 3.69967 |
| chr4 | 147936505 | 147937006 | 884 | Promoter (<=1kb) | 4 | 147909753 | 147936767 | 27015 | 2 | ENSMUST00 | 0 | Plod1 | ENSMUSG0C | false | chr4 | 147936057 | 147937085 | 38.14117 | 119.25307 | 115.13219 | . | . | . | . | . | . |
| chr9 | 119909919 | 119910236 | 884 | Intron (ENSMUST00000036561.6 | 9 | 119894895 | 119926579 | 31685 | 1 | ENSMUST00 | 15025 | Wdr48 | ENSMUSG0C | false | . | . | . | . | . | . | chr9 | 119907328 | 119910777 | 4.65961 | 3.61457 | 2.76599 |
| chr2 | 121354870 | 121355461 | 884 | Promoter (<=1kb) | 2 | 121310561 | 121355330 | 44770 | 2 | ENSMUST00 | 0 | Ppip5k1 | ENSMUSG0C | false | chr2 | 121354397 | 121355435 | 24.07142 | 68.54774 | 65.57966 | chr2 | 121353246 | 121355763 | 3.46336 | 3.25461 | 1.76223 |
| chr5 | 86804031 | 86804574 | 884 | Promoter (<=1kb) | 5 | 86804221 | 86836654 | 32434 | 1 | ENSMUST00 | 0 | Ythdc1 | ENSMUSG0C | false | chr5 | 86803694 | 86805523 | 35.31382 | 118.55977 | 114.46947 | . | . | . | . | . | . |
| chr10 | 128133645 | 128134402 | 884 | Exon (ENSMUST00000092033.2/ | 10 | 128092783 | 128129303 | 36521 | 1 | ENSMUST00 | 40863 | Baz2a | ENSMUSG0C | false | . | . | . | . | . | . | chr10 | 128126743 | 128136296 | 4.86153 | 3.96045 | 2.90995 |
| chr7 | 98851241 | 98851576 | 884 | Intron (ENSMUST00000163913.2 | 7 | 98854732 | 98855095 | 364 | 2 | ENSMUST00 | 3519 | Gm8398 | ENSMUSG0C | false | . | . | . | . | . | . | chr7 | 98850469 | 98851957 | 4.00702 | 3.68889 | 2.17282 |
| chr11 | 51689005 | 51689285 | 883 | Promoter (<=1kb) | 11 | 51685386 | 51688874 | 3489 | 2 | ENSMUST00 | -132 | 0610009B22 | ENSMUSG0C | false | chr11 | 51688104 | 51689576 | 31.24818 | 92.1069 | 88.71768 | . | . | . | . | . | . |
| chr15 | 67045348 | 67045647 | 883 | Distal Intergenic | 15 | 67102875 | 67113992 | 11118 | 2 | ENSMUST00 | 68345 | St3gal1 | ENSMUSG0C | true | . | . | . | . | . | . | chr15 | 67045424 | 67046308 | 2.61658 | 2.72816 | 1.10727 |
| chr16 | 30081771 | 30082261 | 883 | Distal Intergenic | 16 | 30066338 | 30067319 | 982 | 1 | ENSMUST00 | 15434 | Hes1 | ENSMUSG0C | false | . | . | . | . | . | . | chr16 | 30080758 | 30087022 | 7.36233 | 5.17806 | 4.92775 |
| chr11 | 34184216 | 34184603 | 883 | Distal Intergenic | 11 | 34204338 | 34208089 | 3752 | 2 | ENSMUST00 | 23486 | Foxi1 | ENSMUSG0C | false | . | . | . | . | . | . | chr11 | 34177517 | 34186117 | 5.74808 | 4.36831 | 3.63761 |
| chr2 | 159888201 | 159888549 | 883 | Distal Intergenic | 2 | 159858425 | 159859290 | 866 | 2 | ENSMUST00 | -28912 | Gm11446 | ENSMUSG0C | false | . | . | . | . | . | . | chr2 | 159883743 | 159890271 | 4.98798 | 4.05699 | 2.99637 |
| chr11 | 108422387 | 108422708 | 883 | Promoter (<=1kb) | 11 | 108423273 | 108423758 | 486 | 1 | ENSMUST00 | -565 | Gm11667 | ENSMUSG0C | false | . | . | . | . | . | . | . | . | . | . | . | . |
| chr4 | 137349607 | 137349888 | 882 | Intron (ENSMUST00000051477.7 | 4 | 137321762 | 137357699 | 35938 | 2 | ENSMUST00 | 7811 | Cdc42 | ENSMUSG0C | false | . | . | . | . | . | . | chr4 | 137334966 | 137356618 | 5.62758 | 4.35253 | 3.53044 |
| chr1 | 131097599 | 131098034 | 882 | Promoter (<=1kb) | 1 | 131053700 | 131097826 | 44127 | 2 | ENSMUST00 | 0 | Mapkapk2 | ENSMUSG0C | false | chr1 | 131095659 | 131098154 | 28.90341 | 112.84859 | 108.93253 | . | . | . | . | . | . |
| chr2 | 76564564 | 76564857 | 882 | Exon (ENSMUST00000077972.5/ | 2 | 76568120 | 76580028 | 11909 | 1 | ENSMUST00 | -3263 | Osbpl6 | ENSMUSG0C | false | . | . | . | . | . | . | chr2 | 76563488 | 76567561 | 4.40902 | 3.85967 | 2.51429 |
| chr9 | 108479743 | 108480177 | 882 | Promoter (<=1kb) | 9 | 108479862 | 108490530 | 10669 | 1 | ENSMUST00 | 0 | Lamb2 | ENSMUSG0C | false | chr9 | 108478517 | 108480814 | 19.47644 | 60.36021 | 57.50917 | chr9 | 108477974 | 108483462 | 4.94861 | 3.89359 | 3.00507 |
| chr14 | 74915143 | 74915391 | 882 | Intron (ENSMUST00000088970.5 | 14 | 74754674 | 74947877 | 193204 | 2 | ENSMUST00 | 32486 | Lrch1 | ENSMUSG0C | true | . | . | . | . | . | . | chr14 | 74857233 | 74945366 | 6.29465 | 4.64624 | 4.07764 |
| chr12 | 94280920 | 94281302 | 882 | Distal Intergenic | 12 | 93927004 | 93929102 | 2099 | 2 | ENSMUST00 | -351819 | Gm9726 | ENSMUSG0C | false | . | . | . | . | . | . | chr12 | 94279611 | 94284656 | 3.79808 | 3.5006 | 2.0053 |
| chr4 | 40836407 | 40836654 | 882 | Intron (ENSMUST00000030121.7 | 4 | 40850056 | 40850138 | 83 | 2 | ENSMUST00 | 13484 | Mir5123 | ENSMUSG0C | false | . | . | . | . | . | . | chr4 | 40821931 | 40852722 | 6.59841 | 4.80932 | 4.32335 |
| chr11 | 104230431 | 104230747 | 881 | Promoter (<=1kb) | 11 | 104231390 | 104332090 | 100701 | 1 | ENSMUST00 | -643 | Mapt | ENSMUSG0C | true | chr11 | 104230662 | 104231268 | 16.78262 | 53.63071 | 50.86883 | . | . | . | . | . | . |
| chr4 | 144990082 | 144990455 | 881 | Intron (ENSMUST00000036579.8 | 4 | 144974926 | 144983961 | 9036 | 2 | ENSMUST00 | -6122 | Vps13d | ENSMUSG0C | false | . | . | . | . | . | . | chr4 | 144980824 | 144994430 | 5.61677 | 4.30514 | 3.52155 |
| chr4 | 135963520 | 135963830 | 881 | Promoter (<=1kb) | 4 | 135963727 | 135968178 | 4452 | 1 | ENSMUST00 | 0 | Gale | ENSMUSG0C | false | chr4 | 135962895 | 135964814 | 26.1454 | 83.18384 | 79.97624 | chr4 | 135962290 | 135965293 | 5.15033 | 4.12284 | 3.14697 |
| chr12 | 80435919 | 80436877 | 881 | Promoter (<=1kb) | 12 | 80335847 | 80436601 | 100755 | 2 | ENSMUST00 | 0 | Dcaf5 | ENSMUSG0C | false | chr12 | 80434574 | 80437085 | 29.65943 | 90.33628 | 86.98486 | . | . | . | . | . | . |
| chr7 | 4995994 | 4996360 | 881 | Promoter (<=1kb) | 7 | 4983483 | 4996055 | 12573 | 2 | ENSMUST00 | 0 | Zfp579 | ENSMUSG0C | false | chr7 | 4995549 | 4996856 | 19.38504 | 61.02572 | 58.1656 | chr7 | 4994037 | 5000309 | 5.22258 | 4.18877 | 3.19451 |
| chr6 | 139510263 | 139510598 | 881 | Distal Intergenic | 6 | 139493182 | 139501909 | 8728 | 2 | ENSMUST00 | -8355 | Rergl | ENSMUSG0C | false | . | . | . | . | . | . | . | . | . | . | . | . |
| chr17 | 68019436 | 68019801 | 880 | Distal Intergenic | 17 | 67842713 | 68004120 | 161408 | 2 | ENSMUST00 | -15317 | Arhgap28 | ENSMUSG0C | true | . | . | . | . | . | . | chr17 | 68018805 | 68020099 | 3.38836 | 3.11437 | 1.7226 |
| chrX | 50841177 | 50841682 | 880 | Promoter (<=1kb) | 20 | 50841441 | 50880074 | 38634 | 1 | ENSMUST00 | 0 | 2610018G03 | ENSMUSG0C | false | chrX | 50840775 | 50842647 | 20.21941 | 51.82391 | 49.08421 | . | . | . | . | . | . |
| chr11 | 96931637 | 96931948 | 880 | Promoter (<=1kb) | 11 | 96931387 | 96935280 | 3894 | 1 | ENSMUST00 | 251 | Prr15l | ENSMUSG0C | false | . | . | . | . | . | . | chr11 | 96930939 | 96933401 | 5.741 | 4.47904 | 3.61229 |
| chr11 | 116412412 | 116413409 | 880 | Promoter (<=1kb) | 11 | 116336353 | 116413004 | 76652 | 2 | ENSMUST00 | 0 | Rnf157 | ENSMUSG0C | false | chr11 | 116412272 | 116413687 | 34.36326 | 108.53439 | 104.74065 | chr11 | 116411810 | 116413675 | 6.71255 | 4.93125 | 4.4309 |
| chr7 | 78578250 | 78578669 | 880 | Promoter (<=1kb) | 7 | 78192362 | 78578408 | 386047 | 2 | ENSMUST00 | 0 | Ntrk3 | ENSMUSG0C | true | chr7 | 78576814 | 78579420 | 15.45146 | 50.98658 | 48.25883 | . | . | . | . | . | . |
| chr19 | 12719246 | 12719579 | 880 | Promoter (1-2kb) | 19 | 12706055 | 12721203 | 15149 | 2 | ENSMUST00 | 1624 | Gm15962 | ENSMUSG0C | false | . | . | . | . | . | . | chr19 | 12711318 | 12720714 | 4.33765 | 3.73227 | 2.4689 |
| chr11 | 115173851 | 115174293 | 879 | Intron (ENSMUST00000021077.3 | 11 | 115163341 | 115181181 | 17841 | 1 | ENSMUST00 | 10511 | Slc9a3r1 | ENSMUSG0C | false | . | . | . | . | . | . | chr11 | 115165089 | 115178632 | 7.11262 | 4.95304 | 4.75761 |
| chr11 | 95263847 | 95264232 | 879 | Promoter (2-3kb) | 11 | 95261529 | 95269265 | 7737 | 1 | ENSMUST00 | 2319 | Tac4 | ENSMUSG0C | false | . | . | . | . | . | . | chr11 | 95257171 | 95275447 | 6.31854 | 4.58687 | 4.10024 |
| chr5 | 124528487 | 124529153 | 879 | Promoter (2-3kb) | 5 | 124493082 | 124531391 | 38310 | 2 | ENSMUST00 | 2238 | Rilpl1 | ENSMUSG0C | false | chr5 | 124528429 | 124531965 | 33.66875 | 128.30269 | 123.87101 | chr5 | 124523847 | 124533697 | 5.6925 | 4.38176 | 3.58658 |
| chr5 | 142909537 | 142910048 | 879 | Promoter (2-3kb) | 5 | 142903115 | 142906754 | 3640 | 2 | ENSMUST00 | -2784 | Actb | ENSMUSG0C | false | . | . | . | . | . | . | chr5 | 142907369 | 142912953 | 6.83151 | 4.8323 | 4.52174 |
| chr4 | 152086487 | 152087189 | 879 | Promoter (<=1kb) | 4 | 152086858 | 152115390 | 28533 | 1 | ENSMUST00 | 0 | Plekhg5 | ENSMUSG0C | false | chr4 | 152086343 | 152087761 | 34.36702 | 111.64683 | 107.76208 | chr4 | 152086190 | 152088863 | 4.6716 | 3.90229 | 2.75862 |
| chr18 | 6135487 | 6136486 | 879 | Promoter (<=1kb) | 18 | 6024785 | 6135907 | 111123 | 2 | ENSMUST00 | 0 | Arhgap12 | ENSMUSG0C | false | chr18 | 6134705 | 6136106 | 49.92314 | 175.78799 | 169.70897 | . | . | . | . | . | . |
| chr12 | 98233388 | 98233646 | 878 | Intron (ENSMUST00000021390.7 | 12 | 98202304 | 98259459 | 57156 | 2 | ENSMUST00 | 25813 | Galc | ENSMUSG0C | false | . | . | . | . | . | . | chr12 | 98231481 | 98245211 | 5.28965 | 4.20277 | 3.23997 |
| chr15 | 79441684 | 79442285 | 878 | Promoter (<=1kb) | 15 | 79417856 | 79441999 | 24144 | 2 | ENSMUST00 | 0 | Csnk1e | ENSMUSG0C | false | chr15 | 79440865 | 79442525 | 28.53699 | 95.19626 | 91.7378 | chr15 | 79428565 | 79442760 | 6.13618 | 4.60978 | 3.93942 |
| chr3 | 68608769 | 68609238 | 878 | Intron (ENSMUST00000029346.7 | 3 | 68596175 | 68598163 | 1989 | 1 | ENSMUST00 | 12595 | Schip1 | ENSMUSG0C | false | . | . | . | . | . | . | chr3 | 68574597 | 68620372 | 5.61397 | 4.36647 | 3.51459 |
| chr17 | 88529916 | 88530368 | 878 | Promoter (<=1kb) | 17 | 88530118 | 88588367 | 58250 | 1 | ENSMUST00 | 0 | Ppp1r21 | ENSMUSG0C | false | chr17 | 88529505 | 88530968 | 28.91859 | 89.0447 | 85.71584 | chr17 | 88529487 | 88531878 | 3.75088 | 3.51583 | 1.98729 |
| chr3 | 53463159 | 53463923 | 878 | Promoter (<=1kb) | 3 | 53448588 | 53463291 | 14704 | 2 | ENSMUST00 | 0 | Nhlrc3 | ENSMUSG0C | false | chr3 | 53462704 | 53465185 | 39.23734 | 130.83873 | 126.32064 | . | . | . | . | . | . |
| chr1 | 38478495 | 38478772 | 878 | Intron (ENSMUST00000095027.3 | 1 | 38388829 | 38390244 | 1416 | 1 | ENSMUST00 | 89667 | Gm16152 | ENSMUSG0C | false | . | . | . | . | . | . | chr1 | 38477579 | 38479437 | 4.72226 | 3.85685 | 2.80328 |
| chr6 | 88140667 | 88141006 | 878 | Distal Intergenic | 6 | 88104128 | 88104258 | 131 | 2 | ENSMUST00 | -36410 | Mir6376 | ENSMUSG0C | false | . | . | . | . | . | . | chr6 | 88140742 | 88141101 | 3.2962 | 3.17135 | 1.64853 |
| chr3 | 146812378 | 146813315 | 877 | Promoter (<=1kb) | 3 | 146729579 | 146812960 | 83382 | 2 | ENSMUST00 | 0 | Prkacb | ENSMUSG0C | false | chr3 | 146811927 | 146813727 | 38.29964 | 140.16156 | 135.27679 | chr3 | 146810236 | 146812503 | 4.36107 | 3.80659 | 2.48288 |
| chr14 | 57724458 | 57724750 | 877 | 3' UTR | 14 | 57722189 | 57734300 | 12112 | 2 | ENSMUST00 | 9550 | Lats2 | ENSMUSG0C | false | . | . | . | . | . | . | chr14 | 57704250 | 57744175 | 6.91785 | 4.80068 | 4.61004 |
| chr8 | 120276359 | 120276979 | 877 | Intron (ENSMUST00000127664.1 | 8 | 120230536 | 120553634 | 323099 | 1 | ENSMUST00 | 45824 | Gse1 | ENSMUSG0C | false | chr8 | 120276314 | 120277228 | 13.48988 | 31.60083 | 29.11589 | chr8 | 120273860 | 120279682 | 7.5755 | 5.06006 | 5.13676 |
| chr16 | 16213034 | 16213508 | 877 | Promoter (<=1kb) | 16 | 16213318 | 16227287 | 13970 | 1 | ENSMUST00 | 0 | Pkp2 | ENSMUSG0C | false | chr16 | 16212978 | 16214696 | 35.53755 | 114.73917 | 110.76805 | . | . | . | . | . | . |
| chr14 | 120705165 | 120705652 | 877 | Distal Intergenic | 14 | 120749185 | 120750062 | 878 | 1 | ENSMUST00 | -43533 | Gm17613 | ENSMUSG0C | false | . | . | . | . | . | . | chr14 | 120704875 | 120706273 | 2.96719 | 2.95506 | 1.38241 |
| chr10 | 123202058 | 123202343 | 877 | Distal Intergenic | 10 | 123196876 | 123196971 | 96 | 2 | ENSMUST00 | -5088 | Gm27920 | ENSMUSG0C | false | chr10 | 123201979 | 123202456 | 9.63953 | 19.72805 | 17.39678 | chr10 | 123201179 | 123202614 | 2.96241 | 2.95331 | 1.40075 |
| chr12 | 36016602 | 36016978 | 876 | Intron (ENSMUST00000020896.9 | 12 | 36021812 | 36039935 | 18124 | 2 | ENSMUST00 | 22957 | Tspan13 | ENSMUSG0C | false | . | . | . | . | . | . | chr12 | 36012836 | 36017990 | 4.36195 | 3.78173 | 2.49385 |
| chr1 | 44102123 | 44102685 | 876 | Promoter (<=1kb) | 1 | 44086613 | 44102391 | 15779 | 2 | ENSMUST00 | 0 | Tex30 | ENSMUSG0C | false | chr1 | 44101623 | 44102830 | 29.66345 | 100.21 | 96.63225 | . | . | . | . | . | . |
| chr2 | 119742062 | 119742386 | 876 | Promoter (<=1kb) | 2 | 119742337 | 119751263 | 8927 | 1 | ENSMUST00 | 0 | Itpka | ENSMUSG0C | false | chr2 | 119741625 | 119743529 | 33.08674 | 107.33151 | 103.57004 | chr2 | 119740863 | 119745199 | 5.39081 | 4.15988 | 3.37725 |
| chr11 | 120237808 | 120238298 | 876 | Promoter (1-2kb) | 11 | 120236699 | 120292282 | 55584 | 1 | ENSMUST00 | 1110 | Bahcc1 | ENSMUSG0C | false | chr11 | 120232940 | 120238211 | 32.00811 | 98.35596 | 94.82688 | chr11 | 120237752 | 120244869 | 5.41747 | 4.25596 | 3.35787 |
| chr7 | 135479314 | 135479748 | 876 | Distal Intergenic | 7 | 135511466 | 135528654 | 17189 | 2 | ENSMUST00 | 48906 | Clrn3 | ENSMUSG0C | false | . | . | . | . | . | . | chr7 | 135475748 | 135480191 | 4.6542 | 3.90353 | 2.72165 |
| chr16 | 91056551 | 91056847 | 876 | Exon (ENSMUST00000147848.1/ | 16 | 91044656 | 91056677 | 12022 | 1 | ENSMUST00 | 11896 | 4931406G06 | ENSMUSG0C | false | . | . | . | . | . | . | chr16 | 91055848 | 91057249 | 3.33522 | 3.17752 | 1.69815 |
| chr10 | 59741492 | 59741786 | 875 | Promoter (<=1kb) | 10 | 59740774 | 59757174 | 16401 | 1 | ENSMUST00 | 719 | Micu1 | ENSMUSG0C | true | . | . | . | . | . | . | chr10 | 59740710 | 59744639 | 5.66768 | 4.30807 | 3.58646 |
| chr2 | 32881000 | 32881400 | 875 | Intron (ENSMUST00000028135.9 | 2 | 32876114 | 32925254 | 49141 | 1 | ENSMUST00 | 4887 | Fam129b | ENSMUSG0C | false | . | . | . | . | . | . | chr2 | 32877160 | 32918190 | 7.77045 | 5.19056 | 5.2936 |
| chr7 | 80294176 | 80294455 | 875 | Promoter (<=1kb) | 7 | 80294450 | 80316259 | 21810 | 1 | ENSMUST00 | 0 | Prc1 | ENSMUSG0C | false | chr7 | 80294374 | 80294938 | 27.29887 | 88.10642 | 84.80337 | chr7 | 80293719 | 80295226 | 5.16977 | 4.09363 | 3.19473 |
| chr13 | 52530623 | 52530940 | 875 | Promoter (<=1kb) | 13 | 52504375 | 52530836 | 26462 | 2 | ENSMUST00 | 0 | Diras2 | ENSMUSG0C | false | . | . | . | . | . | . | chr13 | 52529872 | 52531033 | 4.57626 | 3.86024 | 2.6563 |
| chr19 | 42752387 | 42753002 | 875 | Promoter (<=1kb) | 19 | 42725858 | 42752775 | 26918 | 2 | ENSMUST00 | 0 | Pyroxd2 | ENSMUSG0C | false | chr19 | 42752018 | 42753504 | 12.91534 | 31.26572 | 28.78545 | chr19 | 42751869 | 42756848 | 4.65033 | 3.76971 | 2.74958 |
| chr1 | 190438879 | 190439264 | 875 | Distal Intergenic | 1 | 190482845 | 190483157 | 313 | 2 | ENSMUST00 | 43893 | Gm23153 | ENSMUSG0C | false | . | . | . | . | . | . | . | . | . | . | . | . |
| chr8 | 121300798 | 121301265 | 875 | Intron (ENSMUST00000127664.1 | 8 | 121256747 | 121356836 | 100090 | 2 | ENSMUST00 | 55571 | Gm26815 | ENSMUSG0C | false | . | . | . | . | . | . | chr8 | 121299982 | 121302568 | 7.20896 | 5.12344 | 4.81006 |
| chr1 | 153332440 | 153333257 | 874 | Promoter (<=1kb) | 1 | 153218922 | 153332786 | 113865 | 2 | ENSMUST00 | 0 | Lamc1 | ENSMUSG0C | false | chr1 | 153331293 | 153333837 | 19.98426 | 92.42262 | 89.02813 | . | . | . | . | . | . |
| chr14 | 8098038 | 8098602 | 874 | Promoter (<=1kb) | 14 | 8098213 | 8165111 | 66899 | 1 | ENSMUST00 | 0 | Pxk | ENSMUSG0C | false | chr14 | 8097842 | 8099268 | 23.3092 | 84.4034 | 81.16875 | chr14 | 8097507 | 8100795 | 3.69272 | 3.44449 | 1.92601 |
| chr9 | 63613216 | 63613516 | 874 | Intron (ENSMUST00000041551.7 | 9 | 63602655 | 63641889 | 39235 | 1 | ENSMUST00 | 10562 | Aagab | ENSMUSG0C | false | . | . | . | . | . | . | chr9 | 63603357 | 63616669 | 6.44347 | 4.69808 | 4.19638 |
| chr19 | 29522009 | 29522608 | 874 | Promoter (<=1kb) | 19 | 29522282 | 29606829 | 84548 | 1 | ENSMUST00 | 0 | C030046E11I | ENSMUSG0C | false | chr19 | 29521423 | 29523832 | 32.8888 | 98.87808 | 95.33504 | . | . | . | . | . | . |
| chr7 | 15980253 | 15980592 | 874 | Intron (ENSMUST00000094821.3 | 7 | 15946977 | 15967567 | 20591 | 2 | ENSMUST00 | -12687 | Ehd2 | ENSMUSG0C | false | . | . | . | . | . | . | chr7 | 15979822 | 15983665 | 6.91297 | 4.87866 | 4.61973 |
| chr6 | 140559885 | 140560181 | 874 | Intron (ENSMUST00000087622.3 | 6 | 140530963 | 140531215 | 253 | 1 | ENSMUST00 | 28923 | Gm25780 | ENSMUSG0C | false | . | . | . | . | . | . | chr6 | 140558264 | 140561876 | 5.36931 | 4.28476 | 3.32365 |
| chr11 | 21571851 | 21572287 | 873 | Promoter (<=1kb) | 11 | 21572235 | 21664385 | 92151 | 1 | ENSMUST00 | 0 | Wdpcp | ENSMUSG0C | false | chr11 | 21570978 | 21572906 | 26.1454 | 83.18384 | 79.97624 | . | . | . | . | . | . |
| chr2 | 73892351 | 73893151 | 873 | Promoter (<=1kb) | 2 | 73816509 | 73892536 | 76028 | 2 | ENSMUST00 | 0 | Atf2 | ENSMUSG0C | false | chr2 | 73891600 | 73892667 | 40.71709 | 132.32185 | 127.74851 | . | . | . | . | . | . |
| chr3 | 98040929 | 98041274 | 873 | Intron (ENSMUST00000079812.6 | 3 | 98013538 | 98150367 | 136830 | 1 | ENSMUST00 | 27392 | Notch2 | ENSMUSG0C | false | . | . | . | . | . | . | chr3 | 98014795 | 98071300 | 5.94878 | 4.48977 | 3.79636 |
| chr10 | 111849300 | 111849776 | 873 | Distal Intergenic | 10 | 111972664 | 111988432 | 15769 | 1 | ENSMUST00 | -122888 | Krr1 | ENSMUSG0C | false | . | . | . | . | . | . | chr10 | 111848667 | 111849873 | 3.23495 | 3.19465 | 1.56761 |
| chr13 | 28502160 | 28502444 | 873 | Intron (ENSMUST00000134787.2 | 13 | 28510744 | 28511232 | 489 | 2 | ENSMUST00 | 8788 | Gm6081 | ENSMUSG0C | false | . | . | . | . | . | . | chr13 | 28502033 | 28502910 | 5.6155 | 4.33078 | 3.57094 |
| chr8 | 121242708 | 121243121 | 873 | Intron (ENSMUST00000127664.1 | 8 | 121198714 | 121201835 | 3122 | 2 | ENSMUST00 | -40874 | 5033426O07 | ENSMUSG0C | false | . | . | . | . | . | . | chr8 | 121236124 | 121247254 | 7.60766 | 5.2125 | 5.14118 |
| chr7 | 25179440 | 25180010 | 872 | Distal Intergenic | 7 | 25120618 | 25159922 | 39305 | 2 | ENSMUST00 | -19519 | Pou2f2 | ENSMUSG0C | false | chr7 | 25178834 | 25180107 | 17.00269 | 41.02908 | 38.42754 | chr7 | 25178428 | 25180099 | 4.17263 | 3.62151 | 2.34355 |
| chr11 | 86357212 | 86357739 | 872 | Promoter (<=1kb) | 11 | 86267033 | 86357602 | 90570 | 2 | ENSMUST00 | 0 | Med13 | ENSMUSG0C | false | chr11 | 86356114 | 86358387 | 25.18546 | 107.13963 | 103.3826 | . | . | . | . | . | . |
| chr5 | 62765996 | 62766513 | 872 | Promoter (<=1kb) | 5 | 62602445 | 62766198 | 163754 | 2 | ENSMUST00 | 0 | Arap2 | ENSMUSG0C | false | chr5 | 62765401 | 62766855 | 24.78212 | 78.86476 | 75.73294 | chr5 | 62762863 | 62766747 | 4.455 | 3.80601 | 2.5777 |
| chr8 | 9977618 | 9978030 | 872 | Promoter (<=1kb) | 8 | 9977707 | 9992155 | 14449 | 1 | ENSMUST00 | 0 | Abhd13 | ENSMUSG0C | false | chr8 | 9976849 | 9978849 | 28.1615 | 93.53628 | 90.11485 | . | . | . | . | . | . |
| chr4 | 133688491 | 133688999 | 872 | Intron (ENSMUST00000105897.4 | 4 | 133668757 | 133672647 | 3891 | 2 | ENSMUST00 | -15845 | Pigv | ENSMUSG0C | false | . | . | . | . | . | . | chr4 | 133686744 | 133692777 | 5.78466 | 4.23388 | 3.6953 |
| chr16 | 97962021 | 97962743 | 872 | Promoter (<=1kb) | 16 | 97947435 | 97962621 | 15187 | 2 | ENSMUST00 | 0 | Zbtb21 | ENSMUSG0C | false | chr16 | 97961578 | 97963139 | 37.68163 | 117.40437 | 113.3432 | chr16 | 97958123 | 97964201 | 4.05813 | 3.63389 | 2.22469 |
| chr11 | 57671851 | 57672181 | 872 | Intron (ENSMUST00000066987.8 | 11 | 57655415 | 57657826 | 2412 | 2 | ENSMUST00 | -14026 | 4933426K07I | ENSMUSG0C | false | . | . | . | . | . | . | chr11 | 57646153 | 57704104 | 7.42247 | 5.08195 | 5.00645 |
| chr5 | 124852346 | 124852672 | 871 | Promoter (<=1kb) | 5 | 124836070 | 124851583 | 15514 | 2 | ENSMUST00 | -764 | Ccdc92 | ENSMUSG0C | true | . | . | . | . | . | . | chr5 | 124839552 | 124861868 | 6.02936 | 4.57753 | 3.85106 |
| chr2 | 173249397 | 173249652 | 871 | Intron (ENSMUST00000036248.7 | 2 | 173228211 | 173234722 | 6512 | 2 | ENSMUST00 | -14676 | Pmepa1 | ENSMUSG0C | true | . | . | . | . | . | . | chr2 | 173223515 | 173275069 | 7.01359 | 4.89215 | 4.68115 |
| chr9 | 116569685 | 116569990 | 871 | Downstream (2-3kb) | 9 | 116577582 | 116822806 | 245225 | 2 | ENSMUST00 | 252816 | Rbms3 | ENSMUSG0C | true | . | . | . | . | . | . | . | . | . | . | . | . |
| chr17 | 44601640 | 44601946 | 871 | Intron (ENSMUST00000162816.2 | 17 | 44641025 | 44641143 | 119 | 2 | ENSMUST00 | 39197 | n-R5s27 | ENSMUSG0C | false | . | . | . | . | . | . | chr17 | 44601844 | 44602227 | 4.02611 | 3.60731 | 2.25427 |
| chr8 | 125341611 | 125342004 | 871 | Distal Intergenic | 8 | 125224400 | 125227323 | 2924 | 2 | ENSMUST00 | -114289 | Gm16237 | ENSMUSG0C | false | . | . | . | . | . | . | chr8 | 125335216 | 125352727 | 6.82037 | 4.91647 | 4.50391 |
| chr17 | 83288075 | 83288521 | 871 | Promoter (<=1kb) | 17 | 83288546 | 83289028 | 483 | 1 | ENSMUST00 | -25 | Rpl21-ps7 | ENSMUSG0C | false | . | . | . | . | . | . | chr17 | 83287263 | 83289778 | 7.57114 | 5.04054 | 5.14387 |
| chr17 | 67996076 | 67996621 | 870 | Intron (ENSMUST00000024840.6 | 17 | 67842713 | 68004120 | 161408 | 2 | ENSMUST00 | 7499 | Arhgap28 | ENSMUSG0C | true | . | . | . | . | . | . | chr17 | 67993413 | 68002591 | 4.56742 | 3.83999 | 2.64221 |
| chr5 | 24425016 | 24425714 | 870 | Promoter (<=1kb) | 5 | 24425232 | 24440950 | 15719 | 1 | ENSMUST00 | 0 | Slc4a2 | ENSMUSG0C | false | chr5 | 24423745 | 24426392 | 21.33535 | 56.153 | 53.36009 | chr5 | 24422685 | 24428786 | 4.83602 | 4.02408 | 2.865 |
| chr2 | 158687853 | 158688144 | 870 | Intron (ENSMUST00000129902.1 | 2 | 158696135 | 158766334 | 70200 | 1 | ENSMUST00 | -7991 | Ppp1r16b | ENSMUSG0C | false | . | . | . | . | . | . | chr2 | 158686612 | 158689720 | 3.31936 | 2.99797 | 1.66875 |
| chr14 | 33696864 | 33697242 | 870 | Distal Intergenic | 14 | 33644495 | 33644601 | 107 | 1 | ENSMUST00 | 52370 | Gm26228 | ENSMUSG0C | false | . | . | . | . | . | . | . | . | . | . | . | . |
| chr10 | 20046098 | 20046407 | 870 | Exon (ENSMUST00000095806.4/ | 10 | 20099525 | 20105437 | 5913 | 1 | ENSMUST00 | -53118 | Map3k5 | ENSMUSG0C | true | . | . | . | . | . | . | chr10 | 20043287 | 20047517 | 4.63903 | 3.83289 | 2.71574 |
| chr8 | 3609101 | 3609378 | 870 | Exon (ENSMUST00000163038.2/ | 8 | 3615529 | 3615855 | 327 | 1 | ENSMUST00 | -6151 | Gm22204 | ENSMUSG0C | false | . | . | . | . | . | . | chr8 | 3608050 | 3610310 | 5.43054 | 4.19336 | 3.40164 |
| chr7 | 99574118 | 99574469 | 869 | Promoter (1-2kb) | 7 | 99573064 | 99588128 | 15065 | 1 | ENSMUST00 | 1055 | Arrb1 | ENSMUSG0C | false | . | . | . | . | . | . | chr7 | 99536482 | 99576026 | 7.32182 | 4.95837 | 4.9327 |
| chr12 | 3856109 | 3856456 | 869 | Intron (ENSMUST00000085826.7 | 12 | 3861021 | 3866162 | 5142 | 1 | ENSMUST00 | -4565 | Dnmt3a | ENSMUSG0C | false | . | . | . | . | . | . | chr12 | 3854212 | 3862098 | 6.77196 | 4.56698 | 4.49547 |
| chr12 | 86421576 | 86421910 | 869 | Promoter (<=1kb) | 12 | 86421644 | 86515522 | 93879 | 1 | ENSMUST00 | 0 | Esrrb | ENSMUSG0C | false | chr12 | 86421349 | 86422647 | 22.05754 | 58.21236 | 55.38818 | chr12 | 86421060 | 86423104 | 5.89799 | 4.43512 | 3.75071 |
| chr4 | 139226503 | 139227202 | 869 | Promoter (<=1kb) | 4 | 139226961 | 139287089 | 60129 | 1 | ENSMUST00 | 0 | Capzb | ENSMUSG0C | false | . | . | . | . | . | . | chr4 | 139223253 | 139239299 | 7.15907 | 4.94882 | 4.80839 |
| chr11 | 101165373 | 101165662 | 869 | Promoter (<=1kb) | 11 | 101162679 | 101165620 | 2942 | 2 | ENSMUST00 | 0 | Plekhh3 | ENSMUSG0C | false | . | . | . | . | . | . | chr11 | 101162175 | 101170220 | 6.22712 | 4.64855 | 4.01342 |
| chr7 | 35753255 | 35753564 | 869 | Promoter (<=1kb) | 7 | 35751319 | 35754355 | 3037 | 2 | ENSMUST00 | 791 | Dpy19l3 | ENSMUSG0C | false | . | . | . | . | . | . | chr7 | 35752993 | 35755046 | 4.29668 | 3.78596 | 2.41652 |
| chrX | 143033148 | 143033425 | 869 | Distal Intergenic | 20 | 143099594 | 143104297 | 4704 | 1 | ENSMUST00 | -66169 | Rgag1 | ENSMUSG0C | false | . | . | . | . | . | . | . | . | . | . | . | . |
| chr11 | 116816726 | 116817183 | 868 | Intron (ENSMUST00000021170.3 | 11 | 116807913 | 116828003 | 20091 | 2 | ENSMUST00 | 10820 | Mxra7 | ENSMUSG0C | false | . | . | . | . | . | . | chr11 | 116810954 | 116829199 | 5.76656 | 4.27097 | 3.65606 |
| chr14 | 66852358 | 66852634 | 868 | Intron (ENSMUST00000022629.8 | 14 | 66802864 | 66868688 | 65825 | 2 | ENSMUST00 | 16054 | Dpysl2 | ENSMUSG0C | true | chr14 | 66851840 | 66852393 | 6.49449 | 11.2705 | 9.07282 | chr14 | 66833125 | 66867182 | 5.87722 | 4.39726 | 3.73623 |
| chr15 | 12718758 | 12719091 | 868 | Intron (ENSMUST00000189469.1 | 15 | 12443331 | 12739924 | 296594 | 2 | ENSMUST00 | 20833 | Pdzd2 | ENSMUSG0C | false | chr15 | 12718736 | 12720598 | 11.22292 | 28.25542 | 25.81138 | . | . | . | . | . | . |
| chr17 | 27131226 | 27131513 | 868 | Promoter (2-3kb) | 17 | 27122668 | 27133623 | 10956 | 2 | ENSMUST00 | 2110 | Uqcc2 | ENSMUSG0C | false | . | . | . | . | . | . | chr17 | 27127363 | 27147146 | 5.99202 | 4.48057 | 3.83593 |
| chr10 | 4128990 | 4129253 | 868 | Intron (ENSMUST00000117291.2 | 10 | 4104313 | 4104469 | 157 | 1 | ENSMUST00 | 24678 | Gm25515 | ENSMUSG0C | false | . | . | . | . | . | . | . | . | . | . | . | . |
| chr2 | 108622822 | 108623162 | 868 | Distal Intergenic | 2 | 108578130 | 108578389 | 260 | 2 | ENSMUST00 | -44434 | Gm13913 | ENSMUSG0C | false | . | . | . | . | . | . | chr2 | 108622504 | 108624040 | 3.43545 | 3.27365 | 1.72721 |
| chr2 | 137116634 | 137117613 | 867 | Promoter (<=1kb) | 2 | 137081456 | 137116644 | 35189 | 2 | ENSMUST00 | 0 | Jag1 | ENSMUSG0C | false | chr2 | 137114213 | 137117889 | 37.86728 | 126.26704 | 121.91066 | . | . | . | . | . | . |
| chr6 | 43642443 | 43642764 | 867 | Intron (ENSMUST00000067888.8 | 6 | 43593043 | 43665881 | 72839 | 2 | ENSMUST00 | 23117 | Tpk1 | ENSMUSG0C | false | . | . | . | . | . | . | chr6 | 43642512 | 43642750 | 3.46636 | 3.32518 | 1.78398 |
| chr15 | 57745267 | 57745924 | 867 | Intron (ENSMUST00000096430.5 | 15 | 57736563 | 57738666 | 2104 | 2 | ENSMUST00 | -6602 | 9330154K18I | ENSMUSG0C | false | . | . | . | . | . | . | chr15 | 57742105 | 57754503 | 5.68542 | 4.44196 | 3.57487 |
| chr12 | 40392917 | 40393248 | 867 | Intron (ENSMUST00000069637.8 | 12 | 40315558 | 40445769 | 130212 | 2 | ENSMUST00 | 52521 | Zfp277 | ENSMUSG0C | false | . | . | . | . | . | . | chr12 | 40378830 | 40393499 | 4.81877 | 3.95373 | 2.86926 |
| chr10 | 69691551 | 69691839 | 867 | Intron (ENSMUST00000182379.2 | 10 | 69706296 | 69710476 | 4181 | 1 | ENSMUST00 | -14457 | Ank3 | ENSMUSG0C | false | . | . | . | . | . | . | chr10 | 69689738 | 69691611 | 3.99759 | 3.56803 | 2.1986 |
| chr9 | 32713828 | 32714132 | 867 | Promoter (2-3kb) | 9 | 32709652 | 32711171 | 1520 | 2 | ENSMUST00 | -2658 | Gm27240 | ENSMUSG0C | false | . | . | . | . | . | . | chr9 | 32697729 | 32732744 | 6.76714 | 4.77458 | 4.47728 |
| chr12 | 55450347 | 55450740 | 866 | Distal Intergenic | 12 | 55489411 | 55492647 | 3237 | 2 | ENSMUST00 | 41907 | Nfkbia | ENSMUSG0C | false | . | . | . | . | . | . | chr12 | 55437151 | 55453280 | 7.38007 | 5.14554 | 4.94833 |
| chr1 | 189918871 | 189919124 | 866 | Intron (ENSMUST00000027897.7 | 1 | 189885046 | 189922253 | 37208 | 2 | ENSMUST00 | 3129 | Smyd2 | ENSMUSG0C | false | . | . | . | . | . | . | chr1 | 189917900 | 189919705 | 3.98069 | 3.54058 | 2.19753 |
| chr17 | 35865598 | 35866855 | 866 | Promoter (<=1kb) | 17 | 35865625 | 35875492 | 9868 | 1 | ENSMUST00 | 0 | Ppp1r18 | ENSMUSG0C | false | chr17 | 35865184 | 35867409 | 19.72513 | 62.50151 | 59.62064 | chr17 | 35864641 | 35867277 | 3.9033 | 3.52343 | 2.11836 |
| chr2 | 129800411 | 129801625 | 866 | Promoter (<=1kb) | 2 | 129800517 | 129828974 | 28458 | 1 | ENSMUST00 | 0 | Stk35 | ENSMUSG0C | false | chr2 | 129800009 | 129802353 | 35.2856 | 127.03558 | 122.65217 | . | . | . | . | . | . |
| chr2 | 160451289 | 160451877 | 866 | Distal Intergenic | 2 | 160363703 | 160367065 | 3363 | 2 | ENSMUST00 | -84225 | Mafb | ENSMUSG0C | true | chr2 | 160451624 | 160452360 | 11.72109 | 25.47406 | 23.06522 | chr2 | 160450283 | 160453067 | 6.2053 | 4.65504 | 3.99045 |
| chr12 | 70692946 | 70693272 | 866 | Distal Intergenic | 12 | 70673646 | 70673746 | 101 | 1 | ENSMUST00 | 19301 | Gm24474 | ENSMUSG0C | false | . | . | . | . | . | . | chr12 | 70691961 | 70693791 | 3.86105 | 3.54319 | 2.07133 |
| chr9 | 80269627 | 80269968 | 866 | Exon (ENSMUST00000035889.9/ | 9 | 80263306 | 80265450 | 2145 | 2 | ENSMUST00 | -4178 | Gm27227 | ENSMUSG0C | false | . | . | . | . | . | . | chr9 | 80269012 | 80270684 | 3.64716 | 3.30023 | 1.92702 |
| chr9 | 114781743 | 114782312 | 865 | Promoter (<=1kb) | 9 | 114756837 | 114781993 | 25157 | 2 | ENSMUST00 | 0 | Cmtm7 | ENSMUSG0C | true | chr9 | 114779700 | 114782404 | 31.82494 | 97.04205 | 93.54184 | chr9 | 114782213 | 114786923 | 4.97492 | 4.01143 | 2.99981 |
| chr11 | 74896471 | 74897280 | 865 | Promoter (<=1kb) | 11 | 74849261 | 74897060 | 47800 | 2 | ENSMUST00 | 0 | Sgsm2 | ENSMUSG0C | false | chr11 | 74896187 | 74898875 | 40.58316 | 133.18515 | 128.57994 | . | . | . | . | . | . |
| chr18 | 36197268 | 36197525 | 865 | Promoter (<=1kb) | 18 | 36017707 | 36197380 | 179674 | 2 | ENSMUST00 | 0 | Nrg2 | ENSMUSG0C | false | chr18 | 36195978 | 36197656 | 12.58327 | 33.30664 | 30.79935 | chr18 | 36195954 | 36198260 | 4.7657 | 3.93779 | 2.84218 |
| chr11 | 106272765 | 106273302 | 865 | Promoter (<=1kb) | 11 | 106263179 | 106272972 | 9794 | 2 | ENSMUST00 | 0 | Smarcd2 | ENSMUSG0C | false | chr11 | 106271426 | 106273693 | 40.4056 | 135.07462 | 130.39536 | chr11 | 106267771 | 106274108 | 5.77787 | 4.43003 | 3.64069 |
| chr10 | 5522905 | 5523327 | 865 | Intron (ENSMUST00000041639.5 | 10 | 5336949 | 5550692 | 213744 | 2 | ENSMUST00 | 27365 | Syne1 | ENSMUSG0C | true | . | . | . | . | . | . | . | . | . | . | . | . |
| chr6 | 92940197 | 92940859 | 865 | Promoter (<=1kb) | 6 | 92940369 | 93272889 | 332521 | 1 | ENSMUST00 | 0 | 9530026P05I | ENSMUSG0C | true | chr6 | 92939881 | 92944204 | 25.90858 | 83.69981 | 80.48042 | . | . | . | . | . | . |
| chr7 | 44748440 | 44748962 | 864 | Promoter (<=1kb) | 7 | 44748485 | 44758105 | 9621 | 1 | ENSMUST00 | 0 | Vrk3 | ENSMUSG0C | false | chr7 | 44748081 | 44749367 | 21.57142 | 58.21323 | 55.38894 | chr7 | 44747714 | 44750298 | 4.15366 | 3.60255 | 2.35267 |
| chr2 | 28192850 | 28193278 | 864 | Promoter (<=1kb) | 2 | 28192992 | 28214425 | 21434 | 1 | ENSMUST00 | 0 | Olfm1 | ENSMUSG0C | false | chr2 | 28192692 | 28193602 | 22.74714 | 65.36804 | 62.44672 | chr2 | 28192710 | 28194006 | 3.25079 | 3.1194 | 1.62189 |
| chr2 | 128943563 | 128944201 | 864 | Promoter (<=1kb) | 2 | 128926268 | 128944077 | 17810 | 2 | ENSMUST00 | 0 | Zc3h8 | ENSMUSG0C | false | chr2 | 128943333 | 128944466 | 18.64736 | 62.11404 | 59.23804 | chr2 | 128943184 | 128943656 | 3.38436 | 3.27524 | 1.66001 |
| chr7 | 80824385 | 80824824 | 864 | Distal Intergenic | 7 | 80711583 | 80803331 | 91749 | 2 | ENSMUST00 | -21055 | Iqgap1 | ENSMUSG0C | false | . | . | . | . | . | . | chr7 | 80823293 | 80826211 | 3.69262 | 3.45449 | 1.98198 |
| chr2 | 34799477 | 34799971 | 864 | Promoter (<=1kb) | 2 | 34777556 | 34799912 | 22357 | 2 | ENSMUST00 | 0 | Rabepk | ENSMUSG0C | false | chr2 | 34799175 | 34800304 | 30.13579 | 99.06311 | 95.51676 | chr2 | 34798058 | 34802092 | 4.67717 | 3.83642 | 2.74909 |
| chr6 | 13285532 | 13285811 | 864 | Distal Intergenic | 6 | 13185613 | 13224965 | 39353 | 2 | ENSMUST00 | -60568 | Vwde | ENSMUSG0C | false | . | . | . | . | . | . | chr6 | 13285195 | 13286356 | 4.12872 | 3.7049 | 2.27585 |
| chr2 | 83644276 | 83645042 | 863 | Promoter (<=1kb) | 2 | 83644435 | 83664622 | 20188 | 1 | ENSMUST00 | 0 | Zc3h15 | ENSMUSG0C | false | chr2 | 83643993 | 83645302 | 31.38972 | 125.99339 | 121.64841 | . | . | . | . | . | . |
| chr9 | 71759908 | 71760278 | 863 | Intron (ENSMUST00000072899.3 | 9 | 71629211 | 71771560 | 142350 | 2 | ENSMUST00 | 11282 | Cgnl1 | ENSMUSG0C | false | . | . | . | . | . | . | chr9 | 71736274 | 71770833 | 5.2398 | 4.14793 | 3.22386 |
| chr8 | 84639208 | 84639941 | 863 | Promoter (<=1kb) | 8 | 84638991 | 84639434 | 444 | 1 | ENSMUST00 | 218 | Cacna1a | ENSMUSG0C | false | chr8 | 84638530 | 84639416 | 15.78968 | 37.54704 | 34.98843 | . | . | . | . | . | . |
| chr15 | 84436364 | 84436635 | 863 | Intron (ENSMUST00000080751.3 | 15 | 84417253 | 84443697 | 26445 | 2 | ENSMUST00 | 7062 | 1810041L15F | ENSMUSG0C | false | . | . | . | . | . | . | chr15 | 84436267 | 84438393 | 3.51703 | 3.36883 | 1.7857 |
| chr3 | 145575951 | 145576417 | 863 | Promoter (<=1kb) | 3 | 145576208 | 145605245 | 29038 | 1 | ENSMUST00 | 0 | Znhit6 | ENSMUSG0C | false | chr3 | 145575887 | 145577178 | 32.87838 | 107.99121 | 104.21945 | chr3 | 145575929 | 145578333 | 5.25448 | 4.18658 | 3.21095 |
| chr2 | 160413133 | 160413387 | 863 | Distal Intergenic | 2 | 160363703 | 160367065 | 3363 | 2 | ENSMUST00 | -46069 | Mafb | ENSMUSG0C | true | . | . | . | . | . | . | chr2 | 160412969 | 160413633 | 4.03355 | 3.49497 | 2.24079 |
| chr18 | 9818522 | 9818881 | 863 | Intron (ENSMUST00000040069.8 | 18 | 9827026 | 9827119 | 94 | 2 | ENSMUST00 | 8238 | Gm23637 | ENSMUSG0C | false | . | . | . | . | . | . | chr18 | 9813990 | 9821255 | 5.03684 | 4.14142 | 3.02798 |
| chr10 | 79858381 | 79858750 | 862 | Promoter (<=1kb) | 10 | 79858772 | 79859387 | 616 | 1 | ENSMUST00 | -22 | Ptbp1 | ENSMUSG0C | false | chr10 | 79854526 | 79858632 | 33.99707 | 114.12061 | 110.16386 | chr10 | 79855944 | 79867013 | 7.93167 | 5.01038 | 5.43402 |
| chr2 | 129198159 | 129198728 | 862 | Promoter (<=1kb) | 2 | 129198764 | 129211616 | 12853 | 1 | ENSMUST00 | -36 | Slc20a1 | ENSMUSG0C | false | chr2 | 129197808 | 129199982 | 29.28796 | 98.53321 | 94.99628 | chr2 | 129196987 | 129198166 | 3.4402 | 3.34197 | 1.70862 |
| chr5 | 35914830 | 35915208 | 862 | Promoter (1-2kb) | 5 | 35913082 | 35996024 | 82943 | 1 | ENSMUST00 | 1749 | Afap1 | ENSMUSG0C | true | . | . | . | . | . | . | chr5 | 35902761 | 35945832 | 6.32118 | 4.58806 | 4.1083 |
| chr3 | 153944129 | 153944801 | 862 | Promoter (<=1kb) | 3 | 153922357 | 153944632 | 22276 | 2 | ENSMUST00 | 0 | Acadm | ENSMUSG0C | false | chr3 | 153943430 | 153945325 | 35.00596 | 114.85688 | 110.88117 | chr3 | 153942037 | 153944263 | 4.48241 | 3.71871 | 2.58572 |
| chr14 | 32900750 | 32901017 | 862 | Intron (ENSMUST00000053175.6 | 14 | 32897719 | 32898729 | 1011 | 1 | ENSMUST00 | 3032 | Gm7734 | ENSMUSG0C | false | . | . | . | . | . | . | chr14 | 32900027 | 32901397 | 4.62549 | 3.92944 | 2.69901 |
| chr5 | 131584167 | 131584717 | 862 | Intron (ENSMUST00000161226.3 | 5 | 131577987 | 131578109 | 123 | 1 | ENSMUST00 | 6181 | Gm27266 | ENSMUSG0C | false | . | . | . | . | . | . | chr5 | 131583629 | 131584983 | 3.43756 | 3.1443 | 1.75871 |
| chr7 | 118539500 | 118539837 | 861 | Intron (ENSMUST00000044195.4 | 7 | 118525062 | 118533356 | 8295 | 2 | ENSMUST00 | -6145 | Coq7 | ENSMUSG0C | false | . | . | . | . | . | . | chr7 | 118536275 | 118546239 | 6.06229 | 4.47014 | 3.88658 |
| chr19 | 25236845 | 25237621 | 861 | Promoter (<=1kb) | 19 | 25236975 | 25434496 | 197522 | 1 | ENSMUST00 | 0 | Kank1 | ENSMUSG0C | true | chr19 | 25236314 | 25239245 | 23.12601 | 77.64799 | 74.53377 | chr19 | 25236144 | 25237180 | 5.29486 | 4.13031 | 3.24295 |
| chr7 | 19822603 | 19822989 | 861 | Promoter (<=1kb) | 7 | 19808462 | 19822770 | 14309 | 2 | ENSMUST00 | 0 | Bcl3 | ENSMUSG0C | false | chr7 | 19822220 | 19823046 | 21.59801 | 56.60135 | 53.80005 | chr7 | 19821732 | 19825440 | 4.97308 | 4.10705 | 2.98414 |
| chr2 | 156732124 | 156732566 | 861 | Intron (ENSMUST00000109567.4 | 2 | 156721492 | 156764363 | 42872 | 1 | ENSMUST00 | 10633 | Dlgap4 | ENSMUSG0C | true | . | . | . | . | . | . | chr2 | 156722290 | 156745258 | 7.00087 | 4.91477 | 4.66073 |
| chr17 | 26948341 | 26948975 | 861 | Promoter (<=1kb) | 17 | 26948056 | 26948119 | 64 | 2 | ENSMUST00 | -223 | Mir3083 | ENSMUSG0C | false | chr17 | 26948318 | 26949056 | 26.65286 | 74.78688 | 71.72144 | chr17 | 26946231 | 26949664 | 3.48984 | 3.31799 | 1.75783 |
| chr14 | 67888177 | 67888500 | 861 | Intron (ENSMUST00000039135.4 | 14 | 67859636 | 67859745 | 110 | 2 | ENSMUST00 | -28433 | Mir6539 | ENSMUSG0C | false | . | . | . | . | . | . | chr14 | 67862320 | 67900048 | 6.11049 | 4.53842 | 3.92537 |
| chr1 | 69429672 | 69429962 | 861 | Distal Intergenic | 1 | 69441056 | 69444084 | 3029 | 1 | ENSMUST00 | -11094 | RP24-280N1 | ENSMUSG0C | false | . | . | . | . | . | . | chr1 | 69429254 | 69431415 | 4.24482 | 3.69717 | 2.41444 |
| chr7 | 28179321 | 28180307 | 860 | Promoter (<=1kb) | 7 | 28179469 | 28187294 | 7826 | 1 | ENSMUST00 | 0 | Dyrk1b | ENSMUSG0C | false | chr7 | 28179405 | 28180796 | 42.42999 | 160.09743 | 154.44568 | . | . | . | . | . | . |
| chr17 | 78218602 | 78218942 | 860 | Intron (ENSMUST00000112498.2 | 17 | 78200248 | 78376592 | 176345 | 1 | ENSMUST00 | 18355 | Crim1 | ENSMUSG0C | true | . | . | . | . | . | . | chr17 | 78202641 | 78231492 | 6.38021 | 4.71597 | 4.13646 |
| chr14 | 57423878 | 57424356 | 860 | Promoter (<=1kb) | 14 | 57424062 | 57517936 | 93875 | 1 | ENSMUST00 | 0 | Ift88 | ENSMUSG0C | false | chr14 | 57423453 | 57424648 | 34.36702 | 111.64683 | 107.76208 | . | . | . | . | . | . |
| chr1 | 64956405 | 64956955 | 860 | Promoter (<=1kb) | 1 | 64785983 | 64956824 | 170842 | 2 | ENSMUST00 | 0 | Plekhm3 | ENSMUSG0C | false | chr1 | 64956081 | 64957136 | 40.83545 | 136.9297 | 132.17467 | chr1 | 64955360 | 64957160 | 3.32067 | 3.22486 | 1.65833 |
| chr1 | 156303386 | 156303853 | 860 | Promoter (<=1kb) | 1 | 156255296 | 156303664 | 48369 | 2 | ENSMUST00 | 0 | Tdrd5 | ENSMUSG0C | true | chr1 | 156299255 | 156304171 | 39.05507 | 126.17973 | 121.82412 | . | . | . | . | . | . |
| chr13 | 42821696 | 42822130 | 860 | Intron (ENSMUST00000128646.2 | 13 | 42782692 | 42788258 | 5567 | 2 | ENSMUST00 | -33439 | Gm15810 | ENSMUSG0C | false | . | . | . | . | . | . | chr13 | 42820630 | 42822930 | 4.80577 | 3.83875 | 2.89523 |
| chr11 | 59208164 | 59208509 | 859 | Promoter (<=1kb) | 11 | 59208321 | 59210738 | 2418 | 1 | ENSMUST00 | 0 | 2310033P09I | ENSMUSG0C | false | chr11 | 59208056 | 59209347 | 35.67729 | 114.98298 | 111.00362 | chr11 | 59207293 | 59210505 | 3.83131 | 3.46737 | 2.08452 |
| chr7 | 131410413 | 131410806 | 859 | Promoter (<=1kb) | 7 | 131410601 | 131446211 | 35611 | 1 | ENSMUST00 | 0 | Acadsb | ENSMUSG0C | false | chr7 | 131409343 | 131411661 | 44.84477 | 157.51604 | 151.94687 | . | . | . | . | . | . |
| chr17 | 35126033 | 35126625 | 859 | Promoter (<=1kb) | 17 | 35126373 | 35126910 | 538 | 1 | ENSMUST00 | 0 | D17H6S53E | ENSMUSG0C | false | chr17 | 35125969 | 35127369 | 30.60524 | 96.90557 | 93.40668 | chr17 | 35125535 | 35130188 | 4.16017 | 3.64157 | 2.31381 |
| chr13 | 107493844 | 107494191 | 859 | Distal Intergenic | 13 | 107413865 | 107414767 | 903 | 2 | ENSMUST00 | -79078 | Apoo-ps | ENSMUSG0C | false | . | . | . | . | . | . | . | . | . | . | . | . |
| chr11 | 102604105 | 102604810 | 859 | Promoter (<=1kb) | 11 | 102604396 | 102608058 | 3663 | 1 | ENSMUST00 | 0 | Fzd2 | ENSMUSG0C | false | chr11 | 102603096 | 102607889 | 33.76198 | 105.43819 | 101.727 | . | . | . | . | . | . |
| chr8 | 126805449 | 126806107 | 859 | Distal Intergenic | 8 | 126944847 | 126944980 | 134 | 2 | ENSMUST00 | 138873 | Gm26397 | ENSMUSG0C | false | . | . | . | . | . | . | chr8 | 126804160 | 126806556 | 6.56769 | 4.74674 | 4.33027 |
| chr8 | 107293107 | 107294618 | 858 | Promoter (<=1kb) | 8 | 107293470 | 107329307 | 35838 | 1 | ENSMUST00 | 0 | Nfat5 | ENSMUSG0C | false | chr8 | 107292711 | 107294946 | 27.18312 | 111.36749 | 107.49936 | . | . | . | . | . | . |
| chr19 | 46499808 | 46500502 | 858 | Promoter (1-2kb) | 19 | 46501648 | 46516444 | 14797 | 1 | ENSMUST00 | -1146 | Trim8 | ENSMUSG0C | false | chr19 | 46499527 | 46499866 | 6.17403 | 14.35629 | 12.10532 | chr19 | 46499125 | 46499819 | 3.4208 | 3.29988 | 1.68435 |
| chr15 | 59168448 | 59168998 | 858 | Distal Intergenic | 15 | 59210908 | 59211720 | 813 | 1 | ENSMUST00 | -41910 | Gm5045 | ENSMUSG0C | false | . | . | . | . | . | . | chr15 | 59167250 | 59173334 | 6.774 | 4.95682 | 4.44708 |
| chr9 | 53383999 | 53384855 | 858 | Promoter (<=1kb) | 9 | 53384023 | 53401867 | 17845 | 1 | ENSMUST00 | 0 | Kdelc2 | ENSMUSG0C | false | chr9 | 53383794 | 53385192 | 31.91637 | 110.38313 | 106.53261 | chr9 | 53384825 | 53385471 | 3.32286 | 3.17987 | 1.6884 |
| chr9 | 119004020 | 119004312 | 858 | Intron (ENSMUST00000073109.6 | 9 | 119031796 | 119031885 | 90 | 1 | ENSMUST00 | -27484 | Mir26a-1 | ENSMUSG0C | false | . | . | . | . | . | . | chr9 | 118927678 | 119010843 | 6.89652 | 4.89961 | 4.57102 |
| chr11 | 87468359 | 87468635 | 858 | Promoter (<=1kb) | 11 | 87467679 | 87469058 | 1380 | 1 | ENSMUST00 | 681 | Gm11489 | ENSMUSG0C | false | . | . | . | . | . | . | chr11 | 87466867 | 87469225 | 5.89145 | 4.43041 | 3.76106 |
| chr16 | 29944017 | 29944305 | 858 | Promoter (2-3kb) | 16 | 29907519 | 29946517 | 38999 | 2 | ENSMUST00 | 2212 | Gm26569 | ENSMUSG0C | false | . | . | . | . | . | . | chr16 | 29943407 | 29947564 | 5.10658 | 4.06635 | 3.12869 |
| chr11 | 87595402 | 87595726 | 857 | Promoter (<=1kb) | 11 | 87595652 | 87601077 | 5426 | 1 | ENSMUST00 | 0 | Mtmr4 | ENSMUSG0C | false | chr11 | 87595219 | 87596051 | 19.02488 | 51.36755 | 48.63442 | chr11 | 87592675 | 87596916 | 6.42731 | 4.6946 | 4.16765 |
| chr15 | 25929772 | 25930253 | 857 | Intron (ENSMUST00000022881.8 | 15 | 25940887 | 25972340 | 31454 | 1 | ENSMUST00 | -10634 | Fam134b | ENSMUSG0C | false | . | . | . | . | . | . | chr15 | 25928258 | 25936828 | 5.85667 | 4.43756 | 3.72128 |
| chr10 | 128471939 | 128472404 | 857 | Intron (ENSMUST00000026433.7 | 10 | 128459236 | 128490173 | 30938 | 1 | ENSMUST00 | 12704 | Smarcc2 | ENSMUSG0C | false | . | . | . | . | . | . | chr10 | 128470761 | 128477757 | 5.27026 | 4.10534 | 3.24837 |
| chr15 | 59647625 | 59648871 | 857 | Promoter (<=1kb) | 15 | 59648350 | 59656550 | 8201 | 1 | ENSMUST00 | 0 | Trib1 | ENSMUSG0C | false | chr15 | 59647395 | 59650262 | 28.1615 | 93.53628 | 90.11485 | chr15 | 59648813 | 59650001 | 4.62384 | 3.86761 | 2.70454 |
| chr10 | 22569399 | 22569807 | 857 | Distal Intergenic | 10 | 22645011 | 22704285 | 59275 | 1 | ENSMUST00 | -75204 | Slc2a12 | ENSMUSG0C | false | . | . | . | . | . | . | chr10 | 22569038 | 22571109 | 3.70394 | 3.38462 | 1.99544 |

| | | | | | | | | | | | | | | | | | | | | | | | | | | |
|---|---|---|---|---|---|---|---|---|---|---|---|---|---|---|---|---|---|---|---|---|---|---|---|---|---|---|
| chrX | 142532761 | 142533019 | 857 | Promoter (<=1kb) | 20 | 142533439 | 142533870 | 432 | 2 | ENSMUST00 | 851 | Gm4995 | ENSMUSG0C | false | . | . | . | . | . | . | . | . | . | . | . | . |
| chr6 | 120782318 | 120782608 | 856 | Intron (ENSMUST00000112682.2 | 6 | 120773768 | 120793982 | 20215 | 1 | ENSMUST00 | 8551 | Slc25a18 | ENSMUSG0C | false | . | . | . | . | . | . | chr6 | 120781119 | 120784596 | 4.25416 | 3.57864 | 2.4295 |
| chr11 | 50376918 | 50377660 | 856 | Promoter (<=1kb) | 11 | 50376990 | 50386528 | 9539 | 1 | ENSMUST00 | 0 | Hnrnph1 | ENSMUSG0C | false | chr11 | 50376642 | 50379439 | 45.10679 | 157.97711 | 152.39525 | . | . | . | . | . | . |
| chr4 | 137301220 | 137301502 | 856 | Distal Intergenic | 4 | 137277489 | 137299726 | 22238 | 1 | ENSMUST00 | 23732 | Wnt4 | ENSMUSG0C | false | . | . | . | . | . | . | chr4 | 137299450 | 137313019 | 6.27354 | 4.63926 | 4.04895 |
| chr8 | 31187161 | 31187709 | 856 | Promoter (<=1kb) | 8 | 31187331 | 31261483 | 74153 | 1 | ENSMUST00 | 0 | Fut10 | ENSMUSG0C | false | chr8 | 31186857 | 31187993 | 22.5292 | 69.37365 | 66.39268 | chr8 | 31186665 | 31188656 | 4.36182 | 3.74306 | 2.48602 |
| chr4 | 106948699 | 106949044 | 856 | Intron (ENSMUST00000150915.2 | 4 | 106955859 | 107048692 | 92834 | 1 | ENSMUST00 | -6815 | Ssbp3 | ENSMUSG0C | false | . | . | . | . | . | . | chr4 | 106945202 | 106965608 | 5.89404 | 4.4035 | 3.75076 |
| chr4 | 148309514 | 148309826 | 856 | Promoter (2-3kb) | 4 | 148306853 | 148316569 | 9717 | 1 | ENSMUST00 | 2662 | Gm13206 | ENSMUSG0C | false | . | . | . | . | . | . | chr4 | 148308840 | 148310329 | 4.23597 | 3.74683 | 2.38261 |
| chr18 | 3669395 | 3669696 | 856 | Distal Intergenic | 18 | 3770741 | 3770804 | 64 | 2 | ENSMUST00 | 101108 | Gm23643 | ENSMUSG0C | false | . | . | . | . | . | . | chr18 | 3664015 | 3670955 | 4.12925 | 3.67983 | 2.28088 |
| chr2 | 31741716 | 31742141 | 855 | Intron (ENSMUST00000142554.2 | 2 | 31759943 | 31804227 | 44285 | 1 | ENSMUST00 | -17802 | Abl1 | ENSMUSG0C | false | . | . | . | . | . | . | chr2 | 31740620 | 31752695 | 6.24045 | 4.61237 | 4.04053 |
| chr2 | 163658245 | 163658702 | 855 | Promoter (<=1kb) | 2 | 163658386 | 163723758 | 65373 | 1 | ENSMUST00 | 0 | Pkig | ENSMUSG0C | false | chr2 | 163658124 | 163659535 | 30.32912 | 88.589 | 85.27629 | chr2 | 163658175 | 163666027 | 5.63736 | 4.28032 | 3.5511 |
| chr10 | 22631609 | 22631871 | 855 | Distal Intergenic | 10 | 22645011 | 22704285 | 59275 | 1 | ENSMUST00 | -13140 | Slc2a12 | ENSMUSG0C | false | . | . | . | . | . | . | chr10 | 22631143 | 22634232 | 3.69144 | 3.35541 | 1.94967 |
| chr19 | 37611913 | 37612245 | 855 | Intron (ENSMUST00000066439.6 | 19 | 37550418 | 37683245 | 132828 | 1 | ENSMUST00 | 61496 | Exoc6 | ENSMUSG0C | false | . | . | . | . | . | . | . | . | . | . | . | . |
| chr14 | 98067387 | 98067667 | 855 | Intron (ENSMUST00000069334.7 | 14 | 98118509 | 98119006 | 498 | 1 | ENSMUST00 | -50842 | Gm16331 | ENSMUSG0C | false | . | . | . | . | . | . | chr14 | 98058294 | 98100007 | 4.23349 | 3.68916 | 2.37626 |
| chr16 | 50822904 | 50823229 | 855 | Distal Intergenic | 16 | 50783478 | 50784065 | 588 | 1 | ENSMUST00 | 39427 | 1700116B05 | ENSMUSG0C | false | . | . | . | . | . | . | chr16 | 50822214 | 50823675 | 4.58345 | 3.89328 | 2.66411 |
| chr11 | 105306566 | 105306872 | 854 | Intron (ENSMUST00000126931.1 | 11 | 105292668 | 105343871 | 51204 | 1 | ENSMUST00 | 13899 | Mrc2 | ENSMUSG0C | true | . | . | . | . | . | . | chr11 | 105306066 | 105319029 | 6.36006 | 4.57632 | 4.15147 |
| chrX | 53370381 | 53370676 | 854 | Promoter (<=1kb) | 20 | 53344598 | 53370502 | 25905 | 2 | ENSMUST00 | 0 | Mospd1 | ENSMUSG0C | false | chrX | 53369514 | 53370565 | 20.21941 | 51.82391 | 49.08421 | . | . | . | . | . | . |
| chr17 | 35979027 | 35980280 | 854 | Promoter (<=1kb) | 17 | 35979851 | 35989462 | 9612 | 1 | ENSMUST00 | 0 | Gnl1 | ENSMUSG0C | false | chr17 | 35978627 | 35981131 | 36.53154 | 123.91184 | 119.64436 | chr17 | 35978591 | 35979041 | 2.78737 | 2.79749 | 1.24041 |
| chr19 | 7172409 | 7172742 | 854 | Intron (ENSMUST00000040261.5 | 19 | 7199540 | 7200464 | 925 | 2 | ENSMUST00 | 27722 | Otub1 | ENSMUSG0C | false | . | . | . | . | . | . | chr19 | 7171429 | 7172890 | 6.23211 | 4.59986 | 4.0702 |
| chr3 | 90417839 | 90418661 | 854 | Intron (ENSMUST00000029542.6 | 3 | 90391380 | 90433636 | 42257 | 2 | ENSMUST00 | 14975 | Ints3 | ENSMUSG0C | false | . | . | . | . | . | . | chr3 | 90415238 | 90423570 | 5.44844 | 4.23884 | 3.38878 |
| chr4 | 6365487 | 6366141 | 854 | Promoter (<=1kb) | 4 | 6365650 | 6396284 | 30635 | 1 | ENSMUST00 | 0 | Sdcbp | ENSMUSG0C | false | chr4 | 6365096 | 6367078 | 35.20524 | 115.20684 | 111.21615 | . | . | . | . | . | . |
| chr12 | 17575894 | 17576429 | 854 | Distal Intergenic | 12 | 17546146 | 17546281 | 136 | 1 | ENSMUST00 | 29749 | Gm22748 | ENSMUSG0C | false | . | . | . | . | . | . | chr12 | 17575224 | 17576833 | 5.88392 | 4.45131 | 3.73201 |
| chr8 | 71974696 | 71975029 | 853 | Intron (ENSMUST00000136878.1 | 8 | 71988483 | 71992915 | 4433 | 2 | ENSMUST00 | 17886 | Cyp4f18 | ENSMUSG0C | false | . | . | . | . | . | . | chr8 | 71974363 | 71975256 | 3.28646 | 3.09536 | 1.65447 |
| chr10 | 13982756 | 13983397 | 853 | Intron (ENSMUST00000187083.1 | 10 | 13967024 | 14079675 | 112652 | 1 | ENSMUST00 | 15733 | Hivep2 | ENSMUSG0C | true | chr10 | 13982367 | 13983535 | 31.63878 | 95.72812 | 92.25946 | chr10 | 13982224 | 13989776 | 4.57573 | 3.78824 | 2.68059 |
| chr14 | 73385139 | 73385583 | 853 | Promoter (<=1kb) | 14 | 73362231 | 73385271 | 23041 | 2 | ENSMUST00 | 0 | Itm2b | ENSMUSG0C | false | chr14 | 73379381 | 73385386 | 38.29964 | 140.16156 | 135.27679 | . | . | . | . | . | . |
| chr4 | 154299682 | 154300318 | 853 | Promoter (<=1kb) | 4 | 154278486 | 154299892 | 21407 | 2 | ENSMUST00 | 0 | Arhgef16 | ENSMUSG0C | true | chr4 | 154295799 | 154300705 | 31.54088 | 108.67455 | 104.87239 | chr4 | 154298019 | 154301857 | 4.67301 | 3.77891 | 2.75618 |
| chr6 | 92943169 | 92943812 | 853 | Promoter (<=1kb) | 6 | 92772699 | 92943492 | 170794 | 2 | ENSMUST00 | 0 | Adamts9 | ENSMUSG0C | false | chr6 | 92939881 | 92944204 | 25.90858 | 83.69981 | 80.48042 | . | . | . | . | . | . |
| chr4 | 83090731 | 83091005 | 853 | Distal Intergenic | 4 | 83078511 | 83078629 | 119 | 1 | ENSMUST00 | 12221 | Gm24884 | ENSMUSG0C | false | . | . | . | . | . | . | chr4 | 83090467 | 83091912 | 3.4684 | 3.2525 | 1.79089 |
| chr3 | 95674222 | 95674665 | 852 | Promoter (2-3kb) | 3 | 95676750 | 95677151 | 402 | 2 | ENSMUST00 | 2486 | Adamtsl4 | ENSMUSG0C | false | . | . | . | . | . | . | chr3 | 95660803 | 95677602 | 6.84699 | 4.91959 | 4.52508 |
| chr19 | 10905305 | 10905699 | 852 | 3' UTR | 19 | 10900282 | 10900993 | 712 | 1 | ENSMUST00 | 5024 | Prpf19 | ENSMUSG0C | false | . | . | . | . | . | . | chr19 | 10903260 | 10910149 | 5.52991 | 4.23684 | 3.45751 |
| chr8 | 127429918 | 127430194 | 852 | Intron (ENSMUST00000162309.2 | 8 | 127447746 | 127474513 | 26768 | 1 | ENSMUST00 | -17552 | Pard3 | ENSMUSG0C | false | . | . | . | . | . | . | chr8 | 127429493 | 127432987 | 3.40678 | 3.19526 | 1.74834 |
| chr3 | 68605118 | 68605358 | 852 | Intron (ENSMUST00000029346.7 | 3 | 68596175 | 68598163 | 1989 | 1 | ENSMUST00 | 8944 | Schip1 | ENSMUSG0C | false | . | . | . | . | . | . | chr3 | 68574597 | 68620372 | 5.61397 | 4.36647 | 3.51459 |
| chr6 | 8495890 | 8496179 | 852 | Promoter (1-2kb) | 6 | 8489933 | 8497979 | 8047 | 2 | ENSMUST00 | 1800 | A430035B10 | ENSMUSG0C | false | . | . | . | . | . | . | chr6 | 8495532 | 8497114 | 5.67118 | 4.34091 | 3.59804 |
| chr13 | 98545928 | 98546451 | 852 | Distal Intergenic | 13 | 98560354 | 98560461 | 108 | 1 | ENSMUST00 | -13903 | Gm22698 | ENSMUSG0C | false | . | . | . | . | . | . | chr13 | 98545309 | 98546901 | 6.31076 | 4.77253 | 4.06072 |
| chr17 | 5849397 | 5849838 | 851 | Intron (ENSMUST00000002436.9 | 17 | 5841328 | 5931033 | 89706 | 1 | ENSMUST00 | 8070 | Snx9 | ENSMUSG0C | false | chr17 | 5849210 | 5849793 | 9.38717 | 20.36066 | 18.02005 | chr17 | 5842208 | 5850409 | 5.53242 | 4.3591 | 3.4424 |
| chr11 | 88047243 | 88047784 | 851 | Promoter (<=1kb) | 11 | 88047373 | 88053755 | 6383 | 1 | ENSMUST00 | 0 | Srsf1 | ENSMUSG0C | false | chr11 | 88046872 | 88048999 | 40.65367 | 138.82761 | 133.98773 | . | . | . | . | . | . |
| chr1 | 82839296 | 82839772 | 851 | Promoter (<=1kb) | 1 | 82839483 | 82901182 | 61700 | 1 | ENSMUST00 | 0 | Agfg1 | ENSMUSG0C | false | chr1 | 82839079 | 82840598 | 25.79552 | 96.83904 | 93.34669 | . | . | . | . | . | . |
| chr3 | 129328293 | 129328637 | 851 | Intron (ENSMUST00000029658.8 | 3 | 129270277 | 129332264 | 61988 | 2 | ENSMUST00 | 3627 | Enpep | ENSMUSG0C | true | chr3 | 129327289 | 129333393 | 25.83068 | 80.74496 | 77.57947 | . | . | . | . | . | . |
| chr17 | 30046629 | 30046915 | 851 | Intron (ENSMUST00000057897.7 | 17 | 30005087 | 30210019 | 204933 | 1 | ENSMUST00 | 41543 | Zfand3 | ENSMUSG0C | false | . | . | . | . | . | . | chr17 | 30040207 | 30049893 | 4.6591 | 3.88122 | 2.74546 |
| chr6 | 71831162 | 71831493 | 851 | Promoter (<=1kb) | 6 | 71831331 | 71875244 | 43914 | 1 | ENSMUST00 | 0 | Immt | ENSMUSG0C | false | chr6 | 71830473 | 71832268 | 39.3713 | 146.56828 | 141.41737 | . | . | . | . | . | . |
| chr19 | 15552779 | 15553241 | 851 | Distal Intergenic | 19 | 15680004 | 15680110 | 107 | 1 | ENSMUST00 | -126763 | Gm24319 | ENSMUSG0C | false | . | . | . | . | . | . | chr19 | 15548063 | 15558324 | 3.96368 | 3.54363 | 2.16221 |
| chr19 | 46367855 | 46368164 | 850 | Promoter (1-2kb) | 19 | 46366734 | 46375252 | 8519 | 1 | ENSMUST00 | 1122 | Tmem180 | ENSMUSG0C | false | . | . | . | . | . | . | . | . | . | . | . | . |
| chr8 | 128358776 | 128359384 | 850 | Promoter (<=1kb) | 8 | 128359073 | 128505462 | 146390 | 1 | ENSMUST00 | 0 | Nrp1 | ENSMUSG0C | false | chr8 | 128356451 | 128361175 | 30.41442 | 103.5799 | 99.9162 | . | . | . | . | . | . |
| chr8 | 72134969 | 72135428 | 850 | Promoter (<=1kb) | 8 | 72134975 | 72144709 | 9735 | 1 | ENSMUST00 | 0 | Tpm4 | ENSMUSG0C | false | chr8 | 72134816 | 72136882 | 41.44235 | 135.77681 | 131.06697 | chr8 | 72132388 | 72135070 | 4.09517 | 3.67812 | 2.26508 |
| chr2 | 179931876 | 179932501 | 850 | Promoter (2-3kb) | 2 | 179922004 | 179929355 | 7352 | 2 | ENSMUST00 | -2522 | Taf4a | ENSMUSG0C | false | . | . | . | . | . | . | chr2 | 179931472 | 179933158 | 3.76485 | 3.31585 | 2.02547 |
| chr4 | 58859303 | 58859605 | 850 | Intron (ENSMUST00000102889.4 | 4 | 58836072 | 58844881 | 8810 | 2 | ENSMUST00 | -14423 | AI314180 | ENSMUSG0C | false | . | . | . | . | . | . | chr4 | 58858516 | 58860678 | 4.11733 | 3.58999 | 2.31807 |
| chr9 | 74791103 | 74791396 | 850 | Distal Intergenic | 9 | 74782070 | 74782212 | 143 | 1 | ENSMUST00 | 9034 | Gm22315 | ENSMUSG0C | false | . | . | . | . | . | . | chr9 | 74789994 | 74791764 | 3.68443 | 3.36483 | 1.96137 |
| chr11 | 101982886 | 101983227 | 849 | Promoter (<=1kb) | 11 | 101971143 | 101983503 | 12361 | 2 | ENSMUST00 | 276 | Dusp3 | ENSMUSG0C | false | chr11 | 101983072 | 101985233 | 23.0285 | 72.05518 | 69.03415 | chr11 | 101974130 | 101983868 | 4.93837 | 3.85129 | 2.98554 |
| chr14 | 16897888 | 16898464 | 849 | Distal Intergenic | 14 | 16430842 | 16575472 | 144631 | 2 | ENSMUST00 | -322417 | Rarb | ENSMUSG0C | false | . | . | . | . | . | . | chr14 | 16896981 | 16899586 | 5.77469 | 4.19969 | 3.67888 |
| chr17 | 45944980 | 45945295 | 849 | Distal Intergenic | 17 | 46018712 | 46021372 | 2661 | 2 | ENSMUST00 | 76077 | Vegfa | ENSMUSG0C | true | . | . | . | . | . | . | chr17 | 45941755 | 45945545 | 5.78054 | 4.32924 | 3.69609 |
| chr19 | 14539082 | 14539421 | 849 | Intron (ENSMUST00000052011.8 | 19 | 14448072 | 14597983 | 149912 | 2 | ENSMUST00 | 58562 | Tle4 | ENSMUSG0C | false | . | . | . | . | . | . | chr19 | 14537127 | 14542277 | 4.38008 | 3.79619 | 2.48568 |
| chr3 | 68573793 | 68574111 | 849 | Promoter (1-2kb) | 3 | 68572245 | 68626464 | 54220 | 1 | ENSMUST00 | 1549 | Schip1 | ENSMUSG0C | false | chr3 | 68572019 | 68576372 | 26.65286 | 74.78688 | 71.72144 | . | . | . | . | . | . |
| chr9 | 70507486 | 70507809 | 849 | Distal Intergenic | 9 | 70425429 | 70503725 | 78297 | 2 | ENSMUST00 | -3762 | Rnf111 | ENSMUSG0C | false | . | . | . | . | . | . | chr9 | 70506132 | 70507660 | 3.06115 | 3.04771 | 1.43386 |
| chr18 | 55663656 | 55664071 | 849 | Distal Intergenic | 18 | 55622636 | 55622920 | 285 | 2 | ENSMUST00 | -40737 | Gm24514 | ENSMUSG0C | false | . | . | . | . | . | . | chr18 | 55663027 | 55664745 | 5.70975 | 4.42549 | 3.57511 |
| chr19 | 53414021 | 53414258 | 848 | Distal Intergenic | 19 | 53379214 | 53390573 | 11360 | 2 | ENSMUST00 | -23449 | Smndc1 | ENSMUSG0C | false | . | . | . | . | . | . | chr19 | 53409099 | 53414773 | 6.07728 | 4.55041 | 3.90583 |
| chr5 | 120648586 | 120648910 | 848 | Promoter (<=1kb) | 5 | 120648812 | 120654902 | 6091 | 1 | ENSMUST00 | 0 | Rasal1 | ENSMUSG0C | false | chr5 | 120648563 | 120649584 | 7.71221 | 14.5447 | 12.28981 | chr5 | 120647200 | 120649809 | 4.97361 | 4.07792 | 2.98001 |
| chr5 | 146876025 | 146876712 | 848 | Distal Intergenic | 5 | 146845071 | 146847726 | 2656 | 1 | ENSMUST00 | 30955 | Rasl11a | ENSMUSG0C | true | . | . | . | . | . | . | chr5 | 146865620 | 146877684 | 5.33365 | 4.16572 | 3.30759 |
| chr9 | 65214685 | 65215656 | 848 | Promoter (<=1kb) | 9 | 65214690 | 65239219 | 24530 | 1 | ENSMUST00 | 0 | Parp16 | ENSMUSG0C | false | chr9 | 65214597 | 65216055 | 28.53699 | 95.19626 | 91.7378 | chr9 | 65214220 | 65219296 | 4.20834 | 3.57428 | 2.38968 |
| chr13 | 59704653 | 59705016 | 848 | Promoter (1-2kb) | 13 | 59700083 | 59706197 | 6115 | 2 | ENSMUST00 | 1181 | Spata31d1a | ENSMUSG0C | false | . | . | . | . | . | . | chr13 | 59704022 | 59705495 | 3.81698 | 3.51733 | 2.02585 |
| chr14 | 32819213 | 32819652 | 848 | Distal Intergenic | 14 | 32833962 | 32836789 | 2828 | 1 | ENSMUST00 | -14310 | Fam170b | ENSMUSG0C | false | . | . | . | . | . | . | chr14 | 32816542 | 32820544 | 4.24344 | 3.48955 | 2.42972 |
| chr1 | 120253673 | 120254083 | 847 | Intron (ENSMUST00000140490.2 | 1 | 120190757 | 120265255 | 74499 | 2 | ENSMUST00 | 11172 | Steap3 | ENSMUSG0C | false | . | . | . | . | . | . | chr1 | 120244209 | 120264315 | 5.23032 | 4.13237 | 3.21241 |
| chr2 | 13573686 | 13574340 | 847 | Promoter (<=1kb) | 2 | 13573927 | 13574738 | 812 | 1 | ENSMUST00 | 0 | Vim | ENSMUSG0C | true | chr2 | 13572823 | 13576831 | 31.81669 | 98.02396 | 94.50304 | . | . | . | . | . | . |
| chr11 | 102189575 | 102189967 | 847 | Promoter (<=1kb) | 11 | 102189620 | 102194081 | 4462 | 1 | ENSMUST00 | 0 | G6pc3 | ENSMUSG0C | false | chr11 | 102189364 | 102190440 | 21.92219 | 60.49891 | 57.64639 | chr11 | 102188849 | 102191275 | 4.39513 | 3.78417 | 2.51606 |
| chr16 | 33036095 | 33036412 | 847 | Intron (ENSMUST00000105610.2 | 16 | 33016950 | 33023937 | 6988 | 2 | ENSMUST00 | -12159 | Iqcg | ENSMUSG0C | false | . | . | . | . | . | . | chr16 | 33036156 | 33037018 | 4.04798 | 3.59006 | 2.26792 |
| chr9 | 63925248 | 63925552 | 847 | Distal Intergenic | 9 | 63954009 | 64016987 | 62979 | 2 | ENSMUST00 | 91435 | Smad6 | ENSMUSG0C | true | . | . | . | . | . | . | chr9 | 63922790 | 63929287 | 7.21154 | 5.06095 | 4.85866 |
| chr4 | 134535316 | 134535687 | 847 | Promoter (<=1kb) | 4 | 134525557 | 134535387 | 9831 | 2 | ENSMUST00 | 0 | Mtfr1l | ENSMUSG0C | false | chr4 | 134533990 | 134535997 | 32.61975 | 142.21701 | 137.24615 | chr4 | 134535673 | 134536855 | 5.45808 | 4.07799 | 3.41485 |
| chr10 | 80766251 | 80766658 | 847 | Promoter (2-3kb) | 10 | 80766859 | 80768987 | 2129 | 2 | ENSMUST00 | 2329 | Gm17151 | ENSMUSG0C | false | . | . | . | . | . | . | chr10 | 80755870 | 80769830 | 6.78362 | 4.72105 | 4.50141 |
| chr9 | 14641789 | 14642122 | 846 | Distal Intergenic | 9 | 14593444 | 14615483 | 22040 | 2 | ENSMUST00 | -26307 | Amotl1 | ENSMUSG0C | false | . | . | . | . | . | . | . | . | . | . | . | . |
| chr7 | 132258772 | 132259106 | 846 | Intron (ENSMUST00000124096.2 | 7 | 132270819 | 132278625 | 7807 | 2 | ENSMUST00 | 19519 | Chst15 | ENSMUSG0C | false | . | . | . | . | . | . | chr7 | 132257715 | 132263177 | 4.97245 | 4.0302 | 2.98893 |
| chr8 | 22942309 | 22942621 | 846 | 3' UTR | 8 | 22913786 | 22923645 | 9860 | 1 | ENSMUST00 | 28524 | Kat6a | ENSMUSG0C | true | . | . | . | . | . | . | chr8 | 22941593 | 22945712 | 4.33102 | 3.72804 | 2.51486 |
| chr4 | 140699746 | 140700057 | 846 | Promoter (<=1kb) | 4 | 140700541 | 140723220 | 22680 | 1 | ENSMUST00 | -484 | Rcc2 | ENSMUSG0C | false | chr4 | 140699593 | 140700679 | 9.72429 | 19.3484 | 17.02254 | chr4 | 140699427 | 140701325 | 3.78124 | 3.27964 | 2.03018 |
| chrX | 134541499 | 134541866 | 846 | Promoter (<=1kb) | 20 | 134537256 | 134541865 | 4610 | 2 | ENSMUST00 | 0 | Timm8a1 | ENSMUSG0C | false | chrX | 134540801 | 134542148 | 18.38128 | 45.59164 | 42.93153 | . | . | . | . | . | . |
| chr18 | 75697349 | 75697816 | 846 | Promoter (<=1kb) | 18 | 75431224 | 75697696 | 266473 | 2 | ENSMUST00 | 0 | Ctif | ENSMUSG0C | false | chr18 | 75695237 | 75698016 | 23.65566 | 74.08845 | 71.03342 | . | . | . | . | . | . |
| chr11 | 82387215 | 82387565 | 845 | Promoter (1-2kb) | 11 | 82388900 | 82446327 | 57428 | 1 | ENSMUST00 | -1335 | Tmem132e | ENSMUSG0C | false | chr11 | 82386895 | 82389905 | 19.27906 | 51.80829 | 49.07 | chr11 | 82386913 | 82390066 | 7.48239 | 5.26205 | 5.02731 |
| chr16 | 85172077 | 85172792 | 845 | Promoter (<=1kb) | 16 | 84954440 | 85173707 | 219268 | 2 | ENSMUST00 | 915 | App | ENSMUSG0C | false | chr16 | 85171787 | 85174244 | 30.94804 | 114.76607 | 110.79114 | chr16 | 85167810 | 85172354 | 5.42256 | 4.17976 | 3.36286 |
| chr15 | 99614847 | 99615188 | 845 | Distal Intergenic | 15 | 99601400 | 99605477 | 4078 | 1 | ENSMUST00 | 13448 | Aqp6 | ENSMUSG0C | false | . | . | . | . | . | . | chr15 | 99614232 | 99615590 | 4.96051 | 4.16896 | 2.95481 |
| chr3 | 89166470 | 89166816 | 845 | Promoter (<=1kb) | 3 | 89167164 | 89170409 | 3246 | 1 | ENSMUST00 | -348 | Clk2 | ENSMUSG0C | false | . | . | . | . | . | . | chr3 | 89163779 | 89168143 | 6.10442 | 4.63703 | 3.89661 |
| chr10 | 14207632 | 14207904 | 845 | Distal Intergenic | 10 | 14233513 | 14233623 | 111 | 1 | ENSMUST00 | -25609 | Gm24343 | ENSMUSG0C | false | . | . | . | . | . | . | . | . | . | . | . | . |
| chr12 | 29029464 | 29029822 | 845 | Distal Intergenic | 12 | 29035180 | 29035294 | 115 | 2 | ENSMUST00 | 5472 | Gm23342 | ENSMUSG0C | false | . | . | . | . | . | . | chr12 | 29028115 | 29030304 | 3.87548 | 3.54546 | 2.06813 |
| chr19 | 47266182 | 47266491 | 845 | 3' UTR | 19 | 47273519 | 47273589 | 71 | 2 | ENSMUST00 | 7098 | Mir6995 | ENSMUSG0C | false | . | . | . | . | . | . | chr19 | 47264187 | 47266853 | 4.1213 | 3.37747 | 2.31459 |
| chr10 | 86287612 | 86287969 | 844 | Intron (ENSMUST00000120638.2 | 10 | 86300372 | 86349506 | 49135 | 1 | ENSMUST00 | -12403 | Timp3 | ENSMUSG0C | false | . | . | . | . | . | . | chr10 | 86287196 | 86291253 | 5.15519 | 4.10271 | 3.1482 |
| chr19 | 5117666 | 5118532 | 844 | Promoter (<=1kb) | 19 | 5107746 | 5118297 | 10552 | 2 | ENSMUST00 | 0 | Klc2 | ENSMUSG0C | false | chr19 | 5117175 | 5118826 | 28.22734 | 101.53345 | 97.92473 | chr19 | 5116044 | 5118904 | 5.75065 | 4.3249 | 3.65419 |
| chr8 | 72547968 | 72548579 | 844 | Promoter (<=1kb) | 8 | 72494558 | 72548310 | 53753 | 2 | ENSMUST00 | 0 | Med26 | ENSMUSG0C | false | chr8 | 72546955 | 72549355 | 36.11848 | 119.98036 | 115.83894 | . | . | . | . | . | . |
| chr16 | 97922197 | 97922779 | 844 | Promoter (<=1kb) | 16 | 97855210 | 97922633 | 67424 | 2 | ENSMUST00 | 0 | C2cd2 | ENSMUSG0C | false | chr16 | 97921637 | 97923023 | 31.80739 | 115.34708 | 111.35392 | . | . | . | . | . | . |
| chr2 | 18672122 | 18673064 | 844 | Promoter (<=1kb) | 2 | 18672384 | 18676230 | 3847 | 1 | ENSMUST00 | 0 | Commd3 | ENSMUSG0C | false | chr2 | 18671836 | 18672324 | 27.40132 | 82.57204 | 79.37107 | chr2 | 18671848 | 18674722 | 3.14612 | 3.08853 | 1.47329 |
| chr2 | 105084035 | 105084350 | 844 | Intron (ENSMUST00000135153.2 | 2 | 105076538 | 105089817 | 13280 | 2 | ENSMUST00 | 5467 | Wt1os | ENSMUSG0C | true | chr2 | 105083047 | 105084091 | 11.1896 | 25.76025 | 23.34976 | chr2 | 105081211 | 105093624 | 5.17671 | 4.11795 | 3.15483 |
| chr7 | 30588522 | 30589042 | 843 | Promoter (<=1kb) | 7 | 30568858 | 30588717 | 19860 | 2 | ENSMUST00 | 0 | Kmt2b | ENSMUSG0C | false | chr7 | 30587480 | 30589984 | 31.70051 | 127.6562 | 123.25787 | . | . | . | . | . | . |
| chr8 | 109868284 | 109869455 | 843 | Promoter (<=1kb) | 8 | 109868542 | 109904971 | 36430 | 1 | ENSMUST00 | 0 | Phlpp2 | ENSMUSG0C | false | chr8 | 109868032 | 109869607 | 27.66025 | 106.38611 | 102.65278 | chr8 | 109867266 | 109868387 | 4.27513 | 3.83054 | 2.39118 |
| chr9 | 122143273 | 122143601 | 843 | Intron (ENSMUST00000120173.2 | 9 | 122160072 | 122166391 | 6320 | 1 | ENSMUST00 | -16471 | Snrk | ENSMUSG0C | false | . | . | . | . | . | . | chr9 | 122139168 | 122143961 | 4.81474 | 3.78854 | 2.88599 |
| chr7 | 80369122 | 80369412 | 843 | Promoter (1-2kb) | 7 | 80349098 | 80371375 | 22278 | 2 | ENSMUST00 | 1963 | Man2a2 | ENSMUSG0C | false | . | . | . | . | . | . | chr7 | 80368144 | 80371674 | 5.59506 | 4.057 | 3.5213 |
| chr7 | 19187949 | 19188214 | 843 | Intron (ENSMUST00000117338.2 | 7 | 19181606 | 19186371 | 4766 | 1 | ENSMUST00 | 6344 | Eml2 | ENSMUSG0C | true | . | . | . | . | . | . | chr7 | 19178043 | 19190895 | 5.91671 | 4.49499 | 3.76502 |
| chr7 | 24902696 | 24903353 | 843 | Promoter (<=1kb) | 7 | 24902912 | 24926592 | 23681 | 1 | ENSMUST00 | 0 | Arhgef1 | ENSMUSG0C | false | chr7 | 24902634 | 24904205 | 26.65955 | 86.95457 | 83.67183 | chr7 | 24902570 | 24903887 | 3.17581 | 3.06549 | 1.52452 |
| chrX | 53045469 | 53045844 | 843 | Promoter (2-3kb) | 20 | 53047997 | 53048078 | 82 | 2 | ENSMUST00 | 2234 | Mir450b | ENSMUSG0C | false | . | . | . | . | . | . | . | . | . | . | . | . |
| chr11 | 8467560 | 8467937 | 842 | Promoter (<=1kb) | 11 | 8451070 | 8468675 | 17606 | 2 | ENSMUST00 | 738 | Tns3 | ENSMUSG0C | false | chr11 | 8467453 | 8469249 | 6.91491 | 17.20307 | 14.90901 | chr11 | 8438250 | 8474887 | 7.10839 | 4.95354 | 4.75045 |
| chr7 | 126428630 | 126429138 | 842 | Promoter (<=1kb) | 7 | 126428759 | 126445901 | 17143 | 1 | ENSMUST00 | 0 | Rabep2 | ENSMUSG0C | false | chr7 | 126428298 | 126429917 | 26.86698 | 95.04005 | 91.5885 | chr7 | 126428306 | 126430309 | 4.69253 | 3.79347 | 2.78825 |
| chr8 | 85036787 | 85037219 | 842 | Promoter (<=1kb) | 8 | 85036915 | 85057583 | 20669 | 1 | ENSMUST00 | 0 | Tnpo2 | ENSMUSG0C | false | chr8 | 85035567 | 85038616 | 22.44583 | 74.56482 | 71.50171 | . | . | . | . | . | . |
| chr6 | 146959998 | 146960360 | 842 | Intron (ENSMUST00000016631.8 | 6 | 146950329 | 146954422 | 4094 | 2 | ENSMUST00 | -5577 | 1700034J05 | ENSMUSG0C | false | . | . | . | . | . | . | chr6 | 146958225 | 146962903 | 6.76492 | 4.98632 | 4.43962 |
| chr16 | 25468679 | 25469006 | 842 | Distal Intergenic | 16 | 25286821 | 25318235 | 31415 | 1 | ENSMUST00 | 181859 | Tprg | ENSMUSG0C | false | . | . | . | . | . | . | chr16 | 25464746 | 25469847 | 3.62992 | 3.29674 | 1.91973 |
| chr8 | 79096960 | 79097309 | 842 | Intron (ENSMUST00000098614.3 | 8 | 79118327 | 79193766 | 75440 | 1 | ENSMUST00 | -21018 | Zfp827 | ENSMUSG0C | false | . | . | . | . | . | . | chr8 | 79096207 | 79098256 | 5.34071 | 4.1647 | 3.32923 |
| chr7 | 109686758 | 109687071 | 841 | Distal Intergenic | 7 | 109703690 | 109712189 | 8500 | 1 | ENSMUST00 | -16619 | Akip1 | ENSMUSG0C | false | . | . | . | . | . | . | . | . | . | . | . | . |
| chr8 | 104867376 | 104867853 | 841 | Promoter (<=1kb) | 8 | 104867488 | 104874080 | 6593 | 1 | ENSMUST00 | 0 | Ces2d-ps | ENSMUSG0C | false | . | . | . | . | . | . | chr8 | 104864625 | 104870535 | 4.84736 | 4.01234 | 2.88046 |
| chr2 | 102185956 | 102186547 | 841 | Promoter (<=1kb) | 2 | 101950203 | 102186358 | 236156 | 2 | ENSMUST00 | 0 | Ldlrad3 | ENSMUSG0C | false | chr2 | 102185400 | 102186830 | 24.78212 | 78.86476 | 75.73294 | chr2 | 102185308 | 102187023 | 3.87136 | 3.53884 | 2.09647 |
| chr3 | 105061455 | 105061879 | 841 | Distal Intergenic | 3 | 105066844 | 105078806 | 11963 | 1 | ENSMUST00 | -4965 | 4930564D02 | ENSMUSG0C | false | . | . | . | . | . | . | . | . | . | . | . | . |
| chr7 | 24486705 | 24487299 | 841 | Intron (ENSMUST00000068023.7 | 7 | 24482023 | 24504539 | 22517 | 1 | ENSMUST00 | 4683 | Cadm4 | ENSMUSG0C | false | . | . | . | . | . | . | chr7 | 24480521 | 24490691 | 5.95135 | 4.53554 | 3.78776 |
| chr7 | 5020236 | 5020738 | 841 | Promoter (<=1kb) | 7 | 5020376 | 5033138 | 12763 | 1 | ENSMUST00 | 0 | Zfp865 | ENSMUSG0C | false | chr7 | 5019648 | 5021260 | 35.56825 | 117.95176 | 113.87694 | . | . | . | . | . | . |
| chr2 | 64097974 | 64098954 | 841 | Promoter (<=1kb) | 2 | 63971507 | 64098012 | 126506 | 2 | ENSMUST00 | 0 | Fign | ENSMUSG0C | true | chr2 | 64096333 | 64099107 | 28.91859 | 89.0447 | 85.71584 | . | . | . | . | . | . |
| chr16 | 91851570 | 91851933 | 840 | Promoter (1-2kb) | 16 | 91853811 | 91863319 | 9509 | 1 | ENSMUST00 | -1878 | Itsn1 | ENSMUSG0C | false | . | . | . | . | . | . | chr16 | 91849364 | 91852438 | 5.57727 | 4.31889 | 3.50137 |
| chr4 | 118134684 | 118135309 | 840 | Promoter (<=1kb) | 4 | 117932154 | 118134890 | 202737 | 2 | ENSMUST00 | 0 | St3gal3 | ENSMUSG0C | true | chr4 | 118132585 | 118135492 | 24.2022 | 107.83455 | 104.06592 | chr4 | 118135121 | 118140157 | 5.86588 | 4.43161 | 3.75583 |
| chr4 | 119865982 | 119866335 | 840 | Intron (ENSMUST00000141605.2 | 4 | 119814495 | 120138045 | 323551 | 1 | ENSMUST00 | 51488 | Hivep3 | ENSMUSG0C | false | . | . | . | . | . | . | chr4 | 119865109 | 119867357 | 4.92082 | 3.89419 | 2.97085 |
| chrX | 12091691 | 12092088 | 840 | Intron (ENSMUST00000043441.7 | 20 | 12036740 | 12080553 | 43814 | 2 | ENSMUST00 | -11139 | Bcor | ENSMUSG0C | false | chrX | 12091578 | 12092219 | 17.92175 | 44.05972 | 41.41886 | . | . | . | . | . | . |
| chr2 | 75633703 | 75634055 | 840 | Promoter (<=1kb) | 2 | 75632638 | 75633382 | 745 | 2 | ENSMUST00 | -322 | Gm13655 | ENSMUSG0C | false | . | . | . | . | . | . | chr2 | 75633073 | 75634846 | 5.33283 | 4.08798 | 3.30541 |
| chr7 | 70643959 | 70644305 | 840 | Distal Intergenic | 7 | 70752416 | 70752500 | 85 | 1 | ENSMUST00 | -108111 | Gm24880 | ENSMUSG0C | false | . | . | . | . | . | . | chr7 | 70643205 | 70644657 | 5.0281 | 4.15186 | 3.03335 |
| chr11 | 57644944 | 57645673 | 839 | Promoter (<=1kb) | 11 | 57645442 | 57787514 | 142073 | 1 | ENSMUST00 | 0 | Galnt10 | ENSMUSG0C | true | chr11 | 57644082 | 57647951 | 34.54477 | 122.48211 | 118.25522 | . | . | . | . | . | . |
| chr3 | 135254822 | 135255170 | 839 | Intron (ENSMUST00000062893.7 | 3 | 135281221 | 135304426 | 23206 | 1 | ENSMUST00 | -26051 | Bdh2 | ENSMUSG0C | false | . | . | . | . | . | . | chr3 | 135253778 | 135257937 | 5.99657 | 4.52312 | 3.83185 |
| chr5 | 31602346 | 31602639 | 839 | Intron (ENSMUST00000142345.2 | 5 | 31598733 | 31622644 | 23912 | 1 | ENSMUST00 | 3614 | Mrpl33 | ENSMUSG0C | false | . | . | . | . | . | . | chr5 | 31601971 | 31602832 | 4.51527 | 3.64471 | 2.64105 |
| chr8 | 86623849 | 86624190 | 839 | Promoter (<=1kb) | 8 | 86624043 | 86716631 | 92589 | 1 | ENSMUST00 | 0 | Lonp2 | ENSMUSG0C | false | chr8 | 86623413 | 86623857 | 22.21283 | 62.70797 | 59.8237 | . | . | . | . | . | . |
| chr15 | 83293915 | 83294277 | 839 | Distal Intergenic | 15 | 83226726 | 83251774 | 25049 | 2 | ENSMUST00 | -42142 | A4galt | ENSMUSG0C | false | . | . | . | . | . | . | chr15 | 83292734 | 83294592 | 3.48937 | 3.25018 | 1.80584 |
| chr7 | 5051451 | 5051881 | 839 | Promoter (<=1kb) | 7 | 5051532 | 5053722 | 2191 | 1 | ENSMUST00 | 0 | Zfp580 | ENSMUSG0C | false | chr7 | 5050922 | 5052722 | 25.90858 | 83.69981 | 80.48042 | chr7 | 5050498 | 5054338 | 6.88179 | 4.81993 | 4.56823 |
| chr11 | 60139290 | 60139625 | 839 | Promoter (<=1kb) | 11 | 60140082 | 60199195 | 59114 | 1 | ENSMUST00 | -457 | Rai1 | ENSMUSG0C | false | chr11 | 60139224 | 60140901 | 27.98104 | 79.82376 | 76.67776 | chr11 | 60138396 | 60141338 | 3.91042 | 3.57283 | 2.09867 |
| chr11 | 116814650 | 116815029 | 838 | Intron (ENSMUST00000021170.3 | 11 | 116807913 | 116828003 | 20091 | 2 | ENSMUST00 | 12974 | Mxra7 | ENSMUSG0C | false | . | . | . | . | . | . | chr11 | 116810954 | 116829199 | 5.76656 | 4.27097 | 3.65606 |
| chr4 | 134755985 | 134756338 | 838 | Intron (ENSMUST00000037828.6 | 4 | 134749416 | 134750850 | 1435 | 2 | ENSMUST00 | -5136 | Ldlrap1 | ENSMUSG0C | false | . | . | . | . | . | . | chr4 | 134750748 | 134766877 | 7.637 | 5.15315 | 5.18044 |
| chr12 | 112595583 | 112595943 | 838 | Intron (ENSMUST00000101029.2 | 12 | 112588784 | 112615556 | 26773 | 1 | ENSMUST00 | 6800 | Inf2 | ENSMUSG0C | false | . | . | . | . | . | . | chr12 | 112585329 | 112600582 | 8.19005 | 5.40781 | 5.63321 |
| chr7 | 130855746 | 130856032 | 838 | Intron (ENSMUST00000124096.2 | 7 | 130865756 | 130913312 | 47557 | 1 | ENSMUST00 | -9724 | Plekha1 | ENSMUSG0C | false | . | . | . | . | . | . | chr7 | 130854861 | 130857242 | 5.53565 | 4.38433 | 3.45522 |
| chr7 | 142025943 | 142026302 | 838 | Intron (ENSMUST00000084418.2 | 7 | 141998661 | 142001441 | 2781 | 1 | ENSMUST00 | 27283 | Brsk2 | ENSMUSG0C | false | chr7 | 142026194 | 142026806 | 10.09794 | 21.03375 | 18.68556 | chr7 | 142014396 | 142028430 | 5.88599 | 4.36974 | 3.76503 |
| chr8 | 111145153 | 111145998 | 838 | Promoter (<=1kb) | 8 | 111128886 | 111145459 | 16574 | 2 | ENSMUST00 | 0 | 9430091E24 | ENSMUSG0C | false | chr8 | 111144711 | 111146229 | 38.42999 | 152.65175 | 147.26898 | . | . | . | . | . | . |
| chr1 | 74761214 | 74761514 | 838 | Distal Intergenic | 1 | 74764798 | 74764890 | 93 | 1 | ENSMUST00 | -3284 | Gm27507 | ENSMUSG0C | false | . | . | . | . | . | . | chr1 | 74760147 | 74763092 | 6.23505 | 4.68861 | 4.00513 |
| chr6 | 114969837 | 114970269 | 837 | Promoter (<=1kb) | 6 | 114862764 | 114969994 | 107231 | 2 | ENSMUST00 | 0 | Vgll4 | ENSMUSG0C | true | chr6 | 114967501 | 114970246 | 26.65955 | 86.95457 | 83.67183 | chr6 | 114966882 | 114970611 | 4.21249 | 3.6549 | 2.39033 |
| chr11 | 4863603 | 4863858 | 837 | Distal Intergenic | 11 | 4873951 | 4894200 | 20250 | 1 | ENSMUST00 | -10093 | Nipsnap1 | ENSMUSG0C | false | . | . | . | . | . | . | chr11 | 4861868 | 4865842 | 3.36617 | 3.25214 | 1.71256 |
| chr13 | 98632302 | 98632624 | 837 | Downstream (2-3kb) | 13 | 98634978 | 98637410 | 2433 | 2 | ENSMUST00 | 4786 | Tmem174 | ENSMUSG0C | false | . | . | . | . | . | . | chr13 | 98631154 | 98632989 | 4.01198 | 3.5026 | 2.22842 |
| chr9 | 43127164 | 43127538 | 837 | Exon (ENSMUST00000034513.7/ | 9 | 43108653 | 43116570 | 7918 | 2 | ENSMUST00 | -10595 | Tmem136 | ENSMUSG0C | false | . | . | . | . | . | . | chr9 | 43126598 | 43131834 | 5.02199 | 4.11654 | 3.02968 |
| chr1 | 38308646 | 38308900 | 837 | Promoter (<=1kb) | 1 | 38307902 | 38311930 | 4029 | 1 | ENSMUST00 | 745 | Gm16151 | ENSMUSG0C | false | . | . | . | . | . | . | chr1 | 38307025 | 38310222 | 5.52473 | 4.379 | 3.44246 |
| chr17 | 6978690 | 6978994 | 837 | Promoter (<=1kb) | 17 | 6978860 | 6998193 | 19334 | 1 | ENSMUST00 | 0 | Rnaset2b | ENSMUSG0C | false | chr17 | 6978383 | 6979605 | 26.18681 | 91.82585 | 88.44535 | chr17 | 6978490 | 6980582 | 4.5926 | 3.83693 | 2.72476 |
| chr8 | 125388693 | 125389199 | 836 | Distal Intergenic | 8 | 125418063 | 125492710 | 74648 | 2 | ENSMUST00 | 103511 | Sipa1l2 | ENSMUSG0C | false | . | . | . | . | . | . | chr8 | 125384422 | 125392083 | 5.56972 | 4.2897 | 3.4878 |
| chr11 | 59506135 | 59506554 | 836 | Promoter (<=1kb) | 11 | 59499544 | 59506220 | 6677 | 2 | ENSMUST00 | 0 | Zkscan17 | ENSMUSG0C | false | chr11 | 59505222 | 59507156 | 33.04283 | 115.53907 | 111.53828 | . | . | . | . | . | . |
| chr12 | 111817063 | 111817397 | 836 | Promoter (2-3kb) | 12 | 111814170 | 111828388 | 14219 | 1 | ENSMUST00 | 2894 | Zfyve21 | ENSMUSG0C | false | . | . | . | . | . | . | chr12 | 111813203 | 111823116 | 5.80855 | 4.38461 | 3.68777 |
| chr19 | 58816753 | 58817210 | 836 | Intron (ENSMUST00000066285.4 | 19 | 58785803 | 58794461 | 8659 | 2 | ENSMUST00 | -22293 | 1700019N19 | ENSMUSG0C | false | . | . | . | . | . | . | chr19 | 58814047 | 58859817 | 7.46659 | 5.12677 | 5.03638 |
| chr4 | 109204435 | 109204871 | 836 | Promoter (2-3kb) | 4 | 109062079 | 109202272 | 140194 | 2 | ENSMUST00 | -2164 | Osbpl9 | ENSMUSG0C | true | . | . | . | . | . | . | chr4 | 109202402 | 109206991 | 6.22791 | 4.58919 | 3.99779 |
| chr16 | 20619918 | 20620217 | 836 | Promoter (1-2kb) | 16 | 20619215 | 20621287 | 2073 | 2 | ENSMUST00 | 1070 | Camk2n2 | ENSMUSG0C | false | chr16 | 20619732 | 20620546 | 13.21926 | 29.21668 | 26.76154 | chr16 | 20619621 | 20622569 | 4.7334 | 3.74519 | 2.81748 |
| chr1 | 135687890 | 135688503 | 836 | Promoter (<=1kb) | 1 | 135606447 | 135688105 | 81659 | 2 | ENSMUST00 | 0 | Nav1 | ENSMUSG0C | false | chr1 | 135687608 | 135688623 | 28.49099 | 81.63235 | 78.4519 | chr1 | 135686800 | 135688948 | 4.13284 | 3.56698 | 2.32732 |
| chr11 | 118425083 | 118425447 | 835 | Promoter (2-3kb) | 11 | 118428203 | 118443843 | 15641 | 1 | ENSMUST00 | -2756 | C1qtnf1 | ENSMUSG0C | false | . | . | . | . | . | . | chr11 | 118411596 | 118448618 | 7.23691 | 5.03811 | 4.84329 |
| chr17 | 21450157 | 21450545 | 835 | Promoter (<=1kb) | 17 | 21450352 | 21465591 | 15240 | 1 | ENSMUST00 | 0 | Zfp51 | ENSMUSG0C | false | chr17 | 21449880 | 21450957 | 24.35436 | 72.61947 | 69.58835 | . | . | . | . | . | . |
| chr1 | 152790902 | 152791658 | 835 | Distal Intergenic | 1 | 152800194 | 152836991 | 36798 | 1 | ENSMUST00 | -8536 | Ncf2 | ENSMUSG0C | true | . | . | . | . | . | . | chr1 | 152788034 | 152800060 | 6.38067 | 4.71623 | 4.13727 |
| chr9 | 103750630 | 103751013 | 835 | Intron (ENSMUST00000189588.1 | 9 | 103752250 | 103761790 | 9541 | 2 | ENSMUST00 | 10777 | Tmem108 | ENSMUSG0C | true | . | . | . | . | . | . | chr9 | 103745931 | 103753624 | 5.24003 | 4.22262 | 3.18902 |
| chr10 | 8785465 | 8785912 | 835 | Intron (ENSMUST00000015449.5 | 10 | 8824011 | 8824147 | 137 | 2 | ENSMUST00 | 38235 | Gm25410 | ENSMUSG0C | false | . | . | . | . | . | . | chr10 | 8785190 | 8788063 | 6.12077 | 4.48297 | 3.96064 |
| chr5 | 32480118 | 32480415 | 835 | Promoter (<=1kb) | 5 | 32479560 | 32479885 | 326 | 2 | ENSMUST00 | -234 | Gm15613 | ENSMUSG0C | false | . | . | . | . | . | . | chr5 | 32471673 | 32481447 | 3.91519 | 3.42294 | 2.15143 |
| chr2 | 52676024 | 52676494 | 834 | Promoter (<=1kb) | 2 | 52675727 | 52676143 | 417 | 2 | ENSMUST00 | 0 | Cacnb4 | ENSMUSG0C | false | chr2 | 52675430 | 52676764 | 19.93088 | 52.12947 | 49.38807 | chr2 | 52674958 | 52677200 | 5.04452 | 4.07467 | 3.04885 |
| chr4 | 21727479 | 21727871 | 834 | Promoter (<=1kb) | 4 | 21727701 | 21750546 | 22846 | 1 | ENSMUST00 | 0 | Ccnc | ENSMUSG0C | false | chr4 | 21727046 | 21728585 | 26.72788 | 101.58999 | 97.97913 | . | . | . | . | . | . |
| chr9 | 63672735 | 63673024 | 834 | Intron (ENSMUST00000034973.4 | 9 | 63654771 | 63666547 | 11777 | 2 | ENSMUST00 | -6189 | Smad3 | ENSMUSG0C | false | . | . | . | . | . | . | chr9 | 63671619 | 63680569 | 5.3648 | 4.16963 | 3.3451 |
| chr12 | 28872326 | 28872624 | 834 | Distal Intergenic | 12 | 28751828 | 28867491 | 115664 | 1 | ENSMUST00 | 120499 | Tssc1 | ENSMUSG0C | false | . | . | . | . | . | . | chr12 | 28872140 | 28876378 | 5.8899 | 4.40225 | 3.76611 |
| chr1 | 153526063 | 153526341 | 834 | Intron (ENSMUST00000041874.8 | 1 | 153548422 | 153549711 | 1290 | 2 | ENSMUST00 | 23370 | Npl | ENSMUSG0C | false | . | . | . | . | . | . | chr1 | 153525248 | 153527202 | 3.31183 | 3.07676 | 1.65846 |
| chr18 | 84015699 | 84016123 | 834 | Exon (ENSMUST00000060303.9/ | 18 | 84012082 | 84086390 | 74309 | 2 | ENSMUST00 | 70267 | Tshz1 | ENSMUSG0C | false | . | . | . | . | . | . | chr18 | 84014205 | 84018386 | 3.72565 | 3.31454 | 1.97496 |
| chr18 | 55133706 | 55134154 | 834 | Distal Intergenic | 18 | 55179840 | 55179919 | 80 | 1 | ENSMUST00 | -45686 | Gm22597 | ENSMUSG0C | false | . | . | . | . | . | . | chr18 | 55130720 | 55134767 | 5.00666 | 4.01828 | 3.01407 |
| chr10 | 75098460 | 75098739 | 833 | Intron (ENSMUST00000164107.1 | 10 | 75060896 | 75184923 | 124028 | 1 | ENSMUST00 | 37565 | Bcr | ENSMUSG0C | false | . | . | . | . | . | . | chr10 | 75096880 | 75101852 | 4.11942 | 3.59347 | 2.31318 |
| chr9 | 51406338 | 51406826 | 833 | Distal Intergenic | 9 | 51289686 | 51328917 | 39232 | 2 | ENSMUST00 | -77422 | 1810046K07 | ENSMUSG0C | false | . | . | . | . | . | . | chr9 | 51404996 | 51411058 | 4.97376 | 4.06092 | 2.98267 |
| chr10 | 34352216 | 34352528 | 833 | Intron (ENSMUST00000105512.2 | 10 | 34389981 | 34397085 | 7105 | 1 | ENSMUST00 | -37453 | Col10a1 | ENSMUSG0C | false | . | . | . | . | . | . | chr10 | 34351539 | 34353794 | 4.10076 | 3.58447 | 2.28824 |
| chr15 | 3995512 | 3996154 | 833 | Promoter (<=1kb) | 15 | 3982036 | 3995752 | 13717 | 2 | ENSMUST00 | 0 | AW549877 | ENSMUSG0C | false | chr15 | 3995190 | 3995797 | 22.60491 | 86.65532 | 83.38163 | . | . | . | . | . | . |
| chr16 | 76372301 | 76373479 | 833 | Promoter (<=1kb) | 16 | 76290862 | 76373049 | 82188 | 2 | ENSMUST00 | 0 | Nrip1 | ENSMUSG0C | false | chr16 | 76370364 | 76374292 | 33.25682 | 103.54333 | 99.8849 | . | . | . | . | . | . |
| chr2 | 61405297 | 61405561 | 833 | Distal Intergenic | 2 | 61492766 | 61492894 | 129 | 1 | ENSMUST00 | -87205 | Gm22338 | ENSMUSG0C | false | . | . | . | . | . | . | chr2 | 61398332 | 61406140 | 5.10401 | 4.1067 | 3.09609 |
| chr1 | 59318991 | 59319479 | 832 | Intron (ENSMUST00000160662.2 | 1 | 59298936 | 59352448 | 53513 | 1 | ENSMUST00 | 20056 | Cdk15 | ENSMUSG0C | false | . | . | . | . | . | . | chr1 | 59314432 | 59322803 | 4.39334 | 3.75254 | 2.52179 |
| chr4 | 53750423 | 53750840 | 832 | Intron (ENSMUST00000128667.2 | 4 | 53779705 | 53788712 | 9008 | 1 | ENSMUST00 | -28865 | Tal2 | ENSMUSG0C | false | . | . | . | . | . | . | chr4 | 53749682 | 53751862 | 4.58443 | 3.91087 | 2.66245 |
| chr7 | 101415232 | 101415532 | 832 | Distal Intergenic | 7 | 101421691 | 101512829 | 91139 | 1 | ENSMUST00 | -6159 | Pde2a | ENSMUSG0C | false | . | . | . | . | . | . | chr7 | 101413160 | 101416221 | 5.64616 | 4.46955 | 3.53528 |
| chr7 | 142095191 | 142095591 | 832 | Promoter (<=1kb) | 7 | 142079490 | 142095272 | 15783 | 2 | ENSMUST00 | 0 | Dusp8 | ENSMUSG0C | false | chr7 | 142094161 | 142096036 | 24.60788 | 97.30984 | 93.80507 | . | . | . | . | . | . |
| chr2 | 129648920 | 129649266 | 832 | Distal Intergenic | 2 | 129620685 | 129628044 | 7360 | 1 | ENSMUST00 | 28236 | Sirpa | ENSMUSG0C | true | . | . | . | . | . | . | chr2 | 129646412 | 129654466 | 5.30193 | 4.20597 | 3.2592 |
| chr5 | 14048880 | 14049275 | 832 | Intron (ENSMUST00000130116.1 | 5 | 14025317 | 14256689 | 231373 | 1 | ENSMUST00 | 23564 | Sema3e | ENSMUSG0C | true | . | . | . | . | . | . | chr5 | 14038084 | 14051077 | 3.98378 | 3.51352 | 2.17931 |
| chr5 | 105782010 | 105782309 | 832 | Intron (ENSMUST00000060531.1 | 5 | 105797321 | 105797713 | 393 | 2 | ENSMUST00 | 15404 | Rps15a-ps5 | ENSMUSG0C | false | . | . | . | . | . | . | chr5 | 105776453 | 105783156 | 4.86662 | 3.80058 | 2.93089 |
| chr3 | 86076557 | 86076864 | 831 | Intron (ENSMUST00000182969.2 | 3 | 86070915 | 86130526 | 59612 | 1 | ENSMUST00 | 5643 | Sh3d19 | ENSMUSG0C | false | chr3 | 86076581 | 86076866 | 5.79018 | 9.39037 | 7.2276 | chr3 | 86074600 | 86079495 | 4.27395 | 3.66881 | 2.45093 |
| chr6 | 87810638 | 87811438 | 831 | Promoter (<=1kb) | 6 | 87788853 | 87811779 | 22927 | 2 | ENSMUST00 | 341 | Rab43 | ENSMUSG0C | false | chr6 | 87810377 | 87812633 | 32.27147 | 102.84008 | 99.19724 | chr6 | 87809262 | 87810841 | 5.20305 | 4.03203 | 3.20761 |
| chr9 | 14802527 | 14802989 | 831 | Exon (ENSMUST00000034405.5/ | 9 | 14797166 | 14803853 | 6688 | 1 | ENSMUST00 | 5362 | Mre11a | ENSMUSG0C | false | . | . | . | . | . | . | chr9 | 14800716 | 14803895 | 4.33927 | 3.71523 | 2.50575 |
| chr8 | 47288286 | 47289588 | 831 | Promoter (<=1kb) | 8 | 47180048 | 47289402 | 109355 | 2 | ENSMUST00 | 0 | Stox2 | ENSMUSG0C | false | chr8 | 47287267 | 47289869 | 24.89402 | 98.85246 | 95.31046 | . | . | . | . | . | . |
| chr9 | 108079075 | 108079449 | 831 | Promoter (<=1kb) | 9 | 108051672 | 108079301 | 27630 | 2 | ENSMUST00 | 0 | Rnf123 | ENSMUSG0C | false | chr9 | 108078441 | 108079690 | 31.16539 | 106.97112 | 103.21663 | chr9 | 108077829 | 108079868 | 5.50262 | 4.33248 | 3.41779 |
| chrX | 13461478 | 13461795 | 831 | Distal Intergenic | 20 | 13466110 | 13489313 | 23204 | 1 | ENSMUST00 | -4315 | Nyx | ENSMUSG0C | false | chrX | 13461343 | 13462282 | 13.32643 | 29.39681 | 26.9387 | . | . | . | . | . | . |
| chr12 | 32504239 | 32504536 | 831 | Distal Intergenic | 12 | 32378704 | 32382943 | 4240 | 1 | ENSMUST00 | 125536 | Ccdc71l | ENSMUSG0C | false | . | . | . | . | . | . | chr12 | 32504033 | 32505881 | 4.59971 | 3.83218 | 2.70701 |
| chr5 | 136083697 | 136084136 | 830 | Promoter (<=1kb) | 5 | 136083916 | 136111860 | 27945 | 1 | ENSMUST00 | 0 | Rasa4 | ENSMUSG0C | false | chr5 | 136083512 | 136084617 | 25.53309 | 82.08177 | 78.89293 | chr5 | 136083376 | 136084928 | 3.68444 | 3.44821 | 1.93219 |
| chr11 | 5196228 | 5196558 | 830 | Intron (ENSMUST00000020662.9 | 11 | 5204833 | 5238448 | 33616 | 2 | ENSMUST00 | 41890 | Kremen1 | ENSMUSG0C | true | . | . | . | . | . | . | chr11 | 5181494 | 5198688 | 5.21662 | 4.12982 | 3.19453 |
| chr11 | 76953624 | 76954072 | 830 | Intron (ENSMUST00000021197.4 | 11 | 76961296 | 76965915 | 4620 | 1 | ENSMUST00 | -7224 | Blmh | ENSMUSG0C | false | . | . | . | . | . | . | . | . | . | . | . | . |
| chr15 | 44671140 | 44671465 | 830 | Downstream (<1kb) | 15 | 44619641 | 44641024 | 21384 | 1 | ENSMUST00 | 51500 | Ebag9 | ENSMUSG0C | false | . | . | . | . | . | . | chr15 | 44669272 | 44673388 | 4.8117 | 3.91147 | 2.88844 |
| chr7 | 70331122 | 70332068 | 830 | Distal Intergenic | 7 | 70351946 | 70360593 | 8648 | 2 | ENSMUST00 | 28525 | Nr2f2 | ENSMUSG0C | false | chr7 | 70330931 | 70332393 | 20.32388 | 57.71489 | 54.90119 | chr7 | 70330943 | 70332691 | 4.49153 | 3.87492 | 2.58151 |
| chr8 | 31032347 | 31032666 | 830 | Distal Intergenic | 8 | 31089471 | 31097047 | 7577 | 1 | ENSMUST00 | -56805 | Dusp26 | ENSMUSG0C | false | chr8 | 31031931 | 31032847 | 14.24974 | 34.23454 | 31.71693 | chr8 | 31032516 | 31033598 | 4.04458 | 3.62631 | 2.25053 |
| chr18 | 67724169 | 67724874 | 829 | Promoter (<=1kb) | 18 | 67665511 | 67724573 | 59063 | 2 | ENSMUST00 | 0 | Ptpn2 | ENSMUSG0C | false | chr18 | 67723607 | 67725123 | 31.01414 | 97.6227 | 94.10706 | chr18 | 67721841 | 67725704 | 3.81613 | 3.48081 | 2.07866 |
| chr19 | 23273600 | 23274017 | 829 | Promoter (<=1kb) | 19 | 23206441 | 23273897 | 67457 | 2 | ENSMUST00 | 0 | Smc5 | ENSMUSG0C | false | chr19 | 23273105 | 23274085 | 25.48472 | 95.2657 | 91.8051 | chr19 | 23272042 | 23273929 | 3.95509 | 3.55895 | 2.19707 |
| chr8 | 22757232 | 22757862 | 829 | Promoter (<=1kb) | 8 | 22757744 | 22782847 | 25104 | 1 | ENSMUST00 | 0 | Plat | ENSMUSG0C | true | chr8 | 22756921 | 22764179 | 35.19295 | 114.13696 | 110.1796 | chr8 | 22756891 | 22757299 | 4.04434 | 3.70093 | 2.19603 |
| chr2 | 167765114 | 167765586 | 829 | Distal Intergenic | 2 | 167777467 | 167782569 | 5103 | 1 | ENSMUST00 | -11881 | Gm14321 | ENSMUSG0C | false | . | . | . | . | . | . | chr2 | 167764213 | 167766418 | 5.97877 | 4.46573 | 3.85358 |
| chr2 | 167800606 | 167800854 | 829 | Distal Intergenic | 2 | 167782053 | 167783486 | 1434 | 1 | ENSMUST00 | 18554 | 9230111E07 | ENSMUSG0C | false | . | . | . | . | . | . | chr2 | 167800118 | 167801527 | 4.08656 | 3.65683 | 2.24364 |
| chr2 | 156974845 | 156975388 | 829 | Intron (ENSMUST00000069600.7 | 2 | 156964158 | 156964293 | 136 | 2 | ENSMUST00 | -10553 | Gm27653 | ENSMUSG0C | false | . | . | . | . | . | . | chr2 | 156972828 | 156978088 | 5.13281 | 4.17527 | 3.15455 |
| chr19 | 47306851 | 47307109 | 829 | Intron (ENSMUST00000081619.3 | 19 | 47273519 | 47273589 | 71 | 2 | ENSMUST00 | -33263 | Mir6995 | ENSMUSG0C | false | . | . | . | . | . | . | chr19 | 47296867 | 47309513 | 5.65055 | 4.24598 | 3.57438 |
| chr15 | 57815838 | 57816243 | 828 | Intron (ENSMUST00000096430.5 | 15 | 57869502 | 57892418 | 22917 | 2 | ENSMUST00 | 76175 | Derl1 | ENSMUSG0C | false | . | . | . | . | . | . | chr15 | 57813470 | 57821267 | 5.16015 | 4.09508 | 3.18154 |
| chr10 | 80398961 | 80399651 | 828 | Promoter (<=1kb) | 10 | 80392539 | 80399550 | 7012 | 2 | ENSMUST00 | 0 | Mbd3 | ENSMUSG0C | false | chr10 | 80398121 | 80399546 | 33.7938 | 119.00099 | 114.88904 | . | . | . | . | . | . |
| chr13 | 21362662 | 21363031 | 828 | Promoter (<=1kb) | 13 | 21362827 | 21369824 | 6998 | 1 | ENSMUST00 | 0 | Zscan12 | ENSMUSG0C | false | chr13 | 21362385 | 21363516 | 25.17393 | 93.69764 | 90.27328 | chr13 | 21362419 | 21364810 | 3.63587 | 3.27994 | 1.90693 |
| chr8 | 90955008 | 90955536 | 828 | Intron (ENSMUST00000048665.6 | 8 | 90907939 | 90908415 | 477 | 2 | ENSMUST00 | -46594 | Gm6658 | ENSMUSG0C | false | chr8 | 90954670 | 90956951 | 14.47545 | 35.30513 | 32.77419 | . | . | . | . | . | . |
| chr3 | 137867460 | 137868147 | 828 | Promoter (<=1kb) | 3 | 137867675 | 137916557 | 48883 | 1 | ENSMUST00 | 0 | Dnajb14 | ENSMUSG0C | false | chr3 | 137867258 | 137868612 | 26.22205 | 82.37943 | 79.18155 | chr3 | 137865283 | 137869163 | 3.98177 | 3.52903 | 2.18873 |
| chr11 | 70750012 | 70750273 | 828 | Exon (ENSMUST00000127333.1/ | 11 | 70748705 | 70755151 | 6447 | 2 | ENSMUST00 | 4878 | Gm12320 | ENSMUSG0C | false | . | . | . | . | . | . | chr11 | 70749987 | 70750350 | 4.80451 | 4.09186 | 2.83618 |
| chr9 | 114528134 | 114528467 | 828 | Intron (ENSMUST00000111816.2 | 9 | 114515697 | 114516424 | 728 | 1 | ENSMUST00 | 12438 | C130032M10 | ENSMUSG0C | false | . | . | . | . | . | . | chr9 | 114525892 | 114529998 | 4.57927 | 3.86564 | 2.65235 |
| chr15 | 25773957 | 25774528 | 827 | Promoter (<=1kb) | 15 | 25773987 | 25781057 | 7071 | 1 | ENSMUST00 | 0 | Myo10 | ENSMUSG0C | false | chr15 | 25773836 | 25774327 | 9.82359 | 19.5136 | 17.18665 | chr15 | 25760036 | 25780626 | 5.60912 | 4.2917 | 3.5207 |
| chr1 | 180966574 | 180966885 | 827 | Intron (ENSMUST00000027800.9 | 1 | 180960067 | 180960841 | 775 | 1 | ENSMUST00 | 6508 | Tmem63a | ENSMUSG0C | false | . | . | . | . | . | . | chr1 | 180965270 | 180969071 | 5.31548 | 4.05219 | 3.30841 |
| chr3 | 88106199 | 88106500 | 827 | Intron (ENSMUST00000071812.5 | 3 | 88082051 | 88121048 | 38998 | 1 | ENSMUST00 | 24149 | Iqgap3 | ENSMUSG0C | false | . | . | . | . | . | . | chr3 | 88104915 | 88107688 | 4.08321 | 3.6231 | 2.29427 |
| chr3 | 135825426 | 135825687 | 827 | Promoter (<=1kb) | 3 | 135825654 | 135859778 | 34125 | 1 | ENSMUST00 | 0 | Slc39a8 | ENSMUSG0C | false | chr3 | 135824972 | 135826805 | 21.33495 | 59.48191 | 56.64483 | chr3 | 135825162 | 135827217 | 3.53168 | 3.19144 | 1.8401 |

| | | | | | | | | | | | | | | | | | | | | | | | | | |
|---|---|---|---|---|---|---|---|---|---|---|---|---|---|---|---|---|---|---|---|---|---|---|---|---|---|
| chr19 | 44144680 | 44144966 | 827 | Promoter (1-2kb) | 19 | 44139266 | 44146446 | 7181 | 2 | ENSMUST00 | 1480 | Bloc1s2 | ENSMUSG0( | false | . | . | . | . | . | . | chr19 | 44143197 | 44146859 | 5.55448 | 4.21889 | 3.50123 |
| chr3 | 96239016 | 96239514 | 827 | Promoter (<=1kb) | 3 | 96238108 | 96239127 | 1020 | 2 | ENSMUST00 | 0 | Hist2h3c2 | ENSMUSG0( | false | chr3 | 96238498 | 96239143 | 21.42557 | 69.98924 | 66.99868 | chr3 | 96236895 | 96239036 | 4.04577 | 3.37246 | 2.23526 |
| chr8 | 110618493 | 110618919 | 826 | Promoter (<=1kb) | 8 | 110618638 | 110720396 | 101759 | 1 | ENSMUST00 | 0 | Vac14 | ENSMUSG0( | false | chr8 | 110617975 | 110619522 | 29.09955 | 105.17428 | 101.47362 | chr8 | 110617790 | 110621349 | 4.64489 | 3.80447 | 2.73746 |
| chr13 | 58453063 | 58453403 | 826 | Distal Intergenic | 13 | 58391142 | 58403343 | 12202 | 2 | ENSMUST00 | -49721 | Hnrnpk | ENSMUSG0( | false | chr13 | 58452745 | 58453502 | 12.53654 | 29.322 | 26.86593 | chr13 | 58452484 | 58453915 | 5.82979 | 4.55293 | 3.6791 |
| chr9 | 121351424 | 121351894 | 826 | Distal Intergenic | 9 | 121366958 | 121474916 | 107959 | 1 | ENSMUST00 | -15064 | Trak1 | ENSMUSG0( | false | . | . | . | . | . | . | chr9 | 121345520 | 121362682 | 6.17533 | 4.48468 | 3.99849 |
| chr14 | 55540156 | 55541060 | 826 | Promoter (<=1kb) | 14 | 55540266 | 55550017 | 9752 | 1 | ENSMUST00 | 0 | Pck2 | ENSMUSG0( | false | chr14 | 55539974 | 55541250 | 14.91789 | 45.43439 | 42.77634 | chr14 | 55539673 | 55541701 | 3.74011 | 3.33781 | 2.00939 |
| chr2 | 158199341 | 158199604 | 826 | Intron (ENSMUST00000103121.4 | 2 | 158192008 | 158195749 | 3742 | 1 | ENSMUST00 | 7334 | 1700060C20I | ENSMUSG0( | false | . | . | . | . | . | . | chr2 | 158198862 | 158200619 | 4.37175 | 3.82595 | 2.50151 |
| chr5 | 20373571 | 20373935 | 826 | Intron (ENSMUST00000088516.4 | 5 | 20264341 | 20264440 | 100 | 1 | ENSMUST00 | 109231 | Gm22755 | ENSMUSG0( | false | . | . | . | . | . | . | chr5 | 20371001 | 20374372 | 4.61088 | 3.86228 | 2.69905 |
| chr9 | 49999940 | 50000319 | 826 | Distal Intergenic | 9 | 49825869 | 49825975 | 107 | 2 | ENSMUST00 | -173966 | Gm25633 | ENSMUSG0( | false | . | . | . | . | . | . | chr9 | 49998981 | 50001457 | 4.42484 | 3.68349 | 2.54375 |
| chr15 | 80233784 | 80234192 | 825 | Promoter (<=1kb) | 15 | 80234080 | 80253366 | 19287 | 1 | ENSMUST00 | 0 | Mief1 | ENSMUSG0( | false | chr15 | 80233528 | 80235026 | 31.20348 | 118.35023 | 114.26361 | chr15 | 80233301 | 80235607 | 4.81374 | 3.96908 | 2.85389 |
| chr5 | 92321656 | 92321939 | 825 | 3' UTR | 5 | 92321331 | 92328079 | 6749 | 2 | ENSMUST00 | 6140 | Cxcl9 | ENSMUSG0( | false | . | . | . | . | . | . | . | . | . | . | . | . |
| chr7 | 119850807 | 119851140 | 825 | Downstream (2-3kb) | 7 | 119844206 | 119845436 | 1231 | 1 | ENSMUST00 | 6602 | 2610020H08I | ENSMUSG0( | false | . | . | . | . | . | . | chr7 | 119849777 | 119852389 | 3.87305 | 3.51474 | 2.07493 |
| chr13 | 34119349 | 34119661 | 825 | Distal Intergenic | 13 | 34127008 | 34130354 | 3347 | 2 | ENSMUST00 | 10693 | Tubb2b | ENSMUSG0( | false | . | . | . | . | . | . | chr13 | 34109058 | 34120340 | 5.26535 | 4.17937 | 3.22481 |
| chr13 | 46268371 | 46268749 | 825 | Distal Intergenic | 13 | 46273721 | 46300115 | 26395 | 1 | ENSMUST00 | -4972 | Stmnd1 | ENSMUSG0( | false | . | . | . | . | . | . | chr13 | 46268084 | 46269364 | 4.52389 | 3.8005 | 2.6399 |
| chr5 | 116448508 | 116449198 | 825 | Promoter (1-2kb) | 5 | 116442432 | 116446844 | 4413 | 2 | ENSMUST00 | -1665 | Srrm4 | ENSMUSG0( | false | . | . | . | . | . | . | . | . | . | . | . | . |
| chr10 | 7077034 | 7077321 | 825 | Distal Intergenic | 10 | 6911543 | 7052478 | 140936 | 2 | ENSMUST00 | -24557 | Ipcef1 | ENSMUSG0( | false | . | . | . | . | . | . | chr10 | 7076489 | 7077599 | 3.88569 | 3.57078 | 2.07521 |
| chr4 | 48042653 | 48043696 | 824 | Promoter (1-2kb) | 4 | 48045153 | 48086445 | 41293 | 1 | ENSMUST00 | -1457 | Nr4a3 | ENSMUSG0( | false | chr4 | 48042390 | 48044094 | 23.12601 | 77.64799 | 74.53377 | . | . | . | . | . | . |
| chr5 | 127617245 | 127617698 | 824 | Promoter (<=1kb) | 5 | 127595664 | 127617416 | 21753 | 2 | ENSMUST00 | 0 | Slc15a4 | ENSMUSG0( | false | chr5 | 127616053 | 127617842 | 37.30079 | 124.19571 | 119.91465 | . | . | . | . | . | . |
| chr6 | 128143358 | 128144039 | 824 | Promoter (<=1kb) | 6 | 127961396 | 128143563 | 182168 | 2 | ENSMUST00 | 0 | Tspan9 | ENSMUSG0( | false | chr6 | 128142694 | 128144365 | 28.51938 | 86.42239 | 83.15316 | chr6 | 128142451 | 128144420 | 3.58849 | 3.3447 | 1.83974 |
| chr2 | 103631284 | 103631810 | 824 | Intron (ENSMUST00000076212.3 | 2 | 103566310 | 103718423 | 152114 | 1 | ENSMUST00 | 64975 | Abtb2 | ENSMUSG0( | false | . | . | . | . | . | . | chr2 | 103621752 | 103634479 | 4.86729 | 3.97695 | 2.90643 |
| chr6 | 88426364 | 88426779 | 824 | Intron (ENSMUST00000165242.1 | 6 | 88257334 | 88446539 | 189206 | 2 | ENSMUST00 | 19760 | Eefsec | ENSMUSG0( | false | . | . | . | . | . | . | chr6 | 88422509 | 88428580 | 5.8759 | 4.49017 | 3.74239 |
| chr13 | 112037776 | 112038162 | 824 | Distal Intergenic | 13 | 112005502 | 112017332 | 11831 | 1 | ENSMUST00 | 32275 | Gm15323 | ENSMUSG0( | false | . | . | . | . | . | . | chr13 | 112036444 | 112038879 | 4.48307 | 3.86335 | 2.58504 |
| chr2 | 160731179 | 160731906 | 823 | Promoter (<=1kb) | 2 | 160731300 | 160762502 | 31203 | 1 | ENSMUST00 | 0 | Plcg1 | ENSMUSG0( | false | chr2 | 160730735 | 160732558 | 28.28183 | 109.60786 | 105.79031 | chr2 | 160729737 | 160731182 | 4.32225 | 3.77291 | 2.45174 |
| chr1 | 16618498 | 16619507 | 823 | Promoter (<=1kb) | 1 | 16540790 | 16619312 | 78523 | 2 | ENSMUST00 | 0 | Ube2w | ENSMUSG0( | false | chr1 | 16617928 | 16619784 | 29.83578 | 117.74437 | 113.67492 | . | . | . | . | . | . |
| chr5 | 99037055 | 99037393 | 823 | Promoter (<=1kb) | 5 | 98929773 | 99037351 | 107579 | 2 | ENSMUST00 | 0 | Prkg2 | ENSMUSG0( | false | chr5 | 99036519 | 99037492 | 17.92175 | 44.05972 | 41.41886 | chr5 | 99036283 | 99037713 | 4.59842 | 3.69973 | 2.70887 |
| chr8 | 62284406 | 62285162 | 823 | Distal Intergenic | 8 | 62057521 | 62123118 | 65598 | 2 | ENSMUST00 | -161289 | Anxa10 | ENSMUSG0( | false | chr8 | 62284049 | 62284795 | 9.06831 | 19.28032 | 16.9548 | chr8 | 62281497 | 62287555 | 4.55194 | 3.84079 | 2.62872 |
| chr5 | 139384560 | 139384908 | 823 | Intron (ENSMUST00000138631.1 | 5 | 139380581 | 139396415 | 15835 | 1 | ENSMUST00 | 3980 | Gpr146 | ENSMUSG0( | false | . | . | . | . | . | . | chr5 | 139381256 | 139395525 | 8.99937 | 5.68291 | 6.29852 |
| chr14 | 61172690 | 61173418 | 823 | Promoter (<=1kb) | 14 | 61172990 | 61195663 | 22674 | 1 | ENSMUST00 | 0 | Sacs | ENSMUSG0( | false | chr14 | 61172490 | 61173715 | 25.4663 | 101.95255 | 98.33054 | chr14 | 61173276 | 61174337 | 4.1615 | 3.74009 | 2.31926 |
| chr1 | 39186780 | 39187090 | 823 | Promoter (<=1kb) | 1 | 39186896 | 39186992 | 97 | 2 | ENSMUST00 | 0 | Mir6349 | ENSMUSG0( | false | . | . | . | . | . | . | chr1 | 39185712 | 39188794 | 5.01008 | 4.11211 | 3.01535 |
| chr3 | 138887910 | 138888193 | 822 | Intron (ENSMUST00000029800.3 | 3 | 138860493 | 138902657 | 42165 | 1 | ENSMUST00 | 27418 | Tspan5 | ENSMUSG0( | true | . | . | . | . | . | . | chr3 | 138887160 | 138888968 | 3.90698 | 3.4702 | 2.14538 |
| chr9 | 116762450 | 116762848 | 822 | Intron (ENSMUST00000111773.4 | 9 | 116577582 | 116822806 | 245225 | 2 | ENSMUST00 | 59958 | Rbms3 | ENSMUSG0( | true | . | . | . | . | . | . | chr9 | 116760630 | 116763722 | 4.62346 | 3.90848 | 2.68517 |
| chr11 | 69603536 | 69603833 | 822 | Promoter (<=1kb) | 11 | 69601358 | 69603613 | 2256 | 2 | ENSMUST00 | 0 | Atp1b2 | ENSMUSG0( | false | . | . | . | . | . | . | chr11 | 69603472 | 69606733 | 4.00382 | 3.53611 | 2.19247 |
| chr18 | 76059089 | 76059744 | 822 | Promoter (<=1kb) | 18 | 76059458 | 76146443 | 86986 | 1 | ENSMUST00 | 0 | Zbtb7c | ENSMUSG0( | true | chr18 | 76059035 | 76063179 | 44.34918 | 154.34741 | 148.89743 | . | . | . | . | . | . |
| chr13 | 60012173 | 60012610 | 822 | Distal Intergenic | 13 | 60174405 | 60177365 | 2961 | 2 | ENSMUST00 | 164755 | Gas1 | ENSMUSG0( | false | . | . | . | . | . | . | chr13 | 60006912 | 60013507 | 4.90407 | 3.99626 | 2.94044 |
| chr3 | 98339774 | 98340159 | 822 | Promoter (<=1kb) | 3 | 98313170 | 98339990 | 26821 | 2 | ENSMUST00 | 0 | Phgdh | ENSMUSG0( | false | chr3 | 98338685 | 98340237 | 19.27906 | 51.80829 | 49.07 | . | . | . | . | . | . |
| chr15 | 73137420 | 73137689 | 822 | Intron (ENSMUST00000044113.1 | 15 | 73090412 | 73094774 | 4363 | 1 | ENSMUST00 | 47009 | Chrac1 | ENSMUSG0( | false | . | . | . | . | . | . | chr15 | 73134756 | 73140961 | 6.23243 | 4.48464 | 4.04089 |
| chr10 | 4429522 | 4430306 | 821 | Promoter (1-2kb) | 10 | 4427243 | 4432295 | 5053 | 2 | ENSMUST00 | 1989 | Rmnd1 | ENSMUSG0( | false | chr10 | 4429163 | 4430557 | 8.3822 | 16.12319 | 13.84377 | chr10 | 4422045 | 4437373 | 5.83291 | 4.40627 | 3.70343 |
| chr11 | 115162776 | 115163438 | 821 | Promoter (<=1kb) | 11 | 115163341 | 115181181 | 17841 | 1 | ENSMUST00 | 0 | Slc9a3r1 | ENSMUSG0( | false | chr11 | 115162609 | 115165876 | 34.68901 | 133.45216 | 128.82967 | chr11 | 115160095 | 115163105 | 5.19967 | 4.25253 | 3.16648 |
| chr17 | 47773653 | 47773975 | 821 | Intron (ENSMUST00000086932.4 | 17 | 47769238 | 47786301 | 17064 | 1 | ENSMUST00 | 4416 | Tfeb | ENSMUSG0( | false | . | . | . | . | . | . | chr17 | 47772764 | 47774408 | 5.43366 | 4.36171 | 3.35242 |
| chrX | 169036457 | 169036763 | 821 | Promoter (<=1kb) | 20 | 169036611 | 169251527 | 214917 | 1 | ENSMUST00 | 0 | Arhgap6 | ENSMUSG0( | false | chrX | 169036407 | 169037107 | 10.56924 | 21.28106 | 18.92771 | . | . | . | . | . | . |
| chr8 | 86787895 | 86788210 | 821 | Distal Intergenic | 8 | 86723940 | 86746006 | 22067 | 2 | ENSMUST00 | -41890 | Siah1a | ENSMUSG0( | false | chr8 | 86787772 | 86788159 | 5.97392 | 9.35129 | 7.18948 | chr8 | 86787203 | 86788979 | 3.68296 | 3.31193 | 1.96458 |
| chr6 | 15752913 | 15753372 | 821 | Intron (ENSMUST00000101663.4 | 6 | 15727799 | 15770349 | 42551 | 1 | ENSMUST00 | 25115 | Mdfic | ENSMUSG0( | true | . | . | . | . | . | . | chr6 | 15748096 | 15764490 | 4.44274 | 3.82749 | 2.54582 |
| chr12 | 21543253 | 21543574 | 821 | Distal Intergenic | 12 | 21417911 | 21419803 | 1893 | 1 | ENSMUST00 | 125343 | Gm4419 | ENSMUSG0( | false | chr12 | 21542953 | 21544077 | 18.64736 | 62.11404 | 59.23804 | . | . | . | . | . | . |
| chr12 | 32715365 | 32715819 | 820 | Distal Intergenic | 12 | 32820335 | 32853369 | 33035 | 1 | ENSMUST00 | -104516 | Nampt | ENSMUSG0( | true | . | . | . | . | . | . | chr12 | 32713700 | 32718759 | 5.2438 | 4.25503 | 3.20864 |
| chr4 | 117581784 | 117582130 | 820 | Promoter (<=1kb) | 4 | 117582687 | 117615582 | 32896 | 1 | ENSMUST00 | -557 | Eri3 | ENSMUSG0( | false | . | . | . | . | . | . | chr4 | 117580895 | 117582709 | 5.4066 | 4.22968 | 3.39796 |
| chr12 | 12941466 | 12942070 | 820 | Promoter (<=1kb) | 12 | 12936096 | 12941914 | 5819 | 2 | ENSMUST00 | 0 | Mycn | ENSMUSG0( | false | chr12 | 12937456 | 12942953 | 27.46248 | 78.93296 | 75.80077 | . | . | . | . | . | . |
| chr5 | 66745606 | 66746062 | 820 | Promoter (<=1kb) | 5 | 66745835 | 67056073 | 310239 | 1 | ENSMUST00 | 0 | Limch1 | ENSMUSG0( | false | chr5 | 66745342 | 66747077 | 20.11728 | 54.87862 | 52.10172 | chr5 | 66745383 | 66746987 | 4.3855 | 3.66529 | 2.533 |
| chr7 | 107691377 | 107691782 | 820 | Intron (ENSMUST00000040056.8 | 7 | 107623971 | 107625161 | 1191 | 2 | ENSMUST00 | -66217 | 5330417H12I | ENSMUSG0( | false | . | . | . | . | . | . | chr7 | 107666979 | 107703704 | 5.18106 | 4.08532 | 3.17145 |
| chr3 | 9799350 | 9799635 | 820 | Intron (ENSMUST00000161949.2 | 3 | 9829620 | 9829728 | 109 | 2 | ENSMUST00 | 30093 | Gm24786 | ENSMUSG0( | false | . | . | . | . | . | . | chr3 | 9778824 | 9802371 | 5.7635 | 4.36373 | 3.65538 |
| chr12 | 87407918 | 87408249 | 819 | Distal Intergenic | 12 | 87305058 | 87388355 | 83298 | 2 | ENSMUST00 | -19564 | Sptlc2 | ENSMUSG0( | false | chr12 | 87407749 | 87408484 | 4.50687 | 8.15551 | 6.02026 | chr12 | 87407543 | 87409740 | 5.72622 | 4.5018 | 3.58534 |
| chr2 | 94103722 | 94104046 | 819 | Intron (ENSMUST00000028619.4 | 2 | 94033522 | 94066764 | 33243 | 2 | ENSMUST00 | -36959 | Hsd17b12 | ENSMUSG0( | false | . | . | . | . | . | . | chr2 | 94102203 | 94104578 | 4.77145 | 4.0094 | 2.81388 |
| chr2 | 30772122 | 30772485 | 819 | Distal Intergenic | 2 | 30795468 | 30796664 | 1197 | 2 | ENSMUST00 | 24179 | 1700001O22 | ENSMUSG0( | false | chr2 | 30771231 | 30772294 | 22.68861 | 66.13821 | 63.20662 | chr2 | 30771226 | 30773975 | 4.38222 | 3.65644 | 2.54422 |
| chr4 | 62566633 | 62566910 | 819 | 5' UTR | 4 | 62559847 | 62659840 | 99994 | 1 | ENSMUST00 | 6787 | Rgs3 | ENSMUSG0( | false | . | . | . | . | . | . | chr4 | 62565172 | 62567705 | 4.32219 | 3.77621 | 2.45766 |
| chr14 | 24779174 | 24779531 | 819 | Distal Intergenic | 14 | 24909414 | 24913440 | 4027 | 1 | ENSMUST00 | -129883 | Gm10398 | ENSMUSG0( | false | chr14 | 24778881 | 24779672 | 11.34984 | 26.03783 | 23.62382 | chr14 | 24778643 | 24780470 | 4.43865 | 3.8885 | 2.535 |
| chr10 | 81408960 | 81409292 | 819 | Promoter (<=1kb) | 10 | 81409575 | 81411579 | 2005 | 1 | ENSMUST00 | -283 | Gm16104 | ENSMUSG0( | false | . | . | . | . | . | . | chr10 | 81392649 | 81426205 | 8.72751 | 5.63067 | 6.0679 |
| chr4 | 54409736 | 54409974 | 819 | Distal Intergenic | 4 | 54468352 | 54468592 | 241 | 1 | ENSMUST00 | -58378 | Gm25687 | ENSMUSG0( | false | . | . | . | . | . | . | chr4 | 54408526 | 54410573 | 3.60793 | 3.32597 | 1.89493 |
| chr14 | 54431372 | 54431908 | 818 | Promoter (<=1kb) | 14 | 54431604 | 54441258 | 9655 | 1 | ENSMUST00 | 0 | Mmp14 | ENSMUSG0( | false | chr14 | 54431085 | 54433050 | 26.19333 | 73.09374 | 70.054 | chr14 | 54430319 | 54434204 | 4.2111 | 3.75246 | 2.34682 |
| chr11 | 47305015 | 47305418 | 818 | Intron (ENSMUST00000109220.2 | 11 | 46978785 | 47379522 | 400738 | 2 | ENSMUST00 | 74104 | Sgcd | ENSMUSG0( | false | . | . | . | . | . | . | . | . | . | . | . | . |
| chr2 | 94069578 | 94069861 | 818 | Promoter (2-3kb) | 2 | 94033522 | 94066764 | 33243 | 2 | ENSMUST00 | -2815 | Hsd17b12 | ENSMUSG0( | false | . | . | . | . | . | . | chr2 | 94069311 | 94070648 | 3.21664 | 3.12297 | 1.58938 |
| chr4 | 109201707 | 109202401 | 818 | Promoter (<=1kb) | 4 | 109061655 | 109202217 | 140563 | 2 | ENSMUST00 | 0 | Osbpl9 | ENSMUSG0( | true | chr4 | 109200480 | 109202667 | 24.86315 | 92.1349 | 88.74321 | . | . | . | . | . | . |
| chr18 | 31902448 | 31902758 | 818 | Exon (ENSMUST00000025264.6/ | 18 | 31909094 | 31911903 | 2810 | 2 | ENSMUST00 | 9145 | Sft2d3 | ENSMUSG0( | false | . | . | . | . | . | . | chr18 | 31902285 | 31902681 | 3.66049 | 3.20598 | 1.93943 |
| chr2 | 59828596 | 59829041 | 818 | Intron (ENSMUST00000112568.2 | 2 | 59815975 | 59816064 | 90 | 2 | ENSMUST00 | -12533 | Gm23842 | ENSMUSG0( | false | . | . | . | . | . | . | chr2 | 59823293 | 59834519 | 7.53663 | 5.1622 | 5.08626 |
| chr12 | 104002749 | 104003356 | 818 | Intron (ENSMUST00000058464.4 | 12 | 103984792 | 103996579 | 11788 | 2 | ENSMUST00 | -6171 | Serpina11 | ENSMUSG0( | false | . | . | . | . | . | . | chr12 | 104001382 | 104003739 | 4.41092 | 3.81538 | 2.51227 |
| chr5 | 129409808 | 129410082 | 817 | Distal Intergenic | 5 | 129501231 | 129571383 | 70153 | 1 | ENSMUST00 | -91149 | Sfswap | ENSMUSG0( | false | . | . | . | . | . | . | chr5 | 129405729 | 129418273 | 5.29114 | 4.12018 | 3.25854 |
| chr7 | 130692764 | 130693434 | 817 | Promoter (<=1kb) | 7 | 130692556 | 130764784 | 72229 | 1 | ENSMUST00 | 209 | Tacc2 | ENSMUSG0( | false | chr7 | 130692522 | 130693037 | 10.88283 | 27.02401 | 24.59589 | chr7 | 130690850 | 130697598 | 6.26132 | 4.62405 | 4.04306 |
| chr7 | 105736374 | 105736893 | 817 | Promoter (<=1kb) | 7 | 105732207 | 105736781 | 4575 | 2 | ENSMUST00 | 0 | Rrp8 | ENSMUSG0( | false | chr7 | 105735728 | 105738289 | 44.26889 | 153.06256 | 147.66058 | . | . | . | . | . | . |
| chr17 | 88626354 | 88626895 | 817 | Promoter (<=1kb) | 17 | 88626555 | 88645724 | 19170 | 1 | ENSMUST00 | 0 | Ston1 | ENSMUSG0( | false | chr17 | 88626424 | 88629225 | 13.60354 | 37.21757 | 34.66072 | chr17 | 88624831 | 88626650 | 4.59272 | 3.91563 | 2.66301 |
| chr11 | 98863452 | 98863935 | 817 | Promoter (<=1kb) | 11 | 98863638 | 98892299 | 28662 | 1 | ENSMUST00 | 0 | Wipf2 | ENSMUSG0( | false | chr11 | 98863105 | 98864606 | 29.20162 | 87.6073 | 84.30928 | chr11 | 98863099 | 98866552 | 3.26189 | 3.1685 | 1.59608 |
| chr6 | 148283510 | 148283780 | 817 | Intron (ENSMUST00000060095.9 | 6 | 148261150 | 148261552 | 403 | 1 | ENSMUST00 | 22361 | Gm15763 | ENSMUSG0( | false | . | . | . | . | . | . | chr6 | 148282474 | 148286757 | 5.12878 | 4.09662 | 3.11127 |
| chr18 | 45937553 | 45937863 | 817 | Distal Intergenic | 18 | 45941007 | 45941133 | 127 | 1 | ENSMUST00 | -3144 | Gm26262 | ENSMUSG0( | false | . | . | . | . | . | . | chr18 | 45936764 | 45938410 | 3.98751 | 3.49678 | 2.20614 |
| chr16 | 45158541 | 45159056 | 816 | Promoter (<=1kb) | 16 | 45158785 | 45188538 | 29754 | 1 | ENSMUST00 | 0 | Atg3 | ENSMUSG0( | false | chr16 | 45157695 | 45160217 | 31.62822 | 118.12274 | 114.03909 | . | . | . | . | . | . |
| chr19 | 10952042 | 10952505 | 816 | Promoter (2-3kb) | 19 | 10941481 | 10949266 | 7786 | 2 | ENSMUST00 | -2777 | Ccdc86 | ENSMUSG0( | false | . | . | . | . | . | . | chr19 | 10946315 | 10954190 | 5.98357 | 4.50647 | 3.81648 |
| chr8 | 119827852 | 119828145 | 816 | Intron (ENSMUST00000168698.1 | 8 | 119809225 | 119840522 | 31298 | 2 | ENSMUST00 | 12377 | Cotl1 | ENSMUSG0( | false | . | . | . | . | . | . | chr8 | 119806261 | 119841342 | 7.1039 | 5.00847 | 4.72992 |
| chr10 | 22926109 | 22926480 | 816 | Distal Intergenic | 10 | 22817279 | 22820128 | 2850 | 2 | ENSMUST00 | -105982 | Tcf21 | ENSMUSG0( | true | chr10 | 22926260 | 22926660 | 5.38833 | 9.07008 | 6.91429 | chr10 | 22925270 | 22927241 | 4.61465 | 3.76695 | 2.71961 |
| chr19 | 46689824 | 46690276 | 816 | Promoter (<=1kb) | 19 | 46689906 | 46703382 | 13477 | 1 | ENSMUST00 | 0 | 2010012O05 | ENSMUSG0( | false | chr19 | 46689613 | 46690621 | 28.49948 | 87.34749 | 84.05397 | chr19 | 46689506 | 46691625 | 5.91694 | 4.50218 | 3.7632 |
| chr17 | 80938086 | 80938450 | 816 | Promoter (<=1kb) | 17 | 80938334 | 80938714 | 381 | 2 | ENSMUST00 | 264 | Gm9386 | ENSMUSG0( | false | . | . | . | . | . | . | chr17 | 80937504 | 80938972 | 4.69523 | 3.67293 | 2.79914 |
| chr2 | 146855667 | 146856093 | 816 | Promoter (<=1kb) | 2 | 146855864 | 146970097 | 114234 | 1 | ENSMUST00 | 0 | Kiz | ENSMUSG0( | false | chr2 | 146855421 | 146856407 | 23.32969 | 96.49238 | 93.00913 | chr2 | 146855021 | 146855679 | 3.75847 | 3.46329 | 1.96813 |
| chr4 | 138496935 | 138497362 | 815 | Distal Intergenic | 4 | 138454314 | 138460123 | 5810 | 1 | ENSMUST00 | 42622 | Camk2n1 | ENSMUSG0( | false | . | . | . | . | . | . | . | . | . | . | . | . |
| chrX | 135993668 | 135993943 | 815 | Promoter (<=1kb) | 20 | 135993824 | 135995355 | 1532 | 1 | ENSMUST00 | 0 | Arxes2 | ENSMUSG0( | false | chrX | 135993395 | 135993990 | 11.02877 | 22.59171 | 20.22099 | . | . | . | . | . | . |
| chr12 | 86973693 | 86973999 | 815 | Distal Intergenic | 12 | 86983381 | 86988676 | 5296 | 2 | ENSMUST00 | 14677 | Zdhhc22 | ENSMUSG0( | false | . | . | . | . | . | . | chr12 | 86971639 | 86975722 | 5.78025 | 4.45948 | 3.65006 |
| chr6 | 51320712 | 51321057 | 815 | Distal Intergenic | 6 | 51269812 | 51269910 | 99 | 2 | ENSMUST00 | -50803 | Mir148a | ENSMUSG0( | false | . | . | . | . | . | . | chr6 | 51320040 | 51321158 | 3.61209 | 3.2997 | 1.89315 |
| chr6 | 71989781 | 71990338 | 815 | Distal Intergenic | 6 | 71961145 | 71963775 | 2631 | 2 | ENSMUST00 | -26007 | Gm26628 | ENSMUSG0( | false | . | . | . | . | . | . | chr6 | 71987940 | 71992521 | 5.97495 | 4.51658 | 3.83046 |
| chr3 | 75677454 | 75677718 | 815 | Distal Intergenic | 3 | 75647446 | 75655731 | 8286 | 2 | ENSMUST00 | -21724 | RP24-105B2: | ENSMUSG0( | false | . | . | . | . | . | . | . | . | . | . | . | . |
| chr17 | 15373874 | 15374174 | 814 | Promoter (<=1kb) | 17 | 15370609 | 15374072 | 3464 | 2 | ENSMUST00 | 0 | Dll1 | ENSMUSG0( | false | . | . | . | . | . | . | chr17 | 15371856 | 15381968 | 5.90641 | 4.43925 | 3.75812 |
| chr3 | 102995755 | 102996178 | 814 | Promoter (<=1kb) | 3 | 102995795 | 103002194 | 6400 | 1 | ENSMUST00 | 0 | Sike1 | ENSMUSG0( | false | chr3 | 102995208 | 102996541 | 32.98514 | 106.12805 | 102.40588 | . | . | . | . | . | . |
| chr8 | 111375088 | 111375552 | 814 | Intron (ENSMUST00000038475.8 | 8 | 111345135 | 111366477 | 21343 | 2 | ENSMUST00 | -8612 | Fa2h | ENSMUSG0( | true | . | . | . | . | . | . | chr8 | 111356598 | 111381726 | 5.44823 | 4.21871 | 3.39859 |
| chr2 | 181287551 | 181288081 | 814 | Promoter (<=1kb) | 2 | 181251449 | 181288035 | 36587 | 2 | ENSMUST00 | 0 | Gmeb2 | ENSMUSG0( | false | chr2 | 181287155 | 181288638 | 28.91859 | 89.0447 | 85.71584 | . | . | . | . | . | . |
| chr14 | 69661369 | 69661677 | 814 | Intron (ENSMUST00000022660.8 | 14 | 69637669 | 69644752 | 7084 | 2 | ENSMUST00 | -16618 | Gm27177 | ENSMUSG0( | false | . | . | . | . | . | . | chr14 | 69659231 | 69662321 | 3.60505 | 3.41623 | 1.84578 |
| chr4 | 134744269 | 134744681 | 814 | Promoter (<=1kb) | 4 | 134741554 | 134743421 | 1868 | 2 | ENSMUST00 | -849 | RP23-132E8. | ENSMUSG0( | false | . | . | . | . | . | . | chr4 | 134743683 | 134746358 | 5.76718 | 4.35803 | 3.67418 |
| chr15 | 82987709 | 82987988 | 814 | Promoter (<=1kb) | 15 | 82982725 | 82987830 | 5106 | 2 | ENSMUST00 | 0 | RP23-223M2 | ENSMUSG0( | false | chr15 | 82987124 | 82987963 | 11.49921 | 31.27799 | 28.79692 | chr15 | 82986791 | 82989401 | 4.42822 | 3.77405 | 2.54648 |
| chr17 | 31855790 | 31856173 | 813 | Promoter (<=1kb) | 17 | 31844248 | 31855792 | 11545 | 2 | ENSMUST00 | 0 | Sik1 | ENSMUSG0( | false | chr17 | 31856014 | 31856714 | 15.62844 | 37.27362 | 34.7164 | chr17 | 31856138 | 31858172 | 6.8028 | 4.9541 | 4.47342 |
| chr5 | 140268683 | 140268979 | 813 | Intron (ENSMUST00000110829.2 | 5 | 140166983 | 140262565 | 95583 | 2 | ENSMUST00 | -6119 | Mad1l1 | ENSMUSG0( | false | . | . | . | . | . | . | chr5 | 140267333 | 140269571 | 3.41364 | 3.23748 | 1.74279 |
| chr4 | 43381657 | 43382404 | 813 | Promoter (<=1kb) | 4 | 43381979 | 43427088 | 45110 | 1 | ENSMUST00 | 0 | Rusc2 | ENSMUSG0( | false | chr4 | 43380841 | 43383763 | 37.94029 | 126.39516 | 122.03232 | . | . | . | . | . | . |
| chr4 | 57300654 | 57300975 | 813 | Promoter (<=1kb) | 4 | 57254917 | 57300915 | 45999 | 2 | ENSMUST00 | 0 | Ptpn3 | ENSMUSG0( | false | chr4 | 57299328 | 57302225 | 26.1063 | 98.41757 | 94.88632 | chr4 | 57294789 | 57301188 | 4.63579 | 3.8844 | 2.72174 |
| chr11 | 103144989 | 103145350 | 813 | Distal Intergenic | 11 | 103133865 | 103138279 | 4415 | 1 | ENSMUST00 | 11125 | Hexim2 | ENSMUSG0( | false | . | . | . | . | . | . | chr11 | 103138329 | 103150034 | 6.87245 | 4.95791 | 4.53389 |
| chr5 | 102813307 | 102813642 | 813 | Intron (ENSMUST00000094559.3 | 5 | 102809049 | 102875860 | 66812 | 1 | ENSMUST00 | 4259 | Arhgap24 | ENSMUSG0( | true | . | . | . | . | . | . | chr5 | 102806003 | 102845305 | 4.78306 | 3.97005 | 2.82906 |
| chr7 | 135588863 | 135589245 | 813 | Intron (ENSMUST00000073961.6 | 7 | 135566162 | 135566282 | 121 | 1 | ENSMUST00 | 22702 | Mir1962 | ENSMUSG0( | false | . | . | . | . | . | . | . | . | . | . | . | . |
| chr5 | 125068562 | 125068959 | 812 | Exon (ENSMUST00000111402.3/ | 5 | 125030800 | 125058400 | 27601 | 2 | ENSMUST00 | -10163 | Ncor2 | ENSMUSG0( | false | . | . | . | . | . | . | . | . | . | . | . | . |
| chr11 | 101162481 | 101163104 | 812 | Promoter (2-3kb) | 11 | 101164167 | 101165324 | 1158 | 2 | ENSMUST00 | 2220 | Plekhh3 | ENSMUSG0( | false | . | . | . | . | . | . | chr11 | 101162175 | 101170220 | 6.22712 | 4.64855 | 4.01342 |
| chr9 | 106821640 | 106822038 | 812 | Promoter (<=1kb) | 9 | 106821874 | 106880443 | 58570 | 1 | ENSMUST00 | 0 | Vprbp | ENSMUSG0( | false | chr9 | 106821212 | 106821746 | 19.27906 | 51.80829 | 49.07 | . | . | . | . | . | . |
| chr17 | 12991681 | 12992735 | 812 | Promoter (<=1kb) | 17 | 12966796 | 12992497 | 25702 | 2 | ENSMUST00 | 0 | Wtap | ENSMUSG0( | false | chr17 | 12991036 | 12993295 | 42.74922 | 148.11885 | 142.90198 | . | . | . | . | . | . |
| chr2 | 167774860 | 167775401 | 812 | Promoter (2-3kb) | 2 | 167777467 | 167782569 | 5103 | 1 | ENSMUST00 | -2066 | Gm14321 | ENSMUSG0( | false | . | . | . | . | . | . | chr2 | 167773374 | 167783003 | 5.39085 | 4.1895 | 3.35037 |
| chr13 | 118376608 | 118377086 | 812 | Distal Intergenic | 13 | 118386630 | 118388124 | 1495 | 1 | ENSMUST00 | -9544 | B430218F22 | ENSMUSG0( | false | . | . | . | . | . | . | . | . | . | . | . | . |
| chr19 | 23056891 | 23057176 | 812 | Intron (ENSMUST00000181623.1 | 19 | 23037390 | 23075853 | 38464 | 2 | ENSMUST00 | 18677 | C330002G04 | ENSMUSG0( | false | . | . | . | . | . | . | . | . | . | . | . | . |
| chr2 | 35581540 | 35582186 | 811 | Promoter (<=1kb) | 2 | 35581060 | 35730992 | 149933 | 1 | ENSMUST00 | 481 | Dab2ip | ENSMUSG0( | false | chr2 | 35581533 | 35583251 | 20.35144 | 58.59763 | 55.77069 | chr2 | 35581146 | 35583413 | 4.17986 | 3.50064 | 2.36858 |
| chr4 | 148806864 | 148807221 | 811 | Promoter (2-3kb) | 4 | 148804465 | 148839746 | 35282 | 1 | ENSMUST00 | 2400 | Casz1 | ENSMUSG0( | true | chr4 | 148806885 | 148807657 | 16.34529 | 41.36735 | 38.7591 | chr4 | 148806488 | 148838873 | 5.49162 | 4.28891 | 3.41701 |
| chr5 | 23435001 | 23435713 | 811 | Promoter (<=1kb) | 5 | 23434449 | 23504235 | 69787 | 1 | ENSMUST00 | 553 | Kmt2e | ENSMUSG0( | false | chr5 | 23431644 | 23437822 | 48.25087 | 161.14223 | 155.45619 | . | . | . | . | . | . |
| chr6 | 136875769 | 136876129 | 811 | Promoter (<=1kb) | 6 | 136872436 | 136875805 | 3370 | 2 | ENSMUST00 | 0 | Mgp | ENSMUSG0( | false | . | . | . | . | . | . | chr6 | 136875748 | 136876097 | 4.45131 | 3.68995 | 2.59469 |
| chr11 | 101712773 | 101713150 | 811 | Distal Intergenic | 11 | 101732919 | 101735696 | 2778 | 1 | ENSMUST00 | -19769 | Dhx8 | ENSMUSG0( | false | . | . | . | . | . | . | chr11 | 101709308 | 101714568 | 4.63527 | 3.87211 | 2.71845 |
| chr13 | 75930954 | 75931506 | 811 | Promoter (<=1kb) | 13 | 75929465 | 75930328 | 864 | 2 | ENSMUST00 | -627 | Gm15528 | ENSMUSG0( | false | . | . | . | . | . | . | chr13 | 75930515 | 75932092 | 2.93018 | 2.92151 | 1.36919 |
| chr2 | 167815059 | 167815424 | 811 | Distal Intergenic | 2 | 167827693 | 167833646 | 5954 | 2 | ENSMUST00 | 18222 | 1200007C13I | ENSMUSG0( | false | . | . | . | . | . | . | chr2 | 167814397 | 167820487 | 6.24217 | 4.56491 | 4.01948 |
| chr1 | 134062518 | 134062805 | 810 | Distal Intergenic | 1 | 134075170 | 134079120 | 3951 | 2 | ENSMUST00 | 16315 | Btg2 | ENSMUSG0( | false | . | . | . | . | . | . | chr1 | 134058485 | 134067448 | 6.65638 | 4.82565 | 4.35972 |
| chr13 | 109236035 | 109236368 | 810 | Intron (ENSMUST00000122041.2 | 13 | 109260654 | 109951364 | 690711 | 1 | ENSMUST00 | -24286 | Pde4d | ENSMUSG0( | false | . | . | . | . | . | . | chr13 | 109235306 | 109236548 | 3.88342 | 3.54124 | 2.08296 |
| chr18 | 61556827 | 61557302 | 810 | Promoter (1-2kb) | 18 | 61555689 | 61587531 | 31843 | 1 | ENSMUST00 | 1139 | Csnk1a1 | ENSMUSG0( | true | chr18 | 61555228 | 61560694 | 42.15588 | 144.81987 | 139.73779 | . | . | . | . | . | . |
| chr6 | 31515644 | 31515980 | 810 | 3' UTR | 6 | 31522992 | 31524311 | 1320 | 2 | ENSMUST00 | 8331 | Podxl | ENSMUSG0( | true | . | . | . | . | . | . | chr6 | 31513503 | 31559838 | 6.27295 | 4.62488 | 4.05806 |
| chr14 | 22174873 | 22175232 | 810 | Intron (ENSMUST00000075639.5 | 14 | 22310165 | 22584439 | 274275 | 1 | ENSMUST00 | -134933 | 1700112E06I | ENSMUSG0( | false | . | . | . | . | . | . | chr14 | 22173884 | 22176588 | 5.1117 | 4.17195 | 3.09387 |
| chr7 | 27929309 | 27929656 | 810 | Promoter (<=1kb) | 7 | 27907392 | 27929430 | 22039 | 2 | ENSMUST00 | 0 | 1700049G17 | ENSMUSG0( | false | chr7 | 27928557 | 27929955 | 26.40393 | 78.95839 | 75.82329 | chr7 | 27926913 | 27930081 | 3.60473 | 3.09091 | 1.89332 |
| chr14 | 25795596 | 25795941 | 810 | Distal Intergenic | 14 | 25809635 | 25809881 | 247 | 2 | ENSMUST00 | 13940 | Gm27185 | ENSMUSG0( | false | . | . | . | . | . | . | . | . | . | . | . | . |
| chr7 | 66450448 | 66450806 | 809 | Distal Intergenic | 7 | 66390892 | 66427517 | 36626 | 2 | ENSMUST00 | -22932 | Aldh1a3 | ENSMUSG0( | false | . | . | . | . | . | . | chr7 | 66438246 | 66451395 | 6.06663 | 4.59192 | 3.88434 |
| chrX | 36796168 | 36796496 | 809 | Promoter (<=1kb) | 20 | 36795654 | 36797643 | 1990 | 1 | ENSMUST00 | 515 | Slc25a5 | ENSMUSG0( | false | chrX | 36795379 | 36796755 | 22.51707 | 59.83231 | 56.98677 | . | . | . | . | . | . |
| chr14 | 57571412 | 57571767 | 809 | Promoter (<=1kb) | 14 | 57549598 | 57571569 | 21972 | 2 | ENSMUST00 | 0 | N6amt2 | ENSMUSG0( | false | chr14 | 57570910 | 57572174 | 31.70772 | 93.8754 | 90.44607 | . | . | . | . | . | . |
| chr1 | 120258744 | 120259104 | 809 | Intron (ENSMUST00000140490.2 | 1 | 120190757 | 120265255 | 74499 | 2 | ENSMUST00 | 6151 | Steap3 | ENSMUSG0( | false | . | . | . | . | . | . | chr1 | 120244209 | 120264315 | 5.23032 | 4.13237 | 3.21241 |
| chr7 | 127210482 | 127210776 | 809 | Promoter (<=1kb) | 7 | 127211608 | 127214298 | 2691 | 1 | ENSMUST00 | -832 | Mylpf | ENSMUSG0( | false | . | . | . | . | . | . | chr7 | 127209269 | 127215944 | 5.73283 | 4.38529 | 3.63334 |
| chr7 | 68687456 | 68687766 | 809 | Distal Intergenic | 7 | 68741511 | 68744906 | 3396 | 2 | ENSMUST00 | 57140 | Arrdc4 | ENSMUSG0( | false | . | . | . | . | . | . | chr7 | 68685426 | 68689298 | 3.84302 | 3.34724 | 2.06938 |
| chr18 | 44441121 | 44441445 | 808 | Intron (ENSMUST00000164666.2 | 18 | 44380500 | 44424969 | 44470 | 1 | ENSMUST00 | 60622 | Dcp2 | ENSMUSG0( | false | . | . | . | . | . | . | chr18 | 44434846 | 44445255 | 6.64362 | 4.65907 | 4.40138 |
| chr5 | 31571035 | 31571426 | 808 | Distal Intergenic | 5 | 31596935 | 31654861 | 57927 | 1 | ENSMUST00 | -25509 | Mrpl33 | ENSMUSG0( | false | . | . | . | . | . | . | chr5 | 31570156 | 31572728 | 5.66703 | 4.47068 | 3.54972 |
| chr5 | 142819438 | 142820073 | 808 | Promoter (1-2kb) | 5 | 142754789 | 142817662 | 62874 | 2 | ENSMUST00 | -1777 | Tnrc18 | ENSMUSG0( | false | chr5 | 142817908 | 142824082 | 29.69128 | 86.53342 | 83.26198 | . | . | . | . | . | . |
| chr3 | 100838996 | 100839323 | 808 | Intron (ENSMUST00000054791.7 | 3 | 100825459 | 100895679 | 70221 | 1 | ENSMUST00 | 13538 | Vtcn1 | ENSMUSG0( | false | . | . | . | . | . | . | chr3 | 100833487 | 100844647 | 4.69269 | 3.92412 | 2.78415 |
| chr7 | 114101442 | 114101789 | 808 | Intron (ENSMUST00000069449.5 | 7 | 114046785 | 114117781 | 70997 | 2 | ENSMUST00 | 15992 | Rras2 | ENSMUSG0( | true | . | . | . | . | . | . | chr7 | 114058507 | 114116129 | 5.64505 | 4.33683 | 3.54844 |
| chr16 | 15084725 | 15085220 | 808 | Distal Intergenic | 16 | 15216859 | 15217328 | 470 | 2 | ENSMUST00 | 132108 | Gm7731 | ENSMUSG0( | false | chr16 | 15084586 | 15085367 | 10.95755 | 25.35628 | 22.94834 | . | . | . | . | . | . |
| chr1 | 155086959 | 155087247 | 808 | Distal Intergenic | 1 | 155063447 | 155083266 | 19820 | 2 | ENSMUST00 | -3694 | Gm9530 | ENSMUSG0( | false | . | . | . | . | . | . | . | . | . | . | . | . |
| chr5 | 64336868 | 64337476 | 807 | Intron (ENSMUST00000043893.7 | 5 | 64349945 | 64350589 | 645 | 1 | ENSMUST00 | -12469 | Tbc1d1 | ENSMUSG0( | false | . | . | . | . | . | . | chr5 | 64333951 | 64340633 | 6.27533 | 4.60721 | 4.05113 |
| chr6 | 145865276 | 145865906 | 807 | Promoter (<=1kb) | 6 | 145858243 | 145865558 | 7316 | 2 | ENSMUST00 | 0 | Bhlhe41 | ENSMUSG0( | false | chr6 | 145864843 | 145866819 | 26.13664 | 77.56537 | 74.45414 | chr6 | 145861038 | 145867407 | 3.647 | 3.29023 | 1.93146 |
| chr5 | 34183220 | 34183494 | 807 | Intron (ENSMUST00000042701.7 | 5 | 34173887 | 34187710 | 13824 | 2 | ENSMUST00 | 4216 | Mxd4 | ENSMUSG0( | false | . | . | . | . | . | . | chr5 | 34179706 | 34186740 | 6.22716 | 4.44968 | 4.05284 |
| chr15 | 75730970 | 75731242 | 807 | Distal Intergenic | 15 | 75746843 | 75747922 | 1080 | 2 | ENSMUST00 | 16680 | Mafa | ENSMUSG0( | false | . | . | . | . | . | . | chr15 | 75729486 | 75733932 | 5.87935 | 4.38557 | 3.73456 |
| chr1 | 55570292 | 55570555 | 807 | Intron (ENSMUST00000042986.9 | 1 | 55443733 | 55445064 | 1332 | 2 | ENSMUST00 | -125229 | Gm6644 | ENSMUSG0( | false | . | . | . | . | . | . | chr1 | 55565857 | 55577601 | 5.51018 | 4.28641 | 3.43047 |
| chr6 | 144103907 | 144104176 | 807 | Intron (ENSMUST00000077160.6 | 6 | 144021908 | 144023778 | 1871 | 1 | ENSMUST00 | 82000 | Sox5os1 | ENSMUSG0( | false | . | . | . | . | . | . | chr6 | 144103132 | 144107582 | 5.73038 | 4.43689 | 3.60871 |
| chr5 | 36292375 | 36292655 | 807 | Intron (ENSMUST00000037370.8 | 5 | 36301570 | 36301628 | 59 | 1 | ENSMUST00 | -8915 | Mir7689 | ENSMUSG0( | false | . | . | . | . | . | . | chr5 | 36292434 | 36293622 | 4.12989 | 3.38929 | 2.32952 |
| chr13 | 55171535 | 55172236 | 806 | Distal Intergenic | 13 | 55156117 | 55156904 | 788 | 1 | ENSMUST00 | 15419 | Fgfr4 | ENSMUSG0( | false | . | . | . | . | . | . | chr13 | 55161802 | 55178663 | 6.64938 | 4.78575 | 4.37276 |
| chr15 | 84356644 | 84357199 | 806 | Distal Intergenic | 15 | 84326119 | 84342978 | 16860 | 1 | ENSMUST00 | 30526 | Parvg | ENSMUSG0( | false | . | . | . | . | . | . | chr15 | 84355589 | 84357349 | 4.47573 | 3.78973 | 2.58407 |
| chr4 | 134766610 | 134767471 | 806 | Promoter (<=1kb) | 4 | 134744735 | 134768024 | 23290 | 2 | ENSMUST00 | 553 | Ldlrap1 | ENSMUSG0( | false | chr4 | 134766368 | 134768617 | 40.29262 | 133.77719 | 129.14264 | chr4 | 134750748 | 134766877 | 7.637 | 5.15315 | 5.18044 |
| chr5 | 135934207 | 135934751 | 806 | Promoter (<=1kb) | 5 | 135908409 | 135934599 | 26191 | 2 | ENSMUST00 | 0 | Ywhag | ENSMUSG0( | false | chr5 | 135933350 | 135935455 | 30.67871 | 107.10089 | 103.34502 | . | . | . | . | . | . |
| chr5 | 93118563 | 93118882 | 806 | Intron (ENSMUST00000074733.7 | 5 | 93093457 | 93174958 | 81502 | 1 | ENSMUST00 | 25107 | Sep-11 | ENSMUSG0( | false | . | . | . | . | . | . | chr5 | 93095018 | 93154711 | 6.59326 | 4.78158 | 4.31782 |
| chr17 | 84291248 | 84291646 | 806 | Intron (ENSMUST00000047524.8 | 17 | 84287641 | 84287747 | 107 | 2 | ENSMUST00 | -3502 | Gm24492 | ENSMUSG0( | false | . | . | . | . | . | . | chr17 | 84287517 | 84294373 | 4.77469 | 3.9692 | 2.81673 |
| chr2 | 53382348 | 53382683 | 806 | Distal Intergenic | 2 | 53397509 | 53398487 | 979 | 2 | ENSMUST00 | 15804 | Gm13501 | ENSMUSG0( | false | . | . | . | . | . | . | chr2 | 53379134 | 53382974 | 4.39508 | 3.80647 | 2.50067 |
| chr10 | 61381838 | 61382183 | 805 | Promoter (1-2kb) | 10 | 61319657 | 61383523 | 63867 | 2 | ENSMUST00 | 1340 | Pald1 | ENSMUSG0( | false | chr10 | 61382020 | 61383994 | 31.64465 | 94.75051 | 91.30649 | chr10 | 61368996 | 61384383 | 5.02563 | 4.04497 | 3.06054 |
| chr14 | 78492639 | 78492979 | 805 | 3' UTR | 14 | 78492246 | 78536808 | 44563 | 2 | ENSMUST00 | 43829 | Akap11 | ENSMUSG0( | true | . | . | . | . | . | . | chr14 | 78492417 | 78494102 | 3.83027 | 3.42716 | 2.06659 |
| chrX | 48623341 | 48623688 | 805 | Promoter (<=1kb) | 20 | 48623413 | 48631036 | 7624 | 1 | ENSMUST00 | 0 | Slc25a14 | ENSMUSG0( | false | chrX | 48623115 | 48624262 | 21.13848 | 54.99947 | 52.22001 | . | . | . | . | . | . |
| chr8 | 60632486 | 60632951 | 805 | Promoter (<=1kb) | 8 | 60632827 | 60674933 | 42107 | 1 | ENSMUST00 | 0 | Mfap3l | ENSMUSG0( | false | chr8 | 60631383 | 60633721 | 32.66734 | 113.81544 | 109.86345 | chr8 | 60631191 | 60633719 | 6.89015 | 4.87244 | 4.60078 |
| chr15 | 81858179 | 81858657 | 805 | Promoter (<=1kb) | 15 | 81848270 | 81858326 | 10057 | 2 | ENSMUST00 | 0 | Tob2 | ENSMUSG0( | false | chr15 | 81855855 | 81859985 | 34.3275 | 112.61823 | 108.70669 | chr15 | 81858434 | 81861484 | 5.30414 | 4.29664 | 3.25085 |
| chr15 | 37791239 | 37792219 | 805 | Promoter (<=1kb) | 15 | 37366175 | 37792000 | 425826 | 2 | ENSMUST00 | 0 | Ncald | ENSMUSG0( | true | chr15 | 37791004 | 37792729 | 32.13701 | 98.5793 | 95.04197 | chr15 | 37786264 | 37791494 | 4.354 | 3.70574 | 2.49785 |
| chr8 | 26779012 | 26779456 | 805 | Distal Intergenic | 8 | 26814593 | 26824423 | 9831 | 2 | ENSMUST00 | 44967 | RP24-502D8. | ENSMUSG0( | false | . | . | . | . | . | . | chr8 | 26778443 | 26781090 | 6.95771 | 5.07118 | 4.60599 |
| chr8 | 4252952 | 4253451 | 804 | Promoter (<=1kb) | 8 | 4253102 | 4256219 | 3118 | 1 | ENSMUST00 | 0 | Snapc2 | ENSMUSG0( | false | chr8 | 4252660 | 4253872 | 19.7569 | 81.42724 | 78.25113 | chr8 | 4252325 | 4254377 | 4.74068 | 3.96865 | 2.79866 |
| chr16 | 17646356 | 17646748 | 804 | Promoter (<=1kb) | 16 | 17646470 | 17649415 | 2946 | 1 | ENSMUST00 | 0 | Ccdc74a | ENSMUSG0( | false | chr16 | 17646147 | 17646967 | 20.61663 | 56.56642 | 53.76645 | chr16 | 17645989 | 17647531 | 4.65599 | 3.9568 | 2.7228 |
| chr12 | 112515857 | 112516128 | 804 | Distal Intergenic | 12 | 112500184 | 112511160 | 10977 | 2 | ENSMUST00 | -4698 | Tmem179 | ENSMUSG0( | false | . | . | . | . | . | . | chr12 | 112515496 | 112518169 | 5.50197 | 4.36858 | 3.41411 |
| chr9 | 44499752 | 44500138 | 804 | Promoter (<=1kb) | 9 | 44499136 | 44510388 | 11253 | 1 | ENSMUST00 | 617 | Bcl9l | ENSMUSG0( | false | . | . | . | . | . | . | chr9 | 44483700 | 44502510 | 6.77697 | 4.85012 | 4.46719 |
| chr6 | 85587463 | 85588344 | 804 | Promoter (<=1kb) | 6 | 85587531 | 85702753 | 115223 | 1 | ENSMUST00 | 0 | Alms1 | ENSMUSG0( | false | chr6 | 85587283 | 85588437 | 19.45739 | 70.46334 | 67.46697 | chr6 | 85588189 | 85588870 | 2.98155 | 3.0572 | 1.38341 |
| chr6 | 88342674 | 88342969 | 804 | Intron (ENSMUST00000165242.1 | 6 | 88405977 | 88406301 | 325 | 1 | ENSMUST00 | -63008 | Gm25178 | ENSMUSG0( | false | . | . | . | . | . | . | chr6 | 88339028 | 88344718 | 7.43935 | 5.14636 | 5.02585 |
| chr16 | 34903941 | 34904235 | 804 | Intron (ENSMUST00000023538.8 | 16 | 34935854 | 34939329 | 3476 | 2 | ENSMUST00 | 35094 | RP23-80O10 | ENSMUSG0( | false | . | . | . | . | . | . | chr16 | 34902265 | 34904558 | 5.06477 | 3.91275 | 3.09683 |
| chr16 | 17605620 | 17605928 | 803 | Distal Intergenic | 16 | 17619354 | 17644828 | 25475 | 1 | ENSMUST00 | -13426 | Smpd4 | ENSMUSG0( | false | . | . | . | . | . | . | chr16 | 17605230 | 17607402 | 3.73914 | 3.35426 | 2.00206 |
| chr14 | 54912649 | 54913266 | 803 | Promoter (<=1kb) | 14 | 54906727 | 54913132 | 6406 | 2 | ENSMUST00 | 0 | Slc22a17 | ENSMUSG0( | false | chr14 | 54912185 | 54913452 | 34.02479 | 107.94543 | 104.17527 | chr14 | 54912066 | 54913778 | 3.93701 | 3.61988 | 2.14945 |
| chr3 | 152193687 | 152194044 | 803 | Promoter (<=1kb) | 3 | 152183955 | 152193803 | 9849 | 2 | ENSMUST00 | 0 | Dnajb4 | ENSMUSG0( | false | chr3 | 152191653 | 152193826 | 25.56571 | 75.64988 | 72.57267 | . | . | . | . | . | . |
| chr16 | 37868104 | 37868802 | 803 | Promoter (<=1kb) | 16 | 37868389 | 37888858 | 20470 | 1 | ENSMUST00 | 0 | Lrrc58 | ENSMUSG0( | false | chr16 | 37867771 | 37870669 | 34.00884 | 130.01486 | 125.52227 | . | . | . | . | . | . |
| chr11 | 32340867 | 32341268 | 803 | Distal Intergenic | 11 | 32347820 | 32428173 | 80354 | 1 | ENSMUST00 | -6552 | Sh3pxd2b | ENSMUSG0( | false | . | . | . | . | . | . | chr11 | 32340628 | 32341590 | 4.79443 | 4.02923 | 2.87427 |
| chr1 | 39749583 | 39749957 | 803 | Distal Intergenic | 1 | 39665306 | 39720997 | 55692 | 2 | ENSMUST00 | -28587 | Rfx8 | ENSMUSG0( | false | . | . | . | . | . | . | chr1 | 39746176 | 39750447 | 5.34297 | 4.23417 | 3.30515 |
| chr10 | 28167134 | 28167504 | 803 | Intron (ENSMUST00000166468.1 | 10 | 28214039 | 28214121 | 83 | 2 | ENSMUST00 | 46617 | Gm22370 | ENSMUSG0( | false | . | . | . | . | . | . | chr10 | 28166663 | 28168217 | 3.68083 | 3.29696 | 1.94915 |
| chr10 | 108535914 | 108536200 | 802 | Intron (ENSMUST00000105276.2 | 10 | 108627286 | 108636964 | 9679 | 2 | ENSMUST00 | 100764 | Syt1 | ENSMUSG0( | false | . | . | . | . | . | . | chr10 | 108534274 | 108538193 | 4.49458 | 3.87119 | 2.58838 |
| chr14 | 77004763 | 77005083 | 802 | Distal Intergenic | 14 | 77034950 | 77036081 | 1132 | 2 | ENSMUST00 | 30998 | Lacc1 | ENSMUSG0( | false | . | . | . | . | . | . | chr14 | 77001969 | 77008459 | 4.51571 | 3.8977 | 2.59777 |
| chr8 | 41026787 | 41027176 | 802 | Intron (ENSMUST00000059115.7 | 8 | 41019523 | 41022902 | 3380 | 2 | ENSMUST00 | -3886 | Mtus1 | ENSMUSG0( | false | . | . | . | . | . | . | chr8 | 41020996 | 41040121 | 5.92676 | 4.49745 | 3.7648 |
| chr13 | 60007810 | 60008125 | 802 | Distal Intergenic | 13 | 60174405 | 60177365 | 2961 | 2 | ENSMUST00 | 169240 | Gas1 | ENSMUSG0( | false | . | . | . | . | . | . | chr13 | 60006912 | 60013507 | 4.90407 | 3.99626 | 2.94044 |
| chr16 | 11253837 | 11254572 | 802 | Promoter (<=1kb) | 16 | 11219292 | 11254270 | 34979 | 2 | ENSMUST00 | 0 | Gspt1 | ENSMUSG0( | false | chr16 | 11252796 | 11255094 | 35.67123 | 127.73884 | 123.33056 | . | . | . | . | . | . |
| chr17 | 4993986 | 4994733 | 802 | Promoter (<=1kb) | 17 | 4994332 | 5097733 | 103402 | 1 | ENSMUST00 | 0 | Arid1b | ENSMUSG0( | false | chr17 | 4993664 | 4997887 | 36.69013 | 115.69205 | 111.68276 | . | . | . | . | . | . |
| chr8 | 26977901 | 26978314 | 802 | Promoter (<=1kb) | 8 | 26977336 | 26981461 | 4126 | 1 | ENSMUST00 | 566 | Zfp703 | ENSMUSG0( | false | chr8 | 26976009 | 26979489 | 27.41053 | 90.23362 | 86.88379 | . | . | . | . | . | . |
| chr11 | 67922051 | 67922455 | 801 | Promoter (<=1kb) | 11 | 67854523 | 67922153 | 67631 | 2 | ENSMUST00 | 0 | Usp43 | ENSMUSG0( | false | chr11 | 67921383 | 67922888 | 33.45771 | 103.89173 | 100.22243 | chr11 | 67921383 | 67923169 | 5.48178 | 4.34015 | 3.44686 |
| chr16 | 4472762 | 4473059 | 801 | Distal Intergenic | 16 | 4493603 | 4523063 | 29461 | 2 | ENSMUST00 | 50004 | Srl | ENSMUSG0( | false | . | . | . | . | . | . | chr16 | 4472553 | 4475716 | 3.65754 | 3.27338 | 1.94367 |
| chr15 | 98533810 | 98534372 | 801 | Promoter (<=1kb) | 15 | 98518821 | 98534244 | 15424 | 2 | ENSMUST00 | 0 | Kansl2 | ENSMUSG0( | false | chr15 | 98532844 | 98534818 | 20.14837 | 78.81087 | 75.68276 | . | . | . | . | . | . |
| chr9 | 72024056 | 72024384 | 801 | Intron (ENSMUST00000183404.2 | 9 | 71882732 | 72000476 | 117745 | 2 | ENSMUST00 | -23581 | Tcf12 | ENSMUSG0( | false | . | . | . | . | . | . | chr9 | 72022847 | 72025920 | 4.29451 | 3.7731 | 2.41427 |
| chr2 | 33575260 | 33575722 | 801 | Intron (ENSMUST00000041730.5 | 2 | 33567156 | 33632283 | 65128 | 2 | ENSMUST00 | 56561 | Lmx1b | ENSMUSG0( | true | . | . | . | . | . | . | chr2 | 33565260 | 33638297 | 6.78481 | 4.81457 | 4.48235 |
| chr14 | 22298397 | 22298754 | 801 | Intron (ENSMUST00000075639.5 | 14 | 22310165 | 22584439 | 274275 | 1 | ENSMUST00 | -11411 | 1700112E06I | ENSMUSG0( | false | . | . | . | . | . | . | chr14 | 22296640 | 22299669 | 4.94954 | 3.96681 | 3.01622 |
| chr3 | 95689778 | 95690231 | 800 | Promoter (1-2kb) | 3 | 95676201 | 95687917 | 11717 | 2 | ENSMUST00 | -1862 | Adamtsl4 | ENSMUSG0( | false | . | . | . | . | . | . | chr3 | 95685856 | 95702888 | 5.06878 | 4.08183 | 3.0659 |
| chr10 | 53751006 | 53751385 | 800 | Promoter (<=1kb) | 10 | 53696997 | 53750909 | 53913 | 2 | ENSMUST00 | -98 | Fam184a | ENSMUSG0( | true | chr10 | 53749886 | 53751730 | 32.25957 | 103.83389 | 100.16508 | chr10 | 53751243 | 53753430 | 4.93021 | 4.02583 | 2.95304 |
| chr18 | 5809322 | 5809671 | 800 | Distal Intergenic | 18 | 5731081 | 5772804 | 41724 | 1 | ENSMUST00 | 78242 | Zeb1 | ENSMUSG0( | false | . | . | . | . | . | . | chr18 | 5807684 | 5810073 | 5.47071 | 4.16293 | 3.42936 |
| chr6 | 88592381 | 88592737 | 800 | Promoter (1-2kb) | 6 | 88586528 | 88590659 | 4132 | 2 | ENSMUST00 | -1723 | Kbtbd12 | ENSMUSG0( | false | . | . | . | . | . | . | chr6 | 88592132 | 88592982 | 3.64274 | 3.47374 | 1.92012 |
| chr17 | 68165488 | 68165749 | 800 | Distal Intergenic | 17 | 68273797 | 68457858 | 184062 | 1 | ENSMUST00 | -108048 | L3mbtl4 | ENSMUSG0( | false | . | . | . | . | . | . | chr17 | 68164538 | 68166576 | 6.23696 | 4.57872 | 4.02364 |
| chr2 | 10372362 | 10372696 | 800 | Promoter (<=1kb) | 2 | 10372436 | 10439137 | 66702 | 1 | ENSMUST00 | 0 | Sfmbt2 | ENSMUSG0( | false | chr2 | 10372318 | 10375667 | 25.56571 | 75.64988 | 72.57267 | chr2 | 10371621 | 10375550 | 3.53833 | 3.3368 | 1.80576 |
| chr19 | 28580323 | 28580745 | 800 | Promoter (2-3kb) | 19 | 28580533 | 28583294 | 2762 | 2 | ENSMUST00 | 2549 | 4933413C19I | ENSMUSG0( | false | . | . | . | . | . | . | . | . | . | . | . | . |
| chr14 | 65261729 | 65262626 | 799 | Promoter (<=1kb) | 14 | 65201026 | 65262463 | 61438 | 2 | ENSMUST00 | 0 | Fzd3 | ENSMUSG0( | false | chr14 | 65261170 | 65263071 | 32.68394 | 98.52523 | 94.99333 | chr14 | 65259845 | 65263295 | 3.94156 | 3.56918 | 2.12897 |
| chr11 | 76449534 | 76450168 | 799 | Intron (ENSMUST00000094012.5 | 11 | 76423048 | 76437220 | 14173 | 2 | ENSMUST00 | -12315 | Abr | ENSMUSG0( | false | . | . | . | . | . | . | chr11 | 76424298 | 76452713 | 6.90719 | 4.90323 | 4.58627 |
| chr15 | 97904503 | 97904894 | 799 | Intron (ENSMUST00000023119.9 | 15 | 97854425 | 97908310 | 53886 | 2 | ENSMUST00 | 3416 | Vdr | ENSMUSG0( | false | . | . | . | . | . | . | chr15 | 97902462 | 97908754 | 5.09257 | 3.97616 | 3.09637 |
| chr19 | 43728968 | 43729256 | 799 | Promoter (2-3kb) | 19 | 43726367 | 43728136 | 1770 | 1 | ENSMUST00 | 2602 | Entpd7 | ENSMUSG0( | false | . | . | . | . | . | . | chr19 | 43728123 | 43730181 | 4.39289 | 3.80516 | 2.51145 |
| chr7 | 64392201 | 64392816 | 799 | Promoter (<=1kb) | 7 | 64376576 | 64392268 | 15693 | 2 | ENSMUST00 | 0 | Mphosph10 | ENSMUSG0( | false | chr7 | 64391514 | 64393386 | 19.35304 | 74.51352 | 71.45322 | . | . | . | . | . | . |
| chr15 | 99018009 | 99018274 | 799 | Distal Intergenic | 15 | 99029891 | 99038105 | 8215 | 1 | ENSMUST00 | -11617 | Tuba1c | ENSMUSG0( | false | . | . | . | . | . | . | chr15 | 99017380 | 99019220 | 3.82524 | 3.54224 | 2.07934 |
| chr8 | 25225652 | 25226065 | 799 | Distal Intergenic | 8 | 25154552 | 25201449 | 46898 | 2 | ENSMUST00 | -24204 | Tacc1 | ENSMUSG0( | false | . | . | . | . | . | . | chr8 | 25224778 | 25226471 | 3.8632 | 3.58325 | 2.05936 |
| chr5 | 148632494 | 148632908 | 798 | Distal Intergenic | 5 | 148505142 | 148552789 | 47648 | 2 | ENSMUST00 | -79706 | Ubl3 | ENSMUSG0( | false | . | . | . | . | . | . | chr5 | 148631497 | 148635604 | 4.97634 | 3.9319 | 3.02677 |
| chrX | 152001661 | 152001963 | 798 | Promoter (<=1kb) | 20 | 152001845 | 152004442 | 2598 | 1 | ENSMUST00 | 0 | Hsd17b10 | ENSMUSG0( | false | chrX | 152001301 | 152001689 | 7.12487 | 12.70834 | 10.47977 | . | . | . | . | . | . |

| | | | | | | | | | | | | | | | | | | | | | | | | | | |
|---|---|---|---|---|---|---|---|---|---|---|---|---|---|---|---|---|---|---|---|---|---|---|---|---|---|---|
| chr14 | 7900508 | 7901047 | 798 | Intron (ENSMUST00000052678.8 | 14 | 7975872 | 7977222 | 1351 | 2 | ENSMUST00 | 76175 | Dnase1l3 | ENSMUSG0C | false | . | . | . | . | . | . | chr14 | 7899745 | 7902015 | 4.92096 | 4.03027 | 2.98767 |
| chr6 | 91754190 | 91754558 | 798 | Intron (ENSMUST00000032185.7 | 6 | 91784730 | 91796244 | 11515 | 2 | ENSMUST00 | 41686 | Grip2 | ENSMUSG0C | false | . | . | . | . | . | . | chr6 | 91751267 | 91755238 | 4.2713 | 3.67063 | 2.44844 |
| chr8 | 40511288 | 40512119 | 798 | Promoter (<=1kb) | 8 | 40511783 | 40551134 | 39352 | 1 | ENSMUST00 | 0 | Vps37a | ENSMUSG0C | false | chr8 | 40510828 | 40512559 | 43.03881 | 155.46181 | 149.96747 | . | . | . | . | . | . |
| chr7 | 127886077 | 127886324 | 798 | 3' UTR | 7 | 127885841 | 127890970 | 5130 | 2 | ENSMUST00 | 4646 | Prss53 | ENSMUSG0C | false | . | . | . | . | . | . | chr7 | 127880545 | 127891013 | 5.86469 | 4.44372 | 3.71216 |
| chr4 | 100170829 | 100171088 | 798 | Intron (ENSMUST00000039630.5 | 4 | 100119621 | 100120841 | 1221 | 1 | ENSMUST00 | 51209 | Gm12701 | ENSMUSG0C | false | . | . | . | . | . | . | chr4 | 100165938 | 100174701 | 4.50598 | 3.80894 | 2.61922 |
| chr9 | 110684050 | 110684330 | 797 | Intron (ENSMUST00000019803.4 | 9 | 110656503 | 110711590 | 55088 | 1 | ENSMUST00 | 27548 | Ccdc12 | ENSMUSG0C | false | . | . | . | . | . | . | chr9 | 110660066 | 110688098 | 6.72087 | 4.76887 | 4.43232 |
| chr13 | 44860295 | 44860679 | 797 | Intron (ENSMUST00000173704.2 | 13 | 44840783 | 44896481 | 55699 | 1 | ENSMUST00 | 19513 | Jarid2 | ENSMUSG0C | false | . | . | . | . | . | . | chr13 | 44854562 | 44868955 | 4.49623 | 3.72677 | 2.61635 |
| chr5 | 114773189 | 114773613 | 797 | Promoter (<=1kb) | 5 | 114727407 | 114773495 | 46089 | 2 | ENSMUST00 | 0 | Git2 | ENSMUSG0C | false | chr5 | 114772533 | 114775921 | 32.0113 | 129.32326 | 124.86235 | . | . | . | . | . | . |
| chr8 | 40797410 | 40797668 | 797 | Distal Intergenic | 8 | 40793273 | 40797303 | 4031 | 1 | ENSMUST00 | 4138 | Adam20 | ENSMUSG0C | false | . | . | . | . | . | . | . | . | . | . | . | . |
| chr14 | 55590834 | 55591737 | 797 | Promoter (<=1kb) | 14 | 55591708 | 55603693 | 11986 | 1 | ENSMUST00 | 0 | Rnf31 | ENSMUSG0C | false | chr14 | 55590291 | 55592805 | 29.54372 | 110.04967 | 106.21311 | . | . | . | . | . | . |
| chr7 | 140675908 | 140676171 | 797 | Distal Intergenic | 7 | 140667289 | 140668248 | 960 | 1 | ENSMUST00 | 8620 | Olfr539 | ENSMUSG0C | false | . | . | . | . | . | . | chr7 | 140676056 | 140676576 | 4.71667 | 3.3688 | 2.81175 |
| chr12 | 88812186 | 88812489 | 797 | Intron (ENSMUST00000167734.2 | 12 | 88795439 | 88853243 | 57805 | 1 | ENSMUST00 | 16748 | Nrxn3 | ENSMUSG0C | false | . | . | . | . | . | . | . | . | . | . | . | . |
| chr3 | 96027908 | 96028188 | 796 | Intron (ENSMUST00000015891.5 | 3 | 96056887 | 96057719 | 833 | 2 | ENSMUST00 | 29531 | Vps45 | ENSMUSG0C | false | . | . | . | . | . | . | chr3 | 96025948 | 96030723 | 4.78459 | 3.92715 | 2.82799 |
| chr14 | 54894018 | 54894689 | 796 | Promoter (<=1kb) | 14 | 54894148 | 54898156 | 4009 | 1 | ENSMUST00 | 0 | Pabpn1 | ENSMUSG0C | false | chr14 | 54894103 | 54895521 | 33.45073 | 115.2328 | 111.2408 | . | . | . | . | . | . |
| chr16 | 14178567 | 14178908 | 796 | Intron (ENSMUST00000023359.7 | 16 | 14175127 | 14188397 | 13271 | 1 | ENSMUST00 | 3441 | Nde1 | ENSMUSG0C | false | . | . | . | . | . | . | chr16 | 14174692 | 14185475 | 6.74955 | 4.90863 | 4.43818 |
| chr9 | 106438154 | 106438567 | 796 | Promoter (<=1kb) | 9 | 106432981 | 106438319 | 5339 | 2 | ENSMUST00 | 0 | Acy1 | ENSMUSG0C | false | chr9 | 106437894 | 106438939 | 15.01947 | 39.77812 | 37.19162 | chr9 | 106437448 | 106439190 | 4.99215 | 4.01961 | 3.00201 |
| chr15 | 97257212 | 97257667 | 796 | Intron (ENSMUST00000059433.6 | 15 | 97244074 | 97247287 | 3214 | 2 | ENSMUST00 | -9926 | Amigo2 | ENSMUSG0C | false | . | . | . | . | . | . | chr15 | 97250601 | 97263586 | 5.27919 | 4.19216 | 3.24693 |
| chr8 | 126746258 | 126746704 | 796 | Distal Intergenic | 8 | 126588296 | 126593986 | 5691 | 2 | ENSMUST00 | -152273 | Irf2bp2 | ENSMUSG0C | false | . | . | . | . | . | . | chr8 | 126744019 | 126748224 | 5.83162 | 4.33447 | 3.7311 |
| chr5 | 123109857 | 123110217 | 796 | Intron (ENSMUST00000161660.2 | 5 | 123099901 | 123115819 | 15919 | 1 | ENSMUST00 | 9957 | Tmem120b | ENSMUSG0C | false | . | . | . | . | . | . | chr5 | 123107529 | 123110953 | 7.96167 | 5.4776 | 5.41543 |
| chr17 | 14960987 | 14961904 | 795 | Promoter (<=1kb) | 17 | 14945009 | 14961273 | 16265 | 2 | ENSMUST00 | 0 | Phf10 | ENSMUSG0C | false | chr17 | 14960475 | 14962497 | 54.36761 | 193.18201 | 186.79927 | chr17 | 14961763 | 14962246 | 4.5986 | 3.66 | 2.70804 |
| chr5 | 31630108 | 31630421 | 795 | Intron (ENSMUST00000142345.2 | 5 | 31624439 | 31640538 | 16100 | 2 | ENSMUST00 | 10117 | Rbks | ENSMUSG0C | false | . | . | . | . | . | . | chr5 | 31627113 | 31634739 | 5.29146 | 4.20461 | 3.2661 |
| chr6 | 124542241 | 124542551 | 795 | Promoter (<=1kb) | 6 | 124530345 | 124542321 | 11977 | 2 | ENSMUST00 | 0 | C1s1 | ENSMUSG0C | false | . | . | . | . | . | . | chr6 | 124542200 | 124543012 | 3.78552 | 3.30211 | 2.039 |
| chr6 | 120978302 | 120978704 | 795 | Intron (ENSMUST00000098457.3 | 6 | 120960012 | 120966118 | 6107 | 2 | ENSMUST00 | -12185 | Mical3 | ENSMUSG0C | false | . | . | . | . | . | . | chr6 | 120976634 | 120979177 | 7.77056 | 5.41626 | 5.26112 |
| chr7 | 49487350 | 49487671 | 795 | Intron (ENSMUST00000183659.2 | 7 | 49450760 | 49464580 | 13821 | 1 | ENSMUST00 | 36591 | Nav2 | ENSMUSG0C | false | . | . | . | . | . | . | . | . | . | . | . | . |
| chr3 | 38254800 | 38255071 | 795 | Distal Intergenic | 3 | 38267836 | 38268084 | 249 | 2 | ENSMUST00 | 13013 | Gm25574 | ENSMUSG0C | false | . | . | . | . | . | . | . | . | . | . | . | . |
| chr15 | 80000589 | 80000992 | 795 | Promoter (1-2kb) | 15 | 79998970 | 79999026 | 57 | 2 | ENSMUST00 | -1564 | Mir7213 | ENSMUSG0C | false | . | . | . | . | . | . | . | . | . | . | . | . |
| chr11 | 76763499 | 76763743 | 794 | Promoter (<=1kb) | 11 | 76726602 | 76763579 | 36978 | 2 | ENSMUST00 | 0 | Gosr1 | ENSMUSG0C | false | chr11 | 76762717 | 76763684 | 29.3377 | 90.74815 | 87.38457 | . | . | . | . | . | . |
| chr15 | 77866635 | 77866988 | 794 | Distal Intergenic | 15 | 77760587 | 77842175 | 81589 | 2 | ENSMUST00 | -24461 | Myh9 | ENSMUSG0C | true | . | . | . | . | . | . | chr15 | 77866259 | 77867811 | 4.78334 | 3.82292 | 2.87755 |
| chr1 | 82253767 | 82254057 | 794 | Intron (ENSMUST00000069799.2 | 1 | 82233112 | 82235933 | 2822 | 1 | ENSMUST00 | 20656 | Gm9747 | ENSMUSG0C | false | . | . | . | . | . | . | chr1 | 82244858 | 82261494 | 5.35178 | 4.22772 | 3.29163 |
| chr3 | 58414971 | 58415701 | 794 | Promoter (<=1kb) | 3 | 58415689 | 58466787 | 51099 | 1 | ENSMUST00 | 0 | Tsc22d2 | ENSMUSG0C | false | chr3 | 58414128 | 58418695 | 40.32275 | 142.69632 | 137.70172 | . | . | . | . | . | . |
| chr1 | 159232102 | 159232548 | 794 | Promoter (<=1kb) | 1 | 159232326 | 159347580 | 115255 | 1 | ENSMUST00 | 0 | Rfwd2 | ENSMUSG0C | false | chr1 | 159231413 | 159233676 | 28.01773 | 86.50358 | 83.23278 | . | . | . | . | . | . |
| chr4 | 152272537 | 152273008 | 794 | Promoter (1-2kb) | 4 | 152274232 | 152285337 | 11106 | 1 | ENSMUST00 | -1224 | Gpr153 | ENSMUSG0C | false | . | . | . | . | . | . | chr4 | 152271972 | 152275112 | 4.94888 | 3.954 | 2.97647 |
| chr11 | 85642576 | 85642989 | 794 | Promoter (1-2kb) | 11 | 85640704 | 85826055 | 185352 | 1 | ENSMUST00 | 1873 | Bcas3 | ENSMUSG0C | false | . | . | . | . | . | . | chr11 | 85639815 | 85643227 | 4.23086 | 3.69506 | 2.37239 |
| chr11 | 86993390 | 86994061 | 793 | Promoter (<=1kb) | 11 | 86936425 | 86993707 | 57283 | 2 | ENSMUST00 | 0 | Ypel2 | ENSMUSG0C | false | chr11 | 86992502 | 86994164 | 33.56525 | 137.72058 | 132.93262 | . | . | . | . | . | . |
| chr11 | 49250359 | 49250962 | 793 | Promoter (<=1kb) | 11 | 49250512 | 49263030 | 12519 | 1 | ENSMUST00 | 0 | Mgat1 | ENSMUSG0C | false | chr11 | 49250056 | 49251442 | 23.28017 | 72.50986 | 69.48009 | chr11 | 49244640 | 49253728 | 6.81123 | 4.86749 | 4.50404 |
| chr19 | 6921393 | 6922140 | 793 | Promoter (<=1kb) | 19 | 6910977 | 6921808 | 10832 | 2 | ENSMUST00 | 0 | Esrra | ENSMUSG0C | false | chr19 | 6920899 | 6922749 | 27.82288 | 84.25431 | 81.02234 | . | . | . | . | . | . |
| chr3 | 95624928 | 95625460 | 793 | Promoter (<=1kb) | 3 | 95624980 | 95630493 | 5514 | 1 | ENSMUST00 | 0 | Ensa | ENSMUSG0C | false | chr3 | 95624921 | 95625958 | 25.16617 | 75.86667 | 72.78426 | . | . | . | . | . | . |
| chrX | 94366705 | 94367138 | 793 | Promoter (<=1kb) | 20 | 94355053 | 94366777 | 11725 | 2 | ENSMUST00 | 0 | 4932442L08F | ENSMUSG0C | false | chrX | 94367045 | 94367623 | 13.78596 | 30.80405 | 28.32676 | . | . | . | . | . | . |
| chr17 | 73283826 | 73284165 | 793 | Distal Intergenic | 17 | 73204150 | 73240371 | 36222 | 1 | ENSMUST00 | 79677 | Lclat1 | ENSMUSG0C | false | . | . | . | . | . | . | . | . | . | . | . | . |
| chr6 | 72355155 | 72355793 | 793 | Promoter (<=1kb) | 6 | 72355447 | 72359762 | 4316 | 1 | ENSMUST00 | 0 | Tmem150a | ENSMUSG0C | false | chr6 | 72354790 | 72356034 | 26.28407 | 85.32412 | 82.07336 | chr6 | 72350510 | 72356490 | 3.55578 | 3.19898 | 1.84912 |
| chr11 | 76854884 | 76855373 | 792 | Distal Intergenic | 11 | 76778424 | 76847018 | 68595 | 2 | ENSMUST00 | -7867 | Cpd | ENSMUSG0C | false | . | . | . | . | . | . | chr11 | 76853455 | 76856274 | 4.38614 | 3.75201 | 2.52636 |
| chr12 | 86076651 | 86076873 | 792 | Promoter (2-3kb) | 12 | 86056744 | 86079041 | 22298 | 2 | ENSMUST00 | 2168 | Tgfb3 | ENSMUSG0C | false | . | . | . | . | . | . | . | . | . | . | . | . |
| chr3 | 94372614 | 94372987 | 792 | Promoter (<=1kb) | 3 | 94372794 | 94397840 | 25047 | 1 | ENSMUST00 | 0 | Rorc | ENSMUSG0C | false | chr3 | 94372718 | 94373954 | 18.84082 | 47.13427 | 44.45482 | chr3 | 94371361 | 94374439 | 5.16649 | 4.17623 | 3.1432 |
| chr6 | 148443981 | 148444511 | 792 | Promoter (<=1kb) | 6 | 148232430 | 148444389 | 211960 | 2 | ENSMUST00 | 0 | Tmtc1 | ENSMUSG0C | true | chr6 | 148441860 | 148445082 | 35.62953 | 109.69414 | 105.87395 | . | . | . | . | . | . |
| chr1 | 138963491 | 138964150 | 792 | Promoter (<=1kb) | 1 | 138963708 | 138966441 | 2734 | 1 | ENSMUST00 | 0 | Dennd1b | ENSMUSG0C | false | chr1 | 138962926 | 138964541 | 32.07243 | 98.46742 | 94.9359 | . | . | . | . | . | . |
| chr6 | 54759876 | 54760234 | 792 | Distal Intergenic | 6 | 54816916 | 54893500 | 76585 | 1 | ENSMUST00 | -56682 | Znrf2 | ENSMUSG0C | false | . | . | . | . | . | . | chr6 | 54759399 | 54760796 | 3.21488 | 3.13163 | 1.52287 |
| chr10 | 80678072 | 80678483 | 792 | Distal Intergenic | 10 | 80681838 | 80687176 | 5339 | 1 | ENSMUST00 | -3355 | Gm26541 | ENSMUSG0C | false | . | . | . | . | . | . | chr10 | 80669195 | 80680494 | 7.16995 | 4.88679 | 4.8213 |
| chr1 | 74236294 | 74236768 | 791 | Promoter (<=1kb) | 1 | 74236497 | 74268209 | 31713 | 1 | ENSMUST00 | 0 | Arpc2 | ENSMUSG0C | false | chr1 | 74235907 | 74238302 | 34.54477 | 122.48211 | 118.25522 | . | . | . | . | . | . |
| chr17 | 27140415 | 27140745 | 791 | Distal Intergenic | 17 | 27122663 | 27133916 | 11254 | 2 | ENSMUST00 | -6500 | Uqcc2 | ENSMUSG0C | false | . | . | . | . | . | . | chr17 | 27127363 | 27147146 | 5.99202 | 4.48057 | 3.83593 |
| chr1 | 39901976 | 39902458 | 791 | Promoter (1-2kb) | 1 | 39900913 | 40026310 | 125398 | 1 | ENSMUST00 | 1064 | Map4k4 | ENSMUSG0C | false | chr1 | 39900215 | 39903023 | 36.76257 | 113.72284 | 109.77938 | chr1 | 39901844 | 39905690 | 5.19093 | 4.08609 | 3.17248 |
| chr1 | 72870992 | 72871349 | 791 | Promoter (2-3kb) | 1 | 72858065 | 72874275 | 16211 | 2 | ENSMUST00 | 2926 | Igfbp5 | ENSMUSG0C | false | chr1 | 72870000 | 72872489 | 10.01484 | 27.17025 | 24.74075 | . | . | . | . | . | . |
| chr1 | 133182362 | 133182678 | 791 | Promoter (1-2kb) | 1 | 133181321 | 133301219 | 119899 | 1 | ENSMUST00 | 1042 | Plekha6 | ENSMUSG0C | false | chr1 | 133182273 | 133182918 | 6.42099 | 15.28918 | 13.02279 | chr1 | 133181212 | 133182924 | 6.82601 | 4.97299 | 4.54123 |
| chrX | 52474255 | 52474542 | 791 | Intron (ENSMUST00000069360.8 | 20 | 52422407 | 52422515 | 109 | 2 | ENSMUST00 | -51741 | Mir717 | ENSMUSG0C | false | . | . | . | . | . | . | . | . | . | . | . | . |
| chr16 | 85137246 | 85137522 | 791 | Intron (ENSMUST00000005406.1 | 16 | 85168849 | 85169001 | 153 | 1 | ENSMUST00 | -31327 | Gm27295 | ENSMUSG0C | false | . | . | . | . | . | . | chr16 | 85126656 | 85166763 | 5.48022 | 4.2963 | 3.40197 |
| chr11 | 100975381 | 100975712 | 790 | Distal Intergenic | 11 | 100956733 | 100970887 | 14155 | 2 | ENSMUST00 | -4495 | Ptrf | ENSMUSG0C | false | . | . | . | . | . | . | chr11 | 100970994 | 100977767 | 4.99847 | 4.08141 | 3.00534 |
| chr5 | 33651982 | 33652762 | 790 | Promoter (<=1kb) | 5 | 33634952 | 33652345 | 17394 | 2 | ENSMUST00 | 0 | Slbp | ENSMUSG0C | false | chr5 | 33651424 | 33652689 | 34.14626 | 110.2223 | 106.38018 | chr5 | 33652746 | 33652992 | 5.91399 | 4.22431 | 3.77918 |
| chr2 | 172983599 | 172983855 | 790 | Promoter (2-3kb) | 2 | 172985877 | 172992053 | 6177 | 1 | ENSMUST00 | -2022 | Spo11 | ENSMUSG0C | false | . | . | . | . | . | . | . | . | . | . | . | . |
| chr11 | 23255380 | 23256122 | 790 | Promoter (<=1kb) | 11 | 23256041 | 23297597 | 41557 | 1 | ENSMUST00 | 0 | Xpo1 | ENSMUSG0C | false | chr11 | 23254787 | 23255543 | 24.45041 | 70.98759 | 67.98131 | . | . | . | . | . | . |
| chr17 | 79074586 | 79074857 | 790 | Intron (ENSMUST00000040789.4 | 17 | 79051906 | 79090243 | 38338 | 1 | ENSMUST00 | 22681 | Qpct | ENSMUSG0C | false | . | . | . | . | . | . | . | . | . | . | . | . |
| chr9 | 116912244 | 116912568 | 790 | Intron (ENSMUST00000111773.4 | 9 | 116582822 | 116822819 | 239998 | 2 | ENSMUST00 | -89426 | Rbms3 | ENSMUSG0C | true | . | . | . | . | . | . | chr9 | 116903777 | 116913504 | 4.73906 | 3.97114 | 2.79562 |
| chr18 | 35873233 | 35874091 | 790 | Distal Intergenic | 18 | 35830389 | 35831093 | 705 | 2 | ENSMUST00 | -42141 | RP24-326K1 | ENSMUSG0C | false | chr18 | 35873085 | 35874233 | 24.03115 | 75.67385 | 72.59326 | chr18 | 35870146 | 35877349 | 5.65833 | 4.25717 | 3.58139 |
| chr3 | 68570721 | 68570952 | 789 | Promoter (1-2kb) | 3 | 68572245 | 68626464 | 54220 | 1 | ENSMUST00 | -1293 | Schip1 | ENSMUSG0C | false | . | . | . | . | . | . | chr3 | 68570245 | 68572599 | 4.19401 | 3.71092 | 2.33294 |
| chr4 | 155668649 | 155669575 | 789 | Promoter (<=1kb) | 4 | 155654682 | 155669198 | 14517 | 2 | ENSMUST00 | 0 | Mib2 | ENSMUSG0C | false | chr4 | 155668118 | 155669937 | 34.54477 | 122.48211 | 118.25522 | . | . | . | . | . | . |
| chr7 | 92799872 | 92800132 | 789 | Intron (ENSMUST00000032879.8 | 7 | 92819905 | 92836647 | 16743 | 1 | ENSMUST00 | -19773 | Rab30 | ENSMUSG0C | true | . | . | . | . | . | . | chr7 | 92798575 | 92801969 | 4.61944 | 3.93173 | 2.7116 |
| chr8 | 11758000 | 11758680 | 789 | Promoter (<=1kb) | 8 | 11758354 | 11835219 | 76866 | 1 | ENSMUST00 | 0 | Arhgef7 | ENSMUSG0C | false | chr8 | 11757637 | 11759504 | 32.33533 | 102.95196 | 99.30833 | chr8 | 11755931 | 11760701 | 3.78715 | 3.47472 | 2.01748 |
| chr9 | 103370674 | 103370976 | 789 | Distal Intergenic | 9 | 103353094 | 103365840 | 12747 | 2 | ENSMUST00 | -4835 | Cdv3 | ENSMUSG0C | false | . | . | . | . | . | . | chr9 | 103368757 | 103372881 | 6.89266 | 4.91327 | 4.56132 |
| chr5 | 114573086 | 114573354 | 789 | Intron (ENSMUST00000043650.7 | 5 | 114568017 | 114594678 | 26662 | 1 | ENSMUST00 | 5070 | Fam222a | ENSMUSG0C | true | chr5 | 114573244 | 114575010 | 11.94783 | 25.26529 | 22.85895 | chr5 | 114571033 | 114614394 | 7.1045 | 4.9325 | 4.74958 |
| chr1 | 163561245 | 163561517 | 789 | Distal Intergenic | 1 | 163607377 | 163610269 | 2893 | 1 | ENSMUST00 | -45860 | RP24-141B1( | ENSMUSG0C | false | . | . | . | . | . | . | chr1 | 163560042 | 163562122 | 4.25635 | 3.68711 | 2.40697 |
| chr13 | 5862358 | 5862627 | 788 | Promoter (<=1kb) | 13 | 5861489 | 5870393 | 8905 | 1 | ENSMUST00 | 870 | Klf6 | ENSMUSG0C | false | chr13 | 5861507 | 5863975 | 32.01914 | 97.37792 | 93.87154 | . | . | . | . | . | . |
| chr5 | 146873184 | 146873586 | 788 | Distal Intergenic | 5 | 146845071 | 146847726 | 2656 | 1 | ENSMUST00 | 28114 | Rasl11a | ENSMUSG0C | true | . | . | . | . | . | . | chr5 | 146865620 | 146877684 | 5.33365 | 4.16572 | 3.30759 |
| chr19 | 10277850 | 10278124 | 788 | Intron (ENSMUST00000039327.5 | 19 | 10253330 | 10256871 | 3542 | 2 | ENSMUST00 | -20980 | Dagla | ENSMUSG0C | false | . | . | . | . | . | . | chr19 | 10270370 | 10305865 | 8.32241 | 5.52593 | 5.72899 |
| chr11 | 69802604 | 69803126 | 788 | Promoter (<=1kb) | 11 | 69796073 | 69801857 | 5785 | 2 | ENSMUST00 | -748 | Fgf11 | ENSMUSG0C | false | chr11 | 69802553 | 69804725 | 12.87622 | 39.27038 | 36.69063 | chr11 | 69799608 | 69805868 | 7.05133 | 4.92123 | 4.73614 |
| chr3 | 122333888 | 122334218 | 788 | Distal Intergenic | 3 | 122293585 | 122293703 | 119 | 2 | ENSMUST00 | -40186 | Mir760 | ENSMUSG0C | false | . | . | . | . | . | . | chr3 | 122333283 | 122335940 | 6.47769 | 4.78261 | 4.22934 |
| chr1 | 181256738 | 181257144 | 788 | Distal Intergenic | 1 | 181241988 | 181242320 | 333 | 2 | ENSMUST00 | -14419 | Rpl35a-ps2 | ENSMUSG0C | false | chr1 | 181255919 | 181257413 | 19.88939 | 55.30079 | 52.51555 | chr1 | 181255532 | 181257983 | 4.33392 | 3.68822 | 2.49216 |
| chr10 | 14212014 | 14212317 | 788 | Distal Intergenic | 10 | 14233513 | 14233623 | 111 | 1 | ENSMUST00 | -21196 | Gm24343 | ENSMUSG0C | false | . | . | . | . | . | . | . | . | . | . | . | . |
| chr8 | 46551644 | 46551992 | 787 | Promoter (<=1kb) | 8 | 46552055 | 46580007 | 27953 | 1 | ENSMUST00 | -63 | Cenpu | ENSMUSG0C | false | chr8 | 46551401 | 46551820 | 18.51447 | 52.87213 | 50.12019 | chr8 | 46551168 | 46552818 | 3.82302 | 3.48395 | 2.06482 |
| chr11 | 32347569 | 32348038 | 787 | Promoter (<=1kb) | 11 | 32347820 | 32428173 | 80354 | 1 | ENSMUST00 | 0 | Sh3pxd2b | ENSMUSG0C | false | chr11 | 32347309 | 32348974 | 24.34554 | 81.76674 | 78.58393 | chr11 | 32347345 | 32350295 | 3.68085 | 3.30451 | 1.96851 |
| chr1 | 54850088 | 54850400 | 787 | Intron (ENSMUST00000179030.2 | 1 | 54766792 | 54884711 | 117920 | 2 | ENSMUST00 | 34311 | Ankrd44 | ENSMUSG0C | true | . | . | . | . | . | . | chr1 | 54848944 | 54859535 | 5.87837 | 4.47531 | 3.73556 |
| chr2 | 160002752 | 160003033 | 787 | Distal Intergenic | 2 | 160016653 | 160040619 | 23967 | 1 | ENSMUST00 | -13620 | 9130015L21F | ENSMUSG0C | false | . | . | . | . | . | . | chr2 | 160001876 | 160004448 | 4.40836 | 3.76657 | 2.52812 |
| chr1 | 151041075 | 151041416 | 787 | Distal Intergenic | 1 | 151062004 | 151062110 | 107 | 2 | ENSMUST00 | 20694 | Gm22756 | ENSMUSG0C | false | . | . | . | . | . | . | . | . | . | . | . | . |
| chr6 | 94625969 | 94626280 | 787 | Intron (ENSMUST00000032105.5 | 6 | 94606364 | 94606415 | 52 | 2 | ENSMUST00 | -19555 | Mir7041 | ENSMUSG0C | false | . | . | . | . | . | . | . | . | . | . | . | . |
| chr4 | 95153904 | 95154319 | 787 | Promoter (2-3kb) | 4 | 95151740 | 95157503 | 5764 | 1 | ENSMUST00 | 2165 | RP23-77I17.4 | ENSMUSG0C | false | . | . | . | . | . | . | chr4 | 95152754 | 95157985 | 3.5377 | 3.22179 | 1.82132 |
| chr2 | 76675005 | 76675449 | 786 | Promoter (<=1kb) | 2 | 76675281 | 76696828 | 21548 | 1 | ENSMUST00 | 0 | Plekha3 | ENSMUSG0C | false | chr2 | 76674806 | 76676198 | 31.01414 | 97.6227 | 94.10706 | chr2 | 76671794 | 76677409 | 4.33168 | 3.69695 | 2.46956 |
| chr19 | 47884325 | 47884612 | 786 | Intron (ENSMUST00000120645.2 | 19 | 47870262 | 47884704 | 14443 | 1 | ENSMUST00 | 14064 | Gsto2 | ENSMUSG0C | false | . | . | . | . | . | . | . | . | . | . | . | . |
| chr1 | 134414995 | 134415754 | 786 | Promoter (<=1kb) | 1 | 134415378 | 134433351 | 17974 | 1 | ENSMUST00 | 0 | Adipor1 | ENSMUSG0C | false | chr1 | 134414933 | 134416522 | 34.4409 | 118.06123 | 113.98219 | . | . | . | . | . | . |
| chr8 | 82863425 | 82864335 | 786 | Promoter (<=1kb) | 8 | 82863356 | 83091271 | 227916 | 1 | ENSMUST00 | 70 | Rnf150 | ENSMUSG0C | true | chr8 | 82863357 | 82865470 | 38.83893 | 127.96716 | 123.54797 | . | . | . | . | . | . |
| chr7 | 24481662 | 24482083 | 786 | Promoter (<=1kb) | 7 | 24482023 | 24504539 | 22517 | 1 | ENSMUST00 | 0 | Cadm4 | ENSMUSG0C | false | chr7 | 24481368 | 24483109 | 24.93112 | 71.82204 | 68.80418 | chr7 | 24480521 | 24490691 | 5.95135 | 4.53554 | 3.78776 |
| chr11 | 117232085 | 117232349 | 786 | Promoter (<=1kb) | 11 | 117232285 | 117362313 | 130029 | 1 | ENSMUST00 | 0 | Sep-09 | ENSMUSG0C | false | chr11 | 117231879 | 117232740 | 7.44452 | 13.65903 | 11.41755 | chr11 | 117231511 | 117233362 | 4.28449 | 3.75542 | 2.40868 |
| chrX | 103616866 | 103617465 | 786 | Promoter (<=1kb) | 20 | 103562897 | 103616954 | 54058 | 2 | ENSMUST00 | 0 | Ftx | ENSMUSG0C | true | chrX | 103616021 | 103616916 | 11.94783 | 25.26529 | 22.85895 | . | . | . | . | . | . |
| chr10 | 53947956 | 53948250 | 785 | Intron (ENSMUST00000003843.9 | 10 | 53907015 | 54075709 | 168695 | 2 | ENSMUST00 | 127459 | Man1a | ENSMUSG0C | false | . | . | . | . | . | . | chr10 | 53947786 | 53948150 | 3.54127 | 3.06039 | 1.84538 |
| chr2 | 91710454 | 91710950 | 785 | Promoter (<=1kb) | 2 | 91710852 | 91712327 | 1476 | 1 | ENSMUST00 | 0 | Harbi1 | ENSMUSG0C | false | chr2 | 91709479 | 91711625 | 38.96697 | 134.74414 | 130.07794 | . | . | . | . | . | . |
| chr9 | 109832525 | 109833165 | 785 | Promoter (<=1kb) | 9 | 109832749 | 109842529 | 9781 | 1 | ENSMUST00 | 0 | Nme6 | ENSMUSG0C | false | chr9 | 109832255 | 109833458 | 23.65566 | 74.08845 | 71.03342 | chr9 | 109832133 | 109833690 | 4.81337 | 3.92971 | 2.90446 |
| chr5 | 105698632 | 105698983 | 785 | Promoter (<=1kb) | 5 | 105699969 | 105815215 | 115247 | 1 | ENSMUST00 | -986 | Lrrc8d | ENSMUSG0C | false | . | . | . | . | . | . | chr5 | 105698293 | 105699788 | 4.87812 | 4.07307 | 2.90278 |
| chr4 | 40887098 | 40887400 | 785 | Distal Intergenic | 4 | 40912741 | 40913355 | 615 | 2 | ENSMUST00 | 25955 | Gm12397 | ENSMUSG0C | false | . | . | . | . | . | . | chr4 | 40882325 | 40890611 | 4.88004 | 4.00478 | 2.90476 |
| chr19 | 47313519 | 47315051 | 785 | Exon (ENSMUST00000081619.3/ | 19 | 47273519 | 47273589 | 71 | 2 | ENSMUST00 | -39931 | Mir6995 | ENSMUSG0C | false | chr19 | 47313175 | 47315281 | 18.90859 | 48.02031 | 45.33304 | chr19 | 47311839 | 47321707 | 6.07643 | 4.48035 | 3.91145 |
| chr4 | 114547725 | 114548133 | 785 | Intron (ENSMUST00000094894.3 | 4 | 114672697 | 114673340 | 644 | 2 | ENSMUST00 | 125207 | RP23-454F1: | ENSMUSG0C | false | . | . | . | . | . | . | chr4 | 114538576 | 114564774 | 5.66596 | 4.26794 | 3.56957 |
| chr10 | 5805451 | 5805793 | 784 | Promoter (<=1kb) | 10 | 5799160 | 5805600 | 6441 | 2 | ENSMUST00 | 0 | Fbxo5 | ENSMUSG0C | false | chr10 | 5804803 | 5805508 | 16.70194 | 60.58573 | 57.73199 | . | . | . | . | . | . |
| chr1 | 82247166 | 82247488 | 784 | Intron (ENSMUST00000069799.2 | 1 | 82233112 | 82235933 | 2822 | 1 | ENSMUST00 | 14055 | Gm9747 | ENSMUSG0C | false | . | . | . | . | . | . | chr1 | 82244858 | 82261494 | 5.35178 | 4.22772 | 3.29163 |
| chr16 | 51986751 | 51987252 | 784 | Distal Intergenic | 16 | 52031549 | 52208047 | 176499 | 1 | ENSMUST00 | -44297 | Cblb | ENSMUSG0C | true | . | . | . | . | . | . | chr16 | 51984722 | 51988455 | 4.93431 | 4.05283 | 2.96223 |
| chr18 | 57242757 | 57243036 | 784 | Exon (ENSMUST00000075770.7/ | 18 | 57142786 | 57292728 | 149943 | 1 | ENSMUST00 | 99972 | Megf10 | ENSMUSG0C | false | . | . | . | . | . | . | chr18 | 57242213 | 57246636 | 3.95459 | 3.58656 | 2.133 |
| chr2 | 143081707 | 143081997 | 784 | 3' UTR | 2 | 143078473 | 143081713 | 3241 | 1 | ENSMUST00 | 3235 | Otor | ENSMUSG0C | false | . | . | . | . | . | . | . | . | . | . | . | . |
| chr5 | 72868344 | 72868769 | 784 | Promoter (<=1kb) | 5 | 72755718 | 72868452 | 112735 | 2 | ENSMUST00 | 0 | Tec | ENSMUSG0C | false | chr5 | 72867770 | 72869007 | 20.65177 | 61.66082 | 58.78991 | chr5 | 72867616 | 72870022 | 3.49464 | 3.16036 | 1.80716 |
| chr9 | 66592139 | 66592844 | 784 | Promoter (<=1kb) | 9 | 66517877 | 66592958 | 75082 | 2 | ENSMUST00 | 114 | Usp3 | ENSMUSG0C | false | chr9 | 66591521 | 66593918 | 35.77904 | 109.9518 | 106.11687 | . | . | . | . | . | . |
| chr3 | 86920375 | 86921015 | 783 | Promoter (<=1kb) | 3 | 86786150 | 86920884 | 134735 | 2 | ENSMUST00 | 0 | Dclk2 | ENSMUSG0C | false | chr3 | 86919574 | 86921174 | 24.82645 | 85.46523 | 82.21117 | chr3 | 86919587 | 86921181 | 3.51324 | 3.26962 | 1.82352 |
| chr3 | 122362912 | 122363210 | 783 | Distal Intergenic | 3 | 122419780 | 122530180 | 110401 | 1 | ENSMUST00 | -56570 | Bcar3 | ENSMUSG0C | true | . | . | . | . | . | . | chr3 | 122358583 | 122365678 | 3.95742 | 3.46571 | 2.18853 |
| chr9 | 63973704 | 63973958 | 783 | Intron (ENSMUST00000041029.5 | 9 | 63954009 | 64016987 | 62979 | 2 | ENSMUST00 | 43029 | Smad6 | ENSMUSG0C | true | . | . | . | . | . | . | chr9 | 63948352 | 64020083 | 7.27873 | 5.05859 | 4.874 |
| chr6 | 71143734 | 71144482 | 783 | Promoter (<=1kb) | 6 | 71128166 | 71144346 | 16181 | 2 | ENSMUST00 | 0 | Thnsl2 | ENSMUSG0C | false | chr6 | 71143314 | 71144570 | 19.90079 | 58.62946 | 55.80003 | chr6 | 71142584 | 71144820 | 5.95693 | 4.53895 | 3.784 |
| chr2 | 61102090 | 61102481 | 783 | Distal Intergenic | 2 | 61030351 | 61030970 | 620 | 2 | ENSMUST00 | -71121 | Gm13579 | ENSMUSG0C | false | . | . | . | . | . | . | chr2 | 61099341 | 61105801 | 4.0096 | 3.59734 | 2.20741 |
| chr4 | 150784639 | 150785198 | 783 | Distal Intergenic | 4 | 150825733 | 150839575 | 13843 | 1 | ENSMUST00 | -40535 | Gm13049 | ENSMUSG0C | false | . | . | . | . | . | . | chr4 | 150784510 | 150786637 | 4.42467 | 3.8393 | 2.5188 |
| chr5 | 74794821 | 74795193 | 783 | Distal Intergenic | 5 | 74731517 | 74734450 | 2934 | 2 | ENSMUST00 | -60372 | Gm15985 | ENSMUSG0C | false | . | . | . | . | . | . | chr5 | 74794732 | 74795052 | 3.29138 | 3.02819 | 1.6326 |
| chr10 | 86331919 | 86332241 | 782 | Intron (ENSMUST00000020234.8 | 10 | 86302854 | 86343981 | 41128 | 1 | ENSMUST00 | 29066 | Timp3 | ENSMUSG0C | false | . | . | . | . | . | . | chr10 | 86303145 | 86347283 | 7.80266 | 5.33584 | 5.30789 |
| chr13 | 63566055 | 63566468 | 782 | Promoter (<=1kb) | 13 | 63511533 | 63565520 | 53988 | 2 | ENSMUST00 | -536 | Ptch1 | ENSMUSG0C | false | chr13 | 63562847 | 63568718 | 33.69019 | 111.49343 | 107.61708 | . | . | . | . | . | . |
| chr15 | 58973033 | 58973354 | 782 | Intron (ENSMUST00000080371.6 | 15 | 58933810 | 58939489 | 5680 | 1 | ENSMUST00 | 39224 | Ndufb9 | ENSMUSG0C | false | . | . | . | . | . | . | chr15 | 58965186 | 59078715 | 7.47325 | 5.12649 | 5.04432 |
| chr2 | 173171459 | 173171748 | 782 | Distal Intergenic | 2 | 173153082 | 173159271 | 6190 | 1 | ENSMUST00 | 18378 | Pck1 | ENSMUSG0C | false | . | . | . | . | . | . | chr2 | 173171318 | 173175002 | 5.00733 | 4.01065 | 3.02514 |
| chr3 | 151856018 | 151856257 | 782 | Distal Intergenic | 3 | 151798918 | 151837630 | 38713 | 2 | ENSMUST00 | -18389 | Ptgfr | ENSMUSG0C | false | . | . | . | . | . | . | chr3 | 151855302 | 151856546 | 4.35187 | 3.75261 | 2.48237 |
| chr4 | 57299852 | 57300651 | 782 | Promoter (<=1kb) | 4 | 57245051 | 57300375 | 55325 | 2 | ENSMUST00 | 0 | Ptpn3 | ENSMUSG0C | false | chr4 | 57299328 | 57302225 | 26.1063 | 98.41757 | 94.88632 | chr4 | 57294789 | 57301188 | 4.63579 | 3.8844 | 2.72174 |
| chr19 | 8989069 | 8989499 | 782 | Promoter (<=1kb) | 19 | 8989284 | 9019195 | 29912 | 1 | ENSMUST00 | 0 | Ahnak | ENSMUSG0C | false | chr19 | 8988723 | 8990788 | 33.41831 | 117.26761 | 113.21259 | chr19 | 8987793 | 8989330 | 6.98554 | 4.85516 | 4.64859 |
| chr17 | 87974817 | 87975768 | 781 | Promoter (<=1kb) | 17 | 87975050 | 87990883 | 15834 | 1 | ENSMUST00 | 0 | Msh6 | ENSMUSG0C | false | chr17 | 87974423 | 87976176 | 27.57146 | 121.15424 | 116.97103 | . | . | . | . | . | . |
| chr11 | 104231075 | 104231534 | 781 | Promoter (<=1kb) | 11 | 104231390 | 104332090 | 100701 | 1 | ENSMUST00 | 0 | Mapt | ENSMUSG0C | true | chr11 | 104231437 | 104234756 | 38.46334 | 130.56975 | 126.05653 | . | . | . | . | . | . |
| chr14 | 46902140 | 46902456 | 781 | Intron (ENSMUST00000022386.9 | 14 | 46884175 | 47105811 | 221637 | 1 | ENSMUST00 | 17966 | Samd4 | ENSMUSG0C | true | . | . | . | . | . | . | chr14 | 46897792 | 46903404 | 5.31812 | 4.11399 | 3.30283 |
| chr19 | 36214592 | 36214866 | 781 | Distal Intergenic | 19 | 36111965 | 36119844 | 7880 | 2 | ENSMUST00 | -94749 | Ankrd1 | ENSMUSG0C | false | . | . | . | . | . | . | chr19 | 36213832 | 36215193 | 2.92726 | 2.93868 | 1.35427 |
| chr4 | 41545977 | 41546273 | 781 | Intron (ENSMUST00000108049.3 | 4 | 41520320 | 41524770 | 4451 | 2 | ENSMUST00 | -21208 | Fam219a | ENSMUSG0C | false | . | . | . | . | . | . | . | . | . | . | . | . |
| chr9 | 67043631 | 67044062 | 781 | Promoter (<=1kb) | 9 | 67022590 | 67043907 | 21318 | 2 | ENSMUST00 | 0 | Tpm1 | ENSMUSG0C | true | chr9 | 67043055 | 67043963 | 27.6601 | 94.57302 | 91.13082 | . | . | . | . | . | . |
| chr16 | 91728559 | 91728874 | 781 | Promoter (<=1kb) | 16 | 91689342 | 91728802 | 39461 | 2 | ENSMUST00 | 0 | Cryzl1 | ENSMUSG0C | false | chr16 | 91728054 | 91729199 | 23.65566 | 74.08845 | 71.03342 | . | . | . | . | . | . |
| chrX | 99216855 | 99217321 | 780 | Distal Intergenic | 20 | 99138131 | 99147230 | 9100 | 1 | ENSMUST00 | 78725 | Efnb1 | ENSMUSG0C | false | chrX | 99216303 | 99217267 | 11.94783 | 25.26529 | 22.85895 | . | . | . | . | . | . |
| chr2 | 37786550 | 37786820 | 780 | Exon (ENSMUST00000050372.7/ | 2 | 37791171 | 37793523 | 2353 | 1 | ENSMUST00 | -4351 | Crb2 | ENSMUSG0C | false | . | . | . | . | . | . | chr2 | 37777133 | 37803419 | 7.19536 | 4.98761 | 4.82001 |
| chr2 | 167062503 | 167063126 | 780 | Promoter (<=1kb) | 2 | 167035793 | 167063015 | 27223 | 2 | ENSMUST00 | 0 | Znfx1 | ENSMUSG0C | false | chr2 | 167061714 | 167063859 | 29.403 | 88.9212 | 85.59992 | . | . | . | . | . | . |
| chr6 | 31217194 | 31217754 | 780 | Promoter (<=1kb) | 6 | 31166524 | 31218346 | 51823 | 2 | ENSMUST00 | 592 | Lincpint | ENSMUSG0C | false | chr6 | 31215613 | 31219210 | 29.2142 | 114.47596 | 110.50954 | . | . | . | . | . | . |
| chr9 | 9238674 | 9239227 | 780 | Promoter (<=1kb) | 9 | 8994329 | 9239101 | 244773 | 2 | ENSMUST00 | 0 | Arhgap42 | ENSMUSG0C | false | chr9 | 9238203 | 9239549 | 14.79283 | 57.11871 | 54.31201 | . | . | . | . | . | . |
| chr8 | 84191976 | 84192340 | 780 | Downstream (<1kb) | 8 | 84192774 | 84197667 | 4894 | 2 | ENSMUST00 | 5327 | Gm26887 | ENSMUSG0C | false | chr8 | 84192248 | 84192996 | 8.63619 | 18.02194 | 15.71366 | chr8 | 84190694 | 84193192 | 4.40739 | 3.61139 | 2.5518 |
| chr6 | 67138387 | 67139065 | 780 | Intron (ENSMUST00000181304.1 | 6 | 67115673 | 67168426 | 52754 | 1 | ENSMUST00 | 22715 | A430010J10F | ENSMUSG0C | false | . | . | . | . | . | . | chr6 | 67123607 | 67146074 | 5.6205 | 4.38535 | 3.51128 |
| chr18 | 44491372 | 44491758 | 779 | Intron (ENSMUST00000164666.2 | 18 | 44380500 | 44424969 | 44470 | 1 | ENSMUST00 | 110873 | Dcp2 | ENSMUSG0C | false | . | . | . | . | . | . | chr18 | 44488874 | 44502831 | 6.07908 | 4.49427 | 3.89377 |
| chr1 | 75236256 | 75236701 | 779 | Promoter (<=1kb) | 1 | 75236406 | 75245688 | 9283 | 1 | ENSMUST00 | 0 | Dnajb2 | ENSMUSG0C | false | chr1 | 75235964 | 75236346 | 8.29801 | 20.47831 | 18.13586 | chr1 | 75234533 | 75238649 | 4.89226 | 4.10199 | 2.9136 |
| chr5 | 107121130 | 107121527 | 779 | Exon (ENSMUST00000031224.9/ | 5 | 107118197 | 107132923 | 14727 | 2 | ENSMUST00 | 11396 | Tgfbr3 | ENSMUSG0C | true | . | . | . | . | . | . | chr5 | 107119899 | 107128040 | 6.2176 | 4.49216 | 4.0381 |
| chr11 | 95009750 | 95010149 | 779 | Promoter (<=1kb) | 11 | 95009879 | 95026087 | 16209 | 1 | ENSMUST00 | 0 | Samd14 | ENSMUSG0C | false | chr11 | 95009394 | 95010240 | 21.15962 | 55.85266 | 53.06144 | chr11 | 95005148 | 95010695 | 4.81833 | 4.0099 | 2.85609 |
| chr7 | 6383166 | 6383532 | 779 | Promoter (<=1kb) | 7 | 6383318 | 6396633 | 13316 | 1 | ENSMUST00 | 0 | Zfp28 | ENSMUSG0C | false | chr7 | 6382653 | 6383974 | 30.41442 | 103.5799 | 99.9162 | chr7 | 6382629 | 6384831 | 4.23592 | 3.81414 | 2.36053 |
| chr14 | 58709886 | 58710191 | 779 | Distal Intergenic | 14 | 58650268 | 58650400 | 133 | 2 | ENSMUST00 | -59487 | Gm25614 | ENSMUSG0C | false | . | . | . | . | . | . | chr14 | 58708949 | 58711065 | 3.80027 | 3.48415 | 2.04571 |
| chr2 | 52950182 | 52950500 | 779 | Intron (ENSMUST00000090952.5 | 2 | 52859723 | 52863516 | 3794 | 2 | ENSMUST00 | -86667 | Gm26603 | ENSMUSG0C | false | . | . | . | . | . | . | chr2 | 52949129 | 52956219 | 4.81718 | 3.92998 | 2.87074 |
| chr19 | 46447922 | 46448286 | 778 | Intron (ENSMUST00000039922.7 | 19 | 46457906 | 46475634 | 17729 | 1 | ENSMUST00 | -9620 | Sufu | ENSMUSG0C | false | . | . | . | . | . | . | chr19 | 46445312 | 46450222 | 5.11208 | 3.84982 | 3.13825 |
| chr6 | 33249123 | 33249525 | 778 | Promoter (<=1kb) | 6 | 33249128 | 33350972 | 101845 | 1 | ENSMUST00 | 0 | Exoc4 | ENSMUSG0C | false | chr6 | 33249030 | 33249796 | 31.37684 | 98.25688 | 94.7338 | chr6 | 33249398 | 33250413 | 3.2736 | 3.20116 | 1.64042 |
| chr8 | 107129229 | 107129502 | 778 | Distal Intergenic | 8 | 107150661 | 107187470 | 36810 | 1 | ENSMUST00 | -21159 | Cyb5b | ENSMUSG0C | false | . | . | . | . | . | . | chr8 | 107127986 | 107130599 | 4.46961 | 3.72636 | 2.58476 |
| chr13 | 48465661 | 48465898 | 778 | Distal Intergenic | 13 | 48489202 | 48506939 | 17738 | 2 | ENSMUST00 | 41041 | Zfp169 | ENSMUSG0C | false | . | . | . | . | . | . | chr13 | 48464452 | 48466553 | 4.23412 | 3.45013 | 2.41637 |
| chr5 | 150384766 | 150385149 | 778 | Intron (ENSMUST00000087204.5 | 5 | 150259930 | 150497752 | 237823 | 1 | ENSMUST00 | 124837 | Fry | ENSMUSG0C | true | . | . | . | . | . | . | chr5 | 150383780 | 150386142 | 5.46322 | 4.19522 | 3.42046 |
| chrX | 99415679 | 99415966 | 778 | Promoter (2-3kb) | 20 | 99409571 | 99418895 | 9325 | 2 | ENSMUST00 | 2929 | Gm14809 | ENSMUSG0C | false | . | . | . | . | . | . | . | . | . | . | . | . |
| chr5 | 74007524 | 74007885 | 778 | Intron (ENSMUST00000119154.2 | 5 | 74014216 | 74029680 | 15465 | 1 | ENSMUST00 | -6331 | Usp46os1 | ENSMUSG0C | false | . | . | . | . | . | . | chr5 | 74001579 | 74008551 | 5.27799 | 4.06988 | 3.27177 |
| chr11 | 76446257 | 76446573 | 777 | Intron (ENSMUST00000094012.5 | 11 | 76423048 | 76437220 | 14173 | 2 | ENSMUST00 | -9038 | Abr | ENSMUSG0C | false | . | . | . | . | . | . | chr11 | 76424298 | 76452713 | 6.90719 | 4.90323 | 4.58627 |
| chr13 | 55190057 | 55190648 | 777 | Distal Intergenic | 13 | 55209782 | 55318325 | 108544 | 1 | ENSMUST00 | -19134 | Nsd1 | ENSMUSG0C | true | chr13 | 55189545 | 55190337 | 9.15642 | 23.76437 | 21.37967 | chr13 | 55189184 | 55191359 | 3.54128 | 3.3909 | 1.79002 |
| chr15 | 34508694 | 34509000 | 777 | Promoter (<=1kb) | 15 | 34509907 | 34512192 | 2286 | 1 | ENSMUST00 | -907 | Pop1 | ENSMUSG0C | false | . | . | . | . | . | . | chr15 | 34508319 | 34517398 | 5.80499 | 4.48832 | 3.66177 |
| chr19 | 3929204 | 3929853 | 777 | Promoter (<=1kb) | 19 | 3913493 | 3929761 | 16269 | 2 | ENSMUST00 | 0 | Aldh3b1 | ENSMUSG0C | false | . | . | . | . | . | . | chr19 | 3928969 | 3931218 | 5.65013 | 4.38621 | 3.52281 |
| chrX | 94234624 | 94235110 | 777 | Promoter (<=1kb) | 20 | 94234927 | 94254063 | 19137 | 1 | ENSMUST00 | 0 | Klhl15 | ENSMUSG0C | false | chrX | 94234167 | 94235882 | 22.05754 | 58.21236 | 55.38818 | . | . | . | . | . | . |
| chr14 | 70584291 | 70584606 | 777 | 3' UTR | 14 | 70577847 | 70582571 | 4725 | 1 | ENSMUST00 | 6445 | Nudt18 | ENSMUSG0C | false | . | . | . | . | . | . | chr14 | 70583047 | 70585331 | 6.34664 | 4.72941 | 4.11177 |
| chr10 | 14452496 | 14452756 | 777 | Intron (ENSMUST00000041168.4 | 10 | 14523078 | 14523545 | 468 | 1 | ENSMUST00 | -70322 | Gm10335 | ENSMUSG0C | false | . | . | . | . | . | . | chr10 | 14452465 | 14452854 | 3.16078 | 3.12867 | 1.48045 |
| chr8 | 111341742 | 111342240 | 776 | Downstream (2-3kb) | 8 | 111311800 | 111338177 | 26378 | 2 | ENSMUST00 | -3566 | Mlkl | ENSMUSG0C | false | . | . | . | . | . | . | chr8 | 111340471 | 111344691 | 3.96639 | 3.50111 | 2.17311 |
| chr2 | 164073969 | 164074391 | 776 | Promoter (<=1kb) | 2 | 164074138 | 164155524 | 81387 | 1 | ENSMUST00 | 0 | Stk4 | ENSMUSG0C | false | chr2 | 164074051 | 164075217 | 42.12929 | 136.96686 | 132.21103 | chr2 | 164072892 | 164078411 | 4.75707 | 4.00898 | 2.7984 |
| chr1 | 160792362 | 160793065 | 776 | Promoter (<=1kb) | 1 | 160256572 | 160792938 | 536367 | 2 | ENSMUST00 | 0 | Rabgap1l | ENSMUSG0C | false | chr1 | 160791788 | 160793644 | 39.29435 | 126.59459 | 122.2257 | . | . | . | . | . | . |
| chr9 | 63968609 | 63968951 | 776 | Intron (ENSMUST00000041029.5 | 9 | 63954009 | 64016987 | 62979 | 2 | ENSMUST00 | 48036 | Smad6 | ENSMUSG0C | true | . | . | . | . | . | . | chr9 | 63948352 | 64020083 | 7.27873 | 5.05859 | 4.874 |
| chr6 | 72512471 | 72513135 | 776 | Promoter (2-3kb) | 6 | 72513644 | 72515290 | 1647 | 2 | ENSMUST00 | 2155 | Sh2d6 | ENSMUSG0C | false | . | . | . | . | . | . | chr6 | 72510186 | 72516424 | 7.38109 | 5.05551 | 4.9697 |
| chr16 | 34490599 | 34490883 | 776 | Intron (ENSMUST00000114961.2 | 16 | 34153467 | 34513944 | 360478 | 2 | ENSMUST00 | 23061 | Kalrn | ENSMUSG0C | false | . | . | . | . | . | . | chr16 | 34490142 | 34491354 | 6.08182 | 4.38084 | 3.92721 |
| chr17 | 65782551 | 65783256 | 776 | Promoter (<=1kb) | 17 | 65782573 | 65803912 | 21340 | 1 | ENSMUST00 | 0 | Ppp4r1 | ENSMUSG0C | false | chr17 | 65782365 | 65784357 | 16.94318 | 73.17311 | 70.13294 | . | . | . | . | . | . |
| chr2 | 6872086 | 6872870 | 775 | Promoter (<=1kb) | 2 | 6546859 | 6872597 | 325739 | 2 | ENSMUST00 | 0 | Celf2 | ENSMUSG0C | false | chr2 | 6871587 | 6873131 | 21.17422 | 79.20749 | 76.06625 | chr2 | 6871865 | 6872953 | 3.41538 | 3.13044 | 1.74298 |
| chr7 | 97081261 | 97082520 | 775 | Promoter (<=1kb) | 7 | 97081751 | 97308940 | 227190 | 1 | ENSMUST00 | 0 | Gab2 | ENSMUSG0C | true | chr7 | 97081213 | 97083356 | 37.54867 | 136.59039 | 131.84451 | chr7 | 97078666 | 97081609 | 4.83743 | 3.99046 | 2.92447 |
| chr7 | 68080615 | 68081113 | 775 | Intron (ENSMUST00000005671.8 | 7 | 67952859 | 68226780 | 273922 | 1 | ENSMUST00 | 127757 | Igf1r | ENSMUSG0C | false | . | . | . | . | . | . | chr7 | 68078670 | 68087520 | 5.45473 | 4.15004 | 3.39634 |
| chr15 | 83171934 | 83172450 | 775 | Promoter (<=1kb) | 15 | 83153494 | 83172592 | 19099 | 2 | ENSMUST00 | 142 | Cyb5r3 | ENSMUSG0C | false | chr15 | 83171457 | 83172969 | 43.84673 | 147.77686 | 142.57693 | chr15 | 83162089 | 83173300 | 7.98675 | 5.40074 | 5.45307 |
| chr13 | 97911036 | 97911401 | 775 | Intron (ENSMUST00000109426.1 | 13 | 97760130 | 97760588 | 459 | 2 | ENSMUST00 | -150449 | Gm10260 | ENSMUSG0C | false | . | . | . | . | . | . | chr13 | 97909477 | 97929349 | 4.81516 | 3.96306 | 2.86901 |
| chrX | 12006880 | 12007479 | 775 | Distal Intergenic | 20 | 11984475 | 11984650 | 176 | 1 | ENSMUST00 | 22406 | Gm14512 | ENSMUSG0C | false | chrX | 12006144 | 12007515 | 12.8669 | 28.00426 | 25.56298 | . | . | . | . | . | . |
| chr16 | 8881859 | 8882139 | 775 | Distal Intergenic | 16 | 8873079 | 8873171 | 93 | 2 | ENSMUST00 | -8689 | Gm22224 | ENSMUSG0C | false | . | . | . | . | . | . | chr16 | 8881909 | 8884143 | 3.58966 | 3.33403 | 1.91244 |
| chr2 | 30093587 | 30094060 | 774 | Promoter (<=1kb) | 2 | 30090944 | 30093648 | 2705 | 2 | ENSMUST00 | 0 | Zdhhc12 | ENSMUSG0C | false | chr2 | 30093188 | 30094333 | 31.51843 | 93.54985 | 90.12806 | chr2 | 30092398 | 30095768 | 4.5559 | 3.80745 | 2.65384 |
| chr11 | 49245236 | 49245567 | 774 | Promoter (<=1kb) | 11 | 49245231 | 49260691 | 15461 | 1 | ENSMUST00 | 6 | Mgat1 | ENSMUSG0C | false | . | . | . | . | . | . | chr11 | 49244640 | 49253728 | 6.81123 | 4.86749 | 4.50404 |
| chr6 | 87811605 | 87812074 | 774 | Promoter (<=1kb) | 6 | 87788853 | 87811779 | 22927 | 2 | ENSMUST00 | 0 | Rab43 | ENSMUSG0C | false | chr6 | 87810377 | 87812633 | 32.27147 | 102.84008 | 99.19724 | . | . | . | . | . | . |
| chr6 | 113492469 | 113492775 | 774 | Promoter (<=1kb) | 6 | 113492019 | 113493335 | 1317 | 1 | ENSMUST00 | 451 | Creld1 | ENSMUSG0C | false | . | . | . | . | . | . | chr6 | 113491965 | 113493110 | 3.71791 | 3.37152 | 1.99257 |
| chr5 | 77378174 | 77378452 | 774 | Intron (ENSMUST00000046746.6 | 5 | 77349240 | 77408045 | 58806 | 2 | ENSMUST00 | 29593 | Igfbp7 | ENSMUSG0C | false | . | . | . | . | . | . | chr5 | 77353137 | 77404361 | 6.51707 | 4.75542 | 4.25399 |
| chrX | 37048630 | 37048893 | 774 | Promoter (<=1kb) | 20 | 37048845 | 37050418 | 1574 | 1 | ENSMUST00 | 0 | Sowahd | ENSMUSG0C | false | chrX | 37047520 | 37049681 | 16.54316 | 39.53096 | 36.94807 | . | . | . | . | . | . |
| chr2 | 142902950 | 142903280 | 774 | Promoter (<=1kb) | 2 | 142902965 | 142903572 | 608 | 1 | ENSMUST00 | 0 | Kif16bos | ENSMUSG0C | false | chr2 | 142902407 | 142903691 | 9.63447 | 24.12454 | 21.73289 | chr2 | 142900386 | 142903706 | 5.23958 | 4.21244 | 3.1947 |
| chr1 | 163929591 | 163930121 | 773 | Promoter (<=1kb) | 1 | 163929765 | 163955126 | 25362 | 1 | ENSMUST00 | 0 | Scyl3 | ENSMUSG0C | false | chr1 | 163929283 | 163930774 | 31.16539 | 106.97112 | 103.21663 | . | . | . | . | . | . |
| chr19 | 58820218 | 58820769 | 773 | Intron (ENSMUST00000066285.4 | 19 | 58785803 | 58794461 | 8659 | 2 | ENSMUST00 | -25758 | 1700019N19 | ENSMUSG0C | false | . | . | . | . | . | . | chr19 | 58814047 | 58859817 | 7.46659 | 5.12677 | 5.03638 |
| chr14 | 65266297 | 65266754 | 773 | Promoter (<=1kb) | 14 | 65266701 | 65321501 | 54801 | 1 | ENSMUST00 | 0 | Fbxo16 | ENSMUSG0C | false | chr14 | 65266032 | 65267025 | 11.73508 | 26.0954 | 23.67916 | chr14 | 65265495 | 65266376 | 4.39405 | 3.84935 | 2.49982 |
| chr5 | 148399723 | 148400142 | 773 | Promoter (<=1kb) | 5 | 148327410 | 148399904 | 72495 | 2 | ENSMUST00 | 0 | Slc7a1 | ENSMUSG0C | false | chr5 | 148398764 | 148400373 | 30.66893 | 90.13895 | 86.79655 | chr5 | 148398697 | 148400532 | 3.58331 | 3.36268 | 1.82755 |
| chr5 | 105699869 | 105700915 | 773 | Promoter (<=1kb) | 5 | 105699969 | 105815215 | 115247 | 1 | ENSMUST00 | 0 | Lrrc8d | ENSMUSG0C | false | chr5 | 105698985 | 105701609 | 37.72209 | 126.01213 | 121.65854 | chr5 | 105700771 | 105705253 | 4.69176 | 3.9581 | 2.74133 |
| chr18 | 69345825 | 69346660 | 773 | Promoter (<=1kb) | 18 | 69346203 | 69683248 | 337046 | 1 | ENSMUST00 | 0 | Tcf4 | ENSMUSG0C | false | chr18 | 69345746 | 69346752 | 33.99591 | 106.86781 | 103.1211 | . | . | . | . | . | . |
| chr7 | 139247392 | 139247789 | 773 | Promoter (<=1kb) | 7 | 139248482 | 139267253 | 18772 | 1 | ENSMUST00 | -693 | Pwwp2b | ENSMUSG0C | false | chr7 | 139247202 | 139249854 | 22.89105 | 88.16154 | 84.85722 | chr7 | 139246761 | 139254099 | 8.53008 | 5.68016 | 5.90094 |
| chr8 | 68880338 | 68881554 | 772 | Promoter (<=1kb) | 8 | 68880491 | 68906934 | 26444 | 1 | ENSMUST00 | 0 | Lpl | ENSMUSG0C | false | chr8 | 68880055 | 68883509 | 28.08037 | 85.65662 | 82.39679 | chr8 | 68879897 | 68880457 | 3.26526 | 3.16727 | 1.56651 |
| chr14 | 26534038 | 26535062 | 772 | Promoter (<=1kb) | 14 | 26413168 | 26534931 | 121764 | 2 | ENSMUST00 | 0 | Slmap | ENSMUSG0C | false | chr14 | 26532901 | 26535352 | 44.72366 | 151.575 | 146.22792 | chr14 | 26525633 | 26534190 | 4.76793 | 3.94538 | 2.82588 |
| chr16 | 14182326 | 14182684 | 772 | Promoter (<=1kb) | 16 | 14183573 | 14190520 | 6948 | 1 | ENSMUST00 | -889 | Nde1 | ENSMUSG0C | false | . | . | . | . | . | . | chr16 | 14174692 | 14185475 | 6.74955 | 4.90863 | 4.43818 |
| chr17 | 68002270 | 68002648 | 772 | Promoter (1-2kb) | 17 | 67842713 | 68004120 | 161408 | 2 | ENSMUST00 | 1472 | Arhgap28 | ENSMUSG0C | true | chr17 | 68001890 | 68004635 | 25.93082 | 85.62936 | 82.37304 | chr17 | 67993413 | 68002591 | 4.56742 | 3.83999 | 2.64221 |
| chr19 | 58818721 | 58819349 | 772 | Intron (ENSMUST00000066285.4 | 19 | 58785803 | 58794461 | 8659 | 2 | ENSMUST00 | -24261 | 1700019N19 | ENSMUSG0C | false | . | . | . | . | . | . | chr19 | 58814047 | 58859817 | 7.46659 | 5.12677 | 5.03638 |
| chr3 | 108840114 | 108840684 | 772 | Promoter (<=1kb) | 3 | 108793176 | 108840511 | 47336 | 2 | ENSMUST00 | 0 | Stxbp3a | ENSMUSG0C | false | chr3 | 108839254 | 108840617 | 22.5292 | 69.37365 | 66.39268 | . | . | . | . | . | . |
| chr2 | 152727139 | 152727466 | 772 | Distal Intergenic | 2 | 152736251 | 152737410 | 1160 | 1 | ENSMUST00 | -8785 | Id1 | ENSMUSG0C | false | . | . | . | . | . | . | chr2 | 152723891 | 152727757 | 5.10737 | 3.93331 | 3.13758 |
| chr7 | 96775069 | 96775293 | 772 | Promoter (2-3kb) | 7 | 96777934 | 96797733 | 19800 | 1 | ENSMUST00 | -2641 | Tenm4 | ENSMUSG0C | false | . | . | . | . | . | . | chr7 | 96774590 | 96775914 | 4.42463 | 3.81828 | 2.5383 |
| chr17 | 35812872 | 35813608 | 771 | Distal Intergenic | 17 | 35821684 | 35822923 | 1240 | 1 | ENSMUST00 | -8076 | Ier3 | ENSMUSG0C | false | . | . | . | . | . | . | chr17 | 35810878 | 35819101 | 6.41913 | 4.79158 | 4.16085 |
| chr11 | 102318689 | 102319962 | 771 | Promoter (<=1kb) | 11 | 102304560 | 102319096 | 14537 | 2 | ENSMUST00 | 0 | Ubtf | ENSMUSG0C | false | chr11 | 102316341 | 102321029 | 40.78419 | 137.94737 | 133.15321 | . | . | . | . | . | . |
| chr9 | 99139623 | 99140581 | 771 | Promoter (<=1kb) | 9 | 99036654 | 99140087 | 103434 | 2 | ENSMUST00 | 0 | Pik3cb | ENSMUSG0C | false | chr9 | 99138922 | 99141004 | 33.41831 | 117.26761 | 113.21259 | . | . | . | . | . | . |
| chr10 | 75837249 | 75837749 | 771 | Promoter (2-3kb) | 10 | 75831845 | 75834881 | 3037 | 2 | ENSMUST00 | -2369 | Gstt2 | ENSMUSG0C | false | chr10 | 75836789 | 75837882 | 18.64736 | 62.11404 | 59.23804 | chr10 | 75833645 | 75838373 | 3.57502 | 3.3152 | 1.87776 |
| chrX | 12135533 | 12136163 | 771 | Intron (ENSMUST00000043441.7 | 20 | 12058981 | 12128400 | 69420 | 2 | ENSMUST00 | -7134 | Bcor | ENSMUSG0C | false | chrX | 12135593 | 12136131 | 9.65017 | 18.71579 | 16.39739 | . | . | . | . | . | . |
| chr17 | 81670072 | 81670450 | 771 | Distal Intergenic | 17 | 81388691 | 81649607 | 260917 | 2 | ENSMUST00 | -20466 | Slc8a1 | ENSMUSG0C | false | chr17 | 81669148 | 81670762 | 27.74132 | 80.34372 | 77.18787 | chr17 | 81669334 | 81670630 | 3.16979 | 3.08148 | 1.51085 |
| chr11 | 117410151 | 117410498 | 771 | Distal Intergenic | 11 | 117414389 | 117415368 | 980 | 1 | ENSMUST00 | -3891 | Gm11729 | ENSMUSG0C | false | . | . | . | . | . | . | chr11 | 117406406 | 117416739 | 6.0329 | 4.55882 | 3.85214 |

| | | | | | | | | | | | | | | | | | | | | | | | | | | |
|---|---|---|---|---|---|---|---|---|---|---|---|---|---|---|---|---|---|---|---|---|---|---|---|---|---|---|
| chr12 | 111377202 | 111378150 | 770 | Promoter (<=1kb) | 12 | 111292972 | 111377718 | 84747 | 2 | ENSMUST00 | 0 | Cdc42bpb | ENSMUSG00 | false | chr12 | 111375721 | 111378772 | 42.53133 | 142.11215 | 137.14622 | . | . | . | . | . | . |
| chr1 | 98073781 | 98074097 | 770 | Intron (ENSMUST00000058762.9 | 1 | 98053990 | 98057883 | 3894 | 2 | ENSMUST00 | -15899 | Pam | ENSMUSG00 | false | . | . | . | . | . | . | chr1 | 98068294 | 98075224 | 4.2597 | 3.67248 | 2.39947 |
| chr1 | 182497716 | 182498100 | 770 | Intron (ENSMUST00000068505.8 | 1 | 182467256 | 182517495 | 50240 | 2 | ENSMUST00 | 19395 | Capn2 | ENSMUSG00 | false | . | . | . | . | . | . | chr1 | 182492178 | 182516499 | 7.84019 | 5.29195 | 5.34945 |
| chr4 | 53430452 | 53430814 | 770 | Distal Intergenic | 4 | 53440413 | 53550200 | 109788 | 1 | ENSMUST00 | -9599 | Slc44a1 | ENSMUSG00 | false | . | . | . | . | . | . | chr4 | 53430405 | 53431010 | 3.69614 | 3.46242 | 1.92526 |
| chr18 | 38906367 | 38906726 | 770 | Intron (ENSMUST00000148850.2 | 18 | 38838673 | 38918771 | 80099 | 2 | ENSMUST00 | 12045 | Fgf1 | ENSMUSG00 | false | . | . | . | . | . | . | chr18 | 38905811 | 38908304 | 3.57846 | 3.39889 | 1.82251 |
| chr9 | 65811510 | 65811773 | 770 | Intron (ENSMUST00000159109.1 | 9 | 65795230 | 65797912 | 2683 | 2 | ENSMUST00 | -13599 | Zfp609 | ENSMUSG00 | false | . | . | . | . | . | . | chr9 | 65810643 | 65815700 | 5.13127 | 4.16567 | 3.0986 |
| chr6 | 112327782 | 112328198 | 770 | Intron (ENSMUST00000032376.5 | 6 | 112285857 | 112288837 | 2981 | 2 | ENSMUST00 | -38946 | 5031434C07I | ENSMUSG00 | false | . | . | . | . | . | . | chr6 | 112327754 | 112328143 | 3.20431 | 2.8903 | 1.56536 |
| chr11 | 54884930 | 54885305 | 769 | Distal Intergenic | 11 | 54902453 | 54909576 | 7124 | 1 | ENSMUST00 | -17148 | Gpx3 | ENSMUSG00 | false | . | . | . | . | . | . | chr11 | 54880911 | 54885790 | 4.35328 | 3.73277 | 2.48101 |
| chr11 | 98780400 | 98780662 | 769 | Distal Intergenic | 11 | 98767932 | 98775333 | 7402 | 2 | ENSMUST00 | -5068 | Nr1d1 | ENSMUSG00 | false | . | . | . | . | . | . | chr11 | 98780022 | 98794752 | 7.02549 | 5.01464 | 4.66092 |
| chr19 | 6241316 | 6241774 | 769 | Promoter (<=1kb) | 19 | 6241668 | 6262335 | 20668 | 1 | ENSMUST00 | 0 | Atg2a | ENSMUSG00 | false | chr19 | 6241598 | 6242537 | 34.90036 | 108.43528 | 104.64344 | chr19 | 6240643 | 6243390 | 4.41442 | 3.85184 | 2.52793 |
| chr6 | 52224565 | 52224842 | 769 | Promoter (1-2kb) | 6 | 52224121 | 52226189 | 2069 | 2 | ENSMUST00 | 1347 | Hoxa9 | ENSMUSG00 | false | chr6 | 52221266 | 52227348 | 34.16929 | 120.7392 | 116.5701 | . | . | . | . | . | . |
| chr7 | 110297148 | 110297787 | 769 | Distal Intergenic | 7 | 110320741 | 110340285 | 19545 | 2 | ENSMUST00 | 42498 | Sbf2 | ENSMUSG00 | true | . | . | . | . | . | . | chr7 | 110295743 | 110298262 | 6.6317 | 4.79008 | 4.36229 |
| chr6 | 117526529 | 117526814 | 769 | Distal Intergenic | 6 | 117559697 | 117568193 | 8497 | 1 | ENSMUST00 | -32883 | Gm9946 | ENSMUSG00 | false | . | . | . | . | . | . | chr6 | 117525715 | 117527547 | 7.47158 | 4.9509 | 5.07423 |
| chr1 | 106629433 | 106629830 | 769 | Intron (ENSMUST00000112751.1 | 1 | 106546496 | 106546631 | 136 | 2 | ENSMUST00 | -82803 | Mir3473f | ENSMUSG00 | false | . | . | . | . | . | . | chr1 | 106627258 | 106640742 | 4.33032 | 3.75351 | 2.45634 |
| chr18 | 44437599 | 44437908 | 768 | Intron (ENSMUST00000164666.2 | 18 | 44380500 | 44424969 | 44470 | 1 | ENSMUST00 | 57100 | Dcp2 | ENSMUSG00 | false | . | . | . | . | . | . | chr18 | 44434846 | 44445255 | 6.64362 | 4.65907 | 4.40138 |
| chr19 | 42526365 | 42526749 | 768 | Intron (ENSMUST00000026188.4 | 19 | 42518822 | 42576100 | 57279 | 1 | ENSMUST00 | 7544 | R3hcc1l | ENSMUSG00 | false | . | . | . | . | . | . | chr19 | 42523799 | 42528396 | 5.00025 | 4.07227 | 3.01528 |
| chrX | 12078571 | 12079103 | 768 | Promoter (1-2kb) | 20 | 12036740 | 12080553 | 43814 | 2 | ENSMUST00 | 1450 | Bcor | ENSMUSG00 | false | chrX | 12076787 | 12081284 | 21.13848 | 54.99947 | 52.22001 | . | . | . | . | . | . |
| chr1 | 59968909 | 59969794 | 768 | Promoter (<=1kb) | 1 | 59970680 | 59981515 | 10836 | 1 | ENSMUST00 | -886 | Fam117b | ENSMUSG00 | true | . | . | . | . | . | . | chr1 | 59964490 | 59970929 | 4.42856 | 3.66443 | 2.5847 |
| chr18 | 84588938 | 84589841 | 768 | Promoter (<=1kb) | 18 | 84207704 | 84589504 | 381801 | 2 | ENSMUST00 | 0 | Zfp407 | ENSMUSG00 | false | chr18 | 84588660 | 84590582 | 34.00884 | 130.01486 | 125.52227 | . | . | . | . | . | . |
| chr14 | 78907316 | 78907614 | 768 | Intron (ENSMUST00000040990.5 | 14 | 78849178 | 79202310 | 353133 | 1 | ENSMUST00 | 58139 | Vwa8 | ENSMUSG00 | false | . | . | . | . | . | . | chr14 | 78906534 | 78909736 | 3.51337 | 3.228 | 1.83077 |
| chr5 | 66365510 | 66365914 | 768 | Intron (ENSMUST00000159786.2 | 5 | 66356918 | 66357185 | 268 | 1 | ENSMUST00 | 8593 | Gm16273 | ENSMUSG00 | false | . | . | . | . | . | . | chr5 | 66352002 | 66369698 | 7.18935 | 5.00706 | 4.81216 |
| chr13 | 13590169 | 13590704 | 767 | Promoter (<=1kb) | 13 | 13590409 | 13777440 | 187032 | 1 | ENSMUST00 | 0 | Lyst | ENSMUSG00 | false | chr13 | 13589771 | 13591179 | 43.4883 | 142.6545 | 137.66066 | . | . | . | . | . | . |
| chr16 | 17489422 | 17489840 | 767 | Promoter (<=1kb) | 16 | 17489611 | 17507485 | 17875 | 1 | ENSMUST00 | 0 | Aifm3 | ENSMUSG00 | false | chr16 | 17489151 | 17490431 | 26.42849 | 77.14894 | 74.04846 | chr16 | 17485729 | 17499611 | 5.1843 | 4.03502 | 3.18262 |
| chr16 | 37809338 | 37809584 | 767 | Intron (ENSMUST00000114763.2 | 16 | 37815706 | 37834426 | 18721 | 1 | ENSMUST00 | -6122 | Fstl1 | ENSMUSG00 | false | . | . | . | . | . | . | chr16 | 37808227 | 37809775 | 4.06507 | 3.56083 | 2.2759 |
| chr17 | 26507977 | 26509004 | 767 | Promoter (<=1kb) | 17 | 26505590 | 26508519 | 2930 | 2 | ENSMUST00 | 0 | Dusp1 | ENSMUSG00 | false | chr17 | 26505964 | 26508522 | 39.08731 | 125.158 | 120.83892 | . | . | . | . | . | . |
| chr8 | 108703462 | 108704964 | 767 | Promoter (<=1kb) | 8 | 108703100 | 108800270 | 97171 | 1 | ENSMUST00 | 363 | Zfhx3 | ENSMUSG00 | false | chr8 | 108703474 | 108705192 | 41.07278 | 140.67711 | 135.76277 | chr8 | 108704809 | 108705039 | 3.8701 | 3.58539 | 2.05955 |
| chr19 | 4191676 | 4192267 | 767 | Promoter (<=1kb) | 19 | 4192158 | 4195419 | 3262 | 1 | ENSMUST00 | 0 | Ppp1ca | ENSMUSG00 | false | chr19 | 4191540 | 4191879 | 5.6323 | 9.47728 | 7.31216 | chr19 | 4190739 | 4193991 | 6.41412 | 4.42752 | 4.20585 |
| chr2 | 74689944 | 74690329 | 767 | Promoter (1-2kb) | 2 | 74691924 | 74695105 | 3182 | 1 | ENSMUST00 | -1595 | Hoxd10 | ENSMUSG00 | false | . | . | . | . | . | . | chr2 | 74686918 | 74693419 | 4.34438 | 3.76944 | 2.4861 |
| chr16 | 18348158 | 18348637 | 766 | Promoter (<=1kb) | 16 | 18348182 | 18407076 | 58895 | 1 | ENSMUST00 | 0 | Arvcf | ENSMUSG00 | true | chr16 | 18347868 | 18349368 | 21.08548 | 68.47759 | 65.51048 | chr16 | 18347547 | 18353348 | 4.32451 | 3.68001 | 2.47557 |
| chr12 | 111485666 | 111486787 | 766 | Promoter (<=1kb) | 12 | 111484611 | 111485823 | 1213 | 2 | ENSMUST00 | 0 | Gm266 | ENSMUSG00 | false | chr12 | 111484468 | 111487284 | 26.28407 | 85.32412 | 82.07336 | chr12 | 111482224 | 111493728 | 5.00155 | 3.95404 | 3.01745 |
| chr15 | 36608082 | 36608675 | 766 | Promoter (<=1kb) | 15 | 36595661 | 36608973 | 13313 | 2 | ENSMUST00 | 298 | Pabpc1 | ENSMUSG00 | false | chr15 | 36607571 | 36609023 | 36.83603 | 116.99731 | 112.9521 | . | . | . | . | . | . |
| chr4 | 88032627 | 88033579 | 766 | Promoter (<=1kb) | 4 | 87769925 | 88033364 | 263440 | 2 | ENSMUST00 | 0 | Mllt3 | ENSMUSG00 | false | chr4 | 88031234 | 88034901 | 31.70772 | 93.8754 | 90.44607 | . | . | . | . | . | . |
| chr10 | 127614812 | 127615379 | 766 | Intron (ENSMUST00000049149.9 | 10 | 127599884 | 127620922 | 21039 | 2 | ENSMUST00 | 5543 | Lrp1 | ENSMUSG00 | false | . | . | . | . | . | . | . | . | . | . | . | . |
| chr8 | 87822155 | 87822519 | 766 | Intron (ENSMUST00000109655.3 | 8 | 87661810 | 87804439 | 142630 | 2 | ENSMUST00 | -17717 | Zfp423 | ENSMUSG00 | true | . | . | . | . | . | . | chr8 | 87821288 | 87824745 | 5.43963 | 4.12087 | 3.41695 |
| chr14 | 45350882 | 45351385 | 766 | Promoter (<=1kb) | 14 | 45351473 | 45373585 | 22113 | 1 | ENSMUST00 | -88 | Styx | ENSMUSG00 | false | chr14 | 45351108 | 45352008 | 33.28429 | 109.73968 | 105.91746 | . | . | . | . | . | . |
| chr4 | 152009465 | 152009930 | 766 | Promoter (<=1kb) | 4 | 152008803 | 152017680 | 8878 | 1 | ENSMUST00 | 663 | Klhl21 | ENSMUSG00 | false | chr4 | 152007699 | 152011153 | 30.9834 | 105.62366 | 101.90845 | chr4 | 152006449 | 152014835 | 5.30205 | 4.17118 | 3.25701 |
| chr17 | 71165685 | 71165978 | 765 | Intron (ENSMUST00000180743.1 | 17 | 71182560 | 71235119 | 52560 | 1 | ENSMUST00 | -16582 | Lpin2 | ENSMUSG00 | false | . | . | . | . | . | . | chr17 | 71157703 | 71180230 | 5.23179 | 4.17367 | 3.19377 |
| chr1 | 33814272 | 33814648 | 765 | Promoter (<=1kb) | 1 | 33761544 | 33814595 | 53052 | 2 | ENSMUST00 | 0 | Zfp451 | ENSMUSG00 | false | chr1 | 33813492 | 33814814 | 34.3512 | 113.70341 | 109.76276 | chr1 | 33810457 | 33814926 | 3.75553 | 3.27946 | 2.01557 |
| chr13 | 60131753 | 60132085 | 765 | Distal Intergenic | 13 | 60174405 | 60177365 | 2961 | 2 | ENSMUST00 | 45280 | Gas1 | ENSMUSG00 | false | . | . | . | . | . | . | chr13 | 60131864 | 60133118 | 4.63241 | 3.80634 | 2.73356 |
| chr9 | 63059183 | 63059535 | 765 | Distal Intergenic | 9 | 63027560 | 63027717 | 158 | 1 | ENSMUST00 | 31624 | Gm24526 | ENSMUSG00 | false | chr9 | 63059114 | 63059654 | 8.3822 | 16.12319 | 13.84377 | chr9 | 63058556 | 63059791 | 6.02332 | 4.52561 | 3.88229 |
| chr4 | 41760296 | 41760649 | 765 | Promoter (<=1kb) | 4 | 41760454 | 41769474 | 9021 | 1 | ENSMUST00 | 0 | Il11ra1 | ENSMUSG00 | false | chr4 | 41760142 | 41761120 | 15.39495 | 41.17039 | 38.56658 | chr4 | 41759912 | 41762981 | 4.66199 | 3.93244 | 2.76293 |
| chrX | 71528761 | 71529072 | 765 | Promoter (2-3kb) | 20 | 71531285 | 71531376 | 92 | 2 | ENSMUST00 | 2304 | Gm23945 | ENSMUSG00 | false | . | . | . | . | . | . | . | . | . | . | . | . |
| chr4 | 124657111 | 124657712 | 765 | Promoter (<=1kb) | 4 | 124528924 | 124657169 | 128246 | 2 | ENSMUST00 | 0 | Gm27247 | ENSMUSG00 | false | chr4 | 124656736 | 124658371 | 25.16654 | 87.04672 | 83.76057 | chr4 | 124656567 | 124660244 | 7.68435 | 5.26787 | 5.21496 |
| chr13 | 18948217 | 18948488 | 764 | Promoter (<=1kb) | 13 | 18948371 | 19150913 | 202543 | 1 | ENSMUST00 | 0 | Amph | ENSMUSG00 | true | chr13 | 18947976 | 18949900 | 35.11596 | 108.8079 | 105.00359 | chr13 | 18947828 | 18949258 | 2.70042 | 2.81322 | 1.16604 |
| chr1 | 172632435 | 172633262 | 764 | Promoter (<=1kb) | 1 | 172630769 | 172632974 | 2206 | 2 | ENSMUST00 | 0 | Dusp23 | ENSMUSG00 | false | chr1 | 172632236 | 172633275 | 24.82645 | 85.46523 | 82.21117 | . | . | . | . | . | . |
| chr5 | 146700318 | 146700966 | 764 | Distal Intergenic | 5 | 146734839 | 146758448 | 23610 | 2 | ENSMUST00 | 57482 | Usp12 | ENSMUSG00 | false | . | . | . | . | . | . | chr5 | 146699668 | 146701542 | 5.47016 | 4.34362 | 3.37679 |
| chr11 | 75139684 | 75139961 | 764 | Intron (ENSMUST00000045281.7 | 11 | 75132435 | 75139369 | 6935 | 1 | ENSMUST00 | 7250 | Smg6 | ENSMUSG00 | false | . | . | . | . | . | . | chr11 | 75136734 | 75141926 | 6.17803 | 4.48241 | 3.99662 |
| chr13 | 44706830 | 44707122 | 764 | Distal Intergenic | 13 | 44729474 | 44731574 | 2101 | 1 | ENSMUST00 | -22352 | Jarid2 | ENSMUSG00 | false | . | . | . | . | . | . | chr13 | 44705169 | 44707763 | 4.94462 | 4.03052 | 2.9764 |
| chr13 | 98358800 | 98359164 | 764 | Distal Intergenic | 13 | 98354242 | 98356703 | 2462 | 1 | ENSMUST00 | 4559 | Foxd1 | ENSMUSG00 | false | chr13 | 98353706 | 98359675 | 27.78601 | 91.88203 | 88.49705 | . | . | . | . | . | . |
| chr12 | 106401365 | 106401691 | 764 | Distal Intergenic | 12 | 106442653 | 106447666 | 5014 | 2 | ENSMUST00 | 45975 | 1700121N20 | ENSMUSG00 | false | . | . | . | . | . | . | chr12 | 106400257 | 106402557 | 4.16712 | 3.56716 | 2.36244 |
| chr2 | 30237490 | 30238158 | 763 | Promoter (<=1kb) | 2 | 30237715 | 30263785 | 26071 | 1 | ENSMUST00 | 0 | Lrrc8a | ENSMUSG00 | false | chr2 | 30236663 | 30238701 | 37.89522 | 119.88892 | 115.74806 | . | . | . | . | . | . |
| chr7 | 66449242 | 66449469 | 763 | Distal Intergenic | 7 | 66390892 | 66427517 | 36626 | 2 | ENSMUST00 | -21726 | Aldh1a3 | ENSMUSG00 | false | . | . | . | . | . | . | chr7 | 66438246 | 66451395 | 6.06663 | 4.59192 | 3.88434 |
| chr10 | 68105220 | 68105597 | 763 | Intron (ENSMUST00000020106.7 | 10 | 68043968 | 68051046 | 7079 | 1 | ENSMUST00 | 61253 | Rtkn2 | ENSMUSG00 | false | . | . | . | . | . | . | chr10 | 68101409 | 68108261 | 4.9792 | 4.05161 | 2.98005 |
| chr13 | 99238772 | 99239298 | 763 | Intron (ENSMUST00000056558.8 | 13 | 99184823 | 99247037 | 62215 | 1 | ENSMUST00 | 53950 | Zfp366 | ENSMUSG00 | false | . | . | . | . | . | . | chr13 | 99235541 | 99241835 | 4.51265 | 3.8659 | 2.60223 |
| chr3 | 105872497 | 105873070 | 763 | Promoter (1-2kb) | 3 | 105870907 | 105924036 | 53130 | 1 | ENSMUST00 | 1591 | Adora3 | ENSMUSG00 | false | . | . | . | . | . | . | chr3 | 105871739 | 105872878 | 5.02086 | 4.14457 | 3.07065 |
| chr3 | 105300791 | 105301185 | 763 | Distal Intergenic | 3 | 105441595 | 105451727 | 10133 | 2 | ENSMUST00 | 150542 | Kcnd3os | ENSMUSG00 | false | . | . | . | . | . | . | . | . | . | . | . | . |
| chr3 | 88729419 | 88729678 | 763 | 3' UTR | 3 | 88722090 | 88722670 | 581 | 1 | ENSMUST00 | 7330 | Gm17146 | ENSMUSG00 | false | . | . | . | . | . | . | chr3 | 88728269 | 88730490 | 4.26038 | 3.63417 | 2.42686 |
| chr2 | 25054243 | 25054748 | 762 | Promoter (<=1kb) | 2 | 25054355 | 25062881 | 8527 | 1 | ENSMUST00 | 0 | Nsmf | ENSMUSG00 | false | chr2 | 25054278 | 25055297 | 29.09699 | 101.16944 | 97.56942 | chr2 | 25053989 | 25055497 | 5.74985 | 4.54731 | 3.61697 |
| chr2 | 152814595 | 152814885 | 762 | Intron (ENSMUST00000007803.6 | 2 | 152781411 | 152828535 | 47125 | 2 | ENSMUST00 | 13650 | Bcl2l1 | ENSMUSG00 | false | . | . | . | . | . | . | chr2 | 152808253 | 152830242 | 5.83553 | 4.3234 | 3.71708 |
| chr14 | 60588511 | 60588834 | 762 | Intron (ENSMUST00000190392.1 | 14 | 60634001 | 60691367 | 57367 | 1 | ENSMUST00 | -45167 | Spata13 | ENSMUSG00 | true | . | . | . | . | . | . | chr14 | 60585347 | 60599096 | 6.17547 | 4.51754 | 3.99543 |
| chr19 | 7482974 | 7483477 | 762 | Promoter (<=1kb) | 19 | 7425901 | 7483281 | 57381 | 2 | ENSMUST00 | 0 | Rtn3 | ENSMUSG00 | false | chr19 | 7481900 | 7483910 | 31.28813 | 116.44205 | 112.41405 | . | . | . | . | . | . |
| chr8 | 61215139 | 61215388 | 762 | Distal Intergenic | 8 | 61224171 | 61396072 | 171902 | 1 | ENSMUST00 | -8783 | Sh3rf1 | ENSMUSG00 | false | . | . | . | . | . | . | chr8 | 61214500 | 61217480 | 3.91198 | 3.5546 | 2.11443 |
| chr4 | 11485637 | 11486114 | 762 | Promoter (<=1kb) | 4 | 11485958 | 11527621 | 41664 | 1 | ENSMUST00 | 0 | 1110037F02I | ENSMUSG00 | false | chr4 | 11485225 | 11487018 | 47.07127 | 162.57588 | 156.84026 | . | . | . | . | . | . |
| chrX | 99975241 | 99975649 | 762 | Promoter (<=1kb) | 20 | 99975606 | 100311299 | 335694 | 1 | ENSMUST00 | 0 | Eda | ENSMUSG00 | false | chrX | 99974986 | 99976711 | 18.84082 | 47.13427 | 44.45482 | . | . | . | . | . | . |
| chr17 | 47975184 | 47975593 | 762 | Distal Intergenic | 17 | 48003379 | 48003572 | 194 | 2 | ENSMUST00 | 27979 | Gm14871 | ENSMUSG00 | false | . | . | . | . | . | . | chr17 | 47973523 | 47976221 | 5.86638 | 4.39509 | 3.7429 |
| chr7 | 28810770 | 28811587 | 761 | Promoter (<=1kb) | 7 | 28810890 | 28822266 | 11377 | 1 | ENSMUST00 | 0 | Hnrnpl | ENSMUSG00 | false | chr7 | 28810797 | 28812960 | 38.55812 | 129.64772 | 125.16568 | . | . | . | . | . | . |
| chr4 | 132922304 | 132922960 | 761 | Promoter (<=1kb) | 4 | 132899213 | 132922558 | 23346 | 2 | ENSMUST00 | 0 | Fam76a | ENSMUSG00 | false | chr4 | 132921541 | 132922574 | 36.7977 | 133.03651 | 128.43381 | . | . | . | . | . | . |
| chr5 | 30772331 | 30772757 | 761 | Intron (ENSMUST00000114729.2 | 5 | 30778130 | 30786934 | 8805 | 1 | ENSMUST00 | -5373 | Dpysl5 | ENSMUSG00 | false | . | . | . | . | . | . | chr5 | 30772715 | 30775704 | 5.52607 | 4.27296 | 3.48816 |
| chr7 | 18873254 | 18873602 | 761 | Distal Intergenic | 7 | 18884690 | 18890438 | 5749 | 1 | ENSMUST00 | -11088 | Pglyrp1 | ENSMUSG00 | false | chr7 | 18872771 | 18873495 | 11.4883 | 23.92001 | 21.53215 | chr7 | 18872583 | 18873376 | 4.65473 | 3.87614 | 2.76407 |
| chr2 | 170158281 | 170158587 | 761 | Distal Intergenic | 2 | 170120719 | 170148103 | 27385 | 2 | ENSMUST00 | -10179 | Zfp217 | ENSMUSG00 | false | . | . | . | . | . | . | chr2 | 170157455 | 170159018 | 5.04661 | 4.16375 | 3.03459 |
| chr11 | 63173598 | 63173950 | 761 | Promoter (<=1kb) | 11 | 63173914 | 63188454 | 14541 | 1 | ENSMUST00 | 0 | B430319H21 | ENSMUSG00 | false | . | . | . | . | . | . | chr11 | 63172118 | 63176091 | 4.6369 | 3.91772 | 2.70825 |
| chr12 | 86793921 | 86794261 | 761 | Distal Intergenic | 12 | 86809957 | 86810258 | 302 | 1 | ENSMUST00 | -15696 | Gm25888 | ENSMUSG00 | false | . | . | . | . | . | . | chr12 | 86789697 | 86796554 | 7.66516 | 5.33848 | 5.17204 |
| chr11 | 76494367 | 76495204 | 760 | Promoter (<=1kb) | 11 | 76458474 | 76495005 | 36532 | 2 | ENSMUST00 | 0 | Abr | ENSMUSG00 | false | chr11 | 76493190 | 76495482 | 27.58279 | 90.54513 | 87.1897 | chr11 | 76485843 | 76495997 | 5.9622 | 4.24487 | 3.82757 |
| chr16 | 85172892 | 85173792 | 760 | Promoter (<=1kb) | 16 | 84954440 | 85173707 | 219268 | 2 | ENSMUST00 | 0 | App | ENSMUSG00 | false | chr16 | 85171787 | 85174244 | 30.94804 | 114.76607 | 110.79114 | . | . | . | . | . | . |
| chr3 | 9896387 | 9896661 | 760 | Distal Intergenic | 3 | 9687479 | 9833679 | 146201 | 2 | ENSMUST00 | -62709 | Pag1 | ENSMUSG00 | true | . | . | . | . | . | . | chr3 | 9895503 | 9897888 | 5.17413 | 3.99997 | 3.1797 |
| chr6 | 61180092 | 61180566 | 760 | Promoter (<=1kb) | 6 | 61180324 | 62382865 | 1202542 | 1 | ENSMUST00 | 0 | Ccser1 | ENSMUSG00 | false | chr6 | 61179826 | 61181514 | 25.53309 | 82.08177 | 78.89293 | . | . | . | . | . | . |
| chrX | 11735081 | 11735396 | 760 | Distal Intergenic | 20 | 11684002 | 11684124 | 123 | 2 | ENSMUST00 | -50958 | Gm14513 | ENSMUSG00 | false | . | . | . | . | . | . | . | . | . | . | . | . |
| chr2 | 8825487 | 8825781 | 760 | Distal Intergenic | 2 | 8633937 | 8634483 | 547 | 1 | ENSMUST00 | 191551 | Gm13255 | ENSMUSG00 | false | . | . | . | . | . | . | . | . | . | . | . | . |
| chr15 | 81487605 | 81487891 | 760 | Distal Intergenic | 15 | 81499084 | 81499548 | 465 | 2 | ENSMUST00 | 11657 | Gm5218 | ENSMUSG00 | false | . | . | . | . | . | . | chr15 | 81487137 | 81488004 | 3.99816 | 3.64193 | 2.20603 |
| chr11 | 98783234 | 98783766 | 759 | Distal Intergenic | 11 | 98767932 | 98775333 | 7402 | 2 | ENSMUST00 | -7902 | Nr1d1 | ENSMUSG00 | false | . | . | . | . | . | . | chr11 | 98780022 | 98794752 | 7.02549 | 5.01464 | 4.66092 |
| chr16 | 21644154 | 21644613 | 759 | 3' UTR | 16 | 21627631 | 21644416 | 16786 | 1 | ENSMUST00 | 16524 | Vps8 | ENSMUSG00 | false | chr16 | 21644103 | 21644448 | 5.90127 | 9.23487 | 7.07661 | chr16 | 21643311 | 21645332 | 6.77387 | 4.95286 | 4.46587 |
| chr11 | 4497463 | 4497777 | 759 | Intron (ENSMUST00000109943.4 | 11 | 4491807 | 4493318 | 1512 | 2 | ENSMUST00 | -4146 | Mtmr3 | ENSMUSG00 | false | . | . | . | . | . | . | chr11 | 4495530 | 4499752 | 5.45059 | 4.2307 | 3.39426 |
| chr2 | 37799075 | 37799431 | 759 | 3' UTR | 2 | 37795169 | 37796616 | 1448 | 1 | ENSMUST00 | 3907 | Crb2 | ENSMUSG00 | false | . | . | . | . | . | . | chr2 | 37777133 | 37803419 | 7.19536 | 4.98761 | 4.82001 |
| chr1 | 79671860 | 79672294 | 759 | Promoter (<=1kb) | 1 | 79606876 | 79671972 | 65097 | 2 | ENSMUST00 | 0 | Ap1s3 | ENSMUSG00 | false | chr1 | 79670988 | 79672518 | 25.56571 | 75.64988 | 72.57267 | chr1 | 79666523 | 79672410 | 4.5349 | 3.86218 | 2.62318 |
| chr7 | 113207065 | 113207941 | 759 | Promoter (<=1kb) | 7 | 113207465 | 113314122 | 106658 | 1 | ENSMUST00 | 0 | Arntl | ENSMUSG00 | false | chr7 | 113206430 | 113207171 | 19.04506 | 49.03148 | 46.32841 | . | . | . | . | . | . |
| chr4 | 82344893 | 82345163 | 759 | Intron (ENSMUST00000050872.9 | 4 | 82439410 | 82439514 | 105 | 1 | ENSMUST00 | -94247 | n-R5s188 | ENSMUSG00 | false | . | . | . | . | . | . | chr4 | 82344940 | 82345386 | 4.26796 | 3.29205 | 2.44199 |
| chr10 | 80621979 | 80622902 | 758 | Promoter (<=1kb) | 10 | 80622780 | 80640771 | 17992 | 1 | ENSMUST00 | 0 | Csnk1g2 | ENSMUSG00 | false | chr10 | 80621560 | 80622431 | 19.04495 | 59.55748 | 56.71581 | . | . | . | . | . | . |
| chr19 | 34921962 | 34922411 | 758 | Promoter (<=1kb) | 19 | 34922358 | 34975731 | 53374 | 1 | ENSMUST00 | 0 | Kif20b | ENSMUSG00 | false | chr19 | 34921589 | 34922788 | 26.61343 | 84.02032 | 80.79139 | chr19 | 34921182 | 34922870 | 3.51877 | 3.31828 | 1.78377 |
| chr11 | 72086688 | 72087135 | 758 | Intron (ENSMUST00000108508.2 | 11 | 72057645 | 72061561 | 3917 | 2 | ENSMUST00 | -25128 | Pitpnm3 | ENSMUSG00 | false | . | . | . | . | . | . | chr11 | 72080141 | 72089941 | 5.61997 | 4.30448 | 3.51854 |
| chr12 | 112488336 | 112488665 | 758 | Promoter (<=1kb) | 12 | 112489448 | 112499927 | 10480 | 1 | ENSMUST00 | -783 | A530016L24I | ENSMUSG00 | false | . | . | . | . | . | . | chr12 | 112488214 | 112488896 | 5.94954 | 4.36818 | 3.81656 |
| chr5 | 37338239 | 37338604 | 758 | Promoter (<=1kb) | 5 | 37338499 | 37425055 | 86557 | 1 | ENSMUST00 | 0 | Evc2 | ENSMUSG00 | false | chr5 | 37337843 | 37338935 | 27.5102 | 81.81912 | 78.63541 | chr5 | 37336340 | 37339292 | 3.80113 | 3.36577 | 2.03207 |
| chr6 | 121004498 | 121004829 | 758 | Promoter (1-2kb) | 6 | 120958343 | 121003153 | 44811 | 2 | ENSMUST00 | -1346 | Mical3 | ENSMUSG00 | false | . | . | . | . | . | . | chr6 | 120994546 | 121005852 | 6.94439 | 4.93247 | 4.60074 |
| chr4 | 114486234 | 114486564 | 758 | Intron (ENSMUST00000094894.3 | 4 | 114406724 | 114615098 | 208375 | 1 | ENSMUST00 | 79511 | Trabd2b | ENSMUSG00 | false | . | . | . | . | . | . | chr4 | 114483011 | 114492191 | 5.92541 | 4.42125 | 3.78407 |
| chr1 | 60657004 | 60657406 | 758 | Distal Intergenic | 1 | 60613698 | 60613803 | 106 | 2 | ENSMUST00 | -43202 | Gm23762 | ENSMUSG00 | false | . | . | . | . | . | . | chr1 | 60655736 | 60657534 | 4.51763 | 3.77894 | 2.66384 |
| chr14 | 47839492 | 47840009 | 757 | Distal Intergenic | 14 | 47729659 | 47736561 | 6903 | 1 | ENSMUST00 | 109834 | Ktn1 | ENSMUSG00 | false | . | . | . | . | . | . | chr14 | 47837556 | 47840903 | 5.37181 | 4.30947 | 3.30593 |
| chr14 | 30974273 | 30974571 | 757 | Exon (ENSMUST00000050171.8/ | 14 | 30951486 | 30987439 | 35954 | 1 | ENSMUST00 | 22788 | Nek4 | ENSMUSG00 | false | . | . | . | . | . | . | chr14 | 30973573 | 30975052 | 4.86597 | 3.79624 | 2.93718 |
| chr18 | 61044806 | 61045186 | 757 | Promoter (<=1kb) | 18 | 61045150 | 61085061 | 39912 | 1 | ENSMUST00 | 0 | Pdgfrb | ENSMUSG00 | false | chr18 | 61045113 | 61046136 | 5.10133 | 8.58512 | 6.43618 | chr18 | 61043544 | 61047108 | 4.09064 | 3.56793 | 2.29529 |
| chr2 | 28401558 | 28401969 | 757 | Distal Intergenic | 2 | 28446800 | 28450506 | 3707 | 1 | ENSMUST00 | -44831 | Ppp1r26 | ENSMUSG00 | false | . | . | . | . | . | . | chr2 | 28401348 | 28402386 | 4.57994 | 3.93624 | 2.64374 |
| chr17 | 24767596 | 24767924 | 757 | Distal Intergenic | 17 | 24753003 | 24758683 | 5681 | 1 | ENSMUST00 | 14594 | Hs3st6 | ENSMUSG00 | false | . | . | . | . | . | . | chr17 | 24755141 | 24772276 | 6.3257 | 4.70291 | 4.09799 |
| chrX | 152342340 | 152342862 | 757 | Promoter (<=1kb) | 20 | 152336852 | 152342425 | 5574 | 2 | ENSMUST00 | 0 | Tspyl2 | ENSMUSG00 | false | chrX | 152341792 | 152343312 | 15.62409 | 36.57069 | 34.02363 | . | . | . | . | . | . |
| chr7 | 37361765 | 37362388 | 757 | Distal Intergenic | 7 | 37299265 | 37304034 | 4770 | 1 | ENSMUST00 | 62501 | RP23-291B1. | ENSMUSG00 | false | chr7 | 37361765 | 37362385 | 7.76973 | 17.46802 | 15.16681 | chr7 | 37359518 | 37363322 | 4.40551 | 3.78336 | 2.55771 |
| chr15 | 81761517 | 81761834 | 756 | Intron (ENSMUST00000109554.1 | 15 | 81744848 | 81796269 | 51422 | 1 | ENSMUST00 | 16670 | Zc3h7b | ENSMUSG00 | false | . | . | . | . | . | . | chr15 | 81757391 | 81763369 | 5.27365 | 4.13537 | 3.24889 |
| chr8 | 80849837 | 80850182 | 756 | Intron (ENSMUST00000034150.8 | 8 | 80764434 | 80880479 | 116046 | 2 | ENSMUST00 | 30297 | Gab1 | ENSMUSG00 | false | . | . | . | . | . | . | chr8 | 80802749 | 80878363 | 5.64706 | 4.31183 | 3.55694 |
| chr9 | 70138580 | 70138971 | 756 | Promoter (2-3kb) | 9 | 70089310 | 70141578 | 52269 | 2 | ENSMUST00 | 2607 | Fam81a | ENSMUSG00 | true | . | . | . | . | . | . | chr9 | 70126965 | 70143919 | 5.28781 | 4.22747 | 3.24757 |
| chr13 | 102898770 | 102899110 | 756 | Intron (ENSMUST00000166726.2 | 13 | 102741969 | 102905786 | 163818 | 2 | ENSMUST00 | 6676 | Mast4 | ENSMUSG00 | true | chr13 | 102898513 | 102899308 | 6.28665 | 10.55282 | 8.36284 | chr13 | 102868528 | 102904155 | 5.63021 | 4.27399 | 3.54307 |
| chr2 | 154613272 | 154613707 | 756 | Promoter (<=1kb) | 2 | 154613297 | 154646821 | 33525 | 1 | ENSMUST00 | 0 | Zfp341 | ENSMUSG00 | false | chr2 | 154612811 | 154614277 | 25.15761 | 80.47005 | 77.31023 | . | . | . | . | . | . |
| chr15 | 31413030 | 31413375 | 756 | Distal Intergenic | 15 | 31446380 | 31446817 | 438 | 2 | ENSMUST00 | 33442 | Gm6361 | ENSMUSG00 | false | . | . | . | . | . | . | chr15 | 31412352 | 31414698 | 5.10299 | 4.05495 | 3.10063 |
| chr2 | 173852326 | 173852826 | 756 | Distal Intergenic | 2 | 173793437 | 173847670 | 54234 | 2 | ENSMUST00 | -4657 | Gm14371 | ENSMUSG00 | false | . | . | . | . | . | . | chr2 | 173847661 | 173860588 | 5.03291 | 4.0817 | 3.03782 |
| chr14 | 70259438 | 70259926 | 755 | Intron (ENSMUST00000078434.6 | 14 | 70217898 | 70289449 | 71552 | 2 | ENSMUST00 | 29523 | Ppp3cc | ENSMUSG00 | false | . | . | . | . | . | . | chr14 | 70257979 | 70260989 | 3.51709 | 3.34661 | 1.80375 |
| chr15 | 31330097 | 31330378 | 755 | Intron (ENSMUST00000110410.3 | 15 | 31293186 | 31367532 | 74347 | 2 | ENSMUST00 | 37154 | Ankrd33b | ENSMUSG00 | false | . | . | . | . | . | . | chr15 | 31315773 | 31357869 | 5.77982 | 4.40135 | 3.65864 |
| chr16 | 32657758 | 32658042 | 755 | Promoter (<=1kb) | 16 | 32657023 | 32668271 | 11249 | 1 | ENSMUST00 | 736 | Tnk2 | ENSMUSG00 | false | . | . | . | . | . | . | chr16 | 32656126 | 32665456 | 6.5689 | 4.78559 | 4.29525 |
| chr2 | 35024504 | 35024785 | 755 | Intron (ENSMUST00000028233.3 | 2 | 34983703 | 35019645 | 35943 | 2 | ENSMUST00 | -4860 | Hc | ENSMUSG00 | false | . | . | . | . | . | . | chr2 | 35024689 | 35025021 | 4.24573 | 3.81882 | 2.36964 |
| chr7 | 120972487 | 120972864 | 755 | Intron (ENSMUST00000140247.1 | 7 | 120957036 | 120981790 | 24755 | 2 | ENSMUST00 | 8926 | Cdr2 | ENSMUSG00 | false | . | . | . | . | . | . | chr7 | 120961858 | 120981649 | 5.33893 | 4.28516 | 3.2901 |
| chr13 | 34734828 | 34735107 | 755 | Promoter (<=1kb) | 13 | 34734850 | 34747613 | 12764 | 1 | ENSMUST00 | 0 | Fam50b | ENSMUSG00 | false | . | . | . | . | . | . | chr13 | 34734558 | 34736219 | 3.70548 | 3.39934 | 1.97817 |
| chr15 | 11160924 | 11161204 | 755 | Intron (ENSMUST00000061318.7 | 15 | 11064790 | 11346867 | 282078 | 1 | ENSMUST00 | 96135 | Adamts12 | ENSMUSG00 | false | . | . | . | . | . | . | chr15 | 11157701 | 11162223 | 4.98014 | 4.07935 | 2.98628 |
| chr5 | 8422662 | 8423006 | 754 | Promoter (<=1kb) | 5 | 8396973 | 8422680 | 25708 | 2 | ENSMUST00 | 0 | Dbf4 | ENSMUSG00 | false | chr5 | 8422049 | 8423680 | 36.4644 | 121.65416 | 117.45652 | chr5 | 8421674 | 8423928 | 3.22548 | 3.13646 | 1.55739 |
| chr11 | 105959444 | 105959723 | 754 | Distal Intergenic | 11 | 105935799 | 105953336 | 17538 | 2 | ENSMUST00 | -6109 | Cyb561 | ENSMUSG00 | false | . | . | . | . | . | . | chr11 | 105958256 | 105960658 | 4.68274 | 3.8473 | 2.75276 |
| chr5 | 74535416 | 74536112 | 754 | Promoter (<=1kb) | 5 | 74535449 | 74598790 | 63342 | 1 | ENSMUST00 | 0 | Fip1l1 | ENSMUSG00 | false | chr5 | 74535063 | 74536477 | 38.0703 | 123.3938 | 119.13641 | . | . | . | . | . | . |
| chr5 | 107151408 | 107151789 | 754 | Intron (ENSMUST00000031224.9 | 5 | 107118197 | 107132923 | 14727 | 2 | ENSMUST00 | -18486 | Tgfbr3 | ENSMUSG00 | true | . | . | . | . | . | . | chr5 | 107143848 | 107172978 | 6.44265 | 4.73295 | 4.19318 |
| chr9 | 58305741 | 58306037 | 754 | Intron (ENSMUST00000061799.8 | 9 | 58287723 | 58313212 | 25490 | 2 | ENSMUST00 | 7175 | Loxl1 | ENSMUSG00 | false | . | . | . | . | . | . | chr9 | 58302831 | 58310544 | 4.56252 | 3.71116 | 2.67437 |
| chr9 | 21463247 | 21463599 | 754 | Intron (ENSMUST00000165766.3 | 9 | 21457107 | 21457570 | 464 | 1 | ENSMUST00 | 6141 | Dnm2 | ENSMUSG00 | false | . | . | . | . | . | . | chr9 | 21460706 | 21464669 | 6.48587 | 4.68389 | 4.26178 |
| chr3 | 89773037 | 89773705 | 754 | Promoter (<=1kb) | 3 | 89773609 | 89783997 | 10389 | 1 | ENSMUST00 | 0 | Ube2q1 | ENSMUSG00 | false | chr3 | 89772697 | 89775534 | 37.85821 | 131.67731 | 127.12434 | chr3 | 89771965 | 89773088 | 3.20876 | 3.1503 | 1.52725 |
| chr10 | 5068236 | 5068639 | 754 | Promoter (<=1kb) | 10 | 5020203 | 5069063 | 48861 | 2 | ENSMUST00 | 424 | Syne1 | ENSMUSG00 | true | . | . | . | . | . | . | chr10 | 5063627 | 5070392 | 5.33842 | 4.22119 | 3.28885 |
| chr2 | 172959405 | 172959665 | 753 | Distal Intergenic | 2 | 172977700 | 172993571 | 15872 | 1 | ENSMUST00 | -18035 | Spo11 | ENSMUSG00 | false | . | . | . | . | . | . | chr2 | 172958080 | 172960043 | 3.22969 | 3.14376 | 1.60456 |
| chr11 | 54336193 | 54336486 | 753 | Promoter (<=1kb) | 11 | 54335497 | 54361539 | 26043 | 1 | ENSMUST00 | 697 | Acsl6 | ENSMUSG00 | false | . | . | . | . | . | . | chr11 | 54335607 | 54336622 | 3.34831 | 2.89076 | 1.69349 |
| chr16 | 92426782 | 92427089 | 753 | Intron (ENSMUST00000060005.9 | 16 | 92391953 | 92400077 | 8125 | 2 | ENSMUST00 | -26706 | Rcan1 | ENSMUSG00 | false | . | . | . | . | . | . | chr16 | 92425267 | 92430211 | 6.11064 | 4.5979 | 3.92954 |
| chr2 | 91730083 | 91730627 | 753 | Promoter (<=1kb) | 2 | 91730134 | 91918849 | 188716 | 1 | ENSMUST00 | 0 | Ambra1 | ENSMUSG00 | false | chr2 | 91729790 | 91731263 | 45.22282 | 155.87894 | 150.36807 | . | . | . | . | . | . |
| chr9 | 15525280 | 15525625 | 753 | Intron (ENSMUST00000045620.4 | 9 | 15520846 | 15545257 | 24412 | 1 | ENSMUST00 | 4435 | Smco4 | ENSMUSG00 | false | chr9 | 15525128 | 15526195 | 8.00962 | 15.03963 | 12.77451 | chr9 | 15520389 | 15531335 | 6.42021 | 4.64905 | 4.18363 |
| chr10 | 86107113 | 86107330 | 753 | Exon (ENSMUST00000131378.1/ | 10 | 86095885 | 86113580 | 17696 | 1 | ENSMUST00 | 11229 | 4930486F22I | ENSMUSG00 | false | . | . | . | . | . | . | chr10 | 86106238 | 86110916 | 5.43863 | 4.21801 | 3.37042 |
| chr1 | 134513806 | 134514134 | 753 | Distal Intergenic | 1 | 134518081 | 134518563 | 483 | 1 | ENSMUST00 | -3947 | Gm15454 | ENSMUSG00 | false | chr1 | 134513225 | 134514120 | 15.77044 | 42.5731 | 39.95169 | chr1 | 134512375 | 134514818 | 3.42133 | 3.2415 | 1.74649 |
| chr13 | 43221350 | 43221639 | 752 | Intron (ENSMUST00000055341.5 | 13 | 43151744 | 43171501 | 19758 | 2 | ENSMUST00 | -49850 | Tbc1d7 | ENSMUSG00 | false | . | . | . | . | . | . | chr13 | 43208585 | 43223004 | 6.08142 | 4.53032 | 3.90259 |
| chrX | 56734581 | 56734823 | 752 | Promoter (2-3kb) | 20 | 56732129 | 56792235 | 60107 | 1 | ENSMUST00 | 2453 | Fhl1 | ENSMUSG00 | false | . | . | . | . | . | . | . | . | . | . | . | . |
| chr17 | 73306513 | 73306826 | 752 | 3' UTR | 17 | 73306464 | 73399296 | 92833 | 2 | ENSMUST00 | 92470 | Capn13 | ENSMUSG00 | false | . | . | . | . | . | . | . | . | . | . | . | . |
| chr4 | 45452645 | 45453173 | 752 | Intron (ENSMUST00000147926.2 | 4 | 45423278 | 45482802 | 59525 | 2 | ENSMUST00 | 29629 | Shb | ENSMUSG00 | false | . | . | . | . | . | . | chr4 | 45451999 | 45455091 | 7.91285 | 5.27298 | 5.45593 |
| chr6 | 47835525 | 47836083 | 752 | Promoter (<=1kb) | 6 | 47835661 | 47843647 | 7987 | 1 | ENSMUST00 | 0 | Zfp398 | ENSMUSG00 | false | chr6 | 47835563 | 47836836 | 41.84113 | 139.79433 | 134.92708 | chr6 | 47835331 | 47838254 | 3.87408 | 3.48275 | 2.12306 |
| chr16 | 95861278 | 95861669 | 752 | Intron (ENSMUST00000136744.2 | 16 | 95831123 | 95898713 | 67591 | 2 | ENSMUST00 | 37044 | 1600002D24I | ENSMUSG00 | false | . | . | . | . | . | . | . | . | . | . | . | . |
| chr16 | 55974426 | 55974840 | 752 | Promoter (<=1kb) | 16 | 55973268 | 55974617 | 1350 | 2 | ENSMUST00 | 0 | RP24-288C12 | ENSMUSG00 | false | chr16 | 55973101 | 55975647 | 44.26367 | 154.19713 | 148.75002 | . | . | . | . | . | . |
| chr10 | 59703378 | 59703673 | 751 | Promoter (<=1kb) | 10 | 59702658 | 59751705 | 49048 | 1 | ENSMUST00 | 721 | Micu1 | ENSMUSG00 | true | chr10 | 59702528 | 59703890 | 22.26925 | 90.51499 | 87.16024 | chr10 | 59701972 | 59705622 | 3.97379 | 3.51895 | 2.202 |
| chr1 | 189867546 | 189867921 | 751 | Intron (ENSMUST00000097442.3 | 1 | 189862659 | 189863433 | 775 | 1 | ENSMUST00 | 4888 | Ptpn14 | ENSMUSG00 | true | . | . | . | . | . | . | chr1 | 189863688 | 189869672 | 7.17559 | 4.95415 | 4.80266 |
| chr4 | 150909491 | 150910056 | 751 | Promoter (<=1kb) | 4 | 150897133 | 150909926 | 12794 | 2 | ENSMUST00 | 0 | Park7 | ENSMUSG00 | false | chr4 | 150909156 | 150910322 | 29.2476 | 106.45832 | 102.71833 | chr4 | 150908005 | 150910648 | 4.53562 | 3.8089 | 2.6432 |
| chr4 | 138481060 | 138481378 | 751 | Distal Intergenic | 4 | 138454314 | 138460123 | 5810 | 1 | ENSMUST00 | 26747 | Camk2n1 | ENSMUSG00 | false | . | . | . | . | . | . | chr4 | 138479434 | 138482434 | 3.99086 | 3.51284 | 2.19925 |
| chr18 | 53793534 | 53793802 | 751 | Distal Intergenic | 18 | 53681724 | 53744547 | 62824 | 2 | ENSMUST00 | -48988 | Cep120 | ENSMUSG00 | false | . | . | . | . | . | . | chr18 | 53792269 | 53794132 | 4.9428 | 4.11433 | 2.94935 |
| chr8 | 111643467 | 111644240 | 751 | Promoter (<=1kb) | 8 | 111643443 | 111671008 | 27566 | 1 | ENSMUST00 | 25 | Zfp1 | ENSMUSG00 | false | chr8 | 111642915 | 111644635 | 29.2476 | 106.45832 | 102.71833 | . | . | . | . | . | . |
| chr12 | 79761360 | 79761623 | 751 | Distal Intergenic | 12 | 79963568 | 79963872 | 305 | 1 | ENSMUST00 | -201945 | RP23-179D3. | ENSMUSG00 | false | . | . | . | . | . | . | chr12 | 79753608 | 79761465 | 5.50685 | 4.27239 | 3.4259 |
| chr6 | 141381403 | 141381894 | 751 | Intron (ENSMUST00000043259.8 | 6 | 141238278 | 141250561 | 12284 | 2 | ENSMUST00 | -130843 | RP23-97L11. | ENSMUSG00 | false | . | . | . | . | . | . | chr6 | 141370327 | 141386627 | 5.21392 | 4.14863 | 3.18933 |
| chr11 | 86935029 | 86935307 | 750 | Downstream (1-2kb) | 11 | 86936425 | 86993707 | 57283 | 2 | ENSMUST00 | 58400 | Ypel2 | ENSMUSG00 | false | . | . | . | . | . | . | chr11 | 86933799 | 86935728 | 3.53415 | 3.06624 | 1.83569 |
| chr8 | 36723750 | 36724022 | 750 | Intron (ENSMUST00000098826.4 | 8 | 36567751 | 36733054 | 165304 | 2 | ENSMUST00 | 9032 | Dlc1 | ENSMUSG00 | true | . | . | . | . | . | . | chr8 | 36707656 | 36733370 | 4.68731 | 3.93374 | 2.7424 |
| chr6 | 127453506 | 127454017 | 750 | Promoter (<=1kb) | 6 | 127453624 | 127472028 | 18405 | 1 | ENSMUST00 | 0 | Parp11 | ENSMUSG00 | false | chr6 | 127453204 | 127454489 | 29.2476 | 106.45832 | 102.71833 | . | . | . | . | . | . |
| chr5 | 33820204 | 33821073 | 750 | Promoter (<=1kb) | 5 | 33820725 | 33897966 | 77242 | 1 | ENSMUST00 | 0 | Whsc1 | ENSMUSG00 | false | chr5 | 33819761 | 33821890 | 35.67123 | 127.73884 | 123.33056 | . | . | . | . | . | . |
| chr8 | 92066007 | 92066314 | 750 | Distal Intergenic | 8 | 92039917 | 92040020 | 104 | 1 | ENSMUST00 | 26091 | Gm24254 | ENSMUSG00 | false | . | . | . | . | . | . | chr8 | 92064727 | 92067058 | 4.23558 | 3.63614 | 2.39963 |
| chr14 | 53329069 | 53329557 | 750 | Distal Intergenic | 14 | 53324632 | 53326237 | 1606 | 1 | ENSMUST00 | 4438 | Trav7n-6 | ENSMUSG00 | false | chr14 | 53328887 | 53329506 | 7.85869 | 22.12435 | 19.76205 | chr14 | 53326867 | 53334472 | 6.19783 | 4.37912 | 4.0048 |
| chr6 | 67134875 | 67135249 | 750 | Intron (ENSMUST00000181304.1 | 6 | 67115673 | 67168426 | 52754 | 1 | ENSMUST00 | 19203 | A430010J10I | ENSMUSG00 | false | . | . | . | . | . | . | chr6 | 67123607 | 67146074 | 5.6205 | 4.38535 | 3.51128 |
| chr8 | 71498847 | 71499328 | 749 | 3' UTR | 8 | 71489459 | 71491411 | 1953 | 1 | ENSMUST00 | 9389 | Gtpbp3 | ENSMUSG00 | false | . | . | . | . | . | . | chr8 | 71495272 | 71500046 | 3.90309 | 3.28772 | 2.14928 |
| chr14 | 25458894 | 25459870 | 749 | Promoter (<=1kb) | 14 | 25459185 | 25538492 | 79308 | 1 | ENSMUST00 | 0 | Zmiz1 | ENSMUSG00 | false | chr14 | 25457181 | 25461653 | 31.54088 | 108.67455 | 104.87239 | . | . | . | . | . | . |
| chr5 | 113771223 | 113771936 | 749 | Promoter (<=1kb) | 5 | 113742446 | 113771649 | 29204 | 2 | ENSMUST00 | 0 | Sart3 | ENSMUSG00 | false | chr5 | 113770528 | 113773818 | 34.91946 | 133.88905 | 129.25287 | . | . | . | . | . | . |
| chr3 | 21934895 | 21935210 | 749 | Distal Intergenic | 3 | 22076652 | 22216594 | 139943 | 1 | ENSMUST00 | -141442 | Tbl1xr1 | ENSMUSG00 | false | . | . | . | . | . | . | chr3 | 21933908 | 21938060 | 4.69098 | 3.94511 | 2.74663 |
| chr3 | 57428768 | 57429240 | 749 | Intron (ENSMUST00000029377.7 | 3 | 57425314 | 57441677 | 16364 | 1 | ENSMUST00 | 3455 | Tm4sf4 | ENSMUSG00 | false | . | . | . | . | . | . | chr3 | 57423717 | 57432176 | 5.2988 | 4.29555 | 3.24175 |
| chrX | 35946475 | 35946737 | 749 | Promoter (<=1kb) | 20 | 35947397 | 36076556 | 129160 | 1 | ENSMUST00 | -660 | Dock11 | ENSMUSG00 | true | . | . | . | . | . | . | . | . | . | . | . | . |
| chr5 | 33228367 | 33228712 | 749 | Distal Intergenic | 5 | 33213518 | 33218238 | 4721 | 2 | ENSMUST00 | -10130 | Spon2 | ENSMUSG00 | false | . | . | . | . | . | . | chr5 | 33221817 | 33232043 | 5.32048 | 4.18629 | 3.28315 |
| chr10 | 45577683 | 45578431 | 749 | Promoter (<=1kb) | 10 | 45577829 | 45712345 | 134517 | 1 | ENSMUST00 | 0 | Hace1 | ENSMUSG00 | false | chr10 | 45577641 | 45578826 | 42.9698 | 143.99599 | 138.94749 | . | . | . | . | . | . |
| chr2 | 127005320 | 127005590 | 748 | Distal Intergenic | 2 | 127008717 | 127037344 | 28628 | 1 | ENSMUST00 | -3127 | Ap4e1 | ENSMUSG00 | false | . | . | . | . | . | . | chr2 | 127005073 | 127010104 | 5.17638 | 4.0967 | 3.18696 |
| chr8 | 83737748 | 83738233 | 748 | Promoter (2-3kb) | 8 | 83723362 | 83741169 | 17808 | 2 | ENSMUST00 | 2936 | Cd97 | ENSMUSG00 | false | . | . | . | . | . | . | . | . | . | . | . | . |
| chr9 | 35116649 | 35117250 | 748 | Promoter (<=1kb) | 9 | 35046579 | 35116810 | 70232 | 2 | ENSMUST00 | 0 | St3gal4 | ENSMUSG00 | false | chr9 | 35116129 | 35117433 | 17.09341 | 55.02552 | 52.24499 | chr9 | 35116075 | 35118239 | 3.93743 | 3.53435 | 2.13929 |
| chr4 | 150570329 | 150570657 | 748 | Promoter (<=1kb) | 4 | 150569657 | 150612069 | 42413 | 1 | ENSMUST00 | 673 | Rere | ENSMUSG00 | false | chr4 | 150568860 | 150571152 | 34.67015 | 117.41676 | 113.35385 | chr4 | 150570369 | 150578877 | 6.14529 | 4.57392 | 3.95861 |
| chr2 | 160620919 | 160621262 | 748 | Promoter (<=1kb) | 2 | 160568379 | 160619971 | 51593 | 2 | ENSMUST00 | -949 | Gm14221 | ENSMUSG00 | false | . | . | . | . | . | . | chr2 | 160619054 | 160622548 | 6.04769 | 4.597 | 3.86282 |
| chr11 | 34160337 | 34160587 | 748 | Distal Intergenic | 11 | 34141827 | 34143421 | 1595 | 1 | ENSMUST00 | 18511 | Gm12121 | ENSMUSG00 | false | . | . | . | . | . | . | chr11 | 34159650 | 34161215 | 4.81561 | 3.88419 | 2.87543 |
| chr13 | 5614870 | 5615202 | 748 | Distal Intergenic | 13 | 5855508 | 5857712 | 2205 | 2 | ENSMUST00 | 242510 | RP23-28J6.1 | ENSMUSG00 | false | . | . | . | . | . | . | chr13 | 5612339 | 5615755 | 5.01508 | 4.12023 | 3.01314 |
| chr11 | 88118809 | 88119106 | 747 | Intron (ENSMUST00000018522.7 | 11 | 88099290 | 88185365 | 86076 | 1 | ENSMUST00 | 19520 | Cuedc1 | ENSMUSG00 | true | . | . | . | . | . | . | chr11 | 88103459 | 88120719 | 6.48176 | 4.71748 | 4.23259 |
| chr19 | 46340890 | 46341321 | 747 | Promoter (<=1kb) | 19 | 46341121 | 46375252 | 34132 | 1 | ENSMUST00 | 0 | Tmem180 | ENSMUSG00 | false | . | . | . | . | . | . | chr19 | 46337591 | 46341950 | 5.94778 | 4.53615 | 3.77912 |
| chr8 | 46740698 | 46741790 | 747 | Promoter (<=1kb) | 8 | 46739745 | 46847458 | 107714 | 1 | ENSMUST00 | 954 | Irf2 | ENSMUSG00 | false | chr8 | 46738444 | 46742094 | 33.45771 | 103.89173 | 100.22243 | . | . | . | . | . | . |
| chr8 | 94601630 | 94602192 | 747 | Promoter (<=1kb) | 8 | 94574943 | 94601726 | 26784 | 2 | ENSMUST00 | 0 | Fam192a | ENSMUSG00 | false | chr8 | 94601927 | 94603405 | 42.91624 | 148.41264 | 143.18958 | . | . | . | . | . | . |
| chr15 | 84654310 | 84654671 | 747 | Distal Intergenic | 15 | 84669620 | 84703673 | 34054 | 1 | ENSMUST00 | -14949 | Prr5 | ENSMUSG00 | false | . | . | . | . | . | . | chr15 | 84654105 | 84654764 | 5.7086 | 4.49569 | 3.60832 |
| chr1 | 64738862 | 64739158 | 747 | Promoter (1-2kb) | 1 | 64730558 | 64737751 | 7194 | 2 | ENSMUST00 | -1112 | Fzd5 | ENSMUSG00 | false | chr1 | 64737999 | 64739348 | 18.1972 | 47.5761 | 44.89159 | chr1 | 64738066 | 64739517 | 4.37992 | 3.84802 | 2.49167 |
| chr7 | 6155049 | 6156412 | 747 | Promoter (<=1kb) | 7 | 6131489 | 6155971 | 24483 | 2 | ENSMUST00 | 0 | Zfp787 | ENSMUSG00 | false | chr7 | 6154864 | 6156928 | 32.66734 | 113.81544 | 109.86345 | . | . | . | . | . | . |
| chr10 | 5515414 | 5515676 | 747 | Intron (ENSMUST00000041639.5 | 10 | 5336949 | 5550692 | 213744 | 2 | ENSMUST00 | 35016 | Syne1 | ENSMUSG00 | true | . | . | . | . | . | . | . | . | . | . | . | . |
| chr1 | 75546114 | 75546637 | 746 | Promoter (<=1kb) | 1 | 75546266 | 75562172 | 15907 | 1 | ENSMUST00 | 0 | Slc4a3 | ENSMUSG00 | false | chr1 | 75545925 | 75546995 | 23.98789 | 70.18118 | 67.18839 | chr1 | 75545900 | 75547041 | 3.41505 | 3.11206 | 1.73373 |
| chr14 | 103070014 | 103070529 | 746 | Promoter (<=1kb) | 14 | 103070216 | 103077579 | 7364 | 1 | ENSMUST00 | 0 | Cln5 | ENSMUSG00 | false | chr14 | 103069501 | 103071221 | 28.91247 | 96.86193 | 93.36482 | . | . | . | . | . | . |
| chr11 | 119547599 | 119548052 | 746 | Promoter (<=1kb) | 11 | 119538719 | 119547753 | 9035 | 2 | ENSMUST00 | 0 | Nptx1 | ENSMUSG00 | false | chr11 | 119546666 | 119548665 | 22.10575 | 73.03297 | 69.99519 | chr11 | 119546007 | 119549713 | 7.10465 | 5.02545 | 4.73159 |
| chr10 | 68138660 | 68138932 | 746 | Intron (ENSMUST00000020106.7 | 10 | 68043968 | 68051046 | 7079 | 1 | ENSMUST00 | 94693 | Rtkn2 | ENSMUSG00 | false | . | . | . | . | . | . | chr10 | 68136755 | 68141633 | 4.96273 | 4.05082 | 2.98182 |
| chr4 | 139130693 | 139131208 | 746 | Promoter (<=1kb) | 4 | 139101814 | 139131113 | 29300 | 2 | ENSMUST00 | 0 | Minos1 | ENSMUSG00 | false | chr4 | 139129787 | 139131399 | 31.16539 | 106.97112 | 103.21663 | . | . | . | . | . | . |
| chr11 | 57983607 | 57983901 | 746 | Distal Intergenic | 11 | 57970244 | 57973134 | 2891 | 2 | ENSMUST00 | -10474 | Gm12249 | ENSMUSG00 | false | . | . | . | . | . | . | chr11 | 57980811 | 57984784 | 5.00394 | 3.96209 | 3.02635 |
| chr17 | 7023428 | 7023741 | 746 | Promoter (2-3kb) | 17 | 7025837 | 7087467 | 61631 | 1 | ENSMUST00 | -2096 | Gm1604b | ENSMUSG00 | false | . | . | . | . | . | . | chr17 | 7022919 | 7025822 | 4.79791 | 3.71476 | 2.86712 |
| chr11 | 75679596 | 75680171 | 745 | Promoter (<=1kb) | 11 | 75679542 | 75700737 | 21196 | 1 | ENSMUST00 | 55 | Crk | ENSMUSG00 | false | chr11 | 75678797 | 75680664 | 41.2653 | 138.78937 | 133.96024 | . | . | . | . | . | . |
| chr11 | 88864403 | 88864801 | 745 | Promoter (<=1kb) | 11 | 88830792 | 88864586 | 33795 | 2 | ENSMUST00 | 0 | Akap1 | ENSMUSG00 | false | chr11 | 88862882 | 88865272 | 29.08755 | 89.33995 | 86.00743 | chr11 | 88860753 | 88865179 | 3.94081 | 3.5221 | 2.14279 |
| chr11 | 8499005 | 8499340 | 745 | Intron (ENSMUST00000020695.7 | 11 | 8451070 | 8468675 | 17606 | 2 | ENSMUST00 | -30331 | Tns3 | ENSMUSG00 | false | chr11 | 8499137 | 8504359 | 19.17126 | 69.02763 | 66.05415 | chr11 | 8476638 | 8500780 | 7.03559 | 4.95227 | 4.67872 |
| chr15 | 76197919 | 76198649 | 745 | Promoter (<=1kb) | 15 | 76170976 | 76198209 | 27234 | 2 | ENSMUST00 | 0 | Plec | ENSMUSG00 | false | chr15 | 76197865 | 76200595 | 17.16829 | 59.16322 | 56.32936 | chr15 | 76192493 | 76209331 | 6.98012 | 4.91238 | 4.64105 |
| chr5 | 72915292 | 72915674 | 745 | Promoter (<=1kb) | 5 | 72914296 | 72914942 | 647 | 2 | ENSMUST00 | -351 | Gm9870 | ENSMUSG00 | false | chr5 | 72914347 | 72917697 | 41.51012 | 139.21701 | 134.36754 | . | . | . | . | . | . |
| chr2 | 17705516 | 17705797 | 745 | Intron (ENSMUST00000028080.6 | 2 | 17343909 | 17731043 | 387135 | 2 | ENSMUST00 | 25246 | Nebl | ENSMUSG00 | true | . | . | . | . | . | . | chr2 | 17675160 | 17718684 | 4.75621 | 3.88836 | 2.82315 |
| chr8 | 115697989 | 115698513 | 745 | Intron (ENSMUST00000069009.6 | 8 | 115705078 | 115707233 | 2156 | 1 | ENSMUST00 | -6565 | Gm15655 | ENSMUSG00 | false | . | . | . | . | . | . | . | . | . | . | . | . |
| chr12 | 102418218 | 102418514 | 744 | Intron (ENSMUST00000110020.2 | 12 | 102394084 | 102423769 | 29686 | 2 | ENSMUST00 | 5255 | Lgmn | ENSMUSG00 | false | . | . | . | . | . | . | . | . | . | . | . | . |
| chr12 | 111782183 | 111782472 | 744 | Promoter (<=1kb) | 12 | 111781947 | 111807844 | 25898 | 1 | ENSMUST00 | 237 | Klc1 | ENSMUSG00 | false | . | . | . | . | . | . | chr12 | 111776671 | 111782879 | 5.48019 | 4.12221 | 3.43546 |
| chr16 | 4907950 | 4908217 | 744 | Intron (ENSMUST00000023159.8 | 16 | 4886100 | 4938296 | 52197 | 1 | ENSMUST00 | 21851 | Mgrn1 | ENSMUSG00 | false | . | . | . | . | . | . | chr16 | 4901148 | 4908946 | 4.52019 | 3.80354 | 2.62131 |
| chr4 | 138216288 | 138216722 | 744 | Promoter (<=1kb) | 4 | 138216296 | 138244683 | 28388 | 1 | ENSMUST00 | 0 | Hp1bp3 | ENSMUSG00 | false | chr4 | 138215850 | 138218184 | 36.7977 | 133.03651 | 128.43381 | . | . | . | . | . | . |
| chr7 | 79578319 | 79579251 | 744 | Distal Intergenic | 7 | 79593369 | 79594924 | 1556 | 2 | ENSMUST00 | 15673 | Rhcg | ENSMUSG00 | false | chr7 | 79578722 | 79579171 | 12.24318 | 32.0257 | 29.53485 | chr7 | 79576878 | 79579493 | 5.82724 | 4.34733 | 3.71437 |
| chr9 | 104741072 | 104741422 | 744 | Intron (ENSMUST00000098443.3 | 9 | 104569787 | 105034544 | 464758 | 1 | ENSMUST00 | 171286 | Cpne4 | ENSMUSG00 | false | . | . | . | . | . | . | . | . | . | . | . | . |

| | | | | | | | | | | | | | | | | | | | | | | | | | | |
|---|---|---|---|---|---|---|---|---|---|---|---|---|---|---|---|---|---|---|---|---|---|---|---|---|---|---|
| chr1 | 182406662 | 182407025 | 744 | Promoter (1-2kb) | 1 | 182408619 | 182409020 | 402 | 2 | ENSMUST00 | 1995 | Gm10517 | ENSMUSG0C | false | . | . | . | . | . | . | chr1 | 182406074 | 182408568 | 4.65619 | 3.95737 | 2.70763 |
| chr15 | 37769691 | 37769966 | 744 | Intron (ENSMUST00000168992.2 | 15 | 37762832 | 37765401 | 2570 | 2 | ENSMUST00 | -4291 | Gm15940 | ENSMUSG0C | false | . | . | . | . | . | . | chr15 | 37765406 | 37771478 | 4.87168 | 3.98023 | 2.91168 |
| chr8 | 111258512 | 111259484 | 743 | Promoter (<=1kb) | 8 | 111154421 | 111259216 | 104796 | 2 | ENSMUST00 | 0 | Glg1 | ENSMUSG0C | false | chr8 | 111258097 | 111259935 | 25.75243 | 103.50989 | 99.85224 | . | . | . | . | . | . |
| chr11 | 54112977 | 54113309 | 743 | Promoter (<=1kb) | 11 | 54113203 | 54114274 | 1072 | 1 | ENSMUST00 | 0 | P4ha2 | ENSMUSG0C | false | . | . | . | . | . | . | chr11 | 54110812 | 54117439 | 8.16018 | 5.41308 | 5.62178 |
| chr11 | 54100772 | 54101187 | 743 | Promoter (<=1kb) | 11 | 54100924 | 54131652 | 30729 | 1 | ENSMUST00 | 0 | P4ha2 | ENSMUSG0C | false | chr11 | 54100609 | 54101866 | 33.91391 | 110.8507 | 106.99079 | chr11 | 54099731 | 54103160 | 5.49345 | 4.15007 | 3.43005 |
| chr10 | 117148335 | 117148764 | 743 | Promoter (<=1kb) | 10 | 117070129 | 117148474 | 78346 | 2 | ENSMUST00 | 0 | Frs2 | ENSMUSG0C | false | chr10 | 117147475 | 117149206 | 41.56779 | 133.79375 | 129.15814 | . | . | . | . | . | . |
| chr10 | 128525792 | 128526248 | 743 | Promoter (<=1kb) | 10 | 128510248 | 128525859 | 15612 | 2 | ENSMUST00 | 0 | Esyt1 | ENSMUSG0C | false | chr10 | 128524149 | 128526021 | 32.63288 | 132.66988 | 128.08362 | . | . | . | . | . | . |
| chr6 | 119448726 | 119449051 | 743 | Promoter (<=1kb) | 6 | 119432531 | 119449386 | 16856 | 2 | ENSMUST00 | 335 | Wnt5b | ENSMUSG0C | false | chr6 | 119448532 | 119449103 | 9.22042 | 18.50693 | 16.19293 | chr6 | 119448074 | 119453219 | 5.22359 | 4.16949 | 3.21432 |
| chr1 | 59574912 | 59575435 | 743 | Intron (ENSMUST00000114243.2 | 1 | 59558008 | 59566636 | 8629 | 1 | ENSMUST00 | 16905 | Gm973 | ENSMUSG0C | false | chr1 | 59574493 | 59575396 | 7.45894 | 16.41436 | 14.12962 | chr1 | 59574320 | 59575762 | 3.97509 | 3.29474 | 2.20992 |
| chr5 | 122158002 | 122158402 | 742 | Promoter (<=1kb) | 5 | 122158278 | 122175273 | 16996 | 1 | ENSMUST00 | 0 | Ppp1cc | ENSMUSG0C | false | chr5 | 122158261 | 122159841 | 27.88725 | 99.90214 | 96.33501 | chr5 | 122156923 | 122158031 | 3.25394 | 3.22097 | 1.5535 |
| chr10 | 36974456 | 36975260 | 742 | Promoter (<=1kb) | 10 | 36974544 | 37001889 | 27346 | 1 | ENSMUST00 | 0 | Hdac2 | ENSMUSG0C | false | chr10 | 36973845 | 36975536 | 32.27911 | 106.92835 | 103.17982 | . | . | . | . | . | . |
| chr9 | 34995920 | 34996264 | 742 | Intron (ENSMUST00000115148.3 | 9 | 35016452 | 35030876 | 14425 | 1 | ENSMUST00 | -20188 | Kirrel3 | ENSMUSG0C | false | . | . | . | . | . | . | chr9 | 34992837 | 34996448 | 5.51987 | 4.32705 | 3.46566 |
| chr13 | 64237381 | 64237648 | 742 | Intron (ENSMUST00000109769.3 | 13 | 64192545 | 64248526 | 55982 | 2 | ENSMUST00 | 10878 | Cdc14b | ENSMUSG0C | false | . | . | . | . | . | . | chr13 | 64233378 | 64237628 | 4.53488 | 3.90319 | 2.62272 |
| chr2 | 130563532 | 130563864 | 742 | Promoter (<=1kb) | 2 | 130563742 | 130565394 | 1653 | 1 | ENSMUST00 | 0 | Mrps26 | ENSMUSG0C | false | chr2 | 130563252 | 130564759 | 32.29185 | 112.09676 | 108.19604 | . | . | . | . | . | . |
| chr6 | 28828657 | 28829021 | 742 | Promoter (1-2kb) | 6 | 28661831 | 28830345 | 168515 | 2 | ENSMUST00 | 1324 | Lrrc4 | ENSMUSG0C | false | . | . | . | . | . | . | chr6 | 28826669 | 28829992 | 4.77862 | 3.87834 | 2.87378 |
| chr16 | 30793136 | 30793426 | 742 | Distal Intergenic | 16 | 30814162 | 30814268 | 107 | 1 | ENSMUST00 | -20736 | Gm26295 | ENSMUSG0C | false | . | . | . | . | . | . | chr16 | 30789222 | 30794139 | 6.35044 | 4.73188 | 4.10754 |
| chr17 | 70755042 | 70755413 | 742 | Promoter (<=1kb) | 17 | 70745329 | 70755582 | 10254 | 2 | ENSMUST00 | 169 | 5031415H12I | ENSMUSG0C | false | chr17 | 70754976 | 70755855 | 15.194 | 36.53499 | 33.98956 | chr17 | 70753352 | 70756203 | 3.73374 | 3.2155 | 2.01619 |
| chr9 | 88547925 | 88548256 | 741 | Promoter (<=1kb) | 9 | 88548020 | 88571061 | 23042 | 1 | ENSMUST00 | 0 | Zfp949 | ENSMUSG0C | false | chr9 | 88547470 | 88549208 | 34.67015 | 117.41676 | 113.35385 | . | . | . | . | . | . |
| chr4 | 101084115 | 101084627 | 741 | Intron (ENSMUST00000106955.1 | 4 | 101069479 | 101103806 | 34328 | 1 | ENSMUST00 | 14637 | Raver2 | ENSMUSG0C | false | . | . | . | . | . | . | chr4 | 101070383 | 101096042 | 5.55154 | 4.29326 | 3.46737 |
| chr8 | 79711601 | 79711862 | 741 | Promoter (<=1kb) | 8 | 79711820 | 79777321 | 65502 | 1 | ENSMUST00 | 0 | Anapc10 | ENSMUSG0C | false | chr8 | 79710841 | 79712537 | 28.22734 | 101.53345 | 97.92473 | . | . | . | . | . | . |
| chr12 | 112516902 | 112517377 | 741 | Distal Intergenic | 12 | 112500184 | 112511160 | 10977 | 2 | ENSMUST00 | -5743 | Tmem179 | ENSMUSG0C | false | . | . | . | . | . | . | chr12 | 112515496 | 112518169 | 5.50197 | 4.36858 | 3.41411 |
| chr15 | 79545853 | 79546445 | 741 | Promoter (<=1kb) | 15 | 79527707 | 79546741 | 19035 | 2 | ENSMUST00 | 296 | Ddx17 | ENSMUSG0C | false | chr15 | 79544943 | 79546662 | 36.46257 | 120.58523 | 116.42456 | . | . | . | . | . | . |
| chr9 | 61370304 | 61370892 | 741 | Promoter (<=1kb) | 9 | 61370878 | 61371627 | 750 | 2 | ENSMUST00 | 735 | Gm10655 | ENSMUSG0C | false | chr9 | 61370729 | 61374567 | 24.24157 | 89.02564 | 85.70093 | chr9 | 61366653 | 61374765 | 6.23582 | 4.63772 | 4.01625 |
| chr10 | 79908502 | 79909095 | 740 | Promoter (<=1kb) | 10 | 79894708 | 79908921 | 14214 | 2 | ENSMUST00 | 0 | Med16 | ENSMUSG0C | false | chr10 | 79908302 | 79909152 | 36.04346 | 118.78738 | 114.68494 | . | . | . | . | . | . |
| chr11 | 96269933 | 96270274 | 740 | Promoter (1-2kb) | 11 | 96271457 | 96276595 | 5139 | 1 | ENSMUST00 | -1183 | Hoxb9 | ENSMUSG0C | false | . | . | . | . | . | . | chr11 | 96269585 | 96272568 | 5.09741 | 3.93611 | 3.1112 |
| chr13 | 100107901 | 100108529 | 740 | Promoter (<=1kb) | 13 | 100107997 | 100114233 | 6237 | 1 | ENSMUST00 | 0 | Serf1 | ENSMUSG0C | false | chr13 | 100107426 | 100108677 | 25.50663 | 88.63404 | 85.31516 | chr13 | 100107334 | 100108617 | 3.96109 | 3.55634 | 2.21519 |
| chr14 | 65127053 | 65127330 | 740 | Distal Intergenic | 14 | 65052060 | 65098106 | 46047 | 2 | ENSMUST00 | -28948 | Extl3 | ENSMUSG0C | false | . | . | . | . | . | . | chr14 | 65125137 | 65127962 | 4.74376 | 3.73094 | 2.8273 |
| chr1 | 152047529 | 152047988 | 740 | Intron (ENSMUST00000187603.1 | 1 | 151893237 | 152055292 | 162056 | 2 | ENSMUST00 | 7304 | 1700025G04 | ENSMUSG0C | true | . | . | . | . | . | . | chr1 | 152043454 | 152055151 | 4.23473 | 3.64901 | 2.41125 |
| chr9 | 122084152 | 122084601 | 740 | Distal Intergenic | 9 | 122117283 | 122137176 | 19894 | 1 | ENSMUST00 | -32682 | Snrk | ENSMUSG0C | false | . | . | . | . | . | . | chr9 | 122083842 | 122086787 | 4.31139 | 3.79965 | 2.49682 |
| chr2 | 17730981 | 17731323 | 740 | Promoter (<=1kb) | 2 | 17343909 | 17731043 | 387135 | 2 | ENSMUST00 | 0 | Nebl | ENSMUSG0C | true | chr2 | 17729605 | 17732624 | 25.17393 | 93.69764 | 90.27328 | . | . | . | . | . | . |
| chr15 | 73716491 | 73716810 | 740 | Distal Intergenic | 15 | 73723145 | 73758766 | 35622 | 1 | ENSMUST00 | -6335 | Ptp4a3 | ENSMUSG0C | false | . | . | . | . | . | . | . | . | . | . | . | . |
| chr14 | 16735097 | 16735673 | 739 | Distal Intergenic | 14 | 16430842 | 16575472 | 144631 | 2 | ENSMUST00 | -159626 | Rarb | ENSMUSG0C | false | . | . | . | . | . | . | chr14 | 16733585 | 16736227 | 5.87914 | 4.422 | 3.77553 |
| chr7 | 67730682 | 67731001 | 739 | 3' UTR | 7 | 67725261 | 67726014 | 754 | 1 | ENSMUST00 | 5422 | Ttc23 | ENSMUSG0C | false | . | . | . | . | . | . | chr7 | 67729777 | 67731053 | 3.80738 | 3.44257 | 2.07388 |
| chr9 | 80172869 | 80173130 | 739 | Intron (ENSMUST00000035889.9 | 9 | 80165227 | 80308097 | 142871 | 1 | ENSMUST00 | 7643 | Myo6 | ENSMUSG0C | true | . | . | . | . | . | . | chr9 | 80167116 | 80198379 | 5.43108 | 4.25219 | 3.37051 |
| chr12 | 13049637 | 13050093 | 739 | Distal Intergenic | 12 | 12936096 | 12941914 | 5819 | 2 | ENSMUST00 | -107724 | Mycn | ENSMUSG0C | false | . | . | . | . | . | . | chr12 | 13047549 | 13049868 | 4.4634 | 3.81848 | 2.58992 |
| chr14 | 29968376 | 29968778 | 739 | Promoter (<=1kb) | 14 | 29968436 | 29973597 | 5162 | 1 | ENSMUST00 | 0 | Selk | ENSMUSG0C | false | chr14 | 29967742 | 29969586 | 33.7938 | 119.00099 | 114.88904 | . | . | . | . | . | . |
| chr10 | 95180840 | 95181194 | 739 | Intron (ENSMUST00000053594.5 | 10 | 95174746 | 95324097 | 149352 | 2 | ENSMUST00 | 142903 | Cradd | ENSMUSG0C | false | . | . | . | . | . | . | chr10 | 95179843 | 95183375 | 4.20064 | 3.64529 | 2.39555 |
| chr2 | 30719023 | 30719416 | 739 | Promoter (<=1kb) | 2 | 30713147 | 30720057 | 6911 | 2 | ENSMUST00 | 641 | Gm14488 | ENSMUSG0C | false | chr2 | 30718167 | 30719397 | 11.02877 | 22.59171 | 20.22099 | chr2 | 30716509 | 30721971 | 7.18867 | 4.77393 | 4.82839 |
| chr1 | 194955728 | 194956142 | 738 | Exon (ENSMUST00000180780.1/ | 1 | 194938821 | 194961292 | 22472 | 1 | ENSMUST00 | 16908 | Cd34 | ENSMUSG0C | false | . | . | . | . | . | . | chr1 | 194955671 | 194956429 | 5.99164 | 4.55125 | 3.82648 |
| chr12 | 108178457 | 108179863 | 738 | Promoter (<=1kb) | 12 | 108179738 | 108203359 | 23622 | 1 | ENSMUST00 | 0 | Ccnk | ENSMUSG0C | false | chr12 | 108179724 | 108180617 | 29.6869 | 105.26892 | 101.56436 | . | . | . | . | . | . |
| chr5 | 103911059 | 103911403 | 738 | Promoter (<=1kb) | 5 | 103862050 | 103911259 | 49210 | 2 | ENSMUST00 | 0 | Klhl8 | ENSMUSG0C | false | chr5 | 103910526 | 103911538 | 40.83545 | 136.9297 | 132.17467 | chr5 | 103909510 | 103911679 | 4.20334 | 3.74312 | 2.33601 |
| chr11 | 50224529 | 50225704 | 738 | Promoter (<=1kb) | 11 | 50225335 | 50235103 | 9769 | 1 | ENSMUST00 | 0 | Mgat4b | ENSMUSG0C | false | chr11 | 50223936 | 50226450 | 22.60491 | 86.65532 | 83.38163 | chr11 | 50220985 | 50224639 | 4.30947 | 3.72961 | 2.43315 |
| chr5 | 24624161 | 24624570 | 738 | Distal Intergenic | 5 | 24645453 | 24648860 | 3408 | 1 | ENSMUST00 | -20883 | 1700022A21I | ENSMUSG0C | false | chr5 | 24624190 | 24624742 | 10.20185 | 22.8781 | 20.50382 | chr5 | 24623705 | 24625243 | 5.78676 | 4.5113 | 3.64045 |
| chr1 | 11033366 | 11033726 | 738 | Intron (ENSMUST00000027056.6 | 1 | 10993518 | 10996601 | 3084 | 1 | ENSMUST00 | 39849 | Prex2 | ENSMUSG0C | false | . | . | . | . | . | . | . | . | . | . | . | . |
| chr17 | 35399679 | 35399985 | 738 | Promoter (<=1kb) | 17 | 35400799 | 35403727 | 2929 | 1 | ENSMUST00 | -814 | Gm8750 | ENSMUSG0C | false | . | . | . | . | . | . | chr17 | 35392097 | 35400320 | 6.23504 | 4.48412 | 4.05111 |
| chr18 | 35836228 | 35836562 | 738 | Intron (ENSMUST00000060722.6 | 18 | 35830389 | 35831093 | 705 | 2 | ENSMUST00 | -5136 | RP24-326K11 | ENSMUSG0C | false | . | . | . | . | . | . | chr18 | 35831800 | 35857612 | 7.70883 | 5.15861 | 5.24645 |
| chr10 | 45066829 | 45067385 | 737 | Promoter (<=1kb) | 10 | 45067206 | 45158997 | 91792 | 1 | ENSMUST00 | 0 | Prep | ENSMUSG0C | false | chr10 | 45066499 | 45067924 | 35.29575 | 125.98199 | 121.63858 | . | . | . | . | . | . |
| chr9 | 65032249 | 65032986 | 737 | Promoter (<=1kb) | 9 | 65032458 | 65082650 | 50193 | 1 | ENSMUST00 | 0 | Dpp8 | ENSMUSG0C | false | chr9 | 65032063 | 65033438 | 35.98183 | 119.73972 | 115.60696 | chr9 | 65031928 | 65034604 | 4.72792 | 3.93879 | 2.82826 |
| chr16 | 24491142 | 24491526 | 737 | Intron (ENSMUST00000038053.7 | 16 | 24448091 | 24992576 | 544486 | 1 | ENSMUST00 | 43052 | Lpp | ENSMUSG0C | false | . | . | . | . | . | . | chr16 | 24489385 | 24492570 | 5.9238 | 4.47158 | 3.8152 |
| chr3 | 85991681 | 85991912 | 737 | Intron (ENSMUST00000182969.2 | 3 | 85993810 | 86002491 | 8682 | 2 | ENSMUST00 | 10579 | Prss48 | ENSMUSG0C | false | . | . | . | . | . | . | chr3 | 85989948 | 85996729 | 5.64841 | 4.2824 | 3.58436 |
| chr7 | 37911759 | 37912020 | 737 | Distal Intergenic | 7 | 37824242 | 37824347 | 106 | 2 | ENSMUST00 | -87413 | Gm22302 | ENSMUSG0C | false | . | . | . | . | . | . | . | . | . | . | . | . |
| chr16 | 56029220 | 56029788 | 737 | Promoter (<=1kb) | 16 | 56015506 | 56029696 | 14191 | 2 | ENSMUST00 | 0 | Pcnp | ENSMUSG0C | false | chr16 | 56028147 | 56030547 | 31.94275 | 111.4613 | 107.58682 | . | . | . | . | . | . |
| chr4 | 150776055 | 150776341 | 737 | Distal Intergenic | 4 | 150825733 | 150839575 | 13843 | 1 | ENSMUST00 | -49392 | Gm13049 | ENSMUSG0C | false | . | . | . | . | . | . | chr4 | 150774614 | 150776551 | 3.08116 | 2.97151 | 1.42976 |
| chr11 | 101230781 | 101231293 | 736 | Distal Intergenic | 11 | 101191115 | 101226463 | 35349 | 2 | ENSMUST00 | -4319 | Ezh1 | ENSMUSG0C | false | . | . | . | . | . | . | chr11 | 101229271 | 101232515 | 5.59162 | 4.37392 | 3.51898 |
| chr5 | 146900442 | 146900761 | 736 | Distal Intergenic | 5 | 146948657 | 146955614 | 6958 | 1 | ENSMUST00 | -47896 | Gtf3a | ENSMUSG0C | false | . | . | . | . | . | . | chr5 | 146900270 | 146902328 | 3.42307 | 3.27845 | 1.70673 |
| chr12 | 3810525 | 3810863 | 736 | Intron (ENSMUST00000173700.1 | 12 | 3807160 | 3913491 | 106332 | 1 | ENSMUST00 | 3366 | Dnmt3a | ENSMUSG0C | false | . | . | . | . | . | . | chr12 | 3808261 | 3828410 | 6.1255 | 4.55618 | 3.94004 |
| chr6 | 72219607 | 72219907 | 736 | Intron (ENSMUST00000042646.7 | 6 | 72206177 | 72235577 | 29401 | 2 | ENSMUST00 | 15670 | Atoh8 | ENSMUSG0C | false | . | . | . | . | . | . | chr6 | 72211696 | 72224884 | 7.46431 | 5.12594 | 5.03432 |
| chr4 | 86777711 | 86778055 | 736 | Promoter (<=1kb) | 4 | 86777843 | 86781752 | 3910 | 1 | ENSMUST00 | 0 | Dennd4c | ENSMUSG0C | false | . | . | . | . | . | . | chr4 | 86775673 | 86778897 | 4.14447 | 3.69517 | 2.30507 |
| chr8 | 125133841 | 125134161 | 736 | Intron (ENSMUST00000074562.5 | 8 | 125087987 | 125102878 | 14892 | 1 | ENSMUST00 | 45855 | Disc1 | ENSMUSG0C | false | . | . | . | . | . | . | chr8 | 125133639 | 125134533 | 6.76821 | 4.97528 | 4.45888 |
| chr19 | 47328579 | 47328865 | 736 | Intron (ENSMUST00000081619.3 | 19 | 47273519 | 47273589 | 71 | 2 | ENSMUST00 | -54991 | Mir6995 | ENSMUSG0C | false | . | . | . | . | . | . | chr19 | 47327826 | 47329356 | 6.59323 | 4.66301 | 4.34728 |
| chr13 | 98420627 | 98420994 | 736 | Distal Intergenic | 13 | 98424109 | 98435065 | 10957 | 2 | ENSMUST00 | 14071 | RP24-76E2.6 | ENSMUSG0C | false | . | . | . | . | . | . | chr13 | 98420526 | 98424983 | 4.88372 | 4.03136 | 2.91585 |
| chr2 | 145856886 | 145857293 | 735 | Intron (ENSMUST00000110005.2 | 2 | 145862644 | 145878974 | 16331 | 1 | ENSMUST00 | -5351 | Rin2 | ENSMUSG0C | true | . | . | . | . | . | . | chr2 | 145849229 | 145860929 | 7.2604 | 5.04351 | 4.86236 |
| chr19 | 46666618 | 46666892 | 735 | Promoter (2-3kb) | 19 | 46668087 | 46669513 | 1427 | 2 | ENSMUST00 | 2621 | Cyp17a1 | ENSMUSG0C | false | . | . | . | . | . | . | chr19 | 46659100 | 46667582 | 5.73382 | 4.29769 | 3.62886 |
| chr19 | 17598642 | 17598960 | 735 | Intron (ENSMUST00000025618.9 | 19 | 17397437 | 17399961 | 2525 | 1 | ENSMUST00 | 201206 | Rfk | ENSMUSG0C | false | . | . | . | . | . | . | chr19 | 17598643 | 17599974 | 2.85398 | 2.86221 | 1.26389 |
| chrX | 42149702 | 42150272 | 735 | Promoter (<=1kb) | 20 | 42150154 | 42211419 | 61266 | 1 | ENSMUST00 | 0 | Stag2 | ENSMUSG0C | false | chrX | 42149209 | 42150570 | 18.84082 | 47.13427 | 44.45482 | . | . | . | . | . | . |
| chr7 | 80803249 | 80803715 | 735 | Promoter (<=1kb) | 7 | 80711583 | 80803331 | 91749 | 2 | ENSMUST00 | 0 | Iqgap1 | ENSMUSG0C | false | chr7 | 80801785 | 80803839 | 36.94879 | 123.57673 | 119.31536 | chr7 | 80803632 | 80803944 | 3.91543 | 3.62741 | 2.09916 |
| chr1 | 3671621 | 3671929 | 735 | Promoter (<=1kb) | 1 | 3214482 | 3671498 | 457017 | 2 | ENSMUST00 | -124 | Xkr4 | ENSMUSG0C | true | . | . | . | . | . | . | . | . | . | . | . | . |
| chr7 | 142019372 | 142019812 | 735 | Intron (ENSMUST00000084418.2 | 7 | 141998661 | 142001441 | 2781 | 1 | ENSMUST00 | 20712 | Brsk2 | ENSMUSG0C | false | chr7 | 142019094 | 142019518 | 4.24176 | 7.36333 | 5.24607 | chr7 | 142014396 | 142028430 | 5.88599 | 4.36974 | 3.76503 |
| chr11 | 28852999 | 28853367 | 734 | Promoter (<=1kb) | 11 | 28853204 | 28926743 | 73540 | 1 | ENSMUST00 | 0 | Efemp1 | ENSMUSG0C | false | chr11 | 28852855 | 28854039 | 19.69817 | 53.33889 | 50.58166 | . | . | . | . | . | . |
| chr1 | 97769773 | 97770202 | 734 | Promoter (<=1kb) | 1 | 97770172 | 97792969 | 22798 | 1 | ENSMUST00 | 0 | Gin1 | ENSMUSG0C | false | chr1 | 97768601 | 97771049 | 39.77878 | 138.39142 | 133.5723 | . | . | . | . | . | . |
| chr9 | 102626082 | 102626715 | 734 | Promoter (<=1kb) | 9 | 102586578 | 102626124 | 39547 | 2 | ENSMUST00 | 0 | Cep63 | ENSMUSG0C | false | chr9 | 102625338 | 102627283 | 32.66734 | 113.81544 | 109.86345 | . | . | . | . | . | . |
| chr11 | 72489621 | 72490247 | 734 | Promoter (<=1kb) | 11 | 72451638 | 72489904 | 38267 | 2 | ENSMUST00 | 0 | Spns2 | ENSMUSG0C | false | chr11 | 72488626 | 72490519 | 31.84599 | 102.09332 | 98.46812 | chr11 | 72488312 | 72490277 | 4.39323 | 3.76556 | 2.54426 |
| chr13 | 115051782 | 115052042 | 734 | Intron (ENSMUST00000061673.7 | 13 | 115088357 | 115090188 | 1832 | 2 | ENSMUST00 | 38146 | Pelo | ENSMUSG0C | false | . | . | . | . | . | . | . | . | . | . | . | . |
| chr6 | 83325667 | 83326443 | 734 | Promoter (<=1kb) | 6 | 83326016 | 83343776 | 17761 | 1 | ENSMUST00 | 0 | Mob1a | ENSMUSG0C | false | chr6 | 83325284 | 83326888 | 25.50501 | 91.48401 | 88.11109 | chr6 | 83325232 | 83328785 | 5.10899 | 3.96805 | 3.13111 |
| chr2 | 156736954 | 156737219 | 734 | Intron (ENSMUST00000109567.4 | 2 | 156721492 | 156764363 | 42872 | 1 | ENSMUST00 | 15463 | Dlgap4 | ENSMUSG0C | true | . | . | . | . | . | . | chr2 | 156722290 | 156745258 | 7.00087 | 4.91477 | 4.66073 |
| chr1 | 122704014 | 122704368 | 734 | Distal Intergenic | 1 | 122810306 | 122810682 | 377 | 1 | ENSMUST00 | -105938 | RP23-148M1 | ENSMUSG0C | false | . | . | . | . | . | . | chr1 | 122702097 | 122706087 | 4.16952 | 3.69872 | 2.32301 |
| chr17 | 79745993 | 79746407 | 733 | Distal Intergenic | 17 | 79706953 | 79715041 | 8089 | 2 | ENSMUST00 | -30953 | Cyp1b1 | ENSMUSG0C | false | . | . | . | . | . | . | chr17 | 79742056 | 79747394 | 4.66385 | 3.93992 | 2.73217 |
| chr5 | 99224664 | 99224998 | 733 | Intron (ENSMUST00000031276.9 | 5 | 99227794 | 99234066 | 6273 | 2 | ENSMUST00 | 9068 | Rasgef1b | ENSMUSG0C | false | . | . | . | . | . | . | chr5 | 99224629 | 99224915 | 4.21901 | 3.36425 | 2.39867 |
| chr6 | 114883233 | 114883677 | 733 | Intron (ENSMUST00000032459.8 | 6 | 114860628 | 114875107 | 14480 | 2 | ENSMUST00 | -8127 | Vgll4 | ENSMUSG0C | true | . | . | . | . | . | . | chr6 | 114853844 | 114919520 | 8.05355 | 5.36011 | 5.51688 |
| chr6 | 54961927 | 54962232 | 733 | Intron (ENSMUST00000168172.1 | 6 | 54923942 | 54971661 | 47720 | 2 | ENSMUST00 | 9429 | Nod1 | ENSMUSG0C | false | . | . | . | . | . | . | chr6 | 54943342 | 54971968 | 5.14308 | 4.10759 | 3.14164 |
| chr2 | 128364234 | 128364494 | 733 | Intron (ENSMUST00000143065.2 | 2 | 128423654 | 128424565 | 912 | 1 | ENSMUST00 | -59160 | Gm14010 | ENSMUSG0C | false | . | . | . | . | . | . | chr2 | 128363915 | 128364802 | 4.81815 | 3.78934 | 2.89085 |
| chr17 | 36930284 | 36930538 | 733 | Intron (ENSMUST00000174380.1 | 17 | 36923792 | 36933432 | 9641 | 1 | ENSMUST00 | 6493 | 1700031A10I | ENSMUSG0C | false | . | . | . | . | . | . | chr17 | 36929814 | 36930552 | 3.00098 | 2.88204 | 1.35514 |
| chr17 | 34600660 | 34601038 | 733 | Promoter (<=1kb) | 17 | 34601151 | 34601576 | 426 | 1 | ENSMUST00 | -113 | Gm20463 | ENSMUSG0C | false | . | . | . | . | . | . | chr17 | 34594616 | 34603372 | 6.22816 | 4.50396 | 4.02393 |
| chr9 | 63132524 | 63133085 | 733 | Promoter (<=1kb) | 9 | 63133283 | 63133389 | 107 | 1 | ENSMUST00 | -198 | Gm25064 | ENSMUSG0C | false | chr9 | 63132093 | 63133795 | 20.65177 | 61.66082 | 58.78991 | chr9 | 63129194 | 63136041 | 4.36968 | 3.72916 | 2.50691 |
| chr11 | 118360537 | 118360861 | 732 | Distal Intergenic | 11 | 118303701 | 118355740 | 52040 | 2 | ENSMUST00 | -4798 | Timp2 | ENSMUSG0C | false | . | . | . | . | . | . | chr11 | 118359762 | 118365457 | 7.10506 | 5.05606 | 4.73517 |
| chr4 | 114908535 | 114909395 | 732 | Promoter (<=1kb) | 4 | 114906282 | 114908873 | 2592 | 2 | ENSMUST00 | 0 | Foxd2 | ENSMUSG0C | false | chr4 | 114904677 | 114911326 | 45.47067 | 151.73354 | 146.38181 | . | . | . | . | . | . |
| chr16 | 49855125 | 49856274 | 732 | Promoter (<=1kb) | 16 | 49855618 | 49915010 | 59393 | 1 | ENSMUST00 | 0 | Cd47 | ENSMUSG0C | false | chr16 | 49854841 | 49857032 | 24.89402 | 98.85246 | 95.31046 | chr16 | 49854263 | 49855144 | 4.0962 | 3.68275 | 2.25107 |
| chr13 | 35707733 | 35708017 | 732 | Intron (ENSMUST00000075220.7 | 13 | 35741313 | 35874058 | 132746 | 1 | ENSMUST00 | -33296 | Cdyl | ENSMUSG0C | true | . | . | . | . | . | . | chr13 | 35706511 | 35709115 | 3.46098 | 3.38601 | 1.72155 |
| chr13 | 80885525 | 80886409 | 732 | Promoter (<=1kb) | 13 | 80886095 | 80894320 | 8226 | 1 | ENSMUST00 | 0 | Arrdc3 | ENSMUSG0C | false | chr13 | 80882761 | 80887175 | 33.47463 | 112.14942 | 108.24687 | . | . | . | . | . | . |
| chr10 | 122712447 | 122712720 | 732 | Intron (ENSMUST00000162853.2 | 10 | 122679315 | 122679415 | 101 | 1 | ENSMUST00 | 33133 | Mir8104 | ENSMUSG0C | false | . | . | . | . | . | . | chr10 | 122701076 | 122720564 | 4.67114 | 3.91157 | 2.74683 |
| chr1 | 186702475 | 186702795 | 732 | Promoter (2-3kb) | 1 | 186705695 | 186707033 | 1339 | 1 | ENSMUST00 | -2900 | RP24-276N2 | ENSMUSG0C | false | chr1 | 186702429 | 186703040 | 9.09886 | 18.30288 | 15.9921 | chr1 | 186700312 | 186704311 | 3.59372 | 3.3045 | 1.86903 |
| chr12 | 81594963 | 81595311 | 731 | Promoter (<=1kb) | 12 | 81573557 | 81595006 | 21450 | 2 | ENSMUST00 | 0 | Med6 | ENSMUSG0C | false | chr12 | 81593966 | 81595618 | 31.90856 | 95.20506 | 91.74643 | . | . | . | . | . | . |
| chr8 | 33599453 | 33600300 | 731 | Promoter (<=1kb) | 8 | 33599621 | 33619794 | 20174 | 1 | ENSMUST00 | 0 | Ppp2cb | ENSMUSG0C | false | chr8 | 33599519 | 33601159 | 31.54088 | 108.67455 | 104.87239 | . | . | . | . | . | . |
| chr14 | 70175845 | 70176271 | 731 | Promoter (<=1kb) | 14 | 70167764 | 70175433 | 7670 | 2 | ENSMUST00 | -413 | Pdlim2 | ENSMUSG0C | true | chr14 | 70173369 | 70177377 | 26.1063 | 98.41757 | 94.88632 | . | . | . | . | . | . |
| chr4 | 133011359 | 133012227 | 731 | Promoter (<=1kb) | 4 | 133011506 | 133077863 | 66358 | 1 | ENSMUST00 | 0 | Ahdc1 | ENSMUSG0C | false | chr4 | 133010598 | 133013973 | 43.65555 | 141.82639 | 136.87535 | . | . | . | . | . | . |
| chr9 | 22071172 | 22071495 | 731 | Promoter (<=1kb) | 9 | 22055460 | 22071345 | 15886 | 2 | ENSMUST00 | 0 | Zfp653 | ENSMUSG0C | false | chr9 | 22070511 | 22071901 | 33.5288 | 108.10788 | 104.32592 | . | . | . | . | . | . |
| chr1 | 74397031 | 74397687 | 731 | Promoter (2-3kb) | 1 | 74394310 | 74394394 | 85 | 1 | ENSMUST00 | 2722 | Mir26b | ENSMUSG0C | false | chr1 | 74397527 | 74398277 | 15.08796 | 37.05233 | 34.49837 | chr1 | 74392453 | 74398560 | 4.69855 | 3.96513 | 2.75686 |
| chr1 | 102875563 | 102875886 | 731 | Distal Intergenic | 1 | 102953415 | 102953548 | 134 | 2 | ENSMUST00 | 77662 | Gm22034 | ENSMUSG0C | false | . | . | . | . | . | . | . | . | . | . | . | . |
| chr9 | 86701865 | 86702123 | 731 | Promoter (1-2kb) | 9 | 86703819 | 86703946 | 128 | 1 | ENSMUST00 | -1696 | Gm22138 | ENSMUSG0C | false | . | . | . | . | . | . | chr9 | 86701383 | 86702599 | 4.472 | 3.6893 | 2.59716 |
| chr6 | 106799921 | 106800274 | 730 | Promoter (<=1kb) | 6 | 106778249 | 106800069 | 21821 | 2 | ENSMUST00 | 0 | Crbn | ENSMUSG0C | false | chr6 | 106799187 | 106800432 | 34.3275 | 112.61823 | 108.70669 | chr6 | 106798218 | 106800458 | 4.81515 | 3.86877 | 2.89302 |
| chr8 | 105471314 | 105471676 | 730 | Promoter (<=1kb) | 8 | 105467493 | 105471526 | 4034 | 2 | ENSMUST00 | 0 | Tppp3 | ENSMUSG0C | false | chr8 | 105471071 | 105471481 | 9.86256 | 23.41135 | 21.03137 | chr8 | 105471137 | 105472840 | 4.16925 | 3.53567 | 2.35406 |
| chr18 | 52942066 | 52942335 | 730 | Distal Intergenic | 18 | 52838010 | 52915312 | 77303 | 1 | ENSMUST00 | 104057 | Sncaip | ENSMUSG0C | true | . | . | . | . | . | . | chr18 | 52938454 | 52942888 | 6.12839 | 4.67692 | 3.93114 |
| chr19 | 30175080 | 30175388 | 730 | Promoter (<=1kb) | 19 | 30098449 | 30175418 | 76970 | 2 | ENSMUST00 | 30 | Gldc | ENSMUSG0C | false | chr19 | 30174568 | 30175616 | 25.55642 | 72.00093 | 68.98035 | chr19 | 30174126 | 30175548 | 5.17645 | 3.64866 | 3.19228 |
| chr2 | 77757656 | 77757971 | 730 | Intron (ENSMUST00000111831.2 | 2 | 77410634 | 77719739 | 309106 | 2 | ENSMUST00 | -37918 | Zfp385b | ENSMUSG0C | true | . | . | . | . | . | . | chr2 | 77749894 | 77759577 | 4.441 | 3.81795 | 2.5402 |
| chr2 | 131262333 | 131263033 | 730 | Promoter (<=1kb) | 2 | 131262495 | 131283437 | 20943 | 1 | ENSMUST00 | 0 | Pank2 | ENSMUSG0C | false | chr2 | 131261792 | 131263701 | 29.50634 | 86.21544 | 82.94717 | . | . | . | . | . | . |
| chr13 | 31915767 | 31916080 | 730 | Intron (ENSMUST00000041859.7 | 13 | 31884225 | 31885224 | 1000 | 1 | ENSMUST00 | 31543 | Gm11380 | ENSMUSG0C | false | . | . | . | . | . | . | chr13 | 31914820 | 31916256 | 4.93822 | 4.10503 | 2.97962 |
| chr1 | 59119724 | 59120573 | 729 | Promoter (<=1kb) | 1 | 59100590 | 59120094 | 19505 | 2 | ENSMUST00 | 0 | Tmem237 | ENSMUSG0C | false | chr1 | 59119469 | 59120710 | 22.31878 | 85.15456 | 81.90936 | chr1 | 59118888 | 59120672 | 3.35163 | 3.28917 | 1.66811 |
| chr7 | 19093798 | 19094374 | 729 | Promoter (<=1kb) | 7 | 19094594 | 19098549 | 3956 | 1 | ENSMUST00 | -220 | Six5 | ENSMUSG0C | false | chr7 | 19092457 | 19093957 | 12.40737 | 26.6269 | 24.20249 | chr7 | 19092010 | 19093879 | 7.70721 | 5.38228 | 5.20778 |
| chr1 | 30873301 | 30874045 | 729 | Promoter (<=1kb) | 1 | 30802339 | 30873581 | 71243 | 2 | ENSMUST00 | 0 | Phf3 | ENSMUSG0C | false | chr1 | 30872553 | 30874500 | 36.88168 | 122.38801 | 118.16834 | . | . | . | . | . | . |
| chr19 | 4333870 | 4334256 | 729 | Promoter (<=1kb) | 19 | 4324906 | 4334001 | 9096 | 2 | ENSMUST00 | 0 | Kdm2a | ENSMUSG0C | false | chr19 | 4332597 | 4334428 | 18.39885 | 52.66533 | 49.91441 | chr19 | 4331351 | 4335335 | 5.94277 | 4.45125 | 3.80268 |
| chr9 | 110756579 | 110756860 | 729 | Distal Intergenic | 9 | 110763646 | 110769804 | 6159 | 1 | ENSMUST00 | -6786 | Myl3 | ENSMUSG0C | false | . | . | . | . | . | . | chr9 | 110754997 | 110760017 | 8.13193 | 5.41736 | 5.56874 |
| chr4 | 117242853 | 117243192 | 729 | Intron (ENSMUST00000165128.2 | 4 | 117213333 | 117251917 | 38585 | 2 | ENSMUST00 | 8725 | Gm1661 | ENSMUSG0C | false | . | . | . | . | . | . | chr4 | 117242141 | 117243941 | 4.71464 | 3.99413 | 2.76592 |
| chr11 | 63827383 | 63827958 | 729 | Distal Intergenic | 11 | 63817019 | 63818300 | 1282 | 2 | ENSMUST00 | -9084 | Gm12288 | ENSMUSG0C | false | . | . | . | . | . | . | . | . | . | . | . | . |
| chr13 | 63371155 | 63371498 | 729 | Intron (ENSMUST00000073029.6 | 13 | 63376761 | 63381045 | 4285 | 2 | ENSMUST00 | 9547 | Gm16132 | ENSMUSG0C | false | . | . | . | . | . | . | chr13 | 63369425 | 63376037 | 5.54057 | 4.3431 | 3.44539 |
| chr19 | 6384196 | 6384483 | 728 | Promoter (<=1kb) | 19 | 6384399 | 6398459 | 14061 | 1 | ENSMUST00 | 0 | Pygm | ENSMUSG0C | false | . | . | . | . | . | . | chr19 | 6382253 | 6385444 | 4.73832 | 3.88448 | 2.81313 |
| chr18 | 46525606 | 46526354 | 728 | Promoter (<=1kb) | 18 | 46501746 | 46525971 | 24226 | 2 | ENSMUST00 | 0 | Fem1c | ENSMUSG0C | false | chr18 | 46523340 | 46525949 | 39.06023 | 122.966 | 118.72274 | . | . | . | . | . | . |
| chr8 | 88696875 | 88697397 | 728 | Promoter (<=1kb) | 8 | 88697028 | 88751945 | 54918 | 1 | ENSMUST00 | 0 | Cyld | ENSMUSG0C | false | chr8 | 88696931 | 88697911 | 27.18312 | 111.36749 | 107.49936 | . | . | . | . | . | . |
| chr9 | 64021729 | 64022384 | 728 | Promoter (<=1kb) | 9 | 63953076 | 64022059 | 68984 | 2 | ENSMUST00 | 0 | Smad6 | ENSMUSG0C | true | chr9 | 64016675 | 64022981 | 24.82645 | 85.46523 | 82.21117 | . | . | . | . | . | . |
| chr3 | 126341530 | 126341831 | 728 | Distal Intergenic | 3 | 126363827 | 126440374 | 76548 | 1 | ENSMUST00 | -21996 | Arsj | ENSMUSG0C | false | . | . | . | . | . | . | chr3 | 126340642 | 126342288 | 5.11504 | 4.20648 | 3.09305 |
| chr17 | 12916176 | 12916596 | 728 | Promoter (<=1kb) | 17 | 12911349 | 12916345 | 4997 | 2 | ENSMUST00 | 0 | Mrpl18 | ENSMUSG0C | false | chr17 | 12916373 | 12917402 | 25.4663 | 101.95255 | 98.33054 | . | . | . | . | . | . |
| chr18 | 4186614 | 4187023 | 728 | Distal Intergenic | 18 | 4198216 | 4198969 | 754 | 2 | ENSMUST00 | 11946 | Gm10557 | ENSMUSG0C | false | . | . | . | . | . | . | . | . | . | . | . | . |
| chr9 | 83304940 | 83305197 | 728 | Distal Intergenic | 9 | 83240491 | 83254540 | 14050 | 2 | ENSMUST00 | -50401 | Gm27216 | ENSMUSG0C | false | . | . | . | . | . | . | chr9 | 83304051 | 83306355 | 4.09191 | 3.5543 | 2.30406 |
| chr12 | 50668856 | 50669201 | 727 | Distal Intergenic | 12 | 50341231 | 50649223 | 307993 | 2 | ENSMUST00 | -19634 | Prkd1 | ENSMUSG0C | false | . | . | . | . | . | . | chr12 | 50668768 | 50669764 | 3.71293 | 3.45356 | 1.93793 |
| chr17 | 71124530 | 71125135 | 727 | Intron (ENSMUST00000180743.1 | 17 | 71182560 | 71235119 | 52560 | 1 | ENSMUST00 | -57425 | Lpin2 | ENSMUSG0C | false | . | . | . | . | . | . | chr17 | 71120006 | 71131073 | 6.06535 | 4.44566 | 3.90237 |
| chr1 | 75142288 | 75143339 | 727 | Promoter (<=1kb) | 1 | 75134554 | 75142638 | 8085 | 2 | ENSMUST00 | 0 | Cnppd1 | ENSMUSG0C | false | chr1 | 75140886 | 75143697 | 38.13901 | 127.82562 | 123.40861 | chr1 | 75143228 | 75144667 | 5.57701 | 4.40593 | 3.50811 |
| chr9 | 118974278 | 118974930 | 727 | Intron (ENSMUST00000073109.6 | 9 | 118929867 | 119033628 | 103762 | 1 | ENSMUST00 | 44412 | Ctdspl | ENSMUSG0C | true | . | . | . | . | . | . | chr9 | 118927678 | 119010843 | 6.89652 | 4.89961 | 4.57102 |
| chr4 | 114903959 | 114904202 | 727 | Promoter (1-2kb) | 4 | 114817161 | 114902859 | 85699 | 2 | ENSMUST00 | -1101 | 9130410C08I | ENSMUSG0C | false | . | . | . | . | . | . | chr4 | 114897050 | 114905478 | 4.35462 | 3.78738 | 2.46946 |
| chr10 | 23181664 | 23181938 | 727 | Intron (ENSMUST00000092665.5 | 10 | 23166865 | 23166971 | 107 | 2 | ENSMUST00 | -14694 | Gm24655 | ENSMUSG0C | false | . | . | . | . | . | . | chr10 | 23179930 | 23182647 | 4.02013 | 3.66417 | 2.20643 |
| chr16 | 30720773 | 30721133 | 727 | Distal Intergenic | 16 | 30814162 | 30814268 | 107 | 1 | ENSMUST00 | -93029 | Gm26295 | ENSMUSG0C | false | . | . | . | . | . | . | chr16 | 30720630 | 30721343 | 3.17044 | 2.99551 | 1.54111 |
| chr16 | 52012986 | 52013474 | 726 | Distal Intergenic | 16 | 52031549 | 52208047 | 176499 | 1 | ENSMUST00 | -18075 | Cblb | ENSMUSG0C | true | . | . | . | . | . | . | chr16 | 52009607 | 52016621 | 4.28782 | 3.76463 | 2.41076 |
| chr16 | 33926633 | 33926981 | 726 | Intron (ENSMUST00000115028.4 | 16 | 33922486 | 33944148 | 21663 | 1 | ENSMUST00 | 4148 | Itgb5 | ENSMUSG0C | true | . | . | . | . | . | . | chr16 | 33911227 | 33952498 | 7.40288 | 5.1092 | 4.9863 |
| chr19 | 36409455 | 36410218 | 726 | Promoter (<=1kb) | 19 | 36409748 | 36456192 | 46445 | 1 | ENSMUST00 | 0 | Pcgf5 | ENSMUSG0C | false | chr19 | 36409130 | 36410665 | 29.69666 | 89.43046 | 86.09753 | chr19 | 36408358 | 36409470 | 3.25431 | 3.11818 | 1.60049 |
| chr9 | 42746029 | 42746264 | 726 | Intron (ENSMUST00000114865.2 | 9 | 42520412 | 42808699 | 288288 | 2 | ENSMUST00 | 62435 | Grik4 | ENSMUSG0C | false | . | . | . | . | . | . | . | . | . | . | . | . |
| chr12 | 72536050 | 72536391 | 726 | Promoter (<=1kb) | 12 | 72536357 | 72580213 | 43857 | 1 | ENSMUST00 | 0 | Pcnxl4 | ENSMUSG0C | false | chr12 | 72535810 | 72536947 | 19.04495 | 59.55748 | 56.71581 | . | . | . | . | . | . |
| chr5 | 54053933 | 54054289 | 726 | Intron (ENSMUST00000117661.3 | 5 | 54074737 | 54104489 | 29753 | 1 | ENSMUST00 | -20448 | Stim2 | ENSMUSG0C | false | . | . | . | . | . | . | chr5 | 54049385 | 54055818 | 3.49069 | 3.18197 | 1.81073 |
| chr6 | 52012071 | 52012657 | 726 | Promoter (<=1kb) | 6 | 51859166 | 52012549 | 153384 | 2 | ENSMUST00 | 0 | Skap2 | ENSMUSG0C | false | chr6 | 52011818 | 52012598 | 23.51194 | 70.23763 | 67.24405 | . | . | . | . | . | . |
| chr12 | 111691337 | 111691867 | 726 | Distal Intergenic | 12 | 111678105 | 111683902 | 5798 | 1 | ENSMUST00 | 13233 | Trmt61a | ENSMUSG0C | false | . | . | . | . | . | . | chr12 | 111690820 | 111692659 | 6.16779 | 4.42629 | 4.01748 |
| chr6 | 29693875 | 29694492 | 725 | Promoter (<=1kb) | 6 | 29694222 | 29718559 | 24338 | 1 | ENSMUST00 | 0 | Tspan33 | ENSMUSG0C | false | chr6 | 29693777 | 29694874 | 29.41006 | 85.09721 | 81.85307 | chr6 | 29693438 | 29694911 | 4.96724 | 3.9875 | 3.00504 |
| chr10 | 59616167 | 59616640 | 725 | Promoter (<=1kb) | 10 | 59602775 | 59616655 | 13881 | 2 | ENSMUST00 | 15 | Mcu | ENSMUSG0C | false | chr10 | 59615385 | 59617197 | 22.31878 | 85.15456 | 81.90936 | . | . | . | . | . | . |
| chr18 | 12168496 | 12168826 | 725 | Promoter (<=1kb) | 18 | 12168717 | 12181116 | 12400 | 1 | ENSMUST00 | 0 | 3110002H16I | ENSMUSG0C | false | chr18 | 12168096 | 12169590 | 25.53309 | 82.08177 | 78.89293 | chr18 | 12167171 | 12170127 | 4.03644 | 3.59967 | 2.19963 |
| chr5 | 76951069 | 76951868 | 725 | Promoter (<=1kb) | 5 | 76913249 | 76951572 | 38324 | 2 | ENSMUST00 | 0 | Ppat | ENSMUSG0C | false | chr5 | 76950025 | 76952599 | 34.14626 | 110.2223 | 106.38018 | . | . | . | . | . | . |
| chr4 | 15265302 | 15266261 | 725 | Promoter (<=1kb) | 4 | 15265831 | 15286753 | 20923 | 1 | ENSMUST00 | 0 | Tmem64 | ENSMUSG0C | false | chr4 | 15265209 | 15266911 | 31.28813 | 116.44205 | 112.41405 | chr4 | 15264923 | 15265346 | 3.60804 | 3.44489 | 1.83905 |
| chr16 | 78235022 | 78235513 | 725 | Distal Intergenic | 16 | 78240258 | 78240388 | 131 | 1 | ENSMUST00 | -4745 | Gm25916 | ENSMUSG0C | false | . | . | . | . | . | . | chr16 | 78233378 | 78237188 | 4.4596 | 3.67453 | 2.5964 |
| chr13 | 48566373 | 48566718 | 725 | Distal Intergenic | 13 | 48572253 | 48572384 | 132 | 2 | ENSMUST00 | 5666 | Gm25232 | ENSMUSG0C | false | . | . | . | . | . | . | chr13 | 48564751 | 48567600 | 4.91388 | 4.10313 | 2.92718 |
| chr4 | 95159856 | 95160168 | 725 | Intron (ENSMUST00000130684.2 | 4 | 95151740 | 95157503 | 5764 | 1 | ENSMUST00 | 8117 | RP23-77I17.4 | ENSMUSG0C | false | . | . | . | . | . | . | chr4 | 95159243 | 95162809 | 4.78211 | 3.95901 | 2.84821 |
| chr1 | 75317309 | 75318109 | 724 | Promoter (<=1kb) | 1 | 75307896 | 75317637 | 9742 | 2 | ENSMUST00 | 0 | Dnpep | ENSMUSG0C | false | chr1 | 75316459 | 75317532 | 29.52499 | 116.10789 | 112.09218 | chr1 | 75318077 | 75318551 | 3.98908 | 3.69419 | 2.19499 |
| chr1 | 143524489 | 143524777 | 724 | Distal Intergenic | 1 | 143633851 | 143636577 | 2727 | 2 | ENSMUST00 | 111800 | Cdc73 | ENSMUSG0C | false | . | . | . | . | . | . | chr1 | 143523120 | 143525330 | 4.11032 | 3.60611 | 2.31651 |
| chr7 | 127967117 | 127968547 | 724 | Promoter (<=1kb) | 7 | 127967457 | 127982368 | 14912 | 1 | ENSMUST00 | 0 | Fus | ENSMUSG0C | false | chr7 | 127966431 | 127969481 | 32.68561 | 114.88856 | 110.91166 | . | . | . | . | . | . |
| chr18 | 76005816 | 76006181 | 724 | Intron (ENSMUST00000058997.8 | 18 | 76059458 | 76146443 | 86986 | 1 | ENSMUST00 | -53277 | Zbtb7c | ENSMUSG0C | true | . | . | . | . | . | . | chr18 | 75999046 | 76019015 | 7.41965 | 5.13063 | 4.98856 |
| chr10 | 128779518 | 128779809 | 724 | Promoter (2-3kb) | 10 | 128776742 | 128777251 | 510 | 2 | ENSMUST00 | -2268 | Rpsa-ps2 | ENSMUSG0C | false | . | . | . | . | . | . | chr10 | 128778481 | 128779736 | 4.67113 | 3.87764 | 2.7284 |
| chr11 | 22512360 | 22512899 | 724 | Promoter (<=1kb) | 11 | 22512088 | 22519234 | 7147 | 1 | ENSMUST00 | 273 | Tmem17 | ENSMUSG0C | false | chr11 | 22512039 | 22513396 | 26.32471 | 106.6389 | 102.89503 | . | . | . | . | . | . |
| chr3 | 90537167 | 90537517 | 724 | Promoter (<=1kb) | 3 | 90537254 | 90543148 | 5895 | 1 | ENSMUST00 | 0 | S100a16 | ENSMUSG0C | false | chr3 | 90537113 | 90538338 | 19.64718 | 50.84783 | 48.12388 | chr3 | 90536771 | 90541456 | 5.23366 | 4.1646 | 3.2227 |
| chr11 | 88113560 | 88113904 | 723 | Intron (ENSMUST00000018522.7 | 11 | 88099290 | 88185365 | 86076 | 1 | ENSMUST00 | 14271 | Cuedc1 | ENSMUSG0C | true | . | . | . | . | . | . | chr11 | 88103459 | 88120719 | 6.48176 | 4.71748 | 4.23259 |
| chr12 | 86078792 | 86079412 | 723 | Promoter (<=1kb) | 12 | 86056744 | 86079041 | 22298 | 2 | ENSMUST00 | 0 | Tgfb3 | ENSMUSG0C | false | chr12 | 86078492 | 86079616 | 24.56002 | 77.54259 | 74.4321 | . | . | . | . | . | . |
| chr9 | 63696634 | 63696953 | 723 | Intron (ENSMUST00000034973.4 | 9 | 63654799 | 63711969 | 57171 | 2 | ENSMUST00 | 15016 | Smad3 | ENSMUSG0C | false | . | . | . | . | . | . | chr9 | 63695180 | 63731540 | 5.72957 | 4.34734 | 3.61571 |
| chr9 | 36974230 | 36974474 | 723 | Intron (ENSMUST00000177218.2 | 9 | 36892478 | 36968490 | 76013 | 2 | ENSMUST00 | -5741 | Pknox2 | ENSMUSG0C | true | . | . | . | . | . | . | chr9 | 36970192 | 36981240 | 5.71459 | 4.43768 | 3.58495 |
| chr13 | 44896353 | 44896863 | 723 | 3' UTR | 13 | 44840783 | 44896481 | 55699 | 1 | ENSMUST00 | 55571 | Jarid2 | ENSMUSG0C | false | . | . | . | . | . | . | chr13 | 44892203 | 44897804 | 4.54964 | 3.85803 | 2.65912 |
| chr4 | 32227911 | 32228164 | 723 | Distal Intergenic | 4 | 32238804 | 32586104 | 347301 | 1 | ENSMUST00 | -10640 | Bach2 | ENSMUSG0C | false | . | . | . | . | . | . | . | . | . | . | . | . |
| chr3 | 103075856 | 103076178 | 723 | Promoter (1-2kb) | 3 | 103074053 | 103088323 | 14271 | 1 | ENSMUST00 | 1804 | Ampd1 | ENSMUSG0C | false | . | . | . | . | . | . | chr3 | 103075729 | 103076567 | 4.05789 | 3.59767 | 2.22282 |
| chr2 | 94263582 | 94263900 | 723 | Promoter (<=1kb) | 2 | 94251879 | 94264739 | 12861 | 2 | ENSMUST00 | 839 | E530001K10I | ENSMUSG0C | false | . | . | . | . | . | . | chr2 | 94263303 | 94266098 | 4.0216 | 3.58773 | 2.23012 |
| chr11 | 79051473 | 79051834 | 722 | Intron (ENSMUST00000018478.5 | 11 | 79045183 | 79057627 | 12445 | 2 | ENSMUST00 | 5793 | Ksr1 | ENSMUSG0C | true | . | . | . | . | . | . | chr11 | 79045596 | 79144720 | 7.09579 | 4.93486 | 4.74337 |
| chr2 | 178327351 | 178327641 | 722 | Promoter (<=1kb) | 2 | 178327556 | 178331766 | 4211 | 1 | ENSMUST00 | 0 | Phactr3 | ENSMUSG0C | false | . | . | . | . | . | . | chr2 | 178326858 | 178327799 | 2.944 | 2.91735 | 1.35473 |
| chr3 | 116396709 | 116396999 | 722 | Intron (ENSMUST00000106491.2 | 3 | 116400186 | 116423024 | 22839 | 2 | ENSMUST00 | 26025 | Cdc14a | ENSMUSG0C | true | . | . | . | . | . | . | chr3 | 116394159 | 116421460 | 5.70692 | 4.42755 | 3.58909 |
| chr7 | 120842669 | 120843833 | 722 | Promoter (<=1kb) | 7 | 120842831 | 120907219 | 64389 | 1 | ENSMUST00 | 0 | Eef2k | ENSMUSG0C | false | chr7 | 120842456 | 120844000 | 22.60491 | 86.65532 | 83.38163 | chr7 | 120843769 | 120845082 | 4.02227 | 3.51726 | 2.20998 |
| chr5 | 53590085 | 53590799 | 722 | Promoter (<=1kb) | 5 | 53590215 | 53657445 | 67231 | 1 | ENSMUST00 | 0 | Rbpj | ENSMUSG0C | false | chr5 | 53588566 | 53591937 | 30.81129 | 113.47932 | 109.54176 | . | . | . | . | . | . |
| chrX | 12159968 | 12160315 | 722 | Promoter (<=1kb) | 20 | 12036743 | 12160355 | 123613 | 2 | ENSMUST00 | 40 | Bcor | ENSMUSG0C | false | chrX | 12159705 | 12160585 | 13.78596 | 30.80405 | 28.32676 | . | . | . | . | . | . |
| chr8 | 90876864 | 90877552 | 722 | Intron (ENSMUST00000048665.6 | 8 | 90907939 | 90908415 | 477 | 2 | ENSMUST00 | 30863 | Gm6658 | ENSMUSG0C | false | chr8 | 90876446 | 90877999 | 24.03115 | 75.67385 | 72.59326 | chr8 | 90875801 | 90878989 | 5.36089 | 4.21626 | 3.30439 |
| chr9 | 98805505 | 98805931 | 722 | Intron (ENSMUST00000186387.1 | 9 | 98748599 | 98820087 | 71489 | 2 | ENSMUST00 | 14156 | RP24-116C1. | ENSMUSG0C | false | . | . | . | . | . | . | chr9 | 98804678 | 98806388 | 3.27212 | 3.18981 | 1.57268 |
| chr1 | 121329377 | 121329707 | 721 | Promoter (1-2kb) | 1 | 121304353 | 121328024 | 23672 | 2 | ENSMUST00 | -1354 | Insig2 | ENSMUSG0C | false | . | . | . | . | . | . | chr1 | 121323223 | 121330049 | 4.29763 | 3.74518 | 2.43067 |
| chr13 | 114867643 | 114867953 | 721 | Intron (ENSMUST00000056117.8 | 13 | 114818905 | 114825230 | 6326 | 1 | ENSMUST00 | 48739 | Mocs2 | ENSMUSG0C | false | . | . | . | . | . | . | chr13 | 114867004 | 114868928 | 3.9251 | 3.60429 | 2.12523 |
| chr15 | 75705414 | 75705928 | 721 | Promoter (1-2kb) | 15 | 75704394 | 75714225 | 9832 | 1 | ENSMUST00 | 1021 | Rhpn1 | ENSMUSG0C | true | chr15 | 75704224 | 75711444 | 34.50201 | 115.01646 | 111.03505 | . | . | . | . | . | . |
| chr19 | 5637369 | 5637957 | 721 | Promoter (<=1kb) | 19 | 5637483 | 5648130 | 10648 | 1 | ENSMUST00 | 0 | Rela | ENSMUSG0C | false | chr19 | 5636775 | 5638669 | 25.79552 | 96.83904 | 93.34669 | . | . | . | . | . | . |
| chr2 | 13574913 | 13575188 | 721 | Promoter (<=1kb) | 2 | 13574322 | 13578604 | 4283 | 1 | ENSMUST00 | 592 | Vim | ENSMUSG0C | true | chr2 | 13572823 | 13576831 | 31.81669 | 98.02396 | 94.50304 | . | . | . | . | . | . |
| chr2 | 68868422 | 68868687 | 721 | Intron (ENSMUST00000028426.3 | 2 | 68861586 | 69108644 | 247059 | 1 | ENSMUST00 | 6837 | Cers6 | ENSMUSG0C | true | . | . | . | . | . | . | chr2 | 68863137 | 68874811 | 5.56542 | 4.31322 | 3.46908 |
| chr4 | 148809218 | 148809493 | 721 | Intron (ENSMUST00000094464.4 | 4 | 148804465 | 148839746 | 35282 | 1 | ENSMUST00 | 4754 | Casz1 | ENSMUSG0C | true | . | . | . | . | . | . | chr4 | 148806488 | 148838873 | 5.49162 | 4.28891 | 3.41701 |
| chr9 | 103111685 | 103112208 | 721 | Promoter (<=1kb) | 9 | 103111787 | 103162657 | 50871 | 1 | ENSMUST00 | 0 | Rab6b | ENSMUSG0C | false | chr9 | 103111207 | 103113371 | 27.57193 | 78.19525 | 75.07539 | . | . | . | . | . | . |
| chr6 | 29718892 | 29719343 | 720 | Distal Intergenic | 6 | 29708028 | 29714822 | 6795 | 1 | ENSMUST00 | 10865 | Tspan33 | ENSMUSG0C | false | chr6 | 29718684 | 29719630 | 8.29801 | 20.47831 | 18.13586 | chr6 | 29716749 | 29719977 | 5.47696 | 4.33258 | 3.40111 |
| chr7 | 35215253 | 35215567 | 720 | Promoter (<=1kb) | 7 | 35200878 | 35215370 | 14493 | 2 | ENSMUST00 | 0 | Lrp3 | ENSMUSG0C | false | chr7 | 35214583 | 35215853 | 25.98482 | 77.30067 | 74.19341 | chr7 | 35214527 | 35215862 | 5.90344 | 4.37761 | 3.74792 |
| chr3 | 98025940 | 98026314 | 720 | Intron (ENSMUST00000079812.6 | 3 | 98013538 | 98150367 | 136830 | 1 | ENSMUST00 | 12403 | Notch2 | ENSMUSG0C | false | . | . | . | . | . | . | chr3 | 98014795 | 98071300 | 5.94878 | 4.48977 | 3.79636 |
| chr6 | 34619594 | 34620247 | 720 | Intron (ENSMUST00000079391.4 | 6 | 34598620 | 34662748 | 64129 | 1 | ENSMUST00 | 20975 | Cald1 | ENSMUSG0C | true | . | . | . | . | . | . | chr6 | 34618764 | 34625785 | 4.63104 | 3.93577 | 2.7056 |
| chrX | 73880683 | 73880965 | 720 | Promoter (<=1kb) | 20 | 73853955 | 73880834 | 26880 | 2 | ENSMUST00 | 0 | L1cam | ENSMUSG0C | false | chrX | 73880391 | 73880858 | 14.2455 | 32.22549 | 29.73087 | . | . | . | . | . | . |
| chr2 | 122348696 | 122349313 | 720 | 3' UTR | 2 | 122349504 | 122354131 | 4628 | 2 | ENSMUST00 | 4818 | Shf | ENSMUSG0C | false | chr2 | 122348388 | 122349553 | 24.82645 | 85.46523 | 82.21117 | chr2 | 122348242 | 122349935 | 4.21524 | 3.41367 | 2.39732 |
| chr18 | 24095207 | 24095552 | 720 | Distal Intergenic | 18 | 24106378 | 24114913 | 8536 | 2 | ENSMUST00 | 19361 | Ino80c | ENSMUSG0C | false | . | . | . | . | . | . | chr18 | 24094865 | 24096963 | 5.22609 | 4.19358 | 3.25162 |
| chr2 | 145841622 | 145841903 | 719 | Intron (ENSMUST00000110005.2 | 2 | 145822289 | 145845087 | 22799 | 1 | ENSMUST00 | 19334 | Rin2 | ENSMUSG0C | true | . | . | . | . | . | . | chr2 | 145827498 | 145846856 | 5.11522 | 4.05923 | 3.13101 |
| chr15 | 10690432 | 10690710 | 719 | 5' UTR | 15 | 10568979 | 10713540 | 144562 | 2 | ENSMUST00 | 22830 | Rai14 | ENSMUSG0C | true | . | . | . | . | . | . | chr15 | 10658484 | 10698797 | 6.0561 | 4.48411 | 3.88941 |
| chr9 | 13558941 | 13559258 | 719 | Distal Intergenic | 9 | 13619990 | 13709388 | 89399 | 1 | ENSMUST00 | -60732 | Maml2 | ENSMUSG0C | false | . | . | . | . | . | . | . | . | . | . | . | . |
| chr12 | 35824663 | 35824995 | 719 | Distal Intergenic | 12 | 35925620 | 35949736 | 24117 | 1 | ENSMUST00 | -100625 | Agr3 | ENSMUSG0C | false | . | . | . | . | . | . | chr12 | 35824152 | 35825166 | 4.15221 | 3.70601 | 2.30022 |
| chrX | 12128312 | 12128937 | 719 | Promoter (<=1kb) | 20 | 12058981 | 12128400 | 69420 | 2 | ENSMUST00 | 0 | Bcor | ENSMUSG0C | false | chrX | 12127816 | 12129066 | 17.92175 | 44.05972 | 41.41886 | . | . | . | . | . | . |
| chr9 | 57076540 | 57077185 | 719 | Promoter (<=1kb) | 9 | 57076628 | 57095403 | 18776 | 1 | ENSMUST00 | 0 | Sin3a | ENSMUSG0C | false | chr9 | 57076296 | 57077594 | 27.34946 | 104.78247 | 101.09169 | . | . | . | . | . | . |
| chr17 | 86937804 | 86938232 | 719 | Distal Intergenic | 17 | 86944109 | 86947887 | 3779 | 2 | ENSMUST00 | 9655 | Atp6v1e2 | ENSMUSG0C | false | chr17 | 86937572 | 86938405 | 10.14579 | 20.57575 | 18.2329 | chr17 | 86937652 | 86938372 | 3.69077 | 3.45057 | 1.91238 |
| chr1 | 40324379 | 40324687 | 719 | Promoter (<=1kb) | 1 | 40324627 | 40365471 | 40845 | 1 | ENSMUST00 | 0 | Il1rl2 | ENSMUSG0C | false | chr1 | 40324485 | 40325998 | 6.89286 | 16.24995 | 13.96717 | chr1 | 40324073 | 40326086 | 4.248 | 3.65506 | 2.38832 |
| chr1 | 132322396 | 132322785 | 718 | Intron (ENSMUST00000072177.8 | 1 | 132327596 | 132330450 | 2855 | 1 | ENSMUST00 | -4811 | Nuak2 | ENSMUSG0C | false | . | . | . | . | . | . | chr1 | 132317064 | 132339308 | 6.74463 | 4.86917 | 4.43585 |
| chr18 | 52945820 | 52946095 | 718 | Distal Intergenic | 18 | 52838010 | 52915312 | 77303 | 1 | ENSMUST00 | 107811 | Sncaip | ENSMUSG0C | true | . | . | . | . | . | . | chr18 | 52944039 | 52948882 | 5.44021 | 4.32749 | 3.36916 |
| chr9 | 65827125 | 65827853 | 718 | Promoter (<=1kb) | 9 | 65692391 | 65827564 | 135174 | 2 | ENSMUST00 | 0 | Zfp609 | ENSMUSG0C | false | chr9 | 65826248 | 65828549 | 30.03893 | 101.89226 | 98.2715 | chr9 | 65817423 | 65827175 | 4.53501 | 3.80778 | 2.62509 |
| chr17 | 36270972 | 36271897 | 718 | Promoter (<=1kb) | 17 | 36258873 | 36271567 | 12695 | 2 | ENSMUST00 | 0 | Trim39 | ENSMUSG0C | false | chr17 | 36270565 | 36272542 | 30.45735 | 121.03082 | 116.85291 | . | . | . | . | . | . |

| | | | | | | | | | | | | | | | | | | | | | | | | | |
|---|---|---|---|---|---|---|---|---|---|---|---|---|---|---|---|---|---|---|---|---|---|---|---|---|---|
| chr3 | 98338999 | 98339347 | 718 | Promoter (<=1kb) | 3 | 98313170 | 98339990 | 26821 | 2 | ENSMUST00 | 643 | Phgdh | ENSMUSG0C | false | chr3 | 98338685 | 98340237 | 19.27906 | 51.80829 | 49.07 | . | . | . | . | . | . |
| chr6 | 54819810 | 54820247 | 718 | Promoter (2-3kb) | 6 | 54817448 | 54885111 | 67664 | 1 | ENSMUST00 | 2363 | Znrf2 | ENSMUSG0C | false | . | . | . | . | . | . | chr6 | 54817953 | 54822750 | 6.26788 | 4.6905 | 4.04182 |
| chr2 | 156717251 | 156717756 | 718 | Promoter (2-3kb) | 2 | 156720572 | 156749091 | 28520 | 1 | ENSMUST00 | -2816 | Dlgap4 | ENSMUSG0C | true | chr2 | 156717214 | 156717930 | 10.10971 | 19.98882 | 17.65231 | chr2 | 156716621 | 156720481 | 7.13343 | 5.10713 | 4.75424 |
| chr2 | 167834647 | 167835033 | 718 | Promoter (1-2kb) | 2 | 167827693 | 167833646 | 5954 | 2 | ENSMUST00 | -1002 | 1200007C13I | ENSMUSG0C | false | . | . | . | . | . | . | chr2 | 167830528 | 167835185 | 5.99375 | 4.5533 | 3.83999 |
| chr10 | 61783775 | 61784100 | 717 | Promoter (<=1kb) | 10 | 61738665 | 61784147 | 45483 | 2 | ENSMUST00 | 47 | H2afy2 | ENSMUSG0C | false | chr10 | 61782647 | 61784301 | 28.56742 | 103.16996 | 99.51957 | chr10 | 61783971 | 61784387 | 3.31767 | 3.22666 | 1.63494 |
| chr3 | 9006763 | 9007016 | 717 | Promoter (2-3kb) | 3 | 8928626 | 9004723 | 76098 | 2 | ENSMUST00 | -2041 | Tpd52 | ENSMUSG0C | false | . | . | . | . | . | . | chr3 | 9004804 | 9008983 | 6.88132 | 5.02165 | 4.53704 |
| chr3 | 138771555 | 138771865 | 717 | Intron (ENSMUST00000029800.3 | 3 | 138778206 | 138902494 | 124289 | 1 | ENSMUST00 | -6341 | Tspan5 | ENSMUSG0C | true | . | . | . | . | . | . | chr3 | 138763693 | 138776582 | 5.61138 | 4.35746 | 3.51014 |
| chr4 | 132843046 | 132843371 | 717 | Promoter (<=1kb) | 4 | 132826929 | 132843169 | 16241 | 2 | ENSMUST00 | 0 | Ppp1r8 | ENSMUSG0C | false | chr4 | 132842194 | 132843679 | 20.74282 | 60.13039 | 57.28273 | chr4 | 132841849 | 132844341 | 4.39588 | 3.77615 | 2.5089 |
| chr5 | 35520647 | 35521194 | 717 | Intron (ENSMUST00000038676.5 | 5 | 35517565 | 35525571 | 8007 | 2 | ENSMUST00 | 4377 | Cpz | ENSMUSG0C | false | . | . | . | . | . | . | . | . | . | . | . | . |
| chr3 | 101437086 | 101437373 | 717 | Intron (ENSMUST00000043983.5 | 3 | 101377125 | 101463060 | 85936 | 1 | ENSMUST00 | 59962 | Igsf3 | ENSMUSG0C | false | . | . | . | . | . | . | . | . | . | . | . | . |
| chr2 | 69722693 | 69723563 | 717 | Promoter (<=1kb) | 2 | 69722805 | 69754012 | 31208 | 1 | ENSMUST00 | 0 | Ppig | ENSMUSG0C | false | chr2 | 69722273 | 69724225 | 44.33739 | 156.62123 | 151.08617 | . | . | . | . | . | . |
| chr11 | 57651228 | 57651741 | 717 | Promoter (1-2kb) | 11 | 57623698 | 57649494 | 25797 | 2 | ENSMUST00 | -1735 | 4933424L21F | ENSMUSG0C | false | . | . | . | . | . | . | chr11 | 57646153 | 57704104 | 7.42247 | 5.08195 | 5.00645 |
| chr2 | 180709785 | 180710439 | 716 | Promoter (<=1kb) | 2 | 180710117 | 180718733 | 8617 | 1 | ENSMUST00 | 0 | Gid8 | ENSMUSG0C | false | chr2 | 180708615 | 180711658 | 35.67123 | 127.73884 | 123.33056 | . | . | . | . | . | . |
| chr4 | 155654408 | 155654799 | 716 | Promoter (1-2kb) | 4 | 155650655 | 155653384 | 2730 | 2 | ENSMUST00 | -1025 | Mmp23 | ENSMUSG0C | false | chr4 | 155651243 | 155655658 | 26.60493 | 74.70471 | 71.64091 | chr4 | 155648087 | 155656351 | 6.88407 | 4.99201 | 4.54195 |
| chr9 | 75592878 | 75593189 | 716 | Intron (ENSMUST00000098552.3 | 9 | 75573816 | 75597651 | 23836 | 2 | ENSMUST00 | 4462 | Tmod2 | ENSMUSG0C | false | . | . | . | . | . | . | chr9 | 75590322 | 75593818 | 3.88341 | 3.38672 | 2.13515 |
| chr2 | 101797549 | 101797911 | 716 | Promoter (<=1kb) | 2 | 101714285 | 101797707 | 83423 | 2 | ENSMUST00 | 0 | Prr5l | ENSMUSG0C | false | chr2 | 101796152 | 101798205 | 29.8354 | 90.64201 | 87.2842 | chr2 | 101796855 | 101798090 | 5.41271 | 4.02607 | 3.38829 |
| chr3 | 84155668 | 84156102 | 716 | Promoter (<=1kb) | 3 | 84087934 | 84155786 | 67853 | 2 | ENSMUST00 | 0 | Mnd1 | ENSMUSG0C | false | chr3 | 84155962 | 84156227 | 5.6323 | 9.47728 | 7.31216 | chr3 | 84155307 | 84156479 | 4.36104 | 3.8205 | 2.46131 |
| chr2 | 161110009 | 161110319 | 716 | Promoter (<=1kb) | 2 | 161008034 | 161109075 | 101042 | 2 | ENSMUST00 | -935 | Chd6 | ENSMUSG0C | false | chr2 | 161109244 | 161110158 | 26.66126 | 88.87019 | 85.5497 | chr2 | 161109445 | 161111487 | 4.40933 | 3.88682 | 2.5295 |
| chr2 | 156737964 | 156738303 | 716 | Intron (ENSMUST00000109567.4 | 2 | 156721492 | 156764363 | 42872 | 1 | ENSMUST00 | 16473 | Dlgap4 | ENSMUSG0C | true | . | . | . | . | . | . | chr2 | 156722290 | 156745258 | 7.00087 | 4.91477 | 4.66073 |
| chr16 | 65814902 | 65815307 | 716 | Promoter (<=1kb) | 16 | 65815633 | 65863057 | 47425 | 1 | ENSMUST00 | -326 | Vgll3 | ENSMUSG0C | false | chr16 | 65814792 | 65816194 | 24.14628 | 82.32003 | 79.12344 | chr16 | 65814784 | 65816158 | 4.37557 | 3.54332 | 2.54445 |
| chr15 | 64301626 | 64301919 | 715 | Intron (ENSMUST00000177371.2 | 15 | 64191537 | 64311689 | 120153 | 2 | ENSMUST00 | 9770 | Asap1 | ENSMUSG0C | false | . | . | . | . | . | . | chr15 | 64295774 | 64304493 | 4.89706 | 3.92154 | 2.95026 |
| chr2 | 39065303 | 39065717 | 715 | Promoter (<=1kb) | 2 | 39016261 | 39065488 | 49228 | 2 | ENSMUST00 | 0 | Golga1 | ENSMUSG0C | false | chr2 | 39064853 | 39066259 | 34.16929 | 120.7392 | 116.5701 | . | . | . | . | . | . |
| chr8 | 33805334 | 33805616 | 715 | Promoter (1-2kb) | 8 | 33784064 | 33806799 | 22736 | 2 | ENSMUST00 | 1183 | Rbpms | ENSMUSG0C | true | . | . | . | . | . | . | chr8 | 33797013 | 33821269 | 7.29084 | 5.02722 | 4.89898 |
| chr11 | 108026541 | 108026811 | 715 | Intron (ENSMUST00000059595.5 | 11 | 107978341 | 108012805 | 34465 | 2 | ENSMUST00 | -13737 | Prkca | ENSMUSG0C | false | . | . | . | . | . | . | chr11 | 108025693 | 108028748 | 4.21076 | 3.59534 | 2.36918 |
| chr11 | 6826240 | 6826489 | 715 | Promoter (<=1kb) | 11 | 6825496 | 6825809 | 314 | 2 | ENSMUST00 | -432 | Gm11982 | ENSMUSG0C | false | . | . | . | . | . | . | chr11 | 6826242 | 6827206 | 3.61654 | 3.22149 | 1.91776 |
| chr4 | 153120421 | 153120754 | 715 | Distal Intergenic | 4 | 153145590 | 153146943 | 1354 | 2 | ENSMUST00 | 26189 | Gm13173 | ENSMUSG0C | false | chr4 | 153118908 | 153120529 | 18.44084 | 48.77526 | 46.0739 | chr4 | 153120491 | 153121459 | 4.78531 | 3.78635 | 2.86266 |
| chr8 | 120538539 | 120539258 | 715 | Promoter (<=1kb) | 8 | 120538696 | 120538754 | 59 | 1 | ENSMUST00 | 0 | Mir7687 | ENSMUSG0C | false | chr8 | 120535456 | 120538941 | 22.15371 | 67.81625 | 64.85734 | chr8 | 120537951 | 120565808 | 7.48404 | 5.10526 | 5.05119 |
| chr15 | 11908421 | 11908867 | 714 | Promoter (2-3kb) | 15 | 11839896 | 11905674 | 65779 | 2 | ENSMUST00 | -2748 | Npr3 | ENSMUSG0C | true | . | . | . | . | . | . | chr15 | 11907654 | 11909880 | 5.64871 | 4.4589 | 3.52516 |
| chr10 | 127339775 | 127340177 | 714 | Promoter (1-2kb) | 10 | 127329889 | 127341589 | 11701 | 2 | ENSMUST00 | 1412 | Gli1 | ENSMUSG0C | false | chr10 | 127339045 | 127340137 | 11.31597 | 24.78923 | 22.38983 | chr10 | 127338758 | 127342756 | 3.59246 | 3.3131 | 1.87872 |
| chr3 | 98246976 | 98247309 | 714 | Distal Intergenic | 3 | 98222156 | 98236748 | 14593 | 1 | ENSMUST00 | 24821 | Reg4 | ENSMUSG0C | false | . | . | . | . | . | . | . | . | . | . | . | . |
| chr6 | 58831282 | 58831982 | 714 | Promoter (<=1kb) | 6 | 58831465 | 58843728 | 12264 | 1 | ENSMUST00 | 0 | Herc3 | ENSMUSG0C | false | chr6 | 58831048 | 58832945 | 41.56779 | 133.79375 | 129.15814 | chr6 | 58830663 | 58833302 | 4.02439 | 3.65185 | 2.20036 |
| chr15 | 32036888 | 32037179 | 714 | Distal Intergenic | 15 | 32177289 | 32178296 | 1008 | 1 | ENSMUST00 | -140110 | Tas2r119 | ENSMUSG0C | false | . | . | . | . | . | . | chr15 | 32033065 | 32044868 | 4.832 | 3.92577 | 2.87384 |
| chr5 | 105731837 | 105732498 | 714 | Promoter (<=1kb) | 5 | 105731990 | 105813639 | 81650 | 1 | ENSMUST00 | 0 | Lrrc8d | ENSMUSG0C | false | chr5 | 105731574 | 105732860 | 22.30049 | 67.20228 | 64.25432 | chr5 | 105730787 | 105733671 | 3.58511 | 3.32623 | 1.87911 |
| chr14 | 121361770 | 121362201 | 714 | Intron (ENSMUST00000079817.7 | 14 | 121286343 | 121379334 | 92992 | 2 | ENSMUST00 | 17133 | Stk24 | ENSMUSG0C | false | . | . | . | . | . | . | chr14 | 121355140 | 121373730 | 4.61394 | 3.83015 | 2.71227 |
| chr1 | 73744063 | 73744366 | 714 | Promoter (<=1kb) | 1 | 73618088 | 73743947 | 125860 | 2 | ENSMUST00 | -117 | RP24-144C1( | ENSMUSG0C | false | . | . | . | . | . | . | chr1 | 73742869 | 73745446 | 5.25482 | 3.9934 | 3.23541 |
| chr11 | 105180913 | 105181610 | 713 | Promoter (<=1kb) | 11 | 105181527 | 105282177 | 100651 | 1 | ENSMUST00 | 0 | Tlk2 | ENSMUSG0C | false | chr11 | 105180525 | 105182894 | 40.64776 | 144.39505 | 139.32506 | chr11 | 105179484 | 105180985 | 4.44591 | 3.75211 | 2.59154 |
| chr14 | 60782568 | 60782791 | 713 | Promoter (1-2kb) | 14 | 60784566 | 60903610 | 119045 | 1 | ENSMUST00 | -1775 | Mipep | ENSMUSG0C | false | . | . | . | . | . | . | chr14 | 60782255 | 60782604 | 2.91507 | 2.98259 | 1.3067 |
| chr7 | 137714323 | 137714634 | 713 | Distal Intergenic | 7 | 137437648 | 137468594 | 30947 | 1 | ENSMUST00 | 276676 | Glrx3 | ENSMUSG0C | false | . | . | . | . | . | . | . | . | . | . | . | . |
| chr7 | 99818121 | 99818683 | 713 | Intron (ENSMUST00000036331.6 | 7 | 99811439 | 99828417 | 16979 | 2 | ENSMUST00 | 9734 | Neu3 | ENSMUSG0C | false | chr7 | 99818089 | 99818903 | 8.08052 | 18.53858 | 16.22368 | chr7 | 99817314 | 99819705 | 4.6839 | 3.91035 | 2.7482 |
| chr5 | 100127127 | 100127586 | 713 | Intron (ENSMUST00000063192.9 | 5 | 100083947 | 100122796 | 38850 | 2 | ENSMUST00 | -4332 | Tmem150c | ENSMUSG0C | true | . | . | . | . | . | . | chr5 | 100107043 | 100130430 | 5.59817 | 4.32599 | 3.51171 |
| chr14 | 76556238 | 76556922 | 713 | Promoter (<=1kb) | 14 | 76532814 | 76556687 | 23874 | 2 | ENSMUST00 | 0 | Serp2 | ENSMUSG0C | false | chr14 | 76556033 | 76557095 | 20.8177 | 63.66574 | 60.7686 | chr14 | 76555703 | 76557002 | 4.32834 | 3.76554 | 2.49873 |
| chr6 | 134189703 | 134190046 | 713 | Intron (ENSMUST00000081028.7 | 6 | 134168879 | 134172007 | 3129 | 2 | ENSMUST00 | -17697 | Gm17088 | ENSMUSG0C | false | . | . | . | . | . | . | chr6 | 134186494 | 134190488 | 5.7551 | 4.46071 | 3.63768 |
| chr12 | 69556127 | 69556525 | 713 | Distal Intergenic | 12 | 69561229 | 69561337 | 109 | 2 | ENSMUST00 | 4812 | Gm23018 | ENSMUSG0C | false | . | . | . | . | . | . | chr12 | 69555906 | 69557204 | 5.71608 | 4.34052 | 3.62711 |
| chr4 | 109225965 | 109226513 | 712 | Distal Intergenic | 4 | 109234485 | 109254251 | 19767 | 1 | ENSMUST00 | -7972 | Calr4 | ENSMUSG0C | false | . | . | . | . | . | . | chr4 | 109224462 | 109228424 | 6.45362 | 4.59713 | 4.24481 |
| chr6 | 124919631 | 124920271 | 712 | Promoter (1-2kb) | 6 | 124913681 | 124917946 | 4266 | 2 | ENSMUST00 | -1686 | Ptms | ENSMUSG0C | false | chr6 | 124919941 | 124920525 | 10.96293 | 22.48196 | 20.11397 | chr6 | 124918423 | 124922012 | 5.01326 | 4.03706 | 3.02451 |
| chr1 | 55746086 | 55746374 | 712 | Intron (ENSMUST00000042986.9 | 1 | 55702025 | 55735227 | 33203 | 1 | ENSMUST00 | 44062 | Plcl1 | ENSMUSG0C | true | . | . | . | . | . | . | chr1 | 55738786 | 55752458 | 4.85141 | 3.98095 | 2.88878 |
| chr5 | 24733288 | 24733559 | 712 | Promoter (2-3kb) | 5 | 24711738 | 24730727 | 18990 | 2 | ENSMUST00 | -2562 | Wdr86 | ENSMUSG0C | false | . | . | . | . | . | . | chr5 | 24723425 | 24734625 | 5.82193 | 4.37932 | 3.69553 |
| chr13 | 51374124 | 51374401 | 712 | Distal Intergenic | 13 | 51408618 | 51422797 | 14180 | 1 | ENSMUST00 | -34217 | S1pr3 | ENSMUSG0C | false | . | . | . | . | . | . | chr13 | 51374036 | 51374786 | 3.33542 | 3.09915 | 1.67462 |
| chr2 | 72266945 | 72267336 | 712 | Distal Intergenic | 2 | 72244965 | 72248975 | 4011 | 2 | ENSMUST00 | -17971 | Rapgef4os3 | ENSMUSG0C | false | . | . | . | . | . | . | chr2 | 72266582 | 72267959 | 5.81547 | 4.53667 | 3.66937 |
| chr17 | 26952449 | 26952772 | 712 | Exon (ENSMUST00000177932.1/ | 17 | 26948056 | 26948119 | 64 | 2 | ENSMUST00 | -4331 | Mir3083 | ENSMUSG0C | false | . | . | . | . | . | . | chr17 | 26950986 | 26954220 | 5.81469 | 4.43814 | 3.67192 |
| chr3 | 85186802 | 85187114 | 712 | Intron (ENSMUST00000154148.2 | 3 | 85317519 | 85332030 | 14512 | 1 | ENSMUST00 | -130405 | RP23-94M23 | ENSMUSG0C | false | chr3 | 85186707 | 85187108 | 8.50221 | 18.80243 | 16.48241 | chr3 | 85186287 | 85188610 | 3.53746 | 3.19063 | 1.85151 |
| chr13 | 110394345 | 110394932 | 711 | Promoter (<=1kb) | 13 | 110395044 | 110400844 | 5801 | 1 | ENSMUST00 | -112 | Plk2 | ENSMUSG0C | false | chr13 | 110394346 | 110396681 | 37.64991 | 125.88531 | 121.55012 | chr13 | 110394160 | 110396575 | 4.81782 | 4.00833 | 2.86688 |
| chr2 | 174089039 | 174089351 | 711 | Intron (ENSMUST00000155000.2 | 2 | 174077066 | 174096967 | 19902 | 1 | ENSMUST00 | 11974 | Stx16 | ENSMUSG0C | false | . | . | . | . | . | . | chr2 | 174073295 | 174092361 | 5.14014 | 4.03613 | 3.13424 |
| chr5 | 103753646 | 103754586 | 711 | Promoter (<=1kb) | 5 | 103754162 | 103855322 | 101161 | 1 | ENSMUST00 | 0 | Aff1 | ENSMUSG0C | false | chr5 | 103752781 | 103755575 | 27.54716 | 98.2761 | 94.74957 | . | . | . | . | . | . |
| chr9 | 72714100 | 72714455 | 711 | Intron (ENSMUST00000034740.9 | 9 | 72739472 | 72748158 | 8687 | 1 | ENSMUST00 | -25017 | Nedd4 | ENSMUSG0C | true | . | . | . | . | . | . | chr9 | 72683575 | 72716820 | 5.72166 | 4.35702 | 3.61419 |
| chr9 | 77754363 | 77754821 | 711 | Promoter (<=1kb) | 9 | 77754535 | 77794485 | 39951 | 1 | ENSMUST00 | 0 | Gclc | ENSMUSG0C | false | chr9 | 77754467 | 77755628 | 35.87559 | 113.23412 | 109.30762 | . | . | . | . | . | . |
| chr8 | 71511649 | 71511953 | 711 | Promoter (<=1kb) | 8 | 71497754 | 71511769 | 14016 | 2 | ENSMUST00 | 0 | Plvap | ENSMUSG0C | false | . | . | . | . | . | . | . | . | . | . | . | . |
| chr4 | 56831334 | 56831710 | 711 | Promoter (1-2kb) | 4 | 56826266 | 56829853 | 3588 | 2 | ENSMUST00 | -1482 | Ctnnal1 | ENSMUSG0C | true | . | . | . | . | . | . | chr4 | 56827266 | 56853148 | 5.48236 | 4.22607 | 3.42275 |
| chr5 | 64581467 | 64581824 | 711 | Distal Intergenic | 5 | 64647633 | 64647749 | 117 | 2 | ENSMUST00 | 65925 | Gm25306 | ENSMUSG0C | false | . | . | . | . | . | . | chr5 | 64578029 | 64582112 | 5.70116 | 4.39721 | 3.6117 |
| chr10 | 121975445 | 121975779 | 710 | Intron (ENSMUST00000020322.6 | 10 | 121780991 | 122047308 | 266318 | 2 | ENSMUST00 | 71529 | Srgap1 | ENSMUSG0C | true | . | . | . | . | . | . | chr10 | 121945243 | 122036389 | 5.74397 | 4.3951 | 3.62974 |
| chr2 | 143545846 | 143546190 | 710 | Promoter (<=1kb) | 2 | 143546156 | 143816285 | 270130 | 1 | ENSMUST00 | 0 | Pcsk2 | ENSMUSG0C | false | chr2 | 143545567 | 143546690 | 15.95428 | 37.82584 | 35.264 | chr2 | 143545345 | 143547090 | 5.48564 | 4.38773 | 3.41297 |
| chr3 | 9033651 | 9033976 | 710 | Distal Intergenic | 3 | 8928626 | 9004723 | 76098 | 2 | ENSMUST00 | -28929 | Tpd52 | ENSMUSG0C | false | . | . | . | . | . | . | chr3 | 9032865 | 9038735 | 5.34195 | 4.17254 | 3.28907 |
| chr6 | 98949596 | 98950096 | 710 | Intron (ENSMUST00000113326.3 | 6 | 98930171 | 98935545 | 5375 | 2 | ENSMUST00 | -14052 | Foxp1 | ENSMUSG0C | false | . | . | . | . | . | . | chr6 | 98946696 | 98951866 | 5.45838 | 4.12852 | 3.41677 |
| chrX | 105763721 | 105763997 | 710 | Promoter (<=1kb) | 20 | 105764279 | 105774939 | 10661 | 1 | ENSMUST00 | -282 | Fgf16 | ENSMUSG0C | false | chrX | 105763531 | 105765408 | 12.40737 | 26.6269 | 24.20249 | . | . | . | . | . | . |
| chr7 | 44815822 | 44816667 | 710 | Promoter (<=1kb) | 7 | 44812257 | 44816658 | 4402 | 2 | ENSMUST00 | 0 | Atf5 | ENSMUSG0C | false | chr7 | 44815266 | 44817335 | 41.911 | 144.38934 | 139.32462 | . | . | . | . | . | . |
| chr3 | 108283894 | 108284624 | 710 | Promoter (<=1kb) | 3 | 108284091 | 108361511 | 77421 | 1 | ENSMUST00 | 0 | Sort1 | ENSMUSG0C | false | chr3 | 108283322 | 108284697 | 35.44477 | 112.48663 | 108.5799 | chr3 | 108282612 | 108285901 | 4.30513 | 3.68197 | 2.42822 |
| chr12 | 86894398 | 86894918 | 710 | Promoter (1-2kb) | 12 | 86891404 | 86892567 | 1164 | 2 | ENSMUST00 | -1832 | Gm26698 | ENSMUSG0C | false | chr12 | 86894492 | 86895224 | 11.02877 | 22.59171 | 20.22099 | chr12 | 86885110 | 86901194 | 5.68305 | 4.41688 | 3.56559 |
| chr4 | 99976436 | 99976732 | 709 | Intron (ENSMUST00000058351.1 | 4 | 99955744 | 99987294 | 31551 | 1 | ENSMUST00 | 20693 | Pgm2 | ENSMUSG0C | false | . | . | . | . | . | . | chr4 | 99968725 | 99977020 | 4.83736 | 3.90531 | 2.88809 |
| chr3 | 127552801 | 127553427 | 709 | Promoter (<=1kb) | 3 | 127536954 | 127553348 | 16395 | 2 | ENSMUST00 | 0 | Larp7 | ENSMUSG0C | false | chr3 | 127552453 | 127553971 | 29.22959 | 88.61999 | 85.30408 | . | . | . | . | . | . |
| chr4 | 43578517 | 43578972 | 709 | Promoter (<=1kb) | 4 | 43566928 | 43578537 | 11610 | 2 | ENSMUST00 | 0 | Gba2 | ENSMUSG0C | false | chr4 | 43577870 | 43579625 | 26.03857 | 105.07202 | 101.37238 | . | . | . | . | . | . |
| chr5 | 67348831 | 67349142 | 709 | Intron (ENSMUST00000113676.2 | 5 | 67332118 | 67353355 | 21238 | 1 | ENSMUST00 | 16714 | Slc30a9 | ENSMUSG0C | false | . | . | . | . | . | . | chr5 | 67348562 | 67349351 | 4.69929 | 4.00386 | 2.74836 |
| chr9 | 77257759 | 77258023 | 709 | Intron (ENSMUST00000185144.2 | 9 | 77249180 | 77252134 | 2955 | 2 | ENSMUST00 | -5626 | Mlip | ENSMUSG0C | false | . | . | . | . | . | . | chr9 | 77257340 | 77259560 | 3.37901 | 3.07834 | 1.72106 |
| chr6 | 50159612 | 50159977 | 709 | Intron (ENSMUST00000166318.2 | 6 | 50180300 | 50196558 | 16259 | 1 | ENSMUST00 | -20323 | Mpp6 | ENSMUSG0C | false | . | . | . | . | . | . | chr6 | 50159031 | 50161700 | 3.77247 | 3.38291 | 2.02279 |
| chr3 | 95194702 | 95195089 | 709 | Promoter (2-3kb) | 3 | 95187589 | 95191898 | 4310 | 2 | ENSMUST00 | -2805 | Gabpb2 | ENSMUSG0C | true | . | . | . | . | . | . | chr3 | 95193311 | 95204022 | 6.01146 | 4.56637 | 3.8326 |
| chr17 | 84365696 | 84366252 | 709 | Intron (ENSMUST00000047524.8 | 17 | 84391429 | 84391535 | 107 | 1 | ENSMUST00 | -25177 | Gm24722 | ENSMUSG0C | false | . | . | . | . | . | . | chr17 | 84361955 | 84374371 | 5.57955 | 4.30181 | 3.49381 |
| chr5 | 136966340 | 136966885 | 708 | Promoter (<=1kb) | 5 | 136966616 | 136975858 | 9243 | 1 | ENSMUST00 | 0 | Cldn15 | ENSMUSG0C | false | . | . | . | . | . | . | chr5 | 136953761 | 136969776 | 6.09241 | 4.43302 | 3.92021 |
| chr10 | 19591528 | 19592182 | 708 | Promoter (<=1kb) | 10 | 19591949 | 19610229 | 18281 | 1 | ENSMUST00 | 0 | Ifngr1 | ENSMUSG0C | false | chr10 | 19591951 | 19594959 | 38.0703 | 123.3938 | 119.13641 | chr10 | 19590567 | 19591687 | 4.39874 | 3.83627 | 2.50362 |
| chr7 | 65933426 | 65933693 | 708 | Intron (ENSMUST00000176199.2 | 7 | 65953012 | 65963193 | 10182 | 1 | ENSMUST00 | -19319 | Pcsk6 | ENSMUSG0C | true | . | . | . | . | . | . | chr7 | 65929189 | 65934196 | 6.32764 | 4.65183 | 4.10713 |
| chr9 | 35045102 | 35045414 | 708 | Downstream (1-2kb) | 9 | 35016452 | 35030876 | 14425 | 1 | ENSMUST00 | 28651 | Kirrel3 | ENSMUSG0C | false | . | . | . | . | . | . | chr9 | 35043483 | 35046427 | 4.2485 | 3.70131 | 2.39402 |
| chr14 | 78710697 | 78710999 | 708 | Intron (ENSMUST00000074729.5 | 14 | 78569609 | 78725089 | 155481 | 2 | ENSMUST00 | 14090 | Dgkh | ENSMUSG0C | false | . | . | . | . | . | . | chr14 | 78703472 | 78719116 | 4.99522 | 4.07558 | 2.9982 |
| chr4 | 134449985 | 134450288 | 708 | Promoter (<=1kb) | 4 | 134448766 | 134450171 | 1406 | 2 | ENSMUST00 | 0 | 1700021N21 | ENSMUSG0C | false | . | . | . | . | . | . | chr4 | 134444756 | 134452233 | 5.18386 | 4.23712 | 3.15068 |
| chr15 | 84302857 | 84303353 | 708 | Intron (ENSMUST00000023072.6 | 15 | 84323209 | 84323352 | 144 | 2 | ENSMUST00 | 19999 | Gm22890 | ENSMUSG0C | false | . | . | . | . | . | . | chr15 | 84297650 | 84306963 | 6.75434 | 4.89978 | 4.45766 |
| chr1 | 39478678 | 39479035 | 707 | Promoter (<=1kb) | 1 | 39371499 | 39478747 | 107249 | 2 | ENSMUST00 | 0 | Tbc1d8 | ENSMUSG0C | false | chr1 | 39477826 | 39479423 | 34.78613 | 113.42423 | 109.48744 | . | . | . | . | . | . |
| chr13 | 40931778 | 40932264 | 707 | Intron (ENSMUST00000110191.3 | 13 | 40917634 | 40960891 | 43258 | 1 | ENSMUST00 | 14145 | Gcnt2 | ENSMUSG0C | false | . | . | . | . | . | . | . | . | . | . | . | . |
| chr1 | 82258294 | 82258609 | 707 | Intron (ENSMUST00000069799.2 | 1 | 82233112 | 82235933 | 2822 | 1 | ENSMUST00 | 25183 | Gm9747 | ENSMUSG0C | false | . | . | . | . | . | . | chr1 | 82244858 | 82261494 | 5.35178 | 4.22772 | 3.29163 |
| chr4 | 57821793 | 57822339 | 707 | Intron (ENSMUST00000098066.3 | 4 | 57829245 | 57855049 | 25805 | 1 | ENSMUST00 | -6906 | Akap2 | ENSMUSG0C | false | . | . | . | . | . | . | chr4 | 57819780 | 57824245 | 5.13956 | 4.0959 | 3.12344 |
| chr11 | 68602472 | 68602801 | 707 | Distal Intergenic | 11 | 68594055 | 68597954 | 3900 | 1 | ENSMUST00 | 8418 | Ccdc42 | ENSMUSG0C | false | . | . | . | . | . | . | chr11 | 68600217 | 68602957 | 4.50566 | 3.83233 | 2.64202 |
| chr15 | 86085932 | 86086262 | 707 | Intron (ENSMUST00000088931.4 | 15 | 86087640 | 86090060 | 2421 | 2 | ENSMUST00 | 3798 | Gm15722 | ENSMUSG0C | false | . | . | . | . | . | . | chr15 | 86074674 | 86092519 | 7.22651 | 5.10744 | 4.82194 |
| chr5 | 7539998 | 7540425 | 707 | Distal Intergenic | 5 | 7399362 | 7400362 | 1001 | 2 | ENSMUST00 | -139637 | Gm4960 | ENSMUSG0C | false | . | . | . | . | . | . | chr5 | 7538094 | 7541801 | 4.0233 | 3.58642 | 2.23455 |
| chr9 | 102252690 | 102253012 | 707 | Intron (ENSMUST00000035129.8 | 9 | 102304924 | 102304975 | 52 | 2 | ENSMUST00 | 51963 | Mir7243 | ENSMUSG0C | false | . | . | . | . | . | . | chr9 | 102247451 | 102256245 | 5.06875 | 3.9593 | 3.08777 |
| chr10 | 85510493 | 85510806 | 706 | Intron (ENSMUST00000105307.2 | 10 | 85598401 | 85660292 | 61892 | 1 | ENSMUST00 | -87595 | Btbd11 | ENSMUSG0C | false | . | . | . | . | . | . | chr10 | 85510469 | 85512380 | 3.75628 | 3.25956 | 2.03953 |
| chrX | 77510596 | 77511250 | 706 | Promoter (<=1kb) | 20 | 77511013 | 77662983 | 151971 | 1 | ENSMUST00 | 0 | Tbl1x | ENSMUSG0C | false | chrX | 77510423 | 77512224 | 23.43614 | 63.09834 | 60.20888 | . | . | . | . | . | . |
| chr1 | 153239842 | 153240112 | 706 | Promoter (<=1kb) | 1 | 153228815 | 153239326 | 10512 | 2 | ENSMUST00 | -517 | Lamc1 | ENSMUSG0C | false | . | . | . | . | . | . | chr1 | 153239708 | 153240965 | 3.36622 | 3.10551 | 1.70524 |
| chr3 | 7428808 | 7429076 | 706 | Intron (ENSMUST00000028999.6 | 3 | 7366604 | 7445366 | 78763 | 1 | ENSMUST00 | 62205 | Pkia | ENSMUSG0C | true | . | . | . | . | . | . | chr3 | 7427815 | 7430766 | 3.68226 | 3.37214 | 1.94365 |
| chr17 | 35516763 | 35517152 | 706 | Promoter (<=1kb) | 17 | 35517100 | 35530996 | 13897 | 1 | ENSMUST00 | 0 | Cchcr1 | ENSMUSG0C | false | chr17 | 35516164 | 35517735 | 25.7338 | 71.40817 | 68.39619 | chr17 | 35515892 | 35517816 | 3.97975 | 3.59259 | 2.17443 |
| chr8 | 25601460 | 25602312 | 706 | Promoter (<=1kb) | 8 | 25601601 | 25677214 | 75614 | 1 | ENSMUST00 | 0 | Whsc1l1 | ENSMUSG0C | false | chr8 | 25601119 | 25603913 | 31.96831 | 119.80809 | 115.67081 | . | . | . | . | . | . |
| chr2 | 157077563 | 157077860 | 706 | Promoter (1-2kb) | 2 | 157015799 | 157079254 | 63456 | 2 | ENSMUST00 | 1394 | Soga1 | ENSMUSG0C | false | chr2 | 157077375 | 157079761 | 27.41053 | 90.23362 | 86.88379 | chr2 | 157074030 | 157079796 | 4.31246 | 3.68722 | 2.46593 |
| chr12 | 86807085 | 86807399 | 706 | Promoter (2-3kb) | 12 | 86809957 | 86810258 | 302 | 1 | ENSMUST00 | -2558 | Gm25888 | ENSMUSG0C | false | . | . | . | . | . | . | chr12 | 86805782 | 86808123 | 5.55102 | 4.30569 | 3.48419 |
| chr3 | 90476150 | 90476520 | 705 | Promoter (<=1kb) | 3 | 90476201 | 90488379 | 12179 | 1 | ENSMUST00 | 0 | Ilf2 | ENSMUSG0C | false | chr3 | 90476084 | 90477052 | 34.00549 | 104.83981 | 101.14426 | chr3 | 90475391 | 90477488 | 3.95328 | 3.48796 | 2.17859 |
| chr19 | 5016170 | 5016564 | 705 | Distal Intergenic | 19 | 5024006 | 5031971 | 7966 | 1 | ENSMUST00 | -7442 | Slc29a2 | ENSMUSG0C | false | . | . | . | . | . | . | chr19 | 5014481 | 5019194 | 5.56102 | 4.37293 | 3.46314 |
| chr4 | 43870521 | 43870805 | 705 | Distal Intergenic | 4 | 43875530 | 43944745 | 69216 | 1 | ENSMUST00 | -4725 | Reck | ENSMUSG0C | false | . | . | . | . | . | . | chr4 | 43870438 | 43870816 | 3.03987 | 3.05891 | 1.46469 |
| chr8 | 23663451 | 23663964 | 705 | Intron (ENSMUST00000131410.2 | 8 | 23669658 | 24063105 | 393448 | 1 | ENSMUST00 | -5694 | Zmat4 | ENSMUSG0C | false | . | . | . | . | . | . | . | . | . | . | . | . |
| chr13 | 20142675 | 20142979 | 705 | Intron (ENSMUST00000181429.2 | 13 | 20111512 | 20342888 | 231377 | 1 | ENSMUST00 | 31164 | Elmo1 | ENSMUSG0C | true | . | . | . | . | . | . | chr13 | 20137737 | 20154822 | 4.88405 | 3.9784 | 2.92438 |
| chr16 | 18683568 | 18683871 | 705 | Distal Intergenic | 16 | 18621811 | 18629938 | 8128 | 2 | ENSMUST00 | -53631 | Sep-05 | ENSMUSG0C | false | . | . | . | . | . | . | chr16 | 18683439 | 18684618 | 4.73584 | 3.97383 | 2.78546 |
| chr6 | 121326449 | 121326963 | 705 | Promoter (<=1kb) | 6 | 121325580 | 121325676 | 97 | 2 | ENSMUST00 | -774 | Gm24855 | ENSMUSG0C | false | . | . | . | . | . | . | chr6 | 121325849 | 121327393 | 5.27972 | 4.18521 | 3.28196 |
| chr16 | 30215825 | 30216136 | 705 | Distal Intergenic | 16 | 30200910 | 30205879 | 4970 | 2 | ENSMUST00 | -9947 | Gm26562 | ENSMUSG0C | false | . | . | . | . | . | . | chr16 | 30215346 | 30216617 | 5.56696 | 4.31306 | 3.50151 |
| chr6 | 128354076 | 128354402 | 704 | Promoter (1-2kb) | 6 | 128334477 | 128355815 | 21339 | 2 | ENSMUST00 | 1413 | Tulp3 | ENSMUSG0C | false | . | . | . | . | . | . | chr6 | 128343079 | 128355029 | 3.93648 | 3.48258 | 2.16297 |
| chr2 | 165311081 | 165311607 | 704 | Promoter (<=1kb) | 2 | 165304694 | 165311642 | 6949 | 2 | ENSMUST00 | 35 | Elmo2 | ENSMUSG0C | false | . | . | . | . | . | . | chr2 | 165307563 | 165325045 | 8.91187 | 5.755 | 6.22321 |
| chr5 | 36581742 | 36582563 | 704 | Promoter (<=1kb) | 5 | 36490604 | 36581803 | 91200 | 2 | ENSMUST00 | 0 | Tbc1d14 | ENSMUSG0C | false | chr5 | 36581265 | 36581929 | 24.40663 | 77.266 | 74.1616 | . | . | . | . | . | . |
| chr17 | 15023980 | 15024890 | 704 | Intron (ENSMUST00000097400.4 | 17 | 15010266 | 15022448 | 12183 | 1 | ENSMUST00 | 13715 | Ermard | ENSMUSG0C | false | chr17 | 15023482 | 15025305 | 45.95321 | 151.4303 | 146.08812 | . | . | . | . | . | . |
| chr7 | 133637340 | 133637727 | 704 | Promoter (<=1kb) | 7 | 133637543 | 133672971 | 35429 | 1 | ENSMUST00 | 0 | Edrf1 | ENSMUSG0C | false | chr7 | 133636998 | 133638678 | 34.33094 | 124.21722 | 119.93477 | . | . | . | . | . | . |
| chr4 | 8690452 | 8691567 | 704 | Promoter (<=1kb) | 4 | 8690470 | 8785591 | 95122 | 1 | ENSMUST00 | 0 | Chd7 | ENSMUSG0C | false | chr4 | 8689673 | 8692342 | 39.81545 | 135.14195 | 130.45284 | . | . | . | . | . | . |
| chr12 | 30567502 | 30567828 | 704 | Distal Intergenic | 12 | 30557914 | 30558120 | 207 | 2 | ENSMUST00 | -9383 | Gm25243 | ENSMUSG0C | false | . | . | . | . | . | . | chr12 | 30566289 | 30569147 | 4.09734 | 3.66876 | 2.25536 |
| chr6 | 141376634 | 141376979 | 704 | Intron (ENSMUST00000043259.8 | 6 | 141238278 | 141250561 | 12284 | 2 | ENSMUST00 | -126074 | RP23-97L11. | ENSMUSG0C | false | . | . | . | . | . | . | chr6 | 141370327 | 141386627 | 5.21392 | 4.14863 | 3.18933 |
| chr2 | 128817580 | 128817944 | 703 | Promoter (<=1kb) | 2 | 128818118 | 128854261 | 36144 | 1 | ENSMUST00 | -174 | Tmem87b | ENSMUSG0C | false | chr2 | 128817330 | 128817853 | 9.86256 | 23.41135 | 21.03137 | . | . | . | . | . | . |
| chr3 | 101923860 | 101924700 | 703 | Promoter (<=1kb) | 3 | 101858930 | 101924449 | 65520 | 2 | ENSMUST00 | 0 | Slc22a15 | ENSMUSG0C | false | chr3 | 101923665 | 101924883 | 36.88168 | 122.38801 | 118.16834 | chr3 | 101923208 | 101924947 | 3.44997 | 3.33428 | 1.72767 |
| chr5 | 43976896 | 43977484 | 703 | Downstream (1-2kb) | 5 | 43978865 | 43981762 | 2898 | 2 | ENSMUST00 | 4278 | Fgfbp1 | ENSMUSG0C | false | . | . | . | . | . | . | chr5 | 43975466 | 43985322 | 3.9051 | 3.48494 | 2.11714 |
| chr10 | 117845102 | 117846351 | 703 | Promoter (<=1kb) | 10 | 117814471 | 117845935 | 31465 | 2 | ENSMUST00 | 0 | Rap1b | ENSMUSG0C | false | chr10 | 117843919 | 117846716 | 31.43325 | 99.35611 | 95.79983 | . | . | . | . | . | . |
| chr7 | 140941521 | 140942036 | 703 | Promoter (<=1kb) | 7 | 140941566 | 140947658 | 6093 | 1 | ENSMUST00 | 0 | Athl1 | ENSMUSG0C | false | chr7 | 140941081 | 140942637 | 15.39495 | 41.17039 | 38.56658 | chr7 | 140939699 | 140943355 | 4.67177 | 3.92406 | 2.73887 |
| chr4 | 88722077 | 88722687 | 703 | Promoter (<=1kb) | 4 | 88718292 | 88722465 | 4174 | 2 | ENSMUST00 | 0 | Klhl9 | ENSMUSG0C | false | chr4 | 88720960 | 88722819 | 33.57447 | 123.878 | 119.61288 | . | . | . | . | . | . |
| chr2 | 159703169 | 159703792 | 703 | Distal Intergenic | 2 | 159777676 | 159778207 | 532 | 2 | ENSMUST00 | 74415 | Gm11445 | ENSMUSG0C | false | . | . | . | . | . | . | chr2 | 159702524 | 159704792 | 4.87913 | 4.07103 | 2.89491 |
| chr4 | 137375134 | 137375481 | 703 | Distal Intergenic | 4 | 137387266 | 137388382 | 1117 | 2 | ENSMUST00 | 12901 | RP23-246F18 | ENSMUSG0C | false | . | . | . | . | . | . | chr4 | 137367417 | 137376426 | 4.18122 | 3.57464 | 2.35005 |
| chr10 | 8181607 | 8181884 | 702 | Distal Intergenic | 10 | 7905653 | 7956230 | 50578 | 2 | ENSMUST00 | -225378 | Tab2 | ENSMUSG0C | true | . | . | . | . | . | . | chr10 | 8180740 | 8182634 | 7.15122 | 5.03252 | 4.82636 |
| chr2 | 34754090 | 34754595 | 702 | Promoter (<=1kb) | 2 | 34676985 | 34754561 | 77577 | 2 | ENSMUST00 | 0 | Gapvd1 | ENSMUSG0C | false | chr2 | 34753418 | 34755238 | 30.26787 | 111.42836 | 107.5545 | . | . | . | . | . | . |
| chr15 | 102089731 | 102090118 | 702 | Exon (ENSMUST00000169681.1/ | 15 | 102102988 | 102116391 | 13404 | 1 | ENSMUST00 | -12870 | Tenc1 | ENSMUSG0C | false | . | . | . | . | . | . | chr15 | 102089015 | 102091634 | 5.36464 | 4.20253 | 3.31404 |
| chr10 | 62843840 | 62844251 | 702 | Intron (ENSMUST00000050826.8 | 10 | 62839097 | 62865304 | 26208 | 2 | ENSMUST00 | 21053 | Tet1 | ENSMUSG0C | false | . | . | . | . | . | . | chr10 | 62841770 | 62845225 | 6.7608 | 4.86152 | 4.43571 |
| chr2 | 17649925 | 17650293 | 702 | Intron (ENSMUST00000028080.6 | 2 | 17343909 | 17731043 | 387135 | 2 | ENSMUST00 | 80750 | Nebl | ENSMUSG0C | true | . | . | . | . | . | . | chr2 | 17644895 | 17651727 | 5.70802 | 4.43304 | 3.5805 |
| chr11 | 57775809 | 57776206 | 702 | Intron (ENSMUST00000066987.8 | 11 | 57764427 | 57764545 | 119 | 1 | ENSMUST00 | 11383 | n-R5s69 | ENSMUSG0C | false | . | . | . | . | . | . | chr11 | 57770983 | 57778876 | 7.36912 | 5.00935 | 4.97771 |
| chr4 | 101041369 | 101041610 | 702 | Distal Intergenic | 4 | 101020298 | 101020696 | 399 | 2 | ENSMUST00 | -20674 | Gm10577 | ENSMUSG0C | false | . | . | . | . | . | . | chr4 | 101039470 | 101042260 | 3.30886 | 3.19175 | 1.62516 |
| chr18 | 35841542 | 35841921 | 702 | Intron (ENSMUST00000060722.6 | 18 | 35830389 | 35831093 | 705 | 2 | ENSMUST00 | -10450 | RP24-326K11 | ENSMUSG0C | false | . | . | . | . | . | . | chr18 | 35831800 | 35857612 | 7.70883 | 5.15861 | 5.24645 |
| chr14 | 46390540 | 46390824 | 701 | Promoter (<=1kb) | 14 | 46383520 | 46390669 | 7150 | 2 | ENSMUST00 | 0 | Bmp4 | ENSMUSG0C | false | . | . | . | . | . | . | chr14 | 46385786 | 46390843 | 3.99631 | 3.58417 | 2.19112 |
| chr15 | 99784691 | 99784964 | 701 | Promoter (<=1kb) | 15 | 99781239 | 99783973 | 2735 | 2 | ENSMUST00 | -719 | Lima1 | ENSMUSG0C | false | . | . | . | . | . | . | chr15 | 99784837 | 99793304 | 6.22394 | 4.62904 | 4.01375 |
| chrX | 42150436 | 42150941 | 701 | Promoter (<=1kb) | 20 | 42150701 | 42277185 | 126485 | 1 | ENSMUST00 | 0 | Stag2 | ENSMUSG0C | false | chrX | 42149209 | 42150570 | 18.84082 | 47.13427 | 44.45482 | . | . | . | . | . | . |
| chr6 | 36777961 | 36778263 | 701 | Intron (ENSMUST00000101534.1 | 6 | 36715663 | 36811361 | 95699 | 2 | ENSMUST00 | 33098 | Ptn | ENSMUSG0C | false | . | . | . | . | . | . | . | . | . | . | . | . |
| chr4 | 126872815 | 126873138 | 701 | Promoter (2-3kb) | 4 | 126868663 | 126870473 | 1811 | 2 | ENSMUST00 | -2343 | Zmym4 | ENSMUSG0C | false | . | . | . | . | . | . | chr4 | 126872747 | 126873179 | 4.34397 | 3.69039 | 2.51105 |
| chr10 | 79535522 | 79535776 | 701 | Promoter (1-2kb) | 10 | 79526430 | 79533787 | 7358 | 2 | ENSMUST00 | -1736 | Ppap2c | ENSMUSG0C | false | . | . | . | . | . | . | . | . | . | . | . | . |
| chr6 | 147718095 | 147718504 | 701 | Promoter (<=1kb) | 6 | 147709982 | 147719451 | 9470 | 2 | ENSMUST00 | 947 | Gm15762 | ENSMUSG0C | false | . | . | . | . | . | . | chr6 | 147718271 | 147719862 | 3.19957 | 3.13742 | 1.56266 |
| chr1 | 62724516 | 62724932 | 701 | Intron (ENSMUST00000126344.1 | 1 | 62714723 | 62718073 | 3351 | 2 | ENSMUST00 | -6444 | RP23-20G7.7 | ENSMUSG0C | false | . | . | . | . | . | . | . | . | . | . | . | . |
| chr17 | 31843969 | 31844272 | 700 | 3' UTR | 17 | 31844248 | 31855792 | 11545 | 2 | ENSMUST00 | 11520 | Sik1 | ENSMUSG0C | false | . | . | . | . | . | . | chr17 | 31842213 | 31847641 | 4.1226 | 3.68821 | 2.28452 |
| chrX | 7574184 | 7574701 | 700 | Promoter (<=1kb) | 20 | 7574492 | 7578415 | 3924 | 1 | ENSMUST00 | 0 | Ppp1r3fos | ENSMUSG0C | false | chrX | 7573333 | 7574848 | 18.38128 | 45.59164 | 42.93153 | . | . | . | . | . | . |
| chr4 | 149679342 | 149679751 | 700 | Intron (ENSMUST00000038859.8 | 4 | 149673439 | 149676093 | 2655 | 2 | ENSMUST00 | -3250 | Pik3cd | ENSMUSG0C | false | . | . | . | . | . | . | chr4 | 149678877 | 149679921 | 3.0224 | 2.95068 | 1.39371 |
| chr8 | 88011463 | 88011774 | 700 | Distal Intergenic | 8 | 87661810 | 87959595 | 297786 | 2 | ENSMUST00 | -51869 | Zfp423 | ENSMUSG0C | true | . | . | . | . | . | . | chr8 | 88009612 | 88012415 | 5.67593 | 4.40788 | 3.60568 |
| chr11 | 116842854 | 116843865 | 700 | Promoter (<=1kb) | 11 | 116837432 | 116843281 | 5850 | 2 | ENSMUST00 | 0 | Jmjd6 | ENSMUSG0C | false | chr11 | 116842041 | 116843458 | 24.82645 | 85.46523 | 82.21117 | . | . | . | . | . | . |
| chr2 | 169538916 | 169539200 | 700 | Distal Intergenic | 2 | 169582845 | 169587745 | 4901 | 2 | ENSMUST00 | 48545 | Gm11011 | ENSMUSG0C | false | . | . | . | . | . | . | chr2 | 169538822 | 169539973 | 3.82338 | 3.55729 | 2.0865 |
| chr10 | 24396097 | 24396393 | 700 | Distal Intergenic | 10 | 24447490 | 24452657 | 5168 | 1 | ENSMUST00 | -51097 | Gm15271 | ENSMUSG0C | false | . | . | . | . | . | . | chr10 | 24396099 | 24396663 | 4.14399 | 3.53845 | 2.34259 |
| chr3 | 69393863 | 69394115 | 700 | Intron (ENSMUST00000029355.7 | 3 | 69431009 | 69432368 | 1360 | 2 | ENSMUST00 | 38253 | Gm17213 | ENSMUSG0C | false | . | . | . | . | . | . | chr3 | 69391821 | 69395444 | 4.50701 | 3.79262 | 2.65411 |
| chr18 | 60824415 | 60824683 | 699 | Intron (ENSMUST00000175934.2 | 18 | 60828067 | 60829027 | 961 | 2 | ENSMUST00 | 4344 | Tcof1 | ENSMUSG0C | false | . | . | . | . | . | . | . | . | . | . | . | . |
| chr3 | 122130671 | 122130959 | 699 | Promoter (1-2kb) | 3 | 122132003 | 122180061 | 48059 | 1 | ENSMUST00 | -1044 | Abca4 | ENSMUSG0C | false | . | . | . | . | . | . | chr3 | 122128691 | 122133156 | 5.55676 | 4.35087 | 3.46867 |
| chr8 | 48425138 | 48425472 | 699 | Intron (ENSMUST00000190840.1 | 8 | 48227682 | 48367363 | 139682 | 2 | ENSMUST00 | -57776 | Tenm3 | ENSMUSG0C | true | chr8 | 48425003 | 48425834 | 10.05864 | 20.96754 | 18.61967 | chr8 | 48424439 | 48426368 | 4.44337 | 3.78743 | 2.55438 |
| chr10 | 121369011 | 121369325 | 699 | Intron (ENSMUST00000040344.6 | 10 | 121365090 | 121397245 | 32156 | 1 | ENSMUST00 | 3922 | Gns | ENSMUSG0C | false | . | . | . | . | . | . | chr10 | 121366145 | 121369544 | 5.43806 | 4.2357 | 3.39368 |
| chr11 | 99126594 | 99127218 | 699 | Distal Intergenic | 11 | 99144196 | 99155077 | 10882 | 2 | ENSMUST00 | 27859 | Ccr7 | ENSMUSG0C | false | . | . | . | . | . | . | . | . | . | . | . | . |
| chr7 | 110539400 | 110539736 | 699 | Intron (ENSMUST00000164759.2 | 7 | 110526792 | 110527121 | 330 | 1 | ENSMUST00 | 12609 | Gm10087 | ENSMUSG0C | false | . | . | . | . | . | . | chr7 | 110537589 | 110541619 | 4.92994 | 4.06433 | 2.93566 |
| chr9 | 117120187 | 117120479 | 699 | Intron (ENSMUST00000111773.4 | 9 | 117065527 | 117065888 | 362 | 1 | ENSMUST00 | 54661 | Gm20397 | ENSMUSG0C | false | . | . | . | . | . | . | chr9 | 117113138 | 117121657 | 4.97914 | 3.95422 | 3.00047 |
| chr12 | 79774901 | 79775275 | 699 | Distal Intergenic | 12 | 79963568 | 79963872 | 305 | 1 | ENSMUST00 | -188293 | RP23-179D3. | ENSMUSG0C | false | . | . | . | . | . | . | chr12 | 79771457 | 79776132 | 5.41323 | 4.28484 | 3.33445 |
| chr10 | 115311343 | 115311937 | 698 | Intron (ENSMUST00000020343.7 | 10 | 115289863 | 115315591 | 25729 | 2 | ENSMUST00 | 3654 | Rab21 | ENSMUSG0C | false | . | . | . | . | . | . | chr10 | 115305250 | 115314291 | 5.86261 | 4.43409 | 3.74616 |
| chr15 | 34237921 | 34238300 | 698 | Promoter (<=1kb) | 15 | 34238026 | 34284295 | 46270 | 1 | ENSMUST00 | 0 | Laptm4b | ENSMUSG0C | false | chr15 | 34237454 | 34239100 | 45.30031 | 150.3008 | 145.00395 | . | . | . | . | . | . |
| chr4 | 62516043 | 62516451 | 698 | Intron (ENSMUST00000137448.1 | 4 | 62509169 | 62519881 | 10713 | 2 | ENSMUST00 | 3430 | Alad | ENSMUSG0C | false | chr4 | 62515264 | 62516219 | 8.2976 | 15.51635 | 13.24704 | chr4 | 62514012 | 62519352 | 6.75426 | 4.89888 | 4.45045 |
| chr2 | 52849042 | 52849338 | 698 | Distal Intergenic | 2 | 52857860 | 53133804 | 275945 | 1 | ENSMUST00 | -8522 | Fmnl2 | ENSMUSG0C | true | . | . | . | . | . | . | chr2 | 52847999 | 52850253 | 4.34516 | 3.67529 | 2.5074 |
| chr10 | 88503709 | 88504185 | 698 | Promoter (<=1kb) | 10 | 88469514 | 88504073 | 34560 | 2 | ENSMUST00 | 0 | Chpt1 | ENSMUSG0C | false | chr10 | 88501897 | 88504448 | 38.2674 | 121.59672 | 117.39927 | chr10 | 88504165 | 88504704 | 3.42011 | 3.30214 | 1.74555 |
| chr4 | 71839359 | 71839805 | 698 | Distal Intergenic | 4 | 71867988 | 71868977 | 990 | 1 | ENSMUST00 | -28183 | Gm11233 | ENSMUSG0C | false | . | . | . | . | . | . | . | . | . | . | . | . |
| chr6 | 100721463 | 100721714 | 698 | Intron (ENSMUST00000032157.7 | 6 | 100726234 | 100726760 | 527 | 1 | ENSMUST00 | -4520 | Gm15576 | ENSMUSG0C | false | . | . | . | . | . | . | chr6 | 100720191 | 100722371 | 3.42477 | 3.15847 | 1.75475 |
| chr18 | 11883486 | 11884066 | 698 | Intron (ENSMUST00000171109.2 | 18 | 11840361 | 11840438 | 78 | 2 | ENSMUST00 | -43049 | Mir1901 | ENSMUSG0C | false | . | . | . | . | . | . | chr18 | 11869945 | 11889578 | 6.23298 | 4.56582 | 4.04247 |
| chr2 | 38350809 | 38351311 | 697 | Promoter (<=1kb) | 2 | 38350759 | 38369733 | 18975 | 1 | ENSMUST00 | 51 | Lhx2 | ENSMUSG0C | false | chr2 | 38350470 | 38352510 | 8.74864 | 23.49345 | 21.11253 | chr2 | 38350410 | 38352653 | 7.68634 | 5.3393 | 5.19643 |
| chr4 | 46489038 | 46489660 | 697 | Promoter (<=1kb) | 4 | 46489248 | 46503632 | 14385 | 1 | ENSMUST00 | 0 | Nans | ENSMUSG0C | false | chr4 | 46488882 | 46490309 | 28.49948 | 87.34749 | 84.05397 | chr4 | 46488969 | 46490179 | 3.20201 | 3.06236 | 1.56991 |
| chr5 | 64343184 | 64343468 | 697 | Intron (ENSMUST00000043893.7 | 5 | 64349945 | 64350589 | 645 | 1 | ENSMUST00 | -6477 | Tbc1d1 | ENSMUSG0C | false | . | . | . | . | . | . | chr5 | 64341943 | 64345940 | 5.86061 | 4.48837 | 3.70318 |
| chr7 | 97842765 | 97843100 | 697 | Promoter (<=1kb) | 7 | 97842935 | 97912381 | 69447 | 1 | ENSMUST00 | 0 | Pak1 | ENSMUSG0C | true | chr7 | 97842956 | 97846713 | 31.85236 | 101.09525 | 97.496 | chr7 | 97841393 | 97842850 | 5.16395 | 4.22984 | 3.14315 |
| chr7 | 137410449 | 137411048 | 697 | Promoter (<=1kb) | 7 | 137375574 | 137410756 | 35183 | 2 | ENSMUST00 | 0 | 9430038I01R | ENSMUSG0C | false | chr7 | 137410007 | 137411686 | 27.78755 | 88.98067 | 85.65775 | . | . | . | . | . | . |
| chr15 | 83148834 | 83149447 | 697 | Promoter (<=1kb) | 15 | 83125978 | 83149331 | 23354 | 2 | ENSMUST00 | 0 | Poldip3 | ENSMUSG0C | false | chr15 | 83148053 | 83149978 | 31.57133 | 109.7552 | 105.93209 | . | . | . | . | . | . |
| chr4 | 100223218 | 100223780 | 697 | Intron (ENSMUST00000039630.5 | 4 | 100119621 | 100120841 | 1221 | 1 | ENSMUST00 | 103598 | Gm12701 | ENSMUSG0C | false | . | . | . | . | . | . | chr4 | 100222070 | 100225444 | 5.66703 | 4.3407 | 3.56148 |
| chr14 | 53721559 | 53722281 | 697 | Distal Intergenic | 14 | 53717015 | 53717289 | 275 | 1 | ENSMUST00 | 4545 | Trav7-6 | ENSMUSG0C | false | . | . | . | . | . | . | chr14 | 53719641 | 53724023 | 4.68452 | 3.93341 | 2.7644 |
| chr11 | 3564498 | 3564843 | 696 | Distal Intergenic | 11 | 3532460 | 3540612 | 8153 | 2 | ENSMUST00 | -23887 | Smtn | ENSMUSG0C | false | . | . | . | . | . | . | chr11 | 3563541 | 3569418 | 5.36923 | 4.11866 | 3.32794 |
| chr2 | 24428283 | 24428573 | 696 | Intron (ENSMUST00000028355.5 | 2 | 24421394 | 24423082 | 1689 | 2 | ENSMUST00 | -5202 | Pax8 | ENSMUSG0C | false | . | . | . | . | . | . | chr2 | 24427779 | 24429002 | 4.83381 | 3.94654 | 2.91679 |
| chr4 | 119239041 | 119239508 | 696 | Promoter (1-2kb) | 4 | 119237942 | 119244149 | 6208 | 1 | ENSMUST00 | 1100 | Lepre1 | ENSMUSG0C | false | . | . | . | . | . | . | chr4 | 119237720 | 119248571 | 6.98267 | 4.77088 | 4.67122 |
| chr11 | 77687075 | 77687696 | 696 | Promoter (<=1kb) | 11 | 77687459 | 77691920 | 4462 | 1 | ENSMUST00 | 0 | Nufip2 | ENSMUSG0C | false | chr11 | 77685741 | 77688196 | 34.54477 | 122.48211 | 118.25522 | . | . | . | . | . | . |
| chr3 | 108769588 | 108769938 | 696 | Intron (ENSMUST00000133931.3 | 3 | 108760214 | 108775002 | 14789 | 1 | ENSMUST00 | 9375 | Aknad1 | ENSMUSG0C | false | . | . | . | . | . | . | chr3 | 108769481 | 108771524 | 3.75878 | 3.41316 | 1.97538 |
| chrX | 133604372 | 133604650 | 696 | Intron (ENSMUST00000167944.2 | 20 | 133587610 | 133685893 | 98284 | 2 | ENSMUST00 | 81243 | Pcdh19 | ENSMUSG0C | false | . | . | . | . | . | . | . | . | . | . | . | . |
| chr4 | 100188301 | 100188615 | 696 | Intron (ENSMUST00000039630.5 | 4 | 100119621 | 100120841 | 1221 | 1 | ENSMUST00 | 68681 | Gm12701 | ENSMUSG0C | false | . | . | . | . | . | . | chr4 | 100186877 | 100189850 | 4.94789 | 3.89694 | 3.00874 |
| chr12 | 9978780 | 9979103 | 696 | Distal Intergenic | 12 | 9865347 | 9865450 | 104 | 2 | ENSMUST00 | -113331 | Gm22845 | ENSMUSG0C | false | . | . | . | . | . | . | chr12 | 9978564 | 9979558 | 4.6222 | 3.97159 | 2.68507 |
| chr1 | 14310114 | 14310509 | 695 | Promoter (<=1kb) | 1 | 14168954 | 14310200 | 141247 | 2 | ENSMUST00 | 0 | Eya1 | ENSMUSG0C | false | chr1 | 14308804 | 14310699 | 19.38504 | 61.02572 | 58.1656 | . | . | . | . | . | . |
| chr7 | 67460024 | 67460321 | 695 | Distal Intergenic | 7 | 67529711 | 67535397 | 5687 | 2 | ENSMUST00 | 75076 | Lrrc28 | ENSMUSG0C | false | . | . | . | . | . | . | chr7 | 67458049 | 67463199 | 4.64009 | 3.8941 | 2.70941 |
| chr6 | 107530098 | 107530395 | 695 | Promoter (<=1kb) | 6 | 107529726 | 107570228 | 40503 | 1 | ENSMUST00 | 373 | Lrrn1 | ENSMUSG0C | false | chr6 | 107529484 | 107530616 | 15.16456 | 35.10921 | 32.57929 | chr6 | 107529564 | 107531737 | 4.13011 | 3.57514 | 2.31508 |
| chr3 | 132856714 | 132857086 | 695 | Distal Intergenic | 3 | 132891176 | 132906359 | 15184 | 2 | ENSMUST00 | 49273 | Npnt | ENSMUSG0C | true | . | . | . | . | . | . | . | . | . | . | . | . |
| chr5 | 138255615 | 138256186 | 695 | Promoter (<=1kb) | 5 | 138255831 | 138259398 | 3568 | 1 | ENSMUST00 | 0 | Lamtor4 | ENSMUSG0C | false | chr5 | 138255010 | 138256431 | 28.666 | 87.63837 | 84.3392 | chr5 | 138255145 | 138256672 | 3.59874 | 3.23723 | 1.90237 |
| chr6 | 47920447 | 47920957 | 695 | Promoter (<=1kb) | 6 | 47920476 | 47932639 | 12164 | 1 | ENSMUST00 | 0 | Zfp212 | ENSMUSG0C | false | chr6 | 47920226 | 47921724 | 42.73649 | 138.01634 | 133.21471 | . | . | . | . | . | . |
| chr15 | 5826957 | 5827217 | 695 | Distal Intergenic | 15 | 5864306 | 5864419 | 114 | 1 | ENSMUST00 | -37089 | Gm27760 | ENSMUSG0C | false | . | . | . | . | . | . | chr15 | 5825129 | 5828365 | 4.13599 | 3.66574 | 2.29193 |
| chr4 | 114576508 | 114576758 | 695 | Intron (ENSMUST00000094894.3 | 4 | 114672697 | 114673340 | 644 | 2 | ENSMUST00 | 96582 | RP23-454F13 | ENSMUSG0C | false | . | . | . | . | . | . | chr4 | 114574979 | 114579790 | 6.8216 | 4.82273 | 4.50379 |
| chr17 | 70528780 | 70529022 | 694 | Intron (ENSMUST00000133983.2 | 17 | 70522162 | 70818302 | 296141 | 1 | ENSMUST00 | 6619 | Dlgap1 | ENSMUSG0C | false | . | . | . | . | . | . | chr17 | 70527571 | 70529779 | 3.20282 | 2.93513 | 1.57331 |
| chr11 | 96861388 | 96861744 | 694 | Downstream (1-2kb) | 11 | 96854131 | 96861203 | 7073 | 1 | ENSMUST00 | 7258 | Copz2 | ENSMUSG0C | false | chr11 | 96861404 | 96862432 | 14.73601 | 38.55974 | 35.98649 | chr11 | 96860440 | 96864502 | 6.8807 | 4.8067 | 4.58039 |
| chr17 | 6006166 | 6006565 | 694 | Promoter (1-2kb) | 17 | 6007580 | 6032929 | 25350 | 1 | ENSMUST00 | -1015 | Synj2 | ENSMUSG0C | false | . | . | . | . | . | . | chr17 | 6005111 | 6007610 | 8.03115 | 5.39042 | 5.49291 |
| chr3 | 90079933 | 90080506 | 694 | Promoter (<=1kb) | 3 | 90080442 | 90087485 | 7044 | 1 | ENSMUST00 | 0 | Tpm3 | ENSMUSG0C | false | chr3 | 90079128 | 90080759 | 41.91343 | 144.39362 | 139.32506 | . | . | . | . | . | . |
| chr5 | 122955191 | 122955539 | 694 | Promoter (2-3kb) | 5 | 122934836 | 122953117 | 18282 | 2 | ENSMUST00 | -2075 | Kdm2b | ENSMUSG0C | false | . | . | . | . | . | . | chr5 | 122954527 | 122957692 | 4.44526 | 3.72161 | 2.56649 |
| chr1 | 179667911 | 179668445 | 694 | Promoter (<=1kb) | 1 | 179668210 | 179687189 | 18980 | 1 | ENSMUST00 | 0 | Sccpdh | ENSMUSG0C | false | chr1 | 179667616 | 179668998 | 24.32174 | 95.77222 | 92.30251 | . | . | . | . | . | . |
| chr1 | 166255616 | 166255873 | 694 | Promoter (1-2kb) | 1 | 166254139 | 166316823 | 62685 | 1 | ENSMUST00 | 1478 | Ildr2 | ENSMUSG0C | true | chr1 | 166253860 | 166256724 | 27.62344 | 81.07606 | 77.90333 | . | . | . | . | . | . |
| chr18 | 68112086 | 68112392 | 694 | Intron (ENSMUST00000063775.3 | 18 | 68260920 | 68260973 | 54 | 2 | ENSMUST00 | 148581 | Mir7219 | ENSMUSG0C | false | . | . | . | . | . | . | . | . | . | . | . | . |
| chr5 | 140812889 | 140813154 | 693 | Intron (ENSMUST00000000153.4 | 5 | 140758408 | 140830431 | 72024 | 2 | ENSMUST00 | 17277 | Gna12 | ENSMUSG0C | false | . | . | . | . | . | . | chr5 | 140787727 | 140831108 | 6.24285 | 4.56075 | 4.04432 |

| | | | | | | | | | | | | | | | | | | | | | | | | | | |
|---|---|---|---|---|---|---|---|---|---|---|---|---|---|---|---|---|---|---|---|---|---|---|---|---|---|---|
| chr13 | 63153427 | 63153679 | 693 | Intron (ENSMUST00000091560.5 | 13 | 63282142 | 63326096 | 43955 | 1 | ENSMUST00 | -128463 | 2010111I01R | ENSMUSG0( | false | . | . | . | . | . | . | . | . | . | . | . | . |
| chr14 | 21751167 | 21751586 | 693 | Promoter (<=1kb) | 14 | 21744874 | 21751181 | 6308 | 2 | ENSMUST00 | 0 | Dusp13 | ENSMUSG0( | false | chr14 | 21750151 | 21752138 | 32.98857 | 124.89146 | 120.57959 | . | . | . | . | . | . |
| chr11 | 120824094 | 120824773 | 693 | Promoter (<=1kb) | 11 | 120805940 | 120824178 | 18239 | 2 | ENSMUST00 | 0 | Fasn | ENSMUSG0( | false | chr11 | 120823409 | 120825174 | 36.7977 | 133.03651 | 128.43381 | chr11 | 120823109 | 120825193 | 4.85423 | 3.9235 | 2.93693 |
| chrX | 75130822 | 75131152 | 693 | Promoter (<=1kb) | 20 | 75109733 | 75131016 | 21284 | 2 | ENSMUST00 | 0 | Mpp1 | ENSMUSG0( | false | chrX | 75129972 | 75131008 | 16.08362 | 38.04472 | 35.47989 | . | . | . | . | . | . |
| chr8 | 33385415 | 33385927 | 693 | Promoter (<=1kb) | 8 | 33234384 | 33385527 | 151144 | 2 | ENSMUST00 | 0 | Wrn | ENSMUSG0( | false | chr8 | 33384518 | 33386849 | 38.4013 | 122.89693 | 118.65718 | . | . | . | . | . | . |
| chr13 | 102842225 | 102842525 | 693 | Intron (ENSMUST00000166726.2 | 13 | 102796913 | 102800921 | 4009 | 2 | ENSMUST00 | -41305 | Mast4 | ENSMUSG0( | true | . | . | . | . | . | . | chr13 | 102834339 | 102867396 | 5.64964 | 4.33081 | 3.54328 |
| chr17 | 87133835 | 87134112 | 693 | Exon (ENSMUST00000041369.6/ | 17 | 87107679 | 87137583 | 29905 | 1 | ENSMUST00 | 26157 | Socs5 | ENSMUSG0( | false | . | . | . | . | . | . | . | . | . | . | . | . |
| chr16 | 52031031 | 52031780 | 692 | Promoter (<=1kb) | 16 | 52031549 | 52208047 | 176499 | 1 | ENSMUST00 | 0 | Cblb | ENSMUSG0( | true | chr16 | 52030893 | 52033595 | 36.7977 | 133.03651 | 128.43381 | . | . | . | . | . | . |
| chr16 | 91646884 | 91647362 | 692 | Promoter (<=1kb) | 16 | 91621186 | 91646952 | 25767 | 2 | ENSMUST00 | 0 | Gart | ENSMUSG0( | false | chr16 | 91646065 | 91649433 | 31.16539 | 106.97112 | 103.21663 | . | . | . | . | . | . |
| chr1 | 37305981 | 37306207 | 692 | Intron (ENSMUST00000146676.1 | 1 | 37300308 | 37399557 | 99250 | 1 | ENSMUST00 | 5674 | Inpp4a | ENSMUSG0( | true | . | . | . | . | . | . | chr1 | 37300992 | 37326505 | 5.58149 | 4.26721 | 3.50707 |
| chr9 | 103999625 | 103999936 | 692 | Promoter (2-3kb) | 9 | 104002544 | 104043811 | 41268 | 1 | ENSMUST00 | -2608 | Nphp3 | ENSMUSG0( | false | . | . | . | . | . | . | . | . | . | . | . | . |
| chr6 | 100127552 | 100128028 | 692 | Distal Intergenic | 6 | 100228566 | 100287441 | 58876 | 2 | ENSMUST00 | 159413 | Rybp | ENSMUSG0( | false | . | . | . | . | . | . | chr6 | 100125670 | 100128898 | 6.78273 | 4.72704 | 4.50739 |
| chr10 | 18022818 | 18023379 | 692 | Promoter (<=1kb) | 10 | 18011263 | 18023252 | 11990 | 2 | ENSMUST00 | 0 | Abracl | ENSMUSG0( | false | chr10 | 18021720 | 18023486 | 32.00811 | 98.35596 | 94.82688 | . | . | . | . | . | . |
| chr13 | 46902952 | 46903196 | 692 | Promoter (<=1kb) | 13 | 46902589 | 46902692 | 104 | 1 | ENSMUST00 | 364 | Gm22960 | ENSMUSG0( | false | . | . | . | . | . | . | chr13 | 46902229 | 46904290 | 3.24695 | 3.12945 | 1.62266 |
| chr14 | 100285870 | 100286320 | 692 | Distal Intergenic | 14 | 100219486 | 100219789 | 304 | 1 | ENSMUST00 | 66385 | Gm16260 | ENSMUSG0( | false | chr14 | 100284160 | 100286947 | 30.32912 | 88.589 | 85.27629 | chr14 | 100284937 | 100287388 | 6.55605 | 4.70889 | 4.33546 |
| chr12 | 87112027 | 87112347 | 691 | Intron (ENSMUST00000037788.4 | 12 | 87097531 | 87102539 | 5009 | 2 | ENSMUST00 | -9489 | Ngb | ENSMUSG0( | false | . | . | . | . | . | . | chr12 | 87110077 | 87113895 | 5.58242 | 4.3245 | 3.48884 |
| chr17 | 26465059 | 26465417 | 691 | Distal Intergenic | 17 | 26506516 | 26507723 | 1208 | 2 | ENSMUST00 | 42306 | Dusp1 | ENSMUSG0( | false | . | . | . | . | . | . | . | . | . | . | . | . |
| chr6 | 11907687 | 11907961 | 691 | Promoter (<=1kb) | 6 | 11907809 | 11933410 | 25602 | 1 | ENSMUST00 | 0 | Phf14 | ENSMUSG0( | false | chr6 | 11906440 | 11908484 | 23.17719 | 89.67316 | 86.33752 | . | . | . | . | . | . |
| chr8 | 105592410 | 105592723 | 691 | Distal Intergenic | 8 | 105605229 | 105622194 | 16966 | 1 | ENSMUST00 | -12506 | Fam65a | ENSMUSG0( | false | . | . | . | . | . | . | . | . | . | . | . | . |
| chr8 | 108714683 | 108715614 | 691 | Promoter (<=1kb) | 8 | 108714644 | 108961630 | 246987 | 1 | ENSMUST00 | 40 | Zfhx3 | ENSMUSG0( | false | chr8 | 108714360 | 108715770 | 30.13579 | 99.06311 | 95.51676 | chr8 | 108713639 | 108718751 | 3.65165 | 3.37374 | 1.93101 |
| chr5 | 135868130 | 135868458 | 691 | Promoter (1-2kb) | 5 | 135866617 | 135868209 | 1593 | 1 | ENSMUST00 | 1514 | Srrm3 | ENSMUSG0( | false | chr5 | 135867751 | 135868589 | 16.08362 | 38.04472 | 35.47989 | chr5 | 135867753 | 135869147 | 3.67225 | 3.23448 | 1.96076 |
| chr7 | 112720134 | 112720631 | 691 | Intron (ENSMUST00000084705.6 | 7 | 112742045 | 112898660 | 156616 | 1 | ENSMUST00 | -21414 | Tead1 | ENSMUSG0( | false | . | . | . | . | . | . | chr7 | 112716220 | 112724700 | 5.77743 | 4.2821 | 3.6747 |
| chr15 | 96284782 | 96285923 | 691 | Promoter (<=1kb) | 15 | 96281499 | 96284495 | 2997 | 2 | ENSMUST00 | -288 | 2610037D02l | ENSMUSG0( | false | chr15 | 96283762 | 96289319 | 34.73729 | 107.12603 | 103.36953 | . | . | . | . | . | . |
| chr15 | 83464131 | 83464871 | 690 | Promoter (<=1kb) | 15 | 83375607 | 83464606 | 89000 | 2 | ENSMUST00 | 0 | Pacsin2 | ENSMUSG0( | false | chr15 | 83463171 | 83465159 | 27.66126 | 83.97217 | 80.74496 | . | . | . | . | . | . |
| chr11 | 67558887 | 67559150 | 690 | Intron (ENSMUST00000108681.2 | 11 | 67547453 | 67585532 | 38080 | 1 | ENSMUST00 | 11435 | Gas7 | ENSMUSG0( | true | . | . | . | . | . | . | chr11 | 67555038 | 67569865 | 4.9616 | 4.01125 | 2.9907 |
| chr2 | 29217997 | 29218308 | 690 | Intron (ENSMUST00000183583.2 | 2 | 29200676 | 29204922 | 4247 | 2 | ENSMUST00 | -13076 | Ntng2 | ENSMUSG0( | true | . | . | . | . | . | . | chr2 | 29214408 | 29229484 | 5.78922 | 4.36589 | 3.66038 |
| chr7 | 107665488 | 107666049 | 690 | Intron (ENSMUST00000040056.8 | 7 | 107623971 | 107625161 | 1191 | 2 | ENSMUST00 | -40328 | 5330417H12l | ENSMUSG0( | false | chr7 | 107664446 | 107667691 | 20.9555 | 57.98476 | 55.16582 | chr7 | 107661425 | 107665686 | 5.62473 | 4.32581 | 3.51662 |
| chr19 | 16872923 | 16873169 | 690 | Promoter (<=1kb) | 19 | 16872316 | 16873830 | 1515 | 2 | ENSMUST00 | 661 | Foxb2 | ENSMUSG0( | false | chr19 | 16872263 | 16873181 | 12.3514 | 27.13773 | 24.70927 | chr19 | 16870641 | 16876124 | 6.25184 | 4.55109 | 4.07521 |
| chr17 | 49482544 | 49482803 | 690 | Intron (ENSMUST00000057610.6 | 17 | 49452284 | 49455435 | 3152 | 1 | ENSMUST00 | 30261 | Mocs1 | ENSMUSG0( | false | . | . | . | . | . | . | chr17 | 49481205 | 49490260 | 5.66436 | 4.31035 | 3.55927 |
| chr3 | 51868875 | 51869217 | 690 | Intron (ENSMUST00000121440.2 | 3 | 51828912 | 51829028 | 117 | 1 | ENSMUST00 | 39964 | n-R5s196 | ENSMUSG0( | false | . | . | . | . | . | . | chr3 | 51864972 | 51875550 | 6.25 | 4.62847 | 4.03397 |
| chr9 | 113296019 | 113296338 | 690 | Distal Intergenic | 9 | 113403475 | 113403771 | 297 | 1 | ENSMUST00 | -107137 | Gm22492 | ENSMUSG0( | false | . | . | . | . | . | . | chr9 | 113295440 | 113297393 | 4.90794 | 3.91065 | 2.97132 |
| chr11 | 93996455 | 93997044 | 689 | Promoter (<=1kb) | 11 | 93996103 | 94126085 | 129983 | 1 | ENSMUST00 | 353 | Spag9 | ENSMUSG0( | false | chr11 | 93994683 | 93997734 | 37.40972 | 147.3676 | 142.18408 | . | . | . | . | . | . |
| chr5 | 64092789 | 64093103 | 689 | Promoter (<=1kb) | 5 | 64092936 | 64128355 | 35420 | 1 | ENSMUST00 | 0 | Pgm1 | ENSMUSG0( | false | chr5 | 64092528 | 64093954 | 25.16617 | 75.86667 | 72.78426 | chr5 | 64092642 | 64095464 | 4.42239 | 3.73784 | 2.54869 |
| chr3 | 75927093 | 75927321 | 689 | Intron (ENSMUST00000117242.2 | 3 | 75894853 | 75902446 | 7594 | 2 | ENSMUST00 | -24648 | Golim4 | ENSMUSG0( | true | . | . | . | . | . | . | chr3 | 75908736 | 75932827 | 4.90077 | 3.96047 | 2.93621 |
| chrX | 20291728 | 20291978 | 689 | Promoter (<=1kb) | 20 | 20105755 | 20291807 | 186053 | 2 | ENSMUST00 | 0 | Slc9a7 | ENSMUSG0( | false | chrX | 20291251 | 20292098 | 21.59801 | 56.60135 | 53.80005 | . | . | . | . | . | . |
| chr7 | 25754544 | 25755099 | 689 | Promoter (<=1kb) | 7 | 25721498 | 25754730 | 33233 | 2 | ENSMUST00 | 0 | Hnrnpul1 | ENSMUSG0( | false | chr7 | 25753297 | 25756211 | 35.36919 | 136.9064 | 132.15405 | . | . | . | . | . | . |
| chr6 | 100288488 | 100288869 | 689 | Promoter (1-2kb) | 6 | 100228566 | 100287441 | 58876 | 2 | ENSMUST00 | -1048 | Rybp | ENSMUSG0( | false | . | . | . | . | . | . | chr6 | 100287772 | 100289184 | 5.57913 | 4.23799 | 3.50814 |
| chr13 | 112357944 | 112358259 | 689 | Intron (ENSMUST00000022275.7 | 13 | 112291121 | 112291273 | 153 | 1 | ENSMUST00 | 66824 | Gm22768 | ENSMUSG0( | false | . | . | . | . | . | . | chr13 | 112357087 | 112358508 | 4.69029 | 4.02632 | 2.73566 |
| chr12 | 79944710 | 79945149 | 689 | Distal Intergenic | 12 | 79963568 | 79963872 | 305 | 1 | ENSMUST00 | -18419 | RP23-179D3. | ENSMUSG0( | false | . | . | . | . | . | . | chr12 | 79942657 | 79955332 | 5.38416 | 4.22706 | 3.33447 |
| chr3 | 87905948 | 87906932 | 688 | Promoter (<=1kb) | 3 | 87906321 | 87916132 | 9812 | 1 | ENSMUST00 | 0 | Hdgf | ENSMUSG0( | false | chr3 | 87905898 | 87907578 | 42.78876 | 142.56003 | 137.56947 | . | . | . | . | . | . |
| chr2 | 93439187 | 93439671 | 688 | Intron (ENSMUST00000099696.2 | 2 | 93419830 | 93452679 | 32850 | 2 | ENSMUST00 | 13008 | Cd82 | ENSMUSG0( | false | . | . | . | . | . | . | chr2 | 93434899 | 93464965 | 6.68053 | 4.76412 | 4.39602 |
| chr9 | 71606575 | 71606940 | 688 | Distal Intergenic | 9 | 71504347 | 71592360 | 88014 | 2 | ENSMUST00 | -14216 | Myzap | ENSMUSG0( | false | . | . | . | . | . | . | chr9 | 71605528 | 71610183 | 5.14246 | 4.05011 | 3.13188 |
| chr5 | 106925813 | 106926114 | 688 | Promoter (<=1kb) | 5 | 106840197 | 106925890 | 85694 | 2 | ENSMUST00 | 0 | Hfm1 | ENSMUSG0( | false | chr5 | 106925543 | 106926395 | 17.00442 | 50.91425 | 48.18763 | . | . | . | . | . | . |
| chr15 | 101977307 | 101977692 | 688 | Distal Intergenic | 15 | 101946004 | 101954287 | 8284 | 2 | ENSMUST00 | -23021 | Krt78 | ENSMUSG0( | false | chr15 | 101976968 | 101977912 | 12.15419 | 27.41675 | 24.98428 | chr15 | 101975790 | 101978730 | 6.00483 | 4.51629 | 3.80634 |
| chr2 | 17647813 | 17648306 | 688 | Intron (ENSMUST00000028080.6 | 2 | 17343909 | 17731043 | 387135 | 2 | ENSMUST00 | 82737 | Nebl | ENSMUSG0( | true | . | . | . | . | . | . | chr2 | 17644895 | 17651727 | 5.70802 | 4.43304 | 3.5805 |
| chr10 | 7473598 | 7473888 | 688 | Promoter (<=1kb) | 10 | 7444873 | 7473477 | 28605 | 2 | ENSMUST00 | -122 | Ulbp1 | ENSMUSG0( | false | . | . | . | . | . | . | chr10 | 7473513 | 7474456 | 3.96946 | 3.49308 | 2.18054 |
| chr4 | 41257818 | 41258137 | 688 | Intron (ENSMUST00000108068.2 | 4 | 41252617 | 41252844 | 228 | 1 | ENSMUST00 | 5202 | Gm12401 | ENSMUSG0( | false | . | . | . | . | . | . | chr4 | 41251196 | 41261743 | 4.92657 | 3.98721 | 2.97941 |
| chr15 | 102245395 | 102245785 | 687 | Promoter (<=1kb) | 15 | 102241943 | 102245231 | 3289 | 2 | ENSMUST00 | -165 | Rarg | ENSMUSG0( | false | chr15 | 102244836 | 102248267 | 30.17592 | 94.17345 | 90.73757 | chr15 | 102241789 | 102251883 | 5.05731 | 4.11666 | 3.0557 |
| chr11 | 59707544 | 59707814 | 687 | Intron (ENSMUST00000138234.1 | 11 | 59727681 | 59776602 | 48922 | 1 | ENSMUST00 | -19867 | Mprip | ENSMUSG0( | false | . | . | . | . | . | . | chr11 | 59704908 | 59728229 | 5.43634 | 4.16695 | 3.39654 |
| chr12 | 80271022 | 80271273 | 687 | Distal Intergenic | 12 | 80167542 | 80260371 | 92830 | 2 | ENSMUST00 | -10652 | Actn1 | ENSMUSG0( | true | . | . | . | . | . | . | chr12 | 80269686 | 80271796 | 5.18831 | 4.07073 | 3.16856 |
| chr10 | 8015010 | 8015404 | 687 | Distal Intergenic | 10 | 7905653 | 7956230 | 50578 | 2 | ENSMUST00 | -58781 | Tab2 | ENSMUSG0( | true | . | . | . | . | . | . | chr10 | 8013399 | 8017130 | 6.28902 | 4.57652 | 4.08766 |
| chr5 | 124327804 | 124328260 | 687 | Promoter (<=1kb) | 5 | 124250959 | 124327972 | 77014 | 2 | ENSMUST00 | 0 | Mphosph9 | ENSMUSG0( | false | chr5 | 124326967 | 124327933 | 30.75439 | 93.21284 | 89.80341 | . | . | . | . | . | . |
| chr11 | 121536018 | 121536239 | 687 | Intron (ENSMUST00000103013.4 | 11 | 121544201 | 121590406 | 46206 | 1 | ENSMUST00 | -7962 | Tbcd | ENSMUSG0( | false | . | . | . | . | . | . | chr11 | 121535748 | 121537102 | 3.91901 | 3.48948 | 2.14477 |
| chr19 | 25610486 | 25610791 | 687 | Promoter (<=1kb) | 19 | 25610537 | 25623920 | 13384 | 1 | ENSMUST00 | 0 | Dmrt3 | ENSMUSG0( | false | chr19 | 25610467 | 25611285 | 14.24974 | 34.23454 | 31.71693 | chr19 | 25609887 | 25611499 | 6.74212 | 4.8843 | 4.4342 |
| chr5 | 129161163 | 129161410 | 687 | Intron (ENSMUST00000056617.8 | 5 | 129128077 | 129128517 | 441 | 2 | ENSMUST00 | -32647 | Rps16-ps2 | ENSMUSG0( | false | . | . | . | . | . | . | chr5 | 129160269 | 129161891 | 3.20826 | 3.16181 | 1.54324 |
| chr2 | 167348689 | 167349765 | 686 | Promoter (<=1kb) | 2 | 167298444 | 167349183 | 50740 | 2 | ENSMUST00 | 0 | B4galt5 | ENSMUSG0( | false | chr2 | 167348132 | 167349979 | 25.53309 | 82.08177 | 78.89293 | chr2 | 167349667 | 167351748 | 4.99383 | 3.99782 | 3.02917 |
| chr10 | 94545127 | 94545553 | 686 | Promoter (<=1kb) | 10 | 94545431 | 94578922 | 33492 | 1 | ENSMUST00 | 0 | Tmcc3 | ENSMUSG0( | false | chr10 | 94545392 | 94546191 | 15.14328 | 38.56274 | 35.98906 | chr10 | 94542791 | 94548485 | 5.10717 | 4.05097 | 3.10342 |
| chr13 | 34874200 | 34874630 | 686 | Promoter (<=1kb) | 13 | 34875494 | 34902877 | 27384 | 1 | ENSMUST00 | -864 | Prpf4b | ENSMUSG0( | false | chr13 | 34873812 | 34876956 | 44.11508 | 143.73827 | 138.70853 | . | . | . | . | . | . |
| chr13 | 74808264 | 74808789 | 686 | Promoter (<=1kb) | 13 | 74694286 | 74807921 | 113636 | 2 | ENSMUST00 | -344 | Cast | ENSMUSG0( | false | chr13 | 74807412 | 74808982 | 23.74947 | 92.7123 | 89.30923 | chr13 | 74808692 | 74809276 | 3.50124 | 3.30053 | 1.81141 |
| chr15 | 59314688 | 59315122 | 686 | Promoter (<=1kb) | 15 | 59315077 | 59328958 | 13882 | 1 | ENSMUST00 | 0 | Sqle | ENSMUSG0( | false | chr15 | 59314322 | 59315989 | 31.51034 | 95.50614 | 92.04281 | . | . | . | . | . | . |
| chr6 | 89358920 | 89359268 | 686 | Promoter (1-2kb) | 6 | 89319015 | 89357717 | 38703 | 2 | ENSMUST00 | -1204 | Plxna1 | ENSMUSG0( | false | . | . | . | . | . | . | chr6 | 89339950 | 89361705 | 7.61354 | 5.20417 | 5.1492 |
| chr4 | 59786584 | 59786995 | 686 | Promoter (2-3kb) | 4 | 59769637 | 59783866 | 14230 | 2 | ENSMUST00 | -2719 | Inip | ENSMUSG0( | false | . | . | . | . | . | . | . | . | . | . | . | . |
| chr5 | 4757633 | 4758319 | 686 | Promoter (<=1kb) | 5 | 4753873 | 4758035 | 4163 | 2 | ENSMUST00 | 0 | Fzd1 | ENSMUSG0( | false | chr5 | 4756381 | 4759090 | 26.52989 | 84.81848 | 81.57964 | . | . | . | . | . | . |
| chr1 | 159386567 | 159386815 | 686 | Distal Intergenic | 1 | 159358509 | 159358903 | 395 | 1 | ENSMUST00 | 28059 | RP23-145H3. | ENSMUSG0( | false | . | . | . | . | . | . | chr1 | 159385051 | 159390744 | 5.42733 | 4.2776 | 3.36708 |
| chr16 | 58502066 | 58502373 | 685 | Promoter (2-3kb) | 16 | 58493743 | 58499341 | 5599 | 2 | ENSMUST00 | -2726 | St3gal6 | ENSMUSG0( | true | . | . | . | . | . | . | chr16 | 58496830 | 58520680 | 5.70067 | 4.33546 | 3.59278 |
| chr19 | 3262192 | 3262536 | 685 | Intron (ENSMUST00000119292.2 | 19 | 3265364 | 3267716 | 2353 | 2 | ENSMUST00 | 5180 | Ighmbp2 | ENSMUSG0( | false | . | . | . | . | . | . | chr19 | 3259601 | 3263900 | 6.21113 | 4.29911 | 4.03454 |
| chr8 | 109608449 | 109608810 | 685 | Promoter (<=1kb) | 8 | 109593248 | 109608673 | 15426 | 2 | ENSMUST00 | 0 | Dhodh | ENSMUSG0( | false | chr8 | 109607763 | 109609294 | 21.58806 | 60.76757 | 57.91108 | . | . | . | . | . | . |
| chr7 | 45922927 | 45923937 | 685 | Promoter (<=1kb) | 7 | 45924072 | 45931607 | 7536 | 1 | ENSMUST00 | -135 | Ccdc114 | ENSMUSG0( | false | chr7 | 45922817 | 45924562 | 19.41141 | 53.64976 | 50.88786 | chr7 | 45915648 | 45924393 | 5.13169 | 3.954 | 3.14707 |
| chr10 | 8513811 | 8514098 | 685 | Intron (ENSMUST00000061601.7 | 10 | 8204753 | 8518825 | 314073 | 2 | ENSMUST00 | 4727 | Ust | ENSMUSG0( | true | . | . | . | . | . | . | chr10 | 8513451 | 8518296 | 4.82778 | 3.94777 | 2.88932 |
| chr8 | 125806531 | 125806791 | 685 | Intron (ENSMUST00000047239.7 | 8 | 125827163 | 125829033 | 1871 | 2 | ENSMUST00 | 22242 | Pcnxl2 | ENSMUSG0( | false | . | . | . | . | . | . | chr8 | 125806235 | 125807167 | 3.77312 | 3.42989 | 2.03989 |
| chr1 | 58284512 | 58284828 | 685 | Intron (ENSMUST00000114366.2 | 1 | 58278326 | 58380259 | 101934 | 1 | ENSMUST00 | 6187 | Aox2 | ENSMUSG0( | false | . | . | . | . | . | . | chr1 | 58284050 | 58285495 | 4.88277 | 4.08969 | 2.90308 |
| chr2 | 113376361 | 113376649 | 685 | Intron (ENSMUST00000099576.3 | 2 | 113384681 | 113385199 | 519 | 1 | ENSMUST00 | -8032 | Gm13967 | ENSMUSG0( | false | . | . | . | . | . | . | . | . | . | . | . | . |
| chr15 | 89075760 | 89076180 | 684 | Promoter (<=1kb) | 15 | 89076013 | 89087077 | 11065 | 1 | ENSMUST00 | 0 | Trabd | ENSMUSG0( | false | chr15 | 89075128 | 89077118 | 35.98183 | 119.73972 | 115.60696 | . | . | . | . | . | . |
| chr13 | 38036905 | 38037299 | 684 | Promoter (<=1kb) | 13 | 38036989 | 38061433 | 24445 | 1 | ENSMUST00 | 0 | Riok1 | ENSMUSG0( | false | chr13 | 38036426 | 38038278 | 39.41127 | 125.71832 | 121.38846 | chr13 | 38036047 | 38040605 | 4.12809 | 3.55154 | 2.31178 |
| chr14 | 34442443 | 34442843 | 684 | Intron (ENSMUST00000049005.9 | 14 | 34414486 | 34460556 | 46071 | 2 | ENSMUST00 | 17713 | Bmpr1a | ENSMUSG0( | true | . | . | . | . | . | . | chr14 | 34439321 | 34450550 | 4.3338 | 3.66849 | 2.47835 |
| chr17 | 79777962 | 79778235 | 684 | Distal Intergenic | 17 | 79706953 | 79715041 | 8089 | 2 | ENSMUST00 | -62922 | Cyp1b1 | ENSMUSG0( | false | . | . | . | . | . | . | chr17 | 79777688 | 79778144 | 3.17318 | 3.12163 | 1.55038 |
| chr3 | 14672019 | 14672337 | 684 | Distal Intergenic | 3 | 14641727 | 14663002 | 21276 | 1 | ENSMUST00 | 30293 | Car13 | ENSMUSG0( | false | . | . | . | . | . | . | chr3 | 14671646 | 14672404 | 4.0285 | 3.38267 | 2.23641 |
| chr9 | 121371555 | 121371892 | 684 | Intron (ENSMUST00000045903.6 | 9 | 121366958 | 121474916 | 107959 | 1 | ENSMUST00 | 4598 | Trak1 | ENSMUSG0( | false | . | . | . | . | . | . | chr9 | 121369443 | 121377566 | 6.26925 | 4.56408 | 4.0918 |
| chr8 | 18594914 | 18595236 | 684 | Promoter (<=1kb) | 8 | 18595131 | 18803189 | 208059 | 1 | ENSMUST00 | 0 | Mcph1 | ENSMUSG0( | false | chr8 | 18594764 | 18596012 | 37.22259 | 125.13314 | 120.81451 | . | . | . | . | . | . |
| chr15 | 95752780 | 95753038 | 684 | Distal Intergenic | 15 | 95775754 | 95776232 | 479 | 1 | ENSMUST00 | -22716 | Gm8849 | ENSMUSG0( | false | . | . | . | . | . | . | chr15 | 95752189 | 95753238 | 4.87677 | 4.08652 | 2.91845 |
| chr6 | 17629094 | 17629447 | 683 | Distal Intergenic | 6 | 17636234 | 17656447 | 20214 | 1 | ENSMUST00 | -6787 | Capza2 | ENSMUSG0( | false | . | . | . | . | . | . | chr6 | 17628658 | 17630493 | 4.19694 | 3.6018 | 2.39365 |
| chr10 | 77259012 | 77259353 | 683 | Promoter (<=1kb) | 10 | 77259300 | 77269586 | 10287 | 1 | ENSMUST00 | 0 | Pofut2 | ENSMUSG0( | false | chr10 | 77258691 | 77260213 | 40.1055 | 130.16986 | 125.67132 | chr10 | 77258533 | 77261105 | 4.43015 | 3.78435 | 2.53269 |
| chr12 | 108792607 | 108793515 | 683 | Promoter (<=1kb) | 12 | 108792973 | 108816632 | 23660 | 1 | ENSMUST00 | 0 | Yy1 | ENSMUSG0( | false | chr12 | 108792910 | 108795425 | 33.7938 | 119.00099 | 114.88904 | . | . | . | . | . | . |
| chr11 | 98970271 | 98970593 | 683 | Exon (ENSMUST00000107474.2/ | 11 | 98960412 | 98974942 | 14531 | 1 | ENSMUST00 | 9860 | Rara | ENSMUSG0( | false | . | . | . | . | . | . | chr11 | 98958785 | 98973618 | 4.55278 | 3.71427 | 2.6612 |
| chr13 | 43615759 | 43616132 | 683 | Promoter (<=1kb) | 13 | 43615983 | 43668066 | 52084 | 1 | ENSMUST00 | 0 | Rnf182 | ENSMUSG0( | false | chr13 | 43615542 | 43616454 | 16.30988 | 55.04095 | 52.25988 | chr13 | 43615729 | 43616643 | 5.35048 | 4.01045 | 3.32858 |
| chr15 | 81856933 | 81857320 | 683 | Promoter (1-2kb) | 15 | 81848270 | 81858326 | 10057 | 2 | ENSMUST00 | 1006 | Tob2 | ENSMUSG0( | false | chr15 | 81855855 | 81859985 | 34.3275 | 112.61823 | 108.70669 | chr15 | 81852673 | 81857140 | 4.32126 | 3.77391 | 2.43283 |
| chr8 | 126594387 | 126594991 | 683 | Promoter (<=1kb) | 8 | 126588296 | 126593986 | 5691 | 2 | ENSMUST00 | -402 | Irf2bp2 | ENSMUSG0( | false | chr8 | 126589312 | 126594549 | 35.95432 | 114.41653 | 110.45183 | chr8 | 126594816 | 126596983 | 8.16677 | 5.62022 | 5.58389 |
| chr12 | 108422828 | 108423550 | 683 | Promoter (<=1kb) | 12 | 108423078 | 108537532 | 114455 | 1 | ENSMUST00 | 0 | Eml1 | ENSMUSG0( | false | chr12 | 108422545 | 108423952 | 42.51921 | 140.97417 | 136.05247 | chr12 | 108422059 | 108433843 | 4.11207 | 3.54569 | 2.29349 |
| chr11 | 96064932 | 96065574 | 682 | Promoter (<=1kb) | 11 | 96047432 | 96065388 | 17957 | 2 | ENSMUST00 | 0 | Ube2z | ENSMUSG0( | false | chr11 | 96064220 | 96065900 | 22.47347 | 97.37944 | 93.87287 | . | . | . | . | . | . |
| chr17 | 47009954 | 47010477 | 682 | Promoter (<=1kb) | 17 | 46928292 | 47010532 | 82241 | 2 | ENSMUST00 | 55 | Ubr2 | ENSMUSG0( | false | chr17 | 47008952 | 47011261 | 28.1615 | 93.53628 | 90.11485 | . | . | . | . | . | . |
| chr3 | 108653720 | 108654210 | 682 | Promoter (<=1kb) | 3 | 108653913 | 108678840 | 24928 | 1 | ENSMUST00 | 0 | Clcc1 | ENSMUSG0( | false | chr3 | 108653419 | 108654919 | 37.70819 | 121.69438 | 117.49508 | . | . | . | . | . | . |
| chr7 | 57590925 | 57591289 | 682 | Promoter (<=1kb) | 7 | 57591162 | 57765781 | 174620 | 1 | ENSMUST00 | 0 | Gabrb3 | ENSMUSG0( | false | chr7 | 57590489 | 57592145 | 19.84988 | 51.19398 | 48.46415 | chr7 | 57590545 | 57592175 | 4.67974 | 3.92902 | 2.76265 |
| chr19 | 36553869 | 36554283 | 682 | Promoter (<=1kb) | 19 | 36554639 | 36621132 | 66494 | 1 | ENSMUST00 | -356 | Hectd2 | ENSMUSG0( | false | chr19 | 36553796 | 36555516 | 42.71282 | 143.54831 | 138.52826 | . | . | . | . | . | . |
| chr2 | 148073133 | 148073411 | 682 | Promoter (1-2kb) | 2 | 148071346 | 148071839 | 494 | 2 | ENSMUST00 | -1295 | Gm14108 | ENSMUSG0( | false | . | . | . | . | . | . | chr2 | 148071881 | 148075123 | 5.12065 | 4.06479 | 3.12109 |
| chr17 | 6816133 | 6816428 | 682 | Intron (ENSMUST00000181895.1 | 17 | 6798390 | 6827990 | 29601 | 2 | ENSMUST00 | 11562 | Gm2885 | ENSMUSG0( | false | . | . | . | . | . | . | chr17 | 6811040 | 6816818 | 5.75765 | 4.4253 | 3.63624 |
| chr9 | 34683012 | 34683354 | 682 | Intron (ENSMUST00000115148.3 | 9 | 34588526 | 34589304 | 779 | 2 | ENSMUST00 | -93709 | Gm26662 | ENSMUSG0( | false | . | . | . | . | . | . | chr9 | 34682241 | 34683694 | 3.19024 | 3.10659 | 1.5662 |
| chr4 | 15957899 | 15958582 | 681 | Promoter (<=1kb) | 4 | 15957925 | 15992589 | 34665 | 1 | ENSMUST00 | 0 | Nbn | ENSMUSG0( | false | chr4 | 15957819 | 15958996 | 35.06866 | 119.18253 | 115.06551 | chr4 | 15957835 | 15959820 | 3.67136 | 3.22339 | 1.9616 |
| chr6 | 98947836 | 98948091 | 681 | Intron (ENSMUST00000113326.3 | 6 | 98930171 | 98935545 | 5375 | 2 | ENSMUST00 | -12292 | Foxp1 | ENSMUSG0( | false | . | . | . | . | . | . | chr6 | 98946696 | 98951866 | 5.45838 | 4.12852 | 3.41677 |
| chrX | 169320270 | 169320638 | 681 | Promoter (<=1kb) | 20 | 169250193 | 169320372 | 70180 | 2 | ENSMUST00 | 0 | Hccs | ENSMUSG0( | false | chrX | 169319768 | 169320406 | 21.59801 | 56.60135 | 53.80005 | . | . | . | . | . | . |
| chr14 | 31019004 | 31019777 | 681 | Promoter (<=1kb) | 14 | 31019138 | 31118901 | 99764 | 1 | ENSMUST00 | 0 | Pbrm1 | ENSMUSG0( | false | chr14 | 31018221 | 31020204 | 25.57207 | 77.50074 | 74.3909 | . | . | . | . | . | . |
| chr18 | 60607712 | 60608250 | 681 | Promoter (1-2kb) | 18 | 60600207 | 60610105 | 9899 | 2 | ENSMUST00 | 1855 | Synpo | ENSMUSG0( | true | chr18 | 60603340 | 60613557 | 27.39171 | 92.13462 | 88.74321 | chr18 | 60593903 | 60608752 | 6.45808 | 4.76452 | 4.19811 |
| chrX | 136033239 | 136033467 | 681 | Promoter (<=1kb) | 20 | 136033367 | 136034946 | 1580 | 1 | ENSMUST00 | 0 | Arxes1 | ENSMUSG0( | false | chrX | 136033350 | 136034096 | 24.81473 | 68.0602 | 65.09722 | . | . | . | . | . | . |
| chr4 | 132421982 | 132422684 | 681 | Promoter (<=1kb) | 4 | 132355923 | 132422489 | 66567 | 2 | ENSMUST00 | 0 | Phactr4 | ENSMUSG0( | true | chr4 | 132421420 | 132422946 | 33.79989 | 120.06606 | 115.92274 | . | . | . | . | . | . |
| chr5 | 73268604 | 73269069 | 681 | Exon (ENSMUST00000188758.1/ | 5 | 73193828 | 73256618 | 62791 | 2 | ENSMUST00 | -11987 | Fryl | ENSMUSG0( | true | . | . | . | . | . | . | chr5 | 73263502 | 73276446 | 5.12469 | 4.16753 | 3.10513 |
| chr7 | 31115631 | 31115977 | 680 | Promoter (<=1kb) | 7 | 31098725 | 31115290 | 16566 | 2 | ENSMUST00 | -342 | Hpn | ENSMUSG0( | false | . | . | . | . | . | . | chr7 | 31114630 | 31116711 | 6.75053 | 4.77712 | 4.44698 |
| chr11 | 52000391 | 52000954 | 680 | Promoter (<=1kb) | 11 | 51985497 | 52000484 | 14988 | 2 | ENSMUST00 | 0 | Ube2b | ENSMUSG0( | false | chr11 | 51998950 | 52000765 | 24.03115 | 75.67385 | 72.59326 | . | . | . | . | . | . |
| chr13 | 75840113 | 75840501 | 680 | Promoter (<=1kb) | 13 | 75839868 | 75850151 | 10284 | 1 | ENSMUST00 | 246 | Glrx | ENSMUSG0( | false | chr13 | 75839868 | 75840968 | 15.53947 | 48.13258 | 45.44172 | chr13 | 75839239 | 75842298 | 4.0166 | 3.50107 | 2.23166 |
| chr1 | 55626298 | 55626521 | 680 | Intron (ENSMUST00000042986.9 | 1 | 55702025 | 55735227 | 33203 | 1 | ENSMUST00 | -75504 | Plcl1 | ENSMUSG0( | true | . | . | . | . | . | . | chr1 | 55625969 | 55634206 | 4.59056 | 3.89106 | 2.67899 |
| chr12 | 73393890 | 73394178 | 680 | Distal Intergenic | 12 | 73364075 | 73371445 | 7371 | 2 | ENSMUST00 | -22446 | D830013O20 | ENSMUSG0( | false | . | . | . | . | . | . | chr12 | 73393428 | 73394495 | 4.63743 | 3.96835 | 2.69529 |
| chr19 | 26914282 | 26914580 | 680 | Distal Intergenic | 19 | 26885925 | 26888639 | 2715 | 1 | ENSMUST00 | 28358 | Gm815 | ENSMUSG0( | false | . | . | . | . | . | . | . | . | . | . | . | . |
| chr4 | 124849576 | 124849887 | 680 | Promoter (<=1kb) | 4 | 124849080 | 124850330 | 1251 | 2 | ENSMUST00 | 443 | 1110065P20l | ENSMUSG0( | false | chr4 | 124849422 | 124852147 | 42.27695 | 136.11826 | 131.39047 | chr4 | 124847029 | 124849942 | 5.89109 | 4.49521 | 3.73465 |
| chr8 | 125348549 | 125348887 | 680 | Distal Intergenic | 8 | 125224400 | 125227323 | 2924 | 2 | ENSMUST00 | -121227 | Gm16237 | ENSMUSG0( | false | . | . | . | . | . | . | chr8 | 125335216 | 125352727 | 6.82037 | 4.91647 | 4.50391 |
| chr11 | 95096761 | 95097319 | 679 | Distal Intergenic | 11 | 95044474 | 95076801 | 32328 | 2 | ENSMUST00 | -19961 | Itga3 | ENSMUSG0( | true | . | . | . | . | . | . | chr11 | 95095874 | 95108125 | 6.99596 | 4.92043 | 4.65315 |
| chr17 | 56008979 | 56009505 | 679 | Promoter (<=1kb) | 17 | 56009201 | 56016790 | 7590 | 1 | ENSMUST00 | 0 | Mpnd | ENSMUSG0( | false | chr17 | 56008496 | 56010059 | 35.41093 | 115.56754 | 111.56183 | . | . | . | . | . | . |
| chr14 | 60676820 | 60677368 | 679 | Intron (ENSMUST00000159169.1 | 14 | 60691975 | 60709569 | 17595 | 1 | ENSMUST00 | -14607 | Spata13 | ENSMUSG0( | true | . | . | . | . | . | . | chr14 | 60673122 | 60679063 | 5.33891 | 4.10601 | 3.32068 |
| chr4 | 123918525 | 123918850 | 679 | Promoter (<=1kb) | 4 | 123919087 | 123920991 | 1905 | 1 | ENSMUST00 | -237 | Rragc | ENSMUSG0( | false | chr4 | 123916660 | 123919069 | 32.3084 | 121.498 | 117.30614 | chr4 | 123918505 | 123921363 | 5.612 | 4.21413 | 3.5291 |
| chr5 | 21005824 | 21006104 | 679 | Intron (ENSMUST00000030556.7 | 5 | 20998453 | 21009606 | 11154 | 2 | ENSMUST00 | 3502 | Ptpn12 | ENSMUSG0( | true | . | . | . | . | . | . | chr5 | 21004220 | 21007801 | 4.17594 | 3.73941 | 2.31445 |
| chr19 | 46612155 | 46612495 | 679 | Intron (ENSMUST00000138302.3 | 19 | 46623401 | 46652522 | 29122 | 1 | ENSMUST00 | -10906 | Wbp1l | ENSMUSG0( | false | . | . | . | . | . | . | chr19 | 46599683 | 46650166 | 6.55215 | 4.7277 | 4.28832 |
| chr16 | 33684255 | 33684851 | 679 | Promoter (<=1kb) | 16 | 33684466 | 33722913 | 38448 | 1 | ENSMUST00 | 0 | Heg1 | ENSMUSG0( | false | chr16 | 33684190 | 33685739 | 34.92444 | 106.42502 | 102.68874 | chr16 | 33684058 | 33685708 | 3.25412 | 3.1767 | 1.55677 |
| chr13 | 12701981 | 12702470 | 679 | Promoter (<=1kb) | 13 | 12702362 | 12702589 | 228 | 1 | ENSMUST00 | 0 | Gm2399 | ENSMUSG0( | false | chr13 | 12702430 | 12703024 | 18.33657 | 60.6815 | 57.82544 | chr13 | 12700774 | 12706233 | 5.06138 | 3.93361 | 3.0952 |
| chr2 | 51287324 | 51287546 | 679 | Intron (ENSMUST00000139497.1 | 2 | 51241831 | 51295945 | 54115 | 1 | ENSMUST00 | 45494 | Gm13497 | ENSMUSG0( | false | . | . | . | . | . | . | chr2 | 51286737 | 51289015 | 4.18868 | 3.66493 | 2.34876 |
| chr18 | 61609637 | 61610162 | 678 | Distal Intergenic | 18 | 61582844 | 61586498 | 3655 | 1 | ENSMUST00 | 26794 | Csnk1a1 | ENSMUSG0( | true | . | . | . | . | . | . | chr18 | 61608642 | 61614659 | 6.67053 | 4.76198 | 4.40969 |
| chr5 | 65536939 | 65537371 | 678 | Promoter (<=1kb) | 5 | 65537233 | 65598988 | 61756 | 1 | ENSMUST00 | 0 | Ube2k | ENSMUSG0( | false | chr5 | 65536471 | 65538620 | 34.36702 | 111.64683 | 107.76208 | . | . | . | . | . | . |
| chr10 | 78591767 | 78592100 | 678 | Promoter (<=1kb) | 10 | 78584503 | 78591964 | 7462 | 2 | ENSMUST00 | 0 | Syde1 | ENSMUSG0( | false | chr10 | 78590821 | 78592452 | 28.40368 | 83.36528 | 80.15265 | chr10 | 78589014 | 78593008 | 3.76476 | 3.41993 | 1.98773 |
| chr8 | 41025676 | 41025972 | 678 | Promoter (2-3kb) | 8 | 41019523 | 41022902 | 3380 | 2 | ENSMUST00 | -2775 | Mtus1 | ENSMUSG0( | false | . | . | . | . | . | . | chr8 | 41020996 | 41040121 | 5.92676 | 4.49745 | 3.7648 |
| chr1 | 79703573 | 79703864 | 678 | 3' UTR | 1 | 79719163 | 79722656 | 3494 | 2 | ENSMUST00 | 18792 | Wdfy1 | ENSMUSG0( | false | . | . | . | . | . | . | chr1 | 79702823 | 79704896 | 4.28554 | 3.64814 | 2.4666 |
| chr3 | 148025809 | 148026063 | 678 | Distal Intergenic | 3 | 148354087 | 148354218 | 132 | 2 | ENSMUST00 | 328155 | Gm23131 | ENSMUSG0( | false | . | . | . | . | . | . | . | . | . | . | . | . |
| chr3 | 37958546 | 37958842 | 678 | Promoter (1-2kb) | 3 | 37959612 | 37960087 | 476 | 2 | ENSMUST00 | 1245 | Gm7799 | ENSMUSG0( | false | . | . | . | . | . | . | chr3 | 37957923 | 37959416 | 5.00244 | 4.06516 | 3.07585 |
| chr5 | 117964291 | 117964628 | 678 | Distal Intergenic | 5 | 117976454 | 117976560 | 107 | 2 | ENSMUST00 | 11932 | Gm25343 | ENSMUSG0( | false | . | . | . | . | . | . | chr5 | 117963900 | 117968142 | 4.51118 | 3.90796 | 2.59775 |
| chr10 | 58636544 | 58637011 | 677 | Intron (ENSMUST00000003312.4 | 10 | 58600789 | 58675654 | 74866 | 2 | ENSMUST00 | 38643 | Edar | ENSMUSG0( | false | . | . | . | . | . | . | chr10 | 58636341 | 58638894 | 4.62756 | 3.89522 | 2.74837 |
| chr16 | 22439247 | 22440168 | 677 | Promoter (<=1kb) | 16 | 22381309 | 22439719 | 58411 | 2 | ENSMUST00 | 0 | Etv5 | ENSMUSG0( | false | chr16 | 22438273 | 22439688 | 28.1615 | 93.53628 | 90.11485 | . | . | . | . | . | . |
| chr9 | 27029086 | 27029350 | 677 | Promoter (<=1kb) | 9 | 27004503 | 27030094 | 25592 | 2 | ENSMUST00 | 744 | Vps26b | ENSMUSG0( | false | chr9 | 27028962 | 27031181 | 42.51875 | 148.84439 | 143.61023 | chr9 | 27025880 | 27029365 | 4.29142 | 3.5259 | 2.45921 |
| chr10 | 78185066 | 78185324 | 677 | Promoter (<=1kb) | 10 | 78170910 | 78185149 | 14240 | 2 | ENSMUST00 | 0 | Pwp2 | ENSMUSG0( | false | chr10 | 78184366 | 78185716 | 23.05105 | 65.89761 | 62.96862 | chr10 | 78184154 | 78186663 | 6.26602 | 4.52088 | 4.0714 |
| chr11 | 116086810 | 116087239 | 677 | Promoter (<=1kb) | 11 | 116078573 | 116086995 | 8423 | 2 | ENSMUST00 | 0 | Wbp2 | ENSMUSG0( | false | chr11 | 116086763 | 116087033 | 20.63265 | 55.76951 | 52.97896 | . | . | . | . | . | . |
| chr9 | 45955042 | 45955654 | 677 | Promoter (<=1kb) | 9 | 45937857 | 45955247 | 17391 | 2 | ENSMUST00 | 0 | Sidt2 | ENSMUSG0( | false | chr9 | 45953182 | 45955922 | 23.32969 | 96.49238 | 93.00913 | chr9 | 45955594 | 45956558 | 3.34827 | 3.14578 | 1.68661 |
| chr6 | 52225115 | 52225466 | 677 | Promoter (<=1kb) | 6 | 52224121 | 52226189 | 2069 | 2 | ENSMUST00 | 723 | Hoxa9 | ENSMUSG0( | false | chr6 | 52221266 | 52227348 | 34.16929 | 120.7392 | 116.5701 | . | . | . | . | . | . |
| chr11 | 19925079 | 19925881 | 677 | Promoter (<=1kb) | 11 | 19924442 | 20022577 | 98136 | 1 | ENSMUST00 | 638 | Spred2 | ENSMUSG0( | true | chr11 | 19923664 | 19926262 | 36.39542 | 121.53264 | 117.33772 | chr11 | 19925807 | 19943090 | 5.3851 | 4.2188 | 3.33929 |
| chr2 | 147012745 | 147013557 | 676 | Promoter (<=1kb) | 2 | 147012996 | 147078000 | 65005 | 1 | ENSMUST00 | 0 | Xrn2 | ENSMUSG0( | false | chr2 | 147012182 | 147014400 | 38.04976 | 128.75285 | 124.30675 | . | . | . | . | . | . |
| chr4 | 32983260 | 32983540 | 676 | Promoter (<=1kb) | 4 | 32983454 | 33022180 | 38727 | 1 | ENSMUST00 | 0 | Rragd | ENSMUSG0( | false | chr4 | 32982548 | 32984479 | 39.28384 | 124.42301 | 120.12974 | . | . | . | . | . | . |
| chr4 | 34614632 | 34615097 | 676 | Promoter (<=1kb) | 4 | 34614957 | 34660167 | 45211 | 1 | ENSMUST00 | 0 | Rars2 | ENSMUSG0( | false | chr4 | 34613891 | 34615909 | 27.97104 | 107.99458 | 104.2211 | . | . | . | . | . | . |
| chr6 | 108828350 | 108828756 | 676 | Promoter (<=1kb) | 6 | 108828641 | 108859356 | 30716 | 1 | ENSMUST00 | 0 | Edem1 | ENSMUSG0( | false | chr6 | 108827844 | 108830493 | 35.02911 | 135.1772 | 130.48503 | . | . | . | . | . | . |
| chr7 | 105810898 | 105811463 | 676 | Promoter (<=1kb) | 7 | 105806658 | 105811099 | 4442 | 2 | ENSMUST00 | 0 | Mrpl17 | ENSMUSG0( | false | chr7 | 105810243 | 105811081 | 36.0064 | 125.12101 | 120.80552 | chr7 | 105809706 | 105812253 | 3.05135 | 2.98901 | 1.45977 |
| chr13 | 46006141 | 46006413 | 676 | Distal Intergenic | 13 | 45965351 | 45966177 | 827 | 1 | ENSMUST00 | 40791 | 5033430I15R | ENSMUSG0( | false | . | . | . | . | . | . | chr13 | 46006007 | 46006658 | 4.90115 | 3.90171 | 2.97147 |
| chr9 | 62270909 | 62271330 | 676 | 3' UTR | 9 | 62270729 | 62282179 | 11451 | 2 | ENSMUST00 | 10849 | Spesp1 | ENSMUSG0( | false | . | . | . | . | . | . | chr9 | 62270261 | 62271640 | 3.48991 | 3.22648 | 1.80315 |
| chr14 | 60116055 | 60116317 | 676 | Intron (ENSMUST00000131670.1 | 14 | 60141063 | 60141410 | 348 | 1 | ENSMUST00 | -24746 | Gm6913 | ENSMUSG0( | false | . | . | . | . | . | . | chr14 | 60115202 | 60116673 | 2.86379 | 2.83351 | 1.31479 |
| chr17 | 15379770 | 15380101 | 675 | Promoter (2-3kb) | 17 | 15374794 | 15376872 | 2079 | 2 | ENSMUST00 | -2899 | Dll1 | ENSMUSG0( | false | chr17 | 15380084 | 15381280 | 17.18351 | 44.2994 | 41.65401 | chr17 | 15371856 | 15381968 | 5.90641 | 4.43925 | 3.75812 |
| chr8 | 85380955 | 85381265 | 675 | Promoter (<=1kb) | 8 | 85324447 | 85380978 | 56532 | 2 | ENSMUST00 | 0 | Mylk3 | ENSMUSG0( | false | . | . | . | . | . | . | chr8 | 85373675 | 85384939 | 5.70069 | 4.34167 | 3.58874 |
| chr10 | 128368543 | 128369157 | 675 | Promoter (<=1kb) | 10 | 128363097 | 128370032 | 6936 | 2 | ENSMUST00 | 875 | Coq10a | ENSMUSG0( | false | chr10 | 128367888 | 128369602 | 28.53699 | 95.19626 | 91.7378 | . | . | . | . | . | . |
| chr6 | 54768499 | 54768749 | 675 | Distal Intergenic | 6 | 54816916 | 54893500 | 76585 | 1 | ENSMUST00 | -48167 | Znrf2 | ENSMUSG0( | false | . | . | . | . | . | . | chr6 | 54767601 | 54771268 | 4.06912 | 3.65749 | 2.2367 |
| chr7 | 27166206 | 27166955 | 675 | Promoter (<=1kb) | 7 | 27158658 | 27166802 | 8145 | 2 | ENSMUST00 | 0 | Egln2 | ENSMUSG0( | false | chr7 | 27165284 | 27167620 | 33.10969 | 106.34647 | 102.61562 | chr7 | 27162888 | 27166285 | 5.20252 | 4.23028 | 3.18303 |
| chr15 | 55806305 | 55806599 | 675 | Intron (ENSMUST00000039769.6 | 15 | 55791938 | 55906310 | 114373 | 2 | ENSMUST00 | 99711 | Sntb1 | ENSMUSG0( | false | . | . | . | . | . | . | . | . | . | . | . | . |
| chr4 | 35525355 | 35525643 | 675 | Distal Intergenic | 4 | 35436214 | 35436781 | 568 | 1 | ENSMUST00 | 89142 | Gm12365 | ENSMUSG0( | false | . | . | . | . | . | . | . | . | . | . | . | . |
| chr17 | 87174223 | 87174906 | 675 | Distal Intergenic | 17 | 87184105 | 87184225 | 121 | 2 | ENSMUST00 | 9319 | Gm22346 | ENSMUSG0( | false | . | . | . | . | . | . | chr17 | 87173521 | 87176844 | 5.34903 | 4.21105 | 3.29772 |
| chr11 | 5260213 | 5260538 | 674 | Promoter (1-2kb) | 11 | 5191552 | 5261558 | 70007 | 2 | ENSMUST00 | 1020 | Kremen1 | ENSMUSG0( | true | chr11 | 5260004 | 5262317 | 38.30962 | 127.04226 | 122.65842 | chr11 | 5235885 | 5260441 | 6.3098 | 4.58865 | 4.10038 |
| chr4 | 126429422 | 126429878 | 674 | Promoter (<=1kb) | 4 | 126332732 | 126429556 | 96825 | 2 | ENSMUST00 | 0 | Ago3 | ENSMUSG0( | false | chr4 | 126428459 | 126430348 | 37.51873 | 124.57824 | 120.27934 | . | . | . | . | . | . |
| chr6 | 89358246 | 89358717 | 674 | Promoter (<=1kb) | 6 | 89319015 | 89357717 | 38703 | 2 | ENSMUST00 | -530 | Plxna1 | ENSMUSG0( | false | . | . | . | . | . | . | chr6 | 89339950 | 89361705 | 7.61354 | 5.20417 | 5.1492 |
| chr4 | 120825345 | 120825920 | 674 | Promoter (<=1kb) | 4 | 120757438 | 120825706 | 68269 | 2 | ENSMUST00 | 0 | Nfyc | ENSMUSG0( | false | chr4 | 120824626 | 120826112 | 36.7977 | 133.03651 | 128.43381 | . | . | . | . | . | . |
| chr11 | 62076548 | 62077000 | 674 | Promoter (<=1kb) | 11 | 62077024 | 62223005 | 145982 | 1 | ENSMUST00 | -24 | Specc1 | ENSMUSG0( | false | chr11 | 62076979 | 62077897 | 30.45704 | 104.66754 | 100.97924 | chr11 | 62076255 | 62078209 | 3.84795 | 3.40317 | 2.10857 |
| chr8 | 84978535 | 84978989 | 674 | Promoter (<=1kb) | 8 | 84976909 | 84978748 | 1840 | 2 | ENSMUST00 | 0 | Junb | ENSMUSG0( | false | chr8 | 84975875 | 84979344 | 20.74539 | 66.97287 | 64.02668 | chr8 | 84974675 | 84986551 | 7.15 | 5.07534 | 4.74925 |
| chr3 | 102744605 | 102744958 | 674 | Intron (ENSMUST00000029451.6 | 3 | 102720231 | 102735417 | 15187 | 2 | ENSMUST00 | -9189 | Tspan2os | ENSMUSG0( | false | . | . | . | . | . | . | chr3 | 102735904 | 102757857 | 6.42245 | 4.63759 | 4.19001 |
| chr13 | 90106187 | 90106448 | 674 | 3' UTR | 13 | 90098981 | 90099114 | 134 | 1 | ENSMUST00 | 7207 | Gm24507 | ENSMUSG0( | false | . | . | . | . | . | . | chr13 | 90105624 | 90107323 | 4.48905 | 3.6491 | 2.59453 |
| chr12 | 71907000 | 71907261 | 674 | Intron (ENSMUST00000085299.2 | 12 | 71846311 | 71847291 | 981 | 2 | ENSMUST00 | -59710 | Gm7985 | ENSMUSG0( | false | . | . | . | . | . | . | chr12 | 71906803 | 71908639 | 3.93754 | 3.43773 | 2.17512 |
| chr10 | 68279913 | 68280293 | 673 | Promoter (1-2kb) | 10 | 68095593 | 68278726 | 183134 | 2 | ENSMUST00 | -1188 | Arid5b | ENSMUSG0( | false | chr10 | 68275857 | 68280673 | 31.01414 | 97.6227 | 94.10706 | chr10 | 68279868 | 68280733 | 3.48139 | 3.33431 | 1.73739 |
| chr12 | 75595293 | 75596560 | 673 | Promoter (<=1kb) | 12 | 75450883 | 75596200 | 145318 | 2 | ENSMUST00 | 0 | Ppp2r5e | ENSMUSG0( | false | chr12 | 75594986 | 75597036 | 36.88168 | 122.38801 | 118.16834 | . | . | . | . | . | . |
| chr2 | 120732143 | 120732463 | 673 | Promoter (<=1kb) | 2 | 120716522 | 120731518 | 14997 | 2 | ENSMUST00 | -626 | Cdan1 | ENSMUSG0( | false | . | . | . | . | . | . | chr2 | 120731791 | 120732712 | 5.08334 | 4.22687 | 3.06454 |
| chr4 | 62519581 | 62520275 | 673 | Promoter (<=1kb) | 4 | 62509169 | 62519881 | 10713 | 2 | ENSMUST00 | 0 | Alad | ENSMUSG0( | false | chr4 | 62518215 | 62520467 | 33.87313 | 113.90014 | 109.94483 | . | . | . | . | . | . |
| chr1 | 178336362 | 178336775 | 673 | Promoter (<=1kb) | 1 | 178335642 | 178336405 | 764 | 2 | ENSMUST00 | 0 | Hnrnpu | ENSMUSG0( | false | chr1 | 178334667 | 178338600 | 62.49637 | 223.48865 | 216.8802 | . | . | . | . | . | . |
| chr4 | 149954904 | 149955421 | 673 | Promoter (<=1kb) | 4 | 149896283 | 149955043 | 58761 | 2 | ENSMUST00 | 0 | Spsb1 | ENSMUSG0( | false | chr4 | 149953980 | 149955073 | 33.1106 | 101.26704 | 97.66567 | chr4 | 149953684 | 149956437 | 4.73248 | 3.96604 | 2.78103 |
| chr17 | 6778927 | 6779211 | 673 | Intron (ENSMUST00000064234.6 | 17 | 6738041 | 6782784 | 44744 | 2 | ENSMUST00 | 3573 | Ezr | ENSMUSG0( | false | chr17 | 6777892 | 6783808 | 43.16833 | 149.98996 | 144.69951 | chr17 | 6754867 | 6782114 | 5.65265 | 4.30593 | 3.559 |
| chr9 | 57542929 | 57543237 | 673 | 3' UTR | 9 | 57531287 | 57537629 | 6343 | 2 | ENSMUST00 | -5301 | Gm26631 | ENSMUSG0( | false | . | . | . | . | . | . | chr9 | 57541237 | 57549125 | 5.67277 | 4.24382 | 3.5915 |
| chr12 | 59130968 | 59131617 | 672 | Promoter (<=1kb) | 12 | 59131022 | 59146872 | 15851 | 1 | ENSMUST00 | 0 | Ctage5 | ENSMUSG0( | false | chr12 | 59128963 | 59132130 | 33.04283 | 115.53907 | 111.53828 | . | . | . | . | . | . |
| chr1 | 6188448 | 6188762 | 672 | Distal Intergenic | 1 | 6206197 | 6234311 | 28115 | 1 | ENSMUST00 | -17435 | Rb1cc1 | ENSMUSG0( | false | . | . | . | . | . | . | chr1 | 6187374 | 6193133 | 4.03131 | 3.62747 | 2.20095 |
| chr9 | 54950485 | 54951020 | 672 | Promoter (<=1kb) | 9 | 54950790 | 54958030 | 7241 | 1 | ENSMUST00 | 0 | Psma4 | ENSMUSG0( | false | chr9 | 54950323 | 54951673 | 25.79552 | 96.83904 | 93.34669 | chr9 | 54949746 | 54952100 | 4.98408 | 3.98383 | 3.04327 |
| chr9 | 60841493 | 60841810 | 672 | Intron (ENSMUST00000050183.6 | 9 | 60821855 | 60838200 | 16346 | 2 | ENSMUST00 | -3294 | Gm9869 | ENSMUSG0( | false | chr9 | 60836859 | 60842659 | 24.93112 | 71.82204 | 68.80418 | chr9 | 60839714 | 60851928 | 5.61355 | 4.39804 | 3.5066 |
| chr3 | 52015680 | 52015961 | 672 | Intron (ENSMUST00000121440.2 | 3 | 51687611 | 52104891 | 417281 | 2 | ENSMUST00 | 88930 | Maml3 | ENSMUSG0( | true | . | . | . | . | . | . | chr3 | 52009386 | 52064052 | 5.58271 | 4.32835 | 3.48679 |
| chr17 | 4946072 | 4946342 | 672 | Distal Intergenic | 17 | 4994332 | 5097733 | 103402 | 1 | ENSMUST00 | -47990 | Arid1b | ENSMUSG0( | false | . | . | . | . | . | . | chr17 | 4944987 | 4948299 | 5.62271 | 4.4199 | 3.52867 |
| chr11 | 95712533 | 95713620 | 672 | Promoter (<=1kb) | 11 | 95712673 | 95772353 | 59681 | 1 | ENSMUST00 | 0 | Zfp652 | ENSMUSG0( | false | chr11 | 95712399 | 95714519 | 37.53511 | 117.15172 | 113.10373 | chr11 | 95712455 | 95716535 | 4.37405 | 3.75193 | 2.48274 |
| chr1 | 86487268 | 86487902 | 672 | Distal Intergenic | 1 | 86473659 | 86474006 | 348 | 1 | ENSMUST00 | 13610 | RP23-403I3.( | ENSMUSG0( | false | chr1 | 86487725 | 86488470 | 30.69271 | 93.10602 | 89.69717 | . | . | . | . | . | . |
| chr14 | 78532445 | 78532699 | 671 | Intron (ENSMUST00000123853.2 | 14 | 78492246 | 78536808 | 44563 | 2 | ENSMUST00 | 4109 | Akap11 | ENSMUSG0( | true | . | . | . | . | . | . | chr14 | 78522737 | 78539051 | 5.22391 | 4.1452 | 3.19494 |
| chr15 | 25775228 | 25775491 | 671 | Promoter (1-2kb) | 15 | 25773987 | 25781057 | 7071 | 1 | ENSMUST00 | 1242 | Myo10 | ENSMUSG0( | false | . | . | . | . | . | . | chr15 | 25760036 | 25780626 | 5.60912 | 4.2917 | 3.5207 |
| chr15 | 59054788 | 59055212 | 671 | Exon (ENSMUST00000080371.6/ | 15 | 58941244 | 59082026 | 140783 | 2 | ENSMUST00 | 26814 | Mtss1 | ENSMUSG0( | true | . | . | . | . | . | . | chr15 | 58965186 | 59078715 | 7.47325 | 5.12649 | 5.04432 |
| chr19 | 20774345 | 20774624 | 671 | Distal Intergenic | 19 | 20692953 | 20727562 | 34610 | 2 | ENSMUST00 | -46784 | Aldh1a7 | ENSMUSG0( | false | . | . | . | . | . | . | . | . | . | . | . | . |
| chr3 | 131110158 | 131110496 | 671 | Promoter (<=1kb) | 3 | 131110471 | 131224356 | 113886 | 1 | ENSMUST00 | 0 | Lef1 | ENSMUSG0( | false | chr3 | 131110142 | 131112789 | 22.5292 | 69.37365 | 66.39268 | chr3 | 131109267 | 131113191 | 5.50748 | 4.3265 | 3.44397 |
| chr4 | 106315802 | 106316477 | 671 | Promoter (<=1kb) | 4 | 106316213 | 106441317 | 125105 | 1 | ENSMUST00 | 0 | Usp24 | ENSMUSG0( | false | chr4 | 106315477 | 106318241 | 33.31129 | 108.75554 | 104.95251 | . | . | . | . | . | . |
| chr11 | 6120614 | 6121092 | 671 | Promoter (1-2kb) | 11 | 6105986 | 6119409 | 13424 | 2 | ENSMUST00 | -1206 | Nudcd3 | ENSMUSG0( | false | . | . | . | . | . | . | chr11 | 6119930 | 6122275 | 4.05856 | 3.55597 | 2.27661 |
| chr14 | 118236916 | 118237238 | 671 | Promoter (<=1kb) | 14 | 118233232 | 118237030 | 3799 | 2 | ENSMUST00 | 0 | Sox21 | ENSMUSG0( | false | chr14 | 118234864 | 118237398 | 19.69817 | 53.33889 | 50.58166 | chr14 | 118234106 | 118237702 | 7.03938 | 4.99832 | 4.69272 |
| chr2 | 29466190 | 29466649 | 671 | Intron (ENSMUST00000028139.5 | 2 | 29459194 | 29459328 | 135 | 1 | ENSMUST00 | 6997 | Gm24976 | ENSMUSG0( | false | . | . | . | . | . | . | chr2 | 29464448 | 29467089 | 4.95472 | 3.93618 | 3.00088 |
| chr11 | 100545481 | 100545804 | 670 | Promoter (<=1kb) | 11 | 100545607 | 100571963 | 26357 | 1 | ENSMUST00 | 0 | Ttc25 | ENSMUSG0( | false | chr11 | 100545213 | 100546009 | 17.00269 | 41.02908 | 38.42754 | chr11 | 100545117 | 100546549 | 4.53262 | 3.84486 | 2.60891 |
| chr9 | 119088262 | 119088814 | 670 | Intron (ENSMUST00000010804.2 | 9 | 119071527 | 119093496 | 21970 | 2 | ENSMUST00 | 4682 | Plcd1 | ENSMUSG0( | false | chr9 | 119087813 | 119089048 | 17.89762 | 44.76895 | 42.12037 | chr9 | 119087543 | 119089820 | 4.96374 | 4.10152 | 2.9651 |
| chr12 | 83991791 | 83992063 | 670 | Intron (ENSMUST00000021649.7 | 12 | 83987861 | 83993873 | 6013 | 1 | ENSMUST00 | 3931 | Acot2 | ENSMUSG0( | false | . | . | . | . | . | . | chr12 | 83986416 | 83992626 | 5.26885 | 4.21659 | 3.23054 |
| chr15 | 66507886 | 66508178 | 670 | Distal Intergenic | 15 | 66379858 | 66500910 | 121053 | 2 | ENSMUST00 | -6977 | Lrrc6 | ENSMUSG0( | false | . | . | . | . | . | . | . | . | . | . | . | . |
| chr17 | 8935589 | 8936605 | 670 | Intron (ENSMUST00000115715.2 | 17 | 8945569 | 8982391 | 36823 | 2 | ENSMUST00 | -8964 | Pde10a | ENSMUSG0( | true | . | . | . | . | . | . | chr17 | 8935246 | 8936702 | 3.82316 | 3.47714 | 2.07286 |
| chr1 | 166378852 | 166379481 | 670 | Promoter (<=1kb) | 1 | 166379097 | 166393621 | 14525 | 1 | ENSMUST00 | 0 | Tada1 | ENSMUSG0( | false | chr1 | 166379287 | 166380486 | 34.34893 | 131.73137 | 127.17394 | chr1 | 166378197 | 166378903 | 4.14773 | 3.7213 | 2.28885 |
| chr2 | 120128629 | 120129095 | 670 | Promoter (<=1kb) | 2 | 120102220 | 120129725 | 27506 | 2 | ENSMUST00 | 630 | Ehd4 | ENSMUSG0( | false | . | . | . | . | . | . | chr2 | 120119955 | 120135886 | 7.40237 | 5.05905 | 4.97897 |
| chr10 | 14545028 | 14545601 | 670 | Promoter (<=1kb) | 10 | 14402585 | 14545036 | 142452 | 2 | ENSMUST00 | 0 | Gpr126 | ENSMUSG0( | false | chr10 | 14544762 | 14545897 | 25.1466 | 74.00616 | 70.95453 | chr10 | 14543907 | 14545545 | 2.85954 | 2.83756 | 1.28617 |

| | | | | | | | | | | | | | | | | | | | | | | | | | | |
|---|---|---|---|---|---|---|---|---|---|---|---|---|---|---|---|---|---|---|---|---|---|---|---|---|---|---|
| chr14 | 120288293 | 120288694 | 669 | Intron (ENSMUST00000088419.7 | 14 | 120275669 | 120431697 | 156029 | 1 | ENSMUST00 | 12625 | Mbnl2 | ENSMUSG0( | false | . | . | . | . | . | . | chr14 | 120277640 | 120316005 | 5.96633 | 4.45953 | 3.81543 |
| chr16 | 62854083 | 62854509 | 669 | Promoter (<=1kb) | 16 | 62854307 | 62929346 | 75040 | 1 | ENSMUST00 | 0 | Pros1 | ENSMUSG0( | false | chr16 | 62853976 | 62855300 | 28.08037 | 85.65662 | 82.39679 | . | . | . | . | . | . |
| chr7 | 68250539 | 68250784 | 669 | Intron (ENSMUST00000032770.2 | 7 | 68236608 | 68264233 | 27626 | 2 | ENSMUST00 | 13449 | Pgpep1l | ENSMUSG0( | false | . | . | . | . | . | . | chr7 | 68249288 | 68251169 | 5.4888 | 4.36093 | 3.40258 |
| chr14 | 61599280 | 61599812 | 669 | Promoter (<=1kb) | 14 | 61599512 | 61605946 | 6435 | 1 | ENSMUST00 | 0 | Trim13 | ENSMUSG0( | false | chr14 | 61599012 | 61600029 | 33.55804 | 103.05054 | 99.40515 | chr14 | 61598741 | 61599298 | 2.90893 | 2.9935 | 1.3443 |
| chr9 | 118972388 | 118972679 | 669 | Intron (ENSMUST00000073109.6 | 9 | 118929867 | 119033628 | 103762 | 1 | ENSMUST00 | 42522 | Ctdspl | ENSMUSG0( | true | . | . | . | . | . | . | chr9 | 118927678 | 119010843 | 6.89652 | 4.89961 | 4.57102 |
| chr16 | 17233379 | 17233884 | 669 | Promoter (<=1kb) | 16 | 17233572 | 17263430 | 29859 | 1 | ENSMUST00 | 0 | Hic2 | ENSMUSG0( | false | chr16 | 17232910 | 17234712 | 35.62953 | 109.69414 | 105.87395 | chr16 | 17232824 | 17235809 | 4.18819 | 3.66086 | 2.38813 |
| chr4 | 65839840 | 65840108 | 669 | Intron (ENSMUST00000084496.2 | 4 | 65605249 | 65613756 | 8508 | 1 | ENSMUST00 | 234592 | Trim32 | ENSMUSG0( | false | . | . | . | . | . | . | chr4 | 65836575 | 65840455 | 4.25045 | 3.67313 | 2.41574 |
| chr13 | 98478048 | 98478617 | 669 | Promoter (<=1kb) | 13 | 98476598 | 98477459 | 862 | 2 | ENSMUST00 | -590 | RP23-369M1 | ENSMUSG0( | false | . | . | . | . | . | . | chr13 | 98474863 | 98480463 | 5.33341 | 4.04738 | 3.29871 |
| chr18 | 80682992 | 80683266 | 668 | Intron (ENSMUST00000167977.2 | 18 | 80606205 | 80708173 | 101969 | 2 | ENSMUST00 | 24907 | Nfatc1 | ENSMUSG0( | false | . | . | . | . | . | . | chr18 | 80682335 | 80685649 | 5.7669 | 4.52846 | 3.62714 |
| chr2 | 84626736 | 84627011 | 668 | Intron (ENSMUST00000099941.3 | 2 | 84624378 | 84638105 | 13728 | 2 | ENSMUST00 | 11094 | Ctnnd1 | ENSMUSG0( | true | . | . | . | . | . | . | chr2 | 84621405 | 84648911 | 6.04389 | 4.54416 | 3.87093 |
| chr7 | 28371408 | 28372000 | 668 | Promoter (<=1kb) | 7 | 28359605 | 28371574 | 11970 | 2 | ENSMUST00 | 0 | Plekhg2 | ENSMUSG0( | true | chr7 | 28370828 | 28372772 | 30.78991 | 105.27288 | 101.56436 | chr7 | 28351351 | 28371775 | 6.79815 | 4.79627 | 4.48731 |
| chr11 | 82951050 | 82951525 | 668 | Promoter (<=1kb) | 11 | 82951349 | 82952170 | 822 | 1 | ENSMUST00 | 0 | Slfn5 | ENSMUSG0( | false | . | . | . | . | . | . | . | . | . | . | . | . |
| chr5 | 122696494 | 122696915 | 668 | Promoter (<=1kb) | 5 | 122697202 | 122697600 | 399 | 2 | ENSMUST00 | 685 | Gm10064 | ENSMUSG0( | false | . | . | . | . | . | . | chr5 | 122696621 | 122701326 | 4.6831 | 3.84502 | 2.79173 |
| chr13 | 21468456 | 21468686 | 668 | Promoter (<=1kb) | 13 | 21467047 | 21468509 | 1463 | 2 | ENSMUST00 | 0 | Nkapl | ENSMUSG0( | false | chr13 | 21467932 | 21469104 | 36.83125 | 118.04518 | 113.96772 | . | . | . | . | . | . |
| chr4 | 132463691 | 132464086 | 668 | Promoter (<=1kb) | 4 | 132458731 | 132463921 | 5191 | 2 | ENSMUST00 | 0 | Med18 | ENSMUSG0( | false | chr4 | 132463229 | 132464465 | 34.71595 | 106.06588 | 102.34386 | . | . | . | . | . | . |
| chr1 | 64690750 | 64691295 | 668 | Promoter (<=1kb) | 1 | 64690822 | 64723946 | 33125 | 1 | ENSMUST00 | 0 | Ccnyl1 | ENSMUSG0( | false | chr1 | 64690907 | 64692184 | 32.87541 | 111.07999 | 107.21675 | . | . | . | . | . | . |
| chr16 | 92094436 | 92094706 | 668 | Promoter (<=1kb) | 16 | 92094568 | 92096490 | 1923 | 1 | ENSMUST00 | 0 | Gm16310 | ENSMUSG0( | false | . | . | . | . | . | . | chr16 | 92093933 | 92095755 | 3.76638 | 3.24876 | 2.0271 |
| chr10 | 59503784 | 59504203 | 667 | Intron (ENSMUST00000020312.7 | 10 | 59448597 | 59458160 | 9564 | 2 | ENSMUST00 | -45625 | Mcu | ENSMUSG0( | false | chr10 | 59503618 | 59504435 | 9.38717 | 20.36066 | 18.02005 | chr10 | 59495881 | 59504668 | 5.58173 | 4.38368 | 3.47182 |
| chr11 | 7160670 | 7160903 | 667 | Promoter (<=1kb) | 11 | 7159882 | 7172879 | 12998 | 1 | ENSMUST00 | 789 | Adcy1 | ENSMUSG0( | true | . | . | . | . | . | . | chr11 | 7147428 | 7161440 | 5.80144 | 4.4368 | 3.66324 |
| chr2 | 57123367 | 57124386 | 667 | Promoter (<=1kb) | 2 | 57106830 | 57124003 | 17174 | 2 | ENSMUST00 | 0 | Nr4a2 | ENSMUSG0( | false | chr2 | 57123281 | 57124500 | 33.66875 | 128.30269 | 123.87101 | . | . | . | . | . | . |
| chr5 | 125056840 | 125057464 | 667 | Promoter (<=1kb) | 5 | 125047802 | 125055988 | 8187 | 2 | ENSMUST00 | -853 | Ncor2 | ENSMUSG0( | false | chr5 | 125055981 | 125058881 | 22.19715 | 59.28558 | 56.44936 | chr5 | 125050765 | 125060996 | 6.95919 | 4.79882 | 4.64527 |
| chr3 | 49757131 | 49757637 | 667 | Promoter (<=1kb) | 3 | 49743296 | 49757316 | 14021 | 2 | ENSMUST00 | 0 | Pcdh18 | ENSMUSG0( | false | chr3 | 49756025 | 49757785 | 23.13664 | 67.80279 | 64.84689 | . | . | . | . | . | . |
| chr1 | 53844761 | 53845057 | 667 | Intron (ENSMUST00000120904.2 | 1 | 53832692 | 53841171 | 8480 | 2 | ENSMUST00 | -3591 | Hecw2 | ENSMUSG0( | false | . | . | . | . | . | . | chr1 | 53844440 | 53846297 | 4.7594 | 4.02524 | 2.82723 |
| chr6 | 72050854 | 72051147 | 667 | Distal Intergenic | 6 | 72097592 | 72154571 | 56980 | 1 | ENSMUST00 | -46445 | St3gal5 | ENSMUSG0( | false | . | . | . | . | . | . | . | . | . | . | . | . |
| chr3 | 153692852 | 153693340 | 667 | Intron (ENSMUST00000064460.5 | 3 | 153708911 | 153709026 | 116 | 1 | ENSMUST00 | -15571 | Gm22206 | ENSMUSG0( | false | . | . | . | . | . | . | chr3 | 153686316 | 153712771 | 6.43925 | 4.66349 | 4.19773 |
| chr8 | 70601194 | 70601513 | 666 | Intron (ENSMUST00000049908.9 | 8 | 70594481 | 70597288 | 2808 | 1 | ENSMUST00 | 6714 | Isyna1 | ENSMUSG0( | false | . | . | . | . | . | . | chr8 | 70592285 | 70603072 | 6.42958 | 4.72399 | 4.17224 |
| chr10 | 39377280 | 39377711 | 666 | Intron (ENSMUST00000148152.2 | 10 | 39369803 | 39564423 | 194621 | 1 | ENSMUST00 | 7478 | Fyn | ENSMUSG0( | true | . | . | . | . | . | . | chr10 | 39371015 | 39390933 | 5.77617 | 4.40934 | 3.64788 |
| chr9 | 37106886 | 37107181 | 666 | Intron (ENSMUST00000039674.6 | 9 | 36890982 | 37083309 | 192328 | 2 | ENSMUST00 | -23578 | Pknox2 | ENSMUSG0( | true | . | . | . | . | . | . | chr9 | 37098428 | 37110906 | 5.53132 | 4.20019 | 3.48047 |
| chr8 | 36528290 | 36528566 | 666 | Distal Intergenic | 8 | 36505489 | 36512593 | 7105 | 1 | ENSMUST00 | 22802 | 6430573F11I | ENSMUSG0( | false | . | . | . | . | . | . | . | . | . | . | . | . |
| chr12 | 76533364 | 76533943 | 666 | Promoter (<=1kb) | 12 | 76533560 | 76579039 | 45480 | 1 | ENSMUST00 | 0 | Plekhg3 | ENSMUSG0( | false | chr12 | 76532849 | 76534984 | 33.45292 | 101.85796 | 98.2438 | chr12 | 76532835 | 76538765 | 4.59558 | 3.84512 | 2.69586 |
| chr7 | 29271834 | 29272458 | 666 | Intron (ENSMUST00000098604.5 | 7 | 29256331 | 29281977 | 25647 | 2 | ENSMUST00 | 9519 | Spint2 | ENSMUSG0( | false | . | . | . | . | . | . | chr7 | 29270378 | 29278106 | 5.53936 | 4.34059 | 3.44839 |
| chr19 | 58903891 | 58904235 | 666 | Distal Intergenic | 19 | 58911673 | 58911759 | 87 | 2 | ENSMUST00 | 7524 | Mir3086 | ENSMUSG0( | false | . | . | . | . | . | . | chr19 | 58902686 | 58904493 | 4.75055 | 3.87798 | 2.8337 |
| chr17 | 66864532 | 66864829 | 666 | Intron (ENSMUST00000037974.8 | 17 | 67026147 | 67026235 | 89 | 2 | ENSMUST00 | 161406 | Mir5709 | ENSMUSG0( | false | . | . | . | . | . | . | chr17 | 66864473 | 66864775 | 3.60487 | 3.29912 | 1.87711 |
| chr12 | 38779378 | 38779688 | 665 | Promoter (<=1kb) | 12 | 38779380 | 38866249 | 86870 | 1 | ENSMUST00 | 0 | Etv1 | ENSMUSG0( | false | chr12 | 38778143 | 38781472 | 25.57207 | 77.50074 | 74.3909 | . | . | . | . | . | . |
| chr19 | 46518468 | 46518854 | 665 | Distal Intergenic | 19 | 46501648 | 46516444 | 14797 | 1 | ENSMUST00 | 16821 | Trim8 | ENSMUSG0( | false | . | . | . | . | . | . | chr19 | 46504684 | 46519488 | 7.5057 | 5.04818 | 5.07578 |
| chr7 | 76369120 | 76369393 | 665 | Intron (ENSMUST00000156166.2 | 7 | 76411062 | 76700123 | 289062 | 1 | ENSMUST00 | -41669 | Agbl1 | ENSMUSG0( | false | . | . | . | . | . | . | . | . | . | . | . | . |
| chr1 | 194793075 | 194793427 | 665 | Intron (ENSMUST00000027952.6 | 1 | 194804710 | 194814227 | 9518 | 1 | ENSMUST00 | -11283 | Plxna2 | ENSMUSG0( | false | . | . | . | . | . | . | chr1 | 194792408 | 194796164 | 5.30049 | 4.0042 | 3.277 |
| chr4 | 129810881 | 129811262 | 665 | Promoter (<=1kb) | 4 | 129811219 | 129839232 | 28014 | 1 | ENSMUST00 | 0 | Ptp4a2 | ENSMUSG0( | false | chr4 | 129810738 | 129811390 | 16.22329 | 39.70485 | 37.12044 | chr4 | 129810716 | 129811523 | 4.03202 | 3.53407 | 2.22831 |
| chr8 | 11497416 | 11497850 | 665 | Promoter (<=1kb) | 8 | 11497506 | 11513287 | 15782 | 1 | ENSMUST00 | 0 | Carkd | ENSMUSG0( | false | chr8 | 11497061 | 11498571 | 35.36919 | 136.9064 | 132.15405 | . | . | . | . | . | . |
| chr2 | 163671249 | 163671530 | 665 | Intron (ENSMUST00000147385.2 | 2 | 163661499 | 163725473 | 63975 | 1 | ENSMUST00 | 9751 | Pkig | ENSMUSG0( | false | . | . | . | . | . | . | chr2 | 163667073 | 163719465 | 6.42152 | 4.6796 | 4.1796 |
| chr13 | 38634959 | 38635217 | 665 | Promoter (<=1kb) | 13 | 38602709 | 38635109 | 32401 | 2 | ENSMUST00 | 0 | Bloc1s5 | ENSMUSG0( | false | chr13 | 38634179 | 38635474 | 33.41831 | 117.26761 | 113.21259 | chr13 | 38632294 | 38635780 | 4.70737 | 4.01539 | 2.74686 |
| chr19 | 45445354 | 45445671 | 665 | Intron (ENSMUST00000065601.6 | 19 | 45450308 | 45451195 | 888 | 2 | ENSMUST00 | 5524 | Gm6807 | ENSMUSG0( | false | . | . | . | . | . | . | chr19 | 45439520 | 45448002 | 4.82555 | 3.94648 | 2.87829 |
| chr7 | 19770377 | 19770789 | 664 | Promoter (<=1kb) | 7 | 19756131 | 19770547 | 14417 | 2 | ENSMUST00 | 0 | Bcam | ENSMUSG0( | false | chr7 | 19768283 | 19771284 | 29.58769 | 108.11006 | 104.32592 | chr7 | 19770687 | 19774598 | 4.10172 | 3.5288 | 2.30412 |
| chr17 | 70596441 | 70596895 | 664 | Intron (ENSMUST00000133983.2 | 17 | 70561787 | 70816093 | 254307 | 1 | ENSMUST00 | 34655 | Dlgap1 | ENSMUSG0( | false | . | . | . | . | . | . | chr17 | 70595676 | 70596902 | 3.00026 | 2.94301 | 1.4135 |
| chr11 | 69632574 | 69633523 | 664 | Promoter (<=1kb) | 11 | 69632990 | 69653297 | 20308 | 1 | ENSMUST00 | 0 | Fxr2 | ENSMUSG0( | false | chr11 | 69632880 | 69633976 | 29.58769 | 108.11006 | 104.32592 | . | . | . | . | . | . |
| chr3 | 81036488 | 81037354 | 664 | Promoter (<=1kb) | 3 | 81037172 | 81214034 | 176863 | 1 | ENSMUST00 | 0 | Pdgfc | ENSMUSG0( | false | chr3 | 81036384 | 81037850 | 20.4053 | 65.4752 | 62.55133 | chr3 | 81036350 | 81037920 | 4.52024 | 3.71011 | 2.63041 |
| chr9 | 70219589 | 70219885 | 664 | Intron (ENSMUST00000034745.7 | 9 | 70207350 | 70400067 | 192718 | 1 | ENSMUST00 | 12240 | Myo1e | ENSMUSG0( | true | . | . | . | . | . | . | chr9 | 70209471 | 70285922 | 6.22404 | 4.59358 | 4.02448 |
| chr4 | 133025363 | 133025710 | 664 | Intron (ENSMUST00000154646.2 | 4 | 133012204 | 133077645 | 65442 | 1 | ENSMUST00 | 13160 | Ahdc1 | ENSMUSG0( | false | . | . | . | . | . | . | chr4 | 133013189 | 133060125 | 8.03751 | 5.40542 | 5.50183 |
| chr8 | 121619311 | 121619764 | 664 | Intron (ENSMUST00000127664.1 | 8 | 121608585 | 121628610 | 20026 | 2 | ENSMUST00 | 8846 | Zcchc14 | ENSMUSG0( | false | . | . | . | . | . | . | chr8 | 121618019 | 121633740 | 6.36546 | 4.59898 | 4.13954 |
| chr18 | 44661947 | 44662662 | 664 | Promoter (<=1kb) | 18 | 44425061 | 44662266 | 237206 | 2 | ENSMUST00 | 0 | Mcc | ENSMUSG0( | true | chr18 | 44659797 | 44662967 | 44.60771 | 154.80135 | 149.33388 | . | . | . | . | . | . |
| chr12 | 80208563 | 80208879 | 663 | Intron (ENSMUST00000021554.9 | 12 | 80168322 | 80260091 | 91770 | 2 | ENSMUST00 | 51212 | Actn1 | ENSMUSG0( | true | . | . | . | . | . | . | chr12 | 80207669 | 80232669 | 6.01537 | 4.51837 | 3.84731 |
| chr7 | 99539308 | 99540113 | 663 | Promoter (<=1kb) | 7 | 99539669 | 99588136 | 48468 | 1 | ENSMUST00 | 0 | Arrb1 | ENSMUSG0( | false | . | . | . | . | . | . | chr7 | 99536482 | 99576026 | 7.32182 | 4.95837 | 4.9327 |
| chr16 | 23352332 | 23352576 | 663 | Intron (ENSMUST00000023601.8 | 16 | 23358316 | 23360348 | 2033 | 1 | ENSMUST00 | -5740 | St6gal1 | ENSMUSG0( | true | . | . | . | . | . | . | chr16 | 23344368 | 23353615 | 5.35656 | 4.16706 | 3.31602 |
| chr4 | 99193873 | 99194347 | 663 | Promoter (<=1kb) | 4 | 99193934 | 99259787 | 65854 | 1 | ENSMUST00 | 0 | Atg4c | ENSMUSG0( | false | chr4 | 99193514 | 99195055 | 25.38893 | 86.51994 | 83.2489 | . | . | . | . | . | . |
| chr6 | 54409257 | 54409641 | 663 | Intron (ENSMUST00000145111.2 | 6 | 54424564 | 54429537 | 4974 | 2 | ENSMUST00 | 19896 | 9130019P16I | ENSMUSG0( | false | . | . | . | . | . | . | chr6 | 54408431 | 54410926 | 5.04308 | 4.04765 | 3.05688 |
| chr13 | 5347977 | 5348278 | 663 | Distal Intergenic | 13 | 5189803 | 5189934 | 132 | 2 | ENSMUST00 | -158044 | Gm24790 | ENSMUSG0( | false | . | . | . | . | . | . | chr13 | 5347495 | 5348476 | 3.51956 | 3.19945 | 1.82853 |
| chr16 | 95824544 | 95824826 | 663 | Distal Intergenic | 16 | 95806029 | 95808317 | 2289 | 2 | ENSMUST00 | -16228 | 2810404F17I | ENSMUSG0( | false | . | . | . | . | . | . | chr16 | 95823405 | 95825755 | 5.96431 | 4.53465 | 3.80959 |
| chr3 | 37907552 | 37908188 | 663 | Distal Intergenic | 3 | 37959612 | 37960087 | 476 | 2 | ENSMUST00 | 51899 | Gm7799 | ENSMUSG0( | false | . | . | . | . | . | . | chr3 | 37906428 | 37909120 | 4.9836 | 4.04855 | 3.03489 |
| chr3 | 153329510 | 153330029 | 663 | Intron (ENSMUST00000064460.5 | 3 | 153313713 | 153313824 | 112 | 1 | ENSMUST00 | 15798 | Gm27405 | ENSMUSG0( | false | . | . | . | . | . | . | . | . | . | . | . | . |
| chr7 | 84397264 | 84397882 | 662 | Intron (ENSMUST00000085077.3 | 7 | 84246281 | 84409959 | 163679 | 2 | ENSMUST00 | 12077 | Arnt2 | ENSMUSG0( | false | . | . | . | . | . | . | chr7 | 84396238 | 84398673 | 5.16877 | 4.07728 | 3.18673 |
| chr6 | 86039032 | 86039293 | 662 | Distal Intergenic | 6 | 86017191 | 86020996 | 3806 | 1 | ENSMUST00 | 21842 | Figla | ENSMUSG0( | false | . | . | . | . | . | . | chr6 | 86038591 | 86039174 | 3.39838 | 3.2452 | 1.68232 |
| chr2 | 5714223 | 5715028 | 662 | Promoter (<=1kb) | 2 | 5293457 | 5714515 | 421059 | 2 | ENSMUST00 | 0 | Camk1d | ENSMUSG0( | false | chr2 | 5713426 | 5715268 | 16.70194 | 60.58573 | 57.73199 | chr2 | 5713189 | 5715491 | 4.9943 | 4.03228 | 3.03155 |
| chr3 | 122989692 | 122990042 | 662 | Distal Intergenic | 3 | 122933601 | 122984447 | 50847 | 2 | ENSMUST00 | -5246 | Usp53 | ENSMUSG0( | true | . | . | . | . | . | . | chr3 | 122985135 | 122990580 | 4.5336 | 3.88052 | 2.61649 |
| chr13 | 30364224 | 30364956 | 662 | Intron (ENSMUST00000066412.7 | 13 | 30336441 | 30382867 | 46427 | 1 | ENSMUST00 | 27784 | Agtr1a | ENSMUSG0( | false | . | . | . | . | . | . | . | . | . | . | . | . |
| chr5 | 135053069 | 135053417 | 662 | Promoter (<=1kb) | 5 | 135053988 | 135054116 | 129 | 2 | ENSMUST00 | 699 | Gm25492 | ENSMUSG0( | false | . | . | . | . | . | . | chr5 | 135050839 | 135057197 | 4.55822 | 3.82811 | 2.68599 |
| chr3 | 8717323 | 8717848 | 662 | Distal Intergenic | 3 | 8679359 | 8679477 | 119 | 2 | ENSMUST00 | -37847 | Gm23670 | ENSMUSG0( | false | chr3 | 8717267 | 8717926 | 7.15905 | 12.35804 | 10.14148 | chr3 | 8712430 | 8718854 | 5.10568 | 4.06101 | 3.11811 |
| chr1 | 177441754 | 177443025 | 662 | Promoter (<=1kb) | 1 | 177429047 | 177443270 | 14224 | 2 | ENSMUST00 | 245 | Gm26801 | ENSMUSG0( | false | chr1 | 177441041 | 177442001 | 38.87692 | 132.39133 | 127.80991 | . | . | . | . | . | . |
| chr8 | 70906356 | 70906680 | 661 | Promoter (<=1kb) | 8 | 70905974 | 70917526 | 11553 | 1 | ENSMUST00 | 383 | Map1s | ENSMUSG0( | false | chr8 | 70905581 | 70906912 | 33.86687 | 104.60014 | 100.91516 | chr8 | 70905175 | 70907910 | 4.62463 | 3.84629 | 2.74195 |
| chr14 | 57740036 | 57740397 | 661 | Intron (ENSMUST00000174694.2 | 14 | 57734180 | 57744654 | 10475 | 2 | ENSMUST00 | 4257 | Lats2 | ENSMUSG0( | false | . | . | . | . | . | . | chr14 | 57704250 | 57744175 | 6.91785 | 4.80068 | 4.61004 |
| chr16 | 26581424 | 26581858 | 661 | Promoter (<=1kb) | 16 | 26581704 | 26725264 | 143561 | 1 | ENSMUST00 | 0 | Il1rap | ENSMUSG0( | false | chr16 | 26581293 | 26582646 | 26.30149 | 81.58109 | 78.4027 | chr16 | 26581164 | 26583039 | 4.55278 | 3.69972 | 2.67993 |
| chr17 | 45599450 | 45599782 | 661 | Promoter (<=1kb) | 17 | 45585200 | 45599603 | 14404 | 2 | ENSMUST00 | 0 | Slc29a1 | ENSMUSG0( | false | chr17 | 45599350 | 45599978 | 14.66885 | 35.63733 | 33.10225 | chr17 | 45591091 | 45600246 | 5.33186 | 4.18113 | 3.28168 |
| chr1 | 60566430 | 60567165 | 661 | Promoter (<=1kb) | 1 | 60482292 | 60566779 | 84488 | 2 | ENSMUST00 | 0 | Raph1 | ENSMUSG0( | false | chr1 | 60565241 | 60567491 | 30.60796 | 113.09483 | 109.17188 | chr1 | 60567161 | 60567772 | 2.97998 | 2.94083 | 1.35499 |
| chr4 | 127168749 | 127169248 | 661 | Promoter (<=1kb) | 4 | 127169204 | 127237022 | 67819 | 1 | ENSMUST00 | 0 | Dlgap3 | ENSMUSG0( | false | chr4 | 127168318 | 127169773 | 22.74714 | 65.36804 | 62.44672 | chr4 | 127168298 | 127170067 | 6.93821 | 4.86699 | 4.60169 |
| chr9 | 90109457 | 90109723 | 661 | Exon (ENSMUST00000185433.1/ | 9 | 90092402 | 90114289 | 21888 | 2 | ENSMUST00 | 4566 | Morf4l1 | ENSMUSG0( | false | . | . | . | . | . | . | chr9 | 90102529 | 90113068 | 5.2252 | 4.20694 | 3.19444 |
| chr13 | 32912703 | 32913008 | 661 | Distal Intergenic | 13 | 32881833 | 32898161 | 16329 | 2 | ENSMUST00 | -14543 | Serpinb1c | ENSMUSG0( | false | . | . | . | . | . | . | chr13 | 32910982 | 32916122 | 4.40915 | 3.69966 | 2.55946 |
| chr17 | 26584521 | 26585044 | 661 | Intron (ENSMUST00000167662.2 | 17 | 26580802 | 26580927 | 126 | 2 | ENSMUST00 | -3595 | Gm22420 | ENSMUSG0( | false | . | . | . | . | . | . | chr17 | 26581058 | 26588125 | 6.16584 | 4.57355 | 3.97658 |
| chr10 | 59702375 | 59702765 | 660 | Promoter (<=1kb) | 10 | 59702477 | 59864132 | 161656 | 1 | ENSMUST00 | 0 | Micu1 | ENSMUSG0( | true | chr10 | 59702528 | 59703890 | 22.26925 | 90.51499 | 87.16024 | chr10 | 59701972 | 59705622 | 3.97379 | 3.51895 | 2.202 |
| chr1 | 131525080 | 131525523 | 660 | Promoter (1-2kb) | 1 | 131285251 | 131527347 | 242097 | 2 | ENSMUST00 | 1824 | Srgap2 | ENSMUSG0( | true | chr1 | 131523158 | 131527435 | 25.99992 | 84.80917 | 81.57088 | . | . | . | . | . | . |
| chr10 | 44529446 | 44529944 | 660 | Promoter (<=1kb) | 10 | 44446717 | 44528501 | 81785 | 2 | ENSMUST00 | -946 | Prdm1 | ENSMUSG0( | false | chr10 | 44528998 | 44530261 | 21.02725 | 63.18829 | 60.29588 | chr10 | 44526893 | 44529919 | 5.41265 | 4.30151 | 3.3347 |
| chr17 | 50508930 | 50509922 | 660 | Promoter (<=1kb) | 17 | 50509547 | 50688493 | 178947 | 1 | ENSMUST00 | 0 | Plcl2 | ENSMUSG0( | false | chr17 | 50508569 | 50510296 | 29.77807 | 93.47606 | 90.05975 | chr17 | 50508129 | 50511680 | 3.64534 | 3.3668 | 1.9063 |
| chr12 | 58268711 | 58269505 | 660 | Promoter (<=1kb) | 12 | 58264720 | 58269258 | 4539 | 2 | ENSMUST00 | 0 | Clec14a | ENSMUSG0( | false | chr12 | 58267268 | 58268926 | 9.13057 | 19.91392 | 17.58092 | chr12 | 58268139 | 58269486 | 4.49185 | 3.92752 | 2.5727 |
| chr12 | 80434981 | 80435384 | 660 | Promoter (1-2kb) | 12 | 80335847 | 80436601 | 100755 | 2 | ENSMUST00 | 1217 | Dcaf5 | ENSMUSG0( | false | chr12 | 80434574 | 80437085 | 29.65943 | 90.33628 | 86.98486 | . | . | . | . | . | . |
| chr8 | 91331505 | 91331832 | 660 | Intron (ENSMUST00000069718.8 | 8 | 91313568 | 91487791 | 174224 | 1 | ENSMUST00 | 17938 | Fto | ENSMUSG0( | false | . | . | . | . | . | . | chr8 | 91326987 | 91333361 | 6.34089 | 4.65362 | 4.10764 |
| chr4 | 153268015 | 153268263 | 660 | Distal Intergenic | 4 | 153218176 | 153218595 | 420 | 1 | ENSMUST00 | 49840 | Gm13174 | ENSMUSG0( | false | . | . | . | . | . | . | chr4 | 153261073 | 153271626 | 5.04729 | 4.02401 | 3.0775 |
| chr2 | 13463228 | 13463608 | 659 | Intron (ENSMUST00000091436.5 | 2 | 13456640 | 13459480 | 2841 | 2 | ENSMUST00 | -3749 | Cubn | ENSMUSG0( | false | . | . | . | . | . | . | chr2 | 13462199 | 13464051 | 4.33651 | 3.79349 | 2.45227 |
| chr2 | 155276393 | 155276997 | 659 | Promoter (<=1kb) | 2 | 155276543 | 155278070 | 1528 | 1 | ENSMUST00 | 0 | Map1lc3a | ENSMUSG0( | false | chr2 | 155276013 | 155277317 | 47.36903 | 159.61537 | 153.98001 | chr2 | 155276050 | 155277658 | 3.46375 | 3.2283 | 1.78199 |
| chr4 | 134682991 | 134683370 | 659 | Intron (ENSMUST00000038628.3 | 4 | 134567889 | 134703767 | 135879 | 2 | ENSMUST00 | 20397 | Man1c1 | ENSMUSG0( | false | . | . | . | . | . | . | chr4 | 134682467 | 134687528 | 5.9319 | 4.46549 | 3.78246 |
| chr14 | 21075806 | 21076299 | 659 | Intron (ENSMUST00000045376.9 | 14 | 21052574 | 21448568 | 395995 | 1 | ENSMUST00 | 23233 | Adk | ENSMUSG0( | false | chr14 | 21075275 | 21076773 | 30.60485 | 90.02879 | 86.68851 | . | . | . | . | . | . |
| chr18 | 62054096 | 62054388 | 659 | Distal Intergenic | 18 | 61953075 | 62015715 | 62641 | 1 | ENSMUST00 | 101022 | Sh3tc2 | ENSMUSG0( | false | . | . | . | . | . | . | chr18 | 62053720 | 62056320 | 4.54152 | 3.76349 | 2.67767 |
| chr4 | 41941967 | 41942366 | 659 | Promoter (<=1kb) | 4 | 41942037 | 41942625 | 589 | 1 | ENSMUST00 | 0 | Gm20878 | ENSMUSG0( | false | . | . | . | . | . | . | . | . | . | . | . | . |
| chr4 | 114902147 | 114902453 | 659 | Promoter (<=1kb) | 4 | 114817161 | 114902859 | 85699 | 2 | ENSMUST00 | 406 | 9130410C08I | ENSMUSG0( | false | chr4 | 114901057 | 114903863 | 19.12836 | 53.15374 | 50.39941 | chr4 | 114897050 | 114905478 | 4.35462 | 3.78738 | 2.46946 |
| chr3 | 9346880 | 9347703 | 659 | Distal Intergenic | 3 | 9402175 | 9402485 | 311 | 1 | ENSMUST00 | -54472 | Gm22604 | ENSMUSG0( | false | chr3 | 9346706 | 9348003 | 10.20265 | 24.60154 | 22.20526 | chr3 | 9346382 | 9349160 | 5.89137 | 4.49563 | 3.76138 |
| chr9 | 119426049 | 119426318 | 659 | Intron (ENSMUST00000035093.9 | 9 | 119440278 | 119444830 | 4553 | 2 | ENSMUST00 | 18512 | 4930516B21I | ENSMUSG0( | false | . | . | . | . | . | . | chr9 | 119424302 | 119426929 | 6.24705 | 4.30112 | 4.06285 |
| chr6 | 18605651 | 18605917 | 658 | Distal Intergenic | 6 | 18366547 | 18514869 | 148323 | 2 | ENSMUST00 | -90783 | Cttnbp2 | ENSMUSG0( | true | . | . | . | . | . | . | chr6 | 18605254 | 18606678 | 3.89284 | 3.59725 | 2.08272 |
| chr10 | 121149700 | 121150005 | 658 | Distal Intergenic | 10 | 121083755 | 121100637 | 16883 | 1 | ENSMUST00 | 65946 | Wif1 | ENSMUSG0( | false | . | . | . | . | . | . | chr10 | 121149523 | 121150065 | 4.34623 | 3.8443 | 2.51057 |
| chr19 | 6399429 | 6399987 | 658 | Promoter (<=1kb) | 19 | 6399456 | 6403633 | 4178 | 1 | ENSMUST00 | 0 | Rasgrp2 | ENSMUSG0( | false | chr19 | 6398550 | 6400677 | 28.26186 | 81.23897 | 78.06556 | chr19 | 6398167 | 6401812 | 5.57956 | 4.20134 | 3.51519 |
| chr4 | 134401192 | 134401529 | 658 | Promoter (1-2kb) | 4 | 134402912 | 134405075 | 2164 | 1 | ENSMUST00 | -1383 | Pafah2 | ENSMUSG0( | false | . | . | . | . | . | . | chr4 | 134401046 | 134403871 | 6.47196 | 4.76737 | 4.20803 |
| chr16 | 76319618 | 76320155 | 658 | Intron (ENSMUST00000121927.2 | 16 | 76294646 | 76323658 | 29013 | 2 | ENSMUST00 | 3503 | Nrip1 | ENSMUSG0( | false | . | . | . | . | . | . | chr16 | 76310676 | 76328402 | 5.18937 | 4.14371 | 3.16272 |
| chr6 | 28943538 | 28943835 | 658 | Distal Intergenic | 6 | 28981634 | 29022165 | 40532 | 1 | ENSMUST00 | -37799 | Gm3294 | ENSMUSG0( | false | . | . | . | . | . | . | chr6 | 28942870 | 28946431 | 3.51364 | 3.25746 | 1.80711 |
| chr11 | 87442874 | 87443552 | 658 | Promoter (<=1kb) | 11 | 87443237 | 87443452 | 216 | 1 | ENSMUST00 | 0 | Rnu3b1 | ENSMUSG0( | false | chr11 | 87442643 | 87443082 | 7.76973 | 17.46802 | 15.16681 | chr11 | 87442309 | 87446291 | 3.31772 | 3.20786 | 1.61008 |
| chr16 | 34924520 | 34925126 | 658 | Intron (ENSMUST00000023538.8 | 16 | 34935854 | 34939329 | 3476 | 2 | ENSMUST00 | 14203 | RP23-80O10 | ENSMUSG0( | false | chr16 | 34920992 | 34925857 | 23.32542 | 76.20397 | 73.11496 | chr16 | 34923663 | 34969430 | 6.94598 | 4.91232 | 4.61654 |
| chr10 | 91568749 | 91568994 | 657 | Distal Intergenic | 10 | 91160417 | 91171582 | 11166 | 2 | ENSMUST00 | -397168 | Tmpo | ENSMUSG0( | false | . | . | . | . | . | . | chr10 | 91567822 | 91570007 | 5.22011 | 4.13421 | 3.25082 |
| chr11 | 82781166 | 82781729 | 657 | Promoter (<=1kb) | 11 | 82781185 | 82800653 | 19469 | 1 | ENSMUST00 | 0 | Lig3 | ENSMUSG0( | false | chr11 | 82780387 | 82782393 | 39.11606 | 129.53703 | 125.06399 | . | . | . | . | . | . |
| chr12 | 55820370 | 55821194 | 657 | Promoter (<=1kb) | 12 | 55602929 | 55821167 | 218239 | 2 | ENSMUST00 | 0 | Ralgapa1 | ENSMUSG0( | false | chr12 | 55819950 | 55821461 | 27.41053 | 90.23362 | 86.88379 | . | . | . | . | . | . |
| chr19 | 40366181 | 40366695 | 657 | Intron (ENSMUST00000099466.4 | 19 | 40295049 | 40402267 | 107219 | 2 | ENSMUST00 | 35572 | Sorbs1 | ENSMUSG0( | false | . | . | . | . | . | . | chr19 | 40365613 | 40366947 | 4.14952 | 3.53684 | 2.32496 |
| chr7 | 120969566 | 120969919 | 657 | Intron (ENSMUST00000140247.1 | 7 | 120957036 | 120981790 | 24755 | 2 | ENSMUST00 | 11871 | Cdr2 | ENSMUSG0( | false | . | . | . | . | . | . | chr7 | 120961858 | 120981649 | 5.33893 | 4.28516 | 3.2901 |
| chr19 | 37176013 | 37176237 | 657 | Promoter (<=1kb) | 19 | 37021879 | 37176063 | 154185 | 2 | ENSMUST00 | 0 | Cpeb3 | ENSMUSG0( | false | chr19 | 37173708 | 37176086 | 24.14628 | 82.32003 | 79.12344 | chr19 | 37175901 | 37178526 | 3.48593 | 3.19866 | 1.78277 |
| chr11 | 60878313 | 60879165 | 657 | Promoter (<=1kb) | 11 | 60864452 | 60879038 | 14587 | 2 | ENSMUST00 | 0 | Tmem11 | ENSMUSG0( | false | chr11 | 60877694 | 60879849 | 33.04283 | 115.53907 | 111.53828 | . | . | . | . | . | . |
| chr1 | 43446665 | 43447221 | 657 | Promoter (<=1kb) | 1 | 43445751 | 43570515 | 124765 | 1 | ENSMUST00 | 915 | Nck2 | ENSMUSG0( | true | chr1 | 43444845 | 43448373 | 43.42055 | 150.43358 | 145.13264 | chr1 | 43446894 | 43509082 | 5.99677 | 4.48008 | 3.83622 |
| chr12 | 86802144 | 86802508 | 657 | Distal Intergenic | 12 | 86809957 | 86810258 | 302 | 1 | ENSMUST00 | -7449 | Gm25888 | ENSMUSG0( | false | chr12 | 86801398 | 86802297 | 8.73111 | 16.23101 | 13.94854 | chr12 | 86797890 | 86804646 | 7.89277 | 5.3352 | 5.36976 |
| chr7 | 38085840 | 38086115 | 656 | Distal Intergenic | 7 | 38098811 | 38099369 | 559 | 2 | ENSMUST00 | 13254 | Ccne1 | ENSMUSG0( | false | chr7 | 38085045 | 38086196 | 11.15939 | 31.88065 | 29.39184 | chr7 | 38084511 | 38088451 | 4.75907 | 3.75227 | 2.82395 |
| chr5 | 17923730 | 17924542 | 656 | Distal Intergenic | 5 | 17781690 | 17888959 | 107270 | 2 | ENSMUST00 | -34772 | Cd36 | ENSMUSG0( | false | . | . | . | . | . | . | chr5 | 17922788 | 17925282 | 4.49689 | 3.74096 | 2.62667 |
| chr1 | 193130095 | 193130422 | 656 | Promoter (<=1kb) | 1 | 193104403 | 193130251 | 25849 | 2 | ENSMUST00 | 0 | Diexf | ENSMUSG0( | false | chr1 | 193129321 | 193130888 | 44.57462 | 145.65463 | 140.53799 | chr1 | 193128092 | 193130830 | 3.87556 | 3.5304 | 2.07432 |
| chr9 | 75441257 | 75441720 | 656 | Promoter (<=1kb) | 9 | 75441524 | 75466432 | 24909 | 1 | ENSMUST00 | 0 | Leo1 | ENSMUSG0( | false | chr9 | 75440979 | 75442254 | 27.49702 | 86.54257 | 83.27029 | chr9 | 75440459 | 75444337 | 3.81626 | 3.5105 | 2.03002 |
| chr6 | 71271449 | 71272070 | 656 | Promoter (<=1kb) | 6 | 71272019 | 71285318 | 13300 | 1 | ENSMUST00 | 0 | Krcc1 | ENSMUSG0( | false | chr6 | 71271644 | 71273367 | 37.45669 | 117.01644 | 112.96991 | . | . | . | . | . | . |
| chr1 | 133434465 | 133434748 | 656 | Distal Intergenic | 1 | 133382303 | 133424377 | 42075 | 2 | ENSMUST00 | -10089 | Sox13 | ENSMUSG0( | false | chr1 | 133434248 | 133434895 | 11.44831 | 25.0134 | 22.61242 | . | . | . | . | . | . |
| chr4 | 54662105 | 54662409 | 656 | Intron (ENSMUST00000127709.1 | 4 | 54669622 | 54670192 | 571 | 2 | ENSMUST00 | 7783 | Gm12477 | ENSMUSG0( | false | . | . | . | . | . | . | . | . | . | . | . | . |
| chr3 | 63179264 | 63179639 | 656 | Distal Intergenic | 3 | 63152217 | 63152358 | 142 | 1 | ENSMUST00 | 27048 | Gm22433 | ENSMUSG0( | false | . | . | . | . | . | . | chr3 | 63177356 | 63180109 | 4.04645 | 3.59034 | 2.26378 |
| chr8 | 36577911 | 36578230 | 655 | Promoter (<=1kb) | 8 | 36572145 | 36577269 | 5125 | 2 | ENSMUST00 | -643 | Dlc1 | ENSMUSG0( | true | . | . | . | . | . | . | chr8 | 36575944 | 36612020 | 5.64773 | 4.3412 | 3.54039 |
| chr10 | 95940478 | 95941350 | 655 | Promoter (<=1kb) | 10 | 95940663 | 96045518 | 104856 | 1 | ENSMUST00 | 0 | Eea1 | ENSMUSG0( | false | chr10 | 95939944 | 95941817 | 30.91854 | 102.46954 | 98.8397 | . | . | . | . | . | . |
| chr10 | 62578255 | 62578630 | 655 | Promoter (<=1kb) | 10 | 62538626 | 62578440 | 39815 | 2 | ENSMUST00 | 0 | 2510003E04I | ENSMUSG0( | false | chr10 | 62577526 | 62578495 | 39.05061 | 143.74966 | 138.71384 | . | . | . | . | . | . |
| chr7 | 28381094 | 28381510 | 655 | Promoter (1-2kb) | 7 | 28376784 | 28379255 | 2472 | 2 | ENSMUST00 | -1840 | Zfp36 | ENSMUSG0( | false | chr7 | 28381008 | 28381445 | 7.01548 | 12.12394 | 9.90998 | chr7 | 28375103 | 28383860 | 6.23801 | 4.67039 | 4.01439 |
| chr5 | 143055612 | 143056072 | 655 | Intron (ENSMUST00000053498.8 | 5 | 142990893 | 143113020 | 122128 | 2 | ENSMUST00 | 56948 | Rnf216 | ENSMUSG0( | false | . | . | . | . | . | . | chr5 | 143046832 | 143061805 | 7.05595 | 4.92101 | 4.70914 |
| chr2 | 20129458 | 20129714 | 655 | Intron (ENSMUST00000114610.2 | 2 | 20037096 | 20037894 | 799 | 2 | ENSMUST00 | -91565 | Gm13360 | ENSMUSG0( | false | . | . | . | . | . | . | chr2 | 20128390 | 20130201 | 4.8078 | 4.02792 | 2.85897 |
| chr4 | 66203028 | 66203317 | 655 | Intron (ENSMUST00000084496.2 | 4 | 66353849 | 66354186 | 338 | 1 | ENSMUST00 | -150532 | Gm11484 | ENSMUSG0( | false | . | . | . | . | . | . | chr4 | 66199366 | 66203835 | 4.35757 | 3.74325 | 2.47897 |
| chr2 | 95761489 | 95761746 | 655 | Distal Intergenic | 2 | 95697917 | 95698036 | 120 | 2 | ENSMUST00 | -63454 | Gm13797 | ENSMUSG0( | false | . | . | . | . | . | . | . | . | . | . | . | . |
| chr1 | 76070752 | 76071206 | 655 | Distal Intergenic | 1 | 76110181 | 76110580 | 400 | 2 | ENSMUST00 | 39374 | RP23-288K1! | ENSMUSG0( | false | . | . | . | . | . | . | chr1 | 76068776 | 76073653 | 3.77789 | 3.44145 | 2.01863 |
| chr11 | 104234647 | 104235031 | 654 | Intron (ENSMUST00000106992.4 | 11 | 104231584 | 104328219 | 96636 | 1 | ENSMUST00 | 3064 | Mapt | ENSMUSG0( | true | chr11 | 104231437 | 104234756 | 38.46334 | 130.56975 | 126.05653 | chr11 | 104233065 | 104238850 | 6.60888 | 4.67556 | 4.3541 |
| chr10 | 76669049 | 76669378 | 654 | Distal Intergenic | 10 | 76595762 | 76623630 | 27869 | 2 | ENSMUST00 | -45420 | Col6a2 | ENSMUSG0( | false | . | . | . | . | . | . | . | . | . | . | . | . |
| chr1 | 152793348 | 152793758 | 654 | Distal Intergenic | 1 | 152800194 | 152836991 | 36798 | 1 | ENSMUST00 | -6436 | Ncf2 | ENSMUSG0( | true | . | . | . | . | . | . | chr1 | 152788034 | 152800060 | 6.38067 | 4.71623 | 4.13727 |
| chr7 | 65082846 | 65083049 | 654 | Intron (ENSMUST00000119118.2 | 7 | 64786690 | 65156416 | 369727 | 2 | ENSMUST00 | 73367 | Fam189a1 | ENSMUSG0( | false | . | . | . | . | . | . | chr7 | 65081993 | 65083529 | 5.36585 | 3.95494 | 3.3478 |
| chr8 | 15052919 | 15053151 | 654 | Distal Intergenic | 8 | 15057653 | 15133541 | 75889 | 1 | ENSMUST00 | -4502 | Myom2 | ENSMUSG0( | true | . | . | . | . | . | . | chr8 | 15052669 | 15059983 | 5.56184 | 4.2692 | 3.4789 |
| chr11 | 63925132 | 63925415 | 654 | Promoter (2-3kb) | 11 | 63885792 | 63922290 | 36499 | 2 | ENSMUST00 | -2843 | Hs3st3b1 | ENSMUSG0( | false | chr11 | 63924926 | 63925627 | 9.86256 | 23.41135 | 21.03137 | chr11 | 63922597 | 63925544 | 5.42872 | 4.29883 | 3.35671 |
| chr10 | 61757988 | 61758293 | 654 | Intron (ENSMUST00000020283.4 | 10 | 61774737 | 61774821 | 85 | 1 | ENSMUST00 | -16444 | Mir5108 | ENSMUSG0( | false | . | . | . | . | . | . | chr10 | 61756253 | 61760614 | 5.95373 | 4.26924 | 3.83028 |
| chr6 | 67102336 | 67103128 | 654 | Distal Intergenic | 6 | 67115673 | 67168426 | 52754 | 1 | ENSMUST00 | -12545 | A430010J10I | ENSMUSG0( | false | . | . | . | . | . | . | chr6 | 67099839 | 67105551 | 5.37506 | 4.25145 | 3.31858 |
| chr1 | 43302214 | 43302553 | 654 | Promoter (<=1kb) | 1 | 43250156 | 43302925 | 52770 | 2 | ENSMUST00 | 372 | RP23-305F2( | ENSMUSG0( | false | . | . | . | . | . | . | chr1 | 43299283 | 43315017 | 5.27004 | 4.116 | 3.25173 |
| chr2 | 121413452 | 121413960 | 653 | Promoter (<=1kb) | 2 | 121413775 | 121438687 | 24913 | 1 | ENSMUST00 | 0 | Pdia3 | ENSMUSG0( | false | chr2 | 121413138 | 121415144 | 37.17318 | 134.81126 | 130.13722 | . | . | . | . | . | . |
| chr1 | 135132487 | 135132787 | 653 | Promoter (<=1kb) | 1 | 135132700 | 135144034 | 11335 | 1 | ENSMUST00 | 0 | Ptpn7 | ENSMUSG0( | false | . | . | . | . | . | . | chr1 | 135126075 | 135132880 | 5.21831 | 4.07933 | 3.22044 |
| chr4 | 43406139 | 43406652 | 653 | Promoter (<=1kb) | 4 | 43406435 | 43427088 | 20654 | 1 | ENSMUST00 | 0 | Rusc2 | ENSMUSG0( | false | chr4 | 43405677 | 43408085 | 15.42443 | 39.05124 | 36.47339 | chr4 | 43382882 | 43406168 | 6.80327 | 4.94439 | 4.48074 |
| chr8 | 41006801 | 41007079 | 653 | Intron (ENSMUST00000059115.7 | 8 | 40998770 | 41000205 | 1436 | 2 | ENSMUST00 | -6597 | Mtus1 | ENSMUSG0( | false | . | . | . | . | . | . | chr8 | 41003261 | 41019903 | 5.33155 | 4.20563 | 3.29685 |
| chr19 | 5662956 | 5663809 | 653 | Promoter (<=1kb) | 19 | 5651185 | 5663707 | 12523 | 2 | ENSMUST00 | 0 | Sipa1 | ENSMUSG0( | false | chr19 | 5662563 | 5664184 | 18.05357 | 46.55767 | 43.88821 | chr19 | 5661394 | 5664763 | 5.89159 | 4.49545 | 3.74751 |
| chr19 | 29014533 | 29014900 | 653 | Intron (ENSMUST00000050148.3 | 19 | 29003408 | 29003842 | 435 | 2 | ENSMUST00 | -10692 | Gm10136 | ENSMUSG0( | false | . | . | . | . | . | . | chr19 | 29014856 | 29017424 | 4.87891 | 4.07337 | 2.94904 |
| chrX | 12917148 | 12917426 | 653 | Promoter (2-3kb) | 20 | 12913910 | 12914592 | 683 | 2 | ENSMUST00 | -2557 | Gm14524 | ENSMUSG0( | false | chrX | 12916693 | 12917211 | 11.02877 | 22.59171 | 20.22099 | . | . | . | . | . | . |
| chr1 | 60655200 | 60655589 | 653 | Distal Intergenic | 1 | 60613698 | 60613803 | 106 | 2 | ENSMUST00 | -41398 | Gm23762 | ENSMUSG0( | false | . | . | . | . | . | . | . | . | . | . | . | . |
| chr1 | 74236878 | 74237272 | 652 | Promoter (<=1kb) | 1 | 74236550 | 74267988 | 31439 | 1 | ENSMUST00 | 329 | Arpc2 | ENSMUSG0( | false | chr1 | 74235907 | 74238302 | 34.54477 | 122.48211 | 118.25522 | . | . | . | . | . | . |
| chr14 | 63321949 | 63322326 | 652 | Distal Intergenic | 14 | 63241102 | 63271692 | 30591 | 2 | ENSMUST00 | -50258 | Gata4 | ENSMUSG0( | false | . | . | . | . | . | . | chr14 | 63320920 | 63323009 | 4.93935 | 3.97228 | 2.97109 |
| chr18 | 44419280 | 44419557 | 652 | 3' UTR | 18 | 44380500 | 44424969 | 44470 | 1 | ENSMUST00 | 38781 | Dcp2 | ENSMUSG0( | false | . | . | . | . | . | . | chr18 | 44417326 | 44422988 | 5.24723 | 4.15903 | 3.21397 |
| chr1 | 179546245 | 179546853 | 652 | Promoter (<=1kb) | 1 | 179528055 | 179546267 | 18213 | 2 | ENSMUST00 | 0 | Tfb2m | ENSMUSG0( | false | chr1 | 179545202 | 179547773 | 26.61085 | 108.21047 | 104.42454 | . | . | . | . | . | . |
| chr2 | 103641159 | 103641408 | 652 | Intron (ENSMUST00000076212.3 | 2 | 103715035 | 103717217 | 2183 | 1 | ENSMUST00 | -73627 | Abtb2 | ENSMUSG0( | false | . | . | . | . | . | . | chr2 | 103637261 | 103646135 | 5.16782 | 4.00674 | 3.1846 |
| chr11 | 80784264 | 80784644 | 652 | Distal Intergenic | 11 | 80583827 | 80780025 | 196199 | 2 | ENSMUST00 | -4240 | Myo1d | ENSMUSG0( | true | . | . | . | . | . | . | chr11 | 80783179 | 80785857 | 3.76677 | 3.46944 | 1.99832 |
| chr6 | 24514930 | 24515247 | 652 | Promoter (<=1kb) | 6 | 24444865 | 24515067 | 70203 | 2 | ENSMUST00 | 0 | Iqub | ENSMUSG0( | false | . | . | . | . | . | . | chr6 | 24514869 | 24515308 | 4.46971 | 3.76262 | 2.56873 |
| chr2 | 120977063 | 120977625 | 652 | Promoter (<=1kb) | 2 | 120977533 | 121007644 | 30112 | 1 | ENSMUST00 | 0 | Tmem62 | ENSMUSG0( | false | chr2 | 120976796 | 120978057 | 31.85236 | 101.09525 | 97.496 | . | . | . | . | . | . |
| chr5 | 150490833 | 150491124 | 652 | Exon (ENSMUST00000087204.5/ | 5 | 150509694 | 150511338 | 1645 | 2 | ENSMUST00 | 20214 | Gm15933 | ENSMUSG0( | false | . | . | . | . | . | . | chr5 | 150488839 | 150496363 | 4.67041 | 3.85736 | 2.7714 |
| chr15 | 76307164 | 76307575 | 651 | Promoter (<=1kb) | 15 | 76296603 | 76307245 | 10643 | 2 | ENSMUST00 | 0 | Oplah | ENSMUSG0( | false | chr15 | 76306973 | 76307820 | 7.90276 | 21.20842 | 18.85721 | chr15 | 76305490 | 76308728 | 3.72027 | 3.4182 | 1.94216 |
| chr16 | 42910802 | 42911176 | 651 | Intron (ENSMUST00000126354.2 | 16 | 42907652 | 43642602 | 734951 | 1 | ENSMUST00 | 3151 | Zbtb20 | ENSMUSG0( | false | chr16 | 42907033 | 42911476 | 14.29631 | 42.77194 | 40.14703 | . | . | . | . | . | . |
| chr19 | 4000250 | 4000743 | 651 | Promoter (<=1kb) | 19 | 4000580 | 4002103 | 1524 | 1 | ENSMUST00 | 0 | Nudt8 | ENSMUSG0( | false | chr19 | 3999790 | 4001275 | 33.91391 | 110.8507 | 106.99079 | chr19 | 3999335 | 4001748 | 5.6504 | 4.48488 | 3.52397 |
| chr18 | 77487903 | 77488141 | 651 | Intron (ENSMUST00000026494.8 | 18 | 77468321 | 77505117 | 36797 | 2 | ENSMUST00 | 16976 | Rnf165 | ENSMUSG0( | false | . | . | . | . | . | . | . | . | . | . | . | . |
| chr3 | 27698734 | 27699047 | 651 | Intron (ENSMUST00000046157.4 | 3 | 27416162 | 27710439 | 294278 | 2 | ENSMUST00 | 11392 | Fndc3b | ENSMUSG0( | true | . | . | . | . | . | . | chr3 | 27676543 | 27709401 | 5.6717 | 4.32901 | 3.57809 |
| chr17 | 62529961 | 62530302 | 651 | Distal Intergenic | 17 | 62456987 | 62457120 | 134 | 2 | ENSMUST00 | -72842 | Gm25800 | ENSMUSG0( | false | . | . | . | . | . | . | chr17 | 62529583 | 62531144 | 4.11635 | 3.62276 | 2.28703 |
| chr4 | 155246620 | 155246980 | 651 | Promoter (2-3kb) | 4 | 155249802 | 155256419 | 6618 | 1 | ENSMUST00 | -2822 | 2610002J02F | ENSMUSG0( | false | . | . | . | . | . | . | chr4 | 155245217 | 155247674 | 5.41926 | 4.12329 | 3.37054 |
| chr11 | 98809545 | 98810185 | 651 | Promoter (<=1kb) | 11 | 98797707 | 98810087 | 12381 | 2 | ENSMUST00 | 0 | Gm12359 | ENSMUSG0( | false | chr11 | 98809154 | 98810753 | 35.62435 | 116.99458 | 112.94973 | . | . | . | . | . | . |
| chr11 | 5505835 | 5506207 | 651 | Distal Intergenic | 11 | 5498840 | 5505699 | 6860 | 1 | ENSMUST00 | 6996 | Gm11963 | ENSMUSG0( | false | chr11 | 5505827 | 5506262 | 6.08969 | 9.87507 | 7.70114 | . | . | . | . | . | . |
| chr7 | 45124315 | 45124606 | 650 | Promoter (<=1kb) | 7 | 45122388 | 45124389 | 2002 | 2 | ENSMUST00 | 0 | Rps11 | ENSMUSG0( | false | chr7 | 45123273 | 45124400 | 28.90751 | 104.81161 | 101.11721 | chr7 | 45120978 | 45125056 | 4.41162 | 3.5835 | 2.557 |
| chr4 | 141303469 | 141303834 | 650 | Promoter (1-2kb) | 4 | 141302407 | 141308734 | 6328 | 1 | ENSMUST00 | 1063 | Epha2 | ENSMUSG0( | false | . | . | . | . | . | . | chr4 | 141302534 | 141316008 | 7.08958 | 4.8653 | 4.73204 |
| chr15 | 97720222 | 97720655 | 650 | Promoter (<=1kb) | 15 | 97711019 | 97721150 | 10132 | 2 | ENSMUST00 | 495 | Endou | ENSMUSG0( | false | . | . | . | . | . | . | chr15 | 97716040 | 97732730 | 6.4081 | 4.59676 | 4.19736 |
| chr18 | 31924357 | 31924658 | 650 | Distal Intergenic | 18 | 31931507 | 31958619 | 27113 | 1 | ENSMUST00 | -6849 | Lims2 | ENSMUSG0( | false | . | . | . | . | . | . | chr18 | 31918598 | 31931008 | 4.89771 | 3.99887 | 2.93822 |
| chr7 | 29309157 | 29309491 | 650 | Promoter (<=1kb) | 7 | 29309513 | 29313135 | 3623 | 1 | ENSMUST00 | -22 | Dpf1 | ENSMUSG0( | false | chr7 | 29309140 | 29309827 | 12.58327 | 33.30664 | 30.79935 | chr7 | 29307040 | 29309901 | 5.79105 | 4.36152 | 3.66111 |
| chr9 | 108249030 | 108249323 | 650 | Intron (ENSMUST00000166905.2 | 9 | 108205958 | 108263736 | 57779 | 2 | ENSMUST00 | 14413 | Dag1 | ENSMUSG0( | true | . | . | . | . | . | . | chr9 | 108210099 | 108251502 | 5.76916 | 4.35802 | 3.65688 |
| chr6 | 120997854 | 120998192 | 650 | Intron (ENSMUST00000098457.3 | 6 | 120931707 | 121003152 | 71446 | 2 | ENSMUST00 | 4960 | Mical3 | ENSMUSG0( | false | . | . | . | . | . | . | chr6 | 120994546 | 121005852 | 6.94439 | 4.93247 | 4.60074 |
| chr1 | 36691307 | 36691662 | 650 | Promoter (<=1kb) | 1 | 36691487 | 36693385 | 1899 | 1 | ENSMUST00 | 0 | Cox5b | ENSMUSG0( | false | chr1 | 36691042 | 36692379 | 24.39013 | 102.54827 | 98.91666 | chr1 | 36690961 | 36692752 | 6.40495 | 4.74181 | 4.21891 |
| chr15 | 75682040 | 75682337 | 649 | Distal Intergenic | 15 | 75657035 | 75678800 | 21766 | 2 | ENSMUST00 | -3241 | Top1mt | ENSMUSG0( | true | chr15 | 75677956 | 75683746 | 14.91789 | 45.43439 | 42.77634 | . | . | . | . | . | . |
| chr9 | 116175093 | 116175582 | 649 | Promoter (<=1kb) | 9 | 116087698 | 116175363 | 87666 | 2 | ENSMUST00 | 0 | Tgfbr2 | ENSMUSG0( | false | chr9 | 116174150 | 116176045 | 29.52499 | 116.10789 | 112.09218 | . | . | . | . | . | . |
| chr14 | 65374853 | 65375250 | 649 | Intron (ENSMUST00000066994.6 | 14 | 65358676 | 65398930 | 40255 | 1 | ENSMUST00 | 16178 | Zfp395 | ENSMUSG0( | false | . | . | . | . | . | . | chr14 | 65365022 | 65390419 | 5.65664 | 4.30592 | 3.56596 |
| chr4 | 108780374 | 108781033 | 649 | Promoter (<=1kb) | 4 | 108637466 | 108780503 | 143038 | 2 | ENSMUST00 | 0 | Zfyve9 | ENSMUSG0( | false | chr4 | 108779751 | 108781324 | 32.64849 | 123.19247 | 118.94151 | chr4 | 108781011 | 108781634 | 2.76843 | 2.71379 | 1.20936 |
| chr3 | 59266983 | 59267294 | 649 | Exon (ENSMUST00000040325.8/ | 3 | 59216271 | 59262831 | 46561 | 2 | ENSMUST00 | -4153 | P2ry12 | ENSMUSG0( | false | . | . | . | . | . | . | chr3 | 59267021 | 59268197 | 4.10974 | 3.58243 | 2.27816 |
| chr17 | 10676745 | 10677031 | 649 | Intron (ENSMUST00000041463.6 | 17 | 10576478 | 10631067 | 54590 | 2 | ENSMUST00 | -45679 | Pacrg | ENSMUSG0( | false | . | . | . | . | . | . | chr17 | 10673669 | 10678435 | 5.44605 | 4.27759 | 3.37178 |
| chr11 | 114851655 | 114851966 | 649 | Promoter (<=1kb) | 11 | 114851842 | 114872617 | 20776 | 1 | ENSMUST00 | 0 | Gprc5c | ENSMUSG0( | false | chr11 | 114850502 | 114853302 | 31.71022 | 94.86349 | 91.41748 | . | . | . | . | . | . |
| chr5 | 135442368 | 135442693 | 649 | Intron (ENSMUST00000060311.8 | 5 | 135376141 | 135394546 | 18406 | 2 | ENSMUST00 | -47823 | Pom121 | ENSMUSG0( | false | . | . | . | . | . | . | chr5 | 135441107 | 135444035 | 5.16195 | 4.03611 | 3.1489 |
| chr11 | 30649066 | 30649397 | 649 | Promoter (<=1kb) | 11 | 30505991 | 30649587 | 143597 | 2 | ENSMUST00 | 190 | Acyp2 | ENSMUSG0( | false | chr11 | 30648808 | 30649813 | 22.60491 | 86.65532 | 83.38163 | . | . | . | . | . | . |
| chr7 | 31111208 | 31111550 | 648 | Promoter (<=1kb) | 7 | 31110566 | 31111009 | 444 | 2 | ENSMUST00 | -200 | Hpn | ENSMUSG0( | false | . | . | . | . | . | . | chr7 | 31108643 | 31112914 | 3.88171 | 3.47007 | 2.1192 |
| chr10 | 12850864 | 12851299 | 648 | Intron (ENSMUST00000076817.4 | 10 | 12382188 | 12861735 | 479548 | 2 | ENSMUST00 | 10436 | Utrn | ENSMUSG0( | true | . | . | . | . | . | . | chr10 | 12848145 | 12859746 | 6.05157 | 4.50322 | 3.87576 |
| chr3 | 89898989 | 89899419 | 648 | Intron (ENSMUST00000029559.6 | 3 | 89869324 | 89913162 | 43839 | 2 | ENSMUST00 | 13743 | Il6ra | ENSMUSG0( | true | . | . | . | . | . | . | chr3 | 89888527 | 89911818 | 6.25567 | 4.6109 | 4.04691 |
| chr8 | 86884702 | 86885388 | 648 | Promoter (<=1kb) | 8 | 86841139 | 86885258 | 44120 | 2 | ENSMUST00 | 0 | N4bp1 | ENSMUSG0( | false | chr8 | 86883811 | 86885338 | 29.52499 | 116.10789 | 112.09218 | . | . | . | . | . | . |
| chr17 | 48816708 | 48816989 | 648 | Distal Intergenic | 17 | 48932379 | 49097588 | 165210 | 1 | ENSMUST00 | -115390 | Lrfn2 | ENSMUSG0( | false | . | . | . | . | . | . | chr17 | 48816186 | 48817100 | 3.29985 | 3.14959 | 1.66255 |
| chr15 | 37779488 | 37779791 | 648 | Intron (ENSMUST00000168992.2 | 15 | 37366175 | 37792000 | 425826 | 2 | ENSMUST00 | 12209 | Ncald | ENSMUSG0( | true | . | . | . | . | . | . | chr15 | 37779288 | 37780636 | 3.89095 | 3.34785 | 2.12582 |
| chr2 | 159631641 | 159632006 | 648 | Distal Intergenic | 2 | 159777676 | 159778207 | 532 | 2 | ENSMUST00 | 146201 | Gm11445 | ENSMUSG0( | false | . | . | . | . | . | . | chr2 | 159630586 | 159632882 | 4.87568 | 4.03084 | 2.94844 |
| chr13 | 29661379 | 29661652 | 648 | Intron (ENSMUST00000140278.2 | 13 | 29732942 | 29733036 | 95 | 1 | ENSMUST00 | -71290 | Gm23729 | ENSMUSG0( | false | . | . | . | . | . | . | chr13 | 29660335 | 29663800 | 3.05033 | 3.03101 | 1.45939 |
| chr9 | 102779429 | 102779792 | 648 | Distal Intergenic | 9 | 102771551 | 102771810 | 260 | 2 | ENSMUST00 | -7620 | RP23-274G2 | ENSMUSG0( | false | . | . | . | . | . | . | chr9 | 102774687 | 102784872 | 7.44631 | 5.21444 | 4.99811 |
| chr18 | 9449678 | 9450356 | 647 | Promoter (<=1kb) | 18 | 9314044 | 9450150 | 136107 | 2 | ENSMUST00 | 0 | Ccny | ENSMUSG0( | false | chr18 | 9448898 | 9450908 | 22.89105 | 88.16154 | 84.85722 | . | . | . | . | . | . |
| chr7 | 46618481 | 46618741 | 647 | Exon (ENSMUST00000033127.5/ | 7 | 46443474 | 46614788 | 171315 | 2 | ENSMUST00 | -3694 | Sergef | ENSMUSG0( | false | . | . | . | . | . | . | chr7 | 46618694 | 46619080 | 3.40804 | 3.30005 | 1.67529 |
| chr4 | 133074097 | 133074403 | 647 | Intron (ENSMUST00000105915.2 | 4 | 133066249 | 133072063 | 5815 | 1 | ENSMUST00 | 7849 | Ahdc1 | ENSMUSG0( | false | . | . | . | . | . | . | chr4 | 133072932 | 133080063 | 5.97164 | 4.45597 | 3.82083 |
| chr10 | 106159658 | 106159946 | 647 | Distal Intergenic | 10 | 106192378 | 106192770 | 393 | 2 | ENSMUST00 | 32824 | Rps15a-ps1 | ENSMUSG0( | false | . | . | . | . | . | . | . | . | . | . | . | . |
| chr11 | 33163030 | 33163396 | 647 | Promoter (<=1kb) | 11 | 33152287 | 33163206 | 10920 | 2 | ENSMUST00 | 0 | Npm1 | ENSMUSG0( | false | chr11 | 33161949 | 33163764 | 18.76905 | 75.7838 | 72.70259 | . | . | . | . | . | . |
| chr10 | 13300265 | 13300564 | 647 | Intron (ENSMUST00000105545.5 | 10 | 13207717 | 13324160 | 116444 | 2 | ENSMUST00 | 23596 | Phactr2 | ENSMUSG0( | false | chr10 | 13300162 | 13300733 | 5.6323 | 9.47728 | 7.31216 | chr10 | 13298126 | 13306713 | 5.43777 | 4.26805 | 3.36071 |
| chr1 | 152669730 | 152670127 | 647 | Intron (ENSMUST00000111859.2 | 1 | 152684026 | 152687501 | 3476 | 2 | ENSMUST00 | 17374 | Gm15479 | ENSMUSG0( | false | . | . | . | . | . | . | chr1 | 152668339 | 152670782 | 3.5528 | 3.35838 | 1.81649 |

| | | | | | | | | | | | | | | | | | | | | | | | | | | |
|---|---|---|---|---|---|---|---|---|---|---|---|---|---|---|---|---|---|---|---|---|---|---|---|---|---|---|
| chr17 | 5756878 | 5757286 | 647 | Distal Intergenic | 17 | 5798657 | 5802893 | 4237 | 1 | ENSMUST00 | -41371 | 3300005D01 | ENSMUSG0( | false | . | . | . | . | . | . | chr17 | 5755286 | 5757077 | 5.30714 | 4.09883 | 3.29051 |
| chr7 | 71188028 | 71188626 | 647 | Distal Intergenic | 7 | 71217049 | 71224409 | 7361 | 2 | ENSMUST00 | -28423 | RP23-257M2 | ENSMUSG0( | false | . | . | . | . | . | . | chr7 | 71187448 | 71189276 | 3.17063 | 3.09105 | 1.56113 |
| chr12 | 35063660 | 35063986 | 646 | Intron (ENSMUST00000048519.1 | 12 | 35047200 | 35134683 | 87484 | 1 | ENSMUST00 | 16461 | Snx13 | ENSMUSG0( | false | . | . | . | . | . | . | chr12 | 35059892 | 35068194 | 4.54516 | 3.79474 | 2.65632 |
| chr14 | 63122298 | 63122834 | 646 | Promoter (<=1kb) | 14 | 63122462 | 63145919 | 23458 | 1 | ENSMUST00 | 0 | Ctsb | ENSMUSG0( | false | chr14 | 63122106 | 63124349 | 28.49948 | 87.34749 | 84.05397 | . | . | . | . | . | . |
| chr18 | 67499186 | 67499492 | 646 | Intron (ENSMUST00000115050.2 | 18 | 67464849 | 67480581 | 15733 | 2 | ENSMUST00 | 34338 | Sfmo1 | ENSMUSG0( | false | . | . | . | . | . | . | chr18 | 67497228 | 67503404 | 5.50174 | 4.28432 | 3.4286 |
| chr5 | 37575197 | 37575506 | 646 | Intron (ENSMUST00000094836.4 | 5 | 37446828 | 37717153 | 270326 | 2 | ENSMUST00 | 141647 | Stk32b | ENSMUSG0( | false | . | . | . | . | . | . | chr5 | 37574280 | 37576488 | 4.72425 | 3.87412 | 2.78944 |
| chr13 | 103741172 | 103741445 | 646 | Downstream (<1kb) | 13 | 103741615 | 103764504 | 22890 | 2 | ENSMUST00 | 23059 | Srek1 | ENSMUSG0( | false | . | . | . | . | . | . | chr13 | 103741280 | 103741696 | 3.86618 | 3.42507 | 2.1161 |
| chr9 | 119444771 | 119445090 | 646 | Promoter (<=1kb) | 9 | 119444923 | 119465517 | 20595 | 1 | ENSMUST00 | 0 | Exog | ENSMUSG0( | false | chr9 | 119444581 | 119445858 | 28.48738 | 97.07422 | 93.57343 | chr9 | 119444666 | 119446600 | 3.71946 | 3.34181 | 1.99602 |
| chr11 | 32407917 | 32408226 | 646 | Promoter (<=1kb) | 11 | 32455370 | 32516687 | 61318 | 1 | ENSMUST00 | -47144 | Ubtd2 | ENSMUSG0( | false | . | . | . | . | . | . | chr11 | 32407459 | 32408595 | 3.78013 | 3.44509 | 2.0157 |
| chr11 | 53342860 | 53343091 | 646 | Promoter (2-3kb) | 11 | 53345290 | 53345677 | 388 | 2 | ENSMUST00 | 2586 | Gm11188 | ENSMUSG0( | false | . | . | . | . | . | . | chr11 | 53341291 | 53346716 | 4.65096 | 3.97609 | 2.7137 |
| chr14 | 25420704 | 25420982 | 645 | Distal Intergenic | 14 | 25459185 | 25538492 | 79308 | 1 | ENSMUST00 | -38203 | Zmiz1 | ENSMUSG0( | false | . | . | . | . | . | . | chr14 | 25420165 | 25423231 | 6.26838 | 4.71268 | 4.02624 |
| chr8 | 128357105 | 128357615 | 645 | Promoter (1-2kb) | 8 | 128359073 | 128505462 | 146390 | 1 | ENSMUST00 | -1458 | Nrp1 | ENSMUSG0( | false | chr8 | 128356451 | 128361175 | 30.41442 | 103.5799 | 99.9162 | chr8 | 128356055 | 128357829 | 4.89727 | 3.98226 | 2.91872 |
| chr6 | 137278838 | 137279136 | 645 | Intron (ENSMUST00000077115.7 | 6 | 137252299 | 137464629 | 212331 | 1 | ENSMUST00 | 26540 | Ptpro | ENSMUSG0( | true | . | . | . | . | . | . | chr6 | 137275709 | 137305131 | 5.12249 | 4.15603 | 3.10285 |
| chr11 | 67535007 | 67535267 | 645 | Intron (ENSMUST00000108681.2 | 11 | 67547154 | 67684974 | 137821 | 1 | ENSMUST00 | -11887 | Gas7 | ENSMUSG0( | true | . | . | . | . | . | . | chr11 | 67521726 | 67547234 | 6.38934 | 4.66296 | 4.16347 |
| chr5 | 108465170 | 108465448 | 645 | Intron (ENSMUST00000112597.2 | 5 | 108461313 | 108486171 | 24859 | 1 | ENSMUST00 | 3858 | Pcgf3 | ENSMUSG0( | false | . | . | . | . | . | . | chr5 | 108461909 | 108466572 | 5.35886 | 4.2105 | 3.29397 |
| chr18 | 80150754 | 80151564 | 645 | Promoter (<=1kb) | 18 | 80126311 | 80151482 | 25172 | 2 | ENSMUST00 | 0 | Adnp2 | ENSMUSG0( | false | chr18 | 80150219 | 80151770 | 22.71493 | 67.06055 | 64.11349 | . | . | . | . | . | . |
| chr12 | 80366015 | 80366486 | 645 | Promoter (<=1kb) | 12 | 80391613 | 80391751 | 139 | 2 | ENSMUST00 | 25265 | Scarna3b | ENSMUSG0( | false | . | . | . | . | . | . | chr12 | 80364452 | 80367653 | 5.52392 | 4.27825 | 3.42268 |
| chrX | 74274898 | 74275144 | 645 | Promoter (<=1kb) | 20 | 74274027 | 74274105 | 79 | 2 | ENSMUST00 | -794 | Mir7091 | ENSMUSG0( | false | . | . | . | . | . | . | . | . | . | . | . | . |
| chr12 | 79859599 | 79859988 | 645 | Distal Intergenic | 12 | 79963568 | 79963872 | 305 | 1 | ENSMUST00 | -103580 | RP23-179D3. | ENSMUSG0( | false | . | . | . | . | . | . | chr12 | 79854046 | 79861014 | 5.11326 | 4.07625 | 3.10953 |
| chr7 | 13037680 | 13038395 | 644 | Promoter (<=1kb) | 7 | 13035120 | 13038275 | 3156 | 2 | ENSMUST00 | 0 | Ube2m | ENSMUSG0( | false | chr7 | 13036853 | 13039040 | 35.07738 | 110.80895 | 106.9512 | . | . | . | . | . | . |
| chr9 | 62343189 | 62344191 | 644 | Promoter (<=1kb) | 9 | 62344063 | 62373596 | 29534 | 1 | ENSMUST00 | 0 | Anp32a | ENSMUSG0( | false | chr9 | 62340409 | 62345269 | 38.55812 | 129.64772 | 125.16568 | . | . | . | . | . | . |
| chr10 | 82629678 | 82630237 | 644 | Promoter (<=1kb) | 10 | 82629828 | 82650324 | 20497 | 1 | ENSMUST00 | 0 | Tdg | ENSMUSG0( | false | chr10 | 82629283 | 82630833 | 18.02578 | 59.25625 | 56.42029 | chr10 | 82628508 | 82631990 | 3.36145 | 3.08732 | 1.69655 |
| chr13 | 95640613 | 95640881 | 644 | Intron (ENSMUST00000068603.6 | 13 | 95601804 | 95618459 | 16656 | 2 | ENSMUST00 | -22155 | F2r | ENSMUSG0( | false | . | . | . | . | . | . | chr13 | 95637625 | 95641811 | 6.80292 | 4.91506 | 4.48347 |
| chr8 | 122283390 | 122283708 | 644 | Promoter (1-2kb) | 8 | 122282141 | 122337251 | 55111 | 1 | ENSMUST00 | 1250 | Zfpm1 | ENSMUSG0( | false | chr8 | 122282038 | 122285122 | 23.88927 | 69.11851 | 66.14216 | chr8 | 122282666 | 122285958 | 5.99671 | 4.57763 | 3.84085 |
| chr6 | 126615691 | 126616236 | 644 | Distal Intergenic | 6 | 126588296 | 126593986 | 5691 | 2 | ENSMUST00 | -21706 | Irf2bp2 | ENSMUSG0( | false | . | . | . | . | . | . | chr6 | 126614520 | 126616494 | 6.00559 | 4.59252 | 3.87504 |
| chr10 | 49131307 | 49131622 | 644 | Intron (ENSMUST00000105487.2 | 10 | 49101128 | 49103931 | 2804 | 2 | ENSMUST00 | -27377 | Grik2 | ENSMUSG0( | false | . | . | . | . | . | . | chr10 | 49130171 | 49131843 | 2.88843 | 2.86247 | 1.28035 |
| chr12 | 15850726 | 15850976 | 644 | Intron (ENSMUST00000180759.1 | 12 | 15817003 | 15888155 | 71153 | 1 | ENSMUST00 | 33724 | Gm5432 | ENSMUSG0( | false | . | . | . | . | . | . | chr12 | 15847592 | 15851469 | 7.16287 | 4.87096 | 4.79181 |
| chr12 | 15713709 | 15713979 | 644 | Distal Intergenic | 12 | 15728097 | 15729095 | 999 | 1 | ENSMUST00 | -14118 | Gm4804 | ENSMUSG0( | false | . | . | . | . | . | . | chr12 | 15710162 | 15714758 | 4.05059 | 3.57762 | 2.23385 |
| chr7 | 45488704 | 45488981 | 643 | Promoter (<=1kb) | 7 | 45472956 | 45488805 | 15850 | 2 | ENSMUST00 | 0 | Dhdh | ENSMUSG0( | false | chr7 | 45488439 | 45489284 | 17.93284 | 48.67312 | 45.97572 | chr7 | 45488148 | 45492330 | 4.69184 | 3.88087 | 2.77913 |
| chr6 | 28423412 | 28424192 | 643 | Promoter (<=1kb) | 6 | 28423560 | 28426602 | 3043 | 1 | ENSMUST00 | 0 | Arf5 | ENSMUSG0( | false | chr6 | 28423162 | 28424631 | 36.809 | 123.33047 | 119.05705 | . | . | . | . | . | . |
| chr13 | 100124628 | 100124967 | 643 | Promoter (<=1kb) | 13 | 100124852 | 100137690 | 12839 | 1 | ENSMUST00 | 0 | Smn1 | ENSMUSG0( | false | chr13 | 100124304 | 100125912 | 42.98468 | 146.27251 | 141.12608 | chr13 | 100124129 | 100126384 | 6.1813 | 4.45943 | 4.00981 |
| chr17 | 63644391 | 63644650 | 643 | Intron (ENSMUST00000097... | 17 | 63057452 | 63500017 | 442566 | 2 | ENSMUST00 | -144375 | Fbxl17 | ENSMUSG0( | false | . | . | . | . | . | . | . | . | . | . | . | . |
| chr3 | 27938584 | 27939589 | 643 | Promoter (<=1kb) | 3 | 27938695 | 28133239 | 194545 | 1 | ENSMUST00 | 0 | Pld1 | ENSMUSG0( | true | chr3 | 27938231 | 27939763 | 24.22294 | 75.10754 | 72.03808 | chr3 | 27939420 | 27944515 | 4.2906 | 3.75366 | 2.41084 |
| chr5 | 125164925 | 125165228 | 643 | Intron (ENSMUST00000111402.3 | 5 | 125017162 | 125179053 | 161892 | 2 | ENSMUST00 | 13825 | Ncor2 | ENSMUSG0( | false | . | . | . | . | . | . | chr5 | 125124978 | 125178451 | 6.70656 | 4.76008 | 4.42049 |
| chr4 | 59330906 | 59331288 | 643 | Promoter (1-2kb) | 4 | 59329210 | 59333022 | 3813 | 1 | ENSMUST00 | 1734 | Susd1 | ENSMUSG0( | false | . | . | . | . | . | . | chr4 | 59330566 | 59331977 | 3.90047 | 3.46992 | 2.13466 |
| chr7 | 114012339 | 114012606 | 643 | Intron (ENSMUST00000046687.1 | 7 | 114046785 | 114117781 | 70997 | 2 | ENSMUST00 | 105175 | Rras2 | ENSMUSG0( | true | . | . | . | . | . | . | chr7 | 114011725 | 114013604 | 4.80962 | 3.93258 | 2.86078 |
| chr14 | 13453903 | 13454155 | 643 | Promoter (<=1kb) | 14 | 13453958 | 13563045 | 109088 | 1 | ENSMUST00 | 0 | Synpr | ENSMUSG0( | false | . | . | . | . | . | . | chr14 | 13453433 | 13454174 | 3.42218 | 3.08864 | 1.74456 |
| chr19 | 46728140 | 46728658 | 642 | Intron (ENSMUST00000003655.7 | 19 | 46707443 | 46741095 | 33653 | 1 | ENSMUST00 | 20698 | As3mt | ENSMUSG0( | false | . | . | . | . | . | . | chr19 | 46726586 | 46729974 | 5.3015 | 4.19832 | 3.26817 |
| chr16 | 32430571 | 32431116 | 642 | Promoter (<=1kb) | 16 | 32430921 | 32474704 | 43784 | 1 | ENSMUST00 | 0 | Pcyt1a | ENSMUSG0( | false | chr16 | 32430226 | 32431794 | 32.98857 | 124.89146 | 120.57959 | . | . | . | . | . | . |
| chr10 | 122047095 | 122047962 | 642 | Promoter (<=1kb) | 10 | 121780991 | 122047308 | 266318 | 2 | ENSMUST00 | 0 | Srgap1 | ENSMUSG0( | true | chr10 | 122044483 | 122048243 | 31.51432 | 94.52584 | 91.08421 | . | . | . | . | . | . |
| chr10 | 110919560 | 110920482 | 642 | Promoter (<=1kb) | 10 | 110920177 | 110939599 | 19423 | 1 | ENSMUST00 | 0 | Csrp2 | ENSMUSG0( | false | chr10 | 110918934 | 110921076 | 35.27562 | 110.11647 | 106.27654 | chr10 | 110918734 | 110919554 | 4.48241 | 3.8018 | 2.57664 |
| chr1 | 118311041 | 118311610 | 642 | Promoter (<=1kb) | 1 | 118294778 | 118311416 | 16639 | 2 | ENSMUST00 | 0 | Tsn | ENSMUSG0( | false | chr1 | 118310257 | 118311782 | 33.47463 | 112.14942 | 108.24687 | . | . | . | . | . | . |
| chr2 | 132029737 | 132030358 | 642 | Promoter (<=1kb) | 2 | 131989415 | 132030001 | 40587 | 2 | ENSMUST00 | 0 | Rassf2 | ENSMUSG0( | false | chr2 | 132028950 | 132030583 | 21.02725 | 63.18829 | 60.29588 | chr2 | 132028350 | 132030731 | 4.13477 | 3.52674 | 2.32143 |
| chr5 | 106696035 | 106696910 | 642 | Promoter (<=1kb) | 5 | 106616739 | 106696830 | 80092 | 2 | ENSMUST00 | 0 | Zfp644 | ENSMUSG0( | false | chr5 | 106695254 | 106697207 | 33.08674 | 107.33151 | 103.57004 | . | . | . | . | . | . |
| chr2 | 152930135 | 152930592 | 642 | Promoter (2-3kb) | 2 | 152931898 | 152933208 | 1311 | 2 | ENSMUST00 | 2616 | Foxs1 | ENSMUSG0( | false | . | . | . | . | . | . | chr2 | 152925247 | 152934463 | 6.47975 | 4.78702 | 4.2189 |
| chr12 | 79904949 | 79905471 | 642 | Distal Intergenic | 12 | 79963568 | 79963872 | 305 | 1 | ENSMUST00 | -58097 | RP23-179D3. | ENSMUSG0( | false | . | . | . | . | . | . | chr12 | 79902506 | 79908120 | 7.24129 | 5.11539 | 4.82473 |
| chr14 | 26913753 | 26914048 | 641 | Intron (ENSMUST00000090337.5 | 14 | 26903243 | 26915258 | 12016 | 1 | ENSMUST00 | 10511 | Asb14 | ENSMUSG0( | false | . | . | . | . | . | . | chr14 | 26912945 | 26914788 | 4.42744 | 3.80766 | 2.58207 |
| chr2 | 4401093 | 4401573 | 641 | Promoter (<=1kb) | 2 | 4401437 | 4431364 | 29928 | 1 | ENSMUST00 | 0 | Frmd4a | ENSMUSG0( | false | chr2 | 4400894 | 4401782 | 22.60412 | 63.38673 | 60.49239 | chr2 | 4400871 | 4402579 | 3.7516 | 3.42883 | 2.02838 |
| chr1 | 170174836 | 170175303 | 641 | Promoter (<=1kb) | 1 | 170141938 | 170174957 | 33020 | 2 | ENSMUST00 | 0 | Uap1 | ENSMUSG0( | false | chr1 | 170174111 | 170175592 | 43.75897 | 147.62399 | 142.42812 | . | . | . | . | . | . |
| chr9 | 21510310 | 21510676 | 641 | Promoter (<=1kb) | 9 | 21507380 | 21510186 | 2807 | 2 | ENSMUST00 | -125 | Tmed1 | ENSMUSG0( | false | chr9 | 21509504 | 21511024 | 24.03115 | 75.67385 | 72.59326 | . | . | . | . | . | . |
| chr14 | 40966716 | 40967367 | 641 | Promoter (<=1kb) | 14 | 40906445 | 40966807 | 60363 | 2 | ENSMUST00 | 0 | Tspan14 | ENSMUSG0( | false | chr14 | 40965844 | 40966967 | 39.05061 | 143.74966 | 138.71384 | . | . | . | . | . | . |
| chr2 | 92058560 | 92058862 | 641 | Promoter (<=1kb) | 2 | 92059489 | 92059369 | 4381 | 2 | ENSMUST00 | 507 | 2900072N19 | ENSMUSG0( | false | chr2 | 92058378 | 92060209 | 25.1466 | 74.00616 | 70.95453 | chr2 | 92058334 | 92060500 | 3.57335 | 3.38199 | 1.83311 |
| chr19 | 47298758 | 47299132 | 641 | Intron (ENSMUST00000081619.3 | 19 | 47273519 | 47273589 | 71 | 2 | ENSMUST00 | -25170 | Mir6995 | ENSMUSG0( | false | . | . | . | . | . | . | chr19 | 47296867 | 47309513 | 5.65055 | 4.24598 | 3.57438 |
| chr14 | 76695799 | 76696263 | 641 | Distal Intergenic | 14 | 76681754 | 76686887 | 5134 | 1 | ENSMUST00 | 14046 | RP23-421C4 | ENSMUSG0( | false | . | . | . | . | . | . | chr14 | 76692388 | 76703896 | 4.91248 | 4.06724 | 2.9299 |
| chr10 | 99011147 | 99011417 | 640 | Intron (ENSMUST00000020107.7 | 10 | 99107036 | 99197988 | 90953 | 1 | ENSMUST00 | -95619 | Poc1b | ENSMUSG0( | false | . | . | . | . | . | . | chr10 | 99010636 | 99012739 | 3.55638 | 3.3779 | 1.81065 |
| chr3 | 107291042 | 107291442 | 640 | Promoter (<=1kb) | 3 | 107291230 | 107312115 | 20886 | 1 | ENSMUST00 | 0 | Slc16a4 | ENSMUSG0( | false | . | . | . | . | . | . | chr3 | 107290234 | 107291802 | 3.97015 | 3.57797 | 2.17284 |
| chr7 | 79202315 | 79202734 | 640 | Distal Intergenic | 7 | 79133769 | 79149060 | 15292 | 2 | ENSMUST00 | -53256 | Mfge8 | ENSMUSG0( | false | . | . | . | . | . | . | . | . | . | . | . | . |
| chr9 | 71741471 | 71741808 | 640 | Intron (ENSMUST00000072899.3 | 9 | 71629211 | 71771560 | 142350 | 2 | ENSMUST00 | 29752 | Cgnl1 | ENSMUSG0( | false | . | . | . | . | . | . | chr9 | 71736274 | 71770833 | 5.2398 | 4.14793 | 3.22386 |
| chr18 | 61911414 | 61911911 | 640 | Promoter (<=1kb) | 18 | 61799395 | 61911824 | 112430 | 2 | ENSMUST00 | 0 | Ablim3 | ENSMUSG0( | false | chr18 | 61910130 | 61911876 | 30.60796 | 113.09483 | 109.17188 | . | . | . | . | . | . |
| chr13 | 93771518 | 93771849 | 640 | Promoter (<=1kb) | 13 | 93771679 | 93943016 | 171338 | 1 | ENSMUST00 | 0 | Arsb | ENSMUSG0( | false | chr13 | 93771425 | 93772881 | 31.07893 | 124.33488 | 120.0426 | chr13 | 93771090 | 93771652 | 4.81333 | 4.09531 | 2.8396 |
| chr11 | 101952740 | 101953343 | 640 | Promoter (<=1kb) | 11 | 101952421 | 101960573 | 8153 | 1 | ENSMUST00 | 320 | 4930417O22 | ENSMUSG0( | false | . | . | . | . | . | . | chr11 | 101953062 | 101953829 | 3.4154 | 3.18413 | 1.73287 |
| chr11 | 88457314 | 88457555 | 640 | Intron (ENSMUST00000107909.2 | 11 | 88432986 | 88433514 | 529 | 2 | ENSMUST00 | -23801 | Gm11510 | ENSMUSG0( | false | . | . | . | . | . | . | chr11 | 88445933 | 88476041 | 6.2627 | 4.6354 | 4.05065 |
| chr4 | 83080399 | 83080725 | 640 | Promoter (1-2kb) | 4 | 83078511 | 83078629 | 119 | 2 | ENSMUST00 | 1889 | Gm24884 | ENSMUSG0( | false | . | . | . | . | . | . | chr4 | 83080073 | 83081204 | 4.12779 | 3.49414 | 2.32145 |
| chr8 | 94789312 | 94789594 | 639 | Distal Intergenic | 8 | 94772225 | 94780022 | 7798 | 1 | ENSMUST00 | 17088 | Cx3cl1 | ENSMUSG0( | false | . | . | . | . | . | . | chr8 | 94787298 | 94790345 | 5.18656 | 4.07326 | 3.17632 |
| chr5 | 120567248 | 120568023 | 639 | Intron (ENSMUST00000046426.8 | 5 | 120534160 | 120588613 | 54454 | 2 | ENSMUST00 | 20590 | Tpcn1 | ENSMUSG0( | false | . | . | . | . | . | . | chr5 | 120558690 | 120580640 | 7.14372 | 4.95764 | 4.77953 |
| chr6 | 88566164 | 88566514 | 639 | Intron (ENSMUST00000184878.2 | 6 | 88586528 | 88590659 | 4132 | 2 | ENSMUST00 | 24145 | Kbtbd12 | ENSMUSG0( | false | . | . | . | . | . | . | chr6 | 88565462 | 88567106 | 4.96487 | 4.05522 | 2.99947 |
| chr17 | 43758695 | 43759138 | 639 | Distal Intergenic | 17 | 43801851 | 44039516 | 237666 | 1 | ENSMUST00 | -42713 | Rcan2 | ENSMUSG0( | false | . | . | . | . | . | . | chr17 | 43756533 | 43759243 | 5.55568 | 4.19034 | 3.4957 |
| chr5 | 114585369 | 114585658 | 639 | Intron (ENSMUST00000043650.7 | 5 | 114568017 | 114594678 | 26662 | 1 | ENSMUST00 | 17353 | Fam222a | ENSMUSG0( | true | . | . | . | . | . | . | chr5 | 114571033 | 114614394 | 7.1045 | 4.9325 | 4.74958 |
| chr15 | 27785275 | 27785927 | 639 | Intron (ENSMUST00000090247.5 | 15 | 27655071 | 27681542 | 26472 | 1 | ENSMUST00 | -103734 | Fam105a | ENSMUSG0( | false | . | . | . | . | . | . | chr15 | 27777264 | 27789305 | 7.24061 | 5.11987 | 4.8476 |
| chr3 | 95016037 | 95016437 | 639 | Promoter (<=1kb) | 3 | 95016249 | 95020138 | 3890 | 1 | ENSMUST00 | 0 | 4930481B07 | ENSMUSG0( | false | chr3 | 95014242 | 95016985 | 30.91854 | 102.46954 | 98.8397 | . | . | . | . | . | . |
| chr9 | 56324503 | 56324892 | 639 | Promoter (<=1kb) | 9 | 56257890 | 56267780 | 9891 | 1 | ENSMUST00 | 66614 | Peak1os | ENSMUSG0( | false | chr9 | 56324531 | 56325132 | 17.64787 | 49.73517 | 47.02245 | . | . | . | . | . | . |
| chr18 | 61665491 | 61665950 | 639 | Promoter (<=1kb) | 18 | 61650974 | 61665537 | 14564 | 2 | ENSMUST00 | 0 | Mir143hg | ENSMUSG0( | false | . | . | . | . | . | . | chr18 | 61664779 | 61666842 | 5.41722 | 4.24945 | 3.35477 |
| chr4 | 143298909 | 143299280 | 638 | Promoter (<=1kb) | 4 | 143268346 | 143299463 | 31118 | 2 | ENSMUST00 | 183 | Pdpn | ENSMUSG0( | true | chr4 | 143297953 | 143299748 | 21.77823 | 66.26614 | 63.32996 | . | . | . | . | . | . |
| chr1 | 155972859 | 155973046 | 638 | Promoter (<=1kb) | 1 | 155844966 | 155972887 | 127922 | 2 | ENSMUST00 | 0 | Cep350 | ENSMUSG0( | false | chr1 | 155972112 | 155973264 | 28.61382 | 119.33878 | 115.2169 | . | . | . | . | . | . |
| chr17 | 45563725 | 45564432 | 638 | Promoter (<=1kb) | 17 | 45563964 | 45567669 | 3706 | 1 | ENSMUST00 | 0 | Slc35b2 | ENSMUSG0( | false | chr17 | 45563840 | 45564964 | 39.39634 | 133.30592 | 128.68834 | chr17 | 45563577 | 45571950 | 4.52176 | 3.76389 | 2.64295 |
| chr10 | 17947313 | 17948190 | 638 | Promoter (<=1kb) | 10 | 17900466 | 17948067 | 47602 | 2 | ENSMUST00 | 0 | Heca | ENSMUSG0( | false | chr10 | 17947067 | 17948455 | 34.79824 | 110.32398 | 106.47989 | . | . | . | . | . | . |
| chr10 | 127516525 | 127516841 | 638 | Promoter (1-2kb) | 10 | 127514939 | 127517152 | 2214 | 1 | ENSMUST00 | 1587 | Ndufa4l2 | ENSMUSG0( | false | . | . | . | . | . | . | chr10 | 127516420 | 127516995 | 4.42467 | 3.8346 | 2.52142 |
| chr14 | 76560874 | 76561191 | 638 | Distal Intergenic | 14 | 76532814 | 76556687 | 23874 | 2 | ENSMUST00 | -4188 | Serp2 | ENSMUSG0( | false | . | . | . | . | . | . | chr14 | 76560547 | 76561640 | 4.43358 | 3.60445 | 2.57203 |
| chr6 | 96044460 | 96044775 | 638 | Distal Intergenic | 6 | 96113154 | 96657198 | 544045 | 1 | ENSMUST00 | -68379 | Fam19a1 | ENSMUSG0( | false | . | . | . | . | . | . | . | . | . | . | . | . |
| chr17 | 26941144 | 26941480 | 638 | Promoter (<=1kb) | 17 | 26941452 | 26970541 | 29090 | 1 | ENSMUST00 | 0 | Syngap1 | ENSMUSG0( | false | . | . | . | . | . | . | chr17 | 26931803 | 26941775 | 5.54742 | 4.33953 | 3.45882 |
| chr1 | 157008540 | 157008843 | 638 | Intron (ENSMUST00000190681.1 | 1 | 156972760 | 157054034 | 81275 | 2 | ENSMUST00 | 45191 | RP24-329K2 | ENSMUSG0( | false | . | . | . | . | . | . | chr1 | 157008415 | 157008693 | 3.59141 | 3.29767 | 1.87303 |
| chr10 | 9881904 | 9882344 | 637 | Intron (ENSMUST00000038213.8 | 10 | 9755547 | 9900683 | 145137 | 2 | ENSMUST00 | 18339 | Stxbp5 | ENSMUSG0( | false | . | . | . | . | . | . | chr10 | 9879365 | 9883912 | 3.89164 | 3.55503 | 2.08267 |
| chr2 | 84714702 | 84715573 | 637 | Promoter (<=1kb) | 2 | 84687970 | 84715180 | 27211 | 2 | ENSMUST00 | 0 | Zdhhc5 | ENSMUSG0( | false | chr2 | 84714015 | 84716660 | 37.13757 | 117.51956 | 113.45509 | . | . | . | . | . | . |
| chr5 | 101954778 | 101955096 | 637 | Intron (ENSMUST00000053177.8 | 5 | 101833323 | 101943954 | 110632 | 2 | ENSMUST00 | -10825 | Wdfy3 | ENSMUSG0( | false | . | . | . | . | . | . | chr5 | 101951013 | 101959268 | 4.37028 | 3.78987 | 2.48323 |
| chr9 | 118928757 | 118929045 | 637 | Promoter (<=1kb) | 9 | 118929867 | 119033628 | 103762 | 1 | ENSMUST00 | -822 | Ctdspl | ENSMUSG0( | true | chr9 | 118925107 | 118929928 | 34.45293 | 106.63475 | 102.89172 | chr9 | 118927678 | 119010483 | 6.89652 | 4.89961 | 4.57102 |
| chr16 | 11908981 | 11909648 | 637 | Promoter (<=1kb) | 16 | 11803721 | 11909423 | 105703 | 2 | ENSMUST00 | 0 | Cpped1 | ENSMUSG0( | false | chr16 | 11908415 | 11909852 | 18.59898 | 66.17558 | 63.2431 | . | . | . | . | . | . |
| chr2 | 117019246 | 117019499 | 637 | Distal Intergenic | 2 | 117055566 | 117056012 | 447 | 1 | ENSMUST00 | -36067 | Gm13981 | ENSMUSG0( | false | . | . | . | . | . | . | chr2 | 117018472 | 117019610 | 2.75334 | 2.79887 | 1.21405 |
| chr12 | 30067849 | 30068134 | 637 | Intron (ENSMUST00000021005.9 | 12 | 30059657 | 30062113 | 2457 | 1 | ENSMUST00 | 8193 | Gm15691 | ENSMUSG0( | false | . | . | . | . | . | . | . | . | . | . | . | . |
| chr4 | 134439254 | 134439558 | 637 | Distal Intergenic | 4 | 134448766 | 134450171 | 1406 | 2 | ENSMUST00 | 10613 | 1700021N21 | ENSMUSG0( | false | . | . | . | . | . | . | chr4 | 134438538 | 134440151 | 5.47299 | 4.2971 | 3.39219 |
| chr5 | 151368411 | 151369025 | 637 | Promoter (<=1kb) | 5 | 151368675 | 151380912 | 12238 | 1 | ENSMUST00 | 0 | 1700028E10 | ENSMUSG0( | false | chr5 | 151367775 | 151368540 | 22.34262 | 62.9334 | 60.04675 | . | . | . | . | . | . |
| chr11 | 76481460 | 76481945 | 636 | Intron (ENSMUST00000094012.5 | 11 | 76464238 | 76493929 | 29692 | 2 | ENSMUST00 | 11984 | Abr | ENSMUSG0( | false | . | . | . | . | . | . | chr11 | 76479251 | 76484606 | 7.02531 | 5.09033 | 4.65222 |
| chr13 | 112403042 | 112403303 | 636 | Distal Intergenic | 13 | 112464070 | 112510086 | 46017 | 1 | ENSMUST00 | -60767 | Il6st | ENSMUSG0( | false | . | . | . | . | . | . | chr13 | 112402242 | 112404991 | 4.56279 | 3.90089 | 2.65969 |
| chr16 | 23107252 | 23107711 | 636 | Promoter (<=1kb) | 16 | 23107444 | 23113947 | 6504 | 1 | ENSMUST00 | 0 | Eif4a2 | ENSMUSG0( | false | chr16 | 23106844 | 23109861 | 28.10168 | 124.31538 | 120.02569 | . | . | . | . | . | . |
| chr1 | 161369261 | 161369521 | 636 | Distal Intergenic | 1 | 161395409 | 161418410 | 23002 | 1 | ENSMUST00 | -25888 | Tnfsf4 | ENSMUSG0( | false | . | . | . | . | . | . | . | . | . | . | . | . |
| chr10 | 75176484 | 75176865 | 636 | Intron (ENSMUST00000164107.1 | 10 | 75212073 | 75312399 | 100327 | 1 | ENSMUST00 | -35208 | Speccl1 | ENSMUSG0( | false | . | . | . | . | . | . | chr10 | 75176177 | 75177163 | 3.45884 | 3.21755 | 1.76959 |
| chr6 | 113623843 | 113624293 | 636 | Promoter (<=1kb) | 6 | 113623959 | 113631633 | 7675 | 1 | ENSMUST00 | 0 | Vhl | ENSMUSG0( | false | chr6 | 113623555 | 113624970 | 22.99841 | 82.87405 | 79.66897 | . | . | . | . | . | . |
| chr2 | 166995655 | 166996429 | 636 | Promoter (<=1kb) | 2 | 166947549 | 166996278 | 48730 | 2 | ENSMUST00 | 0 | Stau1 | ENSMUSG0( | false | chr2 | 166995030 | 166996893 | 41.07278 | 140.67711 | 135.76277 | chr2 | 166987850 | 166995720 | 4.99721 | 4.04688 | 3.01203 |
| chr2 | 14055693 | 14056244 | 636 | Promoter (<=1kb) | 2 | 13975563 | 14056020 | 80458 | 2 | ENSMUST00 | 0 | Ptpla | ENSMUSG0( | false | chr2 | 14054882 | 14056514 | 42.15588 | 144.81987 | 139.73779 | . | . | . | . | . | . |
| chr15 | 53885674 | 53885959 | 636 | Promoter (<=1kb) | 15 | 53884906 | 53886597 | 1692 | 1 | ENSMUST00 | 769 | Gm7489 | ENSMUSG0( | false | . | . | . | . | . | . | chr15 | 53884187 | 53888381 | 4.21608 | 3.7186 | 2.37322 |
| chr8 | 123253840 | 123254291 | 635 | Promoter (<=1kb) | 8 | 123242356 | 123254176 | 11821 | 2 | ENSMUST00 | 0 | Vps9d1 | ENSMUSG0( | false | chr8 | 123253289 | 123255503 | 39.10201 | 127.34201 | 122.95288 | . | . | . | . | . | . |
| chr16 | 18248633 | 18249056 | 635 | Promoter (<=1kb) | 16 | 18239784 | 18248732 | 8949 | 2 | ENSMUST00 | 0 | Ranbp1 | ENSMUSG0( | false | chr16 | 18247796 | 18249956 | 30.60796 | 113.09483 | 109.17188 | . | . | . | . | . | . |
| chr1 | 172447329 | 172447576 | 635 | Distal Intergenic | 1 | 172475358 | 172478575 | 3218 | 1 | ENSMUST00 | -27782 | Slamf9 | ENSMUSG0( | false | . | . | . | . | . | . | chr1 | 172446173 | 172447897 | 3.04786 | 3.04481 | 1.43638 |
| chr4 | 87978037 | 87978320 | 635 | Intron (ENSMUST00000078090.6 | 4 | 87837419 | 88032813 | 195395 | 2 | ENSMUST00 | 54493 | Mllt3 | ENSMUSG0( | false | . | . | . | . | . | . | chr4 | 87975469 | 87980293 | 4.24568 | 3.66385 | 2.41648 |
| chr5 | 137610567 | 137611000 | 635 | Promoter (<=1kb) | 5 | 137607434 | 137610702 | 3269 | 2 | ENSMUST00 | 0 | Pcolce | ENSMUSG0( | false | chr5 | 137610176 | 137611493 | 12.58308 | 29.40204 | 26.94344 | chr5 | 137608490 | 137613729 | 4.66714 | 3.91261 | 2.74881 |
| chr8 | 109705399 | 109706222 | 635 | Promoter (<=1kb) | 8 | 109705549 | 109724932 | 19384 | 1 | ENSMUST00 | 0 | Zfp821 | ENSMUSG0( | false | chr8 | 109705066 | 109706517 | 40.43793 | 136.23427 | 131.50453 | chr8 | 109706165 | 109707093 | 4.24496 | 3.63669 | 2.40523 |
| chr11 | 69599974 | 69600460 | 635 | Promoter (2-3kb) | 11 | 69600666 | 69602741 | 2076 | 2 | ENSMUST00 | 2281 | Atp1b2 | ENSMUSG0( | false | . | . | . | . | . | . | chr11 | 69599270 | 69602398 | 4.93086 | 3.88257 | 2.98001 |
| chr7 | 78623649 | 78623938 | 635 | Intron (ENSMUST00000... | 7 | 78578830 | 78581038 | 2209 | 1 | ENSMUST00 | 44820 | Gm9885 | ENSMUSG0( | false | . | . | . | . | . | . | chr7 | 78613393 | 78625373 | 6.15156 | 4.58262 | 3.94687 |
| chr12 | 33225348 | 33225579 | 635 | Intron (ENSMUST00000077456.7 | 12 | 33193994 | 33228854 | 34861 | 2 | ENSMUST00 | 3275 | Atxn7l1os1 | ENSMUSG0( | false | . | . | . | . | . | . | chr12 | 33224759 | 33227381 | 5.38006 | 4.33991 | 3.30579 |
| chr12 | 3866684 | 3866936 | 634 | Promoter (<=1kb) | 12 | 3866191 | 3876062 | 9872 | 1 | ENSMUST00 | 494 | Dnmt3a | ENSMUSG0( | false | . | . | . | . | . | . | chr12 | 3865258 | 3872731 | 5.39308 | 4.24242 | 3.33177 |
| chr14 | 34449789 | 34450117 | 634 | Intron (ENSMUST00000049005.9 | 14 | 34414486 | 34460556 | 46071 | 2 | ENSMUST00 | 10439 | Bmpr1a | ENSMUSG0( | true | . | . | . | . | . | . | chr14 | 34439321 | 34450550 | 4.3338 | 3.66849 | 2.47835 |
| chr17 | 10840002 | 10840602 | 634 | Promoter (<=1kb) | 17 | 10840384 | 11557234 | 716851 | 1 | ENSMUST00 | 0 | Park2 | ENSMUSG0( | false | chr17 | 10839745 | 10841170 | 33.75878 | 107.48165 | 103.71877 | . | . | . | . | . | . |
| chr9 | 56937171 | 56937494 | 634 | Promoter (<=1kb) | 9 | 56937475 | 56938398 | 924 | 1 | ENSMUST00 | 0 | Imp3 | ENSMUSG0( | false | chr9 | 56937002 | 56938607 | 32.00811 | 98.35596 | 94.82688 | . | . | . | . | . | . |
| chr9 | 75254164 | 75254645 | 634 | Intron (ENSMUST00000036555.6 | 9 | 75232014 | 75305451 | 73438 | 1 | ENSMUST00 | 22151 | Myo5c | ENSMUSG0( | false | . | . | . | . | . | . | chr9 | 75253978 | 75255012 | 2.84001 | 2.85516 | 1.30943 |
| chr7 | 113164728 | 113165003 | 634 | Distal Intergenic | 7 | 113207465 | 113314122 | 106658 | 1 | ENSMUST00 | -42462 | Arntl | ENSMUSG0( | false | . | . | . | . | . | . | . | . | . | . | . | . |
| chr7 | 27837768 | 27838085 | 634 | Distal Intergenic | 7 | 27842855 | 27855086 | 12232 | 1 | ENSMUST00 | -4770 | Zfp59 | ENSMUSG0( | false | chr7 | 27837846 | 27839317 | 21.92587 | 57.98755 | 55.16829 | chr7 | 27836895 | 27837943 | 4.19408 | 3.74372 | 2.33258 |
| chr10 | 24388259 | 24388752 | 634 | Distal Intergenic | 10 | 24447490 | 24452657 | 5168 | 1 | ENSMUST00 | -58738 | Gm15271 | ENSMUSG0( | false | . | . | . | . | . | . | chr10 | 24388007 | 24389279 | 3.59768 | 3.25969 | 1.87782 |
| chr8 | 18917438 | 18917710 | 634 | Distal Intergenic | 8 | 18932989 | 18933091 | 103 | 1 | ENSMUST00 | -15279 | Gm22551 | ENSMUSG0( | false | . | . | . | . | . | . | . | . | . | . | . | . |
| chr5 | 114647636 | 114647997 | 633 | Promoter (2-3kb) | 5 | 114622653 | 114644899 | 22247 | 2 | ENSMUST00 | -2738 | Trpv4 | ENSMUSG0( | false | chr5 | 114647134 | 114648009 | 10.5797 | 24.1096 | 21.71881 | chr5 | 114647099 | 114649465 | 5.4576 | 4.26766 | 3.38086 |
| chr16 | 23250264 | 23250746 | 633 | Intron (ENSMUST00000023601.8 | 16 | 23269629 | 23321179 | 51551 | 1 | ENSMUST00 | -18883 | St6gal1 | ENSMUSG0( | true | . | . | . | . | . | . | chr16 | 23227288 | 23275612 | 5.51887 | 4.26961 | 3.44483 |
| chr18 | 61558473 | 61558743 | 633 | Promoter (2-3kb) | 18 | 61555689 | 61587531 | 31843 | 1 | ENSMUST00 | 2785 | Csnk1a1 | ENSMUSG0( | true | chr18 | 61555228 | 61560694 | 42.15588 | 144.81987 | 139.73779 | chr18 | 61558329 | 61592224 | 4.88929 | 4.02454 | 2.91566 |
| chr15 | 94543118 | 94543621 | 633 | Promoter (<=1kb) | 15 | 94522688 | 94543547 | 20860 | 2 | ENSMUST00 | 0 | Pus7l | ENSMUSG0( | false | chr15 | 94543018 | 94544299 | 39.41886 | 127.89232 | 123.47375 | . | . | . | . | . | . |
| chrX | 106039506 | 106039748 | 633 | Intron (ENSMUST00000113557.2 | 20 | 106027322 | 106124926 | 97605 | 1 | ENSMUST00 | 12185 | Atp7a | ENSMUSG0( | false | . | . | . | . | . | . | . | . | . | . | . | . |
| chr18 | 35045675 | 35045941 | 633 | Distal Intergenic | 18 | 35118888 | 35254773 | 135886 | 1 | ENSMUST00 | -72947 | Ctnna1 | ENSMUSG0( | false | . | . | . | . | . | . | chr18 | 35044479 | 35046499 | 5.36329 | 4.14098 | 3.35294 |
| chr10 | 59114974 | 59115331 | 633 | Intron (ENSMUST00000153031.1 | 10 | 59080034 | 59104357 | 24324 | 2 | ENSMUST00 | 34941 | Sh3rf3 | ENSMUSG0( | false | . | . | . | . | . | . | chr10 | 59113571 | 59118291 | 4.92554 | 4.04794 | 2.96777 |
| chrX | 37052102 | 37052371 | 633 | Distal Intergenic | 20 | 37048845 | 37050418 | 1574 | 2 | ENSMUST00 | 3258 | Sowahd | ENSMUSG0( | false | . | . | . | . | . | . | . | . | . | . | . | . |
| chr2 | 169790529 | 169790983 | 633 | Exon (ENSMUST00000181446.2/ | 2 | 169884893 | 170071816 | 186924 | 2 | ENSMUST00 | -93910 | Tshz2 | ENSMUSG0( | true | chr2 | 169790304 | 169790965 | 10.10971 | 19.98882 | 17.65231 | chr2 | 169789952 | 169793453 | 4.12763 | 3.52202 | 2.32575 |
| chr11 | 98551045 | 98551478 | 632 | Promoter (<=1kb) | 11 | 98551097 | 98558665 | 7569 | 1 | ENSMUST00 | 0 | Zpbp2 | ENSMUSG0( | false | chr11 | 98550566 | 98551756 | 25.97375 | 74.52978 | 71.46892 | chr11 | 98550463 | 98551993 | 4.22482 | 3.61006 | 2.37754 |
| chr14 | 60705038 | 60705356 | 632 | Promoter (1-2kb) | 14 | 60706733 | 60750016 | 43284 | 1 | ENSMUST00 | -1377 | Spata13 | ENSMUSG0( | true | . | . | . | . | . | . | chr14 | 60700874 | 60708424 | 8.33344 | 5.60752 | 5.7266 |
| chr5 | 132190271 | 132190717 | 632 | Intron (ENSMUST00000161226.3 | 5 | 132030909 | 132050409 | 19501 | 2 | ENSMUST00 | -139863 | Auts2 | ENSMUSG0( | true | . | . | . | . | . | . | chr5 | 132185574 | 132192360 | 4.74485 | 3.91337 | 2.84391 |
| chr8 | 11819634 | 11819882 | 632 | Exon (ENSMUST00000033908.7/ | 8 | 11812362 | 11816345 | 3984 | 1 | ENSMUST00 | 7273 | Arhgef7 | ENSMUSG0( | false | . | . | . | . | . | . | chr8 | 11819167 | 11821092 | 4.83671 | 3.84422 | 2.90504 |
| chr9 | 92553448 | 92553690 | 632 | Intron (ENSMUST00000070522.8 | 9 | 92542264 | 92608420 | 66157 | 1 | ENSMUST00 | 11185 | Plod2 | ENSMUSG0( | true | . | . | . | . | . | . | chr9 | 92543215 | 92573465 | 5.80463 | 4.42577 | 3.66882 |
| chr18 | 83907856 | 83908142 | 632 | Distal Intergenic | 18 | 84012082 | 84086390 | 74309 | 1 | ENSMUST00 | 178248 | Tshz1 | ENSMUSG0( | false | . | . | . | . | . | . | chr18 | 83906990 | 83908775 | 4.1667 | 3.63011 | 2.35675 |
| chr8 | 24285017 | 24285426 | 632 | Distal Intergenic | 8 | 24173904 | 24185568 | 11665 | 1 | ENSMUST00 | 111114 | A730045E13 | ENSMUSG0( | false | . | . | . | . | . | . | . | . | . | . | . | . |
| chr6 | 100143794 | 100144161 | 632 | Distal Intergenic | 6 | 100228566 | 100287441 | 58876 | 2 | ENSMUST00 | 143280 | Rybp | ENSMUSG0( | false | . | . | . | . | . | . | chr6 | 100143351 | 100145454 | 3.01265 | 2.97005 | 1.41796 |
| chr3 | 143581993 | 143582289 | 632 | Distal Intergenic | 3 | 143674943 | 143675067 | 125 | 1 | ENSMUST00 | -92654 | Gm24728 | ENSMUSG0( | false | . | . | . | . | . | . | . | . | . | . | . | . |
| chr6 | 115134724 | 115135007 | 631 | Promoter (<=1kb) | 6 | 115134902 | 115276559 | 141658 | 1 | ENSMUST00 | 0 | Syn2 | ENSMUSG0( | false | chr6 | 115134572 | 115135637 | 31.84599 | 102.09332 | 98.46812 | chr6 | 115134443 | 115135741 | 5.4636 | 4.34333 | 3.39514 |
| chr13 | 46929615 | 46930361 | 631 | Promoter (<=1kb) | 13 | 46749087 | 46929718 | 180632 | 2 | ENSMUST00 | 0 | Kif13a | ENSMUSG0( | true | chr13 | 46928802 | 46931023 | 34.78613 | 113.42423 | 109.48744 | . | . | . | . | . | . |
| chr6 | 31398653 | 31399346 | 631 | Promoter (<=1kb) | 6 | 31398735 | 31516811 | 118077 | 1 | ENSMUST00 | 0 | Mkln1 | ENSMUSG0( | false | chr6 | 31398336 | 31399951 | 31.16539 | 106.97112 | 103.21663 | . | . | . | . | . | . |
| chr2 | 49746994 | 49747353 | 631 | Promoter (2-3kb) | 2 | 49744525 | 49757387 | 12863 | 1 | ENSMUST00 | 2470 | Kif5c | ENSMUSG0( | false | . | . | . | . | . | . | chr2 | 49746717 | 49747844 | 4.07154 | 3.56236 | 2.29063 |
| chr4 | 154964125 | 154964531 | 631 | Promoter (<=1kb) | 4 | 154964133 | 154980938 | 16806 | 1 | ENSMUST00 | 0 | Pank4 | ENSMUSG0( | false | chr4 | 154963727 | 154965130 | 31.96453 | 96.29042 | 92.80983 | chr4 | 154963271 | 154965875 | 5.01452 | 4.09461 | 3.03533 |
| chr2 | 135659036 | 135659608 | 631 | Promoter (<=1kb) | 2 | 135659554 | 135742838 | 83285 | 1 | ENSMUST00 | 0 | Plcb4 | ENSMUSG0( | false | chr2 | 135658344 | 135660234 | 29.92778 | 109.76675 | 105.9399 | . | . | . | . | . | . |
| chr11 | 88645218 | 88645664 | 631 | Intron (ENSMUST00000107909.2 | 11 | 88365515 | 88590147 | 224633 | 2 | ENSMUST00 | -55072 | Msi2 | ENSMUSG0( | false | . | . | . | . | . | . | chr11 | 88641230 | 88646639 | 5.43361 | 4.26997 | 3.38044 |
| chr6 | 124740831 | 124741193 | 631 | Promoter (<=1kb) | 6 | 124739186 | 124741065 | 1880 | 2 | ENSMUST00 | 0 | Grcc10 | ENSMUSG0( | false | chr6 | 124740221 | 124741831 | 18.64736 | 62.11404 | 59.23804 | . | . | . | . | . | . |
| chr2 | 23000342 | 23000583 | 631 | Intron (ENSMUST00000126112.2 | 2 | 23019131 | 23019667 | 537 | 2 | ENSMUST00 | 19084 | Gm13357 | ENSMUSG0( | false | . | . | . | . | . | . | chr2 | 22995356 | 23001418 | 5.6009 | 4.27899 | 3.52534 |
| chr1 | 177338733 | 177339052 | 630 | Distal Intergenic | 1 | 177022116 | 177258203 | 236088 | 2 | ENSMUST00 | -80531 | Akt3 | ENSMUSG0( | true | chr1 | 177338537 | 177339308 | 11.56301 | 29.49975 | 27.03959 | chr1 | 177338033 | 177339258 | 2.98471 | 2.9763 | 1.38684 |
| chr3 | 60504864 | 60505426 | 630 | Intron (ENSMUST00000099087.2 | 3 | 60501252 | 60629748 | 128497 | 1 | ENSMUST00 | 3613 | Mbnl1 | ENSMUSG0( | false | . | . | . | . | . | . | chr3 | 60502383 | 60508711 | 4.76066 | 3.9931 | 2.81281 |
| chr8 | 83332109 | 83332675 | 630 | Promoter (<=1kb) | 8 | 83332632 | 83332486 | 19855 | 2 | ENSMUST00 | 0 | Elmod2 | ENSMUSG0( | false | chr8 | 83331567 | 83333039 | 37.63617 | 126.94061 | 122.55849 | chr8 | 83331545 | 83333059 | 5.42755 | 4.27281 | 3.37895 |
| chr10 | 108331819 | 108332330 | 630 | Promoter (<=1kb) | 10 | 108332189 | 108414391 | 82203 | 1 | ENSMUST00 | 0 | Pawr | ENSMUSG0( | false | chr10 | 108331582 | 108333603 | 35.23322 | 112.11895 | 108.21771 | chr10 | 108331233 | 108331942 | 5.1686 | 4.1812 | 3.13036 |
| chr13 | 25055187 | 25055567 | 630 | Promoter (<=1kb) | 13 | 25056004 | 25121523 | 65520 | 1 | ENSMUST00 | -437 | Dcdc2a | ENSMUSG0( | true | chr13 | 25054929 | 25057285 | 27.2424 | 81.35376 | 78.17802 | chr13 | 25054710 | 25055453 | 3.37159 | 3.1988 | 1.69633 |
| chr2 | 34406693 | 34407056 | 630 | Promoter (<=1kb) | 2 | 34406771 | 34444408 | 37638 | 1 | ENSMUST00 | 0 | Mapkap1 | ENSMUSG0( | false | chr2 | 34406362 | 34407501 | 28.53699 | 95.19626 | 91.7378 | chr2 | 34407371 | 34409124 | 3.12059 | 3.02626 | 1.51468 |
| chr18 | 77938206 | 77938567 | 630 | Promoter (<=1kb) | 18 | 77938467 | 78035027 | 96561 | 1 | ENSMUST00 | 0 | Epg5 | ENSMUSG0( | false | chr18 | 77937879 | 77939097 | 46.75885 | 159.71181 | 154.07388 | chr18 | 77937746 | 77942543 | 5.10667 | 3.97993 | 3.10858 |
| chr3 | 149262456 | 149262942 | 630 | Distal Intergenic | 3 | 149221693 | 149225745 | 4053 | 2 | ENSMUST00 | -36712 | Gm10287 | ENSMUSG0( | false | . | . | . | . | . | . | . | . | . | . | . | . |
| chr3 | 153663947 | 153664264 | 630 | Intron (ENSMUST00000064460.5 | 3 | 153621702 | 153621969 | 268 | 1 | ENSMUST00 | 42246 | Gm23285 | ENSMUSG0( | false | . | . | . | . | . | . | chr3 | 153657583 | 153665932 | 6.56941 | 4.57081 | 4.3352 |
| chrX | 100777100 | 100777352 | 629 | Promoter (<=1kb) | 20 | 100777285 | 100818408 | 41124 | 1 | ENSMUST00 | 0 | Dlg3 | ENSMUSG0( | false | chrX | 100776682 | 100777626 | 19.75988 | 50.25065 | 47.53236 | . | . | . | . | . | . |
| chr9 | 107490365 | 107490632 | 629 | Intron (ENSMUST00000010210.7 | 9 | 107505604 | 107513534 | 7931 | 2 | ENSMUST00 | -14972 | Cacna2d2 | ENSMUSG0( | false | . | . | . | . | . | . | . | . | . | . | . | . |
| chr9 | 14627441 | 14627701 | 629 | Distal Intergenic | 9 | 14593444 | 14615483 | 22040 | 2 | ENSMUST00 | -11959 | Amotl1 | ENSMUSG0( | false | . | . | . | . | . | . | . | . | . | . | . | . |
| chr11 | 69088297 | 69088608 | 629 | Promoter (<=1kb) | 11 | 69088490 | 69092384 | 3895 | 1 | ENSMUST00 | 0 | Vamp2 | ENSMUSG0( | false | chr11 | 69088066 | 69090646 | 43.58744 | 151.86528 | 146.50356 | chr11 | 69084778 | 69088340 | 5.05808 | 4.11319 | 3.04009 |
| chr2 | 16394107 | 16394465 | 629 | Intron (ENSMUST00000028081.7 | 2 | 16357117 | 16751857 | 394741 | 1 | ENSMUST00 | 36991 | Plxdc2 | ENSMUSG0( | false | . | . | . | . | . | . | chr2 | 16389376 | 16401429 | 3.88696 | 3.46382 | 2.12715 |
| chr8 | 45658522 | 45658958 | 629 | Promoter (<=1kb) | 8 | 45656870 | 45776882 | 118213 | 2 | ENSMUST00 | 0 | Sorbs2 | ENSMUSG0( | false | chr8 | 45658667 | 45659023 | 4.61021 | 6.59091 | 4.49445 | . | . | . | . | . | . |
| chr12 | 54191978 | 54192337 | 629 | Intron (ENSMUST00000039516.3 | 12 | 54178981 | 54203860 | 24880 | 2 | ENSMUST00 | 15523 | Egln3 | ENSMUSG0( | false | chr12 | 54191776 | 54192456 | 8.39131 | 19.62543 | 17.29633 | chr12 | 54191357 | 54192854 | 3.41072 | 3.24003 | 1.72306 |
| chr11 | 133418223 | 133418596 | 629 | Intron (ENSMUST00000153799.2 | 11 | 133419802 | 133424377 | 42075 | 2 | ENSMUST00 | 5781 | Sox13 | ENSMUSG0( | false | chr11 | 133417877 | 133418807 | 18.02173 | 47.27326 | 44.5908 | chr11 | 133393415 | 133423224 | 5.89736 | 4.38424 | 3.76681 |
| chr2 | 75657911 | 75658208 | 629 | Promoter (1-2kb) | 2 | 75659244 | 75659317 | 74 | 2 | ENSMUST00 | -1036 | Gm24874 | ENSMUSG0( | false | chr2 | 75658153 | 75658982 | 17.13008 | 49.58303 | 46.87408 | chr2 | 75655322 | 75658472 | 3.85349 | 3.43807 | 2.09863 |
| chrX | 74281467 | 74281845 | 628 | Promoter (<=1kb) | 20 | 74273217 | 74281736 | 8520 | 2 | ENSMUST00 | 0 | Dnase1l1 | ENSMUSG0( | true | chrX | 74281767 | 74283502 | 18.84082 | 47.13427 | 44.45482 | . | . | . | . | . | . |
| chr4 | 134355852 | 134356226 | 628 | Downstream (<1kb) | 4 | 134362705 | 134365420 | 2720 | 2 | ENSMUST00 | 9188 | Extl1 | ENSMUSG0( | false | . | . | . | . | . | . | chr4 | 134353684 | 134357410 | 6.42867 | 4.71604 | 4.19315 |
| chr5 | 64234983 | 64235322 | 628 | Intron (ENSMUST00000043893.7 | 5 | 64230299 | 64351482 | 121184 | 1 | ENSMUST00 | 4685 | Tbc1d1 | ENSMUSG0( | false | . | . | . | . | . | . | chr5 | 64231749 | 64248662 | 6.01391 | 4.5094 | 3.84551 |
| chr5 | 66340730 | 66341203 | 628 | Intron (ENSMUST00000159786.2 | 5 | 66301888 | 66337665 | 35778 | 2 | ENSMUST00 | -3066 | Apbb2 | ENSMUSG0( | false | . | . | . | . | . | . | chr5 | 66334637 | 66346684 | 5.95013 | 4.37258 | 3.81282 |
| chr3 | 62379799 | 62380339 | 628 | Intron (ENSMUST00000079300.7 | 3 | 62341101 | 62380991 | 39891 | 2 | ENSMUST00 | 38699 | Arhgef26 | ENSMUSG0( | true | . | . | . | . | . | . | chr3 | 62379074 | 62382110 | 4.29319 | 3.75213 | 2.43025 |
| chr4 | 141536821 | 141537105 | 628 | Promoter (1-2kb) | 4 | 141521803 | 141538253 | 16451 | 2 | ENSMUST00 | 1148 | Spen | ENSMUSG0( | false | chr4 | 141535704 | 141538627 | 38.1391 | 127.82562 | 123.40861 | . | . | . | . | . | . |
| chr10 | 9675087 | 9675537 | 628 | Promoter (<=1kb) | 10 | 9627259 | 9675208 | 47950 | 2 | ENSMUST00 | 0 | Samd5 | ENSMUSG0( | false | chr10 | 9674014 | 9676226 | 24.48636 | 83.88964 | 80.66612 | chr10 | 9673181 | 9678971 | 3.88684 | 3.45899 | 2.10329 |
| chr4 | 57586862 | 57587151 | 628 | Intron (ENSMUST00000102905.2 | 4 | 57568330 | 57854584 | 286255 | 1 | ENSMUST00 | 18533 | Palm2Akap2 | ENSMUSG0( | false | . | . | . | . | . | . | chr4 | 57586116 | 57587514 | 3.21894 | 2.97197 | 1.57754 |
| chr3 | 97297126 | 97297423 | 628 | Promoter (<=1kb) | 3 | 97284908 | 97297778 | 12871 | 2 | ENSMUST00 | 355 | Gm17608 | ENSMUSG0( | false | chr3 | 97296446 | 97298749 | 21.79372 | 61.12524 | 58.26245 | . | . | . | . | . | . |
| chr13 | 109260653 | 109260597 | 627 | Intron (ENSMUST00000... | 13 | 109260654 | 109951364 | 690711 | 1 | ENSMUST00 | -57 | Pde4d | ENSMUSG0( | false | chr13 | 109260086 | 109261287 | 19.17718 | 50.04316 | 47.32813 | . | . | . | . | . | . |
| chr15 | 57829903 | 57830156 | 627 | Intron (ENSMUST00000096430.5 | 15 | 57869502 | 57892418 | 22917 | 2 | ENSMUST00 | 62262 | Derl1 | ENSMUSG0( | false | . | . | . | . | . | . | chr15 | 57828877 | 57831226 | 4.51052 | 3.7887 | 2.61907 |
| chr16 | 59639105 | 59639644 | 627 | Promoter (<=1kb) | 16 | 59612949 | 59639167 | 26219 | 2 | ENSMUST00 | 0 | Arl6 | ENSMUSG0( | false | chr16 | 59638561 | 59639962 | 32.66836 | 102.52041 | 98.88917 | . | . | . | . | . | . |
| chr5 | 114569740 | 114598029 | 627 | Intron (ENSMUST00000043650.7 | 5 | 114568017 | 114594678 | 26662 | 1 | ENSMUST00 | 29674 | Fam222a | ENSMUSG0( | true | . | . | . | . | . | . | chr5 | 114571033 | 114614394 | 7.1045 | 4.9325 | 4.74958 |
| chr14 | 51259757 | 51260052 | 627 | Distal Intergenic | 14 | 51255266 | 51256112 | 847 | 2 | ENSMUST00 | -3646 | Rnase2a | ENSMUSG0( | false | . | . | . | . | . | . | chr14 | 51259960 | 51262675 | 4.08514 | 3.56003 | 2.25602 |
| chr1 | 89588288 | 89588576 | 627 | Intron (ENSMUST00000027521.9 | 1 | 89580236 | 89895277 | 315042 | 1 | ENSMUST00 | 8053 | Agap1 | ENSMUSG0( | false | . | . | . | . | . | . | chr1 | 89564740 | 89589718 | 5.63127 | 4.22514 | 3.56367 |
| chrX | 13242787 | 13243042 | 627 | Intron (ENSMUST00000186931.1 | 20 | 13238327 | 13239553 | 1227 | 1 | ENSMUST00 | 4461 | Gm16481 | ENSMUSG0( | false | chrX | 13241516 | 13243626 | 11.02877 | 22.59171 | 20.22099 | . | . | . | . | . | . |
| chr5 | 73232649 | 73233014 | 627 | Intron (ENSMUST00000101127.6 | 5 | 73210109 | 73212634 | 2526 | 1 | ENSMUST00 | -20016 | Gm17207 | ENSMUSG0( | false | . | . | . | . | . | . | chr5 | 73196545 | 73254445 | 6.29302 | 4.62166 | 4.08669 |
| chr9 | 62783220 | 62783507 | 627 | 3' UTR | 9 | 62804478 | 62808234 | 3757 | 1 | ENSMUST00 | -20971 | Gm26609 | ENSMUSG0( | false | . | . | . | . | . | . | . | . | . | . | . | . |
| chr11 | 63131276 | 63131629 | 626 | Promoter (<=1kb) | 11 | 63131512 | 63159547 | 28036 | 1 | ENSMUST00 | 0 | Pmp22 | ENSMUSG0( | false | chr11 | 63130660 | 63133452 | 20.74539 | 66.97287 | 64.02668 | chr11 | 63129453 | 63136936 | 5.01478 | 3.91056 | 3.06639 |
| chr7 | 142470247 | 142470491 | 626 | Promoter (1-2kb) | 7 | 142471843 | 142494599 | 22757 | 1 | ENSMUST00 | -1352 | Lsp1 | ENSMUSG0( | false | . | . | . | . | . | . | chr7 | 142469601 | 142472837 | 7.10498 | 5.05047 | 4.74174 |
| chr11 | 8579639 | 8579972 | 626 | Intron (ENSMUST00000... | 11 | 8545693 | 8579121 | 33429 | 2 | ENSMUST00 | -519 | Tns3 | ENSMUSG0( | false | . | . | . | . | . | . | chr11 | 8578063 | 8582857 | 6.3882 | 4.68312 | 4.20067 |
| chr9 | 102446085 | 102446454 | 626 | Intron (ENSMUST00000... | 9 | 102505750 | 102546239 | 40490 | 1 | ENSMUST00 | -59296 | Ky | ENSMUSG0( | false | . | . | . | . | . | . | chr9 | 102445467 | 102447247 | 3.06351 | 2.90699 | 1.46111 |
| chr18 | 69517403 | 69517697 | 626 | Intron (ENSMUST00000066717.6 | 18 | 69521538 | 69564687 | 43150 | 1 | ENSMUST00 | -3841 | Tcf4 | ENSMUSG0( | false | . | . | . | . | . | . | chr18 | 69517175 | 69517724 | 6.04634 | 4.34248 | 3.91692 |
| chrX | 11797764 | 11798310 | 626 | Distal Intergenic | 20 | 11850484 | 11850590 | 107 | 1 | ENSMUST00 | -52174 | Gm26314 | ENSMUSG0( | false | . | . | . | . | . | . | . | . | . | . | . | . |
| chr6 | 121325846 | 121326138 | 626 | Promoter (<=1kb) | 6 | 121325580 | 121325676 | 97 | 2 | ENSMUST00 | -171 | Gm24855 | ENSMUSG0( | false | . | . | . | . | . | . | chr6 | 121325849 | 121327393 | 5.27972 | 4.18521 | 3.28196 |
| chr2 | 174049501 | 174049853 | 626 | Promoter (2-3kb) | 2 | 174046619 | 174047273 | 655 | 2 | ENSMUST00 | -2229 | Vamp7-ps | ENSMUSG0( | false | . | . | . | . | . | . | chr2 | 174048519 | 174050410 | 4.72306 | 3.85158 | 2.81522 |
| chrX | 99403435 | 99403680 | 626 | Downstream (1-2kb) | 20 | 99401977 | 99418895 | 9325 | 2 | ENSMUST00 | 15215 | Gm14809 | ENSMUSG0( | false | . | . | . | . | . | . | . | . | . | . | . | . |
| chr17 | 32117738 | 32118205 | 626 | Intron (ENSMUST00000180839.1 | 17 | 32113429 | 32141839 | 28411 | 1 | ENSMUST00 | 4310 | Gm17276 | ENSMUSG0( | false | . | . | . | . | . | . | chr17 | 32113274 | 32122911 | 6.15407 | 4.40853 | 3.96919 |
| chr11 | 84179363 | 84180029 | 625 | Promoter (<=1kb) | 11 | 84179831 | 84214195 | 34365 | 1 | ENSMUST00 | 0 | Acaca | ENSMUSG0( | false | chr11 | 84179012 | 84180621 | 32.09266 | 107.61995 | 103.85494 | . | . | . | . | . | . |
| chr15 | 81465950 | 81466469 | 625 | Promoter (<=1kb) | 15 | 81466316 | 81476369 | 10054 | 1 | ENSMUST00 | 0 | Rbx1 | ENSMUSG0( | false | chr15 | 81465557 | 81466212 | 16.54337 | 42.44649 | 39.82893 | . | . | . | . | . | . |
| chr2 | 27475543 | 27476268 | 625 | Promoter (<=1kb) | 2 | 27445579 | 27475686 | 30108 | 2 | ENSMUST00 | 0 | Brd3 | ENSMUSG0( | false | chr2 | 27474588 | 27476587 | 35.86567 | 122.72874 | 118.49439 | . | . | . | . | . | . |
| chr5 | 125081574 | 125082082 | 625 | Intron (ENSMUST00000111402.3 | 5 | 125030800 | 125058400 | 27601 | 1 | ENSMUST00 | -23175 | Ncor2 | ENSMUSG0( | false | . | . | . | . | . | . | chr5 | 125079756 | 125082652 | 7.59402 | 5.23827 | 5.11893 |
| chr6 | 82490626 | 82491128 | 625 | Intron (ENSMUST00000032122.8 | 6 | 82402475 | 82560103 | 157629 | 1 | ENSMUST00 | 88152 | Tacr1 | ENSMUSG0( | false | . | . | . | . | . | . | chr6 | 82490504 | 82491157 | 2.95239 | 2.96885 | 1.32142 |

| | | | | | | | | | | | | | | | | | | | | | | | | | | |
|---|---|---|---|---|---|---|---|---|---|---|---|---|---|---|---|---|---|---|---|---|---|---|---|---|---|---|
| chr18 | 80363430 | 80363835 | 625 | Promoter (<=1kb) | 18 | 80322944 | 80363650 | 40707 | 2 | ENSMUST00 | 0 | Kcng2 | ENSMUSG0C | false | chr18 | 80361986 | 80364088 | 30.03893 | 101.89226 | 98.2715 | chr18 | 80363365 | 80364690 | 3.80852 | 3.44827 | 2.03841 |
| chr6 | 47805668 | 47805978 | 625 | Promoter (2-3kb) | 6 | 47803576 | 47803652 | 77 | 2 | ENSMUST00 | -2017 | Mir704 | ENSMUSG0C | false | . | . | . | . | . | . | chr6 | 47801958 | 47812228 | 5.59318 | 4.2346 | 3.51569 |
| chr1 | 74587254 | 74587572 | 625 | Promoter (<=1kb) | 1 | 74586896 | 74586972 | 77 | 2 | ENSMUST00 | -283 | Gm25035 | ENSMUSG0C | false | chr1 | 74587078 | 74588189 | 33.1634 | 104.39787 | 100.72088 | chr1 | 74586767 | 74587420 | 4.90215 | 3.83888 | 2.95373 |
| chr10 | 67002524 | 67003070 | 625 | Distal Intergenic | 10 | 66920902 | 66923599 | 2698 | 1 | ENSMUST00 | 81623 | Gm26576 | ENSMUSG0C | false | chr10 | 67002504 | 67005089 | 14.70503 | 33.66069 | 31.14909 | chr10 | 67002337 | 67004855 | 4.24637 | 3.65752 | 2.41375 |
| chr13 | 34572078 | 34572329 | 624 | Distal Intergenic | 13 | 34628868 | 34648568 | 19701 | 2 | ENSMUST00 | 76239 | Pxdc1 | ENSMUSG0C | true | . | . | . | . | . | . | chr13 | 34569305 | 34576256 | 4.71058 | 3.88329 | 2.78127 |
| chr13 | 54302573 | 54303827 | 624 | Distal Intergenic | 13 | 54371349 | 54383917 | 12569 | 1 | ENSMUST00 | -67522 | Cplx2 | ENSMUSG0C | false | chr13 | 54302347 | 54304259 | 33.94791 | 109.87469 | 106.04007 | chr13 | 54302243 | 54302765 | 4.28317 | 3.62913 | 2.43988 |
| chr6 | 30401552 | 30402086 | 624 | Promoter (<=1kb) | 6 | 30401868 | 30453288 | 51421 | 1 | ENSMUST00 | 0 | Klhdc10 | ENSMUSG0C | false | chr6 | 30400851 | 30402755 | 34.04953 | 116.30843 | 112.28568 | . | . | . | . | . | . |
| chr7 | 133765541 | 133765843 | 624 | Intron (ENSMUST00000033290.6 | 7 | 133720946 | 133760269 | 39324 | 2 | ENSMUST00 | -5273 | Dhx32 | ENSMUSG0C | false | . | . | . | . | . | . | chr7 | 133764152 | 133766349 | 3.65546 | 3.08146 | 1.94236 |
| chr1 | 111864745 | 111865198 | 624 | Promoter (<=1kb) | 1 | 111858702 | 111864768 | 6067 | 2 | ENSMUST00 | 0 | Dsel | ENSMUSG0C | false | chr1 | 111864110 | 111865465 | 25.90858 | 83.69981 | 80.48042 | chr1 | 111864214 | 111865712 | 3.36169 | 3.25547 | 1.63799 |
| chr1 | 134556605 | 134557183 | 624 | Promoter (2-3kb) | 1 | 134560171 | 134635285 | 75115 | 1 | ENSMUST00 | -2988 | Kdm5b | ENSMUSG0C | false | chr1 | 134555703 | 134557642 | 19.89052 | 67.91473 | 64.9535 | chr1 | 134555094 | 134557561 | 3.91564 | 3.53609 | 2.12925 |
| chr2 | 75645403 | 75645684 | 624 | Distal Intergenic | 2 | 75632638 | 75633382 | 745 | 2 | ENSMUST00 | -12022 | Gm13655 | ENSMUSG0C | false | . | . | . | . | . | . | chr2 | 75642893 | 75646058 | 5.091 | 4.06002 | 3.09249 |
| chr2 | 114797316 | 114797687 | 624 | Distal Intergenic | 2 | 114854093 | 115043972 | 189880 | 1 | ENSMUST00 | -56406 | Gm13974 | ENSMUSG0C | false | . | . | . | . | . | . | chr2 | 114797127 | 114798346 | 3.88566 | 3.59597 | 2.07308 |
| chr18 | 5162124 | 5162844 | 624 | Promoter (2-3kb) | 18 | 5162880 | 5165731 | 2852 | 2 | ENSMUST00 | 2887 | Gm26682 | ENSMUSG0C | false | chr18 | 5162128 | 5163848 | 21.56542 | 57.37136 | 54.56027 | chr18 | 5161697 | 5163907 | 3.8984 | 3.55869 | 2.09054 |
| chr1 | 158248385 | 158248674 | 623 | Intron (ENSMUST00000004133.8 | 1 | 158245271 | 158356260 | 110990 | 2 | ENSMUST00 | 107586 | Brinp2 | ENSMUSG0C | false | . | . | . | . | . | . | chr1 | 158247946 | 158250762 | 4.52452 | 3.67229 | 2.65247 |
| chr13 | 109440378 | 109440671 | 623 | Promoter (1-2kb) | 13 | 109442183 | 109951364 | 509182 | 1 | ENSMUST00 | -1512 | Pde4d | ENSMUSG0C | false | chr13 | 109440167 | 109441959 | 21.40274 | 64.72344 | 61.81032 | . | . | . | . | . | . |
| chr7 | 145299421 | 145299964 | 623 | Promoter (<=1kb) | 7 | 145300828 | 145309557 | 8730 | 1 | ENSMUST00 | -864 | Mrgprf | ENSMUSG0C | false | chr7 | 145299716 | 145300691 | 11.33197 | 24.23275 | 21.84028 | chr7 | 145298750 | 145300232 | 6.66471 | 4.87953 | 4.35526 |
| chr9 | 95857464 | 95858005 | 623 | Promoter (<=1kb) | 9 | 95857597 | 95868312 | 10716 | 1 | ENSMUST00 | 0 | Atr | ENSMUSG0C | false | chr9 | 95857164 | 95858685 | 29.58769 | 108.11006 | 104.32592 | . | . | . | . | . | . |
| chr1 | 93444807 | 93445600 | 623 | Promoter (<=1kb) | 1 | 93437119 | 93445432 | 8314 | 2 | ENSMUST00 | 0 | Hdlbp | ENSMUSG0C | false | chr1 | 93444164 | 93446315 | 36.7977 | 133.03651 | 128.43381 | . | . | . | . | . | . |
| chr9 | 96983070 | 96983391 | 623 | Promoter (<=1kb) | 9 | 96943485 | 96984195 | 40711 | 2 | ENSMUST00 | 804 | Spsb4 | ENSMUSG0C | false | . | . | . | . | . | . | chr9 | 96975779 | 96994993 | 5.8258 | 4.40484 | 3.6945 |
| chr4 | 83390567 | 83390860 | 623 | Promoter (<=1kb) | 4 | 83378317 | 83390668 | 12352 | 2 | ENSMUST00 | 0 | Gm11413 | ENSMUSG0C | false | . | . | . | . | . | . | chr4 | 83389782 | 83390961 | 3.84421 | 3.43686 | 2.09319 |
| chr8 | 125224470 | 125224757 | 623 | Promoter (2-3kb) | 8 | 125224400 | 125227323 | 2924 | 2 | ENSMUST00 | 2566 | Gm16237 | ENSMUSG0C | false | . | . | . | . | . | . | chr8 | 125223606 | 125224683 | 3.863 | 3.49963 | 2.07083 |
| chr18 | 67583872 | 67584156 | 623 | Distal Intergenic | 18 | 67562387 | 67562851 | 465 | 1 | ENSMUST00 | 21486 | Gm17669 | ENSMUSG0C | false | . | . | . | . | . | . | chr18 | 67581152 | 67586750 | 4.84232 | 3.91546 | 2.89314 |
| chr11 | 58103852 | 58104403 | 622 | Promoter (<=1kb) | 11 | 58104153 | 58118594 | 14442 | 1 | ENSMUST00 | 0 | Cnot8 | ENSMUSG0C | false | chr11 | 58104039 | 58105114 | 38.13901 | 127.82562 | 123.40861 | . | . | . | . | . | . |
| chr13 | 63504407 | 63504764 | 622 | Distal Intergenic | 13 | 63347512 | 63497278 | 149767 | 2 | ENSMUST00 | -7130 | Fancc | ENSMUSG0C | false | . | . | . | . | . | . | chr13 | 63502607 | 63506934 | 4.62602 | 3.91183 | 2.71265 |
| chr13 | 66932778 | 66933549 | 622 | Promoter (<=1kb) | 13 | 66932830 | 66998401 | 65572 | 1 | ENSMUST00 | 0 | Ptdss1 | ENSMUSG0C | false | chr13 | 66932235 | 66934137 | 28.90751 | 104.81161 | 101.11721 | . | . | . | . | . | . |
| chr8 | 25962174 | 25962689 | 622 | Intron (ENSMUST00000037609.6 | 8 | 25944459 | 25976744 | 32286 | 2 | ENSMUST00 | 14055 | Hgsnat | ENSMUSG0C | false | . | . | . | . | . | . | chr8 | 25961491 | 25965675 | 6.4418 | 4.64993 | 4.22382 |
| chr6 | 72244334 | 72244684 | 622 | Distal Intergenic | 6 | 72206177 | 72235577 | 29401 | 2 | ENSMUST00 | -8758 | Atoh8 | ENSMUSG0C | false | . | . | . | . | . | . | chr6 | 72239454 | 72245860 | 6.57411 | 4.7317 | 4.34168 |
| chr5 | 114567175 | 114567497 | 622 | Promoter (<=1kb) | 5 | 114568016 | 114613220 | 45205 | 1 | ENSMUST00 | -519 | Fam222a | ENSMUSG0C | true | chr5 | 114567128 | 114570164 | 43.41453 | 148.15424 | 142.93486 | chr5 | 114566781 | 114568433 | 3.19402 | 3.08382 | 1.50672 |
| chr18 | 35814708 | 35815267 | 622 | Distal Intergenic | 18 | 35829818 | 35861687 | 31870 | 1 | ENSMUST00 | -14551 | Cxxc5 | ENSMUSG0C | false | chr18 | 35814542 | 35815564 | 10.17627 | 21.16558 | 18.81479 | chr18 | 35812838 | 35818671 | 7.77207 | 5.31683 | 5.31473 |
| chr4 | 83525335 | 83525657 | 622 | Promoter (<=1kb) | 4 | 83525545 | 83864670 | 339126 | 1 | ENSMUST00 | 0 | Ccdc171 | ENSMUSG0C | false | chr4 | 83524743 | 83526364 | 41.95543 | 136.66595 | 131.91884 | . | . | . | . | . | . |
| chr5 | 142228799 | 142229149 | 622 | Distal Intergenic | 5 | 142283106 | 142284069 | 964 | 1 | ENSMUST00 | -53957 | Gm26970 | ENSMUSG0C | false | . | . | . | . | . | . | chr5 | 142228406 | 142231122 | 3.89813 | 3.55848 | 2.08709 |
| chr1 | 12764425 | 12764860 | 621 | Intron (ENSMUST00000177608.2 | 1 | 12785443 | 12788818 | 3376 | 1 | ENSMUST00 | -20583 | Sulf1 | ENSMUSG0C | true | . | . | . | . | . | . | chr1 | 12752517 | 12771847 | 5.49437 | 4.27944 | 3.4172 |
| chr4 | 144948079 | 144948352 | 621 | Distal Intergenic | 4 | 144972623 | 144974683 | 2061 | 2 | ENSMUST00 | 26331 | Vps13d | ENSMUSG0C | false | . | . | . | . | . | . | chr4 | 144947700 | 144949709 | 2.9359 | 2.89189 | 1.38181 |
| chr18 | 3382898 | 3383415 | 621 | Promoter (<=1kb) | 18 | 3382988 | 3436374 | 53387 | 1 | ENSMUST00 | 0 | Cul2 | ENSMUSG0C | false | chr18 | 3382779 | 3384086 | 35.61502 | 123.34891 | 119.09213 | . | . | . | . | . | . |
| chr5 | 64294512 | 64294814 | 621 | Intron (ENSMUST00000043893.7 | 5 | 64298530 | 64350919 | 52390 | 1 | ENSMUST00 | -3716 | Tbc1d1 | ENSMUSG0C | false | . | . | . | . | . | . | chr5 | 64273894 | 64301350 | 6.13726 | 4.51007 | 3.95856 |
| chr8 | 70743389 | 70743716 | 621 | Promoter (<=1kb) | 8 | 70743079 | 70750473 | 7395 | 1 | ENSMUST00 | 311 | Pde4c | ENSMUSG0C | false | . | . | . | . | . | . | chr8 | 70740750 | 70748114 | 8.21466 | 5.47394 | 5.63599 |
| chr9 | 53323662 | 53323924 | 621 | Intron (ENSMUST00000051014.1 | 9 | 53341084 | 53368955 | 27872 | 1 | ENSMUST00 | -17160 | Exph5 | ENSMUSG0C | false | . | . | . | . | . | . | chr9 | 53323456 | 53326263 | 4.39417 | 3.619 | 2.54382 |
| chr4 | 45972145 | 45972853 | 621 | Promoter (<=1kb) | 4 | 45972233 | 46034761 | 62529 | 1 | ENSMUST00 | 0 | Tdrd7 | ENSMUSG0C | false | chr4 | 45971834 | 45973478 | 28.74323 | 86.81182 | 83.53605 | . | . | . | . | . | . |
| chr11 | 97050613 | 97050910 | 621 | Promoter (<=1kb) | 11 | 97050628 | 97068904 | 18277 | 1 | ENSMUST00 | 0 | Osbpl7 | ENSMUSG0C | false | chr11 | 97050565 | 97051738 | 21.37974 | 56.22882 | 53.43486 | chr11 | 97050180 | 97052977 | 4.90424 | 3.8571 | 2.97463 |
| chr2 | 92268227 | 92268533 | 621 | Intron (ENSMUST00000044036.9 | 2 | 92233105 | 92233437 | 333 | 1 | ENSMUST00 | 35123 | Gm13786 | ENSMUSG0C | false | . | . | . | . | . | . | chr2 | 92267270 | 92269553 | 3.52445 | 3.08285 | 1.83214 |
| chr17 | 6814673 | 6814975 | 621 | Intron (ENSMUST00000181895.1 | 17 | 6798390 | 6827990 | 29601 | 2 | ENSMUST00 | 13015 | Gm2885 | ENSMUSG0C | false | . | . | . | . | . | . | chr17 | 6811040 | 6816818 | 5.75765 | 4.4253 | 3.63624 |
| chr10 | 39631705 | 39632032 | 620 | Intron (ENSMUST00000019987.6 | 10 | 39641134 | 39645758 | 4625 | 1 | ENSMUST00 | -9102 | Traf3ip2 | ENSMUSG0C | false | . | . | . | . | . | . | chr10 | 39624031 | 39633595 | 5.77874 | 4.37306 | 3.67956 |
| chr13 | 55570923 | 55571171 | 620 | Promoter (<=1kb) | 13 | 55539319 | 55571120 | 31802 | 2 | ENSMUST00 | 0 | Fam193b | ENSMUSG0C | false | chr13 | 55569705 | 55571873 | 27.03867 | 103.18376 | 99.53125 | . | . | . | . | . | . |
| chr5 | 107986819 | 107987475 | 620 | Promoter (<=1kb) | 5 | 107908053 | 107987085 | 79033 | 2 | ENSMUST00 | 0 | Fam69a | ENSMUSG0C | false | chr5 | 107986196 | 107988211 | 27.54716 | 98.2761 | 94.74957 | . | . | . | . | . | . |
| chr4 | 8775031 | 8775418 | 620 | Intron (ENSMUST00000127476.2 | 4 | 8751438 | 8867224 | 115787 | 1 | ENSMUST00 | 23594 | Chd7 | ENSMUSG0C | false | . | . | . | . | . | . | chr4 | 8773919 | 8780117 | 5.69956 | 4.41813 | 3.58628 |
| chr13 | 43365485 | 43365840 | 620 | Distal Intergenic | 13 | 43370710 | 43395202 | 24493 | 1 | ENSMUST00 | -4870 | Sirt5 | ENSMUSG0C | false | chr13 | 43365238 | 43366264 | 26.22066 | 76.7888 | 73.6919 | chr13 | 43364919 | 43367243 | 3.71414 | 3.41676 | 2.00639 |
| chr17 | 84935645 | 84935967 | 620 | Distal Intergenic | 17 | 84956741 | 85023991 | 67251 | 1 | ENSMUST00 | -20774 | Ppm1b | ENSMUSG0C | false | . | . | . | . | . | . | chr17 | 84934594 | 84937369 | 4.75893 | 3.99041 | 2.80661 |
| chr4 | 55385367 | 55385614 | 620 | Exon (ENSMUST00000030134.8/ | 4 | 55392317 | 55392701 | 385 | 2 | ENSMUST00 | 7087 | Gm12516 | ENSMUSG0C | false | . | . | . | . | . | . | chr4 | 55384590 | 55385756 | 3.34005 | 3.26666 | 1.69822 |
| chr17 | 48721879 | 48722241 | 620 | Distal Intergenic | 17 | 48692450 | 48692707 | 258 | 2 | ENSMUST00 | -29173 | Gm23642 | ENSMUSG0C | false | . | . | . | . | . | . | chr17 | 48721520 | 48723583 | 4.16748 | 3.56403 | 2.34312 |
| chr19 | 11121066 | 11121428 | 620 | Promoter (<=1kb) | 19 | 11121510 | 11122425 | 916 | 1 | ENSMUST00 | -82 | RP24-512H1 | ENSMUSG0C | false | chr19 | 11120933 | 11121746 | 19.72513 | 62.50151 | 59.62064 | chr19 | 11121377 | 11121884 | 3.59178 | 3.399 | 1.84004 |
| chr1 | 165844407 | 165845255 | 619 | Intron (ENSMUST00000161971.2 | 1 | 165858216 | 165861555 | 3340 | 1 | ENSMUST00 | -12961 | Cd247 | ENSMUSG0C | false | chr1 | 165844405 | 165845742 | 18.02173 | 47.27326 | 44.5908 | chr1 | 165840591 | 165844460 | 4.11097 | 3.56425 | 2.28175 |
| chr18 | 10609512 | 10610411 | 619 | Promoter (<=1kb) | 18 | 10566611 | 10610129 | 43519 | 2 | ENSMUST00 | 0 | Esco1 | ENSMUSG0C | false | chr18 | 10608994 | 10610706 | 40.83545 | 136.9297 | 132.17467 | . | . | . | . | . | . |
| chr1 | 169531408 | 169531770 | 619 | Promoter (<=1kb) | 1 | 169497934 | 169531464 | 33531 | 2 | ENSMUST00 | 0 | Nuf2 | ENSMUSG0C | false | chr1 | 169531126 | 169531482 | 13.60354 | 37.21757 | 34.66072 | . | . | . | . | . | . |
| chr3 | 86222684 | 86222972 | 619 | Promoter (1-2kb) | 3 | 86224690 | 86782693 | 558004 | 1 | ENSMUST00 | -1718 | Lrba | ENSMUSG0C | false | . | . | . | . | . | . | chr3 | 86222268 | 86223264 | 4.33682 | 3.71521 | 2.48838 |
| chr4 | 74251762 | 74252160 | 619 | Promoter (<=1kb) | 4 | 74251936 | 74405860 | 153925 | 1 | ENSMUST00 | 0 | Kdm4c | ENSMUSG0C | false | chr4 | 74251001 | 74253232 | 32.98857 | 124.89146 | 120.57959 | . | . | . | . | . | . |
| chr1 | 74661624 | 74662102 | 619 | Promoter (<=1kb) | 1 | 74661745 | 74703730 | 41986 | 1 | ENSMUST00 | 0 | Ttll4 | ENSMUSG0C | false | chr1 | 74661680 | 74662863 | 29.52499 | 116.10789 | 112.09218 | . | . | . | . | . | . |
| chr8 | 25596984 | 25597725 | 619 | Promoter (<=1kb) | 8 | 25578492 | 25597487 | 18996 | 2 | ENSMUST00 | 0 | Letm2 | ENSMUSG0C | false | chr8 | 25596721 | 25597950 | 42.0266 | 146.8443 | 141.68535 | chr8 | 25596566 | 25599224 | 4.02404 | 3.62978 | 2.19597 |
| chr10 | 81411481 | 81412012 | 619 | Promoter (1-2kb) | 10 | 81409575 | 81411579 | 2005 | 1 | ENSMUST00 | 1907 | Gm16104 | ENSMUSG0C | false | . | . | . | . | . | . | chr10 | 81392649 | 81426205 | 8.72751 | 5.63067 | 6.0679 |
| chr9 | 104820263 | 104820483 | 619 | Intron (ENSMUST00000098443.3 | 9 | 104994916 | 104995019 | 104 | 1 | ENSMUST00 | -174433 | Gm23391 | ENSMUSG0C | false | . | . | . | . | . | . | . | . | . | . | . | . |
| chr2 | 9878455 | 9878889 | 618 | Promoter (<=1kb) | 2 | 9857078 | 9878600 | 21523 | 2 | ENSMUST00 | 0 | Gata3 | ENSMUSG0C | false | chr2 | 9878010 | 9879779 | 11.56301 | 29.49975 | 27.03959 | chr2 | 9876946 | 9884186 | 5.87686 | 4.47264 | 3.72602 |
| chr10 | 60814821 | 60815121 | 618 | Intron (ENSMUST00000077925.5 | 10 | 60762610 | 60831581 | 68972 | 2 | ENSMUST00 | 16460 | Unc5b | ENSMUSG0C | false | . | . | . | . | . | . | chr10 | 60811545 | 60822537 | 5.55309 | 4.2307 | 3.48027 |
| chr9 | 105121085 | 105121493 | 618 | Distal Intergenic | 9 | 105128903 | 105131312 | 2410 | 2 | ENSMUST00 | 9819 | Nudt16 | ENSMUSG0C | false | . | . | . | . | . | . | . | . | . | . | . | . |
| chr11 | 80779782 | 80780284 | 618 | Promoter (<=1kb) | 11 | 80482126 | 80780025 | 297900 | 2 | ENSMUST00 | 0 | Myo1d | ENSMUSG0C | true | chr11 | 80778230 | 80780044 | 28.91247 | 96.86193 | 93.36482 | . | . | . | . | . | . |
| chr5 | 90234625 | 90234905 | 618 | Intron (ENSMUST00000014421.9 | 5 | 90214724 | 90224001 | 9278 | 2 | ENSMUST00 | -10625 | Cox18 | ENSMUSG0C | false | . | . | . | . | . | . | chr5 | 90233192 | 90242152 | 4.50253 | 3.82528 | 2.59714 |
| chr9 | 106406458 | 106406896 | 618 | Distal Intergenic | 9 | 106429454 | 106431568 | 2115 | 1 | ENSMUST00 | -22558 | Rpl29 | ENSMUSG0C | false | . | . | . | . | . | . | chr9 | 106405125 | 106407317 | 4.64082 | 3.87488 | 2.72717 |
| chr2 | 120102918 | 120103220 | 618 | Intron (ENSMUST00000028755.7 | 2 | 120076214 | 120085772 | 9559 | 2 | ENSMUST00 | -17147 | Sptbn5 | ENSMUSG0C | false | . | . | . | . | . | . | chr2 | 120090578 | 120107323 | 5.14183 | 4.07311 | 3.14279 |
| chr14 | 97877678 | 97877971 | 618 | Intron (ENSMUST00000069334.7 | 14 | 97967829 | 97967991 | 163 | 2 | ENSMUST00 | 90020 | Gm25831 | ENSMUSG0C | false | . | . | . | . | . | . | chr14 | 97877036 | 97880925 | 3.73266 | 3.49018 | 1.95289 |
| chr1 | 7139196 | 7139504 | 618 | Promoter (2-3kb) | 1 | 7136199 | 7136586 | 388 | 2 | ENSMUST00 | -2611 | Gm26983 | ENSMUSG0C | false | chr1 | 7139266 | 7140156 | 6.57187 | 13.44444 | 11.20722 | chr1 | 7136769 | 7142269 | 5.12786 | 4.1036 | 3.11274 |
| chr17 | 33523926 | 33524312 | 617 | Promoter (<=1kb) | 17 | 33524204 | 33553768 | 29565 | 1 | ENSMUST00 | 0 | Adamts10 | ENSMUSG0C | false | chr17 | 33523880 | 33524638 | 21.40274 | 64.72344 | 61.81032 | chr17 | 33523267 | 33525445 | 4.06655 | 3.63168 | 2.25474 |
| chr4 | 143278566 | 143278923 | 617 | Intron (ENSMUST00000030317.8 | 4 | 143268346 | 143299463 | 31118 | 2 | ENSMUST00 | 20540 | Pdpn | ENSMUSG0C | true | . | . | . | . | . | . | chr4 | 143277480 | 143296542 | 5.36947 | 4.22359 | 3.31454 |
| chr9 | 69125378 | 69125876 | 617 | Intron (ENSMUST00000034766.8 | 9 | 69195541 | 69250834 | 55294 | 1 | ENSMUST00 | -69665 | Rora | ENSMUSG0C | false | chr9 | 69125223 | 69126025 | 6.80177 | 13.42905 | 11.19262 | chr9 | 69123940 | 69126708 | 3.56229 | 3.37802 | 1.81547 |
| chr15 | 99221646 | 99222030 | 617 | Promoter (2-3kb) | 15 | 99224976 | 99242817 | 17842 | 1 | ENSMUST00 | -2946 | Kcnh3 | ENSMUSG0C | false | . | . | . | . | . | . | chr15 | 99221254 | 99225886 | 4.91003 | 3.88376 | 2.96688 |
| chr1 | 79668572 | 79668848 | 617 | Intron (ENSMUST00000162342.2 | 1 | 79608845 | 79671965 | 63121 | 2 | ENSMUST00 | 3117 | Ap1s3 | ENSMUSG0C | false | . | . | . | . | . | . | chr1 | 79666523 | 79672410 | 4.5349 | 3.86218 | 2.62318 |
| chr3 | 89962967 | 89963526 | 617 | Promoter (<=1kb) | 3 | 89939481 | 89963508 | 24028 | 2 | ENSMUST00 | 0 | Atp8b2 | ENSMUSG0C | false | chr3 | 89962488 | 89964514 | 29.50911 | 91.04776 | 87.68056 | chr3 | 89958714 | 89964784 | 5.0449 | 3.96997 | 3.08253 |
| chr2 | 167856896 | 167857279 | 617 | Promoter (1-2kb) | 2 | 167858585 | 167862542 | 3958 | 1 | ENSMUST00 | -1306 | Gm14319 | ENSMUSG0C | false | . | . | . | . | . | . | chr2 | 167855418 | 167869855 | 6.99032 | 4.95783 | 4.63791 |
| chr1 | 64041189 | 64041487 | 617 | Promoter (1-2kb) | 1 | 64042438 | 64042501 | 64 | 2 | ENSMUST00 | 1014 | Mir6899 | ENSMUSG0C | false | . | . | . | . | . | . | chr1 | 64034781 | 64045346 | 5.3773 | 4.27595 | 3.31604 |
| chr5 | 111262884 | 111263410 | 617 | Intron (ENSMUST00000040111.9 | 5 | 111274984 | 111275094 | 111 | 2 | ENSMUST00 | 11684 | Gm27602 | ENSMUSG0C | false | . | . | . | . | . | . | chr5 | 111261927 | 111264834 | 5.91099 | 4.52138 | 3.73757 |
| chr16 | 93272644 | 93272982 | 617 | Distal Intergenic | 16 | 93343724 | 93353189 | 9466 | 2 | ENSMUST00 | 80207 | RP24-260I8.2 | ENSMUSG0C | false | . | . | . | . | . | . | . | . | . | . | . | . |
| chr11 | 95079198 | 95079989 | 616 | Promoter (2-3kb) | 11 | 95044474 | 95076801 | 32328 | 2 | ENSMUST00 | -2398 | Itga3 | ENSMUSG0C | true | . | . | . | . | . | . | chr11 | 95077539 | 95082670 | 7.743 | 5.23471 | 5.25962 |
| chr2 | 102936982 | 102937250 | 616 | Distal Intergenic | 2 | 102811141 | 102901665 | 90525 | 2 | ENSMUST00 | -35318 | Cd44 | ENSMUSG0C | false | . | . | . | . | . | . | . | . | . | . | . | . |
| chr11 | 8503717 | 8504015 | 616 | Intron (ENSMUST00000020695.7 | 11 | 8451070 | 8468675 | 17606 | 2 | ENSMUST00 | -35043 | Tns3 | ENSMUSG0C | false | chr11 | 8499137 | 8504359 | 19.17126 | 69.02763 | 66.05415 | . | . | . | . | . | . |
| chr13 | 51573926 | 51574245 | 616 | Distal Intergenic | 13 | 51431041 | 51567084 | 136044 | 2 | ENSMUST00 | -6843 | Shc3 | ENSMUSG0C | false | . | . | . | . | . | . | . | . | . | . | . | . |
| chr13 | 85288889 | 85289738 | 616 | Promoter (<=1kb) | 13 | 85214780 | 85289029 | 74250 | 2 | ENSMUST00 | 0 | Rasa1 | ENSMUSG0C | false | chr13 | 85287580 | 85289440 | 35.06139 | 127.69727 | 123.29834 | . | . | . | . | . | . |
| chr11 | 96826509 | 96826940 | 616 | Promoter (<=1kb) | 11 | 96817599 | 96827719 | 10121 | 2 | ENSMUST00 | 779 | Nfe2l1 | ENSMUSG0C | false | chr11 | 96826093 | 96826719 | 8.34762 | 15.59896 | 13.32767 | chr11 | 96821899 | 96829042 | 7.7991 | 5.17723 | 5.30668 |
| chr10 | 9648350 | 9648714 | 616 | Intron (ENSMUST00000100070.2 | 10 | 9627259 | 9675208 | 47950 | 2 | ENSMUST00 | 26494 | Samd5 | ENSMUSG0C | false | . | . | . | . | . | . | chr10 | 9646052 | 9652313 | 4.83709 | 3.95568 | 2.88396 |
| chr7 | 65734309 | 65734638 | 616 | Distal Intergenic | 7 | 65693467 | 65697879 | 4413 | 1 | ENSMUST00 | 40843 | Tm2d3 | ENSMUSG0C | false | . | . | . | . | . | . | . | . | . | . | . | . |
| chrX | 48314626 | 48315038 | 616 | Distal Intergenic | 20 | 48307043 | 48307143 | 101 | 1 | ENSMUST00 | 7584 | Gm22612 | ENSMUSG0C | false | chrX | 48314253 | 48315184 | 11.94783 | 25.26529 | 22.85895 | . | . | . | . | . | . |
| chr11 | 75587959 | 75588462 | 615 | Promoter (<=1kb) | 11 | 75588097 | 75628804 | 40708 | 1 | ENSMUST00 | 0 | Pitpna | ENSMUSG0C | false | chr11 | 75587816 | 75589321 | 29.2476 | 106.45832 | 102.71833 | chr11 | 75587431 | 75591589 | 4.0768 | 3.61083 | 2.26702 |
| chr15 | 98607671 | 98608196 | 615 | Promoter (<=1kb) | 15 | 98589994 | 98607633 | 17640 | 2 | ENSMUST00 | -39 | Adcy6 | ENSMUSG0C | false | . | . | . | . | . | . | chr15 | 98606507 | 98610285 | 4.43144 | 3.88063 | 2.53251 |
| chr1 | 37299660 | 37300053 | 615 | Promoter (<=1kb) | 1 | 37299865 | 37348512 | 48648 | 1 | ENSMUST00 | 0 | Inpp4a | ENSMUSG0C | true | chr1 | 37299838 | 37301548 | 31.70051 | 127.6562 | 123.25787 | chr1 | 37299104 | 37299760 | 2.87612 | 2.92613 | 1.28614 |
| chr7 | 113820716 | 113821001 | 615 | Intron (ENSMUST00000046687.1 | 7 | 113766174 | 113929876 | 163703 | 1 | ENSMUST00 | 54543 | Spon1 | ENSMUSG0C | false | . | . | . | . | . | . | chr7 | 113817327 | 113823588 | 3.67482 | 3.30434 | 1.93887 |
| chr14 | 57428741 | 57429010 | 615 | Intron (ENSMUST00000122063.2 | 14 | 57424106 | 57517936 | 93831 | 1 | ENSMUST00 | 4636 | Ift88 | ENSMUSG0C | false | . | . | . | . | . | . | chr14 | 57428134 | 57429215 | 3.36268 | 3.28136 | 1.64037 |
| chr8 | 120191152 | 120191433 | 615 | Intron (ENSMUST00000127664.1 | 8 | 120166397 | 120177466 | 11070 | 2 | ENSMUST00 | -13687 | Fam92b | ENSMUSG0C | false | chr8 | 120191035 | 120191417 | 8.80131 | 17.30502 | 15.00832 | chr8 | 120189878 | 120191754 | 5.29039 | 4.20488 | 3.24244 |
| chr15 | 36375459 | 36375868 | 615 | Distal Intergenic | 15 | 36308144 | 36308956 | 813 | 2 | ENSMUST00 | -66504 | Rpl7a-ps3 | ENSMUSG0C | false | chr15 | 36374035 | 36375538 | 16.14863 | 43.24384 | 40.61377 | chr15 | 36371382 | 36376750 | 5.74131 | 4.46347 | 3.60718 |
| chrX | 76100816 | 76101063 | 615 | Distal Intergenic | 20 | 76153331 | 76162605 | 9275 | 2 | ENSMUST00 | 61542 | Gm14710 | ENSMUSG0C | false | . | . | . | . | . | . | . | . | . | . | . | . |
| chr2 | 32846263 | 32846642 | 615 | Promoter (<=1kb) | 2 | 32846186 | 32848168 | 1983 | 1 | ENSMUST00 | 78 | Gm13524 | ENSMUSG0C | false | chr2 | 32845892 | 32847353 | 20.14837 | 78.81087 | 75.68276 | . | . | . | . | . | . |
| chr11 | 108319045 | 108319349 | 614 | Intron (ENSMUST00000059595.5 | 11 | 108343354 | 108395905 | 52552 | 1 | ENSMUST00 | -24005 | Apoh | ENSMUSG0C | false | . | . | . | . | . | . | chr11 | 108317693 | 108319796 | 5.06663 | 4.08732 | 3.10461 |
| chr8 | 22593363 | 22594042 | 614 | Promoter (<=1kb) | 8 | 22577075 | 22593813 | 16739 | 2 | ENSMUST00 | 0 | Vdac3 | ENSMUSG0C | false | chr8 | 22592554 | 22594510 | 36.81325 | 122.26781 | 118.05663 | . | . | . | . | . | . |
| chr17 | 12168562 | 12168985 | 614 | Intron (ENSMUST00000164143.2 | 17 | 12151772 | 12219079 | 67308 | 1 | ENSMUST00 | 16791 | Agpat4 | ENSMUSG0C | true | . | . | . | . | . | . | chr17 | 12154205 | 12176743 | 5.8126 | 4.38993 | 3.68034 |
| chr15 | 38519264 | 38519718 | 614 | Promoter (<=1kb) | 15 | 38487430 | 38519266 | 31837 | 2 | ENSMUST00 | 0 | Azin1 | ENSMUSG0C | false | chr15 | 38517684 | 38520107 | 40.21293 | 133.6382 | 129.0087 | . | . | . | . | . | . |
| chr5 | 124425224 | 124426139 | 614 | Promoter (<=1kb) | 5 | 124368702 | 124425914 | 57213 | 2 | ENSMUST00 | 0 | Sbno1 | ENSMUSG0C | false | chr5 | 124424344 | 124426339 | 32.0113 | 129.32326 | 124.86235 | . | . | . | . | . | . |
| chr2 | 91963362 | 91963872 | 614 | Promoter (<=1kb) | 2 | 91932826 | 91963657 | 30832 | 2 | ENSMUST00 | 0 | Dgkz | ENSMUSG0C | false | chr2 | 91962693 | 91963743 | 29.47202 | 98.86652 | 95.32386 | . | . | . | . | . | . |
| chr8 | 9976152 | 9976526 | 614 | Promoter (<=1kb) | 8 | 9970020 | 9976323 | 6304 | 2 | ENSMUST00 | 0 | Lig4 | ENSMUSG0C | false | chr8 | 9975661 | 9976429 | 27.72203 | 83.13062 | 79.92423 | chr8 | 9974750 | 9977517 | 4.38983 | 3.65035 | 2.53424 |
| chr7 | 139288568 | 139288882 | 614 | Distal Intergenic | 7 | 139248482 | 139267253 | 18772 | 1 | ENSMUST00 | 40087 | Pwwp2b | ENSMUSG0C | false | . | . | . | . | . | . | chr7 | 139288083 | 139289260 | 3.05972 | 3.07162 | 1.46345 |
| chr12 | 71015016 | 71015748 | 614 | Promoter (<=1kb) | 12 | 70991615 | 71015596 | 23982 | 2 | ENSMUST00 | 0 | 3110056K07R | ENSMUSG0C | false | chr12 | 71014262 | 71015695 | 32.60426 | 102.40874 | 98.78054 | . | . | . | . | . | . |
| chr16 | 3884271 | 3884960 | 613 | Promoter (<=1kb) | 16 | 3884619 | 3904780 | 20162 | 1 | ENSMUST00 | 0 | Naa60 | ENSMUSG0C | false | chr16 | 3883780 | 3885605 | 42.07631 | 139.09143 | 134.24335 | . | . | . | . | . | . |
| chr3 | 88455312 | 88455989 | 613 | Promoter (<=1kb) | 3 | 88435959 | 88455739 | 19781 | 2 | ENSMUST00 | 0 | Sema4a | ENSMUSG0C | false | . | . | . | . | . | . | chr3 | 88454614 | 88456784 | 3.86465 | 3.46712 | 2.08753 |
| chr9 | 77508395 | 77508673 | 613 | Intron (ENSMUST00000183873.2 | 9 | 77458090 | 77532444 | 74355 | 2 | ENSMUST00 | 23771 | Lrrc1 | ENSMUSG0C | true | . | . | . | . | . | . | chr9 | 77501306 | 77543222 | 5.57899 | 4.30036 | 3.48961 |
| chr11 | 60537095 | 60537423 | 613 | Promoter (<=1kb) | 11 | 60537978 | 60558512 | 20535 | 1 | ENSMUST00 | -555 | Alkbh5 | ENSMUSG0C | false | chr11 | 60536603 | 60539756 | 32.94511 | 105.03617 | 101.33749 | . | . | . | . | . | . |
| chr2 | 19657413 | 19658029 | 613 | Promoter (<=1kb) | 2 | 19657752 | 19660583 | 2832 | 1 | ENSMUST00 | 0 | Otud1 | ENSMUSG0C | false | chr2 | 19656994 | 19659619 | 33.01748 | 102.11515 | 98.48943 | . | . | . | . | . | . |
| chr7 | 66845784 | 66846132 | 613 | Intron (ENSMUST00000156396.2 | 7 | 66842255 | 67062157 | 219903 | 1 | ENSMUST00 | 3530 | Adamts17 | ENSMUSG0C | false | . | . | . | . | . | . | . | . | . | . | . | . |
| chr8 | 31142620 | 31143113 | 613 | Distal Intergenic | 8 | 31149864 | 31149967 | 104 | 2 | ENSMUST00 | 6854 | Snord13 | ENSMUSG0C | false | . | . | . | . | . | . | chr8 | 31141614 | 31144482 | 3.8348 | 3.45686 | 2.09694 |
| chr13 | 58148588 | 58149045 | 613 | Promoter (<=1kb) | 13 | 58148777 | 58149692 | 916 | 1 | ENSMUST00 | 0 | 1700066J03R | ENSMUSG0C | false | chr13 | 58148364 | 58149139 | 10.47775 | 22.22469 | 19.86031 | chr13 | 58147433 | 58151126 | 4.06165 | 3.6199 | 2.21973 |
| chr2 | 18057562 | 18058071 | 613 | Promoter (<=1kb) | 2 | 18057846 | 18057903 | 58 | 1 | ENSMUST00 | 0 | Mir7655 | ENSMUSG0C | false | chr2 | 18055597 | 18058275 | 30.32912 | 88.589 | 85.27629 | chr2 | 18057975 | 18058495 | 3.65072 | 3.43863 | 1.87948 |
| chr6 | 141380029 | 141380289 | 613 | Intron (ENSMUST00000043259.8 | 6 | 141238278 | 141250561 | 12284 | 2 | ENSMUST00 | -129469 | RP23-97L11. | ENSMUSG0C | false | . | . | . | . | . | . | chr6 | 141370327 | 141386627 | 5.21392 | 4.14863 | 3.18933 |
| chr7 | 30966986 | 30967224 | 612 | Intron (ENSMUST00000001279.9 | 7 | 30957777 | 30973389 | 15613 | 2 | ENSMUST00 | 6165 | Lsr | ENSMUSG0C | false | . | . | . | . | . | . | chr7 | 30963652 | 30967582 | 5.19465 | 4.19131 | 3.16371 |
| chr11 | 54068847 | 54069208 | 612 | Promoter (<=1kb) | 11 | 54054928 | 54069014 | 14087 | 2 | ENSMUST00 | 0 | Pdlim4 | ENSMUSG0C | false | chr11 | 54068682 | 54069337 | 15.08796 | 37.05233 | 34.49837 | chr11 | 54068124 | 54069409 | 6.33281 | 4.76082 | 4.07883 |
| chr16 | 45832452 | 45832730 | 612 | Intron (ENSMUST00000036355.7 | 16 | 45746394 | 45844234 | 97841 | 2 | ENSMUST00 | 11504 | Phldb2 | ENSMUSG0C | false | . | . | . | . | . | . | chr16 | 45828765 | 45843260 | 5.67486 | 4.33105 | 3.58336 |
| chr9 | 96752714 | 96752981 | 612 | Promoter (<=1kb) | 9 | 96682770 | 96752831 | 70062 | 2 | ENSMUST00 | 0 | Zbtb38 | ENSMUSG0C | false | . | . | . | . | . | . | chr9 | 96752101 | 96753966 | 5.28774 | 4.20829 | 3.24482 |
| chr10 | 42277418 | 42278530 | 612 | Promoter (<=1kb) | 10 | 42247431 | 42276755 | 29325 | 2 | ENSMUST00 | -664 | Foxo3 | ENSMUSG0C | false | chr10 | 42273937 | 42278830 | 36.72955 | 143.86444 | 138.82292 | chr10 | 42278404 | 42278775 | 6.44024 | 4.69628 | 4.25024 |
| chr7 | 49360669 | 49360999 | 612 | Intron (ENSMUST00000183659.2 | 7 | 49364721 | 49454642 | 89922 | 1 | ENSMUST00 | -3722 | Nav2 | ENSMUSG0C | false | . | . | . | . | . | . | chr7 | 49359875 | 49361512 | 4.33581 | 3.78484 | 2.45194 |
| chr3 | 97051269 | 97051754 | 612 | 3' UTR | 3 | 97032416 | 97053634 | 21219 | 1 | ENSMUST00 | 18854 | Gja5 | ENSMUSG0C | false | . | . | . | . | . | . | chr3 | 97050689 | 97052471 | 5.23265 | 4.19023 | 3.25086 |
| chr14 | 47921875 | 47922149 | 612 | Distal Intergenic | 14 | 48079375 | 48079752 | 378 | 2 | ENSMUST00 | 157603 | Gm6055 | ENSMUSG0C | false | . | . | . | . | . | . | chr14 | 47921465 | 47922679 | 4.99521 | 4.02022 | 3.04955 |
| chr17 | 85960082 | 85960371 | 612 | Distal Intergenic | 17 | 85796218 | 85824044 | 27827 | 2 | ENSMUST00 | -136039 | RP23-449G1 | ENSMUSG0C | false | . | . | . | . | . | . | chr17 | 85958944 | 85962921 | 4.57652 | 3.82441 | 2.68754 |
| chr1 | 12754619 | 12754910 | 611 | Intron (ENSMUST00000177608.2 | 1 | 12728119 | 12786639 | 58521 | 1 | ENSMUST00 | 26501 | Sulf1 | ENSMUSG0C | true | . | . | . | . | . | . | chr1 | 12752517 | 12771847 | 5.49437 | 4.27944 | 3.4172 |
| chr17 | 26244062 | 26244345 | 611 | Promoter (<=1kb) | 17 | 26211822 | 26244223 | 32402 | 2 | ENSMUST00 | 0 | Itfg3 | ENSMUSG0C | false | chr17 | 26243366 | 26244679 | 18.76905 | 75.7838 | 72.70259 | . | . | . | . | . | . |
| chr2 | 92375200 | 92375544 | 611 | Promoter (<=1kb) | 2 | 92375306 | 92381217 | 5912 | 1 | ENSMUST00 | 0 | Pex16 | ENSMUSG0C | false | chr2 | 92374485 | 92376159 | 31.16539 | 106.97112 | 103.21663 | . | . | . | . | . | . |
| chr8 | 15071216 | 15071677 | 611 | Exon (ENSMUST00000131251.1/ | 8 | 15060190 | 15073840 | 13651 | 1 | ENSMUST00 | 11027 | Myom2 | ENSMUSG0C | true | . | . | . | . | . | . | chr8 | 15062615 | 15075991 | 6.63524 | 4.71645 | 4.35655 |
| chr9 | 32656591 | 32657525 | 611 | Intron (ENSMUST00000183879.1 | 9 | 32636239 | 32651476 | 15238 | 1 | ENSMUST00 | 20353 | Ets1 | ENSMUSG0C | true | . | . | . | . | . | . | chr9 | 32656242 | 32658110 | 6.87778 | 4.79195 | 4.58763 |
| chr2 | 122118088 | 122118497 | 611 | Promoter (<=1kb) | 2 | 122053520 | 122118386 | 64867 | 2 | ENSMUST00 | 0 | Spg11 | ENSMUSG0C | false | chr2 | 122117578 | 122118852 | 39.80928 | 150.74051 | 145.43391 | . | . | . | . | . | . |
| chr7 | 130865652 | 130866343 | 611 | Promoter (<=1kb) | 7 | 130865756 | 130913312 | 47557 | 1 | ENSMUST00 | 0 | Plekha1 | ENSMUSG0C | false | chr7 | 130865745 | 130867110 | 42.98468 | 146.27251 | 141.12608 | . | . | . | . | . | . |
| chr2 | 165906417 | 165906748 | 611 | Promoter (1-2kb) | 2 | 165908051 | 165908175 | 125 | 2 | ENSMUST00 | 1427 | Gm22692 | ENSMUSG0C | false | . | . | . | . | . | . | chr2 | 165904922 | 165909241 | 5.59921 | 4.41708 | 3.48437 |
| chr14 | 76692823 | 76693090 | 611 | Distal Intergenic | 14 | 76681754 | 76686887 | 5134 | 1 | ENSMUST00 | 11070 | RP23-421C24 | ENSMUSG0C | false | . | . | . | . | . | . | chr14 | 76692388 | 76703896 | 4.91248 | 4.06724 | 2.9299 |
| chr14 | 76781690 | 76782036 | 611 | Distal Intergenic | 14 | 76681754 | 76686887 | 5134 | 1 | ENSMUST00 | 99937 | RP23-421C24 | ENSMUSG0C | false | . | . | . | . | . | . | chr14 | 76780483 | 76781875 | 5.05945 | 4.13677 | 3.05162 |
| chr9 | 32648388 | 32648695 | 610 | Intron (ENSMUST00000183879.1 | 9 | 32636239 | 32651476 | 15238 | 1 | ENSMUST00 | 12150 | Ets1 | ENSMUSG0C | true | . | . | . | . | . | . | . | . | . | . | . | . |
| chr19 | 11912106 | 11912791 | 610 | Promoter (<=1kb) | 19 | 11912399 | 11945096 | 32698 | 1 | ENSMUST00 | 0 | Patl1 | ENSMUSG0C | false | chr19 | 11911493 | 11913356 | 34.16929 | 120.7392 | 116.5701 | . | . | . | . | . | . |
| chr10 | 111449014 | 111449316 | 610 | Distal Intergenic | 10 | 111473192 | 111496886 | 23695 | 1 | ENSMUST00 | -23876 | Nap1l1 | ENSMUSG0C | false | . | . | . | . | . | . | chr10 | 111445585 | 111449757 | 3.69234 | 3.44057 | 1.94502 |
| chr1 | 133420322 | 133420864 | 610 | Intron (ENSMUST00000153799.2 | 1 | 133382303 | 133424377 | 42075 | 2 | ENSMUST00 | 3513 | Sox13 | ENSMUSG0C | false | chr1 | 133420822 | 133421787 | 6.02971 | 9.77817 | 7.60541 | chr1 | 133393415 | 133423224 | 5.89736 | 4.38424 | 3.76681 |
| chr16 | 95849010 | 95849284 | 610 | Intron (ENSMUST00000136744.2 | 16 | 95806029 | 95808317 | 2289 | 2 | ENSMUST00 | -40694 | 2810404F17R | ENSMUSG0C | false | . | . | . | . | . | . | . | . | . | . | . | . |
| chr8 | 26983466 | 26983778 | 610 | Distal Intergenic | 8 | 26977336 | 26981461 | 4126 | 1 | ENSMUST00 | 6131 | Zfp703 | ENSMUSG0C | false | . | . | . | . | . | . | chr8 | 26978900 | 26985294 | 4.32877 | 3.61461 | 2.48483 |
| chr16 | 85044841 | 85045196 | 610 | Intron (ENSMUST00000005406.1 | 16 | 85168849 | 85169001 | 153 | 1 | ENSMUST00 | -123653 | Gm27295 | ENSMUSG0C | false | . | . | . | . | . | . | chr16 | 85041565 | 85052812 | 5.14224 | 4.09203 | 3.14604 |
| chr10 | 62204461 | 62204845 | 610 | Intron (ENSMUST00000047883.9 | 10 | 62194167 | 62194227 | 61 | 2 | ENSMUST00 | -10235 | Mir7662 | ENSMUSG0C | false | . | . | . | . | . | . | chr10 | 62185685 | 62229768 | 8.28034 | 5.46771 | 5.70297 |
| chr1 | 62905816 | 62906092 | 610 | Distal Intergenic | 1 | 62821355 | 62832370 | 11016 | 2 | ENSMUST00 | -73447 | RP23-149A5. | ENSMUSG0C | false | . | . | . | . | . | . | chr1 | 62904745 | 62907042 | 4.09421 | 3.66297 | 2.26342 |
| chr11 | 60273141 | 60273494 | 609 | Intron (ENSMUST00000095254.6 | 11 | 60229902 | 60245669 | 15768 | 2 | ENSMUST00 | -27473 | Tom1l2 | ENSMUSG0C | false | . | . | . | . | . | . | chr11 | 60273241 | 60274906 | 4.13569 | 3.63938 | 2.31454 |
| chr14 | 27259445 | 27259739 | 609 | Intron (ENSMUST00000049206.5 | 14 | 27238039 | 27403904 | 165866 | 1 | ENSMUST00 | 21407 | Arhgef3 | ENSMUSG0C | false | . | . | . | . | . | . | chr14 | 27259115 | 27261665 | 3.47634 | 3.25479 | 1.80193 |
| chr3 | 27182769 | 27183027 | 609 | Promoter (<=1kb) | 3 | 27182965 | 27244958 | 61994 | 1 | ENSMUST00 | 0 | Nceh1 | ENSMUSG0C | false | chr3 | 27182540 | 27184235 | 27.6016 | 85.77155 | 82.50918 | chr3 | 27182337 | 27184338 | 3.75728 | 3.49551 | 1.97534 |
| chr3 | 139074433 | 139075702 | 609 | Promoter (<=1kb) | 3 | 138925906 | 139075199 | 149294 | 2 | ENSMUST00 | 0 | Rap1gds1 | ENSMUSG0C | false | chr3 | 139073799 | 139075230 | 42.62012 | 151.29346 | 145.9586 | . | . | . | . | . | . |
| chr5 | 124241011 | 124241390 | 609 | Intron (ENSMUST00000162812.2 | 5 | 124118691 | 124249451 | 130761 | 2 | ENSMUST00 | 8061 | Pitpnm2 | ENSMUSG0C | false | . | . | . | . | . | . | chr5 | 124238588 | 124243039 | 5.88515 | 4.40141 | 3.76533 |
| chr5 | 129415450 | 129415802 | 609 | Distal Intergenic | 5 | 129501231 | 129571383 | 70153 | 1 | ENSMUST00 | -85429 | Sfswap | ENSMUSG0C | false | . | . | . | . | . | . | chr5 | 129405729 | 129418273 | 5.29114 | 4.12018 | 3.25854 |
| chr8 | 33909712 | 33910039 | 609 | Intron (ENSMUST00000053251.6 | 8 | 33863007 | 33891764 | 28758 | 2 | ENSMUST00 | -17949 | Rbpms | ENSMUSG0C | true | . | . | . | . | . | . | chr8 | 33863865 | 33927704 | 6.32022 | 4.67603 | 4.08842 |
| chr9 | 57072752 | 57073887 | 609 | Promoter (<=1kb) | 9 | 57072040 | 57089496 | 17457 | 1 | ENSMUST00 | 713 | Sin3a | ENSMUSG0C | false | chr9 | 57071455 | 57074688 | 31.01414 | 97.6227 | 94.10706 | chr9 | 57073825 | 57075505 | 5.73893 | 4.44541 | 3.6056 |
| chr4 | 8535494 | 8536339 | 609 | Promoter (<=1kb) | 4 | 8535644 | 8607778 | 72135 | 1 | ENSMUST00 | 0 | Rab2a | ENSMUSG0C | false | chr4 | 8535506 | 8537034 | 36.10713 | 116.78466 | 112.7497 | . | . | . | . | . | . |
| chr12 | 70937652 | 70938198 | 608 | Promoter (<=1kb) | 12 | 70937857 | 70964718 | 26862 | 1 | ENSMUST00 | 0 | Actr10 | ENSMUSG0C | false | chr12 | 70937304 | 70938812 | 32.94367 | 134.34938 | 129.69266 | . | . | . | . | . | . |
| chr1 | 75168547 | 75169005 | 608 | Promoter (<=1kb) | 1 | 75168646 | 75170752 | 2107 | 1 | ENSMUST00 | 0 | Zfand2b | ENSMUSG0C | false | chr1 | 75168055 | 75169352 | 30.78991 | 105.27288 | 101.56436 | chr1 | 75167136 | 75169841 | 5.38168 | 4.17846 | 3.37177 |
| chr3 | 63323651 | 63323967 | 608 | Intron (ENSMUST00000029400.1 | 3 | 63295872 | 63382231 | 86360 | 1 | ENSMUST00 | 27780 | Mme | ENSMUSG0C | false | . | . | . | . | . | . | chr3 | 63319672 | 63327568 | 4.32678 | 3.74428 | 2.45114 |
| chr4 | 155173450 | 155173869 | 608 | Intron (ENSMUST00000129314.1 | 4 | 155154075 | 155188296 | 34222 | 2 | ENSMUST00 | 14427 | Ski | ENSMUSG0C | false | . | . | . | . | . | . | chr4 | 155159960 | 155221990 | 6.99931 | 4.87167 | 4.66225 |
| chr5 | 90442350 | 90442701 | 608 | Distal Intergenic | 5 | 90460889 | 90476603 | 15715 | 1 | ENSMUST00 | -18188 | Alb | ENSMUSG0C | false | . | . | . | . | . | . | chr5 | 90441825 | 90444677 | 4.46758 | 3.87924 | 2.57299 |
| chr6 | 52221782 | 52222351 | 608 | Promoter (<=1kb) | 6 | 52216009 | 52221854 | 5846 | 2 | ENSMUST00 | 0 | Hoxa7 | ENSMUSG0C | false | chr6 | 52221266 | 52227348 | 34.16929 | 120.7392 | 116.5701 | chr6 | 52222336 | 52223532 | 4.39861 | 3.82628 | 2.49652 |
| chr3 | 108210103 | 108210724 | 608 | Promoter (<=1kb) | 3 | 108202222 | 108210486 | 8265 | 2 | ENSMUST00 | 0 | Atxn7l2 | ENSMUSG0C | false | chr3 | 108209908 | 108211309 | 19.52531 | 57.12588 | 54.31675 | chr3 | 108209419 | 108211399 | 4.6323 | 3.89844 | 2.71148 |
| chr11 | 54786719 | 54787004 | 608 | Promoter (<=1kb) | 11 | 54740310 | 54787625 | 47316 | 2 | ENSMUST00 | 621 | Cdc42se2 | ENSMUSG0C | true | chr11 | 54785734 | 54788212 | 43.43732 | 157.3205 | 151.76292 | . | . | . | . | . | . |
| chr11 | 94589793 | 94590150 | 608 | Intron (ENSMUST00000103164.3 | 11 | 94570291 | 94601769 | 31479 | 2 | ENSMUST00 | 11619 | Acsf2 | ENSMUSG0C | false | . | . | . | . | . | . | chr11 | 94588503 | 94592861 | 5.27898 | 4.16196 | 3.24456 |
| chr17 | 86610700 | 86610995 | 608 | Intron (ENSMUST00000097275.3 | 17 | 86703298 | 86703407 | 110 | 2 | ENSMUST00 | 92412 | Gm22235 | ENSMUSG0C | false | . | . | . | . | . | . | chr17 | 86610058 | 86615360 | 5.61564 | 4.2317 | 3.54393 |
| chr11 | 87737143 | 87737599 | 607 | Promoter (<=1kb) | 11 | 87737552 | 87743617 | 6066 | 1 | ENSMUST00 | 0 | Supt4a | ENSMUSG0C | false | chr11 | 87736508 | 87737347 | 25.08628 | 74.81181 | 71.74545 | chr11 | 87736288 | 87738772 | 4.33107 | 3.73033 | 2.48005 |
| chr13 | 51100635 | 51101520 | 607 | Promoter (<=1kb) | 13 | 51100880 | 51152546 | 51667 | 1 | ENSMUST00 | 0 | Spin1 | ENSMUSG0C | false | chr13 | 51099595 | 51102016 | 37.76794 | 122.86759 | 118.62959 | . | . | . | . | . | . |
| chr13 | 55599875 | 55600144 | 607 | Promoter (<=1kb) | 13 | 55599896 | 55610443 | 10548 | 1 | ENSMUST00 | 0 | B4galt7 | ENSMUSG0C | false | chr13 | 55599565 | 55600944 | 37.94029 | 126.39516 | 122.03232 | chr13 | 55599296 | 55601891 | 4.16313 | 3.62106 | 2.35973 |
| chr9 | 110002158 | 110002728 | 607 | Intron (ENSMUST00000035055.9 | 9 | 110036742 | 110072705 | 35964 | 1 | ENSMUST00 | -34014 | Map4 | ENSMUSG0C | false | . | . | . | . | . | . | chr9 | 109998891 | 110006603 | 4.86427 | 4.02849 | 2.88824 |
| chr6 | 88724215 | 88724762 | 607 | Promoter (<=1kb) | 6 | 88724412 | 88828360 | 103949 | 1 | ENSMUST00 | 0 | Mgll | ENSMUSG0C | false | chr6 | 88724035 | 88725421 | 27.88725 | 99.90214 | 96.33501 | . | . | . | . | . | . |
| chr5 | 113973100 | 113973352 | 607 | Intron (ENSMUST00000159592.2 | 5 | 113965602 | 113966756 | 1155 | 2 | ENSMUST00 | -6345 | Ssh1 | ENSMUSG0C | false | . | . | . | . | . | . | chr5 | 113973303 | 113977048 | 3.87546 | 3.43264 | 2.11922 |
| chr13 | 76018323 | 76018829 | 607 | Promoter (<=1kb) | 13 | 76001535 | 76018562 | 17028 | 2 | ENSMUST00 | 0 | Rfesd | ENSMUSG0C | false | chr13 | 76017591 | 76019304 | 24.3095 | 71.63969 | 68.62457 | . | . | . | . | . | . |
| chr2 | 32586908 | 32587208 | 607 | Promoter (<=1kb) | 2 | 32587095 | 32599698 | 12604 | 1 | ENSMUST00 | 0 | St6galnac4 | ENSMUSG0C | false | chr2 | 32586689 | 32587897 | 18.57615 | 49.79513 | 47.08208 | chr2 | 32586351 | 32590869 | 4.34468 | 3.69652 | 2.4816 |
| chr3 | 129696108 | 129696554 | 607 | Intron (ENSMUST00000029653.2 | 3 | 129684133 | 129684241 | 109 | 2 | ENSMUST00 | -11868 | Gm24515 | ENSMUSG0C | false | . | . | . | . | . | . | chr3 | 129694610 | 129701042 | 5.50271 | 4.23715 | 3.46885 |
| chr6 | 89634497 | 89634857 | 606 | Distal Intergenic | 6 | 89643988 | 89675529 | 31542 | 1 | ENSMUST00 | -9131 | Txnrd3 | ENSMUSG0C | false | . | . | . | . | . | . | chr6 | 89634682 | 89634927 | 3.33767 | 3.27139 | 1.68085 |
| chr10 | 79681655 | 79682208 | 606 | Promoter (<=1kb) | 10 | 79682195 | 79688394 | 6200 | 1 | ENSMUST00 | 0 | Cdc34 | ENSMUSG0C | false | chr10 | 79682099 | 79683765 | 32.69058 | 104.59052 | 100.90593 | chr10 | 79680842 | 79681673 | 4.73799 | 3.85918 | 2.826 |
| chr14 | 21500890 | 21501386 | 606 | Promoter (<=1kb) | 14 | 21500892 | 21672478 | 171587 | 1 | ENSMUST00 | 0 | Kat6b | ENSMUSG0C | false | chr14 | 21500795 | 21501734 | 17.45443 | 60.55182 | 57.69857 | . | . | . | . | . | . |
| chr17 | 11303173 | 11303458 | 606 | Promoter (<=1kb) | 17 | 11303647 | 11610945 | 307299 | 1 | ENSMUST00 | -189 | Park2 | ENSMUSG0C | false | . | . | . | . | . | . | chr17 | 11301891 | 11304580 | 5.95779 | 4.34413 | 3.8348 |
| chr3 | 129332215 | 129333007 | 606 | Promoter (<=1kb) | 3 | 129269175 | 129332720 | 63546 | 2 | ENSMUST00 | 0 | Enpep | ENSMUSG0C | true | chr3 | 129327289 | 129333393 | 25.83068 | 80.74496 | 77.57947 | . | . | . | . | . | . |
| chr5 | 88764819 | 88765226 | 606 | Promoter (<=1kb) | 5 | 88764996 | 88783281 | 18286 | 1 | ENSMUST00 | 0 | Dck | ENSMUSG0C | false | chr5 | 88764418 | 88765711 | 30.48835 | 90.7988 | 87.43492 | chr5 | 88763765 | 88765645 | 3.61861 | 3.41322 | 1.85934 |
| chr4 | 139091937 | 139092253 | 606 | Promoter (<=1kb) | 4 | 139083207 | 139092106 | 8900 | 2 | ENSMUST00 | 0 | Nbl1 | ENSMUSG0C | false | chr4 | 139091610 | 139093306 | 40.89836 | 130.45236 | 125.94646 | chr4 | 139088999 | 139098811 | 5.83874 | 4.33468 | 3.72707 |
| chr13 | 53531629 | 53531856 | 606 | Distal Intergenic | 13 | 53525703 | 53526059 | 357 | 1 | ENSMUST00 | 5927 | Gm5449 | ENSMUSG0C | false | . | . | . | . | . | . | chr13 | 53531209 | 53532426 | 3.55742 | 3.23524 | 1.87212 |
| chr4 | 114460542 | 114460826 | 606 | Intron (ENSMUST00000094894.3 | 4 | 114406724 | 114615098 | 208375 | 1 | ENSMUST00 | 53819 | Trabd2b | ENSMUSG0C | false | . | . | . | . | . | . | chr4 | 114458782 | 114463222 | 5.76412 | 4.37733 | 3.64386 |
| chr6 | 6299292 | 6299725 | 606 | Promoter (<=1kb) | 6 | 6299421 | 6363892 | 64472 | 1 | ENSMUST00 | 0 | Gm20619 | ENSMUSG0C | false | . | . | . | . | . | . | . | . | . | . | . | . |
| chr7 | 25687706 | 25688169 | 605 | Promoter (<=1kb) | 7 | 25687002 | 25705077 | 18076 | 1 | ENSMUST00 | 705 | Tgfb1 | ENSMUSG0C | false | chr7 | 25686448 | 25689783 | 37.30079 | 124.19571 | 119.91465 | chr7 | 25686345 | 25690723 | 4.57586 | 3.80181 | 2.69028 |
| chr17 | 8923090 | 8923356 | 605 | Promoter (<=1kb) | 17 | 8924143 | 8984288 | 60146 | 1 | ENSMUST00 | -787 | Pde10a | ENSMUSG0C | true | . | . | . | . | . | . | chr17 | 8921749 | 8923627 | 4.01628 | 3.52539 | 2.24375 |
| chr10 | 121476154 | 121476542 | 605 | Promoter (<=1kb) | 10 | 121410350 | 121476250 | 65901 | 2 | ENSMUST00 | 0 | Rassf3 | ENSMUSG0C | false | chr10 | 121476470 | 121476958 | 19.73103 | 51.78704 | 49.0507 | . | . | . | . | . | . |
| chr2 | 151985766 | 151986183 | 605 | Distal Intergenic | 2 | 151969398 | 151980219 | 10822 | 2 | ENSMUST00 | -5548 | Fam110a | ENSMUSG0C | false | . | . | . | . | . | . | chr2 | 151985712 | 151986134 | 3.0286 | 2.96995 | 1.38705 |
| chr4 | 119966790 | 119967126 | 605 | Intron (ENSMUST00000143837.1 | 4 | 120094489 | 120134557 | 40069 | 1 | ENSMUST00 | -127363 | Hivep3 | ENSMUSG0C | false | . | . | . | . | . | . | . | . | . | . | . | . |
| chr8 | 24490997 | 24491288 | 605 | Distal Intergenic | 8 | 24533456 | 24540796 | 7341 | 2 | ENSMUST00 | 49508 | Ido2 | ENSMUSG0C | false | . | . | . | . | . | . | chr8 | 24487687 | 24492698 | 6.78447 | 4.86694 | 4.4711 |
| chr18 | 64646794 | 64647081 | 605 | Intron (ENSMUST00000025482.8 | 18 | 64528979 | 64661000 | 132022 | 2 | ENSMUST00 | 13919 | Atp8b1 | ENSMUSG0C | true | . | . | . | . | . | . | chr18 | 64613324 | 64659141 | 5.21932 | 4.12513 | 3.19621 |
| chr4 | 150572324 | 150572597 | 605 | Promoter (2-3kb) | 4 | 150569657 | 150612069 | 42413 | 1 | ENSMUST00 | 2668 | Rere | ENSMUSG0C | false | . | . | . | . | . | . | chr4 | 150570369 | 150578877 | 6.14529 | 4.57392 | 3.95861 |
| chr10 | 80748645 | 80748956 | 605 | Distal Intergenic | 10 | 80755206 | 80795461 | 40256 | 1 | ENSMUST00 | -6250 | Dot1l | ENSMUSG0C | false | . | . | . | . | . | . | chr10 | 80747739 | 80749290 | 5.83385 | 4.5238 | 3.67561 |
| chr2 | 139678029 | 139678697 | 605 | Promoter (<=1kb) | 2 | 139678178 | 139758580 | 80403 | 1 | ENSMUST00 | 0 | Ism1 | ENSMUSG0C | false | chr2 | 139677037 | 139679067 | 31.13259 | 95.84155 | 92.37124 | chr2 | 139677357 | 139678991 | 4.26013 | 3.60568 | 2.42833 |
| chr7 | 25049735 | 25050001 | 604 | Intron (ENSMUST00000003468.8 | 7 | 25009853 | 25072369 | 62517 | 2 | ENSMUST00 | 22368 | Grik5 | ENSMUSG0C | false | . | . | . | . | . | . | chr7 | 25048845 | 25050650 | 4.88795 | 4.01744 | 2.96193 |
| chr12 | 36002642 | 36002948 | 604 | Intron (ENSMUST00000020898.6 | 12 | 35992907 | 36004087 | 11181 | 1 | ENSMUST00 | 9736 | Agr2 | ENSMUSG0C | false | . | . | . | . | . | . | chr12 | 36001429 | 36003736 | 4.77232 | 3.88266 | 2.8221 |
| chr17 | 29094540 | 29094915 | 604 | Promoter (<=1kb) | 17 | 29093772 | 29100722 | 6951 | 1 | ENSMUST00 | 769 | Cdkn1a | ENSMUSG0C | false | chr17 | 29093483 | 29095648 | 15.53947 | 48.13258 | 45.44172 | chr17 | 29093497 | 29095198 | 6.129 | 4.61254 | 3.95372 |
| chr8 | 75212883 | 75213336 | 604 | Promoter (<=1kb) | 8 | 75213944 | 75224113 | 10170 | 1 | ENSMUST00 | -608 | Rasd2 | ENSMUSG0C | false | chr8 | 75212723 | 75214161 | 25.15761 | 80.47005 | 77.31023 | chr8 | 75211860 | 75214120 | 6.6705 | 4.62075 | 4.40619 |

| | | | | | | | | | | | | | | | | | | | | | | | | | | |
|---|---|---|---|---|---|---|---|---|---|---|---|---|---|---|---|---|---|---|---|---|---|---|---|---|---|---|
| chr7 | 44525115 | 44525451 | 604 | Promoter (<=1kb) | 7 | 44501699 | 44524669 | 22971 | 2 | ENSMUST00 | -447 | Mybpc2 | ENSMUSG0( | false | . | . | . | . | . | . | chr7 | 44524566 | 44526154 | 4.89627 | 4.01152 | 2.90349 |
| chr8 | 84985750 | 84986383 | 604 | Distal Intergenic | 8 | 84990595 | 85003364 | 12770 | 1 | ENSMUST00 | -4212 | Hook2 | ENSMUSG0( | false | chr8 | 84985294 | 84986495 | 23.05105 | 65.89761 | 62.96862 | chr8 | 84974675 | 84986551 | 7.15 | 5.07534 | 4.74925 |
| chr11 | 88542958 | 88543181 | 604 | Intron (ENSMUST00000107909.2 | 11 | 88365515 | 88590147 | 224633 | 2 | ENSMUST00 | 46966 | Msi2 | ENSMUSG0( | false | . | . | . | . | . | . | chr11 | 88537533 | 88544616 | 6.145 | 4.52274 | 3.95223 |
| chr4 | 115709905 | 115710224 | 604 | Promoter (2-3kb) | 4 | 115707251 | 115708303 | 1053 | 1 | ENSMUST00 | 2655 | Gm12849 | ENSMUSG0( | false | . | . | . | . | . | . | chr4 | 115707353 | 115710474 | 3.80958 | 3.35728 | 2.063 |
| chr4 | 114646566 | 114646916 | 604 | Distal Intergenic | 4 | 114672697 | 114673340 | 644 | 2 | ENSMUST00 | 26424 | RP23-454F1: | ENSMUSG0( | false | . | . | . | . | . | . | chr4 | 114642640 | 114648821 | 5.07152 | 4.10947 | 3.07317 |
| chr14 | 25660740 | 25661065 | 603 | Intron (ENSMUST00000162645.2 | 14 | 25656830 | 25658405 | 1576 | 1 | ENSMUST00 | 3911 | Zmiz1 | ENSMUSG0( | false | . | . | . | . | . | . | chr14 | 25660103 | 25661391 | 4.36722 | 3.6525 | 2.51943 |
| chr10 | 28074565 | 28074989 | 603 | Promoter (<=1kb) | 10 | 28074820 | 28597397 | 522578 | 1 | ENSMUST00 | 0 | Ptprk | ENSMUSG0( | false | chr10 | 28073969 | 28076790 | 32.46003 | 103.17023 | 99.51957 | . | . | . | . | . | . |
| chr18 | 36323943 | 36324386 | 603 | Distal Intergenic | 18 | 36345619 | 36366720 | 21102 | 1 | ENSMUST00 | -21233 | Cystm1 | ENSMUSG0( | false | . | . | . | . | . | . | chr18 | 36323785 | 36324556 | 2.92592 | 2.93515 | 1.34488 |
| chr19 | 46645161 | 46645560 | 603 | Intron (ENSMUST00000138302.3 | 19 | 46623401 | 46657385 | 33985 | 1 | ENSMUST00 | 21761 | Wbp1l | ENSMUSG0( | false | . | . | . | . | . | . | chr19 | 46599683 | 46650166 | 6.55215 | 4.7277 | 4.28832 |
| chr15 | 85534273 | 85534640 | 603 | Downstream (<1kb) | 15 | 85510812 | 85511219 | 408 | 1 | ENSMUST00 | 23462 | Gm4825 | ENSMUSG0( | false | . | . | . | . | . | . | chr15 | 85533926 | 85534902 | 4.41604 | 3.59884 | 2.56455 |
| chr7 | 68851934 | 68852215 | 603 | Distal Intergenic | 7 | 68952100 | 68952236 | 137 | 2 | ENSMUST00 | 100021 | Gm23795 | ENSMUSG0( | false | . | . | . | . | . | . | chr7 | 68851673 | 68852436 | 4.41512 | 3.82219 | 2.58167 |
| chr10 | 41707148 | 41707449 | 603 | Promoter (1-2kb) | 10 | 41550608 | 41709297 | 158690 | 2 | ENSMUST00 | 1848 | Ccdc162 | ENSMUSG0( | false | . | . | . | . | . | . | chr10 | 41706976 | 41707445 | 3.59995 | 3.1814 | 1.89392 |
| chr11 | 95444132 | 95444420 | 603 | Intron (ENSMUST00000107724.3 | 11 | 95462546 | 95462721 | 176 | 2 | ENSMUST00 | 18301 | Gm11522 | ENSMUSG0( | false | . | . | . | . | . | . | chr11 | 95438463 | 95446862 | 5.07312 | 3.96274 | 3.09683 |
| chr7 | 82743479 | 82743791 | 603 | Exon (ENSMUST00000144828.1/ | 7 | 82754872 | 82754979 | 108 | 2 | ENSMUST00 | 11188 | n-R5s154 | ENSMUSG0( | false | . | . | . | . | . | . | . | . | . | . | . | . |
| chr6 | 87270079 | 87270337 | 603 | Intron (ENSMUST00000042025.9 | 6 | 87281149 | 87281260 | 112 | 1 | ENSMUST00 | -10812 | n-R5s164 | ENSMUSG0( | false | . | . | . | . | . | . | chr6 | 87269847 | 87271269 | 3.90281 | 3.57557 | 2.16115 |
| chr1 | 51915573 | 51916161 | 602 | Promoter (<=1kb) | 1 | 51749765 | 51916071 | 166307 | 2 | ENSMUST00 | 0 | Myo1b | ENSMUSG0( | false | chr1 | 51914973 | 51916278 | 18.02578 | 59.25625 | 56.42029 | chr1 | 51914945 | 51915650 | 3.35188 | 3.21556 | 1.6853 |
| chr1 | 182509417 | 182509774 | 602 | Intron (ENSMUST00000068505.8 | 1 | 182467256 | 182517495 | 50240 | 2 | ENSMUST00 | 7721 | Capn2 | ENSMUSG0( | false | . | . | . | . | . | . | chr1 | 182492178 | 182516499 | 7.84019 | 5.29195 | 5.34945 |
| chr3 | 146105423 | 146105785 | 602 | Promoter (2-3kb) | 3 | 146040526 | 146108130 | 67605 | 2 | ENSMUST00 | 2345 | Wdr63 | ENSMUSG0( | false | . | . | . | . | . | . | chr3 | 146104593 | 146108507 | 4.61767 | 3.9225 | 2.70507 |
| chr11 | 97485242 | 97485638 | 602 | Intron (ENSMUST00000121799.2 | 11 | 97491144 | 97502398 | 11255 | 1 | ENSMUST00 | -5506 | Arhgap23 | ENSMUSG0( | true | . | . | . | . | . | . | chr11 | 97474352 | 97489987 | 5.93137 | 4.40018 | 3.78292 |
| chr17 | 10086307 | 10086912 | 602 | Distal Intergenic | 17 | 10206471 | 10319361 | 112891 | 2 | ENSMUST00 | 232449 | Qk | ENSMUSG0( | true | . | . | . | . | . | . | chr17 | 10078916 | 10088169 | 4.84297 | 3.88168 | 2.89341 |
| chr7 | 75498179 | 75498547 | 602 | Intron (ENSMUST00000147005.2 | 7 | 75455868 | 75580376 | 124509 | 1 | ENSMUST00 | 42312 | Akap13 | ENSMUSG0( | false | . | . | . | . | . | . | chr7 | 75497206 | 75501500 | 4.64065 | 3.92118 | 2.70755 |
| chr10 | 43977436 | 43977742 | 602 | Intron (ENSMUST00000020017.8 | 10 | 44004154 | 44004701 | 548 | 1 | ENSMUST00 | -26412 | Gm10812 | ENSMUSG0( | false | . | . | . | . | . | . | . | . | . | . | . | . |
| chr4 | 131488348 | 131488669 | 602 | Promoter (1-2kb) | 4 | 131487197 | 131505527 | 18331 | 1 | ENSMUST00 | 1152 | Gm831 | ENSMUSG0( | false | . | . | . | . | . | . | chr4 | 131488351 | 131489644 | 4.66453 | 3.88473 | 2.76991 |
| chr17 | 15140455 | 15140752 | 602 | Distal Intergenic | 17 | 15065615 | 15074752 | 9138 | 1 | ENSMUST00 | 74841 | RP23-464D1( | ENSMUSG0( | false | . | . | . | . | . | . | chr17 | 15139850 | 15140978 | 3.33457 | 3.21875 | 1.65967 |
| chr11 | 119305099 | 119305380 | 601 | Promoter (2-3kb) | 11 | 119307768 | 119345375 | 37608 | 1 | ENSMUST00 | -2388 | Card14 | ENSMUSG0( | false | chr11 | 119304895 | 119305234 | 7.82748 | 15.19303 | 12.92724 | chr11 | 119304168 | 119307174 | 4.88441 | 4.00369 | 2.93715 |
| chr16 | 91648151 | 91648447 | 601 | Promoter (<=1kb) | 16 | 91647885 | 91679221 | 31337 | 1 | ENSMUST00 | 267 | Son | ENSMUSG0( | false | chr16 | 91646065 | 91649433 | 31.16539 | 106.97112 | 103.21663 | . | . | . | . | . | . |
| chr17 | 56935404 | 56935781 | 601 | Promoter (<=1kb) | 17 | 56892612 | 56935388 | 42777 | 2 | ENSMUST00 | -17 | Mllt1 | ENSMUSG0( | false | chr17 | 56934576 | 56935985 | 32.88158 | 104.92503 | 101.22703 | chr17 | 56934431 | 56936004 | 4.08209 | 3.56332 | 2.30159 |
| chr1 | 172432344 | 172432848 | 601 | Distal Intergenic | 1 | 172475358 | 172478575 | 3218 | 1 | ENSMUST00 | -42510 | Slamf9 | ENSMUSG0( | false | . | . | . | . | . | . | chr1 | 172428189 | 172437787 | 6.0765 | 4.59609 | 3.89569 |
| chr3 | 129896611 | 129896901 | 601 | Distal Intergenic | 3 | 129901425 | 129914103 | 12679 | 1 | ENSMUST00 | -4524 | Casp6 | ENSMUSG0( | false | . | . | . | . | . | . | chr3 | 129896066 | 129897068 | 3.82357 | 3.46062 | 2.05905 |
| chr3 | 136609863 | 136610188 | 601 | Distal Intergenic | 3 | 136670124 | 136797838 | 127715 | 1 | ENSMUST00 | -59936 | Ppp3ca | ENSMUSG0( | false | . | . | . | . | . | . | chr3 | 136609554 | 136610817 | 3.74133 | 3.44767 | 2.00704 |
| chr5 | 30848638 | 30849026 | 601 | Intron (ENSMUST00000031058.9 | 5 | 30853799 | 30863358 | 9560 | 1 | ENSMUST00 | -4773 | Mapre3 | ENSMUSG0( | false | . | . | . | . | . | . | chr5 | 30846495 | 30852222 | 5.09006 | 4.09018 | 3.13383 |
| chr16 | 17201333 | 17201637 | 601 | Promoter (<=1kb) | 16 | 17152015 | 17201492 | 49478 | 2 | ENSMUST00 | 0 | Ube2l3 | ENSMUSG0( | false | chr16 | 17200175 | 17201755 | 43.39494 | 141.3761 | 136.44113 | chr16 | 17198507 | 17202116 | 3.99692 | 3.5382 | 2.2084 |
| chr4 | 45506623 | 45507017 | 601 | Intron (ENSMUST00000061986.6 | 4 | 45500485 | 45500552 | 68 | 2 | ENSMUST00 | -6072 | Gm22518 | ENSMUSG0( | false | . | . | . | . | . | . | chr4 | 45486900 | 45516085 | 7.16373 | 5.03069 | 4.78278 |
| chr12 | 26401743 | 26402068 | 601 | Intron (ENSMUST00000020971.7 | 12 | 26414901 | 26415010 | 110 | 2 | ENSMUST00 | 12942 | Mir6538 | ENSMUSG0( | false | . | . | . | . | . | . | . | . | . | . | . | . |
| chr19 | 24556082 | 24556468 | 600 | Promoter (<=1kb) | 19 | 24294794 | 24555872 | 261079 | 2 | ENSMUST00 | -211 | Pip5k1b | ENSMUSG0( | false | chr19 | 24555259 | 24557043 | 29.7561 | 89.53341 | 86.19995 | chr19 | 24555333 | 24557943 | 5.50416 | 4.2393 | 3.44656 |
| chr6 | 142571258 | 142571695 | 600 | Promoter (<=1kb) | 6 | 142564939 | 142571356 | 6418 | 2 | ENSMUST00 | 0 | Kcnj8 | ENSMUSG0( | false | chr6 | 142570923 | 142572394 | 14.24974 | 34.23454 | 31.71693 | chr6 | 142570079 | 142572381 | 4.19747 | 3.54049 | 2.38373 |
| chr9 | 97133141 | 97133414 | 600 | Distal Intergenic | 9 | 97074961 | 97111157 | 36197 | 2 | ENSMUST00 | -21985 | Slc25a36 | ENSMUSG0( | false | . | . | . | . | . | . | chr9 | 97133200 | 97133582 | 3.05563 | 3.01562 | 1.43512 |
| chr7 | 19182060 | 19182549 | 600 | Promoter (<=1kb) | 7 | 19181606 | 19186371 | 4766 | 1 | ENSMUST00 | 455 | Eml2 | ENSMUSG0( | true | chr7 | 19181038 | 19182936 | 28.08037 | 85.65662 | 82.39679 | chr7 | 19178043 | 19190895 | 5.91671 | 4.49499 | 3.76502 |
| chr19 | 11818784 | 11819085 | 600 | Promoter (<=1kb) | 19 | 11775119 | 11818893 | 43775 | 2 | ENSMUST00 | 0 | Stx3 | ENSMUSG0( | false | chr19 | 11818037 | 11820164 | 36.61738 | 114.51753 | 110.55032 | chr19 | 11817174 | 11820525 | 3.92347 | 3.56784 | 2.16012 |
| chr7 | 96672478 | 96672706 | 600 | Intron (ENSMUST00000107162.2 | 7 | 96644172 | 96732280 | 88109 | 1 | ENSMUST00 | 28307 | Tenm4 | ENSMUSG0( | false | . | . | . | . | . | . | chr7 | 96672378 | 96673297 | 4.75453 | 3.96892 | 2.80569 |
| chr3 | 141971571 | 141971871 | 600 | Intron (ENSMUST00000029948.9 | 3 | 141837136 | 141982264 | 145129 | 2 | ENSMUST00 | 10393 | Bmpr1b | ENSMUSG0( | false | . | . | . | . | . | . | chr3 | 141969962 | 141972579 | 3.27863 | 3.05559 | 1.63942 |
| chr12 | 16847405 | 16847788 | 600 | Distal Intergenic | 12 | 16810965 | 16826743 | 15779 | 1 | ENSMUST00 | 36441 | E2f6 | ENSMUSG0( | false | . | . | . | . | . | . | . | . | . | . | . | . |
| chr16 | 17208812 | 17209173 | 600 | Promoter (<=1kb) | 16 | 17208135 | 17213921 | 5787 | 1 | ENSMUST00 | 678 | Rimbp3 | ENSMUSG0( | false | chr16 | 17208423 | 17210710 | 23.61999 | 85.9385 | 82.67356 | chr16 | 17208459 | 17210697 | 6.59345 | 4.79458 | 4.35853 |
| chr14 | 61687107 | 61687402 | 600 | Distal Intergenic | 14 | 61602839 | 61682373 | 79535 | 2 | ENSMUST00 | -4735 | Dleu2 | ENSMUSG0( | false | chr14 | 61686797 | 61687458 | 18.77433 | 54.14365 | 51.37372 | chr14 | 61686362 | 61689518 | 4.21988 | 3.70796 | 2.37131 |
| chr10 | 82763688 | 82764346 | 599 | Promoter (<=1kb) | 10 | 82748701 | 82764144 | 15444 | 2 | ENSMUST00 | 0 | Nfyb | ENSMUSG0( | false | chr10 | 82762672 | 82764547 | 32.64297 | 100.45817 | 96.87489 | chr10 | 82764302 | 82764693 | 5.83271 | 4.5575 | 3.73267 |
| chr17 | 78213140 | 78213570 | 599 | Intron (ENSMUST00000112498.2 | 17 | 78200248 | 78376592 | 176345 | 1 | ENSMUST00 | 12893 | Crim1 | ENSMUSG0( | true | . | . | . | . | . | . | chr17 | 78202641 | 78231492 | 6.38021 | 4.71597 | 4.13646 |
| chr1 | 6076648 | 6076992 | 599 | Distal Intergenic | 1 | 6206197 | 6234311 | 28115 | 1 | ENSMUST00 | -129205 | Rb1cc1 | ENSMUSG0( | false | . | . | . | . | . | . | chr1 | 6075888 | 6077998 | 4.70119 | 3.8822 | 2.80787 |
| chr8 | 70746595 | 70747374 | 599 | Promoter (<=1kb) | 8 | 70747891 | 70750814 | 2924 | 1 | ENSMUST00 | -517 | Pde4c | ENSMUSG0( | false | . | . | . | . | . | . | chr8 | 70740750 | 70748114 | 8.21466 | 5.47394 | 5.63599 |
| chr3 | 96562214 | 96562593 | 599 | Promoter (2-3kb) | 3 | 96560054 | 96560565 | 512 | 1 | ENSMUST00 | 2161 | Txnip | ENSMUSG0( | false | chr3 | 96557994 | 96562539 | 26.0714 | 83.99387 | 80.766 | chr3 | 96561069 | 96567035 | 5.48015 | 4.34024 | 3.394 |
| chr4 | 24966292 | 24966790 | 599 | Promoter (<=1kb) | 4 | 24966407 | 25007300 | 40894 | 1 | ENSMUST00 | 0 | Gpr63 | ENSMUSG0( | false | chr4 | 24966127 | 24967507 | 23.4661 | 79.19912 | 76.05865 | . | . | . | . | . | . |
| chr18 | 33464599 | 33464912 | 599 | Promoter (<=1kb) | 18 | 33464163 | 33472448 | 8286 | 1 | ENSMUST00 | 437 | Gm10549 | ENSMUSG0( | false | chr18 | 33462104 | 33464772 | 29.3377 | 90.74815 | 87.38457 | chr18 | 33464834 | 33465308 | 3.24523 | 3.19337 | 1.54667 |
| chr4 | 41255646 | 41255933 | 599 | Intron (ENSMUST00000108068.2 | 4 | 41252617 | 41252844 | 228 | 1 | ENSMUST00 | 3030 | Gm12401 | ENSMUSG0( | false | . | . | . | . | . | . | chr4 | 41251196 | 41261743 | 4.92657 | 3.98721 | 2.97941 |
| chr12 | 54286940 | 54287256 | 599 | Distal Intergenic | 12 | 54240755 | 54240900 | 146 | 2 | ENSMUST00 | -46041 | Gm24692 | ENSMUSG0( | false | . | . | . | . | . | . | chr12 | 54281533 | 54289226 | 4.50018 | 3.82184 | 2.59526 |
| chr4 | 123282555 | 123282850 | 598 | Promoter (<=1kb) | 4 | 123282825 | 123298925 | 16101 | 1 | ENSMUST00 | 0 | Pabpc4 | ENSMUSG0( | false | chr4 | 123282579 | 123284220 | 24.32174 | 95.77222 | 92.30251 | . | . | . | . | . | . |
| chr5 | 129724938 | 129725517 | 598 | Promoter (<=1kb) | 5 | 129725063 | 129758327 | 33265 | 1 | ENSMUST00 | 0 | Gbas | ENSMUSG0( | false | chr5 | 129724482 | 129726168 | 34.22706 | 108.29755 | 104.50865 | . | . | . | . | . | . |
| chr6 | 87671753 | 87672105 | 598 | Promoter (<=1kb) | 6 | 87628429 | 87672168 | 43740 | 2 | ENSMUST00 | 63 | Aplf | ENSMUSG0( | false | chr6 | 87671311 | 87672242 | 28.69254 | 91.56027 | 88.18607 | chr6 | 87670005 | 87673126 | 4.34392 | 3.68889 | 2.51856 |
| chr11 | 97804974 | 97805297 | 598 | Promoter (<=1kb) | 11 | 97805192 | 97836119 | 30928 | 1 | ENSMUST00 | 0 | Lasp1 | ENSMUSG0( | false | chr11 | 97804698 | 97805247 | 6.05356 | 9.81671 | 7.64356 | chr11 | 97801064 | 97833244 | 8.45203 | 5.49885 | 5.84074 |
| chr11 | 106160007 | 106160427 | 598 | Promoter (<=1kb) | 11 | 106156256 | 106160142 | 3887 | 2 | ENSMUST00 | 0 | Limd2 | ENSMUSG0( | false | chr11 | 106159606 | 106161040 | 26.39899 | 75.26234 | 72.19086 | chr11 | 106159086 | 106161629 | 4.45997 | 3.85708 | 2.56025 |
| chr13 | 32961187 | 32961442 | 598 | Distal Intergenic | 13 | 32965209 | 32979067 | 13859 | 1 | ENSMUST00 | -3767 | Serpinb6b | ENSMUSG0( | false | . | . | . | . | . | . | chr13 | 32957470 | 32971328 | 4.32621 | 3.59198 | 2.48777 |
| chr5 | 150167039 | 150167482 | 598 | Distal Intergenic | 5 | 150080622 | 150082184 | 1563 | 1 | ENSMUST00 | 86418 | Rxfp2 | ENSMUSG0( | false | . | . | . | . | . | . | chr5 | 150164826 | 150168895 | 4.98415 | 4.0526 | 3.01191 |
| chr7 | 66971178 | 66971427 | 598 | Intron (ENSMUST00000148538.2 | 7 | 66909468 | 66969555 | 60088 | 1 | ENSMUST00 | 61711 | Adamts17 | ENSMUSG0( | false | . | . | . | . | . | . | . | . | . | . | . | . |
| chr11 | 96309339 | 96309724 | 598 | Promoter (2-3kb) | 11 | 96291024 | 96306910 | 15887 | 2 | ENSMUST00 | -2430 | Hoxb5os | ENSMUSG0( | false | . | . | . | . | . | . | chr11 | 96308981 | 96316378 | 3.92719 | 3.49798 | 2.14941 |
| chr14 | 76622574 | 76622917 | 598 | Distal Intergenic | 14 | 76672963 | 76687169 | 14207 | 1 | ENSMUST00 | -50046 | RP23-421C2( | ENSMUSG0( | false | . | . | . | . | . | . | chr14 | 76621693 | 76625413 | 5.09533 | 4.18717 | 3.06836 |
| chr8 | 122537187 | 122537455 | 597 | Intron (ENSMUST00000127664.1 | 8 | 122481698 | 122551329 | 69632 | 2 | ENSMUST00 | 13874 | Piezo1 | ENSMUSG0( | false | . | . | . | . | . | . | chr8 | 122535817 | 122552157 | 5.24095 | 4.06992 | 3.22847 |
| chr11 | 8600712 | 8601073 | 597 | Intron (ENSMUST00000020695.7 | 11 | 8545693 | 8579121 | 33429 | 2 | ENSMUST00 | -21592 | Tns3 | ENSMUSG0( | false | . | . | . | . | . | . | . | . | . | . | . | . |
| chr12 | 111166094 | 111166664 | 597 | Promoter (<=1kb) | 12 | 111166370 | 111267153 | 100784 | 1 | ENSMUST00 | 0 | Traf3 | ENSMUSG0( | false | chr12 | 111165749 | 111167378 | 35.38397 | 108.24114 | 104.45254 | chr12 | 111164179 | 111166103 | 4.66567 | 3.95435 | 2.75039 |
| chr17 | 17373942 | 17374262 | 597 | Promoter (<=1kb) | 17 | 17374332 | 17394898 | 20567 | 1 | ENSMUST00 | -70 | Chd1 | ENSMUSG0( | false | chr17 | 17373492 | 17375488 | 25.98482 | 77.30067 | 74.19341 | . | . | . | . | . | . |
| chr1 | 4819594 | 4819879 | 597 | Intron (ENSMUST00000134384.2 | 1 | 4808237 | 4841093 | 32857 | 1 | ENSMUST00 | 11358 | Lypla1 | ENSMUSG0( | false | . | . | . | . | . | . | chr1 | 4818692 | 4822525 | 4.88199 | 4.07844 | 2.90351 |
| chr7 | 122101691 | 122102006 | 597 | Promoter (<=1kb) | 7 | 122085403 | 122101886 | 16484 | 2 | ENSMUST00 | 0 | Ndufab1 | ENSMUSG0( | false | chr7 | 122100946 | 122102271 | 34.36326 | 108.53439 | 104.74065 | chr7 | 122098290 | 122102367 | 5.71206 | 4.36131 | 3.60933 |
| chr5 | 110458978 | 110459242 | 597 | Distal Intergenic | 5 | 110361754 | 110448503 | 86750 | 2 | ENSMUST00 | -10476 | Fbrsl1 | ENSMUSG0( | false | . | . | . | . | . | . | chr5 | 110458161 | 110459676 | 4.48753 | 3.85049 | 2.61781 |
| chr18 | 54985594 | 54986165 | 597 | Exon (ENSMUST00000181829.1/ | 18 | 54888045 | 54990180 | 102136 | 2 | ENSMUST00 | 4015 | Zfp608 | ENSMUSG0( | false | chr18 | 54985294 | 54988807 | 28.53699 | 95.19626 | 91.7378 | chr18 | 54931543 | 54986508 | 4.96857 | 4.00781 | 2.99617 |
| chr9 | 48836046 | 48836604 | 597 | Promoter (<=1kb) | 9 | 48654311 | 48835945 | 181635 | 2 | ENSMUST00 | -102 | Zbtb16 | ENSMUSG0( | false | chr9 | 48834371 | 48836926 | 30.86817 | 91.45322 | 88.08056 | . | . | . | . | . | . |
| chr1 | 178369379 | 178369690 | 597 | Distal Intergenic | 1 | 178363401 | 178363521 | 121 | 2 | ENSMUST00 | -5859 | Gm27458 | ENSMUSG0( | false | chr1 | 178368575 | 178369724 | 12.59008 | 38.01365 | 35.45054 | chr1 | 178368243 | 178370203 | 4.34805 | 3.71336 | 2.52223 |
| chr2 | 33239994 | 33240322 | 596 | Intron (ENSMUST00000004208.5 | 2 | 33216098 | 33230124 | 14027 | 1 | ENSMUST00 | 23897 | Angptl2 | ENSMUSG0( | false | . | . | . | . | . | . | chr2 | 33228177 | 33242280 | 4.41522 | 3.65315 | 2.56656 |
| chr11 | 62281188 | 62281865 | 596 | Promoter (<=1kb) | 11 | 62267224 | 62281375 | 14152 | 2 | ENSMUST00 | 0 | Zswim7 | ENSMUSG0( | false | chr11 | 62280815 | 62282196 | 29.66345 | 100.21 | 96.63225 | . | . | . | . | . | . |
| chr10 | 39395571 | 39395862 | 596 | Intron (ENSMUST00000148152.2 | 10 | 39420009 | 39511859 | 91851 | 1 | ENSMUST00 | -24147 | Fyn | ENSMUSG0( | true | . | . | . | . | . | . | chr10 | 39392220 | 39400863 | 6.30447 | 4.65114 | 4.09998 |
| chr10 | 99681173 | 99681379 | 596 | Distal Intergenic | 10 | 99757705 | 99759658 | 1954 | 2 | ENSMUST00 | 78279 | Csl | ENSMUSG0( | false | . | . | . | . | . | . | chr10 | 99679107 | 99682874 | 3.53255 | 3.27773 | 1.82366 |
| chr5 | 28316776 | 28317563 | 596 | Promoter (<=1kb) | 5 | 28317121 | 28352868 | 35748 | 1 | ENSMUST00 | 0 | Rbm33 | ENSMUSG0( | false | chr5 | 28316314 | 28318412 | 27.39171 | 92.13462 | 88.74321 | . | . | . | . | . | . |
| chr4 | 65532805 | 65533100 | 596 | Intron (ENSMUST00000084496.2 | 4 | 65604986 | 65616238 | 11253 | 1 | ENSMUST00 | -71886 | Trim32 | ENSMUSG0( | false | . | . | . | . | . | . | chr4 | 65531415 | 65533938 | 4.93398 | 3.9332 | 2.97925 |
| chr16 | 34514032 | 34514439 | 596 | Promoter (<=1kb) | 16 | 34152024 | 34514048 | 362025 | 2 | ENSMUST00 | 0 | Kalrn | ENSMUSG0( | false | . | . | . | . | . | . | . | . | . | . | . | . |
| chr4 | 106996838 | 106997065 | 596 | Intron (ENSMUST00000150915.2 | 4 | 107017935 | 107022499 | 4565 | 2 | ENSMUST00 | 25434 | Gm12786 | ENSMUSG0( | false | . | . | . | . | . | . | chr4 | 106990941 | 107007182 | 5.43464 | 4.20931 | 3.38039 |
| chr11 | 106967643 | 106967997 | 596 | Distal Intergenic | 11 | 106972058 | 106973090 | 1033 | 2 | ENSMUST00 | 5093 | Gm11707 | ENSMUSG0( | false | . | . | . | . | . | . | . | . | . | . | . | . |
| chr10 | 111711503 | 111711960 | 596 | Distal Intergenic | 10 | 111586719 | 111594273 | 7555 | 2 | ENSMUST00 | -117231 | 4933440J02F | ENSMUSG0( | false | . | . | . | . | . | . | . | . | . | . | . | . |
| chr7 | 66427369 | 66427839 | 595 | Promoter (<=1kb) | 7 | 66390892 | 66427517 | 36626 | 2 | ENSMUST00 | 0 | Aldh1a3 | ENSMUSG0( | false | chr7 | 66427081 | 66428260 | 22.33599 | 59.52296 | 56.68532 | chr7 | 66427048 | 66428299 | 4.70129 | 3.88138 | 2.82061 |
| chr16 | 14705440 | 14705986 | 595 | Promoter (<=1kb) | 16 | 14705852 | 14709395 | 3544 | 1 | ENSMUST00 | 0 | Snai2 | ENSMUSG0( | false | chr16 | 14704873 | 14707329 | 21.49234 | 58.91379 | 56.0822 | . | . | . | . | . | . |
| chr4 | 151057898 | 151058299 | 595 | Promoter (<=1kb) | 4 | 151047300 | 151057953 | 10654 | 2 | ENSMUST00 | 0 | Vamp3 | ENSMUSG0( | false | chr4 | 151057007 | 151058085 | 42.74922 | 148.11885 | 142.90198 | chr4 | 151054347 | 151058952 | 4.63314 | 3.80747 | 2.72778 |
| chr5 | 32863427 | 32863772 | 595 | Promoter (<=1kb) | 5 | 32863701 | 32887048 | 23348 | 1 | ENSMUST00 | 0 | Depdc5 | ENSMUSG0( | false | chr5 | 32863170 | 32864374 | 20.32388 | 57.71489 | 54.90119 | . | . | . | . | . | . |
| chr12 | 112604035 | 112604370 | 595 | Exon (ENSMUST00000101029.2/ | 12 | 112588784 | 112615556 | 26773 | 1 | ENSMUST00 | 15252 | Inf2 | ENSMUSG0( | false | . | . | . | . | . | . | chr12 | 112602468 | 112606691 | 7.82262 | 5.33288 | 5.31694 |
| chr6 | 51567317 | 51567570 | 595 | Intron (ENSMUST00000049152.9 | 6 | 51544566 | 51589043 | 44478 | 1 | ENSMUST00 | 22752 | Snx10 | ENSMUSG0( | false | . | . | . | . | . | . | chr6 | 51567234 | 51567562 | 5.81446 | 4.55951 | 3.69426 |
| chr2 | 33628316 | 33628720 | 595 | Intron (ENSMUST00000041730.5 | 2 | 33567156 | 33632283 | 65128 | 2 | ENSMUST00 | 3563 | Lmx1b | ENSMUSG0( | true | . | . | . | . | . | . | chr2 | 33565260 | 33638297 | 6.78481 | 4.81457 | 4.48235 |
| chr14 | 21984721 | 21985126 | 595 | 3' UTR | 14 | 21983962 | 21989601 | 5640 | 2 | ENSMUST00 | 4475 | Zfp503 | ENSMUSG0( | false | chr14 | 21984527 | 21987918 | 37.75571 | 118.58805 | 114.49699 | chr14 | 21981011 | 21986083 | 4.65699 | 3.79582 | 2.74477 |
| chr11 | 107348152 | 107348481 | 595 | Promoter (<=1kb) | 11 | 107226257 | 107348130 | 121874 | 2 | ENSMUST00 | -23 | Pitpnc1 | ENSMUSG0( | false | . | . | . | . | . | . | chr11 | 107333685 | 107349983 | 5.09792 | 3.9909 | 3.09753 |
| chr11 | 102214579 | 102214941 | 594 | Intron (ENSMUST00000107151.3 | 11 | 102206628 | 102218928 | 12301 | 2 | ENSMUST00 | 3987 | Hdac5 | ENSMUSG0( | false | . | . | . | . | . | . | chr11 | 102213449 | 102231135 | 7.77054 | 5.31127 | 5.26338 |
| chr18 | 35388106 | 35388386 | 594 | Intron (ENSMUST00000025215.8 | 18 | 35266400 | 35498925 | 232526 | 2 | ENSMUST00 | 110539 | Sil1 | ENSMUSG0( | false | . | . | . | . | . | . | chr18 | 35387265 | 35388876 | 3.66486 | 3.1256 | 1.94966 |
| chr4 | 123718008 | 123718315 | 594 | Promoter (<=1kb) | 4 | 123712710 | 123718186 | 5477 | 2 | ENSMUST00 | 0 | Ndufs5 | ENSMUSG0( | false | chr4 | 123717632 | 123718719 | 27.03504 | 88.5911 | 85.27629 | . | . | . | . | . | . |
| chr8 | 111112073 | 111112356 | 594 | Intron (ENSMUST00000135302.2 | 8 | 111094655 | 111135147 | 40493 | 1 | ENSMUST00 | 17419 | Pdpr | ENSMUSG0( | false | . | . | . | . | . | . | chr8 | 111110632 | 111114038 | 4.19152 | 3.62042 | 2.38349 |
| chr5 | 21248331 | 21248603 | 594 | Intron (ENSMUST00000036031.8 | 5 | 21186267 | 21291701 | 105435 | 1 | ENSMUST00 | 62065 | Gsap | ENSMUSG0( | false | . | . | . | . | . | . | . | . | . | . | . | . |
| chr6 | 148318774 | 148319013 | 594 | Intron (ENSMUST00000060095.9 | 6 | 148354656 | 148355598 | 943 | 1 | ENSMUST00 | -35643 | Rps4l | ENSMUSG0( | false | . | . | . | . | . | . | chr6 | 148308870 | 148320731 | 5.77782 | 4.49565 | 3.63716 |
| chr5 | 73246500 | 73246871 | 594 | Intron (ENSMUST00000101127.6 | 5 | 73119257 | 73256425 | 137169 | 2 | ENSMUST00 | 9554 | Fryl | ENSMUSG0( | true | . | . | . | . | . | . | chr5 | 73196545 | 73254445 | 6.29302 | 4.62166 | 4.08669 |
| chr6 | 38698524 | 38698796 | 594 | Exon (ENSMUST00000161779.2/ | 6 | 38663069 | 38680864 | 17796 | 1 | ENSMUST00 | 35456 | Clec2l | ENSMUSG0( | false | . | . | . | . | . | . | chr6 | 38697150 | 38699956 | 4.76519 | 3.96524 | 2.81256 |
| chr6 | 118808344 | 118808610 | 594 | Intron (ENSMUST00000186889.1 | 6 | 118834900 | 118835000 | 101 | 1 | ENSMUST00 | -26290 | Gm25905 | ENSMUSG0( | false | . | . | . | . | . | . | chr6 | 118807940 | 118809742 | 5.03953 | 3.96835 | 3.07501 |
| chr7 | 99626858 | 99627093 | 594 | Promoter (<=1kb) | 7 | 99624082 | 99627103 | 3022 | 2 | ENSMUST00 | 10 | Gm4980 | ENSMUSG0( | false | chr7 | 99625504 | 99627205 | 14.6238 | 41.22485 | 38.6191 | chr7 | 99625516 | 99627135 | 7.09119 | 4.96505 | 4.71893 |
| chr19 | 45047414 | 45047869 | 593 | Promoter (<=1kb) | 19 | 45047576 | 45056383 | 8808 | 1 | ENSMUST00 | 0 | Sfxn3 | ENSMUSG0( | false | chr19 | 45046467 | 45048099 | 15.37965 | 40.41568 | 37.82122 | chr19 | 45046056 | 45049317 | 4.41989 | 3.70739 | 2.54796 |
| chr6 | 108321701 | 108322055 | 593 | Intron (ENSMUST00000032192.6 | 6 | 108213118 | 108551109 | 337992 | 1 | ENSMUST00 | 108584 | Itpr1 | ENSMUSG0( | false | . | . | . | . | . | . | chr6 | 108320510 | 108322581 | 4.68824 | 3.85537 | 2.78226 |
| chr9 | 51410099 | 51410357 | 593 | Distal Intergenic | 9 | 51289686 | 51328917 | 39232 | 2 | ENSMUST00 | -81183 | 1810046K07I | ENSMUSG0( | false | . | . | . | . | . | . | chr9 | 51404996 | 51411058 | 4.97376 | 4.06092 | 2.98267 |
| chr5 | 21186024 | 21186598 | 593 | Promoter (<=1kb) | 5 | 21186267 | 21291701 | 105435 | 1 | ENSMUST00 | 0 | Gsap | ENSMUSG0( | false | chr5 | 21185878 | 21186941 | 30.6477 | 94.99754 | 91.54929 | chr5 | 21185582 | 21186790 | 3.13633 | 3.09315 | 1.48693 |
| chr4 | 25281514 | 25282013 | 593 | Promoter (<=1kb) | 4 | 25248600 | 25281760 | 33161 | 2 | ENSMUST00 | 0 | Ufl1 | ENSMUSG0( | false | chr4 | 25280968 | 25282172 | 38.87692 | 132.39133 | 127.80991 | chr4 | 25279810 | 25282119 | 4.17517 | 3.67332 | 2.31245 |
| chr16 | 16213989 | 16214419 | 593 | Promoter (<=1kb) | 16 | 16213318 | 16272712 | 59395 | 1 | ENSMUST00 | 672 | Pkp2 | ENSMUSG0( | false | chr16 | 16212978 | 16214696 | 35.53755 | 114.73917 | 110.76805 | chr16 | 16213695 | 16226218 | 5.0735 | 4.12792 | 3.06455 |
| chr4 | 54758133 | 54758494 | 593 | Distal Intergenic | 4 | 54710624 | 54710912 | 289 | 1 | ENSMUST00 | 47510 | Gm24628 | ENSMUSG0( | false | chr4 | 54758124 | 54758708 | 8.58926 | 15.99743 | 13.72071 | chr4 | 54757243 | 54759235 | 4.80673 | 3.97537 | 2.85595 |
| chr4 | 109391743 | 109392163 | 593 | Distal Intergenic | 4 | 109403794 | 109404212 | 419 | 2 | ENSMUST00 | 12049 | Gm12750 | ENSMUSG0( | false | . | . | . | . | . | . | chr4 | 109391349 | 109394447 | 3.77651 | 3.43951 | 2.04858 |
| chr1 | 102862460 | 102862869 | 593 | Distal Intergenic | 1 | 102953415 | 102953548 | 134 | 2 | ENSMUST00 | 90679 | Gm22034 | ENSMUSG0( | false | . | . | . | . | . | . | . | . | . | . | . | . |
| chr17 | 35444811 | 35445216 | 593 | Promoter (<=1kb) | 17 | 35445856 | 35448926 | 3071 | 1 | ENSMUST00 | -640 | Gm8752 | ENSMUSG0( | false | . | . | . | . | . | . | chr17 | 35437568 | 35444857 | 5.8895 | 3.7427 | 3.77509 |
| chr15 | 77911150 | 77911484 | 592 | Distal Intergenic | 15 | 77915047 | 77915700 | 654 | 2 | ENSMUST00 | 4216 | Txn2 | ENSMUSG0( | false | . | . | . | . | . | . | chr15 | 77909803 | 77913581 | 7.31704 | 5.07955 | 4.9193 |
| chr19 | 57017328 | 57017672 | 592 | Distal Intergenic | 19 | 56923540 | 57008228 | 84689 | 2 | ENSMUST00 | -9101 | Afap1l2 | ENSMUSG0( | false | . | . | . | . | . | . | chr19 | 57017473 | 57017943 | 3.06344 | 3.0122 | 1.41468 |
| chr4 | 134950844 | 134951094 | 592 | Distal Intergenic | 4 | 134931010 | 134936078 | 5069 | 1 | ENSMUST00 | 19835 | Syf2 | ENSMUSG0( | false | . | . | . | . | . | . | chr4 | 134950015 | 134951701 | 6.14365 | 4.57725 | 3.94104 |
| chr15 | 77182877 | 77183148 | 592 | Intron (ENSMUST00000048145.6 | 15 | 77078991 | 77153812 | 74822 | 2 | ENSMUST00 | -29066 | Rbfox2 | ENSMUSG0( | false | . | . | . | . | . | . | chr15 | 77182513 | 77188448 | 5.00664 | 3.99889 | 3.01909 |
| chr2 | 20520724 | 20520981 | 592 | Promoter (<=1kb) | 2 | 20519828 | 20810536 | 290709 | 1 | ENSMUST00 | 897 | Etl4 | ENSMUSG0( | true | chr2 | 20519278 | 20521509 | 21.87917 | 60.42461 | 57.57317 | chr2 | 20520657 | 20527429 | 4.7812 | 3.98118 | 2.83529 |
| chr4 | 136595347 | 136595621 | 592 | Intron (ENSMUST00000116273.3 | 4 | 136550848 | 136602308 | 51461 | 2 | ENSMUST00 | 6687 | Kdm1a | ENSMUSG0( | false | . | . | . | . | . | . | chr4 | 136592196 | 136597755 | 4.3058 | 3.69518 | 2.48663 |
| chr9 | 62506458 | 62506761 | 592 | Intron (ENSMUST00000048043.6 | 9 | 62427936 | 62510498 | 82563 | 2 | ENSMUST00 | 3737 | Coro2b | ENSMUSG0( | true | . | . | . | . | . | . | chr9 | 62449515 | 62535966 | 6.98492 | 4.90408 | 4.64549 |
| chr8 | 70564804 | 70565141 | 592 | Intron (ENSMUST00000093454.6 | 8 | 70539675 | 70592858 | 53184 | 1 | ENSMUST00 | 25130 | Ell | ENSMUSG0( | false | . | . | . | . | . | . | chr8 | 70550287 | 70567277 | 5.6244 | 4.31142 | 3.53604 |
| chr2 | 168003453 | 168003759 | 592 | Intron (ENSMUST00000123782.2 | 2 | 168010529 | 168011506 | 978 | 1 | ENSMUST00 | -6770 | Gm14236 | ENSMUSG0( | false | . | . | . | . | . | . | chr2 | 168002556 | 168004420 | 4.9329 | 4.0387 | 2.9499 |
| chr10 | 108531229 | 108531494 | 592 | Intron (ENSMUST00000105276.2 | 10 | 108428067 | 108428160 | 94 | 1 | ENSMUST00 | 103163 | Gm25591 | ENSMUSG0( | false | . | . | . | . | . | . | chr10 | 108530892 | 108533125 | 4.77833 | 4.02337 | 2.81631 |
| chr2 | 166092905 | 166093208 | 591 | Intron (ENSMUST00000109249.3 | 2 | 166074815 | 166078119 | 3305 | 2 | ENSMUST00 | -14787 | Sulf2 | ENSMUSG0( | false | chr2 | 166092157 | 166093106 | 5.905 | 11.42563 | 9.22457 | chr2 | 166085159 | 166097943 | 5.73745 | 4.40556 | 3.61152 |
| chr11 | 97699678 | 97700628 | 591 | Promoter (<=1kb) | 11 | 97689820 | 97700497 | 10678 | 2 | ENSMUST00 | 0 | Pcgf2 | ENSMUSG0( | false | chr11 | 97698478 | 97701285 | 33.7938 | 119.00099 | 114.88904 | chr11 | 97694773 | 97702506 | 4.47077 | 3.79783 | 2.58655 |
| chrX | 140567806 | 140568138 | 591 | Intron (ENSMUST00000112996.3 | 20 | 140543103 | 140548529 | 5427 | 2 | ENSMUST00 | -19278 | Tsc22d3 | ENSMUSG0( | false | . | . | . | . | . | . | . | . | . | . | . | . |
| chrX | 53268997 | 53269327 | 591 | Promoter (<=1kb) | 20 | 53243425 | 53269020 | 25596 | 2 | ENSMUST00 | 0 | Fam122b | ENSMUSG0( | false | chrX | 53268993 | 53269848 | 23.43614 | 63.09834 | 60.20888 | . | . | . | . | . | . |
| chr1 | 166409301 | 166410066 | 591 | Promoter (<=1kb) | 1 | 166384622 | 166409782 | 25161 | 2 | ENSMUST00 | 0 | Pogk | ENSMUSG0( | false | chr1 | 166409016 | 166410272 | 28.31422 | 85.10979 | 81.86543 | . | . | . | . | . | . |
| chr17 | 32536430 | 32536690 | 591 | Promoter (<=1kb) | 17 | 32536558 | 32551798 | 15241 | 1 | ENSMUST00 | 0 | Cyp4f16 | ENSMUSG0( | false | . | . | . | . | . | . | . | . | . | . | . | . |
| chr11 | 48994159 | 48994401 | 591 | Promoter (<=1kb) | 11 | 48987481 | 48994172 | 6692 | 2 | ENSMUST00 | 0 | Tgtp1 | ENSMUSG0( | false | . | . | . | . | . | . | chr11 | 48993396 | 48994693 | 3.99061 | 3.58936 | 2.20141 |
| chr6 | 52172566 | 52172926 | 591 | Promoter (<=1kb) | 6 | 52172881 | 52174059 | 1179 | 1 | ENSMUST00 | 0 | Gm15050 | ENSMUSG0( | false | chr6 | 52172354 | 52173164 | 13.64664 | 30.56956 | 28.09773 | chr6 | 52170880 | 52173229 | 3.81563 | 3.49765 | 2.0493 |
| chr15 | 97927240 | 97927530 | 591 | Distal Intergenic | 15 | 97935285 | 97935595 | 311 | 1 | ENSMUST00 | -7755 | Gm22202 | ENSMUSG0( | false | . | . | . | . | . | . | chr15 | 97924986 | 97928566 | 6.27623 | 4.66463 | 4.05606 |
| chr9 | 89770921 | 89771462 | 591 | Distal Intergenic | 9 | 89759003 | 89759465 | 463 | 1 | ENSMUST00 | 11919 | RP24-340L11 | ENSMUSG0( | false | . | . | . | . | . | . | chr9 | 89768908 | 89779398 | 4.75528 | 3.93987 | 2.80985 |
| chr8 | 3502203 | 3502546 | 590 | Promoter (1-2kb) | 8 | 3500504 | 3506941 | 6438 | 1 | ENSMUST00 | 1700 | Mcoln1 | ENSMUSG0( | false | . | . | . | . | . | . | chr8 | 3499617 | 3504551 | 7.30944 | 5.14934 | 4.90367 |
| chr17 | 28485483 | 28486469 | 590 | Promoter (<=1kb) | 17 | 28399095 | 28486149 | 87055 | 2 | ENSMUST00 | 0 | Fkbp5 | ENSMUSG0( | false | chr17 | 28484845 | 28486181 | 41.78299 | 136.36734 | 131.63638 | chr17 | 28484741 | 28486666 | 4.34938 | 3.65334 | 2.48659 |
| chr3 | 122088402 | 122088763 | 590 | Intron (ENSMUST00000013995.8 | 3 | 122074456 | 122087931 | 13476 | 1 | ENSMUST00 | 13947 | Abca4 | ENSMUSG0( | false | . | . | . | . | . | . | chr3 | 122084432 | 122089037 | 4.64391 | 3.69483 | 2.74239 |
| chr5 | 115547047 | 115547499 | 590 | Promoter (1-2kb) | 5 | 115545304 | 115545775 | 472 | 1 | ENSMUST00 | 1744 | Pxn | ENSMUSG0( | false | . | . | . | . | . | . | chr5 | 115507551 | 115550291 | 7.76238 | 5.17462 | 5.28263 |
| chr7 | 28771862 | 28772177 | 590 | Exon (ENSMUST00000122915.2/ | 7 | 28761769 | 28767512 | 5744 | 2 | ENSMUST00 | -4351 | Nfkbib | ENSMUSG0( | false | . | . | . | . | . | . | chr7 | 28770641 | 28774873 | 5.72688 | 4.37837 | 3.60962 |
| chr9 | 69478400 | 69478665 | 590 | Promoter (<=1kb) | 9 | 69478465 | 69491795 | 13331 | 1 | ENSMUST00 | 0 | Anxa2 | ENSMUSG0( | false | . | . | . | . | . | . | chr9 | 69455103 | 69480655 | 7.58546 | 5.20708 | 5.13718 |
| chr18 | 29703443 | 29704008 | 590 | Distal Intergenic | 18 | 30272747 | 30348126 | 75380 | 1 | ENSMUST00 | -568739 | Pik3c3 | ENSMUSG0( | false | . | . | . | . | . | . | chr18 | 29703706 | 29704784 | 3.26229 | 3.20647 | 1.61418 |
| chr18 | 62150513 | 62150819 | 590 | Distal Intergenic | 18 | 62177817 | 62179959 | 2143 | 2 | ENSMUST00 | 29140 | Adrb2 | ENSMUSG0( | false | . | . | . | . | . | . | chr18 | 62149591 | 62152003 | 4.40668 | 3.80099 | 2.55571 |
| chr4 | 131386944 | 131387249 | 590 | Distal Intergenic | 4 | 131487197 | 131505527 | 18331 | 1 | ENSMUST00 | -99948 | Gm831 | ENSMUSG0( | false | . | . | . | . | . | . | chr4 | 131383399 | 131387394 | 3.53373 | 3.31308 | 1.84011 |
| chr11 | 45535484 | 45535789 | 590 | Distal Intergenic | 11 | 45452399 | 45468504 | 16106 | 2 | ENSMUST00 | -66981 | Gm12162 | ENSMUSG0( | false | . | . | . | . | . | . | . | . | . | . | . | . |
| chr4 | 137356954 | 137357278 | 589 | Promoter (<=1kb) | 4 | 137321762 | 137357699 | 35938 | 2 | ENSMUST00 | 421 | Cdc42 | ENSMUSG0( | false | chr4 | 137356219 | 137358176 | 31.28813 | 116.44205 | 112.41405 | . | . | . | . | . | . |
| chr11 | 23895151 | 23895468 | 589 | Promoter (<=1kb) | 11 | 23862646 | 23895253 | 32608 | 2 | ENSMUST00 | 0 | Papolg | ENSMUSG0( | false | chr11 | 23894200 | 23895678 | 27.34946 | 104.78247 | 101.09169 | . | . | . | . | . | . |
| chr17 | 83213434 | 83213809 | 589 | Promoter (1-2kb) | 17 | 83215292 | 83225070 | 9779 | 1 | ENSMUST00 | -1483 | Pkdcc | ENSMUSG0( | false | chr17 | 83213509 | 83220055 | 37.87741 | 120.9216 | 116.74998 | chr17 | 83212533 | 83214208 | 5.39581 | 4.15799 | 3.38073 |
| chr19 | 53311157 | 53311417 | 589 | Promoter (<=1kb) | 19 | 53310506 | 53375810 | 65305 | 1 | ENSMUST00 | 652 | Mxi1 | ENSMUSG0( | false | chr19 | 53310158 | 53314217 | 35.4076 | 109.31139 | 105.49952 | . | . | . | . | . | . |
| chr14 | 121051798 | 121052451 | 589 | Intron (ENSMUST00000135010.2 | 14 | 121035564 | 121283744 | 248181 | 1 | ENSMUST00 | 16235 | Farp1 | ENSMUSG0( | false | . | . | . | . | . | . | chr14 | 121049104 | 121054112 | 4.4634 | 3.79705 | 2.61369 |
| chr5 | 140565514 | 140565824 | 589 | Promoter (<=1kb) | 5 | 140563119 | 140565071 | 1953 | 2 | ENSMUST00 | -444 | Grifin | ENSMUSG0( | false | . | . | . | . | . | . | chr5 | 140564936 | 140569303 | 4.21005 | 3.64271 | 2.37106 |
| chr10 | 127739251 | 127739923 | 589 | Promoter (<=1kb) | 10 | 127739538 | 127747335 | 7798 | 1 | ENSMUST00 | 0 | Zbtb39 | ENSMUSG0( | false | chr10 | 127739030 | 127740414 | 31.85236 | 101.09525 | 97.496 | chr10 | 127739155 | 127742117 | 3.35156 | 3.09137 | 1.68333 |
| chr7 | 96648803 | 96649107 | 589 | Intron (ENSMUST00000107162.2 | 7 | 96644172 | 96732280 | 88109 | 1 | ENSMUST00 | 4632 | Tenm4 | ENSMUSG0( | false | . | . | . | . | . | . | chr7 | 96647958 | 96649696 | 3.73754 | 3.27443 | 1.99518 |
| chr7 | 19228154 | 19228621 | 589 | Promoter (<=1kb) | 7 | 19228334 | 19247868 | 19535 | 1 | ENSMUST00 | 0 | Opa3 | ENSMUSG0( | false | chr7 | 19228284 | 19228710 | 26.38389 | 80.78929 | 77.61987 | . | . | . | . | . | . |
| chr16 | 8171239 | 8171540 | 589 | Distal Intergenic | 16 | 8203975 | 8204248 | 274 | 1 | ENSMUST00 | -32435 | Gm23476 | ENSMUSG0( | false | . | . | . | . | . | . | . | . | . | . | . | . |
| chr16 | 4640148 | 4640577 | 588 | Promoter (<=1kb) | 16 | 4639989 | 4687292 | 47304 | 1 | ENSMUST00 | 160 | Dnaja3 | ENSMUSG0( | false | chr16 | 4639898 | 4641726 | 32.64888 | 103.50044 | 99.84376 | . | . | . | . | . | . |
| chr2 | 26910501 | 26910781 | 588 | Promoter (<=1kb) | 2 | 26905262 | 26910677 | 5416 | 2 | ENSMUST00 | 0 | Med22 | ENSMUSG0( | false | chr2 | 26909810 | 26912049 | 20.20131 | 69.382 | 66.39919 | . | . | . | . | . | . |
| chr11 | 49232785 | 49233092 | 588 | Distal Intergenic | 11 | 49244191 | 49263022 | 18832 | 1 | ENSMUST00 | -11099 | Mgat1 | ENSMUSG0( | false | . | . | . | . | . | . | chr11 | 49232277 | 49233539 | 3.20292 | 3.11006 | 1.58199 |
| chr12 | 111412765 | 111413461 | 588 | Distal Intergenic | 12 | 111417430 | 111431182 | 13753 | 1 | ENSMUST00 | -3969 | Exoc3l4 | ENSMUSG0( | false | . | . | . | . | . | . | chr12 | 111411393 | 111413852 | 5.10088 | 4.07375 | 3.09391 |
| chr19 | 46478977 | 46479279 | 588 | Intron (ENSMUST00000039922.7 | 19 | 46483097 | 46485982 | 2886 | 1 | ENSMUST00 | -3818 | Sufu | ENSMUSG0( | false | . | . | . | . | . | . | chr19 | 46468103 | 46486697 | 7.92449 | 5.26231 | 5.41796 |
| chr15 | 58822959 | 58823737 | 588 | Promoter (<=1kb) | 15 | 58782269 | 58823530 | 41262 | 2 | ENSMUST00 | 0 | Tmem65 | ENSMUSG0( | false | chr15 | 58822098 | 58824291 | 33.25798 | 110.72714 | 106.87233 | . | . | . | . | . | . |
| chr6 | 66077454 | 66077754 | 588 | Distal Intergenic | 6 | 65989640 | 65989739 | 100 | 2 | ENSMUST00 | -87716 | Gm23625 | ENSMUSG0( | false | . | . | . | . | . | . | chr6 | 66076883 | 66077847 | 3.77044 | 3.34435 | 2.03949 |
| chr5 | 73248375 | 73248662 | 588 | Intron (ENSMUST00000101127.6 | 5 | 73119257 | 73256425 | 137169 | 2 | ENSMUST00 | 7763 | Fryl | ENSMUSG0( | true | . | . | . | . | . | . | chr5 | 73196545 | 73254445 | 6.29302 | 4.62166 | 4.08669 |
| chr3 | 153571167 | 153571504 | 588 | Intron (ENSMUST00000064460.5 | 3 | 153621702 | 153621969 | 268 | 1 | ENSMUST00 | -50198 | Gm22136 | ENSMUSG0( | false | . | . | . | . | . | . | chr3 | 153570549 | 153576267 | 3.53452 | 3.34797 | 1.80763 |
| chr15 | 81523087 | 81523509 | 588 | Distal Intergenic | 15 | 81499084 | 81499548 | 465 | 2 | ENSMUST00 | -23540 | Gm5218 | ENSMUSG0( | false | chr15 | 81522003 | 81524194 | 22.65707 | 70.49107 | 67.49436 | . | . | . | . | . | . |
| chr10 | 4423310 | 4423792 | 587 | Exon (ENSMUST00000132184.1/ | 10 | 4427243 | 4432295 | 5053 | 2 | ENSMUST00 | 8503 | Rmnd1 | ENSMUSG0( | false | . | . | . | . | . | . | chr10 | 4422045 | 4437373 | 5.83291 | 4.40627 | 3.70343 |
| chr13 | 34575336 | 34575652 | 587 | Distal Intergenic | 13 | 34628868 | 34648568 | 19701 | 2 | ENSMUST00 | 72916 | Pxdc1 | ENSMUSG0( | true | . | . | . | . | . | . | chr13 | 34569305 | 34576256 | 4.71058 | 3.88329 | 2.78127 |
| chr5 | 66589210 | 66589527 | 587 | Intron (ENSMUST00000159786.2 | 5 | 66451982 | 66618640 | 166659 | 2 | ENSMUST00 | 29113 | Apbb2 | ENSMUSG0( | false | . | . | . | . | . | . | chr5 | 66558047 | 66617728 | 6.24773 | 4.56399 | 4.04918 |
| chr7 | 120981542 | 120982504 | 587 | Promoter (<=1kb) | 7 | 120957036 | 120981790 | 24755 | 2 | ENSMUST00 | 0 | Cdr2 | ENSMUSG0( | false | chr7 | 120980868 | 120983246 | 36.96683 | 120.40339 | 116.25284 | chr7 | 120961858 | 120981649 | 5.33893 | 4.28516 | 3.2901 |
| chr2 | 27079152 | 27079442 | 587 | Promoter (<=1kb) | 2 | 27079379 | 27108981 | 29603 | 1 | ENSMUST00 | 0 | Adamtsl2 | ENSMUSG0( | false | . | . | . | . | . | . | chr2 | 27077756 | 27080113 | 6.10537 | 4.60349 | 3.89596 |
| chr15 | 99223595 | 99224162 | 587 | Promoter (<=1kb) | 15 | 99224976 | 99242817 | 17842 | 1 | ENSMUST00 | -814 | Kcnh3 | ENSMUSG0( | false | chr15 | 99223127 | 99225653 | 34.11661 | 104.01456 | 100.34361 | chr15 | 99221254 | 99225886 | 4.91003 | 3.88376 | 2.96688 |
| chr6 | 72391109 | 72391651 | 587 | Promoter (<=1kb) | 6 | 72385223 | 72390703 | 5481 | 2 | ENSMUST00 | -407 | Vamp8 | ENSMUSG0( | false | chr6 | 72389183 | 72391497 | 20.5555 | 53.19651 | 50.4419 | chr6 | 72385690 | 72394714 | 4.39484 | 3.72006 | 2.54009 |
| chr1 | 64809026 | 64809407 | 587 | Intron (ENSMUST00000139649.2 | 1 | 64793809 | 64836764 | 42956 | 2 | ENSMUST00 | 27357 | Plekhm3 | ENSMUSG0( | false | . | . | . | . | . | . | chr1 | 64807910 | 64810357 | 4.07953 | 3.70831 | 2.23133 |
| chr10 | 105793815 | 105794081 | 587 | Intron (ENSMUST00000046638.9 | 10 | 105804087 | 105804197 | 111 | 2 | ENSMUST00 | 10116 | n-R5s80 | ENSMUSG0( | false | . | . | . | . | . | . | chr10 | 105793907 | 105794896 | 2.99256 | 2.95804 | 1.42139 |
| chr9 | 102267176 | 102267910 | 587 | Intron (ENSMUST00000035129.8 | 9 | 102304924 | 102304975 | 52 | 2 | ENSMUST00 | 37065 | Mir7243 | ENSMUSG0( | false | chr9 | 102267473 | 102268314 | 13.45719 | 33.53756 | 31.02805 | chr9 | 102258607 | 102274961 | 4.8108 | 3.97089 | 2.85625 |
| chr16 | 90283572 | 90284715 | 586 | Promoter (<=1kb) | 16 | 90228957 | 90284308 | 55352 | 2 | ENSMUST00 | 0 | Scaf4 | ENSMUSG0( | false | chr16 | 90282811 | 90285323 | 29.75837 | 125.79383 | 121.46131 | . | . | . | . | . | . |
| chr3 | 127257563 | 127257892 | 586 | Intron (ENSMUST00000182704.2 | 3 | 126923957 | 127225932 | 301976 | 2 | ENSMUST00 | -31632 | Ank2 | ENSMUSG0( | false | chr3 | 127257496 | 127258209 | 18.84082 | 47.13427 | 44.45482 | . | . | . | . | . | . |
| chr10 | 86933950 | 86934208 | 586 | Intron (ENSMUST00000035288.9 | 10 | 86918662 | 86940640 | 21979 | 2 | ENSMUST00 | 6432 | Stab2 | ENSMUSG0( | false | . | . | . | . | . | . | chr10 | 86933198 | 86934533 | 4.19101 | 3.65737 | 2.40833 |
| chr5 | 76974546 | 76975063 | 586 | Promoter (<=1kb) | 5 | 76974683 | 76999937 | 25255 | 1 | ENSMUST00 | 0 | Srp72 | ENSMUSG0( | false | chr5 | 76974036 | 76975655 | 31.07893 | 124.33488 | 120.0426 | . | . | . | . | . | . |
| chr3 | 37653926 | 37654317 | 586 | Distal Intergenic | 3 | 37640080 | 37644598 | 4519 | 1 | ENSMUST00 | 13847 | Spry1 | ENSMUSG0( | false | . | . | . | . | . | . | chr3 | 37653151 | 37655050 | 4.91027 | 4.00622 | 2.97078 |
| chr11 | 77489539 | 77490128 | 586 | Promoter (<=1kb) | 11 | 77470485 | 77489666 | 19182 | 2 | ENSMUST00 | 0 | Ankrd13b | ENSMUSG0( | false | chr11 | 77489086 | 77490311 | 27.51607 | 79.95555 | 76.80458 | chr11 | 77488828 | 77499060 | 5.56381 | 4.29511 | 3.47349 |
| chr2 | 165034534 | 165035032 | 586 | Promoter (<=1kb) | 2 | 165000357 | 165034867 | 34511 | 2 | ENSMUST00 | 0 | Ncoa5 | ENSMUSG0( | false | chr2 | 165033630 | 165035229 | 29.66345 | 100.21 | 96.63225 | chr2 | 165031140 | 165035422 | 5.92799 | 4.49443 | 3.75369 |
| chr17 | 49539303 | 49539624 | 586 | Intron (ENSMUST00000057610.6 | 17 | 49456022 | 49564337 | 108316 | 2 | ENSMUST00 | 24713 | Daam2 | ENSMUSG0( | true | . | . | . | . | . | . | chr17 | 49511234 | 49548441 | 5.06175 | 4.03075 | 3.07349 |
| chr13 | 45897633 | 45897975 | 586 | Intron (ENSMUST00000091628.4 | 13 | 45549758 | 45964991 | 415234 | 2 | ENSMUST00 | 67016 | Atxn1 | ENSMUSG0( | false | . | . | . | . | . | . | chr13 | 45897011 | 45898862 | 3.09204 | 2.93398 | 1.47583 |
| chr18 | 61368178 | 61368642 | 586 | Intron (ENSMUST00000075299.7 | 18 | 61397813 | 61397925 | 113 | 1 | ENSMUST00 | -29171 | Gm25301 | ENSMUSG0( | false | . | . | . | . | . | . | chr18 | 61366913 | 61371114 | 7.38415 | 5.03346 | 5.00481 |
| chr11 | 23665678 | 23666167 | 585 | Promoter (<=1kb) | 11 | 23665697 | 23731525 | 65829 | 1 | ENSMUST00 | 0 | Pus10 | ENSMUSG0( | false | chr11 | 23664733 | 23667134 | 29.58305 | 102.06358 | 98.44028 | . | . | . | . | . | . |
| chr14 | 67235466 | 67235769 | 585 | Promoter (<=1kb) | 14 | 67234644 | 67428687 | 194044 | 1 | ENSMUST00 | 823 | Ebf2 | ENSMUSG0( | false | . | . | . | . | . | . | chr14 | 67231884 | 67237790 | 5.12034 | 4.11192 | 3.11175 |
| chr19 | 10525175 | 10525936 | 585 | Promoter (<=1kb) | 19 | 10500513 | 10525209 | 24697 | 2 | ENSMUST00 | 0 | Sdhaf2 | ENSMUSG0( | false | chr19 | 10524504 | 10526311 | 34.9887 | 122.22292 | 118.01233 | . | . | . | . | . | . |
| chr6 | 8524867 | 8525093 | 585 | Intron (ENSMUST00000162564.2 | 6 | 8520058 | 8593512 | 73455 | 1 | ENSMUST00 | 4810 | Glcci1 | ENSMUSG0( | false | . | . | . | . | . | . | chr6 | 8522600 | 8527998 | 4.12889 | 3.62209 | 2.29534 |
| chr16 | 96127554 | 96128253 | 585 | Promoter (<=1kb) | 16 | 96120618 | 96127729 | 7112 | 2 | ENSMUST00 | 0 | Hmgn1 | ENSMUSG0( | false | chr16 | 96126477 | 96128557 | 40.93993 | 131.61429 | 127.06391 | . | . | . | . | . | . |
| chr16 | 32332082 | 32332388 | 585 | Promoter (<=1kb) | 16 | 32332252 | 32393747 | 61496 | 1 | ENSMUST00 | 0 | Ubxn7 | ENSMUSG0( | false | chr16 | 32331774 | 32333666 | 37.70819 | 121.69438 | 117.49508 | . | . | . | . | . | . |
| chr1 | 91460482 | 91460807 | 585 | Promoter (1-2kb) | 1 | 91415982 | 91459324 | 43343 | 2 | ENSMUST00 | -1159 | Per2 | ENSMUSG0( | false | . | . | . | . | . | . | chr1 | 91451145 | 91461043 | 6.24965 | 4.62037 | 4.05296 |
| chr11 | 57767885 | 57768222 | 585 | Intron (ENSMUST00000066987.8 | 11 | 57764427 | 57764545 | 119 | 1 | ENSMUST00 | 3459 | n-R5s69 | ENSMUSG0( | false | . | . | . | . | . | . | chr11 | 57744780 | 57769452 | 6.61093 | 4.7685 | 4.34185 |
| chr13 | 21506211 | 21506610 | 585 | Promoter (1-2kb) | 13 | 21504382 | 21505056 | 675 | 2 | ENSMUST00 | -1156 | Zfp389 | ENSMUSG0( | false | chr13 | 21505925 | 21506836 | 28.36989 | 85.2065 | 81.96005 | chr13 | 21505803 | 21506932 | 3.87999 | 3.3988 | 2.12773 |
| chr19 | 36706191 | 36706461 | 585 | Promoter (<=1kb) | 19 | 36614052 | 36705641 | 91590 | 2 | ENSMUST00 | -551 | 1500017E21I | ENSMUSG0( | false | . | . | . | . | . | . | chr19 | 36706340 | 36707152 | 3.65576 | 3.33925 | 1.90163 |
| chr12 | 12262584 | 12262971 | 584 | Promoter (<=1kb) | 12 | 12262139 | 12376359 | 114221 | 1 | ENSMUST00 | 446 | Fam49a | ENSMUSG0( | false | chr12 | 12261863 | 12263016 | 19.89052 | 67.91473 | 64.9535 | chr12 | 12261396 | 12264758 | 5.2038 | 4.07065 | 3.19463 |
| chr12 | 111350088 | 111350510 | 584 | Intron (ENSMUST00000041965.3 | 12 | 111292972 | 111377718 | 84747 | 2 | ENSMUST00 | 27208 | Cdc42bpb | ENSMUSG0( | false | . | . | . | . | . | . | chr12 | 111345052 | 111376732 | 6.23386 | 4.58338 | 4.03546 |
| chr4 | 31963766 | 31964258 | 584 | Promoter (<=1kb) | 4 | 31964097 | 32023467 | 59371 | 1 | ENSMUST00 | 0 | Map3k7 | ENSMUSG0( | false | chr4 | 31963245 | 31965103 | 24.24157 | 89.02564 | 85.70093 | . | . | . | . | . | . |

| | | | | | | | | | | | | | | | | | | | | | | | | | | |
|---|---|---|---|---|---|---|---|---|---|---|---|---|---|---|---|---|---|---|---|---|---|---|---|---|---|---|
| chr4 | 107803735 | 107804002 | 584 | Promoter (<=1kb) | 4 | 107803251 | 107872816 | 69566 | 1 | ENSMUST00 | 485 | Lrp8 | ENSMUSG0C | false | chr4 | 107801651 | 107804211 | 19.04495 | 59.55748 | 56.71581 | chr4 | 107801652 | 107804349 | 5.50588 | 4.26202 | 3.44614 |
| chr7 | 67736422 | 67736687 | 584 | Exon (ENSMUST00000051389.8/ | 7 | 67725261 | 67726014 | 754 | 1 | ENSMUST00 | 11162 | Ttc23 | ENSMUSG0C | false | . | . | . | . | . | . | . | . | . | . | . | . |
| chr4 | 139096219 | 139096611 | 584 | Distal Intergenic | 4 | 139082284 | 139092993 | 10710 | 2 | ENSMUST00 | -3227 | Nbl1 | ENSMUSG0C | false | . | . | . | . | . | . | chr4 | 139088999 | 139098811 | 5.83874 | 4.33468 | 3.72707 |
| chr4 | 150056556 | 150056899 | 584 | Exon (ENSMUST00000134236.1/ | 4 | 150049013 | 150056976 | 7964 | 1 | ENSMUST00 | 7544 | Gm13067 | ENSMUSG0C | false | . | . | . | . | . | . | . | . | . | . | . | . |
| chr8 | 126953971 | 126954241 | 584 | Intron (ENSMUST00000045994.6 | 8 | 126930664 | 126945921 | 15258 | 2 | ENSMUST00 | -8051 | Tomm20 | ENSMUSG0C | false | . | . | . | . | . | . | chr8 | 126952803 | 126955983 | 5.36127 | 4.12735 | 3.33779 |
| chr5 | 95867121 | 95867390 | 584 | Promoter (<=1kb) | 5 | 95867334 | 95867440 | 107 | 1 | ENSMUST00 | 0 | Gm25937 | ENSMUSG0C | false | . | . | . | . | . | . | . | . | . | . | . | . |
| chr8 | 120509266 | 120509581 | 584 | Intron (ENSMUST00000127664.1 | 8 | 120520615 | 120521686 | 1072 | 1 | ENSMUST00 | -11034 | Gm26971 | ENSMUSG0C | false | . | . | . | . | . | . | chr8 | 120497859 | 120511881 | 6.76427 | 4.83633 | 4.46181 |
| chr7 | 19771118 | 19771478 | 583 | Promoter (<=1kb) | 7 | 19765613 | 19771016 | 5404 | 2 | ENSMUST00 | -103 | Bcam | ENSMUSG0C | false | chr7 | 19768283 | 19771284 | 29.58769 | 108.11006 | 104.32592 | chr7 | 19770687 | 19774598 | 4.10172 | 3.5288 | 2.30412 |
| chr15 | 64344826 | 64345400 | 583 | Intron (ENSMUST00000177374.2 | 15 | 64088541 | 64349886 | 261346 | 2 | ENSMUST00 | 4486 | Asap1 | ENSMUSG0C | false | . | . | . | . | . | . | chr15 | 64344269 | 64345793 | 3.31683 | 2.81242 | 1.66003 |
| chr17 | 66757280 | 66757571 | 583 | Intron (ENSMUST00000037974.8 | 17 | 66555252 | 66594621 | 39370 | 2 | ENSMUST00 | -162660 | Themis3 | ENSMUSG0C | false | . | . | . | . | . | . | . | . | . | . | . | . |
| chr1 | 120248325 | 120248711 | 583 | Intron (ENSMUST00000140490.2 | 1 | 120190757 | 120265255 | 74499 | 2 | ENSMUST00 | 16544 | Steap3 | ENSMUSG0C | false | . | . | . | . | . | . | chr1 | 120244209 | 120264315 | 5.23032 | 4.13237 | 3.21241 |
| chr9 | 104001957 | 104002816 | 583 | Promoter (<=1kb) | 9 | 104002544 | 104043811 | 41268 | 1 | ENSMUST00 | 0 | Nphp3 | ENSMUSG0C | false | chr9 | 104001869 | 104002467 | 17.09341 | 55.02552 | 52.24499 | chr9 | 104000807 | 104002058 | 5.27097 | 4.15764 | 3.25134 |
| chr9 | 15715247 | 15715503 | 583 | Intron (ENSMUST00000115588.2 | 9 | 15709739 | 15742384 | 32646 | 1 | ENSMUST00 | 5509 | Slc36a4 | ENSMUSG0C | false | . | . | . | . | . | . | chr9 | 15710631 | 15720191 | 6.1172 | 4.6572 | 3.92515 |
| chr5 | 43982985 | 43983262 | 583 | Promoter (1-2kb) | 5 | 43978865 | 43981762 | 2898 | 2 | ENSMUST00 | -1224 | Fgfbp1 | ENSMUSG0C | false | . | . | . | . | . | . | chr5 | 43975466 | 43985322 | 3.9051 | 3.48494 | 2.11714 |
| chr1 | 74031029 | 74031364 | 583 | Intron (ENSMUST00000169786.2 | 1 | 73993163 | 74037133 | 43971 | 2 | ENSMUST00 | 5769 | Tns1 | ENSMUSG0C | false | . | . | . | . | . | . | chr1 | 74030015 | 74034188 | 4.53486 | 3.7674 | 2.64826 |
| chr11 | 63890300 | 63890638 | 583 | Intron (ENSMUST00000094103.3 | 11 | 63885792 | 63922290 | 36499 | 2 | ENSMUST00 | 31652 | Hs3st3b1 | ENSMUSG0C | false | . | . | . | . | . | . | chr11 | 63886567 | 63903466 | 5.7674 | 4.29978 | 3.66511 |
| chr2 | 168872202 | 168872474 | 583 | Distal Intergenic | 2 | 168882040 | 168882168 | 129 | 2 | ENSMUST00 | 9694 | Gm25214 | ENSMUSG0C | false | . | . | . | . | . | . | chr2 | 168871541 | 168873417 | 5.18439 | 4.14287 | 3.15867 |
| chr7 | 111237837 | 111238138 | 582 | Distal Intergenic | 7 | 111070568 | 111083030 | 12463 | 2 | ENSMUST00 | -154808 | Eif4g2 | ENSMUSG0C | false | . | . | . | . | . | . | . | . | . | . | . | . |
| chr16 | 33062282 | 33062792 | 582 | Promoter (<=1kb) | 16 | 33062521 | 33127695 | 65175 | 1 | ENSMUST00 | 0 | Lmln | ENSMUSG0C | false | chr16 | 33061959 | 33063463 | 30.03893 | 101.89226 | 98.2715 | chr16 | 33061544 | 33063792 | 3.41223 | 3.23207 | 1.74812 |
| chr19 | 56545133 | 56545391 | 582 | Promoter (1-2kb) | 19 | 56529169 | 56547155 | 17987 | 2 | ENSMUST00 | 1764 | Dclre1a | ENSMUSG0C | false | . | . | . | . | . | . | chr19 | 56542255 | 56547600 | 4.83446 | 3.88381 | 2.89984 |
| chr8 | 11327373 | 11327653 | 582 | Intron (ENSMUST00000033899.8 | 8 | 11198423 | 11312826 | 114404 | 2 | ENSMUST00 | -14548 | Col4a1 | ENSMUSG0C | false | . | . | . | . | . | . | chr8 | 11326186 | 11328344 | 4.04351 | 3.55166 | 2.27954 |
| chr15 | 53131440 | 53131697 | 582 | Intron (ENSMUST00000077273.3 | 15 | 53089931 | 53107868 | 17938 | 2 | ENSMUST00 | -23573 | Ext1 | ENSMUSG0C | false | . | . | . | . | . | . | chr15 | 53128914 | 53133791 | 5.07988 | 3.89377 | 3.10401 |
| chr5 | 119370613 | 119371262 | 582 | Distal Intergenic | 5 | 119297245 | 119297361 | 117 | 1 | ENSMUST00 | 73369 | n-R5s175 | ENSMUSG0C | false | . | . | . | . | . | . | . | . | . | . | . | . |
| chr2 | 125279893 | 125280326 | 582 | Downstream (2-3kb) | 2 | 125282688 | 125283978 | 1291 | 2 | ENSMUST00 | 3652 | Gm14003 | ENSMUSG0C | false | . | . | . | . | . | . | . | . | . | . | . | . |
| chr14 | 24216693 | 24216904 | 582 | Intron (ENSMUST00000073687.7 | 14 | 24238172 | 24243391 | 5220 | 2 | ENSMUST00 | 26487 | Gm17105 | ENSMUSG0C | false | . | . | . | . | . | . | chr14 | 24191579 | 24244546 | 6.62149 | 4.76248 | 4.34728 |
| chr4 | 87367826 | 87368363 | 582 | Distal Intergenic | 4 | 87432272 | 87432336 | 65 | 1 | ENSMUST00 | -63909 | Gm23154 | ENSMUSG0C | false | chr4 | 87368252 | 87368868 | 7.14186 | 14.46535 | 12.21151 | chr4 | 87368021 | 87369845 | 4.5459 | 3.73979 | 2.66135 |
| chr1 | 178368871 | 178369280 | 582 | Distal Intergenic | 1 | 178363401 | 178363521 | 121 | 2 | ENSMUST00 | -5351 | Gm27458 | ENSMUSG0C | false | chr1 | 178368575 | 178369724 | 12.59008 | 38.01365 | 35.45054 | chr1 | 178368243 | 178370203 | 4.34805 | 3.71336 | 2.52223 |
| chr17 | 29440083 | 29440413 | 581 | Distal Intergenic | 17 | 29490812 | 29496111 | 5300 | 1 | ENSMUST00 | -50399 | Pim1 | ENSMUSG0C | false | . | . | . | . | . | . | chr17 | 29436890 | 29442166 | 6.63806 | 4.90552 | 4.35203 |
| chr2 | 68861137 | 68862341 | 581 | Promoter (<=1kb) | 2 | 68861441 | 69114282 | 252842 | 1 | ENSMUST00 | 0 | Cers6 | ENSMUSG0C | true | chr2 | 68860798 | 68863593 | 28.56742 | 103.16996 | 99.51957 | chr2 | 68860904 | 68861243 | 4.07615 | 3.69847 | 2.25117 |
| chr8 | 22476405 | 22477348 | 581 | Promoter (<=1kb) | 8 | 22462615 | 22476879 | 14265 | 2 | ENSMUST00 | 0 | Smim19 | ENSMUSG0C | false | chr8 | 22475915 | 22477695 | 19.08793 | 73.09275 | 70.054 | . | . | . | . | . | . |
| chr9 | 44066757 | 44067237 | 581 | Promoter (<=1kb) | 9 | 44067021 | 44095627 | 28607 | 1 | ENSMUST00 | 0 | Usp2 | ENSMUSG0C | false | chr9 | 44066980 | 44067432 | 10.07419 | 29.06419 | 26.60987 | chr9 | 44066569 | 44068210 | 4.09732 | 3.72985 | 2.24579 |
| chr9 | 105131401 | 105131963 | 581 | Promoter (<=1kb) | 9 | 105129332 | 105131805 | 2474 | 2 | ENSMUST00 | 0 | Nudt16 | ENSMUSG0C | false | chr9 | 105131129 | 105132266 | 31.56964 | 96.59996 | 93.11521 | chr9 | 105130438 | 105132330 | 3.86404 | 3.48484 | 2.08937 |
| chr15 | 77153560 | 77154012 | 581 | Promoter (<=1kb) | 15 | 77078991 | 77153812 | 74822 | 2 | ENSMUST00 | 0 | Rbfox2 | ENSMUSG0C | false | chr15 | 77152314 | 77153833 | 28.00574 | 82.6778 | 79.47607 | chr15 | 77153997 | 77155307 | 3.03883 | 3.05165 | 1.42813 |
| chr1 | 55405643 | 55406844 | 581 | Promoter (<=1kb) | 1 | 55405921 | 55754285 | 348365 | 1 | ENSMUST00 | 0 | Plcl1 | ENSMUSG0C | true | chr1 | 55405130 | 55407992 | 37.68014 | 134.63641 | 129.97372 | chr1 | 55404626 | 55405758 | 4.55237 | 3.86683 | 2.662 |
| chr19 | 29647891 | 29648521 | 581 | Promoter (<=1kb) | 19 | 29608214 | 29648415 | 40202 | 2 | ENSMUST00 | 0 | Ermp1 | ENSMUSG0C | false | chr19 | 29647505 | 29648672 | 31.63878 | 95.72812 | 92.25946 | . | . | . | . | . | . |
| chr6 | 135065243 | 135065760 | 581 | Promoter (<=1kb) | 6 | 135065651 | 135084709 | 19059 | 1 | ENSMUST00 | 0 | Gprc5a | ENSMUSG0C | false | chr6 | 135065110 | 135066008 | 26.65286 | 74.78688 | 71.72144 | . | . | . | . | . | . |
| chr8 | 126839177 | 126839998 | 581 | Distal Intergenic | 8 | 126944847 | 126944980 | 134 | 2 | ENSMUST00 | 104982 | Gm26397 | ENSMUSG0C | false | . | . | . | . | . | . | chr8 | 126838034 | 126840386 | 7.40674 | 5.1667 | 4.98361 |
| chr11 | 95105409 | 95105938 | 580 | Distal Intergenic | 11 | 95120119 | 95125296 | 5178 | 1 | ENSMUST00 | -14181 | Dlx3 | ENSMUSG0C | false | chr11 | 95105181 | 95105787 | 6.3356 | 10.63258 | 8.44148 | chr11 | 95095874 | 95108125 | 6.99596 | 4.92043 | 4.65315 |
| chr1 | 150392696 | 150393466 | 580 | Promoter (<=1kb) | 1 | 150392838 | 150449815 | 56978 | 1 | ENSMUST00 | 0 | Tpr | ENSMUSG0C | false | chr1 | 150392105 | 150394544 | 27.34946 | 104.78247 | 101.09169 | . | . | . | . | . | . |
| chr4 | 148513459 | 148513764 | 580 | Intron (ENSMUST00000103221.4 | 4 | 148495183 | 148500460 | 5278 | 2 | ENSMUST00 | -13000 | Angptl7 | ENSMUSG0C | false | . | . | . | . | . | . | chr4 | 148507077 | 148521698 | 5.44417 | 4.08041 | 3.40789 |
| chr9 | 108094509 | 108095013 | 580 | Promoter (<=1kb) | 9 | 108085417 | 108094480 | 9064 | 2 | ENSMUST00 | -30 | Apeh | ENSMUSG0C | false | chr9 | 108093689 | 108095691 | 36.76537 | 115.82222 | 111.81207 | chr9 | 108093434 | 108096226 | 3.85235 | 3.41538 | 2.09572 |
| chr4 | 45184594 | 45184944 | 580 | Promoter (<=1kb) | 4 | 45184875 | 45261900 | 77026 | 1 | ENSMUST00 | 0 | Frmpd1 | ENSMUSG0C | false | chr4 | 45184329 | 45185591 | 30.13579 | 99.06311 | 95.51676 | chr4 | 45183892 | 45185642 | 4.40499 | 3.78816 | 2.53116 |
| chr17 | 23699318 | 23699965 | 580 | Distal Intergenic | 17 | 23705058 | 23706339 | 1282 | 2 | ENSMUST00 | 6374 | 1520401A03I | ENSMUSG0C | false | chr17 | 23699230 | 23700464 | 23.12601 | 77.64799 | 74.53377 | chr17 | 23699158 | 23700704 | 4.65867 | 3.88901 | 2.76451 |
| chr4 | 106955647 | 106955934 | 580 | Promoter (<=1kb) | 4 | 106955859 | 107048692 | 92834 | 1 | ENSMUST00 | 0 | Ssbp3 | ENSMUSG0C | false | . | . | . | . | . | . | chr4 | 106945202 | 106965608 | 5.89404 | 4.4035 | 3.75076 |
| chr1 | 58306417 | 58306715 | 580 | Intron (ENSMUST00000114366.2 | 1 | 58254117 | 58315656 | 61540 | 2 | ENSMUST00 | 8941 | Gm15759 | ENSMUSG0C | false | . | . | . | . | . | . | chr1 | 58304284 | 58306895 | 4.99357 | 4.06757 | 2.98997 |
| chr12 | 71870312 | 71870720 | 580 | Intron (ENSMUST00000085299.2 | 12 | 71846311 | 71847291 | 981 | 2 | ENSMUST00 | -23022 | Gm7985 | ENSMUSG0C | false | . | . | . | . | . | . | chr12 | 71867073 | 71871127 | 3.9038 | 3.45463 | 2.13184 |
| chr6 | 39059463 | 39059966 | 580 | Intron (ENSMUST00000003017.7 | 6 | 39063878 | 39067598 | 3721 | 2 | ENSMUST00 | 7632 | RP23-346B1 | ENSMUSG0C | false | . | . | . | . | . | . | chr6 | 39059277 | 39060761 | 5.04196 | 3.8643 | 3.0482 |
| chrX | 13281581 | 13282102 | 579 | Promoter (<=1kb) | 20 | 13281032 | 13287872 | 6841 | 1 | ENSMUST00 | 550 | Ddx3x | ENSMUSG0C | false | chrX | 13280005 | 13282849 | 28.03146 | 79.91027 | 76.76085 | . | . | . | . | . | . |
| chr7 | 110706701 | 110707073 | 579 | Distal Intergenic | 7 | 110768206 | 110811919 | 43714 | 1 | ENSMUST00 | -61133 | Ampd3 | ENSMUSG0C | false | . | . | . | . | . | . | chr7 | 110706404 | 110707111 | 4.359 | 3.7181 | 2.49674 |
| chr8 | 105965623 | 105966407 | 579 | Promoter (<=1kb) | 8 | 105943590 | 105966044 | 22455 | 2 | ENSMUST00 | 0 | Slc12a4 | ENSMUSG0C | false | chr8 | 105964976 | 105966946 | 29.52499 | 116.10789 | 112.09218 | . | . | . | . | . | . |
| chr15 | 58076230 | 58077086 | 579 | Promoter (<=1kb) | 15 | 58047003 | 58076245 | 29243 | 2 | ENSMUST00 | 0 | Zhx1 | ENSMUSG0C | false | chr15 | 58075166 | 58077523 | 40.5906 | 142.05632 | 137.09286 | . | . | . | . | . | . |
| chr2 | 148865032 | 148865401 | 579 | Promoter (<=1kb) | 2 | 148846904 | 148865038 | 18135 | 2 | ENSMUST00 | 0 | 9230104L09I | ENSMUSG0C | false | . | . | . | . | . | . | chr2 | 148864840 | 148865315 | 3.48868 | 3.07272 | 1.80637 |
| chr7 | 29007659 | 29007910 | 579 | Intron (ENSMUST00000179893.2 | 7 | 28982854 | 29003279 | 20426 | 1 | ENSMUST00 | 24806 | Map4k1 | ENSMUSG0C | false | . | . | . | . | . | . | chr7 | 29007434 | 29008221 | 5.06408 | 4.13024 | 3.06809 |
| chr4 | 32824291 | 32824574 | 579 | Promoter (1-2kb) | 4 | 32821271 | 32822801 | 1531 | 2 | ENSMUST00 | -1491 | Ankrd6 | ENSMUSG0C | true | . | . | . | . | . | . | chr4 | 32823060 | 32825660 | 6.47326 | 4.71868 | 4.26781 |
| chr7 | 107517597 | 107517920 | 579 | Intron (ENSMUST00000130414.2 | 7 | 107504096 | 107548653 | 44558 | 1 | ENSMUST00 | 13502 | Syt9 | ENSMUSG0C | false | . | . | . | . | . | . | chr7 | 107515930 | 107522914 | 4.4782 | 3.78829 | 2.60858 |
| chr4 | 148912953 | 148913233 | 579 | Promoter (1-2kb) | 4 | 148913511 | 148914324 | 814 | 2 | ENSMUST00 | 1091 | Gm15969 | ENSMUSG0C | false | . | . | . | . | . | . | chr4 | 148901819 | 148933146 | 7.00771 | 4.91323 | 4.66351 |
| chr4 | 101356052 | 101357044 | 579 | Promoter (<=1kb) | 4 | 101346531 | 101356248 | 9718 | 2 | ENSMUST00 | 0 | E130102H24I | ENSMUSG0C | false | chr4 | 101354509 | 101357858 | 33.86091 | 106.63318 | 102.89114 | . | . | . | . | . | . |
| chr11 | 84394990 | 84395377 | 578 | Intron (ENSMUST00000103201.2 | 11 | 84442509 | 84449533 | 7025 | 2 | ENSMUST00 | 54156 | Aatf | ENSMUSG0C | false | . | . | . | . | . | . | chr11 | 84395277 | 84395860 | 2.78949 | 2.79494 | 1.23115 |
| chr3 | 98195473 | 98195768 | 578 | Distal Intergenic | 3 | 98222156 | 98236748 | 14593 | 1 | ENSMUST00 | -26388 | Reg4 | ENSMUSG0C | false | . | . | . | . | . | . | chr3 | 98195286 | 98203232 | 4.51299 | 3.82434 | 2.63241 |
| chr6 | 43665857 | 43666374 | 578 | Promoter (<=1kb) | 6 | 43345001 | 43666278 | 321278 | 2 | ENSMUST00 | 0 | Tpk1 | ENSMUSG0C | false | chr6 | 43665308 | 43666768 | 34.22706 | 108.29755 | 104.50865 | chr6 | 43665306 | 43666679 | 3.04764 | 3.01747 | 1.44694 |
| chr8 | 94011940 | 94012951 | 578 | Promoter (<=1kb) | 8 | 93971588 | 94012663 | 41076 | 2 | ENSMUST00 | 0 | Amfr | ENSMUSG0C | false | chr8 | 94011226 | 94013098 | 33.04283 | 115.53907 | 111.53828 | chr8 | 94012796 | 94013559 | 4.92701 | 3.90775 | 2.97957 |
| chr9 | 35026846 | 35027213 | 578 | Intron (ENSMUST00000115148.3 | 9 | 35016452 | 35030876 | 14425 | 1 | ENSMUST00 | 10395 | Kirrel3 | ENSMUSG0C | false | . | . | . | . | . | . | chr9 | 35024970 | 35027974 | 4.8103 | 3.97968 | 2.84919 |
| chr9 | 53339762 | 53340008 | 578 | Promoter (1-2kb) | 9 | 53341084 | 53368955 | 27872 | 1 | ENSMUST00 | -1076 | Exph5 | ENSMUSG0C | false | . | . | . | . | . | . | chr9 | 53339836 | 53341410 | 3.81855 | 3.33382 | 2.0654 |
| chr17 | 49534524 | 49535002 | 578 | Intron (ENSMUST00000057610.6 | 17 | 49456022 | 49564337 | 108316 | 2 | ENSMUST00 | 29335 | Daam2 | ENSMUSG0C | true | . | . | . | . | . | . | chr17 | 49511234 | 49548441 | 5.06175 | 4.03075 | 3.07349 |
| chr11 | 115473771 | 115474131 | 578 | Promoter (1-2kb) | 11 | 115475667 | 115490467 | 14801 | 1 | ENSMUST00 | -1536 | Armc7 | ENSMUSG0C | false | . | . | . | . | . | . | chr11 | 115472068 | 115477549 | 5.34788 | 4.24591 | 3.29751 |
| chr19 | 10770014 | 10770393 | 578 | Intron (ENSMUST00000186596.1 | 19 | 10781915 | 10786039 | 4125 | 1 | ENSMUST00 | -11522 | A430093F15I | ENSMUSG0C | false | . | . | . | . | . | . | chr19 | 10766754 | 10772243 | 5.86305 | 4.46272 | 3.72369 |
| chr4 | 42439252 | 42439679 | 578 | Promoter (<=1kb) | 4 | 42439378 | 42439966 | 589 | 1 | ENSMUST00 | 0 | Gm21586 | ENSMUSG0C | false | . | . | . | . | . | . | . | . | . | . | . | . |
| chr8 | 120504967 | 120505489 | 578 | Intron (ENSMUST00000127664.1 | 8 | 120520615 | 120521686 | 1072 | 1 | ENSMUST00 | -15126 | Gm26971 | ENSMUSG0C | false | . | . | . | . | . | . | chr8 | 120497859 | 120511881 | 6.76427 | 4.83633 | 4.46181 |
| chr10 | 24149122 | 24149698 | 577 | Promoter (<=1kb) | 10 | 24149317 | 24188959 | 39643 | 1 | ENSMUST00 | 0 | Stx7 | ENSMUSG0C | false | chr10 | 24149036 | 24150224 | 27.75693 | 93.7755 | 90.3501 | chr10 | 24148684 | 24154125 | 6.02243 | 4.55226 | 3.84208 |
| chr12 | 15809770 | 15810197 | 577 | Exon (ENSMUST00000020922.7/ | 12 | 15791727 | 15816785 | 25059 | 2 | ENSMUST00 | 6588 | Trib2 | ENSMUSG0C | true | chr12 | 15808022 | 15810153 | 10.31632 | 21.40087 | 19.04606 | chr12 | 15793934 | 15811101 | 7.2046 | 5.07178 | 4.81591 |
| chr3 | 51224408 | 51225099 | 577 | Promoter (<=1kb) | 3 | 51224447 | 51251644 | 27198 | 1 | ENSMUST00 | 0 | Ccrn4l | ENSMUSG0C | false | chr3 | 51223991 | 51225879 | 37.35566 | 118.95506 | 114.85009 | . | . | . | . | . | . |
| chr2 | 116060228 | 116060577 | 577 | Promoter (1-2kb) | 2 | 116049107 | 116058842 | 9736 | 2 | ENSMUST00 | -1387 | Meis2 | ENSMUSG0C | false | chr2 | 116060041 | 116064346 | 29.3377 | 90.74815 | 87.38457 | chr2 | 116058485 | 116061610 | 4.71656 | 3.89534 | 2.78076 |
| chr2 | 59605569 | 59605898 | 577 | Distal Intergenic | 2 | 59612042 | 59845717 | 233676 | 1 | ENSMUST00 | -6144 | Tanc1 | ENSMUSG0C | true | . | . | . | . | . | . | chr2 | 59603840 | 59611869 | 4.71859 | 3.92118 | 2.78796 |
| chr11 | 101126516 | 101126812 | 577 | Promoter (2-3kb) | 11 | 101124500 | 101125828 | 1329 | 1 | ENSMUST00 | 2017 | Tubg1 | ENSMUSG0C | false | . | . | . | . | . | . | . | . | . | . | . | . |
| chr7 | 73390996 | 73391502 | 577 | Promoter (<=1kb) | 7 | 73391160 | 73409451 | 18292 | 1 | ENSMUST00 | 0 | Rgma | ENSMUSG0C | false | chr7 | 73390852 | 73392583 | 21.79372 | 61.12524 | 58.26245 | chr7 | 73389790 | 73392430 | 4.61187 | 3.65475 | 2.71075 |
| chr5 | 74944173 | 74944506 | 577 | Distal Intergenic | 5 | 74949194 | 74966368 | 17175 | 1 | ENSMUST00 | -4688 | Gm6116 | ENSMUSG0C | false | . | . | . | . | . | . | . | . | . | . | . | . |
| chr18 | 11940943 | 11941302 | 577 | Exon (ENSMUST00000171109.2/ | 18 | 11840361 | 11840438 | 78 | 2 | ENSMUST00 | -100506 | Mir1901 | ENSMUSG0C | false | . | . | . | . | . | . | chr18 | 11936954 | 11942682 | 6.83828 | 4.75724 | 4.53728 |
| chr12 | 104935421 | 104935905 | 577 | Intron (ENSMUST00000095439.4 | 12 | 104925501 | 104926142 | 642 | 2 | ENSMUST00 | -9280 | RP24-159O1 | ENSMUSG0C | false | . | . | . | . | . | . | chr12 | 104933545 | 104940268 | 4.94945 | 4.0508 | 2.97944 |
| chr5 | 45857483 | 45857965 | 576 | Promoter (<=1kb) | 5 | 45697181 | 45857615 | 160435 | 2 | ENSMUST00 | 0 | Lcorl | ENSMUSG0C | false | chr5 | 45855601 | 45858324 | 38.87692 | 132.39133 | 127.80991 | . | . | . | . | . | . |
| chr14 | 33923189 | 33923658 | 576 | Promoter (<=1kb) | 14 | 33923587 | 33935280 | 11694 | 1 | ENSMUST00 | 0 | Gdf10 | ENSMUSG0C | false | chr14 | 33923135 | 33924145 | 15.92618 | 39.91766 | 37.3274 | chr14 | 33923199 | 33924209 | 4.24502 | 3.56029 | 2.41531 |
| chr16 | 20381721 | 20381988 | 576 | Intron (ENSMUST00000079158.7 | 16 | 20368877 | 20376642 | 7766 | 2 | ENSMUST00 | -5080 | Abcc5 | ENSMUSG0C | false | . | . | . | . | . | . | chr16 | 20380905 | 20384483 | 4.76945 | 3.75412 | 2.85154 |
| chr17 | 8488951 | 8489237 | 576 | Distal Intergenic | 17 | 8525372 | 8981671 | 456300 | 1 | ENSMUST00 | -36135 | Pde10a | ENSMUSG0C | true | . | . | . | . | . | . | chr17 | 8485107 | 8491054 | 6.22769 | 4.62859 | 4.02201 |
| chr1 | 172514014 | 172514304 | 576 | Distal Intergenic | 1 | 172521044 | 172545870 | 24827 | 1 | ENSMUST00 | -6740 | Ccdc19 | ENSMUSG0C | false | . | . | . | . | . | . | . | . | . | . | . | . |
| chr5 | 35907788 | 35908360 | 576 | Intron (ENSMUST00000064571.5 | 5 | 35913082 | 35996024 | 82943 | 1 | ENSMUST00 | -4722 | Afap1 | ENSMUSG0C | true | . | . | . | . | . | . | chr5 | 35902761 | 35945832 | 6.32118 | 4.58806 | 4.1083 |
| chr8 | 25537208 | 25537702 | 576 | Intron (ENSMUST00000155564.2 | 8 | 25532281 | 25574577 | 42297 | 1 | ENSMUST00 | 4928 | Fgfr1 | ENSMUSG0C | true | . | . | . | . | . | . | chr8 | 25521452 | 25558014 | 5.35885 | 4.21426 | 3.3104 |
| chr1 | 161876418 | 161877068 | 576 | Promoter (<=1kb) | 1 | 161816112 | 161876661 | 60550 | 2 | ENSMUST00 | 0 | Suco | ENSMUSG0C | false | chr1 | 161875523 | 161876835 | 23.46333 | 91.19009 | 87.82124 | . | . | . | . | . | . |
| chr17 | 78417622 | 78418058 | 576 | Promoter (<=1kb) | 17 | 78377885 | 78418131 | 40247 | 2 | ENSMUST00 | 73 | Fez2 | ENSMUSG0C | false | chr17 | 78417090 | 78418392 | 38.58512 | 125.36365 | 121.04008 | chr17 | 78413713 | 78418729 | 4.28243 | 3.65404 | 2.41795 |
| chr11 | 85841948 | 85842279 | 576 | Distal Intergenic | 11 | 85846790 | 85850333 | 3544 | 2 | ENSMUST00 | 8054 | Gm11444 | ENSMUSG0C | false | chr11 | 85841915 | 85842203 | 7.66192 | 14.01863 | 11.77084 | chr11 | 85835908 | 85845824 | 5.08538 | 4.06022 | 3.08415 |
| chr16 | 20516828 | 20517182 | 575 | Promoter (<=1kb) | 16 | 20516982 | 20531650 | 14669 | 1 | ENSMUST00 | 0 | Dvl3 | ENSMUSG0C | false | chr16 | 20516331 | 20518870 | 28.28183 | 109.60786 | 105.79031 | . | . | . | . | . | . |
| chr11 | 5098810 | 5100177 | 575 | Promoter (<=1kb) | 11 | 5069689 | 5099266 | 29578 | 2 | ENSMUST00 | 0 | Ewsr1 | ENSMUSG0C | false | chr11 | 5097235 | 5100631 | 33.83522 | 116.97303 | 112.93475 | . | . | . | . | . | . |
| chr6 | 55676951 | 55677403 | 575 | Downstream (<1kb) | 6 | 55677818 | 55681263 | 3446 | 2 | ENSMUST00 | 3860 | Neurod6 | ENSMUSG0C | false | chr6 | 55676842 | 55677274 | 7.18034 | 12.79967 | 10.57019 | chr6 | 55676834 | 55678926 | 4.83864 | 3.76194 | 2.91193 |
| chr11 | 120261797 | 120262170 | 575 | Intron (ENSMUST00000122148.2 | 11 | 120236699 | 120292282 | 55584 | 1 | ENSMUST00 | 25099 | Bahcc1 | ENSMUSG0C | false | . | . | . | . | . | . | chr11 | 120258803 | 120264750 | 9.27968 | 5.82796 | 6.55297 |
| chr10 | 111460395 | 111460657 | 575 | Distal Intergenic | 10 | 111473192 | 111496886 | 23695 | 1 | ENSMUST00 | -12535 | Nap1l1 | ENSMUSG0C | false | . | . | . | . | . | . | chr10 | 111457974 | 111460598 | 3.23317 | 3.05486 | 1.59246 |
| chr3 | 89960469 | 89960862 | 575 | Promoter (<=1kb) | 3 | 89939481 | 89960629 | 21149 | 2 | ENSMUST00 | 0 | Atp8b2 | ENSMUSG0C | false | chr3 | 89959985 | 89960959 | 33.54585 | 101.01101 | 97.41918 | chr3 | 89958714 | 89964784 | 5.0449 | 3.96997 | 3.08253 |
| chr8 | 119412786 | 119413064 | 575 | Distal Intergenic | 8 | 119394898 | 119411087 | 16190 | 1 | ENSMUST00 | 17889 | Mlycd | ENSMUSG0C | false | . | . | . | . | . | . | . | . | . | . | . | . |
| chr17 | 59170817 | 59171112 | 575 | Distal Intergenic | 17 | 59225723 | 59225843 | 121 | 2 | ENSMUST00 | 54731 | Gm23769 | ENSMUSG0C | false | . | . | . | . | . | . | . | . | . | . | . | . |
| chr7 | 112603528 | 112603877 | 575 | Distal Intergenic | 7 | 112618096 | 112618225 | 130 | 2 | ENSMUST00 | 14348 | Gm24154 | ENSMUSG0C | false | . | . | . | . | . | . | chr7 | 112592016 | 112605817 | 5.40057 | 4.28616 | 3.33023 |
| chr7 | 71230901 | 71231417 | 575 | Distal Intergenic | 7 | 71217049 | 71224409 | 7361 | 1 | ENSMUST00 | 13853 | RP23-257M2 | ENSMUSG0C | false | . | . | . | . | . | . | . | . | . | . | . | . |
| chr11 | 77801142 | 77801552 | 574 | Promoter (<=1kb) | 11 | 77801298 | 77865837 | 64540 | 1 | ENSMUST00 | 0 | Myo18a | ENSMUSG0C | false | chr11 | 77800559 | 77802076 | 16.14593 | 43.98602 | 41.34693 | chr11 | 77799596 | 77817107 | 5.11096 | 4.06241 | 3.11534 |
| chrX | 74368382 | 74368751 | 574 | Promoter (<=1kb) | 20 | 74365718 | 74368673 | 2956 | 2 | ENSMUST00 | 0 | Ubl4 | ENSMUSG0C | false | chrX | 74367926 | 74368772 | 22.97661 | 61.46103 | 58.5941 | . | . | . | . | . | . |
| chr6 | 113483200 | 113483610 | 574 | Promoter (<=1kb) | 6 | 113483297 | 113493338 | 10042 | 1 | ENSMUST00 | 0 | Creld1 | ENSMUSG0C | false | chr6 | 113482373 | 113484218 | 25.31571 | 80.75538 | 77.58801 | . | . | . | . | . | . |
| chr8 | 33731771 | 33732233 | 574 | Promoter (<=1kb) | 8 | 33731833 | 33777164 | 45332 | 1 | ENSMUST00 | 0 | Gtf2e2 | ENSMUSG0C | false | chr8 | 33731482 | 33733039 | 32.31696 | 99.89304 | 96.32935 | . | . | . | . | . | . |
| chr9 | 71771194 | 71771767 | 574 | Promoter (<=1kb) | 9 | 71626509 | 71771602 | 145094 | 2 | ENSMUST00 | 0 | Cgnl1 | ENSMUSG0C | false | chr9 | 71769462 | 71771959 | 21.13367 | 73.82313 | 70.77491 | . | . | . | . | . | . |
| chr9 | 109931421 | 109932157 | 574 | Promoter (<=1kb) | 9 | 109931460 | 110083947 | 152488 | 1 | ENSMUST00 | 0 | Map4 | ENSMUSG0C | false | chr9 | 109931060 | 109933251 | 35.29575 | 125.98199 | 121.63858 | . | . | . | . | . | . |
| chr6 | 5298237 | 5298557 | 574 | Promoter (<=1kb) | 6 | 5264147 | 5298455 | 34309 | 2 | ENSMUST00 | 0 | Pon2 | ENSMUSG0C | false | chr6 | 5297228 | 5298667 | 25.98482 | 77.30067 | 74.19341 | . | . | . | . | . | . |
| chr9 | 96993576 | 96993844 | 574 | Intron (ENSMUST00000055433.4 | 9 | 96943485 | 96984195 | 40711 | 2 | ENSMUST00 | -9382 | Spsb4 | ENSMUSG0C | false | . | . | . | . | . | . | chr9 | 96975779 | 96994993 | 5.8258 | 4.40484 | 3.6945 |
| chr1 | 184831008 | 184831282 | 574 | Exon (ENSMUST00000160982.1/ | 1 | 184813678 | 184827482 | 13805 | 2 | ENSMUST00 | -3527 | 37316 | ENSMUSG0C | false | . | . | . | . | . | . | chr1 | 184822720 | 184831886 | 4.91956 | 3.91689 | 2.97507 |
| chr4 | 152008667 | 152009185 | 574 | Promoter (<=1kb) | 4 | 152008803 | 152017680 | 8878 | 1 | ENSMUST00 | 0 | Klhl21 | ENSMUSG0C | false | chr4 | 152007699 | 152011153 | 30.9834 | 105.62366 | 101.90845 | chr4 | 152006449 | 152014835 | 5.30205 | 4.17118 | 3.25701 |
| chr7 | 68355765 | 68356088 | 574 | Promoter (2-3kb) | 7 | 68338668 | 68353201 | 14534 | 2 | ENSMUST00 | -2565 | Gm16158 | ENSMUSG0C | false | . | . | . | . | . | . | . | . | . | . | . | . |
| chr17 | 5445128 | 5445336 | 573 | Distal Intergenic | 17 | 5432295 | 5440259 | 7965 | 2 | ENSMUST00 | -4870 | Tmem242 | ENSMUSG0C | false | . | . | . | . | . | . | . | . | . | . | . | . |
| chr14 | 67234407 | 67234767 | 573 | Promoter (<=1kb) | 14 | 67234644 | 67428687 | 194044 | 1 | ENSMUST00 | 0 | Ebf2 | ENSMUSG0C | false | chr14 | 67233896 | 67234994 | 8.69047 | 18.11543 | 15.80663 | chr14 | 67231884 | 67237790 | 5.12034 | 4.11192 | 3.11175 |
| chr9 | 59486284 | 59487006 | 573 | Promoter (<=1kb) | 9 | 59388258 | 59486614 | 98357 | 2 | ENSMUST00 | 0 | Arih1 | ENSMUSG0C | false | chr9 | 59484660 | 59487428 | 29.66345 | 100.21 | 96.63225 | . | . | . | . | . | . |
| chr3 | 96696183 | 96696638 | 573 | Promoter (<=1kb) | 3 | 96696384 | 96705962 | 9579 | 1 | ENSMUST00 | 0 | Pias3 | ENSMUSG0C | false | chr3 | 96696049 | 96697777 | 31.07893 | 124.33488 | 120.0426 | chr3 | 96695928 | 96696418 | 4.24489 | 3.56796 | 2.4057 |
| chr6 | 90509120 | 90509467 | 573 | Intron (ENSMUST00000127199.1 | 6 | 90486427 | 90559386 | 72960 | 1 | ENSMUST00 | 22694 | Aldh1l1 | ENSMUSG0C | false | . | . | . | . | . | . | chr6 | 90505671 | 90509951 | 6.56223 | 4.6902 | 4.29382 |
| chr11 | 32410275 | 32410525 | 573 | Intron (ENSMUST00000038753.5 | 11 | 32455370 | 32516687 | 61318 | 1 | ENSMUST00 | -44845 | Ubtd2 | ENSMUSG0C | false | . | . | . | . | . | . | chr11 | 32409945 | 32411455 | 4.88212 | 3.8744 | 2.93023 |
| chr9 | 31043628 | 31043876 | 573 | Intron (ENSMUST00000115222.3 | 9 | 31053296 | 31068755 | 15460 | 1 | ENSMUST00 | -9420 | Zbtb44 | ENSMUSG0C | false | . | . | . | . | . | . | chr9 | 31039987 | 31049325 | 4.23384 | 3.68859 | 2.37688 |
| chr2 | 152398745 | 152399156 | 573 | Promoter (<=1kb) | 2 | 152393611 | 152398063 | 4453 | 2 | ENSMUST00 | -683 | Sox12 | ENSMUSG0C | false | chr2 | 152395767 | 152399538 | 37.89522 | 119.88892 | 115.74806 | chr2 | 152398384 | 152399527 | 4.59345 | 3.79247 | 2.70513 |
| chr16 | 95121433 | 95121735 | 573 | Distal Intergenic | 16 | 95016370 | 95067017 | 50648 | 1 | ENSMUST00 | 105064 | Gm7976 | ENSMUSG0C | false | . | . | . | . | . | . | chr16 | 95119129 | 95122841 | 4.80466 | 3.80795 | 2.8783 |
| chr13 | 15107740 | 15107979 | 573 | Distal Intergenic | 13 | 15459980 | 15461295 | 1316 | 2 | ENSMUST00 | 353316 | Gm26645 | ENSMUSG0C | false | . | . | . | . | . | . | chr13 | 15107030 | 15108336 | 3.59793 | 3.33399 | 1.89801 |
| chr7 | 139434063 | 139434382 | 572 | Intron (ENSMUST00000106098.2 | 7 | 139454054 | 139511476 | 57423 | 1 | ENSMUST00 | -19672 | Inpp5a | ENSMUSG0C | false | . | . | . | . | . | . | chr7 | 139433977 | 139446502 | 4.71083 | 3.92341 | 2.77438 |
| chr2 | 79646027 | 79646341 | 572 | Intron (ENSMUST00000111785.3 | 2 | 79639182 | 79672966 | 33785 | 1 | ENSMUST00 | 6846 | Ssfa2 | ENSMUSG0C | false | . | . | . | . | . | . | chr2 | 79644762 | 79655652 | 5.51555 | 4.19848 | 3.45098 |
| chr3 | 88488714 | 88489277 | 572 | Intron (ENSMUST00000029699.7 | 3 | 88483206 | 88493312 | 10107 | 2 | ENSMUST00 | 4035 | Lmna | ENSMUSG0C | false | . | . | . | . | . | . | chr3 | 88487350 | 88502372 | 7.5952 | 5.18047 | 5.13989 |
| chr4 | 133057344 | 133057692 | 572 | Intron (ENSMUST00000154646.2 | 4 | 133066249 | 133072063 | 5815 | 1 | ENSMUST00 | -8557 | Ahdc1 | ENSMUSG0C | false | . | . | . | . | . | . | chr4 | 133013189 | 133060125 | 8.03751 | 5.40542 | 5.50183 |
| chr1 | 52726331 | 52727522 | 572 | Promoter (<=1kb) | 1 | 52656286 | 52727462 | 71177 | 2 | ENSMUST00 | 0 | Mfsd6 | ENSMUSG0C | false | chr1 | 52726241 | 52727775 | 28.59262 | 111.22587 | 107.35971 | chr1 | 52727416 | 52727947 | 4.33393 | 3.72426 | 2.46 |
| chr16 | 21694136 | 21694974 | 572 | Promoter (<=1kb) | 16 | 21649045 | 21694665 | 45621 | 2 | ENSMUST00 | 0 | 2510009E07I | ENSMUSG0C | false | chr16 | 21693648 | 21695637 | 31.28813 | 116.44205 | 112.41405 | . | . | . | . | . | . |
| chr19 | 31580716 | 31580964 | 572 | Intron (ENSMUST00000065067.7 | 19 | 30568547 | 31664370 | 1095824 | 2 | ENSMUST00 | 83406 | Prkg1 | ENSMUSG0C | true | . | . | . | . | . | . | chr19 | 31580488 | 31581539 | 5.07306 | 3.9762 | 3.08572 |
| chr8 | 35587728 | 35588216 | 572 | Promoter (<=1kb) | 8 | 35587798 | 35679449 | 91652 | 1 | ENSMUST00 | 0 | Mfhas1 | ENSMUSG0C | false | chr8 | 35587571 | 35589216 | 23.47096 | 84.72127 | 81.48416 | chr8 | 35587327 | 35595552 | 4.30177 | 3.67935 | 2.43641 |
| chr5 | 137627896 | 137628230 | 572 | Promoter (<=1kb) | 5 | 137612506 | 137628956 | 16451 | 2 | ENSMUST00 | 726 | Fbxo24 | ENSMUSG0C | false | chr5 | 137627921 | 137628752 | 20.63265 | 55.76951 | 52.97896 | chr5 | 137627385 | 137628010 | 5.42827 | 4.30975 | 3.34168 |
| chr1 | 119309236 | 119309494 | 572 | Intron (ENSMUST00000189894.1 | 1 | 119305346 | 119326833 | 21488 | 2 | ENSMUST00 | 17339 | RP23-400A1 | ENSMUSG0C | false | . | . | . | . | . | . | chr1 | 119308260 | 119309987 | 5.02757 | 4.16184 | 3.02846 |
| chr14 | 31231443 | 31231922 | 571 | Distal Intergenic | 14 | 31217860 | 31230350 | 12491 | 1 | ENSMUST00 | 13584 | Sema3g | ENSMUSG0C | true | . | . | . | . | . | . | chr14 | 31219988 | 31235200 | 8.317 | 5.47646 | 5.7231 |
| chr3 | 142363078 | 142363363 | 571 | Intron (ENSMUST00000029941.1 | 3 | 142302080 | 142391850 | 89771 | 2 | ENSMUST00 | 28487 | Pdlim5 | ENSMUSG0C | false | . | . | . | . | . | . | chr3 | 142361986 | 142364905 | 5.22884 | 4.19983 | 3.18854 |
| chr5 | 52189746 | 52190648 | 571 | Promoter (<=1kb) | 5 | 52150209 | 52190519 | 40311 | 2 | ENSMUST00 | 0 | Dhx15 | ENSMUSG0C | false | chr5 | 52189080 | 52191292 | 31.54088 | 108.67455 | 104.87239 | . | . | . | . | . | . |
| chr9 | 30949738 | 30950029 | 571 | Intron (ENSMUST00000068135.7 | 9 | 30943458 | 30963838 | 20381 | 1 | ENSMUST00 | 6281 | Adamts8 | ENSMUSG0C | false | . | . | . | . | . | . | chr9 | 30948887 | 30951639 | 5.0128 | 4.08778 | 3.03968 |
| chr9 | 115325800 | 115326145 | 571 | Distal Intergenic | 9 | 115242607 | 115310421 | 67815 | 2 | ENSMUST00 | -15380 | Stt3b | ENSMUSG0C | false | . | . | . | . | . | . | . | . | . | . | . | . |
| chr10 | 24682785 | 24683050 | 571 | Intron (ENSMUST00000105520.2 | 10 | 24668486 | 24674258 | 5773 | 2 | ENSMUST00 | -8528 | Enpp1 | ENSMUSG0C | false | . | . | . | . | . | . | chr10 | 24682466 | 24683043 | 2.95267 | 2.95915 | 1.38044 |
| chr10 | 67995414 | 67995712 | 571 | Intron (ENSMUST00000118160.2 | 10 | 67979715 | 68042298 | 62584 | 1 | ENSMUST00 | 15700 | Rtkn2 | ENSMUSG0C | false | . | . | . | . | . | . | chr10 | 67994215 | 67996578 | 3.32201 | 3.23352 | 1.66983 |
| chr2 | 30124379 | 30124792 | 571 | Promoter (<=1kb) | 2 | 30097283 | 30124454 | 27172 | 2 | ENSMUST00 | 0 | Zer1 | ENSMUSG0C | false | chr2 | 30123739 | 30125160 | 19.18445 | 82.69909 | 79.49731 | chr2 | 30122737 | 30125553 | 3.37482 | 3.27422 | 1.67646 |
| chr9 | 121297343 | 121297984 | 571 | Distal Intergenic | 9 | 121102544 | 121277197 | 174654 | 2 | ENSMUST00 | -20147 | Ulk4 | ENSMUSG0C | false | chr9 | 121296988 | 121298937 | 35.62435 | 116.99458 | 112.94973 | chr9 | 121297008 | 121304640 | 5.24869 | 4.22613 | 3.21261 |
| chrX | 7418011 | 7418303 | 571 | Promoter (2-3kb) | 20 | 7411817 | 7421085 | 9269 | 2 | ENSMUST00 | 2782 | 2010204K13I | ENSMUSG0C | false | . | . | . | . | . | . | . | . | . | . | . | . |
| chr4 | 131406725 | 131407246 | 571 | Distal Intergenic | 4 | 131487197 | 131505527 | 18331 | 1 | ENSMUST00 | -79951 | Gm831 | ENSMUSG0C | false | . | . | . | . | . | . | chr4 | 131403590 | 131411016 | 4.73067 | 3.88614 | 2.78676 |
| chr9 | 57870447 | 57870742 | 570 | Distal Intergenic | 9 | 57794687 | 57836793 | 42107 | 2 | ENSMUST00 | -33655 | Arid3b | ENSMUSG0C | false | . | . | . | . | . | . | chr9 | 57863701 | 57872251 | 4.81541 | 3.83026 | 2.89103 |
| chr11 | 96861803 | 96862376 | 570 | Downstream (<1kb) | 11 | 96854131 | 96861203 | 7073 | 1 | ENSMUST00 | 7673 | Copz2 | ENSMUSG0C | false | chr11 | 96861404 | 96862432 | 14.73601 | 38.55974 | 35.98649 | chr11 | 96860440 | 96864502 | 6.8807 | 4.8067 | 4.58039 |
| chr15 | 37136748 | 37137032 | 570 | Distal Intergenic | 15 | 37233036 | 37333647 | 100612 | 1 | ENSMUST00 | -96004 | Grhl2 | ENSMUSG0C | false | . | . | . | . | . | . | chr15 | 37134858 | 37136907 | 4.31234 | 3.50595 | 2.47376 |
| chr1 | 134992282 | 134992605 | 570 | Intron (ENSMUST00000044828.8 | 1 | 134971956 | 134972857 | 902 | 1 | ENSMUST00 | 20327 | Ube2t | ENSMUSG0C | false | . | . | . | . | . | . | . | . | . | . | . | . |
| chr6 | 92460685 | 92461080 | 570 | Intron (ENSMUST00000032093.6 | 6 | 92375384 | 92481392 | 106009 | 2 | ENSMUST00 | 20312 | Prickle2 | ENSMUSG0C | false | . | . | . | . | . | . | chr6 | 92459348 | 92466873 | 4.66008 | 3.93972 | 2.74191 |
| chr7 | 65469660 | 65469985 | 570 | Distal Intergenic | 7 | 65296165 | 65371239 | 75075 | 2 | ENSMUST00 | -98422 | Tjp1 | ENSMUSG0C | true | . | . | . | . | . | . | chr7 | 65466255 | 65471086 | 5.90211 | 4.48768 | 3.77026 |
| chr16 | 78265108 | 78265395 | 570 | Distal Intergenic | 16 | 78240258 | 78240388 | 131 | 1 | ENSMUST00 | 24851 | Gm25916 | ENSMUSG0C | false | chr16 | 78264802 | 78265562 | 11.15024 | 25.69191 | 23.28165 | chr16 | 78264467 | 78266575 | 5.20057 | 4.17619 | 3.20782 |
| chr11 | 69331988 | 69332413 | 570 | Promoter (<=1kb) | 11 | 69329825 | 69332122 | 2298 | 2 | ENSMUST00 | 0 | Kcnab3os | ENSMUSG0C | false | chr11 | 69331855 | 69332661 | 23.24766 | 62.77607 | 59.89146 | chr11 | 69331389 | 69334652 | 5.64109 | 4.43792 | 3.52772 |
| chr1 | 165503182 | 165503501 | 570 | Exon (ENSMUST00000111440.2/ | 1 | 165497999 | 165498249 | 251 | 2 | ENSMUST00 | -4934 | Gm23599 | ENSMUSG0C | false | . | . | . | . | . | . | chr1 | 165502189 | 165505678 | 5.70527 | 4.44067 | 3.59237 |
| chr7 | 118491435 | 118492155 | 570 | Promoter (<=1kb) | 7 | 118485114 | 118491975 | 6862 | 2 | ENSMUST00 | 0 | Itpripl2 | ENSMUSG0C | false | chr7 | 118490421 | 118492334 | 36.04672 | 129.50026 | 125.02953 | . | . | . | . | . | . |
| chr10 | 86021824 | 86022459 | 569 | Promoter (<=1kb) | 10 | 86021972 | 86051873 | 29902 | 1 | ENSMUST00 | 0 | Fbxo7 | ENSMUSG0C | false | chr10 | 86021772 | 86022997 | 21.20881 | 84.61981 | 81.38317 | chr10 | 86021451 | 86024612 | 3.66761 | 3.39744 | 1.91565 |
| chr17 | 56673013 | 56673688 | 569 | Promoter (<=1kb) | 17 | 56673225 | 56711769 | 38545 | 1 | ENSMUST00 | 0 | Ranbp3 | ENSMUSG0C | false | chr17 | 56672778 | 56673047 | 9.74174 | 20.4319 | 18.09061 | . | . | . | . | . | . |
| chr4 | 108882562 | 108882805 | 569 | Intron (ENSMUST00000106650.3 | 4 | 108879200 | 108941185 | 61986 | 1 | ENSMUST00 | 3363 | Rab3b | ENSMUSG0C | true | . | . | . | . | . | . | chr4 | 108880551 | 108888038 | 4.67845 | 3.88915 | 2.75156 |
| chr11 | 30287894 | 30288191 | 569 | Distal Intergenic | 11 | 30216659 | 30268175 | 51517 | 2 | ENSMUST00 | -19720 | Sptbn1 | ENSMUSG0C | false | . | . | . | . | . | . | chr11 | 30287587 | 30292909 | 4.20953 | 3.69116 | 2.33969 |
| chr2 | 35979270 | 35979926 | 569 | Promoter (<=1kb) | 2 | 35751241 | 35979624 | 228384 | 2 | ENSMUST00 | 0 | Ttll11 | ENSMUSG0C | false | chr2 | 35978342 | 35980211 | 34.15937 | 108.17976 | 104.39504 | chr2 | 35977085 | 35980123 | 3.84625 | 3.32141 | 2.09501 |
| chr2 | 155592575 | 155593011 | 569 | Promoter (<=1kb) | 2 | 155563181 | 155592810 | 29630 | 2 | ENSMUST00 | 0 | Gss | ENSMUSG0C | false | chr2 | 155591686 | 155593225 | 22.15371 | 67.81625 | 64.85734 | . | . | . | . | . | . |
| chr9 | 77530455 | 77530761 | 569 | Promoter (1-2kb) | 9 | 77458090 | 77532444 | 74355 | 2 | ENSMUST00 | 1683 | Lrrc1 | ENSMUSG0C | true | . | . | . | . | . | . | chr9 | 77501306 | 77543222 | 5.57899 | 4.30036 | 3.48961 |
| chr15 | 102980805 | 102981180 | 569 | 5' UTR | 15 | 102977032 | 102985099 | 8068 | 1 | ENSMUST00 | 3774 | Hoxc9 | ENSMUSG0C | false | chr15 | 102980785 | 102982803 | 30.41442 | 103.5799 | 99.9162 | chr15 | 102975429 | 102981199 | 5.98646 | 4.4723 | 3.8277 |
| chr16 | 18629787 | 18630162 | 569 | Promoter (<=1kb) | 16 | 18621811 | 18629938 | 8128 | 2 | ENSMUST00 | 0 | Sep-05 | ENSMUSG0C | false | chr16 | 18629329 | 18630303 | 29.41006 | 85.09721 | 81.85307 | chr16 | 18629257 | 18630313 | 3.50537 | 3.09145 | 1.80875 |
| chr1 | 16274866 | 16275129 | 569 | Intron (ENSMUST00000162751.2 | 1 | 16253629 | 16258857 | 5229 | 1 | ENSMUST00 | 21238 | Gm7568 | ENSMUSG0C | false | . | . | . | . | . | . | chr1 | 16274322 | 16275662 | 4.78519 | 3.98471 | 2.84073 |
| chr13 | 72628471 | 72628970 | 568 | Promoter (<=1kb) | 13 | 72628820 | 72634198 | 5379 | 1 | ENSMUST00 | 0 | Irx2 | ENSMUSG0C | false | chr13 | 72628036 | 72629650 | 14.61724 | 33.51261 | 31.00336 | chr13 | 72627431 | 72631986 | 5.20478 | 4.06126 | 3.19751 |
| chr11 | 12464679 | 12465088 | 568 | Promoter (<=1kb) | 11 | 12236608 | 12464804 | 228197 | 2 | ENSMUST00 | 0 | Cobl | ENSMUSG0C | false | chr11 | 12464163 | 12465333 | 29.56773 | 86.32103 | 83.05225 | chr11 | 12464139 | 12465517 | 3.87512 | 3.37105 | 2.12185 |
| chr17 | 67908651 | 67909033 | 568 | Intron (ENSMUST00000024840.6 | 17 | 67845735 | 67901471 | 55737 | 2 | ENSMUST00 | -7181 | Arhgap28 | ENSMUSG0C | true | . | . | . | . | . | . | chr17 | 67892935 | 67914104 | 5.71307 | 4.38398 | 3.59861 |
| chr17 | 64346241 | 64346520 | 568 | Distal Intergenic | 17 | 64281005 | 64331916 | 50912 | 2 | ENSMUST00 | -14326 | Pja2 | ENSMUSG0C | false | . | . | . | . | . | . | chr17 | 64345942 | 64346930 | 3.6184 | 3.38363 | 1.92353 |
| chr9 | 86467013 | 86467446 | 568 | Promoter (<=1kb) | 9 | 86467154 | 86555923 | 88770 | 1 | ENSMUST00 | 0 | Dopey1 | ENSMUSG0C | false | chr9 | 86467107 | 86468011 | 33.41831 | 117.26761 | 113.21259 | chr9 | 86466827 | 86469101 | 3.49652 | 3.33839 | 1.77011 |
| chr1 | 180330272 | 180330540 | 568 | Promoter (<=1kb) | 1 | 180330485 | 180424802 | 94318 | 1 | ENSMUST00 | 0 | Itpkb | ENSMUSG0C | false | chr1 | 180330478 | 180331292 | 12.74236 | 36.28267 | 33.73857 | chr1 | 180330472 | 180333297 | 4.59186 | 3.77988 | 2.71663 |
| chr12 | 58297008 | 58297374 | 568 | Distal Intergenic | 12 | 58264720 | 58269258 | 4539 | 2 | ENSMUST00 | -27751 | Clec14a | ENSMUSG0C | false | . | . | . | . | . | . | . | . | . | . | . | . |
| chrX | 102070348 | 102070671 | 568 | Promoter (<=1kb) | 20 | 102066544 | 102071196 | 4653 | 2 | ENSMUST00 | 525 | Rgag4 | ENSMUSG0C | false | chrX | 102070296 | 102071600 | 13.83063 | 32.8443 | 30.34371 | . | . | . | . | . | . |
| chr6 | 71493659 | 71494645 | 568 | Promoter (<=1kb) | 6 | 71493894 | 71510881 | 16988 | 1 | ENSMUST00 | 0 | Rnf103 | ENSMUSG0C | false | chr6 | 71493411 | 71495597 | 27.20707 | 96.65539 | 93.16586 | . | . | . | . | . | . |
| chr7 | 124868069 | 124868363 | 568 | Distal Intergenic | 7 | 124625670 | 124708935 | 83266 | 2 | ENSMUST00 | -159135 | 3100003L05I | ENSMUSG0C | false | . | . | . | . | . | . | chr7 | 124867201 | 124868724 | 3.46154 | 3.25001 | 1.75711 |
| chr7 | 34233913 | 34234334 | 568 | Promoter (<=1kb) | 7 | 34230420 | 34234075 | 3656 | 2 | ENSMUST00 | 0 | Gm12762 | ENSMUSG0C | false | chr7 | 34233690 | 34234987 | 16.80025 | 47.43613 | 44.75273 | chr7 | 34232817 | 34235157 | 3.83349 | 3.48331 | 2.09725 |
| chr10 | 98493172 | 98493484 | 567 | Distal Intergenic | 10 | 98915152 | 99026143 | 110992 | 1 | ENSMUST00 | -421668 | Atp2b1 | ENSMUSG0C | true | . | . | . | . | . | . | chr10 | 98491771 | 98494725 | 3.86601 | 3.53803 | 2.0676 |
| chr15 | 81810156 | 81810493 | 567 | Promoter (<=1kb) | 15 | 81811046 | 81821256 | 10211 | 1 | ENSMUST00 | -553 | Tef | ENSMUSG0C | false | chr15 | 81810073 | 81812169 | 29.06085 | 90.26336 | 86.91321 | chr15 | 81800930 | 81810375 | 6.79861 | 4.85005 | 4.49384 |
| chr19 | 3686389 | 3686784 | 567 | Promoter (<=1kb) | 19 | 3584828 | 3686556 | 101729 | 2 | ENSMUST00 | 0 | Lrp5 | ENSMUSG0C | false | chr19 | 3685204 | 3686997 | 30.59503 | 95.89513 | 92.42037 | chr19 | 3684060 | 3687204 | 4.26719 | 3.70721 | 2.4061 |
| chr5 | 122372313 | 122372805 | 567 | Promoter (<=1kb) | 5 | 122364580 | 122372364 | 7785 | 2 | ENSMUST00 | 0 | Fam216a | ENSMUSG0C | false | chr5 | 122371498 | 122373147 | 22.99841 | 82.87405 | 79.66897 | chr5 | 122371459 | 122373847 | 4.6526 | 3.87972 | 2.7627 |
| chr5 | 146695709 | 146696060 | 567 | Distal Intergenic | 5 | 146734839 | 146758448 | 23610 | 2 | ENSMUST00 | 62388 | Usp12 | ENSMUSG0C | false | . | . | . | . | . | . | chr5 | 146691248 | 146696389 | 4.5702 | 3.85568 | 2.6568 |
| chr1 | 86440654 | 86440926 | 567 | Distal Intergenic | 1 | 86426329 | 86428052 | 1724 | 1 | ENSMUST00 | 14326 | 1700019O17 | ENSMUSG0C | false | chr1 | 86440745 | 86441363 | 8.58007 | 16.45236 | 14.16747 | chr1 | 86439501 | 86441062 | 4.65536 | 3.96932 | 2.70861 |
| chr7 | 110320287 | 110320620 | 567 | Intron (ENSMUST00000164759.2 | 7 | 110320741 | 110340285 | 19545 | 2 | ENSMUST00 | 19665 | Sbf2 | ENSMUSG0C | true | . | . | . | . | . | . | chr7 | 110317691 | 110322533 | 4.27758 | 3.71535 | 2.44663 |
| chr13 | 44733592 | 44734218 | 567 | Promoter (<=1kb) | 13 | 44734040 | 44874314 | 140275 | 1 | ENSMUST00 | 0 | Jarid2 | ENSMUSG0C | false | chr13 | 44730600 | 44734844 | 29.75681 | 92.45776 | 89.05831 | . | . | . | . | . | . |
| chr6 | 146945016 | 146945323 | 567 | Intron (ENSMUST00000016631.8 | 6 | 146951302 | 146954414 | 3113 | 2 | ENSMUST00 | 9091 | 1700034J05I | ENSMUSG0C | false | . | . | . | . | . | . | chr6 | 146942477 | 146953557 | 4.42545 | 3.77491 | 2.55216 |
| chr4 | 24786279 | 24786536 | 567 | Intron (ENSMUST00000108214.3 | 4 | 24792680 | 24800310 | 7631 | 1 | ENSMUST00 | -6144 | Gm11899 | ENSMUSG0C | false | . | . | . | . | . | . | chr4 | 24785815 | 24787168 | 4.37516 | 3.64782 | 2.50281 |
| chr12 | 110452967 | 110453224 | 566 | Distal Intergenic | 12 | 110485739 | 110583061 | 97323 | 1 | ENSMUST00 | -32515 | Ppp2r5c | ENSMUSG0C | false | . | . | . | . | . | . | . | . | . | . | . | . |
| chr17 | 25084796 | 25085104 | 566 | Intron (ENSMUST00000137386.2 | 17 | 25057702 | 25081181 | 23480 | 2 | ENSMUST00 | -3616 | Tmem204 | ENSMUSG0C | false | . | . | . | . | . | . | chr17 | 25077267 | 25086277 | 7.00235 | 5.00822 | 4.65933 |
| chr5 | 53213273 | 53213642 | 566 | Promoter (<=1kb) | 5 | 53107084 | 53213452 | 106369 | 2 | ENSMUST00 | 0 | Sel1l3 | ENSMUSG0C | false | chr5 | 53212386 | 53214037 | 19.89052 | 67.91473 | 64.9535 | chr5 | 53211936 | 53214334 | 3.93408 | 3.58519 | 2.15829 |
| chr8 | 79374442 | 79374684 | 566 | Intron (ENSMUST00000066091.8 | 8 | 79338395 | 79399518 | 61124 | 2 | ENSMUST00 | 24834 | Smad1 | ENSMUSG0C | false | . | . | . | . | . | . | chr8 | 79373642 | 79375957 | 6.05202 | 4.47493 | 3.89451 |
| chr9 | 77529728 | 77530369 | 566 | Promoter (2-3kb) | 9 | 77458090 | 77532444 | 74355 | 2 | ENSMUST00 | 2075 | Lrrc1 | ENSMUSG0C | true | chr9 | 77529540 | 77529847 | 4.13035 | 5.82909 | 3.75473 | chr9 | 77501306 | 77543222 | 5.57899 | 4.30036 | 3.48961 |
| chr9 | 88346957 | 88347228 | 566 | Intron (ENSMUST00000034992.7 | 9 | 88327633 | 88367844 | 40212 | 1 | ENSMUST00 | 19325 | Nt5e | ENSMUSG0C | false | . | . | . | . | . | . | . | . | . | . | . | . |
| chr16 | 90779828 | 90780137 | 566 | Intron (ENSMUST00000140920.1 | 16 | 90767020 | 90772791 | 5772 | 2 | ENSMUST00 | -7038 | Urb1 | ENSMUSG0C | false | . | . | . | . | . | . | chr16 | 90778899 | 90781112 | 5.37834 | 4.26728 | 3.3759 |
| chr1 | 89575722 | 89576014 | 566 | Intron (ENSMUST00000027521.9 | 1 | 89580236 | 89895277 | 315042 | 1 | ENSMUST00 | -4222 | Agap1 | ENSMUSG0C | false | . | . | . | . | . | . | chr1 | 89564740 | 89589718 | 5.63127 | 4.22514 | 3.56367 |
| chr3 | 37994222 | 37994556 | 566 | Promoter (<=1kb) | 3 | 37995105 | 37995235 | 131 | 2 | ENSMUST00 | 679 | Gm22899 | ENSMUSG0C | false | . | . | . | . | . | . | chr3 | 37993894 | 37994776 | 5.63469 | 4.0597 | 3.562 |
| chr16 | 77618594 | 77618819 | 566 | Intron (ENSMUST00000183099.2 | 16 | 77588578 | 77602094 | 13517 | 2 | ENSMUST00 | -16501 | Gm11146 | ENSMUSG0C | false | . | . | . | . | . | . | chr16 | 77618689 | 77620292 | 3.5829 | 3.3922 | 1.83526 |
| chr11 | 93952411 | 93952688 | 565 | Intron (ENSMUST00000021217.5 | 11 | 93949817 | 93955783 | 5967 | 2 | ENSMUST00 | 3095 | Nme2 | ENSMUSG0C | false | . | . | . | . | . | . | chr11 | 93951624 | 93957429 | 4.20778 | 3.60571 | 2.38129 |
| chr12 | 108221082 | 108221427 | 565 | Intron (ENSMUST00000136175.1 | 12 | 108179738 | 108203359 | 23622 | 1 | ENSMUST00 | 41345 | Ccnk | ENSMUSG0C | false | . | . | . | . | . | . | chr12 | 108216919 | 108227928 | 7.31121 | 5.08617 | 4.91072 |
| chr15 | 64285318 | 64285523 | 565 | Intron (ENSMUST00000177371.2 | 15 | 64191537 | 64311689 | 120153 | 2 | ENSMUST00 | 26166 | Asap1 | ENSMUSG0C | false | . | . | . | . | . | . | chr15 | 64284168 | 64287856 | 4.34547 | 3.58973 | 2.50443 |
| chr2 | 14229139 | 14229412 | 565 | Promoter (<=1kb) | 2 | 14229414 | 14332057 | 102644 | 1 | ENSMUST00 | -2 | Mrc1 | ENSMUSG0C | false | . | . | . | . | . | . | . | . | . | . | . | . |

| | | | | | | | | | | | | | | | | | | | | | | | | | | |
|---|---|---|---|---|---|---|---|---|---|---|---|---|---|---|---|---|---|---|---|---|---|---|---|---|---|---|
| chr4 | 133132143 | 133132487 | 565 | Promoter (1-2kb) | 4 | 133130629 | 133176560 | 45932 | 1 | ENSMUST00 | 1515 | Wasf2 | ENSMUSG0C | true | chr4 | 133129620 | 133132418 | 40.96944 | 136.05835 | 131.33708 | chr4 | 133131598 | 133139920 | 5.43412 | 4.30654 | 3.35263 |
| chr8 | 36697807 | 36698141 | 565 | Intron (ENSMUST00000098826.4 | 8 | 36567751 | 36733054 | 165304 | 2 | ENSMUST00 | 34913 | Dlc1 | ENSMUSG0C | true | . | . | . | . | . | . | chr8 | 36694768 | 36701329 | 4.74882 | 3.93176 | 2.79695 |
| chr8 | 120334880 | 120336098 | 565 | Intron (ENSMUST00000127664.1 | 8 | 120230536 | 120553634 | 323099 | 1 | ENSMUST00 | 104345 | Gse1 | ENSMUSG0C | false | chr8 | 120334886 | 120335828 | 11.2646 | 26.49366 | 24.07236 | chr8 | 120331120 | 120337909 | 6.67651 | 4.80876 | 4.37617 |
| chr2 | 150861883 | 150862185 | 565 | Intron (ENSMUST00000056149.9 | 2 | 150843033 | 150851490 | 8458 | 2 | ENSMUST00 | -10394 | Abhd12 | ENSMUSG0C | false | . | . | . | . | . | . | chr2 | 150861236 | 150862979 | 5.36751 | 4.16501 | 3.35889 |
| chr9 | 78208748 | 78209015 | 565 | Intron (ENSMUST00000034903.5 | 9 | 78191966 | 78209348 | 17383 | 1 | ENSMUST00 | 16783 | Gsta4 | ENSMUSG0C | false | . | . | . | . | . | . | chr9 | 78208033 | 78209347 | 3.56373 | 3.20471 | 1.86067 |
| chr11 | 95032982 | 95033221 | 565 | Promoter (<=1kb) | 11 | 95032759 | 95038283 | 5525 | 1 | ENSMUST00 | 224 | Gm11513 | ENSMUSG0C | false | . | . | . | . | . | . | chr11 | 95031837 | 95049061 | 6.83336 | 4.76219 | 4.52264 |
| chr18 | 76134658 | 76134906 | 565 | Intron (ENSMUST00000058997.8 | 18 | 76170552 | 76170652 | 101 | 1 | ENSMUST00 | -35646 | Mir6358 | ENSMUSG0C | false | . | . | . | . | . | . | chr18 | 76133317 | 76155019 | 6.7969 | 4.75664 | 4.51499 |
| chr13 | 69487913 | 69488218 | 564 | Distal Intergenic | 13 | 69533746 | 69615468 | 81723 | 1 | ENSMUST00 | -45528 | Nsun2 | ENSMUSG0C | false | . | . | . | . | . | . | chr13 | 69487584 | 69489274 | 4.63432 | 3.88794 | 2.70812 |
| chr8 | 121706486 | 121706875 | 564 | Intron (ENSMUST00000127664.1 | 8 | 121729623 | 121789617 | 59995 | 1 | ENSMUST00 | -22748 | Jph3 | ENSMUSG0C | false | . | . | . | . | . | . | chr8 | 121706224 | 121707933 | 4.09715 | 3.56214 | 2.30157 |
| chr9 | 102730059 | 102730333 | 564 | Promoter (<=1kb) | 9 | 102730731 | 102732327 | 1597 | 1 | ENSMUST00 | -398 | Amotl2 | ENSMUSG0C | false | . | . | . | . | . | . | chr9 | 102720260 | 102734975 | 8.68688 | 5.64144 | 6.02662 |
| chr14 | 47418220 | 47418486 | 564 | Promoter (<=1kb) | 14 | 47387779 | 47418407 | 30629 | 2 | ENSMUST00 | 0 | Dlgap5 | ENSMUSG0C | false | chr14 | 47417921 | 47418619 | 15.34636 | 36.79438 | 34.24475 | chr14 | 47417764 | 47418700 | 3.37069 | 3.21435 | 1.71821 |
| chr5 | 64878050 | 64878485 | 564 | Distal Intergenic | 5 | 64924811 | 64932761 | 7951 | 2 | ENSMUST00 | 54276 | Tlr1 | ENSMUSG0C | false | . | . | . | . | . | . | chr5 | 64875130 | 64879652 | 5.60891 | 4.1449 | 3.54722 |
| chr18 | 35304045 | 35304299 | 564 | Intron (ENSMUST00000025215.8 | 18 | 35209011 | 35215024 | 6014 | 2 | ENSMUST00 | -89022 | Lrrtm2 | ENSMUSG0C | false | . | . | . | . | . | . | chr18 | 35302369 | 35304606 | 3.03841 | 2.96197 | 1.44972 |
| chr2 | 120093700 | 120094085 | 564 | Intron (ENSMUST00000028755.7 | 2 | 120076214 | 120085772 | 9559 | 2 | ENSMUST00 | -7929 | Sptbn5 | ENSMUSG0C | false | . | . | . | . | . | . | chr2 | 120090578 | 120107323 | 5.14183 | 4.07311 | 3.14279 |
| chr11 | 18262935 | 18263219 | 564 | Distal Intergenic | 11 | 18206413 | 18206944 | 532 | 2 | ENSMUST00 | -55992 | Gm12019 | ENSMUSG0C | false | . | . | . | . | . | . | chr11 | 18262234 | 18263721 | 5.74604 | 4.3181 | 3.63869 |
| chr2 | 52369999 | 52370321 | 564 | Distal Intergenic | 2 | 52361630 | 52362495 | 866 | 2 | ENSMUST00 | -7505 | Gm13521 | ENSMUSG0C | false | . | . | . | . | . | . | chr2 | 52368596 | 52372932 | 4.18901 | 3.66588 | 2.36008 |
| chr5 | 74792255 | 74792961 | 564 | Distal Intergenic | 5 | 74731517 | 74734450 | 2934 | 2 | ENSMUST00 | -57806 | Gm15985 | ENSMUSG0C | false | . | . | . | . | . | . | chr5 | 74791712 | 74793197 | 2.95389 | 2.93346 | 1.36068 |
| chr17 | 56036473 | 56036838 | 563 | Promoter (<=1kb) | 17 | 56016753 | 56036637 | 19885 | 2 | ENSMUST00 | 0 | Sh3gl1 | ENSMUSG0C | false | chr17 | 56035704 | 56036685 | 31.62822 | 118.12274 | 114.03909 | . | . | . | . | . | . |
| chr16 | 20732855 | 20733351 | 563 | Promoter (<=1kb) | 16 | 20733127 | 20738152 | 5026 | 1 | ENSMUST00 | 0 | Chrd | ENSMUSG0C | false | chr16 | 20732505 | 20734317 | 29.66345 | 100.21 | 96.63225 | chr16 | 20731646 | 20734488 | 4.34418 | 3.84634 | 2.46017 |
| chr10 | 60807790 | 60808183 | 563 | Intron (ENSMUST00000077925.5 | 10 | 60762610 | 60831581 | 68972 | 2 | ENSMUST00 | 23398 | Unc5b | ENSMUSG0C | false | . | . | . | . | . | . | chr10 | 60790706 | 60809046 | 5.64169 | 4.30724 | 3.54777 |
| chr1 | 119648261 | 119648770 | 563 | Promoter (<=1kb) | 1 | 119595074 | 119648617 | 53544 | 2 | ENSMUST00 | 0 | Epb4.1l5 | ENSMUSG0C | false | chr1 | 119646090 | 119649158 | 33.41831 | 117.26761 | 113.21259 | . | . | . | . | . | . |
| chr7 | 102267400 | 102268223 | 563 | Promoter (<=1kb) | 7 | 102267824 | 102436848 | 169025 | 1 | ENSMUST00 | 0 | Stim1 | ENSMUSG0C | false | chr7 | 102267172 | 102268649 | 28.22734 | 101.53345 | 97.92473 | . | . | . | . | . | . |
| chr4 | 134703220 | 134703883 | 563 | Promoter (<=1kb) | 4 | 134567889 | 134703767 | 135879 | 2 | ENSMUST00 | 0 | Man1c1 | ENSMUSG0C | false | chr4 | 134702765 | 134704907 | 29.37731 | 87.91154 | 84.61048 | chr4 | 134702127 | 134705017 | 5.20353 | 4.0496 | 3.1933 |
| chr8 | 87747072 | 87747378 | 563 | Intron (ENSMUST00000165770.3 | 8 | 87783147 | 87803823 | 20677 | 2 | ENSMUST00 | 56445 | Zfp423 | ENSMUSG0C | true | . | . | . | . | . | . | chr8 | 87729485 | 87749127 | 6.31805 | 4.63416 | 4.1004 |
| chr10 | 67342486 | 67342831 | 563 | Distal Intergenic | 10 | 67266689 | 67285305 | 18617 | 2 | ENSMUST00 | -57182 | Nrbf2 | ENSMUSG0C | false | . | . | . | . | . | . | chr10 | 67342103 | 67343511 | 3.78594 | 3.48848 | 1.99767 |
| chr11 | 107277057 | 107277386 | 563 | Intron (ENSMUST00000040380.7 | 11 | 107330708 | 107331058 | 351 | 1 | ENSMUST00 | -53322 | Gm11719 | ENSMUSG0C | false | . | . | . | . | . | . | chr11 | 107276071 | 107280102 | 4.04228 | 3.60796 | 2.23301 |
| chr6 | 146014595 | 146014838 | 563 | Distal Intergenic | 6 | 145951656 | 145957534 | 5879 | 2 | ENSMUST00 | -57062 | Sspnos | ENSMUSG0C | false | . | . | . | . | . | . | chr6 | 146013657 | 146015351 | 3.63501 | 3.26619 | 1.92527 |
| chr4 | 101089518 | 101089816 | 563 | Intron (ENSMUST00000106955.1 | 4 | 101103588 | 101103650 | 63 | 2 | ENSMUST00 | 13834 | Gm22100 | ENSMUSG0C | false | . | . | . | . | . | . | chr4 | 101070383 | 101096042 | 5.55154 | 4.29326 | 3.46737 |
| chr6 | 28606685 | 28606968 | 562 | Intron (ENSMUST00000164915.2 | 6 | 28561817 | 28724971 | 163155 | 1 | ENSMUST00 | 44869 | Snd1 | ENSMUSG0C | false | . | . | . | . | . | . | chr6 | 28602157 | 28607824 | 4.34311 | 3.76479 | 2.47136 |
| chr10 | 8184767 | 8185105 | 562 | Distal Intergenic | 10 | 7905653 | 7956230 | 50578 | 2 | ENSMUST00 | -228538 | Tab2 | ENSMUSG0C | true | . | . | . | . | . | . | chr10 | 8184862 | 8186119 | 3.99825 | 3.52463 | 2.24657 |
| chr19 | 3337012 | 3337326 | 562 | Intron (ENSMUST00000025835.4 | 19 | 3323301 | 3385732 | 62432 | 1 | ENSMUST00 | 13712 | Cpt1a | ENSMUSG0C | false | . | . | . | . | . | . | chr19 | 3325850 | 3346456 | 5.04851 | 4.02752 | 3.06051 |
| chr9 | 114752907 | 114753363 | 562 | Distal Intergenic | 9 | 114731203 | 114749343 | 18141 | 1 | ENSMUST00 | 21705 | Cmtm6 | ENSMUSG0C | false | chr9 | 114753006 | 114753423 | 11.64009 | 27.76484 | 25.32713 | chr9 | 114752296 | 114754520 | 4.15248 | 3.5406 | 2.33784 |
| chr8 | 72354318 | 72354653 | 562 | Intron (ENSMUST00000163643.1 | 8 | 72319033 | 72321656 | 2624 | 1 | ENSMUST00 | 35286 | Klf2 | ENSMUSG0C | false | . | . | . | . | . | . | chr8 | 72352643 | 72356274 | 6.3169 | 4.66576 | 4.13502 |
| chr2 | 121912343 | 121912620 | 562 | Intron (ENSMUST00000110586.4 | 2 | 121898807 | 121933662 | 34856 | 1 | ENSMUST00 | 13537 | Casc4 | ENSMUSG0C | false | . | . | . | . | . | . | chr2 | 121912363 | 121913151 | 4.38783 | 3.46694 | 2.54961 |
| chr9 | 44486393 | 44486659 | 562 | Distal Intergenic | 9 | 44478778 | 44482180 | 3403 | 1 | ENSMUST00 | 7616 | C030014I23R | ENSMUSG0C | false | . | . | . | . | . | . | chr9 | 44483700 | 44502510 | 6.77697 | 4.85012 | 4.46719 |
| chr8 | 105254199 | 105255033 | 562 | Promoter (<=1kb) | 8 | 105252639 | 105255151 | 2513 | 2 | ENSMUST00 | 118 | B3gnt9 | ENSMUSG0C | false | chr8 | 105254023 | 105255259 | 27.03504 | 88.5911 | 85.27629 | chr8 | 105253926 | 105255681 | 3.84569 | 3.48177 | 2.09348 |
| chr12 | 17690280 | 17690967 | 562 | Promoter (<=1kb) | 12 | 17690814 | 17791926 | 101113 | 1 | ENSMUST00 | 0 | Hpcal1 | ENSMUSG0C | false | chr12 | 17690029 | 17692705 | 39.39634 | 133.30592 | 128.68834 | . | . | . | . | . | . |
| chr12 | 110381140 | 110381615 | 562 | Distal Intergenic | 12 | 110279260 | 110280708 | 1449 | 1 | ENSMUST00 | 101881 | Dio3 | ENSMUSG0C | false | . | . | . | . | . | . | chr12 | 110380945 | 110381887 | 3.71474 | 3.40921 | 1.94604 |
| chr11 | 97206962 | 97207249 | 561 | 3' UTR | 11 | 97206908 | 97212462 | 5555 | 2 | ENSMUST00 | 5213 | Npepps | ENSMUSG0C | false | . | . | . | . | . | . | chr11 | 97206659 | 97207811 | 4.75474 | 4.01705 | 2.81604 |
| chr11 | 100850129 | 100850706 | 561 | Promoter (<=1kb) | 11 | 100798667 | 100850538 | 51872 | 2 | ENSMUST00 | 0 | Stat5b | ENSMUSG0C | false | chr11 | 100849708 | 100851276 | 36.25406 | 119.15681 | 115.04134 | . | . | . | . | . | . |
| chr11 | 117780639 | 117781658 | 561 | Promoter (<=1kb) | 11 | 117778742 | 117780944 | 2203 | 2 | ENSMUST00 | 0 | Tmc6 | ENSMUSG0C | false | chr11 | 117778803 | 117781710 | 25.50448 | 78.30592 | 75.18496 | chr11 | 117776172 | 117784117 | 7.16057 | 5.04978 | 4.77595 |
| chr3 | 30514930 | 30515315 | 561 | Intron (ENSMUST00000172754.2 | 3 | 29965456 | 30509822 | 544367 | 2 | ENSMUST00 | -5109 | Mecom | ENSMUSG0C | false | . | . | . | . | . | . | chr3 | 30514311 | 30516163 | 5.54274 | 4.39062 | 3.46091 |
| chr3 | 68572219 | 68572547 | 561 | Promoter (<=1kb) | 3 | 68572245 | 68626464 | 54220 | 1 | ENSMUST00 | 0 | Schip1 | ENSMUSG0C | false | chr3 | 68572019 | 68576372 | 26.65286 | 74.78688 | 71.72144 | chr3 | 68570245 | 68572599 | 4.19401 | 3.71092 | 2.33294 |
| chrX | 140527304 | 140527542 | 561 | Distal Intergenic | 20 | 140539528 | 140542668 | 3141 | 2 | ENSMUST00 | 15126 | Tsc22d3 | ENSMUSG0C | false | . | . | . | . | . | . | . | . | . | . | . | . |
| chr9 | 114785643 | 114786080 | 561 | Distal Intergenic | 9 | 114756837 | 114781993 | 25157 | 2 | ENSMUST00 | -3651 | Cmtm7 | ENSMUSG0C | true | . | . | . | . | . | . | chr9 | 114782213 | 114786923 | 4.97492 | 4.01143 | 2.99981 |
| chr12 | 40430122 | 40430481 | 561 | Intron (ENSMUST00000069637.8 | 12 | 40315558 | 40445769 | 130212 | 2 | ENSMUST00 | 15288 | Zfp277 | ENSMUSG0C | false | . | . | . | . | . | . | chr12 | 40422298 | 40443652 | 4.88409 | 4.00258 | 2.915 |
| chr8 | 65059945 | 65060170 | 561 | Distal Intergenic | 8 | 65075033 | 65077480 | 2448 | 1 | ENSMUST00 | -14863 | Gm10663 | ENSMUSG0C | false | . | . | . | . | . | . | chr8 | 65059200 | 65059948 | 3.78323 | 3.54633 | 1.99175 |
| chr8 | 25543348 | 25543788 | 561 | Intron (ENSMUST00000155564.2 | 8 | 25552678 | 25553560 | 883 | 2 | ENSMUST00 | 9772 | Gm16159 | ENSMUSG0C | false | . | . | . | . | . | . | chr8 | 25521452 | 25558014 | 5.35885 | 4.21426 | 3.3104 |
| chr10 | 39555732 | 39556150 | 561 | Promoter (1-2kb) | 10 | 39556071 | 39558062 | 1992 | 2 | ENSMUST00 | 1912 | Gm16364 | ENSMUSG0C | false | . | . | . | . | . | . | chr10 | 39550959 | 39564232 | 7.35607 | 5.07236 | 4.95132 |
| chr6 | 83775658 | 83776217 | 560 | Promoter (<=1kb) | 6 | 83768885 | 83775813 | 6929 | 2 | ENSMUST00 | 0 | Tex261 | ENSMUSG0C | false | chr6 | 83774759 | 83776337 | 32.94367 | 134.34938 | 129.69266 | chr6 | 83774182 | 83776885 | 4.59104 | 3.72868 | 2.70779 |
| chr12 | 4592571 | 4593557 | 560 | Promoter (<=1kb) | 12 | 4593008 | 4713950 | 120943 | 1 | ENSMUST00 | 0 | Itsn2 | ENSMUSG0C | false | chr12 | 4592382 | 4594262 | 27.54716 | 98.2761 | 94.74957 | chr12 | 4592301 | 4592596 | 2.9261 | 2.88856 | 1.35513 |
| chr1 | 55130974 | 55131569 | 560 | Promoter (<=1kb) | 1 | 55131231 | 55153938 | 22708 | 1 | ENSMUST00 | 0 | Mob4 | ENSMUSG0C | false | chr1 | 55130651 | 55132400 | 44.47867 | 147.74396 | 142.5466 | . | . | . | . | . | . |
| chr5 | 137628880 | 137629300 | 560 | Promoter (<=1kb) | 5 | 137629175 | 137642899 | 13725 | 1 | ENSMUST00 | 0 | Gm20605 | ENSMUSG0C | false | chr5 | 137629072 | 137631603 | 33.41831 | 117.26761 | 113.21259 | . | . | . | . | . | . |
| chr9 | 108236788 | 108237097 | 560 | Intron (ENSMUST00000166905.2 | 9 | 108205958 | 108263736 | 57779 | 2 | ENSMUST00 | 26639 | Dag1 | ENSMUSG0C | true | . | . | . | . | . | . | chr9 | 108210099 | 108251502 | 5.76916 | 4.35802 | 3.65688 |
| chr1 | 64885351 | 64885607 | 560 | Intron (ENSMUST00000139649.2 | 1 | 64860298 | 64928097 | 67800 | 2 | ENSMUST00 | 42490 | Plekhm3 | ENSMUSG0C | false | . | . | . | . | . | . | chr1 | 64883485 | 64886466 | 4.68643 | 3.9587 | 2.7496 |
| chr14 | 13960951 | 13961564 | 560 | Promoter (<=1kb) | 14 | 13949014 | 13961225 | 12212 | 2 | ENSMUST00 | 0 | Thoc7 | ENSMUSG0C | false | chr14 | 13960176 | 13962110 | 32.27147 | 102.84008 | 99.19724 | . | . | . | . | . | . |
| chr12 | 40043077 | 40043317 | 560 | Distal Intergenic | 12 | 40038099 | 40040027 | 1929 | 1 | ENSMUST00 | 4979 | Arl4aos | ENSMUSG0C | false | . | . | . | . | . | . | chr12 | 40042732 | 40045010 | 4.80749 | 3.87548 | 2.86462 |
| chr14 | 97795524 | 97795891 | 560 | Intron (ENSMUST00000069334.7 | 14 | 97967829 | 97967991 | 163 | 2 | ENSMUST00 | 172100 | Gm25831 | ENSMUSG0C | false | . | . | . | . | . | . | chr14 | 97791486 | 97803403 | 3.98924 | 3.5968 | 2.16581 |
| chr8 | 120051128 | 120051389 | 560 | 3' UTR | 8 | 120036194 | 120045287 | 9094 | 2 | ENSMUST00 | -5842 | Gm15684 | ENSMUSG0C | false | . | . | . | . | . | . | chr8 | 120050383 | 120052505 | 4.5779 | 3.75406 | 2.6794 |
| chrX | 75875067 | 75875278 | 559 | Promoter (<=1kb) | 20 | 75785654 | 75875182 | 89529 | 2 | ENSMUST00 | 0 | Pls3 | ENSMUSG0C | true | chrX | 75873307 | 75875446 | 17.92175 | 44.05972 | 41.41886 | . | . | . | . | . | . |
| chr11 | 7064957 | 7065247 | 559 | Promoter (1-2kb) | 11 | 7063489 | 7178506 | 115018 | 1 | ENSMUST00 | 1469 | Adcy1 | ENSMUSG0C | true | chr11 | 7063589 | 7065749 | 23.3092 | 84.4034 | 81.16875 | . | . | . | . | . | . |
| chr15 | 12739677 | 12740209 | 559 | Promoter (<=1kb) | 15 | 12443331 | 12739924 | 296594 | 2 | ENSMUST00 | 0 | Pdzd2 | ENSMUSG0C | false | chr15 | 12738458 | 12740528 | 24.03115 | 75.67385 | 72.59326 | chr15 | 12737558 | 12740976 | 4.75536 | 3.98225 | 2.79909 |
| chr7 | 16816111 | 16816885 | 559 | Promoter (<=1kb) | 7 | 16816317 | 16840927 | 24611 | 1 | ENSMUST00 | 0 | Strn4 | ENSMUSG0C | false | chr7 | 16815903 | 16817616 | 45.52735 | 154.11624 | 148.67334 | . | . | . | . | . | . |
| chr8 | 111094256 | 111094989 | 559 | Promoter (<=1kb) | 8 | 111094630 | 111112579 | 17950 | 1 | ENSMUST00 | 0 | Pdpr | ENSMUSG0C | false | chr8 | 111094098 | 111095587 | 50.54853 | 170.95712 | 164.9677 | chr8 | 111093693 | 111098306 | 4.05127 | 3.49944 | 2.25695 |
| chr9 | 64934056 | 64934385 | 559 | Intron (ENSMUST00000037798.7 | 9 | 64922861 | 64951607 | 28747 | 2 | ENSMUST00 | 17222 | Slc24a1 | ENSMUSG0C | false | . | . | . | . | . | . | chr9 | 64933446 | 64935085 | 5.2635 | 4.24189 | 3.22757 |
| chr4 | 149305584 | 149305911 | 559 | Promoter (1-2kb) | 4 | 149180236 | 149307506 | 127271 | 2 | ENSMUST00 | 1595 | Kif1b | ENSMUSG0C | false | chr4 | 149305850 | 149308579 | 36.82204 | 120.15061 | 116.00636 | chr4 | 149298505 | 149306747 | 5.00289 | 4.11226 | 3.01273 |
| chr4 | 115704853 | 115705181 | 559 | Promoter (2-3kb) | 4 | 115707251 | 115708303 | 1053 | 1 | ENSMUST00 | -2070 | Gm12849 | ENSMUSG0C | false | . | . | . | . | . | . | chr4 | 115704206 | 115705257 | 3.66365 | 3.45417 | 1.90686 |
| chr17 | 23823132 | 23823399 | 559 | Promoter (<=1kb) | 17 | 23823291 | 23823369 | 79 | 1 | ENSMUST00 | 0 | Mir5125 | ENSMUSG0C | false | . | . | . | . | . | . | chr17 | 23820478 | 23824236 | 4.64542 | 3.76984 | 2.74053 |
| chr14 | 120371128 | 120371462 | 559 | Intron (ENSMUST00000088419.7 | 14 | 120384263 | 120388985 | 4723 | 2 | ENSMUST00 | 17523 | Gm26679 | ENSMUSG0C | false | . | . | . | . | . | . | chr14 | 120362275 | 120389547 | 5.43713 | 4.23022 | 3.37672 |
| chr17 | 86800566 | 86800851 | 559 | Intron (ENSMUST00000024954.9 | 17 | 86840030 | 86840716 | 687 | 1 | ENSMUST00 | -39179 | RP24-107B1( | ENSMUSG0C | false | . | . | . | . | . | . | . | . | . | . | . | . |
| chr4 | 129287130 | 129287770 | 558 | Promoter (<=1kb) | 4 | 129287617 | 129308559 | 20943 | 1 | ENSMUST00 | 0 | Sync | ENSMUSG0C | false | chr4 | 129286769 | 129288550 | 28.1615 | 93.53628 | 90.11485 | chr4 | 129284848 | 129289902 | 6.10204 | 4.63545 | 3.89161 |
| chr11 | 4849323 | 4849761 | 558 | Promoter (<=1kb) | 11 | 4765845 | 4849536 | 83692 | 2 | ENSMUST00 | 0 | Nf2 | ENSMUSG0C | false | chr11 | 4849669 | 4850249 | 16.54316 | 39.53096 | 36.94807 | . | . | . | . | . | . |
| chr15 | 51865246 | 51865742 | 558 | Promoter (<=1kb) | 15 | 51786558 | 51865481 | 78924 | 2 | ENSMUST00 | 0 | Eif3h | ENSMUSG0C | false | chr15 | 51864544 | 51865978 | 37.6931 | 122.73721 | 118.50226 | . | . | . | . | . | . |
| chr15 | 91049695 | 91050065 | 558 | Promoter (<=1kb) | 15 | 90933276 | 91049948 | 116673 | 2 | ENSMUST00 | 0 | Kif21a | ENSMUSG0C | false | chr15 | 91048925 | 91050373 | 27.78601 | 91.88203 | 88.49705 | chr15 | 91048590 | 91050575 | 3.25273 | 3.18627 | 1.62226 |
| chr9 | 106202833 | 106203258 | 558 | Promoter (<=1kb) | 9 | 106203108 | 106215389 | 12282 | 1 | ENSMUST00 | 0 | Twf2 | ENSMUSG0C | false | chr9 | 106202779 | 106203827 | 27.11239 | 76.48743 | 73.39479 | chr9 | 106202273 | 106204485 | 5.11385 | 4.22529 | 3.09064 |
| chr11 | 72072380 | 72072728 | 558 | Intron (ENSMUST00000108508.2 | 11 | 72057645 | 72061561 | 3917 | 2 | ENSMUST00 | -10820 | Pitpnm3 | ENSMUSG0C | false | . | . | . | . | . | . | chr11 | 72070648 | 72073649 | 4.57406 | 3.74219 | 2.70205 |
| chr3 | 142169173 | 142169421 | 558 | Promoter (<=1kb) | 3 | 141840367 | 142169213 | 328847 | 2 | ENSMUST00 | 0 | Bmpr1b | ENSMUSG0C | false | chr3 | 142168586 | 142169710 | 27.66126 | 83.97217 | 80.74496 | chr3 | 142168377 | 142169672 | 4.17945 | 3.71043 | 2.32624 |
| chr9 | 64811271 | 64811691 | 558 | Promoter (<=1kb) | 9 | 64811340 | 64919667 | 108328 | 1 | ENSMUST00 | 0 | Dennd4a | ENSMUSG0C | false | chr9 | 64810764 | 64812508 | 33.78609 | 110.6256 | 106.77135 | . | . | . | . | . | . |
| chr4 | 149091382 | 149091644 | 558 | Intron (ENSMUST00000103217.5 | 4 | 149096383 | 149096702 | 320 | 1 | ENSMUST00 | -4739 | Gm9506 | ENSMUSG0C | false | . | . | . | . | . | . | chr4 | 149089749 | 149092669 | 4.53102 | 3.8392 | 2.6367 |
| chr3 | 83401575 | 83401821 | 558 | Distal Intergenic | 3 | 83555363 | 83574122 | 18760 | 2 | ENSMUST00 | 172301 | RP23-24F12. | ENSMUSG0C | false | . | . | . | . | . | . | . | . | . | . | . | . |
| chr17 | 35821472 | 35821890 | 557 | Promoter (<=1kb) | 17 | 35821684 | 35822923 | 1240 | 1 | ENSMUST00 | 0 | Ier3 | ENSMUSG0C | false | chr17 | 35821005 | 35823894 | 34.4409 | 118.06123 | 113.98219 | chr17 | 35820226 | 35824687 | 3.40141 | 3.25998 | 1.6905 |
| chr16 | 55821916 | 55822698 | 557 | Promoter (<=1kb) | 16 | 55811375 | 55822138 | 10764 | 2 | ENSMUST00 | 0 | Nfkbiz | ENSMUSG0C | false | chr16 | 55820908 | 55822916 | 35.61502 | 123.34891 | 119.09213 | . | . | . | . | . | . |
| chr10 | 22676931 | 22677234 | 557 | Intron (ENSMUST00000042261.4 | 10 | 22649008 | 22664690 | 15683 | 1 | ENSMUST00 | 27924 | Slc2a12 | ENSMUSG0C | false | . | . | . | . | . | . | chr10 | 22670879 | 22677260 | 4.29046 | 3.74548 | 2.42265 |
| chr7 | 28919718 | 28920122 | 557 | Promoter (<=1kb) | 7 | 28907014 | 28919221 | 12208 | 2 | ENSMUST00 | -498 | Actn4 | ENSMUSG0C | true | . | . | . | . | . | . | chr7 | 28910727 | 28959574 | 8.49369 | 5.62595 | 5.86532 |
| chr5 | 18360339 | 18360812 | 557 | Promoter (<=1kb) | 5 | 18265135 | 18360413 | 95279 | 2 | ENSMUST00 | 0 | Gnai1 | ENSMUSG0C | false | chr5 | 18358905 | 18360944 | 24.03115 | 75.67385 | 72.59326 | chr5 | 18357849 | 18361057 | 4.35797 | 3.68788 | 2.50849 |
| chr18 | 38797946 | 38798379 | 557 | Intron (ENSMUST00000148850.2 | 18 | 38771496 | 38772347 | 852 | 2 | ENSMUST00 | -25600 | Gm8302 | ENSMUSG0C | false | . | . | . | . | . | . | chr18 | 38796030 | 38799144 | 3.52672 | 3.27767 | 1.85012 |
| chr10 | 24364919 | 24365243 | 557 | Distal Intergenic | 10 | 24447490 | 24452657 | 5168 | 1 | ENSMUST00 | -82247 | Gm15271 | ENSMUSG0C | false | . | . | . | . | . | . | chr10 | 24364096 | 24367834 | 4.28857 | 3.72724 | 2.4501 |
| chr2 | 69264135 | 69264490 | 557 | Intron (ENSMUST00000126240.1 | 2 | 69248307 | 69272491 | 24185 | 1 | ENSMUST00 | 15829 | Gm13613 | ENSMUSG0C | false | . | . | . | . | . | . | chr2 | 69263385 | 69265598 | 3.3495 | 3.2445 | 1.65985 |
| chr17 | 72477045 | 72477379 | 557 | Intron (ENSMUST00000086639.4 | 17 | 72499289 | 72499418 | 130 | 2 | ENSMUST00 | 22039 | Gm24736 | ENSMUSG0C | false | . | . | . | . | . | . | chr17 | 72476588 | 72478745 | 5.4537 | 4.07477 | 3.42383 |
| chr17 | 87202532 | 87202893 | 557 | Distal Intergenic | 17 | 87184105 | 87184225 | 121 | 2 | ENSMUST00 | -18308 | Gm22346 | ENSMUSG0C | false | . | . | . | . | . | . | chr17 | 87202449 | 87204720 | 7.07823 | 5.03967 | 4.70871 |
| chr16 | 50528749 | 50529287 | 557 | Promoter (<=1kb) | 16 | 50528502 | 50558681 | 30180 | 1 | ENSMUST00 | 248 | G730013B05 | ENSMUSG0C | false | chr16 | 50528359 | 50530576 | 39.51976 | 124.83012 | 120.52397 | . | . | . | . | . | . |
| chr11 | 84067071 | 84067365 | 556 | Promoter (<=1kb) | 11 | 84048414 | 84067080 | 18667 | 2 | ENSMUST00 | 0 | Dusp14 | ENSMUSG0C | false | . | . | . | . | . | . | chr11 | 84065469 | 84069880 | 4.51715 | 3.8051 | 2.60899 |
| chr15 | 59062265 | 59062600 | 556 | Intron (ENSMUST00000080371.6 | 15 | 58941244 | 59082026 | 140783 | 2 | ENSMUST00 | 19426 | Mtss1 | ENSMUSG0C | true | . | . | . | . | . | . | chr15 | 58965186 | 59078715 | 7.47325 | 5.12649 | 5.04432 |
| chr16 | 90220520 | 90220779 | 556 | Promoter (<=1kb) | 16 | 90220742 | 90226322 | 5581 | 1 | ENSMUST00 | 0 | Sod1 | ENSMUSG0C | false | chr16 | 90220255 | 90221966 | 29.92778 | 109.76675 | 105.9399 | . | . | . | . | . | . |
| chr11 | 88235244 | 88235878 | 556 | Distal Intergenic | 11 | 88204444 | 88210971 | 6528 | 1 | ENSMUST00 | 30801 | Mrps23 | ENSMUSG0C | false | chr11 | 88235516 | 88236134 | 8.83539 | 19.39511 | 17.06904 | chr11 | 88233690 | 88236557 | 4.41115 | 3.71613 | 2.56478 |
| chr5 | 76079583 | 76079911 | 556 | Distal Intergenic | 5 | 76140271 | 76151515 | 11245 | 1 | ENSMUST00 | -60360 | Srd5a3 | ENSMUSG0C | false | . | . | . | . | . | . | chr5 | 76079307 | 76080153 | 4.11694 | 3.49765 | 2.31003 |
| chr2 | 33636018 | 33636620 | 556 | Intron (ENSMUST00000041730.5 | 2 | 33567156 | 33632283 | 65128 | 2 | ENSMUST00 | -3736 | Lmx1b | ENSMUSG0C | true | chr2 | 33635272 | 33637838 | 12.40737 | 26.6269 | 24.20249 | chr2 | 33565260 | 33638297 | 6.78481 | 4.81457 | 4.48235 |
| chr3 | 89421280 | 89421588 | 556 | Promoter (<=1kb) | 3 | 89421628 | 89429099 | 7472 | 1 | ENSMUST00 | -40 | Shc1 | ENSMUSG0C | false | . | . | . | . | . | . | chr3 | 89419346 | 89423117 | 6.05695 | 4.47955 | 3.88714 |
| chr13 | 97634413 | 97634741 | 556 | Distal Intergenic | 13 | 97735347 | 97743121 | 7775 | 2 | ENSMUST00 | 108380 | 5330416C01I | ENSMUSG0C | false | . | . | . | . | . | . | . | . | . | . | . | . |
| chr3 | 89227383 | 89227732 | 556 | Promoter (<=1kb) | 3 | 89227116 | 89227198 | 83 | 2 | ENSMUST00 | -186 | Mir92b | ENSMUSG0C | false | chr3 | 89227004 | 89227965 | 10.13814 | 22.76719 | 20.39335 | chr3 | 89226945 | 89228348 | 6.12475 | 4.68864 | 3.93233 |
| chr6 | 4441680 | 4441916 | 556 | Distal Intergenic | 6 | 4382878 | 4383204 | 327 | 1 | ENSMUST00 | 58803 | Gm24442 | ENSMUSG0C | false | . | . | . | . | . | . | chr6 | 4440910 | 4443112 | 4.2378 | 3.45126 | 2.41267 |
| chr14 | 34486857 | 34487098 | 556 | Intron (ENSMUST00000049005.9 | 14 | 34501715 | 34502684 | 970 | 1 | ENSMUST00 | -14617 | Gm17110 | ENSMUSG0C | false | . | . | . | . | . | . | chr14 | 34464938 | 34501066 | 4.73779 | 3.87419 | 2.80669 |
| chr12 | 111496535 | 111496810 | 555 | Distal Intergenic | 12 | 111484611 | 111485823 | 1213 | 2 | ENSMUST00 | -10713 | Gm266 | ENSMUSG0C | false | . | . | . | . | . | . | chr12 | 111496444 | 111497522 | 3.27275 | 3.15004 | 1.62928 |
| chr14 | 14076830 | 14077123 | 555 | Intron (ENSMUST00000022257.2 | 14 | 14113910 | 14116550 | 2641 | 2 | ENSMUST00 | 39427 | Psmd6 | ENSMUSG0C | false | . | . | . | . | . | . | chr14 | 14062631 | 14082513 | 5.48806 | 4.32274 | 3.41291 |
| chr16 | 85804120 | 85804944 | 555 | Promoter (1-2kb) | 16 | 85793827 | 85803113 | 9287 | 2 | ENSMUST00 | -1008 | Adamts1 | ENSMUSG0C | false | chr16 | 85803920 | 85804421 | 8.58414 | 21.55966 | 19.20221 | chr16 | 85800084 | 85805307 | 4.30005 | 3.70427 | 2.42091 |
| chr2 | 37443129 | 37443621 | 555 | Promoter (<=1kb) | 2 | 37443279 | 37495589 | 52311 | 1 | ENSMUST00 | 0 | Rabgap1 | ENSMUSG0C | false | chr2 | 37442474 | 37444150 | 37.7199 | 126.00829 | 121.65485 | chr2 | 37442416 | 37444965 | 4.33341 | 3.67452 | 2.49729 |
| chr6 | 55678475 | 55678730 | 555 | Promoter (2-3kb) | 6 | 55677818 | 55681263 | 3446 | 2 | ENSMUST00 | 2533 | Neurod6 | ENSMUSG0C | false | . | . | . | . | . | . | chr6 | 55676834 | 55678926 | 4.83864 | 3.76194 | 2.91193 |
| chr13 | 32802491 | 32802906 | 555 | Promoter (<=1kb) | 13 | 32802457 | 32802849 | 393 | 2 | ENSMUST00 | 0 | D930007J09I | ENSMUSG0C | false | chr13 | 32801439 | 32803311 | 34.54477 | 122.48211 | 118.25522 | chr13 | 32802877 | 32804945 | 4.6464 | 3.74331 | 2.74877 |
| chr12 | 79615986 | 79616334 | 555 | Distal Intergenic | 12 | 79297351 | 79508656 | 211306 | 1 | ENSMUST00 | 318636 | Rad51b | ENSMUSG0C | false | . | . | . | . | . | . | chr12 | 79612895 | 79616940 | 5.18537 | 4.1714 | 3.15883 |
| chr1 | 82634721 | 82635304 | 555 | Promoter (<=1kb) | 1 | 82635013 | 82635110 | 98 | 1 | ENSMUST00 | 0 | n-R5s213 | ENSMUSG0C | false | . | . | . | . | . | . | chr1 | 82623735 | 82642803 | 5.1888 | 4.15508 | 3.1621 |
| chr7 | 27361407 | 27361828 | 555 | Promoter (1-2kb) | 7 | 27363314 | 27363401 | 88 | 1 | ENSMUST00 | -1486 | Gm26043 | ENSMUSG0C | false | chr7 | 27361634 | 27362010 | 11.22292 | 28.25542 | 25.81138 | chr7 | 27360874 | 27362022 | 3.78912 | 3.52141 | 1.99216 |
| chr2 | 5215171 | 5215674 | 555 | Intron (ENSMUST00000027988.7 | 2 | 5137749 | 5230864 | 93116 | 2 | ENSMUST00 | 15190 | Gm26776 | ENSMUSG0C | false | . | . | . | . | . | . | chr2 | 5214866 | 5216480 | 4.4433 | 3.67106 | 2.57942 |
| chr1 | 43130213 | 43130460 | 554 | Intron (ENSMUST00000008280.8 | 1 | 43098710 | 43115947 | 17238 | 1 | ENSMUST00 | 31504 | AI597479 | ENSMUSG0C | false | . | . | . | . | . | . | chr1 | 43130389 | 43131139 | 2.7156 | 2.7068 | 1.18117 |
| chr11 | 78568849 | 78569216 | 554 | Promoter (<=1kb) | 11 | 78567402 | 78568916 | 1515 | 2 | ENSMUST00 | 0 | Nlk | ENSMUSG0C | true | . | . | . | . | . | . | chr11 | 78565466 | 78571591 | 4.72666 | 3.87884 | 2.82952 |
| chr10 | 21160683 | 21161155 | 554 | Promoter (<=1kb) | 10 | 21124936 | 21160905 | 35970 | 2 | ENSMUST00 | 0 | Myb | ENSMUSG0C | false | chr10 | 21158821 | 21161478 | 24.35436 | 72.61947 | 69.58835 | chr10 | 21158270 | 21161787 | 5.73899 | 4.40548 | 3.6165 |
| chr10 | 123196500 | 123197386 | 554 | Promoter (<=1kb) | 10 | 123113244 | 123196935 | 83692 | 2 | ENSMUST00 | 0 | Usp15 | ENSMUSG0C | false | chr10 | 123195574 | 123197730 | 37.27983 | 118.82364 | 114.72062 | . | . | . | . | . | . |
| chr1 | 176813362 | 176814285 | 554 | Promoter (<=1kb) | 1 | 176814660 | 176845497 | 30838 | 1 | ENSMUST00 | -375 | Sdccag8 | ENSMUSG0C | false | chr1 | 176813017 | 176815457 | 33.32866 | 126.59486 | 122.2257 | . | . | . | . | . | . |
| chr9 | 114730981 | 114731464 | 554 | Promoter (<=1kb) | 9 | 114731203 | 114749343 | 18141 | 1 | ENSMUST00 | 0 | Cmtm6 | ENSMUSG0C | false | chr9 | 114730348 | 114732085 | 24.0356 | 94.23969 | 90.80066 | chr9 | 114729006 | 114733084 | 4.39113 | 3.76523 | 2.50971 |
| chr5 | 121386556 | 121386906 | 554 | Promoter (<=1kb) | 5 | 121371725 | 121385632 | 13908 | 2 | ENSMUST00 | -925 | Trafd1 | ENSMUSG0C | false | . | . | . | . | . | . | chr5 | 121386120 | 121387511 | 5.92303 | 4.51611 | 3.77705 |
| chr14 | 56567562 | 56567804 | 554 | Intron (ENSMUST00000065302.7 | 14 | 56526761 | 56571846 | 45086 | 2 | ENSMUST00 | 4042 | Cenpj | ENSMUSG0C | false | . | . | . | . | . | . | . | . | . | . | . | . |
| chr2 | 93791490 | 93791863 | 554 | Promoter (2-3kb) | 2 | 93794570 | 93795080 | 511 | 1 | ENSMUST00 | -2707 | Gm13818 | ENSMUSG0C | false | . | . | . | . | . | . | chr2 | 93791314 | 93792058 | 2.89684 | 2.84181 | 1.3295 |
| chr10 | 12703610 | 12703885 | 554 | Intron (ENSMUST00000076817.4 | 10 | 12793875 | 12793974 | 100 | 2 | ENSMUST00 | 90089 | Gm23044 | ENSMUSG0C | false | . | . | . | . | . | . | chr10 | 12700386 | 12709298 | 4.88951 | 4.01525 | 2.91749 |
| chr6 | 71270689 | 71270972 | 554 | Promoter (<=1kb) | 6 | 71271671 | 71271801 | 131 | 1 | ENSMUST00 | -699 | Mir8112 | ENSMUSG0C | false | chr6 | 71270850 | 71271480 | 17.60262 | 45.78123 | 43.11721 | . | . | . | . | . | . |
| chr16 | 91385417 | 91385825 | 553 | Promoter (1-2kb) | 16 | 91387789 | 91391776 | 3988 | 1 | ENSMUST00 | -1964 | Ifnar2 | ENSMUSG0C | false | . | . | . | . | . | . | chr16 | 91384639 | 91386690 | 3.78714 | 3.46545 | 2.01411 |
| chr17 | 78248878 | 78249270 | 553 | Intron (ENSMUST00000112498.2 | 17 | 78200248 | 78376592 | 176345 | 1 | ENSMUST00 | 48631 | Crim1 | ENSMUSG0C | true | . | . | . | . | . | . | chr17 | 78239579 | 78284617 | 5.22436 | 4.1134 | 3.20479 |
| chr5 | 53607197 | 53607740 | 553 | Intron (ENSMUST00000087360.3 | 5 | 53590574 | 53657445 | 66872 | 1 | ENSMUST00 | 16624 | Rbpj | ENSMUSG0C | false | . | . | . | . | . | . | chr5 | 53596758 | 53625083 | 5.16494 | 4.16876 | 3.14681 |
| chr10 | 128237537 | 128237967 | 553 | Promoter (<=1kb) | 10 | 128238066 | 128252941 | 14876 | 1 | ENSMUST00 | -99 | Timeless | ENSMUSG0C | false | . | . | . | . | . | . | chr10 | 128236496 | 128238664 | 3.63818 | 3.29556 | 1.93286 |
| chr10 | 128096679 | 128097146 | 553 | Intron (ENSMUST00000045621.8 | 10 | 128092783 | 128129303 | 36521 | 1 | ENSMUST00 | 3897 | Baz2a | ENSMUSG0C | false | chr10 | 128096543 | 128098298 | 40.54782 | 137.53284 | 132.75229 | chr10 | 128093379 | 128103131 | 5.08043 | 4.07895 | 3.08556 |
| chr10 | 11280873 | 11281603 | 553 | Promoter (<=1kb) | 10 | 11281330 | 11298052 | 16723 | 1 | ENSMUST00 | 0 | Fbxo30 | ENSMUSG0C | false | chr10 | 11280393 | 11282529 | 34.8365 | 112.46898 | 108.56265 | . | . | . | . | . | . |
| chr3 | 55145053 | 55145569 | 553 | Promoter (2-3kb) | 3 | 55142288 | 55171608 | 29321 | 1 | ENSMUST00 | 2766 | Ccdc169 | ENSMUSG0C | false | . | . | . | . | . | . | chr3 | 55144347 | 55146080 | 3.29774 | 3.11494 | 1.62706 |
| chr3 | 108200436 | 108201003 | 553 | Promoter (<=1kb) | 3 | 108195687 | 108200834 | 5148 | 2 | ENSMUST00 | 0 | Cyb561d1 | ENSMUSG0C | false | chr3 | 108200038 | 108201466 | 20.5121 | 70.85588 | 67.85384 | chr3 | 108199358 | 108202867 | 4.64277 | 3.92158 | 2.73295 |
| chr3 | 153698552 | 153698928 | 553 | Intron (ENSMUST00000064460.5 | 3 | 153708911 | 153709026 | 116 | 1 | ENSMUST00 | -9983 | Gm22206 | ENSMUSG0C | false | . | . | . | . | . | . | chr3 | 153686316 | 153712771 | 6.43925 | 4.66349 | 4.19773 |
| chr4 | 40823207 | 40823506 | 553 | Exon (ENSMUST00000030121.7/ | 4 | 40850056 | 40850138 | 83 | 2 | ENSMUST00 | 26632 | Mir5123 | ENSMUSG0C | false | . | . | . | . | . | . | chr4 | 40821931 | 40852722 | 6.59841 | 4.80932 | 4.32335 |
| chr1 | 128748644 | 128748957 | 553 | Promoter (1-2kb) | 1 | 128750926 | 128751347 | 422 | 1 | ENSMUST00 | -1969 | RP23-424E1 | ENSMUSG0C | false | . | . | . | . | . | . | chr1 | 128748597 | 128750285 | 3.54819 | 3.31635 | 1.86328 |
| chr15 | 77928861 | 77929215 | 552 | Promoter (<=1kb) | 15 | 77915047 | 77929006 | 13960 | 2 | ENSMUST00 | 0 | Txn2 | ENSMUSG0C | false | chr15 | 77928467 | 77929042 | 37.64722 | 120.52259 | 116.3698 | chr15 | 77927421 | 77929830 | 5.3211 | 4.24371 | 3.28236 |
| chr16 | 33169486 | 33169735 | 552 | Distal Intergenic | 16 | 33185071 | 33243311 | 58241 | 1 | ENSMUST00 | -15336 | Osbpl11 | ENSMUSG0C | false | . | . | . | . | . | . | . | . | . | . | . | . |
| chr15 | 98934388 | 98934747 | 552 | Promoter (<=1kb) | 15 | 98931425 | 98934565 | 3141 | 2 | ENSMUST00 | 0 | Tuba1b | ENSMUSG0C | false | chr15 | 98933157 | 98935385 | 28.03146 | 79.91027 | 76.76085 | . | . | . | . | . | . |
| chr3 | 80004967 | 80005372 | 552 | Distal Intergenic | 3 | 79885930 | 79946278 | 60349 | 1 | ENSMUST00 | 119038 | Fam198b | ENSMUSG0C | false | . | . | . | . | . | . | chr3 | 80004619 | 80007354 | 6.39117 | 4.74652 | 4.15183 |
| chr4 | 138044349 | 138044670 | 552 | Intron (ENSMUST00000139759.2 | 4 | 138004752 | 138103394 | 98643 | 1 | ENSMUST00 | 39598 | Eif4g3 | ENSMUSG0C | false | . | . | . | . | . | . | chr4 | 138041910 | 138045462 | 5.01171 | 3.97927 | 3.0261 |
| chr8 | 25100806 | 25101264 | 552 | Promoter (<=1kb) | 8 | 25042944 | 25101771 | 58828 | 2 | ENSMUST00 | 507 | Plekha2 | ENSMUSG0C | true | chr8 | 25099212 | 25103419 | 35.56825 | 117.95176 | 113.87694 | chr8 | 25082618 | 25100838 | 5.3544 | 4.1622 | 3.30614 |
| chr13 | 52980416 | 52981322 | 552 | Promoter (<=1kb) | 13 | 52967209 | 52981073 | 13865 | 2 | ENSMUST00 | 0 | Nfil3 | ENSMUSG0C | false | chr13 | 52979288 | 52982918 | 34.18774 | 104.13715 | 100.46495 | . | . | . | . | . | . |
| chrY | 10642063 | 10642334 | 552 | Promoter (<=1kb) | 21 | 10641257 | 10643107 | 1851 | 2 | ENSMUST00 | 773 | Gm20775 | ENSMUSG0C | false | . | . | . | . | . | . | . | . | . | . | . | . |
| chr11 | 28999241 | 28999538 | 552 | Distal Intergenic | 11 | 29020374 | 29020984 | 611 | 2 | ENSMUST00 | 21446 | Gm12084 | ENSMUSG0C | false | . | . | . | . | . | . | chr11 | 28998228 | 29000483 | 3.95323 | 3.46661 | 2.16376 |
| chr18 | 32693874 | 32694249 | 552 | Intron (ENSMUST00000181632.1 | 18 | 32621920 | 32622056 | 137 | 2 | ENSMUST00 | -71819 | Gm25826 | ENSMUSG0C | false | . | . | . | . | . | . | . | . | . | . | . | . |
| chr10 | 58395416 | 58395703 | 551 | Promoter (1-2kb) | 10 | 58394388 | 58421578 | 27191 | 1 | ENSMUST00 | 1029 | Lims1 | ENSMUSG0C | false | chr10 | 58394347 | 58395666 | 19.69817 | 53.33889 | 50.58166 | chr10 | 58395467 | 58399777 | 3.5284 | 3.22447 | 1.82914 |
| chr10 | 122029835 | 122030100 | 551 | Intron (ENSMUST00000020322.6 | 10 | 121780991 | 122047308 | 266318 | 2 | ENSMUST00 | 17208 | Srgap1 | ENSMUSG0C | true | . | . | . | . | . | . | chr10 | 121945243 | 122036389 | 5.74397 | 4.3951 | 3.62974 |
| chr17 | 43077135 | 43077393 | 551 | Intron (ENSMUST00000024708.4 | 17 | 43016555 | 43089188 | 72634 | 1 | ENSMUST00 | 60581 | Tnfrsf21 | ENSMUSG0C | false | chr17 | 43077267 | 43077841 | 9.63953 | 19.72805 | 17.39678 | chr17 | 43075309 | 43083932 | 4.06236 | 3.6443 | 2.23752 |
| chr6 | 8520458 | 8520788 | 551 | Promoter (<=1kb) | 6 | 8520058 | 8593512 | 73455 | 1 | ENSMUST00 | 401 | Glcci1 | ENSMUSG0C | false | . | . | . | . | . | . | . | . | . | . | . | . |
| chr15 | 78982506 | 78983175 | 551 | Promoter (<=1kb) | 15 | 78983041 | 79005864 | 22824 | 1 | ENSMUST00 | 0 | Triobp | ENSMUSG0C | false | chr15 | 78982378 | 78982895 | 12.50275 | 27.39253 | 24.96176 | chr15 | 78981999 | 78988873 | 5.15157 | 4.09348 | 3.15503 |
| chr3 | 101597223 | 101597548 | 551 | Intron (ENSMUST00000036493.6 | 3 | 101576219 | 101604603 | 28385 | 2 | ENSMUST00 | 7055 | Atp1a1 | ENSMUSG0C | false | . | . | . | . | . | . | chr3 | 101591433 | 101603595 | 6.88213 | 4.93449 | 4.55128 |
| chr6 | 52204411 | 52204758 | 551 | Promoter (<=1kb) | 6 | 52201754 | 52204587 | 2834 | 2 | ENSMUST00 | 0 | Hoxa5 | ENSMUSG0C | false | chr6 | 52203683 | 52204833 | 15.7162 | 39.55585 | 36.97221 | chr6 | 52203796 | 52211563 | 4.96251 | 3.88667 | 3.0102 |
| chrX | 95441369 | 95441610 | 551 | Intron (ENSMUST00000084535.5 | 20 | 95420318 | 95444872 | 24555 | 2 | ENSMUST00 | 3262 | Amer1 | ENSMUSG0C | false | . | . | . | . | . | . | . | . | . | . | . | . |
| chr5 | 135460243 | 135460844 | 551 | Exon (ENSMUST00000060311.8/ | 5 | 135376141 | 135394546 | 18406 | 2 | ENSMUST00 | -65698 | Pom121 | ENSMUSG0C | false | chr5 | 135460134 | 135460848 | 5.05485 | 7.29253 | 5.17558 | chr5 | 135458659 | 135480128 | 5.22779 | 4.14143 | 3.19665 |
| chr6 | 72170831 | 72171076 | 551 | Distal Intergenic | 6 | 72115232 | 72142271 | 27040 | 1 | ENSMUST00 | 55600 | St3gal5 | ENSMUSG0C | false | . | . | . | . | . | . | chr6 | 72169770 | 72171056 | 2.97105 | 2.90024 | 1.38244 |
| chr17 | 13985229 | 13985601 | 551 | Distal Intergenic | 17 | 14000239 | 14001765 | 1527 | 2 | ENSMUST00 | 16164 | RP23-9O15.2 | ENSMUSG0C | false | . | . | . | . | . | . | . | . | . | . | . | . |
| chr15 | 67039394 | 67039928 | 550 | Distal Intergenic | 15 | 66946537 | 66969640 | 23104 | 2 | ENSMUST00 | -69755 | Ndrg1 | ENSMUSG0C | false | chr15 | 67039226 | 67040096 | 23.76298 | 65.38354 | 62.46201 | chr15 | 67038727 | 67040664 | 4.55325 | 3.84107 | 2.68639 |
| chr11 | 80208904 | 80209326 | 550 | Promoter (<=1kb) | 11 | 80209019 | 80266812 | 57794 | 1 | ENSMUST00 | 0 | Rhot1 | ENSMUSG0C | false | chr11 | 80208996 | 80210259 | 31.43325 | 99.35611 | 95.79983 | . | . | . | . | . | . |
| chr17 | 11229502 | 11229732 | 550 | Intron (ENSMUST00000066658.7 | 17 | 11303647 | 11610945 | 307299 | 2 | ENSMUST00 | -73915 | Park2 | ENSMUSG0C | false | . | . | . | . | . | . | chr17 | 11228754 | 11231609 | 4.57671 | 3.75149 | 2.68898 |
| chr2 | 60688938 | 60689427 | 550 | Intron (ENSMUST00000028348.3 | 2 | 60653048 | 60673754 | 20707 | 2 | ENSMUST00 | -15185 | Itgb6 | ENSMUSG0C | false | . | . | . | . | . | . | chr2 | 60688950 | 60689750 | 3.72259 | 3.46964 | 1.98706 |
| chr5 | 52827930 | 52828219 | 550 | Distal Intergenic | 5 | 52834015 | 52867797 | 33783 | 1 | ENSMUST00 | -5796 | Anapc4 | ENSMUSG0C | false | . | . | . | . | . | . | chr5 | 52827361 | 52829616 | 5.24924 | 4.24689 | 3.21783 |
| chr3 | 144534084 | 144534359 | 550 | Intron (ENSMUST00000043325.7 | 3 | 144453963 | 144569881 | 115919 | 2 | ENSMUST00 | 35522 | Hs2st1 | ENSMUSG0C | false | . | . | . | . | . | . | chr3 | 144528524 | 144535848 | 4.74518 | 3.87016 | 2.80068 |
| chr1 | 132355046 | 132355569 | 550 | Downstream (<1kb) | 1 | 132361026 | 132364948 | 3923 | 2 | ENSMUST00 | 9379 | Tmcc2 | ENSMUSG0C | false | . | . | . | . | . | . | chr1 | 132352707 | 132356435 | 8.61007 | 5.59858 | 5.98871 |
| chr18 | 60609190 | 60609665 | 550 | Promoter (<=1kb) | 18 | 60600207 | 60610105 | 9899 | 2 | ENSMUST00 | 440 | Synpo | ENSMUSG0C | true | chr18 | 60603340 | 60613557 | 27.39171 | 92.13462 | 88.74321 | . | . | . | . | . | . |
| chr2 | 119156873 | 119157170 | 550 | Promoter (<=1kb) | 2 | 119137001 | 119157034 | 20034 | 2 | ENSMUST00 | 0 | Rmdn3 | ENSMUSG0C | false | chr2 | 119156068 | 119157543 | 34.3512 | 113.70341 | 109.76276 | . | . | . | . | . | . |
| chr11 | 96350056 | 96350353 | 550 | Promoter (<=1kb) | 11 | 96350525 | 96353658 | 3134 | 1 | ENSMUST00 | -172 | Hoxb2 | ENSMUSG0C | false | . | . | . | . | . | . | . | . | . | . | . | . |
| chr6 | 30460697 | 30460988 | 550 | Promoter (<=1kb) | 6 | 30460985 | 30463313 | 2329 | 1 | ENSMUST00 | 0 | 1700012C14I | ENSMUSG0C | false | . | . | . | . | . | . | chr6 | 30459555 | 30461187 | 4.65505 | 3.85054 | 2.76617 |
| chr9 | 50617113 | 50617563 | 549 | Promoter (<=1kb) | 9 | 50617321 | 50624997 | 7677 | 1 | ENSMUST00 | 0 | Pih1d2 | ENSMUSG0C | false | chr9 | 50616356 | 50618031 | 34.52349 | 109.84589 | 106.01766 | . | . | . | . | . | . |
| chr6 | 121112771 | 121113152 | 549 | Distal Intergenic | 6 | 121007241 | 121081609 | 74369 | 2 | ENSMUST00 | -31163 | Mical3 | ENSMUSG0C | false | . | . | . | . | . | . | chr6 | 121110555 | 121114977 | 5.58267 | 4.27018 | 3.52272 |
| chr11 | 58334833 | 58335291 | 549 | Intron (ENSMUST00000116376.3 | 11 | 58331275 | 58347720 | 16446 | 1 | ENSMUST00 | 3559 | Sh3bp5l | ENSMUSG0C | false | . | . | . | . | . | . | chr11 | 58331188 | 58336230 | 7.18085 | 5.04978 | 4.79472 |
| chr13 | 64312475 | 64312926 | 549 | Promoter (<=1kb) | 13 | 64291836 | 64312710 | 20875 | 2 | ENSMUST00 | 0 | Aaed1 | ENSMUSG0C | false | chr13 | 64311532 | 64312714 | 26.73128 | 81.39996 | 78.22401 | . | . | . | . | . | . |
| chr7 | 28362096 | 28363025 | 549 | Promoter (<=1kb) | 7 | 28359605 | 28363341 | 3737 | 2 | ENSMUST00 | 316 | Plekhg2 | ENSMUSG0C | true | . | . | . | . | . | . | chr7 | 28351351 | 28371775 | 6.79815 | 4.79627 | 4.48731 |
| chr17 | 66446226 | 66446512 | 549 | Promoter (1-2kb) | 17 | 66385746 | 66448369 | 62624 | 2 | ENSMUST00 | 1857 | Mtcl1 | ENSMUSG0C | true | chr17 | 66446496 | 66450714 | 31.54088 | 108.67455 | 104.87239 | chr17 | 66436561 | 66447599 | 5.05611 | 4.05554 | 3.06333 |
| chrX | 157491264 | 157492210 | 549 | Promoter (<=1kb) | 20 | 157445696 | 157492066 | 46371 | 2 | ENSMUST00 | 0 | Sms | ENSMUSG0C | false | chrX | 157490958 | 157492445 | 14.66885 | 35.63733 | 33.10225 | . | . | . | . | . | . |
| chr11 | 102576150 | 102576401 | 549 | Promoter (2-3kb) | 11 | 102576491 | 102579119 | 2629 | 2 | ENSMUST00 | 2718 | Gm11627 | ENSMUSG0C | false | . | . | . | . | . | . | chr11 | 102576085 | 102577098 | 3.60963 | 3.30588 | 1.91106 |
| chr14 | 25473863 | 25474151 | 549 | Intron (ENSMUST00000161766.2 | 14 | 25455737 | 25461128 | 5392 | 2 | ENSMUST00 | -12736 | Zmiz1os1 | ENSMUSG0C | false | . | . | . | . | . | . | chr14 | 25460928 | 25485304 | 5.40667 | 4.23635 | 3.3428 |
| chr10 | 114071633 | 114071942 | 549 | Distal Intergenic | 10 | 114757747 | 114785104 | 27358 | 1 | ENSMUST00 | -685805 | 4930473D10I | ENSMUSG0C | false | chr10 | 114070817 | 114071744 | 10.35471 | 20.92403 | 18.57802 | . | . | . | . | . | . |
| chr6 | 141333363 | 141333602 | 549 | Intron (ENSMUST00000043259.8 | 6 | 141238278 | 141250561 | 12284 | 2 | ENSMUST00 | -82803 | RP23-97L11. | ENSMUSG0C | false | . | . | . | . | . | . | chr6 | 141332152 | 141339237 | 4.48821 | 3.82642 | 2.58244 |
| chr11 | 21033827 | 21034342 | 548 | Distal Intergenic | 11 | 21091291 | 21143005 | 51715 | 1 | ENSMUST00 | -56949 | Peli1 | ENSMUSG0C | false | . | . | . | . | . | . | chr11 | 21033326 | 21034734 | 4.19585 | 3.59778 | 2.39161 |
| chr13 | 100833052 | 100833591 | 548 | Promoter (<=1kb) | 13 | 100802649 | 100833427 | 30779 | 2 | ENSMUST00 | 0 | Slc30a5 | ENSMUSG0C | false | chr13 | 100832248 | 100833839 | 31.96831 | 119.80809 | 115.67081 | . | . | . | . | . | . |
| chr15 | 25826378 | 25826764 | 548 | Distal Intergenic | 15 | 25843264 | 25973687 | 130424 | 1 | ENSMUST00 | -16500 | Fam134b | ENSMUSG0C | false | . | . | . | . | . | . | chr15 | 25824544 | 25829445 | 5.74828 | 4.46732 | 3.62015 |
| chr1 | 53774039 | 53774305 | 548 | Intron (ENSMUST00000027263.8 | 1 | 53761149 | 53785157 | 24009 | 2 | ENSMUST00 | 10852 | Stk17b | ENSMUSG0C | false | . | . | . | . | . | . | chr1 | 53772361 | 53777721 | 3.69619 | 3.44324 | 1.93522 |
| chr6 | 21851914 | 21852488 | 548 | Promoter (<=1kb) | 6 | 21771395 | 21852088 | 80694 | 2 | ENSMUST00 | 0 | Tspan12 | ENSMUSG0C | true | chr6 | 21850969 | 21852979 | 26.18681 | 91.82585 | 88.44535 | . | . | . | . | . | . |
| chr6 | 86770923 | 86771210 | 548 | Intron (ENSMUST00000001187.9 | 6 | 86747471 | 86765910 | 18440 | 2 | ENSMUST00 | -5014 | Anxa4 | ENSMUSG0C | true | . | . | . | . | . | . | chr6 | 86761092 | 86794291 | 5.26625 | 4.09177 | 3.24957 |
| chr7 | 29169349 | 29169722 | 548 | Promoter (<=1kb) | 7 | 29165923 | 29169841 | 3919 | 2 | ENSMUST00 | 119 | Spred3 | ENSMUSG0C | false | chr7 | 29169161 | 29171818 | 19.69817 | 53.33889 | 50.58166 | chr7 | 29168116 | 29172346 | 7.1171 | 5.05011 | 4.73706 |
| chr3 | 134235835 | 134236657 | 548 | Promoter (<=1kb) | 3 | 134236491 | 134262090 | 25600 | 1 | ENSMUST00 | 0 | Cxxc4 | ENSMUSG0C | false | chr3 | 134235553 | 134236537 | 15.50707 | 38.47921 | 35.908 | chr3 | 134235198 | 134237711 | 3.71971 | 3.44201 | 1.95581 |
| chr2 | 18040427 | 18040728 | 548 | Promoter (2-3kb) | 2 | 18025193 | 18037737 | 12545 | 2 | ENSMUST00 | -2691 | A930004D18 | ENSMUSG0C | false | . | . | . | . | . | . | chr2 | 18039586 | 18046995 | 3.8226 | 3.45815 | 2.053 |
| chr5 | 20328922 | 20329197 | 548 | Intron (ENSMUST00000088516.4 | 5 | 20264341 | 20264440 | 100 | 1 | ENSMUST00 | 64582 | Gm22755 | ENSMUSG0C | false | . | . | . | . | . | . | chr5 | 20328071 | 20331637 | 4.7552 | 3.96017 | 2.80488 |
| chr11 | 60086783 | 60087000 | 548 | Distal Intergenic | 11 | 60099547 | 60099673 | 127 | 1 | ENSMUST00 | -12547 | Gm27511 | ENSMUSG0C | false | . | . | . | . | . | . | . | . | . | . | . | . |
| chr7 | 66387476 | 66387844 | 547 | Promoter (<=1kb) | 7 | 66341670 | 66388307 | 46638 | 2 | ENSMUST00 | 463 | Lrrk1 | ENSMUSG0C | false | chr7 | 66387468 | 66389353 | 34.78613 | 113.42423 | 109.48744 | chr7 | 66383007 | 66389812 | 3.14393 | 3.06514 | 1.49318 |
| chr15 | 64312479 | 64313314 | 547 | Promoter (<=1kb) | 15 | 64086857 | 64312673 | 225817 | 2 | ENSMUST00 | 0 | Asap1 | ENSMUSG0C | false | chr15 | 64311537 | 64313724 | 27.46926 | 112.95284 | 109.03484 | chr15 | 64313222 | 64315451 | 3.98543 | 3.61533 | 2.16574 |
| chr8 | 127202244 | 127202534 | 547 | Intron (ENSMUST00000162309.2 | 8 | 127299585 | 127402577 | 102993 | 1 | ENSMUST00 | -97051 | Pard3 | ENSMUSG0C | false | . | . | . | . | . | . | chr8 | 127200419 | 127216139 | 4.67858 | 3.89809 | 2.74264 |
| chr1 | 152066767 | 152067111 | 547 | Intron (ENSMUST00000187603.1 | 1 | 151893237 | 152055292 | 162056 | 2 | ENSMUST00 | -11476 | 1700025G04 | ENSMUSG0C | true | . | . | . | . | . | . | chr1 | 152065155 | 152067582 | 3.80441 | 3.47411 | 2.04868 |
| chr19 | 31659420 | 31659796 | 547 | Intron (ENSMUST00000065067.7 | 19 | 30568547 | 31664370 | 1095824 | 2 | ENSMUST00 | 4574 | Prkg1 | ENSMUSG0C | true | . | . | . | . | . | . | chr19 | 31656063 | 31663727 | 4.06461 | 3.5305 | 2.23999 |
| chr7 | 114071997 | 114072298 | 547 | Intron (ENSMUST00000069449.5 | 7 | 114046785 | 114117781 | 70997 | 2 | ENSMUST00 | 45483 | Rras2 | ENSMUSG0C | true | . | . | . | . | . | . | chr7 | 114058507 | 114116129 | 5.64505 | 4.33683 | 3.54844 |
| chr10 | 91123442 | 91124317 | 547 | Promoter (<=1kb) | 10 | 91116574 | 91124059 | 7486 | 2 | ENSMUST00 | 0 | Slc25a3 | ENSMUSG0C | false | chr10 | 91124194 | 91124535 | 9.86256 | 23.41135 | 21.03137 | . | . | . | . | . | . |

| | | | | | | | | | | | | | | | | | | | | | | | | | | |
|---|---|---|---|---|---|---|---|---|---|---|---|---|---|---|---|---|---|---|---|---|---|---|---|---|---|---|
| chr17 | 84584267 | 84584485 | 547 | Intron (ENSMUST00000047206.5 | 17 | 84624402 | 84624510 | 109 | 1 | ENSMUST00 | -39917 | Gm26066 | ENSMUSG0C | false | . | . | . | . | . | . | chr17 | 84578925 | 84586206 | 5.26213 | 4.07321 | 3.25857 |
| chr17 | 17828556 | 17829086 | 547 | Promoter (1-2kb) | 17 | 17830188 | 17830257 | 70 | 1 | ENSMUST00 | -1102 | Mir99b | ENSMUSG0C | false | chr17 | 17825989 | 17829531 | 29.75681 | 92.45776 | 89.05831 | chr17 | 17828149 | 17841746 | 4.51187 | 3.80845 | 2.61198 |
| chr1 | 91426169 | 91426438 | 547 | Intron (ENSMUST00000069620.9 | 1 | 91412348 | 91414038 | 1691 | 2 | ENSMUST00 | -12132 | Hes6 | ENSMUSG0C | false | . | . | . | . | . | . | chr1 | 91425363 | 91426542 | 4.54437 | 3.88903 | 2.68573 |
| chr3 | 116331002 | 116331265 | 547 | Intron (ENSMUST00000106491.2 | 3 | 116249654 | 116253484 | 3831 | 2 | ENSMUST00 | -77519 | Gpr88 | ENSMUSG0C | false | . | . | . | . | . | . | chr3 | 116319808 | 116393211 | 5.81811 | 4.44924 | 3.68159 |
| chr10 | 120208628 | 120208883 | 546 | Promoter (<=1kb) | 10 | 120208750 | 120224917 | 16168 | 1 | ENSMUST00 | 0 | Tmbim4 | ENSMUSG0C | false | chr10 | 120208618 | 120209998 | 35.34781 | 110.24136 | 106.39787 | . | . | . | . | . | . |
| chr16 | 38902189 | 38902609 | 546 | Promoter (<=1kb) | 16 | 38902316 | 39027159 | 124844 | 1 | ENSMUST00 | 0 | Igsf11 | ENSMUSG0C | false | chr16 | 38902161 | 38902911 | 26.91438 | 86.45937 | 83.18881 | chr16 | 38901918 | 38902661 | 4.32537 | 3.79678 | 2.43619 |
| chr15 | 59167952 | 59168339 | 546 | Distal Intergenic | 15 | 59210908 | 59211720 | 813 | 1 | ENSMUST00 | -42569 | Gm5045 | ENSMUSG0C | false | . | . | . | . | . | . | chr15 | 59167250 | 59173334 | 6.774 | 4.95682 | 4.44708 |
| chr2 | 84727288 | 84727626 | 546 | Promoter (<=1kb) | 2 | 84723122 | 84727350 | 4229 | 2 | ENSMUST00 | 0 | Clp1 | ENSMUSG0C | false | chr2 | 84726571 | 84727937 | 22.94371 | 66.5854 | 63.64468 | . | . | . | . | . | . |
| chr5 | 53589151 | 53589731 | 546 | Promoter (<=1kb) | 5 | 53590215 | 53657445 | 67231 | 1 | ENSMUST00 | -484 | Rbpj | ENSMUSG0C | false | chr5 | 53588566 | 53591937 | 30.81129 | 113.47932 | 109.54176 | . | . | . | . | . | . |
| chr3 | 52267751 | 52268535 | 546 | Promoter (<=1kb) | 3 | 52268337 | 52350109 | 81773 | 1 | ENSMUST00 | 0 | Foxo1 | ENSMUSG0C | false | chr3 | 52267261 | 52270933 | 33.7938 | 119.00099 | 114.88904 | . | . | . | . | . | . |
| chr10 | 13312632 | 13312914 | 546 | Intron (ENSMUST00000105545.5 | 10 | 13207717 | 13324160 | 116444 | 2 | ENSMUST00 | 11246 | Phactr2 | ENSMUSG0C | false | . | . | . | . | . | . | chr10 | 13309818 | 13323388 | 4.90612 | 4.0384 | 2.9333 |
| chr4 | 149073938 | 149074332 | 546 | Intron (ENSMUST00000103217.5 | 4 | 149096383 | 149096702 | 320 | 1 | ENSMUST00 | -22051 | Gm9506 | ENSMUSG0C | false | . | . | . | . | . | . | chr4 | 149067858 | 149076880 | 5.20077 | 4.15256 | 3.17954 |
| chr13 | 102759865 | 102760191 | 546 | Promoter (<=1kb) | 13 | 102760255 | 102761896 | 1642 | 1 | ENSMUST00 | -64 | 1700099I09R | ENSMUSG0C | false | . | . | . | . | . | . | chr13 | 102747137 | 102771216 | 6.28304 | 4.64045 | 4.06151 |
| chr14 | 105767138 | 105767358 | 546 | Distal Intergenic | 14 | 105816908 | 105817000 | 93 | 1 | ENSMUST00 | -49550 | Gm22406 | ENSMUSG0C | false | . | . | . | . | . | . | . | . | . | . | . | . |
| chr18 | 9866205 | 9866533 | 546 | Intron (ENSMUST00000040069.8 | 18 | 9827026 | 9827119 | 94 | 2 | ENSMUST00 | -39087 | Gm23637 | ENSMUSG0C | false | . | . | . | . | . | . | chr18 | 9864894 | 9866756 | 4.19849 | 3.5565 | 2.39726 |
| chr12 | 85227263 | 85227570 | 545 | Promoter (2-3kb) | 12 | 85224526 | 85226493 | 1968 | 1 | ENSMUST00 | 2738 | Eif2b2 | ENSMUSG0C | false | . | . | . | . | . | . | chr12 | 85225589 | 85231278 | 5.34393 | 4.05861 | 3.32517 |
| chr17 | 27135403 | 27135662 | 545 | Promoter (1-2kb) | 17 | 27122663 | 27133916 | 11254 | 2 | ENSMUST00 | -1488 | Uqcc2 | ENSMUSG0C | false | . | . | . | . | . | . | chr17 | 27127363 | 27147146 | 5.99202 | 4.48057 | 3.83593 |
| chr1 | 127773877 | 127774532 | 545 | Promoter (<=1kb) | 1 | 127774164 | 127793589 | 19426 | 1 | ENSMUST00 | 0 | Ccnt2 | ENSMUSG0C | false | chr1 | 127773212 | 127775687 | 37.89522 | 119.88892 | 115.74806 | . | . | . | . | . | . |
| chr19 | 25266229 | 25266543 | 545 | Intron (ENSMUST00000049400.9 | 19 | 25236975 | 25434496 | 197522 | 1 | ENSMUST00 | 29255 | Kank1 | ENSMUSG0C | true | . | . | . | . | . | . | chr19 | 25261572 | 25293097 | 5.29409 | 4.21889 | 3.24888 |
| chr5 | 88675202 | 88676391 | 545 | Promoter (<=1kb) | 5 | 88659448 | 88676171 | 16724 | 2 | ENSMUST00 | 0 | Grsf1 | ENSMUSG0C | false | chr5 | 88674798 | 88676601 | 15.57114 | 68.18018 | 65.21632 | . | . | . | . | . | . |
| chr9 | 44733800 | 44734724 | 545 | Promoter (<=1kb) | 9 | 44686308 | 44735198 | 48891 | 2 | ENSMUST00 | 474 | Phldb1 | ENSMUSG0C | false | chr9 | 44733443 | 44735510 | 29.66345 | 100.21 | 96.63225 | chr9 | 44732060 | 44735669 | 3.74755 | 3.35522 | 1.98328 |
| chr7 | 99953523 | 99953887 | 545 | Intron (ENSMUST00000080817.4 | 7 | 99935278 | 99965734 | 30457 | 2 | ENSMUST00 | 11847 | Rnf169 | ENSMUSG0C | false | . | . | . | . | . | . | chr7 | 99953219 | 99955455 | 5.47856 | 4.36448 | 3.38371 |
| chr18 | 62921941 | 62922369 | 545 | Promoter (<=1kb) | 18 | 62922327 | 62953195 | 30869 | 1 | ENSMUST00 | 0 | Apcdd1 | ENSMUSG0C | false | chr18 | 62921844 | 62923342 | 23.67723 | 66.10762 | 63.1768 | chr18 | 62921519 | 62921971 | 3.48459 | 3.39063 | 1.8128 |
| chr2 | 131452286 | 131452521 | 545 | Promoter (2-3kb) | 2 | 131454128 | 131454640 | 513 | 2 | ENSMUST00 | 2119 | Gm14304 | ENSMUSG0C | false | . | . | . | . | . | . | . | . | . | . | . | . |
| chr3 | 9358090 | 9358518 | 545 | Distal Intergenic | 3 | 9402175 | 9402485 | 311 | 1 | ENSMUST00 | -43657 | Gm22604 | ENSMUSG0C | false | . | . | . | . | . | . | chr3 | 9357880 | 9359121 | 4.48889 | 3.82702 | 2.59138 |
| chr4 | 54425453 | 54425704 | 545 | Distal Intergenic | 4 | 54468352 | 54468592 | 241 | 1 | ENSMUST00 | -42648 | Gm25687 | ENSMUSG0C | false | . | . | . | . | . | . | chr4 | 54422468 | 54426961 | 4.10532 | 3.64374 | 2.28174 |
| chr11 | 101467979 | 101468417 | 544 | Promoter (<=1kb) | 11 | 101468175 | 101471853 | 3679 | 1 | ENSMUST00 | 0 | Rnd2 | ENSMUSG0C | false | chr11 | 101467402 | 101468763 | 25.16654 | 87.04672 | 83.76057 | chr11 | 101463714 | 101469649 | 4.51869 | 3.86017 | 2.61475 |
| chr17 | 69969096 | 69969500 | 544 | Promoter (<=1kb) | 17 | 69969246 | 70819724 | 850479 | 1 | ENSMUST00 | 0 | Dlgap1 | ENSMUSG0C | false | chr17 | 69968953 | 69970217 | 15.39495 | 41.17039 | 38.56658 | chr17 | 69968988 | 69970666 | 5.61683 | 4.37597 | 3.53827 |
| chr12 | 110889134 | 110889658 | 544 | Promoter (<=1kb) | 12 | 110889264 | 110972392 | 83129 | 1 | ENSMUST00 | 0 | Tecpr2 | ENSMUSG0C | false | chr12 | 110888374 | 110890178 | 32.66734 | 113.81544 | 109.86345 | chr12 | 110887996 | 110890571 | 5.72972 | 4.40002 | 3.63583 |
| chr15 | 27466295 | 27466867 | 544 | Promoter (<=1kb) | 15 | 27466677 | 27594909 | 128233 | 1 | ENSMUST00 | 0 | Ank | ENSMUSG0C | false | chr15 | 27465975 | 27467367 | 21.76566 | 71.50774 | 68.49327 | chr15 | 27465412 | 27467509 | 4.17134 | 3.71844 | 2.3432 |
| chr19 | 6276531 | 6276857 | 544 | Promoter (<=1kb) | 19 | 6276725 | 6300096 | 23372 | 1 | ENSMUST00 | 0 | Ehd1 | ENSMUSG0C | false | chr19 | 6276749 | 6279190 | 21.72697 | 98.39969 | 94.87038 | . | . | . | . | . | . |
| chr2 | 68978800 | 68979127 | 544 | Intron (ENSMUST00000028426.3 | 2 | 68959911 | 69051328 | 91418 | 1 | ENSMUST00 | 18890 | Cers6 | ENSMUSG0C | true | . | . | . | . | . | . | chr2 | 68972135 | 68985833 | 4.202 | 3.70826 | 2.34625 |
| chr4 | 148896177 | 148896485 | 544 | Intron (ENSMUST00000094464.4 | 4 | 148901149 | 148932896 | 31748 | 1 | ENSMUST00 | -4664 | Casz1 | ENSMUSG0C | true | . | . | . | . | . | . | chr4 | 148893746 | 148896937 | 5.74911 | 4.24525 | 3.65502 |
| chr7 | 120838814 | 120839064 | 544 | Distal Intergenic | 7 | 120842831 | 120907219 | 64389 | 1 | ENSMUST00 | -3767 | Eef2k | ENSMUSG0C | false | . | . | . | . | . | . | chr7 | 120837696 | 120840063 | 4.93631 | 4.05784 | 2.95967 |
| chr7 | 89041443 | 89041700 | 544 | Distal Intergenic | 7 | 89141856 | 89154300 | 12445 | 2 | ENSMUST00 | 112600 | Tmem135 | ENSMUSG0C | false | . | . | . | . | . | . | chr7 | 89040256 | 89041598 | 4.80589 | 3.88953 | 2.86898 |
| chr15 | 74887717 | 74887961 | 544 | Intron (ENSMUST00000190826.1 | 15 | 74878339 | 74881704 | 3366 | 2 | ENSMUST00 | -6014 | 2010109I03R | ENSMUSG0C | false | . | . | . | . | . | . | . | . | . | . | . | . |
| chr1 | 136128470 | 136128682 | 544 | Promoter (2-3kb) | 1 | 136127719 | 136131171 | 3453 | 2 | ENSMUST00 | 2489 | Gm15850 | ENSMUSG0C | false | . | . | . | . | . | . | . | . | . | . | . | . |
| chr11 | 55417775 | 55418446 | 543 | Promoter (1-2kb) | 11 | 55394500 | 55419898 | 25399 | 2 | ENSMUST00 | 1452 | Sparc | ENSMUSG0C | true | chr11 | 55417325 | 55420160 | 29.3377 | 90.74815 | 87.38457 | . | . | . | . | . | . |
| chr18 | 61087011 | 61087261 | 543 | Distal Intergenic | 18 | 61105572 | 61131134 | 25563 | 1 | ENSMUST00 | -18311 | Csf1r | ENSMUSG0C | false | . | . | . | . | . | . | chr18 | 61086526 | 61088056 | 4.1601 | 3.54945 | 2.36405 |
| chr1 | 164457887 | 164458266 | 543 | Promoter (<=1kb) | 1 | 164437267 | 164458355 | 21089 | 2 | ENSMUST00 | 89 | Atp1b1 | ENSMUSG0C | false | chr1 | 164457085 | 164459055 | 27.73468 | 81.26827 | 78.09463 | chr1 | 164454831 | 164459230 | 4.35089 | 3.68063 | 2.48358 |
| chr7 | 117462674 | 117463016 | 543 | Intron (ENSMUST00000160035.2 | 7 | 117475492 | 117549359 | 73868 | 1 | ENSMUST00 | -12476 | Xylt1 | ENSMUSG0C | false | . | . | . | . | . | . | . | . | . | . | . | . |
| chr16 | 45963158 | 45963456 | 543 | 3' UTR | 16 | 45757142 | 45953598 | 196457 | 2 | ENSMUST00 | -9561 | Phldb2 | ENSMUSG0C | false | . | . | . | . | . | . | chr16 | 45961905 | 45964051 | 6.50635 | 4.86477 | 4.24633 |
| chr9 | 57563015 | 57563312 | 543 | Promoter (2-3kb) | 9 | 57560944 | 57588797 | 27854 | 1 | ENSMUST00 | 2072 | Scamp2 | ENSMUSG0C | false | . | . | . | . | . | . | chr9 | 57560307 | 57566621 | 7.40954 | 5.12519 | 4.97901 |
| chr5 | 41844074 | 41844492 | 543 | Promoter (<=1kb) | 5 | 41787542 | 41844168 | 56627 | 2 | ENSMUST00 | 0 | Bod1l | ENSMUSG0C | false | chr5 | 41842950 | 41845617 | 34.47702 | 107.70252 | 103.93674 | . | . | . | . | . | . |
| chr8 | 121665356 | 121665732 | 543 | Promoter (<=1kb) | 8 | 121666628 | 121666706 | 79 | 2 | ENSMUST00 | 974 | Gm23299 | ENSMUSG0C | false | chr8 | 121665290 | 121665960 | 7.82203 | 16.59716 | 14.30959 | chr8 | 121665075 | 121666018 | 4.4137 | 3.76566 | 2.53342 |
| chrX | 75143206 | 75143503 | 543 | Promoter (<=1kb) | 20 | 75143424 | 75143630 | 207 | 1 | ENSMUST00 | 0 | Gm15378 | ENSMUSG0C | false | chrX | 75143099 | 75143611 | 10.47775 | 22.22469 | 19.86031 | . | . | . | . | . | . |
| chrX | 10719047 | 10719426 | 543 | Promoter (<=1kb) | 20 | 10717473 | 10718710 | 1238 | 2 | ENSMUST00 | -338 | Gm14493 | ENSMUSG0C | false | chrX | 10717005 | 10719476 | 18.84082 | 47.13427 | 44.45482 | . | . | . | . | . | . |
| chr14 | 69474773 | 69475062 | 543 | Intron (ENSMUST00000111062.3 | 14 | 69502856 | 69523605 | 20750 | 1 | ENSMUST00 | -27794 | Gm27175 | ENSMUSG0C | false | . | . | . | . | . | . | . | . | . | . | . | . |
| chr11 | 116110100 | 116110534 | 542 | Promoter (<=1kb) | 11 | 116105752 | 116110237 | 4486 | 2 | ENSMUST00 | 0 | Trim47 | ENSMUSG0C | false | chr11 | 116108781 | 116110664 | 25.48472 | 95.2657 | 91.8051 | chr11 | 116107506 | 116111126 | 4.46337 | 3.8802 | 2.6034 |
| chr14 | 70519835 | 70520513 | 542 | Promoter (<=1kb) | 14 | 70474557 | 70520260 | 45704 | 2 | ENSMUST00 | 0 | Bmp1 | ENSMUSG0C | false | chr14 | 70519471 | 70520854 | 36.04346 | 118.78738 | 114.68494 | chr14 | 70519737 | 70521018 | 3.40014 | 3.32919 | 1.66815 |
| chr16 | 16599587 | 16600203 | 542 | Promoter (<=1kb) | 16 | 16422563 | 16599978 | 177416 | 2 | ENSMUST00 | 0 | Fgd4 | ENSMUSG0C | false | chr16 | 16599022 | 16600812 | 33.0841 | 102.23056 | 98.60399 | chr16 | 16598374 | 16600782 | 3.22774 | 3.15985 | 1.54079 |
| chr11 | 119602722 | 119603087 | 542 | Promoter (<=1kb) | 11 | 119602905 | 119823176 | 220272 | 1 | ENSMUST00 | 0 | Rptor | ENSMUSG0C | false | chr11 | 119602379 | 119604058 | 35.06866 | 119.18253 | 115.06551 | chr11 | 119602246 | 119604702 | 3.9443 | 3.48203 | 2.15679 |
| chr7 | 81454344 | 81454728 | 542 | Promoter (<=1kb) | 7 | 81347026 | 81454758 | 107733 | 2 | ENSMUST00 | 30 | Cpeb1 | ENSMUSG0C | true | chr7 | 81453319 | 81455625 | 30.23563 | 94.27794 | 90.8388 | chr7 | 81449678 | 81454553 | 5.10568 | 4.13331 | 3.08589 |
| chr8 | 13338554 | 13338803 | 542 | Promoter (<=1kb) | 8 | 13339674 | 13377702 | 38029 | 1 | ENSMUST00 | -871 | Tfdp1 | ENSMUSG0C | false | chr8 | 13338544 | 13339623 | 25.55642 | 72.00093 | 68.98035 | chr8 | 13338151 | 13340888 | 4.48694 | 3.78332 | 2.59785 |
| chr11 | 59956791 | 59957151 | 542 | Intron (ENSMUST00000081980.6 | 11 | 59963181 | 59964944 | 1764 | 2 | ENSMUST00 | 7793 | Rasd1 | ENSMUSG0C | false | . | . | . | . | . | . | chr11 | 59956466 | 59958726 | 6.03628 | 4.33223 | 3.89458 |
| chr10 | 24493721 | 24494201 | 542 | Distal Intergenic | 10 | 24514279 | 24514511 | 233 | 2 | ENSMUST00 | 20310 | Gm15272 | ENSMUSG0C | false | . | . | . | . | . | . | chr10 | 24493903 | 24494255 | 3.63838 | 3.45631 | 1.86687 |
| chr12 | 108259745 | 108260272 | 542 | Intron (ENSMUST00000136175.1 | 12 | 108206345 | 108275417 | 69073 | 2 | ENSMUST00 | 15145 | Ccdc85c | ENSMUSG0C | false | . | . | . | . | . | . | chr12 | 108233398 | 108274893 | 6.76084 | 4.78759 | 4.47033 |
| chr11 | 80577719 | 80577965 | 542 | Intron (ENSMUST00000041065.8 | 11 | 80507410 | 80509106 | 1697 | 2 | ENSMUST00 | -68614 | C030013C21I | ENSMUSG0C | false | . | . | . | . | . | . | chr11 | 80576601 | 80579924 | 4.1317 | 3.4848 | 2.30741 |
| chr16 | 49533560 | 49533801 | 542 | Distal Intergenic | 16 | 49424806 | 49425814 | 1009 | 1 | ENSMUST00 | 108755 | Gm6931 | ENSMUSG0C | false | . | . | . | . | . | . | chr16 | 49531864 | 49534399 | 3.36139 | 3.19136 | 1.69682 |
| chr6 | 124712020 | 124712415 | 541 | Promoter (<=1kb) | 6 | 124712336 | 124713068 | 733 | 1 | ENSMUST00 | 0 | Phb2 | ENSMUSG0C | false | chr6 | 124711365 | 124713309 | 36.6511 | 131.68239 | 127.12846 | chr6 | 124708753 | 124714632 | 5.3578 | 4.06637 | 3.31595 |
| chr7 | 139228826 | 139229082 | 541 | 3' UTR | 7 | 139233943 | 139242971 | 9029 | 1 | ENSMUST00 | -4861 | Lrrc27 | ENSMUSG0C | false | . | . | . | . | . | . | chr7 | 139228545 | 139229294 | 5.77812 | 4.32525 | 3.6786 |
| chr12 | 16894168 | 16895671 | 541 | Promoter (<=1kb) | 12 | 16894978 | 16988274 | 93297 | 1 | ENSMUST00 | 0 | Rock2 | ENSMUSG0C | true | chr12 | 16894002 | 16896666 | 32.81923 | 101.77142 | 98.15781 | chr12 | 16893541 | 16894413 | 4.87841 | 4.07896 | 2.89161 |
| chr13 | 58479263 | 58479633 | 541 | Distal Intergenic | 13 | 58391142 | 58403343 | 12202 | 2 | ENSMUST00 | -75921 | Hnrnpk | ENSMUSG0C | false | . | . | . | . | . | . | chr13 | 58474930 | 58480701 | 4.39829 | 3.75615 | 2.53835 |
| chr16 | 20731765 | 20731973 | 541 | Promoter (1-2kb) | 16 | 20725158 | 20730627 | 5470 | 2 | ENSMUST00 | -1139 | Thpo | ENSMUSG0C | false | . | . | . | . | . | . | chr16 | 20731646 | 20734488 | 4.34418 | 3.84634 | 2.46017 |
| chr2 | 157204275 | 157204732 | 541 | Promoter (<=1kb) | 2 | 157145893 | 157204512 | 58620 | 2 | ENSMUST00 | 0 | Rbl1 | ENSMUSG0C | false | chr2 | 157203730 | 157205350 | 33.11161 | 106.34985 | 102.61888 | . | . | . | . | . | . |
| chr9 | 114390492 | 114390853 | 541 | Promoter (<=1kb) | 9 | 114375134 | 114390675 | 15542 | 2 | ENSMUST00 | 0 | Crtap | ENSMUSG0C | false | chr9 | 114389854 | 114391090 | 27.78601 | 91.88203 | 88.49705 | . | . | . | . | . | . |
| chr2 | 91950236 | 91950669 | 541 | Promoter (<=1kb) | 2 | 91932827 | 91950427 | 17601 | 2 | ENSMUST00 | 0 | Dgkz | ENSMUSG0C | false | chr2 | 91950326 | 91950586 | 4.34978 | 5.93396 | 3.85727 | chr2 | 91946924 | 91955908 | 6.11949 | 4.52776 | 3.92628 |
| chr8 | 122282459 | 122283079 | 541 | Promoter (<=1kb) | 8 | 122282141 | 122337251 | 55111 | 1 | ENSMUST00 | 319 | Zfpm1 | ENSMUSG0C | false | chr8 | 122282038 | 122285122 | 23.88927 | 69.11851 | 66.14216 | chr8 | 122282666 | 122285958 | 5.99671 | 4.57763 | 3.84085 |
| chr2 | 160299799 | 160300066 | 541 | Distal Intergenic | 2 | 160311393 | 160313616 | 2224 | 2 | ENSMUST00 | 13550 | Gm826 | ENSMUSG0C | false | . | . | . | . | . | . | chr2 | 160298011 | 160300752 | 4.71186 | 3.84462 | 2.81114 |
| chr12 | 79725913 | 79726326 | 541 | Distal Intergenic | 12 | 79963568 | 79963872 | 305 | 1 | ENSMUST00 | -237242 | RP23-179D3. | ENSMUSG0C | false | . | . | . | . | . | . | chr12 | 79725372 | 79726751 | 3.38713 | 3.21533 | 1.73816 |
| chr8 | 123150716 | 123150971 | 540 | Intron (ENSMUST00000127664.1 | 8 | 123148120 | 123158228 | 10109 | 2 | ENSMUST00 | 7257 | Sult5a1 | ENSMUSG0C | false | . | . | . | . | . | . | chr8 | 123150656 | 123151169 | 3.87133 | 3.60505 | 2.11453 |
| chr2 | 167924893 | 167925494 | 540 | Distal Intergenic | 2 | 167932057 | 167974737 | 42681 | 1 | ENSMUST00 | -6563 | Ptpn1 | ENSMUSG0C | false | chr2 | 167924840 | 167925731 | 9.88648 | 20.14232 | 17.80457 | chr2 | 167924314 | 167926298 | 3.62421 | 3.34029 | 1.89666 |
| chr9 | 102647079 | 102647414 | 540 | Distal Intergenic | 9 | 102628512 | 102629727 | 1216 | 1 | ENSMUST00 | 18568 | Anapc13 | ENSMUSG0C | false | . | . | . | . | . | . | chr9 | 102645417 | 102647945 | 3.65224 | 3.29554 | 1.92892 |
| chr19 | 43675144 | 43675615 | 540 | Promoter (<=1kb) | 19 | 43663822 | 43674881 | 11060 | 2 | ENSMUST00 | -264 | Slc25a28 | ENSMUSG0C | false | chr19 | 43675078 | 43676446 | 28.77697 | 87.83199 | 84.53201 | . | . | . | . | . | . |
| chr17 | 9768394 | 9768607 | 540 | Distal Intergenic | 17 | 9666497 | 9669704 | 3208 | 2 | ENSMUST00 | -98691 | Pabpc6 | ENSMUSG0C | false | . | . | . | . | . | . | . | . | . | . | . | . |
| chr4 | 129406446 | 129406735 | 540 | Distal Intergenic | 4 | 129426269 | 129432529 | 6261 | 2 | ENSMUST00 | 25794 | Zbtb8b | ENSMUSG0C | false | chr4 | 129406148 | 129406871 | 13.74281 | 35.40012 | 32.86803 | chr4 | 129402905 | 129407219 | 3.89135 | 3.50257 | 2.09309 |
| chrX | 51017893 | 51018345 | 540 | Promoter (<=1kb) | 20 | 51003912 | 51018018 | 14107 | 2 | ENSMUST00 | 0 | Rap2c | ENSMUSG0C | false | chrX | 51017427 | 51019512 | 16.08362 | 38.04472 | 35.47989 | . | . | . | . | . | . |
| chr17 | 27392864 | 27393111 | 540 | Distal Intergenic | 17 | 27342460 | 27344825 | 2366 | 1 | ENSMUST00 | 50405 | Gm10505 | ENSMUSG0C | false | . | . | . | . | . | . | chr17 | 27391690 | 27393986 | 4.83968 | 3.91323 | 2.90795 |
| chr2 | 72374113 | 72374378 | 540 | Intron (ENSMUST00000090824.6 | 2 | 72360247 | 72360542 | 296 | 1 | ENSMUST00 | 13867 | Rpl36a-ps4 | ENSMUSG0C | false | . | . | . | . | . | . | chr2 | 72372772 | 72376073 | 4.43193 | 3.62566 | 2.57824 |
| chr10 | 127756141 | 127756506 | 540 | Distal Intergenic | 10 | 127759721 | 127768297 | 8577 | 1 | ENSMUST00 | -3215 | Rdh1 | ENSMUSG0C | false | chr10 | 127756084 | 127756289 | 4.96301 | 7.42671 | 5.30838 | chr10 | 127743127 | 127759693 | 6.17884 | 4.54629 | 3.98922 |
| chr3 | 27637172 | 27637560 | 540 | Intron (ENSMUST00000046157.4 | 3 | 27584900 | 27584995 | 96 | 2 | ENSMUST00 | -52178 | Mir3092 | ENSMUSG0C | false | . | . | . | . | . | . | chr3 | 27630973 | 27640357 | 5.71666 | 4.13593 | 3.62688 |
| chr10 | 127338977 | 127339653 | 539 | Promoter (1-2kb) | 10 | 127329889 | 127341589 | 11701 | 2 | ENSMUST00 | 1936 | Gli1 | ENSMUSG0C | false | chr10 | 127339045 | 127340137 | 11.31597 | 24.78923 | 22.38983 | chr10 | 127338758 | 127342756 | 3.59246 | 3.3131 | 1.87872 |
| chr1 | 74145194 | 74145529 | 539 | Intron (ENSMUST00000080167.5 | 1 | 74153989 | 74160610 | 6622 | 1 | ENSMUST00 | -8460 | Cxcr2 | ENSMUSG0C | false | . | . | . | . | . | . | chr1 | 74144587 | 74147044 | 3.20991 | 3.15901 | 1.59841 |
| chr3 | 8952331 | 8952605 | 539 | Intron (ENSMUST00000091355.6 | 3 | 8931064 | 8962253 | 31190 | 2 | ENSMUST00 | 9648 | Tpd52 | ENSMUSG0C | false | . | . | . | . | . | . | chr3 | 8947714 | 8965965 | 5.38967 | 4.2084 | 3.32941 |
| chr4 | 129223345 | 129223730 | 539 | Exon (ENSMUST00000106051.2/ | 4 | 129225383 | 129227480 | 2098 | 2 | ENSMUST00 | 3750 | C77080 | ENSMUSG0C | false | chr4 | 129223258 | 129223802 | 8.23183 | 15.40765 | 13.13914 | chr4 | 129223195 | 129224042 | 6.7616 | 4.77457 | 4.49006 |
| chr18 | 60729289 | 60729886 | 539 | Distal Intergenic | 18 | 60685978 | 60713389 | 27412 | 2 | ENSMUST00 | -15901 | Ndst1 | ENSMUSG0C | false | . | . | . | . | . | . | chr18 | 60726897 | 60741024 | 7.39668 | 5.03429 | 4.98055 |
| chr7 | 84361142 | 84361437 | 539 | Intron (ENSMUST00000085077.3 | 7 | 84323309 | 84323415 | 107 | 2 | ENSMUST00 | -37728 | Gm25790 | ENSMUSG0C | false | . | . | . | . | . | . | chr7 | 84355508 | 84365417 | 5.93716 | 4.48435 | 3.77591 |
| chr3 | 52576967 | 52577276 | 539 | Distal Intergenic | 3 | 52612835 | 52613836 | 1002 | 1 | ENSMUST00 | -35559 | Gm10293 | ENSMUSG0C | false | . | . | . | . | . | . | chr3 | 52573527 | 52578380 | 3.96078 | 3.42783 | 2.18966 |
| chr2 | 74850044 | 74850315 | 539 | Intron (ENSMUST00000028511.7 | 2 | 74855706 | 74856311 | 606 | 1 | ENSMUST00 | -5391 | Gm14425 | ENSMUSG0C | false | . | . | . | . | . | . | chr2 | 74849105 | 74852751 | 3.63029 | 3.32976 | 1.88637 |
| chr5 | 119400319 | 119400824 | 539 | Distal Intergenic | 5 | 119297245 | 119297361 | 117 | 1 | ENSMUST00 | 103075 | n-R5s175 | ENSMUSG0C | false | . | . | . | . | . | . | . | . | . | . | . | . |
| chr3 | 153690592 | 153690924 | 539 | Intron (ENSMUST00000064460.5 | 3 | 153708911 | 153709026 | 116 | 1 | ENSMUST00 | -17987 | Gm22206 | ENSMUSG0C | false | . | . | . | . | . | . | chr3 | 153686316 | 153712771 | 6.43925 | 4.66349 | 4.19773 |
| chr18 | 61341359 | 61341674 | 539 | Intron (ENSMUST00000075299.7 | 18 | 61397813 | 61397925 | 113 | 1 | ENSMUST00 | -56139 | Gm25301 | ENSMUSG0C | false | . | . | . | . | . | . | chr18 | 61333315 | 61342581 | 5.33079 | 4.15853 | 3.28921 |
| chr6 | 115349433 | 115349700 | 538 | Distal Intergenic | 6 | 115361221 | 115490401 | 129181 | 1 | ENSMUST00 | -11521 | Pparg | ENSMUSG0C | false | . | . | . | . | . | . | chr6 | 115349607 | 115350347 | 2.88283 | 2.79366 | 1.34591 |
| chr7 | 38084843 | 38085344 | 538 | Distal Intergenic | 7 | 38098811 | 38099369 | 559 | 2 | ENSMUST00 | 14025 | Ccne1 | ENSMUSG0C | false | chr7 | 38085045 | 38086196 | 11.15939 | 31.88065 | 29.39184 | chr7 | 38084511 | 38088451 | 4.75907 | 3.75227 | 2.82395 |
| chr15 | 59107457 | 59107729 | 538 | Distal Intergenic | 15 | 58941244 | 59082026 | 140783 | 2 | ENSMUST00 | -25432 | Mtss1 | ENSMUSG0C | true | . | . | . | . | . | . | chr15 | 59105620 | 59110005 | 5.79609 | 4.39139 | 3.65762 |
| chr2 | 83714730 | 83715043 | 538 | Distal Intergenic | 2 | 83724397 | 83756797 | 32401 | 1 | ENSMUST00 | -9354 | Itgav | ENSMUSG0C | false | . | . | . | . | . | . | chr2 | 83713941 | 83715701 | 6.01028 | 4.57137 | 3.82539 |
| chr3 | 100896913 | 100897184 | 538 | Distal Intergenic | 3 | 100922202 | 100931702 | 9501 | 1 | ENSMUST00 | -25018 | Trim45 | ENSMUSG0C | false | . | . | . | . | . | . | chr3 | 100895046 | 100897532 | 3.70003 | 3.36162 | 1.97218 |
| chr2 | 122368617 | 122369327 | 538 | Promoter (<=1kb) | 2 | 122348893 | 122368918 | 20026 | 2 | ENSMUST00 | 0 | Shf | ENSMUSG0C | false | chr2 | 122368187 | 122369883 | 42.75948 | 139.16466 | 134.31596 | chr2 | 122368332 | 122369814 | 4.09324 | 3.61803 | 2.30233 |
| chr9 | 37145827 | 37146131 | 538 | Promoter (<=1kb) | 9 | 36954772 | 37145660 | 190889 | 2 | ENSMUST00 | -168 | Pknox2 | ENSMUSG0C | true | chr9 | 37144737 | 37147348 | 34.92444 | 106.42502 | 102.68874 | . | . | . | . | . | . |
| chr3 | 100826334 | 100826731 | 538 | Promoter (<=1kb) | 3 | 100825459 | 100895679 | 70221 | 1 | ENSMUST00 | 876 | Vtcn1 | ENSMUSG0C | false | chr3 | 100825118 | 100827041 | 15.39495 | 41.17039 | 38.56658 | chr3 | 100826585 | 100832165 | 4.06509 | 3.56252 | 2.26021 |
| chr12 | 73053825 | 73054108 | 538 | Promoter (<=1kb) | 12 | 73043531 | 73053887 | 10357 | 2 | ENSMUST00 | 0 | Six1 | ENSMUSG0C | false | chr12 | 73053350 | 73054533 | 22.15371 | 67.81625 | 64.85734 | chr12 | 73052130 | 73054544 | 3.84894 | 3.51437 | 2.05891 |
| chr18 | 54987656 | 54988169 | 538 | Promoter (2-3kb) | 18 | 54888045 | 54990180 | 102136 | 2 | ENSMUST00 | 2011 | Zfp608 | ENSMUSG0C | false | chr18 | 54985294 | 54988807 | 28.53699 | 95.19626 | 91.7378 | . | . | . | . | . | . |
| chr9 | 104100884 | 104101186 | 538 | Promoter (1-2kb) | 9 | 104084157 | 104102560 | 18404 | 2 | ENSMUST00 | 1374 | Ackr4 | ENSMUSG0C | false | chr9 | 104100927 | 104101613 | 6.38327 | 11.47542 | 9.27082 | chr9 | 104100336 | 104102534 | 4.12157 | 3.52496 | 2.34956 |
| chr11 | 32243627 | 32243937 | 537 | Intron (ENSMUST00000124640.2 | 11 | 32236847 | 32239941 | 3095 | 2 | ENSMUST00 | -3687 | Nprl3 | ENSMUSG0C | false | . | . | . | . | . | . | chr11 | 32242980 | 32244630 | 5.24661 | 4.25725 | 3.19541 |
| chr18 | 52767893 | 52768508 | 537 | Promoter (<=1kb) | 18 | 52767994 | 52915935 | 147942 | 1 | ENSMUST00 | 0 | Sncaip | ENSMUSG0C | true | chr18 | 52767566 | 52770026 | 41.31276 | 141.09892 | 136.17311 | . | . | . | . | . | . |
| chr6 | 36810024 | 36810346 | 537 | Promoter (1-2kb) | 6 | 36715663 | 36811361 | 95699 | 2 | ENSMUST00 | 1015 | Ptn | ENSMUSG0C | false | chr6 | 36809394 | 36810394 | 15.50707 | 38.47921 | 35.908 | chr6 | 36809095 | 36810718 | 3.54322 | 3.28213 | 1.86658 |
| chr13 | 59836033 | 59836446 | 537 | Distal Intergenic | 13 | 59771879 | 59823147 | 51269 | 2 | ENSMUST00 | -12887 | Zcchc6 | ENSMUSG0C | false | . | . | . | . | . | . | chr13 | 59832928 | 59838791 | 4.70155 | 3.94381 | 2.76868 |
| chr4 | 59760297 | 59760666 | 537 | 3' UTR | 4 | 59771297 | 59783808 | 12512 | 2 | ENSMUST00 | 23142 | Inip | ENSMUSG0C | false | . | . | . | . | . | . | chr4 | 59756611 | 59762237 | 4.44224 | 3.62486 | 2.5942 |
| chr18 | 77945790 | 77946050 | 537 | Intron (ENSMUST00000044622.5 | 18 | 77938467 | 78035027 | 96561 | 1 | ENSMUST00 | 7324 | Epg5 | ENSMUSG0C | false | . | . | . | . | . | . | chr18 | 77943925 | 77947807 | 5.01063 | 4.15386 | 3.00201 |
| chr14 | 34215761 | 34216061 | 537 | Distal Intergenic | 14 | 34219595 | 34222371 | 2777 | 1 | ENSMUST00 | -3534 | Syt15 | ENSMUSG0C | false | . | . | . | . | . | . | chr14 | 34214018 | 34216261 | 5.55002 | 4.4174 | 3.46175 |
| chr9 | 114933825 | 114934148 | 537 | Promoter (<=1kb) | 9 | 114899341 | 114933987 | 34647 | 2 | ENSMUST00 | 0 | Gpd1l | ENSMUSG0C | false | chr9 | 114932730 | 114934401 | 30.12659 | 110.14252 | 106.30244 | chr9 | 114931684 | 114934444 | 5.44281 | 4.17389 | 3.40749 |
| chr11 | 79711078 | 79711412 | 537 | Promoter (<=1kb) | 11 | 79711969 | 79712034 | 66 | 1 | ENSMUST00 | -557 | Mir193a | ENSMUSG0C | false | chr11 | 79710752 | 79712376 | 34.76187 | 122.87653 | 118.63768 | . | . | . | . | . | . |
| chr4 | 156084502 | 156084793 | 537 | Distal Intergenic | 4 | 156052715 | 156059802 | 7088 | 2 | ENSMUST00 | -24701 | Gm13648 | ENSMUSG0C | false | . | . | . | . | . | . | chr4 | 156082814 | 156084984 | 5.00531 | 4.02054 | 3.0365 |
| chr9 | 81470236 | 81470466 | 537 | Distal Intergenic | 9 | 81455397 | 81460966 | 5570 | 1 | ENSMUST00 | 14840 | Gm27226 | ENSMUSG0C | false | . | . | . | . | . | . | chr9 | 81469167 | 81471003 | 3.7635 | 3.38289 | 2.02306 |
| chr12 | 76308073 | 76308398 | 536 | Intron (ENSMUST00000021443.5 | 12 | 76324891 | 76334151 | 9261 | 1 | ENSMUST00 | -16493 | Akap5 | ENSMUSG0C | false | . | . | . | . | . | . | chr12 | 76306284 | 76308818 | 4.73953 | 3.74813 | 2.83321 |
| chr14 | 54772852 | 54773239 | 536 | Intron (ENSMUST00000022787.6 | 14 | 54722216 | 54781886 | 59671 | 2 | ENSMUST00 | 8647 | Slc7a8 | ENSMUSG0C | false | . | . | . | . | . | . | . | . | . | . | . | . |
| chr16 | 33829500 | 33830430 | 536 | Promoter (<=1kb) | 16 | 33829665 | 33949146 | 119482 | 1 | ENSMUST00 | 0 | Itgb5 | ENSMUSG0C | true | chr16 | 33828900 | 33834465 | 38.14117 | 119.25307 | 115.13219 | chr16 | 33827896 | 33829519 | 4.07991 | 3.66893 | 2.238 |
| chr1 | 151344225 | 151344877 | 536 | Promoter (<=1kb) | 1 | 151344477 | 151364422 | 19946 | 1 | ENSMUST00 | 0 | Ivns1abp | ENSMUSG0C | false | chr1 | 151343938 | 151345916 | 30.39373 | 108.60838 | 104.8139 | chr1 | 151342994 | 151347059 | 4.14164 | 3.66892 | 2.29971 |
| chr16 | 24367596 | 24367843 | 536 | Distal Intergenic | 16 | 24393350 | 24992576 | 599227 | 1 | ENSMUST00 | -25507 | Lpp | ENSMUSG0C | false | . | . | . | . | . | . | . | . | . | . | . | . |
| chr9 | 43129828 | 43130389 | 536 | Intron (ENSMUST00000034513.7 | 9 | 43108653 | 43116570 | 7918 | 2 | ENSMUST00 | -13259 | Tmem136 | ENSMUSG0C | false | . | . | . | . | . | . | chr9 | 43126598 | 43131834 | 5.02199 | 4.11654 | 3.02968 |
| chr5 | 31697714 | 31698590 | 536 | Promoter (<=1kb) | 5 | 31697970 | 31731448 | 33479 | 1 | ENSMUST00 | 0 | Bre | ENSMUSG0C | false | chr5 | 31697125 | 31698833 | 30.59503 | 95.89513 | 92.42037 | . | . | . | . | . | . |
| chr5 | 133173115 | 133173539 | 536 | Distal Intergenic | 5 | 133475123 | 133475536 | 414 | 1 | ENSMUST00 | -301584 | Gm10051 | ENSMUSG0C | false | . | . | . | . | . | . | chr5 | 133171740 | 133173988 | 4.51386 | 3.80453 | 2.59872 |
| chr12 | 52848454 | 52848712 | 536 | Intron (ENSMUST00000095737.3 | 12 | 52699383 | 53151015 | 451633 | 1 | ENSMUST00 | 149072 | Akap6 | ENSMUSG0C | false | . | . | . | . | . | . | chr12 | 52847879 | 52851010 | 4.61305 | 3.91091 | 2.72789 |
| chr6 | 117820686 | 117820964 | 536 | Distal Intergenic | 6 | 117808720 | 117809220 | 501 | 1 | ENSMUST00 | 11967 | Gm6253 | ENSMUSG0C | false | . | . | . | . | . | . | . | . | . | . | . | . |
| chr4 | 97476782 | 97477192 | 536 | Distal Intergenic | 4 | 97521428 | 97522332 | 905 | 2 | ENSMUST00 | 45140 | Gm12696 | ENSMUSG0C | false | . | . | . | . | . | . | chr4 | 97476055 | 97478070 | 4.51175 | 3.75048 | 2.62732 |
| chr14 | 62850885 | 62851407 | 536 | Intron (ENSMUST00000014691.8 | 14 | 62836908 | 62838207 | 1300 | 2 | ENSMUST00 | -12679 | Gm26536 | ENSMUSG0C | false | . | . | . | . | . | . | chr14 | 62849482 | 62853223 | 4.16667 | 3.48928 | 2.33023 |
| chr14 | 70599136 | 70600040 | 535 | Promoter (<=1kb) | 14 | 70583296 | 70599835 | 16540 | 2 | ENSMUST00 | 0 | Fam160b2 | ENSMUSG0C | false | chr14 | 70598881 | 70600254 | 35.84351 | 110.06287 | 106.22365 | chr14 | 70597977 | 70600490 | 3.77478 | 3.45182 | 1.98545 |
| chr18 | 75386279 | 75386641 | 535 | Intron (ENSMUST00000026999.4 | 18 | 75374914 | 75394001 | 19088 | 1 | ENSMUST00 | 11366 | Smad7 | ENSMUSG0C | false | . | . | . | . | . | . | chr18 | 75369277 | 75392875 | 5.74724 | 4.44386 | 3.62103 |
| chr4 | 35225409 | 35226279 | 535 | Promoter (<=1kb) | 4 | 35191285 | 35226123 | 34839 | 2 | ENSMUST00 | 0 | 3110043O21 | ENSMUSG0C | false | chr4 | 35225022 | 35226833 | 39.50969 | 132.40875 | 127.82707 | . | . | . | . | . | . |
| chr4 | 115052564 | 115052921 | 535 | Distal Intergenic | 4 | 115056426 | 115071755 | 15330 | 1 | ENSMUST00 | -3505 | Tal1 | ENSMUSG0C | false | chr4 | 115052419 | 115053279 | 12.24318 | 32.0257 | 29.53485 | chr4 | 115052160 | 115053322 | 5.32077 | 4.32508 | 3.2508 |
| chr8 | 109586913 | 109587284 | 535 | Distal Intergenic | 8 | 109593247 | 109594051 | 805 | 2 | ENSMUST00 | 6767 | Dhodh | ENSMUSG0C | false | . | . | . | . | . | . | chr8 | 109587117 | 109588099 | 4.31645 | 3.67274 | 2.47118 |
| chr9 | 76594370 | 76594689 | 535 | Distal Intergenic | 9 | 76490054 | 76567116 | 77063 | 2 | ENSMUST00 | -27255 | Fam83b | ENSMUSG0C | false | . | . | . | . | . | . | chr9 | 76592314 | 76595729 | 4.49725 | 3.88091 | 2.5875 |
| chr1 | 182560668 | 182560981 | 535 | Distal Intergenic | 1 | 182565007 | 182606526 | 41520 | 1 | ENSMUST00 | -4026 | Capn8 | ENSMUSG0C | false | . | . | . | . | . | . | chr1 | 182560228 | 182563842 | 4.79607 | 3.89862 | 2.86293 |
| chr14 | 27058601 | 27058862 | 535 | Intron (ENSMUST00000035336.3 | 14 | 27039001 | 27107286 | 68286 | 1 | ENSMUST00 | 19601 | Il17rd | ENSMUSG0C | false | . | . | . | . | . | . | chr14 | 27056393 | 27059807 | 5.10232 | 4.14065 | 3.09017 |
| chr2 | 73386353 | 73386920 | 535 | Promoter (<=1kb) | 2 | 73341506 | 73386572 | 45067 | 2 | ENSMUST00 | 0 | Gpr155 | ENSMUSG0C | false | chr2 | 73386735 | 73387181 | 26.13664 | 77.56537 | 74.45414 | . | . | . | . | . | . |
| chr15 | 74268516 | 74268832 | 535 | Distal Intergenic | 15 | 74093686 | 74094333 | 648 | 2 | ENSMUST00 | -174184 | Gm15387 | ENSMUSG0C | false | . | . | . | . | . | . | chr15 | 74268123 | 74269061 | 4.75062 | 3.97474 | 2.82684 |
| chr4 | 25887993 | 25888278 | 535 | Distal Intergenic | 4 | 25867989 | 25868455 | 467 | 2 | ENSMUST00 | -19539 | Gm11895 | ENSMUSG0C | false | . | . | . | . | . | . | chr4 | 25885708 | 25888346 | 3.90473 | 3.57402 | 2.09164 |
| chr1 | 91398515 | 91399038 | 534 | Promoter (<=1kb) | 1 | 91373861 | 91398700 | 24840 | 2 | ENSMUST00 | 0 | Ilkap | ENSMUSG0C | false | chr1 | 91397475 | 91399330 | 39.97575 | 133.22409 | 128.61581 | . | . | . | . | . | . |
| chr1 | 130715951 | 130716341 | 534 | Promoter (<=1kb) | 1 | 130689183 | 130715814 | 26632 | 2 | ENSMUST00 | -138 | Pfkfb2 | ENSMUSG0C | false | chr1 | 130715318 | 130718138 | 36.42221 | 131.26616 | 126.73016 | . | . | . | . | . | . |
| chr7 | 132930893 | 132931823 | 534 | Promoter (<=1kb) | 7 | 132931142 | 132986391 | 55250 | 1 | ENSMUST00 | 0 | Zranb1 | ENSMUSG0C | false | chr7 | 132930467 | 132932593 | 18.2926 | 68.86671 | 65.89487 | . | . | . | . | . | . |
| chr17 | 10609634 | 10609911 | 534 | Intron (ENSMUST00000041463.6 | 17 | 10576478 | 10631067 | 54590 | 2 | ENSMUST00 | 21156 | Pacrg | ENSMUSG0C | false | . | . | . | . | . | . | chr17 | 10608245 | 10610775 | 5.5053 | 4.2253 | 3.44497 |
| chr19 | 45998170 | 45998637 | 534 | Promoter (<=1kb) | 19 | 45818144 | 45998488 | 180345 | 2 | ENSMUST00 | 0 | 9130011E15I | ENSMUSG0C | false | chr19 | 45997563 | 45998973 | 36.54311 | 113.3445 | 109.41462 | chr19 | 45996044 | 45998913 | 3.98283 | 3.52924 | 2.17316 |
| chrX | 6172915 | 6173226 | 534 | Promoter (<=1kb) | 20 | 6168696 | 6173015 | 4320 | 2 | ENSMUST00 | 0 | Nudt10 | ENSMUSG0C | false | chrX | 6172775 | 6173605 | 10.89686 | 23.4988 | 21.11616 | . | . | . | . | . | . |
| chr13 | 109362446 | 109362718 | 534 | Intron (ENSMUST00000122041.2 | 13 | 109324744 | 109324824 | 81 | 1 | ENSMUST00 | 37703 | Mir582 | ENSMUSG0C | false | . | . | . | . | . | . | chr13 | 109362338 | 109363583 | 3.61463 | 3.33935 | 1.87733 |
| chr6 | 144734867 | 144735104 | 534 | Intron (ENSMUST00000155562.1 | 6 | 144723565 | 144724907 | 1343 | 1 | ENSMUST00 | 11303 | Sox5os5 | ENSMUSG0C | false | . | . | . | . | . | . | chr6 | 144732550 | 144735462 | 4.4224 | 3.73245 | 2.55984 |
| chr7 | 45367061 | 45367622 | 534 | Promoter (<=1kb) | 7 | 45367233 | 45367948 | 716 | 1 | ENSMUST00 | 0 | Mtag2 | ENSMUSG0C | false | chr7 | 45366411 | 45367908 | 23.47016 | 67.50422 | 64.55198 | chr7 | 45365921 | 45368368 | 7.84187 | 5.42146 | 5.31195 |
| chr10 | 20528799 | 20529087 | 534 | Intron (ENSMUST00000020165.8 | 10 | 20518011 | 20519768 | 1758 | 2 | ENSMUST00 | -9032 | Gm17229 | ENSMUSG0C | false | chr10 | 20528587 | 20529334 | 5.68011 | 12.54176 | 10.31921 | chr10 | 20528067 | 20529771 | 4.22416 | 3.7357 | 2.36496 |
| chr5 | 146475308 | 146475639 | 534 | Distal Intergenic | 5 | 146481943 | 146484702 | 2760 | 1 | ENSMUST00 | -6304 | Gm6408 | ENSMUSG0C | false | . | . | . | . | . | . | chr5 | 146474388 | 146476895 | 3.78725 | 3.45261 | 1.99716 |
| chr6 | 115681212 | 115681537 | 533 | Distal Intergenic | 6 | 115630198 | 115676635 | 46438 | 2 | ENSMUST00 | -4578 | Raf1 | ENSMUSG0C | false | . | . | . | . | . | . | chr6 | 115681223 | 115681892 | 5.31497 | 4.13021 | 3.31562 |
| chr14 | 16960013 | 16960612 | 533 | Distal Intergenic | 14 | 16430842 | 16575472 | 144631 | 2 | ENSMUST00 | -384542 | Rarb | ENSMUSG0C | false | . | . | . | . | . | . | chr14 | 16954746 | 16961536 | 4.2934 | 3.68389 | 2.45173 |
| chr4 | 95743031 | 95743266 | 533 | Intron (ENSMUST00000079223.5 | 4 | 95738408 | 95769445 | 31038 | 1 | ENSMUST00 | 4624 | Fggy | ENSMUSG0C | false | . | . | . | . | . | . | chr4 | 95742560 | 95743689 | 4.13647 | 3.72635 | 2.28435 |
| chr7 | 19271546 | 19272088 | 533 | Promoter (<=1kb) | 7 | 19256929 | 19271817 | 14889 | 2 | ENSMUST00 | 0 | Vasp | ENSMUSG0C | false | chr7 | 19269289 | 19272839 | 36.83125 | 118.04518 | 113.96772 | . | . | . | . | . | . |
| chr1 | 183115181 | 183115431 | 533 | Intron (ENSMUST00000003035.6 | 1 | 183086270 | 183147461 | 61192 | 2 | ENSMUST00 | 32030 | Disp1 | ENSMUSG0C | false | . | . | . | . | . | . | chr1 | 183111643 | 183121343 | 6.66914 | 4.87138 | 4.37055 |
| chr13 | 14039191 | 14039645 | 533 | Promoter (<=1kb) | 13 | 13997949 | 14039579 | 41631 | 2 | ENSMUST00 | 0 | Tbce | ENSMUSG0C | false | chr13 | 14038732 | 14040316 | 30.17592 | 94.17345 | 90.73757 | . | . | . | . | . | . |
| chr18 | 35823707 | 35823959 | 533 | Distal Intergenic | 18 | 35829818 | 35861687 | 31870 | 1 | ENSMUST00 | -5859 | Cxxc5 | ENSMUSG0C | false | . | . | . | . | . | . | chr18 | 35821824 | 35824124 | 3.10394 | 3.08835 | 1.49771 |
| chr9 | 96478459 | 96478814 | 533 | Promoter (<=1kb) | 9 | 96470937 | 96478675 | 7739 | 2 | ENSMUST00 | 0 | Rnf7 | ENSMUSG0C | false | chr9 | 96477435 | 96478907 | 30.03893 | 101.89226 | 98.2715 | chr9 | 96473778 | 96478864 | 5.35352 | 4.24499 | 3.30986 |
| chr8 | 70659608 | 70660197 | 533 | Promoter (<=1kb) | 8 | 70646436 | 70659738 | 13303 | 2 | ENSMUST00 | 0 | Pgpep1 | ENSMUSG0C | false | chr8 | 70658804 | 70660437 | 32.16322 | 96.63297 | 93.14677 | chr8 | 70659906 | 70660574 | 6.01536 | 4.46445 | 3.8699 |
| chr3 | 32817305 | 32817741 | 533 | Promoter (<=1kb) | 3 | 32817546 | 32938071 | 120526 | 1 | ENSMUST00 | 0 | Usp13 | ENSMUSG0C | true | chr3 | 32817158 | 32819036 | 36.39278 | 116.22827 | 112.20707 | . | . | . | . | . | . |
| chr8 | 44980081 | 44980448 | 533 | Intron (ENSMUST00000191428.1 | 8 | 44950208 | 45052257 | 102050 | 1 | ENSMUST00 | 29874 | Fat1 | ENSMUSG0C | true | . | . | . | . | . | . | chr8 | 44953863 | 44987890 | 5.49999 | 4.27545 | 3.43184 |
| chr13 | 24831406 | 24832108 | 532 | Promoter (<=1kb) | 13 | 24817948 | 24831517 | 13570 | 2 | ENSMUST00 | 0 | Acot13 | ENSMUSG0C | false | chr13 | 24830659 | 24832491 | 36.809 | 123.33047 | 119.07505 | . | . | . | . | . | . |
| chr8 | 71496571 | 71496825 | 532 | Intron (ENSMUST00000150969.2 | 8 | 71489459 | 71491411 | 1953 | 1 | ENSMUST00 | 7113 | Gtpbp3 | ENSMUSG0C | false | . | . | . | . | . | . | chr8 | 71495272 | 71500046 | 3.90309 | 3.28772 | 2.14928 |
| chr11 | 53707187 | 53707536 | 532 | Promoter (<=1kb) | 11 | 53649519 | 53707319 | 57801 | 2 | ENSMUST00 | 0 | Rad50 | ENSMUSG0C | false | chr11 | 53706462 | 53707792 | 24.55236 | 90.57755 | 87.22021 | . | . | . | . | . | . |
| chr15 | 98295985 | 98296436 | 532 | Promoter (<=1kb) | 15 | 98285588 | 98296130 | 10543 | 2 | ENSMUST00 | 0 | Zfp641 | ENSMUSG0C | false | chr15 | 98295453 | 98296554 | 21.4995 | 59.76749 | 56.92346 | chr15 | 98295236 | 98296367 | 2.96602 | 2.91955 | 1.37526 |
| chr6 | 97312301 | 97312546 | 532 | Promoter (<=1kb) | 6 | 97287082 | 97312380 | 25299 | 2 | ENSMUST00 | 0 | Frmd4b | ENSMUSG0C | false | chr6 | 97311581 | 97312764 | 15.72478 | 37.43704 | 34.87882 | chr6 | 97310781 | 97313395 | 3.41854 | 3.18433 | 1.7373 |
| chr6 | 118069840 | 118070291 | 532 | Intron (ENSMUST00000164960.1 | 6 | 118066385 | 118091546 | 25162 | 1 | ENSMUST00 | 3456 | Rasgef1a | ENSMUSG0C | false | . | . | . | . | . | . | . | . | . | . | . | . |
| chr8 | 13159074 | 13159424 | 532 | Promoter (<=1kb) | 8 | 13159135 | 13175338 | 16204 | 1 | ENSMUST00 | 0 | Lamp1 | ENSMUSG0C | false | chr8 | 13158659 | 13160275 | 41.59011 | 134.93111 | 130.25542 | chr8 | 13158446 | 13162846 | 4.85443 | 4.05072 | 2.89743 |
| chr12 | 40462926 | 40463197 | 532 | Intron (ENSMUST00000037488.6 | 12 | 40446053 | 40846486 | 400434 | 1 | ENSMUST00 | 16874 | Dock4 | ENSMUSG0C | true | . | . | . | . | . | . | chr12 | 40448062 | 40493881 | 5.07226 | 4.00134 | 3.09271 |
| chr2 | 52859554 | 52859853 | 532 | Promoter (1-2kb) | 2 | 52857868 | 53134202 | 276335 | 1 | ENSMUST00 | 1687 | Fmnl2 | ENSMUSG0C | true | chr2 | 52857275 | 52859971 | 40.88443 | 132.61177 | 128.02734 | chr2 | 52859338 | 52863359 | 4.37722 | 3.62844 | 2.51871 |
| chr16 | 90423813 | 90424118 | 532 | Intron (ENSMUST00000065856.6 | 16 | 90386397 | 90499553 | 113157 | 1 | ENSMUST00 | 37417 | Hunk | ENSMUSG0C | false | . | . | . | . | . | . | chr16 | 90424115 | 90424927 | 4.88153 | 4.04041 | 2.95794 |
| chr4 | 123422833 | 123423171 | 532 | Intron (ENSMUST00000097897.5 | 4 | 123435617 | 123435934 | 318 | 1 | ENSMUST00 | -12446 | Gm12924 | ENSMUSG0C | false | . | . | . | . | . | . | chr4 | 123412273 | 123429396 | 5.99663 | 4.50351 | 3.82082 |
| chr12 | 79999327 | 79999603 | 532 | Distal Intergenic | 12 | 79963568 | 79963872 | 305 | 1 | ENSMUST00 | 35760 | RP23-179D3. | ENSMUSG0C | false | . | . | . | . | . | . | chr12 | 79993084 | 80001591 | 4.83553 | 3.97765 | 2.87564 |
| chr14 | 103509612 | 103509839 | 531 | Distal Intergenic | 14 | 103513341 | 103613346 | 100006 | 1 | ENSMUST00 | -3502 | Scel | ENSMUSG0C | true | . | . | . | . | . | . | chr14 | 103509473 | 103510275 | 4.16096 | 3.57291 | 2.31794 |
| chr5 | 32785494 | 32785911 | 531 | Promoter (<=1kb) | 5 | 32736301 | 32785585 | 49285 | 2 | ENSMUST00 | 0 | Pisd | ENSMUSG0C | false | chr5 | 32784971 | 32786245 | 32.88158 | 104.92503 | 101.22703 | chr5 | 32784206 | 32786474 | 3.72962 | 3.37567 | 1.96469 |
| chr1 | 189792534 | 189792852 | 531 | Intron (ENSMUST00000097442.3 | 1 | 189786636 | 189851976 | 65341 | 1 | ENSMUST00 | 5899 | Ptpn14 | ENSMUSG0C | true | . | . | . | . | . | . | chr1 | 189730267 | 189810107 | 6.93269 | 4.87829 | 4.60521 |
| chr4 | 117946988 | 117947278 | 531 | Intron (ENSMUST00000030263.3 | 4 | 117926162 | 117929763 | 3602 | 2 | ENSMUST00 | -17226 | Artn | ENSMUSG0C | false | . | . | . | . | . | . | chr4 | 117946553 | 117949078 | 4.98268 | 4.01452 | 2.99765 |
| chr3 | 121941224 | 121941522 | 531 | Distal Intergenic | 3 | 121953229 | 122016677 | 63449 | 1 | ENSMUST00 | -11707 | Arhgap29 | ENSMUSG0C | false | . | . | . | . | . | . | chr3 | 121940191 | 121942309 | 5.49364 | 4.33531 | 3.46994 |
| chr15 | 101092972 | 101093235 | 531 | Distal Intergenic | 15 | 101115755 | 101120022 | 4268 | 1 | ENSMUST00 | -22520 | Ankrd33 | ENSMUSG0C | false | . | . | . | . | . | . | chr15 | 101092333 | 101093000 | 3.72327 | 3.4135 | 2.00351 |
| chr16 | 20695831 | 20696377 | 531 | Promoter (<=1kb) | 16 | 20696175 | 20701659 | 5485 | 1 | ENSMUST00 | 0 | Fam131a | ENSMUSG0C | false | chr16 | 20695643 | 20696368 | 14.43627 | 33.20709 | 30.70158 | chr16 | 20692677 | 20696663 | 6.75811 | 5.00429 | 4.43548 |
| chr2 | 104070545 | 104070861 | 531 | Promoter (<=1kb) | 2 | 104071010 | 104084957 | 13948 | 1 | ENSMUST00 | -149 | Cd59b | ENSMUSG0C | false | chr2 | 104069617 | 104070899 | 11.90387 | 26.98992 | 24.5636 | chr2 | 104067915 | 104071348 | 4.73792 | 3.68364 | 2.83255 |
| chr2 | 163284899 | 163286040 | 531 | Intron (ENSMUST00000109428.3 | 2 | 163320378 | 163323102 | 2725 | 1 | ENSMUST00 | -34338 | Tox2 | ENSMUSG0C | false | . | . | . | . | . | . | chr2 | 163284077 | 163286098 | 3.6193 | 3.32255 | 1.90744 |
| chr12 | 80132462 | 80132923 | 531 | Promoter (<=1kb) | 12 | 80120548 | 80132844 | 12297 | 2 | ENSMUST00 | 0 | 2310015A10I | ENSMUSG0C | false | chr12 | 80131939 | 80132873 | 15.30398 | 43.94659 | 41.30965 | chr12 | 80116083 | 80133218 | 5.88241 | 4.4251 | 3.73792 |
| chr10 | 27387554 | 27387784 | 531 | Promoter (2-3kb) | 10 | 27389866 | 27397225 | 7360 | 1 | ENSMUST00 | -2082 | RP24-496C19 | ENSMUSG0C | false | . | . | . | . | . | . | . | . | . | . | . | . |
| chr8 | 123813621 | 123813842 | 530 | Intron (ENSMUST00000127664.1 | 8 | 123806068 | 123834485 | 28418 | 1 | ENSMUST00 | 7554 | Rab4a | ENSMUSG0C | false | . | . | . | . | . | . | . | . | . | . | . | . |
| chr12 | 4233966 | 4234426 | 530 | Promoter (<=1kb) | 12 | 4211667 | 4234294 | 22628 | 2 | ENSMUST00 | 0 | Cenpo | ENSMUSG0C | false | chr12 | 4233182 | 4234847 | 21.44446 | 75.31629 | 72.24329 | chr12 | 4231478 | 4235304 | 3.51847 | 3.2737 | 1.82277 |
| chr12 | 87374201 | 87374482 | 530 | Intron (ENSMUST00000021424.4 | 12 | 87346734 | 87388241 | 41508 | 2 | ENSMUST00 | 13759 | Sptlc2 | ENSMUSG0C | false | . | . | . | . | . | . | chr12 | 87367947 | 87387312 | 5.77138 | 4.39496 | 3.64069 |
| chr18 | 60747744 | 60748728 | 530 | Distal Intergenic | 18 | 60774510 | 60778546 | 4037 | 1 | ENSMUST00 | -25782 | Rps14 | ENSMUSG0C | false | chr18 | 60747065 | 60749034 | 29.53108 | 87.21563 | 83.92796 | chr18 | 60742488 | 60750880 | 5.84853 | 4.46265 | 3.69893 |
| chr7 | 55842000 | 55842688 | 530 | Promoter (<=1kb) | 7 | 55842071 | 55930700 | 88630 | 1 | ENSMUST00 | 0 | Cyfip1 | ENSMUSG0C | false | chr7 | 55841802 | 55843400 | 37.68014 | 134.63641 | 129.97372 | chr7 | 55841471 | 55862799 | 4.7417 | 3.9553 | 2.79473 |
| chr9 | 107770883 | 107771164 | 530 | Promoter (<=1kb) | 9 | 107740371 | 107770996 | 30626 | 2 | ENSMUST00 | 0 | Rbm5 | ENSMUSG0C | false | chr9 | 107769446 | 107771471 | 28.90341 | 112.84859 | 108.93253 | . | . | . | . | . | . |
| chr1 | 44119664 | 44120096 | 530 | Promoter (<=1kb) | 1 | 44119968 | 44144770 | 24803 | 1 | ENSMUST00 | 0 | Bivm | ENSMUSG0C | false | chr1 | 44118127 | 44120328 | 19.26894 | 65.0005 | 62.08229 | . | . | . | . | . | . |
| chr2 | 152947147 | 152947459 | 530 | Promoter (2-3kb) | 2 | 152941022 | 152944556 | 3535 | 2 | ENSMUST00 | -2592 | Dusp15 | ENSMUSG0C | false | . | . | . | . | . | . | chr2 | 152946044 | 152955187 | 5.87363 | 4.46622 | 3.72171 |
| chr8 | 11556117 | 11557103 | 530 | Promoter (<=1kb) | 8 | 11556066 | 11563250 | 7185 | 1 | ENSMUST00 | 52 | Ing1 | ENSMUSG0C | false | chr8 | 11554880 | 11558049 | 39.39634 | 133.30592 | 128.68834 | . | . | . | . | . | . |
| chr8 | 93668340 | 93668731 | 530 | Distal Intergenic | 8 | 93499378 | 93535707 | 36330 | 2 | ENSMUST00 | -132634 | Ces5a | ENSMUSG0C | false | . | . | . | . | . | . | chr8 | 93667075 | 93668969 | 6.09324 | 4.55017 | 3.95536 |
| chr10 | 81192523 | 81192938 | 530 | Promoter (1-2kb) | 10 | 81194609 | 81198439 | 3831 | 1 | ENSMUST00 | -1671 | Atcayos | ENSMUSG0C | false | chr10 | 81192017 | 81193242 | 24.3552 | 66.39809 | 63.45943 | chr10 | 81190371 | 81194900 | 5.30511 | 4.26841 | 3.25415 |
| chr1 | 52578191 | 52578577 | 530 | Promoter (1-2kb) | 1 | 52570014 | 52580291 | 10278 | 2 | ENSMUST00 | 1714 | RP24-388H6. | ENSMUSG0C | false | . | . | . | . | . | . | chr1 | 52577889 | 52580731 | 5.30029 | 4.19859 | 3.2445 |
| chr16 | 85979773 | 85980144 | 529 | Distal Intergenic | 16 | 85858170 | 85901125 | 42956 | 2 | ENSMUST00 | -78649 | Adamts5 | ENSMUSG0C | false | . | . | . | . | . | . | . | . | . | . | . | . |

| | | | | | | | | | | | | | | | | | | | | | | | | | | |
|---|---|---|---|---|---|---|---|---|---|---|---|---|---|---|---|---|---|---|---|---|---|---|---|---|---|---|
| chr4 | 152119951 | 152120362 | 529 | Promoter (1-2kb) | 4 | 152117971 | 152119323 | 1353 | 1 | ENSMUST00 | 1981 | Tnfrsf25 | ENSMUSG0C | false | chr4 | 152119592 | 152121333 | 14.333 | 34.37709 | 31.85639 | chr4 | 152118648 | 152122002 | 6.10878 | 4.6144 | 3.91321 |
| chrX | 42147988 | 42148321 | 529 | Promoter (<=1kb) | 20 | 42149317 | 42277185 | 127869 | 1 | ENSMUST00 | -996 | Stag2 | ENSMUSG0C | false | . | . | . | . | . | . | . | . | . | . | . | . |
| chr2 | 150909386 | 150909932 | 529 | Promoter (<=1kb) | 2 | 150909594 | 150931280 | 21687 | 1 | ENSMUST00 | 0 | Gins1 | ENSMUSG0C | false | chr2 | 150909203 | 150910351 | 21.76566 | 71.50774 | 68.49327 | . | . | . | . | . | . |
| chr4 | 119543676 | 119543951 | 529 | Intron (ENSMUST00000176012.2 | 4 | 119540522 | 119573745 | 33224 | 1 | ENSMUST00 | 3155 | Foxj3 | ENSMUSG0C | true | . | . | . | . | . | . | chr4 | 119541733 | 119562342 | 4.43637 | 3.78033 | 2.54926 |
| chr6 | 88655354 | 88655718 | 529 | Distal Intergenic | 6 | 88618617 | 88637950 | 19334 | 2 | ENSMUST00 | -17405 | Kbtbd12 | ENSMUSG0C | false | . | . | . | . | . | . | chr6 | 88654759 | 88656335 | 4.68341 | 3.85477 | 2.76485 |
| chr5 | 100125452 | 100125861 | 529 | Promoter (2-3kb) | 5 | 100083947 | 100122796 | 38850 | 2 | ENSMUST00 | -2657 | Tmem150c | ENSMUSG0C | true | . | . | . | . | . | . | chr5 | 100107043 | 100130430 | 5.59817 | 4.32599 | 3.51171 |
| chr13 | 67754836 | 67755100 | 529 | Promoter (<=1kb) | 13 | 67747800 | 67755134 | 7335 | 2 | ENSMUST00 | 34 | Zfp85 | ENSMUSG0C | false | chr13 | 67754365 | 67755559 | 24.02885 | 69.3614 | 66.38335 | chr13 | 67754283 | 67755849 | 3.68827 | 3.32274 | 1.96173 |
| chr3 | 89164731 | 89165176 | 529 | Promoter (<=1kb) | 3 | 89164795 | 89176921 | 12127 | 1 | ENSMUST00 | 0 | Clk2 | ENSMUSG0C | false | chr3 | 89164269 | 89166387 | 29.45621 | 102.83202 | 99.19408 | chr3 | 89163779 | 89168143 | 6.10442 | 4.63703 | 3.89661 |
| chr2 | 29016197 | 29016469 | 529 | Intron (ENSMUST00000189027.1 | 2 | 28927834 | 29039360 | 111527 | 2 | ENSMUST00 | 22891 | 1700101E01I | ENSMUSG0C | false | . | . | . | . | . | . | . | . | . | . | . | . |
| chr4 | 52438723 | 52439098 | 529 | Promoter (<=1kb) | 4 | 52430285 | 52438965 | 8681 | 2 | ENSMUST00 | 0 | Smc2os | ENSMUSG0C | false | chr4 | 52438330 | 52439908 | 33.7247 | 108.45113 | 104.65779 | . | . | . | . | . | . |
| chr11 | 96993061 | 96993315 | 528 | Distal Intergenic | 11 | 96954219 | 96982959 | 28741 | 2 | ENSMUST00 | -10103 | Sp2 | ENSMUSG0C | false | . | . | . | . | . | . | chr11 | 96984955 | 96998855 | 6.28932 | 4.5897 | 4.08329 |
| chr11 | 76945407 | 76945839 | 528 | Promoter (<=1kb) | 11 | 76945504 | 76987379 | 41876 | 1 | ENSMUST00 | 0 | Blmh | ENSMUSG0C | false | chr11 | 76945142 | 76946667 | 29.48955 | 94.93548 | 91.48817 | chr11 | 76944160 | 76947151 | 4.10471 | 3.58068 | 2.28813 |
| chr16 | 34752038 | 34752366 | 528 | Intron (ENSMUST00000155268.1 | 16 | 34745210 | 34877836 | 132627 | 1 | ENSMUST00 | 6829 | Mylk | ENSMUSG0C | false | . | . | . | . | . | . | chr16 | 34751594 | 34753011 | 4.78689 | 4.02576 | 2.81761 |
| chr17 | 83219944 | 83220400 | 528 | Promoter (<=1kb) | 17 | 83220102 | 83221742 | 1641 | 1 | ENSMUST00 | 0 | Pkdcc | ENSMUSG0C | false | chr17 | 83213509 | 83220055 | 37.87741 | 120.9216 | 116.74998 | chr17 | 83216753 | 83225364 | 6.20423 | 4.53249 | 4.00069 |
| chr1 | 152604175 | 152604514 | 528 | Intron (ENSMUST00000111859.2 | 1 | 152557543 | 152592759 | 35217 | 2 | ENSMUST00 | -11417 | Rgl1 | ENSMUSG0C | false | . | . | . | . | . | . | chr1 | 152596864 | 152604963 | 4.67049 | 3.87984 | 2.74292 |
| chr9 | 99436422 | 99436976 | 528 | Promoter (<=1kb) | 9 | 99387928 | 99436881 | 48954 | 2 | ENSMUST00 | 0 | Mras | ENSMUSG0C | false | chr9 | 99435672 | 99437648 | 30.03704 | 95.90482 | 92.42858 | . | . | . | . | . | . |
| chr8 | 79120858 | 79121113 | 528 | Promoter (<=1kb) | 8 | 79120483 | 79193192 | 72710 | 1 | ENSMUST00 | 376 | Zfp827 | ENSMUSG0C | false | . | . | . | . | . | . | chr8 | 79120536 | 79121080 | 3.18295 | 2.91237 | 1.54262 |
| chr6 | 124636927 | 124637153 | 528 | Promoter (<=1kb) | 6 | 124624625 | 124636085 | 11461 | 2 | ENSMUST00 | -843 | C1s2 | ENSMUSG0C | false | . | . | . | . | . | . | . | . | . | . | . | . |
| chr4 | 114738935 | 114739595 | 528 | Distal Intergenic | 4 | 114680804 | 114687595 | 6792 | 1 | ENSMUST00 | 58132 | Gm12829 | ENSMUSG0C | false | . | . | . | . | . | . | chr4 | 114739077 | 114739457 | 3.35909 | 3.25732 | 1.64184 |
| chr8 | 125337592 | 125337916 | 528 | Distal Intergenic | 8 | 125224400 | 125227323 | 2924 | 2 | ENSMUST00 | -110270 | Gm16237 | ENSMUSG0C | false | . | . | . | . | . | . | chr8 | 125335216 | 125352727 | 6.82037 | 4.91647 | 4.50391 |
| chr8 | 121341887 | 121342158 | 528 | Intron (ENSMUST00000127664.1 | 8 | 121256747 | 121356836 | 100090 | 2 | ENSMUST00 | 14678 | Gm26815 | ENSMUSG0C | false | . | . | . | . | . | . | chr8 | 121340922 | 121349877 | 6.81979 | 4.84985 | 4.53151 |
| chr8 | 33882593 | 33882902 | 528 | Promoter (2-3kb) | 8 | 33884774 | 33885747 | 974 | 2 | ENSMUST00 | 2845 | Gm26978 | ENSMUSG0C | false | . | . | . | . | . | . | chr8 | 33863865 | 33927704 | 6.32022 | 4.67603 | 4.08842 |
| chr9 | 32544745 | 32545064 | 527 | Promoter (1-2kb) | 9 | 32428059 | 32542861 | 114803 | 2 | ENSMUST00 | -1885 | Fli1 | ENSMUSG0C | false | . | . | . | . | . | . | chr9 | 32539978 | 32546053 | 4.89991 | 4.03618 | 2.93455 |
| chr11 | 88820725 | 88821054 | 527 | Distal Intergenic | 11 | 88837012 | 88841540 | 4529 | 2 | ENSMUST00 | 20486 | Akap1 | ENSMUSG0C | false | . | . | . | . | . | . | chr11 | 88820348 | 88821281 | 3.00382 | 2.95523 | 1.39346 |
| chr1 | 72870253 | 72870621 | 527 | Intron (ENSMUST00000027377.8 | 1 | 72858065 | 72874275 | 16211 | 2 | ENSMUST00 | 3654 | Igfbp5 | ENSMUSG0C | false | chr1 | 72870000 | 72872489 | 10.01484 | 27.17025 | 24.74075 | chr1 | 72869237 | 72870646 | 4.23653 | 3.73725 | 2.36821 |
| chr5 | 66338155 | 66338433 | 527 | Promoter (<=1kb) | 5 | 66301888 | 66337665 | 35778 | 2 | ENSMUST00 | -491 | Apbb2 | ENSMUSG0C | false | . | . | . | . | . | . | chr5 | 66334637 | 66346684 | 5.95013 | 4.37258 | 3.81282 |
| chr9 | 69471033 | 69471335 | 527 | Intron (ENSMUST00000034756.9 | 9 | 69478465 | 69491795 | 13331 | 1 | ENSMUST00 | -7130 | Anxa2 | ENSMUSG0C | false | . | . | . | . | . | . | chr9 | 69455103 | 69480655 | 7.58546 | 5.20708 | 5.13718 |
| chr17 | 34589777 | 34590242 | 527 | Promoter (<=1kb) | 17 | 34589806 | 34591754 | 1949 | 1 | ENSMUST00 | 0 | Gpsm3 | ENSMUSG0C | false | chr17 | 34586640 | 34592164 | 28.90751 | 104.81161 | 101.11721 | chr17 | 34586035 | 34591596 | 5.00173 | 4.02952 | 3.00733 |
| chr9 | 52168094 | 52169019 | 527 | Promoter (<=1kb) | 9 | 52111985 | 52168111 | 56127 | 2 | ENSMUST00 | 0 | Zc3h12c | ENSMUSG0C | false | chr9 | 52167458 | 52169267 | 37.30296 | 124.19952 | 119.9176 | chr9 | 52164573 | 52169137 | 3.54382 | 3.31708 | 1.82753 |
| chr6 | 72220377 | 72220716 | 527 | Intron (ENSMUST00000042646.7 | 6 | 72206177 | 72235577 | 29401 | 2 | ENSMUST00 | 14861 | Atoh8 | ENSMUSG0C | false | . | . | . | . | . | . | chr6 | 72211696 | 72224884 | 7.46431 | 5.12594 | 5.03432 |
| chr2 | 32730902 | 32731186 | 527 | Intron (ENSMUST00000074248.5 | 2 | 32727706 | 32755512 | 27807 | 1 | ENSMUST00 | 3197 | Sh2d3c | ENSMUSG0C | false | . | . | . | . | . | . | . | . | . | . | . | . |
| chr18 | 55640094 | 55640368 | 527 | Distal Intergenic | 18 | 55622636 | 55622920 | 285 | 2 | ENSMUST00 | -17175 | Gm24514 | ENSMUSG0C | false | . | . | . | . | . | . | chr18 | 55638386 | 55643134 | 4.73705 | 3.9997 | 2.78615 |
| chr13 | 32681657 | 32681935 | 527 | Distal Intergenic | 13 | 32758202 | 32758310 | 109 | 1 | ENSMUST00 | -76267 | Gm27342 | ENSMUSG0C | false | . | . | . | . | . | . | chr13 | 32678441 | 32682534 | 5.54396 | 4.36904 | 3.43747 |
| chr5 | 45669521 | 45669891 | 526 | Promoter (<=1kb) | 5 | 45669922 | 45700544 | 30623 | 1 | ENSMUST00 | -31 | Ncapg | ENSMUSG0C | false | chr5 | 45668827 | 45670551 | 29.77807 | 93.47606 | 90.05975 | . | . | . | . | . | . |
| chr14 | 16575259 | 16575872 | 526 | Promoter (<=1kb) | 14 | 16430842 | 16575472 | 144631 | 2 | ENSMUST00 | 0 | Rarb | ENSMUSG0C | false | chr14 | 16572513 | 16576178 | 19.18445 | 82.69909 | 79.49731 | . | . | . | . | . | . |
| chr16 | 43531267 | 43531543 | 526 | Intron (ENSMUST00000132881.2 | 16 | 43510308 | 43619137 | 108830 | 1 | ENSMUST00 | 20960 | Zbtb20 | ENSMUSG0C | false | . | . | . | . | . | . | chr16 | 43529498 | 43533262 | 4.15056 | 3.52873 | 2.33958 |
| chr11 | 120242826 | 120243093 | 526 | Intron (ENSMUST00000122148.2 | 11 | 120236699 | 120292282 | 55584 | 1 | ENSMUST00 | 6128 | Bahcc1 | ENSMUSG0C | false | . | . | . | . | . | . | chr11 | 120237752 | 120244869 | 5.41747 | 4.25596 | 3.35787 |
| chr4 | 150454505 | 150454821 | 526 | Intron (ENSMUST00000105682.3 | 4 | 150488393 | 150621962 | 133570 | 1 | ENSMUST00 | -33572 | Rere | ENSMUSG0C | false | . | . | . | . | . | . | chr4 | 150440410 | 150463295 | 5.62337 | 4.29719 | 3.53587 |
| chr6 | 141159510 | 141159792 | 526 | Distal Intergenic | 6 | 141249269 | 141505940 | 256672 | 1 | ENSMUST00 | -89477 | Pde3a | ENSMUSG0C | true | . | . | . | . | . | . | chr6 | 141159335 | 141160789 | 4.19872 | 3.77414 | 2.3462 |
| chr17 | 86444872 | 86445194 | 526 | Intron (ENSMUST00000097275.3 | 17 | 86490562 | 86496112 | 5551 | 1 | ENSMUST00 | -45368 | Prkce | ENSMUSG0C | true | . | . | . | . | . | . | chr17 | 86443786 | 86448490 | 7.30257 | 5.10481 | 4.89887 |
| chr15 | 37460022 | 37460306 | 526 | Promoter (<=1kb) | 15 | 37366218 | 37459360 | 93143 | 2 | ENSMUST00 | -663 | Ncald | ENSMUSG0C | true | chr15 | 37458319 | 37460172 | 15.39495 | 41.17039 | 38.56658 | chr15 | 37456877 | 37461079 | 4.67304 | 3.9677 | 2.72998 |
| chr18 | 69614509 | 69614967 | 526 | Intron (ENSMUST00000066717.6 | 18 | 69599536 | 69651289 | 51754 | 1 | ENSMUST00 | 14974 | Tcf4 | ENSMUSG0C | false | . | . | . | . | . | . | chr18 | 69613659 | 69618744 | 4.6674 | 3.94807 | 2.72945 |
| chr8 | 25439361 | 25439605 | 526 | Distal Intergenic | 8 | 25454273 | 25454704 | 432 | 1 | ENSMUST00 | -14668 | Gm10689 | ENSMUSG0C | false | . | . | . | . | . | . | chr8 | 25435682 | 25441738 | 6.54241 | 4.78324 | 4.28383 |
| chr3 | 100998326 | 100998635 | 526 | Intron (ENSMUST00000147399.3 | 3 | 100993532 | 101022176 | 28645 | 2 | ENSMUST00 | 23541 | Cd101 | ENSMUSG0C | false | . | . | . | . | . | . | chr3 | 100997564 | 100999208 | 4.68957 | 3.89776 | 2.79479 |
| chr13 | 10244305 | 10244712 | 526 | Intron (ENSMUST00000187510.1 | 13 | 10359479 | 10658893 | 299415 | 1 | ENSMUST00 | -114767 | Gm26861 | ENSMUSG0C | false | . | . | . | . | . | . | . | . | . | . | . | . |
| chr11 | 61562751 | 61563039 | 525 | Intron (ENSMUST00000108711.2 | 11 | 61534394 | 61546542 | 12149 | 2 | ENSMUST00 | -16210 | Epn2 | ENSMUSG0C | true | . | . | . | . | . | . | chr11 | 61547634 | 61570806 | 4.88221 | 3.88483 | 2.93676 |
| chr19 | 56548330 | 56548982 | 525 | Promoter (<=1kb) | 19 | 56548261 | 56598846 | 50586 | 1 | ENSMUST00 | 70 | Nhlrc2 | ENSMUSG0C | true | chr19 | 56547547 | 56549649 | 39.62653 | 133.71027 | 129.07751 | . | . | . | . | . | . |
| chr4 | 126255183 | 126255583 | 525 | Promoter (<=1kb) | 4 | 126232167 | 126256277 | 24111 | 2 | ENSMUST00 | 694 | Map7d1 | ENSMUSG0C | false | chr4 | 126254944 | 126257391 | 48.06229 | 188.02159 | 181.71353 | chr4 | 126250228 | 126257655 | 4.99746 | 3.92262 | 3.03857 |
| chr7 | 80198075 | 80198756 | 525 | Promoter (<=1kb) | 7 | 80198538 | 80214065 | 15528 | 1 | ENSMUST00 | 0 | Sema4b | ENSMUSG0C | false | chr7 | 80197944 | 80199464 | 37.58501 | 119.35227 | 115.22867 | chr7 | 80197390 | 80204507 | 5.08167 | 4.14387 | 3.06709 |
| chr4 | 150087120 | 150087657 | 525 | Promoter (<=1kb) | 4 | 150087365 | 150105927 | 18563 | 1 | ENSMUST00 | 0 | Gpr157 | ENSMUSG0C | false | chr4 | 150086980 | 150088393 | 32.14993 | 97.60393 | 94.09416 | chr4 | 150086866 | 150089700 | 4.01429 | 3.64509 | 2.18101 |
| chr5 | 123683551 | 123684502 | 525 | Promoter (<=1kb) | 5 | 123577795 | 123684322 | 106528 | 2 | ENSMUST00 | 0 | Clip1 | ENSMUSG0C | false | chr5 | 123682686 | 123685179 | 39.06023 | 122.966 | 118.72274 | . | . | . | . | . | . |
| chrX | 50012276 | 50012507 | 525 | Promoter (2-3kb) | 20 | 50009306 | 50010235 | 930 | 2 | ENSMUST00 | -2042 | Olfr1323 | ENSMUSG0C | false | . | . | . | . | . | . | . | . | . | . | . | . |
| chr11 | 22981930 | 22982440 | 525 | Promoter (<=1kb) | 11 | 22896136 | 22982382 | 86247 | 2 | ENSMUST00 | 0 | Commd1 | ENSMUSG0C | false | chr11 | 22981480 | 22982775 | 27.91982 | 97.01129 | 93.51138 | . | . | . | . | . | . |
| chr3 | 79213622 | 79213930 | 525 | Distal Intergenic | 3 | 79196374 | 79196477 | 104 | 1 | ENSMUST00 | 17249 | Gm24150 | ENSMUSG0C | false | . | . | . | . | . | . | chr3 | 79202018 | 79214943 | 5.27403 | 4.18459 | 3.23704 |
| chr7 | 64872871 | 64873247 | 525 | Promoter (<=1kb) | 7 | 64867052 | 64872997 | 5946 | 2 | ENSMUST00 | 0 | Ndnl2 | ENSMUSG0C | false | chr7 | 64871935 | 64873471 | 35.61502 | 123.34891 | 119.09213 | . | . | . | . | . | . |
| chr8 | 115038491 | 115038763 | 525 | Intron (ENSMUST00000004756.8 | 8 | 115052814 | 115052913 | 100 | 2 | ENSMUST00 | 14150 | Gm22556 | ENSMUSG0C | false | . | . | . | . | . | . | chr8 | 115037445 | 115040253 | 4.04888 | 3.51677 | 2.26714 |
| chr5 | 140830190 | 140830676 | 524 | Promoter (<=1kb) | 5 | 140758408 | 140830431 | 72024 | 2 | ENSMUST00 | 0 | Gna12 | ENSMUSG0C | false | chr5 | 140829591 | 140830475 | 30.22203 | 93.27019 | 89.85963 | chr5 | 140787727 | 140831108 | 6.24285 | 4.56075 | 4.04432 |
| chr6 | 88924578 | 88924858 | 524 | Distal Intergenic | 6 | 88910158 | 88911778 | 1621 | 1 | ENSMUST00 | 14421 | Tpra1 | ENSMUSG0C | false | . | . | . | . | . | . | chr6 | 88922092 | 88928323 | 9.06661 | 5.7788 | 6.36998 |
| chr4 | 141572014 | 141572317 | 524 | Distal Intergenic | 4 | 141590050 | 141596469 | 6420 | 2 | ENSMUST00 | 24152 | Fblim1 | ENSMUSG0C | false | . | . | . | . | . | . | chr4 | 141568061 | 141575603 | 6.60364 | 4.6991 | 4.34712 |
| chr7 | 43488653 | 43489002 | 524 | Promoter (1-2kb) | 7 | 43472907 | 43490075 | 17169 | 2 | ENSMUST00 | 1073 | Iglon5 | ENSMUSG0C | false | . | . | . | . | . | . | . | . | . | . | . | . |
| chr12 | 30024642 | 30025042 | 524 | Distal Intergenic | 12 | 29999815 | 30002472 | 2658 | 1 | ENSMUST00 | 24828 | Pxdn | ENSMUSG0C | false | . | . | . | . | . | . | chr12 | 30021573 | 30026360 | 4.80877 | 3.9928 | 2.84603 |
| chr15 | 59142625 | 59142986 | 524 | Distal Intergenic | 15 | 58941244 | 59082026 | 140783 | 2 | ENSMUST00 | -60600 | Mtss1 | ENSMUSG0C | true | . | . | . | . | . | . | chr15 | 59140828 | 59146782 | 4.44009 | 3.87565 | 2.53114 |
| chr14 | 121037324 | 121037582 | 524 | Promoter (1-2kb) | 14 | 121035564 | 121283744 | 248181 | 1 | ENSMUST00 | 1761 | Farp1 | ENSMUSG0C | false | chr14 | 121034923 | 121037682 | 40.93993 | 131.61429 | 127.06391 | chr14 | 121036588 | 121040222 | 3.78661 | 3.39253 | 2.03321 |
| chr16 | 26184701 | 26185018 | 524 | Distal Intergenic | 16 | 25959288 | 26105784 | 146497 | 2 | ENSMUST00 | -78918 | Leprel1 | ENSMUSG0C | false | . | . | . | . | . | . | chr16 | 26184488 | 26186872 | 4.36158 | 3.79028 | 2.49054 |
| chr8 | 87746086 | 87746436 | 524 | Intron (ENSMUST00000165770.3 | 8 | 87783147 | 87803823 | 20677 | 2 | ENSMUST00 | 57387 | Zfp423 | ENSMUSG0C | true | . | . | . | . | . | . | chr8 | 87729485 | 87749127 | 6.31805 | 4.63416 | 4.1004 |
| chr7 | 95787754 | 95788121 | 524 | Distal Intergenic | 7 | 95958215 | 95961613 | 3399 | 1 | ENSMUST00 | -170094 | Gm9966 | ENSMUSG0C | false | . | . | . | . | . | . | chr7 | 95784545 | 95792909 | 5.1339 | 4.17838 | 3.10723 |
| chr15 | 86200057 | 86200403 | 524 | Distal Intergenic | 15 | 86212554 | 86212657 | 104 | 2 | ENSMUST00 | 12254 | Gm22818 | ENSMUSG0C | false | . | . | . | . | . | . | chr15 | 86196688 | 86204593 | 6.93026 | 4.73328 | 4.61625 |
| chr12 | 71891515 | 71891755 | 524 | Intron (ENSMUST00000085299.2 | 12 | 71846311 | 71847291 | 981 | 2 | ENSMUST00 | -44225 | Gm7985 | ENSMUSG0C | false | . | . | . | . | . | . | chr12 | 71889934 | 71893643 | 4.03373 | 3.51872 | 2.23556 |
| chr14 | 121611761 | 121612009 | 523 | Intron (ENSMUST00000040700.7 | 14 | 121542039 | 121698417 | 156379 | 2 | ENSMUST00 | 86408 | Dock9 | ENSMUSG0C | false | . | . | . | . | . | . | chr14 | 121611112 | 121612249 | 5.36066 | 4.15904 | 3.3694 |
| chr7 | 78718326 | 78718615 | 523 | Distal Intergenic | 7 | 78775236 | 78783066 | 7831 | 2 | ENSMUST00 | 64451 | Mrpl46 | ENSMUSG0C | false | . | . | . | . | . | . | chr7 | 78715476 | 78719645 | 5.62851 | 4.38751 | 3.5246 |
| chr8 | 110950346 | 110950616 | 523 | Intron (ENSMUST00000117534.2 | 8 | 110940904 | 110957012 | 16109 | 1 | ENSMUST00 | 9443 | St3gal2 | ENSMUSG0C | false | chr8 | 110950094 | 110950489 | 5.86754 | 9.51579 | 7.34688 | chr8 | 110950024 | 110950748 | 3.7897 | 3.45263 | 2.04665 |
| chr5 | 140510219 | 140510510 | 523 | Intron (ENSMUST00000043050.8 | 5 | 140505575 | 140524153 | 18579 | 1 | ENSMUST00 | 4645 | Chst12 | ENSMUSG0C | false | . | . | . | . | . | . | chr5 | 140508836 | 140510861 | 4.14988 | 3.71868 | 2.29452 |
| chr7 | 19181004 | 19181342 | 523 | Promoter (<=1kb) | 7 | 19181159 | 19206482 | 25324 | 1 | ENSMUST00 | 0 | Eml2 | ENSMUSG0C | true | chr7 | 19181038 | 19182936 | 28.08037 | 85.65662 | 82.39679 | chr7 | 19178043 | 19190895 | 5.91671 | 4.49499 | 3.76502 |
| chr6 | 73474609 | 73474888 | 523 | Distal Intergenic | 6 | 73468573 | 73469667 | 1095 | 2 | ENSMUST00 | -4943 | 4931417E11I | ENSMUSG0C | false | . | . | . | . | . | . | chr6 | 73473003 | 73475766 | 3.25767 | 3.10979 | 1.6152 |
| chr7 | 133744770 | 133745060 | 523 | Intron (ENSMUST00000033290.6 | 7 | 133738479 | 133738934 | 456 | 2 | ENSMUST00 | -5837 | Gm15483 | ENSMUSG0C | false | . | . | . | . | . | . | chr7 | 133743183 | 133753176 | 5.21662 | 4.08927 | 3.21065 |
| chr8 | 3209336 | 3209606 | 523 | Intron (ENSMUST00000091291.4 | 8 | 3259704 | 3260641 | 938 | 1 | ENSMUST00 | -50098 | Gm16180 | ENSMUSG0C | false | . | . | . | . | . | . | chr8 | 3208761 | 3212840 | 5.43592 | 4.19508 | 3.38792 |
| chr12 | 86816925 | 86817263 | 523 | Distal Intergenic | 12 | 86809957 | 86810258 | 302 | 1 | ENSMUST00 | 6969 | Gm25888 | ENSMUSG0C | false | . | . | . | . | . | . | chr12 | 86815656 | 86819384 | 5.65895 | 4.35849 | 3.5757 |
| chr7 | 63896368 | 63896664 | 523 | Promoter (<=1kb) | 7 | 63896533 | 63899698 | 3166 | 1 | ENSMUST00 | 0 | Gm27252 | ENSMUSG0C | false | . | . | . | . | . | . | chr7 | 63885397 | 63936279 | 7.51388 | 5.18869 | 5.06896 |
| chr1 | 118481839 | 118482234 | 523 | Promoter (2-3kb) | 1 | 118478604 | 118479591 | 988 | 2 | ENSMUST00 | -2249 | RP24-553C6. | ENSMUSG0C | false | chr1 | 118481424 | 118482973 | 16.48782 | 40.15519 | 37.56212 | chr1 | 118480722 | 118490852 | 5.42823 | 4.04703 | 3.39636 |
| chr11 | 101082340 | 101082746 | 522 | Promoter (<=1kb) | 11 | 101082565 | 101086619 | 4055 | 1 | ENSMUST00 | 0 | Coasy | ENSMUSG0C | false | chr11 | 101081815 | 101083384 | 30.17592 | 94.17345 | 90.73757 | chr11 | 101078620 | 101084538 | 5.0744 | 3.94453 | 3.10504 |
| chr16 | 4077579 | 4078052 | 522 | Promoter (<=1kb) | 16 | 4039971 | 4077827 | 37857 | 2 | ENSMUST00 | 0 | Trap1 | ENSMUSG0C | false | chr16 | 4076892 | 4078331 | 33.7938 | 119.00099 | 114.88904 | . | . | . | . | . | . |
| chr1 | 135793746 | 135794085 | 522 | Intron (ENSMUST00000132795.2 | 1 | 135799402 | 135810989 | 11588 | 1 | ENSMUST00 | -5317 | Tnni1 | ENSMUSG0C | false | chr1 | 135793742 | 135794601 | 4.04026 | 6.18694 | 4.10202 | chr1 | 135793507 | 135794742 | 4.43019 | 3.8812 | 2.52257 |
| chr6 | 145862961 | 145864178 | 522 | Promoter (1-2kb) | 6 | 145863505 | 145865302 | 1798 | 2 | ENSMUST00 | 1124 | Bhlhe41 | ENSMUSG0C | false | chr6 | 145862672 | 145864243 | 22.5292 | 69.37365 | 66.39268 | chr6 | 145861038 | 145867407 | 3.647 | 3.29023 | 1.93146 |
| chr18 | 80637069 | 80637559 | 522 | Intron (ENSMUST00000167977.2 | 18 | 80606205 | 80708173 | 101969 | 2 | ENSMUST00 | 70614 | Nfatc1 | ENSMUSG0C | false | . | . | . | . | . | . | chr18 | 80636225 | 80638726 | 4.98529 | 4.12774 | 2.98118 |
| chr15 | 77314597 | 77314920 | 522 | Distal Intergenic | 15 | 77078991 | 77307053 | 228063 | 2 | ENSMUST00 | -7545 | Rbfox2 | ENSMUSG0C | false | . | . | . | . | . | . | . | . | . | . | . | . |
| chr10 | 79554686 | 79555233 | 522 | Promoter (<=1kb) | 10 | 79540245 | 79555091 | 14847 | 2 | ENSMUST00 | 0 | Mier2 | ENSMUSG0C | false | chr10 | 79554260 | 79555448 | 26.94166 | 91.30816 | 87.93849 | chr10 | 79553593 | 79555625 | 3.62219 | 3.41799 | 1.86952 |
| chr6 | 54326649 | 54327154 | 522 | Promoter (<=1kb) | 6 | 54327012 | 54330200 | 3189 | 1 | ENSMUST00 | 0 | Prr15 | ENSMUSG0C | false | chr6 | 54326642 | 54328336 | 20.06521 | 63.98471 | 61.08233 | chr6 | 54325893 | 54326824 | 3.39282 | 3.30858 | 1.66603 |
| chr6 | 72528394 | 72528723 | 522 | Distal Intergenic | 6 | 72517409 | 72521634 | 4226 | 2 | ENSMUST00 | -6761 | Sh2d6 | ENSMUSG0C | false | . | . | . | . | . | . | chr6 | 72527024 | 72529476 | 4.6066 | 3.78148 | 2.69097 |
| chr17 | 78365002 | 78365300 | 522 | Intron (ENSMUST00000112498.2 | 17 | 78377885 | 78418131 | 40247 | 2 | ENSMUST00 | 52831 | Fez2 | ENSMUSG0C | false | . | . | . | . | . | . | . | . | . | . | . | . |
| chr19 | 36785141 | 36785390 | 522 | Distal Intergenic | 19 | 36731737 | 36736653 | 4917 | 2 | ENSMUST00 | -48489 | Ppp1r3c | ENSMUSG0C | false | . | . | . | . | . | . | chr19 | 36784760 | 36785812 | 2.82775 | 2.61017 | 1.26859 |
| chr2 | 166361184 | 166361455 | 522 | Distal Intergenic | 2 | 166380828 | 166398157 | 17330 | 2 | ENSMUST00 | 36702 | Gm14268 | ENSMUSG0C | false | . | . | . | . | . | . | chr2 | 166359531 | 166362318 | 4.6733 | 4.00112 | 2.73928 |
| chr10 | 75517124 | 75517589 | 521 | Promoter (<=1kb) | 10 | 75517551 | 75537373 | 19823 | 1 | ENSMUST00 | 0 | Snrpd3 | ENSMUSG0C | false | chr10 | 75516392 | 75518670 | 35.98183 | 119.73972 | 115.60696 | . | . | . | . | . | . |
| chr2 | 157201581 | 157201906 | 521 | Promoter (2-3kb) | 2 | 157145893 | 157204512 | 58620 | 2 | ENSMUST00 | 2606 | Rbl1 | ENSMUSG0C | false | . | . | . | . | . | . | chr2 | 157201228 | 157203990 | 3.63944 | 3.38149 | 1.91729 |
| chr8 | 48555832 | 48556183 | 521 | Promoter (<=1kb) | 8 | 48395436 | 48555846 | 160411 | 2 | ENSMUST00 | 0 | Tenm3 | ENSMUSG0C | true | chr8 | 48553777 | 48556246 | 31.85421 | 101.09849 | 97.49914 | . | . | . | . | . | . |
| chr9 | 101038700 | 101039040 | 521 | Distal Intergenic | 9 | 100982032 | 101034898 | 52867 | 2 | ENSMUST00 | -3803 | Pccb | ENSMUSG0C | false | . | . | . | . | . | . | chr9 | 101033076 | 101041068 | 4.72591 | 3.79913 | 2.80177 |
| chr13 | 48625143 | 48625785 | 521 | Promoter (<=1kb) | 13 | 48577873 | 48625573 | 47701 | 2 | ENSMUST00 | 0 | Ptpdc1 | ENSMUSG0C | false | chr13 | 48624908 | 48626279 | 30.60796 | 113.09483 | 109.17188 | chr13 | 48624448 | 48626514 | 3.2538 | 3.1337 | 1.58275 |
| chr6 | 146947259 | 146947521 | 521 | Intron (ENSMUST00000016631.8 | 6 | 146951302 | 146954414 | 3113 | 2 | ENSMUST00 | 6893 | 1700034J05F | ENSMUSG0C | false | . | . | . | . | . | . | chr6 | 146942477 | 146953557 | 4.42545 | 3.77491 | 2.55216 |
| chr4 | 8779391 | 8779628 | 521 | Intron (ENSMUST00000127476.2 | 4 | 8751438 | 8867224 | 115787 | 1 | ENSMUST00 | 27954 | Chd7 | ENSMUSG0C | false | . | . | . | . | . | . | chr4 | 8773919 | 8780117 | 5.69956 | 4.41813 | 3.58628 |
| chr19 | 7143855 | 7144236 | 521 | Intron (ENSMUST00000040261.5 | 19 | 7092014 | 7105729 | 13716 | 2 | ENSMUST00 | -38127 | Flrt1 | ENSMUSG0C | false | chr19 | 7143059 | 7144600 | 15.77044 | 42.5731 | 39.95169 | chr19 | 7127706 | 7162730 | 7.6862 | 5.1249 | 5.24023 |
| chr12 | 70789428 | 70789721 | 521 | Distal Intergenic | 12 | 70825514 | 70902232 | 76719 | 1 | ENSMUST00 | -35793 | Frmd6 | ENSMUSG0C | true | . | . | . | . | . | . | chr12 | 70782534 | 70795474 | 4.35327 | 3.81636 | 2.46879 |
| chr9 | 96436869 | 96437189 | 521 | Promoter (<=1kb) | 9 | 96435784 | 96437434 | 1651 | 2 | ENSMUST00 | 245 | BC043934 | ENSMUSG0C | false | chr9 | 96436926 | 96438702 | 17.10778 | 41.93702 | 39.32339 | chr9 | 96435849 | 96439222 | 5.69412 | 4.37663 | 3.58018 |
| chr7 | 29281200 | 29282093 | 521 | Promoter (<=1kb) | 7 | 29256331 | 29281977 | 25647 | 2 | ENSMUST00 | 0 | Spint2 | ENSMUSG0C | false | chr7 | 29280794 | 29282413 | 23.12601 | 77.64799 | 74.53377 | chr7 | 29279493 | 29282799 | 3.81879 | 3.51816 | 2.03217 |
| chr10 | 83871771 | 83872036 | 521 | Distal Intergenic | 10 | 83722865 | 83762761 | 39897 | 1 | ENSMUST00 | 148907 | 1500009L16F | ENSMUSG0C | false | . | . | . | . | . | . | . | . | . | . | . | . |
| chr6 | 18513476 | 18513779 | 520 | Promoter (<=1kb) | 6 | 18435251 | 18513091 | 77841 | 2 | ENSMUST00 | -386 | Cttnbp2 | ENSMUSG0C | true | chr6 | 18512890 | 18515216 | 24.24157 | 89.02564 | 85.70093 | . | . | . | . | . | . |
| chr15 | 103366512 | 103366920 | 520 | Promoter (<=1kb) | 15 | 103344289 | 103366763 | 22475 | 2 | ENSMUST00 | 0 | Itga5 | ENSMUSG0C | false | chr15 | 103365373 | 103367168 | 35.71972 | 120.33891 | 116.19067 | chr15 | 103365666 | 103367252 | 3.21354 | 3.02818 | 1.59458 |
| chrX | 68678382 | 68678828 | 520 | Promoter (<=1kb) | 20 | 68678541 | 68717963 | 39423 | 1 | ENSMUST00 | 0 | Fmr1 | ENSMUSG0C | false | chrX | 68678124 | 68679977 | 21.37461 | 59.55079 | 56.71094 | . | . | . | . | . | . |
| chr1 | 171251870 | 171252103 | 520 | Promoter (<=1kb) | 1 | 171252116 | 171260637 | 8522 | 1 | ENSMUST00 | -13 | Adamts4 | ENSMUSG0C | false | . | . | . | . | . | . | chr1 | 171247666 | 171252765 | 5.90391 | 4.39144 | 3.74682 |
| chr5 | 125393075 | 125393471 | 520 | Promoter (2-3kb) | 5 | 125385977 | 125390202 | 4226 | 2 | ENSMUST00 | -2874 | Ubc | ENSMUSG0C | false | chr5 | 125393312 | 125394148 | 8.76935 | 16.76631 | 14.47755 | chr5 | 125390597 | 125395558 | 6.07772 | 4.60901 | 3.9043 |
| chr4 | 144983884 | 144984327 | 520 | Promoter (<=1kb) | 4 | 144974926 | 144983961 | 9036 | 2 | ENSMUST00 | 0 | Vps13d | ENSMUSG0C | false | . | . | . | . | . | . | chr4 | 144980824 | 144994430 | 5.61677 | 4.30514 | 3.52155 |
| chr11 | 103101411 | 103101812 | 520 | Promoter (<=1kb) | 11 | 103070304 | 103101658 | 31355 | 2 | ENSMUST00 | 0 | Plcd3 | ENSMUSG0C | false | chr11 | 103100966 | 103102757 | 27.54716 | 98.2761 | 94.74957 | chr11 | 103094623 | 103106775 | 6.90747 | 4.97013 | 4.56297 |
| chr17 | 25539614 | 25539922 | 520 | Distal Intergenic | 17 | 25567988 | 25570214 | 2227 | 2 | ENSMUST00 | 30292 | Sox8 | ENSMUSG0C | false | . | . | . | . | . | . | chr17 | 25536949 | 25544974 | 5.26677 | 4.20031 | 3.22812 |
| chr17 | 72836218 | 72836702 | 520 | Promoter (<=1kb) | 17 | 72836704 | 72851195 | 14492 | 1 | ENSMUST00 | -2 | Ypel5 | ENSMUSG0C | false | chr17 | 72836567 | 72838139 | 37.40972 | 147.3676 | 142.18408 | . | . | . | . | . | . |
| chr6 | 84892956 | 84893584 | 520 | Intron (ENSMUST00000160197.2 | 6 | 84812240 | 84813247 | 1008 | 1 | ENSMUST00 | 80717 | Gm7507 | ENSMUSG0C | false | . | . | . | . | . | . | chr6 | 84891194 | 84895850 | 5.02343 | 4.12314 | 3.03363 |
| chr3 | 85341097 | 85341390 | 520 | Intron (ENSMUST00000154148.2 | 3 | 85360228 | 85361174 | 947 | 2 | ENSMUST00 | 19784 | RP23-94M23 | ENSMUSG0C | false | . | . | . | . | . | . | chr3 | 85340235 | 85341612 | 4.66482 | 3.97033 | 2.72027 |
| chr3 | 83538726 | 83538963 | 520 | Distal Intergenic | 3 | 83555363 | 83574122 | 18760 | 2 | ENSMUST00 | 35159 | RP23-24F12. | ENSMUSG0C | false | . | . | . | . | . | . | . | . | . | . | . | . |
| chr15 | 79402804 | 79403429 | 519 | Promoter (<=1kb) | 15 | 79360684 | 79402919 | 42236 | 2 | ENSMUST00 | 0 | Tmem184b | ENSMUSG0C | false | chr15 | 79402108 | 79404130 | 28.61382 | 119.33878 | 115.2169 | . | . | . | . | . | . |
| chr15 | 76870959 | 76871592 | 519 | Promoter (<=1kb) | 15 | 76852148 | 76871435 | 19288 | 2 | ENSMUST00 | 0 | Zfp251 | ENSMUSG0C | false | chr15 | 76870579 | 76871809 | 32.93275 | 99.9547 | 96.38725 | chr15 | 76870233 | 76872624 | 3.08157 | 2.98514 | 1.44971 |
| chr2 | 132008192 | 132008693 | 519 | Intron (ENSMUST00000028814.9 | 2 | 132000922 | 132021173 | 20252 | 2 | ENSMUST00 | 12480 | Rassf2 | ENSMUSG0C | false | . | . | . | . | . | . | chr2 | 132004737 | 132008762 | 5.71129 | 4.22155 | 3.6412 |
| chr2 | 180042025 | 180042918 | 519 | Promoter (<=1kb) | 2 | 180036374 | 180042395 | 6022 | 2 | ENSMUST00 | 0 | Psma7 | ENSMUSG0C | false | chr2 | 180041455 | 180043558 | 32.63288 | 132.66988 | 128.08362 | . | . | . | . | . | . |
| chr4 | 128730292 | 128730587 | 519 | Promoter (2-3kb) | 4 | 128727982 | 128747132 | 19151 | 1 | ENSMUST00 | 2311 | Phc2 | ENSMUSG0C | false | chr4 | 128727024 | 128730437 | 31.25623 | 96.05631 | 92.57859 | chr4 | 128728570 | 128744451 | 8.13086 | 5.39142 | 5.59738 |
| chr5 | 65537483 | 65537920 | 519 | Promoter (<=1kb) | 5 | 65537352 | 65595047 | 57696 | 1 | ENSMUST00 | 132 | Ube2k | ENSMUSG0C | false | chr5 | 65536471 | 65538620 | 34.36702 | 111.64683 | 107.76208 | . | . | . | . | . | . |
| chr9 | 63967928 | 63968241 | 519 | Intron (ENSMUST00000041029.5 | 9 | 63954009 | 64016987 | 62979 | 2 | ENSMUST00 | 48746 | Smad6 | ENSMUSG0C | true | . | . | . | . | . | . | chr9 | 63948352 | 64020083 | 7.27873 | 5.05859 | 4.874 |
| chr8 | 108697533 | 108697831 | 519 | Distal Intergenic | 8 | 108703100 | 108800270 | 97171 | 1 | ENSMUST00 | -5269 | Zfhx3 | ENSMUSG0C | false | . | . | . | . | . | . | chr8 | 108695243 | 108698085 | 4.08213 | 3.51654 | 2.26937 |
| chr17 | 84512507 | 84512985 | 519 | Promoter (<=1kb) | 17 | 84511895 | 84622142 | 110248 | 1 | ENSMUST00 | 613 | Plekhh2 | ENSMUSG0C | true | chr17 | 84511480 | 84513349 | 27.18312 | 111.36749 | 107.49936 | . | . | . | . | . | . |
| chr9 | 62477080 | 62477430 | 519 | Intron (ENSMUST00000048043.6 | 9 | 62429334 | 62449148 | 19815 | 2 | ENSMUST00 | -27933 | Coro2b | ENSMUSG0C | true | . | . | . | . | . | . | chr9 | 62449515 | 62535966 | 6.98492 | 4.90408 | 4.64549 |
| chr18 | 55491881 | 55492198 | 519 | Distal Intergenic | 18 | 55622636 | 55622920 | 285 | 2 | ENSMUST00 | 130722 | Gm24514 | ENSMUSG0C | false | . | . | . | . | . | . | chr18 | 55489229 | 55492678 | 5.22289 | 4.18788 | 3.18833 |
| chr13 | 8806019 | 8806291 | 518 | Intron (ENSMUST00000054251.7 | 13 | 8835389 | 8837234 | 1846 | 2 | ENSMUST00 | 30943 | Wdr37 | ENSMUSG0C | false | . | . | . | . | . | . | chr13 | 8805586 | 8806954 | 3.15577 | 3.10725 | 1.47942 |
| chr3 | 88409766 | 88410295 | 518 | Promoter (<=1kb) | 3 | 88394143 | 88410316 | 16174 | 2 | ENSMUST00 | 21 | Pmf1 | ENSMUSG0C | false | chr3 | 88409469 | 88410900 | 25.79081 | 75.126 | 72.05579 | chr3 | 88409112 | 88411240 | 5.55893 | 4.38885 | 3.45576 |
| chr7 | 84605794 | 84606336 | 518 | Promoter (<=1kb) | 7 | 84585159 | 84605827 | 20669 | 2 | ENSMUST00 | 0 | Fah | ENSMUSG0C | false | chr7 | 84605357 | 84606814 | 27.90323 | 81.55928 | 78.38181 | chr7 | 84604479 | 84605811 | 3.63589 | 3.35693 | 1.92118 |
| chr15 | 76817794 | 76818346 | 518 | Promoter (<=1kb) | 15 | 76723985 | 76818083 | 94099 | 2 | ENSMUST00 | 0 | Arhgap39 | ENSMUSG0C | false | chr15 | 76816817 | 76818846 | 35.20524 | 115.20684 | 111.21615 | . | . | . | . | . | . |
| chrX | 23692856 | 23693120 | 518 | Promoter (<=1kb) | 20 | 23693051 | 23805990 | 112940 | 1 | ENSMUST00 | 0 | Wdr44 | ENSMUSG0C | false | chrX | 23692413 | 23692873 | 11.4883 | 23.92001 | 21.53215 | . | . | . | . | . | . |
| chr3 | 97776356 | 97776719 | 518 | Intron (ENSMUST00000168438.3 | 3 | 97738854 | 97768009 | 29156 | 2 | ENSMUST00 | -8348 | Pde4dip | ENSMUSG0C | true | . | . | . | . | . | . | chr3 | 97774833 | 97776892 | 3.72704 | 3.38561 | 2.00201 |
| chr18 | 60613348 | 60613798 | 518 | Intron (ENSMUST00000097566.4 | 18 | 60600207 | 60610105 | 9899 | 2 | ENSMUST00 | -3244 | Synpo | ENSMUSG0C | true | chr18 | 60603340 | 60613557 | 27.39171 | 92.13462 | 88.74321 | chr18 | 60613738 | 60622693 | 7.427 | 5.19702 | 4.99228 |
| chr15 | 79962666 | 79962992 | 518 | Intron (ENSMUST00000128931.2 | 15 | 79918449 | 79971118 | 52670 | 2 | ENSMUST00 | 8126 | Cbx7 | ENSMUSG0C | false | . | . | . | . | . | . | chr15 | 79961315 | 79963390 | 4.45542 | 3.91484 | 2.56441 |
| chr5 | 134314224 | 134315066 | 518 | Promoter (<=1kb) | 5 | 134237834 | 134314760 | 76927 | 2 | ENSMUST00 | 0 | Gtf2i | ENSMUSG0C | false | chr5 | 134312869 | 134316109 | 31.91637 | 110.38313 | 106.53261 | . | . | . | . | . | . |
| chr10 | 20502385 | 20502617 | 518 | Intron (ENSMUST00000020165.8 | 10 | 20515327 | 20518164 | 2838 | 2 | ENSMUST00 | 15547 | 4930403O15 | ENSMUSG0C | false | . | . | . | . | . | . | chr10 | 20499070 | 20502586 | 3.65991 | 3.40415 | 1.93161 |
| chr19 | 60995377 | 60995630 | 518 | Intron (ENSMUST00000003313.8 | 19 | 61013417 | 61013545 | 129 | 2 | ENSMUST00 | 17915 | Gm22520 | ENSMUSG0C | false | . | . | . | . | . | . | chr19 | 60992715 | 60996128 | 4.55517 | 3.78005 | 2.65421 |
| chr1 | 88924131 | 88924429 | 518 | Distal Intergenic | 1 | 88932319 | 88933573 | 1255 | 1 | ENSMUST00 | -7890 | RP23-358L5. | ENSMUSG0C | false | . | . | . | . | . | . | chr1 | 88923773 | 88924959 | 5.62208 | 4.41346 | 3.5226 |
| chr10 | 93453241 | 93453495 | 517 | Promoter (<=1kb) | 10 | 93453396 | 93484896 | 31501 | 1 | ENSMUST00 | 0 | Lta4h | ENSMUSG0C | false | chr10 | 93452710 | 93454355 | 24.0356 | 94.23969 | 90.80066 | . | . | . | . | . | . |
| chr13 | 34593034 | 34593351 | 517 | Distal Intergenic | 13 | 34628868 | 34648568 | 19701 | 2 | ENSMUST00 | 55217 | Pxdc1 | ENSMUSG0C | true | . | . | . | . | . | . | chr13 | 34591635 | 34597051 | 4.59359 | 3.89674 | 2.66659 |
| chr13 | 95880591 | 95880865 | 517 | Intron (ENSMUST00000068603.6 | 13 | 95627177 | 95891922 | 264746 | 2 | ENSMUST00 | 11057 | Iqgap2 | ENSMUSG0C | true | . | . | . | . | . | . | chr13 | 95794831 | 95889694 | 5.09372 | 4.07642 | 3.0951 |
| chr19 | 20924000 | 20924360 | 517 | Intron (ENSMUST00000039500.3 | 19 | 20783458 | 20954202 | 170745 | 2 | ENSMUST00 | 29842 | Tmc1 | ENSMUSG0C | false | . | . | . | . | . | . | . | . | . | . | . | . |
| chr5 | 115508756 | 115509492 | 517 | Promoter (1-2kb) | 5 | 115506808 | 115554978 | 48171 | 1 | ENSMUST00 | 1949 | Pxn | ENSMUSG0C | false | chr5 | 115506647 | 115510205 | 34.92026 | 124.22972 | 119.94103 | chr5 | 115507551 | 115550291 | 7.76238 | 5.17462 | 5.28263 |
| chr5 | 140606919 | 140607455 | 517 | Promoter (<=1kb) | 5 | 140607341 | 140615543 | 8203 | 1 | ENSMUST00 | 0 | Lfng | ENSMUSG0C | false | chr5 | 140606439 | 140608441 | 21.17422 | 79.20749 | 76.06625 | chr5 | 140604016 | 140607247 | 5.52811 | 4.37672 | 3.42431 |
| chr7 | 114202911 | 114203240 | 517 | Distal Intergenic | 7 | 114215559 | 114254680 | 39122 | 2 | ENSMUST00 | 51440 | Copb1 | ENSMUSG0C | false | . | . | . | . | . | . | chr7 | 114199145 | 114205819 | 5.60331 | 4.36033 | 3.48713 |
| chr7 | 102242058 | 102242371 | 517 | Intron (ENSMUST00000106923.1 | 7 | 102235903 | 102237654 | 1752 | 1 | ENSMUST00 | 6156 | Pgap2 | ENSMUSG0C | false | . | . | . | . | . | . | chr7 | 102241861 | 102249556 | 6.7928 | 4.84705 | 4.48211 |
| chr16 | 92158969 | 92159465 | 517 | Distal Intergenic | 16 | 92106261 | 92112227 | 5967 | 1 | ENSMUST00 | 52709 | Mrps6 | ENSMUSG0C | false | chr16 | 92158227 | 92159042 | 13.18045 | 29.78312 | 27.32097 | chr16 | 92156045 | 92167743 | 5.46378 | 4.33054 | 3.39881 |
| chr11 | 61929982 | 61930479 | 517 | Promoter (<=1kb) | 11 | 61871307 | 61930252 | 58946 | 2 | ENSMUST00 | 0 | Akap10 | ENSMUSG0C | false | chr11 | 61929327 | 61931046 | 45.10679 | 157.97711 | 152.39525 | . | . | . | . | . | . |
| chr2 | 109960951 | 109961333 | 517 | Intron (ENSMUST00000046548.8 | 2 | 109975542 | 109997652 | 22111 | 1 | ENSMUST00 | -14209 | Lgr4 | ENSMUSG0C | false | . | . | . | . | . | . | chr2 | 109941684 | 109982422 | 4.64613 | 3.80604 | 2.74529 |
| chr6 | 47778348 | 47778612 | 517 | Distal Intergenic | 6 | 47781624 | 47781725 | 102 | 1 | ENSMUST00 | -3012 | Rny3 | ENSMUSG0C | false | chr6 | 47778086 | 47778940 | 4.1696 | 14.1759 | 11.92681 | chr6 | 47778031 | 47780892 | 4.24018 | 3.48371 | 2.39119 |
| chr8 | 23256931 | 23257297 | 516 | Promoter (<=1kb) | 8 | 23241353 | 23257021 | 15669 | 2 | ENSMUST00 | 0 | Golga7 | ENSMUSG0C | false | chr8 | 23256419 | 23257782 | 25.48472 | 95.2657 | 91.8051 | chr8 | 23255225 | 23257942 | 4.81943 | 3.95665 | 2.89826 |
| chr17 | 34627235 | 34627912 | 516 | Promoter (<=1kb) | 17 | 34616662 | 34627435 | 10774 | 2 | ENSMUST00 | 0 | Ppt2 | ENSMUSG0C | false | chr17 | 34626596 | 34628611 | 22.51707 | 59.83231 | 56.98677 | chr17 | 34626584 | 34627250 | 5.00542 | 4.02133 | 3.01636 |
| chr10 | 30600437 | 30600877 | 516 | Promoter (<=1kb) | 10 | 30534225 | 30600749 | 66525 | 2 | ENSMUST00 | 0 | Trmt11 | ENSMUSG0C | false | chr10 | 30599860 | 30601247 | 33.94421 | 102.70458 | 99.07153 | chr10 | 30598679 | 30601392 | 3.6885 | 3.38563 | 1.94237 |
| chr19 | 6046443 | 6047153 | 516 | Promoter (<=1kb) | 19 | 6046576 | 6053712 | 7137 | 1 | ENSMUST00 | 0 | Syvn1 | ENSMUSG0C | false | chr19 | 6045560 | 6048074 | 33.72937 | 104.36225 | 100.686 | . | . | . | . | . | . |
| chr8 | 76902103 | 76902439 | 516 | Promoter (<=1kb) | 8 | 76902288 | 77242996 | 340709 | 1 | ENSMUST00 | 0 | Nr3c2 | ENSMUSG0C | false | chr8 | 76898535 | 76903248 | 21.17422 | 79.20749 | 76.06625 | . | . | . | . | . | . |
| chr3 | 75956383 | 75957221 | 516 | Promoter (<=1kb) | 3 | 75876183 | 75956949 | 80767 | 2 | ENSMUST00 | 0 | Golim4 | ENSMUSG0C | true | chr3 | 75952519 | 75957408 | 32.38164 | 100.00523 | 96.43553 | . | . | . | . | . | . |
| chr11 | 58632981 | 58633341 | 516 | Distal Intergenic | 11 | 58625073 | 58626044 | 972 | 2 | ENSMUST00 | -6938 | Olfr323 | ENSMUSG0C | true | . | . | . | . | . | . | chr11 | 58629570 | 58636373 | 4.35692 | 3.76618 | 2.47571 |
| chr19 | 5490240 | 5490543 | 516 | Promoter (<=1kb) | 19 | 5490485 | 5494029 | 3545 | 1 | ENSMUST00 | 0 | Cfl1 | ENSMUSG0C | false | chr19 | 5490396 | 5492187 | 41.30354 | 154.61249 | 149.14572 | chr19 | 5486437 | 5490291 | 5.06458 | 3.96355 | 3.0584 |
| chr11 | 63889940 | 63890246 | 516 | Intron (ENSMUST00000094103.3 | 11 | 63885792 | 63922290 | 36499 | 2 | ENSMUST00 | 32044 | Hs3st3b1 | ENSMUSG0C | false | . | . | . | . | . | . | chr11 | 63886567 | 63903466 | 5.7674 | 4.29978 | 3.66511 |
| chr13 | 60456225 | 60456511 | 516 | Distal Intergenic | 13 | 60529296 | 60529409 | 114 | 2 | ENSMUST00 | 72898 | Gm25000 | ENSMUSG0C | false | . | . | . | . | . | . | chr13 | 60450423 | 60457848 | 4.93919 | 4.02175 | 2.96511 |
| chr10 | 127411415 | 127411716 | 516 | Intron (ENSMUST00000170164.2 | 10 | 127404284 | 127404392 | 109 | 2 | ENSMUST00 | -7024 | Gm23819 | ENSMUSG0C | false | . | . | . | . | . | . | chr10 | 127410292 | 127414240 | 4.93343 | 3.83661 | 2.98481 |
| chr9 | 114526627 | 114526901 | 516 | Intron (ENSMUST00000111816.2 | 9 | 114515697 | 114516424 | 728 | 1 | ENSMUST00 | 10931 | C130032M1C | ENSMUSG0C | false | . | . | . | . | . | . | chr9 | 114525892 | 114529998 | 4.57927 | 3.86564 | 2.65235 |
| chr15 | 99294816 | 99295399 | 515 | Promoter (<=1kb) | 15 | 99295087 | 99305965 | 10879 | 1 | ENSMUST00 | 0 | Prpf40b | ENSMUSG0C | false | chr15 | 99294285 | 99296025 | 22.03264 | 83.65932 | 80.44393 | . | . | . | . | . | . |
| chr9 | 96611812 | 96612138 | 515 | Intron (ENSMUST00000034984.7 | 9 | 96546282 | 96631536 | 85255 | 2 | ENSMUST00 | 19398 | Rasa2 | ENSMUSG0C | true | . | . | . | . | . | . | chr9 | 96609269 | 96616863 | 5.86264 | 4.45221 | 3.71429 |
| chr12 | 40496546 | 40496892 | 515 | Intron (ENSMUST00000037488.6 | 12 | 40446053 | 40846486 | 400434 | 1 | ENSMUST00 | 50494 | Dock4 | ENSMUSG0C | true | . | . | . | . | . | . | chr12 | 40494912 | 40528013 | 5.05369 | 4.02817 | 3.05886 |
| chr9 | 54560075 | 54560392 | 515 | Promoter (<=1kb) | 9 | 54544794 | 54560218 | 15425 | 2 | ENSMUST00 | 0 | Cib2 | ENSMUSG0C | false | chr9 | 54559677 | 54560474 | 20.09871 | 51.61829 | 48.88371 | chr9 | 54559651 | 54560982 | 3.5387 | 3.22295 | 1.84364 |
| chr13 | 20053278 | 20053538 | 515 | Distal Intergenic | 13 | 20090596 | 20264007 | 173412 | 1 | ENSMUST00 | -37058 | Elmo1 | ENSMUSG0C | true | . | . | . | . | . | . | chr13 | 20044769 | 20055516 | 4.57843 | 3.87081 | 2.66499 |
| chr2 | 70262478 | 70262781 | 515 | Intron (ENSMUST00000060208.5 | 2 | 70096298 | 70183267 | 86970 | 1 | ENSMUST00 | 166181 | Myo3b | ENSMUSG0C | false | . | . | . | . | . | . | chr2 | 70262430 | 70263147 | 4.57218 | 3.70095 | 2.6793 |
| chr8 | 33993050 | 33993300 | 515 | Distal Intergenic | 8 | 34049262 | 34055842 | 6581 | 1 | ENSMUST00 | -55962 | Gm9951 | ENSMUSG0C | false | . | . | . | . | . | . | chr8 | 33992829 | 33993904 | 6.04647 | 4.53285 | 3.87568 |
| chr19 | 46056828 | 46057350 | 515 | Promoter (<=1kb) | 19 | 46056832 | 46061458 | 4627 | 1 | ENSMUST00 | 0 | Pprc1 | ENSMUSG0C | false | chr19 | 46055978 | 46057815 | 37.13757 | 117.51956 | 113.45509 | . | . | . | . | . | . |
| chr7 | 4739995 | 4740363 | 515 | Promoter (<=1kb) | 7 | 4740115 | 4747514 | 7400 | 1 | ENSMUST00 | 0 | Suv420h2 | ENSMUSG0C | false | chr7 | 4740091 | 4741544 | 30.32912 | 88.589 | 85.27629 | . | . | . | . | . | . |
| chr9 | 48760557 | 48760827 | 515 | Intron (ENSMUST00000093852.3 | 9 | 48654311 | 48835945 | 181635 | 2 | ENSMUST00 | 75118 | Zbtb16 | ENSMUSG0C | false | . | . | . | . | . | . | chr9 | 48757527 | 48761450 | 4.85595 | 4.02925 | 2.87236 |
| chr5 | 30232255 | 30232914 | 515 | Promoter (<=1kb) | 5 | 30232581 | 30272012 | 39432 | 1 | ENSMUST00 | 0 | Ept1 | ENSMUSG0C | false | chr5 | 30231760 | 30233605 | 31.25476 | 101.05123 | 97.45738 | . | . | . | . | . | . |
| chr14 | 120374839 | 120375149 | 515 | Intron (ENSMUST00000088419.7 | 14 | 120384263 | 120388985 | 4723 | 2 | ENSMUST00 | 13836 | Gm26679 | ENSMUSG0C | false | . | . | . | . | . | . | chr14 | 120362275 | 120389547 | 5.43713 | 4.23022 | 3.37672 |
| chr11 | 70021162 | 70021697 | 514 | Promoter (2-3kb) | 11 | 70023905 | 70031131 | 7227 | 1 | ENSMUST00 | -2208 | Dlg4 | ENSMUSG0C | false | chr11 | 70020904 | 70021553 | 24.50127 | 70.18111 | 67.18839 | chr11 | 70017824 | 70022124 | 4.03083 | 3.44569 | 2.21537 |
| chr15 | 31367382 | 31367863 | 514 | Promoter (<=1kb) | 15 | 31291479 | 31367759 | 76281 | 2 | ENSMUST00 | 0 | Ankrd33b | ENSMUSG0C | false | chr15 | 31364914 | 31369386 | 33.42846 | 118.3354 | 114.24911 | . | . | . | . | . | . |
| chr2 | 25448255 | 25448599 | 514 | Promoter (2-3kb) | 2 | 25445843 | 25446476 | 634 | 1 | ENSMUST00 | 2413 | Abca2 | ENSMUSG0C | false | . | . | . | . | . | . | chr2 | 25446553 | 25456355 | 6.09426 | 4.46882 | 3.92693 |
| chr5 | 30850210 | 30850494 | 514 | Intron (ENSMUST00000031058.9 | 5 | 30853799 | 30863358 | 9560 | 1 | ENSMUST00 | -3305 | Mapre3 | ENSMUSG0C | false | . | . | . | . | . | . | chr5 | 30846495 | 30852222 | 5.09006 | 4.09018 | 3.13383 |
| chr9 | 59655974 | 59656754 | 514 | Promoter (<=1kb) | 9 | 59656368 | 59679375 | 23008 | 1 | ENSMUST00 | 0 | Pkm | ENSMUSG0C | false | chr9 | 59655687 | 59657702 | 28.91859 | 89.0447 | 85.71584 | . | . | . | . | . | . |
| chr19 | 10240595 | 10240876 | 514 | Promoter (<=1kb) | 19 | 10208272 | 10240748 | 32477 | 2 | ENSMUST00 | 0 | Myrf | ENSMUSG0C | false | chr19 | 10240363 | 10240950 | 13.83063 | 32.8443 | 30.34371 | chr19 | 10240152 | 10240693 | 4.01722 | 3.46363 | 2.23531 |
| chr15 | 97964087 | 97964547 | 514 | Promoter (<=1kb) | 15 | 97964229 | 97970286 | 6058 | 1 | ENSMUST00 | 0 | Tmem106c | ENSMUSG0C | false | chr15 | 97963406 | 97965116 | 29.28796 | 98.53321 | 94.99628 | . | . | . | . | . | . |
| chr12 | 104978955 | 104979374 | 514 | Intron (ENSMUST00000095439.4 | 12 | 104929949 | 104998677 | 68729 | 2 | ENSMUST00 | 19303 | Syne3 | ENSMUSG0C | false | . | . | . | . | . | . | chr12 | 104974830 | 104987771 | 6.02697 | 4.55969 | 3.85063 |
| chr15 | 33250223 | 33250583 | 514 | Intron (ENSMUST00000042167.9 | 15 | 33405206 | 33405939 | 734 | 2 | ENSMUST00 | 155356 | 1700084J12F | ENSMUSG0C | false | . | . | . | . | . | . | chr15 | 33249706 | 33252370 | 3.75675 | 3.48541 | 1.98565 |
| chr14 | 121181841 | 121182090 | 514 | Intron (ENSMUST00000135010.2 | 14 | 121248706 | 121251497 | 2792 | 1 | ENSMUST00 | -66616 | B930095G15 | ENSMUSG0C | false | . | . | . | . | . | . | chr14 | 121180394 | 121182900 | 6.66032 | 4.79565 | 4.39586 |
| chrX | 10734591 | 10734833 | 514 | Distal Intergenic | 20 | 10717473 | 10718710 | 1238 | 2 | ENSMUST00 | -15882 | Gm14493 | ENSMUSG0C | false | . | . | . | . | . | . | . | . | . | . | . | . |
| chr2 | 167819315 | 167819712 | 514 | Distal Intergenic | 2 | 167827693 | 167833646 | 5954 | 2 | ENSMUST00 | 13934 | 1200007C13I | ENSMUSG0C | false | . | . | . | . | . | . | chr2 | 167814397 | 167820487 | 6.24217 | 4.56491 | 4.01948 |
| chr7 | 110985915 | 110986224 | 513 | Distal Intergenic | 7 | 110868270 | 110982461 | 114192 | 2 | ENSMUST00 | -3455 | Mrvi1 | ENSMUSG0C | false | . | . | . | . | . | . | chr7 | 110982186 | 110991297 | 5.7251 | 4.32045 | 3.62307 |
| chr17 | 66519536 | 66519815 | 513 | Promoter (<=1kb) | 17 | 66494512 | 66519670 | 25159 | 2 | ENSMUST00 | 0 | Rab12 | ENSMUSG0C | false | chr17 | 66518531 | 66520403 | 29.58769 | 108.11006 | 104.32592 | . | . | . | . | . | . |
| chr17 | 31790518 | 31790940 | 513 | Distal Intergenic | 17 | 31844248 | 31855792 | 11545 | 2 | ENSMUST00 | 64852 | Sik1 | ENSMUSG0C | false | . | . | . | . | . | . | chr17 | 31789850 | 31791364 | 5.09376 | 4.16074 | 3.13359 |
| chr17 | 25064279 | 25064701 | 513 | Intron (ENSMUST00000137386.2 | 17 | 25057904 | 25078224 | 20321 | 2 | ENSMUST00 | 13523 | Tmem204 | ENSMUSG0C | false | . | . | . | . | . | . | chr17 | 25063881 | 25066178 | 3.40551 | 3.26436 | 1.75018 |
| chr18 | 3510349 | 3510730 | 513 | Promoter (2-3kb) | 18 | 3507957 | 3516404 | 8448 | 1 | ENSMUST00 | 2393 | Bambi | ENSMUSG0C | false | chr18 | 3507865 | 3510534 | 38.97723 | 131.47449 | 126.92584 | chr18 | 3509787 | 3518289 | 4.23988 | 3.50847 | 2.42348 |
| chr5 | 138994821 | 138995202 | 513 | Promoter (<=1kb) | 5 | 138976014 | 138995037 | 19024 | 2 | ENSMUST00 | 0 | Pdgfa | ENSMUSG0C | false | chr5 | 138992846 | 138996442 | 29.41006 | 85.09721 | 81.85307 | . | . | . | . | . | . |
| chr10 | 100487222 | 100487626 | 513 | Promoter (<=1kb) | 10 | 100443902 | 100487350 | 43449 | 2 | ENSMUST00 | 0 | Tmtc3 | ENSMUSG0C | false | chr10 | 100487513 | 100488167 | 19.89052 | 67.91473 | 64.9535 | . | . | . | . | . | . |
| chr15 | 31258335 | 31258633 | 513 | Intron (ENSMUST00000044524.1 | 15 | 31268336 | 31273386 | 5051 | 1 | ENSMUST00 | -9703 | Dap | ENSMUSG0C | false | . | . | . | . | . | . | chr15 | 31255870 | 31259449 | 4.67998 | 3.82382 | 2.77032 |
| chr1 | 126806590 | 126806824 | 513 | Intron (ENSMUST00000094609.4 | 1 | 126527540 | 126830481 | 302942 | 2 | ENSMUST00 | 23657 | Nckap5 | ENSMUSG0C | false | . | . | . | . | . | . | . | . | . | . | . | . |

| | | | | | | | | | | | | | | | | | | | | | | | | | | |
|---|---|---|---|---|---|---|---|---|---|---|---|---|---|---|---|---|---|---|---|---|---|---|---|---|---|---|
| chr10 | 13453428 | 13453666 | 513 | Intron (ENSMUST00000105547.2 | 10 | 13213395 | 13474396 | 261002 | 2 | ENSMUST00 | 20730 | Phactr2 | ENSMUSG0C | false | . | . | . | . | . | . | . | . | . | . | . | . |
| chr10 | 8762033 | 8762595 | 513 | Intron (ENSMUST00000015449.5 | 10 | 8824011 | 8824147 | 137 | 2 | ENSMUST00 | 61552 | Gm25410 | ENSMUSG0C | false | chr10 | 8761269 | 8762607 | 16.01923 | 40.07766 | 37.48569 | chr10 | 8756871 | 8763110 | 4.54887 | 3.87109 | 2.6311 |
| chr3 | 153541443 | 153541730 | 513 | Intron (ENSMUST00000064460.5 | 3 | 153621702 | 153621969 | 268 | 1 | ENSMUST00 | -79972 | Gm22136 | ENSMUSG0C | false | . | . | . | . | . | . | chr3 | 153539516 | 153542283 | 4.0922 | 3.58207 | 2.30981 |
| chr10 | 81168848 | 81169075 | 512 | Promoter (<=1kb) | 10 | 81153266 | 81167923 | 14658 | 2 | ENSMUST00 | -926 | Pias4 | ENSMUSG0C | false | chr10 | 81167982 | 81169233 | 17.64787 | 49.73517 | 47.02245 | . | . | . | . | . | . |
| chr12 | 85937495 | 85937775 | 512 | Promoter (1-2kb) | 12 | 85939389 | 86061893 | 122505 | 1 | ENSMUST00 | -1614 | Ttll5 | ENSMUSG0C | false | . | . | . | . | . | . | chr12 | 85936068 | 85938007 | 3.68052 | 3.42705 | 1.93988 |
| chr14 | 79652643 | 79652999 | 512 | Intron (ENSMUST00000165835.2 | 14 | 79641987 | 79645658 | 3672 | 2 | ENSMUST00 | -6986 | Lect1 | ENSMUSG0C | false | . | . | . | . | . | . | . | . | . | . | . | . |
| chr5 | 116024012 | 116024565 | 512 | Promoter (<=1kb) | 5 | 116013586 | 116024462 | 10877 | 2 | ENSMUST00 | 0 | Prkab1 | ENSMUSG0C | false | chr5 | 116023224 | 116024959 | 35.31382 | 118.55977 | 114.46947 | . | . | . | . | . | . |
| chr6 | 50412189 | 50412433 | 512 | Intron (ENSMUST00000071728.5 | 6 | 50352820 | 50397152 | 44333 | 2 | ENSMUST00 | -15038 | Osbpl3 | ENSMUSG0C | false | . | . | . | . | . | . | chr6 | 50412012 | 50412470 | 4.44029 | 3.69552 | 2.58166 |
| chr8 | 105264458 | 105264809 | 512 | Promoter (<=1kb) | 8 | 105258286 | 105264609 | 6324 | 2 | ENSMUST00 | 0 | Tradd | ENSMUSG0C | false | chr8 | 105263838 | 105265434 | 30.52716 | 100.76356 | 97.17578 | chr8 | 105263274 | 105265318 | 3.15856 | 3.14844 | 1.53544 |
| chr15 | 97247024 | 97248030 | 512 | Promoter (<=1kb) | 15 | 97247107 | 97385689 | 138583 | 1 | ENSMUST00 | 0 | Pced1b | ENSMUSG0C | true | chr15 | 97246325 | 97248785 | 46.1021 | 163.20312 | 157.4384 | . | . | . | . | . | . |
| chr14 | 105258517 | 105258904 | 512 | Promoter (<=1kb) | 14 | 105258573 | 105309298 | 50726 | 1 | ENSMUST00 | 0 | Ndfip2 | ENSMUSG0C | false | chr14 | 105258237 | 105259618 | 37.68163 | 117.40437 | 113.3432 | chr14 | 105258233 | 105258588 | 2.76962 | 2.7875 | 1.1946 |
| chr17 | 79835734 | 79836320 | 512 | Distal Intergenic | 17 | 79848392 | 79896028 | 47637 | 2 | ENSMUST00 | 59708 | Atl2 | ENSMUSG0C | false | . | . | . | . | . | . | chr17 | 79834456 | 79838336 | 4.56637 | 3.93485 | 2.63368 |
| chr9 | 90113689 | 90114439 | 512 | Promoter (<=1kb) | 9 | 90092402 | 90114289 | 21888 | 2 | ENSMUST00 | 0 | Morf4l1 | ENSMUSG0C | false | chr9 | 90112697 | 90115394 | 34.00618 | 119.38669 | 115.26049 | . | . | . | . | . | . |
| chr4 | 151411690 | 151411995 | 512 | Intron (ENSMUST00000169423.3 | 4 | 151376470 | 151379582 | 3113 | 1 | ENSMUST00 | 35221 | 4930589P08I | ENSMUSG0C | false | . | . | . | . | . | . | chr4 | 151410563 | 151412560 | 4.21329 | 3.74776 | 2.35407 |
| chr14 | 22147485 | 22147784 | 512 | Intron (ENSMUST00000075639.5 | 14 | 22021958 | 22022070 | 113 | 2 | ENSMUST00 | -125416 | Gm27532 | ENSMUSG0C | false | chr14 | 22147513 | 22148223 | 8.3822 | 16.12319 | 13.84377 | chr14 | 22143898 | 22155854 | 5.09229 | 4.04853 | 3.09435 |
| chr11 | 32642643 | 32643248 | 511 | Promoter (<=1kb) | 11 | 32642724 | 32746569 | 103846 | 1 | ENSMUST00 | 0 | Fbxw11 | ENSMUSG0C | false | chr11 | 32641800 | 32643992 | 33.41831 | 117.26761 | 113.21259 | . | . | . | . | . | . |
| chr13 | 63431074 | 63431857 | 511 | Promoter (<=1kb) | 13 | 63304709 | 63431745 | 127037 | 2 | ENSMUST00 | 0 | Fancc | ENSMUSG0C | false | chr13 | 63430637 | 63432256 | 32.8888 | 98.87808 | 95.33504 | . | . | . | . | . | . |
| chr17 | 80206662 | 80207476 | 511 | Promoter (<=1kb) | 17 | 80200089 | 80207305 | 7217 | 2 | ENSMUST00 | 0 | Srsf7 | ENSMUSG0C | false | chr17 | 80205681 | 80207946 | 36.04346 | 118.78738 | 114.68494 | . | . | . | . | . | . |
| chr1 | 131251703 | 131252060 | 511 | Downstream (2-3kb) | 1 | 131207654 | 131245258 | 37605 | 2 | ENSMUST00 | -6446 | Rassf5 | ENSMUSG0C | false | . | . | . | . | . | . | . | . | . | . | . | . |
| chr7 | 64839827 | 64840138 | 511 | Intron (ENSMUST00000119118.2 | 7 | 64819765 | 64856397 | 36633 | 2 | ENSMUST00 | 16259 | Fam189a1 | ENSMUSG0C | false | chr7 | 64839718 | 64840408 | 8.21555 | 15.38073 | 13.11249 | chr7 | 64839676 | 64840518 | 2.95443 | 2.92642 | 1.36248 |
| chr8 | 111769296 | 111769558 | 511 | Intron (ENSMUST00000034432.5 | 8 | 111710485 | 111743849 | 33365 | 2 | ENSMUST00 | -25448 | Bcar1 | ENSMUSG0C | false | . | . | . | . | . | . | chr8 | 111768244 | 111772378 | 4.83429 | 3.89484 | 2.8868 |
| chr9 | 44345597 | 44345863 | 511 | Promoter (1-2kb) | 9 | 44336350 | 44344228 | 7879 | 2 | ENSMUST00 | -1370 | Hmbs | ENSMUSG0C | false | . | . | . | . | . | . | chr9 | 44342001 | 44347571 | 4.64457 | 3.84053 | 2.74056 |
| chr1 | 54880070 | 54880327 | 511 | Intron (ENSMUST00000179030.2 | 1 | 54766792 | 54884711 | 117920 | 2 | ENSMUST00 | 4384 | Ankrd44 | ENSMUSG0C | true | . | . | . | . | . | . | chr1 | 54864456 | 54889599 | 5.98614 | 4.56539 | 3.81071 |
| chr4 | 117235952 | 117236263 | 511 | Intron (ENSMUST00000165128.2 | 4 | 117213333 | 117251917 | 38585 | 2 | ENSMUST00 | 15654 | Gm1661 | ENSMUSG0C | false | . | . | . | . | . | . | chr4 | 117235631 | 117236587 | 5.24451 | 4.20969 | 3.24052 |
| chr2 | 68767306 | 68767585 | 511 | Distal Intergenic | 2 | 68716291 | 68717310 | 1020 | 1 | ENSMUST00 | 51016 | 4932414N04 | ENSMUSG0C | false | . | . | . | . | . | . | chr2 | 68765665 | 68768170 | 3.81806 | 3.47174 | 2.06826 |
| chr2 | 159884840 | 159885289 | 511 | Distal Intergenic | 2 | 159858425 | 159859290 | 866 | 2 | ENSMUST00 | -25551 | Gm11446 | ENSMUSG0C | false | . | . | . | . | . | . | chr2 | 159883743 | 159890271 | 4.98798 | 4.05699 | 2.99637 |
| chr10 | 84343516 | 84343764 | 510 | Distal Intergenic | 10 | 84375296 | 84392388 | 17093 | 2 | ENSMUST00 | 48624 | Nuak1 | ENSMUSG0C | false | . | . | . | . | . | . | chr10 | 84338670 | 84343928 | 4.85563 | 4.03051 | 2.89324 |
| chr15 | 44753458 | 44753716 | 510 | Promoter (1-2kb) | 15 | 44671859 | 44752458 | 80600 | 2 | ENSMUST00 | -1001 | Sybu | ENSMUSG0C | false | . | . | . | . | . | . | chr15 | 44753226 | 44753962 | 3.51621 | 3.40819 | 1.75916 |
| chr19 | 4302311 | 4302620 | 510 | Promoter (2-3kb) | 19 | 4292567 | 4305208 | 12642 | 2 | ENSMUST00 | 2588 | Adrbk1 | ENSMUSG0C | false | . | . | . | . | . | . | chr19 | 4293718 | 4307011 | 5.16171 | 4.06783 | 3.17109 |
| chr6 | 31551056 | 31551408 | 510 | Intron (ENSMUST00000026698.7 | 6 | 31519488 | 31563981 | 44494 | 2 | ENSMUST00 | 12573 | Podxl | ENSMUSG0C | true | chr6 | 31549758 | 31552254 | 15.59599 | 36.52319 | 33.97784 | chr6 | 31513503 | 31559838 | 6.27295 | 4.62488 | 4.05806 |
| chr5 | 121584130 | 121584429 | 510 | Intron (ENSMUST00000031411.9 | 5 | 121571472 | 121575127 | 3656 | 2 | ENSMUST00 | -9004 | Aldh2 | ENSMUSG0C | false | . | . | . | . | . | . | chr5 | 121584189 | 121585442 | 5.16409 | 4.24258 | 3.14297 |
| chr9 | 85843249 | 85843951 | 510 | Promoter (<=1kb) | 9 | 85843361 | 85843925 | 565 | 1 | ENSMUST00 | 0 | Tpbg | ENSMUSG0C | false | chr9 | 85841971 | 85844734 | 22.44583 | 74.56482 | 71.50171 | . | . | . | . | . | . |
| chr19 | 22635154 | 22635456 | 510 | Intron (ENSMUST00000074770.6 | 19 | 22692619 | 22772372 | 79754 | 1 | ENSMUST00 | -57163 | Trpm3 | ENSMUSG0C | false | . | . | . | . | . | . | chr19 | 22634698 | 22637818 | 3.43274 | 3.25533 | 1.70895 |
| chr17 | 36042655 | 36042880 | 510 | Promoter (<=1kb) | 17 | 36038407 | 36042690 | 4284 | 2 | ENSMUST00 | 0 | H2-T22 | ENSMUSG0C | false | chr17 | 36041824 | 36043783 | 13.52062 | 44.78815 | 42.13932 | chr17 | 36040880 | 36043900 | 4.46417 | 3.88437 | 2.56293 |
| chr5 | 66150814 | 66151133 | 510 | Promoter (<=1kb) | 5 | 66018602 | 66150898 | 132297 | 2 | ENSMUST00 | 0 | Rbm47 | ENSMUSG0C | false | chr5 | 66150356 | 66151714 | 12.43157 | 35.01519 | 32.48718 | chr5 | 66150291 | 66151792 | 6.64316 | 4.85961 | 4.37022 |
| chr8 | 114671225 | 114671546 | 510 | Intron (ENSMUST00000004756.8 | 8 | 114639546 | 114642114 | 2569 | 2 | ENSMUST00 | -29112 | Gm16117 | ENSMUSG0C | false | . | . | . | . | . | . | chr8 | 114670578 | 114672191 | 3.71628 | 3.41969 | 1.97433 |
| chr5 | 123138435 | 123138863 | 510 | Promoter (<=1kb) | 5 | 123133723 | 123138109 | 4387 | 2 | ENSMUST00 | -327 | AI480526 | ENSMUSG0C | false | chr5 | 123136690 | 123143966 | 31.66066 | 102.77801 | 99.14294 | chr5 | 123131739 | 123139661 | 6.68632 | 4.77641 | 4.39554 |
| chr2 | 172862417 | 172862747 | 510 | Promoter (1-2kb) | 2 | 172864264 | 172864341 | 78 | 2 | ENSMUST00 | 1594 | Gm22773 | ENSMUSG0C | false | chr2 | 172862620 | 172863294 | 11.94783 | 25.26529 | 22.85895 | chr2 | 172861503 | 172865471 | 5.03829 | 4.03624 | 3.05316 |
| chr7 | 101868188 | 101868541 | 509 | Promoter (<=1kb) | 7 | 101863961 | 101868667 | 4707 | 2 | ENSMUST00 | 126 | Folr1 | ENSMUSG0C | false | . | . | . | . | . | . | chr7 | 101866471 | 101873274 | 4.88927 | 3.90902 | 2.93549 |
| chr16 | 22679486 | 22679846 | 509 | Distal Intergenic | 16 | 22469222 | 22657221 | 188000 | 2 | ENSMUST00 | -22266 | Dgkg | ENSMUSG0C | false | . | . | . | . | . | . | chr16 | 22679363 | 22680308 | 4.17121 | 3.38865 | 2.37048 |
| chr14 | 8080203 | 8080545 | 509 | Promoter (<=1kb) | 14 | 8080367 | 8091834 | 11468 | 1 | ENSMUST00 | 0 | Rpp14 | ENSMUSG0C | false | chr14 | 8080276 | 8081204 | 28.91247 | 96.86193 | 93.36482 | chr14 | 8079960 | 8082079 | 3.696 | 3.29133 | 1.97601 |
| chr19 | 23758464 | 23759183 | 509 | Promoter (<=1kb) | 19 | 23758876 | 23949597 | 190722 | 1 | ENSMUST00 | 0 | Apba1 | ENSMUSG0C | false | chr19 | 23757870 | 23759680 | 26.72788 | 101.58999 | 97.97913 | . | . | . | . | . | . |
| chr2 | 44549406 | 44549633 | 509 | Distal Intergenic | 2 | 44565133 | 44571347 | 6215 | 2 | ENSMUST00 | 21714 | Gtdc1 | ENSMUSG0C | false | . | . | . | . | . | . | chr2 | 44548778 | 44550436 | 3.06145 | 3.04343 | 1.45164 |
| chr2 | 64058970 | 64059276 | 509 | Intron (ENSMUST00000131615.3 | 2 | 63980447 | 64097988 | 117542 | 2 | ENSMUST00 | 38712 | Fign | ENSMUSG0C | true | . | . | . | . | . | . | chr2 | 64057522 | 64063924 | 4.28164 | 3.61797 | 2.43443 |
| chr5 | 43573172 | 43573441 | 509 | Intron (ENSMUST00000167522.1 | 5 | 43549930 | 43557769 | 7840 | 2 | ENSMUST00 | -15404 | Gm15865 | ENSMUSG0C | false | chr5 | 43572231 | 43573230 | 7.30631 | 13.42958 | 11.19263 | chr5 | 43560201 | 43596005 | 4.90131 | 4.0072 | 2.93052 |
| chr11 | 75172171 | 75172547 | 509 | Promoter (<=1kb) | 11 | 75172560 | 75177633 | 5074 | 1 | ENSMUST00 | -13 | 1700016P03I | ENSMUSG0C | false | chr11 | 75172287 | 75174081 | 34.14626 | 110.2223 | 106.38018 | chr11 | 75165128 | 75174332 | 5.71251 | 4.38871 | 3.59566 |
| chr3 | 153699078 | 153699382 | 509 | Intron (ENSMUST00000064460.5 | 3 | 153708911 | 153709026 | 116 | 1 | ENSMUST00 | -9529 | Gm22206 | ENSMUSG0C | false | . | . | . | . | . | . | chr3 | 153686316 | 153712771 | 6.43925 | 4.66349 | 4.19773 |
| chr18 | 66510286 | 66510571 | 509 | Distal Intergenic | 18 | 66549935 | 66550045 | 111 | 2 | ENSMUST00 | 39474 | Gm24679 | ENSMUSG0C | false | chr18 | 66510165 | 66510763 | 6.62775 | 15.27588 | 13.00958 | chr18 | 66509464 | 66511703 | 4.95915 | 4.07473 | 2.98048 |
| chr11 | 69122134 | 69122872 | 509 | Promoter (<=1kb) | 11 | 69120054 | 69122589 | 2536 | 2 | ENSMUST00 | 0 | 9130213A22I | ENSMUSG0C | false | chr11 | 69121276 | 69124643 | 27.76267 | 85.10065 | 81.85645 | . | . | . | . | . | . |
| chr1 | 159388568 | 159388825 | 509 | Distal Intergenic | 1 | 159358509 | 159358903 | 395 | 1 | ENSMUST00 | 30060 | RP23-145H3. | ENSMUSG0C | false | . | . | . | . | . | . | chr1 | 159385051 | 159390744 | 5.42733 | 4.2776 | 3.36708 |
| chr16 | 17807332 | 17807734 | 508 | 3' UTR | 16 | 17797282 | 17808287 | 11006 | 1 | ENSMUST00 | 10051 | Scarf2 | ENSMUSG0C | false | chr16 | 17806821 | 17807977 | 21.53151 | 64.09126 | 61.18795 | chr16 | 17806594 | 17808020 | 4.81502 | 3.8603 | 2.88707 |
| chr12 | 110551688 | 110551994 | 508 | Intron (ENSMUST00000084985.4 | 12 | 110601452 | 110666945 | 65494 | 1 | ENSMUST00 | -49458 | Dync1h1 | ENSMUSG0C | false | . | . | . | . | . | . | chr12 | 110549007 | 110557689 | 5.99039 | 4.4389 | 3.83972 |
| chr11 | 116306586 | 116306886 | 508 | Promoter (<=1kb) | 11 | 116288001 | 116306734 | 18734 | 2 | ENSMUST00 | 0 | Exoc7 | ENSMUSG0C | false | chr11 | 116305985 | 116307207 | 21.22555 | 68.73992 | 65.76938 | chr11 | 116305604 | 116307401 | 3.47826 | 3.10086 | 1.78045 |
| chr16 | 92461465 | 92461711 | 508 | Intron (ENSMUST00000060005.9 | 16 | 92391953 | 92466146 | 74194 | 2 | ENSMUST00 | 4435 | Rcan1 | ENSMUSG0C | false | . | . | . | . | . | . | chr16 | 92460879 | 92467402 | 4.41545 | 3.74426 | 2.54063 |
| chr16 | 92697204 | 92697688 | 508 | Promoter (<=1kb) | 16 | 92603656 | 92697328 | 93673 | 2 | ENSMUST00 | 0 | Runx1 | ENSMUSG0C | false | chr16 | 92696643 | 92698812 | 27.94004 | 83.5095 | 80.29546 | chr16 | 92695451 | 92700248 | 5.19566 | 4.17031 | 3.16375 |
| chr5 | 134456542 | 134457020 | 508 | Promoter (<=1kb) | 5 | 134357669 | 134456716 | 99048 | 2 | ENSMUST00 | 0 | Gtf2ird1 | ENSMUSG0C | false | chr5 | 134455170 | 134457151 | 22.8003 | 62.86544 | 59.97937 | chr5 | 134455124 | 134457274 | 3.821 | 3.40331 | 2.07999 |
| chr18 | 61092064 | 61092288 | 508 | Distal Intergenic | 18 | 61105572 | 61131134 | 25563 | 1 | ENSMUST00 | -13284 | Csf1r | ENSMUSG0C | false | . | . | . | . | . | . | chr18 | 61091660 | 61093370 | 3.58285 | 3.37599 | 1.82499 |
| chr5 | 3927913 | 3928692 | 508 | Promoter (<=1kb) | 5 | 3928054 | 3954765 | 26712 | 1 | ENSMUST00 | 0 | Akap9 | ENSMUSG0C | false | chr5 | 3927741 | 3929337 | 29.28796 | 98.53321 | 94.99628 | chr5 | 3927007 | 3927972 | 3.27541 | 3.22199 | 1.57033 |
| chr7 | 107496962 | 107497209 | 508 | Intron (ENSMUST00000130414.2 | 7 | 107504062 | 107548653 | 44592 | 1 | ENSMUST00 | -6853 | Syt9 | ENSMUSG0C | false | . | . | . | . | . | . | chr7 | 107496160 | 107499145 | 5.3088 | 4.18531 | 3.25862 |
| chr14 | 33759606 | 33759939 | 508 | Distal Intergenic | 14 | 33644495 | 33644601 | 107 | 1 | ENSMUST00 | 115112 | Gm26228 | ENSMUSG0C | false | . | . | . | . | . | . | . | . | . | . | . | . |
| chr19 | 8929314 | 8930086 | 508 | Promoter (<=1kb) | 19 | 8929694 | 8941563 | 11870 | 1 | ENSMUST00 | 0 | Eml3 | ENSMUSG0C | false | chr19 | 8929002 | 8930617 | 34.20045 | 107.22294 | 103.46436 | . | . | . | . | . | . |
| chr4 | 114835167 | 114835552 | 508 | Intron (ENSMUST00000137570.1 | 4 | 114844985 | 114845117 | 133 | 2 | ENSMUST00 | 9565 | Gm23230 | ENSMUSG0C | false | . | . | . | . | . | . | chr4 | 114833615 | 114836654 | 4.86379 | 3.84437 | 2.93493 |
| chr2 | 122702420 | 122702990 | 507 | Promoter (<=1kb) | 2 | 122681233 | 122702639 | 21407 | 2 | ENSMUST00 | 0 | Slc30a4 | ENSMUSG0C | false | chr2 | 122701481 | 122703325 | 25.90858 | 83.69981 | 80.48042 | . | . | . | . | . | . |
| chr10 | 7212031 | 7212438 | 507 | Promoter (<=1kb) | 10 | 7119063 | 7212237 | 93175 | 2 | ENSMUST00 | 0 | Cnksr3 | ENSMUSG0C | false | chr10 | 7210033 | 7213358 | 40.23456 | 136.98251 | 132.21786 | . | . | . | . | . | . |
| chr16 | 92465929 | 92466460 | 507 | Promoter (<=1kb) | 16 | 92391953 | 92466146 | 74194 | 2 | ENSMUST00 | 0 | Rcan1 | ENSMUSG0C | false | chr16 | 92465196 | 92466250 | 33.87313 | 113.90014 | 109.94483 | chr16 | 92460879 | 92467402 | 4.41545 | 3.74426 | 2.54063 |
| chr3 | 107680777 | 107681063 | 507 | Intron (ENSMUST00000029490.9 | 3 | 107663118 | 107677767 | 14650 | 2 | ENSMUST00 | -3011 | Ahcyl1 | ENSMUSG0C | false | . | . | . | . | . | . | chr3 | 107679772 | 107682866 | 4.47347 | 3.83422 | 2.5769 |
| chr4 | 125109832 | 125110152 | 507 | Promoter (1-2kb) | 4 | 125107976 | 125109428 | 1453 | 1 | ENSMUST00 | 1857 | Meaf6 | ENSMUSG0C | false | . | . | . | . | . | . | chr4 | 125106469 | 125127334 | 7.64816 | 5.22391 | 5.18478 |
| chr18 | 80712712 | 80713308 | 507 | Promoter (<=1kb) | 18 | 80606205 | 80713071 | 106867 | 2 | ENSMUST00 | 0 | Nfatc1 | ENSMUSG0C | false | chr18 | 80712341 | 80713735 | 25.98482 | 77.30067 | 74.19341 | chr18 | 80711836 | 80715689 | 5.07579 | 4.0937 | 3.06953 |
| chr3 | 118500307 | 118500577 | 507 | Intron (ENSMUST00000180593.1 | 3 | 118562129 | 119432918 | 870790 | 1 | ENSMUST00 | -61552 | Dpyd | ENSMUSG0C | false | . | . | . | . | . | . | chr3 | 118499352 | 118500488 | 4.26546 | 3.60771 | 2.4365 |
| chr4 | 132199069 | 132199347 | 507 | Intron (ENSMUST00000152796.2 | 4 | 132186688 | 132210809 | 24122 | 2 | ENSMUST00 | 11462 | Ythdf2 | ENSMUSG0C | false | . | . | . | . | . | . | chr4 | 132192918 | 132204338 | 4.45686 | 3.69575 | 2.58827 |
| chr14 | 70159344 | 70159613 | 507 | Promoter (<=1kb) | 14 | 70154405 | 70159502 | 5098 | 2 | ENSMUST00 | 0 | 9930012K11I | ENSMUSG0C | false | chr14 | 70158320 | 70159598 | 21.17422 | 79.20749 | 76.06625 | chr14 | 70156516 | 70175803 | 7.12683 | 5 | 4.75261 |
| chr2 | 30601003 | 30601558 | 507 | Intron (ENSMUST00000102853.3 | 2 | 30605217 | 30608217 | 3001 | 1 | ENSMUST00 | -3659 | Cstad | ENSMUSG0C | false | . | . | . | . | . | . | chr2 | 30600055 | 30602050 | 5.57081 | 4.2646 | 3.50515 |
| chr1 | 133471072 | 133471354 | 507 | Distal Intergenic | 1 | 133382303 | 133424377 | 42075 | 2 | ENSMUST00 | -46696 | Sox13 | ENSMUSG0C | false | . | . | . | . | . | . | chr1 | 133468626 | 133471492 | 4.0268 | 3.49346 | 2.23237 |
| chr10 | 19933612 | 19934973 | 507 | Promoter (<=1kb) | 10 | 19934472 | 20142753 | 208282 | 1 | ENSMUST00 | 0 | Map3k5 | ENSMUSG0C | true | chr10 | 19933409 | 19936698 | 31.91637 | 110.38313 | 106.53261 | chr10 | 19933272 | 19933701 | 3.00098 | 2.94904 | 1.35513 |
| chr16 | 18383473 | 18384118 | 506 | Promoter (2-3kb) | 16 | 18380783 | 18406747 | 25965 | 1 | ENSMUST00 | 2691 | Arvcf | ENSMUSG0C | true | . | . | . | . | . | . | chr16 | 18376326 | 18392535 | 8.53722 | 5.66709 | 5.90216 |
| chr3 | 19978548 | 19978958 | 506 | Promoter (1-2kb) | 3 | 19980628 | 19987396 | 6769 | 1 | ENSMUST00 | -1670 | Cp | ENSMUSG0C | false | . | . | . | . | . | . | . | . | . | . | . | . |
| chr1 | 180248788 | 180249096 | 506 | Promoter (<=1kb) | 1 | 180227443 | 180248150 | 20708 | 2 | ENSMUST00 | -639 | Psen2 | ENSMUSG0C | false | . | . | . | . | . | . | . | . | . | . | . | . |
| chr13 | 55513081 | 55513565 | 506 | Promoter (<=1kb) | 13 | 55495795 | 55513438 | 17644 | 2 | ENSMUST00 | 0 | Pdlim7 | ENSMUSG0C | false | chr13 | 55512304 | 55513824 | 23.70827 | 104.82465 | 101.13002 | chr13 | 55510813 | 55514179 | 4.25978 | 3.54681 | 2.42402 |
| chr5 | 72846302 | 72846609 | 506 | Intron (ENSMUST00000071944.7 | 5 | 72755734 | 72868396 | 112663 | 2 | ENSMUST00 | 21787 | Tec | ENSMUSG0C | false | . | . | . | . | . | . | chr5 | 72845283 | 72847474 | 3.9996 | 3.46967 | 2.22189 |
| chr5 | 123523743 | 123524283 | 506 | Promoter (<=1kb) | 5 | 123509765 | 123524143 | 14379 | 2 | ENSMUST00 | 0 | Diablo | ENSMUSG0C | false | chr5 | 123523316 | 123524882 | 37.85821 | 131.67731 | 127.12434 | . | . | . | . | . | . |
| chr6 | 90463481 | 90463886 | 506 | Promoter (<=1kb) | 6 | 90462576 | 90475238 | 12663 | 1 | ENSMUST00 | 906 | Klf15 | ENSMUSG0C | false | chr6 | 90462123 | 90463911 | 27.3492 | 80.60155 | 77.44116 | chr6 | 90462103 | 90464387 | 3.24764 | 3.18149 | 1.56356 |
| chr5 | 124993113 | 124993384 | 506 | Distal Intergenic | 5 | 125003447 | 125012547 | 9101 | 1 | ENSMUST00 | -10063 | Fam101a | ENSMUSG0C | false | . | . | . | . | . | . | . | . | . | . | . | . |
| chr2 | 18028363 | 18029150 | 506 | Promoter (<=1kb) | 2 | 18027926 | 18028760 | 835 | 2 | ENSMUST00 | 0 | A930004D18 | ENSMUSG0C | false | chr2 | 18025808 | 18029357 | 29.56612 | 91.1473 | 87.77953 | . | . | . | . | . | . |
| chr14 | 33751032 | 33751376 | 506 | Distal Intergenic | 14 | 33644495 | 33644601 | 107 | 1 | ENSMUST00 | 106538 | Gm26228 | ENSMUSG0C | false | . | . | . | . | . | . | . | . | . | . | . | . |
| chr3 | 9752091 | 9752554 | 506 | 5' UTR | 3 | 9757607 | 9786142 | 28536 | 1 | ENSMUST00 | -5053 | Gm16337 | ENSMUSG0C | false | . | . | . | . | . | . | chr3 | 9749552 | 9777361 | 5.50618 | 4.27196 | 3.42484 |
| chr9 | 117962136 | 117962416 | 506 | Intron (ENSMUST00000111769.4 | 9 | 117999336 | 117999430 | 95 | 1 | ENSMUST00 | -36920 | Gm22008 | ENSMUSG0C | false | . | . | . | . | . | . | chr9 | 117962192 | 117962829 | 2.62024 | 2.68913 | 1.09925 |
| chr6 | 83417606 | 83417849 | 506 | Intron (ENSMUST00000089622.5 | 6 | 83400996 | 83401102 | 107 | 2 | ENSMUST00 | -16505 | Mir6374 | ENSMUSG0C | false | . | . | . | . | . | . | chr6 | 83405184 | 83419689 | 6.75589 | 4.83654 | 4.47557 |
| chr3 | 88140585 | 88141096 | 505 | Promoter (1-2kb) | 3 | 88142372 | 88172086 | 29715 | 1 | ENSMUST00 | -1276 | Mef2d | ENSMUSG0C | false | . | . | . | . | . | . | chr3 | 88136354 | 88141917 | 6.47517 | 4.6068 | 4.24981 |
| chr11 | 101078982 | 101079245 | 505 | Promoter (<=1kb) | 11 | 101078411 | 101080527 | 2117 | 1 | ENSMUST00 | 572 | Hsd17b1 | ENSMUSG0C | false | . | . | . | . | . | . | chr11 | 101078620 | 101084538 | 5.0744 | 3.94453 | 3.10504 |
| chr16 | 33939775 | 33940043 | 505 | Intron (ENSMUST00000115028.4 | 16 | 33922486 | 33944148 | 21663 | 1 | ENSMUST00 | 17290 | Itgb5 | ENSMUSG0C | true | . | . | . | . | . | . | chr16 | 33911227 | 33952498 | 7.40288 | 5.1092 | 4.9863 |
| chr1 | 14422954 | 14423285 | 505 | Distal Intergenic | 1 | 14168958 | 14310235 | 141278 | 2 | ENSMUST00 | -112720 | Eya1 | ENSMUSG0C | false | . | . | . | . | . | . | chr1 | 14422329 | 14424674 | 4.67044 | 3.84747 | 2.73588 |
| chr4 | 117844524 | 117844839 | 505 | Intron (ENSMUST00000063857.5 | 4 | 117849193 | 117864004 | 14812 | 1 | ENSMUST00 | -4354 | Slc6a9 | ENSMUSG0C | false | . | . | . | . | . | . | chr4 | 117844064 | 117845734 | 5.69112 | 4.28853 | 3.62142 |
| chr8 | 79399016 | 79399615 | 505 | Promoter (<=1kb) | 8 | 79338395 | 79399518 | 61124 | 2 | ENSMUST00 | 0 | Smad1 | ENSMUSG0C | false | chr8 | 79398146 | 79399848 | 34.36702 | 111.64683 | 107.76208 | . | . | . | . | . | . |
| chr2 | 134644034 | 134644289 | 505 | Promoter (<=1kb) | 2 | 134594185 | 134644145 | 49961 | 2 | ENSMUST00 | 0 | Tmx4 | ENSMUSG0C | false | chr2 | 134643280 | 134644493 | 32.68561 | 114.88856 | 110.91166 | chr2 | 134643034 | 134644564 | 3.41001 | 3.26746 | 1.6897 |
| chr2 | 77280157 | 77280831 | 505 | Promoter (<=1kb) | 2 | 77186555 | 77280592 | 94038 | 2 | ENSMUST00 | 0 | Sestd1 | ENSMUSG0C | false | chr2 | 77279943 | 77281168 | 27.03867 | 103.18376 | 99.53125 | chr2 | 77279730 | 77281414 | 3.40709 | 3.2239 | 1.68782 |
| chr10 | 41937225 | 41937525 | 505 | Intron (ENSMUST00000160262.3 | 10 | 41916051 | 41976122 | 60072 | 2 | ENSMUST00 | 38597 | Armc2 | ENSMUSG0C | false | . | . | . | . | . | . | . | . | . | . | . | . |
| chr4 | 141278066 | 141278526 | 505 | Promoter (<=1kb) | 4 | 141278439 | 141279794 | 1356 | 1 | ENSMUST00 | 0 | Gm13056 | ENSMUSG0C | false | . | . | . | . | . | . | chr4 | 141277124 | 141282934 | 7.25702 | 5.14963 | 4.88042 |
| chr18 | 47020725 | 47020954 | 505 | Intron (ENSMUST00000049388.7 | 18 | 46985337 | 46985481 | 145 | 2 | ENSMUST00 | -35245 | Gm22791 | ENSMUSG0C | false | . | . | . | . | . | . | chr18 | 47020041 | 47021508 | 2.77511 | 2.80818 | 1.23359 |
| chr13 | 63337545 | 63337787 | 505 | Intron (ENSMUST00000160278.1 | 13 | 63333210 | 63339956 | 6747 | 1 | ENSMUST00 | 4336 | Gm16133 | ENSMUSG0C | false | . | . | . | . | . | . | chr13 | 63336212 | 63338146 | 6.30534 | 4.72354 | 4.08251 |
| chr11 | 100574822 | 100575189 | 504 | Promoter (<=1kb) | 11 | 100574904 | 100581728 | 6825 | 1 | ENSMUST00 | 0 | Cnp | ENSMUSG0C | false | chr11 | 100574589 | 100575730 | 29.66345 | 100.21 | 96.63225 | chr11 | 100573999 | 100578648 | 4.20396 | 3.70063 | 2.36294 |
| chr17 | 34981643 | 34982215 | 504 | Promoter (<=1kb) | 17 | 34981862 | 34983981 | 2120 | 1 | ENSMUST00 | 0 | Lsm2 | ENSMUSG0C | false | chr17 | 34980890 | 34982714 | 40.37263 | 133.9167 | 129.27921 | chr17 | 34980535 | 34982794 | 4.67382 | 3.86061 | 2.76662 |
| chr16 | 17798454 | 17798797 | 504 | Promoter (1-2kb) | 16 | 17797282 | 17808287 | 11006 | 1 | ENSMUST00 | 1173 | Scarf2 | ENSMUSG0C | false | chr16 | 17798066 | 17798750 | 11.32191 | 27.20368 | 24.77393 | chr16 | 17796984 | 17798959 | 4.83944 | 4.04607 | 2.87295 |
| chr10 | 81378335 | 81378883 | 504 | Promoter (<=1kb) | 10 | 81366222 | 81378516 | 12295 | 2 | ENSMUST00 | 0 | Fzr1 | ENSMUSG0C | false | chr10 | 81377526 | 81378535 | 31.01882 | 98.6291 | 95.09013 | . | . | . | . | . | . |
| chr11 | 106487226 | 106488089 | 504 | Promoter (<=1kb) | 11 | 106394650 | 106487796 | 93147 | 2 | ENSMUST00 | 0 | Ern1 | ENSMUSG0C | false | chr11 | 106486553 | 106488525 | 33.96572 | 122.48407 | 118.2566 | chr11 | 106485011 | 106490016 | 4.65073 | 3.93237 | 2.71964 |
| chr4 | 19560425 | 19560864 | 504 | Intron (ENSMUST00000029885.4 | 4 | 19549575 | 19570083 | 20509 | 2 | ENSMUST00 | 9219 | Cpne3 | ENSMUSG0C | false | . | . | . | . | . | . | chr4 | 19555426 | 19563093 | 4.43444 | 3.77536 | 2.54497 |
| chr9 | 110732682 | 110732957 | 504 | Intron (ENSMUST00000006005.7 | 9 | 110722088 | 110742610 | 20523 | 2 | ENSMUST00 | 9653 | Pth1r | ENSMUSG0C | true | . | . | . | . | . | . | chr9 | 110716295 | 110740574 | 6.49295 | 4.67204 | 4.24267 |
| chr5 | 81476032 | 81476295 | 504 | Intron (ENSMUST00000132375.2 | 5 | 81310025 | 81796634 | 486610 | 1 | ENSMUST00 | 166008 | Lphn3 | ENSMUSG0C | false | . | . | . | . | . | . | chr5 | 81474428 | 81476654 | 3.62152 | 3.36664 | 1.9254 |
| chr14 | 21988326 | 21988573 | 504 | Promoter (1-2kb) | 14 | 21983962 | 21989601 | 5640 | 2 | ENSMUST00 | 1028 | Zfp503 | ENSMUSG0C | false | chr14 | 21988023 | 21988467 | 22.21283 | 62.70797 | 59.8237 | . | . | . | . | . | . |
| chr4 | 45487753 | 45488014 | 504 | Promoter (1-2kb) | 4 | 45447506 | 45489403 | 41898 | 2 | ENSMUST00 | 1389 | Shb | ENSMUSG0C | false | . | . | . | . | . | . | chr4 | 45486900 | 45516085 | 7.16373 | 5.03069 | 4.78278 |
| chr5 | 105708529 | 105708772 | 504 | Intron (ENSMUST00000060531.1 | 5 | 105700866 | 105734238 | 33373 | 1 | ENSMUST00 | 7664 | Lrrc8d | ENSMUSG0C | false | . | . | . | . | . | . | chr5 | 105706707 | 105712616 | 3.71773 | 3.40962 | 1.97711 |
| chr9 | 70848234 | 70848789 | 504 | Intron (ENSMUST00000034731.8 | 9 | 70852576 | 70853418 | 843 | 1 | ENSMUST00 | -3787 | Gm3436 | ENSMUSG0C | false | . | . | . | . | . | . | chr9 | 70847798 | 70849224 | 5.89747 | 4.5995 | 3.74662 |
| chr16 | 18385473 | 18385723 | 503 | Intron (ENSMUST00000115614.3 | 16 | 18380783 | 18406747 | 25965 | 1 | ENSMUST00 | 4691 | Arvcf | ENSMUSG0C | true | . | . | . | . | . | . | chr16 | 18376326 | 18392535 | 8.53722 | 5.66709 | 5.90216 |
| chr11 | 100860136 | 100860578 | 503 | Promoter (<=1kb) | 11 | 100860345 | 100863220 | 2876 | 1 | ENSMUST00 | 0 | Stat5a | ENSMUSG0C | false | chr11 | 100859990 | 100861109 | 30.59503 | 95.89513 | 92.42037 | chr11 | 100859205 | 100861210 | 5.87311 | 4.54401 | 3.71776 |
| chr12 | 78861504 | 78861873 | 503 | Promoter (<=1kb) | 12 | 78842989 | 78861638 | 18650 | 2 | ENSMUST00 | 0 | Atp6v1d | ENSMUSG0C | false | chr12 | 78860566 | 78861640 | 27.20707 | 96.65539 | 93.16586 | . | . | . | . | . | . |
| chr14 | 31250958 | 31251592 | 503 | Promoter (<=1kb) | 14 | 31251450 | 31259944 | 8495 | 1 | ENSMUST00 | 0 | Bap1 | ENSMUSG0C | false | chr14 | 31250459 | 31252425 | 37.17318 | 134.81126 | 130.13722 | . | . | . | . | . | . |
| chr1 | 82200023 | 82200331 | 503 | Distal Intergenic | 1 | 82233112 | 82235933 | 2822 | 1 | ENSMUST00 | -32781 | Gm9747 | ENSMUSG0C | false | . | . | . | . | . | . | chr1 | 82199433 | 82203109 | 5.59685 | 4.46954 | 3.48989 |
| chr17 | 14945180 | 14945442 | 503 | Promoter (1-2kb) | 17 | 14945014 | 14946828 | 1815 | 2 | ENSMUST00 | 1386 | Phf10 | ENSMUSG0C | false | . | . | . | . | . | . | chr17 | 14942333 | 14946656 | 5.02172 | 4.01492 | 3.06428 |
| chr18 | 7868494 | 7870108 | 503 | Promoter (<=1kb) | 18 | 7868832 | 7926505 | 57674 | 1 | ENSMUST00 | 0 | Wac | ENSMUSG0C | false | chr18 | 7867839 | 7871023 | 42.68155 | 140.14148 | 135.26276 | . | . | . | . | . | . |
| chr5 | 116419337 | 116419616 | 503 | Intron (ENSMUST00000036991.4 | 5 | 116408491 | 116422864 | 14374 | 2 | ENSMUST00 | 3248 | Hspb8 | ENSMUSG0C | false | chr5 | 116418999 | 116419736 | 6.75876 | 12.51203 | 10.2897 | chr5 | 116409713 | 116420816 | 4.92389 | 3.90715 | 2.96724 |
| chr14 | 119007326 | 119007791 | 503 | Promoter (<=1kb) | 14 | 118985039 | 119007465 | 22427 | 2 | ENSMUST00 | 0 | Uggt2 | ENSMUSG0C | false | chr14 | 119006416 | 119008128 | 28.91247 | 96.86193 | 93.36482 | . | . | . | . | . | . |
| chrX | 99455675 | 99455877 | 503 | Distal Intergenic | 20 | 99459025 | 99460753 | 1729 | 1 | ENSMUST00 | -3148 | Gm14808 | ENSMUSG0C | false | . | . | . | . | . | . | . | . | . | . | . | . |
| chr17 | 72970959 | 72971296 | 503 | Distal Intergenic | 17 | 72987798 | 72987904 | 107 | 1 | ENSMUST00 | -16502 | Gm23649 | ENSMUSG0C | false | . | . | . | . | . | . | chr17 | 72970900 | 72971294 | 3.31399 | 2.84234 | 1.65874 |
| chr12 | 80046645 | 80047035 | 503 | Distal Intergenic | 12 | 80108034 | 80113000 | 4967 | 1 | ENSMUST00 | -60999 | Gm26669 | ENSMUSG0C | false | . | . | . | . | . | . | chr12 | 80045840 | 80049290 | 5.13307 | 4.09674 | 3.16689 |
| chr10 | 93640822 | 93641408 | 502 | Promoter (<=1kb) | 10 | 93641049 | 93747207 | 106159 | 1 | ENSMUST00 | 0 | Ntn4 | ENSMUSG0C | false | chr10 | 93640662 | 93642462 | 27.78601 | 91.88203 | 88.49705 | chr10 | 93640774 | 93642332 | 4.66034 | 3.90377 | 2.77305 |
| chr12 | 8389544 | 8389786 | 502 | Distal Intergenic | 12 | 8313433 | 8343824 | 30392 | 1 | ENSMUST00 | 76112 | Hs1bp3 | ENSMUSG0C | false | . | . | . | . | . | . | chr12 | 8388849 | 8390530 | 4.49219 | 3.61123 | 2.62147 |
| chr13 | 43232199 | 43232801 | 502 | Intron (ENSMUST00000055341.5 | 13 | 43151744 | 43171501 | 19758 | 2 | ENSMUST00 | -60699 | Tbc1d7 | ENSMUSG0C | false | . | . | . | . | . | . | chr13 | 43225402 | 43235854 | 4.79198 | 3.91033 | 2.85262 |
| chr5 | 149152495 | 149152748 | 502 | Intron (ENSMUST00000180733.1 | 5 | 149155446 | 149156081 | 636 | 2 | ENSMUST00 | 3333 | Gm8615 | ENSMUSG0C | false | . | . | . | . | . | . | . | . | . | . | . | . |
| chr19 | 9074114 | 9074407 | 502 | Intron (ENSMUST00000092955.4 | 19 | 9083636 | 9087958 | 4323 | 2 | ENSMUST00 | 13551 | Scgb1a1 | ENSMUSG0C | false | . | . | . | . | . | . | . | . | . | . | . | . |
| chr17 | 42369886 | 42370161 | 502 | Intron (ENSMUST00000048691.4 | 17 | 42315947 | 42507741 | 191795 | 1 | ENSMUST00 | 53940 | Ptchd4 | ENSMUSG0C | true | . | . | . | . | . | . | chr17 | 42368910 | 42370821 | 4.48901 | 3.89276 | 2.57795 |
| chr5 | 122049741 | 122050285 | 502 | Promoter (<=1kb) | 5 | 121858387 | 122049851 | 191465 | 2 | ENSMUST00 | 0 | Cux2 | ENSMUSG0C | false | chr5 | 122049044 | 122050724 | 36.76257 | 113.72284 | 109.77938 | chr5 | 122048445 | 122051089 | 4.87397 | 3.97334 | 2.92777 |
| chrX | 102071008 | 102071437 | 502 | Promoter (<=1kb) | 20 | 102066544 | 102071196 | 4653 | 2 | ENSMUST00 | 0 | Rgag4 | ENSMUSG0C | false | chrX | 102070296 | 102071600 | 13.83063 | 32.8443 | 30.34371 | . | . | . | . | . | . |
| chr5 | 143548588 | 143548977 | 502 | Promoter (<=1kb) | 5 | 143548736 | 143564525 | 15820 | 1 | ENSMUST00 | 0 | Fam220a | ENSMUSG0C | false | chr5 | 143548668 | 143549562 | 32.03847 | 101.42104 | 97.81718 | chr5 | 143547420 | 143550242 | 5.28155 | 4.24135 | 3.23974 |
| chr5 | 136840349 | 136840636 | 502 | Intron (ENSMUST00000057497.7 | 5 | 136863637 | 136869722 | 6086 | 2 | ENSMUST00 | 29086 | Gm20485 | ENSMUSG0C | false | . | . | . | . | . | . | chr5 | 136840128 | 136841137 | 3.46658 | 3.15212 | 1.78662 |
| chr16 | 57859089 | 57859360 | 502 | Distal Intergenic | 16 | 57884008 | 57884139 | 132 | 2 | ENSMUST00 | 24779 | Gm24039 | ENSMUSG0C | false | . | . | . | . | . | . | chr16 | 57858114 | 57859587 | 4.20327 | 3.65919 | 2.36824 |
| chr17 | 83861544 | 83861905 | 502 | Distal Intergenic | 17 | 83851675 | 83854393 | 2719 | 2 | ENSMUST00 | -7152 | C430042M11 | ENSMUSG0C | false | chr17 | 83861359 | 83863132 | 25.84672 | 90.2271 | 86.88143 | . | . | . | . | . | . |
| chr7 | 28774286 | 28774522 | 501 | Promoter (2-3kb) | 7 | 28776858 | 28777684 | 827 | 1 | ENSMUST00 | -2336 | Sirt2 | ENSMUSG0C | false | . | . | . | . | . | . | chr7 | 28770641 | 28774873 | 5.72688 | 4.37837 | 3.60962 |
| chr13 | 51849276 | 51849666 | 501 | Promoter (2-3kb) | 13 | 51846744 | 51848224 | 1481 | 1 | ENSMUST00 | 2533 | Gadd45g | ENSMUSG0C | false | chr13 | 51849460 | 51849811 | 7.54398 | 13.82373 | 11.57928 | chr13 | 51845166 | 51850273 | 5.3175 | 4.22752 | 3.26442 |
| chr15 | 81744931 | 81745184 | 501 | Promoter (<=1kb) | 15 | 81744848 | 81796269 | 51422 | 1 | ENSMUST00 | 84 | Zc3h7b | ENSMUSG0C | false | chr15 | 81744733 | 81746324 | 28.16151 | 83.89376 | 80.67007 | . | . | . | . | . | . |
| chr16 | 43561309 | 43561668 | 501 | Intron (ENSMUST00000132881.2 | 16 | 43510308 | 43619137 | 108830 | 1 | ENSMUST00 | 51002 | Zbtb20 | ENSMUSG0C | false | . | . | . | . | . | . | chr16 | 43543186 | 43567712 | 5.70766 | 4.3604 | 3.60355 |
| chr19 | 47731182 | 47731984 | 501 | Promoter (<=1kb) | 19 | 47731756 | 47735588 | 3833 | 1 | ENSMUST00 | 0 | Sfr1 | ENSMUSG0C | false | chr19 | 47730652 | 47732360 | 33.5288 | 108.10788 | 104.32592 | . | . | . | . | . | . |
| chr8 | 80852907 | 80853180 | 501 | Intron (ENSMUST00000034150.8 | 8 | 80764434 | 80880479 | 116046 | 2 | ENSMUST00 | 27299 | Gab1 | ENSMUSG0C | false | . | . | . | . | . | . | chr8 | 80802749 | 80878363 | 5.64706 | 4.31183 | 3.55694 |
| chr9 | 92541878 | 92542572 | 501 | Promoter (<=1kb) | 9 | 92542223 | 92608428 | 66206 | 1 | ENSMUST00 | 0 | Plod2 | ENSMUSG0C | true | chr9 | 92541328 | 92546074 | 51.17007 | 179.15596 | 173.00638 | . | . | . | . | . | . |
| chr7 | 145222279 | 145222532 | 501 | Distal Intergenic | 7 | 145253923 | 145283927 | 30005 | 2 | ENSMUST00 | 61395 | Tpcn2 | ENSMUSG0C | false | . | . | . | . | . | . | chr7 | 145221429 | 145223492 | 5.33172 | 4.11521 | 3.29646 |
| chr9 | 69435229 | 69435503 | 501 | Distal Intergenic | 9 | 69450232 | 69451035 | 804 | 2 | ENSMUST00 | 15532 | Gm4978 | ENSMUSG0C | false | . | . | . | . | . | . | . | . | . | . | . | . |
| chr11 | 120518017 | 120518398 | 501 | Distal Intergenic | 11 | 120526855 | 120530554 | 3700 | 2 | ENSMUST00 | 12156 | Gm11789 | ENSMUSG0C | false | . | . | . | . | . | . | chr11 | 120517836 | 120520273 | 5.14815 | 4.17924 | 3.12897 |
| chr17 | 68841043 | 68841264 | 501 | Exon (ENSMUST00000178545.1/ | 17 | 68837136 | 68843138 | 6003 | 1 | ENSMUST00 | 3908 | Tmem200c | ENSMUSG0C | false | chr17 | 68840805 | 68842224 | 21.37974 | 56.22882 | 53.43486 | chr17 | 68840359 | 68842124 | 6.2557 | 4.20782 | 4.07388 |
| chr6 | 129213658 | 129213940 | 501 | Exon (ENSMUST00000181517.2/ | 6 | 129213381 | 129233974 | 20594 | 2 | ENSMUST00 | 20034 | 2310001H17I | ENSMUSG0C | false | . | . | . | . | . | . | . | . | . | . | . | . |
| chr16 | 21246483 | 21246882 | 501 | Downstream (<1kb) | 16 | 21256013 | 21256066 | 54 | 2 | ENSMUST00 | 9184 | Mir7680 | ENSMUSG0C | false | chr16 | 21246024 | 21246848 | 17.27239 | 48.28369 | 45.59018 | chr16 | 21245392 | 21246954 | 5.50426 | 4.31534 | 3.4357 |
| chr10 | 71237940 | 71238250 | 500 | Promoter (<=1kb) | 10 | 71225464 | 71238280 | 12817 | 2 | ENSMUST00 | 30 | Tfam | ENSMUSG0C | false | chr10 | 71237106 | 71238605 | 44.27422 | 151.93042 | 146.56525 | chr10 | 71236526 | 71238894 | 3.31995 | 3.25071 | 1.62649 |
| chr13 | 89740291 | 89740563 | 500 | Promoter (<=1kb) | 13 | 89725434 | 89740397 | 14964 | 2 | ENSMUST00 | 0 | Vcan | ENSMUSG0C | false | chr13 | 89738844 | 89741663 | 25.56571 | 75.64988 | 72.57267 | . | . | . | . | . | . |
| chr3 | 89714874 | 89715238 | 500 | Promoter (<=1kb) | 3 | 89715022 | 89753446 | 38425 | 1 | ENSMUST00 | 0 | Adar | ENSMUSG0C | false | chr3 | 89714676 | 89715826 | 34.46491 | 104.61457 | 100.92722 | chr3 | 89714615 | 89715533 | 3.2263 | 3.21612 | 1.53183 |
| chr7 | 101310446 | 101310973 | 500 | Distal Intergenic | 7 | 101317086 | 101346626 | 29541 | 1 | ENSMUST00 | -6113 | Stard10 | ENSMUSG0C | false | chr7 | 101310088 | 101310911 | 30.56069 | 91.89841 | 88.51308 | chr7 | 101310016 | 101311617 | 4.01105 | 3.65043 | 2.1783 |
| chr8 | 27123182 | 27123400 | 500 | 3' UTR | 8 | 27123832 | 27128632 | 4801 | 2 | ENSMUST00 | 5232 | Brf2 | ENSMUSG0C | false | . | . | . | . | . | . | chr8 | 27122272 | 27124478 | 5.7094 | 4.19742 | 3.61873 |
| chr9 | 21797762 | 21798124 | 500 | Promoter (<=1kb) | 9 | 21766773 | 21798546 | 31774 | 2 | ENSMUST00 | 422 | Kank2 | ENSMUSG0C | false | chr9 | 21797469 | 21799074 | 28.04154 | 116.13699 | 112.1206 | chr9 | 21786534 | 21799409 | 6.22233 | 4.65692 | 4.01382 |
| chr9 | 97110602 | 97111173 | 500 | Promoter (<=1kb) | 9 | 97074961 | 97111157 | 36197 | 2 | ENSMUST00 | 0 | Slc25a36 | ENSMUSG0C | false | chr9 | 97109890 | 97111659 | 30.11944 | 121.38741 | 117.20103 | . | . | . | . | . | . |
| chr3 | 129533446 | 129534172 | 500 | Promoter (1-2kb) | 3 | 129532386 | 129638495 | 106110 | 1 | ENSMUST00 | 1061 | Elovl6 | ENSMUSG0C | false | chr3 | 129532063 | 129534638 | 23.4661 | 79.19912 | 76.05865 | . | . | . | . | . | . |
| chr3 | 122293112 | 122293445 | 500 | Promoter (<=1kb) | 3 | 122255958 | 122293547 | 37590 | 2 | ENSMUST00 | 102 | Gm17494 | ENSMUSG0C | false | chr3 | 122292329 | 122294466 | 35.06866 | 119.18253 | 115.06551 | . | . | . | . | . | . |
| chr13 | 58340742 | 58341002 | 500 | Intron (ENSMUST00000043605.5 | 13 | 58287516 | 58354862 | 67347 | 2 | ENSMUST00 | 13860 | Kif27 | ENSMUSG0C | false | . | . | . | . | . | . | . | . | . | . | . | . |
| chr2 | 82053093 | 82053760 | 500 | Promoter (<=1kb) | 2 | 82053222 | 82259879 | 206658 | 1 | ENSMUST00 | 0 | Zfp804a | ENSMUSG0C | false | chr2 | 82053088 | 82053888 | 12.94928 | 28.76233 | 26.31305 | chr2 | 82053087 | 82054816 | 5.55801 | 4.10693 | 3.48711 |
| chr2 | 32661361 | 32661568 | 500 | Intron (ENSMUST00000009705.8 | 2 | 32652329 | 32652414 | 86 | 1 | ENSMUST00 | 9033 | Mir1954 | ENSMUSG0C | false | . | . | . | . | . | . | . | . | . | . | . | . |
| chrX | 36864152 | 36864532 | 499 | Promoter (<=1kb) | 20 | 36823737 | 36864238 | 40502 | 2 | ENSMUST00 | 0 | C330007P06I | ENSMUSG0C | false | chrX | 36863633 | 36864767 | 13.32643 | 29.39681 | 26.9387 | . | . | . | . | . | . |
| chr18 | 47240594 | 47240860 | 499 | Intron (ENSMUST00000123228.2 | 18 | 47235598 | 47281195 | 45598 | 2 | ENSMUST00 | 40335 | Sema6a | ENSMUSG0C | false | . | . | . | . | . | . | . | . | . | . | . | . |
| chr10 | 82858869 | 82859209 | 499 | Promoter (<=1kb) | 10 | 82859206 | 82897724 | 38519 | 1 | ENSMUST00 | 0 | Txnrd1 | ENSMUSG0C | false | chr10 | 82858525 | 82860999 | 22.90469 | 70.93823 | 67.93287 | . | . | . | . | . | . |
| chr18 | 44439486 | 44439728 | 499 | Intron (ENSMUST00000164666.2 | 18 | 44380500 | 44424969 | 44470 | 1 | ENSMUST00 | 58987 | Dcp2 | ENSMUSG0C | false | . | . | . | . | . | . | chr18 | 44434846 | 44445255 | 6.64362 | 4.65907 | 4.40138 |
| chr19 | 20389798 | 20390135 | 499 | Promoter (<=1kb) | 19 | 20379659 | 20390651 | 10993 | 2 | ENSMUST00 | 516 | Anxa1 | ENSMUSG0C | false | chr19 | 20388965 | 20390989 | 15.39495 | 41.17039 | 38.56658 | . | . | . | . | . | . |
| chr2 | 28225960 | 28226219 | 499 | Intron (ENSMUST00000113920.2 | 2 | 28205659 | 28230736 | 25078 | 1 | ENSMUST00 | 20302 | Olfm1 | ENSMUSG0C | false | . | . | . | . | . | . | . | . | . | . | . | . |
| chr2 | 121579005 | 121579224 | 499 | Intron (ENSMUST00000133898.2 | 2 | 121568947 | 121584012 | 15066 | 2 | ENSMUST00 | 4788 | Frmd5 | ENSMUSG0C | false | . | . | . | . | . | . | . | . | . | . | . | . |
| chr4 | 143287132 | 143287358 | 499 | Intron (ENSMUST00000030317.8 | 4 | 143268346 | 143299463 | 31118 | 2 | ENSMUST00 | 12105 | Pdpn | ENSMUSG0C | true | . | . | . | . | . | . | chr4 | 143277480 | 143296542 | 5.36947 | 4.22359 | 3.31454 |
| chr19 | 23141947 | 23142626 | 499 | Promoter (<=1kb) | 19 | 23141226 | 23166911 | 25686 | 1 | ENSMUST00 | 722 | Klf9 | ENSMUSG0C | false | chr19 | 23140239 | 23143387 | 30.78991 | 105.27288 | 101.56436 | . | . | . | . | . | . |
| chr5 | 103401938 | 103402139 | 499 | Distal Intergenic | 5 | 103425192 | 103598359 | 173168 | 1 | ENSMUST00 | -23053 | Ptpn13 | ENSMUSG0C | false | . | . | . | . | . | . | chr5 | 103401674 | 103402548 | 4.71379 | 3.90338 | 2.80636 |
| chr11 | 114699526 | 114699867 | 499 | Intron (ENSMUST00000045779.5 | 11 | 114675516 | 114690603 | 15088 | 1 | ENSMUST00 | 24011 | Ttyh2 | ENSMUSG0C | false | . | . | . | . | . | . | . | . | . | . | . | . |
| chr11 | 5415688 | 5415893 | 499 | Intron (ENSMUST00000109867.2 | 11 | 5276324 | 5444847 | 168524 | 2 | ENSMUST00 | 28954 | Znrf3 | ENSMUSG0C | false | . | . | . | . | . | . | chr11 | 5410686 | 5427806 | 5.74514 | 4.38198 | 3.63413 |
| chr6 | 120463141 | 120463896 | 498 | Promoter (<=1kb) | 6 | 120463197 | 120483727 | 20531 | 1 | ENSMUST00 | 0 | Il17ra | ENSMUSG0C | false | chr6 | 120463001 | 120464153 | 31.91637 | 110.38313 | 106.53261 | . | . | . | . | . | . |
| chr2 | 33215712 | 33216196 | 498 | Promoter (<=1kb) | 2 | 33216069 | 33247717 | 31649 | 1 | ENSMUST00 | 0 | Angptl2 | ENSMUSG0C | false | chr2 | 33215515 | 33215837 | 6.00779 | 10.46335 | 8.27668 | chr2 | 33215067 | 33215848 | 7.62511 | 5.0171 | 5.19556 |
| chr13 | 95890954 | 95891423 | 498 | Promoter (<=1kb) | 13 | 95627177 | 95891922 | 264746 | 2 | ENSMUST00 | 499 | Iqgap2 | ENSMUSG0C | true | chr13 | 95890627 | 95892671 | 39.90052 | 130.90422 | 126.38404 | . | . | . | . | . | . |
| chr14 | 55578251 | 55578621 | 498 | Promoter (<=1kb) | 14 | 55578360 | 55581528 | 3169 | 1 | ENSMUST00 | 0 | Psme1 | ENSMUSG0C | false | chr14 | 55577811 | 55578944 | 32.21572 | 97.71757 | 94.20073 | chr14 | 55577827 | 55579657 | 3.97787 | 3.46715 | 2.21593 |
| chr3 | 8964034 | 8964389 | 498 | Promoter (<=1kb) | 3 | 8929598 | 8964046 | 34449 | 2 | ENSMUST00 | 0 | Tpd52 | ENSMUSG0C | false | chr3 | 8963869 | 8964497 | 16.66176 | 41.17759 | 38.57329 | chr3 | 8947714 | 8965965 | 5.38967 | 4.2084 | 3.32941 |
| chr5 | 107146156 | 107146478 | 498 | Intron (ENSMUST00000031224.9 | 5 | 107118197 | 107132923 | 14727 | 2 | ENSMUST00 | -13234 | Tgfbr3 | ENSMUSG0C | true | . | . | . | . | . | . | chr5 | 107143848 | 107172978 | 6.44265 | 4.73295 | 4.19318 |
| chr9 | 66697738 | 66697968 | 498 | Distal Intergenic | 9 | 66713686 | 66766845 | 53160 | 1 | ENSMUST00 | -15718 | Car12 | ENSMUSG0C | false | . | . | . | . | . | . | chr9 | 66697683 | 66698268 | 3.50097 | 3.03899 | 1.80209 |
| chr8 | 10924230 | 10924948 | 498 | Exon (ENSMUST00000040608.3/ | 8 | 10924427 | 10928457 | 4031 | 2 | ENSMUST00 | 3509 | 3930402G23 | ENSMUSG0C | false | chr8 | 10924413 | 10925063 | 6.83737 | 14.36054 | 12.10883 | chr8 | 10922915 | 10926903 | 3.94548 | 3.47029 | 2.16665 |
| chr2 | 170520277 | 170520550 | 498 | Distal Intergenic | 2 | 170514890 | 170516633 | 1744 | 1 | ENSMUST00 | 5388 | Pfdn4 | ENSMUSG0C | false | . | . | . | . | . | . | . | . | . | . | . | . |
| chr7 | 99353105 | 99353417 | 498 | Promoter (<=1kb) | 7 | 99345375 | 99353230 | 7856 | 2 | ENSMUST00 | 0 | Serpinh1 | ENSMUSG0C | false | chr7 | 99351184 | 99353540 | 26.6505 | 78.45867 | 75.33303 | chr7 | 99353219 | 99357600 | 5.8752 | 4.49269 | 3.72297 |
| chr8 | 44085090 | 44085339 | 498 | Distal Intergenic | 8 | 44039250 | 44039361 | 112 | 1 | ENSMUST00 | 45841 | Gm24591 | ENSMUSG0C | false | . | . | . | . | . | . | chr8 | 44084334 | 44085628 | 3.40722 | 3.26383 | 1.74792 |
| chr18 | 35930340 | 35930932 | 498 | Distal Intergenic | 18 | 35952824 | 35952907 | 84 | 1 | ENSMUST00 | -21892 | Gm23882 | ENSMUSG0C | false | chr18 | 35930389 | 35930955 | 11.73508 | 26.0954 | 23.67916 | chr18 | 35930143 | 35930959 | 5.28084 | 4.00071 | 3.26033 |
| chr7 | 95269858 | 95270213 | 498 | Distal Intergenic | 7 | 95171864 | 95172157 | 294 | 1 | ENSMUST00 | 97995 | Gm23884 | ENSMUSG0C | false | . | . | . | . | . | . | chr7 | 95268834 | 95270651 | 4.29433 | 3.79912 | 2.41377 |
| chr12 | 112674181 | 112674520 | 497 | Promoter (<=1kb) | 12 | 112653821 | 112674276 | 20456 | 2 | ENSMUST00 | 0 | Akt1 | ENSMUSG0C | false | chr12 | 112673011 | 112674859 | 30.39959 | 93.57949 | 90.15746 | chr12 | 112664706 | 112675798 | 5.0246 | 4.06953 | 3.03809 |
| chr2 | 105398735 | 105399840 | 497 | Promoter (<=1kb) | 2 | 105386291 | 105399319 | 13029 | 2 | ENSMUST00 | 0 | Rcn1 | ENSMUSG0C | false | chr2 | 105398455 | 105399983 | 34.63892 | 109.01322 | 105.20635 | chr2 | 105397721 | 105400427 | 3.31973 | 3.22461 | 1.6635 |
| chr16 | 18343807 | 18344181 | 497 | Promoter (<=1kb) | 16 | 18300827 | 18343927 | 43101 | 2 | ENSMUST00 | 0 | Tango2 | ENSMUSG0C | false | chr16 | 18343048 | 18344368 | 26.49423 | 78.18742 | 75.06905 | chr16 | 18342696 | 18345921 | 4.97292 | 3.86704 | 3.02205 |
| chr14 | 34311278 | 34311573 | 497 | Promoter (<=1kb) | 14 | 34310727 | 34345144 | 34418 | 1 | ENSMUST00 | 552 | Glud1 | ENSMUSG0C | false | chr14 | 34310717 | 34312662 | 35.98183 | 119.73972 | 115.60696 | . | . | . | . | . | . |
| chr4 | 118138902 | 118139278 | 497 | Exon (ENSMUST00000106406.3/ | 4 | 117932158 | 118134914 | 202757 | 2 | ENSMUST00 | -3989 | St3gal3 | ENSMUSG0C | true | . | . | . | . | . | . | chr4 | 118135121 | 118140157 | 5.86588 | 4.43161 | 3.75583 |
| chr5 | 66613785 | 66614087 | 497 | Intron (ENSMUST00000159786.2 | 5 | 66451982 | 66618640 | 166659 | 2 | ENSMUST00 | 4553 | Apbb2 | ENSMUSG0C | false | . | . | . | . | . | . | chr5 | 66558047 | 66617728 | 6.24773 | 4.56399 | 4.04918 |
| chr5 | 53995316 | 53995732 | 497 | Promoter (2-3kb) | 5 | 53998417 | 54121057 | 122641 | 1 | ENSMUST00 | -2685 | Stim2 | ENSMUSG0C | false | chr5 | 53995121 | 53995884 | 14.64398 | 38.39655 | 35.82735 | . | . | . | . | . | . |
| chr2 | 31141443 | 31142153 | 497 | Promoter (<=1kb) | 2 | 31026206 | 31141802 | 115597 | 2 | ENSMUST00 | 0 | Fnbp1 | ENSMUSG0C | false | chr2 | 31140653 | 31142585 | 38.66002 | 128.73949 | 124.2961 | . | . | . | . | . | . |
| chr11 | 54727334 | 54727652 | 497 | Promoter (1-2kb) | 11 | 54728677 | 54734323 | 5647 | 1 | ENSMUST00 | -1025 | Gm12227 | ENSMUSG0C | false | . | . | . | . | . | . | chr11 | 54722256 | 54750513 | 5.8032 | 4.39 | 3.67574 |
| chr8 | 125349359 | 125349715 | 497 | Distal Intergenic | 8 | 125224400 | 125227323 | 2924 | 2 | ENSMUST00 | -122037 | Gm16237 | ENSMUSG0C | false | . | . | . | . | . | . | chr8 | 125335216 | 125352727 | 6.82037 | 4.91647 | 4.50391 |

| | | | | | | | | | | | | | | | | | | | | | | | | | | |
|---|---|---|---|---|---|---|---|---|---|---|---|---|---|---|---|---|---|---|---|---|---|---|---|---|---|---|
| chr17 | 72966100 | 72966316 | 497 | Distal Intergenic | 17 | 72987798 | 72987904 | 107 | 1 | ENSMUST00 | -21482 | Gm23649 | ENSMUSG0C | false | . | . | . | . | . | . | chr17 | 72965314 | 72969391 | 4.61635 | 3.87513 | 2.69667 |
| chr17 | 84367665 | 84367982 | 497 | Intron (ENSMUST00000047524.8 | 17 | 84391429 | 84391535 | 107 | 1 | ENSMUST00 | -23447 | Gm24722 | ENSMUSG0C | false | . | . | . | . | . | . | chr17 | 84361955 | 84374371 | 5.57955 | 4.30181 | 3.49381 |
| chr5 | 123975881 | 123976194 | 496 | Promoter (2-3kb) | 5 | 123973628 | 124003211 | 29584 | 1 | ENSMUST00 | 2254 | Hip1r | ENSMUSG0C | false | . | . | . | . | . | . | chr5 | 123974676 | 123993001 | 6.443 | 4.63774 | 4.21224 |
| chr7 | 30488929 | 30489205 | 496 | Promoter (2-3kb) | 7 | 30486255 | 30486988 | 734 | 1 | ENSMUST00 | 2675 | Nphs1 | ENSMUSG0C | true | . | . | . | . | . | . | chr7 | 30488485 | 30491522 | 3.55097 | 3.09773 | 1.86228 |
| chr6 | 83506804 | 83507193 | 496 | Promoter (<=1kb) | 6 | 83480217 | 83506969 | 26753 | 2 | ENSMUST00 | 0 | Dguok | ENSMUSG0C | false | chr6 | 83506331 | 83507511 | 32.22038 | 106.82387 | 103.07761 | chr6 | 83499827 | 83507611 | 4.23645 | 3.67384 | 2.38781 |
| chr14 | 45329594 | 45329877 | 496 | Promoter (<=1kb) | 14 | 45329824 | 45349071 | 19248 | 1 | ENSMUST00 | 0 | Psmc6 | ENSMUSG0C | false | chr14 | 45329739 | 45330690 | 34.27164 | 115.65593 | 111.648 | chr14 | 45329180 | 45331585 | 4.24393 | 3.69935 | 2.37556 |
| chr19 | 44294734 | 44294994 | 496 | Promoter (1-2kb) | 19 | 44293676 | 44306864 | 13189 | 1 | ENSMUST00 | 1059 | Scd2 | ENSMUSG0C | false | chr19 | 44293182 | 44296580 | 35.73082 | 112.98314 | 109.06473 | . | . | . | . | . | . |
| chr5 | 36484215 | 36484525 | 496 | Promoter (<=1kb) | 5 | 36473670 | 36484285 | 10616 | 2 | ENSMUST00 | 0 | Tada2b | ENSMUSG0C | false | . | . | . | . | . | . | . | . | . | . | . | . |
| chr8 | 107114678 | 107115149 | 496 | Distal Intergenic | 8 | 107075517 | 107096547 | 21031 | 2 | ENSMUST00 | -18132 | Terf2 | ENSMUSG0C | false | chr8 | 107114369 | 107115495 | 12.30394 | 36.76666 | 34.21749 | chr8 | 107112974 | 107120164 | 4.09415 | 3.59087 | 2.26456 |
| chr15 | 75940825 | 75941197 | 496 | Promoter (<=1kb) | 15 | 75940952 | 75949377 | 8426 | 1 | ENSMUST00 | 0 | Zfp623 | ENSMUSG0C | false | chr15 | 75940688 | 75941885 | 28.28183 | 109.60786 | 105.79031 | chr15 | 75940420 | 75944484 | 4.43434 | 3.80843 | 2.53702 |
| chr5 | 65107306 | 65107918 | 496 | Promoter (<=1kb) | 5 | 65107568 | 65168188 | 60621 | 1 | ENSMUST00 | 0 | Klhl5 | ENSMUSG0C | false | chr5 | 65107033 | 65108648 | 34.78613 | 113.42423 | 109.48744 | . | . | . | . | . | . |
| chr5 | 115491818 | 115492283 | 496 | Promoter (<=1kb) | 5 | 115491891 | 115492455 | 565 | 2 | ENSMUST00 | 172 | Gm13836 | ENSMUSG0C | false | chr5 | 115491490 | 115492301 | 10.56924 | 21.28106 | 18.92771 | chr5 | 115490652 | 115493615 | 5.70899 | 4.44985 | 3.58322 |
| chr12 | 86833818 | 86834163 | 496 | Distal Intergenic | 12 | 86809957 | 86810258 | 302 | 1 | ENSMUST00 | 23862 | Gm25888 | ENSMUSG0C | false | . | . | . | . | . | . | chr12 | 86831585 | 86837954 | 7.49129 | 5.26295 | 5.03568 |
| chr1 | 166317085 | 166317384 | 496 | Distal Intergenic | 1 | 166306741 | 166309585 | 2845 | 2 | ENSMUST00 | -7501 | 5330438I03R | ENSMUSG0C | false | . | . | . | . | . | . | chr1 | 166315932 | 166319420 | 3.39903 | 3.28443 | 1.67232 |
| chr9 | 102263458 | 102263688 | 496 | Intron (ENSMUST00000035129.8 | 9 | 102304924 | 102304975 | 52 | 2 | ENSMUST00 | 41287 | Mir7243 | ENSMUSG0C | false | . | . | . | . | . | . | chr9 | 102258607 | 102274961 | 4.8108 | 3.97089 | 2.85625 |
| chr10 | 13109873 | 13110381 | 495 | Intron (ENSMUST00000123135.2 | 10 | 13113465 | 13127341 | 13877 | 1 | ENSMUST00 | -3084 | Plagl1 | ENSMUSG0C | true | . | . | . | . | . | . | chr10 | 13102931 | 13117612 | 5.30894 | 4.20106 | 3.26917 |
| chr13 | 55329711 | 55330228 | 495 | Promoter (<=1kb) | 13 | 55325168 | 55329758 | 4591 | 2 | ENSMUST00 | 0 | Mxd3 | ENSMUSG0C | false | chr13 | 55329368 | 55330650 | 22.89105 | 88.16154 | 84.85722 | chr13 | 55329156 | 55330877 | 7.21907 | 5.14365 | 4.80726 |
| chr14 | 60575472 | 60575790 | 495 | Intron (ENSMUST00000190392.1 | 14 | 60634001 | 60691367 | 57367 | 1 | ENSMUST00 | -58211 | Spata13 | ENSMUSG0C | true | . | . | . | . | . | . | chr14 | 60569890 | 60579408 | 5.07455 | 4.021 | 3.08544 |
| chr11 | 117664954 | 117665189 | 495 | Intron (ENSMUST00000026658.7 | 11 | 117654813 | 117763439 | 108627 | 1 | ENSMUST00 | 10142 | Tnrc6c | ENSMUSG0C | false | . | . | . | . | . | . | chr11 | 117662912 | 117672218 | 4.60401 | 3.89386 | 2.67826 |
| chr18 | 76000719 | 76001184 | 495 | Intron (ENSMUST00000058997.8 | 18 | 76059458 | 76146443 | 86986 | 1 | ENSMUST00 | -58274 | Zbtb7c | ENSMUSG0C | true | . | . | . | . | . | . | chr18 | 75999046 | 76019015 | 7.41965 | 5.13063 | 4.98856 |
| chr10 | 13271796 | 13272080 | 495 | Intron (ENSMUST00000105545.5 | 10 | 13207717 | 13324160 | 116444 | 2 | ENSMUST00 | 52080 | Phactr2 | ENSMUSG0C | false | . | . | . | . | . | . | chr10 | 13268905 | 13274255 | 6.97045 | 4.96959 | 4.60764 |
| chr4 | 144892241 | 144892954 | 495 | Promoter (<=1kb) | 4 | 144892827 | 144927642 | 34816 | 1 | ENSMUST00 | 0 | Dhrs3 | ENSMUSG0C | false | chr4 | 144891998 | 144895332 | 25.47373 | 70.96252 | 67.95692 | chr4 | 144890794 | 144892391 | 5.37423 | 4.13584 | 3.34101 |
| chr16 | 46010324 | 46010690 | 495 | Promoter (<=1kb) | 16 | 45959263 | 46010218 | 50956 | 2 | ENSMUST00 | -107 | Plcxd2 | ENSMUSG0C | false | chr16 | 46009644 | 46011113 | 25.15761 | 80.47005 | 77.31023 | . | . | . | . | . | . |
| chr15 | 86105824 | 86106048 | 495 | Intron (ENSMUST00000088931.4 | 15 | 86087640 | 86090060 | 2421 | 2 | ENSMUST00 | -15765 | Gm15722 | ENSMUSG0C | false | . | . | . | . | . | . | chr15 | 86105499 | 86106755 | 2.71624 | 2.71569 | 1.17423 |
| chr11 | 64464116 | 64464374 | 495 | Intron (ENSMUST00000058652.5 | 11 | 64435156 | 64436653 | 1498 | 2 | ENSMUST00 | -27464 | F930015N05 | ENSMUSG0C | false | . | . | . | . | . | . | chr11 | 64458919 | 64465470 | 5.61991 | 4.31365 | 3.52661 |
| chr18 | 44658901 | 44659206 | 495 | Promoter (2-3kb) | 18 | 44661665 | 44676271 | 14607 | 1 | ENSMUST00 | -2459 | A930012L18I | ENSMUSG0C | false | . | . | . | . | . | . | chr18 | 44646282 | 44660265 | 5.49362 | 4.34447 | 3.41384 |
| chr19 | 55192265 | 55192526 | 495 | Promoter (<=1kb) | 19 | 55192678 | 55192734 | 57 | 1 | ENSMUST00 | -152 | Mir6715 | ENSMUSG0C | false | . | . | . | . | . | . | chr19 | 55191693 | 55192735 | 3.74568 | 3.30284 | 2.02086 |
| chr1 | 62520674 | 62520922 | 495 | Intron (ENSMUST00000075374.4 | 1 | 62534491 | 62535942 | 1452 | 2 | ENSMUST00 | 15020 | RP23-20G7.4 | ENSMUSG0C | false | . | . | . | . | . | . | chr1 | 62518374 | 62523891 | 4.82614 | 3.91958 | 2.86666 |
| chr11 | 101246022 | 101246488 | 494 | Promoter (<=1kb) | 11 | 101246028 | 101248250 | 2223 | 1 | ENSMUST00 | 0 | Ramp2 | ENSMUSG0C | false | chr11 | 101245876 | 101246906 | 23.13664 | 67.80279 | 64.84689 | chr11 | 101244977 | 101247371 | 3.87301 | 3.47814 | 2.08381 |
| chr7 | 141455054 | 141455355 | 494 | Promoter (<=1kb) | 7 | 141455198 | 141460743 | 5546 | 1 | ENSMUST00 | 0 | Pnpla2 | ENSMUSG0C | false | chr7 | 141454899 | 141456677 | 31.04394 | 108.79036 | 104.98655 | . | . | . | . | . | . |
| chr6 | 99619530 | 99619794 | 494 | Intron (ENSMUST00000155466.1 | 6 | 99434782 | 99632376 | 197595 | 2 | ENSMUST00 | 12582 | Gm20696 | ENSMUSG0C | false | . | . | . | . | . | . | . | . | . | . | . | . |
| chr8 | 109778437 | 109779017 | 494 | Promoter (<=1kb) | 8 | 109778554 | 109864204 | 85651 | 1 | ENSMUST00 | 0 | Ap1g1 | ENSMUSG0C | false | chr8 | 109778088 | 109780170 | 43.40503 | 144.75304 | 139.67491 | . | . | . | . | . | . |
| chr17 | 47508001 | 47508287 | 494 | Promoter (2-3kb) | 17 | 47505241 | 47598649 | 93409 | 1 | ENSMUST00 | 2761 | Ccnd3 | ENSMUSG0C | false | . | . | . | . | . | . | chr17 | 47506824 | 47509251 | 4.77877 | 3.89968 | 2.8367 |
| chrX | 48513385 | 48513788 | 494 | Promoter (<=1kb) | 20 | 48474944 | 48513426 | 38483 | 2 | ENSMUST00 | 0 | Aifm1 | ENSMUSG0C | false | chrX | 48512817 | 48514323 | 20.67895 | 53.40692 | 50.64761 | . | . | . | . | . | . |
| chr19 | 3768590 | 3769149 | 494 | Promoter (<=1kb) | 19 | 3768547 | 3796660 | 28114 | 1 | ENSMUST00 | 44 | Suv420h1 | ENSMUSG0C | false | chr19 | 3767123 | 3769600 | 24.98033 | 125.99245 | 121.64841 | . | . | . | . | . | . |
| chr19 | 16773189 | 16773518 | 494 | Intron (ENSMUST00000068156.6 | 19 | 16780547 | 16781103 | 557 | 1 | ENSMUST00 | -7029 | Gm9806 | ENSMUSG0C | false | chr19 | 16772806 | 16773917 | 10.00754 | 19.81923 | 17.48753 | chr19 | 16771363 | 16776645 | 4.45062 | 3.78036 | 2.55285 |
| chrX | 38473599 | 38473992 | 494 | Promoter (2-3kb) | 20 | 38476461 | 38482486 | 6026 | 1 | ENSMUST00 | -2469 | Gm7598 | ENSMUSG0C | false | chrX | 38473221 | 38474198 | 9.65017 | 18.71579 | 16.39739 | . | . | . | . | . | . |
| chr3 | 8713721 | 8713964 | 494 | Distal Intergenic | 3 | 8679359 | 8679477 | 119 | 2 | ENSMUST00 | -34245 | Gm23670 | ENSMUSG0C | false | . | . | . | . | . | . | chr3 | 8712430 | 8718854 | 5.10568 | 4.06101 | 3.11811 |
| chr3 | 83704647 | 83704965 | 494 | Promoter (2-3kb) | 3 | 83702252 | 83702358 | 107 | 1 | ENSMUST00 | 2396 | Gm24584 | ENSMUSG0C | false | . | . | . | . | . | . | . | . | . | . | . | . |
| chr3 | 63543002 | 63543309 | 494 | Distal Intergenic | 3 | 63488993 | 63489098 | 106 | 1 | ENSMUST00 | 54010 | Gm24946 | ENSMUSG0C | false | . | . | . | . | . | . | chr3 | 63539206 | 63546986 | 4.11986 | 3.62901 | 2.28834 |
| chr2 | 27419209 | 27419478 | 493 | Intron (ENSMUST00000185188.2 | 2 | 27262104 | 27426912 | 164809 | 2 | ENSMUST00 | 7434 | Vav2 | ENSMUSG0C | false | . | . | . | . | . | . | chr2 | 27414426 | 27420323 | 3.83549 | 3.35867 | 2.08019 |
| chr10 | 57486218 | 57486461 | 493 | Promoter (<=1kb) | 10 | 57486385 | 57513133 | 26749 | 1 | ENSMUST00 | 0 | Hsf2 | ENSMUSG0C | false | chr10 | 57486338 | 57487441 | 34.82909 | 108.31201 | 104.52283 | . | . | . | . | . | . |
| chr14 | 73117020 | 73117373 | 493 | Distal Intergenic | 14 | 73123037 | 73161986 | 38950 | 1 | ENSMUST00 | -5664 | Rcbtb2 | ENSMUSG0C | false | chr14 | 73116970 | 73117356 | 7.4361 | 14.08204 | 11.83379 | chr14 | 73116830 | 73120495 | 3.1606 | 3.11218 | 1.51106 |
| chr19 | 43731688 | 43731917 | 493 | 3' UTR | 19 | 43726367 | 43728136 | 1770 | 1 | ENSMUST00 | 5322 | Entpd7 | ENSMUSG0C | false | . | . | . | . | . | . | chr19 | 43731195 | 43732382 | 5.69972 | 4.1083 | 3.62057 |
| chr1 | 189879942 | 189880316 | 493 | Promoter (2-3kb) | 1 | 189880642 | 189882372 | 1731 | 2 | ENSMUST00 | 2056 | Smyd2 | ENSMUSG0C | false | . | . | . | . | . | . | chr1 | 189877435 | 189881194 | 6.59888 | 4.8001 | 4.37134 |
| chr9 | 15301540 | 15301811 | 493 | Promoter (<=1kb) | 9 | 15283337 | 15301607 | 18271 | 2 | ENSMUST00 | 0 | 4931406C07I | ENSMUSG0C | false | . | . | . | . | . | . | chr9 | 15301102 | 15303435 | 4.45221 | 3.77335 | 2.59971 |
| chr9 | 121332178 | 121332405 | 493 | Distal Intergenic | 9 | 121366958 | 121474916 | 107959 | 1 | ENSMUST00 | -34553 | Trak1 | ENSMUSG0C | false | . | . | . | . | . | . | chr9 | 121328645 | 121333366 | 5.78279 | 4.39237 | 3.67238 |
| chr4 | 114925320 | 114925597 | 493 | Promoter (<=1kb) | 4 | 114925147 | 114926013 | 867 | 2 | ENSMUST00 | 416 | Foxe3 | ENSMUSG0C | false | chr4 | 114924951 | 114926133 | 20.7012 | 58.37697 | 55.55136 | chr4 | 114924758 | 114926193 | 4.59924 | 3.67572 | 2.71256 |
| chr8 | 127640404 | 127640677 | 493 | Distal Intergenic | 8 | 127597706 | 127597823 | 118 | 1 | ENSMUST00 | 42699 | Gm22968 | ENSMUSG0C | false | . | . | . | . | . | . | chr8 | 127638014 | 127642276 | 4.29978 | 3.69257 | 2.4376 |
| chr11 | 68074343 | 68074835 | 493 | Intron (ENSMUST00000021285.8 | 11 | 68053335 | 68053460 | 126 | 1 | ENSMUST00 | 21009 | Gm26157 | ENSMUSG0C | false | . | . | . | . | . | . | chr11 | 68070889 | 68077090 | 3.99073 | 3.57286 | 2.18212 |
| chr10 | 22148778 | 22149468 | 493 | Promoter (<=1kb) | 10 | 22113050 | 22148819 | 35770 | 2 | ENSMUST00 | 0 | E030030I06R | ENSMUSG0C | false | chr10 | 22148088 | 22150989 | 25.50663 | 88.63404 | 85.31516 | chr10 | 22148184 | 22151783 | 8.15587 | 5.45603 | 5.59505 |
| chr14 | 69256508 | 69256803 | 493 | Promoter (2-3kb) | 14 | 69259146 | 69262037 | 2892 | 1 | ENSMUST00 | -2343 | Gm27222 | ENSMUSG0C | false | . | . | . | . | . | . | chr14 | 69247679 | 69257987 | 5.54859 | 4.38732 | 3.45181 |
| chr14 | 22115142 | 22115484 | 493 | Intron (ENSMUST00000075639.5 | 14 | 22021958 | 22022070 | 113 | 2 | ENSMUST00 | -93073 | Gm27532 | ENSMUSG0C | false | . | . | . | . | . | . | chr14 | 22109984 | 22116660 | 4.10396 | 3.66019 | 2.26221 |
| chr17 | 79731045 | 79731344 | 492 | Distal Intergenic | 17 | 79706953 | 79715041 | 8089 | 2 | ENSMUST00 | -16005 | Cyp1b1 | ENSMUSG0C | false | . | . | . | . | . | . | chr17 | 79726503 | 79733779 | 5.74212 | 4.41435 | 3.61439 |
| chr17 | 83182902 | 83183173 | 492 | Distal Intergenic | 17 | 83215292 | 83225070 | 9779 | 1 | ENSMUST00 | -32119 | Pkdcc | ENSMUSG0C | false | . | . | . | . | . | . | chr17 | 83180936 | 83184029 | 5.7038 | 4.50903 | 3.57371 |
| chr19 | 47886566 | 47887021 | 492 | Intron (ENSMUST00000026048.4 | 19 | 47870262 | 47884704 | 14443 | 1 | ENSMUST00 | 16305 | Gsto2 | ENSMUSG0C | false | . | . | . | . | . | . | chr19 | 47884637 | 47887691 | 3.83516 | 3.42128 | 2.06785 |
| chr2 | 144368831 | 144369373 | 492 | Promoter (<=1kb) | 2 | 144368983 | 144407676 | 38694 | 1 | ENSMUST00 | 0 | Csrp2bp | ENSMUSG0C | false | chr2 | 144368284 | 144369979 | 32.69058 | 104.59052 | 100.90593 | . | . | . | . | . | . |
| chr5 | 76186856 | 76187072 | 492 | Promoter (2-3kb) | 5 | 76183950 | 76196987 | 13038 | 1 | ENSMUST00 | 2907 | Tmem165 | ENSMUSG0C | false | . | . | . | . | . | . | chr5 | 76182437 | 76191091 | 3.91563 | 3.48666 | 2.14013 |
| chr6 | 34684879 | 34685382 | 492 | Intron (ENSMUST00000079391.4 | 6 | 34709442 | 34775473 | 66032 | 1 | ENSMUST00 | -24060 | Cald1 | ENSMUSG0C | true | . | . | . | . | . | . | chr6 | 34680884 | 34687674 | 4.73427 | 3.92076 | 2.79387 |
| chr6 | 112865286 | 112865573 | 492 | Intron (ENSMUST00000088373.5 | 6 | 112795420 | 112829459 | 34040 | 2 | ENSMUST00 | -35828 | Srgap3 | ENSMUSG0C | false | . | . | . | . | . | . | chr6 | 112863878 | 112866566 | 5.95299 | 4.52729 | 3.78409 |
| chr15 | 88751329 | 88752003 | 492 | Promoter (<=1kb) | 15 | 88751660 | 88784516 | 32857 | 1 | ENSMUST00 | 0 | Zbed4 | ENSMUSG0C | false | chr15 | 88750934 | 88752603 | 28.53699 | 95.19626 | 91.7378 | . | . | . | . | . | . |
| chr10 | 29361449 | 29362251 | 492 | Promoter (<=1kb) | 10 | 29346176 | 29362040 | 15865 | 2 | ENSMUST00 | 0 | Rnf146 | ENSMUSG0C | false | chr10 | 29360820 | 29362611 | 25.50663 | 88.63404 | 85.31516 | . | . | . | . | . | . |
| chr8 | 9093374 | 9093907 | 492 | Distal Intergenic | 8 | 8666577 | 8690537 | 23961 | 2 | ENSMUST00 | -402838 | Arglu1 | ENSMUSG0C | false | . | . | . | . | . | . | . | . | . | . | . | . |
| chr17 | 86588563 | 86589256 | 492 | Intron (ENSMUST00000097275.3 | 17 | 86497732 | 86505232 | 7501 | 2 | ENSMUST00 | -83332 | Gm10309 | ENSMUSG0C | false | . | . | . | . | . | . | chr17 | 86585927 | 86589447 | 4.77689 | 3.94623 | 2.8466 |
| chr17 | 10519810 | 10520117 | 492 | Intron (ENSMUST00000161274.1 | 17 | 10496453 | 10553169 | 56717 | 1 | ENSMUST00 | 23358 | Gm16168 | ENSMUSG0C | false | . | . | . | . | . | . | chr17 | 10518495 | 10520877 | 4.24177 | 3.72124 | 2.36599 |
| chr6 | 47772007 | 47772524 | 492 | Promoter (<=1kb) | 6 | 47773329 | 47773429 | 101 | 1 | ENSMUST00 | -805 | Gm24975 | ENSMUSG0C | false | chr6 | 47771783 | 47772516 | 8.30211 | 98.20082 | 94.67821 | . | . | . | . | . | . |
| chr14 | 21523334 | 21523665 | 491 | Intron (ENSMUST00000183201.2 | 14 | 21516751 | 21671206 | 154456 | 1 | ENSMUST00 | 6584 | Kat6b | ENSMUSG0C | false | . | . | . | . | . | . | chr14 | 21519296 | 21529701 | 4.63736 | 3.89681 | 2.70766 |
| chr1 | 6214474 | 6215167 | 491 | Promoter (<=1kb) | 1 | 6214645 | 6276648 | 62004 | 1 | ENSMUST00 | 0 | Rb1cc1 | ENSMUSG0C | false | chr1 | 6213191 | 6216341 | 25.98079 | 111.7719 | 107.88497 | . | . | . | . | . | . |
| chr4 | 139852453 | 139852689 | 491 | Distal Intergenic | 4 | 139737062 | 139833528 | 96467 | 2 | ENSMUST00 | -18926 | Pax7 | ENSMUSG0C | false | . | . | . | . | . | . | chr4 | 139851566 | 139853302 | 4.27189 | 3.7288 | 2.40691 |
| chr6 | 134443628 | 134443878 | 491 | Downstream (2-3kb) | 6 | 134450483 | 134471293 | 20811 | 2 | ENSMUST00 | 27415 | Lrp6 | ENSMUSG0C | false | . | . | . | . | . | . | chr6 | 134443317 | 134444561 | 5.36841 | 4.23535 | 3.30191 |
| chr7 | 19458491 | 19458977 | 491 | Promoter (<=1kb) | 7 | 19426075 | 19458494 | 32420 | 2 | ENSMUST00 | 0 | Mark4 | ENSMUSG0C | false | chr7 | 19457183 | 19459389 | 44.11508 | 143.73827 | 138.70853 | . | . | . | . | . | . |
| chr7 | 46613104 | 46613466 | 491 | Promoter (1-2kb) | 7 | 46443474 | 46614788 | 171315 | 2 | ENSMUST00 | 1322 | Sergef | ENSMUSG0C | false | . | . | . | . | . | . | chr7 | 46612048 | 46613675 | 6.07496 | 4.63353 | 3.89217 |
| chr9 | 72714929 | 72715287 | 491 | Intron (ENSMUST00000034740.9 | 9 | 72739472 | 72748158 | 8687 | 1 | ENSMUST00 | -24185 | Nedd4 | ENSMUSG0C | true | . | . | . | . | . | . | chr9 | 72683575 | 72716820 | 5.72166 | 4.35702 | 3.61419 |
| chr4 | 128891218 | 128891495 | 491 | Intron (ENSMUST00000147852.1 | 4 | 128883588 | 128911328 | 27741 | 1 | ENSMUST00 | 7631 | Trim62 | ENSMUSG0C | false | . | . | . | . | . | . | . | . | . | . | . | . |
| chr2 | 159755807 | 159756020 | 491 | Distal Intergenic | 2 | 159777676 | 159778207 | 532 | 2 | ENSMUST00 | 22187 | Gm11445 | ENSMUSG0C | false | . | . | . | . | . | . | chr2 | 159755027 | 159756996 | 5.18817 | 4.26562 | 3.15016 |
| chr4 | 142776836 | 142777048 | 491 | Distal Intergenic | 4 | 142557390 | 142557836 | 447 | 2 | ENSMUST00 | -219001 | Gm13018 | ENSMUSG0C | false | . | . | . | . | . | . | . | . | . | . | . | . |
| chr10 | 12912053 | 12912529 | 491 | Distal Intergenic | 10 | 12916679 | 12923090 | 6412 | 2 | ENSMUST00 | 10561 | B230208H11 | ENSMUSG0C | false | chr10 | 12911788 | 12912272 | 8.07432 | 16.07958 | 13.80223 | chr10 | 12911483 | 12913618 | 4.91929 | 3.91422 | 2.97582 |
| chr11 | 87462157 | 87462585 | 491 | Promoter (<=1kb) | 11 | 87462286 | 87462500 | 215 | 1 | ENSMUST00 | 0 | Rnu3b4 | ENSMUSG0C | false | chr11 | 87461732 | 87462557 | 19.04495 | 59.55748 | 56.71581 | chr11 | 87461572 | 87462935 | 3.70847 | 3.40135 | 1.93978 |
| chr4 | 129145954 | 129146248 | 490 | Downstream (1-2kb) | 4 | 129136999 | 129144593 | 7595 | 1 | ENSMUST00 | 8956 | Fndc5 | ENSMUSG0C | false | . | . | . | . | . | . | chr4 | 129141119 | 129150343 | 5.31606 | 4.18214 | 3.25946 |
| chr11 | 94473926 | 94474532 | 490 | Promoter (<=1kb) | 11 | 94408391 | 94474100 | 65710 | 2 | ENSMUST00 | 0 | Cacna1g | ENSMUSG0C | false | chr11 | 94472650 | 94474728 | 32.82023 | 98.76002 | 95.21949 | chr11 | 94468980 | 94476124 | 5.80314 | 4.36554 | 3.67462 |
| chr1 | 127803000 | 127803303 | 490 | Exon (ENSMUST00000112570.1/ | 1 | 127796571 | 127808061 | 11491 | 1 | ENSMUST00 | 6430 | Ccnt2 | ENSMUSG0C | false | . | . | . | . | . | . | chr1 | 127802268 | 127803781 | 4.2345 | 3.55869 | 2.40834 |
| chr2 | 16743773 | 16744049 | 490 | Intron (ENSMUST00000028081.7 | 2 | 16638696 | 16751690 | 112995 | 1 | ENSMUST00 | 105078 | Plxdc2 | ENSMUSG0C | false | . | . | . | . | . | . | chr2 | 16743030 | 16746062 | 4.26468 | 3.7012 | 2.39536 |
| chr2 | 45117053 | 45117518 | 490 | Promoter (<=1kb) | 2 | 44983699 | 45117395 | 133697 | 2 | ENSMUST00 | 0 | Zeb2 | ENSMUSG0C | false | chr2 | 45115710 | 45117667 | 28.93884 | 86.19278 | 82.92467 | . | . | . | . | . | . |
| chr5 | 30814542 | 30815188 | 490 | Promoter (<=1kb) | 5 | 30814641 | 30866106 | 51466 | 1 | ENSMUST00 | 0 | Mapre3 | ENSMUSG0C | false | chr5 | 30814249 | 30815503 | 35.71972 | 120.33891 | 116.19067 | . | . | . | . | . | . |
| chr9 | 99145869 | 99146131 | 490 | Distal Intergenic | 9 | 99101350 | 99140621 | 39272 | 2 | ENSMUST00 | -5249 | Pik3cb | ENSMUSG0C | false | . | . | . | . | . | . | chr9 | 99143399 | 99146195 | 4.22559 | 3.68519 | 2.37455 |
| chr2 | 151542292 | 151542699 | 490 | Promoter (<=1kb) | 2 | 151542483 | 151561692 | 19210 | 1 | ENSMUST00 | 0 | Fkbp1a | ENSMUSG0C | false | chr2 | 151541564 | 151543754 | 31.84599 | 102.09332 | 98.46812 | . | . | . | . | . | . |
| chr14 | 78678281 | 78678607 | 490 | Intron (ENSMUST00000074729.5 | 14 | 78569609 | 78725089 | 155481 | 2 | ENSMUST00 | 46482 | Dgkh | ENSMUSG0C | false | . | . | . | . | . | . | chr14 | 78673939 | 78679160 | 5.11458 | 4.17424 | 3.09106 |
| chr14 | 59215341 | 59215558 | 490 | Promoter (<=1kb) | 14 | 59216437 | 59224723 | 8287 | 1 | ENSMUST00 | -879 | Rcbtb1 | ENSMUSG0C | false | . | . | . | . | . | . | chr14 | 59214389 | 59217232 | 3.57096 | 3.29567 | 1.87108 |
| chr3 | 95207315 | 95207585 | 490 | Promoter (1-2kb) | 3 | 95181773 | 95208937 | 27165 | 2 | ENSMUST00 | 1352 | Gabpb2 | ENSMUSG0C | true | . | . | . | . | . | . | chr3 | 95205321 | 95210756 | 5.65965 | 4.33892 | 3.55285 |
| chr18 | 21205008 | 21205422 | 490 | Intron (ENSMUST00000049260.6 | 18 | 21127201 | 21300138 | 172938 | 2 | ENSMUST00 | 94716 | Garem | ENSMUSG0C | false | . | . | . | . | . | . | chr18 | 21202334 | 21207795 | 4.70946 | 3.98529 | 2.76086 |
| chr9 | 65196826 | 65197090 | 490 | Promoter (<=1kb) | 9 | 65196572 | 65196687 | 116 | 2 | ENSMUST00 | -140 | Gm25313 | ENSMUSG0C | false | . | . | . | . | . | . | chr9 | 65194851 | 65201705 | 4.75687 | 3.98918 | 2.80757 |
| chr5 | 136245886 | 136246171 | 489 | Promoter (<=1kb) | 5 | 136218149 | 136244903 | 26755 | 2 | ENSMUST00 | -984 | Sh2b2 | ENSMUSG0C | false | . | . | . | . | . | . | chr5 | 136243626 | 136246018 | 6.17498 | 4.67088 | 3.98284 |
| chr13 | 47039193 | 47039481 | 489 | Intron (ENSMUST00000021806.4 | 13 | 47025179 | 47043124 | 17946 | 2 | ENSMUST00 | 3643 | Tpmt | ENSMUSG0C | false | . | . | . | . | . | . | chr13 | 47038802 | 47040176 | 4.20875 | 3.69242 | 2.37176 |
| chr16 | 11066066 | 11066520 | 489 | Promoter (<=1kb) | 16 | 10963343 | 11066157 | 102815 | 2 | ENSMUST00 | 0 | Litaf | ENSMUSG0C | false | chr16 | 11065844 | 11067095 | 29.69128 | 86.53342 | 83.26198 | chr16 | 11065223 | 11067764 | 4.5885 | 3.90751 | 2.67004 |
| chr5 | 24100448 | 24100661 | 489 | Promoter (<=1kb) | 5 | 24100590 | 24161223 | 60634 | 1 | ENSMUST00 | 0 | Klhl7 | ENSMUSG0C | false | chr5 | 24100027 | 24101627 | 29.60123 | 90.23508 | 86.88511 | . | . | . | . | . | . |
| chr6 | 119848022 | 119848793 | 489 | Promoter (<=1kb) | 6 | 119570796 | 119848150 | 277355 | 2 | ENSMUST00 | 0 | Erc1 | ENSMUSG0C | false | chr6 | 119847023 | 119849337 | 44.11508 | 143.73827 | 138.70853 | . | . | . | . | . | . |
| chr7 | 46033572 | 46033857 | 489 | Promoter (<=1kb) | 7 | 46033698 | 46084212 | 50515 | 1 | ENSMUST00 | 0 | Nomo1 | ENSMUSG0C | false | chr7 | 46033517 | 46034659 | 31.66066 | 102.77801 | 99.14294 | . | . | . | . | . | . |
| chr9 | 63984696 | 63985082 | 489 | Intron (ENSMUST00000041029.5 | 9 | 63954009 | 64016987 | 62979 | 2 | ENSMUST00 | 31905 | Smad6 | ENSMUSG0C | true | . | . | . | . | . | . | chr9 | 63948352 | 64020083 | 7.27873 | 5.05859 | 4.874 |
| chr15 | 31530748 | 31531425 | 489 | Promoter (<=1kb) | 15 | 31455899 | 31531037 | 75139 | 2 | ENSMUST00 | 0 | 38777 | ENSMUSG0C | false | chr15 | 31529514 | 31531708 | 35.41093 | 115.56754 | 111.56183 | . | . | . | . | . | . |
| chr4 | 126468452 | 126468830 | 489 | Promoter (<=1kb) | 4 | 126435012 | 126468583 | 33572 | 2 | ENSMUST00 | 0 | Ago1 | ENSMUSG0C | false | chr4 | 126467729 | 126469026 | 25.16654 | 87.04672 | 83.76057 | . | . | . | . | . | . |
| chr3 | 59005656 | 59006023 | 489 | Promoter (<=1kb) | 3 | 59006978 | 59093788 | 86811 | 1 | ENSMUST00 | -955 | Med12l | ENSMUSG0C | true | chr3 | 59005156 | 59007686 | 27.20707 | 96.65539 | 93.16586 | . | . | . | . | . | . |
| chr3 | 53253487 | 53253763 | 489 | Intron (ENSMUST00000059562.8 | 3 | 53452619 | 53457797 | 5179 | 1 | ENSMUST00 | -198856 | Gm16206 | ENSMUSG0C | false | . | . | . | . | . | . | . | . | . | . | . | . |
| chr4 | 89425917 | 89426159 | 489 | Intron (ENSMUST00000139272.1 | 4 | 89421848 | 89463487 | 41640 | 1 | ENSMUST00 | 4070 | Gm12610 | ENSMUSG0C | false | . | . | . | . | . | . | chr4 | 89425972 | 89426789 | 3.32973 | 2.98544 | 1.66773 |
| chr12 | 86821767 | 86822262 | 489 | Distal Intergenic | 12 | 86809957 | 86810258 | 302 | 1 | ENSMUST00 | 11811 | Gm25888 | ENSMUSG0C | false | . | . | . | . | . | . | chr12 | 86820408 | 86827213 | 6.55613 | 4.76961 | 4.27793 |
| chr16 | 33917899 | 33918190 | 488 | Intron (ENSMUST00000115028.4 | 16 | 33922486 | 33944148 | 21663 | 1 | ENSMUST00 | -4296 | Itgb5 | ENSMUSG0C | true | . | . | . | . | . | . | chr16 | 33911227 | 33952498 | 7.40288 | 5.1092 | 4.9863 |
| chr5 | 67306754 | 67307055 | 488 | Promoter (<=1kb) | 5 | 67306955 | 67355336 | 48382 | 1 | ENSMUST00 | 0 | Slc30a9 | ENSMUSG0C | false | chr5 | 67306320 | 67308135 | 40.56511 | 135.35318 | 130.65489 | . | . | . | . | . | . |
| chrX | 73716206 | 73716783 | 488 | Promoter (<=1kb) | 20 | 73716597 | 73738534 | 21938 | 1 | ENSMUST00 | 0 | Abcd1 | ENSMUSG0C | false | chrX | 73715565 | 73717426 | 13.78596 | 30.80405 | 28.32676 | . | . | . | . | . | . |
| chr14 | 34673744 | 34674335 | 488 | Promoter (<=1kb) | 14 | 34673928 | 34747983 | 74056 | 1 | ENSMUST00 | 0 | Wapal | ENSMUSG0C | false | chr14 | 34673302 | 34675036 | 38.97723 | 131.47449 | 126.92584 | . | . | . | . | . | . |
| chr11 | 5952534 | 5952850 | 488 | Promoter (2-3kb) | 11 | 5900823 | 5950081 | 49259 | 2 | ENSMUST00 | -2454 | Gck | ENSMUSG0C | false | . | . | . | . | . | . | chr11 | 5951444 | 5955358 | 4.62474 | 3.84984 | 2.69795 |
| chr7 | 114109541 | 114109776 | 488 | Intron (ENSMUST00000069449.5 | 7 | 114046785 | 114117781 | 70997 | 2 | ENSMUST00 | 8005 | Rras2 | ENSMUSG0C | true | . | . | . | . | . | . | chr7 | 114058507 | 114116129 | 5.64505 | 4.33683 | 3.54844 |
| chr4 | 142077832 | 142078089 | 488 | Intron (ENSMUST00000036572.3 | 4 | 142083972 | 142088059 | 4088 | 1 | ENSMUST00 | -5883 | Tmem51os1 | ENSMUSG0C | false | . | . | . | . | . | . | chr4 | 142077341 | 142078598 | 4.88962 | 4.10699 | 2.90434 |
| chr11 | 101360593 | 101361012 | 488 | Distal Intergenic | 11 | 101367561 | 101377903 | 10343 | 1 | ENSMUST00 | -6549 | G6pc | ENSMUSG0C | false | . | . | . | . | . | . | chr11 | 101360323 | 101363538 | 4.44384 | 3.76159 | 2.5883 |
| chr2 | 74711180 | 74711400 | 488 | Promoter (<=1kb) | 2 | 74711927 | 74748271 | 36345 | 1 | ENSMUST00 | -527 | Hoxd3 | ENSMUSG0C | false | . | . | . | . | . | . | . | . | . | . | . | . |
| chr4 | 131267699 | 131268052 | 488 | Distal Intergenic | 4 | 131239277 | 131239410 | 134 | 1 | ENSMUST00 | 28423 | Gm25261 | ENSMUSG0C | false | . | . | . | . | . | . | chr4 | 131267576 | 131268121 | 4.47509 | 3.69056 | 2.61371 |
| chr6 | 39264646 | 39264998 | 488 | Promoter (<=1kb) | 6 | 39265844 | 39265942 | 99 | 2 | ENSMUST00 | 944 | Gm26008 | ENSMUSG0C | false | . | . | . | . | . | . | . | . | . | . | . | . |
| chr16 | 21794247 | 21794610 | 488 | Promoter (<=1kb) | 16 | 21794347 | 21806123 | 11777 | 1 | ENSMUST00 | 0 | 1300002E11I | ENSMUSG0C | false | chr16 | 21793830 | 21795312 | 26.61085 | 108.21047 | 104.42454 | . | . | . | . | . | . |
| chr19 | 45302253 | 45302806 | 488 | Promoter (<=1kb) | 19 | 45300463 | 45303034 | 2572 | 2 | ENSMUST00 | 228 | RP23-188J8.: | ENSMUSG0C | false | chr19 | 45300973 | 45302769 | 22.15125 | 59.20708 | 56.37309 | chr19 | 45300529 | 45315943 | 6.02178 | 4.57633 | 3.84298 |
| chr14 | 25421759 | 25422078 | 487 | Distal Intergenic | 14 | 25459185 | 25538492 | 79308 | 1 | ENSMUST00 | -37107 | Zmiz1 | ENSMUSG0C | false | . | . | . | . | . | . | chr14 | 25420165 | 25423231 | 6.26838 | 4.71268 | 4.02624 |
| chr5 | 143625482 | 143625728 | 487 | Promoter (2-3kb) | 5 | 143622534 | 143686526 | 63993 | 1 | ENSMUST00 | 2949 | Cyth3 | ENSMUSG0C | false | . | . | . | . | . | . | chr5 | 143623294 | 143627247 | 5.05306 | 3.88706 | 3.0834 |
| chr14 | 30951216 | 30951557 | 487 | Promoter (<=1kb) | 14 | 30951486 | 30987439 | 35954 | 1 | ENSMUST00 | 0 | Nek4 | ENSMUSG0C | false | chr14 | 30950835 | 30952082 | 29.63113 | 94.2009 | 90.76292 | chr14 | 30950690 | 30952434 | 5.14147 | 4.20193 | 3.1205 |
| chr17 | 24086223 | 24086465 | 487 | Intron (ENSMUST00000102927.4 | 17 | 24088181 | 24093798 | 5618 | 2 | ENSMUST00 | 7333 | Pdpk1 | ENSMUSG0C | false | . | . | . | . | . | . | chr17 | 24082385 | 24089065 | 4.04341 | 3.57122 | 2.25122 |
| chr18 | 5593205 | 5593632 | 487 | Promoter (<=1kb) | 18 | 5593308 | 5641459 | 48152 | 1 | ENSMUST00 | 0 | Zeb1 | ENSMUSG0C | false | chr18 | 5591062 | 5593921 | 37.10487 | 121.71056 | 117.5108 | . | . | . | . | . | . |
| chr19 | 6969585 | 6969928 | 487 | Promoter (<=1kb) | 19 | 6953714 | 6969759 | 16046 | 2 | ENSMUST00 | 0 | Plcb3 | ENSMUSG0C | false | chr19 | 6968628 | 6970505 | 31.85236 | 101.09525 | 97.496 | chr19 | 6967550 | 6971451 | 4.96737 | 4.08907 | 2.97025 |
| chr11 | 119913640 | 119914072 | 487 | Promoter (<=1kb) | 11 | 119913810 | 119919548 | 5739 | 1 | ENSMUST00 | 0 | Chmp6 | ENSMUSG0C | false | chr11 | 119913059 | 119914872 | 28.90751 | 104.81161 | 101.11721 | chr11 | 119912922 | 119917570 | 6.23168 | 4.47351 | 4.04645 |
| chr1 | 172431869 | 172432198 | 487 | Distal Intergenic | 1 | 172475358 | 172478575 | 3218 | 1 | ENSMUST00 | -43160 | Slamf9 | ENSMUSG0C | false | . | . | . | . | . | . | chr1 | 172428189 | 172437787 | 6.0765 | 4.59609 | 3.89569 |
| chr12 | 51848126 | 51848381 | 487 | Distal Intergenic | 12 | 51743722 | 51829536 | 85815 | 2 | ENSMUST00 | -18591 | Hectd1 | ENSMUSG0C | false | . | . | . | . | . | . | . | . | . | . | . | . |
| chr1 | 182763992 | 182764455 | 487 | Promoter (<=1kb) | 1 | 182764012 | 182895232 | 131221 | 1 | ENSMUST00 | 0 | Susd4 | ENSMUSG0C | false | chr1 | 182763639 | 182764659 | 18.77433 | 54.14365 | 51.37372 | chr1 | 182763831 | 182765161 | 4.62809 | 3.85122 | 2.7413 |
| chr1 | 136345456 | 136346450 | 487 | Promoter (<=1kb) | 1 | 136268123 | 136346104 | 77982 | 2 | ENSMUST00 | 0 | Camsap2 | ENSMUSG0C | false | chr1 | 136344371 | 136346807 | 33.04826 | 106.23876 | 102.51532 | . | . | . | . | . | . |
| chr19 | 4332730 | 4333162 | 487 | Promoter (<=1kb) | 19 | 4329191 | 4333919 | 4729 | 2 | ENSMUST00 | 757 | Kdm2a | ENSMUSG0C | false | chr19 | 4332597 | 4334428 | 18.39885 | 52.66533 | 49.91441 | chr19 | 4331351 | 4335335 | 5.94277 | 4.45125 | 3.80268 |
| chr9 | 40730713 | 40731144 | 487 | Intron (ENSMUST00000139577.1 | 9 | 40718906 | 40719986 | 1081 | 2 | ENSMUST00 | -10728 | Gm16095 | ENSMUSG0C | false | chr9 | 40730639 | 40731944 | 17.28391 | 44.47248 | 41.82566 | . | . | . | . | . | . |
| chr7 | 19664827 | 19665506 | 486 | Promoter (<=1kb) | 7 | 19631581 | 19665030 | 33450 | 2 | ENSMUST00 | 0 | Clptm1 | ENSMUSG0C | false | chr7 | 19663940 | 19666077 | 50.52507 | 176.83885 | 170.7393 | . | . | . | . | . | . |
| chr15 | 59080258 | 59080569 | 486 | Promoter (1-2kb) | 15 | 58941244 | 59082026 | 140783 | 2 | ENSMUST00 | 1457 | Mtss1 | ENSMUSG0C | true | chr15 | 59075869 | 59083046 | 40.64776 | 144.39505 | 139.32506 | . | . | . | . | . | . |
| chr1 | 152807611 | 152807836 | 486 | Promoter (<=1kb) | 1 | 152807837 | 152824292 | 16456 | 1 | ENSMUST00 | -1 | Ncf2 | ENSMUSG0C | true | . | . | . | . | . | . | chr1 | 152801274 | 152821507 | 4.847 | 3.96927 | 2.88719 |
| chr5 | 29826721 | 29826968 | 486 | Distal Intergenic | 5 | 29765190 | 29766687 | 1498 | 1 | ENSMUST00 | 61532 | Dnajb6 | ENSMUSG0C | false | . | . | . | . | . | . | chr5 | 29825193 | 29827381 | 3.96914 | 3.59229 | 2.16784 |
| chr2 | 119995747 | 119996165 | 486 | Intron (ENSMUST00000066058.7 | 2 | 120011687 | 120013094 | 1408 | 2 | ENSMUST00 | -15522 | Mapkbp1 | ENSMUSG0C | false | . | . | . | . | . | . | chr2 | 119993061 | 119998247 | 5.22622 | 4.15453 | 3.21809 |
| chr11 | 119083667 | 119083896 | 486 | Promoter (2-3kb) | 11 | 119077573 | 119086221 | 8649 | 2 | ENSMUST00 | 2325 | Cbx4 | ENSMUSG0C | false | . | . | . | . | . | . | chr11 | 119077654 | 119084726 | 7.24094 | 5.11519 | 4.83739 |
| chr9 | 21330255 | 21330514 | 486 | Distal Intergenic | 9 | 21337828 | 21355025 | 17198 | 1 | ENSMUST00 | -7314 | Slc44a2 | ENSMUSG0C | false | . | . | . | . | . | . | chr9 | 21320251 | 21338396 | 4.95977 | 3.98308 | 2.9977 |
| chr4 | 54984641 | 54984895 | 486 | Intron (ENSMUST00000133895.2 | 4 | 55011366 | 55081358 | 69993 | 1 | ENSMUST00 | -26471 | Zfp462 | ENSMUSG0C | true | . | . | . | . | . | . | chr4 | 54964968 | 54991208 | 5.4969 | 4.26974 | 3.41753 |
| chr18 | 3337177 | 3337902 | 486 | Promoter (<=1kb) | 18 | 3266354 | 3337679 | 71326 | 2 | ENSMUST00 | 0 | Crem | ENSMUSG0C | false | chr18 | 3336441 | 3338545 | 35.14787 | 110.9313 | 107.0704 | . | . | . | . | . | . |
| chr3 | 90156972 | 90157423 | 486 | Intron (ENSMUST00000029548.4 | 3 | 90126535 | 90126666 | 132 | 1 | ENSMUST00 | 30438 | Gm23723 | ENSMUSG0C | false | . | . | . | . | . | . | chr3 | 90155667 | 90157759 | 5.10047 | 4.12188 | 3.08018 |
| chrX | 76111162 | 76111381 | 486 | Distal Intergenic | 20 | 76153331 | 76162605 | 9275 | 2 | ENSMUST00 | 51224 | Gm14710 | ENSMUSG0C | false | . | . | . | . | . | . | . | . | . | . | . | . |
| chr3 | 153702511 | 153702782 | 486 | Intron (ENSMUST00000064460.5 | 3 | 153708911 | 153709026 | 116 | 1 | ENSMUST00 | -6129 | Gm22206 | ENSMUSG0C | false | . | . | . | . | . | . | chr3 | 153686316 | 153712771 | 6.43925 | 4.66349 | 4.19773 |
| chr8 | 3994155 | 3994437 | 486 | Promoter (<=1kb) | 8 | 3994628 | 3997505 | 2878 | 1 | ENSMUST00 | -191 | Gm7389 | ENSMUSG0C | false | . | . | . | . | . | . | . | . | . | . | . | . |
| chr7 | 25133234 | 25133492 | 485 | Promoter (<=1kb) | 7 | 25093248 | 25132512 | 39265 | 2 | ENSMUST00 | -723 | Pou2f2 | ENSMUSG0C | false | . | . | . | . | . | . | . | . | . | . | . | . |
| chr13 | 24280931 | 24281244 | 485 | Promoter (<=1kb) | 13 | 24012659 | 24280795 | 268137 | 2 | ENSMUST00 | -137 | Lrrc16a | ENSMUSG0C | false | chr13 | 24280200 | 24281526 | 32.66316 | 99.48935 | 95.93111 | chr13 | 24279814 | 24281558 | 5.0282 | 4.15632 | 3.03521 |
| chr17 | 46055746 | 46056052 | 485 | Distal Intergenic | 17 | 46016996 | 46032377 | 15382 | 2 | ENSMUST00 | -23370 | Vegfa | ENSMUSG0C | true | . | . | . | . | . | . | chr17 | 46048792 | 46077235 | 6.11576 | 4.54762 | 3.92959 |
| chr17 | 80991514 | 80991868 | 485 | Intron (ENSMUST00000025092.4 | 17 | 80944632 | 81001816 | 57185 | 1 | ENSMUST00 | 46883 | Tmem178 | ENSMUSG0C | false | . | . | . | . | . | . | chr17 | 80988419 | 80997814 | 5.61386 | 4.25874 | 3.54679 |
| chr9 | 65630170 | 65630869 | 485 | Promoter (<=1kb) | 9 | 65630582 | 65660505 | 29924 | 1 | ENSMUST00 | 0 | Rbpms2 | ENSMUSG0C | false | chr9 | 65629075 | 65631311 | 18.70486 | 58.09692 | 55.27494 | . | . | . | . | . | . |
| chr12 | 44244523 | 44244738 | 485 | Intron (ENSMUST00000122902.2 | 12 | 44221370 | 44312092 | 90723 | 1 | ENSMUST00 | 23154 | Pnpla8 | ENSMUSG0C | false | . | . | . | . | . | . | chr12 | 44244400 | 44245211 | 3.84384 | 3.47428 | 2.08724 |
| chr19 | 46616552 | 46616827 | 485 | Intron (ENSMUST00000138302.3 | 19 | 46623401 | 46652522 | 29122 | 1 | ENSMUST00 | -6574 | Wbp1l | ENSMUSG0C | false | . | . | . | . | . | . | chr19 | 46599683 | 46650166 | 6.55215 | 4.7277 | 4.28832 |
| chr1 | 87326799 | 87327087 | 485 | Promoter (<=1kb) | 1 | 87326998 | 87355722 | 28725 | 1 | ENSMUST00 | 0 | Gigyf2 | ENSMUSG0C | false | chr1 | 87326615 | 87328066 | 36.52666 | 129.28897 | 124.83229 | chr1 | 87326747 | 87329194 | 4.55731 | 3.95233 | 2.64827 |
| chr19 | 40894441 | 40894804 | 485 | Promoter (<=1kb) | 19 | 40894705 | 40917947 | 23243 | 1 | ENSMUST00 | 0 | Zfp518a | ENSMUSG0C | false | chr19 | 40894179 | 40895684 | 31.54088 | 108.67455 | 104.87239 | . | . | . | . | . | . |
| chr16 | 56083696 | 56083933 | 485 | Intron (ENSMUST00000049128.6 | 16 | 56075409 | 56190011 | 114603 | 1 | ENSMUST00 | 8288 | Senp7 | ENSMUSG0C | false | . | . | . | . | . | . | chr16 | 56079274 | 56085747 | 3.66986 | 3.424 | 1.91089 |
| chr3 | 88913712 | 88914428 | 485 | Promoter (<=1kb) | 3 | 88905107 | 88913931 | 8825 | 2 | ENSMUST00 | 0 | Msto1 | ENSMUSG0C | false | chr3 | 88913350 | 88914844 | 25.84672 | 90.2271 | 86.88143 | . | . | . | . | . | . |
| chr13 | 11828679 | 11828911 | 485 | Intron (ENSMUST00000021750.8 | 13 | 11939558 | 11940543 | 986 | 1 | ENSMUST00 | -110647 | Gm26956 | ENSMUSG0C | false | . | . | . | . | . | . | . | . | . | . | . | . |
| chr10 | 21790121 | 21790413 | 485 | Distal Intergenic | 10 | 21743867 | 21747006 | 3140 | 1 | ENSMUST00 | 46255 | RP23-76C5.1 | ENSMUSG0C | false | . | . | . | . | . | . | chr10 | 21789609 | 21790594 | 4.42661 | 3.84106 | 2.52227 |
| chr19 | 58839779 | 58840104 | 484 | Intron (ENSMUST00000066285.4 | 19 | 58795751 | 58860984 | 65234 | 2 | ENSMUST00 | 20880 | Hspa12a | ENSMUSG0C | true | . | . | . | . | . | . | chr19 | 58814047 | 58859817 | 7.46659 | 5.12677 | 5.03638 |
| chr3 | 122422039 | 122422323 | 484 | Promoter (2-3kb) | 3 | 122419780 | 122530180 | 110401 | 1 | ENSMUST00 | 2260 | Bcar3 | ENSMUSG0C | true | chr3 | 122418997 | 122422602 | 20.02967 | 73.35366 | 70.31202 | chr3 | 122420960 | 122424183 | 4.12678 | 3.41379 | 2.32204 |
| chr4 | 117019270 | 117019621 | 484 | Promoter (<=1kb) | 4 | 117019402 | 117086852 | 67451 | 1 | ENSMUST00 | 0 | Eif2b3 | ENSMUSG0C | false | chr4 | 117018933 | 117020146 | 23.46333 | 91.19009 | 87.82124 | chr4 | 117018841 | 117020766 | 3.59792 | 3.35024 | 1.89117 |
| chr8 | 41278260 | 41278473 | 484 | Intron (ENSMUST00000045218.7 | 8 | 41239759 | 41334086 | 94328 | 1 | ENSMUST00 | 38502 | Pcm1 | ENSMUSG0C | false | . | . | . | . | . | . | . | . | . | . | . | . |
| chr9 | 83440970 | 83441277 | 484 | Promoter (<=1kb) | 9 | 83391364 | 83441127 | 49764 | 2 | ENSMUST00 | 0 | Lca5 | ENSMUSG0C | false | chr9 | 83440355 | 83441682 | 20.65177 | 61.66082 | 58.78991 | . | . | . | . | . | . |
| chr11 | 50138493 | 50138763 | 484 | Intron (ENSMUST00000093138.7 | 11 | 50135427 | 50140568 | 5142 | 1 | ENSMUST00 | 3067 | Tbc1d9b | ENSMUSG0C | false | . | . | . | . | . | . | chr11 | 50137549 | 50140215 | 5.00816 | 3.89784 | 3.04829 |
| chr15 | 98807920 | 98808179 | 484 | Promoter (<=1kb) | 15 | 98803782 | 98807925 | 4144 | 2 | ENSMUST00 | 0 | Ddn | ENSMUSG0C | true | chr15 | 98803793 | 98809569 | 29.66345 | 100.21 | 96.63225 | . | . | . | . | . | . |
| chr12 | 119024094 | 119024424 | 484 | Distal Intergenic | 12 | 118867824 | 118966421 | 98598 | 2 | ENSMUST00 | -57674 | Abcb5 | ENSMUSG0C | false | . | . | . | . | . | . | . | . | . | . | . | . |
| chr1 | 37345581 | 37345929 | 484 | Intron (ENSMUST00000146676.1 | 1 | 37333539 | 37334172 | 634 | 2 | ENSMUST00 | -11410 | 4930439A04I | ENSMUSG0C | false | . | . | . | . | . | . | chr1 | 37329617 | 37358130 | 5.31027 | 4.1114 | 3.2915 |
| chr2 | 72254914 | 72255252 | 484 | Intron (ENSMUST00000090826.6 | 2 | 72244965 | 72248975 | 4011 | 2 | ENSMUST00 | -5940 | Rapgef4os3 | ENSMUSG0C | false | . | . | . | . | . | . | chr2 | 72253306 | 72255856 | 5.20614 | 4.2309 | 3.20005 |
| chr15 | 79839304 | 79839590 | 484 | Distal Intergenic | 15 | 79823896 | 79834688 | 10793 | 2 | ENSMUST00 | -4617 | Cbx6 | ENSMUSG0C | false | . | . | . | . | . | . | chr15 | 79838736 | 79840249 | 4.46583 | 3.61194 | 2.60942 |
| chr16 | 44558187 | 44558643 | 484 | Promoter (<=1kb) | 16 | 44558046 | 44558129 | 84 | 2 | ENSMUST00 | -59 | Mir3081 | ENSMUSG0C | false | chr16 | 44557656 | 44559430 | 26.86698 | 95.04005 | 91.5885 | . | . | . | . | . | . |
| chr12 | 80123371 | 80123870 | 484 | Exon (ENSMUST00000180643.1/ | 12 | 80120548 | 80132844 | 12297 | 2 | ENSMUST00 | 8974 | 2310015A10I | ENSMUSG0C | false | . | . | . | . | . | . | chr12 | 80116083 | 80133218 | 5.88241 | 4.4251 | 3.73792 |
| chr3 | 95667977 | 95668253 | 483 | Distal Intergenic | 3 | 95676750 | 95677151 | 402 | 2 | ENSMUST00 | 8898 | Adamtsl4 | ENSMUSG0C | false | . | . | . | . | . | . | chr3 | 95660803 | 95677602 | 6.84699 | 4.91959 | 4.52508 |
| chr11 | 62551611 | 62551931 | 483 | Promoter (<=1kb) | 11 | 62551676 | 62552853 | 1178 | 1 | ENSMUST00 | 0 | Ubb | ENSMUSG0C | false | chr11 | 62551230 | 62554112 | 22.0147 | 58.97336 | 56.14162 | . | . | . | . | . | . |
| chr11 | 98775215 | 98776072 | 483 | Promoter (<=1kb) | 11 | 98767932 | 98775333 | 7402 | 2 | ENSMUST00 | 0 | Nr1d1 | ENSMUSG0C | false | chr11 | 98774164 | 98775366 | 20.85746 | 56.98322 | 54.17754 | chr11 | 98775631 | 98778417 | 8.10463 | 5.49842 | 5.54643 |
| chr17 | 27141647 | 27142210 | 483 | Downstream (1-2kb) | 17 | 27122663 | 27133916 | 11254 | 2 | ENSMUST00 | -7732 | Uqcc2 | ENSMUSG0C | false | . | . | . | . | . | . | chr17 | 27127363 | 27147146 | 5.99202 | 4.48057 | 3.83593 |
| chr18 | 70622480 | 70622719 | 483 | 3' UTR | 18 | 70617773 | 70625777 | 8005 | 1 | ENSMUST00 | 4708 | Mbd2 | ENSMUSG0C | false | . | . | . | . | . | . | . | . | . | . | . | . |
| chr2 | 35282196 | 35282514 | 483 | Promoter (<=1kb) | 2 | 35282380 | 35307892 | 25513 | 1 | ENSMUST00 | 0 | Gsn | ENSMUSG0C | true | chr2 | 35282148 | 35282846 | 10.51363 | 23.99435 | 21.60407 | chr2 | 35257475 | 35289539 | 6.87726 | 4.86078 | 4.55899 |
| chr2 | 127766919 | 127767268 | 483 | Intron (ENSMUST00000028857.8 | 2 | 127745622 | 127754070 | 8449 | 2 | ENSMUST00 | -12850 | Nphp1 | ENSMUSG0C | false | . | . | . | . | . | . | chr2 | 127767022 | 127767294 | 4.20948 | 3.59871 | 2.37493 |
| chr9 | 21798459 | 21798982 | 483 | Promoter (<=1kb) | 9 | 21766773 | 21798546 | 31774 | 2 | ENSMUST00 | 0 | Kank2 | ENSMUSG0C | false | chr9 | 21797469 | 21799074 | 28.04154 | 116.13699 | 112.1206 | chr9 | 21786534 | 21799409 | 6.22233 | 4.65692 | 4.01382 |
| chr6 | 96027909 | 96028284 | 483 | Distal Intergenic | 6 | 96113154 | 96657198 | 544045 | 1 | ENSMUST00 | -84870 | Fam19a1 | ENSMUSG0C | false | . | . | . | . | . | . | . | . | . | . | . | . |
| chr3 | 90989704 | 90989950 | 483 | Distal Intergenic | 3 | 91071662 | 91075195 | 3534 | 1 | ENSMUST00 | -81712 | 4930529C04I | ENSMUSG0C | false | . | . | . | . | . | . | . | . | . | . | . | . |
| chr3 | 65459728 | 65459958 | 483 | Distal Intergenic | 3 | 65525701 | 65525793 | 93 | 1 | ENSMUST00 | -65743 | Gm22279 | ENSMUSG0C | false | . | . | . | . | . | . | . | . | . | . | . | . |
| chr10 | 90432046 | 90432389 | 483 | Intron (ENSMUST00000183156.2 | 10 | 90450910 | 90451004 | 95 | 1 | ENSMUST00 | -18521 | Gm22602 | ENSMUSG0C | false | . | . | . | . | . | . | chr10 | 90431821 | 90432919 | 3.19043 | 3.16782 | 1.56474 |
| chr19 | 44303643 | 44303853 | 483 | Promoter (<=1kb) | 19 | 44303171 | 44303231 | 61 | 1 | ENSMUST00 | 473 | Mir5114 | ENSMUSG0C | false | . | . | . | . | . | . | chr19 | 44295070 | 44305832 | 5.98031 | 4.42848 | 3.84446 |
| chr7 | 141477930 | 141478355 | 482 | Promoter (1-2kb) | 7 | 141476400 | 141493421 | 17022 | 1 | ENSMUST00 | 1531 | Tspan4 | ENSMUSG0C | false | chr7 | 141477003 | 141478311 | 10.0704 | 19.92358 | 17.59046 | chr7 | 141456108 | 141486560 | 6.62038 | 4.76228 | 4.33989 |
| chr2 | 125858875 | 125859292 | 482 | Promoter (<=1kb) | 2 | 125830304 | 125859139 | 28836 | 2 | ENSMUST00 | 0 | Cops2 | ENSMUSG0C | false | chr2 | 125858259 | 125859691 | 44.9693 | 154.28859 | 148.83946 | . | . | . | . | . | . |
| chr2 | 173581006 | 173581403 | 482 | Promoter (2-3kb) | 2 | 173568666 | 173578365 | 9700 | 2 | ENSMUST00 | -2642 | Ankrd60 | ENSMUSG0C | false | . | . | . | . | . | . | . | . | . | . | . | . |
| chr9 | 49798731 | 49799186 | 482 | Promoter (<=1kb) | 9 | 49517506 | 49798922 | 281417 | 2 | ENSMUST00 | 0 | Ncam1 | ENSMUSG0C | true | chr9 | 49796389 | 49799150 | 29.85632 | 88.73936 | 85.41974 | . | . | . | . | . | . |
| chr15 | 83293155 | 83293506 | 482 | Distal Intergenic | 15 | 83226726 | 83251774 | 25049 | 2 | ENSMUST00 | -41382 | A4galt | ENSMUSG0C | false | . | . | . | . | . | . | chr15 | 83292734 | 83294592 | 3.48937 | 3.25018 | 1.80584 |
| chr9 | 15794402 | 15794678 | 482 | Distal Intergenic | 9 | 15862667 | 15874460 | 11794 | 2 | ENSMUST00 | 79782 | Mtnr1b | ENSMUSG0C | false | . | . | . | . | . | . | chr9 | 15792011 | 15795145 | 5.70064 | 4.41893 | 3.57968 |
| chr4 | 54995417 | 54995684 | 482 | Intron (ENSMUST00000133895.2 | 4 | 55011366 | 55081358 | 69993 | 1 | ENSMUST00 | -15682 | Zfp462 | ENSMUSG0C | true | . | . | . | . | . | . | chr4 | 54992233 | 55006975 | 4.97649 | 3.98236 | 2.99847 |
| chr19 | 24477103 | 24477673 | 482 | Promoter (<=1kb) | 19 | 24476502 | 24477356 | 855 | 2 | ENSMUST00 | 0 | Fam122a | ENSMUSG0C | false | chr19 | 24476142 | 24478155 | 31.88024 | 98.13424 | 94.612 | . | . | . | . | . | . |
| chr7 | 16043497 | 16043775 | 482 | Distal Intergenic | 7 | 16071942 | 16096328 | 24387 | 1 | ENSMUST00 | -28167 | Zfp541 | ENSMUSG0C | false | . | . | . | . | . | . | chr7 | 16025008 | 16047338 | 6.08815 | 4.49198 | 3.92077 |
| chr8 | 121474530 | 121475022 | 482 | Intron (ENSMUST00000127664.1 | 8 | 121434980 | 121435389 | 410 | 1 | ENSMUST00 | 39551 | Gm26784 | ENSMUSG0C | false | . | . | . | . | . | . | chr8 | 121474482 | 121477031 | 3.55199 | 3.37207 | 1.81668 |
| chr8 | 120215473 | 120215778 | 482 | Intron (ENSMUST00000127664.1 | 8 | 120204434 | 120228230 | 23797 | 2 | ENSMUST00 | 12452 | A330074K22 | ENSMUSG0C | false | . | . | . | . | . | . | chr8 | 120214640 | 120217939 | 5.0824 | 4.17837 | 3.06437 |
| chr6 | 93066313 | 93066888 | 482 | Intron (ENSMUST00000181121.2 | 6 | 93137004 | 93137990 | 987 | 2 | ENSMUST00 | 71102 | Gm5313 | ENSMUSG0C | false | chr6 | 93065854 | 93066981 | 8.08052 | 18.53858 | 16.22368 | chr6 | 93061983 | 93070872 | 5.96536 | 4.4407 | 3.83891 |
| chr12 | 79965942 | 79966336 | 482 | Promoter (2-3kb) | 12 | 79963568 | 79963872 | 305 | 1 | ENSMUST00 | 2375 | RP23-179D3. | ENSMUSG0C | false | chr12 | 79965323 | 79966133 | 13.09357 | 30.27553 | 27.8073 | chr12 | 79963528 | 79966903 | 5.11632 | 4.0702 | 3.14196 |

| | | | | | | | | | | | | | | | | | | | | | | | | | | |
|---|---|---|---|---|---|---|---|---|---|---|---|---|---|---|---|---|---|---|---|---|---|---|---|---|---|---|
| chr3 | 51203008 | 51203260 | 481 | Distal Intergenic | 3 | 51224447 | 51251644 | 27198 | 1 | ENSMUST00 | -21187 | Ccrn4l | ENSMUSG0C | false | . | . | . | . | . | . | chr3 | 51201033 | 51204192 | 6.45222 | 4.83018 | 4.17698 |
| chr5 | 124537062 | 124537484 | 481 | Distal Intergenic | 5 | 124540695 | 124550506 | 9812 | 1 | ENSMUST00 | -3211 | Tmed2 | ENSMUSG0C | false | chr5 | 124536865 | 124537314 | 6.9004 | 12.33775 | 10.12162 | chr5 | 124536433 | 124537921 | 4.82409 | 3.97641 | 2.8813 |
| chr6 | 99299273 | 99299559 | 481 | Intron (ENSMUST00000113322.3 | 6 | 99161940 | 99266532 | 104593 | 2 | ENSMUST00 | -32742 | Foxp1 | ENSMUSG0C | false | . | . | . | . | . | . | . | . | . | . | . | . |
| chr5 | 135632537 | 135632908 | 481 | Promoter (<=1kb) | 5 | 135632618 | 135646376 | 13759 | 1 | ENSMUST00 | 0 | Rhbdd2 | ENSMUSG0C | false | chr5 | 135632628 | 135633357 | 31.36125 | 107.32627 | 103.56666 | chr5 | 135632185 | 135633821 | 4.68801 | 4.01848 | 2.73756 |
| chr2 | 109931952 | 109932346 | 481 | Intron (ENSMUST00000046548.8 | 2 | 109917647 | 110014257 | 96611 | 1 | ENSMUST00 | 14306 | Lgr4 | ENSMUSG0C | false | . | . | . | . | . | . | chr2 | 109920715 | 109937278 | 4.86516 | 4.02197 | 2.89791 |
| chr7 | 113247232 | 113247513 | 481 | Intron (ENSMUST00000047321.7 | 7 | 113207465 | 113314122 | 106658 | 1 | ENSMUST00 | 39768 | Arntl | ENSMUSG0C | false | . | . | . | . | . | . | chr7 | 113246286 | 113248300 | 5.80195 | 4.45855 | 3.70568 |
| chr13 | 35503595 | 35503810 | 481 | Distal Intergenic | 13 | 35659862 | 35874064 | 214203 | 1 | ENSMUST00 | -156052 | Cdyl | ENSMUSG0C | true | . | . | . | . | . | . | chr13 | 35501065 | 35504562 | 3.4911 | 3.31509 | 1.80062 |
| chr8 | 122511755 | 122512008 | 481 | Intron (ENSMUST00000127664.1 | 8 | 122520568 | 122520674 | 107 | 2 | ENSMUST00 | 8666 | Gm26497 | ENSMUSG0C | false | . | . | . | . | . | . | chr8 | 122510127 | 122521302 | 5.39156 | 4.16502 | 3.34791 |
| chr17 | 5022084 | 5022326 | 481 | Intron (ENSMUST00000124362.1 | 17 | 5040578 | 5127640 | 87063 | 1 | ENSMUST00 | -18252 | Arid1b | ENSMUSG0C | false | . | . | . | . | . | . | chr17 | 5020739 | 5025977 | 5.86774 | 4.15794 | 3.74996 |
| chr4 | 129033568 | 129033832 | 481 | Distal Intergenic | 4 | 129050097 | 129057983 | 7887 | 2 | ENSMUST00 | 24151 | 1700086P04I | ENSMUSG0C | false | . | . | . | . | . | . | chr4 | 129032304 | 129035991 | 5.85449 | 4.44408 | 3.71528 |
| chr12 | 30561652 | 30561874 | 481 | Distal Intergenic | 12 | 30557914 | 30558120 | 207 | 2 | ENSMUST00 | -3533 | Gm25243 | ENSMUSG0C | false | . | . | . | . | . | . | chr12 | 30561115 | 30561668 | 4.78373 | 4.06501 | 2.81769 |
| chr8 | 121382516 | 121382985 | 481 | Intron (ENSMUST00000127664.1 | 8 | 121256747 | 121356836 | 100090 | 2 | ENSMUST00 | -25681 | Gm26815 | ENSMUSG0C | false | . | . | . | . | . | . | chr8 | 121372330 | 121385684 | 7.83977 | 5.33422 | 5.34287 |
| chr16 | 34966547 | 34966930 | 481 | Intron (ENSMUST00000023538.8 | 16 | 34935854 | 34939329 | 3476 | 2 | ENSMUST00 | -27219 | RP23-80O10 | ENSMUSG0C | false | . | . | . | . | . | . | chr16 | 34923663 | 34969430 | 6.94598 | 4.91232 | 4.61654 |
| chr14 | 73458546 | 73458803 | 480 | Distal Intergenic | 14 | 73510049 | 73518545 | 8497 | 1 | ENSMUST00 | -51246 | Med4 | ENSMUSG0C | false | . | . | . | . | . | . | chr14 | 73454764 | 73459121 | 3.97379 | 3.57554 | 2.16305 |
| chr16 | 32003011 | 32003335 | 480 | Promoter (<=1kb) | 16 | 31962506 | 32003287 | 40782 | 2 | ENSMUST00 | 0 | Senp5 | ENSMUSG0C | false | chr16 | 32002157 | 32003778 | 28.56742 | 103.16996 | 99.51957 | . | . | . | . | . | . |
| chr19 | 7240949 | 7241332 | 480 | Promoter (<=1kb) | 19 | 7225668 | 7241222 | 15555 | 2 | ENSMUST00 | 0 | Naa40 | ENSMUSG0C | false | chr19 | 7240402 | 7241621 | 27.78601 | 91.88203 | 88.49705 | chr19 | 7239533 | 7241659 | 3.73899 | 3.36841 | 2.01247 |
| chr19 | 6233631 | 6234094 | 480 | Promoter (1-2kb) | 19 | 6233937 | 6235843 | 1907 | 2 | ENSMUST00 | 1749 | Ppp2r5b | ENSMUSG0C | false | . | . | . | . | . | . | chr19 | 6233119 | 6235365 | 5.36835 | 4.02057 | 3.33579 |
| chr8 | 121947899 | 121948197 | 480 | Promoter (1-2kb) | 8 | 121949750 | 121997183 | 47434 | 1 | ENSMUST00 | -1553 | Banp | ENSMUSG0C | false | chr8 | 121947767 | 121948704 | 7.13425 | 13.57188 | 11.33167 | chr8 | 121947602 | 121949956 | 4.45765 | 3.71529 | 2.58863 |
| chr3 | 95666202 | 95666432 | 480 | Distal Intergenic | 3 | 95658857 | 95660823 | 1967 | 1 | ENSMUST00 | 7346 | Mcl1 | ENSMUSG0C | false | . | . | . | . | . | . | chr3 | 95660803 | 95677602 | 6.84699 | 4.91959 | 4.52508 |
| chr13 | 37892799 | 37893009 | 480 | Intron (ENSMUST00000122842.2 | 13 | 37888871 | 37951994 | 63124 | 1 | ENSMUST00 | 3929 | Rreb1 | ENSMUSG0C | false | . | . | . | . | . | . | chr13 | 37891842 | 37893472 | 3.8625 | 3.50011 | 2.11561 |
| chr10 | 80165686 | 80166051 | 480 | Promoter (<=1kb) | 10 | 80165985 | 80170316 | 4332 | 1 | ENSMUST00 | 0 | Cirbp | ENSMUSG0C | false | chr10 | 80165265 | 80166711 | 11.91027 | 25.79263 | 23.38204 | chr10 | 80150859 | 80167428 | 9.09735 | 5.82258 | 6.36001 |
| chr13 | 30356492 | 30356706 | 480 | Intron (ENSMUST00000066412.7 | 13 | 30336441 | 30382867 | 46427 | 1 | ENSMUST00 | 20052 | Agtr1a | ENSMUSG0C | false | . | . | . | . | . | . | chr13 | 30355325 | 30357114 | 4.48426 | 3.74781 | 2.60031 |
| chr2 | 38379328 | 38379603 | 480 | Distal Intergenic | 2 | 38359580 | 38363980 | 4401 | 2 | ENSMUST00 | -15349 | Gm13584 | ENSMUSG0C | false | chr2 | 38378964 | 38379669 | 10.96168 | 23.03899 | 20.66252 | chr2 | 38378780 | 38381081 | 5.35169 | 4.07484 | 3.31351 |
| chr13 | 69944719 | 69945025 | 480 | Distal Intergenic | 13 | 70007204 | 70007614 | 411 | 2 | ENSMUST00 | 62589 | Gm6132 | ENSMUSG0C | false | . | . | . | . | . | . | chr13 | 69943118 | 69947833 | 4.94671 | 3.94855 | 2.98156 |
| chr15 | 38483529 | 38483756 | 480 | Distal Intergenic | 15 | 38490473 | 38490547 | 75 | 2 | ENSMUST00 | 6791 | Mir6951 | ENSMUSG0C | false | . | . | . | . | . | . | chr15 | 38483195 | 38484750 | 5.01372 | 4.02929 | 3.04203 |
| chr18 | 35842677 | 35842911 | 480 | Intron (ENSMUST00000060722.6 | 18 | 35830389 | 35831093 | 705 | 2 | ENSMUST00 | -11585 | RP24-326K1: | ENSMUSG0C | false | . | . | . | . | . | . | chr18 | 35831800 | 35857612 | 7.70883 | 5.15861 | 5.24645 |
| chr11 | 30123951 | 30124273 | 479 | Exon (ENSMUST00000006629.8/ | 11 | 30109871 | 30111359 | 1489 | 2 | ENSMUST00 | -12593 | Sptbn1 | ENSMUSG0C | false | . | . | . | . | . | . | . | . | . | . | . | . |
| chr14 | 14612383 | 14612712 | 479 | Distal Intergenic | 14 | 14703025 | 14799943 | 96919 | 1 | ENSMUST00 | -90313 | Slc4a7 | ENSMUSG0C | false | . | . | . | . | . | . | . | . | . | . | . | . |
| chr14 | 79587510 | 79587818 | 479 | Promoter (<=1kb) | 14 | 79587691 | 79629755 | 42065 | 1 | ENSMUST00 | 0 | Sugt1 | ENSMUSG0C | false | chr14 | 79586786 | 79588351 | 25.5656 | 84.01802 | 80.7898 | . | . | . | . | . | . |
| chr4 | 129818603 | 129818888 | 479 | Promoter (1-2kb) | 4 | 129820221 | 129849978 | 29758 | 1 | ENSMUST00 | -1333 | Ptp4a2 | ENSMUSG0C | false | . | . | . | . | . | . | chr4 | 129818600 | 129819771 | 5.74365 | 4.5042 | 3.61498 |
| chr19 | 25243647 | 25243886 | 479 | Intron (ENSMUST00000049400.9 | 19 | 25236975 | 25434496 | 197522 | 1 | ENSMUST00 | 6673 | Kank1 | ENSMUSG0C | true | . | . | . | . | . | . | chr19 | 25239789 | 25260438 | 5.72271 | 4.39219 | 3.6112 |
| chr10 | 117072625 | 117072844 | 479 | 3' UTR | 10 | 117050998 | 117063814 | 12817 | 2 | ENSMUST00 | -8812 | Cct2 | ENSMUSG0C | false | . | . | . | . | . | . | chr10 | 117070579 | 117074194 | 4.8326 | 3.84927 | 2.90208 |
| chrX | 77795787 | 77796086 | 479 | Promoter (<=1kb) | 20 | 77771455 | 77795972 | 24518 | 2 | ENSMUST00 | 0 | Prkx | ENSMUSG0C | false | chrX | 77794773 | 77796146 | 22.51707 | 59.83231 | 56.98677 | . | . | . | . | . | . |
| chr11 | 116418872 | 116419155 | 479 | Distal Intergenic | 11 | 116402841 | 116413032 | 10192 | 2 | ENSMUST00 | -5841 | Rnf157 | ENSMUSG0C | false | . | . | . | . | . | . | chr11 | 116416965 | 116419228 | 6.20979 | 4.72389 | 4.00624 |
| chr7 | 19824079 | 19824504 | 479 | Promoter (1-2kb) | 7 | 19812395 | 19822770 | 10376 | 2 | ENSMUST00 | -1310 | Bcl3 | ENSMUSG0C | false | . | . | . | . | . | . | chr7 | 19821732 | 19825440 | 4.97308 | 4.10705 | 2.98414 |
| chr15 | 91191721 | 91192032 | 479 | Promoter (<=1kb) | 15 | 91145884 | 91191807 | 45924 | 2 | ENSMUST00 | 0 | Abcd2 | ENSMUSG0C | false | chr15 | 91191397 | 91192234 | 10.33961 | 21.99109 | 19.62971 | chr15 | 91191626 | 91192223 | 2.9935 | 2.98749 | 1.37737 |
| chr9 | 48723493 | 48723995 | 479 | Intron (ENSMUST00000093852.3 | 9 | 48654311 | 48835945 | 181635 | 2 | ENSMUST00 | 111950 | Zbtb16 | ENSMUSG0C | false | . | . | . | . | . | . | chr9 | 48721539 | 48724828 | 4.49384 | 3.84728 | 2.57449 |
| chr2 | 168185878 | 168186097 | 479 | Intron (ENSMUST00000057793.5 | 2 | 168178158 | 168178252 | 95 | 2 | ENSMUST00 | -7627 | Gm24561 | ENSMUSG0C | false | . | . | . | . | . | . | chr2 | 168184284 | 168203070 | 5.58311 | 4.24493 | 3.51854 |
| chr5 | 118963203 | 118963425 | 479 | Distal Intergenic | 5 | 119108234 | 119114543 | 6310 | 1 | ENSMUST00 | -144809 | RP24-389L1C | ENSMUSG0C | false | . | . | . | . | . | . | . | . | . | . | . | . |
| chr16 | 6707593 | 6707800 | 478 | Distal Intergenic | 16 | 6809222 | 7409813 | 600592 | 1 | ENSMUST00 | -101422 | Rbfox1 | ENSMUSG0C | false | . | . | . | . | . | . | . | . | . | . | . | . |
| chrX | 9272532 | 9272889 | 478 | Promoter (<=1kb) | 20 | 9272756 | 9313250 | 40495 | 1 | ENSMUST00 | 0 | Xk | ENSMUSG0C | false | chrX | 9272313 | 9273362 | 22.51707 | 59.83231 | 56.98677 | . | . | . | . | . | . |
| chr10 | 10546712 | 10546957 | 478 | Intron (ENSMUST00000019974.3 | 10 | 10545011 | 10558233 | 13223 | 2 | ENSMUST00 | 11276 | Rab32 | ENSMUSG0C | false | chr10 | 10546397 | 10547016 | 9.19064 | 17.46287 | 15.16242 | chr10 | 10546374 | 10547058 | 3.13541 | 3.13759 | 1.47325 |
| chr19 | 53311751 | 53312158 | 478 | Promoter (1-2kb) | 19 | 53310506 | 53375810 | 65305 | 1 | ENSMUST00 | 1246 | Mxi1 | ENSMUSG0C | false | chr19 | 53310158 | 53314217 | 35.4076 | 109.31139 | 105.49952 | . | . | . | . | . | . |
| chr1 | 14431955 | 14432210 | 478 | Distal Intergenic | 1 | 14168958 | 14310235 | 141278 | 2 | ENSMUST00 | -121721 | Eya1 | ENSMUSG0C | false | . | . | . | . | . | . | chr1 | 14431217 | 14433056 | 4.17758 | 3.59818 | 2.35224 |
| chr2 | 52858378 | 52859101 | 478 | Promoter (<=1kb) | 2 | 52857868 | 53134202 | 276335 | 1 | ENSMUST00 | 511 | Fmnl2 | ENSMUSG0C | true | chr2 | 52857275 | 52859971 | 40.88443 | 132.61177 | 128.02734 | . | . | . | . | . | . |
| chr8 | 61933806 | 61934118 | 478 | Intron (ENSMUST00000154398.2 | 8 | 61928089 | 62037701 | 109613 | 1 | ENSMUST00 | 5718 | Ddx60 | ENSMUSG0C | false | . | . | . | . | . | . | chr8 | 61932833 | 61934772 | 4.50552 | 3.79737 | 2.63433 |
| chr14 | 27044299 | 27044623 | 478 | Intron (ENSMUST00000035336.3 | 14 | 27039001 | 27107286 | 68286 | 1 | ENSMUST00 | 5299 | Il17rd | ENSMUSG0C | false | . | . | . | . | . | . | chr14 | 27041643 | 27048842 | 5.16084 | 3.99802 | 3.16031 |
| chr19 | 10695855 | 10696132 | 478 | Intron (ENSMUST00000087951.5 | 19 | 10688815 | 10714419 | 25605 | 1 | ENSMUST00 | 7041 | Vps37c | ENSMUSG0C | false | . | . | . | . | . | . | chr19 | 10693823 | 10703744 | 5.86842 | 4.33637 | 3.75507 |
| chr11 | 5761882 | 5762567 | 478 | Promoter (<=1kb) | 11 | 5713417 | 5762376 | 48960 | 2 | ENSMUST00 | 0 | Urgcp | ENSMUSG0C | false | chr11 | 5761289 | 5762854 | 33.94791 | 109.87469 | 106.04007 | chr11 | 5758473 | 5762853 | 4.20303 | 3.66451 | 2.39807 |
| chr6 | 86849340 | 86849723 | 478 | Promoter (<=1kb) | 6 | 86849517 | 86991864 | 142348 | 1 | ENSMUST00 | 0 | Aak1 | ENSMUSG0C | false | chr6 | 86848422 | 86851621 | 24.48636 | 83.88964 | 80.66612 | . | . | . | . | . | . |
| chr17 | 86572636 | 86572892 | 478 | Intron (ENSMUST00000097275.3 | 17 | 86497732 | 86505232 | 7501 | 2 | ENSMUST00 | -67405 | Gm10309 | ENSMUSG0C | false | . | . | . | . | . | . | chr17 | 86571950 | 86574521 | 4.92186 | 4.00823 | 2.94101 |
| chr15 | 95216208 | 95216617 | 478 | Downstream (2-3kb) | 15 | 95097542 | 95162394 | 64853 | 2 | ENSMUST00 | -53815 | RP24-303N1 | ENSMUSG0C | false | . | . | . | . | . | . | chr15 | 95213810 | 95221642 | 5.84437 | 4.48428 | 3.6948 |
| chr10 | 13868749 | 13869097 | 477 | Promoter (<=1kb) | 10 | 13647054 | 13868980 | 221927 | 2 | ENSMUST00 | 0 | Aig1 | ENSMUSG0C | false | chr10 | 13868363 | 13869834 | 22.15371 | 67.81625 | 64.85734 | chr10 | 13867135 | 13870037 | 3.71167 | 3.33932 | 1.98178 |
| chr9 | 119894571 | 119895127 | 477 | Promoter (<=1kb) | 9 | 119894895 | 119926579 | 31685 | 1 | ENSMUST00 | 0 | Wdr48 | ENSMUSG0C | false | chr9 | 119894501 | 119895905 | 33.83522 | 116.97303 | 112.93475 | . | . | . | . | . | . |
| chr1 | 191317747 | 191318208 | 477 | Promoter (<=1kb) | 1 | 191306789 | 191318194 | 11406 | 2 | ENSMUST00 | 0 | Nenf | ENSMUSG0C | false | chr1 | 191317309 | 191318274 | 38.14117 | 119.25307 | 115.13219 | . | . | . | . | . | . |
| chr2 | 156257098 | 156257361 | 477 | Intron (ENSMUST00000037401.8 | 2 | 156263185 | 156272624 | 9440 | 1 | ENSMUST00 | -5824 | Phf20 | ENSMUSG0C | false | . | . | . | . | . | . | chr2 | 156254965 | 156259335 | 5.1137 | 4.00052 | 3.13456 |
| chr18 | 21299729 | 21300530 | 477 | Promoter (<=1kb) | 18 | 21127201 | 21300138 | 172938 | 2 | ENSMUST00 | 0 | Garem | ENSMUSG0C | false | chr18 | 21299173 | 21300235 | 35.67123 | 127.73884 | 123.33056 | . | . | . | . | . | . |
| chr15 | 80798426 | 80798703 | 477 | Intron (ENSMUST00000067689.7 | 15 | 80711319 | 80941086 | 229768 | 1 | ENSMUST00 | 87108 | Tnrc6b | ENSMUSG0C | false | chr15 | 80797921 | 80799937 | 23.19226 | 64.40237 | 61.49533 | . | . | . | . | . | . |
| chr18 | 62943313 | 62943686 | 477 | Intron (ENSMUST00000096554.5 | 18 | 62922329 | 62952464 | 30136 | 1 | ENSMUST00 | 20985 | Apcdd1 | ENSMUSG0C | false | . | . | . | . | . | . | . | . | . | . | . | . |
| chr2 | 118072762 | 118073124 | 477 | Intron (ENSMUST00000148587.2 | 2 | 118077805 | 118077962 | 158 | 2 | ENSMUST00 | 4838 | Gm25308 | ENSMUSG0C | false | . | . | . | . | . | . | chr2 | 118070959 | 118074915 | 4.18262 | 3.59015 | 2.34681 |
| chr4 | 53952363 | 53952605 | 477 | Distal Intergenic | 4 | 53905696 | 53905820 | 125 | 1 | ENSMUST00 | 46668 | Gm23745 | ENSMUSG0C | false | . | . | . | . | . | . | chr4 | 53948932 | 53953603 | 3.90187 | 3.45261 | 2.1414 |
| chr2 | 75639466 | 75640123 | 477 | Distal Intergenic | 2 | 75632638 | 75633382 | 745 | 2 | ENSMUST00 | -6085 | Gm13655 | ENSMUSG0C | false | chr2 | 75639299 | 75639865 | 13.0727 | 32.19727 | 29.70434 | chr2 | 75639108 | 75641400 | 3.67467 | 3.41747 | 1.92434 |
| chr13 | 28989114 | 28989354 | 477 | Distal Intergenic | 13 | 29014399 | 29015665 | 1267 | 1 | ENSMUST00 | -25045 | A330102I10F | ENSMUSG0C | false | . | . | . | . | . | . | . | . | . | . | . | . |
| chr4 | 148325101 | 148325413 | 477 | Distal Intergenic | 4 | 148306853 | 148316569 | 9717 | 1 | ENSMUST00 | 18249 | Gm13206 | ENSMUSG0C | false | . | . | . | . | . | . | chr4 | 148324659 | 148328387 | 5.58334 | 4.25719 | 3.51337 |
| chr2 | 154615120 | 154615394 | 477 | Promoter (1-2kb) | 2 | 154612937 | 154613437 | 501 | 2 | ENSMUST00 | -1684 | 4930519P11I | ENSMUSG0C | false | . | . | . | . | . | . | chr2 | 154614052 | 154617001 | 3.13056 | 3.06236 | 1.4846 |
| chr13 | 81783543 | 81783973 | 477 | Promoter (<=1kb) | 13 | 81778773 | 81783749 | 4977 | 2 | ENSMUST00 | 0 | Gm17259 | ENSMUSG0C | false | chr13 | 81782643 | 81784311 | 32.48404 | 109.34737 | 105.53431 | . | . | . | . | . | . |
| chr17 | 28622095 | 28622658 | 476 | Promoter (<=1kb) | 17 | 28587648 | 28622521 | 34874 | 2 | ENSMUST00 | 0 | Srpk1 | ENSMUSG0C | false | chr17 | 28621359 | 28622908 | 37.30079 | 124.19571 | 119.91465 | . | . | . | . | . | . |
| chr10 | 21993630 | 21993967 | 476 | Promoter (<=1kb) | 10 | 21993890 | 21999891 | 6002 | 1 | ENSMUST00 | 0 | Sgk1 | ENSMUSG0C | false | chr10 | 21992730 | 21996734 | 36.39542 | 121.53264 | 117.33772 | chr10 | 21991101 | 21993894 | 3.93618 | 3.38892 | 2.16796 |
| chr11 | 57525261 | 57525521 | 476 | Promoter (2-3kb) | 11 | 57522964 | 57529526 | 6563 | 1 | ENSMUST00 | 2298 | Mfap3 | ENSMUSG0C | false | . | . | . | . | . | . | chr11 | 57523233 | 57526409 | 6.15754 | 4.61018 | 3.95451 |
| chr1 | 20890416 | 20890772 | 476 | Promoter (<=1kb) | 1 | 20890606 | 20939650 | 49045 | 1 | ENSMUST00 | 0 | Paqr8 | ENSMUSG0C | false | chr1 | 20890081 | 20891582 | 27.41053 | 90.23362 | 86.88379 | . | . | . | . | . | . |
| chr1 | 75359410 | 75359747 | 476 | Promoter (<=1kb) | 1 | 75360329 | 75368579 | 8251 | 1 | ENSMUST00 | -582 | Des | ENSMUSG0C | false | . | . | . | . | . | . | chr1 | 75356622 | 75362384 | 5.80426 | 4.47779 | 3.65868 |
| chr3 | 36642804 | 36643061 | 476 | Promoter (2-3kb) | 3 | 36638356 | 36640620 | 2265 | 2 | ENSMUST00 | -2185 | Trpc3 | ENSMUSG0C | false | . | . | . | . | . | . | chr3 | 36642800 | 36643263 | 3.38279 | 3.30887 | 1.72286 |
| chr4 | 19569933 | 19570258 | 476 | Promoter (<=1kb) | 4 | 19519254 | 19570108 | 50855 | 2 | ENSMUST00 | 0 | Cpne3 | ENSMUSG0C | false | chr4 | 19569105 | 19570425 | 32.64849 | 123.19247 | 118.94151 | . | . | . | . | . | . |
| chr7 | 100988265 | 100988542 | 476 | Distal Intergenic | 7 | 100997227 | 101011976 | 14750 | 2 | ENSMUST00 | 23434 | P2ry2 | ENSMUSG0C | false | . | . | . | . | . | . | chr7 | 100987831 | 100988649 | 6.13206 | 4.63389 | 3.99331 |
| chr11 | 98960236 | 98960797 | 476 | Promoter (<=1kb) | 11 | 98960412 | 98974942 | 14531 | 1 | ENSMUST00 | 0 | Rara | ENSMUSG0C | false | chr11 | 98959620 | 98961343 | 26.65955 | 86.95457 | 83.67183 | chr11 | 98958785 | 98973618 | 4.55278 | 3.71427 | 2.6612 |
| chr7 | 145199804 | 145200204 | 476 | Distal Intergenic | 7 | 145253923 | 145283927 | 30005 | 2 | ENSMUST00 | 83723 | Tpcn2 | ENSMUSG0C | false | . | . | . | . | . | . | chr7 | 145196929 | 145200615 | 5.05083 | 3.9985 | 3.09965 |
| chr14 | 52104249 | 52104704 | 476 | Promoter (<=1kb) | 14 | 52073380 | 52104028 | 30649 | 2 | ENSMUST00 | -222 | Hnrnpc | ENSMUSG0C | false | chr14 | 52104443 | 52104825 | 4.30255 | 8.94291 | 6.78887 | chr14 | 52104611 | 52107783 | 3.56773 | 3.26365 | 1.84677 |
| chr18 | 46311553 | 46312075 | 476 | Promoter (<=1kb) | 18 | 46282151 | 46311928 | 29778 | 2 | ENSMUST00 | 0 | Ccdc112 | ENSMUSG0C | false | chr18 | 46311332 | 46312310 | 27.88522 | 83.41428 | 80.20149 | chr18 | 46311244 | 46312377 | 3.41011 | 3.26855 | 1.74071 |
| chr1 | 131462421 | 131462691 | 476 | Intron (ENSMUST00000097588.3 | 1 | 131447440 | 131447695 | 256 | 2 | ENSMUST00 | -14727 | RP24-302F18 | ENSMUSG0C | false | . | . | . | . | . | . | chr1 | 131451954 | 131498502 | 5.16635 | 4.13104 | 3.14877 |
| chr10 | 58604776 | 58605355 | 475 | Intron (ENSMUST00000003312.4 | 10 | 58600789 | 58675654 | 74866 | 2 | ENSMUST00 | 70299 | Edar | ENSMUSG0C | false | . | . | . | . | . | . | chr10 | 58604191 | 58605873 | 4.39885 | 3.75442 | 2.57152 |
| chr14 | 87415781 | 87417330 | 475 | Promoter (<=1kb) | 14 | 87416639 | 87514766 | 98128 | 1 | ENSMUST00 | 0 | Tdrd3 | ENSMUSG0C | false | chr14 | 87415190 | 87418026 | 30.39373 | 108.60838 | 104.8139 | . | . | . | . | . | . |
| chr19 | 10038917 | 10039338 | 475 | Promoter (2-3kb) | 19 | 10041548 | 10059671 | 18124 | 1 | ENSMUST00 | -2210 | Fads3 | ENSMUSG0C | false | . | . | . | . | . | . | chr19 | 10036253 | 10050591 | 5.31313 | 4.14637 | 3.29774 |
| chr2 | 25982944 | 25983663 | 475 | Promoter (<=1kb) | 2 | 25926838 | 25983282 | 56445 | 2 | ENSMUST00 | 0 | Camsap1 | ENSMUSG0C | false | chr2 | 25982006 | 25983386 | 26.52689 | 93.43019 | 90.01563 | . | . | . | . | . | . |
| chr6 | 86775169 | 86775442 | 475 | Intron (ENSMUST00000001187.9 | 6 | 86747471 | 86765910 | 18440 | 2 | ENSMUST00 | -9260 | Anxa4 | ENSMUSG0C | true | . | . | . | . | . | . | chr6 | 86761092 | 86794291 | 5.26625 | 4.09177 | 3.24957 |
| chr7 | 28358165 | 28358632 | 475 | Downstream (<1kb) | 7 | 28359605 | 28363341 | 3737 | 2 | ENSMUST00 | 4709 | Plekhg2 | ENSMUSG0C | true | . | . | . | . | . | . | chr7 | 28351351 | 28371775 | 6.79815 | 4.79627 | 4.48731 |
| chr18 | 34776834 | 34777883 | 475 | Promoter (<=1kb) | 18 | 34777008 | 34839370 | 62363 | 1 | ENSMUST00 | 0 | Kdm3b | ENSMUSG0C | false | chr18 | 34776612 | 34778537 | 35.62435 | 116.99458 | 112.94973 | . | . | . | . | . | . |
| chr5 | 149184226 | 149184835 | 475 | Promoter (<=1kb) | 5 | 149184350 | 149215428 | 31079 | 1 | ENSMUST00 | 0 | Uspl1 | ENSMUSG0C | false | chr5 | 149184565 | 149185728 | 27.18497 | 79.384 | 76.2402 | . | . | . | . | . | . |
| chr3 | 52196252 | 52196604 | 475 | Distal Intergenic | 3 | 52268337 | 52350109 | 81773 | 1 | ENSMUST00 | -71733 | Foxo1 | ENSMUSG0C | false | chr3 | 52196002 | 52196469 | 13.83063 | 32.8443 | 30.34371 | chr3 | 52195612 | 52200384 | 5.09884 | 4.11277 | 3.07456 |
| chr11 | 4629040 | 4629290 | 475 | Promoter (<=1kb) | 11 | 4625376 | 4630011 | 4636 | 2 | ENSMUST00 | 721 | Gm11961 | ENSMUSG0C | false | . | . | . | . | . | . | chr11 | 4629058 | 4631520 | 4.22618 | 3.54226 | 2.39735 |
| chr9 | 44107059 | 44107374 | 475 | Promoter (<=1kb) | 9 | 44107245 | 44109184 | 1940 | 1 | ENSMUST00 | 0 | C1qtnf5 | ENSMUSG0C | false | . | . | . | . | . | . | chr9 | 44105712 | 44110614 | 7.52605 | 5.20704 | 5.08185 |
| chr3 | 121321001 | 121321248 | 475 | Intron (ENSMUST00000039442.7 | 3 | 121337692 | 121337959 | 268 | 1 | ENSMUST00 | -16444 | Gm26027 | ENSMUSG0C | false | . | . | . | . | . | . | chr3 | 121320599 | 121321780 | 5.5124 | 4.38153 | 3.41884 |
| chr9 | 79920543 | 79920843 | 475 | Intron (ENSMUST00000093811.5 | 9 | 79913200 | 79913631 | 432 | 1 | ENSMUST00 | 7344 | Gm3211 | ENSMUSG0C | false | . | . | . | . | . | . | chr9 | 79911389 | 79923121 | 5.13391 | 4.11345 | 3.12085 |
| chr10 | 80627653 | 80627988 | 474 | Promoter (1-2kb) | 10 | 80629656 | 80640771 | 11116 | 1 | ENSMUST00 | -1668 | Csnk1g2 | ENSMUSG0C | false | . | . | . | . | . | . | chr10 | 80623344 | 80633861 | 6.51298 | 4.68538 | 4.26272 |
| chr11 | 100610336 | 100610710 | 474 | Promoter (2-3kb) | 11 | 100592964 | 100613342 | 20379 | 2 | ENSMUST00 | 2632 | Dnajc7 | ENSMUSG0C | false | . | . | . | . | . | . | chr11 | 100601454 | 100614332 | 6.03764 | 4.44868 | 3.8868 |
| chr14 | 54577294 | 54578410 | 474 | Promoter (<=1kb) | 14 | 54567469 | 54577661 | 10193 | 2 | ENSMUST00 | 0 | Ajuba | ENSMUSG0C | false | chr14 | 54574109 | 54578573 | 28.90341 | 112.84859 | 108.93253 | chr14 | 54578166 | 54580092 | 6.14347 | 4.64478 | 3.92215 |
| chr18 | 75434596 | 75434892 | 474 | 3' UTR | 18 | 75374914 | 75394001 | 19088 | 1 | ENSMUST00 | 59683 | Smad7 | ENSMUSG0C | false | . | . | . | . | . | . | chr18 | 75433660 | 75435370 | 4.7361 | 3.99373 | 2.80168 |
| chr2 | 71274493 | 71274743 | 474 | 3' UTR | 2 | 71279561 | 71282789 | 3229 | 2 | ENSMUST00 | 8046 | Slc25a12 | ENSMUSG0C | false | chr2 | 71274009 | 71275047 | 11.72773 | 26.68839 | 24.26382 | chr2 | 71273784 | 71276164 | 4.57184 | 3.89277 | 2.66132 |
| chr2 | 119345192 | 119345522 | 474 | Distal Intergenic | 2 | 119351229 | 119354381 | 3153 | 1 | ENSMUST00 | -5707 | Chac1 | ENSMUSG0C | false | . | . | . | . | . | . | chr2 | 119344838 | 119346050 | 3.98635 | 3.5077 | 2.17672 |
| chr3 | 90253990 | 90254743 | 474 | Promoter (<=1kb) | 3 | 90254163 | 90264125 | 9963 | 1 | ENSMUST00 | 0 | Crtc2 | ENSMUSG0C | false | chr3 | 90253884 | 90255447 | 37.22259 | 125.13314 | 120.81451 | . | . | . | . | . | . |
| chr6 | 108282009 | 108282251 | 474 | Intron (ENSMUST00000032192.6 | 6 | 108213118 | 108551109 | 337992 | 1 | ENSMUST00 | 68892 | Itpr1 | ENSMUSG0C | false | . | . | . | . | . | . | chr6 | 108280603 | 108284622 | 4.12843 | 3.48058 | 2.33904 |
| chr6 | 149093498 | 149093770 | 474 | Intron (ENSMUST00000111557.2 | 6 | 149013988 | 149101538 | 87551 | 2 | ENSMUST00 | 7768 | Dennd5b | ENSMUSG0C | false | . | . | . | . | . | . | chr6 | 149085106 | 149097616 | 4.54289 | 3.81059 | 2.63745 |
| chr8 | 110846246 | 110847245 | 474 | Promoter (<=1kb) | 8 | 110846600 | 110881895 | 35296 | 1 | ENSMUST00 | 0 | Cog4 | ENSMUSG0C | false | chr8 | 110845795 | 110847888 | 32.66734 | 113.81544 | 109.86345 | . | . | . | . | . | . |
| chr13 | 93144283 | 93144678 | 474 | Promoter (<=1kb) | 13 | 93040713 | 93144724 | 104012 | 2 | ENSMUST00 | 46 | Cmya5 | ENSMUSG0C | false | chr13 | 93143859 | 93144810 | 23.91008 | 65.63585 | 62.71086 | chr13 | 93142790 | 93145800 | 4.82656 | 3.89807 | 2.91321 |
| chr5 | 134551319 | 134551650 | 474 | Promoter (<=1kb) | 5 | 134489386 | 134552434 | 63049 | 2 | ENSMUST00 | 784 | Clip2 | ENSMUSG0C | false | chr5 | 134550699 | 134552810 | 36.27325 | 129.91212 | 125.42401 | chr5 | 134538830 | 134551710 | 6.68849 | 4.77556 | 4.40881 |
| chr13 | 11099372 | 11099614 | 474 | Distal Intergenic | 13 | 11426779 | 11426898 | 120 | 2 | ENSMUST00 | 327284 | Gm25496 | ENSMUSG0C | false | . | . | . | . | . | . | . | . | . | . | . | . |
| chr11 | 6475896 | 6476295 | 474 | Promoter (<=1kb) | 11 | 6467599 | 6475917 | 8319 | 2 | ENSMUST00 | 0 | Purb | ENSMUSG0C | false | chr11 | 6474409 | 6476644 | 19.88326 | 77.37266 | 74.26472 | . | . | . | . | . | . |
| chr10 | 99106671 | 99107346 | 473 | Promoter (<=1kb) | 10 | 99107036 | 99197988 | 90953 | 1 | ENSMUST00 | 0 | Poc1b | ENSMUSG0C | false | chr10 | 99106403 | 99109000 | 34.18682 | 141.10779 | 136.18079 | . | . | . | . | . | . |
| chr18 | 35022405 | 35022687 | 473 | Distal Intergenic | 18 | 34937414 | 34954357 | 16944 | 2 | ENSMUST00 | -68049 | Hspa9 | ENSMUSG0C | false | . | . | . | . | . | . | chr18 | 35021271 | 35023278 | 5.25281 | 4.18334 | 3.238 |
| chr18 | 60921975 | 60922398 | 473 | Distal Intergenic | 18 | 60925618 | 60988152 | 62535 | 1 | ENSMUST00 | -3220 | Camk2a | ENSMUSG0C | false | . | . | . | . | . | . | chr18 | 60921452 | 60922871 | 5.64828 | 4.47618 | 3.53028 |
| chr3 | 98249865 | 98250166 | 473 | Distal Intergenic | 3 | 98222156 | 98236748 | 14593 | 1 | ENSMUST00 | 27710 | Reg4 | ENSMUSG0C | false | . | . | . | . | . | . | chr3 | 98249535 | 98250158 | 4.0944 | 3.69663 | 2.28151 |
| chr6 | 55567540 | 55567802 | 473 | Distal Intergenic | 6 | 55460236 | 55497428 | 37193 | 1 | ENSMUST00 | 107305 | Adcyap1r1 | ENSMUSG0C | false | . | . | . | . | . | . | chr6 | 55567616 | 55568205 | 3.34614 | 3.17073 | 1.67228 |
| chr1 | 182777135 | 182777439 | 473 | Intron (ENSMUST00000155229.1 | 1 | 182764895 | 182896598 | 131704 | 1 | ENSMUST00 | 12241 | Susd4 | ENSMUSG0C | false | . | . | . | . | . | . | chr1 | 182776908 | 182778253 | 4.3237 | 3.86104 | 2.45013 |
| chr8 | 108708902 | 108709173 | 473 | Intron (ENSMUST00000188994.1 | 8 | 108714644 | 108961630 | 246987 | 1 | ENSMUST00 | -5471 | Zfhx3 | ENSMUSG0C | false | . | . | . | . | . | . | chr8 | 108708274 | 108710085 | 3.94048 | 3.64535 | 2.11581 |
| chr7 | 90089454 | 90089809 | 473 | Distal Intergenic | 7 | 90107477 | 90107968 | 492 | 2 | ENSMUST00 | 18159 | Gm5341 | ENSMUSG0C | false | . | . | . | . | . | . | chr7 | 90089355 | 90092394 | 5.64133 | 4.44876 | 3.52666 |
| chr6 | 32996620 | 32996901 | 473 | Intron (ENSMUST00000066379.5 | 6 | 33038468 | 33043222 | 4755 | 1 | ENSMUST00 | -41567 | Gm14540 | ENSMUSG0C | false | . | . | . | . | . | . | chr6 | 32995158 | 32998321 | 4.517 | 3.80069 | 2.64864 |
| chr4 | 141897748 | 141898037 | 473 | Intron (ENSMUST00000105780.2 | 4 | 141885654 | 141897282 | 11629 | 1 | ENSMUST00 | 12095 | Fhad1os2 | ENSMUSG0C | false | . | . | . | . | . | . | chr4 | 141896610 | 141901092 | 4.66777 | 3.92447 | 2.73167 |
| chr17 | 88260871 | 88261206 | 473 | Distal Intergenic | 17 | 88342388 | 88342482 | 95 | 2 | ENSMUST00 | 81276 | Gm22542 | ENSMUSG0C | false | . | . | . | . | . | . | chr17 | 88255594 | 88263362 | 4.52852 | 3.7778 | 2.62681 |
| chr15 | 5776867 | 5777217 | 473 | Distal Intergenic | 15 | 5864306 | 5864419 | 114 | 1 | ENSMUST00 | -87089 | Gm27760 | ENSMUSG0C | false | . | . | . | . | . | . | . | . | . | . | . | . |
| chr13 | 5608649 | 5608943 | 473 | Distal Intergenic | 13 | 5855508 | 5857712 | 2205 | 2 | ENSMUST00 | 248769 | RP23-28J6.1 | ENSMUSG0C | false | . | . | . | . | . | . | chr13 | 5607667 | 5609247 | 4.12421 | 3.62975 | 2.30951 |
| chr14 | 54426667 | 54427113 | 472 | Promoter (<=1kb) | 14 | 54426909 | 54429750 | 2842 | 1 | ENSMUST00 | 0 | Mrpl52 | ENSMUSG0C | false | chr14 | 54426315 | 54427561 | 38.13901 | 127.82562 | 123.40861 | . | . | . | . | . | . |
| chr12 | 83070963 | 83071198 | 472 | Intron (ENSMUST00000161801.2 | 12 | 83047559 | 83137686 | 90128 | 1 | ENSMUST00 | 23405 | Rgs6 | ENSMUSG0C | false | . | . | . | . | . | . | chr12 | 83069424 | 83074627 | 5.24375 | 4.04679 | 3.22494 |
| chr18 | 58659350 | 58660008 | 472 | Promoter (<=1kb) | 18 | 58659482 | 58679570 | 20089 | 1 | ENSMUST00 | 0 | Isoc1 | ENSMUSG0C | false | chr18 | 58658862 | 58660585 | 31.62822 | 118.12274 | 114.03909 | . | . | . | . | . | . |
| chr4 | 152120539 | 152120915 | 472 | Promoter (2-3kb) | 4 | 152117971 | 152119323 | 1353 | 1 | ENSMUST00 | 2569 | Tnfrsf25 | ENSMUSG0C | false | chr4 | 152119592 | 152121333 | 14.333 | 34.37709 | 31.85639 | chr4 | 152118648 | 152122002 | 6.10878 | 4.6144 | 3.91321 |
| chr4 | 149171606 | 149172017 | 472 | Distal Intergenic | 4 | 149159927 | 149166771 | 6845 | 2 | ENSMUST00 | -4836 | Pgd | ENSMUSG0C | false | . | . | . | . | . | . | chr4 | 149170180 | 149173474 | 4.46561 | 3.70009 | 2.61256 |
| chr7 | 134287664 | 134288103 | 472 | Intron (ENSMUST00000172947.2 | 7 | 134266275 | 134341986 | 75712 | 2 | ENSMUST00 | 53883 | D7Ertd443e | ENSMUSG0C | false | . | . | . | . | . | . | . | . | . | . | . | . |
| chr4 | 118155817 | 118156074 | 472 | Intron (ENSMUST00000106406.3 | 4 | 118149051 | 118161657 | 12607 | 2 | ENSMUST00 | 5583 | Kdm4a | ENSMUSG0C | false | . | . | . | . | . | . | chr4 | 118154585 | 118156722 | 5.48466 | 4.01474 | 3.43636 |
| chr9 | 21329002 | 21329272 | 472 | Distal Intergenic | 9 | 21337828 | 21355025 | 17198 | 1 | ENSMUST00 | -8556 | Slc44a2 | ENSMUSG0C | false | . | . | . | . | . | . | chr9 | 21320251 | 21338396 | 4.95977 | 3.98308 | 2.9977 |
| chr10 | 63497310 | 63497600 | 472 | Intron (ENSMUST00000105441.2 | 10 | 63504037 | 65002833 | 1498797 | 1 | ENSMUST00 | -6437 | Ctnna3 | ENSMUSG0C | false | . | . | . | . | . | . | chr10 | 63495957 | 63498244 | 3.44565 | 3.32844 | 1.73392 |
| chr16 | 78481182 | 78481594 | 472 | Distal Intergenic | 16 | 78423730 | 78423859 | 130 | 2 | ENSMUST00 | -57324 | Gm25038 | ENSMUSG0C | false | . | . | . | . | . | . | chr16 | 78480278 | 78481405 | 3.56268 | 3.32658 | 1.85832 |
| chr2 | 160806361 | 160806667 | 472 | Intron (ENSMUST00000133937.3 | 2 | 160845613 | 160846470 | 858 | 1 | ENSMUST00 | -38946 | Gm11447 | ENSMUSG0C | false | . | . | . | . | . | . | chr2 | 160804125 | 160808352 | 5.0346 | 4.01906 | 3.0928 |
| chr11 | 63020747 | 63021063 | 472 | Distal Intergenic | 11 | 62999115 | 63004134 | 5020 | 2 | ENSMUST00 | -16614 | Cdrt4os1 | ENSMUSG0C | false | . | . | . | . | . | . | chr11 | 63020520 | 63021330 | 3.68111 | 3.28053 | 1.95982 |
| chr12 | 78104487 | 78104698 | 472 | Distal Intergenic | 12 | 78165546 | 78165666 | 121 | 1 | ENSMUST00 | -60848 | Gm24994 | ENSMUSG0C | false | . | . | . | . | . | . | chr12 | 78104087 | 78104772 | 5.00333 | 4.07629 | 3.04015 |
| chr8 | 114709905 | 114710191 | 472 | Promoter (1-2kb) | 8 | 114698439 | 114708006 | 9568 | 2 | ENSMUST00 | -1900 | Gm16118 | ENSMUSG0C | false | . | . | . | . | . | . | chr8 | 114709795 | 114710882 | 3.69101 | 3.26999 | 1.96785 |
| chr1 | 36425106 | 36425363 | 471 | Intron (ENSMUST00000125304.2 | 1 | 36424506 | 36428366 | 3861 | 2 | ENSMUST00 | 3003 | Lman2l | ENSMUSG0C | false | . | . | . | . | . | . | chr1 | 36424774 | 36426225 | 4.33093 | 3.74622 | 2.49723 |
| chr13 | 74807817 | 74808156 | 471 | Promoter (<=1kb) | 13 | 74694286 | 74807921 | 113636 | 2 | ENSMUST00 | 0 | Cast | ENSMUSG0C | false | chr13 | 74807412 | 74808982 | 23.74947 | 92.7123 | 89.30923 | . | . | . | . | . | . |
| chr18 | 34931418 | 34932305 | 471 | Promoter (<=1kb) | 18 | 34902785 | 34932007 | 29223 | 2 | ENSMUST00 | 0 | Etf1 | ENSMUSG0C | false | chr18 | 34930321 | 34933202 | 39.27047 | 129.80659 | 125.32217 | . | . | . | . | . | . |
| chr1 | 182511063 | 182511528 | 471 | Intron (ENSMUST00000068505.8 | 1 | 182467256 | 182517495 | 50240 | 2 | ENSMUST00 | 5967 | Capn2 | ENSMUSG0C | false | . | . | . | . | . | . | chr1 | 182492178 | 182516499 | 7.84019 | 5.29195 | 5.34945 |
| chr3 | 129831020 | 129831484 | 471 | Promoter (<=1kb) | 3 | 129824912 | 129831396 | 6485 | 2 | ENSMUST00 | 0 | Gar1 | ENSMUSG0C | false | chr3 | 129830284 | 129831977 | 48.7104 | 163.09714 | 157.34039 | . | . | . | . | . | . |
| chr7 | 100493772 | 100494237 | 471 | Promoter (<=1kb) | 7 | 100493795 | 100497163 | 3369 | 1 | ENSMUST00 | 0 | Ucp2 | ENSMUSG0C | false | chr7 | 100492519 | 100494857 | 28.20354 | 84.91737 | 81.67689 | chr7 | 100494087 | 100506674 | 4.14782 | 3.62895 | 2.31932 |
| chr9 | 32400627 | 32401047 | 471 | Distal Intergenic | 9 | 32393985 | 32399192 | 5208 | 1 | ENSMUST00 | 6643 | Kcnj1 | ENSMUSG0C | false | . | . | . | . | . | . | . | . | . | . | . | . |
| chr3 | 121609210 | 121609501 | 471 | Distal Intergenic | 3 | 121637145 | 121643485 | 6341 | 2 | ENSMUST00 | 33984 | A730020M07 | ENSMUSG0C | false | . | . | . | . | . | . | chr3 | 121609117 | 121610213 | 3.92331 | 3.43547 | 2.13424 |
| chr14 | 98169462 | 98170101 | 471 | Promoter (<=1kb) | 14 | 97786853 | 98169543 | 382691 | 2 | ENSMUST00 | 0 | Dach1 | ENSMUSG0C | true | chr14 | 98164311 | 98170276 | 41.81742 | 134.22485 | 129.57179 | . | . | . | . | . | . |
| chr14 | 122475122 | 122475586 | 471 | Promoter (<=1kb) | 14 | 122475435 | 122479852 | 4418 | 1 | ENSMUST00 | 0 | Zic2 | ENSMUSG0C | false | chr14 | 122474804 | 122476455 | 15.62409 | 36.57069 | 34.02363 | chr14 | 122473825 | 122477401 | 6.71013 | 4.77809 | 4.41091 |
| chr3 | 128415521 | 128415902 | 471 | Distal Intergenic | 3 | 128540372 | 128540480 | 109 | 1 | ENSMUST00 | -124470 | Gm22293 | ENSMUSG0C | false | . | . | . | . | . | . | chr3 | 128414854 | 128416510 | 5.07501 | 4.18093 | 3.08016 |
| chr2 | 45054098 | 45054438 | 471 | Intron (ENSMUST00000068415.5 | 2 | 45023100 | 45023177 | 78 | 2 | ENSMUST00 | -30922 | Mir5129 | ENSMUSG0C | false | . | . | . | . | . | . | chr2 | 45028593 | 45059388 | 4.87547 | 4.00998 | 2.90178 |
| chr19 | 45293486 | 45293994 | 471 | Distal Intergenic | 19 | 45300463 | 45303034 | 2572 | 2 | ENSMUST00 | 9040 | RP23-188J8.: | ENSMUSG0C | false | chr19 | 45293332 | 45294271 | 12.15419 | 27.41675 | 24.98428 | chr19 | 45292204 | 45298207 | 6.27899 | 4.63491 | 4.0501 |
| chr7 | 46795663 | 46796142 | 470 | Promoter (<=1kb) | 7 | 46796103 | 46823800 | 27698 | 1 | ENSMUST00 | 0 | Gtf2h1 | ENSMUSG0C | false | chr7 | 46794979 | 46796929 | 34.21972 | 118.71816 | 114.6235 | . | . | . | . | . | . |
| chr1 | 12693330 | 12693615 | 470 | Promoter (<=1kb) | 1 | 12692532 | 12852489 | 159958 | 1 | ENSMUST00 | 799 | Sulf1 | ENSMUSG0C | true | chr1 | 12692880 | 12693967 | 22.51707 | 59.83231 | 56.98677 | chr1 | 12693290 | 12699363 | 4.74812 | 3.91894 | 2.8045 |
| chr10 | 41477443 | 41477789 | 470 | Promoter (<=1kb) | 10 | 41477079 | 41478185 | 1107 | 1 | ENSMUST00 | 365 | Mical1 | ENSMUSG0C | false | chr10 | 41476033 | 41477908 | 15.08796 | 37.05233 | 34.49837 | chr10 | 41476070 | 41477765 | 3.74913 | 3.32055 | 2.01213 |
| chr19 | 47559796 | 47560029 | 470 | Distal Intergenic | 19 | 47579678 | 47645246 | 65569 | 1 | ENSMUST00 | -19649 | Slk | ENSMUSG0C | false | . | . | . | . | . | . | chr19 | 47558911 | 47560450 | 5.85239 | 4.54554 | 3.69499 |
| chr7 | 112491745 | 112491963 | 470 | Intron (ENSMUST00000106643.2 | 7 | 112501177 | 112547610 | 46434 | 1 | ENSMUST00 | -9214 | Parva | ENSMUSG0C | true | . | . | . | . | . | . | chr7 | 112487030 | 112537122 | 5.64568 | 4.33316 | 3.55068 |
| chr8 | 13465417 | 13465650 | 470 | 3' UTR | 8 | 13465374 | 13494535 | 29162 | 2 | ENSMUST00 | 28885 | Gas6 | ENSMUSG0C | false | . | . | . | . | . | . | chr8 | 13459474 | 13466027 | 4.90569 | 3.87877 | 2.97234 |
| chr14 | 50930989 | 50931324 | 470 | Promoter (<=1kb) | 14 | 50927215 | 50930856 | 3642 | 2 | ENSMUST00 | -134 | Tmem55b | ENSMUSG0C | false | chr14 | 50931125 | 50931992 | 19.77295 | 52.66101 | 49.91236 | . | . | . | . | . | . |
| chr6 | 50110035 | 50110936 | 470 | Promoter (<=1kb) | 6 | 50110241 | 50198598 | 88358 | 1 | ENSMUST00 | 0 | Mpp6 | ENSMUSG0C | false | chr6 | 50109410 | 50111194 | 30.62272 | 92.00563 | 88.61945 | . | . | . | . | . | . |
| chr5 | 148990880 | 148991333 | 470 | Exon (ENSMUST00000145042.1/ | 5 | 148893775 | 148995147 | 101373 | 2 | ENSMUST00 | 3814 | Katnal1 | ENSMUSG0C | false | . | . | . | . | . | . | chr5 | 148990738 | 148995772 | 3.78992 | 3.4575 | 2.05895 |
| chr1 | 153425022 | 153425286 | 470 | Promoter (<=1kb) | 1 | 153425162 | 153452574 | 27413 | 1 | ENSMUST00 | 0 | Shcbp1l | ENSMUSG0C | false | chr1 | 153424745 | 153425504 | 12.76655 | 31.65917 | 29.17213 | chr1 | 153424070 | 153425487 | 4.98617 | 3.95029 | 3.00566 |
| chr7 | 79715664 | 79716007 | 470 | Promoter (<=1kb) | 7 | 79698098 | 79715720 | 17623 | 2 | ENSMUST00 | 0 | Kif7 | ENSMUSG0C | false | chr7 | 79715217 | 79716305 | 25.50448 | 78.30592 | 75.18496 | . | . | . | . | . | . |
| chr8 | 77764349 | 77764595 | 470 | Promoter (<=1kb) | 8 | 77764359 | 77768198 | 3840 | 1 | ENSMUST00 | 0 | 4933431K23I | ENSMUSG0C | false | . | . | . | . | . | . | chr8 | 77763162 | 77765583 | 3.71448 | 3.3342 | 1.98096 |
| chr4 | 133092378 | 133092737 | 470 | Distal Intergenic | 4 | 133085474 | 133089146 | 3673 | 1 | ENSMUST00 | 6905 | Gm26615 | ENSMUSG0C | false | . | . | . | . | . | . | chr4 | 133082180 | 133096297 | 7.96206 | 5.47868 | 5.42168 |
| chr1 | 136042205 | 136042425 | 470 | Distal Intergenic | 1 | 136050804 | 136051370 | 567 | 1 | ENSMUST00 | -8379 | Ascl5 | ENSMUSG0C | false | . | . | . | . | . | . | chr1 | 136040721 | 136042843 | 4.98766 | 4.02991 | 3.0295 |
| chr11 | 32000313 | 32000690 | 469 | Promoter (<=1kb) | 11 | 32000463 | 32018007 | 17545 | 1 | ENSMUST00 | 0 | Nsg2 | ENSMUSG0C | false | . | . | . | . | . | . | chr11 | 31999534 | 32001827 | 5.38092 | 4.20601 | 3.34401 |
| chr11 | 70844564 | 70845352 | 469 | Promoter (<=1kb) | 11 | 70844778 | 70943105 | 98328 | 1 | ENSMUST00 | 0 | Rabep1 | ENSMUSG0C | false | chr11 | 70844185 | 70845713 | 40.77746 | 134.62177 | 129.96082 | . | . | . | . | . | . |
| chr12 | 98244196 | 98244532 | 469 | Intron (ENSMUST00000021390.7 | 12 | 98202304 | 98259459 | 57156 | 2 | ENSMUST00 | 14927 | Galc | ENSMUSG0C | false | . | . | . | . | . | . | chr12 | 98231481 | 98245211 | 5.28965 | 4.20277 | 3.23997 |
| chr17 | 72922143 | 72922438 | 469 | Promoter (<=1kb) | 17 | 72921188 | 72939345 | 18158 | 1 | ENSMUST00 | 956 | Lbh | ENSMUSG0C | false | . | . | . | . | . | . | chr17 | 72919347 | 72926758 | 5.07529 | 3.99292 | 3.11365 |
| chr3 | 14578563 | 14579271 | 469 | Promoter (<=1kb) | 3 | 14578671 | 14606308 | 27638 | 1 | ENSMUST00 | 0 | E2f5 | ENSMUSG0C | false | chr3 | 14578291 | 14580268 | 31.91637 | 110.38313 | 106.53261 | . | . | . | . | . | . |
| chr3 | 94582291 | 94582750 | 469 | Promoter (<=1kb) | 3 | 94497544 | 94582716 | 85173 | 2 | ENSMUST00 | 0 | Snx27 | ENSMUSG0C | false | chr3 | 94581275 | 94582913 | 27.66025 | 106.38611 | 102.65278 | . | . | . | . | . | . |
| chr8 | 78584928 | 78585178 | 469 | Intron (ENSMUST00000034111.8 | 8 | 78509335 | 78734005 | 224671 | 1 | ENSMUST00 | 75594 | Slc10a7 | ENSMUSG0C | false | . | . | . | . | . | . | chr8 | 78584250 | 78585729 | 3.91218 | 3.3636 | 2.14637 |
| chr11 | 77348113 | 77348438 | 469 | Promoter (<=1kb) | 11 | 77348280 | 77413635 | 65356 | 1 | ENSMUST00 | 0 | Ssh2 | ENSMUSG0C | false | chr11 | 77347879 | 77349056 | 16.70194 | 60.58573 | 57.73199 | . | . | . | . | . | . |
| chr1 | 180850528 | 180851208 | 469 | Promoter (<=1kb) | 1 | 180851151 | 180868097 | 16947 | 1 | ENSMUST00 | 0 | Sde2 | ENSMUSG0C | false | chr1 | 180851100 | 180851884 | 24.64903 | 84.19559 | 80.96455 | . | . | . | . | . | . |
| chr10 | 70151855 | 70152277 | 469 | Intron (ENSMUST00000147545.2 | 10 | 70174509 | 70182206 | 7698 | 1 | ENSMUST00 | -22232 | Ccdc6 | ENSMUSG0C | false | . | . | . | . | . | . | chr10 | 70148720 | 70154033 | 6.46015 | 4.65202 | 4.2548 |
| chr17 | 10216256 | 10216548 | 469 | Exon (ENSMUST00000097414.4/ | 17 | 10206471 | 10319361 | 112891 | 2 | ENSMUST00 | 102813 | Qk | ENSMUSG0C | true | . | . | . | . | . | . | chr17 | 10201700 | 10238338 | 4.5107 | 3.83331 | 2.60572 |
| chr16 | 18451742 | 18452173 | 469 | Promoter (<=1kb) | 16 | 18452039 | 18472249 | 20211 | 1 | ENSMUST00 | 0 | Txnrd2 | ENSMUSG0C | false | . | . | . | . | . | . | chr16 | 18446552 | 18452802 | 5.74275 | 4.36487 | 3.63646 |
| chr1 | 142867272 | 142867625 | 469 | Distal Intergenic | 1 | 142722855 | 142730022 | 7168 | 1 | ENSMUST00 | 144418 | RP23-406B1( | ENSMUSG0C | false | . | . | . | . | . | . | . | . | . | . | . | . |
| chr2 | 27617827 | 27618043 | 468 | Distal Intergenic | 2 | 27676440 | 27741898 | 65459 | 1 | ENSMUST00 | -58397 | Rxra | ENSMUSG0C | false | . | . | . | . | . | . | chr2 | 27616381 | 27618503 | 3.60232 | 3.20618 | 1.8961 |
| chr14 | 70167912 | 70168183 | 468 | Promoter (<=1kb) | 14 | 70164598 | 70167973 | 3376 | 2 | ENSMUST00 | 0 | Pdlim2 | ENSMUSG0C | true | chr14 | 70167652 | 70168043 | 6.00779 | 10.46335 | 8.27668 | chr14 | 70156516 | 70175803 | 7.12683 | 5 | 4.75261 |
| chr5 | 134572475 | 134573043 | 468 | Distal Intergenic | 5 | 134582690 | 134598328 | 15639 | 1 | ENSMUST00 | -9647 | Rfc2 | ENSMUSG0C | false | chr5 | 134571532 | 134572669 | 18.19667 | 46.80325 | 44.13025 | chr5 | 134572357 | 134574085 | 6.85713 | 5.00143 | 4.52705 |
| chr2 | 92603666 | 92603964 | 468 | Intron (ENSMUST00000065797.6 | 2 | 92599707 | 92615250 | 15544 | 1 | ENSMUST00 | 3960 | Chst1 | ENSMUSG0C | true | chr2 | 92599547 | 92603771 | 22.31878 | 85.15456 | 81.90936 | chr2 | 92601677 | 92612313 | 5.96217 | 4.49774 | 3.79786 |
| chr3 | 98194907 | 98195383 | 468 | Distal Intergenic | 3 | 98222156 | 98236748 | 14593 | 1 | ENSMUST00 | -26773 | Reg4 | ENSMUSG0C | false | . | . | . | . | . | . | chr3 | 98195286 | 98203232 | 4.51299 | 3.82434 | 2.63241 |
| chr5 | 108065510 | 108065986 | 468 | Promoter (<=1kb) | 5 | 108065674 | 108107163 | 41490 | 1 | ENSMUST00 | 0 | Mtf2 | ENSMUSG0C | false | chr5 | 108065126 | 108066927 | 35.04076 | 112.82586 | 108.91161 | . | . | . | . | . | . |
| chr2 | 180118964 | 180119555 | 468 | Promoter (<=1kb) | 2 | 180119306 | 180162680 | 43375 | 1 | ENSMUST00 | 0 | Osbpl2 | ENSMUSG0C | false | chr2 | 180118670 | 180119157 | 18.77433 | 54.14365 | 51.37372 | chr2 | 180114077 | 180120696 | 5.8988 | 4.44398 | 3.75334 |
| chr17 | 35916438 | 35916947 | 468 | Promoter (<=1kb) | 17 | 35916566 | 35932283 | 15718 | 1 | ENSMUST00 | 0 | Ppp1r10 | ENSMUSG0C | false | chr17 | 35915629 | 35916728 | 22.78592 | 76.10319 | 73.01487 | . | . | . | . | . | . |
| chr11 | 114044257 | 114044483 | 468 | Intron (ENSMUST00000041627.8 | 11 | 113848562 | 114065975 | 217414 | 2 | ENSMUST00 | 21492 | Sdk2 | ENSMUSG0C | false | . | . | . | . | . | . | chr11 | 114042166 | 114045202 | 6.07085 | 4.58888 | 3.88909 |
| chr9 | 15792726 | 15792970 | 468 | Distal Intergenic | 9 | 15862667 | 15874460 | 11794 | 2 | ENSMUST00 | 81490 | Mtnr1b | ENSMUSG0C | false | . | . | . | . | . | . | chr9 | 15792011 | 15795145 | 5.70064 | 4.41893 | 3.57968 |
| chr1 | 128103205 | 128103893 | 468 | Promoter (<=1kb) | 1 | 128103301 | 128183131 | 79831 | 1 | ENSMUST00 | 0 | R3hdm1 | ENSMUSG0C | false | chr1 | 128102166 | 128104140 | 22.22651 | 95.90443 | 92.42858 | chr1 | 128103884 | 128106436 | 4.71767 | 3.9901 | 2.78908 |
| chr11 | 96284449 | 96284754 | 468 | Promoter (1-2kb) | 11 | 96282725 | 96284693 | 1969 | 1 | ENSMUST00 | 1725 | Hoxb8 | ENSMUSG0C | false | chr11 | 96284031 | 96284998 | 17.35798 | 42.36193 | 39.74529 | chr11 | 96282409 | 96287903 | 4.4578 | 3.82587 | 2.56473 |
| chr17 | 56584720 | 56585192 | 468 | Promoter (<=1kb) | 17 | 56584825 | 56596523 | 11699 | 1 | ENSMUST00 | 0 | Safb | ENSMUSG0C | false | chr17 | 56584951 | 56586211 | 23.85991 | 99.51073 | 95.95212 | . | . | . | . | . | . |
| chr10 | 128492973 | 128493233 | 468 | Promoter (<=1kb) | 10 | 128490860 | 128493875 | 3016 | 2 | ENSMUST00 | 642 | Myl6 | ENSMUSG0C | false | chr10 | 128490711 | 128495254 | 33.41831 | 117.26761 | 113.21259 | . | . | . | . | . | . |
| chr6 | 124662599 | 124663323 | 467 | Promoter (<=1kb) | 6 | 124663027 | 124704370 | 41344 | 1 | ENSMUST00 | 0 | Lpcat3 | ENSMUSG0C | false | chr6 | 124662145 | 124663822 | 27.1957 | 84.10433 | 80.87383 | . | . | . | . | . | . |
| chr9 | 120492511 | 120492813 | 467 | Promoter (<=1kb) | 9 | 120492606 | 120495327 | 2722 | 1 | ENSMUST00 | 0 | Eif1b | ENSMUSG0C | false | chr9 | 120491999 | 120493744 | 43.84554 | 148.90724 | 143.66989 | . | . | . | . | . | . |
| chr18 | 76035107 | 76035766 | 467 | Intron (ENSMUST00000058997.8 | 18 | 76059458 | 76146443 | 86986 | 1 | ENSMUST00 | -23692 | Zbtb7c | ENSMUSG0C | true | chr18 | 76034773 | 76035255 | 13.14203 | 32.98289 | 30.47944 | chr18 | 76032245 | 76037845 | 3.8357 | 3.51533 | 2.06613 |
| chr9 | 56637273 | 56637804 | 467 | Promoter (1-2kb) | 9 | 56619088 | 56635628 | 16541 | 2 | ENSMUST00 | -1646 | Lingo1 | ENSMUSG0C | false | . | . | . | . | . | . | chr9 | 56634857 | 56638086 | 4.71943 | 3.97843 | 2.76558 |
| chr12 | 73002290 | 73002568 | 467 | Distal Intergenic | 12 | 73040015 | 73042815 | 2801 | 2 | ENSMUST00 | 40247 | Six1 | ENSMUSG0C | false | . | . | . | . | . | . | . | . | . | . | . | . |
| chr2 | 170130842 | 170131698 | 467 | Promoter (<=1kb) | 2 | 170108643 | 170131220 | 22578 | 2 | ENSMUST00 | 0 | Zfp217 | ENSMUSG0C | false | chr2 | 170130125 | 170131972 | 30.03893 | 101.89226 | 98.2715 | chr2 | 170131479 | 170135004 | 3.6867 | 3.39433 | 1.9258 |
| chr8 | 25253482 | 25253700 | 467 | Distal Intergenic | 8 | 25154552 | 25201449 | 46898 | 2 | ENSMUST00 | -52034 | Tacc1 | ENSMUSG0C | false | . | . | . | . | . | . | chr8 | 25251758 | 25257009 | 4.24644 | 3.70122 | 2.39253 |
| chr15 | 78406323 | 78406922 | 467 | Promoter (<=1kb) | 15 | 78406712 | 78414002 | 7291 | 1 | ENSMUST00 | 0 | Mpst | ENSMUSG0C | false | chr15 | 78405455 | 78407074 | 31.91637 | 110.38313 | 106.53261 | chr15 | 78406833 | 78407401 | 4.34008 | 3.82512 | 2.45276 |

| | | | | | | | | | | | | | | | | | | | | | | | | | | |
|---|---|---|---|---|---|---|---|---|---|---|---|---|---|---|---|---|---|---|---|---|---|---|---|---|---|---|
| chr6 | 144098151 | 144098407 | 467 | Intron (ENSMUST00000077160.6 | 6 | 144021908 | 144023778 | 1871 | 1 | ENSMUST00 | 76244 | Sox5os1 | ENSMUSG00 | false | . | . | . | . | . | . | chr6 | 144097543 | 144099174 | 3.45048 | 3.24796 | 1.73224 |
| chr15 | 86202654 | 86203085 | 467 | Distal Intergenic | 15 | 86212554 | 86212657 | 104 | 2 | ENSMUST00 | 9572 | Gm22818 | ENSMUSG00 | false | . | . | . | . | . | . | chr15 | 86196688 | 86204593 | 6.93026 | 4.73328 | 4.61625 |
| chr16 | 3452647 | 3453146 | 467 | Distal Intergenic | 16 | 3535485 | 3535611 | 127 | 2 | ENSMUST00 | 82465 | Gm22598 | ENSMUSG00 | false | . | . | . | . | . | . | . | . | . | . | . | . |
| chr6 | 78235922 | 78236207 | 467 | Distal Intergenic | 6 | 78215878 | 78217113 | 1236 | 1 | ENSMUST00 | 20045 | Gm5576 | ENSMUSG00 | false | . | . | . | . | . | . | chr6 | 78234740 | 78236473 | 4.71697 | 3.96848 | 2.82118 |
| chr7 | 39552459 | 39552916 | 467 | Promoter (<=1kb) | 7 | 39552496 | 39556305 | 3810 | 1 | ENSMUST00 | 0 | RP23-402K24 | ENSMUSG00 | false | chr7 | 39552060 | 39553247 | 23.12601 | 77.64799 | 74.53377 | . | . | . | . | . | . |
| chr12 | 80220383 | 80220862 | 466 | Intron (ENSMUST00000021554.9 | 12 | 80168322 | 80260091 | 91770 | 2 | ENSMUST00 | 39229 | Actn1 | ENSMUSG00 | true | . | . | . | . | . | . | chr12 | 80207669 | 80232669 | 6.01537 | 4.51837 | 3.84731 |
| chr5 | 143654981 | 143655466 | 466 | Intron (ENSMUST00000116456.4 | 5 | 143651241 | 143708256 | 57016 | 1 | ENSMUST00 | 3741 | Cyth3 | ENSMUSG00 | false | . | . | . | . | . | . | chr5 | 143651427 | 143662078 | 6.34156 | 4.57429 | 4.14661 |
| chr11 | 8678292 | 8678955 | 466 | Distal Intergenic | 11 | 8431652 | 8664535 | 232884 | 2 | ENSMUST00 | -13758 | Tns3 | ENSMUSG00 | false | . | . | . | . | . | . | chr11 | 8673880 | 8682084 | 6.25879 | 4.6547 | 4.04763 |
| chr12 | 98930120 | 98930360 | 466 | Intron (ENSMUST00000085109.4 | 12 | 98920820 | 98961492 | 40673 | 1 | ENSMUST00 | 9301 | Ttc8 | ENSMUSG00 | false | . | . | . | . | . | . | . | . | . | . | . | . |
| chr18 | 65133179 | 65133414 | 466 | Intron (ENSMUST00000163516.2 | 18 | 65023527 | 65217826 | 194300 | 1 | ENSMUST00 | 109653 | Nedd4l | ENSMUSG00 | false | . | . | . | . | . | . | chr18 | 65132435 | 65138242 | 5.66711 | 4.40632 | 3.55576 |
| chr5 | 124354234 | 124354914 | 466 | Promoter (<=1kb) | 5 | 124345417 | 124354671 | 9255 | 2 | ENSMUST00 | 0 | Cdk2ap1 | ENSMUSG00 | false | chr5 | 124352090 | 124355494 | 28.56742 | 103.16996 | 99.51957 | . | . | . | . | . | . |
| chr7 | 65861849 | 65862229 | 466 | Promoter (<=1kb) | 7 | 65862029 | 66050335 | 188307 | 1 | ENSMUST00 | 0 | Pcsk6 | ENSMUSG00 | true | chr7 | 65861608 | 65863627 | 31.54088 | 108.67455 | 104.87239 | chr7 | 65861140 | 65866313 | 4.55496 | 3.80536 | 2.64963 |
| chr18 | 46092764 | 46093091 | 466 | Distal Intergenic | 18 | 46165302 | 46198818 | 33517 | 2 | ENSMUST00 | 105727 | Trim36 | ENSMUSG00 | false | . | . | . | . | . | . | . | . | . | . | . | . |
| chr2 | 52857564 | 52858259 | 466 | Promoter (<=1kb) | 2 | 52857860 | 53133804 | 275945 | 1 | ENSMUST00 | 0 | Fmnl2 | ENSMUSG00 | true | chr2 | 52857275 | 52859971 | 40.88443 | 132.61177 | 128.02734 | . | . | . | . | . | . |
| chr9 | 117247281 | 117247915 | 466 | Intron (ENSMUST00000111773.4 | 9 | 116943123 | 117251597 | 308475 | 2 | ENSMUST00 | 3682 | Rbms3 | ENSMUSG00 | true | . | . | . | . | . | . | chr9 | 117244709 | 117251011 | 4.98494 | 4.05351 | 2.99079 |
| chr4 | 62622212 | 62622547 | 466 | Promoter (<=1kb) | 4 | 62623531 | 62631320 | 7790 | 1 | ENSMUST00 | -984 | Rgs3 | ENSMUSG00 | false | . | . | . | . | . | . | . | . | . | . | . | . |
| chr8 | 126876546 | 126876865 | 466 | Distal Intergenic | 8 | 126944847 | 126944980 | 134 | 2 | ENSMUST00 | 68115 | Gm26397 | ENSMUSG00 | false | . | . | . | . | . | . | chr8 | 126875667 | 126877243 | 5.03865 | 4.07587 | 3.06828 |
| chr5 | 35948378 | 35948695 | 466 | Intron (ENSMUST00000064571.5 | 5 | 35917897 | 35918201 | 305 | 1 | ENSMUST00 | 30482 | H2af-ps | ENSMUSG00 | false | . | . | . | . | . | . | chr5 | 35946856 | 35958484 | 5.34023 | 4.12733 | 3.32439 |
| chr7 | 114700111 | 114700360 | 466 | Distal Intergenic | 7 | 114705270 | 114706163 | 894 | 2 | ENSMUST00 | 5803 | Gm15500 | ENSMUSG00 | false | . | . | . | . | . | . | chr7 | 114699856 | 114701739 | 5.11854 | 4.04183 | 3.12884 |
| chr8 | 105547028 | 105547365 | 465 | Intron (ENSMUST00000013304.7 | 8 | 105530651 | 105539476 | 8826 | 2 | ENSMUST00 | -7553 | Atp6v0d1 | ENSMUSG00 | false | . | . | . | . | . | . | chr8 | 105536711 | 105553149 | 5.86218 | 4.31178 | 3.75598 |
| chr2 | 18997872 | 18998536 | 465 | Promoter (<=1kb) | 2 | 18998343 | 18999801 | 1459 | 1 | ENSMUST00 | 0 | 4930426L09R | ENSMUSG00 | false | chr2 | 18996496 | 18998878 | 38.46334 | 130.56975 | 126.05653 | . | . | . | . | . | . |
| chr6 | 87913834 | 87914337 | 465 | Promoter (<=1kb) | 6 | 87913935 | 87936629 | 22695 | 1 | ENSMUST00 | 0 | Hmces | ENSMUSG00 | false | chr6 | 87913750 | 87914946 | 32.69058 | 104.59052 | 100.90593 | . | . | . | . | . | . |
| chr8 | 60890202 | 60890488 | 465 | Promoter (<=1kb) | 8 | 60890418 | 60907575 | 17158 | 1 | ENSMUST00 | 0 | 2700029M09 | ENSMUSG00 | false | chr8 | 60889868 | 60891265 | 33.5288 | 108.10788 | 104.32592 | . | . | . | . | . | . |
| chr2 | 173264918 | 173265170 | 465 | Intron (ENSMUST00000036248.7 | 2 | 173228046 | 173276189 | 48144 | 2 | ENSMUST00 | 11019 | Pmepa1 | ENSMUSG00 | true | . | . | . | . | . | . | chr2 | 173223515 | 173275069 | 7.01359 | 4.89215 | 4.68115 |
| chr19 | 37192670 | 37192944 | 465 | Intron (ENSMUST00000180609.1 | 19 | 37024639 | 37198109 | 173471 | 2 | ENSMUST00 | 5165 | Cpeb3 | ENSMUSG00 | false | . | . | . | . | . | . | chr19 | 37192045 | 37193926 | 4.95714 | 4.0592 | 2.9627 |
| chr11 | 119517073 | 119517321 | 465 | Distal Intergenic | 11 | 119501978 | 119507389 | 5412 | 1 | ENSMUST00 | 15096 | Endov | ENSMUSG00 | false | . | . | . | . | . | . | chr11 | 119516287 | 119517615 | 4.39894 | 3.88569 | 2.50084 |
| chr8 | 85026708 | 85027345 | 465 | Promoter (<=1kb) | 8 | 85026833 | 85030285 | 3453 | 1 | ENSMUST00 | 0 | 2310036O22 | ENSMUSG00 | false | chr8 | 85026384 | 85026721 | 9.96269 | 21.3504 | 18.99651 | chr8 | 85023749 | 85026735 | 7.8759 | 5.40813 | 5.35631 |
| chr19 | 16756158 | 16756424 | 465 | Intron (ENSMUST00000068156.6 | 19 | 16780547 | 16781103 | 557 | 1 | ENSMUST00 | -24123 | Gm9806 | ENSMUSG00 | false | . | . | . | . | . | . | chr19 | 16753849 | 16757885 | 4.36724 | 3.76033 | 2.48902 |
| chr12 | 112828946 | 112829790 | 465 | Promoter (<=1kb) | 12 | 112820237 | 112829389 | 9153 | 2 | ENSMUST00 | 0 | Cdca4 | ENSMUSG00 | false | chr12 | 112828574 | 112829497 | 28.22734 | 101.53345 | 97.92473 | . | . | . | . | . | . |
| chr12 | 105109416 | 105109712 | 465 | Distal Intergenic | 12 | 105146014 | 105155009 | 8996 | 1 | ENSMUST00 | -36302 | Tcl1b2 | ENSMUSG00 | false | . | . | . | . | . | . | chr12 | 105108898 | 105110451 | 4.48897 | 3.7652 | 2.62181 |
| chr6 | 32386895 | 32387161 | 465 | Intron (ENSMUST00000115096.3 | 6 | 32451170 | 32454750 | 3581 | 1 | ENSMUST00 | -64009 | Plxna4os2 | ENSMUSG00 | false | . | . | . | . | . | . | chr6 | 32386110 | 32389324 | 4.94933 | 3.90023 | 2.9937 |
| chr2 | 145676774 | 145676995 | 465 | Promoter (1-2kb) | 2 | 145675367 | 145714989 | 39623 | 1 | ENSMUST00 | 1408 | BC039771 | ENSMUSG00 | false | . | . | . | . | . | . | chr2 | 145676010 | 145686155 | 5.31951 | 4.23417 | 3.27212 |
| chr18 | 54453073 | 54453329 | 465 | Distal Intergenic | 18 | 54422295 | 54452840 | 30546 | 1 | ENSMUST00 | 30779 | Redrum | ENSMUSG00 | false | chr18 | 54452503 | 54453674 | 12.1549 | 27.41795 | 24.98545 | . | . | . | . | . | . |
| chr1 | 155167569 | 155167773 | 464 | Intron (ENSMUST00000027743.7 | 1 | 155158703 | 155203517 | 44815 | 1 | ENSMUST00 | 8867 | Stx6 | ENSMUSG00 | false | . | . | . | . | . | . | chr1 | 155166227 | 155168828 | 4.69749 | 3.91927 | 2.78267 |
| chr1 | 162589516 | 162589800 | 464 | Exon (ENSMUST00000132158.2/ | 1 | 162574821 | 162594753 | 19933 | 1 | ENSMUST00 | 14696 | Vamp4 | ENSMUSG00 | true | . | . | . | . | . | . | chr1 | 162587733 | 162591044 | 4.60166 | 3.90294 | 2.67356 |
| chr4 | 102801018 | 102801305 | 464 | Intron (ENSMUST00000097948.3 | 4 | 102760525 | 102973628 | 213104 | 1 | ENSMUST00 | 40494 | Sgip1 | ENSMUSG00 | true | . | . | . | . | . | . | chr4 | 102798049 | 102813929 | 4.80947 | 3.92915 | 2.87079 |
| chr6 | 92905935 | 92906259 | 464 | Intron (ENSMUST00000113438.2 | 6 | 92772702 | 92901441 | 128740 | 2 | ENSMUST00 | -4495 | Adamts9 | ENSMUSG00 | false | . | . | . | . | . | . | chr6 | 92904585 | 92907359 | 3.81453 | 3.31339 | 2.07682 |
| chr18 | 60895348 | 60895971 | 464 | Distal Intergenic | 18 | 60911780 | 60918561 | 6782 | 1 | ENSMUST00 | -15809 | Arsi | ENSMUSG00 | false | . | . | . | . | . | . | chr18 | 60895425 | 60897550 | 4.8659 | 3.93425 | 2.92567 |
| chr2 | 129828144 | 129828391 | 464 | 3' UTR | 2 | 129800517 | 129832287 | 31771 | 1 | ENSMUST00 | 27628 | Stk35 | ENSMUSG00 | false | . | . | . | . | . | . | chr2 | 129812158 | 129831014 | 4.89733 | 3.94874 | 2.94159 |
| chr3 | 108591188 | 108591769 | 464 | Promoter (<=1kb) | 3 | 108591279 | 108645719 | 54441 | 1 | ENSMUST00 | 0 | Wdr47 | ENSMUSG00 | false | chr3 | 108590867 | 108592394 | 33.56525 | 137.72058 | 132.93262 | chr3 | 108590672 | 108592683 | 3.89454 | 3.46747 | 2.09749 |
| chr12 | 76537426 | 76537731 | 464 | Intron (ENSMUST00000075249.4 | 12 | 76533560 | 76579039 | 45480 | 1 | ENSMUST00 | 3867 | Plekhg3 | ENSMUSG00 | false | . | . | . | . | . | . | chr12 | 76532835 | 76538765 | 4.59558 | 3.84512 | 2.69586 |
| chr12 | 104978479 | 104978794 | 464 | Intron (ENSMUST00000095439.4 | 12 | 104929949 | 104998677 | 68729 | 2 | ENSMUST00 | 19883 | Syne3 | ENSMUSG00 | false | . | . | . | . | . | . | chr12 | 104974830 | 104987771 | 6.02697 | 4.55969 | 3.85063 |
| chr2 | 160647957 | 160648364 | 464 | Promoter (1-2kb) | 2 | 160645980 | 160650107 | 4128 | 1 | ENSMUST00 | 1978 | Top1 | ENSMUSG00 | false | chr2 | 160647655 | 160648146 | 8.08052 | 18.53858 | 16.22368 | chr2 | 160647541 | 160654302 | 5.11326 | 4.17537 | 3.09894 |
| chr2 | 91870872 | 91871149 | 464 | Intron (ENSMUST00000099712.4 | 2 | 91920528 | 91920928 | 401 | 2 | ENSMUST00 | 49779 | Gm10805 | ENSMUSG00 | false | . | . | . | . | . | . | chr2 | 91870389 | 91871386 | 3.66594 | 3.4328 | 1.89016 |
| chr17 | 5143344 | 5143617 | 464 | Intron (ENSMUST00000092723.5 | 17 | 5110781 | 5112119 | 1339 | 2 | ENSMUST00 | -31226 | Gm15599 | ENSMUSG00 | false | . | . | . | . | . | . | chr17 | 5138104 | 5145970 | 4.76331 | 3.92754 | 2.84217 |
| chr10 | 93771776 | 93772166 | 464 | Distal Intergenic | 10 | 93778611 | 93779772 | 1162 | 1 | ENSMUST00 | -6445 | Gm3571 | ENSMUSG00 | false | . | . | . | . | . | . | chr10 | 93771667 | 93772564 | 4.12826 | 3.66389 | 2.27826 |
| chr9 | 102763576 | 102763982 | 464 | Distal Intergenic | 9 | 102771551 | 102771810 | 260 | 2 | ENSMUST00 | 7828 | RP23-274G2 | ENSMUSG00 | false | . | . | . | . | . | . | chr9 | 102753752 | 102764393 | 4.95618 | 3.99593 | 2.98018 |
| chr11 | 118354433 | 118354858 | 463 | Promoter (<=1kb) | 11 | 118301069 | 118355506 | 54438 | 2 | ENSMUST00 | 648 | Timp2 | ENSMUSG00 | false | chr11 | 118353231 | 118355882 | 34.66032 | 110.08408 | 106.24455 | chr11 | 118319919 | 118354581 | 6.53665 | 4.67889 | 4.28866 |
| chr19 | 6235732 | 6236029 | 463 | Promoter (<=1kb) | 19 | 6227765 | 6235872 | 8108 | 2 | ENSMUST00 | 0 | Ppp2r5b | ENSMUSG00 | false | chr19 | 6234264 | 6236344 | 43.03529 | 140.75403 | 135.83646 | . | . | . | . | . | . |
| chr8 | 128353722 | 128353941 | 463 | Distal Intergenic | 8 | 128359073 | 128505462 | 146390 | 1 | ENSMUST00 | -5132 | Nrp1 | ENSMUSG00 | false | . | . | . | . | . | . | chr8 | 128352912 | 128354219 | 3.39163 | 3.16034 | 1.69904 |
| chr4 | 45018389 | 45018667 | 463 | Promoter (<=1kb) | 4 | 45018583 | 45033692 | 15110 | 1 | ENSMUST00 | 0 | Polr1e | ENSMUSG00 | false | chr4 | 45018316 | 45019389 | 18.36477 | 56.64417 | 53.84045 | chr4 | 45018020 | 45019926 | 2.98253 | 2.97317 | 1.36241 |
| chr7 | 65885294 | 65885699 | 463 | Intron (ENSMUST00000176199.2 | 7 | 65862432 | 66050340 | 187909 | 1 | ENSMUST00 | 22863 | Pcsk6 | ENSMUSG00 | true | . | . | . | . | . | . | chr7 | 65883866 | 65896928 | 4.99061 | 4.01585 | 3.04032 |
| chr7 | 29441073 | 29441348 | 463 | Intron (ENSMUST00000085809.5 | 7 | 29370788 | 29377745 | 6958 | 2 | ENSMUST00 | -63329 | Sipa1l3 | ENSMUSG00 | false | . | . | . | . | . | . | chr7 | 29436518 | 29441613 | 5.4251 | 4.21976 | 3.40728 |
| chr7 | 24884218 | 24884613 | 463 | Promoter (<=1kb) | 7 | 24884371 | 24889806 | 5436 | 1 | ENSMUST00 | 0 | Rps19 | ENSMUSG00 | false | chr7 | 24884130 | 24885761 | 32.47248 | 106.24805 | 102.52307 | . | . | . | . | . | . |
| chr15 | 81000363 | 81000729 | 463 | Intron (ENSMUST00000042506.9 | 15 | 81007938 | 81009950 | 2013 | 1 | ENSMUST00 | -7209 | Sgsm3 | ENSMUSG00 | false | . | . | . | . | . | . | chr15 | 80999599 | 81004043 | 4.9496 | 4.11318 | 2.96701 |
| chr8 | 128536193 | 128536430 | 463 | Distal Intergenic | 8 | 128650997 | 128651103 | 107 | 1 | ENSMUST00 | -114567 | Gm26002 | ENSMUSG00 | false | . | . | . | . | . | . | chr8 | 128519464 | 128548431 | 5.17648 | 4.1349 | 3.15487 |
| chr2 | 75825662 | 75825879 | 463 | Promoter (2-3kb) | 2 | 75822028 | 75822793 | 766 | 2 | ENSMUST00 | -2870 | Gm13670 | ENSMUSG00 | false | . | . | . | . | . | . | . | . | . | . | . | . |
| chr10 | 24465290 | 24465550 | 463 | Distal Intergenic | 10 | 24447490 | 24452657 | 5168 | 1 | ENSMUST00 | 17801 | Gm15271 | ENSMUSG00 | false | . | . | . | . | . | . | chr10 | 24465254 | 24466368 | 3.36806 | 3.14629 | 1.6981 |
| chr1 | 59488245 | 59488516 | 463 | Exon (ENSMUST00000180778.1/ | 1 | 59488480 | 59492605 | 4126 | 2 | ENSMUST00 | 4089 | Gm26813 | ENSMUSG00 | false | . | . | . | . | . | . | chr1 | 59487236 | 59488801 | 3.59565 | 3.31835 | 1.88953 |
| chr17 | 11481769 | 11482011 | 463 | Intron (ENSMUST00000066658.7 | 17 | 11545505 | 11545798 | 294 | 2 | ENSMUST00 | 63787 | RP23-297J19 | ENSMUSG00 | false | . | . | . | . | . | . | chr17 | 11481290 | 11481953 | 3.80562 | 3.4174 | 2.06433 |
| chr8 | 122576741 | 122577025 | 462 | Promoter (<=1kb) | 8 | 122574637 | 122576907 | 2271 | 2 | ENSMUST00 | 0 | Aprt | ENSMUSG00 | false | chr8 | 122576232 | 122577300 | 22.15371 | 67.81625 | 64.85734 | chr8 | 122575233 | 122577485 | 4.7614 | 3.91019 | 2.82513 |
| chr10 | 7680834 | 7681491 | 462 | Promoter (<=1kb) | 10 | 7644332 | 7681136 | 36805 | 2 | ENSMUST00 | 0 | Pcmt1 | ENSMUSG00 | false | chr10 | 7680225 | 7682548 | 36.25406 | 119.15681 | 115.04134 | . | . | . | . | . | . |
| chr19 | 6340652 | 6341390 | 462 | Promoter (<=1kb) | 19 | 6341135 | 6355612 | 14478 | 1 | ENSMUST00 | 0 | Map4k2 | ENSMUSG00 | false | chr19 | 6340456 | 6341900 | 23.80619 | 80.75649 | 77.58801 | chr19 | 6339224 | 6343379 | 4.09509 | 3.64536 | 2.2597 |
| chr1 | 157507876 | 157508136 | 462 | Promoter (1-2kb) | 1 | 157506810 | 157568425 | 61616 | 1 | ENSMUST00 | 1067 | Sec16b | ENSMUSG00 | false | . | . | . | . | . | . | chr1 | 157506721 | 157508820 | 3.84706 | 3.54432 | 2.07491 |
| chr2 | 152000113 | 152000361 | 462 | Promoter (<=1kb) | 2 | 151999897 | 152009246 | 9350 | 1 | ENSMUST00 | 217 | Slc52a3 | ENSMUSG00 | false | . | . | . | . | . | . | chr2 | 151998261 | 152001333 | 4.96061 | 4.02677 | 2.98118 |
| chr7 | 117470749 | 117471009 | 462 | Intron (ENSMUST00000160035.2 | 7 | 117475492 | 117549359 | 73868 | 1 | ENSMUST00 | -4483 | Xylt1 | ENSMUSG00 | false | . | . | . | . | . | . | chr7 | 117470002 | 117472722 | 4.35468 | 3.66038 | 2.5094 |
| chr7 | 72306578 | 72306865 | 462 | Promoter (<=1kb) | 7 | 72077830 | 72306595 | 228766 | 2 | ENSMUST00 | 0 | Mctp2 | ENSMUSG00 | false | chr7 | 72306035 | 72306916 | 17.04263 | 45.57019 | 42.91153 | chr7 | 72305576 | 72306944 | 4.27823 | 3.39177 | 2.44677 |
| chr15 | 88867064 | 88867351 | 462 | Distal Intergenic | 15 | 88862186 | 88865718 | 3533 | 1 | ENSMUST00 | 4879 | Pim3 | ENSMUSG00 | false | . | . | . | . | . | . | chr15 | 88863272 | 88871136 | 9.58718 | 5.83938 | 6.75823 |
| chr11 | 23633377 | 23633892 | 462 | Promoter (<=1kb) | 11 | 23564961 | 23633549 | 68589 | 2 | ENSMUST00 | 0 | 0610010F05R | ENSMUSG00 | false | chr11 | 23632854 | 23634167 | 39.98587 | 128.8752 | 124.42683 | . | . | . | . | . | . |
| chr14 | 43953445 | 43953769 | 462 | Distal Intergenic | 14 | 43957069 | 43963080 | 6012 | 1 | ENSMUST00 | -3300 | Ang5 | ENSMUSG00 | false | . | . | . | . | . | . | . | . | . | . | . | . |
| chr19 | 40271408 | 40271775 | 462 | Promoter (<=1kb) | 19 | 40221173 | 40271616 | 50444 | 2 | ENSMUST00 | 0 | Pdlim1 | ENSMUSG00 | false | chr19 | 40270513 | 40271964 | 28.83684 | 94.75451 | 91.30991 | . | . | . | . | . | . |
| chr8 | 91536753 | 91536973 | 462 | Intron (ENSMUST00000069718.8 | 8 | 91467685 | 91666946 | 199262 | 1 | ENSMUST00 | 69069 | Fto | ENSMUSG00 | false | . | . | . | . | . | . | chr8 | 91531654 | 91541979 | 5.00584 | 3.9745 | 3.04709 |
| chr11 | 94133376 | 94133736 | 462 | Promoter (<=1kb) | 11 | 94133485 | 94155776 | 22292 | 1 | ENSMUST00 | 0 | B230206L02R | ENSMUSG00 | false | . | . | . | . | . | . | chr11 | 94129611 | 94138962 | 5.93933 | 4.52942 | 3.772 |
| chr12 | 86836584 | 86836987 | 462 | Distal Intergenic | 12 | 86809957 | 86810258 | 302 | 1 | ENSMUST00 | 26628 | Gm25888 | ENSMUSG00 | false | . | . | . | . | . | . | chr12 | 86831585 | 86837954 | 7.49129 | 5.26295 | 5.03568 |
| chr3 | 19953863 | 19954247 | 461 | Promoter (2-3kb) | 3 | 19957054 | 19993138 | 36085 | 1 | ENSMUST00 | -2807 | Cp | ENSMUSG00 | false | . | . | . | . | . | . | . | . | . | . | . | . |
| chr10 | 60002717 | 60003413 | 461 | Promoter (<=1kb) | 10 | 59987409 | 60003135 | 15727 | 2 | ENSMUST00 | 0 | Anapc16 | ENSMUSG00 | false | chr10 | 60001946 | 60003837 | 20.82289 | 72.3363 | 69.31286 | . | . | . | . | . | . |
| chr18 | 13971082 | 13971692 | 461 | Promoter (1-2kb) | 18 | 13687013 | 13972733 | 285721 | 2 | ENSMUST00 | 1041 | Zfp521 | ENSMUSG00 | true | chr18 | 13969413 | 13973642 | 27.20707 | 96.65539 | 93.16586 | . | . | . | . | . | . |
| chr2 | 26604008 | 26604654 | 461 | Promoter (<=1kb) | 2 | 26593057 | 26604211 | 11155 | 2 | ENSMUST00 | 0 | Agpat2 | ENSMUSG00 | false | chr2 | 26603398 | 26604791 | 27.20707 | 96.65539 | 93.16586 | chr2 | 26603020 | 26604927 | 4.20203 | 3.72161 | 2.34955 |
| chr2 | 94406620 | 94406884 | 461 | Promoter (<=1kb) | 2 | 94300767 | 94406679 | 105913 | 2 | ENSMUST00 | 0 | Ttc17 | ENSMUSG00 | false | chr2 | 94405707 | 94407604 | 44.01745 | 150.34283 | 145.04555 | . | . | . | . | . | . |
| chr3 | 88620960 | 88621458 | 461 | Promoter (<=1kb) | 3 | 88621106 | 88648052 | 26947 | 1 | ENSMUST00 | 0 | Arhgef2 | ENSMUSG00 | false | chr3 | 88620710 | 88623767 | 28.41342 | 89.1249 | 85.7944 | . | . | . | . | . | . |
| chr4 | 82859382 | 82859799 | 461 | Promoter (<=1kb) | 4 | 82798738 | 82859673 | 60936 | 2 | ENSMUST00 | 0 | Zdhhc21 | ENSMUSG00 | false | chr4 | 82858685 | 82859999 | 29.92778 | 109.76675 | 105.9399 | chr4 | 82858441 | 82860103 | 3.94331 | 3.43389 | 2.16898 |
| chr8 | 12856025 | 12856386 | 461 | Promoter (<=1kb) | 8 | 12856960 | 12862205 | 5246 | 1 | ENSMUST00 | -574 | Atp11a | ENSMUSG00 | false | . | . | . | . | . | . | chr8 | 12855570 | 12856522 | 3.63295 | 3.34553 | 1.90755 |
| chr7 | 141172493 | 141172949 | 461 | Promoter (<=1kb) | 7 | 141160328 | 141172851 | 12524 | 2 | ENSMUST00 | 0 | Rnh1 | ENSMUSG00 | false | chr7 | 141171961 | 141173303 | 16.88215 | 57.7818 | 54.96725 | chr7 | 141171523 | 141173738 | 4.25764 | 3.70792 | 2.41185 |
| chr3 | 27728214 | 27728479 | 461 | Distal Intergenic | 3 | 27416162 | 27710439 | 294278 | 2 | ENSMUST00 | -17776 | Fndc3b | ENSMUSG00 | true | . | . | . | . | . | . | chr3 | 27727462 | 27730585 | 5.09218 | 4.0151 | 3.09521 |
| chr5 | 150080542 | 150080917 | 461 | Promoter (<=1kb) | 5 | 150080622 | 150082184 | 1563 | 1 | ENSMUST00 | 0 | Rxfp2 | ENSMUSG00 | false | . | . | . | . | . | . | chr5 | 150079739 | 150081310 | 3.95849 | 3.62875 | 2.13101 |
| chr2 | 181487384 | 181488002 | 461 | Distal Intergenic | 2 | 181493206 | 181494980 | 1775 | 1 | ENSMUST00 | -5204 | Abhd16b | ENSMUSG00 | false | . | . | . | . | . | . | chr2 | 181487215 | 181488718 | 4.03516 | 3.56007 | 2.25222 |
| chr6 | 52159237 | 52159620 | 461 | Promoter (<=1kb) | 6 | 52158524 | 52162020 | 3497 | 1 | ENSMUST00 | 714 | Hotairm1 | ENSMUSG00 | false | chr6 | 52157936 | 52159956 | 30.45735 | 121.03082 | 116.85291 | chr6 | 52159328 | 52167512 | 5.47105 | 4.07894 | 3.42628 |
| chr1 | 194976252 | 194976678 | 461 | Promoter (<=1kb) | 1 | 194976375 | 195021493 | 45119 | 1 | ENSMUST00 | 0 | A330023F24R | ENSMUSG00 | false | chr1 | 194975913 | 194977642 | 33.52805 | 106.05383 | 102.33234 | . | . | . | . | . | . |
| chr15 | 67146742 | 67147064 | 460 | Distal Intergenic | 15 | 67102875 | 67113992 | 11118 | 2 | ENSMUST00 | -32751 | St3gal1 | ENSMUSG00 | true | . | . | . | . | . | . | chr15 | 67143105 | 67166416 | 6.05047 | 4.43905 | 3.887 |
| chr1 | 153227984 | 153228227 | 460 | Promoter (<=1kb) | 1 | 153218922 | 153229144 | 10223 | 2 | ENSMUST00 | 917 | Lamc1 | ENSMUSG00 | false | . | . | . | . | . | . | chr1 | 153227473 | 153228569 | 3.66381 | 3.42904 | 1.95577 |
| chr17 | 3597146 | 3597504 | 460 | Distal Intergenic | 17 | 3519263 | 3557713 | 38451 | 2 | ENSMUST00 | -39434 | Tfb1m | ENSMUSG00 | false | . | . | . | . | . | . | chr17 | 3596798 | 3599226 | 4.166 | 3.58578 | 2.36535 |
| chr8 | 10965841 | 10966208 | 460 | Distal Intergenic | 8 | 10924427 | 10928457 | 4031 | 2 | ENSMUST00 | -37385 | 3930402G23 | ENSMUSG00 | false | . | . | . | . | . | . | chr8 | 10964590 | 10967219 | 4.8377 | 4.06247 | 2.86864 |
| chrX | 167332631 | 167332897 | 460 | Promoter (2-3kb) | 20 | 167304931 | 167330558 | 25628 | 2 | ENSMUST00 | -2074 | Tlr7 | ENSMUSG00 | false | . | . | . | . | . | . | . | . | . | . | . | . |
| chr7 | 28911310 | 28911575 | 460 | Exon (ENSMUST00000068045.8/ | 7 | 28901915 | 28907686 | 5772 | 2 | ENSMUST00 | -3625 | Actn4 | ENSMUSG00 | true | . | . | . | . | . | . | chr7 | 28910727 | 28959574 | 8.49369 | 5.62595 | 5.86532 |
| chr18 | 38795783 | 38795993 | 460 | Intron (ENSMUST00000148850.2 | 18 | 38771496 | 38772347 | 852 | 2 | ENSMUST00 | -23437 | Gm8302 | ENSMUSG00 | false | . | . | . | . | . | . | . | . | . | . | . | . |
| chr7 | 96446355 | 96446621 | 460 | Intron (ENSMUST00000107162.2 | 7 | 96435546 | 96435999 | 454 | 2 | ENSMUST00 | -10357 | Rps11-ps5 | ENSMUSG00 | false | . | . | . | . | . | . | chr7 | 96445528 | 96446676 | 3.80828 | 3.45436 | 2.06077 |
| chr18 | 11855506 | 11855943 | 460 | Intron (ENSMUST00000171109.2 | 18 | 11840361 | 11840438 | 78 | 2 | ENSMUST00 | -15069 | Mir1901 | ENSMUSG00 | false | . | . | . | . | . | . | chr18 | 11841487 | 11867262 | 7.10976 | 4.95644 | 4.75629 |
| chr1 | 152702880 | 152703296 | 460 | Intron (ENSMUST00000111859.2 | 1 | 152684026 | 152687501 | 3476 | 2 | ENSMUST00 | -15380 | Gm15479 | ENSMUSG00 | false | . | . | . | . | . | . | chr1 | 152702179 | 152704338 | 4.85934 | 3.95665 | 2.91189 |
| chr6 | 144735468 | 144735767 | 460 | Intron (ENSMUST00000155562.1 | 6 | 144723565 | 144724907 | 1343 | 1 | ENSMUST00 | 11904 | Sox5os5 | ENSMUSG00 | false | . | . | . | . | . | . | . | . | . | . | . | . |
| chr12 | 78144587 | 78144915 | 460 | Distal Intergenic | 12 | 78165546 | 78165666 | 121 | 1 | ENSMUST00 | -20631 | Gm24994 | ENSMUSG00 | false | . | . | . | . | . | . | chr12 | 78144381 | 78145247 | 4.0112 | 3.53322 | 2.22236 |
| chr1 | 131107729 | 131108339 | 460 | Distal Intergenic | 1 | 131102393 | 131102745 | 353 | 2 | ENSMUST00 | -4985 | Gm25717 | ENSMUSG00 | false | . | . | . | . | . | . | chr1 | 131107187 | 131113930 | 4.8831 | 3.79853 | 2.94189 |
| chr1 | 169041042 | 169041417 | 460 | Distal Intergenic | 1 | 169018974 | 169019089 | 116 | 2 | ENSMUST00 | -21954 | Mir6354 | ENSMUSG00 | false | . | . | . | . | . | . | chr1 | 169040684 | 169041469 | 3.91665 | 3.46557 | 2.13736 |
| chr11 | 35979308 | 35979543 | 459 | Promoter (<=1kb) | 11 | 35838400 | 35980527 | 142128 | 2 | ENSMUST00 | 984 | Wwc1 | ENSMUSG00 | false | chr11 | 35979365 | 35981000 | 13.16235 | 40.53663 | 37.94056 | chr11 | 35979136 | 35980883 | 5.04011 | 4.07746 | 3.0548 |
| chr12 | 16865850 | 16866155 | 459 | Distal Intergenic | 12 | 16894978 | 16988274 | 93297 | 1 | ENSMUST00 | -28823 | Rock2 | ENSMUSG00 | true | . | . | . | . | . | . | chr12 | 16864026 | 16867563 | 5.09379 | 4.19219 | 3.07583 |
| chr15 | 101267542 | 101268058 | 459 | Promoter (<=1kb) | 15 | 101266846 | 101274792 | 7947 | 1 | ENSMUST00 | 697 | Nr4a1 | ENSMUSG00 | false | chr15 | 101266077 | 101268132 | 17.60262 | 45.78123 | 43.11721 | chr15 | 101267663 | 101270327 | 3.47023 | 3.29367 | 1.75952 |
| chr2 | 94438027 | 94438292 | 459 | Promoter (<=1kb) | 2 | 94411682 | 94438136 | 26455 | 2 | ENSMUST00 | 0 | Api5 | ENSMUSG00 | false | chr2 | 94436427 | 94438271 | 25.4663 | 101.95255 | 98.33054 | . | . | . | . | . | . |
| chr5 | 137532970 | 137533762 | 459 | Promoter (<=1kb) | 5 | 137483020 | 137533242 | 50223 | 2 | ENSMUST00 | 0 | Epo | ENSMUSG00 | false | chr5 | 137530397 | 137533556 | 27.03504 | 88.5911 | 85.27629 | . | . | . | . | . | . |
| chr8 | 48152914 | 48153173 | 459 | Exon (ENSMUST00000174379.1/ | 8 | 48110211 | 48153233 | 43023 | 1 | ENSMUST00 | 42704 | Dctd | ENSMUSG00 | false | . | . | . | . | . | . | chr8 | 48151273 | 48154362 | 5.80736 | 4.45061 | 3.6749 |
| chr4 | 45341918 | 45342617 | 459 | Promoter (<=1kb) | 4 | 45342101 | 45379759 | 37659 | 1 | ENSMUST00 | 0 | Dcaf10 | ENSMUSG00 | false | chr4 | 45341634 | 45343307 | 27.88725 | 99.90214 | 96.33501 | . | . | . | . | . | . |
| chr4 | 134425135 | 134425454 | 459 | Intron (ENSMUST00000105869.3 | 4 | 134402970 | 134422532 | 19563 | 1 | ENSMUST00 | 22166 | Pafah2 | ENSMUSG00 | false | . | . | . | . | . | . | chr4 | 134424985 | 134425536 | 2.5679 | 2.50567 | 1.05758 |
| chr10 | 68135197 | 68135499 | 459 | Intron (ENSMUST00000020106.7 | 10 | 68043968 | 68051046 | 7079 | 1 | ENSMUST00 | 91230 | Rtkn2 | ENSMUSG00 | false | chr10 | 68134465 | 68136910 | 16.165 | 45.52484 | 42.86639 | chr10 | 68131250 | 68135665 | 4.41564 | 3.75071 | 2.5366 |
| chr15 | 8279392 | 8279807 | 459 | Distal Intergenic | 15 | 8247103 | 8252976 | 5874 | 1 | ENSMUST00 | 32290 | 2410089E03R | ENSMUSG00 | false | chr15 | 8279160 | 8280317 | 18.37877 | 47.11527 | 44.43725 | . | . | . | . | . | . |
| chr5 | 21220661 | 21221008 | 459 | Intron (ENSMUST00000036031.8 | 5 | 21186267 | 21291701 | 105435 | 1 | ENSMUST00 | 34395 | Gsap | ENSMUSG00 | false | . | . | . | . | . | . | chr5 | 21220153 | 21221680 | 3.68302 | 3.35316 | 1.95037 |
| chr12 | 8897156 | 8897468 | 459 | Distal Intergenic | 12 | 8873187 | 8877459 | 4273 | 2 | ENSMUST00 | -19698 | 9930038B18R | ENSMUSG00 | false | . | . | . | . | . | . | chr12 | 8892050 | 8903212 | 6.1858 | 4.65981 | 3.9745 |
| chr16 | 41745150 | 41745373 | 459 | Intron (ENSMUST00000187695.1 | 16 | 41533419 | 41539253 | 5835 | 1 | ENSMUST00 | 211732 | Lsamp | ENSMUSG00 | false | . | . | . | . | . | . | chr16 | 41744911 | 41745979 | 4.81574 | 4.06888 | 2.84399 |
| chr14 | 18820880 | 18821119 | 459 | Intron (ENSMUST00000176555.2 | 14 | 18814169 | 18814303 | 135 | 1 | ENSMUST00 | 6712 | Gm22499 | ENSMUSG00 | false | . | . | . | . | . | . | chr14 | 18820365 | 18821553 | 4.04673 | 3.46982 | 2.26589 |
| chr17 | 42992583 | 42992869 | 458 | Distal Intergenic | 17 | 43016555 | 43089188 | 72634 | 1 | ENSMUST00 | -23686 | Tnfrsf21 | ENSMUSG00 | false | . | . | . | . | . | . | chr17 | 42990241 | 42993592 | 4.60008 | 3.8832 | 2.65987 |
| chr18 | 62977695 | 62978101 | 458 | Promoter (<=1kb) | 18 | 62977836 | 62999450 | 21615 | 1 | ENSMUST00 | 0 | Napg | ENSMUSG00 | false | chr18 | 62977412 | 62979177 | 38.05226 | 120.16034 | 116.01576 | chr18 | 62977097 | 62980532 | 4.44712 | 3.68765 | 2.57664 |
| chr5 | 66586303 | 66586691 | 458 | Intron (ENSMUST00000159786.2 | 5 | 66451982 | 66618640 | 166659 | 2 | ENSMUST00 | 31949 | Apbb2 | ENSMUSG00 | false | . | . | . | . | . | . | chr5 | 66558047 | 66617728 | 6.24773 | 4.56399 | 4.04918 |
| chr2 | 52841144 | 52841591 | 458 | Distal Intergenic | 2 | 52857860 | 53133804 | 275945 | 1 | ENSMUST00 | -16269 | Fmnl2 | ENSMUSG00 | true | . | . | . | . | . | . | chr2 | 52840017 | 52846125 | 4.77462 | 3.96585 | 2.80752 |
| chr10 | 94014286 | 94014492 | 458 | Intron (ENSMUST00000119818.2 | 10 | 93996914 | 94020723 | 23810 | 2 | ENSMUST00 | 6231 | Vezt | ENSMUSG00 | true | . | . | . | . | . | . | chr10 | 94013349 | 94015420 | 4.46333 | 3.80262 | 2.5625 |
| chr10 | 128094640 | 128094934 | 458 | Promoter (1-2kb) | 10 | 128092783 | 128129303 | 36521 | 1 | ENSMUST00 | 1858 | Baz2a | ENSMUSG00 | false | . | . | . | . | . | . | chr10 | 128093379 | 128103131 | 5.08043 | 4.07895 | 3.08556 |
| chr1 | 135765955 | 135766437 | 458 | Promoter (<=1kb) | 1 | 135766119 | 135769136 | 3018 | 1 | ENSMUST00 | 0 | Phlda3 | ENSMUSG00 | false | chr1 | 135765813 | 135767149 | 27.80811 | 86.13519 | 82.86789 | chr1 | 135764697 | 135767254 | 3.80878 | 3.48457 | 2.0575 |
| chr12 | 102354549 | 102354817 | 458 | Promoter (<=1kb) | 12 | 102354745 | 102361507 | 6763 | 1 | ENSMUST00 | 0 | Rin3 | ENSMUSG00 | false | . | . | . | . | . | . | chr12 | 102354330 | 102355014 | 3.65508 | 3.39123 | 1.92421 |
| chr13 | 34111072 | 34111344 | 458 | Distal Intergenic | 13 | 34127008 | 34130354 | 3347 | 2 | ENSMUST00 | 19010 | Tubb2b | ENSMUSG00 | false | . | . | . | . | . | . | chr13 | 34109058 | 34120340 | 5.26535 | 4.17937 | 3.22481 |
| chr3 | 51648258 | 51648620 | 458 | Distal Intergenic | 3 | 51660360 | 51681807 | 21448 | 1 | ENSMUST00 | -11740 | Mgst2 | ENSMUSG00 | false | . | . | . | . | . | . | . | . | . | . | . | . |
| chr11 | 97661610 | 97661939 | 458 | Promoter (1-2kb) | 11 | 97644184 | 97660367 | 16184 | 2 | ENSMUST00 | -1244 | Gm11613 | ENSMUSG00 | false | chr11 | 97661507 | 97664564 | 31.01414 | 97.6227 | 94.10706 | chr11 | 97660731 | 97662935 | 5.22491 | 4.26722 | 3.18868 |
| chr17 | 28875406 | 28875715 | 458 | Intron (ENSMUST00000056866.5 | 17 | 28855136 | 28859503 | 4368 | 2 | ENSMUST00 | -15904 | Pnpla1os | ENSMUSG00 | false | . | . | . | . | . | . | chr17 | 28874228 | 28876520 | 4.9631 | 4.09736 | 2.96113 |
| chr2 | 167828218 | 167828557 | 458 | Exon (ENSMUST00000143649.1/ | 2 | 167827693 | 167833646 | 5954 | 2 | ENSMUST00 | 5089 | 1200007C13R | ENSMUSG00 | false | . | . | . | . | . | . | chr2 | 167826777 | 167828988 | 4.39539 | 3.7382 | 2.55225 |
| chr5 | 20379785 | 20380051 | 458 | Intron (ENSMUST00000088516.4 | 5 | 20264341 | 20264440 | 100 | 1 | ENSMUST00 | 115445 | Gm22755 | ENSMUSG00 | false | . | . | . | . | . | . | chr5 | 20375298 | 20380508 | 3.66481 | 3.28443 | 1.95259 |
| chr1 | 54596078 | 54596468 | 458 | Distal Intergenic | 1 | 54566703 | 54567281 | 579 | 1 | ENSMUST00 | 29376 | RP24-290F15 | ENSMUSG00 | false | . | . | . | . | . | . | chr1 | 54595745 | 54596900 | 4.88389 | 4.07038 | 2.89379 |
| chr15 | 101127255 | 101127532 | 457 | Promoter (<=1kb) | 15 | 101128522 | 101145336 | 16815 | 1 | ENSMUST00 | -990 | Acvrl1 | ENSMUSG00 | false | . | . | . | . | . | . | chr15 | 101127339 | 101129967 | 4.21947 | 3.46857 | 2.39009 |
| chr7 | 100845535 | 100845798 | 457 | Downstream (<1kb) | 7 | 100851112 | 100855429 | 4318 | 2 | ENSMUST00 | 9631 | Relt | ENSMUSG00 | false | . | . | . | . | . | . | chr7 | 100845183 | 100845762 | 4.35564 | 3.63878 | 2.49519 |
| chr16 | 32277328 | 32277817 | 457 | Promoter (<=1kb) | 16 | 32277459 | 32289194 | 11736 | 1 | ENSMUST00 | 0 | Rnf168 | ENSMUSG00 | false | chr16 | 32277025 | 32278537 | 33.94791 | 109.87469 | 106.04007 | . | . | . | . | . | . |
| chr8 | 119666179 | 119666625 | 457 | Promoter (<=1kb) | 8 | 119666523 | 119688222 | 21700 | 1 | ENSMUST00 | 0 | Wfdc1 | ENSMUSG00 | false | . | . | . | . | . | . | chr8 | 119664250 | 119667110 | 6.13153 | 4.55824 | 3.94801 |
| chr17 | 24960630 | 24961038 | 457 | Promoter (<=1kb) | 17 | 24942156 | 24960636 | 18481 | 2 | ENSMUST00 | 0 | Hn1l | ENSMUSG00 | false | chr17 | 24959729 | 24961229 | 32.3084 | 121.498 | 117.30614 | chr17 | 24960983 | 24961577 | 4.15418 | 3.66499 | 2.37283 |
| chr4 | 70391858 | 70392124 | 457 | Intron (ENSMUST00000144099.2 | 4 | 70398886 | 70403541 | 4656 | 2 | ENSMUST00 | 11417 | Cdk5rap2 | ENSMUSG00 | false | . | . | . | . | . | . | chr4 | 70390819 | 70392637 | 4.38609 | 3.58992 | 2.53658 |
| chr3 | 107982777 | 107983076 | 457 | Intron (ENSMUST00000012348.3 | 3 | 107979632 | 107979868 | 237 | 1 | ENSMUST00 | 3146 | Gm12497 | ENSMUSG00 | false | . | . | . | . | . | . | chr3 | 107981942 | 107983168 | 6.38032 | 4.55775 | 4.18777 |
| chr9 | 51765136 | 51765470 | 457 | Promoter (<=1kb) | 9 | 51765337 | 51790249 | 24913 | 1 | ENSMUST00 | 0 | Arhgap20 | ENSMUSG00 | false | chr9 | 51764398 | 51766274 | 28.53699 | 95.19626 | 91.7378 | chr9 | 51764051 | 51767503 | 5.62802 | 4.39324 | 3.52644 |
| chr13 | 38151153 | 38151889 | 457 | Promoter (<=1kb) | 13 | 38151328 | 38198577 | 47250 | 1 | ENSMUST00 | 0 | Dsp | ENSMUSG00 | false | chr13 | 38150957 | 38153283 | 17.89762 | 44.76895 | 42.12037 | chr13 | 38150614 | 38153304 | 4.00697 | 3.57226 | 2.18943 |
| chr6 | 72162195 | 72162512 | 457 | Distal Intergenic | 6 | 72115232 | 72142271 | 27040 | 1 | ENSMUST00 | 46964 | St3gal5 | ENSMUSG00 | false | . | . | . | . | . | . | chr6 | 72159976 | 72163501 | 4.22537 | 3.51631 | 2.394 |
| chr3 | 127837019 | 127837710 | 457 | Promoter (<=1kb) | 3 | 127807264 | 127837492 | 30229 | 2 | ENSMUST00 | 0 | Ap1ar | ENSMUSG00 | false | chr3 | 127836661 | 127838171 | 36.97958 | 118.30276 | 114.21766 | chr3 | 127835453 | 127837037 | 5.0015 | 4.04348 | 3.03127 |
| chr8 | 121376119 | 121376439 | 457 | Intron (ENSMUST00000127664.1 | 8 | 121256747 | 121356836 | 100090 | 2 | ENSMUST00 | -19284 | Gm26815 | ENSMUSG00 | false | . | . | . | . | . | . | chr8 | 121372330 | 121385684 | 7.83977 | 5.33422 | 5.34287 |
| chr9 | 45938902 | 45939183 | 457 | Promoter (<=1kb) | 9 | 45939521 | 45939590 | 70 | 2 | ENSMUST00 | 407 | Mir7087 | ENSMUSG00 | false | . | . | . | . | . | . | chr9 | 45935594 | 45942518 | 5.41154 | 4.18476 | 3.39461 |
| chr3 | 85323357 | 85323660 | 457 | Intron (ENSMUST00000189046.1 | 3 | 85317519 | 85332030 | 14512 | 1 | ENSMUST00 | 5839 | RP23-94M23 | ENSMUSG00 | false | . | . | . | . | . | . | . | . | . | . | . | . |
| chr10 | 94480708 | 94481022 | 456 | Intron (ENSMUST00000132743.1 | 10 | 94514857 | 94590956 | 76100 | 1 | ENSMUST00 | -33835 | Tmcc3 | ENSMUSG00 | false | . | . | . | . | . | . | chr10 | 94477425 | 94481453 | 5.61007 | 4.43652 | 3.49994 |
| chr14 | 60693218 | 60693627 | 456 | Promoter (1-2kb) | 14 | 60691975 | 60709569 | 17595 | 1 | ENSMUST00 | 1244 | Spata13 | ENSMUSG00 | true | . | . | . | . | . | . | chr14 | 60692020 | 60695002 | 7.59295 | 5.30678 | 5.11833 |
| chr15 | 8109062 | 8109409 | 456 | Promoter (<=1kb) | 15 | 8109313 | 8159857 | 50545 | 1 | ENSMUST00 | 0 | Nup155 | ENSMUSG00 | false | chr15 | 8108704 | 8110145 | 30.59503 | 95.89513 | 92.42037 | . | . | . | . | . | . |
| chr16 | 32246832 | 32247234 | 456 | Promoter (<=1kb) | 16 | 32247227 | 32256753 | 9527 | 1 | ENSMUST00 | 0 | Wdr53 | ENSMUSG00 | false | chr16 | 32245965 | 32247998 | 36.97958 | 118.30276 | 114.21766 | . | . | . | . | . | . |
| chr16 | 35490645 | 35490924 | 456 | Promoter (<=1kb) | 16 | 35397313 | 35490873 | 93561 | 2 | ENSMUST00 | 0 | Pdia5 | ENSMUSG00 | false | chr16 | 35489892 | 35491182 | 29.22959 | 88.61999 | 85.30408 | chr16 | 35488992 | 35491412 | 3.53867 | 3.31753 | 1.83616 |
| chr3 | 136570636 | 136570932 | 456 | Distal Intergenic | 3 | 136670124 | 136797838 | 127715 | 1 | ENSMUST00 | -99192 | Ppp3ca | ENSMUSG00 | false | . | . | . | . | . | . | . | . | . | . | . | . |
| chr8 | 120251488 | 120251831 | 456 | Intron (ENSMUST00000127664.1 | 8 | 120230536 | 120553634 | 323099 | 1 | ENSMUST00 | 20953 | Gse1 | ENSMUSG00 | false | . | . | . | . | . | . | chr8 | 120247970 | 120252219 | 6.21079 | 4.4693 | 4.04227 |
| chr8 | 111735637 | 111735945 | 456 | Intron (ENSMUST00000166232.2 | 8 | 111710485 | 111743849 | 33365 | 2 | ENSMUST00 | 7904 | Bcar1 | ENSMUSG00 | false | . | . | . | . | . | . | chr8 | 111710939 | 111747327 | 8.20913 | 5.4583 | 5.64085 |
| chr7 | 101347903 | 101348169 | 456 | Promoter (<=1kb) | 7 | 101348067 | 101411894 | 63828 | 1 | ENSMUST00 | 0 | Arap1 | ENSMUSG00 | false | chr7 | 101347632 | 101348783 | 27.71015 | 85.00858 | 81.76643 | chr7 | 101347372 | 101348996 | 9.38825 | 6.14525 | 6.56721 |
| chr7 | 45921210 | 45921449 | 456 | Promoter (<=1kb) | 7 | 45918023 | 45921426 | 3404 | 2 | ENSMUST00 | 0 | Emp3 | ENSMUSG00 | false | chr7 | 45920389 | 45921478 | 8.87028 | 22.65516 | 20.28383 | chr7 | 45915648 | 45924393 | 5.13169 | 3.954 | 3.14707 |
| chr17 | 62511138 | 62511440 | 456 | Distal Intergenic | 17 | 62456987 | 62457120 | 134 | 2 | ENSMUST00 | -54019 | Gm25800 | ENSMUSG00 | false | . | . | . | . | . | . | chr17 | 62510479 | 62513010 | 4.41834 | 3.68259 | 2.56143 |
| chr2 | 33431021 | 33431441 | 456 | Promoter (<=1kb) | 2 | 33406108 | 33431324 | 25217 | 2 | ENSMUST00 | 0 | Zbtb34 | ENSMUSG00 | false | chr2 | 33430325 | 33431829 | 29.2476 | 106.45832 | 102.71833 | . | . | . | . | . | . |
| chr6 | 86487253 | 86487610 | 456 | Promoter (<=1kb) | 6 | 86483376 | 86488204 | 4829 | 2 | ENSMUST00 | 594 | RP23-281H4. | ENSMUSG00 | false | . | . | . | . | . | . | chr6 | 86483526 | 86489878 | 6.12859 | 4.5529 | 3.93946 |
| chr15 | 67616309 | 67616582 | 456 | Distal Intergenic | 15 | 67226769 | 67377094 | 150326 | 1 | ENSMUST00 | 389541 | RP23-163N4 | ENSMUSG00 | false | . | . | . | . | . | . | chr15 | 67615788 | 67616775 | 3.67875 | 3.37728 | 1.97378 |
| chr9 | 21962442 | 21962736 | 455 | Promoter (<=1kb) | 9 | 21958899 | 21963576 | 4678 | 2 | ENSMUST00 | 840 | Epor | ENSMUSG00 | false | chr9 | 21961366 | 21963078 | 9.18239 | 21.07545 | 18.72512 | chr9 | 21960665 | 21963833 | 4.83894 | 3.98051 | 2.88993 |
| chr11 | 96271065 | 96271355 | 455 | Promoter (<=1kb) | 11 | 96271457 | 96276595 | 5139 | 1 | ENSMUST00 | -102 | Hoxb9 | ENSMUSG00 | false | chr11 | 96270998 | 96272761 | 20.53639 | 56.4273 | 53.62976 | chr11 | 96269585 | 96272568 | 5.09741 | 3.93611 | 3.1112 |
| chr16 | 23259606 | 23259912 | 455 | Intron (ENSMUST00000023601.8 | 16 | 23269629 | 23321179 | 51551 | 1 | ENSMUST00 | -9717 | St6gal1 | ENSMUSG00 | true | . | . | . | . | . | . | chr16 | 23227288 | 23275612 | 5.51887 | 4.26961 | 3.44483 |
| chr4 | 48044613 | 48045760 | 455 | Promoter (<=1kb) | 4 | 48045153 | 48086445 | 41293 | 1 | ENSMUST00 | 0 | Nr4a3 | ENSMUSG00 | false | chr4 | 48044233 | 48048412 | 30.84378 | 98.32127 | 94.79446 | . | . | . | . | . | . |
| chr4 | 102798935 | 102799227 | 455 | Intron (ENSMUST00000097948.3 | 4 | 102760525 | 102973628 | 213104 | 1 | ENSMUST00 | 38411 | Sgip1 | ENSMUSG00 | true | . | . | . | . | . | . | chr4 | 102798049 | 102813929 | 4.80947 | 3.92915 | 2.87079 |
| chr6 | 49822487 | 49822787 | 455 | Promoter (<=1kb) | 6 | 49822710 | 49829507 | 6798 | 1 | ENSMUST00 | 0 | Npy | ENSMUSG00 | false | chr6 | 49822027 | 49823019 | 18.90859 | 48.02031 | 45.33304 | chr6 | 49821860 | 49823846 | 5.70432 | 4.33752 | 3.63705 |
| chr9 | 82974257 | 82974617 | 455 | Promoter (<=1kb) | 9 | 82927158 | 82974268 | 47111 | 2 | ENSMUST00 | 0 | Phip | ENSMUSG00 | false | chr9 | 82972816 | 82976631 | 23.3092 | 84.4034 | 81.16875 | . | . | . | . | . | . |
| chr1 | 91179875 | 91180135 | 455 | Promoter (<=1kb) | 1 | 91180042 | 91206790 | 26749 | 1 | ENSMUST00 | 0 | Ramp1 | ENSMUSG00 | false | chr1 | 91179575 | 91180319 | 18.60216 | 47.49733 | 44.81332 | chr1 | 91179416 | 91182028 | 4.26187 | 3.71909 | 2.39162 |
| chr11 | 80724220 | 80724537 | 455 | Intron (ENSMUST00000041065.8 | 11 | 80673667 | 80780003 | 106337 | 2 | ENSMUST00 | 55466 | Myo1d | ENSMUSG00 | true | . | . | . | . | . | . | chr11 | 80722742 | 80729746 | 5.38488 | 4.11326 | 3.36661 |
| chr2 | 118532849 | 118533391 | 455 | Intron (ENSMUST00000090219.7 | 2 | 118531974 | 118544587 | 12614 | 2 | ENSMUST00 | 11196 | Bmf | ENSMUSG00 | false | . | . | . | . | . | . | chr2 | 118532146 | 118534961 | 4.24094 | 3.5694 | 2.42692 |
| chr9 | 15709596 | 15710512 | 455 | Promoter (<=1kb) | 9 | 15709738 | 15738579 | 28842 | 1 | ENSMUST00 | 0 | Slc36a4 | ENSMUSG00 | false | chr9 | 15709114 | 15710996 | 29.34497 | 88.82044 | 85.50058 | chr9 | 15707792 | 15709622 | 3.35733 | 3.23503 | 1.65041 |
| chr5 | 24724859 | 24725277 | 455 | Intron (ENSMUST00000068693.6 | 5 | 24712265 | 24730680 | 18416 | 2 | ENSMUST00 | 5403 | Wdr86 | ENSMUSG00 | false | . | . | . | . | . | . | chr5 | 24723425 | 24734625 | 5.82193 | 4.37932 | 3.69553 |
| chr19 | 8920245 | 8920718 | 455 | Promoter (<=1kb) | 19 | 8920374 | 8927234 | 6861 | 1 | ENSMUST00 | 0 | B3gat3 | ENSMUSG00 | false | chr19 | 8920094 | 8921667 | 28.49099 | 81.63235 | 78.4519 | . | . | . | . | . | . |
| chr2 | 110173851 | 110174103 | 455 | Distal Intergenic | 2 | 110196889 | 110256131 | 59243 | 1 | ENSMUST00 | -22786 | Gm13936 | ENSMUSG00 | false | . | . | . | . | . | . | chr2 | 110172226 | 110205814 | 5.39899 | 4.25006 | 3.33729 |
| chr7 | 43865450 | 43865650 | 454 | Distal Intergenic | 7 | 43881172 | 43885804 | 4633 | 1 | ENSMUST00 | -15522 | Klk4 | ENSMUSG00 | false | . | . | . | . | . | . | chr7 | 43864411 | 43866084 | 5.18497 | 4.23506 | 3.15057 |
| chr10 | 25449392 | 25449694 | 454 | Intron (ENSMUST00000053748.9 | 10 | 25408368 | 25523519 | 115152 | 1 | ENSMUST00 | 41025 | Epb4.1l2 | ENSMUSG00 | false | . | . | . | . | . | . | chr10 | 25449218 | 25451026 | 4.45218 | 3.813 | 2.557 |
| chr19 | 53409974 | 53410246 | 454 | Distal Intergenic | 19 | 53379214 | 53390573 | 11360 | 2 | ENSMUST00 | -19402 | Smndc1 | ENSMUSG00 | false | . | . | . | . | . | . | chr19 | 53409099 | 53414773 | 6.07728 | 4.55041 | 3.90583 |
| chr19 | 43524172 | 43524867 | 454 | Promoter (<=1kb) | 19 | 43499752 | 43524605 | 24854 | 2 | ENSMUST00 | 0 | Got1 | ENSMUSG00 | false | chr19 | 43523562 | 43525376 | 13.72719 | 56.76396 | 53.95979 | chr19 | 43523310 | 43526485 | 5.50603 | 4.13004 | 3.47386 |
| chr4 | 59547534 | 59547819 | 454 | Promoter (<=1kb) | 4 | 59517473 | 59548737 | 31265 | 2 | ENSMUST00 | 918 | Ptbp3 | ENSMUSG00 | false | chr4 | 59545854 | 59549379 | 31.7391 | 109.03407 | 105.22643 | chr4 | 59537450 | 59547719 | 3.9895 | 3.56933 | 2.1747 |
| chr4 | 137498381 | 137498684 | 454 | Intron (ENSMUST00000030547.9 | 4 | 137468803 | 137570629 | 101827 | 1 | ENSMUST00 | 29579 | Hspg2 | ENSMUSG00 | false | . | . | . | . | . | . | chr4 | 137497674 | 137499182 | 3.29287 | 3.18074 | 1.66282 |
| chr5 | 33220766 | 33221018 | 454 | Promoter (2-3kb) | 5 | 33213518 | 33218238 | 4721 | 2 | ENSMUST00 | -2529 | Spon2 | ENSMUSG00 | false | . | . | . | . | . | . | . | . | . | . | . | . |
| chr6 | 148946043 | 148946357 | 454 | Promoter (<=1kb) | 6 | 148926051 | 148946155 | 20105 | 2 | ENSMUST00 | 0 | Fam60a | ENSMUSG00 | false | chr6 | 148944401 | 148947082 | 36.04346 | 118.78738 | 114.68494 | . | . | . | . | . | . |
| chr9 | 65676348 | 65676682 | 454 | Promoter (<=1kb) | 9 | 65676548 | 65690300 | 13753 | 1 | ENSMUST00 | 0 | Oaz2 | ENSMUSG00 | false | chr9 | 65675887 | 65677358 | 24.35997 | 68.16138 | 65.19754 | chr9 | 65675551 | 65685323 | 4.69201 | 3.84119 | 2.79144 |
| chr17 | 80825612 | 80826346 | 454 | Distal Intergenic | 17 | 80728131 | 80728522 | 392 | 1 | ENSMUST00 | 97482 | Gm9959 | ENSMUSG00 | false | . | . | . | . | . | . | chr17 | 80824776 | 80826826 | 3.52957 | 3.28292 | 1.83567 |
| chr6 | 52165893 | 52166301 | 454 | Promoter (<=1kb) | 6 | 52166518 | 52169564 | 3047 | 1 | ENSMUST00 | -217 | Hoxaas2 | ENSMUSG00 | false | chr6 | 52165559 | 52166190 | 14.1901 | 32.13216 | 29.63978 | chr6 | 52159328 | 52167512 | 5.47105 | 4.07894 | 3.42628 |
| chr3 | 51836156 | 51836667 | 454 | Intron (ENSMUST00000121440.2 | 3 | 51828912 | 51829028 | 117 | 1 | ENSMUST00 | 7245 | n-R5s196 | ENSMUSG00 | false | . | . | . | . | . | . | chr3 | 51831736 | 51847000 | 4.07316 | 3.59293 | 2.2664 |
| chr11 | 108417358 | 108417724 | 454 | Distal Intergenic | 11 | 108423273 | 108423758 | 486 | 1 | ENSMUST00 | -5549 | Gm11667 | ENSMUSG00 | false | . | . | . | . | . | . | . | . | . | . | . | . |
| chr5 | 107303519 | 107303773 | 454 | Promoter (<=1kb) | 5 | 107302946 | 107303047 | 102 | 1 | ENSMUST00 | 574 | Gm22557 | ENSMUSG00 | false | chr5 | 107303238 | 107303864 | 10.05864 | 20.96754 | 18.61967 | chr5 | 107303030 | 107304621 | 4.27183 | 3.64709 | 2.4416 |
| chr12 | 86888414 | 86889089 | 454 | Downstream (2-3kb) | 12 | 86891404 | 86892567 | 1164 | 2 | ENSMUST00 | 3478 | Gm26698 | ENSMUSG00 | false | chr12 | 86888060 | 86889010 | 10.27012 | 20.7831 | 18.43794 | chr12 | 86885110 | 86901194 | 5.68305 | 4.41688 | 3.56559 |
| chr17 | 34632282 | 34632694 | 453 | Promoter (<=1kb) | 17 | 34631404 | 34632397 | 994 | 1 | ENSMUST00 | 879 | Prrt1 | ENSMUSG00 | false | chr17 | 34631792 | 34632576 | 11.36537 | 25.4653 | 23.05655 | chr17 | 34628191 | 34632769 | 5.9403 | 4.50962 | 3.80031 |
| chr6 | 134038808 | 134039096 | 453 | Promoter (2-3kb) | 6 | 134035969 | 134248284 | 212316 | 1 | ENSMUST00 | 2840 | Etv6 | ENSMUSG00 | false | . | . | . | . | . | . | chr6 | 134037014 | 134053440 | 5.9149 | 4.52781 | 3.75986 |
| chrX | 73955085 | 73955323 | 453 | Promoter (2-3kb) | 20 | 73955121 | 73957429 | 2309 | 2 | ENSMUST00 | 2106 | Hcfc1 | ENSMUSG00 | false | . | . | . | . | . | . | . | . | . | . | . | . |
| chr9 | 61373040 | 61373624 | 453 | Promoter (<=1kb) | 9 | 61373219 | 61418486 | 45268 | 1 | ENSMUST00 | 0 | Tle3 | ENSMUSG00 | false | chr9 | 61370729 | 61374567 | 24.24157 | 89.02564 | 85.70093 | chr9 | 61366653 | 61374765 | 6.23582 | 4.63772 | 4.01625 |
| chr1 | 127298641 | 127298911 | 453 | Intron (ENSMUST00000189427.1 | 1 | 127306762 | 127483704 | 176943 | 1 | ENSMUST00 | -7851 | Mgat5 | ENSMUSG00 | true | . | . | . | . | . | . | chr1 | 127297620 | 127306217 | 4.43971 | 3.68092 | 2.58089 |
| chr5 | 33274579 | 33275356 | 453 | Promoter (<=1kb) | 5 | 33247724 | 33275004 | 27281 | 2 | ENSMUST00 | 0 | Ctbp1 | ENSMUSG00 | false | chr5 | 33273539 | 33275535 | 38.84783 | 124.74329 | 120.44118 | . | . | . | . | . | . |
| chr1 | 184033422 | 184034397 | 453 | Promoter (<=1kb) | 1 | 184034381 | 184075636 | 41256 | 1 | ENSMUST00 | 0 | Dusp10 | ENSMUSG00 | false | chr1 | 184033118 | 184034192 | 27.13995 | 99.45934 | 95.90212 | . | . | . | . | . | . |

| | | | | | | | | | | | | | | | | | | | | | | | | | | |
|---|---|---|---|---|---|---|---|---|---|---|---|---|---|---|---|---|---|---|---|---|---|---|---|---|---|---|
| chr17 | 27074035 | 27074344 | 453 | Intron (ENSMUST00000049308.8 | 17 | 27057304 | 27122223 | 64920 | 1 | ENSMUST00 | 16732 | Itpr3 | ENSMUSG0C | false | . | . | . | . | . | . | chr17 | 27056774 | 27076697 | 5.08038 | 3.96808 | 3.08893 |
| chr1 | 73923253 | 73923567 | 453 | Intron (ENSMUST00000169786.2 | 1 | 73925417 | 73928217 | 2801 | 2 | ENSMUST00 | 4650 | Tns1 | ENSMUSG0C | false | . | . | . | . | . | . | . | . | . | . | . | . |
| chr10 | 127751628 | 127751882 | 453 | Promoter (<=1kb) | 10 | 127747276 | 127751732 | 4457 | 2 | ENSMUST00 | 0 | Gpr182 | ENSMUSG0C | false | . | . | . | . | . | . | chr10 | 127743127 | 127759693 | 6.17884 | 4.54629 | 3.98922 |
| chrX | 11837446 | 11837830 | 453 | Distal Intergenic | 20 | 11850484 | 11850590 | 107 | 1 | ENSMUST00 | -12654 | Gm26314 | ENSMUSG0C | false | . | . | . | . | . | . | . | . | . | . | . | . |
| chr7 | 73398409 | 73398815 | 453 | Intron (ENSMUST00000094312.6 | 7 | 73391227 | 73419899 | 28673 | 1 | ENSMUST00 | 7183 | Rgma | ENSMUSG0C | false | . | . | . | . | . | . | chr7 | 73397492 | 73400320 | 4.91984 | 4.07913 | 2.95652 |
| chr4 | 82513136 | 82513396 | 453 | Intron (ENSMUST00000139835.1 | 4 | 82508016 | 82547292 | 39277 | 1 | ENSMUST00 | 5121 | Gm11266 | ENSMUSG0C | false | chr4 | 82513102 | 82513677 | 16.08362 | 38.04472 | 35.47989 | chr4 | 82511629 | 82513800 | 4.90219 | 4.02818 | 2.94353 |
| chr5 | 99447522 | 99447887 | 453 | Intron (ENSMUST00000146396.2 | 5 | 99509944 | 99510682 | 739 | 2 | ENSMUST00 | 62795 | Gm16227 | ENSMUSG0C | false | . | . | . | . | . | . | chr5 | 99442196 | 99449542 | 4.74151 | 3.92904 | 2.79783 |
| chr11 | 101984291 | 101984991 | 452 | Promoter (<=1kb) | 11 | 101971143 | 101984799 | 13657 | 2 | ENSMUST00 | 0 | Dusp3 | ENSMUSG0C | false | chr11 | 101983072 | 101985233 | 23.0285 | 72.05518 | 69.03415 | chr11 | 101984928 | 101988114 | 6.10933 | 4.5538 | 3.96693 |
| chr1 | 171247132 | 171247624 | 452 | Promoter (<=1kb) | 1 | 171234853 | 171247122 | 12270 | 2 | ENSMUST00 | -11 | Ndufs2 | ENSMUSG0C | false | chr1 | 171246458 | 171247900 | 29.86959 | 86.83979 | 83.56236 | . | . | . | . | . | . |
| chr11 | 60811117 | 60811436 | 452 | Promoter (<=1kb) | 11 | 60788104 | 60811265 | 23162 | 2 | ENSMUST00 | 0 | Shmt1 | ENSMUSG0C | false | chr11 | 60810933 | 60811772 | 10.13814 | 22.76719 | 20.39335 | chr11 | 60809624 | 60812145 | 4.63422 | 3.84378 | 2.70843 |
| chr12 | 83328269 | 83328546 | 452 | Intron (ENSMUST00000147469.2 | 12 | 83213745 | 83350895 | 137151 | 2 | ENSMUST00 | 22349 | Dpf3 | ENSMUSG0C | false | . | . | . | . | . | . | chr12 | 83327269 | 83331024 | 4.27118 | 3.73368 | 2.42218 |
| chr18 | 75377235 | 75377805 | 452 | Promoter (2-3kb) | 18 | 75374914 | 75394001 | 19088 | 1 | ENSMUST00 | 2322 | Smad7 | ENSMUSG0C | false | . | . | . | . | . | . | chr18 | 75369277 | 75392875 | 5.74724 | 4.44386 | 3.62103 |
| chr7 | 109960091 | 109960609 | 452 | Promoter (<=1kb) | 7 | 109893780 | 109960400 | 66621 | 2 | ENSMUST00 | 0 | Dennd5a | ENSMUSG0C | false | chr7 | 109959166 | 109960911 | 27.03504 | 88.5911 | 85.27629 | . | . | . | . | . | . |
| chr6 | 30300351 | 30300674 | 452 | Intron (ENSMUST00000102993.4 | 6 | 30214861 | 30304522 | 89662 | 2 | ENSMUST00 | 3848 | Ube2h | ENSMUSG0C | false | chr6 | 30300607 | 30300932 | 4.08106 | 5.99902 | 3.91878 | chr6 | 30296569 | 30303366 | 4.9255 | 4.01255 | 2.94274 |
| chr7 | 27333489 | 27333904 | 452 | Promoter (<=1kb) | 7 | 27305140 | 27333663 | 28524 | 2 | ENSMUST00 | 0 | Ltbp4 | ENSMUSG0C | false | chr7 | 27332275 | 27333899 | 23.50745 | 64.07966 | 61.17685 | chr7 | 27328853 | 27337953 | 4.74964 | 3.95019 | 2.80772 |
| chr8 | 3671989 | 3672295 | 452 | Promoter (<=1kb) | 8 | 3671188 | 3672336 | 1149 | 2 | ENSMUST00 | 41 | Fcor | ENSMUSG0C | false | chr8 | 3671978 | 3672606 | 18.61129 | 49.85614 | 47.14222 | . | . | . | . | . | . |
| chr6 | 117294060 | 117294406 | 452 | Distal Intergenic | 6 | 117213656 | 117214048 | 393 | 2 | ENSMUST00 | -80013 | Rpl28-ps4 | ENSMUSG0C | false | . | . | . | . | . | . | chr6 | 117292305 | 117295611 | 5.09583 | 4.12614 | 3.0804 |
| chr14 | 102199519 | 102199731 | 452 | Distal Intergenic | 14 | 102329530 | 102329661 | 132 | 2 | ENSMUST00 | 129930 | Gm22347 | ENSMUSG0C | false | . | . | . | . | . | . | . | . | . | . | . | . |
| chr12 | 34169069 | 34169329 | 452 | Distal Intergenic | 12 | 34290274 | 34291092 | 819 | 2 | ENSMUST00 | 121763 | Gm18025 | ENSMUSG0C | false | . | . | . | . | . | . | . | . | . | . | . | . |
| chr12 | 108320235 | 108320555 | 452 | Intron (ENSMUST00000021685.6 | 12 | 108313779 | 108328030 | 14252 | 2 | ENSMUST00 | 7475 | 4930478K11I | ENSMUSG0C | false | . | . | . | . | . | . | . | . | . | . | . | . |
| chr1 | 73221860 | 73222357 | 452 | Distal Intergenic | 1 | 73269064 | 73306360 | 37297 | 1 | ENSMUST00 | -46707 | RP24-141F19 | ENSMUSG0C | false | . | . | . | . | . | . | . | . | . | . | . | . |
| chr8 | 122547095 | 122547490 | 451 | Intron (ENSMUST00000127664.1 | 8 | 122481698 | 122551329 | 69632 | 2 | ENSMUST00 | 3839 | Piezo1 | ENSMUSG0C | false | . | . | . | . | . | . | chr8 | 122535817 | 122552157 | 5.24095 | 4.06992 | 3.22847 |
| chr10 | 121878082 | 121878429 | 451 | Intron (ENSMUST00000020322.6 | 10 | 121806640 | 121823133 | 16494 | 2 | ENSMUST00 | -54950 | Srgap1 | ENSMUSG0C | true | . | . | . | . | . | . | chr10 | 121869425 | 121882678 | 5.26441 | 4.14787 | 3.23352 |
| chr11 | 3202312 | 3202922 | 451 | Promoter (<=1kb) | 11 | 3202392 | 3244581 | 42190 | 1 | ENSMUST00 | 0 | Eif4enif1 | ENSMUSG0C | false | chr11 | 3201976 | 3202582 | 5.64303 | 28.58646 | 26.13994 | . | . | . | . | . | . |
| chr18 | 74444379 | 74444649 | 451 | Promoter (1-2kb) | 18 | 74442607 | 74770805 | 328199 | 1 | ENSMUST00 | 1773 | Myo5b | ENSMUSG0C | false | chr18 | 74443774 | 74444853 | 14.61067 | 37.63035 | 35.07038 | chr18 | 74441088 | 74445660 | 3.76081 | 3.28607 | 2.03186 |
| chr3 | 83938914 | 83939185 | 451 | Intron (ENSMUST00000052342.7 | 3 | 83836273 | 83841767 | 5495 | 2 | ENSMUST00 | -97148 | Tlr2 | ENSMUSG0C | false | . | . | . | . | . | . | chr3 | 83938633 | 83939286 | 3.27195 | 3.19584 | 1.61376 |
| chr9 | 69250346 | 69250574 | 451 | Exon (ENSMUST00000132355.1/ | 9 | 69289682 | 69379809 | 90128 | 1 | ENSMUST00 | -39108 | Rora | ENSMUSG0C | false | . | . | . | . | . | . | . | . | . | . | . | . |
| chr9 | 66593060 | 66593293 | 451 | Promoter (<=1kb) | 9 | 66514637 | 66593142 | 78506 | 2 | ENSMUST00 | 0 | Usp3 | ENSMUSG0C | false | chr9 | 66591521 | 66593918 | 35.77904 | 109.9518 | 106.11687 | . | . | . | . | . | . |
| chr13 | 33964454 | 33964870 | 451 | Promoter (<=1kb) | 13 | 33964659 | 33988465 | 23807 | 1 | ENSMUST00 | 0 | Nqo2 | ENSMUSG0C | false | chr13 | 33964344 | 33965896 | 23.91706 | 66.52044 | 63.57986 | . | . | . | . | . | . |
| chr6 | 113377393 | 113377920 | 451 | Promoter (<=1kb) | 6 | 113366025 | 113377883 | 11859 | 2 | ENSMUST00 | 0 | Tada3 | ENSMUSG0C | false | chr6 | 113377011 | 113379381 | 32.64849 | 123.19247 | 118.94151 | . | . | . | . | . | . |
| chr15 | 102624989 | 102625599 | 451 | Promoter (<=1kb) | 15 | 102523843 | 102625067 | 101225 | 2 | ENSMUST00 | 0 | Gm28047 | ENSMUSG0C | false | chr15 | 102624179 | 102626066 | 29.22523 | 92.5029 | 89.10189 | chr15 | 102618999 | 102626125 | 4.24802 | 3.61295 | 2.40981 |
| chr1 | 37864575 | 37865478 | 451 | Promoter (<=1kb) | 1 | 37754572 | 37865079 | 110508 | 2 | ENSMUST00 | 0 | Tsga10 | ENSMUSG0C | false | chr1 | 37864981 | 37865153 | 5.25681 | 8.51884 | 6.37244 | chr1 | 37864064 | 37864693 | 3.47219 | 3.38143 | 1.73706 |
| chr17 | 47140535 | 47140838 | 451 | Promoter (<=1kb) | 17 | 47140875 | 47329536 | 188662 | 1 | ENSMUST00 | -37 | Trerf1 | ENSMUSG0C | false | chr17 | 47139299 | 47142406 | 27.6016 | 85.77155 | 82.50918 | chr17 | 47139017 | 47143146 | 4.41006 | 3.81485 | 2.5073 |
| chr10 | 22731019 | 22731591 | 451 | Promoter (<=1kb) | 10 | 22703879 | 22731447 | 27569 | 2 | ENSMUST00 | 0 | Tbpl1 | ENSMUSG0C | false | chr10 | 22730228 | 22732192 | 39.29051 | 134.21745 | 129.56854 | . | . | . | . | . | . |
| chr2 | 61137312 | 61137538 | 451 | Distal Intergenic | 2 | 61224792 | 61225082 | 291 | 1 | ENSMUST00 | -87254 | Gm13581 | ENSMUSG0C | false | . | . | . | . | . | . | chr2 | 61136622 | 61140725 | 5.9439 | 4.56787 | 3.77059 |
| chr12 | 13598647 | 13598906 | 451 | Distal Intergenic | 12 | 13507862 | 13507968 | 107 | 2 | ENSMUST00 | -90680 | Gm22170 | ENSMUSG0C | false | . | . | . | . | . | . | chr12 | 13598259 | 13599456 | 5.13644 | 4.15155 | 3.14822 |
| chr6 | 134920182 | 134920830 | 450 | Promoter (<=1kb) | 6 | 134920401 | 134925513 | 5113 | 1 | ENSMUST00 | 0 | Cdkn1b | ENSMUSG0C | false | chr6 | 134919565 | 134922632 | 25.50663 | 88.63404 | 85.31516 | . | . | . | . | . | . |
| chr17 | 34660308 | 34660557 | 450 | Distal Intergenic | 17 | 34651627 | 34652103 | 477 | 1 | ENSMUST00 | 8682 | Atf6b | ENSMUSG0C | false | . | . | . | . | . | . | chr17 | 34659116 | 34661951 | 5.72154 | 4.38413 | 3.60456 |
| chr11 | 53636159 | 53636463 | 450 | Promoter (1-2kb) | 11 | 53631324 | 53634702 | 3379 | 2 | ENSMUST00 | -1458 | Il13 | ENSMUSG0C | false | chr11 | 53636100 | 53636628 | 12.15419 | 27.41675 | 24.98428 | chr11 | 53636450 | 53637373 | 3.73698 | 3.46669 | 2.01667 |
| chr16 | 58519763 | 58520111 | 450 | Intron (ENSMUST00000137035.2 | 16 | 58493745 | 58523344 | 29600 | 2 | ENSMUST00 | 3233 | St3gal6 | ENSMUSG0C | true | chr16 | 58518435 | 58519923 | 11.55672 | 26.39465 | 23.97632 | chr16 | 58496830 | 58520680 | 5.70067 | 4.33546 | 3.59278 |
| chr5 | 115119132 | 115119483 | 450 | Promoter (<=1kb) | 5 | 115110299 | 115119287 | 8989 | 2 | ENSMUST00 | 0 | Acads | ENSMUSG0C | false | chr5 | 115118586 | 115119761 | 38.25672 | 133.48087 | 128.85442 | chr5 | 115117256 | 115119855 | 3.5183 | 3.32347 | 1.77397 |
| chr4 | 123343264 | 123343469 | 450 | Promoter (<=1kb) | 4 | 123312645 | 123343252 | 30608 | 2 | ENSMUST00 | -13 | Bmp8a | ENSMUSG0C | false | chr4 | 123342314 | 123343684 | 19.72203 | 59.14077 | 56.30748 | chr4 | 123342077 | 123343751 | 4.73989 | 3.91986 | 2.81092 |
| chr14 | 75753961 | 75754713 | 450 | Promoter (<=1kb) | 14 | 75702351 | 75754493 | 52143 | 2 | ENSMUST00 | 0 | Cog3 | ENSMUSG0C | false | chr14 | 75753380 | 75754982 | 32.0113 | 129.32326 | 124.86235 | chr14 | 75748387 | 75754892 | 4.45254 | 3.77301 | 2.57794 |
| chr13 | 25056837 | 25057397 | 450 | Promoter (<=1kb) | 13 | 25056253 | 25210706 | 154454 | 1 | ENSMUST00 | 585 | Dcdc2a | ENSMUSG0C | true | chr13 | 25054929 | 25057285 | 27.2424 | 81.35376 | 78.17802 | . | . | . | . | . | . |
| chrX | 159817207 | 159817435 | 450 | Exon (ENSMUST00000150433.2/ | 20 | 159840468 | 159975664 | 135197 | 1 | ENSMUST00 | -23033 | Sh3kbp1 | ENSMUSG0C | false | . | . | . | . | . | . | . | . | . | . | . | . |
| chr9 | 31280727 | 31281372 | 450 | Promoter (<=1kb) | 9 | 31280538 | 31330618 | 50081 | 1 | ENSMUST00 | 190 | Prdm10 | ENSMUSG0C | false | chr9 | 31279611 | 31282029 | 30.94804 | 114.76607 | 110.79114 | . | . | . | . | . | . |
| chr5 | 142432071 | 142432389 | 450 | Intron (ENSMUST00000072837.5 | 5 | 142401497 | 142462014 | 60518 | 1 | ENSMUST00 | 30575 | Foxk1 | ENSMUSG0C | false | . | . | . | . | . | . | chr5 | 142423530 | 142434255 | 6.409 | 4.68664 | 4.16411 |
| chr2 | 92551073 | 92551401 | 450 | Distal Intergenic | 2 | 92561269 | 92561680 | 412 | 1 | ENSMUST00 | -9868 | Gm13815 | ENSMUSG0C | false | . | . | . | . | . | . | chr2 | 92543021 | 92555783 | 4.86524 | 4.02879 | 2.88826 |
| chr17 | 10464842 | 10465199 | 450 | Promoter (<=1kb) | 17 | 10464837 | 10473758 | 8922 | 1 | ENSMUST00 | 6 | A230009B12 | ENSMUSG0C | false | . | . | . | . | . | . | chr17 | 10463350 | 10467528 | 4.53143 | 3.66902 | 2.65357 |
| chr14 | 62759744 | 62760593 | 450 | Promoter (<=1kb) | 14 | 62760501 | 62830126 | 69626 | 1 | ENSMUST00 | 0 | 4931440J10F | ENSMUSG0C | false | chr14 | 62759434 | 62761182 | 36.04672 | 129.50026 | 125.02953 | . | . | . | . | . | . |
| chr13 | 45524319 | 45524721 | 449 | Intron (ENSMUST00000000260.7 | 13 | 45513840 | 45545898 | 32059 | 1 | ENSMUST00 | 10480 | Gmpr | ENSMUSG0C | false | . | . | . | . | . | . | chr13 | 45522834 | 45531762 | 4.69665 | 3.835 | 2.77414 |
| chr13 | 55740678 | 55740934 | 449 | Intron (ENSMUST00000021958.5 | 13 | 55727368 | 55776830 | 49463 | 1 | ENSMUST00 | 13311 | Pcbd2 | ENSMUSG0C | false | . | . | . | . | . | . | chr13 | 55739452 | 55743934 | 5.40467 | 4.31375 | 3.33541 |
| chr15 | 34082628 | 34083336 | 449 | Promoter (<=1kb) | 15 | 34082694 | 34103345 | 20652 | 1 | ENSMUST00 | 0 | Mtdh | ENSMUSG0C | false | chr15 | 34081757 | 34084754 | 29.52499 | 116.10789 | 112.09218 | . | . | . | . | . | . |
| chr19 | 10012986 | 10013563 | 449 | Promoter (1-2kb) | 19 | 10015016 | 10027576 | 12561 | 1 | ENSMUST00 | -1453 | Rab3il1 | ENSMUSG0C | false | . | . | . | . | . | . | chr19 | 10011323 | 10014870 | 5.6737 | 4.30843 | 3.56561 |
| chr5 | 125140192 | 125140681 | 449 | Intron (ENSMUST00000111402.3 | 5 | 125017835 | 125124900 | 107066 | 2 | ENSMUST00 | -15293 | Ncor2 | ENSMUSG0C | false | chr5 | 125140272 | 125141740 | 11.71324 | 27.89317 | 25.45456 | chr5 | 125124978 | 125178451 | 6.70656 | 4.76008 | 4.42049 |
| chr8 | 104631231 | 104631547 | 449 | Promoter (<=1kb) | 8 | 104628066 | 104631321 | 3256 | 2 | ENSMUST00 | 0 | Rrad | ENSMUSG0C | false | chr8 | 104630488 | 104631317 | 9.01168 | 19.18248 | 16.85847 | chr8 | 104629601 | 104631637 | 3.71467 | 3.40632 | 1.94614 |
| chr9 | 71750066 | 71750332 | 449 | Intron (ENSMUST00000072899.3 | 9 | 71629211 | 71771560 | 142350 | 2 | ENSMUST00 | 21228 | Cgnl1 | ENSMUSG0C | false | . | . | . | . | . | . | chr9 | 71736274 | 71770833 | 5.2398 | 4.14793 | 3.22386 |
| chr13 | 102899275 | 102899549 | 449 | Intron (ENSMUST00000166726.2 | 13 | 102741969 | 102905786 | 163818 | 2 | ENSMUST00 | 6237 | Mast4 | ENSMUSG0C | true | chr13 | 102898513 | 102899308 | 6.28665 | 10.55282 | 8.36284 | chr13 | 102868528 | 102904155 | 5.63021 | 4.27399 | 3.54307 |
| chr7 | 45628634 | 45629032 | 449 | Promoter (1-2kb) | 7 | 45627488 | 45639093 | 11606 | 1 | ENSMUST00 | 1147 | Rasip1 | ENSMUSG0C | false | chr7 | 45628098 | 45629263 | 23.65566 | 74.08845 | 71.03342 | chr7 | 45628423 | 45629324 | 2.66096 | 2.71893 | 1.14917 |
| chr13 | 34129693 | 34129996 | 449 | Promoter (<=1kb) | 13 | 34127008 | 34130354 | 3347 | 2 | ENSMUST00 | 358 | Tubb2b | ENSMUSG0C | false | chr13 | 34129135 | 34131581 | 24.55417 | 69.38167 | 66.39919 | . | . | . | . | . | . |
| chr6 | 54303941 | 54304270 | 449 | Intron (ENSMUST00000145111.2 | 6 | 54327012 | 54330200 | 3189 | 1 | ENSMUST00 | -22742 | Prr15 | ENSMUSG0C | false | . | . | . | . | . | . | chr6 | 54303044 | 54304590 | 4.41369 | 3.67561 | 2.55581 |
| chr4 | 5782844 | 5783168 | 449 | Intron (ENSMUST00000124256.2 | 4 | 5772879 | 5799946 | 27068 | 1 | ENSMUST00 | 9966 | Fam110b | ENSMUSG0C | false | . | . | . | . | . | . | chr4 | 5782185 | 5785765 | 3.94267 | 3.56755 | 2.14184 |
| chr7 | 4866049 | 4866417 | 449 | Promoter (<=1kb) | 7 | 4844959 | 4866193 | 21235 | 2 | ENSMUST00 | 0 | Isoc2b | ENSMUSG0C | false | chr7 | 4865656 | 4867011 | 15.19686 | 42.27666 | 39.66088 | chr7 | 4862226 | 4867162 | 3.93909 | 3.5309 | 2.16033 |
| chr9 | 32024781 | 32025048 | 449 | Distal Intergenic | 9 | 31936459 | 31936755 | 297 | 2 | ENSMUST00 | -88027 | Gm3756 | ENSMUSG0C | false | . | . | . | . | . | . | . | . | . | . | . | . |
| chr15 | 85653322 | 85654340 | 449 | Distal Intergenic | 15 | 85639747 | 85645934 | 6188 | 1 | ENSMUST00 | 13576 | RP23-64J7.2 | ENSMUSG0C | false | chr15 | 85652927 | 85654833 | 20.20131 | 69.382 | 66.39919 | chr15 | 85652004 | 85661329 | 6.36895 | 4.61573 | 4.152 |
| chr11 | 109439959 | 109440253 | 448 | Promoter (<=1kb) | 11 | 109440156 | 109452813 | 12658 | 1 | ENSMUST00 | 0 | Amz2 | ENSMUSG0C | false | chr11 | 109439719 | 109440803 | 15.85026 | 49.49464 | 46.78799 | . | . | . | . | . | . |
| chr13 | 42100687 | 42100938 | 448 | Intron (ENSMUST00000060148.5 | 13 | 42052021 | 42185026 | 133006 | 1 | ENSMUST00 | 48667 | Hivep1 | ENSMUSG0C | false | . | . | . | . | . | . | chr13 | 42098974 | 42105815 | 5.30539 | 4.14893 | 3.26488 |
| chr15 | 34509782 | 34510159 | 448 | Promoter (<=1kb) | 15 | 34509907 | 34512192 | 2286 | 1 | ENSMUST00 | 0 | Pop1 | ENSMUSG0C | false | . | . | . | . | . | . | chr15 | 34508319 | 34517398 | 5.80499 | 4.48832 | 3.66177 |
| chr10 | 121509843 | 121510250 | 448 | Distal Intergenic | 10 | 121410350 | 121476250 | 65901 | 2 | ENSMUST00 | -33594 | Rassf3 | ENSMUSG0C | false | . | . | . | . | . | . | chr10 | 121509800 | 121510831 | 4.89242 | 4.09176 | 2.90174 |
| chr2 | 154559227 | 154559729 | 448 | Promoter (<=1kb) | 2 | 154547046 | 154558890 | 11845 | 2 | ENSMUST00 | -338 | Necab3 | ENSMUSG0C | false | chr2 | 154557669 | 154559308 | 20.89815 | 57.05354 | 54.24761 | chr2 | 154555502 | 154571286 | 6.03478 | 4.51051 | 3.86945 |
| chr9 | 102834758 | 102835277 | 448 | Promoter (<=1kb) | 9 | 102834917 | 102907542 | 72626 | 1 | ENSMUST00 | 0 | Ryk | ENSMUSG0C | false | chr9 | 102834367 | 102836715 | 30.42596 | 90.69117 | 87.33178 | chr9 | 102833046 | 102850688 | 5.63517 | 4.31932 | 3.53737 |
| chr11 | 69175637 | 69175872 | 448 | Distal Intergenic | 11 | 69156989 | 69169792 | 12804 | 1 | ENSMUST00 | 18649 | Alox12b | ENSMUSG0C | false | . | . | . | . | . | . | chr11 | 69171732 | 69176303 | 5.42866 | 4.24022 | 3.37182 |
| chr9 | 63571297 | 63571548 | 448 | Intron (ENSMUST00000042322.5 | 9 | 63421835 | 63602417 | 180583 | 2 | ENSMUST00 | 30869 | Iqch | ENSMUSG0C | false | . | . | . | . | . | . | chr9 | 63568228 | 63577350 | 4.15727 | 3.62156 | 2.32961 |
| chr5 | 148909462 | 148909768 | 448 | Intron (ENSMUST00000047257.9 | 5 | 148904593 | 148928628 | 24036 | 2 | ENSMUST00 | 18860 | Katnal1 | ENSMUSG0C | false | . | . | . | . | . | . | chr5 | 148907850 | 148910114 | 4.26179 | 3.59882 | 2.44121 |
| chr7 | 127889874 | 127890173 | 448 | Promoter (<=1kb) | 7 | 127885841 | 127890970 | 5130 | 2 | ENSMUST00 | 797 | Prss53 | ENSMUSG0C | false | . | . | . | . | . | . | chr7 | 127880545 | 127891013 | 5.86469 | 4.44372 | 3.71216 |
| chr2 | 132688855 | 132689131 | 448 | Promoter (1-2kb) | 2 | 132690157 | 132750503 | 60347 | 1 | ENSMUST00 | -1026 | 1110034G24 | ENSMUSG0C | false | . | . | . | . | . | . | chr2 | 132683824 | 132693696 | 5.22275 | 4.16804 | 3.18727 |
| chr4 | 55100936 | 55101146 | 448 | Distal Intergenic | 4 | 55116563 | 55116923 | 361 | 1 | ENSMUST00 | -15417 | Gm12515 | ENSMUSG0C | false | . | . | . | . | . | . | chr4 | 55100590 | 55101487 | 4.22785 | 3.74851 | 2.34961 |
| chr4 | 133489022 | 133489327 | 448 | Downstream (1-2kb) | 4 | 133479356 | 133482526 | 3171 | 2 | ENSMUST00 | -6497 | Gm13257 | ENSMUSG0C | false | . | . | . | . | . | . | . | . | . | . | . | . |
| chr19 | 15644903 | 15645196 | 448 | Distal Intergenic | 19 | 15680004 | 15680110 | 107 | 1 | ENSMUST00 | -34808 | Gm24319 | ENSMUSG0C | false | . | . | . | . | . | . | chr19 | 15643565 | 15649571 | 6.62858 | 4.81931 | 4.35258 |
| chr1 | 64532419 | 64533802 | 447 | Promoter (<=1kb) | 1 | 64532645 | 64597695 | 65051 | 1 | ENSMUST00 | 0 | Creb1 | ENSMUSG0C | false | chr1 | 64531819 | 64534191 | 33.1106 | 101.26704 | 97.66567 | . | . | . | . | . | . |
| chr1 | 194619634 | 194620000 | 447 | Promoter (<=1kb) | 1 | 194619829 | 194816869 | 197041 | 1 | ENSMUST00 | 0 | Plxna2 | ENSMUSG0C | false | chr1 | 194618995 | 194621050 | 21.44446 | 75.31629 | 72.24329 | chr1 | 194618894 | 194626082 | 4.03442 | 3.44451 | 2.2464 |
| chr2 | 32626361 | 32626657 | 447 | Promoter (<=1kb) | 2 | 32625437 | 32635054 | 9618 | 1 | ENSMUST00 | 925 | Ak1 | ENSMUSG0C | false | chr2 | 32625128 | 32627071 | 9.13756 | 26.51177 | 24.09012 | chr2 | 32605009 | 32639275 | 6.86759 | 4.92928 | 4.53618 |
| chr8 | 34806992 | 34807602 | 447 | Promoter (<=1kb) | 8 | 34807297 | 34819894 | 12598 | 1 | ENSMUST00 | 0 | Dusp4 | ENSMUSG0C | false | chr8 | 34806492 | 34810291 | 19.52531 | 57.12588 | 54.31675 | chr8 | 34805060 | 34810387 | 5.51025 | 4.28302 | 3.47096 |
| chr11 | 56064696 | 56064971 | 447 | Intron (ENSMUST00000134285.1 | 11 | 56024987 | 56041010 | 16024 | 2 | ENSMUST00 | -23687 | Nmur2 | ENSMUSG0C | false | . | . | . | . | . | . | chr11 | 56064533 | 56065646 | 2.89845 | 2.81374 | 1.32164 |
| chr10 | 128173545 | 128173874 | 447 | Promoter (2-3kb) | 10 | 128129470 | 128176568 | 47099 | 2 | ENSMUST00 | 2694 | Rbms2 | ENSMUSG0C | true | . | . | . | . | . | . | chr10 | 128159882 | 128175743 | 4.81979 | 3.97943 | 2.86388 |
| chr6 | 39203264 | 39203519 | 447 | Intron (ENSMUST00000002305.8 | 6 | 39136623 | 39206789 | 70167 | 2 | ENSMUST00 | 3270 | Kdm7a | ENSMUSG0C | false | . | . | . | . | . | . | chr6 | 39203388 | 39205009 | 5.08162 | 3.91837 | 3.11611 |
| chr17 | 56583528 | 56583801 | 447 | Promoter (<=1kb) | 17 | 56576331 | 56583046 | 6716 | 2 | ENSMUST00 | -483 | Safb2 | ENSMUSG0C | false | chr17 | 56583240 | 56584610 | 39.4261 | 145.54997 | 140.44281 | chr17 | 56580064 | 56583922 | 5.38569 | 4.28979 | 3.30666 |
| chr17 | 27211988 | 27212493 | 447 | Distal Intergenic | 17 | 27189601 | 27204438 | 14838 | 2 | ENSMUST00 | -7551 | Lemd2 | ENSMUSG0C | false | . | . | . | . | . | . | chr17 | 27209279 | 27213622 | 5.47435 | 4.34629 | 3.38789 |
| chr13 | 12461367 | 12461692 | 447 | Promoter (<=1kb) | 13 | 12439418 | 12461493 | 22076 | 2 | ENSMUST00 | 0 | Lgals8 | ENSMUSG0C | false | chr13 | 12460437 | 12462111 | 39.28384 | 124.42301 | 120.12974 | . | . | . | . | . | . |
| chr5 | 115235317 | 115235631 | 447 | Promoter (<=1kb) | 5 | 115235836 | 115240991 | 5156 | 1 | ENSMUST00 | -205 | Pop5 | ENSMUSG0C | false | chr5 | 115235208 | 115236090 | 21.54815 | 66.72034 | 63.77881 | chr5 | 115235057 | 115240113 | 3.84999 | 3.54447 | 2.0518 |
| chr7 | 34313220 | 34313685 | 447 | Promoter (<=1kb) | 7 | 34236707 | 34313551 | 76845 | 2 | ENSMUST00 | 0 | 4931406P16I | ENSMUSG0C | false | chr7 | 34312738 | 34314229 | 36.12563 | 122.12295 | 117.91594 | . | . | . | . | . | . |
| chr8 | 70539216 | 70539851 | 447 | Promoter (<=1kb) | 8 | 70539675 | 70592858 | 53184 | 1 | ENSMUST00 | 0 | Ell | ENSMUSG0C | false | chr8 | 70538131 | 70541044 | 36.17514 | 115.85 | 111.8395 | . | . | . | . | . | . |
| chr19 | 18436394 | 18436708 | 447 | Distal Intergenic | 19 | 18581455 | 18582228 | 774 | 1 | ENSMUST00 | -144747 | Gm8250 | ENSMUSG0C | false | . | . | . | . | . | . | chr19 | 18436066 | 18437549 | 3.97019 | 3.61478 | 2.14462 |
| chr4 | 148914900 | 148915368 | 447 | Promoter (<=1kb) | 4 | 148913511 | 148914324 | 814 | 2 | ENSMUST00 | -577 | Gm15969 | ENSMUSG0C | false | . | . | . | . | . | . | chr4 | 148901819 | 148933146 | 7.00771 | 4.91323 | 4.66351 |
| chr10 | 80939609 | 80940060 | 446 | Distal Intergenic | 10 | 80930091 | 80932204 | 2114 | 1 | ENSMUST00 | 9519 | Gadd45b | ENSMUSG0C | false | chr10 | 80939833 | 80940366 | 9.86256 | 23.41135 | 21.03137 | chr10 | 80936148 | 80942225 | 6.45113 | 4.61033 | 4.23692 |
| chr11 | 93955873 | 93956138 | 446 | Promoter (<=1kb) | 11 | 93949817 | 93955783 | 5967 | 2 | ENSMUST00 | -91 | Nme2 | ENSMUSG0C | false | chr11 | 93954825 | 93956484 | 32.18837 | 99.66983 | 96.10942 | chr11 | 93951624 | 93957429 | 4.20778 | 3.60571 | 2.38129 |
| chr12 | 59011778 | 59012322 | 446 | Promoter (<=1kb) | 12 | 58958383 | 59012017 | 53635 | 2 | ENSMUST00 | 0 | Sec23a | ENSMUSG0C | false | chr12 | 59012234 | 59013989 | 27.18019 | 77.52287 | 74.41271 | . | . | . | . | . | . |
| chr3 | 65927687 | 65928055 | 446 | Distal Intergenic | 3 | 65946782 | 65947998 | 1217 | 2 | ENSMUST00 | 19943 | Ccnl1 | ENSMUSG0C | false | chr3 | 65927064 | 65927955 | 9.22042 | 18.50693 | 16.19293 | chr3 | 65926001 | 65930446 | 5.38488 | 4.15152 | 3.33208 |
| chr11 | 107971760 | 107972004 | 446 | Intron (ENSMUST00000059595.5 | 11 | 107938177 | 107979639 | 41463 | 2 | ENSMUST00 | 7635 | Prkca | ENSMUSG0C | false | . | . | . | . | . | . | chr11 | 107971409 | 107972256 | 4.84573 | 4.11882 | 2.89075 |
| chr15 | 76656682 | 76657121 | 446 | Promoter (<=1kb) | 15 | 76646927 | 76656920 | 9994 | 2 | ENSMUST00 | 0 | Cyhr1 | ENSMUSG0C | false | chr15 | 76655745 | 76657556 | 17.09341 | 55.02552 | 52.24499 | . | . | . | . | . | . |
| chrX | 11418044 | 11418306 | 446 | Distal Intergenic | 20 | 11327822 | 11328151 | 330 | 1 | ENSMUST00 | 90223 | Gm4906 | ENSMUSG0C | false | . | . | . | . | . | . | . | . | . | . | . | . |
| chr2 | 159279017 | 159279320 | 446 | Distal Intergenic | 2 | 159139361 | 159139492 | 132 | 1 | ENSMUST00 | 139657 | Gm25090 | ENSMUSG0C | false | . | . | . | . | . | . | chr2 | 159269480 | 159284862 | 4.11414 | 3.61003 | 2.27932 |
| chr4 | 95051389 | 95052038 | 446 | Promoter (<=1kb) | 4 | 95051534 | 95067422 | 15889 | 1 | ENSMUST00 | 0 | Junos | ENSMUSG0C | false | chr4 | 95048614 | 95052213 | 33.83522 | 116.97303 | 112.93475 | . | . | . | . | . | . |
| chr18 | 75226466 | 75226714 | 446 | Intron (ENSMUST00000039608.7 | 18 | 75182218 | 75182709 | 492 | 2 | ENSMUST00 | -43758 | Gm8807 | ENSMUSG0C | false | . | . | . | . | . | . | chr18 | 75226126 | 75227752 | 4.48583 | 3.74139 | 2.6278 |
| chr4 | 133104085 | 133104309 | 446 | Distal Intergenic | 4 | 133085474 | 133089146 | 3673 | 1 | ENSMUST00 | 18612 | Gm26615 | ENSMUSG0C | false | . | . | . | . | . | . | chr4 | 133100685 | 133104681 | 5.90668 | 4.39897 | 3.76601 |
| chr1 | 152934088 | 152934287 | 446 | Distal Intergenic | 1 | 152949395 | 152950097 | 703 | 2 | ENSMUST00 | 15810 | RP23-258M1 | ENSMUSG0C | false | . | . | . | . | . | . | . | . | . | . | . | . |
| chr12 | 79953384 | 79953659 | 446 | Distal Intergenic | 12 | 79963568 | 79963872 | 305 | 1 | ENSMUST00 | -9909 | RP23-179D3. | ENSMUSG0C | false | . | . | . | . | . | . | chr12 | 79942657 | 79955332 | 5.38416 | 4.22706 | 3.33447 |
| chr1 | 90434782 | 90435331 | 446 | Distal Intergenic | 1 | 90453394 | 90454501 | 1108 | 2 | ENSMUST00 | 19170 | RP24-154L11 | ENSMUSG0C | false | chr1 | 90431359 | 90438623 | 24.40663 | 77.266 | 74.1616 | . | . | . | . | . | . |
| chr15 | 83169519 | 83169748 | 445 | Promoter (<=1kb) | 15 | 83160385 | 83170177 | 9793 | 2 | ENSMUST00 | 429 | Cyb5r3 | ENSMUSG0C | false | . | . | . | . | . | . | chr15 | 83162089 | 83173300 | 7.98675 | 5.40074 | 5.45307 |
| chr12 | 16870463 | 16870830 | 445 | Distal Intergenic | 12 | 16894978 | 16988274 | 93297 | 1 | ENSMUST00 | -24148 | Rock2 | ENSMUSG0C | true | . | . | . | . | . | . | chr12 | 16868702 | 16871132 | 6.43489 | 4.7014 | 4.20933 |
| chr15 | 32496992 | 32497280 | 445 | Intron (ENSMUST00000067458.6 | 15 | 32244813 | 32696341 | 451529 | 1 | ENSMUST00 | 252180 | Sema5a | ENSMUSG0C | true | . | . | . | . | . | . | chr15 | 32493573 | 32498839 | 4.34893 | 3.68215 | 2.48902 |
| chr9 | 63197308 | 63197572 | 445 | Exon (ENSMUST00000034920.9/ | 9 | 63138170 | 63148961 | 10792 | 2 | ENSMUST00 | -48348 | Skor1 | ENSMUSG0C | false | . | . | . | . | . | . | . | . | . | . | . | . |
| chr19 | 10869643 | 10869964 | 445 | Promoter (<=1kb) | 19 | 10857822 | 10869940 | 12119 | 2 | ENSMUST00 | 0 | Tmem132a | ENSMUSG0C | false | chr19 | 10868950 | 10870186 | 18.02173 | 47.27326 | 44.5908 | chr19 | 10868564 | 10870495 | 5.65178 | 4.22897 | 3.57943 |
| chr8 | 71569787 | 71570504 | 445 | Promoter (<=1kb) | 8 | 71568927 | 71586708 | 17782 | 1 | ENSMUST00 | 861 | Slc27a1 | ENSMUSG0C | false | chr8 | 71569284 | 71571112 | 26.22066 | 76.7888 | 73.6919 | chr8 | 71568572 | 71571351 | 3.85889 | 3.42898 | 2.11061 |
| chr9 | 21791813 | 21792219 | 445 | Intron (ENSMUST00000034717.5 | 9 | 21766773 | 21798546 | 31774 | 2 | ENSMUST00 | 6327 | Kank2 | ENSMUSG0C | false | . | . | . | . | . | . | chr9 | 21786534 | 21799409 | 6.22233 | 4.65692 | 4.01382 |
| chr3 | 101612133 | 101612489 | 445 | Distal Intergenic | 3 | 101576219 | 101604603 | 28385 | 2 | ENSMUST00 | -7531 | Atp1a1 | ENSMUSG0C | false | . | . | . | . | . | . | chr3 | 101604927 | 101613489 | 5.50887 | 4.25497 | 3.44478 |
| chr16 | 35031121 | 35031502 | 445 | Intron (ENSMUST00000061156.8 | 16 | 35022421 | 35109175 | 86755 | 1 | ENSMUST00 | 8701 | Ptplb | ENSMUSG0C | false | . | . | . | . | . | . | chr16 | 35030767 | 35034951 | 5.28937 | 4.24744 | 3.26311 |
| chr11 | 6389067 | 6389457 | 445 | Promoter (<=1kb) | 11 | 6389074 | 6406158 | 17085 | 1 | ENSMUST00 | 0 | Zmiz2 | ENSMUSG0C | false | chr11 | 6388725 | 6390322 | 35.96008 | 115.47577 | 111.48098 | . | . | . | . | . | . |
| chr8 | 95331459 | 95332087 | 445 | Promoter (<=1kb) | 8 | 95326137 | 95331950 | 5814 | 2 | ENSMUST00 | 0 | Zfp319 | ENSMUSG0C | false | chr8 | 95330917 | 95332916 | 36.96683 | 120.40339 | 116.25284 | . | . | . | . | . | . |
| chr6 | 86457122 | 86457424 | 445 | Intron (ENSMUST00000050497.8 | 6 | 86464040 | 86469947 | 5908 | 1 | ENSMUST00 | -6616 | C87436 | ENSMUSG0C | false | . | . | . | . | . | . | chr6 | 86456736 | 86458388 | 3.76521 | 3.26634 | 2.01831 |
| chr5 | 73498660 | 73498856 | 445 | Intron (ENSMUST00000087181.6 | 5 | 73492847 | 73520133 | 27287 | 1 | ENSMUST00 | 5814 | Dcun1d4 | ENSMUSG0C | false | . | . | . | . | . | . | . | . | . | . | . | . |
| chr5 | 49043287 | 49043503 | 445 | Intron (ENSMUST00000087395.5 | 5 | 48924116 | 48924491 | 376 | 1 | ENSMUST00 | 119172 | Gm10025 | ENSMUSG0C | false | . | . | . | . | . | . | . | . | . | . | . | . |
| chr1 | 51927326 | 51927563 | 445 | Distal Intergenic | 1 | 51938254 | 51938665 | 412 | 2 | ENSMUST00 | 11102 | RP23-348P7. | ENSMUSG0C | false | . | . | . | . | . | . | . | . | . | . | . | . |
| chr8 | 125488849 | 125489086 | 444 | Intron (ENSMUST00000108775.1 | 8 | 125418063 | 125492710 | 74648 | 2 | ENSMUST00 | 3624 | Sipa1l2 | ENSMUSG0C | false | . | . | . | . | . | . | chr8 | 125488016 | 125490182 | 6.67233 | 4.79284 | 4.38646 |
| chr13 | 29988624 | 29988900 | 444 | Promoter (2-3kb) | 13 | 29906575 | 29985668 | 79094 | 2 | ENSMUST00 | -2957 | E2f3 | ENSMUSG0C | true | . | . | . | . | . | . | chr13 | 29986776 | 29989095 | 4.82825 | 4.07104 | 2.86248 |
| chr16 | 38712753 | 38713499 | 444 | Promoter (<=1kb) | 16 | 38598340 | 38713274 | 114935 | 2 | ENSMUST00 | 0 | Arhgap31 | ENSMUSG0C | false | chr16 | 38711971 | 38713697 | 28.95052 | 83.36136 | 80.14891 | . | . | . | . | . | . |
| chr3 | 121736006 | 121736456 | 444 | Distal Intergenic | 3 | 121723537 | 121735052 | 11516 | 1 | ENSMUST00 | 12470 | F3 | ENSMUSG0C | false | . | . | . | . | . | . | chr3 | 121734745 | 121740674 | 4.74367 | 4.00425 | 2.78613 |
| chr3 | 94371721 | 94372040 | 444 | Promoter (<=1kb) | 3 | 94372794 | 94397840 | 25047 | 1 | ENSMUST00 | -754 | Rorc | ENSMUSG0C | false | . | . | . | . | . | . | chr3 | 94371361 | 94374439 | 5.16649 | 4.17623 | 3.1432 |
| chr3 | 145833982 | 145834364 | 444 | Intron (ENSMUST00000029845.9 | 3 | 145810479 | 145889295 | 78817 | 1 | ENSMUST00 | 23504 | Ddah1 | ENSMUSG0C | false | . | . | . | . | . | . | . | . | . | . | . | . |
| chr4 | 130702521 | 130702783 | 444 | Intron (ENSMUST00000030315.7 | 4 | 130663406 | 130780254 | 116849 | 1 | ENSMUST00 | 39116 | Pum1 | ENSMUSG0C | false | . | . | . | . | . | . | chr4 | 130697929 | 130731647 | 4.40946 | 3.70382 | 2.55396 |
| chr4 | 133240475 | 133240907 | 444 | Promoter (<=1kb) | 4 | 133240818 | 133252929 | 12112 | 1 | ENSMUST00 | 0 | Map3k6 | ENSMUSG0C | false | chr4 | 133240506 | 133242188 | 26.28407 | 85.32412 | 82.07336 | chr4 | 133240235 | 133243355 | 4.7864 | 3.96631 | 2.83351 |
| chr6 | 36781398 | 36781694 | 444 | Intron (ENSMUST00000101534.1 | 6 | 36715663 | 36811361 | 95699 | 2 | ENSMUST00 | 29667 | Ptn | ENSMUSG0C | false | . | . | . | . | . | . | . | . | . | . | . | . |
| chr1 | 182346576 | 182346799 | 444 | Distal Intergenic | 1 | 182313102 | 182341606 | 28505 | 2 | ENSMUST00 | -4971 | Fbxo28 | ENSMUSG0C | false | . | . | . | . | . | . | chr1 | 182346064 | 182347205 | 4.36845 | 3.79079 | 2.49649 |
| chr10 | 120978098 | 120979667 | 444 | Promoter (<=1kb) | 10 | 120923413 | 120979332 | 55920 | 2 | ENSMUST00 | 0 | Lemd3 | ENSMUSG0C | false | chr10 | 120977509 | 120980049 | 38.13901 | 127.82562 | 123.40861 | chr10 | 120976358 | 120978176 | 4.79824 | 4.03188 | 2.85061 |
| chr9 | 109059234 | 109059485 | 444 | Promoter (<=1kb) | 9 | 109057938 | 109059251 | 1314 | 2 | ENSMUST00 | 0 | Trex1 | ENSMUSG0C | false | chr9 | 109058794 | 109059890 | 15.01947 | 39.77812 | 37.19162 | chr9 | 109053192 | 109060472 | 5.12914 | 4.03657 | 3.12589 |
| chr2 | 79496827 | 79497138 | 444 | Distal Intergenic | 2 | 79338569 | 79456785 | 118217 | 2 | ENSMUST00 | -40043 | Cerkl | ENSMUSG0C | false | . | . | . | . | . | . | . | . | . | . | . | . |
| chr6 | 52102759 | 52103003 | 444 | Promoter (<=1kb) | 6 | 52102949 | 52112528 | 9580 | 1 | ENSMUST00 | 0 | Linchoxa1 | ENSMUSG0C | false | . | . | . | . | . | . | chr6 | 52102127 | 52103386 | 5.77086 | 4.34946 | 3.6866 |
| chr6 | 86480304 | 86480803 | 444 | Promoter (<=1kb) | 6 | 86470805 | 86480770 | 9966 | 2 | ENSMUST00 | 0 | A430078I02F | ENSMUSG0C | false | chr6 | 86480441 | 86481097 | 8.42674 | 18.66703 | 16.35069 | chr6 | 86472845 | 86482455 | 6.5006 | 4.7144 | 4.25243 |
| chr9 | 78105098 | 78105375 | 443 | Intron (ENSMUST00000085311.7 | 9 | 78099803 | 78108587 | 8785 | 2 | ENSMUST00 | 3212 | Fbxo9 | ENSMUSG0C | false | . | . | . | . | . | . | chr9 | 78103401 | 78107746 | 5.22062 | 4.17283 | 3.18398 |
| chr7 | 27491813 | 27492073 | 443 | Distal Intergenic | 7 | 27487375 | 27489507 | 2133 | 1 | ENSMUST00 | 4439 | Sertad1 | ENSMUSG0C | false | . | . | . | . | . | . | chr7 | 27488220 | 27512085 | 6.36425 | 4.68802 | 4.12603 |
| chr2 | 116065564 | 116066123 | 443 | Promoter (<=1kb) | 2 | 115863064 | 116065839 | 202776 | 2 | ENSMUST00 | 0 | Meis2 | ENSMUSG0C | false | chr2 | 116066009 | 116067477 | 22.15371 | 67.81625 | 64.85734 | . | . | . | . | . | . |
| chr4 | 109442715 | 109442952 | 443 | Exon (ENSMUST00000064129.8/ | 4 | 109420942 | 109426365 | 5424 | 1 | ENSMUST00 | 21774 | Ttc39a | ENSMUSG0C | false | . | . | . | . | . | . | chr4 | 109441263 | 109443376 | 4.22504 | 3.43659 | 2.39757 |
| chr5 | 142550848 | 142551153 | 443 | Promoter (<=1kb) | 5 | 142484839 | 142551098 | 66260 | 2 | ENSMUST00 | 0 | Radil | ENSMUSG0C | false | chr5 | 142550294 | 142551407 | 24.30838 | 70.74035 | 67.73997 | chr5 | 142550300 | 142551278 | 6.33958 | 4.56007 | 4.1515 |
| chrX | 72987120 | 72987441 | 443 | Promoter (<=1kb) | 20 | 72987339 | 73031543 | 44205 | 1 | ENSMUST00 | 0 | Zfp185 | ENSMUSG0C | false | chrX | 72986865 | 72988251 | 21.13848 | 54.99947 | 52.22001 | . | . | . | . | . | . |
| chr9 | 62980792 | 62981172 | 443 | Promoter (<=1kb) | 9 | 62880077 | 62980879 | 100803 | 2 | ENSMUST00 | 0 | Pias1 | ENSMUSG0C | false | chr9 | 62979447 | 62981327 | 43.18097 | 163.77325 | 157.99008 | chr9 | 62981110 | 62982339 | 3.68377 | 3.4085 | 1.91252 |
| chr19 | 25356676 | 25356900 | 443 | Intron (ENSMUST00000049400.9 | 19 | 25395303 | 25409772 | 14470 | 1 | ENSMUST00 | -38403 | Kank1 | ENSMUSG0C | true | . | . | . | . | . | . | chr19 | 25329142 | 25370790 | 4.73993 | 3.84606 | 2.82412 |
| chr19 | 6398605 | 6398902 | 443 | Promoter (<=1kb) | 19 | 6399340 | 6415210 | 15871 | 1 | ENSMUST00 | -438 | Rasgrp2 | ENSMUSG0C | false | chr19 | 6398550 | 6400677 | 28.26186 | 81.23897 | 78.06556 | chr19 | 6398167 | 6401812 | 5.57956 | 4.20134 | 3.51519 |
| chr17 | 88613131 | 88613385 | 443 | Intron (ENSMUST00000137138.1 | 17 | 88626555 | 88645724 | 19170 | 1 | ENSMUST00 | -13170 | Ston1 | ENSMUSG0C | false | . | . | . | . | . | . | chr17 | 88609322 | 88613984 | 3.90327 | 3.48747 | 2.13151 |
| chr11 | 99200202 | 99200484 | 443 | Distal Intergenic | 11 | 99209992 | 99213443 | 3452 | 2 | ENSMUST00 | 12959 | Smarce1 | ENSMUSG0C | false | . | . | . | . | . | . | . | . | . | . | . | . |
| chr4 | 83323607 | 83324468 | 443 | Promoter (<=1kb) | 4 | 83220300 | 83324255 | 103956 | 2 | ENSMUST00 | 0 | Ttc39b | ENSMUSG0C | false | chr4 | 83323194 | 83324767 | 31.14379 | 104.89771 | 101.20105 | . | . | . | . | . | . |
| chr15 | 80977770 | 80978141 | 443 | Promoter (<=1kb) | 15 | 80977779 | 81006515 | 28737 | 1 | ENSMUST00 | 0 | Sgsm3 | ENSMUSG0C | false | chr15 | 80977670 | 80978584 | 36.76141 | 116.86796 | 112.83147 | chr15 | 80974479 | 80980685 | 3.69096 | 3.44006 | 1.92296 |
| chr4 | 153119618 | 153120053 | 443 | Distal Intergenic | 4 | 153145590 | 153146943 | 1354 | 2 | ENSMUST00 | 26890 | Gm13173 | ENSMUSG0C | false | chr4 | 153118908 | 153120529 | 18.44084 | 48.77526 | 46.0739 | . | . | . | . | . | . |
| chr13 | 41403841 | 41404082 | 443 | Intron (ENSMUST00000021794.7 | 13 | 41359971 | 41370908 | 10938 | 1 | ENSMUST00 | 43871 | Gm26877 | ENSMUSG0C | false | . | . | . | . | . | . | chr13 | 41401406 | 41404342 | 6.38742 | 4.78332 | 4.12974 |
| chr12 | 112976353 | 112977007 | 442 | Promoter (<=1kb) | 12 | 112976482 | 112978930 | 2449 | 1 | ENSMUST00 | 0 | Btbd6 | ENSMUSG0C | false | chr12 | 112976024 | 112977315 | 34.17318 | 106.15081 | 102.42828 | chr12 | 112975529 | 112979185 | 4.09181 | 3.60096 | 2.29866 |
| chr13 | 58127638 | 58128810 | 442 | Promoter (<=1kb) | 13 | 58125879 | 58128556 | 2678 | 2 | ENSMUST00 | 0 | Hnrnpa0 | ENSMUSG0C | false | chr13 | 58126701 | 58129768 | 35.20524 | 115.20684 | 111.21615 | . | . | . | . | . | . |
| chr2 | 35273702 | 35273932 | 442 | Distal Intergenic | 2 | 35279663 | 35293567 | 13905 | 1 | ENSMUST00 | -5731 | Gsn | ENSMUSG0C | true | . | . | . | . | . | . | chr2 | 35257475 | 35289539 | 6.87726 | 4.86078 | 4.55899 |
| chr2 | 26314309 | 26314632 | 442 | Promoter (<=1kb) | 2 | 26315515 | 26348237 | 32723 | 1 | ENSMUST00 | -883 | Gpsm1 | ENSMUSG0C | false | . | . | . | . | . | . | chr2 | 26308853 | 26315234 | 5.20105 | 4.05386 | 3.18987 |
| chr8 | 36594098 | 36594375 | 442 | Intron (ENSMUST00000033923.8 | 8 | 36572145 | 36577269 | 5125 | 2 | ENSMUST00 | -16830 | Dlc1 | ENSMUSG0C | true | . | . | . | . | . | . | chr8 | 36575944 | 36612020 | 5.64773 | 4.3412 | 3.54039 |
| chr4 | 117649417 | 117649990 | 442 | Intron (ENSMUST00000037127.9 | 4 | 117637338 | 117674281 | 36944 | 1 | ENSMUST00 | 12080 | Eri3 | ENSMUSG0C | false | . | . | . | . | . | . | chr4 | 117643452 | 117654917 | 5.66723 | 4.343 | 3.56197 |
| chr8 | 111380402 | 111380594 | 442 | Intron (ENSMUST00000038475.8 | 8 | 111347470 | 111393818 | 46349 | 2 | ENSMUST00 | 13224 | Fa2h | ENSMUSG0C | true | . | . | . | . | . | . | chr8 | 111356598 | 111381726 | 5.44823 | 4.21871 | 3.39859 |
| chr15 | 91571634 | 91571897 | 442 | Promoter (1-2kb) | 15 | 91542225 | 91573194 | 30970 | 2 | ENSMUST00 | 1297 | Slc2a13 | ENSMUSG0C | false | . | . | . | . | . | . | chr15 | 91571678 | 91573875 | 3.66372 | 3.31507 | 1.9461 |
| chr7 | 143822876 | 143823129 | 442 | Promoter (<=1kb) | 7 | 143823145 | 143824215 | 1071 | 1 | ENSMUST00 | -16 | Dhcr7 | ENSMUSG0C | false | chr7 | 143822072 | 143822884 | 21.91892 | 63.05404 | 60.16603 | chr7 | 143818800 | 143823632 | 5.31041 | 4.21003 | 3.25372 |
| chr7 | 45705996 | 45706305 | 442 | Promoter (<=1kb) | 7 | 45705624 | 45710114 | 4491 | 1 | ENSMUST00 | 373 | Dbp | ENSMUSG0C | false | chr7 | 45703819 | 45709044 | 27.54716 | 98.2761 | 94.74957 | chr7 | 45705481 | 45717582 | 6.81134 | 4.85162 | 4.49354 |
| chr16 | 44471208 | 44471461 | 442 | Promoter (<=1kb) | 16 | 44470444 | 44482428 | 11985 | 1 | ENSMUST00 | 765 | Wdr52 | ENSMUSG0C | false | . | . | . | . | . | . | chr16 | 44470705 | 44472128 | 4.92919 | 4.09676 | 2.95594 |
| chr19 | 55628090 | 55628381 | 442 | Distal Intergenic | 19 | 55640589 | 55702204 | 61616 | 2 | ENSMUST00 | 73823 | 4930552P12I | ENSMUSG0C | false | . | . | . | . | . | . | . | . | . | . | . | . |
| chr7 | 63903293 | 63903541 | 442 | Intron (ENSMUST00000063694.8 | 7 | 63897974 | 63899529 | 1556 | 1 | ENSMUST00 | 5320 | Gm27252 | ENSMUSG0C | false | . | . | . | . | . | . | chr7 | 63885397 | 63936279 | 7.51388 | 5.18869 | 5.06896 |
| chr3 | 153301786 | 153302084 | 442 | Intron (ENSMUST00000064460.5 | 3 | 153313713 | 153313824 | 112 | 1 | ENSMUST00 | -11629 | Gm27405 | ENSMUSG0C | false | . | . | . | . | . | . | chr3 | 153299389 | 153305626 | 5.79405 | 4.42861 | 3.6565 |
| chr15 | 82899353 | 82900043 | 442 | Promoter (<=1kb) | 15 | 82899080 | 82905818 | 6739 | 1 | ENSMUST00 | 274 | RP23-223M2 | ENSMUSG0C | false | chr15 | 82898547 | 82900341 | 34.54477 | 122.48211 | 118.25522 | chr15 | 82897974 | 82901584 | 4.00723 | 3.58969 | 2.24102 |
| chr2 | 15054960 | 15055521 | 441 | Promoter (<=1kb) | 2 | 15055282 | 15082456 | 27175 | 1 | ENSMUST00 | 0 | Arl5b | ENSMUSG0C | false | chr2 | 15054450 | 15056235 | 25.84672 | 90.2271 | 86.88143 | . | . | . | . | . | . |
| chr9 | 77917076 | 77917675 | 441 | Promoter (<=1kb) | 9 | 77917364 | 77984519 | 67156 | 1 | ENSMUST00 | 0 | Elovl5 | ENSMUSG0C | false | chr9 | 77916929 | 77918711 | 35.90788 | 118.54927 | 114.46114 | . | . | . | . | . | . |
| chr12 | 86946786 | 86947508 | 441 | Promoter (<=1kb) | 12 | 86947043 | 86962158 | 15116 | 1 | ENSMUST00 | 0 | Cipc | ENSMUSG0C | false | chr12 | 86946143 | 86947252 | 27.08794 | 82.02477 | 78.8401 | . | . | . | . | . | . |
| chr5 | 96313131 | 96313472 | 441 | Distal Intergenic | 5 | 96373955 | 96784728 | 410774 | 1 | ENSMUST00 | -60483 | Fras1 | ENSMUSG0C | false | . | . | . | . | . | . | chr5 | 96312654 | 96315666 | 3.69843 | 3.48828 | 1.9224 |
| chr6 | 67037157 | 67037608 | 441 | Promoter (<=1kb) | 6 | 67035096 | 67037406 | 2311 | 2 | ENSMUST00 | 0 | Gadd45a | ENSMUSG0C | false | chr6 | 67034307 | 67037493 | 35.56825 | 117.95176 | 113.87694 | . | . | . | . | . | . |
| chr12 | 13212151 | 13212392 | 441 | Distal Intergenic | 12 | 13219308 | 13249213 | 29906 | 2 | ENSMUST00 | 36821 | Ddx1 | ENSMUSG0C | false | . | . | . | . | . | . | chr12 | 13211848 | 13212401 | 3.96166 | 3.35478 | 2.18096 |
| chr9 | 90247185 | 90247420 | 441 | Intron (ENSMUST00000041767.8 | 9 | 90241163 | 90255927 | 14765 | 2 | ENSMUST00 | 8507 | Tbc1d2b | ENSMUSG0C | false | . | . | . | . | . | . | . | . | . | . | . | . |
| chr6 | 72234318 | 72234995 | 441 | Promoter (<=1kb) | 6 | 72206177 | 72235577 | 29401 | 2 | ENSMUST00 | 582 | Atoh8 | ENSMUSG0C | false | chr6 | 72233454 | 72235881 | 33.76198 | 105.43819 | 101.727 | . | . | . | . | . | . |
| chr5 | 3516721 | 3516950 | 441 | Intron (ENSMUST00000165117.2 | 5 | 3543844 | 3569810 | 25967 | 1 | ENSMUST00 | -26894 | Fam133b | ENSMUSG0C | false | . | . | . | . | . | . | chr5 | 3514974 | 3517313 | 3.60658 | 3.25429 | 1.89826 |
| chr11 | 117269953 | 117270272 | 441 | Intron (ENSMUST00000093907.5 | 11 | 117266246 | 117362313 | 96068 | 1 | ENSMUST00 | 3708 | Sep-09 | ENSMUSG0C | false | . | . | . | . | . | . | chr11 | 117269100 | 117270548 | 5.25207 | 4.1846 | 3.2161 |
| chr19 | 53364985 | 53365258 | 441 | Intron (ENSMUST00000003870.8 | 19 | 53371566 | 53371766 | 201 | 2 | ENSMUST00 | 6508 | Gm10197 | ENSMUSG0C | false | . | . | . | . | . | . | chr19 | 53364617 | 53367597 | 5.96967 | 4.4782 | 3.84524 |
| chr7 | 64027064 | 64027326 | 441 | Distal Intergenic | 7 | 64057602 | 64062958 | 5357 | 1 | ENSMUST00 | -30276 | Gm20670 | ENSMUSG0C | false | . | . | . | . | . | . | chr7 | 64026253 | 64027735 | 5.87634 | 4.3675 | 3.7581 |
| chr11 | 64442150 | 64442414 | 441 | Intron (ENSMUST00000058652.5 | 11 | 64435156 | 64436653 | 1498 | 2 | ENSMUST00 | -5498 | F930015N05 | ENSMUSG0C | false | . | . | . | . | . | . | chr11 | 64439570 | 64452101 | 5.16119 | 4.1738 | 3.13545 |
| chr14 | 53026768 | 53026990 | 441 | Distal Intergenic | 14 | 53032902 | 53033418 | 517 | 1 | ENSMUST00 | -5912 | Trav13d-3 | ENSMUSG0C | false | . | . | . | . | . | . | . | . | . | . | . | . |
| chr2 | 27438332 | 27438610 | 440 | Distal Intergenic | 2 | 27262104 | 27427033 | 164930 | 2 | ENSMUST00 | -11300 | Vav2 | ENSMUSG0C | false | . | . | . | . | . | . | . | . | . | . | . | . |
| chr10 | 8906789 | 8907028 | 440 | Distal Intergenic | 10 | 8722219 | 8886070 | 163852 | 2 | ENSMUST00 | -20720 | Sash1 | ENSMUSG0C | false | . | . | . | . | . | . | chr10 | 8906384 | 8907363 | 3.03626 | 3.04187 | 1.42837 |
| chr12 | 108220410 | 108220825 | 440 | Intron (ENSMUST00000136175.1 | 12 | 108179738 | 108203359 | 23622 | 1 | ENSMUST00 | 40673 | Ccnk | ENSMUSG0C | false | . | . | . | . | . | . | chr12 | 108216919 | 108227928 | 7.31121 | 5.08617 | 4.91072 |
| chr1 | 189787019 | 189787483 | 440 | Promoter (<=1kb) | 1 | 189786636 | 189851976 | 65341 | 1 | ENSMUST00 | 384 | Ptpn14 | ENSMUSG0C | true | . | . | . | . | . | . | chr1 | 189730267 | 189810107 | 6.93269 | 4.87829 | 4.60521 |
| chr2 | 35200282 | 35201280 | 440 | Promoter (<=1kb) | 2 | 35180205 | 35201120 | 20916 | 2 | ENSMUST00 | 0 | Rab14 | ENSMUSG0C | false | chr2 | 35199491 | 35201101 | 33.7938 | 119.00099 | 114.88904 | . | . | . | . | . | . |
| chr8 | 110919634 | 110920165 | 440 | Promoter (<=1kb) | 8 | 110919922 | 110969823 | 49902 | 1 | ENSMUST00 | 0 | St3gal2 | ENSMUSG0C | false | chr8 | 110919321 | 110921240 | 26.28407 | 85.32412 | 82.07336 | chr8 | 110917856 | 110921888 | 4.4265 | 3.65571 | 2.57146 |
| chr18 | 56896847 | 56897090 | 440 | Intron (ENSMUST00000102912.2 | 18 | 56803669 | 56925515 | 121847 | 2 | ENSMUST00 | 28425 | 37681 | ENSMUSG0C | false | . | . | . | . | . | . | chr18 | 56892786 | 56897353 | 4.70535 | 3.97538 | 2.78548 |
| chr7 | 99441605 | 99441852 | 440 | Intron (ENSMUST00000037528.8 | 7 | 99466004 | 99474022 | 8019 | 1 | ENSMUST00 | -24152 | Klhl35 | ENSMUSG0C | false | . | . | . | . | . | . | chr7 | 99438175 | 99443768 | 8.27229 | 5.37258 | 5.68456 |
| chr8 | 13461202 | 13461447 | 440 | 3' UTR | 8 | 13435459 | 13461452 | 25994 | 1 | ENSMUST00 | 25744 | Tmem255b | ENSMUSG0C | false | . | . | . | . | . | . | chr8 | 13459474 | 13466027 | 4.90569 | 3.87877 | 2.97234 |
| chr1 | 165193910 | 165194613 | 440 | Promoter (<=1kb) | 1 | 165174341 | 165194433 | 20093 | 2 | ENSMUST00 | 0 | Sft2d2 | ENSMUSG0C | false | chr1 | 165193163 | 165195083 | 32.69058 | 104.59052 | 100.90593 | chr1 | 165187356 | 165195202 | 5.29367 | 4.20591 | 3.27431 |
| chr4 | 138938994 | 138939268 | 440 | Distal Intergenic | 4 | 138895379 | 138913945 | 18567 | 2 | ENSMUST00 | -25050 | Otud3 | ENSMUSG0C | false | . | . | . | . | . | . | chr4 | 138938921 | 138939982 | 4.4302 | 3.81407 | 2.58246 |
| chr7 | 25515672 | 25516405 | 440 | 3' UTR | 7 | 25518064 | 25537093 | 19030 | 2 | ENSMUST00 | 20688 | Ceacam2 | ENSMUSG0C | false | . | . | . | . | . | . | chr7 | 25514424 | 25517836 | 6.49736 | 4.26608 | 4.27356 |
| chr4 | 54601746 | 54601979 | 440 | Distal Intergenic | 4 | 54564623 | 54565143 | 521 | 1 | ENSMUST00 | 37124 | Tpt1-ps2 | ENSMUSG0C | false | . | . | . | . | . | . | chr4 | 54600769 | 54602857 | 4.85964 | 3.84127 | 2.90584 |

| | | | | | | | | | | | | | | | | | | | | | | | | | | |
|---|---|---|---|---|---|---|---|---|---|---|---|---|---|---|---|---|---|---|---|---|---|---|---|---|---|---|
| chr10 | 117471691 | 117472028 | 440 | Distal Intergenic | 10 | 117429618 | 117430116 | 499 | 1 | ENSMUST00 | 42074 | Gm9002 | ENSMUSG0C | false | . | . | . | . | . | . | . | . | . | . | . | . |
| chr1 | 90445043 | 90445320 | 440 | Distal Intergenic | 1 | 90453394 | 90454501 | 1108 | 2 | ENSMUST00 | 9181 | RP24-154L11 | ENSMUSG0C | false | . | . | . | . | . | . | chr1 | 90438990 | 90445431 | 5.52456 | 4.20032 | 3.45273 |
| chr18 | 47314606 | 47315003 | 439 | Intron (ENSMUST00000019791.8 | 18 | 47291894 | 47333326 | 41433 | 2 | ENSMUST00 | 18323 | Sema6a | ENSMUSG0C | false | . | . | . | . | . | . | chr18 | 47314264 | 47315553 | 3.88198 | 3.54405 | 2.09858 |
| chr11 | 107668648 | 107668912 | 439 | Promoter (2-3kb) | 11 | 107671052 | 107672994 | 1943 | 1 | ENSMUST00 | -2140 | Helz | ENSMUSG0C | false | . | . | . | . | . | . | chr11 | 107668361 | 107669571 | 4.25923 | 3.42237 | 2.42173 |
| chr12 | 86687759 | 86688254 | 439 | Exon (ENSMUST00000021681.3/ | 12 | 86678700 | 86692091 | 13392 | 1 | ENSMUST00 | 9060 | Vash1 | ENSMUSG0C | false | . | . | . | . | . | . | chr12 | 86685220 | 86694692 | 6.20303 | 4.49357 | 4.02263 |
| chr6 | 92846765 | 92847015 | 439 | Exon (ENSMUST00000133858.1/ | 6 | 92793010 | 92860145 | 67136 | 2 | ENSMUST00 | 13130 | Adamts9 | ENSMUSG0C | false | . | . | . | . | . | . | chr6 | 92839626 | 92847273 | 5.27431 | 4.12091 | 3.24819 |
| chr11 | 114675797 | 114676055 | 439 | Promoter (<=1kb) | 11 | 114675516 | 114690603 | 15088 | 1 | ENSMUST00 | 282 | Ttyh2 | ENSMUSG0C | false | chr11 | 114675270 | 114676405 | 26.40393 | 78.95839 | 75.82329 | chr11 | 114675304 | 114676573 | 3.41202 | 3.27638 | 1.7565 |
| chr8 | 122336402 | 122336708 | 439 | Promoter (2-3kb) | 8 | 122333786 | 122335761 | 1976 | 1 | ENSMUST00 | 2617 | Zfpm1 | ENSMUSG0C | false | chr8 | 122335652 | 122336940 | 20.05275 | 52.33802 | 49.5918 | chr8 | 122335594 | 122337025 | 4.68859 | 3.88161 | 2.75411 |
| chr7 | 112656048 | 112656530 | 439 | Intron (ENSMUST00000146145.2 | 7 | 112679318 | 112899459 | 220142 | 1 | ENSMUST00 | -22788 | Tead1 | ENSMUSG0C | false | . | . | . | . | . | . | chr7 | 112652748 | 112658001 | 4.10876 | 3.60894 | 2.28054 |
| chr1 | 74096819 | 74097031 | 439 | Promoter (<=1kb) | 1 | 73995288 | 74096825 | 101538 | 2 | ENSMUST00 | 0 | Tns1 | ENSMUSG0C | false | . | . | . | . | . | . | . | . | . | . | . | . |
| chr9 | 63074880 | 63075249 | 439 | Distal Intergenic | 9 | 63027560 | 63027717 | 158 | 1 | ENSMUST00 | 47321 | Gm24526 | ENSMUSG0C | false | . | . | . | . | . | . | chr9 | 63069302 | 63075901 | 5.41329 | 4.13744 | 3.37445 |
| chr17 | 82556448 | 82556680 | 439 | Distal Intergenic | 17 | 82539258 | 82539545 | 288 | 1 | ENSMUST00 | 17191 | Gm6594 | ENSMUSG0C | false | . | . | . | . | . | . | . | . | . | . | . | . |
| chr11 | 63545859 | 63546063 | 439 | Distal Intergenic | 11 | 63599632 | 63600923 | 1292 | 2 | ENSMUST00 | 54860 | Gm12287 | ENSMUSG0C | false | . | . | . | . | . | . | chr11 | 63540785 | 63546744 | 4.81426 | 3.84485 | 2.86796 |
| chr17 | 28034598 | 28034852 | 439 | Intron (ENSMUST00000114842.2 | 17 | 27988028 | 28008432 | 20405 | 2 | ENSMUST00 | -26167 | Gm15598 | ENSMUSG0C | false | . | . | . | . | . | . | chr17 | 28029226 | 28043441 | 7.08706 | 4.83268 | 4.74553 |
| chr13 | 44073118 | 44073494 | 439 | Distal Intergenic | 13 | 44121167 | 44125179 | 4013 | 1 | ENSMUST00 | -47673 | Gm5083 | ENSMUSG0C | false | . | . | . | . | . | . | chr13 | 44072111 | 44074608 | 5.25216 | 4.16077 | 3.22818 |
| chr8 | 107792327 | 107792577 | 439 | Distal Intergenic | 8 | 107840957 | 107841057 | 101 | 1 | ENSMUST00 | -48380 | Gm23646 | ENSMUSG0C | false | . | . | . | . | . | . | chr8 | 107790763 | 107793019 | 4.2777 | 3.6654 | 2.44171 |
| chr7 | 71384918 | 71385283 | 439 | Intron (ENSMUST00000190004.1 | 7 | 71370333 | 71379813 | 9481 | 1 | ENSMUST00 | 14586 | RP24-173C11 | ENSMUSG0C | false | . | . | . | . | . | . | chr7 | 71378101 | 71386659 | 5.91662 | 4.56543 | 3.76231 |
| chr16 | 7329309 | 7329537 | 438 | Intron (ENSMUST00000115841.4 | 16 | 7069835 | 7412479 | 342645 | 1 | ENSMUST00 | 259475 | Rbfox1 | ENSMUSG0C | false | . | . | . | . | . | . | chr16 | 7329154 | 7329743 | 3.40248 | 3.30112 | 1.68474 |
| chr16 | 84834864 | 84835108 | 438 | Promoter (<=1kb) | 16 | 84834925 | 84861315 | 26391 | 1 | ENSMUST00 | 0 | Gabpa | ENSMUSG0C | false | chr16 | 84834034 | 84836397 | 49.45498 | 178.53914 | 172.40457 | . | . | . | . | . | . |
| chr11 | 70539960 | 70540409 | 438 | Promoter (<=1kb) | 11 | 70540064 | 70541183 | 1120 | 1 | ENSMUST00 | 0 | Pld2 | ENSMUSG0C | false | chr11 | 70539548 | 70540732 | 18.88512 | 67.59834 | 64.64505 | chr11 | 70539190 | 70540880 | 5.3585 | 4.31403 | 3.2969 |
| chr11 | 95266894 | 95267176 | 438 | Intron (ENSMUST00000021242.4 | 11 | 95261529 | 95269265 | 7737 | 1 | ENSMUST00 | 5366 | Tac4 | ENSMUSG0C | false | . | . | . | . | . | . | chr11 | 95257171 | 95275447 | 6.31854 | 4.58687 | 4.10024 |
| chr5 | 107028988 | 107029237 | 438 | Distal Intergenic | 5 | 106975238 | 106979323 | 4086 | 1 | ENSMUST00 | 53751 | Cdc7 | ENSMUSG0C | false | . | . | . | . | . | . | chr5 | 107027024 | 107030449 | 4.47315 | 3.84585 | 2.57561 |
| chr9 | 44088188 | 44088445 | 438 | Promoter (<=1kb) | 9 | 44087190 | 44092500 | 5311 | 1 | ENSMUST00 | 999 | Usp2 | ENSMUSG0C | false | . | . | . | . | . | . | chr9 | 44083390 | 44090166 | 6.78512 | 4.96584 | 4.45902 |
| chr9 | 21546077 | 21546849 | 438 | Promoter (<=1kb) | 9 | 21546894 | 21589467 | 42574 | 1 | ENSMUST00 | -45 | Carm1 | ENSMUSG0C | false | chr9 | 21545661 | 21546699 | 19.04495 | 59.55748 | 56.71581 | . | . | . | . | . | . |
| chr12 | 86950469 | 86950864 | 438 | Promoter (1-2kb) | 12 | 86952657 | 86962833 | 10177 | 1 | ENSMUST00 | -1793 | Cipc | ENSMUSG0C | false | . | . | . | . | . | . | chr12 | 86948031 | 86951829 | 5.13237 | 4.14882 | 3.11998 |
| chr17 | 26414651 | 26414999 | 438 | Promoter (<=1kb) | 17 | 26414829 | 26441795 | 26967 | 1 | ENSMUST00 | 0 | Neurl1b | ENSMUSG0C | false | chr17 | 26414273 | 26415574 | 30.78865 | 90.34473 | 86.99165 | chr17 | 26414161 | 26415470 | 5.10156 | 4.11627 | 3.09331 |
| chr12 | 108910055 | 108910401 | 438 | Intron (ENSMUST00000167816.2 | 12 | 108894317 | 109028325 | 134009 | 1 | ENSMUST00 | 15739 | Wdr25 | ENSMUSG0C | false | . | . | . | . | . | . | chr12 | 108909156 | 108911461 | 4.76922 | 3.86998 | 2.84121 |
| chr10 | 80345129 | 80345398 | 438 | Promoter (<=1kb) | 10 | 80343006 | 80345238 | 2233 | 2 | ENSMUST00 | 0 | Adamtsl5 | ENSMUSG0C | false | chr10 | 80344776 | 80345635 | 11.02877 | 22.59171 | 20.22099 | chr10 | 80340223 | 80348968 | 6.10026 | 4.58835 | 3.91011 |
| chr1 | 90650863 | 90651328 | 438 | Distal Intergenic | 1 | 90675161 | 90679793 | 4633 | 1 | ENSMUST00 | -23833 | Gm9991 | ENSMUSG0C | false | . | . | . | . | . | . | . | . | . | . | . | . |
| chr4 | 45690679 | 45690876 | 438 | Intron (ENSMUST00000123741.1 | 4 | 45683589 | 45723338 | 39750 | 1 | ENSMUST00 | 7091 | Gm829 | ENSMUSG0C | false | . | . | . | . | . | . | chr4 | 45690408 | 45691328 | 3.30561 | 3.07599 | 1.64896 |
| chr2 | 155629199 | 155629468 | 438 | Promoter (<=1kb) | 2 | 155629060 | 155630384 | 1325 | 1 | ENSMUST00 | 140 | Myh7b | ENSMUSG0C | false | chr2 | 155629170 | 155629637 | 8.50221 | 18.80243 | 16.48241 | chr2 | 155629019 | 155630921 | 4.09839 | 3.57996 | 2.2987 |
| chr4 | 124393638 | 124393871 | 438 | Distal Intergenic | 4 | 124414416 | 124418858 | 4443 | 2 | ENSMUST00 | 24987 | 1700057H15I | ENSMUSG0C | false | . | . | . | . | . | . | chr4 | 124392663 | 124394579 | 4.91672 | 3.98999 | 2.97728 |
| chr6 | 86525711 | 86526316 | 438 | Promoter (<=1kb) | 6 | 86526271 | 86564442 | 38172 | 1 | ENSMUST00 | 0 | 1600020E01I | ENSMUSG0C | false | chr6 | 86523216 | 86527818 | 38.37027 | 120.70945 | 116.54665 | . | . | . | . | . | . |
| chr7 | 99523878 | 99524151 | 437 | Distal Intergenic | 7 | 99535466 | 99606771 | 71306 | 1 | ENSMUST00 | -11315 | Arrb1 | ENSMUSG0C | false | . | . | . | . | . | . | chr7 | 99522439 | 99531882 | 5.37849 | 4.25246 | 3.31489 |
| chr16 | 58512135 | 58512633 | 437 | Intron (ENSMUST00000137035.2 | 16 | 58502861 | 58507898 | 5038 | 2 | ENSMUST00 | -4238 | St3gal6 | ENSMUSG0C | true | chr16 | 58512380 | 58514683 | 5.72276 | 11.47427 | 9.27082 | chr16 | 58496830 | 58520680 | 5.70067 | 4.33546 | 3.59278 |
| chr17 | 8506740 | 8507075 | 437 | Distal Intergenic | 17 | 8525372 | 8981671 | 456300 | 1 | ENSMUST00 | -18297 | Pde10a | ENSMUSG0C | true | . | . | . | . | . | . | chr17 | 8504525 | 8510234 | 7.21779 | 5.10275 | 4.81866 |
| chr17 | 65957317 | 65957668 | 437 | Distal Intergenic | 17 | 65923066 | 65951187 | 28122 | 2 | ENSMUST00 | -6131 | Twsg1 | ENSMUSG0C | false | . | . | . | . | . | . | chr17 | 65956690 | 65958833 | 4.32008 | 3.67536 | 2.48518 |
| chr1 | 189899145 | 189899440 | 437 | Intron (ENSMUST00000027897.7 | 1 | 189880494 | 189889002 | 8509 | 2 | ENSMUST00 | -10144 | Smyd2 | ENSMUSG0C | false | . | . | . | . | . | . | chr1 | 189898312 | 189899741 | 3.57821 | 3.3104 | 1.85524 |
| chrX | 48433632 | 48433883 | 437 | Intron (ENSMUST00000114958.2 | 20 | 48411046 | 48454169 | 43124 | 2 | ENSMUST00 | 20286 | Elf4 | ENSMUSG0C | true | . | . | . | . | . | . | . | . | . | . | . | . |
| chr14 | 70057052 | 70057380 | 437 | Intron (ENSMUST00000022678.3 | 14 | 70077445 | 70080157 | 2713 | 1 | ENSMUST00 | -20065 | Egr3 | ENSMUSG0C | false | . | . | . | . | . | . | chr14 | 70054941 | 70057811 | 4.85052 | 4.01995 | 2.88732 |
| chr2 | 118598106 | 118598335 | 437 | Promoter (<=1kb) | 2 | 118598211 | 118641591 | 43381 | 1 | ENSMUST00 | 0 | Bub1b | ENSMUSG0C | false | chr2 | 118598079 | 118598685 | 22.68498 | 68.76898 | 65.7979 | chr2 | 118597716 | 118598905 | 3.60108 | 3.39518 | 1.91036 |
| chr5 | 121700532 | 121700757 | 437 | Distal Intergenic | 5 | 121711337 | 121749183 | 37847 | 1 | ENSMUST00 | -10580 | Atxn2 | ENSMUSG0C | false | chr5 | 121700387 | 121700861 | 9.20781 | 17.98236 | 15.67656 | chr5 | 121700463 | 121701711 | 3.11036 | 3.06971 | 1.48404 |
| chr2 | 17945582 | 17945865 | 437 | Distal Intergenic | 2 | 17996417 | 17996958 | 542 | 2 | ENSMUST00 | 51093 | H2afb1 | ENSMUSG0C | false | . | . | . | . | . | . | chr2 | 17945598 | 17945925 | 3.54342 | 3.25121 | 1.86057 |
| chr6 | 92041447 | 92041710 | 437 | Intron (ENSMUST00000089334.3 | 6 | 92012394 | 92016117 | 3724 | 2 | ENSMUST00 | -25331 | 4930466I24R | ENSMUSG0C | false | . | . | . | . | . | . | . | . | . | . | . | . |
| chr2 | 26132678 | 26133360 | 437 | Distal Intergenic | 2 | 26138546 | 26139543 | 998 | 1 | ENSMUST00 | -5186 | 1810012K08I | ENSMUSG0C | false | chr2 | 26132583 | 26133600 | 9.51244 | 19.51414 | 17.18665 | chr2 | 26129136 | 26135084 | 6.01114 | 4.43692 | 3.85388 |
| chr5 | 103605678 | 103605979 | 437 | 3' UTR | 5 | 103602587 | 103602879 | 293 | 1 | ENSMUST00 | 3092 | Gm23085 | ENSMUSG0C | false | . | . | . | . | . | . | chr5 | 103602894 | 103607755 | 4.47885 | 3.74726 | 2.61032 |
| chr6 | 52160051 | 52160391 | 437 | Promoter (<=1kb) | 6 | 52161042 | 52161255 | 214 | 1 | ENSMUST00 | -651 | Gm27861 | ENSMUSG0C | false | chr6 | 52160172 | 52161353 | 10.10971 | 19.98882 | 17.65231 | chr6 | 52159328 | 52167512 | 5.47105 | 4.07894 | 3.42628 |
| chr1 | 127959320 | 127959601 | 437 | Promoter (<=1kb) | 1 | 127959516 | 127967452 | 7937 | 1 | ENSMUST00 | 0 | RP24-338G1( | ENSMUSG0C | false | . | . | . | . | . | . | . | . | . | . | . | . |
| chr5 | 136887671 | 136887989 | 436 | Distal Intergenic | 5 | 136757489 | 136883209 | 125721 | 2 | ENSMUST00 | -4463 | Col26a1 | ENSMUSG0C | false | . | . | . | . | . | . | chr5 | 136885321 | 136889075 | 4.822 | 3.84455 | 2.90361 |
| chr11 | 69662406 | 69662826 | 436 | Promoter (<=1kb) | 11 | 69656697 | 69662642 | 5946 | 2 | ENSMUST00 | 0 | Mpdu1 | ENSMUSG0C | false | chr11 | 69661810 | 69663290 | 15.98415 | 46.70646 | 44.03467 | chr11 | 69661142 | 69663902 | 5.79619 | 4.47034 | 3.65344 |
| chr14 | 21523991 | 21524463 | 436 | Intron (ENSMUST00000183201.2 | 14 | 21516751 | 21671206 | 154456 | 1 | ENSMUST00 | 7241 | Kat6b | ENSMUSG0C | false | . | . | . | . | . | . | chr14 | 21519296 | 21529701 | 4.63736 | 3.89681 | 2.70766 |
| chr14 | 20180737 | 20181106 | 436 | Promoter (<=1kb) | 14 | 20140058 | 20181782 | 41725 | 2 | ENSMUST00 | 676 | Kcnk5 | ENSMUSG0C | false | chr14 | 20180394 | 20181858 | 17.08566 | 47.94947 | 45.2625 | chr14 | 20179614 | 20181548 | 5.11281 | 4.09675 | 3.16005 |
| chr7 | 81212960 | 81213628 | 436 | Promoter (<=1kb) | 7 | 81213596 | 81334533 | 120938 | 1 | ENSMUST00 | 0 | Pde8a | ENSMUSG0C | false | chr7 | 81212149 | 81214474 | 27.03504 | 88.5911 | 85.27629 | chr7 | 81211820 | 81214324 | 4.11098 | 3.6083 | 2.28459 |
| chr1 | 52800490 | 52800726 | 436 | Promoter (<=1kb) | 1 | 52797066 | 52800371 | 3306 | 2 | ENSMUST00 | -120 | Inpp1 | ENSMUSG0C | false | . | . | . | . | . | . | chr1 | 52798141 | 52801176 | 4.70521 | 3.90609 | 2.78549 |
| chr6 | 17694069 | 17694777 | 436 | Promoter (<=1kb) | 6 | 17694167 | 17942966 | 248800 | 1 | ENSMUST00 | 0 | St7 | ENSMUSG0C | false | chr6 | 17693660 | 17695108 | 30.87849 | 93.42764 | 90.01563 | . | . | . | . | . | . |
| chr7 | 102317393 | 102317650 | 436 | Intron (ENSMUST00000033289.4 | 7 | 102267824 | 102436848 | 169025 | 1 | ENSMUST00 | 49570 | Stim1 | ENSMUSG0C | false | . | . | . | . | . | . | chr7 | 102316966 | 102318224 | 3.66598 | 3.32847 | 1.95699 |
| chr5 | 76533208 | 76533492 | 436 | Intron (ENSMUST00000087133.5 | 5 | 76529358 | 76570287 | 40930 | 1 | ENSMUST00 | 3851 | Exoc1 | ENSMUSG0C | false | . | . | . | . | . | . | chr5 | 76529877 | 76535256 | 5.4199 | 4.28416 | 3.35627 |
| chr19 | 7416770 | 7417375 | 436 | Promoter (<=1kb) | 19 | 7417625 | 7425904 | 8280 | 1 | ENSMUST00 | -250 | 2700081O15 | ENSMUSG0C | false | chr19 | 7416554 | 7418702 | 27.66286 | 83.97498 | 80.7477 | chr19 | 7416615 | 7417371 | 3.31126 | 3.21391 | 1.60209 |
| chr11 | 88618804 | 88619076 | 436 | Intron (ENSMUST00000107909.2 | 11 | 88365515 | 88590147 | 224633 | 2 | ENSMUST00 | -28658 | Msi2 | ENSMUSG0C | false | . | . | . | . | . | . | chr11 | 88603656 | 88625501 | 5.32676 | 4.10238 | 3.30904 |
| chr2 | 129269997 | 129270223 | 436 | Intron (ENSMUST00000052708.6 | 2 | 129246376 | 129268526 | 22151 | 1 | ENSMUST00 | 23622 | Gm14024 | ENSMUSG0C | false | . | . | . | . | . | . | chr2 | 129269319 | 129270795 | 4.85222 | 3.90638 | 2.93744 |
| chr13 | 93629295 | 93629550 | 436 | Intron (ENSMUST00000099309.4 | 13 | 93625382 | 93628633 | 3252 | 1 | ENSMUST00 | 3914 | Gm15622 | ENSMUSG0C | false | . | . | . | . | . | . | chr13 | 93627939 | 93630769 | 4.96267 | 4.11336 | 3.01005 |
| chr2 | 143712044 | 143712288 | 436 | Intron (ENSMUST00000028905.9 | 2 | 143747583 | 143789186 | 41604 | 2 | ENSMUST00 | 76898 | Pcsk2os2 | ENSMUSG0C | false | . | . | . | . | . | . | chr2 | 143710173 | 143713740 | 4.78244 | 3.98971 | 2.88119 |
| chr14 | 67108953 | 67109253 | 436 | Distal Intergenic | 14 | 67074830 | 67074947 | 118 | 1 | ENSMUST00 | 34124 | Gm27647 | ENSMUSG0C | false | . | . | . | . | . | . | . | . | . | . | . | . |
| chr10 | 34479982 | 34480466 | 435 | Promoter (2-3kb) | 10 | 34483400 | 34611226 | 127827 | 1 | ENSMUST00 | -2934 | Frk | ENSMUSG0C | false | . | . | . | . | . | . | chr10 | 34479877 | 34481455 | 4.7917 | 4.05707 | 2.88397 |
| chr11 | 52098565 | 52099010 | 435 | Promoter (<=1kb) | 11 | 52098681 | 52127778 | 29098 | 1 | ENSMUST00 | 0 | Ppp2ca | ENSMUSG0C | false | chr11 | 52098170 | 52098567 | 10.77139 | 25.03037 | 22.62905 | . | . | . | . | . | . |
| chr17 | 68133624 | 68133889 | 435 | Distal Intergenic | 17 | 67842713 | 68004120 | 161408 | 2 | ENSMUST00 | -129505 | Arhgap28 | ENSMUSG0C | true | . | . | . | . | . | . | chr17 | 68133242 | 68134215 | 3.1738 | 3.02881 | 1.53706 |
| chr4 | 103119116 | 103119825 | 435 | Promoter (<=1kb) | 4 | 103119286 | 103165752 | 46467 | 1 | ENSMUST00 | 0 | Mier1 | ENSMUSG0C | false | chr4 | 103118846 | 103120943 | 35.3795 | 118.67607 | 114.58318 | . | . | . | . | . | . |
| chr6 | 113342027 | 113342546 | 435 | Promoter (1-2kb) | 6 | 113334128 | 113340917 | 6790 | 2 | ENSMUST00 | -1111 | Camk1 | ENSMUSG0C | false | chr6 | 113342483 | 113344245 | 38.58512 | 125.36365 | 121.04008 | chr6 | 113339941 | 113344857 | 6.73316 | 4.71652 | 4.46761 |
| chr14 | 69732403 | 69732877 | 435 | Promoter (<=1kb) | 14 | 69716979 | 69732570 | 15592 | 2 | ENSMUST00 | 0 | Chmp7 | ENSMUSG0C | false | chr14 | 69731620 | 69733869 | 41.18432 | 138.64781 | 133.82259 | . | . | . | . | . | . |
| chr5 | 30907703 | 30908101 | 435 | Promoter (<=1kb) | 5 | 30905885 | 30907788 | 1904 | 2 | ENSMUST00 | 0 | Ost4 | ENSMUSG0C | false | chr5 | 30906784 | 30908449 | 32.69058 | 104.59052 | 100.90593 | chr5 | 30904966 | 30918702 | 6.19394 | 4.49227 | 4.02489 |
| chr15 | 8168905 | 8169337 | 435 | Promoter (<=1kb) | 15 | 8169106 | 8271158 | 102053 | 1 | ENSMUST00 | 0 | 2410089E03I | ENSMUSG0C | false | chr15 | 8168292 | 8169857 | 41.04627 | 146.22916 | 141.09622 | . | . | . | . | . | . |
| chr1 | 166244286 | 166244556 | 435 | Distal Intergenic | 1 | 166201385 | 166238744 | 37360 | 2 | ENSMUST00 | -5543 | Mael | ENSMUSG0C | false | . | . | . | . | . | . | chr1 | 166244229 | 166244835 | 3.84066 | 3.53313 | 2.06212 |
| chr4 | 45462353 | 45462640 | 435 | Intron (ENSMUST00000147926.2 | 4 | 45423278 | 45482802 | 59525 | 2 | ENSMUST00 | 20162 | Shb | ENSMUSG0C | false | . | . | . | . | . | . | chr4 | 45457040 | 45463702 | 5.65864 | 4.22032 | 3.58846 |
| chr14 | 58937447 | 58937774 | 435 | Distal Intergenic | 14 | 58893506 | 58894129 | 624 | 1 | ENSMUST00 | 43942 | Rpl13-ps3 | ENSMUSG0C | false | . | . | . | . | . | . | chr14 | 58936509 | 58939358 | 4.79583 | 4.01575 | 2.85718 |
| chr2 | 92314005 | 92314334 | 435 | Promoter (1-2kb) | 2 | 92312154 | 92312843 | 690 | 1 | ENSMUST00 | 1852 | Gm13785 | ENSMUSG0C | false | . | . | . | . | . | . | chr2 | 92312234 | 92317259 | 4.22313 | 3.57023 | 2.37528 |
| chr3 | 53258533 | 53258885 | 435 | Intron (ENSMUST00000059562.8 | 3 | 53452619 | 53457797 | 5179 | 1 | ENSMUST00 | -193734 | Gm16206 | ENSMUSG0C | false | . | . | . | . | . | . | . | . | . | . | . | . |
| chr11 | 64168630 | 64168884 | 435 | Intron (ENSMUST00000145095.1 | 11 | 64164573 | 64182171 | 17599 | 1 | ENSMUST00 | 4058 | Gm12290 | ENSMUSG0C | false | . | . | . | . | . | . | chr11 | 64164991 | 64169544 | 4.38065 | 3.71133 | 2.547 |
| chr5 | 93005515 | 93005796 | 435 | Distal Intergenic | 5 | 93013032 | 93013139 | 108 | 2 | ENSMUST00 | 7343 | Gm25521 | ENSMUSG0C | false | chr5 | 93005298 | 93006364 | 12.98899 | 30.74239 | 28.2684 | chr5 | 93004108 | 93006581 | 4.09904 | 3.61046 | 2.26218 |
| chr13 | 92326050 | 92326295 | 434 | Intron (ENSMUST00000185852.1 | 13 | 92307497 | 92309457 | 1961 | 2 | ENSMUST00 | -16594 | Msh3 | ENSMUSG0C | false | . | . | . | . | . | . | chr13 | 92324586 | 92328146 | 4.55872 | 3.86213 | 2.63555 |
| chr11 | 80453240 | 80453516 | 434 | Promoter (2-3kb) | 11 | 80456368 | 80471875 | 15508 | 1 | ENSMUST00 | -2852 | Psmd11 | ENSMUSG0C | false | . | . | . | . | . | . | chr11 | 80453090 | 80455171 | 4.2874 | 3.68665 | 2.42417 |
| chr4 | 136181362 | 136181594 | 434 | Promoter (<=1kb) | 4 | 136180783 | 136186807 | 6025 | 1 | ENSMUST00 | 580 | E2f2 | ENSMUSG0C | false | . | . | . | . | . | . | chr4 | 136179975 | 136181453 | 4.13383 | 3.70889 | 2.31338 |
| chr15 | 98881162 | 98881613 | 434 | Promoter (<=1kb) | 15 | 98877778 | 98881404 | 3627 | 2 | ENSMUST00 | 0 | Rhebl1 | ENSMUSG0C | false | chr15 | 98880737 | 98881405 | 24.72749 | 72.3696 | 69.3433 | chr15 | 98880565 | 98881962 | 4.86755 | 3.93036 | 2.92645 |
| chr4 | 102574343 | 102574723 | 434 | Intron (ENSMUST00000106911.2 | 4 | 102570095 | 102605847 | 35753 | 1 | ENSMUST00 | 4249 | Pde4b | ENSMUSG0C | true | chr4 | 102568208 | 102575841 | 28.36989 | 85.2065 | 81.96005 | chr4 | 102573546 | 102597172 | 5.22074 | 4.1871 | 3.19022 |
| chr8 | 83066252 | 83066577 | 434 | Intron (ENSMUST00000078525.5 | 8 | 83165352 | 83272422 | 107071 | 1 | ENSMUST00 | -98775 | Tbc1d9 | ENSMUSG0C | false | . | . | . | . | . | . | chr8 | 83063723 | 83068286 | 4.75927 | 3.98428 | 2.81632 |
| chr19 | 25330613 | 25330823 | 434 | Intron (ENSMUST00000049400.9 | 19 | 25395303 | 25409772 | 14470 | 1 | ENSMUST00 | -64480 | Kank1 | ENSMUSG0C | true | . | . | . | . | . | . | chr19 | 25329142 | 25370790 | 4.73993 | 3.84606 | 2.82412 |
| chr3 | 35753962 | 35754684 | 434 | Promoter (<=1kb) | 3 | 35754134 | 35829688 | 75555 | 1 | ENSMUST00 | 0 | Atp11b | ENSMUSG0C | false | chr3 | 35753595 | 35755263 | 44.74406 | 149.33661 | 144.07773 | . | . | . | . | . | . |
| chr13 | 44715582 | 44715788 | 434 | Distal Intergenic | 13 | 44729474 | 44731574 | 2101 | 1 | ENSMUST00 | -13686 | Jarid2 | ENSMUSG0C | false | . | . | . | . | . | . | chr13 | 44715394 | 44717133 | 4.81544 | 3.97428 | 2.85532 |
| chr19 | 23125640 | 23125906 | 434 | Distal Intergenic | 19 | 23134900 | 23135717 | 818 | 1 | ENSMUST00 | -8994 | 2410080I02R | ENSMUSG0C | false | . | . | . | . | . | . | chr19 | 23125000 | 23126480 | 6.1056 | 4.65699 | 3.89773 |
| chr8 | 61566947 | 61567205 | 434 | Intron (ENSMUST00000121785.2 | 8 | 61515913 | 61591139 | 75227 | 2 | ENSMUST00 | 23934 | Palld | ENSMUSG0C | true | . | . | . | . | . | . | chr8 | 61550887 | 61568090 | 5.03432 | 4.07287 | 3.05874 |
| chr9 | 75523090 | 75523392 | 434 | Intron (ENSMUST00000072232.7 | 9 | 75497796 | 75559657 | 61862 | 2 | ENSMUST00 | 36265 | Tmod3 | ENSMUSG0C | true | . | . | . | . | . | . | . | . | . | . | . | . |
| chr2 | 25167939 | 25168316 | 434 | Distal Intergenic | 2 | 25180758 | 25183339 | 2582 | 1 | ENSMUST00 | -12442 | Nrarp | ENSMUSG0C | false | chr2 | 25167864 | 25168368 | 12.15419 | 27.41675 | 24.98428 | chr2 | 25167776 | 25169934 | 4.01734 | 3.43377 | 2.24159 |
| chr11 | 4291593 | 4292036 | 434 | Distal Intergenic | 11 | 4285460 | 4286939 | 1480 | 1 | ENSMUST00 | 6134 | Gm11958 | ENSMUSG0C | false | . | . | . | . | . | . | chr11 | 4291014 | 4306323 | 6.81252 | 4.86662 | 4.50325 |
| chr8 | 126474454 | 126474753 | 434 | Promoter (<=1kb) | 8 | 126426652 | 126475065 | 48414 | 2 | ENSMUST00 | 312 | Gm17296 | ENSMUSG0C | false | chr8 | 126473965 | 126475285 | 26.91438 | 86.45937 | 83.18881 | chr8 | 126473188 | 126475558 | 4.24538 | 3.72552 | 2.3865 |
| chr15 | 99814390 | 99814765 | 434 | Intron (ENSMUST00000109024.3 | 15 | 99810289 | 99810582 | 294 | 2 | ENSMUST00 | -3809 | Gm4468 | ENSMUSG0C | false | . | . | . | . | . | . | chr15 | 99811513 | 99819053 | 3.94956 | 3.48882 | 2.16316 |
| chr4 | 129136739 | 129137288 | 433 | Promoter (<=1kb) | 4 | 129136999 | 129144593 | 7595 | 1 | ENSMUST00 | 0 | Fndc5 | ENSMUSG0C | false | chr4 | 129136559 | 129137531 | 21.40274 | 64.72344 | 61.81032 | chr4 | 129133723 | 129137736 | 4.59599 | 3.88534 | 2.68145 |
| chr19 | 10182662 | 10183155 | 433 | Promoter (<=1kb) | 19 | 10182888 | 10196870 | 13983 | 1 | ENSMUST00 | 0 | Fads1 | ENSMUSG0C | false | chr19 | 10182386 | 10184244 | 28.56742 | 103.16996 | 99.51957 | . | . | . | . | . | . |
| chr17 | 25273255 | 25274230 | 433 | Promoter (<=1kb) | 17 | 25261921 | 25273383 | 11463 | 2 | ENSMUST00 | 0 | Ube2i | ENSMUSG0C | false | chr17 | 25271619 | 25274673 | 39.52876 | 141.27539 | 136.34271 | . | . | . | . | . | . |
| chr11 | 101118086 | 101118395 | 433 | Promoter (1-2kb) | 11 | 101098000 | 101119765 | 21766 | 2 | ENSMUST00 | 1370 | Fam134c | ENSMUSG0C | false | . | . | . | . | . | . | chr11 | 101109531 | 101119412 | 4.03738 | 3.56069 | 2.2555 |
| chr15 | 94201172 | 94201444 | 433 | Distal Intergenic | 15 | 94270163 | 94404258 | 134096 | 2 | ENSMUST00 | 202814 | Adamts20 | ENSMUSG0C | false | . | . | . | . | . | . | . | . | . | . | . | . |
| chr4 | 119245480 | 119245985 | 433 | Promoter (<=1kb) | 4 | 119245020 | 119248133 | 3114 | 1 | ENSMUST00 | 461 | Lepre1 | ENSMUSG0C | false | . | . | . | . | . | . | chr4 | 119237720 | 119248571 | 6.98267 | 4.77088 | 4.67122 |
| chr7 | 80078860 | 80079297 | 433 | Intron (ENSMUST00000164056.3 | 7 | 80095043 | 80096029 | 987 | 2 | ENSMUST00 | 16732 | Idh2 | ENSMUSG0C | false | . | . | . | . | . | . | chr7 | 80074187 | 80081631 | 6.95437 | 4.98335 | 4.61561 |
| chr9 | 55512009 | 55512395 | 433 | Promoter (<=1kb) | 9 | 55454508 | 55512243 | 57736 | 2 | ENSMUST00 | 0 | Etfa | ENSMUSG0C | false | chr9 | 55511381 | 55512691 | 32.35868 | 122.64178 | 118.41108 | . | . | . | . | . | . |
| chr8 | 94214342 | 94214604 | 433 | Promoter (<=1kb) | 8 | 94214597 | 94314857 | 100261 | 1 | ENSMUST00 | 0 | Nup93 | ENSMUSG0C | false | chr8 | 94214485 | 94215240 | 39.06023 | 122.966 | 118.72274 | chr8 | 94213803 | 94216100 | 4.06186 | 3.4831 | 2.24436 |
| chr2 | 30133131 | 30133426 | 433 | Promoter (<=1kb) | 2 | 30133746 | 30137296 | 3551 | 1 | ENSMUST00 | -320 | Tbc1d13 | ENSMUSG0C | false | chr2 | 30133098 | 30134976 | 35.34781 | 110.24136 | 106.39787 | chr2 | 30132392 | 30135577 | 5.08064 | 4.02768 | 3.09838 |
| chr10 | 42251196 | 42251647 | 433 | Intron (ENSMUST00000105502.2 | 10 | 42197554 | 42258366 | 60813 | 2 | ENSMUST00 | 6719 | Foxo3 | ENSMUSG0C | false | . | . | . | . | . | . | chr10 | 42243354 | 42275609 | 6.05342 | 4.50564 | 3.88797 |
| chr5 | 144223369 | 144223849 | 433 | Promoter (<=1kb) | 5 | 144194442 | 144223615 | 29174 | 2 | ENSMUST00 | 0 | Tecpr1 | ENSMUSG0C | false | chr5 | 144222689 | 144224417 | 23.3092 | 84.4034 | 81.16875 | chr5 | 144220951 | 144225803 | 5.23314 | 4.15965 | 3.19848 |
| chr11 | 63789286 | 63789556 | 433 | Distal Intergenic | 11 | 63817019 | 63818300 | 1282 | 2 | ENSMUST00 | 28744 | Gm12288 | ENSMUSG0C | false | . | . | . | . | . | . | chr11 | 63789114 | 63794351 | 5.46547 | 4.19457 | 3.41907 |
| chr3 | 49278809 | 49279050 | 433 | Distal Intergenic | 3 | 49491351 | 49491478 | 128 | 1 | ENSMUST00 | -212301 | Gm22480 | ENSMUSG0C | false | . | . | . | . | . | . | . | . | . | . | . | . |
| chr17 | 27954923 | 27955323 | 433 | Intron (ENSMUST00000114842.2 | 17 | 27959615 | 27965316 | 5702 | 2 | ENSMUST00 | 9993 | Gm15597 | ENSMUSG0C | false | . | . | . | . | . | . | chr17 | 27952723 | 27958172 | 6.25726 | 4.5441 | 4.03508 |
| chr17 | 34603359 | 34604110 | 432 | Promoter (<=1kb) | 17 | 34601091 | 34603690 | 2600 | 2 | ENSMUST00 | 0 | Rnf5 | ENSMUSG0C | false | chr17 | 34602775 | 34603756 | 20.53639 | 56.4273 | 53.62976 | chr17 | 34594616 | 34603372 | 6.22816 | 4.50396 | 4.02393 |
| chr2 | 91652902 | 91653155 | 432 | Promoter (<=1kb) | 2 | 91653208 | 91672302 | 19095 | 1 | ENSMUST00 | -53 | Arhgap1 | ENSMUSG0C | false | . | . | . | . | . | . | chr2 | 91650770 | 91658145 | 7.66222 | 5.14416 | 5.21273 |
| chr3 | 102204522 | 102204762 | 432 | Promoter (<=1kb) | 3 | 102156701 | 102204693 | 47993 | 2 | ENSMUST00 | 0 | Vangl1 | ENSMUSG0C | false | chr3 | 102203071 | 102204781 | 41.76281 | 135.22986 | 130.53618 | . | . | . | . | . | . |
| chr6 | 87850883 | 87851363 | 432 | Promoter (<=1kb) | 6 | 87842615 | 87851106 | 8492 | 2 | ENSMUST00 | 0 | Cnbp | ENSMUSG0C | false | chr6 | 87849237 | 87851675 | 30.94804 | 114.76607 | 110.79114 | . | . | . | . | . | . |
| chr8 | 36730293 | 36730640 | 432 | Promoter (2-3kb) | 8 | 36567751 | 36733054 | 165304 | 2 | ENSMUST00 | 2414 | Dlc1 | ENSMUSG0C | true | . | . | . | . | . | . | chr8 | 36707656 | 36733370 | 4.68731 | 3.93374 | 2.7424 |
| chr2 | 32137870 | 32138245 | 432 | Distal Intergenic | 2 | 32151082 | 32234537 | 83456 | 1 | ENSMUST00 | -12837 | Prrc2b | ENSMUSG0C | false | chr2 | 32137420 | 32138013 | 18.0267 | 63.35003 | 60.45576 | chr2 | 32137184 | 32138650 | 4.10756 | 3.7152 | 2.25327 |
| chr16 | 94301557 | 94301764 | 432 | Intron (ENSMUST00000163193.2 | 16 | 94129195 | 94313563 | 184369 | 2 | ENSMUST00 | 11799 | Hlcs | ENSMUSG0C | false | . | . | . | . | . | . | chr16 | 94301580 | 94303437 | 4.6378 | 3.90325 | 2.71419 |
| chr18 | 46958766 | 46959203 | 432 | Promoter (<=1kb) | 18 | 46958862 | 47087992 | 129131 | 1 | ENSMUST00 | 0 | Commd10 | ENSMUSG0C | false | chr18 | 46958425 | 46959506 | 29.2476 | 106.45832 | 102.71833 | . | . | . | . | . | . |
| chr18 | 60625373 | 60625659 | 432 | Promoter (<=1kb) | 18 | 60593982 | 60624573 | 30592 | 2 | ENSMUST00 | -801 | Synpo | ENSMUSG0C | true | . | . | . | . | . | . | chr18 | 60624461 | 60629017 | 5.89697 | 4.51569 | 3.75478 |
| chr7 | 127637002 | 127637326 | 432 | Distal Intergenic | 7 | 127608756 | 127615797 | 7042 | 2 | ENSMUST00 | -21206 | Zfp629 | ENSMUSG0C | false | chr7 | 127636649 | 127637757 | 14.64398 | 38.39655 | 35.82735 | chr7 | 127636481 | 127637911 | 5.47361 | 4.28826 | 3.42456 |
| chr7 | 63936298 | 63936744 | 432 | Promoter (<=1kb) | 7 | 63891383 | 63936301 | 44919 | 2 | ENSMUST00 | 0 | Klf13 | ENSMUSG0C | false | chr7 | 63935484 | 63939001 | 41.67902 | 156.4369 | 150.90947 | . | . | . | . | . | . |
| chr9 | 75559409 | 75559814 | 432 | Promoter (<=1kb) | 9 | 75497796 | 75559657 | 61862 | 2 | ENSMUST00 | 0 | Tmod3 | ENSMUSG0C | true | chr9 | 75556512 | 75559968 | 31.38972 | 125.99339 | 121.64841 | chr9 | 75559804 | 75567102 | 4.3979 | 3.82685 | 2.50388 |
| chr2 | 27934216 | 27934509 | 432 | Intron (ENSMUST00000028280.8 | 2 | 27882925 | 27887052 | 4128 | 2 | ENSMUST00 | -47165 | 2810430I11R | ENSMUSG0C | false | . | . | . | . | . | . | chr2 | 27933546 | 27935187 | 4.34562 | 3.55305 | 2.50544 |
| chr8 | 33815422 | 33815732 | 432 | Promoter (2-3kb) | 8 | 33813203 | 33833094 | 19892 | 1 | ENSMUST00 | 2220 | Gm26632 | ENSMUSG0C | false | . | . | . | . | . | . | chr8 | 33797013 | 33821269 | 7.29084 | 5.02722 | 4.89898 |
| chr13 | 52041470 | 52041721 | 432 | Distal Intergenic | 13 | 51973214 | 51976583 | 3370 | 2 | ENSMUST00 | 68257 | Gm26651 | ENSMUSG0C | false | . | . | . | . | . | . | chr13 | 52041382 | 52042553 | 4.78206 | 4.04332 | 2.81703 |
| chr11 | 114795238 | 114795489 | 431 | Promoter (<=1kb) | 11 | 114791217 | 114795873 | 4657 | 2 | ENSMUST00 | 384 | Btbd17 | ENSMUSG0C | false | chr11 | 114795179 | 114795642 | 6.13854 | 9.95394 | 7.77908 | chr11 | 114793751 | 114795931 | 4.71234 | 3.96615 | 2.8049 |
| chr10 | 87492849 | 87493183 | 431 | Promoter (<=1kb) | 10 | 87490819 | 87493660 | 2842 | 2 | ENSMUST00 | 477 | Ascl1 | ENSMUSG0C | false | chr10 | 87492594 | 87493800 | 15.34636 | 36.79438 | 34.24475 | chr10 | 87492340 | 87495350 | 4.15648 | 3.73598 | 2.30472 |
| chr11 | 23770652 | 23771228 | 431 | Promoter (<=1kb) | 11 | 23741514 | 23770970 | 29457 | 2 | ENSMUST00 | 0 | Rel | ENSMUSG0C | false | chr11 | 23769609 | 23771991 | 29.36448 | 91.77025 | 88.3935 | . | . | . | . | . | . |
| chr11 | 46436819 | 46437312 | 431 | Promoter (<=1kb) | 11 | 46436925 | 46440858 | 3934 | 1 | ENSMUST00 | 0 | Med7 | ENSMUSG0C | false | chr11 | 46436468 | 46438045 | 42.64928 | 145.68546 | 140.56778 | . | . | . | . | . | . |
| chr19 | 36834056 | 36835032 | 431 | Promoter (<=1kb) | 19 | 36834232 | 36893477 | 59246 | 1 | ENSMUST00 | 0 | Tnks2 | ENSMUSG0C | false | chr19 | 36833456 | 36836298 | 33.7938 | 119.00099 | 114.88904 | . | . | . | . | . | . |
| chr6 | 108229595 | 108229879 | 431 | Intron (ENSMUST00000032192.6 | 6 | 108213118 | 108551109 | 337992 | 1 | ENSMUST00 | 16478 | Itpr1 | ENSMUSG0C | false | . | . | . | . | . | . | chr6 | 108219855 | 108242489 | 5.57178 | 4.31079 | 3.48186 |
| chr16 | 27388805 | 27389323 | 431 | Promoter (<=1kb) | 16 | 27388869 | 27452218 | 63350 | 1 | ENSMUST00 | 0 | Ccdc50 | ENSMUSG0C | false | chr16 | 27388323 | 27390008 | 29.66345 | 100.21 | 96.63225 | . | . | . | . | . | . |
| chr7 | 45017819 | 45018147 | 431 | Promoter (<=1kb) | 7 | 45017978 | 45021646 | 3669 | 1 | ENSMUST00 | 0 | Rras | ENSMUSG0C | false | chr7 | 45017877 | 45019667 | 28.36989 | 85.2065 | 81.96005 | . | . | . | . | . | . |
| chr9 | 106822460 | 106822732 | 431 | Promoter (<=1kb) | 9 | 106821874 | 106880443 | 58570 | 1 | ENSMUST00 | 587 | Vprbp | ENSMUSG0C | false | chr9 | 106821829 | 106823113 | 31.54088 | 108.67455 | 104.87239 | . | . | . | . | . | . |
| chr1 | 163994494 | 163994953 | 431 | Promoter (<=1kb) | 1 | 163994889 | 163997243 | 2355 | 1 | ENSMUST00 | 0 | Mettl18 | ENSMUSG0C | false | chr1 | 163994206 | 163995669 | 25.99992 | 84.80917 | 81.57088 | chr1 | 163991574 | 163996008 | 5.65475 | 4.33948 | 3.57649 |
| chr19 | 53780931 | 53781200 | 431 | Intron (ENSMUST00000095969.4 | 19 | 53793308 | 53813573 | 20266 | 1 | ENSMUST00 | -12108 | Rbm20 | ENSMUSG0C | false | . | . | . | . | . | . | . | . | . | . | . | . |
| chr13 | 12394926 | 12395239 | 431 | Promoter (<=1kb) | 13 | 12395398 | 12438677 | 43280 | 1 | ENSMUST00 | -159 | Heatr1 | ENSMUSG0C | false | chr13 | 12394669 | 12396294 | 42.33011 | 146.25195 | 141.10883 | . | . | . | . | . | . |
| chr12 | 52604769 | 52605027 | 431 | Promoter (<=1kb) | 12 | 52604485 | 52604591 | 107 | 2 | ENSMUST00 | -179 | Gm23296 | ENSMUSG0C | false | chr12 | 52602865 | 52605494 | 33.05704 | 116.60957 | 112.57934 | . | . | . | . | . | . |
| chr2 | 105057879 | 105058126 | 431 | Distal Intergenic | 2 | 105077895 | 105078107 | 213 | 2 | ENSMUST00 | 19981 | Wt1os | ENSMUSG0C | true | . | . | . | . | . | . | chr2 | 105057120 | 105080197 | 5.15057 | 4.16934 | 3.12788 |
| chr7 | 118275592 | 118275860 | 431 | Distal Intergenic | 7 | 118279861 | 118279965 | 105 | 1 | ENSMUST00 | -4001 | Gm23229 | ENSMUSG0C | false | . | . | . | . | . | . | chr7 | 118274222 | 118279022 | 5.59058 | 4.35566 | 3.51383 |
| chr15 | 99724632 | 99725049 | 431 | Promoter (<=1kb) | 15 | 99715543 | 99724474 | 8932 | 2 | ENSMUST00 | -159 | Gm16537 | ENSMUSG0C | false | . | . | . | . | . | . | chr15 | 99723258 | 99725092 | 6.12716 | 4.24203 | 3.96729 |
| chr10 | 110745040 | 110745521 | 430 | Promoter (<=1kb) | 10 | 110745439 | 110748146 | 2708 | 1 | ENSMUST00 | 0 | E2f7 | ENSMUSG0C | false | chr10 | 110744846 | 110746074 | 26.05581 | 78.34985 | 75.22605 | chr10 | 110743855 | 110746459 | 5.17541 | 4.12717 | 3.18147 |
| chr19 | 6334825 | 6335185 | 430 | Promoter (<=1kb) | 19 | 6334979 | 6340880 | 5902 | 1 | ENSMUST00 | 0 | Men1 | ENSMUSG0C | false | chr19 | 6334640 | 6336004 | 30.00511 | 93.87433 | 90.44607 | chr19 | 6334302 | 6338268 | 4.97357 | 4.0701 | 2.98357 |
| chr1 | 86503858 | 86504572 | 430 | Distal Intergenic | 1 | 86526726 | 86530712 | 3987 | 1 | ENSMUST00 | -22154 | Ptma | ENSMUSG0C | false | chr1 | 86503793 | 86505070 | 13.05315 | 37.5606 | 35.00106 | chr1 | 86499148 | 86507177 | 6.57155 | 4.78752 | 4.28877 |
| chr4 | 123552781 | 123553089 | 430 | Intron (ENSMUST00000097897.5 | 4 | 123349716 | 123544831 | 195116 | 2 | ENSMUST00 | -7951 | Macf1 | ENSMUSG0C | true | . | . | . | . | . | . | chr4 | 123548847 | 123557386 | 5.40924 | 4.22349 | 3.36352 |
| chr9 | 70143442 | 70143698 | 430 | Promoter (<=1kb) | 9 | 70124996 | 70142560 | 17565 | 2 | ENSMUST00 | -883 | Fam81a | ENSMUSG0C | true | . | . | . | . | . | . | chr9 | 70126965 | 70143919 | 5.28781 | 4.22747 | 3.24757 |
| chr12 | 71136885 | 71137218 | 430 | Promoter (<=1kb) | 12 | 71136848 | 71243303 | 106456 | 1 | ENSMUST00 | 38 | 2700049A03I | ENSMUSG0C | false | chr12 | 71135727 | 71138170 | 26.2459 | 113.32486 | 109.39578 | . | . | . | . | . | . |
| chr4 | 125117175 | 125117472 | 430 | Downstream (<1kb) | 4 | 125120094 | 125122520 | 2427 | 2 | ENSMUST00 | 5048 | Zc3h12a | ENSMUSG0C | false | . | . | . | . | . | . | chr4 | 125106469 | 125127334 | 7.64816 | 5.22391 | 5.18478 |
| chr2 | 72117620 | 72117885 | 430 | Intron (ENSMUST00000090826.6 | 2 | 72140711 | 72196261 | 55551 | 1 | ENSMUST00 | -22826 | Rapgef4 | ENSMUSG0C | false | . | . | . | . | . | . | . | . | . | . | . | . |
| chr8 | 83697389 | 83697657 | 430 | Promoter (1-2kb) | 8 | 83670099 | 83699097 | 28999 | 2 | ENSMUST00 | 1440 | Pkn1 | ENSMUSG0C | false | chr8 | 83697311 | 83699766 | 46.69001 | 154.98724 | 149.51399 | chr8 | 83692808 | 83698419 | 7.74578 | 5.12799 | 5.29944 |
| chr4 | 97117118 | 97117461 | 430 | Distal Intergenic | 4 | 97182840 | 97183166 | 327 | 2 | ENSMUST00 | 65705 | Gm10192 | ENSMUSG0C | false | . | . | . | . | . | . | chr4 | 97115811 | 97117835 | 4.45528 | 3.73125 | 2.6052 |
| chr7 | 75369444 | 75369715 | 430 | Distal Intergenic | 7 | 75366055 | 75366264 | 210 | 1 | ENSMUST00 | 3390 | Gm10161 | ENSMUSG0C | false | . | . | . | . | . | . | chr7 | 75369520 | 75370895 | 3.59066 | 3.25161 | 1.88539 |
| chr6 | 54401494 | 54401740 | 430 | Intron (ENSMUST00000145111.2 | 6 | 54424564 | 54429537 | 4974 | 2 | ENSMUST00 | 27797 | 9130019P16I | ENSMUSG0C | false | . | . | . | . | . | . | chr6 | 54391673 | 54403137 | 6.55024 | 4.83886 | 4.27494 |
| chr2 | 158179024 | 158179254 | 430 | Distal Intergenic | 2 | 158192008 | 158195749 | 3742 | 1 | ENSMUST00 | -12754 | 1700060C20I | ENSMUSG0C | false | chr2 | 158179002 | 158179760 | 6.767 | 11.33328 | 9.13518 | chr2 | 158179176 | 158180178 | 5.77331 | 4.30929 | 3.66556 |
| chr17 | 6399128 | 6399560 | 430 | Intron (ENSMUST00000181052.1 | 17 | 6430112 | 6436293 | 6182 | 1 | ENSMUST00 | -30552 | Dynlt1b | ENSMUSG0C | false | . | . | . | . | . | . | chr17 | 6398634 | 6401292 | 8.36031 | 5.19223 | 5.79207 |
| chr12 | 79675583 | 79676237 | 430 | Distal Intergenic | 12 | 79963568 | 79963872 | 305 | 1 | ENSMUST00 | -287331 | RP23-179D3. | ENSMUSG0C | false | . | . | . | . | . | . | chr12 | 79675419 | 79677214 | 4.74811 | 3.97316 | 2.8268 |
| chr14 | 27185892 | 27186214 | 429 | Distal Intergenic | 14 | 27238039 | 27403904 | 165866 | 1 | ENSMUST00 | -51825 | Arhgef3 | ENSMUSG0C | false | . | . | . | . | . | . | . | . | . | . | . | . |
| chr15 | 97773160 | 97773420 | 429 | Distal Intergenic | 15 | 97766859 | 97767972 | 1114 | 2 | ENSMUST00 | -5189 | Rapgef3 | ENSMUSG0C | false | . | . | . | . | . | . | chr15 | 97771183 | 97790304 | 7.43452 | 5.14328 | 5.01036 |
| chr1 | 75382655 | 75383201 | 429 | Promoter (<=1kb) | 1 | 75383556 | 75384975 | 1420 | 1 | ENSMUST00 | -355 | Speg | ENSMUSG0C | true | chr1 | 75381728 | 75383103 | 23.65566 | 74.08845 | 71.03342 | chr1 | 75381009 | 75384668 | 4.13538 | 3.53379 | 2.3139 |
| chr1 | 144004154 | 144004476 | 429 | Promoter (<=1kb) | 1 | 143999338 | 144004161 | 4824 | 2 | ENSMUST00 | 0 | Rgs2 | ENSMUSG0C | false | chr1 | 144003517 | 144005075 | 17.64787 | 49.73517 | 47.02245 | . | . | . | . | . | . |
| chr4 | 135873467 | 135874295 | 429 | Promoter (<=1kb) | 4 | 135870918 | 135873850 | 2933 | 2 | ENSMUST00 | 0 | Pnrc2 | ENSMUSG0C | false | chr4 | 135872022 | 135874455 | 22.68498 | 68.76898 | 65.7979 | chr4 | 135874210 | 135879466 | 7.31253 | 5.17261 | 4.90109 |
| chr7 | 101895977 | 101896301 | 429 | Promoter (<=1kb) | 7 | 101896030 | 101898025 | 1996 | 1 | ENSMUST00 | 0 | Anapc15 | ENSMUSG0C | false | chr7 | 101896043 | 101897252 | 17.00442 | 50.91425 | 48.18763 | . | . | . | . | . | . |
| chr9 | 72985282 | 72985688 | 429 | Promoter (<=1kb) | 9 | 72985429 | 73016340 | 30912 | 1 | ENSMUST00 | 0 | Ccpg1 | ENSMUSG0C | false | chr9 | 72984801 | 72986380 | 44.71377 | 152.69855 | 147.31396 | . | . | . | . | . | . |
| chr10 | 81320371 | 81321282 | 429 | Promoter (<=1kb) | 10 | 81321103 | 81326251 | 5149 | 1 | ENSMUST00 | 0 | Cactin | ENSMUSG0C | false | chr10 | 81320912 | 81322806 | 46.06248 | 156.19411 | 150.67332 | . | . | . | . | . | . |
| chr10 | 44268177 | 44268808 | 429 | Promoter (<=1kb) | 10 | 44268358 | 44364291 | 95934 | 1 | ENSMUST00 | 0 | Atg5 | ENSMUSG0C | false | chr10 | 44267494 | 44269302 | 35.00596 | 114.85688 | 110.88117 | . | . | . | . | . | . |
| chr6 | 125008849 | 125009716 | 429 | Promoter (<=1kb) | 6 | 125009145 | 125036180 | 27036 | 1 | ENSMUST00 | 0 | Zfp384 | ENSMUSG0C | false | chr6 | 125008334 | 125010316 | 26.65955 | 86.95457 | 83.67183 | . | . | . | . | . | . |
| chr1 | 55738913 | 55739203 | 429 | Intron (ENSMUST00000042986.9 | 1 | 55702025 | 55735227 | 33203 | 1 | ENSMUST00 | 36889 | Plcl1 | ENSMUSG0C | true | chr1 | 55737106 | 55739417 | 8.08983 | 23.14769 | 20.77093 | chr1 | 55738786 | 55752458 | 4.85141 | 3.98095 | 2.88878 |
| chr5 | 114091218 | 114091446 | 429 | Promoter (<=1kb) | 5 | 114053767 | 114091380 | 37614 | 2 | ENSMUST00 | 0 | Svop | ENSMUSG0C | false | . | . | . | . | . | . | chr5 | 114091051 | 114091281 | 3.0272 | 3.03353 | 1.43137 |
| chr6 | 67168414 | 67168675 | 429 | Exon (ENSMUST00000181304.1/ | 6 | 67165203 | 67165616 | 414 | 1 | ENSMUST00 | 3212 | Gm27015 | ENSMUSG0C | false | chr6 | 67168288 | 67168896 | 9.52248 | 22.23582 | 19.8701 | chr6 | 67168016 | 67169006 | 4.65725 | 4.00868 | 2.71707 |
| chr3 | 100857372 | 100857698 | 429 | Intron (ENSMUST00000054791.7 | 3 | 100825459 | 100895679 | 70221 | 1 | ENSMUST00 | 31914 | Vtcn1 | ENSMUSG0C | false | . | . | . | . | . | . | chr3 | 100854999 | 100862887 | 5.03234 | 4.00629 | 3.03666 |
| chr8 | 61590896 | 61591493 | 429 | Promoter (<=1kb) | 8 | 61513248 | 61591142 | 77895 | 2 | ENSMUST00 | 0 | Palld | ENSMUSG0C | true | chr8 | 61589311 | 61591302 | 36.69325 | 114.64857 | 110.67819 | chr8 | 61591328 | 61592357 | 3.65079 | 3.3759 | 1.88712 |
| chr3 | 79218846 | 79219166 | 429 | Distal Intergenic | 3 | 79196374 | 79196477 | 104 | 1 | ENSMUST00 | 22473 | Gm24150 | ENSMUSG0C | false | . | . | . | . | . | . | chr3 | 79215948 | 79226268 | 5.05618 | 4.08573 | 3.05849 |
| chr11 | 108424961 | 108425310 | 428 | Promoter (<=1kb) | 11 | 108425192 | 108860615 | 435424 | 1 | ENSMUST00 | 0 | Cep112 | ENSMUSG0C | false | chr11 | 108424572 | 108425870 | 23.65566 | 74.08845 | 71.03342 | chr11 | 108424130 | 108426162 | 4.24172 | 3.67442 | 2.42749 |
| chr12 | 111885960 | 111886396 | 428 | Intron (ENSMUST00000054815.8 | 12 | 111866013 | 111908040 | 42028 | 2 | ENSMUST00 | 21644 | Ppp1r13b | ENSMUSG0C | false | . | . | . | . | . | . | chr12 | 111880166 | 111903803 | 5.80743 | 4.40294 | 3.67276 |
| chr19 | 46476861 | 46477172 | 428 | 3' UTR | 19 | 46483097 | 46485982 | 2886 | 1 | ENSMUST00 | -5925 | Sufu | ENSMUSG0C | false | . | . | . | . | . | . | chr19 | 46468103 | 46486697 | 7.92449 | 5.26231 | 5.41796 |
| chr4 | 125029793 | 125030042 | 428 | Promoter (<=1kb) | 4 | 125029992 | 125055376 | 25385 | 1 | ENSMUST00 | 0 | Gnl2 | ENSMUSG0C | false | chr4 | 125029594 | 125030624 | 42.27695 | 136.11826 | 131.39047 | chr4 | 125028845 | 125031786 | 3.98236 | 3.49891 | 2.2174 |
| chr6 | 136894619 | 136894975 | 428 | Distal Intergenic | 6 | 136872436 | 136875805 | 3370 | 2 | ENSMUST00 | -18815 | Mgp | ENSMUSG0C | false | . | . | . | . | . | . | chr6 | 136894481 | 136895965 | 4.66008 | 3.8735 | 2.78155 |
| chrX | 7670355 | 7670710 | 428 | Promoter (<=1kb) | 20 | 7669330 | 7671313 | 1984 | 2 | ENSMUST00 | 603 | Plp2 | ENSMUSG0C | false | chrX | 7670173 | 7671711 | 20.21941 | 51.82391 | 49.08421 | . | . | . | . | . | . |
| chr9 | 122351514 | 122352088 | 428 | Promoter (<=1kb) | 9 | 122351608 | 122381524 | 29917 | 1 | ENSMUST00 | 0 | Abhd5 | ENSMUSG0C | false | chr9 | 122351245 | 122352433 | 25.18016 | 100.40004 | 96.81754 | . | . | . | . | . | . |
| chr9 | 45138585 | 45138951 | 428 | Promoter (<=1kb) | 9 | 45138437 | 45154152 | 15716 | 1 | ENSMUST00 | 149 | Scn4b | ENSMUSG0C | false | chr9 | 45138152 | 45139055 | 12.58327 | 33.30664 | 30.79935 | chr9 | 45138050 | 45139225 | 6.21487 | 4.6496 | 4.01444 |
| chr9 | 44726536 | 44726997 | 428 | Promoter (<=1kb) | 9 | 44726010 | 44726486 | 477 | 2 | ENSMUST00 | -51 | Phldb1 | ENSMUSG0C | false | . | . | . | . | . | . | chr9 | 44719663 | 44729806 | 5.68645 | 4.29249 | 3.57992 |
| chr1 | 172563003 | 172563613 | 428 | 3' UTR | 1 | 172559906 | 172563716 | 3811 | 1 | ENSMUST00 | 3098 | Vsig8 | ENSMUSG0C | false | chr1 | 172562102 | 172563770 | 21.47392 | 86.08567 | 82.81886 | chr1 | 172561933 | 172563859 | 5.12982 | 4.06493 | 3.14923 |
| chr1 | 54743100 | 54743408 | 428 | Exon (ENSMUST00000179030.2/ | 1 | 54728987 | 54735644 | 6658 | 2 | ENSMUST00 | -7457 | Ankrd44 | ENSMUSG0C | true | . | . | . | . | . | . | chr1 | 54742229 | 54744495 | 5.98304 | 4.59473 | 3.8065 |
| chr5 | 123076186 | 123076518 | 428 | Promoter (<=1kb) | 5 | 123076275 | 123117749 | 41475 | 1 | ENSMUST00 | 0 | Tmem120b | ENSMUSG0C | false | chr5 | 123075871 | 123077894 | 34.00884 | 130.01486 | 125.52227 | chr5 | 123075273 | 123080046 | 4.61242 | 3.90128 | 2.69446 |
| chr14 | 22215844 | 22216191 | 428 | Intron (ENSMUST00000075639.5 | 14 | 22310165 | 22584439 | 274275 | 1 | ENSMUST00 | -93974 | 1700112E06I | ENSMUSG0C | false | . | . | . | . | . | . | chr14 | 22212900 | 22216338 | 4.22296 | 3.54469 | 2.40489 |
| chr4 | 34749314 | 34749568 | 428 | Intron (ENSMUST00000095129.2 | 4 | 34743784 | 34756263 | 12480 | 2 | ENSMUST00 | 6695 | Gm136 | ENSMUSG0C | false | . | . | . | . | . | . | . | . | . | . | . | . |
| chr12 | 17015800 | 17016075 | 428 | Distal Intergenic | 12 | 17003319 | 17011727 | 8409 | 2 | ENSMUST00 | -4074 | 2410004P03I | ENSMUSG0C | false | . | . | . | . | . | . | chr12 | 17014640 | 17018556 | 5.54135 | 4.30287 | 3.45045 |
| chr4 | 123425627 | 123425932 | 428 | Intron (ENSMUST00000097897.5 | 4 | 123435617 | 123435934 | 318 | 1 | ENSMUST00 | -9685 | Gm12924 | ENSMUSG0C | false | . | . | . | . | . | . | chr4 | 123412273 | 123429396 | 5.99663 | 4.50351 | 3.82082 |
| chr5 | 148605208 | 148605425 | 427 | Distal Intergenic | 5 | 148505142 | 148552789 | 47648 | 2 | ENSMUST00 | -52420 | Ubl3 | ENSMUSG0C | false | . | . | . | . | . | . | chr5 | 148604889 | 148606560 | 3.01933 | 2.99583 | 1.4233 |
| chr11 | 45851821 | 45852186 | 427 | Promoter (<=1kb) | 11 | 45852051 | 45910625 | 58575 | 1 | ENSMUST00 | 0 | Clint1 | ENSMUSG0C | false | chr11 | 45851500 | 45852960 | 39.65778 | 126.14424 | 121.78909 | . | . | . | . | . | . |
| chr5 | 146920423 | 146920640 | 427 | Distal Intergenic | 5 | 146948657 | 146955614 | 6958 | 1 | ENSMUST00 | -28017 | Gtf3a | ENSMUSG0C | false | . | . | . | . | . | . | chr5 | 146919863 | 146924296 | 4.88294 | 4.06696 | 2.91401 |
| chr5 | 124111741 | 124112390 | 427 | Promoter (<=1kb) | 5 | 124112297 | 124115483 | 3187 | 1 | ENSMUST00 | 0 | Ogfod2 | ENSMUSG0C | false | chr5 | 124111237 | 124113178 | 36.809 | 123.33047 | 119.07505 | . | . | . | . | . | . |
| chr3 | 30508920 | 30509210 | 427 | Promoter (<=1kb) | 3 | 30236569 | 30509487 | 272919 | 2 | ENSMUST00 | 277 | Mecom | ENSMUSG0C | false | chr3 | 30507549 | 30509587 | 14.66885 | 35.63733 | 33.10225 | chr3 | 30506751 | 30509701 | 5.62817 | 4.3849 | 3.51508 |
| chr3 | 82073755 | 82074027 | 427 | Promoter (<=1kb) | 3 | 82032004 | 82074711 | 42708 | 2 | ENSMUST00 | 684 | Gucy1b3 | ENSMUSG0C | false | chr3 | 82073700 | 82074710 | 10.20265 | 24.60154 | 22.20526 | chr3 | 82072941 | 82074822 | 6.01432 | 4.5244 | 3.86492 |
| chr6 | 136895171 | 136895475 | 427 | Distal Intergenic | 6 | 136872436 | 136875805 | 3370 | 2 | ENSMUST00 | -19367 | Mgp | ENSMUSG0C | false | . | . | . | . | . | . | chr6 | 136894481 | 136895965 | 4.66008 | 3.8735 | 2.78155 |
| chr6 | 85331301 | 85331663 | 427 | Promoter (1-2kb) | 6 | 85318872 | 85332750 | 13879 | 2 | ENSMUST00 | 1087 | Sfxn5 | ENSMUSG0C | false | . | . | . | . | . | . | chr6 | 85331284 | 85332722 | 4.899 | 4.01765 | 2.94858 |

| | | | | | | | | | | | | | | | | | | | | | | | | | | |
|---|---|---|---|---|---|---|---|---|---|---|---|---|---|---|---|---|---|---|---|---|---|---|---|---|---|---|
| chr15 | 99251946 | 99252490 | 427 | Promoter (<=1kb) | 15 | 99242818 | 99251961 | 9144 | 2 | ENSMUST00 | 0 | Mcrs1 | ENSMUSG0C | false | chr15 | 99251025 | 99253132 | 26.22205 | 82.37943 | 79.18155 | chr15 | 99252455 | 99254800 | 3.52233 | 3.35659 | 1.78614 |
| chr19 | 31762592 | 31763057 | 427 | Promoter (1-2kb) | 19 | 31742114 | 31764973 | 22860 | 2 | ENSMUST00 | 1916 | Prkg1 | ENSMUSG0C | true | chr19 | 31762329 | 31765338 | 26.95677 | 77.13882 | 74.03856 | chr19 | 31762049 | 31764545 | 3.9805 | 3.60883 | 2.1536 |
| chr15 | 79984119 | 79984424 | 427 | Distal Intergenic | 15 | 79918417 | 79971119 | 52703 | 2 | ENSMUST00 | -13001 | Cbx7 | ENSMUSG0C | false | . | . | . | . | . | . | chr15 | 79983394 | 79985788 | 4.82306 | 3.90603 | 2.90848 |
| chr10 | 14503829 | 14504266 | 427 | Intron (ENSMUST00000041168.4 | 10 | 14523078 | 14523545 | 468 | 1 | ENSMUST00 | -18812 | Gm10335 | ENSMUSG0C | false | . | . | . | . | . | . | chr10 | 14503031 | 14504416 | 3.07181 | 3.0171 | 1.4772 |
| chr11 | 78833041 | 78833337 | 427 | Intron (ENSMUST00000147875.3 | 11 | 78826629 | 78839275 | 12647 | 1 | ENSMUST00 | 6413 | Lyrm9 | ENSMUSG0C | false | . | . | . | . | . | . | chr11 | 78823496 | 78836454 | 6.00182 | 4.51146 | 3.83406 |
| chr5 | 76928149 | 76928434 | 427 | Promoter (<=1kb) | 5 | 76928368 | 76928845 | 478 | 1 | ENSMUST00 | 0 | 2310040G07 | ENSMUSG0C | false | . | . | . | . | . | . | chr5 | 76927591 | 76930221 | 4.8376 | 3.92712 | 2.88997 |
| chr15 | 80840003 | 80840307 | 427 | Intron (ENSMUST00000067689.7 | 15 | 80940039 | 80940120 | 82 | 1 | ENSMUST00 | -99732 | Mir5113 | ENSMUSG0C | false | . | . | . | . | . | . | chr15 | 80832720 | 80847879 | 5.49721 | 4.14862 | 3.44463 |
| chr13 | 37993942 | 37994300 | 426 | Promoter (<=1kb) | 13 | 37971401 | 37994190 | 22790 | 2 | ENSMUST00 | 0 | Ssr1 | ENSMUSG0C | false | chr13 | 37992943 | 37994742 | 50.25226 | 173.98802 | 167.94014 | . | . | . | . | . | . |
| chr19 | 47579313 | 47579897 | 426 | Promoter (<=1kb) | 19 | 47579678 | 47645246 | 65569 | 1 | ENSMUST00 | 0 | Slk | ENSMUSG0C | false | chr19 | 47578934 | 47580860 | 22.64381 | 71.35664 | 68.34631 | . | . | . | . | . | . |
| chr2 | 119337885 | 119338203 | 426 | Distal Intergenic | 2 | 119328538 | 119332153 | 3616 | 1 | ENSMUST00 | 9348 | Dll4 | ENSMUSG0C | false | chr2 | 119337545 | 119338441 | 23.43614 | 63.09834 | 60.20888 | chr2 | 119337183 | 119338917 | 4.87821 | 4.02501 | 2.91338 |
| chr5 | 24685503 | 24686009 | 426 | Promoter (<=1kb) | 5 | 24685815 | 24710555 | 24741 | 1 | ENSMUST00 | 0 | Nub1 | ENSMUSG0C | false | chr5 | 24685889 | 24687425 | 23.65566 | 74.08845 | 71.03342 | . | . | . | . | . | . |
| chr5 | 92083391 | 92083950 | 426 | Promoter (<=1kb) | 5 | 92052146 | 92083557 | 31412 | 2 | ENSMUST00 | 0 | G3bp2 | ENSMUSG0C | false | chr5 | 92082682 | 92084325 | 32.66734 | 113.81544 | 109.86345 | . | . | . | . | . | . |
| chr6 | 134049242 | 134049470 | 426 | Intron (ENSMUST00000081028.7 | 6 | 134035969 | 134248284 | 212316 | 1 | ENSMUST00 | 13274 | Etv6 | ENSMUSG0C | false | . | . | . | . | . | . | chr6 | 134037014 | 134053440 | 5.9149 | 4.52781 | 3.75986 |
| chr7 | 132841324 | 132841563 | 426 | Intron (ENSMUST00000124096.2 | 7 | 132827781 | 132852379 | 24599 | 2 | ENSMUST00 | 10816 | Mettl10 | ENSMUSG0C | false | . | . | . | . | . | . | chr7 | 132840223 | 132844238 | 4.9076 | 3.97327 | 2.93074 |
| chr9 | 13611131 | 13611392 | 426 | Distal Intergenic | 9 | 13619990 | 13709388 | 89399 | 1 | ENSMUST00 | -8598 | Maml2 | ENSMUSG0C | false | . | . | . | . | . | . | . | . | . | . | . | . |
| chr9 | 63706361 | 63706710 | 426 | Intron (ENSMUST00000034973.4 | 9 | 63654799 | 63711969 | 57171 | 2 | ENSMUST00 | 5259 | Smad3 | ENSMUSG0C | false | . | . | . | . | . | . | chr9 | 63695180 | 63731540 | 5.72957 | 4.34734 | 3.61571 |
| chr15 | 50869535 | 50869877 | 426 | Intron (ENSMUST00000165201.3 | 15 | 50660435 | 50882806 | 222372 | 2 | ENSMUST00 | 12929 | Trps1 | ENSMUSG0C | false | chr15 | 50869392 | 50870337 | 14.84617 | 51.23753 | 48.50555 | chr15 | 50868904 | 50871461 | 4.72515 | 3.89538 | 2.78938 |
| chr17 | 56757430 | 56757798 | 426 | Promoter (<=1kb) | 17 | 56751325 | 56757530 | 6206 | 2 | ENSMUST00 | 0 | Nrtn | ENSMUSG0C | false | chr17 | 56756832 | 56757982 | 23.04863 | 71.19738 | 68.188 | chr17 | 56756785 | 56759234 | 6.10381 | 4.54792 | 3.96627 |
| chr11 | 43773856 | 43774145 | 426 | Downstream (<1kb) | 11 | 43729512 | 43748008 | 18497 | 2 | ENSMUST00 | -25849 | Ttc1 | ENSMUSG0C | false | . | . | . | . | . | . | . | . | . | . | . | . |
| chr3 | 85874073 | 85874321 | 426 | Intron (ENSMUST00000154148.2 | 3 | 85817123 | 85887368 | 70246 | 2 | ENSMUST00 | 13047 | Arfip1 | ENSMUSG0C | false | . | . | . | . | . | . | chr3 | 85873467 | 85874847 | 5.3104 | 4.211 | 3.30362 |
| chr2 | 160066630 | 160067081 | 426 | Distal Intergenic | 2 | 160016653 | 160040619 | 23967 | 1 | ENSMUST00 | 49978 | 9130015L21F | ENSMUSG0C | false | . | . | . | . | . | . | chr2 | 160065057 | 160069010 | 5.33636 | 4.09837 | 3.30668 |
| chr8 | 92227170 | 92227434 | 426 | Distal Intergenic | 8 | 92336668 | 92339250 | 2583 | 2 | ENSMUST00 | 111816 | Gm21817 | ENSMUSG0C | false | . | . | . | . | . | . | chr8 | 92226942 | 92227834 | 4.42253 | 3.67716 | 2.5572 |
| chr8 | 33823868 | 33824195 | 426 | Intron (ENSMUST00000181120.1 | 8 | 33813203 | 33833094 | 19892 | 1 | ENSMUST00 | 10666 | Gm26632 | ENSMUSG0C | false | . | . | . | . | . | . | chr8 | 33822415 | 33859327 | 6.05392 | 4.56707 | 3.87043 |
| chr10 | 86290030 | 86290524 | 425 | Intron (ENSMUST00000120638.2 | 10 | 86300372 | 86349506 | 49135 | 1 | ENSMUST00 | -9848 | Timp3 | ENSMUSG0C | false | . | . | . | . | . | . | chr10 | 86287196 | 86291253 | 5.15519 | 4.10271 | 3.1482 |
| chr11 | 20112393 | 20113067 | 425 | Promoter (<=1kb) | 11 | 20062304 | 20112913 | 50610 | 2 | ENSMUST00 | 0 | Actr2 | ENSMUSG0C | false | chr11 | 20111757 | 20113819 | 36.24025 | 112.82189 | 108.90788 | . | . | . | . | . | . |
| chr3 | 116594690 | 116595069 | 425 | Promoter (<=1kb) | 3 | 116595008 | 116630984 | 35977 | 1 | ENSMUST00 | 0 | Sass6 | ENSMUSG0C | false | chr3 | 116594187 | 116595616 | 35.36919 | 136.9064 | 132.15405 | . | . | . | . | . | . |
| chr4 | 131850337 | 131850573 | 425 | Intron (ENSMUST00000030742.5 | 4 | 131853686 | 131860998 | 7313 | 1 | ENSMUST00 | -3113 | Mecr | ENSMUSG0C | false | . | . | . | . | . | . | chr4 | 131848218 | 131851667 | 5.60333 | 4.20325 | 3.54838 |
| chr7 | 89854795 | 89855187 | 425 | Distal Intergenic | 7 | 89829282 | 89851962 | 22681 | 1 | ENSMUST00 | 25514 | Me3 | ENSMUSG0C | false | . | . | . | . | . | . | chr7 | 89854602 | 89856451 | 4.47575 | 3.81372 | 2.59634 |
| chr12 | 69168405 | 69168694 | 425 | Promoter (<=1kb) | 12 | 69168814 | 69179010 | 10197 | 1 | ENSMUST00 | -120 | Lrr1 | ENSMUSG0C | false | chr12 | 69168116 | 69169221 | 28.49948 | 87.34749 | 84.05397 | chr12 | 69167695 | 69169309 | 4.55536 | 3.59303 | 2.67051 |
| chr11 | 102016719 | 102016934 | 425 | Intron (ENSMUST00000100400.3 | 11 | 102008640 | 102010410 | 1771 | 2 | ENSMUST00 | -6310 | Mpp3 | ENSMUSG0C | false | chr11 | 102015369 | 102016894 | 10.05864 | 20.96754 | 18.61967 | chr11 | 102013085 | 102018629 | 4.41209 | 3.78043 | 2.52457 |
| chr1 | 10039586 | 10039905 | 425 | Promoter (<=1kb) | 1 | 10039767 | 10058872 | 19106 | 1 | ENSMUST00 | 0 | Cspp1 | ENSMUSG0C | false | chr1 | 10037131 | 10040542 | 35.12714 | 133.20123 | 128.59534 | chr1 | 10038935 | 10041270 | 3.83474 | 3.56751 | 2.06372 |
| chr7 | 25619150 | 25619465 | 425 | Promoter (<=1kb) | 7 | 25619414 | 25625551 | 6138 | 1 | ENSMUST00 | 0 | Atp5sl | ENSMUSG0C | false | chr7 | 25618809 | 25620007 | 27.78601 | 91.88203 | 88.49705 | chr7 | 25618872 | 25622316 | 4.14849 | 3.57365 | 2.31963 |
| chr7 | 131371072 | 131371538 | 425 | Promoter (<=1kb) | 7 | 131371141 | 131387839 | 16699 | 1 | ENSMUST00 | 0 | Pstk | ENSMUSG0C | false | chr7 | 131371083 | 131371957 | 23.62199 | 79.492 | 76.34769 | chr7 | 131370803 | 131373147 | 3.26182 | 3.00374 | 1.62925 |
| chr19 | 8914302 | 8914562 | 425 | Exon (ENSMUST00000096246.3/ | 19 | 8920374 | 8927234 | 6861 | 1 | ENSMUST00 | -5812 | B3gat3 | ENSMUSG0C | false | . | . | . | . | . | . | chr19 | 8911015 | 8915692 | 5.21461 | 4.08859 | 3.21453 |
| chr15 | 98952908 | 98953844 | 425 | Promoter (<=1kb) | 15 | 98949841 | 98953551 | 3711 | 2 | ENSMUST00 | 0 | Tuba1a | ENSMUSG0C | false | chr15 | 98952176 | 98954298 | 33.82489 | 107.59698 | 103.83312 | chr15 | 98953723 | 98955481 | 4.37165 | 3.77949 | 2.48087 |
| chr2 | 159299191 | 159299400 | 425 | Distal Intergenic | 2 | 159456170 | 159456457 | 288 | 2 | ENSMUST00 | 157057 | Gm14219 | ENSMUSG0C | false | . | . | . | . | . | . | chr2 | 159296604 | 159299934 | 3.62163 | 3.28852 | 1.90937 |
| chrX | 169997895 | 169998467 | 425 | Intron (ENSMUST00000171433.2 | 20 | 169985801 | 169986939 | 1139 | 2 | ENSMUST00 | -10957 | Gm15247 | ENSMUSG0C | false | . | . | . | . | . | . | . | . | . | . | . | . |
| chr2 | 61396179 | 61396439 | 425 | Distal Intergenic | 2 | 61492766 | 61492894 | 129 | 1 | ENSMUST00 | -96327 | Gm22338 | ENSMUSG0C | false | . | . | . | . | . | . | chr2 | 61395816 | 61397224 | 4.88837 | 4.07364 | 2.92027 |
| chr8 | 121359152 | 121359432 | 425 | Promoter (2-3kb) | 8 | 121256747 | 121356836 | 100090 | 2 | ENSMUST00 | -2317 | Gm26815 | ENSMUSG0C | false | . | . | . | . | . | . | chr8 | 121358623 | 121361301 | 8.88514 | 5.63262 | 6.21498 |
| chr5 | 136021606 | 136021837 | 424 | Promoter (2-3kb) | 5 | 136019558 | 136032864 | 13307 | 1 | ENSMUST00 | 2049 | Dtx2 | ENSMUSG0C | false | . | . | . | . | . | . | chr5 | 136014251 | 136022977 | 5.92015 | 4.36442 | 3.79999 |
| chr11 | 48800010 | 48800382 | 424 | Promoter (<=1kb) | 11 | 48800332 | 48804974 | 4643 | 1 | ENSMUST00 | 0 | Gnb2l1 | ENSMUSG0C | false | chr11 | 48799136 | 48801644 | 21.91694 | 64.77673 | 61.86085 | . | . | . | . | . | . |
| chr11 | 117114666 | 117115634 | 424 | Promoter (<=1kb) | 11 | 117115168 | 117152185 | 37018 | 1 | ENSMUST00 | 0 | Sec14l1 | ENSMUSG0C | false | chr11 | 117114277 | 117117694 | 46.43878 | 156.84802 | 151.30803 | chr11 | 117112932 | 117114786 | 3.43565 | 3.29491 | 1.71201 |
| chr15 | 39112587 | 39112910 | 424 | Promoter (<=1kb) | 15 | 39094191 | 39112716 | 18526 | 2 | ENSMUST00 | 0 | Slc25a32 | ENSMUSG0C | false | chr15 | 39111953 | 39113834 | 29.52499 | 116.10789 | 112.09218 | . | . | . | . | . | . |
| chr3 | 142764977 | 142765263 | 424 | Promoter (<=1kb) | 3 | 142765226 | 142783603 | 18378 | 1 | ENSMUST00 | 0 | Gtf2b | ENSMUSG0C | false | chr3 | 142764529 | 142766473 | 26.41709 | 100.00125 | 96.43182 | . | . | . | . | . | . |
| chr8 | 80738864 | 80739780 | 424 | Promoter (<=1kb) | 8 | 80698507 | 80739497 | 40991 | 2 | ENSMUST00 | 0 | Smarca5 | ENSMUSG0C | false | chr8 | 80738046 | 80740342 | 32.96857 | 104.0584 | 100.38712 | chr8 | 80737467 | 80740332 | 4.70802 | 3.93816 | 2.77678 |
| chr9 | 59667297 | 59667548 | 424 | Intron (ENSMUST00000034834.1 | 9 | 59656368 | 59679375 | 23008 | 1 | ENSMUST00 | 10930 | Pkm | ENSMUSG0C | false | . | . | . | . | . | . | chr9 | 59666557 | 59668237 | 6.40457 | 4.55797 | 4.19672 |
| chr4 | 63403038 | 63403551 | 424 | Promoter (<=1kb) | 4 | 63367125 | 63403354 | 36230 | 2 | ENSMUST00 | 0 | Akna | ENSMUSG0C | false | chr4 | 63402627 | 63403839 | 10.11741 | 21.06654 | 18.71757 | chr4 | 63402504 | 63405386 | 4.14817 | 3.59673 | 2.33545 |
| chr9 | 106788045 | 106788332 | 424 | Promoter (<=1kb) | 9 | 106786759 | 106789176 | 2418 | 2 | ENSMUST00 | 844 | Rad54l2 | ENSMUSG0C | false | chr9 | 106788240 | 106789925 | 26.2459 | 113.32486 | 109.39578 | chr9 | 106783504 | 106790006 | 3.25538 | 3.14663 | 1.56876 |
| chr7 | 143055332 | 143055563 | 424 | Promoter (1-2kb) | 7 | 143032621 | 143053686 | 21066 | 2 | ENSMUST00 | -1647 | R74862 | ENSMUSG0C | false | chr7 | 143052294 | 143056592 | 35.67123 | 127.73884 | 123.33056 | chr7 | 143054510 | 143063668 | 7.18311 | 5.11038 | 4.77801 |
| chr3 | 51778666 | 51779309 | 424 | Intron (ENSMUST00000121440.2 | 3 | 51689437 | 51796535 | 107099 | 2 | ENSMUST00 | 17226 | Maml3 | ENSMUSG0C | true | . | . | . | . | . | . | chr3 | 51767223 | 51782456 | 5.75718 | 4.44357 | 3.63488 |
| chr6 | 97582463 | 97582682 | 424 | Intron (ENSMUST00000113355.3 | 6 | 97572076 | 97572238 | 163 | 2 | ENSMUST00 | -10226 | Gm23510 | ENSMUSG0C | false | . | . | . | . | . | . | chr6 | 97574875 | 97587447 | 5.062 | 4.07112 | 3.05952 |
| chr1 | 59745216 | 59745474 | 424 | Distal Intergenic | 1 | 59763400 | 59879014 | 115615 | 1 | ENSMUST00 | -17926 | Bmpr2 | ENSMUSG0C | true | . | . | . | . | . | . | chr1 | 59744882 | 59745857 | 4.283 | 3.78551 | 2.4042 |
| chr16 | 78219788 | 78220023 | 424 | Distal Intergenic | 16 | 78240258 | 78240388 | 131 | 1 | ENSMUST00 | -20235 | Gm25916 | ENSMUSG0C | false | . | . | . | . | . | . | . | . | . | . | . | . |
| chr11 | 108980474 | 108980725 | 424 | Distal Intergenic | 11 | 109007651 | 109008781 | 1131 | 1 | ENSMUST00 | -26926 | Gm11668 | ENSMUSG0C | false | . | . | . | . | . | . | chr11 | 108980293 | 108981100 | 3.66348 | 3.46296 | 1.9452 |
| chrY | 1074119 | 1074358 | 424 | Distal Intergenic | 21 | 1055764 | 1058868 | 3105 | 2 | ENSMUST00 | -15252 | Tspy-ps | ENSMUSG0C | false | . | . | . | . | . | . | . | . | . | . | . | . |
| chr1 | 194925876 | 194926336 | 423 | Distal Intergenic | 1 | 194938821 | 194961292 | 22472 | 1 | ENSMUST00 | -12485 | Cd34 | ENSMUSG0C | false | . | . | . | . | . | . | . | . | . | . | . | . |
| chr7 | 123377845 | 123378225 | 423 | Promoter (<=1kb) | 7 | 123377982 | 123430358 | 52377 | 1 | ENSMUST00 | 0 | Lcmt1 | ENSMUSG0C | false | chr7 | 123377961 | 123378620 | 31.48209 | 103.47416 | 99.8177 | chr7 | 123377603 | 123379682 | 3.20008 | 3.07874 | 1.57659 |
| chr8 | 95434755 | 95435161 | 423 | Promoter (<=1kb) | 8 | 95420252 | 95434869 | 14618 | 2 | ENSMUST00 | 0 | Gtl3 | ENSMUSG0C | false | chr8 | 95433852 | 95434912 | 42.38816 | 144.10358 | 139.05386 | . | . | . | . | . | . |
| chr9 | 61378418 | 61378627 | 423 | Promoter (2-3kb) | 9 | 61375529 | 61416976 | 41448 | 1 | ENSMUST00 | 2890 | Tle3 | ENSMUSG0C | false | . | . | . | . | . | . | chr9 | 61375865 | 61379180 | 4.0929 | 3.5767 | 2.25879 |
| chr10 | 78574351 | 78574898 | 423 | Promoter (<=1kb) | 10 | 78574500 | 78584498 | 9999 | 1 | ENSMUST00 | 0 | Ilvbl | ENSMUSG0C | false | chr10 | 78574095 | 78575626 | 24.65656 | 75.88111 | 72.79811 | chr10 | 78573794 | 78577008 | 5.86233 | 4.49948 | 3.68948 |
| chr7 | 35555102 | 35555991 | 423 | Promoter (<=1kb) | 7 | 35547521 | 35555194 | 7674 | 2 | ENSMUST00 | 0 | Nudt19 | ENSMUSG0C | false | chr7 | 35554712 | 35556253 | 34.91903 | 109.49904 | 105.68399 | chr7 | 35555934 | 35556589 | 3.89897 | 3.31799 | 2.13481 |
| chr9 | 32227081 | 32227377 | 423 | Promoter (2-3kb) | 9 | 32224592 | 32245983 | 21392 | 1 | ENSMUST00 | 2490 | Arhgap32 | ENSMUSG0C | false | chr9 | 32223834 | 32227622 | 16.52141 | 45.4089 | 42.75185 | . | . | . | . | . | . |
| chr10 | 37140961 | 37141518 | 423 | Promoter (1-2kb) | 10 | 37139914 | 37151315 | 11402 | 1 | ENSMUST00 | 1048 | 5930403N24 | ENSMUSG0C | false | chr10 | 37138147 | 37141657 | 19.27906 | 51.80829 | 49.07 | chr10 | 37136051 | 37141791 | 4.47232 | 3.83588 | 2.60562 |
| chr11 | 59227818 | 59228621 | 423 | Promoter (<=1kb) | 11 | 59211412 | 59228162 | 16751 | 2 | ENSMUST00 | 0 | Arf1 | ENSMUSG0C | false | chr11 | 59226686 | 59228835 | 23.5948 | 97.99914 | 94.47849 | chr11 | 59228533 | 59229230 | 4.18388 | 3.68791 | 2.33794 |
| chr7 | 99980120 | 99980608 | 423 | Promoter (<=1kb) | 7 | 99920257 | 99980458 | 60202 | 2 | ENSMUST00 | 0 | Rnf169 | ENSMUSG0C | false | chr7 | 99979459 | 99980922 | 42.42999 | 160.09743 | 154.44568 | . | . | . | . | . | . |
| chr16 | 9756933 | 9757168 | 423 | Intron (ENSMUST00000115835.2 | 16 | 9577827 | 9992533 | 414707 | 2 | ENSMUST00 | 235365 | Grin2a | ENSMUSG0C | false | . | . | . | . | . | . | chr16 | 9753473 | 9759511 | 5.47538 | 4.31106 | 3.40485 |
| chr7 | 138121131 | 138121397 | 423 | Distal Intergenic | 7 | 138101433 | 138101539 | 107 | 2 | ENSMUST00 | -19593 | Gm25798 | ENSMUSG0C | false | . | . | . | . | . | . | chr7 | 138119026 | 138122562 | 5.3525 | 4.22418 | 3.34817 |
| chr9 | 97011306 | 97011601 | 423 | Intron (ENSMUST00000055433.4 | 9 | 97015966 | 97022043 | 6078 | 1 | ENSMUST00 | -4365 | Gm16010 | ENSMUSG0C | false | . | . | . | . | . | . | chr9 | 97009875 | 97014254 | 4.87082 | 3.81409 | 2.93633 |
| chr5 | 95870564 | 95870856 | 423 | Distal Intergenic | 5 | 95867334 | 95867440 | 107 | 1 | ENSMUST00 | 3231 | Gm25937 | ENSMUSG0C | false | . | . | . | . | . | . | . | . | . | . | . | . |
| chr10 | 37484397 | 37484811 | 423 | Distal Intergenic | 10 | 37337486 | 37379652 | 42167 | 1 | ENSMUST00 | 146912 | Gm26535 | ENSMUSG0C | false | . | . | . | . | . | . | chr10 | 37483637 | 37485292 | 3.6949 | 3.36418 | 1.98222 |
| chr16 | 85047057 | 85047310 | 423 | Intron (ENSMUST00000005406.1 | 16 | 85168849 | 85169001 | 153 | 1 | ENSMUST00 | -121539 | Gm27295 | ENSMUSG0C | false | . | . | . | . | . | . | chr16 | 85041565 | 85052812 | 5.14224 | 4.09203 | 3.14604 |
| chr1 | 159717235 | 159717506 | 422 | Distal Intergenic | 1 | 159737510 | 159924397 | 186888 | 1 | ENSMUST00 | -20004 | Tnr | ENSMUSG0C | false | . | . | . | . | . | . | chr1 | 159717111 | 159717855 | 2.92741 | 2.92435 | 1.35706 |
| chr5 | 146948509 | 146949094 | 422 | Promoter (<=1kb) | 5 | 146948657 | 146955614 | 6958 | 1 | ENSMUST00 | 0 | Gtf3a | ENSMUSG0C | false | chr5 | 146948130 | 146949643 | 30.35121 | 89.59235 | 86.25857 | chr5 | 146947949 | 146950228 | 3.25095 | 3.18449 | 1.55116 |
| chr8 | 46470778 | 46471248 | 422 | Promoter (<=1kb) | 8 | 46471041 | 46536051 | 65011 | 1 | ENSMUST00 | 0 | Acsl1 | ENSMUSG0C | false | chr8 | 46470894 | 46472109 | 40.03016 | 127.86769 | 123.44967 | . | . | . | . | . | . |
| chr14 | 63150300 | 63150667 | 422 | Intron (ENSMUST00000054963.9 | 14 | 63145153 | 63177793 | 32641 | 2 | ENSMUST00 | 27126 | Fdft1 | ENSMUSG0C | false | . | . | . | . | . | . | chr14 | 63148622 | 63154226 | 5.17179 | 4.06034 | 3.1827 |
| chr17 | 67958961 | 67959219 | 422 | Intron (ENSMUST00000024840.6 | 17 | 67845110 | 67950940 | 105831 | 2 | ENSMUST00 | -8022 | Arhgap28 | ENSMUSG0C | true | . | . | . | . | . | . | chr17 | 67956253 | 67965917 | 5.0582 | 4.06136 | 3.07428 |
| chr17 | 25541762 | 25542069 | 422 | Distal Intergenic | 17 | 25567988 | 25570214 | 2227 | 2 | ENSMUST00 | 28145 | Sox8 | ENSMUSG0C | false | . | . | . | . | . | . | chr17 | 25536949 | 25544974 | 5.26677 | 4.20031 | 3.22812 |
| chr4 | 99970110 | 99970453 | 422 | Intron (ENSMUST00000058351.1 | 4 | 99955744 | 99987294 | 31551 | 1 | ENSMUST00 | 14367 | Pgm2 | ENSMUSG0C | false | . | . | . | . | . | . | chr4 | 99968725 | 99977020 | 4.83736 | 3.90531 | 2.88809 |
| chr9 | 14380900 | 14381281 | 422 | Promoter (<=1kb) | 9 | 14353990 | 14381242 | 27253 | 2 | ENSMUST00 | 0 | Endod1 | ENSMUSG0C | false | chr9 | 14380027 | 14381955 | 28.91247 | 96.86193 | 93.36482 | . | . | . | . | . | . |
| chr4 | 14565474 | 14565712 | 422 | Exon (ENSMUST00000042221.8/ | 4 | 14505289 | 14516369 | 11081 | 2 | ENSMUST00 | -49106 | Slc26a7 | ENSMUSG0C | false | . | . | . | . | . | . | chr4 | 14561835 | 14566059 | 4.65819 | 3.96026 | 2.71911 |
| chr12 | 79006822 | 79007485 | 422 | Promoter (<=1kb) | 12 | 78961800 | 79007277 | 45478 | 2 | ENSMUST00 | 0 | Tmem229b | ENSMUSG0C | false | chr12 | 79006402 | 79007629 | 41.35789 | 132.3362 | 127.75856 | chr12 | 79006258 | 79007629 | 5.60481 | 4.36556 | 3.50952 |
| chr3 | 51809888 | 51810135 | 422 | Intron (ENSMUST00000121440.2 | 3 | 51689437 | 51796535 | 107099 | 2 | ENSMUST00 | -13354 | Maml3 | ENSMUSG0C | true | . | . | . | . | . | . | chr3 | 51808592 | 51815795 | 4.43589 | 3.73987 | 2.55618 |
| chr17 | 30468475 | 30468684 | 422 | Intron (ENSMUST00000079924.5 | 17 | 30215524 | 30576287 | 360764 | 2 | ENSMUST00 | 107603 | Btbd9 | ENSMUSG0C | false | . | . | . | . | . | . | chr17 | 30466527 | 30472388 | 5.79007 | 4.41639 | 3.69487 |
| chr9 | 28387007 | 28387238 | 422 | Intron (ENSMUST00000115243.3 | 9 | 28403149 | 28405951 | 2803 | 1 | ENSMUST00 | -15911 | Opcml | ENSMUSG0C | true | . | . | . | . | . | . | chr9 | 28384164 | 28387211 | 4.05454 | 3.58275 | 2.27598 |
| chr8 | 107544810 | 107545114 | 422 | Intron (ENSMUST00000166615.1 | 8 | 107551244 | 107551313 | 70 | 1 | ENSMUST00 | -6130 | Mir140 | ENSMUSG0C | false | . | . | . | . | . | . | chr8 | 107544625 | 107546012 | 5.33493 | 3.9693 | 3.32166 |
| chr12 | 86918587 | 86918854 | 422 | Distal Intergenic | 12 | 86891404 | 86892567 | 1164 | 2 | ENSMUST00 | -26021 | Gm26698 | ENSMUSG0C | false | . | . | . | . | . | . | chr12 | 86917928 | 86919690 | 5.60818 | 4.23622 | 3.53776 |
| chr11 | 69123474 | 69123713 | 422 | Promoter (<=1kb) | 11 | 69120054 | 69122589 | 2536 | 2 | ENSMUST00 | -886 | 9130213A22I | ENSMUSG0C | false | chr11 | 69121276 | 69124643 | 27.76267 | 85.10065 | 81.85645 | . | . | . | . | . | . |
| chr9 | 32556347 | 32556644 | 421 | Distal Intergenic | 9 | 32428059 | 32542861 | 114803 | 2 | ENSMUST00 | -13487 | Fli1 | ENSMUSG0C | false | . | . | . | . | . | . | . | . | . | . | . | . |
| chr10 | 94311766 | 94312226 | 421 | Promoter (<=1kb) | 10 | 94311949 | 94430897 | 118949 | 1 | ENSMUST00 | 0 | Tmcc3 | ENSMUSG0C | false | chr10 | 94311546 | 94312854 | 19.72513 | 62.50151 | 59.62064 | chr10 | 94311462 | 94312566 | 3.69491 | 3.41404 | 1.95827 |
| chr12 | 36042190 | 36042677 | 421 | Promoter (<=1kb) | 12 | 36014557 | 36042489 | 27933 | 2 | ENSMUST00 | 0 | Tspan13 | ENSMUSG0C | false | chr12 | 36041013 | 36042881 | 27.7554 | 114.54269 | 110.5744 | . | . | . | . | . | . |
| chr11 | 79534593 | 79534902 | 421 | Promoter (2-3kb) | 11 | 79536908 | 79570115 | 33208 | 1 | ENSMUST00 | -2006 | Nf1 | ENSMUSG0C | false | . | . | . | . | . | . | chr11 | 79533213 | 79536422 | 5.55159 | 4.38429 | 3.45736 |
| chr13 | 51842808 | 51843086 | 421 | Distal Intergenic | 13 | 51846678 | 51848468 | 1791 | 1 | ENSMUST00 | -3592 | Gadd45g | ENSMUSG0C | false | . | . | . | . | . | . | . | . | . | . | . | . |
| chr5 | 49275463 | 49275666 | 421 | Intron (ENSMUST00000087395.5 | 5 | 48389503 | 49285659 | 896157 | 2 | ENSMUST00 | 9993 | Kcnip4 | ENSMUSG0C | false | . | . | . | . | . | . | chr5 | 49275208 | 49276378 | 2.88516 | 2.90177 | 1.33358 |
| chr18 | 61882075 | 61882415 | 421 | Intron (ENSMUST00000049378.9 | 18 | 61799395 | 61911824 | 112430 | 2 | ENSMUST00 | 29409 | Ablim3 | ENSMUSG0C | false | . | . | . | . | . | . | chr18 | 61877692 | 61898809 | 5.45644 | 4.15279 | 3.39554 |
| chr13 | 81656352 | 81656659 | 421 | Promoter (1-2kb) | 13 | 81657806 | 81671888 | 14083 | 1 | ENSMUST00 | -1147 | Lysmd3 | ENSMUSG0C | false | chr13 | 81656629 | 81659028 | 28.59262 | 111.22587 | 107.35971 | chr13 | 81655875 | 81656995 | 3.62911 | 3.42272 | 1.89308 |
| chr6 | 42692963 | 42693199 | 421 | Promoter (<=1kb) | 6 | 42669942 | 42693087 | 23146 | 2 | ENSMUST00 | 0 | Fam115a | ENSMUSG0C | false | chr6 | 42691997 | 42693318 | 34.52358 | 106.75686 | 103.01088 | . | . | . | . | . | . |
| chr3 | 121646645 | 121646971 | 421 | Distal Intergenic | 3 | 121637145 | 121643485 | 6341 | 2 | ENSMUST00 | -3161 | A730020M07 | ENSMUSG0C | false | . | . | . | . | . | . | chr3 | 121645639 | 121648167 | 4.58042 | 3.87179 | 2.65737 |
| chr1 | 140430571 | 140430869 | 421 | Intron (ENSMUST00000119786.2 | 1 | 140443101 | 140523218 | 80118 | 1 | ENSMUST00 | -12232 | Kcnt2 | ENSMUSG0C | false | . | . | . | . | . | . | . | . | . | . | . | . |
| chr2 | 65334027 | 65334363 | 421 | Intron (ENSMUST00000127623.2 | 2 | 65316430 | 65361401 | 44972 | 2 | ENSMUST00 | 27038 | Slc38a11 | ENSMUSG0C | false | . | . | . | . | . | . | . | . | . | . | . | . |
| chrX | 142499216 | 142499423 | 421 | Distal Intergenic | 20 | 142485322 | 142485447 | 126 | 2 | ENSMUST00 | -13770 | Gm25915 | ENSMUSG0C | false | . | . | . | . | . | . | . | . | . | . | . | . |
| chr5 | 126703851 | 126704288 | 421 | Distal Intergenic | 5 | 126513875 | 126513981 | 107 | 2 | ENSMUST00 | -189871 | Gm24839 | ENSMUSG0C | false | . | . | . | . | . | . | chr5 | 126703207 | 126705284 | 4.0261 | 3.49806 | 2.24342 |
| chr16 | 59600544 | 59601230 | 421 | Promoter (<=1kb) | 16 | 59558013 | 59600886 | 42874 | 2 | ENSMUST00 | 0 | Gm9581 | ENSMUSG0C | false | chr16 | 59600114 | 59601432 | 37.22259 | 125.13314 | 120.81451 | chr16 | 59601121 | 59601563 | 5.88195 | 4.59994 | 3.71819 |
| chr13 | 101465425 | 101465658 | 421 | Distal Intergenic | 13 | 101397670 | 101464160 | 66491 | 1 | ENSMUST00 | 67756 | RP23-170A1S | ENSMUSG0C | false | . | . | . | . | . | . | chr13 | 101464541 | 101465944 | 3.16032 | 3.0557 | 1.53593 |
| chr8 | 125534541 | 125534782 | 420 | Distal Intergenic | 8 | 125418063 | 125492710 | 74648 | 2 | ENSMUST00 | -41832 | Sipa1l2 | ENSMUSG0C | false | . | . | . | . | . | . | chr8 | 125526488 | 125537403 | 5.56127 | 4.26582 | 3.48444 |
| chr6 | 54992741 | 54993104 | 420 | Promoter (<=1kb) | 6 | 54982580 | 54992950 | 10371 | 2 | ENSMUST00 | 0 | Ggct | ENSMUSG0C | false | chr6 | 54992306 | 54993434 | 19.43785 | 59.45512 | 56.6181 | chr6 | 54991477 | 54993579 | 3.90846 | 3.54048 | 2.10768 |
| chr11 | 102406969 | 102407262 | 420 | Promoter (<=1kb) | 11 | 102404609 | 102406955 | 2347 | 2 | ENSMUST00 | -15 | Slc25a39 | ENSMUSG0C | false | chr11 | 102406444 | 102408601 | 32.64849 | 123.19247 | 118.94151 | . | . | . | . | . | . |
| chr10 | 24595203 | 24595930 | 420 | Promoter (<=1kb) | 10 | 24595442 | 24598678 | 3237 | 1 | ENSMUST00 | 0 | Ctgf | ENSMUSG0C | false | chr10 | 24594963 | 24597717 | 25.16654 | 87.04672 | 83.76057 | . | . | . | . | . | . |
| chr6 | 119431985 | 119432369 | 420 | Downstream (<1kb) | 6 | 119353150 | 119417704 | 64555 | 2 | ENSMUST00 | -14282 | Adipor2 | ENSMUSG0C | false | . | . | . | . | . | . | chr6 | 119429592 | 119434115 | 5.87795 | 4.50016 | 3.75512 |
| chr1 | 157472804 | 157473061 | 420 | Intron (ENSMUST00000046743.5 | 1 | 157458580 | 157489060 | 30481 | 1 | ENSMUST00 | 14225 | BC026585 | ENSMUSG0C | false | . | . | . | . | . | . | chr1 | 157470354 | 157473935 | 4.0188 | 3.64281 | 2.19531 |
| chr17 | 88611178 | 88611480 | 420 | Intron (ENSMUST00000137138.1 | 17 | 88597696 | 88645322 | 47627 | 1 | ENSMUST00 | 13483 | Ston1 | ENSMUSG0C | false | . | . | . | . | . | . | chr17 | 88609322 | 88613984 | 3.90327 | 3.48747 | 2.13151 |
| chr4 | 43390962 | 43391252 | 420 | Intron (ENSMUST00000035645.6 | 4 | 43384332 | 43414893 | 30562 | 1 | ENSMUST00 | 6631 | Rusc2 | ENSMUSG0C | false | . | . | . | . | . | . | chr4 | 43382882 | 43406168 | 6.80327 | 4.94439 | 4.48074 |
| chr8 | 87738068 | 87738343 | 420 | Intron (ENSMUST00000165770.3 | 8 | 87783147 | 87803823 | 20677 | 2 | ENSMUST00 | 65480 | Zfp423 | ENSMUSG0C | true | . | . | . | . | . | . | chr8 | 87729485 | 87749127 | 6.31805 | 4.63416 | 4.1004 |
| chr14 | 60251276 | 60251649 | 420 | Promoter (<=1kb) | 14 | 60219204 | 60251507 | 32304 | 2 | ENSMUST00 | 0 | Nupl1 | ENSMUSG0C | false | chr14 | 60250596 | 60252614 | 36.39542 | 121.53264 | 117.33772 | . | . | . | . | . | . |
| chr15 | 36374879 | 36375182 | 420 | Distal Intergenic | 15 | 36308144 | 36308956 | 813 | 2 | ENSMUST00 | -65924 | Rpl7a-ps3 | ENSMUSG0C | false | chr15 | 36374035 | 36375538 | 16.14863 | 43.24384 | 40.61377 | chr15 | 36371382 | 36376750 | 5.74131 | 4.46347 | 3.60718 |
| chr17 | 47909675 | 47910035 | 420 | Intron (ENSMUST00000113265.2 | 17 | 47922378 | 47923429 | 1052 | 1 | ENSMUST00 | -12343 | Gm15556 | ENSMUSG0C | false | chr17 | 47909208 | 47910013 | 20.60796 | 54.90754 | 52.13 | chr17 | 47909016 | 47910501 | 2.9815 | 3.00307 | 1.40039 |
| chr18 | 33659583 | 33659973 | 420 | Distal Intergenic | 18 | 33794892 | 33795986 | 1095 | 1 | ENSMUST00 | -134919 | 2410004N09 | ENSMUSG0C | false | . | . | . | . | . | . | chr18 | 33659377 | 33660366 | 3.35125 | 3.20199 | 1.67949 |
| chr1 | 165589307 | 165589726 | 420 | Downstream (2-3kb) | 1 | 165571850 | 165571976 | 127 | 2 | ENSMUST00 | -17332 | Gm23208 | ENSMUSG0C | false | . | . | . | . | . | . | chr1 | 165587115 | 165592756 | 7.22848 | 5.11851 | 4.82653 |
| chr19 | 47292442 | 47292739 | 420 | Intron (ENSMUST00000081619.3 | 19 | 47273519 | 47273589 | 71 | 2 | ENSMUST00 | -18854 | Mir6995 | ENSMUSG0C | false | . | . | . | . | . | . | chr19 | 47286575 | 47295351 | 6.82895 | 4.98258 | 4.49361 |
| chr2 | 74687354 | 74687910 | 420 | Intron (ENSMUST00000136302.2 | 2 | 74683446 | 74694194 | 10749 | 1 | ENSMUST00 | 3909 | RP23-313J15 | ENSMUSG0C | false | . | . | . | . | . | . | chr2 | 74686918 | 74693419 | 4.34438 | 3.76944 | 2.4861 |
| chr11 | 74837657 | 74837995 | 419 | Promoter (<=1kb) | 11 | 74837468 | 74843775 | 6308 | 1 | ENSMUST00 | 190 | Mnt | ENSMUSG0C | false | chr11 | 74837494 | 74839155 | 33.08631 | 99.21803 | 95.66874 | chr11 | 74832552 | 74837711 | 5.19248 | 3.94315 | 3.20824 |
| chr6 | 29215839 | 29216209 | 419 | Promoter (<=1kb) | 6 | 29206995 | 29215929 | 8935 | 2 | ENSMUST00 | 0 | Impdh1 | ENSMUSG0C | false | chr6 | 29215492 | 29216667 | 30.69271 | 93.10602 | 89.69717 | chr6 | 29215183 | 29216556 | 2.87041 | 2.92456 | 1.30508 |
| chr14 | 52257665 | 52258060 | 419 | Distal Intergenic | 14 | 52261769 | 52269106 | 7338 | 2 | ENSMUST00 | 11046 | Rab2b | ENSMUSG0C | false | chr14 | 52256664 | 52258350 | 36.90527 | 118.17374 | 114.08985 | . | . | . | . | . | . |
| chr1 | 189752785 | 189753038 | 419 | Intron (ENSMUST00000097442.3 | 1 | 189728268 | 189876689 | 148422 | 1 | ENSMUST00 | 24518 | Ptpn14 | ENSMUSG0C | true | . | . | . | . | . | . | chr1 | 189730267 | 189810107 | 6.93269 | 4.87829 | 4.60521 |
| chr2 | 58357691 | 58358102 | 419 | Promoter (<=1kb) | 2 | 58267453 | 58357895 | 90443 | 2 | ENSMUST00 | 0 | Acvr1c | ENSMUSG0C | false | chr2 | 58356830 | 58358149 | 19.30035 | 48.68736 | 45.98855 | chr2 | 58356682 | 58358039 | 4.37234 | 3.5845 | 2.51621 |
| chr5 | 52833864 | 52834254 | 419 | Promoter (<=1kb) | 5 | 52834015 | 52867797 | 33783 | 1 | ENSMUST00 | 0 | Anapc4 | ENSMUSG0C | false | chr5 | 52833457 | 52835017 | 35.31382 | 118.55977 | 114.46947 | . | . | . | . | . | . |
| chr5 | 121545397 | 121546072 | 419 | Promoter (<=1kb) | 5 | 121518613 | 121545482 | 26870 | 2 | ENSMUST00 | 0 | Mapkapk5 | ENSMUSG0C | false | chr5 | 121544415 | 121546654 | 32.29185 | 112.09676 | 108.19604 | . | . | . | . | . | . |
| chr9 | 41156742 | 41157013 | 419 | Promoter (<=1kb) | 9 | 41050554 | 41157502 | 106949 | 2 | ENSMUST00 | 489 | Ubash3b | ENSMUSG0C | false | chr9 | 41156374 | 41158065 | 40.77746 | 134.62177 | 129.96082 | chr9 | 41156097 | 41159183 | 4.64566 | 3.82333 | 2.7567 |
| chr9 | 109968833 | 109969014 | 419 | Intron (ENSMUST00000035055.9 | 9 | 109931864 | 110083947 | 152084 | 1 | ENSMUST00 | 36970 | Map4 | ENSMUSG0C | false | . | . | . | . | . | . | chr9 | 109956210 | 109970265 | 4.89158 | 4.02557 | 2.91459 |
| chr3 | 51752923 | 51753212 | 419 | Intron (ENSMUST00000121440.2 | 3 | 51689437 | 51796535 | 107099 | 2 | ENSMUST00 | 43323 | Maml3 | ENSMUSG0C | true | . | . | . | . | . | . | chr3 | 51741455 | 51754138 | 5.51724 | 4.2147 | 3.45633 |
| chr5 | 149284343 | 149284695 | 419 | Intron (ENSMUST00000071130.3 | 5 | 149281118 | 149281223 | 106 | 2 | ENSMUST00 | -3121 | n-R5s180 | ENSMUSG0C | false | . | . | . | . | . | . | chr5 | 149284094 | 149284798 | 3.51273 | 3.20126 | 1.81739 |
| chrX | 94797488 | 94798318 | 419 | Promoter (<=1kb) | 20 | 94795563 | 94798183 | 2621 | 2 | ENSMUST00 | 0 | Zxda | ENSMUSG0C | false | chrX | 94796903 | 94798710 | 37.68163 | 117.40437 | 113.3432 | . | . | . | . | . | . |
| chr2 | 153494342 | 153494648 | 419 | Promoter (1-2kb) | 2 | 153492790 | 153493841 | 1052 | 1 | ENSMUST00 | 1553 | 4930404H24I | ENSMUSG0C | false | . | . | . | . | . | . | chr2 | 153491609 | 153495011 | 7.33518 | 5.22922 | 4.90791 |
| chr7 | 84749848 | 84750108 | 419 | Intron (ENSMUST00000129059.1 | 7 | 84775703 | 84776020 | 318 | 1 | ENSMUST00 | -25595 | 2610206C17I | ENSMUSG0C | false | chr7 | 84749699 | 84750238 | 12.58308 | 29.40204 | 26.94344 | . | . | . | . | . | . |
| chr2 | 172851849 | 172852140 | 419 | Distal Intergenic | 2 | 172864264 | 172864341 | 78 | 2 | ENSMUST00 | 12201 | Gm22773 | ENSMUSG0C | false | . | . | . | . | . | . | chr2 | 172851923 | 172852419 | 5.07363 | 4.19217 | 3.05546 |
| chr6 | 129232022 | 129232293 | 419 | Promoter (1-2kb) | 6 | 129213381 | 129233974 | 20594 | 2 | ENSMUST00 | 1681 | 2310001H17I | ENSMUSG0C | false | . | . | . | . | . | . | . | . | . | . | . | . |
| chr10 | 61715115 | 61715415 | 418 | Promoter (<=1kb) | 10 | 61715263 | 61738653 | 23391 | 1 | ENSMUST00 | 0 | Aifm2 | ENSMUSG0C | false | chr10 | 61715082 | 61715770 | 27.23813 | 78.54684 | 75.42068 | chr10 | 61714548 | 61716252 | 3.4766 | 3.2868 | 1.80874 |
| chr11 | 49914763 | 49914993 | 418 | Intron (ENSMUST00000093141.6 | 11 | 49902161 | 49980994 | 78834 | 1 | ENSMUST00 | 12603 | Rasgef1c | ENSMUSG0C | false | . | . | . | . | . | . | chr11 | 49914299 | 49915301 | 3.74443 | 3.27685 | 2.00997 |
| chr17 | 71178191 | 71178462 | 418 | Intron (ENSMUST00000180743.1 | 17 | 71182560 | 71235119 | 52560 | 1 | ENSMUST00 | -4098 | Lpin2 | ENSMUSG0C | false | . | . | . | . | . | . | chr17 | 71157703 | 71180230 | 5.23179 | 4.17367 | 3.19377 |
| chr8 | 14888381 | 14888992 | 418 | Promoter (<=1kb) | 8 | 14889004 | 14894786 | 5783 | 1 | ENSMUST00 | -12 | Cln8 | ENSMUSG0C | false | chr8 | 14888012 | 14890072 | 30.5701 | 92.89357 | 89.48846 | chr8 | 14887648 | 14891733 | 4.52554 | 3.80501 | 2.67671 |
| chr2 | 91683032 | 91683314 | 418 | Promoter (<=1kb) | 2 | 91676318 | 91682229 | 5912 | 2 | ENSMUST00 | -804 | Atg13 | ENSMUSG0C | false | . | . | . | . | . | . | chr2 | 91681016 | 91689702 | 5.23827 | 4.0615 | 3.22623 |
| chr3 | 136611746 | 136611969 | 418 | Distal Intergenic | 3 | 136670124 | 136797838 | 127715 | 1 | ENSMUST00 | -58155 | Ppp3ca | ENSMUSG0C | false | . | . | . | . | . | . | . | . | . | . | . | . |
| chr8 | 119558371 | 119558963 | 418 | Promoter (<=1kb) | 8 | 119508156 | 119558761 | 50606 | 2 | ENSMUST00 | 0 | Mbtps1 | ENSMUSG0C | false | chr8 | 119557720 | 119559274 | 30.30193 | 90.47713 | 87.1233 | . | . | . | . | . | . |
| chr1 | 155812748 | 155813258 | 418 | Promoter (<=1kb) | 1 | 155778155 | 155812883 | 34729 | 2 | ENSMUST00 | 0 | Qsox1 | ENSMUSG0C | false | chr1 | 155812138 | 155813540 | 32.69058 | 104.59052 | 100.90593 | chr1 | 155813199 | 155813742 | 3.15678 | 3.13527 | 1.54682 |
| chr9 | 57939959 | 57940291 | 418 | Promoter (<=1kb) | 9 | 57940113 | 57962864 | 22752 | 1 | ENSMUST00 | 0 | Sema7a | ENSMUSG0C | false | chr9 | 57939633 | 57941400 | 28.1615 | 93.53628 | 90.11485 | chr9 | 57939368 | 57941825 | 4.58022 | 3.7506 | 2.69821 |
| chr7 | 114103688 | 114103978 | 418 | Intron (ENSMUST00000069449.5 | 7 | 114046785 | 114117781 | 70997 | 2 | ENSMUST00 | 13803 | Rras2 | ENSMUSG0C | true | chr7 | 114103949 | 114104284 | 8.80466 | 16.8248 | 14.53583 | chr7 | 114058507 | 114116129 | 5.64505 | 4.33683 | 3.54844 |
| chr2 | 32720858 | 32721191 | 418 | Promoter (<=1kb) | 2 | 32721055 | 32754994 | 33940 | 1 | ENSMUST00 | 0 | Sh2d3c | ENSMUSG0C | false | . | . | . | . | . | . | . | . | . | . | . | . |
| chr18 | 44668496 | 44668738 | 418 | Intron (ENSMUST00000181094.1 | 18 | 44425061 | 44662266 | 237206 | 2 | ENSMUST00 | -6231 | Mcc | ENSMUSG0C | true | . | . | . | . | . | . | chr18 | 44667458 | 44669378 | 5.52794 | 4.29309 | 3.46372 |
| chr14 | 58525159 | 58525342 | 418 | Distal Intergenic | 14 | 58650268 | 58650400 | 133 | 2 | ENSMUST00 | 125058 | Gm25614 | ENSMUSG0C | false | . | . | . | . | . | . | . | . | . | . | . | . |
| chr14 | 46388298 | 46388956 | 418 | Promoter (<=1kb) | 14 | 46387519 | 46389282 | 1764 | 1 | ENSMUST00 | 780 | Gm15222 | ENSMUSG0C | false | chr14 | 46386588 | 46388392 | 26.19333 | 73.09374 | 70.054 | chr14 | 46385786 | 46390843 | 3.99631 | 3.58417 | 2.19112 |
| chr7 | 98983693 | 98983886 | 418 | Intron (ENSMUST00000037968.8 | 7 | 98935894 | 98936015 | 122 | 1 | ENSMUST00 | 47800 | Gm23479 | ENSMUSG0C | false | . | . | . | . | . | . | chr7 | 98983147 | 98986492 | 5.33669 | 4.24145 | 3.30537 |
| chr11 | 22858854 | 22859225 | 418 | Promoter (<=1kb) | 11 | 22858931 | 22861264 | 2334 | 1 | ENSMUST00 | 0 | 9130230N09 | ENSMUSG0C | false | chr11 | 22858157 | 22860897 | 30.41442 | 103.5799 | 99.9162 | chr11 | 22858236 | 22858898 | 5.99233 | 4.59791 | 3.81974 |
| chr7 | 110703215 | 110703488 | 418 | Distal Intergenic | 7 | 110626281 | 110639358 | 13078 | 2 | ENSMUST00 | -63858 | RP23-25I3.4 | ENSMUSG0C | false | . | . | . | . | . | . | . | . | . | . | . | . |
| chr3 | 100706659 | 100707435 | 417 | Distal Intergenic | 3 | 100562208 | 100685503 | 123296 | 2 | ENSMUST00 | -21157 | Man1a2 | ENSMUSG0C | true | chr3 | 100705348 | 100707194 | 6.80177 | 13.42905 | 11.19262 | chr3 | 100706343 | 100713148 | 4.75051 | 3.96149 | 2.7997 |
| chr11 | 116004938 | 116005225 | 417 | Exon (ENSMUST00000106461.2/ | 11 | 116008457 | 116012719 | 4263 | 2 | ENSMUST00 | 7494 | Galk1 | ENSMUSG0C | false | chr11 | 116004671 | 116005573 | 10.56684 | 27.64477 | 25.20979 | chr11 | 116004539 | 116005646 | 4.82504 | 3.87045 | 2.87086 |
| chr12 | 111353430 | 111354103 | 417 | Intron (ENSMUST00000041965.3 | 12 | 111292972 | 111377718 | 84747 | 2 | ENSMUST00 | 23615 | Cdc42bpb | ENSMUSG0C | false | . | . | . | . | . | . | chr12 | 111345052 | 111376732 | 6.23386 | 4.58338 | 4.03546 |
| chr13 | 63918010 | 63918468 | 417 | Distal Intergenic | 13 | 63872455 | 63900302 | 27848 | 1 | ENSMUST00 | 45556 | Ercc6l2 | ENSMUSG0C | false | . | . | . | . | . | . | chr13 | 63915196 | 63919976 | 5.59378 | 4.40381 | 3.49158 |
| chr14 | 12198870 | 12199089 | 417 | Intron (ENSMUST00000022264.7 | 14 | 12189943 | 12238563 | 48621 | 1 | ENSMUST00 | 8928 | Ptprg | ENSMUSG0C | false | . | . | . | . | . | . | . | . | . | . | . | . |
| chr16 | 4886139 | 4886360 | 417 | Promoter (<=1kb) | 16 | 4886100 | 4938296 | 52197 | 1 | ENSMUST00 | 40 | Mgrn1 | ENSMUSG0C | false | chr16 | 4886161 | 4886997 | 29.92778 | 109.76675 | 105.9399 | chr16 | 4884949 | 4890671 | 4.43519 | 3.80735 | 2.55461 |
| chr19 | 4148405 | 4148765 | 417 | Promoter (<=1kb) | 19 | 4148619 | 4150504 | 1886 | 1 | ENSMUST00 | 0 | Coro1b | ENSMUSG0C | false | chr19 | 4148113 | 4149802 | 31.91637 | 110.38313 | 106.53261 | chr19 | 4147701 | 4152092 | 7.29509 | 4.99142 | 4.89155 |
| chr4 | 125113545 | 125113774 | 417 | Distal Intergenic | 4 | 125107976 | 125109428 | 1453 | 1 | ENSMUST00 | 5570 | Meaf6 | ENSMUSG0C | false | . | . | . | . | . | . | chr4 | 125106469 | 125127334 | 7.64816 | 5.22391 | 5.18478 |
| chr6 | 86059081 | 86059340 | 417 | Distal Intergenic | 6 | 86078084 | 86119555 | 41472 | 1 | ENSMUST00 | -18744 | Add2 | ENSMUSG0C | false | . | . | . | . | . | . | chr6 | 86057572 | 86060226 | 4.41738 | 3.68017 | 2.55681 |
| chr6 | 137279613 | 137279891 | 417 | Intron (ENSMUST00000077115.7 | 6 | 137252299 | 137464629 | 212331 | 1 | ENSMUST00 | 27315 | Ptpro | ENSMUSG0C | true | . | . | . | . | . | . | chr6 | 137275709 | 137305131 | 5.12249 | 4.15603 | 3.10285 |
| chr18 | 34841346 | 34841569 | 417 | Promoter (<=1kb) | 18 | 34840589 | 34847463 | 6875 | 1 | ENSMUST00 | 758 | Reep2 | ENSMUSG0C | false | . | . | . | . | . | . | chr18 | 34839709 | 34842347 | 5.23259 | 4.05705 | 3.23734 |
| chr5 | 143732169 | 143732662 | 417 | Promoter (<=1kb) | 5 | 143710325 | 143732280 | 21956 | 2 | ENSMUST00 | 0 | Usp42 | ENSMUSG0C | false | chr5 | 143730802 | 143733135 | 30.32912 | 88.589 | 85.27629 | . | . | . | . | . | . |
| chr7 | 29905992 | 29906249 | 417 | Promoter (<=1kb) | 7 | 29893716 | 29906104 | 12389 | 2 | ENSMUST00 | 0 | Zfp27 | ENSMUSG0C | false | chr7 | 29905698 | 29906978 | 20.06521 | 63.98471 | 61.08233 | . | . | . | . | . | . |
| chr1 | 55445737 | 55445942 | 417 | Promoter (<=1kb) | 1 | 55443733 | 55445064 | 1332 | 2 | ENSMUST00 | -674 | Gm6644 | ENSMUSG0C | false | . | . | . | . | . | . | chr1 | 55445058 | 55449958 | 4.55068 | 3.80943 | 2.64653 |
| chr4 | 141274452 | 141274754 | 417 | Distal Intergenic | 4 | 141278439 | 141279794 | 1356 | 1 | ENSMUST00 | -3685 | Gm13056 | ENSMUSG0C | false | . | . | . | . | . | . | chr4 | 141273176 | 141275080 | 3.84347 | 3.44129 | 2.09695 |
| chr2 | 167672187 | 167672414 | 417 | Intron (ENSMUST00000148262.1 | 2 | 167662109 | 167667697 | 5589 | 1 | ENSMUST00 | 10079 | Gm14320 | ENSMUSG0C | false | . | . | . | . | . | . | chr2 | 167671333 | 167673098 | 4.07869 | 3.66935 | 2.24411 |
| chr11 | 23217411 | 23217674 | 416 | Distal Intergenic | 11 | 23256041 | 23297597 | 41557 | 1 | ENSMUST00 | -38367 | Xpo1 | ENSMUSG0C | false | . | . | . | . | . | . | . | . | . | . | . | . |
| chr2 | 45108542 | 45108858 | 416 | Promoter (1-2kb) | 2 | 44988388 | 45110244 | 121857 | 2 | ENSMUST00 | 1386 | Zeb2 | ENSMUSG0C | false | chr2 | 45108545 | 45111853 | 37.2221 | 115.56093 | 111.55734 | . | . | . | . | . | . |
| chr4 | 101144363 | 101144637 | 416 | Intron (ENSMUST00000038463.9 | 4 | 101154249 | 101155119 | 871 | 2 | ENSMUST00 | 10482 | Jak1 | ENSMUSG0C | false | . | . | . | . | . | . | chr4 | 101141547 | 101149273 | 5.88936 | 4.49025 | 3.75587 |
| chr10 | 96615380 | 96615608 | 416 | Promoter (1-2kb) | 10 | 96617001 | 96622813 | 5813 | 1 | ENSMUST00 | -1393 | Btg1 | ENSMUSG0C | false | . | . | . | . | . | . | chr10 | 96614291 | 96616469 | 5.11022 | 4.16996 | 3.0839 |
| chr7 | 89517475 | 89517734 | 416 | Promoter (<=1kb) | 7 | 89507785 | 89517586 | 9802 | 2 | ENSMUST00 | 0 | Prss23 | ENSMUSG0C | false | chr7 | 89517043 | 89518500 | 21.02725 | 63.18829 | 60.29588 | chr7 | 89516785 | 89518441 | 4.16626 | 3.66248 | 2.34081 |
| chr4 | 141098625 | 141098822 | 416 | Intron (ENSMUST00000051907.2 | 4 | 141103985 | 141111546 | 7562 | 1 | ENSMUST00 | -5163 | Spata21 | ENSMUSG0C | false | . | . | . | . | . | . | chr4 | 141096237 | 141098720 | 4.20802 | 3.65724 | 2.37057 |
| chr19 | 58943518 | 58943789 | 416 | Promoter (<=1kb) | 19 | 58943425 | 58971419 | 27995 | 1 | ENSMUST00 | 94 | Eno4 | ENSMUSG0C | false | chr19 | 58942770 | 58944181 | 25.15761 | 80.47005 | 77.31023 | chr19 | 58942687 | 58944311 | 4.24214 | 3.54998 | 2.41079 |
| chr3 | 65442340 | 65442576 | 416 | Distal Intergenic | 3 | 65393337 | 65395184 | 1848 | 1 | ENSMUST00 | 49004 | Gm5540 | ENSMUSG0C | false | . | . | . | . | . | . | chr3 | 65440551 | 65443842 | 4.66122 | 3.82106 | 2.77786 |
| chr8 | 117177068 | 117177582 | 416 | Intron (ENSMUST00000064488.5 | 8 | 117158137 | 117205834 | 47698 | 1 | ENSMUST00 | 18932 | Gan | ENSMUSG0C | false | . | . | . | . | . | . | chr8 | 117172583 | 117179361 | 6.73029 | 4.87581 | 4.42508 |
| chr18 | 48035306 | 48035520 | 416 | Distal Intergenic | 18 | 48045335 | 48048378 | 3044 | 1 | ENSMUST00 | -9815 | Eno1b | ENSMUSG0C | false | . | . | . | . | . | . | chr18 | 48035018 | 48035619 | 3.73428 | 3.56603 | 1.94444 |
| chr7 | 34389406 | 34389873 | 416 | Promoter (<=1kb) | 7 | 34344646 | 34389540 | 44895 | 2 | ENSMUST00 | 0 | Lsm14a | ENSMUSG0C | true | chr7 | 34387343 | 34390805 | 31.16539 | 106.97112 | 103.21663 | . | . | . | . | . | . |
| chr4 | 142097963 | 142098221 | 416 | Distal Intergenic | 4 | 142084856 | 142088077 | 3222 | 1 | ENSMUST00 | 13108 | Tmem51os1 | ENSMUSG0C | false | . | . | . | . | . | . | chr4 | 142097632 | 142098803 | 6.27317 | 4.71722 | 4.0781 |
| chr19 | 16286502 | 16286815 | 416 | Intron (ENSMUST00000025541.5 | 19 | 16277797 | 16278776 | 980 | 1 | ENSMUST00 | 8706 | Gm17068 | ENSMUSG0C | false | . | . | . | . | . | . | chr19 | 16285066 | 16287961 | 3.57171 | 3.24825 | 1.86509 |
| chr4 | 108971022 | 108971315 | 416 | Promoter (<=1kb) | 4 | 108942843 | 108972077 | 29235 | 2 | ENSMUST00 | 762 | 8030443G20 | ENSMUSG0C | false | . | . | . | . | . | . | chr4 | 108969477 | 108974220 | 5.35329 | 4.26775 | 3.30705 |
| chr4 | 149529571 | 149529848 | 416 | Intron (ENSMUST00000030839.7 | 4 | 149537863 | 149537922 | 60 | 1 | ENSMUST00 | -8015 | Mir5616 | ENSMUSG0C | false | . | . | . | . | . | . | chr4 | 149528272 | 149530530 | 5.99824 | 4.47634 | 3.86365 |
| chr12 | 27082748 | 27082945 | 416 | Distal Intergenic | 12 | 27140796 | 27160311 | 19516 | 2 | ENSMUST00 | 77366 | Gm9866 | ENSMUSG0C | false | . | . | . | . | . | . | chr12 | 27082338 | 27083436 | 4.60725 | 3.88171 | 2.70219 |
| chr7 | 110710622 | 110710854 | 415 | Distal Intergenic | 7 | 110768206 | 110811919 | 43714 | 1 | ENSMUST00 | -57352 | Ampd3 | ENSMUSG0C | false | . | . | . | . | . | . | . | . | . | . | . | . |
| chr1 | 43218222 | 43218490 | 415 | Distal Intergenic | 1 | 43123290 | 43196984 | 73695 | 2 | ENSMUST00 | -21239 | Fhl2 | ENSMUSG0C | false | . | . | . | . | . | . | . | . | . | . | . | . |
| chr11 | 23497840 | 23498259 | 415 | Promoter (<=1kb) | 11 | 23487882 | 23497930 | 10049 | 2 | ENSMUST00 | 0 | Ahsa2 | ENSMUSG0C | false | chr11 | 23496813 | 23498021 | 41.10722 | 135.19504 | 130.50221 | . | . | . | . | . | . |
| chr18 | 37835486 | 37835857 | 415 | Intron (ENSMUST00000073447.6 | 18 | 37819546 | 37841873 | 22328 | 1 | ENSMUST00 | 15941 | Pcdhga12 | ENSMUSG0C | false | . | . | . | . | . | . | chr18 | 37835502 | 37836903 | 4.6985 | 3.5825 | 2.80264 |
| chr1 | 152527443 | 152527711 | 415 | Intron (ENSMUST00000111859.2 | 1 | 152551508 | 152553044 | 1537 | 2 | ENSMUST00 | 25333 | Rgl1 | ENSMUSG0C | false | . | . | . | . | . | . | . | . | . | . | . | . |
| chr2 | 32908426 | 32908722 | 415 | Promoter (2-3kb) | 2 | 32910908 | 32912587 | 1680 | 1 | ENSMUST00 | -2186 | Fam129b | ENSMUSG0C | false | . | . | . | . | . | . | chr2 | 32877160 | 32918190 | 7.77045 | 5.19056 | 5.2936 |
| chr9 | 107748696 | 107748949 | 415 | Promoter (<=1kb) | 9 | 107745194 | 107749911 | 4718 | 2 | ENSMUST00 | 962 | Rbm5 | ENSMUSG0C | false | . | . | . | . | . | . | chr9 | 107747010 | 107750847 | 4.08582 | 3.58886 | 2.25749 |
| chr13 | 59971513 | 59971750 | 415 | Distal Intergenic | 13 | 59771879 | 59823147 | 51269 | 2 | ENSMUST00 | -148367 | Zcchc6 | ENSMUSG0C | false | chr13 | 59971201 | 59972007 | 6.75876 | 12.51203 | 10.2897 | chr13 | 59971138 | 59972283 | 4.44171 | 3.79562 | 2.58648 |
| chrX | 53269462 | 53269921 | 415 | Promoter (<=1kb) | 20 | 53243415 | 53269805 | 26391 | 2 | ENSMUST00 | 0 | Fam122b | ENSMUSG0C | false | chrX | 53268993 | 53269848 | 23.43614 | 63.09834 | 60.20888 | . | . | . | . | . | . |
| chr1 | 127232619 | 127232855 | 415 | Intron (ENSMUST00000189427.1 | 1 | 127205028 | 127398792 | 193765 | 1 | ENSMUST00 | 27592 | Mgat5 | ENSMUSG0C | true | . | . | . | . | . | . | chr1 | 127226370 | 127236809 | 5.7385 | 4.45909 | 3.61218 |
| chr1 | 166230399 | 166230654 | 415 | Intron (ENSMUST00000038782.3 | 1 | 166201385 | 166238744 | 37360 | 2 | ENSMUST00 | 8090 | Mael | ENSMUSG0C | false | . | . | . | . | . | . | chr1 | 166229168 | 166233313 | 3.32861 | 3.14313 | 1.68225 |

| | | | | | | | | | | | | | | | | | | | | | | | | | | |
|---|---|---|---|---|---|---|---|---|---|---|---|---|---|---|---|---|---|---|---|---|---|---|---|---|---|---|
| chr19 | 40685661 | 40685907 | 415 | Intron (ENSMUST00000134063.2 | 19 | 40681667 | 40723193 | 41527 | 1 | ENSMUST00 | 3995 | Entpd1 | ENSMUSG0C | false | . | . | . | . | . | . | . | . | . | . | . | . |
| chr13 | 107469571 | 107469904 | 415 | Distal Intergenic | 13 | 107413865 | 107414767 | 903 | 2 | ENSMUST00 | -54805 | Apoo-ps | ENSMUSG0C | false | chr13 | 107469727 | 107470454 | 11.44606 | 24.42448 | 22.03097 | chr13 | 107469341 | 107471153 | 3.83119 | 3.32217 | 2.08034 |
| chr12 | 78639988 | 78640350 | 415 | Intron (ENSMUST00000052472.4 | 12 | 78691535 | 78734513 | 42979 | 1 | ENSMUST00 | -51185 | Fam71d | ENSMUSG0C | false | . | . | . | . | . | . | chr12 | 78639371 | 78640893 | 5.66804 | 4.45746 | 3.57153 |
| chr2 | 127482532 | 127482837 | 415 | Promoter (<=1kb) | 2 | 127458130 | 127482499 | 24370 | 2 | ENSMUST00 | -34 | Kcnip3 | ENSMUSG0C | false | chr2 | 127481915 | 127483499 | 22.33355 | 63.77912 | 60.87984 | chr2 | 127481856 | 127484884 | 4.75547 | 3.87461 | 2.84902 |
| chr13 | 112100824 | 112101312 | 415 | Distal Intergenic | 13 | 112005502 | 112017332 | 11831 | 1 | ENSMUST00 | 95323 | Gm15323 | ENSMUSG0C | false | chr13 | 112100438 | 112101481 | 13.89301 | 35.6666 | 33.12951 | . | . | . | . | . | . |
| chr16 | 25058901 | 25059948 | 415 | Promoter (<=1kb) | 16 | 25059639 | 25069058 | 9420 | 1 | ENSMUST00 | 0 | A230028O05 | ENSMUSG0C | false | chr16 | 25058670 | 25059871 | 21.46036 | 80.68572 | 77.52129 | chr16 | 25058468 | 25060207 | 5.90202 | 4.5232 | 3.75314 |
| chr4 | 141310376 | 141310595 | 414 | Promoter (1-2kb) | 4 | 141308894 | 141310958 | 2065 | 1 | ENSMUST00 | 1483 | Epha2 | ENSMUSG0C | false | . | . | . | . | . | . | chr4 | 141302534 | 141316008 | 7.08958 | 4.8653 | 4.73204 |
| chr14 | 25587708 | 25588046 | 414 | Intron (ENSMUST00000162645.2 | 14 | 25607357 | 25643657 | 36301 | 1 | ENSMUST00 | -19311 | Zmiz1 | ENSMUSG0C | false | . | . | . | . | . | . | chr14 | 25585435 | 25590318 | 5.95302 | 4.46379 | 3.80715 |
| chr14 | 70351232 | 70351578 | 414 | Promoter (<=1kb) | 14 | 70303469 | 70351425 | 47957 | 2 | ENSMUST00 | 0 | Slc39a14 | ENSMUSG0C | false | chr14 | 70350670 | 70351850 | 35.38397 | 108.24114 | 104.45254 | chr14 | 70350473 | 70352235 | 4.56468 | 3.82854 | 2.65305 |
| chr15 | 75716502 | 75716768 | 414 | Distal Intergenic | 15 | 75704394 | 75714225 | 9832 | 1 | ENSMUST00 | 12109 | Rhpn1 | ENSMUSG0C | true | . | . | . | . | . | . | chr15 | 75708646 | 75721294 | 7.3449 | 5.11049 | 4.92697 |
| chr15 | 99457747 | 99458057 | 414 | Promoter (<=1kb) | 15 | 99422035 | 99457748 | 35714 | 2 | ENSMUST00 | 0 | Nckap5l | ENSMUSG0C | false | chr15 | 99456565 | 99458437 | 33.29458 | 104.62617 | 100.93848 | chr15 | 99455697 | 99458712 | 3.47458 | 3.25781 | 1.75757 |
| chr3 | 57445717 | 57445973 | 414 | Distal Intergenic | 3 | 57425314 | 57441677 | 16364 | 1 | ENSMUST00 | 20404 | Tm4sf4 | ENSMUSG0C | false | . | . | . | . | . | . | chr3 | 57444897 | 57448591 | 4.68557 | 3.81748 | 2.7892 |
| chr4 | 6190877 | 6191213 | 414 | Promoter (<=1kb) | 4 | 6191098 | 6221688 | 30591 | 1 | ENSMUST00 | 0 | Ubxn2b | ENSMUSG0C | false | chr4 | 6190582 | 6191771 | 44.10828 | 149.36552 | 144.10564 | . | . | . | . | . | . |
| chr6 | 108782907 | 108783210 | 414 | Promoter (<=1kb) | 6 | 108783059 | 108823723 | 40665 | 1 | ENSMUST00 | 0 | Arl8b | ENSMUSG0C | true | chr6 | 108783011 | 108784750 | 48.77102 | 166.70177 | 160.83063 | . | . | . | . | . | . |
| chr15 | 76723773 | 76724046 | 414 | Promoter (<=1kb) | 15 | 76721466 | 76723845 | 2380 | 2 | ENSMUST00 | 0 | Lrrc24 | ENSMUSG0C | false | chr15 | 76722485 | 76725119 | 19.69817 | 53.33889 | 50.58166 | chr15 | 76722042 | 76725994 | 5.97827 | 4.47626 | 3.81427 |
| chr4 | 138972683 | 138973332 | 414 | Promoter (<=1kb) | 4 | 138972888 | 138990720 | 17833 | 1 | ENSMUST00 | 0 | Tmco4 | ENSMUSG0C | false | chr4 | 138972203 | 138973837 | 26.86698 | 95.04005 | 91.5885 | . | . | . | . | . | . |
| chr17 | 84185686 | 84186301 | 414 | Promoter (1-2kb) | 17 | 84183931 | 84187947 | 4017 | 2 | ENSMUST00 | 1646 | Zfp36l2 | ENSMUSG0C | false | chr17 | 84184922 | 84188829 | 24.14628 | 82.32003 | 79.12344 | chr17 | 84185089 | 84187043 | 4.25783 | 3.76851 | 2.38031 |
| chr7 | 145037059 | 145037305 | 414 | Distal Intergenic | 7 | 144929931 | 144939869 | 9939 | 2 | ENSMUST00 | -97191 | Ccnd1 | ENSMUSG0C | false | . | . | . | . | . | . | . | . | . | . | . | . |
| chr16 | 44494577 | 44494851 | 414 | Intron (ENSMUST00000023370.5 | 16 | 44470444 | 44482428 | 11985 | 1 | ENSMUST00 | 24134 | Wdr52 | ENSMUSG0C | false | . | . | . | . | . | . | chr16 | 44492743 | 44496360 | 6.31318 | 4.76529 | 4.0799 |
| chr12 | 25768750 | 25769003 | 414 | Distal Intergenic | 12 | 25687801 | 25687906 | 106 | 1 | ENSMUST00 | 80950 | Gm22527 | ENSMUSG0C | false | . | . | . | . | . | . | chr12 | 25768527 | 25768862 | 3.95234 | 3.49652 | 2.16737 |
| chr13 | 37636423 | 37636645 | 414 | Distal Intergenic | 13 | 37508940 | 37512074 | 3135 | 1 | ENSMUST00 | 127484 | RP23-21N15 | ENSMUSG0C | false | . | . | . | . | . | . | . | . | . | . | . | . |
| chr1 | 47051195 | 47051427 | 414 | Distal Intergenic | 1 | 47067788 | 47068025 | 238 | 1 | ENSMUST00 | -16361 | RP23-69F3.1 | ENSMUSG0C | false | . | . | . | . | . | . | chr1 | 47049093 | 47051266 | 2.9599 | 2.88149 | 1.35513 |
| chr6 | 115658768 | 115659005 | 413 | Intron (ENSMUST00000112949.2 | 6 | 115618569 | 115675964 | 57396 | 2 | ENSMUST00 | 16959 | Raf1 | ENSMUSG0C | false | . | . | . | . | . | . | chr6 | 115655794 | 115662536 | 4.67401 | 3.8093 | 2.77744 |
| chr13 | 58408717 | 58408978 | 413 | Exon (ENSMUST00000042450.8/ | 13 | 58391142 | 58403343 | 12202 | 2 | ENSMUST00 | -5375 | Hnrnpk | ENSMUSG0C | false | . | . | . | . | . | . | . | . | . | . | . | . |
| chr16 | 37665585 | 37665810 | 413 | Distal Intergenic | 16 | 37647170 | 37654453 | 7284 | 2 | ENSMUST00 | -11133 | Ndufb4 | ENSMUSG0C | false | . | . | . | . | . | . | chr16 | 37663595 | 37666617 | 4.61887 | 3.90723 | 2.70222 |
| chr17 | 43030373 | 43030688 | 413 | Intron (ENSMUST00000024708.4 | 17 | 43016555 | 43089188 | 72634 | 1 | ENSMUST00 | 13819 | Tnfrsf21 | ENSMUSG0C | false | . | . | . | . | . | . | chr17 | 43017914 | 43036611 | 4.69484 | 3.87934 | 2.77439 |
| chr17 | 26502962 | 26503233 | 413 | Downstream (2-3kb) | 17 | 26506516 | 26507723 | 1208 | 2 | ENSMUST00 | 4490 | Dusp1 | ENSMUSG0C | false | . | . | . | . | . | . | chr17 | 26502045 | 26505674 | 6.04575 | 4.53758 | 3.87264 |
| chr1 | 165459953 | 165460705 | 413 | Promoter (<=1kb) | 1 | 165329511 | 165460465 | 130955 | 2 | ENSMUST00 | 0 | Dcaf6 | ENSMUSG0C | false | chr1 | 165459002 | 165461051 | 38.74602 | 137.65443 | 132.86864 | . | . | . | . | . | . |
| chr2 | 4504510 | 4504735 | 413 | Intron (ENSMUST00000177457.2 | 2 | 4473194 | 4498103 | 24910 | 1 | ENSMUST00 | 31317 | Frmd4a | ENSMUSG0C | false | . | . | . | . | . | . | chr2 | 4503811 | 4504934 | 3.6561 | 3.46737 | 1.90146 |
| chr17 | 49562529 | 49562915 | 413 | Promoter (1-2kb) | 17 | 49456022 | 49564337 | 108316 | 2 | ENSMUST00 | 1422 | Daam2 | ENSMUSG0C | true | chr17 | 49562125 | 49564376 | 19.02488 | 51.36755 | 48.63442 | chr17 | 49553664 | 49563281 | 6.14765 | 4.61155 | 3.94797 |
| chr14 | 51229851 | 51230120 | 413 | Distal Intergenic | 14 | 51255266 | 51256112 | 847 | 2 | ENSMUST00 | 25992 | Rnase2a | ENSMUSG0C | false | . | . | . | . | . | . | chr14 | 51229582 | 51230070 | 2.97111 | 2.95732 | 1.35513 |
| chr1 | 54830331 | 54830573 | 413 | Intron (ENSMUST00000179030.2 | 1 | 54762368 | 54813443 | 51076 | 2 | ENSMUST00 | -16889 | Ankrd44 | ENSMUSG0C | true | . | . | . | . | . | . | chr1 | 54821851 | 54833551 | 6.37475 | 4.70533 | 4.12466 |
| chr8 | 91360615 | 91360872 | 413 | Intron (ENSMUST00000069718.8 | 8 | 91387736 | 91409840 | 22105 | 1 | ENSMUST00 | -26864 | Fto | ENSMUSG0C | false | . | . | . | . | . | . | chr8 | 91337651 | 91367410 | 5.62588 | 4.32828 | 3.53466 |
| chr9 | 20581327 | 20581591 | 413 | Promoter (<=1kb) | 9 | 20581525 | 20595194 | 13670 | 1 | ENSMUST00 | 0 | Zfp846 | ENSMUSG0C | false | chr9 | 20581018 | 20582070 | 18.31284 | 64.75944 | 61.84419 | chr9 | 20581135 | 20582542 | 3.8646 | 3.57584 | 2.05644 |
| chr16 | 34347589 | 34347952 | 413 | Intron (ENSMUST00000076810.6 | 16 | 34357268 | 34361799 | 4532 | 2 | ENSMUST00 | 13847 | Kalrn | ENSMUSG0C | false | . | . | . | . | . | . | . | . | . | . | . | . |
| chr10 | 128832713 | 128832969 | 413 | Intron (ENSMUST00000105230.2 | 10 | 128821771 | 128877638 | 55868 | 1 | ENSMUST00 | 10943 | Sarnp | ENSMUSG0C | false | . | . | . | . | . | . | chr10 | 128829105 | 128833258 | 3.22119 | 3.13416 | 1.5711 |
| chr11 | 119565756 | 119566004 | 413 | Intron (ENSMUST00000135630.1 | 11 | 119548630 | 119569046 | 20417 | 1 | ENSMUST00 | 17127 | Gm11762 | ENSMUSG0C | false | . | . | . | . | . | . | chr11 | 119565829 | 119566057 | 4.33921 | 3.84959 | 2.49332 |
| chr8 | 104395640 | 104396019 | 413 | Promoter (<=1kb) | 8 | 104348191 | 104395807 | 47617 | 2 | ENSMUST00 | 0 | Cmtm4 | ENSMUSG0C | false | chr8 | 104394767 | 104396288 | 33.10969 | 106.34647 | 102.61562 | . | . | . | . | . | . |
| chr15 | 38420770 | 38421034 | 413 | Distal Intergenic | 15 | 38384981 | 38386300 | 1320 | 1 | ENSMUST00 | 35790 | Gm26863 | ENSMUSG0C | false | . | . | . | . | . | . | chr15 | 38419041 | 38421526 | 6.47817 | 4.55053 | 4.25073 |
| chr16 | 18388817 | 18389139 | 412 | Intron (ENSMUST00000115614.3 | 16 | 18392588 | 18407076 | 14489 | 1 | ENSMUST00 | -3449 | Arvcf | ENSMUSG0C | true | . | . | . | . | . | . | chr16 | 18376326 | 18392535 | 8.53722 | 5.66709 | 5.90216 |
| chr7 | 111413859 | 111414126 | 412 | Distal Intergenic | 7 | 111070568 | 111083030 | 12463 | 2 | ENSMUST00 | -330830 | Eif4g2 | ENSMUSG0C | false | . | . | . | . | . | . | . | . | . | . | . | . |
| chr9 | 108597479 | 108597733 | 412 | Promoter (<=1kb) | 9 | 108578826 | 108597600 | 18775 | 2 | ENSMUST00 | 0 | P4htm | ENSMUSG0C | false | chr9 | 108596863 | 108598133 | 28.53732 | 88.37692 | 85.06884 | chr9 | 108596599 | 108602820 | 4.71757 | 3.8751 | 2.78944 |
| chr10 | 84487691 | 84487964 | 412 | Promoter (<=1kb) | 10 | 84488293 | 84497435 | 9143 | 1 | ENSMUST00 | -329 | 4930463O16 | ENSMUSG0C | false | . | . | . | . | . | . | chr10 | 84485926 | 84487783 | 3.10208 | 3.05326 | 1.43527 |
| chr4 | 81442199 | 81442677 | 412 | Promoter (<=1kb) | 4 | 81278500 | 81442805 | 164306 | 2 | ENSMUST00 | 128 | Mpdz | ENSMUSG0C | false | chr4 | 81442015 | 81443104 | 25.56571 | 75.64988 | 72.57267 | chr4 | 81442051 | 81442578 | 3.52642 | 3.31709 | 1.85058 |
| chr7 | 126502265 | 126502539 | 412 | Promoter (<=1kb) | 7 | 126491715 | 126502414 | 10700 | 2 | ENSMUST00 | 0 | Atxn2l | ENSMUSG0C | false | chr7 | 126501677 | 126504111 | 42.82515 | 147.12178 | 141.95673 | . | . | . | . | . | . |
| chr10 | 79914919 | 79915203 | 412 | Promoter (1-2kb) | 10 | 79910053 | 79916930 | 6878 | 2 | ENSMUST00 | 1727 | R3hdm4 | ENSMUSG0C | false | . | . | . | . | . | . | chr10 | 79909270 | 79922043 | 5.7269 | 4.35016 | 3.60803 |
| chr5 | 29568973 | 29569535 | 412 | Promoter (<=1kb) | 5 | 29569242 | 29676077 | 106836 | 1 | ENSMUST00 | 0 | Ube3c | ENSMUSG0C | false | chr5 | 29568783 | 29569055 | 9.74174 | 20.4319 | 18.09061 | . | . | . | . | . | . |
| chr4 | 35112207 | 35112432 | 412 | Intron (ENSMUST00000102975.4 | 4 | 35152056 | 35154005 | 1950 | 1 | ENSMUST00 | -39624 | Ifnk | ENSMUSG0C | false | . | . | . | . | . | . | chr4 | 35112247 | 35112515 | 3.52343 | 3.35314 | 1.82935 |
| chr19 | 24999199 | 24999655 | 412 | Promoter (<=1kb) | 19 | 24999529 | 25202432 | 202904 | 1 | ENSMUST00 | 0 | Dock8 | ENSMUSG0C | false | chr19 | 24998983 | 25000509 | 31.43241 | 100.35837 | 96.77719 | chr19 | 24999092 | 25000896 | 5.32403 | 4.18245 | 3.33873 |
| chr11 | 98740364 | 98740738 | 412 | Promoter (<=1kb) | 11 | 98740638 | 98753379 | 12742 | 1 | ENSMUST00 | 0 | Thra | ENSMUSG0C | false | chr11 | 98740244 | 98743094 | 32.66734 | 113.81544 | 109.86345 | chr11 | 98739896 | 98756634 | 7.38853 | 5.10697 | 4.96875 |
| chr4 | 55013758 | 55014189 | 412 | Promoter (2-3kb) | 4 | 55011480 | 55081330 | 69851 | 1 | ENSMUST00 | 2279 | Zfp462 | ENSMUSG0C | true | . | . | . | . | . | . | chr4 | 55012881 | 55015436 | 3.40582 | 3.04348 | 1.73265 |
| chrX | 11821832 | 11822058 | 412 | Distal Intergenic | 20 | 11850484 | 11850590 | 107 | 1 | ENSMUST00 | -28426 | Gm26314 | ENSMUSG0C | false | . | . | . | . | . | . | . | . | . | . | . | . |
| chr6 | 121146874 | 121147200 | 412 | Distal Intergenic | 6 | 121183667 | 121196185 | 12519 | 1 | ENSMUST00 | -36467 | Pex26 | ENSMUSG0C | false | . | . | . | . | . | . | chr6 | 121143778 | 121148146 | 5.47647 | 4.19044 | 3.41087 |
| chr2 | 105063346 | 105063691 | 412 | Distal Intergenic | 2 | 105077895 | 105078107 | 213 | 2 | ENSMUST00 | 14416 | Wt1os | ENSMUSG0C | true | . | . | . | . | . | . | chr2 | 105057120 | 105080197 | 5.15057 | 4.16934 | 3.12788 |
| chr4 | 9202856 | 9203088 | 412 | Distal Intergenic | 4 | 9210003 | 9210108 | 106 | 1 | ENSMUST00 | -6915 | Gm23423 | ENSMUSG0C | false | . | . | . | . | . | . | chr4 | 9202505 | 9203765 | 4.03246 | 3.59105 | 2.2406 |
| chr5 | 144271847 | 144272091 | 412 | Intron (ENSMUST00000155491.2 | 5 | 144277061 | 144280513 | 3453 | 1 | ENSMUST00 | -4970 | Gm15708 | ENSMUSG0C | false | . | . | . | . | . | . | chr5 | 144256424 | 144275062 | 6.51227 | 4.68639 | 4.25571 |
| chr15 | 101183235 | 101183476 | 411 | Intron (ENSMUST00000000544.1 | 15 | 101174067 | 101213679 | 39613 | 1 | ENSMUST00 | 9169 | Acvr1b | ENSMUSG0C | false | . | . | . | . | . | . | chr15 | 101174616 | 101193068 | 5.45533 | 4.23935 | 3.38802 |
| chr7 | 27396216 | 27396436 | 411 | Promoter (<=1kb) | 7 | 27356383 | 27396198 | 39816 | 2 | ENSMUST00 | -19 | Sptbn4 | ENSMUSG0C | false | . | . | . | . | . | . | chr7 | 27396228 | 27397609 | 4.31104 | 3.53764 | 2.48943 |
| chr11 | 96971990 | 96972307 | 411 | Intron (ENSMUST00000107626.2 | 11 | 96953473 | 96968665 | 15193 | 2 | ENSMUST00 | -3326 | Sp2 | ENSMUSG0C | false | . | . | . | . | . | . | chr11 | 96963311 | 96976922 | 7.31488 | 5.11145 | 4.90474 |
| chr18 | 40219340 | 40219545 | 411 | Promoter (<=1kb) | 18 | 40204865 | 40219399 | 14535 | 2 | ENSMUST00 | 0 | Yipf5 | ENSMUSG0C | false | chr18 | 40218507 | 40219431 | 28.22734 | 101.53345 | 97.92473 | . | . | . | . | . | . |
| chr19 | 7268045 | 7268443 | 411 | Promoter (<=1kb) | 19 | 7268301 | 7275225 | 6925 | 1 | ENSMUST00 | 0 | Rcor2 | ENSMUSG0C | false | chr19 | 7267778 | 7269372 | 23.47016 | 67.50422 | 64.55198 | chr19 | 7266091 | 7270232 | 5.81658 | 4.28552 | 3.71613 |
| chr1 | 164898986 | 164899427 | 411 | Distal Intergenic | 1 | 164931644 | 164935527 | 3884 | 2 | ENSMUST00 | 36100 | Xcl1 | ENSMUSG0C | false | chr1 | 164898967 | 164899567 | 22.51707 | 59.83231 | 56.98677 | chr1 | 164898511 | 164900110 | 4.29219 | 3.74558 | 2.4339 |
| chr2 | 154564171 | 154564442 | 411 | Exon (ENSMUST00000103145.5/ | 2 | 154559634 | 154569720 | 10087 | 2 | ENSMUST00 | 5278 | E2f1 | ENSMUSG0C | false | chr2 | 154562854 | 154564520 | 10.47775 | 22.22469 | 19.86031 | chr2 | 154555502 | 154571286 | 6.03478 | 4.51051 | 3.86945 |
| chr10 | 81570819 | 81571093 | 411 | Downstream (1-2kb) | 10 | 81574561 | 81590466 | 15906 | 1 | ENSMUST00 | -3468 | Tle2 | ENSMUSG0C | false | chr10 | 81570656 | 81572272 | 28.7177 | 82.96152 | 79.75617 | chr10 | 81568638 | 81573670 | 5.32097 | 4.17822 | 3.27113 |
| chr17 | 66207796 | 66208084 | 411 | Distal Intergenic | 17 | 66123520 | 66152167 | 28648 | 1 | ENSMUST00 | 84277 | Ddx11 | ENSMUSG0C | false | . | . | . | . | . | . | chr17 | 66207456 | 66209749 | 4.01745 | 3.498 | 2.23347 |
| chr1 | 36709816 | 36710188 | 411 | Promoter (<=1kb) | 1 | 36698116 | 36710008 | 11893 | 2 | ENSMUST00 | 0 | Actr1b | ENSMUSG0C | false | chr1 | 36709036 | 36710077 | 40.83545 | 136.9297 | 132.17467 | . | . | . | . | . | . |
| chr6 | 59024443 | 59024729 | 411 | Promoter (<=1kb) | 6 | 58932090 | 59024549 | 92460 | 2 | ENSMUST00 | 0 | Fam13a | ENSMUSG0C | false | chr6 | 59024080 | 59024888 | 16.7644 | 42.82802 | 40.20103 | chr6 | 59023731 | 59024945 | 3.78625 | 3.50813 | 2.0038 |
| chr10 | 75939927 | 75940187 | 411 | Promoter (<=1kb) | 10 | 75937723 | 75940672 | 2950 | 2 | ENSMUST00 | 485 | Gm867 | ENSMUSG0C | false | . | . | . | . | . | . | chr10 | 75934956 | 75940928 | 4.36053 | 3.69776 | 2.49095 |
| chr5 | 72921303 | 72921579 | 411 | Intron (ENSMUST00000144843.2 | 5 | 72914296 | 72914942 | 647 | 2 | ENSMUST00 | -6362 | Gm9870 | ENSMUSG0C | false | . | . | . | . | . | . | chr5 | 72915763 | 72933978 | 5.15676 | 4.0985 | 3.1473 |
| chr11 | 117605869 | 117606146 | 411 | Distal Intergenic | 11 | 117611247 | 117613857 | 2611 | 1 | ENSMUST00 | -5101 | 2900041M22 | ENSMUSG0C | false | . | . | . | . | . | . | chr11 | 117605634 | 117607034 | 4.67024 | 3.73759 | 2.7515 |
| chr1 | 43827597 | 43828088 | 411 | Promoter (<=1kb) | 1 | 43746966 | 43827800 | 80835 | 2 | ENSMUST00 | 0 | Uxs1 | ENSMUSG0C | false | chr1 | 43826816 | 43828457 | 28.89995 | 120.94616 | 116.77392 | . | . | . | . | . | . |
| chr10 | 76997950 | 76998196 | 411 | Distal Intergenic | 10 | 77011995 | 77022214 | 10220 | 1 | ENSMUST00 | -13799 | Gm10787 | ENSMUSG0C | false | . | . | . | . | . | . | chr10 | 76997694 | 76998368 | 4.36129 | 3.65374 | 2.50711 |
| chr3 | 50115655 | 50115934 | 411 | Distal Intergenic | 3 | 50259932 | 50275128 | 15197 | 1 | ENSMUST00 | -143998 | RP24-377J3. | ENSMUSG0C | false | . | . | . | . | . | . | chr3 | 50115570 | 50115959 | 2.9477 | 2.90066 | 1.35513 |
| chr7 | 141131118 | 141131421 | 410 | Promoter (<=1kb) | 7 | 141131286 | 141157021 | 25736 | 1 | ENSMUST00 | 0 | Ptdss2 | ENSMUSG0C | false | chr7 | 141130877 | 141132105 | 47.4287 | 160.87634 | 155.20065 | chr7 | 141130583 | 141132360 | 5.09134 | 3.95426 | 3.11892 |
| chr3 | 56183178 | 56184013 | 410 | Promoter (<=1kb) | 3 | 55625205 | 56183701 | 558497 | 2 | ENSMUST00 | 0 | Nbea | ENSMUSG0C | false | chr3 | 56182369 | 56184233 | 24.32174 | 95.77222 | 92.30251 | . | . | . | . | . | . |
| chr3 | 116007291 | 116007662 | 410 | Promoter (<=1kb) | 3 | 116007462 | 116029016 | 21555 | 1 | ENSMUST00 | 0 | Extl2 | ENSMUSG0C | false | chr3 | 116006495 | 116008826 | 23.3092 | 84.4034 | 81.16875 | . | . | . | . | . | . |
| chr5 | 92278019 | 92278316 | 410 | Promoter (<=1kb) | 5 | 92257659 | 92278170 | 20512 | 2 | ENSMUST00 | 0 | Naaa | ENSMUSG0C | false | chr5 | 92277532 | 92278322 | 31.57443 | 95.61693 | 92.15154 | . | . | . | . | . | . |
| chr19 | 25304563 | 25304838 | 410 | Intron (ENSMUST00000049400.9 | 19 | 25236975 | 25434496 | 197522 | 1 | ENSMUST00 | 67589 | Kank1 | ENSMUSG0C | true | . | . | . | . | . | . | chr19 | 25294157 | 25326429 | 5.37785 | 4.23317 | 3.31781 |
| chr9 | 66945906 | 66946270 | 410 | Promoter (<=1kb) | 9 | 66946086 | 66949512 | 3427 | 1 | ENSMUST00 | 0 | Rps27l | ENSMUSG0C | false | chr9 | 66945369 | 66946939 | 33.73597 | 126.30199 | 121.9427 | . | . | . | . | . | . |
| chr2 | 33627025 | 33627241 | 410 | Intron (ENSMUST00000041730.5 | 2 | 33567156 | 33632283 | 65128 | 2 | ENSMUST00 | 5042 | Lmx1b | ENSMUSG0C | true | . | . | . | . | . | . | chr2 | 33565260 | 33638297 | 6.78481 | 4.81457 | 4.48235 |
| chr10 | 17471416 | 17471661 | 410 | Distal Intergenic | 10 | 17723228 | 17725673 | 2446 | 1 | ENSMUST00 | -251567 | Cited2 | ENSMUSG0C | false | . | . | . | . | . | . | chr10 | 17470740 | 17472975 | 3.98191 | 3.57313 | 2.16724 |
| chrX | 152144036 | 152144495 | 410 | Promoter (<=1kb) | 20 | 152144268 | 152225235 | 80968 | 1 | ENSMUST00 | 0 | Iqsec2 | ENSMUSG0C | false | chrX | 152143983 | 152145711 | 21.13848 | 54.99947 | 52.22001 | . | . | . | . | . | . |
| chr12 | 102302916 | 102303227 | 410 | Intron (ENSMUST00000056950.8 | 12 | 102283641 | 102390855 | 107215 | 1 | ENSMUST00 | 19276 | Rin3 | ENSMUSG0C | false | . | . | . | . | . | . | chr12 | 102301484 | 102309684 | 6.64516 | 4.78421 | 4.36044 |
| chr16 | 64851494 | 64852694 | 410 | Promoter (<=1kb) | 16 | 64851996 | 64859507 | 7512 | 1 | ENSMUST00 | 0 | Cggbp1 | ENSMUSG0C | false | chr16 | 64850498 | 64851808 | 42.34764 | 140.67596 | 135.76277 | . | . | . | . | . | . |
| chr8 | 103288517 | 103288754 | 410 | Distal Intergenic | 8 | 102864779 | 102865853 | 1075 | 2 | ENSMUST00 | -422665 | Gm8730 | ENSMUSG0C | false | . | . | . | . | . | . | chr8 | 103288175 | 103293261 | 5.7653 | 4.41075 | 3.63503 |
| chr7 | 30000659 | 30000872 | 410 | Intron (ENSMUST00000148442.2 | 7 | 29987773 | 30023647 | 35875 | 1 | ENSMUST00 | 12887 | Zfp568 | ENSMUSG0C | false | . | . | . | . | . | . | chr7 | 29999979 | 30001495 | 3.87344 | 3.56166 | 2.12821 |
| chr5 | 72040136 | 72040537 | 410 | Intron (ENSMUST00000031122.7 | 5 | 72032718 | 72033488 | 771 | 2 | ENSMUST00 | -6649 | Gm15617 | ENSMUSG0C | false | . | . | . | . | . | . | chr5 | 72038906 | 72040858 | 3.65032 | 3.34882 | 1.92291 |
| chr2 | 167853455 | 167853864 | 410 | Distal Intergenic | 2 | 167858585 | 167862542 | 3958 | 1 | ENSMUST00 | -4721 | Gm14319 | ENSMUSG0C | false | . | . | . | . | . | . | chr2 | 167850474 | 167854412 | 5.03048 | 3.96235 | 3.05598 |
| chr10 | 107924104 | 107924382 | 410 | Distal Intergenic | 10 | 107762223 | 107912134 | 149912 | 2 | ENSMUST00 | -11971 | Otogl | ENSMUSG0C | false | . | . | . | . | . | . | chr10 | 107924003 | 107924506 | 4.32359 | 3.80715 | 2.46633 |
| chr10 | 121857705 | 121857959 | 409 | Intron (ENSMUST00000020322.6 | 10 | 121806640 | 121823133 | 16494 | 2 | ENSMUST00 | -34573 | Srgap1 | ENSMUSG0C | true | . | . | . | . | . | . | chr10 | 121853826 | 121867005 | 4.67918 | 3.79658 | 2.7761 |
| chr16 | 20535380 | 20536149 | 409 | Promoter (<=1kb) | 16 | 20535478 | 20544056 | 8579 | 1 | ENSMUST00 | 0 | Ap2m1 | ENSMUSG0C | false | chr16 | 20535449 | 20536335 | 43.19602 | 139.91907 | 135.04468 | . | . | . | . | . | . |
| chr5 | 73729674 | 73730003 | 409 | Distal Intergenic | 5 | 73657535 | 73679482 | 21948 | 1 | ENSMUST00 | 72140 | Spata18 | ENSMUSG0C | false | . | . | . | . | . | . | . | . | . | . | . | . |
| chr6 | 97614905 | 97615225 | 409 | Promoter (2-3kb) | 6 | 97350933 | 97617529 | 266597 | 2 | ENSMUST00 | 2304 | Frmd4b | ENSMUSG0C | false | . | . | . | . | . | . | chr6 | 97612810 | 97617843 | 4.53399 | 3.713 | 2.65016 |
| chr6 | 89362444 | 89362760 | 409 | Promoter (<=1kb) | 6 | 89316316 | 89362613 | 46298 | 2 | ENSMUST00 | 0 | Plxna1 | ENSMUSG0C | false | chr6 | 89362742 | 89363328 | 17.45018 | 43.2582 | 40.62776 | . | . | . | . | . | . |
| chr8 | 11405774 | 11406099 | 409 | Promoter (<=1kb) | 8 | 11404908 | 11421254 | 16347 | 1 | ENSMUST00 | 867 | Col4a2 | ENSMUSG0C | false | . | . | . | . | . | . | chr8 | 11405665 | 11406765 | 3.4816 | 3.28242 | 1.79796 |
| chr8 | 33892248 | 33892522 | 409 | Promoter (<=1kb) | 8 | 33863007 | 33891764 | 28758 | 2 | ENSMUST00 | -485 | Rbpms | ENSMUSG0C | true | . | . | . | . | . | . | chr8 | 33863865 | 33927704 | 6.32022 | 4.67603 | 4.08842 |
| chr9 | 43210227 | 43210645 | 409 | Promoter (<=1kb) | 9 | 43123939 | 43210369 | 86431 | 2 | ENSMUST00 | 0 | Pou2f3 | ENSMUSG0C | false | chr9 | 43210225 | 43210775 | 11.08881 | 23.82316 | 21.43778 | chr9 | 43210308 | 43211451 | 4.20498 | 3.50789 | 2.37492 |
| chr5 | 103393102 | 103393563 | 409 | Distal Intergenic | 5 | 103425192 | 103598359 | 173168 | 1 | ENSMUST00 | -31629 | Ptpn13 | ENSMUSG0C | false | . | . | . | . | . | . | chr5 | 103392235 | 103395296 | 3.60455 | 3.30834 | 1.90131 |
| chr3 | 40949456 | 40949998 | 409 | Promoter (<=1kb) | 3 | 40950631 | 40977793 | 27163 | 1 | ENSMUST00 | -633 | Larp1b | ENSMUSG0C | false | chr3 | 40949721 | 40951569 | 28.91247 | 96.86193 | 93.36482 | . | . | . | . | . | . |
| chr10 | 22873196 | 22873452 | 409 | Distal Intergenic | 10 | 22817279 | 22820128 | 2850 | 2 | ENSMUST00 | -53069 | Tcf21 | ENSMUSG0C | true | . | . | . | . | . | . | chr10 | 22866220 | 22874539 | 4.6225 | 3.87067 | 2.69769 |
| chr12 | 21160611 | 21160882 | 409 | Intron (ENSMUST00000174530.1 | 12 | 21111954 | 21268179 | 156226 | 1 | ENSMUST00 | 48658 | Asap2 | ENSMUSG0C | false | . | . | . | . | . | . | chr12 | 21148547 | 21176379 | 6.51645 | 4.71535 | 4.25897 |
| chrX | 94122260 | 94122721 | 409 | Promoter (<=1kb) | 20 | 94074632 | 94122900 | 48269 | 2 | ENSMUST00 | 179 | Zfx | ENSMUSG0C | false | chrX | 94121286 | 94123843 | 19.75988 | 50.25065 | 47.53236 | . | . | . | . | . | . |
| chr4 | 148920600 | 148920816 | 409 | Intron (ENSMUST00000094464.4 | 4 | 148913511 | 148914324 | 814 | 2 | ENSMUST00 | -6277 | Gm15969 | ENSMUSG0C | false | . | . | . | . | . | . | chr4 | 148901819 | 148933146 | 7.00771 | 4.91323 | 4.66351 |
| chr18 | 3666496 | 3666819 | 409 | Distal Intergenic | 18 | 3770741 | 3770804 | 64 | 2 | ENSMUST00 | 103985 | Gm23643 | ENSMUSG0C | false | . | . | . | . | . | . | chr18 | 3664015 | 3670955 | 4.12925 | 3.67983 | 2.28088 |
| chr2 | 84743568 | 84743910 | 409 | Promoter (<=1kb) | 2 | 84742169 | 84743655 | 1487 | 2 | ENSMUST00 | 0 | Gm19426 | ENSMUSG0C | false | chr2 | 84742264 | 84744578 | 30.59503 | 95.89513 | 92.42037 | . | . | . | . | . | . |
| chr18 | 68117543 | 68117827 | 409 | Intron (ENSMUST00000063775.3 | 18 | 68260920 | 68260973 | 54 | 2 | ENSMUST00 | 143146 | Mir7219 | ENSMUSG0C | false | . | . | . | . | . | . | chr18 | 68117175 | 68119651 | 4.3261 | 3.73932 | 2.46289 |
| chr10 | 115205441 | 115205727 | 408 | Intron (ENSMUST00000020339.8 | 10 | 115078641 | 115185022 | 106382 | 2 | ENSMUST00 | -20420 | Tph2 | ENSMUSG0C | false | . | . | . | . | . | . | . | . | . | . | . | . |
| chr12 | 16853642 | 16853871 | 408 | Distal Intergenic | 12 | 16894978 | 16988274 | 93297 | 1 | ENSMUST00 | -41107 | Rock2 | ENSMUSG0C | true | . | . | . | . | . | . | chr12 | 16848168 | 16854626 | 5.81138 | 4.51483 | 3.66527 |
| chr18 | 60526071 | 60526634 | 408 | Promoter (<=1kb) | 18 | 60526221 | 60558762 | 32542 | 1 | ENSMUST00 | 0 | Dctn4 | ENSMUSG0C | false | chr18 | 60525701 | 60527274 | 30.84885 | 99.33006 | 95.77879 | . | . | . | . | . | . |
| chr10 | 126935644 | 126936009 | 408 | Distal Intergenic | 10 | 126887249 | 126901371 | 14123 | 2 | ENSMUST00 | -34274 | Xrcc6bp1 | ENSMUSG0C | false | . | . | . | . | . | . | chr10 | 126935236 | 126936469 | 9.72017 | 6.18388 | 6.85522 |
| chr2 | 60865868 | 60866113 | 408 | Intron (ENSMUST00000028347.7 | 2 | 60751953 | 60881438 | 129486 | 2 | ENSMUST00 | 15325 | Rbms1 | ENSMUSG0C | true | . | . | . | . | . | . | chr2 | 60861855 | 60880641 | 6.87834 | 4.85657 | 4.57159 |
| chr4 | 123917161 | 123918093 | 408 | Promoter (<=1kb) | 4 | 123917446 | 123936997 | 19552 | 1 | ENSMUST00 | 0 | Rragc | ENSMUSG0C | false | chr4 | 123916660 | 123919069 | 32.3084 | 121.498 | 117.30614 | . | . | . | . | . | . |
| chr3 | 101524550 | 101524786 | 408 | Distal Intergenic | 3 | 101576316 | 101577560 | 1245 | 2 | ENSMUST00 | 52774 | Atp1a1 | ENSMUSG0C | false | . | . | . | . | . | . | . | . | . | . | . | . |
| chr5 | 103374235 | 103374503 | 408 | Distal Intergenic | 5 | 103425192 | 103598359 | 173168 | 1 | ENSMUST00 | -50689 | Ptpn13 | ENSMUSG0C | false | . | . | . | . | . | . | chr5 | 103373752 | 103374567 | 4.4171 | 3.89416 | 2.50687 |
| chr14 | 21994634 | 21994834 | 408 | Distal Intergenic | 14 | 21983962 | 21989601 | 5640 | 2 | ENSMUST00 | -5034 | Zfp503 | ENSMUSG0C | false | chr14 | 21993562 | 21996021 | 31.01414 | 97.6227 | 94.10706 | . | . | . | . | . | . |
| chr15 | 57723765 | 57724069 | 408 | Intron (ENSMUST00000096430.5 | 15 | 57736563 | 57738666 | 2104 | 2 | ENSMUST00 | 14597 | 9330154K18I | ENSMUSG0C | false | . | . | . | . | . | . | chr15 | 57721101 | 57725779 | 3.55347 | 3.26003 | 1.86235 |
| chr3 | 67515066 | 67515678 | 408 | Promoter (<=1kb) | 3 | 67478886 | 67515523 | 36638 | 2 | ENSMUST00 | 0 | Rarres1 | ENSMUSG0C | false | chr3 | 67514339 | 67515861 | 26.65955 | 86.95457 | 83.67183 | chr3 | 67515612 | 67515970 | 3.4621 | 3.264 | 1.79167 |
| chr3 | 106721704 | 106722377 | 408 | Promoter (<=1kb) | 3 | 106721728 | 106732211 | 10484 | 1 | ENSMUST00 | 0 | Lrif1 | ENSMUSG0C | false | chr3 | 106721230 | 106723023 | 32.07243 | 98.46742 | 94.9359 | . | . | . | . | . | . |
| chr14 | 66818585 | 66818901 | 408 | Intron (ENSMUST00000022629.8 | 14 | 66784529 | 66793758 | 9230 | 1 | ENSMUST00 | 34057 | Gm10032 | ENSMUSG0C | false | . | . | . | . | . | . | chr14 | 66815115 | 66830778 | 7.77502 | 5.24019 | 5.30996 |
| chr10 | 63496644 | 63497006 | 408 | Intron (ENSMUST00000105441.2 | 10 | 63504037 | 65002833 | 1498797 | 1 | ENSMUST00 | -7031 | Ctnna3 | ENSMUSG0C | false | chr10 | 63496503 | 63497215 | 6.42188 | 10.41091 | 8.23098 | chr10 | 63495957 | 63498244 | 3.44565 | 3.32844 | 1.73392 |
| chr6 | 121145882 | 121146162 | 408 | Distal Intergenic | 6 | 121183667 | 121196185 | 12519 | 1 | ENSMUST00 | -37505 | Pex26 | ENSMUSG0C | false | . | . | . | . | . | . | chr6 | 121143778 | 121148146 | 5.47647 | 4.19044 | 3.41087 |
| chr2 | 39154857 | 39155337 | 408 | Intron (ENSMUST00000038874.6 | 2 | 39161251 | 39161989 | 739 | 2 | ENSMUST00 | 6652 | Gm16523 | ENSMUSG0C | false | . | . | . | . | . | . | chr2 | 39149319 | 39156055 | 5.39771 | 4.23651 | 3.33948 |
| chr10 | 12792086 | 12792354 | 408 | Promoter (1-2kb) | 10 | 12793875 | 12793974 | 100 | 2 | ENSMUST00 | 1620 | Gm23044 | ENSMUSG0C | false | . | . | . | . | . | . | chr10 | 12790606 | 12799534 | 5.35568 | 4.24048 | 3.29665 |
| chr5 | 25532723 | 25532980 | 408 | Promoter (2-3kb) | 5 | 25530013 | 25531466 | 1454 | 1 | ENSMUST00 | 2711 | RP23-405O1 | ENSMUSG0C | false | . | . | . | . | . | . | chr5 | 25530644 | 25533143 | 4.13994 | 3.5679 | 2.30637 |
| chr5 | 143624015 | 143624279 | 407 | Promoter (1-2kb) | 5 | 143622534 | 143686526 | 63993 | 1 | ENSMUST00 | 1482 | Cyth3 | ENSMUSG0C | false | chr5 | 143622170 | 143624398 | 40.66609 | 142.19028 | 137.22005 | chr5 | 143623294 | 143627247 | 5.05306 | 3.88706 | 3.0834 |
| chr1 | 36844990 | 36845325 | 407 | Intron (ENSMUST00000027290.6 | 1 | 36818660 | 36824781 | 6122 | 2 | ENSMUST00 | -20210 | Tmem131 | ENSMUSG0C | false | . | . | . | . | . | . | chr1 | 36839564 | 36847738 | 4.82478 | 3.86744 | 2.886 |
| chr1 | 133800956 | 133801380 | 407 | Promoter (<=1kb) | 1 | 133699457 | 133801041 | 101585 | 2 | ENSMUST00 | 0 | Atp2b4 | ENSMUSG0C | false | chr1 | 133800021 | 133801596 | 17.10778 | 41.93702 | 39.32339 | chr1 | 133799317 | 133802174 | 4.5737 | 3.86022 | 2.66153 |
| chr6 | 142507813 | 142508155 | 407 | Promoter (<=1kb) | 6 | 142490249 | 142507957 | 17709 | 2 | ENSMUST00 | 0 | Ldhb | ENSMUSG0C | false | chr6 | 142507058 | 142508304 | 14.73601 | 38.55974 | 35.98649 | chr6 | 142505920 | 142508471 | 4.32485 | 3.68555 | 2.45254 |
| chr6 | 149309260 | 149309810 | 407 | Promoter (<=1kb) | 6 | 149309414 | 149326671 | 17258 | 1 | ENSMUST00 | 0 | 2810474O19 | ENSMUSG0C | false | chr6 | 149308882 | 149310424 | 28.89701 | 129.08754 | 124.63428 | . | . | . | . | . | . |
| chr1 | 4785442 | 4785874 | 407 | Promoter (<=1kb) | 1 | 4773206 | 4785710 | 12505 | 2 | ENSMUST00 | 0 | Mrpl15 | ENSMUSG0C | false | chr1 | 4784947 | 4786386 | 29.58769 | 108.11006 | 104.32592 | . | . | . | . | . | . |
| chr8 | 123113780 | 123114166 | 407 | Intron (ENSMUST00000127664.1 | 8 | 123117374 | 123135180 | 17807 | 1 | ENSMUST00 | -3208 | Cpne7 | ENSMUSG0C | false | . | . | . | . | . | . | . | . | . | . | . | . |
| chr18 | 15064674 | 15064980 | 407 | Promoter (1-2kb) | 18 | 14968685 | 15063582 | 94898 | 2 | ENSMUST00 | -1093 | Kctd1 | ENSMUSG0C | false | chr18 | 15062362 | 15065257 | 28.1615 | 93.53628 | 90.11485 | chr18 | 15062478 | 15066597 | 4.85515 | 3.85397 | 2.93151 |
| chr5 | 121836533 | 121837283 | 407 | Promoter (<=1kb) | 5 | 121815488 | 121836825 | 21338 | 2 | ENSMUST00 | 0 | Sh2b3 | ENSMUSG0C | false | chr5 | 121835543 | 121838063 | 34.66576 | 107.00249 | 103.24738 | chr5 | 121837267 | 121839323 | 5.65924 | 4.22393 | 3.56804 |
| chr4 | 120287563 | 120288148 | 407 | Promoter (<=1kb) | 4 | 120267079 | 120287349 | 20271 | 2 | ENSMUST00 | -215 | Foxo6 | ENSMUSG0C | false | chr4 | 120286553 | 120288405 | 30.17592 | 94.17345 | 90.73757 | chr4 | 120286572 | 120290219 | 4.15182 | 3.67163 | 2.31183 |
| chr9 | 119409255 | 119409679 | 407 | Intron (ENSMUST00000035093.9 | 9 | 119402501 | 119433501 | 31001 | 1 | ENSMUST00 | 6755 | Acvr2b | ENSMUSG0C | false | . | . | . | . | . | . | chr9 | 119407292 | 119411802 | 5.25494 | 4.02994 | 3.24537 |
| chr5 | 134539973 | 134540272 | 407 | Intron (ENSMUST00000036999.6 | 5 | 134489386 | 134552434 | 63049 | 2 | ENSMUST00 | 12162 | Clip2 | ENSMUSG0C | false | . | . | . | . | . | . | chr5 | 134538830 | 134551710 | 6.68849 | 4.77556 | 4.40881 |
| chr7 | 144943262 | 144943559 | 407 | Distal Intergenic | 7 | 144929931 | 144939869 | 9939 | 2 | ENSMUST00 | -3394 | Ccnd1 | ENSMUSG0C | false | chr7 | 144941140 | 144943434 | 27.88725 | 99.90214 | 96.33501 | chr7 | 144938154 | 144943823 | 5.32689 | 4.2503 | 3.27022 |
| chr5 | 73152240 | 73152467 | 407 | Intron (ENSMUST00000101127.6 | 5 | 73021211 | 73148136 | 126926 | 2 | ENSMUST00 | -4105 | Fryl | ENSMUSG0C | true | . | . | . | . | . | . | chr5 | 73148451 | 73194976 | 5.88301 | 4.44395 | 3.74315 |
| chr4 | 101292942 | 101293546 | 407 | Intron (ENSMUST00000131864.1 | 4 | 101284221 | 101285203 | 983 | 1 | ENSMUST00 | 8722 | Gm12801 | ENSMUSG0C | false | chr4 | 101292040 | 101293747 | 23.60507 | 64.24684 | 61.34294 | chr4 | 101291052 | 101294245 | 4.50505 | 3.85025 | 2.60632 |
| chr18 | 82870141 | 82870513 | 407 | Exon (ENSMUST00000181345.1/ | 18 | 82855897 | 82859162 | 3266 | 1 | ENSMUST00 | 14245 | 4930445N18 | ENSMUSG0C | false | chr18 | 82869862 | 82870434 | 9.88648 | 20.14232 | 17.80457 | chr18 | 82867704 | 82870337 | 4.92089 | 3.98798 | 2.96174 |
| chr16 | 34927015 | 34927401 | 407 | Intron (ENSMUST00000023538.8 | 16 | 34935854 | 34939329 | 3476 | 2 | ENSMUST00 | 11928 | RP23-80O10 | ENSMUSG0C | false | . | . | . | . | . | . | chr16 | 34923663 | 34969430 | 6.94598 | 4.91232 | 4.61654 |
| chr19 | 46470513 | 46470927 | 406 | Intron (ENSMUST00000039922.7 | 19 | 46483097 | 46485982 | 2886 | 1 | ENSMUST00 | -12170 | Sufu | ENSMUSG0C | false | . | . | . | . | . | . | chr19 | 46468103 | 46486697 | 7.92449 | 5.26231 | 5.41796 |
| chr1 | 87510320 | 87510647 | 406 | Promoter (<=1kb) | 1 | 87476834 | 87510362 | 33529 | 2 | ENSMUST00 | 0 | Ngef | ENSMUSG0C | false | chr1 | 87509293 | 87510649 | 23.28017 | 72.50986 | 69.48009 | chr1 | 87508932 | 87511417 | 5.46366 | 4.23354 | 3.38653 |
| chr1 | 175692612 | 175692995 | 406 | Promoter (<=1kb) | 1 | 175662421 | 175692776 | 30356 | 2 | ENSMUST00 | 0 | Opn3 | ENSMUSG0C | false | chr1 | 175691698 | 175692850 | 22.89105 | 88.16154 | 84.85722 | . | . | . | . | . | . |
| chr4 | 33031234 | 33031970 | 406 | Promoter (<=1kb) | 4 | 33031416 | 33052363 | 20948 | 1 | ENSMUST00 | 0 | Ube2j1 | ENSMUSG0C | false | chr4 | 33030273 | 33032302 | 28.1615 | 93.53628 | 90.11485 | . | . | . | . | . | . |
| chr4 | 130174722 | 130174997 | 406 | Promoter (<=1kb) | 4 | 130164454 | 130174802 | 10349 | 2 | ENSMUST00 | 0 | Tinagl1 | ENSMUSG0C | false | chr4 | 130174513 | 130175098 | 10.29113 | 20.81812 | 18.4725 | chr4 | 130172313 | 130175844 | 4.68522 | 3.93643 | 2.73692 |
| chr7 | 88430094 | 88430431 | 406 | Promoter (<=1kb) | 7 | 88430273 | 88491466 | 61194 | 1 | ENSMUST00 | 0 | Rab38 | ENSMUSG0C | false | chr7 | 88429918 | 88431011 | 21.92219 | 60.49891 | 57.64639 | . | . | . | . | . | . |
| chr9 | 63997506 | 63997798 | 406 | Intron (ENSMUST00000041029.5 | 9 | 63954009 | 64016987 | 62979 | 2 | ENSMUST00 | 19189 | Smad6 | ENSMUSG0C | true | . | . | . | . | . | . | chr9 | 63948352 | 64020083 | 7.27873 | 5.05859 | 4.874 |
| chr15 | 100468740 | 100469181 | 406 | Promoter (<=1kb) | 15 | 100469034 | 100479252 | 10219 | 1 | ENSMUST00 | 0 | Letmd1 | ENSMUSG0C | false | chr15 | 100468181 | 100469981 | 36.76257 | 113.72284 | 109.77938 | chr15 | 100467383 | 100470985 | 4.63747 | 3.91689 | 2.75023 |
| chr2 | 129626270 | 129626485 | 406 | Intron (ENSMUST00000161620.2 | 2 | 129620685 | 129628044 | 7360 | 1 | ENSMUST00 | 5586 | Sirpa | ENSMUSG0C | true | . | . | . | . | . | . | chr2 | 129621436 | 129635375 | 5.74226 | 4.37432 | 3.64032 |
| chr15 | 5244126 | 5244923 | 406 | Promoter (<=1kb) | 15 | 5233399 | 5244172 | 10774 | 2 | ENSMUST00 | 0 | Ptger4 | ENSMUSG0C | false | chr15 | 5239457 | 5245131 | 38.68621 | 127.7006 | 123.29971 | . | . | . | . | . | . |
| chr16 | 25465492 | 25465744 | 406 | Distal Intergenic | 16 | 25286821 | 25318235 | 31415 | 1 | ENSMUST00 | 178672 | Tprg | ENSMUSG0C | false | . | . | . | . | . | . | chr16 | 25464746 | 25469847 | 3.62992 | 3.29674 | 1.91973 |
| chr11 | 117949312 | 117950055 | 406 | Distal Intergenic | 11 | 117966079 | 117969186 | 3108 | 2 | ENSMUST00 | 19131 | Socs3 | ENSMUSG0C | false | chr11 | 117949418 | 117950173 | 12.99241 | 30.10301 | 27.6364 | chr11 | 117949439 | 117952468 | 5.44541 | 4.26334 | 3.37136 |
| chr17 | 36025956 | 36026236 | 406 | Intron (ENSMUST00000173900.1 | 17 | 36005695 | 36020560 | 14866 | 2 | ENSMUST00 | -5397 | H2-T24 | ENSMUSG0C | false | . | . | . | . | . | . | chr17 | 36019995 | 36026641 | 6.63467 | 4.74288 | 4.36704 |
| chr1 | 74000251 | 74000559 | 406 | Promoter (1-2kb) | 1 | 73992205 | 74001567 | 9363 | 2 | ENSMUST00 | 1008 | Tns1 | ENSMUSG0C | false | . | . | . | . | . | . | chr1 | 73999408 | 74002515 | 4.00759 | 3.54768 | 2.2425 |
| chr17 | 27826003 | 27826429 | 406 | Distal Intergenic | 17 | 27751232 | 27820558 | 69327 | 2 | ENSMUST00 | -5446 | D17Wsu92e | ENSMUSG0C | false | chr17 | 27826256 | 27826938 | 15.69042 | 38.08285 | 35.51791 | chr17 | 27825647 | 27827658 | 4.42096 | 3.71372 | 2.57905 |
| chr19 | 60755803 | 60756375 | 406 | Promoter (<=1kb) | 19 | 60755987 | 60759913 | 3927 | 1 | ENSMUST00 | 0 | Nanos1 | ENSMUSG0C | false | chr19 | 60755532 | 60757412 | 28.1615 | 93.53628 | 90.11485 | chr19 | 60755262 | 60758355 | 3.98377 | 3.46427 | 2.22199 |
| chr17 | 71598692 | 71598939 | 406 | Promoter (<=1kb) | 17 | 71557027 | 71598755 | 41729 | 2 | ENSMUST00 | 0 | Trmt61b | ENSMUSG0C | false | chr17 | 71597789 | 71599268 | 25.16654 | 87.04672 | 83.76057 | . | . | . | . | . | . |
| chr10 | 85957713 | 85957958 | 405 | Promoter (<=1kb) | 10 | 85938637 | 85957823 | 19187 | 2 | ENSMUST00 | 0 | Rtcb | ENSMUSG0C | false | chr10 | 85956815 | 85958197 | 40.75532 | 136.78966 | 132.04059 | . | . | . | . | . | . |
| chr11 | 106364840 | 106365126 | 405 | Promoter (<=1kb) | 11 | 106365472 | 106377114 | 11643 | 1 | ENSMUST00 | -346 | 2310007L24F | ENSMUSG0C | false | . | . | . | . | . | . | chr11 | 106364074 | 106366705 | 6.01822 | 4.49039 | 3.85011 |
| chr10 | 83648409 | 83649001 | 405 | Promoter (<=1kb) | 10 | 83600033 | 83648738 | 48706 | 2 | ENSMUST00 | 0 | Appl2 | ENSMUSG0C | false | chr10 | 83647819 | 83649464 | 34.00537 | 102.80987 | 99.17244 | chr10 | 83647309 | 83649476 | 4.4831 | 3.79959 | 2.60159 |
| chr14 | 21623763 | 21624127 | 405 | Promoter (1-2kb) | 14 | 21622542 | 21634589 | 12048 | 1 | ENSMUST00 | 1222 | Kat6b | ENSMUSG0C | false | . | . | . | . | . | . | chr14 | 21623489 | 21624156 | 4.81466 | 4.05176 | 2.83931 |
| chr17 | 80726911 | 80727152 | 405 | Promoter (<=1kb) | 17 | 80580512 | 80728093 | 147582 | 2 | ENSMUST00 | 941 | Map4k3 | ENSMUSG0C | false | chr17 | 80726713 | 80728878 | 38.77008 | 154.42084 | 148.9682 | chr17 | 80713848 | 80726989 | 4.98201 | 3.9926 | 3.00553 |
| chr1 | 16687876 | 16688114 | 405 | Promoter (<=1kb) | 1 | 16688051 | 16691947 | 3897 | 1 | ENSMUST00 | 0 | Ly96 | ENSMUSG0C | false | chr1 | 16687810 | 16689283 | 23.85253 | 69.94444 | 66.95615 | chr1 | 16687690 | 16688173 | 4.91362 | 4.05642 | 2.94522 |
| chr2 | 121611961 | 121612186 | 405 | Intron (ENSMUST00000110593.3 | 2 | 121545529 | 121586433 | 40905 | 2 | ENSMUST00 | -25529 | Frmd5 | ENSMUSG0C | false | . | . | . | . | . | . | . | . | . | . | . | . |
| chr2 | 120170142 | 120170464 | 405 | Exon (ENSMUST00000136845.2/ | 2 | 120089175 | 120154606 | 65432 | 2 | ENSMUST00 | -15537 | Ehd4 | ENSMUSG0C | false | . | . | . | . | . | . | chr2 | 120169905 | 120170875 | 7.18142 | 5.09885 | 4.87546 |
| chr2 | 148451664 | 148451883 | 405 | Distal Intergenic | 2 | 148436640 | 148443563 | 6924 | 2 | ENSMUST00 | -8102 | Cd93 | ENSMUSG0C | false | . | . | . | . | . | . | chr2 | 148450928 | 148452385 | 3.39922 | 3.18897 | 1.73583 |
| chr5 | 117364448 | 117364699 | 405 | Promoter (<=1kb) | 5 | 117363522 | 117377807 | 14286 | 1 | ENSMUST00 | 927 | Wsb2 | ENSMUSG0C | false | chr5 | 117364677 | 117366041 | 7.9533 | 20.3031 | 17.96454 | chr5 | 117358665 | 117372620 | 6.58746 | 4.86077 | 4.29355 |
| chr10 | 20312864 | 20313157 | 405 | Promoter (<=1kb) | 10 | 20312537 | 20340263 | 27727 | 1 | ENSMUST00 | 328 | Bclaf1 | ENSMUSG0C | false | chr10 | 20311327 | 20313751 | 35.2856 | 127.03558 | 122.65217 | . | . | . | . | . | . |
| chr8 | 31031771 | 31032019 | 405 | Distal Intergenic | 8 | 31089471 | 31097047 | 7577 | 1 | ENSMUST00 | -57452 | Dusp26 | ENSMUSG0C | false | chr8 | 31031931 | 31032847 | 14.24974 | 34.23454 | 31.71693 | . | . | . | . | . | . |
| chr10 | 8460116 | 8460333 | 405 | Intron (ENSMUST00000061601.7 | 10 | 8204753 | 8518825 | 314073 | 2 | ENSMUST00 | 58492 | Ust | ENSMUSG0C | true | . | . | . | . | . | . | chr10 | 8459821 | 8462435 | 5.39647 | 4.36286 | 3.31932 |
| chr16 | 28851468 | 28851850 | 405 | Intron (ENSMUST00000100023.1 | 16 | 28826176 | 28929698 | 103523 | 2 | ENSMUST00 | 77848 | Mb21d2 | ENSMUSG0C | false | chr16 | 28850748 | 28852000 | 13.78596 | 30.80405 | 28.32676 | . | . | . | . | . | . |
| chr3 | 142917679 | 142917944 | 405 | Distal Intergenic | 3 | 142882495 | 142894644 | 12150 | 1 | ENSMUST00 | 35185 | 9530052C20I | ENSMUSG0C | false | . | . | . | . | . | . | chr3 | 142916611 | 142919722 | 4.07942 | 3.53967 | 2.27488 |
| chr6 | 93218019 | 93218304 | 405 | Intron (ENSMUST00000181121.2 | 6 | 93137004 | 93137990 | 987 | 2 | ENSMUST00 | -80030 | Gm5313 | ENSMUSG0C | false | . | . | . | . | . | . | chr6 | 93217150 | 93219095 | 5.76781 | 4.29714 | 3.6736 |
| chr6 | 141251252 | 141251468 | 405 | Promoter (<=1kb) | 6 | 141238278 | 141250561 | 12284 | 2 | ENSMUST00 | -692 | RP23-97L11. | ENSMUSG0C | false | chr6 | 141249145 | 141251922 | 34.07522 | 104.96031 | 101.26198 | chr6 | 141251076 | 141262548 | 4.7705 | 3.8909 | 2.85315 |
| chr1 | 154785006 | 154785226 | 404 | Intron (ENSMUST00000188965.1 | 1 | 154481075 | 154816941 | 335867 | 2 | ENSMUST00 | 31715 | Cacna1e | ENSMUSG0C | false | . | . | . | . | . | . | chr1 | 154784771 | 154786214 | 5.30973 | 4.19735 | 3.26513 |
| chr11 | 75893283 | 75893522 | 404 | Intron (ENSMUST00000121287.2 | 11 | 75832137 | 75911062 | 78926 | 2 | ENSMUST00 | 17540 | Rph3al | ENSMUSG0C | false | . | . | . | . | . | . | chr11 | 75892886 | 75894517 | 6.01319 | 4.62662 | 3.82655 |
| chr17 | 31296430 | 31296809 | 404 | Promoter (<=1kb) | 17 | 31296219 | 31350696 | 54478 | 1 | ENSMUST00 | 212 | Slc37a1 | ENSMUSG0C | false | chr17 | 31295863 | 31297198 | 25.56571 | 75.64988 | 72.57267 | chr17 | 31295768 | 31297866 | 3.88674 | 3.49974 | 2.13633 |
| chr4 | 53713903 | 53714478 | 404 | Promoter (<=1kb) | 4 | 53713998 | 53765785 | 51788 | 1 | ENSMUST00 | 0 | Fktn | ENSMUSG0C | false | chr4 | 53713868 | 53714963 | 30.94804 | 114.76607 | 110.79114 | chr4 | 53713629 | 53716128 | 4.12149 | 3.6299 | 2.28147 |
| chr4 | 44072353 | 44072852 | 404 | Promoter (<=1kb) | 4 | 44036826 | 44072778 | 35953 | 2 | ENSMUST00 | 0 | Gne | ENSMUSG0C | false | chr4 | 44071698 | 44073354 | 36.12563 | 122.12295 | 117.91594 | chr4 | 44069446 | 44073518 | 3.56735 | 3.32854 | 1.8435 |
| chr8 | 36692351 | 36692541 | 404 | Intron (ENSMUST00000098826.4 | 8 | 36567751 | 36733054 | 165304 | 2 | ENSMUST00 | 40513 | Dlc1 | ENSMUSG0C | true | . | . | . | . | . | . | chr8 | 36689162 | 36693241 | 3.98479 | 3.60699 | 2.16075 |
| chr12 | 80463012 | 80463592 | 404 | Promoter (<=1kb) | 12 | 80463095 | 80498135 | 35041 | 1 | ENSMUST00 | 0 | Exd2 | ENSMUSG0C | false | chr12 | 80462742 | 80464239 | 37.2221 | 115.56093 | 111.55734 | . | . | . | . | . | . |
| chr4 | 116877052 | 116878060 | 404 | Promoter (<=1kb) | 4 | 116877376 | 116989264 | 111889 | 1 | ENSMUST00 | 0 | Zswim5 | ENSMUSG0C | false | chr4 | 116876655 | 116878267 | 35.07738 | 110.80895 | 106.9512 | . | . | . | . | . | . |
| chr5 | 123394338 | 123395108 | 404 | Promoter (<=1kb) | 5 | 123394798 | 123446896 | 52099 | 1 | ENSMUST00 | 0 | Mlxip | ENSMUSG0C | false | chr5 | 123394150 | 123396353 | 24.65524 | 104.07409 | 100.40247 | chr5 | 123393827 | 123394381 | 3.37409 | 3.17808 | 1.67977 |
| chr11 | 95515575 | 95515790 | 404 | Promoter (1-2kb) | 11 | 95509845 | 95514570 | 4726 | 2 | ENSMUST00 | -1006 | Nxph3 | ENSMUSG0C | false | chr11 | 95512463 | 95515642 | 20.31581 | 74.8081 | 71.74177 | chr11 | 95514812 | 95517733 | 7.19932 | 5.16539 | 4.79467 |
| chr5 | 123881885 | 123882193 | 404 | Promoter (1-2kb) | 5 | 123876736 | 123880020 | 3285 | 2 | ENSMUST00 | -1866 | Hcar1 | ENSMUSG0C | false | . | . | . | . | . | . | . | . | . | . | . | . |
| chr1 | 156040658 | 156041027 | 404 | Promoter (<=1kb) | 1 | 156040890 | 156051669 | 10780 | 1 | ENSMUST00 | 0 | Tor1aip2 | ENSMUSG0C | false | chr1 | 156040804 | 156041431 | 18.02468 | 55.19937 | 52.41551 | chr1 | 156040208 | 156047758 | 4.78046 | 3.98339 | 2.81997 |
| chr7 | 19561770 | 19562601 | 404 | Promoter (<=1kb) | 7 | 19530967 | 19562398 | 31432 | 2 | ENSMUST00 | 0 | Ppp1r37 | ENSMUSG0C | false | chr7 | 19561027 | 19563379 | 26.17789 | 120.04698 | 115.90433 | . | . | . | . | . | . |
| chr8 | 70506287 | 70506798 | 404 | Promoter (<=1kb) | 8 | 70504292 | 70506721 | 2430 | 2 | ENSMUST00 | 0 | 2810428I15R | ENSMUSG0C | false | chr8 | 70506017 | 70506799 | 21.77823 | 66.26614 | 63.32996 | chr8 | 70500932 | 70507284 | 5.18595 | 4.03112 | 3.19085 |
| chr8 | 121604968 | 121605548 | 404 | Promoter (<=1kb) | 8 | 121600350 | 121604514 | 4165 | 2 | ENSMUST00 | -455 | Zcchc14 | ENSMUSG0C | false | . | . | . | . | . | . | chr8 | 121602612 | 121606344 | 5.77038 | 4.15543 | 3.68262 |
| chr18 | 49979308 | 49979964 | 404 | Promoter (<=1kb) | 18 | 49979427 | 50091001 | 111575 | 1 | ENSMUST00 | 0 | Tnfaip8 | ENSMUSG0C | false | chr18 | 49978679 | 49980374 | 33.75878 | 107.48165 | 103.71877 | . | . | . | . | . | . |
| chr8 | 44941976 | 44942241 | 404 | Intron (ENSMUST00000191428.1 | 8 | 44935447 | 45052257 | 116811 | 1 | ENSMUST00 | 6530 | Fat1 | ENSMUSG0C | true | . | . | . | . | . | . | chr8 | 44939815 | 44952756 | 5.42131 | 4.25024 | 3.3828 |
| chr1 | 172207157 | 172207452 | 403 | Promoter (<=1kb) | 1 | 172199775 | 172206804 | 7030 | 2 | ENSMUST00 | -354 | Pea15a | ENSMUSG0C | false | . | . | . | . | . | . | chr1 | 172200936 | 172207651 | 6.63015 | 4.77931 | 4.3486 |
| chr10 | 68444442 | 68444738 | 403 | Intron (ENSMUST00000020103.3 | 10 | 68436280 | 68525767 | 89488 | 2 | ENSMUST00 | 81029 | 1700040L02F | ENSMUSG0C | false | . | . | . | . | . | . | . | . | . | . | . | . |
| chr11 | 23771306 | 23771546 | 403 | Promoter (<=1kb) | 11 | 23741514 | 23770970 | 29457 | 2 | ENSMUST00 | -337 | Rel | ENSMUSG0C | false | chr11 | 23769609 | 23771991 | 29.36448 | 91.77025 | 88.3935 | . | . | . | . | . | . |

| | | | | | | | | | | | | | | | | | | | | | | | | | | |
|---|---|---|---|---|---|---|---|---|---|---|---|---|---|---|---|---|---|---|---|---|---|---|---|---|---|---|
| chr12 | 12262012 | 12262492 | 403 | Promoter (<=1kb) | 12 | 12262139 | 12376359 | 114221 | 1 | ENSMUST00 | 0 | Fam49a | ENSMUSG0C | false | chr12 | 12261863 | 12263016 | 19.89052 | 67.91473 | 64.9535 | chr12 | 12261396 | 12264758 | 5.2038 | 4.07065 | 3.19463 |
| chr10 | 128921437 | 128921695 | 403 | Promoter (1-2kb) | 10 | 128918420 | 128922888 | 4469 | 2 | ENSMUST00 | 1193 | Rdh5 | ENSMUSG0C | false | . | . | . | . | . | . | chr10 | 128919550 | 128924490 | 5.64648 | 4.33088 | 3.54938 |
| chr2 | 132846451 | 132847117 | 403 | Promoter (<=1kb) | 2 | 132846666 | 132866785 | 20120 | 1 | ENSMUST00 | 0 | Crls1 | ENSMUSG0C | false | chr2 | 132846572 | 132848126 | 35.56825 | 117.95176 | 113.87694 | chr2 | 132846290 | 132846606 | 3.9499 | 3.47103 | 2.14482 |
| chr5 | 103387439 | 103387721 | 403 | Distal Intergenic | 5 | 103425192 | 103598359 | 173168 | 1 | ENSMUST00 | -37471 | Ptpn13 | ENSMUSG0C | false | . | . | . | . | . | . | chr5 | 103387607 | 103388496 | 2.73041 | 2.82053 | 1.20874 |
| chr9 | 35439021 | 35439426 | 403 | Intron (ENSMUST00000151682.2 | 9 | 35452076 | 35507651 | 55576 | 1 | ENSMUST00 | -12650 | Cdon | ENSMUSG0C | false | . | . | . | . | . | . | chr9 | 35437995 | 35440379 | 4.6397 | 3.79706 | 2.7528 |
| chr7 | 112299782 | 112300209 | 403 | Intron (ENSMUST00000050149.6 | 7 | 112316029 | 112347491 | 31463 | 1 | ENSMUST00 | -15820 | Mical2 | ENSMUSG0C | false | . | . | . | . | . | . | chr7 | 112293818 | 112300553 | 6.12041 | 4.56532 | 3.92522 |
| chr10 | 121739819 | 121740088 | 403 | Promoter (<=1kb) | 10 | 121739937 | 121752865 | 12929 | 1 | ENSMUST00 | 0 | BC048403 | ENSMUSG0C | false | chr10 | 121739395 | 121740639 | 35.13972 | 124.62851 | 120.32814 | chr10 | 121739361 | 121740919 | 4.19076 | 3.62404 | 2.35969 |
| chr2 | 140395164 | 140395508 | 403 | Promoter (<=1kb) | 2 | 140395309 | 142392966 | 1997658 | 1 | ENSMUST00 | 0 | Macrod2 | ENSMUSG0C | false | chr2 | 140394864 | 140396429 | 26.74277 | 88.06109 | 84.75893 | chr2 | 140394864 | 140396730 | 3.38595 | 3.22792 | 1.72352 |
| chr2 | 92424997 | 92425267 | 403 | Exon (ENSMUST00000125488.1/ | 2 | 92426904 | 92432392 | 5489 | 2 | ENSMUST00 | 7125 | Cry2 | ENSMUSG0C | false | . | . | . | . | . | . | chr2 | 92421995 | 92433198 | 7.705 | 5.3487 | 5.2209 |
| chr16 | 77618988 | 77619271 | 403 | Intron (ENSMUST00000183099.2 | 16 | 77588578 | 77602094 | 13517 | 2 | ENSMUST00 | -16895 | Gm11146 | ENSMUSG0C | false | . | . | . | . | . | . | chr16 | 77618689 | 77620292 | 3.5829 | 3.3922 | 1.83526 |
| chr17 | 47973800 | 47974505 | 403 | Distal Intergenic | 17 | 48003379 | 48003572 | 194 | 2 | ENSMUST00 | 29067 | Gm14871 | ENSMUSG0C | false | chr17 | 47973525 | 47974624 | 15.92618 | 39.91766 | 37.3274 | chr17 | 47973523 | 47976221 | 5.86638 | 4.39509 | 3.7429 |
| chr18 | 50069826 | 50070207 | 403 | Intron (ENSMUST00000148989.2 | 18 | 50052299 | 50053844 | 1546 | 2 | ENSMUST00 | -15983 | C030005K06I | ENSMUSG0C | false | . | . | . | . | . | . | chr18 | 50054967 | 50071969 | 5.75206 | 4.39775 | 3.62765 |
| chr1 | 23255917 | 23256482 | 403 | Distal Intergenic | 1 | 23245211 | 23245331 | 121 | 1 | ENSMUST00 | 10707 | Gm24176 | ENSMUSG0C | false | chr1 | 23256193 | 23259443 | 33.26644 | 105.59763 | 101.88331 | chr1 | 23255105 | 23255975 | 4.18648 | 3.68431 | 2.38816 |
| chr13 | 44024619 | 44024943 | 403 | Distal Intergenic | 13 | 44121167 | 44125179 | 4013 | 1 | ENSMUST00 | -96224 | Gm5083 | ENSMUSG0C | false | . | . | . | . | . | . | chr13 | 44024117 | 44026966 | 4.12624 | 3.62755 | 2.28723 |
| chr12 | 79838202 | 79838492 | 403 | Distal Intergenic | 12 | 79963568 | 79963872 | 305 | 1 | ENSMUST00 | -125076 | RP23-179D3. | ENSMUSG0C | false | . | . | . | . | . | . | chr12 | 79837421 | 79839476 | 5.18729 | 3.93671 | 3.18617 |
| chr15 | 80012058 | 80012632 | 402 | Promoter (2-3kb) | 15 | 79995900 | 80014808 | 18909 | 2 | ENSMUST00 | 2176 | Pdgfb | ENSMUSG0C | false | chr15 | 80012006 | 80015223 | 33.97523 | 103.77077 | 100.10409 | chr15 | 80009666 | 80015472 | 5.02785 | 4.1066 | 3.0255 |
| chr2 | 164441170 | 164441479 | 402 | Promoter (1-2kb) | 2 | 164424247 | 164443186 | 18940 | 2 | ENSMUST00 | 1707 | Sdc4 | ENSMUSG0C | true | chr2 | 164438320 | 164443697 | 20.02967 | 73.35366 | 70.31202 | . | . | . | . | . | . |
| chr1 | 36860355 | 36860872 | 402 | Intron (ENSMUST00000027290.6 | 1 | 36818660 | 36824781 | 6122 | 2 | ENSMUST00 | -35575 | Tmem131 | ENSMUSG0C | false | . | . | . | . | . | . | chr1 | 36858902 | 36862488 | 5.67821 | 4.18363 | 3.60011 |
| chr4 | 75278131 | 75278441 | 402 | Promoter (<=1kb) | 4 | 75277354 | 75278305 | 952 | 2 | ENSMUST00 | 0 | Tmem261 | ENSMUSG0C | false | chr4 | 75277417 | 75278561 | 26.1316 | 74.80193 | 71.73632 | . | . | . | . | . | . |
| chr7 | 65419070 | 65419350 | 402 | Distal Intergenic | 7 | 65296165 | 65371239 | 75075 | 2 | ENSMUST00 | -47832 | Tjp1 | ENSMUSG0C | true | . | . | . | . | . | . | chr7 | 65409463 | 65433128 | 6.06932 | 4.54697 | 3.8847 |
| chr9 | 62341810 | 62343091 | 402 | Promoter (<=1kb) | 9 | 62341902 | 62374718 | 32817 | 1 | ENSMUST00 | 0 | Anp32a | ENSMUSG0C | false | chr9 | 62340409 | 62345269 | 38.55812 | 129.64772 | 125.16568 | . | . | . | . | . | . |
| chr9 | 58308419 | 58308715 | 402 | Intron (ENSMUST00000061799.8 | 9 | 58287723 | 58313212 | 25490 | 2 | ENSMUST00 | 4497 | Loxl1 | ENSMUSG0C | false | . | . | . | . | . | . | chr9 | 58302831 | 58310544 | 4.56252 | 3.71116 | 2.67437 |
| chr8 | 111541874 | 111542362 | 402 | Promoter (<=1kb) | 8 | 111541259 | 111621664 | 80406 | 1 | ENSMUST00 | 616 | Znrf1 | ENSMUSG0C | false | . | . | . | . | . | . | chr8 | 111537768 | 111552011 | 5.57979 | 4.33787 | 3.48276 |
| chr4 | 136206088 | 136206636 | 402 | Promoter (<=1kb) | 4 | 136206365 | 136245216 | 38852 | 1 | ENSMUST00 | 0 | Asap3 | ENSMUSG0C | false | chr4 | 136205955 | 136207125 | 29.14328 | 85.59041 | 82.33509 | chr4 | 136205771 | 136208529 | 4.62227 | 3.60277 | 2.74134 |
| chr5 | 134639064 | 134639591 | 402 | Promoter (<=1kb) | 5 | 134619880 | 134639348 | 19469 | 2 | ENSMUST00 | 0 | Eif4h | ENSMUSG0C | false | chr5 | 134637596 | 134639924 | 37.7199 | 126.00829 | 121.65485 | . | . | . | . | . | . |
| chr1 | 90203737 | 90204042 | 402 | Promoter (<=1kb) | 1 | 90203980 | 90216751 | 12772 | 1 | ENSMUST00 | 0 | Ackr3 | ENSMUSG0C | false | chr1 | 90203977 | 90206221 | 15.92618 | 39.91766 | 37.3274 | chr1 | 90202500 | 90212099 | 3.28128 | 3.13332 | 1.63821 |
| chr1 | 106242135 | 106242360 | 402 | Intron (ENSMUST00000061047.6 | 1 | 106281425 | 106287422 | 5998 | 1 | ENSMUST00 | -39065 | Phlpp1 | ENSMUSG0C | false | . | . | . | . | . | . | chr1 | 106240384 | 106245684 | 5.95652 | 4.50217 | 3.79421 |
| chr7 | 127345363 | 127346412 | 402 | Promoter (<=1kb) | 7 | 127342797 | 127345314 | 2518 | 2 | ENSMUST00 | -50 | Zfp768 | ENSMUSG0C | false | chr7 | 127344643 | 127346973 | 28.22734 | 101.53345 | 97.92473 | . | . | . | . | . | . |
| chr8 | 82858675 | 82858914 | 402 | Distal Intergenic | 8 | 82863356 | 83091271 | 227916 | 1 | ENSMUST00 | -4442 | Rnf150 | ENSMUSG0C | true | . | . | . | . | . | . | chr8 | 82858133 | 82859787 | 4.82369 | 4.04906 | 2.86175 |
| chr12 | 104985848 | 104986162 | 402 | Intron (ENSMUST00000095439.4 | 12 | 104929949 | 104998677 | 68729 | 2 | ENSMUST00 | 12515 | Syne3 | ENSMUSG0C | false | . | . | . | . | . | . | chr12 | 104974830 | 104987771 | 6.02697 | 4.55969 | 3.85063 |
| chr1 | 84883948 | 84884213 | 402 | Intron (ENSMUST00000097672.3 | 1 | 84869151 | 84900487 | 31337 | 1 | ENSMUST00 | 14798 | Fbxo36 | ENSMUSG0C | false | . | . | . | . | . | . | chr1 | 84883793 | 84884388 | 3.11133 | 3.01304 | 1.51565 |
| chr9 | 102996250 | 102996641 | 402 | Intron (ENSMUST00000185215.1 | 9 | 102991895 | 102992216 | 322 | 1 | ENSMUST00 | 4356 | Gm25842 | ENSMUSG0C | false | . | . | . | . | . | . | chr9 | 102991889 | 103008204 | 5.47516 | 4.26994 | 3.39839 |
| chr9 | 71345674 | 71346065 | 401 | Distal Intergenic | 9 | 71215789 | 71296243 | 80455 | 1 | ENSMUST00 | 129886 | Aldh1a2 | ENSMUSG0C | false | . | . | . | . | . | . | . | . | . | . | . | . |
| chr17 | 34655532 | 34656005 | 401 | Distal Intergenic | 17 | 34651627 | 34652103 | 477 | 1 | ENSMUST00 | 3906 | Atf6b | ENSMUSG0C | false | . | . | . | . | . | . | chr17 | 34654434 | 34657679 | 4.55416 | 3.89933 | 2.63159 |
| chr17 | 46160785 | 46161101 | 401 | Promoter (<=1kb) | 17 | 46161032 | 46169368 | 8337 | 1 | ENSMUST00 | 0 | Gtpbp2 | ENSMUSG0C | false | chr17 | 46159755 | 46161963 | 39.51976 | 124.83012 | 120.52397 | . | . | . | . | . | . |
| chr4 | 148804153 | 148804497 | 401 | Promoter (<=1kb) | 4 | 148804429 | 148945129 | 140701 | 1 | ENSMUST00 | 0 | Casz1 | ENSMUSG0C | true | chr4 | 148803997 | 148806784 | 38.29964 | 140.16156 | 135.27679 | chr4 | 148803702 | 148804250 | 4.3582 | 3.67235 | 2.49285 |
| chr5 | 65335322 | 65335888 | 401 | Promoter (<=1kb) | 5 | 65261852 | 65335670 | 73819 | 2 | ENSMUST00 | 0 | Rfc1 | ENSMUSG0C | false | chr5 | 65334890 | 65336115 | 38.03245 | 121.19012 | 117.00504 | chr5 | 65334289 | 65336489 | 3.6019 | 3.3743 | 1.90259 |
| chr6 | 121245648 | 121245997 | 401 | Promoter (<=1kb) | 6 | 121245906 | 121270916 | 25011 | 1 | ENSMUST00 | 0 | Usp18 | ENSMUSG0C | false | chr6 | 121245119 | 121246418 | 23.81181 | 65.46733 | 62.54538 | . | . | . | . | . | . |
| chr9 | 71467376 | 71467733 | 401 | Distal Intergenic | 9 | 71478471 | 71484958 | 6488 | 2 | ENSMUST00 | 17225 | Polr2m | ENSMUSG0C | false | . | . | . | . | . | . | chr9 | 71466318 | 71468500 | 4.47065 | 3.84444 | 2.56323 |
| chr9 | 119967731 | 119967999 | 401 | 3' UTR | 9 | 119961618 | 119966169 | 4552 | 1 | ENSMUST00 | 6114 | Ttc21a | ENSMUSG0C | false | . | . | . | . | . | . | chr9 | 119967128 | 119967743 | 3.17747 | 3.14284 | 1.56787 |
| chr4 | 98806394 | 98806738 | 401 | Intron (ENSMUST00000102790.3 | 4 | 98754898 | 98817537 | 62640 | 2 | ENSMUST00 | 10799 | Kank4 | ENSMUSG0C | false | . | . | . | . | . | . | . | . | . | . | . | . |
| chr2 | 33616357 | 33616728 | 401 | Intron (ENSMUST00000041730.5 | 2 | 33567156 | 33632283 | 65128 | 2 | ENSMUST00 | 15555 | Lmx1b | ENSMUSG0C | true | chr2 | 33616045 | 33617221 | 5.56731 | 11.5554 | 9.34776 | chr2 | 33565260 | 33638297 | 6.78481 | 4.81457 | 4.48235 |
| chr11 | 97438936 | 97439176 | 401 | Promoter (2-3kb) | 11 | 97436285 | 97444228 | 7944 | 1 | ENSMUST00 | 2652 | Arhgap23 | ENSMUSG0C | true | . | . | . | . | . | . | chr11 | 97431804 | 97440135 | 9.81823 | 6.18842 | 6.94298 |
| chr1 | 120074219 | 120074617 | 401 | Promoter (<=1kb) | 1 | 120067377 | 120074074 | 6698 | 2 | ENSMUST00 | -146 | Tmem37 | ENSMUSG0C | false | chr1 | 120072898 | 120074442 | 18.85995 | 50.28701 | 47.56678 | chr1 | 120072417 | 120075840 | 5.62407 | 4.252 | 3.56349 |
| chr15 | 36892892 | 36893126 | 401 | Distal Intergenic | 15 | 36870501 | 36879816 | 9316 | 2 | ENSMUST00 | -13077 | Gm10384 | ENSMUSG0C | false | . | . | . | . | . | . | chr15 | 36892121 | 36894159 | 3.42626 | 3.17657 | 1.73295 |
| chr7 | 80457329 | 80457625 | 401 | Intron (ENSMUST00000170315.1 | 7 | 80429210 | 80430740 | 1531 | 1 | ENSMUST00 | 28120 | Gm15544 | ENSMUSG0C | false | . | . | . | . | . | . | chr7 | 80456026 | 80457958 | 2.80125 | 2.83557 | 1.24522 |
| chr15 | 73732029 | 73732694 | 401 | Promoter (2-3kb) | 15 | 73729164 | 73729290 | 127 | 1 | ENSMUST00 | 2866 | Gm22106 | ENSMUSG0C | false | . | . | . | . | . | . | chr15 | 73729948 | 73733332 | 4.85267 | 3.92399 | 2.89329 |
| chr18 | 64807596 | 64807838 | 401 | Distal Intergenic | 18 | 64786329 | 64786428 | 100 | 2 | ENSMUST00 | -21169 | Gm24504 | ENSMUSG0C | false | . | . | . | . | . | . | . | . | . | . | . | . |
| chr10 | 110807535 | 110807800 | 401 | Intron (ENSMUST00000188776.1 | 10 | 110796173 | 110796300 | 128 | 1 | ENSMUST00 | 11363 | Gm23698 | ENSMUSG0C | false | . | . | . | . | . | . | chr10 | 110803361 | 110808639 | 4.77259 | 3.87002 | 2.82892 |
| chr3 | 65658753 | 65659558 | 401 | Promoter (<=1kb) | 3 | 65659288 | 65659426 | 139 | 1 | ENSMUST00 | 0 | Mir8120 | ENSMUSG0C | false | chr3 | 65658315 | 65659888 | 31.89884 | 96.17711 | 92.69861 | . | . | . | . | . | . |
| chr8 | 84687745 | 84688078 | 400 | Promoter (<=1kb) | 8 | 84670483 | 84687862 | 17380 | 2 | ENSMUST00 | 0 | Nacc1 | ENSMUSG0C | false | chr8 | 84686292 | 84688224 | 30.14656 | 119.38538 | 115.2599 | . | . | . | . | . | . |
| chr16 | 89960734 | 89961130 | 400 | Promoter (<=1kb) | 16 | 89787111 | 89960827 | 173717 | 2 | ENSMUST00 | 0 | Tiam1 | ENSMUSG0C | false | chr16 | 89960264 | 89960876 | 12.40737 | 26.6269 | 24.20249 | chr16 | 89959477 | 89962284 | 4.51657 | 3.84983 | 2.61607 |
| chr15 | 103214887 | 103215533 | 400 | Promoter (<=1kb) | 15 | 103196687 | 103215352 | 18666 | 2 | ENSMUST00 | 0 | Cbx5 | ENSMUSG0C | false | chr15 | 103214892 | 103216001 | 12.92336 | 34.59913 | 32.07519 | chr15 | 103213987 | 103216744 | 4.06078 | 3.66084 | 2.22595 |
| chr11 | 96988401 | 96988740 | 400 | Distal Intergenic | 11 | 96954219 | 96982959 | 28741 | 2 | ENSMUST00 | -5443 | Sp2 | ENSMUSG0C | false | chr11 | 96988127 | 96988449 | 4.97263 | 8.68936 | 6.53912 | chr11 | 96984955 | 96998855 | 6.28932 | 4.5897 | 4.08329 |
| chr11 | 49847328 | 49847574 | 400 | Promoter (<=1kb) | 11 | 49846893 | 49873397 | 26505 | 1 | ENSMUST00 | 436 | Mapk9 | ENSMUSG0C | false | chr11 | 49846235 | 49847738 | 33.04283 | 115.53907 | 111.53828 | . | . | . | . | . | . |
| chr11 | 69125448 | 69125675 | 400 | Promoter (<=1kb) | 11 | 69125896 | 69149115 | 23220 | 1 | ENSMUST00 | -221 | Aloxe3 | ENSMUSG0C | false | chr11 | 69124925 | 69126298 | 16.16105 | 50.86513 | 48.14017 | chr11 | 69124511 | 69126544 | 3.60281 | 3.40145 | 1.86943 |
| chr13 | 98316326 | 98317133 | 400 | Promoter (<=1kb) | 13 | 98309896 | 98316967 | 7072 | 2 | ENSMUST00 | 0 | Btf3 | ENSMUSG0C | false | chr13 | 98315862 | 98317835 | 29.83578 | 117.74437 | 113.67492 | . | . | . | . | . | . |
| chr17 | 31777104 | 31777427 | 400 | Distal Intergenic | 17 | 31844248 | 31855792 | 11545 | 2 | ENSMUST00 | 78365 | Sik1 | ENSMUSG0C | false | . | . | . | . | . | . | chr17 | 31776737 | 31778634 | 4.73225 | 3.83111 | 2.82663 |
| chr2 | 45109284 | 45110254 | 400 | Promoter (<=1kb) | 2 | 44988388 | 45110244 | 121857 | 2 | ENSMUST00 | 0 | Zeb2 | ENSMUSG0C | false | chr2 | 45108545 | 45111853 | 37.2221 | 115.56093 | 111.55734 | . | . | . | . | . | . |
| chr4 | 123370112 | 123370321 | 400 | Promoter (<=1kb) | 4 | 123349633 | 123369931 | 20299 | 2 | ENSMUST00 | -182 | Macf1 | ENSMUSG0C | true | . | . | . | . | . | . | chr4 | 123369856 | 123370617 | 4.51278 | 3.53589 | 2.63499 |
| chr5 | 110802173 | 110802439 | 400 | Intron (ENSMUST00000031490.6 | 5 | 110784488 | 110810097 | 25610 | 2 | ENSMUST00 | 7658 | Ulk1 | ENSMUSG0C | false | . | . | . | . | . | . | chr5 | 110797106 | 110809391 | 6.92459 | 4.77628 | 4.6086 |
| chr12 | 112554785 | 112555136 | 400 | Distal Intergenic | 12 | 112588784 | 112615556 | 26773 | 1 | ENSMUST00 | -33648 | Inf2 | ENSMUSG0C | false | . | . | . | . | . | . | chr12 | 112552233 | 112556761 | 4.66336 | 3.79449 | 2.7644 |
| chr3 | 89251341 | 89251860 | 400 | Promoter (2-3kb) | 3 | 89248867 | 89249722 | 856 | 1 | ENSMUST00 | 2475 | Krtcap2 | ENSMUSG0C | false | . | . | . | . | . | . | chr3 | 89250573 | 89253531 | 4.93424 | 3.94571 | 2.97227 |
| chr11 | 60139687 | 60140099 | 400 | Promoter (<=1kb) | 11 | 60140082 | 60199195 | 59114 | 1 | ENSMUST00 | 0 | Rai1 | ENSMUSG0C | false | chr11 | 60139224 | 60140901 | 27.98104 | 79.82376 | 76.67776 | chr11 | 60138396 | 60141338 | 3.91042 | 3.57283 | 2.09867 |
| chr5 | 124030345 | 124030593 | 400 | Promoter (1-2kb) | 5 | 124004641 | 124032260 | 27620 | 2 | ENSMUST00 | 1667 | Vps37b | ENSMUSG0C | false | . | . | . | . | . | . | chr5 | 124024488 | 124032679 | 7.6835 | 5.26739 | 5.19795 |
| chr2 | 32661687 | 32662041 | 400 | Intron (ENSMUST00000009705.8 | 2 | 32652329 | 32652414 | 86 | 1 | ENSMUST00 | 9359 | Mir1954 | ENSMUSG0C | false | . | . | . | . | . | . | . | . | . | . | . | . |
| chr3 | 108146056 | 108146343 | 399 | Promoter (<=1kb) | 3 | 108107280 | 108146146 | 38867 | 2 | ENSMUST00 | 0 | Gnai3 | ENSMUSG0C | false | chr3 | 108145284 | 108146677 | 29.61888 | 117.31387 | 113.25776 | chr3 | 108141029 | 108147867 | 4.96005 | 4.03426 | 2.9789 |
| chr10 | 75932289 | 75932605 | 399 | Promoter (<=1kb) | 10 | 75923222 | 75932475 | 9254 | 2 | ENSMUST00 | 0 | Mmp11 | ENSMUSG0C | false | chr10 | 75932124 | 75932953 | 12.0178 | 35.52963 | 32.99667 | chr10 | 75929929 | 75933151 | 5.39687 | 4.27468 | 3.36518 |
| chr10 | 80039557 | 80039982 | 399 | Promoter (<=1kb) | 10 | 80035949 | 80039710 | 3762 | 2 | ENSMUST00 | 0 | Polr2e | ENSMUSG0C | false | chr10 | 80038985 | 80039798 | 30.19165 | 95.18787 | 91.73401 | chr10 | 80031609 | 80040449 | 6.23886 | 4.55556 | 4.03957 |
| chr11 | 61619782 | 61620102 | 399 | Distal Intergenic | 11 | 61653265 | 61672784 | 19520 | 1 | ENSMUST00 | -33163 | Grap | ENSMUSG0C | false | chr11 | 61618794 | 61620874 | 12.76342 | 27.83059 | 25.39249 | chr11 | 61610870 | 61621664 | 5.06575 | 4.05211 | 3.07721 |
| chr2 | 74578796 | 74579271 | 399 | Promoter (<=1kb) | 2 | 74520291 | 74578948 | 58658 | 2 | ENSMUST00 | 0 | Lnp | ENSMUSG0C | false | chr2 | 74577860 | 74579489 | 26.32471 | 106.6389 | 102.89503 | . | . | . | . | . | . |
| chr15 | 80133045 | 80133542 | 399 | Promoter (<=1kb) | 15 | 80133154 | 80161701 | 28548 | 1 | ENSMUST00 | 0 | Tab1 | ENSMUSG0C | false | chr15 | 80132695 | 80134176 | 24.78212 | 78.86476 | 75.73294 | chr15 | 80132452 | 80135076 | 5.60141 | 4.37527 | 3.50624 |
| chr18 | 7910403 | 7910661 | 399 | Promoter (<=1kb) | 18 | 7910601 | 7916387 | 5787 | 1 | ENSMUST00 | 0 | Wac | ENSMUSG0C | false | . | . | . | . | . | . | chr18 | 7906408 | 7914385 | 3.75861 | 3.39961 | 1.99304 |
| chr19 | 55106870 | 55107072 | 399 | Distal Intergenic | 19 | 55069734 | 55099451 | 29718 | 2 | ENSMUST00 | -7420 | Gpam | ENSMUSG0C | false | . | . | . | . | . | . | chr19 | 55105863 | 55107788 | 6.02905 | 4.43378 | 3.88519 |
| chr1 | 98095389 | 98095789 | 399 | Promoter (<=1kb) | 1 | 97821094 | 98095621 | 274528 | 2 | ENSMUST00 | 0 | Pam | ENSMUSG0C | false | chr1 | 98094581 | 98096244 | 36.30304 | 111.89011 | 107.99921 | . | . | . | . | . | . |
| chr6 | 113326600 | 113327118 | 399 | Promoter (<=1kb) | 6 | 113326972 | 113334188 | 7217 | 1 | ENSMUST00 | 0 | Ogg1 | ENSMUSG0C | false | chr6 | 113326915 | 113327680 | 14.02077 | 44.3905 | 41.74443 | . | . | . | . | . | . |
| chr9 | 83582850 | 83583120 | 399 | Intron (ENSMUST00000188241.1 | 9 | 83576831 | 83597524 | 20694 | 1 | ENSMUST00 | 6020 | Sh3bgrl2 | ENSMUSG0C | true | . | . | . | . | . | . | chr9 | 83578741 | 83594009 | 5.16719 | 4.03655 | 3.17944 |
| chr9 | 113741040 | 113741634 | 399 | Distal Intergenic | 9 | 113651744 | 113708259 | 56516 | 2 | ENSMUST00 | -32782 | Pdcd6ip | ENSMUSG0C | false | chr9 | 113740556 | 113742818 | 39.80159 | 147.3544 | 142.17488 | . | . | . | . | . | . |
| chr9 | 119725340 | 119725631 | 399 | Distal Intergenic | 9 | 119608456 | 119719032 | 110577 | 2 | ENSMUST00 | -6309 | Scn10a | ENSMUSG0C | false | . | . | . | . | . | . | . | . | . | . | . | . |
| chr5 | 77400225 | 77400510 | 399 | Intron (ENSMUST00000046746.6 | 5 | 77349240 | 77408045 | 58806 | 2 | ENSMUST00 | 7535 | Igfbp7 | ENSMUSG0C | false | . | . | . | . | . | . | chr5 | 77353137 | 77404361 | 6.51707 | 4.75542 | 4.25399 |
| chr7 | 3290212 | 3290613 | 399 | Promoter (<=1kb) | 7 | 3290556 | 3299345 | 8790 | 1 | ENSMUST00 | 0 | Myadm | ENSMUSG0C | false | chr7 | 3289024 | 3290357 | 13.70367 | 30.66556 | 28.1932 | . | . | . | . | . | . |
| chr11 | 57696159 | 57696376 | 399 | Intron (ENSMUST00000066987.8 | 11 | 57655415 | 57657826 | 2412 | 2 | ENSMUST00 | -38334 | 4933426K07I | ENSMUSG0C | false | . | . | . | . | . | . | chr11 | 57646153 | 57704104 | 7.42247 | 5.08195 | 5.00645 |
| chr1 | 89556474 | 89556754 | 399 | Intron (ENSMUST00000027521.9 | 1 | 89563360 | 89563474 | 115 | 1 | ENSMUST00 | -6606 | Gm25180 | ENSMUSG0C | false | . | . | . | . | . | . | chr1 | 89553718 | 89560143 | 7.51776 | 5.25618 | 5.0628 |
| chr18 | 82854779 | 82855004 | 399 | Promoter (<=1kb) | 18 | 82855894 | 82870604 | 14711 | 1 | ENSMUST00 | -890 | 4930445N18 | ENSMUSG0C | false | . | . | . | . | . | . | chr18 | 82854604 | 82856624 | 3.96784 | 3.52576 | 2.19508 |
| chr11 | 77607320 | 77608166 | 398 | Promoter (<=1kb) | 11 | 77529162 | 77607815 | 78654 | 2 | ENSMUST00 | 0 | Taok1 | ENSMUSG0C | false | chr11 | 77606424 | 77608552 | 31.91637 | 110.38313 | 106.53261 | . | . | . | . | . | . |
| chr13 | 43191746 | 43192195 | 398 | Distal Intergenic | 13 | 43151744 | 43171501 | 19758 | 2 | ENSMUST00 | -20246 | Tbc1d7 | ENSMUSG0C | false | . | . | . | . | . | . | chr13 | 43191139 | 43192316 | 4.66507 | 3.73675 | 2.76897 |
| chr15 | 102517969 | 102518877 | 398 | Promoter (<=1kb) | 15 | 102518192 | 102523674 | 5483 | 1 | ENSMUST00 | 0 | Tarbp2 | ENSMUSG0C | false | chr15 | 102518102 | 102519426 | 22.78592 | 76.10319 | 73.01487 | . | . | . | . | . | . |
| chr17 | 33915812 | 33916299 | 398 | Promoter (<=1kb) | 17 | 33915899 | 33920415 | 4517 | 1 | ENSMUST00 | 0 | Tapbp | ENSMUSG0C | false | chr17 | 33916115 | 33917028 | 17.46222 | 42.53877 | 39.91948 | chr17 | 33915195 | 33919300 | 5.2192 | 4.01698 | 3.2267 |
| chrX | 99191511 | 99191789 | 398 | Distal Intergenic | 20 | 99138131 | 99147230 | 9100 | 1 | ENSMUST00 | 53381 | Efnb1 | ENSMUSG0C | false | . | . | . | . | . | . | . | . | . | . | . | . |
| chr2 | 104098627 | 104098852 | 398 | Promoter (2-3kb) | 2 | 104095840 | 104114263 | 18424 | 1 | ENSMUST00 | 2788 | Cd59a | ENSMUSG0C | true | chr2 | 104098304 | 104098985 | 8.08967 | 17.56143 | 15.25973 | chr2 | 104096873 | 104114534 | 4.7333 | 3.92676 | 2.79529 |
| chr12 | 99469659 | 99469917 | 398 | Distal Intergenic | 12 | 99388439 | 99450111 | 61673 | 2 | ENSMUST00 | -19549 | Foxn3 | ENSMUSG0C | false | . | . | . | . | . | . | . | . | . | . | . | . |
| chr9 | 46073305 | 46073569 | 398 | Intron (ENSMUST00000126865.2 | 9 | 46123237 | 46202885 | 79649 | 1 | ENSMUST00 | -49668 | Sik3 | ENSMUSG0C | false | . | . | . | . | . | . | chr9 | 46065832 | 46078010 | 4.88141 | 3.97476 | 2.92323 |
| chr14 | 47406134 | 47406320 | 398 | Intron (ENSMUST00000043296.4 | 14 | 47398497 | 47411666 | 13170 | 2 | ENSMUST00 | 5346 | Dlgap5 | ENSMUSG0C | false | . | . | . | . | . | . | chr14 | 47405944 | 47407116 | 3.51369 | 3.20318 | 1.80091 |
| chr2 | 33130218 | 33131442 | 398 | Promoter (<=1kb) | 2 | 33054158 | 33131436 | 77279 | 2 | ENSMUST00 | 0 | Garnl3 | ENSMUSG0C | false | chr2 | 33129759 | 33132001 | 37.04683 | 119.47973 | 115.35273 | chr2 | 33129635 | 33130416 | 6.08972 | 4.56942 | 3.94427 |
| chr5 | 143112466 | 143113162 | 398 | Promoter (<=1kb) | 5 | 142990893 | 143113020 | 122128 | 2 | ENSMUST00 | 0 | Rnf216 | ENSMUSG0C | false | chr5 | 143111903 | 143113458 | 29.34276 | 99.62727 | 96.06711 | chr5 | 143100232 | 143113412 | 4.82716 | 3.92416 | 2.88218 |
| chr16 | 18453870 | 18454191 | 398 | Promoter (1-2kb) | 16 | 18452045 | 18472886 | 20842 | 1 | ENSMUST00 | 1826 | Txnrd2 | ENSMUSG0C | false | . | . | . | . | . | . | . | . | . | . | . | . |
| chrX | 101274023 | 101274269 | 398 | Promoter (<=1kb) | 20 | 101274030 | 101297465 | 23436 | 1 | ENSMUST00 | 0 | Med12 | ENSMUSG0C | false | chrX | 101273700 | 101274973 | 16.14593 | 43.98602 | 41.34693 | . | . | . | . | . | . |
| chr7 | 96965155 | 96965382 | 398 | Intron (ENSMUST00000044466.6 | 7 | 96947910 | 96951886 | 3977 | 2 | ENSMUST00 | -13270 | C230038L03F | ENSMUSG0C | false | . | . | . | . | . | . | chr7 | 96952896 | 96991589 | 4.87493 | 3.9704 | 2.91243 |
| chr8 | 72160801 | 72161090 | 398 | Promoter (<=1kb) | 8 | 72147601 | 72160855 | 13255 | 2 | ENSMUST00 | 0 | Gm16091 | ENSMUSG0C | false | chr8 | 72160295 | 72160929 | 20.27628 | 60.14116 | 57.29097 | . | . | . | . | . | . |
| chr10 | 83859628 | 83859985 | 398 | Distal Intergenic | 10 | 83722865 | 83762761 | 39897 | 1 | ENSMUST00 | 136764 | 1500009L16F | ENSMUSG0C | false | . | . | . | . | . | . | . | . | . | . | . | . |
| chr17 | 10481762 | 10482001 | 398 | Intron (ENSMUST00000159602.1 | 17 | 10496446 | 10569396 | 72951 | 1 | ENSMUST00 | -14445 | Gm16168 | ENSMUSG0C | false | . | . | . | . | . | . | chr17 | 10472916 | 10482475 | 5.53103 | 4.25636 | 3.45832 |
| chr4 | 148066578 | 148066860 | 398 | Promoter (1-2kb) | 4 | 148068031 | 148068492 | 462 | 2 | ENSMUST00 | 1632 | Gm13202 | ENSMUSG0C | false | . | . | . | . | . | . | . | . | . | . | . | . |
| chr12 | 111663278 | 111663565 | 397 | Distal Intergenic | 12 | 111669355 | 111672338 | 2984 | 2 | ENSMUST00 | 8773 | Ckb | ENSMUSG0C | false | . | . | . | . | . | . | chr12 | 111662983 | 111665961 | 5.6742 | 4.24256 | 3.59899 |
| chr12 | 85824506 | 85825217 | 397 | Promoter (<=1kb) | 12 | 85824659 | 85857405 | 32747 | 1 | ENSMUST00 | 0 | Ttll5 | ENSMUSG0C | false | chr12 | 85823896 | 85825603 | 24.89402 | 98.85246 | 95.31046 | chr12 | 85823594 | 85825711 | 5.5576 | 4.2702 | 3.48906 |
| chr16 | 32913921 | 32914580 | 397 | Promoter (<=1kb) | 16 | 32914100 | 33015647 | 101548 | 1 | ENSMUST00 | 0 | Lrch3 | ENSMUSG0C | false | chr16 | 32913705 | 32915472 | 35.22365 | 121.58154 | 117.38489 | . | . | . | . | . | . |
| chr5 | 90890955 | 90891256 | 397 | Promoter (<=1kb) | 5 | 90891241 | 90893115 | 1875 | 1 | ENSMUST00 | 0 | Cxcl1 | ENSMUSG0C | false | chr5 | 90890807 | 90891600 | 20.42746 | 52.97813 | 50.22504 | chr5 | 90890713 | 90893350 | 3.86829 | 3.55076 | 2.0801 |
| chr8 | 11797144 | 11797438 | 397 | Intron (ENSMUST00000033908.7 | 8 | 11782641 | 11835219 | 52579 | 1 | ENSMUST00 | 14504 | Arhgef7 | ENSMUSG0C | false | . | . | . | . | . | . | chr8 | 11794930 | 11798601 | 4.72114 | 3.94725 | 2.82239 |
| chr9 | 114805119 | 114805721 | 397 | Intron (ENSMUST00000047013.3 | 9 | 114756837 | 114781993 | 25157 | 2 | ENSMUST00 | -23127 | Cmtm7 | ENSMUSG0C | true | . | . | . | . | . | . | chr9 | 114801208 | 114806677 | 5.65143 | 4.12227 | 3.58214 |
| chr19 | 25374855 | 25375157 | 397 | Intron (ENSMUST00000049400.9 | 19 | 25395303 | 25409772 | 14470 | 1 | ENSMUST00 | -20146 | Kank1 | ENSMUSG0C | true | . | . | . | . | . | . | chr19 | 25372205 | 25390774 | 5.07753 | 4.06725 | 3.07529 |
| chr8 | 105796188 | 105796476 | 397 | Intron (ENSMUST00000041400.5 | 8 | 105768308 | 105827350 | 59043 | 2 | ENSMUST00 | 30874 | Ranbp10 | ENSMUSG0C | false | . | . | . | . | . | . | chr8 | 105787963 | 105815970 | 5.80473 | 4.39186 | 3.67716 |
| chr2 | 154500797 | 154501061 | 397 | Intron (ENSMUST00000109725.2 | 2 | 154510536 | 154539356 | 28821 | 1 | ENSMUST00 | -9475 | Cbfa2t2 | ENSMUSG0C | false | . | . | . | . | . | . | chr2 | 154498493 | 154505686 | 4.96722 | 4.07088 | 2.99062 |
| chr5 | 28071839 | 28072126 | 397 | Promoter (<=1kb) | 5 | 28071363 | 28078662 | 7300 | 1 | ENSMUST00 | 477 | Insig1 | ENSMUSG0C | false | chr5 | 28071277 | 28072315 | 39.37477 | 126.73389 | 122.35824 | chr5 | 28069872 | 28073371 | 3.7566 | 3.35457 | 1.99974 |
| chr11 | 77097010 | 77097237 | 397 | Intron (ENSMUST00000108400.2 | 11 | 77089944 | 77103062 | 13119 | 2 | ENSMUST00 | 5825 | Efcab5 | ENSMUSG0C | false | . | . | . | . | . | . | . | . | . | . | . | . |
| chr18 | 14782979 | 14783569 | 397 | Promoter (<=1kb) | 18 | 14783245 | 14900359 | 117115 | 1 | ENSMUST00 | 0 | Taf4b | ENSMUSG0C | false | chr18 | 14782562 | 14784378 | 32.27147 | 102.84008 | 99.19724 | . | . | . | . | . | . |
| chr11 | 97684776 | 97685079 | 397 | Promoter (<=1kb) | 11 | 97685826 | 97688621 | 2796 | 1 | ENSMUST00 | -747 | Cisd3 | ENSMUSG0C | false | . | . | . | . | . | . | chr11 | 97683689 | 97687456 | 6.46404 | 4.74717 | 4.22185 |
| chr13 | 44817799 | 44818044 | 397 | Intron (ENSMUST00000174068.2 | 13 | 44838165 | 44838283 | 119 | 2 | ENSMUST00 | 20239 | Gm22213 | ENSMUSG0C | false | . | . | . | . | . | . | chr13 | 44734396 | 44823919 | 5.05049 | 4.0328 | 3.07079 |
| chr9 | 107438890 | 107439242 | 397 | Intron (ENSMUST00000010210.7 | 9 | 107443950 | 107445031 | 1082 | 2 | ENSMUST00 | 5789 | Gm17118 | ENSMUSG0C | false | . | . | . | . | . | . | . | . | . | . | . | . |
| chr10 | 108494043 | 108494354 | 397 | Distal Intergenic | 10 | 108428067 | 108428160 | 94 | 1 | ENSMUST00 | 65977 | Gm25591 | ENSMUSG0C | false | . | . | . | . | . | . | . | . | . | . | . | . |
| chr7 | 39556549 | 39556787 | 397 | Intron (ENSMUST00000188003.1 | 7 | 39552525 | 39565210 | 12686 | 1 | ENSMUST00 | 4025 | RP23-402K2 | ENSMUSG0C | false | . | . | . | . | . | . | . | . | . | . | . | . |
| chr10 | 78104747 | 78105365 | 396 | Distal Intergenic | 10 | 78069360 | 78079525 | 10166 | 1 | ENSMUST00 | 35388 | Icosl | ENSMUSG0C | false | . | . | . | . | . | . | . | . | . | . | . | . |
| chr15 | 75921229 | 75921664 | 396 | Promoter (<=1kb) | 15 | 75916463 | 75921560 | 5098 | 2 | ENSMUST00 | 0 | Pycrl | ENSMUSG0C | false | chr15 | 75920962 | 75921570 | 28.50037 | 84.48055 | 81.24493 | chr15 | 75920066 | 75924880 | 5.20598 | 4.11154 | 3.21735 |
| chr18 | 34926830 | 34927076 | 396 | Intron (ENSMUST00000025218.7 | 18 | 34931834 | 34931941 | 108 | 2 | ENSMUST00 | 4865 | Etf1 | ENSMUSG0C | false | . | . | . | . | . | . | chr18 | 34913989 | 34930918 | 5.3617 | 4.17498 | 3.32231 |
| chr19 | 56516933 | 56517146 | 396 | Distal Intergenic | 19 | 56529582 | 56537382 | 7801 | 2 | ENSMUST00 | 20236 | Dclre1a | ENSMUSG0C | false | . | . | . | . | . | . | chr19 | 56516347 | 56517847 | 3.66743 | 3.43543 | 1.91084 |
| chr11 | 121434764 | 121435046 | 396 | Promoter (<=1kb) | 11 | 121434913 | 121450491 | 15579 | 1 | ENSMUST00 | 0 | Fn3k | ENSMUSG0C | false | chr11 | 121434257 | 121435431 | 18.79251 | 47.82232 | 45.13606 | chr11 | 121433646 | 121435844 | 5.40849 | 3.9851 | 3.37481 |
| chr7 | 78846944 | 78847361 | 396 | Promoter (<=1kb) | 7 | 78827474 | 78847236 | 19763 | 2 | ENSMUST00 | 0 | Det1 | ENSMUSG0C | false | chr7 | 78846364 | 78847461 | 32.64849 | 123.19247 | 118.94151 | . | . | . | . | . | . |
| chr8 | 36610610 | 36610868 | 396 | Intron (ENSMUST00000033923.8 | 8 | 36593231 | 36613874 | 20644 | 2 | ENSMUST00 | 3006 | Dlc1 | ENSMUSG0C | true | . | . | . | . | . | . | chr8 | 36575944 | 36612020 | 5.64773 | 4.3412 | 3.54039 |
| chr8 | 36721775 | 36721983 | 396 | Intron (ENSMUST00000098826.4 | 8 | 36567751 | 36733054 | 165304 | 2 | ENSMUST00 | 11071 | Dlc1 | ENSMUSG0C | true | . | . | . | . | . | . | chr8 | 36707656 | 36733370 | 4.68731 | 3.93374 | 2.7424 |
| chr9 | 70223223 | 70223441 | 396 | Intron (ENSMUST00000034745.7 | 9 | 70207350 | 70400067 | 192718 | 1 | ENSMUST00 | 15874 | Myo1e | ENSMUSG0C | true | . | . | . | . | . | . | chr9 | 70209471 | 70285922 | 6.22404 | 4.59358 | 4.02448 |
| chr9 | 57130590 | 57130877 | 396 | Promoter (<=1kb) | 9 | 57130690 | 57134155 | 3466 | 1 | ENSMUST00 | 0 | Man2c1 | ENSMUSG0C | false | chr9 | 57130158 | 57131721 | 37.5719 | 132.25589 | 127.68574 | . | . | . | . | . | . |
| chr11 | 98836623 | 98837182 | 396 | Promoter (<=1kb) | 11 | 98836785 | 98853697 | 16913 | 1 | ENSMUST00 | 0 | Rapgefl1 | ENSMUSG0C | false | chr11 | 98836489 | 98837525 | 30.59503 | 95.89513 | 92.42037 | chr11 | 98835982 | 98837946 | 6.14087 | 4.50514 | 3.97859 |
| chr8 | 105605052 | 105605590 | 396 | Promoter (<=1kb) | 8 | 105605229 | 105622194 | 16966 | 1 | ENSMUST00 | 0 | Fam65a | ENSMUSG0C | false | chr8 | 105600584 | 105607833 | 28.7777 | 83.06458 | 79.8585 | chr8 | 105598855 | 105624293 | 5.6449 | 4.36488 | 3.54309 |
| chr1 | 135430280 | 135430671 | 396 | Promoter (<=1kb) | 1 | 135382312 | 135430499 | 48188 | 2 | ENSMUST00 | 0 | Ipo9 | ENSMUSG0C | false | chr1 | 135429257 | 135430977 | 26.52689 | 93.43019 | 90.01563 | chr1 | 135427951 | 135431094 | 4.76832 | 4.0173 | 2.81633 |
| chr1 | 54825779 | 54826081 | 396 | Intron (ENSMUST00000179030.2 | 1 | 54762368 | 54813443 | 51076 | 2 | ENSMUST00 | -12337 | Ankrd44 | ENSMUSG0C | true | . | . | . | . | . | . | chr1 | 54821851 | 54833551 | 6.37475 | 4.70533 | 4.12466 |
| chr10 | 19964199 | 19964622 | 396 | Intron (ENSMUST00000095806.4 | 10 | 19951055 | 20023699 | 72645 | 1 | ENSMUST00 | 13145 | Map3k5 | ENSMUSG0C | true | . | . | . | . | . | . | chr10 | 19952351 | 19978792 | 5.69092 | 4.38418 | 3.5805 |
| chr4 | 126141047 | 126141403 | 396 | Distal Intergenic | 4 | 126145979 | 126147735 | 1757 | 1 | ENSMUST00 | -4576 | Gm12946 | ENSMUSG0C | false | . | . | . | . | . | . | chr4 | 126139743 | 126142440 | 7.01043 | 4.99644 | 4.6336 |
| chr13 | 41372948 | 41373182 | 396 | Intron (ENSMUST00000021794.7 | 13 | 41359971 | 41370908 | 10938 | 1 | ENSMUST00 | 12978 | Gm26877 | ENSMUSG0C | false | . | . | . | . | . | . | chr13 | 41372897 | 41373150 | 2.62544 | 2.72725 | 1.11466 |
| chr17 | 85432804 | 85433025 | 396 | Intron (ENSMUST00000095188.5 | 17 | 85498633 | 85499008 | 376 | 1 | ENSMUST00 | -65608 | Rpl31-ps16 | ENSMUSG0C | false | . | . | . | . | . | . | chr17 | 85432256 | 85433690 | 4.65944 | 3.80977 | 2.76206 |
| chrX | 74282540 | 74282792 | 395 | Promoter (<=1kb) | 20 | 74282697 | 74290151 | 7455 | 1 | ENSMUST00 | 0 | Taz | ENSMUSG0C | false | chrX | 74281767 | 74283502 | 18.84082 | 47.13427 | 44.45482 | . | . | . | . | . | . |
| chr9 | 108569661 | 108569980 | 395 | Promoter (<=1kb) | 9 | 108569892 | 108572771 | 2880 | 1 | ENSMUST00 | 0 | Dalrd3 | ENSMUSG0C | false | chr9 | 108569535 | 108570677 | 18.82282 | 71.67798 | 68.66222 | chr9 | 108567861 | 108571025 | 5.72605 | 4.3938 | 3.64898 |
| chr10 | 111539934 | 111540292 | 395 | Distal Intergenic | 10 | 111507696 | 111508165 | 470 | 1 | ENSMUST00 | 32239 | Phlda1 | ENSMUSG0C | false | . | . | . | . | . | . | chr10 | 111537069 | 111540404 | 6.18723 | 4.57319 | 3.99271 |
| chr16 | 4964117 | 4964631 | 395 | Promoter (<=1kb) | 16 | 4941415 | 4964237 | 22823 | 2 | ENSMUST00 | 0 | Anks3 | ENSMUSG0C | false | chr16 | 4963351 | 4965084 | 37.45829 | 123.39867 | 119.14091 | chr16 | 4962238 | 4965423 | 4.55384 | 3.79717 | 2.65235 |
| chr16 | 91044136 | 91044793 | 395 | Promoter (<=1kb) | 16 | 91014037 | 91044543 | 30507 | 2 | ENSMUST00 | 0 | Paxbp1 | ENSMUSG0C | false | chr16 | 91043343 | 91045245 | 36.04672 | 129.50026 | 125.02953 | . | . | . | . | . | . |
| chr2 | 122028188 | 122028626 | 395 | Promoter (<=1kb) | 2 | 122028546 | 122056598 | 28053 | 1 | ENSMUST00 | 0 | Eif3j1 | ENSMUSG0C | false | chr2 | 122027707 | 122028413 | 28.00751 | 87.44511 | 84.14996 | . | . | . | . | . | . |
| chr5 | 24687760 | 24687991 | 395 | Promoter (1-2kb) | 5 | 24685815 | 24710555 | 24741 | 1 | ENSMUST00 | 1946 | Nub1 | ENSMUSG0C | false | . | . | . | . | . | . | chr5 | 24686523 | 24688998 | 5.70522 | 4.32303 | 3.6074 |
| chr7 | 133171712 | 133171908 | 395 | Distal Intergenic | 7 | 133025702 | 133124354 | 98653 | 2 | ENSMUST00 | -47359 | Ctbp2 | ENSMUSG0C | false | . | . | . | . | . | . | . | . | . | . | . | . |
| chr10 | 22570967 | 22571270 | 395 | Distal Intergenic | 10 | 22645011 | 22704285 | 59275 | 1 | ENSMUST00 | -73741 | Slc2a12 | ENSMUSG0C | false | . | . | . | . | . | . | chr10 | 22569038 | 22571109 | 3.70394 | 3.38462 | 1.99544 |
| chr17 | 84502822 | 84503164 | 395 | Distal Intergenic | 17 | 84511895 | 84622142 | 110248 | 1 | ENSMUST00 | -8731 | Plekhh2 | ENSMUSG0C | true | chr17 | 84502135 | 84504110 | 11.32191 | 27.20368 | 24.77393 | chr17 | 84497894 | 84511794 | 4.6554 | 3.81889 | 2.73035 |
| chr17 | 66297630 | 66297841 | 395 | Distal Intergenic | 17 | 66343946 | 66346545 | 2600 | 2 | ENSMUST00 | 48704 | Mtcl1 | ENSMUSG0C | true | . | . | . | . | . | . | . | . | . | . | . | . |
| chr7 | 143825456 | 143825914 | 395 | Promoter (2-3kb) | 7 | 143823167 | 143848409 | 25243 | 1 | ENSMUST00 | 2290 | Dhcr7 | ENSMUSG0C | false | . | . | . | . | . | . | chr7 | 143825088 | 143827843 | 5.59602 | 4.38894 | 3.51135 |
| chr4 | 155694238 | 155694536 | 395 | Promoter (<=1kb) | 4 | 155694342 | 155696481 | 2140 | 1 | ENSMUST00 | 0 | B930041F14 | ENSMUSG0C | false | chr4 | 155693984 | 155695391 | 26.18681 | 91.82585 | 88.44535 | chr4 | 155693802 | 155695492 | 4.8937 | 4.12241 | 2.89862 |
| chr1 | 72254671 | 72254986 | 395 | Promoter (<=1kb) | 1 | 72255008 | 72255198 | 191 | 1 | ENSMUST00 | -22 | Gm25939 | ENSMUSG0C | false | chr1 | 72254269 | 72255424 | 22.21283 | 62.70797 | 59.8237 | chr1 | 72254950 | 72256745 | 3.57594 | 3.24604 | 1.87931 |
| chrY | 90772239 | 90772804 | 395 | Distal Intergenic | 21 | 90784738 | 90816439 | 31702 | 1 | ENSMUST00 | -11934 | Erdr1 | ENSMUSG0C | false | . | . | . | . | . | . | . | . | . | . | . | . |
| chr8 | 120510993 | 120511242 | 395 | Intron (ENSMUST00000127664.1 | 8 | 120520615 | 120521686 | 1072 | 1 | ENSMUST00 | -9373 | Gm26971 | ENSMUSG0C | false | . | . | . | . | . | . | chr8 | 120497859 | 120511881 | 6.76427 | 4.83633 | 4.46181 |
| chr18 | 76119659 | 76120288 | 395 | Intron (ENSMUST00000058997.8 | 18 | 76170552 | 76170652 | 101 | 1 | ENSMUST00 | -50264 | Mir6358 | ENSMUSG0C | false | . | . | . | . | . | . | chr18 | 76061445 | 76132293 | 6.70015 | 4.79833 | 4.4158 |
| chr1 | 62858135 | 62858359 | 395 | Distal Intergenic | 1 | 62821355 | 62832370 | 11016 | 2 | ENSMUST00 | -25766 | RP23-149A5. | ENSMUSG0C | false | . | . | . | . | . | . | chr1 | 62857305 | 62862928 | 5.80429 | 4.51093 | 3.66956 |
| chr18 | 34624154 | 34624799 | 394 | Promoter (<=1kb) | 18 | 34598615 | 34624601 | 25987 | 2 | ENSMUST00 | 0 | Brd8 | ENSMUSG0C | false | chr18 | 34623373 | 34625328 | 34.46491 | 104.61457 | 100.92722 | . | . | . | . | . | . |
| chr11 | 100738185 | 100738501 | 394 | Promoter (<=1kb) | 11 | 100715009 | 100738215 | 23207 | 2 | ENSMUST00 | 0 | Rab5c | ENSMUSG0C | false | chr11 | 100738405 | 100738878 | 10.17572 | 22.26956 | 19.90332 | . | . | . | . | . | . |
| chr19 | 53330131 | 53330451 | 394 | Promoter (<=1kb) | 19 | 53330446 | 53375810 | 45365 | 1 | ENSMUST00 | 0 | Mxi1 | ENSMUSG0C | false | chr19 | 53329381 | 53330780 | 33.94791 | 109.87469 | 106.04007 | chr19 | 53330413 | 53330766 | 2.93418 | 2.83544 | 1.35513 |
| chr19 | 46356726 | 46357095 | 394 | Promoter (<=1kb) | 19 | 46356880 | 46375252 | 18373 | 1 | ENSMUST00 | 0 | Tmem180 | ENSMUSG0C | false | chr19 | 46356699 | 46358297 | 28.48018 | 82.55319 | 79.35308 | chr19 | 46356343 | 46359337 | 3.73595 | 3.20855 | 2.00706 |
| chr8 | 123982775 | 123983173 | 394 | Promoter (<=1kb) | 8 | 123952459 | 123983122 | 30664 | 2 | ENSMUST00 | 0 | Abcb10 | ENSMUSG0C | false | chr8 | 123982163 | 123983624 | 33.7938 | 119.00099 | 114.88904 | chr8 | 123982440 | 123984046 | 3.53233 | 3.32575 | 1.80163 |
| chr9 | 103376747 | 103376993 | 394 | Distal Intergenic | 9 | 103353094 | 103365840 | 12747 | 2 | ENSMUST00 | -10908 | Cdv3 | ENSMUSG0C | false | . | . | . | . | . | . | chr9 | 103375369 | 103379120 | 5.44758 | 4.13243 | 3.4216 |
| chr10 | 111377829 | 111378255 | 394 | Distal Intergenic | 10 | 111298679 | 111301736 | 3058 | 1 | ENSMUST00 | 79151 | Bbs10 | ENSMUSG0C | false | . | . | . | . | . | . | chr10 | 111372873 | 111379701 | 4.78915 | 4.01987 | 2.82418 |
| chr9 | 57606147 | 57606387 | 394 | Promoter (<=1kb) | 9 | 57600019 | 57606281 | 6263 | 2 | ENSMUST00 | 0 | Cplx3 | ENSMUSG0C | false | . | . | . | . | . | . | chr9 | 57606175 | 57606986 | 3.02085 | 2.67432 | 1.42219 |
| chr6 | 121488269 | 121488626 | 394 | Distal Intergenic | 6 | 121404801 | 121473678 | 68878 | 2 | ENSMUST00 | -14592 | Iqsec3 | ENSMUSG0C | false | . | . | . | . | . | . | . | . | . | . | . | . |
| chr3 | 89093283 | 89093511 | 394 | Promoter (<=1kb) | 3 | 89083981 | 89093363 | 9383 | 2 | ENSMUST00 | 0 | Rusc1 | ENSMUSG0C | false | . | . | . | . | . | . | chr3 | 89087499 | 89093721 | 3.97851 | 3.49946 | 2.17472 |
| chr1 | 136138175 | 136138417 | 394 | Intron (ENSMUST00000075164.5 | 1 | 136131454 | 136149993 | 18540 | 1 | ENSMUST00 | 6722 | Kif21b | ENSMUSG0C | false | . | . | . | . | . | . | chr1 | 136136450 | 136141742 | 3.35814 | 3.22403 | 1.65544 |
| chr8 | 3492949 | 3493249 | 394 | Promoter (<=1kb) | 8 | 3493138 | 3497208 | 4071 | 1 | ENSMUST00 | 0 | Zfp358 | ENSMUSG0C | false | chr8 | 3492673 | 3494001 | 26.72788 | 101.58999 | 97.97913 | chr8 | 3492637 | 3494824 | 6.38513 | 4.71355 | 4.17954 |
| chr16 | 90403541 | 90403784 | 394 | Intron (ENSMUST00000065856.6 | 16 | 90386397 | 90499553 | 113157 | 1 | ENSMUST00 | 17145 | Hunk | ENSMUSG0C | false | . | . | . | . | . | . | chr16 | 90400784 | 90404150 | 4.52588 | 3.79392 | 2.63265 |
| chr19 | 31106220 | 31106478 | 394 | Intron (ENSMUST00000065067.7 | 19 | 31082841 | 31086590 | 3750 | 1 | ENSMUST00 | 23380 | Cstf2t | ENSMUSG0C | false | . | . | . | . | . | . | chr19 | 31105834 | 31107256 | 3.84421 | 3.5325 | 2.08278 |
| chr6 | 32986561 | 32986856 | 394 | Intron (ENSMUST00000066379.5 | 6 | 33038468 | 33043222 | 4755 | 1 | ENSMUST00 | -51612 | Gm14540 | ENSMUSG0C | false | . | . | . | . | . | . | chr6 | 32982645 | 32986965 | 4.1184 | 3.60909 | 2.28152 |
| chr8 | 124231209 | 124231764 | 394 | Promoter (<=1kb) | 8 | 124231391 | 124345722 | 114332 | 1 | ENSMUST00 | 0 | Galnt2 | ENSMUSG0C | false | chr8 | 124230883 | 124232575 | 38.18876 | 121.46067 | 117.27263 | . | . | . | . | . | . |
| chr5 | 18694795 | 18695077 | 394 | Distal Intergenic | 5 | 18623742 | 18626379 | 2638 | 2 | ENSMUST00 | -68417 | Gm17037 | ENSMUSG0C | false | . | . | . | . | . | . | . | . | . | . | . | . |
| chr5 | 92830839 | 92831137 | 394 | Intron (ENSMUST00000113055.3 | 5 | 92804220 | 92809488 | 5269 | 2 | ENSMUST00 | -21352 | 4932430I15R | ENSMUSG0C | false | . | . | . | . | . | . | chr5 | 92813516 | 92845987 | 5.7053 | 4.3919 | 3.59815 |
| chr16 | 18382019 | 18382251 | 393 | Promoter (1-2kb) | 16 | 18380783 | 18406747 | 25965 | 1 | ENSMUST00 | 1237 | Arvcf | ENSMUSG0C | true | . | . | . | . | . | . | chr16 | 18376326 | 18392535 | 8.53722 | 5.66709 | 5.90216 |
| chr11 | 87086746 | 87086998 | 393 | Promoter (<=1kb) | 11 | 87077732 | 87086774 | 9043 | 2 | ENSMUST00 | 0 | Smg8 | ENSMUSG0C | false | chr11 | 87085908 | 87087315 | 37.4597 | 129.87831 | 125.39164 | . | . | . | . | . | . |
| chr12 | 29957048 | 29957358 | 393 | Intron (ENSMUST00000122328.2 | 12 | 29938050 | 29989546 | 51497 | 1 | ENSMUST00 | 18999 | Pxdn | ENSMUSG0C | false | . | . | . | . | . | . | chr12 | 29956473 | 29958310 | 4.67419 | 3.87374 | 2.76954 |
| chr5 | 138754839 | 138755107 | 393 | Promoter (<=1kb) | 5 | 138755083 | 138809381 | 54299 | 1 | ENSMUST00 | 0 | Fam20c | ENSMUSG0C | false | chr5 | 138754087 | 138756196 | 28.06191 | 82.77493 | 79.57206 | chr5 | 138753866 | 138756351 | 4.3192 | 3.63182 | 2.46278 |
| chr3 | 98198068 | 98198486 | 393 | Distal Intergenic | 3 | 98222156 | 98236748 | 14593 | 1 | ENSMUST00 | -23670 | Reg4 | ENSMUSG0C | false | . | . | . | . | . | . | chr3 | 98195286 | 98203232 | 4.51299 | 3.82434 | 2.63241 |
| chrX | 163887302 | 163887498 | 393 | Distal Intergenic | 20 | 163909017 | 163933666 | 24650 | 1 | ENSMUST00 | -21519 | Ap1s2 | ENSMUSG0C | false | . | . | . | . | . | . | . | . | . | . | . | . |
| chr8 | 25074847 | 25075086 | 393 | Intron (ENSMUST00000064883.7 | 8 | 25062433 | 25091346 | 28914 | 2 | ENSMUST00 | 16260 | Plekha2 | ENSMUSG0C | true | . | . | . | . | . | . | chr8 | 25067309 | 25079381 | 5.19206 | 4.11384 | 3.1814 |
| chr10 | 79874283 | 79874760 | 393 | Promoter (<=1kb) | 10 | 79860475 | 79874570 | 14096 | 2 | ENSMUST00 | 0 | BC005764 | ENSMUSG0C | false | chr10 | 79874058 | 79874856 | 24.3552 | 66.39809 | 63.45943 | chr10 | 79873870 | 79874832 | 3.91852 | 3.52884 | 2.13557 |
| chr15 | 80255597 | 80256102 | 393 | Promoter (<=1kb) | 15 | 80255184 | 80257540 | 2357 | 1 | ENSMUST00 | 414 | Atf4 | ENSMUSG0C | false | chr15 | 80255173 | 80257302 | 45.68299 | 161.30347 | 155.60802 | . | . | . | . | . | . |
| chr9 | 43116013 | 43116658 | 393 | Promoter (<=1kb) | 9 | 43108653 | 43116570 | 7918 | 2 | ENSMUST00 | 0 | Tmem136 | ENSMUSG0C | false | chr9 | 43115803 | 43116749 | 29.81577 | 89.63673 | 86.30187 | chr9 | 43113999 | 43116918 | 4.91258 | 4.01078 | 2.97285 |
| chr1 | 167054116 | 167054341 | 393 | Intron (ENSMUST00000165874.2 | 1 | 167001589 | 167091009 | 89421 | 1 | ENSMUST00 | 52528 | Fam78b | ENSMUSG0C | true | . | . | . | . | . | . | chr1 | 167042661 | 167060137 | 5.59987 | 4.30177 | 3.51451 |
| chr9 | 108648808 | 108649523 | 393 | Promoter (<=1kb) | 9 | 108602942 | 108649380 | 46439 | 2 | ENSMUST00 | 0 | Arih2 | ENSMUSG0C | false | chr9 | 108648312 | 108650207 | 41.61197 | 136.07095 | 131.34828 | . | . | . | . | . | . |
| chr18 | 65144300 | 65144573 | 393 | Intron (ENSMUST00000163516.2 | 18 | 65248861 | 65248926 | 66 | 1 | ENSMUST00 | -104288 | Mir122a | ENSMUSG0C | false | . | . | . | . | . | . | chr18 | 65142111 | 65145595 | 4.23892 | 3.65516 | 2.42586 |
| chr3 | 149233157 | 149233438 | 393 | Distal Intergenic | 3 | 149221693 | 149225745 | 4053 | 2 | ENSMUST00 | -7413 | Gm10287 | ENSMUSG0C | false | . | . | . | . | . | . | . | . | . | . | . | . |
| chr7 | 16937843 | 16938108 | 393 | Distal Intergenic | 7 | 16944682 | 16948828 | 4147 | 1 | ENSMUST00 | -6574 | Pnmal2 | ENSMUSG0C | false | . | . | . | . | . | . | chr7 | 16937974 | 16950184 | 4.88741 | 4.01233 | 2.92708 |
| chr14 | 54876206 | 54877322 | 393 | Promoter (<=1kb) | 14 | 54875597 | 54877538 | 1942 | 2 | ENSMUST00 | 216 | Ppp1r3e | ENSMUSG0C | false | chr14 | 54875917 | 54877905 | 36.53154 | 123.91184 | 119.64436 | chr14 | 54875596 | 54876241 | 4.73413 | 3.98586 | 2.82915 |
| chr10 | 71028031 | 71028593 | 393 | Intron (ENSMUST00000143791.2 | 10 | 71082876 | 71083651 | 776 | 1 | ENSMUST00 | -54283 | Gm15647 | ENSMUSG0C | false | . | . | . | . | . | . | chr10 | 71023281 | 71029421 | 3.96647 | 3.49689 | 2.18756 |
| chr18 | 33862084 | 33862376 | 393 | Intron (ENSMUST00000025234.5 | 18 | 33795164 | 33795295 | 132 | 1 | ENSMUST00 | 66921 | Gm23639 | ENSMUSG0C | false | . | . | . | . | . | . | chr18 | 33861621 | 33864746 | 5.36645 | 4.25909 | 3.30026 |
| chr11 | 113698152 | 113698485 | 392 | 3' UTR | 11 | 113698172 | 113708967 | 10796 | 2 | ENSMUST00 | 10482 | Cpsf4l | ENSMUSG0C | false | . | . | . | . | . | . | chr11 | 113697883 | 113699032 | 4.06533 | 3.68895 | 2.23519 |
| chr2 | 29944721 | 29944991 | 392 | Promoter (1-2kb) | 2 | 29942725 | 29944764 | 2040 | 1 | ENSMUST00 | 1997 | Gle1 | ENSMUSG0C | false | . | . | . | . | . | . | chr2 | 29943809 | 29945128 | 2.78432 | 2.8023 | 1.23131 |
| chr11 | 104596465 | 104596734 | 392 | Distal Intergenic | 11 | 104608000 | 104670476 | 62477 | 1 | ENSMUST00 | -11266 | Itgb3 | ENSMUSG0C | false | . | . | . | . | . | . | . | . | . | . | . | . |
| chr13 | 43212230 | 43212493 | 392 | Intron (ENSMUST00000055341.5 | 13 | 43151744 | 43171501 | 19758 | 2 | ENSMUST00 | -40730 | Tbc1d7 | ENSMUSG0C | false | . | . | . | . | . | . | chr13 | 43208585 | 43223004 | 6.08142 | 4.53032 | 3.90259 |

| | | | | | | | | | | | | | | | | | | | | | | | | | | |
|---|---|---|---|---|---|---|---|---|---|---|---|---|---|---|---|---|---|---|---|---|---|---|---|---|---|---|
| chr14 | 60741649 | 60741902 | 392 | Intron (ENSMUST00000160973.2 | 14 | 60732906 | 60764549 | 31644 | 1 | ENSMUST00 | 8744 | Spata13 | ENSMUSG0C | true | . | . | . | . | . | . | chr14 | 60731957 | 60745391 | 6.28488 | 4.60271 | 4.07482 |
| chr14 | 118132593 | 118132911 | 392 | Promoter (<=1kb) | 14 | 118111911 | 118132765 | 20855 | 2 | ENSMUST00 | 0 | Tgds | ENSMUSG0C | false | chr14 | 118131916 | 118133153 | 30.08928 | 92.05882 | 88.67145 | chr14 | 118130635 | 118133376 | 4.48441 | 3.88216 | 2.56914 |
| chr15 | 11906158 | 11906509 | 392 | Promoter (<=1kb) | 15 | 11839896 | 11905674 | 65779 | 2 | ENSMUST00 | -485 | Npr3 | ENSMUSG0C | true | chr15 | 11902214 | 11907460 | 38.7176 | 123.44448 | 119.18625 | . | . | . | . | . | . |
| chr18 | 75405729 | 75405976 | 392 | Distal Intergenic | 18 | 75374914 | 75394001 | 19088 | 1 | ENSMUST00 | 30816 | Smad7 | ENSMUSG0C | false | . | . | . | . | . | . | chr18 | 75404389 | 75406415 | 7.13292 | 4.92703 | 4.80646 |
| chr5 | 138138736 | 138138977 | 392 | Downstream (<1kb) | 5 | 138144667 | 138152899 | 8233 | 2 | ENSMUST00 | 13922 | Zfp113 | ENSMUSG0C | false | . | . | . | . | . | . | chr5 | 138138175 | 138139446 | 5.50054 | 4.2749 | 3.43779 |
| chr12 | 112678621 | 112679392 | 392 | Promoter (<=1kb) | 12 | 112678828 | 112682747 | 3920 | 1 | ENSMUST00 | 0 | Zbtb42 | ENSMUSG0C | false | chr12 | 112678302 | 112679730 | 36.69013 | 115.69205 | 111.68276 | chr12 | 112677723 | 112680351 | 4.26712 | 3.6295 | 2.42363 |
| chr13 | 43480925 | 43481793 | 392 | Promoter (<=1kb) | 13 | 43402673 | 43480973 | 78301 | 2 | ENSMUST00 | 0 | Ranbp9 | ENSMUSG0C | false | chr13 | 43479386 | 43482222 | 37.59926 | 118.31785 | 114.23224 | . | . | . | . | . | . |
| chrX | 12136700 | 12136931 | 392 | Intron (ENSMUST00000043441.7 | 20 | 12058981 | 12128400 | 69420 | 2 | ENSMUST00 | -8301 | Bcor | ENSMUSG0C | false | chrX | 12136360 | 12136985 | 5.44843 | 8.50832 | 6.36429 | . | . | . | . | . | . |
| chr7 | 16052778 | 16053105 | 392 | Distal Intergenic | 7 | 16071942 | 16096328 | 24387 | 1 | ENSMUST00 | -18837 | Zfp541 | ENSMUSG0C | false | chr7 | 16052049 | 16053426 | 14.6071 | 44.0986 | 41.45673 | chr7 | 16051766 | 16053447 | 4.22753 | 3.6663 | 2.37863 |
| chr2 | 9182320 | 9182593 | 392 | Distal Intergenic | 2 | 9189515 | 9197477 | 7963 | 1 | ENSMUST00 | -6922 | 1700061F12I | ENSMUSG0C | false | . | . | . | . | . | . | chr2 | 9182186 | 9182606 | 4.90493 | 4.05605 | 2.92593 |
| chr11 | 63029896 | 63030100 | 392 | Distal Intergenic | 11 | 62999115 | 63004134 | 5020 | 2 | ENSMUST00 | -25763 | Cdrt4os1 | ENSMUSG0C | false | . | . | . | . | . | . | chr11 | 63029485 | 63030322 | 4.57242 | 3.65698 | 2.68274 |
| chr1 | 15636490 | 15636775 | 392 | Intron (ENSMUST00000175681.2 | 1 | 15685935 | 15686046 | 112 | 2 | ENSMUST00 | 49271 | Gm25227 | ENSMUSG0C | false | . | . | . | . | . | . | chr1 | 15636020 | 15638340 | 4.38534 | 3.65671 | 2.53034 |
| chr6 | 127149287 | 127149794 | 392 | Promoter (<=1kb) | 6 | 127149389 | 127212119 | 62731 | 1 | ENSMUST00 | 0 | 9330179D12I | ENSMUSG0C | false | chr6 | 127149169 | 127151101 | 16.34529 | 41.36735 | 38.7591 | chr6 | 127148760 | 127155682 | 5.84503 | 4.49672 | 3.70608 |
| chr14 | 76731605 | 76731992 | 392 | Distal Intergenic | 14 | 76681754 | 76686887 | 5134 | 1 | ENSMUST00 | 49852 | RP23-421C24 | ENSMUSG0C | false | . | . | . | . | . | . | chr14 | 76731162 | 76732148 | 3.58374 | 3.19439 | 1.88358 |
| chr10 | 63176445 | 63176674 | 391 | Intron (ENSMUST00000095580.2 | 10 | 63115795 | 63203952 | 88158 | 2 | ENSMUST00 | 27278 | Mypn | ENSMUSG0C | false | . | . | . | . | . | . | . | . | . | . | . | . |
| chr15 | 4796682 | 4797130 | 391 | Exon (ENSMUST00000162585.2/ | 15 | 4765945 | 4791164 | 25220 | 1 | ENSMUST00 | 30738 | C6 | ENSMUSG0C | false | . | . | . | . | . | . | chr15 | 4795646 | 4798528 | 4.16065 | 3.64941 | 2.31393 |
| chr16 | 50753617 | 50753879 | 391 | Distal Intergenic | 16 | 50719294 | 50732773 | 13480 | 2 | ENSMUST00 | -20845 | 5330426P16I | ENSMUSG0C | false | . | . | . | . | . | . | chr16 | 50751317 | 50756974 | 5.63491 | 4.41224 | 3.52706 |
| chr16 | 32970463 | 32970706 | 391 | Intron (ENSMUST00000023491.7 | 16 | 32979397 | 32998578 | 19182 | 1 | ENSMUST00 | -8691 | Lrch3 | ENSMUSG0C | false | . | . | . | . | . | . | chr16 | 32970094 | 32971379 | 3.77278 | 3.53936 | 1.97836 |
| chr18 | 32067621 | 32068239 | 391 | Promoter (<=1kb) | 18 | 32067734 | 32104331 | 36598 | 1 | ENSMUST00 | 0 | Iws1 | ENSMUSG0C | false | chr18 | 32067200 | 32068691 | 28.90751 | 104.81161 | 101.11721 | . | . | . | . | . | . |
| chr17 | 35134878 | 35135713 | 391 | Promoter (<=1kb) | 17 | 35135178 | 35147305 | 12128 | 1 | ENSMUST00 | 0 | Bag6 | ENSMUSG0C | false | chr17 | 35134562 | 35136490 | 30.17592 | 94.17345 | 90.73757 | . | . | . | . | . | . |
| chr19 | 38498310 | 38498571 | 391 | Intron (ENSMUST00000181994.1 | 19 | 38481121 | 38783430 | 302310 | 1 | ENSMUST00 | 17190 | Plce1 | ENSMUSG0C | true | . | . | . | . | . | . | chr19 | 38495787 | 38511149 | 5.1247 | 4.07357 | 3.12523 |
| chr5 | 122988263 | 122988724 | 391 | Promoter (<=1kb) | 5 | 122870681 | 122988533 | 117853 | 2 | ENSMUST00 | 0 | Kdm2b | ENSMUSG0C | false | chr5 | 122987524 | 122989984 | 17.09341 | 55.02552 | 52.24499 | chr5 | 122985809 | 122988300 | 5.28924 | 4.2616 | 3.26899 |
| chr9 | 27128643 | 27128905 | 391 | Intron (ENSMUST00000034472.9 | 9 | 27100668 | 27155418 | 54751 | 2 | ENSMUST00 | 26513 | Jam3 | ENSMUSG0C | false | . | . | . | . | . | . | . | . | . | . | . | . |
| chr18 | 38686933 | 38687159 | 391 | Intron (ENSMUST00000127448.2 | 18 | 38676718 | 39110534 | 433817 | 1 | ENSMUST00 | 10216 | Arhgap26 | ENSMUSG0C | false | . | . | . | . | . | . | chr18 | 38684676 | 38687532 | 3.92089 | 3.57477 | 2.11344 |
| chr13 | 9276447 | 9277009 | 391 | Promoter (<=1kb) | 13 | 9276528 | 9668928 | 392401 | 1 | ENSMUST00 | 0 | Dip2c | ENSMUSG0C | false | chr13 | 9276148 | 9278135 | 38.7176 | 123.44448 | 119.18625 | . | . | . | . | . | . |
| chr1 | 133693130 | 133693339 | 391 | Distal Intergenic | 1 | 133679091 | 133690108 | 11018 | 2 | ENSMUST00 | -3023 | Lax1 | ENSMUSG0C | false | . | . | . | . | . | . | chr1 | 133689861 | 133697286 | 7.10741 | 4.96538 | 4.74647 |
| chr8 | 84984564 | 84984975 | 391 | Distal Intergenic | 8 | 84990595 | 85003364 | 12770 | 1 | ENSMUST00 | -5620 | Hook2 | ENSMUSG0C | false | . | . | . | . | . | . | chr8 | 84974675 | 84986551 | 7.15 | 5.07534 | 4.74925 |
| chr12 | 40441000 | 40441285 | 391 | Intron (ENSMUST00000069637.8 | 12 | 40315558 | 40445769 | 130212 | 2 | ENSMUST00 | 4484 | Zfp277 | ENSMUSG0C | false | . | . | . | . | . | . | chr12 | 40422298 | 40443652 | 4.88409 | 4.00258 | 2.915 |
| chr8 | 61584807 | 61585178 | 391 | Intron (ENSMUST00000121785.2 | 8 | 61515913 | 61591139 | 75227 | 2 | ENSMUST00 | 5961 | Palld | ENSMUSG0C | true | . | . | . | . | . | . | chr8 | 61569315 | 61589884 | 5.62358 | 4.37337 | 3.52305 |
| chr12 | 21183315 | 21183635 | 391 | Intron (ENSMUST00000174530.1 | 12 | 21240825 | 21247323 | 6499 | 2 | ENSMUST00 | 63688 | Itgb1bp1 | ENSMUSG0C | false | . | . | . | . | . | . | chr12 | 21178850 | 21190099 | 5.11522 | 4.01562 | 3.13353 |
| chr5 | 85612688 | 85612987 | 391 | Distal Intergenic | 5 | 85829627 | 85829787 | 161 | 1 | ENSMUST00 | -216640 | Gm24626 | ENSMUSG0C | false | . | . | . | . | . | . | . | . | . | . | . | . |
| chr8 | 87708076 | 87708371 | 391 | Intron (ENSMUST00000165770.3 | 8 | 87636657 | 87637013 | 357 | 1 | ENSMUST00 | 71420 | Gm24841 | ENSMUSG0C | false | . | . | . | . | . | . | chr8 | 87706758 | 87709336 | 4.48364 | 3.82447 | 2.58327 |
| chr13 | 37514217 | 37514681 | 391 | Distal Intergenic | 13 | 37508940 | 37512074 | 3135 | 1 | ENSMUST00 | 5278 | RP23-21N15 | ENSMUSG0C | false | . | . | . | . | . | . | chr13 | 37505795 | 37515330 | 6.33539 | 4.55559 | 4.12407 |
| chr5 | 138160933 | 138161202 | 390 | Promoter (<=1kb) | 5 | 138161071 | 138164646 | 3576 | 1 | ENSMUST00 | 0 | Cops6 | ENSMUSG0C | false | chr5 | 138160735 | 138161817 | 21.77823 | 66.26614 | 63.32996 | chr5 | 138160417 | 138162204 | 5.40635 | 4.17485 | 3.36072 |
| chr10 | 79780150 | 79780413 | 390 | Promoter (<=1kb) | 10 | 79780581 | 79782607 | 2027 | 1 | ENSMUST00 | -168 | Fstl3 | ENSMUSG0C | false | chr10 | 79780286 | 79781351 | 18.84082 | 47.13427 | 44.45482 | chr10 | 79777026 | 79781482 | 5.85357 | 4.52993 | 3.6966 |
| chr13 | 64432172 | 64432707 | 390 | Promoter (<=1kb) | 13 | 64432553 | 64439720 | 7168 | 1 | ENSMUST00 | 0 | Cdk20 | ENSMUSG0C | false | chr13 | 64431923 | 64433090 | 26.19333 | 73.09374 | 70.054 | chr13 | 64431443 | 64433124 | 5.16253 | 4.188 | 3.19647 |
| chr19 | 47698001 | 47698235 | 390 | Distal Intergenic | 19 | 47646344 | 47692094 | 45751 | 2 | ENSMUST00 | -5908 | Col17a1 | ENSMUSG0C | false | . | . | . | . | . | . | chr19 | 47697960 | 47702263 | 4.14895 | 3.61656 | 2.32554 |
| chr6 | 116337825 | 116338275 | 390 | Promoter (<=1kb) | 6 | 116338024 | 116409540 | 71517 | 1 | ENSMUST00 | 0 | 39508 | ENSMUSG0C | false | chr6 | 116337597 | 116338886 | 33.41831 | 117.26761 | 113.21259 | chr6 | 116337547 | 116339458 | 3.42373 | 3.28742 | 1.71033 |
| chr1 | 65281026 | 65281269 | 390 | Promoter (<=1kb) | 1 | 65282056 | 65345341 | 63286 | 1 | ENSMUST00 | -787 | Pth2r | ENSMUSG0C | false | . | . | . | . | . | . | chr1 | 65280461 | 65281944 | 4.89991 | 4.0195 | 2.92565 |
| chr7 | 29512035 | 29512224 | 390 | Intron (ENSMUST00000181975.2 | 7 | 29516771 | 29518555 | 1785 | 2 | ENSMUST00 | 6331 | Sipa1l3 | ENSMUSG0C | false | . | . | . | . | . | . | chr7 | 29510878 | 29515001 | 4.93221 | 3.90673 | 2.99896 |
| chr3 | 116419896 | 116420152 | 390 | Promoter (2-3kb) | 3 | 116400186 | 116423024 | 22839 | 2 | ENSMUST00 | 2872 | Cdc14a | ENSMUSG0C | true | chr3 | 116419661 | 116421046 | 10.89686 | 23.4988 | 21.11616 | chr3 | 116394159 | 116421460 | 5.70692 | 4.42755 | 3.58909 |
| chr2 | 11603096 | 11603377 | 390 | Promoter (<=1kb) | 2 | 11585439 | 11603235 | 17797 | 2 | ENSMUST00 | 0 | Rbm17 | ENSMUSG0C | false | chr2 | 11602268 | 11603483 | 28.69254 | 91.56027 | 88.18607 | . | . | . | . | . | . |
| chr10 | 26373954 | 26374496 | 390 | Promoter (<=1kb) | 10 | 26274468 | 26373956 | 99489 | 2 | ENSMUST00 | 0 | L3mbtl3 | ENSMUSG0C | true | chr10 | 26372988 | 26376464 | 34.78613 | 113.42423 | 109.48744 | . | . | . | . | . | . |
| chr6 | 108574022 | 108574469 | 390 | Downstream (2-3kb) | 6 | 108563445 | 108565693 | 2249 | 2 | ENSMUST00 | -8330 | Gm16433 | ENSMUSG0C | false | . | . | . | . | . | . | chr6 | 108569788 | 108574684 | 5.11474 | 4.14126 | 3.10691 |
| chr7 | 122670906 | 122671133 | 390 | Promoter (<=1kb) | 7 | 122671391 | 122768421 | 97031 | 1 | ENSMUST00 | -258 | Cacng3 | ENSMUSG0C | false | . | . | . | . | . | . | . | . | . | . | . | . |
| chr16 | 33676238 | 33676707 | 390 | Distal Intergenic | 16 | 33684466 | 33722913 | 38448 | 1 | ENSMUST00 | -7759 | Heg1 | ENSMUSG0C | false | . | . | . | . | . | . | chr16 | 33675640 | 33678302 | 5.44421 | 4.34759 | 3.43396 |
| chr3 | 60126972 | 60127479 | 390 | Promoter (<=1kb) | 3 | 60127641 | 60127747 | 107 | 2 | ENSMUST00 | 268 | Gm24382 | ENSMUSG0C | false | . | . | . | . | . | . | chr3 | 60120518 | 60128464 | 4.1733 | 3.66476 | 2.33216 |
| chr4 | 54718144 | 54718493 | 390 | Distal Intergenic | 4 | 54710624 | 54710912 | 289 | 1 | ENSMUST00 | 7521 | Gm24628 | ENSMUSG0C | false | . | . | . | . | . | . | chr4 | 54717573 | 54718740 | 3.10298 | 3.00917 | 1.48933 |
| chr2 | 71966840 | 71967282 | 390 | Distal Intergenic | 2 | 71956053 | 71957368 | 1316 | 1 | ENSMUST00 | 10788 | Gm13746 | ENSMUSG0C | false | . | . | . | . | . | . | chr2 | 71966744 | 71966994 | 3.8132 | 3.58986 | 2.02454 |
| chr14 | 100750123 | 100750514 | 390 | Distal Intergenic | 14 | 100760168 | 100760268 | 101 | 2 | ENSMUST00 | 9754 | Gm23538 | ENSMUSG0C | false | chr14 | 100749988 | 100750735 | 10.25583 | 21.2993 | 18.94588 | . | . | . | . | . | . |
| chr1 | 89573625 | 89573823 | 390 | Intron (ENSMUST00000027521.9 | 1 | 89568765 | 89576602 | 7838 | 1 | ENSMUST00 | 4861 | RP23-205B8. | ENSMUSG0C | false | . | . | . | . | . | . | chr1 | 89564740 | 89589718 | 5.63127 | 4.22514 | 3.56367 |
| chr17 | 28261978 | 28262268 | 389 | Intron (ENSMUST00000002320.9 | 17 | 28271708 | 28273163 | 1456 | 1 | ENSMUST00 | -9440 | Ppard | ENSMUSG0C | false | . | . | . | . | . | . | chr17 | 28259989 | 28267005 | 3.86487 | 3.41614 | 2.11509 |
| chr17 | 87265839 | 87266178 | 389 | Promoter (<=1kb) | 17 | 87254658 | 87265935 | 11278 | 2 | ENSMUST00 | 0 | Mcfd2 | ENSMUSG0C | false | chr17 | 87265120 | 87266517 | 35.00596 | 114.85688 | 110.88117 | . | . | . | . | . | . |
| chr19 | 21106381 | 21106667 | 389 | Distal Intergenic | 19 | 20783458 | 20954202 | 170745 | 2 | ENSMUST00 | -152180 | Tmc1 | ENSMUSG0C | false | . | . | . | . | . | . | chr19 | 21104721 | 21109391 | 4.4041 | 3.71096 | 2.54536 |
| chr7 | 130233836 | 130234383 | 389 | Intron (ENSMUST00000124096.2 | 7 | 130199807 | 130228756 | 28950 | 2 | ENSMUST00 | -5081 | Fgfr2 | ENSMUSG0C | false | . | . | . | . | . | . | chr7 | 130228840 | 130264807 | 4.60924 | 3.83033 | 2.69727 |
| chr8 | 94771985 | 94772489 | 389 | Promoter (<=1kb) | 8 | 94772009 | 94778945 | 6937 | 1 | ENSMUST00 | 0 | Cx3cl1 | ENSMUSG0C | false | chr8 | 94771750 | 94772906 | 17.09341 | 55.02552 | 52.24499 | chr8 | 94770747 | 94774171 | 4.2759 | 3.75021 | 2.41539 |
| chr7 | 100916512 | 100916846 | 389 | Intron (ENSMUST00000107032.1 | 7 | 100869746 | 100932161 | 62416 | 2 | ENSMUST00 | 15315 | Arhgef17 | ENSMUSG0C | false | . | . | . | . | . | . | chr7 | 100901870 | 100926579 | 7.14639 | 4.97053 | 4.78972 |
| chr10 | 26804378 | 26804634 | 389 | Intron (ENSMUST00000176060.2 | 10 | 26822602 | 26850001 | 27400 | 1 | ENSMUST00 | -17968 | Arhgap18 | ENSMUSG0C | false | . | . | . | . | . | . | chr10 | 26803676 | 26808321 | 5.63777 | 4.31219 | 3.534 |
| chr17 | 86185630 | 86185928 | 389 | Intron (ENSMUST00000097275.3 | 17 | 86168174 | 86655415 | 487242 | 1 | ENSMUST00 | 17457 | Prkce | ENSMUSG0C | true | . | . | . | . | . | . | chr17 | 86181776 | 86189028 | 4.85954 | 3.97288 | 2.89005 |
| chr11 | 49217595 | 49217859 | 389 | 3' UTR | 11 | 49214549 | 49218809 | 4261 | 1 | ENSMUST00 | 3047 | Zfp62 | ENSMUSG0C | false | . | . | . | . | . | . | chr11 | 49217052 | 49218134 | 4.58506 | 3.79282 | 2.71083 |
| chr7 | 49450313 | 49450636 | 389 | Promoter (<=1kb) | 7 | 49450760 | 49464580 | 13821 | 1 | ENSMUST00 | -124 | Nav2 | ENSMUSG0C | false | chr7 | 49450235 | 49451151 | 7.81205 | 13.8349 | 11.58824 | chr7 | 49446529 | 49453829 | 5.00391 | 3.99232 | 3.02197 |
| chr7 | 141092381 | 141092649 | 389 | Promoter (2-3kb) | 7 | 141089861 | 141090509 | 649 | 1 | ENSMUST00 | 2521 | Pkp3 | ENSMUSG0C | false | . | . | . | . | . | . | chr7 | 141092541 | 141093011 | 5.1295 | 4.01252 | 3.14782 |
| chr1 | 86154568 | 86154888 | 389 | Promoter (<=1kb) | 1 | 86154780 | 86246326 | 91547 | 1 | ENSMUST00 | 0 | Armc9 | ENSMUSG0C | false | chr1 | 86154131 | 86155428 | 32.75675 | 107.77596 | 104.00922 | chr1 | 86153866 | 86155605 | 6.12065 | 4.58895 | 3.97969 |
| chr1 | 87913589 | 87913791 | 389 | Intron (ENSMUST00000027517.8 | 1 | 87917898 | 87921971 | 4074 | 1 | ENSMUST00 | -4107 | Dgkd | ENSMUSG0C | false | . | . | . | . | . | . | chr1 | 87912458 | 87914578 | 3.71167 | 3.37687 | 1.97467 |
| chr14 | 44009729 | 44009985 | 389 | Distal Intergenic | 14 | 44001664 | 44006418 | 4755 | 2 | ENSMUST00 | -3312 | Ang6 | ENSMUSG0C | false | . | . | . | . | . | . | . | . | . | . | . | . |
| chr4 | 148922335 | 148922605 | 389 | Intron (ENSMUST00000094464.4 | 4 | 148913511 | 148914324 | 814 | 2 | ENSMUST00 | -8012 | Gm15969 | ENSMUSG0C | false | . | . | . | . | . | . | chr4 | 148901819 | 148933146 | 7.00771 | 4.91323 | 4.66351 |
| chr5 | 132705259 | 132705493 | 389 | Distal Intergenic | 5 | 132801678 | 132801956 | 279 | 1 | ENSMUST00 | -96185 | Gm25722 | ENSMUSG0C | false | . | . | . | . | . | . | . | . | . | . | . | . |
| chr5 | 92112790 | 92113013 | 389 | Distal Intergenic | 5 | 92118177 | 92118283 | 107 | 2 | ENSMUST00 | 5270 | Gm24931 | ENSMUSG0C | false | chr5 | 92112124 | 92113154 | 9.65017 | 18.71579 | 16.39739 | chr5 | 92112324 | 92113259 | 3.81724 | 3.49948 | 2.07673 |
| chr14 | 60486703 | 60486937 | 389 | Promoter (1-2kb) | 14 | 60485208 | 60599031 | 113824 | 1 | ENSMUST00 | 1496 | RP24-127N1 | ENSMUSG0C | false | . | . | . | . | . | . | chr14 | 60485211 | 60487404 | 5.5341 | 4.42585 | 3.43099 |
| chr11 | 61653055 | 61653326 | 388 | Promoter (<=1kb) | 11 | 61653265 | 61672784 | 19520 | 1 | ENSMUST00 | 0 | Grap | ENSMUSG0C | false | . | . | . | . | . | . | . | . | . | . | . | . |
| chr19 | 10101424 | 10102114 | 388 | Promoter (<=1kb) | 19 | 10064164 | 10101503 | 37340 | 2 | ENSMUST00 | 0 | Fads2 | ENSMUSG0C | false | chr19 | 10100545 | 10102386 | 22.83944 | 61.22662 | 58.36318 | chr19 | 10100640 | 10102614 | 4.58078 | 3.96348 | 2.68335 |
| chr7 | 74060364 | 74060757 | 388 | Distal Intergenic | 7 | 73939119 | 74013682 | 74564 | 2 | ENSMUST00 | -46683 | St8sia2 | ENSMUSG0C | false | . | . | . | . | . | . | chr7 | 74055877 | 74067021 | 4.21683 | 3.65298 | 2.3751 |
| chr15 | 59197728 | 59197975 | 388 | Distal Intergenic | 15 | 59210908 | 59211720 | 813 | 1 | ENSMUST00 | -12933 | Gm5045 | ENSMUSG0C | false | . | . | . | . | . | . | chr15 | 59193661 | 59205577 | 5.53199 | 4.27632 | 3.45834 |
| chr2 | 49836079 | 49836316 | 388 | Intron (ENSMUST00000028103.7 | 2 | 49840090 | 49942537 | 102448 | 1 | ENSMUST00 | -3774 | Lypd6b | ENSMUSG0C | false | . | . | . | . | . | . | . | . | . | . | . | . |
| chr5 | 147721460 | 147721698 | 388 | Intron (ENSMUST00000031653.6 | 5 | 147635159 | 147725988 | 90830 | 2 | ENSMUST00 | -4290 | Flt1 | ENSMUSG0C | false | . | . | . | . | . | . | . | . | . | . | . | . |
| chr6 | 24325636 | 24325843 | 388 | Distal Intergenic | 6 | 24088283 | 24168092 | 79810 | 2 | ENSMUST00 | -157545 | Slc13a1 | ENSMUSG0C | false | . | . | . | . | . | . | chr6 | 24325189 | 24326500 | 4.75588 | 3.89866 | 2.84741 |
| chr6 | 34686427 | 34686679 | 388 | Exon (ENSMUST00000155714.2/ | 6 | 34709442 | 34775473 | 66032 | 1 | ENSMUST00 | -22763 | Cald1 | ENSMUSG0C | true | . | . | . | . | . | . | chr6 | 34680884 | 34687674 | 4.73427 | 3.92076 | 2.79387 |
| chr8 | 36613752 | 36614182 | 388 | Promoter (<=1kb) | 8 | 36567751 | 36613943 | 46193 | 2 | ENSMUST00 | 0 | Dlc1 | ENSMUSG0C | true | chr8 | 36610903 | 36614533 | 20.9437 | 83.15932 | 79.95254 | . | . | . | . | . | . |
| chr12 | 40613101 | 40613333 | 388 | Intron (ENSMUST00000037488.6 | 12 | 40446053 | 40846486 | 400434 | 1 | ENSMUST00 | 167049 | Dock4 | ENSMUSG0C | true | . | . | . | . | . | . | chr12 | 40606881 | 40620640 | 3.9863 | 3.53232 | 2.19655 |
| chrX | 12083741 | 12083961 | 388 | Intron (ENSMUST00000043441.7 | 20 | 12036740 | 12080553 | 43814 | 2 | ENSMUST00 | -3189 | Bcor | ENSMUSG0C | false | . | . | . | . | . | . | . | . | . | . | . | . |
| chr15 | 81937412 | 81937902 | 388 | Promoter (<=1kb) | 15 | 81936982 | 81946869 | 9888 | 1 | ENSMUST00 | 431 | Csdc2 | ENSMUSG0C | false | chr15 | 81936283 | 81937895 | 13.91144 | 32.98249 | 30.47944 | chr15 | 81935029 | 81939977 | 4.75257 | 3.93487 | 2.80237 |
| chr6 | 89035114 | 89035349 | 388 | Intron (ENSMUST00000168837.3 | 6 | 89096110 | 89107834 | 11725 | 1 | ENSMUST00 | -60761 | 4933427D06I | ENSMUSG0C | false | . | . | . | . | . | . | chr6 | 89033650 | 89037786 | 4.66592 | 3.88673 | 2.74934 |
| chr5 | 76911872 | 76912077 | 388 | Downstream (1-2kb) | 5 | 76875935 | 76905514 | 29580 | 2 | ENSMUST00 | -6359 | Aasdh | ENSMUSG0C | false | . | . | . | . | . | . | chr5 | 76911250 | 76913252 | 4.79492 | 3.90541 | 2.86922 |
| chr7 | 143837785 | 143838256 | 388 | Promoter (2-3kb) | 7 | 143840929 | 143843299 | 2371 | 2 | ENSMUST00 | -2673 | Dhcr7 | ENSMUSG0C | false | . | . | . | . | . | . | chr7 | 143830649 | 143839948 | 5.20091 | 4.06122 | 3.19483 |
| chr2 | 102856200 | 102856419 | 388 | Intron (ENSMUST00000005218.9 | 2 | 102897960 | 102898066 | 107 | 1 | ENSMUST00 | -41541 | Gm23731 | ENSMUSG0C | false | . | . | . | . | . | . | chr2 | 102855803 | 102856543 | 2.80137 | 2.88132 | 1.2618 |
| chr4 | 136321218 | 136321475 | 388 | Intron (ENSMUST00000131671.2 | 4 | 136311685 | 136316423 | 4739 | 1 | ENSMUST00 | 9534 | Hnrnpr | ENSMUSG0C | false | . | . | . | . | . | . | chr4 | 136319766 | 136321896 | 5.84815 | 4.46521 | 3.69679 |
| chr10 | 69579609 | 69579966 | 388 | Intron (ENSMUST00000182379.2 | 10 | 69534222 | 70015633 | 481412 | 1 | ENSMUST00 | 45388 | Ank3 | ENSMUSG0C | false | . | . | . | . | . | . | . | . | . | . | . | . |
| chr2 | 91236989 | 91237317 | 388 | Promoter (<=1kb) | 2 | 91237146 | 91238352 | 1207 | 1 | ENSMUST00 | 0 | A330069E16 | ENSMUSG0C | false | chr2 | 91236355 | 91237807 | 24.40663 | 77.266 | 74.1616 | . | . | . | . | . | . |
| chr7 | 66439157 | 66439636 | 387 | Distal Intergenic | 7 | 66390892 | 66427517 | 36626 | 2 | ENSMUST00 | -11641 | Aldh1a3 | ENSMUSG0C | false | . | . | . | . | . | . | chr7 | 66438246 | 66451395 | 6.06663 | 4.59192 | 3.88434 |
| chr7 | 16923841 | 16924393 | 387 | Promoter (<=1kb) | 7 | 16915379 | 16924114 | 8736 | 2 | ENSMUST00 | 0 | Calm3 | ENSMUSG0C | false | chr7 | 16921011 | 16924086 | 41.49189 | 142.53105 | 137.54099 | . | . | . | . | . | . |
| chr19 | 56619043 | 56619263 | 387 | Distal Intergenic | 19 | 56548261 | 56598846 | 50586 | 1 | ENSMUST00 | 70783 | Nhlrc2 | ENSMUSG0C | true | . | . | . | . | . | . | chr19 | 56617978 | 56620301 | 3.43513 | 3.21322 | 1.75838 |
| chr4 | 155223018 | 155223537 | 387 | Promoter (<=1kb) | 4 | 155154075 | 155222592 | 68518 | 2 | ENSMUST00 | -427 | Ski | ENSMUSG0C | false | chr4 | 155223453 | 155223772 | 14.64398 | 38.39655 | 35.82735 | . | . | . | . | . | . |
| chr9 | 49060645 | 49060886 | 387 | Exon (ENSMUST00000034803.8/ | 9 | 49055581 | 49078772 | 23192 | 1 | ENSMUST00 | 5065 | Zw10 | ENSMUSG0C | false | . | . | . | . | . | . | chr9 | 49059704 | 49061389 | 3.77579 | 3.45046 | 2.00882 |
| chr9 | 103004472 | 103004653 | 387 | Intron (ENSMUST00000185215.1 | 9 | 103008002 | 103090451 | 82450 | 1 | ENSMUST00 | -3349 | Slco2a1 | ENSMUSG0C | true | . | . | . | . | . | . | chr9 | 102991889 | 103008204 | 5.47516 | 4.26994 | 3.39839 |
| chr5 | 111200963 | 111201165 | 387 | Intron (ENSMUST00000040111.9 | 5 | 111225458 | 111229415 | 3958 | 1 | ENSMUST00 | -24293 | Ttc28 | ENSMUSG0C | false | chr5 | 111200913 | 111201103 | 3.37938 | 4.21603 | 2.20503 | chr5 | 111200058 | 111202014 | 5.60516 | 4.23471 | 3.51841 |
| chr11 | 80772669 | 80772884 | 387 | Intron (ENSMUST00000041065.8 | 11 | 80673667 | 80780003 | 106337 | 2 | ENSMUST00 | 7119 | Myo1d | ENSMUSG0C | true | . | . | . | . | . | . | chr11 | 80758294 | 80778239 | 6.06128 | 4.46131 | 3.88566 |
| chr12 | 40499572 | 40499918 | 387 | Intron (ENSMUST00000037488.6 | 12 | 40446053 | 40846486 | 400434 | 1 | ENSMUST00 | 53520 | Dock4 | ENSMUSG0C | true | . | . | . | . | . | . | chr12 | 40494912 | 40528013 | 5.05369 | 4.02817 | 3.05886 |
| chr3 | 123267158 | 123267626 | 387 | Promoter (<=1kb) | 3 | 123267496 | 123365635 | 98140 | 1 | ENSMUST00 | 0 | Sec24d | ENSMUSG0C | false | chr3 | 123266862 | 123268531 | 24.14628 | 82.32003 | 79.12344 | . | . | . | . | . | . |
| chr13 | 101767990 | 101768532 | 387 | Promoter (<=1kb) | 13 | 101680563 | 101768217 | 87655 | 2 | ENSMUST00 | 0 | Pik3r1 | ENSMUSG0C | false | chr13 | 101766190 | 101769111 | 27.30634 | 119.5799 | 115.45193 | . | . | . | . | . | . |
| chr15 | 89147105 | 89147287 | 387 | Promoter (2-3kb) | 15 | 89142486 | 89149606 | 7121 | 2 | ENSMUST00 | 2319 | Mapk11 | ENSMUSG0C | false | . | . | . | . | . | . | chr15 | 89145732 | 89150483 | 5.03705 | 4.13995 | 3.0374 |
| chr14 | 36911868 | 36912106 | 387 | Promoter (2-3kb) | 14 | 36879096 | 36909181 | 30086 | 2 | ENSMUST00 | -2688 | Ccser2 | ENSMUSG0C | false | . | . | . | . | . | . | chr14 | 36906395 | 36918566 | 3.99854 | 3.5267 | 2.20501 |
| chr6 | 117292807 | 117293033 | 387 | Distal Intergenic | 6 | 117213656 | 117214048 | 393 | 2 | ENSMUST00 | -78760 | Rpl28-ps4 | ENSMUSG0C | false | . | . | . | . | . | . | chr6 | 117292305 | 117295611 | 5.09583 | 4.12614 | 3.0804 |
| chr11 | 59347943 | 59348211 | 387 | Distal Intergenic | 11 | 59337051 | 59337146 | 96 | 2 | ENSMUST00 | -10798 | Gm24756 | ENSMUSG0C | false | . | . | . | . | . | . | chr11 | 59347548 | 59348448 | 6.55647 | 4.63528 | 4.32105 |
| chr14 | 76254875 | 76255115 | 387 | Distal Intergenic | 14 | 76236688 | 76237353 | 666 | 2 | ENSMUST00 | -17523 | Rps2-ps6 | ENSMUSG0C | false | . | . | . | . | . | . | chr14 | 76254575 | 76259568 | 4.86247 | 3.98568 | 2.92539 |
| chr18 | 13328540 | 13328767 | 387 | Distal Intergenic | 18 | 13209702 | 13209808 | 107 | 2 | ENSMUST00 | -118733 | Gm22251 | ENSMUSG0C | false | . | . | . | . | . | . | chr18 | 13325173 | 13328781 | 4.64263 | 3.85051 | 2.72366 |
| chr1 | 194159181 | 194159440 | 387 | Distal Intergenic | 1 | 193923749 | 193923985 | 237 | 2 | ENSMUST00 | -235197 | RP24-404L18 | ENSMUSG0C | false | . | . | . | . | . | . | chr1 | 194158632 | 194160031 | 2.87786 | 2.9118 | 1.33339 |
| chr1 | 74295514 | 74296105 | 386 | Promoter (<=1kb) | 1 | 74288251 | 74295695 | 7445 | 2 | ENSMUST00 | 0 | Tmbim1 | ENSMUSG0C | false | chr1 | 74295970 | 74296658 | 11.16678 | 24.53592 | 22.14134 | chr1 | 74285402 | 74303246 | 7.11676 | 4.92079 | 4.75747 |
| chr7 | 16918919 | 16919171 | 386 | Promoter (1-2kb) | 7 | 16915937 | 16917203 | 1267 | 2 | ENSMUST00 | -1717 | Calm3 | ENSMUSG0C | false | . | . | . | . | . | . | chr7 | 16913262 | 16922976 | 7.95697 | 5.3563 | 5.42533 |
| chr11 | 3575324 | 3575582 | 386 | Distal Intergenic | 11 | 3532460 | 3540612 | 8153 | 2 | ENSMUST00 | -34713 | Smtn | ENSMUSG0C | false | . | . | . | . | . | . | chr11 | 3571707 | 3576040 | 6.68511 | 4.75587 | 4.39333 |
| chr13 | 117025314 | 117026017 | 386 | Promoter (<=1kb) | 13 | 116854824 | 117025516 | 170693 | 2 | ENSMUST00 | 0 | Parp8 | ENSMUSG0C | false | chr13 | 117023675 | 117025980 | 31.54088 | 108.67455 | 104.87239 | . | . | . | . | . | . |
| chr9 | 123529617 | 123529916 | 386 | Promoter (<=1kb) | 9 | 123529882 | 123592598 | 62717 | 1 | ENSMUST00 | 0 | Sacm1l | ENSMUSG0C | false | chr9 | 123529641 | 123531067 | 36.53584 | 119.65029 | 115.52113 | . | . | . | . | . | . |
| chr1 | 92020577 | 92020827 | 386 | Intron (ENSMUST00000097644.3 | 1 | 92003670 | 92006860 | 3191 | 2 | ENSMUST00 | -13718 | Hdac4 | ENSMUSG0C | false | . | . | . | . | . | . | chr1 | 92020013 | 92021617 | 2.96185 | 2.97768 | 1.40235 |
| chr3 | 36475680 | 36476014 | 386 | Promoter (<=1kb) | 3 | 36448925 | 36475843 | 26919 | 2 | ENSMUST00 | 0 | Anxa5 | ENSMUSG0C | false | chr3 | 36474815 | 36475713 | 28.08037 | 85.65662 | 82.39679 | . | . | . | . | . | . |
| chr9 | 102841865 | 102842078 | 386 | Intron (ENSMUST00000175883.2 | 9 | 102845833 | 102861897 | 16065 | 1 | ENSMUST00 | -3755 | Ryk | ENSMUSG0C | false | . | . | . | . | . | . | chr9 | 102833046 | 102850688 | 5.63517 | 4.31932 | 3.53737 |
| chr10 | 75211877 | 75212503 | 386 | Promoter (<=1kb) | 10 | 75212073 | 75312399 | 100327 | 1 | ENSMUST00 | 0 | Specc1l | ENSMUSG0C | false | chr10 | 75211632 | 75212747 | 26.97976 | 80.89639 | 77.72532 | . | . | . | . | . | . |
| chr2 | 29716759 | 29717160 | 386 | Intron (ENSMUST00000095087.5 | 2 | 29730593 | 29735848 | 5256 | 1 | ENSMUST00 | -13433 | Rapgef1 | ENSMUSG0C | false | . | . | . | . | . | . | chr2 | 29715709 | 29718850 | 5.30682 | 4.01857 | 3.29979 |
| chr1 | 136625236 | 136625650 | 386 | Promoter (<=1kb) | 1 | 136624901 | 136630053 | 5153 | 1 | ENSMUST00 | 336 | Zfp281 | ENSMUSG0C | false | chr1 | 136624228 | 136626150 | 33.94791 | 109.87469 | 106.04007 | chr1 | 136625633 | 136629427 | 3.49675 | 3.12532 | 1.81248 |
| chr8 | 90551818 | 90552129 | 386 | Distal Intergenic | 8 | 90247040 | 90348343 | 101304 | 2 | ENSMUST00 | -203476 | Tox3 | ENSMUSG0C | false | . | . | . | . | . | . | chr8 | 90550475 | 90552636 | 5.27908 | 4.22606 | 3.22107 |
| chr7 | 100955359 | 100955607 | 386 | Intron (ENSMUST00000060174.4 | 7 | 100937638 | 100964391 | 26754 | 2 | ENSMUST00 | 8784 | P2ry6 | ENSMUSG0C | false | . | . | . | . | . | . | chr7 | 100955206 | 100955630 | 4.33869 | 3.70502 | 2.49209 |
| chr9 | 60843548 | 60843761 | 386 | Intron (ENSMUST00000050183.6 | 9 | 60821855 | 60838200 | 16346 | 2 | ENSMUST00 | -5349 | Gm9869 | ENSMUSG0C | false | . | . | . | . | . | . | chr9 | 60839714 | 60851928 | 5.61355 | 4.39804 | 3.5066 |
| chr5 | 117318873 | 117319384 | 386 | Promoter (<=1kb) | 5 | 117319266 | 117355005 | 35740 | 1 | ENSMUST00 | 0 | Vsig10 | ENSMUSG0C | false | chr5 | 117318562 | 117320661 | 29.66345 | 100.21 | 96.63225 | . | . | . | . | . | . |
| chr5 | 20809329 | 20809576 | 386 | Intron (ENSMUST00000118174.2 | 5 | 20806090 | 20806464 | 375 | 1 | ENSMUST00 | 3240 | Rpl31-ps5 | ENSMUSG0C | false | . | . | . | . | . | . | . | . | . | . | . | . |
| chr4 | 142316434 | 142316662 | 386 | Intron (ENSMUST00000125582.1 | 4 | 142307765 | 142346475 | 38711 | 2 | ENSMUST00 | 29813 | Gm13052 | ENSMUSG0C | false | . | . | . | . | . | . | chr4 | 142316319 | 142316878 | 3.81156 | 3.49338 | 2.07158 |
| chr3 | 126698947 | 126699229 | 386 | Intron (ENSMUST00000106402.2 | 3 | 126667329 | 126669032 | 1704 | 2 | ENSMUST00 | -29916 | Gm15551 | ENSMUSG0C | false | . | . | . | . | . | . | chr3 | 126691383 | 126703472 | 4.8505 | 3.97802 | 2.88932 |
| chr9 | 37008124 | 37008386 | 386 | Intron (ENSMUST00000177218.2 | 9 | 37015758 | 37020671 | 4914 | 1 | ENSMUST00 | -7372 | Gm3896 | ENSMUSG0C | false | . | . | . | . | . | . | chr9 | 37001731 | 37010197 | 5.19895 | 4.15664 | 3.17524 |
| chr5 | 143527528 | 143528242 | 385 | Promoter (<=1kb) | 5 | 143505478 | 143528036 | 22559 | 2 | ENSMUST00 | 0 | Rac1 | ENSMUSG0C | false | chr5 | 143526334 | 143528725 | 33.65816 | 114.56059 | 110.59126 | . | . | . | . | . | . |
| chr13 | 34814643 | 34814922 | 385 | Distal Intergenic | 13 | 34875494 | 34902877 | 27384 | 1 | ENSMUST00 | -60572 | Prpf4b | ENSMUSG0C | false | . | . | . | . | . | . | chr13 | 34814325 | 34815234 | 5.72139 | 4.38312 | 3.5951 |
| chr7 | 140116362 | 140116611 | 385 | Promoter (<=1kb) | 7 | 140105710 | 140116476 | 10767 | 2 | ENSMUST00 | 0 | Echs1 | ENSMUSG0C | false | chr7 | 140115662 | 140117006 | 27.78755 | 88.98067 | 85.65775 | . | . | . | . | . | . |
| chr3 | 136697701 | 136697944 | 385 | Intron (ENSMUST00000124206.1 | 3 | 136670770 | 136935779 | 265010 | 1 | ENSMUST00 | 26932 | Ppp3ca | ENSMUSG0C | false | . | . | . | . | . | . | chr3 | 136696619 | 136698748 | 4.42038 | 3.68693 | 2.54473 |
| chr9 | 67046076 | 67046401 | 385 | Promoter (2-3kb) | 9 | 67022593 | 67043962 | 21370 | 2 | ENSMUST00 | -2115 | Tpm1 | ENSMUSG0C | true | . | . | . | . | . | . | chr9 | 67044218 | 67047652 | 5.48647 | 4.39358 | 3.39737 |
| chr9 | 64281373 | 64281916 | 385 | Promoter (<=1kb) | 9 | 64281607 | 64304792 | 23186 | 1 | ENSMUST00 | 0 | Tipin | ENSMUSG0C | false | chr9 | 64281205 | 64282333 | 33.0268 | 99.11563 | 95.56912 | . | . | . | . | . | . |
| chr13 | 23739510 | 23739866 | 385 | Promoter (<=1kb) | 13 | 23738808 | 23740367 | 1560 | 1 | ENSMUST00 | 703 | Hist1h1c | ENSMUSG0C | false | chr13 | 23738302 | 23740845 | 31.6659 | 164.62325 | 158.82181 | . | . | . | . | . | . |
| chr5 | 32990348 | 32990620 | 385 | Intron (ENSMUST00000087897.5 | 5 | 32967719 | 32974176 | 6458 | 1 | ENSMUST00 | 22630 | Depdc5 | ENSMUSG0C | false | . | . | . | . | . | . | chr5 | 32989461 | 32996528 | 5.20315 | 4.09317 | 3.21514 |
| chr13 | 47185764 | 47186134 | 385 | Intron (ENSMUST00000068891.5 | 13 | 47194003 | 47247991 | 53989 | 1 | ENSMUST00 | -7869 | Rnf144b | ENSMUSG0C | true | . | . | . | . | . | . | chr13 | 47183764 | 47187186 | 4.71376 | 3.8619 | 2.80112 |
| chr6 | 144318358 | 144318787 | 385 | Intron (ENSMUST00000141102.2 | 6 | 144154829 | 144209568 | 54740 | 2 | ENSMUST00 | -108791 | Sox5 | ENSMUSG0C | true | . | . | . | . | . | . | chr6 | 144318192 | 144320787 | 4.69233 | 3.89751 | 2.798 |
| chr2 | 32734929 | 32735369 | 385 | Promoter (<=1kb) | 2 | 32735281 | 32741321 | 6041 | 1 | ENSMUST00 | 0 | Sh2d3c | ENSMUSG0C | false | . | . | . | . | . | . | . | . | . | . | . | . |
| chr19 | 25171400 | 25171759 | 385 | Exon (ENSMUST00000025831.6/ | 19 | 25166085 | 25166216 | 132 | 2 | ENSMUST00 | -5185 | Gm24252 | ENSMUSG0C | false | . | . | . | . | . | . | chr19 | 25162630 | 25181812 | 5.3445 | 4.22539 | 3.29608 |
| chr8 | 44949968 | 44950200 | 385 | Promoter (<=1kb) | 8 | 44950208 | 45052257 | 102050 | 1 | ENSMUST00 | -8 | Fat1 | ENSMUSG0C | true | . | . | . | . | . | . | chr8 | 44939815 | 44952756 | 5.42131 | 4.25024 | 3.3828 |
| chr5 | 126714776 | 126715011 | 385 | Distal Intergenic | 5 | 126513875 | 126513981 | 107 | 2 | ENSMUST00 | -200796 | Gm24839 | ENSMUSG0C | false | . | . | . | . | . | . | chr5 | 126714323 | 126715138 | 6.28195 | 4.79502 | 4.04648 |
| chr11 | 57983979 | 57984242 | 385 | Distal Intergenic | 11 | 57970244 | 57973134 | 2891 | 2 | ENSMUST00 | -10846 | Gm12249 | ENSMUSG0C | false | . | . | . | . | . | . | chr11 | 57980811 | 57984784 | 5.00394 | 3.96209 | 3.02635 |
| chr4 | 131260589 | 131260889 | 385 | Distal Intergenic | 4 | 131239277 | 131239410 | 134 | 1 | ENSMUST00 | 21313 | Gm25261 | ENSMUSG0C | false | . | . | . | . | . | . | chr4 | 131258846 | 131261997 | 5.70489 | 4.1926 | 3.61984 |
| chr8 | 22647163 | 22647399 | 385 | Promoter (2-3kb) | 8 | 22644861 | 22646286 | 1426 | 1 | ENSMUST00 | 2303 | Gm26909 | ENSMUSG0C | false | . | . | . | . | . | . | . | . | . | . | . | . |
| chr2 | 30242556 | 30242907 | 385 | Promoter (1-2kb) | 2 | 30240567 | 30282148 | 41582 | 1 | ENSMUST00 | 1990 | Gm28035 | ENSMUSG0C | false | . | . | . | . | . | . | . | . | . | . | . | . |
| chr13 | 5592553 | 5592816 | 385 | Distal Intergenic | 13 | 5855508 | 5857712 | 2205 | 2 | ENSMUST00 | 264896 | RP23-28J6.1 | ENSMUSG0C | false | . | . | . | . | . | . | . | . | . | . | . | . |
| chr5 | 147051129 | 147051383 | 384 | Intron (ENSMUST00000016664.7 | 5 | 147016653 | 147076586 | 59932 | 2 | ENSMUST00 | 25203 | Lnx2 | ENSMUSG0C | false | chr5 | 147051077 | 147052037 | 5.97761 | 10.04734 | 7.87135 | chr5 | 147046705 | 147052738 | 5.83351 | 4.47539 | 3.70334 |
| chr16 | 52433282 | 52433563 | 384 | Intron (ENSMUST00000023312.8 | 16 | 52251467 | 52452465 | 200999 | 2 | ENSMUST00 | 18902 | Alcam | ENSMUSG0C | true | . | . | . | . | . | . | chr16 | 52428200 | 52436533 | 4.42754 | 3.67314 | 2.55046 |
| chr16 | 96145039 | 96145505 | 384 | Promoter (<=1kb) | 16 | 96145407 | 96146764 | 1358 | 1 | ENSMUST00 | 0 | Wrb | ENSMUSG0C | false | chr16 | 96145333 | 96146075 | 27.54716 | 98.2761 | 94.74957 | chr16 | 96144970 | 96146871 | 4.47913 | 3.7945 | 2.58207 |
| chr2 | 27886127 | 27886484 | 384 | Promoter (<=1kb) | 2 | 27886425 | 28039514 | 153090 | 1 | ENSMUST00 | 0 | Col5a1 | ENSMUSG0C | false | chr2 | 27885967 | 27887278 | 26.60951 | 79.31685 | 76.1741 | chr2 | 27885611 | 27888553 | 4.29162 | 3.66787 | 2.46333 |
| chr4 | 83412666 | 83412959 | 384 | Distal Intergenic | 4 | 83417724 | 83453340 | 35617 | 1 | ENSMUST00 | -4765 | Snapc3 | ENSMUSG0C | false | . | . | . | . | . | . | chr4 | 83411883 | 83413297 | 4.02764 | 3.5913 | 2.21029 |
| chr4 | 134853930 | 134854441 | 384 | Promoter (<=1kb) | 4 | 134802759 | 134853345 | 50587 | 2 | ENSMUST00 | -586 | Tmem57 | ENSMUSG0C | false | chr4 | 134852051 | 134854505 | 36.04346 | 118.78738 | 114.68494 | . | . | . | . | . | . |
| chr9 | 57765849 | 57766158 | 384 | Promoter (<=1kb) | 9 | 57750711 | 57765860 | 15150 | 2 | ENSMUST00 | 0 | Clk3 | ENSMUSG0C | false | chr9 | 57763942 | 57766341 | 37.17318 | 134.81126 | 130.13722 | chr9 | 57765990 | 57766475 | 4.12814 | 3.61922 | 2.34453 |
| chr8 | 71668109 | 71668356 | 384 | Promoter (<=1kb) | 8 | 71660346 | 71668082 | 7737 | 2 | ENSMUST00 | -28 | Unc13a | ENSMUSG0C | false | . | . | . | . | . | . | . | . | . | . | . | . |
| chr11 | 101713367 | 101713667 | 384 | Distal Intergenic | 11 | 101732919 | 101735696 | 2778 | 1 | ENSMUST00 | -19252 | Dhx8 | ENSMUSG0C | false | . | . | . | . | . | . | chr11 | 101709308 | 101714568 | 4.63527 | 3.87211 | 2.71845 |
| chr7 | 80864858 | 80865243 | 384 | Promoter (1-2kb) | 7 | 80862900 | 80875098 | 12199 | 1 | ENSMUST00 | 1959 | Zscan2 | ENSMUSG0C | false | . | . | . | . | . | . | . | . | . | . | . | . |
| chr6 | 59426051 | 59426701 | 384 | Promoter (<=1kb) | 6 | 59352461 | 59426290 | 73830 | 2 | ENSMUST00 | 0 | Gprin3 | ENSMUSG0C | false | chr6 | 59425318 | 59426818 | 30.59503 | 95.89513 | 92.42037 | chr6 | 59424013 | 59427111 | 2.80721 | 2.84187 | 1.25478 |
| chr3 | 103968168 | 103968765 | 384 | Promoter (<=1kb) | 3 | 103968258 | 104007488 | 39231 | 1 | ENSMUST00 | 0 | Phtf1 | ENSMUSG0C | false | chr3 | 103967695 | 103969065 | 28.56742 | 103.16996 | 99.51957 | . | . | . | . | . | . |
| chr11 | 98409465 | 98409736 | 384 | Promoter (2-3kb) | 11 | 98412470 | 98437716 | 25247 | 1 | ENSMUST00 | -2734 | Erbb2 | ENSMUSG0C | false | . | . | . | . | . | . | chr11 | 98408782 | 98410315 | 4.65991 | 3.55495 | 2.76348 |
| chr12 | 9174420 | 9174678 | 384 | Distal Intergenic | 12 | 9029997 | 9036394 | 6398 | 1 | ENSMUST00 | 144424 | Ttc32 | ENSMUSG0C | false | . | . | . | . | . | . | chr12 | 9172890 | 9177711 | 4.77444 | 3.89476 | 2.82995 |
| chr15 | 60824754 | 60825172 | 384 | Promoter (<=1kb) | 15 | 60818994 | 60825080 | 6087 | 2 | ENSMUST00 | 0 | Fam84b | ENSMUSG0C | false | chr15 | 60822622 | 60825350 | 25.17393 | 93.69764 | 90.27328 | . | . | . | . | . | . |
| chr2 | 163202908 | 163203314 | 384 | Promoter (<=1kb) | 2 | 163203125 | 163324170 | 121046 | 1 | ENSMUST00 | 0 | Tox2 | ENSMUSG0C | false | chr2 | 163202708 | 163203711 | 13.16235 | 40.53663 | 37.94056 | chr2 | 163202750 | 163205601 | 5.03462 | 4.01166 | 3.06637 |
| chr6 | 148971728 | 148971993 | 384 | Distal Intergenic | 6 | 148967316 | 148967437 | 122 | 1 | ENSMUST00 | 4413 | Gm25539 | ENSMUSG0C | false | . | . | . | . | . | . | chr6 | 148971275 | 148972349 | 3.70852 | 3.47763 | 1.935 |
| chr7 | 64050270 | 64050571 | 384 | Distal Intergenic | 7 | 64057602 | 64062958 | 5357 | 1 | ENSMUST00 | -7031 | Gm20670 | ENSMUSG0C | false | . | . | . | . | . | . | chr7 | 64047583 | 64051824 | 7.77647 | 5.37371 | 5.28037 |
| chr6 | 47757718 | 47758094 | 384 | Distal Intergenic | 6 | 47773329 | 47773429 | 101 | 1 | ENSMUST00 | -15235 | Gm24975 | ENSMUSG0C | false | chr6 | 47757752 | 47758418 | 14.34657 | 63.50294 | 60.60785 | . | . | . | . | . | . |
| chr5 | 124010839 | 124011425 | 383 | Exon (ENSMUST00000040967.7/ | 5 | 124000599 | 124001674 | 1076 | 1 | ENSMUST00 | 10241 | Hip1r | ENSMUSG0C | false | . | . | . | . | . | . | chr5 | 124009872 | 124020161 | 4.58057 | 3.77289 | 2.6871 |
| chr4 | 136143574 | 136143843 | 383 | Promoter (<=1kb) | 4 | 136143864 | 136144716 | 853 | 1 | ENSMUST00 | -21 | Id3 | ENSMUSG0C | false | chr4 | 136142962 | 136145178 | 19.27017 | 55.84607 | 53.05495 | chr4 | 136142629 | 136145211 | 5.36542 | 4.15748 | 3.32061 |
| chr8 | 123342347 | 123342606 | 383 | Intron (ENSMUST00000127664.1 | 8 | 123332713 | 123369515 | 36803 | 1 | ENSMUST00 | 9635 | Spire2 | ENSMUSG0C | false | . | . | . | . | . | . | chr8 | 123342485 | 123342805 | 4.12759 | 3.44317 | 2.32565 |
| chr10 | 122008582 | 122009099 | 383 | Intron (ENSMUST00000020322.6 | 10 | 121780991 | 122047308 | 266318 | 2 | ENSMUST00 | 38209 | Srgap1 | ENSMUSG0C | true | . | . | . | . | . | . | chr10 | 121945243 | 122036389 | 5.74397 | 4.3951 | 3.62974 |
| chr16 | 33929236 | 33929683 | 383 | Intron (ENSMUST00000115028.4 | 16 | 33922486 | 33944148 | 21663 | 1 | ENSMUST00 | 6751 | Itgb5 | ENSMUSG0C | true | . | . | . | . | . | . | chr16 | 33911227 | 33952498 | 7.40288 | 5.1092 | 4.9863 |
| chr19 | 14502272 | 14502650 | 383 | Intron (ENSMUST00000052011.8 | 19 | 14448072 | 14597983 | 149912 | 2 | ENSMUST00 | 95333 | Tle4 | ENSMUSG0C | false | . | . | . | . | . | . | chr19 | 14496852 | 14509645 | 5.46229 | 4.19323 | 3.40627 |
| chr5 | 24601569 | 24602644 | 383 | Promoter (<=1kb) | 5 | 24590818 | 24601976 | 11159 | 2 | ENSMUST00 | 0 | Smarcd3 | ENSMUSG0C | false | chr5 | 24601323 | 24603193 | 16.88215 | 57.7818 | 54.96725 | chr5 | 24601136 | 24606675 | 7.02969 | 4.92989 | 4.67941 |
| chr9 | 50835413 | 50835682 | 383 | Promoter (<=1kb) | 9 | 50835571 | 50843091 | 7521 | 1 | ENSMUST00 | 0 | Alg9 | ENSMUSG0C | false | . | . | . | . | . | . | chr9 | 50835090 | 50835900 | 4.15274 | 3.69792 | 2.31121 |
| chr5 | 77394263 | 77394750 | 383 | Intron (ENSMUST00000046746.6 | 5 | 77349240 | 77408045 | 58806 | 2 | ENSMUST00 | 13295 | Igfbp7 | ENSMUSG0C | false | . | . | . | . | . | . | chr5 | 77353137 | 77404361 | 6.51707 | 4.75542 | 4.25399 |
| chrX | 142902736 | 142902981 | 383 | Intron (ENSMUST00000041317.2 | 20 | 142851146 | 142966728 | 115583 | 2 | ENSMUST00 | 63747 | Ammecr1 | ENSMUSG0C | false | . | . | . | . | . | . | . | . | . | . | . | . |
| chr19 | 12476578 | 12476974 | 383 | Intron (ENSMUST00000045521.7 | 19 | 12460779 | 12465284 | 4506 | 1 | ENSMUST00 | 15800 | Mpeg1 | ENSMUSG0C | false | . | . | . | . | . | . | chr19 | 12475291 | 12477507 | 4.4595 | 3.60276 | 2.59019 |
| chr8 | 103481552 | 103481759 | 383 | Distal Intergenic | 8 | 102864779 | 102865853 | 1075 | 2 | ENSMUST00 | -615700 | Gm8730 | ENSMUSG0C | false | . | . | . | . | . | . | chr8 | 103480970 | 103482577 | 4.00715 | 3.59072 | 2.24329 |
| chr6 | 43932550 | 43932762 | 383 | Distal Intergenic | 6 | 43887194 | 43887324 | 131 | 1 | ENSMUST00 | 45357 | Gm22939 | ENSMUSG0C | false | . | . | . | . | . | . | chr6 | 43931777 | 43933509 | 3.89427 | 3.57103 | 2.11693 |
| chr2 | 7242185 | 7242416 | 383 | Intron (ENSMUST00000002176.7 | 2 | 7350097 | 7350193 | 97 | 1 | ENSMUST00 | -107681 | Gm24340 | ENSMUSG0C | false | . | . | . | . | . | . | . | . | . | . | . | . |
| chr16 | 26035078 | 26035347 | 383 | Intron (ENSMUST00000039990.5 | 16 | 26031810 | 26040870 | 9061 | 2 | ENSMUST00 | 5523 | AU015336 | ENSMUSG0C | false | . | . | . | . | . | . | chr16 | 26035252 | 26056185 | 6.03158 | 4.46161 | 3.87334 |
| chr5 | 118424748 | 118425032 | 383 | Distal Intergenic | 5 | 118418727 | 118418818 | 92 | 2 | ENSMUST00 | -5931 | Gm26455 | ENSMUSG0C | false | . | . | . | . | . | . | chr5 | 118424215 | 118425560 | 3.47384 | 3.33567 | 1.73304 |
| chr4 | 133093484 | 133093717 | 383 | Distal Intergenic | 4 | 133085474 | 133089146 | 3673 | 1 | ENSMUST00 | 8011 | Gm26615 | ENSMUSG0C | false | . | . | . | . | . | . | chr4 | 133082180 | 133096297 | 7.96206 | 5.47868 | 5.42168 |
| chr7 | 126565736 | 126566028 | 383 | Promoter (<=1kb) | 7 | 126565919 | 126566380 | 462 | 1 | ENSMUST00 | 0 | G730046D07 | ENSMUSG0C | false | chr7 | 126565521 | 126567070 | 31.54088 | 108.67455 | 104.87239 | chr7 | 126564903 | 126565779 | 4.11473 | 3.69586 | 2.26361 |
| chr1 | 90463404 | 90463700 | 383 | Distal Intergenic | 1 | 90453394 | 90454501 | 1108 | 2 | ENSMUST00 | -8904 | RP24-154L11 | ENSMUSG0C | false | . | . | . | . | . | . | chr1 | 90461309 | 90477894 | 5.37806 | 4.21863 | 3.31987 |
| chr12 | 32061113 | 32061467 | 382 | Promoter (<=1kb) | 12 | 31958476 | 32061235 | 102760 | 2 | ENSMUST00 | 0 | Prkar2b | ENSMUSG0C | false | chr12 | 32060375 | 32061738 | 23.37426 | 74.48241 | 71.42245 | chr12 | 32059848 | 32061834 | 5.17804 | 3.91933 | 3.1876 |
| chr12 | 87388120 | 87388524 | 382 | Promoter (<=1kb) | 12 | 87305058 | 87388355 | 83298 | 2 | ENSMUST00 | 0 | Sptlc2 | ENSMUSG0C | false | chr12 | 87386808 | 87388351 | 37.92415 | 138.37384 | 133.55817 | . | . | . | . | . | . |
| chr13 | 51777129 | 51777420 | 382 | Promoter (<=1kb) | 13 | 51725390 | 51777111 | 51722 | 2 | ENSMUST00 | -19 | Sema4d | ENSMUSG0C | false | . | . | . | . | . | . | chr13 | 51777084 | 51777929 | 4.36003 | 3.65968 | 2.51985 |
| chr18 | 52951194 | 52951399 | 382 | Distal Intergenic | 18 | 52838010 | 52915312 | 77303 | 1 | ENSMUST00 | 113185 | Sncaip | ENSMUSG0C | true | . | . | . | . | . | . | chr18 | 52950572 | 52952080 | 3.45239 | 3.23588 | 1.76688 |
| chrX | 151343819 | 151344267 | 382 | Promoter (<=1kb) | 20 | 151344175 | 151471842 | 127668 | 1 | ENSMUST00 | 0 | Fam120c | ENSMUSG0C | false | chrX | 151343760 | 151345169 | 23.43614 | 63.09834 | 60.20888 | . | . | . | . | . | . |
| chr4 | 40143203 | 40143791 | 382 | Promoter (<=1kb) | 4 | 40143081 | 40198338 | 55258 | 1 | ENSMUST00 | 123 | Aco1 | ENSMUSG0C | false | chr4 | 40142530 | 40144718 | 34.3275 | 112.61823 | 108.70669 | . | . | . | . | . | . |
| chr4 | 134327222 | 134327591 | 382 | Intron (ENSMUST00000105875.2 | 4 | 134319988 | 134329628 | 9641 | 1 | ENSMUST00 | 7235 | Trim63 | ENSMUSG0C | false | . | . | . | . | . | . | chr4 | 134326497 | 134329268 | 5.45912 | 4.30304 | 3.39778 |
| chr5 | 65019847 | 65020153 | 382 | Intron (ENSMUST00000031080.9 | 5 | 64972425 | 65005891 | 33467 | 1 | ENSMUST00 | 47423 | Fam114a1 | ENSMUSG0C | false | . | . | . | . | . | . | chr5 | 65019418 | 65022572 | 7.26089 | 5.21682 | 4.84068 |
| chr5 | 72855692 | 72855947 | 382 | Intron (ENSMUST00000071944.7 | 5 | 72755734 | 72868396 | 112663 | 2 | ENSMUST00 | 12449 | Tec | ENSMUSG0C | false | . | . | . | . | . | . | . | . | . | . | . | . |
| chr8 | 25087826 | 25088141 | 382 | Intron (ENSMUST00000064883.7 | 8 | 25062433 | 25091346 | 28914 | 2 | ENSMUST00 | 3205 | Plekha2 | ENSMUSG0C | true | . | . | . | . | . | . | chr8 | 25082618 | 25100838 | 5.3544 | 4.1622 | 3.30614 |
| chr8 | 111731776 | 111732323 | 382 | Intron (ENSMUST00000166232.2 | 8 | 111710485 | 111743849 | 33365 | 2 | ENSMUST00 | 11526 | Bcar1 | ENSMUSG0C | false | chr8 | 111730354 | 111732664 | 27.54716 | 98.2761 | 94.74957 | chr8 | 111710939 | 111747327 | 8.20913 | 5.4583 | 5.64085 |
| chr4 | 119684349 | 119684570 | 382 | Distal Intergenic | 4 | 119656607 | 119658954 | 2348 | 2 | ENSMUST00 | -25396 | Guca2b | ENSMUSG0C | false | . | . | . | . | . | . | chr4 | 119682921 | 119684541 | 3.51328 | 3.31167 | 1.77955 |
| chr4 | 117645898 | 117646251 | 382 | Intron (ENSMUST00000037127.9 | 4 | 117637338 | 117674281 | 36944 | 1 | ENSMUST00 | 8561 | Eri3 | ENSMUSG0C | false | . | . | . | . | . | . | chr4 | 117643452 | 117654917 | 5.66723 | 4.343 | 3.56197 |

| | | | | | | | | | | | | | | | | | | | | | | | | | | |
|---|---|---|---|---|---|---|---|---|---|---|---|---|---|---|---|---|---|---|---|---|---|---|---|---|---|---|
| chr5 | 139388539 | 139388859 | 382 | Promoter (<=1kb) | 5 | 139389785 | 139396414 | 6630 | 1 | ENSMUST00 | -926 | Gpr146 | ENSMUSG0C | false | . | . | . | . | . | . | chr5 | 139381256 | 139395525 | 8.99937 | 5.68291 | 6.29852 |
| chr16 | 42517451 | 42517679 | 382 | Distal Intergenic | 16 | 42248442 | 42340651 | 92210 | 2 | ENSMUST00 | -176801 | Gap43 | ENSMUSG0C | false | . | . | . | . | . | . | chr16 | 42515803 | 42517827 | 4.3861 | 3.76327 | 2.55295 |
| chr18 | 21185339 | 21185546 | 382 | Intron (ENSMUST00000049260.6 | 18 | 21076667 | 21077110 | 444 | 2 | ENSMUST00 | -108230 | Gm6378 | ENSMUSG0C | false | . | . | . | . | . | . | chr18 | 21184280 | 21187575 | 5.4355 | 4.23033 | 3.3806 |
| chr6 | 144711870 | 144712130 | 382 | Intron (ENSMUST00000141102.2 | 6 | 144720911 | 144759883 | 38973 | 1 | ENSMUST00 | -8781 | Sox5os4 | ENSMUSG0C | false | . | . | . | . | . | . | chr6 | 144706059 | 144712669 | 4.36393 | 3.70966 | 2.49703 |
| chr5 | 64680013 | 64680289 | 382 | Distal Intergenic | 5 | 64695850 | 64698520 | 2671 | 2 | ENSMUST00 | 18231 | Gm26761 | ENSMUSG0C | false | . | . | . | . | . | . | . | . | . | . | . | . |
| chr1 | 61527179 | 61527393 | 382 | Distal Intergenic | 1 | 61548122 | 61548511 | 390 | 1 | ENSMUST00 | -20729 | RP23-280M7 | ENSMUSG0C | false | . | . | . | . | . | . | chr1 | 61526896 | 61533551 | 4.85222 | 3.97276 | 2.89832 |
| chr5 | 136888503 | 136888915 | 381 | Distal Intergenic | 5 | 136757489 | 136883209 | 125721 | 2 | ENSMUST00 | -5295 | Col26a1 | ENSMUSG0C | false | . | . | . | . | . | . | chr5 | 136885321 | 136889075 | 4.822 | 3.84455 | 2.90361 |
| chr9 | 21858177 | 21858441 | 381 | Distal Intergenic | 9 | 21800184 | 21852635 | 52452 | 2 | ENSMUST00 | -5543 | Dock6 | ENSMUSG0C | false | . | . | . | . | . | . | chr9 | 21857640 | 21859055 | 3.46752 | 3.3899 | 1.75362 |
| chr9 | 56192100 | 56192370 | 381 | Distal Intergenic | 9 | 56135885 | 56161070 | 25186 | 2 | ENSMUST00 | -31031 | Tspan3 | ENSMUSG0C | false | . | . | . | . | . | . | chr9 | 56191561 | 56192857 | 4.3024 | 3.68696 | 2.45308 |
| chr7 | 63816065 | 63816312 | 381 | Distal Intergenic | 7 | 63754712 | 63757925 | 3214 | 1 | ENSMUST00 | 61354 | Otud7a | ENSMUSG0C | false | . | . | . | . | . | . | . | . | . | . | . | . |
| chr9 | 63896500 | 63896809 | 381 | Distal Intergenic | 9 | 63954009 | 64016987 | 62979 | 2 | ENSMUST00 | 120178 | Smad6 | ENSMUSG0C | true | . | . | . | . | . | . | chr9 | 63894722 | 63897655 | 5.97667 | 4.36657 | 3.81609 |
| chr3 | 37639742 | 37640016 | 381 | Promoter (<=1kb) | 3 | 37639947 | 37644598 | 4652 | 1 | ENSMUST00 | 0 | Spry1 | ENSMUSG0C | false | chr3 | 37639368 | 37642407 | 23.85991 | 99.51073 | 95.95212 | chr3 | 37639237 | 37639764 | 3.79355 | 3.4993 | 1.99462 |
| chr2 | 165874766 | 165875189 | 381 | Promoter (<=1kb) | 2 | 165827129 | 165875777 | 48649 | 2 | ENSMUST00 | 588 | Zmynd8 | ENSMUSG0C | false | . | . | . | . | . | . | chr2 | 165851451 | 165889342 | 6.61553 | 4.6931 | 4.35892 |
| chr4 | 126967551 | 126968436 | 381 | Promoter (<=1kb) | 4 | 126861939 | 126968135 | 106197 | 2 | ENSMUST00 | 0 | Zmym4 | ENSMUSG0C | false | chr4 | 126967024 | 126968587 | 24.89402 | 98.85246 | 95.31046 | . | . | . | . | . | . |
| chr9 | 24503030 | 24503629 | 381 | Promoter (<=1kb) | 9 | 24411778 | 24503140 | 91363 | 2 | ENSMUST00 | 0 | Dpy19l1 | ENSMUSG0C | false | chr9 | 24502352 | 24504092 | 20.74539 | 66.97287 | 64.02668 | . | . | . | . | . | . |
| chr17 | 27203170 | 27203443 | 381 | Promoter (<=1kb) | 17 | 27189601 | 27204438 | 14838 | 2 | ENSMUST00 | 995 | Lemd2 | ENSMUSG0C | false | chr17 | 27202960 | 27204531 | 33.7938 | 119.00099 | 114.88904 | chr17 | 27191319 | 27203697 | 8.84294 | 5.47024 | 6.1733 |
| chr18 | 83954623 | 83955118 | 381 | Distal Intergenic | 18 | 84012082 | 84086390 | 74309 | 2 | ENSMUST00 | 131272 | Tshz1 | ENSMUSG0C | false | . | . | . | . | . | . | chr18 | 83954616 | 83956633 | 4.5346 | 3.89014 | 2.6372 |
| chr19 | 5059411 | 5059693 | 381 | Distal Intergenic | 19 | 5068078 | 5070637 | 2560 | 1 | ENSMUST00 | -8385 | Cd248 | ENSMUSG0C | false | . | . | . | . | . | . | chr19 | 5057884 | 5062125 | 5.35627 | 4.08701 | 3.34292 |
| chr12 | 16850367 | 16850673 | 381 | Distal Intergenic | 12 | 16810965 | 16826743 | 15779 | 1 | ENSMUST00 | 39403 | E2f6 | ENSMUSG0C | false | . | . | . | . | . | . | chr12 | 16848168 | 16854626 | 5.81138 | 4.51483 | 3.66527 |
| chr11 | 22871680 | 22872081 | 381 | Promoter (2-3kb) | 11 | 22868901 | 22869004 | 104 | 1 | ENSMUST00 | 2780 | Gm24917 | ENSMUSG0C | false | . | . | . | . | . | . | chr11 | 22871696 | 22873248 | 3.55161 | 3.2923 | 1.82362 |
| chr6 | 144670525 | 144670764 | 381 | Promoter (1-2kb) | 6 | 144670427 | 144672720 | 2294 | 2 | ENSMUST00 | 1956 | Sox5it | ENSMUSG0C | false | . | . | . | . | . | . | chr6 | 144669585 | 144672014 | 3.26048 | 3.16427 | 1.61875 |
| chr1 | 134099062 | 134099511 | 381 | Distal Intergenic | 1 | 134109368 | 134109799 | 432 | 1 | ENSMUST00 | -9857 | Gm8618 | ENSMUSG0C | false | chr1 | 134099435 | 134099888 | 10.36119 | 22.58756 | 20.21866 | chr1 | 134095503 | 134101645 | 7.68348 | 5.30363 | 5.20617 |
| chr11 | 105510767 | 105511060 | 381 | Distal Intergenic | 11 | 105401652 | 105456735 | 55084 | 2 | ENSMUST00 | -54033 | 40238 | ENSMUSG0C | false | . | . | . | . | . | . | chr11 | 105508005 | 105511620 | 4.37401 | 3.68617 | 2.52223 |
| chr13 | 97390621 | 97390843 | 381 | Distal Intergenic | 13 | 97434970 | 97497007 | 62038 | 1 | ENSMUST00 | -44127 | Lincenc1 | ENSMUSG0C | false | . | . | . | . | . | . | chr13 | 97388892 | 97391211 | 3.75713 | 3.46594 | 1.96891 |
| chr6 | 111892443 | 111892703 | 381 | Distal Intergenic | 6 | 111992110 | 111992424 | 315 | 1 | ENSMUST00 | -99407 | Gm26362 | ENSMUSG0C | false | . | . | . | . | . | . | chr6 | 111891785 | 111892857 | 3.29315 | 3.10991 | 1.62903 |
| chr1 | 181841960 | 181842742 | 380 | Promoter (<=1kb) | 1 | 181815316 | 181842401 | 27086 | 2 | ENSMUST00 | 0 | Lbr | ENSMUSG0C | false | chr1 | 181841117 | 181843213 | 29.28796 | 98.53321 | 94.99628 | chr1 | 181840293 | 181842010 | 3.01401 | 2.99209 | 1.41041 |
| chr11 | 117119677 | 117120047 | 380 | Intron (ENSMUST00000127090.2 | 11 | 117115252 | 117128763 | 13512 | 1 | ENSMUST00 | 4426 | Sec14l1 | ENSMUSG0C | false | . | . | . | . | . | . | chr11 | 117116362 | 117143393 | 7.76107 | 5.12248 | 5.29175 |
| chr13 | 75320265 | 75320595 | 380 | Distal Intergenic | 13 | 75089916 | 75106603 | 16688 | 1 | ENSMUST00 | 230350 | Pcsk1 | ENSMUSG0C | false | . | . | . | . | . | . | . | . | . | . | . | . |
| chr15 | 32442816 | 32443230 | 380 | Intron (ENSMUST00000067458.6 | 15 | 32244813 | 32696341 | 451529 | 1 | ENSMUST00 | 198004 | Sema5a | ENSMUSG0C | true | . | . | . | . | . | . | chr15 | 32436609 | 32448232 | 4.8943 | 3.99707 | 2.94397 |
| chr18 | 76929871 | 76930244 | 380 | Promoter (<=1kb) | 18 | 76930126 | 76971449 | 41324 | 1 | ENSMUST00 | 0 | Hdhd2 | ENSMUSG0C | false | chr18 | 76929562 | 76931051 | 41.94252 | 141.08568 | 136.16306 | . | . | . | . | . | . |
| chr1 | 143946997 | 143947220 | 380 | Distal Intergenic | 1 | 144001883 | 144004032 | 2150 | 2 | ENSMUST00 | 56812 | Rgs2 | ENSMUSG0C | false | . | . | . | . | . | . | chr1 | 143946689 | 143948068 | 3.86854 | 3.5611 | 2.12128 |
| chr1 | 183147367 | 183147692 | 380 | Promoter (<=1kb) | 1 | 183086270 | 183147461 | 61192 | 2 | ENSMUST00 | 0 | Disp1 | ENSMUSG0C | false | chr1 | 183146159 | 183147549 | 15.17612 | 37.20362 | 34.64814 | chr1 | 183138001 | 183155887 | 4.39977 | 3.65871 | 2.54516 |
| chr7 | 112498703 | 112498949 | 380 | Promoter (2-3kb) | 7 | 112501177 | 112547610 | 46434 | 1 | ENSMUST00 | -2228 | Parva | ENSMUSG0C | true | . | . | . | . | . | . | chr7 | 112487030 | 112537122 | 5.64568 | 4.33316 | 3.55068 |
| chr13 | 32962105 | 32962448 | 380 | Promoter (2-3kb) | 13 | 32965209 | 32979067 | 13859 | 1 | ENSMUST00 | -2761 | Serpinb6b | ENSMUSG0C | false | . | . | . | . | . | . | chr13 | 32957470 | 32971328 | 4.32621 | 3.59198 | 2.48777 |
| chr18 | 76111614 | 76111833 | 380 | Intron (ENSMUST00000058997.8 | 18 | 76059458 | 76146443 | 86986 | 1 | ENSMUST00 | 52157 | Zbtb7c | ENSMUSG0C | true | . | . | . | . | . | . | chr18 | 76061445 | 76132293 | 6.70015 | 4.79833 | 4.4158 |
| chr12 | 78227091 | 78227686 | 380 | Promoter (<=1kb) | 12 | 78226655 | 78684767 | 458113 | 1 | ENSMUST00 | 437 | Gphn | ENSMUSG0C | false | chr12 | 78225441 | 78228015 | 33.69441 | 105.32094 | 101.61211 | . | . | . | . | . | . |
| chr7 | 100950580 | 100950854 | 380 | Intron (ENSMUST00000060174.4 | 7 | 100937638 | 100964391 | 26754 | 2 | ENSMUST00 | 13537 | P2ry6 | ENSMUSG0C | false | . | . | . | . | . | . | chr7 | 100948011 | 100950928 | 4.64651 | 3.83748 | 2.74968 |
| chr3 | 108739888 | 108740173 | 380 | Promoter (<=1kb) | 3 | 108739658 | 108782296 | 42639 | 1 | ENSMUST00 | 231 | Aknad1 | ENSMUSG0C | false | . | . | . | . | . | . | chr3 | 108738933 | 108740883 | 3.99831 | 3.53199 | 2.22767 |
| chr3 | 108186103 | 108186684 | 380 | Promoter (<=1kb) | 3 | 108186335 | 108191776 | 5442 | 1 | ENSMUST00 | 0 | Amigo1 | ENSMUSG0C | false | chr3 | 108185509 | 108187025 | 29.22734 | 90.55506 | 87.19949 | chr3 | 108185550 | 108190724 | 4.72302 | 3.94917 | 2.78536 |
| chr3 | 142065164 | 142065414 | 380 | Intron (ENSMUST00000098568.2 | 3 | 141837136 | 141982264 | 145129 | 2 | ENSMUST00 | -82901 | Bmpr1b | ENSMUSG0C | false | . | . | . | . | . | . | chr3 | 142063277 | 142066227 | 4.57154 | 3.86243 | 2.64319 |
| chr10 | 122984895 | 122985137 | 380 | Promoter (<=1kb) | 10 | 122985634 | 122985723 | 90 | 2 | ENSMUST00 | 586 | Mirlet7i | ENSMUSG0C | false | chr10 | 122984594 | 122987014 | 36.09495 | 113.61411 | 109.67428 | . | . | . | . | . | . |
| chr3 | 116210671 | 116210941 | 380 | Distal Intergenic | 3 | 116249654 | 116253484 | 3831 | 2 | ENSMUST00 | 42543 | Gpr88 | ENSMUSG0C | false | . | . | . | . | . | . | . | . | . | . | . | . |
| chr8 | 87235714 | 87235985 | 380 | Intron (ENSMUST00000183344.1 | 8 | 87203026 | 87293504 | 90479 | 1 | ENSMUST00 | 32689 | Gm27169 | ENSMUSG0C | false | . | . | . | . | . | . | chr8 | 87235330 | 87236262 | 4.05271 | 3.43517 | 2.26166 |
| chr8 | 120548659 | 120548913 | 380 | Intron (ENSMUST00000127664.1 | 8 | 120538696 | 120538754 | 59 | 1 | ENSMUST00 | 9964 | Mir7687 | ENSMUSG0C | false | . | . | . | . | . | . | chr8 | 120537951 | 120565808 | 7.48404 | 5.10526 | 5.05119 |
| chr7 | 25278882 | 25279368 | 379 | Promoter (<=1kb) | 7 | 25279641 | 25284932 | 5292 | 1 | ENSMUST00 | -273 | Cic | ENSMUSG0C | false | chr7 | 25278343 | 25279876 | 25.53309 | 82.08177 | 78.89293 | chr7 | 25269284 | 25281823 | 5.87867 | 4.36785 | 3.75349 |
| chr6 | 92174465 | 92174727 | 379 | Promoter (1-2kb) | 6 | 92174838 | 92175902 | 1065 | 2 | ENSMUST00 | 1175 | Mrps25 | ENSMUSG0C | false | . | . | . | . | . | . | chr6 | 92173429 | 92177313 | 6.8074 | 4.84073 | 4.49534 |
| chr13 | 48168451 | 48168681 | 379 | Intron (ENSMUST00000180921.1 | 13 | 48261427 | 48264036 | 2610 | 1 | ENSMUST00 | -92746 | Id4 | ENSMUSG0C | false | . | . | . | . | . | . | . | . | . | . | . | . |
| chr17 | 66564181 | 66564530 | 379 | Intron (ENSMUST00000024914.2 | 17 | 66555252 | 66594621 | 39370 | 2 | ENSMUST00 | 30091 | Themis3 | ENSMUSG0C | false | . | . | . | . | . | . | chr17 | 66564209 | 66564549 | 2.92436 | 2.85801 | 1.37707 |
| chr8 | 70680153 | 70680360 | 379 | Promoter (2-3kb) | 8 | 70677665 | 70678116 | 452 | 1 | ENSMUST00 | 2489 | Lsm4 | ENSMUSG0C | false | . | . | . | . | . | . | chr8 | 70678490 | 70683293 | 6.70585 | 4.76208 | 4.42699 |
| chrX | 101637211 | 101637442 | 379 | Promoter (2-3kb) | 20 | 101640060 | 101682940 | 42881 | 1 | ENSMUST00 | -2618 | Ogt | ENSMUSG0C | false | . | . | . | . | . | . | . | . | . | . | . | . |
| chr5 | 81509024 | 81509325 | 379 | Intron (ENSMUST00000132375.2 | 5 | 81645989 | 81688522 | 42534 | 1 | ENSMUST00 | -136664 | Lphn3 | ENSMUSG0C | false | . | . | . | . | . | . | chr5 | 81508688 | 81510949 | 3.27766 | 3.19493 | 1.60391 |
| chr10 | 26860006 | 26860238 | 379 | Intron (ENSMUST00000176060.2 | 10 | 26863342 | 26918458 | 55117 | 1 | ENSMUST00 | -3104 | Arhgap18 | ENSMUSG0C | false | . | . | . | . | . | . | chr10 | 26857976 | 26867351 | 4.83934 | 3.98113 | 2.87017 |
| chr13 | 113180313 | 113180625 | 379 | Promoter (<=1kb) | 13 | 113171638 | 113180897 | 9260 | 2 | ENSMUST00 | 272 | Gzmk | ENSMUSG0C | false | . | . | . | . | . | . | chr13 | 113176806 | 113183700 | 4.79857 | 4.02053 | 2.8354 |
| chr11 | 20616162 | 20616406 | 379 | Intron (ENSMUST00000093292.5 | 11 | 20631979 | 20653021 | 21043 | 1 | ENSMUST00 | -15573 | Sertad2 | ENSMUSG0C | true | . | . | . | . | . | . | chr11 | 20613119 | 20617578 | 4.88871 | 4.05396 | 2.91585 |
| chr8 | 91523322 | 91523573 | 379 | Exon (ENSMUST00000128081.2/ | 8 | 91467685 | 91666946 | 199262 | 1 | ENSMUST00 | 55638 | Fto | ENSMUSG0C | false | . | . | . | . | . | . | chr8 | 91521033 | 91528017 | 5.83853 | 4.42897 | 3.70928 |
| chr11 | 60106325 | 60106907 | 379 | Promoter (1-2kb) | 11 | 60105177 | 60185431 | 80255 | 1 | ENSMUST00 | 1149 | Rai1 | ENSMUSG0C | false | chr11 | 60104773 | 60106757 | 36.76944 | 114.78011 | 110.80492 | chr11 | 60106353 | 60117543 | 5.28818 | 4.15483 | 3.25623 |
| chr11 | 57773634 | 57774156 | 379 | Intron (ENSMUST00000066987.8 | 11 | 57764427 | 57764545 | 119 | 1 | ENSMUST00 | 9208 | n-R5s69 | ENSMUSG0C | false | . | . | . | . | . | . | chr11 | 57770983 | 57778876 | 7.36912 | 5.00935 | 4.97771 |
| chr2 | 156775226 | 156775479 | 379 | Promoter (<=1kb) | 2 | 156775420 | 156781658 | 6239 | 1 | ENSMUST00 | 0 | Myl9 | ENSMUSG0C | false | chr2 | 156775091 | 156775680 | 7.17313 | 13.20777 | 10.97558 | chr2 | 156774761 | 156776209 | 4.58511 | 3.82445 | 2.69354 |
| chr9 | 90226440 | 90226830 | 379 | Intron (ENSMUST00000041767.8 | 9 | 90221295 | 90221625 | 331 | 2 | ENSMUST00 | -4816 | Gm16200 | ENSMUSG0C | false | . | . | . | . | . | . | . | . | . | . | . | . |
| chr16 | 35200826 | 35201165 | 379 | Intron (ENSMUST00000114913.1 | 16 | 35234796 | 35234923 | 128 | 2 | ENSMUST00 | 33758 | Gm25179 | ENSMUSG0C | false | . | . | . | . | . | . | chr16 | 35198472 | 35207433 | 6.25906 | 4.64811 | 4.02553 |
| chr6 | 18682063 | 18682282 | 379 | Distal Intergenic | 6 | 18727292 | 18727398 | 107 | 1 | ENSMUST00 | -45010 | Gm23648 | ENSMUSG0C | false | . | . | . | . | . | . | chr6 | 18681924 | 18682552 | 5.1351 | 4.17998 | 3.10885 |
| chr16 | 55966212 | 55966480 | 379 | Promoter (<=1kb) | 16 | 55966275 | 55971435 | 5161 | 1 | ENSMUST00 | 0 | Rpl24 | ENSMUSG0C | false | chr16 | 55965761 | 55967413 | 18.02578 | 59.25625 | 56.42029 | . | . | . | . | . | . |
| chr13 | 70578073 | 70578307 | 379 | Distal Intergenic | 13 | 70573280 | 70576105 | 2826 | 1 | ENSMUST00 | 4794 | RP23-376D8. | ENSMUSG0C | false | . | . | . | . | . | . | . | . | . | . | . | . |
| chr2 | 38339167 | 38339455 | 378 | Promoter (<=1kb) | 2 | 38339281 | 38369199 | 29919 | 1 | ENSMUST00 | 0 | Lhx2 | ENSMUSG0C | false | chr2 | 38339114 | 38340086 | 19.88939 | 55.30079 | 52.51555 | chr2 | 38338698 | 38341125 | 7.49614 | 5.06479 | 5.06339 |
| chr8 | 71981129 | 71981336 | 378 | Intron (ENSMUST00000136878.1 | 8 | 71988483 | 71992915 | 4433 | 2 | ENSMUST00 | 11579 | Cyp4f18 | ENSMUSG0C | false | . | . | . | . | . | . | chr8 | 71980993 | 71981688 | 5.1874 | 4.1766 | 3.21112 |
| chr10 | 79989954 | 79990247 | 378 | Promoter (1-2kb) | 10 | 79988623 | 79994684 | 6062 | 1 | ENSMUST00 | 1332 | Cnn2 | ENSMUSG0C | false | chr10 | 79988286 | 79990465 | 33.5288 | 108.10788 | 104.32592 | chr10 | 79985659 | 79991351 | 6.40037 | 4.65973 | 4.17147 |
| chr16 | 4419749 | 4420814 | 378 | Promoter (<=1kb) | 16 | 4287529 | 4419808 | 132280 | 2 | ENSMUST00 | 0 | Adcy9 | ENSMUSG0C | false | chr16 | 4419243 | 4421071 | 31.91637 | 110.38313 | 106.53261 | chr16 | 4420792 | 4421336 | 3.88093 | 3.34641 | 2.11582 |
| chr7 | 66331828 | 66332120 | 378 | Intron (ENSMUST00000015277.8 | 7 | 66341500 | 66343265 | 1766 | 2 | ENSMUST00 | 11145 | Lrrk1 | ENSMUSG0C | false | . | . | . | . | . | . | chr7 | 66330932 | 66333871 | 3.00779 | 2.95222 | 1.42348 |
| chr11 | 42044839 | 42045072 | 378 | Distal Intergenic | 11 | 41910203 | 42000857 | 90655 | 2 | ENSMUST00 | -43983 | Gabrg2 | ENSMUSG0C | false | . | . | . | . | . | . | chr11 | 42044825 | 42045161 | 3.46409 | 3.32547 | 1.72957 |
| chr4 | 136763492 | 136763702 | 378 | Intron (ENSMUST00000105845.3 | 4 | 136661463 | 136694040 | 32578 | 2 | ENSMUST00 | -69453 | Ephb2 | ENSMUSG0C | false | . | . | . | . | . | . | chr4 | 136763414 | 136764003 | 3.66469 | 3.32404 | 1.94347 |
| chr4 | 132329343 | 132329679 | 378 | Promoter (<=1kb) | 4 | 132311763 | 132329489 | 17727 | 2 | ENSMUST00 | 0 | Trnau1ap | ENSMUSG0C | false | chr4 | 132328653 | 132330129 | 29.92778 | 109.76675 | 105.9399 | . | . | . | . | . | . |
| chr5 | 66603734 | 66604072 | 378 | Intron (ENSMUST00000159786.2 | 5 | 66451982 | 66618640 | 166659 | 2 | ENSMUST00 | 14568 | Apbb2 | ENSMUSG0C | false | . | . | . | . | . | . | chr5 | 66558047 | 66617728 | 6.24773 | 4.56399 | 4.04918 |
| chr10 | 81343203 | 81343539 | 378 | Promoter (<=1kb) | 10 | 81337762 | 81343266 | 5505 | 2 | ENSMUST00 | 0 | Gipc3 | ENSMUSG0C | false | chr10 | 81342803 | 81343699 | 24.80288 | 69.81013 | 66.8223 | chr10 | 81339953 | 81351319 | 5.28841 | 4.20398 | 3.24758 |
| chr10 | 41887191 | 41887657 | 378 | Promoter (<=1kb) | 10 | 41887439 | 41908424 | 20986 | 1 | ENSMUST00 | 0 | Sesn1 | ENSMUSG0C | false | chr10 | 41886943 | 41888211 | 28.91247 | 96.86193 | 93.36482 | chr10 | 41886776 | 41889286 | 4.27698 | 3.77888 | 2.41165 |
| chr16 | 10366729 | 10367022 | 378 | Intron (ENSMUST00000043415.7 | 16 | 10361254 | 10395448 | 34195 | 2 | ENSMUST00 | 28426 | Tekt5 | ENSMUSG0C | false | . | . | . | . | . | . | chr16 | 10366088 | 10367740 | 5.88423 | 4.44058 | 3.72343 |
| chr18 | 82692840 | 82693060 | 378 | Promoter (<=1kb) | 18 | 82593593 | 82692883 | 99291 | 2 | ENSMUST00 | 0 | Zfp236 | ENSMUSG0C | false | chr18 | 82691477 | 82693572 | 23.88927 | 69.11851 | 66.14216 | . | . | . | . | . | . |
| chr4 | 152273845 | 152274353 | 378 | Promoter (<=1kb) | 4 | 152274232 | 152285337 | 11106 | 1 | ENSMUST00 | 0 | Gpr153 | ENSMUSG0C | false | chr4 | 152273869 | 152274536 | 15.50707 | 38.47921 | 35.908 | chr4 | 152271972 | 152275112 | 4.94888 | 3.954 | 2.97647 |
| chr9 | 37544493 | 37544716 | 378 | Promoter (1-2kb) | 9 | 37544954 | 37546138 | 1185 | 2 | ENSMUST00 | 1422 | Nrgn | ENSMUSG0C | false | chr9 | 37544354 | 37545220 | 10.51363 | 23.99435 | 21.60407 | chr9 | 37544181 | 37545633 | 5.22636 | 4.22789 | 3.18289 |
| chr14 | 64440536 | 64441073 | 378 | 5' UTR | 14 | 64122623 | 64455903 | 333281 | 2 | ENSMUST00 | 14830 | Msra | ENSMUSG0C | false | chr14 | 64440031 | 64441524 | 24.55236 | 90.57755 | 87.22021 | . | . | . | . | . | . |
| chr10 | 117746232 | 117746557 | 378 | Promoter (<=1kb) | 10 | 117733679 | 117746358 | 12680 | 2 | ENSMUST00 | 0 | Slc35e3 | ENSMUSG0C | false | chr10 | 117745534 | 117746867 | 27.29887 | 88.10642 | 84.80337 | chr10 | 117745089 | 117747957 | 3.59248 | 3.27041 | 1.90125 |
| chr10 | 126986634 | 126987126 | 378 | Intron (ENSMUST00000105256.3 | 10 | 126978717 | 126999975 | 21259 | 1 | ENSMUST00 | 7918 | Ctdsp2 | ENSMUSG0C | false | . | . | . | . | . | . | chr10 | 126980927 | 126992543 | 7.08562 | 4.77321 | 4.73941 |
| chr5 | 107222481 | 107222771 | 378 | Intron (ENSMUST00000031224.9 | 5 | 107241623 | 107242367 | 745 | 1 | ENSMUST00 | -18852 | Gm8145 | ENSMUSG0C | false | . | . | . | . | . | . | chr5 | 107211338 | 107245281 | 5.12218 | 4.09923 | 3.11878 |
| chr6 | 86526352 | 86527429 | 378 | Promoter (<=1kb) | 6 | 86527401 | 86564442 | 37042 | 1 | ENSMUST00 | 0 | 1600020E01I | ENSMUSG0C | false | chr6 | 86523216 | 86527818 | 38.37027 | 120.70945 | 116.54665 | . | . | . | . | . | . |
| chr11 | 79339363 | 79340091 | 377 | Promoter (<=1kb) | 11 | 79339693 | 79345186 | 5494 | 1 | ENSMUST00 | 0 | Nf1 | ENSMUSG0C | false | chr11 | 79338660 | 79340697 | 35.18842 | 108.93304 | 105.12779 | . | . | . | . | . | . |
| chr16 | 33879595 | 33879869 | 377 | Intron (ENSMUST00000115028.4 | 16 | 33885026 | 33910585 | 25560 | 1 | ENSMUST00 | -5157 | Itgb5 | ENSMUSG0C | true | . | . | . | . | . | . | chr16 | 33868993 | 33898226 | 6.80171 | 4.94523 | 4.47886 |
| chr11 | 88234117 | 88234386 | 377 | Distal Intergenic | 11 | 88204444 | 88210971 | 6528 | 1 | ENSMUST00 | 29674 | Mrps23 | ENSMUSG0C | false | . | . | . | . | . | . | chr11 | 88233690 | 88236557 | 4.41115 | 3.71613 | 2.56478 |
| chr2 | 144011259 | 144011646 | 377 | Promoter (<=1kb) | 2 | 143947395 | 144011263 | 63869 | 2 | ENSMUST00 | 0 | Rrbp1 | ENSMUSG0C | false | chr2 | 144009741 | 144011296 | 37.3746 | 124.32532 | 120.03539 | . | . | . | . | . | . |
| chr7 | 70337778 | 70337991 | 377 | Distal Intergenic | 7 | 70351946 | 70360593 | 8648 | 2 | ENSMUST00 | 22602 | Nr2f2 | ENSMUSG0C | false | chr7 | 70337629 | 70338074 | 8.73111 | 16.23101 | 13.94854 | chr7 | 70337082 | 70338342 | 2.96249 | 2.84585 | 1.37479 |
| chr7 | 89483851 | 89484075 | 377 | Distal Intergenic | 7 | 89510007 | 89511368 | 1362 | 1 | ENSMUST00 | -25932 | Prss23os | ENSMUSG0C | false | . | . | . | . | . | . | chr7 | 89480307 | 89485104 | 4.42544 | 3.8149 | 2.52302 |
| chrX | 167330461 | 167330686 | 377 | Promoter (<=1kb) | 20 | 167304929 | 167330528 | 25600 | 2 | ENSMUST00 | 0 | Tlr7 | ENSMUSG0C | false | . | . | . | . | . | . | . | . | . | . | . | . |
| chr10 | 81424225 | 81424496 | 377 | Promoter (2-3kb) | 10 | 81399138 | 81427114 | 27977 | 2 | ENSMUST00 | 2618 | Nfic | ENSMUSG0C | false | chr10 | 81424144 | 81425098 | 7.82203 | 16.59716 | 14.30959 | chr10 | 81392649 | 81426205 | 8.72751 | 5.63067 | 6.0679 |
| chr7 | 112762713 | 112762970 | 377 | Intron (ENSMUST00000084705.6 | 7 | 112759546 | 112842374 | 82829 | 1 | ENSMUST00 | 3168 | Tead1 | ENSMUSG0C | false | . | . | . | . | . | . | chr7 | 112739113 | 112766200 | 5.84261 | 4.33795 | 3.71566 |
| chr19 | 5060772 | 5061093 | 377 | Distal Intergenic | 19 | 5068078 | 5070637 | 2560 | 1 | ENSMUST00 | -6985 | Cd248 | ENSMUSG0C | false | . | . | . | . | . | . | chr19 | 5057884 | 5062125 | 5.35627 | 4.08701 | 3.34292 |
| chr14 | 116925156 | 116925576 | 377 | Promoter (<=1kb) | 14 | 116925297 | 117979529 | 1054233 | 1 | ENSMUST00 | 0 | Gpc6 | ENSMUSG0C | false | chr14 | 116925098 | 116927440 | 24.50127 | 70.18111 | 67.18839 | chr14 | 116925486 | 116928341 | 3.36847 | 3.10967 | 1.70841 |
| chr11 | 101627359 | 101627684 | 377 | Promoter (<=1kb) | 11 | 101627366 | 101627556 | 191 | 1 | ENSMUST00 | 0 | Gm23849 | ENSMUSG0C | false | chr11 | 101627645 | 101628686 | 25.16654 | 87.04672 | 83.76057 | . | . | . | . | . | . |
| chr3 | 105800662 | 105801468 | 377 | Promoter (<=1kb) | 3 | 105727267 | 105801330 | 74064 | 2 | ENSMUST00 | 0 | Rap1a | ENSMUSG0C | false | chr3 | 105799997 | 105801764 | 28.9509 | 90.07051 | 86.72839 | . | . | . | . | . | . |
| chr10 | 80264691 | 80265221 | 377 | Promoter (<=1kb) | 10 | 80264984 | 80288408 | 23425 | 1 | ENSMUST00 | 0 | Dazap1 | ENSMUSG0C | false | chr10 | 80264951 | 80266229 | 38.23518 | 126.91195 | 122.53338 | chr10 | 80259637 | 80275268 | 5.67236 | 4.28519 | 3.5841 |
| chr3 | 53525788 | 53526035 | 377 | Exon (ENSMUST00000091137.4/ | 3 | 53512002 | 53512126 | 125 | 1 | ENSMUST00 | 13787 | Gm23675 | ENSMUSG0C | false | . | . | . | . | . | . | . | . | . | . | . | . |
| chr5 | 146471672 | 146471905 | 377 | 3' UTR | 5 | 146481943 | 146484702 | 2760 | 1 | ENSMUST00 | -10038 | Gm6408 | ENSMUSG0C | false | . | . | . | . | . | . | chr5 | 146470762 | 146472219 | 4.76069 | 4.00812 | 2.81772 |
| chr17 | 15993765 | 15994003 | 377 | Distal Intergenic | 17 | 15837549 | 15837655 | 107 | 2 | ENSMUST00 | -156111 | Gm25598 | ENSMUSG0C | false | . | . | . | . | . | . | chr17 | 15993445 | 15995149 | 4.36753 | 3.72931 | 2.52649 |
| chr14 | 61866142 | 61866474 | 377 | Distal Intergenic | 14 | 61602839 | 61682373 | 79535 | 2 | ENSMUST00 | -183770 | Dleu2 | ENSMUSG0C | false | . | . | . | . | . | . | chr14 | 61866073 | 61866454 | 3.26104 | 3.11529 | 1.61527 |
| chr18 | 54971297 | 54971576 | 377 | Intron (ENSMUST00000064763.5 | 18 | 54981435 | 54987689 | 6255 | 1 | ENSMUST00 | -9859 | Gm26742 | ENSMUSG0C | false | . | . | . | . | . | . | chr18 | 54931543 | 54986508 | 4.96857 | 4.00781 | 2.99617 |
| chr5 | 114707701 | 114707881 | 376 | Promoter (<=1kb) | 5 | 114707760 | 114722327 | 14568 | 1 | ENSMUST00 | 0 | Tchp | ENSMUSG0C | false | chr5 | 114707703 | 114708580 | 28.53699 | 95.19626 | 91.7378 | chr5 | 114707363 | 114708929 | 7.03659 | 5.04842 | 4.64939 |
| chr16 | 8671513 | 8671983 | 376 | Promoter (<=1kb) | 16 | 8658587 | 8672153 | 13567 | 2 | ENSMUST00 | 170 | Carhsp1 | ENSMUSG0C | false | chr16 | 8671377 | 8672582 | 35.06866 | 119.18253 | 115.06551 | chr16 | 8663381 | 8672619 | 4.87856 | 3.94015 | 2.92794 |
| chr15 | 97821322 | 97821564 | 376 | Intron (ENSMUST00000079838.8 | 15 | 97792688 | 97814863 | 22176 | 2 | ENSMUST00 | -6460 | Hdac7 | ENSMUSG0C | false | . | . | . | . | . | . | chr15 | 97819806 | 97833530 | 5.44469 | 4.22025 | 3.37668 |
| chr18 | 57977499 | 57977706 | 376 | Distal Intergenic | 18 | 57878678 | 57946821 | 68144 | 1 | ENSMUST00 | 98822 | Slc12a2 | ENSMUSG0C | false | . | . | . | . | . | . | chr18 | 57977104 | 57978066 | 4.14355 | 3.68761 | 2.28789 |
| chr1 | 172501709 | 172501980 | 376 | Promoter (<=1kb) | 1 | 172501252 | 172506733 | 5482 | 1 | ENSMUST00 | 458 | Tagln2 | ENSMUSG0C | false | chr1 | 172499800 | 172502632 | 28.23176 | 83.06847 | 79.86218 | chr1 | 172501063 | 172503936 | 5.77421 | 4.5263 | 3.63388 |
| chr2 | 84798734 | 84798934 | 376 | Promoter (<=1kb) | 2 | 84798828 | 84810176 | 11349 | 1 | ENSMUST00 | 0 | Ube2l6 | ENSMUSG0C | false | chr2 | 84798207 | 84799862 | 24.81473 | 68.0602 | 65.09722 | . | . | . | . | . | . |
| chr4 | 116075350 | 116075988 | 376 | Promoter (<=1kb) | 4 | 116075463 | 116083056 | 7594 | 1 | ENSMUST00 | 0 | Lrrc41 | ENSMUSG0C | false | chr4 | 116074263 | 116077277 | 38.36185 | 128.21687 | 123.78947 | . | . | . | . | . | . |
| chr6 | 134589496 | 134589752 | 376 | Distal Intergenic | 6 | 134511284 | 134566965 | 55682 | 2 | ENSMUST00 | -22532 | Lrp6 | ENSMUSG0C | false | . | . | . | . | . | . | chr6 | 134588434 | 134589938 | 2.91013 | 2.92546 | 1.33903 |
| chr6 | 146501965 | 146502513 | 376 | Promoter (<=1kb) | 6 | 146108299 | 146502099 | 393801 | 2 | ENSMUST00 | 0 | Itpr2 | ENSMUSG0C | false | chr6 | 146501332 | 146502915 | 36.87877 | 123.45338 | 119.19501 | . | . | . | . | . | . |
| chr7 | 133098434 | 133098681 | 376 | Intron (ENSMUST00000124096.2 | 7 | 133025720 | 133111791 | 86072 | 2 | ENSMUST00 | 13110 | Ctbp2 | ENSMUSG0C | false | . | . | . | . | . | . | chr7 | 133097458 | 133106487 | 5.4045 | 4.25576 | 3.34215 |
| chr9 | 56864887 | 56865161 | 376 | Promoter (<=1kb) | 9 | 56865104 | 56899870 | 34767 | 1 | ENSMUST00 | 0 | Cspg4 | ENSMUSG0C | false | chr9 | 56864988 | 56865426 | 5.25681 | 8.51884 | 6.37244 | chr9 | 56859778 | 56865049 | 5.56524 | 4.35295 | 3.47288 |
| chr6 | 91960189 | 91960581 | 376 | Distal Intergenic | 6 | 91978878 | 92076004 | 97127 | 1 | ENSMUST00 | -18297 | Fgd5 | ENSMUSG0C | false | . | . | . | . | . | . | chr6 | 91959159 | 91963347 | 4.23505 | 3.68567 | 2.39153 |
| chr8 | 122943488 | 122943716 | 376 | Intron (ENSMUST00000127664.1 | 8 | 122884151 | 122915987 | 31837 | 2 | ENSMUST00 | -27502 | Ankrd11 | ENSMUSG0C | false | . | . | . | . | . | . | chr8 | 122908962 | 122946959 | 5.2818 | 4.08553 | 3.26755 |
| chr7 | 112239074 | 112239351 | 376 | Intron (ENSMUST00000050149.6 | 7 | 112225890 | 112348368 | 122479 | 1 | ENSMUST00 | 13185 | Mical2 | ENSMUSG0C | false | . | . | . | . | . | . | chr7 | 112238355 | 112239623 | 7.15927 | 4.54234 | 4.81361 |
| chr3 | 87150666 | 87150996 | 376 | Intron (ENSMUST00000159976.2 | 3 | 87084628 | 87174550 | 89923 | 2 | ENSMUST00 | 23554 | Kirrel | ENSMUSG0C | true | . | . | . | . | . | . | chr3 | 87139743 | 87171483 | 6.81014 | 4.88372 | 4.49361 |
| chr15 | 80710863 | 80711520 | 376 | Promoter (<=1kb) | 15 | 80711319 | 80941086 | 229768 | 1 | ENSMUST00 | 0 | Tnrc6b | ENSMUSG0C | false | chr15 | 80710752 | 80712434 | 41.30354 | 154.61249 | 149.14572 | . | . | . | . | . | . |
| chr15 | 94629023 | 94629371 | 376 | Promoter (<=1kb) | 15 | 94629185 | 95096097 | 466913 | 1 | ENSMUST00 | 0 | Tmem117 | ENSMUSG0C | false | chr15 | 94628841 | 94629705 | 15.98761 | 37.88226 | 35.32005 | . | . | . | . | . | . |
| chr16 | 51287107 | 51287360 | 376 | Distal Intergenic | 16 | 51362452 | 51363832 | 1381 | 2 | ENSMUST00 | 76472 | Gm6767 | ENSMUSG0C | false | . | . | . | . | . | . | chr16 | 51286927 | 51287978 | 4.81014 | 3.92237 | 2.85995 |
| chr15 | 77203474 | 77203705 | 376 | Intron (ENSMUST00000048145.6 | 15 | 77242406 | 77242551 | 146 | 2 | ENSMUST00 | 38846 | Gm24056 | ENSMUSG0C | false | . | . | . | . | . | . | chr15 | 77200992 | 77206317 | 6.95383 | 5.03221 | 4.58681 |
| chr15 | 84305480 | 84305734 | 376 | Intron (ENSMUST00000023072.6 | 15 | 84323209 | 84323352 | 144 | 2 | ENSMUST00 | 17618 | Gm22890 | ENSMUSG0C | false | . | . | . | . | . | . | chr15 | 84297650 | 84306963 | 6.75434 | 4.89978 | 4.45766 |
| chr5 | 140808104 | 140808386 | 375 | Intron (ENSMUST00000000153.4 | 5 | 140758408 | 140830431 | 72024 | 2 | ENSMUST00 | 22045 | Gna12 | ENSMUSG0C | false | . | . | . | . | . | . | chr5 | 140787727 | 140831108 | 6.24285 | 4.56075 | 4.04432 |
| chr11 | 77776161 | 77776418 | 375 | Promoter (<=1kb) | 11 | 77777265 | 77865980 | 88716 | 1 | ENSMUST00 | -847 | Myo18a | ENSMUSG0C | false | . | . | . | . | . | . | . | . | . | . | . | . |
| chr11 | 23256930 | 23257640 | 375 | Promoter (<=1kb) | 11 | 23256589 | 23267714 | 11126 | 1 | ENSMUST00 | 342 | Xpo1 | ENSMUSG0C | false | chr11 | 23255903 | 23258180 | 38.01358 | 126.5237 | 122.15934 | chr11 | 23257537 | 23261648 | 4.19815 | 3.68155 | 2.35175 |
| chr11 | 29172692 | 29173081 | 375 | Promoter (<=1kb) | 11 | 29172890 | 29209641 | 36752 | 1 | ENSMUST00 | 0 | Smek2 | ENSMUSG0C | false | chr11 | 29171793 | 29174277 | 42.09286 | 140.23274 | 135.34074 | . | . | . | . | . | . |
| chr18 | 52769228 | 52769489 | 375 | Promoter (<=1kb) | 18 | 52768732 | 52838040 | 69309 | 1 | ENSMUST00 | 497 | Sncaip | ENSMUSG0C | true | chr18 | 52767566 | 52770026 | 41.31276 | 141.09892 | 136.17311 | chr18 | 52769397 | 52783745 | 4.44145 | 3.73481 | 2.56927 |
| chr19 | 47067489 | 47067900 | 375 | Promoter (<=1kb) | 19 | 47067748 | 47083479 | 15732 | 1 | ENSMUST00 | 0 | Taf5 | ENSMUSG0C | false | chr19 | 47067164 | 47068977 | 36.42221 | 131.26616 | 126.73016 | . | . | . | . | . | . |
| chr4 | 126256190 | 126256565 | 375 | Promoter (<=1kb) | 4 | 126232167 | 126256277 | 24111 | 2 | ENSMUST00 | 0 | Map7d1 | ENSMUSG0C | false | chr4 | 126254944 | 126257391 | 48.06229 | 188.02159 | 181.71353 | chr4 | 126250228 | 126257655 | 4.99746 | 3.92262 | 3.03857 |
| chr6 | 121272340 | 121272721 | 375 | Distal Intergenic | 6 | 121245906 | 121270916 | 25011 | 1 | ENSMUST00 | 26435 | Usp18 | ENSMUSG0C | false | chr6 | 121272080 | 121273002 | 18.77433 | 54.14365 | 51.37372 | . | . | . | . | . | . |
| chr6 | 143152555 | 143152801 | 375 | Distal Intergenic | 6 | 143167230 | 143208542 | 41313 | 1 | ENSMUST00 | -14429 | Etnk1 | ENSMUSG0C | false | . | . | . | . | . | . | chr6 | 143152296 | 143152734 | 3.05858 | 2.99172 | 1.43933 |
| chr2 | 26021504 | 26021882 | 375 | Promoter (<=1kb) | 2 | 25998543 | 26021747 | 23205 | 2 | ENSMUST00 | 0 | Ubac1 | ENSMUSG0C | false | chr2 | 26020809 | 26022200 | 31.19586 | 98.93996 | 95.39634 | . | . | . | . | . | . |
| chr17 | 79355541 | 79355993 | 375 | Promoter (<=1kb) | 17 | 79334025 | 79355091 | 21067 | 2 | ENSMUST00 | -451 | Cdc42ep3 | ENSMUSG0C | false | chr17 | 79353864 | 79356423 | 40.11768 | 137.88335 | 133.09009 | . | . | . | . | . | . |
| chr15 | 99096980 | 99097286 | 375 | Promoter (2-3kb) | 15 | 99099412 | 99104737 | 5326 | 1 | ENSMUST00 | -2126 | Dnajc22 | ENSMUSG0C | false | chr15 | 99096554 | 99098767 | 9.01289 | 21.84568 | 19.48558 | chr15 | 99093546 | 99099596 | 5.88288 | 4.52988 | 3.73597 |
| chr4 | 59783662 | 59783966 | 375 | Promoter (<=1kb) | 4 | 59769637 | 59783866 | 14230 | 2 | ENSMUST00 | 0 | Inip | ENSMUSG0C | false | chr4 | 59782788 | 59783948 | 40.29262 | 133.77719 | 129.14264 | . | . | . | . | . | . |
| chr15 | 33342345 | 33342677 | 375 | Intron (ENSMUST00000042167.9 | 15 | 33405206 | 33405939 | 734 | 2 | ENSMUST00 | 63262 | 1700084J12F | ENSMUSG0C | false | . | . | . | . | . | . | chr15 | 33340716 | 33343621 | 3.51173 | 3.33977 | 1.77506 |
| chr2 | 132284803 | 132285077 | 375 | Promoter (<=1kb) | 2 | 132285176 | 132295573 | 10398 | 1 | ENSMUST00 | -99 | Cds2 | ENSMUSG0C | false | . | . | . | . | . | . | chr2 | 132276320 | 132287879 | 4.58762 | 3.81537 | 2.69582 |
| chr1 | 52388229 | 52388422 | 375 | Distal Intergenic | 1 | 52444436 | 52444544 | 109 | 2 | ENSMUST00 | 56122 | Gm23460 | ENSMUSG0C | false | . | . | . | . | . | . | . | . | . | . | . | . |
| chr11 | 80523132 | 80523458 | 375 | Intron (ENSMUST00000041065.8 | 11 | 80507410 | 80509106 | 1697 | 2 | ENSMUST00 | -14027 | C030013C21I | ENSMUSG0C | false | . | . | . | . | . | . | chr11 | 80521891 | 80525746 | 5.79583 | 4.35194 | 3.7035 |
| chr15 | 86185301 | 86186127 | 375 | Promoter (<=1kb) | 15 | 86185728 | 86187152 | 1425 | 1 | ENSMUST00 | 0 | Gm15569 | ENSMUSG0C | false | chr15 | 86184618 | 86186620 | 30.60796 | 113.09483 | 109.17188 | chr15 | 86185990 | 86192464 | 8.70941 | 5.50823 | 6.04495 |
| chr9 | 63087023 | 63087233 | 375 | Distal Intergenic | 9 | 63133283 | 63133389 | 107 | 1 | ENSMUST00 | -46050 | Gm25064 | ENSMUSG0C | false | . | . | . | . | . | . | chr9 | 63086558 | 63088842 | 4.32075 | 3.77487 | 2.44355 |
| chr12 | 99563358 | 99563807 | 375 | Promoter (<=1kb) | 12 | 99563842 | 99620038 | 56198 | 1 | ENSMUST00 | -34 | Gm26839 | ENSMUSG0C | false | chr12 | 99561991 | 99564031 | 33.04283 | 115.53907 | 111.53828 | . | . | . | . | . | . |
| chr15 | 59120039 | 59120267 | 374 | Distal Intergenic | 15 | 58941244 | 59082026 | 140783 | 2 | ENSMUST00 | -38014 | Mtss1 | ENSMUSG0C | true | . | . | . | . | . | . | chr15 | 59118386 | 59123267 | 4.46297 | 3.74289 | 2.58134 |
| chr1 | 92178361 | 92178789 | 374 | Promoter (1-2kb) | 1 | 92029955 | 92180297 | 150343 | 2 | ENSMUST00 | 1508 | Hdac4 | ENSMUSG0C | false | chr1 | 92178778 | 92180363 | 21.79372 | 61.12524 | 58.26245 | chr1 | 92177024 | 92179782 | 6.0152 | 4.45666 | 3.84635 |
| chr2 | 26493515 | 26493821 | 374 | Intron (ENSMUST00000028288.4 | 2 | 26457903 | 26503822 | 45920 | 2 | ENSMUST00 | 10001 | Notch1 | ENSMUSG0C | false | . | . | . | . | . | . | . | . | . | . | . | . |
| chr6 | 21949594 | 21950113 | 374 | Promoter (<=1kb) | 6 | 21949649 | 21974530 | 24882 | 1 | ENSMUST00 | 0 | Ing3 | ENSMUSG0C | false | chr6 | 21949556 | 21950660 | 34.76894 | 112.35082 | 108.44631 | . | . | . | . | . | . |
| chr7 | 65155917 | 65156157 | 374 | Promoter (<=1kb) | 7 | 64786690 | 65156416 | 369727 | 2 | ENSMUST00 | 259 | Fam189a1 | ENSMUSG0C | false | chr7 | 65155442 | 65156992 | 24.72749 | 72.3696 | 69.3433 | chr7 | 65155492 | 65156960 | 4.91813 | 3.98055 | 2.96941 |
| chrX | 103821887 | 103822199 | 374 | Promoter (<=1kb) | 20 | 103697414 | 103821965 | 124552 | 2 | ENSMUST00 | 0 | Slc16a2 | ENSMUSG0C | false | chrX | 103820678 | 103822759 | 14.2455 | 32.22549 | 29.73087 | . | . | . | . | . | . |
| chr17 | 26451937 | 26452327 | 374 | Distal Intergenic | 17 | 26414965 | 26446349 | 31385 | 1 | ENSMUST00 | 36973 | Neurl1b | ENSMUSG0C | false | . | . | . | . | . | . | chr17 | 26451215 | 26452814 | 4.18104 | 3.47268 | 2.36407 |
| chrX | 51205683 | 51206046 | 374 | Promoter (<=1kb) | 20 | 51117269 | 51205832 | 88564 | 2 | ENSMUST00 | 0 | Mbnl3 | ENSMUSG0C | false | chrX | 51204522 | 51206287 | 19.75988 | 50.25065 | 47.53236 | . | . | . | . | . | . |
| chr17 | 6685135 | 6685803 | 374 | Intron (ENSMUST00000115763.4 | 17 | 6689072 | 6738042 | 48971 | 1 | ENSMUST00 | -3269 | Sytl3 | ENSMUSG0C | false | . | . | . | . | . | . | chr17 | 6683520 | 6686540 | 9.55911 | 5.51229 | 6.8442 |
| chr11 | 3913771 | 3914035 | 374 | Promoter (<=1kb) | 11 | 3907021 | 3914664 | 7644 | 2 | ENSMUST00 | 629 | Slc35e4 | ENSMUSG0C | false | chr11 | 3912589 | 3914725 | 24.51421 | 78.37994 | 75.25539 | chr11 | 3912558 | 3914820 | 5.529 | 4.16978 | 3.47676 |
| chr15 | 102470338 | 102470604 | 374 | Promoter (<=1kb) | 15 | 102470632 | 102500059 | 29428 | 1 | ENSMUST00 | -28 | Pcbp2 | ENSMUSG0C | false | chr15 | 102470489 | 102471870 | 22.78592 | 76.10319 | 73.01487 | . | . | . | . | . | . |
| chr12 | 54862526 | 54863104 | 374 | Promoter (<=1kb) | 12 | 54858817 | 54862877 | 4061 | 2 | ENSMUST00 | 0 | Cfl2 | ENSMUSG0C | false | chr12 | 54861687 | 54863386 | 34.16929 | 120.7392 | 116.5701 | . | . | . | . | . | . |
| chr6 | 100170810 | 100171129 | 374 | Distal Intergenic | 6 | 100228566 | 100287441 | 58876 | 2 | ENSMUST00 | 116312 | Rybp | ENSMUSG0C | false | . | . | . | . | . | . | chr6 | 100169895 | 100172561 | 6.80218 | 4.98719 | 4.46701 |
| chr2 | 105069351 | 105069633 | 374 | Distal Intergenic | 2 | 105077895 | 105078107 | 213 | 2 | ENSMUST00 | 8474 | Wt1os | ENSMUSG0C | true | . | . | . | . | . | . | chr2 | 105057120 | 105080197 | 5.15057 | 4.16934 | 3.12788 |
| chr7 | 124573766 | 124574012 | 374 | Distal Intergenic | 7 | 124476541 | 124494301 | 17761 | 2 | ENSMUST00 | -79466 | Gm15339 | ENSMUSG0C | false | . | . | . | . | . | . | . | . | . | . | . | . |
| chr1 | 7397932 | 7398360 | 374 | Promoter (<=1kb) | 1 | 7349406 | 7397801 | 48396 | 2 | ENSMUST00 | -132 | Gm26901 | ENSMUSG0C | false | chr1 | 7398087 | 7398800 | 29.41006 | 85.09721 | 81.85307 | . | . | . | . | . | . |
| chr14 | 46941916 | 46942145 | 374 | Intron (ENSMUST00000022386.9 | 14 | 46954830 | 46954928 | 99 | 2 | ENSMUST00 | 12783 | Mir378c | ENSMUSG0C | false | . | . | . | . | . | . | chr14 | 46941019 | 46943569 | 3.80294 | 3.33735 | 2.06286 |
| chr8 | 61288791 | 61289051 | 374 | Promoter (<=1kb) | 8 | 61281522 | 61288872 | 7351 | 2 | ENSMUST00 | 0 | RP23-5F21.1 | ENSMUSG0C | false | . | . | . | . | . | . | chr8 | 61281009 | 61293339 | 5.05439 | 4.05325 | 3.0617 |
| chr12 | 80023576 | 80023795 | 374 | Distal Intergenic | 12 | 79963568 | 79963872 | 305 | 1 | ENSMUST00 | 60009 | RP23-179D3. | ENSMUSG0C | false | . | . | . | . | . | . | chr12 | 80020894 | 80023973 | 4.65589 | 3.9516 | 2.7178 |
| chr11 | 94928824 | 94929219 | 373 | Distal Intergenic | 11 | 94936224 | 94953042 | 16819 | 1 | ENSMUST00 | -7005 | Col1a1 | ENSMUSG0C | false | . | . | . | . | . | . | chr11 | 94926406 | 94931022 | 5.04668 | 3.94408 | 3.08549 |
| chr15 | 67145922 | 67146188 | 373 | Distal Intergenic | 15 | 67102875 | 67113992 | 11118 | 2 | ENSMUST00 | -31931 | St3gal1 | ENSMUSG0C | true | . | . | . | . | . | . | chr15 | 67143105 | 67166416 | 6.05047 | 4.43905 | 3.887 |
| chr10 | 59951816 | 59952210 | 373 | Promoter (<=1kb) | 10 | 59949675 | 59951770 | 2096 | 2 | ENSMUST00 | -47 | Ddit4 | ENSMUSG0C | false | chr10 | 59950517 | 59952770 | 25.75243 | 103.50989 | 99.85224 | chr10 | 59949666 | 59953527 | 4.31885 | 3.68721 | 2.49347 |
| chr15 | 59100953 | 59101338 | 373 | Distal Intergenic | 15 | 58941244 | 59082026 | 140783 | 2 | ENSMUST00 | -18928 | Mtss1 | ENSMUSG0C | true | . | . | . | . | . | . | . | . | . | . | . | . |
| chr15 | 73425996 | 73426224 | 373 | Promoter (2-3kb) | 15 | 73215557 | 73423191 | 207635 | 2 | ENSMUST00 | -2806 | Ptk2 | ENSMUSG0C | false | . | . | . | . | . | . | chr15 | 73423923 | 73426772 | 3.74803 | 3.37619 | 1.98742 |
| chr19 | 21821864 | 21822139 | 373 | Intron (ENSMUST00000025663.6 | 19 | 21778342 | 21858327 | 79986 | 1 | ENSMUST00 | 43523 | Tmem2 | ENSMUSG0C | false | . | . | . | . | . | . | chr19 | 21821944 | 21822383 | 3.43944 | 3.31937 | 1.74095 |
| chr3 | 129322960 | 129323308 | 373 | Intron (ENSMUST00000029658.8 | 3 | 129270277 | 129332264 | 61988 | 2 | ENSMUST00 | 8956 | Enpep | ENSMUSG0C | true | . | . | . | . | . | . | chr3 | 129303627 | 129327892 | 5.35019 | 4.2164 | 3.29511 |
| chr5 | 107184523 | 107184769 | 373 | Intron (ENSMUST00000031224.9 | 5 | 107109281 | 107192367 | 83087 | 2 | ENSMUST00 | 7598 | Tgfbr3 | ENSMUSG0C | true | . | . | . | . | . | . | chr5 | 107183014 | 107186515 | 5.34937 | 4.07827 | 3.33495 |
| chr3 | 53211317 | 53211565 | 373 | Intron (ENSMUST00000059562.8 | 3 | 53041583 | 53261679 | 220097 | 1 | ENSMUST00 | 169735 | Lhfp | ENSMUSG0C | false | . | . | . | . | . | . | chr3 | 53210933 | 53212659 | 3.70338 | 3.29967 | 2.00126 |
| chr12 | 21163641 | 21164025 | 373 | Intron (ENSMUST00000174530.1 | 12 | 21111954 | 21268179 | 156226 | 1 | ENSMUST00 | 51688 | Asap2 | ENSMUSG0C | false | . | . | . | . | . | . | chr12 | 21148547 | 21176379 | 6.51645 | 4.71535 | 4.25897 |
| chr15 | 102073583 | 102074210 | 373 | Promoter (<=1kb) | 15 | 102073773 | 102097172 | 23400 | 1 | ENSMUST00 | 0 | Eif4b | ENSMUSG0C | false | chr15 | 102073251 | 102073636 | 9.22042 | 18.50693 | 16.19293 | . | . | . | . | . | . |
| chr2 | 11367320 | 11367715 | 373 | Distal Intergenic | 2 | 11352514 | 11352769 | 256 | 1 | ENSMUST00 | 14807 | Gm13295 | ENSMUSG0C | false | . | . | . | . | . | . | . | . | . | . | . | . |
| chr11 | 31807464 | 31807767 | 373 | Intron (ENSMUST00000129425.2 | 11 | 31790113 | 31824473 | 34361 | 2 | ENSMUST00 | 16706 | D630024D03 | ENSMUSG0C | false | . | . | . | . | . | . | chr11 | 31807104 | 31810464 | 5.47128 | 4.18915 | 3.42525 |
| chr11 | 78435796 | 78436058 | 373 | Distal Intergenic | 11 | 78444107 | 78444901 | 795 | 1 | ENSMUST00 | -8049 | Gm11195 | ENSMUSG0C | false | . | . | . | . | . | . | chr11 | 78435902 | 78436733 | 5.23005 | 4.19451 | 3.23446 |
| chr6 | 118845901 | 118846159 | 373 | Intron (ENSMUST00000186889.1 | 6 | 118834900 | 118835000 | 101 | 1 | ENSMUST00 | 11002 | Gm25905 | ENSMUSG0C | false | . | . | . | . | . | . | chr6 | 118843658 | 118848402 | 5.8475 | 4.40619 | 3.70753 |
| chr9 | 37248351 | 37248558 | 373 | Intron (ENSMUST00000115068.3 | 9 | 37252922 | 37253365 | 444 | 1 | ENSMUST00 | -4364 | Gm10177 | ENSMUSG0C | false | . | . | . | . | . | . | chr9 | 37247052 | 37249515 | 6.04849 | 4.68071 | 3.86379 |
| chr7 | 25832091 | 25832329 | 373 | Distal Intergenic | 7 | 25828587 | 25828877 | 291 | 2 | ENSMUST00 | -3215 | Gm4613 | ENSMUSG0C | false | . | . | . | . | . | . | chr7 | 25831547 | 25832553 | 5.10024 | 4.13395 | 3.08923 |
| chr3 | 94404033 | 94404263 | 373 | Promoter (1-2kb) | 3 | 94392971 | 94402848 | 9878 | 2 | ENSMUST00 | -1186 | Gm17179 | ENSMUSG0C | false | . | . | . | . | . | . | chr3 | 94403221 | 94406496 | 6.08922 | 4.53013 | 3.93833 |
| chr18 | 61360564 | 61360848 | 373 | Intron (ENSMUST00000075299.7 | 18 | 61397813 | 61397925 | 113 | 1 | ENSMUST00 | -36965 | Gm25301 | ENSMUSG0C | false | . | . | . | . | . | . | chr18 | 61359328 | 61363669 | 5.31584 | 4.26737 | 3.26768 |
| chr8 | 121286800 | 121287135 | 373 | Intron (ENSMUST00000127664.1 | 8 | 121256747 | 121356836 | 100090 | 2 | ENSMUST00 | 69701 | Gm26815 | ENSMUSG0C | false | . | . | . | . | . | . | chr8 | 121285086 | 121289147 | 5.87486 | 4.23382 | 3.74695 |
| chr2 | 145863050 | 145863270 | 372 | Promoter (<=1kb) | 2 | 145862644 | 145878974 | 16331 | 1 | ENSMUST00 | 407 | Rin2 | ENSMUSG0C | true | chr2 | 145860076 | 145863605 | 23.70866 | 76.9101 | 73.81126 | chr2 | 145862482 | 145865970 | 5.39781 | 4.29406 | 3.37842 |
| chr7 | 130972887 | 130973151 | 372 | Intron (ENSMUST00000150717.2 | 7 | 130981670 | 130982443 | 774 | 1 | ENSMUST00 | -8519 | Htra1 | ENSMUSG0C | false | . | . | . | . | . | . | chr7 | 130956209 | 130988327 | 6.29742 | 4.64648 | 4.07297 |
| chr6 | 48566481 | 48566796 | 372 | Intron (ENSMUST00000114545.2 | 6 | 48569698 | 48572670 | 2973 | 2 | ENSMUST00 | 5874 | Rarres2 | ENSMUSG0C | false | . | . | . | . | . | . | chr6 | 48565716 | 48567175 | 4.63542 | 3.97426 | 2.6952 |
| chr10 | 13553057 | 13553291 | 372 | Promoter (<=1kb) | 10 | 13523842 | 13553142 | 29301 | 2 | ENSMUST00 | 0 | Pex3 | ENSMUSG0C | false | chr10 | 13552480 | 13554033 | 25.48472 | 95.2657 | 91.8051 | . | . | . | . | . | . |
| chr11 | 116852261 | 116852611 | 372 | Promoter (<=1kb) | 11 | 116852440 | 116874712 | 22273 | 1 | ENSMUST00 | 0 | Mfsd11 | ENSMUSG0C | false | chr11 | 116851455 | 116854638 | 40.75607 | 149.08708 | 143.84373 | . | . | . | . | . | . |
| chr14 | 70551979 | 70552267 | 372 | Promoter (<=1kb) | 14 | 70552212 | 70554719 | 2508 | 1 | ENSMUST00 | 0 | Hr | ENSMUSG0C | false | . | . | . | . | . | . | chr14 | 70551984 | 70555696 | 6.15727 | 4.33196 | 3.98723 |
| chr1 | 75345185 | 75345440 | 372 | Distal Intergenic | 1 | 75360329 | 75368579 | 8251 | 1 | ENSMUST00 | -14889 | Des | ENSMUSG0C | false | . | . | . | . | . | . | chr1 | 75340961 | 75345934 | 4.949 | 3.97142 | 2.9865 |
| chr1 | 165633289 | 165633568 | 372 | Promoter (<=1kb) | 1 | 165592181 | 165634541 | 42361 | 2 | ENSMUST00 | 973 | Mpzl1 | ENSMUSG0C | false | chr1 | 165633299 | 165635197 | 21.23866 | 90.05283 | 86.71227 | chr1 | 165629759 | 165635444 | 3.56725 | 3.34446 | 1.82568 |
| chr18 | 39194982 | 39195229 | 372 | Intron (ENSMUST00000097593.3 | 18 | 39227368 | 39372511 | 145144 | 1 | ENSMUST00 | -32139 | Arhgap26 | ENSMUSG0C | false | . | . | . | . | . | . | chr18 | 39193471 | 39197393 | 4.91054 | 4.00067 | 2.92834 |
| chr5 | 65199508 | 65199891 | 372 | Promoter (<=1kb) | 5 | 65199696 | 65260415 | 60720 | 1 | ENSMUST00 | 0 | Wdr19 | ENSMUSG0C | false | chr5 | 65199640 | 65200386 | 23.34944 | 75.34173 | 72.26852 | . | . | . | . | . | . |
| chr3 | 97732789 | 97732978 | 372 | Intron (ENSMUST00000168438.3 | 3 | 97740817 | 97767931 | 27115 | 2 | ENSMUST00 | 34953 | Pde4dip | ENSMUSG0C | true | . | . | . | . | . | . | chr3 | 97729211 | 97741784 | 3.96932 | 3.48187 | 2.18916 |
| chr16 | 4003451 | 4003944 | 372 | Promoter (<=1kb) | 16 | 4001003 | 4003770 | 2768 | 2 | ENSMUST00 | 0 | Slx4 | ENSMUSG0C | false | chr16 | 4003174 | 4004370 | 18.33657 | 60.6815 | 57.82544 | chr16 | 4002504 | 4004478 | 4.00489 | 3.50722 | 2.21869 |
| chrX | 38189653 | 38190073 | 372 | Promoter (<=1kb) | 20 | 38189793 | 38197046 | 7254 | 1 | ENSMUST00 | 0 | Zbtb33 | ENSMUSG0C | false | chrX | 38189347 | 38190847 | 21.59801 | 56.60135 | 53.80005 | . | . | . | . | . | . |
| chr15 | 79951951 | 79952208 | 372 | Intron (ENSMUST00000128931.2 | 15 | 79918148 | 79933899 | 15752 | 2 | ENSMUST00 | -18053 | Cbx7 | ENSMUSG0C | false | . | . | . | . | . | . | chr15 | 79949890 | 79954213 | 7.95672 | 5.38582 | 5.41593 |

| | | | | | | | | | | | | | | | | | | | | | | | | | | |
|---|---|---|---|---|---|---|---|---|---|---|---|---|---|---|---|---|---|---|---|---|---|---|---|---|---|---|
| chr7 | 66955393 | 66955713 | 372 | Intron (ENSMUST00000156396.2 | 7 | 66909468 | 66969555 | 60088 | 1 | ENSMUST00 | 45926 | Adamts17 | ENSMUSG0( | false | . | . | . | . | . | . | . | . | . | . | . | . |
| chr10 | 117048479 | 117048800 | 372 | Downstream (2-3kb) | 10 | 117045341 | 117046768 | 1428 | 1 | ENSMUST00 | 3139 | Lrrc10 | ENSMUSG0( | false | . | . | . | . | . | . | chr10 | 117045739 | 117050616 | 4.20138 | 3.5215 | 2.382 |
| chr10 | 80752062 | 80752346 | 372 | Promoter (2-3kb) | 10 | 80755206 | 80795461 | 40256 | 1 | ENSMUST00 | -2860 | Dot1l | ENSMUSG0( | false | . | . | . | . | . | . | chr10 | 80750612 | 80754824 | 4.13572 | 3.6561 | 2.30135 |
| chr2 | 68080026 | 68080643 | 372 | Promoter (1-2kb) | 2 | 68081791 | 68082329 | 539 | 1 | ENSMUST00 | -1148 | Gm13607 | ENSMUSG0( | false | . | . | . | . | . | . | . | . | . | . | . | . |
| chr16 | 58041575 | 58041875 | 372 | Distal Intergenic | 16 | 57884008 | 57884139 | 132 | 2 | ENSMUST00 | -157437 | Gm24039 | ENSMUSG0( | false | . | . | . | . | . | . | chr16 | 58040395 | 58042495 | 3.17139 | 3.0694 | 1.50205 |
| chr15 | 5802614 | 5803052 | 372 | Distal Intergenic | 15 | 5864306 | 5864419 | 114 | 1 | ENSMUST00 | -61254 | Gm27760 | ENSMUSG0( | false | . | . | . | . | . | . | chr15 | 5801820 | 5803604 | 4.47498 | 3.73107 | 2.60899 |
| chr11 | 95982532 | 95982802 | 371 | Promoter (<=1kb) | 11 | 95973144 | 95982594 | 9451 | 2 | ENSMUST00 | 0 | Igf2bp1 | ENSMUSG0( | false | chr11 | 95981866 | 95983014 | 12.39106 | 30.348 | 27.8772 | chr11 | 95980123 | 95983650 | 3.78542 | 3.29226 | 2.04651 |
| chr11 | 95834384 | 95834636 | 371 | Promoter (1-2kb) | 11 | 95832723 | 95835890 | 3168 | 2 | ENSMUST00 | 1254 | Abi3 | ENSMUSG0( | false | . | . | . | . | . | . | chr11 | 95829550 | 95838805 | 5.29848 | 4.1616 | 3.26577 |
| chr11 | 3863791 | 3864066 | 371 | Promoter (<=1kb) | 11 | 3703731 | 3863903 | 160173 | 2 | ENSMUST00 | 0 | Osbp2 | ENSMUSG0( | false | chr11 | 3863478 | 3864305 | 18.44084 | 48.77526 | 46.0739 | chr11 | 3862373 | 3864559 | 3.97366 | 3.5266 | 2.17832 |
| chr19 | 9088536 | 9088755 | 371 | Promoter (<=1kb) | 19 | 9083636 | 9087958 | 4323 | 2 | ENSMUST00 | -579 | Scgb1a1 | ENSMUSG0( | false | . | . | . | . | . | . | . | . | . | . | . | . |
| chr2 | 69240849 | 69241125 | 371 | Promoter (1-2kb) | 2 | 69238310 | 69239078 | 769 | 2 | ENSMUST00 | -1772 | Abcb11 | ENSMUSG0( | false | . | . | . | . | . | . | chr2 | 69239319 | 69241315 | 4.55564 | 3.6536 | 2.6765 |
| chr2 | 93808143 | 93808417 | 371 | Intron (ENSMUST00000184931.2 | 2 | 93762561 | 93795989 | 33429 | 2 | ENSMUST00 | -12155 | Ext2 | ENSMUSG0( | true | . | . | . | . | . | . | chr2 | 93805378 | 93812725 | 6.22701 | 4.60265 | 4.02277 |
| chr9 | 107981614 | 107981927 | 371 | Promoter (<=1kb) | 9 | 107980890 | 107983199 | 2310 | 1 | ENSMUST00 | 725 | Uba7 | ENSMUSG0( | false | . | . | . | . | . | . | . | . | . | . | . | . |
| chr19 | 21652975 | 21653755 | 371 | Promoter (<=1kb) | 19 | 21597312 | 21652976 | 55665 | 2 | ENSMUST00 | 0 | 1110059E24I | ENSMUSG0( | false | chr19 | 21651908 | 21654841 | 33.41831 | 117.26761 | 113.21259 | . | . | . | . | . | . |
| chr8 | 86745392 | 86746083 | 371 | Promoter (<=1kb) | 8 | 86723940 | 86746006 | 22067 | 2 | ENSMUST00 | 0 | Siah1a | ENSMUSG0( | false | chr8 | 86744704 | 86746872 | 29.21782 | 100.39505 | 96.81345 | . | . | . | . | . | . |
| chr1 | 38545646 | 38545895 | 371 | Intron (ENSMUST00000095027.3 | 1 | 38177326 | 38627185 | 449860 | 2 | ENSMUST00 | 81290 | Aff3 | ENSMUSG0( | true | . | . | . | . | . | . | chr1 | 38544410 | 38551371 | 5.22999 | 4.16319 | 3.21526 |
| chr6 | 147091318 | 147091737 | 371 | Promoter (<=1kb) | 6 | 147091379 | 147112778 | 21400 | 1 | ENSMUST00 | 0 | Klhl42 | ENSMUSG0( | false | chr6 | 147090907 | 147092605 | 24.55236 | 90.57755 | 87.22021 | . | . | . | . | . | . |
| chr11 | 114852009 | 114852318 | 371 | Promoter (<=1kb) | 11 | 114852319 | 114863897 | 11579 | 1 | ENSMUST00 | -1 | Gprc5c | ENSMUSG0( | false | chr11 | 114850502 | 114853302 | 31.71022 | 94.86349 | 91.41748 | . | . | . | . | . | . |
| chr5 | 135919044 | 135919276 | 371 | Intron (ENSMUST00000055808.5 | 5 | 135930761 | 135934550 | 3790 | 2 | ENSMUST00 | 15274 | Ywhag | ENSMUSG0( | false | . | . | . | . | . | . | chr5 | 135911165 | 135921283 | 4.85161 | 3.88824 | 2.92073 |
| chr12 | 69241688 | 69242173 | 371 | Promoter (<=1kb) | 12 | 69241832 | 69284632 | 42801 | 1 | ENSMUST00 | 0 | Klhdc1 | ENSMUSG0( | false | chr12 | 69241385 | 69242592 | 36.9999 | 115.1778 | 111.19392 | chr12 | 69241174 | 69241843 | 3.87506 | 3.57168 | 2.12418 |
| chr1 | 39762073 | 39762299 | 371 | Distal Intergenic | 1 | 39665306 | 39720997 | 55692 | 2 | ENSMUST00 | -41077 | Rfx8 | ENSMUSG0( | false | chr1 | 39761731 | 39762300 | 12.68821 | 30.86953 | 28.39186 | chr1 | 39759438 | 39762551 | 4.33134 | 3.78958 | 2.45114 |
| chr16 | 8514409 | 8514674 | 371 | Promoter (<=1kb) | 16 | 8513485 | 8619570 | 106086 | 1 | ENSMUST00 | 925 | Abat | ENSMUSG0( | false | chr16 | 8513058 | 8514438 | 27.40733 | 77.91289 | 74.79585 | chr16 | 8513054 | 8516114 | 4.31331 | 3.7593 | 2.49116 |
| chr12 | 17725174 | 17725488 | 371 | Intron (ENSMUST00000071858.3 | 12 | 17690814 | 17791926 | 101113 | 1 | ENSMUST00 | 34361 | Hpcal1 | ENSMUSG0( | false | . | . | . | . | . | . | chr12 | 17723568 | 17729175 | 5.4309 | 4.07721 | 3.3921 |
| chr3 | 153550014 | 153550242 | 371 | Intron (ENSMUST00000064460.5 | 3 | 153621702 | 153621969 | 268 | 1 | ENSMUST00 | -71460 | Gm22136 | ENSMUSG0( | false | . | . | . | . | . | . | chr3 | 153543231 | 153554124 | 4.38082 | 3.74714 | 2.51003 |
| chr15 | 68508339 | 68508525 | 371 | Distal Intergenic | 15 | 68340983 | 68363176 | 22194 | 2 | ENSMUST00 | -145164 | Gm20732 | ENSMUSG0( | false | . | . | . | . | . | . | chr15 | 68507438 | 68509221 | 3.80682 | 3.50538 | 2.01439 |
| chr10 | 39899017 | 39899319 | 371 | Promoter (<=1kb) | 10 | 39890018 | 39899238 | 9221 | 2 | ENSMUST00 | 0 | AA474331 | ENSMUSG0( | false | chr10 | 39898529 | 39899580 | 28.9509 | 90.07051 | 86.72839 | chr10 | 39898443 | 39899656 | 3.47074 | 3.2936 | 1.72942 |
| chr11 | 100399474 | 100399655 | 370 | Promoter (1-2kb) | 11 | 100383315 | 100397763 | 14449 | 2 | ENSMUST00 | -1712 | Jup | ENSMUSG0( | false | . | . | . | . | . | . | chr11 | 100398067 | 100400413 | 4.9091 | 4.05794 | 2.92372 |
| chr8 | 84913237 | 84913513 | 370 | Promoter (<=1kb) | 8 | 84913391 | 84922915 | 9525 | 1 | ENSMUST00 | 0 | Dnase2a | ENSMUSG0( | false | chr8 | 84912839 | 84913660 | 19.22969 | 50.92535 | 48.19825 | chr8 | 84912777 | 84913852 | 3.33994 | 3.14202 | 1.66002 |
| chr13 | 69482042 | 69482407 | 370 | Distal Intergenic | 13 | 69533746 | 69615468 | 81723 | 1 | ENSMUST00 | -51339 | Nsun2 | ENSMUSG0( | false | chr13 | 69481744 | 69482790 | 11.2646 | 26.49366 | 24.07236 | chr13 | 69480935 | 69483242 | 4.14373 | 3.61906 | 2.30774 |
| chr15 | 76189905 | 76190177 | 370 | Promoter (1-2kb) | 15 | 76175380 | 76188043 | 12664 | 2 | ENSMUST00 | -1863 | Plec | ENSMUSG0( | false | . | . | . | . | . | . | chr15 | 76189097 | 76191015 | 5.91643 | 4.50779 | 3.75268 |
| chr1 | 91540457 | 91540796 | 370 | Promoter (<=1kb) | 1 | 91540544 | 91556088 | 15545 | 1 | ENSMUST00 | 0 | Asb1 | ENSMUSG0( | false | chr1 | 91539734 | 91541534 | 37.70819 | 121.69438 | 117.49508 | chr1 | 91539333 | 91541759 | 4.19895 | 3.64786 | 2.36113 |
| chr2 | 122238234 | 122238491 | 370 | Intron (ENSMUST00000110551.3 | 2 | 122234749 | 122265340 | 30592 | 1 | ENSMUST00 | 3486 | Sord | ENSMUSG0( | false | . | . | . | . | . | . | chr2 | 122233807 | 122239647 | 6.06461 | 4.59249 | 3.88635 |
| chr9 | 23223308 | 23223620 | 370 | Promoter (<=1kb) | 9 | 23223076 | 23485202 | 262127 | 1 | ENSMUST00 | 233 | Bmper | ENSMUSG0( | false | chr9 | 23222397 | 23223906 | 31.56046 | 98.57728 | 95.04015 | chr9 | 23221947 | 23224065 | 3.50174 | 3.29353 | 1.81269 |
| chr2 | 66256493 | 66256944 | 370 | Promoter (<=1kb) | 2 | 66184327 | 66256617 | 72291 | 2 | ENSMUST00 | 0 | Ttc21b | ENSMUSG0( | false | chr2 | 66256085 | 66257392 | 28.31422 | 85.10979 | 81.86543 | . | . | . | . | . | . |
| chr1 | 170521664 | 170521914 | 370 | Intron (ENSMUST00000160456.2 | 1 | 170327214 | 170557110 | 229897 | 2 | ENSMUST00 | 35196 | Nos1ap | ENSMUSG0( | false | . | . | . | . | . | . | chr1 | 170515636 | 170524080 | 5.30484 | 4.19024 | 3.27564 |
| chr4 | 57739105 | 57739396 | 370 | Promoter (2-3kb) | 4 | 57742095 | 57896982 | 154888 | 1 | ENSMUST00 | -2699 | Akap2 | ENSMUSG0( | false | . | . | . | . | . | . | chr4 | 57738339 | 57740025 | 4.23438 | 3.60674 | 2.39311 |
| chr2 | 165886389 | 165886707 | 370 | Promoter (1-2kb) | 2 | 165785519 | 165884874 | 99356 | 2 | ENSMUST00 | -1516 | Zmynd8 | ENSMUSG0( | false | . | . | . | . | . | . | chr2 | 165851451 | 165889342 | 6.61553 | 4.6931 | 4.35892 |
| chr14 | 26980323 | 26980805 | 370 | Distal Intergenic | 14 | 26918988 | 26971232 | 52245 | 2 | ENSMUST00 | -9092 | Appl1 | ENSMUSG0( | false | . | . | . | . | . | . | chr14 | 26979823 | 26980912 | 4.41611 | 3.76424 | 2.55411 |
| chr6 | 72387610 | 72387889 | 370 | Promoter (2-3kb) | 6 | 72385228 | 72389911 | 4684 | 2 | ENSMUST00 | 2022 | Vamp8 | ENSMUSG0( | false | . | . | . | . | . | . | chr6 | 72385690 | 72394714 | 4.39484 | 3.72006 | 2.54009 |
| chr5 | 43456120 | 43456394 | 370 | Distal Intergenic | 5 | 43515569 | 43616585 | 101017 | 1 | ENSMUST00 | -59175 | C1qtnf7 | ENSMUSG0( | true | . | . | . | . | . | . | chr5 | 43454125 | 43458196 | 4.54715 | 3.87103 | 2.63681 |
| chr9 | 57503464 | 57503721 | 370 | Promoter (<=1kb) | 9 | 57504102 | 57505447 | 1346 | 1 | ENSMUST00 | -381 | Rpp25 | ENSMUSG0( | false | chr9 | 57501755 | 57504739 | 20.27628 | 60.14116 | 57.29097 | chr9 | 57500796 | 57505928 | 4.78679 | 4.00087 | 2.83045 |
| chr15 | 103304290 | 103304633 | 370 | Distal Intergenic | 15 | 103308236 | 103310438 | 2203 | 2 | ENSMUST00 | 5805 | Gpr84 | ENSMUSG0( | false | chr15 | 103304505 | 103305880 | 24.3552 | 66.39809 | 63.45943 | chr15 | 103298885 | 103310553 | 5.16019 | 4.10407 | 3.16269 |
| chr3 | 123001068 | 123001391 | 370 | Distal Intergenic | 3 | 123015259 | 123015368 | 110 | 2 | ENSMUST00 | 13977 | Gm22066 | ENSMUSG0( | false | . | . | . | . | . | . | chr3 | 122998598 | 123002901 | 3.91269 | 3.52677 | 2.10616 |
| chrX | 169106398 | 169106660 | 370 | Promoter (<=1kb) | 20 | 169037674 | 169106536 | 68863 | 2 | ENSMUST00 | 0 | Gm15261 | ENSMUSG0( | false | . | . | . | . | . | . | . | . | . | . | . | . |
| chr17 | 87367500 | 87367820 | 370 | Intron (ENSMUST00000041110.6 | 17 | 87362239 | 87364462 | 2224 | 2 | ENSMUST00 | -3039 | 0610012D04I | ENSMUSG0( | false | . | . | . | . | . | . | chr17 | 87364501 | 87371587 | 7.75503 | 5.20981 | 5.26833 |
| chr10 | 92125710 | 92125927 | 370 | Intron (ENSMUST00000182033.2 | 10 | 92133222 | 92133347 | 126 | 2 | ENSMUST00 | 7420 | Gm27582 | ENSMUSG0( | false | . | . | . | . | . | . | . | . | . | . | . | . |
| chr7 | 110953225 | 110953437 | 369 | Intron (ENSMUST00000005751.7 | 7 | 110868272 | 110946187 | 77916 | 2 | ENSMUST00 | -7039 | Mrvi1 | ENSMUSG0( | false | . | . | . | . | . | . | . | . | . | . | . | . |
| chr14 | 25606682 | 25606999 | 369 | Promoter (<=1kb) | 14 | 25607357 | 25643657 | 36301 | 1 | ENSMUST00 | -358 | Zmiz1 | ENSMUSG0( | false | chr14 | 25606620 | 25608499 | 15.16532 | 49.65084 | 46.94072 | chr14 | 25606027 | 25610598 | 4.72292 | 3.88282 | 2.81242 |
| chr1 | 156310572 | 156310946 | 369 | Promoter (<=1kb) | 1 | 156310727 | 156328035 | 17309 | 1 | ENSMUST00 | 0 | Nphs2 | ENSMUSG0( | true | chr1 | 156309730 | 156319004 | 28.81932 | 90.81086 | 87.44578 | . | . | . | . | . | . |
| chr4 | 118037545 | 118037817 | 369 | Intron (ENSMUST00000030263.3 | 4 | 117940354 | 118031882 | 91529 | 2 | ENSMUST00 | -5664 | St3gal3 | ENSMUSG0( | true | . | . | . | . | . | . | chr4 | 118036457 | 118045566 | 4.0353 | 3.47805 | 2.24897 |
| chr7 | 128246816 | 128247288 | 369 | Promoter (<=1kb) | 7 | 128246871 | 128253712 | 6842 | 1 | ENSMUST00 | 0 | Tgfb1i1 | ENSMUSG0( | false | chr7 | 128246452 | 128247624 | 22.90469 | 70.93823 | 67.93287 | chr7 | 128245755 | 128248385 | 3.83675 | 3.4231 | 2.05771 |
| chr8 | 120449661 | 120449903 | 369 | Intron (ENSMUST00000127664.1 | 8 | 120488447 | 120581390 | 92944 | 1 | ENSMUST00 | -38544 | Gse1 | ENSMUSG0( | false | . | . | . | . | . | . | chr8 | 120448683 | 120450891 | 5.62445 | 4.3356 | 3.53594 |
| chr9 | 57547369 | 57547676 | 369 | Promoter (1-2kb) | 9 | 57548590 | 57549672 | 1083 | 2 | ENSMUST00 | 1996 | Mpi | ENSMUSG0( | false | . | . | . | . | . | . | chr9 | 57541237 | 57549125 | 5.67277 | 4.24382 | 3.5915 |
| chr9 | 65140926 | 65141207 | 369 | Promoter (<=1kb) | 9 | 65141154 | 65185857 | 44704 | 1 | ENSMUST00 | 0 | Igdcc3 | ENSMUSG0( | false | chr9 | 65140715 | 65142580 | 25.86957 | 74.34999 | 71.29122 | . | . | . | . | . | . |
| chrX | 152147456 | 152147662 | 369 | Intron (ENSMUST00000168786.2 | 20 | 152144272 | 152225236 | 80965 | 1 | ENSMUST00 | 3185 | Iqsec2 | ENSMUSG0( | false | . | . | . | . | . | . | . | . | . | . | . | . |
| chr13 | 99355368 | 99355733 | 369 | Intron (ENSMUST00000052249.5 | 13 | 99344786 | 99415561 | 70776 | 1 | ENSMUST00 | 10583 | Mrps27 | ENSMUSG0( | false | . | . | . | . | . | . | . | . | . | . | . | . |
| chr12 | 21111528 | 21111964 | 369 | Promoter (<=1kb) | 12 | 21111748 | 21270171 | 158424 | 1 | ENSMUST00 | 0 | Asap2 | ENSMUSG0( | false | chr12 | 21111284 | 21113355 | 41.65482 | 133.94409 | 129.3063 | . | . | . | . | . | . |
| chr7 | 66927848 | 66928061 | 369 | Intron (ENSMUST00000156396.2 | 7 | 66909468 | 66969555 | 60088 | 1 | ENSMUST00 | 18381 | Adamts17 | ENSMUSG0( | false | . | . | . | . | . | . | . | . | . | . | . | . |
| chr15 | 102659566 | 102659788 | 369 | Distal Intergenic | 15 | 102664739 | 102667743 | 3005 | 2 | ENSMUST00 | 7955 | Atp5g2 | ENSMUSG0( | false | . | . | . | . | . | . | chr15 | 102655445 | 102661031 | 4.34433 | 3.73001 | 2.48191 |
| chr17 | 49500393 | 49500681 | 369 | Intron (ENSMUST00000057610.6 | 17 | 49452284 | 49455435 | 3152 | 1 | ENSMUST00 | 48110 | Mocs1 | ENSMUSG0( | false | . | . | . | . | . | . | chr17 | 49499397 | 49507267 | 4.19929 | 3.58358 | 2.38859 |
| chr3 | 144586910 | 144587255 | 369 | Promoter (<=1kb) | 3 | 144586825 | 144586952 | 128 | 2 | ENSMUST00 | 0 | Gm24406 | ENSMUSG0( | false | . | . | . | . | . | . | chr3 | 144583802 | 144589715 | 5.5211 | 4.30146 | 3.44364 |
| chr6 | 112244049 | 112244291 | 369 | Distal Intergenic | 6 | 112234163 | 112234262 | 100 | 2 | ENSMUST00 | -9788 | n-R5s165 | ENSMUSG0( | false | . | . | . | . | . | . | . | . | . | . | . | . |
| chr9 | 102992865 | 102993175 | 369 | Promoter (<=1kb) | 9 | 102991895 | 102992216 | 322 | 1 | ENSMUST00 | 971 | Gm25842 | ENSMUSG0( | false | . | . | . | . | . | . | chr9 | 102991889 | 103008204 | 5.47516 | 4.26994 | 3.39839 |
| chr4 | 126140404 | 126140741 | 369 | 3' UTR | 4 | 126145979 | 126147735 | 1757 | 1 | ENSMUST00 | -5238 | Gm12946 | ENSMUSG0( | false | . | . | . | . | . | . | chr4 | 126139743 | 126142440 | 7.01043 | 4.99644 | 4.6336 |
| chr10 | 24599376 | 24600027 | 369 | Promoter (2-3kb) | 10 | 24595249 | 24597108 | 1860 | 2 | ENSMUST00 | -2269 | Gm15270 | ENSMUSG0( | false | . | . | . | . | . | . | chr10 | 24597031 | 24599930 | 4.39111 | 3.6267 | 2.53425 |
| chr10 | 124423545 | 124423819 | 369 | Distal Intergenic | 10 | 124455059 | 124455317 | 259 | 1 | ENSMUST00 | -31240 | Gm23777 | ENSMUSG0( | false | . | . | . | . | . | . | chr10 | 124422835 | 124424375 | 3.47718 | 3.31906 | 1.80236 |
| chr10 | 93262285 | 93262478 | 368 | Intron (ENSMUST00000008542.6 | 10 | 93249323 | 93310811 | 61489 | 2 | ENSMUST00 | 48333 | Elk3 | ENSMUSG0( | false | . | . | . | . | . | . | chr10 | 93261484 | 93262680 | 5.54661 | 4.17821 | 3.48839 |
| chr5 | 114603344 | 114603552 | 368 | Intron (ENSMUST00000043650.7 | 5 | 114626744 | 114630638 | 3895 | 2 | ENSMUST00 | 27086 | Trpv4 | ENSMUSG0( | false | . | . | . | . | . | . | chr5 | 114571033 | 114614394 | 7.1045 | 4.9325 | 4.74958 |
| chr10 | 59859219 | 59859530 | 368 | Promoter (1-2kb) | 10 | 59861518 | 59863403 | 1886 | 1 | ENSMUST00 | -1988 | Micu1 | ENSMUSG0( | true | . | . | . | . | . | . | chr10 | 59857833 | 59865240 | 4.40218 | 3.74218 | 2.52608 |
| chr15 | 11906634 | 11906956 | 368 | Promoter (<=1kb) | 15 | 11839896 | 11905674 | 65779 | 2 | ENSMUST00 | -961 | Npr3 | ENSMUSG0( | true | chr15 | 11902214 | 11907460 | 38.7176 | 123.44448 | 119.18625 | . | . | . | . | . | . |
| chr16 | 17069912 | 17070359 | 368 | Promoter (<=1kb) | 16 | 17070158 | 17086341 | 16184 | 1 | ENSMUST00 | 0 | Ypel1 | ENSMUSG0( | false | chr16 | 17069675 | 17070569 | 21.42557 | 69.98924 | 66.99868 | chr16 | 17068756 | 17071207 | 3.10475 | 3.04476 | 1.48261 |
| chr6 | 90983526 | 90983759 | 368 | Distal Intergenic | 6 | 91013317 | 91017429 | 4113 | 2 | ENSMUST00 | 33670 | Nup210 | ENSMUSG0( | false | . | . | . | . | . | . | chr6 | 90982644 | 90988187 | 5.21445 | 4.07964 | 3.20504 |
| chr15 | 88850380 | 88850710 | 368 | Distal Intergenic | 15 | 88862186 | 88865718 | 3533 | 1 | ENSMUST00 | -11476 | Pim3 | ENSMUSG0( | false | . | . | . | . | . | . | chr15 | 88850349 | 88851601 | 3.39854 | 3.22511 | 1.74654 |
| chr8 | 69974173 | 69974529 | 368 | Promoter (<=1kb) | 8 | 69907069 | 69974383 | 67315 | 2 | ENSMUST00 | 0 | Gatad2a | ENSMUSG0( | false | chr8 | 69972761 | 69974894 | 43.06936 | 146.42058 | 141.27246 | . | . | . | . | . | . |
| chr10 | 22578734 | 22578962 | 368 | Distal Intergenic | 10 | 22645011 | 22704285 | 59275 | 1 | ENSMUST00 | -66049 | Slc2a12 | ENSMUSG0( | false | . | . | . | . | . | . | chr10 | 22578730 | 22579064 | 3.26938 | 3.10838 | 1.62242 |
| chr7 | 144553631 | 144553928 | 368 | Promoter (<=1kb) | 7 | 144476758 | 144553729 | 76972 | 2 | ENSMUST00 | 0 | Ppfia1 | ENSMUSG0( | false | chr7 | 144552932 | 144554372 | 32.47248 | 106.24805 | 102.52307 | chr7 | 144550289 | 144554604 | 3.85277 | 3.34655 | 2.10842 |
| chr7 | 68682324 | 68682609 | 368 | Distal Intergenic | 7 | 68741511 | 68744906 | 3396 | 2 | ENSMUST00 | 62297 | Arrdc4 | ENSMUSG0( | false | . | . | . | . | . | . | chr7 | 68681789 | 68684213 | 5.4606 | 4.32818 | 3.38781 |
| chr4 | 45459490 | 45459715 | 368 | Intron (ENSMUST00000147926.2 | 4 | 45423278 | 45482802 | 59525 | 2 | ENSMUST00 | 23087 | Shb | ENSMUSG0( | false | . | . | . | . | . | . | chr4 | 45457040 | 45463702 | 5.65864 | 4.22032 | 3.58846 |
| chr8 | 119649614 | 119649834 | 368 | Distal Intergenic | 8 | 119623854 | 119635680 | 11827 | 2 | ENSMUST00 | -13935 | Kcng4 | ENSMUSG0( | false | . | . | . | . | . | . | chr8 | 119649334 | 119652098 | 3.6622 | 3.27726 | 1.93865 |
| chr10 | 84533754 | 84534273 | 368 | Promoter (<=1kb) | 10 | 84526305 | 84533888 | 7584 | 2 | ENSMUST00 | 0 | Ckap4 | ENSMUSG0( | false | chr10 | 84533060 | 84534572 | 29.02424 | 85.38521 | 82.13421 | chr10 | 84532105 | 84534498 | 3.00673 | 2.98686 | 1.38213 |
| chr19 | 28436509 | 28436686 | 368 | Intron (ENSMUST00000162022.2 | 19 | 28262973 | 28540420 | 277448 | 2 | ENSMUST00 | 103734 | Glis3 | ENSMUSG0( | false | . | . | . | . | . | . | chr19 | 28436141 | 28437230 | 3.46274 | 3.073 | 1.77176 |
| chr15 | 75795470 | 75795928 | 368 | Intron (ENSMUST00000100538.2 | 15 | 75759859 | 75759989 | 131 | 1 | ENSMUST00 | 35612 | Gm23747 | ENSMUSG0( | false | . | . | . | . | . | . | chr15 | 75794484 | 75796829 | 5.67267 | 4.25937 | 3.59229 |
| chr11 | 104259454 | 104259722 | 368 | Promoter (<=1kb) | 11 | 104250902 | 104259361 | 8460 | 2 | ENSMUST00 | -94 | Gm11659 | ENSMUSG0( | false | . | . | . | . | . | . | chr11 | 104242646 | 104282246 | 6.16611 | 4.45267 | 3.98703 |
| chr6 | 145725840 | 145726043 | 368 | Distal Intergenic | 6 | 145738693 | 145746273 | 7581 | 2 | ENSMUST00 | 20230 | Gm15704 | ENSMUSG0( | false | . | . | . | . | . | . | chr6 | 145725966 | 145726945 | 3.94815 | 3.43796 | 2.1705 |
| chr5 | 103592738 | 103593107 | 368 | Intron (ENSMUST00000048957.9 | 5 | 103602587 | 103602879 | 293 | 1 | ENSMUST00 | -9480 | Gm23085 | ENSMUSG0( | false | . | . | . | . | . | . | chr5 | 103591220 | 103600143 | 5.06031 | 4.03902 | 3.07298 |
| chr12 | 23189564 | 23189839 | 368 | Distal Intergenic | 12 | 23199780 | 23199889 | 110 | 1 | ENSMUST00 | -9941 | Gm26140 | ENSMUSG0( | false | chr12 | 23189096 | 23189872 | 9.01168 | 19.18248 | 16.85847 | chr12 | 23188748 | 23189754 | 3.6492 | 3.42345 | 1.92218 |
| chr7 | 136152459 | 136152678 | 368 | Distal Intergenic | 7 | 135952798 | 135953783 | 986 | 1 | ENSMUST00 | 199662 | Gm9341 | ENSMUSG0( | false | . | . | . | . | . | . | . | . | . | . | . | . |
| chr8 | 72268314 | 72268626 | 367 | Distal Intergenic | 8 | 72252935 | 72256912 | 3978 | 1 | ENSMUST00 | 15380 | Ap1m1 | ENSMUSG0( | false | . | . | . | . | . | . | . | . | . | . | . | . |
| chr10 | 79861127 | 79861418 | 367 | Promoter (<=1kb) | 10 | 79861738 | 79863050 | 1313 | 1 | ENSMUST00 | -320 | Ptbp1 | ENSMUSG0( | false | . | . | . | . | . | . | chr10 | 79855944 | 79867013 | 7.93167 | 5.01038 | 5.43402 |
| chr4 | 53111785 | 53112081 | 367 | Intron (ENSMUST00000030010.3 | 4 | 53030787 | 53159895 | 129109 | 2 | ENSMUST00 | 47814 | Abca1 | ENSMUSG0( | false | . | . | . | . | . | . | chr4 | 53110201 | 53112663 | 3.42492 | 3.1259 | 1.7519 |
| chr11 | 76027621 | 76028050 | 367 | Promoter (<=1kb) | 11 | 76019194 | 76027782 | 8589 | 2 | ENSMUST00 | 0 | Fam101b | ENSMUSG0( | false | chr11 | 76026815 | 76028460 | 32.00811 | 98.35596 | 94.82688 | chr11 | 76026169 | 76028496 | 5.21191 | 4.24137 | 3.16398 |
| chr13 | 53377301 | 53377569 | 367 | Promoter (<=1kb) | 13 | 53332716 | 53377365 | 44650 | 2 | ENSMUST00 | 0 | Sptlc1 | ENSMUSG0( | false | chr13 | 53376577 | 53377871 | 40.99673 | 133.90312 | 129.26613 | chr13 | 53375503 | 53377715 | 5.01279 | 4.12366 | 3.02341 |
| chr3 | 14533690 | 14534037 | 367 | Promoter (<=1kb) | 3 | 14533788 | 14572658 | 38871 | 1 | ENSMUST00 | 0 | Lrrcc1 | ENSMUSG0( | false | chr3 | 14533323 | 14534756 | 15.2551 | 59.70792 | 56.86497 | chr3 | 14533393 | 14535769 | 3.52161 | 3.34299 | 1.78624 |
| chr4 | 119356333 | 119356715 | 367 | Promoter (2-3kb) | 4 | 119352247 | 119359587 | 7341 | 2 | ENSMUST00 | 2872 | Ccdc30 | ENSMUSG0( | false | . | . | . | . | . | . | chr4 | 119356267 | 119356907 | 6.6342 | 4.93375 | 4.34384 |
| chr5 | 21061643 | 21061895 | 367 | Distal Intergenic | 5 | 20986645 | 21055911 | 69267 | 2 | ENSMUST00 | -5733 | Ptpn12 | ENSMUSG0( | true | . | . | . | . | . | . | chr5 | 21060987 | 21062595 | 3.78444 | 3.49458 | 1.98543 |
| chr5 | 21785113 | 21785354 | 367 | Promoter (<=1kb) | 5 | 21785283 | 21803787 | 18505 | 1 | ENSMUST00 | 0 | Psmc2 | ENSMUSG0( | false | chr5 | 21784084 | 21785577 | 42.47133 | 144.2491 | 139.19682 | . | . | . | . | . | . |
| chr5 | 103788964 | 103789155 | 367 | Intron (ENSMUST00000126335.1 | 5 | 103784359 | 103811595 | 27237 | 1 | ENSMUST00 | 4606 | Aff1 | ENSMUSG0( | false | . | . | . | . | . | . | chr5 | 103765592 | 103792229 | 6.51088 | 4.73748 | 4.26006 |
| chr5 | 103629497 | 103629775 | 367 | Promoter (<=1kb) | 5 | 103605711 | 103629403 | 23693 | 2 | ENSMUST00 | -95 | Slc10a6 | ENSMUSG0( | false | . | . | . | . | . | . | chr5 | 103627004 | 103630959 | 5.1666 | 4.08182 | 3.15645 |
| chr6 | 87535371 | 87535630 | 367 | Promoter (2-3kb) | 6 | 87459405 | 87533259 | 73855 | 2 | ENSMUST00 | -2113 | Arhgap25 | ENSMUSG0( | false | . | . | . | . | . | . | chr6 | 87535330 | 87536039 | 3.61162 | 3.12158 | 1.90427 |
| chr5 | 52429202 | 52429397 | 367 | Intron (ENSMUST00000059428.5 | 5 | 52374651 | 52471543 | 96893 | 2 | ENSMUST00 | 42146 | Ccdc149 | ENSMUSG0( | false | . | . | . | . | . | . | chr5 | 52428864 | 52430170 | 3.58417 | 3.32236 | 1.89014 |
| chr4 | 153956906 | 153957600 | 367 | Promoter (<=1kb) | 4 | 153957237 | 153961918 | 4682 | 1 | ENSMUST00 | 0 | A430005L14I | ENSMUSG0( | false | chr4 | 153956707 | 153958174 | 38.14117 | 119.25307 | 115.13219 | chr4 | 153956491 | 153959308 | 4.71705 | 3.7724 | 2.81569 |
| chr9 | 96437931 | 96438341 | 367 | Promoter (<=1kb) | 9 | 96435783 | 96438138 | 2356 | 2 | ENSMUST00 | 0 | BC043934 | ENSMUSG0( | false | chr9 | 96436926 | 96438702 | 17.10778 | 41.93702 | 39.32339 | chr9 | 96435849 | 96439222 | 5.69412 | 4.37663 | 3.58018 |
| chr3 | 32836683 | 32837104 | 367 | Intron (ENSMUST00000072312.6 | 3 | 32817711 | 32915033 | 97323 | 1 | ENSMUST00 | 18973 | Usp13 | ENSMUSG0( | true | . | . | . | . | . | . | . | . | . | . | . | . |
| chr14 | 54863790 | 54864277 | 367 | Promoter (<=1kb) | 14 | 54853787 | 54864158 | 10372 | 2 | ENSMUST00 | 0 | Homez | ENSMUSG0( | false | chr14 | 54863467 | 54864678 | 37.54867 | 136.59039 | 131.84451 | chr14 | 54862341 | 54864820 | 3.40264 | 3.29311 | 1.72532 |
| chr13 | 97928281 | 97928651 | 367 | Intron (ENSMUST00000109426.1 | 13 | 97760130 | 97760588 | 459 | 2 | ENSMUST00 | -167694 | Gm10260 | ENSMUSG0( | false | . | . | . | . | . | . | chr13 | 97909477 | 97929349 | 4.81516 | 3.96306 | 2.86901 |
| chr11 | 46029555 | 46029855 | 367 | Distal Intergenic | 11 | 46036440 | 46036781 | 342 | 1 | ENSMUST00 | -6585 | Gm12165 | ENSMUSG0( | false | . | . | . | . | . | . | . | . | . | . | . | . |
| chr16 | 30185479 | 30185747 | 367 | Distal Intergenic | 16 | 30195071 | 30200299 | 5229 | 1 | ENSMUST00 | -9324 | RP23-11P15. | ENSMUSG0( | false | . | . | . | . | . | . | chr16 | 30184555 | 30189184 | 7.55656 | 5.24836 | 5.09841 |
| chr4 | 136147812 | 136148181 | 366 | Promoter (2-3kb) | 4 | 136144974 | 136145755 | 782 | 1 | ENSMUST00 | 2839 | Id3 | ENSMUSG0( | false | chr4 | 136147444 | 136148224 | 14.46613 | 33.25751 | 30.75175 | chr4 | 136146751 | 136148293 | 4.0621 | 3.59404 | 2.2575 |
| chr2 | 164430712 | 164431008 | 366 | Intron (ENSMUST00000017153.3 | 2 | 164424247 | 164443186 | 18940 | 2 | ENSMUST00 | 12178 | Sdc4 | ENSMUSG0( | true | chr2 | 164430677 | 164431038 | 5.71153 | 8.93077 | 6.77673 | chr2 | 164415904 | 164437244 | 5.93196 | 4.473 | 3.77292 |
| chr2 | 165316801 | 165317148 | 366 | Promoter (<=1kb) | 2 | 165291117 | 165316903 | 25787 | 2 | ENSMUST00 | 0 | Elmo2 | ENSMUSG0( | false | . | . | . | . | . | . | chr2 | 165307563 | 165325045 | 8.91187 | 5.755 | 6.22321 |
| chr2 | 164515713 | 164516072 | 366 | Distal Intergenic | 2 | 164499916 | 164501639 | 1724 | 1 | ENSMUST00 | 15798 | Pigt | ENSMUSG0( | false | . | . | . | . | . | . | chr2 | 164509177 | 164517266 | 5.6289 | 4.32047 | 3.52645 |
| chr10 | 99203723 | 99204212 | 366 | Distal Intergenic | 10 | 99192892 | 99196850 | 3959 | 1 | ENSMUST00 | 10832 | Poc1b | ENSMUSG0( | false | chr10 | 99203639 | 99204461 | 10.50614 | 21.17611 | 18.82507 | chr10 | 99199695 | 99205008 | 4.87834 | 4.03183 | 2.90004 |
| chr15 | 76511792 | 76512200 | 366 | Promoter (<=1kb) | 15 | 76502015 | 76511951 | 9937 | 2 | ENSMUST00 | 0 | Dgat1 | ENSMUSG0( | false | chr15 | 76512105 | 76512529 | 10.79468 | 28.07545 | 25.63352 | . | . | . | . | . | . |
| chr7 | 112453971 | 112454228 | 366 | Intron (ENSMUST00000106643.2 | 7 | 112427905 | 112552929 | 125025 | 1 | ENSMUST00 | 26067 | Parva | ENSMUSG0( | true | . | . | . | . | . | . | chr7 | 112434747 | 112475955 | 6.44046 | 4.72904 | 4.19131 |
| chr7 | 105621085 | 105621369 | 366 | Intron (ENSMUST00000106791.2 | 7 | 105617031 | 105618026 | 996 | 2 | ENSMUST00 | -3060 | Trim3 | ENSMUSG0( | false | . | . | . | . | . | . | chr7 | 105618845 | 105625752 | 4.54007 | 3.8111 | 2.64709 |
| chr4 | 135565058 | 135565607 | 366 | Intron (ENSMUST00000105855.1 | 4 | 135541888 | 135573630 | 31743 | 2 | ENSMUST00 | 8023 | Grhl3 | ENSMUSG0( | false | . | . | . | . | . | . | . | . | . | . | . | . |
| chr6 | 34780403 | 34781019 | 366 | Promoter (<=1kb) | 6 | 34780432 | 34857932 | 77501 | 1 | ENSMUST00 | 0 | Agbl3 | ENSMUSG0( | false | chr6 | 34780162 | 34781417 | 28.06786 | 91.41865 | 88.0463 | . | . | . | . | . | . |
| chr15 | 26895174 | 26895753 | 366 | Promoter (<=1kb) | 15 | 26543084 | 26895049 | 351966 | 2 | ENSMUST00 | -126 | Fbxl7 | ENSMUSG0( | false | chr15 | 26894129 | 26896386 | 33.07612 | 110.40385 | 106.55179 | . | . | . | . | . | . |
| chr10 | 102528049 | 102528294 | 366 | Intron (ENSMUST00000055355.4 | 10 | 102512222 | 102546560 | 34339 | 1 | ENSMUST00 | 15828 | Rassf9 | ENSMUSG0( | false | . | . | . | . | . | . | chr10 | 102528028 | 102529628 | 4.30354 | 3.50422 | 2.46901 |
| chr11 | 62646067 | 62646285 | 366 | Promoter (1-2kb) | 11 | 62605823 | 62648040 | 42218 | 2 | ENSMUST00 | 1755 | Lrrc75a | ENSMUSG0( | false | . | . | . | . | . | . | chr11 | 62645300 | 62649776 | 3.45477 | 3.20498 | 1.78642 |
| chr13 | 100744062 | 100744722 | 366 | Promoter (<=1kb) | 13 | 100735955 | 100744659 | 8705 | 2 | ENSMUST00 | 0 | Mrps36 | ENSMUSG0( | false | chr13 | 100743765 | 100745190 | 44.3943 | 146.46883 | 141.31972 | . | . | . | . | . | . |
| chr19 | 8897960 | 8898297 | 366 | Promoter (<=1kb) | 19 | 8898090 | 8916663 | 18574 | 1 | ENSMUST00 | 0 | Ganab | ENSMUSG0( | false | chr19 | 8897342 | 8899234 | 32.29185 | 112.09676 | 108.19604 | . | . | . | . | . | . |
| chrX | 12840336 | 12840629 | 366 | Distal Intergenic | 20 | 12870269 | 12870362 | 94 | 2 | ENSMUST00 | 29733 | Gm25063 | ENSMUSG0( | false | chrX | 12839896 | 12840655 | 9.19064 | 17.46287 | 15.16242 | . | . | . | . | . | . |
| chr4 | 114904929 | 114905669 | 366 | Promoter (2-3kb) | 4 | 114817161 | 114902859 | 85699 | 2 | ENSMUST00 | -2071 | 9130410C08I | ENSMUSG0( | false | chr4 | 114904677 | 114911326 | 45.47067 | 151.73354 | 146.38181 | chr4 | 114897050 | 114905478 | 4.35462 | 3.78738 | 2.46946 |
| chr16 | 95769668 | 95769977 | 366 | Distal Intergenic | 16 | 95806029 | 95808317 | 2289 | 2 | ENSMUST00 | 38340 | 2810404F17I | ENSMUSG0( | false | . | . | . | . | . | . | . | . | . | . | . | . |
| chr9 | 40614170 | 40614439 | 366 | Distal Intergenic | 9 | 40670613 | 40671014 | 402 | 1 | ENSMUST00 | -56174 | Gm21915 | ENSMUSG0( | false | . | . | . | . | . | . | chr9 | 40612387 | 40618093 | 4.45996 | 3.66496 | 2.59375 |
| chr18 | 78549327 | 78549690 | 366 | Distal Intergenic | 18 | 78533458 | 78533564 | 107 | 1 | ENSMUST00 | 15870 | Gm26430 | ENSMUSG0( | false | . | . | . | . | . | . | . | . | . | . | . | . |
| chr12 | 16198870 | 16199144 | 366 | Distal Intergenic | 12 | 15817003 | 15888155 | 71153 | 1 | ENSMUST00 | 381868 | Gm5432 | ENSMUSG0( | false | . | . | . | . | . | . | chr12 | 16198451 | 16199608 | 4.69398 | 3.62563 | 2.79103 |
| chr15 | 101147934 | 101148218 | 365 | Distal Intergenic | 15 | 101137600 | 101142624 | 5025 | 1 | ENSMUST00 | 10335 | Acvrl1 | ENSMUSG0( | false | . | . | . | . | . | . | . | . | . | . | . | . |
| chr12 | 104864974 | 104865273 | 365 | Promoter (<=1kb) | 12 | 104763114 | 104865076 | 101963 | 2 | ENSMUST00 | 0 | Clmn | ENSMUSG0( | false | chr12 | 104864253 | 104865506 | 20.60195 | 76.26862 | 73.17842 | chr12 | 104863571 | 104865888 | 3.90875 | 3.5026 | 2.1583 |
| chr15 | 58994364 | 58994610 | 365 | Intron (ENSMUST00000080371.6 | 15 | 58933810 | 58939489 | 5680 | 1 | ENSMUST00 | 60555 | Ndufb9 | ENSMUSG0( | false | . | . | . | . | . | . | chr15 | 58965186 | 59078715 | 7.47325 | 5.12649 | 5.04432 |
| chr17 | 46027432 | 46027825 | 365 | Promoter (2-3kb) | 17 | 46018606 | 46024881 | 6276 | 2 | ENSMUST00 | -2552 | Vegfa | ENSMUSG0( | true | chr17 | 46023564 | 46033809 | 32.66734 | 113.81544 | 109.86345 | . | . | . | . | . | . |
| chr2 | 28514628 | 28514918 | 365 | Promoter (1-2kb) | 2 | 28513326 | 28553081 | 39756 | 1 | ENSMUST00 | 1303 | Ralgds | ENSMUSG0( | false | . | . | . | . | . | . | chr2 | 28512883 | 28514877 | 3.65863 | 3.26341 | 1.92829 |
| chr3 | 35932268 | 35933207 | 365 | Promoter (<=1kb) | 3 | 35892105 | 35932334 | 40230 | 2 | ENSMUST00 | 0 | Dcun1d1 | ENSMUSG0( | false | chr3 | 35931692 | 35933728 | 28.1615 | 93.53628 | 90.11485 | . | . | . | . | . | . |
| chr5 | 24577071 | 24577813 | 365 | Promoter (<=1kb) | 5 | 24565341 | 24577467 | 12127 | 2 | ENSMUST00 | 0 | Abcf2 | ENSMUSG0( | false | chr5 | 24576525 | 24578236 | 30.41422 | 89.70082 | 86.36422 | . | . | . | . | . | . |
| chr9 | 63399095 | 63399478 | 365 | Promoter (<=1kb) | 9 | 63394447 | 63399241 | 4795 | 2 | ENSMUST00 | 0 | 2300009A05I | ENSMUSG0( | false | chr9 | 63398366 | 63399901 | 21.20881 | 84.61981 | 81.38317 | chr9 | 63395634 | 63400384 | 4.37014 | 3.80719 | 2.48769 |
| chr10 | 95144511 | 95144755 | 365 | Distal Intergenic | 10 | 95174746 | 95324097 | 149352 | 2 | ENSMUST00 | 179342 | Cradd | ENSMUSG0( | false | . | . | . | . | . | . | chr10 | 95142298 | 95147251 | 5.59095 | 4.28663 | 3.51913 |
| chr3 | 95228571 | 95229052 | 365 | Promoter (<=1kb) | 3 | 95228732 | 95236409 | 7678 | 1 | ENSMUST00 | 0 | Cdc42se1 | ENSMUSG0( | false | chr3 | 95227430 | 95229926 | 20.14837 | 78.81087 | 75.68276 | . | . | . | . | . | . |
| chr3 | 107333850 | 107334142 | 365 | Promoter (<=1kb) | 3 | 107326110 | 107333289 | 7180 | 2 | ENSMUST00 | -562 | Rbm15 | ENSMUSG0( | false | chr3 | 107333310 | 107334618 | 21.91509 | 66.5122 | 63.57177 | . | . | . | . | . | . |
| chr10 | 42023248 | 42023543 | 365 | Distal Intergenic | 10 | 41916807 | 42018442 | 101636 | 2 | ENSMUST00 | -4807 | Armc2 | ENSMUSG0( | false | . | . | . | . | . | . | . | . | . | . | . | . |
| chr7 | 16047811 | 16048373 | 365 | Distal Intergenic | 7 | 16071942 | 16096328 | 24387 | 1 | ENSMUST00 | -23569 | Zfp541 | ENSMUSG0( | false | chr7 | 16046337 | 16049003 | 27.82288 | 84.25431 | 81.02234 | . | . | . | . | . | . |
| chr2 | 93776239 | 93776615 | 365 | Intron (ENSMUST00000184931.2 | 2 | 93794570 | 93795080 | 511 | 1 | ENSMUST00 | -17955 | Gm13818 | ENSMUSG0( | false | . | . | . | . | . | . | chr2 | 93770070 | 93782161 | 5.86588 | 4.4453 | 3.71814 |
| chrX | 167038571 | 167038814 | 365 | Distal Intergenic | 20 | 167028902 | 167029436 | 535 | 2 | ENSMUST00 | -9136 | Gm8814 | ENSMUSG0( | false | . | . | . | . | . | . | . | . | . | . | . | . |
| chr11 | 88382348 | 88382610 | 365 | Intron (ENSMUST00000107909.2 | 11 | 88356291 | 88363947 | 7657 | 1 | ENSMUST00 | 26058 | Gm15893 | ENSMUSG0( | false | . | . | . | . | . | . | chr11 | 88380987 | 88383820 | 4.87197 | 4.04748 | 2.90971 |
| chr13 | 45455357 | 45455642 | 365 | Distal Intergenic | 13 | 45486224 | 45486330 | 107 | 2 | ENSMUST00 | 30688 | Gm23387 | ENSMUSG0( | false | . | . | . | . | . | . | chr13 | 45453237 | 45457856 | 4.83005 | 4.04045 | 2.8659 |
| chr5 | 3505111 | 3505379 | 365 | Intron (ENSMUST00000165117.2 | 5 | 3473071 | 3473178 | 108 | 2 | ENSMUST00 | -31934 | Gm17590 | ENSMUSG0( | false | . | . | . | . | . | . | chr5 | 3504319 | 3505565 | 3.82796 | 3.47606 | 2.06634 |
| chr3 | 79286550 | 79287501 | 365 | Distal Intergenic | 3 | 79345376 | 79464129 | 118754 | 1 | ENSMUST00 | -57875 | Gm17359 | ENSMUSG0( | false | chr3 | 79284979 | 79287963 | 29.92778 | 109.76675 | 105.9399 | . | . | . | . | . | . |
| chr9 | 96696217 | 96696465 | 365 | Intron (ENSMUST00000152594.2 | 9 | 96705828 | 96711449 | 5622 | 1 | ENSMUST00 | -9363 | RP24-217M7 | ENSMUSG0( | false | . | . | . | . | . | . | chr9 | 96692956 | 96697178 | 3.52483 | 3.34469 | 1.81624 |
| chr8 | 22537148 | 22537474 | 364 | Intron (ENSMUST00000067786.7 | 8 | 22577075 | 22593813 | 16739 | 2 | ENSMUST00 | 56339 | Vdac3 | ENSMUSG0( | false | . | . | . | . | . | . | chr8 | 22536560 | 22537864 | 3.07353 | 2.9749 | 1.4241 |
| chr10 | 7589600 | 7590123 | 364 | Promoter (<=1kb) | 10 | 7589800 | 7608567 | 18768 | 1 | ENSMUST00 | 0 | Lrp11 | ENSMUSG0( | false | chr10 | 7589305 | 7590761 | 30.3748 | 101.49462 | 97.88963 | . | . | . | . | . | . |
| chr15 | 98606176 | 98606412 | 364 | Promoter (1-2kb) | 15 | 98589994 | 98607633 | 17640 | 2 | ENSMUST00 | 1221 | Adcy6 | ENSMUSG0( | false | . | . | . | . | . | . | . | . | . | . | . | . |
| chr17 | 56325858 | 56326606 | 364 | Promoter (<=1kb) | 17 | 56326062 | 56356056 | 29995 | 1 | ENSMUST00 | 0 | Kdm4b | ENSMUSG0( | false | chr17 | 56325969 | 56327059 | 27.66126 | 83.97217 | 80.74496 | . | . | . | . | . | . |
| chr3 | 133310019 | 133310759 | 364 | Promoter (<=1kb) | 3 | 133310110 | 133378078 | 67969 | 1 | ENSMUST00 | 0 | Ppa2 | ENSMUSG0( | false | chr3 | 133309986 | 133311194 | 28.28183 | 109.60786 | 105.79031 | . | . | . | . | . | . |
| chr4 | 41505491 | 41505688 | 364 | Promoter (2-3kb) | 4 | 41505009 | 41507754 | 2746 | 2 | ENSMUST00 | 2066 | 1110017D15I | ENSMUSG0( | false | . | . | . | . | . | . | chr4 | 41504779 | 41507701 | 5.135 | 4.20167 | 3.11487 |
| chr7 | 98309002 | 98309656 | 364 | Promoter (<=1kb) | 7 | 98213658 | 98309528 | 95871 | 2 | ENSMUST00 | 0 | Acer3 | ENSMUSG0( | false | chr7 | 98308665 | 98310071 | 25.90858 | 83.69981 | 80.48042 | . | . | . | . | . | . |
| chrX | 73967565 | 73967854 | 364 | Promoter (1-2kb) | 20 | 73942792 | 73966357 | 23566 | 2 | ENSMUST00 | -1209 | Hcfc1 | ENSMUSG0( | false | chrX | 73965220 | 73968256 | 24.81473 | 68.0602 | 65.09722 | . | . | . | . | . | . |
| chr1 | 89090015 | 89090393 | 364 | Intron (ENSMUST00000066279.5 | 1 | 89070453 | 89112860 | 42408 | 1 | ENSMUST00 | 19563 | Sh3bp4 | ENSMUSG0( | false | . | . | . | . | . | . | chr1 | 89071619 | 89098621 | 6.06397 | 4.51902 | 3.881 |
| chr9 | 96987203 | 96987538 | 364 | Intron (ENSMUST00000055433.4 | 9 | 96943485 | 96984195 | 40711 | 2 | ENSMUST00 | -3009 | Spsb4 | ENSMUSG0( | false | . | . | . | . | . | . | chr9 | 96975779 | 96994993 | 5.8258 | 4.40484 | 3.6945 |
| chr17 | 26710052 | 26710502 | 364 | Distal Intergenic | 17 | 26715650 | 26776635 | 60986 | 1 | ENSMUST00 | -5148 | Crebrf | ENSMUSG0( | false | . | . | . | . | . | . | chr17 | 26707095 | 26711677 | 5.75587 | 4.28483 | 3.66024 |
| chr1 | 133025053 | 133025489 | 364 | Promoter (<=1kb) | 1 | 132959484 | 133025338 | 65855 | 2 | ENSMUST00 | 0 | Mdm4 | ENSMUSG0( | false | chr1 | 133024460 | 133026209 | 28.90751 | 104.81161 | 101.11721 | . | . | . | . | . | . |
| chr7 | 141047258 | 141047468 | 364 | Distal Intergenic | 7 | 141061274 | 141072119 | 10846 | 1 | ENSMUST00 | -13806 | B4galnt4 | ENSMUSG0( | false | . | . | . | . | . | . | chr7 | 141046795 | 141048933 | 3.50285 | 3.33276 | 1.77277 |
| chr3 | 51868312 | 51868610 | 364 | Intron (ENSMUST00000121440.2 | 3 | 51828912 | 51829028 | 117 | 1 | ENSMUST00 | 39401 | n-R5s196 | ENSMUSG0( | false | . | . | . | . | . | . | chr3 | 51864972 | 51875550 | 6.25 | 4.62847 | 4.03397 |
| chr2 | 38658237 | 38658758 | 364 | Distal Intergenic | 2 | 38671158 | 38687651 | 16494 | 1 | ENSMUST00 | -12400 | Gpr144-ps | ENSMUSG0( | false | chr2 | 38658123 | 38658664 | 12.79009 | 27.87535 | 25.43694 | chr2 | 38658042 | 38658849 | 4.27517 | 3.49092 | 2.43273 |
| chr1 | 83713397 | 83713781 | 364 | Intron (ENSMUST00000189405.1 | 1 | 83795912 | 83796032 | 121 | 1 | ENSMUST00 | -82131 | Gm25754 | ENSMUSG0( | false | . | . | . | . | . | . | . | . | . | . | . | . |
| chr15 | 62392445 | 62392699 | 364 | Distal Intergenic | 15 | 62222603 | 62242505 | 19903 | 1 | ENSMUST00 | 169843 | Pvt1 | ENSMUSG0( | false | . | . | . | . | . | . | . | . | . | . | . | . |
| chr19 | 45918461 | 45918658 | 364 | Intron (ENSMUST00000045396.7 | 19 | 45998137 | 46013386 | 15250 | 1 | ENSMUST00 | -79479 | 4930505N22 | ENSMUSG0( | false | . | . | . | . | . | . | chr19 | 45916903 | 45920732 | 5.3038 | 4.17016 | 3.28303 |
| chr14 | 105619451 | 105619852 | 364 | Distal Intergenic | 14 | 105623642 | 105623770 | 129 | 2 | ENSMUST00 | 3918 | Mir6390 | ENSMUSG0( | false | . | . | . | . | . | . | . | . | . | . | . | . |
| chr11 | 87436146 | 87436376 | 364 | Intron (ENSMUST00000060835.6 | 11 | 87443237 | 87443452 | 216 | 1 | ENSMUST00 | -6861 | Rnu3b1 | ENSMUSG0( | false | . | . | . | . | . | . | chr11 | 87436059 | 87436922 | 3.28226 | 3.18478 | 1.59734 |
| chr1 | 62928041 | 62928306 | 364 | Distal Intergenic | 1 | 62821355 | 62832370 | 11016 | 2 | ENSMUST00 | -95672 | RP23-149A5. | ENSMUSG0( | false | . | . | . | . | . | . | chr1 | 62926621 | 62929567 | 4.22749 | 3.65922 | 2.3783 |
| chr10 | 78069144 | 78069420 | 363 | Promoter (<=1kb) | 10 | 78069360 | 78079525 | 10166 | 1 | ENSMUST00 | 0 | Icosl | ENSMUSG0( | false | chr10 | 78068651 | 78070259 | 26.46939 | 80.00455 | 76.85218 | chr10 | 78068826 | 78070211 | 4.36833 | 3.90414 | 2.46798 |
| chr11 | 61579039 | 61579811 | 363 | Promoter (<=1kb) | 11 | 61517249 | 61579669 | 62421 | 2 | ENSMUST00 | 0 | Epn2 | ENSMUSG0( | true | chr11 | 61577769 | 61579775 | 35.61502 | 123.34891 | 119.09213 | . | . | . | . | . | . |
| chr11 | 88150176 | 88150413 | 363 | Intron (ENSMUST00000018522.7 | 11 | 88162261 | 88194140 | 31880 | 1 | ENSMUST00 | -11848 | Cuedc1 | ENSMUSG0( | true | . | . | . | . | . | . | chr11 | 88146778 | 88183132 | 6.97604 | 4.89368 | 4.62779 |
| chr12 | 91588109 | 91588577 | 363 | Promoter (1-2kb) | 12 | 91586625 | 91589639 | 3015 | 2 | ENSMUST00 | 1062 | Gtf2a1 | ENSMUSG0( | false | chr12 | 91587910 | 91591057 | 26.41709 | 100.00125 | 96.43182 | chr12 | 91587017 | 91588233 | 4.4858 | 3.81795 | 2.5948 |
| chr12 | 73794197 | 73794446 | 363 | Distal Intergenic | 12 | 73901375 | 73947530 | 46156 | 1 | ENSMUST00 | -106929 | Hif1a | ENSMUSG0( | false | . | . | . | . | . | . | chr12 | 73793910 | 73795515 | 5.64286 | 3.99099 | 3.58114 |
| chr15 | 11898923 | 11899123 | 363 | Intron (ENSMUST00000066529.3 | 15 | 11839896 | 11905674 | 65779 | 2 | ENSMUST00 | 6551 | Npr3 | ENSMUSG0( | true | . | . | . | . | . | . | chr15 | 11893760 | 11902915 | 4.91242 | 4.01511 | 2.94512 |
| chr17 | 45557451 | 45557651 | 363 | Promoter (1-2kb) | 17 | 45555716 | 45563169 | 7454 | 1 | ENSMUST00 | 1736 | Nfkbie | ENSMUSG0( | false | . | . | . | . | . | . | chr17 | 45555051 | 45558483 | 6.04677 | 4.65587 | 3.86605 |
| chr17 | 46069516 | 46069926 | 363 | Distal Intergenic | 17 | 46016996 | 46032377 | 15382 | 2 | ENSMUST00 | -37140 | Vegfa | ENSMUSG0( | true | . | . | . | . | . | . | chr17 | 46048792 | 46077235 | 6.11576 | 4.54762 | 3.92959 |
| chr17 | 78246518 | 78246924 | 363 | Intron (ENSMUST00000112498.2 | 17 | 78200248 | 78376592 | 176345 | 1 | ENSMUST00 | 46271 | Crim1 | ENSMUSG0( | true | . | . | . | . | . | . | chr17 | 78239579 | 78284617 | 5.22436 | 4.1134 | 3.20479 |
| chr1 | 75506309 | 75506589 | 363 | Promoter (<=1kb) | 1 | 75485825 | 75506361 | 20537 | 2 | ENSMUST00 | 0 | Obsl1 | ENSMUSG0( | false | chr1 | 75505204 | 75506553 | 24.55236 | 90.57755 | 87.22021 | chr1 | 75504966 | 75506568 | 3.59493 | 3.44057 | 1.8457 |
| chr2 | 11501292 | 11501635 | 363 | Promoter (<=1kb) | 2 | 11493903 | 11501296 | 7394 | 2 | ENSMUST00 | 0 | Pfkfb3 | ENSMUSG0( | false | chr2 | 11500699 | 11503640 | 31.43507 | 99.35931 | 95.8029 | chr2 | 11499274 | 11501299 | 5.1877 | 4.23502 | 3.14541 |
| chr4 | 102638311 | 102638557 | 363 | Distal Intergenic | 4 | 102589843 | 102603441 | 13599 | 1 | ENSMUST00 | 48469 | Pde4b | ENSMUSG0( | true | . | . | . | . | . | . | chr4 | 102637493 | 102641859 | 6.17662 | 4.64515 | 3.9647 |
| chr5 | 125166034 | 125166266 | 363 | Intron (ENSMUST00000111402.3 | 5 | 125017162 | 125179053 | 161892 | 2 | ENSMUST00 | 12787 | Ncor2 | ENSMUSG0( | false | . | . | . | . | . | . | chr5 | 125124978 | 125178451 | 6.70656 | 4.76008 | 4.42049 |
| chr5 | 114443933 | 114444214 | 363 | Promoter (<=1kb) | 5 | 114431034 | 114444059 | 13026 | 2 | ENSMUST00 | 0 | Mmab | ENSMUSG0( | false | chr5 | 114443548 | 114444071 | 19.89052 | 67.91473 | 64.9535 | chr5 | 114442458 | 114445646 | 3.62445 | 3.29222 | 1.92494 |
| chr18 | 83929945 | 83930139 | 363 | Distal Intergenic | 18 | 84012082 | 84086390 | 74309 | 2 | ENSMUST00 | 156251 | Tshz1 | ENSMUSG0( | false | . | . | . | . | . | . | chr18 | 83928334 | 83930423 | 3.98099 | 3.49443 | 2.21665 |
| chr13 | 42576237 | 42576590 | 363 | Distal Intergenic | 13 | 42680623 | 43138524 | 457902 | 1 | ENSMUST00 | -104033 | Phactr1 | ENSMUSG0( | false | . | . | . | . | . | . | chr13 | 42575977 | 42576351 | 4.63998 | 3.89802 | 2.71727 |
| chr11 | 101417293 | 101417557 | 363 | Promoter (<=1kb) | 11 | 101410808 | 101417384 | 6577 | 2 | ENSMUST00 | 0 | Aarsd1 | ENSMUSG0( | false | chr11 | 101416477 | 101417754 | 22.68762 | 81.35053 | 78.1749 | chr11 | 101416213 | 101418084 | 3.41529 | 3.30491 | 1.72566 |
| chr13 | 28483068 | 28483309 | 363 | Promoter (2-3kb) | 13 | 28460778 | 28480841 | 20064 | 2 | ENSMUST00 | -2228 | 2610307P16I | ENSMUSG0( | false | . | . | . | . | . | . | . | . | . | . | . | . |
| chr1 | 152677658 | 152677982 | 363 | Intron (ENSMUST00000111859.2 | 1 | 152684026 | 152687501 | 3476 | 2 | ENSMUST00 | 9519 | Gm15479 | ENSMUSG0( | false | . | . | . | . | . | . | chr1 | 152676620 | 152678812 | 5.11656 | 4.08047 | 3.14493 |
| chr10 | 12737999 | 12738316 | 363 | Exon (ENSMUST00000076817.4/ | 10 | 12793875 | 12793974 | 100 | 2 | ENSMUST00 | 55658 | Gm23044 | ENSMUSG0( | false | . | . | . | . | . | . | chr10 | 12736694 | 12739352 | 6.98187 | 5.06168 | 4.62005 |
| chr1 | 64052432 | 64052697 | 363 | Intron (ENSMUST00000114086.2 | 1 | 64042438 | 64042501 | 64 | 2 | ENSMUST00 | -9932 | Mir6899 | ENSMUSG0( | false | . | . | . | . | . | . | chr1 | 64049784 | 64053841 | 6.14264 | 4.54916 | 3.96547 |
| chr5 | 135887719 | 135887977 | 362 | Promoter (<=1kb) | 5 | 135887919 | 135889563 | 1645 | 1 | ENSMUST00 | 0 | Hspb1 | ENSMUSG0( | false | chr5 | 135887419 | 135888188 | 15.62409 | 36.57069 | 34.02363 | chr5 | 135887579 | 135890810 | 4.19739 | 3.73367 | 2.33372 |
| chr3 | 138501025 | 138501212 | 362 | Distal Intergenic | 3 | 138458960 | 138489382 | 30423 | 2 | ENSMUST00 | -11644 | Metap1 | ENSMUSG0( | false | . | . | . | . | . | . | . | . | . | . | . | . |
| chr1 | 172206651 | 172206935 | 362 | Promoter (<=1kb) | 1 | 172196728 | 172206804 | 10077 | 2 | ENSMUST00 | 0 | Pea15a | ENSMUSG0( | false | chr1 | 172205664 | 172207142 | 27.11239 | 76.48743 | 73.39479 | chr1 | 172200936 | 172207651 | 6.63015 | 4.77931 | 4.3486 |
| chrX | 36874194 | 36874491 | 362 | Promoter (<=1kb) | 20 | 36874358 | 36884215 | 9858 | 1 | ENSMUST00 | 0 | Ube2a | ENSMUSG0( | false | chrX | 36873986 | 36875275 | 22.90469 | 70.93823 | 67.93287 | . | . | . | . | . | . |
| chr14 | 20885219 | 20885473 | 362 | Distal Intergenic | 14 | 20929433 | 21033673 | 104241 | 1 | ENSMUST00 | -43960 | Vcl | ENSMUSG0( | false | . | . | . | . | . | . | chr14 | 20885293 | 20886061 | 4.91222 | 3.93816 | 2.9522 |

| | | | | | | | | | | | | | | | | | | | | | | | | | | |
|---|---|---|---|---|---|---|---|---|---|---|---|---|---|---|---|---|---|---|---|---|---|---|---|---|---|---|
| chr17 | 71480627 | 71480851 | 362 | Distal Intergenic | 17 | 71344489 | 71475343 | 130855 | 2 | ENSMUST00 | -5285 | Smchd1 | ENSMUSG00 | false | . | . | . | . | . | . | chr17 | 71479529 | 71481294 | 3.78268 | 3.45559 | 2.06343 |
| chr17 | 24454623 | 24454958 | 362 | Promoter (<=1kb) | 17 | 24443792 | 24455311 | 11520 | 2 | ENSMUST00 | 353 | E4f1 | ENSMUSG00 | false | chr17 | 24454317 | 24455384 | 34.16929 | 120.7392 | 116.5701 | . | . | . | . | . | . |
| chr1 | 52817448 | 52817908 | 362 | Promoter (<=1kb) | 1 | 52785427 | 52817688 | 32262 | 2 | ENSMUST00 | 0 | Inpp1 | ENSMUSG00 | false | chr1 | 52816600 | 52818107 | 32.90413 | 103.94604 | 100.27567 | . | . | . | . | . | . |
| chr3 | 131494058 | 131494266 | 362 | Distal Intergenic | 3 | 131564768 | 131643670 | 78903 | 1 | ENSMUST00 | -70502 | Papss1 | ENSMUSG00 | false | . | . | . | . | . | . | chr3 | 131493287 | 131494655 | 3.9521 | 3.41494 | 2.16433 |
| chr4 | 107889699 | 107890214 | 362 | Promoter (<=1kb) | 4 | 107889813 | 107899384 | 9572 | 1 | ENSMUST00 | 0 | 0610037L13 | ENSMUSG00 | false | chr4 | 107889275 | 107890655 | 14.91789 | 45.43439 | 42.77634 | chr4 | 107888953 | 107891144 | 3.53619 | 3.22998 | 1.85622 |
| chr8 | 27174598 | 27174851 | 362 | Promoter (<=1kb) | 8 | 27138773 | 27174646 | 35874 | 2 | ENSMUST00 | 0 | Rab11fip1 | ENSMUSG00 | false | chr8 | 27173868 | 27175786 | 38.03245 | 121.19012 | 117.00504 | chr8 | 27172077 | 27177912 | 4.742 | 3.87663 | 2.84361 |
| chr9 | 57589291 | 57589859 | 362 | Promoter (<=1kb) | 9 | 57589452 | 57596233 | 6782 | 1 | ENSMUST00 | 0 | Ulk3 | ENSMUSG00 | false | chr9 | 57589080 | 57590210 | 30.78991 | 105.27288 | 101.56436 | chr9 | 57588773 | 57590443 | 3.75455 | 3.42623 | 2.03872 |
| chr6 | 67075002 | 67075198 | 362 | Intron (ENSMUST00000130657.2 | 6 | 67036897 | 67037457 | 561 | 2 | ENSMUST00 | -37546 | Gadd45a | ENSMUSG00 | false | . | . | . | . | . | . | chr6 | 67069252 | 67079338 | 6.07124 | 4.48472 | 3.90237 |
| chr17 | 49559604 | 49559833 | 362 | Intron (ENSMUST00000057610.6 | 17 | 49456022 | 49564337 | 108316 | 2 | ENSMUST00 | 4504 | Daam2 | ENSMUSG00 | true | . | . | . | . | . | . | chr17 | 49553664 | 49563281 | 6.14765 | 4.61155 | 3.94797 |
| chr1 | 44567529 | 44567776 | 362 | Intron (ENSMUST00000074525.4 | 1 | 44552597 | 44708632 | 156036 | 1 | ENSMUST00 | 14933 | Gulp1 | ENSMUSG00 | true | . | . | . | . | . | . | chr1 | 44564703 | 44568481 | 4.0088 | 3.63286 | 2.17868 |
| chr9 | 36797213 | 36797532 | 362 | Promoter (<=1kb) | 9 | 36779159 | 36797273 | 18115 | 2 | ENSMUST00 | 0 | Ei24 | ENSMUSG00 | false | chr9 | 36796370 | 36797563 | 43.93277 | 149.05946 | 143.81693 | chr9 | 36793274 | 36797816 | 4.3424 | 3.78926 | 2.45401 |
| chr14 | 22213706 | 22213992 | 362 | Intron (ENSMUST00000075639.5 | 14 | 22310165 | 22584439 | 274275 | 1 | ENSMUST00 | -96173 | 1700112E06 | ENSMUSG00 | false | . | . | . | . | . | . | chr14 | 22212900 | 22216338 | 4.22296 | 3.54469 | 2.40489 |
| chr2 | 93013976 | 93014339 | 362 | Promoter (<=1kb) | 2 | 93009973 | 93013910 | 3938 | 2 | ENSMUST00 | -67 | Prdm11 | ENSMUSG00 | false | chr2 | 93014233 | 93015383 | 23.43614 | 63.09834 | 60.20888 | chr2 | 93013792 | 93015901 | 3.91582 | 3.4512 | 2.13459 |
| chr2 | 16169298 | 16169518 | 362 | Distal Intergenic | 2 | 16270488 | 16270794 | 307 | 2 | ENSMUST00 | 101276 | Gm13319 | ENSMUSG00 | false | . | . | . | . | . | . | . | . | . | . | . | . |
| chr6 | 54475977 | 54476184 | 362 | Intron (ENSMUST00000126637.2 | 6 | 54472002 | 54485360 | 13359 | 1 | ENSMUST00 | 3976 | Wipf3 | ENSMUSG00 | true | . | . | . | . | . | . | chr6 | 54456883 | 54482420 | 5.37713 | 4.1745 | 3.32864 |
| chr9 | 77547017 | 77547335 | 362 | Promoter (1-2kb) | 9 | 77545202 | 77546467 | 1266 | 1 | ENSMUST00 | 1816 | Gm19569 | ENSMUSG00 | false | . | . | . | . | . | . | chr9 | 77545346 | 77549514 | 6.11128 | 4.58585 | 3.90858 |
| chr2 | 145777853 | 145778038 | 361 | Distal Intergenic | 2 | 145784734 | 145822302 | 37569 | 1 | ENSMUST00 | -6696 | Rin2 | ENSMUSG00 | true | . | . | . | . | . | . | chr2 | 145776882 | 145778488 | 3.73476 | 3.4255 | 1.98571 |
| chr11 | 8587336 | 8587576 | 361 | Intron (ENSMUST00000020695.7 | 11 | 8545693 | 8579121 | 33429 | 2 | ENSMUST00 | -8216 | Tns3 | ENSMUSG00 | false | . | . | . | . | . | . | chr11 | 8583919 | 8594376 | 5.3263 | 4.09262 | 3.31573 |
| chr11 | 7065630 | 7065836 | 361 | Promoter (2-3kb) | 11 | 7063489 | 7178506 | 115018 | 1 | ENSMUST00 | 2142 | Adcy1 | ENSMUSG00 | true | chr11 | 7063589 | 7065749 | 23.3092 | 84.4034 | 81.16875 | . | . | . | . | . | . |
| chr19 | 36818011 | 36818456 | 361 | Distal Intergenic | 19 | 36834232 | 36893477 | 59246 | 1 | ENSMUST00 | -15776 | Tnks2 | ENSMUSG00 | false | chr19 | 36817962 | 36818941 | 13.73236 | 30.71385 | 28.24041 | chr19 | 36817824 | 36818850 | 3.7299 | 3.51071 | 1.94329 |
| chr19 | 38641187 | 38641470 | 361 | Intron (ENSMUST00000182481.2 | 19 | 38583368 | 38652007 | 68640 | 1 | ENSMUST00 | 57820 | Plce1 | ENSMUSG00 | true | . | . | . | . | . | . | chr19 | 38638925 | 38642252 | 3.65753 | 3.3782 | 1.95121 |
| chrX | 81071088 | 81071362 | 361 | Promoter (<=1kb) | 20 | 81070716 | 81097872 | 27157 | 1 | ENSMUST00 | 373 | Tmem47 | ENSMUSG00 | false | chrX | 81070309 | 81071644 | 20.67895 | 53.40692 | 50.64761 | . | . | . | . | . | . |
| chr4 | 134930678 | 134930982 | 361 | Promoter (<=1kb) | 4 | 134930898 | 134937548 | 6651 | 1 | ENSMUST00 | 0 | Syf2 | ENSMUSG00 | false | chr4 | 134930934 | 134931820 | 45.49427 | 162.13052 | 156.40686 | chr4 | 134924825 | 134933087 | 5.30601 | 4.17978 | 3.26976 |
| chr9 | 5308615 | 5308859 | 361 | Promoter (<=1kb) | 9 | 5308828 | 5323713 | 14886 | 1 | ENSMUST00 | 0 | Casp4 | ENSMUSG00 | false | . | . | . | . | . | . | . | . | . | . | . | . |
| chr3 | 31012827 | 31013093 | 361 | Intron (ENSMUST00000108249.3 | 3 | 31005193 | 31034513 | 29321 | 1 | ENSMUST00 | 7635 | Prkci | ENSMUSG00 | false | . | . | . | . | . | . | chr3 | 30996848 | 31027341 | 5.11782 | 4.10598 | 3.11002 |
| chr8 | 110217679 | 110218004 | 361 | Promoter (<=1kb) | 8 | 110217989 | 110224486 | 6498 | 1 | ENSMUST00 | 0 | Cmtr2 | ENSMUSG00 | false | chr8 | 110217355 | 110217761 | 8.21885 | 16.32505 | 14.04127 | chr8 | 110217163 | 110218795 | 5.09937 | 4.20982 | 3.07625 |
| chr6 | 72519896 | 72520293 | 361 | Promoter (<=1kb) | 6 | 72515327 | 72519977 | 4651 | 2 | ENSMUST00 | 0 | Sh2d6 | ENSMUSG00 | false | . | . | . | . | . | . | chr6 | 72518099 | 72520608 | 4.68498 | 3.97638 | 2.7464 |
| chr11 | 115475424 | 115475720 | 361 | Promoter (<=1kb) | 11 | 115475667 | 115490467 | 14801 | 1 | ENSMUST00 | 0 | Armc7 | ENSMUSG00 | false | chr11 | 115475189 | 115476342 | 33.96634 | 105.79253 | 102.076 | chr11 | 115472068 | 115477549 | 5.34788 | 4.24591 | 3.29751 |
| chr8 | 126788649 | 126788974 | 361 | Distal Intergenic | 8 | 126944847 | 126944980 | 134 | 2 | ENSMUST00 | 156006 | Gm26397 | ENSMUSG00 | false | . | . | . | . | . | . | chr8 | 126788204 | 126789639 | 5.23112 | 4.12944 | 3.24521 |
| chrX | 159371894 | 159372297 | 361 | Promoter (<=1kb) | 20 | 159372178 | 159376558 | 4381 | 1 | ENSMUST00 | 0 | Eif1ax | ENSMUSG00 | false | chrX | 159371820 | 159372033 | 8.27158 | 15.0213 | 12.75664 | . | . | . | . | . | . |
| chr4 | 144533500 | 144533701 | 361 | Distal Intergenic | 4 | 144526411 | 144527042 | 632 | 2 | ENSMUST00 | -6459 | Gm13121 | ENSMUSG00 | false | . | . | . | . | . | . | . | . | . | . | . | . |
| chr4 | 109173404 | 109173609 | 361 | Intron (ENSMUST00000160774.2 | 4 | 109160924 | 109161197 | 274 | 2 | ENSMUST00 | -12208 | Gm23743 | ENSMUSG00 | false | . | . | . | . | . | . | chr4 | 109172476 | 109177683 | 4.66267 | 3.69415 | 2.77594 |
| chr19 | 44314543 | 44315214 | 361 | Distal Intergenic | 19 | 44303171 | 44303231 | 61 | 1 | ENSMUST00 | 11373 | Mir5114 | ENSMUSG00 | false | chr19 | 44314370 | 44315405 | 12.58327 | 33.30664 | 30.79935 | chr19 | 44310172 | 44316564 | 5.73307 | 4.39799 | 3.62835 |
| chr8 | 92253067 | 92253278 | 361 | Distal Intergenic | 8 | 92336668 | 92339250 | 2583 | 2 | ENSMUST00 | 85972 | Gm21817 | ENSMUSG00 | false | . | . | . | . | . | . | chr8 | 92251699 | 92253810 | 4.53494 | 3.90002 | 2.61595 |
| chr13 | 78656821 | 78657017 | 361 | Distal Intergenic | 13 | 78372987 | 78379134 | 6148 | 2 | ENSMUST00 | -277688 | Gm6317 | ENSMUSG00 | false | . | . | . | . | . | . | . | . | . | . | . | . |
| chr12 | 15649361 | 15649587 | 361 | Distal Intergenic | 12 | 15728097 | 15729095 | 999 | 1 | ENSMUST00 | -78510 | Gm4804 | ENSMUSG00 | false | . | . | . | . | . | . | chr12 | 15643700 | 15661713 | 4.77631 | 3.96574 | 2.83123 |
| chr7 | 71467437 | 71467668 | 361 | Distal Intergenic | 7 | 71370333 | 71379813 | 9481 | 1 | ENSMUST00 | 97105 | RP24-173C1 | ENSMUSG00 | false | . | . | . | . | . | . | chr7 | 71463195 | 71468768 | 4.4454 | 3.87005 | 2.53324 |
| chr11 | 115187279 | 115187595 | 360 | Promoter (<=1kb) | 11 | 115182832 | 115187332 | 4501 | 2 | ENSMUST00 | 0 | Nat9 | ENSMUSG00 | false | chr11 | 115186754 | 115187994 | 26.28407 | 85.32412 | 82.07336 | chr11 | 115186254 | 115189000 | 4.40797 | 3.76104 | 2.513 |
| chr12 | 31499494 | 31500175 | 360 | Promoter (<=1kb) | 12 | 31484829 | 31499616 | 14788 | 2 | ENSMUST00 | 0 | Cbll1 | ENSMUSG00 | false | chr12 | 31498404 | 31500658 | 33.06624 | 113.49754 | 109.55873 | . | . | . | . | . | . |
| chr19 | 55742676 | 55743268 | 360 | Promoter (<=1kb) | 19 | 55742845 | 55910681 | 167837 | 1 | ENSMUST00 | 0 | Tcf7l2 | ENSMUSG00 | false | chr19 | 55741009 | 55745168 | 33.32866 | 126.59486 | 122.2257 | . | . | . | . | . | . |
| chr5 | 125154936 | 125155276 | 360 | Intron (ENSMUST00000111402.3 | 5 | 125017162 | 125179053 | 161892 | 2 | ENSMUST00 | 23777 | Ncor2 | ENSMUSG00 | false | . | . | . | . | . | . | chr5 | 125124978 | 125178451 | 6.70656 | 4.76008 | 4.42049 |
| chr5 | 145083755 | 145084155 | 360 | Promoter (<=1kb) | 5 | 145083830 | 145108761 | 24932 | 1 | ENSMUST00 | 0 | Arpc1a | ENSMUSG00 | false | chr5 | 145083439 | 145084861 | 37.26972 | 119.86725 | 115.72688 | chr5 | 145083246 | 145087102 | 3.31265 | 3.18149 | 1.66714 |
| chr6 | 118066825 | 118067061 | 360 | Promoter (<=1kb) | 6 | 118066385 | 118091546 | 25162 | 1 | ENSMUST00 | 441 | Rasgef1a | ENSMUSG00 | false | chr6 | 118065871 | 118067211 | 28.80981 | 88.85442 | 85.53428 | chr6 | 118066880 | 118067233 | 5.28806 | 4.19117 | 3.25021 |
| chr10 | 12965870 | 12966077 | 360 | Promoter (1-2kb) | 10 | 12939983 | 12964259 | 24277 | 2 | ENSMUST00 | -1612 | Stx11 | ENSMUSG00 | false | . | . | . | . | . | . | chr10 | 12964698 | 12965925 | 3.41496 | 3.16722 | 1.74459 |
| chr10 | 127030644 | 127030972 | 360 | Promoter (<=1kb) | 10 | 127011572 | 127030803 | 19232 | 2 | ENSMUST00 | 0 | Tsfm | ENSMUSG00 | false | chr10 | 127030208 | 127030924 | 27.88725 | 99.90214 | 96.33501 | . | . | . | . | . | . |
| chr11 | 103141119 | 103141353 | 360 | Distal Intergenic | 11 | 103133865 | 103138279 | 4415 | 1 | ENSMUST00 | 7255 | Hexim2 | ENSMUSG00 | false | . | . | . | . | . | . | chr11 | 103138329 | 103150034 | 6.87245 | 4.95791 | 4.53389 |
| chr5 | 144768436 | 144768758 | 360 | Promoter (<=1kb) | 5 | 144768544 | 144777128 | 8585 | 1 | ENSMUST00 | 0 | Trrap | ENSMUSG00 | false | chr5 | 144767881 | 144769477 | 32.31418 | 113.17245 | 109.24831 | . | . | . | . | . | . |
| chr10 | 70158435 | 70158644 | 360 | Intron (ENSMUST00000147545.2 | 10 | 70174509 | 70182206 | 7698 | 1 | ENSMUST00 | -15865 | Ccdc6 | ENSMUSG00 | false | . | . | . | . | . | . | chr10 | 70157764 | 70160392 | 6.52181 | 4.85974 | 4.24262 |
| chr5 | 36796594 | 36796974 | 360 | Promoter (<=1kb) | 5 | 36794867 | 36796772 | 1906 | 2 | ENSMUST00 | 0 | Mrfap1 | ENSMUSG00 | false | chr5 | 36795370 | 36797310 | 32.27147 | 102.84008 | 99.19724 | chr5 | 36794910 | 36797265 | 4.3313 | 3.67231 | 2.46797 |
| chrX | 94724551 | 94725222 | 360 | Promoter (<=1kb) | 20 | 94724569 | 94730187 | 5619 | 1 | ENSMUST00 | 0 | Zxdb | ENSMUSG00 | false | . | . | . | . | . | . | . | . | . | . | . | . |
| chr8 | 25438164 | 25438543 | 360 | Distal Intergenic | 8 | 25454273 | 25454704 | 432 | 1 | ENSMUST00 | -15730 | Gm10689 | ENSMUSG00 | false | . | . | . | . | . | . | chr8 | 25435682 | 25441738 | 6.54241 | 4.78324 | 4.28383 |
| chr16 | 11568224 | 11568505 | 360 | Intron (ENSMUST00000180792.2 | 16 | 11494821 | 11496306 | 1486 | 2 | ENSMUST00 | -71919 | Gm15897 | ENSMUSG00 | false | . | . | . | . | . | . | chr16 | 11565128 | 11569221 | 5.38779 | 4.10608 | 3.36493 |
| chr7 | 4876928 | 4877150 | 360 | Promoter (<=1kb) | 7 | 4877053 | 4895717 | 18665 | 1 | ENSMUST00 | 0 | Isoc2a | ENSMUSG00 | false | chr7 | 4876914 | 4877704 | 19.85205 | 54.41815 | 51.64575 | chr7 | 4876421 | 4878565 | 4.70897 | 3.92586 | 2.75932 |
| chr2 | 75757415 | 75757679 | 360 | Distal Intergenic | 2 | 75777188 | 75782306 | 5119 | 1 | ENSMUST00 | -19509 | Gm13657 | ENSMUSG00 | false | . | . | . | . | . | . | chr2 | 75756402 | 75757941 | 4.09619 | 3.64214 | 2.31582 |
| chr2 | 151536136 | 151536614 | 360 | Distal Intergenic | 2 | 151540731 | 151542348 | 1618 | 2 | ENSMUST00 | 5734 | Gm14167 | ENSMUSG00 | false | . | . | . | . | . | . | chr2 | 151535859 | 151537625 | 3.73189 | 3.44556 | 1.98316 |
| chr11 | 22589789 | 22590139 | 360 | Distal Intergenic | 11 | 22600335 | 22610879 | 10545 | 2 | ENSMUST00 | 20740 | Gm26829 | ENSMUSG00 | false | . | . | . | . | . | . | . | . | . | . | . | . |
| chr5 | 143758134 | 143758415 | 360 | Promoter (<=1kb) | 5 | 143758325 | 143766974 | 8650 | 1 | ENSMUST00 | 0 | D130017N08 | ENSMUSG00 | false | chr5 | 143757857 | 143758937 | 15.50707 | 38.47921 | 35.908 | . | . | . | . | . | . |
| chr1 | 12644676 | 12644952 | 360 | Distal Intergenic | 1 | 12655867 | 12656553 | 687 | 1 | ENSMUST00 | -10915 | Gm6216 | ENSMUSG00 | false | . | . | . | . | . | . | . | . | . | . | . | . |
| chr7 | 131161932 | 131162181 | 359 | Intron (ENSMUST00000124096.2 | 7 | 131174402 | 131250945 | 76544 | 1 | ENSMUST00 | -12221 | 5430419D17 | ENSMUSG00 | false | . | . | . | . | . | . | chr7 | 131160681 | 131162687 | 3.61711 | 3.42506 | 1.87925 |
| chr10 | 18708936 | 18709143 | 359 | Intron (ENSMUST00000019999.5 | 10 | 18588011 | 18743758 | 155748 | 2 | ENSMUST00 | 34615 | D10Bwg137 | ENSMUSG00 | false | . | . | . | . | . | . | chr10 | 18707554 | 18712829 | 3.67702 | 3.43058 | 1.91637 |
| chr12 | 70093612 | 70093803 | 359 | Intron (ENSMUST00000085314.4 | 12 | 70011435 | 70102833 | 91399 | 2 | ENSMUST00 | 9030 | Nin | ENSMUSG00 | false | . | . | . | . | . | . | chr12 | 70092948 | 70095543 | 3.96657 | 3.44449 | 2.19825 |
| chr14 | 19738524 | 19738810 | 359 | Distal Intergenic | 14 | 19751257 | 19811787 | 60531 | 1 | ENSMUST00 | -12447 | Nid2 | ENSMUSG00 | false | . | . | . | . | . | . | chr14 | 19737868 | 19738974 | 4.66828 | 3.83729 | 2.74744 |
| chr18 | 61096444 | 61096693 | 359 | Distal Intergenic | 18 | 61105572 | 61131134 | 25563 | 1 | ENSMUST00 | -8879 | Csf1r | ENSMUSG00 | false | chr18 | 61096192 | 61096968 | 8.27158 | 15.0213 | 12.75664 | chr18 | 61094767 | 61097367 | 4.34494 | 3.80371 | 2.45858 |
| chr4 | 108833229 | 108833825 | 359 | Promoter (<=1kb) | 4 | 108814295 | 108833614 | 19320 | 2 | ENSMUST00 | 0 | Btf3l4 | ENSMUSG00 | false | chr4 | 108832647 | 108835582 | 31.28813 | 116.44205 | 112.41405 | . | . | . | . | . | . |
| chr4 | 123404088 | 123404304 | 359 | Intron (ENSMUST00000097897.5 | 4 | 123397275 | 123407515 | 10241 | 2 | ENSMUST00 | 3211 | Macf1 | ENSMUSG00 | true | . | . | . | . | . | . | chr4 | 123397088 | 123411313 | 6.71434 | 4.80088 | 4.42637 |
| chr8 | 11379690 | 11379972 | 359 | Intron (ENSMUST00000033899.8 | 8 | 11404908 | 11421254 | 16347 | 1 | ENSMUST00 | -24936 | Col4a2 | ENSMUSG00 | false | . | . | . | . | . | . | . | . | . | . | . | . |
| chr17 | 5643347 | 5643602 | 359 | Intron (ENSMUST00000089185.4 | 17 | 5492600 | 5753891 | 261292 | 1 | ENSMUST00 | 150748 | Zdhhc14 | ENSMUSG00 | true | . | . | . | . | . | . | chr17 | 5642711 | 5644622 | 3.295 | 3.07932 | 1.64908 |
| chr19 | 56821744 | 56821983 | 359 | Promoter (<=1kb) | 19 | 56787481 | 56822190 | 34710 | 2 | ENSMUST00 | 207 | A630007B06 | ENSMUSG00 | false | chr19 | 56821099 | 56822749 | 44.61662 | 165.21439 | 159.37868 | . | . | . | . | . | . |
| chr8 | 40972697 | 40972925 | 359 | Intron (ENSMUST00000034004.9 | 8 | 40998770 | 41000205 | 1436 | 2 | ENSMUST00 | 27280 | Mtus1 | ENSMUSG00 | false | . | . | . | . | . | . | . | . | . | . | . | . |
| chr16 | 31111821 | 31112058 | 359 | Intron (ENSMUST00000058033.7 | 16 | 30955627 | 31081432 | 125806 | 2 | ENSMUST00 | -30390 | Xxylt1 | ENSMUSG00 | false | . | . | . | . | . | . | chr16 | 31111821 | 31112522 | 3.26288 | 3.15514 | 1.57925 |
| chr9 | 45092951 | 45093197 | 359 | Intron (ENSMUST00000050020.6 | 9 | 45079183 | 45108530 | 29348 | 1 | ENSMUST00 | 13769 | Amica1 | ENSMUSG00 | false | . | . | . | . | . | . | . | . | . | . | . | . |
| chr9 | 109816434 | 109816771 | 359 | Promoter (<=1kb) | 9 | 109816571 | 109826633 | 10063 | 1 | ENSMUST00 | 0 | Spink8 | ENSMUSG00 | false | . | . | . | . | . | . | . | . | . | . | . | . |
| chr3 | 51774920 | 51775156 | 359 | Intron (ENSMUST00000121440.2 | 3 | 51689437 | 51796535 | 107099 | 2 | ENSMUST00 | 21379 | Maml3 | ENSMUSG00 | true | . | . | . | . | . | . | chr3 | 51767223 | 51782456 | 5.75718 | 4.44357 | 3.63488 |
| chr17 | 28052315 | 28052675 | 359 | Exon (ENSMUST00000114842.2/ | 17 | 28066797 | 28068476 | 1680 | 2 | ENSMUST00 | 15801 | Tcp11 | ENSMUSG00 | false | . | . | . | . | . | . | chr17 | 28050256 | 28053615 | 6.74781 | 4.87761 | 4.47038 |
| chr2 | 61344867 | 61345468 | 359 | Distal Intergenic | 2 | 61280278 | 61280409 | 132 | 1 | ENSMUST00 | 64590 | Gm26422 | ENSMUSG00 | false | . | . | . | . | . | . | chr2 | 61344859 | 61347837 | 3.27854 | 3.11583 | 1.62388 |
| chr2 | 119873158 | 119873459 | 359 | Distal Intergenic | 2 | 119878069 | 119878403 | 335 | 2 | ENSMUST00 | 4944 | Gm13998 | ENSMUSG00 | false | chr2 | 119872141 | 119873656 | 30.17592 | 94.17345 | 90.73757 | chr2 | 119872041 | 119873826 | 3.32933 | 3.21221 | 1.62121 |
| chr2 | 148073705 | 148073945 | 359 | Promoter (1-2kb) | 2 | 148071346 | 148071839 | 494 | 2 | ENSMUST00 | -1867 | Gm14108 | ENSMUSG00 | false | . | . | . | . | . | . | chr2 | 148071881 | 148075123 | 5.12065 | 4.06479 | 3.12109 |
| chr4 | 101285449 | 101285675 | 359 | Promoter (1-2kb) | 4 | 101284221 | 101285203 | 983 | 1 | ENSMUST00 | 1229 | Gm12801 | ENSMUSG00 | false | . | . | . | . | . | . | chr4 | 101284816 | 101286271 | 4.26942 | 3.76266 | 2.45398 |
| chr16 | 11008436 | 11008771 | 359 | Promoter (<=1kb) | 16 | 11008898 | 11015186 | 6289 | 1 | ENSMUST00 | -127 | Gm4262 | ENSMUSG00 | false | chr16 | 11008077 | 11008765 | 9.02267 | 17.18543 | 14.89151 | chr16 | 11006018 | 11010119 | 5.87028 | 4.53825 | 3.72697 |
| chr11 | 83941951 | 83942225 | 358 | Promoter (<=1kb) | 11 | 83942062 | 83963088 | 21027 | 1 | ENSMUST00 | 0 | Ddx52 | ENSMUSG00 | false | chr11 | 83941557 | 83942838 | 17.78121 | 70.23115 | 67.23819 | chr11 | 83941465 | 83943418 | 4.05364 | 3.62735 | 2.26891 |
| chr11 | 72207409 | 72207681 | 358 | Promoter (<=1kb) | 11 | 72207547 | 72210015 | 2469 | 1 | ENSMUST00 | 0 | Txndc17 | ENSMUSG00 | false | chr11 | 72206093 | 72208424 | 25.84672 | 90.2271 | 86.88143 | . | . | . | . | . | . |
| chr13 | 68680250 | 68680715 | 358 | Intron (ENSMUST00000022013.7 | 13 | 68597439 | 68614231 | 16793 | 1 | ENSMUST00 | 82812 | 1700001L19 | ENSMUSG00 | false | . | . | . | . | . | . | chr13 | 68679955 | 68681780 | 4.52804 | 3.73206 | 2.6563 |
| chr16 | 30388588 | 30388979 | 358 | Promoter (<=1kb) | 16 | 30385893 | 30388837 | 2945 | 2 | ENSMUST00 | 0 | Atp13a3 | ENSMUSG00 | false | chr16 | 30387757 | 30389792 | 27.29656 | 83.33401 | 80.12321 | . | . | . | . | . | . |
| chr16 | 87495734 | 87496091 | 358 | Promoter (<=1kb) | 16 | 87483326 | 87495873 | 12548 | 2 | ENSMUST00 | 0 | Cct8 | ENSMUSG00 | false | chr16 | 87495038 | 87496706 | 37.34335 | 127.50397 | 123.10989 | . | . | . | . | . | . |
| chr17 | 25919618 | 25920654 | 358 | Promoter (<=1kb) | 17 | 25882114 | 25919642 | 37529 | 2 | ENSMUST00 | 0 | Rab40c | ENSMUSG00 | false | chr17 | 25918691 | 25919790 | 39.46206 | 129.05257 | 124.59962 | . | . | . | . | . | . |
| chr1 | 143952949 | 143953183 | 358 | Distal Intergenic | 1 | 144001883 | 144004032 | 2150 | 2 | ENSMUST00 | 50849 | Rgs2 | ENSMUSG00 | false | . | . | . | . | . | . | . | . | . | . | . | . |
| chr2 | 112265493 | 112266063 | 358 | Promoter (<=1kb) | 2 | 112265825 | 112359127 | 93303 | 1 | ENSMUST00 | 0 | Slc12a6 | ENSMUSG00 | false | chr2 | 112265293 | 112267278 | 33.41831 | 117.26761 | 113.21259 | chr2 | 112265124 | 112270238 | 4.03999 | 3.6413 | 2.20283 |
| chr18 | 38928696 | 38929114 | 358 | Promoter (<=1kb) | 18 | 38858568 | 38929157 | 70590 | 2 | ENSMUST00 | 43 | Fgf1 | ENSMUSG00 | false | . | . | . | . | . | . | . | . | . | . | . | . |
| chr2 | 20609340 | 20609562 | 358 | Intron (ENSMUST00000114610.2 | 2 | 20559935 | 20709635 | 149701 | 1 | ENSMUST00 | 49406 | Etl4 | ENSMUSG00 | true | . | . | . | . | . | . | chr2 | 20607513 | 20612109 | 4.92916 | 3.92302 | 2.99672 |
| chr7 | 145129985 | 145130221 | 358 | Distal Intergenic | 7 | 145253923 | 145283927 | 30005 | 2 | ENSMUST00 | 153706 | Tpcn2 | ENSMUSG00 | false | . | . | . | . | . | . | chr7 | 145130050 | 145130433 | 3.26881 | 3.19901 | 1.63162 |
| chr14 | 56587678 | 56587960 | 358 | Exon (ENSMUST00000161553.1/ | 14 | 56575619 | 56659794 | 84176 | 1 | ENSMUST00 | 12060 | Parp4 | ENSMUSG00 | true | . | . | . | . | . | . | chr14 | 56576330 | 56589138 | 4.82597 | 3.96005 | 2.86782 |
| chr1 | 106716881 | 106717112 | 358 | Promoter (2-3kb) | 1 | 106538178 | 106714274 | 176097 | 2 | ENSMUST00 | -2608 | Bcl2 | ENSMUSG00 | false | . | . | . | . | . | . | chr1 | 106714557 | 106720254 | 4.66173 | 3.83764 | 2.74504 |
| chr8 | 71476355 | 71476961 | 358 | 3' UTR | 8 | 71470411 | 71476098 | 5688 | 1 | ENSMUST00 | 5945 | Dda1 | ENSMUSG00 | false | chr8 | 71476070 | 71477063 | 18.70486 | 58.09692 | 55.27494 | chr8 | 71475391 | 71477623 | 4.08039 | 3.52882 | 2.2665 |
| chr15 | 95842659 | 95842895 | 358 | Intron (ENSMUST00000071874.6 | 15 | 95824459 | 95830072 | 5614 | 2 | ENSMUST00 | -12588 | Gm17546 | ENSMUSG00 | false | . | . | . | . | . | . | chr15 | 95842210 | 95848596 | 5.24923 | 4.16785 | 3.21628 |
| chr10 | 128344982 | 128345203 | 358 | Intron (ENSMUST00000005826.7 | 10 | 128349241 | 128349375 | 135 | 1 | ENSMUST00 | -4038 | Gm23182 | ENSMUSG00 | false | chr10 | 128345191 | 128345427 | 5.97392 | 9.35129 | 7.18948 | chr10 | 128338322 | 128347776 | 3.98634 | 3.51075 | 2.20546 |
| chr15 | 32239716 | 32239924 | 358 | Promoter (1-2kb) | 15 | 32241853 | 32241923 | 71 | 2 | ENSMUST00 | 1999 | Snord123 | ENSMUSG00 | false | . | . | . | . | . | . | chr15 | 32239025 | 32241508 | 3.67813 | 3.18535 | 1.95927 |
| chr8 | 114703580 | 114703793 | 358 | Intron (ENSMUST00000004756.8 | 8 | 114698439 | 114708006 | 9568 | 2 | ENSMUST00 | 4213 | Gm16118 | ENSMUSG00 | false | . | . | . | . | . | . | chr8 | 114702990 | 114704337 | 3.05811 | 3.01264 | 1.45817 |
| chr10 | 88383401 | 88383595 | 358 | Promoter (<=1kb) | 10 | 88379244 | 88383711 | 4468 | 2 | ENSMUST00 | 116 | Gm26546 | ENSMUSG00 | false | . | . | . | . | . | . | chr10 | 88382921 | 88385846 | 3.29417 | 3.10733 | 1.63885 |
| chr14 | 70191844 | 70192379 | 358 | Exon (ENSMUST00000022682.5/ | 14 | 70205324 | 70217659 | 12336 | 1 | ENSMUST00 | -12945 | Gm26908 | ENSMUSG00 | false | chr14 | 70191374 | 70192726 | 19.61815 | 75.94017 | 72.85583 | chr14 | 70190541 | 70199927 | 4.55845 | 3.79328 | 2.66728 |
| chr12 | 104904225 | 104904440 | 358 | Distal Intergenic | 12 | 104925501 | 104926142 | 642 | 2 | ENSMUST00 | 21702 | RP24-159O1 | ENSMUSG00 | false | . | . | . | . | . | . | chr12 | 104903887 | 104904951 | 5.71697 | 4.38529 | 3.64885 |
| chr12 | 79998512 | 79999079 | 358 | Distal Intergenic | 12 | 79963568 | 79963872 | 305 | 1 | ENSMUST00 | 34945 | RP23-179D3. | ENSMUSG00 | false | . | . | . | . | . | . | chr12 | 79993084 | 80001591 | 4.83553 | 3.97765 | 2.87564 |
| chr8 | 122664406 | 122664643 | 357 | Intron (ENSMUST00000127664.1 | 8 | 122629463 | 122678072 | 48610 | 2 | ENSMUST00 | 13429 | Cbfa2t3 | ENSMUSG00 | false | . | . | . | . | . | . | chr8 | 122648562 | 122676943 | 7.87697 | 5.30313 | 5.37575 |
| chr8 | 123805698 | 123806060 | 357 | Promoter (<=1kb) | 8 | 123805985 | 123835287 | 29303 | 1 | ENSMUST00 | 0 | Rab4a | ENSMUSG00 | false | chr8 | 123805532 | 123806981 | 23.93078 | 87.47927 | 84.18233 | chr8 | 123805549 | 123807934 | 4.16939 | 3.71669 | 2.32598 |
| chr1 | 84838546 | 84838973 | 357 | Promoter (<=1kb) | 1 | 84794231 | 84838731 | 44501 | 2 | ENSMUST00 | 0 | Trip12 | ENSMUSG00 | false | chr1 | 84837968 | 84840939 | 30.78991 | 105.27288 | 101.56436 | . | . | . | . | . | . |
| chr2 | 120970383 | 120970816 | 357 | Promoter (<=1kb) | 2 | 120860269 | 120970715 | 110447 | 2 | ENSMUST00 | 0 | Ubr1 | ENSMUSG00 | false | chr2 | 120969913 | 120971260 | 28.53699 | 95.19626 | 91.7378 | . | . | . | . | . | . |
| chr2 | 117090105 | 117090376 | 357 | Distal Intergenic | 2 | 117121374 | 117182279 | 60906 | 1 | ENSMUST00 | -30998 | Spred1 | ENSMUSG00 | false | . | . | . | . | . | . | chr2 | 117089644 | 117091017 | 4.32197 | 3.74483 | 2.47094 |
| chr4 | 76366717 | 76367009 | 357 | Intron (ENSMUST00000107289.3 | 4 | 75944958 | 76344243 | 399286 | 2 | ENSMUST00 | -22475 | Ptprd | ENSMUSG00 | true | . | . | . | . | . | . | chr4 | 76343707 | 76371478 | 4.49023 | 3.82181 | 2.58441 |
| chr7 | 80687692 | 80687897 | 357 | Promoter (<=1kb) | 7 | 80586627 | 80688877 | 102251 | 2 | ENSMUST00 | 980 | Crtc3 | ENSMUSG00 | false | chr7 | 80687626 | 80689456 | 30.90191 | 98.42355 | 94.89227 | chr7 | 80679765 | 80688014 | 5.47936 | 4.26578 | 3.4104 |
| chr7 | 100908036 | 100908361 | 357 | Intron (ENSMUST00000107032.1 | 7 | 100869746 | 100932161 | 62416 | 2 | ENSMUST00 | 23800 | Arhgef17 | ENSMUSG00 | false | chr7 | 100908338 | 100908675 | 4.42908 | 6.05774 | 3.97659 | chr7 | 100901870 | 100926579 | 7.14639 | 4.97053 | 4.78972 |
| chr6 | 88446413 | 88446639 | 357 | Promoter (<=1kb) | 6 | 88257334 | 88446539 | 189206 | 2 | ENSMUST00 | 0 | Eefsec | ENSMUSG00 | false | chr6 | 88445685 | 88446905 | 40.23456 | 136.98251 | 132.21786 | chr6 | 88445596 | 88446788 | 3.69432 | 3.41464 | 1.96803 |
| chr10 | 117089719 | 117089983 | 357 | Intron (ENSMUST00000020381.3 | 10 | 117050998 | 117063814 | 12817 | 2 | ENSMUST00 | -25906 | Cct2 | ENSMUSG00 | false | . | . | . | . | . | . | chr10 | 117086815 | 117095074 | 5.71886 | 4.41367 | 3.58491 |
| chr6 | 83439396 | 83439718 | 357 | Promoter (1-2kb) | 6 | 83362373 | 83441675 | 79303 | 2 | ENSMUST00 | 1957 | Tet3 | ENSMUSG00 | true | chr6 | 83438843 | 83440773 | 9.86256 | 23.41135 | 21.03137 | chr6 | 83421103 | 83440609 | 6.41331 | 4.64298 | 4.1723 |
| chr9 | 96158565 | 96158818 | 357 | Intron (ENSMUST00000085217.6 | 9 | 96170916 | 96173465 | 2550 | 1 | ENSMUST00 | -12098 | Gk5 | ENSMUSG00 | false | . | . | . | . | . | . | . | . | . | . | . | . |
| chrX | 137049484 | 137049705 | 357 | Promoter (<=1kb) | 20 | 137049594 | 137082503 | 32910 | 1 | ENSMUST00 | 0 | Fam199x | ENSMUSG00 | false | chrX | 137049118 | 137050290 | 21.13848 | 54.99947 | 52.22001 | . | . | . | . | . | . |
| chr7 | 143008646 | 143008921 | 357 | Promoter (<=1kb) | 7 | 142988969 | 143008033 | 19065 | 2 | ENSMUST00 | -614 | Tspan32os | ENSMUSG00 | false | . | . | . | . | . | . | chr7 | 143004528 | 143022688 | 7.71992 | 5.26439 | 5.23131 |
| chr8 | 61397172 | 61397559 | 357 | Distal Intergenic | 8 | 61364019 | 61364483 | 465 | 2 | ENSMUST00 | -32690 | Gm10015 | ENSMUSG00 | false | . | . | . | . | . | . | chr8 | 61396733 | 61397921 | 3.10283 | 3.05248 | 1.49905 |
| chr5 | 105776850 | 105777352 | 357 | Intron (ENSMUST00000060531.1 | 5 | 105797321 | 105797713 | 393 | 2 | ENSMUST00 | 20361 | Rps15a-ps5 | ENSMUSG00 | false | . | . | . | . | . | . | chr5 | 105776453 | 105783156 | 4.86662 | 3.80058 | 2.93089 |
| chr1 | 87879662 | 87880038 | 357 | Promoter (<=1kb) | 1 | 87880435 | 87915744 | 35310 | 1 | ENSMUST00 | -397 | Dgkd | ENSMUSG00 | false | . | . | . | . | . | . | chr1 | 87878000 | 87881940 | 4.76819 | 3.85677 | 2.8369 |
| chr13 | 60340176 | 60340462 | 357 | Distal Intergenic | 13 | 60293672 | 60293772 | 101 | 1 | ENSMUST00 | 46505 | Gm24999 | ENSMUSG00 | false | . | . | . | . | . | . | chr13 | 60338549 | 60344628 | 4.38308 | 3.79503 | 2.48565 |
| chr2 | 121308855 | 121309462 | 357 | 3' UTR | 2 | 121293703 | 121296008 | 2306 | 2 | ENSMUST00 | -12848 | Gm14978 | ENSMUSG00 | false | . | . | . | . | . | . | chr2 | 121307126 | 121310501 | 4.14149 | 3.44807 | 2.33126 |
| chr18 | 35847904 | 35848412 | 357 | Intron (ENSMUST00000060722.6 | 18 | 35830389 | 35831093 | 705 | 2 | ENSMUST00 | -16812 | RP24-326K1 | ENSMUSG00 | false | chr18 | 35847706 | 35848814 | 10.51363 | 23.99435 | 21.60407 | chr18 | 35831800 | 35857612 | 7.70883 | 5.15861 | 5.24645 |
| chr13 | 5604225 | 5604493 | 357 | Distal Intergenic | 13 | 5855508 | 5857712 | 2205 | 2 | ENSMUST00 | 253219 | RP23-28J6.1 | ENSMUSG00 | false | . | . | . | . | . | . | chr13 | 5602685 | 5605930 | 3.87241 | 3.47298 | 2.08213 |
| chr5 | 38441432 | 38441870 | 356 | Promoter (<=1kb) | 5 | 38415440 | 38440829 | 25390 | 2 | ENSMUST00 | -604 | Slc2a9 | ENSMUSG00 | false | . | . | . | . | . | . | chr5 | 38439809 | 38442320 | 4.45705 | 3.77489 | 2.59918 |
| chr5 | 135064020 | 135064307 | 356 | Promoter (<=1kb) | 5 | 135052959 | 135064078 | 11120 | 2 | ENSMUST00 | 0 | Wbscr22 | ENSMUSG00 | false | chr5 | 135063491 | 135065117 | 20.9917 | 88.60235 | 85.28741 | . | . | . | . | . | . |
| chr10 | 9900578 | 9901400 | 356 | Promoter (<=1kb) | 10 | 9755547 | 9900683 | 145137 | 2 | ENSMUST00 | 0 | Stxbp5 | ENSMUSG00 | false | chr10 | 9899546 | 9902121 | 26.03857 | 105.07202 | 101.37238 | . | . | . | . | . | . |
| chr11 | 8539953 | 8540367 | 356 | Intron (ENSMUST00000020695.7 | 11 | 8537539 | 8545351 | 7813 | 2 | ENSMUST00 | 4984 | Tns3 | ENSMUSG00 | false | . | . | . | . | . | . | chr11 | 8538859 | 8546017 | 6.35703 | 4.56721 | 4.16167 |
| chr11 | 102761402 | 102761767 | 356 | Promoter (<=1kb) | 11 | 102761439 | 102780262 | 18824 | 1 | ENSMUST00 | 0 | Adam11 | ENSMUSG00 | false | chr11 | 102761199 | 102762462 | 25.7338 | 71.40817 | 68.39619 | chr11 | 102761398 | 102763210 | 5.82924 | 4.34519 | 3.71591 |
| chr17 | 45933600 | 45933844 | 356 | Distal Intergenic | 17 | 46018712 | 46021372 | 2661 | 2 | ENSMUST00 | 87528 | Vegfa | ENSMUSG00 | true | . | . | . | . | . | . | chr17 | 45928605 | 45934413 | 4.63767 | 3.72884 | 2.75421 |
| chr2 | 77770400 | 77770636 | 356 | Intron (ENSMUST00000111831.2 | 2 | 77411100 | 77816816 | 405717 | 2 | ENSMUST00 | 46180 | Zfp385b | ENSMUSG00 | true | . | . | . | . | . | . | chr2 | 77767132 | 77782197 | 4.901 | 4.03491 | 2.92527 |
| chr5 | 77364782 | 77365057 | 356 | Intron (ENSMUST00000046746.6 | 5 | 77340368 | 77342936 | 2569 | 1 | ENSMUST00 | 24415 | Polr2b | ENSMUSG00 | false | . | . | . | . | . | . | chr5 | 77353137 | 77404361 | 6.51707 | 4.75542 | 4.25399 |
| chr9 | 63665954 | 63666227 | 356 | Promoter (<=1kb) | 9 | 63654771 | 63666547 | 11777 | 2 | ENSMUST00 | 320 | Smad3 | ENSMUSG00 | false | . | . | . | . | . | . | chr9 | 63662341 | 63669966 | 7.25169 | 5.00736 | 4.88788 |
| chr4 | 116734385 | 116734735 | 356 | Intron (ENSMUST00000124628.2 | 4 | 116741683 | 116804102 | 62420 | 1 | ENSMUST00 | -6948 | Tesk2 | ENSMUSG00 | false | . | . | . | . | . | . | chr4 | 116731176 | 116741002 | 4.98596 | 4.04488 | 2.98621 |
| chr13 | 43453306 | 43453520 | 356 | Intron (ENSMUST00000144326.2 | 13 | 43402673 | 43480973 | 78301 | 2 | ENSMUST00 | 27453 | Ranbp9 | ENSMUSG00 | false | . | . | . | . | . | . | chr13 | 43441880 | 43457436 | 4.46486 | 3.6925 | 2.595 |
| chr11 | 48816967 | 48817318 | 356 | Promoter (<=1kb) | 11 | 48809097 | 48816953 | 7857 | 2 | ENSMUST00 | -15 | Trim41 | ENSMUSG00 | false | chr11 | 48815920 | 48818558 | 26.65955 | 86.95457 | 83.67183 | . | . | . | . | . | . |
| chr12 | 102308440 | 102308712 | 356 | Intron (ENSMUST00000056950.8 | 12 | 102283641 | 102390855 | 107215 | 1 | ENSMUST00 | 24800 | Rin3 | ENSMUSG00 | false | . | . | . | . | . | . | chr12 | 102301484 | 102309684 | 6.64516 | 4.78421 | 4.36044 |
| chr1 | 90710673 | 90710914 | 356 | Distal Intergenic | 1 | 90675161 | 90679793 | 4633 | 1 | ENSMUST00 | 35513 | Gm9991 | ENSMUSG00 | false | . | . | . | . | . | . | . | . | . | . | . | . |
| chr2 | 25196648 | 25196884 | 356 | Promoter (<=1kb) | 2 | 25195057 | 25196759 | 1703 | 2 | ENSMUST00 | 0 | Tor4a | ENSMUSG00 | false | chr2 | 25196158 | 25196963 | 7.14186 | 14.46535 | 12.21151 | chr2 | 25193209 | 25197932 | 3.91699 | 3.59343 | 2.10252 |
| chr4 | 124569428 | 124569756 | 356 | Promoter (<=1kb) | 4 | 124569190 | 124574645 | 5456 | 1 | ENSMUST00 | 239 | 4933407E24 | ENSMUSG00 | false | . | . | . | . | . | . | chr4 | 124568013 | 124576752 | 4.65263 | 3.69297 | 2.75743 |
| chr2 | 163489917 | 163490103 | 356 | Promoter (2-3kb) | 2 | 163492318 | 163502612 | 10295 | 1 | ENSMUST00 | -2215 | R3hdml | ENSMUSG00 | false | . | . | . | . | . | . | chr2 | 163489306 | 163495730 | 4.41102 | 3.63186 | 2.55954 |
| chr4 | 115708753 | 115709283 | 356 | Promoter (1-2kb) | 4 | 115707251 | 115708303 | 1053 | 1 | ENSMUST00 | 1503 | Gm12849 | ENSMUSG00 | false | . | . | . | . | . | . | chr4 | 115707353 | 115710474 | 3.80958 | 3.35728 | 2.063 |
| chr9 | 67157753 | 67157964 | 356 | Distal Intergenic | 9 | 67104208 | 67104355 | 148 | 1 | ENSMUST00 | 53546 | Gm22145 | ENSMUSG00 | false | . | . | . | . | . | . | chr9 | 67156689 | 67159818 | 3.68956 | 3.47155 | 1.91284 |
| chr17 | 83916629 | 83916958 | 356 | Distal Intergenic | 17 | 83947588 | 83947696 | 109 | 1 | ENSMUST00 | -30630 | Gm23905 | ENSMUSG00 | false | . | . | . | . | . | . | chr17 | 83913091 | 83919286 | 6.62106 | 4.7113 | 4.34904 |
| chr4 | 40844217 | 40844476 | 356 | Intron (ENSMUST00000030121.7 | 4 | 40850056 | 40850138 | 83 | 2 | ENSMUST00 | 5662 | Mir5123 | ENSMUSG00 | false | . | . | . | . | . | . | chr4 | 40821931 | 40852722 | 6.59841 | 4.80932 | 4.32335 |
| chr9 | 21291121 | 21291709 | 356 | Promoter (<=1kb) | 9 | 21288463 | 21291209 | 2747 | 2 | ENSMUST00 | 0 | Cdkn2d | ENSMUSG00 | false | chr9 | 21290440 | 21291318 | 31.66066 | 102.77801 | 99.14294 | chr9 | 21285555 | 21291921 | 5.06285 | 4.15318 | 3.06441 |
| chr2 | 160739948 | 160740237 | 355 | Promoter (<=1kb) | 2 | 160740987 | 160762506 | 21520 | 1 | ENSMUST00 | -750 | Plcg1 | ENSMUSG00 | false | . | . | . | . | . | . | chr2 | 160732227 | 160746274 | 4.65715 | 3.84763 | 2.74112 |
| chr14 | 16364535 | 16365366 | 355 | Promoter (<=1kb) | 14 | 16365179 | 16430787 | 65609 | 1 | ENSMUST00 | 0 | Top2b | ENSMUSG00 | false | chr14 | 16363963 | 16366999 | 34.91417 | 125.28625 | 120.96531 | . | . | . | . | . | . |
| chr11 | 20039835 | 20040083 | 355 | Distal Intergenic | 11 | 20078686 | 20112867 | 34182 | 2 | ENSMUST00 | 72784 | Actr2 | ENSMUSG00 | false | . | . | . | . | . | . | chr11 | 20033774 | 20044254 | 5.74342 | 4.41024 | 3.62353 |
| chr11 | 115564077 | 115564583 | 355 | Promoter (<=1kb) | 11 | 115564434 | 115583985 | 19552 | 1 | ENSMUST00 | 0 | Nup85 | ENSMUSG00 | false | chr11 | 115563642 | 115565044 | 24.48636 | 83.88964 | 80.66612 | chr11 | 115563719 | 115565460 | 4.55412 | 3.92327 | 2.63013 |
| chr3 | 51202423 | 51202611 | 355 | Distal Intergenic | 3 | 51224447 | 51251644 | 27198 | 1 | ENSMUST00 | -21836 | Ccrn4l | ENSMUSG00 | false | . | . | . | . | . | . | chr3 | 51201033 | 51204192 | 6.45222 | 4.83018 | 4.17698 |
| chr2 | 132577782 | 132578371 | 355 | Promoter (<=1kb) | 2 | 132529082 | 132578178 | 49097 | 2 | ENSMUST00 | 0 | Gpcpd1 | ENSMUSG00 | true | chr2 | 132575235 | 132578510 | 31.25476 | 101.05123 | 97.45738 | . | . | . | . | . | . |
| chr7 | 29453349 | 29453697 | 355 | Intron (ENSMUST00000085809.5 | 7 | 29384256 | 29505452 | 121197 | 2 | ENSMUST00 | 51755 | Sipa1l3 | ENSMUSG00 | false | . | . | . | . | . | . | chr7 | 29451630 | 29455276 | 5.78646 | 4.41831 | 3.66885 |
| chr1 | 127205947 | 127206344 | 355 | Promoter (<=1kb) | 1 | 127205028 | 127398792 | 193765 | 1 | ENSMUST00 | 920 | Mgat5 | ENSMUSG00 | true | chr1 | 127204578 | 127207504 | 33.22886 | 104.51182 | 100.83223 | . | . | . | . | . | . |
| chr11 | 93968261 | 93968559 | 355 | Promoter (<=1kb) | 11 | 93956979 | 93968521 | 11543 | 2 | ENSMUST00 | 0 | Nme1 | ENSMUSG00 | false | chr11 | 93967612 | 93968449 | 24.84317 | 70.77231 | 67.77174 | chr11 | 93966360 | 93969999 | 4.89241 | 4.04593 | 2.89918 |
| chr1 | 57868620 | 57869009 | 355 | Intron (ENSMUST00000172287.2 | 1 | 57845571 | 57946965 | 101395 | 1 | ENSMUST00 | 23050 | Spats2l | ENSMUSG00 | false | . | . | . | . | . | . | chr1 | 57866940 | 57870483 | 4.22504 | 3.41377 | 2.40461 |
| chr4 | 32916028 | 32916370 | 355 | Intron (ENSMUST00000035719.5 | 4 | 32804035 | 32923455 | 119421 | 2 | ENSMUST00 | 7085 | Ankrd6 | ENSMUSG00 | true | . | . | . | . | . | . | chr4 | 32914595 | 32917196 | 6.22268 | 4.67704 | 4.06258 |
| chr9 | 37488828 | 37489334 | 355 | Promoter (<=1kb) | 9 | 37489321 | 37524148 | 34828 | 1 | ENSMUST00 | 0 | Msantd2 | ENSMUSG00 | false | chr9 | 37488179 | 37490348 | 44.60098 | 167.51549 | 161.60988 | . | . | . | . | . | . |
| chr8 | 82870939 | 82871259 | 355 | Intron (ENSMUST00000078525.5 | 8 | 82863356 | 83091271 | 227916 | 1 | ENSMUST00 | 7584 | Rnf150 | ENSMUSG00 | true | . | . | . | . | . | . | chr8 | 82865363 | 82877746 | 4.58463 | 3.79894 | 2.66584 |
| chr3 | 89430129 | 89430757 | 355 | Promoter (<=1kb) | 3 | 89430214 | 89432743 | 2530 | 1 | ENSMUST00 | 0 | Pygo2 | ENSMUSG00 | false | chr3 | 89429914 | 89431194 | 29.71419 | 101.30199 | 97.70005 | chr3 | 89429775 | 89432259 | 3.85074 | 3.28849 | 2.09586 |
| chr5 | 93136931 | 93137246 | 355 | Intron (ENSMUST00000074733.7 | 5 | 93093457 | 93174958 | 81502 | 1 | ENSMUST00 | 43475 | Sep-11 | ENSMUSG00 | false | . | . | . | . | . | . | chr5 | 93095018 | 93154711 | 6.59326 | 4.78158 | 4.31782 |
| chr12 | 86823971 | 86824347 | 355 | Distal Intergenic | 12 | 86809957 | 86810258 | 302 | 1 | ENSMUST00 | 14015 | Gm25888 | ENSMUSG00 | false | . | . | . | . | . | . | chr12 | 86820408 | 86827213 | 6.55613 | 4.76961 | 4.27793 |
| chr15 | 25680628 | 25680885 | 355 | Promoter (<=1kb) | 15 | 25681798 | 25682125 | 328 | 1 | ENSMUST00 | -913 | Gm2862 | ENSMUSG00 | false | . | . | . | . | . | . | chr15 | 25676232 | 25685645 | 5.10402 | 4.00285 | 3.13785 |
| chr15 | 57712408 | 57712661 | 355 | Intron (ENSMUST00000096430.5 | 15 | 57708114 | 57708927 | 814 | 2 | ENSMUST00 | -3482 | Gm16006 | ENSMUSG00 | false | . | . | . | . | . | . | chr15 | 57695806 | 57714739 | 5.09262 | 4.12618 | 3.0824 |
| chr12 | 22030691 | 22031006 | 355 | Distal Intergenic | 12 | 22040913 | 22041022 | 110 | 1 | ENSMUST00 | -9907 | Gm22908 | ENSMUSG00 | false | . | . | . | . | . | . | . | . | . | . | . | . |
| chrY | 90765351 | 90765840 | 355 | Distal Intergenic | 21 | 90754513 | 90754821 | 309 | 2 | ENSMUST00 | -10531 | Gm21860 | ENSMUSG00 | false | . | . | . | . | . | . | . | . | . | . | . | . |
| chr1 | 133863631 | 133863865 | 355 | Distal Intergenic | 1 | 133835189 | 133838372 | 3184 | 1 | ENSMUST00 | 28443 | RP23-18J18. | ENSMUSG00 | false | . | . | . | . | . | . | chr1 | 133863632 | 133864072 | 3.5081 | 3.23709 | 1.79449 |
| chr7 | 34202434 | 34202732 | 354 | Exon (ENSMUST00000038027.4/ | 7 | 34184499 | 34196661 | 12163 | 2 | ENSMUST00 | -5774 | Pdcd2l | ENSMUSG00 | false | . | . | . | . | . | . | chr7 | 34201107 | 34204044 | 5.47026 | 4.2242 | 3.40227 |
| chr11 | 8444789 | 8445054 | 354 | Exon (ENSMUST00000020695.7/ | 11 | 8451070 | 8468675 | 17606 | 2 | ENSMUST00 | 23621 | Tns3 | ENSMUSG00 | false | . | . | . | . | . | . | chr11 | 8438250 | 8474887 | 7.10839 | 4.95354 | 4.75045 |
| chr12 | 113013863 | 113014416 | 354 | Promoter (<=1kb) | 12 | 113014508 | 113074401 | 59894 | 1 | ENSMUST00 | -92 | Pacs2 | ENSMUSG00 | true | chr12 | 113013711 | 113014344 | 22.29138 | 63.70553 | 60.80725 | chr12 | 113012974 | 113013930 | 5.03806 | 3.9758 | 3.05708 |
| chr13 | 46742635 | 46742904 | 354 | Distal Intergenic | 13 | 46679905 | 46727940 | 48036 | 2 | ENSMUST00 | -14696 | Nup153 | ENSMUSG00 | false | . | . | . | . | . | . | chr13 | 46742337 | 46745736 | 6.13228 | 4.58343 | 3.93818 |
| chr17 | 79165247 | 79165476 | 354 | Distal Intergenic | 17 | 79051906 | 79090243 | 38338 | 1 | ENSMUST00 | 113342 | Qpct | ENSMUSG00 | false | . | . | . | . | . | . | chr17 | 79163950 | 79165559 | 3.16651 | 3.05439 | 1.54561 |
| chr4 | 155839514 | 155839925 | 354 | Promoter (<=1kb) | 4 | 155839680 | 155842130 | 2451 | 1 | ENSMUST00 | 0 | Mxra8 | ENSMUSG00 | false | chr4 | 155839481 | 155841295 | 13.67473 | 40.1468 | 37.55384 | chr4 | 155837961 | 155842164 | 5.80082 | 4.37605 | 3.67392 |
| chr5 | 30775046 | 30775306 | 354 | Promoter (2-3kb) | 5 | 30778130 | 30786934 | 8805 | 1 | ENSMUST00 | -2824 | Dpysl5 | ENSMUSG00 | false | . | . | . | . | . | . | chr5 | 30772715 | 30775704 | 5.52607 | 4.27296 | 3.48816 |
| chr9 | 111118052 | 111118668 | 354 | Promoter (<=1kb) | 9 | 111118111 | 111225668 | 107558 | 1 | ENSMUST00 | 0 | Lrrfip2 | ENSMUSG00 | false | chr9 | 111117612 | 111119605 | 33.04283 | 115.53907 | 111.53828 | . | . | . | . | . | . |
| chr2 | 103572442 | 103572739 | 354 | Intron (ENSMUST00000076212.3 | 2 | 103566310 | 103718423 | 152114 | 1 | ENSMUST00 | 6133 | Abtb2 | ENSMUSG00 | false | . | . | . | . | . | . | chr2 | 103567791 | 103586817 | 4.63826 | 3.84919 | 2.72485 |
| chr7 | 98156040 | 98156300 | 354 | Intron (ENSMUST00000040971.8 | 7 | 98145516 | 98162210 | 16695 | 2 | ENSMUST00 | 5910 | Capn5 | ENSMUSG00 | false | . | . | . | . | . | . | . | . | . | . | . | . |
| chr18 | 34854495 | 34854766 | 354 | Distal Intergenic | 18 | 34859823 | 34864984 | 5162 | 1 | ENSMUST00 | -5057 | Egr1 | ENSMUSG00 | false | . | . | . | . | . | . | chr18 | 34854282 | 34856996 | 5.66667 | 4.31088 | 3.55875 |
| chr4 | 14563120 | 14563415 | 354 | Intron (ENSMUST00000042221.8 | 4 | 14505289 | 14516369 | 11081 | 2 | ENSMUST00 | -46752 | Slc26a7 | ENSMUSG00 | false | . | . | . | . | . | . | chr4 | 14561835 | 14566059 | 4.65819 | 3.96026 | 2.71911 |
| chr1 | 21008911 | 21009276 | 354 | Intron (ENSMUST00000037998.5 | 1 | 21004313 | 21043656 | 39344 | 2 | ENSMUST00 | 34380 | Tram2 | ENSMUSG00 | false | . | . | . | . | . | . | chr1 | 21007403 | 21016311 | 6.38947 | 4.64139 | 4.16543 |
| chr14 | 31930456 | 31930856 | 354 | Distal Intergenic | 14 | 31906528 | 31929879 | 23352 | 1 | ENSMUST00 | 23929 | D830044D21 | ENSMUSG00 | false | . | . | . | . | . | . | chr14 | 31929401 | 31936104 | 4.99921 | 3.98187 | 3.04329 |
| chr4 | 12136070 | 12136313 | 354 | Distal Intergenic | 4 | 12140264 | 12144688 | 4425 | 1 | ENSMUST00 | -3951 | Rbm12b1 | ENSMUSG00 | false | chr4 | 12136061 | 12137120 | 14.31595 | 48.63377 | 45.93701 | chr4 | 12135546 | 12137003 | 3.57243 | 3.02605 | 1.88209 |
| chr7 | 80025652 | 80025979 | 354 | Promoter (<=1kb) | 7 | 80026228 | 80092751 | 66524 | 1 | ENSMUST00 | -249 | Zfp710 | ENSMUSG00 | false | chr7 | 80024655 | 80028372 | 33.9579 | 107.82887 | 104.06033 | chr7 | 80023473 | 80025921 | 3.47992 | 3.28125 | 1.77065 |
| chr18 | 38241052 | 38241240 | 354 | Intron (ENSMUST00000181871.1 | 18 | 38244828 | 38245382 | 555 | 1 | ENSMUST00 | -3588 | Gm10268 | ENSMUSG00 | false | . | . | . | . | . | . | chr18 | 38238635 | 38241845 | 4.62307 | 3.82198 | 2.71514 |
| chr11 | 120194815 | 120195060 | 354 | Distal Intergenic | 11 | 120187707 | 120189852 | 2146 | 2 | ENSMUST00 | -4964 | 2810410L24 | ENSMUSG00 | false | . | . | . | . | . | . | chr11 | 120194289 | 120195340 | 5.02824 | 4.14293 | 3.02384 |
| chr3 | 159885242 | 159885665 | 354 | Intron (ENSMUST00000068952.5 | 3 | 159916255 | 159916361 | 107 | 2 | ENSMUST00 | 30696 | Gm23941 | ENSMUSG00 | false | . | . | . | . | . | . | chr3 | 159882555 | 159887252 | 5.25564 | 3.99629 | 3.25229 |
| chr2 | 93892001 | 93892263 | 354 | Distal Intergenic | 2 | 93898042 | 93898133 | 92 | 1 | ENSMUST00 | -5779 | Gm23487 | ENSMUSG00 | false | . | . | . | . | . | . | . | . | . | . | . | . |
| chr11 | 63494235 | 63494573 | 354 | Distal Intergenic | 11 | 63413169 | 63417888 | 4720 | 1 | ENSMUST00 | 81067 | B130011K05 | ENSMUSG00 | false | . | . | . | . | . | . | chr11 | 63493865 | 63497633 | 3.63527 | 3.36618 | 1.87651 |
| chr13 | 5822678 | 5822932 | 354 | Distal Intergenic | 13 | 5855508 | 5857712 | 2205 | 2 | ENSMUST00 | 34780 | RP23-28J6.1 | ENSMUSG00 | false | . | . | . | . | . | . | chr13 | 5820517 | 5827722 | 4.31021 | 3.73598 | 2.44429 |
| chr7 | 99523027 | 99523371 | 353 | Distal Intergenic | 7 | 99535466 | 99606771 | 71306 | 1 | ENSMUST00 | -12095 | Arrb1 | ENSMUSG00 | false | . | . | . | . | . | . | chr7 | 99522439 | 99531882 | 5.37849 | 4.25246 | 3.31489 |
| chr12 | 25163447 | 25163759 | 353 | Distal Intergenic | 12 | 25093801 | 25096092 | 2292 | 2 | ENSMUST00 | -67356 | Id2 | ENSMUSG00 | false | . | . | . | . | . | . | chr12 | 25163654 | 25164191 | 3.06899 | 3.02632 | 1.42037 |
| chr19 | 46471657 | 46471998 | 353 | Intron (ENSMUST00000039922.7 | 19 | 46483097 | 46485982 | 2886 | 1 | ENSMUST00 | -11099 | Sufu | ENSMUSG00 | false | . | . | . | . | . | . | chr19 | 46468103 | 46486697 | 7.92449 | 5.26231 | 5.41796 |
| chr2 | 26315334 | 26315728 | 353 | Promoter (<=1kb) | 2 | 26315515 | 26348237 | 32723 | 1 | ENSMUST00 | 0 | Gpsm1 | ENSMUSG00 | false | chr2 | 26315294 | 26316499 | 32.27147 | 102.84008 | 99.19724 | . | . | . | . | . | . |
| chr2 | 93976408 | 93976830 | 353 | Intron (ENSMUST00000183110.1 | 2 | 93980034 | 93983575 | 3542 | 1 | ENSMUST00 | -3204 | 4921507L20 | ENSMUSG00 | false | chr2 | 93976270 | 93976934 | 11.2646 | 26.49366 | 24.07236 | chr2 | 93975707 | 93977715 | 4.07775 | 3.5062 | 2.28819 |
| chr3 | 9833490 | 9833856 | 353 | Promoter (<=1kb) | 3 | 9687479 | 9833679 | 146201 | 2 | ENSMUST00 | 0 | Pag1 | ENSMUSG00 | true | chr3 | 9830234 | 9834375 | 30.45735 | 121.03082 | 116.85291 | . | . | . | . | . | . |
| chr4 | 128654606 | 128654914 | 353 | Promoter (<=1kb) | 4 | 128654702 | 128752881 | 98180 | 1 | ENSMUST00 | 0 | Phc2 | ENSMUSG00 | false | chr4 | 128654558 | 128655651 | 18.02817 | 71.61055 | 68.59548 | . | . | . | . | . | . |
| chr5 | 107195696 | 107196097 | 353 | Intron (ENSMUST00000031224.9 | 5 | 107109281 | 107192367 | 83087 | 2 | ENSMUST00 | -3330 | Tgfbr3 | ENSMUSG00 | true | . | . | . | . | . | . | chr5 | 107189975 | 107206509 | 4.95135 | 4.04289 | 2.96667 |

| | | | | | | | | | | | | | | | | | | | | | | | | | | |
|---|---|---|---|---|---|---|---|---|---|---|---|---|---|---|---|---|---|---|---|---|---|---|---|---|---|---|
| chr6 | 137065520 | 137065733 | 353 | Intron (ENSMUST00000032347.5 | 6 | 137056099 | 137169685 | 113587 | 2 | ENSMUST00 | 103952 | Rerg | ENSMUSG0C | false | . | . | . | . | . | . | . | . | . | . | . | . |
| chr9 | 110624329 | 110624619 | 353 | Promoter (<=1kb) | 9 | 110621137 | 110624393 | 3257 | 2 | ENSMUST00 | 0 | Nradd | ENSMUSG0C | false | chr9 | 110624023 | 110625607 | 15.7376 | 52.33027 | 49.58585 | chr9 | 110623760 | 110626234 | 6.08485 | 4.59413 | 3.88504 |
| chr1 | 59900726 | 59900992 | 353 | Distal Intergenic | 1 | 59913006 | 59985346 | 72341 | 1 | ENSMUST00 | -12014 | Fam117b | ENSMUSG0C | true | . | . | . | . | . | . | chr1 | 59899114 | 59902971 | 4.43258 | 3.77456 | 2.55304 |
| chr7 | 49175701 | 49175951 | 353 | Intron (ENSMUST00000183659.2 | 7 | 49246189 | 49409643 | 163455 | 1 | ENSMUST00 | -70238 | Nav2 | ENSMUSG0C | false | . | . | . | . | . | . | chr7 | 49173121 | 49176990 | 5.11703 | 4.10573 | 3.09981 |
| chr4 | 62360371 | 62360662 | 353 | Promoter (<=1kb) | 4 | 62300342 | 62360470 | 60129 | 2 | ENSMUST00 | 0 | Fkbp15 | ENSMUSG0C | false | chr4 | 62359649 | 62361552 | 36.67561 | 120.95901 | 116.78313 | . | . | . | . | . | . |
| chr6 | 118479144 | 118479635 | 353 | Promoter (<=1kb) | 6 | 118461950 | 118479320 | 17371 | 2 | ENSMUST00 | 0 | Zfp9 | ENSMUSG0C | false | chr6 | 118478769 | 118479810 | 29.86959 | 86.83979 | 83.56236 | chr6 | 118478187 | 118479699 | 3.20611 | 3.15748 | 1.51599 |
| chr19 | 6061040 | 6061295 | 353 | Promoter (<=1kb) | 19 | 6061192 | 6062472 | 1281 | 1 | ENSMUST00 | 0 | Znhit2 | ENSMUSG0C | false | chr19 | 6060656 | 6062281 | 37.16048 | 123.94915 | 119.68073 | chr19 | 6060497 | 6063005 | 7.27297 | 5.12921 | 4.87048 |
| chr11 | 45670249 | 45670803 | 353 | Distal Intergenic | 11 | 45639383 | 45639514 | 132 | 1 | ENSMUST00 | 30867 | Gm22284 | ENSMUSG0C | false | . | . | . | . | . | . | . | . | . | . | . | . |
| chr11 | 54744248 | 54744609 | 353 | Promoter (2-3kb) | 11 | 54746357 | 54746824 | 468 | 2 | ENSMUST00 | 2215 | Gm12229 | ENSMUSG0C | false | . | . | . | . | . | . | chr11 | 54722256 | 54750513 | 5.8032 | 4.39 | 3.67574 |
| chr2 | 115385357 | 115385680 | 353 | Distal Intergenic | 2 | 115344385 | 115346968 | 2584 | 1 | ENSMUST00 | 40973 | Gm13976 | ENSMUSG0C | false | . | . | . | . | . | . | chr2 | 115384975 | 115386138 | 3.69404 | 3.37423 | 1.93066 |
| chr2 | 45061597 | 45061949 | 353 | Intron (ENSMUST00000068415.5 | 2 | 45023100 | 45023177 | 78 | 2 | ENSMUST00 | -38421 | Mir5129 | ENSMUSG0C | false | . | . | . | . | . | . | chr2 | 45060607 | 45073593 | 4.65688 | 3.82601 | 2.75469 |
| chr8 | 68207202 | 68207430 | 353 | Distal Intergenic | 8 | 68218173 | 68228413 | 10241 | 2 | ENSMUST00 | 20983 | Gm26659 | ENSMUSG0C | false | . | . | . | . | . | . | . | . | . | . | . | . |
| chr13 | 78697938 | 78698209 | 353 | Distal Intergenic | 13 | 78372987 | 78379134 | 6148 | 2 | ENSMUST00 | -318805 | Gm6317 | ENSMUSG0C | false | . | . | . | . | . | . | chr13 | 78697933 | 78698236 | 3.75539 | 2.93748 | 2.02299 |
| chr11 | 97315481 | 97315938 | 352 | Promoter (<=1kb) | 11 | 97315716 | 97329920 | 14205 | 1 | ENSMUST00 | 0 | Mrpl45 | ENSMUSG0C | false | chr11 | 97315303 | 97315622 | 8.68527 | 17.11081 | 14.8173 | chr11 | 97315147 | 97316788 | 4.37811 | 3.75658 | 2.48551 |
| chr19 | 18758963 | 18759271 | 352 | Intron (ENSMUST00000040489.7 | 19 | 18749983 | 18892510 | 142528 | 1 | ENSMUST00 | 8981 | Trpm6 | ENSMUSG0C | false | . | . | . | . | . | . | chr19 | 18758531 | 18759256 | 3.07584 | 3.078 | 1.4927 |
| chr9 | 109097945 | 109098263 | 352 | Promoter (<=1kb) | 9 | 109093844 | 109097796 | 3953 | 2 | ENSMUST00 | -150 | Gm7628 | ENSMUSG0C | false | chr9 | 109094798 | 109099441 | 38.03245 | 121.19012 | 117.00504 | chr9 | 109096591 | 109107997 | 6.47539 | 4.76075 | 4.21763 |
| chr7 | 116442981 | 116443559 | 352 | Promoter (<=1kb) | 7 | 116337276 | 116443458 | 106183 | 2 | ENSMUST00 | 0 | Pik3c2a | ENSMUSG0C | false | chr7 | 116442665 | 116444451 | 32.27911 | 106.92835 | 103.17982 | chr7 | 116439880 | 116445274 | 4.66141 | 3.96823 | 2.71734 |
| chr13 | 68578002 | 68578242 | 352 | Intron (ENSMUST00000045827.4 | 13 | 68560780 | 68582121 | 21342 | 2 | ENSMUST00 | 3879 | Mtrr | ENSMUSG0C | false | . | . | . | . | . | . | . | . | . | . | . | . |
| chr5 | 30863964 | 30864176 | 352 | Intron (ENSMUST00000031058.9 | 5 | 30869579 | 30879191 | 9613 | 1 | ENSMUST00 | -5403 | Tmem214 | ENSMUSG0C | false | . | . | . | . | . | . | chr5 | 30862900 | 30864585 | 5.19143 | 3.88299 | 3.19532 |
| chrX | 12087401 | 12087749 | 352 | Intron (ENSMUST00000043441.7 | 20 | 12036740 | 12080553 | 43814 | 2 | ENSMUST00 | -6849 | Bcor | ENSMUSG0C | false | chrX | 12087071 | 12088799 | 10.20265 | 24.60154 | 22.20526 | . | . | . | . | . | . |
| chr17 | 31386042 | 31386304 | 352 | Promoter (<=1kb) | 17 | 31386234 | 31476310 | 90077 | 1 | ENSMUST00 | 0 | Pde9a | ENSMUSG0C | false | chr17 | 31386037 | 31386819 | 14.30691 | 45.69287 | 43.03132 | chr17 | 31384502 | 31386906 | 4.48898 | 3.77971 | 2.5884 |
| chr1 | 93235569 | 93235856 | 352 | Promoter (<=1kb) | 1 | 93235841 | 93301065 | 65225 | 1 | ENSMUST00 | 0 | Sned1 | ENSMUSG0C | false | chr1 | 93235486 | 93236791 | 34.20045 | 107.22294 | 103.46436 | chr1 | 93235148 | 93236858 | 3.63186 | 3.39824 | 1.8834 |
| chr15 | 36403846 | 36404083 | 352 | Distal Intergenic | 15 | 36477668 | 36496791 | 19124 | 2 | ENSMUST00 | 92708 | Ankrd46 | ENSMUSG0C | false | . | . | . | . | . | . | . | . | . | . | . | . |
| chr16 | 34352682 | 34353008 | 352 | Intron (ENSMUST00000076810.6 | 16 | 34357268 | 34361799 | 4532 | 2 | ENSMUST00 | 8791 | Kalrn | ENSMUSG0C | false | . | . | . | . | . | . | chr16 | 34352175 | 34353256 | 3.90072 | 3.42206 | 2.13269 |
| chr17 | 47295777 | 47296160 | 352 | Promoter (1-2kb) | 17 | 47294069 | 47359463 | 65395 | 1 | ENSMUST00 | 1709 | Trerf1 | ENSMUSG0C | false | . | . | . | . | . | . | . | . | . | . | . | . |
| chr12 | 77626906 | 77627098 | 352 | Distal Intergenic | 12 | 77814680 | 77814809 | 130 | 2 | ENSMUST00 | 187711 | Gm24070 | ENSMUSG0C | false | . | . | . | . | . | . | chr12 | 77626321 | 77627222 | 3.96584 | 3.46542 | 2.17693 |
| chr14 | 32827234 | 32827539 | 352 | Distal Intergenic | 14 | 32833962 | 32836789 | 2828 | 1 | ENSMUST00 | -6423 | Fam170b | ENSMUSG0C | false | . | . | . | . | . | . | chr14 | 32821466 | 32828973 | 8.34134 | 5.48292 | 5.74355 |
| chr2 | 75618809 | 75619119 | 352 | Distal Intergenic | 2 | 75632638 | 75633382 | 745 | 2 | ENSMUST00 | 14263 | Gm13655 | ENSMUSG0C | false | . | . | . | . | . | . | chr2 | 75617759 | 75619855 | 4.69454 | 3.92942 | 2.80142 |
| chr8 | 128466983 | 128467355 | 352 | Intron (ENSMUST00000026917.8 | 8 | 128359241 | 128359320 | 80 | 1 | ENSMUST00 | 107743 | Mir1903 | ENSMUSG0C | false | . | . | . | . | . | . | chr8 | 128463254 | 128470568 | 3.75612 | 3.2535 | 2.02615 |
| chr16 | 16542994 | 16543246 | 352 | Promoter (<=1kb) | 16 | 16542323 | 16542742 | 420 | 1 | ENSMUST00 | 672 | Gm16139 | ENSMUSG0C | false | . | . | . | . | . | . | chr16 | 16542168 | 16544863 | 5.65278 | 4.37091 | 3.5895 |
| chr10 | 61759964 | 61760234 | 352 | Intron (ENSMUST00000020283.4 | 10 | 61774737 | 61774821 | 85 | 1 | ENSMUST00 | -14503 | Mir5108 | ENSMUSG0C | false | . | . | . | . | . | . | chr10 | 61756253 | 61760614 | 5.95373 | 4.26924 | 3.83028 |
| chr11 | 83291555 | 83291875 | 352 | Promoter (<=1kb) | 11 | 83291699 | 83294631 | 2933 | 1 | ENSMUST00 | 0 | AA465934 | ENSMUSG0C | false | chr11 | 83291080 | 83292571 | 25.68869 | 73.13053 | 70.09039 | . | . | . | . | . | . |
| chr17 | 66411593 | 66411876 | 352 | Promoter (2-3kb) | 17 | 66413631 | 66413984 | 354 | 2 | ENSMUST00 | 2108 | Gm4705 | ENSMUSG0C | false | . | . | . | . | . | . | chr17 | 66401889 | 66429482 | 6.16935 | 4.5489 | 3.97689 |
| chr17 | 8562247 | 8562488 | 352 | Intron (ENSMUST00000115715.2 | 17 | 8566399 | 8566869 | 471 | 2 | ENSMUST00 | 4381 | Gm17087 | ENSMUSG0C | false | . | . | . | . | . | . | chr17 | 8561715 | 8563121 | 4.00211 | 3.49148 | 2.23106 |
| chrX | 13262191 | 13262427 | 352 | Distal Intergenic | 20 | 13234684 | 13247736 | 13053 | 2 | ENSMUST00 | -14456 | RP23-158L1 | ENSMUSG0C | false | . | . | . | . | . | . | . | . | . | . | . | . |
| chr5 | 9627342 | 9627603 | 351 | Intron (ENSMUST00000004076.3 | 5 | 9485236 | 9725352 | 240117 | 2 | ENSMUST00 | 97749 | Grm3 | ENSMUSG0C | false | . | . | . | . | . | . | chr5 | 9627090 | 9628689 | 4.8321 | 4.04814 | 2.87533 |
| chr3 | 87985459 | 87985855 | 351 | Downstream (1-2kb) | 3 | 87971129 | 87980451 | 9323 | 1 | ENSMUST00 | 14331 | Nes | ENSMUSG0C | false | . | . | . | . | . | . | chr3 | 87973300 | 87988519 | 8.41459 | 5.52768 | 5.81173 |
| chr10 | 80260810 | 80261174 | 351 | Promoter (<=1kb) | 10 | 80258151 | 80261012 | 2862 | 2 | ENSMUST00 | 0 | Gamt | ENSMUSG0C | false | chr10 | 80260517 | 80262231 | 19.37025 | 50.37409 | 47.65327 | chr10 | 80259637 | 80275268 | 5.67236 | 4.28519 | 3.5841 |
| chr15 | 11862282 | 11862617 | 351 | Intron (ENSMUST00000066529.3 | 15 | 11839896 | 11905674 | 65779 | 2 | ENSMUST00 | 43057 | Npr3 | ENSMUSG0C | true | . | . | . | . | . | . | chr15 | 11851382 | 11874908 | 5.13432 | 4.1471 | 3.11197 |
| chr15 | 32244632 | 32244925 | 351 | Promoter (<=1kb) | 15 | 32244813 | 32696341 | 451529 | 1 | ENSMUST00 | 0 | Sema5a | ENSMUSG0C | true | chr15 | 32243266 | 32247730 | 21.73259 | 92.96881 | 89.56307 | . | . | . | . | . | . |
| chr15 | 25603480 | 25603734 | 351 | Distal Intergenic | 15 | 25622525 | 25813673 | 191149 | 1 | ENSMUST00 | -18791 | Myo10 | ENSMUSG0C | false | . | . | . | . | . | . | chr15 | 25603355 | 25603821 | 3.28645 | 3.21384 | 1.59669 |
| chr9 | 78045294 | 78045532 | 351 | Distal Intergenic | 9 | 78051958 | 78065624 | 13667 | 1 | ENSMUST00 | -6426 | Gcm1 | ENSMUSG0C | false | . | . | . | . | . | . | chr9 | 78044561 | 78047289 | 4.80808 | 4.02519 | 2.8599 |
| chr17 | 87270604 | 87270856 | 351 | Distal Intergenic | 17 | 87254658 | 87265935 | 11278 | 2 | ENSMUST00 | -4670 | Mcfd2 | ENSMUSG0C | false | . | . | . | . | . | . | chr17 | 87270123 | 87271820 | 4.92681 | 3.96377 | 2.98632 |
| chr8 | 127157236 | 127157488 | 351 | Intron (ENSMUST00000162309.2 | 8 | 127073240 | 127426684 | 353445 | 1 | ENSMUST00 | 83997 | Pard3 | ENSMUSG0C | false | . | . | . | . | . | . | chr8 | 127156957 | 127158191 | 4.62467 | 3.77391 | 2.7289 |
| chr5 | 34949223 | 34949775 | 351 | Promoter (<=1kb) | 5 | 34949445 | 35033595 | 84151 | 1 | ENSMUST00 | 0 | Rgs12 | ENSMUSG0C | false | chr5 | 34948690 | 34949989 | 28.1615 | 93.53628 | 90.11485 | chr5 | 34948619 | 34949956 | 2.89881 | 2.84459 | 1.29834 |
| chr8 | 13105376 | 13105889 | 351 | Promoter (<=1kb) | 8 | 13105621 | 13147940 | 42320 | 1 | ENSMUST00 | 0 | Cul4a | ENSMUSG0C | false | chr8 | 13104180 | 13106753 | 40.01227 | 134.38658 | 129.72945 | . | . | . | . | . | . |
| chr10 | 117071126 | 117071541 | 351 | 3' UTR | 10 | 117050998 | 117063814 | 12817 | 2 | ENSMUST00 | -7313 | Cct2 | ENSMUSG0C | false | . | . | . | . | . | . | chr10 | 117070579 | 117074194 | 4.8326 | 3.84927 | 2.90208 |
| chr1 | 89088170 | 89088449 | 351 | Intron (ENSMUST00000066279.5 | 1 | 89070453 | 89112860 | 42408 | 1 | ENSMUST00 | 17718 | Sh3bp4 | ENSMUSG0C | false | . | . | . | . | . | . | chr1 | 89071619 | 89098621 | 6.06397 | 4.51902 | 3.881 |
| chr8 | 47243496 | 47244033 | 351 | Intron (ENSMUST00000079195.5 | 8 | 47180048 | 47289402 | 109355 | 2 | ENSMUST00 | 45369 | Stox2 | ENSMUSG0C | false | chr8 | 47242647 | 47244577 | 14.20671 | 34.84183 | 32.31654 | chr8 | 47241723 | 47245095 | 3.45087 | 3.29689 | 1.72654 |
| chr6 | 145210844 | 145211241 | 351 | Promoter (<=1kb) | 6 | 145211134 | 145215791 | 4658 | 1 | ENSMUST00 | 0 | Lyrm5 | ENSMUSG0C | false | chr6 | 145210438 | 145210938 | 10.58711 | 29.50236 | 27.04178 | . | . | . | . | . | . |
| chr3 | 88592316 | 88592812 | 351 | Distal Intergenic | 3 | 88579671 | 88588397 | 8727 | 1 | ENSMUST00 | 12646 | Ssr2 | ENSMUSG0C | false | chr3 | 88592125 | 88592728 | 11.16317 | 23.37655 | 20.99793 | chr3 | 88591756 | 88592933 | 5.49561 | 4.15039 | 3.45522 |
| chr14 | 52019688 | 52020255 | 351 | Promoter (<=1kb) | 14 | 52006086 | 52019713 | 13628 | 2 | ENSMUST00 | 0 | Zfp219 | ENSMUSG0C | false | chr14 | 52019298 | 52021224 | 33.10969 | 106.34647 | 102.61562 | . | . | . | . | . | . |
| chr16 | 49933137 | 49933340 | 351 | Distal Intergenic | 16 | 49867968 | 49911081 | 43114 | 1 | ENSMUST00 | 65170 | Cd47 | ENSMUSG0C | false | . | . | . | . | . | . | chr16 | 49932887 | 49933327 | 4.09604 | 3.6726 | 2.28768 |
| chr5 | 93104881 | 93105209 | 351 | Intron (ENSMUST00000074733.7 | 5 | 93093457 | 93174958 | 81502 | 1 | ENSMUST00 | 11425 | Sep-11 | ENSMUSG0C | false | . | . | . | . | . | . | chr5 | 93095018 | 93154711 | 6.59326 | 4.78158 | 4.31782 |
| chr6 | 51880799 | 51881017 | 351 | Intron (ENSMUST00000078214.5 | 6 | 51859166 | 52012549 | 153384 | 2 | ENSMUST00 | 131532 | Skap2 | ENSMUSG0C | false | . | . | . | . | . | . | chr6 | 51880315 | 51881358 | 3.91924 | 3.37792 | 2.15237 |
| chr3 | 27608384 | 27608584 | 351 | Intron (ENSMUST00000046157.4 | 3 | 27584900 | 27584995 | 96 | 2 | ENSMUST00 | -23390 | Mir3092 | ENSMUSG0C | false | . | . | . | . | . | . | chr3 | 27606689 | 27609428 | 6.03495 | 4.43485 | 3.9035 |
| chr2 | 52958932 | 52959501 | 351 | Intron (ENSMUST00000090952.5 | 2 | 52859723 | 52863516 | 3794 | 2 | ENSMUST00 | -95417 | Gm26603 | ENSMUSG0C | false | . | . | . | . | . | . | chr2 | 52957581 | 52964204 | 5.57716 | 4.2199 | 3.5098 |
| chr7 | 44378117 | 44378551 | 350 | Intron (ENSMUST00000150034.1 | 7 | 44360043 | 44375030 | 14988 | 2 | ENSMUST00 | -3088 | 1700008O03 | ENSMUSG0C | false | . | . | . | . | . | . | chr7 | 44377155 | 44379053 | 5.30218 | 4.24896 | 3.24387 |
| chr11 | 76461326 | 76461611 | 350 | Exon (ENSMUST00000094012.5/ | 11 | 76454544 | 76468298 | 13755 | 2 | ENSMUST00 | 6687 | Abr | ENSMUSG0C | false | . | . | . | . | . | . | chr11 | 76460441 | 76470304 | 6.56354 | 4.63779 | 4.31962 |
| chr10 | 86368859 | 86369131 | 350 | Intron (ENSMUST00000120638.2 | 10 | 86302854 | 86343981 | 41128 | 1 | ENSMUST00 | 66006 | Timp3 | ENSMUSG0C | false | . | . | . | . | . | . | chr10 | 86368821 | 86369579 | 4.54567 | 3.85505 | 2.67231 |
| chr11 | 9011050 | 9011384 | 350 | Promoter (<=1kb) | 11 | 8993137 | 9011160 | 18024 | 2 | ENSMUST00 | 0 | Hus1 | ENSMUSG0C | false | chr11 | 9010305 | 9011731 | 27.39171 | 92.13462 | 88.74321 | . | . | . | . | . | . |
| chr16 | 72808092 | 72808318 | 350 | Intron (ENSMUST00000023600.7 | 16 | 72663149 | 73046095 | 382947 | 1 | ENSMUST00 | 144944 | Robo1 | ENSMUSG0C | false | chr16 | 72807879 | 72808982 | 8.01187 | 19.41161 | 17.08515 | . | . | . | . | . | . |
| chr3 | 99940802 | 99941017 | 350 | Intron (ENSMUST00000127518.1 | 3 | 99885417 | 100143322 | 257906 | 1 | ENSMUST00 | 55386 | Spag17 | ENSMUSG0C | false | . | . | . | . | . | . | chr3 | 99940629 | 99941219 | 3.20758 | 3.18237 | 1.57743 |
| chr5 | 134688420 | 134689175 | 350 | Promoter (<=1kb) | 5 | 134656039 | 134688590 | 32552 | 2 | ENSMUST00 | 0 | Limk1 | ENSMUSG0C | false | chr5 | 134687933 | 134689442 | 25.1466 | 74.00616 | 70.95453 | . | . | . | . | . | . |
| chr6 | 87838590 | 87838999 | 350 | Promoter (<=1kb) | 6 | 87818447 | 87838759 | 20313 | 2 | ENSMUST00 | 0 | Isy1 | ENSMUSG0C | false | chr6 | 87837982 | 87839537 | 33.28429 | 109.73968 | 105.91746 | . | . | . | . | . | . |
| chr2 | 103706287 | 103706599 | 350 | Intron (ENSMUST00000076212.3 | 2 | 103715035 | 103717217 | 2183 | 1 | ENSMUST00 | -8436 | Abtb2 | ENSMUSG0C | false | . | . | . | . | . | . | chr2 | 103705305 | 103708147 | 6.9412 | 4.98849 | 4.63796 |
| chr10 | 111237366 | 111237621 | 350 | Intron (ENSMUST00000095310.1 | 10 | 111298679 | 111301736 | 3058 | 1 | ENSMUST00 | -61058 | Bbs10 | ENSMUSG0C | false | . | . | . | . | . | . | chr10 | 111232705 | 111239147 | 4.92603 | 3.95685 | 2.97209 |
| chr15 | 55072096 | 55072314 | 350 | Promoter (<=1kb) | 15 | 55015129 | 55072152 | 57024 | 2 | ENSMUST00 | 0 | Taf2 | ENSMUSG0C | false | chr15 | 55071338 | 55072780 | 41.2653 | 138.78937 | 133.96024 | chr15 | 55071100 | 55072838 | 4.82447 | 4.06953 | 2.85391 |
| chr18 | 34840451 | 34840736 | 350 | Promoter (<=1kb) | 18 | 34840589 | 34847463 | 6875 | 1 | ENSMUST00 | 0 | Reep2 | ENSMUSG0C | false | chr18 | 34840320 | 34841334 | 24.35436 | 72.61947 | 69.58835 | chr18 | 34839709 | 34842347 | 5.23259 | 4.05705 | 3.23734 |
| chr5 | 142405329 | 142405825 | 350 | Intron (ENSMUST00000072837.5 | 5 | 142401497 | 142462014 | 60518 | 1 | ENSMUST00 | 3833 | Foxk1 | ENSMUSG0C | false | . | . | . | . | . | . | chr5 | 142402291 | 142422557 | 5.92639 | 4.38004 | 3.78969 |
| chr14 | 18893386 | 18893999 | 350 | Promoter (<=1kb) | 14 | 18573577 | 18893776 | 320200 | 2 | ENSMUST00 | 0 | Ube2e2 | ENSMUSG0C | false | chr14 | 18892782 | 18894436 | 18.02173 | 47.27326 | 44.5908 | . | . | . | . | . | . |
| chr3 | 108562104 | 108562644 | 350 | Promoter (<=1kb) | 3 | 108556425 | 108562170 | 5746 | 2 | ENSMUST00 | 0 | Tmem167b | ENSMUSG0C | false | chr3 | 108561517 | 108563451 | 35.20524 | 115.20684 | 111.21615 | . | . | . | . | . | . |
| chr2 | 75208799 | 75209319 | 350 | Intron (ENSMUST00000186411.1 | 2 | 75192277 | 75192759 | 483 | 1 | ENSMUST00 | 16523 | Gm13653 | ENSMUSG0C | false | chr2 | 75208826 | 75209471 | 9.64628 | 19.2185 | 16.89346 | chr2 | 75208472 | 75209959 | 4.85439 | 3.9671 | 2.93705 |
| chr15 | 75841866 | 75842095 | 350 | Promoter (<=1kb) | 15 | 75754432 | 75841915 | 87484 | 2 | ENSMUST00 | 0 | Zc3h3 | ENSMUSG0C | false | chr15 | 75841123 | 75842571 | 35.56825 | 117.95176 | 113.87694 | . | . | . | . | . | . |
| chr16 | 32547759 | 32548065 | 350 | Intron (ENSMUST00000160832.1 | 16 | 32591301 | 32591909 | 609 | 2 | ENSMUST00 | 43844 | Gm5405 | ENSMUSG0C | false | . | . | . | . | . | . | chr16 | 32547175 | 32548323 | 4.71992 | 3.88984 | 2.80671 |
| chr12 | 108269153 | 108269443 | 350 | Intron (ENSMUST00000136175.1 | 12 | 108206345 | 108275417 | 69073 | 2 | ENSMUST00 | 5974 | Ccdc85c | ENSMUSG0C | false | . | . | . | . | . | . | chr12 | 108233398 | 108274893 | 6.76084 | 4.78759 | 4.47033 |
| chr17 | 78521249 | 78521511 | 350 | Intron (ENSMUST00000024880.9 | 17 | 78533903 | 78534013 | 111 | 2 | ENSMUST00 | 12502 | Gm22562 | ENSMUSG0C | false | . | . | . | . | . | . | chr17 | 78519699 | 78522529 | 3.32464 | 3.20313 | 1.64858 |
| chr7 | 135597930 | 135598212 | 350 | Intron (ENSMUST00000073961.6 | 7 | 135566162 | 135566282 | 121 | 1 | ENSMUST00 | 31769 | Mir1962 | ENSMUSG0C | false | . | . | . | . | . | . | . | . | . | . | . | . |
| chr2 | 172901874 | 172902093 | 350 | Intron (ENSMUST00000009143.7 | 2 | 172864264 | 172864341 | 78 | 2 | ENSMUST00 | -37534 | Gm22773 | ENSMUSG0C | false | chr2 | 172901585 | 172901973 | 8.27158 | 15.0213 | 12.75664 | chr2 | 172899319 | 172903119 | 5.23548 | 3.97477 | 3.23651 |
| chr11 | 79002674 | 79002914 | 349 | Distal Intergenic | 11 | 78962974 | 78984946 | 21973 | 2 | ENSMUST00 | -17729 | Lgals9 | ENSMUSG0C | false | . | . | . | . | . | . | chr11 | 79002413 | 79004093 | 5.26006 | 4.15271 | 3.22665 |
| chr18 | 36735016 | 36735288 | 349 | Promoter (<=1kb) | 18 | 36735068 | 36742391 | 7324 | 1 | ENSMUST00 | 0 | Tmco6 | ENSMUSG0C | false | chr18 | 36734573 | 36735835 | 31.31841 | 96.16425 | 92.68594 | chr18 | 36733915 | 36736094 | 4.35446 | 3.77728 | 2.46439 |
| chr12 | 32605416 | 32605683 | 349 | Distal Intergenic | 12 | 32820335 | 32853369 | 33035 | 1 | ENSMUST00 | -214652 | Nampt | ENSMUSG0C | true | . | . | . | . | . | . | chr12 | 32604311 | 32607595 | 5.19097 | 4.16045 | 3.14599 |
| chr3 | 51197048 | 51197527 | 349 | Distal Intergenic | 3 | 51224447 | 51251644 | 27198 | 1 | ENSMUST00 | -26920 | Ccrn4l | ENSMUSG0C | false | . | . | . | . | . | . | chr3 | 51195656 | 51197695 | 3.01714 | 2.97652 | 1.42461 |
| chr17 | 67955786 | 67956140 | 349 | Intron (ENSMUST00000024840.6 | 17 | 67845110 | 67950940 | 105831 | 2 | ENSMUST00 | -4847 | Arhgap28 | ENSMUSG0C | true | chr17 | 67952886 | 67956202 | 20.45667 | 53.83482 | 51.06929 | . | . | . | . | . | . |
| chr19 | 9053153 | 9053378 | 349 | Intron (ENSMUST00000092955.4 | 19 | 9083636 | 9087958 | 4323 | 2 | ENSMUST00 | 34580 | Scgb1a1 | ENSMUSG0C | false | . | . | . | . | . | . | chr19 | 9052994 | 9053851 | 4.61556 | 3.96946 | 2.70562 |
| chr1 | 79858449 | 79858839 | 349 | Promoter (<=1kb) | 1 | 79794197 | 79858696 | 64500 | 2 | ENSMUST00 | 0 | Serpine2 | ENSMUSG0C | false | chr1 | 79857873 | 79859155 | 24.50127 | 70.18111 | 67.18839 | chr1 | 79858154 | 79859177 | 4.894 | 3.89049 | 2.9455 |
| chrX | 99215210 | 99215452 | 349 | Distal Intergenic | 20 | 99138131 | 99147230 | 9100 | 1 | ENSMUST00 | 77080 | Efnb1 | ENSMUSG0C | false | . | . | . | . | . | . | . | . | . | . | . | . |
| chrX | 101332634 | 101332839 | 349 | Distal Intergenic | 20 | 101314391 | 101318151 | 3761 | 1 | ENSMUST00 | 18244 | Nlgn3 | ENSMUSG0C | false | . | . | . | . | . | . | . | . | . | . | . | . |
| chr8 | 27085391 | 27086106 | 349 | Promoter (<=1kb) | 8 | 27085583 | 27123435 | 37853 | 1 | ENSMUST00 | 0 | Gpr124 | ENSMUSG0C | false | chr8 | 27085211 | 27087724 | 25.48472 | 95.2657 | 91.8051 | chr8 | 27083119 | 27091776 | 4.52211 | 3.74431 | 2.66045 |
| chr4 | 45819752 | 45819968 | 349 | Intron (ENSMUST00000044297.6 | 4 | 45813007 | 45816363 | 3357 | 2 | ENSMUST00 | -3390 | Igfbpl1 | ENSMUSG0C | false | chr4 | 45819270 | 45820069 | 10.13814 | 22.76719 | 20.39335 | chr4 | 45819257 | 45820107 | 4.60459 | 3.87169 | 2.69924 |
| chr2 | 44951904 | 44952141 | 349 | Distal Intergenic | 2 | 44635264 | 44927657 | 292394 | 2 | ENSMUST00 | -24248 | Gtdc1 | ENSMUSG0C | false | . | . | . | . | . | . | chr2 | 44950418 | 44953463 | 5.08094 | 4.0784 | 3.0727 |
| chr17 | 43767866 | 43768178 | 349 | Distal Intergenic | 17 | 43801851 | 44039516 | 237666 | 1 | ENSMUST00 | -33673 | Rcan2 | ENSMUSG0C | false | . | . | . | . | . | . | . | . | . | . | . | . |
| chr12 | 104687014 | 104687196 | 349 | Downstream (<1kb) | 12 | 104687742 | 104751952 | 64211 | 2 | ENSMUST00 | 64756 | Dicer1 | ENSMUSG0C | false | . | . | . | . | . | . | chr12 | 104686864 | 104687955 | 3.53007 | 3.36594 | 1.78566 |
| chr1 | 135105080 | 135105342 | 349 | Promoter (<=1kb) | 1 | 134983301 | 135105276 | 121976 | 2 | ENSMUST00 | 0 | Lgr6 | ENSMUSG0C | false | chr1 | 135104968 | 135105547 | 7.81205 | 13.8349 | 11.58824 | chr1 | 135104781 | 135106103 | 5.83954 | 4.36055 | 3.72709 |
| chr10 | 95162945 | 95163271 | 349 | Distal Intergenic | 10 | 95174746 | 95324097 | 149352 | 2 | ENSMUST00 | 160826 | Cradd | ENSMUSG0C | false | . | . | . | . | . | . | chr10 | 95161637 | 95165733 | 4.69044 | 3.9319 | 2.75703 |
| chr4 | 83052116 | 83052345 | 349 | Promoter (<=1kb) | 4 | 82897920 | 83052167 | 154248 | 2 | ENSMUST00 | 0 | Frem1 | ENSMUSG0C | false | chr4 | 83051315 | 83052311 | 9.27429 | 18.59722 | 16.28163 | . | . | . | . | . | . |
| chr9 | 37030661 | 37030878 | 349 | Intron (ENSMUST00000177218.2 | 9 | 37026399 | 37026507 | 109 | 1 | ENSMUST00 | 4263 | n-R5s82 | ENSMUSG0C | false | . | . | . | . | . | . | chr9 | 37021034 | 37031907 | 6.40408 | 4.65381 | 4.17785 |
| chr11 | 59777367 | 59777604 | 349 | Promoter (2-3kb) | 11 | 59775286 | 59780321 | 5036 | 2 | ENSMUST00 | 2717 | Gm12264 | ENSMUSG0C | false | . | . | . | . | . | . | chr11 | 59775352 | 59780895 | 5.11217 | 3.97205 | 3.11627 |
| chr1 | 60615077 | 60615431 | 349 | Promoter (1-2kb) | 1 | 60613698 | 60613803 | 106 | 2 | ENSMUST00 | -1275 | Gm23762 | ENSMUSG0C | false | . | . | . | . | . | . | chr1 | 60614639 | 60615524 | 3.55348 | 3.34564 | 1.85422 |
| chr10 | 119424130 | 119424458 | 349 | Intron (ENSMUST00000185238.1 | 10 | 119413444 | 119426864 | 13421 | 1 | ENSMUST00 | 10687 | RP24-528H2 | ENSMUSG0C | false | . | . | . | . | . | . | . | . | . | . | . | . |
| chr8 | 120990973 | 120991276 | 349 | Intron (ENSMUST00000127664.1 | 8 | 121002927 | 121003190 | 264 | 1 | ENSMUST00 | -11651 | RP23-80D9.2 | ENSMUSG0C | false | . | . | . | . | . | . | . | . | . | . | . | . |
| chr5 | 114380365 | 114380850 | 348 | Promoter (<=1kb) | 5 | 114363567 | 114380515 | 16949 | 2 | ENSMUST00 | 0 | Kctd10 | ENSMUSG0C | false | chr5 | 114379830 | 114381806 | 25.90858 | 83.69981 | 80.48042 | . | . | . | . | . | . |
| chr6 | 88924975 | 88925250 | 348 | Distal Intergenic | 6 | 88910158 | 88911778 | 1621 | 1 | ENSMUST00 | 14818 | Tpra1 | ENSMUSG0C | false | . | . | . | . | . | . | chr6 | 88922092 | 88928323 | 9.06661 | 5.7788 | 6.36998 |
| chr11 | 90369528 | 90369859 | 348 | Intron (ENSMUST00000004051.7 | 11 | 90340827 | 90381935 | 41109 | 2 | ENSMUST00 | 12076 | Hlf | ENSMUSG0C | false | . | . | . | . | . | . | chr11 | 90365274 | 90371966 | 5.21089 | 4.09868 | 3.19361 |
| chr11 | 3409058 | 3409459 | 348 | Promoter (<=1kb) | 11 | 3344256 | 3409174 | 64919 | 2 | ENSMUST00 | 0 | Limk2 | ENSMUSG0C | false | chr11 | 3408169 | 3409742 | 29.66345 | 100.21 | 96.63225 | chr11 | 3407553 | 3409857 | 3.92393 | 3.35063 | 2.16257 |
| chr14 | 118937571 | 118937853 | 348 | Promoter (<=1kb) | 14 | 118937932 | 118981702 | 43771 | 1 | ENSMUST00 | -79 | Dnajc3 | ENSMUSG0C | false | chr14 | 118937104 | 118938956 | 41.07278 | 140.67711 | 135.76277 | . | . | . | . | . | . |
| chr17 | 71228057 | 71228587 | 348 | Promoter (2-3kb) | 17 | 71231433 | 71238085 | 6653 | 1 | ENSMUST00 | -2846 | Lpin2 | ENSMUSG0C | false | . | . | . | . | . | . | chr17 | 71186127 | 71229089 | 5.51375 | 4.29183 | 3.43736 |
| chr19 | 17393844 | 17394349 | 348 | Promoter (<=1kb) | 19 | 17394043 | 17401349 | 7307 | 1 | ENSMUST00 | 0 | Rfk | ENSMUSG0C | false | chr19 | 17393476 | 17395235 | 37.48252 | 127.75053 | 123.3422 | . | . | . | . | . | . |
| chr19 | 57003303 | 57003851 | 348 | Promoter (2-3kb) | 19 | 56930098 | 57001046 | 70949 | 2 | ENSMUST00 | -2258 | Afap1l2 | ENSMUSG0C | false | . | . | . | . | . | . | . | . | . | . | . | . |
| chr1 | 15805900 | 15806206 | 348 | Promoter (<=1kb) | 1 | 15805664 | 15843470 | 37807 | 1 | ENSMUST00 | 237 | Terf1 | ENSMUSG0C | false | chr1 | 15805135 | 15806856 | 24.89402 | 98.85246 | 95.31046 | . | . | . | . | . | . |
| chr6 | 102523166 | 102523379 | 348 | Distal Intergenic | 6 | 102163306 | 102464667 | 301362 | 2 | ENSMUST00 | -58500 | Cntn3 | ENSMUSG0C | false | . | . | . | . | . | . | . | . | . | . | . | . |
| chr18 | 80479040 | 80479399 | 348 | Distal Intergenic | 18 | 80407959 | 80469695 | 61737 | 2 | ENSMUST00 | -9346 | Ctdp1 | ENSMUSG0C | false | . | . | . | . | . | . | chr18 | 80474858 | 80481285 | 5.64513 | 4.38222 | 3.53512 |
| chr14 | 65326662 | 65326877 | 348 | Distal Intergenic | 14 | 65358676 | 65398930 | 40255 | 1 | ENSMUST00 | -31799 | Zfp395 | ENSMUSG0C | false | . | . | . | . | . | . | chr14 | 65326304 | 65328021 | 3.54843 | 3.23094 | 1.83705 |
| chr17 | 65964383 | 65964576 | 348 | Downstream (2-3kb) | 17 | 65968013 | 65972394 | 4382 | 2 | ENSMUST00 | 7818 | Ankrd12 | ENSMUSG0C | false | . | . | . | . | . | . | chr17 | 65963360 | 65965614 | 3.36122 | 3.20396 | 1.70057 |
| chr5 | 90223789 | 90224291 | 348 | Promoter (<=1kb) | 5 | 90214724 | 90223972 | 9249 | 2 | ENSMUST00 | 0 | Cox18 | ENSMUSG0C | false | chr5 | 90223166 | 90224759 | 43.04933 | 143.01288 | 138.00682 | . | . | . | . | . | . |
| chr2 | 71031466 | 71031694 | 348 | Intron (ENSMUST00000100037.3 | 2 | 70993345 | 71018378 | 25034 | 2 | ENSMUST00 | -13089 | Mettl8 | ENSMUSG0C | false | . | . | . | . | . | . | chr2 | 71030550 | 71032190 | 4.28416 | 3.66587 | 2.45403 |
| chr8 | 84976145 | 84976779 | 348 | Promoter (1-2kb) | 8 | 84976909 | 84978748 | 1840 | 2 | ENSMUST00 | 1969 | Junb | ENSMUSG0C | false | chr8 | 84975875 | 84979344 | 20.74539 | 66.97287 | 64.02668 | chr8 | 84974675 | 84986551 | 7.15 | 5.07534 | 4.74925 |
| chr19 | 31468493 | 31468731 | 348 | Intron (ENSMUST00000065067.7 | 19 | 30568547 | 31664370 | 1095824 | 2 | ENSMUST00 | 195639 | Prkg1 | ENSMUSG0C | true | . | . | . | . | . | . | chr19 | 31466298 | 31469803 | 3.3002 | 3.16637 | 1.63441 |
| chr10 | 4434380 | 4434613 | 348 | Promoter (<=1kb) | 10 | 4434581 | 4440469 | 5889 | 1 | ENSMUST00 | 0 | 1700052N19 | ENSMUSG0C | false | . | . | . | . | . | . | chr10 | 4422045 | 4437373 | 5.83291 | 4.40627 | 3.70343 |
| chr10 | 13323802 | 13324339 | 348 | Promoter (<=1kb) | 10 | 13207717 | 13324160 | 116444 | 2 | ENSMUST00 | 0 | Phactr2 | ENSMUSG0C | false | chr10 | 13323002 | 13324565 | 20.82289 | 72.3363 | 69.31286 | . | . | . | . | . | . |
| chr11 | 101442004 | 101442419 | 348 | Promoter (<=1kb) | 11 | 101442298 | 101445529 | 3232 | 1 | ENSMUST00 | 0 | Rpl27 | ENSMUSG0C | false | chr11 | 101442399 | 101443270 | 19.74353 | 71.90538 | 68.88615 | . | . | . | . | . | . |
| chr1 | 184845717 | 184846143 | 348 | Promoter (<=1kb) | 1 | 184813068 | 184845881 | 32814 | 2 | ENSMUST00 | 0 | 37316 | ENSMUSG0C | false | chr1 | 184844456 | 184846746 | 38.68621 | 127.7006 | 123.29971 | . | . | . | . | . | . |
| chr4 | 148970263 | 148970557 | 348 | Exon (ENSMUST00000103217.5/ | 4 | 148960360 | 148966998 | 6639 | 1 | ENSMUST00 | 9904 | Gm17029 | ENSMUSG0C | false | . | . | . | . | . | . | chr4 | 148968291 | 148974289 | 7.92118 | 5.40621 | 5.39072 |
| chrX | 7319131 | 7319412 | 347 | Promoter (<=1kb) | 20 | 7159469 | 7319233 | 159765 | 2 | ENSMUST00 | 0 | Clcn5 | ENSMUSG0C | false | chrX | 7318483 | 7319382 | 28.49099 | 81.63235 | 78.4519 | . | . | . | . | . | . |
| chr10 | 95394748 | 95394971 | 347 | Promoter (1-2kb) | 10 | 95387667 | 95392967 | 5301 | 2 | ENSMUST00 | -1782 | Socs2 | ENSMUSG0C | false | chr10 | 95394672 | 95395470 | 10.58711 | 29.50236 | 27.04178 | . | . | . | . | . | . |
| chr11 | 116198783 | 116199434 | 347 | Promoter (<=1kb) | 11 | 116171888 | 116198801 | 26914 | 2 | ENSMUST00 | 0 | Acox1 | ENSMUSG0C | false | chr11 | 116198099 | 116199882 | 27.08421 | 81.07839 | 77.90557 | . | . | . | . | . | . |
| chr14 | 79486651 | 79486998 | 347 | Intron (ENSMUST00000040131.6 | 14 | 79459937 | 79481268 | 21332 | 2 | ENSMUST00 | -5384 | Wbp4 | ENSMUSG0C | false | chr14 | 79486710 | 79487981 | 19.90079 | 58.62946 | 55.80003 | chr14 | 79482839 | 79490765 | 4.91206 | 4.05656 | 2.95254 |
| chr15 | 27913044 | 27913268 | 347 | Intron (ENSMUST00000090247.5 | 15 | 27730652 | 28025848 | 295197 | 2 | ENSMUST00 | 112580 | Trio | ENSMUSG0C | false | . | . | . | . | . | . | chr15 | 27912832 | 27914426 | 4.22732 | 3.3292 | 2.40557 |
| chr16 | 91597588 | 91597838 | 347 | Promoter (<=1kb) | 16 | 91574503 | 91597800 | 23298 | 2 | ENSMUST00 | 0 | Tmem50b | ENSMUSG0C | false | chr16 | 91596520 | 91597737 | 25.17393 | 93.69764 | 90.27328 | chr16 | 91596486 | 91598420 | 3.59694 | 3.39144 | 1.83425 |
| chr2 | 181365137 | 181365525 | 347 | Promoter (<=1kb) | 2 | 181365405 | 181380782 | 15378 | 1 | ENSMUST00 | 0 | Zgpat | ENSMUSG0C | false | chr2 | 181364391 | 181366453 | 31.83799 | 103.0912 | 99.44543 | . | . | . | . | . | . |
| chr4 | 129110112 | 129110519 | 347 | Exon (ENSMUST00000106064.4/ | 4 | 129106869 | 129110544 | 3676 | 1 | ENSMUST00 | 3244 | Tmem54 | ENSMUSG0C | false | . | . | . | . | . | . | chr4 | 129106734 | 129114119 | 7.73893 | 5.31745 | 5.25102 |
| chr5 | 124482923 | 124483590 | 347 | Promoter (<=1kb) | 5 | 124483134 | 124491124 | 7991 | 1 | ENSMUST00 | 0 | Snrnp35 | ENSMUSG0C | false | chr5 | 124482408 | 124484009 | 23.68814 | 67.0026 | 64.0563 | . | . | . | . | . | . |
| chr5 | 92387525 | 92387845 | 347 | Promoter (<=1kb) | 5 | 92387763 | 92392725 | 4963 | 1 | ENSMUST00 | 0 | Art3 | ENSMUSG0C | false | . | . | . | . | . | . | . | . | . | . | . | . |
| chr11 | 98938228 | 98938447 | 347 | Promoter (<=1kb) | 11 | 98938328 | 98950361 | 12034 | 1 | ENSMUST00 | 0 | Rara | ENSMUSG0C | false | chr11 | 98936314 | 98938742 | 41.35789 | 132.3362 | 127.75856 | chr11 | 98928668 | 98957642 | 6.70032 | 4.70048 | 4.41905 |
| chr4 | 11253658 | 11254214 | 347 | Promoter (<=1kb) | 4 | 11199158 | 11254258 | 55101 | 2 | ENSMUST00 | 44 | Ints8 | ENSMUSG0C | false | chr4 | 11253039 | 11254812 | 22.00414 | 89.03333 | 85.70822 | . | . | . | . | . | . |
| chr4 | 138469041 | 138469423 | 347 | Distal Intergenic | 4 | 138454314 | 138460123 | 5810 | 1 | ENSMUST00 | 14728 | Camk2n1 | ENSMUSG0C | false | . | . | . | . | . | . | chr4 | 138468565 | 138471372 | 4.85657 | 4.0786 | 2.88086 |
| chr5 | 137651069 | 137651289 | 347 | 3' UTR | 5 | 137643032 | 137645714 | 2683 | 2 | ENSMUST00 | -5356 | Irs3 | ENSMUSG0C | false | . | . | . | . | . | . | chr5 | 137649328 | 137653192 | 7.00301 | 4.9223 | 4.68019 |
| chr7 | 145197116 | 145197396 | 347 | Distal Intergenic | 7 | 145253923 | 145283927 | 30005 | 2 | ENSMUST00 | 86531 | Tpcn2 | ENSMUSG0C | false | . | . | . | . | . | . | chr7 | 145196929 | 145200615 | 5.05083 | 3.9985 | 3.09965 |
| chr5 | 34513755 | 34514236 | 347 | Promoter (<=1kb) | 5 | 34496087 | 34513963 | 17877 | 2 | ENSMUST00 | 0 | Tnip2 | ENSMUSG0C | false | chr5 | 34512942 | 34514474 | 32.16725 | 95.65015 | 92.18271 | chr5 | 34511842 | 34514882 | 3.97068 | 3.44712 | 2.19923 |
| chr6 | 115730469 | 115730691 | 347 | Exon (ENSMUST00000112946.2/ | 6 | 115729141 | 115734951 | 5811 | 2 | ENSMUST00 | 4260 | Tmem40 | ENSMUSG0C | false | chr6 | 115730374 | 115730769 | 6.65711 | 11.15507 | 8.95905 | chr6 | 115730223 | 115731251 | 4.22523 | 3.59198 | 2.40287 |
| chr5 | 88554031 | 88554340 | 347 | Promoter (<=1kb) | 5 | 88554483 | 88556083 | 1601 | 1 | ENSMUST00 | -143 | Utp3 | ENSMUSG0C | false | chr5 | 88553392 | 88554295 | 21.46036 | 80.68572 | 77.52129 | . | . | . | . | . | . |
| chr6 | 37619010 | 37619290 | 347 | Distal Intergenic | 6 | 37580335 | 37580427 | 93 | 1 | ENSMUST00 | 38676 | Gm25062 | ENSMUSG0C | false | . | . | . | . | . | . | chr6 | 37613519 | 37619590 | 4.74098 | 3.99613 | 2.78613 |
| chr19 | 26355690 | 26355915 | 347 | Distal Intergenic | 19 | 26260369 | 26260493 | 125 | 2 | ENSMUST00 | -95198 | Gm25176 | ENSMUSG0C | false | . | . | . | . | . | . | chr19 | 26351132 | 26356191 | 4.5733 | 3.82867 | 2.65437 |
| chr7 | 143407876 | 143408334 | 347 | Intron (ENSMUST00000009689.5 | 7 | 143447605 | 143447665 | 61 | 1 | ENSMUST00 | -39271 | Gm27901 | ENSMUSG0C | false | . | . | . | . | . | . | chr7 | 143407161 | 143409878 | 4.98044 | 4.06372 | 2.99644 |
| chr1 | 125663245 | 125663545 | 347 | Distal Intergenic | 1 | 125667988 | 125668616 | 629 | 2 | ENSMUST00 | 5071 | RP24-97E12. | ENSMUSG0C | false | . | . | . | . | . | . | . | . | . | . | . | . |
| chr17 | 34587247 | 34587787 | 346 | Promoter (<=1kb) | 17 | 34587041 | 34587818 | 778 | 1 | ENSMUST00 | 207 | Notch4 | ENSMUSG0C | false | chr17 | 34586640 | 34592164 | 28.90751 | 104.81161 | 101.11721 | chr17 | 34586035 | 34591596 | 5.00173 | 4.02952 | 3.00733 |
| chr11 | 20200893 | 20201500 | 346 | Promoter (<=1kb) | 11 | 20201432 | 20226856 | 25425 | 1 | ENSMUST00 | 0 | Rab1 | ENSMUSG0C | false | chr11 | 20200618 | 20201079 | 20.53639 | 56.4273 | 53.62976 | . | . | . | . | . | . |
| chr10 | 77314850 | 77315123 | 346 | Intron (ENSMUST00000020496.8 | 10 | 77325715 | 77340176 | 14462 | 2 | ENSMUST00 | 25053 | Adarb1 | ENSMUSG0C | false | . | . | . | . | . | . | chr10 | 77314652 | 77315772 | 4.02773 | 3.65632 | 2.2086 |
| chr15 | 97840047 | 97840448 | 346 | Intron (ENSMUST00000116408.3 | 15 | 97792709 | 97844164 | 51456 | 2 | ENSMUST00 | 3716 | Hdac7 | ENSMUSG0C | false | . | . | . | . | . | . | chr15 | 97835434 | 97841080 | 5.51907 | 4.35222 | 3.43438 |
| chr17 | 32753679 | 32753947 | 346 | Distal Intergenic | 17 | 32773252 | 32787547 | 14296 | 2 | ENSMUST00 | 33600 | Zfp871 | ENSMUSG0C | false | . | . | . | . | . | . | . | . | . | . | . | . |
| chr19 | 10605299 | 10605848 | 346 | Promoter (<=1kb) | 19 | 10605625 | 10629813 | 24189 | 1 | ENSMUST00 | 0 | Ddb1 | ENSMUSG0C | false | chr19 | 10603045 | 10606602 | 41.22328 | 133.19821 | 128.59248 | . | . | . | . | . | . |
| chr11 | 121388141 | 121388613 | 346 | Promoter (<=1kb) | 11 | 121356125 | 121388251 | 32127 | 2 | ENSMUST00 | 0 | Rab40b | ENSMUSG0C | false | chr11 | 121387488 | 121388799 | 34.36702 | 111.64683 | 107.76208 | chr11 | 121387273 | 121389131 | 4.12093 | 3.44228 | 2.32 |
| chr5 | 145167032 | 145167338 | 346 | Promoter (<=1kb) | 5 | 145147514 | 145167108 | 19595 | 2 | ENSMUST00 | 0 | Ptcd1 | ENSMUSG0C | false | chr5 | 145166174 | 145168160 | 29.52499 | 116.10789 | 112.09218 | . | . | . | . | . | . |
| chrX | 74328691 | 74329160 | 346 | Promoter (<=1kb) | 20 | 74329066 | 74332243 | 3178 | 1 | ENSMUST00 | 0 | Plxna3 | ENSMUSG0C | false | chrX | 74328470 | 74329456 | 11.4883 | 23.92001 | 21.53215 | . | . | . | . | . | . |
| chr12 | 40448123 | 40448432 | 346 | Promoter (2-3kb) | 12 | 40446053 | 40846486 | 400434 | 1 | ENSMUST00 | 2071 | Dock4 | ENSMUSG0C | true | chr12 | 40444767 | 40448342 | 40.65367 | 138.82761 | 133.98773 | chr12 | 40448062 | 40493881 | 5.07226 | 4.00134 | 3.09271 |
| chr10 | 43221006 | 43221567 | 346 | Promoter (<=1kb) | 10 | 43221486 | 43405395 | 183910 | 1 | ENSMUST00 | 0 | Pdss2 | ENSMUSG0C | false | chr10 | 43220403 | 43222284 | 25.50663 | 88.63404 | 85.31516 | . | . | . | . | . | . |
| chr19 | 32266986 | 32267207 | 346 | Intron (ENSMUST00000099514.4 | 19 | 32160299 | 32276926 | 116628 | 2 | ENSMUST00 | 9719 | Sgms1 | ENSMUSG0C | false | . | . | . | . | . | . | chr19 | 32265512 | 32283023 | 5.2656 | 4.10074 | 3.25195 |
| chr9 | 103656118 | 103656394 | 346 | Intron (ENSMUST00000049452.9 | 9 | 103482947 | 103692807 | 209861 | 2 | ENSMUST00 | 36413 | Tmem108 | ENSMUSG0C | true | . | . | . | . | . | . | chr9 | 103650371 | 103666133 | 5.65063 | 4.36287 | 3.53959 |
| chr14 | 79426335 | 79426780 | 346 | Promoter (<=1kb) | 14 | 79426511 | 79431039 | 4529 | 1 | ENSMUST00 | 0 | Kbtbd7 | ENSMUSG0C | false | chr14 | 79426350 | 79427272 | 21.28758 | 100.96285 | 97.37373 | chr14 | 79426062 | 79428080 | 6.08891 | 4.24009 | 3.9478 |
| chr15 | 98868712 | 98868988 | 346 | Promoter (2-3kb) | 15 | 98831669 | 98871183 | 39515 | 2 | ENSMUST00 | 2195 | Kmt2d | ENSMUSG0C | false | chr15 | 98868485 | 98872894 | 47.14861 | 169.72867 | 163.76503 | chr15 | 98866239 | 98870766 | 5.47992 | 4.27424 | 3.40843 |
| chr12 | 17881600 | 17882182 | 346 | Distal Intergenic | 12 | 17924294 | 17925668 | 1375 | 1 | ENSMUST00 | -42112 | B430203G13 | ENSMUSG0C | false | chr12 | 17881272 | 17883012 | 27.54716 | 98.2761 | 94.74957 | chr12 | 17880832 | 17881678 | 3.66289 | 3.29033 | 1.94237 |
| chr7 | 52005728 | 52006486 | 346 | Promoter (<=1kb) | 7 | 51997165 | 52006018 | 8854 | 2 | ENSMUST00 | 0 | Svip | ENSMUSG0C | false | chr7 | 52005354 | 52006907 | 24.14628 | 82.32003 | 79.12344 | . | . | . | . | . | . |
| chr2 | 157170228 | 157170606 | 346 | Promoter (1-2kb) | 2 | 157168820 | 157169179 | 360 | 1 | ENSMUST00 | 1409 | Gm14277 | ENSMUSG0C | false | . | . | . | . | . | . | chr2 | 157169520 | 157171891 | 4.30337 | 3.74852 | 2.42457 |
| chr8 | 110772428 | 110772895 | 346 | Intron (ENSMUST00000150680.1 | 8 | 110754992 | 110758970 | 3979 | 1 | ENSMUST00 | 17437 | Gm15894 | ENSMUSG0C | false | . | . | . | . | . | . | chr8 | 110771936 | 110776315 | 5.81817 | 4.50309 | 3.66655 |
| chr13 | 81130706 | 81130926 | 346 | Intron (ENSMUST00000128120.1 | 13 | 81065477 | 81065560 | 84 | 1 | ENSMUST00 | 65230 | Gm25241 | ENSMUSG0C | false | . | . | . | . | . | . | chr13 | 81128690 | 81133234 | 5.13344 | 4.16391 | 3.11287 |
| chr9 | 63109158 | 63109617 | 346 | Distal Intergenic | 9 | 63133283 | 63133389 | 107 | 1 | ENSMUST00 | -23666 | Gm25064 | ENSMUSG0C | false | chr9 | 63108809 | 63109798 | 17.34451 | 52.33425 | 49.58816 | chr9 | 63108661 | 63110251 | 5.86601 | 4.5651 | 3.72634 |
| chr14 | 53331017 | 53331206 | 346 | Distal Intergenic | 14 | 53337122 | 53337638 | 517 | 1 | ENSMUST00 | -5916 | Trav13n-3 | ENSMUSG0C | false | . | . | . | . | . | . | chr14 | 53326867 | 53334472 | 6.19783 | 4.37912 | 4.0048 |
| chr8 | 105577341 | 105577527 | 345 | Distal Intergenic | 8 | 105566700 | 105568298 | 1599 | 2 | ENSMUST00 | -9044 | Agrp | ENSMUSG0C | true | . | . | . | . | . | . | chr8 | 105571509 | 105577963 | 4.48434 | 3.72744 | 2.59946 |
| chr2 | 70661307 | 70661706 | 345 | Promoter (<=1kb) | 2 | 70661576 | 70691725 | 30150 | 1 | ENSMUST00 | 0 | Gorasp2 | ENSMUSG0C | false | chr2 | 70660882 | 70662957 | 40.46964 | 137.39561 | 132.61725 | . | . | . | . | . | . |
| chr11 | 97744563 | 97745372 | 345 | Promoter (<=1kb) | 11 | 97715157 | 97744704 | 29548 | 2 | ENSMUST00 | 0 | Pip4k2b | ENSMUSG0C | false | chr11 | 97743491 | 97745609 | 38.13901 | 127.82562 | 123.40861 | . | . | . | . | . | . |
| chr1 | 16334546 | 16335055 | 345 | Intron (ENSMUST00000162751.2 | 1 | 16344198 | 16375395 | 31198 | 2 | ENSMUST00 | 40340 | Stau2 | ENSMUSG0C | false | . | . | . | . | . | . | chr1 | 16334197 | 16336428 | 4.82613 | 3.9304 | 2.8968 |
| chr3 | 88623116 | 88623386 | 345 | Promoter (<=1kb) | 3 | 88623354 | 88633617 | 10264 | 1 | ENSMUST00 | 0 | Arhgef2 | ENSMUSG0C | false | chr3 | 88620710 | 88623767 | 28.41342 | 89.1249 | 85.7944 | chr3 | 88621743 | 88631594 | 4.65392 | 3.86367 | 2.72435 |
| chr4 | 120177341 | 120177596 | 345 | Distal Intergenic | 4 | 120161206 | 120167360 | 6155 | 1 | ENSMUST00 | 16136 | Edn2 | ENSMUSG0C | false | . | . | . | . | . | . | chr4 | 120176074 | 120178737 | 4.64576 | 3.82674 | 2.73912 |
| chr6 | 40325227 | 40325653 | 345 | Promoter (<=1kb) | 6 | 40325478 | 40396762 | 71285 | 1 | ENSMUST00 | 0 | Agk | ENSMUSG0C | false | chr6 | 40324965 | 40326179 | 24.02885 | 69.3614 | 66.38335 | . | . | . | . | . | . |
| chr6 | 119498639 | 119499108 | 345 | Intron (ENSMUST00000178696.2 | 6 | 119479668 | 119482630 | 2963 | 1 | ENSMUST00 | 18972 | Fbxl14 | ENSMUSG0C | false | . | . | . | . | . | . | chr6 | 119498158 | 119500722 | 4.60649 | 3.79762 | 2.71627 |
| chr9 | 95066853 | 95067335 | 345 | Intron (ENSMUST00000033463.9 | 9 | 95121686 | 95124551 | 2866 | 1 | ENSMUST00 | -54351 | Slc9a9 | ENSMUSG0C | false | . | . | . | . | . | . | chr9 | 95066388 | 95068153 | 4.15817 | 3.46912 | 2.35543 |
| chr9 | 31210971 | 31211758 | 345 | Promoter (<=1kb) | 9 | 31149557 | 31211815 | 62259 | 2 | ENSMUST00 | 57 | Aplp2 | ENSMUSG0C | false | chr9 | 31210441 | 31212694 | 35.75768 | 125.74667 | 121.41521 | . | . | . | . | . | . |
| chr6 | 90711513 | 90711786 | 345 | Intron (ENSMUST00000101153.4 | 6 | 90659598 | 90716529 | 56932 | 2 | ENSMUST00 | 4743 | Iqsec1 | ENSMUSG0C | false | . | . | . | . | . | . | chr6 | 90695436 | 90737892 | 6.93129 | 4.85273 | 4.61297 |
| chr12 | 12972843 | 12973289 | 345 | Distal Intergenic | 12 | 12936096 | 12941914 | 5819 | 2 | ENSMUST00 | -30930 | Mycn | ENSMUSG0C | false | . | . | . | . | . | . | chr12 | 12972330 | 12976013 | 5.97919 | 4.52073 | 3.86248 |
| chr16 | 10784999 | 10786247 | 345 | Promoter (<=1kb) | 16 | 10783808 | 10785536 | 1729 | 2 | ENSMUST00 | 0 | Socs1 | ENSMUSG0C | false | chr16 | 10784188 | 10786523 | 46.52121 | 165.10674 | 159.2771 | chr16 | 10786155 | 10787838 | 3.82525 | 3.47054 | 2.03393 |
| chr18 | 62113575 | 62114020 | 345 | Distal Intergenic | 18 | 62177817 | 62179959 | 2143 | 2 | ENSMUST00 | 65939 | Adrb2 | ENSMUSG0C | false | . | . | . | . | . | . | . | . | . | . | . | . |
| chr5 | 37405078 | 37405458 | 345 | Promoter (<=1kb) | 5 | 37404384 | 37404863 | 480 | 2 | ENSMUST00 | -216 | Gm15732 | ENSMUSG0C | false | . | . | . | . | . | . | chr5 | 37404047 | 37408204 | 3.97525 | 3.51327 | 2.18657 |
| chr14 | 117267897 | 117268164 | 345 | Intron (ENSMUST00000078849.5 | 14 | 116925531 | 117975406 | 1049876 | 1 | ENSMUST00 | 342367 | Gpc6 | ENSMUSG0C | false | . | . | . | . | . | . | chr14 | 117267739 | 117268784 | 3.59993 | 3.23999 | 1.87134 |
| chr13 | 51644899 | 51645156 | 345 | Promoter (<=1kb) | 13 | 51645232 | 51650660 | 5429 | 1 | ENSMUST00 | -76 | Cks2 | ENSMUSG0C | false | chr13 | 51644323 | 51646300 | 20.60195 | 76.26862 | 73.17842 | chr13 | 51644286 | 51646206 | 4.39585 | 3.7608 | 2.52476 |
| chr5 | 107250621 | 107250915 | 345 | Intron (ENSMUST00000031224.9 | 5 | 107241623 | 107242367 | 745 | 1 | ENSMUST00 | 8999 | Gm8145 | ENSMUSG0C | false | . | . | . | . | . | . | chr5 | 107250320 | 107254987 | 5.52178 | 4.3563 | 3.42599 |
| chr4 | 137887951 | 137888276 | 345 | Intron (ENSMUST00000102518.4 | 4 | 137882370 | 137883937 | 1568 | 2 | ENSMUST00 | -4015 | Gm13012 | ENSMUSG0C | false | . | . | . | . | . | . | chr4 | 137887880 | 137888836 | 5.31168 | 4.11511 | 3.28403 |
| chr4 | 149868011 | 149868289 | 345 | Distal Intergenic | 4 | 149904006 | 149908149 | 4144 | 1 | ENSMUST00 | -35717 | Gm13070 | ENSMUSG0C | false | . | . | . | . | . | . | chr4 | 149863103 | 149868831 | 4.3298 | 3.68834 | 2.48071 |
| chr10 | 12705821 | 12706421 | 345 | Intron (ENSMUST00000076817.4 | 10 | 12793875 | 12793974 | 100 | 2 | ENSMUST00 | 87553 | Gm23044 | ENSMUSG0C | false | . | . | . | . | . | . | chr10 | 12700386 | 12709298 | 4.88951 | 4.01525 | 2.91749 |
| chr15 | 80003655 | 80003967 | 345 | Intron (ENSMUST00000000500.6 | 15 | 79998970 | 79999026 | 57 | 2 | ENSMUST00 | -4630 | Mir7213 | ENSMUSG0C | false | . | . | . | . | . | . | chr15 | 80003121 | 80008590 | 4.67086 | 3.88569 | 2.76659 |
| chr15 | 100599786 | 100600308 | 344 | Promoter (<=1kb) | 15 | 100575318 | 100599984 | 24667 | 2 | ENSMUST00 | 0 | Pou6f1 | ENSMUSG0C | false | chr15 | 100598613 | 100600666 | 39.82221 | 128.59171 | 124.15125 | chr15 | 100600194 | 100601176 | 5.02908 | 4.1598 | 3.0146 |
| chr7 | 143513914 | 143514358 | 344 | 3' UTR | 7 | 143501545 | 143502541 | 997 | 2 | ENSMUST00 | -11374 | Phlda2 | ENSMUSG0C | false | . | . | . | . | . | . | chr7 | 143513229 | 143515593 | 4.90891 | 3.91044 | 2.95511 |
| chr11 | 110347132 | 110347429 | 344 | Distal Intergenic | 11 | 110325496 | 110337716 | 12221 | 2 | ENSMUST00 | -9417 | Abca5 | ENSMUSG0C | false | . | . | . | . | . | . | . | . | . | . | . | . |
| chr10 | 59920887 | 59921443 | 344 | Distal Intergenic | 10 | 59949675 | 59951770 | 2096 | 2 | ENSMUST00 | 30327 | Ddit4 | ENSMUSG0C | false | chr10 | 59920707 | 59921675 | 9.7287 | 26.0223 | 23.6086 | chr10 | 59918664 | 59922070 | 6.15988 | 4.45838 | 4.00029 |
| chr14 | 26637864 | 26638186 | 344 | Promoter (<=1kb) | 14 | 26638074 | 26657258 | 19185 | 1 | ENSMUST00 | 0 | Arf4 | ENSMUSG0C | false | chr14 | 26637196 | 26637927 | 14.18495 | 33.44911 | 30.9408 | . | . | . | . | . | . |
| chr15 | 79616154 | 79616399 | 344 | Intron (ENSMUST00000109648.3 | 15 | 79561500 | 79605084 | 43585 | 2 | ENSMUST00 | -11071 | Dmc1 | ENSMUSG0C | false | . | . | . | . | . | . | . | . | . | . | . | . |

| | | | | | | | | | | | | | | | | | | | | | | | | | | |
|---|---|---|---|---|---|---|---|---|---|---|---|---|---|---|---|---|---|---|---|---|---|---|---|---|---|---|
| chrX | 77511538 | 77512049 | 344 | Promoter (<=1kb) | 20 | 77511214 | 77659743 | 148530 | 1 | ENSMUST00 | 325 | Tbl1x | ENSMUSG00 | false | chrX | 77510423 | 77512224 | 23.43614 | 63.09834 | 60.20888 | . | . | . | . | . | . |
| chr6 | 103272287 | 103272547 | 344 | Distal Intergenic | 6 | 103510876 | 103733035 | 222160 | 1 | ENSMUST00 | -238329 | Chl1 | ENSMUSG00 | false | . | . | . | . | . | . | chr6 | 103271196 | 103273915 | 3.70924 | 3.37744 | 1.95418 |
| chr8 | 23208271 | 23208594 | 344 | Promoter (<=1kb) | 8 | 23172946 | 23208453 | 35508 | 2 | ENSMUST00 | 0 | Agpat6 | ENSMUSG00 | false | chr8 | 23207317 | 23209189 | 22.31878 | 85.15456 | 81.90936 | . | . | . | . | . | . |
| chr9 | 121719074 | 121719761 | 344 | Promoter (<=1kb) | 9 | 121719169 | 121742718 | 23550 | 1 | ENSMUST00 | 0 | Nktr | ENSMUSG00 | true | chr9 | 121718570 | 121720727 | 32.69058 | 104.59052 | 100.90593 | . | . | . | . | . | . |
| chr1 | 151974908 | 151975180 | 344 | Intron (ENSMUST00000187603.1 | 1 | 151892903 | 151921227 | 28325 | 2 | ENSMUST00 | -53682 | 1700025G04 | ENSMUSG00 | true | . | . | . | . | . | . | chr1 | 151972781 | 151980078 | 5.57591 | 4.3299 | 3.48688 |
| chr19 | 45013909 | 45014198 | 344 | Promoter (<=1kb) | 19 | 45015176 | 45027104 | 11929 | 1 | ENSMUST00 | -978 | Lzts2 | ENSMUSG00 | false | . | . | . | . | . | . | chr19 | 45012490 | 45017626 | 4.84179 | 3.90507 | 2.90417 |
| chr17 | 72836852 | 72837652 | 344 | Promoter (<=1kb) | 17 | 72836704 | 72851195 | 14492 | 1 | ENSMUST00 | 149 | Ypel5 | ENSMUSG00 | false | chr17 | 72836567 | 72838139 | 37.40972 | 147.3676 | 142.18408 | . | . | . | . | . | . |
| chr9 | 114844015 | 114844564 | 344 | Promoter (<=1kb) | 9 | 114789345 | 114844152 | 54808 | 2 | ENSMUST00 | 0 | Cmtm8 | ENSMUSG00 | false | chr9 | 114842973 | 114844728 | 37.34461 | 119.99735 | 115.85562 | . | . | . | . | . | . |
| chr15 | 100489252 | 100489566 | 344 | Exon (ENSMUST00000061457.5/ | 15 | 100479570 | 100495239 | 15670 | 2 | ENSMUST00 | 5673 | Csrnp2 | ENSMUSG00 | false | . | . | . | . | . | . | chr15 | 100486390 | 100491572 | 5.41408 | 4.26528 | 3.35969 |
| chr7 | 19572978 | 19573549 | 344 | Promoter (<=1kb) | 7 | 19564949 | 19573343 | 8395 | 2 | ENSMUST00 | 0 | Gemin7 | ENSMUSG00 | false | chr7 | 19572424 | 19574065 | 30.32912 | 88.589 | 85.27629 | chr7 | 19570365 | 19574181 | 6.23861 | 4.63142 | 4.02771 |
| chrX | 167208415 | 167208636 | 344 | Promoter (<=1kb) | 20 | 167207292 | 167208827 | 1536 | 2 | ENSMUST00 | 191 | Tmsb4x | ENSMUSG00 | false | chrX | 167206266 | 167210665 | 18.38128 | 45.59164 | 42.93153 | . | . | . | . | . | . |
| chr19 | 31424309 | 31424624 | 344 | Intron (ENSMUST00000065067.7 | 19 | 30568547 | 31664370 | 1095824 | 2 | ENSMUST00 | 239746 | Prkg1 | ENSMUSG00 | true | . | . | . | . | . | . | . | . | . | . | . | . |
| chr7 | 127244303 | 127244638 | 344 | Promoter (<=1kb) | 7 | 127244526 | 127254803 | 10278 | 1 | ENSMUST00 | 0 | Zfp771 | ENSMUSG00 | false | chr7 | 127243936 | 127245261 | 34.6346 | 112.11573 | 108.21484 | chr7 | 127243747 | 127246061 | 3.64317 | 3.3013 | 1.93563 |
| chr19 | 22800485 | 22800749 | 344 | Intron (ENSMUST00000074770.6 | 19 | 22750600 | 22750679 | 80 | 1 | ENSMUST00 | 49886 | Mir204 | ENSMUSG00 | false | . | . | . | . | . | . | chr19 | 22800068 | 22801029 | 3.56505 | 3.33286 | 1.87347 |
| chr17 | 44637289 | 44637528 | 344 | Intron (ENSMUST00000162816.2 | 17 | 44641025 | 44641143 | 119 | 2 | ENSMUST00 | 3615 | n-R5s27 | ENSMUSG00 | false | . | . | . | . | . | . | chr17 | 44636712 | 44637714 | 3.55597 | 3.24153 | 1.84974 |
| chr4 | 54340213 | 54340411 | 344 | Distal Intergenic | 4 | 54235670 | 54235876 | 207 | 2 | ENSMUST00 | -104338 | Gm12469 | ENSMUSG00 | false | . | . | . | . | . | . | chr4 | 54336484 | 54341744 | 5.26949 | 4.2658 | 3.22204 |
| chr10 | 68625473 | 68625747 | 343 | Distal Intergenic | 10 | 68430953 | 68541896 | 110944 | 2 | ENSMUST00 | -83578 | 1700040L02F | ENSMUSG00 | false | . | . | . | . | . | . | chr10 | 68623248 | 68627234 | 5.31525 | 4.27204 | 3.27061 |
| chr10 | 63023325 | 63024379 | 343 | Promoter (<=1kb) | 10 | 63014664 | 63023894 | 9231 | 2 | ENSMUST00 | 0 | Hnrnph3 | ENSMUSG00 | false | chr10 | 63022070 | 63025156 | 42.74922 | 148.11885 | 142.90198 | . | . | . | . | . | . |
| chr10 | 62792189 | 62792518 | 343 | Promoter (<=1kb) | 10 | 62743928 | 62792368 | 48441 | 2 | ENSMUST00 | 0 | Ccar1 | ENSMUSG00 | false | chr10 | 62791319 | 62793219 | 36.54057 | 115.43316 | 111.4393 | . | . | . | . | . | . |
| chr12 | 73752867 | 73753090 | 343 | Intron (ENSMUST00000021527.9 | 12 | 73901375 | 73947530 | 46156 | 1 | ENSMUST00 | -148285 | Hif1a | ENSMUSG00 | false | . | . | . | . | . | . | chr12 | 73751719 | 73756344 | 3.92998 | 3.48537 | 2.1414 |
| chr13 | 56702893 | 56703199 | 343 | Promoter (<=1kb) | 13 | 56703010 | 56742377 | 39368 | 1 | ENSMUST00 | 0 | Smad5 | ENSMUSG00 | false | chr13 | 56702432 | 56704287 | 34.46491 | 104.61457 | 100.92722 | . | . | . | . | . | . |
| chr14 | 65837717 | 65838015 | 343 | Promoter (<=1kb) | 14 | 65837302 | 65866604 | 29303 | 1 | ENSMUST00 | 416 | Ccdc25 | ENSMUSG00 | false | chr14 | 65836958 | 65838434 | 31.38972 | 125.99339 | 121.64841 | . | . | . | . | . | . |
| chr1 | 91095866 | 91096178 | 343 | Intron (ENSMUST00000189505.1 | 1 | 91111214 | 91116921 | 5708 | 1 | ENSMUST00 | -15036 | Lrrfip1 | ENSMUSG00 | true | . | . | . | . | . | . | chr1 | 91055222 | 91105818 | 7.56473 | 5.21453 | 5.11657 |
| chr6 | 113237576 | 113237908 | 343 | Promoter (<=1kb) | 6 | 113237843 | 113281202 | 43360 | 1 | ENSMUST00 | 0 | Mtmr14 | ENSMUSG00 | false | chr6 | 113237289 | 113238881 | 41.35843 | 143.41548 | 138.39708 | chr6 | 113237082 | 113239742 | 4.28873 | 3.50826 | 2.46119 |
| chr7 | 114564225 | 114564495 | 343 | Promoter (1-2kb) | 7 | 114550166 | 114562972 | 12807 | 2 | ENSMUST00 | -1254 | Cyp2r1 | ENSMUSG00 | false | . | . | . | . | . | . | chr7 | 114562267 | 114565374 | 3.86463 | 3.55045 | 2.0589 |
| chr9 | 27029910 | 27030400 | 343 | Promoter (<=1kb) | 9 | 27004503 | 27030094 | 25592 | 2 | ENSMUST00 | 0 | Vps26b | ENSMUSG00 | false | chr9 | 27028962 | 27031181 | 42.51875 | 148.84439 | 143.61023 | . | . | . | . | . | . |
| chr9 | 103008251 | 103008528 | 343 | Promoter (<=1kb) | 9 | 103008499 | 103096002 | 87504 | 1 | ENSMUST00 | 0 | Slco2a1 | ENSMUSG00 | true | chr9 | 103007566 | 103008928 | 30.2403 | 90.37072 | 87.01736 | . | . | . | . | . | . |
| chr8 | 126734636 | 126735103 | 343 | Distal Intergenic | 8 | 126588296 | 126593986 | 5691 | 2 | ENSMUST00 | -140651 | Irf2bp2 | ENSMUSG00 | false | . | . | . | . | . | . | chr8 | 126733289 | 126737272 | 7.19033 | 5.00384 | 4.80902 |
| chr11 | 98785802 | 98786017 | 343 | Distal Intergenic | 11 | 98795516 | 98807859 | 12344 | 1 | ENSMUST00 | -9499 | Msl1 | ENSMUSG00 | false | . | . | . | . | . | . | chr11 | 98780022 | 98794752 | 7.02549 | 5.01464 | 4.66092 |
| chr18 | 69344189 | 69344633 | 343 | Promoter (<=1kb) | 18 | 69344492 | 69565506 | 221015 | 1 | ENSMUST00 | 0 | Tcf4 | ENSMUSG00 | false | chr18 | 69343931 | 69345637 | 23.09106 | 69.49222 | 66.50756 | chr18 | 69343638 | 69345358 | 3.2489 | 3.19933 | 1.55274 |
| chr4 | 106847888 | 106848246 | 343 | Promoter (<=1kb) | 4 | 106841761 | 106847920 | 6160 | 2 | ENSMUST00 | 0 | Gm12746 | ENSMUSG00 | false | . | . | . | . | . | . | chr4 | 106846953 | 106849468 | 6.9846 | 4.73761 | 4.66476 |
| chr2 | 173278936 | 173279298 | 343 | Promoter (2-3kb) | 2 | 173276698 | 173278428 | 1731 | 1 | ENSMUST00 | 2239 | Pmepa1os | ENSMUSG00 | false | . | . | . | . | . | . | chr2 | 173276948 | 173281914 | 5.70628 | 4.41535 | 3.58938 |
| chr9 | 113342575 | 113343068 | 343 | Distal Intergenic | 9 | 113403475 | 113403771 | 297 | 1 | ENSMUST00 | -60407 | Gm22492 | ENSMUSG00 | false | . | . | . | . | . | . | chr9 | 113339166 | 113343758 | 4.62012 | 3.83176 | 2.72203 |
| chr2 | 125672832 | 125673492 | 343 | Promoter (<=1kb) | 2 | 125673095 | 125674785 | 1691 | 1 | ENSMUST00 | 0 | Eid1 | ENSMUSG00 | false | chr2 | 125672595 | 125674020 | 30.59503 | 95.89513 | 92.42037 | . | . | . | . | . | . |
| chr12 | 32890469 | 32890650 | 343 | Distal Intergenic | 12 | 32924957 | 32953783 | 28827 | 2 | ENSMUST00 | 63133 | 4933406C10I | ENSMUSG00 | false | . | . | . | . | . | . | . | . | . | . | . | . |
| chr2 | 5200309 | 5200620 | 343 | Intron (ENSMUST00000027988.7 | 2 | 5137749 | 5230864 | 93116 | 2 | ENSMUST00 | 30244 | Gm26776 | ENSMUSG00 | false | . | . | . | . | . | . | chr2 | 5196720 | 5202535 | 4.35636 | 3.76045 | 2.46313 |
| chr3 | 85378792 | 85379035 | 343 | Intron (ENSMUST00000154148.2 | 3 | 85360228 | 85361174 | 947 | 2 | ENSMUST00 | -17619 | RP23-94M23 | ENSMUSG00 | false | . | . | . | . | . | . | chr3 | 85377576 | 85380465 | 4.83943 | 4.02641 | 2.88675 |
| chr17 | 14011837 | 14012148 | 343 | Distal Intergenic | 17 | 14000239 | 14001765 | 1527 | 2 | ENSMUST00 | -10073 | RP23-9O15.2 | ENSMUSG00 | false | . | . | . | . | . | . | . | . | . | . | . | . |
| chr1 | 43173485 | 43173780 | 342 | Intron (ENSMUST00000185893.1 | 1 | 43123074 | 43163961 | 40888 | 2 | ENSMUST00 | -9525 | Fhl2 | ENSMUSG00 | false | . | . | . | . | . | . | chr1 | 43173360 | 43173837 | 2.85526 | 2.86504 | 1.29965 |
| chr7 | 99565223 | 99565607 | 342 | Intron (ENSMUST00000098266.3 | 7 | 99573064 | 99588128 | 15065 | 1 | ENSMUST00 | -7457 | Arrb1 | ENSMUSG00 | false | . | . | . | . | . | . | chr7 | 99536482 | 99576026 | 7.32182 | 4.95837 | 4.9327 |
| chr10 | 84388362 | 84388683 | 342 | Intron (ENSMUST00000020220.9 | 10 | 84375296 | 84392388 | 17093 | 2 | ENSMUST00 | 3705 | Nuak1 | ENSMUSG00 | false | . | . | . | . | . | . | chr10 | 84386221 | 84399643 | 5.21453 | 4.13999 | 3.18346 |
| chr11 | 73018766 | 73019056 | 342 | Promoter (<=1kb) | 11 | 73019008 | 73042062 | 23055 | 1 | ENSMUST00 | 0 | Camkk1 | ENSMUSG00 | false | chr11 | 73018603 | 73019752 | 23.28017 | 72.50986 | 69.48009 | chr11 | 73018430 | 73020084 | 4.23042 | 3.76706 | 2.36132 |
| chr16 | 4913695 | 4913941 | 342 | Exon (ENSMUST00000023159.8/ | 16 | 4939111 | 4941020 | 1910 | 1 | ENSMUST00 | -25170 | Nudt16l1 | ENSMUSG00 | false | . | . | . | . | . | . | chr16 | 4913089 | 4914448 | 6.18901 | 4.53096 | 4.0019 |
| chr16 | 43573554 | 43573901 | 342 | Intron (ENSMUST00000151244.2 | 16 | 43620877 | 43623955 | 3079 | 1 | ENSMUST00 | -46976 | Zbtb20 | ENSMUSG00 | false | . | . | . | . | . | . | chr16 | 43569396 | 43579622 | 5.24946 | 4.20024 | 3.21225 |
| chr17 | 71475286 | 71475644 | 342 | Promoter (<=1kb) | 17 | 71344489 | 71475343 | 130855 | 2 | ENSMUST00 | 0 | Smchd1 | ENSMUSG00 | false | chr17 | 71474211 | 71476282 | 40.80059 | 132.46654 | 127.88422 | . | . | . | . | . | . |
| chr1 | 120265191 | 120265512 | 342 | Promoter (<=1kb) | 1 | 120190757 | 120265255 | 74499 | 2 | ENSMUST00 | 0 | Steap3 | ENSMUSG00 | false | chr1 | 120263582 | 120265737 | 29.35304 | 95.67908 | 92.21078 | chr1 | 120265500 | 120267419 | 4.24208 | 3.72811 | 2.42983 |
| chr2 | 102381241 | 102381433 | 342 | Intron (ENSMUST00000102573.2 | 2 | 102304494 | 102400257 | 95764 | 2 | ENSMUST00 | 18824 | Trim44 | ENSMUSG00 | false | . | . | . | . | . | . | chr2 | 102378602 | 102385369 | 4.82589 | 3.90454 | 2.88799 |
| chr2 | 157278939 | 157279486 | 342 | Promoter (<=1kb) | 2 | 157279017 | 157289561 | 10545 | 1 | ENSMUST00 | 0 | Rpn2 | ENSMUSG00 | false | chr2 | 157278838 | 157280074 | 34.92026 | 124.22972 | 119.94103 | . | . | . | . | . | . |
| chr1 | 183120361 | 183120583 | 342 | Intron (ENSMUST00000003035.6 | 1 | 183086270 | 183147461 | 61192 | 2 | ENSMUST00 | 26878 | Disp1 | ENSMUSG00 | false | chr1 | 183120557 | 183120860 | 5.76737 | 9.70104 | 7.52928 | chr1 | 183111643 | 183121343 | 6.66914 | 4.87138 | 4.37055 |
| chrX | 7657261 | 7657655 | 342 | Promoter (<=1kb) | 20 | 7657472 | 7661128 | 3657 | 1 | ENSMUST00 | 0 | Prickle3 | ENSMUSG00 | false | chrX | 7657113 | 7658374 | 15.62409 | 36.57069 | 34.02363 | . | . | . | . | . | . |
| chr8 | 95084778 | 95085093 | 342 | Intron (ENSMUST00000034239.7 | 8 | 95081201 | 95099874 | 18674 | 1 | ENSMUST00 | 3578 | Katnb1 | ENSMUSG00 | false | . | . | . | . | . | . | chr8 | 95081517 | 95086253 | 5.77799 | 4.48414 | 3.64759 |
| chr2 | 103632293 | 103632546 | 342 | Intron (ENSMUST00000076212.3 | 2 | 103566310 | 103718423 | 152114 | 1 | ENSMUST00 | 65984 | Abtb2 | ENSMUSG00 | false | . | . | . | . | . | . | chr2 | 103621752 | 103634479 | 4.86729 | 3.97695 | 2.90643 |
| chr9 | 121857441 | 121858144 | 342 | Promoter (<=1kb) | 9 | 121848560 | 121858000 | 9441 | 2 | ENSMUST00 | 0 | Higd1a | ENSMUSG00 | false | chr9 | 121856654 | 121858574 | 32.87005 | 117.31267 | 113.25658 | . | . | . | . | . | . |
| chr2 | 5746512 | 5746937 | 342 | Distal Intergenic | 2 | 5293457 | 5714515 | 421059 | 2 | ENSMUST00 | -31998 | Camk1d | ENSMUSG00 | false | chr2 | 5746390 | 5746840 | 12.0809 | 26.07937 | 23.66483 | chr2 | 5745672 | 5747567 | 3.80628 | 3.46881 | 2.04292 |
| chr5 | 74348597 | 74349120 | 342 | Intron (ENSMUST00000072857.7 | 5 | 74356566 | 74358415 | 1850 | 2 | ENSMUST00 | 9295 | Gm15982 | ENSMUSG00 | false | . | . | . | . | . | . | chr5 | 74348278 | 74349372 | 4.03133 | 3.33613 | 2.24539 |
| chr4 | 57295235 | 57295516 | 342 | Intron (ENSMUST00000075637.5 | 4 | 57300096 | 57307305 | 7210 | 1 | ENSMUST00 | -4580 | Gm12536 | ENSMUSG00 | false | . | . | . | . | . | . | chr4 | 57294789 | 57301188 | 4.63579 | 3.8844 | 2.72174 |
| chr4 | 53219315 | 53219678 | 342 | Promoter (1-2kb) | 4 | 53214557 | 53217857 | 3301 | 2 | ENSMUST00 | -1459 | 4930412L05F | ENSMUSG00 | false | . | . | . | . | . | . | chr4 | 53217207 | 53221444 | 7.22057 | 5.07507 | 4.83453 |
| chr17 | 83312337 | 83312799 | 342 | Distal Intergenic | 17 | 83288546 | 83289028 | 483 | 1 | ENSMUST00 | 23792 | Rpl21-ps7 | ENSMUSG00 | false | . | . | . | . | . | . | chr17 | 83311635 | 83313428 | 4.64766 | 3.96327 | 2.70983 |
| chr18 | 37172350 | 37172576 | 342 | Intron (ENSMUST00000070797.5 | 18 | 37175596 | 37178506 | 2911 | 2 | ENSMUST00 | 5930 | Gm10544 | ENSMUSG00 | false | . | . | . | . | . | . | chr18 | 37172319 | 37172850 | 3.21099 | 3.07268 | 1.51982 |
| chr1 | 119525959 | 119526427 | 342 | Promoter (<=1kb) | 1 | 119520090 | 119526359 | 6270 | 2 | ENSMUST00 | 0 | Gm27184 | ENSMUSG00 | false | chr1 | 119525551 | 119526807 | 30.03893 | 101.89226 | 98.2715 | chr1 | 119525242 | 119528011 | 5.49691 | 4.22768 | 3.4334 |
| chr15 | 102246145 | 102246623 | 341 | Promoter (<=1kb) | 15 | 102234940 | 102246500 | 11561 | 2 | ENSMUST00 | 0 | Rarg | ENSMUSG00 | false | chr15 | 102244836 | 102248267 | 30.17592 | 94.17345 | 90.73757 | chr15 | 102241789 | 102251883 | 5.05731 | 4.11666 | 3.0557 |
| chr1 | 134083017 | 134083376 | 341 | Distal Intergenic | 1 | 134075170 | 134079120 | 3951 | 2 | ENSMUST00 | -3898 | Btg2 | ENSMUSG00 | false | . | . | . | . | . | . | chr1 | 134079534 | 134084201 | 5.79764 | 4.37454 | 3.66417 |
| chr11 | 109995708 | 109996016 | 341 | Promoter (<=1kb) | 11 | 109932190 | 109995845 | 63656 | 2 | ENSMUST00 | 0 | Abca8b | ENSMUSG00 | false | chr11 | 109995458 | 109996216 | 11.28637 | 28.99159 | 26.53998 | chr11 | 109995182 | 109996222 | 4.52753 | 3.9226 | 2.60339 |
| chr16 | 16302739 | 16303067 | 341 | Promoter (<=1kb) | 16 | 16302965 | 16308942 | 5978 | 1 | ENSMUST00 | 0 | Yars2 | ENSMUSG00 | false | chr16 | 16302532 | 16303955 | 41.2653 | 138.78937 | 133.96024 | . | . | . | . | . | . |
| chr5 | 139149728 | 139150333 | 341 | Promoter (<=1kb) | 5 | 139108511 | 139150001 | 41491 | 2 | ENSMUST00 | 0 | Prkar1b | ENSMUSG00 | false | chr5 | 139149210 | 139149956 | 10.20448 | 21.76199 | 19.40286 | . | . | . | . | . | . |
| chr4 | 102883267 | 102883469 | 341 | Exon (ENSMUST00000142430.2/ | 4 | 102870564 | 102970603 | 100040 | 1 | ENSMUST00 | 12704 | Sgip1 | ENSMUSG00 | true | . | . | . | . | . | . | chr4 | 102882823 | 102887128 | 4.0975 | 3.64078 | 2.29202 |
| chr5 | 66620229 | 66620762 | 341 | Promoter (1-2kb) | 5 | 66298861 | 66618828 | 319968 | 2 | ENSMUST00 | -1402 | Apbb2 | ENSMUSG00 | false | chr5 | 66619941 | 66620884 | 26.33979 | 77.91899 | 74.80188 | chr5 | 66618990 | 66621424 | 6.79745 | 5.02559 | 4.46868 |
| chr5 | 131839139 | 131839458 | 341 | Intron (ENSMUST00000161226.3 | 5 | 131439080 | 132050337 | 611258 | 2 | ENSMUST00 | 210879 | Auts2 | ENSMUSG00 | true | . | . | . | . | . | . | chr5 | 131837773 | 131840213 | 5.04198 | 4.0695 | 3.08218 |
| chr8 | 48337589 | 48337813 | 341 | Intron (ENSMUST00000129821.2 | 8 | 48227682 | 48367363 | 139682 | 2 | ENSMUST00 | 29550 | Tenm3 | ENSMUSG00 | true | . | . | . | . | . | . | . | . | . | . | . | . |
| chr2 | 37780421 | 37780691 | 341 | Intron (ENSMUST00000050372.7 | 2 | 37776249 | 37799103 | 22855 | 1 | ENSMUST00 | 4173 | Crb2 | ENSMUSG00 | false | . | . | . | . | . | . | chr2 | 37777133 | 37803419 | 7.19536 | 4.98761 | 4.82001 |
| chr8 | 79639023 | 79639637 | 341 | Promoter (<=1kb) | 8 | 79639618 | 79677724 | 38107 | 1 | ENSMUST00 | 0 | Otud4 | ENSMUSG00 | false | chr8 | 79638419 | 79640775 | 37.25272 | 138.2439 | 133.43781 | . | . | . | . | . | . |
| chr6 | 144759167 | 144759447 | 341 | Intron (ENSMUST00000155562.1 | 6 | 144104452 | 144781919 | 677468 | 2 | ENSMUST00 | 22472 | Sox5 | ENSMUSG00 | true | . | . | . | . | . | . | chr6 | 144758737 | 144760275 | 5.84873 | 4.43163 | 3.70274 |
| chr13 | 94788796 | 94789446 | 341 | Promoter (<=1kb) | 13 | 94788943 | 94842899 | 53957 | 1 | ENSMUST00 | 0 | Tbca | ENSMUSG00 | false | chr13 | 94788578 | 94789846 | 37.19513 | 119.7376 | 115.60696 | chr13 | 94788674 | 94790498 | 3.77287 | 3.45577 | 2.02984 |
| chr15 | 7198832 | 7199099 | 341 | Distal Intergenic | 15 | 7212191 | 7223069 | 10879 | 2 | ENSMUST00 | 23970 | Egflam | ENSMUSG00 | false | . | . | . | . | . | . | chr15 | 7197761 | 7199461 | 5.05748 | 4.08356 | 3.05375 |
| chr5 | 139382841 | 139383217 | 341 | Promoter (2-3kb) | 5 | 139380581 | 139396415 | 15835 | 1 | ENSMUST00 | 2261 | Gpr146 | ENSMUSG00 | false | chr5 | 139380120 | 139383385 | 24.55236 | 90.57755 | 87.22021 | chr5 | 139381256 | 139395525 | 8.99937 | 5.68291 | 6.29852 |
| chr4 | 106972743 | 106973014 | 341 | Intron (ENSMUST00000150915.2 | 4 | 106966426 | 107031067 | 64642 | 1 | ENSMUST00 | 6318 | Ssbp3 | ENSMUSG00 | false | . | . | . | . | . | . | chr4 | 106968395 | 106974041 | 5.32502 | 4.11636 | 3.30677 |
| chr10 | 4107676 | 4108031 | 341 | Intron (ENSMUST00000117291.2 | 10 | 4104313 | 4104469 | 157 | 1 | ENSMUST00 | 3364 | Gm25515 | ENSMUSG00 | false | . | . | . | . | . | . | chr10 | 4105669 | 4108607 | 5.02974 | 4.06391 | 3.02794 |
| chr11 | 44367253 | 44367483 | 341 | Distal Intergenic | 11 | 44387115 | 44387274 | 160 | 2 | ENSMUST00 | 19791 | Gm12155 | ENSMUSG00 | false | . | . | . | . | . | . | . | . | . | . | . | . |
| chrX | 135990785 | 135990995 | 341 | Promoter (2-3kb) | 20 | 135988547 | 135989266 | 720 | 1 | ENSMUST00 | 2239 | Gm15016 | ENSMUSG00 | false | . | . | . | . | . | . | . | . | . | . | . | . |
| chr12 | 5186897 | 5187127 | 341 | Intron (ENSMUST00000020958.8 | 12 | 4999266 | 4999368 | 103 | 2 | ENSMUST00 | -187530 | Gm23167 | ENSMUSG00 | false | . | . | . | . | . | . | chr12 | 5185725 | 5189130 | 5.82568 | 4.49661 | 3.67837 |
| chr6 | 140148236 | 140148576 | 341 | Distal Intergenic | 6 | 140166535 | 140166645 | 111 | 2 | ENSMUST00 | 18069 | Gm24174 | ENSMUSG00 | false | . | . | . | . | . | . | chr6 | 140147471 | 140148769 | 3.68522 | 3.16454 | 1.96542 |
| chr7 | 71411597 | 71411819 | 341 | Distal Intergenic | 7 | 71370333 | 71379813 | 9481 | 1 | ENSMUST00 | 41265 | RP24-173C1 | ENSMUSG00 | false | chr7 | 71411271 | 71412350 | 13.32643 | 29.39681 | 26.9387 | chr7 | 71408173 | 71411949 | 5.13174 | 4.08589 | 3.11505 |
| chr3 | 100682821 | 100683114 | 340 | Promoter (1-2kb) | 3 | 100620033 | 100684613 | 64581 | 2 | ENSMUST00 | 1499 | Man1a2 | ENSMUSG00 | true | chr3 | 100680989 | 100686556 | 30.41442 | 103.5799 | 99.9162 | . | . | . | . | . | . |
| chr10 | 20554857 | 20555072 | 340 | Intron (ENSMUST00000020165.8 | 10 | 20407087 | 20548470 | 141384 | 2 | ENSMUST00 | -6388 | Pde7b | ENSMUSG00 | false | . | . | . | . | . | . | . | . | . | . | . | . |
| chr14 | 31001324 | 31002071 | 340 | Promoter (<=1kb) | 14 | 31001393 | 31012441 | 11049 | 1 | ENSMUST00 | 0 | Glt8d1 | ENSMUSG00 | false | chr14 | 31000426 | 31002352 | 33.04283 | 115.53907 | 111.53828 | . | . | . | . | . | . |
| chr17 | 47484234 | 47484631 | 340 | Distal Intergenic | 17 | 47488050 | 47502287 | 14238 | 2 | ENSMUST00 | 17656 | Taf8 | ENSMUSG00 | false | . | . | . | . | . | . | chr17 | 47483789 | 47485864 | 4.32642 | 3.6465 | 2.49295 |
| chr17 | 73845714 | 73846109 | 340 | Distal Intergenic | 17 | 73804841 | 73832093 | 27253 | 1 | ENSMUST00 | 40874 | Ehd3 | ENSMUSG00 | false | . | . | . | . | . | . | . | . | . | . | . | . |
| chrX | 167382634 | 167382911 | 340 | Promoter (<=1kb) | 20 | 167346322 | 167382749 | 36428 | 2 | ENSMUST00 | 0 | Prps2 | ENSMUSG00 | false | chrX | 167381725 | 167383017 | 18.84082 | 47.13427 | 44.45482 | . | . | . | . | . | . |
| chr5 | 92309722 | 92310165 | 340 | Promoter (<=1kb) | 5 | 92284010 | 92310024 | 26015 | 2 | ENSMUST00 | 0 | Sdad1 | ENSMUSG00 | false | chr5 | 92309245 | 92310576 | 25.62849 | 83.19241 | 79.98436 | . | . | . | . | . | . |
| chr5 | 117976636 | 117976935 | 340 | Promoter (<=1kb) | 5 | 117976770 | 118010191 | 33422 | 1 | ENSMUST00 | 0 | Fbxo21 | ENSMUSG00 | false | chr5 | 117976342 | 117977742 | 34.49762 | 142.80731 | 137.81027 | chr5 | 117975985 | 117978599 | 5.5059 | 4.3586 | 3.42256 |
| chr12 | 76875180 | 76875395 | 340 | Intron (ENSMUST00000125842.2 | 12 | 76887058 | 76910246 | 23189 | 1 | ENSMUST00 | -11663 | Fntb | ENSMUSG00 | false | . | . | . | . | . | . | chr12 | 76874034 | 76875372 | 3.0085 | 3.03674 | 1.39842 |
| chrX | 106027126 | 106027351 | 340 | Promoter (<=1kb) | 20 | 106027276 | 106124917 | 97642 | 1 | ENSMUST00 | 0 | Atp7a | ENSMUSG00 | false | chrX | 106027255 | 106028197 | 17.00269 | 41.02908 | 38.42754 | . | . | . | . | . | . |
| chr4 | 14826302 | 14826761 | 340 | Promoter (<=1kb) | 4 | 14809498 | 14826587 | 17090 | 2 | ENSMUST00 | 0 | Otud6b | ENSMUSG00 | false | chr4 | 14825500 | 14827022 | 24.31009 | 69.8498 | 66.86187 | . | . | . | . | . | . |
| chr6 | 118128204 | 118128459 | 340 | Promoter (1-2kb) | 6 | 118107452 | 118129620 | 22169 | 2 | ENSMUST00 | 1161 | Csgalnact2 | ENSMUSG00 | false | . | . | . | . | . | . | . | . | . | . | . | . |
| chr9 | 6313896 | 6314129 | 340 | Intron (ENSMUST00000168039.2 | 9 | 6265028 | 6266547 | 1520 | 2 | ENSMUST00 | -47350 | Ddi1 | ENSMUSG00 | false | . | . | . | . | . | . | . | . | . | . | . | . |
| chr3 | 107314648 | 107314862 | 340 | Distal Intergenic | 3 | 107326110 | 107333289 | 7180 | 2 | ENSMUST00 | 18427 | Rbm15 | ENSMUSG00 | false | . | . | . | . | . | . | chr3 | 107310153 | 107317382 | 4.20648 | 3.66271 | 2.36734 |
| chr11 | 97400355 | 97400628 | 340 | Distal Intergenic | 11 | 97415533 | 97502402 | 86870 | 1 | ENSMUST00 | -14905 | Arhgap23 | ENSMUSG00 | true | . | . | . | . | . | . | chr11 | 97400274 | 97400978 | 4.40213 | 3.81989 | 2.56326 |
| chr12 | 69437181 | 69437409 | 340 | Distal Intergenic | 12 | 69371835 | 69372458 | 624 | 2 | ENSMUST00 | -64724 | Gm9887 | ENSMUSG00 | false | . | . | . | . | . | . | chr12 | 69436684 | 69444561 | 5.15172 | 4.15476 | 3.12711 |
| chr5 | 129260833 | 129261041 | 340 | Distal Intergenic | 5 | 129128077 | 129128517 | 441 | 2 | ENSMUST00 | -132317 | Rps16-ps2 | ENSMUSG00 | false | . | . | . | . | . | . | . | . | . | . | . | . |
| chr10 | 12530780 | 12531003 | 340 | Intron (ENSMUST00000076817.4 | 10 | 12368623 | 12368832 | 210 | 1 | ENSMUST00 | 162158 | Gm25139 | ENSMUSG00 | false | . | . | . | . | . | . | chr10 | 12530563 | 12531943 | 4.43177 | 3.86768 | 2.53606 |
| chr7 | 25835206 | 25835869 | 340 | Distal Intergenic | 7 | 25828587 | 25828877 | 291 | 2 | ENSMUST00 | -6330 | Gm4613 | ENSMUSG00 | false | . | . | . | . | . | . | chr7 | 25835034 | 25836175 | 4.99559 | 4.1404 | 3.02681 |
| chr10 | 111595932 | 111596131 | 340 | Promoter (1-2kb) | 10 | 111586719 | 111594273 | 7555 | 2 | ENSMUST00 | -1660 | 4933440J02F | ENSMUSG00 | false | . | . | . | . | . | . | chr10 | 111595564 | 111596260 | 5.86157 | 4.31397 | 3.76177 |
| chr9 | 34689415 | 34689600 | 340 | Intron (ENSMUST00000115148.3 | 9 | 34588526 | 34589304 | 779 | 2 | ENSMUST00 | -100112 | Gm26662 | ENSMUSG00 | false | . | . | . | . | . | . | chr9 | 34688749 | 34690636 | 4.35138 | 3.6219 | 2.4975 |
| chr11 | 50236583 | 50237018 | 340 | Promoter (<=1kb) | 11 | 50237009 | 50288839 | 51831 | 1 | ENSMUST00 | 0 | Gm26542 | ENSMUSG00 | false | chr11 | 50236154 | 50237650 | 26.61085 | 108.21047 | 104.42454 | chr11 | 50235920 | 50238486 | 4.62795 | 3.77419 | 2.73646 |
| chr12 | 50629895 | 50630234 | 339 | Intron (ENSMUST00000002765.7 | 12 | 50341231 | 50649223 | 307993 | 2 | ENSMUST00 | 18989 | Prkd1 | ENSMUSG00 | false | . | . | . | . | . | . | chr12 | 50628911 | 50630085 | 4.75931 | 4.04183 | 2.80083 |
| chr10 | 111164445 | 111164917 | 339 | Promoter (<=1kb) | 10 | 111164802 | 111297247 | 132446 | 1 | ENSMUST00 | 0 | Osbpl8 | ENSMUSG00 | false | chr10 | 111164281 | 111166238 | 36.53815 | 117.53558 | 113.47088 | . | . | . | . | . | . |
| chr10 | 120112635 | 120113092 | 339 | Promoter (<=1kb) | 10 | 120083608 | 120112987 | 29380 | 2 | ENSMUST00 | 0 | Helb | ENSMUSG00 | false | chr10 | 120112319 | 120113648 | 27.51607 | 79.95555 | 76.80458 | . | . | . | . | . | . |
| chr15 | 58214936 | 58215418 | 339 | Promoter (<=1kb) | 15 | 58175879 | 58214892 | 39014 | 2 | ENSMUST00 | -45 | Fbxo32 | ENSMUSG00 | false | chr15 | 58213813 | 58215815 | 29.14352 | 87.50663 | 84.2095 | chr15 | 58212631 | 58216083 | 4.59828 | 3.94214 | 2.66723 |
| chr16 | 37615509 | 37615783 | 339 | Intron (ENSMUST00000160847.1 | 16 | 37621849 | 37632020 | 10172 | 1 | ENSMUST00 | -6066 | Hgd | ENSMUSG00 | false | . | . | . | . | . | . | chr16 | 37614959 | 37618754 | 4.96558 | 4.04706 | 2.98024 |
| chr17 | 67904340 | 67904612 | 339 | Promoter (2-3kb) | 17 | 67845735 | 67901471 | 55737 | 2 | ENSMUST00 | -2870 | Arhgap28 | ENSMUSG00 | true | . | . | . | . | . | . | chr17 | 67892935 | 67914104 | 5.71307 | 4.38398 | 3.59861 |
| chr18 | 13815159 | 13815364 | 339 | Intron (ENSMUST00000025288.7 | 18 | 13687013 | 13972733 | 285721 | 2 | ENSMUST00 | 157369 | Zfp521 | ENSMUSG00 | true | . | . | . | . | . | . | chr18 | 13813194 | 13816052 | 4.29835 | 3.68495 | 2.46993 |
| chr19 | 53653980 | 53654338 | 339 | Distal Intergenic | 19 | 53643358 | 53644808 | 1451 | 1 | ENSMUST00 | 10623 | Smc3 | ENSMUSG00 | false | . | . | . | . | . | . | chr19 | 53653888 | 53654738 | 3.82177 | 3.49032 | 2.02569 |
| chr2 | 5895119 | 5895626 | 339 | Promoter (<=1kb) | 2 | 5870987 | 5895432 | 24446 | 2 | ENSMUST00 | 0 | Sec61a2 | ENSMUSG00 | false | chr2 | 5894821 | 5895828 | 31.81669 | 98.02396 | 94.50304 | . | . | . | . | . | . |
| chr1 | 165667362 | 165667686 | 339 | Intron (ENSMUST00000040357.9 | 1 | 165592181 | 165634541 | 42361 | 2 | ENSMUST00 | -32822 | Mpzl1 | ENSMUSG00 | false | . | . | . | . | . | . | chr1 | 165663194 | 165707027 | 5.93252 | 4.47383 | 3.77552 |
| chrX | 20704415 | 20704740 | 339 | Promoter (<=1kb) | 20 | 20703960 | 20717685 | 13726 | 1 | ENSMUST00 | 456 | Usp11 | ENSMUSG00 | false | chrX | 20703768 | 20705244 | 12.58327 | 33.30664 | 30.79935 | . | . | . | . | . | . |
| chr2 | 135658533 | 135658780 | 339 | Promoter (<=1kb) | 2 | 135659011 | 135742751 | 83741 | 1 | ENSMUST00 | -231 | Plcb4 | ENSMUSG00 | false | chr2 | 135658344 | 135660234 | 29.92778 | 109.76675 | 105.9399 | . | . | . | . | . | . |
| chr1 | 136131123 | 136131513 | 339 | Promoter (<=1kb) | 1 | 136131389 | 136177998 | 46610 | 1 | ENSMUST00 | 0 | Kif21b | ENSMUSG00 | false | chr1 | 136130929 | 136131999 | 22.44583 | 74.56482 | 71.50171 | chr1 | 136130113 | 136133251 | 3.95004 | 3.63828 | 2.1239 |
| chr4 | 11007290 | 11007943 | 339 | Promoter (<=1kb) | 4 | 10988662 | 11007676 | 19015 | 2 | ENSMUST00 | 0 | Plekhf2 | ENSMUSG00 | false | chr4 | 11005934 | 11008487 | 38.50526 | 122.00793 | 117.8042 | . | . | . | . | . | . |
| chr10 | 78124190 | 78124561 | 339 | Distal Intergenic | 10 | 78162067 | 78169768 | 7702 | 2 | ENSMUST00 | 45207 | D10Jhu81e | ENSMUSG00 | false | chr10 | 78124273 | 78124732 | 4.66184 | 7.82715 | 5.69732 | chr10 | 78122907 | 78125702 | 4.51385 | 3.643 | 2.64751 |
| chr4 | 129357842 | 129358108 | 339 | Exon (ENSMUST00000030610.2/ | 4 | 129343940 | 129346930 | 2991 | 1 | ENSMUST00 | 13903 | Zbtb8os | ENSMUSG00 | false | chr4 | 129356376 | 129357908 | 14.70503 | 33.66069 | 31.14909 | chr4 | 129350486 | 129375787 | 4.55883 | 3.85552 | 2.64801 |
| chr7 | 139272281 | 139272464 | 339 | Distal Intergenic | 7 | 139248482 | 139267253 | 18772 | 1 | ENSMUST00 | 23800 | Pwwp2b | ENSMUSG00 | false | . | . | . | . | . | . | chr7 | 139270937 | 139275472 | 8.52846 | 5.56821 | 5.90402 |
| chr8 | 22687872 | 22688141 | 339 | Promoter (1-2kb) | 8 | 22686265 | 22695646 | 9382 | 1 | ENSMUST00 | 1608 | Gm15346 | ENSMUSG00 | false | . | . | . | . | . | . | chr8 | 22686451 | 22699233 | 6.12277 | 4.57796 | 3.92686 |
| chr11 | 18712208 | 18712460 | 339 | Promoter (1-2kb) | 11 | 18595248 | 18710222 | 114975 | 2 | ENSMUST00 | -1987 | Gm12023 | ENSMUSG00 | false | . | . | . | . | . | . | chr11 | 18712128 | 18712565 | 3.67173 | 3.27901 | 1.93859 |
| chr13 | 103211498 | 103211767 | 339 | Intron (ENSMUST00000166726.2 | 13 | 103159795 | 103164069 | 4275 | 2 | ENSMUST00 | -47430 | Gm6211 | ENSMUSG00 | false | . | . | . | . | . | . | . | . | . | . | . | . |
| chr16 | 29734956 | 29735226 | 339 | Distal Intergenic | 16 | 29811058 | 29837814 | 26757 | 1 | ENSMUST00 | -75832 | Gm26834 | ENSMUSG00 | false | . | . | . | . | . | . | chr16 | 29734294 | 29735810 | 3.98774 | 3.39378 | 2.21004 |
| chr5 | 137293365 | 137293665 | 339 | Promoter (<=1kb) | 5 | 137293780 | 137293876 | 97 | 1 | ENSMUST00 | -115 | Mir8116 | ENSMUSG00 | false | chr5 | 137293142 | 137295178 | 30.78865 | 90.34473 | 86.99165 | chr5 | 137293109 | 137295664 | 7.26578 | 5.11561 | 4.89474 |
| chr2 | 167690629 | 167691299 | 338 | Promoter (<=1kb) | 2 | 167691177 | 167729172 | 37996 | 1 | ENSMUST00 | 0 | A530013C23 | ENSMUSG00 | false | chr2 | 167688507 | 167691530 | 36.76257 | 113.72284 | 109.77938 | chr2 | 167691144 | 167694518 | 4.76166 | 3.87486 | 2.8261 |
| chr10 | 115241874 | 115242084 | 338 | Intron (ENSMUST00000020339.8 | 10 | 115197877 | 115251493 | 53617 | 2 | ENSMUST00 | 9409 | Tbc1d15 | ENSMUSG00 | false | . | . | . | . | . | . | chr10 | 115241369 | 115250913 | 4.81613 | 4.0347 | 2.85181 |
| chr15 | 44628301 | 44628515 | 338 | Exon (ENSMUST00000022964.7/ | 15 | 44619641 | 44641024 | 21384 | 1 | ENSMUST00 | 8661 | Ebag9 | ENSMUSG00 | false | . | . | . | . | . | . | chr15 | 44627493 | 44628961 | 3.48362 | 3.37434 | 1.7397 |
| chr19 | 17797829 | 17798043 | 338 | Intron (ENSMUST00000025618.9 | 19 | 17432832 | 17837632 | 404801 | 2 | ENSMUST00 | 39589 | Pcsk5 | ENSMUSG00 | true | . | . | . | . | . | . | chr19 | 17796432 | 17798717 | 4.24138 | 3.6271 | 2.41496 |
| chr4 | 155624565 | 155624925 | 338 | Promoter (<=1kb) | 4 | 155624854 | 155630340 | 5487 | 1 | ENSMUST00 | 0 | Cdk11b | ENSMUSG00 | false | chr4 | 155623921 | 155625589 | 31.16539 | 106.97112 | 103.21663 | chr4 | 155623632 | 155626506 | 6.55562 | 4.72454 | 4.33546 |
| chr5 | 129586056 | 129586299 | 338 | Promoter (1-2kb) | 5 | 129584169 | 129608875 | 24707 | 1 | ENSMUST00 | 1888 | Mmp17 | ENSMUSG00 | false | chr5 | 129583568 | 129586214 | 36.24269 | 113.86979 | 109.91653 | chr5 | 129583379 | 129586671 | 4.85672 | 4.00401 | 2.89177 |
| chr6 | 33773015 | 33773244 | 338 | Intron (ENSMUST00000052266.9 | 6 | 33701434 | 33862160 | 160727 | 1 | ENSMUST00 | 71582 | Exoc4 | ENSMUSG00 | false | . | . | . | . | . | . | . | . | . | . | . | . |
| chr6 | 87042477 | 87042841 | 338 | Promoter (<=1kb) | 6 | 87042846 | 87055424 | 12579 | 1 | ENSMUST00 | -5 | Gfpt1 | ENSMUSG00 | false | chr6 | 87042081 | 87043758 | 26.72788 | 101.58999 | 97.97913 | . | . | . | . | . | . |
| chr7 | 35388732 | 35389034 | 338 | Intron (ENSMUST00000032705.7 | 7 | 35382565 | 35385210 | 2646 | 1 | ENSMUST00 | 6168 | Rhpn2 | ENSMUSG00 | false | . | . | . | . | . | . | chr7 | 35388378 | 35391555 | 5.24347 | 4.02438 | 3.25059 |
| chr9 | 82800421 | 82800631 | 338 | Distal Intergenic | 9 | 82829540 | 82847687 | 18148 | 1 | ENSMUST00 | -28909 | Irak1bp1 | ENSMUSG00 | false | . | . | . | . | . | . | . | . | . | . | . | . |
| chr11 | 114684493 | 114684780 | 338 | Intron (ENSMUST00000045779.5 | 11 | 114675516 | 114690603 | 15088 | 1 | ENSMUST00 | 8978 | Ttyh2 | ENSMUSG00 | false | . | . | . | . | . | . | chr11 | 114684432 | 114685244 | 4.00603 | 3.58811 | 2.24038 |
| chr5 | 34466179 | 34466494 | 338 | Promoter (<=1kb) | 5 | 34466129 | 34477689 | 11561 | 1 | ENSMUST00 | 51 | Fam193a | ENSMUSG00 | false | . | . | . | . | . | . | chr5 | 34465382 | 34469742 | 3.9432 | 3.59158 | 2.13788 |
| chr18 | 34861925 | 34862207 | 338 | Promoter (<=1kb) | 18 | 34861207 | 34864956 | 3750 | 1 | ENSMUST00 | 719 | Egr1 | ENSMUSG00 | false | chr18 | 34859236 | 34862643 | 33.41831 | 117.26761 | 113.21259 | chr18 | 34858199 | 34863782 | 4.89469 | 4.00673 | 2.91424 |
| chr11 | 95365725 | 95366069 | 338 | Intron (ENSMUST00000037502.6 | 11 | 95375467 | 95377585 | 2119 | 1 | ENSMUST00 | -9398 | Fam117a | ENSMUSG00 | false | . | . | . | . | . | . | chr11 | 95364434 | 95372547 | 6.52296 | 4.667 | 4.26701 |
| chrX | 159809643 | 159809828 | 338 | Intron (ENSMUST00000112456.3 | 20 | 159840468 | 159975664 | 135197 | 1 | ENSMUST00 | -30640 | Sh3kbp1 | ENSMUSG00 | false | . | . | . | . | . | . | . | . | . | . | . | . |
| chr11 | 120231296 | 120231728 | 338 | Promoter (<=1kb) | 11 | 120229802 | 120231585 | 1784 | 2 | ENSMUST00 | 0 | 2900052L18F | ENSMUSG00 | false | chr11 | 120229779 | 120231698 | 35.98183 | 119.73972 | 115.60696 | chr11 | 120231721 | 120232770 | 11.0773 | 6.71912 | 7.9603 |
| chr2 | 74715881 | 74716126 | 338 | Promoter (<=1kb) | 2 | 74710193 | 74715996 | 5804 | 2 | ENSMUST00 | 0 | Hoxd3os1 | ENSMUSG00 | false | chr2 | 74715539 | 74716285 | 7.14815 | 15.37828 | 13.11012 | chr2 | 74713292 | 74716754 | 5.00916 | 4.13191 | 3.01287 |
| chr6 | 72076127 | 72076688 | 338 | Distal Intergenic | 6 | 72097592 | 72154571 | 56980 | 1 | ENSMUST00 | -20904 | St3gal5 | ENSMUSG00 | false | . | . | . | . | . | . | chr6 | 72075855 | 72077995 | 4.75064 | 3.99388 | 2.83376 |
| chr11 | 4697732 | 4697997 | 338 | Distal Intergenic | 11 | 4701982 | 4704053 | 2072 | 2 | ENSMUST00 | 6056 | Uqcr10 | ENSMUSG00 | false | . | . | . | . | . | . | chr11 | 4697323 | 4698271 | 3.48775 | 3.2625 | 1.8134 |
| chr12 | 76403568 | 76404374 | 338 | Promoter (<=1kb) | 12 | 76404176 | 76406934 | 2759 | 1 | ENSMUST00 | 0 | Hspa2 | ENSMUSG00 | false | chr12 | 76403128 | 76406376 | 31.07551 | 97.73014 | 94.2132 | . | . | . | . | . | . |
| chr17 | 5001046 | 5001307 | 338 | Intron (ENSMUST00000124362.1 | 17 | 4996429 | 5098186 | 101758 | 1 | ENSMUST00 | 4618 | Arid1b | ENSMUSG00 | false | . | . | . | . | . | . | chr17 | 4997270 | 5019445 | 5.48495 | 4.15675 | 3.43424 |
| chr4 | 11975244 | 11975609 | 338 | Intron (ENSMUST00000154194.1 | 4 | 11980162 | 11981661 | 1500 | 1 | ENSMUST00 | -4553 | 1700123M08 | ENSMUSG00 | false | . | . | . | . | . | . | chr4 | 11974214 | 11976665 | 5.84877 | 4.53422 | 3.69662 |
| chr5 | 8056203 | 8056704 | 337 | Promoter (<=1kb) | 5 | 8056527 | 8069151 | 12625 | 1 | ENSMUST00 | 0 | Sri | ENSMUSG00 | false | chr5 | 8055753 | 8057709 | 30.78865 | 90.34473 | 86.99165 | . | . | . | . | . | . |
| chr19 | 32755896 | 32756609 | 337 | Promoter (<=1kb) | 19 | 32757497 | 32826160 | 68664 | 1 | ENSMUST00 | -888 | Pten | ENSMUSG00 | false | chr19 | 32755517 | 32759967 | 38.34636 | 121.73327 | 117.53308 | . | . | . | . | . | . |
| chr14 | 12132299 | 12132549 | 337 | Intron (ENSMUST00000022264.7 | 14 | 12091160 | 12122132 | 30973 | 1 | ENSMUST00 | 41140 | Ptprg | ENSMUSG00 | false | . | . | . | . | . | . | . | . | . | . | . | . |
| chr11 | 120672860 | 120673464 | 337 | Promoter (<=1kb) | 11 | 120672973 | 120674346 | 1374 | 1 | ENSMUST00 | 0 | Aspscr1 | ENSMUSG00 | false | chr11 | 120672681 | 120674029 | 29.3377 | 90.74815 | 87.38457 | . | . | . | . | . | . |
| chrX | 52987875 | 52988302 | 337 | Promoter (<=1kb) | 20 | 52988137 | 53021659 | 33523 | 1 | ENSMUST00 | 0 | Hprt | ENSMUSG00 | false | chrX | 52987728 | 52989091 | 21.59801 | 56.60135 | 53.80005 | . | . | . | . | . | . |
| chr2 | 144545960 | 144546274 | 337 | Intron (ENSMUST00000028915.5 | 2 | 144543821 | 144550790 | 6970 | 2 | ENSMUST00 | 4516 | Rbbp9 | ENSMUSG00 | false | . | . | . | . | . | . | chr2 | 144544550 | 144548141 | 6.05889 | 4.60451 | 3.87024 |
| chr3 | 9110798 | 9111040 | 337 | Distal Intergenic | 3 | 8928626 | 9004723 | 76098 | 2 | ENSMUST00 | -106076 | Tpd52 | ENSMUSG00 | false | chr3 | 9110969 | 9111240 | 4.69649 | 6.99982 | 4.89137 | chr3 | 9109447 | 9112114 | 5.15216 | 4.01034 | 3.17919 |
| chr3 | 58525789 | 58526057 | 337 | Promoter (<=1kb) | 3 | 58521969 | 58525873 | 3905 | 2 | ENSMUST00 | 0 | Serp1 | ENSMUSG00 | false | chr3 | 58524418 | 58526697 | 43.8714 | 181.39459 | 175.20454 | . | . | . | . | . | . |
| chr4 | 135987047 | 135987588 | 337 | Promoter (<=1kb) | 4 | 135975460 | 135987264 | 11805 | 2 | ENSMUST00 | 0 | Pithd1 | ENSMUSG00 | false | chr4 | 135986291 | 135987789 | 26.28407 | 85.32412 | 82.07336 | chr4 | 135985127 | 135987675 | 4.63725 | 3.74914 | 2.74075 |
| chr4 | 134852760 | 134853406 | 337 | Promoter (<=1kb) | 4 | 134802759 | 134853345 | 50587 | 2 | ENSMUST00 | 0 | Tmem57 | ENSMUSG00 | false | chr4 | 134852051 | 134854505 | 36.04346 | 118.78738 | 114.68494 | . | . | . | . | . | . |
| chr5 | 123344719 | 123345709 | 337 | Promoter (<=1kb) | 5 | 123344500 | 123360278 | 15779 | 1 | ENSMUST00 | 220 | Bcl7a | ENSMUSG00 | false | chr5 | 123343837 | 123346514 | 34.14626 | 110.2223 | 106.38018 | . | . | . | . | . | . |
| chr5 | 122354132 | 122354492 | 337 | Promoter (<=1kb) | 5 | 122354369 | 122364984 | 10616 | 1 | ENSMUST00 | 0 | Vps29 | ENSMUSG00 | false | chr5 | 122353779 | 122355233 | 32.11694 | 110.747 | 106.89115 | . | . | . | . | . | . |
| chr7 | 128417794 | 128418572 | 337 | Promoter (<=1kb) | 7 | 128373621 | 128418173 | 44553 | 2 | ENSMUST00 | 0 | Rgs10 | ENSMUSG00 | false | chr7 | 128417382 | 128418878 | 27.29656 | 83.33401 | 80.12321 | chr7 | 128417312 | 128419277 | 4.01732 | 3.59431 | 2.21817 |
| chr8 | 13677129 | 13677803 | 337 | Promoter (<=1kb) | 8 | 13566950 | 13677603 | 110654 | 2 | ENSMUST00 | 0 | Rasa3 | ENSMUSG00 | false | chr8 | 13676275 | 13678189 | 36.39105 | 113.08218 | 109.16298 | . | . | . | . | . | . |
| chr7 | 99405612 | 99405992 | 337 | Intron (ENSMUST00000037528.8 | 7 | 99381549 | 99460983 | 79435 | 1 | ENSMUST00 | 24064 | Gdpd5 | ENSMUSG00 | true | . | . | . | . | . | . | chr7 | 99399797 | 99417330 | 7.01691 | 4.96429 | 4.66748 |
| chr5 | 100415910 | 100416472 | 337 | Promoter (<=1kb) | 5 | 100361649 | 100416234 | 54586 | 2 | ENSMUST00 | 0 | Sec31a | ENSMUSG00 | false | chr5 | 100415482 | 100416869 | 34.60421 | 120.46815 | 116.3158 | . | . | . | . | . | . |
| chr19 | 45016211 | 45017057 | 337 | Promoter (1-2kb) | 19 | 45015210 | 45027104 | 11895 | 1 | ENSMUST00 | 1002 | Lzts2 | ENSMUSG00 | false | chr19 | 45014893 | 45017754 | 22.05754 | 58.21236 | 55.38818 | chr19 | 45012490 | 45017626 | 4.84179 | 3.90507 | 2.90417 |
| chr12 | 21138512 | 21138789 | 337 | Intron (ENSMUST00000174530.1 | 12 | 21111954 | 21268179 | 156226 | 1 | ENSMUST00 | 26559 | Asap2 | ENSMUSG00 | false | . | . | . | . | . | . | chr12 | 21122843 | 21147439 | 6.77508 | 4.82373 | 4.46417 |
| chr7 | 75835323 | 75835507 | 337 | Distal Intergenic | 7 | 75848338 | 75874130 | 25793 | 1 | ENSMUST00 | -12831 | Klhl25 | ENSMUSG00 | false | . | . | . | . | . | . | . | . | . | . | . | . |
| chr3 | 116291429 | 116291675 | 337 | Intron (ENSMUST00000106491.2 | 3 | 116249654 | 116253484 | 3831 | 2 | ENSMUST00 | -37946 | Gpr88 | ENSMUSG00 | false | . | . | . | . | . | . | chr3 | 116290475 | 116294463 | 4.52794 | 3.8676 | 2.61391 |
| chr10 | 11149360 | 11149936 | 337 | Promoter (<=1kb) | 10 | 11149427 | 11188150 | 38724 | 1 | ENSMUST00 | 0 | Shprh | ENSMUSG00 | false | chr10 | 11148950 | 11150431 | 35.29575 | 125.98199 | 121.63858 | . | . | . | . | . | . |
| chr18 | 77732634 | 77733116 | 337 | Promoter (<=1kb) | 18 | 77729681 | 77733470 | 3790 | 2 | ENSMUST00 | 354 | A330094K24 | ENSMUSG00 | false | chr18 | 77732034 | 77733389 | 10.14765 | 21.66544 | 19.30706 | chr18 | 77731377 | 77733851 | 4.82693 | 4.04298 | 2.84945 |
| chr7 | 45370101 | 45370705 | 336 | Promoter (<=1kb) | 7 | 45367891 | 45370583 | 2693 | 2 | ENSMUST00 | 0 | Lin7b | ENSMUSG00 | false | chr7 | 45369930 | 45371040 | 17.19387 | 43.56631 | 40.93357 | chr7 | 45369742 | 45371023 | 5.99417 | 4.6318 | 3.81918 |
| chr15 | 85336209 | 85336909 | 336 | Promoter (<=1kb) | 15 | 85336381 | 85463836 | 127456 | 1 | ENSMUST00 | 0 | Atxn10 | ENSMUSG00 | true | chr15 | 85336032 | 85337422 | 31.19535 | 97.93977 | 94.4203 | . | . | . | . | . | . |
| chr13 | 93499520 | 93500041 | 336 | Promoter (<=1kb) | 13 | 93430097 | 93499808 | 69712 | 2 | ENSMUST00 | 0 | Jmy | ENSMUSG00 | false | chr13 | 93497758 | 93500424 | 37.95786 | 125.34732 | 121.02549 | . | . | . | . | . | . |
| chr18 | 13886340 | 13886632 | 336 | Intron (ENSMUST00000025288.7 | 18 | 13687013 | 13972733 | 285721 | 2 | ENSMUST00 | 86101 | Zfp521 | ENSMUSG00 | true | . | . | . | . | . | . | chr18 | 13884925 | 13898147 | 5.50345 | 4.32012 | 3.42059 |
| chr1 | 40222623 | 40222973 | 336 | Promoter (2-3kb) | 1 | 40225080 | 40316177 | 91098 | 1 | ENSMUST00 | -2107 | Il1r1 | ENSMUSG00 | false | . | . | . | . | . | . | chr1 | 40221760 | 40225801 | 4.10563 | 3.65212 | 2.26792 |
| chr4 | 129377678 | 129378165 | 336 | Promoter (<=1kb) | 4 | 129353628 | 129378116 | 24489 | 2 | ENSMUST00 | 0 | Zbtb8a | ENSMUSG00 | true | chr4 | 129376396 | 129378534 | 18.2599 | 77.14165 | 74.04132 | chr4 | 129378069 | 129387044 | 4.79047 | 3.93835 | 2.85504 |
| chr5 | 125140862 | 125141576 | 336 | Intron (ENSMUST00000111402.3 | 5 | 125017835 | 125124900 | 107066 | 2 | ENSMUST00 | -15963 | Ncor2 | ENSMUSG00 | false | chr5 | 125140272 | 125141740 | 11.71324 | 27.89317 | 25.45456 | chr5 | 125124978 | 125178451 | 6.70656 | 4.76008 | 4.42049 |
| chrX | 99243033 | 99243219 | 336 | Distal Intergenic | 20 | 99138131 | 99147230 | 9100 | 1 | ENSMUST00 | 104903 | Efnb1 | ENSMUSG00 | false | . | . | . | . | . | . | . | . | . | . | . | . |
| chr8 | 123859375 | 123859731 | 336 | Promoter (<=1kb) | 8 | 123840831 | 123859530 | 18700 | 2 | ENSMUST00 | 0 | Ccsap | ENSMUSG00 | false | chr8 | 123858622 | 123860243 | 34.45293 | 106.63475 | 102.89172 | chr8 | 123858428 | 123859805 | 5.18631 | 4.0218 | 3.17163 |
| chr5 | 103389101 | 103389294 | 336 | Distal Intergenic | 5 | 103425192 | 103598359 | 173168 | 1 | ENSMUST00 | -35898 | Ptpn13 | ENSMUSG00 | false | . | . | . | . | . | . | . | . | . | . | . | . |
| chr9 | 53307931 | 53308209 | 336 | Intron (ENSMUST00000051014.1 | 9 | 53301670 | 53377514 | 75845 | 1 | ENSMUST00 | 6262 | Exph5 | ENSMUSG00 | false | . | . | . | . | . | . | chr9 | 53307727 | 53308720 | 3.66372 | 3.41629 | 1.89321 |
| chr11 | 98954411 | 98954753 | 336 | Intron (ENSMUST00000107474.2 | 11 | 98960412 | 98974942 | 14531 | 1 | ENSMUST00 | -5659 | Rara | ENSMUSG00 | false | . | . | . | . | . | . | chr11 | 98928668 | 98957642 | 6.70032 | 4.70048 | 4.41905 |
| chr5 | 121618550 | 121618950 | 336 | Promoter (<=1kb) | 5 | 121598284 | 121618938 | 20655 | 2 | ENSMUST00 | 0 | Acad12 | ENSMUSG00 | false | chr5 | 121618332 | 121619845 | 29.08755 | 89.33995 | 86.00743 | . | . | . | . | . | . |
| chr10 | 95217822 | 95218237 | 336 | Intron (ENSMUST00000053594.5 | 10 | 95174746 | 95324097 | 149352 | 2 | ENSMUST00 | 105860 | Cradd | ENSMUSG00 | false | . | . | . | . | . | . | chr10 | 95217715 | 95219394 | 5.72451 | 4.38435 | 3.60575 |
| chr2 | 109917584 | 109917858 | 336 | Promoter (<=1kb) | 2 | 109917647 | 110014257 | 96611 | 1 | ENSMUST00 | 0 | Lgr4 | ENSMUSG00 | false | chr2 | 109917200 | 109921050 | 28.59262 | 111.22587 | 107.35971 | . | . | . | . | . | . |
| chr11 | 96687022 | 96687238 | 336 | Intron (ENSMUST00000103154.5 | 11 | 96708235 | 96713202 | 4968 | 1 | ENSMUST00 | -20997 | Skap1 | ENSMUSG00 | false | . | . | . | . | . | . | . | . | . | . | . | . |
| chr19 | 24251923 | 24252167 | 336 | Distal Intergenic | 19 | 24261862 | 24278171 | 16310 | 2 | ENSMUST00 | 26004 | Fxn | ENSMUSG00 | false | . | . | . | . | . | . | . | . | . | . | . | . |
| chr9 | 20607146 | 20607722 | 336 | Promoter (<=1kb) | 9 | 20607481 | 20617271 | 9791 | 1 | ENSMUST00 | 0 | Fbxl12os | ENSMUSG00 | false | chr9 | 20606949 | 20608085 | 32.4768 | 102.1866 | 98.56064 | . | . | . | . | . | . |
| chr19 | 25168894 | 25169151 | 336 | Promoter (2-3kb) | 19 | 25166085 | 25166216 | 132 | 2 | ENSMUST00 | -2679 | Gm24252 | ENSMUSG00 | false | . | . | . | . | . | . | chr19 | 25162630 | 25181812 | 5.3445 | 4.22539 | 3.29608 |
| chr8 | 79113600 | 79113844 | 336 | Intron (ENSMUST00000098614.3 | 8 | 79118327 | 79193766 | 75440 | 1 | ENSMUST00 | -4483 | Zfp827 | ENSMUSG00 | false | . | . | . | . | . | . | chr8 | 79111391 | 79116090 | 5.16683 | 3.99061 | 3.14481 |
| chr2 | 159353459 | 159353747 | 336 | Distal Intergenic | 2 | 159456170 | 159456457 | 288 | 2 | ENSMUST00 | 102710 | Gm14219 | ENSMUSG00 | false | . | . | . | . | . | . | chr2 | 159353426 | 159358106 | 4.96229 | 3.96932 | 2.99075 |
| chr8 | 88082185 | 88082415 | 336 | Intron (ENSMUST00000181500.1 | 8 | 88044898 | 88118607 | 73710 | 2 | ENSMUST00 | 36192 | Gm2716 | ENSMUSG00 | false | . | . | . | . | . | . | chr8 | 88081760 | 88082957 | 3.56091 | 3.17253 | 1.86265 |
| chr8 | 3991367 | 3991654 | 336 | Promoter (2-3kb) | 8 | 3994628 | 3997505 | 2878 | 1 | ENSMUST00 | -2974 | Gm7389 | ENSMUSG00 | false | . | . | . | . | . | . | chr8 | 3991050 | 3992838 | 4.22712 | 3.59426 | 2.41753 |
| chr14 | 21031933 | 21032485 | 335 | Promoter (2-3kb) | 14 | 21031459 | 21034566 | 3108 | 2 | ENSMUST00 | 2081 | Ap3m1 | ENSMUSG00 | false | . | . | . | . | . | . | . | . | . | . | . | . |
| chr18 | 68084660 | 68084916 | 335 | Intron (ENSMUST00000063775.3 | 18 | 67933257 | 68255549 | 322293 | 1 | ENSMUST00 | 151404 | Ldlrad4 | ENSMUSG00 | false | . | . | . | . | . | . | chr18 | 68084426 | 68085380 | 3.65498 | 3.28561 | 1.90382 |

| | | | | | | | | | | | | | | | | | | | | | | | | | | |
|---|---|---|---|---|---|---|---|---|---|---|---|---|---|---|---|---|---|---|---|---|---|---|---|---|---|---|
| chr18 | 68058442 | 68058702 | 335 | Intron (ENSMUST00000063775.3 | 18 | 67933257 | 68255549 | 322293 | 1 | ENSMUST00 | 125186 | Ldlrad4 | ENSMUSG00 | false | chr18 | 68057848 | 68058647 | 10.30097 | 28.33038 | 25.88568 | chr18 | 68057251 | 68060499 | 3.88822 | 3.53367 | 2.0861 |
| chr19 | 53310308 | 53310859 | 335 | Promoter (<=1kb) | 19 | 53310506 | 53375810 | 65305 | 1 | ENSMUST00 | 0 | Mxi1 | ENSMUSG00 | false | chr19 | 53310158 | 53314217 | 35.4076 | 109.31139 | 105.49952 | chr19 | 53309637 | 53310613 | 5.43718 | 4.2978 | 3.40102 |
| chr3 | 96635337 | 96635834 | 335 | Promoter (<=1kb) | 3 | 96635376 | 96645366 | 9991 | 1 | ENSMUST00 | 0 | Pex11b | ENSMUSG00 | false | chr3 | 96635047 | 96636574 | 31.61155 | 112.91588 | 108.99915 | chr3 | 96634176 | 96637139 | 4.4252 | 3.73972 | 2.57099 |
| chr4 | 48046945 | 48047345 | 335 | Promoter (1-2kb) | 4 | 48045231 | 48050165 | 4935 | 1 | ENSMUST00 | 1715 | Nr4a3 | ENSMUSG00 | false | chr4 | 48044233 | 48048412 | 30.84378 | 98.32127 | 94.79446 | chr4 | 48046055 | 48048268 | 5.18325 | 4.15882 | 3.15309 |
| chr9 | 14276140 | 14276900 | 335 | Promoter (<=1kb) | 9 | 14276301 | 14326138 | 49838 | 1 | ENSMUST00 | 0 | Sesn3 | ENSMUSG00 | false | chr9 | 14276184 | 14277840 | 33.55804 | 103.05054 | 99.40515 | . | . | . | . | . | . |
| chr9 | 107872389 | 107872989 | 335 | Promoter (<=1kb) | 9 | 107773559 | 107872724 | 99166 | 2 | ENSMUST00 | 0 | Rbm6 | ENSMUSG00 | false | chr9 | 107871754 | 107873634 | 32.75367 | 101.65766 | 98.0462 | . | . | . | . | . | . |
| chr8 | 106210818 | 106211179 | 335 | Promoter (<=1kb) | 8 | 106200438 | 106210933 | 10496 | 2 | ENSMUST00 | 0 | Slc7a6os | ENSMUSG00 | false | chr8 | 106209902 | 106211779 | 31.91637 | 110.38313 | 106.53261 | chr8 | 106207883 | 106212081 | 5.57922 | 4.3257 | 3.48605 |
| chr13 | 70637624 | 70638235 | 335 | Promoter (<=1kb) | 13 | 70588689 | 70637634 | 48946 | 2 | ENSMUST00 | 0 | BC018507 | ENSMUSG00 | false | chr13 | 70636576 | 70638601 | 30.41442 | 103.5799 | 99.9162 | . | . | . | . | . | . |
| chr10 | 96697809 | 96698136 | 335 | Distal Intergenic | 10 | 96617001 | 96622813 | 5813 | 1 | ENSMUST00 | 80809 | Btg1 | ENSMUSG00 | false | . | . | . | . | . | . | chr10 | 96696982 | 96698787 | 6.19154 | 4.50847 | 4.03319 |
| chr2 | 129821231 | 129821466 | 335 | Intron (ENSMUST00000165413.3 | 2 | 129800517 | 129832287 | 31771 | 1 | ENSMUST00 | 20715 | Stk35 | ENSMUSG00 | false | . | . | . | . | . | . | chr2 | 129812158 | 129831014 | 4.89733 | 3.94874 | 2.94159 |
| chr5 | 143315027 | 143315473 | 335 | Promoter (<=1kb) | 5 | 143301195 | 143315064 | 13870 | 2 | ENSMUST00 | 0 | E130309D02 | ENSMUSG00 | false | chr5 | 143314251 | 143315661 | 36.04346 | 118.78738 | 114.68494 | chr5 | 143313354 | 143315832 | 5.31547 | 4.2253 | 3.26823 |
| chr9 | 108227639 | 108227954 | 335 | Intron (ENSMUST00000166905.2 | 9 | 108205958 | 108263736 | 57779 | 2 | ENSMUST00 | 35782 | Dag1 | ENSMUSG00 | true | . | . | . | . | . | . | chr9 | 108210099 | 108251502 | 5.76916 | 4.35802 | 3.65688 |
| chr1 | 38753329 | 38753664 | 335 | Distal Intergenic | 1 | 38794509 | 38821215 | 26707 | 2 | ENSMUST00 | 67551 | Lonrf2 | ENSMUSG00 | false | . | . | . | . | . | . | . | . | . | . | . | . |
| chr4 | 126150235 | 126150579 | 335 | Promoter (1-2kb) | 4 | 126148532 | 126149865 | 1334 | 1 | ENSMUST00 | 1704 | Eva1b | ENSMUSG00 | false | chr4 | 126147596 | 126150575 | 31.96693 | 102.30585 | 98.67843 | chr4 | 126145657 | 126153251 | 4.91678 | 3.70661 | 2.96666 |
| chr3 | 122773553 | 122773750 | 335 | Intron (ENSMUST00000066728.5 | 3 | 122729158 | 122859374 | 130217 | 1 | ENSMUST00 | 44396 | Pde5a | ENSMUSG00 | false | . | . | . | . | . | . | . | . | . | . | . | . |
| chr10 | 11343274 | 11344015 | 335 | Promoter (<=1kb) | 10 | 11343404 | 11459644 | 116241 | 1 | ENSMUST00 | 0 | Epm2a | ENSMUSG00 | false | chr10 | 11342945 | 11344338 | 23.04863 | 71.19738 | 68.188 | . | . | . | . | . | . |
| chr18 | 7692890 | 7693339 | 335 | Distal Intergenic | 18 | 7347962 | 7626863 | 278902 | 2 | ENSMUST00 | -66028 | Mpp7 | ENSMUSG00 | false | chr18 | 7692569 | 7693676 | 16.95696 | 43.908 | 41.27272 | chr18 | 7692422 | 7693749 | 3.79838 | 3.55 | 1.99982 |
| chr1 | 120557116 | 120557715 | 335 | Distal Intergenic | 1 | 120602418 | 120607992 | 5575 | 1 | ENSMUST00 | -44703 | En1 | ENSMUSG00 | false | . | . | . | . | . | . | chr1 | 120555379 | 120559086 | 5.23275 | 4.1913 | 3.21542 |
| chr2 | 117432160 | 117432479 | 335 | Distal Intergenic | 2 | 117414392 | 117417697 | 3306 | 2 | ENSMUST00 | -14464 | Gm13982 | ENSMUSG00 | false | . | . | . | . | . | . | chr2 | 117432133 | 117433127 | 4.75705 | 3.78273 | 2.83936 |
| chr2 | 92747478 | 92747881 | 335 | Intron (ENSMUST00000152230.1 | 2 | 92742230 | 92759208 | 16979 | 1 | ENSMUST00 | 5249 | Gm13816 | ENSMUSG00 | false | . | . | . | . | . | . | chr2 | 92746675 | 92748472 | 5.04438 | 4.00899 | 3.0597 |
| chr4 | 118427885 | 118428370 | 334 | Promoter (<=1kb) | 4 | 118428093 | 118432953 | 4861 | 1 | ENSMUST00 | 0 | Elovl1 | ENSMUSG00 | false | chr4 | 118427573 | 118429131 | 37.6931 | 122.73721 | 118.50226 | . | . | . | . | . | . |
| chr1 | 36368674 | 36369325 | 334 | Promoter (<=1kb) | 1 | 36335730 | 36369157 | 33428 | 2 | ENSMUST00 | 0 | Kansl3 | ENSMUSG00 | true | chr1 | 36367800 | 36369502 | 24.39013 | 102.54827 | 98.91666 | chr1 | 36369283 | 36370175 | 3.39601 | 3.10098 | 1.69746 |
| chr1 | 152356752 | 152357101 | 334 | Distal Intergenic | 1 | 152370838 | 152372082 | 1245 | 2 | ENSMUST00 | 14981 | Tsen15 | ENSMUSG00 | false | . | . | . | . | . | . | chr1 | 152356123 | 152357600 | 5.75479 | 4.30863 | 3.6516 |
| chr15 | 84855975 | 84856515 | 334 | Promoter (<=1kb) | 15 | 84785384 | 84856049 | 70666 | 2 | ENSMUST00 | 0 | Phf21b | ENSMUSG00 | false | chr15 | 84852984 | 84856941 | 17.45443 | 60.55182 | 57.69857 | chr15 | 84852969 | 84856987 | 4.65188 | 3.77993 | 2.75722 |
| chr10 | 60814396 | 60814710 | 334 | Intron (ENSMUST00000077925.5 | 10 | 60762610 | 60831581 | 68972 | 2 | ENSMUST00 | 16871 | Unc5b | ENSMUSG00 | false | . | . | . | . | . | . | chr10 | 60811545 | 60822537 | 5.55309 | 4.2307 | 3.48027 |
| chr11 | 30986193 | 30986502 | 334 | Promoter (<=1kb) | 11 | 30976707 | 30986350 | 9644 | 2 | ENSMUST00 | 0 | Chac2 | ENSMUSG00 | false | chr11 | 30985860 | 30987022 | 25.92862 | 72.63998 | 69.60807 | chr11 | 30985107 | 30987277 | 4.76486 | 3.98392 | 2.8205 |
| chr13 | 59585034 | 59585441 | 334 | Promoter (<=1kb) | 13 | 59585259 | 59599790 | 14532 | 1 | ENSMUST00 | 0 | Naa35 | ENSMUSG00 | false | chr13 | 59584388 | 59585130 | 14.64398 | 38.39655 | 35.82735 | . | . | . | . | . | . |
| chr13 | 95222921 | 95223340 | 334 | Promoter (<=1kb) | 13 | 95024505 | 95223066 | 198562 | 2 | ENSMUST00 | 0 | Pde8b | ENSMUSG00 | false | chr13 | 95222477 | 95223520 | 17.46222 | 42.53877 | 39.91948 | chr13 | 95222480 | 95223197 | 4.03558 | 3.32696 | 2.24973 |
| chrX | 155280759 | 155281152 | 334 | Intron (ENSMUST00000026324.9 | 20 | 155293843 | 155296913 | 3071 | 1 | ENSMUST00 | -12691 | Acot9 | ENSMUSG00 | false | . | . | . | . | . | . | . | . | . | . | . | . |
| chr3 | 103809323 | 103809675 | 334 | Promoter (<=1kb) | 3 | 103800605 | 103809378 | 8774 | 2 | ENSMUST00 | 0 | Dclre1b | ENSMUSG00 | false | chr3 | 103809519 | 103810080 | 34.14524 | 105.08128 | 101.38137 | . | . | . | . | . | . |
| chr4 | 43442206 | 43442741 | 334 | Promoter (<=1kb) | 4 | 43441939 | 43448064 | 6126 | 1 | ENSMUST00 | 268 | Tesk1 | ENSMUSG00 | false | chr4 | 43442217 | 43444119 | 26.86698 | 95.04005 | 91.5885 | . | . | . | . | . | . |
| chr6 | 143099762 | 143100313 | 334 | Promoter (<=1kb) | 6 | 143010920 | 143100107 | 89188 | 2 | ENSMUST00 | 0 | C2cd5 | ENSMUSG00 | false | chr6 | 143099152 | 143100613 | 37.54867 | 136.59039 | 131.84451 | chr6 | 143098087 | 143100971 | 3.82925 | 3.48017 | 2.09156 |
| chr7 | 35365178 | 35365484 | 334 | Intron (ENSMUST00000032705.7 | 7 | 35379282 | 35381622 | 2341 | 1 | ENSMUST00 | -13798 | Rhpn2 | ENSMUSG00 | false | . | . | . | . | . | . | chr7 | 35364516 | 35366217 | 4.7624 | 3.91433 | 2.86312 |
| chr7 | 45570045 | 45570609 | 334 | Promoter (<=1kb) | 7 | 45570153 | 45589710 | 19558 | 1 | ENSMUST00 | 0 | Bcat2 | ENSMUSG00 | false | chr7 | 45569998 | 45571389 | 30.07331 | 91.05476 | 87.68732 | chr7 | 45566728 | 45575570 | 5.46963 | 4.20393 | 3.40899 |
| chr8 | 107096107 | 107096682 | 334 | Promoter (<=1kb) | 8 | 107069400 | 107096545 | 27146 | 2 | ENSMUST00 | 0 | Terf2 | ENSMUSG00 | false | chr8 | 107095512 | 107097189 | 23.42648 | 72.77338 | 69.73964 | . | . | . | . | . | . |
| chr17 | 43789297 | 43789517 | 334 | Distal Intergenic | 17 | 43801851 | 44039516 | 237666 | 1 | ENSMUST00 | -12334 | Rcan2 | ENSMUSG00 | false | . | . | . | . | . | . | . | . | . | . | . | . |
| chr14 | 55092201 | 55092561 | 334 | Promoter (<=1kb) | 14 | 55060262 | 55092324 | 32063 | 2 | ENSMUST00 | 0 | Zfhx2 | ENSMUSG00 | false | chr14 | 55091378 | 55092748 | 20.32388 | 57.71489 | 54.90119 | chr14 | 55091313 | 55092691 | 3.28895 | 3.24618 | 1.65756 |
| chr11 | 31671677 | 31671986 | 334 | Promoter (<=1kb) | 11 | 31665181 | 31671885 | 6705 | 2 | ENSMUST00 | 0 | Bod1 | ENSMUSG00 | false | chr11 | 31671232 | 31672519 | 34.79824 | 110.32398 | 106.47989 | . | . | . | . | . | . |
| chr7 | 116038599 | 116038870 | 334 | Promoter (<=1kb) | 7 | 115471344 | 116038744 | 567401 | 2 | ENSMUST00 | 0 | Sox6 | ENSMUSG00 | false | . | . | . | . | . | . | chr7 | 116037583 | 116040756 | 5.37088 | 4.07398 | 3.3296 |
| chr10 | 69738664 | 69738914 | 334 | Intron (ENSMUST00000182194.2 | 10 | 69706478 | 69822635 | 116158 | 1 | ENSMUST00 | 32187 | Ank3 | ENSMUSG00 | false | . | . | . | . | . | . | chr10 | 69738503 | 69739322 | 3.15277 | 2.72518 | 1.53516 |
| chr13 | 119631617 | 119632084 | 334 | Intron (ENSMUST00000099241.2 | 13 | 119623819 | 119654354 | 30536 | 1 | ENSMUST00 | 7799 | Ccl28 | ENSMUSG00 | false | chr13 | 119631546 | 119632308 | 15.64407 | 45.32187 | 42.668 | . | . | . | . | . | . |
| chr12 | 36926391 | 36926663 | 334 | Distal Intergenic | 12 | 36816205 | 36816278 | 74 | 1 | ENSMUST00 | 110187 | Mir5099 | ENSMUSG00 | false | . | . | . | . | . | . | chr12 | 36926143 | 36927302 | 3.89512 | 3.45188 | 2.11033 |
| chr7 | 24399389 | 24400005 | 333 | Promoter (<=1kb) | 7 | 24399619 | 24408273 | 8655 | 1 | ENSMUST00 | 0 | Smg9 | ENSMUSG00 | false | chr7 | 24399214 | 24400413 | 24.16179 | 66.94102 | 63.99775 | . | . | . | . | . | . |
| chr12 | 111813950 | 111814749 | 333 | Promoter (<=1kb) | 12 | 111814170 | 111828388 | 14219 | 1 | ENSMUST00 | 0 | Zfyve21 | ENSMUSG00 | false | chr12 | 111814116 | 111814909 | 33.07612 | 110.40385 | 106.55179 | chr12 | 111813203 | 111823116 | 5.80855 | 4.38461 | 3.68777 |
| chr14 | 57744443 | 57744873 | 333 | Promoter (<=1kb) | 14 | 57734180 | 57744654 | 10475 | 2 | ENSMUST00 | 0 | Lats2 | ENSMUSG00 | false | chr14 | 57743310 | 57746899 | 34.76187 | 122.87653 | 118.63768 | . | . | . | . | . | . |
| chr16 | 92399975 | 92400350 | 333 | Promoter (<=1kb) | 16 | 92391953 | 92400077 | 8125 | 2 | ENSMUST00 | 0 | Rcan1 | ENSMUSG00 | false | chr16 | 92399134 | 92400236 | 15.97138 | 43.67439 | 41.04113 | chr16 | 92400332 | 92403993 | 7.0854 | 4.9845 | 4.70773 |
| chr7 | 141193864 | 141194220 | 333 | Promoter (<=1kb) | 7 | 141189105 | 141193960 | 4856 | 2 | ENSMUST00 | 0 | Hras | ENSMUSG00 | false | chr7 | 141192831 | 141195274 | 22.06604 | 78.32127 | 75.1984 | . | . | . | . | . | . |
| chr8 | 67870436 | 67870722 | 333 | Intron (ENSMUST00000038959.1 | 8 | 67878278 | 67904120 | 25843 | 2 | ENSMUST00 | 33398 | Psd3 | ENSMUSG00 | true | . | . | . | . | . | . | chr8 | 67870538 | 67877087 | 3.98545 | 3.45277 | 2.21737 |
| chr5 | 47983004 | 47983263 | 333 | Promoter (<=1kb) | 5 | 47983155 | 47990332 | 7178 | 1 | ENSMUST00 | 0 | Slit2 | ENSMUSG00 | false | chr5 | 47982701 | 47985505 | 19.43846 | 52.88828 | 50.13554 | chr5 | 47982868 | 47986636 | 5.08586 | 3.8792 | 3.11285 |
| chr18 | 80633593 | 80634080 | 333 | Intron (ENSMUST00000167977.2 | 18 | 80606205 | 80708173 | 101969 | 2 | ENSMUST00 | 74093 | Nfatc1 | ENSMUSG00 | false | . | . | . | . | . | . | chr18 | 80632924 | 80635177 | 6.60201 | 4.81632 | 4.32739 |
| chr6 | 91889295 | 91889530 | 333 | Promoter (2-3kb) | 6 | 91892347 | 91895293 | 2947 | 1 | ENSMUST00 | -2817 | Ccdc174 | ENSMUSG00 | false | . | . | . | . | . | . | chr6 | 91887867 | 91890472 | 3.07839 | 2.99063 | 1.49025 |
| chr5 | 19228123 | 19228479 | 333 | Promoter (1-2kb) | 5 | 19227046 | 20702903 | 1475858 | 1 | ENSMUST00 | 1078 | Magi2 | ENSMUSG00 | true | chr5 | 19223861 | 19232405 | 34.78864 | 106.1911 | 102.46782 | . | . | . | . | . | . |
| chr10 | 128091466 | 128091977 | 333 | Promoter (<=1kb) | 10 | 128092783 | 128129303 | 36521 | 1 | ENSMUST00 | -806 | Baz2a | ENSMUSG00 | false | chr10 | 128091204 | 128094095 | 33.28429 | 109.73968 | 105.91746 | chr10 | 128090498 | 128092091 | 4.71862 | 3.81521 | 2.79631 |
| chrX | 162643484 | 162643775 | 333 | Promoter (<=1kb) | 20 | 162411954 | 162643601 | 231648 | 2 | ENSMUST00 | 0 | Reps2 | ENSMUSG00 | false | chrX | 162643170 | 162643946 | 20.67895 | 53.40692 | 50.64761 | . | . | . | . | . | . |
| chr2 | 50110152 | 50110565 | 333 | Intron (ENSMUST00000112712.4 | 2 | 50128369 | 50193563 | 65195 | 1 | ENSMUST00 | -17804 | Lypd6 | ENSMUSG00 | false | . | . | . | . | . | . | chr2 | 50109251 | 50111092 | 3.6132 | 3.35253 | 1.86513 |
| chr6 | 34865115 | 34865489 | 333 | Promoter (1-2kb) | 6 | 34863146 | 34874946 | 11801 | 1 | ENSMUST00 | 1970 | Tmem140 | ENSMUSG00 | false | . | . | . | . | . | . | chr6 | 34863195 | 34879034 | 5.53817 | 4.34013 | 3.44642 |
| chr11 | 85578248 | 85578991 | 333 | Promoter (2-3kb) | 11 | 85581042 | 85603044 | 22003 | 1 | ENSMUST00 | -2051 | Bcas3 | ENSMUSG00 | false | . | . | . | . | . | . | chr11 | 85570899 | 85579415 | 4.61039 | 3.87464 | 2.69094 |
| chr16 | 33056316 | 33056821 | 333 | Promoter (<=1kb) | 16 | 33056453 | 33060189 | 3737 | 1 | ENSMUST00 | 0 | Rpl35a | ENSMUSG00 | false | chr16 | 33055841 | 33057426 | 27.07707 | 105.22646 | 101.52403 | . | . | . | . | . | . |
| chr7 | 133748302 | 133748558 | 333 | Intron (ENSMUST00000033290.6 | 7 | 133738479 | 133738934 | 456 | 2 | ENSMUST00 | -9369 | Gm15483 | ENSMUSG00 | false | . | . | . | . | . | . | chr7 | 133743183 | 133753176 | 5.21662 | 4.08927 | 3.21065 |
| chr12 | 108443292 | 108443710 | 333 | Promoter (2-3kb) | 12 | 108440530 | 108441270 | 741 | 2 | ENSMUST00 | -2023 | Gm15636 | ENSMUSG00 | false | chr12 | 108443541 | 108443858 | 6.263 | 10.51426 | 8.32665 | chr12 | 108434869 | 108445741 | 5.03088 | 3.96016 | 3.06008 |
| chr8 | 31272666 | 31273182 | 333 | Distal Intergenic | 8 | 31214886 | 31215351 | 466 | 2 | ENSMUST00 | -57316 | Gm8051 | ENSMUSG00 | false | . | . | . | . | . | . | chr8 | 31272801 | 31273510 | 3.61014 | 3.40469 | 1.87012 |
| chr4 | 12490157 | 12490405 | 333 | Distal Intergenic | 4 | 12271399 | 12314583 | 43185 | 1 | ENSMUST00 | 218759 | Gm11846 | ENSMUSG00 | false | . | . | . | . | . | . | chr4 | 12489611 | 12492934 | 3.37407 | 3.23783 | 1.69022 |
| chrX | 169991079 | 169991591 | 333 | Intron (ENSMUST00000171433.2 | 20 | 169985801 | 169986939 | 1139 | 2 | ENSMUST00 | -4141 | Gm15247 | ENSMUSG00 | false | . | . | . | . | . | . | . | . | . | . | . | . |
| chr19 | 55593434 | 55593622 | 333 | Intron (ENSMUST00000095950.2 | 19 | 55640589 | 55702204 | 61616 | 2 | ENSMUST00 | 108582 | 4930552P12I | ENSMUSG00 | false | . | . | . | . | . | . | . | . | . | . | . | . |
| chr3 | 153895668 | 153895873 | 332 | Intron (ENSMUST00000005630.5 | 3 | 153857269 | 153904731 | 47463 | 2 | ENSMUST00 | 8858 | Msh4 | ENSMUSG00 | false | . | . | . | . | . | . | chr3 | 153895382 | 153896306 | 3.91296 | 3.50567 | 2.12745 |
| chr17 | 29445187 | 29445387 | 332 | Distal Intergenic | 17 | 29490812 | 29496111 | 5300 | 1 | ENSMUST00 | -45425 | Pim1 | ENSMUSG00 | false | . | . | . | . | . | . | chr17 | 29443894 | 29452487 | 4.93685 | 4.01842 | 2.95062 |
| chr19 | 36843866 | 36844118 | 332 | Intron (ENSMUST00000025729.6 | 19 | 36849371 | 36855012 | 5642 | 1 | ENSMUST00 | -5253 | Tnks2 | ENSMUSG00 | false | . | . | . | . | . | . | chr19 | 36835417 | 36846536 | 4.71753 | 3.92952 | 2.78616 |
| chr1 | 152819616 | 152819948 | 332 | Intron (ENSMUST00000186568.1 | 1 | 152823569 | 152824484 | 916 | 1 | ENSMUST00 | -3621 | Ncf2 | ENSMUSG00 | true | . | . | . | . | . | . | chr1 | 152801274 | 152821507 | 4.847 | 3.96927 | 2.88719 |
| chr2 | 79598920 | 79599146 | 332 | Distal Intergenic | 2 | 79635352 | 79672966 | 37615 | 1 | ENSMUST00 | -36206 | Ssfa2 | ENSMUSG00 | false | . | . | . | . | . | . | chr2 | 79597578 | 79605221 | 5.34068 | 4.26617 | 3.27267 |
| chr2 | 103797419 | 103797844 | 332 | Promoter (<=1kb) | 2 | 103762941 | 103797649 | 34709 | 2 | ENSMUST00 | 0 | Caprin1 | ENSMUSG00 | false | chr2 | 103795571 | 103798285 | 34.87302 | 114.62315 | 110.65308 | . | . | . | . | . | . |
| chr5 | 64360509 | 64360810 | 332 | Distal Intergenic | 5 | 64349945 | 64350589 | 645 | 1 | ENSMUST00 | 10565 | Tbc1d1 | ENSMUSG00 | false | . | . | . | . | . | . | chr5 | 64357689 | 64361652 | 5.99462 | 4.35746 | 3.86792 |
| chr6 | 49263820 | 49264166 | 332 | Promoter (<=1kb) | 6 | 49243921 | 49264052 | 20132 | 2 | ENSMUST00 | 0 | Tra2a | ENSMUSG00 | false | chr6 | 49262330 | 49264348 | 38.83361 | 140.02353 | 135.14601 | . | . | . | . | . | . |
| chr7 | 64287346 | 64287713 | 332 | Promoter (<=1kb) | 7 | 64287653 | 64303172 | 15520 | 1 | ENSMUST00 | 0 | Mtmr10 | ENSMUSG00 | false | chr7 | 64287101 | 64288643 | 31.61155 | 112.91588 | 108.99915 | . | . | . | . | . | . |
| chr4 | 59315479 | 59315794 | 332 | Promoter (<=1kb) | 4 | 59314683 | 59315966 | 1284 | 2 | ENSMUST00 | 172 | Susd1 | ENSMUSG00 | false | . | . | . | . | . | . | chr4 | 59314795 | 59316255 | 4.86183 | 3.87844 | 2.92015 |
| chr1 | 164248733 | 164249233 | 332 | Promoter (<=1kb) | 1 | 164249046 | 164265385 | 16340 | 1 | ENSMUST00 | 0 | Slc19a2 | ENSMUSG00 | false | chr1 | 164248386 | 164249924 | 31.48209 | 103.47416 | 99.8177 | chr1 | 164247897 | 164251110 | 4.80821 | 3.91888 | 2.904 |
| chr18 | 84041189 | 84041400 | 332 | Intron (ENSMUST00000060303.9 | 18 | 84012082 | 84086390 | 74309 | 2 | ENSMUST00 | 44990 | Tshz1 | ENSMUSG00 | false | . | . | . | . | . | . | chr18 | 84039971 | 84053779 | 4.96682 | 4.00459 | 2.98491 |
| chr18 | 64109488 | 64109795 | 332 | Distal Intergenic | 18 | 64254359 | 64276146 | 21788 | 1 | ENSMUST00 | -144564 | St8sia3 | ENSMUSG00 | false | . | . | . | . | . | . | . | . | . | . | . | . |
| chr9 | 37021755 | 37021958 | 332 | Intron (ENSMUST00000177218.2 | 9 | 37026399 | 37026507 | 109 | 1 | ENSMUST00 | -4441 | n-R5s82 | ENSMUSG00 | false | . | . | . | . | . | . | chr9 | 37021034 | 37031907 | 6.40408 | 4.65381 | 4.17785 |
| chrX | 13347088 | 13347337 | 332 | Distal Intergenic | 20 | 13326023 | 13327019 | 997 | 2 | ENSMUST00 | -20070 | Gm7129 | ENSMUSG00 | false | chrX | 13346676 | 13347645 | 12.40737 | 26.6269 | 24.20249 | . | . | . | . | . | . |
| chr4 | 120290159 | 120290424 | 332 | Promoter (<=1kb) | 4 | 120291350 | 120303892 | 12543 | 1 | ENSMUST00 | -926 | Foxo6os | ENSMUSG00 | false | . | . | . | . | . | . | chr4 | 120286572 | 120290219 | 4.15182 | 3.67163 | 2.31183 |
| chr2 | 120850163 | 120850689 | 332 | Promoter (<=1kb) | 2 | 120732816 | 120850406 | 117591 | 2 | ENSMUST00 | 0 | Ttbk2 | ENSMUSG00 | false | chr2 | 120849487 | 120850506 | 42.17744 | 140.37996 | 135.48283 | . | . | . | . | . | . |
| chr17 | 47629471 | 47630099 | 332 | Promoter (<=1kb) | 17 | 47630690 | 47686738 | 56049 | 1 | ENSMUST00 | -591 | Usp49 | ENSMUSG00 | false | chr17 | 47628770 | 47630664 | 32.69058 | 104.59052 | 100.90593 | . | . | . | . | . | . |
| chr1 | 160053959 | 160054283 | 332 | Intron (ENSMUST00000168250.2 | 1 | 160044447 | 160071509 | 27063 | 1 | ENSMUST00 | 9513 | 4930523C07I | ENSMUSG00 | false | . | . | . | . | . | . | chr1 | 160043756 | 160061722 | 5.00419 | 4.03176 | 3.02332 |
| chr7 | 63911812 | 63912192 | 332 | Intron (ENSMUST00000063694.8 | 7 | 63916857 | 63920539 | 3683 | 1 | ENSMUST00 | -4665 | E030018B13 | ENSMUSG00 | false | . | . | . | . | . | . | chr7 | 63885397 | 63936279 | 7.51388 | 5.18869 | 5.06896 |
| chr9 | 102690045 | 102690382 | 332 | Promoter (2-3kb) | 9 | 102692885 | 102693834 | 950 | 1 | ENSMUST00 | -2503 | RP24-221A1 | ENSMUSG00 | false | chr9 | 102689705 | 102690990 | 12.9904 | 42.26624 | 39.65078 | . | . | . | . | . | . |
| chr1 | 73224294 | 73224688 | 332 | Distal Intergenic | 1 | 73269064 | 73306360 | 37297 | 1 | ENSMUST00 | -44376 | RP24-141F1 | ENSMUSG00 | false | . | . | . | . | . | . | . | . | . | . | . | . |
| chr15 | 102405243 | 102406008 | 331 | Promoter (<=1kb) | 15 | 102406143 | 102408235 | 2093 | 1 | ENSMUST00 | -135 | Sp1 | ENSMUSG00 | false | chr15 | 102404801 | 102405657 | 11.80999 | 32.51257 | 30.01635 | . | . | . | . | . | . |
| chr2 | 91196584 | 91196836 | 331 | Promoter (1-2kb) | 2 | 91184092 | 91195151 | 11060 | 2 | ENSMUST00 | -1434 | Nr1h3 | ENSMUSG00 | false | . | . | . | . | . | . | . | . | . | . | . | . |
| chr7 | 97083843 | 97084108 | 331 | Promoter (2-3kb) | 7 | 97081751 | 97308940 | 227190 | 1 | ENSMUST00 | 2093 | Gab2 | ENSMUSG00 | true | . | . | . | . | . | . | chr7 | 97082937 | 97085999 | 4.33316 | 3.67721 | 2.47772 |
| chr11 | 22990357 | 22990798 | 331 | Promoter (<=1kb) | 11 | 22990519 | 23003780 | 13262 | 1 | ENSMUST00 | 0 | Cct4 | ENSMUSG00 | false | chr11 | 22990063 | 22991717 | 20.88808 | 77.73511 | 74.61916 | . | . | . | . | . | . |
| chr14 | 60582729 | 60582938 | 331 | Intron (ENSMUST00000190392.1 | 14 | 60634001 | 60691367 | 57367 | 1 | ENSMUST00 | -51063 | Spata13 | ENSMUSG00 | true | . | . | . | . | . | . | chr14 | 60582610 | 60583069 | 3.3982 | 3.30951 | 1.66706 |
| chr15 | 64382528 | 64383034 | 331 | Promoter (<=1kb) | 15 | 64088539 | 64382919 | 294381 | 2 | ENSMUST00 | 0 | Asap1 | ENSMUSG00 | false | chr15 | 64381829 | 64383394 | 29.41006 | 85.09721 | 81.85307 | chr15 | 64382940 | 64384947 | 4.88721 | 4.02955 | 2.97126 |
| chr16 | 52008533 | 52008825 | 331 | Distal Intergenic | 16 | 52031549 | 52208047 | 176499 | 1 | ENSMUST00 | -22724 | Cblb | ENSMUSG00 | true | . | . | . | . | . | . | chr16 | 52006772 | 52008620 | 4.44137 | 3.85659 | 2.53731 |
| chr15 | 98608979 | 98609699 | 331 | Promoter (1-2kb) | 15 | 98589994 | 98607633 | 17640 | 2 | ENSMUST00 | -1347 | Adcy6 | ENSMUSG00 | false | chr15 | 98608565 | 98610228 | 30.78991 | 105.27288 | 101.56436 | chr15 | 98606507 | 98610285 | 4.43144 | 3.88063 | 2.53251 |
| chr2 | 26409012 | 26409345 | 331 | Promoter (<=1kb) | 2 | 26396249 | 26409200 | 12952 | 2 | ENSMUST00 | 0 | Inpp5e | ENSMUSG00 | false | chr2 | 26408352 | 26409246 | 29.21465 | 104.38775 | 100.71102 | chr2 | 26406424 | 26411650 | 6.13376 | 4.51137 | 3.97268 |
| chr5 | 146694267 | 146694457 | 331 | Distal Intergenic | 5 | 146734839 | 146758448 | 23610 | 2 | ENSMUST00 | 63991 | Usp12 | ENSMUSG00 | false | . | . | . | . | . | . | chr5 | 146691248 | 146696389 | 4.5702 | 3.85568 | 2.6568 |
| chrX | 20637067 | 20637289 | 331 | Intron (ENSMUST00000115375.2 | 20 | 20620098 | 20650684 | 30587 | 1 | ENSMUST00 | 16970 | Rbm10 | ENSMUSG00 | false | . | . | . | . | . | . | . | . | . | . | . | . |
| chr9 | 42761036 | 42761294 | 331 | Intron (ENSMUST00000114865.2 | 9 | 42520412 | 42808699 | 288288 | 2 | ENSMUST00 | 47405 | Grik4 | ENSMUSG00 | false | . | . | . | . | . | . | chr9 | 42760968 | 42761955 | 7.38923 | 4.95766 | 5.00635 |
| chr18 | 80476032 | 80476403 | 331 | Distal Intergenic | 18 | 80407959 | 80469695 | 61737 | 2 | ENSMUST00 | -6338 | Ctdp1 | ENSMUSG00 | false | . | . | . | . | . | . | chr18 | 80474858 | 80481285 | 5.64513 | 4.38222 | 3.53512 |
| chr7 | 120830420 | 120830891 | 331 | Distal Intergenic | 7 | 120842831 | 120907219 | 64389 | 1 | ENSMUST00 | -11940 | Eef2k | ENSMUSG00 | false | chr7 | 120830240 | 120831642 | 34.78613 | 113.42423 | 109.48744 | chr7 | 120830223 | 120832426 | 3.75346 | 3.35944 | 1.991 |
| chr13 | 84222137 | 84222371 | 331 | Promoter (<=1kb) | 13 | 84222314 | 84295870 | 73557 | 1 | ENSMUST00 | 0 | Tmem161b | ENSMUSG00 | false | chr13 | 84221590 | 84223059 | 25.53352 | 72.8631 | 69.82863 | . | . | . | . | . | . |
| chr14 | 21981540 | 21981810 | 331 | Downstream (2-3kb) | 14 | 21983962 | 21989601 | 5640 | 2 | ENSMUST00 | 7791 | Zfp503 | ENSMUSG00 | false | . | . | . | . | . | . | chr14 | 21981011 | 21986083 | 4.65699 | 3.79582 | 2.74477 |
| chr12 | 52603304 | 52603680 | 331 | Promoter (<=1kb) | 12 | 52603788 | 52603894 | 107 | 1 | ENSMUST00 | -108 | Gm24859 | ENSMUSG00 | false | chr12 | 52602865 | 52605494 | 33.05704 | 116.60957 | 112.57934 | . | . | . | . | . | . |
| chrX | 104474392 | 104474587 | 331 | Distal Intergenic | 20 | 104482818 | 104507490 | 24673 | 1 | ENSMUST00 | -8231 | Uprt | ENSMUSG00 | false | . | . | . | . | . | . | . | . | . | . | . | . |
| chr6 | 72685240 | 72685489 | 331 | Intron (ENSMUST00000069536.6 | 6 | 72699214 | 72705090 | 5877 | 1 | ENSMUST00 | -13725 | Gm15402 | ENSMUSG00 | false | . | . | . | . | . | . | chr6 | 72684563 | 72686078 | 4.90806 | 4.02319 | 2.97756 |
| chr11 | 49088392 | 49088655 | 331 | Promoter (<=1kb) | 11 | 49088234 | 49088543 | 310 | 2 | ENSMUST00 | 0 | Gm12188 | ENSMUSG00 | false | . | . | . | . | . | . | chr11 | 49086710 | 49089112 | 3.43131 | 3.28878 | 1.71524 |
| chr5 | 93005203 | 93005438 | 331 | Distal Intergenic | 5 | 93013032 | 93013139 | 108 | 2 | ENSMUST00 | 7701 | Gm25521 | ENSMUSG00 | false | chr5 | 93005298 | 93006364 | 12.98899 | 30.74239 | 28.2684 | chr5 | 93004108 | 93006581 | 4.09904 | 3.61046 | 2.26218 |
| chr1 | 155059575 | 155059869 | 331 | Promoter (2-3kb) | 1 | 155011332 | 155057179 | 45848 | 2 | ENSMUST00 | -2397 | RP23-76L21. | ENSMUSG00 | false | . | . | . | . | . | . | chr1 | 155059632 | 155060515 | 2.87515 | 2.78412 | 1.33004 |
| chr11 | 60416906 | 60417236 | 330 | Promoter (<=1kb) | 11 | 60417145 | 60450927 | 33783 | 1 | ENSMUST00 | 0 | Gid4 | ENSMUSG00 | false | chr11 | 60416404 | 60418696 | 32.29185 | 112.09676 | 108.19604 | . | . | . | . | . | . |
| chr10 | 68208725 | 68208943 | 330 | Intron (ENSMUST00000020106.7 | 10 | 68095593 | 68278726 | 183134 | 2 | ENSMUST00 | 69783 | Arid5b | ENSMUSG00 | false | . | . | . | . | . | . | chr10 | 68207933 | 68210076 | 5.08582 | 4.00461 | 3.09752 |
| chr14 | 79631031 | 79631394 | 330 | Distal Intergenic | 14 | 79637816 | 79642195 | 4380 | 2 | ENSMUST00 | 10801 | Lect1 | ENSMUSG00 | false | . | . | . | . | . | . | . | . | . | . | . | . |
| chr16 | 92428261 | 92428504 | 330 | Intron (ENSMUST00000060005.9 | 16 | 92391953 | 92400077 | 8125 | 2 | ENSMUST00 | -28185 | Rcan1 | ENSMUSG00 | false | . | . | . | . | . | . | chr16 | 92425267 | 92430211 | 6.11064 | 4.5979 | 3.92954 |
| chr1 | 37997934 | 37998222 | 330 | Promoter (<=1kb) | 1 | 37998010 | 38055579 | 57570 | 1 | ENSMUST00 | 0 | Eif5b | ENSMUSG00 | false | chr1 | 37996353 | 37999132 | 38.63617 | 126.53137 | 122.16663 | . | . | . | . | . | . |
| chr2 | 93334401 | 93334713 | 330 | Promoter (<=1kb) | 2 | 93201760 | 93334487 | 132728 | 2 | ENSMUST00 | 0 | Tspan18 | ENSMUSG00 | false | chr2 | 93333540 | 93334966 | 28.59991 | 96.29427 | 92.81311 | chr2 | 93333386 | 93335021 | 3.30736 | 3.20101 | 1.60412 |
| chr5 | 124094832 | 124095089 | 330 | Promoter (<=1kb) | 5 | 124087943 | 124095786 | 7844 | 2 | ENSMUST00 | 697 | Abcb9 | ENSMUSG00 | false | chr5 | 124095050 | 124095851 | 28.97134 | 97.95801 | 94.43773 | chr5 | 124091809 | 124096473 | 5.50921 | 4.17361 | 3.46115 |
| chr6 | 137142867 | 137143231 | 330 | Intron (ENSMUST00000032347.5 | 6 | 137056099 | 137169685 | 113587 | 2 | ENSMUST00 | 26454 | Rerg | ENSMUSG00 | false | . | . | . | . | . | . | . | . | . | . | . | . |
| chr7 | 89839848 | 89840060 | 330 | Intron (ENSMUST00000032856.7 | 7 | 89829282 | 89851962 | 22681 | 1 | ENSMUST00 | 10567 | Me3 | ENSMUSG00 | false | . | . | . | . | . | . | chr7 | 89837393 | 89843944 | 5.63494 | 4.37723 | 3.53051 |
| chr9 | 102843607 | 102844007 | 330 | Promoter (1-2kb) | 9 | 102845833 | 102861897 | 16065 | 1 | ENSMUST00 | -1826 | Ryk | ENSMUSG00 | false | . | . | . | . | . | . | chr9 | 102833046 | 102850688 | 5.63517 | 4.31932 | 3.53737 |
| chr15 | 96360696 | 96360943 | 330 | Intron (ENSMUST00000096250.4 | 15 | 96356840 | 96359783 | 2944 | 1 | ENSMUST00 | 3857 | Arid2 | ENSMUSG00 | false | . | . | . | . | . | . | chr15 | 96360105 | 96361838 | 4.27759 | 3.57614 | 2.45167 |
| chr5 | 103435891 | 103436157 | 330 | Intron (ENSMUST00000048957.9 | 5 | 103425192 | 103598359 | 173168 | 1 | ENSMUST00 | 10700 | Ptpn13 | ENSMUSG00 | false | . | . | . | . | . | . | chr5 | 103435354 | 103440723 | 4.34219 | 3.75689 | 2.46138 |
| chr16 | 11175980 | 11176537 | 330 | Promoter (<=1kb) | 16 | 11136592 | 11176393 | 39802 | 2 | ENSMUST00 | 0 | Zc3h7a | ENSMUSG00 | false | chr16 | 11174891 | 11177068 | 42.33011 | 146.25195 | 141.10883 | . | . | . | . | . | . |
| chr14 | 55085612 | 55085935 | 330 | Intron (ENSMUST00000183822.1 | 14 | 55074569 | 55080452 | 5884 | 2 | ENSMUST00 | -5161 | Zfhx2 | ENSMUSG00 | false | . | . | . | . | . | . | chr14 | 55085425 | 55086689 | 4.69106 | 3.64576 | 2.78337 |
| chr15 | 102203435 | 102204301 | 330 | Promoter (<=1kb) | 15 | 102203648 | 102215605 | 11958 | 1 | ENSMUST00 | 0 | Zfp740 | ENSMUSG00 | false | chr15 | 102202622 | 102204939 | 30.68508 | 100.04181 | 96.47092 | . | . | . | . | . | . |
| chr19 | 36918828 | 36919278 | 330 | Promoter (<=1kb) | 19 | 36917550 | 36919615 | 2066 | 2 | ENSMUST00 | 337 | Fgfbp3 | ENSMUSG00 | false | chr19 | 36918249 | 36919478 | 30.17592 | 94.17345 | 90.73757 | chr19 | 36918051 | 36920020 | 3.89098 | 3.38735 | 2.1202 |
| chr13 | 95616964 | 95617434 | 330 | Promoter (1-2kb) | 13 | 95601804 | 95618459 | 16656 | 2 | ENSMUST00 | 1025 | F2r | ENSMUSG00 | false | chr13 | 95616110 | 95618852 | 30.42596 | 90.69117 | 87.33178 | chr13 | 95610650 | 95618067 | 4.64293 | 3.76201 | 2.74776 |
| chr12 | 21373369 | 21373930 | 330 | Promoter (<=1kb) | 12 | 21323509 | 21373593 | 50085 | 2 | ENSMUST00 | 0 | Adam17 | ENSMUSG00 | false | chr12 | 21372741 | 21374479 | 29.77807 | 93.47606 | 90.05975 | chr12 | 21372152 | 21375040 | 3.34957 | 3.21177 | 1.64595 |
| chr12 | 69378487 | 69378703 | 330 | Distal Intergenic | 12 | 69371835 | 69372458 | 624 | 2 | ENSMUST00 | -6030 | Gm9887 | ENSMUSG00 | false | . | . | . | . | . | . | chr12 | 69373982 | 69379784 | 3.93811 | 3.5697 | 2.12584 |
| chr18 | 82534286 | 82534685 | 330 | Intron (ENSMUST00000091789.5 | 18 | 82523544 | 82524021 | 478 | 2 | ENSMUST00 | -10266 | Rpl21-ps8 | ENSMUSG00 | false | . | . | . | . | . | . | . | . | . | . | . | . |
| chr13 | 46548149 | 46548395 | 330 | Intron (ENSMUST00000021802.9 | 13 | 46518684 | 46518761 | 78 | 1 | ENSMUST00 | 29466 | Gm24915 | ENSMUSG00 | false | . | . | . | . | . | . | chr13 | 46547436 | 46548971 | 4.27485 | 3.74167 | 2.4076 |
| chr6 | 67123508 | 67123768 | 330 | Intron (ENSMUST00000181304.1 | 6 | 67115673 | 67168426 | 52754 | 1 | ENSMUST00 | 7836 | A430010J10I | ENSMUSG00 | false | . | . | . | . | . | . | chr6 | 67123607 | 67146074 | 5.6205 | 4.38535 | 3.51128 |
| chr11 | 98051815 | 98052125 | 329 | Promoter (1-2kb) | 11 | 98036623 | 98053462 | 16840 | 2 | ENSMUST00 | 1337 | Stac2 | ENSMUSG00 | false | chr11 | 98051992 | 98053609 | 18.51447 | 52.87213 | 50.12019 | chr11 | 98051686 | 98053723 | 4.08379 | 3.44401 | 2.28578 |
| chr11 | 12036541 | 12036928 | 329 | Promoter (<=1kb) | 11 | 11930508 | 12037401 | 106894 | 2 | ENSMUST00 | 473 | Grb10 | ENSMUSG00 | false | chr11 | 12035826 | 12038123 | 26.00477 | 73.67448 | 70.62826 | chr11 | 12034418 | 12038326 | 3.27373 | 3.08101 | 1.6369 |
| chr17 | 78206508 | 78206768 | 329 | Intron (ENSMUST00000112498.2 | 17 | 78200248 | 78376592 | 176345 | 1 | ENSMUST00 | 6261 | Crim1 | ENSMUSG00 | true | . | . | . | . | . | . | chr17 | 78202641 | 78231492 | 6.38021 | 4.71597 | 4.13646 |
| chr17 | 64628839 | 64629083 | 329 | Intron (ENSMUST00000086723.4 | 17 | 64600736 | 64755110 | 154375 | 1 | ENSMUST00 | 28104 | Man2a1 | ENSMUSG00 | false | . | . | . | . | . | . | chr17 | 64628811 | 64629294 | 3.06288 | 2.98944 | 1.45447 |
| chr17 | 24886483 | 24886820 | 329 | Promoter (<=1kb) | 17 | 24886643 | 24892149 | 5507 | 1 | ENSMUST00 | 0 | Spsb3 | ENSMUSG00 | false | chr17 | 24886605 | 24887644 | 38.25637 | 125.86897 | 121.53396 | . | . | . | . | . | . |
| chr2 | 71367513 | 71367845 | 329 | Promoter (<=1kb) | 2 | 71271063 | 71367749 | 96687 | 2 | ENSMUST00 | 0 | Slc25a12 | ENSMUSG00 | false | chr2 | 71366842 | 71368486 | 22.15371 | 67.81625 | 64.85734 | . | . | . | . | . | . |
| chr3 | 102734396 | 102735045 | 329 | Promoter (<=1kb) | 3 | 102734529 | 102772292 | 37764 | 1 | ENSMUST00 | 0 | Tspan2 | ENSMUSG00 | false | chr3 | 102734254 | 102736336 | 41.24086 | 137.63742 | 132.85336 | chr3 | 102733783 | 102734587 | 5.10639 | 4.18753 | 3.09634 |
| chr4 | 136021512 | 136021967 | 329 | Promoter (<=1kb) | 4 | 136003368 | 136021763 | 18396 | 2 | ENSMUST00 | 0 | Tceb3 | ENSMUSG00 | false | chr4 | 136020557 | 136021737 | 44.95315 | 147.43549 | 142.24948 | . | . | . | . | . | . |
| chr6 | 99666479 | 99667025 | 329 | Promoter (<=1kb) | 6 | 99257484 | 99666797 | 409314 | 2 | ENSMUST00 | 0 | Gm20696 | ENSMUSG00 | false | chr6 | 99665745 | 99667267 | 38.27806 | 124.82941 | 120.52397 | . | . | . | . | . | . |
| chr6 | 115994761 | 115995473 | 329 | Promoter (<=1kb) | 6 | 115954811 | 115995005 | 40195 | 2 | ENSMUST00 | 0 | Plxnd1 | ENSMUSG00 | false | chr6 | 115992161 | 115995454 | 32.66836 | 102.52041 | 98.88917 | chr6 | 115980470 | 115999580 | 5.17804 | 4.10687 | 3.16948 |
| chr2 | 20510283 | 20510537 | 329 | Promoter (<=1kb) | 2 | 20510047 | 20810524 | 300478 | 1 | ENSMUST00 | 237 | Etl4 | ENSMUSG00 | true | . | . | . | . | . | . | chr2 | 20508122 | 20510806 | 3.03403 | 2.97542 | 1.39057 |
| chr10 | 22653828 | 22654065 | 329 | Intron (ENSMUST00000042261.4 | 10 | 22649008 | 22664690 | 15683 | 2 | ENSMUST00 | 4821 | Slc2a12 | ENSMUSG00 | false | . | . | . | . | . | . | chr10 | 22652851 | 22655129 | 5.21893 | 4.11907 | 3.19197 |
| chr3 | 49944771 | 49945006 | 329 | Distal Intergenic | 3 | 49743296 | 49757316 | 14021 | 2 | ENSMUST00 | -187456 | Pcdh18 | ENSMUSG00 | false | . | . | . | . | . | . | . | . | . | . | . | . |
| chr7 | 83891727 | 83892379 | 329 | Promoter (<=1kb) | 7 | 83892000 | 83901532 | 9533 | 1 | ENSMUST00 | 0 | Mesdc2 | ENSMUSG00 | false | chr7 | 83891929 | 83893016 | 28.91859 | 89.0447 | 85.71584 | . | . | . | . | . | . |
| chr9 | 65252076 | 65252251 | 329 | Distal Intergenic | 9 | 65265180 | 65280605 | 15426 | 1 | ENSMUST00 | -12929 | Cilp | ENSMUSG00 | false | . | . | . | . | . | . | chr9 | 65251607 | 65252224 | 2.95247 | 2.95601 | 1.38703 |
| chr18 | 36320092 | 36320532 | 329 | Distal Intergenic | 18 | 36345619 | 36366720 | 21102 | 1 | ENSMUST00 | -25087 | Cystm1 | ENSMUSG00 | false | chr18 | 36319794 | 36320996 | 27.1142 | 89.69608 | 86.35963 | chr18 | 36318761 | 36321156 | 5.16295 | 4.03887 | 3.18719 |
| chr15 | 57801193 | 57801545 | 329 | Intron (ENSMUST00000096430.5 | 15 | 57736563 | 57738666 | 2104 | 2 | ENSMUST00 | -62528 | 9330154K18I | ENSMUSG00 | false | . | . | . | . | . | . | chr15 | 57801135 | 57803252 | 3.91426 | 3.22904 | 2.158 |
| chr7 | 145251667 | 145251869 | 329 | Downstream (2-3kb) | 7 | 145253923 | 145283927 | 30005 | 2 | ENSMUST00 | 32058 | Tpcn2 | ENSMUSG00 | false | . | . | . | . | . | . | chr7 | 145250570 | 145253714 | 5.12618 | 4.07282 | 3.1468 |
| chr1 | 60343227 | 60343619 | 329 | Promoter (<=1kb) | 1 | 60343323 | 60372707 | 29385 | 1 | ENSMUST00 | 0 | Cyp20a1 | ENSMUSG00 | false | chr1 | 60342766 | 60344158 | 24.40663 | 77.266 | 74.1616 | chr1 | 60342750 | 60344421 | 3.59405 | 3.38245 | 1.88903 |
| chr8 | 9167342 | 9167549 | 329 | Distal Intergenic | 8 | 9568132 | 9568249 | 118 | 2 | ENSMUST00 | 400700 | n-R5s95 | ENSMUSG00 | false | . | . | . | . | . | . | . | . | . | . | . | . |
| chr2 | 72383379 | 72383643 | 329 | Intron (ENSMUST00000090824.6 | 2 | 72360247 | 72360542 | 296 | 1 | ENSMUST00 | 23133 | Rpl36a-ps4 | ENSMUSG00 | false | . | . | . | . | . | . | chr2 | 72382311 | 72385045 | 5.23105 | 4.03841 | 3.21946 |
| chrX | 101260823 | 101261126 | 329 | Promoter (2-3kb) | 20 | 101261376 | 101263975 | 2600 | 2 | ENSMUST00 | 2849 | Gm614 | ENSMUSG00 | false | . | . | . | . | . | . | . | . | . | . | . | . |
| chr5 | 148569260 | 148569782 | 328 | Distal Intergenic | 5 | 148505142 | 148552789 | 47648 | 2 | ENSMUST00 | -16472 | Ubl3 | ENSMUSG00 | false | . | . | . | . | . | . | chr5 | 148568728 | 148569712 | 3.41776 | 3.26843 | 1.69235 |
| chr17 | 5872616 | 5872836 | 328 | Intron (ENSMUST00000002436.9 | 17 | 5841328 | 5931033 | 89706 | 1 | ENSMUST00 | 31289 | Snx9 | ENSMUSG00 | false | . | . | . | . | . | . | chr17 | 5867189 | 5879266 | 5.9622 | 4.44173 | 3.80171 |
| chr1 | 43196123 | 43196518 | 328 | Promoter (<=1kb) | 1 | 43131764 | 43196761 | 64998 | 2 | ENSMUST00 | 243 | Fhl2 | ENSMUSG00 | false | . | . | . | . | . | . | . | . | . | . | . | . |
| chr11 | 69410406 | 69411456 | 328 | Promoter (2-3kb) | 11 | 69398508 | 69413675 | 15168 | 2 | ENSMUST00 | 2219 | Kdm6b | ENSMUSG00 | false | chr11 | 69408142 | 69412281 | 27.88725 | 99.90214 | 96.33501 | chr11 | 69410823 | 69417315 | 7.23301 | 5.12321 | 4.82613 |
| chr13 | 55536457 | 55536836 | 328 | Promoter (<=1kb) | 13 | 55530410 | 55536658 | 6249 | 2 | ENSMUST00 | 0 | Ddx41 | ENSMUSG00 | false | chr13 | 55535668 | 55537037 | 29.85283 | 94.59187 | 91.14945 | chr13 | 55534615 | 55537341 | 5.06613 | 4.00303 | 3.104 |
| chr19 | 30030223 | 30030895 | 328 | Promoter (<=1kb) | 19 | 30030513 | 30058812 | 28300 | 1 | ENSMUST00 | 0 | Uhrf2 | ENSMUSG00 | false | chr19 | 30029826 | 30031769 | 38.76242 | 127.83364 | 123.41602 | . | . | . | . | . | . |
| chr2 | 14073490 | 14074403 | 328 | Promoter (<=1kb) | 2 | 14074100 | 14148338 | 74239 | 1 | ENSMUST00 | 0 | Stam | ENSMUSG00 | false | chr2 | 14073137 | 14075226 | 27.66025 | 106.38611 | 102.65278 | . | . | . | . | . | . |
| chr6 | 89368867 | 89369056 | 328 | Distal Intergenic | 6 | 89316316 | 89362613 | 46298 | 2 | ENSMUST00 | -6255 | Plxna1 | ENSMUSG00 | false | . | . | . | . | . | . | . | . | . | . | . | . |
| chr9 | 96363501 | 96364006 | 328 | Promoter (<=1kb) | 9 | 96332655 | 96364442 | 31788 | 2 | ENSMUST00 | 436 | Atp1b3 | ENSMUSG00 | false | chr9 | 96362708 | 96364402 | 29.69666 | 89.43046 | 86.09753 | . | . | . | . | . | . |
| chr9 | 108403147 | 108403378 | 328 | Downstream (<1kb) | 9 | 108392834 | 108397763 | 4930 | 1 | ENSMUST00 | 10314 | 1700102P08I | ENSMUSG00 | false | . | . | . | . | . | . | . | . | . | . | . | . |
| chr5 | 131308372 | 131308586 | 328 | Promoter (<=1kb) | 5 | 131306850 | 131308078 | 1229 | 2 | ENSMUST00 | -295 | Wbscr17 | ENSMUSG00 | false | . | . | . | . | . | . | . | . | . | . | . | . |
| chr18 | 14682413 | 14683237 | 328 | Promoter (<=1kb) | 18 | 14624198 | 14682766 | 58569 | 2 | ENSMUST00 | 0 | Ss18 | ENSMUSG00 | false | chr18 | 14681165 | 14683465 | 36.04672 | 129.50026 | 125.02953 | . | . | . | . | . | . |
| chr1 | 185816437 | 185816696 | 328 | Distal Intergenic | 1 | 186087731 | 186117310 | 29580 | 2 | ENSMUST00 | 300614 | Lyplal1 | ENSMUSG00 | false | . | . | . | . | . | . | chr1 | 185815376 | 185816527 | 3.41411 | 3.15098 | 1.75212 |
| chr16 | 23224805 | 23225483 | 328 | Promoter (<=1kb) | 16 | 23220656 | 23225368 | 4713 | 2 | ENSMUST00 | 0 | BC106179 | ENSMUSG00 | false | chr16 | 23224518 | 23227486 | 26.72788 | 101.58999 | 97.97913 | chr16 | 23223566 | 23225144 | 3.02066 | 3.03305 | 1.43135 |
| chr6 | 83033365 | 83033705 | 328 | Promoter (<=1kb) | 6 | 83030934 | 83033471 | 2538 | 2 | ENSMUST00 | 0 | Dok1 | ENSMUSG00 | false | chr6 | 83032551 | 83034569 | 30.73327 | 90.24956 | 86.89949 | . | . | . | . | . | . |
| chr5 | 88565261 | 88565873 | 328 | Distal Intergenic | 5 | 88554483 | 88556083 | 1601 | 1 | ENSMUST00 | 10779 | Utp3 | ENSMUSG00 | false | chr5 | 88564232 | 88566691 | 28.91247 | 96.86193 | 93.36482 | . | . | . | . | . | . |
| chr14 | 52014330 | 52015044 | 328 | Promoter (1-2kb) | 14 | 52016865 | 52019787 | 2923 | 1 | ENSMUST00 | -1821 | Tmem253 | ENSMUSG00 | false | chr14 | 52013893 | 52015600 | 30.32912 | 88.589 | 85.27629 | . | . | . | . | . | . |
| chr14 | 54191862 | 54192141 | 328 | Promoter (<=1kb) | 14 | 54192303 | 54192361 | 59 | 1 | ENSMUST00 | -162 | Traj27 | ENSMUSG00 | false | . | . | . | . | . | . | chr14 | 54191241 | 54192605 | 3.11235 | 3.01356 | 1.48278 |
| chrX | 123103261 | 123103998 | 328 | Promoter (<=1kb) | 20 | 123103495 | 123145651 | 42157 | 1 | ENSMUST00 | 0 | 3110007F17I | ENSMUSG00 | false | chrX | 123102881 | 123104946 | 19.30035 | 48.68736 | 45.98855 | . | . | . | . | . | . |
| chrX | 103913833 | 103914037 | 328 | Distal Intergenic | 20 | 103925536 | 103926984 | 1449 | 2 | ENSMUST00 | 12947 | Gm9166 | ENSMUSG00 | false | . | . | . | . | . | . | . | . | . | . | . | . |
| chr15 | 72875555 | 72875799 | 328 | Intron (ENSMUST00000023276.9 | 15 | 72913357 | 72913560 | 204 | 1 | ENSMUST00 | -37558 | Gm3150 | ENSMUSG00 | false | . | . | . | . | . | . | chr15 | 72871481 | 72876459 | 4.31483 | 3.63205 | 2.46941 |
| chr1 | 167295890 | 167296192 | 328 | Distal Intergenic | 1 | 167290125 | 167290225 | 101 | 1 | ENSMUST00 | 5766 | Mir6347 | ENSMUSG00 | false | . | . | . | . | . | . | chr1 | 167295599 | 167296957 | 3.95444 | 3.56477 | 2.15932 |
| chr6 | 4600796 | 4601575 | 327 | Promoter (<=1kb) | 6 | 4600840 | 4643381 | 42542 | 1 | ENSMUST00 | 0 | Casd1 | ENSMUSG00 | false | chr6 | 4600584 | 4601859 | 36.88168 | 122.38801 | 118.16834 | chr6 | 4600342 | 4601814 | 4.11164 | 3.74454 | 2.25496 |
| chr10 | 59978761 | 59979239 | 327 | Distal Intergenic | 10 | 59988404 | 59990383 | 1980 | 2 | ENSMUST00 | 11144 | Anapc16 | ENSMUSG00 | false | . | . | . | . | . | . | chr10 | 59970305 | 59981054 | 7.15674 | 4.95725 | 4.79406 |
| chr11 | 7076299 | 7076646 | 327 | Intron (ENSMUST00000020706.4 | 11 | 7063489 | 7178506 | 115018 | 1 | ENSMUST00 | 12811 | Adcy1 | ENSMUSG00 | true | . | . | . | . | . | . | chr11 | 7065998 | 7086778 | 4.86984 | 3.96898 | 2.91077 |
| chr13 | 43218840 | 43219093 | 327 | Intron (ENSMUST00000055341.5 | 13 | 43151744 | 43171501 | 19758 | 2 | ENSMUST00 | -47340 | Tbc1d7 | ENSMUSG00 | false | . | . | . | . | . | . | chr13 | 43208585 | 43223004 | 6.08142 | 4.53032 | 3.90259 |
| chr13 | 63263366 | 63263622 | 327 | Intron (ENSMUST00000091560.5 | 13 | 63282142 | 63326096 | 43955 | 1 | ENSMUST00 | -18520 | 2010111I01R | ENSMUSG00 | false | . | . | . | . | . | . | chr13 | 63252739 | 63272327 | 6.37956 | 4.73666 | 4.1425 |
| chr3 | 135606617 | 135606847 | 327 | Promoter (1-2kb) | 3 | 135584657 | 135608270 | 23614 | 2 | ENSMUST00 | 1423 | Nfkb1 | ENSMUSG00 | false | . | . | . | . | . | . | chr3 | 135601632 | 135607878 | 6.30926 | 4.60341 | 4.09794 |
| chr6 | 91972279 | 91972633 | 327 | Distal Intergenic | 6 | 91978878 | 92076004 | 97127 | 1 | ENSMUST00 | -6245 | Fgd5 | ENSMUSG00 | false | . | . | . | . | . | . | chr6 | 91971411 | 91973719 | 4.00563 | 3.52592 | 2.24219 |
| chr1 | 55618310 | 55618697 | 327 | Intron (ENSMUST00000042986.9 | 1 | 55702025 | 55735227 | 33203 | 1 | ENSMUST00 | -83328 | Plcl1 | ENSMUSG00 | true | . | . | . | . | . | . | chr1 | 55616806 | 55624269 | 5.67386 | 4.38362 | 3.56181 |
| chr17 | 56162259 | 56162540 | 327 | Promoter (<=1kb) | 17 | 56162477 | 56173955 | 11479 | 1 | ENSMUST00 | 0 | Tnfaip8l1 | ENSMUSG00 | false | chr17 | 56162034 | 56162807 | 7.48194 | 15.52168 | 13.25109 | chr17 | 56161867 | 56162873 | 5.79673 | 4.40533 | 3.70346 |
| chr11 | 37062962 | 37063163 | 327 | Intron (ENSMUST00000102801.2 | 11 | 36139480 | 36944166 | 804687 | 2 | ENSMUST00 | -118797 | Tenm2 | ENSMUSG00 | false | . | . | . | . | . | . | chr11 | 37062891 | 37063132 | 3.86767 | 3.59657 | 2.08563 |
| chr4 | 148151619 | 148152231 | 327 | Promoter (<=1kb) | 4 | 148145716 | 148151908 | 6193 | 2 | ENSMUST00 | 0 | Fbxo6 | ENSMUSG00 | false | chr4 | 148150673 | 148152220 | 31.96453 | 96.29042 | 92.80983 | chr4 | 148148886 | 148152963 | 5.22456 | 4.18047 | 3.19373 |
| chr17 | 12195232 | 12195649 | 327 | Intron (ENSMUST00000164143.2 | 17 | 12189921 | 12190456 | 536 | 1 | ENSMUST00 | 5312 | Tdgf1-ps2 | ENSMUSG00 | false | . | . | . | . | . | . | chr17 | 12189838 | 12208021 | 6.13066 | 4.60409 | 3.94082 |
| chr2 | 153161157 | 153161455 | 327 | Promoter (<=1kb) | 2 | 153161303 | 153204862 | 43560 | 1 | ENSMUST00 | 0 | Tm9sf4 | ENSMUSG00 | false | chr2 | 153160857 | 153162449 | 21.13367 | 73.82313 | 70.77491 | . | . | . | . | . | . |
| chr15 | 36374166 | 36374509 | 327 | Distal Intergenic | 15 | 36308144 | 36308956 | 813 | 2 | ENSMUST00 | -65211 | Rpl7a-ps3 | ENSMUSG00 | false | chr15 | 36374035 | 36375538 | 16.14863 | 43.24384 | 40.61377 | chr15 | 36371382 | 36376750 | 5.74131 | 4.46347 | 3.60718 |
| chr18 | 53944200 | 53944429 | 327 | Intron (ENSMUST00000069597.6 | 18 | 53984321 | 53985328 | 1008 | 1 | ENSMUST00 | -39892 | Gm5507 | ENSMUSG00 | false | . | . | . | . | . | . | chr18 | 53942655 | 53945080 | 4.06116 | 3.65213 | 2.22104 |
| chr11 | 19307544 | 19307802 | 327 | Distal Intergenic | 11 | 19360926 | 19361069 | 144 | 2 | ENSMUST00 | 53267 | Gm12025 | ENSMUSG00 | false | . | . | . | . | . | . | chr11 | 19306827 | 19309425 | 3.81568 | 3.35813 | 2.06556 |
| chr4 | 53384918 | 53385237 | 327 | Distal Intergenic | 4 | 53401948 | 53404030 | 2083 | 2 | ENSMUST00 | 18793 | Gm12496 | ENSMUSG00 | false | . | . | . | . | . | . | chr4 | 53384570 | 53385578 | 4.84236 | 4.01658 | 2.87682 |
| chr2 | 27328753 | 27328975 | 327 | Promoter (2-3kb) | 2 | 27326704 | 27331297 | 4594 | 1 | ENSMUST00 | 2050 | AA645442 | ENSMUSG00 | false | . | . | . | . | . | . | . | . | . | . | . | . |
| chr18 | 9844062 | 9844298 | 327 | Intron (ENSMUST00000040069.8 | 18 | 9827026 | 9827119 | 94 | 2 | ENSMUST00 | -16944 | Gm23637 | ENSMUSG00 | false | . | . | . | . | . | . | chr18 | 9842580 | 9846507 | 5.07276 | 4.1781 | 3.05948 |
| chr2 | 168070275 | 168070698 | 327 | Distal Intergenic | 2 | 168064820 | 168064897 | 78 | 2 | ENSMUST00 | -5379 | Gm24327 | ENSMUSG00 | false | . | . | . | . | . | . | chr2 | 168063587 | 168080595 | 6.53927 | 4.82888 | 4.26326 |
| chr16 | 85133265 | 85133851 | 327 | Intron (ENSMUST00000005406.1 | 16 | 85168849 | 85169001 | 153 | 1 | ENSMUST00 | -34998 | Gm27295 | ENSMUSG00 | false | . | . | . | . | . | . | chr16 | 85126656 | 85166763 | 5.48022 | 4.2963 | 3.40197 |

| | | | | | | | | | | | | | | | | | | | | | | | | | | |
|---|---|---|---|---|---|---|---|---|---|---|---|---|---|---|---|---|---|---|---|---|---|---|---|---|---|---|
| chr8 | 13209883 | 13210125 | 327 | Distal Intergenic | 8 | 13200820 | 13207707 | 6888 | 1 | ENSMUST00 | 9064 | RP23-185G9 | ENSMUSG00 | false | . | . | . | . | . | . | chr8 | 13208705 | 13211559 | 4.95412 | 3.93735 | 2.99787 |
| chr11 | 117849066 | 117849369 | 326 | Promoter (<=1kb) | 11 | 117849251 | 117855732 | 6482 | 1 | ENSMUST00 | 0 | Birc5 | ENSMUSG00 | false | chr11 | 117848353 | 117849591 | 24.78212 | 78.86476 | 75.73294 | chr11 | 117848347 | 117849854 | 4.57767 | 3.9707 | 2.64904 |
| chr10 | 4481654 | 4481871 | 326 | Promoter (<=1kb) | 10 | 4482502 | 4562231 | 79730 | 1 | ENSMUST00 | -631 | Ccdc170 | ENSMUSG00 | false | . | . | . | . | . | . | chr10 | 4480086 | 4482966 | 3.74845 | 3.50043 | 1.97181 |
| chr12 | 5374121 | 5374513 | 326 | Promoter (1-2kb) | 12 | 5077472 | 5375682 | 298211 | 2 | ENSMUST00 | 1169 | Klhl29 | ENSMUSG00 | true | chr12 | 5373404 | 5376248 | 26.18681 | 91.82585 | 88.44535 | chr12 | 5372978 | 5376323 | 3.31501 | 3.17659 | 1.67432 |
| chr14 | 20831549 | 20831825 | 326 | Distal Intergenic | 14 | 20836662 | 20843388 | 6727 | 1 | ENSMUST00 | -4837 | Plau | ENSMUSG00 | false | . | . | . | . | . | . | chr14 | 20830942 | 20832148 | 4.1134 | 3.57485 | 2.30733 |
| chr15 | 28133576 | 28133852 | 326 | Distal Intergenic | 15 | 28203752 | 28472045 | 268294 | 1 | ENSMUST00 | -69900 | Dnah5 | ENSMUSG00 | false | . | . | . | . | . | . | chr15 | 28133161 | 28134509 | 4.40511 | 3.78365 | 2.55652 |
| chr15 | 68201092 | 68201343 | 326 | Intron (ENSMUST00000162054.2 | 15 | 68084321 | 68258787 | 174467 | 2 | ENSMUST00 | 57444 | Zfat | ENSMUSG00 | false | . | . | . | . | . | . | chr15 | 68199773 | 68201566 | 3.53169 | 3.29301 | 1.85058 |
| chr17 | 45931103 | 45931562 | 326 | Distal Intergenic | 17 | 46018712 | 46021372 | 2661 | 2 | ENSMUST00 | 89810 | Vegfa | ENSMUSG00 | true | . | . | . | . | . | . | chr17 | 45928605 | 45934413 | 4.63767 | 3.72884 | 2.75421 |
| chr19 | 38485493 | 38485828 | 326 | Intron (ENSMUST00000181994.1 | 19 | 38481121 | 38783430 | 302310 | 1 | ENSMUST00 | 4373 | Plce1 | ENSMUSG00 | true | . | . | . | . | . | . | chr19 | 38483400 | 38489979 | 4.74928 | 3.83992 | 2.83904 |
| chr1 | 163779448 | 163780000 | 326 | Promoter (<=1kb) | 1 | 163779583 | 163917109 | 137527 | 1 | ENSMUST00 | 0 | Kifap3 | ENSMUSG00 | false | chr1 | 163779596 | 163780335 | 31.91637 | 110.38313 | 106.53261 | chr1 | 163779037 | 163780356 | 3.58889 | 3.40551 | 1.85465 |
| chr2 | 180966981 | 180967293 | 326 | Promoter (<=1kb) | 2 | 180967225 | 180982458 | 15234 | 1 | ENSMUST00 | 0 | Arfgap1 | ENSMUSG00 | false | chr2 | 180966622 | 180968168 | 31.69034 | 97.80461 | 94.28741 | chr2 | 180966283 | 180969972 | 5.17373 | 4.221 | 3.1597 |
| chr3 | 102366112 | 102366523 | 326 | Distal Intergenic | 3 | 102469928 | 102521013 | 51086 | 1 | ENSMUST00 | -103405 | Ngf | ENSMUSG00 | false | . | . | . | . | . | . | . | . | . | . | . | . |
| chr6 | 89554138 | 89554397 | 326 | Intron (ENSMUST00000032172.9 | 6 | 89383146 | 89595652 | 212507 | 2 | ENSMUST00 | 41255 | Chchd6 | ENSMUSG00 | false | . | . | . | . | . | . | chr6 | 89551829 | 89557049 | 4.41772 | 3.81106 | 2.52097 |
| chrX | 73966255 | 73966508 | 326 | Promoter (<=1kb) | 20 | 73942792 | 73966357 | 23566 | 2 | ENSMUST00 | 0 | Hcfc1 | ENSMUSG00 | false | chrX | 73965220 | 73968256 | 24.81473 | 68.0602 | 65.09722 | . | . | . | . | . | . |
| chr9 | 112207775 | 112208022 | 326 | Intron (ENSMUST00000159432.2 | 9 | 112116220 | 112217261 | 101042 | 2 | ENSMUST00 | 9239 | Arpp21 | ENSMUSG00 | false | . | . | . | . | . | . | chr9 | 112207829 | 112208335 | 3.91275 | 3.5068 | 2.15255 |
| chr8 | 94549967 | 94550252 | 326 | Intron (ENSMUST00000048653.3 | 8 | 94562650 | 94570114 | 7465 | 1 | ENSMUST00 | -12398 | Cpne2 | ENSMUSG00 | false | . | . | . | . | . | . | . | . | . | . | . | . |
| chr8 | 3676335 | 3676599 | 326 | Promoter (<=1kb) | 8 | 3676299 | 3681255 | 4957 | 1 | ENSMUST00 | 37 | Trappc5 | ENSMUSG00 | false | chr8 | 3676017 | 3677566 | 31.91637 | 110.38313 | 106.53261 | . | . | . | . | . | . |
| chr3 | 67490216 | 67490412 | 326 | Intron (ENSMUST00000178314.2 | 3 | 67478886 | 67515523 | 36638 | 2 | ENSMUST00 | 25111 | Rarres1 | ENSMUSG00 | false | . | . | . | . | . | . | chr3 | 67489799 | 67490820 | 3.89498 | 3.43218 | 2.13468 |
| chr14 | 66809309 | 66809809 | 326 | Intron (ENSMUST00000022629.8 | 14 | 66784529 | 66793758 | 9230 | 1 | ENSMUST00 | 24781 | Gm10032 | ENSMUSG00 | false | . | . | . | . | . | . | chr14 | 66809094 | 66811856 | 4.77025 | 3.8128 | 2.85526 |
| chr3 | 79519069 | 79519397 | 326 | Intron (ENSMUST00000076136.5 | 3 | 79504228 | 79515252 | 11025 | 2 | ENSMUST00 | -3818 | Fnip2 | ENSMUSG00 | false | . | . | . | . | . | . | chr3 | 79518892 | 79520188 | 4.1493 | 3.51552 | 2.33588 |
| chr2 | 84670394 | 84670884 | 326 | Promoter (<=1kb) | 2 | 84669215 | 84670770 | 1556 | 2 | ENSMUST00 | 0 | 2700094K13I | ENSMUSG00 | false | chr2 | 84669973 | 84671112 | 28.91247 | 96.86193 | 93.36482 | chr2 | 84669788 | 84671646 | 4.77645 | 3.86325 | 2.85662 |
| chr4 | 153133230 | 153133482 | 326 | Distal Intergenic | 4 | 153145590 | 153146943 | 1354 | 2 | ENSMUST00 | 13461 | Gm13173 | ENSMUSG00 | false | . | . | . | . | . | . | chr4 | 153127760 | 153135799 | 5.054 | 4.04265 | 3.06739 |
| chr13 | 107551621 | 107551857 | 326 | Distal Intergenic | 13 | 107687397 | 107692162 | 4766 | 1 | ENSMUST00 | -135540 | RP23-125L8. | ENSMUSG00 | false | . | . | . | . | . | . | chr13 | 107551068 | 107551869 | 3.945 | 3.48714 | 2.15123 |
| chr3 | 95985366 | 95985694 | 325 | Downstream (2-3kb) | 3 | 95988819 | 95995668 | 6850 | 2 | ENSMUST00 | 9974 | Plekho1 | ENSMUSG00 | true | chr3 | 95985612 | 95986225 | 4.50584 | 6.69248 | 4.59165 | chr3 | 95984661 | 95993242 | 6.4304 | 4.71231 | 4.18378 |
| chr3 | 95699642 | 95699941 | 325 | Distal Intergenic | 3 | 95676201 | 95687917 | 11717 | 2 | ENSMUST00 | -11726 | Adamtsl4 | ENSMUSG00 | false | . | . | . | . | . | . | chr3 | 95685856 | 95702888 | 5.06878 | 4.08183 | 3.0659 |
| chr10 | 68442997 | 68443190 | 325 | Intron (ENSMUST00000020103.3 | 10 | 68436280 | 68525767 | 89488 | 2 | ENSMUST00 | 82577 | 1700040L02F | ENSMUSG00 | false | . | . | . | . | . | . | chr10 | 68442759 | 68444184 | 5.88944 | 4.53394 | 3.72038 |
| chr14 | 57889721 | 57890644 | 325 | Promoter (<=1kb) | 14 | 57832702 | 57890262 | 57561 | 2 | ENSMUST00 | 0 | Zdhhc20 | ENSMUSG00 | false | chr14 | 57889078 | 57890767 | 43.19602 | 139.91907 | 135.04468 | chr14 | 57886950 | 57890960 | 3.63603 | 3.3775 | 1.87705 |
| chr16 | 43363642 | 43364215 | 325 | Promoter (<=1kb) | 16 | 43363873 | 43609520 | 245648 | 1 | ENSMUST00 | 0 | Zbtb20 | ENSMUSG00 | false | . | . | . | . | . | . | chr16 | 43359177 | 43365677 | 4.92703 | 4.0399 | 2.94152 |
| chr16 | 94536937 | 94537472 | 325 | Distal Intergenic | 16 | 94497783 | 94526830 | 29048 | 2 | ENSMUST00 | -10108 | Dscr3 | ENSMUSG00 | false | . | . | . | . | . | . | chr16 | 94535920 | 94537708 | 5.54134 | 4.23361 | 3.47706 |
| chr18 | 5819737 | 5820145 | 325 | Distal Intergenic | 18 | 5731081 | 5772804 | 41724 | 1 | ENSMUST00 | 88657 | Zeb1 | ENSMUSG00 | false | chr18 | 5819564 | 5820389 | 14.24974 | 34.23454 | 31.71693 | chr18 | 5818136 | 5820555 | 4.28947 | 3.75198 | 2.41779 |
| chr1 | 165617193 | 165617542 | 325 | Intron (ENSMUST00000068705.7 | 1 | 165592181 | 165634541 | 42361 | 2 | ENSMUST00 | 16999 | Mpzl1 | ENSMUSG00 | false | . | . | . | . | . | . | chr1 | 165615531 | 165619626 | 4.96655 | 4.01111 | 2.99301 |
| chr2 | 155133456 | 155134008 | 325 | Promoter (<=1kb) | 2 | 155133509 | 155139289 | 5781 | 1 | ENSMUST00 | 0 | Itch | ENSMUSG00 | false | chr2 | 155133033 | 155134461 | 32.31418 | 113.17245 | 109.24831 | . | . | . | . | . | . |
| chr5 | 122643706 | 122643995 | 325 | Promoter (<=1kb) | 5 | 122643911 | 122684289 | 40379 | 1 | ENSMUST00 | 0 | P2rx7 | ENSMUSG00 | false | . | . | . | . | . | . | . | . | . | . | . | . |
| chr4 | 121017004 | 121017453 | 325 | Promoter (<=1kb) | 4 | 120968317 | 121017247 | 48931 | 2 | ENSMUST00 | 0 | Smap2 | ENSMUSG00 | false | chr4 | 121016006 | 121017644 | 46.06248 | 156.19411 | 150.67332 | chr4 | 121017407 | 121017672 | 2.86408 | 2.85454 | 1.32218 |
| chrX | 129749906 | 129750386 | 325 | Promoter (<=1kb) | 20 | 129749976 | 130465834 | 715859 | 1 | ENSMUST00 | 0 | Diap2 | ENSMUSG00 | false | chrX | 129749389 | 129750968 | 21.59801 | 56.60135 | 53.80005 | . | . | . | . | . | . |
| chr10 | 122746367 | 122746580 | 325 | Intron (ENSMUST00000162853.2 | 10 | 122757198 | 122811471 | 54274 | 1 | ENSMUST00 | -10618 | Ppm1h | ENSMUSG00 | false | . | . | . | . | . | . | chr10 | 122745820 | 122747329 | 3.78115 | 3.38617 | 2.02587 |
| chr11 | 95343173 | 95343618 | 325 | Intron (ENSMUST00000037502.6 | 11 | 95339962 | 95375087 | 35126 | 1 | ENSMUST00 | 3212 | Fam117a | ENSMUSG00 | false | . | . | . | . | . | . | chr11 | 95342290 | 95345340 | 5.34084 | 4.17953 | 3.29757 |
| chr10 | 17722090 | 17722583 | 325 | Promoter (<=1kb) | 10 | 17723228 | 17725673 | 2446 | 1 | ENSMUST00 | -645 | Cited2 | ENSMUSG00 | false | chr10 | 17721687 | 17725607 | 33.41831 | 117.26761 | 113.21259 | . | . | . | . | . | . |
| chr17 | 27406884 | 27407200 | 325 | Distal Intergenic | 17 | 27342460 | 27344825 | 2366 | 1 | ENSMUST00 | 64425 | Gm10505 | ENSMUSG00 | false | . | . | . | . | . | . | chr17 | 27405718 | 27410499 | 6.06906 | 4.38016 | 3.91867 |
| chr4 | 154025730 | 154026182 | 325 | Promoter (<=1kb) | 4 | 154020588 | 154025912 | 5325 | 2 | ENSMUST00 | 0 | Smim1 | ENSMUSG00 | false | chr4 | 154024913 | 154026694 | 30.60524 | 96.90557 | 93.40668 | chr4 | 154021005 | 154028094 | 7.34568 | 5.07579 | 4.95328 |
| chr2 | 72673602 | 72674142 | 325 | Distal Intergenic | 2 | 72703510 | 72729468 | 25959 | 1 | ENSMUST00 | -29368 | 8430437L04F | ENSMUSG00 | false | . | . | . | . | . | . | chr2 | 72672965 | 72675132 | 3.38344 | 3.15544 | 1.728 |
| chr12 | 32378543 | 32379017 | 325 | Promoter (<=1kb) | 12 | 32378704 | 32382943 | 4240 | 1 | ENSMUST00 | 0 | Ccdc71l | ENSMUSG00 | false | chr12 | 32378503 | 32381087 | 22.06604 | 78.32127 | 75.1984 | . | . | . | . | . | . |
| chr16 | 25011145 | 25011421 | 325 | Distal Intergenic | 16 | 25059639 | 25069058 | 9420 | 1 | ENSMUST00 | -48218 | A230028O05 | ENSMUSG00 | false | . | . | . | . | . | . | . | . | . | . | . | . |
| chr10 | 14060355 | 14060558 | 325 | Intron (ENSMUST00000187083.1 | 10 | 13967024 | 14007405 | 40382 | 2 | ENSMUST00 | -52951 | Gm26835 | ENSMUSG00 | false | . | . | . | . | . | . | chr10 | 14054380 | 14062954 | 5.60088 | 4.29997 | 3.513 |
| chr14 | 120333885 | 120334198 | 325 | Intron (ENSMUST00000088419.7 | 14 | 120384263 | 120388985 | 4723 | 2 | ENSMUST00 | 54787 | Gm26679 | ENSMUSG00 | false | . | . | . | . | . | . | chr14 | 120329884 | 120339572 | 6.16737 | 4.46838 | 4.00867 |
| chr2 | 13750172 | 13750524 | 324 | Intron (ENSMUST00000003509.8 | 2 | 13657154 | 13793293 | 136140 | 2 | ENSMUST00 | 42769 | St8sia6 | ENSMUSG00 | false | . | . | . | . | . | . | chr2 | 13749531 | 13750738 | 3.7314 | 3.42435 | 2.00425 |
| chr2 | 165320665 | 165321051 | 324 | Promoter (<=1kb) | 2 | 165288031 | 165320826 | 32796 | 2 | ENSMUST00 | 0 | Elmo2 | ENSMUSG00 | false | . | . | . | . | . | . | chr2 | 165307563 | 165325045 | 8.91187 | 5.755 | 6.22321 |
| chr11 | 60417404 | 60417859 | 324 | Promoter (<=1kb) | 11 | 60417262 | 60445277 | 28016 | 1 | ENSMUST00 | 143 | Gid4 | ENSMUSG00 | false | chr11 | 60416404 | 60418696 | 32.29185 | 112.09676 | 108.19604 | . | . | . | . | . | . |
| chr12 | 100127885 | 100128229 | 324 | Intron (ENSMUST00000021596.7 | 12 | 100110186 | 100123364 | 13179 | 1 | ENSMUST00 | 17700 | Psmc1 | ENSMUSG00 | false | . | . | . | . | . | . | chr12 | 100126788 | 100129574 | 7.24765 | 5.11573 | 4.862 |
| chr17 | 43081356 | 43081590 | 324 | Intron (ENSMUST00000024708.4 | 17 | 43016555 | 43089188 | 72634 | 1 | ENSMUST00 | 64802 | Tnfrsf21 | ENSMUSG00 | false | . | . | . | . | . | . | chr17 | 43075309 | 43083932 | 4.06236 | 3.6443 | 2.23752 |
| chr1 | 36939801 | 36940109 | 324 | Promoter (<=1kb) | 1 | 36792194 | 36939527 | 147334 | 2 | ENSMUST00 | -275 | Tmem131 | ENSMUSG00 | false | chr1 | 36938405 | 36940540 | 37.17318 | 134.81126 | 130.13722 | . | . | . | . | . | . |
| chr1 | 34023432 | 34023693 | 324 | Intron (ENSMUST00000182513.2 | 1 | 34011825 | 34308661 | 296837 | 1 | ENSMUST00 | 11608 | Dst | ENSMUSG00 | true | . | . | . | . | . | . | chr1 | 34020714 | 34026603 | 4.63881 | 3.8982 | 2.69619 |
| chr4 | 34550386 | 34550696 | 324 | Promoter (<=1kb) | 4 | 34550937 | 34566908 | 15972 | 1 | ENSMUST00 | -241 | Akirin2 | ENSMUSG00 | false | chr4 | 34549956 | 34552534 | 33.03282 | 126.03245 | 121.67878 | . | . | . | . | . | . |
| chr4 | 119415456 | 119415668 | 324 | Promoter (<=1kb) | 4 | 119323841 | 119415521 | 91681 | 2 | ENSMUST00 | 0 | Ccdc30 | ENSMUSG00 | false | chr4 | 119414935 | 119415914 | 18.70486 | 58.09692 | 55.27494 | chr4 | 119414703 | 119415948 | 3.29916 | 3.14254 | 1.65957 |
| chr4 | 155189729 | 155189978 | 324 | Promoter (1-2kb) | 4 | 155154075 | 155188296 | 34222 | 2 | ENSMUST00 | -1434 | Ski | ENSMUSG00 | false | . | . | . | . | . | . | chr4 | 155159960 | 155221990 | 6.99931 | 4.87167 | 4.66225 |
| chr7 | 68128938 | 68129633 | 324 | Intron (ENSMUST00000005671.8 | 7 | 68236608 | 68264233 | 27626 | 2 | ENSMUST00 | 134600 | Pgpep1l | ENSMUSG00 | false | chr7 | 68128511 | 68130116 | 18.02578 | 59.25625 | 56.42029 | . | . | . | . | . | . |
| chr9 | 66090618 | 66090878 | 324 | Promoter (<=1kb) | 9 | 66089124 | 66091347 | 2224 | 2 | ENSMUST00 | 469 | Snx1 | ENSMUSG00 | false | . | . | . | . | . | . | chr9 | 66090378 | 66094572 | 5.17902 | 4.04243 | 3.19238 |
| chr18 | 20896217 | 20896582 | 324 | Promoter (<=1kb) | 18 | 20846145 | 20896078 | 49934 | 2 | ENSMUST00 | -140 | Trappc8 | ENSMUSG00 | false | chr18 | 20895173 | 20897177 | 32.86763 | 120.45166 | 116.30058 | . | . | . | . | . | . |
| chr9 | 79966005 | 79966326 | 324 | Intron (ENSMUST00000093811.5 | 9 | 79820661 | 79962033 | 141373 | 2 | ENSMUST00 | -3973 | Filip1 | ENSMUSG00 | false | . | . | . | . | . | . | chr9 | 79964843 | 79967276 | 3.96395 | 3.49573 | 2.20882 |
| chr16 | 35022206 | 35022598 | 324 | Promoter (<=1kb) | 16 | 35022421 | 35109175 | 86755 | 1 | ENSMUST00 | 0 | Ptplb | ENSMUSG00 | false | chr16 | 35022017 | 35023359 | 27.7554 | 114.54269 | 110.5744 | chr16 | 35022035 | 35024053 | 3.23683 | 3.1235 | 1.56443 |
| chr6 | 125378156 | 125378632 | 324 | Promoter (2-3kb) | 6 | 125362660 | 125380793 | 18134 | 2 | ENSMUST00 | 2161 | Plekhg6 | ENSMUSG00 | false | . | . | . | . | . | . | chr6 | 125377085 | 125381156 | 7.16203 | 4.89207 | 4.82497 |
| chr1 | 63964504 | 63964772 | 324 | Promoter (<=1kb) | 1 | 63964727 | 63968999 | 4273 | 1 | ENSMUST00 | 0 | Gm13749 | ENSMUSG00 | false | . | . | . | . | . | . | chr1 | 63963757 | 63964957 | 4.0895 | 3.6779 | 2.2523 |
| chr7 | 115706278 | 115706489 | 324 | Intron (ENSMUST00000072804.5 | 7 | 115474475 | 115824707 | 350233 | 2 | ENSMUST00 | 118218 | Sox6 | ENSMUSG00 | false | . | . | . | . | . | . | chr7 | 115705592 | 115707231 | 3.80342 | 3.37267 | 2.05706 |
| chr5 | 150149507 | 150149742 | 324 | Distal Intergenic | 5 | 150080622 | 150082184 | 1563 | 1 | ENSMUST00 | 68886 | Rxfp2 | ENSMUSG00 | false | . | . | . | . | . | . | chr5 | 150148830 | 150161159 | 5.72506 | 4.43285 | 3.60287 |
| chr3 | 51713298 | 51713615 | 324 | Intron (ENSMUST00000121440.2 | 3 | 51693771 | 51695090 | 1320 | 1 | ENSMUST00 | 19528 | Gm10729 | ENSMUSG00 | false | . | . | . | . | . | . | chr3 | 51705501 | 51729803 | 5.39003 | 4.2072 | 3.33836 |
| chr3 | 156814699 | 156814975 | 324 | Promoter (1-2kb) | 3 | 156816037 | 156816511 | 475 | 1 | ENSMUST00 | -1062 | Gm15578 | ENSMUSG00 | false | . | . | . | . | . | . | chr3 | 156814456 | 156815515 | 3.73675 | 3.48512 | 2.00066 |
| chr13 | 31309232 | 31309442 | 324 | Intron (ENSMUST00000144541.1 | 13 | 31299628 | 31396579 | 96952 | 1 | ENSMUST00 | 9605 | Gm11376 | ENSMUSG00 | false | . | . | . | . | . | . | chr13 | 31308600 | 31310158 | 4.0181 | 3.48975 | 2.22474 |
| chr6 | 128355258 | 128356001 | 323 | Promoter (<=1kb) | 6 | 128321140 | 128355851 | 34712 | 2 | ENSMUST00 | 0 | Tulp3 | ENSMUSG00 | false | chr6 | 128354529 | 128356336 | 28.1183 | 82.87242 | 79.66893 | . | . | . | . | . | . |
| chr7 | 16348776 | 16349026 | 323 | Intron (ENSMUST00000094815.3 | 7 | 16313517 | 16317697 | 4181 | 1 | ENSMUST00 | 35260 | Bbc3 | ENSMUSG00 | false | . | . | . | . | . | . | chr7 | 16348291 | 16350172 | 3.82066 | 3.52395 | 2.03161 |
| chr10 | 127653380 | 127653742 | 323 | Promoter (<=1kb) | 10 | 127653079 | 127655015 | 1937 | 1 | ENSMUST00 | 302 | Stat6 | ENSMUSG00 | false | . | . | . | . | . | . | chr10 | 127652196 | 127654838 | 5.64464 | 4.19339 | 3.56983 |
| chr11 | 69980581 | 69981111 | 323 | Promoter (<=1kb) | 11 | 69980035 | 69980696 | 662 | 2 | ENSMUST00 | 0 | Elp5 | ENSMUSG00 | false | chr11 | 69979551 | 69981936 | 34.36702 | 111.64683 | 107.76208 | . | . | . | . | . | . |
| chr12 | 84996080 | 84996874 | 323 | Promoter (<=1kb) | 12 | 84996321 | 85070492 | 74172 | 1 | ENSMUST00 | 0 | Ylpm1 | ENSMUSG00 | false | chr12 | 84996250 | 84997380 | 39.12038 | 124.14072 | 119.86815 | . | . | . | . | . | . |
| chr19 | 5424660 | 5425271 | 323 | Promoter (<=1kb) | 19 | 5422806 | 5424759 | 1954 | 2 | ENSMUST00 | 0 | Drap1 | ENSMUSG00 | false | chr19 | 5423832 | 5426468 | 29.11804 | 94.27461 | 90.83557 | . | . | . | . | . | . |
| chr1 | 75124780 | 75125330 | 323 | Promoter (<=1kb) | 1 | 75125136 | 75125200 | 65 | 2 | ENSMUST00 | 0 | Nhej1 | ENSMUSG00 | false | chr1 | 75124518 | 75125711 | 20.60796 | 54.90754 | 52.13 | chr1 | 75124181 | 75125967 | 4.57384 | 3.9242 | 2.66433 |
| chr1 | 178530535 | 178530754 | 323 | Promoter (1-2kb) | 1 | 178529125 | 178932730 | 403606 | 1 | ENSMUST00 | 1411 | Kif26b | ENSMUSG00 | false | chr1 | 178527358 | 178530919 | 20.79993 | 62.77878 | 59.89386 | chr1 | 178527430 | 178531162 | 6.05657 | 4.63626 | 3.88095 |
| chr4 | 152325666 | 152326056 | 323 | Promoter (<=1kb) | 4 | 152325742 | 152332476 | 6735 | 1 | ENSMUST00 | 0 | Rpl22 | ENSMUSG00 | false | chr4 | 152325753 | 152326885 | 50.54853 | 170.95712 | 164.9677 | . | . | . | . | . | . |
| chr5 | 140388980 | 140389439 | 323 | Promoter (<=1kb) | 5 | 140340304 | 140389251 | 48948 | 2 | ENSMUST00 | 0 | Snx8 | ENSMUSG00 | false | chr5 | 140388513 | 140389686 | 34.73729 | 107.12603 | 103.36953 | chr5 | 140381298 | 140389755 | 4.4341 | 3.83696 | 2.53558 |
| chr7 | 66689478 | 66690212 | 323 | Promoter (<=1kb) | 7 | 66644565 | 66689596 | 45032 | 2 | ENSMUST00 | 0 | Asb7 | ENSMUSG00 | false | chr7 | 66688676 | 66690647 | 31.82494 | 97.04205 | 93.54184 | chr7 | 66690150 | 66690878 | 3.63752 | 3.47499 | 1.86513 |
| chr4 | 118290736 | 118291112 | 323 | Promoter (<=1kb) | 4 | 118208213 | 118291405 | 83193 | 2 | ENSMUST00 | 293 | Ptprf | ENSMUSG00 | false | chr4 | 118290285 | 118291695 | 25.56571 | 75.64988 | 72.57267 | . | . | . | . | . | . |
| chr11 | 101424840 | 101425201 | 323 | Promoter (<=1kb) | 11 | 101425085 | 101435673 | 10589 | 1 | ENSMUST00 | 0 | Rundc1 | ENSMUSG00 | false | chr11 | 101424134 | 101425924 | 30.60796 | 113.09483 | 109.17188 | chr11 | 101423983 | 101427225 | 4.44266 | 3.8334 | 2.58514 |
| chr18 | 39213743 | 39213992 | 323 | Intron (ENSMUST00000097593.3 | 18 | 39227368 | 39372511 | 145144 | 1 | ENSMUST00 | -13376 | Arhgap26 | ENSMUSG00 | false | . | . | . | . | . | . | chr18 | 39213565 | 39213976 | 3.10277 | 3.09557 | 1.51278 |
| chr6 | 148946435 | 148946813 | 323 | Promoter (<=1kb) | 6 | 148921035 | 148946467 | 25433 | 2 | ENSMUST00 | 0 | Fam60a | ENSMUSG00 | false | chr6 | 148944401 | 148947082 | 36.04346 | 118.78738 | 114.68494 | . | . | . | . | . | . |
| chr2 | 91984469 | 91984862 | 323 | Intron (ENSMUST00000028663.4 | 2 | 91932824 | 91975864 | 43041 | 2 | ENSMUST00 | -8606 | Dgkz | ENSMUSG00 | false | . | . | . | . | . | . | chr2 | 91983325 | 91984739 | 4.8008 | 3.95199 | 2.84962 |
| chr2 | 17703760 | 17704045 | 323 | Intron (ENSMUST00000028080.6 | 2 | 17343909 | 17731043 | 387135 | 2 | ENSMUST00 | 26998 | Nebl | ENSMUSG00 | true | . | . | . | . | . | . | chr2 | 17675160 | 17718684 | 4.75621 | 3.88836 | 2.82315 |
| chr10 | 42030999 | 42031281 | 323 | Distal Intergenic | 10 | 41916807 | 42018442 | 101636 | 2 | ENSMUST00 | -12558 | Armc2 | ENSMUSG00 | false | . | . | . | . | . | . | chr10 | 42030870 | 42031711 | 6.00563 | 4.56548 | 3.82927 |
| chr6 | 100163226 | 100163597 | 323 | Distal Intergenic | 6 | 100228566 | 100287441 | 58876 | 2 | ENSMUST00 | 123844 | Rybp | ENSMUSG00 | false | chr6 | 100163172 | 100163502 | 8.21885 | 16.32505 | 14.04127 | chr6 | 100162903 | 100164715 | 6.08938 | 4.6874 | 3.89326 |
| chr6 | 86343817 | 86344198 | 323 | Distal Intergenic | 6 | 86334031 | 86334240 | 210 | 1 | ENSMUST00 | 9787 | Gm10443 | ENSMUSG00 | false | . | . | . | . | . | . | chr6 | 86342876 | 86347309 | 5.03053 | 3.95011 | 3.06041 |
| chr5 | 64314351 | 64314539 | 323 | Intron (ENSMUST00000043893.7 | 5 | 64320256 | 64320754 | 499 | 1 | ENSMUST00 | -5717 | Gm6044 | ENSMUSG00 | false | . | . | . | . | . | . | chr5 | 64312182 | 64328219 | 6.3028 | 4.70024 | 4.07007 |
| chr2 | 180837357 | 180837689 | 323 | Promoter (<=1kb) | 2 | 180837838 | 180837955 | 118 | 1 | ENSMUST00 | -149 | Gm22842 | ENSMUSG00 | false | . | . | . | . | . | . | chr2 | 180837164 | 180838016 | 3.14542 | 3.12275 | 1.50326 |
| chr11 | 78536096 | 78536582 | 322 | Promoter (<=1kb) | 11 | 78536361 | 78541737 | 5377 | 1 | ENSMUST00 | 0 | Ift20 | ENSMUSG00 | false | chr11 | 78535381 | 78536261 | 15.85026 | 49.49464 | 46.78799 | . | . | . | . | . | . |
| chr5 | 38561349 | 38561941 | 322 | Promoter (<=1kb) | 5 | 38526814 | 38561666 | 34853 | 2 | ENSMUST00 | 0 | Wdr1 | ENSMUSG00 | false | chr5 | 38560225 | 38562421 | 31.96831 | 119.80809 | 115.67081 | . | . | . | . | . | . |
| chr11 | 118363027 | 118363530 | 322 | Distal Intergenic | 11 | 118303701 | 118355740 | 52040 | 2 | ENSMUST00 | -7288 | Timp2 | ENSMUSG00 | false | . | . | . | . | . | . | chr11 | 118359762 | 118365457 | 7.10506 | 5.05606 | 4.73517 |
| chr10 | 41450158 | 41450537 | 322 | Promoter (<=1kb) | 10 | 41450396 | 41465571 | 15176 | 1 | ENSMUST00 | 0 | Zbtb24 | ENSMUSG00 | false | chr10 | 41449557 | 41451045 | 38.76865 | 124.60608 | 120.30663 | . | . | . | . | . | . |
| chr11 | 90002811 | 90003083 | 322 | Promoter (<=1kb) | 11 | 89982665 | 90002894 | 20230 | 2 | ENSMUST00 | 0 | Pctp | ENSMUSG00 | false | chr11 | 90002186 | 90003289 | 24.30838 | 70.74035 | 67.73997 | chr11 | 90002287 | 90003241 | 3.48621 | 3.36535 | 1.75159 |
| chr14 | 73454177 | 73454625 | 322 | Distal Intergenic | 14 | 73510049 | 73518545 | 8497 | 1 | ENSMUST00 | -55424 | Med4 | ENSMUSG00 | false | . | . | . | . | . | . | . | . | . | . | . | . |
| chr17 | 23676627 | 23676969 | 322 | Promoter (<=1kb) | 17 | 23675447 | 23677449 | 2003 | 2 | ENSMUST00 | 480 | Tnfrsf12a | ENSMUSG00 | false | chr17 | 23676112 | 23677653 | 26.28407 | 85.32412 | 82.07336 | chr17 | 23673161 | 23678044 | 6.00185 | 4.61543 | 3.82218 |
| chr18 | 32377045 | 32377415 | 322 | Promoter (<=1kb) | 18 | 32377217 | 32435736 | 58520 | 1 | ENSMUST00 | 0 | Bin1 | ENSMUSG00 | false | chr18 | 32376592 | 32378381 | 31.58442 | 93.66337 | 90.24092 | . | . | . | . | . | . |
| chr18 | 67932909 | 67933633 | 322 | Promoter (<=1kb) | 18 | 67933257 | 68255549 | 322293 | 1 | ENSMUST00 | 0 | Ldlrad4 | ENSMUSG00 | false | chr18 | 67932777 | 67934639 | 36.04346 | 118.78738 | 114.68494 | chr18 | 67932035 | 67934630 | 3.93263 | 3.46698 | 2.16447 |
| chr1 | 65179047 | 65179336 | 322 | Promoter (<=1kb) | 1 | 65158616 | 65179181 | 20566 | 2 | ENSMUST00 | 0 | Idh1 | ENSMUSG00 | false | chr1 | 65177109 | 65180019 | 28.91859 | 89.0447 | 85.71584 | . | . | . | . | . | . |
| chr3 | 57470999 | 57471323 | 322 | Intron (ENSMUST00000029380.8 | 3 | 57425314 | 57441677 | 16364 | 1 | ENSMUST00 | 45686 | Tm4sf4 | ENSMUSG00 | false | chr3 | 57470852 | 57471327 | 6.12159 | 11.42045 | 9.22029 | chr3 | 57469907 | 57474203 | 5.35164 | 4.23409 | 3.30612 |
| chr5 | 103377841 | 103378245 | 322 | Distal Intergenic | 5 | 103425192 | 103598359 | 173168 | 1 | ENSMUST00 | -46947 | Ptpn13 | ENSMUSG00 | false | . | . | . | . | . | . | chr5 | 103376057 | 103378714 | 5.86262 | 4.48493 | 3.69682 |
| chr1 | 127213752 | 127214184 | 322 | Intron (ENSMUST00000189427.1 | 1 | 127205028 | 127398792 | 193765 | 1 | ENSMUST00 | 8725 | Mgat5 | ENSMUSG00 | true | . | . | . | . | . | . | chr1 | 127206353 | 127222843 | 5.20875 | 4.0815 | 3.19162 |
| chr17 | 24752795 | 24753039 | 322 | Promoter (<=1kb) | 17 | 24753003 | 24758683 | 5681 | 1 | ENSMUST00 | 0 | Hs3st6 | ENSMUSG00 | false | chr17 | 24752281 | 24755341 | 33.1785 | 101.38432 | 97.78168 | . | . | . | . | . | . |
| chr5 | 65157821 | 65158262 | 322 | Promoter (1-2kb) | 5 | 65156070 | 65168142 | 12073 | 1 | ENSMUST00 | 1752 | Klhl5 | ENSMUSG00 | false | chr5 | 65157462 | 65158077 | 5.9709 | 9.68309 | 7.51161 | chr5 | 65157555 | 65158250 | 3.48892 | 3.25422 | 1.80407 |
| chr11 | 85492221 | 85492607 | 322 | Promoter (<=1kb) | 11 | 85492568 | 85495914 | 3347 | 1 | ENSMUST00 | 0 | Bcas3 | ENSMUSG00 | false | . | . | . | . | . | . | . | . | . | . | . | . |
| chr17 | 24251123 | 24251654 | 322 | Promoter (<=1kb) | 17 | 24223232 | 24251409 | 28178 | 2 | ENSMUST00 | 0 | Ccnf | ENSMUSG00 | false | chr17 | 24250691 | 24252009 | 25.00426 | 73.75784 | 70.7114 | chr17 | 24250106 | 24252129 | 3.53263 | 3.30107 | 1.84759 |
| chr6 | 31620896 | 31621209 | 322 | Intron (ENSMUST00000124919.2 | 6 | 31605289 | 31743314 | 138026 | 1 | ENSMUST00 | 15608 | Gm13847 | ENSMUSG00 | false | . | . | . | . | . | . | chr6 | 31612785 | 31643065 | 5.56305 | 4.31748 | 3.47573 |
| chr17 | 33732703 | 33732926 | 322 | Distal Intergenic | 17 | 33729509 | 33729584 | 76 | 1 | ENSMUST00 | 3195 | Gm22943 | ENSMUSG00 | false | . | . | . | . | . | . | chr17 | 33732164 | 33733758 | 3.95268 | 3.55569 | 2.15689 |
| chr16 | 30236350 | 30236628 | 322 | Distal Intergenic | 16 | 30200910 | 30205879 | 4970 | 2 | ENSMUST00 | -30472 | Gm26562 | ENSMUSG00 | false | chr16 | 30236118 | 30236495 | 7.30631 | 13.42958 | 11.19263 | . | . | . | . | . | . |
| chr19 | 23028416 | 23028669 | 322 | Distal Intergenic | 19 | 23037390 | 23075853 | 38464 | 2 | ENSMUST00 | 47184 | C330002G04 | ENSMUSG00 | false | . | . | . | . | . | . | . | . | . | . | . | . |
| chr13 | 98440668 | 98440917 | 322 | Distal Intergenic | 13 | 98424109 | 98435065 | 10957 | 2 | ENSMUST00 | -5604 | RP24-76E2.6 | ENSMUSG00 | false | . | . | . | . | . | . | chr13 | 98440202 | 98441668 | 3.66646 | 3.31118 | 1.93286 |
| chr11 | 69067686 | 69067927 | 321 | Promoter (2-3kb) | 11 | 69070806 | 69073030 | 2225 | 1 | ENSMUST00 | -2879 | Tmem107 | ENSMUSG00 | false | . | . | . | . | . | . | chr11 | 69066211 | 69068630 | 7.30405 | 5.06242 | 4.90746 |
| chr15 | 7726970 | 7727206 | 321 | Distal Intergenic | 15 | 7811011 | 7837575 | 26565 | 1 | ENSMUST00 | -83805 | Gdnf | ENSMUSG00 | false | . | . | . | . | . | . | chr15 | 7726724 | 7727647 | 2.94843 | 2.83695 | 1.35344 |
| chr5 | 137288050 | 137288355 | 321 | Promoter (<=1kb) | 5 | 137288277 | 137294466 | 6190 | 1 | ENSMUST00 | 0 | Ache | ENSMUSG00 | false | chr5 | 137287934 | 137288703 | 11.56301 | 29.49975 | 27.03959 | chr5 | 137287075 | 137289323 | 5.82274 | 4.55922 | 3.67036 |
| chr5 | 140606448 | 140606727 | 321 | Promoter (<=1kb) | 5 | 140607341 | 140615543 | 8203 | 1 | ENSMUST00 | -614 | Lfng | ENSMUSG00 | false | chr5 | 140606439 | 140608441 | 21.17422 | 79.20749 | 76.06625 | chr5 | 140604016 | 140607247 | 5.52811 | 4.37672 | 3.42431 |
| chr6 | 115863336 | 115863719 | 321 | Intron (ENSMUST00000112925.2 | 6 | 115853645 | 115875297 | 21653 | 1 | ENSMUST00 | 9692 | Ift122 | ENSMUSG00 | false | . | . | . | . | . | . | chr6 | 115862331 | 115864587 | 4.89458 | 4.0163 | 2.9257 |
| chr7 | 92769111 | 92769350 | 321 | Intron (ENSMUST00000032879.8 | 7 | 92741603 | 92837117 | 95515 | 1 | ENSMUST00 | 27509 | Rab30 | ENSMUSG00 | true | . | . | . | . | . | . | chr7 | 92764576 | 92769835 | 3.35455 | 3.14281 | 1.6895 |
| chr9 | 55149052 | 55150029 | 321 | Promoter (<=1kb) | 9 | 55149395 | 55206399 | 57005 | 1 | ENSMUST00 | 0 | Ube2q2 | ENSMUSG00 | false | chr9 | 55148832 | 55150580 | 30.82847 | 106.35825 | 102.62659 | chr9 | 55148740 | 55149137 | 3.18289 | 3.19714 | 1.49815 |
| chr9 | 67840183 | 67840649 | 321 | Promoter (<=1kb) | 9 | 67840396 | 67995634 | 155239 | 1 | ENSMUST00 | 0 | Vps13c | ENSMUSG00 | false | chr9 | 67839796 | 67841312 | 36.42221 | 131.26616 | 126.73016 | . | . | . | . | . | . |
| chr11 | 80810140 | 80810493 | 321 | Promoter (<=1kb) | 11 | 80810175 | 80822033 | 11859 | 1 | ENSMUST00 | 0 | Tmem98 | ENSMUSG00 | false | chr11 | 80809976 | 80811357 | 20.27628 | 60.14116 | 57.29097 | chr11 | 80808590 | 80810333 | 3.62352 | 3.32235 | 1.90864 |
| chr10 | 68125786 | 68126207 | 321 | Intron (ENSMUST00000020106.7 | 10 | 68043968 | 68051046 | 7079 | 1 | ENSMUST00 | 81819 | Rtkn2 | ENSMUSG00 | false | . | . | . | . | . | . | chr10 | 68124928 | 68129398 | 3.5431 | 3.34166 | 1.83799 |
| chr4 | 70534725 | 70535242 | 321 | Promoter (<=1kb) | 4 | 70427065 | 70534928 | 107864 | 2 | ENSMUST00 | 0 | Megf9 | ENSMUSG00 | false | chr4 | 70533328 | 70535844 | 36.46257 | 120.58523 | 116.42456 | . | . | . | . | . | . |
| chr17 | 43147340 | 43147630 | 321 | Distal Intergenic | 17 | 43270329 | 43324737 | 54409 | 1 | ENSMUST00 | -122699 | Gpr110 | ENSMUSG00 | false | . | . | . | . | . | . | chr17 | 43146554 | 43148633 | 4.86238 | 3.92665 | 2.91451 |
| chr6 | 47830256 | 47830689 | 321 | Promoter (<=1kb) | 6 | 47821658 | 47830568 | 8911 | 2 | ENSMUST00 | 0 | Zfp786 | ENSMUSG00 | false | chr6 | 47830002 | 47830883 | 34.74108 | 114.39095 | 110.42815 | chr6 | 47829706 | 47831026 | 4.83291 | 4.08324 | 2.85829 |
| chr8 | 94516266 | 94516498 | 321 | Exon (ENSMUST00000053085.6/ | 8 | 94521431 | 94523509 | 2079 | 1 | ENSMUST00 | -4933 | Nlrc5 | ENSMUSG00 | false | . | . | . | . | . | . | . | . | . | . | . | . |
| chr7 | 70065494 | 70065953 | 321 | Distal Intergenic | 7 | 69957776 | 69957904 | 129 | 2 | ENSMUST00 | -107591 | Gm24120 | ENSMUSG00 | false | . | . | . | . | . | . | chr7 | 70065494 | 70068605 | 4.26383 | 3.68388 | 2.40565 |
| chr11 | 106501290 | 106501667 | 321 | Promoter (<=1kb) | 11 | 106501245 | 106501377 | 133 | 1 | ENSMUST00 | 46 | Gm22711 | ENSMUSG00 | false | chr11 | 106500875 | 106501735 | 38.14117 | 119.25307 | 115.13219 | chr11 | 106501603 | 106501864 | 4.4883 | 3.6498 | 2.61022 |
| chr14 | 87255464 | 87255745 | 321 | Distal Intergenic | 14 | 87346245 | 87346387 | 143 | 1 | ENSMUST00 | -90500 | Gm23278 | ENSMUSG00 | false | . | . | . | . | . | . | chr14 | 87253607 | 87256977 | 3.79403 | 3.45802 | 2.05291 |
| chr14 | 97588178 | 97588533 | 321 | Distal Intergenic | 14 | 97336495 | 97336632 | 138 | 2 | ENSMUST00 | -251547 | Gm23276 | ENSMUSG00 | false | chr14 | 97588264 | 97588675 | 4.42115 | 6.83029 | 4.72578 | chr14 | 97587130 | 97589021 | 4.58164 | 3.82448 | 2.69644 |
| chr18 | 67577631 | 67577947 | 321 | Distal Intergenic | 18 | 67562387 | 67562851 | 465 | 1 | ENSMUST00 | 15245 | Gm17669 | ENSMUSG00 | false | . | . | . | . | . | . | chr18 | 67575475 | 67579988 | 6.90057 | 4.98929 | 4.56165 |
| chr18 | 55695762 | 55696049 | 321 | Distal Intergenic | 18 | 55745251 | 55746228 | 978 | 2 | ENSMUST00 | 50179 | Gm26959 | ENSMUSG00 | false | . | . | . | . | . | . | chr18 | 55693097 | 55696396 | 4.5344 | 3.88448 | 2.65659 |
| chr9 | 89776031 | 89776334 | 321 | Distal Intergenic | 9 | 89759003 | 89759465 | 463 | 1 | ENSMUST00 | 17029 | RP24-340L11 | ENSMUSG00 | false | . | . | . | . | . | . | chr9 | 89768908 | 89779398 | 4.75528 | 3.93987 | 2.80985 |
| chr13 | 19395473 | 19395999 | 320 | Promoter (<=1kb) | 13 | 19357677 | 19395752 | 38076 | 2 | ENSMUST00 | 0 | Stard3nl | ENSMUSG00 | false | chr13 | 19394935 | 19396398 | 36.04346 | 118.78738 | 114.68494 | . | . | . | . | . | . |
| chr1 | 12691909 | 12692168 | 320 | Promoter (<=1kb) | 1 | 12692277 | 12861192 | 168916 | 1 | ENSMUST00 | -109 | Sulf1 | ENSMUSG00 | true | chr1 | 12691712 | 12692719 | 17.92175 | 44.05972 | 41.41886 | chr1 | 12689349 | 12692023 | 4.3571 | 3.78338 | 2.48252 |
| chr11 | 44518697 | 44519538 | 320 | Promoter (<=1kb) | 11 | 44518964 | 44565520 | 46557 | 1 | ENSMUST00 | 0 | Rnf145 | ENSMUSG00 | false | chr11 | 44518262 | 44520961 | 40.23456 | 136.98251 | 132.21786 | chr11 | 44518156 | 44522498 | 3.70908 | 3.34851 | 1.98894 |
| chr12 | 80945751 | 80946243 | 320 | Promoter (<=1kb) | 12 | 80946376 | 80950497 | 4122 | 1 | ENSMUST00 | -133 | Srsf5 | ENSMUSG00 | false | chr12 | 80944805 | 80947053 | 36.26418 | 124.50906 | 120.21295 | . | . | . | . | . | . |
| chr17 | 29683227 | 29683444 | 320 | Promoter (1-2kb) | 17 | 29682149 | 29690095 | 7947 | 1 | ENSMUST00 | 1079 | Cmtr1 | ENSMUSG00 | false | . | . | . | . | . | . | chr17 | 29682908 | 29683590 | 3.7861 | 3.49853 | 2.0545 |
| chr2 | 31019734 | 31019968 | 320 | Promoter (<=1kb) | 2 | 31019819 | 31021381 | 1563 | 1 | ENSMUST00 | 0 | Usp20 | ENSMUSG00 | false | . | . | . | . | . | . | chr2 | 31018637 | 31021079 | 4.66014 | 3.58712 | 2.75751 |
| chr2 | 128125698 | 128126204 | 320 | Promoter (<=1kb) | 2 | 128126038 | 128130015 | 3978 | 1 | ENSMUST00 | 0 | Bcl2l11 | ENSMUSG00 | false | chr2 | 128125321 | 128128418 | 32.64849 | 123.19247 | 118.94151 | chr2 | 128122465 | 128127189 | 4.22997 | 3.65307 | 2.38221 |
| chr6 | 97405494 | 97405766 | 320 | Intron (ENSMUST00000113355.3 | 6 | 97323658 | 97396736 | 73079 | 2 | ENSMUST00 | -8759 | Frmd4b | ENSMUSG00 | false | . | . | . | . | . | . | chr6 | 97404992 | 97406311 | 4.98435 | 4.04962 | 2.9845 |
| chr7 | 66109132 | 66109669 | 320 | Promoter (<=1kb) | 7 | 66109515 | 66173789 | 64275 | 1 | ENSMUST00 | 0 | Chsy1 | ENSMUSG00 | false | chr7 | 66108901 | 66110654 | 27.49702 | 86.54257 | 83.27029 | chr7 | 66106375 | 66113379 | 4.42596 | 3.75674 | 2.53397 |
| chr10 | 108335152 | 108335401 | 320 | Promoter (2-3kb) | 10 | 108332189 | 108414391 | 82203 | 1 | ENSMUST00 | 2964 | Pawr | ENSMUSG00 | false | . | . | . | . | . | . | chr10 | 108334141 | 108343133 | 4.77514 | 3.97163 | 2.82359 |
| chr1 | 89069973 | 89071406 | 320 | Promoter (<=1kb) | 1 | 89070415 | 89155068 | 84654 | 1 | ENSMUST00 | 0 | Sh3bp4 | ENSMUSG00 | false | chr1 | 89069458 | 89069983 | 9.44256 | 24.88688 | 22.48677 | . | . | . | . | . | . |
| chr4 | 86882419 | 86882651 | 320 | Intron (ENSMUST00000045224.8 | 4 | 86874414 | 86932491 | 58078 | 1 | ENSMUST00 | 8006 | Acer2 | ENSMUSG00 | false | . | . | . | . | . | . | chr4 | 86881704 | 86884128 | 3.54565 | 3.27819 | 1.84593 |
| chr14 | 76899364 | 76899692 | 320 | Distal Intergenic | 14 | 77034950 | 77036081 | 1132 | 2 | ENSMUST00 | 136389 | Lacc1 | ENSMUSG00 | false | . | . | . | . | . | . | chr14 | 76898777 | 76900911 | 5.19331 | 4.24653 | 3.15027 |
| chr7 | 25076888 | 25077570 | 320 | Promoter (<=1kb) | 7 | 25077205 | 25083492 | 6288 | 1 | ENSMUST00 | 0 | Zfp574 | ENSMUSG00 | false | chr7 | 25077166 | 25078317 | 37.30079 | 124.19571 | 119.91465 | . | . | . | . | . | . |
| chr3 | 123093328 | 123093617 | 320 | Intron (ENSMUST00000139160.2 | 3 | 123076519 | 123112810 | 36292 | 2 | ENSMUST00 | 19193 | Synpo2 | ENSMUSG00 | false | . | . | . | . | . | . | chr3 | 123088474 | 123094788 | 4.57971 | 3.83586 | 2.65793 |
| chr8 | 68281655 | 68281834 | 320 | Intron (ENSMUST00000066594.3 | 8 | 68276567 | 68347694 | 71128 | 1 | ENSMUST00 | 5089 | Sh2d4a | ENSMUSG00 | true | . | . | . | . | . | . | chr8 | 68277725 | 68299894 | 4.63858 | 3.82888 | 2.72658 |
| chr8 | 72318637 | 72319109 | 320 | Promoter (<=1kb) | 8 | 72319033 | 72321656 | 2624 | 1 | ENSMUST00 | 0 | Klf2 | ENSMUSG00 | false | chr8 | 72318583 | 72321527 | 16.96705 | 61.94942 | 59.07605 | chr8 | 72317912 | 72320075 | 3.41972 | 3.1996 | 1.7168 |
| chr5 | 13982747 | 13982947 | 320 | Distal Intergenic | 5 | 14025276 | 14254314 | 229039 | 1 | ENSMUST00 | -42329 | Sema3e | ENSMUSG00 | true | . | . | . | . | . | . | chr5 | 13982519 | 13984353 | 4.08511 | 3.60549 | 2.30484 |
| chr1 | 43467743 | 43468002 | 320 | Intron (ENSMUST00000187435.1 | 1 | 43457273 | 43462222 | 4950 | 1 | ENSMUST00 | 10471 | Nck2 | ENSMUSG00 | true | . | . | . | . | . | . | chr1 | 43446894 | 43509082 | 5.99677 | 4.48008 | 3.83622 |
| chr11 | 63920605 | 63921278 | 320 | Promoter (1-2kb) | 11 | 63885792 | 63922290 | 36499 | 2 | ENSMUST00 | 1012 | Hs3st3b1 | ENSMUSG00 | false | chr11 | 63918689 | 63922332 | 41.51012 | 139.21701 | 134.36754 | . | . | . | . | . | . |
| chr2 | 79081473 | 79081729 | 320 | Distal Intergenic | 2 | 79038533 | 79038734 | 202 | 2 | ENSMUST00 | -42740 | Gm14469 | ENSMUSG00 | false | . | . | . | . | . | . | . | . | . | . | . | . |
| chr1 | 43181067 | 43181375 | 320 | Intron (ENSMUST00000185893.1 | 1 | 43189156 | 43189638 | 483 | 1 | ENSMUST00 | -7781 | Gm8210 | ENSMUSG00 | false | . | . | . | . | . | . | . | . | . | . | . | . |
| chr15 | 95526918 | 95527233 | 319 | Promoter (1-2kb) | 15 | 95219933 | 95528252 | 308320 | 2 | ENSMUST00 | 1019 | Nell2 | ENSMUSG00 | true | chr15 | 95526988 | 95528909 | 23.05105 | 65.89761 | 62.96862 | chr15 | 95526608 | 95528583 | 3.51491 | 3.28216 | 1.81492 |
| chr16 | 26248836 | 26249164 | 319 | Distal Intergenic | 16 | 26356642 | 26371841 | 15200 | 2 | ENSMUST00 | 122677 | Cldn1 | ENSMUSG00 | false | . | . | . | . | . | . | chr16 | 26248000 | 26249523 | 5.09539 | 4.18966 | 3.09391 |
| chr16 | 56716835 | 56717566 | 319 | Promoter (<=1kb) | 16 | 56690332 | 56717450 | 27119 | 2 | ENSMUST00 | 0 | Tfg | ENSMUSG00 | false | chr16 | 56716231 | 56717497 | 30.89292 | 96.41597 | 92.93404 | . | . | . | . | . | . |
| chr16 | 84835280 | 84835832 | 319 | Promoter (<=1kb) | 16 | 84827871 | 84835503 | 7633 | 2 | ENSMUST00 | 0 | Atp5j | ENSMUSG00 | false | chr16 | 84834034 | 84836397 | 49.45498 | 178.53914 | 172.40457 | . | . | . | . | . | . |
| chr18 | 79257112 | 79257557 | 319 | Distal Intergenic | 18 | 78750380 | 79109391 | 359012 | 2 | ENSMUST00 | -147722 | Setbp1 | ENSMUSG00 | false | . | . | . | . | . | . | chr18 | 79257144 | 79258127 | 3.46032 | 3.28564 | 1.74829 |
| chr1 | 51478168 | 51478654 | 319 | Promoter (<=1kb) | 1 | 51465862 | 51478399 | 12538 | 2 | ENSMUST00 | 0 | Nabp1 | ENSMUSG00 | false | chr1 | 51477533 | 51478508 | 30.44846 | 92.68264 | 89.28044 | . | . | . | . | . | . |
| chr2 | 32624402 | 32624681 | 319 | Promoter (<=1kb) | 2 | 32625437 | 32635054 | 9618 | 1 | ENSMUST00 | -756 | Ak1 | ENSMUSG00 | false | . | . | . | . | . | . | chr2 | 32605009 | 32639275 | 6.86759 | 4.92928 | 4.53618 |
| chr2 | 24408144 | 24408391 | 319 | Promoter (2-3kb) | 2 | 24405629 | 24407007 | 1379 | 1 | ENSMUST00 | 2516 | Psd4 | ENSMUSG00 | false | . | . | . | . | . | . | chr2 | 24407880 | 24409844 | 4.74512 | 3.91569 | 2.81509 |
| chr7 | 4925622 | 4925802 | 319 | Promoter (<=1kb) | 7 | 4925844 | 4944821 | 18978 | 1 | ENSMUST00 | -42 | Ssc5d | ENSMUSG00 | false | . | . | . | . | . | . | chr7 | 4920462 | 4926188 | 4.73104 | 3.83915 | 2.79979 |
| chr9 | 119334397 | 119334734 | 319 | Intron (ENSMUST00000170400.3 | 9 | 119339676 | 119342752 | 3077 | 1 | ENSMUST00 | -4942 | Acaa1a | ENSMUSG00 | false | . | . | . | . | . | . | chr9 | 119334291 | 119334900 | 3.56242 | 3.09461 | 1.86794 |
| chr11 | 54530388 | 54530678 | 319 | Intron (ENSMUST00000102743.4 | 11 | 54522911 | 54561596 | 38686 | 1 | ENSMUST00 | 7478 | Rapgef6 | ENSMUSG00 | true | . | . | . | . | . | . | chr11 | 54523692 | 54544299 | 4.68081 | 3.90811 | 2.75331 |
| chr11 | 97831954 | 97832284 | 319 | Promoter (<=1kb) | 11 | 97833183 | 97836205 | 3023 | 1 | ENSMUST00 | -899 | Lasp1 | ENSMUSG00 | false | . | . | . | . | . | . | chr11 | 97801064 | 97833244 | 8.45203 | 5.49885 | 5.84074 |
| chr2 | 32362920 | 32363455 | 319 | Promoter (<=1kb) | 2 | 32363010 | 32367407 | 4398 | 1 | ENSMUST00 | 0 | Ciz1 | ENSMUSG00 | false | chr2 | 32362481 | 32363897 | 30.17592 | 94.17345 | 90.73757 | chr2 | 32360623 | 32364173 | 4.73103 | 3.92594 | 2.82173 |
| chr10 | 17511476 | 17511708 | 319 | Distal Intergenic | 10 | 17723228 | 17725673 | 2446 | 1 | ENSMUST00 | -211520 | Cited2 | ENSMUSG00 | false | . | . | . | . | . | . | . | . | . | . | . | . |
| chr10 | 117810668 | 117810908 | 319 | Distal Intergenic | 10 | 117750643 | 117792705 | 42063 | 2 | ENSMUST00 | -17964 | Nup107 | ENSMUSG00 | false | . | . | . | . | . | . | chr10 | 117810094 | 117812350 | 5.62435 | 4.22864 | 3.55721 |
| chr12 | 84148199 | 84148604 | 319 | Promoter (<=1kb) | 12 | 84146131 | 84148489 | 2359 | 2 | ENSMUST00 | 0 | Pnma1 | ENSMUSG00 | false | chr12 | 84148042 | 84148694 | 25.12339 | 69.47189 | 66.48775 | . | . | . | . | . | . |
| chr15 | 37007041 | 37007931 | 319 | Promoter (<=1kb) | 15 | 36997031 | 37007402 | 10372 | 2 | ENSMUST00 | 0 | Zfp706 | ENSMUSG00 | false | chr15 | 37005733 | 37008177 | 30.03893 | 101.89226 | 98.2715 | . | . | . | . | . | . |
| chr18 | 35919020 | 35919351 | 319 | Distal Intergenic | 18 | 35952824 | 35952907 | 84 | 1 | ENSMUST00 | -33473 | Gm23882 | ENSMUSG00 | false | chr18 | 35918794 | 35920648 | 19.69154 | 51.71931 | 48.98378 | chr18 | 35918346 | 35921757 | 4.57948 | 3.8339 | 2.68759 |
| chrX | 146164727 | 146164906 | 319 | Distal Intergenic | 20 | 146175942 | 146176577 | 636 | 1 | ENSMUST00 | -11036 | Hmgb1-ps2 | ENSMUSG00 | false | . | . | . | . | . | . | . | . | . | . | . | . |
| chr13 | 29631484 | 29631731 | 319 | Intron (ENSMUST00000140278.2 | 13 | 29732942 | 29733036 | 95 | 1 | ENSMUST00 | -101211 | Gm23729 | ENSMUSG00 | false | . | . | . | . | . | . | chr13 | 29631459 | 29632040 | 3.88713 | 3.57201 | 2.08583 |
| chr18 | 13251522 | 13251754 | 319 | Distal Intergenic | 18 | 13209702 | 13209808 | 107 | 2 | ENSMUST00 | -41715 | Gm22251 | ENSMUSG00 | false | . | . | . | . | . | . | chr18 | 13251072 | 13251985 | 4.6991 | 3.95175 | 2.79114 |
| chr9 | 32871227 | 32871520 | 319 | Distal Intergenic | 9 | 32904937 | 32928966 | 24030 | 2 | ENSMUST00 | 57446 | Gm27162 | ENSMUSG00 | false | . | . | . | . | . | . | . | . | . | . | . | . |
| chr10 | 79857530 | 79857836 | 318 | Promoter (<=1kb) | 10 | 79858772 | 79859387 | 616 | 1 | ENSMUST00 | -936 | Ptbp1 | ENSMUSG00 | false | chr10 | 79854526 | 79858632 | 33.99707 | 114.12061 | 110.16386 | chr10 | 79855944 | 79867013 | 7.93167 | 5.01038 | 5.43402 |
| chr17 | 57032064 | 57032375 | 318 | Promoter (<=1kb) | 17 | 57021058 | 57031507 | 10450 | 2 | ENSMUST00 | -558 | Khsrp | ENSMUSG00 | false | chr17 | 57032151 | 57032660 | 13.41152 | 31.46699 | 28.98322 | . | . | . | . | . | . |
| chr3 | 107760063 | 107760643 | 318 | Promoter (<=1kb) | 3 | 107741048 | 107760236 | 19189 | 2 | ENSMUST00 | 0 | Csf1 | ENSMUSG00 | false | chr3 | 107759461 | 107760773 | 27.1916 | 82.20596 | 79.01564 | chr3 | 107758037 | 107760756 | 3.32312 | 3.20124 | 1.64053 |
| chr6 | 34172117 | 34172471 | 318 | Intron (ENSMUST00000126498.1 | 6 | 34158085 | 34176970 | 18886 | 2 | ENSMUST00 | 4499 | Slc35b4 | ENSMUSG00 | false | . | . | . | . | . | . | chr6 | 34170482 | 34177467 | 5.2089 | 4.17803 | 3.18044 |
| chr11 | 19019546 | 19020012 | 318 | Promoter (<=1kb) | 11 | 18879817 | 19018985 | 139169 | 2 | ENSMUST00 | -562 | Meis1 | ENSMUSG00 | false | chr11 | 19018184 | 19021149 | 22.06604 | 78.32127 | 75.1984 | . | . | . | . | . | . |
| chr13 | 68581905 | 68582281 | 318 | Promoter (<=1kb) | 13 | 68582248 | 68592279 | 10032 | 1 | ENSMUST00 | 0 | Fastkd3 | ENSMUSG00 | false | chr13 | 68581239 | 68582933 | 31.54088 | 108.67455 | 104.87239 | . | . | . | . | . | . |
| chr14 | 30479467 | 30479754 | 318 | Promoter (<=1kb) | 14 | 30479565 | 30527056 | 47492 | 1 | ENSMUST00 | 0 | Dcp1a | ENSMUSG00 | false | chr14 | 30479073 | 30480785 | 21.73903 | 87.55687 | 84.25908 | . | . | . | . | . | . |
| chr14 | 70949704 | 70949968 | 318 | Intron (ENSMUST00000022699.8 | 14 | 70890130 | 70979838 | 89709 | 1 | ENSMUST00 | 59575 | Gfra2 | ENSMUSG00 | false | . | . | . | . | . | . | . | . | . | . | . | . |
| chr14 | 103778239 | 103778504 | 318 | Distal Intergenic | 14 | 103814626 | 103843696 | 29071 | 2 | ENSMUST00 | 65192 | Ednrb | ENSMUSG00 | false | chr14 | 103778182 | 103778698 | 10.08475 | 20.47385 | 18.13181 | chr14 | 103777808 | 103778962 | 3.5303 | 3.38569 | 1.77972 |
| chr1 | 40266489 | 40266805 | 318 | Promoter (<=1kb) | 1 | 40266586 | 40316198 | 49613 | 1 | ENSMUST00 | 0 | Il1r1 | ENSMUSG00 | false | chr1 | 40266288 | 40267027 | 23.56964 | 65.0516 | 62.13324 | chr1 | 40266105 | 40267594 | 3.21998 | 3.14257 | 1.58555 |
| chr7 | 120917622 | 120918005 | 318 | Promoter (<=1kb) | 7 | 120917744 | 120947432 | 29689 | 1 | ENSMUST00 | 0 | Polr3e | ENSMUSG00 | false | chr7 | 120917145 | 120918777 | 41.2835 | 132.20779 | 127.63918 | . | . | . | . | . | . |
| chr8 | 27106852 | 27107212 | 318 | Intron (ENSMUST00000178514.2 | 8 | 27114133 | 27118834 | 4702 | 1 | ENSMUST00 | -6921 | Gpr124 | ENSMUSG00 | false | . | . | . | . | . | . | . | . | . | . | . | . |
| chr4 | 83250693 | 83250930 | 318 | Intron (ENSMUST00000102823.4 | 4 | 83247613 | 83266039 | 18427 | 2 | ENSMUST00 | 15109 | Ttc39b | ENSMUSG00 | false | . | . | . | . | . | . | chr4 | 83250588 | 83252444 | 3.13021 | 3.07243 | 1.50959 |
| chr18 | 63708509 | 63708799 | 318 | Promoter (<=1kb) | 18 | 63708695 | 63989760 | 281066 | 1 | ENSMUST00 | 0 | Wdr7 | ENSMUSG00 | false | chr18 | 63708304 | 63709486 | 32.97563 | 101.03377 | 97.44133 | chr18 | 63708781 | 63709742 | 5.22945 | 4.22725 | 3.18885 |
| chr13 | 45001491 | 45002285 | 318 | Promoter (<=1kb) | 13 | 44922079 | 45002096 | 80018 | 2 | ENSMUST00 | 0 | Dtnbp1 | ENSMUSG00 | false | chr13 | 45001036 | 45002726 | 29.2476 | 106.45832 | 102.71833 | . | . | . | . | . | . |
| chr15 | 36960878 | 36961067 | 318 | Distal Intergenic | 15 | 36997031 | 37007402 | 10372 | 2 | ENSMUST00 | 46335 | Zfp706 | ENSMUSG00 | false | . | . | . | . | . | . | chr15 | 36959870 | 36961581 | 4.72463 | 3.95821 | 2.79607 |
| chr19 | 18821848 | 18822171 | 318 | Intron (ENSMUST00000040489.7 | 19 | 18776746 | 18776961 | 216 | 2 | ENSMUST00 | -44888 | Gm24190 | ENSMUSG00 | false | . | . | . | . | . | . | chr19 | 18821210 | 18822434 | 4.64784 | 3.95575 | 2.75566 |
| chr17 | 30188683 | 30189014 | 318 | Intron (ENSMUST00000057897.7 | 17 | 30311079 | 30311151 | 73 | 1 | ENSMUST00 | -122065 | Gm25049 | ENSMUSG00 | false | . | . | . | . | . | . | chr17 | 30177119 | 30208101 | 5.95143 | 4.39921 | 3.81342 |
| chr11 | 22584746 | 22584976 | 318 | Distal Intergenic | 11 | 22600335 | 22610879 | 10545 | 2 | ENSMUST00 | 25903 | Gm26829 | ENSMUSG00 | false | . | . | . | . | . | . | chr11 | 22584102 | 22585126 | 5.65622 | 4.41076 | 3.5445 |

| | | | | | | | | | | | | | | | | | | | | | | | | | | |
|---|---|---|---|---|---|---|---|---|---|---|---|---|---|---|---|---|---|---|---|---|---|---|---|---|---|---|
| chr1 | 39846010 | 39846346 | 318 | Promoter (<=1kb) | 1 | 39839175 | 39845699 | 6525 | 2 | ENSMUST00 | -312 | RP23-469H1: | ENSMUSG0( | false | | | | | | | | | | | | |
| chr6 | 141371870 | 141372100 | 318 | Intron (ENSMUST00000043259.8 | 6 | 141238278 | 141250561 | 12284 | 2 | ENSMUST00 | -121310 | RP23-97L11. | ENSMUSG0( | false | | | | | | | chr6 | 141370327 | 141386627 | 5.21392 | 4.14863 | 3.18933 |
| chr6 | 115480629 | 115481062 | 317 | Intron (ENSMUST00000171644.2 | 6 | 115422066 | 115490401 | 68336 | 1 | ENSMUST00 | 58564 | Pparg | ENSMUSG0( | false | | | | | | | | | | | | |
| chr6 | 52234461 | 52235158 | 317 | Promoter (<=1kb) | 6 | 52231197 | 52234939 | 3743 | 2 | ENSMUST00 | 0 | Hoxa10 | ENSMUSG0( | false | chr6 | 52227601 | 52235632 | 41.69514 | 140.65352 | 135.74945 | chr6 | 52234919 | 52241057 | 4.85457 | 4.03897 | 2.87932 |
| chr3 | 103790904 | 103791578 | 317 | Promoter (<=1kb) | 3 | 103739815 | 103791563 | 51749 | 2 | ENSMUST00 | 0 | Hipk1 | ENSMUSG0( | false | chr3 | 103789748 | 103791963 | 31.16539 | 106.97112 | 103.21663 | | | | | | |
| chr11 | 106216730 | 106217194 | 317 | Promoter (<=1kb) | 11 | 106216926 | 106249139 | 32214 | 1 | ENSMUST00 | 0 | Ddx42 | ENSMUSG0( | false | chr11 | 106215465 | 106217940 | 40.65367 | 138.82761 | 133.98773 | | | | | | |
| chr12 | 83317771 | 83318030 | 317 | Intron (ENSMUST00000147469.2 | 12 | 83260493 | 83294428 | 33936 | 2 | ENSMUST00 | -23344 | Dpf3 | ENSMUSG0( | false | | | | | | | chr12 | 83317552 | 83318603 | 4.40962 | 3.56685 | 2.54617 |
| chr14 | 30888346 | 30888654 | 317 | Promoter (1-2kb) | 14 | 30886521 | 30901846 | 15326 | 1 | ENSMUST00 | 1826 | Itih4 | ENSMUSG0( | false | | | | | | | chr14 | 30887231 | 30888556 | 4.03437 | 3.55443 | 2.22122 |
| chr15 | 12824587 | 12825029 | 317 | Promoter (<=1kb) | 15 | 12824815 | 12935290 | 110476 | 1 | ENSMUST00 | 0 | Drosha | ENSMUSG0( | false | chr15 | 12823969 | 12825597 | 30.60796 | 113.09483 | 109.17188 | chr15 | 12823582 | 12826224 | 4.69736 | 3.99366 | 2.77209 |
| chr16 | 94580900 | 94581173 | 317 | Intron (ENSMUST00000119878.2 | 16 | 94570907 | 94695517 | 124611 | 1 | ENSMUST00 | 9994 | Dyrk1a | ENSMUSG0( | false | | | | | | | chr16 | 94571320 | 94615960 | 5.49747 | 4.23365 | 3.43578 |
| chr9 | 59544572 | 59544834 | 317 | Intron (ENSMUST00000026262.6 | 9 | 59539667 | 59565105 | 25439 | 1 | ENSMUST00 | 4906 | Hexa | ENSMUSG0( | false | | | | | | | chr9 | 59542778 | 59545329 | 4.13886 | 3.65901 | 2.34176 |
| chr3 | 88058128 | 88058655 | 317 | Promoter (<=1kb) | 3 | 88056523 | 88058495 | 1973 | 2 | ENSMUST00 | 0 | Apoa1bp | ENSMUSG0( | false | chr3 | 88057578 | 88058938 | 29.71419 | 101.30199 | 97.70005 | chr3 | 88057259 | 88059148 | 5.96725 | 4.47554 | 3.80123 |
| chr6 | 43310156 | 43310383 | 317 | Promoter (<=1kb) | 6 | 43303674 | 43309554 | 5881 | 2 | ENSMUST00 | -603 | Nobox | ENSMUSG0( | false | | | | | | | chr6 | 43309438 | 43311583 | 3.49775 | 3.27718 | 1.82126 |
| chr9 | 78448031 | 78448350 | 317 | Promoter (<=1kb) | 9 | 78448208 | 78474152 | 25945 | 1 | ENSMUST00 | 0 | Mto1 | ENSMUSG0( | false | chr9 | 78447873 | 78449175 | 39.62653 | 133.71027 | 129.07751 | chr9 | 78447644 | 78449711 | 5.19651 | 4.18569 | 3.1697 |
| chr2 | 33607492 | 33607863 | 317 | Intron (ENSMUST00000041730.5 | 2 | 33567156 | 33632283 | 65128 | 2 | ENSMUST00 | 24420 | Lmx1b | ENSMUSG0( | true | | | | | | | chr2 | 33565260 | 33638297 | 6.78481 | 4.81457 | 4.48235 |
| chr16 | 92173286 | 92173589 | 317 | Distal Intergenic | 16 | 92106261 | 92112227 | 5967 | 1 | ENSMUST00 | 67026 | Mrps6 | ENSMUSG0( | false | | | | | | | chr16 | 92170108 | 92180343 | 5.69391 | 4.37461 | 3.59071 |
| chr1 | 130731817 | 130732062 | 317 | Promoter (<=1kb) | 1 | 130731976 | 130744622 | 12647 | 1 | ENSMUST00 | 0 | AA986860 | ENSMUSG0( | false | chr1 | 130731948 | 130732360 | 11.40015 | 24.34736 | 21.95423 | chr1 | 130732013 | 130736173 | 5.51371 | 4.2295 | 3.45563 |
| chr14 | 57056374 | 57056798 | 317 | Promoter (1-2kb) | 14 | 57035608 | 57058030 | 22423 | 2 | ENSMUST00 | 1232 | Gja3 | ENSMUSG0( | false | chr14 | 57056396 | 57058454 | 32.01914 | 97.37792 | 93.87154 | chr14 | 57053149 | 57056474 | 4.05021 | 3.5021 | 2.24184 |
| chr14 | 32861172 | 32861425 | 317 | Intron (ENSMUST00000053175.6 | 14 | 32856789 | 32939487 | 82699 | 1 | ENSMUST00 | 4384 | Vstm4 | ENSMUSG0( | false | | | | | | | chr14 | 32858959 | 32861977 | 6.14822 | 4.66947 | 3.93387 |
| chr1 | 171287836 | 171288145 | 317 | Promoter (<=1kb) | 1 | 171281945 | 171287991 | 6047 | 2 | ENSMUST00 | 0 | Usp21 | ENSMUSG0( | false | chr1 | 171287191 | 171288223 | 33.38818 | 124.5882 | 120.28912 | chr1 | 171285329 | 171288647 | 4.98608 | 4.04168 | 3.03001 |
| chr9 | 110633910 | 110634389 | 317 | Promoter (<=1kb) | 9 | 110629570 | 110634449 | 4880 | 2 | ENSMUST00 | 60 | Nbeal2 | ENSMUSG0( | false | | | | | | | chr9 | 110632191 | 110635511 | 6.98922 | 4.76714 | 4.66266 |
| chr2 | 132723884 | 132724126 | 317 | Intron (ENSMUST00000061891.5 | 2 | 132717106 | 132717208 | 103 | 1 | ENSMUST00 | 6779 | Gm22245 | ENSMUSG0( | false | | | | | | | chr2 | 132724103 | 132725781 | 4.29432 | 3.69979 | 2.44193 |
| chr18 | 73536174 | 73536677 | 317 | Promoter (2-3kb) | 18 | 73533403 | 73533550 | 148 | 2 | ENSMUST00 | -2625 | n-R5-8s1 | ENSMUSG0( | false | | | | | | | chr18 | 73536427 | 73537004 | 3.82563 | 3.36551 | 2.07391 |
| chr17 | 35000817 | 35001697 | 316 | Promoter (<=1kb) | 17 | 35000987 | 35016322 | 15336 | 1 | ENSMUST00 | 0 | Vars | ENSMUSG0( | false | chr17 | 35000585 | 35002024 | 26.72788 | 101.58999 | 97.97913 | chr17 | 34999793 | 35002592 | 5.36325 | 4.09482 | 3.33862 |
| chr12 | 24574376 | 24574575 | 316 | Promoter (2-3kb) | 12 | 24572287 | 24617387 | 45101 | 1 | ENSMUST00 | 2090 | Grhl1 | ENSMUSG0( | false | | | | | | | chr12 | 24571597 | 24576131 | 4.40813 | 3.73639 | 2.52624 |
| chr15 | 98777938 | 98778152 | 316 | Promoter (<=1kb) | 15 | 98771753 | 98778150 | 6398 | 2 | ENSMUST00 | 0 | Wnt10b | ENSMUSG0( | false | | | | | | | chr15 | 98775676 | 98778436 | 4.71083 | 3.81658 | 2.80395 |
| chr19 | 45874991 | 45875310 | 316 | Intron (ENSMUST00000045396.7 | 19 | 45792346 | 45816061 | 23716 | 2 | ENSMUST00 | -58931 | Kcnip2 | ENSMUSG0( | false | | | | | | | chr19 | 45872918 | 45876647 | 4.52597 | 3.85941 | 2.63692 |
| chr7 | 140125472 | 140125775 | 316 | Promoter (<=1kb) | 7 | 140125657 | 140134334 | 8678 | 1 | ENSMUST00 | 0 | Paox | ENSMUSG0( | false | chr7 | 140125241 | 140126186 | 34.83228 | 119.81895 | 115.68042 | chr7 | 140125008 | 140126292 | 4.19046 | 3.64845 | 2.35013 |
| chr1 | 189752387 | 189752693 | 316 | Intron (ENSMUST00000097442.3 | 1 | 189728268 | 189876689 | 148422 | 1 | ENSMUST00 | 24120 | Ptpn14 | ENSMUSG0( | true | | | | | | | chr1 | 189730267 | 189810107 | 6.93269 | 4.87829 | 4.60521 |
| chr2 | 74826017 | 74826500 | 316 | Promoter (<=1kb) | 2 | 74825830 | 74868814 | 42985 | 1 | ENSMUST00 | 188 | Mtx2 | ENSMUSG0( | false | chr2 | 74825369 | 74826551 | 32.64849 | 123.19247 | 118.94151 | | | | | | |
| chr3 | 132683692 | 132684322 | 316 | Promoter (<=1kb) | 3 | 132660498 | 132683879 | 23382 | 2 | ENSMUST00 | 0 | Aimp1 | ENSMUSG0( | false | chr3 | 132682978 | 132685271 | 32.67316 | 105.57997 | 101.8659 | | | | | | |
| chr6 | 119902538 | 119903120 | 316 | Promoter (<=1kb) | 6 | 119902698 | 119915335 | 12638 | 1 | ENSMUST00 | 0 | Rad52 | ENSMUSG0( | false | chr6 | 119902226 | 119903655 | 38.55812 | 129.64772 | 125.16568 | chr6 | 119901878 | 119910158 | 4.28788 | 3.7252 | 2.42516 |
| chr9 | 69092105 | 69092547 | 316 | Intron (ENSMUST00000034766.8 | 9 | 69195541 | 69250834 | 55294 | 1 | ENSMUST00 | -102994 | Rora | ENSMUSG0( | false | | | | | | | chr9 | 69091054 | 69093502 | 6.32528 | 4.66528 | 4.13989 |
| chr6 | 91562935 | 91563176 | 316 | Distal Intergenic | 6 | 91516047 | 91517481 | 1435 | 1 | ENSMUST00 | 46889 | Lsm3 | ENSMUSG0( | false | | | | | | | chr6 | 91561703 | 91573814 | 5.57785 | 4.24835 | 3.49935 |
| chr15 | 103177576 | 103177876 | 316 | Distal Intergenic | 15 | 103153307 | 103158002 | 4696 | 2 | ENSMUST00 | -19575 | Smug1 | ENSMUSG0( | false | | | | | | | chr15 | 103176992 | 103179383 | 4.02794 | 3.61227 | 2.21156 |
| chr11 | 75139025 | 75139348 | 316 | 3' UTR | 11 | 75132435 | 75139369 | 6935 | 1 | ENSMUST00 | 6591 | Smg6 | ENSMUSG0( | false | | | | | | | chr11 | 75136734 | 75141926 | 6.17803 | 4.48241 | 3.99662 |
| chr2 | 5793822 | 5794024 | 316 | Downstream (<1kb) | 2 | 5794665 | 5798598 | 3934 | 2 | ENSMUST00 | 4574 | Cdc123 | ENSMUSG0( | false | | | | | | | | | | | | |
| chr17 | 24672177 | 24672504 | 316 | Promoter (2-3kb) | 17 | 24669752 | 24682015 | 12264 | 1 | ENSMUST00 | 2426 | Zfp598 | ENSMUSG0( | false | | | | | | | chr17 | 24668886 | 24676794 | 7.81405 | 5.36667 | 5.31609 |
| chr15 | 99030124 | 99030486 | 316 | Promoter (<=1kb) | 15 | 99029891 | 99038105 | 8215 | 1 | ENSMUST00 | 234 | Tuba1c | ENSMUSG0( | false | chr15 | 99029288 | 99031062 | 27.71015 | 85.00858 | 81.76643 | chr15 | 99029075 | 99031519 | 4.25563 | 3.71636 | 2.40423 |
| chr4 | 119175701 | 119175931 | 316 | Promoter (1-2kb) | 4 | 119169513 | 119174195 | 4683 | 2 | ENSMUST00 | -1507 | Zfp691 | ENSMUSG0( | false | | | | | | | chr4 | 119174446 | 119176102 | 4.16588 | 3.69192 | 2.30932 |
| chr11 | 95824092 | 95824576 | 316 | Promoter (<=1kb) | 11 | 95824499 | 95830701 | 6203 | 1 | ENSMUST00 | 0 | Phospho1 | ENSMUSG0( | false | chr11 | 95823751 | 95825261 | 27.71015 | 85.00858 | 81.76643 | chr11 | 95823663 | 95825242 | 4.34345 | 3.80682 | 2.49752 |
| chr18 | 5404548 | 5404771 | 316 | Distal Intergenic | 18 | 5210031 | 5334439 | 124409 | 2 | ENSMUST00 | -70110 | Zfp438 | ENSMUSG0( | false | | | | | | | | | | | | |
| chr11 | 52167981 | 52168315 | 316 | Intron (ENSMUST00000181262.1 | 11 | 52155378 | 52156283 | 906 | 2 | ENSMUST00 | -11699 | Olfr1372-ps1 | ENSMUSG0( | false | chr11 | 52167719 | 52168683 | 29.93585 | 89.84451 | 86.50629 | chr11 | 52166041 | 52168637 | 4.03372 | 3.61712 | 2.23828 |
| chr8 | 31657740 | 31658051 | 316 | Distal Intergenic | 8 | 31661331 | 31661431 | 101 | 2 | ENSMUST00 | 3380 | Gm24184 | ENSMUSG0( | false | | | | | | | | | | | | |
| chr3 | 131802939 | 131803207 | 316 | Distal Intergenic | 3 | 131918259 | 131918353 | 95 | 1 | ENSMUST00 | -115052 | Gm26103 | ENSMUSG0( | false | | | | | | | | | | | | |
| chr12 | 113140081 | 113140604 | 315 | Promoter (<=1kb) | 12 | 113140236 | 113145504 | 5269 | 1 | ENSMUST00 | 0 | Crip2 | ENSMUSG0( | false | chr12 | 113139865 | 113141336 | 23.3092 | 84.4034 | 81.16875 | chr12 | 113139363 | 113149705 | 6.62613 | 4.83812 | 4.33223 |
| chr12 | 72664678 | 72665279 | 315 | Promoter (<=1kb) | 12 | 72650356 | 72664828 | 14473 | 2 | ENSMUST00 | 0 | Dhrs7 | ENSMUSG0( | false | chr12 | 72663805 | 72665580 | 26.19333 | 73.09374 | 70.054 | | | | | | |
| chr17 | 29451339 | 29451674 | 315 | Distal Intergenic | 17 | 29490812 | 29496111 | 5300 | 1 | ENSMUST00 | -39138 | Pim1 | ENSMUSG0( | false | chr17 | 29451035 | 29451518 | 9.94526 | 25.28477 | 22.8772 | chr17 | 29443894 | 29452487 | 4.93685 | 4.01842 | 2.95062 |
| chr2 | 31572377 | 31572985 | 315 | Promoter (<=1kb) | 2 | 31572651 | 31617526 | 44876 | 1 | ENSMUST00 | 0 | Fubp3 | ENSMUSG0( | false | chr2 | 31571910 | 31573549 | 27.18312 | 111.36749 | 107.49936 | chr2 | 31571924 | 31575035 | 3.53305 | 3.29702 | 1.85008 |
| chrX | 7899275 | 7899544 | 315 | Promoter (<=1kb) | 20 | 7899357 | 7908351 | 8995 | 1 | ENSMUST00 | 0 | Timm17b | ENSMUSG0( | false | chrX | 7898746 | 7899988 | 17.46222 | 42.53877 | 39.91948 | | | | | | |
| chr9 | 107635146 | 107635555 | 315 | Promoter (<=1kb) | 9 | 107614138 | 107635342 | 21205 | 2 | ENSMUST00 | 0 | Gnai2 | ENSMUSG0( | false | chr9 | 107632708 | 107635411 | 27.71015 | 85.00858 | 81.76643 | | | | | | |
| chr11 | 26210209 | 26210576 | 315 | Promoter (<=1kb) | 11 | 25617668 | 26210569 | 592902 | 2 | ENSMUST00 | 0 | 5730522E02I | ENSMUSG0( | false | chr11 | 26209894 | 26211068 | 26.07877 | 74.71087 | 71.64691 | | | | | | |
| chr1 | 55732698 | 55733222 | 315 | Intron (ENSMUST00000042986.9 | 1 | 55702025 | 55735227 | 33203 | 1 | ENSMUST00 | 30674 | Plcl1 | ENSMUSG0( | true | | | | | | | chr1 | 55730782 | 55736479 | 4.72182 | 3.89251 | 2.79474 |
| chr9 | 32409381 | 32409667 | 315 | Distal Intergenic | 9 | 32393985 | 32399192 | 5208 | 1 | ENSMUST00 | 15397 | Kcnj1 | ENSMUSG0( | false | chr9 | 32409074 | 32409671 | 11.39974 | 34.91658 | 32.39005 | chr9 | 32408277 | 32409928 | 5.41112 | 4.16133 | 3.3729 |
| chr4 | 45408677 | 45409091 | 315 | Promoter (<=1kb) | 4 | 45395923 | 45408761 | 12839 | 2 | ENSMUST00 | 0 | Slc25a51 | ENSMUSG0( | false | chr4 | 45407590 | 45409701 | 31.25232 | 111.22263 | 107.35811 | | | | | | |
| chr2 | 153267575 | 153267862 | 315 | 3' UTR | 2 | 153257572 | 153268063 | 10492 | 1 | ENSMUST00 | 10004 | Pofut1 | ENSMUSG0( | true | | | | | | | chr2 | 153267389 | 153267961 | 4.42906 | 3.48929 | 2.56835 |
| chr13 | 92593150 | 92593400 | 315 | Distal Intergenic | 13 | 92574631 | 92576232 | 1602 | 2 | ENSMUST00 | -16919 | Spz1 | ENSMUSG0( | false | | | | | | | chr13 | 92592427 | 92595642 | 4.40973 | 3.63511 | 2.57033 |
| chr17 | 15825801 | 15826815 | 315 | Promoter (<=1kb) | 17 | 15806253 | 15826586 | 20334 | 2 | ENSMUST00 | 0 | Rgmb | ENSMUSG0( | false | chr17 | 15824815 | 15827519 | 37.37859 | 116.88169 | 112.84478 | | | | | | |
| chr15 | 78836170 | 78836535 | 315 | Distal Intergenic | 15 | 78842647 | 78850902 | 8256 | 1 | ENSMUST00 | -6112 | Cdc42ep1 | ENSMUSG0( | false | | | | | | | chr15 | 78830112 | 78842533 | 8.19317 | 5.56805 | 5.60698 |
| chr5 | 100115846 | 100116179 | 315 | Intron (ENSMUST00000063192.9 | 5 | 100083947 | 100122796 | 38850 | 2 | ENSMUST00 | 6617 | Tmem150c | ENSMUSG0( | true | | | | | | | chr5 | 100107043 | 100130430 | 5.59817 | 4.32599 | 3.51171 |
| chr12 | 69462044 | 69462407 | 315 | Distal Intergenic | 12 | 69371835 | 69372458 | 624 | 2 | ENSMUST00 | -89587 | Gm9887 | ENSMUSG0( | false | chr12 | 69461850 | 69462301 | 8.2222 | 14.94027 | 12.68024 | chr12 | 69456245 | 69463797 | 5.86899 | 4.48095 | 3.71706 |
| chr13 | 110943427 | 110943662 | 315 | Distal Intergenic | 13 | 111255013 | 111257749 | 2737 | 1 | ENSMUST00 | -311351 | Actbl2 | ENSMUSG0( | false | | | | | | | chr13 | 110943048 | 110944463 | 3.30169 | 3.14468 | 1.63298 |
| chr3 | 86551306 | 86551651 | 315 | Intron (ENSMUST00000107635.1 | 3 | 86545581 | 86548283 | 2703 | 2 | ENSMUST00 | -3024 | Mab21l2 | ENSMUSG0( | false | | | | | | | chr3 | 86549838 | 86552087 | 4.5347 | 3.89433 | 2.62667 |
| chr1 | 92850320 | 92850528 | 315 | Promoter (<=1kb) | 1 | 92850378 | 92850443 | 66 | 1 | ENSMUST00 | 0 | Mir149 | ENSMUSG0( | false | chr1 | 92850025 | 92850464 | 6.83755 | 11.44763 | 9.24644 | chr1 | 92844008 | 92853336 | 4.9948 | 4.07551 | 3.01307 |
| chr10 | 19999736 | 20000003 | 315 | Intron (ENSMUST00000095806.4 | 10 | 19951055 | 20023699 | 72645 | 1 | ENSMUST00 | 48682 | Map3k5 | ENSMUSG0( | true | | | | | | | chr10 | 19996646 | 20008131 | 4.6971 | 3.8756 | 2.77918 |
| chrX | 77414222 | 77414476 | 315 | Distal Intergenic | 20 | 77456586 | 77457228 | 643 | 1 | ENSMUST00 | -42110 | Gm5937 | ENSMUSG0( | false | | | | | | | | | | | | |
| chr6 | 115601728 | 115602111 | 314 | Promoter (<=1kb) | 6 | 115601938 | 115618670 | 16733 | 1 | ENSMUST00 | 0 | Mkrn2 | ENSMUSG0( | false | chr6 | 115601705 | 115602741 | 34.84454 | 128.36943 | 123.93644 | chr6 | 115601567 | 115603543 | 3.63295 | 3.30659 | 1.93386 |
| chr10 | 78069570 | 78069981 | 314 | Promoter (<=1kb) | 10 | 78069360 | 78079525 | 10166 | 1 | ENSMUST00 | 211 | Icosl | ENSMUSG0( | false | chr10 | 78068651 | 78070259 | 26.46939 | 80.00455 | 76.85218 | chr10 | 78068826 | 78070211 | 4.36833 | 3.90414 | 2.46798 |
| chrX | 74393095 | 74393408 | 314 | Promoter (<=1kb) | 20 | 74393290 | 74433993 | 40704 | 1 | ENSMUST00 | 0 | Ikbkg | ENSMUSG0( | false | chrX | 74392559 | 74393958 | 14.2455 | 32.22549 | 29.73087 | | | | | | |
| chr6 | 124423283 | 124423514 | 314 | Distal Intergenic | 6 | 124396817 | 124415067 | 18251 | 2 | ENSMUST00 | -8217 | Pex5 | ENSMUSG0( | false | | | | | | | | | | | | |
| chr11 | 76846867 | 76847267 | 314 | Promoter (<=1kb) | 11 | 76778424 | 76847018 | 68595 | 2 | ENSMUST00 | 0 | Cpd | ENSMUSG0( | false | chr11 | 76845296 | 76847512 | 42.05941 | 137.95251 | 133.15681 | | | | | | |
| chr16 | 17530810 | 17531213 | 314 | Promoter (<=1kb) | 16 | 17527982 | 17531136 | 3155 | 2 | ENSMUST00 | 0 | Thap7 | ENSMUSG0( | false | chr16 | 17530073 | 17531784 | 19.72513 | 62.50151 | 59.62064 | chr16 | 17525320 | 17532033 | 4.80793 | 3.90711 | 2.85739 |
| chr16 | 91484899 | 91485335 | 314 | Promoter (<=1kb) | 16 | 91485238 | 91507441 | 22204 | 1 | ENSMUST00 | 0 | Ifnar1 | ENSMUSG0( | false | chr16 | 91484460 | 91486279 | 31.40916 | 110.4849 | 106.632 | | | | | | |
| chr18 | 46741753 | 46742498 | 314 | Promoter (<=1kb) | 18 | 46741917 | 46790826 | 48910 | 1 | ENSMUST00 | 0 | Ap3s1 | ENSMUSG0( | false | chr18 | 46740400 | 46743200 | 23.21435 | 101.83275 | 98.21896 | | | | | | |
| chr2 | 25365196 | 25365780 | 314 | Promoter (<=1kb) | 2 | 25359889 | 25365682 | 5794 | 2 | ENSMUST00 | 0 | Uap1l1 | ENSMUSG0( | false | chr2 | 25364801 | 25365954 | 33.30314 | 106.68535 | 102.93993 | chr2 | 25362750 | 25366265 | 6.16994 | 4.63103 | 3.98027 |
| chr3 | 85573822 | 85574320 | 314 | Promoter (<=1kb) | 3 | 85574119 | 85646579 | 72461 | 1 | ENSMUST00 | 0 | Pet112 | ENSMUSG0( | false | chr3 | 85574092 | 85574815 | 24.82645 | 85.46523 | 82.21117 | | | | | | |
| chr9 | 62828716 | 62829077 | 314 | Distal Intergenic | 9 | 62838785 | 62852006 | 13222 | 1 | ENSMUST00 | -9708 | Cln6 | ENSMUSG0( | false | chr9 | 62828063 | 62828925 | 8.76935 | 16.76631 | 14.47755 | chr9 | 62828185 | 62830264 | 3.95035 | 3.52699 | 2.1544 |
| chr11 | 69580196 | 69580499 | 314 | Promoter (<=1kb) | 11 | 69577708 | 69580255 | 2548 | 2 | ENSMUST00 | 0 | Wrap53 | ENSMUSG0( | false | chr11 | 69579501 | 69582098 | 31.85236 | 101.09525 | 97.496 | | | | | | |
| chr19 | 5848084 | 5848605 | 314 | Promoter (<=1kb) | 19 | 5848508 | 5851372 | 2865 | 1 | ENSMUST00 | 0 | Frmd8os | ENSMUSG0( | false | chr19 | 5846849 | 5849375 | 7.82203 | 16.59716 | 14.30959 | chr19 | 5846278 | 5852772 | 9.4084 | 5.8562 | 6.60958 |
| chr8 | 124434353 | 124434732 | 314 | Promoter (<=1kb) | 8 | 124369053 | 124434338 | 65286 | 2 | ENSMUST00 | -16 | Pgbd5 | ENSMUSG0( | false | chr8 | 124433676 | 124434584 | 18.34506 | 46.29042 | 43.62275 | | | | | | |
| chr1 | 183439143 | 183439347 | 314 | Distal Intergenic | 1 | 183418503 | 183437053 | 18551 | 1 | ENSMUST00 | 20641 | Hhipl2 | ENSMUSG0( | false | chr1 | 183438856 | 183439355 | 18.67167 | 50.75271 | 48.02938 | | | | | | |
| chr4 | 149822322 | 149822517 | 314 | Distal Intergenic | 4 | 149817732 | 149817840 | 109 | 1 | ENSMUST00 | 4591 | n-R5s193 | ENSMUSG0( | false | | | | | | | chr4 | 149820569 | 149823210 | 5.53517 | 4.35229 | 3.4375 |
| chr12 | 9029912 | 9030587 | 314 | Promoter (<=1kb) | 12 | 9029997 | 9036394 | 6398 | 1 | ENSMUST00 | 0 | Ttc32 | ENSMUSG0( | false | chr12 | 9029631 | 9031447 | 31.01414 | 97.6227 | 94.10706 | | | | | | |
| chr5 | 46213155 | 46213354 | 314 | Distal Intergenic | 5 | 46414344 | 46414616 | 273 | 1 | ENSMUST00 | -200990 | Gm7931 | ENSMUSG0( | false | | | | | | | chr5 | 46212679 | 46213461 | 4.00369 | 3.68092 | 2.16751 |
| chr5 | 67832697 | 67832893 | 314 | Intron (ENSMUST00000037380.9 | 5 | 67847066 | 67909013 | 61948 | 1 | ENSMUST00 | -14173 | Gm15477 | ENSMUSG0( | false | | | | | | | chr5 | 67829717 | 67835039 | 4.35036 | 3.77032 | 2.46971 |
| chr1 | 176668063 | 176668284 | 314 | Distal Intergenic | 1 | 176700617 | 176700724 | 108 | 1 | ENSMUST00 | -32333 | Gm26104 | ENSMUSG0( | false | | | | | | | | | | | | |
| chr10 | 100286794 | 100287000 | 314 | Distal Intergenic | 10 | 100306571 | 100307185 | 615 | 2 | ENSMUST00 | 20185 | Gm9476 | ENSMUSG0( | false | | | | | | | | | | | | |
| chr17 | 35838124 | 35838430 | 313 | Promoter (<=1kb) | 17 | 35833921 | 35838306 | 4386 | 2 | ENSMUST00 | 0 | Tubb5 | ENSMUSG0( | false | chr17 | 35836718 | 35838317 | 21.62913 | 60.83906 | 57.9818 | | | | | | |
| chr7 | 28388250 | 28388660 | 313 | Intron (ENSMUST00000003536.8 | 7 | 28386146 | 28392708 | 6563 | 2 | ENSMUST00 | 4048 | Med29 | ENSMUSG0( | false | | | | | | | chr7 | 28386416 | 28389258 | 5.59227 | 4.35178 | 3.47291 |
| chr2 | 155885115 | 155885582 | 313 | Intron (ENSMUST00000136933.2 | 2 | 155887119 | 155910604 | 23486 | 2 | ENSMUST00 | 25022 | Uqcc1 | ENSMUSG0( | false | | | | | | | chr2 | 155868791 | 155890222 | 5.22346 | 4.19096 | 3.19144 |
| chr13 | 114838620 | 114838976 | 313 | Intron (ENSMUST00000056117.8 | 13 | 114818905 | 114825230 | 6326 | 1 | ENSMUST00 | 19716 | Mocs2 | ENSMUSG0( | false | | | | | | | chr13 | 114836770 | 114842005 | 5.38285 | 4.19314 | 3.33615 |
| chr16 | 23300123 | 23300386 | 313 | Intron (ENSMUST00000023601.8 | 16 | 23290470 | 23358595 | 68126 | 1 | ENSMUST00 | 9654 | St6gal1 | ENSMUSG0( | true | | | | | | | chr16 | 23277163 | 23310148 | 4.85522 | 3.95632 | 2.90417 |
| chr17 | 29428335 | 29428639 | 313 | Distal Intergenic | 17 | 29376501 | 29378452 | 1952 | 1 | ENSMUST00 | 51835 | Fgd2 | ENSMUSG0( | false | | | | | | | | | | | | |
| chr14 | 118706165 | 118706620 | 313 | Promoter (<=1kb) | 14 | 118482692 | 118706219 | 223528 | 2 | ENSMUST00 | 0 | Abcc4 | ENSMUSG0( | false | chr14 | 118705611 | 118706713 | 21.79372 | 61.12524 | 58.26245 | chr14 | 118705593 | 118706811 | 6.40271 | 4.73947 | 4.15964 |
| chr8 | 70234481 | 70234875 | 313 | Promoter (<=1kb) | 8 | 70234617 | 70242651 | 8035 | 1 | ENSMUST00 | 0 | Sugp2 | ENSMUSG0( | false | chr8 | 70233576 | 70235234 | 23.2161 | 80.55299 | 77.39275 | | | | | | |
| chr5 | 9161414 | 9161935 | 313 | Promoter (<=1kb) | 5 | 9118801 | 9161776 | 42976 | 2 | ENSMUST00 | 0 | Dmtf1 | ENSMUSG0( | false | chr5 | 9160571 | 9162609 | 38.14117 | 119.25307 | 115.13219 | | | | | | |
| chrX | 134756451 | 134756632 | 313 | Promoter (<=1kb) | 20 | 134756595 | 134761425 | 4831 | 1 | ENSMUST00 | 0 | Armcx3 | ENSMUSG0( | false | chrX | 134756578 | 134757470 | 19.30035 | 48.68736 | 45.98855 | | | | | | |
| chr7 | 97446586 | 97446810 | 313 | Distal Intergenic | 7 | 97451334 | 97457779 | 6446 | 1 | ENSMUST00 | -4524 | Kctd14 | ENSMUSG0( | false | | | | | | | chr7 | 97446311 | 97447440 | 4.45327 | 3.82856 | 2.58812 |
| chr13 | 58970725 | 58970913 | 313 | Intron (ENSMUST00000079828.5 | 13 | 58807900 | 59133970 | 326071 | 1 | ENSMUST00 | 162826 | Ntrk2 | ENSMUSG0( | false | | | | | | | | | | | | |
| chr10 | 117376197 | 117376964 | 313 | Promoter (<=1kb) | 10 | 117349042 | 117376929 | 27888 | 2 | ENSMUST00 | 0 | Cpsf6 | ENSMUSG0( | false | chr10 | 117375634 | 117377721 | 31.38507 | 94.30289 | 90.86349 | | | | | | |
| chr10 | 75591611 | 75591870 | 313 | Promoter (<=1kb) | 10 | 75590142 | 75590653 | 512 | 2 | ENSMUST00 | -959 | Gm9985 | ENSMUSG0( | false | | | | | | | chr10 | 75585895 | 75593802 | 6.23311 | 4.61744 | 4.02118 |
| chr17 | 79895469 | 79896310 | 313 | Promoter (<=1kb) | 17 | 79848392 | 79896028 | 47637 | 2 | ENSMUST00 | 0 | Atl2 | ENSMUSG0( | false | chr17 | 79894559 | 79896570 | 32.67316 | 105.57997 | 101.8659 | chr17 | 79896276 | 79896608 | 5.68137 | 4.4204 | 3.61292 |
| chr18 | 73905070 | 73905417 | 313 | Distal Intergenic | 18 | 73863672 | 73871980 | 8309 | 1 | ENSMUST00 | 41399 | Mro | ENSMUSG0( | false | | | | | | | chr18 | 73903825 | 73905596 | 4.39521 | 3.84579 | 2.5103 |
| chr19 | 8880507 | 8881032 | 313 | Promoter (<=1kb) | 19 | 8880016 | 8880933 | 918 | 2 | ENSMUST00 | 0 | AI462493 | ENSMUSG0( | false | chr19 | 8879994 | 8881359 | 28.70854 | 85.79402 | 82.53136 | chr19 | 8878949 | 8881392 | 4.78404 | 4.03421 | 2.82388 |
| chr1 | 158732913 | 158733456 | 313 | Intron (ENSMUST00000159861.3 | 1 | 158765085 | 158814469 | 49385 | 2 | ENSMUST00 | 81013 | Pappa2 | ENSMUSG0( | false | | | | | | | chr1 | 158732280 | 158734502 | 4.41094 | 3.84021 | 2.52502 |
| chr2 | 79081844 | 79082185 | 313 | Distal Intergenic | 2 | 79038533 | 79038734 | 202 | 2 | ENSMUST00 | -43111 | Gm14469 | ENSMUSG0( | false | | | | | | | | | | | | |
| chr13 | 44229080 | 44229332 | 313 | Distal Intergenic | 13 | 44121294 | 44125179 | 3886 | 1 | ENSMUST00 | 107787 | Gm5083 | ENSMUSG0( | false | chr13 | 44228751 | 44229291 | 6.37612 | 11.07527 | 8.87976 | chr13 | 44226456 | 44236587 | 5.70374 | 4.41191 | 3.58124 |
| chr7 | 93697603 | 93697976 | 313 | Distal Intergenic | 7 | 94197210 | 94197461 | 252 | 1 | ENSMUST00 | -499234 | Gm5899 | ENSMUSG0( | false | | | | | | | | | | | | |
| chr17 | 33843002 | 33843512 | 312 | Promoter (<=1kb) | 17 | 33843091 | 33849774 | 6684 | 1 | ENSMUST00 | 0 | Cd320 | ENSMUSG0( | false | chr17 | 33842822 | 33843840 | 27.41053 | 90.23362 | 86.88379 | chr17 | 33839123 | 33844713 | 4.29447 | 3.64469 | 2.46277 |
| chr11 | 53520683 | 53521016 | 312 | Promoter (<=1kb) | 11 | 53519927 | 53541847 | 21921 | 1 | ENSMUST00 | 757 | Sep-08 | ENSMUSG0( | false | chr11 | 53519402 | 53521185 | 40.11361 | 128.01187 | 123.59241 | chr11 | 53519096 | 53521911 | 4.4349 | 3.84194 | 2.57204 |
| chr10 | 58383310 | 58383577 | 312 | Intron (ENSMUST00000020078.8 | 10 | 58394373 | 58421783 | 27411 | 1 | ENSMUST00 | -10796 | Lims1 | ENSMUSG0( | false | | | | | | | chr10 | 58381582 | 58387543 | 5.37473 | 4.21579 | 3.3328 |
| chr10 | 88458448 | 88458640 | 312 | Promoter (<=1kb) | 10 | 88459569 | 88473235 | 13667 | 1 | ENSMUST00 | -929 | Sycp3 | ENSMUSG0( | false | | | | | | | chr10 | 88458303 | 88459666 | 3.05559 | 2.95792 | 1.39796 |
| chr13 | 73603848 | 73604109 | 312 | Promoter (<=1kb) | 13 | 73604002 | 73620639 | 16638 | 1 | ENSMUST00 | 0 | Clptm1l | ENSMUSG0( | false | chr13 | 73603140 | 73605160 | 30.26787 | 111.42836 | 107.5545 | chr13 | 73603042 | 73606018 | 4.69638 | 3.82125 | 2.77881 |
| chr15 | 6662078 | 6662270 | 312 | Promoter (1-2kb) | 15 | 6660405 | 6663251 | 2847 | 1 | ENSMUST00 | 1674 | Fyb | ENSMUSG0( | false | | | | | | | chr15 | 6661750 | 6662585 | 3.95668 | 3.49294 | 2.16388 |
| chr18 | 75378679 | 75379049 | 312 | Intron (ENSMUST00000026999.4 | 18 | 75374914 | 75394001 | 19088 | 1 | ENSMUST00 | 3766 | Smad7 | ENSMUSG0( | false | | | | | | | chr18 | 75369277 | 75392875 | 5.74724 | 4.44386 | 3.62103 |
| chr5 | 123344238 | 123344612 | 312 | Promoter (<=1kb) | 5 | 123344448 | 123374082 | 29635 | 1 | ENSMUST00 | 0 | Bcl7a | ENSMUSG0( | false | chr5 | 123343837 | 123346514 | 34.14626 | 110.2223 | 106.38018 | | | | | | |
| chr7 | 101394185 | 101394471 | 312 | Promoter (<=1kb) | 7 | 101394368 | 101412586 | 18219 | 1 | ENSMUST00 | 0 | Arap1 | ENSMUSG0( | false | | | | | | | chr7 | 101391376 | 101394892 | 6.29135 | 4.66495 | 4.06264 |
| chr10 | 108332539 | 108333427 | 312 | Promoter (<=1kb) | 10 | 108332189 | 108414391 | 82203 | 1 | ENSMUST00 | 351 | Pawr | ENSMUSG0( | false | chr10 | 108331582 | 108333603 | 35.23322 | 112.11895 | 108.21771 | | | | | | |
| chr6 | 40435953 | 40436322 | 312 | Promoter (<=1kb) | 6 | 40401375 | 40436135 | 34761 | 2 | ENSMUST00 | 0 | E330009J07F | ENSMUSG0( | false | chr6 | 40434915 | 40436523 | 27.18312 | 111.36749 | 107.49936 | | | | | | |
| chr14 | 45497094 | 45497342 | 312 | Intron (ENSMUST00000045905.7 | 14 | 45469314 | 45477861 | 8548 | 2 | ENSMUST00 | -19234 | Fermt2 | ENSMUSG0( | true | | | | | | | chr14 | 45489964 | 45528839 | 6.12493 | 4.54024 | 3.93691 |
| chr7 | 112320329 | 112320654 | 312 | Exon (ENSMUST00000050149.6/ | 7 | 112316029 | 112347491 | 31463 | 1 | ENSMUST00 | 4301 | Mical2 | ENSMUSG0( | false | | | | | | | chr7 | 112315227 | 112322046 | 5.14867 | 4.13411 | 3.15663 |
| chr5 | 142814994 | 142815429 | 312 | Promoter (1-2kb) | 5 | 142754796 | 142817387 | 62592 | 2 | ENSMUST00 | 1958 | Tnrc18 | ENSMUSG0( | false | chr5 | 142813148 | 142817670 | 35.38397 | 108.24114 | 104.45254 | | | | | | |
| chr3 | 52103911 | 52104405 | 312 | Promoter (<=1kb) | 3 | 51687611 | 52104891 | 417281 | 2 | ENSMUST00 | 486 | Maml3 | ENSMUSG0( | true | chr3 | 52102288 | 52105222 | 37.14626 | 116.48058 | 112.45192 | | | | | | |
| chr1 | 93825167 | 93825615 | 312 | Promoter (<=1kb) | 1 | 93825240 | 93852348 | 27109 | 1 | ENSMUST00 | 0 | D2hgdh | ENSMUSG0( | false | chr1 | 93824772 | 93826075 | 20.65177 | 61.66082 | 58.78991 | chr1 | 93824618 | 93826620 | 4.9927 | 4.06462 | 3.00741 |
| chr11 | 64548117 | 64548323 | 312 | Distal Intergenic | 11 | 64590495 | 64590640 | 146 | 2 | ENSMUST00 | 42317 | Gm24275 | ENSMUSG0( | false | | | | | | | chr11 | 64539605 | 64548358 | 5.34347 | 4.23728 | 3.29791 |
| chr17 | 28032259 | 28032530 | 312 | Intron (ENSMUST00000114842.2 | 17 | 27988028 | 28008432 | 20405 | 2 | ENSMUST00 | -23828 | Gm15598 | ENSMUSG0( | false | | | | | | | chr17 | 28029226 | 28043441 | 7.08706 | 4.83268 | 4.74553 |
| chr19 | 4220892 | 4221310 | 312 | Exon (ENSMUST00000132305.1/ | 19 | 4215402 | 4215488 | 87 | 2 | ENSMUST00 | -5405 | Gm26115 | ENSMUSG0( | false | | | | | | | chr19 | 4219961 | 4222780 | 6.64534 | 4.76361 | 4.37575 |
| chr16 | 52071895 | 52072308 | 312 | Intron (ENSMUST00000114471.1 | 16 | 52105015 | 52105147 | 133 | 2 | ENSMUST00 | 32839 | Gm23513 | ENSMUSG0( | false | | | | | | | chr16 | 52047691 | 52073501 | 5.26548 | 4.17513 | 3.22843 |
| chr17 | 8565353 | 8565579 | 312 | Promoter (1-2kb) | 17 | 8566399 | 8566869 | 471 | 2 | ENSMUST00 | 1290 | Gm17087 | ENSMUSG0( | false | | | | | | | chr17 | 8564707 | 8567009 | 3.7332 | 3.44911 | 2.01534 |
| chr19 | 6057725 | 6058096 | 311 | Promoter (<=1kb) | 19 | 6053630 | 6057751 | 4122 | 2 | ENSMUST00 | 0 | Mrpl49 | ENSMUSG0( | false | chr19 | 6057225 | 6058915 | 31.14379 | 104.89771 | 101.20105 | chr19 | 6055698 | 6059221 | 5.22484 | 4.10346 | 3.21382 |
| chr5 | 31048084 | 31048351 | 311 | Promoter (<=1kb) | 5 | 31048312 | 31052769 | 4458 | 1 | ENSMUST00 | 0 | Atraid | ENSMUSG0( | false | chr5 | 31047525 | 31049639 | 30.44846 | 92.68264 | 89.28044 | | | | | | |
| chr3 | 95993025 | 95993235 | 311 | Promoter (2-3kb) | 3 | 95988819 | 95995668 | 6850 | 2 | ENSMUST00 | 2433 | Plekho1 | ENSMUSG0( | true | chr3 | 95992078 | 95997273 | 32.06657 | 104.51031 | 100.83098 | chr3 | 95984661 | 95993242 | 6.4304 | 4.71231 | 4.18378 |
| chr11 | 70700319 | 70700858 | 311 | Promoter (<=1kb) | 11 | 70700548 | 70707568 | 7021 | 1 | ENSMUST00 | 0 | Kif1c | ENSMUSG0( | false | chr11 | 70699759 | 70701628 | 31.96831 | 119.80809 | 115.67081 | | | | | | |
| chr14 | 60590173 | 60590567 | 311 | Intron (ENSMUST00000190392.1 | 14 | 60634001 | 60691367 | 57367 | 1 | ENSMUST00 | -43434 | Spata13 | ENSMUSG0( | true | | | | | | | chr14 | 60585347 | 60599096 | 6.17547 | 4.51754 | 3.99543 |
| chr17 | 67901195 | 67901441 | 311 | Promoter (<=1kb) | 17 | 67845735 | 67901471 | 55737 | 2 | ENSMUST00 | 30 | Arhgap28 | ENSMUSG0( | true | | | | | | | chr17 | 67892935 | 67914104 | 5.71307 | 4.38398 | 3.59861 |
| chr1 | 37540761 | 37541226 | 311 | Promoter (<=1kb) | 1 | 37439340 | 37541016 | 101677 | 2 | ENSMUST00 | 0 | Mgat4a | ENSMUSG0( | true | chr1 | 37539932 | 37541579 | 29.36448 | 91.77025 | 88.3935 | | | | | | |
| chr4 | 115101757 | 115101993 | 311 | Distal Intergenic | 4 | 115090932 | 115093864 | 2933 | 1 | ENSMUST00 | 10826 | Pdzk1ip1 | ENSMUSG0( | false | | | | | | | chr4 | 115101952 | 115104077 | 3.68461 | 3.44793 | 1.97994 |
| chr11 | 84828975 | 84829276 | 311 | Promoter (<=1kb) | 11 | 84821992 | 84828994 | 7003 | 2 | ENSMUST00 | 0 | Dhrs11 | ENSMUSG0( | false | chr11 | 84828291 | 84829675 | 37.30079 | 124.19571 | 119.91465 | chr11 | 84826370 | 84830121 | 3.47367 | 3.22383 | 1.78724 |
| chr10 | 84917083 | 84918118 | 311 | Promoter (<=1kb) | 10 | 84917616 | 85016446 | 98831 | 1 | ENSMUST00 | 0 | Ric8b | ENSMUSG0( | false | chr10 | 84916703 | 84918437 | 32.88158 | 104.92503 | 101.22703 | chr10 | 84918013 | 84920241 | 4.11503 | 3.69892 | 2.26775 |
| chr18 | 34859615 | 34860057 | 311 | Promoter (<=1kb) | 18 | 34859823 | 34864984 | 5162 | 1 | ENSMUST00 | 0 | Egr1 | ENSMUSG0( | false | chr18 | 34859236 | 34862643 | 33.41831 | 117.26761 | 113.21259 | chr18 | 34858199 | 34863782 | 4.89469 | 4.00673 | 2.91424 |
| chr10 | 53414841 | 53415119 | 311 | Distal Intergenic | 10 | 53278081 | 53379851 | 101771 | 2 | ENSMUST00 | -34991 | Cep85l | ENSMUSG0( | true | | | | | | | chr10 | 53407459 | 53424022 | 5.24828 | 4.19397 | 3.20837 |
| chr14 | 20734219 | 20734772 | 311 | Downstream (<1kb) | 14 | 20723739 | 20730327 | 6589 | 2 | ENSMUST00 | -3893 | Ndst2 | ENSMUSG0( | false | chr14 | 20733546 | 20735008 | 31.07893 | 124.33488 | 120.0426 | | | | | | |
| chr13 | 101757690 | 101758046 | 311 | 5' UTR | 13 | 101755571 | 101768213 | 12643 | 2 | ENSMUST00 | 10167 | Pik3r1 | ENSMUSG0( | false | | | | | | | chr13 | 101748779 | 101766882 | 5.55016 | 4.25775 | 3.4729 |
| chr4 | 106650872 | 106651124 | 311 | Promoter (<=1kb) | 4 | 106651069 | 106653539 | 2471 | 1 | ENSMUST00 | 0 | Pars2 | ENSMUSG0( | false | chr4 | 106651039 | 106651672 | 21.87981 | 58.7423 | 55.91244 | chr4 | 106649637 | 106652500 | 5.3602 | 4.00044 | 3.33642 |
| chr8 | 115754772 | 115755177 | 311 | Distal Intergenic | 8 | 115701435 | 115707794 | 6360 | 2 | ENSMUST00 | -46979 | Maf | ENSMUSG0( | false | | | | | | | | | | | | |
| chr8 | 61582612 | 61582923 | 311 | Intron (ENSMUST00000121785.2 | 8 | 61515913 | 61591139 | 75227 | 2 | ENSMUST00 | 8216 | Palld | ENSMUSG0( | true | | | | | | | chr8 | 61569315 | 61589884 | 5.62358 | 4.37337 | 3.52305 |
| chr2 | 38926015 | 38926568 | 311 | Promoter (<=1kb) | 2 | 38723374 | 38926217 | 202844 | 2 | ENSMUST00 | 0 | Nr6a1 | ENSMUSG0( | false | chr2 | 38925035 | 38926515 | 35.56825 | 117.95176 | 113.87694 | | | | | | |
| chr12 | 3169835 | 3170386 | 311 | Distal Intergenic | 12 | 3235504 | 3241410 | 5907 | 1 | ENSMUST00 | -65118 | Rab10os | ENSMUSG0( | false | | | | | | | chr12 | 3167677 | 3171335 | 3.16385 | 3.08339 | 1.4861 |
| chr14 | 62855811 | 62856149 | 311 | Intron (ENSMUST00000014691.8 | 14 | 62865542 | 62865651 | 110 | 1 | ENSMUST00 | -9393 | Mir6541 | ENSMUSG0( | false | | | | | | | chr14 | 62855129 | 62859378 | 5.08747 | 4.22052 | 3.06735 |
| chr15 | 101181608 | 101181876 | 310 | Intron (ENSMUST00000000544.1 | 15 | 101174067 | 101213679 | 39613 | 1 | ENSMUST00 | 7542 | Acvr1b | ENSMUSG0( | false | | | | | | | chr15 | 101174616 | 101193068 | 5.45533 | 4.23935 | 3.38802 |
| chr2 | 172937739 | 172938030 | 310 | Promoter (2-3kb) | 2 | 172916118 | 172940092 | 23975 | 2 | ENSMUST00 | 2062 | Bmp7 | ENSMUSG0( | false | | | | | | | chr2 | 172924796 | 172941177 | 5.52242 | 4.25198 | 3.44947 |
| chr14 | 70241661 | 70242065 | 310 | Intron (ENSMUST00000078434.6 | 14 | 70180468 | 70207637 | 27170 | 2 | ENSMUST00 | -34025 | Sorbs3 | ENSMUSG0( | false | | | | | | | chr14 | 70241103 | 70243666 | 3.67242 | 3.4214 | 1.90187 |
| chr15 | 77873632 | 77874162 | 310 | Distal Intergenic | 15 | 77760587 | 77842175 | 81589 | 2 | ENSMUST00 | -31458 | Myh9 | ENSMUSG0( | true | | | | | | | chr15 | 77872327 | 77874662 | 4.90598 | 3.90272 | 2.96392 |
| chr19 | 53049511 | 53049785 | 310 | Distal Intergenic | 19 | 52991183 | 53040214 | 49032 | 2 | ENSMUST00 | -9298 | Xpnpep1 | ENSMUSG0( | false | | | | | | | | | | | | |
| chr2 | 120731164 | 120731723 | 310 | Promoter (<=1kb) | 2 | 120716154 | 120731517 | 15364 | 2 | ENSMUST00 | 0 | Cdan1 | ENSMUSG0( | false | chr2 | 120729861 | 120732120 | 36.0064 | 125.12101 | 120.80552 | | | | | | |
| chr6 | 33891956 | 33892496 | 310 | Intron (ENSMUST00000052266.9 | 6 | 33862100 | 33874181 | 12082 | 1 | ENSMUST00 | 29857 | Exoc4 | ENSMUSG0( | false | | | | | | | chr6 | 33890561 | 33893038 | 4.25907 | 3.61017 | 2.43546 |
| chr7 | 67681152 | 67681388 | 310 | Promoter (1-2kb) | 7 | 67679296 | 67726052 | 46757 | 1 | ENSMUST00 | 1857 | Ttc23 | ENSMUSG0( | false | | | | | | | chr7 | 67679722 | 67681984 | 4.074 | 3.54991 | 2.26893 |
| chr8 | 120563863 | 120564357 | 310 | Intron (ENSMUST00000127664.1 | 8 | 120571281 | 120573272 | 1992 | 1 | ENSMUST00 | -6924 | Gse1 | ENSMUSG0( | false | | | | | | | chr8 | 120537951 | 120565808 | 7.48404 | 5.10526 | 5.05119 |
| chr10 | 123076043 | 123076624 | 310 | Promoter (<=1kb) | 10 | 122992061 | 123076505 | 84445 | 2 | ENSMUST00 | 0 | Mon2 | ENSMUSG0( | false | chr10 | 123075268 | 123077129 | 34.74927 | 111.27614 | 107.40929 | | | | | | |
| chr18 | 39187879 | 39188229 | 310 | Intron (ENSMUST00000097593.3 | 18 | 39227368 | 39372511 | 145144 | 1 | ENSMUST00 | -39139 | Arhgap26 | ENSMUSG0( | false | | | | | | | chr18 | 39187112 | 39189636 | 6.19093 | 4.64886 | 4.03647 |
| chr10 | 22604868 | 22605096 | 310 | Distal Intergenic | 10 | 22645011 | 22704285 | 59275 | 1 | ENSMUST00 | -39915 | Slc2a12 | ENSMUSG0( | false | | | | | | | chr10 | 22604319 | 22605710 | 5.41163 | 4.30081 | 3.33826 |
| chr10 | 127621468 | 127622263 | 310 | Promoter (<=1kb) | 10 | 127587627 | 127621148 | 33522 | 2 | ENSMUST00 | -321 | Lrp1 | ENSMUSG0( | false | chr10 | 127621282 | 127622474 | 25.56571 | 75.64988 | 72.57267 | chr10 | 127615771 | 127622467 | 4.42783 | 3.68176 | 2.56096 |
| chr15 | 36173754 | 36174127 | 310 | Promoter (<=1kb) | 15 | 36174010 | 36177007 | 2998 | 1 | ENSMUST00 | 0 | Polr2k | ENSMUSG0( | false | chr15 | 36173499 | 36174698 | 28.06191 | 82.77493 | 79.57206 | chr15 | 36173202 | 36175423 | 5.112 | 4.17662 | 3.09328 |
| chr12 | 21285803 | 21286345 | 310 | Promoter (<=1kb) | 12 | 21286297 | 21315056 | 28760 | 1 | ENSMUST00 | 0 | Cpsf3 | ENSMUSG0( | false | chr12 | 21285266 | 21287173 | 38.15516 | 131.11487 | 126.58785 | chr12 | 21285063 | 21285823 | 4.47756 | 3.8969 | 2.56049 |
| chr17 | 30575868 | 30576338 | 310 | Promoter (<=1kb) | 17 | 30215524 | 30576287 | 360764 | 2 | ENSMUST00 | 0 | Btbd9 | ENSMUSG0( | false | chr17 | 30575193 | 30577241 | 27.78601 | 91.88203 | 88.49705 | | | | | | |
| chr2 | 160346622 | 160346897 | 310 | Distal Intergenic | 2 | 160311744 | 160334162 | 22419 | 2 | ENSMUST00 | -12461 | Gm826 | ENSMUSG0( | false | | | | | | | chr2 | 160345410 | 160356586 | 4.87436 | 3.97596 | 2.90772 |
| chr4 | 52461400 | 52461647 | 310 | Intron (ENSMUST00000102915.4 | 4 | 52443915 | 52444172 | 258 | 2 | ENSMUST00 | -17229 | Tmc4-ps | ENSMUSG0( | false | | | | | | | | | | | | |
| chrX | 166728178 | 166728388 | 310 | Distal Intergenic | 20 | 166662025 | 166689812 | 27788 | 1 | ENSMUST00 | 66154 | Gm1720 | ENSMUSG0( | false | | | | | | | | | | | | |
| chr18 | 75788510 | 75788871 | 310 | Distal Intergenic | 18 | 75781032 | 75782303 | 1272 | 1 | ENSMUST00 | 7479 | 1700034B16I | ENSMUSG0( | false | | | | | | | chr18 | 75788257 | 75789891 | 4.53751 | 3.76525 | 2.64049 |
| chr8 | 26887106 | 26887863 | 310 | Distal Intergenic | 8 | 26814593 | 26824423 | 9831 | 2 | ENSMUST00 | -62684 | RP24-502D8. | ENSMUSG0( | false | | | | | | | chr8 | 26886385 | 26888430 | 5.75484 | 4.30426 | 3.6506 |
| chr14 | 61360215 | 61360529 | 309 | Promoter (<=1kb) | 14 | 61339763 | 61360445 | 20683 | 2 | ENSMUST00 | 0 | Ebpl | ENSMUSG0( | false | chr14 | 61359907 | 61360988 | 26.39107 | 78.00815 | 74.89062 | chr14 | 61360135 | 61361252 | 5.75908 | 4.40381 | 3.69867 |
| chr14 | 79633284 | 79633473 | 309 | Distal Intergenic | 14 | 79637816 | 79642195 | 4380 | 2 | ENSMUST00 | 8722 | Lect1 | ENSMUSG0( | false | | | | | | | | | | | | |
| chr15 | 77820294 | 77821094 | 309 | Intron (ENSMUST00000016771.7 | 15 | 77794652 | 77813231 | 18580 | 2 | ENSMUST00 | -7064 | Myh9 | ENSMUSG0( | true | | | | | | | chr15 | 77781877 | 77840076 | 8.49833 | 5.61339 | 5.87304 |
| chr1 | 52232294 | 52233515 | 309 | Promoter (<=1kb) | 1 | 52163448 | 52233223 | 69776 | 2 | ENSMUST00 | 0 | Gls | ENSMUSG0( | false | chr1 | 52231711 | 52233894 | 37.91348 | 118.86035 | 114.75604 | | | | | | |
| chr2 | 131186728 | 131187192 | 309 | Promoter (<=1kb) | 2 | 131186949 | 131198495 | 11547 | 1 | ENSMUST00 | 0 | Cdc25b | ENSMUSG0( | false | chr2 | 131186305 | 131187478 | 14.00287 | 32.47356 | 29.97816 | chr2 | 131186438 | 131187611 | 3.4179 | 3.27015 | 1.75402 |
| chr3 | 7369856 | 7370128 | 309 | Intron (ENSMUST00000028999.6 | 3 | 7366604 | 7445366 | 78763 | 1 | ENSMUST00 | 3253 | Pkia | ENSMUSG0( | true | | | | | | | chr3 | 7367305 | 7376983 | 4.50721 | 3.84318 | 2.59382 |
| chr3 | 87765140 | 87765347 | 309 | Promoter (2-3kb) | 3 | 87749098 | 87762214 | 13117 | 2 | ENSMUST00 | -2927 | Pear1 | ENSMUSG0( | false | | | | | | | chr3 | 87763394 | 87765267 | 4.19285 | 3.65031 | 2.35304 |
| chr4 | 149445171 | 149445673 | 309 | Distal Intergenic | 4 | 149449687 | 149453424 | 3738 | 2 | ENSMUST00 | 7751 | Rbp7 | ENSMUSG0( | false | chr4 | 149445062 | 149446178 | 7.48194 | 15.52168 | 13.25109 | chr4 | 149444993 | 149446394 | 4.16482 | 3.60739 | 2.34365 |
| chr9 | 75314064 | 75314586 | 309 | Promoter (2-3kb) | 9 | 75311395 | 75344964 | 33570 | 1 | ENSMUST00 | 2670 | Gnb5 | ENSMUSG0( | false | chr9 | 75313818 | 75314989 | 27.6016 | 85.77155 | 82.50918 | | | | | | |
| chr17 | 88466625 | 88466829 | 309 | Intron (ENSMUST00000112238.3 | 17 | 88440775 | 88487251 | 46477 | 1 | ENSMUST00 | 25851 | Foxn2 | ENSMUSG0( | false | | | | | | | chr17 | 88462698 | 88467147 | 4.61432 | 3.84565 | 2.70156 |
| chr9 | 37099988 | 37100268 | 309 | Intron (ENSMUST00000039674.6 | 9 | 36890982 | 37083309 | 192328 | 2 | ENSMUST00 | -16680 | Pknox2 | ENSMUSG0( | true | | | | | | | chr9 | 37098428 | 37110906 | 5.53132 | 4.20019 | 3.48047 |
| chr1 | 136624543 | 136625108 | 309 | Promoter (<=1kb) | 1 | 136624901 | 136630053 | 5153 | 1 | ENSMUST00 | 0 | Zfp281 | ENSMUSG0( | false | chr1 | 136624228 | 136626150 | 33.94791 | 109.87469 | 106.04007 | | | | | | |
| chr6 | 86357015 | 86357373 | 309 | Distal Intergenic | 6 | 86365683 | 86370057 | 4375 | 1 | ENSMUST00 | -8310 | Fam136a | ENSMUSG0( | false | | | | | | | chr6 | 86356929 | 86365074 | 4.66248 | 3.70767 | 2.75361 |
| chr15 | 83009057 | 83009342 | 309 | Intron (ENSMUST00000109503.4 | 15 | 82997721 | 83023138 | 25418 | 2 | ENSMUST00 | 13796 | Nfam1 | ENSMUSG0( | false | | | | | | | chr15 | 83009031 | 83009425 | 3.01134 | 3.06395 | 1.41742 |
| chr10 | 90618052 | 90618265 | 309 | Intron (ENSMUST00000183156.2 | 10 | 90576992 | 90969820 | 392829 | 1 | ENSMUST00 | 41061 | Anks1b | ENSMUSG0( | true | | | | | | | chr10 | 90617601 | 90619307 | 3.13868 | 2.9645 | 1.51841 |
| chr5 | 96911929 | 96912153 | 309 | Distal Intergenic | 5 | 96907632 | 96907825 | 194 | 1 | ENSMUST00 | 4298 | Gm24623 | ENSMUSG0( | false | chr5 | 96911519 | 96912088 | 7.48194 | 15.52168 | 13.25109 | chr5 | 96911502 | 96913313 | 3.21852 | 3.09399 | 1.53272 |
| chr7 | 145046524 | 145046742 | 309 | Distal Intergenic | 7 | 144929931 | 144939869 | 9939 | 2 | ENSMUST00 | -106656 | Ccnd1 | ENSMUSG0( | false | | | | | | | | | | | | |
| chr7 | 102687021 | 102687447 | 309 | Promoter (<=1kb) | 7 | 102677580 | 102687326 | 9747 | 2 | ENSMUST00 | 0 | Trim68 | ENSMUSG0( | false | chr7 | 102686722 | 102687967 | 19.45739 | 70.46334 | 67.46697 | chr7 | 102685623 | 102688123 | 3.19603 | 3.12095 | 1.50959 |
| chr11 | 68022893 | 68023193 | 309 | Intron (ENSMUST00000021285.8 | 11 | 68035038 | 68035710 | 673 | 2 | ENSMUST00 | 12517 | Gm12303 | ENSMUSG0( | false | | | | | | | chr11 | 68022476 | 68024266 | 4.66804 | 3.74548 | 2.7712 |
| chrX | 99317396 | 99317687 | 309 | Distal Intergenic | 20 | 99409571 | 99418895 | 9325 | 2 | ENSMUST00 | 101208 | Gm14809 | ENSMUSG0( | false | | | | | | | | | | | | |
| chr17 | 24645401 | 24645791 | 308 | Promoter (<=1kb) | 17 | 24639282 | 24644957 | 5676 | 2 | ENSMUST00 | -445 | Slc9a3r2 | ENSMUSG0( | false | chr17 | 24642925 | 24645980 | 11.05033 | 23.18759 | 20.81059 | chr17 | 24630730 | 24649025 | 8.70874 | 5.59711 | 6.05272 |
| chr12 | 101893382 | 101893582 | 308 | Promoter (<=1kb) | 12 | 101837546 | 101894316 | 56771 | 2 | ENSMUST00 | 734 | Trip11 | ENSMUSG0( | false | | | | | | | chr12 | 101893359 | 101893777 | 4.52722 | 3.90391 | 2.5988 |
| chr14 | 57392429 | 57392673 | 308 | Intron (ENSMUST00000022517.7 | 14 | 57275034 | 57398483 | 123450 | 2 | ENSMUST00 | 5810 | Cryl1 | ENSMUSG0( | false | | | | | | | chr14 | 57391629 | 57393483 | 4.0984 | 3.55962 | 2.29666 |
| chr15 | 79804532 | 79804901 | 308 | Promoter (<=1kb) | 15 | 79786352 | 79804709 | 18358 | 2 | ENSMUST00 | 0 | Nptxr | ENSMUSG0( | false | chr15 | 79803764 | 79805110 | 24.93112 | 71.82204 | 68.80418 | chr15 | 79803686 | 79805115 | 6.24238 | 4.74524 | 4.0085 |
| chr16 | 30510532 | 30510886 | 308 | Downstream (<1kb) | 16 | 30513970 | 30540909 | 26940 | 2 | ENSMUST00 | 30023 | Tmem44 | ENSMUSG0( | false | chr16 | 30510208 | 30511499 | 19.52531 | 57.12588 | 54.31675 | | | | | | |

| | | | | | | | | | | | | | | | | | | | | | | | | | | |
|---|---|---|---|---|---|---|---|---|---|---|---|---|---|---|---|---|---|---|---|---|---|---|---|---|---|---|
| chr19 | 6127844 | 6128401 | 308 | Promoter (<=1kb) | 19 | 6119399 | 6128304 | 8906 | 2 | ENSMUST00 | 0 | Snx15 | ENSMUSG0C | false | chr19 | 6127537 | 6128305 | 18.72985 | 51.651 | 48.91581 | chr19 | 6125067 | 6128746 | 3.41889 | 3.21653 | 1.75911 |
| chr6 | 12749148 | 12749564 | 308 | Promoter (<=1kb) | 6 | 12311610 | 12749253 | 437644 | 2 | ENSMUST00 | 0 | Thsd7a | ENSMUSG0C | true | chr6 | 12745333 | 12750108 | 39.82066 | 130.7653 | 126.24818 | . | . | . | . | . | . |
| chr4 | 57844963 | 57845249 | 308 | Promoter (<=1kb) | 4 | 57845247 | 57896982 | 51736 | 1 | ENSMUST00 | 0 | Akap2 | ENSMUSG0C | false | chr4 | 57845075 | 57848262 | 24.35436 | 72.61947 | 69.58835 | . | . | . | . | . | . |
| chr9 | 116863726 | 116864015 | 308 | Intron (ENSMUST00000111773.4 | 9 | 116582822 | 116822819 | 239998 | 2 | ENSMUST00 | -40908 | Rbms3 | ENSMUSG0C | true | . | . | . | . | . | . | chr9 | 116861631 | 116869998 | 4.54253 | 3.878 | 2.63257 |
| chr5 | 21353610 | 21353918 | 308 | Intron (ENSMUST00000115245.2 | 5 | 21372642 | 21378374 | 5733 | 1 | ENSMUST00 | -18724 | Fgl2 | ENSMUSG0C | false | . | . | . | . | . | . | chr5 | 21353162 | 21354525 | 4.17191 | 3.67164 | 2.35342 |
| chr14 | 37135153 | 37135646 | 308 | Promoter (<=1kb) | 14 | 37120444 | 37135322 | 14879 | 2 | ENSMUST00 | 0 | Ghitm | ENSMUSG0C | false | chr14 | 37134341 | 37136021 | 33.47463 | 112.14942 | 108.24687 | . | . | . | . | . | . |
| chr4 | 138434492 | 138434784 | 308 | Promoter (<=1kb) | 4 | 138434671 | 138442265 | 7595 | 1 | ENSMUST00 | 0 | Mul1 | ENSMUSG0C | false | chr4 | 138434165 | 138435568 | 32.16322 | 96.63297 | 93.14677 | chr4 | 138434223 | 138437878 | 4.78241 | 3.82988 | 2.84605 |
| chr11 | 20248987 | 20249712 | 308 | Promoter (<=1kb) | 11 | 20227037 | 20249418 | 22382 | 2 | ENSMUST00 | 0 | Cep68 | ENSMUSG0C | false | chr11 | 20248487 | 20250058 | 38.46334 | 130.56975 | 126.05653 | chr11 | 20244795 | 20250083 | 3.82323 | 3.49825 | 2.06457 |
| chr4 | 149737585 | 149738181 | 308 | Promoter (<=1kb) | 4 | 149715375 | 149738044 | 22670 | 2 | ENSMUST00 | 0 | Tmem201 | ENSMUSG0C | false | chr4 | 149737028 | 149738543 | 36.39388 | 118.3417 | 114.25511 | . | . | . | . | . | . |
| chr2 | 161787255 | 161787570 | 308 | Intron (ENSMUST00000109443.2 | 2 | 161698009 | 161800155 | 102147 | 2 | ENSMUST00 | 12585 | Ptprt | ENSMUSG0C | false | . | . | . | . | . | . | chr2 | 161786412 | 161789280 | 5.3941 | 4.3494 | 3.3206 |
| chr14 | 74871301 | 74871675 | 308 | Intron (ENSMUST00000088970.5 | 14 | 74754674 | 74947877 | 193204 | 2 | ENSMUST00 | 76202 | Lrch1 | ENSMUSG0C | true | . | . | . | . | . | . | chr14 | 74857233 | 74945366 | 6.29465 | 4.64624 | 4.07764 |
| chr6 | 83054421 | 83054848 | 308 | Promoter (<=1kb) | 6 | 83054653 | 83057680 | 3028 | 1 | ENSMUST00 | 0 | Aup1 | ENSMUSG0C | false | chr6 | 83053281 | 83056016 | 41.35843 | 143.41548 | 138.39708 | . | . | . | . | . | . |
| chr11 | 107027702 | 107028376 | 308 | Promoter (<=1kb) | 11 | 107028118 | 107030438 | 2321 | 1 | ENSMUST00 | 0 | 1810010H24I | ENSMUSG0C | false | chr11 | 107027494 | 107028820 | 20.02967 | 73.35366 | 70.31202 | chr11 | 107024089 | 107029891 | 4.48121 | 3.86241 | 2.56711 |
| chr4 | 129394538 | 129394839 | 308 | Distal Intergenic | 4 | 129379844 | 129380179 | 336 | 1 | ENSMUST00 | 14695 | Gm12980 | ENSMUSG0C | false | . | . | . | . | . | . | chr4 | 129390716 | 129394877 | 3.63378 | 3.31345 | 1.93208 |
| chr2 | 50631344 | 50631619 | 308 | Distal Intergenic | 2 | 50657656 | 50658067 | 412 | 2 | ENSMUST00 | 26448 | Gm13484 | ENSMUSG0C | false | . | . | . | . | . | . | chr2 | 50631207 | 50631975 | 3.9289 | 3.65393 | 2.12591 |
| chr9 | 71215366 | 71216031 | 307 | Promoter (<=1kb) | 9 | 71215789 | 71296243 | 80455 | 1 | ENSMUST00 | 0 | Aldh1a2 | ENSMUSG0C | false | chr9 | 71215273 | 71217229 | 25.90858 | 83.69981 | 80.48042 | chr9 | 71215323 | 71217289 | 4.52473 | 3.85691 | 2.62799 |
| chr6 | 17636761 | 17637167 | 307 | Promoter (<=1kb) | 6 | 17636983 | 17666972 | 29990 | 1 | ENSMUST00 | 0 | Capza2 | ENSMUSG0C | false | chr6 | 17636619 | 17636840 | 6.61735 | 11.47251 | 9.2704 | . | . | . | . | . | . |
| chr1 | 143739441 | 143739800 | 307 | Promoter (<=1kb) | 1 | 143739557 | 143742739 | 3183 | 1 | ENSMUST00 | 0 | Glrx2 | ENSMUSG0C | false | chr1 | 143738928 | 143740484 | 26.97976 | 80.89639 | 77.72532 | . | . | . | . | . | . |
| chr11 | 94074630 | 94074971 | 307 | Promoter (2-3kb) | 11 | 94071663 | 94094229 | 22567 | 1 | ENSMUST00 | 2968 | Spag9 | ENSMUSG0C | false | . | . | . | . | . | . | chr11 | 94073149 | 94075030 | 4.22075 | 3.4137 | 2.41556 |
| chr15 | 32676571 | 32676946 | 307 | Intron (ENSMUST00000067458.6 | 15 | 32920723 | 33034730 | 114008 | 1 | ENSMUST00 | -243777 | Sdc2 | ENSMUSG0C | false | . | . | . | . | . | . | chr15 | 32676598 | 32677475 | 2.54151 | 2.61784 | 1.0436 |
| chr17 | 8283459 | 8284107 | 307 | Promoter (<=1kb) | 17 | 8283762 | 8297661 | 13900 | 1 | ENSMUST00 | 0 | Mpc1 | ENSMUSG0C | false | chr17 | 8282625 | 8284867 | 41.81742 | 134.22485 | 129.57179 | . | . | . | . | . | . |
| chr3 | 89391737 | 89392215 | 307 | Promoter (<=1kb) | 3 | 89377644 | 89392069 | 14426 | 2 | ENSMUST00 | 0 | Zbtb7b | ENSMUSG0C | false | chr3 | 89391380 | 89396724 | 30.32912 | 88.589 | 85.27629 | chr3 | 89387156 | 89392657 | 4.07018 | 3.58009 | 2.2478 |
| chr7 | 65424622 | 65424919 | 307 | Distal Intergenic | 7 | 65296165 | 65371239 | 75075 | 2 | ENSMUST00 | -53384 | Tjp1 | ENSMUSG0C | true | . | . | . | . | . | . | chr7 | 65409463 | 65433128 | 6.06932 | 4.54697 | 3.8847 |
| chr9 | 63715455 | 63715744 | 307 | Intron (ENSMUST00000034973.4 | 9 | 63654799 | 63711969 | 57171 | 2 | ENSMUST00 | -3487 | Smad3 | ENSMUSG0C | false | . | . | . | . | . | . | chr9 | 63695180 | 63731540 | 5.72957 | 4.34734 | 3.61571 |
| chr8 | 46152073 | 46152699 | 307 | Promoter (<=1kb) | 8 | 46033261 | 46152159 | 118899 | 2 | ENSMUST00 | 0 | Snx25 | ENSMUSG0C | false | chr8 | 46151196 | 46153054 | 30.59681 | 95.89824 | 92.42339 | . | . | . | . | . | . |
| chr8 | 123949180 | 123949600 | 307 | Promoter (<=1kb) | 8 | 123897123 | 123949265 | 52143 | 2 | ENSMUST00 | 0 | Nup133 | ENSMUSG0C | false | chr8 | 123948488 | 123950292 | 36.7977 | 133.03651 | 128.43381 | . | . | . | . | . | . |
| chrX | 142228296 | 142228711 | 307 | Promoter (<=1kb) | 20 | 142228703 | 142239692 | 10990 | 1 | ENSMUST00 | 0 | Nxt2 | ENSMUSG0C | false | chrX | 142227929 | 142228954 | 14.70503 | 33.66069 | 31.14909 | . | . | . | . | . | . |
| chr18 | 21205747 | 21206051 | 307 | Intron (ENSMUST00000049260.6 | 18 | 21127201 | 21300138 | 172938 | 2 | ENSMUST00 | 94087 | Garem | ENSMUSG0C | false | . | . | . | . | . | . | chr18 | 21202334 | 21207795 | 4.70946 | 3.98529 | 2.76086 |
| chr10 | 86808489 | 86808690 | 307 | Intron (ENSMUST00000099396.2 | 10 | 86779005 | 86838389 | 59385 | 1 | ENSMUST00 | 29485 | Nt5dc3 | ENSMUSG0C | false | . | . | . | . | . | . | chr10 | 86807739 | 86810002 | 4.39868 | 3.68099 | 2.55287 |
| chr10 | 23754226 | 23754543 | 307 | Distal Intergenic | 10 | 23785347 | 23785475 | 129 | 2 | ENSMUST00 | 30932 | Snora33 | ENSMUSG0C | false | . | . | . | . | . | . | chr10 | 23752881 | 23757038 | 3.70769 | 3.28211 | 1.97317 |
| chr17 | 69417283 | 69417633 | 307 | Promoter (<=1kb) | 17 | 69417402 | 69421632 | 4231 | 1 | ENSMUST00 | 0 | C030034I22R | ENSMUSG0C | false | chr17 | 69416699 | 69418386 | 28.91247 | 96.86193 | 93.36482 | . | . | . | . | . | . |
| chr5 | 67429453 | 67429687 | 307 | Promoter (1-2kb) | 5 | 67392147 | 67427799 | 35653 | 2 | ENSMUST00 | -1655 | Bend4 | ENSMUSG0C | false | . | . | . | . | . | . | . | . | . | . | . | . |
| chr16 | 52121106 | 52121328 | 307 | Intron (ENSMUST00000114471.1 | 16 | 52105015 | 52105147 | 133 | 2 | ENSMUST00 | -15960 | Gm23513 | ENSMUSG0C | false | . | . | . | . | . | . | chr16 | 52119455 | 52121445 | 5.16287 | 4.07365 | 3.1884 |
| chr5 | 16370823 | 16371075 | 307 | 3' UTR | 5 | 16508165 | 16509183 | 1019 | 2 | ENSMUST00 | 138108 | Gm8984 | ENSMUSG0C | false | . | . | . | . | . | . | chr5 | 16370610 | 16370945 | 4.39743 | 3.76213 | 2.51681 |
| chr1 | 62799327 | 62799623 | 307 | Intron (ENSMUST00000075144.6 | 1 | 62821355 | 62832370 | 11016 | 2 | ENSMUST00 | 32747 | RP23-149A5. | ENSMUSG0C | false | . | . | . | . | . | . | chr1 | 62794449 | 62800989 | 3.79616 | 3.40869 | 2.02444 |
| chrX | 37191092 | 37191316 | 306 | Promoter (<=1kb) | 20 | 37187588 | 37191163 | 3576 | 2 | ENSMUST00 | 0 | Ndufa1 | ENSMUSG0C | false | chrX | 37190506 | 37192515 | 18.84082 | 47.13427 | 44.45482 | . | . | . | . | . | . |
| chr11 | 77442682 | 77443075 | 306 | Intron (ENSMUST00000037912.6 | 11 | 77462411 | 77470484 | 8074 | 1 | ENSMUST00 | -19336 | Coro6 | ENSMUSG0C | false | . | . | . | . | . | . | chr11 | 77441460 | 77445660 | 3.32201 | 3.19331 | 1.62988 |
| chr17 | 4073507 | 4073772 | 306 | Distal Intergenic | 17 | 3635240 | 3696261 | 61022 | 2 | ENSMUST00 | -377247 | Nox3 | ENSMUSG0C | false | . | . | . | . | . | . | . | . | . | . | . | . |
| chr2 | 4918793 | 4919133 | 306 | Promoter (<=1kb) | 2 | 4919019 | 4938730 | 19712 | 1 | ENSMUST00 | 0 | Phyh | ENSMUSG0C | false | chr2 | 4918699 | 4919626 | 26.03857 | 105.07202 | 101.37238 | chr2 | 4917155 | 4919739 | 4.74673 | 3.93549 | 2.80974 |
| chr4 | 118048028 | 118048286 | 306 | Intron (ENSMUST00000030263.3 | 4 | 117940354 | 118031882 | 91529 | 2 | ENSMUST00 | -16147 | St3gal3 | ENSMUSG0C | true | . | . | . | . | . | . | chr4 | 118046938 | 118052090 | 4.08715 | 3.64437 | 2.26859 |
| chr7 | 137437433 | 137437935 | 306 | Promoter (<=1kb) | 7 | 137437648 | 137468594 | 30947 | 1 | ENSMUST00 | 0 | Glrx3 | ENSMUSG0C | false | chr7 | 137437318 | 137438870 | 34.74108 | 114.39095 | 110.42815 | chr7 | 137436953 | 137437457 | 3.68737 | 3.46057 | 1.95725 |
| chrX | 52153561 | 52153750 | 306 | Intron (ENSMUST00000033450.2 | 20 | 52053021 | 52165252 | 112232 | 2 | ENSMUST00 | 11502 | Gpc4 | ENSMUSG0C | false | . | . | . | . | . | . | . | . | . | . | . | . |
| chr12 | 76837288 | 76837879 | 306 | Promoter (<=1kb) | 12 | 76837414 | 76921412 | 83999 | 1 | ENSMUST00 | 0 | Fntb | ENSMUSG0C | false | chr12 | 76837212 | 76838449 | 18.33657 | 60.6815 | 57.82544 | chr12 | 76836923 | 76838329 | 3.17008 | 3.01969 | 1.54137 |
| chr15 | 76671470 | 76671909 | 306 | Promoter (<=1kb) | 15 | 76671615 | 76694919 | 23305 | 1 | ENSMUST00 | 0 | Ppp1r16a | ENSMUSG0C | false | chr15 | 76671572 | 76672462 | 31.91637 | 110.38313 | 106.53261 | chr15 | 76671087 | 76673219 | 5.24661 | 4.24559 | 3.22365 |
| chr10 | 117094950 | 117095355 | 306 | Intron (ENSMUST00000020381.3 | 10 | 117050998 | 117063814 | 12817 | 2 | ENSMUST00 | -31137 | Cct2 | ENSMUSG0C | false | . | . | . | . | . | . | chr10 | 117086815 | 117095074 | 5.71886 | 4.41367 | 3.58491 |
| chr11 | 77513070 | 77513571 | 306 | Promoter (<=1kb) | 11 | 77508099 | 77513370 | 5272 | 2 | ENSMUST00 | 0 | Trp53i13 | ENSMUSG0C | false | chr11 | 77512623 | 77514094 | 23.80619 | 80.75649 | 77.58801 | chr11 | 77512407 | 77515867 | 5.13544 | 4.05533 | 3.12826 |
| chr5 | 134313489 | 134314212 | 306 | Promoter (<=1kb) | 5 | 134286909 | 134314378 | 27470 | 2 | ENSMUST00 | 166 | Gtf2i | ENSMUSG0C | false | chr5 | 134312869 | 134316109 | 31.91637 | 110.38313 | 106.53261 | chr5 | 134311650 | 134313588 | 4.01533 | 3.63157 | 2.18958 |
| chr2 | 138256437 | 138257198 | 306 | Promoter (<=1kb) | 2 | 138256565 | 138287422 | 30858 | 1 | ENSMUST00 | 0 | Btbd3 | ENSMUSG0C | false | chr2 | 138256276 | 138258341 | 37.49432 | 126.6904 | 122.3202 | . | . | . | . | . | . |
| chr9 | 61368594 | 61368954 | 306 | Promoter (2-3kb) | 9 | 61370878 | 61371627 | 750 | 2 | ENSMUST00 | 2673 | Gm10655 | ENSMUSG0C | false | . | . | . | . | . | . | chr9 | 61366653 | 61374765 | 6.23582 | 4.63772 | 4.01625 |
| chr5 | 23675820 | 23676226 | 306 | Promoter (1-2kb) | 5 | 23674289 | 23674381 | 93 | 2 | ENSMUST00 | -1440 | Gm24009 | ENSMUSG0C | false | chr5 | 23674847 | 23676543 | 36.12563 | 122.12295 | 117.91594 | . | . | . | . | . | . |
| chr8 | 12354666 | 12354870 | 306 | Promoter (1-2kb) | 8 | 12356153 | 12356469 | 317 | 1 | ENSMUST00 | -1283 | Gm22207 | ENSMUSG0C | false | . | . | . | . | . | . | chr8 | 12354445 | 12355325 | 4.57633 | 3.89056 | 2.69184 |
| chr17 | 67133757 | 67134030 | 306 | Intron (ENSMUST00000037974.8 | 17 | 67026147 | 67026235 | 89 | 2 | ENSMUST00 | -107523 | Mir5709 | ENSMUSG0C | false | . | . | . | . | . | . | . | . | . | . | . | . |
| chrX | 108731853 | 108732064 | 306 | Distal Intergenic | 20 | 108664004 | 108664891 | 888 | 2 | ENSMUST00 | -66963 | Gm379 | ENSMUSG0C | false | . | . | . | . | . | . | . | . | . | . | . | . |
| chr5 | 64806053 | 64806397 | 306 | Promoter (1-2kb) | 5 | 64709722 | 64804117 | 94396 | 2 | ENSMUST00 | -1937 | Gm20033 | ENSMUSG0C | false | chr5 | 64802420 | 64806226 | 29.58769 | 108.11006 | 104.32592 | chr5 | 64804440 | 64817688 | 5.05462 | 4.05864 | 3.05357 |
| chr17 | 10360844 | 10361084 | 306 | Distal Intergenic | 17 | 10324602 | 10329307 | 4706 | 1 | ENSMUST00 | 36243 | RP24-162O2 | ENSMUSG0C | false | . | . | . | . | . | . | chr17 | 10355565 | 10365097 | 4.72976 | 3.90325 | 2.7952 |
| chr14 | 46787790 | 46788501 | 305 | Promoter (<=1kb) | 14 | 46775582 | 46788411 | 12830 | 2 | ENSMUST00 | 0 | Cnih1 | ENSMUSG0C | false | chr14 | 46787422 | 46788855 | 41.44478 | 148.06744 | 142.8609 | chr14 | 46784664 | 46789409 | 4.03126 | 3.55964 | 2.20107 |
| chr10 | 78009188 | 78009832 | 305 | Promoter (<=1kb) | 10 | 77986947 | 78009807 | 22861 | 2 | ENSMUST00 | 0 | Pfkl | ENSMUSG0C | false | chr10 | 78008931 | 78010334 | 35.67123 | 127.73884 | 123.33056 | chr10 | 78008065 | 78010377 | 4.48754 | 3.83139 | 2.58522 |
| chr12 | 86726331 | 86726665 | 305 | Promoter (<=1kb) | 12 | 86700502 | 86726460 | 25959 | 2 | ENSMUST00 | 0 | Angel1 | ENSMUSG0C | false | chr12 | 86725747 | 86726552 | 33.04844 | 119.73734 | 115.60696 | chr12 | 86725046 | 86726998 | 3.36309 | 3.27711 | 1.71824 |
| chr18 | 79026904 | 79027201 | 305 | Intron (ENSMUST00000025430.9 | 18 | 78750380 | 79109391 | 359012 | 2 | ENSMUST00 | 82190 | Setbp1 | ENSMUSG0C | false | . | . | . | . | . | . | chr18 | 79026097 | 79027981 | 4.74246 | 4.0507 | 2.78158 |
| chr2 | 101678289 | 101678548 | 305 | Promoter (<=1kb) | 2 | 101678429 | 101686235 | 7807 | 1 | ENSMUST00 | 0 | Traf6 | ENSMUSG0C | false | chr2 | 101678047 | 101679533 | 24.24157 | 89.02564 | 85.70093 | chr2 | 101677498 | 101678353 | 4.12533 | 3.67753 | 2.33687 |
| chr4 | 127021753 | 127022507 | 305 | Promoter (<=1kb) | 4 | 127021324 | 127031236 | 9913 | 1 | ENSMUST00 | 430 | Sfpq | ENSMUSG0C | false | chr4 | 127021239 | 127023612 | 36.04672 | 129.50026 | 125.02953 | . | . | . | . | . | . |
| chr9 | 21615790 | 21616530 | 305 | Promoter (<=1kb) | 9 | 21616169 | 21704217 | 88049 | 1 | ENSMUST00 | 0 | Smarca4 | ENSMUSG0C | false | chr9 | 21616122 | 21617444 | 33.7938 | 119.00099 | 114.88904 | . | . | . | . | . | . |
| chr3 | 94863269 | 94863489 | 305 | Promoter (<=1kb) | 3 | 94863728 | 94864550 | 823 | 1 | ENSMUST00 | -239 | Pogz | ENSMUSG0C | false | . | . | . | . | . | . | chr3 | 94862427 | 94863809 | 3.71452 | 3.37076 | 1.9711 |
| chr8 | 10910467 | 10910757 | 305 | Distal Intergenic | 8 | 10924427 | 10928457 | 4031 | 2 | ENSMUST00 | 17700 | 3930402G23 | ENSMUSG0C | false | . | . | . | . | . | . | . | . | . | . | . | . |
| chr4 | 70536556 | 70536850 | 305 | Promoter (1-2kb) | 4 | 70461826 | 70534995 | 73170 | 2 | ENSMUST00 | -1562 | Megf9 | ENSMUSG0C | false | . | . | . | . | . | . | chr4 | 70535476 | 70537261 | 5.40623 | 4.27142 | 3.34578 |
| chr10 | 130412196 | 130412488 | 305 | Promoter (<=1kb) | 10 | 130412499 | 130413312 | 814 | 1 | ENSMUST00 | -11 | Gm9770 | ENSMUSG0C | false | chr10 | 130412221 | 130412640 | 20.82289 | 72.3363 | 69.31286 | . | . | . | . | . | . |
| chr17 | 66352200 | 66352627 | 305 | Intron (ENSMUST00000086693.6 | 17 | 66343946 | 66346545 | 2600 | 2 | ENSMUST00 | -5656 | Mtcl1 | ENSMUSG0C | true | . | . | . | . | . | . | chr17 | 66349054 | 66377196 | 6.56105 | 4.76301 | 4.29736 |
| chr3 | 121390923 | 121391132 | 305 | Distal Intergenic | 3 | 121426541 | 121458205 | 31665 | 1 | ENSMUST00 | -35409 | Cnn3 | ENSMUSG0C | false | . | . | . | . | . | . | chr3 | 121390034 | 121391228 | 3.94169 | 3.58472 | 2.12032 |
| chr5 | 150263746 | 150263940 | 305 | Intron (ENSMUST00000087204.5 | 5 | 150259930 | 150497752 | 237823 | 1 | ENSMUST00 | 3817 | Fry | ENSMUSG0C | true | . | . | . | . | . | . | chr5 | 150258249 | 150264947 | 5.29784 | 4.15974 | 3.29734 |
| chr1 | 39367633 | 39367950 | 305 | Promoter (<=1kb) | 1 | 39367851 | 39371908 | 4058 | 1 | ENSMUST00 | 0 | Rpl31 | ENSMUSG0C | false | chr1 | 39367167 | 39368903 | 23.4661 | 79.19912 | 76.05865 | . | . | . | . | . | . |
| chr3 | 138613235 | 138613529 | 305 | Distal Intergenic | 3 | 138526415 | 138526494 | 80 | 1 | ENSMUST00 | 86821 | Mir1956 | ENSMUSG0C | false | . | . | . | . | . | . | chr3 | 138612689 | 138613495 | 2.93479 | 2.77473 | 1.35369 |
| chr8 | 73162021 | 73162335 | 305 | Intron (ENSMUST00000004497.9 | 8 | 73175119 | 73175258 | 140 | 2 | ENSMUST00 | 12923 | Gm22532 | ENSMUSG0C | false | . | . | . | . | . | . | chr8 | 73160245 | 73165634 | 4.96341 | 4.05335 | 2.98462 |
| chr16 | 96757124 | 96757350 | 305 | Intron (ENSMUST00000056102.7 | 16 | 96467606 | 96525793 | 58188 | 1 | ENSMUST00 | 289519 | Pcp4 | ENSMUSG0C | false | . | . | . | . | . | . | chr16 | 96757159 | 96758588 | 3.78436 | 3.30537 | 2.04389 |
| chr10 | 5185129 | 5185383 | 305 | Distal Intergenic | 10 | 5069192 | 5071975 | 2784 | 1 | ENSMUST00 | 115938 | Gm10097 | ENSMUSG0C | false | . | . | . | . | . | . | chr10 | 5184699 | 5195639 | 5.38954 | 4.23968 | 3.32126 |
| chr8 | 121246324 | 121246577 | 305 | Intron (ENSMUST00000127664.1 | 8 | 121198714 | 121201835 | 3122 | 2 | ENSMUST00 | -44490 | 5033426O07 | ENSMUSG0C | false | . | . | . | . | . | . | chr8 | 121236124 | 121247254 | 7.60766 | 5.2125 | 5.14118 |
| chr8 | 122443496 | 122443888 | 304 | Promoter (<=1kb) | 8 | 122433596 | 122443422 | 9827 | 2 | ENSMUST00 | -75 | Mvd | ENSMUSG0C | false | chr8 | 122442663 | 122443547 | 18.39885 | 52.66533 | 49.91441 | . | . | . | . | . | . |
| chr11 | 116021073 | 116021356 | 304 | Downstream (<1kb) | 11 | 116023509 | 116024473 | 965 | 2 | ENSMUST00 | 3117 | H3f3b | ENSMUSG0C | false | . | . | . | . | . | . | chr11 | 116020551 | 116023719 | 5.57175 | 4.15554 | 3.50082 |
| chr11 | 79047401 | 79047669 | 304 | Exon (ENSMUST00000018478.5/ | 11 | 79045183 | 79057627 | 12445 | 2 | ENSMUST00 | 9958 | Ksr1 | ENSMUSG0C | true | . | . | . | . | . | . | chr11 | 79045596 | 79144720 | 7.09579 | 4.93486 | 4.74337 |
| chr12 | 29938380 | 29938818 | 304 | Promoter (<=1kb) | 12 | 29938050 | 29989546 | 51497 | 1 | ENSMUST00 | 331 | Pxdn | ENSMUSG0C | false | chr12 | 29937557 | 29939537 | 47.33181 | 157.24483 | 151.69102 | chr12 | 29936541 | 29947539 | 4.38488 | 3.75724 | 2.49928 |
| chr15 | 81803057 | 81803377 | 304 | Promoter (<=1kb) | 15 | 81802826 | 81805911 | 3086 | 1 | ENSMUST00 | 232 | Tef | ENSMUSG0C | false | chr15 | 81802642 | 81804673 | 20.25921 | 52.6909 | 49.93975 | chr15 | 81800930 | 81810375 | 6.79861 | 4.85005 | 4.49384 |
| chr17 | 29347676 | 29348046 | 304 | Promoter (<=1kb) | 17 | 29332072 | 29347904 | 15833 | 2 | ENSMUST00 | 0 | Mtch1 | ENSMUSG0C | false | chr17 | 29346218 | 29348377 | 26.13664 | 77.56537 | 74.45414 | . | . | . | . | . | . |
| chr1 | 67041834 | 67042154 | 304 | Promoter (2-3kb) | 1 | 67000517 | 67038872 | 38356 | 2 | ENSMUST00 | -2963 | Lancl1 | ENSMUSG0C | false | . | . | . | . | . | . | chr1 | 67041499 | 67042983 | 6.03525 | 4.53012 | 3.86058 |
| chr1 | 24099878 | 24100575 | 304 | Promoter (<=1kb) | 1 | 24011245 | 24100194 | 88950 | 2 | ENSMUST00 | 0 | Fam135a | ENSMUSG0C | false | chr1 | 24099193 | 24100931 | 25.75243 | 103.50989 | 99.85224 | . | . | . | . | . | . |
| chr2 | 12923786 | 12924171 | 304 | Promoter (<=1kb) | 2 | 12924041 | 12980404 | 56364 | 1 | ENSMUST00 | 0 | Pter | ENSMUSG0C | false | chr2 | 12923666 | 12924791 | 11.90309 | 30.75663 | 28.28125 | chr2 | 12923419 | 12925214 | 3.79423 | 3.33771 | 2.04922 |
| chr2 | 119742724 | 119743081 | 304 | Promoter (<=1kb) | 2 | 119742337 | 119751263 | 8927 | 1 | ENSMUST00 | 388 | Itpka | ENSMUSG0C | false | chr2 | 119741625 | 119743529 | 33.08674 | 107.33151 | 103.57004 | chr2 | 119740863 | 119745199 | 5.39081 | 4.15988 | 3.37725 |
| chr5 | 96802334 | 96802591 | 304 | Intron (ENSMUST00000031447.7 | 5 | 96793385 | 96845968 | 52584 | 1 | ENSMUST00 | 8950 | Anxa3 | ENSMUSG0C | true | . | . | . | . | . | . | chr5 | 96794753 | 96812235 | 5.5921 | 4.32447 | 3.49382 |
| chr7 | 137750910 | 137751190 | 304 | Distal Intergenic | 7 | 137437648 | 137468594 | 30947 | 1 | ENSMUST00 | 313263 | Glrx3 | ENSMUSG0C | false | . | . | . | . | . | . | . | . | . | . | . | . |
| chr10 | 121625864 | 121626707 | 304 | Promoter (<=1kb) | 10 | 121587380 | 121626316 | 38937 | 2 | ENSMUST00 | 0 | Xpot | ENSMUSG0C | false | chr10 | 121625236 | 121627023 | 39.13739 | 139.46275 | 134.60301 | . | . | . | . | . | . |
| chr13 | 47125607 | 47125878 | 304 | Promoter (2-3kb) | 13 | 47122720 | 47247991 | 125272 | 1 | ENSMUST00 | 2888 | Rnf144b | ENSMUSG0C | true | . | . | . | . | . | . | chr13 | 47121911 | 47127413 | 4.95977 | 3.98421 | 3.01017 |
| chr2 | 69897106 | 69897678 | 304 | Promoter (<=1kb) | 2 | 69897246 | 70024013 | 126768 | 1 | ENSMUST00 | 0 | Ubr3 | ENSMUSG0C | false | chr2 | 69896922 | 69899033 | 34.54477 | 122.48211 | 118.25522 | . | . | . | . | . | . |
| chr11 | 80377886 | 80378144 | 304 | Promoter (<=1kb) | 11 | 80360492 | 80378015 | 17524 | 2 | ENSMUST00 | 0 | 5730455P16I | ENSMUSG0C | false | chr11 | 80377345 | 80378452 | 36.88899 | 121.33285 | 117.14651 | chr11 | 80377161 | 80378606 | 4.9193 | 4.1152 | 2.93783 |
| chr8 | 61574692 | 61574939 | 304 | Intron (ENSMUST00000121785.2 | 8 | 61515913 | 61591139 | 75227 | 2 | ENSMUST00 | 16200 | Palld | ENSMUSG0C | true | . | . | . | . | . | . | chr8 | 61569315 | 61589884 | 5.62358 | 4.37337 | 3.52305 |
| chr1 | 23117471 | 23117772 | 304 | Intron (ENSMUST00000191045.1 | 1 | 23100473 | 23102253 | 1781 | 2 | ENSMUST00 | -15219 | 4933415F23I | ENSMUSG0C | false | . | . | . | . | . | . | . | . | . | . | . | . |
| chr13 | 29704332 | 29704669 | 304 | Intron (ENSMUST00000140278.2 | 13 | 29732942 | 29733036 | 95 | 1 | ENSMUST00 | -28273 | Gm23729 | ENSMUSG0C | false | . | . | . | . | . | . | chr13 | 29697735 | 29704646 | 4.65988 | 3.89142 | 2.72494 |
| chr11 | 102408459 | 102408746 | 303 | Promoter (<=1kb) | 11 | 102404854 | 102407946 | 3093 | 2 | ENSMUST00 | -514 | Slc25a39 | ENSMUSG0C | false | chr11 | 102406444 | 102408601 | 32.64849 | 123.19247 | 118.94151 | chr11 | 102407921 | 102409731 | 4.81374 | 4.01023 | 2.85015 |
| chr11 | 76027001 | 76027439 | 303 | Promoter (<=1kb) | 11 | 76019194 | 76027782 | 8589 | 2 | ENSMUST00 | 343 | Fam101b | ENSMUSG0C | false | chr11 | 76026815 | 76028460 | 32.00811 | 98.35596 | 94.82688 | chr11 | 76026169 | 76028496 | 5.21191 | 4.24137 | 3.16398 |
| chr12 | 98627957 | 98628362 | 303 | Promoter (<=1kb) | 12 | 98628157 | 98669815 | 41659 | 1 | ENSMUST00 | 0 | Spata7 | ENSMUSG0C | false | chr12 | 98627633 | 98628884 | 24.03115 | 75.67385 | 72.59326 | chr12 | 98627320 | 98628717 | 3.45672 | 3.2204 | 1.77987 |
| chr19 | 26706980 | 26707252 | 303 | Intron (ENSMUST00000025862.8 | 19 | 26716115 | 26771753 | 55639 | 1 | ENSMUST00 | -8863 | Smarca2 | ENSMUSG0C | false | . | . | . | . | . | . | chr19 | 26705288 | 26709198 | 4.17392 | 3.68555 | 2.3409 |
| chr8 | 25510057 | 25510316 | 303 | Distal Intergenic | 8 | 25513654 | 25555568 | 41915 | 1 | ENSMUST00 | -3338 | Fgfr1 | ENSMUSG0C | true | . | . | . | . | . | . | chr8 | 25509883 | 25510496 | 3.09444 | 3.01497 | 1.45584 |
| chr9 | 27026336 | 27026626 | 303 | Intron (ENSMUST00000034470.9 | 9 | 27004503 | 27030094 | 25592 | 2 | ENSMUST00 | 3468 | Vps26b | ENSMUSG0C | false | . | . | . | . | . | . | chr9 | 27025880 | 27029365 | 4.29142 | 3.5259 | 2.45921 |
| chr9 | 58583104 | 58583522 | 303 | Promoter (<=1kb) | 9 | 58582444 | 58643668 | 61225 | 1 | ENSMUST00 | 661 | Nptn | ENSMUSG0C | false | chr9 | 58582230 | 58583686 | 39.20496 | 131.87447 | 127.31225 | chr9 | 58583444 | 58595403 | 4.26417 | 3.64376 | 2.40869 |
| chr17 | 5552411 | 5552639 | 303 | Intron (ENSMUST00000089185.4 | 17 | 5492600 | 5753891 | 261292 | 1 | ENSMUST00 | 59812 | Zdhhc14 | ENSMUSG0C | true | . | . | . | . | . | . | chr17 | 5549454 | 5554385 | 5.39068 | 4.24528 | 3.35705 |
| chr11 | 80735084 | 80735319 | 303 | Intron (ENSMUST00000041065.8 | 11 | 80673667 | 80780003 | 106337 | 2 | ENSMUST00 | 44684 | Myo1d | ENSMUSG0C | true | . | . | . | . | . | . | chr11 | 80730876 | 80736610 | 5.73884 | 4.35877 | 3.63747 |
| chr4 | 133041694 | 133042386 | 303 | Promoter (2-3kb) | 4 | 133039493 | 133077242 | 37750 | 1 | ENSMUST00 | 2202 | Ahdc1 | ENSMUSG0C | false | . | . | . | . | . | . | chr4 | 133013189 | 133060125 | 8.03751 | 5.40542 | 5.50183 |
| chr1 | 163403596 | 163404000 | 303 | Promoter (<=1kb) | 1 | 163384908 | 163403669 | 18762 | 2 | ENSMUST00 | 0 | Gorab | ENSMUSG0C | false | chr1 | 163402860 | 163404270 | 21.79372 | 61.12524 | 58.26245 | chr1 | 163402050 | 163404342 | 4.40811 | 3.67911 | 2.55897 |
| chr7 | 27447738 | 27448135 | 303 | Promoter (<=1kb) | 7 | 27447978 | 27466144 | 18167 | 1 | ENSMUST00 | 0 | Blvrb | ENSMUSG0C | false | chr7 | 27447900 | 27448417 | 23.34944 | 75.34173 | 72.26852 | chr7 | 27446146 | 27449888 | 5.57918 | 4.3431 | 3.4976 |
| chr8 | 90828552 | 90828962 | 303 | Promoter (<=1kb) | 8 | 90828835 | 91054508 | 225674 | 1 | ENSMUST00 | 0 | Chd9 | ENSMUSG0C | false | chr8 | 90828109 | 90829753 | 24.99264 | 71.03027 | 68.02251 | chr8 | 90828089 | 90831159 | 5.48929 | 4.37584 | 3.40909 |
| chr2 | 161109174 | 161109501 | 303 | Promoter (<=1kb) | 2 | 161008034 | 161109075 | 101042 | 2 | ENSMUST00 | -100 | Chd6 | ENSMUSG0C | false | chr2 | 161109244 | 161110158 | 26.66126 | 88.87019 | 85.5497 | chr2 | 161109445 | 161111487 | 4.40933 | 3.88682 | 2.5295 |
| chr18 | 44785272 | 44785524 | 303 | Intron (ENSMUST00000089874.5 | 18 | 44425061 | 44812182 | 387122 | 2 | ENSMUST00 | 26658 | Mcc | ENSMUSG0C | true | . | . | . | . | . | . | . | . | . | . | . | . |
| chr2 | 160174374 | 160174941 | 303 | Distal Intergenic | 2 | 160311393 | 160313616 | 2224 | 2 | ENSMUST00 | 138675 | Gm826 | ENSMUSG0C | false | . | . | . | . | . | . | chr2 | 160172180 | 160175717 | 4.51414 | 3.79665 | 2.65101 |
| chr2 | 61396797 | 61397006 | 303 | Distal Intergenic | 2 | 61492766 | 61492894 | 129 | 1 | ENSMUST00 | -95760 | Gm22338 | ENSMUSG0C | false | . | . | . | . | . | . | chr2 | 61395816 | 61397224 | 4.88837 | 4.07364 | 2.92027 |
| chr5 | 137630687 | 137631081 | 303 | Promoter (<=1kb) | 5 | 137630130 | 137640180 | 10051 | 1 | ENSMUST00 | 558 | Lrch4 | ENSMUSG0C | false | chr5 | 137629072 | 137631603 | 33.41831 | 117.26761 | 113.21259 | chr5 | 137630189 | 137634654 | 7.31247 | 5.1581 | 4.90713 |
| chr13 | 97297648 | 97297943 | 303 | Distal Intergenic | 13 | 97292019 | 97292108 | 90 | 1 | ENSMUST00 | 5630 | Gm22007 | ENSMUSG0C | false | . | . | . | . | . | . | . | . | . | . | . | . |
| chr18 | 35850799 | 35851185 | 303 | Intron (ENSMUST00000060722.6 | 18 | 35830389 | 35831093 | 705 | 2 | ENSMUST00 | -19707 | RP24-326K1 | ENSMUSG0C | false | . | . | . | . | . | . | chr18 | 35831800 | 35857612 | 7.70883 | 5.15861 | 5.24645 |
| chr4 | 136247849 | 136248365 | 302 | Promoter (<=1kb) | 4 | 136247952 | 136274898 | 26947 | 1 | ENSMUST00 | 0 | Tcea3 | ENSMUSG0C | false | chr4 | 136247308 | 136248988 | 31.24818 | 92.1069 | 88.71768 | chr4 | 136246390 | 136250396 | 5.33059 | 4.23099 | 3.28527 |
| chr7 | 44304844 | 44305463 | 302 | Promoter (1-2kb) | 7 | 44303770 | 44306959 | 3190 | 2 | ENSMUST00 | 1496 | Clec11a | ENSMUSG0C | false | chr7 | 44304267 | 44305249 | 18.02749 | 67.47041 | 64.5209 | . | . | . | . | . | . |
| chr5 | 143635371 | 143635759 | 302 | Promoter (<=1kb) | 5 | 143636351 | 143692554 | 56204 | 1 | ENSMUST00 | -592 | Cyth3 | ENSMUSG0C | false | chr5 | 143635357 | 143635806 | 5.48026 | 8.55952 | 6.41235 | chr5 | 143628431 | 143643057 | 5.82289 | 4.44189 | 3.69355 |
| chr14 | 16256267 | 16256476 | 302 | Intron (ENSMUST00000022310.6 | 14 | 16238656 | 16249808 | 11153 | 2 | ENSMUST00 | -6460 | Oxsm | ENSMUSG0C | false | . | . | . | . | . | . | chr14 | 16255361 | 16256889 | 5.0348 | 4.10918 | 3.05195 |
| chr2 | 172440282 | 172440625 | 302 | Promoter (<=1kb) | 2 | 172440556 | 172469908 | 29353 | 1 | ENSMUST00 | 0 | Rtfdc1 | ENSMUSG0C | false | chr2 | 172439724 | 172441491 | 23.17719 | 89.67316 | 86.33752 | chr2 | 172439733 | 172442067 | 4.46218 | 3.80825 | 2.60184 |
| chr7 | 100919675 | 100920168 | 302 | Intron (ENSMUST00000107032.1 | 7 | 100869746 | 100932161 | 62416 | 2 | ENSMUST00 | 11993 | Arhgef17 | ENSMUSG0C | false | . | . | . | . | . | . | chr7 | 100901870 | 100926579 | 7.14639 | 4.97053 | 4.78972 |
| chr5 | 62751832 | 62752018 | 302 | Intron (ENSMUST00000076623.5 | 5 | 62749296 | 62765656 | 16361 | 2 | ENSMUST00 | 13638 | Arap2 | ENSMUSG0C | false | . | . | . | . | . | . | chr5 | 62751555 | 62753279 | 4.58617 | 3.93852 | 2.6653 |
| chr1 | 185898568 | 185898792 | 302 | Distal Intergenic | 1 | 186087731 | 186117310 | 29580 | 2 | ENSMUST00 | 218518 | Lyplal1 | ENSMUSG0C | false | . | . | . | . | . | . | . | . | . | . | . | . |
| chr12 | 116405023 | 116405364 | 302 | Promoter (<=1kb) | 12 | 116405402 | 116463531 | 58130 | 1 | ENSMUST00 | -38 | Ncapg2 | ENSMUSG0C | false | chr12 | 116404853 | 116405992 | 25.50663 | 88.63404 | 85.31516 | chr12 | 116404664 | 116406242 | 5.97492 | 4.60533 | 3.79762 |
| chr10 | 37231974 | 37232248 | 302 | Distal Intergenic | 10 | 37139914 | 37151315 | 11402 | 1 | ENSMUST00 | 92061 | 5930403N24 | ENSMUSG0C | false | . | . | . | . | . | . | chr10 | 37231334 | 37232712 | 3.05179 | 3.04376 | 1.40207 |
| chr5 | 44261289 | 44261809 | 302 | Distal Intergenic | 5 | 44175164 | 44226606 | 51443 | 2 | ENSMUST00 | -34684 | Tapt1 | ENSMUSG0C | false | . | . | . | . | . | . | . | . | . | . | . | . |
| chr5 | 137645605 | 137646047 | 302 | Promoter (<=1kb) | 5 | 137643032 | 137645714 | 2683 | 2 | ENSMUST00 | 0 | Irs3 | ENSMUSG0C | false | chr5 | 137644938 | 137646119 | 15.65496 | 40.89989 | 38.30132 | chr5 | 137642779 | 137646719 | 4.58973 | 3.85457 | 2.71996 |
| chr8 | 84659933 | 84660451 | 302 | Promoter (2-3kb) | 8 | 84661331 | 84662852 | 1522 | 2 | ENSMUST00 | 2401 | Ier2 | ENSMUSG0C | false | chr8 | 84659950 | 84663752 | 33.52805 | 106.05383 | 102.33234 | chr8 | 84659817 | 84660793 | 6.22441 | 4.42466 | 4.03762 |
| chr9 | 99242938 | 99243701 | 302 | Promoter (<=1kb) | 9 | 99243367 | 99263762 | 20396 | 1 | ENSMUST00 | 0 | Cep70 | ENSMUSG0C | false | chr9 | 99242651 | 99244141 | 42.16172 | 139.23978 | 134.38916 | . | . | . | . | . | . |
| chr4 | 101037225 | 101037477 | 302 | Distal Intergenic | 4 | 101020298 | 101020696 | 399 | 2 | ENSMUST00 | -16530 | Gm10577 | ENSMUSG0C | false | . | . | . | . | . | . | chr4 | 101036389 | 101037869 | 4.4606 | 3.86694 | 2.55743 |
| chr11 | 94601680 | 94602162 | 302 | Promoter (<=1kb) | 11 | 94557102 | 94601871 | 44770 | 2 | ENSMUST00 | 0 | Acsf2 | ENSMUSG0C | false | chr11 | 94601426 | 94602381 | 19.64801 | 57.34576 | 54.5348 | chr11 | 94601014 | 94602800 | 4.33472 | 3.77167 | 2.45255 |
| chr4 | 123624522 | 123624996 | 302 | Intron (ENSMUST00000097897.5 | 4 | 123630295 | 123630703 | 409 | 1 | ENSMUST00 | -5299 | Gm12926 | ENSMUSG0C | false | . | . | . | . | . | . | chr4 | 123587116 | 123630412 | 5.80173 | 4.46538 | 3.66108 |
| chr11 | 103884782 | 103885089 | 302 | Intron (ENSMUST00000103075.5 | 11 | 103879915 | 103880243 | 329 | 2 | ENSMUST00 | -4540 | Gm8738 | ENSMUSG0C | false | . | . | . | . | . | . | chr11 | 103882205 | 103888074 | 5.46826 | 4.08776 | 3.42706 |
| chr13 | 107587818 | 107588055 | 302 | Distal Intergenic | 13 | 107687397 | 107692162 | 4766 | 1 | ENSMUST00 | -99342 | RP23-125L8. | ENSMUSG0C | false | . | . | . | . | . | . | chr13 | 107587054 | 107588104 | 4.32861 | 3.76909 | 2.51646 |
| chr16 | 91801542 | 91801776 | 301 | Promoter (2-3kb) | 16 | 91804622 | 91816917 | 12296 | 1 | ENSMUST00 | -2846 | Itsn1 | ENSMUSG0C | false | . | . | . | . | . | . | . | . | . | . | . | . |
| chr17 | 8506183 | 8506411 | 301 | Distal Intergenic | 17 | 8525372 | 8981671 | 456300 | 1 | ENSMUST00 | -18961 | Pde10a | ENSMUSG0C | true | chr17 | 8505790 | 8506275 | 7.45894 | 16.41436 | 14.12962 | chr17 | 8504525 | 8510234 | 7.21779 | 5.10275 | 4.81866 |
| chr7 | 140881699 | 140882516 | 301 | Promoter (<=1kb) | 7 | 140863670 | 140881869 | 18200 | 2 | ENSMUST00 | 0 | Sirt3 | ENSMUSG0C | false | chr7 | 140881281 | 140883193 | 28.04154 | 116.13699 | 112.1206 | . | . | . | . | . | . |
| chr3 | 96727444 | 96727973 | 301 | Promoter (<=1kb) | 3 | 96727611 | 96791155 | 63545 | 1 | ENSMUST00 | 0 | Rnf115 | ENSMUSG0C | false | chr3 | 96726884 | 96728769 | 34.74108 | 114.39095 | 110.42815 | . | . | . | . | . | . |
| chr6 | 91728990 | 91729285 | 301 | Intron (ENSMUST00000032185.7 | 6 | 91684067 | 91759063 | 74997 | 1 | ENSMUST00 | 44924 | Slc6a6 | ENSMUSG0C | false | . | . | . | . | . | . | chr6 | 91726179 | 91731827 | 5.90933 | 4.2 | 3.79661 |
| chr7 | 80134560 | 80134756 | 301 | Distal Intergenic | 7 | 80094846 | 80115392 | 20547 | 2 | ENSMUST00 | -19169 | Idh2 | ENSMUSG0C | false | . | . | . | . | . | . | chr7 | 80132335 | 80135432 | 6.30904 | 4.54613 | 4.12205 |
| chrX | 74085010 | 74085689 | 301 | Promoter (<=1kb) | 20 | 74026592 | 74085638 | 59047 | 2 | ENSMUST00 | 0 | Mecp2 | ENSMUSG0C | false | chrX | 74084207 | 74085691 | 26.65286 | 74.78688 | 71.72144 | . | . | . | . | . | . |
| chr8 | 120425062 | 120426057 | 301 | Intron (ENSMUST00000127664.1 | 8 | 120488447 | 120581390 | 92944 | 1 | ENSMUST00 | -62390 | Gse1 | ENSMUSG0C | false | . | . | . | . | . | . | chr8 | 120415621 | 120427536 | 6.84739 | 4.85598 | 4.53347 |
| chr9 | 74952825 | 74953680 | 301 | Promoter (<=1kb) | 9 | 74953053 | 75032468 | 79416 | 1 | ENSMUST00 | 0 | Fam214a | ENSMUSG0C | false | chr9 | 74952562 | 74953836 | 37.5617 | 145.4381 | 140.34033 | . | . | . | . | . | . |
| chr10 | 22637800 | 22638095 | 301 | Distal Intergenic | 10 | 22645011 | 22704285 | 59275 | 1 | ENSMUST00 | -6916 | Slc2a12 | ENSMUSG0C | false | . | . | . | . | . | . | chr10 | 22637309 | 22639518 | 4.08689 | 3.67237 | 2.23998 |
| chr5 | 81474581 | 81475061 | 301 | Intron (ENSMUST00000132375.2 | 5 | 81310025 | 81796634 | 486610 | 1 | ENSMUST00 | 164557 | Lphn3 | ENSMUSG0C | false | . | . | . | . | . | . | chr5 | 81474428 | 81476654 | 3.62152 | 3.36664 | 1.9254 |
| chr17 | 43156146 | 43156435 | 301 | Distal Intergenic | 17 | 43270329 | 43324737 | 54409 | 1 | ENSMUST00 | -113894 | Gpr110 | ENSMUSG0C | false | . | . | . | . | . | . | chr17 | 43155358 | 43159256 | 5.96873 | 4.44672 | 3.85454 |
| chr9 | 32142913 | 32143143 | 301 | Intron (ENSMUST00000174641.2 | 9 | 32129570 | 32181966 | 52397 | 1 | ENSMUST00 | 13344 | Arhgap32 | ENSMUSG0C | false | . | . | . | . | . | . | . | . | . | . | . | . |
| chr18 | 84086334 | 84086958 | 301 | Promoter (<=1kb) | 18 | 84011627 | 84086404 | 74778 | 2 | ENSMUST00 | 0 | Tshz1 | ENSMUSG0C | false | chr18 | 84084710 | 84090178 | 35.29575 | 125.98199 | 121.63858 | . | . | . | . | . | . |
| chr9 | 66863477 | 66863781 | 301 | Exon (ENSMUST00000041139.7/ | 9 | 66814994 | 66834726 | 19733 | 2 | ENSMUST00 | -28752 | Aph1c | ENSMUSG0C | false | . | . | . | . | . | . | chr9 | 66860939 | 66865039 | 5.30744 | 4.13827 | 3.30598 |
| chr5 | 34520866 | 34521145 | 301 | Distal Intergenic | 5 | 34525838 | 34551700 | 25863 | 1 | ENSMUST00 | -4693 | Sh3bp2 | ENSMUSG0C | false | . | . | . | . | . | . | . | . | . | . | . | . |
| chr4 | 149820615 | 149820921 | 301 | Promoter (2-3kb) | 4 | 149817732 | 149817840 | 109 | 1 | ENSMUST00 | 2884 | n-R5s193 | ENSMUSG0C | false | . | . | . | . | . | . | chr4 | 149820569 | 149823210 | 5.53517 | 4.35229 | 3.4375 |
| chr6 | 72958090 | 72958896 | 301 | Promoter (<=1kb) | 6 | 72957347 | 72958488 | 1142 | 2 | ENSMUST00 | 0 | Tmsb10 | ENSMUSG0C | false | chr6 | 72956611 | 72959041 | 38.01312 | 122.22395 | 118.01301 | . | . | . | . | . | . |
| chr2 | 124747742 | 124747961 | 301 | Distal Intergenic | 2 | 124708655 | 124711083 | 2429 | 1 | ENSMUST00 | 39088 | Gm13994 | ENSMUSG0C | false | . | . | . | . | . | . | chr2 | 124747186 | 124748047 | 4.72858 | 3.90271 | 2.81181 |
| chr1 | 12690010 | 12690280 | 300 | Promoter (1-2kb) | 1 | 12692277 | 12861192 | 168916 | 1 | ENSMUST00 | -1997 | Sulf1 | ENSMUSG0C | true | . | . | . | . | . | . | chr1 | 12689349 | 12692023 | 4.3571 | 3.78338 | 2.48252 |
| chr10 | 19062470 | 19062796 | 300 | Distal Intergenic | 10 | 19008266 | 19015657 | 7392 | 2 | ENSMUST00 | -46814 | Tnfaip3 | ENSMUSG0C | false | . | . | . | . | . | . | chr10 | 19061791 | 19063280 | 4.21436 | 3.73982 | 2.35477 |
| chr13 | 63272518 | 63272997 | 300 | Intron (ENSMUST00000091560.5 | 13 | 63282142 | 63326096 | 43955 | 1 | ENSMUST00 | -9145 | 2010111I01R | ENSMUSG0C | false | chr13 | 63271624 | 63282991 | 26.57451 | 76.48061 | 73.3894 | . | . | . | . | . | . |
| chr15 | 99398418 | 99398721 | 300 | Promoter (2-3kb) | 15 | 99401085 | 99402102 | 1018 | 1 | ENSMUST00 | -2364 | Tmbim6 | ENSMUSG0C | false | . | . | . | . | . | . | chr15 | 99393778 | 99402130 | 6.4261 | 4.61526 | 4.21071 |
| chr18 | 77064923 | 77065944 | 300 | Promoter (<=1kb) | 18 | 77065208 | 77155708 | 90501 | 1 | ENSMUST00 | 0 | Pias2 | ENSMUSG0C | false | chr18 | 77064771 | 77066818 | 35.78724 | 111.00061 | 107.13931 | . | . | . | . | . | . |
| chr5 | 134770930 | 134771341 | 300 | Distal Intergenic | 5 | 134703781 | 134747241 | 43461 | 2 | ENSMUST00 | -23690 | Eln | ENSMUSG0C | false | . | . | . | . | . | . | chr5 | 134770712 | 134774141 | 6.42644 | 4.70561 | 4.20353 |
| chr6 | 145763780 | 145763997 | 300 | Intron (ENSMUST00000111704.2 | 6 | 145750571 | 145815188 | 64618 | 1 | ENSMUST00 | 13210 | Rassf8 | ENSMUSG0C | true | . | . | . | . | . | . | chr6 | 145748529 | 145783801 | 6.70363 | 4.81765 | 4.41144 |
| chr7 | 132349082 | 132349394 | 300 | Intron (ENSMUST00000124096.2 | 7 | 132235780 | 132317228 | 81449 | 2 | ENSMUST00 | -31855 | Chst15 | ENSMUSG0C | false | . | . | . | . | . | . | chr7 | 132348648 | 132349873 | 3.71707 | 3.38167 | 2.00796 |
| chr9 | 56152153 | 56152370 | 300 | Intron (ENSMUST00000034876.8 | 9 | 56135885 | 56161070 | 25186 | 2 | ENSMUST00 | 8700 | Tspan3 | ENSMUSG0C | false | . | . | . | . | . | . | chr9 | 56151595 | 56161546 | 4.944 | 4.05437 | 2.99412 |
| chr5 | 3342125 | 3342699 | 300 | Promoter (1-2kb) | 5 | 3343893 | 3523218 | 179326 | 1 | ENSMUST00 | -1194 | Cdk6 | ENSMUSG0C | false | chr5 | 3341034 | 3344969 | 32.55001 | 105.36325 | 101.65399 | chr5 | 3341133 | 3344421 | 4.3677 | 3.73533 | 2.50877 |
| chr12 | 9584796 | 9585108 | 300 | Distal Intergenic | 12 | 9574442 | 9581499 | 7058 | 1 | ENSMUST00 | 10355 | Osr1 | ENSMUSG0C | false | chr12 | 9584583 | 9585248 | 8.16709 | 15.30057 | 13.0337 | . | . | . | . | . | . |
| chr3 | 96601010 | 96601711 | 300 | Promoter (<=1kb) | 3 | 96601133 | 96625352 | 24220 | 1 | ENSMUST00 | 0 | Lix1l | ENSMUSG0C | false | chr3 | 96600614 | 96602004 | 29.3377 | 90.74815 | 87.38457 | chr3 | 96600137 | 96605853 | 5.74186 | 4.36179 | 3.64522 |
| chr12 | 108471105 | 108471644 | 300 | Promoter (<=1kb) | 12 | 108471853 | 108539564 | 67712 | 1 | ENSMUST00 | -209 | Eml1 | ENSMUSG0C | false | . | . | . | . | . | . | chr12 | 108470736 | 108472206 | 4.18885 | 3.60023 | 2.34013 |
| chr4 | 136462366 | 136462853 | 300 | Promoter (<=1kb) | 4 | 136462250 | 136463977 | 1728 | 1 | ENSMUST00 | 117 | 6030445D17I | ENSMUSG0C | false | chr4 | 136462373 | 136463335 | 31.64465 | 94.75051 | 91.30649 | . | . | . | . | . | . |
| chr7 | 126487038 | 126487330 | 300 | Promoter (<=1kb) | 7 | 126487355 | 126490730 | 3376 | 1 | ENSMUST00 | -25 | Tufm | ENSMUSG0C | false | chr7 | 126486726 | 126488354 | 31.31841 | 96.16425 | 92.68594 | chr7 | 126486623 | 126488410 | 4.09576 | 3.50574 | 2.28952 |
| chr2 | 73529258 | 73529792 | 300 | Promoter (<=1kb) | 2 | 73429610 | 73529734 | 100125 | 2 | ENSMUST00 | 0 | Wipf1 | ENSMUSG0C | false | chr2 | 73528721 | 73530153 | 20.31581 | 74.8081 | 71.74177 | . | . | . | . | . | . |
| chr3 | 153686871 | 153688059 | 300 | Intron (ENSMUST00000064460.5 | 3 | 153708911 | 153709026 | 116 | 1 | ENSMUST00 | -20852 | Gm22206 | ENSMUSG0C | false | . | . | . | . | . | . | chr3 | 153686316 | 153712771 | 6.43925 | 4.66349 | 4.19773 |
| chr15 | 80842560 | 80842798 | 300 | Intron (ENSMUST00000067689.7 | 15 | 80940039 | 80940120 | 82 | 1 | ENSMUST00 | -97241 | Mir5113 | ENSMUSG0C | false | . | . | . | . | . | . | chr15 | 80832720 | 80847879 | 5.49721 | 4.14862 | 3.44463 |
| chr13 | 94284979 | 94285806 | 299 | Promoter (<=1kb) | 13 | 94201310 | 94285829 | 84520 | 2 | ENSMUST00 | 23 | Scamp1 | ENSMUSG0C | false | chr13 | 94284370 | 94286213 | 34.92026 | 124.22972 | 119.94103 | . | . | . | . | . | . |
| chr1 | 65178198 | 65178887 | 299 | Promoter (<=1kb) | 1 | 65158616 | 65179181 | 20566 | 2 | ENSMUST00 | 294 | Idh1 | ENSMUSG0C | false | chr1 | 65177109 | 65180019 | 28.91859 | 89.0447 | 85.71584 | . | . | . | . | . | . |
| chr1 | 134962349 | 134962665 | 299 | Promoter (<=1kb) | 1 | 134962565 | 134974162 | 11598 | 1 | ENSMUST00 | 0 | Ube2t | ENSMUSG0C | false | chr1 | 134962060 | 134963089 | 26.44925 | 85.62254 | 82.36654 | chr1 | 134962006 | 134963348 | 4.25172 | 3.52816 | 2.43842 |
| chr3 | 129369566 | 129370051 | 299 | Distal Intergenic | 3 | 129298594 | 129332720 | 34127 | 2 | ENSMUST00 | -36847 | Enpep | ENSMUSG0C | true | . | . | . | . | . | . | chr3 | 129365437 | 129373366 | 4.57359 | 3.87373 | 2.65814 |
| chr7 | 112444518 | 112444937 | 299 | Intron (ENSMUST00000106643.2 | 7 | 112427905 | 112552929 | 125025 | 1 | ENSMUST00 | 16614 | Parva | ENSMUSG0C | true | . | . | . | . | . | . | chr7 | 112434747 | 112475955 | 6.44046 | 4.72904 | 4.19131 |
| chr9 | 43318676 | 43318939 | 299 | Intron (ENSMUST00000034511.5 | 9 | 43310763 | 43336125 | 25363 | 1 | ENSMUST00 | 7914 | Trim29 | ENSMUSG0C | false | . | . | . | . | . | . | chr9 | 43318373 | 43319844 | 5.50569 | 4.2756 | 3.44997 |
| chr1 | 7100596 | 7100918 | 299 | Intron (ENSMUST00000061280.1 | 1 | 7089166 | 7172122 | 82957 | 1 | ENSMUST00 | 11431 | Pcmtd1 | ENSMUSG0C | true | . | . | . | . | . | . | chr1 | 7098260 | 7109458 | 4.35067 | 3.6967 | 2.50301 |
| chr12 | 21165806 | 21166447 | 299 | Intron (ENSMUST00000174530.1 | 12 | 21111954 | 21268179 | 156226 | 1 | ENSMUST00 | 53853 | Asap2 | ENSMUSG0C | false | . | . | . | . | . | . | chr12 | 21148547 | 21176379 | 6.51645 | 4.71535 | 4.25897 |
| chr18 | 69593192 | 69593578 | 299 | Promoter (<=1kb) | 18 | 69593417 | 69683252 | 89836 | 1 | ENSMUST00 | 0 | Tcf4 | ENSMUSG0C | false | chr18 | 69592339 | 69594600 | 15.30398 | 43.94659 | 41.30965 | . | . | . | . | . | . |
| chrX | 166931156 | 166931375 | 299 | Distal Intergenic | 20 | 166935469 | 166935915 | 447 | 1 | ENSMUST00 | -4094 | Gm15231 | ENSMUSG0C | false | . | . | . | . | . | . | . | . | . | . | . | . |
| chr6 | 34453401 | 34453678 | 299 | Distal Intergenic | 6 | 34443523 | 34445436 | 1914 | 2 | ENSMUST00 | -7966 | Gm13857 | ENSMUSG0C | false | . | . | . | . | . | . | chr6 | 34452891 | 34454471 | 5.83308 | 4.46275 | 3.70218 |
| chr4 | 62732864 | 62733128 | 299 | Distal Intergenic | 4 | 62723516 | 62727537 | 4022 | 2 | ENSMUST00 | -5328 | Gm11211 | ENSMUSG0C | false | . | . | . | . | . | . | chr4 | 62726795 | 62739050 | 7.64763 | 5.19923 | 5.18438 |
| chr10 | 24367016 | 24367318 | 299 | Distal Intergenic | 10 | 24447490 | 24452657 | 5168 | 1 | ENSMUST00 | -80172 | Gm15271 | ENSMUSG0C | false | . | . | . | . | . | . | chr10 | 24364096 | 24367834 | 4.28857 | 3.72724 | 2.4501 |
| chr13 | 25808909 | 25809184 | 299 | Distal Intergenic | 13 | 25921033 | 25930941 | 9909 | 2 | ENSMUST00 | 121757 | Gm11351 | ENSMUSG0C | false | . | . | . | . | . | . | chr13 | 25807480 | 25809831 | 3.83644 | 3.42083 | 2.0939 |
| chrX | 101126703 | 101126970 | 299 | Intron (ENSMUST00000156473.2 | 20 | 101141433 | 101141552 | 120 | 2 | ENSMUST00 | 14582 | Gm24061 | ENSMUSG0C | false | . | . | . | . | . | . | . | . | . | . | . | . |
| chr1 | 133609822 | 133610342 | 299 | Promoter (<=1kb) | 1 | 133603871 | 133610291 | 6421 | 2 | ENSMUST00 | 0 | Snrpe | ENSMUSG0C | false | chr1 | 133609373 | 133610411 | 25.61729 | 89.79346 | 86.4557 | . | . | . | . | . | . |
| chr17 | 35353471 | 35354130 | 299 | Distal Intergenic | 17 | 35358989 | 35359972 | 984 | 1 | ENSMUST00 | -4859 | H2-Q3 | ENSMUSG0C | false | . | . | . | . | . | . | . | . | . | . | . | . |
| chr7 | 63896774 | 63897240 | 299 | Promoter (<=1kb) | 7 | 63896533 | 63899698 | 3166 | 1 | ENSMUST00 | 242 | Gm27252 | ENSMUSG0C | false | . | . | . | . | . | . | chr7 | 63885397 | 63936279 | 7.51388 | 5.18869 | 5.06896 |
| chr1 | 134060796 | 134061479 | 298 | Distal Intergenic | 1 | 134075170 | 134079120 | 3951 | 2 | ENSMUST00 | 17641 | Btg2 | ENSMUSG0C | false | chr1 | 134060670 | 134061705 | 14.47252 | 34.61563 | 32.09162 | chr1 | 134058485 | 134067448 | 6.65638 | 4.82565 | 4.35972 |
| chr15 | 34511903 | 34512389 | 298 | Promoter (1-2kb) | 15 | 34514170 | 34518018 | 3849 | 1 | ENSMUST00 | -1781 | Pop1 | ENSMUSG0C | false | . | . | . | . | . | . | chr15 | 34508319 | 34517398 | 5.80499 | 4.48832 | 3.66177 |
| chr16 | 32608737 | 32609235 | 298 | Promoter (<=1kb) | 16 | 32608920 | 32632794 | 23875 | 1 | ENSMUST00 | 0 | Tfrc | ENSMUSG0C | false | chr16 | 32608478 | 32609750 | 39.74135 | 132.8143 | 128.22414 | . | . | . | . | . | . |
| chr16 | 72662970 | 72663738 | 298 | Promoter (<=1kb) | 16 | 72663149 | 73046095 | 382947 | 1 | ENSMUST00 | 0 | Robo1 | ENSMUSG0C | false | chr16 | 72662877 | 72665011 | 28.00751 | 87.44511 | 84.14996 | . | . | . | . | . | . |
| chr1 | 13646425 | 13646642 | 298 | Intron (ENSMUST00000027071.6 | 1 | 13629995 | 13638411 | 8417 | 2 | ENSMUST00 | -8015 | Lactb2 | ENSMUSG0C | false | . | . | . | . | . | . | chr1 | 13646097 | 13646787 | 2.87058 | 2.91138 | 1.31781 |
| chr4 | 76451186 | 76451526 | 298 | Promoter (1-2kb) | 4 | 75941238 | 76449980 | 508743 | 2 | ENSMUST00 | -1207 | Ptprd | ENSMUSG0C | true | chr4 | 76450773 | 76451286 | 8.63619 | 18.02194 | 15.71366 | chr4 | 76450883 | 76452522 | 4.63665 | 3.96911 | 2.71169 |
| chr7 | 144581640 | 144582187 | 298 | Promoter (<=1kb) | 7 | 144577318 | 144582463 | 5146 | 2 | ENSMUST00 | 276 | Fadd | ENSMUSG0C | false | chr7 | 144581347 | 144582972 | 34.38402 | 121.12925 | 116.94628 | chr7 | 144580870 | 144581856 | 5.58984 | 4.1907 | 3.52014 |

| | | | | | | | | | | | | | | | | | | | | | | | | | | |
|---|---|---|---|---|---|---|---|---|---|---|---|---|---|---|---|---|---|---|---|---|---|---|---|---|---|---|
| chr8 | 119910631 | 119911392 | 298 | Promoter (<=1kb) | 8 | 119910814 | 119957560 | 46747 | 1 | ENSMUST00 | 0 | Usp10 | ENSMUSG0C | false | chr8 | 119910112 | 119911760 | 20.20131 | 69.382 | 66.39919 | chr8 | 119911246 | 119915223 | 4.4436 | 3.76443 | 2.55371 |
| chr9 | 57637590 | 57638189 | 298 | Intron (ENSMUST00000034863.6 | 9 | 57626647 | 57645653 | 19007 | 2 | ENSMUST00 | 7464 | Csk | ENSMUSG0C | false | . | . | . | . | . | . | chr9 | 57632762 | 57645064 | 6.57255 | 4.74185 | 4.30478 |
| chr10 | 83347547 | 83347873 | 298 | Distal Intergenic | 10 | 83230848 | 83337882 | 107035 | 2 | ENSMUST00 | -9666 | Slc41a2 | ENSMUSG0C | true | . | . | . | . | . | . | chr10 | 83346049 | 83348940 | 4.65571 | 3.85652 | 2.75851 |
| chr13 | 25077355 | 25077658 | 298 | Intron (ENSMUST00000036932.9 | 13 | 25096347 | 25119406 | 23060 | 1 | ENSMUST00 | -18689 | Dcdc2a | ENSMUSG0C | true | . | . | . | . | . | . | chr13 | 25068613 | 25089235 | 4.80481 | 3.93018 | 2.85848 |
| chr1 | 59912660 | 59913934 | 298 | Promoter (<=1kb) | 1 | 59913006 | 59985346 | 72341 | 1 | ENSMUST00 | 0 | Fam117b | ENSMUSG0C | true | chr1 | 59912260 | 59915684 | 34.92026 | 124.22972 | 119.94103 | . | . | . | . | . | . |
| chr6 | 141118294 | 141118604 | 298 | Distal Intergenic | 6 | 141249269 | 141505940 | 256672 | 1 | ENSMUST00 | -130665 | Pde3a | ENSMUSG0C | true | . | . | . | . | . | . | . | . | . | . | . | . |
| chr4 | 127076983 | 127077557 | 298 | Promoter (<=1kb) | 4 | 127077383 | 127124370 | 46988 | 1 | ENSMUST00 | 0 | Zmym6 | ENSMUSG0C | false | chr4 | 127076728 | 127077872 | 23.54255 | 69.40097 | 66.41734 | . | . | . | . | . | . |
| chr9 | 118939470 | 118939983 | 298 | Intron (ENSMUST00000073109.6 | 9 | 118929867 | 119033628 | 103762 | 1 | ENSMUST00 | 9604 | Ctdspl | ENSMUSG0C | true | . | . | . | . | . | . | chr9 | 118927678 | 119010843 | 6.89652 | 4.89961 | 4.57102 |
| chr3 | 51839179 | 51839711 | 298 | Intron (ENSMUST00000121440.2 | 3 | 51828912 | 51829028 | 117 | 1 | ENSMUST00 | 10268 | n-R5s196 | ENSMUSG0C | false | . | . | . | . | . | . | chr3 | 51831736 | 51847000 | 4.07316 | 3.59293 | 2.2664 |
| chr13 | 6193373 | 6193934 | 298 | Distal Intergenic | 13 | 5997455 | 5997551 | 97 | 2 | ENSMUST00 | -195823 | Gm26259 | ENSMUSG0C | false | . | . | . | . | . | . | chr13 | 6192501 | 6194786 | 5.1606 | 4.16876 | 3.13601 |
| chr1 | 64088055 | 64088550 | 298 | Intron (ENSMUST00000149820.1 | 1 | 64078654 | 64090782 | 12129 | 1 | ENSMUST00 | 9402 | Gm13748 | ENSMUSG0C | false | . | . | . | . | . | . | chr1 | 64074337 | 64110188 | 4.96577 | 4.0468 | 2.98015 |
| chrX | 123497870 | 123498317 | 297 | Distal Intergenic | 20 | 123483317 | 123483814 | 498 | 2 | ENSMUST00 | -14057 | Gm17412 | ENSMUSG0C | false | . | . | . | . | . | . | . | . | . | . | . | . |
| chr18 | 5186968 | 5187607 | 297 | Distal Intergenic | 18 | 5162880 | 5165731 | 2852 | 2 | ENSMUST00 | -21238 | Gm26682 | ENSMUSG0C | false | . | . | . | . | . | . | chr18 | 5183663 | 5187429 | 6.5798 | 4.77195 | 4.30572 |
| chr11 | 108342714 | 108343415 | 182 | Promoter (<=1kb) | 11 | 108343354 | 108395905 | 52552 | 1 | ENSMUST00 | 0 | Apoh | ENSMUSG0C | false | chr11 | 108342513 | 108344600 | 38.27806 | 124.82941 | 120.52397 | chr11 | 108341583 | 108342723 | 4.01394 | 3.62274 | 2.20918 |
| chr11 | 108268914 | 108269226 | 182 | Intron (ENSMUST00000059595.5 | 11 | 108343354 | 108395905 | 52552 | 1 | ENSMUST00 | -74128 | Apoh | ENSMUSG0C | false | . | . | . | . | . | . | chr11 | 108267109 | 108271570 | 6.00913 | 4.36645 | 3.86724 |
| chr4 | 120405284 | 120405788 | 182 | Promoter (<=1kb) | 4 | 120405307 | 120530184 | 124878 | 1 | ENSMUST00 | 0 | Scmh1 | ENSMUSG0C | false | chr4 | 120404779 | 120406505 | 32.64849 | 123.19247 | 118.94151 | . | . | . | . | . | . |
| chr11 | 85832579 | 85833101 | 182 | Promoter (<=1kb) | 11 | 85832551 | 85841948 | 9398 | 1 | ENSMUST00 | 29 | Tbx2 | ENSMUSG0C | false | chr11 | 85832779 | 85833988 | 15.23487 | 37.30433 | 34.74706 | chr11 | 85829587 | 85833766 | 5.80049 | 4.32881 | 3.6947 |
| chr6 | 127150962 | 127151144 | 182 | Promoter (<=1kb) | 6 | 127125779 | 127151048 | 25270 | 2 | ENSMUST00 | 0 | Ccnd2 | ENSMUSG0C | false | chr6 | 127149169 | 127151101 | 16.34529 | 41.36735 | 38.7591 | chr6 | 127148760 | 127155682 | 5.84503 | 4.49672 | 3.70608 |
| chr13 | 45507308 | 45507571 | 182 | Promoter (<=1kb) | 13 | 45507444 | 45546382 | 38939 | 1 | ENSMUST00 | 0 | Gmpr | ENSMUSG0C | false | chr13 | 45507198 | 45508834 | 27.71015 | 85.00858 | 81.76643 | chr13 | 45506699 | 45509680 | 3.66321 | 3.42715 | 1.9571 |
| chr4 | 129556436 | 129556718 | 182 | Promoter (<=1kb) | 4 | 129555476 | 129556459 | 984 | 2 | ENSMUST00 | 0 | Lck | ENSMUSG0C | false | chr4 | 129555572 | 129557250 | 15.77044 | 42.5731 | 39.95169 | chr4 | 129555580 | 129557321 | 6.84933 | 4.95909 | 4.53576 |
| chr11 | 60352790 | 60353246 | 182 | Promoter (<=1kb) | 11 | 60226714 | 60352905 | 126192 | 2 | ENSMUST00 | 0 | Tom1l2 | ENSMUSG0C | false | chr11 | 60351504 | 60353003 | 23.93078 | 87.47927 | 84.18233 | . | . | . | . | . | . |
| chr3 | 105864201 | 105864391 | 182 | Distal Intergenic | 3 | 105870858 | 105908353 | 37496 | 1 | ENSMUST00 | -6467 | Adora3 | ENSMUSG0C | false | . | . | . | . | . | . | . | . | . | . | . | . |
| chr11 | 77766811 | 77767140 | 182 | Promoter (<=1kb) | 11 | 77766736 | 77865980 | 99245 | 1 | ENSMUST00 | 76 | Myo18a | ENSMUSG0C | false | chr11 | 77766313 | 77767112 | 13.57127 | 32.39957 | 29.9044 | chr11 | 77762694 | 77767023 | 4.9493 | 3.85185 | 2.99435 |
| chr11 | 77781254 | 77781498 | 182 | Intron (ENSMUST00000108375.3 | 11 | 77777316 | 77864848 | 87533 | 1 | ENSMUST00 | 3939 | Myo18a | ENSMUSG0C | false | . | . | . | . | . | . | chr11 | 77780682 | 77787654 | 5.96582 | 4.26257 | 3.83545 |
| chr11 | 77515031 | 77515290 | 182 | Promoter (<=1kb) | 11 | 77515121 | 77520045 | 4925 | 1 | ENSMUST00 | 0 | Abhd15 | ENSMUSG0C | false | chr11 | 77514569 | 77516040 | 23.47016 | 67.50422 | 64.55198 | chr11 | 77512407 | 77515867 | 5.13544 | 4.05533 | 3.12826 |
| chr17 | 53655133 | 53655401 | 182 | Intron (ENSMUST00000000724.9 | 17 | 53659238 | 53663649 | 4412 | 1 | ENSMUST00 | -3837 | Kat2b | ENSMUSG0C | false | . | . | . | . | . | . | chr17 | 53652574 | 53657149 | 4.63259 | 3.86546 | 2.72831 |
| chr11 | 75268133 | 75268333 | 182 | Distal Intergenic | 11 | 75301305 | 75342188 | 40884 | 2 | ENSMUST00 | 73855 | Rpa1 | ENSMUSG0C | false | . | . | . | . | . | . | . | . | . | . | . | . |
| chrX | 13280338 | 13280599 | 182 | Promoter (<=1kb) | 20 | 13280970 | 13294052 | 13083 | 1 | ENSMUST00 | -371 | Ddx3x | ENSMUSG0C | false | chrX | 13280005 | 13282849 | 28.03146 | 79.91027 | 76.76085 | . | . | . | . | . | . |
| chr6 | 52190988 | 52191255 | 182 | Promoter (<=1kb) | 6 | 52189671 | 52191753 | 2083 | 2 | ENSMUST00 | 498 | Hoxa4 | ENSMUSG0C | false | chr6 | 52190914 | 52191800 | 29.59164 | 87.32004 | 84.03093 | chr6 | 52188874 | 52193165 | 4.2446 | 3.64299 | 2.41132 |
| chr14 | 54432283 | 54432552 | 182 | Promoter (<=1kb) | 14 | 54431604 | 54441258 | 9655 | 1 | ENSMUST00 | 680 | Mmp14 | ENSMUSG0C | false | chr14 | 54431085 | 54433050 | 26.19333 | 73.09374 | 70.054 | chr14 | 54430319 | 54434204 | 4.2111 | 3.75246 | 2.34682 |
| chr3 | 90509043 | 90509236 | 182 | Promoter (<=1kb) | 3 | 90503131 | 90509022 | 5892 | 2 | ENSMUST00 | -22 | Chtop | ENSMUSG0C | false | chr3 | 90508236 | 90509564 | 39.54591 | 131.3782 | 126.83877 | . | . | . | . | . | . |
| chr11 | 106387727 | 106388005 | 182 | Promoter (<=1kb) | 11 | 106378806 | 106388072 | 9267 | 2 | ENSMUST00 | 67 | Icam2 | ENSMUSG0C | false | . | . | . | . | . | . | . | . | . | . | . | . |
| chr11 | 106382612 | 106382904 | 182 | Promoter (<=1kb) | 11 | 106377656 | 106382781 | 5126 | 2 | ENSMUST00 | 0 | Icam2 | ENSMUSG0C | false | . | . | . | . | . | . | . | . | . | . | . | . |
| chr11 | 61615995 | 61616317 | 182 | Distal Intergenic | 11 | 61517250 | 61579687 | 62438 | 2 | ENSMUST00 | -36309 | Epn2 | ENSMUSG0C | true | . | . | . | . | . | . | chr11 | 61610870 | 61621664 | 5.06575 | 4.05211 | 3.07721 |
| chr4 | 136469960 | 136470661 | 182 | Promoter (<=1kb) | 4 | 136469850 | 136540667 | 70818 | 1 | ENSMUST00 | 111 | Luzp1 | ENSMUSG0C | false | chr4 | 136469495 | 136471700 | 34.47702 | 107.70252 | 103.93674 | . | . | . | . | . | . |
| chr4 | 136469034 | 136469242 | 182 | Promoter (<=1kb) | 4 | 136469761 | 136549315 | 79555 | 1 | ENSMUST00 | -519 | Luzp1 | ENSMUSG0C | false | . | . | . | . | . | . | chr4 | 136468830 | 136469667 | 4.36581 | 3.81648 | 2.4931 |
| chr10 | 76961883 | 76962288 | 182 | Promoter (<=1kb) | 10 | 76761860 | 76961887 | 200028 | 2 | ENSMUST00 | 0 | Pcbp3 | ENSMUSG0C | false | chr10 | 76961345 | 76962056 | 20.06521 | 63.98471 | 61.08233 | . | . | . | . | . | . |
| chrX | 20848640 | 20848928 | 182 | Promoter (<=1kb) | 20 | 20848589 | 20853450 | 4862 | 1 | ENSMUST00 | 52 | Araf | ENSMUSG0C | true | chrX | 20848525 | 20849511 | 17.92175 | 44.05972 | 41.41886 | . | . | . | . | . | . |
| chrX | 20961930 | 20962184 | 182 | Promoter (<=1kb) | 20 | 20941722 | 20961991 | 20270 | 2 | ENSMUST00 | 0 | Uxt | ENSMUSG0C | false | chrX | 20961230 | 20962646 | 14.70503 | 33.66069 | 31.14909 | . | . | . | . | . | . |
| chr12 | 100199266 | 100199559 | 182 | Promoter (<=1kb) | 12 | 100199435 | 100209806 | 10372 | 1 | ENSMUST00 | 0 | Calm1 | ENSMUSG0C | false | chr12 | 100198919 | 100201212 | 21.48562 | 91.50835 | 88.13527 | . | . | . | . | . | . |
| chr10 | 78297823 | 78298335 | 182 | Promoter (<=1kb) | 10 | 78271563 | 78298198 | 26636 | 2 | ENSMUST00 | 0 | Agpat3 | ENSMUSG0C | false | . | . | . | . | . | . | chr10 | 78294895 | 78301060 | 6.73612 | 4.81929 | 4.42476 |
| chr17 | 56218758 | 56218967 | 182 | Promoter (<=1kb) | 17 | 56186682 | 56218889 | 32208 | 2 | ENSMUST00 | 0 | Dpp9 | ENSMUSG0C | false | chr17 | 56217823 | 56219558 | 28.12215 | 95.42207 | 91.9593 | chr17 | 56211598 | 56220063 | 6.93474 | 4.88891 | 4.6356 |
| chr15 | 102399528 | 102399894 | 182 | Distal Intergenic | 15 | 102406143 | 102408235 | 2093 | 1 | ENSMUST00 | -6249 | Sp1 | ENSMUSG0C | false | chr15 | 102399060 | 102400347 | 15.47449 | 37.71435 | 35.15295 | chr15 | 102396804 | 102401498 | 5.3749 | 4.27518 | 3.30533 |
| chr15 | 102326045 | 102326904 | 182 | Promoter (<=1kb) | 15 | 102326116 | 102331873 | 5758 | 1 | ENSMUST00 | 0 | Pfdn5 | ENSMUSG0C | false | chr15 | 102325753 | 102327357 | 28.48738 | 97.07422 | 93.57343 | . | . | . | . | . | . |
| chr11 | 101465002 | 101465331 | 182 | Promoter (<=1kb) | 11 | 101464999 | 101471853 | 6855 | 1 | ENSMUST00 | 4 | Rnd2 | ENSMUSG0C | false | chr11 | 101464825 | 101466732 | 41.77201 | 145.26822 | 140.17451 | chr11 | 101463714 | 101469649 | 4.51869 | 3.86017 | 2.61475 |
| chr4 | 129148397 | 129148627 | 182 | 3' UTR | 4 | 129136999 | 129144593 | 7595 | 1 | ENSMUST00 | 11399 | Fndc5 | ENSMUSG0C | false | . | . | . | . | . | . | chr4 | 129141119 | 129150343 | 5.31606 | 4.18214 | 3.25946 |
| chr2 | 164769572 | 164769950 | 182 | Promoter (<=1kb) | 2 | 164769898 | 164772907 | 3010 | 1 | ENSMUST00 | 0 | Ube2c | ENSMUSG0C | false | chr2 | 164769302 | 164770309 | 26.40393 | 78.95839 | 75.82329 | chr2 | 164769297 | 164770373 | 4.81591 | 3.98955 | 2.89752 |
| chr3 | 88150520 | 88150786 | 182 | Intron (ENSMUST00000001455.7 | 3 | 88142574 | 88156003 | 13430 | 1 | ENSMUST00 | 7947 | Mef2d | ENSMUSG0C | false | . | . | . | . | . | . | chr3 | 88143086 | 88154835 | 7.43337 | 5.03451 | 5.02118 |
| chr3 | 88139132 | 88139557 | 182 | Promoter (2-3kb) | 3 | 88142372 | 88172086 | 29715 | 1 | ENSMUST00 | -2815 | Mef2d | ENSMUSG0C | false | chr3 | 88138472 | 88139892 | 22.69969 | 67.91293 | 64.95291 | chr3 | 88136354 | 88141917 | 6.47517 | 4.6068 | 4.24981 |
| chr6 | 28479892 | 28480394 | 182 | Promoter (<=1kb) | 6 | 28480348 | 28877723 | 397376 | 1 | ENSMUST00 | 0 | Snd1 | ENSMUSG0C | false | chr6 | 28479479 | 28480083 | 24.84317 | 70.77231 | 67.77174 | . | . | . | . | . | . |
| chr6 | 28848876 | 28849095 | 182 | Promoter (<=1kb) | 6 | 28848907 | 28880308 | 31402 | 1 | ENSMUST00 | 0 | Snd1 | ENSMUSG0C | false | . | . | . | . | . | . | chr6 | 28838725 | 28849885 | 5.40567 | 4.29116 | 3.35017 |
| chr6 | 28705515 | 28705733 | 182 | Promoter (<=1kb) | 6 | 28705720 | 28724981 | 19262 | 1 | ENSMUST00 | 0 | Snd1 | ENSMUSG0C | false | . | . | . | . | . | . | chr6 | 28705021 | 28711323 | 5.33556 | 4.27576 | 3.27874 |
| chr10 | 77109284 | 77109484 | 182 | Intron (ENSMUST00000081654.7 | 10 | 77052181 | 77113676 | 61496 | 2 | ENSMUST00 | 4192 | Col18a1 | ENSMUSG0C | false | . | . | . | . | . | . | chr10 | 77096587 | 77112087 | 5.00583 | 4.02872 | 3.02421 |
| chr8 | 123373520 | 123373937 | 182 | Promoter (<=1kb) | 8 | 123373824 | 123401892 | 28069 | 1 | ENSMUST00 | 0 | Tcf25 | ENSMUSG0C | false | chr8 | 123373094 | 123374909 | 40.23391 | 129.30441 | 124.84723 | . | . | . | . | . | . |
| chr11 | 101891517 | 101891778 | 182 | Promoter (2-3kb) | 11 | 101877510 | 101894374 | 16865 | 2 | ENSMUST00 | 2596 | Meox1 | ENSMUSG0C | false | . | . | . | . | . | . | chr11 | 101891400 | 101892029 | 4.24544 | 3.69674 | 2.41449 |
| chr11 | 94937390 | 94937722 | 182 | Promoter (<=1kb) | 11 | 94937686 | 94938103 | 418 | 1 | ENSMUST00 | 0 | Col1a1 | ENSMUSG0C | false | chr11 | 94937285 | 94937862 | 11.02877 | 22.59171 | 20.22099 | chr11 | 94936490 | 94938939 | 5.5761 | 4.23463 | 3.50064 |
| chr11 | 95088883 | 95089171 | 182 | Distal Intergenic | 11 | 95044474 | 95076801 | 32328 | 2 | ENSMUST00 | -12083 | Itga3 | ENSMUSG0C | true | . | . | . | . | . | . | chr11 | 95083817 | 95090459 | 4.77961 | 3.94636 | 2.81919 |
| chr6 | 128344396 | 128344594 | 182 | Intron (ENSMUST00000001562.6 | 6 | 128334477 | 128355815 | 21339 | 2 | ENSMUST00 | 11221 | Tulp3 | ENSMUSG0C | false | . | . | . | . | . | . | chr6 | 128343079 | 128355029 | 3.93648 | 3.48258 | 2.16297 |
| chr11 | 100396071 | 100396608 | 182 | Promoter (1-2kb) | 11 | 100370619 | 100397749 | 27131 | 2 | ENSMUST00 | 1141 | Jup | ENSMUSG0C | false | chr11 | 100394893 | 100397863 | 34.54477 | 122.48211 | 118.25522 | chr11 | 100384756 | 100396489 | 9.22555 | 5.81405 | 6.48761 |
| chr11 | 100415423 | 100415778 | 182 | Promoter (<=1kb) | 11 | 100415697 | 100424824 | 9128 | 1 | ENSMUST00 | 0 | Fkbp10 | ENSMUSG0C | false | . | . | . | . | . | . | chr11 | 100413310 | 100415965 | 3.87681 | 3.54621 | 2.06891 |
| chr12 | 40223156 | 40223537 | 182 | Promoter (<=1kb) | 12 | 40201567 | 40223189 | 21623 | 2 | ENSMUST00 | 0 | Ifrd1 | ENSMUSG0C | false | chr12 | 40223472 | 40224030 | 12.39106 | 30.348 | 27.8772 | . | . | . | . | . | . |
| chr15 | 102990449 | 102990988 | 182 | Promoter (<=1kb) | 15 | 102990607 | 102993821 | 3215 | 1 | ENSMUST00 | 0 | Hoxc8 | ENSMUSG0C | false | chr15 | 102990385 | 102993111 | 31.24818 | 92.1069 | 88.71768 | chr15 | 102986473 | 102990693 | 4.48121 | 3.82974 | 2.59026 |
| chr15 | 102994287 | 102994564 | 182 | Promoter (2-3kb) | 15 | 102991889 | 102994111 | 2223 | 1 | ENSMUST00 | 2399 | Hoxc8 | ENSMUSG0C | false | . | . | . | . | . | . | chr15 | 102993412 | 102997788 | 5.35742 | 4.3108 | 3.28377 |
| chr5 | 148608325 | 148608529 | 182 | Distal Intergenic | 5 | 148505142 | 148552789 | 47648 | 2 | ENSMUST00 | -55537 | Ubl3 | ENSMUSG0C | false | . | . | . | . | . | . | chr5 | 148607994 | 148609062 | 4.78501 | 3.84002 | 2.86292 |
| chr11 | 101058785 | 101059010 | 182 | Intron (ENSMUST00000103110.4 | 11 | 101070012 | 101077672 | 7661 | 1 | ENSMUST00 | -11002 | Naglu | ENSMUSG0C | false | . | . | . | . | . | . | chr11 | 101054317 | 101067609 | 9.20102 | 5.81671 | 6.45648 |
| chr11 | 101064051 | 101064312 | 182 | Distal Intergenic | 11 | 101070012 | 101077672 | 7661 | 1 | ENSMUST00 | -5700 | Naglu | ENSMUSG0C | false | . | . | . | . | . | . | chr11 | 101054317 | 101067609 | 9.20102 | 5.81671 | 6.45648 |
| chr11 | 101069962 | 101070215 | 182 | Promoter (<=1kb) | 11 | 101070012 | 101077672 | 7661 | 1 | ENSMUST00 | 0 | Naglu | ENSMUSG0C | false | chr11 | 101069577 | 101070879 | 35.20524 | 115.20684 | 111.21615 | chr11 | 101069392 | 101071435 | 4.97623 | 4.06431 | 3.03269 |
| chr9 | 18292037 | 18292286 | 182 | Promoter (<=1kb) | 9 | 18292267 | 18314000 | 21734 | 1 | ENSMUST00 | 0 | Chordc1 | ENSMUSG0C | false | chr9 | 18291639 | 18293431 | 17.23216 | 63.31984 | 60.4264 | . | . | . | . | . | . |
| chr7 | 30193811 | 30194259 | 182 | Promoter (<=1kb) | 7 | 30190049 | 30194150 | 4102 | 2 | ENSMUST00 | 0 | Capns1 | ENSMUSG0C | false | chr7 | 30193627 | 30195415 | 21.20881 | 84.61981 | 81.38317 | . | . | . | . | . | . |
| chr5 | 143329142 | 143329342 | 182 | Promoter (<=1kb) | 5 | 143316489 | 143329256 | 12768 | 2 | ENSMUST00 | 0 | Zdhhc4 | ENSMUSG0C | false | chr5 | 143328488 | 143330078 | 24.78633 | 81.65401 | 78.47337 | chr5 | 143327803 | 143330070 | 4.42362 | 3.65759 | 2.55829 |
| chr2 | 127006010 | 127006268 | 182 | Promoter (2-3kb) | 2 | 127008717 | 127037344 | 28628 | 1 | ENSMUST00 | -2449 | Ap4e1 | ENSMUSG0C | false | . | . | . | . | . | . | chr2 | 127005073 | 127010104 | 5.17638 | 4.0967 | 3.18696 |
| chr11 | 78245593 | 78246147 | 182 | Promoter (<=1kb) | 11 | 78206746 | 78245987 | 39242 | 2 | ENSMUST00 | 0 | Supt6 | ENSMUSG0C | false | chr11 | 78244811 | 78246806 | 28.59991 | 96.29427 | 92.81311 | . | . | . | . | . | . |
| chr7 | 38107347 | 38107848 | 182 | Promoter (<=1kb) | 7 | 38097984 | 38107534 | 9551 | 2 | ENSMUST00 | 0 | Ccne1 | ENSMUSG0C | false | chr7 | 38107709 | 38108320 | 18.95815 | 63.55374 | 60.65727 | chr7 | 38106772 | 38108265 | 4.90599 | 4.01118 | 2.98211 |
| chr7 | 16313622 | 16314311 | 182 | Promoter (<=1kb) | 7 | 16313517 | 16317697 | 4181 | 1 | ENSMUST00 | 106 | Bbc3 | ENSMUSG0C | false | chr7 | 16313234 | 16314485 | 34.81759 | 111.39526 | 107.52535 | chr7 | 16309137 | 16315495 | 5.75225 | 4.42325 | 3.62489 |
| chr2 | 91194974 | 91195304 | 182 | Promoter (<=1kb) | 2 | 91184061 | 91195025 | 10965 | 2 | ENSMUST00 | 0 | Nr1h3 | ENSMUSG0C | false | chr2 | 91194699 | 91195264 | 10.47775 | 22.22469 | 19.86031 | . | . | . | . | . | . |
| chr3 | 104788602 | 104789092 | 182 | Promoter (<=1kb) | 3 | 104789011 | 104794459 | 5449 | 1 | ENSMUST00 | 0 | Rhoc | ENSMUSG0C | false | chr3 | 104788434 | 104790298 | 29.54326 | 90.13423 | 86.79193 | chr3 | 104788261 | 104791231 | 5.41717 | 3.99308 | 3.38207 |
| chr17 | 28254942 | 28255156 | 182 | Intron (ENSMUST00000002320.9 | 17 | 28271708 | 28273163 | 1456 | 1 | ENSMUST00 | -16552 | Ppard | ENSMUSG0C | false | . | . | . | . | . | . | chr17 | 28251377 | 28258644 | 5.73452 | 4.39083 | 3.62025 |
| chr14 | 55635516 | 55635808 | 182 | Promoter (<=1kb) | 14 | 55625400 | 55635684 | 10285 | 2 | ENSMUST00 | 0 | Ipo4 | ENSMUSG0C | false | chr14 | 55634877 | 55636107 | 30.81035 | 92.32966 | 88.93555 | chr14 | 55634537 | 55635978 | 4.96921 | 4.1047 | 2.98005 |
| chr14 | 55745575 | 55745816 | 182 | Promoter (<=1kb) | 14 | 55739020 | 55745599 | 6580 | 2 | ENSMUST00 | 0 | Dhrs1 | ENSMUSG0C | false | chr14 | 55745042 | 55746841 | 36.76537 | 115.82222 | 111.81207 | chr14 | 55743738 | 55747345 | 6.12391 | 4.46048 | 3.96696 |
| chr17 | 24645016 | 24645303 | 182 | Promoter (<=1kb) | 17 | 24639282 | 24644957 | 5676 | 2 | ENSMUST00 | -60 | Slc9a3r2 | ENSMUSG0C | false | chr17 | 24642925 | 24645980 | 11.05033 | 23.18759 | 20.81059 | chr17 | 24630730 | 24649025 | 8.70874 | 5.59711 | 6.05272 |
| chr15 | 76080536 | 76081014 | 182 | Promoter (<=1kb) | 15 | 76070184 | 76080946 | 10763 | 2 | ENSMUST00 | 0 | Puf60 | ENSMUSG0C | false | chr15 | 76079693 | 76081549 | 22.68762 | 81.35053 | 78.1749 | . | . | . | . | . | . |
| chr2 | 32288206 | 32288426 | 182 | Promoter (<=1kb) | 2 | 32288253 | 32307921 | 19669 | 1 | ENSMUST00 | 0 | Golga2 | ENSMUSG0C | false | chr2 | 32287238 | 32289480 | 31.80086 | 99.99902 | 96.43139 | . | . | . | . | . | . |
| chr7 | 25707544 | 25708382 | 182 | Downstream (2-3kb) | 7 | 25712935 | 25716106 | 3172 | 2 | ENSMUST00 | 7724 | Ccdc97 | ENSMUSG0C | false | . | . | . | . | . | . | chr7 | 25704659 | 25712354 | 4.6492 | 3.76801 | 2.75693 |
| chr12 | 50647591 | 50647995 | 182 | Promoter (1-2kb) | 12 | 50341231 | 50649223 | 307993 | 2 | ENSMUST00 | 1228 | Prkd1 | ENSMUSG0C | false | chr12 | 50647570 | 50649686 | 30.41442 | 103.5799 | 99.9162 | chr12 | 50645591 | 50649902 | 3.80439 | 3.39333 | 2.04575 |
| chr2 | 76647958 | 76648538 | 182 | Promoter (<=1kb) | 2 | 76629898 | 76648015 | 18118 | 2 | ENSMUST00 | 0 | Prkra | ENSMUSG0C | false | chr2 | 76646657 | 76648968 | 31.85236 | 101.09525 | 97.496 | . | . | . | . | . | . |
| chr5 | 135186710 | 135187349 | 182 | Promoter (<=1kb) | 5 | 135187264 | 135246129 | 58866 | 1 | ENSMUST00 | 0 | Baz1b | ENSMUSG0C | false | chr5 | 135186308 | 135188508 | 35.20524 | 115.20684 | 111.21615 | . | . | . | . | . | . |
| chr7 | 45866019 | 45866720 | 182 | Promoter (<=1kb) | 7 | 45831883 | 45867129 | 35247 | 2 | ENSMUST00 | 409 | Grin2d | ENSMUSG0C | false | chr7 | 45865939 | 45866990 | 26.65955 | 86.95457 | 83.67183 | chr7 | 45866137 | 45866981 | 4.54507 | 3.8993 | 2.64606 |
| chr11 | 60727014 | 60727444 | 182 | Promoter (<=1kb) | 11 | 60714143 | 60727263 | 13121 | 2 | ENSMUST00 | 0 | Flii | ENSMUSG0C | false | chr11 | 60726256 | 60729003 | 34.63892 | 109.01322 | 105.20635 | . | . | . | . | . | . |
| chr6 | 88898726 | 88898932 | 182 | Promoter (<=1kb) | 6 | 88883475 | 88898780 | 15306 | 2 | ENSMUST00 | 0 | Mcm2 | ENSMUSG0C | false | chr6 | 88898170 | 88898753 | 18.02173 | 47.27326 | 44.5908 | chr6 | 88898158 | 88907962 | 5.34589 | 4.21337 | 3.29833 |
| chr12 | 31265111 | 31265391 | 182 | Promoter (<=1kb) | 12 | 31265234 | 31329493 | 64260 | 1 | ENSMUST00 | 0 | Lamb1 | ENSMUSG0C | false | chr12 | 31264923 | 31266287 | 17.53869 | 44.15654 | 41.51399 | chr12 | 31264572 | 31269502 | 3.99338 | 3.54192 | 2.21735 |
| chr8 | 4275865 | 4276089 | 182 | Promoter (<=1kb) | 8 | 4259731 | 4275885 | 16155 | 2 | ENSMUST00 | 0 | Timm44 | ENSMUSG0C | false | chr8 | 4275117 | 4276486 | 29.36448 | 91.77025 | 88.3935 | chr8 | 4273499 | 4276634 | 4.70508 | 3.96379 | 2.74805 |
| chr7 | 141562011 | 141562427 | 182 | Promoter (<=1kb) | 7 | 141562180 | 141633011 | 70832 | 1 | ENSMUST00 | 0 | Ap2a2 | ENSMUSG0C | false | chr7 | 141561658 | 141563249 | 30.94804 | 114.76607 | 110.79114 | . | . | . | . | . | . |
| chr7 | 19628473 | 19629017 | 182 | Promoter (<=1kb) | 7 | 19616375 | 19628784 | 12410 | 2 | ENSMUST00 | 0 | Relb | ENSMUSG0C | false | chr7 | 19628045 | 19629963 | 39.02359 | 127.20569 | 122.81842 | . | . | . | . | . | . |
| chr12 | 31950275 | 31950812 | 182 | Promoter (<=1kb) | 12 | 31926254 | 31950535 | 24282 | 2 | ENSMUST00 | 0 | Hbp1 | ENSMUSG0C | false | chr12 | 31948692 | 31950317 | 42.96212 | 142.86137 | 137.8616 | . | . | . | . | . | . |
| chr10 | 80142195 | 80142442 | 182 | Promoter (<=1kb) | 10 | 80142315 | 80145818 | 3504 | 1 | ENSMUST00 | 0 | Atp5d | ENSMUSG0C | false | chr10 | 80141816 | 80143104 | 25.7338 | 71.40817 | 68.39619 | chr10 | 80140751 | 80148113 | 6.54123 | 4.83987 | 4.26386 |
| chr9 | 45984440 | 45984995 | 182 | Promoter (<=1kb) | 9 | 45965311 | 45984871 | 19561 | 2 | ENSMUST00 | 0 | Pafah1b2 | ENSMUSG0C | false | chr9 | 45983749 | 45985474 | 26.41709 | 100.00125 | 96.43182 | chr9 | 45976663 | 45985562 | 4.05405 | 3.63163 | 2.24436 |
| chr1 | 39474586 | 39474797 | 182 | Intron (ENSMUST00000054462.5 | 1 | 39371499 | 39478747 | 107249 | 2 | ENSMUST00 | 3950 | Tbc1d8 | ENSMUSG0C | false | . | . | . | . | . | . | chr1 | 39474278 | 39475091 | 3.28618 | 3.13702 | 1.62804 |
| chr10 | 58446699 | 58446963 | 182 | Promoter (<=1kb) | 10 | 58446852 | 58494155 | 47304 | 1 | ENSMUST00 | 0 | Ranbp2 | ENSMUSG0C | false | chr10 | 58446089 | 58447838 | 41.81742 | 134.22485 | 129.57179 | . | . | . | . | . | . |
| chr16 | 20498757 | 20499066 | 182 | Promoter (<=1kb) | 16 | 20498817 | 20509955 | 11139 | 1 | ENSMUST00 | 0 | Eif2b5 | ENSMUSG0C | false | chr16 | 20498071 | 20499669 | 25.48472 | 95.2657 | 91.8051 | chr16 | 20497033 | 20500365 | 3.92641 | 3.37638 | 2.18038 |
| chr7 | 45700598 | 45700854 | 182 | Promoter (<=1kb) | 7 | 45701016 | 45703477 | 2462 | 1 | ENSMUST00 | -162 | Car11 | ENSMUSG0C | false | chr7 | 45699729 | 45701036 | 16.66176 | 41.17759 | 38.57329 | chr7 | 45699898 | 45701543 | 5.44459 | 4.36442 | 3.34685 |
| chr10 | 80701812 | 80702382 | 182 | Promoter (<=1kb) | 10 | 80685256 | 80701820 | 16565 | 2 | ENSMUST00 | 0 | Mob3a | ENSMUSG0C | false | chr10 | 80701379 | 80702000 | 28.96167 | 93.99567 | 90.56499 | . | . | . | . | . | . |
| chr15 | 98662778 | 98663066 | 182 | Promoter (<=1kb) | 15 | 98645134 | 98662889 | 17756 | 2 | ENSMUST00 | 0 | Ddx23 | ENSMUSG0C | false | chr15 | 98662100 | 98663226 | 34.14626 | 110.2223 | 106.38018 | . | . | . | . | . | . |
| chr7 | 25072560 | 25073028 | 182 | Promoter (<=1kb) | 7 | 25009853 | 25072369 | 62517 | 2 | ENSMUST00 | -192 | Grik5 | ENSMUSG0C | false | chr7 | 25072030 | 25073077 | 22.97661 | 61.46103 | 58.5941 | chr7 | 25071953 | 25073636 | 4.1809 | 3.61076 | 2.3354 |
| chr7 | 28311913 | 28312194 | 182 | Promoter (<=1kb) | 7 | 28305516 | 28312072 | 6557 | 2 | ENSMUST00 | 0 | Timm50 | ENSMUSG0C | false | chr7 | 28311136 | 28312482 | 42.74922 | 148.11885 | 142.90198 | chr7 | 28310602 | 28312618 | 3.61305 | 3.31346 | 1.90409 |
| chr14 | 70474171 | 70474411 | 182 | Downstream (<1kb) | 14 | 70458523 | 70463340 | 4818 | 1 | ENSMUST00 | 15649 | Phyhip | ENSMUSG0C | false | . | . | . | . | . | . | chr14 | 70470535 | 70475401 | 5.45533 | 4.17788 | 3.42554 |
| chr17 | 35701436 | 35701675 | 182 | Promoter (<=1kb) | 17 | 35685306 | 35701964 | 16659 | 2 | ENSMUST00 | 289 | Ddr1 | ENSMUSG0C | false | chr17 | 35699302 | 35701573 | 21.75525 | 76.8157 | 73.7174 | chr17 | 35696430 | 35706756 | 5.35332 | 4.19448 | 3.30476 |
| chr7 | 19327105 | 19327388 | 182 | Distal Intergenic | 7 | 19302721 | 19310045 | 7325 | 2 | ENSMUST00 | -17061 | Fosb | ENSMUSG0C | false | chr7 | 19326495 | 19327541 | 9.28568 | 20.18442 | 17.84641 | chr7 | 19326443 | 19328062 | 5.82901 | 4.29291 | 3.72172 |
| chr8 | 70282712 | 70283081 | 182 | Promoter (<=1kb) | 8 | 70282827 | 70286530 | 3704 | 1 | ENSMUST00 | 0 | Homer3 | ENSMUSG0C | false | chr8 | 70282389 | 70283850 | 29.28796 | 98.53321 | 94.99628 | . | . | . | . | . | . |
| chr5 | 8996942 | 8997502 | 182 | Promoter (<=1kb) | 5 | 8966033 | 8997324 | 31292 | 2 | ENSMUST00 | 0 | Crot | ENSMUSG0C | false | chr5 | 8996006 | 8997859 | 25.37642 | 79.93365 | 76.78373 | . | . | . | . | . | . |
| chr4 | 129672441 | 129672985 | 182 | Promoter (<=1kb) | 4 | 129643980 | 129672767 | 28788 | 2 | ENSMUST00 | 0 | Kpna6 | ENSMUSG0C | false | chr4 | 129671587 | 129673549 | 41.4024 | 145.73851 | 140.61644 | . | . | . | . | . | . |
| chr11 | 90390702 | 90391090 | 182 | Promoter (<=1kb) | 11 | 90336536 | 90390895 | 54360 | 2 | ENSMUST00 | 0 | Hlf | ENSMUSG0C | false | chr11 | 90388817 | 90391517 | 18.02336 | 51.19578 | 48.46415 | chr11 | 90388455 | 90392196 | 4.31586 | 3.75682 | 2.45428 |
| chr11 | 90382452 | 90382690 | 182 | Promoter (<=1kb) | 11 | 90340827 | 90381935 | 41109 | 2 | ENSMUST00 | -518 | Hlf | ENSMUSG0C | false | . | . | . | . | . | . | chr11 | 90381657 | 90384253 | 3.82277 | 3.52622 | 2.04408 |
| chr13 | 91520485 | 91520712 | 182 | Intron (ENSMUST00000156547.2 | 13 | 91539343 | 91657776 | 118434 | 1 | ENSMUST00 | -18631 | Ssbp2 | ENSMUSG0C | true | . | . | . | . | . | . | chr13 | 91517753 | 91523504 | 4.36033 | 3.64539 | 2.50015 |
| chr11 | 100951058 | 100951294 | 182 | Distal Intergenic | 11 | 100886810 | 100939540 | 52731 | 2 | ENSMUST00 | -11519 | Stat3 | ENSMUSG0C | false | . | . | . | . | . | . | chr11 | 100950390 | 100955692 | 5.1087 | 4.1694 | 3.09166 |
| chr11 | 100912466 | 100912655 | 182 | Intron (ENSMUST00000103114.2 | 11 | 100905656 | 100908224 | 2569 | 2 | ENSMUST00 | -4243 | Stat3 | ENSMUSG0C | false | . | . | . | . | . | . | chr11 | 100911718 | 100915085 | 6.71838 | 4.88236 | 4.39897 |
| chr7 | 27591392 | 27592097 | 182 | Promoter (<=1kb) | 7 | 27591552 | 27640826 | 49275 | 1 | ENSMUST00 | 0 | Akt2 | ENSMUSG0C | false | chr7 | 27591673 | 27593253 | 29.58769 | 108.11006 | 104.32592 | . | . | . | . | . | . |
| chr2 | 72311721 | 72312120 | 182 | Intron (ENSMUST00000090824.6 | 2 | 72297901 | 72355622 | 57722 | 1 | ENSMUST00 | 13821 | Zak | ENSMUSG0C | false | . | . | . | . | . | . | chr2 | 72307524 | 72323444 | 4.91051 | 3.9112 | 2.97144 |
| chr6 | 124759353 | 124759541 | 182 | Promoter (2-3kb) | 6 | 124742544 | 124756524 | 13981 | 2 | ENSMUST00 | -2830 | Atn1 | ENSMUSG0C | false | . | . | . | . | . | . | chr6 | 124757014 | 124759681 | 6.18502 | 4.6504 | 3.98377 |
| chr6 | 29467627 | 29467843 | 182 | Promoter (<=1kb) | 6 | 29467718 | 29470512 | 2795 | 1 | ENSMUST00 | 0 | Atp6v1f | ENSMUSG0C | false | chr6 | 29467299 | 29468653 | 24.72749 | 72.3696 | 69.3433 | . | . | . | . | . | . |
| chr6 | 52714119 | 52714532 | 182 | Promoter (<=1kb) | 6 | 52714219 | 52733255 | 19037 | 1 | ENSMUST00 | 0 | Tax1bp1 | ENSMUSG0C | false | chr6 | 52713330 | 52714872 | 27.7554 | 114.54269 | 110.5744 | . | . | . | . | . | . |
| chr8 | 3500328 | 3500625 | 182 | Promoter (<=1kb) | 8 | 3500457 | 3515232 | 14776 | 1 | ENSMUST00 | 0 | Mcoln1 | ENSMUSG0C | false | chr8 | 3499872 | 3501326 | 29.63113 | 94.2009 | 90.76292 | chr8 | 3499617 | 3504551 | 7.30944 | 5.14934 | 4.90367 |
| chr7 | 43443362 | 43443720 | 182 | Promoter (<=1kb) | 7 | 43444108 | 43457800 | 13693 | 1 | ENSMUST00 | -388 | Etfb | ENSMUSG0C | false | chr7 | 43442913 | 43444736 | 24.35436 | 72.61947 | 69.58835 | chr7 | 43442747 | 43443447 | 4.29835 | 3.62205 | 2.45094 |
| chr6 | 55402503 | 55402776 | 182 | Distal Intergenic | 6 | 55376295 | 55388530 | 12236 | 1 | ENSMUST00 | 26209 | Ghrhr | ENSMUSG0C | false | . | . | . | . | . | . | . | . | . | . | . | . |
| chr1 | 33719850 | 33720245 | 182 | Promoter (<=1kb) | 1 | 33719887 | 33726891 | 7005 | 1 | ENSMUST00 | 0 | Rab23 | ENSMUSG0C | false | chr1 | 33719716 | 33720760 | 25.15761 | 80.47005 | 77.31023 | chr1 | 33719673 | 33721214 | 3.87673 | 3.57454 | 2.06485 |
| chr12 | 85205192 | 85205398 | 182 | Distal Intergenic | 12 | 85219481 | 85226628 | 7148 | 1 | ENSMUST00 | -14083 | Eif2b2 | ENSMUSG0C | false | . | . | . | . | . | . | . | . | . | . | . | . |
| chr11 | 61620381 | 61620679 | 182 | Distal Intergenic | 11 | 61653265 | 61672784 | 19520 | 1 | ENSMUST00 | -32586 | Grap | ENSMUSG0C | false | chr11 | 61618794 | 61620874 | 12.76342 | 27.83059 | 25.39249 | chr11 | 61610870 | 61621664 | 5.06575 | 4.05211 | 3.07721 |
| chr3 | 87930120 | 87930601 | 182 | Promoter (<=1kb) | 3 | 87922601 | 87930127 | 7527 | 2 | ENSMUST00 | 0 | Rrnad1 | ENSMUSG0C | false | chr3 | 87929380 | 87931180 | 34.56782 | 111.9988 | 108.10535 | . | . | . | . | . | . |
| chr9 | 64253150 | 64253766 | 182 | Promoter (<=1kb) | 9 | 64185770 | 64253631 | 67862 | 2 | ENSMUST00 | 0 | Map2k1 | ENSMUSG0C | false | chr9 | 64252324 | 64253690 | 33.7938 | 119.00099 | 114.88904 | . | . | . | . | . | . |
| chr12 | 72085117 | 72086109 | 182 | Promoter (<=1kb) | 12 | 72085589 | 72094867 | 9279 | 1 | ENSMUST00 | 0 | Jkamp | ENSMUSG0C | false | chr12 | 72085706 | 72087055 | 34.3275 | 112.61823 | 108.70669 | chr12 | 72082188 | 72085236 | 3.38963 | 3.17266 | 1.71248 |
| chr15 | 66961589 | 66961905 | 182 | Intron (ENSMUST00000005256.7 | 15 | 66943079 | 66967925 | 24847 | 2 | ENSMUST00 | 6020 | Ndrg1 | ENSMUSG0C | false | . | . | . | . | . | . | chr15 | 66959859 | 66970307 | 5.98016 | 4.46549 | 3.81257 |
| chr11 | 69681907 | 69682254 | 182 | Promoter (<=1kb) | 11 | 69673121 | 69682084 | 8964 | 2 | ENSMUST00 | 0 | Senp3 | ENSMUSG0C | false | chr11 | 69681078 | 69682590 | 32.44658 | 100.11784 | 96.54536 | chr11 | 69680256 | 69687418 | 8.8727 | 5.82026 | 6.17215 |
| chr17 | 88004917 | 88005113 | 182 | Intron (ENSMUST00000130379.2 | 17 | 87991418 | 88008696 | 17279 | 2 | ENSMUST00 | 3583 | Fbxo11 | ENSMUSG0C | false | . | . | . | . | . | . | chr17 | 88003553 | 88005754 | 4.30766 | 3.67106 | 2.48556 |
| chr13 | 13504017 | 13504279 | 182 | Intron (ENSMUST00000005532.7 | 13 | 13437602 | 13512275 | 74674 | 1 | ENSMUST00 | 66416 | Nid1 | ENSMUSG0C | false | . | . | . | . | . | . | . | . | . | . | . | . |
| chr11 | 59662227 | 59662753 | 182 | Promoter (<=1kb) | 11 | 59662495 | 59780860 | 118366 | 1 | ENSMUST00 | 0 | Mprip | ENSMUSG0C | false | chr11 | 59661712 | 59664872 | 16.78262 | 53.63071 | 50.86883 | . | . | . | . | . | . |
| chr7 | 25268302 | 25268625 | 182 | Promoter (<=1kb) | 7 | 25268387 | 25271272 | 2886 | 1 | ENSMUST00 | 0 | Cic | ENSMUSG0C | false | chr7 | 25268350 | 25269389 | 32.41473 | 98.06107 | 94.53979 | . | . | . | . | . | . |
| chr1 | 88008286 | 88008546 | 182 | Promoter (<=1kb) | 1 | 87946443 | 88008534 | 62092 | 2 | ENSMUST00 | 0 | Usp40 | ENSMUSG0C | false | chr1 | 88007807 | 88008854 | 33.08631 | 99.21803 | 95.66874 | chr1 | 88007290 | 88009151 | 3.67577 | 3.41295 | 1.9038 |
| chr7 | 67952935 | 67953409 | 182 | Promoter (<=1kb) | 7 | 67952859 | 68226780 | 273922 | 1 | ENSMUST00 | 77 | Igf1r | ENSMUSG0C | false | chr7 | 67950399 | 67954090 | 38.55812 | 129.64772 | 125.16568 | chr7 | 67953281 | 67983240 | 5.79319 | 4.4206 | 3.65863 |
| chr7 | 67968524 | 67968808 | 182 | Intron (ENSMUST00000005671.8 | 7 | 67952859 | 68226780 | 273922 | 1 | ENSMUST00 | 15666 | Igf1r | ENSMUSG0C | false | . | . | . | . | . | . | chr7 | 67953281 | 67983240 | 5.79319 | 4.4206 | 3.65863 |
| chr8 | 3279437 | 3279739 | 182 | Promoter (<=1kb) | 8 | 3150922 | 3279617 | 128696 | 2 | ENSMUST00 | 0 | Insr | ENSMUSG0C | false | chr8 | 3278295 | 3280203 | 31.95414 | 97.26555 | 93.76166 | . | . | . | . | . | . |
| chr7 | 111122076 | 111122542 | 182 | Distal Intergenic | 7 | 111070568 | 111083030 | 12463 | 2 | ENSMUST00 | -39047 | Eif4g2 | ENSMUSG0C | false | chr7 | 111121782 | 111123426 | 26.85422 | 77.8849 | 74.7682 | . | . | . | . | . | . |
| chr7 | 110941984 | 110942282 | 182 | Intron (ENSMUST00000005751.7 | 7 | 110868272 | 110946187 | 77916 | 2 | ENSMUST00 | 3905 | Mrvi1 | ENSMUSG0C | false | . | . | . | . | . | . | chr7 | 110941590 | 110942876 | 3.83617 | 3.51836 | 2.09872 |
| chr7 | 110982372 | 110982700 | 182 | Promoter (<=1kb) | 7 | 110868270 | 110982461 | 114192 | 2 | ENSMUST00 | 0 | Mrvi1 | ENSMUSG0C | false | . | . | . | . | . | . | chr7 | 110982186 | 110991297 | 5.7251 | 4.32045 | 3.62307 |
| chr7 | 80997780 | 80998030 | 182 | Intron (ENSMUST00000107353.2 | 7 | 80993681 | 81045164 | 51484 | 1 | ENSMUST00 | 4100 | Zfp592 | ENSMUSG0C | false | . | . | . | . | . | . | chr7 | 80994427 | 81011007 | 5.53832 | 4.32158 | 3.44803 |
| chr1 | 165836545 | 165836843 | 182 | Intron (ENSMUST00000161971.2 | 1 | 165858216 | 165861555 | 3340 | 1 | ENSMUST00 | -21373 | Cd247 | ENSMUSG0C | false | . | . | . | . | . | . | chr1 | 165833325 | 165839586 | 4.76919 | 3.8939 | 2.83602 |
| chr2 | 155882953 | 155883162 | 182 | Intron (ENSMUST00000136933.2 | 2 | 155887119 | 155910604 | 23486 | 2 | ENSMUST00 | 27442 | Uqcc1 | ENSMUSG0C | false | . | . | . | . | . | . | chr2 | 155868791 | 155890222 | 5.22346 | 4.19096 | 3.19144 |
| chr2 | 155918114 | 155918510 | 182 | Intron (ENSMUST00000136933.2 | 2 | 155887119 | 155910604 | 23486 | 2 | ENSMUST00 | -7511 | Uqcc1 | ENSMUSG0C | false | . | . | . | . | . | . | chr2 | 155914466 | 155919478 | 5.32661 | 4.17258 | 3.29527 |
| chr2 | 155863118 | 155863343 | 182 | Intron (ENSMUST00000136933.2 | 2 | 155848184 | 155851692 | 3509 | 2 | ENSMUST00 | -11427 | Uqcc1 | ENSMUSG0C | false | . | . | . | . | . | . | chr2 | 155860492 | 155867272 | 5.81598 | 4.2461 | 3.69487 |
| chr17 | 28952203 | 28952766 | 182 | Promoter (<=1kb) | 17 | 28952436 | 28969549 | 17114 | 1 | ENSMUST00 | 0 | Kctd20 | ENSMUSG0C | false | chr17 | 28951668 | 28953760 | 28.45235 | 93.08366 | 89.67602 | . | . | . | . | . | . |
| chr19 | 4283074 | 4283376 | 182 | Promoter (<=1kb) | 19 | 4270180 | 4283135 | 12956 | 2 | ENSMUST00 | 0 | Ankrd13d | ENSMUSG0C | false | chr19 | 4282703 | 4283507 | 25.18016 | 100.40004 | 96.81754 | chr19 | 4282579 | 4283963 | 5.64794 | 4.40429 | 3.53247 |
| chr7 | 16119882 | 16120141 | 182 | Promoter (<=1kb) | 7 | 16119895 | 16121524 | 1630 | 1 | ENSMUST00 | 0 | Kptn | ENSMUSG0C | false | chr7 | 16119437 | 16120524 | 28.91859 | 89.0447 | 85.71584 | chr7 | 16119319 | 16120963 | 4.04092 | 3.69245 | 2.2039 |
| chr16 | 17451779 | 17452140 | 182 | Promoter (<=1kb) | 16 | 17451987 | 17486255 | 34269 | 1 | ENSMUST00 | 0 | Crkl | ENSMUSG0C | false | chr16 | 17451315 | 17453886 | 34.00884 | 130.01486 | 125.52227 | . | . | . | . | . | . |
| chr5 | 136051597 | 136052026 | 182 | Promoter (2-3kb) | 5 | 136054492 | 136064324 | 9833 | 1 | ENSMUST00 | -2466 | Upk3bl | ENSMUSG0C | false | . | . | . | . | . | . | chr5 | 136050355 | 136053748 | 6.35695 | 4.76018 | 4.11724 |
| chr13 | 29859249 | 29859591 | 182 | Distal Intergenic | 13 | 29325301 | 29855674 | 530374 | 2 | ENSMUST00 | -3576 | Cdkal1 | ENSMUSG0C | false | . | . | . | . | . | . | . | . | . | . | . | . |
| chr13 | 29582011 | 29582340 | 182 | Intron (ENSMUST00000140278.2 | 13 | 29443776 | 29517454 | 73679 | 2 | ENSMUST00 | -64558 | Cdkal1 | ENSMUSG0C | false | . | . | . | . | . | . | chr13 | 29581720 | 29583474 | 6.32492 | 4.6438 | 4.1397 |
| chr7 | 130949593 | 130949817 | 182 | Intron (ENSMUST00000150717.2 | 7 | 130943489 | 130985660 | 42172 | 1 | ENSMUST00 | 6105 | Htra1 | ENSMUSG0C | false | . | . | . | . | . | . | chr7 | 130934926 | 130953774 | 6.08548 | 4.53495 | 3.90785 |
| chr9 | 21927349 | 21927762 | 182 | Promoter (<=1kb) | 9 | 21927471 | 21935872 | 8402 | 1 | ENSMUST00 | 0 | Ccdc159 | ENSMUSG0C | false | chr9 | 21926813 | 21927406 | 14.75456 | 35.78426 | 33.24598 | chr9 | 21926014 | 21928120 | 4.74443 | 3.70188 | 2.84119 |
| chr5 | 88720721 | 88721231 | 182 | Promoter (<=1kb) | 5 | 88720855 | 88747350 | 26496 | 1 | ENSMUST00 | 0 | Mob1b | ENSMUSG0C | false | chr5 | 88720019 | 88722250 | 31.38972 | 125.99339 | 121.64841 | . | . | . | . | . | . |
| chr14 | 50785341 | 50785642 | 182 | Promoter (<=1kb) | 14 | 50765415 | 50785518 | 20104 | 2 | ENSMUST00 | 0 | Ttc5 | ENSMUSG0C | false | chr14 | 50784823 | 50785681 | 31.62822 | 118.12274 | 114.03909 | chr14 | 50783693 | 50785856 | 3.59414 | 3.28458 | 1.88913 |
| chr8 | 122662170 | 122662528 | 182 | Intron (ENSMUST00000127664.1 | 8 | 122629463 | 122678072 | 48610 | 2 | ENSMUST00 | 15544 | Cbfa2t3 | ENSMUSG0C | false | . | . | . | . | . | . | chr8 | 122648562 | 122676943 | 7.87697 | 5.30313 | 5.37575 |
| chr8 | 122639766 | 122639984 | 182 | Intron (ENSMUST00000127664.1 | 8 | 122639018 | 122646466 | 7449 | 2 | ENSMUST00 | 6482 | Cbfa2t3 | ENSMUSG0C | false | . | . | . | . | . | . | chr8 | 122635146 | 122641135 | 6.79662 | 4.86734 | 4.50965 |
| chr8 | 122677091 | 122677343 | 182 | Promoter (<=1kb) | 8 | 122629463 | 122678072 | 48610 | 2 | ENSMUST00 | 729 | Cbfa2t3 | ENSMUSG0C | false | chr8 | 122676506 | 122679041 | 35.62435 | 116.99458 | 112.94973 | . | . | . | . | . | . |
| chr15 | 79030528 | 79030925 | 182 | Promoter (<=1kb) | 15 | 79030874 | 79038348 | 7475 | 1 | ENSMUST00 | 0 | Gcat | ENSMUSG0C | false | chr15 | 79027416 | 79030756 | 27.03504 | 88.5911 | 85.27629 | chr15 | 79028839 | 79031311 | 5.33413 | 4.20473 | 3.28922 |
| chr4 | 141301932 | 141302385 | 182 | Promoter (<=1kb) | 4 | 141302407 | 141308734 | 6328 | 1 | ENSMUST00 | -22 | Epha2 | ENSMUSG0C | false | chr4 | 141301143 | 141303243 | 41.2653 | 138.78937 | 133.96024 | . | . | . | . | . | . |
| chr9 | 108490539 | 108490935 | 182 | Promoter (<=1kb) | 9 | 108490676 | 108501697 | 11022 | 1 | ENSMUST00 | 0 | Usp19 | ENSMUSG0C | false | chr9 | 108490636 | 108491934 | 29.92778 | 109.76675 | 105.9399 | . | . | . | . | . | . |
| chr4 | 137367910 | 137368136 | 182 | Distal Intergenic | 4 | 137319696 | 137357720 | 38025 | 2 | ENSMUST00 | -10191 | Cdc42 | ENSMUSG0C | false | . | . | . | . | . | . | chr4 | 137367417 | 137376426 | 4.18122 | 3.57464 | 2.35005 |
| chr2 | 166112987 | 166113227 | 182 | Intron (ENSMUST00000109249.3 | 2 | 166074815 | 166078119 | 3305 | 2 | ENSMUST00 | -34869 | Sulf2 | ENSMUSG0C | false | . | . | . | . | . | . | chr2 | 166111370 | 166117038 | 4.57128 | 3.82866 | 2.65452 |
| chr6 | 83569146 | 83569387 | 182 | Intron (ENSMUST00000068054.7 | 6 | 83543206 | 83572504 | 29299 | 2 | ENSMUST00 | 3117 | Stambp | ENSMUSG0C | false | . | . | . | . | . | . | chr6 | 83566781 | 83573087 | 4.86122 | 3.78814 | 2.93222 |
| chr11 | 100408872 | 100409114 | 182 | Promoter (2-3kb) | 11 | 100409366 | 100411874 | 2509 | 2 | ENSMUST00 | 2760 | Leprel4 | ENSMUSG0C | false | . | . | . | . | . | . | chr11 | 100407887 | 100409947 | 5.38575 | 4.3145 | 3.31572 |
| chr9 | 120933330 | 120933612 | 182 | Promoter (<=1kb) | 9 | 120933400 | 120960507 | 27108 | 1 | ENSMUST00 | 0 | Ctnnb1 | ENSMUSG0C | false | chr9 | 120933357 | 120935851 | 27.83859 | 82.38849 | 79.1904 | . | . | . | . | . | . |
| chr9 | 120963205 | 120963425 | 182 | Intron (ENSMUST00000170406.2 | 9 | 120956867 | 120957824 | 958 | 1 | ENSMUST00 | 6339 | Ctnnb1 | ENSMUSG0C | false | . | . | . | . | . | . | chr9 | 120957401 | 120965230 | 7.3009 | 5.00727 | 4.91838 |
| chr9 | 120934347 | 120934634 | 182 | Promoter (<=1kb) | 9 | 120933818 | 120951293 | 17476 | 1 | ENSMUST00 | 530 | Ctnnb1 | ENSMUSG0C | false | chr9 | 120933357 | 120935851 | 27.83859 | 82.38849 | 79.1904 | chr9 | 120934463 | 120950300 | 5.02026 | 3.91783 | 3.06038 |
| chr17 | 34931180 | 34931468 | 182 | Promoter (<=1kb) | 17 | 34931253 | 34935921 | 4669 | 1 | ENSMUST00 | 0 | Neu1 | ENSMUSG0C | false | chr17 | 34931249 | 34932414 | 25.90299 | 79.94019 | 76.78951 | chr17 | 34930771 | 34935446 | 5.8546 | 4.41264 | 3.721 |
| chr17 | 28313302 | 28313538 | 182 | Promoter (<=1kb) | 17 | 28313530 | 28326567 | 13038 | 1 | ENSMUST00 | 0 | Fance | ENSMUSG0C | false | chr17 | 28313508 | 28314368 | 38.25637 | 125.86897 | 121.53396 | chr17 | 28312915 | 28315421 | 4.27987 | 3.68258 | 2.44436 |
| chr2 | 152830492 | 152830763 | 182 | Promoter (<=1kb) | 2 | 152780672 | 152830666 | 49995 | 2 | ENSMUST00 | 0 | Bcl2l1 | ENSMUSG0C | false | chr2 | 152829469 | 152831763 | 41.35843 | 143.41548 | 138.39708 | . | . | . | . | . | . |
| chr2 | 152816301 | 152816495 | 182 | Intron (ENSMUST00000007803.6 | 2 | 152781411 | 152828535 | 47125 | 2 | ENSMUST00 | 12040 | Bcl2l1 | ENSMUSG0C | false | . | . | . | . | . | . | chr2 | 152808253 | 152830242 | 5.83553 | 4.3234 | 3.71708 |
| chr2 | 152831221 | 152831703 | 182 | Promoter (<=1kb) | 2 | 152829757 | 152831710 | 1954 | 2 | ENSMUST00 | 7 | Bcl2l1 | ENSMUSG0C | false | chr2 | 152829469 | 152831763 | 41.35843 | 143.41548 | 138.39708 | . | . | . | . | . | . |
| chr17 | 57031017 | 57031749 | 182 | Promoter (<=1kb) | 17 | 57021058 | 57031507 | 10450 | 2 | ENSMUST00 | 0 | Khsrp | ENSMUSG0C | false | chr17 | 57030151 | 57031593 | 31.76828 | 95.95177 | 92.47445 | chr17 | 57028736 | 57031130 | 5.44389 | 4.01519 | 3.40772 |
| chr11 | 50378411 | 50378630 | 182 | Promoter (<=1kb) | 11 | 50378301 | 50385923 | 7623 | 1 | ENSMUST00 | 111 | Hnrnph1 | ENSMUSG0C | false | chr11 | 50376642 | 50379439 | 45.10679 | 157.97711 | 152.39525 | . | . | . | . | . | . |
| chr4 | 136140411 | 136140727 | 182 | Promoter (2-3kb) | 4 | 136143497 | 136145392 | 1896 | 1 | ENSMUST00 | -2770 | Id3 | ENSMUSG0C | false | chr4 | 136140348 | 136140740 | 3.70442 | 6.08744 | 4.00582 | chr4 | 136139844 | 136141145 | 4.32057 | 3.79388 | 2.4373 |
| chr4 | 133729570 | 133729861 | 182 | Intron (ENSMUST00000105897.4 | 4 | 133720779 | 133723879 | 3101 | 2 | ENSMUST00 | -5692 | Arid1a | ENSMUSG0C | false | . | . | . | . | . | . | chr4 | 133723677 | 133736326 | 5.40007 | 4.1324 | 3.3715 |
| chr7 | 142387881 | 142388076 | 182 | Promoter (<=1kb) | 7 | 142375910 | 142388038 | 12129 | 2 | ENSMUST00 | 0 | Ctsd | ENSMUSG0C | false | chr7 | 142386717 | 142388441 | 33.35327 | 102.69651 | 99.06359 | chr7 | 142385787 | 142388278 | 4.39298 | 3.77139 | 2.51009 |
| chr18 | 56562429 | 56562645 | 182 | Promoter (<=1kb) | 18 | 56562443 | 56587189 | 24747 | 1 | ENSMUST00 | 0 | Phax | ENSMUSG0C | false | chr18 | 56561767 | 56563208 | 18.02336 | 51.19578 | 48.46415 | chr18 | 56561700 | 56563622 | 3.55744 | 3.31936 | 1.86304 |
| chr3 | 100705394 | 100705709 | 182 | Distal Intergenic | 3 | 100562208 | 100685503 | 123296 | 2 | ENSMUST00 | -19892 | Man1a2 | ENSMUSG0C | true | chr3 | 100705348 | 100707194 | 6.80177 | 13.42905 | 11.19262 | . | . | . | . | . | . |
| chr12 | 85288366 | 85288708 | 182 | Promoter (<=1kb) | 12 | 85272547 | 85288438 | 15892 | 2 | ENSMUST00 | 0 | Acyp1 | ENSMUSG0C | false | chr12 | 85287825 | 85289333 | 19.06563 | 48.28789 | 45.59428 | chr12 | 85287483 | 85292663 | 4.73802 | 4.00512 | 2.78959 |
| chr11 | 102222033 | 102222314 | 182 | Intron (ENSMUST00000107151.3 | 11 | 102207031 | 102218929 | 11899 | 2 | ENSMUST00 | -3105 | Hdac5 | ENSMUSG0C | false | . | . | . | . | . | . | chr11 | 102213449 | 102231135 | 7.77054 | 5.31127 | 5.26338 |
| chr11 | 102229906 | 102230344 | 182 | Promoter (<=1kb) | 11 | 102194432 | 102230166 | 35735 | 2 | ENSMUST00 | 0 | Hdac5 | ENSMUSG0C | false | chr11 | 102229308 | 102230763 | 28.56742 | 103.16996 | 99.51957 | chr11 | 102213449 | 102231135 | 7.77054 | 5.31127 | 5.26338 |
| chr2 | 172954321 | 172954535 | 182 | Distal Intergenic | 2 | 172868012 | 172940321 | 72310 | 2 | ENSMUST00 | -14001 | Bmp7 | ENSMUSG0C | false | . | . | . | . | . | . | chr2 | 172953818 | 172955166 | 5.16523 | 4.16875 | 3.13356 |
| chr2 | 172906198 | 172906460 | 182 | Intron (ENSMUST00000009143.7 | 2 | 172916118 | 172940092 | 23975 | 2 | ENSMUST00 | 33632 | Bmp7 | ENSMUSG0C | false | . | . | . | . | . | . | chr2 | 172904237 | 172908799 | 6.43181 | 4.74985 | 4.16523 |
| chr5 | 115300895 | 115301339 | 182 | Promoter (<=1kb) | 5 | 115297110 | 115300999 | 3890 | 2 | ENSMUST00 | 0 | Dynll1 | ENSMUSG0C | false | chr5 | 115299554 | 115300971 | 33.47463 | 112.14942 | 108.24687 | . | . | . | . | . | . |
| chr10 | 77581565 | 77582127 | 182 | Promoter (<=1kb) | 10 | 77581720 | 77598732 | 17013 | 1 | ENSMUST00 | 0 | Pttg1ip | ENSMUSG0C | false | chr10 | 77581193 | 77581597 | 9.65017 | 18.71579 | 16.39739 | chr10 | 77580829 | 77594032 | 8.74443 | 5.72701 | 6.06549 |
| chr13 | 98890286 | 98890960 | 182 | Promoter (<=1kb) | 13 | 98849223 | 98890998 | 41776 | 2 | ENSMUST00 | 38 | Tnpo1 | ENSMUSG0C | false | chr13 | 98889912 | 98891282 | 27.24215 | 82.29425 | 79.10038 | chr13 | 98887643 | 98898885 | 4.47967 | 3.8014 | 2.58678 |
| chr13 | 98926213 | 98926817 | 182 | Promoter (<=1kb) | 13 | 98839019 | 98926384 | 87366 | 2 | ENSMUST00 | 0 | Tnpo1 | ENSMUSG0C | false | chr13 | 98924893 | 98927111 | 50.0437 | 183.17024 | 176.94609 | . | . | . | . | . | . |
| chr2 | 118479635 | 118479911 | 182 | Promoter (<=1kb) | 2 | 118475850 | 118479711 | 3862 | 2 | ENSMUST00 | 0 | Srp14 | ENSMUSG0C | false | chr2 | 118478601 | 118480175 | 25.48472 | 95.2657 | 91.8051 | . | . | . | . | . | . |
| chr7 | 31033872 | 31034080 | 182 | 3' UTR | 7 | 31032732 | 31039046 | 6315 | 2 | ENSMUST00 | 4966 | Fxyd5 | ENSMUSG0C | false | . | . | . | . | . | . | . | . | . | . | . | . |
| chr15 | 100569420 | 100569719 | 182 | Distal Intergenic | 15 | 100575321 | 100580231 | 4911 | 2 | ENSMUST00 | 10512 | Pou6f1 | ENSMUSG0C | false | . | . | . | . | . | . | chr15 | 100568426 | 100570546 | 4.40625 | 3.79398 | 2.53259 |

| | | | | | | | | | | | | | | | | | | | | | | | | | | |
|---|---|---|---|---|---|---|---|---|---|---|---|---|---|---|---|---|---|---|---|---|---|---|---|---|---|---|
| chr1 | 132326851 | 132327077 | 182 | Promoter (<=1kb) | 1 | 132327596 | 132330450 | 2855 | 1 | ENSMUST00 | -519 | Nuak2 | ENSMUSG0( | false | . | . | . | . | . | . | chr1 | 132317064 | 132339308 | 6.74463 | 4.86917 | 4.43585 |
| chr4 | 140961190 | 140961433 | 182 | Promoter (<=1kb) | 4 | 140961203 | 140979193 | 17991 | 1 | ENSMUST00 | 0 | Sdhb | ENSMUSG0( | false | chr4 | 140961260 | 140962243 | 41.75335 | 144.11185 | 139.06203 | chr4 | 140958427 | 140964559 | 4.93225 | 3.97851 | 2.99313 |
| chr9 | 107533858 | 107534142 | 182 | Promoter (<=1kb) | 9 | 107533945 | 107538656 | 4712 | 1 | ENSMUST00 | 0 | Tmem115 | ENSMUSG0( | false | chr9 | 107533968 | 107536510 | 25.84672 | 90.2271 | 86.88143 | chr9 | 107533484 | 107537392 | 6.33705 | 4.57243 | 4.13006 |
| chr9 | 107555190 | 107555448 | 182 | Promoter (<=1kb) | 9 | 107554656 | 107562263 | 7608 | 1 | ENSMUST00 | 535 | Rassf1 | ENSMUSG0( | false | chr9 | 107554606 | 107556283 | 26.52689 | 93.43019 | 90.01563 | chr9 | 107555352 | 107558233 | 5.76852 | 4.26742 | 3.65541 |
| chr9 | 107551379 | 107551663 | 182 | Promoter (<=1kb) | 9 | 107551555 | 107562263 | 10709 | 1 | ENSMUST00 | 0 | Rassf1 | ENSMUSG0( | false | chr9 | 107551259 | 107552030 | 16.34529 | 41.36735 | 38.7591 | chr9 | 107550899 | 107554394 | 4.41992 | 3.73757 | 2.53811 |
| chr9 | 107554340 | 107554663 | 182 | Promoter (<=1kb) | 9 | 107554656 | 107562263 | 7608 | 1 | ENSMUST00 | 0 | Rassf1 | ENSMUSG0( | false | chr9 | 107553604 | 107554526 | 16.01923 | 40.07766 | 37.48569 | chr9 | 107550899 | 107554394 | 4.41992 | 3.73757 | 2.53811 |
| chr11 | 94499638 | 94500104 | 182 | Promoter (<=1kb) | 11 | 94489599 | 94499974 | 10376 | 2 | ENSMUST00 | 0 | Epn3 | ENSMUSG0( | false | chr11 | 94495773 | 94500513 | 28.91859 | 89.0447 | 85.71584 | . | . | . | . | . | . |
| chr11 | 61437547 | 61437782 | 182 | Distal Intergenic | 11 | 61452263 | 61453919 | 1657 | 2 | ENSMUST00 | 16137 | Rnf112 | ENSMUSG0( | false | . | . | . | . | . | . | . | . | . | . | . | . |
| chr3 | 88296543 | 88296935 | 182 | Promoter (<=1kb) | 3 | 88282760 | 88296838 | 14079 | 2 | ENSMUST00 | 0 | Tsacc | ENSMUSG0( | false | chr3 | 88296175 | 88296929 | 15.59995 | 42.26899 | 39.65329 | . | . | . | . | . | . |
| chr12 | 83632018 | 83632484 | 182 | Promoter (<=1kb) | 12 | 83632252 | 83683123 | 50872 | 1 | ENSMUST00 | 0 | Rbm25 | ENSMUSG0( | false | chr12 | 83631427 | 83632065 | 12.9904 | 42.26624 | 39.65078 | . | . | . | . | . | . |
| chr1 | 180256224 | 180256469 | 182 | Promoter (<=1kb) | 1 | 180227004 | 180256300 | 29297 | 2 | ENSMUST00 | 0 | Psen2 | ENSMUSG0( | false | chr1 | 180255386 | 180256935 | 32.3084 | 121.498 | 117.30614 | . | . | . | . | . | . |
| chr19 | 10176916 | 10177161 | 182 | Distal Intergenic | 19 | 10182888 | 10196870 | 13983 | 1 | ENSMUST00 | -5727 | Fads1 | ENSMUSG0( | false | . | . | . | . | . | . | chr19 | 10176590 | 10178442 | 4.55282 | 3.90486 | 2.63119 |
| chr11 | 46064947 | 46065351 | 182 | Exon (ENSMUST00000011400.7/ | 11 | 46056058 | 46065082 | 9025 | 1 | ENSMUST00 | 8890 | Adam19 | ENSMUSG0( | false | . | . | . | . | . | . | . | . | . | . | . | . |
| chr5 | 107868902 | 107869423 | 182 | Promoter (<=1kb) | 5 | 107788287 | 107869168 | 80882 | 2 | ENSMUST00 | 0 | Evi5 | ENSMUSG0( | false | chr5 | 107868027 | 107869684 | 40.31728 | 129.44856 | 124.98527 | . | . | . | . | . | . |
| chr5 | 114690829 | 114691217 | 182 | Promoter (<=1kb) | 5 | 114669403 | 114690984 | 21582 | 2 | ENSMUST00 | 0 | Gltp | ENSMUSG0( | false | chr5 | 114690140 | 114691487 | 30.59503 | 95.89513 | 92.42037 | chr5 | 114691207 | 114691573 | 5.64782 | 3.78134 | 3.57611 |
| chr14 | 18270755 | 18271281 | 182 | Promoter (<=1kb) | 14 | 18267823 | 18271067 | 3245 | 2 | ENSMUST00 | 0 | Rpl15 | ENSMUSG0( | false | chr14 | 18269737 | 18271010 | 28.53699 | 95.19626 | 91.7378 | . | . | . | . | . | . |
| chr6 | 29575450 | 29575665 | 182 | Intron (ENSMUST00000115251.2 | 6 | 29578552 | 29582386 | 3835 | 2 | ENSMUST00 | 6721 | Tnpo3 | ENSMUSG0( | false | . | . | . | . | . | . | chr6 | 29575429 | 29576117 | 3.39767 | 3.2867 | 1.67169 |
| chr17 | 56474906 | 56475111 | 182 | Promoter (1-2kb) | 17 | 56422237 | 56476369 | 54133 | 2 | ENSMUST00 | 1258 | Ptprs | ENSMUSG0( | false | chr17 | 56474360 | 56476713 | 21.20328 | 55.92731 | 53.13585 | chr17 | 56472349 | 56476690 | 7.48518 | 5.21435 | 5.04173 |
| chr17 | 56467500 | 56467803 | 182 | Intron (ENSMUST00000067538.5 | 17 | 56412431 | 56462216 | 49786 | 2 | ENSMUST00 | -5285 | Ptprs | ENSMUSG0( | false | . | . | . | . | . | . | chr17 | 56466795 | 56471317 | 4.55921 | 3.79034 | 2.64724 |
| chr9 | 121759751 | 121760601 | 182 | Promoter (<=1kb) | 9 | 121760033 | 121771742 | 11710 | 1 | ENSMUST00 | 0 | Zfp651 | ENSMUSG0( | false | chr9 | 121759121 | 121761008 | 32.73014 | 99.60502 | 96.04542 | chr9 | 121758798 | 121762224 | 4.25498 | 3.62486 | 2.42707 |
| chr2 | 25211239 | 25211550 | 182 | Promoter (<=1kb) | 2 | 25199712 | 25211489 | 11778 | 2 | ENSMUST00 | 0 | Nelfb | ENSMUSG0( | false | chr2 | 25210115 | 25211701 | 31.85236 | 101.09525 | 97.496 | chr2 | 25207650 | 25212040 | 4.56251 | 3.59642 | 2.68585 |
| chr11 | 58354455 | 58354806 | 182 | Distal Intergenic | 11 | 58345312 | 58346950 | 1639 | 1 | ENSMUST00 | 9144 | Sh3bp5l | ENSMUSG0( | false | . | . | . | . | . | . | chr11 | 58354418 | 58355825 | 4.28083 | 3.77774 | 2.45675 |
| chr15 | 67130556 | 67130773 | 182 | Distal Intergenic | 15 | 67102875 | 67113992 | 11118 | 2 | ENSMUST00 | -16565 | St3gal1 | ENSMUSG0( | true | . | . | . | . | . | . | chr15 | 67124446 | 67141856 | 4.85469 | 4.00506 | 2.89973 |
| chr10 | 79984101 | 79984717 | 182 | Promoter (<=1kb) | 10 | 79977118 | 79984330 | 7213 | 2 | ENSMUST00 | 0 | Tmem259 | ENSMUSG0( | false | chr10 | 79983154 | 79984959 | 30.81035 | 92.32966 | 88.93555 | . | . | . | . | . | . |
| chr13 | 58066525 | 58066867 | 182 | Intron (ENSMUST00000160860.2 | 13 | 58004957 | 58102428 | 97472 | 2 | ENSMUST00 | 35561 | Klhl3 | ENSMUSG0( | false | . | . | . | . | . | . | chr13 | 58064550 | 58070843 | 5.88033 | 4.46811 | 3.7345 |
| chr16 | 4606345 | 4606778 | 182 | Promoter (1-2kb) | 16 | 4608627 | 4615942 | 7316 | 1 | ENSMUST00 | -1849 | Glis2 | ENSMUSG0( | false | . | . | . | . | . | . | chr16 | 4591906 | 4609359 | 7.50331 | 5.19962 | 5.05636 |
| chr10 | 70966684 | 70966960 | 182 | Intron (ENSMUST00000143791.2 | 10 | 70934502 | 70948489 | 13988 | 2 | ENSMUST00 | -18196 | Bicc1 | ENSMUSG0( | false | chr10 | 70966931 | 70967191 | 4.42115 | 6.83029 | 4.72578 | chr10 | 70965109 | 70967880 | 5.96903 | 4.50713 | 3.8026 |
| chr2 | 128687312 | 128687537 | 182 | Promoter (<=1kb) | 2 | 128610104 | 128687391 | 77288 | 2 | ENSMUST00 | 0 | Anapc1 | ENSMUSG0( | false | chr2 | 128686446 | 128687786 | 33.41831 | 117.26761 | 113.21259 | chr2 | 128685327 | 128688035 | 4.1007 | 3.53217 | 2.28932 |
| chr2 | 128715444 | 128715948 | 182 | Intron (ENSMUST00000014505.4 | 2 | 128698956 | 128802894 | 103939 | 1 | ENSMUST00 | 16489 | Mertk | ENSMUSG0( | true | . | . | . | . | . | . | chr2 | 128701787 | 128720514 | 6.13295 | 4.54405 | 3.94372 |
| chr2 | 128698060 | 128698384 | 182 | Promoter (<=1kb) | 2 | 128698956 | 128802894 | 103939 | 1 | ENSMUST00 | -572 | Mertk | ENSMUSG0( | true | chr2 | 128697969 | 128703570 | 36.46257 | 120.58523 | 116.42456 | chr2 | 128691254 | 128698190 | 4.49449 | 3.79066 | 2.60905 |
| chr14 | 63368521 | 63368799 | 182 | Distal Intergenic | 14 | 63372837 | 63417187 | 44351 | 2 | ENSMUST00 | 48388 | Blk | ENSMUSG0( | false | . | . | . | . | . | . | chr14 | 63367809 | 63369426 | 5.08666 | 3.96887 | 3.1039 |
| chr4 | 151165448 | 151165752 | 182 | Intron (ENSMUST00000169423.3 | 4 | 151086039 | 151139584 | 53546 | 2 | ENSMUST00 | -25865 | Camta1 | ENSMUSG0( | false | . | . | . | . | . | . | chr4 | 151162762 | 151183610 | 6.13162 | 4.48987 | 3.9543 |
| chr4 | 151179097 | 151179385 | 182 | Intron (ENSMUST00000169423.3 | 4 | 151086039 | 151139584 | 53546 | 2 | ENSMUST00 | -39514 | Camta1 | ENSMUSG0( | false | . | . | . | . | . | . | chr4 | 151162762 | 151183610 | 6.13162 | 4.48987 | 3.9543 |
| chr3 | 107746018 | 107746257 | 182 | Intron (ENSMUST00000118593.2 | 3 | 107748542 | 107750741 | 2200 | 2 | ENSMUST00 | 4484 | Csf1 | ENSMUSG0( | false | . | . | . | . | . | . | chr3 | 107745803 | 107747270 | 4.62971 | 3.74017 | 2.73338 |
| chr13 | 92311925 | 92312218 | 182 | Promoter (2-3kb) | 13 | 92307497 | 92309457 | 1961 | 2 | ENSMUST00 | -2469 | Msh3 | ENSMUSG0( | false | . | . | . | . | . | . | chr13 | 92311818 | 92313740 | 5.03884 | 4.13883 | 3.05184 |
| chr2 | 26933407 | 26933764 | 182 | Promoter (<=1kb) | 2 | 26924970 | 26933781 | 8812 | 2 | ENSMUST00 | 17 | Surf4 | ENSMUSG0( | false | chr2 | 26932319 | 26934235 | 42.33011 | 146.25195 | 141.10883 | . | . | . | . | . | . |
| chr8 | 110997708 | 110997975 | 182 | Promoter (<=1kb) | 8 | 110974996 | 110997800 | 22805 | 2 | ENSMUST00 | 0 | Ddx19a | ENSMUSG0( | false | chr8 | 110997039 | 110998471 | 18.27513 | 72.99583 | 69.9605 | . | . | . | . | . | . |
| chr2 | 25455638 | 25455856 | 182 | Promoter (<=1kb) | 2 | 25456838 | 25458676 | 1839 | 1 | ENSMUST00 | -982 | Clic3 | ENSMUSG0( | true | chr2 | 25455822 | 25461073 | 24.56002 | 77.54259 | 74.4321 | chr2 | 25446553 | 25456355 | 6.09426 | 4.46882 | 3.92693 |
| chr7 | 66405577 | 66405852 | 182 | Promoter (<=1kb) | 7 | 66399895 | 66406002 | 6108 | 2 | ENSMUST00 | 150 | Aldh1a3 | ENSMUSG0( | false | . | . | . | . | . | . | . | . | . | . | . | . |
| chr12 | 80230088 | 80230344 | 182 | Intron (ENSMUST00000021554.9 | 12 | 80168322 | 80260091 | 91770 | 2 | ENSMUST00 | 29747 | Actn1 | ENSMUSG0( | true | . | . | . | . | . | . | chr12 | 80207669 | 80232669 | 6.01537 | 4.51837 | 3.84731 |
| chr12 | 80218226 | 80218502 | 182 | Intron (ENSMUST00000021554.9 | 12 | 80168322 | 80260091 | 91770 | 2 | ENSMUST00 | 41589 | Actn1 | ENSMUSG0( | true | . | . | . | . | . | . | chr12 | 80207669 | 80232669 | 6.01537 | 4.51837 | 3.84731 |
| chr19 | 46075814 | 46076052 | 182 | Promoter (<=1kb) | 19 | 46075863 | 46085528 | 9666 | 1 | ENSMUST00 | 0 | Nolc1 | ENSMUSG0( | false | chr19 | 46075397 | 46076916 | 27.74132 | 80.34372 | 77.18787 | chr19 | 46075057 | 46078097 | 4.67749 | 3.7326 | 2.77974 |
| chr10 | 7118533 | 7118735 | 182 | Downstream (<1kb) | 10 | 7119063 | 7135506 | 16444 | 2 | ENSMUST00 | 16771 | Cnksr3 | ENSMUSG0( | false | . | . | . | . | . | . | chr10 | 7118068 | 7119226 | 3.12093 | 3.02805 | 1.50378 |
| chrX | 71556606 | 71557056 | 182 | Promoter (<=1kb) | 20 | 71556946 | 71559273 | 2328 | 1 | ENSMUST00 | 0 | Hmgb3 | ENSMUSG0( | false | chrX | 71555539 | 71557518 | 25.7338 | 71.40817 | 68.39619 | . | . | . | . | . | . |
| chr4 | 53115857 | 53116055 | 182 | Intron (ENSMUST00000030010.3 | 4 | 53030787 | 53159895 | 129109 | 2 | ENSMUST00 | 43840 | Abca1 | ENSMUSG0( | false | . | . | . | . | . | . | chr4 | 53114826 | 53118669 | 4.22918 | 3.68391 | 2.39273 |
| chr10 | 80940669 | 80941056 | 182 | Distal Intergenic | 10 | 80930091 | 80932204 | 2114 | 1 | ENSMUST00 | 10579 | Gadd45b | ENSMUSG0( | false | . | . | . | . | . | . | chr10 | 80936148 | 80942225 | 6.45113 | 4.61033 | 4.23692 |
| chr13 | 43727051 | 43727353 | 182 | Distal Intergenic | 13 | 43785107 | 43803130 | 18024 | 1 | ENSMUST00 | -57754 | Cd83 | ENSMUSG0( | false | . | . | . | . | . | . | chr13 | 43726588 | 43727520 | 4.24522 | 3.47226 | 2.41835 |
| chr2 | 27009765 | 27010091 | 182 | Promoter (<=1kb) | 2 | 27009926 | 27016849 | 6924 | 1 | ENSMUST00 | 0 | Cacfd1 | ENSMUSG0( | false | chr2 | 27009437 | 27010847 | 32.66734 | 113.81544 | 109.86345 | chr2 | 27007563 | 27011439 | 4.79428 | 3.85674 | 2.88621 |
| chr3 | 95434336 | 95434613 | 182 | Promoter (<=1kb) | 3 | 95434388 | 95491622 | 57235 | 1 | ENSMUST00 | 0 | Arnt | ENSMUSG0( | false | chr3 | 95433809 | 95435481 | 28.91247 | 96.86193 | 93.36482 | . | . | . | . | . | . |
| chr8 | 68930148 | 68930376 | 182 | Distal Intergenic | 8 | 68890744 | 68895819 | 5076 | 1 | ENSMUST00 | 39405 | Lpl | ENSMUSG0( | false | . | . | . | . | . | . | chr8 | 68923527 | 68931359 | 4.50778 | 3.74958 | 2.61471 |
| chr2 | 180228661 | 180228944 | 182 | Promoter (2-3kb) | 2 | 180176373 | 180225859 | 49487 | 2 | ENSMUST00 | -2803 | Lama5 | ENSMUSG0( | true | chr2 | 180227200 | 180228692 | 9.76265 | 21.55577 | 19.19873 | chr2 | 180225908 | 180236447 | 7.12665 | 5.06061 | 4.74583 |
| chr9 | 40800667 | 40801133 | 182 | Promoter (<=1kb) | 9 | 40800984 | 40805200 | 4217 | 1 | ENSMUST00 | 0 | Hspa8 | ENSMUSG0( | false | chr9 | 40799626 | 40800806 | 21.96151 | 62.27057 | 59.39301 | . | . | . | . | . | . |
| chr17 | 6073706 | 6073917 | 182 | Intron (ENSMUST00000097432.4 | 17 | 6044066 | 6079727 | 35662 | 2 | ENSMUST00 | 5810 | Serac1 | ENSMUSG0( | false | . | . | . | . | . | . | chr17 | 6072820 | 6074291 | 6.48339 | 4.83118 | 4.21802 |
| chr17 | 6079641 | 6079903 | 182 | Promoter (<=1kb) | 17 | 6042196 | 6079741 | 37546 | 2 | ENSMUST00 | 0 | Serac1 | ENSMUSG0( | false | chr17 | 6079019 | 6080497 | 38.82705 | 122.56369 | 118.33408 | . | . | . | . | . | . |
| chr3 | 96044538 | 96045020 | 182 | Intron (ENSMUST00000015891.5 | 3 | 96056887 | 96057719 | 833 | 2 | ENSMUST00 | 12699 | Vps45 | ENSMUSG0( | false | . | . | . | . | . | . | . | . | . | . | . | . |
| chr3 | 95929565 | 95930213 | 182 | Promoter (<=1kb) | 3 | 95929582 | 95945318 | 15737 | 1 | ENSMUST00 | 0 | Anp32e | ENSMUSG0( | false | chr3 | 95929228 | 95930742 | 39.61801 | 127.15503 | 122.77008 | chr3 | 95930115 | 95932922 | 3.58446 | 3.41165 | 1.8255 |
| chr10 | 7965080 | 7965302 | 182 | Distal Intergenic | 10 | 7905653 | 7956230 | 50578 | 2 | ENSMUST00 | -8851 | Tab2 | ENSMUSG0( | true | . | . | . | . | . | . | chr10 | 7962889 | 7966091 | 5.26626 | 4.12835 | 3.25921 |
| chr6 | 137558923 | 137559214 | 182 | Intron (ENSMUST00000058210.7 | 6 | 137477668 | 137571017 | 93350 | 2 | ENSMUST00 | 11803 | Eps8 | ENSMUSG0( | false | . | . | . | . | . | . | chr6 | 137557484 | 137559802 | 2.90855 | 2.92957 | 1.30626 |
| chr3 | 95702095 | 95702461 | 182 | Distal Intergenic | 3 | 95676201 | 95687917 | 11717 | 2 | ENSMUST00 | -14179 | Adamtsl4 | ENSMUSG0( | false | chr3 | 95701798 | 95702505 | 9.84369 | 19.54702 | 17.21913 | chr3 | 95685856 | 95702888 | 5.06878 | 4.08183 | 3.0659 |
| chr5 | 45856575 | 45856756 | 182 | Promoter (<=1kb) | 5 | 45700450 | 45856552 | 156103 | 2 | ENSMUST00 | -24 | Lcorl | ENSMUSG0( | false | chr5 | 45855601 | 45858324 | 38.87692 | 132.39133 | 127.80991 | . | . | . | . | . | . |
| chr13 | 56134658 | 56135066 | 182 | Promoter (<=1kb) | 13 | 56104441 | 56134825 | 30385 | 2 | ENSMUST00 | 0 | H2afy | ENSMUSG0( | false | chr13 | 56134430 | 56136012 | 34.00884 | 130.01486 | 125.52227 | chr13 | 56132562 | 56136152 | 4.04555 | 3.54219 | 2.25967 |
| chr7 | 98816224 | 98816447 | 182 | Distal Intergenic | 7 | 98835112 | 98846587 | 11476 | 1 | ENSMUST00 | -18665 | Wnt11 | ENSMUSG0( | false | chr7 | 98814693 | 98816760 | 27.06219 | 100.28806 | 96.70959 | . | . | . | . | . | . |
| chr7 | 139234726 | 139235065 | 182 | Promoter (<=1kb) | 7 | 139233943 | 139242971 | 9029 | 1 | ENSMUST00 | 784 | Lrrc27 | ENSMUSG0( | false | . | . | . | . | . | . | chr7 | 139233932 | 139235767 | 4.24404 | 3.7709 | 2.42945 |
| chr7 | 139213086 | 139213410 | 182 | Promoter (<=1kb) | 7 | 139213272 | 139229008 | 15737 | 1 | ENSMUST00 | 0 | Lrrc27 | ENSMUSG0( | false | chr7 | 139212604 | 139213790 | 25.4311 | 72.68642 | 69.65319 | . | . | . | . | . | . |
| chr5 | 151108692 | 151108887 | 182 | Intron (ENSMUST00000062015.9 | 5 | 151063593 | 151095747 | 32155 | 2 | ENSMUST00 | -12946 | Stard13 | ENSMUSG0( | false | chr5 | 151108315 | 151108823 | 8.80131 | 17.30502 | 15.00832 | . | . | . | . | . | . |
| chr2 | 105138980 | 105139204 | 182 | Intron (ENSMUST00000143043.2 | 2 | 105130883 | 105173616 | 42734 | 1 | ENSMUST00 | 8098 | Wt1 | ENSMUSG0( | true | . | . | . | . | . | . | chr2 | 105133597 | 105164717 | 4.85658 | 3.95538 | 2.90242 |
| chr2 | 105130314 | 105130988 | 182 | Promoter (<=1kb) | 2 | 105130800 | 105133950 | 3151 | 1 | ENSMUST00 | 0 | Wt1 | ENSMUSG0( | true | chr2 | 105122726 | 105134202 | 33.52805 | 106.05383 | 102.33234 | . | . | . | . | . | . |
| chr2 | 105127131 | 105127461 | 182 | Promoter (<=1kb) | 2 | 105127210 | 105172681 | 45472 | 1 | ENSMUST00 | 0 | Wt1 | ENSMUSG0( | true | chr2 | 105122726 | 105134202 | 33.52805 | 106.05383 | 102.33234 | . | . | . | . | . | . |
| chr1 | 194938786 | 194939053 | 182 | Promoter (<=1kb) | 1 | 194938821 | 194961292 | 22472 | 1 | ENSMUST00 | 0 | Cd34 | ENSMUSG0( | false | chr1 | 194938763 | 194939554 | 10.95946 | 24.18275 | 21.79051 | chr1 | 194938703 | 194939211 | 3.93916 | 3.4008 | 2.16504 |
| chr5 | 146923651 | 146923857 | 182 | Distal Intergenic | 5 | 146948657 | 146955614 | 6958 | 1 | ENSMUST00 | -24800 | Gtf3a | ENSMUSG0( | false | . | . | . | . | . | . | chr5 | 146919863 | 146924296 | 4.88294 | 4.06696 | 2.91401 |
| chr5 | 147076469 | 147076932 | 182 | Promoter (<=1kb) | 5 | 147016655 | 147076586 | 59932 | 2 | ENSMUST00 | 0 | Lnx2 | ENSMUSG0( | false | chr5 | 147075302 | 147078299 | 32.66836 | 102.52041 | 98.88917 | . | . | . | . | . | . |
| chr15 | 84923166 | 84923397 | 182 | Promoter (<=1kb) | 15 | 84923428 | 84942955 | 19528 | 1 | ENSMUST00 | -31 | Nup50 | ENSMUSG0( | false | chr15 | 84922584 | 84924126 | 20.67859 | 81.70428 | 78.522 | . | . | . | . | . | . |
| chr11 | 115420418 | 115420655 | 182 | Promoter (<=1kb) | 11 | 115420312 | 115422429 | 2118 | 1 | ENSMUST00 | 107 | Kctd2 | ENSMUSG0( | false | chr11 | 115419229 | 115421314 | 42.16172 | 139.23978 | 134.38916 | . | . | . | . | . | . |
| chr2 | 164366898 | 164367135 | 182 | Intron (ENSMUST00000167427.1 | 2 | 164354070 | 164356507 | 2438 | 2 | ENSMUST00 | -10392 | Slpi | ENSMUSG0( | false | . | . | . | . | . | . | chr2 | 164365965 | 164368814 | 5.81203 | 4.50267 | 3.65337 |
| chr2 | 164440459 | 164440638 | 182 | Promoter (2-3kb) | 2 | 164424247 | 164443186 | 18940 | 2 | ENSMUST00 | 2548 | Sdc4 | ENSMUSG0( | true | chr2 | 164438320 | 164443697 | 20.02967 | 73.35366 | 70.31202 | . | . | . | . | . | . |
| chr11 | 100440984 | 100441313 | 182 | Promoter (<=1kb) | 11 | 100422321 | 100441089 | 18769 | 2 | ENSMUST00 | 0 | Nt5c3b | ENSMUSG0( | false | chr11 | 100440528 | 100441216 | 33.10969 | 106.34647 | 102.61562 | . | . | . | . | . | . |
| chr11 | 78697071 | 78697612 | 182 | Promoter (<=1kb) | 11 | 78567168 | 78697373 | 130206 | 2 | ENSMUST00 | 0 | Nlk | ENSMUSG0( | true | chr11 | 78696349 | 78697538 | 36.17194 | 114.79427 | 110.81874 | . | . | . | . | . | . |
| chr11 | 80383131 | 80383522 | 182 | Promoter (<=1kb) | 11 | 80383279 | 80396245 | 12967 | 1 | ENSMUST00 | 0 | Zfp207 | ENSMUSG0( | false | chr11 | 80382405 | 80385417 | 24.86315 | 92.1349 | 88.74321 | . | . | . | . | . | . |
| chr11 | 118428065 | 118428621 | 182 | Promoter (<=1kb) | 11 | 118428203 | 118443843 | 15641 | 1 | ENSMUST00 | 0 | C1qtnf1 | ENSMUSG0( | false | chr11 | 118427431 | 118430359 | 22.74714 | 65.36804 | 62.44672 | chr11 | 118411596 | 118448618 | 7.23691 | 5.03811 | 4.84329 |
| chr11 | 118427062 | 118427469 | 182 | Promoter (<=1kb) | 11 | 118428203 | 118443843 | 15641 | 1 | ENSMUST00 | -734 | C1qtnf1 | ENSMUSG0( | false | chr11 | 118427431 | 118430359 | 22.74714 | 65.36804 | 62.44672 | chr11 | 118411596 | 118448618 | 7.23691 | 5.03811 | 4.84329 |
| chr11 | 77907087 | 77907320 | 182 | Distal Intergenic | 11 | 77880615 | 77894096 | 13482 | 2 | ENSMUST00 | -12992 | Pipox | ENSMUSG0( | false | . | . | . | . | . | . | chr11 | 77906193 | 77907370 | 5.31905 | 4.23213 | 3.27274 |
| chr11 | 77893890 | 77894386 | 182 | Promoter (<=1kb) | 11 | 77880615 | 77894096 | 13482 | 2 | ENSMUST00 | 0 | Pipox | ENSMUSG0( | false | . | . | . | . | . | . | chr11 | 77893363 | 77897732 | 4.7599 | 4.01036 | 2.81137 |
| chr11 | 118348960 | 118349147 | 182 | Intron (ENSMUST00000017610.4 | 11 | 118301069 | 118355506 | 54438 | 2 | ENSMUST00 | 6359 | Timp2 | ENSMUSG0( | false | . | . | . | . | . | . | chr11 | 118319919 | 118354581 | 6.53665 | 4.67889 | 4.28866 |
| chr8 | 122376393 | 122376947 | 182 | Promoter (<=1kb) | 8 | 122376609 | 122417359 | 40751 | 1 | ENSMUST00 | 0 | Zc3h18 | ENSMUSG0( | false | chr8 | 122376081 | 122377570 | 28.41342 | 89.1249 | 85.7944 | chr8 | 122375128 | 122380558 | 3.60707 | 3.27107 | 1.91049 |
| chr14 | 16365487 | 16366028 | 182 | Promoter (<=1kb) | 14 | 16365179 | 16430787 | 65609 | 1 | ENSMUST00 | 309 | Top2b | ENSMUSG0( | false | chr14 | 16363963 | 16366999 | 34.91417 | 125.28625 | 120.96531 | . | . | . | . | . | . |
| chr14 | 16578063 | 16578351 | 182 | Promoter (2-3kb) | 14 | 16430842 | 16575472 | 144631 | 2 | ENSMUST00 | -2592 | Rarb | ENSMUSG0( | false | . | . | . | . | . | . | chr14 | 16576099 | 16578520 | 5.1848 | 4.07653 | 3.18327 |
| chr14 | 16516117 | 16516385 | 182 | Intron (ENSMUST00000063750.6 | 14 | 16430842 | 16575472 | 144631 | 2 | ENSMUST00 | 59087 | Rarb | ENSMUSG0( | false | chr14 | 16515883 | 16516149 | 6.38327 | 11.47542 | 9.27082 | chr14 | 16513332 | 16520797 | 5.11377 | 4.06158 | 3.14269 |
| chr14 | 16519668 | 16519942 | 182 | Intron (ENSMUST00000063750.6 | 14 | 16430842 | 16575472 | 144631 | 2 | ENSMUST00 | 55530 | Rarb | ENSMUSG0( | false | . | . | . | . | . | . | chr14 | 16513332 | 16520797 | 5.11377 | 4.06158 | 3.14269 |
| chr11 | 80080779 | 80081145 | 182 | Promoter (<=1kb) | 11 | 80046493 | 80080938 | 34446 | 2 | ENSMUST00 | 0 | Crlf3 | ENSMUSG0( | true | chr11 | 80079550 | 80081430 | 31.38972 | 125.99339 | 121.64841 | . | . | . | . | . | . |
| chr11 | 78531038 | 78531278 | 182 | Intron (ENSMUST00000017759.3 | 11 | 78522850 | 78536271 | 13422 | 2 | ENSMUST00 | 4993 | Tnfaip1 | ENSMUSG0( | false | . | . | . | . | . | . | chr11 | 78530172 | 78532036 | 3.02254 | 3.01899 | 1.38382 |
| chr2 | 165326561 | 165327203 | 182 | Promoter (<=1kb) | 2 | 165315672 | 165326479 | 10808 | 2 | ENSMUST00 | -83 | Elmo2 | ENSMUSG0( | false | chr2 | 165326644 | 165327257 | 20.44396 | 54.62578 | 51.85264 | chr2 | 165326956 | 165327929 | 5.04752 | 4.09871 | 3.0734 |
| chr2 | 165325595 | 165325822 | 182 | Promoter (<=1kb) | 2 | 165289155 | 165326479 | 37325 | 2 | ENSMUST00 | 657 | Elmo2 | ENSMUSG0( | false | chr2 | 165324516 | 165326561 | 32.66734 | 113.81544 | 109.86345 | . | . | . | . | . | . |
| chr2 | 165321629 | 165321849 | 182 | Promoter (<=1kb) | 2 | 165288031 | 165320826 | 32796 | 2 | ENSMUST00 | -804 | Elmo2 | ENSMUSG0( | false | . | . | . | . | . | . | chr2 | 165307563 | 165325045 | 8.91187 | 5.755 | 6.22321 |
| chr2 | 163644654 | 163645270 | 182 | Promoter (<=1kb) | 2 | 163623272 | 163645131 | 21860 | 2 | ENSMUST00 | 0 | Serinc3 | ENSMUSG0( | false | chr2 | 163642816 | 163645495 | 37.54867 | 136.59039 | 131.84451 | . | . | . | . | . | . |
| chr2 | 164513494 | 164513793 | 182 | Distal Intergenic | 2 | 164499916 | 164501639 | 1724 | 1 | ENSMUST00 | 13579 | Pigt | ENSMUSG0( | false | . | . | . | . | . | . | chr2 | 164509177 | 164517266 | 5.6289 | 4.32047 | 3.52645 |
| chr2 | 164500371 | 164500662 | 182 | Promoter (<=1kb) | 2 | 164499916 | 164501639 | 1724 | 1 | ENSMUST00 | 456 | Pigt | ENSMUSG0( | false | . | . | . | . | . | . | chr2 | 164496710 | 164500875 | 5.3608 | 4.17074 | 3.31633 |
| chr11 | 101784431 | 101784729 | 182 | Promoter (<=1kb) | 11 | 101773862 | 101784491 | 10630 | 2 | ENSMUST00 | 0 | Etv4 | ENSMUSG0( | false | chr11 | 101784049 | 101785755 | 23.80275 | 64.58516 | 61.676 | chr11 | 101783730 | 101786247 | 4.40284 | 3.79612 | 2.50277 |
| chr13 | 73763468 | 73763791 | 182 | Promoter (<=1kb) | 13 | 73763697 | 73816743 | 53047 | 1 | ENSMUST00 | 0 | Slc12a7 | ENSMUSG0( | false | chr13 | 73763388 | 73764824 | 24.55417 | 69.38167 | 66.39919 | chr13 | 73763189 | 73765277 | 3.6005 | 3.37467 | 1.90059 |
| chr11 | 75655594 | 75656211 | 182 | Promoter (<=1kb) | 11 | 75655873 | 75673908 | 18036 | 1 | ENSMUST00 | 0 | Myo1c | ENSMUSG0( | false | chr11 | 75651407 | 75657466 | 25.17393 | 93.69764 | 90.27328 | chr11 | 75652271 | 75655636 | 6.31319 | 4.65216 | 4.07889 |
| chr11 | 75679115 | 75679457 | 182 | Promoter (<=1kb) | 11 | 75679259 | 75706065 | 26807 | 1 | ENSMUST00 | 0 | Crk | ENSMUSG0( | false | chr11 | 75678797 | 75680664 | 41.2653 | 138.78937 | 133.96024 | . | . | . | . | . | . |
| chr11 | 100622764 | 100623120 | 182 | Promoter (<=1kb) | 11 | 100622876 | 100627607 | 4732 | 1 | ENSMUST00 | 0 | Nkiras2 | ENSMUSG0( | false | chr11 | 100622779 | 100623594 | 18.2599 | 77.14165 | 74.04132 | . | . | . | . | . | . |
| chr12 | 110510292 | 110510518 | 182 | Intron (ENSMUST00000084985.4 | 12 | 110485739 | 110583061 | 97323 | 1 | ENSMUST00 | 24554 | Ppp2r5c | ENSMUSG0( | false | . | . | . | . | . | . | chr12 | 110492860 | 110513828 | 5.53082 | 4.27464 | 3.44454 |
| chr12 | 110476726 | 110476912 | 182 | Distal Intergenic | 12 | 110485739 | 110583061 | 97323 | 1 | ENSMUST00 | -8827 | Ppp2r5c | ENSMUSG0( | false | . | . | . | . | . | . | chr12 | 110475835 | 110477598 | 4.41501 | 3.73478 | 2.57454 |
| chr5 | 143633043 | 143633251 | 182 | Promoter (<=1kb) | 5 | 143634066 | 143692652 | 58587 | 1 | ENSMUST00 | -815 | Cyth3 | ENSMUSG0( | false | . | . | . | . | . | . | chr5 | 143628431 | 143643057 | 5.82289 | 4.44189 | 3.69355 |
| chr5 | 143657096 | 143657333 | 182 | Intron (ENSMUST00000116456.4 | 5 | 143651241 | 143708256 | 57016 | 1 | ENSMUST00 | 5856 | Cyth3 | ENSMUSG0( | false | . | . | . | . | . | . | chr5 | 143651427 | 143662078 | 6.34156 | 4.57429 | 4.14661 |
| chr5 | 139793463 | 139793666 | 182 | Promoter (1-2kb) | 5 | 139791534 | 139799500 | 7967 | 1 | ENSMUST00 | 1930 | Mafk | ENSMUSG0( | false | chr5 | 139791432 | 139793535 | 32.3084 | 121.498 | 117.30614 | chr5 | 139792309 | 139799489 | 7.14368 | 5.03259 | 4.78629 |
| chr10 | 128589634 | 128589910 | 182 | Promoter (<=1kb) | 10 | 128567523 | 128589652 | 22130 | 2 | ENSMUST00 | 0 | Erbb3 | ENSMUSG0( | false | chr10 | 128589190 | 128590713 | 25.50663 | 88.63404 | 85.31516 | chr10 | 128589323 | 128590850 | 3.1703 | 3.03605 | 1.54074 |
| chr11 | 98358223 | 98358735 | 182 | Promoter (<=1kb) | 11 | 98358368 | 98381099 | 22732 | 1 | ENSMUST00 | 0 | Stard3 | ENSMUSG0( | false | chr11 | 98357889 | 98359357 | 19.14982 | 55.63055 | 52.84086 | chr11 | 98355645 | 98364253 | 5.76527 | 4.38786 | 3.64659 |
| chr1 | 143777006 | 143777411 | 182 | Promoter (<=1kb) | 1 | 143777272 | 143807305 | 30034 | 1 | ENSMUST00 | 0 | Uchl5 | ENSMUSG0( | false | chr1 | 143776236 | 143778292 | 27.97104 | 107.99458 | 104.2211 | . | . | . | . | . | . |
| chr11 | 70653493 | 70654206 | 182 | Promoter (<=1kb) | 11 | 70652622 | 70654598 | 1977 | 2 | ENSMUST00 | 392 | Pfn1 | ENSMUSG0( | false | chr11 | 70652869 | 70656609 | 41.07278 | 140.67711 | 135.76277 | chr11 | 70647853 | 70653572 | 5.12137 | 4.04412 | 3.13154 |
| chr2 | 164070631 | 164071388 | 182 | Promoter (<=1kb) | 2 | 164053540 | 164071102 | 17563 | 2 | ENSMUST00 | 0 | Tomm34 | ENSMUSG0( | false | chr2 | 164070106 | 164071514 | 50.11335 | 172.5629 | 166.54207 | chr2 | 164071302 | 164071818 | 3.93795 | 3.60864 | 2.1426 |
| chr11 | 79059750 | 79060162 | 182 | Promoter (<=1kb) | 11 | 79044920 | 79059875 | 14956 | 2 | ENSMUST00 | 0 | Ksr1 | ENSMUSG0( | true | . | . | . | . | . | . | chr11 | 79045596 | 79144720 | 7.09579 | 4.93486 | 4.74337 |
| chr11 | 79146414 | 79146616 | 182 | Promoter (<=1kb) | 11 | 79026707 | 79146407 | 119701 | 2 | ENSMUST00 | -8 | Ksr1 | ENSMUSG0( | true | chr11 | 79143897 | 79146881 | 38.4896 | 132.80048 | 128.21092 | . | . | . | . | . | . |
| chr11 | 79083152 | 79083393 | 182 | Intron (ENSMUST00000018478.5 | 11 | 79044920 | 79059875 | 14956 | 2 | ENSMUST00 | -23278 | Ksr1 | ENSMUSG0( | true | . | . | . | . | . | . | chr11 | 79045596 | 79144720 | 7.09579 | 4.93486 | 4.74337 |
| chr11 | 106999514 | 106999734 | 182 | Promoter (<=1kb) | 11 | 106988629 | 106999541 | 10913 | 2 | ENSMUST00 | 0 | Kpna2 | ENSMUSG0( | false | chr11 | 106998544 | 106999580 | 36.03628 | 116.66104 | 112.62998 | chr11 | 106997249 | 107000191 | 4.37561 | 3.79383 | 2.49556 |
| chr11 | 88099731 | 88100051 | 182 | Promoter (<=1kb) | 11 | 88099290 | 88185365 | 86076 | 1 | ENSMUST00 | 442 | Cuedc1 | ENSMUSG0( | true | chr11 | 88098491 | 88100196 | 39.74135 | 132.8143 | 128.22414 | chr11 | 88099933 | 88101241 | 4.84832 | 4.05533 | 2.87122 |
| chr11 | 88162078 | 88162350 | 182 | Promoter (<=1kb) | 11 | 88162261 | 88194140 | 31880 | 1 | ENSMUST00 | 0 | Cuedc1 | ENSMUSG0( | true | chr11 | 88161702 | 88162230 | 7.43959 | 17.32416 | 15.02694 | chr11 | 88146778 | 88183132 | 6.97604 | 4.89368 | 4.62779 |
| chr11 | 88132004 | 88132291 | 182 | Intron (ENSMUST00000018522.7 | 11 | 88162261 | 88194140 | 31880 | 1 | ENSMUST00 | -29970 | Cuedc1 | ENSMUSG0( | true | . | . | . | . | . | . | chr11 | 88130558 | 88134123 | 4.74153 | 3.86104 | 2.80616 |
| chr11 | 95834929 | 95835410 | 182 | Promoter (<=1kb) | 11 | 95832723 | 95835890 | 3168 | 2 | ENSMUST00 | 480 | Abi3 | ENSMUSG0( | false | . | . | . | . | . | . | chr11 | 95829550 | 95838805 | 5.29848 | 4.1616 | 3.26577 |
| chr11 | 84819249 | 84819605 | 182 | Promoter (<=1kb) | 11 | 84813061 | 84819515 | 6455 | 2 | ENSMUST00 | 0 | Mrm1 | ENSMUSG0( | false | chr11 | 84818196 | 84819934 | 27.85705 | 92.98368 | 89.57738 | . | . | . | . | . | . |
| chr11 | 104234164 | 104234405 | 182 | Promoter (2-3kb) | 11 | 104231584 | 104328219 | 96636 | 1 | ENSMUST00 | 2581 | Mapt | ENSMUSG0( | true | chr11 | 104231437 | 104234756 | 38.46334 | 130.56975 | 126.05653 | chr11 | 104233065 | 104238850 | 6.60888 | 4.67556 | 4.3541 |
| chr11 | 104469152 | 104469542 | 182 | Promoter (<=1kb) | 11 | 104333501 | 104468861 | 135361 | 2 | ENSMUST00 | -292 | Kansl1 | ENSMUSG0( | false | chr11 | 104468564 | 104469932 | 30.41442 | 103.5799 | 99.9162 | . | . | . | . | . | . |
| chr1 | 51914966 | 51915253 | 182 | Promoter (<=1kb) | 1 | 51810681 | 51915912 | 105232 | 2 | ENSMUST00 | 659 | Myo1b | ENSMUSG0( | false | chr1 | 51914973 | 51916278 | 18.02578 | 59.25625 | 56.42029 | chr1 | 51914945 | 51915650 | 3.35188 | 3.21556 | 1.6853 |
| chr1 | 51822081 | 51822316 | 182 | Promoter (1-2kb) | 1 | 51799536 | 51820546 | 21011 | 2 | ENSMUST00 | -1536 | Myo1b | ENSMUSG0( | false | . | . | . | . | . | . | chr1 | 51820030 | 51822945 | 3.36877 | 3.07405 | 1.72036 |
| chr11 | 71019585 | 71019970 | 182 | Promoter (<=1kb) | 11 | 71009933 | 71019841 | 9909 | 2 | ENSMUST00 | 0 | Derl2 | ENSMUSG0( | false | chr11 | 71019547 | 71020536 | 21.54815 | 66.72034 | 63.77881 | . | . | . | . | . | . |
| chr11 | 69420109 | 69420462 | 182 | Downstream (<1kb) | 11 | 69398508 | 69413675 | 15168 | 2 | ENSMUST00 | -6435 | Kdm6b | ENSMUSG0( | false | chr11 | 69419049 | 69420647 | 17.64787 | 49.73517 | 47.02245 | chr11 | 69419407 | 69421881 | 5.4946 | 4.19627 | 3.42749 |
| chr11 | 62538728 | 62539113 | 182 | Promoter (<=1kb) | 11 | 62533887 | 62539003 | 5117 | 2 | ENSMUST00 | 0 | Cenpv | ENSMUSG0( | false | chr11 | 62538296 | 62539629 | 29.25995 | 87.70835 | 84.40896 | chr11 | 62537698 | 62539665 | 3.18195 | 3.0196 | 1.55081 |
| chr11 | 97698777 | 97699417 | 182 | Promoter (<=1kb) | 11 | 97688823 | 97699384 | 10562 | 2 | ENSMUST00 | 0 | Pcgf2 | ENSMUSG0( | false | chr11 | 97698478 | 97701285 | 33.7938 | 119.00099 | 114.88904 | chr11 | 97694773 | 97702506 | 4.47077 | 3.79783 | 2.58655 |
| chr11 | 97733663 | 97733914 | 182 | Intron (ENSMUST00000018691.8 | 11 | 97720396 | 97726743 | 6348 | 2 | ENSMUST00 | -6921 | Pip4k2b | ENSMUSG0( | false | . | . | . | . | . | . | chr11 | 97732967 | 97738104 | 4.38191 | 3.80976 | 2.49294 |
| chr11 | 55428192 | 55428608 | 182 | Distal Intergenic | 11 | 55394500 | 55420080 | 25581 | 2 | ENSMUST00 | -8113 | Sparc | ENSMUSG0( | true | . | . | . | . | . | . | chr11 | 55426310 | 55429682 | 5.1823 | 4.17012 | 3.15798 |
| chr11 | 55410427 | 55410689 | 182 | Intron (ENSMUST00000108858.2 | 11 | 55395520 | 55402165 | 6646 | 2 | ENSMUST00 | -8263 | Sparc | ENSMUSG0( | true | . | . | . | . | . | . | chr11 | 55405757 | 55417699 | 6.44715 | 4.69413 | 4.21932 |
| chr11 | 96789035 | 96789554 | 182 | Promoter (<=1kb) | 11 | 96789127 | 96803969 | 14843 | 1 | ENSMUST00 | 0 | Cbx1 | ENSMUSG0( | false | chr11 | 96788747 | 96790143 | 29.73511 | 88.5301 | 85.22105 | chr11 | 96788512 | 96791001 | 4.70967 | 3.87715 | 2.82251 |
| chr11 | 84513402 | 84513661 | 182 | Promoter (<=1kb) | 11 | 84422855 | 84513522 | 90668 | 2 | ENSMUST00 | 0 | Aatf | ENSMUSG0( | false | chr11 | 84512655 | 84514216 | 28.22734 | 101.53345 | 97.92473 | . | . | . | . | . | . |
| chr12 | 110591187 | 110591490 | 182 | Distal Intergenic | 12 | 110601452 | 110666945 | 65494 | 1 | ENSMUST00 | -9962 | Dync1h1 | ENSMUSG0( | false | chr12 | 110590657 | 110592416 | 31.1999 | 108.05416 | 104.27962 | chr12 | 110590265 | 110592649 | 5.13247 | 4.15576 | 3.11453 |
| chr12 | 110552095 | 110552487 | 182 | Intron (ENSMUST00000084985.4 | 12 | 110601452 | 110666945 | 65494 | 1 | ENSMUST00 | -48965 | Dync1h1 | ENSMUSG0( | false | . | . | . | . | . | . | chr12 | 110549007 | 110557689 | 5.99039 | 4.4389 | 3.83972 |
| chr11 | 68967633 | 68968097 | 182 | Promoter (<=1kb) | 11 | 68968131 | 68972516 | 4386 | 1 | ENSMUST00 | -34 | Slc25a35 | ENSMUSG0( | false | chr11 | 68967300 | 68968611 | 18.88512 | 67.59834 | 64.64505 | chr11 | 68966301 | 68969994 | 5.04313 | 4.14481 | 3.02479 |
| chr11 | 35769457 | 35770175 | 182 | Promoter (<=1kb) | 11 | 35769484 | 35791285 | 21802 | 1 | ENSMUST00 | 0 | Pank3 | ENSMUSG0( | false | chr11 | 35769023 | 35770839 | 27.03504 | 88.5911 | 85.27629 | . | . | . | . | . | . |
| chr11 | 53769764 | 53770114 | 182 | Promoter (<=1kb) | 11 | 53770014 | 53777323 | 7310 | 1 | ENSMUST00 | 0 | Irf1 | ENSMUSG0( | false | chr11 | 53769560 | 53771265 | 28.90751 | 104.81161 | 101.11721 | chr11 | 53768436 | 53769839 | 4.8183 | 3.97626 | 2.86434 |
| chr11 | 70410024 | 70410252 | 182 | Promoter (<=1kb) | 11 | 70392883 | 70410028 | 17146 | 2 | ENSMUST00 | 0 | Pelp1 | ENSMUSG0( | false | chr11 | 70409194 | 70410542 | 36.04672 | 129.50026 | 125.02953 | chr11 | 70408721 | 70410718 | 4.3747 | 3.59667 | 2.52275 |
| chr11 | 60913418 | 60913834 | 182 | Promoter (<=1kb) | 11 | 60902246 | 60913812 | 11567 | 2 | ENSMUST00 | 0 | Gm16515 | ENSMUSG0( | false | chr11 | 60911793 | 60914427 | 37.32329 | 122.0922 | 117.88763 | chr11 | 60898871 | 60915700 | 5.92882 | 4.41351 | 3.79632 |
| chr5 | 33024515 | 33024804 | 182 | Promoter (2-3kb) | 5 | 33021791 | 33026981 | 5191 | 1 | ENSMUST00 | 2725 | Ywhah | ENSMUSG0( | false | . | . | . | . | . | . | chr5 | 33020191 | 33028393 | 5.43632 | 4.2855 | 3.37828 |
| chr6 | 34176593 | 34177209 | 182 | Promoter (<=1kb) | 6 | 34153380 | 34177111 | 23732 | 2 | ENSMUST00 | 0 | Slc35b4 | ENSMUSG0( | false | chr6 | 34176058 | 34177139 | 28.90751 | 104.81161 | 101.11721 | chr6 | 34170482 | 34177467 | 5.2089 | 4.17803 | 3.18044 |
| chr9 | 21918134 | 21918403 | 182 | Promoter (<=1kb) | 9 | 21907491 | 21918192 | 10702 | 2 | ENSMUST00 | 0 | Rab3d | ENSMUSG0( | false | chr9 | 21917580 | 21918741 | 24.48636 | 83.88964 | 80.66612 | chr9 | 21917513 | 21918876 | 3.23264 | 3.17276 | 1.60191 |
| chr7 | 141437490 | 141437774 | 182 | Promoter (<=1kb) | 7 | 141429749 | 141437631 | 7883 | 2 | ENSMUST00 | 0 | Slc25a22 | ENSMUSG0( | false | chr7 | 141437009 | 141437898 | 19.17732 | 54.04958 | 51.28253 | chr7 | 141432253 | 141438158 | 5.83688 | 4.42084 | 3.69359 |
| chr8 | 70600540 | 70600931 | 182 | Intron (ENSMUST00000049908.9 | 8 | 70594481 | 70597288 | 2808 | 1 | ENSMUST00 | 6060 | Isyna1 | ENSMUSG0( | false | . | . | . | . | . | . | chr8 | 70592285 | 70603072 | 6.42958 | 4.72399 | 4.17224 |
| chr8 | 70575027 | 70575234 | 182 | Intron (ENSMUST00000093454.6 | 8 | 70594481 | 70597288 | 2808 | 1 | ENSMUST00 | -19247 | Isyna1 | ENSMUSG0( | false | . | . | . | . | . | . | chr8 | 70574618 | 70576077 | 5.57937 | 4.2893 | 3.52492 |
| chr7 | 16444848 | 16445068 | 182 | Distal Intergenic | 7 | 16452779 | 16455489 | 2711 | 1 | ENSMUST00 | -7711 | Tmem160 | ENSMUSG0( | false | . | . | . | . | . | . | chr7 | 16443897 | 16445981 | 3.85346 | 3.51383 | 2.06543 |
| chr11 | 44563005 | 44563296 | 182 | Promoter (2-3kb) | 11 | 44560019 | 44561977 | 1959 | 1 | ENSMUST00 | 2987 | Rnf145 | ENSMUSG0( | false | . | . | . | . | . | . | . | . | . | . | . | . |
| chr17 | 35241303 | 35241992 | 182 | Promoter (<=1kb) | 17 | 35241746 | 35253707 | 11962 | 1 | ENSMUST00 | 0 | Ddx39b | ENSMUSG0( | false | chr17 | 35241108 | 35241702 | 9.54222 | 21.71909 | 19.36044 | . | . | . | . | . | . |
| chr8 | 83656970 | 83657179 | 182 | Intron (ENSMUST00000019577.8 | 8 | 83652678 | 83664788 | 12111 | 1 | ENSMUST00 | 4293 | Gipc1 | ENSMUSG0( | false | . | . | . | . | . | . | . | . | . | . | . | . |
| chr11 | 78178259 | 78178768 | 182 | Promoter (<=1kb) | 11 | 78178651 | 78181909 | 3259 | 1 | ENSMUST00 | 0 | Tlcd1 | ENSMUSG0( | false | chr11 | 78176183 | 78179336 | 13.57108 | 58.53591 | 55.70979 | chr11 | 78176113 | 78179448 | 5.11362 | 4.02761 | 3.12991 |
| chr17 | 57250472 | 57250700 | 182 | Promoter (1-2kb) | 17 | 57249451 | 57263697 | 14247 | 1 | ENSMUST00 | 1022 | Trip10 | ENSMUSG0( | false | chr17 | 57247969 | 57250927 | 29.63113 | 94.2009 | 90.76292 | chr17 | 57248321 | 57254242 | 4.6037 | 3.90098 | 2.69294 |
| chr5 | 138171946 | 138172212 | 182 | Promoter (<=1kb) | 5 | 138172002 | 138178708 | 6707 | 1 | ENSMUST00 | 0 | Ap4m1 | ENSMUSG0( | false | chr5 | 138170353 | 138172591 | 27.24215 | 82.29425 | 79.10038 | chr5 | 138170112 | 138173019 | 4.035 | 3.50486 | 2.25711 |
| chr10 | 79935207 | 79935452 | 182 | Intron (ENSMUST00000019708.6 | 10 | 79930424 | 79931218 | 795 | 1 | ENSMUST00 | 4784 | Arid3a | ENSMUSG0( | false | . | . | . | . | . | . | chr10 | 79931436 | 79938436 | 8.97089 | 5.80763 | 6.24553 |
| chr10 | 79937257 | 79937449 | 182 | Intron (ENSMUST00000019708.6 | 10 | 79930424 | 79931218 | 795 | 1 | ENSMUST00 | 6834 | Arid3a | ENSMUSG0( | false | chr10 | 79936883 | 79937621 | 17.45018 | 43.2582 | 40.62776 | chr10 | 79931436 | 79938436 | 8.97089 | 5.80763 | 6.24553 |
| chr6 | 5496440 | 5496632 | 182 | Promoter (<=1kb) | 6 | 5483351 | 5496309 | 12959 | 2 | ENSMUST00 | -132 | Pdk4 | ENSMUSG0( | false | chr6 | 5495847 | 5497159 | 23.05105 | 65.89761 | 62.96862 | chr6 | 5495242 | 5497047 | 3.21388 | 3.1949 | 1.52209 |
| chr17 | 56199898 | 56200186 | 182 | Intron (ENSMUST00000038794.4 | 17 | 56175744 | 56183920 | 8177 | 2 | ENSMUST00 | -15979 | D17Wsu104e | ENSMUSG0( | false | chr17 | 56199916 | 56200182 | 21.65864 | 58.36299 | 55.53803 | . | . | . | . | . | . |
| chr10 | 4430882 | 4431248 | 182 | Promoter (1-2kb) | 10 | 4427243 | 4432295 | 5053 | 2 | ENSMUST00 | 1047 | Rmnd1 | ENSMUSG0( | false | . | . | . | . | . | . | chr10 | 4422045 | 4437373 | 5.83291 | 4.40627 | 3.70343 |
| chr10 | 4472096 | 4472328 | 182 | Distal Intergenic | 10 | 4482502 | 4562231 | 79730 | 1 | ENSMUST00 | -10174 | Ccdc170 | ENSMUSG0( | false | . | . | . | . | . | . | . | . | . | . | . | . |
| chr10 | 34019418 | 34019921 | 182 | Promoter (<=1kb) | 10 | 33996555 | 34019624 | 23070 | 2 | ENSMUST00 | 0 | Rwdd1 | ENSMUSG0( | false | chr10 | 34018380 | 34019734 | 36.88168 | 122.38801 | 118.16834 | . | . | . | . | . | . |
| chr10 | 30618320 | 30618650 | 182 | Promoter (<=1kb) | 10 | 30608141 | 30618479 | 10339 | 2 | ENSMUST00 | 0 | Hint3 | ENSMUSG0( | false | chr10 | 30617599 | 30619020 | 30.60796 | 113.09483 | 109.17188 | . | . | . | . | . | . |
| chr10 | 7725834 | 7726281 | 182 | Promoter (<=1kb) | 10 | 7726000 | 7763149 | 37150 | 1 | ENSMUST00 | 0 | Katna1 | ENSMUSG0( | false | chr10 | 7725607 | 7726872 | 27.78601 | 91.88203 | 88.49705 | chr10 | 7725640 | 7729873 | 4.83468 | 4.03745 | 2.86022 |
| chr10 | 42501835 | 42502371 | 182 | Promoter (<=1kb) | 10 | 42502030 | 42534787 | 32758 | 1 | ENSMUST00 | 0 | Snx3 | ENSMUSG0( | false | chr10 | 42501601 | 42503518 | 47.82016 | 161.55559 | 155.85097 | . | . | . | . | . | . |
| chr10 | 13714949 | 13715289 | 182 | Intron (ENSMUST00000162610.2 | 10 | 13653588 | 13844286 | 190699 | 2 | ENSMUST00 | 128997 | Aig1 | ENSMUSG0( | false | chr10 | 13714661 | 13715258 | 5.28342 | 9.57744 | 7.40675 | chr10 | 13714419 | 13716480 | 4.19322 | 3.48418 | 2.37337 |
| chr10 | 13869450 | 13869766 | 182 | Promoter (<=1kb) | 10 | 13647054 | 13868980 | 221927 | 2 | ENSMUST00 | -471 | Aig1 | ENSMUSG0( | false | chr10 | 13868363 | 13869834 | 22.15371 | 67.81625 | 64.85734 | chr10 | 13867135 | 13870037 | 3.71167 | 3.33932 | 1.98178 |
| chr10 | 41490100 | 41490564 | 182 | Promoter (<=1kb) | 10 | 41485642 | 41490330 | 4689 | 2 | ENSMUST00 | 0 | Smpd2 | ENSMUSG0( | false | chr10 | 41489428 | 41491002 | 32.66734 | 113.81544 | 109.86345 | . | . | . | . | . | . |
| chr10 | 41476175 | 41476617 | 182 | Promoter (<=1kb) | 10 | 41476314 | 41487030 | 10717 | 1 | ENSMUST00 | 0 | Mical1 | ENSMUSG0( | false | chr10 | 41476033 | 41477908 | 15.08796 | 37.05233 | 34.49837 | chr10 | 41476070 | 41477765 | 3.74913 | 3.32055 | 2.01213 |
| chr10 | 39646616 | 39646972 | 182 | Promoter (<=1kb) | 10 | 39646714 | 39654823 | 8110 | 1 | ENSMUST00 | 0 | Traf3ip2 | ENSMUSG0( | false | . | . | . | . | . | . | chr10 | 39646601 | 39647098 | 3.05159 | 2.82867 | 1.44917 |
| chr10 | 39629859 | 39630338 | 182 | Intron (ENSMUST00000019987.6 | 10 | 39641134 | 39645758 | 4625 | 1 | ENSMUST00 | -10796 | Traf3ip2 | ENSMUSG0( | false | chr10 | 39629946 | 39630283 | 4.54943 | 6.49485 | 4.40153 | chr10 | 39624031 | 39633595 | 5.77874 | 4.37306 | 3.67956 |
| chr10 | 39488326 | 39488602 | 182 | Intron (ENSMUST00000148152.2 | 10 | 39511739 | 39564422 | 52684 | 1 | ENSMUST00 | -23137 | Fyn | ENSMUSG0( | true | . | . | . | . | . | . | chr10 | 39483418 | 39514959 | 6.19365 | 4.56605 | 3.99401 |
| chr10 | 39467388 | 39467643 | 182 | Intron (ENSMUST00000148152.2 | 10 | 39454915 | 39564355 | 109441 | 1 | ENSMUST00 | 12474 | Fyn | ENSMUSG0( | true | . | . | . | . | . | . | chr10 | 39448300 | 39473913 | 6.46111 | 4.64933 | 4.22352 |
| chr10 | 39471658 | 39472331 | 182 | Intron (ENSMUST00000148152.2 | 10 | 39454915 | 39564355 | 109441 | 1 | ENSMUST00 | 16744 | Fyn | ENSMUSG0( | true | . | . | . | . | . | . | chr10 | 39448300 | 39473913 | 6.46111 | 4.64933 | 4.22352 |
| chr10 | 18844945 | 18845234 | 182 | Promoter (<=1kb) | 10 | 18845071 | 18857073 | 12003 | 1 | ENSMUST00 | 0 | Perp | ENSMUSG0( | false | chr10 | 18844766 | 18845515 | 21.40274 | 64.72344 | 61.81032 | chr10 | 18844922 | 18845574 | 3.56137 | 3.15254 | 1.85201 |
| chr10 | 53872095 | 53872408 | 182 | Distal Intergenic | 10 | 53696997 | 53750909 | 53913 | 2 | ENSMUST00 | -121187 | Fam184a | ENSMUSG0( | true | . | . | . | . | . | . | chr10 | 53871438 | 53873464 | 4.60895 | 3.89593 | 2.67429 |
| chr10 | 57532030 | 57532449 | 182 | Promoter (<=1kb) | 10 | 57523035 | 57532428 | 9394 | 2 | ENSMUST00 | 0 | Serinc1 | ENSMUSG0( | false | chr10 | 57531168 | 57532829 | 34.54477 | 122.48211 | 118.25522 | . | . | . | . | . | . |
| chr10 | 29313152 | 29313589 | 182 | Promoter (<=1kb) | 10 | 29313166 | 29345560 | 32395 | 1 | ENSMUST00 | 0 | Echdc1 | ENSMUSG0( | false | chr10 | 29312648 | 29314111 | 31.19535 | 97.93977 | 94.4203 | . | . | . | . | . | . |
| chr10 | 27735644 | 27735900 | 182 | Distal Intergenic | 10 | 27422392 | 27619758 | 197367 | 2 | ENSMUST00 | -115887 | Lama2 | ENSMUSG0( | false | . | . | . | . | . | . | . | . | . | . | . | . |
| chr10 | 59221328 | 59222139 | 182 | Promoter (<=1kb) | 10 | 59164113 | 59221847 | 57735 | 2 | ENSMUST00 | 0 | Sep-10 | ENSMUSG0( | false | chr10 | 59220670 | 59223467 | 27.66025 | 106.38611 | 102.65278 | . | . | . | . | . | . |
| chr10 | 58323215 | 58323800 | 182 | Promoter (<=1kb) | 10 | 58323466 | 58424691 | 101226 | 1 | ENSMUST00 | 0 | Lims1 | ENSMUSG0( | false | chr10 | 58322741 | 58323283 | 7.15345 | 16.30454 | 14.02114 | . | . | . | . | . | . |
| chr10 | 72654669 | 72654899 | 182 | Promoter (<=1kb) | 10 | 72654845 | 72674964 | 20120 | 1 | ENSMUST00 | 0 | Zwint | ENSMUSG0( | false | chr10 | 72654498 | 72655923 | 31.83343 | 96.06423 | 92.58627 | chr10 | 72654006 | 72657545 | 6.08676 | 4.606 | 3.87739 |
| chr10 | 71285001 | 71285437 | 182 | Promoter (<=1kb) | 10 | 71254980 | 71285262 | 30283 | 2 | ENSMUST00 | 0 | Ube2d1 | ENSMUSG0( | false | chr10 | 71284153 | 71285709 | 34.16929 | 120.7392 | 116.5701 | . | . | . | . | . | . |
| chr10 | 98930143 | 98930421 | 182 | Intron (ENSMUST00000020107.7 | 10 | 98915152 | 99026143 | 110992 | 1 | ENSMUST00 | 14992 | Atp2b1 | ENSMUSG0( | true | . | . | . | . | . | . | chr10 | 98924769 | 98936268 | 4.32918 | 3.69852 | 2.48767 |
| chr10 | 69212946 | 69213321 | 182 | Promoter (<=1kb) | 10 | 69213084 | 69266190 | 53107 | 1 | ENSMUST00 | 0 | Rhobtb1 | ENSMUSG0( | false | chr10 | 69211966 | 69213907 | 26.86698 | 95.04005 | 91.5885 | . | . | . | . | . | . |
| chr10 | 68277038 | 68277335 | 182 | Promoter (1-2kb) | 10 | 68095593 | 68278726 | 183134 | 2 | ENSMUST00 | 1391 | Arid5b | ENSMUSG0( | false | chr10 | 68275857 | 68280673 | 31.01414 | 97.6227 | 94.10706 | . | . | . | . | . | . |
| chr10 | 68192795 | 68193116 | 182 | Intron (ENSMUST00000020106.7 | 10 | 68095593 | 68278726 | 183134 | 2 | ENSMUST00 | 85610 | Arid5b | ENSMUSG0( | false | . | . | . | . | . | . | chr10 | 68191635 | 68193809 | 3.35252 | 3.24653 | 1.63403 |
| chr10 | 99062699 | 99062897 | 182 | Distal Intergenic | 10 | 99107036 | 99197988 | 90953 | 1 | ENSMUST00 | -44139 | Poc1b | ENSMUSG0( | false | . | . | . | . | . | . | chr10 | 99062095 | 99064568 | 5.42532 | 4.29898 | 3.3479 |
| chr10 | 99961647 | 99961838 | 182 | Distal Intergenic | 10 | 100015630 | 100100413 | 84784 | 1 | ENSMUST00 | -53792 | Kitl | ENSMUSG0( | false | . | . | . | . | . | . | . | . | . | . | . | . |
| chr10 | 21923663 | 21924033 | 182 | Intron (ENSMUST00000120509.2 | 10 | 21928470 | 21990402 | 61933 | 1 | ENSMUST00 | -4437 | Sgk1 | ENSMUSG0( | false | . | . | . | . | . | . | chr10 | 21922739 | 21924324 | 3.58921 | 3.27814 | 1.89733 |
| chr10 | 25362480 | 25362739 | 182 | Promoter (2-3kb) | 10 | 25359798 | 25523519 | 163722 | 1 | ENSMUST00 | 2683 | Epb4.1l2 | ENSMUSG0( | false | . | . | . | . | . | . | chr10 | 25360776 | 25363877 | 4.62066 | 3.92548 | 2.6912 |
| chr10 | 92722233 | 92722629 | 182 | Promoter (<=1kb) | 10 | 92684746 | 92722420 | 37675 | 2 | ENSMUST00 | 0 | Nedd1 | ENSMUSG0( | false | chr10 | 92721740 | 92722810 | 37.0612 | 128.08391 | 123.66181 | chr10 | 92721675 | 92723104 | 3.13429 | 3.02642 | 1.46374 |
| chr10 | 19526439 | 19526742 | 182 | Distal Intergenic | 10 | 19591949 | 19610229 | 18281 | 1 | ENSMUST00 | -65207 | Ifngr1 | ENSMUSG0( | false | . | . | . | . | . | . | . | . | . | . | . | . |
| chr10 | 19508264 | 19508606 | 182 | Distal Intergenic | 10 | 19591949 | 19610229 | 18281 | 1 | ENSMUST00 | -83343 | Ifngr1 | ENSMUSG0( | false | . | . | . | . | . | . | . | . | . | . | . | . |
| chr10 | 94505286 | 94505565 | 182 | Intron (ENSMUST00000132743.1 | 10 | 94514857 | 94590956 | 76100 | 1 | ENSMUST00 | -9292 | Tmcc3 | ENSMUSG0( | false | chr10 | 94505525 | 94505774 | 5.05485 | 7.29253 | 5.17558 | chr10 | 94505385 | 94507246 | 3.84005 | 3.40678 | 2.09134 |
| chr10 | 94479630 | 94479899 | 182 | Intron (ENSMUST00000132743.1 | 10 | 94514857 | 94590956 | 76100 | 1 | ENSMUST00 | -34958 | Tmcc3 | ENSMUSG0( | false | . | . | . | . | . | . | chr10 | 94477425 | 94481453 | 5.61007 | 4.43652 | 3.49994 |
| chr10 | 95392150 | 95392447 | 182 | Promoter (<=1kb) | 10 | 95387667 | 95392967 | 5301 | 2 | ENSMUST00 | 520 | Socs2 | ENSMUSG0( | false | . | . | . | . | . | . | chr10 | 95391484 | 95394454 | 3.62473 | 3.37345 | 1.87327 |

| | | | | | | | | | | | | | | | | | | | | | | | | | | |
|---|---|---|---|---|---|---|---|---|---|---|---|---|---|---|---|---|---|---|---|---|---|---|---|---|---|---|
| chr10 | 85185146 | 85185503 | 182 | Promoter (<=1kb) | 10 | 85131702 | 85185054 | 53353 | 2 | ENSMUST00 | -93 | Cry1 | ENSMUSG0C | false | chr10 | 85183451 | 85186083 | 25.18546 | 107.13963 | 103.3826 | . | . | . | . | . | . |
| chr10 | 86302774 | 86303013 | 182 | Promoter (<=1kb) | 10 | 86302854 | 86343981 | 41128 | 1 | ENSMUST00 | 0 | Timp3 | ENSMUSG0C | false | chr10 | 86300140 | 86302829 | 30.26787 | 111.42836 | 107.5545 | . | . | . | . | . | . |
| chr10 | 86705431 | 86705958 | 182 | Promoter (<=1kb) | 10 | 86690840 | 86705509 | 14670 | 2 | ENSMUST00 | 0 | Hsp90b1 | ENSMUSG0C | false | chr10 | 86705710 | 86706864 | 21.916 | 65.643 | 62.71761 | . | . | . | . | . | . |
| chr10 | 61680197 | 61680696 | 182 | Promoter (<=1kb) | 10 | 61680310 | 61693295 | 12986 | 1 | ENSMUST00 | 0 | Sar1a | ENSMUSG0C | false | chr10 | 61679667 | 61681197 | 29.3377 | 90.74815 | 87.38457 | . | . | . | . | . | . |
| chr10 | 61147482 | 61147790 | 182 | Promoter (<=1kb) | 10 | 61098642 | 61147644 | 49003 | 2 | ENSMUST00 | 0 | Sgpl1 | ENSMUSG0C | false | chr10 | 61145939 | 61148106 | 34.36702 | 111.64683 | 107.76208 | . | . | . | . | . | . |
| chr10 | 61081026 | 61081234 | 182 | Distal Intergenic | 10 | 61089343 | 61094328 | 4986 | 1 | ENSMUST00 | -8109 | Pcbd1 | ENSMUSG0C | false | . | . | . | . | . | . | chr10 | 61080295 | 61081142 | 4.66864 | 3.99583 | 2.7265 |
| chr10 | 60816925 | 60817180 | 182 | Intron (ENSMUST00000077925.5 | 10 | 60762610 | 60831581 | 68972 | 2 | ENSMUST00 | 14401 | Unc5b | ENSMUSG0C | false | . | . | . | . | . | . | chr10 | 60811545 | 60822537 | 5.55309 | 4.2307 | 3.48027 |
| chr10 | 125325734 | 125325920 | 182 | Intron (ENSMUST00000063318.3 | 10 | 125226464 | 125328963 | 102500 | 2 | ENSMUST00 | 3043 | Slc16a7 | ENSMUSG0C | false | . | . | . | . | . | . | chr10 | 125324986 | 125326331 | 3.09554 | 3.10769 | 1.50452 |
| chr10 | 125966016 | 125966337 | 182 | Promoter (<=1kb) | 10 | 125966219 | 126015359 | 49141 | 1 | ENSMUST00 | 0 | Lrig3 | ENSMUSG0C | false | chr10 | 125965700 | 125967874 | 29.28796 | 98.53321 | 94.99628 | . | . | . | . | . | . |
| chr10 | 121993386 | 121993600 | 182 | Intron (ENSMUST00000020322.6 | 10 | 121780991 | 122047308 | 266318 | 2 | ENSMUST00 | 53708 | Srgap1 | ENSMUSG0C | true | . | . | . | . | . | . | chr10 | 121945243 | 122036389 | 5.74397 | 4.3951 | 3.62974 |
| chr10 | 115251503 | 115251724 | 182 | Promoter (<=1kb) | 10 | 115197877 | 115251493 | 53617 | 2 | ENSMUST00 | -11 | Tbc1d15 | ENSMUSG0C | false | chr10 | 115250494 | 115252670 | 40.9965 | 143.89526 | 138.84959 | . | . | . | . | . | . |
| chr11 | 20332745 | 20333375 | 182 | Promoter (<=1kb) | 11 | 20302180 | 20332713 | 30534 | 2 | ENSMUST00 | -33 | Slc1a4 | ENSMUSG0C | false | chr11 | 20331759 | 20333631 | 28.79974 | 86.91003 | 83.63136 | chr11 | 20331503 | 20333950 | 4.39516 | 3.71038 | 2.51618 |
| chr10 | 116301316 | 116301509 | 182 | Promoter (<=1kb) | 10 | 116301374 | 116389538 | 88165 | 1 | ENSMUST00 | 0 | Ptprb | ENSMUSG0C | false | chr10 | 116301308 | 116302192 | 7.96309 | 14.9624 | 12.70017 | chr10 | 116300481 | 116301770 | 4.54644 | 3.83629 | 2.66146 |
| chr10 | 80226275 | 80226748 | 182 | Promoter (<=1kb) | 10 | 80226434 | 80243903 | 17470 | 1 | ENSMUST00 | 0 | Mum1 | ENSMUSG0C | false | chr10 | 80225979 | 80227278 | 25.84672 | 90.2271 | 86.88143 | chr10 | 80226050 | 80228763 | 4.03176 | 3.63655 | 2.20393 |
| chr10 | 80433557 | 80433847 | 182 | Promoter (<=1kb) | 10 | 80409514 | 80433637 | 24124 | 2 | ENSMUST00 | 0 | Tcf3 | ENSMUSG0C | false | chr10 | 80432209 | 80434035 | 41.61197 | 136.07095 | 131.34828 | . | . | . | . | . | . |
| chr11 | 12536480 | 12536765 | 182 | Distal Intergenic | 11 | 12236678 | 12464960 | 228283 | 2 | ENSMUST00 | -71521 | Cobl | ENSMUSG0C | false | . | . | . | . | . | . | chr11 | 12531938 | 12538351 | 4.20628 | 3.585 | 2.37047 |
| chr10 | 75037074 | 75037293 | 182 | Promoter (<=1kb) | 10 | 75037084 | 75054103 | 17020 | 1 | ENSMUST00 | 0 | Rab36 | ENSMUSG0C | false | chr10 | 75036771 | 75037449 | 9.76265 | 21.55577 | 19.19873 | chr10 | 75037217 | 75037536 | 3.31152 | 3.14377 | 1.65276 |
| chr10 | 117709973 | 117710938 | 182 | Promoter (<=1kb) | 10 | 117688888 | 117710758 | 21871 | 2 | ENSMUST00 | 0 | Mdm2 | ENSMUSG0C | false | chr10 | 117709420 | 117711415 | 35.2856 | 127.03558 | 122.65217 | . | . | . | . | . | . |
| chr10 | 110918867 | 110919193 | 182 | Promoter (<=1kb) | 10 | 110920177 | 110939599 | 19423 | 1 | ENSMUST00 | -984 | Csrp2 | ENSMUSG0C | false | chr10 | 110918934 | 110921076 | 35.27562 | 110.11647 | 106.27654 | chr10 | 110918734 | 110919574 | 4.48241 | 3.8018 | 2.57664 |
| chr10 | 80797841 | 80798350 | 182 | Promoter (<=1kb) | 10 | 80798198 | 80804924 | 6727 | 1 | ENSMUST00 | 0 | Sf3a2 | ENSMUSG0C | false | chr10 | 80797167 | 80799728 | 24.60788 | 97.30984 | 93.80507 | . | . | . | . | . | . |
| chr10 | 80900766 | 80901208 | 182 | Promoter (<=1kb) | 10 | 80899450 | 80900969 | 1520 | 2 | ENSMUST00 | 0 | Timm13 | ENSMUSG0C | false | chr10 | 80899740 | 80900929 | 26.03857 | 105.07202 | 101.37238 | chr10 | 80899166 | 80901694 | 4.66098 | 3.90798 | 2.72875 |
| chr4 | 144949562 | 144949868 | 182 | Distal Intergenic | 4 | 144972623 | 144974683 | 2061 | 2 | ENSMUST00 | 24815 | Vps13d | ENSMUSG0C | false | . | . | . | . | . | . | chr4 | 144947700 | 144949709 | 2.9359 | 2.89189 | 1.38181 |
| chr10 | 81346636 | 81346992 | 182 | Promoter (<=1kb) | 10 | 81346052 | 81346730 | 679 | 2 | ENSMUST00 | 0 | Hmg20b | ENSMUSG0C | false | chr10 | 81346895 | 81347134 | 7.96309 | 14.9624 | 12.70017 | chr10 | 81339953 | 81351319 | 5.28841 | 4.20398 | 3.24758 |
| chr10 | 77417686 | 77418037 | 182 | Promoter (<=1kb) | 10 | 77325338 | 77417854 | 92517 | 2 | ENSMUST00 | 0 | Adarb1 | ENSMUSG0C | false | chr10 | 77416895 | 77418831 | 28.12373 | 86.68959 | 83.41467 | chr10 | 77416518 | 77418791 | 6.01625 | 4.52948 | 3.83448 |
| chr11 | 32676926 | 32677140 | 182 | Intron (ENSMUST00000109366.2 | 11 | 32642904 | 32720574 | 77671 | 1 | ENSMUST00 | 34023 | Fbxw11 | ENSMUSG0C | false | . | . | . | . | . | . | chr11 | 32670392 | 32685981 | 4.84811 | 3.97997 | 2.89823 |
| chr11 | 32538461 | 32538704 | 182 | Intron (ENSMUST00000102821.3 | 11 | 32533305 | 32624587 | 91283 | 1 | ENSMUST00 | 5157 | Stk10 | ENSMUSG0C | false | . | . | . | . | . | . | . | . | . | . | . | . |
| chr11 | 30131684 | 30131967 | 182 | Intron (ENSMUST00000006629.8 | 11 | 30109871 | 30111359 | 1489 | 2 | ENSMUST00 | -20326 | Sptbn1 | ENSMUSG0C | false | . | . | . | . | . | . | . | . | . | . | . | . |
| chr11 | 30291287 | 30291493 | 182 | Distal Intergenic | 11 | 30216659 | 30268175 | 51517 | 2 | ENSMUST00 | -23113 | Sptbn1 | ENSMUSG0C | false | . | . | . | . | . | . | chr11 | 30287587 | 30292909 | 4.20953 | 3.69116 | 2.33969 |
| chr11 | 30326267 | 30326468 | 182 | Distal Intergenic | 11 | 30216659 | 30268175 | 51517 | 2 | ENSMUST00 | -58093 | Sptbn1 | ENSMUSG0C | false | . | . | . | . | . | . | . | . | . | . | . | . |
| chr11 | 30197883 | 30198208 | 182 | Promoter (<=1kb) | 11 | 30106785 | 30198257 | 91473 | 2 | ENSMUST00 | 49 | Sptbn1 | ENSMUSG0C | false | chr11 | 30197643 | 30198889 | 20.53639 | 56.4273 | 53.62976 | chr11 | 30194584 | 30200966 | 3.79922 | 3.47993 | 2.01627 |
| chr11 | 54028073 | 54028383 | 182 | Promoter (<=1kb) | 11 | 53983123 | 54028090 | 44968 | 2 | ENSMUST00 | 0 | Slc22a4 | ENSMUSG0C | false | chr11 | 54027815 | 54028717 | 13.94198 | 39.24413 | 36.66504 | chr11 | 54027732 | 54029145 | 4.1339 | 3.49806 | 2.32595 |
| chr11 | 46291156 | 46291412 | 182 | Intron (ENSMUST00000165599.3 | 11 | 46196071 | 46270669 | 74599 | 2 | ENSMUST00 | -20488 | Cyfip2 | ENSMUSG0C | false | . | . | . | . | . | . | chr11 | 46290894 | 46291790 | 3.40791 | 3.09996 | 1.72999 |
| chr11 | 51584430 | 51584782 | 182 | Promoter (<=1kb) | 11 | 51584757 | 51603254 | 18498 | 1 | ENSMUST00 | 0 | Phykpl | ENSMUSG0C | false | chr11 | 51583944 | 51585416 | 28.56742 | 103.16996 | 99.51957 | chr11 | 51583104 | 51586617 | 3.96976 | 3.453 | 2.19621 |
| chr11 | 53279760 | 53280123 | 182 | Intron (ENSMUST00000020630.7 | 11 | 53262880 | 53267241 | 4362 | 2 | ENSMUST00 | -12520 | Hspa4 | ENSMUSG0C | false | . | . | . | . | . | . | chr11 | 53277056 | 53281698 | 3.86967 | 3.48046 | 2.09446 |
| chr11 | 50024827 | 50025285 | 182 | Promoter (<=1kb) | 11 | 50025346 | 50104758 | 79413 | 1 | ENSMUST00 | -61 | Rnf130 | ENSMUSG0C | false | chr11 | 50024682 | 50026322 | 32.66734 | 113.81544 | 109.86345 | . | . | . | . | . | . |
| chr11 | 50036882 | 50037140 | 182 | Intron (ENSMUST00000054684.8 | 11 | 50025348 | 50104753 | 79406 | 1 | ENSMUST00 | 11535 | Rnf130 | ENSMUSG0C | false | . | . | . | . | . | . | chr11 | 50033034 | 50039318 | 4.11228 | 3.67096 | 2.27366 |
| chr11 | 51263001 | 51263220 | 182 | Promoter (<=1kb) | 11 | 51263114 | 51281764 | 18651 | 1 | ENSMUST00 | 0 | Clk4 | ENSMUSG0C | false | chr11 | 51262442 | 51264681 | 30.60796 | 113.09483 | 109.17188 | . | . | . | . | . | . |
| chr11 | 51857742 | 51858116 | 182 | Promoter (<=1kb) | 11 | 51813455 | 51857653 | 44199 | 2 | ENSMUST00 | -90 | Jade2 | ENSMUSG0C | false | chr11 | 51855275 | 51859109 | 33.55804 | 103.05054 | 99.40515 | . | . | . | . | . | . |
| chr11 | 51858371 | 51859095 | 182 | Promoter (<=1kb) | 11 | 51813455 | 51857653 | 44199 | 2 | ENSMUST00 | -719 | Jade2 | ENSMUSG0C | false | chr11 | 51855275 | 51859109 | 33.55804 | 103.05054 | 99.40515 | . | . | . | . | . | . |
| chr11 | 52004053 | 52004541 | 182 | Promoter (<=1kb) | 11 | 52004221 | 52033802 | 29582 | 1 | ENSMUST00 | 0 | Cdkl3 | ENSMUSG0C | false | chr11 | 52003475 | 52004117 | 17.28391 | 44.47248 | 41.82566 | chr11 | 52000966 | 52005906 | 4.50276 | 3.85227 | 2.59917 |
| chr11 | 5297248 | 5297567 | 182 | Intron (ENSMUST00000109867.2 | 11 | 5191552 | 5261558 | 70007 | 2 | ENSMUST00 | -35691 | Kremen1 | ENSMUSG0C | true | . | . | . | . | . | . | chr11 | 5295333 | 5298636 | 5.85542 | 4.37827 | 3.74912 |
| chr11 | 54955884 | 54956153 | 182 | Promoter (<=1kb) | 11 | 54910785 | 54956122 | 45338 | 2 | ENSMUST00 | 0 | Tnip1 | ENSMUSG0C | false | chr11 | 54955487 | 54956498 | 33.41831 | 117.26761 | 113.21259 | . | . | . | . | . | . |
| chr7 | 27860361 | 27860664 | 182 | Promoter (1-2kb) | 7 | 27862557 | 27879567 | 17011 | 1 | ENSMUST00 | -1893 | Zfp607 | ENSMUSG0C | false | chr7 | 27859993 | 27861362 | 20.27628 | 60.14116 | 57.29097 | chr7 | 27858891 | 27862600 | 3.91111 | 3.41005 | 2.15579 |
| chr11 | 8457928 | 8458209 | 182 | Intron (ENSMUST00000020695.7 | 11 | 8451070 | 8468675 | 17606 | 2 | ENSMUST00 | 10466 | Tns3 | ENSMUSG0C | false | . | . | . | . | . | . | chr11 | 8438250 | 8474887 | 7.10839 | 4.95354 | 4.75045 |
| chr11 | 8446118 | 8446347 | 182 | Intron (ENSMUST00000020695.7 | 11 | 8451070 | 8468675 | 17606 | 2 | ENSMUST00 | 22328 | Tns3 | ENSMUSG0C | false | . | . | . | . | . | . | chr11 | 8438250 | 8474887 | 7.10839 | 4.95354 | 4.75045 |
| chr11 | 8443629 | 8444065 | 182 | Exon (ENSMUST00000020695.7/ | 11 | 8451070 | 8468675 | 17606 | 2 | ENSMUST00 | 24610 | Tns3 | ENSMUSG0C | false | . | . | . | . | . | . | chr11 | 8438250 | 8474887 | 7.10839 | 4.95354 | 4.75045 |
| chr11 | 8525667 | 8526008 | 182 | Intron (ENSMUST00000020695.7 | 11 | 8537539 | 8545351 | 7813 | 2 | ENSMUST00 | 19343 | Tns3 | ENSMUSG0C | false | . | . | . | . | . | . | chr11 | 8522913 | 8527168 | 5.22155 | 3.92632 | 3.2343 |
| chr1 | 134066255 | 134066447 | 182 | Distal Intergenic | 1 | 134075170 | 134079120 | 3951 | 2 | ENSMUST00 | 12673 | Btg2 | ENSMUSG0C | false | . | . | . | . | . | . | chr1 | 134058485 | 134067448 | 6.65638 | 4.82565 | 4.35972 |
| chr11 | 7101756 | 7102260 | 182 | Intron (ENSMUST00000020706.4 | 11 | 7063489 | 7178506 | 115018 | 1 | ENSMUST00 | 38268 | Adcy1 | ENSMUSG0C | true | . | . | . | . | . | . | chr11 | 7095593 | 7110246 | 5.98026 | 4.46357 | 3.82741 |
| chr11 | 59191761 | 59192123 | 182 | Promoter (<=1kb) | 11 | 59183875 | 59191952 | 8078 | 2 | ENSMUST00 | 0 | Guk1 | ENSMUSG0C | false | chr11 | 59191203 | 59192601 | 32.62678 | 97.43105 | 93.92233 | chr11 | 59186978 | 59192740 | 4.28508 | 3.71734 | 2.41498 |
| chr11 | 3289507 | 3290303 | 182 | Promoter (<=1kb) | 11 | 3289993 | 3295429 | 5437 | 1 | ENSMUST00 | 0 | Patz1 | ENSMUSG0C | false | chr11 | 3288485 | 3291480 | 34.92444 | 106.42502 | 102.68874 | . | . | . | . | . | . |
| chr11 | 3292107 | 3292327 | 182 | Promoter (<=1kb) | 11 | 3292195 | 3294668 | 2474 | 1 | ENSMUST00 | 0 | Patz1 | ENSMUSG0C | false | . | . | . | . | . | . | chr11 | 3290999 | 3294526 | 5.47377 | 4.06255 | 3.42082 |
| chr11 | 28812722 | 28812903 | 182 | Intron (ENSMUST00000138164.1 | 11 | 28853204 | 28926743 | 73540 | 1 | ENSMUST00 | -40301 | Efemp1 | ENSMUSG0C | false | . | . | . | . | . | . | chr11 | 28811965 | 28816045 | 4.09984 | 3.68608 | 2.26904 |
| chr11 | 87579442 | 87579734 | 182 | Promoter (<=1kb) | 11 | 87578928 | 87589274 | 10347 | 1 | ENSMUST00 | 515 | Sep-04 | ENSMUSG0C | false | . | . | . | . | . | . | chr11 | 87577600 | 87579497 | 5.23962 | 4.02541 | 3.23451 |
| chr11 | 57627999 | 57628191 | 182 | Intron (ENSMUST00000142486.1 | 11 | 57645442 | 57787514 | 142073 | 1 | ENSMUST00 | -17251 | Galnt10 | ENSMUSG0C | true | . | . | . | . | . | . | chr11 | 57625503 | 57628742 | 4.3087 | 3.68467 | 2.44199 |
| chr11 | 60699587 | 60699908 | 182 | Promoter (<=1kb) | 11 | 60699723 | 60714186 | 14464 | 1 | ENSMUST00 | 0 | Llgl1 | ENSMUSG0C | false | chr11 | 60698688 | 60700396 | 34.81759 | 111.39526 | 107.52535 | chr11 | 60698585 | 60702033 | 4.65245 | 3.76135 | 2.76304 |
| chr12 | 33181852 | 33182074 | 182 | Intron (ENSMUST00000077456.7 | 12 | 33147805 | 33328849 | 181045 | 1 | ENSMUST00 | 34048 | Atxn7l1 | ENSMUSG0C | false | . | . | . | . | . | . | chr12 | 33172634 | 33186886 | 4.76751 | 3.94578 | 2.83396 |
| chr12 | 33302813 | 33303250 | 182 | Promoter (<=1kb) | 12 | 33302515 | 33368273 | 65759 | 1 | ENSMUST00 | 299 | Atxn7l1 | ENSMUSG0C | false | chr12 | 33302526 | 33303732 | 21.79225 | 59.43074 | 56.59393 | . | . | . | . | . | . |
| chr12 | 17266407 | 17266692 | 182 | Promoter (<=1kb) | 12 | 17266545 | 17281810 | 15266 | 1 | ENSMUST00 | 0 | Pdia6 | ENSMUSG0C | false | chr12 | 17266519 | 17267741 | 33.04283 | 115.53907 | 111.53828 | . | . | . | . | . | . |
| chr12 | 13269033 | 13269236 | 182 | Promoter (<=1kb) | 12 | 13269127 | 13583810 | 314684 | 1 | ENSMUST00 | 0 | Nbas | ENSMUSG0C | false | chr12 | 13268801 | 13270170 | 26.32471 | 106.6389 | 102.89503 | chr12 | 13268553 | 13270414 | 4.02687 | 3.49821 | 2.23229 |
| chr12 | 8913748 | 8913952 | 182 | Distal Intergenic | 12 | 8921664 | 8938742 | 17079 | 1 | ENSMUST00 | -7712 | Laptm4a | ENSMUSG0C | true | . | . | . | . | . | . | chr12 | 8911996 | 8918641 | 6.86212 | 4.89812 | 4.53488 |
| chr12 | 35046849 | 35047992 | 182 | Promoter (<=1kb) | 12 | 35047189 | 35147469 | 100281 | 1 | ENSMUST00 | 0 | Snx13 | ENSMUSG0C | false | chr12 | 35046630 | 35048578 | 37.47034 | 122.34885 | 118.13707 | . | . | . | . | . | . |
| chr12 | 16589808 | 16590229 | 182 | Promoter (<=1kb) | 12 | 16535669 | 16589770 | 54102 | 2 | ENSMUST00 | -39 | Lpin1 | ENSMUSG0C | false | chr12 | 16588648 | 16589858 | 32.29185 | 112.09676 | 108.19604 | chr12 | 16582080 | 16592266 | 4.5687 | 3.8842 | 2.65368 |
| chr12 | 8521435 | 8521709 | 182 | Distal Intergenic | 12 | 8529345 | 8539545 | 10201 | 2 | ENSMUST00 | 17836 | Slc7a15 | ENSMUSG0C | false | . | . | . | . | . | . | . | . | . | . | . | . |
| chr12 | 10390724 | 10391123 | 182 | Promoter (<=1kb) | 12 | 10390780 | 10395562 | 4783 | 1 | ENSMUST00 | 0 | Rdh14 | ENSMUSG0C | false | chr12 | 10390672 | 10391830 | 20.9437 | 83.15932 | 79.95254 | . | . | . | . | . | . |
| chr12 | 10525041 | 10525234 | 182 | Distal Intergenic | 12 | 10390780 | 10395562 | 4783 | 1 | ENSMUST00 | 134262 | Rdh14 | ENSMUSG0C | false | . | . | . | . | . | . | chr12 | 10524421 | 10525681 | 3.5699 | 3.3475 | 1.88684 |
| chr11 | 112986007 | 112986203 | 182 | Distal Intergenic | 11 | 112825179 | 112832469 | 7291 | 1 | ENSMUST00 | 160829 | 4933434M16 | ENSMUSG0C | false | . | . | . | . | . | . | . | . | . | . | . | . |
| chr12 | 5213363 | 5213724 | 182 | Intron (ENSMUST00000020958.8 | 12 | 5077472 | 5375682 | 298211 | 2 | ENSMUST00 | 161958 | Klhl29 | ENSMUSG0C | true | . | . | . | . | . | . | chr12 | 5210811 | 5216549 | 5.72225 | 4.09005 | 3.64171 |
| chr12 | 5250175 | 5250376 | 182 | Intron (ENSMUST00000020958.8 | 12 | 5077472 | 5375682 | 298211 | 2 | ENSMUST00 | 125306 | Klhl29 | ENSMUSG0C | true | . | . | . | . | . | . | chr12 | 5242191 | 5258451 | 5.65112 | 4.34582 | 3.54796 |
| chr12 | 4907570 | 4907757 | 182 | Promoter (<=1kb) | 12 | 4879032 | 4907705 | 28674 | 2 | ENSMUST00 | 0 | Ubxn2a | ENSMUSG0C | false | chr12 | 4906536 | 4908244 | 36.39542 | 121.53264 | 117.33772 | . | . | . | . | . | . |
| chr12 | 26469734 | 26470005 | 182 | Promoter (<=1kb) | 12 | 26469204 | 26479835 | 10632 | 1 | ENSMUST00 | 531 | Cmpk2 | ENSMUSG0C | false | chr12 | 26468972 | 26470438 | 38.87692 | 132.39133 | 127.80991 | chr12 | 26469963 | 26471616 | 3.23561 | 3.17797 | 1.6088 |
| chr12 | 31654526 | 31654899 | 182 | Promoter (<=1kb) | 12 | 31640055 | 31654826 | 14772 | 2 | ENSMUST00 | 0 | Dus4l | ENSMUSG0C | false | chr12 | 31654187 | 31655804 | 38.97723 | 131.47449 | 126.92584 | . | . | . | . | . | . |
| chr12 | 3871033 | 3871321 | 182 | Intron (ENSMUST00000172719.2 | 12 | 3866191 | 3876062 | 9872 | 1 | ENSMUST00 | 4843 | Dnmt3a | ENSMUSG0C | false | . | . | . | . | . | . | chr12 | 3865258 | 3872731 | 5.39308 | 4.24242 | 3.33177 |
| chr12 | 29937922 | 29938237 | 182 | Promoter (<=1kb) | 12 | 29938036 | 30017658 | 79623 | 1 | ENSMUST00 | 0 | Pxdn | ENSMUSG0C | false | chr12 | 29937557 | 29939537 | 47.33181 | 157.24483 | 151.69102 | chr12 | 29936541 | 29947539 | 4.38488 | 3.75724 | 2.49928 |
| chr11 | 83473039 | 83473429 | 182 | Promoter (<=1kb) | 11 | 83473086 | 83489213 | 16128 | 1 | ENSMUST00 | 0 | Taf15 | ENSMUSG0C | false | chr11 | 83472283 | 83474074 | 23.61999 | 85.9385 | 82.67356 | . | . | . | . | . | . |
| chr11 | 105967833 | 105968126 | 182 | Promoter (<=1kb) | 11 | 105967945 | 105989964 | 22020 | 1 | ENSMUST00 | 0 | Ace | ENSMUSG0C | false | chr11 | 105967580 | 105968446 | 20.72239 | 59.25234 | 56.41743 | chr11 | 105967288 | 105968712 | 7.79042 | 5.32035 | 5.31632 |
| chr11 | 105126111 | 105126363 | 182 | Promoter (<=1kb) | 11 | 105126425 | 105140394 | 13970 | 1 | ENSMUST00 | -62 | Mettl2 | ENSMUSG0C | false | chr11 | 105125994 | 105126324 | 17.92175 | 44.05972 | 41.41886 | . | . | . | . | . | . |
| chr11 | 105294178 | 105294441 | 182 | Promoter (1-2kb) | 11 | 105292668 | 105343871 | 51204 | 1 | ENSMUST00 | 1511 | Mrc2 | ENSMUSG0C | true | chr11 | 105293126 | 105294450 | 18.57313 | 46.67894 | 44.00877 | chr11 | 105293909 | 105305115 | 5.5399 | 4.23471 | 3.46028 |
| chr11 | 105312925 | 105313561 | 182 | Intron (ENSMUST00000126931.1 | 11 | 105329052 | 105332299 | 3248 | 1 | ENSMUST00 | -15491 | Mrc2 | ENSMUSG0C | true | . | . | . | . | . | . | chr11 | 105306066 | 105319029 | 6.36006 | 4.57632 | 4.15147 |
| chr11 | 81153513 | 81153766 | 182 | Promoter (<=1kb) | 11 | 80880169 | 81153321 | 273153 | 2 | ENSMUST00 | -193 | Asic2 | ENSMUSG0C | true | chr11 | 81151425 | 81153765 | 25.90858 | 83.69981 | 80.48042 | . | . | . | . | . | . |
| chr11 | 80184101 | 80184377 | 182 | Promoter (<=1kb) | 11 | 80184223 | 80190011 | 5789 | 1 | ENSMUST00 | 0 | Rnf135 | ENSMUSG0C | false | chr11 | 80183321 | 80184415 | 23.34944 | 75.34173 | 72.26852 | chr11 | 80182939 | 80185042 | 3.56632 | 3.39317 | 1.83406 |
| chr11 | 106716260 | 106716527 | 182 | Promoter (<=1kb) | 11 | 106654218 | 106715394 | 61177 | 2 | ENSMUST00 | -867 | Pecam1 | ENSMUSG0C | false | . | . | . | . | . | . | chr11 | 106716125 | 106716939 | 3.22969 | 2.88101 | 1.59055 |
| chr11 | 108583805 | 108583993 | 182 | Intron (ENSMUST00000150863.3 | 11 | 108587077 | 108757737 | 170661 | 1 | ENSMUST00 | -3084 | Cep112 | ENSMUSG0C | false | . | . | . | . | . | . | chr11 | 108583889 | 108585059 | 3.26636 | 3.20231 | 1.57019 |
| chr11 | 115603583 | 115604209 | 182 | Promoter (<=1kb) | 11 | 115584255 | 115604051 | 19797 | 2 | ENSMUST00 | 0 | Gga3 | ENSMUSG0C | false | chr11 | 115602764 | 115604989 | 42.74922 | 148.11885 | 142.90198 | . | . | . | . | . | . |
| chr11 | 74650372 | 74650582 | 182 | Promoter (<=1kb) | 11 | 74649830 | 74657339 | 7510 | 1 | ENSMUST00 | 543 | Cluh | ENSMUSG0C | false | chr11 | 74648841 | 74650947 | 35.67123 | 127.73884 | 123.33056 | chr11 | 74648886 | 74653397 | 4.70613 | 3.95834 | 2.75929 |
| chr11 | 115612566 | 115613041 | 182 | Promoter (<=1kb) | 11 | 115607918 | 115612887 | 4970 | 2 | ENSMUST00 | 0 | Mif4gd | ENSMUSG0C | false | chr11 | 115611872 | 115613594 | 40.85493 | 131.46736 | 126.92584 | chr11 | 115611527 | 115616014 | 4.75349 | 3.90772 | 2.82178 |
| chr11 | 116012662 | 116012863 | 182 | Promoter (<=1kb) | 11 | 116008457 | 116012719 | 4263 | 2 | ENSMUST00 | 0 | Galk1 | ENSMUSG0C | false | chr11 | 116012188 | 116013235 | 21.37461 | 59.55079 | 56.71094 | chr11 | 116011820 | 116013241 | 3.1966 | 2.97941 | 1.57183 |
| chr11 | 116168030 | 116168327 | 182 | Promoter (<=1kb) | 11 | 116142285 | 116168153 | 25869 | 2 | ENSMUST00 | 0 | Fbf1 | ENSMUSG0C | false | chr11 | 116167477 | 116168650 | 22.24324 | 59.3644 | 56.5279 | chr11 | 116167025 | 116169019 | 4.61433 | 3.9693 | 2.67586 |
| chr11 | 115823547 | 115824130 | 182 | Promoter (<=1kb) | 11 | 115824049 | 115855780 | 31732 | 1 | ENSMUST00 | 0 | Llgl2 | ENSMUSG0C | false | chr11 | 115823561 | 115824840 | 29.2476 | 106.45832 | 102.71833 | chr11 | 115823381 | 115825618 | 4.38468 | 3.834 | 2.48208 |
| chr11 | 73048040 | 73048396 | 182 | Promoter (<=1kb) | 11 | 73047873 | 73056737 | 8865 | 1 | ENSMUST00 | 168 | 1200014J11F | ENSMUSG0C | false | chr11 | 73047761 | 73048865 | 24.12502 | 101.02714 | 97.43493 | . | . | . | . | . | . |
| chr11 | 116581112 | 116581683 | 182 | Promoter (<=1kb) | 11 | 116537740 | 116581447 | 43708 | 2 | ENSMUST00 | 0 | Ube2o | ENSMUSG0C | false | chr11 | 116580541 | 116581616 | 25.48472 | 95.2657 | 91.8051 | chr11 | 116580016 | 116581701 | 3.57542 | 3.18493 | 1.85995 |
| chr11 | 70237479 | 70238053 | 182 | Promoter (<=1kb) | 11 | 70235206 | 70237853 | 2648 | 2 | ENSMUST00 | 0 | 0610010K14I | ENSMUSG0C | false | chr11 | 70237200 | 70240661 | 38.20206 | 124.69703 | 120.39601 | chr11 | 70236886 | 70239229 | 4.7163 | 3.95495 | 2.76883 |
| chr11 | 76791373 | 76791617 | 182 | Promoter (<=1kb) | 11 | 76786145 | 76791022 | 4878 | 2 | ENSMUST00 | -352 | Cpd | ENSMUSG0C | false | . | . | . | . | . | . | chr11 | 76791438 | 76792108 | 4.51055 | 3.88282 | 2.60797 |
| chr11 | 76355545 | 76355736 | 182 | Intron (ENSMUST00000021204.3 | 11 | 76358803 | 76399068 | 40266 | 2 | ENSMUST00 | 43332 | Nxn | ENSMUSG0C | true | . | . | . | . | . | . | chr11 | 76353940 | 76397781 | 6.09271 | 4.53696 | 3.91295 |
| chr11 | 76324860 | 76325105 | 182 | Intron (ENSMUST00000021204.3 | 11 | 76358803 | 76399068 | 40266 | 2 | ENSMUST00 | 73963 | Nxn | ENSMUSG0C | true | . | . | . | . | . | . | chr11 | 76321722 | 76329132 | 6.38123 | 4.7269 | 4.13711 |
| chr11 | 76390745 | 76390994 | 182 | Intron (ENSMUST00000021204.3 | 11 | 76358803 | 76399068 | 40266 | 2 | ENSMUST00 | 8074 | Nxn | ENSMUSG0C | true | . | . | . | . | . | . | chr11 | 76353940 | 76397781 | 6.09271 | 4.53696 | 3.91295 |
| chr11 | 76380073 | 76380525 | 182 | Intron (ENSMUST00000021204.3 | 11 | 76358803 | 76399068 | 40266 | 2 | ENSMUST00 | 18543 | Nxn | ENSMUSG0C | true | . | . | . | . | . | . | chr11 | 76353940 | 76397781 | 6.09271 | 4.53696 | 3.91295 |
| chr11 | 94323391 | 94323629 | 182 | Promoter (1-2kb) | 11 | 94291139 | 94321988 | 30850 | 2 | ENSMUST00 | -1404 | Luc7l3 | ENSMUSG0C | false | . | . | . | . | . | . | chr11 | 94322570 | 94327227 | 5.14346 | 4.13085 | 3.14416 |
| chr11 | 96271839 | 96272035 | 182 | Promoter (<=1kb) | 11 | 96271523 | 96276595 | 5073 | 1 | ENSMUST00 | 317 | Hoxb9 | ENSMUSG0C | false | chr11 | 96270998 | 96272761 | 20.53639 | 56.4273 | 53.62976 | chr11 | 96269585 | 96272568 | 5.09741 | 3.93611 | 3.1112 |
| chr11 | 96777542 | 96777818 | 182 | Promoter (<=1kb) | 11 | 96767877 | 96777559 | 9683 | 2 | ENSMUST00 | 0 | Snx11 | ENSMUSG0C | false | chr11 | 96777059 | 96777646 | 24.03115 | 75.67385 | 72.59326 | chr11 | 96775419 | 96778355 | 3.62842 | 3.36837 | 1.93168 |
| chr11 | 69098590 | 69098936 | 182 | Promoter (<=1kb) | 11 | 69098948 | 69109960 | 11013 | 1 | ENSMUST00 | -12 | Per1 | ENSMUSG0C | true | chr11 | 69097223 | 69098719 | 11.28697 | 24.15705 | 21.76529 | chr11 | 69097810 | 69105920 | 6.46779 | 4.59054 | 4.2103 |
| chr11 | 69094925 | 69095336 | 182 | Promoter (<=1kb) | 11 | 69095217 | 69106458 | 11242 | 1 | ENSMUST00 | 0 | Per1 | ENSMUSG0C | true | chr11 | 69095226 | 69097021 | 30.07331 | 91.05476 | 87.68732 | . | . | . | . | . | . |
| chr11 | 68386791 | 68387277 | 182 | Promoter (<=1kb) | 11 | 68209364 | 68386826 | 177463 | 2 | ENSMUST00 | 0 | Ntn1 | ENSMUSG0C | false | chr11 | 68384591 | 68387483 | 22.10575 | 73.03297 | 69.99519 | chr11 | 68384510 | 68387569 | 5.19498 | 3.94343 | 3.20337 |
| chr11 | 102309036 | 102309316 | 182 | Promoter (<=1kb) | 11 | 102308530 | 102309440 | 911 | 2 | ENSMUST00 | 124 | Ubtf | ENSMUSG0C | false | chr11 | 102309181 | 102309720 | 11.55672 | 26.39465 | 23.97632 | chr11 | 102308313 | 102311528 | 7.43571 | 5.10275 | 5.04611 |
| chr11 | 103016922 | 103017222 | 182 | Promoter (<=1kb) | 11 | 102994047 | 103017168 | 23122 | 2 | ENSMUST00 | 0 | Dcakd | ENSMUSG0C | false | chr11 | 103016419 | 103017187 | 30.28657 | 98.33401 | 94.80609 | . | . | . | . | . | . |
| chr11 | 103267335 | 103267895 | 182 | Promoter (<=1kb) | 11 | 103219762 | 103267472 | 47711 | 2 | ENSMUST00 | 0 | Map3k14 | ENSMUSG0C | false | chr11 | 103266683 | 103268189 | 28.22734 | 101.53345 | 97.92473 | chr11 | 103265600 | 103268313 | 4.1413 | 3.66067 | 2.29885 |
| chr12 | 98197599 | 98197801 | 182 | Distal Intergenic | 12 | 98202304 | 98259459 | 57156 | 2 | ENSMUST00 | 61658 | Galc | ENSMUSG0C | false | . | . | . | . | . | . | chr12 | 98196932 | 98199409 | 5.31653 | 3.97646 | 3.2995 |
| chr12 | 98249990 | 98250240 | 182 | Intron (ENSMUST00000021390.7 | 12 | 98202304 | 98259459 | 57156 | 2 | ENSMUST00 | 9219 | Galc | ENSMUSG0C | false | . | . | . | . | . | . | chr12 | 98249310 | 98251433 | 3.5663 | 3.28558 | 1.84541 |
| chr16 | 20717564 | 20718108 | 182 | Promoter (<=1kb) | 16 | 20717665 | 20722267 | 4603 | 1 | ENSMUST00 | 0 | Polr2h | ENSMUSG0C | false | chr16 | 20715653 | 20718713 | 17.53424 | 68.85774 | 65.88623 | chr16 | 20714689 | 20719050 | 5.50285 | 4.33961 | 3.41363 |
| chr12 | 87147375 | 87147819 | 182 | Promoter (<=1kb) | 12 | 87147707 | 87164721 | 17015 | 1 | ENSMUST00 | 0 | Gstz1 | ENSMUSG0C | false | chr12 | 87146697 | 87148637 | 42.6403 | 153.60713 | 148.18196 | . | . | . | . | . | . |
| chr12 | 87381432 | 87381769 | 182 | Intron (ENSMUST00000021424.4 | 12 | 87346734 | 87388241 | 41508 | 2 | ENSMUST00 | 6472 | Sptlc2 | ENSMUSG0C | false | . | . | . | . | . | . | chr12 | 87367947 | 87387312 | 5.77138 | 4.39496 | 3.64069 |
| chr12 | 104864611 | 104864944 | 182 | Promoter (<=1kb) | 12 | 104763114 | 104865076 | 101963 | 2 | ENSMUST00 | 132 | Clmn | ENSMUSG0C | false | chr12 | 104864253 | 104865506 | 20.60195 | 76.26862 | 73.17842 | chr12 | 104863571 | 104865888 | 3.90875 | 3.5026 | 2.1583 |
| chr12 | 79172170 | 79172834 | 182 | Promoter (<=1kb) | 12 | 79156017 | 79172667 | 16651 | 2 | ENSMUST00 | 0 | Vti1b | ENSMUSG0C | false | chr12 | 79171474 | 79173188 | 35.51886 | 140.45512 | 135.55692 | . | . | . | . | . | . |
| chr13 | 8866635 | 8866881 | 182 | Intron (ENSMUST00000054251.7 | 13 | 8852669 | 8870979 | 18311 | 2 | ENSMUST00 | 4098 | Wdr37 | ENSMUSG0C | false | . | . | . | . | . | . | chr13 | 8861181 | 8870213 | 5.45675 | 4.30296 | 3.39541 |
| chr12 | 117761760 | 117762053 | 182 | Distal Intergenic | 12 | 117804289 | 117870047 | 65759 | 1 | ENSMUST00 | -42236 | Cdca7l | ENSMUSG0C | false | . | . | . | . | . | . | chr12 | 117760605 | 117763615 | 3.46919 | 3.30497 | 1.76033 |
| chr12 | 101958159 | 101958407 | 182 | Promoter (<=1kb) | 12 | 101918901 | 101958246 | 39346 | 2 | ENSMUST00 | 0 | Atxn3 | ENSMUSG0C | false | chr12 | 101957223 | 101958769 | 34.34893 | 131.73137 | 127.17394 | . | . | . | . | . | . |
| chr12 | 83444553 | 83444760 | 182 | Intron (ENSMUST00000178756.2 | 12 | 83269315 | 83439920 | 170606 | 2 | ENSMUST00 | -4634 | Dpf3 | ENSMUSG0C | false | . | . | . | . | . | . | chr12 | 83444145 | 83447069 | 3.19547 | 3.09768 | 1.54242 |
| chr12 | 83770417 | 83770595 | 182 | Exon (ENSMUST00000121733.2/ | 12 | 83763662 | 83781132 | 17471 | 1 | ENSMUST00 | 6756 | Papln | ENSMUSG0C | false | . | . | . | . | . | . | chr12 | 83768670 | 83771926 | 5.12696 | 3.81255 | 3.14148 |
| chr12 | 83921683 | 83922094 | 182 | Promoter (<=1kb) | 12 | 83794034 | 83921934 | 127901 | 2 | ENSMUST00 | 0 | Numb | ENSMUSG0C | false | chr12 | 83920493 | 83921919 | 36.04672 | 129.50026 | 125.02953 | . | . | . | . | . | . |
| chr12 | 84408667 | 84409097 | 182 | Promoter (<=1kb) | 12 | 84373875 | 84409029 | 35155 | 2 | ENSMUST00 | 0 | Entpd5 | ENSMUSG0C | false | chr12 | 84407946 | 84409513 | 36.04346 | 118.78738 | 114.68494 | chr12 | 84407993 | 84409696 | 3.23924 | 3.12296 | 1.62014 |
| chr12 | 85474435 | 85474780 | 182 | Promoter (<=1kb) | 12 | 85474691 | 85476007 | 1317 | 1 | ENSMUST00 | 0 | Fos | ENSMUSG0C | false | chr12 | 85472966 | 85475150 | 24.03115 | 75.67385 | 72.59326 | chr12 | 85471270 | 85476604 | 4.14697 | 3.66678 | 2.30093 |
| chr12 | 85813645 | 85814019 | 182 | Downstream (1-2kb) | 12 | 85819332 | 85822250 | 2919 | 2 | ENSMUST00 | 8231 | 0610007P14I | ENSMUSG0C | false | . | . | . | . | . | . | chr12 | 85813516 | 85815190 | 4.78207 | 3.83615 | 2.84673 |
| chr12 | 86692931 | 86693146 | 182 | Distal Intergenic | 12 | 86702355 | 86705964 | 3610 | 2 | ENSMUST00 | 12818 | Angel1 | ENSMUSG0C | false | . | . | . | . | . | . | chr12 | 86685220 | 86694692 | 6.20303 | 4.49357 | 4.02263 |
| chr12 | 108209374 | 108209818 | 182 | Intron (ENSMUST00000136175.1 | 12 | 108179738 | 108203359 | 23622 | 1 | ENSMUST00 | 29637 | Ccnk | ENSMUSG0C | false | . | . | . | . | . | . | chr12 | 108207980 | 108215965 | 7.17873 | 5.06814 | 4.78346 |
| chr12 | 108209014 | 108209233 | 182 | Intron (ENSMUST00000136175.1 | 12 | 108179738 | 108203359 | 23622 | 1 | ENSMUST00 | 29277 | Ccnk | ENSMUSG0C | false | . | . | . | . | . | . | chr12 | 108207980 | 108215965 | 7.17873 | 5.06814 | 4.78346 |
| chr12 | 108338986 | 108339179 | 182 | Intron (ENSMUST00000021684.4 | 12 | 108334377 | 108362231 | 27855 | 1 | ENSMUST00 | 4610 | Cyp46a1 | ENSMUSG0C | false | . | . | . | . | . | . | chr12 | 108338793 | 108339290 | 4.17248 | 3.75394 | 2.30808 |
| chr12 | 108304070 | 108304344 | 182 | Promoter (1-2kb) | 12 | 108306270 | 108328300 | 22031 | 1 | ENSMUST00 | -1926 | Hhipl1 | ENSMUSG0C | false | . | . | . | . | . | . | chr12 | 108300429 | 108307899 | 5.251 | 4.15318 | 3.21807 |
| chr12 | 108805614 | 108805862 | 182 | Promoter (<=1kb) | 12 | 108806239 | 108815564 | 9326 | 1 | ENSMUST00 | -377 | Yy1 | ENSMUSG0C | false | . | . | . | . | . | . | chr12 | 108795103 | 108814854 | 5.27563 | 4.12358 | 3.24997 |
| chr12 | 108832386 | 108832597 | 182 | Promoter (<=1kb) | 12 | 108827224 | 108831617 | 4394 | 2 | ENSMUST00 | -770 | Slc25a29 | ENSMUSG0C | false | chr12 | 108832249 | 108833572 | 12.97578 | 28.80697 | 26.35731 | chr12 | 108830624 | 108835287 | 6.71004 | 4.82515 | 4.4293 |
| chr12 | 108880646 | 108880909 | 182 | Exon (ENSMUST00000160974.1/ | 12 | 108865871 | 108875281 | 9411 | 2 | ENSMUST00 | -5366 | Wars | ENSMUSG0C | false | . | . | . | . | . | . | chr12 | 108877969 | 108888605 | 5.39827 | 4.24317 | 3.3389 |
| chr12 | 108878571 | 108878897 | 182 | Intron (ENSMUST00000161410.2 | 12 | 108865871 | 108875281 | 9411 | 2 | ENSMUST00 | -3291 | Wars | ENSMUSG0C | false | . | . | . | . | . | . | chr12 | 108877969 | 108888605 | 5.39827 | 4.24317 | 3.3389 |
| chr12 | 110696198 | 110696754 | 182 | Promoter (<=1kb) | 12 | 110690605 | 110696395 | 5791 | 2 | ENSMUST00 | 0 | Hsp90aa1 | ENSMUSG0C | false | chr12 | 110695420 | 110696379 | 37.40972 | 147.3676 | 142.18408 | . | . | . | . | . | . |
| chr12 | 110695839 | 110696114 | 182 | Promoter (<=1kb) | 12 | 110695286 | 110695951 | 666 | 2 | ENSMUST00 | 0 | Hsp90aa1 | ENSMUSG0C | false | chr12 | 110695420 | 110696379 | 37.40972 | 147.3676 | 142.18408 | . | . | . | . | . | . |
| chr12 | 111538592 | 111538907 | 182 | Promoter (<=1kb) | 12 | 111538853 | 111544923 | 6071 | 1 | ENSMUST00 | 0 | Eif5 | ENSMUSG0C | false | chr12 | 111538117 | 111539637 | 32.3084 | 121.498 | 117.30614 | . | . | . | . | . | . |
| chr12 | 111896865 | 111897223 | 182 | Intron (ENSMUST00000054815.8 | 12 | 111866013 | 111908040 | 42028 | 2 | ENSMUST00 | 10817 | Ppp1r13b | ENSMUSG0C | false | . | . | . | . | . | . | chr12 | 111880166 | 111903803 | 5.80743 | 4.40294 | 3.67276 |
| chr12 | 111819954 | 111820290 | 182 | Intron (ENSMUST00000021714.7 | 12 | 111814170 | 111828388 | 14219 | 1 | ENSMUST00 | 5785 | Zfyve21 | ENSMUSG0C | false | . | . | . | . | . | . | chr12 | 111813203 | 111823116 | 5.80855 | 4.38461 | 3.68777 |
| chr12 | 111759699 | 111760086 | 182 | Promoter (<=1kb) | 12 | 111759013 | 111806727 | 47715 | 1 | ENSMUST00 | 687 | Klc1 | ENSMUSG0C | false | chr12 | 111758148 | 111760750 | 28.51907 | 101.09029 | 97.496 | chr12 | 111759928 | 111772632 | 6.01486 | 4.51861 | 3.83756 |
| chr12 | 111746155 | 111746373 | 182 | Intron (ENSMUST00000160366.2 | 12 | 111758849 | 111806775 | 47927 | 1 | ENSMUST00 | -12476 | Klc1 | ENSMUSG0C | false | . | . | . | . | . | . | chr12 | 111743231 | 111747984 | 5.31732 | 3.94321 | 3.29765 |
| chr13 | 24960885 | 24961121 | 182 | Intron (ENSMUST00000021773.6 | 13 | 24955854 | 24962818 | 6965 | 1 | ENSMUST00 | 5032 | Gpld1 | ENSMUSG0C | false | . | . | . | . | . | . | chr13 | 24947308 | 24961124 | 4.97023 | 4.07319 | 2.98257 |
| chr13 | 30973785 | 30974195 | 182 | Promoter (<=1kb) | 13 | 30813919 | 30974047 | 160129 | 2 | ENSMUST00 | 0 | Exoc2 | ENSMUSG0C | true | chr13 | 30973358 | 30974593 | 36.14217 | 127.51552 | 123.12013 | . | . | . | . | . | . |
| chr13 | 41358783 | 41359222 | 182 | Promoter (<=1kb) | 13 | 41309916 | 41359001 | 49086 | 2 | ENSMUST00 | 0 | Nedd9 | ENSMUSG0C | false | chr13 | 41355780 | 41359729 | 22.15371 | 67.81625 | 64.85734 | . | . | . | . | . | . |
| chr13 | 34002211 | 34002504 | 182 | Promoter (<=1kb) | 13 | 34002363 | 34033677 | 31315 | 1 | ENSMUST00 | 0 | Ripk1 | ENSMUSG0C | false | chr13 | 34001889 | 34003343 | 38.25702 | 141.19 | 136.26103 | . | . | . | . | . | . |
| chr13 | 34994002 | 34994354 | 182 | Promoter (<=1kb) | 13 | 34977748 | 34994144 | 16397 | 2 | ENSMUST00 | 0 | Eci2 | ENSMUSG0C | false | chr13 | 34992948 | 34994858 | 22.15371 | 67.81625 | 64.85734 | . | . | . | . | . | . |
| chr13 | 35058191 | 35058448 | 182 | Distal Intergenic | 13 | 34986106 | 35027094 | 40989 | 2 | ENSMUST00 | -31098 | Eci2 | ENSMUSG0C | false | . | . | . | . | . | . | chr13 | 35058029 | 35059284 | 3.8316 | 3.40364 | 2.08194 |
| chr13 | 51588130 | 51588401 | 182 | Distal Intergenic | 13 | 51431041 | 51567084 | 136044 | 2 | ENSMUST00 | -21047 | Shc3 | ENSMUSG0C | false | . | . | . | . | . | . | . | . | . | . | . | . |
| chr13 | 63236354 | 63236767 | 182 | Intron (ENSMUST00000091560.5 | 13 | 63282142 | 63326096 | 43955 | 1 | ENSMUST00 | -45375 | 2010111I01R | ENSMUSG0C | false | . | . | . | . | . | . | chr13 | 63235330 | 63236376 | 2.83583 | 2.70396 | 1.27388 |
| chr13 | 63176505 | 63176732 | 182 | Intron (ENSMUST00000091560.5 | 13 | 63282142 | 63326096 | 43955 | 1 | ENSMUST00 | -105410 | 2010111I01R | ENSMUSG0C | false | . | . | . | . | . | . | chr13 | 63174733 | 63177604 | 4.6697 | 3.85326 | 2.77017 |
| chr13 | 63256251 | 63256471 | 182 | Intron (ENSMUST00000091560.5 | 13 | 63282142 | 63326096 | 43955 | 1 | ENSMUST00 | -25671 | 2010111I01R | ENSMUSG0C | false | chr13 | 63255485 | 63256554 | 7.14815 | 15.37828 | 13.11012 | chr13 | 63252739 | 63272327 | 6.37956 | 4.73666 | 4.1425 |
| chr13 | 53257323 | 53257569 | 182 | Intron (ENSMUST00000130235.2 | 13 | 53131960 | 53275658 | 143699 | 2 | ENSMUST00 | 18089 | Ror2 | ENSMUSG0C | false | . | . | . | . | . | . | . | . | . | . | . | . |
| chr13 | 63567776 | 63568173 | 182 | Promoter (2-3kb) | 13 | 63511533 | 63565520 | 53988 | 2 | ENSMUST00 | -2257 | Ptch1 | ENSMUSG0C | false | chr13 | 63562847 | 63568718 | 33.69019 | 111.49343 | 107.61708 | . | . | . | . | . | . |
| chr13 | 63553793 | 63553996 | 182 | Intron (ENSMUST00000021921.5 | 13 | 63511533 | 63565520 | 53988 | 2 | ENSMUST00 | 11524 | Ptch1 | ENSMUSG0C | false | . | . | . | . | . | . | chr13 | 63548517 | 63562687 | 5.58712 | 4.19239 | 3.52613 |
| chr13 | 54101153 | 54101346 | 182 | Intron (ENSMUST00000021930.8 | 13 | 54071845 | 54108346 | 36502 | 1 | ENSMUST00 | 29309 | Sfxn1 | ENSMUSG0C | false | . | . | . | . | . | . | chr13 | 54100779 | 54101941 | 4.46997 | 3.5335 | 2.60313 |
| chr13 | 66905130 | 66905480 | 182 | Promoter (<=1kb) | 13 | 66900617 | 66905378 | 4762 | 2 | ENSMUST00 | 0 | Uqcrb | ENSMUSG0C | false | chr13 | 66904045 | 66905759 | 29.2476 | 106.45832 | 102.71833 | . | . | . | . | . | . |
| chr13 | 58394334 | 58394538 | 182 | Promoter (<=1kb) | 13 | 58391960 | 58394785 | 2826 | 2 | ENSMUST00 | 247 | Hnrnpk | ENSMUSG0C | false | . | . | . | . | . | . | chr13 | 58393966 | 58398296 | 3.13784 | 2.92956 | 1.52285 |
| chr13 | 85189244 | 85189551 | 182 | Promoter (<=1kb) | 13 | 85189408 | 85213723 | 24316 | 1 | ENSMUST00 | 0 | Ccnh | ENSMUSG0C | false | chr13 | 85189018 | 85190300 | 41.67628 | 135.08018 | 130.40039 | . | . | . | . | . | . |
| chr13 | 59407258 | 59407498 | 182 | Distal Intergenic | 13 | 59449537 | 59450228 | 692 | 2 | ENSMUST00 | 42730 | Agtpbp1 | ENSMUSG0C | false | . | . | . | . | . | . | . | . | . | . | . | . |
| chr13 | 60686012 | 60686301 | 182 | Intron (ENSMUST00000077453.6 | 13 | 60602211 | 60763185 | 160975 | 1 | ENSMUST00 | 83802 | Dapk1 | ENSMUSG0C | false | . | . | . | . | . | . | chr13 | 60683506 | 60688289 | 3.94744 | 3.4739 | 2.16956 |
| chr13 | 60674882 | 60675100 | 182 | Intron (ENSMUST00000077453.6 | 13 | 60602211 | 60763185 | 160975 | 1 | ENSMUST00 | 72672 | Dapk1 | ENSMUSG0C | false | . | . | . | . | . | . | chr13 | 60674440 | 60675363 | 3.98418 | 3.64502 | 2.20762 |
| chr13 | 74062202 | 74062610 | 182 | Promoter (<=1kb) | 13 | 74036495 | 74062285 | 25791 | 2 | ENSMUST00 | 0 | Cep72 | ENSMUSG0C | false | chr13 | 74061625 | 74062582 | 30.78991 | 105.27288 | 101.56436 | . | . | . | . | . | . |
| chr13 | 74009268 | 74009551 | 182 | Promoter (<=1kb) | 13 | 74009407 | 74031371 | 21965 | 1 | ENSMUST00 | 0 | Tppp | ENSMUSG0C | false | chr13 | 74008885 | 74010801 | 24.40663 | 77.266 | 74.1616 | . | . | . | . | . | . |
| chr13 | 74640132 | 74640386 | 182 | Promoter (<=1kb) | 13 | 74639872 | 74691875 | 52004 | 1 | ENSMUST00 | 261 | Erap1 | ENSMUSG0C | false | chr13 | 74639074 | 74640621 | 41.07278 | 140.67711 | 135.76277 | . | . | . | . | . | . |
| chr13 | 73467108 | 73467369 | 182 | Promoter (<=1kb) | 13 | 73467197 | 73516422 | 49226 | 1 | ENSMUST00 | 0 | Lpcat1 | ENSMUSG0C | false | chr13 | 73466857 | 73468175 | 31.69034 | 97.80461 | 94.28741 | chr13 | 73466338 | 73469558 | 3.1845 | 3.13298 | 1.57061 |
| chr13 | 95779550 | 95779829 | 182 | Intron (ENSMUST00000068603.6 | 13 | 95696853 | 95702739 | 5887 | 1 | ENSMUST00 | 82698 | F2rl2 | ENSMUSG0C | false | . | . | . | . | . | . | chr13 | 95772302 | 95783884 | 6.26953 | 4.56663 | 4.0844 |
| chr13 | 95879098 | 95879308 | 182 | Intron (ENSMUST00000068603.6 | 13 | 95627177 | 95891922 | 264746 | 2 | ENSMUST00 | 12614 | Iqgap2 | ENSMUSG0C | true | . | . | . | . | . | . | chr13 | 95794831 | 95889694 | 5.09372 | 4.07642 | 3.0951 |
| chr13 | 95826291 | 95826648 | 182 | Intron (ENSMUST00000068603.6 | 13 | 95627177 | 95891922 | 264746 | 2 | ENSMUST00 | 65274 | Iqgap2 | ENSMUSG0C | true | . | . | . | . | . | . | chr13 | 95794831 | 95889694 | 5.09372 | 4.07642 | 3.0951 |
| chr13 | 95514883 | 95515096 | 182 | Intron (ENSMUST00000022185.8 | 13 | 95511732 | 95525240 | 13509 | 2 | ENSMUST00 | 10144 | F2rl1 | ENSMUSG0C | false | . | . | . | . | . | . | chr13 | 95514784 | 95515273 | 3.36668 | 3.28747 | 1.6609 |
| chr13 | 94358834 | 94359240 | 182 | Promoter (<=1kb) | 13 | 94358960 | 94566316 | 207357 | 1 | ENSMUST00 | 0 | Ap3b1 | ENSMUSG0C | false | chr13 | 94357953 | 94359860 | 37.5443 | 122.47784 | 118.25522 | . | . | . | . | . | . |
| chr13 | 106946894 | 106947206 | 182 | Promoter (<=1kb) | 13 | 106947129 | 106960224 | 13096 | 1 | ENSMUST00 | 0 | Dimt1 | ENSMUSG0C | false | chr13 | 106946275 | 106947899 | 37.82652 | 124.04217 | 119.77048 | . | . | . | . | . | . |
| chr13 | 108599677 | 108599944 | 182 | Intron (ENSMUST00000133929.2 | 13 | 108654177 | 109951474 | 1297298 | 1 | ENSMUST00 | -54233 | Pde4d | ENSMUSG0C | false | . | . | . | . | . | . | . | . | . | . | . | . |
| chr13 | 104109484 | 104109908 | 182 | Promoter (<=1kb) | 13 | 104023439 | 104109614 | 86176 | 2 | ENSMUST00 | 0 | Nln | ENSMUSG0C | false | chr13 | 104108731 | 104110687 | 32.3084 | 121.498 | 117.30614 | chr13 | 104108552 | 104111010 | 3.31729 | 3.21958 | 1.67211 |
| chr13 | 117024265 | 117025022 | 182 | Promoter (<=1kb) | 13 | 116854824 | 117025516 | 170693 | 2 | ENSMUST00 | 494 | Parp8 | ENSMUSG0C | false | chr13 | 117023675 | 117025980 | 31.54088 | 108.67455 | 104.87239 | chr13 | 117023387 | 117024309 | 2.96546 | 2.9302 | 1.33049 |
| chr14 | 11553341 | 11553624 | 182 | Promoter (<=1kb) | 14 | 11553532 | 12242041 | 688510 | 1 | ENSMUST00 | 0 | Ptprg | ENSMUSG0C | false | chr14 | 11551784 | 11556756 | 21.44446 | 75.31629 | 72.24329 | . | . | . | . | . | . |
| chr13 | 112439361 | 112439737 | 182 | Distal Intergenic | 13 | 112464070 | 112510086 | 46017 | 1 | ENSMUST00 | -24333 | Il6st | ENSMUSG0C | false | chr13 | 112439437 | 112439617 | 5.35832 | 8.05648 | 5.92315 | chr13 | 112438022 | 112440699 | 5.04558 | 3.81413 | 3.07753 |
| chr13 | 114387733 | 114388193 | 182 | Promoter (<=1kb) | 13 | 114287795 | 114388094 | 100300 | 2 | ENSMUST00 | 0 | Ndufs4 | ENSMUSG0C | false | chr13 | 114387066 | 114388440 | 37.59926 | 118.31785 | 114.23224 | chr13 | 114388191 | 114388631 | 4.5533 | 3.69119 | 2.67014 |
| chr13 | 114343133 | 114343390 | 182 | Intron (ENSMUST00000022286.6 | 13 | 114287795 | 114388094 | 100300 | 2 | ENSMUST00 | 44704 | Ndufs4 | ENSMUSG0C | false | . | . | . | . | . | . | chr13 | 114342800 | 114345567 | 4.48882 | 3.84585 | 2.58788 |
| chr13 | 114158191 | 114158407 | 182 | Distal Intergenic | 13 | 114287795 | 114388094 | 100300 | 2 | ENSMUST00 | 229687 | Ndufs4 | ENSMUSG0C | false | . | . | . | . | . | . | . | . | . | . | . | . |
| chr14 | 21553064 | 21553289 | 182 | Intron (ENSMUST00000183201.2 | 14 | 21516751 | 21671206 | 154456 | 1 | ENSMUST00 | 36314 | Kat6b | ENSMUSG0C | false | . | . | . | . | . | . | chr14 | 21544903 | 21565964 | 5.23527 | 4.17113 | 3.20783 |
| chr14 | 17660636 | 17661067 | 182 | Promoter (<=1kb) | 14 | 17660960 | 18036559 | 375600 | 1 | ENSMUST00 | 0 | Thrb | ENSMUSG0C | false | chr14 | 17659199 | 17661639 | 35.20524 | 115.20684 | 111.21615 | . | . | . | . | . | . |
| chr14 | 24197927 | 24198218 | 182 | Intron (ENSMUST00000073687.7 | 14 | 24133953 | 24177789 | 43837 | 2 | ENSMUST00 | -20139 | Dlg5 | ENSMUSG0C | true | . | . | . | . | . | . | chr14 | 24191579 | 24244546 | 6.62149 | 4.76248 | 4.34728 |
| chr14 | 16249170 | 16249617 | 182 | Promoter (<=1kb) | 14 | 16249280 | 16311926 | 62647 | 1 | ENSMUST00 | 0 | Ngly1 | ENSMUSG0C | false | chr14 | 16248824 | 16250539 | 28.53699 | 95.19626 | 91.7378 | . | . | . | . | . | . |
| chr14 | 41013709 | 41014048 | 182 | Promoter (<=1kb) | 14 | 40993740 | 41013775 | 20036 | 2 | ENSMUST00 | 0 | Fam213a | ENSMUSG0C | false | chr14 | 41013064 | 41014319 | 17.74057 | 61.94745 | 59.07436 | chr14 | 41006611 | 41014538 | 4.04691 | 3.52092 | 2.2655 |
| chr14 | 34509124 | 34509298 | 182 | Downstream (2-3kb) | 14 | 34434462 | 34503341 | 68880 | 2 | ENSMUST00 | -5784 | Bmpr1a | ENSMUSG0C | true | . | . | . | . | . | . | chr14 | 34508079 | 34509644 | 4.78233 | 3.95195 | 2.88493 |
| chr14 | 34548547 | 34548885 | 182 | Intron (ENSMUST00000090040.5 | 14 | 34511619 | 34522801 | 11183 | 2 | ENSMUST00 | -25747 | 9230112D13I | ENSMUSG0C | false | chr14 | 34548457 | 34549653 | 37.0612 | 128.08391 | 123.66181 | chr14 | 34547537 | 34550058 | 3.36884 | 3.27518 | 1.66231 |
| chr14 | 19762141 | 19762360 | 182 | Intron (ENSMUST00000022340.3 | 14 | 19751257 | 19811787 | 60531 | 1 | ENSMUST00 | 10885 | Nid2 | ENSMUSG0C | false | . | . | . | . | . | . | chr14 | 19758989 | 19771108 | 5.58176 | 4.33118 | 3.48417 |
| chr14 | 20388840 | 20389095 | 182 | Promoter (<=1kb) | 14 | 20384638 | 20388910 | 4273 | 2 | ENSMUST00 | 0 | Dnajc9 | ENSMUSG0C | false | chr14 | 20388139 | 20389535 | 34.98938 | 113.78057 | 109.83514 | chr14 | 20387840 | 20389573 | 5.36632 | 4.32812 | 3.30517 |
| chr14 | 20964358 | 20964546 | 182 | Intron (ENSMUST00000022369.7 | 14 | 20929433 | 21033673 | 104241 | 1 | ENSMUST00 | 34926 | Vcl | ENSMUSG0C | false | . | . | . | . | . | . | chr14 | 20962606 | 20965995 | 4.51987 | 3.87926 | 2.60492 |
| chr14 | 20930942 | 20931118 | 182 | Promoter (1-2kb) | 14 | 20929433 | 21033673 | 104241 | 1 | ENSMUST00 | 1510 | Vcl | ENSMUSG0C | false | . | . | . | . | . | . | chr14 | 20930127 | 20934058 | 3.20405 | 3.10837 | 1.53176 |
| chr14 | 20914461 | 20914671 | 182 | Distal Intergenic | 14 | 20929433 | 21033673 | 104241 | 1 | ENSMUST00 | -14762 | Vcl | ENSMUSG0C | false | . | . | . | . | . | . | chr14 | 20914217 | 20914724 | 2.82819 | 2.70804 | 1.25914 |
| chr14 | 45318427 | 45318812 | 182 | Promoter (<=1kb) | 14 | 45283092 | 45318572 | 35481 | 2 | ENSMUST00 | 0 | Ero1l | ENSMUSG0C | false | chr14 | 45317289 | 45319128 | 39.25979 | 127.61615 | 123.21935 | . | . | . | . | . | . |
| chr14 | 46884958 | 46885184 | 182 | Promoter (<=1kb) | 14 | 46884175 | 47105811 | 221637 | 1 | ENSMUST00 | 784 | Samd4 | ENSMUSG0C | true | chr14 | 46882071 | 46885731 | 34.78613 | 113.42423 | 109.48744 | chr14 | 46881114 | 46887653 | 4.95941 | 4.04007 | 2.96585 |
| chr14 | 47888133 | 47888408 | 182 | Distal Intergenic | 14 | 47729659 | 47736561 | 6903 | 1 | ENSMUST00 | 158475 | Ktn1 | ENSMUSG0C | false | . | . | . | . | . | . | chr14 | 47887039 | 47893837 | 4.36041 | 3.66154 | 2.50717 |
| chr14 | 48134432 | 48134692 | 182 | Intron (ENSMUST00000073150.4 | 14 | 48120869 | 48260883 | 140015 | 1 | ENSMUST00 | 13564 | Peli2 | ENSMUSG0C | false | chr14 | 48133026 | 48134696 | 7.96309 | 14.9624 | 12.70017 | chr14 | 48122053 | 48141561 | 6.29 | 4.61313 | 4.06953 |
| chr14 | 26764889 | 26765189 | 182 | Intron (ENSMUST00000022433.5 | 14 | 26771029 | 26773772 | 2744 | 1 | ENSMUST00 | -5840 | Dnah12 | ENSMUSG0C | false | . | . | . | . | . | . | . | . | . | . | . | . |
| chr14 | 31348623 | 31349016 | 182 | Intron (ENSMUST00000022451.8 | 14 | 31336707 | 31371984 | 35278 | 1 | ENSMUST00 | 11917 | Capn7 | ENSMUSG0C | false | . | . | . | . | . | . | chr14 | 31346919 | 31349448 | 4.19001 | 3.58231 | 2.36101 |
| chr14 | 27238044 | 27238348 | 182 | Promoter (<=1kb) | 14 | 27238039 | 27403904 | 165866 | 1 | ENSMUST00 | 6 | Arhgef3 | ENSMUSG0C | false | chr14 | 27237740 | 27238500 | 7.16187 | 18.18295 | 15.87379 | chr14 | 27237556 | 27238533 | 4.02204 | 3.546 | 2.23392 |

| | | | | | | | | | | | | | | | | | | | | | | | | | | |
|---|---|---|---|---|---|---|---|---|---|---|---|---|---|---|---|---|---|---|---|---|---|---|---|---|---|---|
| chr14 | 31206587 | 31206837 | 182 | Promoter (<=1kb) | 14 | 31170930 | 31206801 | 35872 | 2 | ENSMUST00 | 0 | Nisch | ENSMUSG0( | false | chr14 | 31205406 | 31207302 | 24.87197 | 72.62122 | 69.58942 | . | . | . | . | . | . |
| chr14 | 61556616 | 61557283 | 182 | Promoter (<=1kb) | 14 | 61531993 | 61556886 | 24894 | 2 | ENSMUST00 | 0 | Spryd7 | ENSMUSG0( | false | chr14 | 61556004 | 61557669 | 38.15516 | 131.11487 | 126.58785 | . | . | . | . | . | . |
| chr14 | 56777558 | 56777801 | 182 | Promoter (<=1kb) | 14 | 56748858 | 56777927 | 29070 | 2 | ENSMUST00 | 126 | Pspc1 | ENSMUSG0( | false | chr14 | 56776691 | 56779122 | 31.08358 | 101.75523 | 98.14187 | . | . | . | . | . | . |
| chr14 | 30621395 | 30621595 | 182 | Intron (ENSMUST00000112210.4 | 14 | 30600262 | 30626172 | 25911 | 2 | ENSMUST00 | 4577 | Prkcd | ENSMUSG0( | false | . | . | . | . | . | . | chr14 | 30619049 | 30622602 | 4.50388 | 3.83668 | 2.60886 |
| chr14 | 57664956 | 57665271 | 182 | Promoter (<=1kb) | 14 | 57596990 | 57664992 | 68003 | 2 | ENSMUST00 | 0 | Xpo4 | ENSMUSG0( | false | chr14 | 57663945 | 57666077 | 25.17393 | 93.69764 | 90.27328 | . | . | . | . | . | . |
| chr14 | 57735957 | 57736273 | 182 | Promoter (1-2kb) | 14 | 57722189 | 57734300 | 12112 | 2 | ENSMUST00 | -1658 | Lats2 | ENSMUSG0( | false | . | . | . | . | . | . | chr14 | 57704250 | 57744175 | 6.91785 | 4.80068 | 4.61004 |
| chr14 | 57798041 | 57798219 | 182 | Promoter (<=1kb) | 14 | 57798180 | 57804980 | 6801 | 1 | ENSMUST00 | 0 | Sap18 | ENSMUSG0( | false | chr14 | 57797680 | 57798077 | 13.59473 | 31.77968 | 29.29185 | . | . | . | . | . | . |
| chr14 | 57826136 | 57826638 | 182 | Promoter (<=1kb) | 14 | 57806561 | 57826163 | 19603 | 2 | ENSMUST00 | 0 | Ska3 | ENSMUSG0( | false | chr14 | 57825764 | 57827010 | 29.58769 | 108.11006 | 104.32592 | chr14 | 57825546 | 57828186 | 6.4789 | 4.82128 | 4.19467 |
| chr14 | 64949726 | 64950242 | 182 | Promoter (<=1kb) | 14 | 64811600 | 64949850 | 138251 | 2 | ENSMUST00 | 0 | Hmbox1 | ENSMUSG0( | false | chr14 | 64948455 | 64950969 | 28.1615 | 93.53628 | 90.11485 | . | . | . | . | . | . |
| chr14 | 57997194 | 57997413 | 182 | Promoter (1-2kb) | 14 | 57916280 | 57999262 | 82983 | 2 | ENSMUST00 | 1849 | Micu2 | ENSMUSG0( | true | . | . | . | . | . | . | chr14 | 57993689 | 57999995 | 3.84376 | 3.4543 | 2.10473 |
| chr14 | 58074901 | 58075091 | 182 | Promoter (<=1kb) | 14 | 58075115 | 58110174 | 35060 | 1 | ENSMUST00 | -24 | Fgf9 | ENSMUSG0( | false | chr14 | 58073800 | 58076799 | 22.5292 | 69.37365 | 66.39268 | chr14 | 58069813 | 58076920 | 5.9191 | 4.53596 | 3.79262 |
| chr14 | 60738612 | 60738867 | 182 | Intron (ENSMUST00000160973.2 | 14 | 60732906 | 60764549 | 31644 | 1 | ENSMUST00 | 5707 | Spata13 | ENSMUSG0( | true | . | . | . | . | . | . | chr14 | 60731957 | 60745391 | 6.28488 | 4.60271 | 4.07482 |
| chr14 | 29721968 | 29722151 | 182 | Promoter (<=1kb) | 14 | 28904943 | 29721864 | 816922 | 2 | ENSMUST00 | -105 | Cacna2d3 | ENSMUSG0( | false | chr14 | 29721447 | 29723107 | 25.98482 | 77.30067 | 74.19341 | chr14 | 29721362 | 29723201 | 5.44896 | 4.27033 | 3.40106 |
| chr14 | 60784542 | 60784854 | 182 | Promoter (<=1kb) | 14 | 60784566 | 60903610 | 119045 | 1 | ENSMUST00 | 0 | Mipep | ENSMUSG0( | false | chr14 | 60784529 | 60785401 | 38.05226 | 120.16034 | 116.01576 | chr14 | 60784195 | 60785424 | 3.42905 | 3.31507 | 1.71208 |
| chr14 | 74831209 | 74831507 | 182 | Intron (ENSMUST00000088970.5 | 14 | 74738506 | 74749755 | 11250 | 1 | ENSMUST00 | 92704 | Esd | ENSMUSG0( | false | . | . | . | . | . | . | chr14 | 74819121 | 74836959 | 5.2109 | 4.16836 | 3.18482 |
| chr14 | 78537704 | 78537957 | 182 | Promoter (<=1kb) | 14 | 78492246 | 78536860 | 44615 | 2 | ENSMUST00 | -845 | Akap11 | ENSMUSG0( | true | . | . | . | . | . | . | chr14 | 78522737 | 78539051 | 5.22391 | 4.1452 | 3.19494 |
| chr14 | 79390300 | 79390836 | 182 | Promoter (<=1kb) | 14 | 79334507 | 79390778 | 56272 | 2 | ENSMUST00 | 0 | Naa16 | ENSMUSG0( | false | chr14 | 79389734 | 79391248 | 36.53154 | 123.91184 | 119.64436 | . | . | . | . | . | . |
| chr14 | 65592984 | 65593221 | 182 | Promoter (<=1kb) | 14 | 65530446 | 65593112 | 62667 | 2 | ENSMUST00 | 0 | Elp3 | ENSMUSG0( | false | chr14 | 65592319 | 65593487 | 27.39617 | 87.32104 | 84.03119 | chr14 | 65591901 | 65593557 | 5.25753 | 4.01478 | 3.2514 |
| chr14 | 66111226 | 66111450 | 182 | Intron (ENSMUST00000070515.1 | 14 | 66084377 | 66124500 | 40124 | 2 | ENSMUST00 | 13050 | Ephx2 | ENSMUSG0( | false | . | . | . | . | . | . | chr14 | 66110650 | 66112816 | 4.48848 | 3.78716 | 2.62693 |
| chr14 | 69805293 | 69805867 | 182 | Promoter (<=1kb) | 14 | 69784990 | 69805530 | 20541 | 2 | ENSMUST00 | 0 | Rhobtb2 | ENSMUSG0( | false | chr14 | 69804386 | 69806027 | 28.53919 | 82.65468 | 79.45325 | . | . | . | . | . | . |
| chr14 | 70288977 | 70289526 | 182 | Promoter (<=1kb) | 14 | 70217898 | 70289449 | 71552 | 2 | ENSMUST00 | 0 | Ppp3cc | ENSMUSG0( | false | chr14 | 70288628 | 70289902 | 34.43176 | 108.65345 | 104.85863 | . | . | . | . | . | . |
| chr14 | 70889925 | 70890254 | 182 | Promoter (<=1kb) | 14 | 70890130 | 70979838 | 89709 | 1 | ENSMUST00 | 0 | Gfra2 | ENSMUSG0( | false | chr14 | 70889621 | 70890952 | 9.01289 | 21.84568 | 19.48558 | chr14 | 70888617 | 70891453 | 4.78245 | 3.94008 | 2.85537 |
| chr14 | 73372055 | 73372549 | 182 | Intron (ENSMUST00000022704.7 | 14 | 73362231 | 73385271 | 23041 | 2 | ENSMUST00 | 12722 | Itm2b | ENSMUSG0( | false | chr14 | 73372177 | 73372414 | 4.99839 | 7.78073 | 5.65212 | chr14 | 73365895 | 73382079 | 6.04681 | 4.50984 | 3.87534 |
| chr14 | 73552664 | 73552878 | 182 | Promoter (<=1kb) | 14 | 73552665 | 73596142 | 43478 | 1 | ENSMUST00 | 0 | Sucla2 | ENSMUSG0( | false | chr14 | 73552312 | 73553765 | 35.98183 | 119.73972 | 115.60696 | chr14 | 73550753 | 73555854 | 4.02279 | 3.53105 | 2.20838 |
| chr14 | 120288980 | 120289488 | 182 | Intron (ENSMUST00000088419.7 | 14 | 120275669 | 120431697 | 156029 | 1 | ENSMUST00 | 13312 | Mbnl2 | ENSMUSG0( | false | . | . | . | . | . | . | chr14 | 120277640 | 120316005 | 5.96633 | 4.45953 | 3.81543 |
| chr15 | 6934812 | 6935003 | 182 | Distal Intergenic | 15 | 6815037 | 6874969 | 59933 | 2 | ENSMUST00 | -59844 | Osmr | ENSMUSG0( | false | . | . | . | . | . | . | chr15 | 6933792 | 6935616 | 4.73048 | 3.90441 | 2.78065 |
| chr15 | 6683322 | 6683569 | 182 | Distal Intergenic | 15 | 6660405 | 6663251 | 2847 | 1 | ENSMUST00 | 22918 | Fyb | ENSMUSG0( | false | . | . | . | . | . | . | chr15 | 6682845 | 6683921 | 5.181 | 4.031 | 3.19232 |
| chr15 | 6385547 | 6385900 | 182 | Promoter (<=1kb) | 15 | 6386598 | 6440705 | 54108 | 1 | ENSMUST00 | -698 | Dab2 | ENSMUSG0( | false | . | . | . | . | . | . | chr15 | 6383263 | 6386496 | 4.13234 | 3.62074 | 2.30439 |
| chr14 | 54570083 | 54570303 | 182 | Exon (ENSMUST00000054487.8/ | 14 | 54567469 | 54577661 | 10193 | 2 | ENSMUST00 | 7358 | Ajuba | ENSMUSG0( | false | . | . | . | . | . | . | chr14 | 54568405 | 54575512 | 5.48983 | 4.2726 | 3.41523 |
| chr15 | 3974774 | 3975048 | 182 | Promoter (1-2kb) | 15 | 3969038 | 3972860 | 3823 | 2 | ENSMUST00 | -1915 | Fbxo4 | ENSMUSG0( | true | . | . | . | . | . | . | chr15 | 3975000 | 3978565 | 4.16374 | 3.67485 | 2.31344 |
| chr15 | 4026399 | 4026780 | 182 | Promoter (<=1kb) | 15 | 4026684 | 4153963 | 127280 | 1 | ENSMUST00 | 0 | Oxct1 | ENSMUSG0( | false | chr15 | 4025971 | 4028201 | 33.65816 | 114.56059 | 110.59126 | . | . | . | . | . | . |
| chr15 | 12718198 | 12718439 | 182 | Intron (ENSMUST00000189469.1 | 15 | 12443331 | 12739924 | 296594 | 2 | ENSMUST00 | 21485 | Pdzd2 | ENSMUSG0( | false | . | . | . | . | . | . | chr15 | 12717304 | 12718642 | 4.97517 | 4.10274 | 2.9831 |
| chr15 | 11883268 | 11883533 | 182 | Exon (ENSMUST00000066529.3/ | 15 | 11839896 | 11905674 | 65779 | 2 | ENSMUST00 | 22141 | Npr3 | ENSMUSG0( | true | . | . | . | . | . | . | chr15 | 11881241 | 11889185 | 5.24561 | 4.23363 | 3.20362 |
| chr15 | 32433040 | 32433292 | 182 | Intron (ENSMUST00000067458.6 | 15 | 32244813 | 32696341 | 451529 | 1 | ENSMUST00 | 188228 | Sema5a | ENSMUSG0( | true | . | . | . | . | . | . | chr15 | 32425857 | 32434592 | 5.25149 | 4.17269 | 3.22521 |
| chr15 | 32505924 | 32506288 | 182 | Intron (ENSMUST00000067458.6 | 15 | 32244813 | 32696341 | 451529 | 1 | ENSMUST00 | 261112 | Sema5a | ENSMUSG0( | true | . | . | . | . | . | . | chr15 | 32499761 | 32540421 | 4.87542 | 4.01677 | 2.90332 |
| chr15 | 32537109 | 32537393 | 182 | Intron (ENSMUST00000067458.6 | 15 | 32244813 | 32696341 | 451529 | 1 | ENSMUST00 | 292297 | Sema5a | ENSMUSG0( | true | . | . | . | . | . | . | chr15 | 32499761 | 32540421 | 4.87542 | 4.01677 | 2.90332 |
| chr15 | 10661807 | 10662288 | 182 | Intron (ENSMUST00000090339.4 | 15 | 10568979 | 10713540 | 144562 | 2 | ENSMUST00 | 51252 | Rai14 | ENSMUSG0( | true | . | . | . | . | . | . | chr15 | 10658484 | 10698797 | 6.0561 | 4.48411 | 3.88941 |
| chr15 | 34046432 | 34046652 | 182 | Distal Intergenic | 15 | 34082694 | 34103345 | 20652 | 1 | ENSMUST00 | -36042 | Mtdh | ENSMUSG0( | false | . | . | . | . | . | . | . | . | . | . | . | . |
| chr15 | 32998529 | 32998728 | 182 | Intron (ENSMUST00000022871.5 | 15 | 32920774 | 32921305 | 532 | 1 | ENSMUST00 | 77756 | Sdc2 | ENSMUSG0( | false | . | . | . | . | . | . | chr15 | 32997685 | 33000925 | 4.14561 | 3.60371 | 2.35273 |
| chr15 | 27525377 | 27525650 | 182 | Intron (ENSMUST00000022875.6 | 15 | 27571449 | 27593163 | 21715 | 1 | ENSMUST00 | -45799 | Ank | ENSMUSG0( | false | . | . | . | . | . | . | . | . | . | . | . | . |
| chr15 | 25926287 | 25926548 | 182 | Intron (ENSMUST00000022881.8 | 15 | 25940887 | 25972340 | 31454 | 1 | ENSMUST00 | -14339 | Fam134b | ENSMUSG0( | false | . | . | . | . | . | . | chr15 | 25918964 | 25927270 | 5.03918 | 4.09395 | 3.04714 |
| chr15 | 25843107 | 25843740 | 182 | Promoter (<=1kb) | 15 | 25843264 | 25973687 | 130424 | 1 | ENSMUST00 | 0 | Fam134b | ENSMUSG0( | false | chr15 | 25842720 | 25844970 | 41.07278 | 140.67711 | 135.76277 | . | . | . | . | . | . |
| chr15 | 25783114 | 25783300 | 182 | Promoter (2-3kb) | 15 | 25780190 | 25781368 | 1179 | 1 | ENSMUST00 | 2925 | Myo10 | ENSMUSG0( | false | . | . | . | . | . | . | chr15 | 25781620 | 25786320 | 4.97313 | 4.06246 | 2.98415 |
| chr15 | 25807498 | 25807773 | 182 | Exon (ENSMUST00000110457.2/ | 15 | 25780190 | 25781368 | 1179 | 1 | ENSMUST00 | 27309 | Myo10 | ENSMUSG0( | false | . | . | . | . | . | . | chr15 | 25805586 | 25810070 | 5.51807 | 4.36356 | 3.42397 |
| chr15 | 39943494 | 39944191 | 182 | Promoter (<=1kb) | 15 | 39870603 | 39943788 | 73186 | 2 | ENSMUST00 | 0 | Lrp12 | ENSMUSG0( | false | chr15 | 39942756 | 39944821 | 31.49598 | 116.8353 | 112.79929 | . | . | . | . | . | . |
| chr15 | 51991080 | 51991467 | 182 | Promoter (<=1kb) | 15 | 51962605 | 51991760 | 29156 | 2 | ENSMUST00 | 293 | Rad21 | ENSMUSG0( | true | chr15 | 51989574 | 51991790 | 32.98857 | 124.89146 | 120.57959 | . | . | . | . | . | . |
| chr15 | 35326504 | 35326761 | 182 | Distal Intergenic | 15 | 35296098 | 35303304 | 7207 | 1 | ENSMUST00 | 30407 | Osr2 | ENSMUSG0( | false | chr15 | 35326442 | 35327177 | 13.48988 | 31.60083 | 29.11589 | chr15 | 35326570 | 35327133 | 4.06921 | 3.63141 | 2.29139 |
| chr15 | 35295924 | 35296188 | 182 | Promoter (<=1kb) | 15 | 35296098 | 35303304 | 7207 | 1 | ENSMUST00 | 0 | Osr2 | ENSMUSG0( | false | chr15 | 35295490 | 35297547 | 28.24444 | 85.94312 | 82.67722 | chr15 | 35292873 | 35297748 | 5.6906 | 4.42763 | 3.58601 |
| chr15 | 68259572 | 68259811 | 182 | Promoter (<=1kb) | 15 | 68083766 | 68258856 | 175091 | 2 | ENSMUST00 | -717 | Zfat | ENSMUSG0( | false | chr15 | 68257894 | 68259920 | 25.57207 | 77.50074 | 74.3909 | chr15 | 68259708 | 68264615 | 3.7026 | 3.36162 | 1.94402 |
| chr15 | 43282623 | 43282970 | 182 | Promoter (<=1kb) | 15 | 43250040 | 43282736 | 32697 | 2 | ENSMUST00 | 0 | Eif3e | ENSMUSG0( | false | chr15 | 43281778 | 43283136 | 33.73597 | 126.30199 | 121.9427 | . | . | . | . | . | . |
| chr15 | 44619380 | 44619724 | 182 | Promoter (<=1kb) | 15 | 44619641 | 44641024 | 21384 | 1 | ENSMUST00 | 0 | Ebag9 | ENSMUSG0( | false | chr15 | 44619120 | 44620360 | 38.47989 | 123.03305 | 118.78763 | chr15 | 44618450 | 44623380 | 3.68052 | 3.44681 | 1.90725 |
| chr15 | 59092093 | 59092362 | 182 | Distal Intergenic | 15 | 58941244 | 59082026 | 140783 | 2 | ENSMUST00 | -10068 | Mtss1 | ENSMUSG0( | true | . | . | . | . | . | . | chr15 | 59082267 | 59100762 | 5.7793 | 4.47653 | 3.63859 |
| chr15 | 58995197 | 58995485 | 182 | Intron (ENSMUST00000080371.6 | 15 | 58933810 | 58939489 | 5680 | 1 | ENSMUST00 | 61388 | Ndufb9 | ENSMUSG0( | false | . | . | . | . | . | . | chr15 | 58965186 | 59078715 | 7.47325 | 5.12649 | 5.04432 |
| chr15 | 58141275 | 58141572 | 182 | Promoter (<=1kb) | 15 | 58141436 | 58158652 | 17217 | 1 | ENSMUST00 | 0 | Wdyhv1 | ENSMUSG0( | false | chr15 | 58140916 | 58142176 | 30.60796 | 113.09483 | 109.17188 | . | . | . | . | . | . |
| chr15 | 58078339 | 58078544 | 182 | Promoter (1-2kb) | 15 | 58068660 | 58076541 | 7882 | 2 | ENSMUST00 | -1799 | Zhx1 | ENSMUSG0( | false | . | . | . | . | . | . | chr15 | 58077164 | 58078932 | 3.87181 | 3.49962 | 2.08636 |
| chr15 | 85581560 | 85581847 | 182 | Promoter (<=1kb) | 15 | 85535439 | 85581821 | 46383 | 2 | ENSMUST00 | 0 | Wnt7b | ENSMUSG0( | false | chr15 | 85580363 | 85582124 | 16.14593 | 43.98602 | 41.34693 | chr15 | 85580251 | 85583390 | 5.27783 | 4.10154 | 3.27147 |
| chr15 | 81808840 | 81809111 | 182 | Promoter (1-2kb) | 15 | 81811046 | 81821256 | 10211 | 1 | ENSMUST00 | -1935 | Tef | ENSMUSG0( | false | . | . | . | . | . | . | chr15 | 81800930 | 81810375 | 6.79861 | 4.85005 | 4.49384 |
| chr15 | 85036954 | 85037230 | 182 | Promoter (<=1kb) | 15 | 85037062 | 85062828 | 25767 | 1 | ENSMUST00 | 0 | Fam118a | ENSMUSG0( | false | chr15 | 85036337 | 85037731 | 33.10969 | 106.34647 | 102.61562 | chr15 | 85035923 | 85038696 | 3.76592 | 3.48389 | 1.97741 |
| chr15 | 77804800 | 77805144 | 182 | Intron (ENSMUST00000016771.7 | 15 | 77794652 | 77813231 | 18580 | 2 | ENSMUST00 | 8087 | Myh9 | ENSMUSG0( | true | . | . | . | . | . | . | chr15 | 77781877 | 77840076 | 8.49833 | 5.61339 | 5.87304 |
| chr15 | 82345797 | 82346255 | 182 | Promoter (<=1kb) | 15 | 82345954 | 82349090 | 3137 | 1 | ENSMUST00 | 0 | Smdt1 | ENSMUSG0( | false | chr15 | 82345970 | 82347455 | 33.5288 | 108.10788 | 104.32592 | . | . | . | . | . | . |
| chr15 | 96709540 | 96710282 | 182 | Distal Intergenic | 15 | 96687394 | 96699698 | 12305 | 2 | ENSMUST00 | -9843 | Slc38a2 | ENSMUSG0( | false | chr15 | 96709302 | 96710655 | 26.25447 | 78.69737 | 75.57 | chr15 | 96709144 | 96710923 | 3.51407 | 3.33727 | 1.78803 |
| chr15 | 97774005 | 97774911 | 182 | Distal Intergenic | 15 | 97766859 | 97767972 | 1114 | 2 | ENSMUST00 | -6034 | Rapgef3 | ENSMUSG0( | false | . | . | . | . | . | . | chr15 | 97771183 | 97790304 | 7.43452 | 5.14328 | 5.01036 |
| chr15 | 97772617 | 97772834 | 182 | Distal Intergenic | 15 | 97766859 | 97767972 | 1114 | 2 | ENSMUST00 | -4646 | Rapgef3 | ENSMUSG0( | false | . | . | . | . | . | . | chr15 | 97771183 | 97790304 | 7.43452 | 5.14328 | 5.01036 |
| chr15 | 97837754 | 97837963 | 182 | Intron (ENSMUST00000116408.3 | 15 | 97792681 | 97831767 | 39087 | 2 | ENSMUST00 | -5988 | Hdac7 | ENSMUSG0( | false | . | . | . | . | . | . | chr15 | 97835434 | 97841080 | 5.51907 | 4.35222 | 3.43438 |
| chr15 | 97892578 | 97892796 | 182 | Intron (ENSMUST00000023119.9 | 15 | 97876776 | 97896813 | 20038 | 2 | ENSMUST00 | 4017 | Vdr | ENSMUSG0( | false | . | . | . | . | . | . | chr15 | 97890193 | 97894481 | 3.62044 | 3.26013 | 1.90833 |
| chr16 | 8830938 | 8831174 | 182 | Promoter (<=1kb) | 16 | 8830100 | 8858916 | 28817 | 1 | ENSMUST00 | 839 | 1810013L24R | ENSMUSG0( | false | chr16 | 8830049 | 8831883 | 32.61975 | 142.21701 | 137.24615 | . | . | . | . | . | . |
| chr16 | 23993404 | 23993892 | 182 | Distal Intergenic | 16 | 23965052 | 23988852 | 23801 | 2 | ENSMUST00 | -4553 | Bcl6 | ENSMUSG0( | false | . | . | . | . | . | . | chr16 | 23989690 | 23996301 | 4.07154 | 3.65227 | 2.23755 |
| chr16 | 25549627 | 25549908 | 182 | Distal Intergenic | 16 | 25683763 | 25821318 | 137556 | 1 | ENSMUST00 | -133855 | Trp63 | ENSMUSG0( | false | . | . | . | . | . | . | . | . | . | . | . | . |
| chr16 | 30065060 | 30065416 | 182 | Promoter (<=1kb) | 16 | 30065340 | 30067796 | 2457 | 1 | ENSMUST00 | 0 | Hes1 | ENSMUSG0( | false | chr16 | 30063120 | 30067742 | 24.1047 | 77.63509 | 74.52327 | . | . | . | . | . | . |
| chr16 | 30066833 | 30067297 | 182 | Promoter (<=1kb) | 16 | 30066338 | 30067319 | 982 | 1 | ENSMUST00 | 496 | Hes1 | ENSMUSG0( | false | chr16 | 30063120 | 30067742 | 24.1047 | 77.63509 | 74.52327 | chr16 | 30067043 | 30072068 | 6.09156 | 4.6344 | 3.88906 |
| chr16 | 30499406 | 30499616 | 182 | Distal Intergenic | 16 | 30513970 | 30540909 | 26940 | 2 | ENSMUST00 | 41293 | Tmem44 | ENSMUSG0( | false | . | . | . | . | . | . | chr16 | 30499213 | 30499879 | 6.79 | 4.99774 | 4.49751 |
| chr16 | 5255844 | 5256206 | 182 | Promoter (<=1kb) | 16 | 5244152 | 5255983 | 11832 | 2 | ENSMUST00 | 0 | Fam86 | ENSMUSG0( | false | chr16 | 5254553 | 5256620 | 29.06336 | 92.21697 | 88.82447 | chr16 | 5253239 | 5256784 | 4.59203 | 3.86124 | 2.68801 |
| chr16 | 13189454 | 13189810 | 182 | Distal Intergenic | 16 | 13132391 | 13144491 | 12101 | 1 | ENSMUST00 | 57064 | Ercc4 | ENSMUSG0( | false | . | . | . | . | . | . | chr16 | 13188422 | 13190534 | 4.59735 | 3.75293 | 2.70347 |
| chr15 | 76351291 | 76351706 | 182 | Promoter (<=1kb) | 15 | 76351294 | 76354372 | 3079 | 1 | ENSMUST00 | 0 | Maf1 | ENSMUSG0( | false | chr15 | 76349778 | 76352229 | 28.59262 | 111.22587 | 107.35971 | . | . | . | . | . | . |
| chr15 | 76477212 | 76477493 | 182 | Promoter (<=1kb) | 15 | 76477445 | 76500972 | 23528 | 1 | ENSMUST00 | 0 | Hsf1 | ENSMUSG0( | false | chr15 | 76476684 | 76478589 | 39.54136 | 129.19044 | 124.73553 | . | . | . | . | . | . |
| chr15 | 76538657 | 76538958 | 182 | Promoter (<=1kb) | 15 | 76535730 | 76538746 | 3017 | 2 | ENSMUST00 | 0 | Fbxl6 | ENSMUSG0( | false | chr15 | 76537923 | 76539844 | 27.20494 | 88.89825 | 85.57716 | chr15 | 76536854 | 76540588 | 6.72736 | 4.87802 | 4.42236 |
| chr15 | 76069396 | 76069950 | 182 | Promoter (<=1kb) | 15 | 76047158 | 76069754 | 22597 | 2 | ENSMUST00 | 0 | Scrib | ENSMUSG0( | false | chr15 | 76068809 | 76069842 | 27.13995 | 99.45934 | 95.90212 | chr15 | 76067246 | 76070913 | 5.68831 | 4.29984 | 3.62222 |
| chr15 | 75703350 | 75703608 | 182 | Promoter (<=1kb) | 15 | 75704280 | 75714408 | 10129 | 1 | ENSMUST00 | -672 | Rhpn1 | ENSMUSG0( | true | chr15 | 75703357 | 75704139 | 26.40393 | 78.95839 | 75.82329 | . | . | . | . | . | . |
| chr16 | 55973538 | 55973764 | 182 | Promoter (<=1kb) | 16 | 55973883 | 56008913 | 35031 | 1 | ENSMUST00 | -119 | Zbtb11 | ENSMUSG0( | false | chr16 | 55973101 | 55975647 | 44.26367 | 154.19713 | 148.75002 | . | . | . | . | . | . |
| chr15 | 89429784 | 89430017 | 182 | Promoter (<=1kb) | 15 | 89426360 | 89429860 | 3501 | 2 | ENSMUST00 | 0 | Chkb | ENSMUSG0( | false | chr15 | 89428885 | 89430631 | 40.43883 | 128.57333 | 124.13428 | . | . | . | . | . | . |
| chr16 | 52452064 | 52452412 | 182 | Promoter (<=1kb) | 16 | 52251467 | 52452465 | 200999 | 2 | ENSMUST00 | 53 | Alcam | ENSMUSG0( | true | chr16 | 52450004 | 52454370 | 27.24215 | 82.29425 | 79.10038 | . | . | . | . | . | . |
| chr16 | 51915717 | 51915918 | 182 | Distal Intergenic | 16 | 52031549 | 52208047 | 176499 | 1 | ENSMUST00 | -115631 | Cblb | ENSMUSG0( | true | . | . | . | . | . | . | chr16 | 51915431 | 51917899 | 3.71214 | 3.42083 | 1.95763 |
| chr16 | 52060282 | 52060529 | 182 | Intron (ENSMUST00000114471.1 | 16 | 52031549 | 52208047 | 176499 | 1 | ENSMUST00 | 28734 | Cblb | ENSMUSG0( | true | . | . | . | . | . | . | chr16 | 52047691 | 52073501 | 5.26548 | 4.17513 | 3.22843 |
| chr16 | 50431384 | 50432468 | 182 | Promoter (<=1kb) | 16 | 50191844 | 50432390 | 240547 | 2 | ENSMUST00 | 0 | Bbx | ENSMUSG0( | false | chr16 | 50429049 | 50432934 | 35.20524 | 115.20684 | 111.21615 | . | . | . | . | . | . |
| chr16 | 50431094 | 50431349 | 182 | Promoter (<=1kb) | 16 | 50425961 | 50432309 | 6349 | 2 | ENSMUST00 | 960 | Bbx | ENSMUSG0( | false | chr16 | 50429049 | 50432934 | 35.20524 | 115.20684 | 111.21615 | . | . | . | . | . | . |
| chr16 | 45102779 | 45102964 | 182 | Intron (ENSMUST00000099498.3 | 16 | 45094053 | 45127924 | 33872 | 1 | ENSMUST00 | 8727 | Ccdc80 | ENSMUSG0( | false | . | . | . | . | . | . | chr16 | 45098204 | 45106638 | 4.5684 | 3.92095 | 2.64446 |
| chr16 | 15637700 | 15638035 | 182 | Promoter (<=1kb) | 16 | 15637866 | 15684749 | 46884 | 1 | ENSMUST00 | 0 | Prkdc | ENSMUSG0( | false | chr16 | 15636693 | 15638435 | 25.25706 | 81.58189 | 78.40311 | . | . | . | . | . | . |
| chr16 | 13668042 | 13668423 | 182 | Promoter (<=1kb) | 16 | 13537964 | 13668170 | 130207 | 2 | ENSMUST00 | 0 | Parn | ENSMUSG0( | false | chr16 | 13667412 | 13668408 | 18.82282 | 71.67798 | 68.66222 | chr16 | 13667614 | 13668767 | 5.11925 | 4.12449 | 3.11449 |
| chr16 | 70396351 | 70396602 | 182 | Intron (ENSMUST00000163832.2 | 16 | 70314134 | 70569714 | 255581 | 1 | ENSMUST00 | 82218 | Gbe1 | ENSMUSG0( | false | . | . | . | . | . | . | chr16 | 70396061 | 70397046 | 2.94843 | 2.99364 | 1.37678 |
| chr16 | 42897219 | 42897441 | 182 | Intron (ENSMUST00000126354.2 | 16 | 42907392 | 43072468 | 165077 | 1 | ENSMUST00 | -9951 | Zbtb20 | ENSMUSG0( | false | . | . | . | . | . | . | chr16 | 42896901 | 42899356 | 3.62146 | 3.42768 | 1.85675 |
| chr16 | 42907421 | 42907761 | 182 | Promoter (<=1kb) | 16 | 42907542 | 43569781 | 662240 | 1 | ENSMUST00 | 0 | Zbtb20 | ENSMUSG0( | false | chr16 | 42907033 | 42911476 | 14.29631 | 42.77194 | 40.14703 | . | . | . | . | . | . |
| chr16 | 18289111 | 18289662 | 182 | Promoter (<=1kb) | 16 | 18253948 | 18289246 | 35299 | 2 | ENSMUST00 | 0 | Dgcr8 | ENSMUSG0( | false | chr16 | 18288146 | 18289955 | 30.67334 | 114.24463 | 110.28603 | . | . | . | . | . | . |
| chr16 | 57231408 | 57231624 | 182 | Promoter (<=1kb) | 16 | 57168866 | 57231466 | 62601 | 2 | ENSMUST00 | 0 | Tbc1d23 | ENSMUSG0( | false | chr16 | 57230448 | 57231979 | 34.17132 | 121.80145 | 117.60049 | . | . | . | . | . | . |
| chr16 | 56667105 | 56667388 | 182 | Intron (ENSMUST00000048471.8 | 16 | 56690968 | 56704463 | 13496 | 2 | ENSMUST00 | 37075 | Tfg | ENSMUSG0( | false | . | . | . | . | . | . | . | . | . | . | . | . |
| chr16 | 17496501 | 17496926 | 182 | Promoter (2-3kb) | 16 | 17499699 | 17507481 | 7783 | 1 | ENSMUST00 | -2773 | Aifm3 | ENSMUSG0( | false | . | . | . | . | . | . | chr16 | 17485729 | 17499611 | 5.1843 | 4.03502 | 3.18262 |
| chr16 | 17497072 | 17497260 | 182 | Promoter (2-3kb) | 16 | 17499699 | 17507481 | 7783 | 1 | ENSMUST00 | -2439 | Aifm3 | ENSMUSG0( | false | . | . | . | . | . | . | chr16 | 17485729 | 17499611 | 5.1843 | 4.03502 | 3.18262 |
| chr16 | 16870628 | 16871022 | 182 | Promoter (<=1kb) | 16 | 16870736 | 16885849 | 15114 | 1 | ENSMUST00 | 0 | Top3b | ENSMUSG0( | false | chr16 | 16870250 | 16871600 | 34.74108 | 114.39095 | 110.42815 | . | . | . | . | . | . |
| chr16 | 38400073 | 38400259 | 182 | Intron (ENSMUST00000002926.6 | 16 | 38371998 | 38374137 | 2140 | 1 | ENSMUST00 | 28076 | Popdc2 | ENSMUSG0( | false | chr16 | 38399626 | 38400302 | 7.14815 | 15.37828 | 13.11012 | chr16 | 38399120 | 38400267 | 4.23035 | 3.43922 | 2.40939 |
| chr16 | 38399577 | 38399843 | 182 | Intron (ENSMUST00000002926.6 | 16 | 38371998 | 38374137 | 2140 | 1 | ENSMUST00 | 27580 | Popdc2 | ENSMUSG0( | false | chr16 | 38399626 | 38400302 | 7.14815 | 15.37828 | 13.11012 | chr16 | 38399120 | 38400267 | 4.23035 | 3.43922 | 2.40939 |
| chr16 | 37539590 | 37539857 | 182 | Promoter (<=1kb) | 16 | 37509790 | 37539789 | 30000 | 2 | ENSMUST00 | 0 | Gtf2e1 | ENSMUSG0( | false | chr16 | 37539029 | 37540718 | 30.17592 | 94.17345 | 90.73757 | chr16 | 37538079 | 37541163 | 4.9737 | 4.11839 | 2.98435 |
| chr16 | 35177842 | 35178071 | 182 | Intron (ENSMUST00000114913.1 | 16 | 35154877 | 35305737 | 150861 | 1 | ENSMUST00 | 22966 | Adcy5 | ENSMUSG0( | true | . | . | . | . | . | . | chr16 | 35176357 | 35187367 | 5.48942 | 4.28454 | 3.41207 |
| chr16 | 35769166 | 35769551 | 182 | Promoter (<=1kb) | 16 | 35694062 | 35769356 | 75295 | 2 | ENSMUST00 | 0 | Dirc2 | ENSMUSG0( | false | chr16 | 35768281 | 35771151 | 24.60788 | 97.30984 | 93.80507 | chr16 | 35767992 | 35771593 | 3.89547 | 3.46454 | 2.13759 |
| chr16 | 21947249 | 21947652 | 182 | Promoter (<=1kb) | 16 | 21934327 | 21947552 | 13226 | 2 | ENSMUST00 | 0 | Tmem41a | ENSMUSG0( | false | chr16 | 21946729 | 21947835 | 25.79552 | 96.83904 | 93.34669 | chr16 | 21945881 | 21947906 | 4.52724 | 3.80739 | 2.66373 |
| chr16 | 22265326 | 22265578 | 182 | Promoter (<=1kb) | 16 | 22246462 | 22265829 | 19368 | 2 | ENSMUST00 | 251 | Tra2b | ENSMUSG0( | false | chr16 | 22263924 | 22266550 | 36.04937 | 140.37729 | 135.48044 | . | . | . | . | . | . |
| chr16 | 78376131 | 78376393 | 182 | Promoter (<=1kb) | 16 | 78359860 | 78376810 | 16951 | 2 | ENSMUST00 | 417 | Btg3 | ENSMUSG0( | false | chr16 | 78375631 | 78377323 | 25.75243 | 103.50989 | 99.85224 | . | . | . | . | . | . |
| chr16 | 23127670 | 23127995 | 182 | Promoter (<=1kb) | 16 | 23113943 | 23127737 | 13795 | 2 | ENSMUST00 | 0 | Rfc4 | ENSMUSG0( | false | chr16 | 23127320 | 23128515 | 25.48472 | 95.2657 | 91.8051 | chr16 | 23126728 | 23129060 | 4.47886 | 3.80087 | 2.59071 |
| chr16 | 72717491 | 72717706 | 182 | Intron (ENSMUST00000023600.7 | 16 | 72663149 | 73046095 | 382947 | 1 | ENSMUST00 | 54343 | Robo1 | ENSMUSG0( | false | . | . | . | . | . | . | chr16 | 72717425 | 72718155 | 4.28281 | 3.63092 | 2.43706 |
| chr16 | 23349157 | 23349333 | 182 | Intron (ENSMUST00000023601.8 | 16 | 23358316 | 23360348 | 2033 | 1 | ENSMUST00 | -8983 | St6gal1 | ENSMUSG0( | true | . | . | . | . | . | . | chr16 | 23344368 | 23353615 | 5.35656 | 4.16706 | 3.31602 |
| chr16 | 85756168 | 85756449 | 182 | Distal Intergenic | 16 | 85798231 | 85799638 | 1408 | 2 | ENSMUST00 | 43189 | Adamts1 | ENSMUSG0( | false | . | . | . | . | . | . | chr16 | 85755334 | 85756944 | 3.71271 | 3.44594 | 1.95342 |
| chr16 | 85803179 | 85803391 | 182 | Promoter (<=1kb) | 16 | 85793827 | 85803113 | 9287 | 2 | ENSMUST00 | -67 | Adamts1 | ENSMUSG0( | false | chr16 | 85800671 | 85803715 | 26.05581 | 78.34985 | 75.22605 | chr16 | 85800084 | 85805307 | 4.30005 | 3.70427 | 2.42091 |
| chr16 | 36770242 | 36770470 | 182 | Promoter (2-3kb) | 16 | 36772071 | 36773278 | 1208 | 2 | ENSMUST00 | 2808 | Slc15a2 | ENSMUSG0( | false | . | . | . | . | . | . | . | . | . | . | . | . |
| chr16 | 36640130 | 36640329 | 182 | Promoter (2-3kb) | 16 | 36620713 | 36642805 | 22093 | 2 | ENSMUST00 | 2476 | Cd86 | ENSMUSG0( | false | . | . | . | . | . | . | chr16 | 36639655 | 36641079 | 3.80729 | 3.42569 | 2.06859 |
| chr16 | 35992662 | 35992877 | 182 | Intron (ENSMUST00000172991.1 | 16 | 36000187 | 36033908 | 33722 | 1 | ENSMUST00 | -7310 | Kpna1 | ENSMUSG0( | false | . | . | . | . | . | . | chr16 | 35991968 | 35994964 | 5.1828 | 4.06272 | 3.19191 |
| chr16 | 62846817 | 62847152 | 182 | Promoter (<=1kb) | 16 | 62793685 | 62847040 | 53356 | 2 | ENSMUST00 | 0 | Arl13b | ENSMUSG0( | false | chr16 | 62845942 | 62847074 | 52.11642 | 180.80226 | 174.62196 | chr16 | 62845315 | 62847523 | 3.65827 | 3.41118 | 1.88304 |
| chr16 | 62874129 | 62874422 | 182 | Intron (ENSMUST00000023629.8 | 16 | 62854307 | 62929346 | 75040 | 1 | ENSMUST00 | 19823 | Pros1 | ENSMUSG0( | false | . | . | . | . | . | . | chr16 | 62866329 | 62875410 | 4.59164 | 3.90036 | 2.66788 |
| chr16 | 93883675 | 93883975 | 182 | Promoter (<=1kb) | 16 | 93883901 | 93906115 | 22215 | 1 | ENSMUST00 | 0 | Chaf1b | ENSMUSG0( | false | chr16 | 93883326 | 93884864 | 30.44295 | 97.61457 | 94.10351 | . | . | . | . | . | . |
| chr16 | 93711550 | 93712228 | 182 | Promoter (<=1kb) | 16 | 93711907 | 93810585 | 98679 | 1 | ENSMUST00 | 0 | Dopey2 | ENSMUSG0( | false | chr16 | 93711890 | 93712743 | 34.54477 | 122.48211 | 118.25522 | chr16 | 93712223 | 93714534 | 4.08957 | 3.6332 | 2.25397 |
| chr16 | 93682798 | 93683255 | 182 | Promoter (<=1kb) | 16 | 93683215 | 93690990 | 7776 | 1 | ENSMUST00 | 0 | Cbr3 | ENSMUSG0( | false | chr16 | 93682573 | 93684551 | 20.76252 | 61.01094 | 58.15269 | chr16 | 93682402 | 93684537 | 5.08052 | 4.14861 | 3.06303 |
| chr16 | 92415730 | 92415991 | 182 | Intron (ENSMUST00000060005.9 | 16 | 92391953 | 92400077 | 8125 | 2 | ENSMUST00 | -15654 | Rcan1 | ENSMUSG0( | false | . | . | . | . | . | . | chr16 | 92415129 | 92416512 | 5.12964 | 4.17261 | 3.09669 |
| chr16 | 92696829 | 92697031 | 182 | Promoter (<=1kb) | 16 | 92686989 | 92697325 | 10337 | 2 | ENSMUST00 | 294 | Runx1 | ENSMUSG0( | false | chr16 | 92696643 | 92698812 | 27.94004 | 83.5095 | 80.29546 | chr16 | 92695451 | 92700248 | 5.19566 | 4.17031 | 3.16375 |
| chr16 | 91647571 | 91648018 | 182 | Promoter (<=1kb) | 16 | 91647879 | 91665708 | 17830 | 1 | ENSMUST00 | 0 | Son | ENSMUSG0( | false | chr16 | 91646065 | 91649433 | 31.16539 | 106.97112 | 103.21663 | . | . | . | . | . | . |
| chr16 | 91399486 | 91399716 | 182 | Intron (ENSMUST00000023693.8 | 16 | 91406164 | 91425822 | 19659 | 1 | ENSMUST00 | -6448 | Il10rb | ENSMUSG0( | false | . | . | . | . | . | . | chr16 | 91399479 | 91399930 | 2.92753 | 2.93274 | 1.36135 |
| chr16 | 91406082 | 91406386 | 182 | Promoter (<=1kb) | 16 | 91406164 | 91425822 | 19659 | 1 | ENSMUST00 | 0 | Il10rb | ENSMUSG0( | false | chr16 | 91405640 | 91407210 | 26.18681 | 91.82585 | 88.44535 | chr16 | 91405562 | 91408156 | 3.71117 | 3.35173 | 1.98203 |
| chr16 | 90934847 | 90935099 | 182 | Promoter (<=1kb) | 16 | 90925970 | 90934927 | 8958 | 2 | ENSMUST00 | 0 | 1110004E09R | ENSMUSG0( | false | chr16 | 90933837 | 90935564 | 25.48472 | 95.2657 | 91.8051 | . | . | . | . | . | . |
| chr15 | 98917548 | 98918255 | 182 | Promoter (<=1kb) | 15 | 98903921 | 98918231 | 14311 | 2 | ENSMUST00 | 0 | Lmbr1l | ENSMUSG0( | false | chr15 | 98917164 | 98918235 | 35.95432 | 114.41653 | 110.45183 | . | . | . | . | . | . |
| chr15 | 99396898 | 99397326 | 182 | Intron (ENSMUST00000023749.9 | 15 | 99393610 | 99408610 | 15001 | 1 | ENSMUST00 | 3289 | Tmbim6 | ENSMUSG0( | false | . | . | . | . | . | . | chr15 | 99393778 | 99402130 | 6.4261 | 4.61526 | 4.21071 |
| chr15 | 100422822 | 100423413 | 182 | Promoter (<=1kb) | 15 | 100387898 | 100423055 | 35158 | 2 | ENSMUST00 | 0 | Slc11a2 | ENSMUSG0( | false | chr15 | 100422204 | 100424245 | 39.70201 | 131.65079 | 127.10004 | chr15 | 100417554 | 100425131 | 6.25151 | 4.61613 | 4.04938 |
| chr15 | 100923484 | 100923668 | 182 | Intron (ENSMUST00000082209.7 | 15 | 100933429 | 100940798 | 7370 | 1 | ENSMUST00 | -9761 | Scn8a | ENSMUSG0( | false | . | . | . | . | . | . | . | . | . | . | . | . |
| chr15 | 101262541 | 101262857 | 182 | Distal Intergenic | 15 | 101266846 | 101274792 | 7947 | 1 | ENSMUST00 | -3989 | Nr4a1 | ENSMUSG0( | false | . | . | . | . | . | . | . | . | . | . | . | . |
| chr15 | 102721783 | 102722354 | 182 | Promoter (<=1kb) | 15 | 102706779 | 102722178 | 15400 | 2 | ENSMUST00 | 0 | Calcoco1 | ENSMUSG0( | false | chr15 | 102721189 | 102722153 | 23.12601 | 77.64799 | 74.53377 | chr15 | 102720354 | 102723786 | 3.72435 | 3.47928 | 1.94971 |
| chr3 | 51220873 | 51221112 | 182 | Distal Intergenic | 3 | 51224447 | 51251644 | 27198 | 1 | ENSMUST00 | -3335 | Ccrn4l | ENSMUSG0( | false | . | . | . | . | . | . | chr3 | 51212148 | 51224324 | 4.29558 | 3.75668 | 2.42602 |
| chr6 | 24527501 | 24527857 | 182 | Promoter (<=1kb) | 6 | 24518666 | 24527570 | 8905 | 2 | ENSMUST00 | 0 | Ndufa5 | ENSMUSG0( | false | chr6 | 24526536 | 24528331 | 28.80981 | 88.85442 | 85.53428 | chr6 | 24526474 | 24534302 | 5.36778 | 4.25265 | 3.31628 |
| chr5 | 123907048 | 123907294 | 182 | Promoter (<=1kb) | 5 | 123907175 | 123928832 | 21658 | 1 | ENSMUST00 | 0 | Denr | ENSMUSG0( | false | chr5 | 123907208 | 123908092 | 26.72788 | 101.58999 | 97.97913 | chr5 | 123906744 | 123909035 | 4.89302 | 3.93732 | 2.94393 |
| chr16 | 30257732 | 30257935 | 182 | 3' UTR | 16 | 30256378 | 30267532 | 11155 | 2 | ENSMUST00 | 9597 | Cpn2 | ENSMUSG0( | false | . | . | . | . | . | . | chr16 | 30257559 | 30258096 | 3.68995 | 3.38843 | 1.9571 |
| chr6 | 124857055 | 124857721 | 182 | Promoter (<=1kb) | 6 | 124841206 | 124857729 | 16524 | 2 | ENSMUST00 | 8 | Leprel2 | ENSMUSG0( | false | chr6 | 124856897 | 124857761 | 19.39874 | 49.63541 | 46.92579 | . | . | . | . | . | . |
| chr2 | 84774798 | 84775243 | 182 | Promoter (<=1kb) | 2 | 84773131 | 84775128 | 1998 | 2 | ENSMUST00 | 0 | Serping1 | ENSMUSG0( | false | . | . | . | . | . | . | chr2 | 84773836 | 84775831 | 5.2778 | 4.15422 | 3.26847 |
| chr8 | 4324864 | 4325304 | 182 | Promoter (<=1kb) | 8 | 4325210 | 4359998 | 34789 | 1 | ENSMUST00 | 0 | Ccl25 | ENSMUSG0( | false | chr8 | 4323747 | 4325926 | 35.82539 | 116.29276 | 112.27101 | chr8 | 4311994 | 4326296 | 5.54778 | 4.25812 | 3.46123 |
| chr2 | 126151933 | 126152279 | 182 | Promoter (<=1kb) | 2 | 126152141 | 126165276 | 13136 | 1 | ENSMUST00 | 0 | Dtwd1 | ENSMUSG0( | false | chr2 | 126151549 | 126152953 | 28.22734 | 101.53345 | 97.92473 | . | . | . | . | . | . |
| chr8 | 119660856 | 119661520 | 182 | Distal Intergenic | 8 | 119666523 | 119688222 | 21700 | 1 | ENSMUST00 | -5003 | Wfdc1 | ENSMUSG0( | false | . | . | . | . | . | . | chr8 | 119659713 | 119662363 | 8.19394 | 5.62527 | 5.61282 |
| chr16 | 97535045 | 97535310 | 182 | Promoter (<=1kb) | 16 | 97535308 | 97560339 | 25032 | 1 | ENSMUST00 | 0 | Mx2 | ENSMUSG0( | false | chr16 | 97534754 | 97536023 | 11.22292 | 28.25542 | 25.81138 | chr16 | 97534378 | 97539123 | 7.43346 | 5.26162 | 5.00178 |
| chr16 | 97536128 | 97536381 | 182 | Promoter (<=1kb) | 16 | 97536083 | 97560899 | 24817 | 1 | ENSMUST00 | 46 | Mx2 | ENSMUSG0( | false | . | . | . | . | . | . | chr16 | 97534378 | 97539123 | 7.43346 | 5.26162 | 5.00178 |
| chr5 | 137314050 | 137314305 | 182 | Promoter (<=1kb) | 5 | 137309899 | 137314241 | 4343 | 2 | ENSMUST00 | 0 | Trip6 | ENSMUSG0( | false | chr5 | 137312954 | 137314692 | 24.30838 | 70.74035 | 67.73997 | chr5 | 137311496 | 137315031 | 4.90485 | 4.00152 | 2.93486 |
| chr5 | 24481859 | 24482169 | 182 | Intron (ENSMUST00000024123.4 | 5 | 24452177 | 24502047 | 49871 | 1 | ENSMUST00 | 29683 | Agap3 | ENSMUSG0( | false | . | . | . | . | . | . | chr5 | 24481028 | 24486934 | 6.61014 | 4.77469 | 4.33237 |
| chr14 | 55823289 | 55823639 | 182 | Promoter (1-2kb) | 14 | 55824795 | 55833943 | 9149 | 1 | ENSMUST00 | -1156 | Nfatc4 | ENSMUSG0( | false | chr14 | 55822836 | 55824209 | 22.21283 | 62.70797 | 59.8237 | chr14 | 55822511 | 55827107 | 5.07816 | 4.14481 | 3.0616 |
| chr9 | 106453612 | 106453924 | 182 | Promoter (<=1kb) | 9 | 106453845 | 106454836 | 992 | 1 | ENSMUST00 | 0 | Pcbp4 | ENSMUSG0( | false | chr9 | 106453266 | 106454683 | 27.57193 | 78.19525 | 75.07539 | chr9 | 106452714 | 106456807 | 4.65098 | 3.94386 | 2.7101 |
| chr8 | 104641651 | 104641840 | 182 | Promoter (<=1kb) | 8 | 104641807 | 104658587 | 16781 | 1 | ENSMUST00 | 0 | 1700082M22 | ENSMUSG0( | false | chr8 | 104640850 | 104642429 | 39.02359 | 127.20569 | 122.81842 | . | . | . | . | . | . |
| chr17 | 3678175 | 3678389 | 182 | Intron (ENSMUST00000115800.1 | 17 | 3635240 | 3696261 | 61022 | 2 | ENSMUST00 | 17872 | Nox3 | ENSMUSG0( | false | . | . | . | . | . | . | chr17 | 3677813 | 3679347 | 3.48821 | 3.28801 | 1.79383 |
| chr17 | 7169698 | 7170047 | 182 | Promoter (<=1kb) | 17 | 7170115 | 7303313 | 133199 | 1 | ENSMUST00 | -68 | Rps6ka2 | ENSMUSG0( | false | chr17 | 7169205 | 7169866 | 16.96414 | 42.42906 | 39.81211 | . | . | . | . | . | . |
| chr17 | 11950006 | 11950183 | 182 | Intron (ENSMUST00000191124.1 | 17 | 12029916 | 12063358 | 33443 | 1 | ENSMUST00 | -79733 | Park2 | ENSMUSG0( | false | . | . | . | . | . | . | chr17 | 11949587 | 11950568 | 3.54925 | 3.19744 | 1.8512 |
| chr17 | 11963052 | 11963233 | 182 | Intron (ENSMUST00000191124.1 | 17 | 12029916 | 12063358 | 33443 | 1 | ENSMUST00 | -66683 | Park2 | ENSMUSG0( | false | . | . | . | . | . | . | chr17 | 11962721 | 11963625 | 3.70665 | 3.28944 | 1.97595 |
| chr17 | 8818900 | 8819198 | 182 | Intron (ENSMUST00000115715.2 | 17 | 8801779 | 8986645 | 184867 | 1 | ENSMUST00 | 17122 | Pde10a | ENSMUSG0( | true | . | . | . | . | . | . | chr17 | 8817580 | 8819489 | 4.92873 | 4.0649 | 3.00503 |
| chr17 | 8846539 | 8846821 | 182 | Intron (ENSMUST00000115715.2 | 17 | 8849985 | 8982400 | 132416 | 1 | ENSMUST00 | -3164 | Pde10a | ENSMUSG0( | true | . | . | . | . | . | . | . | . | . | . | . | . |
| chr17 | 23740107 | 23740515 | 182 | Promoter (<=1kb) | 17 | 23736186 | 23740330 | 4145 | 2 | ENSMUST00 | 0 | Paqr4 | ENSMUSG0( | false | chr17 | 23739513 | 23741047 | 29.22523 | 92.5029 | 89.10189 | chr17 | 23739155 | 23743063 | 4.49337 | 3.79493 | 2.61097 |
| chr17 | 43077693 | 43077905 | 182 | Intron (ENSMUST00000024708.4 | 17 | 43016555 | 43089188 | 72634 | 1 | ENSMUST00 | 61139 | Tnfrsf21 | ENSMUSG0( | false | chr17 | 43077267 | 43077841 | 9.63953 | 19.72805 | 17.39678 | chr17 | 43075309 | 43083932 | 4.06236 | 3.6443 | 2.23752 |
| chr17 | 46032250 | 46032576 | 182 | Promoter (<=1kb) | 17 | 46016993 | 46032369 | 15377 | 2 | ENSMUST00 | 0 | Vegfa | ENSMUSG0( | true | chr17 | 46023564 | 46033809 | 32.66734 | 113.81544 | 109.86345 | . | . | . | . | . | . |
| chr17 | 46084241 | 46084583 | 182 | Distal Intergenic | 17 | 46110986 | 46128910 | 17925 | 1 | ENSMUST00 | -26403 | Mrps18a | ENSMUSG0( | false | chr17 | 46084246 | 46084634 | 6.10183 | 9.89467 | 7.72061 | chr17 | 46081746 | 46085243 | 5.14134 | 4.1226 | 3.129 |
| chr17 | 46612758 | 46613146 | 182 | Intron (ENSMUST00000181301.1 | 17 | 46564451 | 46629504 | 65054 | 2 | ENSMUST00 | 16358 | Ptk7 | ENSMUSG0( | false | . | . | . | . | . | . | chr17 | 46605355 | 46631878 | 5.60661 | 4.26306 | 3.52431 |
| chr17 | 46628725 | 46628944 | 182 | Promoter (<=1kb) | 17 | 46564451 | 46629504 | 65054 | 2 | ENSMUST00 | 560 | Ptk7 | ENSMUSG0( | false | chr17 | 46628274 | 46630084 | 29.66345 | 100.21 | 96.63225 | chr17 | 46605355 | 46631878 | 5.60661 | 4.26306 | 3.52431 |
| chr17 | 46751706 | 46752366 | 182 | Promoter (<=1kb) | 17 | 46735705 | 46752214 | 16510 | 2 | ENSMUST00 | 0 | Cnpy3 | ENSMUSG0( | false | chr17 | 46751480 | 46752624 | 20.5121 | 70.85588 | 67.85384 | chr17 | 46750587 | 46751711 | 3.33505 | 3.12334 | 1.6723 |
| chr17 | 48409568 | 48409792 | 182 | Promoter (<=1kb) | 17 | 48386885 | 48409765 | 22881 | 2 | ENSMUST00 | 0 | Nfya | ENSMUSG0( | false | chr17 | 48408332 | 48409762 | 41.49189 | 142.53105 | 137.54099 | . | . | . | . | . | . |
| chr17 | 29330843 | 29331175 | 182 | Promoter (2-3kb) | 17 | 29332114 | 29334005 | 1892 | 2 | ENSMUST00 | 2830 | Mtch1 | ENSMUSG0( | false | . | . | . | . | . | . | chr17 | 29329048 | 29335325 | 5.42072 | 4.20084 | 3.36531 |
| chr17 | 29398045 | 29398383 | 182 | Distal Intergenic | 17 | 29376501 | 29378452 | 1952 | 1 | ENSMUST00 | 21545 | Fgd2 | ENSMUSG0( | false | . | . | . | . | . | . | chr17 | 29396996 | 29399392 | 4.27884 | 3.786 | 2.45157 |
| chr17 | 29450759 | 29451212 | 182 | Distal Intergenic | 17 | 29490812 | 29496111 | 5300 | 1 | ENSMUST00 | -39600 | Pim1 | ENSMUSG0( | false | chr17 | 29451035 | 29451518 | 9.94526 | 25.28477 | 22.8772 | chr17 | 29443894 | 29452487 | 4.93685 | 4.01842 | 2.95062 |
| chr17 | 31855218 | 31855472 | 182 | Promoter (<=1kb) | 17 | 31844248 | 31855792 | 11545 | 2 | ENSMUST00 | 320 | Sik1 | ENSMUSG0( | false | chr17 | 31854734 | 31855826 | 30.94804 | 114.76607 | 110.79114 | . | . | . | . | . | . |
| chr17 | 67907640 | 67907897 | 182 | Intron (ENSMUST00000024840.6 | 17 | 67845735 | 67901471 | 55737 | 2 | ENSMUST00 | -6170 | Arhgap28 | ENSMUSG0( | true | . | . | . | . | . | . | chr17 | 67892935 | 67914104 | 5.71307 | 4.38398 | 3.59861 |
| chr17 | 32321167 | 32321414 | 182 | Promoter (<=1kb) | 17 | 32303676 | 32321153 | 17478 | 2 | ENSMUST00 | -15 | Akap8 | ENSMUSG0( | false | chr17 | 32319923 | 32322060 | 28.91247 | 96.86193 | 93.36482 | . | . | . | . | . | . |
| chr17 | 32364350 | 32364607 | 182 | Promoter (<=1kb) | 17 | 32361675 | 32364463 | 2789 | 2 | ENSMUST00 | 0 | Wiz | ENSMUSG0( | false | chr17 | 32363372 | 32365041 | 37.73948 | 119.61958 | 115.49072 | chr17 | 32360420 | 32365678 | 4.55703 | 3.85553 | 2.65108 |
| chr17 | 71206582 | 71206787 | 182 | Promoter (1-2kb) | 17 | 71204676 | 71248242 | 43567 | 1 | ENSMUST00 | 1907 | Lpin2 | ENSMUSG0( | false | . | . | . | . | . | . | chr17 | 71186127 | 71229089 | 5.51375 | 4.29183 | 3.43736 |
| chr17 | 73808546 | 73808789 | 182 | Intron (ENSMUST00000024860.7 | 17 | 73804841 | 73832093 | 27253 | 1 | ENSMUST00 | 3706 | Ehd3 | ENSMUSG0( | false | chr17 | 73808367 | 73808715 | 4.04026 | 6.18694 | 4.10202 | chr17 | 73807995 | 73808758 | 4.69511 | 4.04282 | 2.74054 |
| chr17 | 78225608 | 78225858 | 182 | Intron (ENSMUST00000112498.2 | 17 | 78200248 | 78376592 | 176345 | 1 | ENSMUST00 | 25361 | Crim1 | ENSMUSG0( | true | . | . | . | . | . | . | chr17 | 78202641 | 78231492 | 6.38021 | 4.71597 | 4.13646 |
| chr17 | 79781770 | 79781965 | 182 | Distal Intergenic | 17 | 79706953 | 79715041 | 8089 | 2 | ENSMUST00 | -66730 | Cyp1b1 | ENSMUSG0( | false | . | . | . | . | . | . | . | . | . | . | . | . |
| chr17 | 80060791 | 80061004 | 182 | Promoter (1-2kb) | 17 | 80045049 | 80062135 | 17087 | 2 | ENSMUST00 | 1131 | Hnrnpll | ENSMUSG0( | false | chr17 | 80060963 | 80063060 | 39.4261 | 145.54997 | 140.44281 | chr17 | 80058717 | 80061162 | 3.78737 | 3.44429 | 2.05196 |
| chr17 | 65950813 | 65951252 | 182 | Promoter (<=1kb) | 17 | 65923066 | 65951187 | 28122 | 2 | ENSMUST00 | 0 | Twsg1 | ENSMUSG0( | false | chr17 | 65950415 | 65951253 | 30.60583 | 111.04385 | 107.18215 | chr17 | 65949903 | 65951658 | 4.55032 | 3.81363 | 2.68316 |
| chr17 | 85985708 | 85985889 | 182 | Intron (ENSMUST00000095187.3 | 17 | 85984665 | 86145175 | 160511 | 2 | ENSMUST00 | 159286 | Srbd1 | ENSMUSG0( | false | . | . | . | . | . | . | chr17 | 85985115 | 85986853 | 4.04836 | 3.60152 | 2.23581 |
| chr17 | 86754815 | 86755126 | 182 | Promoter (<=1kb) | 17 | 86753907 | 86833410 | 79504 | 1 | ENSMUST00 | 909 | Epas1 | ENSMUSG0( | false | chr17 | 86752839 | 86756072 | 28.88092 | 86.09254 | 82.82554 | chr17 | 86754755 | 86758321 | 4.1837 | 3.6793 | 2.33155 |
| chr17 | 24936819 | 24937135 | 182 | Promoter (<=1kb) | 17 | 24892153 | 24936938 | 44786 | 2 | ENSMUST00 | 0 | Mapk8ip3 | ENSMUSG0( | false | chr17 | 24935707 | 24937390 | 28.8526 | 98.7319 | 95.19158 | . | . | . | . | . | . |
| chr17 | 25082059 | 25082539 | 182 | Promoter (<=1kb) | 17 | 25057702 | 25081181 | 23480 | 2 | ENSMUST00 | -879 | Tmem204 | ENSMUSG0( | false | chr17 | 25081688 | 25083565 | 15.01947 | 39.77812 | 37.19162 | chr17 | 25077267 | 25086277 | 7.00235 | 5.00822 | 4.65933 |
| chr17 | 25081190 | 25081462 | 182 | Promoter (<=1kb) | 17 | 25057702 | 25081181 | 23480 | 2 | ENSMUST00 | -10 | Tmem204 | ENSMUSG0( | false | . | . | . | . | . | . | chr17 | 25077267 | 25086277 | 7.00235 | 5.00822 | 4.65933 |
| chr17 | 26095238 | 26095622 | 182 | Promoter (<=1kb) | 17 | 26091734 | 26095508 | 3775 | 2 | ENSMUST00 | 0 | Nme4 | ENSMUSG0( | false | chr17 | 26095049 | 26095767 | 21.91997 | 62.19815 | 59.32085 | chr17 | 26095146 | 26095880 | 4.22097 | 3.56132 | 2.37072 |
| chr17 | 26781352 | 26781569 | 182 | Promoter (<=1kb) | 17 | 26781158 | 26792425 | 11268 | 1 | ENSMUST00 | 195 | Bnip1 | ENSMUSG0( | false | chr17 | 26780865 | 26781966 | 32.99268 | 103.08465 | 99.4389 | chr17 | 26780617 | 26785656 | 4.73293 | 3.84498 | 2.82106 |
| chr17 | 56289945 | 56290304 | 182 | Promoter (<=1kb) | 17 | 56278962 | 56290511 | 11550 | 2 | ENSMUST00 | 207 | Plin3 | ENSMUSG0( | false | chr17 | 56289551 | 56291172 | 9.94526 | 25.28477 | 22.8772 | chr17 | 56287767 | 56291695 | 4.39609 | 3.74014 | 2.51749 |
| chr17 | 56289650 | 56289943 | 182 | Promoter (<=1kb) | 17 | 56278962 | 56290511 | 11550 | 2 | ENSMUST00 | 568 | Plin3 | ENSMUSG0( | false | chr17 | 56289551 | 56291172 | 9.94526 | 25.28477 | 22.8772 | chr17 | 56287767 | 56291695 | 4.39609 | 3.74014 | 2.51749 |
| chr17 | 27130299 | 27130587 | 182 | Intron (ENSMUST00000025045.9 | 17 | 27122668 | 27133623 | 10956 | 2 | ENSMUST00 | 3036 | Uqcc2 | ENSMUSG0( | false | . | . | . | . | . | . | chr17 | 27127363 | 27147146 | 5.99202 | 4.48057 | 3.83593 |
| chr6 | 28135001 | 28135284 | 182 | Promoter (<=1kb) | 6 | 27934377 | 28135178 | 200802 | 2 | ENSMUST00 | 0 | Grm8 | ENSMUSG0( | false | chr6 | 28134897 | 28136135 | 14.96389 | 42.58086 | 39.95799 | chr6 | 28133682 | 28136032 | 5.3495 | 4.28459 | 3.284 |
| chr17 | 27623385 | 27623856 | 182 | Promoter (<=1kb) | 17 | 27579382 | 27623465 | 44084 | 2 | ENSMUST00 | 0 | Nudt3 | ENSMUSG0( | false | chr17 | 27622163 | 27624145 | 39.98587 | 128.8752 | 124.42683 | chr17 | 27623831 | 27626291 | 3.96391 | 3.46175 | 2.17801 |
| chr17 | 27839905 | 27840205 | 182 | Promoter (<=1kb) | 17 | 27839974 | 27851964 | 11991 | 1 | ENSMUST00 | 0 | Snrpc | ENSMUSG0( | false | chr17 | 27839197 | 27840871 | 31.62822 | 118.12274 | 114.03909 | . | . | . | . | . | . |
| chr17 | 27907614 | 27907845 | 182 | Promoter (<=1kb) | 17 | 27901128 | 27907724 | 6597 | 2 | ENSMUST00 | 0 | Taf11 | ENSMUSG0( | false | chr17 | 27906960 | 27910842 | 24.39013 | 102.54827 | 98.91666 | chr17 | 27906430 | 27908798 | 4.0785 | 3.54669 | 2.27111 |
| chr17 | 28007215 | 28007571 | 182 | Promoter (<=1kb) | 17 | 28007585 | 28062599 | 55015 | 1 | ENSMUST00 | -14 | Anks1 | ENSMUSG0( | false | chr17 | 28007125 | 28009684 | 10.52322 | 21.7476 | 19.38849 | chr17 | 28005688 | 28028167 | 8.71897 | 5.48322 | 6.09085 |
| chr17 | 28177187 | 28177457 | 182 | Promoter (<=1kb) | 17 | 28177207 | 28205886 | 28680 | 1 | ENSMUST00 | 0 | Zfp523 | ENSMUSG0( | false | chr17 | 28176587 | 28177973 | 31.54088 | 108.67455 | 104.87239 | . | . | . | . | . | . |
| chr18 | 4866435 | 4866646 | 182 | Distal Intergenic | 18 | 4920540 | 5119291 | 198752 | 1 | ENSMUST00 | -53894 | Svil | ENSMUSG0( | false | . | . | . | . | . | . | . | . | . | . | . | . |
| chr18 | 5591591 | 5592059 | 182 | Promoter (<=1kb) | 18 | 5591860 | 5775467 | 183608 | 1 | ENSMUST00 | 0 | Zeb1 | ENSMUSG0( | false | chr18 | 5591062 | 5593921 | 37.10487 | 121.71056 | 117.5108 | . | . | . | . | . | . |
| chr18 | 6516117 | 6516545 | 182 | Promoter (<=1kb) | 18 | 6435951 | 6516108 | 80158 | 2 | ENSMUST00 | -10 | Epc1 | ENSMUSG0( | false | chr18 | 6514144 | 6516887 | 36.809 | 123.33047 | 119.07505 | . | . | . | . | . | . |
| chr18 | 6490791 | 6491180 | 182 | Promoter (<=1kb) | 18 | 6435953 | 6490856 | 54904 | 2 | ENSMUST00 | 0 | Epc1 | ENSMUSG0( | false | chr18 | 6489051 | 6491620 | 35.33424 | 109.18479 | 105.37405 | . | . | . | . | . | . |
| chr17 | 84639976 | 84640261 | 182 | Intron (ENSMUST00000025101.8 | 17 | 84626496 | 84655558 | 29063 | 1 | ENSMUST00 | 13481 | Dync2li1 | ENSMUSG0( | false | . | . | . | . | . | . | . | . | . | . | . | . |
| chr18 | 31634550 | 31635169 | 182 | Promoter (<=1kb) | 18 | 31634383 | 31723061 | 88679 | 1 | ENSMUST00 | 168 | Sap130 | ENSMUSG0( | false | chr18 | 31633629 | 31635801 | 38.86769 | 126.9345 | 122.55549 | . | . | . | . | . | . |

| | | | | | | | | | | | | | | | | | | | | | | | | | | |
|---|---|---|---|---|---|---|---|---|---|---|---|---|---|---|---|---|---|---|---|---|---|---|---|---|---|---|
| chr18 | 24603691 | 24603980 | 182 | Promoter (<=1kb) | 18 | 24579881 | 24603817 | 23937 | 2 | ENSMUST00 | 0 | Slc39a6 | ENSMUSG0C | false | chr18 | 24602821 | 24604600 | 36.39542 | 121.53264 | 117.33772 | . | . | . | . | . | . |
| chr18 | 9957794 | 9958294 | 182 | Promoter (<=1kb) | 18 | 9958180 | 9995484 | 37305 | 1 | ENSMUST00 | 0 | Thoc1 | ENSMUSG0C | false | chr18 | 9957507 | 9958955 | 23.46333 | 91.19009 | 87.82124 | . | . | . | . | . | . |
| chr18 | 10181730 | 10182401 | 182 | Promoter (<=1kb) | 18 | 10064401 | 10181792 | 117392 | 2 | ENSMUST00 | 0 | Rock1 | ENSMUSG0C | false | chr18 | 10180415 | 10182911 | 39.80159 | 147.3544 | 142.17488 | . | . | . | . | . | . |
| chr17 | 32787911 | 32788408 | 182 | Promoter (<=1kb) | 17 | 32771236 | 32788287 | 17052 | 2 | ENSMUST00 | 0 | Zfp871 | ENSMUSG0C | false | chr17 | 32786896 | 32788757 | 34.0961 | 113.25239 | 109.32404 | . | . | . | . | . | . |
| chr17 | 33919323 | 33919678 | 182 | Promoter (<=1kb) | 17 | 33919332 | 33929288 | 9957 | 1 | ENSMUST00 | 0 | Tapbp | ENSMUSG0C | false | chr17 | 33919171 | 33920968 | 26.18681 | 91.82585 | 88.44535 | . | . | . | . | . | . |
| chr17 | 34024592 | 34025041 | 182 | Promoter (<=1kb) | 17 | 34020792 | 34024680 | 3889 | 2 | ENSMUST00 | 0 | Ring1 | ENSMUSG0C | false | chr17 | 34024085 | 34025483 | 38.34636 | 121.73327 | 117.53308 | chr17 | 34023682 | 34026109 | 5.46449 | 4.26468 | 3.42702 |
| chr17 | 34119298 | 34119610 | 182 | Promoter (<=1kb) | 17 | 34112030 | 34120411 | 8382 | 2 | ENSMUST00 | 801 | Brd2 | ENSMUSG0C | false | chr17 | 34116580 | 34119365 | 28.28183 | 109.60786 | 105.79031 | . | . | . | . | . | . |
| chr18 | 36018151 | 36018389 | 182 | Exon (ENSMUST00000115712.2/ | 18 | 35987019 | 36006084 | 19066 | 1 | ENSMUST00 | 31133 | Psd2 | ENSMUSG0C | false | chr18 | 36017700 | 36019090 | 18.27513 | 72.99583 | 69.9605 | . | . | . | . | . | . |
| chr18 | 35383552 | 35383921 | 182 | Intron (ENSMUST00000025215.8 | 18 | 35266400 | 35498925 | 232526 | 2 | ENSMUST00 | 115004 | Sil1 | ENSMUSG0C | false | . | . | . | . | . | . | chr18 | 35382703 | 35386338 | 3.46513 | 3.1609 | 1.78491 |
| chr18 | 35498741 | 35499105 | 182 | Promoter (<=1kb) | 18 | 35266400 | 35498925 | 232526 | 2 | ENSMUST00 | 0 | Sil1 | ENSMUSG0C | false | chr18 | 35498166 | 35499383 | 36.46257 | 120.58523 | 116.42456 | chr18 | 35496774 | 35499212 | 6.83536 | 4.94897 | 4.52558 |
| chr18 | 33213903 | 33214155 | 182 | Promoter (<=1kb) | 18 | 33199355 | 33213862 | 14508 | 2 | ENSMUST00 | -42 | Stard4 | ENSMUSG0C | false | chr18 | 33213183 | 33214422 | 25.15761 | 80.47005 | 77.31023 | . | . | . | . | . | . |
| chr18 | 32172351 | 32172752 | 182 | Intron (ENSMUST00000096575.3 | 18 | 32163089 | 32236751 | 73663 | 1 | ENSMUST00 | 9263 | Map3k2 | ENSMUSG0C | false | . | . | . | . | . | . | chr18 | 32169492 | 32178009 | 5.19521 | 4.09241 | 3.19205 |
| chr18 | 12643456 | 12643847 | 182 | Promoter (<=1kb) | 18 | 12643344 | 12737049 | 93706 | 1 | ENSMUST00 | 113 | Ttc39c | ENSMUSG0C | false | chr18 | 12643179 | 12644524 | 24.82645 | 85.46523 | 82.21117 | chr18 | 12643758 | 12644422 | 5.17542 | 4.29125 | 3.15144 |
| chr18 | 38418741 | 38419307 | 182 | Promoter (<=1kb) | 18 | 38418975 | 38464399 | 45425 | 1 | ENSMUST00 | 0 | Ndfip1 | ENSMUSG0C | false | chr18 | 38418542 | 38420768 | 41.33371 | 134.48721 | 129.82906 | . | . | . | . | . | . |
| chr18 | 38277883 | 38278168 | 182 | Exon (ENSMUST00000025311.5/ | 18 | 38267092 | 38284401 | 17310 | 2 | ENSMUST00 | 6233 | Pcdh12 | ENSMUSG0C | false | . | . | . | . | . | . | chr18 | 38274249 | 38279359 | 5.63879 | 4.38394 | 3.54879 |
| chr18 | 36560428 | 36560938 | 182 | Promoter (<=1kb) | 18 | 36560611 | 36658908 | 98298 | 1 | ENSMUST00 | 0 | Ankhd1 | ENSMUSG0C | false | chr18 | 36560556 | 36561910 | 42.07631 | 139.09143 | 134.24335 | . | . | . | . | . | . |
| chr18 | 42511276 | 42511643 | 182 | Promoter (<=1kb) | 18 | 42511510 | 42575551 | 64042 | 1 | ENSMUST00 | 0 | Tcerg1 | ENSMUSG0C | false | chr18 | 42511414 | 42512593 | 39.97929 | 126.69925 | 122.32521 | . | . | . | . | . | . |
| chr18 | 49755254 | 49755890 | 182 | Promoter (<=1kb) | 18 | 49696145 | 49755601 | 59457 | 2 | ENSMUST00 | 0 | Dtwd2 | ENSMUSG0C | false | chr18 | 49755061 | 49756385 | 31.96831 | 119.80809 | 115.67081 | . | . | . | . | . | . |
| chr18 | 67464684 | 67465131 | 182 | Promoter (<=1kb) | 18 | 67464849 | 67480581 | 15733 | 1 | ENSMUST00 | 0 | Slmo1 | ENSMUSG0C | false | chr18 | 67464224 | 67465553 | 32.48404 | 109.34737 | 105.53431 | chr18 | 67464223 | 67466709 | 5.83535 | 4.49712 | 3.72947 |
| chr18 | 53245454 | 53246060 | 182 | Promoter (<=1kb) | 18 | 53245662 | 53390823 | 145162 | 1 | ENSMUST00 | 0 | Snx24 | ENSMUSG0C | false | chr18 | 53245095 | 53246206 | 30.59503 | 95.89513 | 92.42037 | . | . | . | . | . | . |
| chr18 | 79009146 | 79009466 | 182 | Intron (ENSMUST00000025430.9 | 18 | 78750380 | 79109391 | 359012 | 2 | ENSMUST00 | 99925 | Setbp1 | ENSMUSG0C | false | . | . | . | . | . | . | chr18 | 79007044 | 79009937 | 3.67533 | 3.36663 | 1.96421 |
| chr18 | 73815400 | 73815847 | 182 | Promoter (<=1kb) | 18 | 73770040 | 73815392 | 45353 | 2 | ENSMUST00 | -9 | Me2 | ENSMUSG0C | false | chr18 | 73814766 | 73816183 | 32.44109 | 108.24224 | 104.45363 | chr18 | 73814336 | 73816754 | 4.51383 | 3.85796 | 2.60806 |
| chr18 | 74267955 | 74268541 | 182 | Promoter (<=1kb) | 18 | 74268272 | 74282685 | 14414 | 1 | ENSMUST00 | 0 | Mbd1 | ENSMUSG0C | false | chr18 | 74267664 | 74269096 | 23.80619 | 80.75649 | 77.58801 | . | . | . | . | . | . |
| chr18 | 64566016 | 64566240 | 182 | Intron (ENSMUST00000025482.8 | 18 | 64499665 | 64516557 | 16893 | 2 | ENSMUST00 | -49460 | Nars | ENSMUSG0C | false | . | . | . | . | . | . | chr18 | 64564456 | 64568047 | 7.21287 | 5.09643 | 4.82155 |
| chr18 | 56708838 | 56709081 | 182 | Promoter (1-2kb) | 18 | 56707813 | 56753424 | 45612 | 1 | ENSMUST00 | 1026 | Lmnb1 | ENSMUSG0C | false | chr18 | 56707031 | 56709440 | 34.17132 | 121.80145 | 117.60049 | chr18 | 56708716 | 56710745 | 3.3482 | 3.21297 | 1.67145 |
| chr18 | 58555553 | 58555955 | 182 | Promoter (<=1kb) | 18 | 58556257 | 58612773 | 56517 | 1 | ENSMUST00 | -302 | Slc27a6 | ENSMUSG0C | false | . | . | . | . | . | . | chr18 | 58555426 | 58556955 | 3.44062 | 3.24989 | 1.75693 |
| chr18 | 90510092 | 90510455 | 182 | Promoter (<=1kb) | 18 | 90510154 | 90543267 | 33114 | 1 | ENSMUST00 | 0 | Tmx3 | ENSMUSG0C | false | chr18 | 90509491 | 90510945 | 39.97575 | 133.22411 | 128.61581 | . | . | . | . | . | . |
| chr18 | 61176971 | 61177248 | 182 | Promoter (<=1kb) | 18 | 61131279 | 61177050 | 45772 | 2 | ENSMUST00 | 0 | Hmgxb3 | ENSMUSG0C | false | chr18 | 61176314 | 61177815 | 34.3811 | 132.86635 | 128.27481 | . | . | . | . | . | . |
| chr19 | 14506813 | 14507020 | 182 | Intron (ENSMUST00000052011.8 | 19 | 14448072 | 14597983 | 149912 | 2 | ENSMUST00 | 90963 | Tle4 | ENSMUSG0C | false | . | . | . | . | . | . | chr19 | 14496852 | 14509645 | 5.46229 | 4.19323 | 3.40627 |
| chr18 | 84685477 | 84685760 | 182 | Promoter (<=1kb) | 18 | 84667470 | 84685633 | 18164 | 2 | ENSMUST00 | 0 | Cndp2 | ENSMUSG0C | false | chr18 | 84684779 | 84686215 | 22.89105 | 88.16154 | 84.85722 | chr18 | 84682942 | 84686268 | 3.37017 | 3.25417 | 1.6687 |
| chr19 | 10895175 | 10895425 | 182 | Promoter (<=1kb) | 19 | 10895231 | 10905532 | 10302 | 1 | ENSMUST00 | 0 | Prpf19 | ENSMUSG0C | false | chr19 | 10895211 | 10896661 | 26.52689 | 93.43019 | 90.01563 | chr19 | 10894632 | 10897803 | 5.6691 | 4.3837 | 3.56358 |
| chr19 | 21274190 | 21274623 | 182 | Promoter (<=1kb) | 19 | 21273246 | 21277038 | 3793 | 1 | ENSMUST00 | 945 | Zfand5 | ENSMUSG0C | false | chr19 | 21271332 | 21275517 | 33.89795 | 105.67399 | 101.95845 | . | . | . | . | . | . |
| chr19 | 7494165 | 7495027 | 182 | Promoter (<=1kb) | 19 | 7494438 | 7538608 | 44171 | 1 | ENSMUST00 | 0 | Atl3 | ENSMUSG0C | false | chr19 | 7493707 | 7495706 | 35.36919 | 136.9064 | 132.15405 | . | . | . | . | . | . |
| chr19 | 6306141 | 6306658 | 182 | Promoter (<=1kb) | 19 | 6306456 | 6325652 | 19197 | 1 | ENSMUST00 | 0 | Cdc42bpg | ENSMUSG0C | false | chr19 | 6306058 | 6307381 | 29.47202 | 98.86652 | 95.32386 | chr19 | 6304627 | 6310527 | 4.28153 | 3.54914 | 2.43346 |
| chr19 | 28990305 | 28990561 | 182 | Promoter (<=1kb) | 19 | 28990494 | 29017569 | 27076 | 1 | ENSMUST00 | 0 | Cdc37l1 | ENSMUSG0C | false | chr19 | 28989833 | 28991359 | 30.41442 | 103.5799 | 99.9162 | . | . | . | . | . | . |
| chr19 | 29047740 | 29048089 | 182 | Promoter (<=1kb) | 19 | 29020833 | 29047961 | 27129 | 2 | ENSMUST00 | 0 | Ak3 | ENSMUSG0C | false | chr19 | 29046371 | 29048801 | 36.17514 | 115.85 | 111.8395 | . | . | . | . | . | . |
| chr19 | 6084866 | 6085061 | 182 | Promoter (<=1kb) | 19 | 6084983 | 6091777 | 6795 | 1 | ENSMUST00 | 0 | Cdca5 | ENSMUSG0C | false | chr19 | 6084044 | 6085628 | 17.04805 | 44.82024 | 42.17082 | chr19 | 6083481 | 6085993 | 6.94562 | 4.89914 | 4.59827 |
| chr19 | 29940277 | 29940488 | 182 | Intron (ENSMUST00000120388.3 | 19 | 29945790 | 29960704 | 14915 | 1 | ENSMUST00 | -5302 | Il33 | ENSMUSG0C | false | . | . | . | . | . | . | . | . | . | . | . | . |
| chr19 | 36835177 | 36835362 | 182 | Promoter (<=1kb) | 19 | 36834404 | 36855937 | 21534 | 1 | ENSMUST00 | 774 | Tnks2 | ENSMUSG0C | false | chr19 | 36833456 | 36836298 | 33.7938 | 119.00099 | 114.88904 | . | . | . | . | . | . |
| chr19 | 4130182 | 4130472 | 182 | Promoter (2-3kb) | 19 | 4127572 | 4131257 | 3686 | 1 | ENSMUST00 | 2611 | Tmem134 | ENSMUSG0C | false | . | . | . | . | . | . | chr19 | 4126408 | 4133141 | 4.36291 | 3.575 | 2.49929 |
| chr19 | 4125785 | 4126034 | 182 | Promoter (<=1kb) | 19 | 4125934 | 4132285 | 6352 | 1 | ENSMUST00 | 0 | Tmem134 | ENSMUSG0C | false | chr19 | 4125868 | 4127236 | 30.78991 | 105.27288 | 101.56436 | . | . | . | . | . | . |
| chr19 | 4229577 | 4230086 | 182 | Promoter (1-2kb) | 19 | 4231899 | 4233631 | 1733 | 1 | ENSMUST00 | -1813 | Pold4 | ENSMUSG0C | false | . | . | . | . | . | . | chr19 | 4228207 | 4233506 | 8.46056 | 5.68198 | 5.82099 |
| chr19 | 5273061 | 5273373 | 182 | Promoter (<=1kb) | 19 | 5133688 | 5273119 | 139432 | 2 | ENSMUST00 | 0 | Pacs1 | ENSMUSG0C | true | chr19 | 5271956 | 5273622 | 41.4024 | 145.73851 | 140.61644 | chr19 | 5269809 | 5274576 | 4.35528 | 3.7387 | 2.47376 |
| chr19 | 24961521 | 24961897 | 182 | Promoter (<=1kb) | 19 | 24919916 | 24961616 | 41701 | 2 | ENSMUST00 | 0 | Cbwd1 | ENSMUSG0C | false | chr19 | 24960896 | 24961679 | 20.20131 | 69.382 | 66.39919 | . | . | . | . | . | . |
| chr19 | 32107910 | 32108301 | 182 | Distal Intergenic | 19 | 31984654 | 32061469 | 76816 | 2 | ENSMUST00 | -46442 | Asah2 | ENSMUSG0C | false | chr19 | 32107593 | 32108438 | 35.41093 | 115.56754 | 111.56183 | . | . | . | . | . | . |
| chr19 | 5023921 | 5024176 | 182 | Promoter (<=1kb) | 19 | 5024006 | 5031971 | 7966 | 1 | ENSMUST00 | 0 | Slc29a2 | ENSMUSG0C | false | chr19 | 5023330 | 5024767 | 37.0612 | 128.08391 | 123.66181 | chr19 | 5020815 | 5024806 | 4.14069 | 3.62648 | 2.28954 |
| chr19 | 5488232 | 5488483 | 182 | Promoter (<=1kb) | 19 | 5482832 | 5488402 | 5571 | 2 | ENSMUST00 | 0 | Mus81 | ENSMUSG0C | false | chr19 | 5487532 | 5489063 | 16.80025 | 47.43613 | 44.75273 | chr19 | 5486437 | 5490291 | 5.06458 | 3.96355 | 3.0584 |
| chr19 | 5489085 | 5489352 | 182 | Promoter (<=1kb) | 19 | 5482832 | 5488402 | 5571 | 2 | ENSMUST00 | -684 | Mus81 | ENSMUSG0C | false | chr19 | 5489205 | 5489624 | 14.24369 | 37.68267 | 35.12186 | chr19 | 5486437 | 5490291 | 5.06458 | 3.96355 | 3.0584 |
| chr19 | 5473777 | 5474018 | 182 | Promoter (<=1kb) | 19 | 5473973 | 5477578 | 3606 | 1 | ENSMUST00 | 0 | Efemp2 | ENSMUSG0C | false | chr19 | 5473603 | 5475286 | 20.14468 | 62.43501 | 59.55626 | chr19 | 5473231 | 5475447 | 3.84953 | 3.58392 | 2.05184 |
| chr19 | 5474480 | 5475012 | 182 | Promoter (<=1kb) | 19 | 5474688 | 5481853 | 7166 | 1 | ENSMUST00 | 0 | Efemp2 | ENSMUSG0C | false | chr19 | 5473603 | 5475286 | 20.14468 | 62.43501 | 59.55626 | chr19 | 5473231 | 5475447 | 3.84953 | 3.58392 | 2.05184 |
| chr19 | 3665673 | 3665918 | 182 | Intron (ENSMUST00000025856.1 | 19 | 3584828 | 3686556 | 101729 | 2 | ENSMUST00 | 20638 | Lrp5 | ENSMUSG0C | false | . | . | . | . | . | . | chr19 | 3665159 | 3668073 | 4.54409 | 3.88728 | 2.66405 |
| chr19 | 3634938 | 3635163 | 182 | Intron (ENSMUST00000025856.1 | 19 | 3584836 | 3615792 | 30957 | 2 | ENSMUST00 | -19147 | Lrp5 | ENSMUSG0C | false | . | . | . | . | . | . | . | . | . | . | . | . |
| chr19 | 26605727 | 26605967 | 182 | Promoter (<=1kb) | 19 | 26605119 | 26778275 | 173157 | 1 | ENSMUST00 | 609 | Smarca2 | ENSMUSG0C | false | chr19 | 26603929 | 26608671 | 23.80619 | 80.75649 | 77.58801 | . | . | . | . | . | . |
| chr19 | 5767404 | 5767695 | 182 | Intron (ENSMUST00000025890.8 | 19 | 5758427 | 5771401 | 12975 | 2 | ENSMUST00 | 3706 | Scyl1 | ENSMUSG0C | false | . | . | . | . | . | . | chr19 | 5766114 | 5772182 | 5.63678 | 4.25073 | 3.56165 |
| chr19 | 5771338 | 5771540 | 182 | Promoter (<=1kb) | 19 | 5758427 | 5771401 | 12975 | 2 | ENSMUST00 | 0 | Scyl1 | ENSMUSG0C | false | chr19 | 5770509 | 5771473 | 32.3084 | 121.498 | 117.30614 | chr19 | 5766114 | 5772182 | 5.63678 | 4.25073 | 3.56165 |
| chr19 | 6140809 | 6141267 | 182 | Promoter (<=1kb) | 19 | 6134374 | 6141140 | 6767 | 2 | ENSMUST00 | 0 | Arl2 | ENSMUSG0C | false | chr19 | 6140301 | 6141783 | 30.94897 | 95.52228 | 92.05737 | . | . | . | . | . | . |
| chr19 | 6840558 | 6840781 | 182 | Promoter (<=1kb) | 19 | 6829085 | 6840601 | 11517 | 2 | ENSMUST00 | 0 | Rps6ka4 | ENSMUSG0C | false | chr19 | 6839670 | 6841212 | 34.16929 | 120.7392 | 116.5701 | chr19 | 6839295 | 6841504 | 4.92915 | 4.08747 | 3.00024 |
| chr19 | 6987308 | 6987933 | 182 | Promoter (<=1kb) | 19 | 6982473 | 6987629 | 5157 | 2 | ENSMUST00 | 0 | Vegfb | ENSMUSG0C | false | chr19 | 6986900 | 6988151 | 33.89127 | 107.71272 | 103.94643 | chr19 | 6986619 | 6988373 | 4.7714 | 3.85366 | 2.86175 |
| chr19 | 7267182 | 7267436 | 182 | Promoter (<=1kb) | 19 | 7267325 | 7270144 | 2820 | 1 | ENSMUST00 | 0 | Rcor2 | ENSMUSG0C | false | . | . | . | . | . | . | chr19 | 7266091 | 7270232 | 5.81658 | 4.28552 | 3.71613 |
| chr19 | 37434591 | 37434822 | 182 | Promoter (<=1kb) | 19 | 37434810 | 37440731 | 5922 | 1 | ENSMUST00 | 0 | Hhex | ENSMUSG0C | false | chr19 | 37434061 | 37436006 | 18.61129 | 49.85614 | 47.14222 | chr19 | 37433933 | 37435410 | 4.1337 | 3.45854 | 2.32786 |
| chr19 | 37813403 | 37813659 | 182 | Distal Intergenic | 19 | 37697808 | 37701528 | 3721 | 1 | ENSMUST00 | 115596 | Cyp26a1 | ENSMUSG0C | false | . | . | . | . | . | . | chr19 | 37810499 | 37816117 | 5.83071 | 4.48483 | 3.6923 |
| chr19 | 60811486 | 60811922 | 182 | Promoter (<=1kb) | 19 | 60811585 | 60830864 | 19280 | 1 | ENSMUST00 | 0 | Fam45a | ENSMUSG0C | false | chr19 | 60811097 | 60812456 | 24.14628 | 82.32003 | 79.12344 | chr19 | 60810890 | 60813495 | 3.7012 | 3.38821 | 1.93802 |
| chr19 | 60874349 | 60874634 | 182 | Promoter (<=1kb) | 19 | 60864051 | 60874556 | 10506 | 2 | ENSMUST00 | 0 | Prdx3 | ENSMUSG0C | false | chr19 | 60874075 | 60874992 | 17.4042 | 56.42813 | 53.62976 | chr19 | 60870881 | 60875062 | 4.28747 | 3.68047 | 2.4328 |
| chr19 | 38560178 | 38560393 | 182 | Intron (ENSMUST00000182481.2 | 19 | 38583368 | 38652007 | 68640 | 1 | ENSMUST00 | -22975 | Plce1 | ENSMUSG0C | true | . | . | . | . | . | . | chr19 | 38551907 | 38564462 | 3.84478 | 3.45977 | 2.09455 |
| chr19 | 40432124 | 40432401 | 182 | Distal Intergenic | 19 | 40295049 | 40402267 | 107219 | 2 | ENSMUST00 | -29858 | Sorbs1 | ENSMUSG0C | false | . | . | . | . | . | . | . | . | . | . | . | . |
| chr19 | 40513479 | 40513960 | 182 | Distal Intergenic | 19 | 40553427 | 40554741 | 1315 | 2 | ENSMUST00 | 40781 | Aldh18a1 | ENSMUSG0C | false | chr19 | 40512533 | 40514324 | 24.24157 | 89.02564 | 85.70093 | . | . | . | . | . | . |
| chr19 | 53390126 | 53390714 | 182 | Promoter (<=1kb) | 19 | 53379214 | 53390573 | 11360 | 2 | ENSMUST00 | 0 | Smndc1 | ENSMUSG0C | false | chr19 | 53389502 | 53391452 | 41.911 | 144.38934 | 139.32462 | . | . | . | . | . | . |
| chr19 | 53038471 | 53038719 | 182 | Promoter (<=1kb) | 19 | 52943417 | 53038553 | 95137 | 2 | ENSMUST00 | 0 | Xpnpep1 | ENSMUSG0C | false | chr19 | 53037782 | 53039160 | 30.94804 | 114.76607 | 110.79114 | chr19 | 53037126 | 53042423 | 5.46211 | 4.19899 | 3.43522 |
| chr19 | 46523859 | 46524144 | 182 | Distal Intergenic | 19 | 46501648 | 46516444 | 14797 | 1 | ENSMUST00 | 22212 | Trim8 | ENSMUSG0C | false | chr19 | 46523878 | 46524562 | 6.466 | 12.01425 | 9.80126 | chr19 | 46521335 | 46535388 | 7.6256 | 5.19094 | 5.16609 |
| chr19 | 46500668 | 46500996 | 182 | Promoter (<=1kb) | 19 | 46501648 | 46516444 | 14797 | 1 | ENSMUST00 | -652 | Trim8 | ENSMUSG0C | false | chr19 | 46500135 | 46506359 | 34.21972 | 118.71816 | 114.6235 | . | . | . | . | . | . |
| chr19 | 46524628 | 46524838 | 182 | Distal Intergenic | 19 | 46501648 | 46516444 | 14797 | 1 | ENSMUST00 | 22981 | Trim8 | ENSMUSG0C | false | . | . | . | . | . | . | chr19 | 46521335 | 46535388 | 7.6256 | 5.19094 | 5.16609 |
| chr19 | 46531483 | 46531842 | 182 | Intron (ENSMUST00000026009.8 | 19 | 46501648 | 46516444 | 14797 | 1 | ENSMUST00 | 29836 | Trim8 | ENSMUSG0C | false | . | . | . | . | . | . | chr19 | 46521335 | 46535388 | 7.6256 | 5.19094 | 5.16609 |
| chr19 | 47699875 | 47700105 | 182 | Distal Intergenic | 19 | 47646344 | 47692094 | 45751 | 2 | ENSMUST00 | -7782 | Col17a1 | ENSMUSG0C | false | . | . | . | . | . | . | chr19 | 47697960 | 47702263 | 4.14895 | 3.61656 | 2.32554 |
| chr19 | 56944326 | 56944599 | 182 | Intron (ENSMUST00000118800.2 | 19 | 56920398 | 56928553 | 8156 | 2 | ENSMUST00 | -15774 | Afap1l2 | ENSMUSG0C | false | . | . | . | . | . | . | . | . | . | . | . | . |
| chr19 | 56908371 | 56908632 | 182 | Intron (ENSMUST00000026068.7 | 19 | 56920398 | 56928553 | 8156 | 2 | ENSMUST00 | 19921 | Afap1l2 | ENSMUSG0C | false | . | . | . | . | . | . | . | . | . | . | . | . |
| chr19 | 56977584 | 56977832 | 182 | Intron (ENSMUST00000118800.2 | 19 | 56930098 | 57001046 | 70949 | 2 | ENSMUST00 | 23214 | Afap1l2 | ENSMUSG0C | false | . | . | . | . | . | . | chr19 | 56976918 | 56977864 | 2.98594 | 3.03906 | 1.39572 |
| chr19 | 57103520 | 57103770 | 182 | Intron (ENSMUST00000079360.6 | 19 | 57036479 | 57118913 | 82435 | 2 | ENSMUST00 | 15143 | Ablim1 | ENSMUSG0C | false | . | . | . | . | . | . | . | . | . | . | . | . |
| chr19 | 58879667 | 58879874 | 182 | Distal Intergenic | 19 | 58795751 | 58860984 | 65234 | 2 | ENSMUST00 | -18684 | Hspa12a | ENSMUSG0C | true | . | . | . | . | . | . | chr19 | 58878512 | 58880046 | 3.70132 | 3.48969 | 1.93584 |
| chr19 | 58829665 | 58829929 | 182 | Intron (ENSMUST00000066285.4 | 19 | 58795751 | 58860984 | 65234 | 2 | ENSMUST00 | 31055 | Hspa12a | ENSMUSG0C | true | . | . | . | . | . | . | chr19 | 58814047 | 58859817 | 7.46659 | 5.12677 | 5.03638 |
| chr7 | 81904176 | 81904361 | 182 | Intron (ENSMUST00000107305.2 | 7 | 81897811 | 81934023 | 36213 | 2 | ENSMUST00 | 29662 | Hdgfrp3 | ENSMUSG0C | false | . | . | . | . | . | . | chr7 | 81903621 | 81904539 | 3.88633 | 3.49181 | 2.1523 |
| chr11 | 120581575 | 120581859 | 182 | Promoter (<=1kb) | 11 | 120578104 | 120581624 | 3521 | 2 | ENSMUST00 | 0 | Arhgdia | ENSMUSG0C | false | chr11 | 120579222 | 120582581 | 31.43241 | 100.35837 | 96.77719 | chr11 | 120581803 | 120582772 | 3.54382 | 3.38989 | 1.79214 |
| chr7 | 97480834 | 97481211 | 182 | Promoter (<=1kb) | 7 | 97480956 | 97541397 | 60442 | 1 | ENSMUST00 | 0 | Ints4 | ENSMUSG0C | false | chr7 | 97480918 | 97481811 | 21.57688 | 72.03428 | 69.01365 | chr7 | 97480287 | 97483366 | 5.29937 | 4.29542 | 3.24619 |
| chr7 | 142060942 | 142061194 | 182 | Promoter (<=1kb) | 7 | 142008553 | 142061034 | 52482 | 2 | ENSMUST00 | 0 | Mob2 | ENSMUSG0C | true | chr7 | 142059991 | 142061607 | 28.91247 | 96.86193 | 93.36482 | chr7 | 142055004 | 142061788 | 6.37873 | 4.67403 | 4.16006 |
| chr11 | 121204339 | 121204779 | 182 | Promoter (<=1kb) | 11 | 121177591 | 121204711 | 27121 | 2 | ENSMUST00 | 0 | Ogfod3 | ENSMUSG0C | false | chr11 | 121203972 | 121205344 | 34.00537 | 102.80987 | 99.17244 | chr11 | 121203789 | 121205511 | 5.4031 | 4.34202 | 3.33454 |
| chr19 | 43525108 | 43525387 | 182 | Promoter (<=1kb) | 19 | 43499752 | 43524605 | 24854 | 2 | ENSMUST00 | -504 | Got1 | ENSMUSG0C | false | chr19 | 43523562 | 43525376 | 13.72719 | 56.76396 | 53.95979 | chr19 | 43523310 | 43526485 | 5.50603 | 4.13004 | 3.47386 |
| chr19 | 44135538 | 44135859 | 182 | Promoter (<=1kb) | 19 | 44108644 | 44135876 | 27233 | 2 | ENSMUST00 | 17 | Cwf19l1 | ENSMUSG0C | false | chr19 | 44134936 | 44135949 | 24.78212 | 78.86476 | 75.73294 | . | . | . | . | . | . |
| chr2 | 52623571 | 52623800 | 182 | Intron (ENSMUST00000178799.2 | 2 | 52619754 | 52620185 | 432 | 1 | ENSMUST00 | 3818 | Bloc1s2-ps | ENSMUSG0C | false | . | . | . | . | . | . | chr2 | 52622888 | 52624616 | 4.71656 | 3.74891 | 2.79748 |
| chr19 | 46045941 | 46046339 | 182 | Promoter (<=1kb) | 19 | 46032593 | 46045214 | 12622 | 2 | ENSMUST00 | -728 | Ldb1 | ENSMUSG0C | false | chr19 | 46043949 | 46046177 | 34.10395 | 106.03094 | 102.3112 | chr19 | 46045644 | 46047114 | 3.8061 | 3.50605 | 2.01095 |
| chr19 | 46305593 | 46305973 | 182 | Promoter (<=1kb) | 19 | 46305695 | 46312090 | 6396 | 1 | ENSMUST00 | 0 | Nfkb2 | ENSMUSG0C | false | chr19 | 46304097 | 46306268 | 27.29606 | 82.38838 | 79.1904 | chr19 | 46302648 | 46306957 | 4.47557 | 3.86215 | 2.56811 |
| chr19 | 46328750 | 46328980 | 182 | Promoter (<=1kb) | 19 | 46328187 | 46330445 | 2259 | 1 | ENSMUST00 | 564 | Fbxl15 | ENSMUSG0C | false | chr19 | 46327814 | 46329602 | 36.46257 | 120.58523 | 116.42456 | chr19 | 46326626 | 46330389 | 6.33395 | 4.67328 | 4.11438 |
| chr9 | 123478535 | 123478802 | 182 | Promoter (<=1kb) | 9 | 123478701 | 123521548 | 42848 | 1 | ENSMUST00 | 0 | Limd1 | ENSMUSG0C | false | chr9 | 123478684 | 123479821 | 24.86315 | 92.1349 | 88.74321 | chr9 | 123477828 | 123484562 | 4.86734 | 3.85268 | 2.9337 |
| chrX | 151803002 | 151803229 | 182 | Promoter (<=1kb) | 20 | 151803243 | 151873033 | 69791 | 1 | ENSMUST00 | -14 | Huwe1 | ENSMUSG0C | false | chrX | 151802550 | 151803098 | 11.94783 | 25.26529 | 22.85895 | . | . | . | . | . | . |
| chr14 | 7900032 | 7900289 | 182 | Intron (ENSMUST00000052678.8 | 14 | 7975872 | 7977222 | 1351 | 2 | ENSMUST00 | 76933 | Dnase1l3 | ENSMUSG0C | false | . | . | . | . | . | . | chr14 | 7899745 | 7902015 | 4.92096 | 4.03027 | 2.98767 |
| chr12 | 119237963 | 119238368 | 182 | Promoter (<=1kb) | 12 | 119201463 | 119238770 | 37308 | 2 | ENSMUST00 | 402 | Itgb8 | ENSMUSG0C | false | chr12 | 119236888 | 119239435 | 23.00904 | 72.91611 | 69.8813 | . | . | . | . | . | . |
| chr12 | 119206839 | 119207099 | 182 | Intron (ENSMUST00000026360.8 | 12 | 119201463 | 119238770 | 37308 | 2 | ENSMUST00 | 31671 | Itgb8 | ENSMUSG0C | false | . | . | . | . | . | . | chr12 | 119205654 | 119222572 | 4.19208 | 3.68012 | 2.33746 |
| chr12 | 119218002 | 119218236 | 182 | Intron (ENSMUST00000026360.8 | 12 | 119201463 | 119238770 | 37308 | 2 | ENSMUST00 | 20534 | Itgb8 | ENSMUSG0C | false | . | . | . | . | . | . | chr12 | 119205654 | 119222572 | 4.19208 | 3.68012 | 2.33746 |
| chrX | 152232701 | 152232953 | 182 | Promoter (<=1kb) | 20 | 152233020 | 152273768 | 40749 | 1 | ENSMUST00 | -67 | Kdm5c | ENSMUSG0C | false | . | . | . | . | . | . | . | . | . | . | . | . |
| chr10 | 128889033 | 128889440 | 182 | Promoter (2-3kb) | 10 | 128882295 | 128891718 | 9424 | 2 | ENSMUST00 | 2278 | Gdf11 | ENSMUSG0C | false | . | . | . | . | . | . | . | . | . | . | . | . |
| chr10 | 128565929 | 128566412 | 182 | Promoter (<=1kb) | 10 | 128557766 | 128565934 | 8169 | 2 | ENSMUST00 | 0 | Pa2g4 | ENSMUSG0C | false | chr10 | 128565011 | 128566823 | 31.66066 | 102.77801 | 99.14294 | . | . | . | . | . | . |
| chr10 | 127669195 | 127669466 | 182 | Promoter (<=1kb) | 10 | 127664984 | 127668568 | 3585 | 2 | ENSMUST00 | -628 | Nab2 | ENSMUSG0C | false | . | . | . | . | . | . | chr10 | 127668820 | 127678463 | 4.93617 | 4.00247 | 2.9603 |
| chr10 | 127380451 | 127380697 | 182 | Promoter (<=1kb) | 10 | 127380662 | 127499360 | 118699 | 1 | ENSMUST00 | 0 | R3hdm2 | ENSMUSG0C | false | chr10 | 127380110 | 127381752 | 33.54585 | 101.01101 | 97.41918 | . | . | . | . | . | . |
| chr10 | 127266215 | 127266521 | 182 | Promoter (<=1kb) | 10 | 127266262 | 127281959 | 15698 | 1 | ENSMUST00 | 0 | Dctn2 | ENSMUSG0C | false | chr10 | 127266048 | 127267362 | 34.91417 | 125.28625 | 120.96531 | chr10 | 127265797 | 127269225 | 5.88685 | 4.45699 | 3.75228 |
| chr4 | 106678820 | 106679097 | 182 | Promoter (<=1kb) | 4 | 106662256 | 106678944 | 16689 | 2 | ENSMUST00 | 0 | Ttc4 | ENSMUSG0C | false | chr4 | 106678141 | 106679264 | 29.58769 | 108.11006 | 104.32592 | chr4 | 106677829 | 106679707 | 5.62144 | 4.3202 | 3.51054 |
| chr10 | 127081519 | 127081774 | 182 | Promoter (2-3kb) | 10 | 127078907 | 127093169 | 14263 | 1 | ENSMUST00 | 2613 | Agap2 | ENSMUSG0C | false | chr10 | 127081232 | 127081777 | 9.19064 | 17.46287 | 15.16242 | chr10 | 127078565 | 127082404 | 6.18229 | 4.56655 | 3.98455 |
| chr18 | 77565250 | 77565482 | 182 | Promoter (<=1kb) | 18 | 77458685 | 77565147 | 106463 | 2 | ENSMUST00 | -104 | Rnf165 | ENSMUSG0C | false | chr18 | 77564720 | 77566297 | 19.30035 | 48.68736 | 45.98855 | chr18 | 77564981 | 77566250 | 3.74122 | 3.43655 | 1.96575 |
| chr18 | 77773906 | 77774310 | 182 | Promoter (<=1kb) | 18 | 77773956 | 77782866 | 8911 | 1 | ENSMUST00 | 0 | Atp5a1 | ENSMUSG0C | false | chr18 | 77773316 | 77775134 | 35.29575 | 125.98199 | 121.63858 | . | . | . | . | . | . |
| chr7 | 140036310 | 140036733 | 182 | Promoter (<=1kb) | 7 | 140036391 | 140040605 | 4215 | 1 | ENSMUST00 | 0 | Zfp511 | ENSMUSG0C | false | chr7 | 140035620 | 140037192 | 37.86004 | 121.9582 | 117.75514 | chr7 | 140034172 | 140037529 | 4.22514 | 3.73355 | 2.36517 |
| chr7 | 141447436 | 141447733 | 182 | Promoter (<=1kb) | 7 | 141447631 | 141448616 | 986 | 1 | ENSMUST00 | 0 | Rplp2 | ENSMUSG0C | false | chr7 | 141446958 | 141448497 | 29.33348 | 103.60687 | 99.9429 | . | . | . | . | . | . |
| chr7 | 141475075 | 141475442 | 182 | Promoter (<=1kb) | 7 | 141475240 | 141493427 | 18188 | 1 | ENSMUST00 | 0 | Tspan4 | ENSMUSG0C | false | chr7 | 141474160 | 141476599 | 25.18546 | 107.13963 | 103.3826 | chr7 | 141456108 | 141486560 | 6.62038 | 4.76228 | 4.33989 |
| chr7 | 141519082 | 141519270 | 182 | Intron (ENSMUST00000153191.2 | 7 | 141511775 | 141526467 | 14693 | 2 | ENSMUST00 | 7197 | Chid1 | ENSMUSG0C | false | . | . | . | . | . | . | chr7 | 141518425 | 141521037 | 4.85075 | 3.94392 | 2.92706 |
| chr14 | 121108484 | 121108775 | 182 | Intron (ENSMUST00000135010.2 | 14 | 121102079 | 121238659 | 136581 | 1 | ENSMUST00 | 6406 | Farp1 | ENSMUSG0C | false | . | . | . | . | . | . | chr14 | 121103510 | 121111640 | 5.60892 | 4.31508 | 3.50533 |
| chr14 | 121098997 | 121099387 | 182 | Promoter (2-3kb) | 14 | 121102079 | 121238659 | 136581 | 1 | ENSMUST00 | -2692 | Farp1 | ENSMUSG0C | false | . | . | . | . | . | . | chr14 | 121096048 | 121101856 | 5.65796 | 4.33676 | 3.54371 |
| chr14 | 121062191 | 121062411 | 182 | Intron (ENSMUST00000135010.2 | 14 | 121035564 | 121283744 | 248181 | 1 | ENSMUST00 | 26628 | Farp1 | ENSMUSG0C | false | . | . | . | . | . | . | chr14 | 121058413 | 121066025 | 5.3821 | 4.1988 | 3.34196 |
| chr11 | 117654090 | 117654320 | 182 | Promoter (<=1kb) | 11 | 117654289 | 117763437 | 109149 | 1 | ENSMUST00 | 0 | Tnrc6c | ENSMUSG0C | false | chr11 | 117653596 | 117656293 | 35.34781 | 110.24136 | 106.39787 | . | . | . | . | . | . |
| chr11 | 118728090 | 118728338 | 182 | Intron (ENSMUST00000117731.2 | 11 | 118489764 | 118761041 | 271278 | 2 | ENSMUST00 | 32703 | Rbfox3 | ENSMUSG0C | false | . | . | . | . | . | . | . | . | . | . | . | . |
| chr11 | 119299676 | 119300043 | 182 | Promoter (<=1kb) | 11 | 119292841 | 119300014 | 7174 | 2 | ENSMUST00 | 0 | Eif4a3 | ENSMUSG0C | false | chr11 | 119298749 | 119300721 | 25.45057 | 108.67924 | 104.87682 | . | . | . | . | . | . |
| chr11 | 119772571 | 119773046 | 182 | Intron (ENSMUST00000125583.2 | 11 | 119798773 | 119846955 | 48183 | 1 | ENSMUST00 | -25727 | Rptor | ENSMUSG0C | false | . | . | . | . | . | . | chr11 | 119769610 | 119775681 | 5.86948 | 4.43812 | 3.71402 |
| chr11 | 119677217 | 119677424 | 182 | Intron (ENSMUST00000125583.2 | 11 | 119655944 | 119865810 | 209867 | 1 | ENSMUST00 | 21274 | Rptor | ENSMUSG0C | false | . | . | . | . | . | . | chr11 | 119675334 | 119679416 | 3.92555 | 3.36849 | 2.15053 |
| chr7 | 81313690 | 81313878 | 182 | Intron (ENSMUST00000026672.7 | 7 | 81306693 | 81319440 | 12748 | 1 | ENSMUST00 | 6998 | Pde8a | ENSMUSG0C | false | . | . | . | . | . | . | . | . | . | . | . | . |
| chr16 | 87558691 | 87558911 | 182 | Intron (ENSMUST00000026700.7 | 16 | 87553330 | 87595336 | 42007 | 1 | ENSMUST00 | 5362 | Map3k7cl | ENSMUSG0C | false | . | . | . | . | . | . | . | . | . | . | . | . |
| chr9 | 109074020 | 109074209 | 182 | Promoter (<=1kb) | 9 | 109057933 | 109074071 | 16139 | 2 | ENSMUST00 | 0 | Atrip | ENSMUSG0C | false | chr9 | 109073136 | 109074690 | 28.49948 | 87.34749 | 84.05397 | chr9 | 109072846 | 109075044 | 4.63986 | 3.79994 | 2.73495 |
| chrX | 81070558 | 81070833 | 182 | Promoter (<=1kb) | 20 | 81070698 | 81094776 | 24079 | 1 | ENSMUST00 | 0 | Tmem47 | ENSMUSG0C | false | chrX | 81070309 | 81071644 | 20.67895 | 53.40692 | 50.64761 | . | . | . | . | . | . |
| chr17 | 25844373 | 25844723 | 182 | Promoter (<=1kb) | 17 | 25838840 | 25844463 | 5624 | 2 | ENSMUST00 | 0 | Rhot2 | ENSMUSG0C | false | chr17 | 25843814 | 25844907 | 30.42596 | 90.69117 | 87.33178 | chr17 | 25841531 | 25845607 | 4.92524 | 3.92025 | 2.98804 |
| chr5 | 29951542 | 29951778 | 182 | Distal Intergenic | 5 | 30013161 | 30019968 | 6808 | 1 | ENSMUST00 | -61383 | Il6 | ENSMUSG0C | false | . | . | . | . | . | . | . | . | . | . | . | . |
| chr3 | 41574798 | 41575030 | 182 | Intron (ENSMUST00000026865.9 | 3 | 41564880 | 41616862 | 51983 | 1 | ENSMUST00 | 9919 | Jade1 | ENSMUSG0C | false | . | . | . | . | . | . | chr3 | 41573000 | 41576866 | 4.45013 | 3.68831 | 2.59245 |
| chr2 | 10048574 | 10048833 | 182 | Promoter (<=1kb) | 2 | 9914552 | 10048596 | 134045 | 2 | ENSMUST00 | 0 | Taf3 | ENSMUSG0C | false | chr2 | 10047371 | 10049480 | 36.77131 | 134.07788 | 129.4352 | . | . | . | . | . | . |
| chr9 | 123112983 | 123113306 | 182 | Promoter (<=1kb) | 9 | 123113215 | 123136129 | 22915 | 1 | ENSMUST00 | 0 | Exosc7 | ENSMUSG0C | false | chr9 | 123112407 | 123114045 | 27.97104 | 107.99458 | 104.2211 | . | . | . | . | . | . |
| chr7 | 74458809 | 74459108 | 182 | Intron (ENSMUST00000098371.3 | 7 | 74359654 | 74504335 | 144682 | 2 | ENSMUST00 | 45227 | Slco3a1 | ENSMUSG0C | false | . | . | . | . | . | . | . | . | . | . | . | . |
| chr4 | 99969294 | 99969635 | 182 | Intron (ENSMUST00000058351.1 | 4 | 99955744 | 99987294 | 31551 | 1 | ENSMUST00 | 13551 | Pgm2 | ENSMUSG0C | false | . | . | . | . | . | . | chr4 | 99968725 | 99977020 | 4.83736 | 3.90531 | 2.88809 |
| chr8 | 128679911 | 128680198 | 182 | Distal Intergenic | 8 | 128685654 | 128733200 | 47547 | 1 | ENSMUST00 | -5456 | Itgb1 | ENSMUSG0C | true | . | . | . | . | . | . | chr8 | 128679197 | 128680734 | 5.83485 | 4.48531 | 3.68626 |
| chr8 | 127129014 | 127129280 | 182 | Intron (ENSMUST00000162309.2 | 8 | 127073240 | 127426684 | 353445 | 1 | ENSMUST00 | 55775 | Pard3 | ENSMUSG0C | false | . | . | . | . | . | . | chr8 | 127128644 | 127139305 | 4.20446 | 3.66177 | 2.36227 |
| chr5 | 139015159 | 139015377 | 182 | Downstream (1-2kb) | 5 | 139018666 | 139023810 | 5145 | 2 | ENSMUST00 | 8433 | Prkar1b | ENSMUSG0C | false | . | . | . | . | . | . | chr5 | 139015009 | 139015462 | 3.93849 | 3.47854 | 2.17316 |
| chr5 | 138995417 | 138996038 | 182 | Promoter (<=1kb) | 5 | 138988441 | 138996001 | 7561 | 2 | ENSMUST00 | 0 | Pdgfa | ENSMUSG0C | false | chr5 | 138992846 | 138996442 | 29.41006 | 85.09721 | 81.85307 | chr5 | 138995540 | 139000004 | 4.38678 | 3.74869 | 2.49984 |
| chr18 | 75385133 | 75385378 | 182 | Intron (ENSMUST00000026999.4 | 18 | 75374914 | 75394001 | 19088 | 1 | ENSMUST00 | 10220 | Smad7 | ENSMUSG0C | false | . | . | . | . | . | . | chr18 | 75369277 | 75392875 | 5.74724 | 4.44386 | 3.62103 |
| chr1 | 4807729 | 4808199 | 182 | Promoter (<=1kb) | 1 | 4807788 | 4848410 | 40623 | 1 | ENSMUST00 | 0 | Lypla1 | ENSMUSG0C | false | chr1 | 4807230 | 4808912 | 34.78613 | 113.42423 | 109.48744 | . | . | . | . | . | . |
| chr1 | 10038081 | 10038391 | 182 | Promoter (<=1kb) | 1 | 10024601 | 10038127 | 13527 | 2 | ENSMUST00 | 0 | Cops5 | ENSMUSG0C | false | chr1 | 10037131 | 10040542 | 35.12714 | 133.20123 | 128.59534 | . | . | . | . | . | . |
| chr1 | 16518857 | 16519490 | 182 | Promoter (<=1kb) | 1 | 16228674 | 16519306 | 290633 | 2 | ENSMUST00 | 0 | Stau2 | ENSMUSG0C | false | chr1 | 16518543 | 16520190 | 31.24818 | 92.1069 | 88.71768 | . | . | . | . | . | . |
| chr15 | 59245993 | 59246201 | 182 | Distal Intergenic | 15 | 59210908 | 59211720 | 813 | 1 | ENSMUST00 | 35086 | Gm5045 | ENSMUSG0C | false | . | . | . | . | . | . | chr15 | 59246031 | 59246317 | 4.05491 | 3.21214 | 2.26354 |
| chr1 | 13589770 | 13590112 | 182 | Promoter (<=1kb) | 1 | 13564698 | 13589910 | 25213 | 2 | ENSMUST00 | 0 | Tram1 | ENSMUSG0C | false | chr1 | 13588480 | 13590440 | 31.85236 | 101.09525 | 97.496 | . | . | . | . | . | . |
| chr1 | 13644233 | 13644441 | 182 | Intron (ENSMUST00000027071.6 | 1 | 13629995 | 13638411 | 8417 | 2 | ENSMUST00 | -5823 | Lactb2 | ENSMUSG0C | false | . | . | . | . | . | . | chr1 | 13643550 | 13645065 | 4.46982 | 3.79776 | 2.61839 |
| chr1 | 55363425 | 55363602 | 182 | Promoter (<=1kb) | 1 | 55346257 | 55363469 | 17213 | 2 | ENSMUST00 | 0 | Boll | ENSMUSG0C | false | chr1 | 55363020 | 55363973 | 17.92175 | 44.05972 | 41.41886 | chr1 | 55363159 | 55363888 | 3.82479 | 3.49567 | 2.09639 |
| chr1 | 67038656 | 67039054 | 182 | Promoter (<=1kb) | 1 | 67000517 | 67038865 | 38349 | 2 | ENSMUST00 | 0 | Lancl1 | ENSMUSG0C | false | chr1 | 67038121 | 67039355 | 33.04283 | 115.53907 | 111.53828 | chr1 | 67037366 | 67039579 | 5.06542 | 3.87733 | 3.08721 |
| chr1 | 58973433 | 58974007 | 182 | Promoter (<=1kb) | 1 | 58973522 | 58981475 | 7954 | 1 | ENSMUST00 | 0 | Stradb | ENSMUSG0C | false | chr1 | 58972637 | 58974577 | 36.809 | 123.33047 | 119.07505 | . | . | . | . | . | . |
| chr1 | 43694796 | 43695051 | 182 | Distal Intergenic | 1 | 43730602 | 43742578 | 11977 | 1 | ENSMUST00 | -35551 | 1500015O10 | ENSMUSG0C | false | . | . | . | . | . | . | chr1 | 43694384 | 43695440 | 5.53323 | 4.23885 | 3.46654 |
| chr1 | 40681069 | 40681667 | 182 | Promoter (<=1kb) | 1 | 40681712 | 40768885 | 87174 | 1 | ENSMUST00 | -45 | Slc9a2 | ENSMUSG0C | false | chr1 | 40680316 | 40682232 | 28.7777 | 83.06458 | 79.8585 | chr1 | 40680269 | 40682307 | 4.73179 | 3.86624 | 2.83282 |
| chr1 | 37429514 | 37430263 | 182 | Promoter (<=1kb) | 1 | 37430170 | 37439124 | 8955 | 1 | ENSMUST00 | 0 | Unc50 | ENSMUSG0C | false | chr1 | 37429187 | 37431192 | 19.50994 | 80.00812 | 76.85559 | chr1 | 37428768 | 37429672 | 4.55693 | 3.81446 | 2.68767 |
| chr1 | 34025918 | 34026127 | 182 | Intron (ENSMUST00000182513.2 | 1 | 34011825 | 34308661 | 296837 | 1 | ENSMUST00 | 14094 | Dst | ENSMUSG0C | true | . | . | . | . | . | . | chr1 | 34020714 | 34026603 | 4.63881 | 3.8982 | 2.69619 |
| chr1 | 34051224 | 34051576 | 182 | Intron (ENSMUST00000182513.2 | 1 | 34089292 | 34119153 | 29862 | 1 | ENSMUST00 | -37716 | Dst | ENSMUSG0C | true | . | . | . | . | . | . | chr1 | 34048127 | 34055658 | 4.48528 | 3.80645 | 2.61101 |
| chr1 | 74587674 | 74588490 | 182 | Promoter (<=1kb) | 1 | 74565127 | 74588028 | 22902 | 2 | ENSMUST00 | 0 | Zfp142 | ENSMUSG0C | false | chr1 | 74587078 | 74588189 | 33.1634 | 104.39787 | 100.72088 | . | . | . | . | . | . |
| chr1 | 23921647 | 23922504 | 182 | Promoter (<=1kb) | 1 | 23845625 | 23922317 | 76693 | 2 | ENSMUST00 | 0 | Smap1 | ENSMUSG0C | false | chr1 | 23921556 | 23922782 | 32.29185 | 112.09676 | 108.19604 | chr1 | 23915404 | 23922686 | 4.32915 | 3.70478 | 2.46464 |
| chr1 | 74714199 | 74714458 | 182 | Promoter (<=1kb) | 1 | 74713574 | 74737892 | 24319 | 1 | ENSMUST00 | 626 | Cyp27a1 | ENSMUSG0C | true | chr1 | 74712846 | 74714683 | 26.66126 | 88.87019 | 85.5497 | chr1 | 74714232 | 74717018 | 5.21045 | 4.16823 | 3.24082 |
| chr1 | 74332306 | 74332674 | 182 | Promoter (<=1kb) | 1 | 74332610 | 74353686 | 21077 | 1 | ENSMUST00 | 0 | Pnkd | ENSMUSG0C | false | chr1 | 74332151 | 74333246 | 21.20057 | 58.40949 | 55.58382 | chr1 | 74331414 | 74333595 | 4.76722 | 3.8164 | 2.842 |
| chr1 | 71102944 | 71103238 | 182 | Promoter (<=1kb) | 1 | 71027498 | 71103146 | 75649 | 2 | ENSMUST00 | 0 | Bard1 | ENSMUSG0C | false | chr1 | 71102039 | 71103659 | 34.27164 | 115.65593 | 111.648 | . | . | . | . | . | . |
| chr1 | 75386107 | 75386427 | 182 | Promoter (<=1kb) | 1 | 75385610 | 75432304 | 46695 | 1 | ENSMUST00 | 498 | Speg | ENSMUSG0C | true | chr1 | 75384399 | 75387322 | 24.11245 | 66.85627 | 63.91418 | chr1 | 75386249 | 75400648 | 7.0728 | 5.02664 | 4.70473 |
| chr1 | 75361828 | 75362055 | 182 | Promoter (<=1kb) | 1 | 75362030 | 75363638 | 1609 | 1 | ENSMUST00 | 0 | Des | ENSMUSG0C | false | . | . | . | . | . | . | chr1 | 75356622 | 75362384 | 5.80426 | 4.47779 | 3.65868 |
| chr1 | 75344323 | 75344734 | 182 | Distal Intergenic | 1 | 75360329 | 75368579 | 8251 | 1 | ENSMUST00 | -15595 | Des | ENSMUSG0C | false | . | . | . | . | . | . | chr1 | 75340961 | 75345934 | 4.949 | 3.97142 | 2.9865 |
| chr1 | 84839244 | 84839746 | 182 | Promoter (<=1kb) | 1 | 84721189 | 84839304 | 118116 | 2 | ENSMUST00 | 0 | Trip12 | ENSMUSG0C | false | chr1 | 84837968 | 84840939 | 30.78991 | 105.27288 | 101.56436 | . | . | . | . | . | . |
| chr1 | 85894466 | 85894651 | 182 | Promoter (<=1kb) | 1 | 85894619 | 85908359 | 13741 | 1 | ENSMUST00 | 0 | Itm2c | ENSMUSG0C | false | chr1 | 85894103 | 85895855 | 28.49948 | 87.34749 | 84.05397 | chr1 | 85893777 | 85895969 | 3.89056 | 3.56088 | 2.12253 |
| chr1 | 86064343 | 86064844 | 182 | Promoter (<=1kb) | 1 | 86064387 | 86070776 | 6390 | 1 | ENSMUST00 | 0 | Psmd1 | ENSMUSG0C | false | chr1 | 86064354 | 86065837 | 46.03907 | 157.30389 | 151.74776 | . | . | . | . | . | . |
| chr1 | 86528046 | 86528765 | 182 | Promoter (1-2kb) | 1 | 86526807 | 86530209 | 3403 | 1 | ENSMUST00 | 1240 | Ptma | ENSMUSG0C | false | chr1 | 86525464 | 86529269 | 41.35843 | 143.41548 | 138.39708 | . | . | . | . | . | . |
| chr1 | 86582223 | 86582623 | 182 | Promoter (<=1kb) | 1 | 86542994 | 86582547 | 39554 | 2 | ENSMUST00 | 0 | Pde6d | ENSMUSG0C | false | chr1 | 86581764 | 86583368 | 36.96067 | 121.45832 | 117.27072 | chr1 | 86581127 | 86583314 | 3.49305 | 3.32577 | 1.77726 |
| chr1 | 92980959 | 92981342 | 182 | Promoter (2-3kb) | 1 | 92978922 | 92986391 | 7470 | 1 | ENSMUST00 | 2038 | Gpr35 | ENSMUSG0C | false | . | . | . | . | . | . | . | . | . | . | . | . |
| chr1 | 93686705 | 93687022 | 182 | Promoter (<=1kb) | 1 | 93686263 | 93694070 | 7808 | 1 | ENSMUST00 | 443 | Bok | ENSMUSG0C | false | chr1 | 93685234 | 93686792 | 23.13664 | 67.80279 | 64.84689 | chr1 | 93685377 | 93687202 | 3.48118 | 3.17875 | 1.7988 |
| chr1 | 93754391 | 93755311 | 182 | Promoter (<=1kb) | 1 | 93701935 | 93754780 | 52846 | 2 | ENSMUST00 | 0 | Thap4 | ENSMUSG0C | false | chr1 | 93753617 | 93755649 | 47.77854 | 170.84077 | 164.85675 | . | . | . | . | . | . |
| chr1 | 93803860 | 93804402 | 182 | Promoter (<=1kb) | 1 | 93803965 | 93822095 | 18131 | 1 | ENSMUST00 | 0 | Ing5 | ENSMUSG0C | false | chr1 | 93803311 | 93804724 | 31.91277 | 108.3221 | 104.53201 | chr1 | 93801156 | 93805057 | 4.71121 | 3.88113 | 2.8127 |
| chr1 | 87755796 | 87756137 | 182 | Promoter (<=1kb) | 1 | 87755870 | 87771398 | 15529 | 1 | ENSMUST00 | 0 | Atg16l1 | ENSMUSG0C | false | chr1 | 87755464 | 87757081 | 41.67902 | 156.4369 | 150.90947 | chr1 | 87755535 | 87758908 | 3.41421 | 3.17799 | 1.74042 |
| chr1 | 91054944 | 91055127 | 182 | Promoter (1-2kb) | 1 | 91053586 | 91128287 | 74702 | 1 | ENSMUST00 | 1359 | Lrrfip1 | ENSMUSG0C | true | chr1 | 91051607 | 91058713 | 31.85236 | 101.09525 | 97.496 | . | . | . | . | . | . |
| chr1 | 91351390 | 91351606 | 182 | Promoter (<=1kb) | 1 | 91351016 | 91362416 | 11401 | 1 | ENSMUST00 | 375 | Klhl30 | ENSMUSG0C | false | . | . | . | . | . | . | chr1 | 91350502 | 91353461 | 5.92886 | 4.38143 | 3.8118 |
| chr1 | 92127807 | 92127980 | 182 | Intron (ENSMUST00000097644.3 | 1 | 92030050 | 92113331 | 83282 | 2 | ENSMUST00 | -14477 | Hdac4 | ENSMUSG0C | false | . | . | . | . | . | . | chr1 | 92127225 | 92132642 | 5.04853 | 3.91786 | 3.06445 |
| chr1 | 92130532 | 92130820 | 182 | Intron (ENSMUST00000097644.3 | 1 | 92030050 | 92113331 | 83282 | 2 | ENSMUST00 | -17202 | Hdac4 | ENSMUSG0C | false | . | . | . | . | . | . | chr1 | 92127225 | 92132642 | 5.04853 | 3.91786 | 3.06445 |
| chr1 | 98079707 | 98079940 | 182 | Intron (ENSMUST00000058762.9 | 1 | 97821094 | 98095621 | 274528 | 2 | ENSMUST00 | 15681 | Pam | ENSMUSG0C | false | . | . | . | . | . | . | chr1 | 98076522 | 98088542 | 5.45317 | 4.29285 | 3.37862 |
| chr1 | 125676914 | 125677143 | 182 | Promoter (<=1kb) | 1 | 125676995 | 125873862 | 196868 | 1 | ENSMUST00 | 0 | Gpr39 | ENSMUSG0C | false | . | . | . | . | . | . | chr1 | 125676795 | 125677280 | 5.52486 | 4.35165 | 3.48324 |
| chr1 | 128416959 | 128417479 | 182 | Promoter (<=1kb) | 1 | 128363707 | 128417359 | 53653 | 2 | ENSMUST00 | 0 | Dars | ENSMUSG0C | false | chr1 | 128416488 | 128417732 | 25.88745 | 125.26566 | 120.94565 | . | . | . | . | . | . |
| chr1 | 143633103 | 143633427 | 182 | Intron (ENSMUST00000018337.8 | 1 | 143633851 | 143636577 | 2727 | 2 | ENSMUST00 | 3150 | Cdc73 | ENSMUSG0C | false | . | . | . | . | . | . | chr1 | 143631518 | 143633771 | 4.66077 | 3.91353 | 2.71923 |
| chr1 | 119638757 | 119638999 | 182 | Intron (ENSMUST00000052404.7 | 1 | 119617655 | 119634000 | 16346 | 2 | ENSMUST00 | -4758 | Epb4.1l5 | ENSMUSG0C | false | . | . | . | . | . | . | chr1 | 119619960 | 119646490 | 4.72133 | 3.93355 | 2.78285 |
| chr1 | 120229004 | 120229238 | 182 | Intron (ENSMUST00000140490.2 | 1 | 120190757 | 120265255 | 74499 | 2 | ENSMUST00 | 36017 | Steap3 | ENSMUSG0C | false | . | . | . | . | . | . | chr1 | 120220892 | 120232141 | 6.47152 | 4.73253 | 4.21633 |
| chr1 | 138619446 | 138619892 | 182 | Promoter (<=1kb) | 1 | 138483837 | 138619749 | 135913 | 2 | ENSMUST00 | 0 | Nek7 | ENSMUSG0C | false | chr1 | 138617343 | 138620436 | 36.04672 | 129.50026 | 125.02953 | . | . | . | . | . | . |
| chr1 | 138618008 | 138618212 | 182 | Promoter (1-2kb) | 1 | 138485707 | 138619674 | 133968 | 2 | ENSMUST00 | 1462 | Nek7 | ENSMUSG0C | false | chr1 | 138617343 | 138620436 | 36.04672 | 129.50026 | 125.02953 | chr1 | 138617200 | 138619038 | 5.15956 | 4.08899 | 3.16942 |
| chr1 | 130462414 | 130463096 | 182 | Promoter (<=1kb) | 1 | 130439027 | 130462744 | 23718 | 2 | ENSMUST00 | 0 | Cd55 | ENSMUSG0C | false | chr1 | 130461445 | 130463349 | 24.78212 | 78.86476 | 75.73294 | . | . | . | . | . | . |
| chr1 | 135743921 | 135744209 | 182 | Intron (ENSMUST00000097561.3 | 1 | 135729147 | 135752232 | 23086 | 1 | ENSMUST00 | 14775 | Csrp1 | ENSMUSG0C | false | . | . | . | . | . | . | . | . | . | . | . | . |
| chr1 | 131525961 | 131526243 | 182 | Promoter (1-2kb) | 1 | 131285251 | 131527347 | 242097 | 2 | ENSMUST00 | 1104 | Srgap2 | ENSMUSG0C | true | chr1 | 131523158 | 131527435 | 25.99992 | 84.80917 | 81.57088 | . | . | . | . | . | . |
| chr1 | 131378382 | 131378605 | 182 | Intron (ENSMUST00000097588.3 | 1 | 131296641 | 131364064 | 67424 | 2 | ENSMUST00 | -14319 | Srgap2 | ENSMUSG0C | true | . | . | . | . | . | . | chr1 | 131377737 | 131386854 | 5.13189 | 4.0644 | 3.1348 |
| chr1 | 131238991 | 131239268 | 182 | Intron (ENSMUST00000027688.9 | 1 | 131176410 | 131245258 | 68849 | 2 | ENSMUST00 | 5990 | Rassf5 | ENSMUSG0C | false | . | . | . | . | . | . | . | . | . | . | . | . |
| chr1 | 132007460 | 132008413 | 182 | Promoter (<=1kb) | 1 | 132007607 | 132026448 | 18842 | 1 | ENSMUST00 | 0 | Elk4 | ENSMUSG0C | false | chr1 | 132006980 | 132009267 | 31.54088 | 108.67455 | 104.87239 | . | . | . | . | . | . |
| chr1 | 153900494 | 153900911 | 182 | Promoter (<=1kb) | 1 | 153900606 | 153907140 | 6535 | 1 | ENSMUST00 | 0 | Glul | ENSMUSG0C | false | chr1 | 153899280 | 153901286 | 40.89836 | 130.45236 | 125.94646 | . | . | . | . | . | . |
| chr1 | 152817874 | 152818170 | 182 | Intron (ENSMUST00000186568.1 | 1 | 152823569 | 152824484 | 916 | 1 | ENSMUST00 | -5399 | Ncf2 | ENSMUSG0C | true | . | . | . | . | . | . | chr1 | 152801274 | 152821507 | 4.847 | 3.96927 | 2.88719 |
| chr1 | 152594843 | 152595072 | 182 | Promoter (2-3kb) | 1 | 152557543 | 152592759 | 35217 | 2 | ENSMUST00 | -2085 | Rgl1 | ENSMUSG0C | false | . | . | . | . | . | . | chr1 | 152587354 | 152595118 | 4.48145 | 3.81233 | 2.58473 |
| chr1 | 152624565 | 152624886 | 182 | Promoter (<=1kb) | 1 | 152557543 | 152625029 | 67487 | 2 | ENSMUST00 | 143 | Rgl1 | ENSMUSG0C | false | chr1 | 152623682 | 152625635 | 24.03115 | 75.67385 | 72.59326 | . | . | . | . | . | . |
| chr1 | 152555984 | 152556544 | 182 | Promoter (2-3kb) | 1 | 152551508 | 152553044 | 1537 | 2 | ENSMUST00 | -2941 | Rgl1 | ENSMUSG0C | false | . | . | . | . | . | . | chr1 | 152554068 | 152559430 | 6.27115 | 4.74329 | 4.04038 |
| chr1 | 152531573 | 152531937 | 182 | Exon (ENSMUST00000111859.2/ | 1 | 152551508 | 152553044 | 1537 | 2 | ENSMUST00 | 21107 | Rgl1 | ENSMUSG0C | false | . | . | . | . | . | . | . | . | . | . | . | . |
| chr1 | 151677370 | 151677659 | 182 | Promoter (<=1kb) | 1 | 151677167 | 151719342 | 42176 | 1 | ENSMUST00 | 204 | Fam129a | ENSMUSG0C | false | . | . | . | . | . | . | chr1 | 151676392 | 151678352 | 3.28648 | 3.22998 | 1.586 |
| chr1 | 178187465 | 178187941 | 182 | Promoter (<=1kb) | 1 | 178187532 | 178252598 | 65067 | 1 | ENSMUST00 | 0 | Desi2 | ENSMUSG0C | false | chr1 | 178187528 | 178188717 | 46.18486 | 178.62439 | 172.4859 | . | . | . | . | . | . |
| chr1 | 182496931 | 182497169 | 182 | Intron (ENSMUST00000068505.8 | 1 | 182467256 | 182517495 | 50240 | 2 | ENSMUST00 | 20326 | Capn2 | ENSMUSG0C | false | . | . | . | . | . | . | chr1 | 182492178 | 182516499 | 7.84019 | 5.29195 | 5.34945 |
| chr1 | 182125036 | 182125350 | 182 | Promoter (<=1kb) | 1 | 182124750 | 182132407 | 7658 | 1 | ENSMUST00 | 287 | Srp9 | ENSMUSG0C | false | chr1 | 182124638 | 182125968 | 32.27147 | 102.84008 | 99.19724 | . | . | . | . | . | . |
| chr1 | 172514906 | 172515134 | 182 | Distal Intergenic | 1 | 172521044 | 172545870 | 24827 | 1 | ENSMUST00 | -5910 | Ccdc19 | ENSMUSG0C | false | . | . | . | . | . | . | . | . | . | . | . | . |

| | | | | | | | | | | | | | | | | | | | | | | | | | | |
|---|---|---|---|---|---|---|---|---|---|---|---|---|---|---|---|---|---|---|---|---|---|---|---|---|---|---|
| chr1 | 172501040 | 172501268 | 182 | Promoter (<=1kb) | 1 | 172501252 | 172506733 | 5482 | 1 | ENSMUST00 | 0 | Tagln2 | ENSMUSG0C | false | chr1 | 172499800 | 172502632 | 28.23176 | 83.06847 | 79.86218 | chr1 | 172501063 | 172503936 | 5.77421 | 4.5263 | 3.63388 |
| chr1 | 165461028 | 165461323 | 182 | Promoter (<=1kb) | 1 | 165461037 | 165481214 | 20178 | 1 | ENSMUST00 | 0 | Mpc2 | ENSMUSG0C | false | chr1 | 165461194 | 165462209 | 27.49702 | 86.54257 | 83.27029 | . | . | . | . | . | . |
| chr1 | 189756846 | 189757227 | 182 | Intron (ENSMUST00000097442.3 | 1 | 189728268 | 189876689 | 148422 | 1 | ENSMUST00 | 28579 | Ptpn14 | ENSMUSG0C | true | . | . | . | . | . | . | chr1 | 189730267 | 189810107 | 6.93269 | 4.87829 | 4.60521 |
| chr1 | 185362845 | 185363203 | 182 | Promoter (<=1kb) | 1 | 185363095 | 185428355 | 65261 | 1 | ENSMUST00 | 0 | Eprs | ENSMUSG0C | false | chr1 | 185362319 | 185364185 | 40.55256 | 150.97545 | 145.64915 | . | . | . | . | . | . |
| chr1 | 185091874 | 185092084 | 182 | Distal Intergenic | 1 | 184896443 | 184999549 | 103107 | 2 | ENSMUST00 | -92326 | Mark1 | ENSMUSG0C | false | . | . | . | . | . | . | . | . | . | . | . | . |
| chr1 | 191855071 | 191855377 | 182 | Distal Intergenic | 1 | 191827148 | 191829988 | 2841 | 1 | ENSMUST00 | 27924 | Nek2 | ENSMUSG0C | false | . | . | . | . | . | . | chr1 | 191854353 | 191856164 | 4.5858 | 3.91795 | 2.68646 |
| chr1 | 191462221 | 191462479 | 182 | Distal Intergenic | 1 | 191351981 | 191397041 | 45061 | 2 | ENSMUST00 | -65181 | Ppp2r5a | ENSMUSG0C | false | . | . | . | . | . | . | chr1 | 191457318 | 191471879 | 4.86599 | 4.03205 | 2.90022 |
| chr1 | 191444540 | 191444717 | 182 | Distal Intergenic | 1 | 191351981 | 191397041 | 45061 | 2 | ENSMUST00 | -47500 | Ppp2r5a | ENSMUSG0C | false | . | . | . | . | . | . | chr1 | 191443645 | 191445803 | 5.6789 | 4.43248 | 3.56297 |
| chr1 | 191340380 | 191340630 | 182 | Intron (ENSMUST00000027940.5 | 1 | 191325912 | 191350914 | 25003 | 1 | ENSMUST00 | 14469 | Tmem206 | ENSMUSG0C | false | . | . | . | . | . | . | chr1 | 191339806 | 191340860 | 6.17785 | 4.67841 | 3.96366 |
| chr2 | 3474930 | 3475288 | 182 | Promoter (<=1kb) | 2 | 3455815 | 3475031 | 19217 | 2 | ENSMUST00 | 0 | Suv39h2 | ENSMUSG0C | false | chr2 | 3474048 | 3475781 | 40.43883 | 128.57333 | 124.13428 | . | . | . | . | . | . |
| chr2 | 4501727 | 4501961 | 182 | Intron (ENSMUST00000177457.2 | 2 | 4473194 | 4498103 | 24910 | 1 | ENSMUST00 | 28534 | Frmd4a | ENSMUSG0C | false | . | . | . | . | . | . | chr2 | 4501044 | 4502271 | 4.22632 | 3.76978 | 2.35624 |
| chr1 | 170867619 | 170867897 | 182 | Promoter (<=1kb) | 1 | 170704674 | 170867771 | 163098 | 2 | ENSMUST00 | 0 | Atf6 | ENSMUSG0C | false | chr1 | 170866958 | 170869093 | 39.45224 | 138.91832 | 134.07426 | . | . | . | . | . | . |
| chr2 | 5063786 | 5064127 | 182 | Promoter (<=1kb) | 2 | 5020642 | 5064045 | 43404 | 2 | ENSMUST00 | 0 | Optn | ENSMUSG0C | true | chr2 | 5061506 | 5064203 | 37.26972 | 119.86725 | 115.72688 | . | . | . | . | . | . |
| chr2 | 5058157 | 5058371 | 182 | Intron (ENSMUST00000114996.2 | 2 | 5020644 | 5063938 | 43295 | 2 | ENSMUST00 | 5567 | Optn | ENSMUSG0C | true | . | . | . | . | . | . | chr2 | 5041736 | 5062961 | 5.08721 | 4.02119 | 3.09837 |
| chr1 | 167349968 | 167350348 | 182 | Promoter (<=1kb) | 1 | 167349991 | 167368530 | 18540 | 1 | ENSMUST00 | 0 | Aldh9a1 | ENSMUSG0C | false | chr1 | 167349624 | 167350909 | 25.84672 | 90.2271 | 86.88143 | . | . | . | . | . | . |
| chr2 | 13384884 | 13385217 | 182 | Intron (ENSMUST00000091436.5 | 2 | 13445021 | 13455624 | 10604 | 2 | ENSMUST00 | 70407 | Cubn | ENSMUSG0C | false | . | . | . | . | . | . | chr2 | 13383881 | 13386383 | 5.33592 | 4.27318 | 3.27845 |
| chr2 | 13410011 | 13410190 | 182 | Intron (ENSMUST00000091436.5 | 2 | 13445021 | 13455624 | 10604 | 2 | ENSMUST00 | 45434 | Cubn | ENSMUSG0C | false | . | . | . | . | . | . | chr2 | 13409573 | 13411165 | 4.62996 | 3.88842 | 2.72023 |
| chr2 | 18996938 | 18997170 | 182 | Promoter (<=1kb) | 2 | 18842255 | 18998126 | 155872 | 2 | ENSMUST00 | 956 | Pip4k2a | ENSMUSG0C | false | chr2 | 18996496 | 18998878 | 38.46334 | 130.56975 | 126.05653 | chr2 | 18995091 | 18997086 | 3.76347 | 3.48584 | 1.98793 |
| chr2 | 18392395 | 18392977 | 182 | Promoter (<=1kb) | 2 | 18206333 | 18392811 | 186479 | 2 | ENSMUST00 | 0 | Dnajc1 | ENSMUSG0C | false | chr2 | 18391673 | 18393132 | 31.62822 | 118.12274 | 114.03909 | . | . | . | . | . | . |
| chr2 | 16399057 | 16399287 | 182 | Intron (ENSMUST00000028081.7 | 2 | 16357117 | 16751857 | 394741 | 1 | ENSMUST00 | 41941 | Plxdc2 | ENSMUSG0C | false | . | . | . | . | . | . | chr2 | 16389376 | 16401429 | 3.88696 | 3.46382 | 2.12715 |
| chr2 | 49838724 | 49839024 | 182 | Promoter (1-2kb) | 2 | 49840090 | 49942537 | 102448 | 1 | ENSMUST00 | -1066 | Lypd6b | ENSMUSG0C | false | . | . | . | . | . | . | chr2 | 49836358 | 49847151 | 4.90555 | 4.05449 | 2.92293 |
| chr2 | 49861160 | 49861404 | 182 | Intron (ENSMUST00000028103.7 | 2 | 49840090 | 49942537 | 102448 | 1 | ENSMUST00 | 21071 | Lypd6b | ENSMUSG0C | false | . | . | . | . | . | . | chr2 | 49860697 | 49861642 | 3.67278 | 3.44027 | 1.90344 |
| chr2 | 50296548 | 50296847 | 182 | Promoter (<=1kb) | 2 | 50279881 | 50296695 | 16815 | 2 | ENSMUST00 | 0 | Mmadhc | ENSMUSG0C | false | chr2 | 50295404 | 50297372 | 26.1692 | 76.69956 | 73.60538 | . | . | . | . | . | . |
| chr2 | 11501950 | 11502260 | 182 | Promoter (<=1kb) | 2 | 11471430 | 11502025 | 30596 | 2 | ENSMUST00 | 0 | Pfkfb3 | ENSMUSG0C | false | chr2 | 11500699 | 11503640 | 31.43507 | 99.35931 | 95.8029 | . | . | . | . | . | . |
| chr2 | 23155928 | 23156166 | 182 | Promoter (<=1kb) | 2 | 23115606 | 23156024 | 40419 | 2 | ENSMUST00 | 0 | Mastl | ENSMUSG0C | false | chr2 | 23155399 | 23157490 | 28.67964 | 89.59386 | 86.26003 | . | . | . | . | . | . |
| chr2 | 32875510 | 32876191 | 182 | Promoter (<=1kb) | 2 | 32876114 | 32925254 | 49141 | 1 | ENSMUST00 | 0 | Fam129b | ENSMUSG0C | false | chr2 | 32875288 | 32879031 | 32.3084 | 121.498 | 117.30614 | chr2 | 32869147 | 32875861 | 5.74163 | 4.37142 | 3.62068 |
| chr2 | 32598584 | 32598814 | 182 | Promoter (<=1kb) | 2 | 32599709 | 32615124 | 15416 | 1 | ENSMUST00 | -895 | St6galnac6 | ENSMUSG0C | true | . | . | . | . | . | . | chr2 | 32598512 | 32598899 | 3.00551 | 2.98534 | 1.3713 |
| chr2 | 28640826 | 28641359 | 182 | Promoter (<=1kb) | 2 | 28641228 | 28672507 | 31280 | 1 | ENSMUST00 | 0 | Tsc1 | ENSMUSG0C | false | chr2 | 28640640 | 28640883 | 8.08052 | 18.53858 | 16.22368 | chr2 | 28640321 | 28642908 | 3.68235 | 3.39602 | 1.91678 |
| chr2 | 28650041 | 28650232 | 182 | Intron (ENSMUST00000156857.2 | 2 | 28658567 | 28691167 | 32601 | 1 | ENSMUST00 | -8335 | Tsc1 | ENSMUSG0C | false | . | . | . | . | . | . | chr2 | 28648758 | 28651761 | 3.65232 | 3.25879 | 1.93627 |
| chr2 | 32465362 | 32465600 | 182 | Distal Intergenic | 2 | 32414490 | 32451445 | 36956 | 2 | ENSMUST00 | -13918 | Slc25a25 | ENSMUSG0C | false | . | . | . | . | . | . | chr2 | 32463809 | 32466367 | 6.31936 | 4.60617 | 4.12814 |
| chr2 | 32312321 | 32312605 | 182 | Promoter (<=1kb) | 2 | 32309485 | 32312265 | 2781 | 2 | ENSMUST00 | -57 | Dnm1 | ENSMUSG0C | false | chr2 | 32311860 | 32312859 | 24.07142 | 68.54774 | 65.57966 | chr2 | 32311185 | 32312975 | 6.59467 | 4.52431 | 4.34609 |
| chr2 | 58516112 | 58516371 | 182 | Promoter (<=1kb) | 2 | 58388644 | 58516272 | 127629 | 2 | ENSMUST00 | 0 | Acvr1 | ENSMUSG0C | false | chr2 | 58516147 | 58516393 | 5.86754 | 9.51579 | 7.34688 | chr2 | 58513168 | 58518712 | 4.38962 | 3.79661 | 2.50474 |
| chr2 | 31759763 | 31760058 | 182 | Promoter (<=1kb) | 2 | 31759943 | 31804227 | 44285 | 1 | ENSMUST00 | 0 | Abl1 | ENSMUSG0C | false | chr2 | 31759357 | 31761932 | 31.85236 | 101.09525 | 97.496 | . | . | . | . | . | . |
| chr2 | 31769726 | 31770127 | 182 | Intron (ENSMUST00000142554.2 | 2 | 31760173 | 31779024 | 18852 | 1 | ENSMUST00 | 9554 | Abl1 | ENSMUSG0C | false | . | . | . | . | . | . | chr2 | 31760789 | 31776841 | 6.98405 | 4.98586 | 4.62933 |
| chr2 | 30392710 | 30393196 | 182 | Promoter (<=1kb) | 2 | 30392454 | 30397813 | 5360 | 1 | ENSMUST00 | 257 | Dolpp1 | ENSMUSG0C | false | chr2 | 30391708 | 30393784 | 32.69058 | 104.59052 | 100.90593 | chr2 | 30393171 | 30395281 | 4.50255 | 3.83436 | 2.61373 |
| chr2 | 30807595 | 30807857 | 182 | Promoter (<=1kb) | 2 | 30807826 | 30823033 | 15208 | 1 | ENSMUST00 | 0 | Ntmt1 | ENSMUSG0C | false | chr2 | 30807144 | 30807622 | 17.92175 | 44.05972 | 41.41886 | chr2 | 30806776 | 30808989 | 4.43015 | 3.87715 | 2.52753 |
| chr2 | 30358536 | 30358778 | 182 | Promoter (<=1kb) | 2 | 30344809 | 30359240 | 14432 | 2 | ENSMUST00 | 462 | Sh3glb2 | ENSMUSG0C | false | chr2 | 30358513 | 30359650 | 31.48209 | 103.47416 | 99.8177 | chr2 | 30355332 | 30359891 | 5.91362 | 4.50021 | 3.75375 |
| chr2 | 45067113 | 45067543 | 182 | Intron (ENSMUST00000068415.5 | 2 | 44988388 | 45110244 | 121857 | 2 | ENSMUST00 | 42701 | Zeb2 | ENSMUSG0C | false | . | . | . | . | . | . | chr2 | 45060607 | 45073593 | 4.65688 | 3.82601 | 2.75469 |
| chr2 | 35267838 | 35268410 | 182 | Distal Intergenic | 2 | 35279663 | 35293567 | 13905 | 1 | ENSMUST00 | -11253 | Gsn | ENSMUSG0C | true | . | . | . | . | . | . | chr2 | 35257475 | 35289539 | 6.87726 | 4.86078 | 4.55899 |
| chr2 | 35336992 | 35337292 | 182 | Promoter (<=1kb) | 2 | 35313986 | 35336976 | 22991 | 2 | ENSMUST00 | -17 | Stom | ENSMUSG0C | false | chr2 | 35336294 | 35337626 | 29.66345 | 100.21 | 96.63225 | chr2 | 35335537 | 35337849 | 4.3255 | 3.73064 | 2.45714 |
| chr2 | 35582625 | 35582935 | 182 | Promoter (<=1kb) | 2 | 35582840 | 35730992 | 148153 | 1 | ENSMUST00 | 0 | Dab2ip | ENSMUSG0C | false | chr2 | 35581533 | 35583251 | 20.35144 | 58.59763 | 55.77069 | chr2 | 35581146 | 35583413 | 4.17986 | 3.50064 | 2.36858 |
| chr2 | 26525692 | 26525895 | 182 | Distal Intergenic | 2 | 26471276 | 26516663 | 45388 | 2 | ENSMUST00 | -9030 | Notch1 | ENSMUSG0C | false | . | . | . | . | . | . | . | . | . | . | . | . |
| chr19 | 58822476 | 58823046 | 182 | Intron (ENSMUST00000066285.4 | 19 | 58785803 | 58794461 | 8659 | 2 | ENSMUST00 | -28016 | 1700019N19 | ENSMUSG0C | false | . | . | . | . | . | . | chr19 | 58814047 | 58859817 | 7.46659 | 5.12677 | 5.03638 |
| chr2 | 26098707 | 26098925 | 182 | Intron (ENSMUST00000028300.5 | 2 | 26055535 | 26092177 | 36643 | 2 | ENSMUST00 | -6531 | Nacc2 | ENSMUSG0C | false | . | . | . | . | . | . | chr2 | 26090412 | 26100336 | 5.43617 | 4.26207 | 3.38086 |
| chr2 | 26091455 | 26092232 | 182 | Promoter (<=1kb) | 2 | 26055535 | 26092177 | 36643 | 2 | ENSMUST00 | 0 | Nacc2 | ENSMUSG0C | false | chr2 | 26091165 | 26092272 | 18.70405 | 48.44756 | 45.75203 | chr2 | 26090412 | 26100336 | 5.43617 | 4.26207 | 3.38086 |
| chr2 | 53191633 | 53192423 | 182 | Promoter (<=1kb) | 2 | 53191726 | 53217534 | 25809 | 1 | ENSMUST00 | 0 | Arl6ip6 | ENSMUSG0C | false | chr2 | 53190411 | 53192920 | 29.52499 | 116.10789 | 112.09218 | . | . | . | . | . | . |
| chr2 | 60908275 | 60908540 | 182 | Intron (ENSMUST00000028347.7 | 2 | 60842297 | 60889030 | 46734 | 2 | ENSMUST00 | -19246 | Rbms1 | ENSMUSG0C | true | . | . | . | . | . | . | chr2 | 60890567 | 60932725 | 5.60936 | 4.28041 | 3.52606 |
| chr2 | 60756443 | 60756751 | 182 | Exon (ENSMUST00000028347.7/ | 2 | 60753560 | 60762321 | 8762 | 2 | ENSMUST00 | 5570 | Rbms1 | ENSMUSG0C | true | . | . | . | . | . | . | chr2 | 60756001 | 60759166 | 4.15499 | 3.62905 | 2.3537 |
| chr2 | 60815704 | 60815885 | 182 | Intron (ENSMUST00000028347.7 | 2 | 60755416 | 60780941 | 25526 | 2 | ENSMUST00 | -34764 | Rbms1 | ENSMUSG0C | true | . | . | . | . | . | . | chr2 | 60798942 | 60834915 | 5.34112 | 4.16576 | 3.30137 |
| chr2 | 59882184 | 59882778 | 182 | Promoter (<=1kb) | 2 | 59852364 | 59882584 | 30221 | 2 | ENSMUST00 | 0 | Wdsub1 | ENSMUSG0C | false | chr2 | 59881834 | 59882942 | 43.58161 | 145.05977 | 139.97232 | chr2 | 59881416 | 59883355 | 3.17207 | 3.04611 | 1.53915 |
| chr2 | 59160665 | 59160897 | 182 | Promoter (<=1kb) | 2 | 59160850 | 59355204 | 194355 | 1 | ENSMUST00 | 0 | Pkp4 | ENSMUSG0C | false | chr2 | 59160059 | 59162590 | 28.51907 | 101.09029 | 97.496 | . | . | . | . | . | . |
| chr2 | 59161307 | 59162088 | 182 | Promoter (<=1kb) | 2 | 59161927 | 59215031 | 53105 | 1 | ENSMUST00 | 0 | Pkp4 | ENSMUSG0C | false | chr2 | 59160059 | 59162590 | 28.51907 | 101.09029 | 97.496 | chr2 | 59161984 | 59164579 | 3.45993 | 3.28562 | 1.73074 |
| chr2 | 79638866 | 79639213 | 182 | Promoter (<=1kb) | 2 | 79639182 | 79672966 | 33785 | 1 | ENSMUST00 | 0 | Ssfa2 | ENSMUSG0C | false | chr2 | 79635281 | 79639748 | 36.04672 | 129.50026 | 125.02953 | chr2 | 79636998 | 79643476 | 4.38951 | 3.81687 | 2.49348 |
| chr2 | 71211611 | 71212101 | 182 | Promoter (<=1kb) | 2 | 71211706 | 71263298 | 51593 | 1 | ENSMUST00 | 0 | Dync1i2 | ENSMUSG0C | false | chr2 | 71211651 | 71212884 | 39.20194 | 124.28158 | 119.99248 | . | . | . | . | . | . |
| chr2 | 84810990 | 84811275 | 182 | 3' UTR | 2 | 84806229 | 84809234 | 3006 | 1 | ENSMUST00 | 4762 | Ube2l6 | ENSMUSG0C | false | chr2 | 84811070 | 84811502 | 12.39106 | 30.348 | 27.8772 | chr2 | 84810645 | 84811487 | 3.53806 | 3.37317 | 1.78021 |
| chr2 | 71786806 | 71787090 | 182 | Promoter (<=1kb) | 2 | 71786929 | 71820943 | 34015 | 1 | ENSMUST00 | 0 | Itga6 | ENSMUSG0C | false | chr2 | 71786545 | 71788373 | 31.91637 | 110.38313 | 106.53261 | chr2 | 71786536 | 71788062 | 3.7797 | 3.4796 | 2.00396 |
| chr2 | 106974365 | 106974599 | 182 | Promoter (<=1kb) | 2 | 106962527 | 106974424 | 11898 | 2 | ENSMUST00 | 0 | Arl14ep | ENSMUSG0C | false | chr2 | 106973606 | 106974761 | 26.6505 | 78.45867 | 75.33303 | chr2 | 106973065 | 106974784 | 4.27265 | 3.80927 | 2.39771 |
| chr2 | 105904505 | 105904748 | 182 | Promoter (<=1kb) | 2 | 105701027 | 105904564 | 203538 | 2 | ENSMUST00 | 0 | Elp4 | ENSMUSG0C | false | chr2 | 105903837 | 105905507 | 24.55236 | 90.57755 | 87.22021 | . | . | . | . | . | . |
| chr2 | 103796660 | 103797284 | 182 | Promoter (<=1kb) | 2 | 103765018 | 103797104 | 32087 | 2 | ENSMUST00 | 0 | Caprin1 | ENSMUSG0C | false | chr2 | 103795571 | 103798285 | 34.87302 | 114.62315 | 110.65308 | . | . | . | . | . | . |
| chr2 | 103761148 | 103761390 | 182 | Promoter (<=1kb) | 2 | 103721256 | 103761270 | 40015 | 2 | ENSMUST00 | 0 | Nat10 | ENSMUSG0C | false | chr2 | 103760437 | 103761988 | 31.40916 | 110.4849 | 106.632 | chr2 | 103759944 | 103762055 | 4.16892 | 3.72529 | 2.31549 |
| chr2 | 94111237 | 94111486 | 182 | Intron (ENSMUST00000028619.4 | 2 | 94033522 | 94066764 | 33243 | 2 | ENSMUST00 | -44474 | Hsd17b12 | ENSMUSG0C | false | . | . | . | . | . | . | chr2 | 94110928 | 94111992 | 4.18728 | 3.57345 | 2.36544 |
| chr2 | 125866090 | 125866321 | 182 | Promoter (<=1kb) | 2 | 125866107 | 125929570 | 63464 | 1 | ENSMUST00 | 0 | Galk2 | ENSMUSG0C | false | chr2 | 125866000 | 125867056 | 25.17599 | 69.56199 | 66.57701 | . | . | . | . | . | . |
| chr2 | 93458813 | 93459111 | 182 | Promoter (2-3kb) | 2 | 93421827 | 93461912 | 40086 | 2 | ENSMUST00 | 2801 | Cd82 | ENSMUSG0C | false | . | . | . | . | . | . | chr2 | 93434899 | 93464965 | 6.68053 | 4.76412 | 4.39602 |
| chr2 | 92601369 | 92601593 | 182 | Promoter (1-2kb) | 2 | 92599707 | 92615250 | 15544 | 1 | ENSMUST00 | 1663 | Chst1 | ENSMUSG0C | true | chr2 | 92599547 | 92603771 | 22.31878 | 85.15456 | 81.90936 | . | . | . | . | . | . |
| chr2 | 92659630 | 92659882 | 182 | Distal Intergenic | 2 | 92599707 | 92615250 | 15544 | 1 | ENSMUST00 | 59924 | Chst1 | ENSMUSG0C | true | . | . | . | . | . | . | chr2 | 92659016 | 92660290 | 3.80473 | 3.46038 | 2.07451 |
| chr2 | 92009606 | 92009812 | 182 | Intron (ENSMUST00000028663.4 | 2 | 91982328 | 92024502 | 42175 | 2 | ENSMUST00 | 14690 | Creb3l1 | ENSMUSG0C | false | . | . | . | . | . | . | chr2 | 92006719 | 92018935 | 6.953 | 4.90744 | 4.61859 |
| chr2 | 92012779 | 92013087 | 182 | Intron (ENSMUST00000028663.4 | 2 | 91982328 | 92024502 | 42175 | 2 | ENSMUST00 | 11415 | Creb3l1 | ENSMUSG0C | false | . | . | . | . | . | . | chr2 | 92006719 | 92018935 | 6.953 | 4.90744 | 4.61859 |
| chr2 | 121506596 | 121506795 | 182 | Promoter (<=1kb) | 2 | 121506723 | 121543404 | 36682 | 1 | ENSMUST00 | 0 | Wdr76 | ENSMUSG0C | false | chr2 | 121505826 | 121507746 | 32.98857 | 124.89146 | 120.57959 | . | . | . | . | . | . |
| chr2 | 91650063 | 91650377 | 182 | Promoter (<=1kb) | 2 | 91650118 | 91672326 | 22209 | 1 | ENSMUST00 | 0 | Arhgap1 | ENSMUSG0C | false | chr2 | 91648776 | 91651859 | 28.06786 | 91.41865 | 88.0463 | . | . | . | . | . | . |
| chr2 | 91656208 | 91656446 | 182 | Intron (ENSMUST00000111329.2 | 2 | 91653208 | 91672302 | 19095 | 1 | ENSMUST00 | 3001 | Arhgap1 | ENSMUSG0C | false | . | . | . | . | . | . | chr2 | 91650770 | 91658145 | 7.66222 | 5.14416 | 5.21273 |
| chr2 | 91265014 | 91265428 | 182 | Promoter (<=1kb) | 2 | 91265293 | 91276931 | 11639 | 1 | ENSMUST00 | 0 | Arfgap2 | ENSMUSG0C | false | chr2 | 91264604 | 91266283 | 22.68762 | 81.35053 | 78.1749 | . | . | . | . | . | . |
| chr2 | 121171052 | 121171289 | 182 | Promoter (<=1kb) | 2 | 121171168 | 121198770 | 27603 | 1 | ENSMUST00 | 0 | Tubgcp4 | ENSMUSG0C | false | chr2 | 121169593 | 121171927 | 33.52805 | 106.05383 | 102.33234 | . | . | . | . | . | . |
| chr2 | 136679756 | 136680208 | 182 | Distal Intergenic | 2 | 136713453 | 136782428 | 68976 | 1 | ENSMUST00 | -33245 | Snap25 | ENSMUSG0C | false | chr2 | 136678336 | 136680011 | 21.76566 | 71.50774 | 68.49327 | chr2 | 136677840 | 136679972 | 5.06206 | 4.18367 | 3.04713 |
| chr2 | 90846956 | 90847288 | 182 | Promoter (<=1kb) | 2 | 90847155 | 90866810 | 19656 | 1 | ENSMUST00 | 0 | Mtch2 | ENSMUSG0C | false | chr2 | 90846506 | 90848212 | 35.02417 | 111.75519 | 107.8695 | . | . | . | . | . | . |
| chr2 | 120609267 | 120609580 | 182 | Promoter (<=1kb) | 2 | 120609383 | 120619906 | 10524 | 1 | ENSMUST00 | 0 | Haus2 | ENSMUSG0C | false | chr2 | 120608534 | 120610256 | 26.18681 | 91.82585 | 88.44535 | chr2 | 120607906 | 120610603 | 4.10409 | 3.56843 | 2.29713 |
| chr2 | 120567479 | 120567974 | 182 | Promoter (<=1kb) | 2 | 120567671 | 120601255 | 33585 | 1 | ENSMUST00 | 0 | Snap23 | ENSMUSG0C | false | chr2 | 120566919 | 120568583 | 17.6846 | 53.76268 | 50.99834 | . | . | . | . | . | . |
| chr2 | 119757945 | 119758621 | 182 | Promoter (<=1kb) | 2 | 119751320 | 119758525 | 7206 | 2 | ENSMUST00 | 0 | Ltk | ENSMUSG0C | false | chr2 | 119757645 | 119759259 | 34.98938 | 113.78057 | 109.83514 | chr2 | 119757391 | 119759373 | 4.85253 | 3.83206 | 2.92321 |
| chr2 | 131996433 | 131996671 | 182 | Exon (ENSMUST00000028814.9/ | 2 | 132000922 | 132021173 | 20252 | 2 | ENSMUST00 | 24502 | Rassf2 | ENSMUSG0C | false | . | . | . | . | . | . | chr2 | 131995801 | 131998546 | 3.84517 | 3.55547 | 2.05559 |
| chr2 | 126707159 | 126707445 | 182 | Promoter (<=1kb) | 2 | 126707328 | 126759248 | 51921 | 1 | ENSMUST00 | 0 | Usp8 | ENSMUSG0C | false | chr2 | 126706747 | 126707944 | 26.1063 | 98.41757 | 94.88632 | . | . | . | . | . | . |
| chr2 | 128863858 | 128864065 | 182 | Promoter (<=1kb) | 2 | 128863932 | 128897030 | 33099 | 1 | ENSMUST00 | 0 | Fbln7 | ENSMUSG0C | false | chr2 | 128863594 | 128864410 | 18.39457 | 51.06108 | 48.33263 | chr2 | 128863762 | 128864464 | 3.28376 | 3.25595 | 1.62833 |
| chr2 | 130178717 | 130179448 | 182 | Promoter (<=1kb) | 2 | 130171414 | 130179303 | 7890 | 2 | ENSMUST00 | 0 | Snrpb | ENSMUSG0C | false | chr2 | 130177981 | 130180043 | 37.18052 | 130.46529 | 125.95659 | . | . | . | . | . | . |
| chr2 | 130284311 | 130284620 | 182 | Promoter (<=1kb) | 2 | 130279309 | 130284547 | 5239 | 2 | ENSMUST00 | 0 | Idh3b | ENSMUSG0C | false | chr2 | 130283581 | 130285094 | 34.74108 | 114.39095 | 110.42815 | chr2 | 130282852 | 130285268 | 4.03238 | 3.64409 | 2.19439 |
| chr2 | 144009469 | 144009884 | 182 | Promoter (1-2kb) | 2 | 143947395 | 144011263 | 63869 | 2 | ENSMUST00 | 1379 | Rrbp1 | ENSMUSG0C | false | chr2 | 144009741 | 144011296 | 37.3746 | 124.32532 | 120.03539 | chr2 | 144001586 | 144010150 | 4.4582 | 3.69929 | 2.57041 |
| chr2 | 153009276 | 153009469 | 182 | 3' UTR | 2 | 153009470 | 153012490 | 3021 | 2 | ENSMUST00 | 3021 | Pdrg1 | ENSMUSG0C | false | . | . | . | . | . | . | chr2 | 153008550 | 153010422 | 4.26142 | 3.47903 | 2.43324 |
| chr2 | 153649257 | 153649724 | 182 | Promoter (<=1kb) | 2 | 153649450 | 153687730 | 38281 | 1 | ENSMUST00 | 0 | Dnmt3b | ENSMUSG0C | false | chr2 | 153648932 | 153650679 | 29.66345 | 100.21 | 96.63225 | chr2 | 153648750 | 153650754 | 4.79224 | 3.89846 | 2.86691 |
| chr2 | 154407456 | 154408189 | 182 | Promoter (<=1kb) | 2 | 154376313 | 154408081 | 31769 | 2 | ENSMUST00 | 0 | Snta1 | ENSMUSG0C | false | chr2 | 154406634 | 154408534 | 31.70772 | 93.8754 | 90.44607 | . | . | . | . | . | . |
| chr2 | 154569889 | 154570239 | 182 | Promoter (<=1kb) | 2 | 154559407 | 154569892 | 10486 | 2 | ENSMUST00 | 0 | E2f1 | ENSMUSG0C | false | chr2 | 154568805 | 154570548 | 28.7177 | 82.96152 | 79.75617 | chr2 | 154555502 | 154571286 | 6.03478 | 4.51051 | 3.86945 |
| chr3 | 8948942 | 8949127 | 182 | Intron (ENSMUST00000091355.6 | 3 | 8931064 | 8962253 | 31190 | 2 | ENSMUST00 | 13126 | Tpd52 | ENSMUSG0C | false | . | . | . | . | . | . | chr3 | 8947714 | 8965965 | 5.38967 | 4.2084 | 3.32941 |
| chr2 | 173026958 | 173027181 | 182 | Intron (ENSMUST00000173393.2 | 2 | 173023050 | 173034700 | 11651 | 1 | ENSMUST00 | 3909 | Rbm38 | ENSMUSG0C | false | . | . | . | . | . | . | chr2 | 173018630 | 173028303 | 6.68949 | 4.89157 | 4.38552 |
| chr2 | 174343929 | 174344362 | 182 | Promoter (<=1kb) | 2 | 174344650 | 174345728 | 1079 | 1 | ENSMUST00 | -288 | Gnas | ENSMUSG0C | true | . | . | . | . | . | . | chr2 | 174331748 | 174347496 | 6.56852 | 4.7785 | 4.29829 |
| chr2 | 167956653 | 167956950 | 182 | Intron (ENSMUST00000147210.2 | 2 | 167951048 | 167965067 | 14020 | 1 | ENSMUST00 | 5606 | Ptpn1 | ENSMUSG0C | false | . | . | . | . | . | . | chr2 | 167933270 | 167963827 | 6.4817 | 4.58258 | 4.24812 |
| chr2 | 167949302 | 167949562 | 182 | Promoter (1-2kb) | 2 | 167951048 | 167965067 | 14020 | 1 | ENSMUST00 | -1486 | Ptpn1 | ENSMUSG0C | false | . | . | . | . | . | . | chr2 | 167933270 | 167963827 | 6.4817 | 4.58258 | 4.24812 |
| chr2 | 168955152 | 168955661 | 182 | Promoter (<=1kb) | 2 | 168893331 | 168955587 | 62257 | 2 | ENSMUST00 | 0 | Zfp64 | ENSMUSG0C | false | chr2 | 168954334 | 168956001 | 30.13579 | 99.06311 | 95.51676 | chr2 | 168954005 | 168955851 | 4.33868 | 3.72481 | 2.49881 |
| chr3 | 14653511 | 14653764 | 182 | Intron (ENSMUST00000029071.8 | 3 | 14641727 | 14663002 | 21276 | 1 | ENSMUST00 | 11785 | Car13 | ENSMUSG0C | false | . | . | . | . | . | . | chr3 | 14642905 | 14655474 | 4.82781 | 3.95203 | 2.88721 |
| chr2 | 180582637 | 180583065 | 182 | Promoter (<=1kb) | 2 | 180582712 | 180586287 | 3576 | 1 | ENSMUST00 | 0 | Mrgbp | ENSMUSG0C | false | chr2 | 180582646 | 180583426 | 25.45057 | 108.67924 | 104.87682 | chr2 | 180582299 | 180583642 | 4.67993 | 3.85844 | 2.79582 |
| chr2 | 156179414 | 156179693 | 182 | Promoter (<=1kb) | 2 | 156147246 | 156179580 | 32335 | 2 | ENSMUST00 | 0 | Rbm39 | ENSMUSG0C | false | chr2 | 156178007 | 156180253 | 34.16929 | 120.7392 | 116.5701 | . | . | . | . | . | . |
| chr2 | 157135122 | 157135359 | 182 | Promoter (<=1kb) | 2 | 157097533 | 157135265 | 37733 | 2 | ENSMUST00 | 0 | Samhd1 | ENSMUSG0C | false | chr2 | 157133813 | 157135626 | 22.03264 | 83.65932 | 80.44393 | chr2 | 157129981 | 157136107 | 4.39307 | 3.81834 | 2.49612 |
| chr3 | 32405166 | 32405401 | 182 | Intron (ENSMUST00000108243.2 | 3 | 32397671 | 32466107 | 68437 | 1 | ENSMUST00 | 7496 | Pik3ca | ENSMUSG0C | false | . | . | . | . | . | . | chr3 | 32401518 | 32410064 | 5.45009 | 4.2562 | 3.39377 |
| chr3 | 32708339 | 32708795 | 182 | Promoter (<=1kb) | 3 | 32708546 | 32726968 | 18423 | 1 | ENSMUST00 | 0 | Actl6a | ENSMUSG0C | false | chr3 | 32708011 | 32708374 | 13.2347 | 31.16438 | 28.68482 | . | . | . | . | . | . |
| chr3 | 32736953 | 32737152 | 182 | Promoter (<=1kb) | 3 | 32736990 | 32751566 | 14577 | 1 | ENSMUST00 | 0 | Ndufb5 | ENSMUSG0C | false | chr3 | 32737005 | 32737938 | 30.22771 | 102.23431 | 98.60725 | chr3 | 32736075 | 32739879 | 4.17297 | 3.52331 | 2.36424 |
| chr3 | 33799672 | 33800361 | 182 | Promoter (<=1kb) | 3 | 33799955 | 33814860 | 14906 | 1 | ENSMUST00 | 0 | Ttc14 | ENSMUSG0C | false | chr3 | 33799366 | 33800882 | 39.62653 | 133.71027 | 129.07751 | chr3 | 33799234 | 33799678 | 4.77517 | 4.04371 | 2.80875 |
| chr3 | 30508296 | 30508574 | 182 | Promoter (<=1kb) | 3 | 30236569 | 30509487 | 272919 | 2 | ENSMUST00 | 913 | Mecom | ENSMUSG0C | false | chr3 | 30507549 | 30509587 | 14.66885 | 35.63733 | 33.10225 | chr3 | 30506751 | 30509701 | 5.62817 | 4.3849 | 3.51508 |
| chr3 | 29951082 | 29951380 | 182 | 3' UTR | 3 | 29952716 | 29997663 | 44948 | 2 | ENSMUST00 | 46283 | Mecom | ENSMUSG0C | false | . | . | . | . | . | . | . | . | . | . | . | . |
| chr3 | 36476525 | 36476887 | 182 | Promoter (<=1kb) | 3 | 36475937 | 36482299 | 6363 | 1 | ENSMUST00 | 589 | 1810062G17 | ENSMUSG0C | false | chr3 | 36475860 | 36477050 | 26.38351 | 74.32498 | 71.26698 | chr3 | 36476431 | 36477881 | 3.58383 | 3.36033 | 1.826 |
| chr3 | 53863663 | 53864180 | 182 | Promoter (<=1kb) | 3 | 53855212 | 53863830 | 8619 | 2 | ENSMUST00 | 0 | Ufm1 | ENSMUSG0C | false | chr3 | 53862964 | 53863991 | 42.23227 | 138.25212 | 133.44542 | . | . | . | . | . | . |
| chr3 | 60563633 | 60563861 | 182 | Intron (ENSMUST00000099087.2 | 3 | 60501252 | 60629748 | 128497 | 1 | ENSMUST00 | 62382 | Mbnl1 | ENSMUSG0C | false | . | . | . | . | . | . | chr3 | 60558656 | 60566617 | 4.34049 | 3.63722 | 2.50486 |
| chr3 | 68676885 | 68677084 | 182 | Distal Intergenic | 3 | 68690644 | 68698547 | 7904 | 1 | ENSMUST00 | -13560 | Il12a | ENSMUSG0C | false | . | . | . | . | . | . | chr3 | 68675981 | 68677502 | 3.54205 | 3.39641 | 1.81562 |
| chr3 | 68606716 | 68607032 | 182 | Intron (ENSMUST00000029346.7 | 3 | 68596175 | 68598163 | 1989 | 1 | ENSMUST00 | 10542 | Schip1 | ENSMUSG0C | false | . | . | . | . | . | . | chr3 | 68574597 | 68620372 | 5.61397 | 4.36647 | 3.51459 |
| chr3 | 69721821 | 69722412 | 182 | Promoter (<=1kb) | 3 | 69721985 | 69724461 | 2477 | 1 | ENSMUST00 | 0 | Nmd3 | ENSMUSG0C | false | chr3 | 69721465 | 69723120 | 34.22706 | 108.29755 | 104.50865 | . | . | . | . | . | . |
| chr3 | 70102894 | 70103111 | 182 | Distal Intergenic | 3 | 70007613 | 70028708 | 21096 | 1 | ENSMUST00 | 95282 | Otol1 | ENSMUSG0C | false | . | . | . | . | . | . | chr3 | 70102314 | 70104149 | 4.50617 | 3.78192 | 2.61106 |
| chr3 | 57572717 | 57572916 | 182 | Promoter (2-3kb) | 3 | 57455649 | 57575828 | 120180 | 2 | ENSMUST00 | 2912 | Wwtr1 | ENSMUSG0C | false | . | . | . | . | . | . | chr3 | 57570735 | 57574583 | 4.8265 | 3.86007 | 2.89702 |
| chr3 | 63964536 | 63964828 | 182 | Promoter (<=1kb) | 3 | 63933507 | 63964666 | 31160 | 2 | ENSMUST00 | 0 | Slc33a1 | ENSMUSG0C | false | chr3 | 63962339 | 63965122 | 33.96572 | 122.48407 | 118.2566 | . | . | . | . | . | . |
| chr3 | 65929499 | 65929770 | 182 | Distal Intergenic | 3 | 65946782 | 65947998 | 1217 | 2 | ENSMUST00 | 18228 | Ccnl1 | ENSMUSG0C | false | . | . | . | . | . | . | chr3 | 65926001 | 65930446 | 5.38488 | 4.15152 | 3.33208 |
| chr3 | 103058192 | 103058441 | 182 | Promoter (<=1kb) | 3 | 103058339 | 103067914 | 9576 | 1 | ENSMUST00 | 0 | Nras | ENSMUSG0C | true | chr3 | 103057815 | 103059145 | 36.809 | 123.33047 | 119.07505 | . | . | . | . | . | . |
| chr3 | 102152951 | 102153159 | 182 | Distal Intergenic | 3 | 102144051 | 102145585 | 1535 | 1 | ENSMUST00 | 8901 | Casq2 | ENSMUSG0C | false | . | . | . | . | . | . | chr3 | 102152076 | 102154358 | 4.80732 | 3.99358 | 2.87432 |
| chr3 | 98024381 | 98024762 | 182 | Intron (ENSMUST00000079812.6 | 3 | 98013538 | 98150367 | 136830 | 1 | ENSMUST00 | 10844 | Notch2 | ENSMUSG0C | false | . | . | . | . | . | . | chr3 | 98014795 | 98071300 | 5.94878 | 4.48977 | 3.79636 |
| chr3 | 98013815 | 98014240 | 182 | Promoter (<=1kb) | 3 | 98013538 | 98150367 | 136830 | 1 | ENSMUST00 | 278 | Notch2 | ENSMUSG0C | false | chr3 | 98012900 | 98014942 | 35.22365 | 121.58154 | 117.38489 | . | . | . | . | . | . |
| chr3 | 107714500 | 107714732 | 182 | Distal Intergenic | 3 | 107663120 | 107696560 | 33441 | 2 | ENSMUST00 | -17941 | Ahcyl1 | ENSMUSG0C | false | . | . | . | . | . | . | chr3 | 107712381 | 107715538 | 4.67141 | 3.813 | 2.76803 |
| chr3 | 107696164 | 107696706 | 182 | Promoter (<=1kb) | 3 | 107663120 | 107696560 | 33441 | 2 | ENSMUST00 | 0 | Ahcyl1 | ENSMUSG0C | false | chr3 | 107695484 | 107696643 | 36.67291 | 122.02113 | 117.81725 | . | . | . | . | . | . |
| chr3 | 105687430 | 105687697 | 182 | Promoter (<=1kb) | 3 | 105678270 | 105687574 | 9305 | 2 | ENSMUST00 | 0 | Ddx20 | ENSMUSG0C | false | chr3 | 105686766 | 105688240 | 32.64849 | 123.19247 | 118.94151 | . | . | . | . | . | . |
| chr3 | 93520386 | 93520610 | 182 | Promoter (<=1kb) | 3 | 93520488 | 93526287 | 5800 | 1 | ENSMUST00 | 0 | S100a11 | ENSMUSG0C | false | chr3 | 93520371 | 93523048 | 29.97852 | 88.95019 | 85.62817 | chr3 | 93520128 | 93524973 | 5.6407 | 4.38912 | 3.52992 |
| chr3 | 93014502 | 93014854 | 182 | Promoter (<=1kb) | 3 | 93014519 | 93015669 | 1151 | 2 | ENSMUST00 | 815 | Crct1 | ENSMUSG0C | false | chr3 | 93014414 | 93014905 | 12.8954 | 29.30041 | 26.84444 | chr3 | 93014160 | 93014902 | 3.281 | 2.90369 | 1.62888 |
| chr3 | 90464516 | 90464776 | 182 | Promoter (1-2kb) | 3 | 90450591 | 90465866 | 15276 | 2 | ENSMUST00 | 1090 | Npr1 | ENSMUSG0C | false | chr3 | 90462429 | 90466691 | 46.53382 | 157.01303 | 151.46753 | . | . | . | . | . | . |
| chr3 | 89913058 | 89913365 | 182 | Promoter (<=1kb) | 3 | 89869324 | 89913162 | 43839 | 2 | ENSMUST00 | 0 | Il6ra | ENSMUSG0C | true | chr3 | 89912092 | 89913983 | 31.16539 | 106.97112 | 103.21663 | . | . | . | . | . | . |
| chr3 | 114031853 | 114032052 | 182 | Promoter (1-2kb) | 3 | 114030540 | 114220718 | 190179 | 1 | ENSMUST00 | 1314 | Col11a1 | ENSMUSG0C | false | . | . | . | . | . | . | chr3 | 114030118 | 114032263 | 4.50916 | 3.84721 | 2.64421 |
| chr3 | 123446762 | 123447133 | 182 | Promoter (<=1kb) | 3 | 123446913 | 123506602 | 59690 | 1 | ENSMUST00 | 0 | Prss12 | ENSMUSG0C | false | chr3 | 123446595 | 123447803 | 28.00751 | 87.44511 | 84.14996 | chr3 | 123446370 | 123448826 | 3.85205 | 3.41691 | 2.06909 |
| chr3 | 131111988 | 131112563 | 182 | Promoter (<=1kb) | 3 | 131112797 | 131117387 | 4591 | 1 | ENSMUST00 | -234 | Lef1 | ENSMUSG0C | false | chr3 | 131110142 | 131112789 | 22.5292 | 69.37365 | 66.39268 | chr3 | 131109267 | 131113191 | 5.50748 | 4.3265 | 3.44397 |
| chr3 | 129878399 | 129878723 | 182 | Promoter (<=1kb) | 3 | 129878622 | 129895821 | 17200 | 1 | ENSMUST00 | 0 | Pla2g12a | ENSMUSG0C | false | chr3 | 129878062 | 129879464 | 36.12563 | 122.12295 | 117.91594 | . | . | . | . | . | . |
| chr3 | 129769590 | 129769911 | 182 | Distal Intergenic | 3 | 129677575 | 129755322 | 77748 | 2 | ENSMUST00 | -14269 | Egf | ENSMUSG0C | false | . | . | . | . | . | . | chr3 | 129766906 | 129771147 | 5.1145 | 4.00179 | 3.12888 |
| chr3 | 129313299 | 129313581 | 182 | Intron (ENSMUST00000029658.8 | 3 | 129276764 | 129309034 | 32271 | 2 | ENSMUST00 | -4266 | Enpep | ENSMUSG0C | true | . | . | . | . | . | . | chr3 | 129303627 | 129327892 | 5.35019 | 4.2164 | 3.29511 |
| chr3 | 131489969 | 131490504 | 182 | Distal Intergenic | 3 | 131564768 | 131643670 | 78903 | 1 | ENSMUST00 | -74264 | Papss1 | ENSMUSG0C | false | chr3 | 131489555 | 131490997 | 21.54279 | 65.84157 | 62.9152 | chr3 | 131489608 | 131491228 | 6.01991 | 4.59933 | 3.8277 |
| chr1 | 21961477 | 21961749 | 182 | Promoter (<=1kb) | 1 | 21398403 | 21961581 | 563179 | 2 | ENSMUST00 | 0 | Kcnq5 | ENSMUSG0C | false | chr1 | 21960440 | 21962504 | 16.02374 | 53.68177 | 50.91932 | chr1 | 21960522 | 21963173 | 4.13536 | 3.59427 | 2.33272 |
| chr3 | 88621646 | 88621940 | 182 | Promoter (<=1kb) | 3 | 88621555 | 88646929 | 25375 | 1 | ENSMUST00 | 92 | Arhgef2 | ENSMUSG0C | false | chr3 | 88620710 | 88623767 | 28.41342 | 89.1249 | 85.7944 | chr3 | 88621743 | 88631594 | 4.65392 | 3.86367 | 2.72435 |
| chr3 | 88552798 | 88553019 | 182 | Promoter (<=1kb) | 3 | 88549820 | 88552859 | 3040 | 2 | ENSMUST00 | 0 | Lamtor2 | ENSMUSG0C | false | chr3 | 88552107 | 88553013 | 30.75439 | 93.21284 | 89.80341 | . | . | . | . | . | . |
| chr3 | 87763952 | 87764343 | 182 | Promoter (1-2kb) | 3 | 87749098 | 87762214 | 13117 | 2 | ENSMUST00 | -1739 | Pear1 | ENSMUSG0C | false | chr3 | 87763938 | 87764321 | 8.73111 | 16.23101 | 13.94854 | chr3 | 87763394 | 87765267 | 4.19285 | 3.65031 | 2.35304 |
| chr3 | 86852887 | 86853101 | 182 | Intron (ENSMUST00000029719.8 | 3 | 86786150 | 86920884 | 134735 | 2 | ENSMUST00 | 67783 | Dclk2 | ENSMUSG0C | false | . | . | . | . | . | . | chr3 | 86846839 | 86853593 | 3.95845 | 3.57179 | 2.16593 |
| chr3 | 86038642 | 86038886 | 182 | Promoter (<=1kb) | 3 | 86038877 | 86083294 | 44418 | 1 | ENSMUST00 | 0 | Sh3d19 | ENSMUSG0C | false | . | . | . | . | . | . | chr3 | 86038138 | 86041689 | 5.84822 | 4.4832 | 3.73307 |
| chr3 | 85971809 | 85972070 | 182 | Promoter (<=1kb) | 3 | 85971109 | 86083538 | 112430 | 1 | ENSMUST00 | 701 | Sh3d19 | ENSMUSG0C | false | chr3 | 85970752 | 85972621 | 36.31701 | 113.99837 | 110.04225 | . | . | . | . | . | . |
| chr3 | 96593916 | 96594418 | 182 | Promoter (<=1kb) | 3 | 96577872 | 96594181 | 16310 | 2 | ENSMUST00 | 0 | Polr3gl | ENSMUSG0C | false | chr3 | 96593397 | 96594507 | 21.91997 | 62.19815 | 59.32085 | . | . | . | . | . | . |
| chr3 | 123385688 | 123386069 | 182 | Promoter (<=1kb) | 3 | 123364645 | 123385948 | 21304 | 2 | ENSMUST00 | 0 | Mettl14 | ENSMUSG0C | false | chr3 | 123385144 | 123386655 | 31.70772 | 93.8754 | 90.44607 | . | . | . | . | . | . |
| chr3 | 95106694 | 95107062 | 182 | Promoter (<=1kb) | 3 | 95058530 | 95106875 | 48346 | 2 | ENSMUST00 | 0 | Pip5k1a | ENSMUSG0C | true | chr3 | 95105324 | 95107428 | 28.22848 | 94.63612 | 91.19299 | . | . | . | . | . | . |
| chr3 | 121171333 | 121171836 | 182 | Promoter (<=1kb) | 3 | 121155400 | 121171695 | 16296 | 2 | ENSMUST00 | 0 | Rwdd3 | ENSMUSG0C | false | chr3 | 121171056 | 121172163 | 22.64381 | 71.35664 | 68.34631 | chr3 | 121171105 | 121172460 | 4.07414 | 3.50472 | 2.28857 |
| chr3 | 138753291 | 138753727 | 182 | Intron (ENSMUST00000029800.3 | 3 | 138742769 | 138894484 | 151716 | 1 | ENSMUST00 | 10523 | Tspan5 | ENSMUSG0C | true | . | . | . | . | . | . | chr3 | 138743273 | 138755777 | 5.55338 | 4.30175 | 3.46487 |
| chr3 | 135485463 | 135485747 | 182 | Promoter (<=1kb) | 3 | 135485611 | 135555566 | 69956 | 1 | ENSMUST00 | 0 | Manba | ENSMUSG0C | false | chr3 | 135485185 | 135486284 | 32.8888 | 98.87808 | 95.33504 | . | . | . | . | . | . |
| chr3 | 159840203 | 159840677 | 182 | Promoter (<=1kb) | 3 | 159839737 | 159935833 | 96097 | 1 | ENSMUST00 | 467 | Wls | ENSMUSG0C | false | chr3 | 159839398 | 159841336 | 24.71438 | 74.15884 | 71.1036 | . | . | . | . | . | . |
| chr3 | 157534133 | 157534725 | 182 | Promoter (<=1kb) | 3 | 157534160 | 157548390 | 14231 | 1 | ENSMUST00 | 0 | Zranb2 | ENSMUSG0C | false | chr3 | 157533711 | 157535409 | 26.81296 | 97.85361 | 94.33582 | . | . | . | . | . | . |
| chr3 | 148892961 | 148893179 | 182 | Intron (ENSMUST00000106128.2 | 3 | 148815586 | 148954635 | 139050 | 2 | ENSMUST00 | 61456 | Lphn2 | ENSMUSG0C | false | . | . | . | . | . | . | chr3 | 148884697 | 148902761 | 4.68075 | 3.92083 | 2.75759 |
| chr3 | 145758787 | 145759183 | 182 | Promoter (<=1kb) | 3 | 145758675 | 145894277 | 135603 | 1 | ENSMUST00 | 113 | Ddah1 | ENSMUSG0C | false | chr3 | 145758375 | 145759815 | 28.33848 | 93.85651 | 90.42933 | chr3 | 145758206 | 145760257 | 5.47326 | 4.30929 | 3.40299 |
| chr3 | 145728359 | 145728610 | 182 | Distal Intergenic | 3 | 145758675 | 145894277 | 135603 | 1 | ENSMUST00 | -30065 | Ddah1 | ENSMUSG0C | false | . | . | . | . | . | . | chr3 | 145728346 | 145730104 | 4.30419 | 3.67711 | 2.47637 |
| chr3 | 154596733 | 154597201 | 182 | Promoter (<=1kb) | 3 | 154597197 | 154611556 | 14360 | 1 | ENSMUST00 | 0 | Cryz | ENSMUSG0C | false | chr3 | 154596276 | 154597735 | 36.88168 | 122.38801 | 118.16834 | . | . | . | . | . | . |
| chr4 | 9669055 | 9669450 | 182 | Promoter (<=1kb) | 4 | 9449079 | 9669162 | 220084 | 2 | ENSMUST00 | 0 | Asph | ENSMUSG0C | false | chr4 | 9668334 | 9669818 | 35.98183 | 119.73972 | 115.60696 | chr4 | 9667826 | 9669885 | 3.99333 | 3.45109 | 2.21683 |
| chr4 | 15945229 | 15945542 | 182 | Promoter (<=1kb) | 4 | 15917240 | 15945507 | 28268 | 2 | ENSMUST00 | 0 | Decr1 | ENSMUSG0C | false | chr4 | 15944626 | 15945984 | 27.39617 | 87.32104 | 84.03119 | . | . | . | . | . | . |
| chr4 | 17853272 | 17853493 | 182 | Promoter (<=1kb) | 4 | 17853458 | 18119145 | 265688 | 1 | ENSMUST00 | 0 | Mmp16 | ENSMUSG0C | false | chr4 | 17852871 | 17854099 | 21.52556 | 63.2201 | 60.32722 | chr4 | 17853328 | 17853991 | 3.46693 | 3.35012 | 1.76963 |
| chr4 | 19574934 | 19575293 | 182 | Promoter (<=1kb) | 4 | 19575162 | 19598654 | 23493 | 1 | ENSMUST00 | 0 | Rmdn1 | ENSMUSG0C | false | chr4 | 19574627 | 19575981 | 34.3275 | 112.61823 | 108.70669 | . | . | . | . | . | . |
| chr4 | 21848370 | 21848580 | 182 | Promoter (<=1kb) | 4 | 21848081 | 21862123 | 14043 | 1 | ENSMUST00 | 290 | Pnisr | ENSMUSG0C | false | chr4 | 21847275 | 21849107 | 39.70409 | 136.04816 | 131.32747 | . | . | . | . | . | . |
| chr3 | 145118656 | 145118891 | 182 | Promoter (<=1kb) | 3 | 145118600 | 145153915 | 35316 | 1 | ENSMUST00 | 57 | Odf2l | ENSMUSG0C | false | chr3 | 145118077 | 145119514 | 20.82289 | 72.3363 | 69.31286 | chr3 | 145116849 | 145120125 | 4.03643 | 3.5929 | 2.22585 |
| chr3 | 142618745 | 142618984 | 182 | Promoter (1-2kb) | 3 | 142620602 | 142638008 | 17407 | 1 | ENSMUST00 | -1618 | Gbp2 | ENSMUSG0C | false | . | . | . | . | . | . | chr3 | 142618545 | 142621601 | 4.06897 | 3.59973 | 2.25628 |
| chr3 | 142394533 | 142394913 | 182 | Promoter (<=1kb) | 3 | 142239586 | 142395696 | 156111 | 2 | ENSMUST00 | 783 | Pdlim5 | ENSMUSG0C | false | chr3 | 142393844 | 142396226 | 35.56825 | 117.95176 | 113.87694 | . | . | . | . | . | . |
| chr3 | 142348792 | 142349040 | 182 | Intron (ENSMUST00000029941.1 | 3 | 142302080 | 142391850 | 89771 | 2 | ENSMUST00 | 42810 | Pdlim5 | ENSMUSG0C | false | . | . | . | . | . | . | chr3 | 142342654 | 142360103 | 4.39932 | 3.72752 | 2.52416 |
| chr3 | 142395280 | 142395791 | 182 | Promoter (<=1kb) | 3 | 142239586 | 142395696 | 156111 | 2 | ENSMUST00 | 0 | Pdlim5 | ENSMUSG0C | false | chr3 | 142393844 | 142396226 | 35.56825 | 117.95176 | 113.87694 | . | . | . | . | . | . |
| chr4 | 46553837 | 46554034 | 182 | Intron (ENSMUST00000030021.8 | 4 | 46541316 | 46542802 | 1487 | 2 | ENSMUST00 | -11036 | Coro2a | ENSMUSG0C | false | . | . | . | . | . | . | . | . | . | . | . | . |
| chr4 | 48473350 | 48473587 | 182 | Promoter (<=1kb) | 4 | 48430858 | 48473459 | 42602 | 2 | ENSMUST00 | 0 | Tex10 | ENSMUSG0C | false | chr4 | 48472261 | 48473732 | 32.49479 | 112.46503 | 108.55889 | . | . | . | . | . | . |
| chr4 | 48585315 | 48585798 | 182 | Promoter (<=1kb) | 4 | 48585365 | 48662085 | 76721 | 1 | ENSMUST00 | 0 | Tmeff1 | ENSMUSG0C | false | chr4 | 48584659 | 48586353 | 24.03115 | 75.67385 | 72.59326 | chr4 | 48584849 | 48586653 | 5.18318 | 4.19996 | 3.17741 |
| chr4 | 66388729 | 66388950 | 182 | Intron (ENSMUST00000084496.2 | 4 | 65380803 | 66404483 | 1023681 | 2 | ENSMUST00 | 15533 | Astn2 | ENSMUSG0C | true | . | . | . | . | . | . | chr4 | 66387210 | 66390107 | 3.99902 | 3.61784 | 2.18293 |
| chr4 | 62518485 | 62519000 | 182 | Promoter (<=1kb) | 4 | 62509169 | 62519881 | 10713 | 2 | ENSMUST00 | 881 | Alad | ENSMUSG0C | false | chr4 | 62518215 | 62520467 | 33.87313 | 113.90014 | 109.94483 | chr4 | 62514012 | 62519352 | 6.75426 | 4.89888 | 4.45045 |
| chr4 | 62514983 | 62515365 | 182 | Intron (ENSMUST00000137448.1 | 4 | 62509169 | 62519881 | 10713 | 2 | ENSMUST00 | 4516 | Alad | ENSMUSG0C | false | chr4 | 62515264 | 62516219 | 8.2976 | 15.51635 | 13.24704 | chr4 | 62514012 | 62519352 | 6.75426 | 4.89888 | 4.45045 |
| chr4 | 76327311 | 76327642 | 182 | Promoter (2-3kb) | 4 | 76140358 | 76325288 | 184931 | 2 | ENSMUST00 | -2024 | Ptprd | ENSMUSG0C | true | chr4 | 76327299 | 76327958 | 8.45881 | 17.21912 | 14.92473 | chr4 | 76323260 | 76332890 | 4.03079 | 3.5236 | 2.22355 |
| chr4 | 53450876 | 53451119 | 182 | Intron (ENSMUST00000107646.3 | 4 | 53440684 | 53563450 | 122767 | 1 | ENSMUST00 | 10193 | Slc44a1 | ENSMUSG0C | false | . | . | . | . | . | . | chr4 | 53449540 | 53452110 | 4.05659 | 3.45789 | 2.27426 |
| chr4 | 40853856 | 40854373 | 182 | Promoter (<=1kb) | 4 | 40804602 | 40854005 | 49404 | 2 | ENSMUST00 | 0 | B4galt1 | ENSMUSG0C | true | chr4 | 40851570 | 40854097 | 37.91348 | 118.86035 | 114.75604 | . | . | . | . | . | . |
| chr4 | 41314699 | 41315012 | 182 | Promoter (<=1kb) | 4 | 41291300 | 41314889 | 23590 | 2 | ENSMUST00 | 0 | Dcaf12 | ENSMUSG0C | false | chr4 | 41313584 | 41315385 | 37.0612 | 128.08391 | 123.66181 | . | . | . | . | . | . |
| chr4 | 41569278 | 41569817 | 182 | Promoter (<=1kb) | 4 | 41517691 | 41569509 | 51819 | 2 | ENSMUST00 | 0 | Fam219a | ENSMUSG0C | false | chr4 | 41568168 | 41569606 | 34.11661 | 104.01456 | 100.34361 | chr4 | 41563319 | 41570177 | 5.29575 | 4.11611 | 3.25298 |
| chr4 | 43558359 | 43558581 | 182 | Promoter (<=1kb) | 4 | 43555106 | 43558460 | 3355 | 2 | ENSMUST00 | 0 | Tln1 | ENSMUSG0C | false | chr4 | 43558329 | 43559317 | 6.00779 | 10.46335 | 8.27668 | chr4 | 43556567 | 43561682 | 7.2451 | 5.00392 | 4.84927 |
| chr4 | 86534209 | 86534445 | 182 | Intron (ENSMUST00000151481.2 | 4 | 86444641 | 86539117 | 94477 | 2 | ENSMUST00 | 4672 | Fam154a | ENSMUSG0C | false | . | . | . | . | . | . | chr4 | 86532837 | 86535151 | 5.83243 | 4.50475 | 3.71514 |
| chr4 | 87934205 | 87934496 | 182 | Intron (ENSMUST00000078090.6 | 4 | 87832781 | 87874542 | 41762 | 2 | ENSMUST00 | -59664 | Mllt3 | ENSMUSG0C | false | . | . | . | . | . | . | chr4 | 87933223 | 87936886 | 6.33341 | 4.75754 | 4.09872 |
| chr4 | 102639957 | 102640257 | 182 | Distal Intergenic | 4 | 102589843 | 102603441 | 13599 | 1 | ENSMUST00 | 50115 | Pde4b | ENSMUSG0C | true | . | . | . | . | . | . | chr4 | 102637493 | 102641859 | 6.17662 | 4.64515 | 3.9647 |
| chr4 | 101420011 | 101420495 | 182 | Promoter (<=1kb) | 4 | 101420497 | 101463680 | 43184 | 1 | ENSMUST00 | -2 | Ak4 | ENSMUSG0C | false | chr4 | 101419025 | 101420579 | 33.10969 | 106.34647 | 102.61562 | chr4 | 101420112 | 101420785 | 3.30385 | 3.11841 | 1.6572 |
| chr4 | 101264637 | 101265241 | 182 | Promoter (<=1kb) | 4 | 101181912 | 101265223 | 83312 | 2 | ENSMUST00 | 0 | Jak1 | ENSMUSG0C | false | chr4 | 101263825 | 101265786 | 39.02359 | 127.20569 | 122.81842 | . | . | . | . | . | . |
| chr4 | 101254035 | 101254294 | 182 | Intron (ENSMUST00000102781.4 | 4 | 101181912 | 101265223 | 83312 | 2 | ENSMUST00 | 10929 | Jak1 | ENSMUSG0C | false | . | . | . | . | . | . | chr4 | 101243772 | 101264376 | 5.46033 | 4.21492 | 3.39754 |
| chr4 | 101185312 | 101185580 | 182 | Promoter (<=1kb) | 4 | 101179591 | 101185425 | 5835 | 2 | ENSMUST00 | 0 | Jak1 | ENSMUSG0C | false | . | . | . | . | . | . | . | . | . | . | . | . |
| chr4 | 100992481 | 100992726 | 182 | Promoter (1-2kb) | 4 | 100994611 | 101003061 | 8451 | 1 | ENSMUST00 | -1885 | Cachd1 | ENSMUSG0C | false | . | . | . | . | . | . | chr4 | 100990036 | 100996788 | 4.59397 | 3.79686 | 2.68268 |
| chr4 | 100865448 | 100865690 | 182 | Intron (ENSMUST00000030257.9 | 4 | 100776675 | 101029220 | 252546 | 1 | ENSMUST00 | 88774 | Cachd1 | ENSMUSG0C | false | . | . | . | . | . | . | chr4 | 100863785 | 100867033 | 4.29863 | 3.76952 | 2.43386 |
| chr4 | 100777641 | 100778035 | 182 | Promoter (<=1kb) | 4 | 100776675 | 101029220 | 252546 | 1 | ENSMUST00 | 967 | Cachd1 | ENSMUSG0C | false | chr4 | 100776119 | 100778615 | 28.83684 | 94.75451 | 91.30991 | chr4 | 100777967 | 100779974 | 3.55423 | 3.40887 | 1.8082 |
| chr4 | 111414979 | 111415395 | 182 | Promoter (<=1kb) | 4 | 111415006 | 111460298 | 45293 | 1 | ENSMUST00 | 0 | Bend5 | ENSMUSG0C | false | chr4 | 111414260 | 111415782 | 35.20524 | 115.20684 | 111.21615 | chr4 | 111414088 | 111417075 | 3.75716 | 3.33992 | 2.02228 |
| chr4 | 99033052 | 99033320 | 182 | Promoter (<=1kb) | 4 | 99033179 | 99037959 | 4781 | 1 | ENSMUST00 | 0 | Angptl3 | ENSMUSG0C | false | . | . | . | . | . | . | chr4 | 99032106 | 99034372 | 4.29194 | 3.80546 | 2.40976 |
| chr4 | 97868608 | 97869191 | 182 | Intron (ENSMUST00000152023.2 | 4 | 97880119 | 98065355 | 185237 | 1 | ENSMUST00 | -10928 | Nfia | ENSMUSG0C | false | chr4 | 97868386 | 97869737 | 11.74122 | 27.32396 | 24.89351 | chr4 | 97866265 | 97870412 | 4.08618 | 3.54965 | 2.28233 |
| chr4 | 97990526 | 97990738 | 182 | Intron (ENSMUST00000152023.2 | 4 | 97911033 | 98042595 | 131563 | 1 | ENSMUST00 | 79494 | Nfia | ENSMUSG0C | false | . | . | . | . | . | . | chr4 | 97988906 | 97993591 | 4.11434 | 3.63182 | 2.28144 |
| chr4 | 130629444 | 130629699 | 182 | Distal Intergenic | 4 | 130663321 | 130781564 | 118244 | 1 | ENSMUST00 | -33622 | Pum1 | ENSMUSG0C | false | chr4 | 130629533 | 130629813 | 7.81205 | 13.8349 | 11.58824 | . | . | . | . | . | . |
| chr4 | 108619837 | 108620061 | 182 | Promoter (<=1kb) | 4 | 108619940 | 108634123 | 14184 | 1 | ENSMUST00 | 0 | Cc2d1b | ENSMUSG0C | false | chr4 | 108619225 | 108620804 | 33.25048 | 115.91603 | 111.90475 | chr4 | 108618687 | 108621017 | 4.86705 | 3.89876 | 2.92237 |
| chr4 | 108071340 | 108071525 | 182 | Promoter (<=1kb) | 4 | 108044378 | 108071363 | 26986 | 2 | ENSMUST00 | 0 | Scp2 | ENSMUSG0C | false | chr4 | 108070413 | 108071809 | 17.96111 | 52.67509 | 49.92405 | chr4 | 108068357 | 108072272 | 3.37568 | 3.16186 | 1.69525 |
| chr4 | 120924949 | 120925179 | 182 | Promoter (<=1kb) | 4 | 120921196 | 120925005 | 3810 | 2 | ENSMUST00 | 0 | Exo5 | ENSMUSG0C | false | chr4 | 120924201 | 120925562 | 25.82118 | 72.45564 | 69.42885 | chr4 | 120924071 | 120925911 | 4.46025 | 3.76801 | 2.61928 |
| chr4 | 120569861 | 120570355 | 182 | Promoter (<=1kb) | 4 | 120539868 | 120570276 | 30409 | 2 | ENSMUST00 | 0 | Ctps | ENSMUSG0C | false | chr4 | 120569090 | 120570929 | 22.60491 | 86.65532 | 83.38163 | . | . | . | . | . | . |
| chr4 | 119977821 | 119978051 | 182 | Intron (ENSMUST00000143837.1 | 4 | 120094489 | 120134557 | 40069 | 1 | ENSMUST00 | -116438 | Hivep3 | ENSMUSG0C | false | . | . | . | . | . | . | . | . | . | . | . | . |
| chr4 | 120082036 | 120082374 | 182 | Intron (ENSMUST00000106307.3 | 4 | 120094489 | 120134557 | 40069 | 1 | ENSMUST00 | -12115 | Hivep3 | ENSMUSG0C | false | . | . | . | . | . | . | chr4 | 120080308 | 120082813 | 4.50108 | 3.80371 | 2.64199 |
| chr4 | 120032327 | 120032858 | 182 | Intron (ENSMUST00000106307.3 | 4 | 120094489 | 120134557 | 40069 | 1 | ENSMUST00 | -61631 | Hivep3 | ENSMUSG0C | false | chr4 | 120032346 | 120033678 | 23.88927 | 69.11851 | 66.14216 | chr4 | 120032328 | 120034133 | 5.051 | 4.15045 | 3.06122 |
| chr4 | 123904826 | 123905326 | 182 | Promoter (<=1kb) | 4 | 123904832 | 123912269 | 7438 | 1 | ENSMUST00 | 0 | Mycbp | ENSMUSG0C | false | chr4 | 123904277 | 123905645 | 33.7938 | 119.00099 | 114.88904 | chr4 | 123904299 | 123906905 | 3.68666 | 3.42934 | 1.94229 |

|||||||||||||||||||||||||||
|---|---|---|---|---|---|---|---|---|---|---|---|---|---|---|---|---|---|---|---|---|---|---|---|---|---|---|
|chr4|123572484|123572830|182|Intron (ENSMUST00000097897.5|4|123538880|123564694|25815|2|ENSMUST00|-7791|Macf1|ENSMUSG0C|true|chr4|123572504|123573089|6.00779|10.46335|8.27668|chr4|123558735|123586068|5.19694|4.11261|3.17674|
|chr4|123043772|123044118|182|Exon (ENSMUST00000102649.3/|4|123048756|123054203|5448|1|ENSMUST00|-4638|Trit1|ENSMUSG0C|false|.|.|.|.|.|.|chr4|123041465|123048354|6.50436|4.7749|4.23787|
|chr4|135972432|135972730|182|Promoter (<=1kb)|4|135968224|135972626|4403|2|ENSMUST00|0|Lypla2|ENSMUSG0C|false|chr4|135971920|135973034|35.75768|125.74667|121.41521|chr4|135970686|135973478|4.17679|3.64304|2.36254|
|chr4|135876082|135876265|182|Promoter (2-3kb)|4|135870918|135873850|2933|2|ENSMUST00|-2233|Pnrc2|ENSMUSG0C|false|chr4|135876153|135877006|10.96168|23.03899|20.66252|chr4|135874210|135879466|7.31253|5.17261|4.90109|
|chr4|116994365|116994579|182|Promoter (<=1kb)|4|116989965|116994413|4449|2|ENSMUST00|0|Urod|ENSMUSG0C|false|chr4|116993601|116994502|22.8003|62.86544|59.97937|chr4|116992876|116996990|5.81634|4.52882|3.66042|
|chr4|116150293|116150697|182|Promoter (<=1kb)|4|116150518|116159844|9327|1|ENSMUST00|0|Pomgnt1|ENSMUSG0C|false|chr4|116149764|116151273|29.75681|92.45776|89.05831|chr4|116149659|116152435|3.98362|3.51592|2.17893|
|chr4|116144455|116144741|182|Promoter (<=1kb)|4|116132406|116144609|12204|2|ENSMUST00|0|Lurap1|ENSMUSG0C|false|chr4|116143868|116144863|17.45443|60.55182|57.69857|chr4|116143576|116145225|3.11084|3.08982|1.51881|
|chr4|115839062|115839316|182|Promoter (<=1kb)|4|115839198|115879250|40053|1|ENSMUST00|0|Mknk1|ENSMUSG0C|false|chr4|115838693|115839945|38.43098|125.09557|120.78131|chr4|115838623|115841752|5.15726|4.16531|3.13421|
|chr4|115103564|115103802|182|Downstream (2-3kb)|4|115090932|115093864|2933|1|ENSMUST00|12633|Pdzk1ip1|ENSMUSG0C|false|.|.|.|.|.|.|chr4|115101952|115104077|3.68461|3.44793|1.97994|
|chr4|139192388|139193152|182|Promoter (<=1kb)|4|139192899|139291818|98920|1|ENSMUST00|0|Capzb|ENSMUSG0C|false|chr4|139192025|139192789|22.63194|64.29881|61.39256|.|.|.|.|.|.|
|chr4|138555591|138555801|182|Distal Intergenic|4|138571046|138581083|10038|2|ENSMUST00|25282|Vwa5b1|ENSMUSG0C|false|chr4|138555526|138555911|6.77105|11.72451|9.51382|chr4|138554304|138555896|6.35918|4.52112|4.1625|
|chr4|138304626|138304977|182|Promoter (<=1kb)|4|138304730|138312628|7899|1|ENSMUST00|0|Ddost|ENSMUSG0C|false|chr4|138304060|138304627|18.34506|46.29042|43.62275|chr4|138303953|138306078|4.28228|3.51166|2.44542|
|chr4|137993603|137994018|182|Promoter (<=1kb)|4|137993484|138058035|64552|1|ENSMUST00|120|Eif4g3|ENSMUSG0C|false|chr4|137992884|137995121|29.28796|98.53321|94.99628|.|.|.|.|.|.|
|chr4|137483944|137484187|182|Intron (ENSMUST00000030547.9|4|137468803|137570629|101827|1|ENSMUST00|15142|Hspg2|ENSMUSG0C|false|.|.|.|.|.|.|chr4|137484181|137487113|4.37241|3.84074|2.48161|
|chr4|130168360|130168727|182|Promoter (<=1kb)|4|130166977|130169484|2508|2|ENSMUST00|757|Tinagl1|ENSMUSG0C|false|.|.|.|.|.|.|chr4|130167523|130170285|4.83497|3.86522|2.89943|
|chr4|129121015|129121294|182|Promoter (<=1kb)|4|129111576|129121291|9716|2|ENSMUST00|0|Hpca|ENSMUSG0C|false|chr4|129120896|129121613|19.75988|50.25065|47.53236|chr4|129120382|129121766|4.91658|3.81047|2.97246|
|chr4|129011810|129012037|182|Distal Intergenic|4|128993353|129008229|14877|1|ENSMUST00|18458|Ak2|ENSMUSG0C|false|.|.|.|.|.|.|.|.|.|.|.|.|
|chr4|129057991|129058740|182|Promoter (<=1kb)|4|129058271|129084526|26256|1|ENSMUST00|0|Rnf19b|ENSMUSG0C|false|chr4|129058051|129059547|26.72788|101.58999|97.97913|.|.|.|.|.|.|
|chr4|128805781|128806612|182|Promoter (<=1kb)|4|128773088|128806045|32958|2|ENSMUST00|0|Zfp362|ENSMUSG0C|false|chr4|128805149|128807225|28.03146|79.91027|76.76085|.|.|.|.|.|.|
|chr4|134468870|134469157|182|Promoter (<=1kb)|4|134468409|134473664|5256|1|ENSMUST00|462|Stmn1|ENSMUSG0C|false|chr4|134467498|134469579|34.25306|109.37451|105.56091|chr4|134469049|134469845|2.6606|2.77252|1.14055|
|chr4|134460206|134460449|182|Distal Intergenic|4|134468320|134473843|5524|1|ENSMUST00|-7871|Stmn1|ENSMUSG0C|false|.|.|.|.|.|.|.|.|.|.|.|.|
|chr4|134128283|134128947|182|Promoter (<=1kb)|4|134127406|134128789|1384|2|ENSMUST00|0|Sh3bgrl3|ENSMUSG0C|false|chr4|134128046|134129392|30.26787|111.42836|107.5545|chr4|134127087|134129640|3.08093|2.9745|1.46214|
|chr4|126242129|126242334|182|Promoter (2-3kb)|4|126232167|126240111|7945|2|ENSMUST00|-2019|Map7d1|ENSMUSG0C|false|.|.|.|.|.|.|chr4|126238132|126243427|4.67672|3.84797|2.76909|
|chr4|133277561|133277968|182|Promoter (<=1kb)|4|133266045|133277792|11748|2|ENSMUST00|0|Tmem222|ENSMUSG0C|false|chr4|133276832|133278394|26.28407|85.32412|82.07336|.|.|.|.|.|.|
|chr4|133241128|133241482|182|Promoter (<=1kb)|4|133240818|133252929|12112|1|ENSMUST00|311|Map3k6|ENSMUSG0C|false|chr4|133240506|133242188|26.28407|85.32412|82.07336|chr4|133240235|133243355|4.7864|3.96631|2.83351|
|chr4|132515671|132515888|182|Distal Intergenic|4|132492032|132510501|18470|2|ENSMUST00|-5171|Sesn2|ENSMUSG0C|false|.|.|.|.|.|.|chr4|132513633|132516273|5.14354|4.16547|3.12732|
|chr4|131939479|131939698|182|Promoter (2-3kb)|4|131923413|131937387|13975|2|ENSMUST00|-2093|Epb4.1|ENSMUSG0C|false|.|.|.|.|.|.|chr4|131939165|131940029|2.83143|2.87765|1.27419|
|chr5|5380066|5380779|182|Promoter (<=1kb)|5|4803391|5380197|576807|2|ENSMUST00|0|Cdk14|ENSMUSG0C|false|chr5|5379620|5381052|25.34239|74.34718|71.28911|.|.|.|.|.|.|
|chr5|5357479|5357697|182|Intron (ENSMUST00000030763.7|5|5131876|5380193|248318|2|ENSMUST00|22496|Cdk14|ENSMUSG0C|false|.|.|.|.|.|.|.|.|.|.|.|.|
|chr4|152177761|152178221|182|Promoter (<=1kb)|4|152178134|152271852|93719|1|ENSMUST00|0|Acot7|ENSMUSG0C|false|chr4|152177486|152178864|26.65955|86.95457|83.67183|chr4|152177538|152179235|5.69924|4.28629|3.62815|
|chr4|152150498|152150786|182|Promoter (1-2kb)|4|152120331|152152207|31877|2|ENSMUST00|1421|Espn|ENSMUSG0C|false|.|.|.|.|.|.|.|.|.|.|.|.|
|chr4|149415628|149416162|182|Intron (ENSMUST00000103212.4|4|149329676|149425905|96230|2|ENSMUST00|9743|Ube4b|ENSMUSG0C|false|.|.|.|.|.|.|chr4|149414712|149425607|5.07523|3.96443|3.10543|
|chr4|149400816|149401178|182|Promoter (1-2kb)|4|149398512|149399378|867|2|ENSMUST00|-1439|Ube4b|ENSMUSG0C|false|.|.|.|.|.|.|chr4|149390699|149406335|4.99887|3.95109|3.0296|
|chr4|150914308|150914755|182|Promoter (<=1kb)|4|150897133|150914437|17305|2|ENSMUST00|0|Park7|ENSMUSG0C|false|chr4|150913674|150915272|27.40481|79.76363|76.61784|chr4|150913459|150915867|4.13743|3.66|2.316|
|chr4|150902021|150902308|182|Promoter (<=1kb)|4|150897384|150901835|4452|2|ENSMUST00|-187|Park7|ENSMUSG0C|false|.|.|.|.|.|.|chr4|150901186|150905355|4.96689|4.03795|3.02715|
|chr4|149099221|149099932|182|Promoter (<=1kb)|4|148960535|149099876|139342|2|ENSMUST00|0|Pex14|ENSMUSG0C|false|chr4|149098516|149100205|45.26388|159.40776|153.7765|chr4|149094679|149100402|4.30823|3.73955|2.44038|
|chr4|148805773|148806472|182|Promoter (1-2kb)|4|148804465|148839746|35282|1|ENSMUST00|1309|Casz1|ENSMUSG0C|true|chr4|148803997|148806784|38.29964|140.16156|135.27679|.|.|.|.|.|.|
|chr4|148878902|148879259|182|Intron (ENSMUST00000094464.4|4|148889381|148891936|2556|1|ENSMUST00|-10122|Casz1|ENSMUSG0C|true|.|.|.|.|.|.|chr4|148876805|148882838|7.02287|4.88609|4.67766|
|chr4|148804723|148805200|182|Promoter (<=1kb)|4|148804465|148839746|35282|1|ENSMUST00|259|Casz1|ENSMUSG0C|true|chr4|148803997|148806784|38.29964|140.16156|135.27679|.|.|.|.|.|.|
|chr4|149989154|149989432|182|Intron (ENSMUST00000084117.7|4|149995848|150001265|5418|2|ENSMUST00|11833|H6pd|ENSMUSG0C|false|.|.|.|.|.|.|chr4|149985077|150009610|6.9066|4.88029|4.57705|
|chr4|148156098|148156332|182|Promoter (2-3kb)|4|148153537|148158599|5063|2|ENSMUST00|2267|Fbxo44|ENSMUSG0C|false|.|.|.|.|.|.|chr4|148154210|148157639|5.27444|4.19088|3.22587|
|chr5|23434262|23434516|182|Promoter (<=1kb)|5|23434441|23504235|69795|1|ENSMUST00|0|Kmt2e|ENSMUSG0C|false|chr5|23431644|23437822|48.25087|161.14223|155.45619|.|.|.|.|.|.|
|chr4|154139941|154140183|182|Promoter (<=1kb)|4|154056253|154140208|83956|2|ENSMUST00|25|Trp73|ENSMUSG0C|false|chr4|154139476|154141532|24.30838|70.74035|67.73997|chr4|154139239|154143166|6.40665|4.55048|4.18956|
|chr4|154142244|154142618|182|Promoter (<=1kb)|4|154142372|154156816|14445|1|ENSMUST00|0|Wrap73|ENSMUSG0C|false|chr4|154141650|154143092|26.28407|85.32412|82.07336|chr4|154139239|154143166|6.40665|4.55048|4.18956|
|chr4|155086223|155086538|182|Promoter (<=1kb)|4|155074110|155086371|12262|2|ENSMUST00|0|Rer1|ENSMUSG0C|false|chr4|155085258|155086435|20.109|88.33971|85.03291|.|.|.|.|.|.|
|chr4|155145942|155146205|182|Distal Intergenic|4|155110690|155128778|18089|1|ENSMUST00|35253|Morn1|ENSMUSG0C|false|.|.|.|.|.|.|chr4|155144041|155153706|6.53431|4.75254|4.26776|
|chr4|155302075|155302447|182|Promoter (<=1kb)|4|155294278|155301171|6894|2|ENSMUST00|-905|Prkcz|ENSMUSG0C|false|.|.|.|.|.|.|chr4|155301428|155302868|6.80302|4.92995|4.49631|
|chr5|35004021|35004248|182|Intron (ENSMUST00000030984.8|5|34999184|35039635|40452|1|ENSMUST00|4838|Rgs12|ENSMUSG0C|false|.|.|.|.|.|.|chr5|35003219|35005717|4.03287|3.47966|2.25309|
|chr5|34565727|34566062|182|Distal Intergenic|5|34573664|34632308|58645|1|ENSMUST00|-7602|Add1|ENSMUSG0C|false|.|.|.|.|.|.|chr5|34565056|34566889|4.56541|3.88556|2.66924|
|chr5|38039070|38039378|182|Promoter (<=1kb)|5|38039233|38136817|97585|1|ENSMUST00|0|Stx18|ENSMUSG0C|false|chr5|38038465|38040150|30.17592|94.17345|90.73757|.|.|.|.|.|.|
|chr5|41707805|41708548|182|Promoter (<=1kb)|5|41624976|41708155|83180|2|ENSMUST00|0|Rab28|ENSMUSG0C|false|chr5|41706910|41708875|45.58365|150.79124|145.48296|.|.|.|.|.|.|
|chr5|32346309|32346865|182|Intron (ENSMUST00000101377.3|5|32353569|32366520|12952|1|ENSMUST00|-6704|Plb1|ENSMUSG0C|false|chr5|32346060|32346376|7.12487|12.70834|10.47977|chr5|32345292|32347541|5.35296|4.19919|3.30033|
|chr5|31179825|31180252|182|Promoter (<=1kb)|5|31156005|31180132|24128|2|ENSMUST00|0|Gtf3c2|ENSMUSG0C|false|chr5|31179015|31180907|29.71786|90.43782|87.08427|chr5|31178159|31181078|4.80429|3.9289|2.8571|
|chr5|31192670|31193387|182|Promoter (<=1kb)|5|31187559|31193139|5581|2|ENSMUST00|0|Eif2b4|ENSMUSG0C|false|chr5|31192419|31194275|33.45073|115.2328|111.2408|chr5|31191719|31192724|5.61552|4.40384|3.49296|
|chr5|73362820|73363092|182|Distal Intergenic|5|73332128|73338947|6820|2|ENSMUST00|-23874|Ociad2|ENSMUSG0C|false|.|.|.|.|.|.|.|.|.|.|.|.|
|chr5|73647422|73647934|182|Promoter (<=1kb)|5|73632749|73647713|14965|2|ENSMUST00|0|Sgcb|ENSMUSG0C|false|chr5|73646992|73648042|32.25957|103.83389|100.16508|chr5|73646663|73648066|3.39581|3.27643|1.6818|
|chr5|30916031|30916444|182|Promoter (2-3kb)|5|30913739|30919703|5965|1|ENSMUST00|2293|Emilin1|ENSMUSG0C|false|.|.|.|.|.|.|chr5|30904966|30918702|6.19394|4.49227|4.02489|
|chr5|36695780|36696195|182|Promoter (<=1kb)|5|36600504|36695969|95466|2|ENSMUST00|0|D5Ertd579e|ENSMUSG0C|false|chr5|36694931|36696494|23.12601|77.64799|74.53377|.|.|.|.|.|.|
|chr5|36530477|36530693|182|Promoter (<=1kb)|5|36490605|36530623|40019|2|ENSMUST00|0|Tbc1d14|ENSMUSG0C|false|chr5|36529746|36530852|28.08037|85.65662|82.39679|chr5|36528092|36531734|4.272|3.60522|2.44492|
|chr5|65415567|65415943|182|Intron (ENSMUST00000117542.2|5|65413224|65422660|9437|2|ENSMUST00|6717|Ugdh|ENSMUSG0C|false|.|.|.|.|.|.|chr5|65415378|65416500|4.45528|3.71021|2.5974|
|chr5|65706983|65707221|182|Distal Intergenic|5|65615260|65697856|82597|2|ENSMUST00|-9128|Pds5a|ENSMUSG0C|false|.|.|.|.|.|.|chr5|65706376|65707669|3.40812|3.15651|1.73476|
|chr5|66527163|66527393|182|Intron (ENSMUST00000159786.2|5|66452261|66514837|62577|2|ENSMUST00|-12327|Apbb2|ENSMUSG0C|false|.|.|.|.|.|.|chr5|66502348|66557053|5.30285|4.18237|3.2769|
|chr5|66406136|66406438|182|Intron (ENSMUST00000159786.2|5|66364974|66451629|86656|2|ENSMUST00|45191|Apbb2|ENSMUSG0C|false|.|.|.|.|.|.|chr5|66404491|66409180|6.02158|4.52811|3.8468|
|chr5|66543808|66544049|182|Promoter (2-3kb)|5|66452267|66546957|94691|2|ENSMUST00|2908|Apbb2|ENSMUSG0C|false|.|.|.|.|.|.|chr5|66502348|66557053|5.30285|4.18237|3.2769|
|chr5|66545462|66545709|182|Promoter (1-2kb)|5|66452267|66546957|94691|2|ENSMUST00|1248|Apbb2|ENSMUSG0C|false|.|.|.|.|.|.|chr5|66502348|66557053|5.30285|4.18237|3.2769|
|chr5|72821115|72821301|182|Intron (ENSMUST00000071944.7|5|72755738|72786020|30283|2|ENSMUST00|-35096|Tec|ENSMUSG0C|false|.|.|.|.|.|.|chr5|72819807|72821762|4.08116|3.63609|2.2445|
|chr5|75046919|75047116|182|Promoter (2-3kb)|5|75001026|75044774|43749|2|ENSMUST00|-2146|Chic2|ENSMUSG0C|false|.|.|.|.|.|.|.|.|.|.|.|.|
|chr5|76140461|76140781|182|Promoter (<=1kb)|5|76140700|76149755|9056|1|ENSMUST00|0|Srd5a3|ENSMUSG0C|false|chr5|76139804|76141310|28.53732|88.37692|85.06884|.|.|.|.|.|.|
|chr5|108312593|108312909|182|Promoter (<=1kb)|5|108312609|108348233|35625|1|ENSMUST00|0|Pigg|ENSMUSG0C|false|chr5|108312006|108313586|33.32351|103.65904|99.99326|.|.|.|.|.|.|
|chr5|107597418|107597871|182|Promoter (<=1kb)|5|107548967|107597639|48673|2|ENSMUST00|0|Glmn|ENSMUSG0C|false|chr5|107597049|107598312|26.52689|93.43019|90.01563|.|.|.|.|.|.|
|chr5|107081405|107081594|182|Distal Intergenic|5|107118197|107132923|14727|2|ENSMUST00|51329|Tgfbr3|ENSMUSG0C|true|.|.|.|.|.|.|chr5|107079090|107082975|3.48278|3.27962|1.7633|
|chr5|107200899|107201203|182|Intron (ENSMUST00000031224.9|5|107109281|107192367|83087|2|ENSMUST00|-8533|Tgfbr3|ENSMUSG0C|true|.|.|.|.|.|.|chr5|107189975|107206509|4.95135|4.04289|2.96667|
|chr5|105876194|105876621|182|Promoter (<=1kb)|5|105876565|105914780|38216|1|ENSMUST00|0|Zfp326|ENSMUSG0C|false|chr5|105875939|105878057|20.9917|88.60235|85.28741|.|.|.|.|.|.|
|chr5|103836450|103836677|182|Intron (ENSMUST00000054979.4|5|103870310|103874369|4060|2|ENSMUST00|37692|Klhl8|ENSMUSG0C|false|.|.|.|.|.|.|chr5|103827430|103841060|6.32258|4.7151|4.08994|
|chr5|103835673|103835891|182|Intron (ENSMUST00000054979.4|5|103870310|103874369|4060|2|ENSMUST00|38478|Klhl8|ENSMUSG0C|false|.|.|.|.|.|.|chr5|103827430|103841060|6.32258|4.7151|4.08994|
|chr5|103778457|103778737|182|Intron (ENSMUST00000126335.1|5|103784359|103811595|27237|1|ENSMUST00|-5622|Aff1|ENSMUSG0C|false|.|.|.|.|.|.|chr5|103765592|103792229|6.51088|4.73748|4.26006|
|chr5|100846006|100846379|182|Promoter (<=1kb)|5|100846130|100899099|52970|1|ENSMUST00|0|Agpat9|ENSMUSG0C|false|chr5|100845079|100847111|30.42596|90.69117|87.33178|.|.|.|.|.|.|
|chr5|100845486|100845813|182|Promoter (<=1kb)|5|100845713|100846810|1098|1|ENSMUST00|0|Agpat9|ENSMUSG0C|false|chr5|100845079|100847111|30.42596|90.69117|87.33178|.|.|.|.|.|.|
|chr5|100038420|100038735|182|Promoter (<=1kb)|5|100034721|100038869|4149|2|ENSMUST00|134|Hnrnpdl|ENSMUSG0C|false|chr5|100036831|100040800|37.7199|126.00829|121.65485|.|.|.|.|.|.|
|chr5|100038782|100039201|182|Promoter (<=1kb)|5|100034721|100038869|4149|2|ENSMUST00|0|Hnrnpdl|ENSMUSG0C|false|chr5|100036831|100040800|37.7199|126.00829|121.65485|.|.|.|.|.|.|
|chr5|92809040|92809347|182|Promoter (<=1kb)|5|92809378|92965318|155941|1|ENSMUST00|-31|Shroom3|ENSMUSG0C|true|chr5|92808748|92813129|21.42557|69.98924|66.99868|.|.|.|.|.|.|
|chr5|93267339|93267694|182|Promoter (<=1kb)|5|93267589|93268454|866|1|ENSMUST00|0|Ccng2|ENSMUSG0C|false|chr5|93267238|93268862|34.68901|133.45216|128.82967|.|.|.|.|.|.|
|chr5|93268032|93268386|182|Promoter (<=1kb)|5|93268314|93276224|7911|1|ENSMUST00|0|Ccng2|ENSMUSG0C|false|chr5|93267238|93268862|34.68901|133.45216|128.82967|.|.|.|.|.|.|
|chr5|124578989|124579252|182|Promoter (<=1kb)|5|124579140|124597680|18541|1|ENSMUST00|0|Gtf2h3|ENSMUSG0C|false|chr5|124578504|124579886|22.60491|86.65532|83.38163|chr5|124577917|124581071|4.76076|3.80779|2.84853|
|chr5|124552659|124552904|182|Promoter (<=1kb)|5|124552864|124569660|16797|1|ENSMUST00|0|Ddx55|ENSMUSG0C|false|chr5|124551849|124553504|39.97929|126.69925|122.32521|chr5|124551670|124553488|5.17372|4.09429|3.17319|
|chr5|124540604|124540862|182|Promoter (<=1kb)|5|124540695|124550506|9812|1|ENSMUST00|0|Tmed2|ENSMUSG0C|false|chr5|124540331|124541863|21.13367|73.82313|70.77491|.|.|.|.|.|.|
|chr5|124509782|124510071|182|Intron (ENSMUST00000062153.7|5|124493082|124531391|38310|2|ENSMUST00|21320|Rilpl1|ENSMUSG0C|false|.|.|.|.|.|.|chr5|124509539|124511959|4.87435|4.09135|2.89051|
|chr5|123748903|123749750|182|Promoter (<=1kb)|5|123749726|123821593|71868|1|ENSMUST00|0|Kntc1|ENSMUSG0C|false|chr5|123749665|123750060|16.76261|41.34956|38.74308|chr5|123749747|123750212|4.93426|4.16197|2.94884|
|chr5|92505464|92505804|182|Promoter (<=1kb)|5|92443873|92505608|61736|2|ENSMUST00|0|Scarb2|ENSMUSG0C|false|chr5|92504372|92506287|28.10168|124.31538|120.02569|chr5|92505777|92506700|4.08177|3.6548|2.28451|
|chr5|129019705|129020514|182|Promoter (<=1kb)|5|129020069|129024323|4255|1|ENSMUST00|0|Ran|ENSMUSG0C|false|chr5|129019359|129021396|34.18682|141.10779|136.18079|.|.|.|.|.|.|
|chr5|129787076|129787400|182|Promoter (<=1kb)|5|129765558|129787204|21647|2|ENSMUST00|0|Psph|ENSMUSG0C|false|chr5|129786551|129788333|35.31687|113.30753|109.37892|.|.|.|.|.|.|
|chr5|121452255|121452622|182|Promoter (<=1kb)|5|121452575|121494445|41871|1|ENSMUST00|0|Tmem116|ENSMUSG0C|false|chr5|121451584|121453273|30.03893|101.89226|98.2715|.|.|.|.|.|.|
|chr5|122621492|122621720|182|Distal Intergenic|5|122550204|122614518|64315|2|ENSMUST00|-6975|Ift81|ENSMUSG0C|false|.|.|.|.|.|.|chr5|122619549|122623735|4.4334|3.78869|2.54699|
|chr5|122770940|122771224|182|Intron (ENSMUST00000111668.3|5|122731173|122779410|48238|2|ENSMUST00|8186|Camkk2|ENSMUSG0C|false|.|.|.|.|.|.|.|.|.|.|.|.|
|chr5|122989013|122989471|182|Promoter (<=1kb)|5|122870668|122989099|118432|2|ENSMUST00|0|Kdm2b|ENSMUSG0C|false|chr5|122987524|122989984|17.09341|55.02552|52.24499|.|.|.|.|.|.|
|chr5|123014795|123015680|182|Promoter (<=1kb)|5|123007579|123015080|7502|2|ENSMUST00|0|Kdm2b|ENSMUSG0C|false|chr5|123014659|123016346|32.27147|102.84008|99.19724|chr5|123013785|123014820|4.16977|3.77565|2.30553|
|chr5|125143625|125143980|182|Intron (ENSMUST00000111402.3|5|125017835|125124900|107066|2|ENSMUST00|-18726|Ncor2|ENSMUSG0C|false|.|.|.|.|.|.|chr5|125124978|125178451|6.70656|4.76008|4.42049|
|chr5|125217851|125218182|182|Distal Intergenic|5|125017153|125179219|162067|2|ENSMUST00|-38633|Ncor2|ENSMUSG0C|false|.|.|.|.|.|.|chr5|125217473|125218451|4.14455|3.47928|2.33755|
|chr5|125053143|125053540|182|Promoter (2-3kb)|5|125047802|125055988|8187|2|ENSMUST00|2448|Ncor2|ENSMUSG0C|false|.|.|.|.|.|.|chr5|125050765|125060996|6.95919|4.79882|4.64527|
|chr5|125150904|125151271|182|Intron (ENSMUST00000111402.3|5|125017835|125124900|107066|2|ENSMUST00|-26005|Ncor2|ENSMUSG0C|false|.|.|.|.|.|.|chr5|125124978|125178451|6.70656|4.76008|4.42049|
|chr5|125475680|125475906|182|Promoter (<=1kb)|5|125475814|125517412|41599|1|ENSMUST00|0|Aacs|ENSMUSG0C|false|chr5|125475256|125476543|27.88725|99.90214|96.33501|.|.|.|.|.|.|
|chr5|96793287|96793778|182|Promoter (<=1kb)|5|96793385|96845968|52584|1|ENSMUST00|0|Anxa3|ENSMUSG0C|true|chr5|96792343|96794781|28.49099|81.63235|78.4519|.|.|.|.|.|.|
|chr5|110230818|110231401|182|Promoter (<=1kb)|5|110231004|110256651|25648|1|ENSMUST00|0|Ankle2|ENSMUSG0C|false|chr5|110230299|110231981|34.92026|124.22972|119.94103|.|.|.|.|.|.|
|chr5|115523947|115524306|182|Intron (ENSMUST00000067268.9|5|115520305|115555982|35678|1|ENSMUST00|3643|Pxn|ENSMUSG0C|false|.|.|.|.|.|.|chr5|115507551|115550291|7.76238|5.17462|5.28263|
|chr6|17079296|17079572|182|Intron (ENSMUST00000115467.5|6|17065224|17105828|40605|1|ENSMUST00|14073|Tes|ENSMUSG0C|false|.|.|.|.|.|.|chr6|17079245|17080115|3.87137|3.5347|2.09041|
|chr6|15282040|15282380|182|Intron (ENSMUST00000115477.2|6|15254283|15286770|32488|1|ENSMUST00|27758|Foxp2|ENSMUSG0C|false|.|.|.|.|.|.|.|.|.|.|.|.|
|chr5|114100182|114100710|182|Promoter (<=1kb)|5|114100333|114123506|23174|1|ENSMUST00|0|Usp30|ENSMUSG0C|false|chr5|114099830|114101172|31.89884|96.17711|92.69861|chr5|114099758|114101552|3.79057|3.38431|2.05636|
|chr5|145116422|145116736|182|Promoter (2-3kb)|5|145114316|145122635|8320|1|ENSMUST00|2107|Arpc1b|ENSMUSG0C|false|.|.|.|.|.|.|chr5|145113465|145118752|4.56678|3.84193|2.6654|
|chr5|145130225|145130467|182|3' UTR|5|145127223|145128088|866|1|ENSMUST00|3003|Arpc1b|ENSMUSG0C|false|.|.|.|.|.|.|chr5|145128945|145131198|4.61541|3.63109|2.72954|
|chr5|146220976|146221504|182|Promoter (<=1kb)|5|146211676|146221418|9743|2|ENSMUST00|0|Rnf6|ENSMUSG0C|false|chr5|146220447|146221984|33.94988|109.87815|106.04343|.|.|.|.|.|.|
|chr5|146762669|146763025|182|Intron (ENSMUST00000085614.5|5|146740431|146758456|18026|2|ENSMUST00|-4214|Usp12|ENSMUSG0C|false|.|.|.|.|.|.|chr5|146761783|146767841|3.67138|3.3407|1.94814|
|chr5|146867456|146867921|182|Distal Intergenic|5|146845071|146847726|2656|1|ENSMUST00|22386|Rasl11a|ENSMUSG0C|true|.|.|.|.|.|.|chr5|146865620|146877684|5.33365|4.16572|3.30759|
|chr5|147519707|147519911|182|Promoter (<=1kb)|5|147519687|147548496|28810|1|ENSMUST00|21|Pan3|ENSMUSG0C|false|.|.|.|.|.|.|chr5|147516948|147520150|2.9446|2.94071|1.35719|
|chr5|147459889|147460107|182|Intron (ENSMUST00000031651.9|5|147453289|147548496|95208|1|ENSMUST00|6601|Pan3|ENSMUSG0C|false|.|.|.|.|.|.|chr5|147458526|147463399|4.09626|3.49906|2.27578|
|chr5|150665613|150665913|182|Promoter (<=1kb)|5|150635972|150665612|29641|2|ENSMUST00|-2|N4bp2l2|ENSMUSG0C|false|chr5|150664539|150665702|35.97658|118.66996|114.57709|.|.|.|.|.|.|
|chr6|21949232|21949495|182|Promoter (<=1kb)|6|21949571|21976038|26468|1|ENSMUST00|-76|Ing3|ENSMUSG0C|false|chr6|21948939|21949473|18.02173|47.27326|44.5908|.|.|.|.|.|.|
|chr6|25809210|25809442|182|Promoter (<=1kb)|6|25743737|25809226|65490|2|ENSMUST00|0|Pot1a|ENSMUSG0C|false|chr6|25808383|25809801|19.45739|70.46334|67.46697|.|.|.|.|.|.|
|chr5|135168258|135168788|182|Promoter (<=1kb)|5|135168283|135181855|13573|1|ENSMUST00|0|Bcl7b|ENSMUSG0C|false|chr5|135167981|135169578|27.24215|82.29425|79.10038|chr5|135167909|135170511|4.07754|3.65485|2.27853|
|chr6|24543077|24543322|182|Intron (ENSMUST00000031696.4|6|24556072|24573164|17093|1|ENSMUST00|-12750|Asb15|ENSMUSG0C|true|chr6|24543105|24545291|14.6238|41.22485|38.6191|chr6|24540933|24543169|4.54464|3.92618|2.61377|
|chr6|47594216|47594442|182|Promoter (<=1kb)|6|47530274|47595030|64757|2|ENSMUST00|588|Ezh2|ENSMUSG0C|false|chr6|47593688|47596080|34.34893|131.73137|127.17394|.|.|.|.|.|.|
|chr5|136406294|136406518|182|Intron (ENSMUST00000148082.1|5|136274070|136373348|99279|2|ENSMUST00|-32947|Cux1|ENSMUSG0C|false|.|.|.|.|.|.|.|.|.|.|.|.|
|chr5|136566908|136567731|182|Promoter (<=1kb)|5|136248135|136567248|319114|2|ENSMUST00|0|Cux1|ENSMUSG0C|false|chr5|136565019|136569262|36.46257|120.58523|116.42456|.|.|.|.|.|.|
|chr5|137348727|137348970|182|Promoter (1-2kb)|5|137350109|137374528|24420|1|ENSMUST00|-1139|Ephb4|ENSMUSG0C|false|.|.|.|.|.|.|chr5|137347819|137351010|4.1561|3.65922|2.31468|
|chr5|137530899|137531450|182|Promoter (<=1kb)|5|137528503|137531297|2795|2|ENSMUST00|0|Gnb2|ENSMUSG0C|false|chr5|137530397|137533556|27.03504|88.5911|85.27629|chr5|137529479|137532031|4.39789|3.61512|2.52536|
|chr5|137684501|137685064|182|Promoter (<=1kb)|5|137650483|137684726|34244|2|ENSMUST00|0|Agfg2|ENSMUSG0C|false|chr5|137683658|137684817|20.31581|74.8081|71.74177|.|.|.|.|.|.|
|chr5|138170983|138171278|182|Promoter (<=1kb)|5|138168456|138171266|2811|2|ENSMUST00|0|Mcm7|ENSMUSG0C|false|chr5|138170353|138172591|27.24215|82.29425|79.10038|chr5|138170112|138173019|4.035|3.50486|2.25711|
|chr6|6881931|6882130|182|Promoter (<=1kb)|6|6877805|6882068|4264|2|ENSMUST00|0|Dlx5|ENSMUSG0C|false|chr6|6881365|6882228|14.10153|31.98288|29.49352|chr6|6879390|6882102|4.98249|3.91401|3.0307|
|chr6|34610364|34610866|182|Intron (ENSMUST00000079391.4|6|34598620|34662748|64129|1|ENSMUST00|11745|Cald1|ENSMUSG0C|true|.|.|.|.|.|.|chr6|34600355|34614741|4.22277|3.68306|2.37293|
|chr6|33823706|33824132|182|Intron (ENSMUST00000052266.9|6|33862100|33874181|12082|1|ENSMUST00|-37968|Exoc4|ENSMUSG0C|false|.|.|.|.|.|.|chr6|33823076|33824999|5.30044|4.28112|3.24005|
|chr6|30509652|30509881|182|Promoter (<=1kb)|6|30479053|30509704|30652|2|ENSMUST00|0|Tmem209|ENSMUSG0C|false|chr6|30509099|30510160|31.25476|101.05123|97.45738|.|.|.|.|.|.|
|chr6|48488329|48488683|182|Promoter (<=1kb)|6|48488042|48501250|13209|1|ENSMUST00|288|Sspo|ENSMUSG0C|false|.|.|.|.|.|.|.|.|.|.|.|.|
|chr6|58596449|58596687|182|Promoter (<=1kb)|6|58596671|58692451|95781|1|ENSMUST00|0|Abcg2|ENSMUSG0C|false|chr6|58596029|58596768|7.01212|23.94238|21.55444|chr6|58596152|58597223|3.84163|3.18016|2.08995|
|chr6|49073697|49073909|182|Promoter (<=1kb)|6|49073795|49084717|10923|1|ENSMUST00|0|Malsu1|ENSMUSG0C|false|chr6|49073330|49074698|33.52805|106.05383|102.33234|.|.|.|.|.|.|
|chr6|38354183|38354471|182|Promoter (<=1kb)|6|38305286|38354273|48988|2|ENSMUST00|0|Zc3hav1|ENSMUSG0C|false|chr6|38353230|38355708|31.01414|97.6227|94.10706|.|.|.|.|.|.|
|chr6|37871329|37871712|182|Promoter (<=1kb)|6|37870811|37966296|95486|1|ENSMUST00|519|Trim24|ENSMUSG0C|false|chr6|37870693|37872083|30.26787|111.42836|107.5545|.|.|.|.|.|.|
|chr6|41026066|41026278|182|Distal Intergenic|6|41030268|41035509|5242|2|ENSMUST00|9231|2210010C04I|ENSMUSG0C|false|.|.|.|.|.|.|.|.|.|.|.|.|
|chr6|40471304|40471570|182|Promoter (<=1kb)|6|40471352|40481823|10472|1|ENSMUST00|0|Ssbp1|ENSMUSG0C|false|chr6|40470723|40472224|28.28183|109.60786|105.79031|.|.|.|.|.|.|
|chr6|39810543|39810791|182|Promoter (<=1kb)|6|39802192|39810885|8694|2|ENSMUST00|94|Mrps33|ENSMUSG0C|false|chr6|39810232|39811219|18.02578|59.25625|56.42029|.|.|.|.|.|.|
|chr6|65042501|65042813|182|Promoter (<=1kb)|6|65042667|65116050|73384|1|ENSMUST00|0|Smarcad1|ENSMUSG0C|false|chr6|65042284|65043527|32.90413|103.94604|100.27567|.|.|.|.|.|.|
|chr6|39419870|39420464|182|Promoter (<=1kb)|6|39397804|39420462|22659|2|ENSMUST00|0|Mkrn1|ENSMUSG0C|false|chr6|39419347|39420676|32.98857|124.89146|120.57959|.|.|.|.|.|.|
|chr6|86758004|86758220|182|Promoter (2-3kb)|6|86737728|86760737|23010|2|ENSMUST00|2517|Anxa4|ENSMUSG0C|true|.|.|.|.|.|.|chr6|86756577|86760107|4.98978|4.12031|3.04624|
|chr6|86070762|86071142|182|Distal Intergenic|6|86078084|86119555|41472|1|ENSMUST00|-6942|Add2|ENSMUSG0C|false|.|.|.|.|.|.|.|.|.|.|.|.|
|chr6|85961884|85962123|182|Promoter (<=1kb)|6|85954962|85961667|6706|2|ENSMUST00|-218|Dusp11|ENSMUSG0C|false|chr6|85960339|85962539|34.78613|113.42423|109.48744|.|.|.|.|.|.|
|chr6|85961257|85961502|182|Promoter (<=1kb)|6|85955064|85961607|6544|2|ENSMUST00|105|Dusp11|ENSMUSG0C|false|chr6|85960339|85962539|34.78613|113.42423|109.48744|.|.|.|.|.|.|
|chr6|92414607|92414827|182|Intron (ENSMUST00000032093.6|6|92375384|92481392|106009|2|ENSMUST00|66565|Prickle2|ENSMUSG0C|false|.|.|.|.|.|.|chr6|92414292|92416053|2.88246|2.92967|1.30867|
|chr6|92464810|92465133|182|Intron (ENSMUST00000032093.6|6|92375384|92481392|106009|2|ENSMUST00|16259|Prickle2|ENSMUSG0C|false|.|.|.|.|.|.|chr6|92459348|92466873|4.66008|3.93972|2.74191|
|chr6|94647662|94647907|182|Intron (ENSMUST00000032105.5|6|94626450|94655525|29076|2|ENSMUST00|7618|Lrig1|ENSMUSG0C|false|.|.|.|.|.|.|chr6|94647579|94651502|3.37294|3.13323|1.70352|
|chr6|83007699|83007937|182|Intron (ENSMUST00000113980.3|6|83028251|83029823|1573|1|ENSMUST00|-20314|M1ap|ENSMUSG0C|false|.|.|.|.|.|.|.|.|.|.|.|.|
|chr6|82652568|82653164|182|Promoter (<=1kb)|6|82621852|82652870|31019|2|ENSMUST00|0|Pole4|ENSMUSG0C|false|chr6|82652157|82653322|30.45735|121.03082|116.85291|.|.|.|.|.|.|
|chr6|87887764|87887999|182|Promoter (<=1kb)|6|87887814|87913595|25782|1|ENSMUST00|0|Copg1|ENSMUSG0C|false|chr6|87887849|87888619|31.7393|99.89149|96.32799|chr6|87886956|87889190|4.54626|3.92911|2.64917|
|chr6|99096131|99096356|182|Promoter (<=1kb)|6|99009933|99096205|86273|2|ENSMUST00|0|Foxp1|ENSMUSG0C|false|chr6|99094864|99096288|7.48194|15.52168|13.25109|chr6|99089995|99102481|4.84431|4.00533|2.88029|
|chr6|98955688|98955934|182|Intron (ENSMUST00000113326.3|6|98930171|98935545|5375|2|ENSMUST00|-20144|Foxp1|ENSMUSG0C|false|.|.|.|.|.|.|.|.|.|.|.|.|
|chr6|100704611|100704840|182|Promoter (<=1kb)|6|100704734|100805081|100348|1|ENSMUST00|0|Gxylt2|ENSMUSG0C|false|chr6|100704452|100706031|22.81947|62.04271|59.16861|chr6|100704720|100707186|3.5451|3.29975|1.85359|
|chr6|88820842|88821154|182|Intron (ENSMUST00000113585.3|6|88835915|88841935|6021|2|ENSMUST00|20781|Abtb1|ENSMUSG0C|false|.|.|.|.|.|.|chr6|88819627|88821263|4.30519|3.77849|2.46106|
|chr6|89347836|89348092|182|Intron (ENSMUST00000163139.2|6|89319015|89357717|38703|2|ENSMUST00|9625|Plxna1|ENSMUSG0C|false|.|.|.|.|.|.|chr6|89339950|89361705|7.61354|5.20417|5.1492|
|chr6|89456951|89457205|182|Intron (ENSMUST00000032172.9|6|89316316|89362613|46298|2|ENSMUST00|-94339|Plxna1|ENSMUSG0C|false|.|.|.|.|.|.|.|.|.|.|.|.|
|chr6|90508244|90508514|182|Intron (ENSMUST00000127199.1|6|90486427|90559386|72960|1|ENSMUST00|21818|Aldh1l1|ENSMUSG0C|false|.|.|.|.|.|.|chr6|90505671|90509951|6.56223|4.6902|4.29382|
|chr6|90604652|90605066|182|Promoter (<=1kb)|6|90604725|90646412|41688|1|ENSMUST00|0|Slc41a3|ENSMUSG0C|false|chr6|90604548|90605461|23.87381|72.67255|69.64001|chr6|90605000|90607465|3.25167|3.202|1.58366|
|chr6|90621584|90621776|182|Promoter (2-3kb)|6|90619246|90646412|27167|1|ENSMUST00|2339|Slc41a3|ENSMUSG0C|false|.|.|.|.|.|.|.|.|.|.|.|.|
|chr6|90988551|90988825|182|Distal Intergenic|6|91013317|91017429|4113|2|ENSMUST00|28604|Nup210|ENSMUSG0C|false|chr6|90987314|90989227|27.66126|83.97217|80.74496|.|.|.|.|.|.|
|chr6|91515721|91516014|182|Promoter (<=1kb)|6|91489305|91515884|26580|2|ENSMUST00|0|Xpc|ENSMUSG0C|false|chr6|91515061|91516824|27.1142|89.69608|86.35963|.|.|.|.|.|.|
|chr6|91685240|91685759|182|Promoter (1-2kb)|6|91684067|91759063|74997|1|ENSMUST00|1174|Slc6a6|ENSMUSG0C|false|chr6|91683631|91686902|37.7199|126.00829|121.65485|chr6|91685018|91701814|5.17334|4.04439|3.18665|
|chr6|108237651|108237953|182|Intron (ENSMUST00000032192.6|6|108213118|108551109|337992|1|ENSMUST00|24534|Itpr1|ENSMUSG0C|false|.|.|.|.|.|.|chr6|108219855|108242489|5.57178|4.31079|3.48186|
|chr6|108234664|108234879|182|Intron (ENSMUST00000032192.6|6|108213118|108551109|337992|1|ENSMUST00|21547|Itpr1|ENSMUSG0C|false|.|.|.|.|.|.|chr6|108219855|108242489|5.57178|4.31079|3.48186|
|chr6|115971213|115971478|182|Promoter (1-2kb)|6|115966963|115969426|2464|2|ENSMUST00|-1788|Plxnd1|ENSMUSG0C|false|.|.|.|.|.|.|chr6|115971456|115972298|3.8635|3.55924|2.11477|
|chr6|115991630|115991845|182|Intron (ENSMUST00000015511.9|6|115954811|115995005|40195|2|ENSMUST00|3160|Plxnd1|ENSMUSG0C|false|chr6|115991058|115992036|7.88522|15.75676|13.48224|chr6|115980470|115999580|5.17804|4.10687|3.16948|
|chr6|129533007|129533427|182|Promoter (<=1kb)|6|129533200|129542331|9132|1|ENSMUST00|0|Gabarapl1|ENSMUSG0C|false|chr6|129532540|129534683|30.78991|105.27288|101.56436|.|.|.|.|.|.|
|chr6|134048702|134048915|182|Intron (ENSMUST00000081028.7|6|134035969|134248284|212316|1|ENSMUST00|12734|Etv6|ENSMUSG0C|false|.|.|.|.|.|.|chr6|134037014|134053440|5.9149|4.52781|3.75986|
|chr6|134515462|134515658|182|Intron (ENSMUST00000032322.9|6|134450483|134471293|20811|2|ENSMUST00|-44170|Lrp6|ENSMUSG0C|false|.|.|.|.|.|.|chr6|134509605|134529849|4.94608|3.98494|2.97789|
|chr6|135038648|135038848|182|Distal Intergenic|6|135019000|135021178|2179|1|ENSMUST00|19649|Ddx47|ENSMUSG0C|false|.|.|.|.|.|.|.|.|.|.|.|.|
|chr6|135472278|135472583|182|Distal Intergenic|6|135367493|135383173|15681|1|ENSMUST00|104786|Emp1|ENSMUSG0C|false|.|.|.|.|.|.|chr6|135472053|135473453|5.16874|4.24936|3.16647|
|chr6|139861306|139861548|182|3' UTR|6|139843648|139969284|125637|1|ENSMUST00|17659|Pik3c2g|ENSMUSG0C|false|.|.|.|.|.|.|chr6|139860240|139861475|3.52965|3.30266|1.81771|
|chr6|140424930|140425363|182|Promoter (<=1kb)|6|140424099|140594906|170808|1|ENSMUST00|832|Plekha5|ENSMUSG0C|true|chr6|140423533|140426226|36.12563|122.12295|117.91594|.|.|.|.|.|.|
|chr6|141395923|141396187|182|Intron (ENSMUST00000043259.8|6|141524386|141569720|45335|1|ENSMUST00|-128199|Slco1c1|ENSMUSG0C|false|.|.|.|.|.|.|chr6|141387816|141400666|4.81448|4.05067|2.84093|
|chr6|145768739|145769049|182|Intron (ENSMUST00000111704.2|6|145750571|145815188|64618|1|ENSMUST00|18169|Rassf8|ENSMUSG0C|true|.|.|.|.|.|.|chr6|145748529|145783801|6.70363|4.81765|4.41144|
|chr6|145024143|145024401|182|Intron (ENSMUST00000032402.6|6|145015676|145032843|17168|2|ENSMUST00|8442|Bcat1|ENSMUSG0C|false|.|.|.|.|.|.|.|.|.|.|.|.|
|chr6|144927680|144927881|182|Distal Intergenic|6|145015676|145032843|17168|2|ENSMUST00|104962|Bcat1|ENSMUSG0C|false|.|.|.|.|.|.|chr6|144927723|144928082|3.10932|3.10942|1.47449|
|chr6|143167650|143167968|182|Promoter (<=1kb)|6|143167230|143208542|41313|1|ENSMUST00|421|Etnk1|ENSMUSG0C|false|chr6|143166144|143168273|38.97723|131.47449|126.92584|.|.|.|.|.|.|
|chr6|113471347|113471550|182|Promoter (<=1kb)|6|113471427|113483140|11714|1|ENSMUST00|0|Il17rc|ENSMUSG0C|false|chr6|113471067|113472016|18.84007|53.45216|50.69242|chr6|113468959|113472716|4.62328|3.87023|2.71371|
|chr6|148896009|148896475|182|Promoter (<=1kb)|6|148842492|148896237|53746|2|ENSMUST00|0|Caprin2|ENSMUSG0C|false|chr6|148895238|148896628|31.87132|99.11874|95.57203|.|.|.|.|.|.|
|chr6|114643018|114643238|182|Promoter (<=1kb)|6|114643097|114860614|217518|1|ENSMUST00|0|Atg7|ENSMUSG0C|false|chr6|114642620|114644013|31.70772|93.8754|90.44607|chr6|114642558|114644579|4.68723|3.99456|2.75022|
|chr6|114916646|114916852|182|Intron (ENSMUST00000032459.8|6|114861424|114921821|60398|2|ENSMUST00|4969|Vgll4|ENSMUSG0C|true|.|.|.|.|.|.|chr6|114853844|114919520|8.05355|5.36011|5.51688|
|chr6|114888914|114889261|182|Intron (ENSMUST00000032459.8|6|114860628|114875107|14480|2|ENSMUST00|-13808|Vgll4|ENSMUSG0C|true|.|.|.|.|.|.|chr6|114853844|114919520|8.05355|5.36011|5.51688|
|chr6|114912776|114913043|182|Intron (ENSMUST00000032459.8|6|114861424|114921821|60398|2|ENSMUST00|8778|Vgll4|ENSMUSG0C|true|.|.|.|.|.|.|chr6|114853844|114919520|8.05355|5.36011|5.51688|
|chr6|128764193|128764478|182|Distal Intergenic|6|128780043|128788486|8444|2|ENSMUST00|24008|Klrb1c|ENSMUSG0C|false|chr6|128764324|128764808|8.08052|18.53858|16.22368|.|.|.|.|.|.|
|chr6|125331016|125331240|182|Exon (ENSMUST00000176655.2/|6|125321521|125344612|23092|1|ENSMUST00|9496|Scnn1a|ENSMUSG0C|false|chr6|125330708|125331390|11.39974|34.91658|32.39005|chr6|125330782|125331975|3.80806|3.53097|2.03908|
|chr6|126939573|126940002|182|Promoter (<=1kb)|6|126923050|126939587|16538|2|ENSMUST00|0|Rad51ap1|ENSMUSG0C|false|chr6|126939092|126940812|28.00751|87.44511|84.14996|chr6|126939312|126941132|4.92309|3.94106|2.97004|
|chr6|128438529|128438782|182|Promoter (<=1kb)|6|128429735|128438677|8943|2|ENSMUST00|0|Fkbp4|ENSMUSG0C|false|chr6|128437487|128439261|36.27325|129.91212|125.42401|.|.|.|.|.|.|
|chr7|6331217|6331438|182|Promoter (<=1kb)|7|6315660|6331285|15626|2|ENSMUST00|0|Zfp583|ENSMUSG0C|false|chr7|6330784|6331810|18.70486|58.09692|55.27494|chr7|6330320|6331730|3.80694|3.36815|2.06967|
|chr7|55850122|55850455|182|Intron (ENSMUST00000168271.3|7|55842255|55930471|88217|1|ENSMUST00|7868|Cyfip1|ENSMUSG0C|false|.|.|.|.|.|.|chr7|55841471|55862799|4.7417|3.9553|2.79473|
|chr7|56050103|56050404|182|Promoter (<=1kb)|7|56050155|56231794|181640|1|ENSMUST00|0|Herc2|ENSMUSG0C|false|chr7|56049558|56051192|25.56571|75.64988|72.57267|.|.|.|.|.|.|
|chr7|43579531|43579837|182|Promoter (<=1kb)|7|43579610|43593235|13626|1|ENSMUST00|0|Zfp719|ENSMUSG0C|false|chr7|43579211|43580522|27.41053|90.23362|86.88379|chr7|43579006|43581706|4.69305|3.98469|2.73998|
|chr7|51861888|51862241|182|Promoter (<=1kb)|7|51862015|51994975|132961|1|ENSMUST00|0|Gas2|ENSMUSG0C|false|chr7|51861279|51862822|25.78858|76.95798|73.85856|chr7|51861295|51863047|5.53185|4.04795|3.4781|
|chr7|65932250|65932482|182|Intron (ENSMUST00000176199.2|7|65953012|65963193|10182|1|ENSMUST00|-20530|Pcsk6|ENSMUSG0C|true|.|.|.|.|.|.|chr7|65929189|65934196|6.32764|4.65183|4.10713|
|chr7|65387156|65387350|182|Distal Intergenic|7|65296165|65371239|75075|2|ENSMUST00|-15918|Tjp1|ENSMUSG0C|true|.|.|.|.|.|.|chr7|65385226|65389130|6.48767|4.73299|4.25669|

| | | | | | | | | | | | | | | | | | | | | | | | | | | |
|---|---|---|---|---|---|---|---|---|---|---|---|---|---|---|---|---|---|---|---|---|---|---|---|---|---|---|
| chr7 | 65417282 | 65417599 | 182 | Distal Intergenic | 7 | 65296165 | 65371239 | 75075 | 2 | ENSMUST00 | -46044 | Tjp1 | ENSMUSG0C | true | . | . | . | . | . | . | chr7 | 65409463 | 65433128 | 6.06932 | 4.54697 | 3.8847 |
| chr7 | 64160708 | 64160997 | 182 | Intron (ENSMUST00000085222.6 | 7 | 64153928 | 64269132 | 115205 | 1 | ENSMUST00 | 6781 | Trpm1 | ENSMUSG0C | false | . | . | . | . | . | . | chr7 | 64158668 | 64162494 | 4.08646 | 3.49224 | 2.29226 |
| chr7 | 80681280 | 80681489 | 182 | Intron (ENSMUST00000122255.2 | 7 | 80586627 | 80688877 | 102251 | 2 | ENSMUST00 | 7388 | Crtc3 | ENSMUSG0C | false | . | . | . | . | . | . | chr7 | 80679765 | 80688014 | 5.47936 | 4.26578 | 3.4104 |
| chr7 | 80121187 | 80121504 | 182 | Distal Intergenic | 7 | 80094846 | 80115392 | 20547 | 2 | ENSMUST00 | -5796 | Idh2 | ENSMUSG0C | false | . | . | . | . | . | . | chr7 | 80120639 | 80123945 | 5.92421 | 4.52192 | 3.77874 |
| chr7 | 79583255 | 79583547 | 182 | Distal Intergenic | 7 | 79593369 | 79594924 | 1556 | 2 | ENSMUST00 | 11377 | Rhcg | ENSMUSG0C | false | . | . | . | . | . | . | chr7 | 79582196 | 79584968 | 3.68427 | 3.25928 | 1.9655 |
| chr7 | 70346072 | 70346495 | 182 | Distal Intergenic | 7 | 70351946 | 70360593 | 8648 | 2 | ENSMUST00 | 14098 | Nr2f2 | ENSMUSG0C | false | chr7 | 70345833 | 70346633 | 16.93343 | 41.6405 | 39.02962 | chr7 | 70342111 | 70350630 | 5.07918 | 4.15019 | 3.06419 |
| chr7 | 70369381 | 70369760 | 182 | Promoter (2-3kb) | 7 | 70354439 | 70366735 | 12297 | 2 | ENSMUST00 | -2647 | Nr2f2 | ENSMUSG0C | false | chr7 | 70369313 | 70370722 | 21.3896 | 57.0703 | 54.26409 | chr7 | 70369479 | 70370899 | 2.97914 | 2.86691 | 1.38556 |
| chr7 | 67372437 | 67373524 | 182 | Promoter (<=1kb) | 7 | 67231163 | 67372858 | 141696 | 2 | ENSMUST00 | 0 | Mef2a | ENSMUSG0C | false | chr7 | 67371358 | 67373918 | 33.94791 | 109.87469 | 106.04007 | chr7 | 67373519 | 67374144 | 3.13148 | 3.0855 | 1.47828 |
| chr7 | 29238157 | 29238516 | 182 | Promoter (<=1kb) | 7 | 29238329 | 29247529 | 9201 | 1 | ENSMUST00 | 0 | Yif1b | ENSMUSG0C | false | chr7 | 29238327 | 29238999 | 16.91761 | 46.11281 | 43.44656 | chr7 | 29236563 | 29239872 | 3.80381 | 3.49004 | 2.03667 |
| chr7 | 29180483 | 29180734 | 182 | Promoter (<=1kb) | 7 | 29174188 | 29180552 | 6365 | 2 | ENSMUST00 | 0 | Psmd8 | ENSMUSG0C | false | chr7 | 29179346 | 29180586 | 29.50911 | 91.04776 | 87.68056 | . | . | . | . | . | . |
| chr7 | 28587704 | 28588009 | 182 | Intron (ENSMUST00000040531.8 | 7 | 28569299 | 28598086 | 28788 | 2 | ENSMUST00 | 10077 | Pak4 | ENSMUSG0C | false | . | . | . | . | . | . | chr7 | 28584075 | 28589478 | 8.07384 | 5.35319 | 5.52158 |
| chr7 | 28598122 | 28598319 | 182 | Promoter (<=1kb) | 7 | 28399549 | 28598144 | 198596 | 2 | ENSMUST00 | 0 | Pak4 | ENSMUSG0C | false | chr7 | 28596642 | 28598238 | 21.07794 | 58.1971 | 55.37426 | chr7 | 28590479 | 28598779 | 4.7239 | 3.86249 | 2.79695 |
| chr7 | 79125779 | 79126091 | 182 | Intron (ENSMUST00000032827.7 | 7 | 79117027 | 79131018 | 13992 | 2 | ENSMUST00 | 4927 | Hapln3 | ENSMUSG0C | false | . | . | . | . | . | . | chr7 | 79125317 | 79127444 | 4.24043 | 3.73578 | 2.37379 |
| chr7 | 79130961 | 79131242 | 182 | Promoter (<=1kb) | 7 | 79117027 | 79131018 | 13992 | 2 | ENSMUST00 | 0 | Hapln3 | ENSMUSG0C | false | . | . | . | . | . | . | chr7 | 79130142 | 79132933 | 4.35974 | 3.67056 | 2.51303 |
| chr7 | 90030295 | 90030691 | 182 | Distal Intergenic | 7 | 89954656 | 89980983 | 26328 | 2 | ENSMUST00 | -49313 | Eed | ENSMUSG0C | false | . | . | . | . | . | . | chr7 | 90029068 | 90030948 | 3.51 | 3.26999 | 1.82659 |
| chr7 | 92800353 | 92800535 | 182 | Intron (ENSMUST00000032879.8 | 7 | 92819905 | 92836647 | 16743 | 1 | ENSMUST00 | -19370 | Rab30 | ENSMUSG0C | true | . | . | . | . | . | . | chr7 | 92798575 | 92801969 | 4.61944 | 3.93173 | 2.7116 |
| chr7 | 92741508 | 92741743 | 182 | Promoter (<=1kb) | 7 | 92741603 | 92837117 | 95515 | 1 | ENSMUST00 | 0 | Rab30 | ENSMUSG0C | true | chr7 | 92740196 | 92743123 | 27.03504 | 88.5911 | 85.27629 | . | . | . | . | . | . |
| chr7 | 92741896 | 92742131 | 182 | Promoter (<=1kb) | 7 | 92741603 | 92837117 | 95515 | 1 | ENSMUST00 | 294 | Rab30 | ENSMUSG0C | true | chr7 | 92740196 | 92743123 | 27.03504 | 88.5911 | 85.27629 | . | . | . | . | . | . |
| chr7 | 101414812 | 101415078 | 182 | Distal Intergenic | 7 | 101421691 | 101512829 | 91139 | 1 | ENSMUST00 | -6613 | Pde2a | ENSMUSG0C | false | . | . | . | . | . | . | chr7 | 101413160 | 101416221 | 5.64616 | 4.46955 | 3.53528 |
| chr7 | 114585051 | 114585228 | 182 | Distal Intergenic | 7 | 114550166 | 114562972 | 12807 | 2 | ENSMUST00 | -22080 | Cyp2r1 | ENSMUSG0C | false | . | . | . | . | . | . | chr7 | 114580734 | 114585793 | 4.94026 | 4.04895 | 2.95084 |
| chr7 | 114416216 | 114416750 | 182 | Promoter (<=1kb) | 7 | 114415281 | 114539251 | 123971 | 1 | ENSMUST00 | 936 | Pde3b | ENSMUSG0C | false | chr7 | 114414652 | 114417189 | 26.18681 | 91.82585 | 88.44535 | chr7 | 114416495 | 114418375 | 3.71596 | 3.44861 | 1.96705 |
| chr7 | 101320966 | 101321320 | 182 | Promoter (<=1kb) | 7 | 101321113 | 101346308 | 25196 | 1 | ENSMUST00 | 0 | Stard10 | ENSMUSG0C | false | chr7 | 101320580 | 101321978 | 23.87381 | 72.67255 | 69.64001 | chr7 | 101320561 | 101323782 | 4.03379 | 3.60603 | 2.23446 |
| chr7 | 100121310 | 100121692 | 182 | Promoter (<=1kb) | 7 | 100082111 | 100121565 | 39455 | 2 | ENSMUST00 | 0 | Pold3 | ENSMUSG0C | false | chr7 | 100120657 | 100122085 | 46.23398 | 165.7672 | 159.91937 | . | . | . | . | . | . |
| chr7 | 125472100 | 125472614 | 182 | Exon (ENSMUST00000033006.8/ | 7 | 125471167 | 125478559 | 7393 | 2 | ENSMUST00 | 5945 | Nsmce1 | ENSMUSG0C | false | . | . | . | . | . | . | chr7 | 125470936 | 125474354 | 4.07645 | 3.67094 | 2.28354 |
| chr7 | 114254333 | 114254844 | 182 | Promoter (<=1kb) | 7 | 114215559 | 114254680 | 39122 | 2 | ENSMUST00 | 0 | Copb1 | ENSMUSG0C | false | chr7 | 114253808 | 114255033 | 24.69612 | 110.86208 | 107.00209 | . | . | . | . | . | . |
| chr7 | 112479256 | 112479489 | 182 | Intron (ENSMUST00000106643.2 | 7 | 112501177 | 112547610 | 46434 | 1 | ENSMUST00 | -21688 | Parva | ENSMUSG0C | true | . | . | . | . | . | . | chr7 | 112477579 | 112486053 | 5.9433 | 4.42533 | 3.78015 |
| chr7 | 112428217 | 112428426 | 182 | Promoter (<=1kb) | 7 | 112427905 | 112552929 | 125025 | 1 | ENSMUST00 | 313 | Parva | ENSMUSG0C | true | chr7 | 112427013 | 112429707 | 31.16539 | 106.97112 | 103.21663 | . | . | . | . | . | . |
| chr7 | 112496215 | 112496412 | 182 | Intron (ENSMUST00000106643.2 | 7 | 112501177 | 112547610 | 46434 | 1 | ENSMUST00 | -4765 | Parva | ENSMUSG0C | true | . | . | . | . | . | . | chr7 | 112487030 | 112537122 | 5.64568 | 4.33316 | 3.55068 |
| chr7 | 112405165 | 112405366 | 182 | Promoter (1-2kb) | 7 | 112407108 | 112413053 | 5946 | 1 | ENSMUST00 | -1742 | Micalcl | ENSMUSG0C | false | . | . | . | . | . | . | chr7 | 112403199 | 112406515 | 5.44368 | 4.32878 | 3.37289 |
| chr7 | 112148018 | 112148324 | 182 | Intron (ENSMUST00000033036.5 | 7 | 112116019 | 112159057 | 43039 | 2 | ENSMUST00 | 10733 | Dkk3 | ENSMUSG0C | false | . | . | . | . | . | . | chr7 | 112147161 | 112148260 | 4.46122 | 3.73921 | 2.60101 |
| chr7 | 122969546 | 122970066 | 182 | Promoter (<=1kb) | 7 | 122970564 | 123002557 | 31994 | 1 | ENSMUST00 | -498 | Rbbp6 | ENSMUSG0C | false | chr7 | 122969333 | 122971749 | 35.07281 | 114.97435 | 110.99648 | . | . | . | . | . | . |
| chr7 | 110879871 | 110880323 | 182 | Intron (ENSMUST00000127935.2 | 7 | 110850607 | 110862953 | 12347 | 2 | ENSMUST00 | -16919 | Lyve1 | ENSMUSG0C | false | . | . | . | . | . | . | . | . | . | . | . | . |
| chr7 | 110844314 | 110844514 | 182 | Promoter (<=1kb) | 7 | 110813531 | 110844457 | 30927 | 2 | ENSMUST00 | 0 | Rnf141 | ENSMUSG0C | false | chr7 | 110843466 | 110844572 | 36.39222 | 115.17619 | 111.19315 | chr7 | 110842615 | 110844737 | 3.93304 | 3.58676 | 2.16146 |
| chr7 | 127841404 | 127841878 | 182 | Promoter (<=1kb) | 7 | 127841764 | 127849019 | 7256 | 1 | ENSMUST00 | 0 | Stx4a | ENSMUSG0C | false | chr7 | 127840892 | 127842835 | 28.35378 | 86.13386 | 82.86658 | chr7 | 127840746 | 127841448 | 3.94716 | 3.61201 | 2.124 |
| chr7 | 127470994 | 127471937 | 182 | Promoter (<=1kb) | 7 | 127471019 | 127473854 | 2836 | 1 | ENSMUST00 | 0 | Prr14 | ENSMUSG0C | false | chr7 | 127470082 | 127472628 | 27.78601 | 91.88203 | 88.49705 | . | . | . | . | . | . |
| chr7 | 46639597 | 46639844 | 182 | Promoter (<=1kb) | 7 | 46443168 | 46639807 | 196640 | 2 | ENSMUST00 | 0 | Sergef | ENSMUSG0C | false | chr7 | 46639184 | 46640418 | 25.16654 | 87.04672 | 83.76057 | chr7 | 46639005 | 46641113 | 3.78274 | 3.48018 | 2.01663 |
| chr7 | 101905743 | 101906200 | 182 | Promoter (<=1kb) | 7 | 101905837 | 101911901 | 6065 | 1 | ENSMUST00 | 0 | Lamtor1 | ENSMUSG0C | false | chr7 | 101905830 | 101906676 | 28.90751 | 104.81161 | 101.11721 | chr7 | 101905133 | 101910546 | 5.38251 | 4.15439 | 3.35869 |
| chr7 | 128461067 | 128461503 | 182 | Promoter (<=1kb) | 7 | 128441403 | 128461209 | 19807 | 2 | ENSMUST00 | 0 | Tial1 | ENSMUSG0C | false | chr7 | 128460002 | 128461770 | 30.19165 | 95.18787 | 91.73401 | . | . | . | . | . | . |
| chr7 | 130519390 | 130519944 | 182 | Promoter (<=1kb) | 7 | 130391526 | 130519530 | 128005 | 2 | ENSMUST00 | 0 | Ate1 | ENSMUSG0C | false | chr7 | 130518559 | 130520450 | 33.0841 | 102.23056 | 98.60399 | . | . | . | . | . | . |
| chr7 | 120634949 | 120635247 | 182 | Promoter (<=1kb) | 7 | 120635176 | 120659478 | 24303 | 1 | ENSMUST00 | 0 | Uqcrc2 | ENSMUSG0C | false | chr7 | 120634318 | 120636330 | 39.37288 | 136.56555 | 131.82602 | . | . | . | . | . | . |
| chr7 | 119895603 | 119895872 | 182 | Promoter (<=1kb) | 7 | 119895861 | 119916754 | 20894 | 1 | ENSMUST00 | 0 | Lyrm1 | ENSMUSG0C | false | chr7 | 119894701 | 119896756 | 21.02725 | 63.18829 | 60.29588 | . | . | . | . | . | . |
| chr7 | 132232969 | 132233241 | 182 | Intron (ENSMUST00000124096.2 | 7 | 132270819 | 132278625 | 7807 | 2 | ENSMUST00 | 45384 | Chst15 | ENSMUSG0C | false | . | . | . | . | . | . | . | . | . | . | . | . |
| chr7 | 132307825 | 132308267 | 182 | Intron (ENSMUST00000124096.2 | 7 | 132235780 | 132316714 | 80935 | 2 | ENSMUST00 | 8447 | Chst15 | ENSMUSG0C | false | . | . | . | . | . | . | . | . | . | . | . | . |
| chr7 | 132316254 | 132317021 | 182 | Promoter (<=1kb) | 7 | 132235780 | 132316714 | 80935 | 2 | ENSMUST00 | 0 | Chst15 | ENSMUSG0C | false | chr7 | 132315520 | 132317720 | 20.25082 | 84.28156 | 81.04926 | . | . | . | . | . | . |
| chr7 | 132389432 | 132389623 | 182 | Intron (ENSMUST00000124096.2 | 7 | 132235780 | 132317228 | 81449 | 2 | ENSMUST00 | -72205 | Chst15 | ENSMUSG0C | false | . | . | . | . | . | . | . | . | . | . | . | . |
| chr7 | 132747283 | 132747551 | 182 | Intron (ENSMUST00000124096.2 | 7 | 132713365 | 132776916 | 63552 | 2 | ENSMUST00 | 29365 | Fam53b | ENSMUSG0C | false | . | . | . | . | . | . | . | . | . | . | . | . |
| chr7 | 132813566 | 132814014 | 182 | Promoter (<=1kb) | 7 | 132712088 | 132813886 | 101799 | 2 | ENSMUST00 | 0 | Fam53b | ENSMUSG0C | false | chr7 | 132811861 | 132814347 | 24.22294 | 75.10754 | 72.03808 | . | . | . | . | . | . |
| chr7 | 132726615 | 132726855 | 182 | Intron (ENSMUST00000124096.2 | 7 | 132713365 | 132776916 | 63552 | 2 | ENSMUST00 | 50061 | Fam53b | ENSMUSG0C | false | . | . | . | . | . | . | chr7 | 132722805 | 132729612 | 4.35419 | 3.59271 | 2.51349 |
| chr7 | 132775419 | 132775607 | 182 | Promoter (1-2kb) | 7 | 132713365 | 132776916 | 63552 | 2 | ENSMUST00 | 1309 | Fam53b | ENSMUSG0C | false | . | . | . | . | . | . | chr7 | 132775295 | 132776705 | 3.68371 | 3.39104 | 1.96331 |
| chr7 | 133776635 | 133776937 | 182 | Promoter (<=1kb) | 7 | 133720942 | 133776803 | 55862 | 2 | ENSMUST00 | 0 | Dhx32 | ENSMUSG0C | false | chr7 | 133775865 | 133777162 | 26.70311 | 93.76232 | 90.33739 | chr7 | 133774191 | 133777235 | 4.43927 | 3.87218 | 2.53305 |
| chr7 | 102223039 | 102223515 | 182 | Promoter (<=1kb) | 7 | 102223045 | 102237226 | 14182 | 1 | ENSMUST00 | 0 | Pgap2 | ENSMUSG0C | false | chr7 | 102222429 | 102223527 | 27.66126 | 83.97217 | 80.74496 | chr7 | 102222557 | 102224468 | 3.46955 | 3.20141 | 1.76451 |
| chrX | 13070567 | 13071028 | 182 | Promoter (<=1kb) | 20 | 13071498 | 13123449 | 51952 | 1 | ENSMUST00 | -470 | Usp9x | ENSMUSG0C | false | chrX | 13070252 | 13073605 | 25.27427 | 69.73028 | 66.74294 | . | . | . | . | . | . |
| chrX | 13574402 | 13574608 | 182 | Intron (ENSMUST00000156096.2 | 20 | 13533340 | 13559331 | 25992 | 2 | ENSMUST00 | -15072 | Cask | ENSMUSG0C | false | . | . | . | . | . | . | . | . | . | . | . | . |
| chr7 | 109672152 | 109672370 | 182 | Distal Intergenic | 7 | 109703690 | 109712189 | 8500 | 1 | ENSMUST00 | -31320 | Akip1 | ENSMUSG0C | false | chr7 | 109671273 | 109672671 | 20.88808 | 77.73511 | 74.61916 | chr7 | 109669958 | 109673113 | 4.43113 | 3.79156 | 2.56081 |
| chr7 | 109576705 | 109577075 | 182 | Intron (ENSMUST00000079282.2 | 7 | 109523920 | 109616568 | 92649 | 2 | ENSMUST00 | 39493 | St5 | ENSMUSG0C | false | . | . | . | . | . | . | chr7 | 109574563 | 109578210 | 4.32346 | 3.74777 | 2.46892 |
| chr7 | 109628968 | 109629294 | 182 | Distal Intergenic | 7 | 109523911 | 109617147 | 93237 | 2 | ENSMUST00 | -11822 | St5 | ENSMUSG0C | false | . | . | . | . | . | . | chr7 | 109627739 | 109630692 | 4.97021 | 4.00109 | 3.03977 |
| chrX | 20617726 | 20618071 | 182 | Promoter (<=1kb) | 20 | 20617783 | 20650901 | 33119 | 1 | ENSMUST00 | 0 | Rbm10 | ENSMUSG0C | false | chrX | 20617730 | 20618826 | 24.3552 | 66.39809 | 63.45943 | . | . | . | . | . | . |
| chr7 | 144770235 | 144770450 | 182 | Distal Intergenic | 7 | 144650503 | 144751974 | 101472 | 2 | ENSMUST00 | -18262 | Ano1 | ENSMUSG0C | false | . | . | . | . | . | . | . | . | . | . | . | . |
| chr7 | 144449725 | 144449926 | 182 | Promoter (<=1kb) | 7 | 144438786 | 144449774 | 10989 | 2 | ENSMUST00 | 0 | Cttn | ENSMUSG0C | false | . | . | . | . | . | . | chr7 | 144449756 | 144450730 | 6.48739 | 4.39986 | 4.25873 |
| chrX | 48450425 | 48450604 | 182 | Intron (ENSMUST00000114958.2 | 20 | 48411046 | 48454169 | 43124 | 2 | ENSMUST00 | 3565 | Elf4 | ENSMUSG0C | true | . | . | . | . | . | . | . | . | . | . | . | . |
| chrX | 7789910 | 7790105 | 182 | Promoter (<=1kb) | 20 | 7789994 | 7812483 | 22490 | 1 | ENSMUST00 | 0 | Gripap1 | ENSMUSG0C | false | chrX | 7789795 | 7790688 | 16.54316 | 39.53096 | 36.94807 | . | . | . | . | . | . |
| chrX | 10216352 | 10216641 | 182 | Promoter (<=1kb) | 20 | 10158216 | 10216453 | 58238 | 2 | ENSMUST00 | 0 | Rpgr | ENSMUSG0C | false | chrX | 10215717 | 10216848 | 18.84082 | 47.13427 | 44.45482 | . | . | . | . | . | . |
| chrX | 96095891 | 96096132 | 182 | Promoter (<=1kb) | 20 | 96096042 | 96168552 | 72511 | 1 | ENSMUST00 | 0 | Msn | ENSMUSG0C | false | chrX | 96095680 | 96097279 | 20.67895 | 53.40692 | 50.64761 | . | . | . | . | . | . |
| chrX | 134059121 | 134059606 | 182 | Promoter (<=1kb) | 20 | 134059187 | 134086046 | 26860 | 1 | ENSMUST00 | 0 | Cstf2 | ENSMUSG0C | false | chrX | 134058855 | 134059973 | 18.84082 | 47.13427 | 44.45482 | . | . | . | . | . | . |
| chrX | 103356325 | 103356606 | 182 | Promoter (<=1kb) | 20 | 103356476 | 103396092 | 39617 | 1 | ENSMUST00 | 0 | Chic1 | ENSMUSG0C | false | chrX | 103356107 | 103357299 | 18.38128 | 45.59164 | 42.93153 | . | . | . | . | . | . |
| chrX | 163908882 | 163909326 | 182 | Promoter (<=1kb) | 20 | 163909017 | 163933666 | 24650 | 1 | ENSMUST00 | 0 | Ap1s2 | ENSMUSG0C | false | chrX | 163908206 | 163908912 | 16.08362 | 38.04472 | 35.47989 | . | . | . | . | . | . |
| chr8 | 13676777 | 13677037 | 182 | Promoter (<=1kb) | 8 | 13595810 | 13677451 | 81642 | 2 | ENSMUST00 | 414 | Rasa3 | ENSMUSG0C | false | chr8 | 13676275 | 13678189 | 36.39105 | 113.08218 | 109.16298 | chr8 | 13675870 | 13676805 | 3.34678 | 3.18377 | 1.64588 |
| chr8 | 13600121 | 13600365 | 182 | Intron (ENSMUST00000117551.2 | 8 | 13595809 | 13606108 | 10300 | 2 | ENSMUST00 | 5743 | Rasa3 | ENSMUSG0C | false | . | . | . | . | . | . | chr8 | 13600199 | 13601793 | 3.53609 | 3.20606 | 1.83944 |
| chr8 | 13648462 | 13648789 | 182 | Intron (ENSMUST00000117551.2 | 8 | 13595810 | 13677451 | 81642 | 2 | ENSMUST00 | 28662 | Rasa3 | ENSMUSG0C | false | . | . | . | . | . | . | chr8 | 13647613 | 13649538 | 4.33751 | 3.82005 | 2.44817 |
| chr8 | 18846194 | 18846577 | 182 | Promoter (<=1kb) | 8 | 18846277 | 18891361 | 45085 | 1 | ENSMUST00 | 0 | Agpat5 | ENSMUSG0C | false | chr8 | 18845670 | 18847232 | 29.99031 | 99.80121 | 96.2382 | . | . | . | . | . | . |
| chr8 | 27089130 | 27089317 | 182 | Intron (ENSMUST00000178514.2 | 8 | 27085598 | 27123436 | 37839 | 1 | ENSMUST00 | 3533 | Gpr124 | ENSMUSG0C | false | . | . | . | . | . | . | chr8 | 27083119 | 27091776 | 4.52211 | 3.74431 | 2.66045 |
| chr8 | 11234112 | 11234377 | 182 | Intron (ENSMUST00000033898.9 | 8 | 11212401 | 11216801 | 4401 | 2 | ENSMUST00 | -17312 | Col4a1 | ENSMUSG0C | false | . | . | . | . | . | . | . | . | . | . | . | . |
| chr8 | 11270880 | 11271169 | 182 | Intron (ENSMUST00000033898.9 | 8 | 11312805 | 11449287 | 136483 | 1 | ENSMUST00 | -41636 | Col4a2 | ENSMUSG0C | false | . | . | . | . | . | . | . | . | . | . | . | . |
| chr8 | 11635709 | 11635967 | 182 | Promoter (<=1kb) | 8 | 11611583 | 11635754 | 24172 | 2 | ENSMUST00 | 0 | Ankrd10 | ENSMUSG0C | false | chr8 | 11634291 | 11636731 | 43.41453 | 148.15424 | 142.93486 | . | . | . | . | . | . |
| chr8 | 11783383 | 11783590 | 182 | Promoter (<=1kb) | 8 | 11782641 | 11835219 | 52579 | 1 | ENSMUST00 | 743 | Arhgef7 | ENSMUSG0C | false | . | . | . | . | . | . | chr8 | 11782632 | 11786218 | 4.55476 | 3.85609 | 2.6779 |
| chr8 | 36581190 | 36581477 | 182 | Intron (ENSMUST00000033923.8 | 8 | 36572145 | 36577269 | 5125 | 2 | ENSMUST00 | -3922 | Dlc1 | ENSMUSG0C | true | . | . | . | . | . | . | chr8 | 36575944 | 36612020 | 5.64773 | 4.3412 | 3.54039 |
| chr8 | 36585558 | 36585791 | 182 | Intron (ENSMUST00000033923.8 | 8 | 36572145 | 36577269 | 5125 | 2 | ENSMUST00 | -8290 | Dlc1 | ENSMUSG0C | true | . | . | . | . | . | . | chr8 | 36575944 | 36612020 | 5.64773 | 4.3412 | 3.54039 |
| chr8 | 36745972 | 36746434 | 182 | Intron (ENSMUST00000163663.2 | 8 | 36567751 | 36733054 | 165304 | 2 | ENSMUST00 | -12919 | Dlc1 | ENSMUSG0C | true | . | . | . | . | . | . | . | . | . | . | . | . |
| chr8 | 35495385 | 35495626 | 182 | Promoter (<=1kb) | 8 | 35465265 | 35495533 | 30269 | 2 | ENSMUST00 | 0 | Eri1 | ENSMUSG0C | false | chr8 | 35494364 | 35496178 | 36.66269 | 126.29414 | 121.93508 | . | . | . | . | . | . |
| chr8 | 22974619 | 22974855 | 182 | Promoter (<=1kb) | 8 | 22974844 | 23149933 | 175090 | 1 | ENSMUST00 | 0 | Ank1 | ENSMUSG0C | false | chr8 | 22974162 | 22975667 | 22.21283 | 62.70797 | 59.8237 | chr8 | 22974121 | 22976043 | 5.13089 | 4.02553 | 3.13402 |
| chr8 | 23352418 | 23352740 | 182 | Distal Intergenic | 8 | 23411502 | 23449632 | 38131 | 1 | ENSMUST00 | -58762 | Sfrp1 | ENSMUSG0C | false | . | . | . | . | . | . | chr8 | 23352577 | 23353420 | 3.62256 | 3.27379 | 1.91225 |
| chr8 | 25016566 | 25017188 | 182 | Promoter (<=1kb) | 8 | 24949611 | 25016922 | 67312 | 2 | ENSMUST00 | 0 | Adam9 | ENSMUSG0C | false | chr8 | 25015978 | 25017012 | 34.16929 | 120.7392 | 116.5701 | . | . | . | . | . | . |
| chr8 | 25070570 | 25071110 | 182 | Intron (ENSMUST00000064883.7 | 8 | 25062433 | 25091346 | 28914 | 2 | ENSMUST00 | 20236 | Plekha2 | ENSMUSG0C | true | . | . | . | . | . | . | chr8 | 25067309 | 25079381 | 5.19206 | 4.11384 | 3.1814 |
| chr8 | 25521476 | 25521691 | 182 | Promoter (1-2kb) | 8 | 25519837 | 25555729 | 35893 | 1 | ENSMUST00 | 1640 | Fgfr1 | ENSMUSG0C | true | . | . | . | . | . | . | chr8 | 25521452 | 25558014 | 5.35885 | 4.21426 | 3.3104 |
| chr8 | 33914916 | 33915187 | 182 | Intron (ENSMUST00000053251.6 | 8 | 33803939 | 33929805 | 125867 | 2 | ENSMUST00 | 14618 | Rbpms | ENSMUSG0C | true | . | . | . | . | . | . | chr8 | 33863865 | 33927704 | 6.32022 | 4.67603 | 4.08842 |
| chr8 | 33976569 | 33976745 | 182 | Distal Intergenic | 8 | 33782643 | 33929863 | 147221 | 2 | ENSMUST00 | -46707 | Rbpms | ENSMUSG0C | true | . | . | . | . | . | . | . | . | . | . | . | . |
| chr8 | 33786448 | 33786707 | 182 | Intron (ENSMUST00000053251.6 | 8 | 33784072 | 33797214 | 13143 | 2 | ENSMUST00 | 10507 | Rbpms | ENSMUSG0C | true | . | . | . | . | . | . | chr8 | 33784277 | 33787664 | 5.5958 | 4.22668 | 3.52901 |
| chr8 | 33807772 | 33808113 | 182 | Promoter (<=1kb) | 8 | 33784061 | 33806836 | 22776 | 2 | ENSMUST00 | -937 | Rbpms | ENSMUSG0C | true | . | . | . | . | . | . | chr8 | 33797013 | 33821269 | 7.29084 | 5.02722 | 4.89898 |
| chr8 | 64880889 | 64881126 | 182 | Distal Intergenic | 8 | 64739863 | 64849818 | 109956 | 2 | ENSMUST00 | -31072 | Klhl2 | ENSMUSG0C | false | . | . | . | . | . | . | . | . | . | . | . | . |
| chr8 | 57652106 | 57652831 | 182 | Promoter (<=1kb) | 8 | 57524700 | 57653003 | 128304 | 2 | ENSMUST00 | 172 | Galnt7 | ENSMUSG0C | false | chr8 | 57651425 | 57653684 | 28.22734 | 101.53345 | 97.92473 | . | . | . | . | . | . |
| chr8 | 45689719 | 45690016 | 182 | Promoter (1-2kb) | 8 | 45688337 | 45759101 | 70765 | 1 | ENSMUST00 | 1383 | Sorbs2 | ENSMUSG0C | false | . | . | . | . | . | . | . | . | . | . | . | . |
| chr8 | 46617163 | 46617540 | 182 | Promoter (<=1kb) | 8 | 46617449 | 46638706 | 21258 | 1 | ENSMUST00 | 0 | Casp3 | ENSMUSG0C | false | chr8 | 46616644 | 46617278 | 21.42557 | 69.98924 | 66.99868 | . | . | . | . | . | . |
| chr8 | 61225663 | 61225873 | 182 | Promoter (1-2kb) | 8 | 61224171 | 61396072 | 171902 | 1 | ENSMUST00 | 1493 | Sh3rf1 | ENSMUSG0C | false | chr8 | 61225383 | 61226302 | 5.28342 | 9.57744 | 7.40675 | chr8 | 61225708 | 61227995 | 3.70003 | 3.32417 | 1.98205 |
| chr8 | 60993089 | 60993575 | 182 | Promoter (<=1kb) | 8 | 60993195 | 61131339 | 138145 | 1 | ENSMUST00 | 0 | Nek1 | ENSMUSG0C | true | chr8 | 60992445 | 60994609 | 33.5288 | 108.10788 | 104.32592 | . | . | . | . | . | . |
| chr8 | 87472478 | 87472852 | 182 | Promoter (<=1kb) | 8 | 87468854 | 87472592 | 3739 | 2 | ENSMUST00 | 0 | Cbln1 | ENSMUSG0C | false | chr8 | 87472162 | 87473695 | 20.4053 | 65.4752 | 62.55133 | chr8 | 87468771 | 87473905 | 5.79353 | 4.39234 | 3.65572 |
| chr8 | 88361830 | 88362259 | 182 | Promoter (<=1kb) | 8 | 88331059 | 88362194 | 31136 | 2 | ENSMUST00 | 0 | Brd7 | ENSMUSG0C | false | chr8 | 88360901 | 88363027 | 43.4858 | 141.53311 | 136.59532 | . | . | . | . | . | . |
| chr8 | 85554762 | 85555408 | 182 | Promoter (<=1kb) | 8 | 85537640 | 85555271 | 17632 | 2 | ENSMUST00 | 0 | Dnaja2 | ENSMUSG0C | false | chr8 | 85554247 | 85556254 | 36.81759 | 121.2078 | 117.02259 | . | . | . | . | . | . |
| chr8 | 85840681 | 85841095 | 182 | Promoter (<=1kb) | 8 | 85717557 | 85840949 | 123393 | 2 | ENSMUST00 | 0 | Itfg1 | ENSMUSG0C | false | chr8 | 85839917 | 85841768 | 34.04953 | 116.30843 | 112.28568 | . | . | . | . | . | . |
| chr8 | 82773917 | 82774274 | 182 | Promoter (<=1kb) | 8 | 82763621 | 82774160 | 10540 | 2 | ENSMUST00 | 0 | Zfp330 | ENSMUSG0C | false | chr8 | 82773478 | 82774728 | 22.68762 | 81.35053 | 78.1749 | . | . | . | . | . | . |
| chr8 | 109692725 | 109693425 | 182 | Promoter (<=1kb) | 8 | 109671322 | 109693294 | 21973 | 2 | ENSMUST00 | 0 | Ist1 | ENSMUSG0C | false | chr8 | 109691717 | 109693283 | 27.7677 | 107.59161 | 103.82806 | . | . | . | . | . | . |
| chr8 | 110808516 | 110808817 | 182 | Promoter (2-3kb) | 8 | 110755525 | 110805924 | 50400 | 2 | ENSMUST00 | -2593 | Il34 | ENSMUSG0C | false | . | . | . | . | . | . | chr8 | 110803880 | 110810225 | 4.5376 | 3.71366 | 2.65908 |
| chr8 | 95131116 | 95131503 | 182 | Intron (ENSMUST00000034240.8 | 8 | 95099832 | 95142540 | 42709 | 2 | ENSMUST00 | 11037 | Kifc3 | ENSMUSG0C | false | . | . | . | . | . | . | . | . | . | . | . | . |
| chr8 | 72136440 | 72136827 | 182 | Promoter (1-2kb) | 8 | 72135259 | 72153142 | 17884 | 1 | ENSMUST00 | 1182 | Tpm4 | ENSMUSG0C | false | chr8 | 72134816 | 72136882 | 41.44235 | 135.77681 | 131.06697 | chr8 | 72136645 | 72138280 | 5.37089 | 4.33128 | 3.3064 |
| chr8 | 120494891 | 120495254 | 182 | Intron (ENSMUST00000127664.1 | 8 | 120488447 | 120581390 | 92944 | 1 | ENSMUST00 | 6445 | Gse1 | ENSMUSG0C | false | . | . | . | . | . | . | chr8 | 120489514 | 120496763 | 8.44988 | 5.58728 | 5.83793 |
| chr8 | 120370013 | 120370228 | 182 | Intron (ENSMUST00000127664.1 | 8 | 120488447 | 120581390 | 92944 | 1 | ENSMUST00 | -118219 | Gse1 | ENSMUSG0C | false | . | . | . | . | . | . | chr8 | 120360953 | 120384219 | 7.36608 | 4.94236 | 4.97938 |
| chr8 | 120264983 | 120265331 | 182 | Intron (ENSMUST00000127664.1 | 8 | 120230536 | 120553634 | 323099 | 1 | ENSMUST00 | 34448 | Gse1 | ENSMUSG0C | false | chr8 | 120264577 | 120265125 | 7.13425 | 13.57188 | 11.33167 | chr8 | 120263198 | 120270766 | 5.71876 | 4.35069 | 3.62232 |
| chr8 | 120427109 | 120427307 | 182 | Intron (ENSMUST00000127664.1 | 8 | 120488447 | 120581390 | 92944 | 1 | ENSMUST00 | -61140 | Gse1 | ENSMUSG0C | false | . | . | . | . | . | . | chr8 | 120415621 | 120427536 | 6.84739 | 4.85598 | 4.53347 |
| chr8 | 120489110 | 120489570 | 182 | Promoter (<=1kb) | 8 | 120488447 | 120581390 | 92944 | 1 | ENSMUST00 | 664 | Gse1 | ENSMUSG0C | false | chr8 | 120486986 | 120490241 | 34.92444 | 106.42502 | 102.68874 | chr8 | 120489514 | 120496763 | 8.44988 | 5.58728 | 5.83793 |
| chr8 | 120504142 | 120504535 | 182 | Intron (ENSMUST00000127664.1 | 8 | 120488447 | 120581390 | 92944 | 1 | ENSMUST00 | 15696 | Gse1 | ENSMUSG0C | false | . | . | . | . | . | . | chr8 | 120497859 | 120511881 | 6.76427 | 4.83633 | 4.46181 |
| chr8 | 120067949 | 120068186 | 182 | Intron (ENSMUST00000127664.1 | 8 | 120082403 | 120086835 | 4433 | 2 | ENSMUST00 | 18649 | Zdhhc7 | ENSMUSG0C | false | . | . | . | . | . | . | chr8 | 120067680 | 120068875 | 8.04652 | 5.59221 | 5.48001 |
| chr8 | 119840419 | 119840854 | 182 | Promoter (<=1kb) | 8 | 119809225 | 119840522 | 31298 | 2 | ENSMUST00 | 0 | Cotl1 | ENSMUSG0C | false | chr8 | 119838819 | 119841220 | 27.03867 | 103.18376 | 99.53125 | chr8 | 119806261 | 119841342 | 7.1039 | 5.00847 | 4.72992 |
| chr8 | 70752388 | 70752744 | 182 | Promoter (1-2kb) | 8 | 70754679 | 70758677 | 3999 | 1 | ENSMUST00 | -1935 | Rab3a | ENSMUSG0C | false | . | . | . | . | . | . | chr8 | 70749419 | 70756647 | 5.88874 | 4.48997 | 3.73674 |
| chr8 | 104170428 | 104170663 | 182 | Promoter (<=1kb) | 8 | 104170513 | 104219092 | 48580 | 1 | ENSMUST00 | 0 | Bean1 | ENSMUSG0C | false | chr8 | 104170268 | 104171332 | 23.47016 | 67.50422 | 64.55198 | chr8 | 104170203 | 104171365 | 3.75689 | 3.36563 | 2.02127 |
| chr8 | 104341146 | 104341555 | 182 | Promoter (<=1kb) | 8 | 104340594 | 104347672 | 7079 | 1 | ENSMUST00 | 553 | Cmtm3 | ENSMUSG0C | false | chr8 | 104340179 | 104341479 | 24.91517 | 69.11501 | 66.14118 | chr8 | 104339769 | 104341914 | 3.91904 | 3.5879 | 2.16318 |
| chr8 | 104534232 | 104534712 | 182 | Promoter (<=1kb) | 8 | 104511028 | 104534636 | 23609 | 2 | ENSMUST00 | 0 | Nae1 | ENSMUSG0C | false | chr8 | 104533603 | 104535957 | 26.52689 | 93.43019 | 90.01563 | . | . | . | . | . | . |
| chr8 | 105225072 | 105225363 | 182 | Promoter (<=1kb) | 8 | 105225145 | 105253053 | 27909 | 1 | ENSMUST00 | 0 | D230025D16 | ENSMUSG0C | false | chr8 | 105224498 | 105225991 | 36.69013 | 115.69205 | 111.68276 | chr8 | 105224348 | 105228044 | 4.73248 | 3.95066 | 2.8182 |
| chr8 | 105937013 | 105937321 | 182 | Promoter (1-2kb) | 8 | 105935728 | 105938392 | 2665 | 2 | ENSMUST00 | 1071 | Psmb10 | ENSMUSG0C | false | chr8 | 105936542 | 105938522 | 28.22734 | 101.53345 | 97.92473 | . | . | . | . | . | . |
| chr8 | 106337904 | 106338272 | 182 | Promoter (<=1kb) | 8 | 106252548 | 106337988 | 85441 | 2 | ENSMUST00 | 0 | Smpd3 | ENSMUSG0C | false | chr8 | 106337553 | 106338028 | 11.23051 | 24.64421 | 22.24776 | chr8 | 106336593 | 106338473 | 5.85319 | 4.56058 | 3.75746 |
| chr8 | 107157737 | 107157966 | 182 | Intron (ENSMUST00000034400.3 | 8 | 107150661 | 107187470 | 36810 | 1 | ENSMUST00 | 7077 | Cyb5b | ENSMUSG0C | false | . | . | . | . | . | . | chr8 | 107157034 | 107158886 | 4.76385 | 3.88172 | 2.8436 |
| chr9 | 14784496 | 14784968 | 182 | Promoter (<=1kb) | 9 | 14784654 | 14834646 | 49993 | 1 | ENSMUST00 | 0 | Mre11a | ENSMUSG0C | false | chr9 | 14783999 | 14785418 | 33.04283 | 115.53907 | 111.53828 | . | . | . | . | . | . |
| chr8 | 111992286 | 111992523 | 182 | Promoter (<=1kb) | 8 | 111966908 | 111992302 | 25395 | 2 | ENSMUST00 | 0 | Adat1 | ENSMUSG0C | false | chr8 | 111991651 | 111992564 | 17.04032 | 66.12922 | 63.19824 | chr8 | 111989897 | 111992984 | 4.52719 | 3.81973 | 2.65618 |
| chr8 | 111939965 | 111940177 | 182 | Promoter (<=1kb) | 8 | 111940703 | 111953612 | 12910 | 1 | ENSMUST00 | -526 | Gabarapl2 | ENSMUSG0C | false | chr8 | 111939834 | 111941588 | 37.1464 | 154.63618 | 149.16937 | chr8 | 111939605 | 111940290 | 4.51492 | 3.80444 | 2.60848 |
| chr8 | 111743712 | 111743993 | 182 | Promoter (<=1kb) | 8 | 111710485 | 111743849 | 33365 | 2 | ENSMUST00 | 0 | Bcar1 | ENSMUSG0C | false | chr8 | 111742412 | 111744224 | 40.35223 | 130.59746 | 126.08324 | chr8 | 111710939 | 111747327 | 8.20913 | 5.4583 | 5.64085 |
| chr8 | 124721953 | 124722234 | 182 | Promoter (<=1kb) | 8 | 124722139 | 124734123 | 11985 | 1 | ENSMUST00 | 0 | Arv1 | ENSMUSG0C | false | chr8 | 124721276 | 124722857 | 33.41831 | 117.26761 | 113.21259 | . | . | . | . | . | . |
| chr8 | 124654402 | 124654704 | 182 | Intron (ENSMUST00000034464.6 | 8 | 124635766 | 124663369 | 27604 | 2 | ENSMUST00 | 8665 | 2310022B05 | ENSMUSG0C | false | . | . | . | . | . | . | chr8 | 124641308 | 124664635 | 6.64691 | 4.65387 | 4.38894 |
| chr9 | 31203786 | 31204101 | 182 | Intron (ENSMUST00000072634.8 | 9 | 31149557 | 31211815 | 62259 | 2 | ENSMUST00 | 7714 | Aplp2 | ENSMUSG0C | false | . | . | . | . | . | . | chr9 | 31201279 | 31210333 | 5.34556 | 4.22255 | 3.30008 |
| chr9 | 7872802 | 7873256 | 182 | Promoter (<=1kb) | 9 | 7848699 | 7873024 | 24326 | 2 | ENSMUST00 | 0 | Birc3 | ENSMUSG0C | false | chr9 | 7872437 | 7873684 | 25.53309 | 82.08177 | 78.89293 | . | . | . | . | . | . |
| chr9 | 7868008 | 7868289 | 182 | Intron (ENSMUST00000013949.9 | 9 | 7866193 | 7872857 | 6665 | 2 | ENSMUST00 | 4568 | Birc3 | ENSMUSG0C | false | . | . | . | . | . | . | . | . | . | . | . | . |
| chr9 | 44084775 | 44085002 | 182 | Promoter (<=1kb) | 9 | 44084950 | 44093926 | 8977 | 1 | ENSMUST00 | 0 | Usp2 | ENSMUSG0C | false | chr9 | 44084344 | 44085603 | 25.53309 | 82.08177 | 78.89293 | chr9 | 44083390 | 44090166 | 6.78512 | 4.96584 | 4.45902 |
| chr9 | 42988095 | 42988329 | 182 | Intron (ENSMUST00000165665.2 | 9 | 42518135 | 42944493 | 426359 | 2 | ENSMUST00 | -43603 | Grik4 | ENSMUSG0C | false | . | . | . | . | . | . | chr9 | 42987839 | 42988939 | 3.98507 | 3.44483 | 2.1947 |
| chr9 | 42264146 | 42264629 | 182 | Promoter (<=1kb) | 9 | 42254177 | 42264300 | 10124 | 2 | ENSMUST00 | 0 | Sc5d | ENSMUSG0C | false | chr9 | 42263561 | 42264865 | 21.77823 | 66.26614 | 63.32996 | chr9 | 42263302 | 42264906 | 3.64559 | 3.36819 | 1.88002 |
| chr9 | 50566389 | 50566587 | 182 | Distal Intergenic | 9 | 50557148 | 50561268 | 4121 | 2 | ENSMUST00 | -5122 | Tex12 | ENSMUSG0C | false | . | . | . | . | . | . | chr9 | 50564938 | 50567248 | 3.39032 | 3.13238 | 1.71803 |
| chr9 | 46282887 | 46283153 | 182 | Promoter (<=1kb) | 9 | 46283008 | 46294099 | 11092 | 1 | ENSMUST00 | 0 | Bud13 | ENSMUSG0C | false | chr9 | 46282506 | 46283758 | 39.20496 | 131.87447 | 127.31225 | . | . | . | . | . | . |
| chr9 | 46273099 | 46273302 | 182 | Promoter (<=1kb) | 9 | 46273197 | 46281489 | 8293 | 1 | ENSMUST00 | 0 | Zpr1 | ENSMUSG0C | false | chr9 | 46272634 | 46274141 | 36.46257 | 120.58523 | 116.42456 | chr9 | 46272179 | 46275218 | 3.435 | 3.18991 | 1.76628 |
| chr9 | 21131090 | 21131481 | 182 | Distal Intergenic | 9 | 21104069 | 21127346 | 23278 | 2 | ENSMUST00 | -3745 | Tyk2 | ENSMUSG0C | false | chr9 | 21130643 | 21131837 | 22.90469 | 70.93823 | 67.93287 | chr9 | 21129968 | 21131629 | 5.80353 | 4.42259 | 3.65977 |
| chr9 | 21723408 | 21723671 | 182 | Promoter (<=1kb) | 9 | 21723576 | 21749916 | 26341 | 1 | ENSMUST00 | 0 | Ldlr | ENSMUSG0C | false | chr9 | 21722916 | 21724671 | 17.71499 | 57.83842 | 55.02271 | chr9 | 21722801 | 21725254 | 4.27179 | 3.64547 | 2.44224 |
| chr9 | 21869458 | 21869655 | 182 | Distal Intergenic | 9 | 21800184 | 21852635 | 52452 | 2 | ENSMUST00 | -16824 | Dock6 | ENSMUSG0C | false | . | . | . | . | . | . | . | . | . | . | . | . |
| chr9 | 69467731 | 69467941 | 182 | Intron (ENSMUST00000034756.9 | 9 | 69478465 | 69491795 | 13331 | 1 | ENSMUST00 | -10524 | Anxa2 | ENSMUSG0C | false | . | . | . | . | . | . | chr9 | 69455103 | 69480655 | 7.58546 | 5.20708 | 5.13718 |
| chr9 | 69477377 | 69477685 | 182 | Promoter (<=1kb) | 9 | 69478465 | 69491795 | 13331 | 1 | ENSMUST00 | -780 | Anxa2 | ENSMUSG0C | false | . | . | . | . | . | . | chr9 | 69455103 | 69480655 | 7.58546 | 5.20708 | 5.13718 |
| chr9 | 69455303 | 69455578 | 182 | Promoter (1-2kb) | 9 | 69454066 | 69483859 | 29794 | 1 | ENSMUST00 | 1238 | Anxa2 | ENSMUSG0C | false | chr9 | 69453593 | 69455744 | 36.10714 | 116.78466 | 112.7497 | chr9 | 69455103 | 69480655 | 7.58546 | 5.20708 | 5.13718 |
| chr9 | 71760548 | 71760963 | 182 | Intron (ENSMUST00000072899.3 | 9 | 71629211 | 71771560 | 142350 | 2 | ENSMUST00 | 10597 | Cgnl1 | ENSMUSG0C | false | . | . | . | . | . | . | chr9 | 71736274 | 71770833 | 5.2398 | 4.14793 | 3.22386 |
| chr9 | 69301738 | 69301982 | 182 | Intron (ENSMUST00000034766.8 | 9 | 69289682 | 69379809 | 90128 | 1 | ENSMUST00 | 12057 | Rora | ENSMUSG0C | false | . | . | . | . | . | . | chr9 | 69291208 | 69308799 | 5.17084 | 4.06697 | 3.16692 |
| chr9 | 68873775 | 68873987 | 182 | Intron (ENSMUST00000034766.8 | 9 | 68655303 | 68664180 | 8878 | 1 | ENSMUST00 | 218473 | Rora | ENSMUSG0C | false | . | . | . | . | . | . | . | . | . | . | . | . |
| chr9 | 62340817 | 62341471 | 182 | Promoter (<=1kb) | 9 | 62341293 | 62377701 | 36409 | 1 | ENSMUST00 | 0 | Anp32a | ENSMUSG0C | false | chr9 | 62340409 | 62345269 | 38.55812 | 129.64772 | 125.16568 | . | . | . | . | . | . |
| chr9 | 62122575 | 62122764 | 182 | Promoter (<=1kb) | 9 | 62057248 | 62122610 | 65363 | 2 | ENSMUST00 | 0 | Glce | ENSMUSG0C | false | chr9 | 62121399 | 62123449 | 23.3092 | 84.4034 | 81.16875 | . | . | . | . | . | . |
| chr9 | 83600655 | 83600890 | 182 | 3' UTR | 9 | 83576831 | 83597524 | 20694 | 1 | ENSMUST00 | 23825 | Sh3bgrl2 | ENSMUSG0C | true | . | . | . | . | . | . | chr9 | 83598641 | 83606741 | 6.13876 | 4.64085 | 3.94135 |
| chr9 | 83585757 | 83585960 | 182 | Intron (ENSMUST00000188241.1 | 9 | 83576831 | 83597524 | 20694 | 1 | ENSMUST00 | 8927 | Sh3bgrl2 | ENSMUSG0C | true | . | . | . | . | . | . | chr9 | 83578741 | 83594009 | 5.16719 | 4.03655 | 3.17944 |
| chr9 | 54863577 | 54864389 | 182 | Promoter (<=1kb) | 9 | 54863755 | 54912534 | 48780 | 1 | ENSMUST00 | 0 | Ireb2 | ENSMUSG0C | false | chr9 | 54863122 | 54864695 | 41.07278 | 140.67711 | 135.76277 | . | . | . | . | . | . |
| chr9 | 55326567 | 55326988 | 182 | Promoter (<=1kb) | 9 | 55221301 | 55326749 | 105449 | 2 | ENSMUST00 | 0 | Nrg4 | ENSMUSG0C | false | chr9 | 55326284 | 55327538 | 31.25655 | 95.06709 | 91.61534 | chr9 | 55325915 | 55327615 | 4.97661 | 4.11442 | 2.98828 |
| chr9 | 57765001 | 57765645 | 182 | Promoter (<=1kb) | 9 | 57750711 | 57765860 | 15150 | 2 | ENSMUST00 | 215 | Clk3 | ENSMUSG0C | false | chr9 | 57763942 | 57766341 | 37.17318 | 134.81126 | 130.13722 | . | . | . | . | . | . |
| chr9 | 58253063 | 58253269 | 182 | Promoter (<=1kb) | 9 | 58253164 | 58262523 | 9360 | 1 | ENSMUST00 | 0 | Stoml1 | ENSMUSG0C | false | chr9 | 58253191 | 58253958 | 42.22279 | 143.81401 | 138.77701 | chr9 | 58252556 | 58255789 | 4.55946 | 3.87216 | 2.64575 |
| chr9 | 58273886 | 58274095 | 182 | Distal Intergenic | 9 | 58253164 | 58262523 | 9360 | 1 | ENSMUST00 | 20723 | Stoml1 | ENSMUSG0C | false | . | . | . | . | . | . | chr9 | 58271896 | 58276504 | 4.48501 | 3.8046 | 2.61422 |
| chr9 | 77750554 | 77751081 | 182 | Distal Intergenic | 9 | 77754535 | 77794485 | 39951 | 1 | ENSMUST00 | -3454 | Gclc | ENSMUSG0C | false | chr9 | 77750140 | 77751518 | 25.84672 | 90.2271 | 86.88143 | . | . | . | . | . | . |
| chr9 | 89704874 | 89705175 | 182 | Promoter (<=1kb) | 9 | 89699205 | 89705068 | 5864 | 2 | ENSMUST00 | 0 | Tmed3 | ENSMUSG0C | false | chr9 | 89704222 | 89705572 | 28.8526 | 98.7319 | 95.19158 | chr9 | 89703100 | 89706714 | 3.82434 | 3.4178 | 2.07973 |
| chr9 | 90047957 | 90048307 | 182 | Distal Intergenic | 9 | 90054152 | 90065422 | 11271 | 1 | ENSMUST00 | -5845 | Ctsh | ENSMUSG0C | false | . | . | . | . | . | . | chr9 | 90046598 | 90049431 | 4.32634 | 3.77038 | 2.49638 |
| chr9 | 67035898 | 67036184 | 182 | Exon (ENSMUST00000113696.2/ | 9 | 67029920 | 67032825 | 2906 | 2 | ENSMUST00 | -3074 | Tpm1 | ENSMUSG0C | true | . | . | . | . | . | . | chr9 | 67035130 | 67042899 | 5.46761 | 4.24053 | 3.41337 |
| chr9 | 92250077 | 92250765 | 182 | Promoter (<=1kb) | 9 | 92250342 | 92271975 | 21634 | 1 | ENSMUST00 | 0 | Plscr1 | ENSMUSG0C | false | chr9 | 92249860 | 92252220 | 39.33916 | 127.75399 | 123.34521 | . | . | . | . | . | . |
| chr9 | 66975151 | 66975565 | 182 | Promoter (<=1kb) | 9 | 66955393 | 66975484 | 20092 | 2 | ENSMUST00 | 0 | Lactb | ENSMUSG0C | false | chr9 | 66974580 | 66975855 | 32.3084 | 121.498 | 117.30614 | . | . | . | . | . | . |
| chr9 | 92559732 | 92559994 | 182 | Intron (ENSMUST00000070522.8 | 9 | 92573588 | 92591684 | 18097 | 1 | ENSMUST00 | -13594 | Plod2 | ENSMUSG0C | true | . | . | . | . | . | . | chr9 | 92543215 | 92573465 | 5.80463 | 4.42577 | 3.66882 |
| chr9 | 66251232 | 66251740 | 182 | Intron (ENSMUST00000034944.3 | 9 | 66268374 | 66271715 | 3342 | 1 | ENSMUST00 | -16634 | Dapk2 | ENSMUSG0C | false | . | . | . | . | . | . | . | . | . | . | . | . |
| chr9 | 66276952 | 66277270 | 182 | Distal Intergenic | 9 | 66268374 | 66271715 | 3342 | 1 | ENSMUST00 | 8579 | Dapk2 | ENSMUSG0C | false | . | . | . | . | . | . | chr9 | 66274995 | 66277364 | 3.89587 | 3.52469 | 2.14733 |
| chr9 | 65460825 | 65461260 | 182 | Promoter (<=1kb) | 9 | 65460937 | 65488470 | 27534 | 1 | ENSMUST00 | 0 | Spg21 | ENSMUSG0C | false | chr9 | 65460642 | 65461561 | 32.50483 | 115.59841 | 111.59217 | . | . | . | . | . | . |
| chr9 | 63677678 | 63678259 | 182 | Intron (ENSMUST00000034973.4 | 9 | 63654771 | 63666547 | 11777 | 2 | ENSMUST00 | -11132 | Smad3 | ENSMUSG0C | false | . | . | . | . | . | . | chr9 | 63671619 | 63680569 | 5.3648 | 4.16963 | 3.3451 |
| chr9 | 95511746 | 95512189 | 182 | Promoter (<=1kb) | 9 | 95456898 | 95511894 | 54997 | 2 | ENSMUST00 | 0 | U2surp | ENSMUSG0C | false | chr9 | 95510814 | 95512717 | 32.87838 | 107.99121 | 104.21945 | . | . | . | . | . | . |
| chr9 | 96364152 | 96364712 | 182 | Promoter (<=1kb) | 9 | 96332655 | 96364442 | 31788 | 2 | ENSMUST00 | 0 | Atp1b3 | ENSMUSG0C | false | chr9 | 96364501 | 96365263 | 20.65177 | 61.66082 | 58.78991 | . | . | . | . | . | . |
| chr9 | 114802193 | 114802443 | 182 | Intron (ENSMUST00000047013.3 | 9 | 114756837 | 114781993 | 25157 | 2 | ENSMUST00 | -20201 | Cmtm7 | ENSMUSG0C | true | . | . | . | . | . | . | chr9 | 114801208 | 114806677 | 5.65143 | 4.12227 | 3.58214 |
| chr9 | 115310846 | 115311037 | 182 | Promoter (<=1kb) | 9 | 115242607 | 115310421 | 67815 | 2 | ENSMUST00 | -426 | Stt3b | ENSMUSG0C | false | chr9 | 115307644 | 115311336 | 30.14656 | 119.38538 | 115.2599 | . | . | . | . | . | . |
| chr9 | 116234258 | 116234525 | 182 | Distal Intergenic | 9 | 116087698 | 116175363 | 87666 | 2 | ENSMUST00 | -58896 | Tgfbr2 | ENSMUSG0C | false | . | . | . | . | . | . | chr9 | 116227870 | 116235420 | 4.58647 | 3.82659 | 2.67853 |
| chr9 | 116324377 | 116324732 | 182 | Distal Intergenic | 9 | 116087698 | 116175363 | 87666 | 2 | ENSMUST00 | -149015 | Tgfbr2 | ENSMUSG0C | false | . | . | . | . | . | . | . | . | . | . | . | . |
| chr9 | 116401886 | 116402106 | 182 | Distal Intergenic | 9 | 116087698 | 116175363 | 87666 | 2 | ENSMUST00 | -226524 | Tgfbr2 | ENSMUSG0C | false | . | . | . | . | . | . | . | . | . | . | . | . |
| chr9 | 98296439 | 98296862 | 182 | Promoter (<=1kb) | 9 | 98296583 | 98411434 | 114852 | 1 | ENSMUST00 | 0 | Nmnat3 | ENSMUSG0C | false | chr9 | 98296463 | 98297381 | 24.06788 | 78.48502 | 75.35915 | . | . | . | . | . | . |
| chr9 | 99568621 | 99569037 | 182 | Promoter (<=1kb) | 9 | 99478372 | 99568899 | 90528 | 2 | ENSMUST00 | 0 | Armc8 | ENSMUSG0C | false | chr9 | 99567865 | 99569427 | 30.76814 | 122.68065 | 118.44827 | . | . | . | . | . | . |
| chr9 | 100545618 | 100545927 | 182 | Promoter (<=1kb) | 9 | 100497800 | 100545731 | 47932 | 2 | ENSMUST00 | 0 | Nck1 | ENSMUSG0C | false | chr9 | 100544788 | 100547004 | 29.2476 | 106.45832 | 102.71833 | . | . | . | . | . | . |
| chr9 | 109875262 | 109875478 | 182 | Promoter (<=1kb) | 9 | 109875579 | 109893888 | 18310 | 1 | ENSMUST00 | -101 | Cdc25a | ENSMUSG0C | false | chr9 | 109875378 | 109876629 | 28.59262 | 111.22587 | 107.35971 | . | . | . | . | . | . |
| chr9 | 110333236 | 110333725 | 182 | Promoter (<=1kb) | 9 | 110333288 | 110373081 | 39794 | 1 | ENSMUST00 | 0 | Scap | ENSMUSG0C | false | chr9 | 110333216 | 110334287 | 28.91859 | 89.0447 | 85.71584 | chr9 | 110332906 | 110339014 | 3.65846 | 3.40453 | 1.90737 |
| chr9 | 111213576 | 111213835 | 182 | Intron (ENSMUST00000035078.7 | 9 | 111229870 | 111230948 | 1079 | 2 | ENSMUST00 | 17113 | Mlh1 | ENSMUSG0C | false | . | . | . | . | . | . | chr9 | 111213753 | 111215720 | 3.07133 | 3.06632 | 1.43043 |
| chr9 | 111179027 | 111179350 | 182 | Intron (ENSMUST00000035078.7 | 9 | 111229870 | 111230948 | 1079 | 2 | ENSMUST00 | 51598 | Mlh1 | ENSMUSG0C | false | . | . | . | . | . | . | chr9 | 111178449 | 111180265 | 4.36053 | 3.6998 | 2.50652 |
| chr9 | 120127555 | 120127750 | 182 | Promoter (<=1kb) | 9 | 120127766 | 120132369 | 4604 | 1 | ENSMUST00 | -16 | Rpsa | ENSMUSG0C | false | chr9 | 120127025 | 120127564 | 14.15711 | 33.4017 | 30.89395 | . | . | . | . | . | . |
| chr9 | 120110263 | 120110550 | 182 | Promoter (<=1kb) | 9 | 120110374 | 120124504 | 14131 | 1 | ENSMUST00 | 0 | Slc25a38 | ENSMUSG0C | false | chr9 | 120110272 | 120111312 | 40.89836 | 130.45236 | 125.94646 | . | . | . | . | . | . |
| chr9 | 121710226 | 121710476 | 182 | Promoter (<=1kb) | 9 | 121710389 | 121712921 | 2533 | 1 | ENSMUST00 | 0 | Deb1 | ENSMUSG0C | false | chr9 | 121710227 | 121711199 | 37.18052 | 130.46529 | 125.95659 | chr9 | 121703875 | 121712716 | 6.06469 | 4.507 | 3.89251 |
| chr9 | 102731090 | 102731276 | 182 | Promoter (<=1kb) | 9 | 102730731 | 102732327 | 1597 | 1 | ENSMUST00 | 360 | Amotl2 | ENSMUSG0C | false | . | . | . | . | . | . | chr9 | 102720260 | 102734975 | 8.68688 | 5.64144 | 6.02662 |
| chr9 | 102977863 | 102978289 | 182 | Distal Intergenic | 9 | 102988712 | 103020775 | 32064 | 1 | ENSMUST00 | -10423 | Slco2a1 | ENSMUSG0C | true | . | . | . | . | . | . | . | . | . | . | . | . |
| chr9 | 103078263 | 103078643 | 182 | Intron (ENSMUST00000035148.7 | 9 | 103111787 | 103162657 | 50871 | 1 | ENSMUST00 | -33144 | Rab6b | ENSMUSG0C | false | . | . | . | . | . | . | . | . | . | . | . | . |
| chr9 | 103230611 | 103230904 | 182 | Promoter (<=1kb) | 9 | 103208795 | 103230444 | 21650 | 2 | ENSMUST00 | -168 | Trf | ENSMUSG0C | false | . | . | . | . | . | . | . | . | . | . | . | . |
| chr9 | 107628805 | 107629002 | 182 | Intron (ENSMUST00000055704.6 | 9 | 107614138 | 107635342 | 21205 | 2 | ENSMUST00 | 6340 | Gnai2 | ENSMUSG0C | false | . | . | . | . | . | . | chr9 | 107617961 | 107633388 | 7.13311 | 4.9094 | 4.77996 |
| chr9 | 105395250 | 105395469 | 182 | Promoter (<=1kb) | 9 | 105395392 | 105405758 | 10367 | 1 | ENSMUST00 | 0 | Aste1 | ENSMUSG0C | false | chr9 | 105394719 | 105396315 | 27.41053 | 90.23362 | 86.88379 | chr9 | 105394705 | 105396092 | 4.20162 | 3.64512 | 2.39804 |
| chr9 | 107735993 | 107736384 | 182 | Distal Intergenic | 9 | 107743333 | 107744234 | 902 | 2 | ENSMUST00 | 7850 | Rbm5 | ENSMUSG0C | false | chr9 | 107735725 | 107736006 | 5.97392 | 9.35129 | 7.18948 | chr9 | 107734148 | 107736914 | 6.94684 | 4.94491 | 4.65981 |
| chr9 | 107982726 | 107983026 | 182 | Promoter (<=1kb) | 9 | 107983025 | 107984053 | 1029 | 1 | ENSMUST00 | 0 | Uba7 | ENSMUSG0C | false | . | . | . | . | . | . | . | . | . | . | . | . |
| chr9 | 108280196 | 108280476 | 182 | Distal Intergenic | 9 | 108290448 | 108296498 | 6051 | 2 | ENSMUST00 | -9972 | Nicn1 | ENSMUSG0C | false | . | . | . | . | . | . | chr9 | 108279075 | 108281445 | 3.76417 | 3.403 | 2.0186 |
| chr4 | 124704916 | 124705395 | 182 | Intron (ENSMUST00000106199.4 | 4 | 124700720 | 124708611 | 7892 | 1 | ENSMUST00 | 4197 | Fhl3 | ENSMUSG0C | false | . | . | . | . | . | . | . | . | . | . | . | . |
| chr1 | 151900910 | 151901347 | 182 | Intron (ENSMUST00000187603.1 | 1 | 151892903 | 151921227 | 28325 | 2 | ENSMUST00 | 19880 | 1700025G04 | ENSMUSG0C | true | . | . | . | . | . | . | chr1 | 151896806 | 151901695 | 5.38261 | 4.34371 | 3.31168 |
| chr18 | 65437587 | 65437828 | 182 | Exon (ENSMUST00000049248.5/ | 18 | 65430963 | 65478823 | 47861 | 1 | ENSMUST00 | 6625 | Malt1 | ENSMUSG0C | true | . | . | . | . | . | . | chr18 | 65432205 | 65440925 | 4.98691 | 4.11378 | 2.99843 |
| chr2 | 103970053 | 103970390 | 182 | Promoter (<=1kb) | 2 | 103970129 | 103981855 | 11727 | 1 | ENSMUST00 | 0 | Lmo2 | ENSMUSG0C | false | chr2 | 103969199 | 103971275 | 25.10597 | 76.67836 | 73.5843 | chr2 | 103969703 | 103970858 | 3.81706 | 3.37082 | 2.0545 |
| chr2 | 103624892 | 103625130 | 182 | Intron (ENSMUST00000076212.3 | 2 | 103566310 | 103718423 | 152114 | 1 | ENSMUST00 | 58583 | Abtb2 | ENSMUSG0C | false | . | . | . | . | . | . | chr2 | 103621752 | 103634479 | 4.86729 | 3.97695 | 2.90643 |
| chr5 | 120586877 | 120587305 | 182 | Promoter (1-2kb) | 5 | 120534160 | 120588613 | 54454 | 2 | ENSMUST00 | 1308 | Tpcn1 | ENSMUSG0C | false | chr5 | 120586812 | 120588712 | 29.2476 | 106.45832 | 102.71833 | chr5 | 120581611 | 120588072 | 5.52017 | 4.24074 | 3.45534 |

| | | | | | | | | | | | | | | | | | | | | | | | | | | |
|---|---|---|---|---|---|---|---|---|---|---|---|---|---|---|---|---|---|---|---|---|---|---|---|---|---|---|
| chr5 | 120588688 | 120588933 | 182 | Promoter (<=1kb) | 5 | 120534160 | 120588613 | 54454 | 2 | ENSMUST00 | -76 | Tpcn1 | ENSMUSG0C | false | chr5 | 120586812 | 120588712 | 29.2476 | 106.45832 | 102.71833 | . | . | . | . | . | . |
| chr7 | 125707646 | 125708237 | 182 | Promoter (<=1kb) | 7 | 125707915 | 125761832 | 53918 | 1 | ENSMUST00 | 0 | D430042O09 | ENSMUSG0C | false | chr7 | 125706384 | 125708627 | 38.04976 | 128.75285 | 124.30675 | . | . | . | . | . | . |
| chr2 | 101750888 | 101751214 | 182 | Intron (ENSMUST00000171088.2 | 2 | 101717250 | 101797650 | 80401 | 2 | ENSMUST00 | 46436 | Prr5l | ENSMUSG0C | false | . | . | . | . | . | . | chr2 | 101750438 | 101752738 | 3.85645 | 3.38429 | 2.09336 |
| chr4 | 121021936 | 121022307 | 182 | Distal Intergenic | 4 | 120968317 | 121017247 | 48931 | 2 | ENSMUST00 | -4690 | Smap2 | ENSMUSG0C | false | . | . | . | . | . | . | chr4 | 121021403 | 121022220 | 3.6911 | 3.31379 | 1.94319 |
| chr1 | 78657664 | 78658493 | 182 | Promoter (<=1kb) | 1 | 78657825 | 78707743 | 49919 | 1 | ENSMUST00 | 0 | Acsl3 | ENSMUSG0C | false | chr1 | 78657393 | 78659137 | 35.29575 | 125.98199 | 121.63858 | chr1 | 78658375 | 78659120 | 3.56292 | 3.39067 | 1.81361 |
| chr5 | 117977149 | 117977364 | 182 | Promoter (<=1kb) | 5 | 117976770 | 118010191 | 33422 | 1 | ENSMUST00 | 380 | Fbxo21 | ENSMUSG0C | false | chr5 | 117976342 | 117977742 | 34.49762 | 142.80731 | 137.81027 | chr5 | 117975985 | 117978599 | 5.5059 | 4.3586 | 3.42256 |
| chr9 | 56871250 | 56871519 | 182 | Intron (ENSMUST00000035661.5 | 9 | 56865104 | 56899870 | 34767 | 1 | ENSMUST00 | 6147 | Cspg4 | ENSMUSG0C | false | . | . | . | . | . | . | chr9 | 56869458 | 56871598 | 5.83101 | 4.39874 | 3.72193 |
| chr16 | 75592777 | 75593070 | 182 | Promoter (<=1kb) | 16 | 75592844 | 75601695 | 8852 | 1 | ENSMUST00 | 0 | Rbm11 | ENSMUSG0C | false | chr16 | 75592585 | 75594022 | 31.85236 | 101.09525 | 97.496 | chr16 | 75592528 | 75593986 | 3.29446 | 3.20073 | 1.58598 |
| chr18 | 80714867 | 80715104 | 182 | Promoter (1-2kb) | 18 | 80606205 | 80713071 | 106867 | 2 | ENSMUST00 | -1797 | Nfatc1 | ENSMUSG0C | false | chr18 | 80714491 | 80715784 | 14.82622 | 34.53743 | 32.01512 | chr18 | 80711836 | 80715689 | 5.07579 | 4.0937 | 3.06953 |
| chr1 | 75292797 | 75293159 | 182 | Distal Intergenic | 1 | 75272199 | 75278415 | 6217 | 2 | ENSMUST00 | -14383 | Resp18 | ENSMUSG0C | false | . | . | . | . | . | . | chr1 | 75289827 | 75294939 | 5.35548 | 4.22327 | 3.31093 |
| chr15 | 98108314 | 98108713 | 182 | Promoter (<=1kb) | 15 | 98108471 | 98132447 | 23977 | 1 | ENSMUST00 | 0 | Pfkm | ENSMUSG0C | false | chr15 | 98108058 | 98109151 | 26.03697 | 72.82576 | 69.79147 | chr15 | 98108034 | 98109328 | 3.68482 | 3.4471 | 1.96389 |
| chr15 | 78934857 | 78935261 | 182 | Promoter (<=1kb) | 15 | 78934933 | 78943634 | 8702 | 1 | ENSMUST00 | 0 | Nol12 | ENSMUSG0C | false | chr15 | 78934764 | 78935809 | 32.62334 | 101.43139 | 97.82731 | chr15 | 78934343 | 78936538 | 4.08551 | 3.51136 | 2.27064 |
| chr13 | 64129246 | 64129563 | 182 | Promoter (<=1kb) | 13 | 64096310 | 64129330 | 33021 | 2 | ENSMUST00 | 0 | Slc35d2 | ENSMUSG0C | false | chr13 | 64128692 | 64129858 | 19.57973 | 66.4542 | 63.51413 | chr13 | 64128715 | 64130359 | 3.80953 | 3.47594 | 2.04913 |
| chr13 | 63990058 | 63990334 | 182 | Distal Intergenic | 13 | 64058274 | 64089186 | 30913 | 2 | ENSMUST00 | 98852 | Hsd17b3 | ENSMUSG0C | false | . | . | . | . | . | . | . | . | . | . | . | . |
| chr16 | 45742780 | 45743212 | 182 | Promoter (<=1kb) | 16 | 45729725 | 45742955 | 13231 | 2 | ENSMUST00 | 0 | Abhd10 | ENSMUSG0C | false | chr16 | 45742277 | 45743436 | 18.02749 | 67.47041 | 64.5209 | chr16 | 45741571 | 45744810 | 3.88492 | 3.52308 | 2.08764 |
| chr1 | 75141465 | 75141916 | 182 | Promoter (<=1kb) | 1 | 75135768 | 75141691 | 5924 | 2 | ENSMUST00 | 0 | Cnppd1 | ENSMUSG0C | false | chr1 | 75140886 | 75143697 | 38.13901 | 127.82562 | 123.40861 | . | . | . | . | . | . |
| chr15 | 78790239 | 78790681 | 182 | Promoter (<=1kb) | 15 | 78787411 | 78790515 | 3105 | 2 | ENSMUST00 | 0 | Card10 | ENSMUSG0C | true | . | . | . | . | . | . | chr15 | 78770626 | 78800649 | 7.36271 | 5.05378 | 4.95398 |
| chr6 | 88759918 | 88760265 | 182 | Intron (ENSMUST00000113585.3 | 6 | 88724828 | 88828360 | 103533 | 1 | ENSMUST00 | 35091 | Mgll | ENSMUSG0C | false | . | . | . | . | . | . | chr6 | 88758421 | 88760631 | 3.42327 | 3.18226 | 1.75337 |
| chr6 | 88761846 | 88762023 | 182 | Intron (ENSMUST00000113585.3 | 6 | 88724828 | 88828360 | 103533 | 1 | ENSMUST00 | 37019 | Mgll | ENSMUSG0C | false | . | . | . | . | . | . | chr6 | 88761792 | 88762332 | 4.43086 | 3.8147 | 2.59318 |
| chr5 | 111229705 | 111229970 | 182 | Intron (ENSMUST00000040111.9 | 5 | 111233388 | 111238464 | 5077 | 1 | ENSMUST00 | -3418 | Ttc28 | ENSMUSG0C | false | . | . | . | . | . | . | chr5 | 111227053 | 111231630 | 6.63109 | 4.54688 | 4.37946 |
| chr5 | 111139316 | 111139503 | 182 | Intron (ENSMUST00000040111.9 | 5 | 111225458 | 111229415 | 3958 | 1 | ENSMUST00 | -85955 | Ttc28 | ENSMUSG0C | false | . | . | . | . | . | . | chr5 | 111137002 | 111140665 | 3.23852 | 3.14403 | 1.61079 |
| chr5 | 111205791 | 111206082 | 182 | Intron (ENSMUST00000040111.9 | 5 | 111225458 | 111229415 | 3958 | 1 | ENSMUST00 | -19376 | Ttc28 | ENSMUSG0C | false | . | . | . | . | . | . | chr5 | 111203210 | 111208761 | 4.08653 | 3.54857 | 2.2972 |
| chr9 | 21426108 | 21426312 | 182 | Promoter (<=1kb) | 9 | 21425244 | 21507754 | 82511 | 1 | ENSMUST00 | 865 | Dnm2 | ENSMUSG0C | false | chr9 | 21424870 | 21426244 | 32.3084 | 121.498 | 117.30614 | chr9 | 21425583 | 21440915 | 6.54133 | 4.65697 | 4.31065 |
| chr9 | 95407142 | 95407534 | 182 | Promoter (<=1kb) | 9 | 95399292 | 95406722 | 7431 | 2 | ENSMUST00 | -421 | Chst2 | ENSMUSG0C | false | chr9 | 95405075 | 95408037 | 26.703 | 78.54972 | 75.42343 | chr9 | 95404750 | 95408450 | 4.85791 | 3.91044 | 2.90355 |
| chr10 | 75200765 | 75201145 | 182 | Distal Intergenic | 10 | 75212073 | 75312399 | 100327 | 1 | ENSMUST00 | -10928 | Specc1l | ENSMUSG0C | false | . | . | . | . | . | . | . | . | . | . | . | . |
| chr10 | 75241001 | 75241209 | 182 | Exon (ENSMUST00000105421.3/ | 10 | 75229855 | 75312399 | 82545 | 1 | ENSMUST00 | 11147 | Specc1l | ENSMUSG0C | false | . | . | . | . | . | . | chr10 | 75239344 | 75242323 | 6.33386 | 4.76511 | 4.08758 |
| chr17 | 47688050 | 47688275 | 182 | Promoter (<=1kb) | 17 | 47686646 | 47688183 | 1538 | 2 | ENSMUST00 | 0 | Tomm6 | ENSMUSG0C | false | chr17 | 47687260 | 47689501 | 34.78613 | 113.42423 | 109.48744 | chr17 | 47686192 | 47689517 | 5.23678 | 4.09637 | 3.20214 |
| chr3 | 116361248 | 116361453 | 182 | Intron (ENSMUST00000106491.2 | 3 | 116400186 | 116423024 | 22839 | 2 | ENSMUST00 | 61571 | Cdc14a | ENSMUSG0C | true | . | . | . | . | . | . | chr3 | 116319808 | 116393211 | 5.81811 | 4.44924 | 3.68159 |
| chr6 | 43265433 | 43265873 | 182 | Promoter (<=1kb) | 6 | 43265582 | 43289320 | 23739 | 1 | ENSMUST00 | 0 | Arhgef5 | ENSMUSG0C | false | chr6 | 43265083 | 43266670 | 27.66126 | 83.97217 | 80.74496 | . | . | . | . | . | . |
| chr8 | 111537025 | 111537272 | 182 | Promoter (<=1kb) | 8 | 111537120 | 111622973 | 85854 | 1 | ENSMUST00 | 0 | Znrf1 | ENSMUSG0C | false | chr8 | 111536021 | 111538448 | 42.22279 | 143.81401 | 138.77701 | . | . | . | . | . | . |
| chr9 | 80158852 | 80159073 | 182 | Distal Intergenic | 9 | 80165031 | 80311729 | 146699 | 1 | ENSMUST00 | -5958 | Myo6 | ENSMUSG0C | true | . | . | . | . | . | . | chr9 | 80157427 | 80160962 | 4.62742 | 3.79838 | 2.70841 |
| chr8 | 111393623 | 111394024 | 182 | Promoter (<=1kb) | 8 | 111345141 | 111393824 | 48684 | 2 | ENSMUST00 | 0 | Fa2h | ENSMUSG0C | true | chr8 | 111392792 | 111394218 | 21.75525 | 76.8157 | 73.7174 | chr8 | 111390850 | 111394204 | 4.46985 | 3.74648 | 2.58644 |
| chr16 | 22067495 | 22067794 | 182 | Intron (ENSMUST00000100052.5 | 16 | 22061366 | 22063697 | 2332 | 2 | ENSMUST00 | -3799 | Igf2bp2 | ENSMUSG0C | false | . | . | . | . | . | . | . | . | . | . | . | . |
| chr8 | 111300132 | 111300447 | 182 | Promoter (<=1kb) | 8 | 111270944 | 111300222 | 29279 | 2 | ENSMUST00 | 0 | Rfwd3 | ENSMUSG0C | false | chr8 | 111299240 | 111300356 | 40.48558 | 151.98326 | 146.61765 | . | . | . | . | . | . |
| chr16 | 21422991 | 21423624 | 182 | Promoter (<=1kb) | 16 | 21423118 | 21579247 | 156130 | 1 | ENSMUST00 | 0 | Vps8 | ENSMUSG0C | false | chr16 | 21422763 | 21423824 | 40.17707 | 149.16289 | 143.90889 | . | . | . | . | . | . |
| chr7 | 100495749 | 100496036 | 182 | Promoter (<=1kb) | 7 | 100496079 | 100498256 | 2178 | 1 | ENSMUST00 | -43 | Ucp2 | ENSMUSG0C | false | chr7 | 100495153 | 100495958 | 6.38327 | 11.47542 | 9.27082 | chr7 | 100494087 | 100506674 | 4.14782 | 3.62895 | 2.31932 |
| chr1 | 155558040 | 155558621 | 182 | Promoter (<=1kb) | 1 | 155558120 | 155572288 | 14169 | 1 | ENSMUST00 | 0 | Acbd6 | ENSMUSG0C | false | chr1 | 155557631 | 155559183 | 42.50686 | 139.83865 | 134.96973 | chr1 | 155557592 | 155560826 | 4.53142 | 3.84981 | 2.64735 |
| chr1 | 155244157 | 155244609 | 182 | Promoter (<=1kb) | 1 | 155212471 | 155244444 | 31974 | 2 | ENSMUST00 | 0 | BC034090 | ENSMUSG0C | false | chr1 | 155243730 | 155245165 | 31.84599 | 102.09332 | 98.46812 | chr1 | 155243581 | 155245355 | 4.05572 | 3.60945 | 2.26946 |
| chr1 | 155237375 | 155237642 | 182 | Intron (ENSMUST00000186156.1 | 1 | 155224911 | 155240906 | 15996 | 2 | ENSMUST00 | 3264 | BC034090 | ENSMUSG0C | false | . | . | . | . | . | . | chr1 | 155232733 | 155238771 | 5.23157 | 4.18441 | 3.18545 |
| chr15 | 76710511 | 76710724 | 182 | Promoter (<=1kb) | 15 | 76710623 | 76713570 | 2948 | 1 | ENSMUST00 | 0 | Lrrc14 | ENSMUSG0C | false | chr15 | 76710078 | 76711301 | 31.92468 | 100.21517 | 96.63724 | chr15 | 76709960 | 76713178 | 5.02817 | 4.01202 | 3.03619 |
| chr8 | 110719460 | 110719687 | 182 | Promoter (1-2kb) | 8 | 110721476 | 110741400 | 19925 | 1 | ENSMUST00 | -1789 | Mtss1l | ENSMUSG0C | false | . | . | . | . | . | . | . | . | . | . | . | . |
| chr8 | 110721296 | 110721709 | 182 | Promoter (<=1kb) | 8 | 110721476 | 110741400 | 19925 | 1 | ENSMUST00 | 0 | Mtss1l | ENSMUSG0C | false | chr8 | 110721059 | 110721907 | 13.52062 | 44.78815 | 42.13932 | . | . | . | . | . | . |
| chr19 | 6521539 | 6521732 | 182 | Intron (ENSMUST00000113462.2 | 19 | 6517205 | 6532296 | 15092 | 1 | ENSMUST00 | 4335 | Nrxn2 | ENSMUSG0C | false | . | . | . | . | . | . | chr19 | 6521294 | 6522525 | 3.634 | 3.24262 | 1.92225 |
| chr6 | 84385730 | 84386032 | 182 | Distal Intergenic | 6 | 84210484 | 84211041 | 558 | 1 | ENSMUST00 | 175247 | Dysf | ENSMUSG0C | false | . | . | . | . | . | . | chr6 | 84385830 | 84386715 | 2.96902 | 2.95319 | 1.37154 |
| chr1 | 5083043 | 5083566 | 182 | Promoter (<=1kb) | 1 | 5083173 | 5162549 | 79377 | 1 | ENSMUST00 | 0 | Atp6v1h | ENSMUSG0C | false | chr1 | 5082795 | 5083891 | 31.91637 | 110.38313 | 106.53261 | chr1 | 5082924 | 5086684 | 4.35362 | 3.77158 | 2.48259 |
| chr1 | 4857473 | 4858117 | 182 | Promoter (<=1kb) | 1 | 4857814 | 4897905 | 40092 | 1 | ENSMUST00 | 0 | Tcea1 | ENSMUSG0C | false | chr1 | 4857039 | 4858858 | 36.84594 | 114.91216 | 110.93493 | . | . | . | . | . | . |
| chr17 | 88628892 | 88629219 | 182 | Promoter (1-2kb) | 17 | 88627364 | 88637634 | 10271 | 1 | ENSMUST00 | 1529 | Ston1 | ENSMUSG0C | false | chr17 | 88626424 | 88629225 | 13.60354 | 37.21757 | 34.66072 | chr17 | 88627674 | 88642305 | 5.51401 | 4.25963 | 3.44192 |
| chr17 | 88629299 | 88629610 | 182 | Promoter (1-2kb) | 17 | 88627364 | 88637634 | 10271 | 1 | ENSMUST00 | 1936 | Ston1 | ENSMUSG0C | false | . | . | . | . | . | . | chr17 | 88627674 | 88642305 | 5.51401 | 4.25963 | 3.44192 |
| chr17 | 88598614 | 88599004 | 182 | Promoter (<=1kb) | 17 | 88597696 | 88645322 | 47627 | 1 | ENSMUST00 | 919 | Ston1 | ENSMUSG0C | false | chr17 | 88597214 | 88598641 | 21.40274 | 64.72344 | 61.81032 | chr17 | 88595166 | 88604773 | 4.12794 | 3.59375 | 2.29889 |
| chr3 | 82144903 | 82145202 | 182 | Promoter (<=1kb) | 3 | 82092428 | 82145877 | 53450 | 2 | ENSMUST00 | 675 | Gucy1a3 | ENSMUSG0C | false | chr3 | 82144916 | 82145857 | 9.64628 | 19.2185 | 16.89346 | chr3 | 82144721 | 82145770 | 5.76111 | 4.43566 | 3.63347 |
| chr3 | 82152516 | 82152774 | 182 | Distal Intergenic | 3 | 82092428 | 82145877 | 53450 | 2 | ENSMUST00 | -6640 | Gucy1a3 | ENSMUSG0C | false | . | . | . | . | . | . | . | . | . | . | . | . |
| chr2 | 119897484 | 119897754 | 182 | Promoter (<=1kb) | 2 | 119897305 | 119960794 | 63490 | 1 | ENSMUST00 | 180 | Mga | ENSMUSG0C | false | chr2 | 119896566 | 119898614 | 13.51324 | 34.99091 | 32.46387 | chr2 | 119896548 | 119901453 | 4.29045 | 3.62202 | 2.45668 |
| chr11 | 17159045 | 17159469 | 182 | Promoter (<=1kb) | 11 | 17159263 | 17200375 | 41113 | 1 | ENSMUST00 | 0 | Ppp3r1 | ENSMUSG0C | false | chr11 | 17158587 | 17160727 | 37.92415 | 138.37384 | 133.55817 | chr11 | 17158835 | 17159063 | 3.2181 | 3.23256 | 1.52539 |
| chr2 | 85047930 | 85048646 | 182 | Promoter (<=1kb) | 2 | 85048022 | 85052134 | 4113 | 1 | ENSMUST00 | 0 | Tnks1bp1 | ENSMUSG0C | false | chr2 | 85047255 | 85048940 | 30.35223 | 94.4818 | 91.04041 | . | . | . | . | . | . |
| chr4 | 116720816 | 116721271 | 182 | Promoter (<=1kb) | 4 | 116720948 | 116804242 | 83295 | 1 | ENSMUST00 | 0 | Tesk2 | ENSMUSG0C | false | chr4 | 116720168 | 116722603 | 32.98857 | 124.89146 | 120.57959 | . | . | . | . | . | . |
| chr18 | 80261609 | 80261833 | 182 | Promoter (1-2kb) | 18 | 80262840 | 80263485 | 646 | 1 | ENSMUST00 | -1007 | Pqlc1 | ENSMUSG0C | false | . | . | . | . | . | . | chr18 | 80261559 | 80263853 | 4.21156 | 3.50951 | 2.38894 |
| chr10 | 117094536 | 117094834 | 182 | Intron (ENSMUST00000020381.3 | 10 | 117050998 | 117063814 | 12817 | 2 | ENSMUST00 | -30723 | Cct2 | ENSMUSG0C | false | . | . | . | . | . | . | chr10 | 117086815 | 117095074 | 5.71886 | 4.41367 | 3.58491 |
| chr4 | 116557392 | 116557736 | 182 | Promoter (<=1kb) | 4 | 116557658 | 116593882 | 36225 | 1 | ENSMUST00 | 0 | Gpbp1l1 | ENSMUSG0C | false | chr4 | 116555293 | 116559172 | 37.92415 | 138.37384 | 133.55817 | . | . | . | . | . | . |
| chr16 | 38550167 | 38550366 | 182 | Promoter (<=1kb) | 16 | 38525137 | 38550258 | 25122 | 2 | ENSMUST00 | 0 | Poglut1 | ENSMUSG0C | false | chr16 | 38549386 | 38550580 | 37.68163 | 117.40437 | 113.3432 | chr16 | 38549151 | 38550717 | 3.65527 | 3.42271 | 1.881 |
| chr5 | 130110730 | 130110942 | 182 | Intron (ENSMUST00000118993.2 | 5 | 130093589 | 130102050 | 8462 | 1 | ENSMUST00 | 17142 | Tpst1 | ENSMUSG0C | false | . | . | . | . | . | . | chr5 | 130110355 | 130111807 | 4.27288 | 3.69701 | 2.42452 |
| chr12 | 86932276 | 86932525 | 182 | Distal Intergenic | 12 | 86947043 | 86962158 | 15116 | 1 | ENSMUST00 | -14518 | Cipc | ENSMUSG0C | false | . | . | . | . | . | . | chr12 | 86931420 | 86933864 | 4.43386 | 3.66285 | 2.55663 |
| chr16 | 37869314 | 37869546 | 182 | Promoter (<=1kb) | 16 | 37868389 | 37888858 | 20470 | 1 | ENSMUST00 | 926 | Lrrc58 | ENSMUSG0C | false | chr16 | 37867771 | 37870669 | 34.00884 | 130.01486 | 125.52227 | . | . | . | . | . | . |
| chr16 | 37871467 | 37871769 | 182 | Intron (ENSMUST00000078717.6 | 16 | 37868389 | 37888858 | 20470 | 1 | ENSMUST00 | 3079 | Lrrc58 | ENSMUSG0C | false | . | . | . | . | . | . | chr16 | 37870054 | 37882292 | 5.93789 | 4.46579 | 3.79659 |
| chr17 | 47521672 | 47521954 | 182 | Intron (ENSMUST00000182129.2 | 17 | 47505241 | 47598649 | 93409 | 1 | ENSMUST00 | 16432 | Ccnd3 | ENSMUSG0C | false | . | . | . | . | . | . | chr17 | 47521584 | 47523652 | 3.45261 | 3.17128 | 1.74463 |
| chr17 | 47593237 | 47593591 | 182 | Promoter (<=1kb) | 17 | 47593436 | 47598174 | 4739 | 1 | ENSMUST00 | 0 | Ccnd3 | ENSMUSG0C | false | chr17 | 47592999 | 47595153 | 45.95321 | 151.4303 | 146.08812 | . | . | . | . | . | . |
| chr12 | 86872189 | 86872598 | 182 | Distal Intergenic | 12 | 86880703 | 86884814 | 4112 | 2 | ENSMUST00 | 12216 | Irf2bpl | ENSMUSG0C | false | . | . | . | . | . | . | chr12 | 86871591 | 86874830 | 6.00169 | 4.57238 | 3.81491 |
| chr11 | 103953981 | 103954201 | 182 | Promoter (<=1kb) | 11 | 103821782 | 103954056 | 132275 | 2 | ENSMUST00 | 0 | Nsf | ENSMUSG0C | true | chr11 | 103951021 | 103954455 | 30.26787 | 111.42836 | 107.5545 | . | . | . | . | . | . |
| chr2 | 119288464 | 119288820 | 182 | Promoter (<=1kb) | 2 | 119288740 | 119295487 | 6748 | 1 | ENSMUST00 | 0 | Vps18 | ENSMUSG0C | false | chr2 | 119288276 | 119289498 | 30.02262 | 88.06211 | 84.75982 | chr2 | 119287858 | 119290609 | 4.06803 | 3.6083 | 2.23251 |
| chr9 | 53431368 | 53431568 | 182 | Intron (ENSMUST00000005262.1 | 9 | 53440105 | 53442560 | 2456 | 2 | ENSMUST00 | 10992 | Atm | ENSMUSG0C | false | . | . | . | . | . | . | . | . | . | . | . | . |
| chr9 | 53517671 | 53517947 | 182 | Intron (ENSMUST00000118282.2 | 9 | 53527194 | 53536608 | 9415 | 2 | ENSMUST00 | 18661 | Atm | ENSMUSG0C | false | . | . | . | . | . | . | chr9 | 53514120 | 53518429 | 4.87902 | 3.88407 | 2.93002 |
| chr11 | 103363390 | 103363709 | 182 | Promoter (<=1kb) | 11 | 103331498 | 103363692 | 32195 | 2 | ENSMUST00 | 0 | Arhgap27 | ENSMUSG0C | false | chr11 | 103362467 | 103363725 | 17.46863 | 44.03677 | 41.39737 | chr11 | 103359206 | 103364527 | 5.2545 | 4.2405 | 3.21907 |
| chr17 | 5599034 | 5599361 | 182 | Intron (ENSMUST00000089185.4 | 17 | 5492600 | 5753891 | 261292 | 1 | ENSMUST00 | 106435 | Zdhhc14 | ENSMUSG0C | true | . | . | . | . | . | . | chr17 | 5587797 | 5599687 | 5.93548 | 4.48203 | 3.77591 |
| chr12 | 85698156 | 85698566 | 182 | Intron (ENSMUST00000040536.5 | 12 | 85686669 | 85709087 | 22419 | 1 | ENSMUST00 | 11488 | Batf | ENSMUSG0C | false | chr12 | 85697928 | 85698747 | 15.42443 | 39.05124 | 36.47339 | . | . | . | . | . | . |
| chr11 | 86358196 | 86358522 | 182 | Promoter (<=1kb) | 11 | 86267033 | 86357602 | 90570 | 2 | ENSMUST00 | -595 | Med13 | ENSMUSG0C | false | chr11 | 86356114 | 86358387 | 25.18546 | 107.13963 | 103.3826 | . | . | . | . | . | . |
| chr9 | 37328754 | 37329105 | 182 | Intron (ENSMUST00000037275.5 | 9 | 37275835 | 37348392 | 72558 | 2 | ENSMUST00 | 19287 | Ccdc15 | ENSMUSG0C | false | . | . | . | . | . | . | chr9 | 37328046 | 37329863 | 5.96426 | 4.50945 | 3.79613 |
| chr6 | 90716250 | 90716639 | 182 | Promoter (<=1kb) | 6 | 90659598 | 90716529 | 56932 | 2 | ENSMUST00 | 0 | Iqsec1 | ENSMUSG0C | false | chr6 | 90715816 | 90716965 | 24.82645 | 85.46523 | 82.21117 | chr6 | 90695436 | 90737892 | 6.93129 | 4.85273 | 4.61297 |
| chr6 | 90660834 | 90661060 | 182 | 3' UTR | 6 | 90662812 | 90667999 | 5188 | 2 | ENSMUST00 | 6939 | Iqsec1 | ENSMUSG0C | false | . | . | . | . | . | . | chr6 | 90658675 | 90665095 | 6.23415 | 4.63243 | 4.01533 |
| chr3 | 69044593 | 69045047 | 182 | Promoter (<=1kb) | 3 | 69035288 | 69044755 | 9468 | 2 | ENSMUST00 | 0 | Trim59 | ENSMUSG0C | false | chr3 | 69043976 | 69045799 | 34.38775 | 109.60938 | 105.79044 | . | . | . | . | . | . |
| chr14 | 67715744 | 67716602 | 182 | Promoter (<=1kb) | 14 | 67715937 | 67739661 | 23725 | 1 | ENSMUST00 | 0 | Kctd9 | ENSMUSG0C | false | chr14 | 67714976 | 67715747 | 29.22958 | 88.61999 | 85.30408 | . | . | . | . | . | . |
| chr11 | 116093837 | 116094070 | 182 | Distal Intergenic | 11 | 116081829 | 116086995 | 5167 | 2 | ENSMUST00 | -6843 | Wbp2 | ENSMUSG0C | false | . | . | . | . | . | . | chr11 | 116093669 | 116094361 | 5.69942 | 4.45557 | 3.59412 |
| chr19 | 10604724 | 10604968 | 182 | Promoter (<=1kb) | 19 | 10592200 | 10604258 | 12059 | 2 | ENSMUST00 | -467 | Dak | ENSMUSG0C | false | chr19 | 10603045 | 10606602 | 41.22328 | 133.19821 | 128.59248 | chr19 | 10601470 | 10605052 | 4.9759 | 3.96533 | 3.02446 |
| chr6 | 112333994 | 112334241 | 182 | Distal Intergenic | 6 | 112369275 | 112375720 | 6446 | 2 | ENSMUST00 | 41479 | Ssu2 | ENSMUSG0C | false | . | . | . | . | . | . | . | . | . | . | . | . |
| chr8 | 117332387 | 117332640 | 182 | Intron (ENSMUST00000166750.2 | 8 | 117349170 | 117461503 | 112334 | 1 | ENSMUST00 | -16530 | Cmip | ENSMUSG0C | false | . | . | . | . | . | . | chr8 | 117331134 | 117336079 | 6.7378 | 4.89636 | 4.42057 |
| chr11 | 4264162 | 4264413 | 182 | Promoter (2-3kb) | 11 | 4266796 | 4268249 | 1454 | 1 | ENSMUST00 | -2383 | Lif | ENSMUSG0C | false | . | . | . | . | . | . | chr11 | 4260098 | 4268486 | 5.73067 | 4.42918 | 3.60325 |
| chr10 | 84622241 | 84622474 | 182 | Promoter (<=1kb) | 10 | 84622437 | 84727177 | 104741 | 1 | ENSMUST00 | 0 | Polr3b | ENSMUSG0C | false | chr10 | 84621752 | 84623247 | 22.29294 | 68.06667 | 65.1033 | chr10 | 84621586 | 84623718 | 5.22736 | 4.21269 | 3.21556 |
| chr12 | 73113043 | 73113316 | 182 | Promoter (<=1kb) | 12 | 73099609 | 73113425 | 13817 | 2 | ENSMUST00 | 109 | Six4 | ENSMUSG0C | false | chr12 | 73111613 | 73113805 | 18.2926 | 68.86671 | 65.89487 | chr12 | 73110875 | 73113168 | 6.06701 | 4.5763 | 3.89898 |
| chr18 | 53176203 | 53176462 | 182 | Promoter (<=1kb) | 18 | 53176365 | 53220860 | 44496 | 1 | ENSMUST00 | 0 | Snx2 | ENSMUSG0C | false | chr18 | 53175802 | 53177208 | 35.47494 | 134.93686 | 130.2605 | . | . | . | . | . | . |
| chr1 | 89927249 | 89927533 | 182 | Promoter (1-2kb) | 1 | 89927966 | 89929185 | 1220 | 2 | ENSMUST00 | 1652 | Gbx2 | ENSMUSG0C | false | chr1 | 89927016 | 89927617 | 11.90309 | 30.75663 | 28.28125 | chr1 | 89927074 | 89927992 | 4.03028 | 3.54383 | 2.23602 |
| chr8 | 74993520 | 74994128 | 182 | Promoter (<=1kb) | 8 | 74993679 | 74999798 | 6120 | 1 | ENSMUST00 | 0 | Hmgxb4 | ENSMUSG0C | true | chr8 | 74993050 | 74994752 | 29.28796 | 98.53321 | 94.99628 | . | . | . | . | . | . |
| chr8 | 75018346 | 75018597 | 182 | Promoter (2-3kb) | 8 | 75016222 | 75031913 | 15692 | 1 | ENSMUST00 | 2125 | Hmgxb4 | ENSMUSG0C | true | . | . | . | . | . | . | chr8 | 75017226 | 75019828 | 3.77536 | 3.32138 | 2.01818 |
| chr14 | 65376654 | 65376829 | 182 | Intron (ENSMUST00000066994.6 | 14 | 65358676 | 65398930 | 40255 | 1 | ENSMUST00 | 17979 | Zfp395 | ENSMUSG0C | false | . | . | . | . | . | . | chr14 | 65365022 | 65390419 | 5.65664 | 4.30592 | 3.56596 |
| chr5 | 103389534 | 103389818 | 182 | Distal Intergenic | 5 | 103425192 | 103598359 | 173168 | 1 | ENSMUST00 | -35374 | Ptpn13 | ENSMUSG0C | false | . | . | . | . | . | . | . | . | . | . | . | . |
| chr5 | 103337276 | 103337512 | 182 | Distal Intergenic | 5 | 103425192 | 103598359 | 173168 | 1 | ENSMUST00 | -87680 | Ptpn13 | ENSMUSG0C | false | chr5 | 103336102 | 103337433 | 8.80131 | 17.30502 | 15.00832 | chr5 | 103336069 | 103337817 | 3.59794 | 3.35796 | 1.83972 |
| chr10 | 122912901 | 122913390 | 182 | Intron (ENSMUST00000161487.2 | 10 | 122895368 | 122941449 | 46082 | 1 | ENSMUST00 | 17534 | Ppm1h | ENSMUSG0C | false | . | . | . | . | . | . | chr10 | 122912613 | 122913575 | 6.90397 | 5.0277 | 4.59685 |
| chr10 | 122678642 | 122679614 | 182 | Promoter (<=1kb) | 10 | 122678762 | 122880333 | 201572 | 1 | ENSMUST00 | 0 | Ppm1h | ENSMUSG0C | false | chr10 | 122678407 | 122680961 | 35.8325 | 117.35963 | 113.30182 | . | . | . | . | . | . |
| chr10 | 122818609 | 122818812 | 182 | Intron (ENSMUST00000162853.2 | 10 | 122811447 | 122886293 | 74847 | 1 | ENSMUST00 | 7163 | Ppm1h | ENSMUSG0C | false | . | . | . | . | . | . | chr10 | 122817595 | 122819107 | 3.85529 | 3.36725 | 2.09229 |
| chr10 | 122096783 | 122097428 | 182 | Promoter (<=1kb) | 10 | 122078114 | 122097371 | 19258 | 2 | ENSMUST00 | 0 | Tmem5 | ENSMUSG0C | false | chr10 | 122096005 | 122097350 | 29.66345 | 100.21 | 96.63225 | . | . | . | . | . | . |
| chr4 | 108300679 | 108301218 | 182 | Promoter (<=1kb) | 4 | 108229724 | 108301096 | 71373 | 2 | ENSMUST00 | 0 | Zyg11b | ENSMUSG0C | false | chr4 | 108300058 | 108301693 | 34.54477 | 122.48211 | 118.25522 | . | . | . | . | . | . |
| chr8 | 84567740 | 84567926 | 182 | Exon (ENSMUST00000121390.2/ | 8 | 84571797 | 84580462 | 8666 | 1 | ENSMUST00 | -3871 | Cacna1a | ENSMUSG0C | false | . | . | . | . | . | . | chr8 | 84567772 | 84568458 | 3.87121 | 3.57676 | 2.05532 |
| chr8 | 84414363 | 84414724 | 182 | Promoter (<=1kb) | 8 | 84415364 | 84640246 | 224883 | 1 | ENSMUST00 | -640 | Cacna1a | ENSMUSG0C | false | chr8 | 84414241 | 84415968 | 20.27628 | 60.14116 | 57.29097 | chr8 | 84414464 | 84416306 | 5.1587 | 3.95274 | 3.15369 |
| chr10 | 82629082 | 82629345 | 182 | Promoter (<=1kb) | 10 | 82629828 | 82650324 | 20497 | 1 | ENSMUST00 | -483 | Tdg | ENSMUSG0C | false | chr10 | 82629283 | 82630833 | 18.02578 | 59.25625 | 56.42029 | chr10 | 82628508 | 82631990 | 3.36145 | 3.08732 | 1.69655 |
| chr13 | 55483155 | 55483360 | 182 | Promoter (<=1kb) | 13 | 55482280 | 55483969 | 1690 | 2 | ENSMUST00 | 609 | Dbn1 | ENSMUSG0C | false | . | . | . | . | . | . | chr13 | 55482428 | 55483943 | 3.53673 | 3.30589 | 1.81903 |
| chr17 | 24414536 | 24415014 | 182 | Promoter (<=1kb) | 17 | 24414675 | 24425895 | 11221 | 1 | ENSMUST00 | 0 | Rnps1 | ENSMUSG0C | false | chr17 | 24413610 | 24415692 | 28.53699 | 95.19626 | 91.7378 | . | . | . | . | . | . |
| chr9 | 107709592 | 107710152 | 182 | Promoter (<=1kb) | 9 | 107681502 | 107710475 | 28974 | 2 | ENSMUST00 | 323 | Sema3f | ENSMUSG0C | false | chr9 | 107707503 | 107710482 | 34.42823 | 105.56979 | 101.85753 | chr9 | 107693726 | 107710535 | 7.184 | 5.04064 | 4.79532 |
| chr11 | 102442030 | 102442305 | 182 | Promoter (2-3kb) | 11 | 102438748 | 102439777 | 1030 | 2 | ENSMUST00 | -2254 | Fam171a2 | ENSMUSG0C | false | . | . | . | . | . | . | chr11 | 102441257 | 102442959 | 4.70803 | 3.93334 | 2.75442 |
| chr11 | 102445513 | 102445805 | 182 | Promoter (1-2kb) | 11 | 102440233 | 102447594 | 7362 | 2 | ENSMUST00 | 1789 | Fam171a2 | ENSMUSG0C | false | . | . | . | . | . | . | chr11 | 102445028 | 102447787 | 5.54764 | 4.31769 | 3.46753 |
| chr11 | 114679303 | 114679523 | 182 | Intron (ENSMUST00000045779.5 | 11 | 114675516 | 114690603 | 15088 | 1 | ENSMUST00 | 3788 | Ttyh2 | ENSMUSG0C | false | . | . | . | . | . | . | . | . | . | . | . | . |
| chr5 | 96164109 | 96164367 | 182 | Promoter (<=1kb) | 5 | 96131097 | 96164171 | 33075 | 2 | ENSMUST00 | 0 | Cnot6l | ENSMUSG0C | false | . | . | . | . | . | . | chr5 | 96163214 | 96164183 | 4.47367 | 3.8739 | 2.56511 |
| chr14 | 78724675 | 78725112 | 182 | Promoter (<=1kb) | 14 | 78569609 | 78725089 | 155481 | 2 | ENSMUST00 | 0 | Dgkh | ENSMUSG0C | false | chr14 | 78724500 | 78725859 | 15.7376 | 52.33027 | 49.58585 | chr14 | 78723971 | 78726906 | 4.1608 | 3.54553 | 2.32388 |
| chr14 | 78775914 | 78776208 | 182 | Distal Intergenic | 14 | 78569609 | 78725089 | 155481 | 2 | ENSMUST00 | -50826 | Dgkh | ENSMUSG0C | false | . | . | . | . | . | . | chr14 | 78775000 | 78776889 | 3.92404 | 3.50609 | 2.16943 |
| chr2 | 69129546 | 69129786 | 182 | Distal Intergenic | 2 | 69135800 | 69189330 | 53531 | 1 | ENSMUST00 | -6014 | Nostrin | ENSMUSG0C | false | . | . | . | . | . | . | chr2 | 69128242 | 69129775 | 4.5602 | 3.85353 | 2.69193 |
| chr2 | 69053736 | 69053926 | 182 | Intron (ENSMUST00000028426.3 | 2 | 69135800 | 69189330 | 53531 | 1 | ENSMUST00 | -81874 | Nostrin | ENSMUSG0C | false | . | . | . | . | . | . | chr2 | 69051725 | 69055298 | 4.31463 | 3.76123 | 2.44237 |
| chr6 | 83794848 | 83795287 | 182 | Promoter (<=1kb) | 6 | 83794982 | 83802556 | 7575 | 1 | ENSMUST00 | 0 | Nagk | ENSMUSG0C | false | chr6 | 83794348 | 83795957 | 35.16326 | 111.99725 | 108.10535 | chr6 | 83794081 | 83796782 | 3.8567 | 3.47451 | 2.06922 |
| chr13 | 103271380 | 103271574 | 182 | Intron (ENSMUST00000166726.2 | 13 | 102732486 | 103334192 | 601707 | 2 | ENSMUST00 | 62618 | Mast4 | ENSMUSG0C | true | . | . | . | . | . | . | . | . | . | . | . | . |
| chr13 | 103270941 | 103271201 | 182 | Intron (ENSMUST00000166726.2 | 13 | 102732486 | 103334192 | 601707 | 2 | ENSMUST00 | 62991 | Mast4 | ENSMUSG0C | true | . | . | . | . | . | . | . | . | . | . | . | . |
| chr10 | 81600721 | 81601005 | 182 | Promoter (<=1kb) | 10 | 81590904 | 81600875 | 9972 | 2 | ENSMUST00 | 0 | Tle6 | ENSMUSG0C | false | . | . | . | . | . | . | chr10 | 81600180 | 81603284 | 5.55517 | 4.36004 | 3.46327 |
| chr4 | 107618006 | 107618238 | 182 | Exon (ENSMUST00000125573.1/ | 4 | 107615000 | 107632393 | 17394 | 1 | ENSMUST00 | 3007 | Glis1 | ENSMUSG0C | false | . | . | . | . | . | . | . | . | . | . | . | . |
| chr6 | 83711009 | 83711266 | 182 | Promoter (<=1kb) | 6 | 83711264 | 83738313 | 27050 | 1 | ENSMUST00 | 0 | Vax2 | ENSMUSG0C | false | chr6 | 83710552 | 83711584 | 17.30849 | 43.01685 | 40.38835 | chr6 | 83710123 | 83712423 | 6.18165 | 4.62412 | 4.01968 |
| chr2 | 67565420 | 67565972 | 182 | Promoter (<=1kb) | 2 | 67565871 | 67810894 | 245024 | 1 | ENSMUST00 | 0 | B3galt1 | ENSMUSG0C | false | chr2 | 67565176 | 67566948 | 18.77433 | 54.14365 | 51.37372 | chr2 | 67564945 | 67567572 | 5.22415 | 4.19739 | 3.21045 |
| chr14 | 77036626 | 77037057 | 182 | Promoter (<=1kb) | 14 | 77036772 | 77112204 | 75433 | 1 | ENSMUST00 | 0 | Ccdc122 | ENSMUSG0C | false | chr14 | 77036071 | 77037177 | 30.07331 | 91.05476 | 87.68732 | . | . | . | . | . | . |
| chr8 | 71610899 | 71611221 | 182 | Promoter (<=1kb) | 8 | 71611024 | 71624909 | 13886 | 1 | ENSMUST00 | 0 | Colgalt1 | ENSMUSG0C | false | chr8 | 71611041 | 71612143 | 38.18876 | 121.46067 | 117.27263 | . | . | . | . | . | . |
| chr6 | 83427467 | 83427863 | 182 | Intron (ENSMUST00000089622.5 | 6 | 83362373 | 83441675 | 79303 | 2 | ENSMUST00 | 13812 | Tet3 | ENSMUSG0C | true | . | . | . | . | . | . | chr6 | 83421103 | 83440609 | 6.41331 | 4.64298 | 4.1723 |
| chr17 | 70990430 | 70990666 | 182 | Promoter (<=1kb) | 17 | 70976936 | 70990530 | 13595 | 2 | ENSMUST00 | 0 | Myl12b | ENSMUSG0C | false | chr17 | 70989464 | 70991264 | 29.99031 | 99.80121 | 96.2382 | chr17 | 70983359 | 71000483 | 5.96068 | 4.55086 | 3.78929 |
| chr17 | 70990744 | 70991129 | 182 | Promoter (<=1kb) | 17 | 70973920 | 70990787 | 16868 | 2 | ENSMUST00 | 0 | Myl12b | ENSMUSG0C | false | chr17 | 70989464 | 70991264 | 29.99031 | 99.80121 | 96.2382 | chr17 | 70983359 | 71000483 | 5.96068 | 4.55086 | 3.78929 |
| chr9 | 45944573 | 45944866 | 182 | Promoter (<=1kb) | 9 | 45942346 | 45944631 | 2286 | 2 | ENSMUST00 | 0 | Sidt2 | ENSMUSG0C | false | . | . | . | . | . | . | chr9 | 45943650 | 45947822 | 6.38099 | 4.51987 | 4.19123 |
| chr9 | 45966450 | 45966657 | 182 | 3' UTR | 9 | 45937857 | 45955247 | 17391 | 2 | ENSMUST00 | -11204 | Sidt2 | ENSMUSG0C | false | . | . | . | . | . | . | chr9 | 45963529 | 45968445 | 6.00453 | 4.42254 | 3.85296 |
| chr10 | 81266572 | 81266868 | 182 | Promoter (<=1kb) | 10 | 81265430 | 81266924 | 1495 | 2 | ENSMUST00 | 56 | Mrpl54 | ENSMUSG0C | false | chr10 | 81266206 | 81267114 | 25.16617 | 75.86667 | 72.78426 | chr10 | 81265703 | 81269340 | 4.66593 | 3.88425 | 2.77737 |
| chr11 | 83964291 | 83964712 | 182 | Promoter (<=1kb) | 11 | 83964428 | 84040954 | 76527 | 1 | ENSMUST00 | 0 | Synrg | ENSMUSG0C | false | chr11 | 83963960 | 83965228 | 31.56964 | 96.59996 | 93.11521 | chr11 | 83963621 | 83969402 | 3.81256 | 3.49752 | 2.05604 |
| chr7 | 35118977 | 35119344 | 182 | Promoter (<=1kb) | 7 | 35119293 | 35121926 | 2634 | 1 | ENSMUST00 | 0 | Cebpa | ENSMUSG0C | false | chr7 | 35117750 | 35120763 | 21.02725 | 63.18829 | 60.29588 | . | . | . | . | . | . |
| chr14 | 75010985 | 75011257 | 182 | Distal Intergenic | 14 | 75016027 | 75052532 | 36506 | 1 | ENSMUST00 | -4770 | 5031414D18 | ENSMUSG0C | false | . | . | . | . | . | . | chr14 | 75010515 | 75011832 | 4.33624 | 3.78421 | 2.46354 |
| chr10 | 81182864 | 81183176 | 182 | Promoter (<=1kb) | 10 | 81183007 | 81193197 | 10191 | 1 | ENSMUST00 | 0 | Dapk3 | ENSMUSG0C | false | chr10 | 81182941 | 81183458 | 16.88215 | 57.7818 | 54.96725 | chr10 | 81180614 | 81187537 | 5.21602 | 4.06527 | 3.22051 |
| chr4 | 103313584 | 103314149 | 182 | Promoter (<=1kb) | 4 | 103313812 | 103366443 | 52632 | 1 | ENSMUST00 | 0 | Oma1 | ENSMUSG0C | false | chr4 | 103313729 | 103314546 | 17.49727 | 64.69686 | 61.78638 | chr4 | 103313413 | 103314381 | 3.04468 | 3.05804 | 1.38941 |
| chr4 | 103321004 | 103321283 | 182 | Intron (ENSMUST00000035780.3 | 4 | 103324484 | 103328889 | 4406 | 1 | ENSMUST00 | -3201 | Oma1 | ENSMUSG0C | false | . | . | . | . | . | . | chr4 | 103320376 | 103322124 | 4.46741 | 3.77115 | 2.57873 |
| chr12 | 54162769 | 54163147 | 182 | Distal Intergenic | 12 | 54178981 | 54203860 | 24880 | 2 | ENSMUST00 | 40713 | Egln3 | ENSMUSG0C | false | chr12 | 54162447 | 54163232 | 8.29801 | 20.47831 | 18.13586 | chr12 | 54162530 | 54163499 | 4.00307 | 3.63303 | 2.17219 |
| chr16 | 58458526 | 58458742 | 182 | Promoter (1-2kb) | 16 | 58460199 | 58465827 | 5629 | 1 | ENSMUST00 | -1457 | Dcbld2 | ENSMUSG0C | false | . | . | . | . | . | . | chr16 | 58458372 | 58460878 | 4.18849 | 3.61495 | 2.37027 |
| chr2 | 59785273 | 59785541 | 182 | Exon (ENSMUST00000112568.2/ | 2 | 59779711 | 59785499 | 5789 | 1 | ENSMUST00 | 5563 | Tanc1 | ENSMUSG0C | true | . | . | . | . | . | . | chr2 | 59785448 | 59786416 | 2.78293 | 2.83604 | 1.23104 |
| chr2 | 59626522 | 59626736 | 182 | Intron (ENSMUST00000112568.2 | 2 | 59630063 | 59647005 | 16943 | 1 | ENSMUST00 | -3327 | Tanc1 | ENSMUSG0C | true | . | . | . | . | . | . | chr2 | 59625432 | 59634695 | 6.09177 | 4.47351 | 3.93532 |
| chr19 | 10474524 | 10474751 | 182 | Exon (ENSMUST00000038842.3/ | 19 | 10474257 | 10482897 | 8641 | 2 | ENSMUST00 | 8146 | Ppp1r32 | ENSMUSG0C | false | . | . | . | . | . | . | . | . | . | . | . | . |
| chr7 | 5080113 | 5080365 | 182 | Promoter (<=1kb) | 7 | 5080235 | 5098178 | 17944 | 1 | ENSMUST00 | 0 | Epn1 | ENSMUSG0C | false | chr7 | 5079795 | 5081124 | 31.96831 | 119.80809 | 115.67081 | chr7 | 5079040 | 5089985 | 4.0894 | 3.60068 | 2.27543 |
| chr4 | 101647181 | 101648041 | 182 | Promoter (<=1kb) | 4 | 101647718 | 101659364 | 11647 | 1 | ENSMUST00 | 0 | Leprot | ENSMUSG0C | false | chr4 | 101647700 | 101649031 | 32.98514 | 106.12805 | 102.40588 | . | . | . | . | . | . |
| chr7 | 99858833 | 99859156 | 182 | Promoter (<=1kb) | 7 | 99837569 | 99858883 | 21315 | 2 | ENSMUST00 | 0 | Spcs2 | ENSMUSG0C | false | chr7 | 99857968 | 99859477 | 27.6016 | 85.77155 | 82.50918 | chr7 | 99852448 | 99859525 | 4.35269 | 3.7333 | 2.48612 |
| chr6 | 97148772 | 97149126 | 182 | Promoter (<=1kb) | 6 | 97111972 | 97148883 | 36912 | 2 | ENSMUST00 | 0 | Eogt | ENSMUSG0C | false | chr6 | 97148008 | 97149516 | 31.62822 | 118.12274 | 114.03909 | chr6 | 97147589 | 97149924 | 5.46177 | 4.29858 | 3.42327 |
| chr13 | 59971233 | 59971430 | 182 | Distal Intergenic | 13 | 59771879 | 59823147 | 51269 | 2 | ENSMUST00 | -148087 | Zcchc6 | ENSMUSG0C | false | chr13 | 59971201 | 59972007 | 6.75876 | 12.51203 | 10.2897 | chr13 | 59971138 | 59972283 | 4.44171 | 3.79562 | 2.58648 |
| chr13 | 59822916 | 59823407 | 182 | Promoter (<=1kb) | 13 | 59771879 | 59823147 | 51269 | 2 | ENSMUST00 | 0 | Zcchc6 | ENSMUSG0C | false | chr13 | 59822193 | 59823730 | 50.54853 | 170.95712 | 164.9677 | . | . | . | . | . | . |
| chr13 | 59875759 | 59876095 | 182 | Distal Intergenic | 13 | 59771879 | 59823147 | 51269 | 2 | ENSMUST00 | -52613 | Zcchc6 | ENSMUSG0C | false | chr13 | 59876075 | 59878826 | 17.64787 | 49.73517 | 47.02245 | . | . | . | . | . | . |
| chr2 | 163705972 | 163706272 | 182 | Intron (ENSMUST00000147385.2 | 2 | 163694038 | 163726152 | 32115 | 1 | ENSMUST00 | 11935 | Pkig | ENSMUSG0C | false | . | . | . | . | . | . | chr2 | 163667073 | 163719465 | 6.42152 | 4.6796 | 4.1796 |
| chr2 | 163675062 | 163675341 | 182 | Intron (ENSMUST00000147385.2 | 2 | 163661499 | 163725473 | 63975 | 1 | ENSMUST00 | 13564 | Pkig | ENSMUSG0C | false | . | . | . | . | . | . | chr2 | 163667073 | 163719465 | 6.42152 | 4.6796 | 4.1796 |
| chr19 | 56705370 | 56705603 | 182 | Distal Intergenic | 19 | 56722372 | 56724862 | 2491 | 1 | ENSMUST00 | -16769 | Adrb1 | ENSMUSG0C | false | . | . | . | . | . | . | chr19 | 56705293 | 56705633 | 3.02068 | 2.7776 | 1.43533 |
| chr2 | 38990404 | 38990775 | 182 | Distal Intergenic | 2 | 38997476 | 39006168 | 8693 | 1 | ENSMUST00 | -6701 | Wdr38 | ENSMUSG0C | false | . | . | . | . | . | . | chr2 | 38990205 | 38991244 | 4.29203 | 3.53655 | 2.4652 |
| chr4 | 100095924 | 100096422 | 182 | Promoter (<=1kb) | 4 | 100095791 | 100444765 | 348975 | 1 | ENSMUST00 | 134 | Ror1 | ENSMUSG0C | false | chr4 | 100095760 | 100097247 | 34.72914 | 110.20382 | 106.3632 | . | . | . | . | . | . |
| chr12 | 59219228 | 59219552 | 182 | Promoter (<=1kb) | 12 | 59200655 | 59219725 | 19071 | 2 | ENSMUST00 | 173 | Fbxo33 | ENSMUSG0C | false | chr12 | 59217895 | 59220396 | 38.67513 | 141.95352 | 136.99454 | . | . | . | . | . | . |
| chr7 | 99139542 | 99139804 | 182 | Promoter (1-2kb) | 7 | 98886743 | 99141144 | 254402 | 2 | ENSMUST00 | 1340 | Uvrag | ENSMUSG0C | false | chr7 | 99139179 | 99141230 | 40.8303 | 133.61433 | 128.98549 | chr7 | 99127487 | 99140202 | 5.46741 | 4.21908 | 3.40558 |
| chr7 | 99130494 | 99130693 | 182 | Intron (ENSMUST00000037968.8 | 7 | 98886743 | 99141144 | 254402 | 2 | ENSMUST00 | 10451 | Uvrag | ENSMUSG0C | false | . | . | . | . | . | . | chr7 | 99127487 | 99140202 | 5.46741 | 4.21908 | 3.40558 |
| chr16 | 55838631 | 55838931 | 182 | Promoter (<=1kb) | 16 | 55811375 | 55838899 | 27525 | 2 | ENSMUST00 | 0 | Nfkbiz | ENSMUSG0C | false | chr16 | 55838318 | 55839457 | 29.47077 | 87.11163 | 83.82504 | chr16 | 55837769 | 55839638 | 3.29079 | 3.13577 | 1.65342 |
| chr19 | 3708186 | 3708679 | 182 | Promoter (<=1kb) | 19 | 3708333 | 3717881 | 9549 | 1 | ENSMUST00 | 0 | 1810055G02 | ENSMUSG0C | false | chr19 | 3707741 | 3709052 | 38.03245 | 121.19012 | 117.00504 | chr19 | 3707181 | 3709698 | 4.1542 | 3.54832 | 2.35149 |
| chr6 | 100670472 | 100670863 | 182 | Promoter (<=1kb) | 6 | 100662839 | 100671133 | 8295 | 2 | ENSMUST00 | 270 | Shq1 | ENSMUSG0C | false | chr6 | 100670213 | 100671482 | 26.89699 | 109.78668 | 105.95901 | . | . | . | . | . | . |
| chr2 | 38099274 | 38099543 | 182 | Intron (ENSMUST00000102787.4 | 2 | 38021266 | 38166081 | 144816 | 2 | ENSMUST00 | 66538 | Dennd1a | ENSMUSG0C | false | . | . | . | . | . | . | chr2 | 38099413 | 38100748 | 4.21821 | 3.62801 | 2.40846 |
| chr7 | 98655647 | 98655932 | 182 | Promoter (<=1kb) | 7 | 98590615 | 98656569 | 65955 | 2 | ENSMUST00 | 637 | 2210018M11 | ENSMUSG0C | false | chr7 | 98655540 | 98657644 | 39.24815 | 133.0453 | 128.44171 | chr7 | 98653773 | 98655773 | 3.8943 | 3.58459 | 2.09454 |
| chr11 | 80775301 | 80775520 | 182 | Intron (ENSMUST00000041065.8 | 11 | 80673667 | 80780003 | 106337 | 2 | ENSMUST00 | 4483 | Myo1d | ENSMUSG0C | true | . | . | . | . | . | . | chr11 | 80758294 | 80778239 | 6.06128 | 4.46131 | 3.88566 |
| chr10 | 87016126 | 87016363 | 182 | Intron (ENSMUST00000162837.1 | 10 | 86841198 | 87008025 | 166828 | 2 | ENSMUST00 | -8102 | Stab2 | ENSMUSG0C | false | . | . | . | . | . | . | chr10 | 87015230 | 87016576 | 4.66001 | 3.67989 | 2.76289 |
| chr16 | 33581670 | 33582172 | 182 | Intron (ENSMUST00000121925.2 | 16 | 33596562 | 33664135 | 67574 | 1 | ENSMUST00 | -14390 | Slc12a8 | ENSMUSG0C | false | . | . | . | . | . | . | . | . | . | . | . | . |
| chr2 | 29252869 | 29253149 | 182 | Promoter (<=1kb) | 2 | 29203822 | 29253005 | 49184 | 2 | ENSMUST00 | 0 | Ntng2 | ENSMUSG0C | true | chr2 | 29249202 | 29253779 | 38.46334 | 130.56975 | 126.05653 | . | . | . | . | . | . |
| chr10 | 85916906 | 85917209 | 182 | Promoter (<=1kb) | 10 | 85891970 | 85916945 | 24976 | 2 | ENSMUST00 | 0 | Prdm4 | ENSMUSG0C | false | chr10 | 85916118 | 85917629 | 39.1345 | 126.3175 | 121.95772 | chr10 | 85915493 | 85917889 | 4.32818 | 3.75781 | 2.47143 |
| chr11 | 100320231 | 100320641 | 182 | Promoter (<=1kb) | 11 | 100320518 | 100321349 | 832 | 1 | ENSMUST00 | 0 | Eif1 | ENSMUSG0C | false | chr11 | 100319937 | 100321297 | 31.8806 | 105.19856 | 101.49664 | . | . | . | . | . | . |
| chr9 | 102449497 | 102449706 | 182 | Distal Intergenic | 9 | 102505750 | 102546239 | 40490 | 1 | ENSMUST00 | -56044 | Ky | ENSMUSG0C | false | . | . | . | . | . | . | chr9 | 102449118 | 102449995 | 2.77568 | 2.87585 | 1.24139 |
| chr10 | 84967570 | 84967941 | 182 | Intron (ENSMUST00000038523.8 | 10 | 84917656 | 85016446 | 98791 | 1 | ENSMUST00 | 49915 | Ric8b | ENSMUSG0C | false | . | . | . | . | . | . | chr10 | 84963136 | 84972104 | 5.87946 | 4.4866 | 3.72979 |
| chr10 | 80156787 | 80157026 | 182 | 3' UTR | 10 | 80151238 | 80153696 | 2459 | 1 | ENSMUST00 | 5550 | Midn | ENSMUSG0C | false | . | . | . | . | . | . | chr10 | 80150859 | 80167428 | 9.09735 | 5.82258 | 6.36001 |
| chr7 | 97579508 | 97579967 | 182 | Promoter (<=1kb) | 7 | 97579896 | 97692782 | 112887 | 1 | ENSMUST00 | 0 | Rsf1 | ENSMUSG0C | false | chr7 | 97578774 | 97581491 | 38.42999 | 152.65175 | 147.26898 | . | . | . | . | . | . |
| chr4 | 44756489 | 44756763 | 182 | Promoter (<=1kb) | 4 | 44756568 | 44932203 | 175636 | 1 | ENSMUST00 | 0 | Zcchc7 | ENSMUSG0C | false | chr4 | 44755579 | 44757654 | 40.18469 | 140.22169 | 135.33289 | . | . | . | . | . | . |
| chr19 | 10326157 | 10326462 | 182 | Distal Intergenic | 19 | 10245265 | 10304877 | 59613 | 2 | ENSMUST00 | -21281 | Dagla | ENSMUSG0C | false | . | . | . | . | . | . | chr19 | 10322178 | 10329326 | 5.27385 | 4.17732 | 3.23885 |
| chr19 | 10294632 | 10294933 | 182 | Intron (ENSMUST00000039327.5 | 19 | 10245268 | 10304857 | 59590 | 2 | ENSMUST00 | 9924 | Dagla | ENSMUSG0C | false | . | . | . | . | . | . | chr19 | 10270370 | 10305865 | 8.32241 | 5.52593 | 5.72899 |
| chr10 | 111339444 | 111339738 | 182 | Distal Intergenic | 10 | 111298679 | 111301736 | 3058 | 1 | ENSMUST00 | 40766 | Bbs10 | ENSMUSG0C | false | . | . | . | . | . | . | chr10 | 111335890 | 111343265 | 5.40426 | 4.1855 | 3.34231 |
| chr10 | 111287475 | 111287650 | 182 | Intron (ENSMUST00000095310.1 | 10 | 111298679 | 111301736 | 3058 | 1 | ENSMUST00 | -11029 | Bbs10 | ENSMUSG0C | false | . | . | . | . | . | . | chr10 | 111287294 | 111287569 | 3.86864 | 3.46156 | 2.06175 |
| chr15 | 88875628 | 88875978 | 182 | Distal Intergenic | 15 | 88862186 | 88865718 | 3533 | 1 | ENSMUST00 | 13443 | Pim3 | ENSMUSG0C | false | . | . | . | . | . | . | chr15 | 88875156 | 88876576 | 6.52728 | 4.67111 | 4.26869 |
| chr15 | 88867540 | 88867845 | 182 | Distal Intergenic | 15 | 88862186 | 88865718 | 3533 | 1 | ENSMUST00 | 5355 | Pim3 | ENSMUSG0C | false | . | . | . | . | . | . | chr15 | 88863272 | 88871136 | 9.58718 | 5.83938 | 6.75823 |
| chr13 | 81691961 | 81692173 | 182 | Intron (ENSMUST00000048993.6 | 13 | 81673880 | 81687285 | 13406 | 2 | ENSMUST00 | -4677 | Polr3g | ENSMUSG0C | false | chr13 | 81691806 | 81692046 | 8.36866 | 16.57842 | 14.29226 | chr13 | 81690603 | 81692515 | 2.91509 | 2.86941 | 1.35156 |
| chr13 | 81686342 | 81686840 | 182 | Promoter (<=1kb) | 13 | 81673880 | 81687285 | 13406 | 2 | ENSMUST00 | 445 | Polr3g | ENSMUSG0C | false | . | . | . | . | . | . | chr13 | 81685069 | 81687312 | 3.48138 | 3.25144 | 1.78999 |
| chr2 | 160872517 | 160873296 | 182 | Promoter (<=1kb) | 2 | 160748708 | 160872998 | 124291 | 2 | ENSMUST00 | 0 | Zhx3 | ENSMUSG0C | false | chr2 | 160872139 | 160873587 | 31.96831 | 119.80809 | 115.67081 | . | . | . | . | . | . |
| chr5 | 86172453 | 86173025 | 182 | Promoter (<=1kb) | 5 | 86110720 | 86172803 | 62084 | 2 | ENSMUST00 | 0 | Uba6 | ENSMUSG0C | false | chr5 | 86171817 | 86173491 | 41.49189 | 142.53105 | 137.54099 | chr5 | 86171427 | 86173526 | 4.66901 | 3.93236 | 2.72719 |
| chr9 | 37139633 | 37139825 | 182 | Promoter (2-3kb) | 9 | 36936439 | 37137567 | 201129 | 2 | ENSMUST00 | -2067 | Pknox2 | ENSMUSG0C | true | . | . | . | . | . | . | chr9 | 37137127 | 37144958 | 5.79298 | 4.47095 | 3.65934 |
| chr4 | 43405090 | 43405312 | 182 | Promoter (1-2kb) | 4 | 43406435 | 43427088 | 20654 | 1 | ENSMUST00 | -1123 | Rusc2 | ENSMUSG0C | false | . | . | . | . | . | . | chr4 | 43382882 | 43406168 | 6.80327 | 4.94439 | 4.48074 |
| chr11 | 54437960 | 54438284 | 182 | Promoter (<=1kb) | 11 | 54438199 | 54518235 | 80037 | 1 | ENSMUST00 | 0 | Fnip1 | ENSMUSG0C | false | chr11 | 54437572 | 54439590 | 33.94791 | 109.87469 | 106.04007 | . | . | . | . | . | . |
| chr4 | 43045745 | 43046193 | 182 | Promoter (<=1kb) | 4 | 43032414 | 43045797 | 13384 | 2 | ENSMUST00 | 0 | Fam214b | ENSMUSG0C | false | chr4 | 43045142 | 43046346 | 19.14982 | 55.63055 | 52.84086 | . | . | . | . | . | . |
| chr13 | 24617840 | 24618113 | 182 | Intron (ENSMUST00000110383.2 | 13 | 24614627 | 24705897 | 91271 | 1 | ENSMUST00 | 3214 | Fam65b | ENSMUSG0C | false | . | . | . | . | . | . | chr13 | 24616471 | 24619638 | 4.21231 | 3.64673 | 2.36172 |
| chr13 | 24565971 | 24566353 | 182 | Distal Intergenic | 13 | 24582189 | 24733816 | 151628 | 1 | ENSMUST00 | -15836 | Fam65b | ENSMUSG0C | false | . | . | . | . | . | . | chr13 | 24564838 | 24567964 | 5.75538 | 4.49221 | 3.62154 |
| chr15 | 84987708 | 84988208 | 182 | Promoter (<=1kb) | 15 | 84943936 | 84988000 | 44065 | 2 | ENSMUST00 | 0 | 5031439G07 | ENSMUSG0C | false | chr15 | 84987053 | 84988586 | 24.55236 | 90.57755 | 87.22021 | chr15 | 84985840 | 84988951 | 4.64287 | 3.81903 | 2.70534 |
| chr2 | 53070776 | 53070984 | 182 | Intron (ENSMUST00000090952.5 | 2 | 53114115 | 53133804 | 19690 | 1 | ENSMUST00 | -43131 | Fmnl2 | ENSMUSG0C | true | . | . | . | . | . | . | chr2 | 53069122 | 53071178 | 3.80822 | 3.38507 | 2.07197 |
| chr2 | 53095005 | 53095271 | 182 | Intron (ENSMUST00000090952.5 | 2 | 53114115 | 53133804 | 19690 | 1 | ENSMUST00 | -18844 | Fmnl2 | ENSMUSG0C | true | . | . | . | . | . | . | chr2 | 53094285 | 53096316 | 6.25344 | 4.63005 | 4.02606 |
| chr4 | 41755583 | 41755841 | 182 | Promoter (<=1kb) | 4 | 41755758 | 41757480 | 1723 | 1 | ENSMUST00 | 0 | Galt | ENSMUSG0C | false | chr4 | 41754961 | 41756541 | 23.93078 | 87.47927 | 84.18233 | . | . | . | . | . | . |
| chr2 | 26628292 | 26628503 | 182 | Promoter (<=1kb) | 2 | 26628457 | 26636497 | 8041 | 1 | ENSMUST00 | 0 | Fam69b | ENSMUSG0C | false | chr2 | 26627864 | 26628866 | 21.79372 | 61.12524 | 58.26245 | . | . | . | . | . | . |
| chr15 | 93275096 | 93275556 | 182 | Promoter (<=1kb) | 15 | 93239742 | 93275161 | 35420 | 2 | ENSMUST00 | 0 | Gxylt1 | ENSMUSG0C | false | chr15 | 93274138 | 93275225 | 35.67123 | 127.73884 | 123.33056 | . | . | . | . | . | . |
| chr1 | 89092652 | 89092840 | 182 | Intron (ENSMUST00000066279.5 | 1 | 89070453 | 89112860 | 42408 | 1 | ENSMUST00 | 22200 | Sh3bp4 | ENSMUSG0C | false | . | . | . | . | . | . | chr1 | 89071619 | 89098621 | 6.06397 | 4.51902 | 3.881 |
| chr17 | 36958185 | 36958665 | 182 | Promoter (<=1kb) | 17 | 36958592 | 36965622 | 7031 | 1 | ENSMUST00 | 0 | Znrd1as | ENSMUSG0C | false | chr17 | 36957679 | 36959243 | 19.17126 | 69.02763 | 66.05415 | chr17 | 36956323 | 36959716 | 5.61048 | 4.33555 | 3.51521 |
| chr18 | 15063911 | 15064416 | 182 | Promoter (<=1kb) | 18 | 14968685 | 15063582 | 94898 | 2 | ENSMUST00 | -330 | Kctd1 | ENSMUSG0C | false | chr18 | 15062362 | 15065257 | 28.1615 | 93.53628 | 90.11485 | chr18 | 15062478 | 15066597 | 4.85515 | 3.85397 | 2.93151 |
| chr4 | 41179640 | 41179830 | 182 | Intron (ENSMUST00000040008.3 | 4 | 41182920 | 41190823 | 7904 | 1 | ENSMUST00 | -3090 | Ube2r2 | ENSMUSG0C | false | . | . | . | . | . | . | chr4 | 41178048 | 41181548 | 3.56505 | 3.13895 | 1.85578 |

| | | | | | | | | | | | | | | | | | | | | | | | | | | |
|---|---|---|---|---|---|---|---|---|---|---|---|---|---|---|---|---|---|---|---|---|---|---|---|---|---|---|
| chr1 | 88277478 | 88277782 | 182 | Promoter (<=1kb) | 1 | 88277661 | 88308073 | 30413 | 1 | ENSMUST00 | 0 | Trpm8 | ENSMUSG0C | false | chr1 | 88276574 | 88278131 | 25.44072 | 266.78333 | 259.59549 | chr1 | 88275882 | 88278450 | 7.48282 | 3.31096 | 5.068 |
| chr5 | 77402895 | 77403175 | 182 | Intron (ENSMUST00000046746.6 | 5 | 77349240 | 77408045 | 58806 | 2 | ENSMUST00 | 4870 | Igfbp7 | ENSMUSG0C | false | . | . | . | . | . | . | chr5 | 77353137 | 77404361 | 6.51707 | 4.75542 | 4.25399 |
| chr15 | 91673009 | 91673278 | 182 | Promoter (<=1kb) | 15 | 91673175 | 91816120 | 142946 | 1 | ENSMUST00 | 0 | Lrrk2 | ENSMUSG0C | false | chr15 | 91672736 | 91674386 | 17.61183 | 48.10911 | 45.41927 | chr15 | 91672920 | 91675488 | 3.85528 | 3.51076 | 2.05536 |
| chr15 | 91573087 | 91573441 | 182 | Promoter (<=1kb) | 15 | 91267696 | 91573261 | 305566 | 2 | ENSMUST00 | 0 | Slc2a13 | ENSMUSG0C | false | chr15 | 91572223 | 91573629 | 34.56959 | 108.89287 | 105.0881 | chr15 | 91571678 | 91573875 | 3.66372 | 3.31507 | 1.9461 |
| chr3 | 59330146 | 59330399 | 182 | Exon (ENSMUST00000039419.6/ | 3 | 59316735 | 59344256 | 27522 | 2 | ENSMUST00 | 13857 | Igsf10 | ENSMUSG0C | false | . | . | . | . | . | . | chr3 | 59330073 | 59331184 | 4.37555 | 3.69823 | 2.54158 |
| chr17 | 79615007 | 79615534 | 182 | Promoter (<=1kb) | 17 | 79614900 | 79682152 | 67253 | 1 | ENSMUST00 | 108 | Rmdn2 | ENSMUSG0C | false | chr17 | 79614607 | 79615964 | 27.34946 | 104.78247 | 101.09169 | chr17 | 79614042 | 79615733 | 3.72035 | 3.25596 | 1.98371 |
| chr6 | 67073770 | 67074020 | 182 | Exon (ENSMUST00000149889.1/ | 6 | 67036897 | 67037457 | 561 | 2 | ENSMUST00 | -36314 | Gadd45a | ENSMUSG0C | false | . | . | . | . | . | . | chr6 | 67069252 | 67079338 | 6.07124 | 4.48472 | 3.90237 |
| chr11 | 51756387 | 51756678 | 182 | Promoter (<=1kb) | 11 | 51692264 | 51756618 | 64355 | 2 | ENSMUST00 | 0 | Sec24a | ENSMUSG0C | false | chr11 | 51755431 | 51757268 | 40.33899 | 132.76065 | 128.17128 | chr11 | 51753763 | 51757273 | 4.34117 | 3.67066 | 2.50377 |
| chr9 | 26803644 | 26803942 | 182 | Promoter (1-2kb) | 9 | 26765203 | 26802585 | 37383 | 2 | ENSMUST00 | -1060 | Glb1l2 | ENSMUSG0C | false | . | . | . | . | . | . | . | . | . | . | . | . |
| chr7 | 34230231 | 34230538 | 182 | Promoter (<=1kb) | 7 | 34202122 | 34230281 | 28160 | 2 | ENSMUST00 | 0 | Gpi1 | ENSMUSG0C | false | chr7 | 34229243 | 34230723 | 19.74353 | 71.90538 | 68.88615 | chr7 | 34226191 | 34231057 | 4.94131 | 3.87684 | 2.98759 |
| chr18 | 39196635 | 39196870 | 182 | Intron (ENSMUST00000097593.3 | 18 | 39227368 | 39372511 | 145144 | 1 | ENSMUST00 | -30498 | Arhgap26 | ENSMUSG0C | false | . | . | . | . | . | . | chr18 | 39193471 | 39197393 | 4.91054 | 4.00067 | 2.92834 |
| chr15 | 89315240 | 89315555 | 182 | Promoter (<=1kb) | 15 | 89288236 | 89315311 | 27076 | 2 | ENSMUST00 | 0 | Sbf1 | ENSMUSG0C | false | chr15 | 89314018 | 89315977 | 41.19707 | 132.05855 | 127.49266 | . | . | . | . | . | . |
| chr17 | 79442743 | 79442991 | 182 | Distal Intergenic | 17 | 79334025 | 79355091 | 21067 | 2 | ENSMUST00 | -87653 | Cdc42ep3 | ENSMUSG0C | false | . | . | . | . | . | . | . | . | . | . | . | . |
| chr17 | 79354906 | 79355312 | 182 | Promoter (<=1kb) | 17 | 79334025 | 79355091 | 21067 | 2 | ENSMUST00 | 0 | Cdc42ep3 | ENSMUSG0C | false | chr17 | 79353864 | 79356423 | 40.11768 | 137.88335 | 133.09009 | . | . | . | . | . | . |
| chr17 | 14992595 | 14993541 | 182 | Downstream (2-3kb) | 17 | 14978864 | 14991577 | 12714 | 1 | ENSMUST00 | 13732 | Ermard | ENSMUSG0C | false | chr17 | 14992064 | 14992914 | 51.00806 | 172.93202 | 166.90143 | . | . | . | . | . | . |
| chr17 | 15010012 | 15010367 | 182 | Promoter (<=1kb) | 17 | 15010266 | 15022448 | 12183 | 1 | ENSMUST00 | 0 | Ermard | ENSMUSG0C | false | chr17 | 15009455 | 15011012 | 43.8213 | 195.4646 | 189.06622 | chr17 | 15009541 | 15011647 | 4.75185 | 3.47166 | 2.83337 |
| chr1 | 86434104 | 86434732 | 182 | Distal Intergenic | 1 | 86426329 | 86428052 | 1724 | 1 | ENSMUST00 | 7776 | 1700019O17 | ENSMUSG0C | false | chr1 | 86433982 | 86435081 | 20.5121 | 70.85588 | 67.85384 | chr1 | 86432854 | 86438129 | 4.97301 | 4.04235 | 2.9885 |
| chr18 | 38938215 | 38938611 | 182 | Intron (ENSMUST00000148850.2 | 18 | 38838675 | 38929404 | 90730 | 2 | ENSMUST00 | -8812 | Fgf1 | ENSMUSG0C | false | . | . | . | . | . | . | . | . | . | . | . | . |
| chr5 | 140571691 | 140572032 | 182 | Distal Intergenic | 5 | 140563119 | 140565071 | 1953 | 2 | ENSMUST00 | -6621 | Grifin | ENSMUSG0C | false | . | . | . | . | . | . | . | . | . | . | . | . |
| chr2 | 25332512 | 25332743 | 182 | Promoter (<=1kb) | 2 | 25332729 | 25352212 | 19484 | 1 | ENSMUST00 | 0 | Man1b1 | ENSMUSG0C | false | chr2 | 25331751 | 25333538 | 24.55236 | 90.57755 | 87.22021 | chr2 | 25331422 | 25334399 | 6.04247 | 4.54554 | 3.86639 |
| chr12 | 28612489 | 28612749 | 182 | Intron (ENSMUST00000036136.7 | 12 | 28594173 | 28623290 | 29118 | 2 | ENSMUST00 | 10541 | Colec11 | ENSMUSG0C | false | . | . | . | . | . | . | chr12 | 28612321 | 28613035 | 4.33742 | 3.55477 | 2.47025 |
| chr6 | 42714742 | 42714924 | 182 | Distal Intergenic | 6 | 42709097 | 42710072 | 976 | 2 | ENSMUST00 | -4671 | Fam115a | ENSMUSG0C | false | . | . | . | . | . | . | . | . | . | . | . | . |
| chr5 | 139724948 | 139725234 | 182 | Intron (ENSMUST00000044642.1 | 5 | 139706696 | 139732834 | 26139 | 2 | ENSMUST00 | 7600 | Micall2 | ENSMUSG0C | false | . | . | . | . | . | . | chr5 | 139722629 | 139726712 | 5.64092 | 4.23845 | 3.58902 |
| chr9 | 119322842 | 119323094 | 182 | Promoter (<=1kb) | 9 | 119238432 | 119322427 | 83996 | 2 | ENSMUST00 | -416 | Oxsr1 | ENSMUSG0C | false | chr9 | 119321480 | 119323666 | 23.93078 | 87.47927 | 84.18233 | . | . | . | . | . | . |
| chr2 | 25224856 | 25225516 | 182 | Promoter (<=1kb) | 2 | 25222160 | 25224702 | 2543 | 2 | ENSMUST00 | -155 | Tubb4b | ENSMUSG0C | false | chr2 | 25224949 | 25225717 | 16.72886 | 41.29202 | 38.68602 | chr2 | 25225427 | 25226092 | 3.7981 | 3.41112 | 2.04669 |
| chr2 | 25224196 | 25224520 | 182 | Promoter (<=1kb) | 2 | 25223523 | 25224654 | 1132 | 2 | ENSMUST00 | 134 | Tubb4b | ENSMUSG0C | false | chr2 | 25223627 | 25224712 | 34.54477 | 122.48211 | 118.25522 | . | . | . | . | . | . |
| chr17 | 26090155 | 26090451 | 182 | Promoter (<=1kb) | 17 | 26081211 | 26090300 | 9090 | 2 | ENSMUST00 | 0 | Decr2 | ENSMUSG0C | false | chr17 | 26089549 | 26090527 | 23.16072 | 66.96479 | 64.02055 | chr17 | 26089430 | 26090990 | 3.56065 | 3.34266 | 1.87151 |
| chr9 | 64017049 | 64017408 | 182 | Promoter (<=1kb) | 9 | 63954009 | 64016987 | 62979 | 2 | ENSMUST00 | -63 | Smad6 | ENSMUSG0C | true | chr9 | 64016675 | 64022981 | 24.82645 | 85.46523 | 82.21117 | chr9 | 63948352 | 64020083 | 7.27873 | 5.05859 | 4.874 |
| chr9 | 63928453 | 63928738 | 182 | Distal Intergenic | 9 | 63954009 | 64016987 | 62979 | 2 | ENSMUST00 | 88249 | Smad6 | ENSMUSG0C | true | . | . | . | . | . | . | chr9 | 63922790 | 63929287 | 7.21154 | 5.06095 | 4.85866 |
| chr9 | 63952630 | 63952824 | 182 | Downstream (<1kb) | 9 | 63954009 | 64016987 | 62979 | 2 | ENSMUST00 | 64163 | Smad6 | ENSMUSG0C | true | . | . | . | . | . | . | chr9 | 63948352 | 64020083 | 7.27873 | 5.05859 | 4.874 |
| chr9 | 63923299 | 63923524 | 182 | Distal Intergenic | 9 | 63954009 | 64016987 | 62979 | 2 | ENSMUST00 | 93463 | Smad6 | ENSMUSG0C | true | . | . | . | . | . | . | chr9 | 63922790 | 63929287 | 7.21154 | 5.06095 | 4.85866 |
| chr9 | 63927650 | 63928037 | 182 | Distal Intergenic | 9 | 63954009 | 64016987 | 62979 | 2 | ENSMUST00 | 88950 | Smad6 | ENSMUSG0C | true | chr9 | 63927416 | 63928107 | 11.16678 | 24.53592 | 22.14134 | chr9 | 63922790 | 63929287 | 7.21154 | 5.06095 | 4.85866 |
| chr3 | 145946643 | 145946974 | 182 | Distal Intergenic | 3 | 145939086 | 145944198 | 5113 | 1 | ENSMUST00 | 7558 | 2410004B18 | ENSMUSG0C | false | . | . | . | . | . | . | chr3 | 145937374 | 145952538 | 5.27117 | 4.192 | 3.23206 |
| chr3 | 145945018 | 145945332 | 182 | Distal Intergenic | 3 | 145939086 | 145944198 | 5113 | 1 | ENSMUST00 | 5933 | 2410004B18 | ENSMUSG0C | false | . | . | . | . | . | . | chr3 | 145937374 | 145952538 | 5.27117 | 4.192 | 3.23206 |
| chr18 | 74778852 | 74779187 | 182 | Promoter (<=1kb) | 18 | 74779197 | 74806207 | 27011 | 1 | ENSMUST00 | -10 | Acaa2 | ENSMUSG0C | false | chr18 | 74778092 | 74780533 | 27.39617 | 87.32104 | 84.03119 | . | . | . | . | . | . |
| chr7 | 30534886 | 30535199 | 182 | Promoter (<=1kb) | 7 | 30522226 | 30534180 | 11955 | 2 | ENSMUST00 | -707 | Arhgap33 | ENSMUSG0C | false | chr7 | 30534310 | 30535717 | 30.32912 | 88.589 | 85.27629 | chr7 | 30534482 | 30535677 | 3.75907 | 3.39357 | 2.00569 |
| chr3 | 62433722 | 62434138 | 182 | Intron (ENSMUST00000079300.7 | 3 | 62419668 | 62460212 | 40545 | 1 | ENSMUST00 | 14055 | Arhgef26 | ENSMUSG0C | true | . | . | . | . | . | . | chr3 | 62430774 | 62435924 | 4.18342 | 3.6888 | 2.33197 |
| chr3 | 61364340 | 61364749 | 182 | Promoter (<=1kb) | 3 | 61364507 | 61368703 | 4197 | 1 | ENSMUST00 | 0 | Rap2b | ENSMUSG0C | false | chr3 | 61362950 | 61366087 | 27.03867 | 103.18376 | 99.53125 | . | . | . | . | . | . |
| chr3 | 61363637 | 61364087 | 182 | Promoter (<=1kb) | 3 | 61364507 | 61368703 | 4197 | 1 | ENSMUST00 | -420 | Rap2b | ENSMUSG0C | false | chr3 | 61362950 | 61366087 | 27.03867 | 103.18376 | 99.53125 | . | . | . | . | . | . |
| chr18 | 9210762 | 9211210 | 182 | Promoter (1-2kb) | 18 | 9212856 | 9216201 | 3346 | 1 | ENSMUST00 | -1646 | Fzd8 | ENSMUSG0C | false | chr18 | 9210613 | 9214860 | 27.78601 | 91.88203 | 88.49705 | chr18 | 9210168 | 9215430 | 5.43327 | 4.13916 | 3.36349 |
| chr18 | 9212052 | 9212437 | 182 | Promoter (<=1kb) | 18 | 9212856 | 9216201 | 3346 | 1 | ENSMUST00 | -419 | Fzd8 | ENSMUSG0C | false | chr18 | 9210613 | 9214860 | 27.78601 | 91.88203 | 88.49705 | chr18 | 9210168 | 9215430 | 5.43327 | 4.13916 | 3.36349 |
| chr17 | 87306867 | 87307068 | 182 | Promoter (2-3kb) | 17 | 87309332 | 87330077 | 20746 | 1 | ENSMUST00 | -2264 | Ttc7 | ENSMUSG0C | false | . | . | . | . | . | . | chr17 | 87306252 | 87307309 | 6.45583 | 4.60359 | 4.24458 |
| chr17 | 87282700 | 87283301 | 182 | Promoter (<=1kb) | 17 | 87282886 | 87381769 | 98884 | 1 | ENSMUST00 | 0 | Ttc7 | ENSMUSG0C | false | chr17 | 87281464 | 87284423 | 37.82652 | 124.04217 | 119.77048 | . | . | . | . | . | . |
| chr10 | 62725828 | 62726232 | 182 | Promoter (<=1kb) | 10 | 62659043 | 62726128 | 67086 | 2 | ENSMUST00 | 0 | Stox1 | ENSMUSG0C | false | chr10 | 62725516 | 62726603 | 34.00537 | 102.80987 | 99.17244 | chr10 | 62724980 | 62726725 | 4.22476 | 3.55411 | 2.40169 |
| chr8 | 85432471 | 85433043 | 182 | Promoter (<=1kb) | 8 | 85408759 | 85432841 | 24083 | 2 | ENSMUST00 | 0 | 4921524J17F | ENSMUSG0C | false | chr8 | 85430895 | 85433408 | 28.29403 | 89.8816 | 86.54237 | . | . | . | . | . | . |
| chr17 | 3557771 | 3558071 | 182 | Promoter (<=1kb) | 17 | 3519263 | 3557713 | 38451 | 2 | ENSMUST00 | -59 | Tfb1m | ENSMUSG0C | false | chr17 | 3556940 | 3557897 | 27.29656 | 83.33401 | 80.12321 | . | . | . | . | . | . |
| chr10 | 62329788 | 62330025 | 182 | Promoter (2-3kb) | 10 | 62296093 | 62327767 | 31675 | 2 | ENSMUST00 | -2022 | Hk1 | ENSMUSG0C | false | . | . | . | . | . | . | chr10 | 62329327 | 62339742 | 4.9923 | 4.02342 | 3.00782 |
| chr10 | 62218903 | 62219145 | 182 | Intron (ENSMUST00000047883.9 | 10 | 62185396 | 62231218 | 45823 | 2 | ENSMUST00 | 12073 | Tspan15 | ENSMUSG0C | false | . | . | . | . | . | . | chr10 | 62185685 | 62229768 | 8.28034 | 5.46771 | 5.70297 |
| chr10 | 62222826 | 62223032 | 182 | Intron (ENSMUST00000047883.9 | 10 | 62185396 | 62231218 | 45823 | 2 | ENSMUST00 | 8186 | Tspan15 | ENSMUSG0C | false | . | . | . | . | . | . | chr10 | 62185685 | 62229768 | 8.28034 | 5.46771 | 5.70297 |
| chr1 | 119446254 | 119446717 | 182 | Distal Intergenic | 1 | 119415465 | 119422248 | 6784 | 2 | ENSMUST00 | -24007 | Inhbb | ENSMUSG0C | false | . | . | . | . | . | . | chr1 | 119445773 | 119447267 | 6.06775 | 4.53555 | 3.93483 |
| chr3 | 144688307 | 144688491 | 182 | Downstream (<1kb) | 3 | 144688844 | 144720323 | 31480 | 2 | ENSMUST00 | 31832 | Sh3glb1 | ENSMUSG0C | false | . | . | . | . | . | . | chr3 | 144681472 | 144689973 | 5.50423 | 4.31451 | 3.423 |
| chr19 | 44407683 | 44407908 | 182 | Promoter (<=1kb) | 19 | 44394451 | 44407709 | 13259 | 2 | ENSMUST00 | 0 | Scd1 | ENSMUSG0C | false | chr19 | 44407790 | 44408746 | 12.99241 | 30.10301 | 27.6364 | chr19 | 44407248 | 44408680 | 4.19511 | 3.66125 | 2.34415 |
| chr8 | 84104442 | 84104933 | 182 | Promoter (<=1kb) | 8 | 84097075 | 84104773 | 7699 | 2 | ENSMUST00 | 0 | Dcaf15 | ENSMUSG0C | false | chr8 | 84103873 | 84104844 | 21.13848 | 54.99947 | 52.22001 | chr8 | 84102725 | 84105731 | 4.73976 | 3.94023 | 2.83827 |
| chr12 | 13249062 | 13249288 | 182 | Promoter (<=1kb) | 12 | 13219308 | 13249213 | 29906 | 2 | ENSMUST00 | 0 | Ddx1 | ENSMUSG0C | false | chr12 | 13248352 | 13249753 | 29.35304 | 95.67908 | 92.21078 | chr12 | 13247507 | 13250002 | 3.58112 | 3.34412 | 1.86735 |
| chr12 | 12940114 | 12940557 | 182 | Promoter (<=1kb) | 12 | 12936096 | 12940616 | 4521 | 2 | ENSMUST00 | 59 | Mycn | ENSMUSG0C | false | chr12 | 12937456 | 12942953 | 27.46248 | 78.93296 | 75.80077 | chr12 | 12937234 | 12941381 | 4.36203 | 3.60348 | 2.51894 |
| chr10 | 61424521 | 61424786 | 182 | 3' UTR | 10 | 61417972 | 61425338 | 7367 | 1 | ENSMUST00 | 6550 | Nodal | ENSMUSG0C | false | . | . | . | . | . | . | chr10 | 61424090 | 61428083 | 6.39114 | 4.69537 | 4.15569 |
| chr6 | 40435191 | 40435840 | 182 | Promoter (<=1kb) | 6 | 40407500 | 40436133 | 28634 | 2 | ENSMUST00 | 293 | E330009J07F | ENSMUSG0C | false | chr6 | 40434915 | 40436523 | 27.18312 | 111.36749 | 107.49936 | chr6 | 40434431 | 40435196 | 4.83459 | 4.04013 | 2.88801 |
| chr15 | 101287294 | 101287568 | 182 | Promoter (2-3kb) | 15 | 101284301 | 101290934 | 6634 | 1 | ENSMUST00 | 2994 | Atg101 | ENSMUSG0C | false | . | . | . | . | . | . | chr15 | 101284730 | 101290185 | 8.19197 | 5.36025 | 5.65555 |
| chr3 | 37640831 | 37641138 | 182 | Promoter (<=1kb) | 3 | 37640080 | 37644598 | 4519 | 1 | ENSMUST00 | 752 | Spry1 | ENSMUSG0C | false | chr3 | 37639368 | 37642407 | 23.85991 | 99.51073 | 95.95212 | . | . | . | . | . | . |
| chr5 | 34187991 | 34188254 | 182 | Promoter (<=1kb) | 5 | 34176579 | 34187720 | 11142 | 2 | ENSMUST00 | -272 | Mxd4 | ENSMUSG0C | false | chr5 | 34188018 | 34188670 | 22.15371 | 67.81625 | 64.85734 | . | . | . | . | . | . |
| chr11 | 58302898 | 58303156 | 182 | Distal Intergenic | 11 | 58307069 | 58314613 | 7545 | 1 | ENSMUST00 | -3913 | Zfp692 | ENSMUSG0C | false | chr11 | 58301401 | 58303425 | 14.58106 | 49.93175 | 47.21736 | . | . | . | . | . | . |
| chr9 | 63602427 | 63602783 | 182 | Promoter (<=1kb) | 9 | 63602655 | 63641889 | 39235 | 1 | ENSMUST00 | 0 | Aagab | ENSMUSG0C | false | chr9 | 63602084 | 63603798 | 45.26388 | 159.40776 | 153.7765 | . | . | . | . | . | . |
| chr9 | 63618195 | 63618396 | 182 | Intron (ENSMUST00000041551.7 | 9 | 63602655 | 63641889 | 39235 | 1 | ENSMUST00 | 15541 | Aagab | ENSMUSG0C | false | . | . | . | . | . | . | chr9 | 63617899 | 63618249 | 3.37279 | 3.30162 | 1.64746 |
| chr3 | 36862982 | 36863354 | 182 | Promoter (<=1kb) | 3 | 36863106 | 37053033 | 189928 | 1 | ENSMUST00 | 0 | 4932438A13 | ENSMUSG0C | false | chr3 | 36862581 | 36863954 | 27.34946 | 104.78247 | 101.09169 | . | . | . | . | . | . |
| chr4 | 134704234 | 134704524 | 182 | Promoter (<=1kb) | 4 | 134561690 | 134704290 | 142601 | 2 | ENSMUST00 | 0 | Man1c1 | ENSMUSG0C | false | chr4 | 134702765 | 134704907 | 29.37731 | 87.91154 | 84.61048 | chr4 | 134702127 | 134705017 | 5.20353 | 4.0496 | 3.1933 |
| chr5 | 33695354 | 33695703 | 182 | Distal Intergenic | 5 | 33669569 | 33672201 | 2633 | 1 | ENSMUST00 | 25786 | Tacc3 | ENSMUSG0C | false | chr5 | 33695092 | 33696139 | 20.27628 | 60.14116 | 57.29097 | chr5 | 33694889 | 33696611 | 4.27433 | 3.70154 | 2.43607 |
| chr5 | 137307393 | 137307818 | 182 | Promoter (<=1kb) | 5 | 137295704 | 137307674 | 11971 | 2 | ENSMUST00 | 0 | Srrt | ENSMUSG0C | false | chr5 | 137306616 | 137308486 | 34.74108 | 114.39095 | 110.42815 | . | . | . | . | . | . |
| chr1 | 192779113 | 192779387 | 182 | Distal Intergenic | 1 | 192512828 | 192771213 | 258386 | 2 | ENSMUST00 | -7901 | Hhat | ENSMUSG0C | false | . | . | . | . | . | . | chr1 | 192775490 | 192780726 | 4.58756 | 3.78752 | 2.66597 |
| chr1 | 192759456 | 192759679 | 182 | Exon (ENSMUST00000044190.6/ | 1 | 192543149 | 192771060 | 227912 | 2 | ENSMUST00 | 11381 | Hhat | ENSMUSG0C | false | . | . | . | . | . | . | chr1 | 192759021 | 192760617 | 5.18018 | 4.2043 | 3.15504 |
| chr5 | 33225121 | 33225344 | 182 | Distal Intergenic | 5 | 33213518 | 33218238 | 4721 | 2 | ENSMUST00 | -6884 | Spon2 | ENSMUSG0C | false | . | . | . | . | . | . | chr5 | 33221817 | 33232043 | 5.32048 | 4.18629 | 3.28315 |
| chr9 | 90264685 | 90264873 | 182 | Intron (ENSMUST00000041767.8 | 9 | 90202027 | 90270769 | 68743 | 2 | ENSMUST00 | 5896 | Tbc1d2b | ENSMUSG0C | false | . | . | . | . | . | . | chr9 | 90264105 | 90265003 | 6.00031 | 4.60277 | 3.81288 |
| chr9 | 90269064 | 90269339 | 182 | Promoter (1-2kb) | 9 | 90202027 | 90270769 | 68743 | 2 | ENSMUST00 | 1430 | Tbc1d2b | ENSMUSG0C | false | . | . | . | . | . | . | chr9 | 90269262 | 90270189 | 3.8506 | 3.38297 | 2.10037 |
| chr9 | 90260623 | 90260879 | 182 | Intron (ENSMUST00000041767.8 | 9 | 90241163 | 90255927 | 14765 | 2 | ENSMUST00 | -4697 | Tbc1d2b | ENSMUSG0C | false | . | . | . | . | . | . | chr9 | 90258932 | 90261177 | 4.87342 | 4.03529 | 2.92395 |
| chr5 | 137072187 | 137072476 | 182 | Promoter (<=1kb) | 5 | 137061504 | 137072272 | 10769 | 2 | ENSMUST00 | 0 | Serpine1 | ENSMUSG0C | false | . | . | . | . | . | . | chr5 | 137071594 | 137074126 | 4.74285 | 3.8439 | 2.82946 |
| chr5 | 137032024 | 137032296 | 182 | Promoter (1-2kb) | 5 | 137030701 | 137031407 | 707 | 1 | ENSMUST00 | 1324 | Vgf | ENSMUSG0C | false | chr5 | 137031245 | 137033291 | 15.50707 | 38.47921 | 35.908 | chr5 | 137029894 | 137033165 | 7.20686 | 4.82218 | 4.83474 |
| chr12 | 113155863 | 113156317 | 182 | Promoter (<=1kb) | 12 | 113156421 | 113165456 | 9036 | 1 | ENSMUST00 | -104 | 4930427A07I | ENSMUSG0C | false | chr12 | 113155178 | 113156098 | 16.71423 | 46.51484 | 43.84613 | chr12 | 113154877 | 113156741 | 4.21205 | 3.55531 | 2.38958 |
| chr12 | 113155464 | 113155707 | 182 | Promoter (<=1kb) | 12 | 113156421 | 113165456 | 9036 | 1 | ENSMUST00 | -714 | 4930427A07I | ENSMUSG0C | false | chr12 | 113155178 | 113156098 | 16.71423 | 46.51484 | 43.84613 | chr12 | 113154877 | 113156741 | 4.21205 | 3.55531 | 2.38958 |
| chr12 | 3426537 | 3426875 | 182 | Promoter (<=1kb) | 12 | 3426857 | 3506847 | 79991 | 1 | ENSMUST00 | 0 | Asxl2 | ENSMUSG0C | false | chr12 | 3425990 | 3428222 | 44.33166 | 157.76141 | 152.18738 | . | . | . | . | . | . |
| chr1 | 34801609 | 34801929 | 182 | Promoter (<=1kb) | 1 | 34801704 | 34813309 | 11606 | 1 | ENSMUST00 | 0 | Arhgef4 | ENSMUSG0C | false | chr1 | 34801511 | 34802534 | 24.14628 | 82.32003 | 79.12344 | chr1 | 34801170 | 34802503 | 6.62408 | 4.89966 | 4.35781 |
| chr14 | 47276823 | 47277298 | 182 | Promoter (<=1kb) | 14 | 47240944 | 47276832 | 35889 | 2 | ENSMUST00 | 0 | Wdhd1 | ENSMUSG0C | false | chr14 | 47276211 | 47277770 | 31.54088 | 108.67455 | 104.87239 | . | . | . | . | . | . |
| chr10 | 20311879 | 20312498 | 182 | Promoter (<=1kb) | 10 | 20312469 | 20342501 | 30033 | 1 | ENSMUST00 | 0 | Bclaf1 | ENSMUSG0C | false | chr10 | 20311327 | 20313751 | 35.2856 | 127.03558 | 122.65217 | . | . | . | . | . | . |
| chr14 | 46670769 | 46670951 | 182 | Distal Intergenic | 14 | 46760541 | 46771525 | 10985 | 1 | ENSMUST00 | -89590 | Cdkn3 | ENSMUSG0C | false | . | . | . | . | . | . | chr14 | 46670122 | 46671358 | 3.83367 | 3.37853 | 2.08899 |
| chr3 | 30956749 | 30956925 | 182 | Intron (ENSMUST00000064718.6 | 3 | 30961103 | 30965935 | 4833 | 2 | ENSMUST00 | 9010 | Phc3 | ENSMUSG0C | false | . | . | . | . | . | . | chr3 | 30955706 | 30960693 | 3.38873 | 3.21991 | 1.72748 |
| chr5 | 67662302 | 67662590 | 182 | Promoter (2-3kb) | 5 | 67622680 | 67660034 | 37355 | 2 | ENSMUST00 | -2269 | Atp8a1 | ENSMUSG0C | true | . | . | . | . | . | . | chr5 | 67662062 | 67663129 | 3.44338 | 3.32471 | 1.76635 |
| chr4 | 133017143 | 133017387 | 182 | Intron (ENSMUST00000154646.2 | 4 | 133012204 | 133077645 | 65442 | 1 | ENSMUST00 | 4940 | Ahdc1 | ENSMUSG0C | false | . | . | . | . | . | . | chr4 | 133013189 | 133060125 | 8.03751 | 5.40542 | 5.50183 |
| chr2 | 130642737 | 130642979 | 182 | Promoter (<=1kb) | 2 | 130632842 | 130642803 | 9962 | 2 | ENSMUST00 | 0 | Lzts3 | ENSMUSG0C | false | chr2 | 130642161 | 130643208 | 26.39899 | 75.26234 | 72.19086 | chr2 | 130641282 | 130643207 | 4.65429 | 3.91755 | 2.7135 |
| chr14 | 45477817 | 45478063 | 182 | Promoter (<=1kb) | 14 | 45469314 | 45477861 | 8548 | 2 | ENSMUST00 | 0 | Fermt2 | ENSMUSG0C | true | . | . | . | . | . | . | chr14 | 45470113 | 45484905 | 5.01031 | 4.09563 | 3.01362 |
| chr12 | 5416731 | 5416989 | 182 | Distal Intergenic | 12 | 5411641 | 5416631 | 4991 | 1 | ENSMUST00 | 5091 | 2810032G03 | ENSMUSG0C | false | . | . | . | . | . | . | chr12 | 5415800 | 5417313 | 4.13334 | 3.49198 | 2.32531 |
| chr9 | 78480827 | 78481143 | 182 | Promoter (<=1kb) | 9 | 78479915 | 78480841 | 927 | 2 | ENSMUST00 | 0 | Eef1a1 | ENSMUSG0C | false | chr9 | 78480141 | 78481717 | 35.02911 | 135.1772 | 130.48503 | . | . | . | . | . | . |
| chr11 | 78115330 | 78115782 | 182 | Promoter (<=1kb) | 11 | 78115606 | 78156700 | 41095 | 1 | ENSMUST00 | 0 | Fam222b | ENSMUSG0C | false | chr11 | 78114982 | 78116222 | 23.53563 | 65.86355 | 62.93705 | chr11 | 78114707 | 78116504 | 4.54859 | 3.87804 | 2.67224 |
| chr9 | 99629412 | 99630073 | 182 | Promoter (<=1kb) | 9 | 99629496 | 99637796 | 8301 | 1 | ENSMUST00 | 0 | Dzip1l | ENSMUSG0C | false | chr9 | 99629371 | 99630536 | 21.77823 | 66.26614 | 63.32996 | chr9 | 99628926 | 99632578 | 3.34757 | 3.15166 | 1.68433 |
| chr12 | 111712991 | 111713357 | 182 | Promoter (<=1kb) | 12 | 111713261 | 111734220 | 20960 | 1 | ENSMUST00 | 0 | Apopt1 | ENSMUSG0C | false | chr12 | 111712442 | 111714088 | 37.91852 | 123.12976 | 118.88376 | . | . | . | . | . | . |
| chr6 | 57824563 | 57825080 | 182 | Promoter (<=1kb) | 6 | 57774709 | 57824640 | 49932 | 2 | ENSMUST00 | 0 | Vopp1 | ENSMUSG0C | false | chr6 | 57824019 | 57825531 | 36.12563 | 122.12295 | 117.91594 | . | . | . | . | . | . |
| chr9 | 13827700 | 13828072 | 182 | Promoter (<=1kb) | 9 | 13827716 | 13846541 | 18826 | 1 | ENSMUST00 | 0 | Fam76b | ENSMUSG0C | false | chr9 | 13827687 | 13828673 | 30.25803 | 109.37234 | 105.55905 | . | . | . | . | . | . |
| chr3 | 40950140 | 40950410 | 182 | Promoter (<=1kb) | 3 | 40950631 | 40977793 | 27163 | 1 | ENSMUST00 | -221 | Larp1b | ENSMUSG0C | false | chr3 | 40949721 | 40951569 | 28.91247 | 96.86193 | 93.36482 | . | . | . | . | . | . |
| chr3 | 40893919 | 40894329 | 182 | Promoter (<=1kb) | 3 | 40894145 | 40938138 | 43994 | 1 | ENSMUST00 | 0 | 3110057O12 | ENSMUSG0C | false | chr3 | 40893351 | 40895033 | 26.74277 | 88.06109 | 84.75893 | chr3 | 40892630 | 40895332 | 4.58126 | 3.83072 | 2.70693 |
| chr2 | 158139695 | 158139935 | 182 | Promoter (<=1kb) | 2 | 158134322 | 158139736 | 5415 | 2 | ENSMUST00 | 0 | Tgm2 | ENSMUSG0C | false | . | . | . | . | . | . | chr2 | 158138948 | 158141373 | 8.01704 | 5.50791 | 5.47459 |
| chr19 | 18631836 | 18632052 | 182 | Promoter (<=1kb) | 19 | 18631950 | 18636385 | 4436 | 1 | ENSMUST00 | 0 | Nmrk1 | ENSMUSG0C | false | chr19 | 18630087 | 18632373 | 31.38972 | 125.99339 | 121.64841 | . | . | . | . | . | . |
| chr10 | 67912365 | 67912699 | 182 | Promoter (<=1kb) | 10 | 67886103 | 67912662 | 26560 | 2 | ENSMUST00 | 0 | Zfp365 | ENSMUSG0C | false | chr10 | 67911967 | 67912778 | 21.08548 | 68.47759 | 65.51048 | chr10 | 67911634 | 67912718 | 4.04301 | 3.53507 | 2.24868 |
| chr10 | 67056461 | 67056718 | 182 | Intron (ENSMUST00000020023.7 | 10 | 67096125 | 67191748 | 95624 | 1 | ENSMUST00 | -39407 | Jmjd1c | ENSMUSG0C | false | . | . | . | . | . | . | chr10 | 67055207 | 67059305 | 4.18071 | 3.45776 | 2.37763 |
| chr3 | 49854386 | 49854664 | 182 | Distal Intergenic | 3 | 49743296 | 49757316 | 14021 | 2 | ENSMUST00 | -97071 | Pcdh18 | ENSMUSG0C | false | . | . | . | . | . | . | . | . | . | . | . | . |
| chr12 | 111062425 | 111062661 | 182 | Intron (ENSMUST00000084968.8 | 12 | 111039798 | 111113384 | 73587 | 1 | ENSMUST00 | 22628 | Rcor1 | ENSMUSG0C | false | . | . | . | . | . | . | chr12 | 111058346 | 111063236 | 3.38464 | 3.24799 | 1.71349 |
| chr2 | 129614695 | 129615043 | 182 | Intron (ENSMUST00000153491.2 | 2 | 129620685 | 129628044 | 7360 | 1 | ENSMUST00 | -5642 | Sirpa | ENSMUSG0C | true | . | . | . | . | . | . | chr2 | 129613761 | 129620104 | 6.112 | 4.56754 | 3.9269 |
| chr2 | 129633479 | 129633703 | 182 | Distal Intergenic | 2 | 129620685 | 129628044 | 7360 | 1 | ENSMUST00 | 12795 | Sirpa | ENSMUSG0C | true | . | . | . | . | . | . | chr2 | 129621436 | 129635375 | 5.74226 | 4.37432 | 3.64032 |
| chr2 | 129627749 | 129627939 | 182 | Intron (ENSMUST00000161620.2 | 2 | 129620685 | 129628044 | 7360 | 1 | ENSMUST00 | 7065 | Sirpa | ENSMUSG0C | true | . | . | . | . | . | . | chr2 | 129621436 | 129635375 | 5.74226 | 4.37432 | 3.64032 |
| chr12 | 110978577 | 110979322 | 182 | Promoter (<=1kb) | 12 | 110975353 | 110979040 | 3688 | 2 | ENSMUST00 | 0 | Ankrd9 | ENSMUSG0C | false | chr12 | 110978291 | 110979566 | 28.53699 | 95.19626 | 91.7378 | chr12 | 110977974 | 110979557 | 4.01223 | 3.34835 | 2.22479 |
| chr5 | 125441412 | 125441714 | 182 | Promoter (<=1kb) | 5 | 125441568 | 125460872 | 19305 | 1 | ENSMUST00 | 0 | Bri3bp | ENSMUSG0C | false | chr5 | 125441176 | 125442448 | 38.14117 | 119.25307 | 115.13219 | chr5 | 125441219 | 125442746 | 3.81758 | 3.51584 | 2.02028 |
| chr13 | 49371537 | 49371844 | 182 | Intron (ENSMUST00000048544.7 | 13 | 49341549 | 49387025 | 45477 | 1 | ENSMUST00 | 29989 | Bicd2 | ENSMUSG0C | false | . | . | . | . | . | . | chr13 | 49360131 | 49378531 | 5.26531 | 4.14063 | 3.2473 |
| chr11 | 99230374 | 99230759 | 182 | Promoter (<=1kb) | 11 | 99219629 | 99230445 | 10817 | 2 | ENSMUST00 | 0 | Smarce1 | ENSMUSG0C | false | chr11 | 99229408 | 99231335 | 31.62822 | 118.12274 | 114.03909 | . | . | . | . | . | . |
| chr5 | 125267062 | 125267302 | 182 | Distal Intergenic | 5 | 125277787 | 125284159 | 6373 | 2 | ENSMUST00 | 16857 | Scarb1 | ENSMUSG0C | false | . | . | . | . | . | . | chr5 | 125267069 | 125268583 | 5.7806 | 4.49527 | 3.63896 |
| chr11 | 99147823 | 99148049 | 182 | Intron (ENSMUST00000103134.3 | 11 | 99144196 | 99155077 | 10882 | 2 | ENSMUST00 | 7028 | Ccr7 | ENSMUSG0C | false | . | . | . | . | . | . | . | . | . | . | . | . |
| chr12 | 110737766 | 110738033 | 182 | Promoter (<=1kb) | 12 | 110737949 | 110795028 | 57080 | 1 | ENSMUST00 | 0 | Wdr20 | ENSMUSG0C | false | chr12 | 110737110 | 110739434 | 33.38818 | 124.5882 | 120.28912 | . | . | . | . | . | . |
| chr5 | 124960545 | 124960776 | 182 | Distal Intergenic | 5 | 125003447 | 125012547 | 9101 | 1 | ENSMUST00 | -42671 | Fam101a | ENSMUSG0C | false | . | . | . | . | . | . | chr5 | 124960209 | 124961127 | 3.9145 | 3.47137 | 2.14974 |
| chr5 | 125003282 | 125003501 | 182 | Promoter (<=1kb) | 5 | 125003447 | 125012547 | 9101 | 1 | ENSMUST00 | 0 | Fam101a | ENSMUSG0C | false | chr5 | 125003372 | 125003903 | 7.72573 | 18.36004 | 16.04866 | chr5 | 125003072 | 125004610 | 3.90854 | 3.51281 | 2.15377 |
| chr9 | 107340477 | 107340827 | 182 | Promoter (<=1kb) | 9 | 107340602 | 107349683 | 9082 | 1 | ENSMUST00 | 0 | 6430571L13F | ENSMUSG0C | false | chr9 | 107340165 | 107341059 | 17.09341 | 55.02552 | 52.24499 | . | . | . | . | . | . |
| chr6 | 55650016 | 55650363 | 182 | Distal Intergenic | 6 | 55677818 | 55681263 | 3446 | 2 | ENSMUST00 | 30900 | Neurod6 | ENSMUSG0C | false | . | . | . | . | . | . | chr6 | 55648400 | 55651115 | 4.67151 | 3.88099 | 2.76616 |
| chr11 | 98936204 | 98936456 | 182 | Promoter (1-2kb) | 11 | 98937698 | 98974162 | 36465 | 1 | ENSMUST00 | -1242 | Rara | ENSMUSG0C | false | chr11 | 98936314 | 98938742 | 41.35789 | 132.3362 | 127.75856 | chr11 | 98928668 | 98957642 | 6.70032 | 4.70048 | 4.41905 |
| chr11 | 98965826 | 98965999 | 182 | Intron (ENSMUST00000107474.2 | 11 | 98960412 | 98974942 | 14531 | 1 | ENSMUST00 | 5415 | Rara | ENSMUSG0C | false | . | . | . | . | . | . | chr11 | 98958785 | 98973618 | 4.55278 | 3.71427 | 2.6612 |
| chr3 | 135307626 | 135307799 | 182 | Promoter (<=1kb) | 3 | 135307700 | 135342535 | 34836 | 1 | ENSMUST00 | 0 | Slc9b2 | ENSMUSG0C | false | chr3 | 135307288 | 135307791 | 10.89613 | 22.929 | 20.55399 | . | . | . | . | . | . |
| chr11 | 98839835 | 98840184 | 182 | Intron (ENSMUST00000107479.2 | 11 | 98836785 | 98853697 | 16913 | 1 | ENSMUST00 | 3051 | Rapgefl1 | ENSMUSG0C | false | . | . | . | . | . | . | chr11 | 98839442 | 98840325 | 3.10256 | 3.07584 | 1.47907 |
| chr6 | 55241259 | 55241493 | 182 | Intron (ENSMUST00000053094.7 | 6 | 55203383 | 55320222 | 116840 | 1 | ENSMUST00 | 37877 | Fam188b | ENSMUSG0C | false | . | . | . | . | . | . | . | . | . | . | . | . |
| chr13 | 48842306 | 48842528 | 182 | Intron (ENSMUST00000035540.7 | 13 | 48801750 | 48870885 | 69136 | 2 | ENSMUST00 | 28357 | Phf2 | ENSMUSG0C | false | . | . | . | . | . | . | chr13 | 48840650 | 48856638 | 4.24545 | 3.54861 | 2.41708 |
| chr11 | 76253823 | 76254101 | 182 | Distal Intergenic | 11 | 76249212 | 76250508 | 1297 | 1 | ENSMUST00 | 4612 | Rnmtl1 | ENSMUSG0C | false | . | . | . | . | . | . | chr11 | 76252612 | 76258538 | 5.07624 | 3.99195 | 3.10276 |
| chr16 | 10542863 | 10543216 | 182 | Promoter (<=1kb) | 16 | 10502531 | 10543054 | 40524 | 2 | ENSMUST00 | 0 | Dexi | ENSMUSG0C | false | chr16 | 10541909 | 10543316 | 26.86698 | 95.04005 | 91.5885 | chr16 | 10539283 | 10547437 | 4.45024 | 3.72545 | 2.5691 |
| chr9 | 119102238 | 119102421 | 182 | Promoter (<=1kb) | 9 | 119102478 | 119125295 | 22818 | 1 | ENSMUST00 | -57 | Dlec1 | ENSMUSG0C | false | chr9 | 119102059 | 119103232 | 24.06788 | 78.48502 | 75.35915 | chr9 | 119100561 | 119103393 | 4.73205 | 4.0101 | 2.78489 |
| chr4 | 84884242 | 84884548 | 182 | Promoter (<=1kb) | 4 | 84884309 | 85131921 | 247613 | 1 | ENSMUST00 | 0 | Cntln | ENSMUSG0C | false | chr4 | 84883849 | 84885458 | 30.77379 | 96.20782 | 92.72798 | . | . | . | . | . | . |
| chr13 | 47043376 | 47043703 | 182 | Promoter (<=1kb) | 13 | 47043499 | 47084600 | 41102 | 1 | ENSMUST00 | 0 | Kdm1b | ENSMUSG0C | false | chr13 | 47043467 | 47044803 | 32.66734 | 113.81544 | 109.86345 | . | . | . | . | . | . |
| chr2 | 156278628 | 156278860 | 182 | Intron (ENSMUST00000037401.8 | 2 | 156282631 | 156292528 | 9898 | 1 | ENSMUST00 | -3771 | Phf20 | ENSMUSG0C | false | . | . | . | . | . | . | chr2 | 156276344 | 156279552 | 6.61434 | 4.90082 | 4.31329 |
| chr2 | 156298491 | 156298922 | 182 | Promoter (2-3kb) | 2 | 156296405 | 156304692 | 8288 | 1 | ENSMUST00 | 2087 | Phf20 | ENSMUSG0C | false | . | . | . | . | . | . | chr2 | 156296883 | 156299991 | 6.14036 | 4.63131 | 3.94331 |
| chr5 | 24751827 | 24752361 | 182 | Promoter (<=1kb) | 5 | 24751003 | 24751324 | 322 | 2 | ENSMUST00 | -504 | Crygn | ENSMUSG0C | false | chr5 | 24750232 | 24752112 | 18.39885 | 52.66533 | 49.91441 | chr5 | 24749817 | 24752596 | 3.96722 | 3.59791 | 2.16982 |
| chr8 | 47354817 | 47355044 | 182 | Promoter (2-3kb) | 8 | 47180048 | 47352348 | 172301 | 2 | ENSMUST00 | -2470 | Stox2 | ENSMUSG0C | false | . | . | . | . | . | . | chr8 | 47354358 | 47355682 | 4.97914 | 3.96655 | 3.02595 |
| chr9 | 122187146 | 122187777 | 182 | Intron (ENSMUST00000042546.2 | 9 | 122166744 | 122167475 | 732 | 1 | ENSMUST00 | 20403 | Snrk | ENSMUSG0C | false | . | . | . | . | . | . | . | . | . | . | . | . |
| chr13 | 45976323 | 45976625 | 182 | Distal Intergenic | 13 | 45965351 | 45966177 | 827 | 1 | ENSMUST00 | 10973 | 5033430I15R | ENSMUSG0C | false | . | . | . | . | . | . | . | . | . | . | . | . |
| chr7 | 113768638 | 113768880 | 182 | Promoter (2-3kb) | 7 | 113766174 | 113929876 | 163703 | 1 | ENSMUST00 | 2465 | Spon1 | ENSMUSG0C | false | . | . | . | . | . | . | chr7 | 113765597 | 113769620 | 4.68296 | 3.94314 | 2.74544 |
| chr16 | 26105318 | 26105970 | 182 | Promoter (<=1kb) | 16 | 25959288 | 26105784 | 146497 | 2 | ENSMUST00 | 0 | Leprel1 | ENSMUSG0C | false | chr16 | 26102904 | 26106511 | 34.21972 | 118.71816 | 114.6235 | . | . | . | . | . | . |
| chr16 | 26164244 | 26164435 | 182 | Distal Intergenic | 16 | 25959288 | 26105784 | 146497 | 2 | ENSMUST00 | -58461 | Leprel1 | ENSMUSG0C | false | . | . | . | . | . | . | chr16 | 26162973 | 26165141 | 3.71901 | 3.37646 | 1.98757 |
| chr3 | 97767765 | 97768383 | 182 | Promoter (<=1kb) | 3 | 97738854 | 97768009 | 29156 | 2 | ENSMUST00 | 0 | Pde4dip | ENSMUSG0C | true | chr3 | 97763573 | 97768572 | 31.25476 | 101.05123 | 97.45738 | chr3 | 97768345 | 97768945 | 4.72476 | 4.03852 | 2.79626 |
| chr3 | 97868162 | 97868476 | 182 | Promoter (<=1kb) | 3 | 97746693 | 97868248 | 121556 | 2 | ENSMUST00 | 0 | Pde4dip | ENSMUSG0C | true | chr3 | 97867118 | 97868651 | 25.90858 | 83.69981 | 80.48042 | chr3 | 97865926 | 97868623 | 3.35558 | 3.16658 | 1.6975 |
| chr8 | 46941170 | 46941448 | 182 | Distal Intergenic | 8 | 46986925 | 47030714 | 43790 | 1 | ENSMUST00 | -45477 | Enpp6 | ENSMUSG0C | false | . | . | . | . | . | . | chr8 | 46940834 | 46943045 | 7.10776 | 5.00701 | 4.73513 |
| chr13 | 45389400 | 45389945 | 182 | Promoter (<=1kb) | 13 | 45389742 | 45412019 | 22278 | 1 | ENSMUST00 | 0 | Mylip | ENSMUSG0C | false | chr13 | 45389204 | 45392361 | 36.81325 | 122.26781 | 118.05663 | . | . | . | . | . | . |
| chr10 | 43489554 | 43489857 | 182 | Intron (ENSMUST00000040147.7 | 10 | 43479140 | 43515396 | 36257 | 1 | ENSMUST00 | 10415 | Bend3 | ENSMUSG0C | false | . | . | . | . | . | . | chr10 | 43488051 | 43491444 | 6.08389 | 4.66697 | 3.89456 |
| chr1 | 171437686 | 171438018 | 182 | Promoter (<=1kb) | 1 | 171437579 | 171460846 | 23268 | 1 | ENSMUST00 | 108 | F11r | ENSMUSG0C | false | chr1 | 171437150 | 171440146 | 21.08548 | 68.47759 | 65.51048 | . | . | . | . | . | . |
| chr10 | 43252264 | 43252695 | 182 | Intron (ENSMUST00000160823.2 | 10 | 43221736 | 43346116 | 124381 | 1 | ENSMUST00 | 30529 | Pdss2 | ENSMUSG0C | false | . | . | . | . | . | . | chr10 | 43250837 | 43253804 | 4.98911 | 4.09945 | 3.00958 |
| chr11 | 98338832 | 98339050 | 182 | Distal Intergenic | 11 | 98325415 | 98329648 | 4234 | 2 | ENSMUST00 | -9185 | Neurod2 | ENSMUSG0C | false | chr11 | 98338360 | 98339625 | 24.81473 | 68.0602 | 65.09722 | chr11 | 98338284 | 98339968 | 4.36231 | 3.52212 | 2.50841 |
| chr7 | 45333466 | 45333932 | 182 | Promoter (<=1kb) | 7 | 45303155 | 45333780 | 30626 | 2 | ENSMUST00 | 0 | Trpm4 | ENSMUSG0C | false | chr7 | 45332526 | 45333989 | 25.50663 | 88.63404 | 85.31516 | chr7 | 45332151 | 45334090 | 4.85422 | 3.72964 | 2.90984 |
| chr11 | 75152736 | 75153051 | 182 | Intron (ENSMUST00000045281.7 | 11 | 75159633 | 75163182 | 3550 | 1 | ENSMUST00 | -6582 | Smg6 | ENSMUSG0C | false | . | . | . | . | . | . | . | . | . | . | . | . |
| chr7 | 45192107 | 45192332 | 182 | Promoter (2-3kb) | 7 | 45183928 | 45190066 | 6139 | 2 | ENSMUST00 | -2042 | Ccdc155 | ENSMUSG0C | false | . | . | . | . | . | . | chr7 | 45191294 | 45192562 | 5.90574 | 4.34822 | 3.77554 |
| chr1 | 56972563 | 56973557 | 182 | Promoter (<=1kb) | 1 | 56793981 | 56972502 | 178522 | 2 | ENSMUST00 | -62 | Satb2 | ENSMUSG0C | false | chr1 | 56971937 | 56973997 | 33.25446 | 136.03296 | 131.3158 | chr1 | 56969693 | 56972732 | 4.78049 | 3.80518 | 2.86658 |
| chr1 | 55612947 | 55613224 | 182 | Intron (ENSMUST00000042986.9 | 1 | 55702025 | 55735227 | 33203 | 1 | ENSMUST00 | -88801 | Plcl1 | ENSMUSG0C | true | . | . | . | . | . | . | chr1 | 55608366 | 55615053 | 4.52589 | 3.89067 | 2.60304 |
| chr1 | 55664858 | 55665302 | 182 | Intron (ENSMUST00000042986.9 | 1 | 55702025 | 55735227 | 33203 | 1 | ENSMUST00 | -36723 | Plcl1 | ENSMUSG0C | true | . | . | . | . | . | . | chr1 | 55664304 | 55669510 | 3.61029 | 3.34318 | 1.88584 |
| chr1 | 55585611 | 55585788 | 182 | Intron (ENSMUST00000042986.9 | 1 | 55702025 | 55735227 | 33203 | 1 | ENSMUST00 | -116237 | Plcl1 | ENSMUSG0C | true | . | . | . | . | . | . | chr1 | 55584373 | 55589373 | 4.24125 | 3.71771 | 2.38874 |
| chr11 | 97806149 | 97806423 | 182 | Promoter (<=1kb) | 11 | 97806310 | 97806900 | 591 | 1 | ENSMUST00 | 0 | Lasp1 | ENSMUSG0C | false | . | . | . | . | . | . | chr11 | 97801064 | 97833244 | 8.45203 | 5.49885 | 5.84074 |
| chr11 | 97825244 | 97825502 | 182 | Intron (ENSMUST00000043843.6 | 11 | 97833183 | 97836205 | 3023 | 1 | ENSMUST00 | -7681 | Lasp1 | ENSMUSG0C | false | . | . | . | . | . | . | chr11 | 97801064 | 97833244 | 8.45203 | 5.49885 | 5.84074 |
| chr4 | 88094486 | 88094758 | 182 | Promoter (<=1kb) | 4 | 88094629 | 88121812 | 27184 | 1 | ENSMUST00 | 0 | Focad | ENSMUSG0C | false | chr4 | 88094020 | 88095311 | 34.27164 | 115.65593 | 111.648 | chr4 | 88093831 | 88095056 | 4.38539 | 3.78244 | 2.49125 |
| chr13 | 32338486 | 32338920 | 182 | Promoter (<=1kb) | 13 | 31819579 | 32338712 | 519134 | 2 | ENSMUST00 | 0 | Gmds | ENSMUSG0C | true | chr13 | 32337977 | 32339540 | 25.4663 | 101.95255 | 98.33054 | . | . | . | . | . | . |
| chr7 | 45019246 | 45019516 | 182 | Promoter (<=1kb) | 7 | 45019470 | 45020405 | 936 | 1 | ENSMUST00 | 0 | Rras | ENSMUSG0C | false | chr7 | 45017877 | 45019667 | 28.36989 | 85.2065 | 81.96005 | chr7 | 45018306 | 45019972 | 7.21749 | 5.17377 | 4.81389 |
| chr7 | 45017364 | 45017642 | 182 | Promoter (<=1kb) | 7 | 45017978 | 45021646 | 3669 | 1 | ENSMUST00 | -336 | Rras | ENSMUSG0C | false | chr7 | 45016468 | 45017564 | 21.04643 | 56.48158 | 53.68221 | . | . | . | . | . | . |
| chr6 | 124863646 | 124863920 | 182 | Promoter (<=1kb) | 6 | 124858444 | 124863917 | 5474 | 2 | ENSMUST00 | 0 | Gpr162 | ENSMUSG0C | false | chr6 | 124863267 | 124864173 | 16.14089 | 38.84904 | 36.27268 | chr6 | 124863230 | 124864425 | 3.92179 | 3.59959 | 2.09963 |
| chr8 | 13785524 | 13785960 | 182 | Promoter (<=1kb) | 8 | 13785615 | 13798538 | 12924 | 1 | ENSMUST00 | 0 | Upf3a | ENSMUSG0C | false | chr8 | 13785055 | 13786498 | 23.4661 | 79.19912 | 76.05865 | . | . | . | . | . | . |
| chr2 | 173234804 | 173235047 | 182 | Promoter (<=1kb) | 2 | 173228211 | 173234722 | 6512 | 2 | ENSMUST00 | -83 | Pmepa1 | ENSMUSG0C | true | . | . | . | . | . | . | chr2 | 173223515 | 173275069 | 7.01359 | 4.89215 | 4.68115 |
| chr7 | 45015960 | 45016448 | 182 | Promoter (<=1kb) | 7 | 45002948 | 45016363 | 13416 | 2 | ENSMUST00 | 0 | Scaf1 | ENSMUSG0C | false | chr7 | 45015107 | 45016269 | 37.22259 | 125.13314 | 120.81451 | . | . | . | . | . | . |
| chr4 | 62502253 | 62502494 | 182 | Promoter (<=1kb) | 4 | 62499008 | 62502276 | 3269 | 2 | ENSMUST00 | 0 | Hdhd3 | ENSMUSG0C | false | chr4 | 62501735 | 62502741 | 26.18681 | 91.82585 | 88.44535 | chr4 | 62501435 | 62503178 | 3.66132 | 3.3504 | 1.91351 |
| chr6 | 124805259 | 124805889 | 182 | Promoter (2-3kb) | 6 | 124808661 | 124810619 | 1959 | 1 | ENSMUST00 | -2772 | Spsb2 | ENSMUSG0C | false | chr6 | 124805331 | 124806580 | 17.28391 | 44.47248 | 41.82566 | chr6 | 124804618 | 124808507 | 5.49826 | 4.36598 | 3.4161 |
| chr6 | 39557286 | 39558009 | 182 | Promoter (<=1kb) | 6 | 39462378 | 39557867 | 95490 | 2 | ENSMUST00 | 0 | Dennd2a | ENSMUSG0C | false | chr6 | 39557112 | 39558181 | 27.74132 | 80.34372 | 77.18787 | chr6 | 39557115 | 39558192 | 4.39837 | 3.82494 | 2.50213 |
| chr13 | 30484481 | 30484732 | 182 | Distal Intergenic | 13 | 30540308 | 30545362 | 5055 | 2 | ENSMUST00 | 60630 | Uqcrfs1 | ENSMUSG0C | false | . | . | . | . | . | . | . | . | . | . | . | . |
| chr13 | 30545244 | 30545519 | 182 | Promoter (<=1kb) | 13 | 30540308 | 30545362 | 5055 | 2 | ENSMUST00 | 0 | Uqcrfs1 | ENSMUSG0C | false | chr13 | 30544221 | 30545909 | 42.1856 | 147.12523 | 141.95732 | . | . | . | . | . | . |
| chr1 | 170450270 | 170450501 | 182 | Intron (ENSMUST00000160456.2 | 1 | 170338209 | 170454914 | 116706 | 2 | ENSMUST00 | 4413 | Nos1ap | ENSMUSG0C | false | . | . | . | . | . | . | chr1 | 170444264 | 170462009 | 5.07205 | 3.99273 | 3.09536 |
| chr1 | 170589626 | 170590043 | 182 | Promoter (<=1kb) | 1 | 170317459 | 170589861 | 272403 | 2 | ENSMUST00 | 0 | Nos1ap | ENSMUSG0C | false | chr1 | 170588872 | 170590414 | 37.74925 | 129.3071 | 124.84747 | . | . | . | . | . | . |
| chr3 | 96105199 | 96105532 | 182 | Promoter (<=1kb) | 3 | 96104548 | 96141553 | 37006 | 1 | ENSMUST00 | 652 | Otud7b | ENSMUSG0C | true | chr3 | 96103895 | 96107098 | 33.65816 | 114.56059 | 110.59126 | chr3 | 96105433 | 96123323 | 4.88166 | 4.00172 | 2.91968 |
| chr7 | 44866295 | 44866865 | 182 | Promoter (<=1kb) | 7 | 44863068 | 44866774 | 3707 | 2 | ENSMUST00 | 0 | Ptov1 | ENSMUSG0C | false | . | . | . | . | . | . | chr7 | 44860730 | 44868940 | 6.16567 | 4.38467 | 3.98867 |
| chr8 | 13200388 | 13200729 | 182 | Promoter (<=1kb) | 8 | 13176869 | 13200624 | 23756 | 2 | ENSMUST00 | 0 | Grtp1 | ENSMUSG0C | false | chr8 | 13200086 | 13201411 | 23.54255 | 69.40097 | 66.41734 | . | . | . | . | . | . |
| chr11 | 97150008 | 97150256 | 182 | Promoter (<=1kb) | 11 | 97136177 | 97150046 | 13870 | 2 | ENSMUST00 | 0 | Tbkbp1 | ENSMUSG0C | false | chr11 | 97149875 | 97150039 | 4.57782 | 7.38195 | 5.26448 | chr11 | 97149976 | 97152332 | 5.15955 | 4.20764 | 3.14497 |
| chr13 | 44630899 | 44631179 | 182 | Distal Intergenic | 13 | 44729474 | 44731574 | 2101 | 1 | ENSMUST00 | -98295 | Jarid2 | ENSMUSG0C | false | . | . | . | . | . | . | chr13 | 44621314 | 44631701 | 5.57294 | 4.3396 | 3.4838 |
| chr13 | 44630193 | 44630412 | 182 | Distal Intergenic | 13 | 44729474 | 44731574 | 2101 | 1 | ENSMUST00 | -99062 | Jarid2 | ENSMUSG0C | false | . | . | . | . | . | . | chr13 | 44621314 | 44631701 | 5.57294 | 4.3396 | 3.4838 |
| chr3 | 95905149 | 95905832 | 182 | Promoter (<=1kb) | 3 | 95897768 | 95904691 | 6924 | 2 | ENSMUST00 | -459 | Car14 | ENSMUSG0C | false | chr3 | 95905129 | 95905921 | 8.8423 | 19.93079 | 17.59637 | chr3 | 95901722 | 95907425 | 5.6994 | 4.40757 | 3.58314 |
| chr6 | 124490744 | 124490956 | 182 | Promoter (2-3kb) | 6 | 124493113 | 124510643 | 17531 | 1 | ENSMUST00 | -2157 | C1rl | ENSMUSG0C | false | . | . | . | . | . | . | . | . | . | . | . | . |
| chr8 | 13104769 | 13105094 | 182 | Promoter (<=1kb) | 8 | 13079896 | 13104831 | 24936 | 2 | ENSMUST00 | 0 | Pcid2 | ENSMUSG0C | false | chr8 | 13104180 | 13106753 | 40.01227 | 134.38658 | 129.72945 | chr8 | 13103522 | 13105267 | 4.87722 | 3.9288 | 2.93465 |
| chr17 | 46650125 | 46650626 | 182 | Promoter (<=1kb) | 17 | 46650337 | 46664364 | 14028 | 1 | ENSMUST00 | 0 | Cul7 | ENSMUSG0C | false | chr17 | 46649997 | 46651405 | 36.111 | 117.84728 | 113.77644 | chr17 | 46649685 | 46652126 | 5.96056 | 4.46845 | 3.84791 |
| chr13 | 43503018 | 43503224 | 182 | Distal Intergenic | 13 | 43402673 | 43480973 | 78301 | 2 | ENSMUST00 | -22046 | Ranbp9 | ENSMUSG0C | false | . | . | . | . | . | . | chr13 | 43502496 | 43504465 | 3.8401 | 3.43343 | 2.08025 |
| chr10 | 53379493 | 53379984 | 182 | Promoter (<=1kb) | 10 | 53278081 | 53379851 | 101771 | 2 | ENSMUST00 | 0 | Cep85l | ENSMUSG0C | true | chr10 | 53376609 | 53380307 | 36.04672 | 129.50026 | 125.02953 | . | . | . | . | . | . |
| chr4 | 58943636 | 58943947 | 182 | Promoter (<=1kb) | 4 | 58943628 | 58958355 | 14728 | 1 | ENSMUST00 | 9 | Zkscan16 | ENSMUSG0C | false | chr4 | 58943316 | 58944136 | 29.4107 | 87.008 | 83.72497 | chr4 | 58943202 | 58944348 | 4.27568 | 3.58459 | 2.43515 |
| chr6 | 37442173 | 37442642 | 182 | Promoter (<=1kb) | 6 | 37327255 | 37442146 | 114892 | 2 | ENSMUST00 | -28 | Creb3l2 | ENSMUSG0C | true | chr6 | 37439782 | 37442252 | 26.03857 | 105.07202 | 101.37238 | . | . | . | . | . | . |
| chr6 | 37427496 | 37427686 | 182 | Intron (ENSMUST00000041093.5 | 6 | 37327255 | 37442146 | 114892 | 2 | ENSMUST00 | 14460 | Creb3l2 | ENSMUSG0C | true | . | . | . | . | . | . | chr6 | 37373151 | 37440158 | 6.00246 | 4.49234 | 3.83883 |
| chr9 | 66350231 | 66350555 | 182 | Promoter (<=1kb) | 9 | 66350450 | 66374070 | 23621 | 1 | ENSMUST00 | 0 | Herc1 | ENSMUSG0C | false | chr9 | 66349776 | 66351501 | 45.47067 | 151.73354 | 146.38181 | . | . | . | . | . | . |
| chr13 | 41000924 | 41001436 | 182 | Promoter (<=1kb) | 13 | 41001023 | 41013015 | 11993 | 1 | ENSMUST00 | 0 | Pak1ip1 | ENSMUSG0C | false | chr13 | 41000191 | 41002089 | 28.32768 | 117.73571 | 113.66808 | chr13 | 41000190 | 41005644 | 4.08783 | 3.53884 | 2.29204 |
| chr5 | 145191409 | 145191689 | 182 | Promoter (<=1kb) | 5 | 145183706 | 145191592 | 7887 | 2 | ENSMUST00 | 0 | Atp5j2 | ENSMUSG0C | false | chr5 | 145190630 | 145191996 | 31.13118 | 94.84997 | 91.40416 | chr5 | 145188209 | 145192073 | 5.05712 | 3.94764 | 3.09065 |
| chr2 | 34614199 | 34614504 | 182 | Intron (ENSMUST00000124443.2 | 2 | 34619582 | 34623919 | 4338 | 1 | ENSMUST00 | -5078 | Mapkap1 | ENSMUSG0C | false | . | . | . | . | . | . | chr2 | 34613669 | 34615356 | 3.55587 | 3.2953 | 1.82147 |
| chr3 | 95282859 | 95283519 | 182 | Promoter (<=1kb) | 3 | 95282944 | 95296166 | 13223 | 1 | ENSMUST00 | 0 | Fam63a | ENSMUSG0C | false | chr3 | 95281057 | 95284014 | 34.78613 | 113.42423 | 109.48744 | chr3 | 95283433 | 95285689 | 4.01105 | 3.63611 | 2.20953 |
| chr4 | 57810502 | 57810754 | 182 | Intron (ENSMUST00000098066.3 | 4 | 57829245 | 57855049 | 25805 | 1 | ENSMUST00 | -18491 | Akap2 | ENSMUSG0C | false | . | . | . | . | . | . | chr4 | 57809325 | 57810841 | 4.19674 | 3.73449 | 2.35391 |
| chr6 | 35177425 | 35177746 | 182 | Promoter (<=1kb) | 6 | 35177430 | 35247449 | 70020 | 1 | ENSMUST00 | 0 | Nup205 | ENSMUSG0C | false | chr6 | 35177153 | 35178417 | 29.66345 | 100.21 | 96.63225 | . | . | . | . | . | . |
| chr4 | 57256860 | 57257066 | 182 | Intron (ENSMUST00000075637.5 | 4 | 57218347 | 57222054 | 3708 | 2 | ENSMUST00 | -34807 | Ptpn3 | ENSMUSG0C | false | . | . | . | . | . | . | chr4 | 57256079 | 57260126 | 4.37335 | 3.7161 | 2.51884 |
| chr2 | 33585000 | 33585334 | 182 | Intron (ENSMUST00000041730.5 | 2 | 33567156 | 33632283 | 65128 | 2 | ENSMUST00 | 46949 | Lmx1b | ENSMUSG0C | true | . | . | . | . | . | . | chr2 | 33565260 | 33638297 | 6.78481 | 4.81457 | 4.48235 |
| chr3 | 95203141 | 95203390 | 182 | 3' UTR | 3 | 95181773 | 95208937 | 27165 | 2 | ENSMUST00 | 5547 | Gabpb2 | ENSMUSG0C | true | . | . | . | . | . | . | chr3 | 95193311 | 95204022 | 6.01146 | 4.56637 | 3.8326 |
| chr9 | 119052131 | 119052459 | 182 | Promoter (<=1kb) | 9 | 119052778 | 119058055 | 5278 | 1 | ENSMUST00 | -319 | Vill | ENSMUSG0C | false | . | . | . | . | . | . | chr9 | 119051377 | 119053614 | 5.22234 | 4.04957 | 3.21543 |
| chr11 | 74522732 | 74522916 | 182 | Intron (ENSMUST00000102521.2 | 11 | 74437043 | 74590124 | 153082 | 2 | ENSMUST00 | 67208 | Rap1gap2 | ENSMUSG0C | false | chr11 | 74521980 | 74523361 | 18.72985 | 51.651 | 48.91581 | chr11 | 74521529 | 74523816 | 4.50589 | 3.81873 | 2.64229 |

| | | | | | | | | | | | | | | | | | | | | | | | | | | |
|---|---|---|---|---|---|---|---|---|---|---|---|---|---|---|---|---|---|---|---|---|---|---|---|---|---|---|
| chr4 | 56862600 | 56862826 | 182 | Promoter (2-3kb) | 4 | 56840746 | 56865154 | 24409 | 2 | ENSMUST00 | 2328 | Ctnnal1 | ENSMUSG0C | true | chr4 | 56862824 | 56865430 | 41.48112 | 133.644 | 129.01402 | chr4 | 56854119 | 56864552 | 4.03652 | 3.57633 | 2.22068 |
| chr4 | 56864372 | 56864598 | 182 | Promoter (<=1kb) | 4 | 56840746 | 56865154 | 24409 | 2 | ENSMUST00 | 556 | Ctnnal1 | ENSMUSG0C | true | chr4 | 56862824 | 56865430 | 41.48112 | 133.644 | 129.01402 | chr4 | 56854119 | 56864552 | 4.03652 | 3.57633 | 2.22068 |
| chr4 | 56865203 | 56865527 | 182 | Promoter (<=1kb) | 4 | 56810935 | 56865188 | 54254 | 2 | ENSMUST00 | -16 | Ctnnal1 | ENSMUSG0C | true | chr4 | 56862824 | 56865430 | 41.48112 | 133.644 | 129.01402 | chr4 | 56865487 | 56869170 | 5.47346 | 4.30542 | 3.36427 |
| chr4 | 56858762 | 56859289 | 182 | Intron (ENSMUST00000045142.9 | 4 | 56840746 | 56865154 | 24409 | 2 | ENSMUST00 | 5865 | Ctnnal1 | ENSMUSG0C | true | | | | | | | chr4 | 56854119 | 56864552 | 4.03652 | 3.57633 | 2.22068 |
| chr5 | 30862894 | 30863110 | 182 | Exon (ENSMUST00000146459.1/ | 5 | 30869579 | 30879191 | 9613 | 1 | ENSMUST00 | -6469 | Tmem214 | ENSMUSG0C | false | | | | | | | chr5 | 30862900 | 30864585 | 5.19143 | 3.88299 | 3.19532 |
| chr2 | 33165130 | 33165418 | 182 | Promoter (2-3kb) | 2 | 33137636 | 33168222 | 30587 | 2 | ENSMUST00 | 2804 | Ralgps1 | ENSMUSG0C | false | | | | | | | chr2 | 33154669 | 33171007 | 7.40171 | 5.13174 | 4.97591 |
| chr2 | 142901391 | 142901739 | 182 | Promoter (<=1kb) | 2 | 142617474 | 142901531 | 284058 | 2 | ENSMUST00 | 0 | Kif16b | ENSMUSG0C | false | chr2 | 142900680 | 142902004 | 37.5443 | 122.47784 | 118.25522 | chr2 | 142900386 | 142903706 | 5.23958 | 4.21244 | 3.1947 |
| chr5 | 144316651 | 144317054 | 182 | Intron (ENSMUST00000155491.2 | 5 | 144281990 | 144288907 | 6918 | 1 | ENSMUST00 | -27745 | Baiap2l1 | ENSMUSG0C | false | | | | | | | chr5 | 144315523 | 144319209 | 3.63661 | 3.24059 | 1.92432 |
| chr1 | 105993817 | 105994087 | 182 | Intron (ENSMUST00000118196.2 | 1 | 105990680 | 106034067 | 43379 | 1 | ENSMUST00 | 3129 | Zcchc2 | ENSMUSG0C | false | chr1 | 105993233 | 105994297 | 33.06624 | 113.49754 | 109.55873 | chr1 | 105991697 | 106000156 | 4.75264 | 3.97886 | 2.80533 |
| chr8 | 10899620 | 10899862 | 182 | Distal Intergenic | 8 | 10924427 | 10928457 | 4031 | 2 | ENSMUST00 | 28595 | 3930402G23 | ENSMUSG0C | false | chr8 | 10899661 | 10900229 | 11.20895 | 23.45317 | 21.07269 | chr8 | 10899146 | 10900515 | 4.59118 | 3.82964 | 2.67588 |
| chr17 | 44777003 | 44777738 | 182 | Promoter (<=1kb) | 17 | 44777152 | 45119290 | 342139 | 1 | ENSMUST00 | 0 | Supt3 | ENSMUSG0C | false | chr17 | 44776597 | 44778038 | 33.25048 | 115.91603 | 111.90475 | | | | | | |
| chr11 | 95038141 | 95038379 | 182 | Promoter (2-3kb) | 11 | 95029654 | 95040558 | 10905 | 2 | ENSMUST00 | 2179 | Pdk2 | ENSMUSG0C | false | | | | | | | chr11 | 95031837 | 95049061 | 6.83336 | 4.76219 | 4.52264 |
| chr11 | 95041304 | 95041513 | 182 | Promoter (<=1kb) | 11 | 95026258 | 95041354 | 15097 | 2 | ENSMUST00 | 0 | Pdk2 | ENSMUSG0C | false | chr11 | 95039880 | 95041440 | 40.45295 | 134.0567 | 129.41484 | chr11 | 95031837 | 95049061 | 6.83336 | 4.76219 | 4.52264 |
| chr2 | 180273193 | 180273602 | 182 | Promoter (<=1kb) | 2 | 180258540 | 180273496 | 14957 | 2 | ENSMUST00 | 0 | Cables2 | ENSMUSG0C | false | chr2 | 180272453 | 180273457 | 34.04953 | 116.30843 | 112.28568 | chr2 | 180271344 | 180276255 | 4.85323 | 4.07719 | 2.88156 |
| chr3 | 152713990 | 152714175 | 182 | Promoter (<=1kb) | 3 | 152714100 | 152788898 | 74799 | 1 | ENSMUST00 | 0 | Pigk | ENSMUSG0C | false | chr3 | 152713711 | 152714806 | 29.75837 | 125.79383 | 121.46131 | chr3 | 152713733 | 152715351 | 5.13818 | 4.24405 | 3.11262 |
| chr2 | 180115375 | 180115634 | 182 | Distal Intergenic | 2 | 180119306 | 180162680 | 43375 | 1 | ENSMUST00 | -3672 | Osbpl2 | ENSMUSG0C | false | | | | | | | chr2 | 180114077 | 180120696 | 5.8988 | 4.44398 | 3.75334 |
| chr14 | 21962559 | 21962944 | 182 | Promoter (<=1kb) | 14 | 21983962 | 21989601 | 5640 | 2 | ENSMUST00 | 26657 | Zfp503 | ENSMUSG0C | false | chr14 | 21962606 | 21963094 | 4.50584 | 6.69248 | 4.59165 | chr14 | 21962260 | 21963277 | 3.98081 | 3.61133 | 2.21132 |
| chr14 | 21995491 | 21995779 | 182 | Promoter (<=1kb) | 14 | 21983962 | 21989601 | 5640 | 2 | ENSMUST00 | -5891 | Zfp503 | ENSMUSG0C | false | chr14 | 21993562 | 21996021 | 31.01414 | 97.6227 | 94.10706 | | | | | | |
| chr13 | 37910968 | 37911160 | 182 | Intron (ENSMUST00000110238.3 | 13 | 37888871 | 37951994 | 63124 | 1 | ENSMUST00 | 22098 | Rreb1 | ENSMUSG0C | false | | | | | | | chr13 | 37907864 | 37913535 | 5.86085 | 4.40073 | 3.75871 |
| chr11 | 94548422 | 94548668 | 182 | Promoter (<=1kb) | 11 | 94544635 | 94548602 | 3968 | 2 | ENSMUST00 | 0 | Rsad1 | ENSMUSG0C | false | chr11 | 94548415 | 94549345 | 23.12601 | 77.64799 | 74.53377 | chr11 | 94548094 | 94549728 | 6.01374 | 4.45106 | 3.86837 |
| chr15 | 31539715 | 31539915 | 182 | Promoter (<=1kb) | 15 | 31455899 | 31531037 | 75139 | 2 | ENSMUST00 | -8679 | 38777 | ENSMUSG0C | false | | | | | | | chr15 | 31539046 | 31544069 | 5.25379 | 4.14739 | 3.21602 |
| chr3 | 152265560 | 152265993 | 182 | Promoter (<=1kb) | 3 | 152236986 | 152266320 | 29335 | 2 | ENSMUST00 | 327 | Nexn | ENSMUSG0C | false | chr3 | 152264840 | 152266333 | 20.4053 | 65.4752 | 62.55133 | chr3 | 152261393 | 152266449 | 3.59081 | 3.37956 | 1.84594 |
| chr9 | 118149956 | 118150245 | 182 | Promoter (<=1kb) | 9 | 118064529 | 118150196 | 85668 | 2 | ENSMUST00 | 0 | Cmc1 | ENSMUSG0C | false | chr9 | 118149159 | 118150425 | 40.53126 | 139.72299 | 134.85652 | | | | | | |
| chr5 | 53633965 | 53634269 | 182 | Intron (ENSMUST00000087360.3 | 5 | 53590574 | 53657445 | 66872 | 1 | ENSMUST00 | 43392 | Rbpj | ENSMUSG0C | false | | | | | | | chr5 | 53631067 | 53638773 | 5.07176 | 4.05082 | 3.09551 |
| chr2 | 32381748 | 32382063 | 182 | Promoter (<=1kb) | 2 | 32377097 | 32381862 | 4766 | 2 | ENSMUST00 | 0 | 1110008P14R | ENSMUSG0C | false | chr2 | 32381188 | 32382428 | 27.58279 | 90.54513 | 87.1897 | chr2 | 32380564 | 32382711 | 4.11421 | 3.66191 | 2.2743 |
| chr14 | 21082891 | 21083097 | 182 | Intron (ENSMUST00000045376.9 | 14 | 21052574 | 21448568 | 395995 | 1 | ENSMUST00 | 30318 | Adk | ENSMUSG0C | false | | | | | | | chr14 | 21082275 | 21083864 | 4.98141 | 4.06312 | 2.99148 |
| chr8 | 105496786 | 105497149 | 182 | Distal Intergenic | 8 | 105472481 | 105484350 | 11870 | 2 | ENSMUST00 | -12437 | Zdhhc1 | ENSMUSG0C | false | chr8 | 105496272 | 105497231 | 38.14666 | 123.52654 | 119.2661 | chr8 | 105493645 | 105497412 | 4.30992 | 3.56255 | 2.48068 |
| chr3 | 28805279 | 28805566 | 182 | Promoter (<=1kb) | 3 | 28805436 | 28807352 | 1917 | 1 | ENSMUST00 | 0 | Rpl22l1 | ENSMUSG0C | false | chr3 | 28804965 | 28806327 | 34.27164 | 115.65593 | 111.648 | | | | | | |
| chr11 | 121451795 | 121452252 | 182 | Promoter (<=1kb) | 11 | 121451949 | 121617164 | 165216 | 1 | ENSMUST00 | 0 | Tbcd | ENSMUSG0C | false | chr11 | 121451320 | 121452786 | 25.84672 | 90.2271 | 86.88143 | chr11 | 121451286 | 121453521 | 4.36496 | 3.73039 | 2.52308 |
| chr3 | 123267819 | 123268205 | 182 | Promoter (<=1kb) | 3 | 123267496 | 123365635 | 98140 | 1 | ENSMUST00 | 324 | Sec24d | ENSMUSG0C | false | chr3 | 123266862 | 123268531 | 24.14628 | 82.32003 | 79.12344 | chr3 | 123268142 | 123268660 | 4.61929 | 4.00551 | 2.70702 |
| chr2 | 32249100 | 32249350 | 182 | Promoter (1-2kb) | 2 | 32247573 | 32251771 | 4199 | 1 | ENSMUST00 | 1528 | Pomt1 | ENSMUSG0C | false | | | | | | | chr2 | 32248460 | 32249504 | 3.86031 | 3.36191 | 2.09827 |
| chr2 | 174110091 | 174110551 | 182 | Promoter (<=1kb) | 2 | 174110349 | 174123070 | 12722 | 1 | ENSMUST00 | 0 | Npepl1 | ENSMUSG0C | false | chr2 | 174109904 | 174111766 | 29.25679 | 89.63524 | 86.30125 | chr2 | 174109729 | 174116112 | 5.48827 | 4.29987 | 3.41377 |
| chr4 | 70709099 | 70709424 | 182 | Distal Intergenic | 4 | 70461826 | 70534995 | 73170 | 2 | ENSMUST00 | -174105 | Megf9 | ENSMUSG0C | false | | | | | | | | | | | | |
| chr4 | 70509642 | 70509978 | 182 | Intron (ENSMUST00000107359.3 | 4 | 70427065 | 70534928 | 107864 | 2 | ENSMUST00 | 24950 | Megf9 | ENSMUSG0C | false | | | | | | | chr4 | 70509070 | 70513155 | 4.17468 | 3.63411 | 2.32637 |
| chr1 | 185203788 | 185204129 | 182 | Promoter (<=1kb) | 1 | 185204168 | 185286743 | 82576 | 1 | ENSMUST00 | -39 | Rab3gap2 | ENSMUSG0C | false | chr1 | 185203116 | 185205017 | 23.70827 | 104.82465 | 101.13002 | | | | | | |
| chr8 | 31111546 | 31111960 | 182 | Promoter (<=1kb) | 8 | 31111820 | 31131482 | 19663 | 1 | ENSMUST00 | 0 | Rnf122 | ENSMUSG0C | false | chr8 | 31111227 | 31112953 | 27.20707 | 96.65539 | 93.16586 | | | | | | |
| chr17 | 78835315 | 78835671 | 182 | Promoter (<=1kb) | 17 | 78752906 | 78835381 | 82476 | 2 | ENSMUST00 | 0 | Heatr5b | ENSMUSG0C | false | chr17 | 78834676 | 78836245 | 39.02359 | 127.20569 | 122.81842 | | | | | | |
| chr15 | 12204908 | 12205173 | 182 | Promoter (<=1kb) | 15 | 12205028 | 12272240 | 67213 | 1 | ENSMUST00 | 0 | Mtmr12 | ENSMUSG0C | false | chr15 | 12204978 | 12206098 | 35.8325 | 117.35963 | 113.30182 | | | | | | |
| chr15 | 12257810 | 12258011 | 182 | Intron (ENSMUST00000038172.9 | 15 | 12223354 | 12245052 | 21699 | 1 | ENSMUST00 | 34457 | Mtmr12 | ENSMUSG0C | false | | | | | | | chr15 | 12257507 | 12258598 | 3.086 | 3.06112 | 1.46152 |
| chr2 | 167421492 | 167422036 | 182 | Promoter (<=1kb) | 2 | 167421712 | 167477000 | 55289 | 1 | ENSMUST00 | 0 | Slc9a8 | ENSMUSG0C | false | chr2 | 167420993 | 167422465 | 24.86315 | 92.1349 | 88.74321 | chr2 | 167420660 | 167422853 | 4.22416 | 3.64759 | 2.38613 |
| chr5 | 36988924 | 36989311 | 182 | Promoter (<=1kb) | 5 | 36966104 | 36988982 | 22879 | 2 | ENSMUST00 | 0 | Wfs1 | ENSMUSG0C | false | chr5 | 36988297 | 36989768 | 27.49702 | 86.54257 | 83.27029 | chr5 | 36987850 | 36989930 | 3.9134 | 3.56754 | 2.10179 |
| chr5 | 142815962 | 142816753 | 182 | Promoter (<=1kb) | 5 | 142754796 | 142817387 | 62592 | 2 | ENSMUST00 | 634 | Tnrc18 | ENSMUSG0C | false | chr5 | 142813148 | 142817670 | 35.38397 | 108.24114 | 104.45254 | | | | | | |
| chr5 | 142822836 | 142823780 | 182 | Distal Intergenic | 5 | 142754789 | 142817662 | 62874 | 2 | ENSMUST00 | -5175 | Tnrc18 | ENSMUSG0C | false | chr5 | 142817908 | 142824082 | 29.69128 | 86.53342 | 83.26198 | chr5 | 142821951 | 142824303 | 4.24728 | 3.72552 | 2.37961 |
| chr10 | 34207085 | 34207344 | 182 | Promoter (<=1kb) | 10 | 34151394 | 34207551 | 56158 | 2 | ENSMUST00 | 207 | Dse | ENSMUSG0C | false | chr10 | 34206373 | 34207924 | 41.12505 | 143.00313 | 137.99791 | | | | | | |
| chr10 | 34193187 | 34193403 | 182 | Intron (ENSMUST00000048010.7 | 10 | 34151394 | 34207551 | 56158 | 2 | ENSMUST00 | 14148 | Dse | ENSMUSG0C | false | | | | | | | chr10 | 34192601 | 34195266 | 5.66117 | 4.28654 | 3.56968 |
| chr2 | 167058282 | 167058486 | 182 | Promoter (1-2kb) | 2 | 167044032 | 167060425 | 16394 | 2 | ENSMUST00 | 1939 | Znfx1 | ENSMUSG0C | false | | | | | | | chr2 | 167058296 | 167061973 | 3.95041 | 3.39064 | 2.17129 |
| chr17 | 35551987 | 35552257 | 182 | Promoter (<=1kb) | 17 | 35552128 | 35557180 | 5053 | 1 | ENSMUST00 | 0 | Cdsn | ENSMUSG0C | false | | | | | | | chr17 | 35546287 | 35552496 | 5.78427 | 4.44046 | 3.65751 |
| chr18 | 64682555 | 64682866 | 182 | Distal Intergenic | 18 | 64528979 | 64661000 | 132022 | 2 | ENSMUST00 | -21556 | Atp8b1 | ENSMUSG0C | true | | | | | | | chr18 | 64681798 | 64685128 | 4.24698 | 3.65242 | 2.43695 |
| chr9 | 49797757 | 49798282 | 182 | Promoter (<=1kb) | 9 | 49504361 | 49798729 | 294369 | 2 | ENSMUST00 | 447 | Ncam1 | ENSMUSG0C | true | chr9 | 49796389 | 49799150 | 29.85632 | 88.73936 | 85.41974 | | | | | | |
| chr14 | 20348123 | 20348407 | 182 | Promoter (<=1kb) | 14 | 20348162 | 20368563 | 20402 | 1 | ENSMUST00 | 0 | Fam149b | ENSMUSG0C | false | chr14 | 20347149 | 20349125 | 33.41831 | 117.26761 | 113.21259 | | | | | | |
| chr17 | 43801741 | 43801974 | 182 | Promoter (<=1kb) | 17 | 43801851 | 44039516 | 237666 | 1 | ENSMUST00 | 0 | Rcan2 | ENSMUSG0C | false | chr17 | 43801261 | 43802494 | 20.74539 | 66.97287 | 64.02668 | chr17 | 43800825 | 43802620 | 3.76601 | 3.45731 | 2.00726 |
| chr2 | 166713597 | 166714109 | 182 | Promoter (<=1kb) | 2 | 166566342 | 166713832 | 147491 | 2 | ENSMUST00 | 0 | Prex1 | ENSMUSG0C | false | chr2 | 166712878 | 166714083 | 28.53699 | 95.19626 | 91.7378 | chr2 | 166712875 | 166714644 | 3.74954 | 3.21503 | 2.00338 |
| chr5 | 142451743 | 142452056 | 182 | Intron (ENSMUST00000072837.5 | 5 | 142463931 | 142478715 | 14785 | 1 | ENSMUST00 | -11875 | Ap5z1 | ENSMUSG0C | false | | | | | | | chr5 | 142451310 | 142453222 | 3.50126 | 3.29223 | 1.79127 |
| chr8 | 31033278 | 31033578 | 182 | Distal Intergenic | 8 | 31089471 | 31097047 | 7577 | 1 | ENSMUST00 | -55893 | Dusp26 | ENSMUSG0C | false | chr8 | 31033002 | 31033577 | 9.76265 | 21.55577 | 19.19873 | chr8 | 31032516 | 31033598 | 4.04458 | 3.62631 | 2.25053 |
| chr2 | 165887020 | 165887363 | 182 | Promoter (2-3kb) | 2 | 165785519 | 165884874 | 99356 | 2 | ENSMUST00 | -2147 | Zmynd8 | ENSMUSG0C | false | | | | | | | chr2 | 165851451 | 165889342 | 6.61553 | 4.6931 | 4.35892 |
| chr16 | 92057949 | 92058315 | 182 | Promoter (<=1kb) | 16 | 92058270 | 92112227 | 53958 | 1 | ENSMUST00 | 0 | Mrps6 | ENSMUSG0C | false | chr16 | 92057556 | 92059912 | 26.61085 | 108.21047 | 104.42454 | | | | | | |
| chr5 | 44587431 | 44587653 | 182 | Intron (ENSMUST00000070748.5 | 5 | 44472133 | 44799707 | 327575 | 1 | ENSMUST00 | 212054 | Ldb2 | ENSMUSG0C | false | | | | | | | | | | | | |
| chr2 | 30048745 | 30048996 | 182 | Promoter (<=1kb) | 2 | 30031546 | 30048881 | 17336 | 2 | ENSMUST00 | 0 | Wdr34 | ENSMUSG0C | false | chr2 | 30048435 | 30049494 | 22.5292 | 69.37365 | 66.39268 | chr2 | 30047919 | 30050108 | 3.49005 | 3.27427 | 1.81929 |
| chr3 | 122634857 | 122635576 | 182 | Distal Intergenic | 3 | 122538719 | 122619667 | 80949 | 2 | ENSMUST00 | -15191 | Fnbp1l | ENSMUSG0C | true | chr3 | 122635432 | 122635639 | 6.38327 | 11.47542 | 9.27082 | chr3 | 122628629 | 122639499 | 4.55949 | 3.79906 | 2.66137 |
| chr3 | 122632288 | 122632507 | 182 | Distal Intergenic | 3 | 122538719 | 122619667 | 80949 | 2 | ENSMUST00 | -12622 | Fnbp1l | ENSMUSG0C | true | | | | | | | chr3 | 122628629 | 122639499 | 4.55949 | 3.79906 | 2.66137 |
| chr3 | 122590337 | 122590748 | 182 | Intron (ENSMUST00000162409.2 | 3 | 122542105 | 122619553 | 77449 | 2 | ENSMUST00 | 28805 | Fnbp1l | ENSMUSG0C | true | | | | | | | chr3 | 122585238 | 122617925 | 4.82785 | 4.00989 | 2.86632 |
| chr11 | 120304944 | 120305487 | 182 | Distal Intergenic | 11 | 120289793 | 120292167 | 2375 | 1 | ENSMUST00 | 15152 | Bahcc1 | ENSMUSG0C | false | chr11 | 120304949 | 120305940 | 22.23551 | 60.19247 | 57.34209 | chr11 | 120304915 | 120308233 | 4.76282 | 3.79866 | 2.85595 |
| chr11 | 120281261 | 120281539 | 182 | Intron (ENSMUST00000122148.2 | 11 | 120289793 | 120292167 | 2375 | 1 | ENSMUST00 | -8254 | Bahcc1 | ENSMUSG0C | false | | | | | | | | | | | | |
| chr11 | 120086711 | 120086995 | 182 | Promoter (<=1kb) | 11 | 120064430 | 120086827 | 22398 | 2 | ENSMUST00 | 0 | Cep131 | ENSMUSG0C | false | chr11 | 120086093 | 120086892 | 25.50663 | 88.63404 | 85.31516 | chr11 | 120085344 | 120087126 | 5.33067 | 4.5521 | 3.2717 |
| chr5 | 43232814 | 43233370 | 182 | Promoter (<=1kb) | 5 | 43233463 | 43289724 | 56262 | 1 | ENSMUST00 | -93 | Cpeb2 | ENSMUSG0C | false | chr5 | 43232302 | 43236237 | 40.20312 | 135.82269 | 131.11238 | | | | | | |
| chr4 | 46664229 | 46664453 | 182 | 3' UTR | 4 | 46632580 | 46650209 | 17630 | 2 | ENSMUST00 | -14021 | Tbc1d2 | ENSMUSG0C | false | chr4 | 46664438 | 46664986 | 17.92175 | 44.05972 | 41.41886 | chr4 | 46664047 | 46665630 | 5.77403 | 4.4657 | 3.70294 |
| chr10 | 128498708 | 128499025 | 182 | Promoter (<=1kb) | 10 | 128494157 | 128498685 | 4529 | 2 | ENSMUST00 | -24 | Myl6b | ENSMUSG0C | false | | | | | | | chr10 | 128494763 | 128498723 | 6.35318 | 4.70636 | 4.12163 |
| chr10 | 128496383 | 128496757 | 182 | Promoter (1-2kb) | 10 | 128494157 | 128498685 | 4529 | 2 | ENSMUST00 | 1928 | Myl6b | ENSMUSG0C | false | | | | | | | chr10 | 128494763 | 128498723 | 6.35318 | 4.70636 | 4.12163 |
| chr4 | 150447471 | 150447909 | 182 | Intron (ENSMUST00000105682.3 | 4 | 150488393 | 150621962 | 133570 | 1 | ENSMUST00 | -40484 | Rere | ENSMUSG0C | false | | | | | | | chr4 | 150440410 | 150463295 | 5.62337 | 4.29719 | 3.53587 |
| chr4 | 150406587 | 150406771 | 182 | Promoter (<=1kb) | 4 | 150406394 | 150470381 | 63988 | 1 | ENSMUST00 | 194 | Rere | ENSMUSG0C | false | | | | | | | chr4 | 150389021 | 150411738 | 5.13627 | 4.01964 | 3.14914 |
| chr5 | 21501720 | 21501993 | 182 | Intron (ENSMUST00000051358.5 | 5 | 21543527 | 21575900 | 32374 | 1 | ENSMUST00 | -41534 | Lrrc17 | ENSMUSG0C | false | | | | | | | chr5 | 21501280 | 21503719 | 4.22556 | 3.68184 | 2.37621 |
| chr5 | 135744077 | 135744320 | 182 | Promoter (<=1kb) | 5 | 135735485 | 135744271 | 8787 | 2 | ENSMUST00 | 0 | Tmem120a | ENSMUSG0C | false | chr5 | 135743398 | 135744879 | 32.31418 | 113.17245 | 109.24831 | chr5 | 135742838 | 135744985 | 4.65779 | 3.8088 | 2.75075 |
| chr5 | 21383704 | 21384029 | 182 | Intron (ENSMUST00000115245.2 | 5 | 21372642 | 21378374 | 5733 | 1 | ENSMUST00 | 11063 | Fgl2 | ENSMUSG0C | false | | | | | | | chr5 | 21382114 | 21384797 | 4.90899 | 4.09939 | 2.91908 |
| chr10 | 17722918 | 17723368 | 182 | Promoter (<=1kb) | 10 | 17723228 | 17725673 | 2446 | 1 | ENSMUST00 | 0 | Cited2 | ENSMUSG0C | false | chr10 | 17721687 | 17725607 | 31.41831 | 117.26761 | 113.21259 | | | | | | |
| chr10 | 17562889 | 17563190 | 182 | Distal Intergenic | 10 | 17723228 | 17725673 | 2446 | 1 | ENSMUST00 | -160038 | Cited2 | ENSMUSG0C | false | | | | | | | chr10 | 17558936 | 17563847 | 4.67516 | 3.75075 | 2.78175 |
| chr4 | 149678099 | 149678352 | 182 | Promoter (2-3kb) | 4 | 149673439 | 149676093 | 2655 | 2 | ENSMUST00 | -2007 | Pik3cd | ENSMUSG0C | false | | | | | | | | | | | | |
| chr4 | 149697888 | 149698061 | 182 | Promoter (<=1kb) | 4 | 149660184 | 149698552 | 38369 | 2 | ENSMUST00 | 491 | Pik3cd | ENSMUSG0C | false | chr4 | 149698048 | 149698982 | 18.02336 | 51.19578 | 48.46415 | chr4 | 149696010 | 149698960 | 3.53169 | 3.32761 | 1.84418 |
| chr15 | 5262729 | 5262996 | 182 | Distal Intergenic | 15 | 5233399 | 5244187 | 10789 | 2 | ENSMUST00 | -18543 | Ptger4 | ENSMUSG0C | false | | | | | | | chr15 | 5258589 | 5263270 | 5.08372 | 4.13888 | 3.08264 |
| chr15 | 5262182 | 5262435 | 182 | Distal Intergenic | 15 | 5233399 | 5244187 | 10789 | 2 | ENSMUST00 | -17996 | Ptger4 | ENSMUSG0C | false | chr15 | 5262135 | 5262599 | 7.48194 | 15.52168 | 13.25109 | chr15 | 5258589 | 5263270 | 5.08372 | 4.13888 | 3.08264 |
| chr2 | 135722610 | 135722906 | 182 | Intron (ENSMUST00000144931.1 | 2 | 135741830 | 135910484 | 168655 | 1 | ENSMUST00 | -18924 | Plcb4 | ENSMUSG0C | false | | | | | | | chr2 | 135722177 | 135723789 | 3.62903 | 3.31804 | 1.89221 |
| chr9 | 108240020 | 108240232 | 182 | Intron (ENSMUST00000166905.2 | 9 | 108205958 | 108263736 | 57779 | 2 | ENSMUST00 | 23504 | Dag1 | ENSMUSG0C | true | | | | | | | chr9 | 108210099 | 108251502 | 5.76916 | 4.35802 | 3.65688 |
| chr16 | 90743631 | 90743871 | 182 | Intron (ENSMUST00000133973.2 | 16 | 90738324 | 90749785 | 11462 | 1 | ENSMUST00 | 5308 | Mrap | ENSMUSG0C | false | | | | | | | | | | | | |
| chr5 | 135533497 | 135533710 | 182 | Intron (ENSMUST00000060311.8 | 5 | 135406523 | 135544122 | 138600 | 2 | ENSMUST00 | 11412 | Hip1 | ENSMUSG0C | true | | | | | | | chr5 | 135531281 | 135535104 | 4.51579 | 3.77818 | 2.61688 |
| chr4 | 34882516 | 34883161 | 182 | Promoter (<=1kb) | 4 | 34803113 | 34882930 | 79818 | 2 | ENSMUST00 | 0 | Zfp292 | ENSMUSG0C | false | chr4 | 34882101 | 34883867 | 28.91247 | 96.86193 | 93.36482 | | | | | | |
| chr5 | 19232296 | 19232494 | 182 | Intron (ENSMUST00000088516.4 | 5 | 19227046 | 20702903 | 1475858 | 1 | ENSMUST00 | 5251 | Magi2 | ENSMUSG0C | true | chr5 | 19223861 | 19232465 | 34.78864 | 106.1911 | 102.46782 | chr5 | 19231939 | 19248475 | 4.30115 | 3.72983 | 2.43444 |
| chr10 | 128179926 | 128180351 | 182 | Promoter (<=1kb) | 10 | 128129470 | 128180297 | 50828 | 2 | ENSMUST00 | 0 | Rbms2 | ENSMUSG0C | true | chr10 | 128179852 | 128180477 | 7.45894 | 16.41436 | 14.12962 | chr10 | 128177251 | 128186172 | 4.50393 | 3.86903 | 2.59287 |
| chr4 | 33261295 | 33261623 | 182 | Intron (ENSMUST00000137589.1 | 4 | 33245423 | 33248510 | 3088 | 2 | ENSMUST00 | -12786 | Pnrc1 | ENSMUSG0C | false | | | | | | | chr4 | 33260130 | 33262096 | 5.71449 | 4.48298 | 3.59831 |
| chr3 | 144506272 | 144506512 | 182 | Intron (ENSMUST00000043325.7 | 3 | 144453963 | 144569881 | 115919 | 2 | ENSMUST00 | 63369 | Hs2st1 | ENSMUSG0C | false | | | | | | | chr3 | 144500981 | 144508673 | 5.4449 | 4.19664 | 3.38898 |
| chr7 | 131362537 | 131363121 | 182 | Promoter (<=1kb) | 7 | 131342718 | 131362698 | 19981 | 2 | ENSMUST00 | 0 | 2310057M21R | ENSMUSG0C | false | chr7 | 131361788 | 131363493 | 35.82539 | 116.29276 | 112.27101 | | | | | | |
| chr9 | 57560825 | 57561337 | 182 | Promoter (<=1kb) | 9 | 57560944 | 57588797 | 27854 | 1 | ENSMUST00 | 0 | Scamp2 | ENSMUSG0C | false | chr9 | 57560513 | 57561714 | 24.16644 | 68.71214 | 65.74169 | chr9 | 57560307 | 57566621 | 7.40954 | 5.12519 | 4.97901 |
| chr8 | 3863763 | 3864008 | 182 | Distal Intergenic | 8 | 3847965 | 3854309 | 6345 | 2 | ENSMUST00 | -9455 | Cd209e | ENSMUSG0C | false | | | | | | | | | | | | |
| chr7 | 45920449 | 45920704 | 182 | Promoter (<=1kb) | 7 | 45918023 | 45920849 | 2827 | 2 | ENSMUST00 | 145 | Emp3 | ENSMUSG0C | false | chr7 | 45920389 | 45921478 | 8.87028 | 22.65516 | 20.28383 | chr7 | 45915648 | 45924393 | 5.13169 | 3.954 | 3.14707 |
| chr10 | 127563904 | 127564176 | 182 | Intron (ENSMUST00000049149.9 | 10 | 127546395 | 127549917 | 3523 | 2 | ENSMUST00 | -13988 | Lrp1 | ENSMUSG0C | false | | | | | | | chr10 | 127564150 | 127564729 | 8.16576 | 5.41449 | 5.622 |
| chr17 | 49554959 | 49555360 | 182 | Intron (ENSMUST00000057610.6 | 17 | 49456022 | 49564337 | 108316 | 2 | ENSMUST00 | 8977 | Daam2 | ENSMUSG0C | true | | | | | | | chr17 | 49553664 | 49563281 | 6.14765 | 4.61155 | 3.94797 |
| chr3 | 8923371 | 8923980 | 182 | Promoter (<=1kb) | 3 | 8802146 | 8923918 | 121773 | 2 | ENSMUST00 | 0 | Mrps28 | ENSMUSG0C | false | chr3 | 8922912 | 8924319 | 31.61691 | 99.67752 | 96.11594 | | | | | | |
| chr4 | 32387914 | 32388240 | 182 | Intron (ENSMUST00000108180.3 | 4 | 32417435 | 32586108 | 168674 | 1 | ENSMUST00 | -29195 | Bach2 | ENSMUSG0C | false | | | | | | | chr4 | 32386584 | 32390421 | 3.7493 | 3.42774 | 2.00584 |
| chr5 | 3341103 | 3341606 | 182 | Promoter (2-3kb) | 5 | 3343893 | 3523218 | 179326 | 1 | ENSMUST00 | -2287 | Cdk6 | ENSMUSG0C | false | chr5 | 3341034 | 3344969 | 32.55001 | 105.36325 | 101.65399 | chr5 | 3341133 | 3344421 | 4.3677 | 3.73533 | 2.50877 |
| chr17 | 46546241 | 46546466 | 182 | Promoter (<=1kb) | 17 | 46500605 | 46546388 | 45784 | 2 | ENSMUST00 | 0 | Cul9 | ENSMUSG0C | false | chr17 | 46545596 | 46546716 | 27.03504 | 88.5911 | 85.27629 | chr17 | 46545023 | 46548108 | 4.11298 | 3.63528 | 2.28309 |
| chr3 | 109027449 | 109027883 | 182 | Distal Intergenic | 3 | 108970997 | 109027667 | 56611 | 2 | ENSMUST00 | 0 | Fam102b | ENSMUSG0C | false | chr3 | 109026385 | 109028112 | 19.38504 | 61.02572 | 58.1656 | | | | | | |
| chrX | 12158757 | 12159443 | 182 | Promoter (<=1kb) | 20 | 12036743 | 12160355 | 123613 | 2 | ENSMUST00 | 912 | Bcor | ENSMUSG0C | false | chrX | 12158400 | 12159041 | 7.35251 | 12.67314 | 10.44663 | | | | | | |
| chr7 | 27674290 | 27674741 | 182 | Promoter (<=1kb) | 7 | 27656375 | 27674594 | 18220 | 2 | ENSMUST00 | 0 | Map3k10 | ENSMUSG0C | false | chr7 | 27673368 | 27675137 | 22.06604 | 78.32127 | 75.1984 | chr7 | 27671908 | 27675344 | 4.11023 | 3.50702 | 2.31117 |
| chr1 | 160917819 | 160918049 | 182 | Intron (ENSMUST00000161609.2 | 1 | 160906418 | 160974978 | 68561 | 1 | ENSMUST00 | 11402 | Rc3h1 | ENSMUSG0C | false | | | | | | | chr1 | 160912997 | 160919191 | 5.67815 | 4.30447 | 3.61318 |
| chr19 | 32243376 | 32243588 | 182 | Intron (ENSMUST00000099514.4 | 19 | 32218757 | 32248009 | 29253 | 2 | ENSMUST00 | 4421 | Sgms1 | ENSMUSG0C | false | | | | | | | chr19 | 32238839 | 32243880 | 6.18778 | 4.61868 | 3.97593 |
| chr19 | 32291978 | 32292190 | 182 | Promoter (2-3kb) | 19 | 32220587 | 32289579 | 68993 | 2 | ENSMUST00 | -2400 | Sgms1 | ENSMUSG0C | false | | | | | | | chr19 | 32284373 | 32299675 | 4.90937 | 3.97268 | 2.94183 |
| chr19 | 32304977 | 32305318 | 182 | Intron (ENSMUST00000099514.4 | 19 | 32220587 | 32289579 | 68993 | 2 | ENSMUST00 | -15399 | Sgms1 | ENSMUSG0C | false | | | | | | | chr19 | 32301122 | 32307156 | 5.60323 | 4.40652 | 3.49303 |
| chr11 | 107124467 | 107124770 | 182 | Intron (ENSMUST00000057892.5 | 11 | 107033081 | 107131922 | 98842 | 2 | ENSMUST00 | 7152 | Bptf | ENSMUSG0C | false | | | | | | | chr11 | 107113194 | 107126929 | 4.9512 | 4.02953 | 2.97886 |
| chr11 | 72301462 | 72301701 | 182 | Promoter (<=1kb) | 11 | 72301629 | 72309061 | 7433 | 1 | ENSMUST00 | 0 | Xaf1 | ENSMUSG0C | false | chr11 | 72301385 | 72302721 | 15.08796 | 37.05233 | 34.49837 | chr11 | 72300867 | 72304472 | 4.7758 | 3.96598 | 2.82634 |
| chr9 | 65826675 | 65826910 | 182 | Promoter (<=1kb) | 9 | 65692391 | 65827564 | 135174 | 2 | ENSMUST00 | 654 | Zfp609 | ENSMUSG0C | false | chr9 | 65826248 | 65828549 | 30.03893 | 101.89226 | 98.2715 | chr9 | 65817423 | 65827175 | 4.53501 | 3.80778 | 2.62509 |
| chr7 | 19695234 | 19695493 | 182 | Promoter (2-3kb) | 7 | 19689484 | 19692658 | 3175 | 2 | ENSMUST00 | -2577 | Apoc1 | ENSMUSG0C | false | | | | | | | | | | | | |
| chr19 | 36925852 | 36926529 | 182 | Promoter (<=1kb) | 19 | 36926070 | 37012752 | 86674 | 1 | ENSMUST00 | 0 | Btaf1 | ENSMUSG0C | false | chr19 | 36925411 | 36928022 | 31.66066 | 102.77801 | 99.14294 | | | | | | |
| chr4 | 142223916 | 142224159 | 182 | Intron (ENSMUST00000036476.4 | 4 | 142141541 | 142210206 | 68666 | 2 | ENSMUST00 | -13711 | Kazn | ENSMUSG0C | false | | | | | | | chr4 | 142218898 | 142232146 | 6.00295 | 4.43269 | 3.85028 |
| chr4 | 142225799 | 142225990 | 182 | Intron (ENSMUST00000036476.4 | 4 | 142139240 | 142239401 | 100162 | 2 | ENSMUST00 | 13411 | Kazn | ENSMUSG0C | false | | | | | | | chr4 | 142218898 | 142232146 | 6.00295 | 4.43269 | 3.85028 |
| chr11 | 71004300 | 71004574 | 182 | Promoter (<=1kb) | 11 | 70984091 | 71004437 | 20347 | 2 | ENSMUST00 | 0 | Dhx33 | ENSMUSG0C | false | chr11 | 71003453 | 71005291 | 22.31878 | 85.15456 | 81.90936 | chr11 | 71001559 | 71008095 | 4.5289 | 3.85489 | 2.65528 |
| chr10 | 3898166 | 3898380 | 182 | Intron (ENSMUST00000120274.2 | 10 | 3903582 | 3919362 | 15781 | 1 | ENSMUST00 | -5202 | Plekhg1 | ENSMUSG0C | true | | | | | | | chr10 | 3882895 | 3909570 | 5.33531 | 4.14005 | 3.29397 |
| chr9 | 65681018 | 65681202 | 182 | Intron (ENSMUST00000128928.2 | 9 | 65676591 | 65688538 | 11948 | 1 | ENSMUST00 | 4428 | Oaz2 | ENSMUSG0C | false | | | | | | | chr9 | 65675551 | 65685323 | 4.69201 | 3.84119 | 2.79144 |
| chr9 | 65671516 | 65671880 | 182 | Promoter (1-2kb) | 9 | 65669712 | 65687825 | 18114 | 1 | ENSMUST00 | 1805 | Oaz2 | ENSMUSG0C | false | | | | | | | | | | | | |
| chr10 | 4028828 | 4029084 | 182 | Intron (ENSMUST00000117291.2 | 10 | 4032280 | 4048951 | 16672 | 1 | ENSMUST00 | -3196 | Mthfd1l | ENSMUSG0C | false | | | | | | | | | | | | |
| chr2 | 91182871 | 91183098 | 182 | Promoter (<=1kb) | 2 | 91137360 | 91182888 | 45529 | 2 | ENSMUST00 | 0 | Madd | ENSMUSG0C | false | chr2 | 91182039 | 91183580 | 24.65524 | 104.07409 | 100.40247 | | | | | | |
| chr4 | 141790507 | 141791035 | 182 | Promoter (<=1kb) | 4 | 141760189 | 141790891 | 30703 | 2 | ENSMUST00 | 0 | Dnajc16 | ENSMUSG0C | false | chr4 | 141789947 | 141791329 | 31.70051 | 127.6562 | 123.25787 | chr4 | 141789605 | 141793271 | 4.57361 | 3.8656 | 2.66603 |
| chr11 | 32349345 | 32349699 | 182 | Promoter (1-2kb) | 11 | 32347840 | 32390300 | 42461 | 1 | ENSMUST00 | 1506 | Sh3pxd2b | ENSMUSG0C | false | | | | | | | chr11 | 32347345 | 32350295 | 3.68085 | 3.30451 | 1.96851 |
| chr7 | 19399815 | 19400007 | 182 | Promoter (<=1kb) | 7 | 19394440 | 19399878 | 5439 | 2 | ENSMUST00 | 0 | Klc3 | ENSMUSG0C | false | | | | | | | chr7 | 19399481 | 19400393 | 4.31912 | 3.77042 | 2.44958 |
| chr14 | 27070514 | 27070993 | 182 | Intron (ENSMUST00000035336.3 | 14 | 27039001 | 27107286 | 68286 | 1 | ENSMUST00 | 31514 | Il17rd | ENSMUSG0C | false | | | | | | | chr14 | 27068032 | 27074912 | 5.10086 | 4.02367 | 3.09953 |
| chr1 | 165724586 | 165724892 | 182 | Distal Intergenic | 1 | 165648945 | 165708094 | 59150 | 2 | ENSMUST00 | -16493 | Rcsd1 | ENSMUSG0C | true | | | | | | | chr1 | 165723018 | 165727515 | 4.62584 | 3.45829 | 3.52846 |
| chr1 | 165701284 | 165701530 | 182 | Intron (ENSMUST00000040357.5 | 1 | 165648945 | 165708094 | 59150 | 2 | ENSMUST00 | 6564 | Rcsd1 | ENSMUSG0C | true | | | | | | | chr1 | 165663194 | 165707027 | 5.93252 | 4.47383 | 3.77552 |
| chr9 | 8134049 | 8134539 | 182 | Promoter (<=1kb) | 9 | 8076633 | 8134294 | 57662 | 2 | ENSMUST00 | 0 | AK129341 | ENSMUSG0C | false | chr9 | 8133662 | 8134918 | 26.18681 | 91.82585 | 88.44535 | | | | | | |
| chr3 | 106788631 | 106788932 | 182 | Promoter (1-2kb) | 3 | 106758861 | 106790149 | 31289 | 2 | ENSMUST00 | 1217 | Cd53 | ENSMUSG0C | false | | | | | | | | | | | | |
| chr4 | 129618868 | 129619274 | 182 | Promoter (<=1kb) | 4 | 129615126 | 129619099 | 3974 | 2 | ENSMUST00 | 0 | Iqcc | ENSMUSG0C | false | chr4 | 129618457 | 129619553 | 21.19411 | 66.07429 | 63.14441 | chr4 | 129617394 | 129623366 | 7.42581 | 5.02559 | 5.00857 |
| chr7 | 19094617 | 19094841 | 182 | Promoter (<=1kb) | 7 | 19094594 | 19098549 | 3956 | 1 | ENSMUST00 | 24 | Six5 | ENSMUSG0C | false | chr7 | 19094120 | 19096125 | 21.40274 | 64.72344 | 61.81032 | | | | | | |
| chr1 | 165247149 | 165247368 | 182 | Distal Intergenic | 1 | 165214922 | 165236943 | 22022 | 2 | ENSMUST00 | -10207 | Tiprl | ENSMUSG0C | false | | | | | | | | | | | | |
| chr1 | 165236956 | 165237251 | 182 | Promoter (<=1kb) | 1 | 165214922 | 165236943 | 22022 | 2 | ENSMUST00 | -14 | Tiprl | ENSMUSG0C | false | chr1 | 165235872 | 165237874 | 22.53436 | 92.00179 | 88.61587 | | | | | | |
| chr1 | 165198902 | 165199156 | 182 | Distal Intergenic | 1 | 165174341 | 165194433 | 20093 | 2 | ENSMUST00 | -4470 | Sft2d2 | ENSMUSG0C | false | | | | | | | | | | | | |
| chr17 | 84515430 | 84515659 | 182 | Intron (ENSMUST00000047206.5 | 17 | 84511895 | 84622142 | 110248 | 1 | ENSMUST00 | 3536 | Plekhh2 | ENSMUSG0C | true | | | | | | | chr17 | 84513757 | 84526219 | 4.51864 | 3.79359 | 2.6341 |
| chr4 | 129461287 | 129461608 | 182 | Promoter (<=1kb) | 4 | 129461581 | 129488498 | 26918 | 1 | ENSMUST00 | 0 | Bsdc1 | ENSMUSG0C | false | chr4 | 129461576 | 129462666 | 34.78613 | 113.42423 | 109.48744 | chr4 | 129460856 | 129464569 | 3.97952 | 3.5011 | 2.17206 |
| chr4 | 141060376 | 141060856 | 182 | Promoter (<=1kb) | 4 | 141016637 | 141060550 | 43914 | 2 | ENSMUST00 | 0 | Crocc | ENSMUSG0C | false | chr4 | 141059952 | 141061435 | 15.55856 | 58.09642 | 55.27494 | | | | | | |
| chr19 | 45659404 | 45660184 | 182 | Promoter (<=1kb) | 19 | 45578254 | 45660312 | 82059 | 2 | ENSMUST00 | 128 | Fbxw4 | ENSMUSG0C | false | chr19 | 45659211 | 45660899 | 31.48209 | 103.47416 | 99.8177 | chr19 | 45658545 | 45659480 | 4.47431 | 3.60538 | 2.61171 |
| chr4 | 129184390 | 129184727 | 182 | Promoter (1-2kb) | 4 | 129149372 | 129182530 | 33159 | 2 | ENSMUST00 | -1861 | S100pbp | ENSMUSG0C | true | | | | | | | chr4 | 129183108 | 129188971 | 3.74369 | 3.42065 | 1.97895 |
| chr7 | 18840437 | 18840774 | 182 | Promoter (<=1kb) | 7 | 18839966 | 18858654 | 18689 | 1 | ENSMUST00 | 472 | Mill2 | ENSMUSG0C | true | chr7 | 18839929 | 18840777 | 17.4338 | 43.97722 | 41.34015 | chr7 | 18839515 | 18841690 | 3.24572 | 3.16358 | 1.56811 |
| chrX | 159711173 | 159711358 | 182 | Promoter (2-3kb) | 20 | 159708601 | 159974167 | 265567 | 1 | ENSMUST00 | 2573 | Sh3kbp1 | ENSMUSG0C | false | | | | | | | | | | | | |
| chrX | 159640075 | 159640325 | 182 | Intron (ENSMUST00000112456.3 | 20 | 159627541 | 159973169 | 345629 | 1 | ENSMUST00 | 12535 | Sh3kbp1 | ENSMUSG0C | false | | | | | | | | | | | | |
| chrX | 159627151 | 159627330 | 182 | Promoter (<=1kb) | 20 | 159627272 | 159978069 | 350798 | 1 | ENSMUST00 | 0 | Sh3kbp1 | ENSMUSG0C | false | chrX | 159626654 | 159628266 | 20.67895 | 53.40692 | 50.64761 | | | | | | |
| chr3 | 132880840 | 132881364 | 182 | Downstream (<1kb) | 3 | 132891176 | 132906359 | 15184 | 2 | ENSMUST00 | 24995 | Npnt | ENSMUSG0C | true | | | | | | | chr3 | 132880303 | 132885324 | 5.40591 | 4.23547 | 3.34377 |
| chr3 | 132857829 | 132858105 | 182 | Distal Intergenic | 3 | 132891176 | 132906359 | 15184 | 2 | ENSMUST00 | 48254 | Npnt | ENSMUSG0C | true | | | | | | | chr3 | 132857929 | 132860116 | 2.96706 | 2.91364 | 1.35513 |
| chr4 | 128884026 | 128884240 | 182 | Promoter (<=1kb) | 4 | 128883588 | 128911328 | 27741 | 1 | ENSMUST00 | 439 | Trim62 | ENSMUSG0C | false | chr4 | 128883298 | 128884769 | 28.48738 | 97.07422 | 93.57343 | chr4 | 128883147 | 128884887 | 3.62856 | 3.34723 | 1.89535 |
| chr4 | 139574379 | 139574688 | 182 | Promoter (<=1kb) | 4 | 139574720 | 139620382 | 45663 | 1 | ENSMUST00 | -32 | Iffo2 | ENSMUSG0C | false | chr4 | 139574658 | 139577815 | 38.13901 | 127.82562 | 123.40861 | | | | | | |
| chr4 | 139581597 | 139581862 | 182 | Intron (ENSMUST00000123827.3 | 4 | 139574720 | 139620382 | 45663 | 1 | ENSMUST00 | 6878 | Iffo2 | ENSMUSG0C | false | chr4 | 139580247 | 139582371 | 12.5733 | 28.75278 | 26.30366 | chr4 | 139576299 | 139606726 | 8.92638 | 5.72021 | 6.23508 |
| chr17 | 71616022 | 71616284 | 182 | Promoter (<=1kb) | 17 | 71616215 | 71659031 | 42817 | 1 | ENSMUST00 | 0 | Wdr43 | ENSMUSG0C | false | chr17 | 71615666 | 71617723 | 39.80159 | 147.3544 | 142.17488 | | | | | | |
| chr4 | 19708803 | 19709295 | 182 | Promoter (<=1kb) | 4 | 19608303 | 19708993 | 100691 | 2 | ENSMUST00 | 0 | Wwp1 | ENSMUSG0C | false | chr4 | 19708046 | 19709648 | 28.22734 | 101.53345 | 97.92473 | | | | | | |
| chr13 | 20145905 | 20146084 | 182 | Intron (ENSMUST00000181429.2 | 13 | 20111512 | 20342888 | 231377 | 1 | ENSMUST00 | 34394 | Elmo1 | ENSMUSG0C | true | | | | | | | chr13 | 20137737 | 20154822 | 4.88405 | 3.9784 | 2.92438 |
| chr16 | 20140999 | 20141423 | 182 | Promoter (<=1kb) | 16 | 20141063 | 20232573 | 91511 | 1 | ENSMUST00 | 0 | Yeats2 | ENSMUSG0C | false | chr16 | 20140561 | 20141914 | 37.26972 | 119.86725 | 115.72688 | | | | | | |
| chr18 | 11392634 | 11393036 | 182 | Distal Intergenic | 18 | 11633276 | 11743207 | 109932 | 1 | ENSMUST00 | -240240 | Rbbp8 | ENSMUSG0C | false | | | | | | | chr18 | 11392498 | 11393536 | 3.89424 | 3.54726 | 2.12479 |
| chr17 | 43249694 | 43250031 | 182 | Distal Intergenic | 17 | 43270329 | 43324737 | 54409 | 1 | ENSMUST00 | -20298 | Gpr110 | ENSMUSG0C | false | | | | | | | | | | | | |
| chr17 | 43147920 | 43148270 | 182 | Distal Intergenic | 17 | 43270329 | 43324737 | 54409 | 1 | ENSMUST00 | -122059 | Gpr110 | ENSMUSG0C | false | | | | | | | chr17 | 43146554 | 43148633 | 4.86238 | 3.92665 | 2.91451 |
| chr5 | 148994649 | 148994990 | 182 | Intron (ENSMUST00000085546.3 | 5 | 148893375 | 148995147 | 101373 | 2 | ENSMUST00 | 157 | Katnal1 | ENSMUSG0C | false | chr5 | 148994297 | 148995572 | 23.23877 | 64.48248 | 61.57465 | chr5 | 148990738 | 148995772 | 3.78992 | 3.4575 | 2.05895 |
| chr5 | 148928695 | 148928983 | 182 | Promoter (<=1kb) | 5 | 148885726 | 148929500 | 43275 | 2 | ENSMUST00 | 517 | Katnal1 | ENSMUSG0C | false | chr5 | 148928039 | 148929495 | 40.26965 | 130.45439 | 125.94711 | chr5 | 148926977 | 148929738 | 4.23962 | 3.7432 | 2.38934 |
| chr5 | 148923538 | 148923792 | 182 | Intron (ENSMUST00000047257.5 | 5 | 148904593 | 148928628 | 24036 | 2 | ENSMUST00 | 4836 | Katnal1 | ENSMUSG0C | false | chr5 | 148923479 | 148924297 | 13.64828 | 32.53182 | 30.03459 | chr5 | 148923123 | 148924439 | 3.44597 | 3.1823 | 1.7623 |
| chr1 | 54438897 | 54439130 | 182 | Promoter (<=1kb) | 1 | 54396004 | 54438971 | 42968 | 2 | ENSMUST00 | 0 | Gtf3c3 | ENSMUSG0C | false | chr1 | 54438271 | 54439517 | 22.06604 | 78.32127 | 75.1984 | chr1 | 54437016 | 54439488 | 3.62534 | 3.4205 | 1.88059 |
| chr5 | 148464445 | 148464699 | 182 | Distal Intergenic | 5 | 148527410 | 148599904 | 72495 | 1 | ENSMUST00 | -62542 | Slc7a1 | ENSMUSG0C | false | | | | | | | | | | | | |
| chr7 | 92668653 | 92668896 | 182 | Promoter (1-2kb) | 7 | 92643712 | 92669933 | 26222 | 2 | ENSMUST00 | 1037 | Pcf11 | ENSMUSG0C | false | chr7 | 92667825 | 92669960 | 39.85937 | 131.92529 | 127.36071 | chr7 | 92665253 | 92668745 | 4.79914 | 3.93626 | 2.85256 |
| chr11 | 69579277 | 69579693 | 182 | Promoter (<=1kb) | 11 | 69561759 | 69579324 | 17566 | 2 | ENSMUST00 | 0 | Wrap53 | ENSMUSG0C | false | chr11 | 69579501 | 69582098 | 31.85236 | 101.09525 | 97.496 | | | | | | |
| chr19 | 27429854 | 27430124 | 182 | Promoter (<=1kb) | 19 | 27388702 | 27429820 | 41119 | 2 | ENSMUST00 | -35 | D19Wbg1357 | ENSMUSG0C | false | chr19 | 27428956 | 27431040 | 32.66734 | 113.81544 | 109.86345 | | | | | | |
| chr7 | 16295467 | 16295756 | 182 | Distal Intergenic | 7 | 16282755 | 16286795 | 4041 | 2 | ENSMUST00 | -8673 | Ccdc9 | ENSMUSG0C | false | chr7 | 16294945 | 16295821 | 21.49234 | 58.91379 | 56.0822 | chr7 | 16295229 | 16298235 | 6.20923 | 4.6252 | 3.99375 |
| chr7 | 16294929 | 16295123 | 182 | Distal Intergenic | 7 | 16282755 | 16286795 | 4041 | 2 | ENSMUST00 | -8135 | Ccdc9 | ENSMUSG0C | false | chr7 | 16294945 | 16295821 | 21.49234 | 58.91379 | 56.0822 | | | | | | |
| chr16 | 18733202 | 18733463 | 182 | Distal Intergenic | 16 | 18776847 | 18778260 | 1414 | 1 | ENSMUST00 | -43384 | Cldn5 | ENSMUSG0C | true | chr16 | 18732913 | 18733740 | 10.47775 | 22.22469 | 19.86031 | chr16 | 18732618 | 18733830 | 4.02517 | 3.60677 | 2.25252 |
| chr6 | 15792753 | 15793065 | 182 | Intron (ENSMUST00000101663.4 | 6 | 15727799 | 15770349 | 42551 | 1 | ENSMUST00 | 64955 | Mdfic | ENSMUSG0C | true | | | | | | | chr6 | 15787856 | 15795075 | 6.51926 | 4.79406 | 4.25431 |
| chr12 | 104713911 | 104714129 | 182 | Intron (ENSMUST00000041987.6 | 12 | 104687742 | 104751952 | 64211 | 2 | ENSMUST00 | 37823 | Dicer1 | ENSMUSG0C | false | | | | | | | chr12 | 104712662 | 104716204 | 5.60676 | 4.39762 | 3.50553 |
| chr13 | 101696836 | 101697079 | 182 | Promoter (2-3kb) | 13 | 101689040 | 101694600 | 5561 | 2 | ENSMUST00 | -2237 | Pik3r1 | ENSMUSG0C | false | | | | | | | chr13 | 101690067 | 101698367 | 4.61928 | 3.86189 | 2.69344 |
| chr13 | 101719388 | 101719613 | 182 | Intron (ENSMUST00000055518.7 | 13 | 101701449 | 101702104 | 656 | 2 | ENSMUST00 | -17285 | Pik3r1 | ENSMUSG0C | false | | | | | | | chr13 | 101715036 | 101723721 | 4.94025 | 4.05283 | 2.99666 |
| chr1 | 52844724 | 52845010 | 182 | Promoter (<=1kb) | 1 | 52844929 | 52848693 | 3765 | 1 | ENSMUST00 | 0 | Hibch | ENSMUSG0C | false | chr1 | 52844480 | 52844797 | 12.58327 | 33.30664 | 30.79935 | chr1 | 52844108 | 52846226 | 4.34574 | 3.73717 | 2.50184 |
| chr1 | 52722758 | 52722940 | 182 | Intron (ENSMUST00000156876.2 | 1 | 52675623 | 52727358 | 51736 | 1 | ENSMUST00 | 4418 | Mfsd6 | ENSMUSG0C | false | | | | | | | chr1 | 52722399 | 52724580 | 4.38276 | 3.81521 | 2.51058 |
| chr9 | 96119199 | 96119428 | 182 | Promoter (<=1kb) | 9 | 96119362 | 96180190 | 60829 | 1 | ENSMUST00 | 0 | Gk5 | ENSMUSG0C | false | chr9 | 96118703 | 96120086 | 45.10679 | 157.97711 | 152.39525 | chr9 | 96118635 | 96120302 | 3.88646 | 3.62753 | 2.10924 |
| chr9 | 32032096 | 32032520 | 182 | Distal Intergenic | 9 | 32116136 | 32268446 | 152311 | 1 | ENSMUST00 | -83616 | Arhgap32 | ENSMUSG0C | false | chr9 | 32031756 | 32033342 | 28.03146 | 79.91027 | 76.76085 | | | | | | |
| chr18 | 63348163 | 63348771 | 182 | Intron (ENSMUST00000047480.7 | 18 | 63059718 | 63386457 | 326740 | 2 | ENSMUST00 | 37686 | Piezo2 | ENSMUSG0C | false | | | | | | | | | | | | |
| chr7 | 14622941 | 14623209 | 182 | Promoter (<=1kb) | 7 | 14530652 | 14622479 | 91828 | 2 | ENSMUST00 | -463 | Nlrp5-ps | ENSMUSG0C | false | chr7 | 14622078 | 14623016 | 18.30549 | 46.98978 | 44.31301 | chr7 | 14622391 | 14623541 | 2.84289 | 2.93433 | 1.30376 |
| chr11 | 113748862 | 113749135 | 182 | Promoter (2-3kb) | 11 | 113729305 | 113751154 | 21850 | 2 | ENSMUST00 | 2019 | Cdc42ep4 | ENSMUSG0C | false | | | | | | | chr11 | 113747048 | 113750021 | 3.19795 | 3.06178 | 1.57205 |
| chr5 | 115732583 | 115732957 | 182 | Promoter (<=1kb) | 5 | 115648175 | 115731621 | 83447 | 2 | ENSMUST00 | -963 | Ccdc64 | ENSMUSG0C | false | chr5 | 115731087 | 115732982 | 19.52531 | 57.12588 | 54.31675 | chr5 | 115730797 | 115733021 | 5.3037 | 4.09041 | 3.29064 |
| chr14 | 122912933 | 122913295 | 182 | Promoter (<=1kb) | 14 | 122816223 | 122913120 | 96898 | 2 | ENSMUST00 | 0 | Ggact | ENSMUSG0C | false | chr14 | 122912508 | 122913604 | 31.43325 | 99.35611 | 95.79983 | | | | | | |
| chr14 | 122534042 | 122534406 | 182 | Promoter (<=1kb) | 14 | 122534324 | 122582977 | 48654 | 1 | ENSMUST00 | 0 | Pcca | ENSMUSG0C | false | chr14 | 122533621 | 122534902 | 28.36989 | 85.2065 | 81.96005 | chr14 | 122533477 | 122535111 | 3.08819 | 3.06592 | 1.43711 |
| chr11 | 113560244 | 113560469 | 182 | Intron (ENSMUST00000017539.4 | 11 | 113244853 | 113565756 | 320904 | 1 | ENSMUST00 | 5287 | Slc39a11 | ENSMUSG0C | false | | | | | | | chr11 | 113557859 | 113560931 | 4.15802 | 3.55589 | 2.39039 |
| chr14 | 32994985 | 32995199 | 182 | Promoter (<=1kb) | 14 | 32995265 | 33008459 | 13195 | 1 | ENSMUST00 | -66 | Lrrc18 | ENSMUSG0C | false | | | | | | | chr14 | 32994449 | 32995864 | 5.4916 | 4.27913 | 3.44503 |
| chr14 | 32946157 | 32946358 | 182 | Distal Intergenic | 14 | 32991382 | 33015292 | 23911 | 1 | ENSMUST00 | -45024 | Lrrc18 | ENSMUSG0C | false | | | | | | | chr14 | 32945937 | 32960190 | 5.28697 | 4.21543 | 3.24118 |
| chr12 | 102757417 | 102758152 | 182 | Promoter (<=1kb) | 12 | 102757975 | 102777701 | 19727 | 1 | ENSMUST00 | 0 | Ubr7 | ENSMUSG0C | false | chr12 | 102756811 | 102759183 | 29.58769 | 108.11006 | 104.32592 | | | | | | |
| chr9 | 62491614 | 62491846 | 182 | Intron (ENSMUST00000048043.6 | 9 | 62427936 | 62510498 | 82563 | 2 | ENSMUST00 | 18652 | Coro2b | ENSMUSG0C | true | | | | | | | chr9 | 62449515 | 62535966 | 6.98492 | 4.90408 | 4.64549 |
| chr3 | 87158072 | 87158317 | 182 | Intron (ENSMUST00000159976.2 | 3 | 87084628 | 87174550 | 89923 | 2 | ENSMUST00 | 16233 | Kirrel | ENSMUSG0C | true | | | | | | | chr3 | 87139743 | 87171483 | 6.81014 | 4.88372 | 4.49361 |
| chr3 | 87163297 | 87163584 | 182 | Intron (ENSMUST00000159976.2 | 3 | 87084628 | 87174550 | 89923 | 2 | ENSMUST00 | 10966 | Kirrel | ENSMUSG0C | true | | | | | | | chr3 | 87139743 | 87171483 | 6.81014 | 4.88372 | 4.49361 |
| chr9 | 31688189 | 31688459 | 182 | Distal Intergenic | 9 | 31426199 | 31464238 | 38040 | 2 | ENSMUST00 | -223952 | Tmem45b | ENSMUSG0C | false | chr9 | 31688049 | 31688624 | 7.82203 | 16.59716 | 14.30959 | chr9 | 31686931 | 31688986 | 4.16543 | 3.55476 | 2.35681 |
| chr6 | 141053980 | 141054355 | 182 | Distal Intergenic | 6 | 141249269 | 141505940 | 256672 | 1 | ENSMUST00 | -194914 | Pde3a | ENSMUSG0C | true | | | | | | | | | | | | |
| chr13 | 98236447 | 98236674 | 182 | Distal Intergenic | 13 | 98246845 | 98262992 | 16148 | 2 | ENSMUST00 | 26318 | Utp15 | ENSMUSG0C | false | | | | | | | chr13 | 98236065 | 98236511 | 2.94142 | 2.94053 | 1.36184 |
| chr1 | 133237080 | 133237340 | 182 | Intron (ENSMUST00000187285.1 | 1 | 133246097 | 133303435 | 57339 | 1 | ENSMUST00 | -8757 | Plekha6 | ENSMUSG0C | false | | | | | | | chr1 | 133236575 | 133241534 | 5.4642 | 4.18784 | 3.40705 |
| chr1 | 133168468 | 133168833 | 182 | Intron (ENSMUST00000153291.1 | 1 | 133181321 | 133301219 | 119899 | 1 | ENSMUST00 | -12488 | Plekha6 | ENSMUSG0C | false | | | | | | | chr1 | 133167271 | 133169127 | 4.77947 | 3.88191 | 2.86967 |
| chr1 | 43933579 | 43933954 | 182 | Promoter (<=1kb) | 1 | 43933646 | 43949405 | 15759 | 1 | ENSMUST00 | 0 | Tpp2 | ENSMUSG0C | false | chr1 | 43933120 | 43935537 | 42.07631 | 139.09143 | 134.24335 | | | | | | |
| chr18 | 77758780 | 77758977 | 182 | Intron (ENSMUST00000048192.7 | 18 | 77757567 | 77767780 | 10214 | 2 | ENSMUST00 | 8803 | Haus1 | ENSMUSG0C | false | | | | | | | chr18 | 77758073 | 77759740 | 4.38243 | 3.81917 | 2.49728 |
| chr15 | 5116290 | 5116578 | 182 | Promoter (<=1kb) | 15 | 5116613 | 5119140 | 2528 | 1 | ENSMUST00 | -35 | Rpl37 | ENSMUSG0C | false | chr15 | 5115562 | 5117636 | 28.5703 | 92.31807 | 88.92419 | | | | | | |
| chr1 | 43092310 | 43092907 | 182 | Promoter (<=1kb) | 1 | 43092588 | 43093835 | 1248 | 1 | ENSMUST00 | 0 | 8430432A02 | ENSMUSG0C | false | chr1 | 43091387 | 43093290 | 32.03847 | 101.42104 | 97.81718 | | | | | | |
| chr11 | 109611662 | 109612192 | 182 | Promoter (<=1kb) | 11 | 109611464 | 109612192 | 729 | 2 | ENSMUST00 | 0 | Wipi1 | ENSMUSG0C | false | | | | | | | chr11 | 109612157 | 109612846 | 3.9233 | 3.60854 | 2.10621 |
| chr11 | 109610858 | 109611483 | 182 | Promoter (<=1kb) | 11 | 109573331 | 109611432 | 38102 | 2 | ENSMUST00 | 0 | Wipi1 | ENSMUSG0C | false | chr11 | 109610494 | 109611486 | 27.97104 | 107.99458 | 104.2211 | | | | | | |

| | | | | | | | | | | | | | | | | | | | | | | | | | | |
|---|---|---|---|---|---|---|---|---|---|---|---|---|---|---|---|---|---|---|---|---|---|---|---|---|---|---|
| chr1 | 42952961 | 42953288 | 182 | Promoter (<=1kb) | 1 | 42952963 | 43035456 | 82494 | 1 | ENSMUST00 | 0 | Gpr45 | ENSMUSG0( | false | chr1 | 42952733 | 42953876 | 21.13848 | 54.99947 | 52.22001 | chr1 | 42952984 | 42953782 | 5.09867 | 3.78256 | 3.12231 |
| chr3 | 94413042 | 94413435 | 182 | Promoter (<=1kb) | 3 | 94413296 | 94432633 | 19338 | 1 | ENSMUST00 | 0 | Tdrkh | ENSMUSG0( | false | chr3 | 94412473 | 94414007 | 19.72513 | 62.50151 | 59.62064 | chr3 | 94412278 | 94414601 | 4.26606 | 3.67353 | 2.403 |
| chr18 | 31759772 | 31760144 | 182 | Promoter (<=1kb) | 18 | 31759824 | 31784071 | 24248 | 1 | ENSMUST00 | 0 | Ammecr1l | ENSMUSG0( | false | chr18 | 31759113 | 31761179 | 42.48137 | 137.57564 | 132.7934 | . | . | . | . | . | . |
| chr5 | 114585784 | 114586165 | 182 | Intron (ENSMUST00000043650.7 | 5 | 114568017 | 114594678 | 26662 | 1 | ENSMUST00 | 17768 | Fam222a | ENSMUSG0( | true | . | . | . | . | . | . | chr5 | 114571033 | 114614394 | 7.1045 | 4.9325 | 4.74958 |
| chr5 | 114567747 | 114568154 | 182 | Promoter (<=1kb) | 5 | 114568016 | 114613220 | 45205 | 1 | ENSMUST00 | 0 | Fam222a | ENSMUSG0( | true | chr5 | 114567128 | 114570164 | 43.41453 | 148.15424 | 142.93486 | chr5 | 114566781 | 114568433 | 3.19402 | 3.08382 | 1.50672 |
| chr5 | 114591180 | 114591586 | 182 | Intron (ENSMUST00000043650.7 | 5 | 114568017 | 114594678 | 26662 | 1 | ENSMUST00 | 23164 | Fam222a | ENSMUSG0( | true | . | . | . | . | . | . | chr5 | 114571033 | 114614394 | 7.1045 | 4.9325 | 4.74958 |
| chr2 | 71055582 | 71055863 | 182 | Promoter (<=1kb) | 2 | 71055744 | 71095858 | 40115 | 1 | ENSMUST00 | 0 | Dcaf17 | ENSMUSG0( | false | chr2 | 71054773 | 71056569 | 40.23456 | 136.98251 | 132.21786 | chr2 | 71053912 | 71057105 | 3.55622 | 3.29132 | 1.83446 |
| chr2 | 70802729 | 70803040 | 182 | Intron (ENSMUST00000135128.2 | 2 | 70750994 | 70825228 | 74235 | 2 | ENSMUST00 | 22188 | Tlk1 | ENSMUSG0( | false | . | . | . | . | . | . | chr2 | 70801426 | 70803778 | 5.44744 | 4.17898 | 3.40857 |
| chr3 | 101377012 | 101377185 | 182 | Promoter (<=1kb) | 3 | 101377125 | 101463060 | 85936 | 1 | ENSMUST00 | 0 | Igsf3 | ENSMUSG0( | false | chr3 | 101376757 | 101378196 | 26.30149 | 81.58109 | 78.4027 | . | . | . | . | . | . |
| chr7 | 129514667 | 129514980 | 182 | Distal Intergenic | 7 | 129591863 | 129635738 | 43876 | 1 | ENSMUST00 | -76883 | Wdr11 | ENSMUSG0( | false | . | . | . | . | . | . | . | . | . | . | . | . |
| chr12 | 116047807 | 116048117 | 182 | Promoter (<=1kb) | 12 | 116047829 | 116051625 | 3797 | 1 | ENSMUST00 | 0 | Zfp386 | ENSMUSG0( | false | chr12 | 116047153 | 116049091 | 30.52716 | 100.76356 | 97.17578 | . | . | . | . | . | . |
| chr6 | 117906765 | 117907210 | 182 | Promoter (<=1kb) | 6 | 117906790 | 117923798 | 17009 | 1 | ENSMUST00 | 0 | Hnrnpf | ENSMUSG0( | false | chr6 | 117906713 | 117908804 | 44.72366 | 151.575 | 146.22792 | . | . | . | . | . | . |
| chr9 | 107985714 | 107986143 | 182 | Promoter (<=1kb) | 9 | 107984227 | 107985879 | 1653 | 2 | ENSMUST00 | 0 | Fam212a | ENSMUSG0( | false | chr9 | 107984961 | 107986547 | 27.03504 | 88.5911 | 85.27629 | chr9 | 107984326 | 107986831 | 3.179 | 3.1386 | 1.55862 |
| chr15 | 81936332 | 81936799 | 182 | Promoter (<=1kb) | 15 | 81936759 | 81950941 | 14183 | 1 | ENSMUST00 | 0 | Csdc2 | ENSMUSG0( | false | chr15 | 81936283 | 81937895 | 13.91144 | 32.98249 | 30.47944 | chr15 | 81935029 | 81939977 | 4.75257 | 3.93487 | 2.80237 |
| chr1 | 132298446 | 132299187 | 182 | Promoter (<=1kb) | 1 | 132298626 | 132307357 | 8732 | 1 | ENSMUST00 | 0 | Klhdc8a | ENSMUSG0( | false | chr1 | 132298350 | 132299550 | 19.74353 | 71.90538 | 68.88615 | chr1 | 132297839 | 132299518 | 3.47448 | 3.2271 | 1.80909 |
| chr4 | 155777108 | 155777398 | 182 | Promoter (2-3kb) | 4 | 155768149 | 155774698 | 6550 | 2 | ENSMUST00 | -2411 | Vwa1 | ENSMUSG0( | false | . | . | . | . | . | . | chr4 | 155772708 | 155782290 | 6.11654 | 4.55391 | 3.92825 |
| chr1 | 134555674 | 134556099 | 182 | Distal Intergenic | 1 | 134560171 | 134635285 | 75115 | 1 | ENSMUST00 | -4072 | Kdm5b | ENSMUSG0( | false | chr1 | 134555703 | 134557642 | 19.89052 | 67.91473 | 64.9535 | chr1 | 134555094 | 134557561 | 3.91564 | 3.53609 | 2.12925 |
| chr1 | 33757478 | 33757920 | 182 | Promoter (<=1kb) | 1 | 33745484 | 33757795 | 12312 | 2 | ENSMUST00 | 0 | Bag2 | ENSMUSG0( | false | chr1 | 33757109 | 33758128 | 29.63113 | 94.2009 | 90.76292 | . | . | . | . | . | . |
| chr13 | 115090138 | 115090393 | 182 | Promoter (<=1kb) | 13 | 115088357 | 115090188 | 1832 | 2 | ENSMUST00 | 0 | Pelo | ENSMUSG0( | false | chr13 | 115088957 | 115091242 | 22.30817 | 62.01665 | 59.14301 | . | . | . | . | . | . |
| chr13 | 115054705 | 115055012 | 182 | Intron (ENSMUST00000061673.7 | 13 | 115088357 | 115090188 | 1832 | 2 | ENSMUST00 | 35176 | Pelo | ENSMUSG0( | false | chr13 | 115054456 | 115055124 | 6.75876 | 12.51203 | 10.2897 | . | . | . | . | . | . |
| chr15 | 81181348 | 81181557 | 182 | Intron (ENSMUST00000134469.2 | 15 | 81045133 | 81190522 | 145390 | 2 | ENSMUST00 | 8965 | Mkl1 | ENSMUSG0( | false | . | . | . | . | . | . | chr15 | 81178114 | 81182466 | 4.31673 | 3.46826 | 2.49255 |
| chr9 | 48495208 | 48495443 | 182 | Promoter (<=1kb) | 9 | 48495345 | 48495964 | 620 | 1 | ENSMUST00 | 0 | Gm5617 | ENSMUSG0( | false | chr9 | 48494451 | 48496186 | 36.53257 | 122.84278 | 118.60545 | . | . | . | . | . | . |
| chr1 | 134361705 | 134362089 | 182 | Promoter (<=1kb) | 1 | 134346097 | 134361979 | 15883 | 2 | ENSMUST00 | 0 | Tmem183a | ENSMUSG0( | false | chr1 | 134361026 | 134362594 | 36.10714 | 116.78466 | 112.7497 | . | . | . | . | . | . |
| chr13 | 113616115 | 113616317 | 182 | Promoter (2-3kb) | 13 | 113592180 | 113618564 | 26385 | 2 | ENSMUST00 | 2247 | Snx18 | ENSMUSG0( | false | chr13 | 113614178 | 113618647 | 32.67316 | 105.57997 | 101.8659 | chr13 | 113595158 | 113617709 | 5.59629 | 4.30467 | 3.50767 |
| chr2 | 76369938 | 76370148 | 182 | Promoter (<=1kb) | 2 | 76369984 | 76383768 | 13785 | 1 | ENSMUST00 | 0 | Rbm45 | ENSMUSG0( | false | chr2 | 76369918 | 76370856 | 33.65816 | 114.56059 | 110.59126 | . | . | . | . | . | . |
| chr2 | 75832099 | 75832649 | 182 | Promoter (<=1kb) | 2 | 75832177 | 75931350 | 99174 | 1 | ENSMUST00 | 0 | Agps | ENSMUSG0( | false | chr2 | 75831720 | 75833180 | 21.73903 | 87.55687 | 84.25908 | chr2 | 75831689 | 75834108 | 4.53886 | 3.92525 | 2.62888 |
| chr9 | 75436038 | 75436272 | 182 | Distal Intergenic | 9 | 75441524 | 75466432 | 24909 | 1 | ENSMUST00 | -5252 | Leo1 | ENSMUSG0( | false | . | . | . | . | . | . | chr9 | 75434049 | 75437217 | 5.50176 | 4.33886 | 3.42079 |
| chr7 | 127208213 | 127208775 | 182 | Promoter (<=1kb) | 7 | 127197465 | 127208468 | 11004 | 2 | ENSMUST00 | 0 | Tbc1d10b | ENSMUSG0( | false | chr7 | 127206978 | 127208714 | 27.03867 | 103.18376 | 99.53125 | . | . | . | . | . | . |
| chr4 | 126533204 | 126533582 | 182 | Promoter (<=1kb) | 4 | 126489541 | 126533472 | 43932 | 2 | ENSMUST00 | 0 | Ago4 | ENSMUSG0( | false | chr4 | 126532569 | 126534044 | 23.17719 | 89.67316 | 86.33752 | . | . | . | . | . | . |
| chr6 | 6955855 | 6956305 | 182 | Promoter (<=1kb) | 6 | 6956004 | 7042979 | 86976 | 1 | ENSMUST00 | 0 | Acn9 | ENSMUSG0( | false | chr6 | 6955350 | 6956778 | 25.27427 | 69.73028 | 66.74294 | . | . | . | . | . | . |
| chr11 | 61175370 | 61175627 | 182 | Promoter (<=1kb) | 11 | 61151785 | 61175055 | 23271 | 2 | ENSMUST00 | -316 | Usp22 | ENSMUSG0( | false | . | . | . | . | . | . | . | . | . | . | . | . |
| chr11 | 61022158 | 61022593 | 182 | Promoter (<=1kb) | 11 | 61022564 | 61071131 | 48568 | 1 | ENSMUST00 | 0 | Kcnj12 | ENSMUSG0( | false | chr11 | 61021904 | 61023783 | 19.61815 | 75.94017 | 72.85583 | chr11 | 61021835 | 61022972 | 3.01092 | 2.94417 | 1.41918 |
| chr9 | 57076141 | 57076406 | 182 | Promoter (<=1kb) | 9 | 57076376 | 57128364 | 51989 | 1 | ENSMUST00 | 0 | Sin3a | ENSMUSG0( | false | chr9 | 57076296 | 57077594 | 27.34946 | 104.78247 | 101.09169 | . | . | . | . | . | . |
| chr5 | 121838761 | 121839157 | 182 | Promoter (1-2kb) | 5 | 121815648 | 121836859 | 21212 | 2 | ENSMUST00 | -1903 | Sh2b3 | ENSMUSG0( | false | chr5 | 121838190 | 121839252 | 18.42046 | 46.41892 | 43.75084 | chr5 | 121837267 | 121839323 | 5.65924 | 4.22393 | 3.56804 |
| chr5 | 121837509 | 121837863 | 182 | Promoter (<=1kb) | 5 | 121815648 | 121836859 | 21212 | 2 | ENSMUST00 | -651 | Sh2b3 | ENSMUSG0( | false | chr5 | 121835543 | 121838063 | 34.66576 | 107.00249 | 103.24738 | chr5 | 121837267 | 121839323 | 5.65924 | 4.22393 | 3.56804 |
| chr15 | 79359896 | 79360297 | 182 | Downstream (<1kb) | 15 | 79348061 | 79359072 | 11012 | 1 | ENSMUST00 | 11836 | Maff | ENSMUSG0( | false | . | . | . | . | . | . | chr15 | 79357019 | 79362227 | 4.03994 | 3.43272 | 2.25642 |
| chr15 | 79347827 | 79348435 | 182 | Promoter (<=1kb) | 15 | 79348061 | 79359072 | 11012 | 1 | ENSMUST00 | 0 | Maff | ENSMUSG0( | false | chr15 | 79347265 | 79348573 | 32.68175 | 111.76729 | 107.88066 | chr15 | 79347113 | 79349118 | 3.71452 | 3.23574 | 1.97575 |
| chr15 | 79328302 | 79328542 | 182 | Promoter (<=1kb) | 15 | 79308854 | 79328390 | 19537 | 2 | ENSMUST00 | 0 | Pla2g6 | ENSMUSG0( | false | chr15 | 79327467 | 79328736 | 19.14982 | 55.63055 | 52.84086 | chr15 | 79322453 | 79328725 | 4.68 | 3.859 | 2.75171 |
| chr7 | 126776786 | 126777155 | 182 | Promoter (<=1kb) | 7 | 126776955 | 126780514 | 3560 | 1 | ENSMUST00 | 0 | Ypel3 | ENSMUSG0( | false | chr7 | 126776355 | 126777937 | 31.45397 | 113.65508 | 109.71495 | chr7 | 126774405 | 126781721 | 5.58029 | 4.28527 | 3.5405 |
| chr18 | 21201842 | 21202067 | 182 | Intron (ENSMUST00000049260.6 | 18 | 21127201 | 21300138 | 172938 | 2 | ENSMUST00 | 98071 | Garem | ENSMUSG0( | false | . | . | . | . | . | . | . | . | . | . | . | . |
| chr1 | 153543095 | 153543310 | 182 | Intron (ENSMUST00000041874.8 | 1 | 153548422 | 153549711 | 1290 | 2 | ENSMUST00 | 6401 | Npl | ENSMUSG0( | false | . | . | . | . | . | . | chr1 | 153539148 | 153550339 | 5.26804 | 4.15937 | 3.23842 |
| chr19 | 47549285 | 47549498 | 182 | Distal Intergenic | 19 | 47524707 | 47537507 | 12801 | 2 | ENSMUST00 | -11779 | Obfc1 | ENSMUSG0( | false | . | . | . | . | . | . | chr19 | 47548672 | 47550070 | 5.04098 | 4.15961 | 3.04432 |
| chr2 | 152725389 | 152725658 | 182 | Distal Intergenic | 2 | 152736251 | 152737410 | 1160 | 1 | ENSMUST00 | -10593 | Id1 | ENSMUSG0( | false | . | . | . | . | . | . | chr2 | 152723891 | 152727757 | 5.10737 | 3.93331 | 3.13758 |
| chr7 | 97788343 | 97789214 | 182 | Distal Intergenic | 7 | 97726379 | 97738247 | 11869 | 2 | ENSMUST00 | -50097 | Aqp11 | ENSMUSG0( | false | chr7 | 97788076 | 97789484 | 38.29027 | 120.57137 | 116.41801 | . | . | . | . | . | . |
| chr2 | 167543921 | 167544346 | 182 | Distal Intergenic | 2 | 167538195 | 167542814 | 4620 | 1 | ENSMUST00 | 5727 | Snai1 | ENSMUSG0( | false | . | . | . | . | . | . | chr2 | 167541955 | 167544771 | 7.69518 | 5.21237 | 5.24384 |
| chr8 | 75033554 | 75034044 | 182 | Promoter (<=1kb) | 8 | 75033686 | 75070121 | 36436 | 1 | ENSMUST00 | 0 | Tom1 | ENSMUSG0( | false | chr8 | 75033611 | 75034936 | 29.09105 | 93.24493 | 89.8345 | . | . | . | . | . | . |
| chr14 | 101021591 | 101021779 | 182 | Distal Intergenic | 14 | 101197690 | 101200069 | 2380 | 2 | ENSMUST00 | 178290 | Prr30 | ENSMUSG0( | false | . | . | . | . | . | . | . | . | . | . | . | . |
| chr18 | 10414765 | 10414996 | 182 | Intron (ENSMUST00000173356.2 | 18 | 10458225 | 10538289 | 80065 | 1 | ENSMUST00 | -43229 | Greb1l | ENSMUSG0( | false | . | . | . | . | . | . | chr18 | 10413420 | 10414999 | 3.23488 | 3.05509 | 1.58843 |
| chr15 | 7241015 | 7241214 | 182 | Intron (ENSMUST00000058593.4 | 15 | 7253511 | 7254768 | 1258 | 2 | ENSMUST00 | 13554 | Egflam | ENSMUSG0( | false | . | . | . | . | . | . | chr15 | 7240637 | 7241762 | 4.04261 | 3.60302 | 2.24325 |
| chr7 | 126283094 | 126283354 | 182 | Intron (ENSMUST00000056028.9 | 7 | 126272619 | 126294999 | 22381 | 1 | ENSMUST00 | 10476 | Sbk1 | ENSMUSG0( | false | . | . | . | . | . | . | chr7 | 126282178 | 126286434 | 4.60436 | 3.79502 | 2.72929 |
| chr4 | 35099940 | 35100274 | 182 | Intron (ENSMUST00000102975.4 | 4 | 35152056 | 35154005 | 1950 | 1 | ENSMUST00 | -51782 | Ifnk | ENSMUSG0( | false | chr4 | 35099818 | 35100578 | 12.92336 | 34.59913 | 32.07519 | chr4 | 35099244 | 35100562 | 4.47934 | 3.70385 | 2.60936 |
| chr7 | 16909975 | 16910286 | 182 | 3' UTR | 7 | 16906514 | 16910900 | 4387 | 1 | ENSMUST00 | 3462 | Ptgir | ENSMUSG0( | false | . | . | . | . | . | . | chr7 | 16900106 | 16910944 | 5.45499 | 4.23801 | 3.39476 |
| chr5 | 122136905 | 122137095 | 182 | Promoter (<=1kb) | 5 | 122108040 | 122138062 | 30023 | 2 | ENSMUST00 | 967 | Ccdc63 | ENSMUSG0( | false | . | . | . | . | . | . | chr5 | 122135320 | 122137818 | 5.7733 | 4.46559 | 3.63807 |
| chr16 | 44347341 | 44347788 | 182 | Promoter (<=1kb) | 16 | 44347414 | 44370768 | 23355 | 1 | ENSMUST00 | 0 | Spice1 | ENSMUSG0( | false | chr16 | 44346842 | 44348178 | 21.75525 | 76.8157 | 73.7174 | . | . | . | . | . | . |
| chr18 | 60596140 | 60596471 | 182 | Exon (ENSMUST00000097566.4/ | 18 | 60593991 | 60600497 | 6507 | 2 | ENSMUST00 | 4026 | Synpo | ENSMUSG0( | true | chr18 | 60596246 | 60596759 | 10.52322 | 21.7476 | 19.38849 | chr18 | 60593903 | 60608752 | 6.45808 | 4.76452 | 4.19811 |
| chr18 | 60651920 | 60652330 | 182 | Intron (ENSMUST00000137894.1 | 18 | 60600209 | 60648319 | 48111 | 2 | ENSMUST00 | -3602 | Synpo | ENSMUSG0( | true | . | . | . | . | . | . | chr18 | 60651799 | 60652445 | 4.04647 | 3.53769 | 2.22366 |
| chr11 | 75168152 | 75168371 | 182 | Promoter (<=1kb) | 11 | 75167153 | 75168146 | 994 | 2 | ENSMUST00 | -7 | Hic1 | ENSMUSG0( | false | chr11 | 75168275 | 75169318 | 17.76238 | 66.08042 | 63.15025 | chr11 | 75165128 | 75174332 | 5.71251 | 4.38871 | 3.59566 |
| chr4 | 116821454 | 116821894 | 182 | Promoter (<=1kb) | 4 | 116819904 | 116821707 | 1804 | 2 | ENSMUST00 | 0 | Hpdl | ENSMUSG0( | false | chr4 | 116820878 | 116821929 | 20.9555 | 57.98476 | 55.16582 | chr4 | 116821041 | 116822210 | 3.62345 | 3.16773 | 1.90909 |
| chr6 | 57691931 | 57692209 | 182 | Promoter (<=1kb) | 6 | 57686576 | 57692078 | 5503 | 2 | ENSMUST00 | 0 | Pyurf | ENSMUSG0( | false | chr6 | 57691329 | 57692533 | 24.65524 | 104.07409 | 100.40247 | . | . | . | . | . | . |
| chr13 | 76003272 | 76003529 | 182 | Intron (ENSMUST00000050997.1 | 13 | 76001535 | 76018562 | 17028 | 2 | ENSMUST00 | 15033 | Rfesd | ENSMUSG0( | false | . | . | . | . | . | . | chr13 | 76002882 | 76004160 | 4.33641 | 3.78411 | 2.5062 |
| chr6 | 52208750 | 52209180 | 182 | Promoter (<=1kb) | 6 | 52206288 | 52208722 | 2435 | 2 | ENSMUST00 | -29 | Hoxa6 | ENSMUSG0( | false | chr6 | 52208245 | 52210010 | 10.58711 | 29.50236 | 27.04178 | chr6 | 52203796 | 52211563 | 4.96251 | 3.88667 | 3.0102 |
| chr4 | 133670372 | 133670608 | 182 | Promoter (2-3kb) | 4 | 133660387 | 133672647 | 12261 | 2 | ENSMUST00 | 2039 | Pigv | ENSMUSG0( | false | chr4 | 133670509 | 133673626 | 34.67549 | 114.27542 | 110.31616 | chr4 | 133666195 | 133677441 | 5.95677 | 4.52886 | 3.78779 |
| chr11 | 103144666 | 103144892 | 182 | Distal Intergenic | 11 | 103133865 | 103138279 | 4415 | 1 | ENSMUST00 | 10802 | Hexim2 | ENSMUSG0( | false | . | . | . | . | . | . | chr11 | 103138329 | 103150034 | 6.87245 | 4.95791 | 4.53389 |
| chr11 | 103132742 | 103133082 | 182 | Promoter (<=1kb) | 11 | 103133314 | 103139191 | 5878 | 1 | ENSMUST00 | -232 | Hexim2 | ENSMUSG0( | false | chr11 | 103131357 | 103132961 | 39.90052 | 130.90422 | 126.38404 | . | . | . | . | . | . |
| chr4 | 137594059 | 137594341 | 182 | Promoter (<=1kb) | 4 | 137594140 | 137658531 | 64392 | 1 | ENSMUST00 | 0 | Usp48 | ENSMUSG0( | false | chr4 | 137593688 | 137595481 | 33.47463 | 112.14942 | 108.24687 | . | . | . | . | . | . |
| chr4 | 137610439 | 137610701 | 182 | Exon (ENSMUST00000105840.2/ | 4 | 137618471 | 137625291 | 6821 | 1 | ENSMUST00 | -7770 | Usp48 | ENSMUSG0( | false | . | . | . | . | . | . | chr4 | 137609125 | 137610531 | 3.85195 | 3.54675 | 2.07915 |
| chr1 | 182670666 | 182670841 | 182 | Distal Intergenic | 1 | 182747126 | 182749180 | 2055 | 2 | ENSMUST00 | 78339 | Ccdc185 | ENSMUSG0( | false | . | . | . | . | . | . | chr1 | 182669635 | 182671287 | 3.21888 | 3.13802 | 1.57447 |
| chrX | 161717260 | 161717522 | 182 | Promoter (<=1kb) | 20 | 161717627 | 161779496 | 61870 | 1 | ENSMUST00 | -105 | Rai2 | ENSMUSG0( | false | chrX | 161716902 | 161718697 | 18.02336 | 51.19578 | 48.46415 | . | . | . | . | . | . |
| chr19 | 50678563 | 50678938 | 182 | Promoter (<=1kb) | 19 | 50143301 | 50678646 | 535346 | 2 | ENSMUST00 | 0 | Sorcs1 | ENSMUSG0( | false | chr19 | 50677906 | 50679093 | 20.34299 | 52.83396 | 50.08234 | chr19 | 50677998 | 50678888 | 5.75215 | 4.43755 | 3.68242 |
| chr17 | 29888095 | 29888373 | 182 | Promoter (<=1kb) | 17 | 29827958 | 29887882 | 59925 | 2 | ENSMUST00 | -214 | Mdga1 | ENSMUSG0( | false | chr17 | 29886537 | 29888883 | 22.5292 | 69.37365 | 66.39268 | chr17 | 29886337 | 29890016 | 6.22314 | 4.58529 | 4.05182 |
| chr19 | 60212602 | 60212808 | 182 | Exon (ENSMUST00000065286.1/ | 19 | 60226334 | 60227678 | 1345 | 1 | ENSMUST00 | -13526 | 4933412A08 | ENSMUSG0( | false | . | . | . | . | . | . | chr19 | 60209053 | 60220176 | 4.2874 | 3.68168 | 2.43595 |
| chr8 | 90348077 | 90348618 | 182 | Promoter (<=1kb) | 8 | 90247040 | 90348343 | 101304 | 2 | ENSMUST00 | 0 | Tox3 | ENSMUSG0( | false | chr8 | 90347017 | 90349010 | 26.78979 | 82.44355 | 79.24416 | chr8 | 90346476 | 90349964 | 4.81693 | 4.00968 | 2.87842 |
| chr5 | 121183814 | 121184056 | 182 | Intron (ENSMUST00000100770.3 | 5 | 121130535 | 121191367 | 60833 | 2 | ENSMUST00 | 7311 | Ptpn11 | ENSMUSG0( | false | . | . | . | . | . | . | chr5 | 121179014 | 121190676 | 5.96142 | 4.50958 | 3.80604 |
| chr5 | 121190828 | 121191571 | 182 | Promoter (<=1kb) | 5 | 121130533 | 121191397 | 60865 | 2 | ENSMUST00 | 0 | Ptpn11 | ENSMUSG0( | false | chr5 | 121190294 | 121191978 | 36.25188 | 118.09364 | 114.014 | . | . | . | . | . | . |
| chr4 | 127061040 | 127061317 | 182 | Promoter (<=1kb) | 4 | 127047094 | 127061152 | 14059 | 2 | ENSMUST00 | 0 | Zmym1 | ENSMUSG0( | false | chr4 | 127060456 | 127061634 | 32.00811 | 98.35596 | 94.82688 | chr4 | 127060108 | 127061675 | 5.08607 | 4.12447 | 3.07149 |
| chr5 | 101933396 | 101933730 | 182 | Promoter (2-3kb) | 5 | 101922817 | 101930853 | 8037 | 2 | ENSMUST00 | -2544 | Wdfy3 | ENSMUSG0( | false | . | . | . | . | . | . | chr5 | 101932371 | 101933836 | 4.52314 | 3.76465 | 2.64814 |
| chr5 | 101978135 | 101978317 | 182 | Intron (ENSMUST00000053177.8 | 5 | 101836158 | 101981558 | 145401 | 2 | ENSMUST00 | 3241 | Wdfy3 | ENSMUSG0( | false | . | . | . | . | . | . | chr5 | 101978024 | 101979375 | 3.45219 | 3.14572 | 1.77328 |
| chr18 | 36240007 | 36240213 | 182 | Distal Intergenic | 18 | 36281097 | 36301521 | 20425 | 1 | ENSMUST00 | -40884 | Pura | ENSMUSG0( | false | . | . | . | . | . | . | chr18 | 36239423 | 36240491 | 5.42166 | 4.35363 | 3.34208 |
| chr2 | 113440695 | 113440968 | 182 | Promoter (<=1kb) | 2 | 113441064 | 113709855 | 268792 | 1 | ENSMUST00 | -96 | Fmn1 | ENSMUSG0( | false | chr2 | 113440160 | 113441633 | 16.14593 | 43.98602 | 41.34693 | chr2 | 113439638 | 113445640 | 4.21191 | 3.72328 | 2.35219 |
| chr1 | 92680827 | 92681051 | 182 | Distal Intergenic | 1 | 92644218 | 92648841 | 4624 | 2 | ENSMUST00 | -31987 | Otos | ENSMUSG0( | false | . | . | . | . | . | . | chr1 | 92680001 | 92684688 | 6.40462 | 4.67491 | 4.16393 |
| chr11 | 94243142 | 94243528 | 182 | Promoter (<=1kb) | 11 | 94235956 | 94242707 | 6752 | 2 | ENSMUST00 | -436 | Wfikkn2 | ENSMUSG0( | false | chr11 | 94242877 | 94243767 | 8.16212 | 17.69099 | 15.38708 | chr11 | 94242522 | 94244888 | 5.60247 | 4.27771 | 3.53253 |
| chr9 | 100470942 | 100471123 | 182 | Promoter (<=1kb) | 9 | 100459077 | 100470609 | 11533 | 2 | ENSMUST00 | -334 | Il20rb | ENSMUSG0( | false | . | . | . | . | . | . | chr9 | 100470864 | 100471627 | 5.08303 | 4.21172 | 3.06501 |
| chr18 | 12941602 | 12942019 | 182 | Promoter (<=1kb) | 18 | 12755314 | 12941841 | 186528 | 2 | ENSMUST00 | 0 | Osbpl1a | ENSMUSG0( | false | chr18 | 12941134 | 12942141 | 44.57462 | 145.65463 | 140.53799 | . | . | . | . | . | . |
| chr5 | 114127570 | 114127856 | 182 | Promoter (<=1kb) | 5 | 114123926 | 114128176 | 4251 | 2 | ENSMUST00 | 320 | Alkbh2 | ENSMUSG0( | false | chr5 | 114127243 | 114128311 | 25.76091 | 81.55552 | 78.37826 | . | . | . | . | . | . |
| chrX | 137038117 | 137038390 | 182 | Promoter (<=1kb) | 20 | 136981116 | 137038300 | 57185 | 2 | ENSMUST00 | 0 | Slc25a53 | ENSMUSG0( | false | chrX | 137037403 | 137038705 | 17.92175 | 44.05972 | 41.41886 | . | . | . | . | . | . |
| chr16 | 87354174 | 87354404 | 182 | Promoter (<=1kb) | 16 | 87354185 | 87368742 | 14558 | 1 | ENSMUST00 | 0 | N6amt1 | ENSMUSG0( | false | chr16 | 87353948 | 87354949 | 37.09717 | 122.76616 | 118.52905 | chr16 | 87353636 | 87356917 | 4.1583 | 3.60481 | 2.33606 |
| chr12 | 102286828 | 102287206 | 182 | Intron (ENSMUST00000056950.8 | 12 | 102283641 | 102390855 | 107215 | 1 | ENSMUST00 | 3188 | Rin3 | ENSMUSG0( | false | . | . | . | . | . | . | chr12 | 102282553 | 102299505 | 4.61003 | 3.86052 | 2.6933 |
| chr6 | 31197783 | 31198047 | 182 | Exon (ENSMUST00000115107.2/ | 6 | 31166524 | 31218346 | 51823 | 2 | ENSMUST00 | 20299 | Lincpint | ENSMUSG0( | false | . | . | . | . | . | . | chr6 | 31172537 | 31202809 | 5.30919 | 4.15007 | 3.26883 |
| chr3 | 121640652 | 121640891 | 182 | Promoter (2-3kb) | 3 | 121637145 | 121643485 | 6341 | 2 | ENSMUST00 | 2594 | A730020M07 | ENSMUSG0( | false | . | . | . | . | . | . | . | . | . | . | . | . |
| chr10 | 120764187 | 120764386 | 182 | Promoter (1-2kb) | 10 | 120765774 | 120778886 | 13113 | 1 | ENSMUST00 | -1388 | 4921513I03R | ENSMUSG0( | false | . | . | . | . | . | . | . | . | . | . | . | . |
| chr12 | 71309602 | 71309927 | 182 | Promoter (<=1kb) | 12 | 71309884 | 71320107 | 10224 | 1 | ENSMUST00 | 0 | Dact1 | ENSMUSG0( | false | chr12 | 71309332 | 71310789 | 20.5121 | 70.85588 | 67.85384 | chr12 | 71309434 | 71311799 | 3.50682 | 3.17858 | 1.82499 |
| chr8 | 72571148 | 72571516 | 182 | Promoter (<=1kb) | 8 | 72565188 | 72571088 | 5901 | 2 | ENSMUST00 | -61 | Smim7 | ENSMUSG0( | false | chr8 | 72570135 | 72572933 | 22.15371 | 67.81625 | 64.85734 | . | . | . | . | . | . |
| chr10 | 99380175 | 99380475 | 182 | Intron (ENSMUST00000189035.1 | 10 | 99420522 | 99423047 | 2526 | 1 | ENSMUST00 | -40047 | B530045E10 | ENSMUSG0( | false | . | . | . | . | . | . | . | . | . | . | . | . |
| chr2 | 168078435 | 168078731 | 182 | Promoter (2-3kb) | 2 | 168081004 | 168101203 | 20200 | 1 | ENSMUST00 | -2273 | Pard6b | ENSMUSG0( | false | . | . | . | . | . | . | chr2 | 168063587 | 168080595 | 6.53927 | 4.82888 | 4.26326 |
| chr17 | 50624985 | 50625162 | 182 | Intron (ENSMUST00000043938.6 | 17 | 50698525 | 50699880 | 1356 | 1 | ENSMUST00 | -73363 | Gm7334 | ENSMUSG0( | false | . | . | . | . | . | . | chr17 | 50623168 | 50627324 | 6.11363 | 4.44066 | 3.94765 |
| chr18 | 76052795 | 76053009 | 182 | Intron (ENSMUST00000058997.8 | 18 | 76059458 | 76146443 | 86986 | 1 | ENSMUST00 | -6449 | Zbtb7c | ENSMUSG0( | true | . | . | . | . | . | . | chr18 | 76050774 | 76058938 | 6.53532 | 4.72087 | 4.27307 |
| chr18 | 76078461 | 76078646 | 182 | Intron (ENSMUST00000058997.8 | 18 | 76059458 | 76146443 | 86986 | 1 | ENSMUST00 | 19004 | Zbtb7c | ENSMUSG0( | true | . | . | . | . | . | . | chr18 | 76061445 | 76132293 | 6.70015 | 4.79833 | 4.4158 |
| chr18 | 75874337 | 75874632 | 182 | Intron (ENSMUST00000058997.8 | 18 | 75820178 | 76148561 | 328384 | 1 | ENSMUST00 | 54160 | Zbtb7c | ENSMUSG0( | true | . | . | . | . | . | . | chr18 | 75873296 | 75875508 | 4.67256 | 3.97514 | 2.72241 |
| chr5 | 4595088 | 4595273 | 182 | Distal Intergenic | 5 | 4753873 | 4758035 | 4163 | 2 | ENSMUST00 | 162762 | Fzd1 | ENSMUSG0( | false | . | . | . | . | . | . | . | . | . | . | . | . |
| chr9 | 51278409 | 51278875 | 182 | Promoter (<=1kb) | 9 | 51278513 | 51284210 | 5698 | 1 | ENSMUST00 | 0 | 2010007H06 | ENSMUSG0( | false | chr9 | 51278067 | 51279570 | 26.44925 | 85.62254 | 82.36654 | chr9 | 51277482 | 51279701 | 3.84213 | 3.36473 | 2.08855 |
| chr11 | 43528439 | 43529087 | 182 | Promoter (<=1kb) | 11 | 43528784 | 43586997 | 58214 | 1 | ENSMUST00 | 0 | Ccnjl | ENSMUSG0( | false | chr11 | 43528218 | 43529735 | 30.91854 | 102.46954 | 98.8397 | chr11 | 43528200 | 43530406 | 3.49707 | 3.33368 | 1.77001 |
| chr7 | 28383083 | 28383473 | 182 | Downstream (2-3kb) | 7 | 28376784 | 28379255 | 2472 | 2 | ENSMUST00 | -3829 | Zfp36 | ENSMUSG0( | false | . | . | . | . | . | . | chr7 | 28375103 | 28383860 | 6.23801 | 4.67039 | 4.01439 |
| chr9 | 110611492 | 110611729 | 182 | Intron (ENSMUST00000153838.3 | 9 | 110602174 | 110618148 | 15975 | 1 | ENSMUST00 | 9319 | Setd2 | ENSMUSG0( | false | . | . | . | . | . | . | chr9 | 110604849 | 110618387 | 5.08369 | 4.06361 | 3.09341 |
| chr4 | 45529880 | 45531229 | 182 | Promoter (<=1kb) | 4 | 45423278 | 45530330 | 107053 | 2 | ENSMUST00 | 0 | Shb | ENSMUSG0( | false | chr4 | 45528278 | 45531712 | 31.40916 | 110.4849 | 106.632 | chr4 | 45530927 | 45532459 | 3.72102 | 3.38037 | 2.00084 |
| chr4 | 45489801 | 45490164 | 182 | Promoter (<=1kb) | 4 | 45458129 | 45489794 | 31666 | 2 | ENSMUST00 | -8 | Shb | ENSMUSG0( | false | . | . | . | . | . | . | chr4 | 45486900 | 45516085 | 7.16373 | 5.03069 | 4.78278 |
| chr3 | 38484666 | 38485255 | 182 | Promoter (<=1kb) | 3 | 38449261 | 38484844 | 35584 | 2 | ENSMUST00 | 0 | Ankrd50 | ENSMUSG0( | false | chr3 | 38485055 | 38485573 | 31.01414 | 97.6227 | 94.10706 | . | . | . | . | . | . |
| chr7 | 6171944 | 6172562 | 182 | Promoter (<=1kb) | 7 | 6172226 | 6183951 | 11726 | 1 | ENSMUST00 | 0 | Zfp444 | ENSMUSG0( | false | chr7 | 6171255 | 6173219 | 22.90469 | 70.93823 | 67.93287 | . | . | . | . | . | . |
| chr13 | 64152014 | 64152767 | 182 | Promoter (<=1kb) | 13 | 64135403 | 64152429 | 17027 | 2 | ENSMUST00 | 0 | Zfp367 | ENSMUSG0( | false | chr13 | 64151825 | 64153338 | 35.31382 | 118.55977 | 114.46947 | chr13 | 64151346 | 64152258 | 5.25485 | 4.30295 | 3.2136 |
| chr11 | 43682520 | 43682885 | 182 | Promoter (<=1kb) | 11 | 43682548 | 43684054 | 1507 | 1 | ENSMUST00 | 0 | Pwwp2a | ENSMUSG0( | false | chr11 | 43681948 | 43683166 | 31.19374 | 94.95832 | 91.51084 | . | . | . | . | . | . |
| chr11 | 43681561 | 43682305 | 182 | Promoter (<=1kb) | 11 | 43681998 | 43707191 | 25194 | 1 | ENSMUST00 | 0 | Pwwp2a | ENSMUSG0( | false | chr11 | 43681095 | 43681749 | 17.64787 | 49.73517 | 47.02245 | . | . | . | . | . | . |
| chr7 | 97332093 | 97332616 | 182 | Promoter (<=1kb) | 7 | 97332327 | 97350213 | 17887 | 1 | ENSMUST00 | 0 | Kctd21 | ENSMUSG0( | false | chr7 | 97331532 | 97333065 | 36.6152 | 115.56235 | 111.55841 | chr7 | 97330997 | 97334609 | 4.63952 | 3.87645 | 2.70315 |
| chr9 | 74156805 | 74157260 | 182 | Exon (ENSMUST00000055879.8/ | 9 | 74110404 | 74113679 | 3276 | 1 | ENSMUST00 | 46402 | Wdr72 | ENSMUSG0( | false | . | . | . | . | . | . | . | . | . | . | . | . |
| chr1 | 64779978 | 64780215 | 182 | Distal Intergenic | 1 | 64730558 | 64737751 | 7194 | 2 | ENSMUST00 | -42228 | Fzd5 | ENSMUSG0( | false | . | . | . | . | . | . | chr1 | 64779654 | 64780460 | 2.72999 | 2.76879 | 1.20067 |
| chr11 | 101155705 | 101155912 | 182 | Promoter (<=1kb) | 11 | 101155907 | 101161787 | 5881 | 1 | ENSMUST00 | 0 | Tubg2 | ENSMUSG0( | false | chr11 | 101155759 | 101156260 | 12.24318 | 32.0257 | 29.53485 | chr11 | 101155369 | 101156262 | 3.76293 | 3.34639 | 2.02731 |
| chr17 | 86166455 | 86167016 | 182 | Promoter (<=1kb) | 17 | 86167785 | 86657919 | 490135 | 1 | ENSMUST00 | -769 | Prkce | ENSMUSG0( | true | chr17 | 86166332 | 86169593 | 32.75195 | 98.6424 | 95.10313 | . | . | . | . | . | . |
| chr19 | 4615475 | 4616007 | 182 | Promoter (<=1kb) | 19 | 4611785 | 4615667 | 3883 | 2 | ENSMUST00 | 0 | Lrfn4 | ENSMUSG0( | false | chr19 | 4614686 | 4616300 | 28.6362 | 83.76622 | 80.54601 | chr19 | 4614855 | 4617378 | 4.00718 | 3.61262 | 2.17637 |
| chr17 | 83631570 | 83632226 | 182 | Promoter (<=1kb) | 17 | 83585957 | 83631895 | 45939 | 2 | ENSMUST00 | 0 | Kcng3 | ENSMUSG0( | false | chr17 | 83631115 | 83633458 | 24.78212 | 78.86476 | 75.73294 | chr17 | 83630677 | 83633722 | 5.65253 | 4.32635 | 3.55513 |
| chr18 | 61526768 | 61527007 | 182 | Intron (ENSMUST00000171629.1 | 18 | 61493794 | 61536536 | 42743 | 2 | ENSMUST00 | 9529 | Arhgef37 | ENSMUSG0( | false | . | . | . | . | . | . | . | . | . | . | . | . |
| chrX | 152764247 | 152764524 | 182 | Intron (ENSMUST00000101141.3 | 20 | 152609509 | 152769461 | 159953 | 2 | ENSMUST00 | 4937 | Shroom2 | ENSMUSG0( | false | . | . | . | . | . | . | . | . | . | . | . | . |
| chr10 | 80159145 | 80159393 | 182 | Distal Intergenic | 10 | 80165985 | 80170316 | 4332 | 1 | ENSMUST00 | -6592 | Cirbp | ENSMUSG0( | false | chr10 | 80159097 | 80159881 | 9.74399 | 19.38118 | 17.0552 | chr10 | 80150859 | 80167428 | 9.09735 | 5.82258 | 6.36001 |
| chr1 | 9748317 | 9748643 | 182 | Promoter (<=1kb) | 1 | 9718622 | 9748382 | 29761 | 2 | ENSMUST00 | 0 | Vcpip1 | ENSMUSG0( | false | chr1 | 9746906 | 9748499 | 37.63617 | 126.94061 | 122.55849 | . | . | . | . | . | . |
| chr14 | 70577600 | 70577829 | 182 | Promoter (<=1kb) | 14 | 70577847 | 70582571 | 4725 | 1 | ENSMUST00 | -18 | Nudt18 | ENSMUSG0( | false | chr14 | 70577252 | 70578497 | 39.37288 | 136.56555 | 131.82602 | . | . | . | . | . | . |
| chr1 | 36017252 | 36017479 | 182 | Distal Intergenic | 1 | 36068400 | 36106446 | 38047 | 1 | ENSMUST00 | -50921 | Hs6st1 | ENSMUSG0( | false | . | . | . | . | . | . | chr1 | 36012244 | 36019068 | 4.80731 | 3.9122 | 2.86313 |
| chr7 | 127421860 | 127422134 | 182 | Promoter (<=1kb) | 7 | 127418967 | 127422027 | 3061 | 2 | ENSMUST00 | 0 | Zfp688 | ENSMUSG0( | false | chr7 | 127421244 | 127422700 | 27.91253 | 86.3188 | 83.05016 | chr7 | 127421356 | 127423441 | 4.09904 | 3.31895 | 2.29692 |
| chr7 | 127426437 | 127426705 | 182 | Distal Intergenic | 7 | 127418969 | 127422068 | 3100 | 2 | ENSMUST00 | -4370 | Zfp688 | ENSMUSG0( | false | . | . | . | . | . | . | . | . | . | . | . | . |
| chr4 | 119174018 | 119174320 | 182 | Promoter (<=1kb) | 4 | 119169513 | 119174195 | 4683 | 2 | ENSMUST00 | 0 | Zfp691 | ENSMUSG0( | false | chr4 | 119173076 | 119174292 | 28.91859 | 89.0447 | 85.71584 | . | . | . | . | . | . |
| chr11 | 75251217 | 75251505 | 182 | Intron (ENSMUST00000102514.3 | 11 | 75193783 | 75267769 | 73987 | 1 | ENSMUST00 | 57435 | Rtn4rl1 | ENSMUSG0( | false | . | . | . | . | . | . | . | . | . | . | . | . |
| chr11 | 75212246 | 75212543 | 182 | Intron (ENSMUST00000102514.3 | 11 | 75193783 | 75267769 | 73987 | 1 | ENSMUST00 | 18464 | Rtn4rl1 | ENSMUSG0( | false | . | . | . | . | . | . | chr11 | 75211683 | 75215082 | 4.64264 | 3.7272 | 2.72584 |
| chr2 | 6126240 | 6126488 | 182 | Intron (ENSMUST00000054254.6 | 2 | 6101190 | 6130139 | 28950 | 2 | ENSMUST00 | 3651 | Proser2 | ENSMUSG0( | false | . | . | . | . | . | . | chr2 | 6125226 | 6127099 | 4.20499 | 3.5096 | 2.38706 |
| chr8 | 87891911 | 87892135 | 182 | Intron (ENSMUST00000109655.3 | 8 | 87661812 | 87895735 | 233924 | 2 | ENSMUST00 | 3600 | Zfp423 | ENSMUSG0( | true | . | . | . | . | . | . | chr8 | 87891982 | 87896514 | 4.79857 | 3.93021 | 2.87804 |
| chr8 | 87957951 | 87958377 | 182 | Promoter (1-2kb) | 8 | 87661810 | 87959595 | 297786 | 2 | ENSMUST00 | 1218 | Zfp423 | ENSMUSG0( | true | chr8 | 87957648 | 87961172 | 32.66734 | 113.81544 | 109.86345 | chr8 | 87942134 | 87958259 | 5.08353 | 3.95987 | 3.11076 |
| chr8 | 87794653 | 87794861 | 182 | Intron (ENSMUST00000165770.3 | 8 | 87783147 | 87803823 | 20677 | 2 | ENSMUST00 | 8962 | Zfp423 | ENSMUSG0( | true | . | . | . | . | . | . | chr8 | 87794189 | 87796678 | 4.32732 | 3.7058 | 2.47398 |
| chr8 | 87745134 | 87745394 | 182 | Intron (ENSMUST00000165770.3 | 8 | 87783147 | 87803823 | 20677 | 2 | ENSMUST00 | 58429 | Zfp423 | ENSMUSG0( | true | . | . | . | . | . | . | chr8 | 87729485 | 87749127 | 6.31805 | 4.63416 | 4.1004 |
| chr5 | 137741593 | 137741909 | 182 | Promoter (<=1kb) | 5 | 137730913 | 137741607 | 10695 | 2 | ENSMUST00 | 0 | Nyap1 | ENSMUSG0( | false | chr5 | 137740937 | 137742017 | 17.34451 | 52.33425 | 49.58816 | chr5 | 137740899 | 137742013 | 6.07252 | 4.54701 | 3.92694 |
| chr2 | 164460841 | 164461347 | 182 | Promoter (<=1kb) | 2 | 164460971 | 164465508 | 4538 | 1 | ENSMUST00 | 0 | Sys1 | ENSMUSG0( | false | chr2 | 164460919 | 164462087 | 31.43241 | 100.35837 | 96.77719 | . | . | . | . | . | . |
| chr11 | 102948560 | 102948983 | 182 | Promoter (1-2kb) | 11 | 102939264 | 102946688 | 7425 | 2 | ENSMUST00 | -1873 | C1ql1 | ENSMUSG0( | false | chr11 | 102948140 | 102949075 | 18.59898 | 66.17558 | 63.2431 | chr11 | 102945019 | 102949394 | 5.6668 | 4.35893 | 3.56122 |
| chr3 | 104864361 | 104864548 | 182 | Promoter (<=1kb) | 3 | 104864503 | 104926860 | 62358 | 1 | ENSMUST00 | 0 | St7l | ENSMUSG0( | false | chr3 | 104863463 | 104866019 | 34.16929 | 120.7392 | 116.5701 | . | . | . | . | . | . |
| chr3 | 104864961 | 104865150 | 182 | Promoter (<=1kb) | 3 | 104865071 | 104927894 | 62824 | 1 | ENSMUST00 | 0 | St7l | ENSMUSG0( | false | chr3 | 104863463 | 104866019 | 34.16929 | 120.7392 | 116.5701 | . | . | . | . | . | . |
| chr7 | 127233028 | 127233366 | 182 | Promoter (<=1kb) | 7 | 127233061 | 127238179 | 5119 | 1 | ENSMUST00 | 0 | Zfp553 | ENSMUSG0( | false | chr7 | 127232200 | 127233944 | 22.89105 | 88.16154 | 84.85722 | . | . | . | . | . | . |
| chr8 | 40975729 | 40975982 | 182 | Intron (ENSMUST00000034004.9 | 8 | 40998770 | 41000205 | 1436 | 2 | ENSMUST00 | 24223 | Mtus1 | ENSMUSG0( | false | . | . | . | . | . | . | chr8 | 40975804 | 40976186 | 5.00563 | 4.11266 | 3.0037 |
| chr7 | 116308248 | 116308592 | 182 | Promoter (<=1kb) | 7 | 116123493 | 116308376 | 184884 | 2 | ENSMUST00 | 0 | Plekha7 | ENSMUSG0( | false | chr7 | 116307045 | 116309157 | 35.20524 | 115.20684 | 111.21615 | chr7 | 116306793 | 116309625 | 4.02643 | 3.60414 | 2.20051 |
| chr19 | 5929617 | 5929864 | 182 | Distal Intergenic | 19 | 5917556 | 5924816 | 7261 | 2 | ENSMUST00 | -4802 | Cdc42ep2 | ENSMUSG0( | false | . | . | . | . | . | . | chr19 | 5927850 | 5930766 | 4.06173 | 3.38733 | 2.25342 |
| chr11 | 19963802 | 19964107 | 182 | Intron (ENSMUST00000093299.7 | 11 | 19954570 | 20018133 | 63564 | 1 | ENSMUST00 | 9233 | Spred2 | ENSMUSG0( | true | . | . | . | . | . | . | chr11 | 19944374 | 19999086 | 6.36759 | 4.68062 | 4.13462 |
| chr11 | 19969709 | 19969895 | 182 | Intron (ENSMUST00000093299.7 | 11 | 19954570 | 20018133 | 63564 | 1 | ENSMUST00 | 15140 | Spred2 | ENSMUSG0( | true | . | . | . | . | . | . | chr11 | 19944374 | 19999086 | 6.36759 | 4.68062 | 4.13462 |
| chr10 | 22913174 | 22913452 | 182 | Distal Intergenic | 10 | 22817279 | 22820128 | 2850 | 2 | ENSMUST00 | -93047 | Tcf21 | ENSMUSG0( | true | . | . | . | . | . | . | . | . | . | . | . | . |
| chr4 | 146891756 | 146892001 | 182 | Intron (ENSMUST00000181199.1 | 4 | 146892406 | 146909590 | 17185 | 2 | ENSMUST00 | 17589 | Gm21411 | ENSMUSG0( | false | . | . | . | . | . | . | . | . | . | . | . | . |
| chr6 | 29985178 | 29985503 | 182 | Promoter (<=1kb) | 6 | 29985329 | 29993530 | 8202 | 1 | ENSMUST00 | 0 | 1700023L04R | ENSMUSG0( | false | chr6 | 29984813 | 29985641 | 16.52141 | 45.4089 | 42.75185 | chr6 | 29984784 | 29985799 | 3.16984 | 3.10748 | 1.50038 |
| chr5 | 52471284 | 52471685 | 182 | Promoter (<=1kb) | 5 | 52374651 | 52471543 | 96893 | 2 | ENSMUST00 | 0 | Ccdc149 | ENSMUSG0( | false | chr5 | 52470936 | 52471906 | 25.7338 | 71.40817 | 68.39619 | chr5 | 52471146 | 52471903 | 3.24886 | 3.17097 | 1.57544 |
| chr10 | 95145630 | 95146028 | 182 | Distal Intergenic | 10 | 95174746 | 95324097 | 149352 | 2 | ENSMUST00 | 178069 | Cradd | ENSMUSG0( | false | . | . | . | . | . | . | chr10 | 95142298 | 95147251 | 5.59095 | 4.28663 | 3.51913 |
| chr10 | 95323501 | 95323748 | 182 | Promoter (<=1kb) | 10 | 95174746 | 95324097 | 149352 | 2 | ENSMUST00 | 349 | Cradd | ENSMUSG0( | false | chr10 | 95323306 | 95324150 | 24.73908 | 82.49783 | 79.2979 | . | . | . | . | . | . |
| chr11 | 68637919 | 68638133 | 182 | Distal Intergenic | 11 | 68594055 | 68597954 | 3900 | 1 | ENSMUST00 | 43865 | Ccdc42 | ENSMUSG0( | false | . | . | . | . | . | . | chr11 | 68637953 | 68639022 | 2.86876 | 2.91439 | 1.309 |
| chr11 | 68623775 | 68624065 | 182 | Distal Intergenic | 11 | 68594055 | 68597954 | 3900 | 1 | ENSMUST00 | 29721 | Ccdc42 | ENSMUSG0( | false | chr11 | 68623493 | 68623933 | 8.94881 | 17.06333 | 14.77052 | . | . | . | . | . | . |
| chr8 | 11555699 | 11556029 | 182 | Promoter (<=1kb) | 8 | 11556066 | 11563250 | 7185 | 1 | ENSMUST00 | -37 | Ing1 | ENSMUSG0( | false | chr8 | 11554880 | 11558049 | 39.39634 | 133.30592 | 128.68834 | . | . | . | . | . | . |
| chr16 | 20672579 | 20673202 | 182 | Promoter (<=1kb) | 16 | 20672723 | 20692884 | 20162 | 1 | ENSMUST00 | 0 | Eif4g1 | ENSMUSG0( | false | chr16 | 20672338 | 20674211 | 39.51976 | 124.83012 | 120.52397 | . | . | . | . | . | . |
| chr13 | 104287737 | 104287974 | 182 | Promoter (<=1kb) | 13 | 104287873 | 104494763 | 206891 | 1 | ENSMUST00 | 0 | Adamts6 | ENSMUSG0( | false | chr13 | 104285383 | 104288917 | 31.32084 | 94.19205 | 90.75528 | . | . | . | . | . | . |
| chr17 | 9764620 | 9764925 | 182 | Distal Intergenic | 17 | 9666497 | 9669704 | 3208 | 2 | ENSMUST00 | -94917 | Pabpc6 | ENSMUSG0( | false | chr17 | 9764667 | 9765360 | 16.06223 | 39.4301 | 36.8496 | . | . | . | . | . | . |
| chr9 | 78617203 | 78617457 | 182 | Promoter (1-2kb) | 9 | 78615763 | 78618089 | 2327 | 1 | ENSMUST00 | 1441 | Cd109 | ENSMUSG0( | false | . | . | . | . | . | . | chr9 | 78614436 | 78618392 | 5.64883 | 4.40591 | 3.51997 |
| chr11 | 49200179 | 49200408 | 182 | Promoter (2-3kb) | 11 | 49203292 | 49218553 | 15262 | 1 | ENSMUST00 | -2884 | Zfp62 | ENSMUSG0( | false | . | . | . | . | . | . | chr11 | 49199134 | 49202381 | 3.93913 | 3.60937 | 2.1219 |
| chr4 | 41503091 | 41503368 | 182 | Promoter (<=1kb) | 4 | 41495604 | 41503076 | 7473 | 2 | ENSMUST00 | -16 | AI464131 | ENSMUSG0( | false | chr4 | 41502473 | 41503143 | 14.96389 | 42.58086 | 39.95799 | chr4 | 41502097 | 41503825 | 5.45376 | 4.17076 | 3.41977 |
| chr17 | 57060198 | 57060661 | 182 | Promoter (<=1kb) | 17 | 57043711 | 57059863 | 16153 | 2 | ENSMUST00 | -336 | Slc25a23 | ENSMUSG0( | false | chr17 | 57059228 | 57061163 | 17.49727 | 64.69686 | 61.78638 | chr17 | 57058859 | 57063211 | 4.63146 | 3.8307 | 2.72869 |
| chr7 | 109518942 | 109519338 | 182 | Promoter (<=1kb) | 7 | 109519147 | 109522367 | 3221 | 1 | ENSMUST00 | 0 | Rpl27a | ENSMUSG0( | false | chr7 | 109518375 | 109520221 | 36.91116 | 131.06699 | 126.54268 | . | . | . | . | . | . |
| chr19 | 16780316 | 16780640 | 182 | Promoter (<=1kb) | 19 | 16780547 | 16781103 | 557 | 1 | ENSMUST00 | 0 | Gm9806 | ENSMUSG0( | false | chr19 | 16779373 | 16781820 | 31.70051 | 127.6562 | 123.25787 | . | . | . | . | . | . |
| chr4 | 138464718 | 138465105 | 182 | Distal Intergenic | 4 | 138454314 | 138460123 | 5810 | 1 | ENSMUST00 | 10405 | Camk2n1 | ENSMUSG0( | false | . | . | . | . | . | . | chr4 | 138463360 | 138465484 | 5.99771 | 4.45783 | 3.84992 |
| chrX | 104199890 | 104200321 | 182 | Promoter (<=1kb) | 20 | 104077435 | 104201117 | 123683 | 2 | ENSMUST00 | 796 | C77370 | ENSMUSG0( | false | chrX | 104199573 | 104201696 | 20.21941 | 51.82391 | 49.08421 | . | . | . | . | . | . |
| chr2 | 181675589 | 181675837 | 182 | Distal Intergenic | 2 | 181669836 | 181671640 | 1805 | 2 | ENSMUST00 | -3950 | Sox18 | ENSMUSG0( | false | . | . | . | . | . | . | chr2 | 181670327 | 181676580 | 3.99625 | 3.33699 | 2.21903 |
| chr11 | 43470159 | 43470408 | 182 | Distal Intergenic | 11 | 43474276 | 43491505 | 17230 | 1 | ENSMUST00 | -3868 | C1qtnf2 | ENSMUSG0( | false | . | . | . | . | . | . | . | . | . | . | . | . |
| chr16 | 30692529 | 30693070 | 182 | Distal Intergenic | 16 | 30599723 | 30602797 | 3075 | 1 | ENSMUST00 | 92807 | Fam43a | ENSMUSG0( | false | . | . | . | . | . | . | chr16 | 30691145 | 30692703 | 4.72215 | 4.00507 | 2.76653 |
| chr5 | 115134625 | 115135109 | 182 | Promoter (<=1kb) | 5 | 115122550 | 115134975 | 12426 | 2 | ENSMUST00 | 0 | Unc119b | ENSMUSG0( | false | chr5 | 115134185 | 115135180 | 35.95432 | 114.41653 | 110.45183 | chr5 | 115133212 | 115135295 | 3.81408 | 3.4266 | 2.08804 |
| chr11 | 121686832 | 121687052 | 182 | Distal Intergenic | 11 | 121627268 | 121673153 | 45886 | 2 | ENSMUST00 | -13680 | B3gntl1 | ENSMUSG0( | false | . | . | . | . | . | . | . | . | . | . | . | . |
| chr5 | 143269435 | 143270101 | 182 | Promoter (<=1kb) | 5 | 143249697 | 143270022 | 20326 | 2 | ENSMUST00 | 0 | Zfp316 | ENSMUSG0( | false | chr5 | 143269182 | 143270604 | 28.91247 | 96.86193 | 93.36482 | chr5 | 143268974 | 143270802 | 3.80386 | 3.50767 | 2.02619 |
| chr17 | 27555200 | 27555615 | 182 | Promoter (1-2kb) | 17 | 27556620 | 27563674 | 7055 | 1 | ENSMUST00 | -1005 | Hmga1 | ENSMUSG0( | false | chr17 | 27555151 | 27557949 | 32.14993 | 97.60393 | 94.09416 | chr17 | 27554969 | 27556354 | 5.6026 | 4.43863 | 3.49058 |
| chr6 | 138672689 | 138673003 | 182 | Distal Intergenic | 6 | 138657095 | 138658444 | 1350 | 2 | ENSMUST00 | -14246 | Igbp1b | ENSMUSG0( | false | . | . | . | . | . | . | chr6 | 138671679 | 138673767 | 4.5458 | 3.77986 | 2.6976 |
| chr3 | 38885499 | 38885863 | 182 | Promoter (1-2kb) | 3 | 38886940 | 39011985 | 125046 | 1 | ENSMUST00 | -1077 | Fat4 | ENSMUSG0( | false | chr3 | 38884547 | 38888528 | 37.05418 | 118.43227 | 114.34519 | . | . | . | . | . | . |
| chr15 | 76009357 | 76009650 | 182 | Promoter (<=1kb) | 15 | 76001093 | 76009498 | 8406 | 2 | ENSMUST00 | 0 | Fam83h | ENSMUSG0( | false | chr15 | 76008984 | 76010319 | 41.50429 | 134.78256 | 130.11549 | chr15 | 76009086 | 76010228 | 3.59208 | 3.26568 | 1.886 |
| chr8 | 35376413 | 35376696 | 182 | Promoter (<=1kb) | 8 | 35375741 | 35388124 | 12384 | 1 | ENSMUST00 | 673 | Ppp1r3b | ENSMUSG0( | false | chr8 | 35375405 | 35376980 | 12.76655 | 31.65917 | 29.17213 | chr8 | 35375774 | 35377272 | 2.83249 | 2.84747 | 1.28493 |
| chr19 | 12475533 | 12475734 | 182 | Intron (ENSMUST00000045521.7 | 19 | 12460779 | 12465284 | 4506 | 1 | ENSMUST00 | 14755 | Mpeg1 | ENSMUSG0( | false | . | . | . | . | . | . | chr19 | 12475291 | 12477507 | 4.4595 | 3.60276 | 2.59019 |
| chr6 | 34877850 | 34878233 | 182 | Promoter (<=1kb) | 6 | 34871771 | 34878021 | 6251 | 2 | ENSMUST00 | 0 | 3110062M04 | ENSMUSG0( | false | chr6 | 34877381 | 34878524 | 27.03504 | 88.5911 | 85.27629 | chr6 | 34863195 | 34879034 | 5.53817 | 4.34013 | 3.44642 |
| chr10 | 75560034 | 75560270 | 182 | Promoter (<=1kb) | 10 | 75550125 | 75560330 | 10206 | 2 | ENSMUST00 | 60 | Lrrc75b | ENSMUSG0( | false | chr10 | 75559770 | 75560540 | 14.6071 | 44.0986 | 41.45673 | chr10 | 75559181 | 75560809 | 5.42721 | 4.26856 | 3.40622 |
| chr5 | 72203133 | 72203694 | 182 | Promoter (<=1kb) | 5 | 72203329 | 72230475 | 27147 | 1 | ENSMUST00 | 0 | Atp10d | ENSMUSG0( | false | chr5 | 72203003 | 72204225 | 21.91422 | 67.38415 | 64.43526 | chr5 | 72201458 | 72204813 | 5.06198 | 4.14522 | 3.04611 |
| chr6 | 39573731 | 39574187 | 182 | Promoter (<=1kb) | 6 | 39573873 | 39588769 | 14897 | 1 | ENSMUST00 | 0 | Adck2 | ENSMUSG0( | false | chr6 | 39573285 | 39574769 | 36.42221 | 131.26616 | 126.73016 | chr6 | 39573243 | 39575412 | 4.75328 | 3.98118 | 2.82819 |
| chr18 | 83815211 | 83815501 | 182 | Distal Intergenic | 18 | 84012082 | 84086390 | 74309 | 2 | ENSMUST00 | 270889 | Tshz1 | ENSMUSG0( | false | . | . | . | . | . | . | . | . | . | . | . | . |
| chr9 | 96972053 | 96972300 | 182 | Promoter (<=1kb) | 9 | 96943972 | 96971975 | 28004 | 2 | ENSMUST00 | -79 | Spsb4 | ENSMUSG0( | false | . | . | . | . | . | . | chr9 | 96971657 | 96974342 | 5.54331 | 4.16362 | 3.49322 |
| chr9 | 96993087 | 96993323 | 182 | Intron (ENSMUST00000055433.4 | 9 | 96943485 | 96984195 | 40711 | 2 | ENSMUST00 | -8893 | Spsb4 | ENSMUSG0( | false | . | . | . | . | . | . | chr9 | 96975779 | 96994993 | 5.8258 | 4.40484 | 3.6945 |
| chr15 | 75616371 | 75616731 | 182 | Promoter (<=1kb) | 15 | 75616679 | 75625300 | 8622 | 1 | ENSMUST00 | 0 | Zfp41 | ENSMUSG0( | false | chr15 | 75615707 | 75617294 | 17.71499 | 57.83842 | 55.02271 | chr15 | 75615615 | 75617342 | 5.21478 | 4.18872 | 3.17278 |
| chr19 | 18712951 | 18713376 | 182 | Promoter (<=1kb) | 19 | 18713193 | 18718428 | 5236 | 1 | ENSMUST00 | 0 | D030056L22 | ENSMUSG0( | false | chr19 | 18712461 | 18714024 | 28.1615 | 93.53628 | 90.11485 | . | . | . | . | . | . |
| chr6 | 135323425 | 135323692 | 182 | Distal Intergenic | 6 | 135309784 | 135310347 | 564 | 2 | ENSMUST00 | -13079 | Pbp2 | ENSMUSG0( | false | . | . | . | . | . | . | chr6 | 135323247 | 135323970 | 4.71379 | 3.89517 | 2.80331 |
| chr3 | 16183618 | 16183867 | 182 | Promoter (<=1kb) | 3 | 16183183 | 16217035 | 33853 | 1 | ENSMUST00 | 436 | Ythdf3 | ENSMUSG0( | false | chr3 | 16182684 | 16184590 | 38.11044 | 121.32513 | 117.13898 | . | . | . | . | . | . |
| chr2 | 30600319 | 30600630 | 182 | Intron (ENSMUST00000102853.3 | 2 | 30605217 | 30608217 | 3001 | 1 | ENSMUST00 | -4587 | Cstad | ENSMUSG0( | false | . | . | . | . | . | . | chr2 | 30600055 | 30602050 | 5.57081 | 4.2646 | 3.50515 |
| chr2 | 30613760 | 30613970 | 182 | Distal Intergenic | 2 | 30605217 | 30608217 | 3001 | 1 | ENSMUST00 | 8544 | Cstad | ENSMUSG0( | false | . | . | . | . | . | . | chr2 | 30613479 | 30614237 | 3.34307 | 3.15994 | 1.68913 |
| chr8 | 116943279 | 116943502 | 182 | Promoter (<=1kb) | 8 | 116943393 | 116960445 | 17053 | 1 | ENSMUST00 | 0 | Atmin | ENSMUSG0( | false | chr8 | 116943321 | 116944464 | 29.10187 | 109.2007 | 105.38981 | . | . | . | . | . | . |
| chr9 | 118963214 | 118963481 | 182 | Intron (ENSMUST00000073109.6 | 9 | 118929867 | 119033628 | 103762 | 1 | ENSMUST00 | 33348 | Ctdspl | ENSMUSG0( | true | . | . | . | . | . | . | chr9 | 118927678 | 119010843 | 6.89652 | 4.89961 | 4.57102 |
| chr9 | 118949724 | 118949900 | 182 | Intron (ENSMUST00000073109.6 | 9 | 118929867 | 119033628 | 103762 | 1 | ENSMUST00 | 19858 | Ctdspl | ENSMUSG0( | true | . | . | . | . | . | . | chr9 | 118927678 | 119010843 | 6.89652 | 4.89961 | 4.57102 |
| chr6 | 35330903 | 35331166 | 182 | Distal Intergenic | 6 | 35312746 | 35326141 | 13396 | 2 | ENSMUST00 | -4763 | Fam180a | ENSMUSG0( | false | . | . | . | . | . | . | chr6 | 35328656 | 35334639 | 4.39332 | 3.7234 | 2.52503 |
| chr8 | 14779303 | 14779522 | 182 | Promoter (1-2kb) | 8 | 14777773 | 14847398 | 69626 | 1 | ENSMUST00 | 1531 | Dlgap2 | ENSMUSG0( | false | . | . | . | . | . | . | . | . | . | . | . | . |
| chr12 | 65172700 | 65172906 | 182 | Promoter (<=1kb) | 12 | 65132734 | 65172604 | 39871 | 2 | ENSMUST00 | -97 | Mis18bp1 | ENSMUSG0( | false | chr12 | 65172151 | 65173446 | 20.31581 | 74.8081 | 71.74177 | . | . | . | . | . | . |

| | | | | | | | | | | | | | | | | | | | | | | | | | | |
|---|---|---|---|---|---|---|---|---|---|---|---|---|---|---|---|---|---|---|---|---|---|---|---|---|---|---|
| chr13 | 54611041 | 54611734 | 182 | Promoter (<=1kb) | 13 | 54592401 | 54611344 | 18944 | 2 | ENSMUST00 | 0 | Cltb | ENSMUSG0C | false | chr13 | 54610388 | 54611912 | 38.6007 | 121.10697 | 116.92481 | chr13 | 54604859 | 54612332 | 4.18969 | 3.66231 | 2.3575 |
| chr7 | 109986029 | 109986572 | 182 | Promoter (<=1kb) | 7 | 109972187 | 109986451 | 14265 | 2 | ENSMUST00 | 0 | Tmem41b | ENSMUSG0C | false | chr7 | 109985350 | 109987042 | 25.17393 | 93.69764 | 90.27328 | . | . | . | . | . | . |
| chr15 | 75747798 | 75748583 | 182 | Promoter (<=1kb) | 15 | 75746843 | 75747922 | 1080 | 2 | ENSMUST00 | 0 | Mafa | ENSMUSG0C | false | chr15 | 75745907 | 75748880 | 24.03115 | 75.67385 | 72.59326 | chr15 | 75745856 | 75749278 | 6.53429 | 4.75316 | 4.26674 |
| chr10 | 56164520 | 56164740 | 182 | Intron (ENSMUST00000099739.3 | 10 | 56106917 | 56116880 | 9964 | 1 | ENSMUST00 | 57604 | Msl3l2 | ENSMUSG0C | false | . | . | . | . | . | . | chr10 | 56163967 | 56166316 | 4.09311 | 3.61856 | 2.28213 |
| chr4 | 117126184 | 117126382 | 182 | Promoter (<=1kb) | 4 | 117126794 | 117128336 | 1543 | 1 | ENSMUST00 | -412 | Tctex1d4 | ENSMUSG0C | false | . | . | . | . | . | . | chr4 | 117124494 | 117127382 | 6.63787 | 4.73349 | 4.35283 |
| chr19 | 46598893 | 46599182 | 182 | Promoter (<=1kb) | 19 | 46599084 | 46657389 | 58306 | 1 | ENSMUST00 | 0 | Wbp1l | ENSMUSG0C | false | chr19 | 46599005 | 46600299 | 26.65955 | 86.95457 | 83.67183 | . | . | . | . | . | . |
| chr11 | 64396475 | 64396784 | 182 | Distal Intergenic | 11 | 64435332 | 64522841 | 87510 | 1 | ENSMUST00 | -38548 | Hs3st3a1 | ENSMUSG0C | true | . | . | . | . | . | . | chr11 | 64395439 | 64396689 | 3.32813 | 3.227 | 1.63275 |
| chr9 | 60648832 | 60649045 | 182 | Promoter (<=1kb) | 9 | 60610270 | 60649710 | 39441 | 2 | ENSMUST00 | 665 | Lrrc49 | ENSMUSG0C | true | chr9 | 60648251 | 60649883 | 12.58327 | 33.30664 | 30.79935 | chr9 | 60643773 | 60650965 | 3.96179 | 3.59314 | 2.14865 |
| chr4 | 151995083 | 151995910 | 182 | Promoter (<=1kb) | 4 | 151989633 | 151996258 | 6626 | 2 | ENSMUST00 | 348 | Phf13 | ENSMUSG0C | false | chr4 | 151994455 | 151997533 | 32.98857 | 124.89146 | 120.57959 | . | . | . | . | . | . |
| chr17 | 17448695 | 17448915 | 182 | Promoter (<=1kb) | 17 | 17448820 | 17459387 | 10568 | 1 | ENSMUST00 | 0 | Lix1 | ENSMUSG0C | false | chr17 | 17448655 | 17449642 | 10.20265 | 24.60154 | 22.20526 | chr17 | 17448661 | 17449889 | 4.66667 | 3.79559 | 2.76682 |
| chr19 | 7142686 | 7142880 | 182 | Intron (ENSMUST00000040261.5 | 19 | 7092014 | 7105729 | 13716 | 2 | ENSMUST00 | -36958 | Flrt1 | ENSMUSG0C | false | . | . | . | . | . | . | chr19 | 7127706 | 7162730 | 7.6862 | 5.1249 | 5.24023 |
| chr19 | 7122324 | 7122609 | 182 | Intron (ENSMUST00000040261.5 | 19 | 7092014 | 7105729 | 13716 | 2 | ENSMUST00 | -16596 | Flrt1 | ENSMUSG0C | false | . | . | . | . | . | . | chr19 | 7120136 | 7123619 | 8.00868 | 5.27125 | 5.51846 |
| chr1 | 93267357 | 93267556 | 182 | Intron (ENSMUST00000062202.8 | 1 | 93274431 | 93282317 | 7887 | 1 | ENSMUST00 | -6875 | Sned1 | ENSMUSG0C | false | . | . | . | . | . | . | chr1 | 93267177 | 93267751 | 3.04814 | 3.03414 | 1.46977 |
| chr15 | 83223644 | 83223837 | 182 | Downstream (2-3kb) | 15 | 83226726 | 83251731 | 25006 | 2 | ENSMUST00 | 27894 | A4galt | ENSMUSG0C | false | . | . | . | . | . | . | chr15 | 83222754 | 83224600 | 4.16204 | 3.66093 | 2.3567 |
| chr18 | 10052357 | 10052870 | 182 | Distal Intergenic | 18 | 10021704 | 10045119 | 23416 | 2 | ENSMUST00 | -7239 | Usp14 | ENSMUSG0C | true | chr18 | 10051557 | 10053003 | 23.5743 | 68.56921 | 65.60024 | chr18 | 10052721 | 10053089 | 3.35567 | 3.21736 | 1.70517 |
| chr15 | 80771984 | 80772261 | 182 | Intron (ENSMUST00000067689.7 | 15 | 80711319 | 80941086 | 229768 | 1 | ENSMUST00 | 60666 | Tnrc6b | ENSMUSG0C | false | . | . | . | . | . | . | chr15 | 80769539 | 80772948 | 4.51599 | 3.81076 | 2.60434 |
| chr2 | 70337250 | 70337464 | 182 | Intron (ENSMUST00000060208.5 | 2 | 70458982 | 70459560 | 579 | 1 | ENSMUST00 | -121518 | Rpl9-ps7 | ENSMUSG0C | false | . | . | . | . | . | . | chr2 | 70336894 | 70339313 | 6.6141 | 4.87694 | 4.32257 |
| chr11 | 97502505 | 97502826 | 182 | Promoter (2-3kb) | 11 | 97490783 | 97500340 | 9558 | 2 | ENSMUST00 | -2166 | 4933428G20 | ENSMUSG0C | false | . | . | . | . | . | . | chr11 | 97500597 | 97503780 | 4.48772 | 3.78379 | 2.61862 |
| chr18 | 24152163 | 24152371 | 182 | Distal Intergenic | 18 | 24104757 | 24121953 | 17197 | 2 | ENSMUST00 | -30211 | Ino80c | ENSMUSG0C | false | . | . | . | . | . | . | chr18 | 24151287 | 24153407 | 5.04223 | 4.07602 | 3.10274 |
| chr7 | 96212682 | 96213071 | 182 | Promoter (1-2kb) | 7 | 96211671 | 96911093 | 699423 | 1 | ENSMUST00 | 1012 | Tenm4 | ENSMUSG0C | false | chr7 | 96212520 | 96213657 | 18.84082 | 47.13427 | 44.45482 | chr7 | 96209771 | 96213085 | 4.07405 | 3.53581 | 2.27113 |
| chr7 | 96704072 | 96704336 | 182 | Exon (ENSMUST00000107162.2/ | 7 | 96644172 | 96732280 | 88109 | 1 | ENSMUST00 | 59901 | Tenm4 | ENSMUSG0C | false | . | . | . | . | . | . | . | . | . | . | . | . |
| chr19 | 40660280 | 40660535 | 182 | Promoter (<=1kb) | 19 | 40660493 | 40723097 | 62605 | 1 | ENSMUST00 | 0 | Entpd1 | ENSMUSG0C | false | chr19 | 40659504 | 40660819 | 10.85587 | 23.4294 | 21.04904 | chr19 | 40658829 | 40660618 | 3.60875 | 3.39122 | 1.86275 |
| chr15 | 98871385 | 98872324 | 182 | Promoter (<=1kb) | 15 | 98831672 | 98871204 | 39533 | 2 | ENSMUST00 | -182 | Kmt2d | ENSMUSG0C | false | chr15 | 98868485 | 98872894 | 47.14861 | 169.72867 | 163.76503 | . | . | . | . | . | . |
| chr7 | 128717989 | 128718313 | 182 | Intron (ENSMUST00000057557.8 | 7 | 128712611 | 128714858 | 2248 | 2 | ENSMUST00 | -3132 | Mcmbp | ENSMUSG0C | false | . | . | . | . | . | . | . | . | . | . | . | . |
| chr15 | 98148378 | 98148661 | 182 | Promoter (2-3kb) | 15 | 98135714 | 98145706 | 9993 | 2 | ENSMUST00 | -2673 | Asb8 | ENSMUSG0C | false | . | . | . | . | . | . | . | . | . | . | . | . |
| chr7 | 141462457 | 141462756 | 182 | Promoter (<=1kb) | 7 | 141463111 | 141464729 | 1619 | 1 | ENSMUST00 | -355 | Efcab4a | ENSMUSG0C | false | chr7 | 141461557 | 141462588 | 12.72529 | 41.01831 | 38.41836 | chr7 | 141456108 | 141486560 | 6.62038 | 4.76228 | 4.33989 |
| chr12 | 70853074 | 70853281 | 182 | Intron (ENSMUST00000057859.7 | 12 | 70825514 | 70902232 | 76719 | 1 | ENSMUST00 | 27561 | Frmd6 | ENSMUSG0C | true | . | . | . | . | . | . | chr12 | 70826659 | 70861086 | 4.82543 | 3.92114 | 2.87602 |
| chr12 | 70858498 | 70858749 | 182 | Intron (ENSMUST00000057859.7 | 12 | 70825514 | 70902232 | 76719 | 1 | ENSMUST00 | 32985 | Frmd6 | ENSMUSG0C | true | . | . | . | . | . | . | chr12 | 70826659 | 70861086 | 4.82543 | 3.92114 | 2.87602 |
| chr12 | 70826703 | 70827003 | 182 | Promoter (1-2kb) | 12 | 70825514 | 70902232 | 76719 | 1 | ENSMUST00 | 1190 | Frmd6 | ENSMUSG0C | true | chr12 | 70825459 | 70828098 | 39.00716 | 125.01923 | 120.70579 | chr12 | 70826659 | 70861086 | 4.82543 | 3.92114 | 2.87602 |
| chr13 | 95612886 | 95613139 | 182 | Intron (ENSMUST00000059193.5 | 13 | 95601804 | 95618459 | 16656 | 2 | ENSMUST00 | 5320 | F2r | ENSMUSG0C | false | . | . | . | . | . | . | chr13 | 95610650 | 95618067 | 4.64293 | 3.76201 | 2.74776 |
| chr2 | 83812496 | 83812805 | 182 | Promoter (<=1kb) | 2 | 83812636 | 83883486 | 70851 | 1 | ENSMUST00 | 0 | Fam171b | ENSMUSG0C | false | chr2 | 83812364 | 83813184 | 23.16072 | 66.96479 | 64.02055 | chr2 | 83812281 | 83813078 | 4.4779 | 3.69176 | 2.60509 |
| chr11 | 120902447 | 120902699 | 182 | Intron (ENSMUST00000056781.4 | 11 | 120873792 | 120885911 | 12120 | 2 | ENSMUST00 | -16537 | Ccdc57 | ENSMUSG0C | false | chr11 | 120901830 | 120902741 | 7.46892 | 13.27344 | 11.04042 | chr11 | 120901637 | 120903309 | 3.28588 | 3.09636 | 1.65046 |
| chr2 | 163481536 | 163481729 | 182 | Distal Intergenic | 2 | 163466379 | 163472629 | 6251 | 2 | ENSMUST00 | -8908 | Fitm2 | ENSMUSG0C | false | chr2 | 163481508 | 163482317 | 11.94783 | 25.26529 | 22.85895 | chr2 | 163480709 | 163482521 | 5.2109 | 4.14313 | 3.18483 |
| chr8 | 4259105 | 4259375 | 182 | Promoter (<=1kb) | 8 | 4257648 | 4259274 | 1627 | 2 | ENSMUST00 | 0 | Ctxn1 | ENSMUSG0C | false | chr8 | 4258335 | 4259772 | 15.85026 | 49.49464 | 46.78799 | chr8 | 4258292 | 4260018 | 7.12633 | 5.06268 | 4.7359 |
| chr2 | 119541210 | 119541403 | 182 | Promoter (<=1kb) | 2 | 119516511 | 119541513 | 25003 | 2 | ENSMUST00 | 110 | Exd1 | ENSMUSG0C | false | . | . | . | . | . | . | . | . | . | . | . | . |
| chr6 | 128362522 | 128362880 | 182 | Promoter (<=1kb) | 6 | 128357000 | 128362812 | 5813 | 2 | ENSMUST00 | 0 | Rhno1 | ENSMUSG0C | false | chr6 | 128361968 | 128363462 | 22.15647 | 106.74507 | 102.9996 | chr6 | 128361723 | 128363593 | 6.29021 | 4.67788 | 4.06607 |
| chr7 | 145206192 | 145206601 | 182 | Distal Intergenic | 7 | 145253923 | 145283927 | 30005 | 2 | ENSMUST00 | 77326 | Tpcn2 | ENSMUSG0C | false | chr7 | 145206041 | 145206738 | 7.90289 | 16.26095 | 13.97799 | chr7 | 145205749 | 145208181 | 6.26922 | 4.48757 | 4.07663 |
| chr10 | 70159833 | 70160226 | 182 | Intron (ENSMUST00000147545.2 | 10 | 70174509 | 70182206 | 7698 | 1 | ENSMUST00 | -14283 | Ccdc6 | ENSMUSG0C | false | . | . | . | . | . | . | chr10 | 70157764 | 70160392 | 6.52181 | 4.85974 | 4.24262 |
| chr10 | 70167722 | 70167931 | 182 | Intron (ENSMUST00000147545.2 | 10 | 70174509 | 70182206 | 7698 | 1 | ENSMUST00 | -6578 | Ccdc6 | ENSMUSG0C | false | . | . | . | . | . | . | chr10 | 70161380 | 70168358 | 5.98338 | 4.50782 | 3.80721 |
| chr1 | 38767004 | 38767230 | 182 | Distal Intergenic | 1 | 38794509 | 38821215 | 26707 | 2 | ENSMUST00 | 53985 | Lonrf2 | ENSMUSG0C | false | . | . | . | . | . | . | chr1 | 38766767 | 38767552 | 6.16982 | 4.65411 | 3.95412 |
| chr17 | 62961610 | 62961823 | 182 | Distal Intergenic | 17 | 62604184 | 62881317 | 277134 | 2 | ENSMUST00 | -80294 | Efna5 | ENSMUSG0C | false | . | . | . | . | . | . | chr17 | 62961164 | 62963938 | 2.91956 | 2.88993 | 1.34888 |
| chr12 | 75450096 | 75450362 | 182 | Downstream (<1kb) | 12 | 75411720 | 75416781 | 5062 | 2 | ENSMUST00 | -33316 | Gphb5 | ENSMUSG0C | false | chr12 | 75449665 | 75450394 | 6.80124 | 12.58378 | 10.36022 | chr12 | 75448651 | 75450755 | 5.85731 | 4.42597 | 3.76347 |
| chr12 | 75369419 | 75369807 | 182 | Intron (ENSMUST00000118602.2 | 12 | 75411720 | 75416781 | 5062 | 2 | ENSMUST00 | 46974 | Gphb5 | ENSMUSG0C | false | . | . | . | . | . | . | . | . | . | . | . | . |
| chr4 | 5783899 | 5784237 | 182 | Intron (ENSMUST00000124256.2 | 4 | 5772879 | 5799946 | 27068 | 1 | ENSMUST00 | 11021 | Fam110b | ENSMUSG0C | false | . | . | . | . | . | . | chr4 | 5782185 | 5785765 | 3.94267 | 3.56755 | 2.14184 |
| chr4 | 74013167 | 74013760 | 182 | Promoter (<=1kb) | 4 | 74013442 | 74202214 | 188773 | 1 | ENSMUST00 | 0 | Frmd3 | ENSMUSG0C | false | chr4 | 74013134 | 74014153 | 15.84839 | 38.35189 | 35.78466 | chr4 | 74012569 | 74014658 | 3.41534 | 3.15899 | 1.75001 |
| chr19 | 40895032 | 40895352 | 182 | Promoter (<=1kb) | 19 | 40894705 | 40917947 | 23243 | 1 | ENSMUST00 | 328 | Zfp518a | ENSMUSG0C | false | chr19 | 40894179 | 40895684 | 31.54088 | 108.67455 | 104.87239 | . | . | . | . | . | . |
| chr9 | 40269589 | 40269800 | 182 | Promoter (<=1kb) | 9 | 40269687 | 40288407 | 18721 | 1 | ENSMUST00 | 0 | Scn3b | ENSMUSG0C | false | . | . | . | . | . | . | chr9 | 40268046 | 40269980 | 5.66255 | 4.04038 | 3.5682 |
| chr5 | 124446539 | 124446814 | 182 | Promoter (<=1kb) | 5 | 124446704 | 124457118 | 10415 | 1 | ENSMUST00 | 0 | Setd8 | ENSMUSG0C | false | chr5 | 124444651 | 124446796 | 30.58814 | 94.89368 | 91.44746 | chr5 | 124446227 | 124457002 | 6.32983 | 4.7089 | 4.09735 |
| chr5 | 124445205 | 124446211 | 182 | Promoter (<=1kb) | 5 | 124445493 | 124460883 | 15391 | 1 | ENSMUST00 | 0 | Setd8 | ENSMUSG0C | false | chr5 | 124444651 | 124446796 | 30.58814 | 94.89368 | 91.44746 | chr5 | 124444273 | 124445243 | 4.59476 | 3.96071 | 2.6772 |
| chr11 | 36508564 | 36508829 | 182 | Intron (ENSMUST00000102801.2 | 11 | 36163719 | 36495139 | 331421 | 2 | ENSMUST00 | -13426 | Tenm2 | ENSMUSG0C | false | . | . | . | . | . | . | chr11 | 36508657 | 36510522 | 3.13779 | 3.1214 | 1.52634 |
| chr11 | 36509700 | 36509955 | 182 | Intron (ENSMUST00000102801.2 | 11 | 36163719 | 36495139 | 331421 | 2 | ENSMUST00 | -14562 | Tenm2 | ENSMUSG0C | false | . | . | . | . | . | . | chr11 | 36508657 | 36510522 | 3.13779 | 3.1214 | 1.52634 |
| chr7 | 30095049 | 30095279 | 182 | Promoter (<=1kb) | 7 | 30095162 | 30100118 | 4957 | 1 | ENSMUST00 | 0 | Zfp260 | ENSMUSG0C | false | chr7 | 30094381 | 30095835 | 26.6505 | 78.45867 | 75.33303 | . | . | . | . | . | . |
| chr11 | 53470409 | 53470654 | 182 | Promoter (<=1kb) | 11 | 53469749 | 53470479 | 731 | 2 | ENSMUST00 | 0 | Gm9837 | ENSMUSG0C | false | . | . | . | . | . | . | chr11 | 53470022 | 53471265 | 3.92082 | 3.48639 | 2.14486 |
| chr4 | 12087841 | 12088093 | 182 | Promoter (<=1kb) | 4 | 12039355 | 12087957 | 48603 | 2 | ENSMUST00 | 0 | Tmem67 | ENSMUSG0C | false | chr4 | 12087306 | 12088431 | 19.30515 | 51.05533 | 48.32729 | chr4 | 12087403 | 12089222 | 5.56961 | 4.28986 | 3.52471 |
| chr7 | 47002903 | 47003165 | 182 | Intron (ENSMUST00000061639.7 | 7 | 46990396 | 47008411 | 18016 | 2 | ENSMUST00 | 5246 | Spty2d1 | ENSMUSG0C | false | . | . | . | . | . | . | chr7 | 47000137 | 47007497 | 5.98469 | 4.53303 | 3.82111 |
| chr5 | 123675208 | 123675567 | 182 | Intron (ENSMUST00000111564.2 | 5 | 123617706 | 123666690 | 48985 | 2 | ENSMUST00 | -8519 | Clip1 | ENSMUSG0C | false | . | . | . | . | . | . | chr5 | 123656964 | 123683302 | 5.99044 | 4.55496 | 3.80916 |
| chr7 | 127876521 | 127876901 | 182 | Promoter (<=1kb) | 7 | 127865170 | 127876828 | 11659 | 2 | ENSMUST00 | 0 | Zfp668 | ENSMUSG0C | false | chr7 | 127875160 | 127877572 | 30.25803 | 109.37234 | 105.55905 | . | . | . | . | . | . |
| chr10 | 76032139 | 76032760 | 182 | Promoter (<=1kb) | 10 | 76032654 | 76043234 | 10581 | 1 | ENSMUST00 | 0 | Zfp280b | ENSMUSG0C | false | chr10 | 76031985 | 76033514 | 33.54585 | 101.01101 | 97.41918 | chr10 | 76031167 | 76034461 | 3.34023 | 3.16477 | 1.65321 |
| chr11 | 20631357 | 20632351 | 182 | Promoter (<=1kb) | 11 | 20631979 | 20653021 | 21043 | 1 | ENSMUST00 | 0 | Sertad2 | ENSMUSG0C | true | chr11 | 20631080 | 20633425 | 34.92026 | 124.22972 | 119.94103 | . | . | . | . | . | . |
| chr11 | 20543576 | 20544078 | 182 | Promoter (<=1kb) | 11 | 20543334 | 20653021 | 109688 | 1 | ENSMUST00 | 243 | Sertad2 | ENSMUSG0C | true | chr11 | 20543047 | 20544428 | 26.61085 | 108.21047 | 104.42454 | chr11 | 20544024 | 20546590 | 3.49809 | 3.40157 | 1.76367 |
| chr11 | 97434923 | 97435353 | 182 | Promoter (<=1kb) | 11 | 97436285 | 97444228 | 7944 | 1 | ENSMUST00 | -932 | Arhgap23 | ENSMUSG0C | true | chr11 | 97434949 | 97435291 | 8.1714 | 16.72606 | 14.43788 | chr11 | 97431804 | 97440135 | 9.81823 | 6.18842 | 6.94298 |
| chr11 | 97415846 | 97416114 | 182 | Promoter (<=1kb) | 11 | 97415533 | 97502402 | 86870 | 1 | ENSMUST00 | 314 | Arhgap23 | ENSMUSG0C | true | chr11 | 97415033 | 97416334 | 26.7 | 83.22852 | 80.01951 | chr11 | 97414588 | 97417653 | 6.47594 | 4.82039 | 4.21762 |
| chr11 | 97435876 | 97436117 | 182 | Promoter (<=1kb) | 11 | 97436285 | 97444228 | 7944 | 1 | ENSMUST00 | -168 | Arhgap23 | ENSMUSG0C | true | . | . | . | . | . | . | chr11 | 97431804 | 97440135 | 9.81823 | 6.18842 | 6.94298 |
| chr1 | 88702057 | 88702590 | 182 | Promoter (<=1kb) | 1 | 88698225 | 88702221 | 3997 | 2 | ENSMUST00 | 0 | Arl4c | ENSMUSG0C | false | chr1 | 88698405 | 88702774 | 23.74947 | 92.7123 | 89.30923 | chr1 | 88702440 | 88702985 | 3.93913 | 3.63546 | 2.12035 |
| chr12 | 69576022 | 69576292 | 182 | Downstream (1-2kb) | 12 | 69577628 | 69583028 | 5401 | 2 | ENSMUST00 | 6736 | Vcpkmt | ENSMUSG0C | false | . | . | . | . | . | . | chr12 | 69573229 | 69581639 | 5.27043 | 4.12296 | 3.23088 |
| chr5 | 74161607 | 74161952 | 182 | Distal Intergenic | 5 | 74195288 | 74199481 | 4194 | 1 | ENSMUST00 | -33336 | Rasl11b | ENSMUSG0C | false | . | . | . | . | . | . | chr5 | 74159254 | 74162088 | 4.23771 | 3.55646 | 2.41124 |
| chr2 | 92459239 | 92459878 | 182 | Promoter (<=1kb) | 2 | 92452764 | 92459794 | 7031 | 2 | ENSMUST00 | 0 | Slc35c1 | ENSMUSG0C | false | chr2 | 92458898 | 92460809 | 27.41053 | 90.23362 | 86.88379 | chr2 | 92455932 | 92459269 | 4.7473 | 3.86344 | 2.8315 |
| chr6 | 28833179 | 28833732 | 182 | Promoter (1-2kb) | 6 | 28828133 | 28831747 | 3615 | 2 | ENSMUST00 | -1433 | Lrrc4 | ENSMUSG0C | false | chr6 | 28831812 | 28833824 | 33.66105 | 104.24397 | 100.57021 | chr6 | 28830957 | 28833786 | 3.57503 | 3.3004 | 1.88157 |
| chr6 | 28816055 | 28816309 | 182 | Intron (ENSMUST00000164915.2 | 6 | 28661831 | 28830345 | 168515 | 2 | ENSMUST00 | 14036 | Lrrc4 | ENSMUSG0C | false | . | . | . | . | . | . | chr6 | 28814408 | 28822515 | 6.50428 | 4.75801 | 4.26658 |
| chrX | 51018802 | 51018999 | 182 | Promoter (<=1kb) | 20 | 51003912 | 51018018 | 14107 | 2 | ENSMUST00 | -785 | Rap2c | ENSMUSG0C | false | chrX | 51017427 | 51019512 | 16.08362 | 38.04472 | 35.47989 | . | . | . | . | . | . |
| chr3 | 98374525 | 98374787 | 182 | Distal Intergenic | 3 | 98382481 | 98431948 | 49468 | 1 | ENSMUST00 | -7694 | Zfp697 | ENSMUSG0C | false | . | . | . | . | . | . | chr3 | 98373850 | 98375485 | 3.7783 | 3.40617 | 2.0168 |
| chr3 | 88425083 | 88425267 | 182 | Promoter (<=1kb) | 3 | 88410498 | 88425139 | 14642 | 2 | ENSMUST00 | 0 | Slc25a44 | ENSMUSG0C | false | chr3 | 88424399 | 88425681 | 28.50037 | 84.48055 | 81.24493 | . | . | . | . | . | . |
| chr2 | 109932450 | 109932704 | 182 | Intron (ENSMUST00000046548.8 | 2 | 109917647 | 110014257 | 96611 | 1 | ENSMUST00 | 14804 | Lgr4 | ENSMUSG0C | false | . | . | . | . | . | . | chr2 | 109920715 | 109937278 | 4.86516 | 4.02197 | 2.89791 |
| chr1 | 172376189 | 172376562 | 182 | Promoter (<=1kb) | 1 | 172376531 | 172384099 | 7569 | 1 | ENSMUST00 | 0 | Pigm | ENSMUSG0C | false | chr1 | 172375815 | 172377599 | 34.33094 | 124.21722 | 119.93477 | chr1 | 172375571 | 172380228 | 4.61053 | 3.88089 | 2.69733 |
| chr11 | 67024973 | 67025227 | 182 | Promoter (<=1kb) | 11 | 67025154 | 67034790 | 9637 | 1 | ENSMUST00 | 0 | Tmem220 | ENSMUSG0C | false | chr11 | 67025097 | 67025690 | 21.91791 | 63.91367 | 61.01354 | chr11 | 67024251 | 67025832 | 3.55906 | 3.38335 | 1.8057 |
| chr16 | 76532234 | 76532582 | 182 | Distal Intergenic | 16 | 76403209 | 76403712 | 504 | 2 | ENSMUST00 | -128523 | Gm9843 | ENSMUSG0C | false | . | . | . | . | . | . | . | . | . | . | . | . |
| chr14 | 47366944 | 47367157 | 182 | Promoter (<=1kb) | 14 | 47367751 | 47380418 | 12668 | 1 | ENSMUST00 | -594 | Lgals3 | ENSMUSG0C | false | . | . | . | . | . | . | chr14 | 47366580 | 47367254 | 4.13735 | 3.59075 | 2.34038 |
| chr4 | 63734015 | 63734215 | 182 | Intron (ENSMUST00000062246.6 | 4 | 63727084 | 63745113 | 18030 | 2 | ENSMUST00 | 10898 | Tnfsf15 | ENSMUSG0C | false | . | . | . | . | . | . | chr4 | 63733970 | 63735062 | 5.37486 | 4.35931 | 3.3177 |
| chr4 | 63744999 | 63745286 | 182 | Promoter (<=1kb) | 4 | 63727084 | 63745113 | 18030 | 2 | ENSMUST00 | 0 | Tnfsf15 | ENSMUSG0C | false | chr4 | 63744609 | 63745629 | 15.01947 | 39.77812 | 37.19162 | chr4 | 63737893 | 63745853 | 5.56462 | 4.30016 | 3.48505 |
| chr4 | 63785394 | 63785716 | 182 | Distal Intergenic | 4 | 63727084 | 63745113 | 18030 | 2 | ENSMUST00 | -40282 | Tnfsf15 | ENSMUSG0C | false | . | . | . | . | . | . | chr4 | 63785333 | 63785721 | 3.96267 | 3.60079 | 2.15335 |
| chr9 | 35267565 | 35267837 | 182 | Promoter (<=1kb) | 9 | 35216965 | 35267780 | 50816 | 2 | ENSMUST00 | 0 | Fam118b | ENSMUSG0C | false | chr9 | 35266797 | 35268694 | 27.78601 | 91.88203 | 88.49705 | . | . | . | . | . | . |
| chr9 | 66511643 | 66512013 | 182 | Promoter (2-3kb) | 9 | 66508459 | 66514593 | 6135 | 2 | ENSMUST00 | 2580 | Fbxl22 | ENSMUSG0C | false | chr9 | 66511174 | 66512787 | 26.28407 | 85.32412 | 82.07336 | . | . | . | . | . | . |
| chr11 | 97841174 | 97841409 | 182 | Promoter (<=1kb) | 11 | 97841293 | 97844099 | 2807 | 1 | ENSMUST00 | 0 | B230217C12 | ENSMUSG0C | false | chr11 | 97840756 | 97841660 | 10.7597 | 23.84104 | 21.45527 | chr11 | 97839128 | 97841643 | 3.54395 | 3.21602 | 1.82871 |
| chr3 | 127930008 | 127930222 | 182 | Distal Intergenic | 3 | 127869688 | 127896323 | 26636 | 2 | ENSMUST00 | -33686 | 5730508B09 | ENSMUSG0C | false | . | . | . | . | . | . | chr3 | 127929850 | 127930891 | 3.60357 | 3.28126 | 1.90122 |
| chr9 | 35570049 | 35570464 | 182 | Promoter (<=1kb) | 9 | 35560820 | 35570398 | 9579 | 2 | ENSMUST00 | 0 | Hyls1 | ENSMUSG0C | false | chr9 | 35569470 | 35570915 | 30.17592 | 94.17345 | 90.73757 | chr9 | 35569209 | 35570691 | 3.05306 | 2.86956 | 1.4423 |
| chr4 | 139131882 | 139132108 | 182 | Promoter (<=1kb) | 4 | 139101814 | 139131113 | 29300 | 2 | ENSMUST00 | -770 | Minos1 | ENSMUSG0C | false | . | . | . | . | . | . | chr4 | 139131555 | 139132114 | 6.3061 | 4.70558 | 4.13437 |
| chr9 | 65550945 | 65551321 | 182 | Distal Intergenic | 9 | 65554386 | 65580040 | 25655 | 2 | ENSMUST00 | 28719 | Plekho2 | ENSMUSG0C | false | . | . | . | . | . | . | . | . | . | . | . | . |
| chr6 | 49319138 | 49319510 | 182 | Promoter (<=1kb) | 6 | 49319274 | 49341582 | 22309 | 1 | ENSMUST00 | 0 | Ccdc126 | ENSMUSG0C | false | chr6 | 49319068 | 49320144 | 22.99841 | 82.87405 | 79.66897 | . | . | . | . | . | . |
| chr8 | 45999737 | 46000252 | 182 | Promoter (<=1kb) | 8 | 45996902 | 45999867 | 2966 | 2 | ENSMUST00 | 0 | Ankrd37 | ENSMUSG0C | false | chr8 | 45999323 | 46000786 | 21.1594 | 63.42564 | 60.53069 | . | . | . | . | . | . |
| chr8 | 125592363 | 125592630 | 182 | Distal Intergenic | 8 | 125669818 | 125673359 | 3542 | 1 | ENSMUST00 | -77188 | Map10 | ENSMUSG0C | false | . | . | . | . | . | . | . | . | . | . | . | . |
| chr8 | 125604913 | 125605473 | 182 | Distal Intergenic | 8 | 125669818 | 125673359 | 3542 | 1 | ENSMUST00 | -64345 | Map10 | ENSMUSG0C | false | . | . | . | . | . | . | chr8 | 125604152 | 125606262 | 5.80946 | 4.48749 | 3.66281 |
| chr18 | 5334719 | 5334994 | 182 | Promoter (<=1kb) | 18 | 5210031 | 5334439 | 124409 | 2 | ENSMUST00 | -281 | Zfp438 | ENSMUSG0C | false | chr18 | 5333642 | 5334839 | 18.02749 | 67.47041 | 64.5209 | . | . | . | . | . | . |
| chr11 | 108064472 | 108064675 | 182 | Intron (ENSMUST00000059595.5 | 11 | 107978341 | 108012805 | 34465 | 2 | ENSMUST00 | -51668 | Prkca | ENSMUSG0C | false | . | . | . | . | . | . | . | . | . | . | . | . |
| chr4 | 134551990 | 134552302 | 182 | Promoter (<=1kb) | 4 | 134537892 | 134552166 | 14275 | 2 | ENSMUST00 | 0 | Sepn1 | ENSMUSG0C | false | chr4 | 134551500 | 134552505 | 18.84082 | 47.13427 | 44.45482 | chr4 | 134551168 | 134552708 | 3.31975 | 3.15372 | 1.67263 |
| chr7 | 107574011 | 107574660 | 182 | Intron (ENSMUST00000120990.1 | 7 | 107567446 | 107591094 | 23649 | 1 | ENSMUST00 | 6566 | Olfml1 | ENSMUSG0C | false | . | . | . | . | . | . | chr7 | 107573869 | 107576080 | 3.17803 | 3.14128 | 1.52129 |
| chr11 | 114877125 | 114877346 | 182 | Distal Intergenic | 11 | 114864492 | 114870607 | 6116 | 1 | ENSMUST00 | 12634 | Gprc5c | ENSMUSG0C | false | . | . | . | . | . | . | chr11 | 114872046 | 114877510 | 5.24149 | 4.10614 | 3.22308 |
| chr11 | 114850648 | 114851094 | 182 | Promoter (<=1kb) | 11 | 114851152 | 114863733 | 12582 | 1 | ENSMUST00 | -58 | Gprc5c | ENSMUSG0C | false | chr11 | 114850502 | 114853302 | 31.71022 | 94.86349 | 91.41748 | . | . | . | . | . | . |
| chr3 | 100797531 | 100797752 | 182 | Distal Intergenic | 3 | 100825459 | 100895679 | 70221 | 1 | ENSMUST00 | -27707 | Vtcn1 | ENSMUSG0C | false | . | . | . | . | . | . | chr3 | 100794605 | 100798870 | 5.16708 | 4.05467 | 3.17162 |
| chr2 | 134785964 | 134786212 | 182 | Promoter (<=1kb) | 2 | 134786159 | 135475258 | 689100 | 1 | ENSMUST00 | 0 | Plcb1 | ENSMUSG0C | false | chr2 | 134786019 | 134787187 | 16.88215 | 57.7818 | 54.96725 | chr2 | 134785855 | 134786886 | 4.45266 | 3.74575 | 2.60073 |
| chrX | 166510296 | 166510615 | 182 | Promoter (<=1kb) | 20 | 166499812 | 166510478 | 10667 | 2 | ENSMUST00 | 0 | Tceanc | ENSMUSG0C | false | chrX | 166509715 | 166510472 | 21.59801 | 56.60135 | 53.80005 | . | . | . | . | . | . |
| chr5 | 143731414 | 143731806 | 182 | Promoter (<=1kb) | 5 | 143710325 | 143732280 | 21956 | 2 | ENSMUST00 | 474 | Usp42 | ENSMUSG0C | false | chr5 | 143730802 | 143733135 | 30.32912 | 88.589 | 85.27629 | . | . | . | . | . | . |
| chr2 | 127792124 | 127792616 | 182 | Promoter (<=1kb) | 2 | 127791388 | 127792488 | 1101 | 2 | ENSMUST00 | 0 | 1500011K16 | ENSMUSG0C | false | chr2 | 127791824 | 127792813 | 22.05754 | 58.21236 | 55.38818 | . | . | . | . | . | . |
| chr1 | 64899877 | 64900058 | 182 | Intron (ENSMUST00000139649.2 | 1 | 64860298 | 64928097 | 67800 | 2 | ENSMUST00 | 28039 | Plekhm3 | ENSMUSG0C | false | . | . | . | . | . | . | chr1 | 64898946 | 64900091 | 3.64719 | 3.41122 | 1.88674 |
| chr18 | 38145649 | 38145862 | 182 | Distal Intergenic | 18 | 38188888 | 38203163 | 14276 | 2 | ENSMUST00 | 57301 | Pcdh1 | ENSMUSG0C | false | . | . | . | . | . | . | chr18 | 38144395 | 38145702 | 4.56061 | 3.8535 | 2.63275 |
| chr18 | 38174350 | 38174685 | 182 | Distal Intergenic | 18 | 38188888 | 38203163 | 14276 | 2 | ENSMUST00 | 28478 | Pcdh1 | ENSMUSG0C | false | . | . | . | . | . | . | chr18 | 38173904 | 38175924 | 3.64412 | 3.32517 | 1.93547 |
| chr18 | 38185883 | 38186154 | 182 | 3' UTR | 18 | 38188888 | 38203163 | 14276 | 2 | ENSMUST00 | 17009 | Pcdh1 | ENSMUSG0C | false | chr18 | 38185112 | 38186528 | 20.38514 | 52.9059 | 50.15308 | . | . | . | . | . | . |
| chr7 | 127137828 | 127138145 | 182 | Promoter (<=1kb) | 7 | 127133702 | 127137823 | 4122 | 2 | ENSMUST00 | -6 | Spn | ENSMUSG0C | false | . | . | . | . | . | . | chr7 | 127137152 | 127137960 | 4.97163 | 4.11079 | 3.01357 |
| chr8 | 126736536 | 126736821 | 182 | Distal Intergenic | 8 | 126588296 | 126593986 | 5691 | 2 | ENSMUST00 | -142551 | Irf2bp2 | ENSMUSG0C | false | . | . | . | . | . | . | chr8 | 126733289 | 126737272 | 7.19033 | 5.00384 | 4.80902 |
| chr11 | 110815213 | 110815463 | 182 | Distal Intergenic | 11 | 110968033 | 111027965 | 59933 | 1 | ENSMUST00 | -152570 | Kcnj16 | ENSMUSG0C | false | . | . | . | . | . | . | . | . | . | . | . | . |
| chr5 | 73490974 | 73491705 | 182 | Promoter (<=1kb) | 5 | 73491026 | 73560794 | 69769 | 1 | ENSMUST00 | 0 | Dcun1d4 | ENSMUSG0C | false | chr5 | 73490170 | 73492849 | 32.27147 | 102.84008 | 99.19724 | . | . | . | . | . | . |
| chr5 | 73545689 | 73545957 | 182 | Intron (ENSMUST00000087181.6 | 5 | 73510868 | 73534691 | 23824 | 1 | ENSMUST00 | 34822 | Dcun1d4 | ENSMUSG0C | false | . | . | . | . | . | . | chr5 | 73545423 | 73546074 | 4.61581 | 3.82554 | 2.72551 |
| chr4 | 65679111 | 65679326 | 182 | Intron (ENSMUST00000084496.2 | 4 | 65605249 | 65613756 | 8508 | 1 | ENSMUST00 | 73863 | Trim32 | ENSMUSG0C | false | . | . | . | . | . | . | chr4 | 65678677 | 65680294 | 4.08638 | 3.55756 | 2.26973 |
| chr2 | 69885581 | 69885917 | 182 | Promoter (<=1kb) | 2 | 69871198 | 69885615 | 14418 | 2 | ENSMUST00 | 0 | Mettl5 | ENSMUSG0C | false | chr2 | 69884960 | 69886184 | 20.06521 | 63.98471 | 61.08233 | . | . | . | . | . | . |
| chr16 | 92301208 | 92301538 | 182 | Promoter (<=1kb) | 16 | 92301286 | 92313041 | 11756 | 1 | ENSMUST00 | 0 | Smim11 | ENSMUSG0C | false | chr16 | 92300602 | 92302068 | 26.6505 | 78.45867 | 75.33303 | chr16 | 92300286 | 92303536 | 4.10043 | 3.64578 | 2.30542 |
| chr9 | 60837384 | 60837621 | 182 | Promoter (<=1kb) | 9 | 60821855 | 60838200 | 16346 | 2 | ENSMUST00 | 579 | Gm9869 | ENSMUSG0C | false | chr9 | 60836859 | 60842659 | 24.93112 | 71.82204 | 68.80418 | . | . | . | . | . | . |
| chr9 | 60846974 | 60847388 | 182 | Intron (ENSMUST00000050183.6 | 9 | 60821855 | 60838200 | 16346 | 2 | ENSMUST00 | -8775 | Gm9869 | ENSMUSG0C | false | . | . | . | . | . | . | chr9 | 60839714 | 60851928 | 5.61355 | 4.39804 | 3.5066 |
| chr17 | 27635228 | 27635516 | 182 | Promoter (<=1kb) | 17 | 27630422 | 27635276 | 4855 | 2 | ENSMUST00 | 0 | Rps10 | ENSMUSG0C | false | chr17 | 27634180 | 27635882 | 30.62272 | 92.00563 | 88.61945 | chr17 | 27631062 | 27637618 | 4.84233 | 3.8179 | 2.90957 |
| chr2 | 48813575 | 48814182 | 182 | Promoter (<=1kb) | 2 | 48814109 | 48903269 | 89161 | 1 | ENSMUST00 | 0 | Acvr2a | ENSMUSG0C | false | chr2 | 48813188 | 48815735 | 32.23851 | 100.76218 | 97.17527 | . | . | . | . | . | . |
| chr5 | 144184914 | 144185148 | 182 | 3' UTR | 5 | 144190286 | 144194441 | 4156 | 1 | ENSMUST00 | -5138 | Bhlha15 | ENSMUSG0C | false | . | . | . | . | . | . | chr5 | 144181835 | 144185686 | 5.69891 | 4.32402 | 3.61705 |
| chr7 | 143009104 | 143009415 | 182 | Promoter (1-2kb) | 7 | 142988969 | 143008033 | 19065 | 2 | ENSMUST00 | -1072 | Tspan32os | ENSMUSG0C | false | . | . | . | . | . | . | chr7 | 143004528 | 143022688 | 7.71992 | 5.26439 | 5.23131 |
| chr7 | 4658313 | 4658612 | 182 | Promoter (<=1kb) | 7 | 4631501 | 4658950 | 27450 | 2 | ENSMUST00 | 338 | Ppp6r1 | ENSMUSG0C | false | chr7 | 4657404 | 4659752 | 39.54591 | 131.3782 | 126.83877 | . | . | . | . | . | . |
| chr7 | 4658902 | 4659296 | 182 | Promoter (<=1kb) | 7 | 4631501 | 4658950 | 27450 | 2 | ENSMUST00 | 0 | Ppp6r1 | ENSMUSG0C | false | chr7 | 4657404 | 4659752 | 39.54591 | 131.3782 | 126.83877 | . | . | . | . | . | . |
| chr11 | 54787140 | 54787407 | 182 | Promoter (<=1kb) | 11 | 54740310 | 54787625 | 47316 | 2 | ENSMUST00 | 218 | Cdc42se2 | ENSMUSG0C | true | chr11 | 54785734 | 54788212 | 43.43732 | 157.3205 | 151.76292 | . | . | . | . | . | . |
| chr1 | 54925888 | 54926595 | 182 | Promoter (<=1kb) | 1 | 54645340 | 54926387 | 281048 | 2 | ENSMUST00 | 0 | Ankrd44 | ENSMUSG0C | true | chr1 | 54925084 | 54926821 | 33.08674 | 107.33151 | 103.57004 | chr1 | 54926542 | 54927069 | 5.97917 | 4.64153 | 3.82214 |
| chr2 | 74711812 | 74712175 | 182 | Promoter (<=1kb) | 2 | 74703248 | 74712054 | 8807 | 2 | ENSMUST00 | 0 | Hoxd3os1 | ENSMUSG0C | false | chr2 | 74711584 | 74713681 | 43.85173 | 143.28317 | 138.27043 | . | . | . | . | . | . |
| chr17 | 6746047 | 6746320 | 182 | Intron (ENSMUST00000064234.6 | 17 | 6740837 | 6742821 | 1985 | 2 | ENSMUST00 | -3227 | Ezr | ENSMUSG0C | false | . | . | . | . | . | . | chr17 | 6734583 | 6749633 | 6.00896 | 4.44798 | 3.8635 |
| chr17 | 6782697 | 6783565 | 182 | Promoter (<=1kb) | 17 | 6738041 | 6782784 | 44744 | 2 | ENSMUST00 | 0 | Ezr | ENSMUSG0C | false | chr17 | 6777892 | 6783808 | 43.16833 | 149.98996 | 144.69951 | . | . | . | . | . | . |
| chr2 | 26964233 | 26964698 | 182 | Promoter (<=1kb) | 2 | 26953563 | 26964386 | 10824 | 2 | ENSMUST00 | 0 | Rexo4 | ENSMUSG0C | false | chr2 | 26963536 | 26965076 | 29.2476 | 106.45832 | 102.71833 | chr2 | 26962087 | 26965131 | 4.42554 | 3.70003 | 2.55324 |
| chr16 | 44101432 | 44101673 | 182 | Exon (ENSMUST00000114666.3/ | 16 | 44097175 | 44138980 | 41806 | 2 | ENSMUST00 | 37307 | Atp6v1a | ENSMUSG0C | false | . | . | . | . | . | . | chr16 | 44101173 | 44102302 | 3.61842 | 3.23893 | 1.89712 |
| chr13 | 98697419 | 98697679 | 182 | Promoter (2-3kb) | 13 | 98686235 | 98694834 | 8600 | 2 | ENSMUST00 | -2586 | Tmem171 | ENSMUSG0C | false | . | . | . | . | . | . | chr13 | 98696816 | 98698103 | 2.77368 | 2.83049 | 1.22416 |
| chr8 | 72474753 | 72475343 | 182 | Promoter (<=1kb) | 8 | 72460489 | 72475233 | 14745 | 2 | ENSMUST00 | 0 | Cherp | ENSMUSG0C | false | chr8 | 72474318 | 72476077 | 33.09821 | 104.28432 | 100.60858 | . | . | . | . | . | . |
| chr7 | 49449824 | 49450027 | 182 | Promoter (<=1kb) | 7 | 49450760 | 49464580 | 13821 | 1 | ENSMUST00 | -733 | Nav2 | ENSMUSG0C | false | . | . | . | . | . | . | chr7 | 49446529 | 49453829 | 5.00391 | 3.99232 | 3.02197 |
| chr7 | 49475446 | 49475633 | 182 | Intron (ENSMUST00000183659.2 | 7 | 49450760 | 49464580 | 13821 | 1 | ENSMUST00 | 24687 | Nav2 | ENSMUSG0C | false | . | . | . | . | . | . | chr7 | 49475024 | 49478318 | 4.82944 | 4.0116 | 2.86901 |
| chr7 | 49132865 | 49133082 | 182 | Intron (ENSMUST00000183659.2 | 7 | 49246189 | 49409643 | 163455 | 1 | ENSMUST00 | -113107 | Nav2 | ENSMUSG0C | false | . | . | . | . | . | . | . | . | . | . | . | . |
| chr7 | 49280527 | 49280780 | 182 | Intron (ENSMUST00000183659.2 | 7 | 49304138 | 49411406 | 107269 | 1 | ENSMUST00 | -23358 | Nav2 | ENSMUSG0C | false | . | . | . | . | . | . | chr7 | 49279823 | 49281872 | 4.44421 | 3.74118 | 2.56199 |
| chr7 | 49521438 | 49521653 | 182 | Intron (ENSMUST00000183659.2 | 7 | 49548192 | 49556996 | 8805 | 1 | ENSMUST00 | -26539 | Nav2 | ENSMUSG0C | false | . | . | . | . | . | . | chr7 | 49520693 | 49523191 | 4.96489 | 4.064 | 2.98742 |
| chr7 | 49321175 | 49321380 | 182 | Intron (ENSMUST00000183659.2 | 7 | 49304254 | 49305917 | 1664 | 1 | ENSMUST00 | 16922 | Nav2 | ENSMUSG0C | false | . | . | . | . | . | . | chr7 | 49320259 | 49321459 | 4.10806 | 3.4911 | 2.30366 |
| chr2 | 30286305 | 30286512 | 182 | Promoter (<=1kb) | 2 | 30286397 | 30307732 | 21336 | 1 | ENSMUST00 | 0 | Nup188 | ENSMUSG0C | false | chr2 | 30285584 | 30287487 | 34.78613 | 113.42423 | 109.48744 | . | . | . | . | . | . |
| chr8 | 117172887 | 117173121 | 182 | Intron (ENSMUST00000064488.5 | 8 | 117158137 | 117205834 | 47698 | 1 | ENSMUST00 | 14751 | Gan | ENSMUSG0C | false | . | . | . | . | . | . | chr8 | 117172583 | 117179361 | 6.73029 | 4.87581 | 4.42508 |
| chr15 | 90679272 | 90679625 | 182 | Promoter (<=1kb) | 15 | 90487482 | 90679371 | 191890 | 2 | ENSMUST00 | 0 | Cpne8 | ENSMUSG0C | false | chr15 | 90678118 | 90680514 | 25.16654 | 87.04672 | 83.76057 | chr15 | 90677941 | 90680555 | 3.37083 | 3.25226 | 1.65398 |
| chr6 | 72510723 | 72511063 | 182 | Downstream (2-3kb) | 6 | 72513644 | 72515290 | 1647 | 2 | ENSMUST00 | 4227 | Sh2d6 | ENSMUSG0C | false | . | . | . | . | . | . | chr6 | 72510186 | 72516424 | 7.38109 | 5.05551 | 4.9697 |
| chr6 | 72514068 | 72514275 | 182 | Promoter (1-2kb) | 6 | 72513644 | 72515290 | 1647 | 2 | ENSMUST00 | 1015 | Sh2d6 | ENSMUSG0C | false | . | . | . | . | . | . | chr6 | 72510186 | 72516424 | 7.38109 | 5.05551 | 4.9697 |
| chr9 | 120577128 | 120577462 | 182 | Promoter (<=1kb) | 9 | 120577346 | 120578072 | 727 | 1 | ENSMUST00 | 0 | 5830454E08 | ENSMUSG0C | false | chr9 | 120576838 | 120577979 | 22.37683 | 79.83291 | 76.68538 | chr9 | 120576533 | 120578356 | 5.20142 | 4.23175 | 3.16252 |
| chr12 | 69447847 | 69448140 | 182 | Distal Intergenic | 12 | 69371835 | 69372458 | 624 | 2 | ENSMUST00 | -75390 | Gm9887 | ENSMUSG0C | false | . | . | . | . | . | . | chr12 | 69445497 | 69452349 | 6.03809 | 4.48697 | 3.87006 |
| chr12 | 69424899 | 69425248 | 182 | Distal Intergenic | 12 | 69371835 | 69372458 | 624 | 2 | ENSMUST00 | -52442 | Gm9887 | ENSMUSG0C | false | . | . | . | . | . | . | chr12 | 69417044 | 69426977 | 5.04673 | 4.12135 | 3.05328 |
| chr17 | 29268744 | 29269167 | 182 | Promoter (<=1kb) | 17 | 29268788 | 29302881 | 34094 | 1 | ENSMUST00 | 0 | BC004004 | ENSMUSG0C | false | chr17 | 29268447 | 29269771 | 19.35304 | 74.51352 | 71.45322 | chr17 | 29267954 | 29271766 | 3.98432 | 3.52849 | 2.18847 |
| chr9 | 114758490 | 114758675 | 182 | Intron (ENSMUST00000035009.9 | 9 | 114780100 | 114782257 | 2158 | 1 | ENSMUST00 | -21425 | Gm9888 | ENSMUSG0C | false | . | . | . | . | . | . | chr9 | 114758262 | 114780983 | 6.81555 | 4.89533 | 4.50006 |
| chr6 | 73248400 | 73248771 | 182 | Promoter (<=1kb) | 6 | 73248652 | 73264273 | 15622 | 1 | ENSMUST00 | 0 | Suclg1 | ENSMUSG0C | false | chr6 | 73247964 | 73249716 | 30.78991 | 105.27288 | 101.56436 | . | . | . | . | . | . |
| chr8 | 84979954 | 84980465 | 182 | Promoter (1-2kb) | 8 | 84976909 | 84978748 | 1840 | 2 | ENSMUST00 | -1207 | Junb | ENSMUSG0C | false | chr8 | 84979560 | 84980539 | 14.16908 | 34.09626 | 31.58084 | chr8 | 84974675 | 84986551 | 7.15 | 5.07534 | 4.74925 |
| chr11 | 98786463 | 98787108 | 182 | Distal Intergenic | 11 | 98795516 | 98807859 | 12344 | 1 | ENSMUST00 | -8408 | Msl1 | ENSMUSG0C | false | . | . | . | . | . | . | chr11 | 98780022 | 98794752 | 7.02549 | 5.01464 | 4.66092 |
| chr16 | 56075260 | 56076006 | 182 | Promoter (<=1kb) | 16 | 56075409 | 56126544 | 51136 | 1 | ENSMUST00 | 0 | Senp7 | ENSMUSG0C | false | chr16 | 56075165 | 56076480 | 34.64357 | 105.94113 | 102.22192 | . | . | . | . | . | . |
| chr19 | 31764252 | 31764483 | 182 | Promoter (<=1kb) | 19 | 31742114 | 31764973 | 22860 | 2 | ENSMUST00 | 490 | Prkg1 | ENSMUSG0C | true | chr19 | 31762329 | 31765338 | 26.95677 | 77.13882 | 74.03856 | chr19 | 31762049 | 31764545 | 3.9805 | 3.60883 | 2.1536 |
| chr18 | 75695257 | 75695512 | 182 | Promoter (2-3kb) | 18 | 75431224 | 75697696 | 266473 | 2 | ENSMUST00 | 2184 | Ctif | ENSMUSG0C | false | chr18 | 75695237 | 75698016 | 23.65566 | 74.08845 | 71.03342 | chr18 | 75694176 | 75696498 | 4.60551 | 3.8793 | 2.68165 |
| chr13 | 60057701 | 60057987 | 182 | Distal Intergenic | 13 | 60174405 | 60177365 | 2961 | 2 | ENSMUST00 | 119378 | Gas1 | ENSMUSG0C | false | . | . | . | . | . | . | chr13 | 60057459 | 60058653 | 4.41101 | 3.89771 | 2.50833 |
| chr6 | 53615836 | 53616096 | 182 | Intron (ENSMUST00000047450.7 | 6 | 53573384 | 53696007 | 122624 | 1 | ENSMUST00 | 42453 | Creb5 | ENSMUSG0C | false | . | . | . | . | . | . | chr6 | 53615239 | 53616087 | 2.81179 | 2.91682 | 1.26378 |
| chr9 | 8003711 | 8004281 | 182 | Promoter (<=1kb) | 9 | 7973796 | 8003757 | 29962 | 2 | ENSMUST00 | 0 | Yap1 | ENSMUSG0C | false | chr9 | 8002688 | 8005179 | 35.84351 | 110.06287 | 106.22365 | . | . | . | . | . | . |
| chr7 | 80387893 | 80388128 | 182 | Promoter (<=1kb) | 7 | 80377758 | 80387946 | 10189 | 2 | ENSMUST00 | 0 | Fes | ENSMUSG0C | false | chr7 | 80387019 | 80388177 | 7.51422 | 13.34765 | 11.11425 | chr7 | 80386474 | 80388442 | 5.21552 | 4.03402 | 3.22878 |
| chr5 | 4192249 | 4192450 | 182 | Promoter (<=1kb) | 5 | 4192367 | 4197651 | 5285 | 1 | ENSMUST00 | 0 | Mterf1b | ENSMUSG0C | false | chr5 | 4191986 | 4193509 | 43.81247 | 194.26276 | 187.87305 | chr5 | 4192051 | 4193570 | 3.29619 | 3.19035 | 1.59041 |
| chr9 | 53248036 | 53248260 | 182 | Promoter (<=1kb) | 9 | 53098454 | 53248112 | 149659 | 2 | ENSMUST00 | 0 | Ddx10 | ENSMUSG0C | false | chr9 | 53247237 | 53248682 | 38.55812 | 129.64772 | 125.16568 | . | . | . | . | . | . |
| chr7 | 27178747 | 27179062 | 182 | Promoter (<=1kb) | 7 | 27168424 | 27178896 | 10473 | 2 | ENSMUST00 | 0 | Rab4b | ENSMUSG0C | false | chr7 | 27178237 | 27179485 | 29.79559 | 88.63454 | 85.31541 | . | . | . | . | . | . |
| chr9 | 37555180 | 37555454 | 182 | Promoter (2-3kb) | 9 | 37544492 | 37552904 | 8413 | 2 | ENSMUST00 | -2277 | Nrgn | ENSMUSG0C | false | . | . | . | . | . | . | . | . | . | . | . | . |
| chr17 | 28691162 | 28691503 | 182 | Promoter (<=1kb) | 17 | 28691342 | 28741164 | 49823 | 1 | ENSMUST00 | 0 | Mapk14 | ENSMUSG0C | false | chr17 | 28690674 | 28692802 | 43.58744 | 151.86528 | 146.50356 | . | . | . | . | . | . |
| chr18 | 69667173 | 69667446 | 182 | Intron (ENSMUST00000066717.6 | 18 | 69679699 | 69684528 | 4830 | 1 | ENSMUST00 | -12253 | Tcf4 | ENSMUSG0C | false | . | . | . | . | . | . | chr18 | 69665126 | 69667715 | 3.89112 | 3.49655 | 2.13685 |
| chr4 | 109043665 | 109043997 | 182 | Promoter (2-3kb) | 4 | 109046310 | 109050900 | 4591 | 1 | ENSMUST00 | -2313 | Nrd1 | ENSMUSG0C | false | . | . | . | . | . | . | chr4 | 109043109 | 109045879 | 3.98998 | 3.50945 | 2.22692 |
| chr2 | 130295007 | 130295719 | 182 | Promoter (<=1kb) | 2 | 130295169 | 130370481 | 75313 | 1 | ENSMUST00 | 0 | Ebf4 | ENSMUSG0C | false | chr2 | 130294730 | 130296281 | 15.45146 | 50.98658 | 48.25883 | chr2 | 130293549 | 130296921 | 5.46342 | 4.23521 | 3.41383 |
| chr15 | 100636813 | 100637159 | 182 | Promoter (<=1kb) | 15 | 100621342 | 100636855 | 15514 | 2 | ENSMUST00 | 0 | Smagp | ENSMUSG0C | false | chr15 | 100636116 | 100637647 | 36.76257 | 113.72284 | 109.77938 | chr15 | 100635361 | 100637620 | 3.85169 | 3.47786 | 2.08285 |
| chr2 | 125469487 | 125469693 | 182 | Intron (ENSMUST00000028633.7 | 2 | 125505089 | 125507881 | 2793 | 1 | ENSMUST00 | -35396 | Gm9913 | ENSMUSG0C | false | . | . | . | . | . | . | . | . | . | . | . | . |
| chr19 | 47432465 | 47432717 | 182 | Intron (ENSMUST00000081619.3 | 19 | 47260174 | 47464411 | 204238 | 2 | ENSMUST00 | 31694 | Sh3pxd2a | ENSMUSG0C | false | . | . | . | . | . | . | . | . | . | . | . | . |
| chr2 | 17363847 | 17364037 | 182 | Promoter (2-3kb) | 2 | 17348694 | 17361086 | 12393 | 2 | ENSMUST00 | -2762 | Nebl | ENSMUSG0C | true | . | . | . | . | . | . | chr2 | 17362870 | 17366947 | 5.09212 | 3.96546 | 3.12404 |
| chr7 | 25379630 | 25379891 | 182 | 3' UTR | 7 | 25384805 | 25385573 | 769 | 1 | ENSMUST00 | -4914 | 4732471J01R | ENSMUSG0C | false | chr7 | 25379383 | 25380204 | 11.64009 | 27.76484 | 25.32713 | chr7 | 25376454 | 25381332 | 4.0129 | 3.53444 | 2.24503 |
| chr9 | 106367678 | 106367981 | 182 | Promoter (<=1kb) | 9 | 106368632 | 106375724 | 7093 | 1 | ENSMUST00 | -651 | Dusp7 | ENSMUSG0C | false | chr9 | 106367929 | 106370134 | 27.97104 | 107.99458 | 104.2211 | . | . | . | . | . | . |
| chr11 | 75976850 | 75977037 | 182 | Distal Intergenic | 11 | 75999912 | 76003569 | 3658 | 1 | ENSMUST00 | -22875 | 1700016K19 | ENSMUSG0C | false | . | . | . | . | . | . | . | . | . | . | . | . |
| chr3 | 126596059 | 126596431 | 182 | Distal Intergenic | 3 | 126560302 | 126684688 | 88387 | 1 | ENSMUST00 | 0 | Camk2d | ENSMUSG0C | false | chr3 | 126595914 | 126598682 | 36.46498 | 119.52629 | 115.39857 | . | . | . | . | . | . |
| chr4 | 129640665 | 129641250 | 182 | Promoter (<=1kb) | 4 | 129626078 | 129641065 | 14988 | 2 | ENSMUST00 | 0 | Txlna | ENSMUSG0C | false | chr4 | 129639863 | 129641496 | 19.52531 | 57.12588 | 54.31675 | . | . | . | . | . | . |
| chr6 | 72438945 | 72439217 | 182 | Promoter (<=1kb) | 6 | [illegible] | [illegible] | 6760 | 1 | ENSMUST00 | 341 | Mat2a | ENSMUSG0C | false | chr6 | 72437281 | 72440865 | 28.40054 | 134.09712 | 129.45372 | . | . | . | . | . | . |
| chr15 | 76659749 | 76660296 | 182 | Promoter (<=1kb) | 15 | 76643395 | 76660061 | 16667 | 2 | ENSMUST00 | 0 | Cyhr1 | ENSMUSG0C | false | chr15 | 76659480 | 76661269 | 13.94362 | 38.54297 | 35.97016 | chr15 | 76657269 | 76661601 | 5.18146 | 4.11399 | 3.17215 |
| chr3 | 121427649 | 121427843 | 182 | Promoter (1-2kb) | 3 | 121426541 | 121458205 | 31665 | 1 | ENSMUST00 | 1109 | Cnn3 | ENSMUSG0C | false | chr3 | 121425851 | 121428534 | 43.84918 | 145.52415 | 140.42462 | . | . | . | . | . | . |
| chr1 | 180510373 | 180510685 | 182 | Distal Intergenic | 1 | 180438320 | 180483504 | 45176 | 1 | ENSMUST00 | -26870 | 6330403A02 | ENSMUSG0C | false | . | . | . | . | . | . | chr1 | 180507395 | 180511832 | 5.07172 | 4.07804 | 3.0995 |
| chr3 | 122729597 | 122730594 | 182 | Intron (ENSMUST00000075632.7 | 3 | 122729158 | 122859374 | 130217 | 1 | ENSMUST00 | 440 | Pde5a | ENSMUSG0C | false | chr3 | 122728814 | 122731114 | 33.08674 | 107.33151 | 103.57004 | chr3 | 122729944 | 122730785 | 2.68071 | 2.65351 | 1.16719 |
| chr4 | 93334922 | 93335529 | 182 | Promoter (<=1kb) | 4 | 93334138 | 93335511 | 1374 | 2 | ENSMUST00 | 0 | Tusc1 | ENSMUSG0C | false | chr4 | 93334516 | 93335970 | 36.04672 | 129.50026 | 125.02953 | . | . | . | . | . | . |
| chr10 | 86771726 | 86771973 | 182 | Intron ([illegible] | 10 | 86779005 | 86838389 | 59385 | 1 | ENSMUST00 | -7032 | Nt5dc3 | ENSMUSG0C | false | . | . | . | . | . | . | chr10 | 86771240 | 86772426 | 3.43745 | 3.29154 | 1.76858 |
| chr11 | 93884969 | 93885360 | 182 | Promoter (<=1kb) | 11 | 93859243 | 93885766 | 26524 | 2 | ENSMUST00 | 406 | Utp18 | ENSMUSG0C | false | chr11 | 93884540 | 93887391 | 41.19049 | 135.33968 | 130.64169 | chr11 | 93884485 | 93884986 | 7.07098 | 5.14667 | 4.68641 |
| chr15 | 9135114 | 9135294 | 182 | Intron (ENSMUST00000096482.4 | 15 | 9127048 | 9140352 | 13305 | 2 | ENSMUST00 | 5058 | Skp2 | ENSMUSG0C | false | . | . | . | . | . | . | chr15 | 9134256 | 9135946 | 3.26048 | 3.03489 | 1.61447 |
| chr12 | 104965370 | 104965555 | 182 | Intron (ENSMUST00000095439.4 | 12 | 104929949 | 104998677 | 68729 | 2 | ENSMUST00 | 33122 | Syne3 | ENSMUSG0C | false | . | . | . | . | . | . | chr12 | 104960454 | 104967194 | 6.25608 | 4.13377 | 3.08121 |
| chr19 | 38223981 | 38224233 | 182 | Promoter (<=1kb) | 19 | 38188481 | 38224138 | 35658 | 2 | ENSMUST00 | 0 | Fra10ac1 | ENSMUSG0C | false | chr19 | 38223425 | 38224996 | 23.61999 | 85.9385 | 82.67356 | . | . | . | . | . | . |
| chr13 | 41662206 | 41662758 | 182 | Distal Intergenic | 13 | 41655697 | 41656524 | 828 | 1 | ENSMUST00 | 6510 | Gm5082 | ENSMUSG0C | false | . | . | . | . | . | . | chr13 | 41658087 | 41664664 | 5.64312 | 4.43444 | 3.52317 |
| chr15 | 7070486 | 7070690 | 182 | Distal Intergenic | 15 | 7129559 | 7191743 | 62185 | 1 | ENSMUST00 | -58869 | Lifr | ENSMUSG0C | false | . | . | . | . | . | . | . | . | . | . | . | . |
| chr15 | 83349886 | 83350392 | 182 | Promoter (<=1kb) | 15 | 83299740 | 83350247 | 50508 | 2 | ENSMUST00 | 0 | Arfgap3 | ENSMUSG0C | false | chr15 | 83349550 | 83350723 | 19.61815 | 75.94017 | 72.85583 | chr15 | 83348799 | 83349951 | 5.66828 | 4.41492 | 3.61518 |
| chr8 | 105437313 | 105437614 | 182 | Intron (ENSMUST00000139900.2 | 8 | 105427640 | 105464084 | 36445 | 1 | ENSMUST00 | 9674 | Lrrc36 | ENSMUSG0C | false | . | . | . | . | . | . | chr8 | 105436098 | 105438102 | 4.43887 | 3.76167 | 2.58171 |
| chr12 | 8511034 | 8511253 | 182 | Distal Intergenic | 12 | 8497763 | 8499985 | 2223 | 1 | ENSMUST00 | -11050 | Rhob | ENSMUSG0C | false | chr12 | 8510984 | 8511787 | 6.58118 | 14.31666 | 12.06619 | chr12 | 8509731 | 8512412 | 5.11283 | 4.02341 | 3.13197 |
| chr10 | 81560338 | 81560580 | 182 | Promoter (<=1kb) | 10 | 81559494 | 81566362 | 6869 | 1 | ENSMUST00 | 845 | Aes | ENSMUSG0C | false | chr10 | 81559124 | 81561616 | 23.12601 | 77.64799 | 74.53377 | . | . | . | . | . | . |
| chr2 | 173795228 | 173795611 | 182 | Intron (ENSMUST00000138925.1 | 2 | 173768200 | 173778125 | 9926 | 1 | ENSMUST00 | 27029 | Vapb | ENSMUSG0C | false | . | . | . | . | . | . | chr2 | 173788713 | 173798959 | 4.81417 | 3.91068 | 2.86471 |
| chr17 | 73072281 | 73072570 | 182 | Distal Intergenic | 17 | 73107985 | 73161991 | 54007 | 1 | ENSMUST00 | -35415 | Lclat1 | ENSMUSG0C | false | . | . | . | . | . | . | chr17 | 73071267 | 73073107 | 3.79601 | 3.42915 | 2.07243 |
| chr6 | 117495016 | 117495289 | 182 | Distal Intergenic | 6 | 117559697 | 117568193 | 8497 | 1 | ENSMUST00 | -64408 | Gm9946 | ENSMUSG0C | false | . | . | . | . | . | . | chr6 | 117492016 | 117499530 | 5.51915 | 4.32118 | 3.4422 |
| chr6 | 117407505 | 117407848 | 182 | Distal Intergenic | 6 | 117559697 | 117568193 | 8497 | 1 | ENSMUST00 | -151849 | Gm9946 | ENSMUSG0C | false | . | . | . | . | . | . | chr6 | 117406854 | 117409614 | 3.55968 | 3.20235 | 1.85945 |

| | | | | | | | | | | | | | | | | | | | | | | | | | | |
|---|---|---|---|---|---|---|---|---|---|---|---|---|---|---|---|---|---|---|---|---|---|---|---|---|---|---|
| chr5 | 34526106 | 34526453 | 182 | Promoter (<=1kb) | 5 | 34525846 | 34563625 | 37780 | 1 | ENSMUST00 | 261 | Sh3bp2 | ENSMUSG0C | false | chr5 | 34525486 | 34526710 | 24.81473 | 68.0602 | 65.09722 | chr5 | 34525252 | 34527508 | 4.75815 | 3.96988 | 2.7987 |
| chr7 | 133963600 | 133963968 | 182 | Promoter (2-3kb) | 7 | 133886196 | 133961265 | 75070 | 2 | ENSMUST00 | -2336 | Adam12 | ENSMUSG0C | false | . | . | . | . | . | . | chr7 | 133963443 | 133964232 | 4.20865 | 3.75995 | 2.36036 |
| chr8 | 34064298 | 34064523 | 182 | Distal Intergenic | 8 | 34054622 | 34057046 | 2425 | 1 | ENSMUST00 | 9677 | Gm9951 | ENSMUSG0C | false | . | . | . | . | . | . | . | . | . | . | . | . |
| chr5 | 151412980 | 151413206 | 182 | Intron (ENSMUST00000186838.1 | 5 | 151367720 | 151428210 | 60491 | 2 | ENSMUST00 | 15004 | D730045B01 | ENSMUSG0C | false | . | . | . | . | . | . | chr5 | 151411739 | 151414029 | 3.98746 | 3.52656 | 2.19278 |
| chr4 | 53631795 | 53632075 | 182 | Promoter (<=1kb) | 4 | 53631716 | 53659678 | 27963 | 1 | ENSMUST00 | 80 | Fsd1l | ENSMUSG0C | false | chr4 | 53631039 | 53632373 | 32.38209 | 104.04922 | 100.37821 | chr4 | 53631039 | 53632182 | 3.46978 | 3.33971 | 1.72576 |
| chr8 | 45122284 | 45122530 | 182 | Distal Intergenic | 8 | 45069374 | 45087356 | 17983 | 1 | ENSMUST00 | 52911 | Mtnr1a | ENSMUSG0C | false | . | . | . | . | . | . | . | . | . | . | . | . |
| chr9 | 110476508 | 110476863 | 182 | Promoter (<=1kb) | 9 | 110425928 | 110476590 | 50663 | 2 | ENSMUST00 | 0 | Klhl18 | ENSMUSG0C | false | chr9 | 110475529 | 110477398 | 23.80619 | 80.75649 | 77.58801 | . | . | . | . | . | . |
| chr7 | 28966040 | 28966297 | 182 | Distal Intergenic | 7 | 28893248 | 28962340 | 69093 | 2 | ENSMUST00 | -3701 | Actn4 | ENSMUSG0C | true | . | . | . | . | . | . | chr7 | 28962675 | 28973922 | 7.08725 | 4.96018 | 4.72253 |
| chr7 | 28911690 | 28912019 | 182 | Intron (ENSMUST00000068045.8 | 7 | 28901915 | 28907686 | 5772 | 2 | ENSMUST00 | -4005 | Actn4 | ENSMUSG0C | true | . | . | . | . | . | . | chr7 | 28910727 | 28959574 | 8.49369 | 5.62595 | 5.86532 |
| chr7 | 28958474 | 28958725 | 182 | Intron (ENSMUST00000068045.8 | 7 | 28898620 | 28962223 | 63604 | 2 | ENSMUST00 | 3498 | Actn4 | ENSMUSG0C | true | . | . | . | . | . | . | chr7 | 28910727 | 28959574 | 8.49369 | 5.62595 | 5.86532 |
| chr9 | 123259889 | 123260275 | 182 | Promoter (<=1kb) | 9 | 123259058 | 123260789 | 1732 | 2 | ENSMUST00 | 514 | Tmem158 | ENSMUSG0C | false | chr9 | 123259525 | 123261205 | 20.4053 | 65.4752 | 62.55133 | chr9 | 123259434 | 123261224 | 4.51486 | 3.80384 | 2.61046 |
| chr7 | 6286195 | 6286563 | 182 | Promoter (<=1kb) | 7 | 6286579 | 6307883 | 21305 | 1 | ENSMUST00 | -16 | Zfp667 | ENSMUSG0C | false | chr7 | 6285947 | 6287260 | 24.3552 | 66.39809 | 63.45943 | chr7 | 6285931 | 6287364 | 3.52063 | 3.25913 | 1.83209 |
| chr15 | 76925410 | 76925765 | 182 | Distal Intergenic | 15 | 76910851 | 76918010 | 7160 | 2 | ENSMUST00 | -7401 | Zfp647 | ENSMUSG0C | false | chr15 | 76924945 | 76926154 | 25.16654 | 87.04672 | 83.76057 | chr15 | 76924210 | 76926433 | 4.10021 | 3.65213 | 2.30067 |
| chr10 | 81407557 | 81407808 | 182 | Promoter (<=1kb) | 10 | 81400769 | 81407641 | 6873 | 2 | ENSMUST00 | 0 | Nfic | ENSMUSG0C | false | . | . | . | . | . | . | chr10 | 81392649 | 81426205 | 8.72751 | 5.63067 | 6.0679 |
| chr10 | 81431671 | 81432014 | 182 | Promoter (<=1kb) | 10 | 81400953 | 81431005 | 30053 | 2 | ENSMUST00 | -667 | Nfic | ENSMUSG0C | false | . | . | . | . | . | . | chr10 | 81427223 | 81436730 | 7.9801 | 5.39294 | 5.46678 |
| chr13 | 17694611 | 17695300 | 182 | Promoter (<=1kb) | 13 | 16857855 | 17694732 | 836878 | 2 | ENSMUST00 | 0 | Sugct | ENSMUSG0C | false | chr13 | 17694330 | 17694872 | 25.90858 | 83.69981 | 80.48042 | . | . | . | . | . | . |
| chr5 | 36796035 | 36796364 | 182 | Promoter (<=1kb) | 5 | 36794867 | 36796772 | 1906 | 2 | ENSMUST00 | 408 | Mrfap1 | ENSMUSG0C | false | chr5 | 36795370 | 36797310 | 32.27147 | 102.84008 | 99.19724 | chr5 | 36794910 | 36797265 | 4.3313 | 3.67231 | 2.46797 |
| chr7 | 112660404 | 112660655 | 182 | Intron (ENSMUST00000146145.2 | 7 | 112679318 | 112899459 | 220142 | 1 | ENSMUST00 | -18663 | Tead1 | ENSMUSG0C | false | . | . | . | . | . | . | chr7 | 112659009 | 112665328 | 4.94469 | 4.01538 | 2.96709 |
| chr7 | 112663231 | 112663622 | 182 | Intron (ENSMUST00000146145.2 | 7 | 112679318 | 112899459 | 220142 | 1 | ENSMUST00 | -15696 | Tead1 | ENSMUSG0C | false | chr7 | 112662557 | 112663376 | 10.11741 | 21.06654 | 18.71757 | chr7 | 112659009 | 112665328 | 4.94469 | 4.01538 | 2.96709 |
| chr7 | 112717680 | 112717923 | 182 | Intron (ENSMUST00000084705.6 | 7 | 112742045 | 112898660 | 156616 | 1 | ENSMUST00 | -24122 | Tead1 | ENSMUSG0C | false | . | . | . | . | . | . | chr7 | 112716220 | 112724700 | 5.77743 | 4.2821 | 3.6747 |
| chr1 | 74025312 | 74025626 | 182 | Intron (ENSMUST00000169786.2 | 1 | 73990374 | 74016422 | 26049 | 2 | ENSMUST00 | -8891 | Tns1 | ENSMUSG0C | false | . | . | . | . | . | . | chr1 | 74023768 | 74026067 | 3.53515 | 3.29983 | 1.86529 |
| chr1 | 73932566 | 73932838 | 182 | Promoter (1-2kb) | 1 | 73925417 | 73930811 | 5395 | 2 | ENSMUST00 | -1756 | Tns1 | ENSMUSG0C | false | . | . | . | . | . | . | chr1 | 73927832 | 73933110 | 4.94644 | 3.95251 | 2.98269 |
| chr1 | 74008688 | 74008948 | 182 | Intron (ENSMUST00000169786.2 | 1 | 73992205 | 74001567 | 9363 | 2 | ENSMUST00 | -7122 | Tns1 | ENSMUSG0C | false | . | . | . | . | . | . | chr1 | 74006078 | 74010176 | 3.31651 | 3.21932 | 1.61985 |
| chr1 | 73902725 | 73902960 | 182 | Distal Intergenic | 1 | 73925417 | 73928217 | 2801 | 2 | ENSMUST00 | 25257 | Tns1 | ENSMUSG0C | false | . | . | . | . | . | . | chr1 | 73902016 | 73903067 | 3.7807 | 3.53465 | 2.02164 |
| chr1 | 74026669 | 74026901 | 182 | Intron (ENSMUST00000169786.2 | 1 | 73993163 | 74037133 | 43971 | 2 | ENSMUST00 | 10232 | Tns1 | ENSMUSG0C | false | . | . | . | . | . | . | . | . | . | . | . | . |
| chr1 | 73905196 | 73905590 | 182 | Distal Intergenic | 1 | 73925417 | 73928217 | 2801 | 2 | ENSMUST00 | 22627 | Tns1 | ENSMUSG0C | false | . | . | . | . | . | . | . | . | . | . | . | . |
| chr1 | 74034874 | 74035065 | 182 | Promoter (2-3kb) | 1 | 73993163 | 74037133 | 43971 | 2 | ENSMUST00 | 2068 | Tns1 | ENSMUSG0C | false | . | . | . | . | . | . | . | . | . | . | . | . |
| chr6 | 89034350 | 89034666 | 182 | Intron (ENSMUST00000168837.3 | 6 | 89096110 | 89107834 | 11725 | 1 | ENSMUST00 | -61444 | 4933427D06I | ENSMUSG0C | false | . | . | . | . | . | . | chr6 | 89033650 | 89037786 | 4.66592 | 3.88673 | 2.74934 |
| chr10 | 117353153 | 117353442 | 182 | Promoter (2-3kb) | 10 | 117355526 | 117356056 | 531 | 2 | ENSMUST00 | 2614 | Cpsf6 | ENSMUSG0C | false | . | . | . | . | . | . | chr10 | 117350614 | 117354234 | 3.10317 | 3.08384 | 1.44255 |
| chr14 | 25708624 | 25708857 | 182 | Downstream (2-3kb) | 14 | 25711644 | 25719792 | 8149 | 2 | ENSMUST00 | 10935 | Zcchc24 | ENSMUSG0C | false | . | . | . | . | . | . | . | . | . | . | . | . |
| chr14 | 25768853 | 25769455 | 182 | Promoter (<=1kb) | 14 | 25711642 | 25769039 | 57398 | 2 | ENSMUST00 | 0 | Zcchc24 | ENSMUSG0C | false | chr14 | 25767764 | 25769441 | 33.08631 | 99.21803 | 95.66874 | chr14 | 25767571 | 25769404 | 5.01396 | 3.93512 | 3.05098 |
| chr8 | 70523003 | 70523320 | 182 | Promoter (<=1kb) | 8 | 70508272 | 70523163 | 14892 | 2 | ENSMUST00 | 0 | Kxd1 | ENSMUSG0C | false | chr8 | 70522446 | 70523697 | 31.80867 | 99.00969 | 95.46461 | chr8 | 70522176 | 70523760 | 3.54285 | 3.11925 | 1.85521 |
| chr2 | 31371571 | 31371900 | 182 | Intron (ENSMUST00000113532.3 | 2 | 31400277 | 31405814 | 5538 | 1 | ENSMUST00 | -28377 | Hmcn2 | ENSMUSG0C | false | . | . | . | . | . | . | chr2 | 31371105 | 31372187 | 4.57374 | 3.92074 | 2.65919 |
| chr14 | 98167253 | 98167487 | 182 | Promoter (2-3kb) | 14 | 97786853 | 98169543 | 382691 | 2 | ENSMUST00 | 2056 | Dach1 | ENSMUSG0C | true | chr14 | 98164311 | 98170276 | 41.81742 | 134.22485 | 129.57179 | . | . | . | . | . | . |
| chr7 | 75848158 | 75848423 | 182 | Promoter (<=1kb) | 7 | 75848338 | 75874130 | 25793 | 1 | ENSMUST00 | 0 | Klhl25 | ENSMUSG0C | false | chr7 | 75847679 | 75849583 | 41.07278 | 140.67711 | 135.76277 | . | . | . | . | . | . |
| chrX | 52613721 | 52614061 | 182 | Promoter (<=1kb) | 20 | 52272426 | 52613921 | 341496 | 2 | ENSMUST00 | 0 | Gpc3 | ENSMUSG0C | true | chrX | 52612296 | 52614474 | 16.08362 | 38.04472 | 35.47989 | . | . | . | . | . | . |
| chr11 | 72796084 | 72796381 | 182 | Promoter (<=1kb) | 11 | 72796226 | 72851380 | 55155 | 1 | ENSMUST00 | 0 | Zzef1 | ENSMUSG0C | true | chr11 | 72795110 | 72797378 | 36.66269 | 126.29414 | 121.93508 | . | . | . | . | . | . |
| chr7 | 114181640 | 114181879 | 182 | Distal Intergenic | 7 | 114046785 | 114117781 | 70997 | 2 | ENSMUST00 | -63860 | Rras2 | ENSMUSG0C | true | . | . | . | . | . | . | chr7 | 114180313 | 114185262 | 4.90433 | 4.01205 | 2.93003 |
| chr6 | 72655322 | 72655691 | 182 | Intron (ENSMUST00000069536.6 | 6 | 72648815 | 72671404 | 22590 | 2 | ENSMUST00 | 15713 | Tcf7l1 | ENSMUSG0C | false | chr6 | 72655030 | 72655884 | 14.08946 | 35.32397 | 32.79257 | chr6 | 72652506 | 72662237 | 5.35583 | 4.22796 | 3.3027 |
| chr12 | 40381960 | 40382201 | 182 | Intron (ENSMUST00000069637.8 | 12 | 40315558 | 40445769 | 130212 | 2 | ENSMUST00 | 63568 | Zfp277 | ENSMUSG0C | false | chr12 | 40381556 | 40381977 | 4.97263 | 8.68936 | 6.53912 | chr12 | 40378830 | 40393499 | 4.81877 | 3.95373 | 2.86926 |
| chr5 | 76912180 | 76912404 | 182 | Downstream (<1kb) | 5 | 76875935 | 76905514 | 29580 | 2 | ENSMUST00 | -6667 | Aasdh | ENSMUSG0C | false | . | . | . | . | . | . | chr5 | 76911250 | 76913252 | 4.79492 | 3.90541 | 2.86922 |
| chr8 | 91425352 | 91425577 | 182 | Intron (ENSMUST00000069718.8 | 8 | 91390555 | 91409464 | 18910 | 1 | ENSMUST00 | 34798 | Fto | ENSMUSG0C | false | . | . | . | . | . | . | chr8 | 91423893 | 91427928 | 4.19619 | 3.71364 | 2.32909 |
| chr8 | 91350734 | 91350957 | 182 | Intron (ENSMUST00000069718.8 | 8 | 91387736 | 91409840 | 22105 | 1 | ENSMUST00 | -36779 | Fto | ENSMUSG0C | false | . | . | . | . | . | . | chr8 | 91337651 | 91367410 | 5.62588 | 4.32828 | 3.53466 |
| chr2 | 112454879 | 112455125 | 182 | Promoter (<=1kb) | 2 | 112454997 | 112472162 | 17166 | 1 | ENSMUST00 | 0 | Emc7 | ENSMUSG0C | false | chr2 | 112454162 | 112455814 | 34.00549 | 104.83981 | 101.14426 | . | . | . | . | . | . |
| chr1 | 82286554 | 82287042 | 182 | 3' UTR | 1 | 82233101 | 82291416 | 58316 | 2 | ENSMUST00 | 4374 | Irs1 | ENSMUSG0C | false | chr1 | 82285600 | 82287741 | 10.47576 | 22.7833 | 20.40945 | . | . | . | . | . | . |
| chr11 | 94718204 | 94718473 | 182 | Distal Intergenic | 11 | 94694498 | 94704499 | 10002 | 2 | ENSMUST00 | -13706 | Gm11541 | ENSMUSG0C | false | . | . | . | . | . | . | chr11 | 94715033 | 94719556 | 5.32725 | 4.25479 | 3.2741 |
| chr15 | 27783466 | 27783902 | 182 | Intron (ENSMUST00000090247.5 | 15 | 27655071 | 27681542 | 26472 | 2 | ENSMUST00 | -101925 | Fam105a | ENSMUSG0C | false | . | . | . | . | . | . | chr15 | 27777264 | 27789305 | 7.24061 | 5.11987 | 4.8476 |
| chr5 | 140419211 | 140419458 | 182 | Promoter (<=1kb) | 5 | 140419305 | 140443358 | 24054 | 1 | ENSMUST00 | 0 | Eif3b | ENSMUSG0C | false | chr5 | 140419171 | 140420345 | 38.94548 | 127.06984 | 122.68537 | . | . | . | . | . | . |
| chr10 | 86743415 | 86743729 | 182 | Intron (ENSMUST00000075632.7 | 10 | 86731754 | 86732409 | 656 | 2 | ENSMUST00 | -11007 | Fabp3-ps1 | ENSMUSG0C | false | . | . | . | . | . | . | . | . | . | . | . | . |
| chr7 | 13278368 | 13278864 | 182 | Promoter (<=1kb) | 7 | 13278826 | 13311425 | 32600 | 1 | ENSMUST00 | 0 | Lig1 | ENSMUSG0C | false | chr7 | 13277932 | 13279594 | 19.89052 | 67.91473 | 64.9535 | . | . | . | . | . | . |
| chr6 | 52177056 | 52177363 | 182 | Promoter (<=1kb) | 6 | 52177498 | 52180851 | 3354 | 1 | ENSMUST00 | -135 | 5730596B20I | ENSMUSG0C | false | chr6 | 52176947 | 52177703 | 7.48194 | 15.52168 | 13.25109 | chr6 | 52174986 | 52186369 | 4.8401 | 4.02656 | 2.87141 |
| chr19 | 5067800 | 5068192 | 182 | Promoter (<=1kb) | 19 | 5068078 | 5070637 | 2560 | 1 | ENSMUST00 | 0 | Cd248 | ENSMUSG0C | false | chr19 | 5067956 | 5068600 | 18.15775 | 45.21172 | 42.55974 | chr19 | 5067164 | 5068809 | 3.85346 | 3.2892 | 2.09892 |
| chr5 | 142367342 | 142367765 | 182 | Distal Intergenic | 5 | 142401497 | 142462014 | 60518 | 1 | ENSMUST00 | -33732 | Foxk1 | ENSMUSG0C | false | chr5 | 142366791 | 142368168 | 22.61859 | 73.09582 | 70.05593 | . | . | . | . | . | . |
| chr5 | 142427862 | 142428093 | 182 | Intron (ENSMUST00000072837.5 | 5 | 142401497 | 142462014 | 60518 | 1 | ENSMUST00 | 26366 | Foxk1 | ENSMUSG0C | false | . | . | . | . | . | . | chr5 | 142423530 | 142434255 | 6.409 | 4.68664 | 4.16411 |
| chr5 | 142401353 | 142401741 | 182 | Promoter (<=1kb) | 5 | 142401497 | 142462014 | 60518 | 1 | ENSMUST00 | 0 | Foxk1 | ENSMUSG0C | false | chr5 | 142400816 | 142402701 | 34.78613 | 113.42423 | 109.48744 | . | . | . | . | . | . |
| chr17 | 65772196 | 65772808 | 182 | Promoter (<=1kb) | 17 | 65651726 | 65772752 | 121027 | 2 | ENSMUST00 | 0 | Rab31 | ENSMUSG0C | false | chr17 | 65771973 | 65773014 | 27.66126 | 83.97217 | 80.74496 | chr17 | 65771110 | 65773717 | 3.19156 | 3.03064 | 1.54921 |
| chr11 | 3648979 | 3649233 | 182 | Promoter (<=1kb) | 11 | 3646840 | 3649042 | 2203 | 2 | ENSMUST00 | 0 | Tug1 | ENSMUSG0C | false | chr11 | 3647807 | 3650043 | 35.56825 | 117.95176 | 113.87694 | . | . | . | . | . | . |
| chr7 | 43562082 | 43562452 | 182 | Promoter (<=1kb) | 7 | 43562256 | 43575461 | 13206 | 1 | ENSMUST00 | 0 | Zfp658 | ENSMUSG0C | false | chr7 | 43561859 | 43563041 | 21.76566 | 71.50774 | 68.49327 | . | . | . | . | . | . |
| chr19 | 6974847 | 6975270 | 182 | Promoter (<=1kb) | 19 | 6975048 | 6977324 | 2277 | 1 | ENSMUST00 | 0 | Ppp1r14b | ENSMUSG0C | false | chr19 | 6974159 | 6976424 | 25.84672 | 90.2271 | 86.88143 | . | . | . | . | . | . |
| chr1 | 155098282 | 155098809 | 182 | Promoter (<=1kb) | 1 | 155096361 | 155099636 | 3276 | 2 | ENSMUST00 | 827 | Ier5 | ENSMUSG0C | false | chr1 | 155097721 | 155099716 | 34.1261 | 124.90291 | 120.59063 | chr1 | 155097851 | 155100038 | 3.51631 | 3.25977 | 1.81589 |
| chr8 | 125012760 | 125013121 | 182 | Promoter (<=1kb) | 8 | 125012997 | 125034192 | 21196 | 1 | ENSMUST00 | 0 | Tsnax | ENSMUSG0C | false | chr8 | 125012518 | 125013834 | 30.14656 | 119.38538 | 115.2599 | chr8 | 125012680 | 125014182 | 3.80677 | 3.53406 | 2.02264 |
| chr1 | 44584879 | 44585073 | 182 | Intron (ENSMUST00000074525.4 | 1 | 44552597 | 44708632 | 156036 | 1 | ENSMUST00 | 32283 | Gulp1 | ENSMUSG0C | true | . | . | . | . | . | . | chr1 | 44583660 | 44586527 | 5.26874 | 4.13444 | 3.26857 |
| chr1 | 44551839 | 44552701 | 182 | Promoter (<=1kb) | 1 | 44552597 | 44708632 | 156036 | 1 | ENSMUST00 | 0 | Gulp1 | ENSMUSG0C | true | chr1 | 44551388 | 44552745 | 35.15467 | 116.16882 | 112.14974 | . | . | . | . | . | . |
| chr13 | 12565754 | 12566042 | 182 | Promoter (<=1kb) | 13 | 12565883 | 12609526 | 43644 | 1 | ENSMUST00 | 0 | Ero1lb | ENSMUSG0C | false | chr13 | 12565502 | 12566770 | 27.03867 | 103.18376 | 99.53125 | chr13 | 12565526 | 12566833 | 6.4251 | 4.68403 | 4.22801 |
| chr3 | 96997772 | 96998076 | 182 | Distal Intergenic | 3 | 97032416 | 97050873 | 18458 | 1 | ENSMUST00 | -34340 | Gja5 | ENSMUSG0C | false | . | . | . | . | . | . | . | . | . | . | . | . |
| chr1 | 39687298 | 39687591 | 182 | Intron (ENSMUST00000151913.1 | 1 | 39680932 | 39691871 | 10940 | 2 | ENSMUST00 | 4280 | Rfx8 | ENSMUSG0C | false | . | . | . | . | . | . | . | . | . | . | . | . |
| chr1 | 39746523 | 39746731 | 182 | Distal Intergenic | 1 | 39665306 | 39720997 | 55692 | 2 | ENSMUST00 | -25527 | Rfx8 | ENSMUSG0C | false | . | . | . | . | . | . | chr1 | 39746176 | 39750447 | 5.34297 | 4.23417 | 3.30515 |
| chr9 | 7836901 | 7837172 | 182 | Promoter (<=1kb) | 9 | 7818227 | 7837057 | 18831 | 2 | ENSMUST00 | 0 | Birc2 | ENSMUSG0C | false | chr9 | 7836210 | 7838078 | 17.49727 | 64.69686 | 61.78638 | . | . | . | . | . | . |
| chr4 | 107314199 | 107314479 | 182 | Promoter (<=1kb) | 4 | 107314363 | 107359822 | 45460 | 1 | ENSMUST00 | 0 | Yipf1 | ENSMUSG0C | false | chr4 | 107313597 | 107314233 | 10.54274 | 25.8059 | 23.39399 | chr4 | 107313492 | 107323582 | 6.11384 | 4.58418 | 3.92071 |
| chr12 | 16810850 | 16811314 | 182 | Promoter (<=1kb) | 12 | 16810965 | 16826743 | 15779 | 1 | ENSMUST00 | 0 | E2f6 | ENSMUSG0C | false | chr12 | 16810615 | 16811935 | 39.45224 | 138.91832 | 134.07426 | chr12 | 16810560 | 16812339 | 3.47691 | 3.34742 | 1.77428 |
| chr19 | 44146223 | 44146462 | 182 | Promoter (<=1kb) | 19 | 44139266 | 44146446 | 7181 | 2 | ENSMUST00 | 0 | Bloc1s2 | ENSMUSG0C | false | chr19 | 44145674 | 44146784 | 24.24157 | 89.02564 | 85.70093 | chr19 | 44143197 | 44146859 | 5.55448 | 4.21889 | 3.50123 |
| chr18 | 46615290 | 46615509 | 182 | Distal Intergenic | 18 | 46597708 | 46609171 | 11464 | 1 | ENSMUST00 | 17583 | Eif1a | ENSMUSG0C | false | . | . | . | . | . | . | chr18 | 46615133 | 46615968 | 3.92821 | 3.60991 | 2.13059 |
| chr18 | 46621545 | 46621759 | 182 | Distal Intergenic | 18 | 46597708 | 46609171 | 11464 | 1 | ENSMUST00 | 23838 | Eif1a | ENSMUSG0C | false | . | . | . | . | . | . | chr18 | 46620634 | 46622247 | 4.66671 | 3.85876 | 2.74326 |
| chr6 | 112273500 | 112273830 | 182 | Promoter (<=1kb) | 6 | 112273758 | 112330425 | 56668 | 1 | ENSMUST00 | 0 | Lmcd1 | ENSMUSG0C | false | chr6 | 112273304 | 112274523 | 11.15821 | 25.11025 | 22.70824 | chr6 | 112273242 | 112276142 | 5.0923 | 4.15644 | 3.07356 |
| chr7 | 82869756 | 82870039 | 182 | Promoter (2-3kb) | 7 | 82867333 | 82871563 | 4231 | 1 | ENSMUST00 | 2424 | Mex3b | ENSMUSG0C | false | chr7 | 82869396 | 82870335 | 11.82498 | 29.97654 | 27.5129 | chr7 | 82868322 | 82870254 | 6.05655 | 4.46636 | 3.88837 |
| chr5 | 98329168 | 98329426 | 182 | Promoter (<=1kb) | 5 | 98329354 | 98801910 | 472557 | 1 | ENSMUST00 | 0 | 1700007G11 | ENSMUSG0C | false | chr5 | 98329057 | 98329625 | 18.70486 | 58.09692 | 55.27494 | . | . | . | . | . | . |
| chr14 | 66824801 | 66824993 | 182 | Intron (ENSMUST00000022629.8 | 14 | 66784529 | 66793758 | 9230 | 1 | ENSMUST00 | 40273 | Gm10032 | ENSMUSG0C | false | . | . | . | . | . | . | chr14 | 66815115 | 66830778 | 7.77502 | 5.24019 | 5.30996 |
| chr2 | 14910594 | 14910807 | 182 | Intron (ENSMUST00000114723.3 | 2 | 14873993 | 14963667 | 89675 | 1 | ENSMUST00 | 36602 | Cacnb2 | ENSMUSG0C | false | . | . | . | . | . | . | chr2 | 14909209 | 14912201 | 3.26663 | 3.16732 | 1.59089 |
| chr19 | 16168687 | 16169208 | 182 | Intron (ENSMUST00000025541.5 | 19 | 16164312 | 16164805 | 494 | 2 | ENSMUST00 | -3883 | E030024N20 | ENSMUSG0C | false | . | . | . | . | . | . | chr19 | 16167099 | 16171547 | 6.67313 | 4.81612 | 4.38185 |
| chr5 | 93138845 | 93139118 | 182 | Intron (ENSMUST00000074733.7 | 5 | 93093457 | 93174958 | 81502 | 1 | ENSMUST00 | 45389 | Sep-11 | ENSMUSG0C | false | . | . | . | . | . | . | chr5 | 93095018 | 93154711 | 6.59326 | 4.78158 | 4.31782 |
| chr5 | 93113617 | 93113921 | 182 | Intron (ENSMUST00000074733.7 | 5 | 93093457 | 93174958 | 81502 | 1 | ENSMUST00 | 20161 | Sep-11 | ENSMUSG0C | false | . | . | . | . | . | . | chr5 | 93095018 | 93154711 | 6.59326 | 4.78158 | 4.31782 |
| chr5 | 93108757 | 93109003 | 182 | Intron (ENSMUST00000074733.7 | 5 | 93093457 | 93174958 | 81502 | 1 | ENSMUST00 | 15301 | Sep-11 | ENSMUSG0C | false | . | . | . | . | . | . | chr5 | 93095018 | 93154711 | 6.59326 | 4.78158 | 4.31782 |
| chr5 | 93130106 | 93130366 | 182 | Intron (ENSMUST00000074733.7 | 5 | 93093457 | 93174958 | 81502 | 1 | ENSMUST00 | 36650 | Sep-11 | ENSMUSG0C | false | . | . | . | . | . | . | chr5 | 93095018 | 93154711 | 6.59326 | 4.78158 | 4.31782 |
| chr12 | 108393776 | 108393949 | 182 | Intron (ENSMUST00000138456.2 | 12 | 108410542 | 108539575 | 129034 | 1 | ENSMUST00 | -16593 | Eml1 | ENSMUSG0C | false | . | . | . | . | . | . | chr12 | 108391550 | 108395350 | 4.13346 | 3.63494 | 2.32961 |
| chr3 | 108017816 | 108018076 | 182 | Promoter (<=1kb) | 3 | 108012255 | 108017973 | 5719 | 2 | ENSMUST00 | 0 | Gstm1 | ENSMUSG0C | false | chr3 | 108017241 | 108018783 | 10.13814 | 22.76719 | 20.39335 | chr3 | 108017378 | 108018938 | 4.0493 | 3.64417 | 2.21063 |
| chr7 | 66963578 | 66963853 | 182 | Intron (ENSMUST00000156396.2 | 7 | 66909468 | 66969555 | 60088 | 1 | ENSMUST00 | 54111 | Adamts17 | ENSMUSG0C | false | . | . | . | . | . | . | . | . | . | . | . | . |
| chr7 | 66859380 | 66859596 | 182 | Intron (ENSMUST00000156396.2 | 7 | 66849707 | 66871825 | 22119 | 1 | ENSMUST00 | 9674 | Adamts17 | ENSMUSG0C | false | . | . | . | . | . | . | . | . | . | . | . | . |
| chr2 | 92183115 | 92183430 | 182 | Promoter (<=1kb) | 2 | 92184106 | 92361829 | 177724 | 1 | ENSMUST00 | -676 | Phf21a | ENSMUSG0C | false | chr2 | 92182752 | 92186025 | 28.03146 | 79.91027 | 76.76085 | . | . | . | . | . | . |
| chr1 | 37996897 | 37997329 | 182 | Promoter (<=1kb) | 1 | 37983867 | 37997208 | 13342 | 2 | ENSMUST00 | 0 | Txndc9 | ENSMUSG0C | false | chr1 | 37996353 | 37999132 | 38.63617 | 126.53137 | 122.16663 | . | . | . | . | . | . |
| chr9 | 63356500 | 63356716 | 182 | Intron (ENSMUST00000034920.9 | 9 | 63163769 | 63377902 | 214134 | 2 | ENSMUST00 | 21186 | Map2k5 | ENSMUSG0C | false | . | . | . | . | . | . | chr9 | 63355688 | 63357333 | 3.62184 | 3.35299 | 1.88191 |
| chr9 | 63273731 | 63273986 | 182 | Intron (ENSMUST00000034920.9 | 9 | 63163769 | 63377902 | 214134 | 2 | ENSMUST00 | 103916 | Map2k5 | ENSMUSG0C | false | . | . | . | . | . | . | chr9 | 63273565 | 63274371 | 6.72738 | 4.60999 | 4.452 |
| chr18 | 10103241 | 10103510 | 182 | Intron (ENSMUST00000067947.5 | 18 | 10145453 | 10145965 | 513 | 2 | ENSMUST00 | 42455 | Gm2531 | ENSMUSG0C | false | . | . | . | . | . | . | . | . | . | . | . | . |
| chr15 | 83100044 | 83100320 | 182 | Promoter (<=1kb) | 15 | 83100204 | 83116674 | 16471 | 1 | ENSMUST00 | 0 | Serhl | ENSMUSG0C | false | chr15 | 83099604 | 83100808 | 18.33134 | 50.1574 | 47.44131 | chr15 | 83099498 | 83100953 | 4.68538 | 3.92802 | 2.76096 |
| chr10 | 90575435 | 90575644 | 182 | Promoter (<=1kb) | 10 | 90575727 | 90973300 | 397574 | 1 | ENSMUST00 | -83 | Anks1b | ENSMUSG0C | true | . | . | . | . | . | . | chr10 | 90575142 | 90576361 | 3.00253 | 2.95727 | 1.40166 |
| chr2 | 11777582 | 11777858 | 182 | Promoter (<=1kb) | 2 | 11742573 | 11777582 | 35010 | 2 | ENSMUST00 | -1 | Fbxo18 | ENSMUSG0C | false | chr2 | 11776733 | 11779311 | 36.89429 | 120.27677 | 116.13108 | . | . | . | . | . | . |
| chr14 | 36921390 | 36921565 | 182 | Promoter (1-2kb) | 14 | 36879359 | 36919535 | 40177 | 2 | ENSMUST00 | -1856 | Ccser2 | ENSMUSG0C | false | . | . | . | . | . | . | chr14 | 36919621 | 36923846 | 5.56295 | 4.16578 | 3.50382 |
| chr14 | 36968618 | 36968944 | 182 | Promoter (<=1kb) | 14 | 36874936 | 36968687 | 93752 | 2 | ENSMUST00 | 0 | Ccser2 | ENSMUSG0C | false | chr14 | 36967727 | 36969299 | 38.99286 | 128.2356 | 123.80784 | chr14 | 36965546 | 36971612 | 4.04539 | 3.42379 | 2.26179 |
| chr2 | 132262991 | 132263305 | 182 | Promoter (<=1kb) | 2 | 132263148 | 132296193 | 33046 | 1 | ENSMUST00 | 0 | Cds2 | ENSMUSG0C | false | chr2 | 132263073 | 132264530 | 32.32209 | 130.99449 | 126.47215 | . | . | . | . | . | . |
| chr15 | 103258353 | 103258701 | 182 | Promoter (<=1kb) | 15 | 103249349 | 103258403 | 9055 | 2 | ENSMUST00 | 0 | Nfe2 | ENSMUSG0C | false | chr15 | 103258367 | 103258880 | 7.35251 | 12.67314 | 10.44663 | chr15 | 103257779 | 103258889 | 3.82997 | 3.31922 | 2.0869 |
| chr18 | 82910698 | 82911180 | 182 | Promoter (<=1kb) | 18 | 82910879 | 83005314 | 94436 | 1 | ENSMUST00 | 0 | Zfp516 | ENSMUSG0C | false | chr18 | 82909434 | 82915656 | 29.53108 | 87.21563 | 83.92796 | chr18 | 82910930 | 82912602 | 3.96326 | 3.62245 | 2.16872 |
| chr13 | 67081261 | 67081666 | 182 | Promoter (<=1kb) | 13 | 67069398 | 67081512 | 12115 | 2 | ENSMUST00 | 0 | Zfp708 | ENSMUSG0C | false | chr13 | 67080753 | 67082022 | 27.9498 | 82.58102 | 79.37986 | chr13 | 67080071 | 67082367 | 4.78445 | 4.04497 | 2.81742 |
| chr16 | 9825140 | 9825381 | 182 | Intron (ENSMUST00000115835.2 | 16 | 9577827 | 9992533 | 414707 | 2 | ENSMUST00 | 167152 | Grin2a | ENSMUSG0C | false | . | . | . | . | . | . | . | . | . | . | . | . |
| chr7 | 143548643 | 143549198 | 182 | Promoter (<=1kb) | 7 | 143513579 | 143549090 | 35512 | 2 | ENSMUST00 | 0 | Nap1l4 | ENSMUSG0C | false | chr7 | 143548070 | 143549784 | 45.42985 | 166.67271 | 160.80592 | chr7 | 143540031 | 143550074 | 4.56108 | 3.80762 | 2.66455 |
| chr17 | 22866907 | 22867382 | 182 | Promoter (<=1kb) | 17 | 22846697 | 22867134 | 20438 | 2 | ENSMUST00 | 0 | Zfp945 | ENSMUSG0C | false | chr17 | 22865878 | 22867248 | 28.7777 | 83.06458 | 79.8585 | . | . | . | . | . | . |
| chr6 | 51905753 | 51906026 | 182 | Intron (ENSMUST00000078214.5 | 6 | 51859166 | 52012549 | 153384 | 2 | ENSMUST00 | 106523 | Skap2 | ENSMUSG0C | false | . | . | . | . | . | . | chr6 | 51904885 | 51906420 | 5.07396 | 4.06136 | 3.10789 |
| chr11 | 117384940 | 117385237 | 182 | Distal Intergenic | 11 | 117359360 | 117360866 | 1507 | 1 | ENSMUST00 | 25581 | Sep-09 | ENSMUSG0C | false | chr11 | 117385100 | 117385455 | 6.43345 | 10.42957 | 8.24651 | chr11 | 117384316 | 117387448 | 6.24823 | 4.41622 | 4.08168 |
| chr11 | 117312552 | 117312921 | 182 | Intron (ENSMUST00000093907.5 | 11 | 117308154 | 117352212 | 44059 | 1 | ENSMUST00 | 4399 | Sep-09 | ENSMUSG0C | false | . | . | . | . | . | . | chr11 | 117307956 | 117318257 | 4.37981 | 3.73647 | 2.50647 |
| chr11 | 117316837 | 117317254 | 182 | Intron (ENSMUST00000093907.5 | 11 | 117308154 | 117352212 | 44059 | 1 | ENSMUST00 | 8684 | Sep-09 | ENSMUSG0C | false | chr11 | 117316756 | 117317778 | 5.94284 | 10.35463 | 8.1753 | chr11 | 117307956 | 117318257 | 4.37981 | 3.73647 | 2.50647 |
| chr7 | 112023394 | 112023926 | 182 | Promoter (<=1kb) | 7 | 112023506 | 112111386 | 87881 | 1 | ENSMUST00 | 0 | Usp47 | ENSMUSG0C | false | chr7 | 112022814 | 112025018 | 37.48252 | 127.75053 | 123.3422 | . | . | . | . | . | . |
| chr7 | 16406134 | 16406410 | 182 | Intron (ENSMUST00000098789.3 | 7 | 16401196 | 16437694 | 36499 | 1 | ENSMUST00 | 4939 | Zc3h4 | ENSMUSG0C | false | . | . | . | . | . | . | chr7 | 16402625 | 16410024 | 6.51218 | 4.79197 | 4.24167 |
| chr11 | 69395789 | 69396127 | 182 | Promoter (<=1kb) | 11 | 69395814 | 69396680 | 867 | 1 | ENSMUST00 | 0 | Naa38 | ENSMUSG0C | false | chr11 | 69394552 | 69396853 | 24.07142 | 68.54774 | 65.57966 | . | . | . | . | . | . |
| chr5 | 77099671 | 77099989 | 182 | Intron (ENSMUST00000148568.1 | 5 | 77086988 | 77095277 | 8290 | 2 | ENSMUST00 | -4395 | Hopx | ENSMUSG0C | false | . | . | . | . | . | . | . | . | . | . | . | . |
| chr5 | 122702297 | 122702525 | 182 | Distal Intergenic | 5 | 122697202 | 122697600 | 399 | 2 | ENSMUST00 | -4698 | Gm10064 | ENSMUSG0C | false | . | . | . | . | . | . | . | . | . | . | . | . |
| chr19 | 24280306 | 24280641 | 182 | Promoter (<=1kb) | 19 | 24261453 | 24280605 | 19153 | 2 | ENSMUST00 | 0 | Fxn | ENSMUSG0C | false | chr19 | 24279685 | 24281055 | 25.1466 | 74.00616 | 70.95453 | chr19 | 24279342 | 24281199 | 4.77924 | 3.90961 | 2.83248 |
| chr11 | 85615462 | 85615768 | 182 | Intron (ENSMUST00000108061.2 | 11 | 85640704 | 85826055 | 185352 | 1 | ENSMUST00 | -24936 | Bcas3 | ENSMUSG0C | false | . | . | . | . | . | . | . | . | . | . | . | . |
| chr11 | 93886801 | 93887017 | 182 | Promoter (<=1kb) | 11 | 93886880 | 93930278 | 43399 | 1 | ENSMUST00 | 0 | Mbtd1 | ENSMUSG0C | false | chr11 | 93884540 | 93887391 | 41.19049 | 135.33968 | 130.64169 | chr11 | 93886707 | 93889928 | 4.48394 | 3.78631 | 2.59385 |
| chr4 | 10905394 | 10905699 | 182 | Distal Intergenic | 4 | 10879539 | 10891778 | 12240 | 1 | ENSMUST00 | 25856 | 2610301B20I | ENSMUSG0C | false | . | . | . | . | . | . | chr4 | 10905311 | 10905600 | 4.60462 | 3.63504 | 2.70702 |
| chr6 | 56796792 | 56797110 | 182 | Promoter (<=1kb) | 6 | 56789523 | 56796747 | 7225 | 2 | ENSMUST00 | -46 | Kbtbd2 | ENSMUSG0C | false | chr6 | 56796286 | 56798543 | 30.35223 | 94.4818 | 91.04041 | . | . | . | . | . | . |
| chr9 | 43094007 | 43094366 | 182 | Intron (ENSMUST00000165665.2 | 9 | 42963843 | 43105486 | 141644 | 2 | ENSMUST00 | 11120 | Arhgef12 | ENSMUSG0C | true | . | . | . | . | . | . | chr9 | 43060024 | 43103547 | 5.90513 | 4.47373 | 3.75735 |
| chr11 | 69581141 | 69581682 | 182 | Promoter (<=1kb) | 11 | 69580405 | 69591873 | 11469 | 1 | ENSMUST00 | 737 | Trp53 | ENSMUSG0C | false | chr11 | 69579501 | 69582098 | 31.85236 | 101.09525 | 97.496 | . | . | . | . | . | . |
| chr5 | 43960007 | 43960310 | 182 | Distal Intergenic | 5 | 43943902 | 43944483 | 582 | 2 | ENSMUST00 | -15525 | Gm5292 | ENSMUSG0C | false | . | . | . | . | . | . | chr5 | 43957762 | 43962221 | 3.59478 | 3.3816 | 1.836 |
| chr5 | 43949015 | 43949212 | 182 | Distal Intergenic | 5 | 43943902 | 43944483 | 582 | 2 | ENSMUST00 | -4533 | Gm5292 | ENSMUSG0C | false | chr5 | 43947943 | 43949058 | 12.40737 | 26.6269 | 24.20249 | chr5 | 43947617 | 43949965 | 3.46477 | 3.2764 | 1.76252 |
| chr6 | 117841078 | 117841598 | 182 | Promoter (<=1kb) | 6 | 117841180 | 117844518 | 3339 | 1 | ENSMUST00 | 0 | Zfp637 | ENSMUSG0C | false | chr6 | 117840572 | 117842150 | 37.70819 | 121.69438 | 117.49508 | . | . | . | . | . | . |
| chr11 | 107577580 | 107577800 | 182 | Intron (ENSMUST00000141730.2 | 11 | 107531757 | 107548373 | 16617 | 2 | ENSMUST00 | -29208 | A830035A12 | ENSMUSG0C | false | . | . | . | . | . | . | chr11 | 107576584 | 107579379 | 5.00531 | 3.93642 | 3.07147 |
| chr17 | 35822996 | 35823429 | 182 | Promoter (<=1kb) | 17 | 35823230 | 35832791 | 9562 | 1 | ENSMUST00 | 0 | Flot1 | ENSMUSG0C | false | chr17 | 35821005 | 35823894 | 34.4409 | 118.06123 | 113.98219 | chr17 | 35820226 | 35824687 | 3.40141 | 3.25998 | 1.6905 |
| chr9 | 110781286 | 110781628 | 182 | Distal Intergenic | 9 | 110769001 | 110769794 | 794 | 1 | ENSMUST00 | 12286 | Myl3 | ENSMUSG0C | false | chr9 | 110780486 | 110782085 | 15.01947 | 39.77812 | 37.19162 | chr9 | 110772879 | 110782615 | 6.7891 | 4.9169 | 4.4717 |
| chr17 | 35861198 | 35861649 | 182 | Promoter (<=1kb) | 17 | 35861318 | 35865402 | 4085 | 1 | ENSMUST00 | 0 | Nrm | ENSMUSG0C | false | chr17 | 35860920 | 35861831 | 39.86176 | 129.74701 | 125.26318 | chr17 | 35860741 | 35862099 | 4.52688 | 3.85743 | 2.67087 |
| chr4 | 62634160 | 62634424 | 182 | Promoter (<=1kb) | 4 | 62634378 | 62697928 | 63551 | 1 | ENSMUST00 | 0 | Rgs3 | ENSMUSG0C | false | . | . | . | . | . | . | chr4 | 62633973 | 62636466 | 3.90803 | 3.58649 | 2.11306 |
| chr3 | 41742274 | 41742567 | 182 | Promoter (<=1kb) | 3 | 41626717 | 41742514 | 115798 | 2 | ENSMUST00 | 0 | Sclt1 | ENSMUSG0C | false | chr3 | 41741443 | 41743295 | 24.81506 | 73.42724 | 70.38435 | . | . | . | . | . | . |
| chr7 | 44952604 | 44953009 | 182 | Exon (ENSMUST00000120929.2/ | 7 | 44943261 | 44958032 | 14772 | 1 | ENSMUST00 | 9344 | Tsks | ENSMUSG0C | false | chr7 | 44951540 | 44952944 | 18.33657 | 60.6815 | 57.82544 | chr7 | 44951512 | 44953755 | 3.48863 | 3.16304 | 1.79666 |
| chr3 | 141491622 | 141492079 | 182 | Intron (ENSMUST00000106236.3 | 3 | 141465671 | 141830538 | 364868 | 1 | ENSMUST00 | 25952 | Unc5c | ENSMUSG0C | true | . | . | . | . | . | . | chr3 | 141489640 | 141499892 | 4.31369 | 3.67642 | 2.44983 |
| chr12 | 70974592 | 70975077 | 182 | Promoter (<=1kb) | 12 | 70974621 | 70996347 | 21727 | 1 | ENSMUST00 | 0 | Psma3 | ENSMUSG0C | false | chr12 | 70974273 | 70975788 | 42.50686 | 139.83865 | 134.96973 | . | . | . | . | . | . |
| chr4 | 54966161 | 54966568 | 182 | Intron (ENSMUST00000133895.2 | 4 | 54947976 | 55083563 | 135588 | 1 | ENSMUST00 | 18186 | Zfp462 | ENSMUSG0C | true | chr4 | 54966022 | 54966999 | 6.74139 | 13.32281 | 11.08953 | chr4 | 54964968 | 54991208 | 5.4969 | 4.26974 | 3.41753 |
| chr2 | 121866785 | 121867163 | 182 | Promoter (<=1kb) | 2 | 121866970 | 121936205 | 69236 | 1 | ENSMUST00 | 0 | Casc4 | ENSMUSG0C | false | chr2 | 121866208 | 121868007 | 30.41442 | 103.5799 | 99.9162 | chr2 | 121866183 | 121869214 | 3.87419 | 3.50064 | 2.07794 |
| chr9 | 44478602 | 44478918 | 182 | Promoter (<=1kb) | 9 | 44478778 | 44482180 | 3403 | 1 | ENSMUST00 | 0 | C030014I23R | ENSMUSG0C | false | chr9 | 44478008 | 44480080 | 23.05105 | 65.89761 | 62.96862 | chr9 | 44477636 | 44480742 | 4.26784 | 3.72227 | 2.41582 |
| chr8 | 69625113 | 69625490 | 182 | Promoter (<=1kb) | 8 | 69618869 | 69625545 | 6677 | 2 | ENSMUST00 | 55 | Zfp868 | ENSMUSG0C | false | chr8 | 69624639 | 69625923 | 26.91438 | 86.45937 | 83.18881 | . | . | . | . | . | . |
| chr18 | 38296617 | 38297304 | 182 | Promoter (<=1kb) | 18 | 38296635 | 38317847 | 21213 | 1 | ENSMUST00 | 0 | Rnf14 | ENSMUSG0C | false | chr18 | 38296359 | 38297841 | 28.59262 | 111.22587 | 107.35971 | chr18 | 38295947 | 38301658 | 5.0746 | 3.9538 | 3.07792 |
| chr14 | 61037814 | 61038120 | 182 | Promoter (<=1kb) | 14 | 60963834 | 61037987 | 74154 | 2 | ENSMUST00 | 0 | Tnfrsf19 | ENSMUSG0C | false | chr14 | 61037400 | 61038207 | 19.17126 | 69.02763 | 66.05415 | . | . | . | . | . | . |
| chr8 | 47413479 | 47413962 | 182 | Promoter (<=1kb) | 8 | 47413129 | 47415359 | 2231 | 1 | ENSMUST00 | 351 | Gm10083 | ENSMUSG0C | false | chr8 | 47412486 | 47414325 | 25.4393 | 79.11702 | 75.98065 | chr8 | 47411851 | 47414270 | 4.88572 | 3.88095 | 2.9457 |
| chr7 | 38538163 | 38538419 | 182 | Distal Intergenic | 7 | 38518139 | 38528193 | 10055 | 2 | ENSMUST00 | -9971 | Gm5591 | ENSMUSG0C | false | . | . | . | . | . | . | . | . | . | . | . | . |
| chr9 | 51075894 | 51076121 | 182 | Promoter (<=1kb) | 9 | 51056780 | 51077094 | 20315 | 2 | ENSMUST00 | 973 | Layn | ENSMUSG0C | false | . | . | . | . | . | . | chr9 | 51075652 | 51079387 | 3.4741 | 3.29042 | 1.7771 |
| chr14 | 51779219 | 51779520 | 182 | Distal Intergenic | 14 | 51763878 | 51773590 | 9713 | 2 | ENSMUST00 | -5630 | Ang4 | ENSMUSG0C | false | . | . | . | . | . | . | chr14 | 51778670 | 51780251 | 5.0273 | 4.0957 | 3.06223 |
| chr16 | 14159248 | 14159533 | 182 | Promoter (<=1kb) | 16 | 14109166 | 14159274 | 50109 | 2 | ENSMUST00 | 0 | Marf1 | ENSMUSG0C | false | . | . | . | . | . | . | . | . | . | . | . | . |
| chr12 | 79028971 | 79029371 | 182 | Promoter (<=1kb) | 12 | 79029163 | 79081648 | 52486 | 1 | ENSMUST00 | 0 | Plekhh1 | ENSMUSG0C | false | chr12 | 79028669 | 79030046 | 33.9579 | 107.82887 | 104.06033 | chr12 | 79028208 | 79030063 | 3.68211 | 3.3969 | 1.98064 |
| chr7 | 110561922 | 110562175 | 182 | Intron (ENSMUST00000164759.2 | 7 | 110526792 | 110527121 | 330 | 1 | ENSMUST00 | 35131 | Gm10087 | ENSMUSG0C | false | . | . | . | . | . | . | chr7 | 110559199 | 110562964 | 4.01744 | 3.54673 | 2.19937 |
| chr10 | 63429830 | 63430199 | 182 | Promoter (<=1kb) | 10 | 63430098 | 65002965 | 1572868 | 1 | ENSMUST00 | 0 | Ctnna3 | ENSMUSG0C | false | chr10 | 63428590 | 63430584 | 35.2856 | 127.03558 | 122.65217 | . | . | . | . | . | . |
| chr3 | 52007576 | 52007923 | 182 | Intron (ENSMUST00000121440.2 | 3 | 51687611 | 52104891 | 417281 | 2 | ENSMUST00 | 96968 | Maml3 | ENSMUSG0C | true | chr3 | 52007630 | 52007854 | 8.63619 | 18.02194 | 15.71366 | chr3 | 52005145 | 52008316 | 4.97561 | 4.08859 | 2.9881 |
| chr3 | 51749678 | 51750293 | 182 | Intron (ENSMUST00000121440.2 | 3 | 51689437 | 51796535 | 107099 | 2 | ENSMUST00 | 46242 | Maml3 | ENSMUSG0C | true | . | . | . | . | . | . | chr3 | 51741455 | 51754138 | 5.51724 | 4.2147 | 3.45633 |
| chr3 | 51752185 | 51752442 | 182 | Intron (ENSMUST00000121440.2 | 3 | 51689437 | 51796535 | 107099 | 2 | ENSMUST00 | 44093 | Maml3 | ENSMUSG0C | true | . | . | . | . | . | . | chr3 | 51741455 | 51754138 | 5.51724 | 4.2147 | 3.45633 |
| chr3 | 52046960 | 52047225 | 182 | Intron (ENSMUST00000121440.2 | 3 | 51687611 | 52104891 | 417281 | 2 | ENSMUST00 | 57666 | Maml3 | ENSMUSG0C | true | . | . | . | . | . | . | chr3 | 52009386 | 52064052 | 5.58271 | 4.32835 | 3.48679 |
| chr3 | 52046408 | 52046652 | 182 | Intron (ENSMUST00000121440.2 | 3 | 51687611 | 52104891 | 417281 | 2 | ENSMUST00 | 58239 | Maml3 | ENSMUSG0C | true | chr3 | 52046256 | 52046934 | 8.61023 | 17.47713 | 15.17558 | chr3 | 52009386 | 52064052 | 5.58271 | 4.32835 | 3.48679 |
| chr3 | 52074898 | 52075091 | 182 | Intron (ENSMUST00000121440.2 | 3 | 51687611 | 52104891 | 417281 | 2 | ENSMUST00 | 29800 | Maml3 | ENSMUSG0C | true | . | . | . | . | . | . | chr3 | 52065361 | 52076212 | 4.4072 | 3.71398 | 2.5587 |
| chr3 | 52055148 | 52055596 | 182 | Intron (ENSMUST00000121440.2 | 3 | 51687611 | 52104891 | 417281 | 2 | ENSMUST00 | 49295 | Maml3 | ENSMUSG0C | true | . | . | . | . | . | . | chr3 | 52009386 | 52064052 | 5.58271 | 4.32835 | 3.48679 |
| chr3 | 79484412 | 79484734 | 182 | Intron (ENSMUST00000076136.5 | 3 | 79459593 | 79500522 | 40930 | 2 | ENSMUST00 | 15788 | Fnip2 | ENSMUSG0C | false | . | . | . | . | . | . | . | . | . | . | . | . |
| chr5 | 117241124 | 117241369 | 182 | Promoter (<=1kb) | 5 | 117240667 | 117273869 | 33203 | 1 | ENSMUST00 | 458 | Taok3 | ENSMUSG0C | false | chr5 | 117240148 | 117241406 | 12.5733 | 28.75278 | 26.30366 | chr5 | 117241259 | 117243329 | 3.78503 | 3.40832 | 2.01219 |
| chr11 | 120151184 | 120151389 | 182 | Promoter (<=1kb) | 11 | 120103960 | 120151346 | 47387 | 2 | ENSMUST00 | 0 | Slc38a10 | ENSMUSG0C | false | chr11 | 120149746 | 120151354 | 26.86698 | 95.04005 | 91.5885 | . | . | . | . | . | . |
| chr6 | 117168543 | 117168818 | 182 | Promoter (<=1kb) | 6 | 117168574 | 117181366 | 12793 | 1 | ENSMUST00 | 0 | Cxcl12 | ENSMUSG0C | false | chr6 | 117167809 | 117170507 | 34.00537 | 102.80987 | 99.17244 | . | . | . | . | . | . |
| chr8 | 121652610 | 121652974 | 182 | Promoter (<=1kb) | 8 | 121598703 | 121652901 | 54199 | 2 | ENSMUST00 | 0 | Zcchc14 | ENSMUSG0C | false | chr8 | 121652509 | 121653931 | 28.00751 | 87.44511 | 84.14996 | . | . | . | . | . | . |
| chr10 | 61313813 | 61314048 | 182 | Distal Intergenic | 10 | 61309830 | 61310276 | 447 | 1 | ENSMUST00 | 3984 | Rpl27a-ps1 | ENSMUSG0C | false | . | . | . | . | . | . | chr10 | 61312220 | 61315624 | 5.20834 | 4.09549 | 3.19373 |
| chr5 | 43515650 | 43515925 | 182 | Promoter (<=1kb) | 5 | 43515588 | 43618817 | 103230 | 1 | ENSMUST00 | 63 | C1qtnf7 | ENSMUSG0C | true | chr5 | 43515384 | 43518831 | 23.48881 | 65.78277 | 62.85688 | . | . | . | . | . | . |
| chr12 | 52855014 | 52855462 | 182 | Intron (ENSMUST00000095737.3 | 12 | 52699383 | 53151015 | 451633 | 1 | ENSMUST00 | 155632 | Akap6 | ENSMUSG0C | false | . | . | . | . | . | . | chr12 | 52853999 | 52855472 | 4.79907 | 3.98306 | 2.8461 |
| chr17 | 42840843 | 42841110 | 182 | Intron (ENSMUST00000024709.7 | 17 | 42792951 | 42876424 | 83474 | 2 | ENSMUST00 | 35314 | Cd2ap | ENSMUSG0C | true | . | . | . | . | . | . | chr17 | 42834080 | 42845253 | 4.73325 | 3.91855 | 2.78884 |
| chr17 | 42875105 | 42875407 | 182 | Promoter (1-2kb) | 17 | 42792951 | 42876424 | 83474 | 2 | ENSMUST00 | 1017 | Cd2ap | ENSMUSG0C | true | chr17 | 42874504 | 42877282 | 43.42055 | 150.43358 | 145.13264 | . | . | . | . | . | . |
| chr11 | 98348074 | 98348376 | 182 | Promoter (<=1kb) | 11 | 98348404 | 98357796 | 9393 | 1 | ENSMUST00 | -28 | Ppp1r1b | ENSMUSG0C | false | chr11 | 98348228 | 98349138 | 9.86256 | 23.41135 | 21.03137 | chr11 | 98348122 | 98349097 | 10.2863 | 6.58771 | 7.30942 |
| chr15 | 53160138 | 53160390 | 182 | Intron (ENSMUST00000077273.3 | 15 | 53083146 | 53196255 | 113110 | 2 | ENSMUST00 | 35865 | Ext1 | ENSMUSG0C | false | . | . | . | . | . | . | chr15 | 53155580 | 53161791 | 4.74684 | 3.83611 | 2.83507 |
| chr15 | 53188155 | 53188422 | 182 | Intron (ENSMUST00000077273.3 | 15 | 53083146 | 53196255 | 113110 | 2 | ENSMUST00 | 7833 | Ext1 | ENSMUSG0C | false | . | . | . | . | . | . | chr15 | 53187230 | 53188880 | 4.83468 | 3.85765 | 2.90953 |
| chr16 | 34045581 | 34045789 | 182 | Intron (ENSMUST00000114973.2 | 16 | 33974258 | 34095987 | 121730 | 2 | ENSMUST00 | 50198 | Kalrn | ENSMUSG0C | false | . | . | . | . | . | . | . | . | . | . | . | . |
| chr5 | 123930577 | 123930894 | 182 | Promoter (<=1kb) | 5 | 123930689 | 123936645 | 5957 | 1 | ENSMUST00 | 0 | Ccdc62 | ENSMUSG0C | false | chr5 | 123930269 | 123931336 | 25.15761 | 80.47005 | 77.31023 | . | . | . | . | . | . |
| chr5 | 143180726 | 143181013 | 182 | Promoter (<=1kb) | 5 | 143172186 | 143180775 | 8590 | 2 | ENSMUST00 | 0 | Rbak | ENSMUSG0C | false | chr5 | 143179842 | 143181208 | 21.7465 | 82.16968 | 78.97962 | . | . | . | . | . | . |
| chr12 | 29353860 | 29354175 | 182 | Distal Intergenic | 12 | 29528384 | 29923209 | 394826 | 1 | ENSMUST00 | -174209 | Myt1l | ENSMUSG0C | false | . | . | . | . | . | . | . | . | . | . | . | . |
| chr15 | 38176455 | 38176663 | 182 | Distal Intergenic | 15 | 38219203 | 38226735 | 7533 | 1 | ENSMUST00 | -42540 | Odf1 | ENSMUSG0C | false | . | . | . | . | . | . | . | . | . | . | . | . |
| chr10 | 23787174 | 23787491 | 182 | Promoter (<=1kb) | 10 | 23785211 | 23787195 | 1985 | 2 | ENSMUST00 | 0 | Rps12 | ENSMUSG0C | false | chr10 | 23786112 | 23788131 | 25.4663 | 101.95255 | 98.33054 | . | . | . | . | . | . |
| chr1 | 159099220 | 159099456 | 182 | Distal Intergenic | 1 | 159168157 | 159168642 | 486 | 2 | ENSMUST00 | 69186 | Gm4953 | ENSMUSG0C | false | . | . | . | . | . | . | chr1 | 159098417 | 159099503 | 4.35797 | 3.73933 | 2.49379 |
| chr17 | 23599024 | 23599291 | 182 | Promoter (<=1kb) | 17 | 23575852 | 23599487 | 23636 | 2 | ENSMUST00 | 196 | Zfp13 | ENSMUSG0C | false | chr17 | 23598594 | 23599797 | 22.51707 | 59.83231 | 56.98677 | chr17 | 23598278 | 23600356 | 4.13133 | 3.67229 | 2.28861 |
| chrX | 106186784 | 106187056 | 182 | Promoter (<=1kb) | 20 | 106187100 | 106203699 | 16600 | 1 | ENSMUST00 | -44 | Pgk1 | ENSMUSG0C | false | chrX | 106186491 | 106186788 | 11.02877 | 22.59171 | 20.22099 | . | . | . | . | . | . |
| chr10 | 39171235 | 39171504 | 182 | Distal Intergenic | 10 | 39150971 | 39163794 | 12824 | 2 | ENSMUST00 | -7442 | Wisp3 | ENSMUSG0C | false | chr10 | 39171283 | 39172006 | 15.85026 | 49.49464 | 46.78799 | chr10 | 39165020 | 39174077 | 5.50767 | 4.30295 | 3.42682 |
| chr17 | 10268722 | 10269112 | 182 | Intron (ENSMUST00000097414.4 | 17 | 10206471 | 10319361 | 112891 | 2 | ENSMUST00 | 50249 | Qk | ENSMUSG0C | true | . | . | . | . | . | . | chr17 | 10262681 | 10274077 | 4.10711 | 3.65276 | 2.26384 |
| chr17 | 55944917 | 55945312 | 182 | Promoter (<=1kb) | 17 | 55938381 | 55945259 | 6879 | 2 | ENSMUST00 | 0 | Zfp119b | ENSMUSG0C | false | chr17 | 55944460 | 55945971 | 40.55256 | 150.97545 | 145.64915 | . | . | . | . | . | . |
| chr11 | 60177396 | 60177708 | 182 | Promoter (1-2kb) | 11 | 60175879 | 60199197 | 23319 | 1 | ENSMUST00 | 1518 | Rai1 | ENSMUSG0C | false | chr11 | 60175495 | 60177680 | 20.5121 | 70.85588 | 67.85384 | chr11 | 60176890 | 60178980 | 8.09767 | 5.33041 | 5.55498 |
| chr1 | 181150850 | 181151121 | 182 | Promoter (<=1kb) | 1 | 181150931 | 181168994 | 18064 | 1 | ENSMUST00 | 0 | Cnih4 | ENSMUSG0C | false | chr1 | 181150788 | 181151670 | 28.22734 | 101.53345 | 97.92473 | chr1 | 181150493 | 181151953 | 3.16295 | 3.04573 | 1.53012 |
| chr6 | 57702936 | 57703457 | 182 | Promoter (<=1kb) | 6 | 57703051 | 57737972 | 34922 | 1 | ENSMUST00 | 0 | Lancl2 | ENSMUSG0C | false | chr6 | 57702226 | 57703887 | 35.20524 | 115.20684 | 111.21615 | . | . | . | . | . | . |
| chr18 | 50011517 | 50011790 | 182 | Intron (ENSMUST00000148989.2 | 18 | 50019567 | 50090794 | 71228 | 1 | ENSMUST00 | -7777 | Tnfaip8 | ENSMUSG0C | false | . | . | . | . | . | . | . | . | . | . | . | . |
| chr5 | 108619251 | 108619489 | 182 | Intron (ENSMUST00000046603.9 | 5 | 108580079 | 108625091 | 45013 | 2 | ENSMUST00 | 5602 | Gak | ENSMUSG0C | false | . | . | . | . | . | . | chr5 | 108619307 | 108620211 | 3.42628 | 3.19445 | 1.755 |
| chr5 | 121507232 | 121507412 | 182 | Intron (ENSMUST00000111795.2 | 5 | 121500098 | 121503435 | 3338 | 2 | ENSMUST00 | -3798 | Adam1b | ENSMUSG0C | false | . | . | . | . | . | . | chr5 | 121506450 | 121507958 | 3.99606 | 3.35394 | 2.22345 |
| chr12 | 24680768 | 24681066 | 182 | Promoter (<=1kb) | 12 | 24665953 | 24681023 | 15071 | 2 | ENSMUST00 | 0 | Cys1 | ENSMUSG0C | false | chr12 | 24679777 | 24682165 | 34.00537 | 102.80987 | 99.17244 | chr12 | 24680915 | 24683067 | 3.63656 | 3.29087 | 1.89507 |
| chr17 | 27562269 | 27562464 | 182 | Exon (ENSMUST00000117600.2/ | 17 | 27564682 | 27565667 | 986 | 2 | ENSMUST00 | 3203 | AI413582 | ENSMUSG0C | false | . | . | . | . | . | . | chr17 | 27561495 | 27563286 | 3.4394 | 3.16843 | 1.76461 |
| chr4 | 89137340 | 89137579 | 182 | Promoter (<=1kb) | 4 | 89137122 | 89181081 | 43960 | 1 | ENSMUST00 | 219 | Mtap | ENSMUSG0C | false | chr4 | 89136849 | 89138351 | 33.50033 | 109.08844 | 105.27855 | . | . | . | . | . | . |
| chr4 | 89138013 | 89138331 | 182 | Promoter (<=1kb) | 4 | 89137122 | 89181081 | 43960 | 1 | ENSMUST00 | 892 | Mtap | ENSMUSG0C | false | chr4 | 89136849 | 89138351 | 33.50033 | 109.08844 | 105.27855 | . | . | . | . | . | . |
| chr18 | 21177503 | 21177717 | 182 | Intron (ENSMUST00000049260.6 | 18 | 21076667 | 21077110 | 444 | 2 | ENSMUST00 | -100394 | Gm6378 | ENSMUSG0C | false | . | . | . | . | . | . | chr18 | 21176472 | 21181524 | 4.5098 | 3.77789 | 2.649 |
| chr18 | 21128087 | 21128325 | 182 | 3' UTR | 18 | 21076667 | 21077110 | 444 | 2 | ENSMUST00 | -50978 | Gm6378 | ENSMUSG0C | false | . | . | . | . | . | . | chr18 | 21126098 | 21129897 | 4.85872 | 4.01393 | 2.89413 |
| chr6 | 22045229 | 22045500 | 182 | Intron (ENSMUST00000137437.2 | 6 | 22025943 | 22085138 | 59196 | 1 | ENSMUST00 | 19287 | Cped1 | ENSMUSG0C | false | . | . | . | . | . | . | chr6 | 22044754 | 22045532 | 3.17288 | 3.01554 | 1.531 |
| chr5 | 93150373 | 93150672 | 182 | Intron (ENSMUST00000074733.7 | 5 | 93187453 | 93188220 | 768 | 2 | ENSMUST00 | 37548 | Ccni | ENSMUSG0C | true | . | . | . | . | . | . | chr5 | 93095018 | 93154711 | 6.59326 | 4.78158 | 4.31782 |
| chr8 | 47673750 | 47674206 | 182 | Promoter (<=1kb) | 8 | 47667178 | 47673689 | 6512 | 2 | ENSMUST00 | -62 | Ing2 | ENSMUSG0C | false | chr8 | 47673181 | 47676469 | 40.65367 | 138.82761 | 133.98773 | . | . | . | . | . | . |
| chr18 | 5446608 | 5446967 | 182 | Distal Intergenic | 18 | 5518415 | 5521109 | 2695 | 2 | ENSMUST00 | 74142 | Gm10125 | ENSMUSG0C | false | . | . | . | . | . | . | . | . | . | . | . | . |
| chr3 | 51415766 | 51416037 | 182 | Promoter (<=1kb) | 3 | 51416016 | 51475985 | 59970 | 1 | ENSMUST00 | 0 | Naa15 | ENSMUSG0C | false | chr3 | 51415428 | 51417724 | 34.91417 | 125.28625 | 120.96531 | . | . | . | . | . | . |

| | | | | | | | | | | | | | | | | | | | | | | | | | | |
|---|---|---|---|---|---|---|---|---|---|---|---|---|---|---|---|---|---|---|---|---|---|---|---|---|---|---|
| chr9 | 44498726 | 44499119 | 182 | Promoter (<=1kb) | 9 | 44499136 | 44510388 | 11253 | 1 | ENSMUST00 | -17 | Bcl9l | ENSMUSG0C | false | chr9 | 44498133 | 44499362 | 10.53897 | 22.32804 | 19.96106 | chr9 | 44483700 | 44502510 | 6.77697 | 4.85012 | 4.46719 |
| chr17 | 22165658 | 22165951 | 182 | Promoter (<=1kb) | 17 | 22144359 | 22166190 | 21832 | 2 | ENSMUST00 | 239 | Zfp947 | ENSMUSG0C | false | chr17 | 22165296 | 22166172 | 20.05708 | 59.74671 | 56.90333 | chr17 | 22164916 | 22166687 | 3.02037 | 2.98694 | 1.39028 |
| chr14 | 22166012 | 22166267 | 182 | Intron (ENSMUST00000075639.5 | 14 | 22310165 | 22584439 | 274275 | 1 | ENSMUST00 | -143898 | 1700112E06I | ENSMUSG0C | false | . | . | . | . | . | . | chr14 | 22164591 | 22169694 | 6.26653 | 4.63848 | 4.05874 |
| chr14 | 22175915 | 22176158 | 182 | Intron (ENSMUST00000075639.5 | 14 | 22310165 | 22584439 | 274275 | 1 | ENSMUST00 | -134007 | 1700112E06I | ENSMUSG0C | false | . | . | . | . | . | . | chr14 | 22173884 | 22176588 | 5.1117 | 4.17195 | 3.09387 |
| chr7 | 45395481 | 45396022 | 182 | Promoter (<=1kb) | 7 | 45376462 | 45395692 | 19231 | 2 | ENSMUST00 | 0 | Snrnp70 | ENSMUSG0C | false | chr7 | 45394890 | 45395811 | 40.88443 | 132.61177 | 128.02734 | . | . | . | . | . | . |
| chr4 | 150236750 | 150237323 | 182 | Promoter (<=1kb) | 4 | 150236952 | 150248879 | 11928 | 1 | ENSMUST00 | 0 | Eno1 | ENSMUSG0C | false | chr4 | 150236391 | 150238373 | 33.32866 | 126.59486 | 122.2257 | . | . | . | . | . | . |
| chr8 | 94917524 | 94918111 | 182 | Promoter (<=1kb) | 8 | 94902869 | 94918098 | 15230 | 2 | ENSMUST00 | 0 | Ccdc102a | ENSMUSG0C | false | chr8 | 94917111 | 94918353 | 21.42557 | 69.98924 | 66.99868 | chr8 | 94917115 | 94918356 | 4.13794 | 3.61415 | 2.34752 |
| chrX | 108834169 | 108834422 | 182 | Promoter (<=1kb) | 20 | 108737016 | 108834348 | 97333 | 2 | ENSMUST00 | 0 | Brwd3 | ENSMUSG0C | false | chrX | 108832866 | 108834360 | 17.00269 | 41.02908 | 38.42754 | . | . | . | . | . | . |
| chr10 | 31608882 | 31609840 | 182 | Promoter (<=1kb) | 10 | 31493193 | 31609184 | 115992 | 2 | ENSMUST00 | 0 | Rnf217 | ENSMUSG0C | false | chr10 | 31608164 | 31610207 | 37.7199 | 126.00829 | 121.65485 | chr10 | 31606070 | 31608928 | 4.35828 | 3.78102 | 2.48252 |
| chr7 | 4167109 | 4167492 | 182 | Promoter (2-3kb) | 7 | 4151260 | 4164860 | 13601 | 2 | ENSMUST00 | -2250 | Cdc42ep5 | ENSMUSG0C | false | . | . | . | . | . | . | . | . | . | . | . | . |
| chr14 | 48203768 | 48204219 | 182 | Intron (ENSMUST00000073150.4 | 14 | 48198716 | 48228413 | 29698 | 1 | ENSMUST00 | 5053 | Gm6498 | ENSMUSG0C | false | . | . | . | . | . | . | chr14 | 48197046 | 48208841 | 5.12168 | 4.05556 | 3.13469 |
| chr7 | 34985581 | 34985851 | 182 | Intron (ENSMUST00000075068.8 | 7 | 34969510 | 34980794 | 11285 | 1 | ENSMUST00 | 16072 | Pepd | ENSMUSG0C | false | . | . | . | . | . | . | chr7 | 34983638 | 34987677 | 6.30038 | 4.69301 | 4.07806 |
| chr3 | 96240348 | 96240609 | 182 | Promoter (<=1kb) | 3 | 96239749 | 96240276 | 528 | 2 | ENSMUST00 | -73 | Hist2h2aa1 | ENSMUSG0C | false | chr3 | 96240485 | 96240743 | 8.8423 | 19.93079 | 17.59637 | . | . | . | . | . | . |
| chr2 | 38926651 | 38926904 | 182 | Promoter (<=1kb) | 2 | 38740543 | 38926275 | 185733 | 2 | ENSMUST00 | -377 | Nr6a1 | ENSMUSG0C | false | chr2 | 38926655 | 38927318 | 20.41515 | 53.76371 | 50.99926 | . | . | . | . | . | . |
| chr18 | 73861798 | 73862019 | 182 | Promoter (1-2kb) | 18 | 73863631 | 73870209 | 6579 | 1 | ENSMUST00 | -1612 | Mro | ENSMUSG0C | false | . | . | . | . | . | . | chr18 | 73861669 | 73862048 | 2.95418 | 2.87764 | 1.35513 |
| chr18 | 73839991 | 73840203 | 182 | Distal Intergenic | 18 | 73859385 | 73878730 | 19346 | 1 | ENSMUST00 | -19182 | Mro | ENSMUSG0C | false | . | . | . | . | . | . | . | . | . | . | . | . |
| chr17 | 47323013 | 47323193 | 182 | Intron (ENSMUST00000188947.1 | 17 | 47294069 | 47359463 | 65395 | 1 | ENSMUST00 | 28945 | Trerf1 | ENSMUSG0C | false | . | . | . | . | . | . | . | . | . | . | . | . |
| chr17 | 47174696 | 47174931 | 182 | Intron (ENSMUST00000188947.1 | 17 | 47141216 | 47318790 | 177575 | 1 | ENSMUST00 | 33481 | Trerf1 | ENSMUSG0C | false | chr17 | 47174701 | 47175281 | 14.58643 | 34.13128 | 31.61576 | . | . | . | . | . | . |
| chr17 | 47295311 | 47295613 | 182 | Promoter (1-2kb) | 17 | 47294069 | 47359463 | 65395 | 1 | ENSMUST00 | 1243 | Trerf1 | ENSMUSG0C | false | . | . | . | . | . | . | . | . | . | . | . | . |
| chr17 | 47140946 | 47141329 | 182 | Promoter (<=1kb) | 17 | 47141216 | 47318790 | 177575 | 1 | ENSMUST00 | 0 | Trerf1 | ENSMUSG0C | false | chr17 | 47139299 | 47142406 | 27.6016 | 85.77155 | 82.50918 | chr17 | 47139017 | 47143146 | 4.41006 | 3.81485 | 2.5073 |
| chr13 | 67859198 | 67859443 | 182 | Distal Intergenic | 13 | 67863326 | 67863923 | 598 | 1 | ENSMUST00 | -3883 | BC048507 | ENSMUSG0C | false | . | . | . | . | . | . | chr13 | 67857058 | 67864527 | 4.45316 | 3.8043 | 2.549 |
| chr7 | 24507211 | 24507707 | 182 | Promoter (<=1kb) | 7 | 24507130 | 24515680 | 8551 | 1 | ENSMUST00 | 82 | Zfp428 | ENSMUSG0C | false | chr7 | 24506592 | 24508247 | 39.51976 | 124.83012 | 120.52397 | . | . | . | . | . | . |
| chr5 | 21264830 | 21265279 | 182 | Intron (ENSMUST00000036031.8 | 5 | 21319023 | 21322746 | 3724 | 2 | ENSMUST00 | 57467 | Ccdc146 | ENSMUSG0C | false | . | . | . | . | . | . | chr5 | 21264614 | 21265978 | 3.84181 | 3.38745 | 2.09094 |
| chr1 | 118388829 | 118389438 | 182 | Promoter (<=1kb) | 1 | 118389058 | 118609432 | 220375 | 1 | ENSMUST00 | 0 | Clasp1 | ENSMUSG0C | false | chr1 | 118387964 | 118390405 | 29.66345 | 100.21 | 96.63225 | . | . | . | . | . | . |
| chr15 | 3270410 | 3270781 | 182 | Promoter (<=1kb) | 15 | 3270767 | 3280508 | 9742 | 1 | ENSMUST00 | 0 | Sepp1 | ENSMUSG0C | false | chr15 | 3270402 | 3272276 | 19.52687 | 54.66541 | 51.89166 | . | . | . | . | . | . |
| chr2 | 132742108 | 132742299 | 182 | Intron (ENSMUST00000061891.5 | 2 | 132717106 | 132717208 | 103 | 1 | ENSMUST00 | 25003 | Gm22245 | ENSMUSG0C | false | . | . | . | . | . | . | chr2 | 132741970 | 132742564 | 4.0419 | 3.40073 | 2.25036 |
| chr14 | 33687526 | 33687798 | 182 | Distal Intergenic | 14 | 33644495 | 33644601 | 107 | 1 | ENSMUST00 | 43032 | Gm26228 | ENSMUSG0C | false | . | . | . | . | . | . | . | . | . | . | . | . |
| chr2 | 38484633 | 38484876 | 182 | Distal Intergenic | 2 | 38480338 | 38480445 | 108 | 1 | ENSMUST00 | 4296 | Gm25081 | ENSMUSG0C | false | . | . | . | . | . | . | . | . | . | . | . | . |
| chr8 | 47764590 | 47764826 | 182 | Distal Intergenic | 8 | 47719246 | 47719369 | 124 | 1 | ENSMUST00 | 45345 | n-R5s96 | ENSMUSG0C | false | chr8 | 47763766 | 47764916 | 6.89298 | 11.53747 | 9.33025 | chr8 | 47762950 | 47764955 | 3.9038 | 3.48015 | 2.15657 |
| chr5 | 96917747 | 96918013 | 182 | Distal Intergenic | 5 | 96907632 | 96907825 | 194 | 1 | ENSMUST00 | 10116 | Gm24623 | ENSMUSG0C | false | . | . | . | . | . | . | chr5 | 96916423 | 96918601 | 4.94258 | 3.85789 | 3.00394 |
| chr11 | 5250813 | 5251068 | 182 | Promoter (<=1kb) | 11 | 5251850 | 5251956 | 107 | 1 | ENSMUST00 | -782 | Gm25142 | ENSMUSG0C | false | . | . | . | . | . | . | chr11 | 5235885 | 5260441 | 6.3098 | 4.58865 | 4.10038 |
| chr16 | 87644243 | 87644534 | 182 | Distal Intergenic | 16 | 87621014 | 87621120 | 107 | 2 | ENSMUST00 | -23124 | Gm22808 | ENSMUSG0C | false | . | . | . | . | . | . | chr16 | 87643359 | 87645741 | 4.64765 | 3.83883 | 2.76069 |
| chr11 | 20859083 | 20859580 | 182 | Distal Intergenic | 11 | 20847753 | 20847881 | 129 | 2 | ENSMUST00 | -11203 | Gm22807 | ENSMUSG0C | false | . | . | . | . | . | . | chr11 | 20858251 | 20860793 | 3.69424 | 3.51711 | 1.91287 |
| chr11 | 101641777 | 101642082 | 182 | Promoter (<=1kb) | 11 | 101641748 | 101641938 | 191 | 1 | ENSMUST00 | 30 | Gm24950 | ENSMUSG0C | false | . | . | . | . | . | . | chr11 | 101640831 | 101647012 | 4.11995 | 3.68866 | 2.28525 |
| chr5 | 149320823 | 149321092 | 182 | Distal Intergenic | 5 | 149281118 | 149281223 | 106 | 2 | ENSMUST00 | -39601 | n-R5s180 | ENSMUSG0C | false | . | . | . | . | . | . | chr5 | 149320265 | 149321110 | 5.73613 | 4.03104 | 3.63681 |
| chr12 | 13145523 | 13145826 | 182 | Distal Intergenic | 12 | 13163629 | 13163763 | 135 | 2 | ENSMUST00 | 17937 | Gm24208 | ENSMUSG0C | false | . | . | . | . | . | . | chr12 | 13145093 | 13146546 | 3.82015 | 3.48294 | 2.02427 |
| chr12 | 13177699 | 13177917 | 182 | Distal Intergenic | 12 | 13163629 | 13163763 | 135 | 2 | ENSMUST00 | -13937 | Gm24208 | ENSMUSG0C | false | . | . | . | . | . | . | . | . | . | . | . | . |
| chr3 | 88879408 | 88879632 | 182 | Intron (ENSMUST00000107498.3 | 3 | 88862919 | 88863080 | 162 | 1 | ENSMUST00 | 16490 | Gm23054 | ENSMUSG0C | false | . | . | . | . | . | . | chr3 | 88878907 | 88879833 | 4.67421 | 3.74198 | 2.76892 |
| chr4 | 49251043 | 49251250 | 182 | Intron (ENSMUST00000076670.2 | 4 | 49289298 | 49289403 | 106 | 1 | ENSMUST00 | -38048 | Gm22685 | ENSMUSG0C | false | . | . | . | . | . | . | chr4 | 49250271 | 49254046 | 4.84306 | 3.98037 | 2.88173 |
| chr9 | 63035481 | 63035756 | 182 | Distal Intergenic | 9 | 63027560 | 63027717 | 158 | 1 | ENSMUST00 | 7922 | Gm24526 | ENSMUSG0C | false | . | . | . | . | . | . | chr9 | 63035033 | 63035866 | 4.45755 | 3.83059 | 2.54873 |
| chr3 | 144643327 | 144643781 | 182 | Distal Intergenic | 3 | 144586825 | 144586952 | 128 | 2 | ENSMUST00 | -56376 | Gm24406 | ENSMUSG0C | false | . | . | . | . | . | . | chr3 | 144642467 | 144644851 | 3.96283 | 3.44681 | 2.18857 |
| chr9 | 78175337 | 78175716 | 182 | Promoter (<=1kb) | 9 | 78175303 | 78175633 | 331 | 2 | ENSMUST00 | 0 | Rn7sk | ENSMUSG0C | false | . | . | . | . | . | . | . | . | . | . | . | . |
| chr4 | 25661294 | 25661487 | 182 | Intron (ENSMUST00000084770.4 | 4 | 25566660 | 25566822 | 163 | 2 | ENSMUST00 | -94473 | Gm24977 | ENSMUSG0C | false | . | . | . | . | . | . | chr4 | 25661360 | 25662041 | 3.2587 | 3.1392 | 1.60941 |
| chr15 | 58816272 | 58816453 | 182 | Intron (ENSMUST00000072113.5 | 15 | 58819323 | 58819478 | 156 | 2 | ENSMUST00 | 3025 | Gm24041 | ENSMUSG0C | false | . | . | . | . | . | . | chr15 | 58813332 | 58822800 | 5.28733 | 4.15484 | 3.25305 |
| chr3 | 51859288 | 51859541 | 182 | Intron (ENSMUST00000121440.2 | 3 | 51828912 | 51829028 | 117 | 1 | ENSMUST00 | 30377 | n-R5s196 | ENSMUSG0C | false | . | . | . | . | . | . | chr3 | 51859194 | 51859631 | 4.49283 | 3.73891 | 2.62327 |
| chr3 | 51869306 | 51869523 | 182 | Intron (ENSMUST00000121440.2 | 3 | 51828912 | 51829028 | 117 | 1 | ENSMUST00 | 40395 | n-R5s196 | ENSMUSG0C | false | . | . | . | . | . | . | chr3 | 51864972 | 51875550 | 6.25 | 4.62847 | 4.03397 |
| chr8 | 115466704 | 115466923 | 182 | Distal Intergenic | 8 | 115591170 | 115591273 | 104 | 2 | ENSMUST00 | 124350 | Gm23677 | ENSMUSG0C | false | . | . | . | . | . | . | . | . | . | . | . | . |
| chr11 | 65608922 | 65609282 | 182 | Distal Intergenic | 11 | 65626394 | 65626503 | 110 | 1 | ENSMUST00 | -17112 | Gm26128 | ENSMUSG0C | false | . | . | . | . | . | . | . | . | . | . | . | . |
| chr6 | 135636689 | 135636922 | 182 | Distal Intergenic | 6 | 135584192 | 135584306 | 115 | 1 | ENSMUST00 | 52498 | Gm25136 | ENSMUSG0C | false | . | . | . | . | . | . | . | . | . | . | . | . |
| chr11 | 101645504 | 101645931 | 182 | Promoter (<=1kb) | 11 | 101645740 | 101645930 | 191 | 2 | ENSMUST00 | 0 | Gm23971 | ENSMUSG0C | false | . | . | . | . | . | . | chr11 | 101640831 | 101647012 | 4.11995 | 3.68866 | 2.28525 |
| chr17 | 84585249 | 84585566 | 182 | Intron (ENSMUST00000047206.5 | 17 | 84624402 | 84624510 | 109 | 1 | ENSMUST00 | -38836 | Gm26066 | ENSMUSG0C | false | . | . | . | . | . | . | chr17 | 84578925 | 84586206 | 5.26213 | 4.07321 | 3.25857 |
| chr11 | 104254340 | 104254578 | 182 | Intron (ENSMUST00000106992.4 | 11 | 104251012 | 104251144 | 133 | 2 | ENSMUST00 | -3197 | Gm22743 | ENSMUSG0C | false | . | . | . | . | . | . | chr11 | 104242646 | 104282246 | 6.16611 | 4.45267 | 3.98703 |
| chr11 | 104247226 | 104247530 | 182 | Intron (ENSMUST00000106992.4 | 11 | 104251012 | 104251144 | 133 | 2 | ENSMUST00 | 3614 | Gm22743 | ENSMUSG0C | false | . | . | . | . | . | . | chr11 | 104242646 | 104282246 | 6.16611 | 4.45267 | 3.98703 |
| chr7 | 19472292 | 19472674 | 182 | Distal Intergenic | 7 | 19460797 | 19460935 | 139 | 2 | ENSMUST00 | -11358 | Gm23099 | ENSMUSG0C | false | . | . | . | . | . | . | . | . | . | . | . | . |
| chr18 | 65150703 | 65150899 | 182 | Intron (ENSMUST00000163516.2 | 18 | 65248861 | 65248926 | 66 | 1 | ENSMUST00 | -97962 | Mir122a | ENSMUSG0C | false | . | . | . | . | . | . | chr18 | 65149599 | 65151406 | 6.97989 | 4.85243 | 4.67904 |
| chr1 | 23271489 | 23271687 | 182 | Promoter (<=1kb) | 1 | 23272264 | 23272345 | 82 | 1 | ENSMUST00 | -577 | Mir30a | ENSMUSG0C | false | chr1 | 23271151 | 23276131 | 19.17732 | 54.04958 | 51.28253 | . | . | . | . | . | . |
| chr1 | 74396596 | 74396875 | 182 | Promoter (2-3kb) | 1 | 74394310 | 74394394 | 85 | 1 | ENSMUST00 | 2287 | Mir26b | ENSMUSG0C | false | . | . | . | . | . | . | chr1 | 74392453 | 74398560 | 4.69855 | 3.96513 | 2.75686 |
| chr9 | 119005390 | 119005590 | 182 | Intron (ENSMUST00000073109.6 | 9 | 119031796 | 119031885 | 90 | 1 | ENSMUST00 | -26206 | Mir26a-1 | ENSMUSG0C | false | . | . | . | . | . | . | chr9 | 118927678 | 119010843 | 6.89652 | 4.89961 | 4.57102 |
| chr1 | 162225851 | 162226106 | 182 | Promoter (2-3kb) | 1 | 162223368 | 162223477 | 110 | 1 | ENSMUST00 | 2484 | Mir214 | ENSMUSG0C | false | . | . | . | . | . | . | . | . | . | . | . | . |
| chr19 | 29128336 | 29128519 | 182 | Intron (ENSMUST00000064393.5 | 19 | 29135279 | 29135375 | 97 | 1 | ENSMUST00 | -6760 | Mir101b | ENSMUSG0C | false | . | . | . | . | . | . | . | . | . | . | . | . |
| chr7 | 99553537 | 99553821 | 182 | Promoter (1-2kb) | 7 | 99552269 | 99552363 | 95 | 1 | ENSMUST00 | 1269 | Mir326 | ENSMUSG0C | false | . | . | . | . | . | . | chr7 | 99536482 | 99576026 | 7.32182 | 4.95837 | 4.9327 |
| chr5 | 139364701 | 139365187 | 182 | Intron (ENSMUST00000066052.7 | 5 | 139369650 | 139369745 | 96 | 2 | ENSMUST00 | 4558 | Mir339 | ENSMUSG0C | false | . | . | . | . | . | . | chr5 | 139364092 | 139366758 | 7.94465 | 5.46117 | 5.39437 |
| chr8 | 126928857 | 126929117 | 182 | Downstream (1-2kb) | 8 | 126944847 | 126944980 | 134 | 2 | ENSMUST00 | 15863 | Gm26397 | ENSMUSG0C | false | . | . | . | . | . | . | chr8 | 126925616 | 126933815 | 4.49681 | 3.6256 | 2.63517 |
| chr8 | 126856395 | 126857088 | 182 | Distal Intergenic | 8 | 126944847 | 126944980 | 134 | 2 | ENSMUST00 | 87892 | Gm26397 | ENSMUSG0C | false | . | . | . | . | . | . | chr8 | 126855989 | 126856437 | 3.56996 | 3.41783 | 1.84742 |
| chr8 | 126853015 | 126853352 | 182 | Distal Intergenic | 8 | 126944847 | 126944980 | 134 | 2 | ENSMUST00 | 91628 | Gm26397 | ENSMUSG0C | false | . | . | . | . | . | . | chr8 | 126847776 | 126853210 | 4.5877 | 3.85968 | 2.679 |
| chr8 | 126833639 | 126833958 | 182 | Distal Intergenic | 8 | 126944847 | 126944980 | 134 | 2 | ENSMUST00 | 111022 | Gm26397 | ENSMUSG0C | false | . | . | . | . | . | . | chr8 | 126824457 | 126835290 | 5.5694 | 4.27648 | 3.48091 |
| chr8 | 126824738 | 126824962 | 182 | Distal Intergenic | 8 | 126944847 | 126944980 | 134 | 2 | ENSMUST00 | 120018 | Gm26397 | ENSMUSG0C | false | . | . | . | . | . | . | chr8 | 126824457 | 126835290 | 5.5694 | 4.27648 | 3.48091 |
| chr10 | 108389997 | 108390207 | 182 | Intron (ENSMUST00000095313.3 | 10 | 108363665 | 108363798 | 134 | 1 | ENSMUST00 | 26333 | Gm23105 | ENSMUSG0C | false | . | . | . | . | . | . | chr10 | 108383715 | 108391831 | 4.99674 | 4.04583 | 3.02244 |
| chr10 | 8756739 | 8757140 | 182 | Exon (ENSMUST00000015449.5/ | 10 | 8824011 | 8824147 | 137 | 2 | ENSMUST00 | 67007 | Gm25410 | ENSMUSG0C | false | chr10 | 8755557 | 8757768 | 8.50221 | 18.80243 | 16.48241 | chr10 | 8756871 | 8763110 | 4.54887 | 3.87109 | 2.6311 |
| chr10 | 47474398 | 47474589 | 182 | Distal Intergenic | 10 | 46838346 | 46838507 | 162 | 2 | ENSMUST00 | -635892 | Gm25650 | ENSMUSG0C | false | . | . | . | . | . | . | . | . | . | . | . | . |
| chr7 | 130098598 | 130098874 | 182 | Distal Intergenic | 7 | 130035094 | 130035203 | 110 | 1 | ENSMUST00 | 63505 | Gm23847 | ENSMUSG0C | false | . | . | . | . | . | . | chr7 | 130098300 | 130099432 | 4.73655 | 4.02132 | 2.80864 |
| chr10 | 4132627 | 4132953 | 182 | Intron (ENSMUST00000117291.2 | 10 | 4104313 | 4104469 | 157 | 1 | ENSMUST00 | 28315 | Gm25515 | ENSMUSG0C | false | . | . | . | . | . | . | . | . | . | . | . | . |
| chr2 | 63701984 | 63702217 | 182 | Distal Intergenic | 2 | 63432088 | 63432194 | 107 | 2 | ENSMUST00 | -269791 | Gm23503 | ENSMUSG0C | false | chr2 | 63701903 | 63703074 | 8.50137 | 15.8526 | 13.57707 | chr2 | 63701955 | 63703908 | 4.28156 | 3.79438 | 2.40494 |
| chr3 | 86107157 | 86107395 | 182 | Exon (ENSMUST00000182666.2/ | 3 | 86100534 | 86100655 | 122 | 1 | ENSMUST00 | 6624 | Gm25188 | ENSMUSG0C | false | . | . | . | . | . | . | chr3 | 86106054 | 86108034 | 4.2092 | 3.54995 | 2.37563 |
| chr8 | 25275772 | 25276140 | 182 | Distal Intergenic | 8 | 25154552 | 25201449 | 46898 | 2 | ENSMUST00 | -74324 | Tacc1 | ENSMUSG0C | false | . | . | . | . | . | . | chr8 | 25274758 | 25277183 | 5.90896 | 4.54281 | 3.75849 |
| chr4 | 144893336 | 144893909 | 182 | Promoter (<=1kb) | 4 | 144893219 | 144927639 | 34421 | 1 | ENSMUST00 | 118 | Dhrs3 | ENSMUSG0C | false | chr4 | 144891998 | 144895332 | 25.47373 | 70.96252 | 67.95692 | chr4 | 144893840 | 144898334 | 3.97359 | 3.51387 | 2.17115 |
| chr4 | 144930360 | 144930697 | 182 | Distal Intergenic | 4 | 144918879 | 144919972 | 1094 | 1 | ENSMUST00 | 11482 | Dhrs3 | ENSMUSG0C | false | . | . | . | . | . | . | chr4 | 144929102 | 144931190 | 6.53574 | 4.74081 | 4.26509 |
| chr4 | 139380410 | 139380767 | 182 | Promoter (<=1kb) | 4 | 139380659 | 139489530 | 108872 | 1 | ENSMUST00 | 0 | Ubr4 | ENSMUSG0C | false | chr4 | 139379645 | 139381792 | 38.86769 | 126.9345 | 122.55549 | . | . | . | . | . | . |
| chr4 | 132396332 | 132396618 | 182 | Promoter (1-2kb) | 4 | 132377461 | 132398199 | 20739 | 2 | ENSMUST00 | 1581 | Phactr4 | ENSMUSG0C | true | chr4 | 132394676 | 132398208 | 24.76026 | 74.23955 | 71.18308 | chr4 | 132379835 | 132397156 | 5.13856 | 4.06122 | 3.15072 |
| chr4 | 132398104 | 132398444 | 182 | Promoter (<=1kb) | 4 | 132358249 | 132398211 | 39963 | 2 | ENSMUST00 | 0 | Phactr4 | ENSMUSG0C | true | chr4 | 132394676 | 132398208 | 24.76026 | 74.23955 | 71.18308 | chr4 | 132398431 | 132412921 | 4.68778 | 3.92949 | 2.75307 |
| chr4 | 62348010 | 62348279 | 182 | Intron (ENSMUST00000084527.4 | 4 | 62344137 | 62360469 | 16333 | 2 | ENSMUST00 | 12190 | Fkbp15 | ENSMUSG0C | false | . | . | . | . | . | . | chr4 | 62348052 | 62349367 | 4.60395 | 3.51711 | 2.71851 |
| chr4 | 62293027 | 62293267 | 182 | Promoter (1-2kb) | 4 | 62291547 | 62297477 | 5931 | 1 | ENSMUST00 | 1481 | Slc31a2 | ENSMUSG0C | false | . | . | . | . | . | . | chr4 | 62287890 | 62304836 | 6.16439 | 4.4648 | 3.98538 |
| chr4 | 62286984 | 62287238 | 182 | Promoter (<=1kb) | 4 | 62286449 | 62298411 | 11963 | 1 | ENSMUST00 | 536 | Slc31a2 | ENSMUSG0C | false | chr4 | 62285924 | 62287996 | 33.56007 | 105.08768 | 101.38747 | . | . | . | . | . | . |
| chr4 | 4793097 | 4793371 | 182 | Promoter (<=1kb) | 4 | 4762484 | 4793355 | 30872 | 2 | ENSMUST00 | 0 | Impad1 | ENSMUSG0C | false | chr4 | 4792040 | 4793755 | 39.97575 | 133.22411 | 128.61581 | . | . | . | . | . | . |
| chr9 | 108578644 | 108578905 | 182 | Promoter (<=1kb) | 9 | 108572313 | 108578670 | 6358 | 2 | ENSMUST00 | 0 | Wdr6 | ENSMUSG0C | false | chr9 | 108578036 | 108579239 | 26.5582 | 79.22746 | 76.08501 | chr9 | 108577463 | 108579704 | 4.9109 | 4.06359 | 2.92127 |
| chr7 | 81850244 | 81850482 | 182 | Distal Intergenic | 7 | 81855533 | 81855794 | 262 | 2 | ENSMUST00 | 5312 | Gm10160 | ENSMUSG0C | false | . | . | . | . | . | . | chr7 | 81849920 | 81852312 | 3.89172 | 3.42178 | 2.13772 |
| chr9 | 89211141 | 89211548 | 182 | Promoter (<=1kb) | 9 | 89211157 | 89226458 | 15302 | 1 | ENSMUST00 | 0 | Mthfs | ENSMUSG0C | false | chr9 | 89211168 | 89211897 | 24.45906 | 68.33184 | 65.3672 | . | . | . | . | . | . |
| chr12 | 56438476 | 56438706 | 182 | Distal Intergenic | 12 | 56373593 | 56373823 | 231 | 2 | ENSMUST00 | -64654 | Gm6265 | ENSMUSG0C | false | . | . | . | . | . | . | . | . | . | . | . | . |
| chr7 | 36095515 | 36095825 | 182 | Distal Intergenic | 7 | 36118274 | 36118828 | 555 | 1 | ENSMUST00 | -22449 | Rpl17-ps9 | ENSMUSG0C | false | . | . | . | . | . | . | . | . | . | . | . | . |
| chr7 | 35988250 | 35988733 | 182 | Distal Intergenic | 7 | 36118274 | 36118828 | 555 | 1 | ENSMUST00 | -129541 | Rpl17-ps9 | ENSMUSG0C | false | . | . | . | . | . | . | chr7 | 35988072 | 35989682 | 5.16371 | 4.08384 | 3.16297 |
| chr12 | 8969913 | 8970140 | 182 | Intron (ENSMUST00000020899.4 | 12 | 8973892 | 9028847 | 54956 | 1 | ENSMUST00 | -3752 | Wdr35 | ENSMUSG0C | false | . | . | . | . | . | . | chr12 | 8969500 | 8970807 | 4.16302 | 3.7595 | 2.30726 |
| chr9 | 25048295 | 25048686 | 182 | Distal Intergenic | 9 | 25089422 | 25089490 | 69 | 1 | ENSMUST00 | -40736 | Gm10181 | ENSMUSG0C | false | . | . | . | . | . | . | . | . | . | . | . | . |
| chr9 | 22085228 | 22085561 | 182 | Promoter (<=1kb) | 9 | 22072246 | 22085402 | 13157 | 2 | ENSMUST00 | 0 | Ecsit | ENSMUSG0C | false | chr9 | 22084690 | 22086122 | 25.15761 | 80.47005 | 77.31023 | chr9 | 22083597 | 22086895 | 4.27847 | 3.75537 | 2.44583 |
| chrX | 166781514 | 166781687 | 182 | Distal Intergenic | 20 | 166854211 | 166855011 | 801 | 1 | ENSMUST00 | -72524 | Rpl7a-ps11 | ENSMUSG0C | false | . | . | . | . | . | . | . | . | . | . | . | . |
| chr17 | 46247969 | 46248334 | 182 | Promoter (<=1kb) | 17 | 46243920 | 46248054 | 4135 | 2 | ENSMUST00 | 0 | Polr1c | ENSMUSG0C | false | chr17 | 46247345 | 46249077 | 38.16745 | 122.49164 | 118.26377 | . | . | . | . | . | . |
| chr5 | 72652474 | 72652741 | 182 | Intron (ENSMUST00000087212.3 | 5 | 72647796 | 72671078 | 23283 | 1 | ENSMUST00 | 4679 | Nipal1 | ENSMUSG0C | false | . | . | . | . | . | . | . | . | . | . | . | . |
| chr17 | 33092474 | 33092723 | 182 | Promoter (<=1kb) | 17 | 33092458 | 33105058 | 12601 | 1 | ENSMUST00 | 17 | Zfp563 | ENSMUSG0C | false | chr17 | 33092120 | 33093457 | 20.67151 | 72.03884 | 69.0182 | . | . | . | . | . | . |
| chr5 | 24525958 | 24526444 | 182 | Promoter (<=1kb) | 5 | 24502757 | 24527202 | 24446 | 2 | ENSMUST00 | 758 | Gbx1 | ENSMUSG0C | false | chr5 | 24525602 | 24527538 | 25.56571 | 75.64988 | 72.57267 | chr5 | 24524809 | 24526030 | 4.33552 | 3.78926 | 2.46082 |
| chr2 | 157566162 | 157566425 | 182 | Promoter (<=1kb) | 2 | 157556362 | 157566337 | 9976 | 2 | ENSMUST00 | 0 | Blcap | ENSMUSG0C | false | chr2 | 157565403 | 157567312 | 41.4024 | 145.73851 | 140.61644 | . | . | . | . | . | . |
| chr2 | 156777416 | 156777936 | 182 | Promoter (<=1kb) | 2 | 156778517 | 156781164 | 2648 | 1 | ENSMUST00 | -581 | Myl9 | ENSMUSG0C | false | . | . | . | . | . | . | . | . | . | . | . | . |
| chr6 | 121141379 | 121141650 | 182 | Distal Intergenic | 6 | 121183667 | 121196185 | 12519 | 1 | ENSMUST00 | -42017 | Pex26 | ENSMUSG0C | false | . | . | . | . | . | . | chr6 | 121140632 | 121142501 | 4.39009 | 3.76092 | 2.50977 |
| chr14 | 74877344 | 74877783 | 182 | Intron (ENSMUST00000088970.5 | 14 | 74754674 | 74947877 | 193204 | 2 | ENSMUST00 | 70094 | Lrch1 | ENSMUSG0C | true | . | . | . | . | . | . | chr14 | 74857233 | 74945366 | 6.29465 | 4.64624 | 4.07764 |
| chr14 | 74936492 | 74936700 | 182 | Intron (ENSMUST00000088970.5 | 14 | 74754674 | 74947877 | 193204 | 2 | ENSMUST00 | 11177 | Lrch1 | ENSMUSG0C | true | . | . | . | . | . | . | chr14 | 74857233 | 74945366 | 6.29465 | 4.64624 | 4.07764 |
| chr14 | 74867563 | 74867910 | 182 | Intron (ENSMUST00000088970.5 | 14 | 74754674 | 74947877 | 193204 | 2 | ENSMUST00 | 79967 | Lrch1 | ENSMUSG0C | true | . | . | . | . | . | . | chr14 | 74857233 | 74945366 | 6.29465 | 4.64624 | 4.07764 |
| chr2 | 131210208 | 131210550 | 182 | Promoter (<=1kb) | 2 | 131210370 | 131213002 | 2633 | 1 | ENSMUST00 | 0 | Ap5s1 | ENSMUSG0C | false | chr2 | 131210000 | 131211274 | 31.91637 | 110.38313 | 106.53261 | . | . | . | . | . | . |
| chr10 | 20505507 | 20505730 | 182 | Intron (ENSMUST00000020165.8 | 10 | 20515327 | 20518164 | 2838 | 2 | ENSMUST00 | 12434 | 4930403O15 | ENSMUSG0C | false | . | . | . | . | . | . | . | . | . | . | . | . |
| chr2 | 92431309 | 92431857 | 182 | Promoter (<=1kb) | 2 | 92426904 | 92432392 | 5489 | 2 | ENSMUST00 | 535 | Cry2 | ENSMUSG0C | false | chr2 | 92431256 | 92434116 | 30.13579 | 99.06311 | 95.51676 | chr2 | 92421995 | 92433198 | 7.705 | 5.3487 | 5.2209 |
| chr2 | 92442705 | 92443002 | 182 | Distal Intergenic | 2 | 92403646 | 92434043 | 30398 | 2 | ENSMUST00 | -8663 | Cry2 | ENSMUSG0C | false | . | . | . | . | . | . | chr2 | 92442484 | 92443052 | 5.04926 | 3.84892 | 3.08475 |
| chr3 | 103020289 | 103020607 | 182 | Promoter (<=1kb) | 3 | 103020546 | 103058189 | 37644 | 1 | ENSMUST00 | 0 | Csde1 | ENSMUSG0C | false | chr3 | 103019827 | 103021910 | 28.56742 | 103.16996 | 99.51957 | . | . | . | . | . | . |
| chr3 | 94932905 | 94933238 | 182 | Promoter (<=1kb) | 3 | 94933056 | 94944745 | 11690 | 1 | ENSMUST00 | 0 | Selenbp1 | ENSMUSG0C | false | . | . | . | . | . | . | chr3 | 94930042 | 94935789 | 5.46862 | 4.30057 | 3.39851 |
| chr2 | 29827106 | 29827356 | 182 | Promoter (<=1kb) | 2 | 29827342 | 29843374 | 16033 | 1 | ENSMUST00 | 0 | Urm1 | ENSMUSG0C | false | chr2 | 29826439 | 29828177 | 30.41442 | 103.5799 | 99.9162 | . | . | . | . | . | . |
| chr3 | 19310897 | 19311732 | 182 | Promoter (<=1kb) | 3 | 19223108 | 19311322 | 88215 | 2 | ENSMUST00 | 0 | Pde7a | ENSMUSG0C | false | chr3 | 19310411 | 19312172 | 41.65439 | 138.35785 | 133.54871 | . | . | . | . | . | . |
| chr13 | 97919613 | 97919895 | 182 | Intron (ENSMUST00000109426.1 | 13 | 97760130 | 97760588 | 459 | 2 | ENSMUST00 | -159026 | Gm10260 | ENSMUSG0C | false | . | . | . | . | . | . | chr13 | 97909477 | 97929349 | 4.81516 | 3.96306 | 2.86901 |
| chr13 | 81352342 | 81352578 | 182 | Intron (ENSMUST00000095585.5 | 13 | 81095068 | 81343315 | 248248 | 2 | ENSMUST00 | -9028 | Gpr98 | ENSMUSG0C | false | . | . | . | . | . | . | . | . | . | . | . | . |
| chr13 | 55760920 | 55761156 | 182 | Intron (ENSMUST00000021958.5 | 13 | 55775940 | 55784479 | 8540 | 2 | ENSMUST00 | 23323 | 4930451E10I | ENSMUSG0C | false | chr13 | 55760823 | 55761194 | 5.43604 | 8.18012 | 6.04458 | chr13 | 55759365 | 55764010 | 3.5108 | 3.28016 | 1.79677 |
| chr13 | 55756636 | 55756840 | 182 | Intron (ENSMUST00000021958.5 | 13 | 55775940 | 55784479 | 8540 | 2 | ENSMUST00 | 27639 | 4930451E10I | ENSMUSG0C | false | . | . | . | . | . | . | chr13 | 55755814 | 55757391 | 3.4595 | 3.1527 | 1.78528 |
| chr13 | 23551504 | 23551826 | 182 | Promoter (<=1kb) | 13 | 23551258 | 23551648 | 391 | 2 | ENSMUST00 | 0 | Hist1h4f | ENSMUSG0C | false | chr13 | 23551125 | 23552074 | 13.05315 | 37.5606 | 35.00106 | chr13 | 23550665 | 23552415 | 3.21921 | 3.16102 | 1.52627 |
| chr10 | 37078586 | 37079001 | 182 | Distal Intergenic | 10 | 37133375 | 37138920 | 5546 | 2 | ENSMUST00 | 59919 | Marcks | ENSMUSG0C | false | . | . | . | . | . | . | . | . | . | . | . | . |
| chr17 | 4994876 | 4995127 | 182 | Promoter (<=1kb) | 17 | 4994939 | 5343505 | 348567 | 1 | ENSMUST00 | 0 | Arid1b | ENSMUSG0C | false | chr17 | 4993664 | 4997887 | 36.69013 | 115.69205 | 111.68276 | . | . | . | . | . | . |
| chr11 | 88686062 | 88686400 | 182 | 3' UTR | 11 | 88343889 | 88718101 | 374213 | 2 | ENSMUST00 | 31701 | Msi2 | ENSMUSG0C | false | . | . | . | . | . | . | chr11 | 88673697 | 88698174 | 6.07352 | 4.42058 | 3.91072 |
| chr11 | 88552915 | 88553234 | 182 | Intron (ENSMUST00000107909.2 | 11 | 88365515 | 88590147 | 224633 | 2 | ENSMUST00 | 36913 | Msi2 | ENSMUSG0C | false | . | . | . | . | . | . | chr11 | 88545893 | 88565759 | 5.157 | 4.05043 | 3.16526 |
| chr19 | 8992748 | 8993087 | 182 | Intron (ENSMUST00000092956.2 | 19 | 8989284 | 9076919 | 87636 | 1 | ENSMUST00 | 3465 | Ahnak | ENSMUSG0C | false | . | . | . | . | . | . | chr19 | 8990337 | 9001009 | 6.29078 | 4.5762 | 4.06914 |
| chr11 | 58214790 | 58215057 | 182 | Promoter (<=1kb) | 11 | 58215028 | 58222779 | 7752 | 1 | ENSMUST00 | 0 | Irgm2 | ENSMUSG0C | false | chr11 | 58214493 | 58215673 | 23.28017 | 72.50986 | 69.48009 | . | . | . | . | . | . |
| chr8 | 105255088 | 105255328 | 182 | Promoter (<=1kb) | 8 | 105252638 | 105255153 | 2516 | 2 | ENSMUST00 | 0 | B3gnt9 | ENSMUSG0C | false | chr8 | 105254023 | 105255259 | 27.03504 | 88.5911 | 85.27629 | chr8 | 105253926 | 105255681 | 3.84569 | 3.48177 | 2.09348 |
| chr8 | 44934979 | 44935500 | 182 | Promoter (<=1kb) | 8 | 44935447 | 45052257 | 116811 | 1 | ENSMUST00 | 0 | Fat1 | ENSMUSG0C | true | chr8 | 44934665 | 44936566 | 26.19333 | 73.09374 | 70.054 | . | . | . | . | . | . |
| chr8 | 44805500 | 44805809 | 182 | Distal Intergenic | 8 | 44935447 | 45052257 | 116811 | 1 | ENSMUST00 | -129638 | Fat1 | ENSMUSG0C | true | . | . | . | . | . | . | . | . | . | . | . | . |
| chr8 | 44977900 | 44978142 | 182 | Intron (ENSMUST00000191428.1 | 8 | 44950208 | 45052257 | 102050 | 1 | ENSMUST00 | 27693 | Fat1 | ENSMUSG0C | true | . | . | . | . | . | . | chr8 | 44953863 | 44987890 | 5.49999 | 4.27545 | 3.43184 |
| chr16 | 33008865 | 33009283 | 182 | Exon (ENSMUST00000135193.3/ | 16 | 33001903 | 33002021 | 119 | 1 | ENSMUST00 | 6963 | n-R5s33 | ENSMUSG0C | false | . | . | . | . | . | . | chr16 | 33007066 | 33008877 | 4.69125 | 3.75735 | 2.78788 |
| chr9 | 119181235 | 119181437 | 182 | Intron (ENSMUST00000093775.6 | 9 | 119169668 | 119173652 | 3985 | 2 | ENSMUST00 | -7584 | Slc22a14 | ENSMUSG0C | false | . | . | . | . | . | . | chr9 | 119179771 | 119182458 | 4.55118 | 3.88937 | 2.63029 |
| chr9 | 35127221 | 35127486 | 182 | Intron (ENSMUST00000093873.3 | 9 | 35118195 | 35126897 | 8703 | 1 | ENSMUST00 | 9027 | 4930581F22I | ENSMUSG0C | false | . | . | . | . | . | . | . | . | . | . | . | . |
| chr11 | 63922190 | 63922674 | 182 | Promoter (<=1kb) | 11 | 63885792 | 63922290 | 36499 | 2 | ENSMUST00 | 0 | Hs3st3b1 | ENSMUSG0C | false | chr11 | 63918689 | 63922332 | 41.51012 | 139.21701 | 134.36754 | chr11 | 63922597 | 63925544 | 5.42872 | 4.29883 | 3.35671 |
| chr7 | 102065049 | 102065561 | 182 | Promoter (<=1kb) | 7 | 102019137 | 102065158 | 46022 | 2 | ENSMUST00 | 0 | Rnf121 | ENSMUSG0C | false | chr7 | 102064411 | 102066178 | 18.88512 | 67.59834 | 64.64505 | . | . | . | . | . | . |
| chr7 | 73391649 | 73392045 | 182 | Promoter (<=1kb) | 7 | 73391227 | 73419899 | 28673 | 1 | ENSMUST00 | 423 | Rgma | ENSMUSG0C | false | chr7 | 73390852 | 73392583 | 21.79372 | 61.12524 | 58.26245 | chr7 | 73389790 | 73392430 | 4.61187 | 3.65475 | 2.71075 |
| chr7 | 45162167 | 45162428 | 182 | Promoter (1-2kb) | 7 | 45163921 | 45176138 | 12218 | 1 | ENSMUST00 | -1493 | Slc17a7 | ENSMUSG0C | false | chr7 | 45161878 | 45162477 | 12.92288 | 28.71786 | 26.27042 | chr7 | 45161619 | 45162779 | 5.83533 | 4.54254 | 3.68007 |
| chr1 | 133382725 | 133382927 | 182 | Promoter (2-3kb) | 1 | 133383978 | 133385526 | 1549 | 2 | ENSMUST00 | 2599 | Sox13 | ENSMUSG0C | false | . | . | . | . | . | . | chr1 | 133382342 | 133383491 | 3.62173 | 3.35141 | 1.88138 |
| chr7 | 28277949 | 28278280 | 182 | Promoter (<=1kb) | 7 | 28277706 | 28280129 | 2424 | 1 | ENSMUST00 | 244 | Eid2b | ENSMUSG0C | false | chr7 | 28277660 | 28278372 | 30.41442 | 103.5799 | 99.9162 | chr7 | 28277974 | 28278619 | 3.67679 | 3.48667 | 1.90537 |
| chr4 | 132074019 | 132074232 | 182 | Promoter (<=1kb) | 4 | 132074697 | 132076992 | 2296 | 1 | ENSMUST00 | -465 | Gm10300 | ENSMUSG0C | false | chr4 | 132074095 | 132075741 | 43.58744 | 151.86528 | 146.50356 | chr4 | 132073635 | 132074332 | 4.13788 | 3.68042 | 2.28228 |
| chr5 | 73236868 | 73237067 | 182 | Intron (ENSMUST00000101127.6 | 5 | 73119257 | 73256425 | 137169 | 2 | ENSMUST00 | 19358 | Fryl | ENSMUSG0C | true | . | . | . | . | . | . | chr5 | 73196545 | 73254445 | 6.29302 | 4.62166 | 4.08669 |
| chr5 | 73255571 | 73256371 | 182 | Promoter (<=1kb) | 5 | 73119257 | 73256425 | 137169 | 2 | ENSMUST00 | 54 | Fryl | ENSMUSG0C | true | chr5 | 73254020 | 73256650 | 42.59918 | 148.98648 | 143.74643 | . | . | . | . | . | . |
| chr1 | 87853038 | 87853429 | 182 | Promoter (<=1kb) | 1 | 87853287 | 87944490 | 91204 | 1 | ENSMUST00 | 0 | Dgkd | ENSMUSG0C | false | chr1 | 87852728 | 87854508 | 30.45704 | 104.66754 | 100.97924 | . | . | . | . | . | . |
| chr7 | 16018461 | 16018720 | 182 | Distal Intergenic | 7 | 15971262 | 15999495 | 28234 | 2 | ENSMUST00 | -18967 | Gltscr1 | ENSMUSG0C | false | . | . | . | . | . | . | chr7 | 16008939 | 16024059 | 5.6506 | 4.26598 | 3.56534 |
| chr7 | 16005175 | 16005372 | 182 | Distal Intergenic | 7 | 15971262 | 15999495 | 28234 | 2 | ENSMUST00 | -5681 | Gltscr1 | ENSMUSG0C | false | . | . | . | . | . | . | chr7 | 15999873 | 16007086 | 5.38556 | 4.2466 | 3.32251 |
| chr4 | 114304679 | 114304910 | 182 | Distal Intergenic | 4 | 114406724 | 114615098 | 208375 | 1 | ENSMUST00 | -101814 | Trabd2b | ENSMUSG0C | false | . | . | . | . | . | . | chr4 | 114303193 | 114308525 | 5.01086 | 4.08211 | 3.01047 |
| chr1 | 43077573 | 43077788 | 182 | Intron (ENSMUST00000095014.2 | 1 | 43059891 | 43092447 | 32557 | 2 | ENSMUST00 | 14659 | Tgfbrap1 | ENSMUSG0C | false | . | . | . | . | . | . | chr1 | 43075544 | 43081053 | 4.91233 | 3.89321 | 2.94525 |
| chr4 | 58995226 | 58995632 | 182 | Promoter (<=1kb) | 4 | 58995114 | 59022683 | 27570 | 1 | ENSMUST00 | 113 | Dnajc25 | ENSMUSG0C | false | . | . | . | . | . | . | chr4 | 58994753 | 58995706 | 3.24701 | 3.16666 | 1.58473 |
| chr4 | 43972181 | 43972491 | 182 | Intron (ENSMUST00000030202.8 | 4 | 43983483 | 43985423 | 1941 | 1 | ENSMUST00 | -10992 | Ccin | ENSMUSG0C | false | . | . | . | . | . | . | chr4 | 43972177 | 43972734 | 3.87988 | 3.53012 | 2.07399 |
| chr17 | 86507670 | 86507891 | 182 | Promoter (2-3kb) | 17 | 86497732 | 86505232 | 7501 | 2 | ENSMUST00 | -2439 | Gm10309 | ENSMUSG0C | false | . | . | . | . | . | . | . | . | . | . | . | . |
| chr10 | 128058743 | 128059506 | 182 | Promoter (<=1kb) | 10 | 128058954 | 128077272 | 18319 | 1 | ENSMUST00 | 0 | Ptges3 | ENSMUSG0C | false | chr10 | 128058154 | 128060485 | 33.52805 | 106.05383 | 102.33234 | . | . | . | . | . | . |
| chr17 | 46253962 | 46254342 | 182 | Promoter (<=1kb) | 17 | 46254165 | 46257316 | 3152 | 1 | ENSMUST00 | 0 | Lrrc73 | ENSMUSG0C | false | chr17 | 46253492 | 46254618 | 26.78267 | 75.92174 | 72.83789 | chr17 | 46253116 | 46254776 | 4.26127 | 3.64077 | 2.43351 |
| chr17 | 29032352 | 29032639 | 182 | Promoter (<=1kb) | 17 | 29032673 | 29042243 | 9571 | 1 | ENSMUST00 | -34 | Srsf3 | ENSMUSG0C | false | chr17 | 29031776 | 29035290 | 39.67387 | 146.00111 | 140.87399 | . | . | . | . | . | . |
| chr14 | 79160501 | 79160765 | 182 | Intron (ENSMUST00000040990.5 | 14 | 79212355 | 79223876 | 11522 | 2 | ENSMUST00 | 63111 | Zfp957 | ENSMUSG0C | false | . | . | . | . | . | . | chr14 | 79154001 | 79161573 | 4.37416 | 3.79583 | 2.49942 |
| chr17 | 22423933 | 22424537 | 182 | Promoter (<=1kb) | 17 | 22424222 | 22447416 | 23195 | 1 | ENSMUST00 | 0 | Zfp946 | ENSMUSG0C | false | chr17 | 22423609 | 22425123 | 22.71493 | 67.06055 | 64.11349 | chr17 | 22423589 | 22425165 | 2.97841 | 2.98862 | 1.39763 |
| chr12 | 17063876 | 17064068 | 182 | Distal Intergenic | 12 | 17003319 | 17011727 | 8409 | 2 | ENSMUST00 | -52150 | 2410004P03I | ENSMUSG0C | false | . | . | . | . | . | . | chr12 | 17061085 | 17064225 | 6.62006 | 4.90177 | 4.32889 |
| chr16 | 15887019 | 15887314 | 182 | Promoter (<=1kb) | 16 | 15887286 | 15888466 | 1181 | 1 | ENSMUST00 | 0 | Cebpd | ENSMUSG0C | false | chr16 | 15886653 | 15888845 | 34.74108 | 114.39095 | 110.42815 | . | . | . | . | . | . |
| chr19 | 8942085 | 8942569 | 182 | Promoter (<=1kb) | 19 | 8942493 | 8945873 | 3381 | 1 | ENSMUST00 | 0 | Mta2 | ENSMUSG0C | false | chr19 | 8941466 | 8943320 | 33.7938 | 119.00099 | 114.88904 | . | . | . | . | . | . |
| chr19 | 8830593 | 8830905 | 182 | Exon (ENSMUST00000096753.3/ | 19 | 8837467 | 8848683 | 11217 | 1 | ENSMUST00 | -6562 | Bscl2 | ENSMUSG0C | false | . | . | . | . | . | . | chr19 | 8828114 | 8833308 | 5.31614 | 4.28934 | 3.28488 |
| chr19 | 8819098 | 8819417 | 182 | Promoter (<=1kb) | 19 | 8819401 | 8834142 | 14742 | 1 | ENSMUST00 | 0 | Hnrnpul2 | ENSMUSG0C | false | chr19 | 8818123 | 8819393 | 18.02468 | 55.19937 | 52.41551 | . | . | . | . | . | . |
| chrX | 95026581 | 95026864 | 182 | Promoter (<=1kb) | 20 | 95022510 | 95026682 | 4173 | 2 | ENSMUST00 | 0 | Spin4 | ENSMUSG0C | false | chrX | 95025694 | 95027004 | 17.92175 | 44.05972 | 41.41886 | . | . | . | . | . | . |
| chr15 | 57697055 | 57697291 | 182 | Promoter (1-2kb) | 15 | 57695729 | 57821110 | 125382 | 1 | ENSMUST00 | 1327 | Zhx2 | ENSMUSG0C | true | . | . | . | . | . | . | chr15 | 57695806 | 57714739 | 5.09262 | 4.12618 | 3.0824 |
| chr8 | 20424754 | 20424945 | 182 | Promoter (<=1kb) | 8 | 20385782 | 20424814 | 39033 | 2 | ENSMUST00 | 0 | 2610005L07I | ENSMUSG0C | false | chr8 | 20424027 | 20425282 | 27.72379 | 160.87808 | 155.20201 | chr8 | 20423243 | 20425478 | 7.43915 | 3.36614 | 5.03504 |
| chr18 | 35277744 | 35277970 | 182 | Intron (ENSMUST00000025215.8 | 18 | 35209011 | 35215024 | 6014 | 2 | ENSMUST00 | -62721 | Lrrtm2 | ENSMUSG0C | false | . | . | . | . | . | . | chr18 | 35270353 | 35281972 | 4.90959 | 3.90299 | 2.98034 |
| chr18 | 14937934 | 14938155 | 182 | Distal Intergenic | 18 | 15011246 | 15011692 | 447 | 1 | ENSMUST00 | -73091 | Gm6304 | ENSMUSG0C | false | . | . | . | . | . | . | chr18 | 14934962 | 14938659 | 4.44122 | 3.66591 | 2.57945 |
| chr15 | 66577462 | 66578110 | 182 | Promoter (<=1kb) | 15 | 66577572 | 66645255 | 67684 | 1 | ENSMUST00 | 0 | Phf20l1 | ENSMUSG0C | false | chr15 | 66577170 | 66578557 | 37.91348 | 118.86035 | 114.75604 | . | . | . | . | . | . |
| chr14 | 25141989 | 25142209 | 182 | Distal Intergenic | 14 | 24909414 | 24913440 | 4027 | 1 | ENSMUST00 | 232576 | Gm10398 | ENSMUSG0C | false | . | . | . | . | . | . | chr14 | 25141455 | 25142477 | 4.64353 | 3.75128 | 2.74373 |
| chr6 | 100102186 | 100102422 | 182 | Distal Intergenic | 6 | 100228566 | 100287441 | 58876 | 2 | ENSMUST00 | 185019 | Rybp | ENSMUSG0C | false | . | . | . | . | . | . | chr6 | 100102013 | 100103196 | 4.85816 | 4.00126 | 2.89317 |
| chr5 | 74922259 | 74922579 | 182 | Distal Intergenic | 5 | 74949194 | 74966368 | 17175 | 1 | ENSMUST00 | -26615 | Gm6116 | ENSMUSG0C | false | . | . | . | . | . | . | . | . | . | . | . | . |
| chr6 | 72643262 | 72643444 | 182 | Intron (ENSMUST00000069536.6 | 6 | 72636253 | 72637717 | 1465 | 1 | ENSMUST00 | 7010 | Gm15401 | ENSMUSG0C | false | . | . | . | . | . | . | chr6 | 72641932 | 72644077 | 4.30527 | 3.74146 | 2.43194 |
| chr6 | 72371376 | 72371871 | 182 | Intron (ENSMUST00000074231.3 | 6 | 72368049 | 72380468 | 12420 | 2 | ENSMUST00 | 8597 | Vamp5 | ENSMUSG0C | true | . | . | . | . | . | . | chr6 | 72369039 | 72384082 | 5.14581 | 4.03784 | 3.16124 |
| chr6 | 54431817 | 54432107 | 182 | Promoter (1-2kb) | 6 | 54269681 | 54430221 | 160541 | 2 | ENSMUST00 | -1597 | 9130019P16I | ENSMUSG0C | false | chr6 | 54429012 | 54432013 | 22.68762 | 81.35053 | 78.1749 | chr6 | 54431186 | 54455170 | 5.36229 | 4.20786 | 3.31695 |
| chr5 | 23410192 | 23410523 | 182 | Intron (ENSMUST00000185943.1 | 5 | 23422404 | 23433309 | 10906 | 2 | ENSMUST00 | 22786 | 5031425E22I | ENSMUSG0C | false | . | . | . | . | . | . | chr5 | 23407933 | 23417945 | 6.48738 | 4.78205 | 4.22896 |
| chr5 | 23418457 | 23418675 | 182 | Intron (ENSMUST00000185943.1 | 5 | 23422404 | 23433309 | 10906 | 2 | ENSMUST00 | 14634 | 5031425E22I | ENSMUSG0C | false | chr5 | 23417333 | 23419520 | 18.55771 | 70.26926 | 67.27464 | . | . | . | . | . | . |
| chr6 | 35252614 | 35252858 | 182 | Promoter (<=1kb) | 6 | 35252654 | 35263496 | 10843 | 1 | ENSMUST00 | 0 | 1810058I24R | ENSMUSG0C | false | chr6 | 35252601 | 35253613 | 27.18312 | 111.36749 | 107.49936 | . | . | . | . | . | . |
| chr17 | 8310950 | 8311149 | 182 | Promoter (<=1kb) | 17 | 8311102 | 8327442 | 16341 | 1 | ENSMUST00 | 0 | Sft2d1 | ENSMUSG0C | false | chr17 | 8310558 | 8311862 | 39.02457 | 132.65163 | 128.06667 | . | . | . | . | . | . |
| chr1 | 134955827 | 134956193 | 182 | Promoter (<=1kb) | 1 | 134754658 | 134955940 | 201283 | 2 | ENSMUST00 | 0 | Ppp1r12b | ENSMUSG0C | false | chr1 | 134954938 | 134956311 | 37.85821 | 131.67731 | 127.12434 | chr1 | 134954528 | 134956381 | 3.99698 | 3.46512 | 2.2124 |
| chr18 | 53904319 | 53904611 | 182 | Intron (ENSMUST00000069597.6 | 18 | 53862113 | 53955684 | 93572 | 1 | ENSMUST00 | 42207 | Csnk1g3 | ENSMUSG0C | false | . | . | . | . | . | . | chr18 | 53899459 | 53905488 | 5.44692 | 4.27473 | 3.35509 |
| chr18 | 6764909 | 6765476 | 182 | Promoter (<=1kb) | 18 | 6765205 | 6790231 | 25027 | 1 | ENSMUST00 | 0 | Rab18 | ENSMUSG0C | false | chr18 | 6764641 | 6766162 | 33.30314 | 106.68535 | 102.93993 | . | . | . | . | . | . |
| chr1 | 55525372 | 55525773 | 182 | Intron (ENSMUST00000042986.9 | 1 | 55443733 | 55445064 | 1332 | 2 | ENSMUST00 | -80309 | Gm6644 | ENSMUSG0C | false | . | . | . | . | . | . | chr1 | 55521380 | 55534511 | 4.64334 | 3.78945 | 2.73695 |
| chr1 | 55528922 | 55529245 | 182 | Intron (ENSMUST00000042986.9 | 1 | 55443733 | 55445064 | 1332 | 2 | ENSMUST00 | -83859 | Gm6644 | ENSMUSG0C | false | . | . | . | . | . | . | chr1 | 55521380 | 55534511 | 4.64334 | 3.78945 | 2.73695 |
| chr1 | 55495335 | 55495628 | 182 | Intron (ENSMUST00000042986.9 | 1 | 55443733 | 55445064 | 1332 | 2 | ENSMUST00 | -50272 | Gm6644 | ENSMUSG0C | false | . | . | . | . | . | . | . | . | . | . | . | . |
| chr1 | 55521664 | 55521960 | 182 | Intron (ENSMUST00000042986.9 | 1 | 55443733 | 55445064 | 1332 | 2 | ENSMUST00 | -76601 | Gm6644 | ENSMUSG0C | false | . | . | . | . | . | . | chr1 | 55521380 | 55534511 | 4.64334 | 3.78945 | 2.73695 |
| chr1 | 55513011 | 55513292 | 182 | Intron (ENSMUST00000042986.9 | 1 | 55443733 | 55445064 | 1332 | 2 | ENSMUST00 | -67948 | Gm6644 | ENSMUSG0C | false | . | . | . | . | . | . | chr1 | 55512970 | 55516588 | 4.68678 | 3.80902 | 2.79522 |
| chr4 | 155780872 | 155781297 | 182 | Downstream (<1kb) | 4 | 155781591 | 155785874 | 4284 | 2 | ENSMUST00 | 4577 | Tmem88b | ENSMUSG0C | false | . | . | . | . | . | . | chr4 | 155772708 | 155782290 | 6.11654 | 4.55391 | 3.92825 |
| chr4 | 155238367 | 155238550 | 182 | Distal Intergenic | 4 | 155249802 | 155256419 | 6618 | 1 | ENSMUST00 | -11252 | 2610002J02R | ENSMUSG0C | false | . | . | . | . | . | . | chr4 | 155232833 | 155238793 | 8.44933 | 5.67419 | 5.82221 |
| chr4 | 126162691 | 126163050 | 182 | Promoter (<=1kb) | 4 | 126150602 | 126162665 | 12064 | 2 | ENSMUST00 | -27 | Sh3d21 | ENSMUSG0C | false | chr4 | 126162443 | 126163467 | 29.17286 | 86.59738 | 83.32484 | chr4 | 126162371 | 126164086 | 3.52645 | 3.30446 | 1.83872 |
| chr7 | 142211917 | 142212175 | 182 | Promoter (<=1kb) | 7 | 142211859 | 142212590 | 732 | 1 | ENSMUST00 | 59 | Gm7579 | ENSMUSG0C | false | . | . | . | . | . | . | . | . | . | . | . | . |
| chr7 | 126482459 | 126482764 | 182 | Distal Intergenic | 7 | 126487355 | 126490730 | 3376 | 1 | ENSMUST00 | -4591 | Tufm | ENSMUSG0C | false | chr7 | 126481885 | 126482537 | 7.14815 | 15.37828 | 13.11012 | chr7 | 126482212 | 126483413 | 4.18345 | 3.7549 | 2.32255 |
| chr7 | 101836687 | 101837139 | 182 | Promoter (<=1kb) | 7 | 101836751 | 101837363 | 613 | 1 | ENSMUST00 | 0 | Gm10602 | ENSMUSG0C | false | chr7 | 101835409 | 101838921 | 30.94804 | 114.76607 | 110.79114 | . | . | . | . | . | . |
| chr9 | 114368135 | 114368398 | 182 | Promoter (<=1kb) | 9 | 114368303 | 114377229 | 8927 | 1 | ENSMUST00 | 0 | 4930520O04 | ENSMUSG0C | false | chr9 | 114367864 | 114368718 | 19.04358 | 50.60438 | 47.88221 | chr9 | 114367911 | 114368836 | 4.4319 | 3.80268 | 2.57684 |
| chr7 | 68320261 | 68320498 | 182 | Intron (ENSMUST00000125685.2 | 7 | 68331124 | 68353725 | 22602 | 1 | ENSMUST00 | -10626 | Fam169b | ENSMUSG0C | false | . | . | . | . | . | . | . | . | . | . | . | . |
| chr7 | 68275644 | 68275955 | 182 | Promoter (<=1kb) | 7 | 68275995 | 68353646 | 77652 | 1 | ENSMUST00 | -40 | Fam169b | ENSMUSG0C | false | chr7 | 68275563 | 68276607 | 19.85205 | 54.41815 | 51.64575 | chr7 | 68274907 | 68277496 | 4.20642 | 3.65058 | 2.35589 |
| chr7 | 45413633 | 45413866 | 182 | Promoter (<=1kb) | 7 | 45413695 | 45417178 | 3484 | 1 | ENSMUST00 | 0 | Ntf5 | ENSMUSG0C | false | chr7 | 45413339 | 45414538 | 26.6505 | 78.45867 | 75.33303 | chr7 | 45412960 | 45416613 | 5.41903 | 4.17003 | 3.40026 |
| chr7 | 45128694 | 45129245 | 182 | Promoter (<=1kb) | 7 | 45125558 | 45128748 | 3191 | 2 | ENSMUST00 | 0 | Rpl13a | ENSMUSG0C | false | chr7 | 45128991 | 45129861 | 13.89301 | 35.6666 | 33.12951 | . | . | . | . | . | . |
| chr7 | 45136182 | 45136481 | 182 | Promoter (<=1kb) | 7 | 45125922 | 45136231 | 10310 | 2 | ENSMUST00 | 0 | Rpl13a | ENSMUSG0C | false | chr7 | 45135798 | 45136549 | 26.05581 | 78.34985 | 75.22605 | chr7 | 45134927 | 45136776 | 4.57199 | 3.88887 | 2.706 |
| chr9 | 61337143 | 61337444 | 182 | Distal Intergenic | 9 | 61370878 | 61371627 | 750 | 2 | ENSMUST00 | 34183 | Gm10655 | ENSMUSG0C | false | . | . | . | . | . | . | chr9 | 61336781 | 61337764 | 4.79244 | 3.94455 | 2.86537 |
| chr7 | 25610179 | 25610420 | 182 | Distal Intergenic | 7 | 25614621 | 25615892 | 1272 | 2 | ENSMUST00 | 5472 | Erich4 | ENSMUSG0C | false | . | . | . | . | . | . | . | . | . | . | . | . |

| | | | | | | | | | | | | | | | | | | | | | | | | | |
|---|---|---|---|---|---|---|---|---|---|---|---|---|---|---|---|---|---|---|---|---|---|---|---|---|---|
| chr7 | 24316480 | 24316851 | 182 | Promoter (<=1kb) | 7 | 24301704 | 24316605 | 14902 | 2 | ENSMUST00 | 0 | Zfp94 | ENSMUSG0C | false | chr7 | 24316135 | 24317143 | 20.02586 | 58.85371 | 56.02288 | chr7 | 24315738 | 24317358 | 3.58268 | 3.22531 | 1.88054 |
| chr3 | 105877376 | 105877817 | 182 | Intron (ENSMUST00000164730.1 | 3 | 105870907 | 105924036 | 53130 | 1 | ENSMUST00 | 6470 | Adora3 | ENSMUSG0C | false | . | . | . | . | . | . | . | . | . | . | . | . |
| chr3 | 96555050 | 96555387 | 182 | Promoter (2-3kb) | 3 | 96555765 | 96557815 | 2051 | 2 | ENSMUST00 | 2428 | Gm15441 | ENSMUSG0C | false | . | . | . | . | . | . | chr3 | 96551412 | 96557179 | 4.83722 | 4.04794 | 2.86193 |
| chr3 | 96564749 | 96565028 | 182 | Promoter (1-2kb) | 3 | 96555788 | 96566801 | 11014 | 2 | ENSMUST00 | 1773 | Gm15441 | ENSMUSG0C | false | . | . | . | . | . | . | chr3 | 96561069 | 96567035 | 5.48015 | 4.34024 | 3.394 |
| chr7 | 4915114 | 4915501 | 182 | Promoter (<=1kb) | 7 | 4915217 | 4922003 | 6787 | 1 | ENSMUST00 | 0 | Zfp628 | ENSMUSG0C | false | chr7 | 4914514 | 4916194 | 30.17592 | 94.17345 | 90.73757 | . | . | . | . | . | . |
| chr8 | 25465719 | 25465983 | 182 | Distal Intergenic | 8 | 25454273 | 25454704 | 432 | 1 | ENSMUST00 | 11447 | Gm10689 | ENSMUSG0C | false | chr8 | 25465846 | 25466220 | 11.64009 | 27.76484 | 25.32713 | chr8 | 25462225 | 25466639 | 4.97289 | 3.99886 | 3.01441 |
| chr3 | 88526355 | 88526660 | 182 | Distal Intergenic | 3 | 88532395 | 88541396 | 9002 | 1 | ENSMUST00 | -5735 | Mex3a | ENSMUSG0C | false | . | . | . | . | . | . | chr3 | 88526143 | 88527178 | 4.69383 | 3.9384 | 2.77979 |
| chr3 | 88531584 | 88531951 | 182 | Promoter (<=1kb) | 3 | 88532395 | 88541396 | 9002 | 1 | ENSMUST00 | -444 | Mex3a | ENSMUSG0C | false | chr3 | 88531122 | 88532849 | 32.44658 | 100.11784 | 96.54536 | chr3 | 88530483 | 88533230 | 4.01692 | 3.605 | 2.18493 |
| chr3 | 51708709 | 51709210 | 182 | Intron (ENSMUST00000121440.2 | 3 | 51693771 | 51695090 | 1320 | 1 | ENSMUST00 | 14939 | Gm10729 | ENSMUSG0C | false | chr3 | 51708796 | 51709325 | 8.11812 | 15.68191 | 13.40967 | chr3 | 51705501 | 51729803 | 5.39003 | 4.2072 | 3.33836 |
| chr2 | 163229152 | 163229430 | 182 | Intron (ENSMUST00000109428.3 | 2 | 163225454 | 163323102 | 97649 | 1 | ENSMUST00 | 3699 | Tox2 | ENSMUSG0C | false | . | . | . | . | . | . | . | . | . | . | . | . |
| chr2 | 152962166 | 152962450 | 182 | Promoter (<=1kb) | 2 | 152962485 | 153007340 | 44856 | 1 | ENSMUST00 | -35 | Ttll9 | ENSMUSG0C | false | chr2 | 152962141 | 152962790 | 16.10786 | 38.79296 | 36.21762 | chr2 | 152961789 | 152962378 | 3.4043 | 3.26595 | 1.67505 |
| chr3 | 17616759 | 17617003 | 182 | Distal Intergenic | 3 | 17783514 | 17786898 | 3385 | 2 | ENSMUST00 | 169895 | Gm10742 | ENSMUSG0C | false | . | . | . | . | . | . | chr3 | 17616685 | 17617299 | 3.11648 | 2.92505 | 1.50047 |
| chr2 | 144165408 | 144165760 | 182 | Distal Intergenic | 2 | 144174315 | 144179086 | 4772 | 2 | ENSMUST00 | 13326 | Gm5535 | ENSMUSG0C | false | chr2 | 144164562 | 144166242 | 23.0285 | 72.05518 | 69.03415 | chr2 | 144164052 | 144166701 | 4.22189 | 3.64064 | 2.42937 |
| chr2 | 132647523 | 132647774 | 182 | Promoter (1-2kb) | 2 | 132645881 | 132652999 | 7119 | 1 | ENSMUST00 | 1643 | AU019990 | ENSMUSG0C | false | . | . | . | . | . | . | chr2 | 132645606 | 132648470 | 4.60592 | 3.97308 | 2.67712 |
| chr9 | 124312621 | 124312956 | 182 | Promoter (<=1kb) | 9 | 124291804 | 124312696 | 20893 | 2 | ENSMUST00 | 0 | 2010315B03 | ENSMUSG0C | false | chr9 | 124311612 | 124312922 | 25.27427 | 69.73028 | 66.74294 | . | . | . | . | . | . |
| chr2 | 120091085 | 120091297 | 182 | 3' UTR | 2 | 120076214 | 120085772 | 9559 | 2 | ENSMUST00 | -5314 | Sptbn5 | ENSMUSG0C | false | . | . | . | . | . | . | chr2 | 120090578 | 120107323 | 5.14183 | 4.07311 | 3.14279 |
| chr2 | 118926366 | 118926662 | 182 | Promoter (<=1kb) | 2 | 118926496 | 118928585 | 2090 | 1 | ENSMUST00 | 0 | Chst14 | ENSMUSG0C | false | chr2 | 118925995 | 118927451 | 38.29964 | 140.16156 | 135.27679 | chr2 | 118925573 | 118927772 | 5.82056 | 4.3478 | 3.72242 |
| chr19 | 23687106 | 23687580 | 182 | Promoter (<=1kb) | 19 | 23687400 | 23721129 | 33730 | 1 | ENSMUST00 | 0 | Ptar1 | ENSMUSG0C | false | chr19 | 23686825 | 23688717 | 25.17802 | 89.90526 | 86.56598 | . | . | . | . | . | . |
| chr10 | 67281952 | 67282175 | 182 | Intron (ENSMUST00000077839.7 | 10 | 67267841 | 67285180 | 17340 | 2 | ENSMUST00 | 3005 | Nrbf2 | ENSMUSG0C | false | . | . | . | . | . | . | chr10 | 67275859 | 67284544 | 5.03837 | 4.06125 | 3.05103 |
| chr2 | 93014684 | 93015023 | 182 | Promoter (<=1kb) | 2 | 93009973 | 93013910 | 3938 | 2 | ENSMUST00 | -775 | Prdm11 | ENSMUSG0C | false | chr2 | 93014233 | 93015383 | 23.43614 | 63.09834 | 60.20888 | chr2 | 93013792 | 93015901 | 3.91582 | 3.4512 | 2.13459 |
| chr13 | 17993301 | 17993548 | 182 | Promoter (<=1kb) | 13 | 17986640 | 17993351 | 6712 | 2 | ENSMUST00 | 0 | Yae1d1 | ENSMUSG0C | false | chr13 | 17992320 | 17993900 | 30.17592 | 94.17345 | 90.73757 | . | . | . | . | . | . |
| chr10 | 40302246 | 40302692 | 182 | Promoter (<=1kb) | 10 | 40287458 | 40302188 | 14731 | 2 | ENSMUST00 | -59 | Amd1 | ENSMUSG0C | false | chr10 | 40300536 | 40303093 | 35.20524 | 115.20684 | 111.21615 | . | . | . | . | . | . |
| chr10 | 40301447 | 40301674 | 182 | Promoter (<=1kb) | 10 | 40287458 | 40302188 | 14731 | 2 | ENSMUST00 | 514 | Amd1 | ENSMUSG0C | false | chr10 | 40300536 | 40303093 | 35.20524 | 115.20684 | 111.21615 | . | . | . | . | . | . |
| chr2 | 79543498 | 79543800 | 182 | Distal Intergenic | 2 | 79338569 | 79456785 | 118217 | 2 | ENSMUST00 | -86714 | Cerkl | ENSMUSG0C | false | . | . | . | . | . | . | chr2 | 79543244 | 79544727 | 3.19753 | 3.03299 | 1.57467 |
| chr2 | 79510988 | 79511221 | 182 | Distal Intergenic | 2 | 79338569 | 79456785 | 118217 | 2 | ENSMUST00 | -54204 | Cerkl | ENSMUSG0C | false | . | . | . | . | . | . | . | . | . | . | . | . |
| chr2 | 75224576 | 75224834 | 182 | Intron (ENSMUST00000186411.1 | 2 | 75192277 | 75192759 | 483 | 1 | ENSMUST00 | 32300 | Gm13653 | ENSMUSG0C | false | . | . | . | . | . | . | chr2 | 75224472 | 75225359 | 4.14402 | 3.68953 | 2.32398 |
| chrY | 10643734 | 10643931 | 182 | Promoter (<=1kb) | 21 | 10641257 | 10643107 | 1851 | 2 | ENSMUST00 | -628 | Gm20775 | ENSMUSG0C | false | chrY | 10643624 | 10643797 | 5.97392 | 9.35129 | 7.18948 | . | . | . | . | . | . |
| chr11 | 120185263 | 120185552 | 182 | Exon (ENSMUST00000155681.1/ | 11 | 120185400 | 120188751 | 3352 | 2 | ENSMUST00 | 3199 | 2810410L24 | ENSMUSG0C | false | chr11 | 120184503 | 120186230 | 13.67473 | 40.1468 | 37.55384 | chr11 | 120182684 | 120187708 | 6.97187 | 4.88035 | 4.63971 |
| chr15 | 101293041 | 101293506 | 182 | Promoter (<=1kb) | 15 | 101293212 | 101296615 | 3404 | 1 | ENSMUST00 | 0 | 6030408B16 | ENSMUSG0C | false | . | . | . | . | . | . | chr15 | 101291483 | 101296758 | 5.6438 | 4.08858 | 3.5627 |
| chr11 | 116648368 | 116648715 | 182 | Intron (ENSMUST00000021166.5 | 11 | 116653534 | 116668092 | 14559 | 1 | ENSMUST00 | -4819 | Gm11744 | ENSMUSG0C | false | . | . | . | . | . | . | . | . | . | . | . | . |
| chr2 | 31142263 | 31142484 | 182 | Promoter (<=1kb) | 2 | 31026207 | 31142008 | 115802 | 2 | ENSMUST00 | -256 | Fnbp1 | ENSMUSG0C | false | chr2 | 31140653 | 31142585 | 38.66002 | 128.73949 | 124.2961 | chr2 | 31142238 | 31143170 | 5.75057 | 4.41945 | 3.67368 |
| chr2 | 31106979 | 31107192 | 182 | Promoter (1-2kb) | 2 | 31096091 | 31105450 | 9360 | 2 | ENSMUST00 | -1530 | Fnbp1 | ENSMUSG0C | false | . | . | . | . | . | . | chr2 | 31094325 | 31113159 | 5.94479 | 4.49985 | 3.7863 |
| chr11 | 96413645 | 96413938 | 182 | Distal Intergenic | 11 | 96392112 | 96392666 | 555 | 1 | ENSMUST00 | 21534 | Gm11531 | ENSMUSG0C | false | . | . | . | . | . | . | chr11 | 96413229 | 96414277 | 3.5425 | 3.28537 | 1.84617 |
| chr11 | 96351846 | 96352062 | 182 | Promoter (<=1kb) | 11 | 96351631 | 96354012 | 2382 | 1 | ENSMUST00 | 216 | Hoxb2 | ENSMUSG0C | false | chr11 | 96351914 | 96353821 | 12.30394 | 36.76666 | 34.21749 | chr11 | 96351920 | 96353494 | 4.56363 | 3.90607 | 2.63737 |
| chr3 | 58576883 | 58577267 | 182 | Promoter (<=1kb) | 3 | 58576698 | 58588812 | 12115 | 1 | ENSMUST00 | 186 | Selt | ENSMUSG0C | false | chr3 | 58576362 | 58577992 | 21.44446 | 75.31629 | 72.24329 | . | . | . | . | . | . |
| chr11 | 3514651 | 3514860 | 182 | Promoter (<=1kb) | 11 | 3514684 | 3517348 | 2665 | 1 | ENSMUST00 | 0 | Selm | ENSMUSG0C | false | chr11 | 3514172 | 3515438 | 13.94362 | 38.54297 | 35.97016 | . | . | . | . | . | . |
| chr11 | 3517224 | 3517431 | 182 | Promoter (2-3kb) | 11 | 3514867 | 3517351 | 2485 | 1 | ENSMUST00 | 2358 | Selm | ENSMUSG0C | false | chr11 | 3517048 | 3517826 | 7.68513 | 13.62737 | 11.38679 | chr11 | 3516385 | 3518288 | 3.91725 | 3.53849 | 2.16476 |
| chr10 | 80051937 | 80052357 | 182 | Promoter (1-2kb) | 10 | 80053488 | 80056439 | 2952 | 1 | ENSMUST00 | -1131 | Gpx4 | ENSMUSG0C | false | . | . | . | . | . | . | chr10 | 80046083 | 80053338 | 6.68412 | 4.69614 | 4.42765 |
| chr8 | 128512739 | 128512997 | 182 | Distal Intergenic | 8 | 128650997 | 128651103 | 107 | 1 | ENSMUST00 | -138000 | Gm26002 | ENSMUSG0C | false | . | . | . | . | . | . | chr8 | 128511642 | 128513750 | 4.01522 | 3.43565 | 2.22498 |
| chr8 | 128544475 | 128544830 | 182 | Distal Intergenic | 8 | 128650997 | 128651103 | 107 | 1 | ENSMUST00 | -106167 | Gm26002 | ENSMUSG0C | false | . | . | . | . | . | . | chr8 | 128519464 | 128548431 | 5.17648 | 4.1349 | 3.15487 |
| chrX | 53046078 | 53046290 | 182 | Promoter (1-2kb) | 20 | 53047997 | 53048078 | 82 | 2 | ENSMUST00 | 1788 | Mir450b | ENSMUSG0C | false | . | . | . | . | . | . | . | . | . | . | . | . |
| chr2 | 27411790 | 27412051 | 182 | Intron (ENSMUST00000185188.2 | 2 | 27401852 | 27401951 | 100 | 1 | ENSMUST00 | 9939 | Gm24049 | ENSMUSG0C | false | . | . | . | . | . | . | chr2 | 27411152 | 27412943 | 5.00269 | 4.16743 | 2.99587 |
| chr9 | 67181885 | 67182152 | 182 | Distal Intergenic | 9 | 67236651 | 67236735 | 85 | 2 | ENSMUST00 | 54583 | Mir190a | ENSMUSG0C | false | . | . | . | . | . | . | chr9 | 67181578 | 67182767 | 4.99355 | 4.09373 | 3.00004 |
| chr13 | 28958574 | 28958935 | 182 | Distal Intergenic | 13 | 28948919 | 28953713 | 4795 | 2 | ENSMUST00 | -4862 | Sox4 | ENSMUSG0C | false | chr13 | 28957933 | 28960010 | 27.63319 | 79.22645 | 76.08477 | . | . | . | . | . | . |
| chr13 | 28951203 | 28951544 | 182 | Promoter (2-3kb) | 13 | 28948919 | 28953713 | 4795 | 2 | ENSMUST00 | 2169 | Sox4 | ENSMUSG0C | false | chr13 | 28949862 | 28954154 | 30.8734 | 92.43849 | 89.0432 | chr13 | 28950712 | 28953349 | 3.40198 | 3.22275 | 1.7076 |
| chr11 | 118342688 | 118342923 | 182 | Promoter (<=1kb) | 11 | 118332360 | 118342500 | 10141 | 2 | ENSMUST00 | -189 | BC100451 | ENSMUSG0C | false | . | . | . | . | . | . | chr11 | 118319919 | 118354581 | 6.53665 | 4.67889 | 4.28866 |
| chr11 | 118317303 | 118317533 | 182 | Exon (ENSMUST00000017610.4/ | 11 | 118332415 | 118334016 | 1602 | 2 | ENSMUST00 | 16483 | BC100451 | ENSMUSG0C | false | . | . | . | . | . | . | chr11 | 118316643 | 118318771 | 4.02422 | 3.66503 | 2.19005 |
| chr14 | 54216830 | 54217076 | 182 | Promoter (<=1kb) | 14 | 54217292 | 54217357 | 66 | 1 | ENSMUST00 | -216 | Traj3 | ENSMUSG0C | false | . | . | . | . | . | . | chr14 | 54215806 | 54217860 | 4.41282 | 3.73265 | 2.56623 |
| chr11 | 57727580 | 57727832 | 182 | Intron (ENSMUST00000066987.8 | 11 | 57741300 | 57741402 | 103 | 1 | ENSMUST00 | -13468 | Gm24582 | ENSMUSG0C | false | . | . | . | . | . | . | chr11 | 57724712 | 57734543 | 6.29874 | 4.67914 | 4.07864 |
| chr9 | 111562804 | 111562998 | 182 | Intron (ENSMUST00000161995.1 | 9 | 111613898 | 111614015 | 118 | 2 | ENSMUST00 | 51017 | n-R5s90 | ENSMUSG0C | false | . | . | . | . | . | . | . | . | . | . | . | . |
| chr11 | 83339844 | 83340041 | 182 | Intron (ENSMUST00000018875.7 | 11 | 83366870 | 83367001 | 132 | 2 | ENSMUST00 | 26960 | Gm23951 | ENSMUSG0C | false | . | . | . | . | . | . | chr11 | 83337613 | 83339988 | 3.31509 | 3.11686 | 1.65939 |
| chr6 | 39007325 | 39007597 | 182 | Intron (ENSMUST00000003017.7 | 6 | 39013666 | 39013791 | 126 | 1 | ENSMUST00 | -6069 | Gm23010 | ENSMUSG0C | false | . | . | . | . | . | . | chr6 | 39004756 | 39008368 | 4.60603 | 3.82528 | 2.71154 |
| chr1 | 13738279 | 13738496 | 182 | Distal Intergenic | 1 | 13774334 | 13774464 | 131 | 2 | ENSMUST00 | 35968 | Gm26273 | ENSMUSG0C | false | . | . | . | . | . | . | chr1 | 13737381 | 13742832 | 4.42 | 3.67834 | 2.55838 |
| chr8 | 7601846 | 7602190 | 182 | Distal Intergenic | 8 | 7638764 | 7638892 | 129 | 1 | ENSMUST00 | -36574 | Gm25169 | ENSMUSG0C | false | . | . | . | . | . | . | . | . | . | . | . | . |
| chr8 | 23284034 | 23284348 | 182 | Distal Intergenic | 8 | 23290033 | 23290164 | 132 | 2 | ENSMUST00 | 5816 | Gm24335 | ENSMUSG0C | false | . | . | . | . | . | . | . | . | . | . | . | . |
| chr4 | 53951986 | 53952287 | 182 | Distal Intergenic | 4 | 53905696 | 53905820 | 125 | 1 | ENSMUST00 | 46291 | Gm23745 | ENSMUSG0C | false | . | . | . | . | . | . | chr4 | 53948932 | 53953603 | 3.90187 | 3.45261 | 2.1414 |
| chr15 | 94955904 | 94956089 | 182 | Intron (ENSMUST00000080141.4 | 15 | 94987007 | 94987138 | 132 | 2 | ENSMUST00 | 31049 | Gm23129 | ENSMUSG0C | false | . | . | . | . | . | . | . | . | . | . | . | . |
| chr17 | 33760196 | 33760485 | 182 | Promoter (<=1kb) | 17 | 33746750 | 33760306 | 13557 | 2 | ENSMUST00 | 0 | Rab11b | ENSMUSG0C | false | chr17 | 33759353 | 33760455 | 28.28183 | 109.60786 | 105.79031 | . | . | . | . | . | . |
| chr16 | 78260258 | 78260570 | 182 | Distal Intergenic | 16 | 78240258 | 78240388 | 131 | 1 | ENSMUST00 | 20001 | Gm25916 | ENSMUSG0C | false | . | . | . | . | . | . | chr16 | 78259806 | 78261218 | 4.08564 | 3.62833 | 2.2997 |
| chr3 | 67568240 | 67568466 | 182 | Distal Intergenic | 3 | 67573160 | 67573291 | 132 | 2 | ENSMUST00 | 4825 | Gm25919 | ENSMUSG0C | false | . | . | . | . | . | . | chr3 | 67567523 | 67568272 | 3.12979 | 3.07185 | 1.46452 |
| chr15 | 69303689 | 69303977 | 182 | Distal Intergenic | 15 | 69451015 | 69451146 | 132 | 2 | ENSMUST00 | 147169 | Gm24232 | ENSMUSG0C | false | . | . | . | . | . | . | . | . | . | . | . | . |
| chr2 | 181659665 | 181659978 | 182 | Promoter (2-3kb) | 2 | 181662306 | 181662412 | 107 | 1 | ENSMUST00 | -2328 | Gm25879 | ENSMUSG0C | false | . | . | . | . | . | . | chr2 | 181656347 | 181660213 | 4.90313 | 4.0137 | 2.96333 |
| chr7 | 109561664 | 109562279 | 182 | Intron (ENSMUST00000079282.2 | 7 | 109521280 | 109521409 | 130 | 1 | ENSMUST00 | 40385 | Gm24888 | ENSMUSG0C | false | . | . | . | . | . | . | chr7 | 109558924 | 109563977 | 4.69759 | 3.86892 | 2.79221 |
| chr11 | 74908614 | 74908937 | 182 | Promoter (2-3kb) | 11 | 74906109 | 74906199 | 91 | 1 | ENSMUST00 | 2506 | Gm22771 | ENSMUSG0C | false | . | . | . | . | . | . | chr11 | 74906804 | 74909835 | 4.11539 | 3.65194 | 2.27976 |
| chr3 | 38015792 | 38016017 | 182 | Distal Intergenic | 3 | 37995105 | 37995235 | 131 | 2 | ENSMUST00 | -20558 | Gm22899 | ENSMUSG0C | false | . | . | . | . | . | . | chr3 | 38013969 | 38017570 | 4.88229 | 4.08788 | 2.92086 |
| chr12 | 73209242 | 73209618 | 182 | Intron (ENSMUST00000187549.1 | 12 | 73145774 | 73145905 | 132 | 2 | ENSMUST00 | -63338 | Gm23910 | ENSMUSG0C | false | . | . | . | . | . | . | chr12 | 73208704 | 73211456 | 3.98001 | 3.44027 | 2.2195 |
| chr18 | 82947677 | 82947903 | 182 | Intron (ENSMUST00000171238.2 | 18 | 82976323 | 82976414 | 92 | 1 | ENSMUST00 | -28420 | Gm26258 | ENSMUSG0C | false | . | . | . | . | . | . | chr18 | 82942535 | 82949651 | 4.25334 | 3.56915 | 2.42349 |
| chr2 | 167622454 | 167622638 | 182 | Intron (ENSMUST00000109207.4 | 2 | 167626315 | 167626403 | 89 | 2 | ENSMUST00 | 3765 | Gm23135 | ENSMUSG0C | false | . | . | . | . | . | . | chr2 | 167616687 | 167624077 | 5.13013 | 4.05974 | 3.12778 |
| chr11 | 48826328 | 48826830 | 182 | Promoter (<=1kb) | 11 | 48825346 | 48826655 | 1310 | 2 | ENSMUST00 | 0 | Gm12184 | ENSMUSG0C | false | chr11 | 48825828 | 48827167 | 20.5724 | 58.98804 | 56.15619 | . | . | . | . | . | . |
| chr6 | 53760690 | 53760963 | 182 | Distal Intergenic | 6 | 53819975 | 53820040 | 66 | 1 | ENSMUST00 | -59012 | Gm16499 | ENSMUSG0C | false | . | . | . | . | . | . | . | . | . | . | . | . |
| chr13 | 98367581 | 98368017 | 182 | Distal Intergenic | 13 | 98354242 | 98356703 | 2462 | 1 | ENSMUST00 | 13340 | Foxd1 | ENSMUSG0C | false | . | . | . | . | . | . | chr13 | 98366713 | 98370947 | 4.92348 | 3.98982 | 2.95701 |
| chr4 | 156234549 | 156234971 | 182 | Promoter (<=1kb) | 4 | 156229044 | 156234857 | 5814 | 2 | ENSMUST00 | 0 | Klhl17 | ENSMUSG0C | false | chr4 | 156233883 | 156235233 | 17.04032 | 66.12922 | 63.19824 | chr4 | 156233813 | 156235528 | 4.67407 | 3.92711 | 2.73135 |
| chr4 | 134517038 | 134517318 | 182 | Intron (ENSMUST00000105866.2 | 4 | 134510999 | 134523927 | 12929 | 1 | ENSMUST00 | 6040 | Aunip | ENSMUSG0C | false | . | . | . | . | . | . | chr4 | 134510323 | 134518262 | 5.68606 | 4.35573 | 3.59434 |
| chr4 | 131481198 | 131481433 | 182 | Distal Intergenic | 4 | 131487197 | 131505527 | 18331 | 1 | ENSMUST00 | -5764 | Gm831 | ENSMUSG0C | false | . | . | . | . | . | . | . | . | . | . | . | . |
| chr4 | 124843417 | 124843607 | 182 | Intron (ENSMUST00000106193.2 | 4 | 124849080 | 124850330 | 1251 | 2 | ENSMUST00 | 6723 | 1110065P20 | ENSMUSG0C | false | . | . | . | . | . | . | chr4 | 124842715 | 124844296 | 4.39371 | 3.7956 | 2.50403 |
| chr15 | 95823916 | 95824141 | 182 | Intron (ENSMUST00000071874.6 | 15 | 95824459 | 95830072 | 5614 | 2 | ENSMUST00 | 5931 | Gm17546 | ENSMUSG0C | false | . | . | . | . | . | . | chr15 | 95822757 | 95825389 | 3.72158 | 3.3401 | 1.9944 |
| chr11 | 101315815 | 101316362 | 182 | Promoter (<=1kb) | 11 | 101316213 | 101320008 | 3796 | 1 | ENSMUST00 | 0 | Psme3 | ENSMUSG0C | false | chr11 | 101315586 | 101317002 | 34.92026 | 124.22972 | 119.94103 | chr11 | 101315365 | 101318696 | 4.96377 | 4.17814 | 2.97531 |
| chr7 | 66365992 | 66366220 | 182 | Promoter (<=1kb) | 7 | 66365905 | 66386309 | 20405 | 1 | ENSMUST00 | 88 | Gm10974 | ENSMUSG0C | false | . | . | . | . | . | . | chr7 | 66364515 | 66372493 | 5.22527 | 4.14692 | 3.19405 |
| chr7 | 65693302 | 65693558 | 182 | Promoter (<=1kb) | 7 | 65693417 | 65701902 | 8486 | 1 | ENSMUST00 | 0 | Tm2d3 | ENSMUSG0C | false | chr7 | 65692880 | 65694150 | 35.8325 | 117.35963 | 113.30182 | . | . | . | . | . | . |
| chr3 | 88798952 | 88799196 | 182 | Intron (ENSMUST00000135913.1 | 3 | 88775216 | 88825587 | 50372 | 1 | ENSMUST00 | 23737 | 5830417I10R | ENSMUSG0C | false | . | . | . | . | . | . | . | . | . | . | . | . |
| chr15 | 96055315 | 96055769 | 182 | Intron (ENSMUST00000100271.3 | 15 | 96051616 | 96078134 | 26519 | 2 | ENSMUST00 | 22365 | D030018L15 | ENSMUSG0C | false | . | . | . | . | . | . | chr15 | 96054907 | 96057007 | 4.66694 | 3.96012 | 2.72621 |
| chr13 | 97546250 | 97546485 | 182 | Distal Intergenic | 13 | 97482955 | 97489700 | 6746 | 1 | ENSMUST00 | 63296 | Lincenc1 | ENSMUSG0C | false | . | . | . | . | . | . | chr13 | 97545926 | 97548907 | 3.1867 | 3.12648 | 1.50253 |
| chr5 | 104508356 | 104508640 | 182 | Promoter (<=1kb) | 5 | 104508352 | 104522383 | 14032 | 1 | ENSMUST00 | 5 | BC005561 | ENSMUSG0C | false | chr5 | 104507986 | 104509227 | 28.91859 | 89.0447 | 85.71584 | . | . | . | . | . | . |
| chr1 | 182908348 | 182908632 | 182 | Distal Intergenic | 1 | 182954788 | 182976044 | 21257 | 1 | ENSMUST00 | -46156 | Tlr5 | ENSMUSG0C | false | . | . | . | . | . | . | chr1 | 182908075 | 182909090 | 4.82822 | 4.07374 | 2.88998 |
| chr2 | 105091151 | 105091388 | 182 | Promoter (<=1kb) | 2 | 105092018 | 105093760 | 1743 | 1 | ENSMUST00 | -630 | Gm11060 | ENSMUSG0C | false | . | . | . | . | . | . | chr2 | 105081211 | 105093624 | 5.17671 | 4.11795 | 3.15483 |
| chr12 | 107638378 | 107638680 | 182 | Distal Intergenic | 12 | 107613000 | 107613840 | 841 | 1 | ENSMUST00 | 25379 | Gm9006 | ENSMUSG0C | false | . | . | . | . | . | . | chr12 | 107638060 | 107639022 | 3.20646 | 3.17335 | 1.58632 |
| chr9 | 73039442 | 73039910 | 182 | Promoter (<=1kb) | 9 | 73013711 | 73039699 | 25989 | 2 | ENSMUST00 | 0 | Pigb | ENSMUSG0C | false | chr9 | 73039068 | 73040618 | 30.41442 | 103.5799 | 99.9162 | chr9 | 73038574 | 73040809 | 4.62032 | 3.84132 | 2.71163 |
| chr6 | 88045224 | 88045406 | 182 | Promoter (<=1kb) | 6 | 87999106 | 88045270 | 46165 | 2 | ENSMUST00 | 0 | Rab7 | ENSMUSG0C | false | chr6 | 88044096 | 88045316 | 34.36702 | 111.64683 | 107.76208 | . | . | . | . | . | . |
| chr6 | 72986878 | 72987119 | 182 | Distal Intergenic | 6 | 72957361 | 72958748 | 1388 | 2 | ENSMUST00 | -28131 | Tmsb10 | ENSMUSG0C | false | . | . | . | . | . | . | chr6 | 72984886 | 72987837 | 4.55933 | 3.83687 | 2.67777 |
| chr5 | 9100598 | 9101323 | 182 | Promoter (<=1kb) | 5 | 9100737 | 9118983 | 18247 | 1 | ENSMUST00 | 0 | Tmem243 | ENSMUSG0C | false | chr5 | 9100104 | 9101841 | 34.16929 | 120.7392 | 116.5701 | . | . | . | . | . | . |
| chr3 | 85485620 | 85485822 | 182 | Intron (ENSMUST00000154148.2 | 3 | 85467377 | 85467498 | 122 | 1 | ENSMUST00 | 18244 | Mir466k | ENSMUSG0C | false | . | . | . | . | . | . | chr3 | 85484761 | 85487913 | 3.3847 | 3.24839 | 1.67641 |
| chr15 | 37549281 | 37549502 | 182 | Intron (ENSMUST00000168992.2 | 15 | 37603415 | 37603534 | 120 | 2 | ENSMUST00 | 54032 | Gm24833 | ENSMUSG0C | false | . | . | . | . | . | . | chr15 | 37549091 | 37550395 | 4.1803 | 3.69864 | 2.35416 |
| chr11 | 64541529 | 64541741 | 182 | Distal Intergenic | 11 | 64590495 | 64590640 | 146 | 2 | ENSMUST00 | 48899 | Gm24275 | ENSMUSG0C | false | . | . | . | . | . | . | chr11 | 64539605 | 64548358 | 5.34347 | 4.23728 | 3.29791 |
| chr18 | 35919574 | 35920077 | 182 | Distal Intergenic | 18 | 35952824 | 35952907 | 84 | 1 | ENSMUST00 | -32747 | Gm23882 | ENSMUSG0C | false | chr18 | 35918794 | 35920648 | 19.69154 | 51.71931 | 48.98378 | chr18 | 35918346 | 35921757 | 4.57948 | 3.8339 | 2.68759 |
| chr1 | 165021290 | 165021487 | 182 | Promoter (2-3kb) | 1 | 165024345 | 165024456 | 112 | 1 | ENSMUST00 | -2858 | Gm26042 | ENSMUSG0C | false | . | . | . | . | . | . | . | . | . | . | . | . |
| chr13 | 71362454 | 71362670 | 182 | Distal Intergenic | 13 | 71107089 | 71107209 | 121 | 1 | ENSMUST00 | 255366 | Mir466f-4 | ENSMUSG0C | false | . | . | . | . | . | . | . | . | . | . | . | . |
| chr13 | 60464817 | 60465079 | 182 | Distal Intergenic | 13 | 60529296 | 60529409 | 114 | 2 | ENSMUST00 | 64330 | Gm25000 | ENSMUSG0C | false | . | . | . | . | . | . | chr13 | 60464211 | 60468871 | 5.68489 | 4.41885 | 3.55497 |
| chr13 | 60513758 | 60514160 | 182 | Distal Intergenic | 13 | 60529296 | 60529409 | 114 | 2 | ENSMUST00 | 15249 | Gm25000 | ENSMUSG0C | false | . | . | . | . | . | . | . | . | . | . | . | . |
| chr13 | 60262591 | 60262830 | 182 | Distal Intergenic | 13 | 60293672 | 60293772 | 101 | 1 | ENSMUST00 | -30842 | Gm24999 | ENSMUSG0C | false | . | . | . | . | . | . | chr13 | 60261613 | 60263980 | 4.88347 | 3.97943 | 2.91472 |
| chr11 | 76523448 | 76523681 | 182 | Intron (ENSMUST00000072740.7 | 11 | 76514069 | 76514571 | 503 | 1 | ENSMUST00 | 9380 | Gm12341 | ENSMUSG0C | false | . | . | . | . | . | . | chr11 | 76522111 | 76524929 | 4.13657 | 3.43385 | 2.32156 |
| chr19 | 18421540 | 18421752 | 182 | Distal Intergenic | 19 | 18581455 | 18582228 | 774 | 1 | ENSMUST00 | -159703 | Gm8250 | ENSMUSG0C | false | . | . | . | . | . | . | chr19 | 18419648 | 18422209 | 3.97672 | 3.53662 | 2.17944 |
| chr19 | 18599011 | 18599336 | 182 | Intron (ENSMUST00000025631.6 | 19 | 18581455 | 18582228 | 774 | 1 | ENSMUST00 | 17557 | Gm8250 | ENSMUSG0C | false | . | . | . | . | . | . | chr19 | 18596963 | 18611084 | 4.82912 | 4.00871 | 2.86974 |
| chr2 | 12053286 | 12053660 | 182 | Distal Intergenic | 2 | 12082612 | 12084003 | 1392 | 2 | ENSMUST00 | 30343 | Gm13310 | ENSMUSG0C | false | . | . | . | . | . | . | chr2 | 12052987 | 12053702 | 2.88054 | 2.77489 | 1.31408 |
| chr4 | 46304809 | 46305156 | 182 | Intron (ENSMUST00000123933.2 | 4 | 46327296 | 46327765 | 470 | 1 | ENSMUST00 | -22140 | Gm12447 | ENSMUSG0C | false | chr4 | 46304404 | 46305687 | 31.85236 | 101.09525 | 97.496 | . | . | . | . | . | . |
| chr9 | 116100549 | 116100826 | 182 | Intron (ENSMUST00000035014.6 | 9 | 116040593 | 116041064 | 472 | 2 | ENSMUST00 | -59486 | Gm9385 | ENSMUSG0C | false | . | . | . | . | . | . | chr9 | 116095463 | 116105255 | 8.03235 | 5.43993 | 5.4794 |
| chr5 | 130192324 | 130192614 | 182 | Intron (ENSMUST00000119797.2 | 5 | 130188395 | 130188796 | 402 | 1 | ENSMUST00 | 3930 | Gm15920 | ENSMUSG0C | false | . | . | . | . | . | . | chr5 | 130172335 | 130207681 | 5.43982 | 4.22227 | 3.37739 |
| chr4 | 29134015 | 29134233 | 182 | Distal Intergenic | 4 | 29125946 | 29126124 | 179 | 1 | ENSMUST00 | 8070 | Gm11918 | ENSMUSG0C | false | chr4 | 29133371 | 29134016 | 9.88648 | 20.14232 | 17.80457 | chr4 | 29133371 | 29135776 | 3.87509 | 3.55966 | 2.08924 |
| chr4 | 100351945 | 100352350 | 182 | Intron (ENSMUST00000039630.5 | 4 | 100361362 | 100362059 | 698 | 2 | ENSMUST00 | 9709 | Gm12706 | ENSMUSG0C | false | . | . | . | . | . | . | chr4 | 100347301 | 100354535 | 6.63157 | 4.80816 | 4.33798 |
| chr11 | 45240720 | 45240993 | 182 | Distal Intergenic | 11 | 45280594 | 45280954 | 361 | 2 | ENSMUST00 | 39961 | Gm12161 | ENSMUSG0C | false | . | . | . | . | . | . | . | . | . | . | . | . |
| chr4 | 149048710 | 149048992 | 182 | Intron (ENSMUST00000103217.5 | 4 | 149096383 | 149096702 | 320 | 1 | ENSMUST00 | -47391 | Gm9506 | ENSMUSG0C | false | . | . | . | . | . | . | chr4 | 149044122 | 149052366 | 7.01115 | 4.98336 | 4.65021 |
| chr2 | 59562664 | 59562903 | 182 | Promoter (1-2kb) | 2 | 59564268 | 59564684 | 417 | 2 | ENSMUST00 | 1781 | Gm13552 | ENSMUSG0C | false | . | . | . | . | . | . | . | . | . | . | . | . |
| chr2 | 93758389 | 93758697 | 182 | Intron (ENSMUST00000184931.2 | 2 | 93794570 | 93795080 | 511 | 1 | ENSMUST00 | -35873 | Gm13818 | ENSMUSG0C | false | . | . | . | . | . | . | chr2 | 93757685 | 93760236 | 4.90066 | 3.87491 | 2.97166 |
| chr16 | 51305106 | 51305414 | 182 | Distal Intergenic | 16 | 51362452 | 51363832 | 1381 | 2 | ENSMUST00 | 58418 | Gm6767 | ENSMUSG0C | false | . | . | . | . | . | . | chr16 | 51304950 | 51305805 | 4.69874 | 4.0009 | 2.7462 |
| chr11 | 63863843 | 63864067 | 182 | Distal Intergenic | 11 | 63846183 | 63846818 | 636 | 2 | ENSMUST00 | -17026 | Hmgb1-ps3 | ENSMUSG0C | false | . | . | . | . | . | . | chr11 | 63861623 | 63864289 | 3.98164 | 3.632 | 2.17845 |
| chr4 | 123434971 | 123435283 | 182 | Promoter (<=1kb) | 4 | 123435617 | 123435934 | 318 | 1 | ENSMUST00 | -334 | Gm12924 | ENSMUSG0C | false | . | . | . | . | . | . | chr4 | 123432183 | 123439235 | 5.18997 | 4.08191 | 3.17757 |
| chr15 | 97784213 | 97784855 | 182 | Promoter (<=1kb) | 15 | 97784355 | 97792692 | 8338 | 1 | ENSMUST00 | 0 | Slc48a1 | ENSMUSG0C | false | chr15 | 97784332 | 97785489 | 42.74922 | 148.11885 | 142.90198 | chr15 | 97771183 | 97790304 | 7.43452 | 5.14328 | 5.01036 |
| chr11 | 8797510 | 8797836 | 182 | Distal Intergenic | 11 | 8805952 | 8806848 | 897 | 2 | ENSMUST00 | 9012 | Gm11990 | ENSMUSG0C | false | . | . | . | . | . | . | . | . | . | . | . | . |
| chrX | 108445051 | 108445367 | 182 | Promoter (1-2kb) | 20 | 108446389 | 108449268 | 2880 | 1 | ENSMUST00 | -1022 | Gm9218 | ENSMUSG0C | false | . | . | . | . | . | . | . | . | . | . | . | . |
| chr11 | 63308847 | 63309048 | 182 | Distal Intergenic | 11 | 63312310 | 63313311 | 1002 | 2 | ENSMUST00 | 4263 | Gm12286 | ENSMUSG0C | false | . | . | . | . | . | . | chr11 | 63307751 | 63309914 | 3.85549 | 3.501 | 2.11759 |
| chr16 | 94602817 | 94603038 | 182 | Intron (ENSMUST00000119878.2 | 16 | 94620377 | 94620587 | 211 | 1 | ENSMUST00 | -17339 | Gm15971 | ENSMUSG0C | false | . | . | . | . | . | . | chr16 | 94571320 | 94615960 | 5.49747 | 4.23365 | 3.43578 |
| chr2 | 173992070 | 173992406 | 182 | Distal Intergenic | 2 | 173984482 | 173984691 | 210 | 2 | ENSMUST00 | -7380 | Gm14337 | ENSMUSG0C | false | . | . | . | . | . | . | chr2 | 173991174 | 173993442 | 5.14357 | 4.10249 | 3.17196 |
| chr11 | 64715416 | 64715618 | 182 | Distal Intergenic | 11 | 64759717 | 64760027 | 311 | 2 | ENSMUST00 | 44409 | Gm12291 | ENSMUSG0C | false | . | . | . | . | . | . | chr11 | 64714531 | 64716056 | 4.35237 | 3.66734 | 2.49615 |
| chr11 | 64708248 | 64708593 | 182 | Distal Intergenic | 11 | 64759717 | 64760027 | 311 | 2 | ENSMUST00 | 51434 | Gm12291 | ENSMUSG0C | false | . | . | . | . | . | . | chr11 | 64706948 | 64713410 | 4.30838 | 3.74761 | 2.44128 |
| chr11 | 64726810 | 64727027 | 182 | Distal Intergenic | 11 | 64759717 | 64760027 | 311 | 2 | ENSMUST00 | 33000 | Gm12291 | ENSMUSG0C | false | . | . | . | . | . | . | chr11 | 64726167 | 64728278 | 3.11756 | 3.07064 | 1.5175 |
| chr2 | 83740552 | 83740960 | 182 | Exon (ENSMUST00000155521.1/ | 2 | 83744484 | 83745736 | 1253 | 2 | ENSMUST00 | 4776 | Dnmt3l-ps1 | ENSMUSG0C | false | . | . | . | . | . | . | chr2 | 83738044 | 83745203 | 5.80522 | 4.36226 | 3.6965 |
| chr11 | 19828006 | 19828269 | 182 | Distal Intergenic | 11 | 19783652 | 19784483 | 832 | 1 | ENSMUST00 | 44355 | Gm12029 | ENSMUSG0C | false | . | . | . | . | . | . | chr11 | 19827692 | 19831074 | 3.14358 | 3.08329 | 1.47727 |
| chr6 | 149445916 | 149446114 | 182 | Intron (ENSMUST00000086829.5 | 6 | 149433780 | 149434473 | 694 | 2 | ENSMUST00 | -11444 | Gm15785 | ENSMUSG0C | false | . | . | . | . | . | . | chr6 | 149446013 | 149446390 | 2.97357 | 2.98746 | 1.3784 |
| chr5 | 150409584 | 150409826 | 182 | Intron (ENSMUST00000087204.5 | 5 | 150509694 | 150511338 | 1645 | 2 | ENSMUST00 | 101512 | Gm15933 | ENSMUSG0C | false | . | . | . | . | . | . | chr5 | 150408365 | 150411632 | 5.34997 | 4.21823 | 3.29677 |
| chr5 | 150466136 | 150466525 | 182 | Exon (ENSMUST00000087204.5/ | 5 | 150509694 | 150511338 | 1645 | 2 | ENSMUST00 | 44813 | Gm15933 | ENSMUSG0C | false | . | . | . | . | . | . | chr5 | 150464615 | 150467435 | 7.08694 | 5.02005 | 4.70994 |
| chr4 | 8647093 | 8647766 | 182 | Distal Intergenic | 4 | 8672633 | 8673582 | 950 | 2 | ENSMUST00 | 25816 | Gm11809 | ENSMUSG0C | false | chr4 | 8646552 | 8648739 | 24.03115 | 75.67385 | 72.59326 | . | . | . | . | . | . |
| chr1 | 31791824 | 31792152 | 182 | Distal Intergenic | 1 | 31736057 | 31736891 | 835 | 1 | ENSMUST00 | 55768 | Gm7910 | ENSMUSG0C | false | . | . | . | . | . | . | . | . | . | . | . | . |
| chr4 | 155505266 | 155505474 | 182 | Promoter (2-3kb) | 4 | 155502316 | 155502706 | 391 | 2 | ENSMUST00 | -2561 | Gm13171 | ENSMUSG0C | false | . | . | . | . | . | . | chr4 | 155497429 | 155517972 | 5.62103 | 4.35714 | 3.51731 |
| chr17 | 48005138 | 48005336 | 182 | Promoter (1-2kb) | 17 | 48003379 | 48003572 | 194 | 2 | ENSMUST00 | -1567 | Gm14871 | ENSMUSG0C | false | . | . | . | . | . | . | . | . | . | . | . | . |
| chr17 | 47976090 | 47976465 | 182 | Distal Intergenic | 17 | 48003379 | 48003572 | 194 | 2 | ENSMUST00 | 27107 | Gm14871 | ENSMUSG0C | false | chr17 | 47975786 | 47976500 | 12.3514 | 27.13773 | 24.70927 | chr17 | 47973523 | 47976221 | 5.86638 | 4.39509 | 3.7429 |
| chr11 | 117939782 | 117940073 | 182 | Distal Intergenic | 11 | 117921905 | 117922328 | 424 | 1 | ENSMUST00 | 17878 | Gm11726 | ENSMUSG0C | false | . | . | . | . | . | . | . | . | . | . | . | . |
| chr16 | 11537708 | 11537904 | 182 | Intron (ENSMUST00000180792.2 | 16 | 11494821 | 11496306 | 1486 | 2 | ENSMUST00 | -41403 | Gm15897 | ENSMUSG0C | false | . | . | . | . | . | . | chr16 | 11536875 | 11538744 | 5.34737 | 4.24032 | 3.30212 |
| chr13 | 25115572 | 25115865 | 182 | Intron (ENSMUST00000036932.9 | 13 | 25130747 | 25131064 | 318 | 2 | ENSMUST00 | 15199 | Gm11348 | ENSMUSG0C | false | chr13 | 25115467 | 25116122 | 10.05864 | 20.96754 | 18.61967 | chr13 | 25115051 | 25116470 | 5.52927 | 4.33868 | 3.43135 |
| chr7 | 130320080 | 130320322 | 182 | Intron (ENSMUST00000124096.2 | 7 | 130352038 | 130354362 | 2325 | 2 | ENSMUST00 | 34040 | Gm5903 | ENSMUSG0C | false | . | . | . | . | . | . | . | . | . | . | . | . |
| chr11 | 22636849 | 22637204 | 182 | Distal Intergenic | 11 | 22626375 | 22626664 | 290 | 2 | ENSMUST00 | -10186 | Gm12053 | ENSMUSG0C | false | . | . | . | . | . | . | chr11 | 22635852 | 22640892 | 6.051 | 4.54017 | 3.86945 |
| chr13 | 31911579 | 31911807 | 182 | Intron (ENSMUST00000041859.7 | 13 | 31884225 | 31885224 | 1000 | 1 | ENSMUST00 | 27355 | Gm11380 | ENSMUSG0C | false | . | . | . | . | . | . | chr13 | 31911749 | 31912165 | 5.42885 | 4.35494 | 3.34855 |
| chr13 | 31993494 | 31993716 | 182 | Intron (ENSMUST00000041859.7 | 13 | 31884225 | 31885224 | 1000 | 1 | ENSMUST00 | 109270 | Gm11380 | ENSMUSG0C | false | . | . | . | . | . | . | chr13 | 31990566 | 31994385 | 5.04334 | 4.1016 | 3.04615 |
| chr11 | 61974376 | 61974655 | 182 | Promoter (<=1kb) | 11 | 61973966 | 61974350 | 385 | 2 | ENSMUST00 | -27 | Gm12274 | ENSMUSG0C | false | . | . | . | . | . | . | chr11 | 61973202 | 61974583 | 5.77182 | 4.46716 | 3.65535 |
| chr4 | 15550538 | 15550750 | 182 | Distal Intergenic | 4 | 15531657 | 15532111 | 455 | 2 | ENSMUST00 | -18428 | Gm11859 | ENSMUSG0C | false | . | . | . | . | . | . | chr4 | 15550067 | 15552760 | 3.27032 | 3.15859 | 1.58918 |
| chr17 | 8698342 | 8698549 | 182 | Intron (ENSMUST00000115715.2 | 17 | 8765372 | 8765927 | 556 | 2 | ENSMUST00 | 67378 | Gm15426 | ENSMUSG0C | false | . | . | . | . | . | . | chr17 | 8697762 | 8702488 | 3.56825 | 3.38569 | 1.82101 |
| chr4 | 84407584 | 84407828 | 182 | Intron (ENSMUST00000102820.3 | 4 | 84428818 | 84429124 | 307 | 2 | ENSMUST00 | 21296 | Gm12421 | ENSMUSG0C | false | . | . | . | . | . | . | . | . | . | . | . | . |
| chr4 | 9027890 | 9028116 | 182 | Distal Intergenic | 4 | 8941259 | 8941656 | 398 | 1 | ENSMUST00 | 86632 | Rps18-ps2 | ENSMUSG0C | false | chr4 | 9027573 | 9028722 | 8.93054 | 17.03311 | 14.7407 | chr4 | 9027455 | 9029178 | 4.52949 | 3.84646 | 2.6319 |
| chr13 | 30396523 | 30396836 | 182 | Downstream (2-3kb) | 13 | 30399617 | 30399934 | 318 | 2 | ENSMUST00 | 3098 | Gm11369 | ENSMUSG0C | false | . | . | . | . | . | . | . | . | . | . | . | . |
| chr11 | 68121324 | 68121518 | 182 | Intron (ENSMUST00000021285.8 | 11 | 68095580 | 68095964 | 385 | 2 | ENSMUST00 | -25361 | Gm12304 | ENSMUSG0C | false | . | . | . | . | . | . | chr11 | 68121221 | 68121544 | 3.5829 | 3.04223 | 1.85742 |
| chr11 | 65020876 | 65021100 | 182 | Intron (ENSMUST00000047463.9 | 11 | 65037052 | 65037541 | 490 | 2 | ENSMUST00 | 16441 | Gm12294 | ENSMUSG0C | false | . | . | . | . | . | . | chr11 | 65020037 | 65021489 | 4.96581 | 3.99911 | 3.01297 |
| chr6 | 145340619 | 145340945 | 182 | Distal Intergenic | 6 | 145315613 | 145316608 | 996 | 2 | ENSMUST00 | -24012 | Gm15707 | ENSMUSG0C | false | chr6 | 145339893 | 145340769 | 14.96389 | 42.58086 | 39.95799 | . | . | . | . | . | . |
| chr4 | 89152262 | 89152507 | 182 | Intron (ENSMUST00000058030.7 | 4 | 89161699 | 89162191 | 493 | 2 | ENSMUST00 | 9684 | Gm12607 | ENSMUSG0C | false | . | . | . | . | . | . | . | . | . | . | . | . |
| chr2 | 170287411 | 170287799 | 182 | Promoter (2-3kb) | 2 | 170285035 | 170285880 | 846 | 1 | ENSMUST00 | 2377 | Gm14270 | ENSMUSG0C | false | chr2 | 170287072 | 170287903 | 16.76111 | 40.61943 | 38.02256 | chr2 | 170286445 | 170288689 | 2.94281 | 2.92825 | 1.38316 |
| chr2 | 170258478 | 170258712 | 182 | Distal Intergenic | 2 | 170285035 | 170285880 | 846 | 1 | ENSMUST00 | -26323 | Gm14270 | ENSMUSG0C | false | . | . | . | . | . | . | . | . | . | . | . | . |
| chr3 | 94821303 | 94821726 | 182 | Distal Intergenic | 3 | 94832977 | 94833805 | 829 | 1 | ENSMUST00 | -11251 | Gm15263 | ENSMUSG0C | false | chr3 | 94820977 | 94822042 | 21.42557 | 69.98924 | 66.99868 | . | . | . | . | . | . |
| chr11 | 22763032 | 22763308 | 182 | Intron (ENSMUST00000189990.1 | 11 | 22758599 | 22760025 | 1427 | 2 | ENSMUST00 | -3008 | Gm12057 | ENSMUSG0C | false | . | . | . | . | . | . | . | . | . | . | . | . |
| chr5 | 103620376 | 103620596 | 182 | Intron (ENSMUST00000031263.1 | 5 | 103624471 | 103625046 | 576 | 1 | ENSMUST00 | -3875 | Gm15844 | ENSMUSG0C | false | . | . | . | . | . | . | chr5 | 103617128 | 103621618 | 4.46422 | 3.80493 | 2.5717 |
| chr15 | 37417347 | 37417588 | 182 | Intron (ENSMUST00000168992.2 | 15 | 37421248 | 37422694 | 1447 | 1 | ENSMUST00 | -3660 | Gm8668 | ENSMUSG0C | false | . | . | . | . | . | . | chr15 | 37415977 | 37419194 | 4.14939 | 3.51106 | 2.34462 |
| chr2 | 8728136 | 8728359 | 182 | Distal Intergenic | 2 | 8633937 | 8634483 | 547 | 1 | ENSMUST00 | 94200 | Gm13255 | ENSMUSG0C | false | . | . | . | . | . | . | . | . | . | . | . | . |
| chr11 | 88460449 | 88460643 | 182 | Intron (ENSMUST00000107909.2 | 11 | 88432986 | 88433514 | 529 | 2 | ENSMUST00 | -26936 | Gm11510 | ENSMUSG0C | false | . | . | . | . | . | . | chr11 | 88445933 | 88476041 | 6.2627 | 4.6354 | 4.05065 |
| chr14 | 78644055 | 78644297 | 182 | Intron (ENSMUST00000074729.5 | 14 | 78603710 | 78603831 | 122 | 1 | ENSMUST00 | 40346 | Gm26251 | ENSMUSG0C | false | . | . | . | . | . | . | chr14 | 78639208 | 78651275 | 4.93549 | 3.97381 | 2.99528 |
| chr5 | 119246990 | 119247451 | 182 | Distal Intergenic | 5 | 119297245 | 119297361 | 117 | 1 | ENSMUST00 | -49794 | n-R5s175 | ENSMUSG0C | false | . | . | . | . | . | . | . | . | . | . | . | . |
| chr5 | 119356238 | 119356420 | 182 | Distal Intergenic | 5 | 119297245 | 119297361 | 117 | 1 | ENSMUST00 | 58994 | n-R5s175 | ENSMUSG0C | false | . | . | . | . | . | . | . | . | . | . | . | . |
| chr6 | 88363689 | 88363881 | 182 | Intron (ENSMUST00000165242.1 | 6 | 88405977 | 88406301 | 325 | 1 | ENSMUST00 | -42096 | Gm25178 | ENSMUSG0C | false | . | . | . | . | . | . | chr6 | 88363025 | 88364743 | 3.1582 | 2.98896 | 1.51174 |
| chr4 | 82266118 | 82266582 | 182 | Distal Intergenic | 4 | 82439410 | 82439514 | 105 | 1 | ENSMUST00 | -172828 | n-R5s188 | ENSMUSG0C | false | . | . | . | . | . | . | . | . | . | . | . | . |
| chr3 | 83610427 | 83610658 | 182 | Distal Intergenic | 3 | 83634577 | 83634874 | 298 | 2 | ENSMUST00 | 24216 | Gm22342 | ENSMUSG0C | false | . | . | . | . | . | . | . | . | . | . | . | . |
| chr19 | 43939896 | 43940404 | 182 | Distal Intergenic | 19 | 43932316 | 43932432 | 117 | 1 | ENSMUST00 | 7581 | n-R5s21 | ENSMUSG0C | false | chr19 | 43939281 | 43940742 | 26.21985 | 74.04404 | 70.99222 | chr19 | 43939233 | 43940726 | 3.70106 | 3.38684 | 1.97864 |
| chr3 | 60123785 | 60124113 | 182 | Distal Intergenic | 3 | 60127641 | 60127747 | 107 | 2 | ENSMUST00 | 3634 | Gm24382 | ENSMUSG0C | false | . | . | . | . | . | . | chr3 | 60120518 | 60128464 | 4.1733 | 3.66476 | 2.33216 |
| chr3 | 104597140 | 104597416 | 182 | Distal Intergenic | 3 | 104601158 | 104601239 | 82 | 2 | ENSMUST00 | 3823 | Gm26091 | ENSMUSG0C | false | . | . | . | . | . | . | chr3 | 104596927 | 104597577 | 2.78338 | 2.85188 | 1.22471 |
| chr3 | 121342081 | 121342277 | 182 | Intron (ENSMUST00000039442.7 | 3 | 121337692 | 121337959 | 268 | 1 | ENSMUST00 | 4390 | Gm26027 | ENSMUSG0C | false | . | . | . | . | . | . | chr3 | 121337943 | 121343113 | 3.99072 | 3.52601 | 2.18837 |
| chr9 | 106008296 | 106008532 | 182 | Intron (ENSMUST00000121963.1 | 9 | 106001718 | 106001798 | 81 | 1 | ENSMUST00 | 6579 | Gm24631 | ENSMUSG0C | false | . | . | . | . | . | . | . | . | . | . | . | . |
| chr3 | 88536462 | 88537002 | 182 | Promoter (<=1kb) | 3 | 88536301 | 88536382 | 82 | 2 | ENSMUST00 | -81 | Mir1905 | ENSMUSG0C | false | chr3 | 88536138 | 88537358 | 18.95331 | 81.30177 | 78.12804 | chr3 | 88536082 | 88537473 | 4.18782 | 3.62893 | 2.36506 |
| chr9 | 8325389 | 8325603 | 182 | Distal Intergenic | 9 | 8246786 | 8246881 | 96 | 1 | ENSMUST00 | 78604 | Mir1899 | ENSMUSG0C | false | . | . | . | . | . | . | . | . | . | . | . | . |
| chr4 | 41636297 | 41636756 | 182 | Promoter (2-3kb) | 4 | 41639378 | 41641416 | 2039 | 1 | ENSMUST00 | -2622 | Gm12406 | ENSMUSG0C | false | . | . | . | . | . | . | . | . | . | . | . | . |
| chr11 | 95631543 | 95631743 | 182 | Distal Intergenic | 11 | 95622660 | 95629836 | 7177 | 1 | ENSMUST00 | 8884 | Gm11528 | ENSMUSG0C | false | . | . | . | . | . | . | chr11 | 95630766 | 95631998 | 2.84059 | 2.83908 | 1.27112 |
| chr16 | 91461810 | 91462061 | 182 | Distal Intergenic | 16 | 91465104 | 91470121 | 5018 | 1 | ENSMUST00 | -3043 | A930006K02 | ENSMUSG0C | false | . | . | . | . | . | . | chr16 | 91461134 | 91466910 | 4.53361 | 3.81817 | 2.62159 |
| chr4 | 95278764 | 95279136 | 182 | Distal Intergenic | 4 | 95193362 | 95206248 | 12887 | 1 | ENSMUST00 | 85403 | Gm12708 | ENSMUSG0C | false | . | . | . | . | . | . | chr4 | 95277868 | 95279910 | 5.50005 | 4.36049 | 3.42056 |
| chr8 | 45610763 | 45611052 | 182 | Intron (ENSMUST00000067065.8 | 8 | 45587912 | 45596744 | 8833 | 2 | ENSMUST00 | -14020 | Gm16351 | ENSMUSG0C | false | . | . | . | . | . | . | chr8 | 45610489 | 45612056 | 3.16308 | 3.09583 | 1.55015 |
| chr11 | 79704269 | 79704656 | 182 | Promoter (<=1kb) | 11 | 79704630 | 79705420 | 791 | 1 | ENSMUST00 | 0 | Gm11205 | ENSMUSG0C | false | chr11 | 79703487 | 79704556 | 28.91247 | 96.86193 | 93.36482 | . | . | . | . | . | . |
| chr1 | 93210421 | 93210662 | 182 | Distal Intergenic | 1 | 93215899 | 93231072 | 15174 | 1 | ENSMUST00 | -5237 | E030010N08 | ENSMUSG0C | false | . | . | . | . | . | . | chr1 | 93210088 | 93211973 | 5.50157 | 4.34556 | 3.43814 |
| chrX | 36807802 | 36808064 | 182 | Distal Intergenic | 20 | 36814771 | 36817200 | 2430 | 1 | ENSMUST00 | -6707 | Gm14549 | ENSMUSG0C | false | chrX | 36807277 | 36808144 | 11.4883 | 23.92001 | 21.53215 | . | . | . | . | . | . |
| chr14 | 100284766 | 100285170 | 182 | Distal Intergenic | 14 | 100219486 | 100219789 | 304 | 1 | ENSMUST00 | 65281 | Gm16260 | ENSMUSG0C | false | chr14 | 100284160 | 100286947 | 30.32912 | 88.589 | 85.27629 | chr14 | 100284937 | 100287388 | 6.55605 | 4.70889 | 4.33546 |
| chrX | 99285705 | 99285939 | 182 | Distal Intergenic | 20 | 99409571 | 99418895 | 9325 | 2 | ENSMUST00 | 132956 | Gm14809 | ENSMUSG0C | false | . | . | . | . | . | . | . | . | . | . | . | . |
| chr11 | 119833242 | 119833617 | 182 | Promoter (<=1kb) | 11 | 119820812 | 119834365 | 13554 | 2 | ENSMUST00 | 748 | Rptoros | ENSMUSG0C | false | . | . | . | . | . | . | chr11 | 119831481 | 119836988 | 5.44333 | 4.10671 | 3.40297 |
| chr2 | 33187625 | 33187922 | 182 | Intron (ENSMUST00000042615.7 | 2 | 33177902 | 33181881 | 3980 | 1 | ENSMUST00 | 9724 | Gm13528 | ENSMUSG0C | false | . | . | . | . | . | . | chr2 | 33175823 | 33189808 | 7.09799 | 4.90853 | 4.74291 |
| chr2 | 44147626 | 44147943 | 182 | Intron (ENSMUST00000112824.2 | 2 | 44059238 | 44065324 | 6087 | 2 | ENSMUST00 | -82303 | Arhgap15os | ENSMUSG0C | false | . | . | . | . | . | . | . | . | . | . | . | . |
| chr11 | 16800610 | 16800888 | 182 | Intron (ENSMUST00000102884.4 | 11 | 16815431 | 16830702 | 15272 | 2 | ENSMUST00 | 29814 | Egfros | ENSMUSG0C | false | . | . | . | . | . | . | . | . | . | . | . | . |
| chr4 | 150758921 | 150759194 | 182 | Distal Intergenic | 4 | 150825733 | 150839575 | 13843 | 1 | ENSMUST00 | -66539 | Gm13049 | ENSMUSG0C | false | . | . | . | . | . | . | chr4 | 150757809 | 150759700 | 5.10694 | 4.10195 | 3.14093 |
| chr3 | 53284581 | 53284933 | 182 | Distal Intergenic | 3 | 53452619 | 53457797 | 5179 | 1 | ENSMUST00 | -167686 | Gm16206 | ENSMUSG0C | false | . | . | . | . | . | . | . | . | . | . | . | . |
| chr1 | 172550583 | 172550886 | 182 | Promoter (<=1kb) | 1 | 172538133 | 172550669 | 12537 | 2 | ENSMUST00 | 0 | 4933439K11 | ENSMUSG0C | false | chr1 | 172550086 | 172550983 | 16.54316 | 39.53096 | 36.94807 | chr1 | 172549319 | 172551328 | 4.91126 | 4.01745 | 2.94867 |
| chr2 | 3749046 | 3749304 | 182 | Intron (ENSMUST00000115052.3 | 2 | 3752986 | 3755177 | 2192 | 2 | ENSMUST00 | 5873 | Gm13185 | ENSMUSG0C | false | chr2 | 3749239 | 3749869 | 9.69585 | 19.82268 | 17.49052 | chr2 | 3748520 | 3753877 | 3.62264 | 3.31703 | 1.91049 |
| chr4 | 84019528 | 84019985 | 182 | Distal Intergenic | 4 | 84042039 | 84044879 | 2841 | 2 | ENSMUST00 | 24894 | 6030471H07 | ENSMUSG0C | false | . | . | . | . | . | . | chr4 | 84018335 | 84020917 | 5.45387 | 4.23509 | 3.42635 |
| chr7 | 84689578 | 84689868 | 182 | Promoter (<=1kb) | 7 | 84689640 | 84776549 | 86910 | 1 | ENSMUST00 | 0 | 2610206C17 | ENSMUSG0C | false | chr7 | 84688896 | 84690552 | 29.50911 | 91.04776 | 87.68056 | . | . | . | . | . | . |
| chr2 | 165855651 | 165856234 | 182 | Promoter (1-2kb) | 2 | 165857684 | 165859034 | 1351 | 1 | ENSMUST00 | -1450 | BC046401 | ENSMUSG0C | false | . | . | . | . | . | . | chr2 | 165851451 | 165889342 | 6.61553 | 4.6931 | 4.35892 |
| chr2 | 30545179 | 30545400 | 182 | Distal Intergenic | 2 | 30534729 | 30539022 | 4294 | 1 | ENSMUST00 | 10451 | Gm14487 | ENSMUSG0C | false | chr2 | 30544951 | 30545539 | 9.32368 | 22.97803 | 20.60213 | chr2 | 30544532 | 30545882 | 4.85915 | 4.01128 | 2.90762 |
| chr13 | 42916417 | 42916725 | 182 | Intron (ENSMUST00000128646.2 | 13 | 42868338 | 42869226 | 889 | 2 | ENSMUST00 | -47192 | Gm15812 | ENSMUSG0C | false | chr13 | 42916148 | 42917655 | 22.27177 | 70.67676 | 67.6792 | chr13 | 42916055 | 42918264 | 5.35453 | 4.26257 | 3.28981 |

| | | | | | | | | | | | | | | | | | | | | | | | | | | |
|---|---|---|---|---|---|---|---|---|---|---|---|---|---|---|---|---|---|---|---|---|---|---|---|---|---|---|
| chr10 | 105578418 | 105578619 | 182 | Intron (ENSMUST00000161364.2 | 10 | 105574559 | 105583874 | 9316 | 1 | ENSMUST00 | 3860 | Gm15663 | ENSMUSG0C | false | . | . | . | . | . | . | chr10 | 105578075 | 105579348 | 5.14117 | 4.16235 | 3.16088 |
| chr17 | 28882377 | 28882572 | 182 | Intron (ENSMUST00000056866.5 | 17 | 28905265 | 28906771 | 1507 | 1 | ENSMUST00 | -22693 | 1700030A11 | ENSMUSG0C | false | . | . | . | . | . | . | chr17 | 28881963 | 28882785 | 4.03961 | 3.46173 | 2.26893 |
| chr6 | 34669024 | 34669274 | 182 | Intron (ENSMUST00000079391.4 | 6 | 34651354 | 34653894 | 2541 | 2 | ENSMUST00 | -15131 | Gm13861 | ENSMUSG0C | false | . | . | . | . | . | . | chr6 | 34666478 | 34672346 | 4.53955 | 3.76821 | 2.66652 |
| chr4 | 116597506 | 116597755 | 182 | Promoter (<=1kb) | 4 | 116589733 | 116597630 | 7898 | 2 | ENSMUST00 | 0 | C530005A16 | ENSMUSG0C | false | chr4 | 116596656 | 116598461 | 24.40663 | 77.266 | 74.1616 | chr4 | 116596320 | 116599190 | 3.45479 | 3.22552 | 1.75481 |
| chr2 | 167851526 | 167851935 | 182 | Distal Intergenic | 2 | 167858585 | 167862542 | 3958 | 1 | ENSMUST00 | -6650 | Gm14319 | ENSMUSG0C | false | . | . | . | . | . | . | chr2 | 167850474 | 167854412 | 5.03048 | 3.96235 | 3.05598 |
| chr2 | 102038556 | 102038789 | 182 | Promoter (<=1kb) | 2 | 102039629 | 102066725 | 27097 | 1 | ENSMUST00 | -840 | Gm13919 | ENSMUSG0C | false | . | . | . | . | . | . | . | . | . | . | . | . |
| chr1 | 61385045 | 61385235 | 182 | Intron (ENSMUST00000141000.1 | 1 | 61378432 | 61395904 | 17473 | 1 | ENSMUST00 | 6614 | 9530026F06 | ENSMUSG0C | false | . | . | . | . | . | . | chr1 | 61384733 | 61388389 | 3.54539 | 3.38388 | 1.80969 |
| chr1 | 61378278 | 61378522 | 182 | Promoter (<=1kb) | 1 | 61378432 | 61395904 | 17473 | 1 | ENSMUST00 | 0 | 9530026F06 | ENSMUSG0C | false | . | . | . | . | . | . | chr1 | 61376362 | 61380536 | 4.64842 | 3.89817 | 2.72461 |
| chr7 | 96981033 | 96981259 | 182 | Intron (ENSMUST00000044466.6 | 7 | 96947910 | 96951886 | 3977 | 2 | ENSMUST00 | -29148 | C230038L03 | ENSMUSG0C | false | . | . | . | . | . | . | chr7 | 96952896 | 96991589 | 4.87493 | 3.9704 | 2.91243 |
| chr7 | 96985844 | 96986201 | 182 | Intron (ENSMUST00000044466.6 | 7 | 96947910 | 96951886 | 3977 | 2 | ENSMUST00 | -33959 | C230038L03 | ENSMUSG0C | false | . | . | . | . | . | . | chr7 | 96952896 | 96991589 | 4.87493 | 3.9704 | 2.91243 |
| chr8 | 11173217 | 11173558 | 182 | Distal Intergenic | 8 | 11187755 | 11204503 | 16749 | 1 | ENSMUST00 | -14197 | Gm15418 | ENSMUSG0C | false | . | . | . | . | . | . | chr8 | 11169588 | 11174102 | 5.12745 | 4.19846 | 3.11143 |
| chr4 | 107016408 | 107016642 | 182 | Intron (ENSMUST00000150915.2 | 4 | 107017935 | 107022499 | 4565 | 2 | ENSMUST00 | 5857 | Gm12786 | ENSMUSG0C | false | . | . | . | . | . | . | chr4 | 107009465 | 107020061 | 6.67324 | 4.69024 | 4.42102 |
| chr4 | 151302380 | 151302634 | 182 | Intron (ENSMUST00000169423.3 | 4 | 151376470 | 151379582 | 3113 | 1 | ENSMUST00 | -73836 | 4930589P08 | ENSMUSG0C | false | . | . | . | . | . | . | chr4 | 151300466 | 151304389 | 7.3711 | 5.1333 | 4.95551 |
| chr11 | 101881751 | 101882004 | 182 | Intron (ENSMUST00000057054.7 | 11 | 101886237 | 101891206 | 4970 | 1 | ENSMUST00 | -4233 | Gm11551 | ENSMUSG0C | false | . | . | . | . | . | . | chr11 | 101881513 | 101882112 | 3.91624 | 3.56195 | 2.10505 |
| chr2 | 167565833 | 167566040 | 182 | Intron (ENSMUST00000127006.1 | 2 | 167553314 | 167607845 | 54532 | 1 | ENSMUST00 | 12520 | Gm11476 | ENSMUSG0C | false | . | . | . | . | . | . | . | . | . | . | . | . |
| chr2 | 167559530 | 167559714 | 182 | Intron (ENSMUST00000127006.1 | 2 | 167553314 | 167607845 | 54532 | 1 | ENSMUST00 | 6217 | Gm11476 | ENSMUSG0C | false | . | . | . | . | . | . | . | . | . | . | . | . |
| chr1 | 75370389 | 75370694 | 182 | Promoter (<=1kb) | 1 | 75368272 | 75370451 | 2180 | 2 | ENSMUST00 | 0 | Gm15179 | ENSMUSG0C | false | . | . | . | . | . | . | chr1 | 75369321 | 75372101 | 7.41167 | 5.1468 | 4.97977 |
| chr6 | 145713237 | 145713440 | 182 | Distal Intergenic | 6 | 145738693 | 145746273 | 7581 | 2 | ENSMUST00 | 32833 | Gm15704 | ENSMUSG0C | false | . | . | . | . | . | . | chr6 | 145712653 | 145713870 | 4.03646 | 3.34946 | 2.24941 |
| chr4 | 131920732 | 131921027 | 182 | Promoter (<=1kb) | 4 | 131899478 | 131920029 | 20552 | 2 | ENSMUST00 | -704 | Gm12992 | ENSMUSG0C | false | chr4 | 131920283 | 131922489 | 21.79372 | 61.12524 | 58.26245 | chr4 | 131920243 | 131922772 | 5.42572 | 3.9801 | 3.38953 |
| chr2 | 33440147 | 33440426 | 182 | Distal Intergenic | 2 | 33446802 | 33449207 | 2406 | 1 | ENSMUST00 | -6376 | Gm13536 | ENSMUSG0C | false | . | . | . | . | . | . | . | . | . | . | . | . |
| chr11 | 75021929 | 75022156 | 182 | Intron (ENSMUST00000045281.7 | 11 | 75036476 | 75039348 | 2873 | 2 | ENSMUST00 | 17192 | Gm12333 | ENSMUSG0C | false | . | . | . | . | . | . | . | . | . | . | . | . |
| chr11 | 31816222 | 31816426 | 182 | Intron (ENSMUST00000129425.2 | 11 | 31790113 | 31824473 | 34361 | 2 | ENSMUST00 | 8047 | D630024D03 | ENSMUSG0C | false | . | . | . | . | . | . | . | . | . | . | . | . |
| chr14 | 75834530 | 75835064 | 182 | Distal Intergenic | 14 | 75843379 | 75844964 | 1586 | 2 | ENSMUST00 | 9900 | Gm4285 | ENSMUSG0C | false | chr14 | 75834423 | 75835535 | 20.92485 | 54.63506 | 51.8618 | chr14 | 75832212 | 75838305 | 5.41657 | 4.276 | 3.35538 |
| chr4 | 98790112 | 98790351 | 182 | Intron (ENSMUST00000102790.3 | 4 | 98793768 | 98798900 | 5133 | 1 | ENSMUST00 | -3417 | Kank4os | ENSMUSG0C | false | . | . | . | . | . | . | chr4 | 98789928 | 98790967 | 3.39606 | 3.11842 | 1.72209 |
| chr4 | 114682342 | 114682571 | 182 | Promoter (1-2kb) | 4 | 114680804 | 114687595 | 6792 | 1 | ENSMUST00 | 1539 | Gm12829 | ENSMUSG0C | false | . | . | . | . | . | . | chr4 | 114677087 | 114691330 | 5.31309 | 4.19527 | 3.28131 |
| chr14 | 11984155 | 11984446 | 182 | Promoter (2-3kb) | 14 | 11965447 | 11981223 | 15777 | 2 | ENSMUST00 | -2933 | Gm3848 | ENSMUSG0C | false | . | . | . | . | . | . | . | . | . | . | . | . |
| chr14 | 11811270 | 11811658 | 182 | Intron (ENSMUST00000022264.7 | 14 | 11980198 | 11981208 | 1011 | 2 | ENSMUST00 | 169550 | Gm3848 | ENSMUSG0C | false | . | . | . | . | . | . | . | . | . | . | . | . |
| chr7 | 124593472 | 124593714 | 182 | Distal Intergenic | 7 | 124476541 | 124494301 | 17761 | 2 | ENSMUST00 | -99172 | Gm15339 | ENSMUSG0C | false | . | . | . | . | . | . | . | . | . | . | . | . |
| chr13 | 28531705 | 28531951 | 182 | Intron (ENSMUST00000148885.2 | 13 | 28460034 | 28521108 | 61075 | 2 | ENSMUST00 | -10598 | 2610307P16 | ENSMUSG0C | false | . | . | . | . | . | . | . | . | . | . | . | . |
| chr11 | 98357611 | 98357947 | 182 | Promoter (<=1kb) | 11 | 98350718 | 98358283 | 7566 | 2 | ENSMUST00 | 336 | 1700003D09 | ENSMUSG0C | false | chr11 | 98357889 | 98359357 | 19.14982 | 55.63055 | 52.84086 | chr11 | 98355645 | 98364253 | 5.76527 | 4.38786 | 3.64659 |
| chrX | 167063711 | 167063937 | 182 | Intron (ENSMUST00000133867.2 | 20 | 167063552 | 167068389 | 4838 | 2 | ENSMUST00 | 4452 | Gm8817 | ENSMUSG0C | false | . | . | . | . | . | . | . | . | . | . | . | . |
| chr9 | 40723320 | 40723515 | 182 | Intron (ENSMUST00000139577.1 | 9 | 40718906 | 40719986 | 1081 | 2 | ENSMUST00 | -3335 | Gm16095 | ENSMUSG0C | false | . | . | . | . | . | . | chr9 | 40722650 | 40724634 | 5.05695 | 4.13649 | 3.04547 |
| chr4 | 146537328 | 146537750 | 182 | Exon (ENSMUST00000105735.3/ | 4 | 146519039 | 146546873 | 27835 | 2 | ENSMUST00 | 9123 | 1700095A21 | ENSMUSG0C | false | . | . | . | . | . | . | . | . | . | . | . | . |
| chr8 | 3211706 | 3211911 | 182 | Intron (ENSMUST00000091291.4 | 8 | 3259704 | 3260641 | 938 | 1 | ENSMUST00 | -47793 | Gm16180 | ENSMUSG0C | false | . | . | . | . | . | . | chr8 | 3208761 | 3212840 | 5.43592 | 4.19508 | 3.38792 |
| chr8 | 3220969 | 3221350 | 182 | Intron (ENSMUST00000091291.4 | 8 | 3259704 | 3260641 | 938 | 1 | ENSMUST00 | -38354 | Gm16180 | ENSMUSG0C | false | chr8 | 3220969 | 3221812 | 17.6846 | 53.76268 | 50.99834 | chr8 | 3219942 | 3222910 | 4.33234 | 3.7568 | 2.44296 |
| chr16 | 33949109 | 33949305 | 182 | 3' UTR | 16 | 33966239 | 33967312 | 1074 | 1 | ENSMUST00 | -16934 | Gm15829 | ENSMUSG0C | false | . | . | . | . | . | . | chr16 | 33911227 | 33952498 | 7.40288 | 5.1092 | 4.9863 |
| chr13 | 112002899 | 112003273 | 182 | Promoter (2-3kb) | 13 | 112005502 | 112017332 | 11831 | 1 | ENSMUST00 | -2229 | Gm15323 | ENSMUSG0C | false | . | . | . | . | . | . | chr13 | 112001965 | 112004080 | 4.95112 | 4.0437 | 2.97813 |
| chr4 | 147674992 | 147675318 | 182 | Exon (ENSMUST00000117638.1/ | 4 | 147668518 | 147670854 | 2337 | 2 | ENSMUST00 | -4139 | Gm16211 | ENSMUSG0C | false | . | . | . | . | . | . | . | . | . | . | . | . |
| chr10 | 24479438 | 24479631 | 182 | Distal Intergenic | 10 | 24447490 | 24452657 | 5168 | 1 | ENSMUST00 | 31949 | Gm15271 | ENSMUSG0C | false | . | . | . | . | . | . | chr10 | 24478946 | 24480493 | 3.65887 | 3.40677 | 1.91809 |
| chr11 | 85713618 | 85713863 | 182 | Intron (ENSMUST00000108061.2 | 11 | 85702200 | 85719744 | 17545 | 2 | ENSMUST00 | 5881 | Bcas3os1 | ENSMUSG0C | false | . | . | . | . | . | . | . | . | . | . | . | . |
| chr11 | 102628657 | 102628896 | 182 | Distal Intergenic | 11 | 102619796 | 102624416 | 4621 | 2 | ENSMUST00 | -4242 | 2810433D01 | ENSMUSG0C | false | . | . | . | . | . | . | chr11 | 102621435 | 102629837 | 4.81998 | 3.90168 | 2.88673 |
| chr2 | 116076066 | 116076293 | 182 | Intron (ENSMUST00000152412.1 | 2 | 116067968 | 116085910 | 17943 | 1 | ENSMUST00 | 8099 | G630016G05 | ENSMUSG0C | false | chr2 | 116075411 | 116076396 | 18.19994 | 52.30853 | 49.5647 | chr2 | 116074680 | 116076218 | 3.92253 | 3.41484 | 2.14488 |
| chr11 | 62495201 | 62495724 | 182 | Intron (ENSMUST00000014389.5 | 11 | 62482797 | 62501704 | 18908 | 1 | ENSMUST00 | 12405 | Gm12278 | ENSMUSG0C | false | . | . | . | . | . | . | chr11 | 62494403 | 62496808 | 6.31672 | 4.47095 | 4.11911 |
| chr18 | 74505447 | 74505650 | 182 | Intron (ENSMUST00000125882.2 | 18 | 74525253 | 74531106 | 5854 | 2 | ENSMUST00 | 25456 | 1700120E14 | ENSMUSG0C | false | . | . | . | . | . | . | chr18 | 74504326 | 74506638 | 5.96401 | 4.46476 | 3.80574 |
| chr5 | 73153963 | 73154167 | 182 | Promoter (2-3kb) | 5 | 73157014 | 73157880 | 867 | 1 | ENSMUST00 | -2847 | 1700071G01 | ENSMUSG0C | false | . | . | . | . | . | . | chr5 | 73148451 | 73194976 | 5.88301 | 4.44395 | 3.74315 |
| chr4 | 100166812 | 100167127 | 182 | Intron (ENSMUST00000039630.5 | 4 | 100119621 | 100120841 | 1221 | 1 | ENSMUST00 | 47192 | Gm12701 | ENSMUSG0C | false | . | . | . | . | . | . | chr4 | 100165938 | 100174701 | 4.50598 | 3.80894 | 2.61922 |
| chr11 | 97546849 | 97547233 | 182 | Intron (ENSMUST00000107596.3 | 11 | 97566711 | 97576186 | 9476 | 1 | ENSMUST00 | -19478 | Gm11612 | ENSMUSG0C | false | chr11 | 97546418 | 97547365 | 15.50707 | 38.47921 | 35.908 | chr11 | 97546339 | 97547503 | 8.17375 | 5.36841 | 5.66523 |
| chr16 | 4781586 | 4781841 | 182 | Promoter (2-3kb) | 16 | 4777743 | 4779454 | 1712 | 2 | ENSMUST00 | -2133 | Gm15835 | ENSMUSG0C | false | . | . | . | . | . | . | chr16 | 4776742 | 4789496 | 4.39738 | 3.71911 | 2.52689 |
| chr13 | 111725444 | 111725890 | 182 | Promoter (<=1kb) | 13 | 111725517 | 111735124 | 9608 | 1 | ENSMUST00 | 0 | Gm15287 | ENSMUSG0C | false | chr13 | 111724511 | 111726223 | 25.16654 | 87.04672 | 83.76057 | . | . | . | . | . | . |
| chr2 | 70509191 | 70509452 | 182 | Promoter (<=1kb) | 2 | 70503433 | 70509199 | 5767 | 2 | ENSMUST00 | 0 | Erich2os | ENSMUSG0C | false | chr2 | 70508762 | 70509795 | 17.18351 | 44.2994 | 41.65401 | chr2 | 70508565 | 70509757 | 3.76844 | 3.43846 | 2.04455 |
| chr11 | 119126475 | 119126695 | 182 | Intron (ENSMUST00000144096.1 | 11 | 119123011 | 119138421 | 15411 | 2 | ENSMUST00 | 11726 | Gm11755 | ENSMUSG0C | false | . | . | . | . | . | . | chr11 | 119126043 | 119128960 | 6.47423 | 4.66106 | 4.2144 |
| chr4 | 117985312 | 117985554 | 182 | Intron (ENSMUST00000146982.1 | 4 | 117975036 | 117993972 | 18937 | 1 | ENSMUST00 | 10277 | 9530034E10 | ENSMUSG0C | false | . | . | . | . | . | . | chr4 | 117972459 | 117988550 | 5.57621 | 4.24526 | 3.52237 |
| chr3 | 96219922 | 96220309 | 182 | Promoter (<=1kb) | 3 | 96209627 | 96220168 | 10542 | 2 | ENSMUST00 | 0 | Gm15444 | ENSMUSG0C | false | chr3 | 96219035 | 96222442 | 35.13211 | 109.86813 | 106.03947 | . | . | . | . | . | . |
| chr11 | 117394619 | 117394910 | 182 | Distal Intergenic | 11 | 117414389 | 117415368 | 980 | 1 | ENSMUST00 | -19479 | Gm11729 | ENSMUSG0C | false | . | . | . | . | . | . | chr11 | 117393965 | 117395787 | 5.60463 | 4.31457 | 3.56383 |
| chr2 | 60117395 | 60117644 | 182 | Intron (ENSMUST00000112550.2 | 2 | 60125096 | 60125819 | 724 | 1 | ENSMUST00 | -7452 | Gm13620 | ENSMUSG0C | false | . | . | . | . | . | . | chr2 | 60113393 | 60124063 | 4.90918 | 3.93506 | 2.95446 |
| chr2 | 60117978 | 60118212 | 182 | Intron (ENSMUST00000112550.2 | 2 | 60125096 | 60125819 | 724 | 1 | ENSMUST00 | -6884 | Gm13620 | ENSMUSG0C | false | . | . | . | . | . | . | chr2 | 60113393 | 60124063 | 4.90918 | 3.93506 | 2.95446 |
| chr4 | 148948036 | 148948294 | 182 | Promoter (<=1kb) | 4 | 148947492 | 148948914 | 1423 | 2 | ENSMUST00 | 620 | Gm13205 | ENSMUSG0C | false | . | . | . | . | . | . | chr4 | 148943504 | 148957694 | 6.65935 | 4.74149 | 4.37423 |
| chr7 | 65481945 | 65482193 | 182 | Distal Intergenic | 7 | 65582031 | 65618869 | 36839 | 1 | ENSMUST00 | -99838 | BC046251 | ENSMUSG0C | false | . | . | . | . | . | . | chr7 | 65481395 | 65487238 | 5.07365 | 4.02733 | 3.07685 |
| chr2 | 27324276 | 27324557 | 182 | Promoter (2-3kb) | 2 | 27326704 | 27331297 | 4594 | 1 | ENSMUST00 | -2147 | AA645442 | ENSMUSG0C | false | . | . | . | . | . | . | chr2 | 27323689 | 27324972 | 3.8664 | 3.51102 | 2.08899 |
| chr6 | 32378670 | 32378877 | 182 | Intron (ENSMUST00000115096.3 | 6 | 32451170 | 32454750 | 3581 | 1 | ENSMUST00 | -72293 | Plxna4os2 | ENSMUSG0C | false | . | . | . | . | . | . | . | . | . | . | . | . |
| chr3 | 126667517 | 126667794 | 182 | Promoter (1-2kb) | 3 | 126667329 | 126669032 | 1704 | 2 | ENSMUST00 | 1238 | Gm15551 | ENSMUSG0C | false | . | . | . | . | . | . | chr3 | 126666998 | 126670167 | 5.50406 | 4.20102 | 3.45409 |
| chr8 | 125299233 | 125299505 | 182 | Distal Intergenic | 8 | 125224400 | 125227323 | 2924 | 2 | ENSMUST00 | -71911 | Gm16237 | ENSMUSG0C | false | . | . | . | . | . | . | chr8 | 125299258 | 125300295 | 2.66484 | 2.64234 | 1.15184 |
| chr5 | 123397903 | 123398131 | 182 | Promoter (<=1kb) | 5 | 123393649 | 123397957 | 4309 | 2 | ENSMUST00 | 0 | Gm15747 | ENSMUSG0C | false | chr5 | 123397696 | 123398378 | 7.54473 | 13.39761 | 11.16265 | chr5 | 123395911 | 123400888 | 4.80474 | 3.87842 | 2.88571 |
| chr3 | 156929087 | 156929300 | 182 | Intron (ENSMUST00000074015.5 | 3 | 156903863 | 156920780 | 16918 | 2 | ENSMUST00 | -8308 | Gm15577 | ENSMUSG0C | false | . | . | . | . | . | . | chr3 | 156928412 | 156929769 | 3.36574 | 3.04676 | 1.70241 |
| chr3 | 102743358 | 102743562 | 182 | Intron (ENSMUST00000029451.6 | 3 | 102720231 | 102735417 | 15187 | 2 | ENSMUST00 | -7942 | Tspan2os | ENSMUSG0C | false | . | . | . | . | . | . | chr3 | 102735904 | 102757857 | 6.42245 | 4.63759 | 4.19001 |
| chr6 | 32487228 | 32487539 | 182 | Intron (ENSMUST00000115096.3 | 6 | 32511073 | 32515576 | 4504 | 1 | ENSMUST00 | -23534 | Plxna4os1 | ENSMUSG0C | false | . | . | . | . | . | . | chr6 | 32487114 | 32488078 | 3.40669 | 3.13575 | 1.74675 |
| chr4 | 148915890 | 148916206 | 182 | Promoter (1-2kb) | 4 | 148913511 | 148914324 | 814 | 2 | ENSMUST00 | -1567 | Gm15969 | ENSMUSG0C | false | . | . | . | . | . | . | chr4 | 148901819 | 148933146 | 7.00771 | 4.91323 | 4.66351 |
| chr4 | 149889889 | 149890170 | 182 | Distal Intergenic | 4 | 149904006 | 149908149 | 4144 | 1 | ENSMUST00 | -13836 | Gm13070 | ENSMUSG0C | false | . | . | . | . | . | . | chr4 | 149889628 | 149890679 | 4.37275 | 3.86579 | 2.48791 |
| chr2 | 75788916 | 75789168 | 182 | Distal Intergenic | 2 | 75777188 | 75782306 | 5119 | 1 | ENSMUST00 | 11729 | Gm13657 | ENSMUSG0C | false | . | . | . | . | . | . | . | . | . | . | . | . |
| chr2 | 75792497 | 75792787 | 182 | Distal Intergenic | 2 | 75777188 | 75782306 | 5119 | 1 | ENSMUST00 | 15310 | Gm13657 | ENSMUSG0C | false | . | . | . | . | . | . | . | . | . | . | . | . |
| chr2 | 21141607 | 21142162 | 182 | Promoter (<=1kb) | 2 | 21044808 | 21141954 | 97147 | 2 | ENSMUST00 | 0 | Gm13377 | ENSMUSG0C | false | chr2 | 21141410 | 21142657 | 20.9555 | 57.98476 | 55.16582 | chr2 | 21141746 | 21142740 | 4.87637 | 3.71924 | 2.94036 |
| chr6 | 145175836 | 145176105 | 182 | Exon (ENSMUST00000111727.2/ | 6 | 145182275 | 145187630 | 5356 | 1 | ENSMUST00 | -6170 | Gm15543 | ENSMUSG0C | false | . | . | . | . | . | . | chr6 | 145173875 | 145176214 | 4.62733 | 3.91615 | 2.71166 |
| chr7 | 58844156 | 58844469 | 182 | Distal Intergenic | 7 | 58836793 | 58837938 | 1146 | 2 | ENSMUST00 | -6219 | Gm6226 | ENSMUSG0C | false | . | . | . | . | . | . | chr7 | 58841028 | 58844965 | 5.03896 | 4.01061 | 3.06597 |
| chr11 | 57677689 | 57677948 | 182 | Intron (ENSMUST00000066987.8 | 11 | 57655415 | 57657826 | 2412 | 2 | ENSMUST00 | -19864 | 4933426K07 | ENSMUSG0C | false | . | . | . | . | . | . | chr11 | 57646153 | 57704104 | 7.42247 | 5.08195 | 5.00645 |
| chr11 | 57684961 | 57685240 | 182 | Intron (ENSMUST00000066987.8 | 11 | 57655415 | 57657826 | 2412 | 2 | ENSMUST00 | -27136 | 4933426K07 | ENSMUSG0C | false | . | . | . | . | . | . | chr11 | 57646153 | 57704104 | 7.42247 | 5.08195 | 5.00645 |
| chr11 | 57660826 | 57661099 | 182 | Intron (ENSMUST00000066987.8 | 11 | 57655415 | 57657826 | 2412 | 2 | ENSMUST00 | -3001 | 4933426K07 | ENSMUSG0C | false | . | . | . | . | . | . | chr11 | 57646153 | 57704104 | 7.42247 | 5.08195 | 5.00645 |
| chr4 | 106839013 | 106839259 | 182 | Downstream (2-3kb) | 4 | 106841761 | 106847920 | 6160 | 2 | ENSMUST00 | 8661 | Gm12746 | ENSMUSG0C | false | chr4 | 106839087 | 106840723 | 27.09851 | 84.88239 | 81.64215 | chr4 | 106837700 | 106843971 | 4.90694 | 4.01968 | 2.94365 |
| chr15 | 37433351 | 37433634 | 182 | Promoter (<=1kb) | 15 | 37427397 | 37433414 | 6018 | 2 | ENSMUST00 | 0 | Gm15941 | ENSMUSG0C | false | . | . | . | . | . | . | chr15 | 37431122 | 37434915 | 6.16538 | 4.65218 | 3.95735 |
| chr15 | 37432523 | 37432904 | 182 | Promoter (<=1kb) | 15 | 37421134 | 37432664 | 11531 | 2 | ENSMUST00 | 0 | Gm15941 | ENSMUSG0C | false | . | . | . | . | . | . | chr15 | 37431122 | 37434915 | 6.16538 | 4.65218 | 3.95735 |
| chr2 | 92055343 | 92055665 | 182 | Exon (ENSMUST00000123840.1/ | 2 | 92054989 | 92059369 | 4381 | 2 | ENSMUST00 | 3704 | 2900072N19 | ENSMUSG0C | false | chr2 | 92055270 | 92056117 | 9.7287 | 26.0223 | 23.6086 | . | . | . | . | . | . |
| chr18 | 26005690 | 26006190 | 182 | Distal Intergenic | 18 | 25823581 | 25824357 | 777 | 2 | ENSMUST00 | -181334 | Gm7847 | ENSMUSG0C | false | . | . | . | . | . | . | . | . | . | . | . | . |
| chr13 | 78210140 | 78210551 | 182 | Intron (ENSMUST00000145613.2 | 13 | 78202787 | 78279391 | 76605 | 1 | ENSMUST00 | 7354 | A830082K12 | ENSMUSG0C | false | chr13 | 78209762 | 78210902 | 8.63619 | 18.02194 | 15.71366 | chr13 | 78209605 | 78211050 | 5.23326 | 4.01965 | 3.21806 |
| chr13 | 78199013 | 78199533 | 182 | Promoter (<=1kb) | 13 | 78199738 | 78200515 | 778 | 1 | ENSMUST00 | -205 | A830082K12 | ENSMUSG0C | false | chr13 | 78197737 | 78199944 | 25.05667 | 70.24663 | 67.25255 | chr13 | 78197868 | 78204489 | 4.55487 | 3.83516 | 2.65489 |
| chr2 | 152654517 | 152654801 | 182 | Promoter (<=1kb) | 2 | 152652883 | 152654330 | 1448 | 2 | ENSMUST00 | -188 | 1700030C14 | ENSMUSG0C | false | . | . | . | . | . | . | . | . | . | . | . | . |
| chr6 | 144491857 | 144492278 | 182 | Intron (ENSMUST00000156443.1 | 6 | 144481225 | 144501416 | 20192 | 1 | ENSMUST00 | 10633 | Sox5os2 | ENSMUSG0C | false | . | . | . | . | . | . | chr6 | 144490642 | 144492940 | 4.33759 | 3.76965 | 2.51017 |
| chr7 | 110908620 | 110908831 | 182 | Intron (ENSMUST00000127935.2 | 7 | 110904985 | 110905851 | 867 | 1 | ENSMUST00 | 3636 | Gm16336 | ENSMUSG0C | false | . | . | . | . | . | . | . | . | . | . | . | . |
| chr16 | 5254453 | 5254693 | 182 | Promoter (<=1kb) | 16 | 5254786 | 5289621 | 34836 | 1 | ENSMUST00 | -93 | Gm15983 | ENSMUSG0C | false | chr16 | 5254553 | 5256620 | 29.06336 | 92.21697 | 88.82447 | chr16 | 5253239 | 5256784 | 4.59203 | 3.86124 | 2.68801 |
| chr2 | 21035492 | 21035711 | 182 | Distal Intergenic | 2 | 21097099 | 21120659 | 23561 | 1 | ENSMUST00 | -61388 | Gm13376 | ENSMUSG0C | false | . | . | . | . | . | . | chr2 | 21033559 | 21038134 | 3.93044 | 3.54193 | 2.16761 |
| chr5 | 74001841 | 74002140 | 182 | 3' UTR | 5 | 74014216 | 74029680 | 15465 | 1 | ENSMUST00 | -12076 | Usp46os1 | ENSMUSG0C | false | . | . | . | . | . | . | chr5 | 74001579 | 74008551 | 5.27799 | 4.06988 | 3.27177 |
| chr5 | 67842346 | 67842601 | 182 | Intron (ENSMUST00000037380.9 | 5 | 67847066 | 67909013 | 61948 | 1 | ENSMUST00 | -4465 | Gm15477 | ENSMUSG0C | false | . | . | . | . | . | . | chr5 | 67840654 | 67846460 | 5.77871 | 4.36447 | 3.69241 |
| chr2 | 30677052 | 30677403 | 182 | Promoter (<=1kb) | 2 | 30658613 | 30678011 | 19399 | 2 | ENSMUST00 | 608 | Gm14486 | ENSMUSG0C | false | . | . | . | . | . | . | chr2 | 30676525 | 30678218 | 5.66002 | 4.08031 | 3.59549 |
| chr15 | 97765567 | 97765826 | 182 | Promoter (<=1kb) | 15 | 97766549 | 97768713 | 2165 | 1 | ENSMUST00 | -723 | Rapgef3os2 | ENSMUSG0C | false | chr15 | 97764504 | 97768671 | 37.30079 | 124.19571 | 119.91465 | chr15 | 97747182 | 97765592 | 5.8605 | 4.39118 | 3.71878 |
| chr3 | 90447892 | 90448122 | 182 | Downstream (2-3kb) | 3 | 90451583 | 90457095 | 5513 | 1 | ENSMUST00 | -3461 | Gm16048 | ENSMUSG0C | false | . | . | . | . | . | . | chr3 | 90444443 | 90452742 | 6.22784 | 4.66702 | 4.02057 |
| chr4 | 82530984 | 82531246 | 182 | Intron (ENSMUST00000139835.1 | 4 | 82508016 | 82547292 | 39277 | 1 | ENSMUST00 | 22969 | Gm11266 | ENSMUSG0C | false | . | . | . | . | . | . | . | . | . | . | . | . |
| chr2 | 124848429 | 124848720 | 182 | Distal Intergenic | 2 | 124708655 | 124711083 | 2429 | 1 | ENSMUST00 | 139775 | Gm13994 | ENSMUSG0C | false | . | . | . | . | . | . | . | . | . | . | . | . |
| chr3 | 37937434 | 37937675 | 182 | Distal Intergenic | 3 | 37959612 | 37960087 | 476 | 2 | ENSMUST00 | 22412 | Gm7799 | ENSMUSG0C | false | . | . | . | . | . | . | chr3 | 37933256 | 37939361 | 5.79607 | 4.42106 | 3.66786 |
| chr4 | 142179600 | 142179909 | 182 | Promoter (<=1kb) | 4 | 142180602 | 142184139 | 3538 | 1 | ENSMUST00 | -693 | Gm13062 | ENSMUSG0C | false | . | . | . | . | . | . | chr4 | 142179296 | 142181188 | 5.76053 | 4.33202 | 3.66783 |
| chr14 | 25488220 | 25488512 | 182 | Intron (ENSMUST00000161766.2 | 14 | 25455737 | 25461128 | 5392 | 2 | ENSMUST00 | -27093 | Zmiz1os1 | ENSMUSG0C | false | . | . | . | . | . | . | chr14 | 25486554 | 25489968 | 6.20956 | 4.60644 | 4.00958 |
| chr18 | 33794715 | 33794979 | 182 | Promoter (<=1kb) | 18 | 33794892 | 33795986 | 1095 | 1 | ENSMUST00 | 0 | 2410004N09 | ENSMUSG0C | false | chr18 | 33794397 | 33795757 | 34.92026 | 124.22972 | 119.94103 | . | . | . | . | . | . |
| chr18 | 33741816 | 33742024 | 182 | Distal Intergenic | 18 | 33794892 | 33795986 | 1095 | 1 | ENSMUST00 | -52868 | 2410004N09 | ENSMUSG0C | false | . | . | . | . | . | . | chr18 | 33741166 | 33742472 | 3.36529 | 3.27296 | 1.69573 |
| chr2 | 145675271 | 145675777 | 182 | Promoter (<=1kb) | 2 | 145675367 | 145714989 | 39623 | 1 | ENSMUST00 | 0 | BC039771 | ENSMUSG0C | false | chr2 | 145674554 | 145676544 | 26.52689 | 93.43019 | 90.01563 | . | . | . | . | . | . |
| chr18 | 56774325 | 56774791 | 182 | Intron (ENSMUST00000128240.1 | 18 | 56769717 | 56801688 | 31972 | 1 | ENSMUST00 | 4609 | Gm15345 | ENSMUSG0C | false | chr18 | 56774403 | 56775071 | 5.36929 | 8.07392 | 5.94005 | chr18 | 56774460 | 56775114 | 3.87508 | 3.33244 | 2.10672 |
| chr18 | 56754970 | 56755236 | 182 | Distal Intergenic | 18 | 56769717 | 56801688 | 31972 | 1 | ENSMUST00 | -14481 | Gm15345 | ENSMUSG0C | false | chr18 | 56754441 | 56755308 | 7.72573 | 18.36004 | 16.04866 | chr18 | 56753533 | 56757612 | 3.84422 | 3.46396 | 2.09571 |
| chr16 | 30959839 | 30960100 | 182 | Intron (ENSMUST00000055389.7 | 16 | 30969393 | 30973518 | 4126 | 2 | ENSMUST00 | 13418 | Gm15742 | ENSMUSG0C | false | . | . | . | . | . | . | chr16 | 30959851 | 30970783 | 4.98026 | 4.05336 | 2.99614 |
| chr11 | 64180249 | 64180478 | 182 | Exon (ENSMUST00000145095.1/ | 11 | 64164573 | 64182171 | 17599 | 1 | ENSMUST00 | 15677 | Gm12290 | ENSMUSG0C | false | . | . | . | . | . | . | chr11 | 64179597 | 64181738 | 4.7057 | 3.84436 | 2.7994 |
| chr10 | 83836373 | 83836644 | 182 | Distal Intergenic | 10 | 83722865 | 83762761 | 39897 | 1 | ENSMUST00 | 113509 | 1500009L16 | ENSMUSG0C | false | . | . | . | . | . | . | . | . | . | . | . | . |
| chr11 | 107376299 | 107376647 | 182 | Intron (ENSMUST00000103064.4 | 11 | 107386983 | 107388971 | 1989 | 1 | ENSMUST00 | -10336 | Gm11712 | ENSMUSG0C | false | . | . | . | . | . | . | chr11 | 107374940 | 107381526 | 5.71552 | 4.41302 | 3.58844 |
| chr11 | 107378567 | 107378819 | 182 | Intron (ENSMUST00000103064.4 | 11 | 107386983 | 107388971 | 1989 | 1 | ENSMUST00 | -8164 | Gm11712 | ENSMUSG0C | false | . | . | . | . | . | . | chr11 | 107374940 | 107381526 | 5.71552 | 4.41302 | 3.58844 |
| chr17 | 28930784 | 28931069 | 182 | Intron (ENSMUST00000148464.1 | 17 | 28934421 | 28935078 | 658 | 1 | ENSMUST00 | -3352 | Gm16191 | ENSMUSG0C | false | . | . | . | . | . | . | chr17 | 28929891 | 28931763 | 3.85975 | 3.52949 | 2.06739 |
| chr17 | 86670690 | 86671169 | 182 | Distal Intergenic | 17 | 86703298 | 86703407 | 110 | 2 | ENSMUST00 | 32238 | Gm22235 | ENSMUSG0C | false | . | . | . | . | . | . | . | . | . | . | . | . |
| chr17 | 86711925 | 86712161 | 182 | Distal Intergenic | 17 | 86703298 | 86703407 | 110 | 2 | ENSMUST00 | -8519 | Gm22235 | ENSMUSG0C | false | . | . | . | . | . | . | chr17 | 86711937 | 86713784 | 3.18742 | 3.13019 | 1.50297 |
| chr17 | 86617881 | 86618188 | 182 | Intron (ENSMUST00000097275.3 | 17 | 86703298 | 86703407 | 110 | 2 | ENSMUST00 | 85219 | Gm22235 | ENSMUSG0C | false | . | . | . | . | . | . | chr17 | 86616853 | 86621360 | 5.8599 | 4.38553 | 3.73139 |
| chr17 | 86611772 | 86612048 | 182 | Intron (ENSMUST00000097275.3 | 17 | 86703298 | 86703407 | 110 | 2 | ENSMUST00 | 91359 | Gm22235 | ENSMUSG0C | false | . | . | . | . | . | . | chr17 | 86610058 | 86615360 | 5.61564 | 4.2317 | 3.54393 |
| chr4 | 34425786 | 34426040 | 182 | Distal Intergenic | 4 | 34434296 | 34434401 | 106 | 1 | ENSMUST00 | -8256 | Gm23466 | ENSMUSG0C | false | . | . | . | . | . | . | . | . | . | . | . | . |
| chr3 | 100735643 | 100735907 | 182 | Promoter (2-3kb) | 3 | 100738452 | 100738713 | 262 | 2 | ENSMUST00 | 2806 | Gm23465 | ENSMUSG0C | false | . | . | . | . | . | . | chr3 | 100734767 | 100736545 | 3.97181 | 3.47992 | 2.1968 |
| chr10 | 70483786 | 70484027 | 182 | Intron (ENSMUST00000105436.3 | 10 | 70501192 | 70501503 | 312 | 2 | ENSMUST00 | 17476 | Gm22320 | ENSMUSG0C | false | . | . | . | . | . | . | chr10 | 70483150 | 70485810 | 4.17553 | 3.5308 | 2.35529 |
| chr3 | 132835122 | 132835435 | 182 | Intron (ENSMUST00000169172.2 | 3 | 132787225 | 132787335 | 111 | 2 | ENSMUST00 | -47788 | Gm22322 | ENSMUSG0C | false | . | . | . | . | . | . | . | . | . | . | . | . |
| chr6 | 30352853 | 30353102 | 182 | Distal Intergenic | 6 | 30347371 | 30347466 | 96 | 2 | ENSMUST00 | -5388 | Gm25625 | ENSMUSG0C | false | . | . | . | . | . | . | chr6 | 30352472 | 30353968 | 5.73359 | 4.44105 | 3.65722 |
| chr6 | 47681001 | 47681842 | 182 | Distal Intergenic | 6 | 47637143 | 47637219 | 77 | 1 | ENSMUST00 | 43859 | Gm25283 | ENSMUSG0C | false | . | . | . | . | . | . | . | . | . | . | . | . |
| chr6 | 47663704 | 47664155 | 182 | Distal Intergenic | 6 | 47637143 | 47637219 | 77 | 1 | ENSMUST00 | 26562 | Gm25283 | ENSMUSG0C | false | chr6 | 47663853 | 47664155 | 22.15371 | 67.81625 | 64.85734 | . | . | . | . | . | . |
| chr6 | 47655057 | 47655516 | 182 | Distal Intergenic | 6 | 47637143 | 47637219 | 77 | 1 | ENSMUST00 | 17915 | Gm25283 | ENSMUSG0C | false | chr6 | 47654721 | 47655533 | 15.44916 | 124.7347 | 120.43309 | chr6 | 47654282 | 47656331 | 10.90277 | 4.33282 | 7.99539 |
| chr6 | 47676660 | 47677152 | 182 | Distal Intergenic | 6 | 47637143 | 47637219 | 77 | 1 | ENSMUST00 | 39518 | Gm25283 | ENSMUSG0C | false | . | . | . | . | . | . | . | . | . | . | . | . |
| chr6 | 47672338 | 47673128 | 182 | Distal Intergenic | 6 | 47637143 | 47637219 | 77 | 1 | ENSMUST00 | 35196 | Gm25283 | ENSMUSG0C | false | chr6 | 47672087 | 47673247 | 11.30501 | 147.47621 | 142.28978 | chr6 | 47671883 | 47673500 | 10.26163 | 3.78615 | 7.35578 |
| chr6 | 47685301 | 47685764 | 182 | Distal Intergenic | 6 | 47637143 | 47637219 | 77 | 1 | ENSMUST00 | 48159 | Gm25283 | ENSMUSG0C | false | . | . | . | . | . | . | . | . | . | . | . | . |
| chr6 | 47668033 | 47668503 | 182 | Distal Intergenic | 6 | 47637143 | 47637219 | 77 | 1 | ENSMUST00 | 30891 | Gm25283 | ENSMUSG0C | false | chr6 | 47668287 | 47668554 | 16.43574 | 91.94095 | 88.5556 | chr6 | 47668334 | 47669306 | 6.54991 | 3.85602 | 4.29221 |
| chr6 | 47659344 | 47660136 | 182 | Distal Intergenic | 6 | 47637143 | 47637219 | 77 | 1 | ENSMUST00 | 22202 | Gm25283 | ENSMUSG0C | false | . | . | . | . | . | . | . | . | . | . | . | . |
| chr7 | 118277656 | 118277885 | 182 | Promoter (1-2kb) | 7 | 118279861 | 118279965 | 105 | 1 | ENSMUST00 | -1976 | Gm23229 | ENSMUSG0C | false | chr7 | 118277147 | 118277690 | 10.54274 | 25.8059 | 23.39399 | chr7 | 118274222 | 118279022 | 5.59058 | 4.35566 | 3.51383 |
| chr4 | 62879878 | 62880083 | 182 | Distal Intergenic | 4 | 62939341 | 62939474 | 134 | 1 | ENSMUST00 | -59258 | Gm25391 | ENSMUSG0C | false | . | . | . | . | . | . | chr4 | 62878464 | 62879969 | 3.51867 | 3.22245 | 1.81692 |
| chr4 | 127278620 | 127278889 | 182 | Distal Intergenic | 4 | 127284731 | 127284833 | 103 | 2 | ENSMUST00 | 5944 | Gm22221 | ENSMUSG0C | false | . | . | . | . | . | . | . | . | . | . | . | . |
| chr11 | 119387320 | 119387513 | 182 | Promoter (2-3kb) | 11 | 119390472 | 119390561 | 90 | 1 | ENSMUST00 | -2959 | Mir1932 | ENSMUSG0C | false | . | . | . | . | . | . | chr11 | 119386374 | 119396745 | 5.43205 | 4.26989 | 3.36389 |
| chr8 | 3803120 | 3803372 | 182 | Promoter (<=1kb) | 8 | 3803069 | 3803358 | 290 | 2 | ENSMUST00 | 0 | Rprl3 | ENSMUSG0C | false | . | . | . | . | . | . | . | . | . | . | . | . |
| chr4 | 131261289 | 131261570 | 182 | Distal Intergenic | 4 | 131239277 | 131239410 | 134 | 1 | ENSMUST00 | 22013 | Gm25261 | ENSMUSG0C | false | . | . | . | . | . | . | chr4 | 131258846 | 131261997 | 5.70489 | 4.1926 | 3.61984 |
| chr4 | 131173801 | 131174028 | 182 | Distal Intergenic | 4 | 131239277 | 131239410 | 134 | 1 | ENSMUST00 | -65249 | Gm25261 | ENSMUSG0C | false | chr4 | 131173889 | 131174066 | 5.05485 | 7.29253 | 5.17558 | chr4 | 131173511 | 131174506 | 4.83502 | 3.97675 | 2.91231 |
| chr5 | 90892774 | 90893003 | 182 | Promoter (<=1kb) | 5 | 90893380 | 90893496 | 117 | 1 | ENSMUST00 | -377 | Gm22816 | ENSMUSG0C | false | . | . | . | . | . | . | chr5 | 90890713 | 90893350 | 3.86829 | 3.55076 | 2.0801 |
| chr5 | 132928838 | 132929018 | 182 | Distal Intergenic | 5 | 132801678 | 132801956 | 279 | 1 | ENSMUST00 | 127161 | Gm25722 | ENSMUSG0C | false | . | . | . | . | . | . | chr5 | 132927590 | 132930738 | 3.27414 | 3.16581 | 1.60252 |
| chr7 | 135598526 | 135598771 | 182 | Intron (ENSMUST00000073961.6 | 7 | 135566162 | 135566282 | 121 | 1 | ENSMUST00 | 32365 | Mir1962 | ENSMUSG0C | false | . | . | . | . | . | . | . | . | . | . | . | . |
| chr3 | 108554629 | 108554905 | 182 | Promoter (<=1kb) | 3 | 108554338 | 108554751 | 414 | 2 | ENSMUST00 | 0 | Scarna2 | ENSMUSG0C | false | chr3 | 108554052 | 108554913 | 27.78601 | 91.88203 | 88.49705 | . | . | . | . | . | . |
| chr17 | 83935712 | 83935974 | 182 | Distal Intergenic | 17 | 83947588 | 83947696 | 109 | 1 | ENSMUST00 | -11614 | Gm23905 | ENSMUSG0C | false | . | . | . | . | . | . | chr17 | 83935321 | 83943115 | 7.64209 | 5.27918 | 5.16816 |
| chr17 | 83954504 | 83954781 | 182 | Distal Intergenic | 17 | 83947588 | 83947696 | 109 | 1 | ENSMUST00 | 6917 | Gm23905 | ENSMUSG0C | false | . | . | . | . | . | . | chr17 | 83952763 | 83964037 | 8.60917 | 5.64195 | 5.95295 |
| chr12 | 31895948 | 31896172 | 182 | Intron (ENSMUST00000036862.4 | 12 | 31858458 | 31858592 | 135 | 1 | ENSMUST00 | 37491 | Gm25913 | ENSMUSG0C | false | . | . | . | . | . | . | chr12 | 31895315 | 31896699 | 3.61625 | 3.3445 | 1.9048 |
| chr15 | 72688601 | 72688834 | 182 | Intron (ENSMUST00000023276.9 | 15 | 72721321 | 72721387 | 67 | 2 | ENSMUST00 | 32553 | Gm28020 | ENSMUSG0C | false | . | . | . | . | . | . | chr15 | 72686038 | 72689611 | 5.42595 | 4.11861 | 3.39229 |
| chr15 | 72774791 | 72775000 | 182 | Intron (ENSMUST00000023276.9 | 15 | 72721321 | 72721387 | 67 | 2 | ENSMUST00 | -53405 | Gm28020 | ENSMUSG0C | false | . | . | . | . | . | . | chr15 | 72772332 | 72775331 | 4.37032 | 3.67134 | 2.53137 |
| chr15 | 72773796 | 72774044 | 182 | Intron (ENSMUST00000023276.9 | 15 | 72721321 | 72721387 | 67 | 2 | ENSMUST00 | -52410 | Gm28020 | ENSMUSG0C | false | . | . | . | . | . | . | chr15 | 72772332 | 72775331 | 4.37032 | 3.67134 | 2.53137 |
| chr3 | 138526938 | 138527546 | 182 | Promoter (<=1kb) | 3 | 138526415 | 138526494 | 80 | 1 | ENSMUST00 | 524 | Mir1956 | ENSMUSG0C | false | chr3 | 138526131 | 138527087 | 25.49012 | 74.6041 | 71.54089 | . | . | . | . | . | . |
| chr12 | 54256879 | 54257091 | 182 | Distal Intergenic | 12 | 54240755 | 54240900 | 146 | 2 | ENSMUST00 | -15980 | Gm24692 | ENSMUSG0C | false | . | . | . | . | . | . | chr12 | 54256532 | 54258234 | 5.10618 | 4.14897 | 3.08011 |
| chr12 | 54265980 | 54266207 | 182 | Distal Intergenic | 12 | 54240755 | 54240900 | 146 | 2 | ENSMUST00 | -25081 | Gm24692 | ENSMUSG0C | false | . | . | . | . | . | . | chr12 | 54259658 | 54266087 | 5.59493 | 4.35111 | 3.49425 |
| chr9 | 44396855 | 44397139 | 182 | Promoter (<=1kb) | 9 | 44395797 | 44395915 | 119 | 2 | ENSMUST00 | -941 | Gm26306 | ENSMUSG0C | false | chr9 | 44396250 | 44397700 | 17.10421 | 70.31819 | 67.3235 | chr9 | 44395975 | 44399288 | 3.95836 | 3.52211 | 2.19331 |
| chr17 | 83274044 | 83274295 | 182 | Distal Intergenic | 17 | 83269971 | 83270105 | 135 | 1 | ENSMUST00 | 4074 | Gm24240 | ENSMUSG0C | false | . | . | . | . | . | . | chr17 | 83272007 | 83278538 | 5.76726 | 4.43266 | 3.64322 |
| chr10 | 17367184 | 17367395 | 182 | Distal Intergenic | 10 | 17116804 | 17116900 | 97 | 1 | ENSMUST00 | 250381 | Gm25382 | ENSMUSG0C | false | . | . | . | . | . | . | chr10 | 17365343 | 17367652 | 3.18472 | 2.8961 | 1.55843 |
| chr10 | 17411421 | 17411665 | 182 | Distal Intergenic | 10 | 17116804 | 17116900 | 97 | 1 | ENSMUST00 | 294618 | Gm25382 | ENSMUSG0C | false | . | . | . | . | . | . | . | . | . | . | . | . |
| chr1 | 184854092 | 184854368 | 182 | Distal Intergenic | 1 | 184858138 | 184858412 | 275 | 2 | ENSMUST00 | 4044 | Gm23780 | ENSMUSG0C | false | . | . | . | . | . | . | chr1 | 184846562 | 184855069 | 5.5375 | 4.29731 | 3.4563 |
| chr8 | 127624836 | 127625037 | 182 | Distal Intergenic | 8 | 127597706 | 127597823 | 118 | 1 | ENSMUST00 | 27131 | Gm22968 | ENSMUSG0C | false | . | . | . | . | . | . | chr8 | 127624250 | 127625180 | 3.29317 | 3.23041 | 1.64701 |
| chr8 | 127858129 | 127858370 | 182 | Distal Intergenic | 8 | 127597706 | 127597823 | 118 | 1 | ENSMUST00 | 260424 | Gm22968 | ENSMUSG0C | false | . | . | . | . | . | . | chr8 | 127854533 | 127858409 | 4.46793 | 3.7993 | 2.57626 |
| chr8 | 127675413 | 127675683 | 182 | Distal Intergenic | 8 | 127597706 | 127597823 | 118 | 1 | ENSMUST00 | 77708 | Gm22968 | ENSMUSG0C | false | . | . | . | . | . | . | chr8 | 127673630 | 127677018 | 5.03984 | 4.09361 | 3.04381 |
| chr8 | 127849838 | 127850181 | 182 | Distal Intergenic | 8 | 127597706 | 127597823 | 118 | 1 | ENSMUST00 | 252133 | Gm22968 | ENSMUSG0C | false | . | . | . | . | . | . | . | . | . | . | . | . |
| chr19 | 18819213 | 18819440 | 182 | Intron (ENSMUST00000040489.7 | 19 | 18776746 | 18776961 | 216 | 2 | ENSMUST00 | -42253 | Gm24190 | ENSMUSG0C | false | . | . | . | . | . | . | chr19 | 18818709 | 18820118 | 4.30452 | 3.75638 | 2.43635 |
| chr6 | 93648608 | 93648885 | 182 | Distal Intergenic | 6 | 93509573 | 93509702 | 130 | 1 | ENSMUST00 | 139036 | Gm22312 | ENSMUSG0C | false | . | . | . | . | . | . | . | . | . | . | . | . |
| chr6 | 93462914 | 93463267 | 182 | Distal Intergenic | 6 | 93509573 | 93509702 | 130 | 1 | ENSMUST00 | -46306 | Gm22312 | ENSMUSG0C | false | . | . | . | . | . | . | . | . | . | . | . | . |
| chr10 | 14978243 | 14978684 | 182 | Distal Intergenic | 10 | 14984670 | 14984799 | 130 | 1 | ENSMUST00 | -5986 | Gm23892 | ENSMUSG0C | false | . | . | . | . | . | . | chr10 | 14977867 | 14978719 | 2.98348 | 2.94797 | 1.36275 |
| chr4 | 45500394 | 45500582 | 182 | Promoter (<=1kb) | 4 | 45500485 | 45500552 | 68 | 2 | ENSMUST00 | 0 | Gm22518 | ENSMUSG0C | false | . | . | . | . | . | . | chr4 | 45486900 | 45516085 | 7.16373 | 5.03069 | 4.78278 |
| chr19 | 60999564 | 61000052 | 182 | Intron (ENSMUST00000003313.8 | 19 | 61013417 | 61013545 | 129 | 2 | ENSMUST00 | 13493 | Gm22520 | ENSMUSG0C | false | . | . | . | . | . | . | chr19 | 60997276 | 61001132 | 5.39691 | 4.33144 | 3.34485 |
| chr19 | 5433596 | 5433835 | 182 | Promoter (2-3kb) | 19 | 5435900 | 5435965 | 66 | 1 | ENSMUST00 | -2065 | Gm25432 | ENSMUSG0C | false | chr19 | 5433448 | 5433907 | 8.3822 | 16.12319 | 13.84377 | chr19 | 5432781 | 5434371 | 5.26915 | 4.16119 | 3.26333 |
| chr1 | 9822849 | 9823094 | 182 | Intron (ENSMUST00000166384.2 | 1 | 9834498 | 9834602 | 105 | 2 | ENSMUST00 | 11508 | Gm22607 | ENSMUSG0C | false | . | . | . | . | . | . | chr1 | 9821367 | 9824738 | 6.25682 | 4.62326 | 4.06263 |
| chr13 | 82258482 | 82258876 | 182 | Distal Intergenic | 13 | 82050751 | 82050881 | 131 | 1 | ENSMUST00 | 207732 | Gm24295 | ENSMUSG0C | false | . | . | . | . | . | . | . | . | . | . | . | . |
| chr16 | 74093169 | 74093369 | 182 | Intron (ENSMUST00000117785.2 | 16 | 74058899 | 74059033 | 135 | 2 | ENSMUST00 | -34137 | Gm22163 | ENSMUSG0C | false | . | . | . | . | . | . | chr16 | 74090760 | 74094201 | 4.77226 | 3.99214 | 2.80493 |
| chr1 | 186478618 | 186478888 | 182 | Distal Intergenic | 1 | 186676742 | 186676876 | 135 | 1 | ENSMUST00 | -197854 | Gm26169 | ENSMUSG0C | false | . | . | . | . | . | . | . | . | . | . | . | . |
| chr6 | 93849534 | 93849798 | 182 | Intron (ENSMUST00000093769.5 | 6 | 93890615 | 93890750 | 136 | 1 | ENSMUST00 | -40817 | Gm22840 | ENSMUSG0C | false | . | . | . | . | . | . | . | . | . | . | . | . |
| chr12 | 80574597 | 80574931 | 182 | Intron (ENSMUST00000021558.6 | 12 | 80622830 | 80622933 | 104 | 2 | ENSMUST00 | 48002 | Gm23817 | ENSMUSG0C | false | . | . | . | . | . | . | chr12 | 80574596 | 80575774 | 6.2777 | 4.61134 | 4.09103 |
| chr13 | 43864402 | 43864723 | 182 | Distal Intergenic | 13 | 43872802 | 43872909 | 108 | 1 | ENSMUST00 | -8079 | Gm23117 | ENSMUSG0C | false | chr13 | 43864248 | 43864590 | 8.8433 | 18.37784 | 16.06619 | chr13 | 43864129 | 43865741 | 5.12202 | 4.20733 | 3.10266 |
| chr6 | 118839479 | 118839793 | 182 | Intron (ENSMUST00000186889.1 | 6 | 118834900 | 118835000 | 101 | 1 | ENSMUST00 | 4580 | Gm25905 | ENSMUSG0C | false | . | . | . | . | . | . | chr6 | 118838856 | 118839519 | 3.22155 | 3.08944 | 1.58574 |
| chr6 | 118864658 | 118865012 | 182 | Intron (ENSMUST00000186889.1 | 6 | 118834900 | 118835000 | 101 | 1 | ENSMUST00 | 29759 | Gm25905 | ENSMUSG0C | false | . | . | . | . | . | . | . | . | . | . | . | . |
| chr11 | 72803681 | 72803967 | 182 | Intron (ENSMUST00000150551.2 | 11 | 72800287 | 72800478 | 192 | 2 | ENSMUST00 | -3204 | Gm22075 | ENSMUSG0C | false | . | . | . | . | . | . | chr11 | 72796653 | 72812302 | 6.21579 | 4.59023 | 4.00723 |
| chr15 | 97945305 | 97945787 | 182 | Distal Intergenic | 15 | 97935285 | 97935595 | 311 | 1 | ENSMUST00 | 10021 | Gm22202 | ENSMUSG0C | false | . | . | . | . | . | . | chr15 | 97945004 | 97946228 | 6.12741 | 4.57177 | 3.96252 |
| chr15 | 97925775 | 97926009 | 182 | Distal Intergenic | 15 | 97935285 | 97935595 | 311 | 1 | ENSMUST00 | -9276 | Gm22202 | ENSMUSG0C | false | . | . | . | . | . | . | chr15 | 97924986 | 97928566 | 6.27623 | 4.66463 | 4.05606 |
| chr3 | 153689920 | 153690248 | 182 | Intron (ENSMUST00000064460.5 | 3 | 153708911 | 153709026 | 116 | 1 | ENSMUST00 | -18663 | Gm22206 | ENSMUSG0C | false | . | . | . | . | . | . | chr3 | 153686316 | 153712771 | 6.43925 | 4.66349 | 4.19773 |
| chr16 | 87050029 | 87050344 | 182 | Distal Intergenic | 16 | 86845470 | 86845594 | 125 | 1 | ENSMUST00 | 204560 | Gm25715 | ENSMUSG0C | false | . | . | . | . | . | . | . | . | . | . | . | . |
| chr11 | 57962463 | 57962914 | 182 | Distal Intergenic | 11 | 57969342 | 57969612 | 271 | 1 | ENSMUST00 | -6428 | Gm26025 | ENSMUSG0C | false | chr11 | 57962276 | 57963391 | 27.40132 | 82.57204 | 79.37107 | chr11 | 57961875 | 57963469 | 3.24715 | 3.15891 | 1.55959 |
| chr7 | 73662720 | 73663006 | 182 | Promoter (<=1kb) | 7 | 73662048 | 73662300 | 253 | 1 | ENSMUST00 | 673 | Gm23020 | ENSMUSG0C | false | . | . | . | . | . | . | . | . | . | . | . | . |
| chr7 | 79313510 | 79313801 | 182 | Intron (ENSMUST00000037315.7 | 7 | 79352488 | 79352745 | 258 | 2 | ENSMUST00 | 38944 | Gm24224 | ENSMUSG0C | false | . | . | . | . | . | . | chr7 | 79311920 | 79316435 | 5.38863 | 4.18655 | 3.35018 |
| chr7 | 79318388 | 79318662 | 182 | Intron (ENSMUST00000037315.7 | 7 | 79352488 | 79352745 | 258 | 2 | ENSMUST00 | 34083 | Gm24224 | ENSMUSG0C | false | . | . | . | . | . | . | . | . | . | . | . | . |
| chr11 | 89186894 | 89187137 | 182 | Distal Intergenic | 11 | 89136505 | 89136766 | 262 | 2 | ENSMUST00 | -50129 | Gm24734 | ENSMUSG0C | false | chr11 | 89186667 | 89187483 | 11.17602 | 23.97021 | 21.58181 | chr11 | 89186020 | 89187560 | 4.23766 | 3.47855 | 2.41442 |
| chr17 | 72478133 | 72478336 | 182 | Intron (ENSMUST00000086639.4 | 17 | 72499289 | 72499418 | 130 | 2 | ENSMUST00 | 21082 | Gm24736 | ENSMUSG0C | false | . | . | . | . | . | . | chr17 | 72476588 | 72478745 | 5.4537 | 4.07477 | 3.42383 |
| chr13 | 99047338 | 99047529 | 182 | Distal Intergenic | 13 | 99040398 | 99040650 | 253 | 2 | ENSMUST00 | -6689 | Gm24733 | ENSMUSG0C | false | . | . | . | . | . | . | chr13 | 99046944 | 99047433 | 3.08306 | 3.0949 | 1.469 |
| chr16 | 8163982 | 8164309 | 182 | Distal Intergenic | 16 | 8203975 | 8204248 | 274 | 1 | ENSMUST00 | -39666 | Gm23476 | ENSMUSG0C | false | . | . | . | . | . | . | . | . | . | . | . | . |
| chr16 | 8178893 | 8179351 | 182 | Distal Intergenic | 16 | 8203975 | 8204248 | 274 | 1 | ENSMUST00 | -24624 | Gm23476 | ENSMUSG0C | false | . | . | . | . | . | . | . | . | . | . | . | . |
| chr17 | 63111041 | 63111250 | 182 | Intron (ENSMUST00000024761.7 | 17 | 63096890 | 63097000 | 111 | 2 | ENSMUST00 | -14042 | Gm25348 | ENSMUSG0C | false | . | . | . | . | . | . | chr17 | 63110799 | 63111913 | 3.62121 | 3.37018 | 1.91875 |
| chr2 | 104205620 | 104205843 | 182 | Intron (ENSMUST00000141159.2 | 2 | 104228112 | 104228238 | 127 | 2 | ENSMUST00 | 22395 | Gm22501 | ENSMUSG0C | false | . | . | . | . | . | . | chr2 | 104205122 | 104205982 | 3.30956 | 3.03539 | 1.64046 |
| chr17 | 10411243 | 10411424 | 182 | Intron (ENSMUST00000041463.6 | 17 | 10456216 | 10474123 | 17908 | 1 | ENSMUST00 | -44792 | A230009B12 | ENSMUSG0C | false | . | . | . | . | . | . | chr17 | 10410257 | 10411705 | 4.06384 | 3.50321 | 2.25548 |
| chr6 | 85936884 | 85937382 | 182 | Distal Intergenic | 6 | 85932681 | 85933379 | 699 | 2 | ENSMUST00 | -3506 | Nat8b | ENSMUSG0C | false | chr6 | 85936553 | 85937879 | 21.0928 | 59.0607 | 56.22787 | chr6 | 85936111 | 85937850 | 3.00115 | 2.97707 | 1.4196 |
| chr11 | 69685720 | 69686005 | 182 | Promoter (<=1kb) | 11 | 69682932 | 69685784 | 2853 | 2 | ENSMUST00 | 0 | Tnfsf13 | ENSMUSG0C | false | . | . | . | . | . | . | chr11 | 69680256 | 69687418 | 8.8727 | 5.82026 | 6.17215 |
| chr15 | 86094801 | 86095023 | 182 | Intron (ENSMUST00000088931.4 | 15 | 86087640 | 86090060 | 2421 | 2 | ENSMUST00 | -4742 | Gm15722 | ENSMUSG0C | false | . | . | . | . | . | . | chr15 | 86093572 | 86096550 | 6.54504 | 4.84352 | 4.26819 |
| chr17 | 87331015 | 87331347 | 182 | Promoter (<=1kb) | 17 | 87329764 | 87332052 | 2289 | 2 | ENSMUST00 | 705 | Gm15978 | ENSMUSG0C | false | . | . | . | . | . | . | chr17 | 87325598 | 87334451 | 7.19659 | 4.99755 | 4.81063 |
| chr2 | 30470713 | 30470956 | 182 | Downstream (<1kb) | 2 | 30471537 | 30474219 | 2683 | 2 | ENSMUST00 | 3263 | Ier5l | ENSMUSG0C | false | chr2 | 30469735 | 30471289 | 20.26563 | 55.95697 | 53.16479 | chr2 | 30469252 | 30473405 | 4.8209 | 4.06708 | 2.84851 |
| chr1 | 164199569 | 164199749 | 182 | Intron (ENSMUST00000086040.5 | 1 | 164166579 | 164166933 | 355 | 1 | ENSMUST00 | 32991 | Gm16588 | ENSMUSG0C | false | . | . | . | . | . | . | chr1 | 164199590 | 164200260 | 3.8369 | 3.40278 | 2.09512 |
| chr13 | 44085700 | 44086007 | 182 | Distal Intergenic | 13 | 44121167 | 44125179 | 4013 | 1 | ENSMUST00 | -35160 | Gm5083 | ENSMUSG0C | false | . | . | . | . | . | . | chr13 | 44085323 | 44086822 | 3.98593 | 3.60332 | 2.17107 |

| | | | | | | | | | | | | | | | | | | | | | | | | | | |
|---|---|---|---|---|---|---|---|---|---|---|---|---|---|---|---|---|---|---|---|---|---|---|---|---|---|---|
| chr8 | 98996070 | 98996278 | 182 | Distal Intergenic | 8 | 99011886 | 99032740 | 20855 | 1 | ENSMUST00 | -15608 | Gm15679 | ENSMUSG0( | false | . | . | . | . | . | . | . | . | . | . | . | . |
| chr1 | 58341646 | 58341947 | 182 | Intron (ENSMUST00000114366.2 | 1 | 58254117 | 58315656 | 61540 | 2 | ENSMUST00 | -25991 | Gm15759 | ENSMUSG0( | false | . | . | . | . | . | . | chr1 | 58338368 | 58342831 | 5.64738 | 4.2411 | 3.58443 |
| chr8 | 71908729 | 71909103 | 182 | Promoter (<=1kb) | 8 | 71908905 | 71913587 | 4683 | 1 | ENSMUST00 | 0 | Zfp882 | ENSMUSG0( | false | chr8 | 71908250 | 71909533 | 27.78601 | 91.88203 | 88.49705 | . | . | . | . | . | . |
| chr6 | 8259307 | 8259522 | 182 | Promoter (<=1kb) | 6 | 8259350 | 8427138 | 167789 | 1 | ENSMUST00 | 0 | Gm16039 | ENSMUSG0( | false | chr6 | 8258438 | 8260172 | 36.10317 | 115.72481 | 111.71513 | . | . | . | . | . | . |
| chr17 | 10527824 | 10528135 | 182 | Intron (ENSMUST00000161274.1 | 17 | 10496453 | 10553169 | 56717 | 1 | ENSMUST00 | 31372 | Gm16168 | ENSMUSG0( | false | . | . | . | . | . | . | chr17 | 10526632 | 10533668 | 4.66575 | 3.85507 | 2.75675 |
| chr10 | 86831225 | 86831629 | 182 | Intron (ENSMUST00000099396.2 | 10 | 86839094 | 86874954 | 35861 | 1 | ENSMUST00 | -7465 | Gm16268 | ENSMUSG0( | false | . | . | . | . | . | . | . | . | . | . | . | . |
| chr8 | 69008097 | 69008346 | 182 | Distal Intergenic | 8 | 69044956 | 69063422 | 18467 | 1 | ENSMUST00 | -36610 | Gm15716 | ENSMUSG0( | false | . | . | . | . | . | . | . | . | . | . | . | . |
| chr8 | 14961550 | 14961757 | 182 | Promoter (<=1kb) | 8 | 14961066 | 14961586 | 521 | 2 | ENSMUST00 | 0 | Gm16350 | ENSMUSG0( | false | . | . | . | . | . | . | chr8 | 14960786 | 14962773 | 4.76816 | 3.96097 | 2.8455 |
| chr8 | 127544238 | 127544412 | 182 | Intron (ENSMUST00000162309.2 | 8 | 127561435 | 127561869 | 435 | 1 | ENSMUST00 | -17023 | Gm6921 | ENSMUSG0( | false | . | . | . | . | . | . | chr8 | 127541083 | 127545342 | 4.55577 | 3.71778 | 2.67228 |
| chr1 | 132375426 | 132375643 | 182 | Intron (ENSMUST00000045473.1 | 1 | 132381129 | 132382509 | 1381 | 1 | ENSMUST00 | -5486 | Gm15849 | ENSMUSG0( | false | . | . | . | . | . | . | chr1 | 132369807 | 132377989 | 6.45268 | 4.60038 | 4.22615 |
| chr5 | 123135532 | 123135815 | 182 | Promoter (2-3kb) | 5 | 123133723 | 123138109 | 4387 | 2 | ENSMUST00 | 2294 | AI480526 | ENSMUSG0( | false | chr5 | 123135355 | 123136555 | 12.76676 | 28.45456 | 26.00909 | chr5 | 123131739 | 123139661 | 6.68632 | 4.77641 | 4.39554 |
| chr9 | 104063218 | 104063726 | 182 | Promoter (<=1kb) | 9 | 104063377 | 104127463 | 64087 | 1 | ENSMUST00 | 0 | Acad11 | ENSMUSG0( | false | chr9 | 104061947 | 104064222 | 37.85821 | 131.67731 | 127.12434 | . | . | . | . | . | . |
| chr16 | 26047844 | 26048172 | 182 | Intron (ENSMUST00000039990.5 | 16 | 26031810 | 26040870 | 9061 | 2 | ENSMUST00 | -6975 | AU015336 | ENSMUSG0( | false | . | . | . | . | . | . | chr16 | 26035252 | 26056185 | 6.03158 | 4.46161 | 3.87334 |
| chr10 | 71025226 | 71025520 | 182 | Intron (ENSMUST00000143791.2 | 10 | 71082876 | 71083651 | 776 | 1 | ENSMUST00 | -57356 | Gm15647 | ENSMUSG0( | false | chr10 | 71024507 | 71025449 | 9.33584 | 19.216 | 16.89141 | chr10 | 71023281 | 71029421 | 3.96647 | 3.49689 | 2.18756 |
| chr10 | 128923010 | 128923616 | 182 | Promoter (<=1kb) | 10 | 128917882 | 128923524 | 5643 | 2 | ENSMUST00 | 0 | Bloc1s1 | ENSMUSG0( | false | chr10 | 128922352 | 128924170 | 31.96831 | 119.80809 | 115.67081 | chr10 | 128919550 | 128924490 | 5.64648 | 4.33088 | 3.54938 |
| chr1 | 162037950 | 162038238 | 182 | Intron (ENSMUST00000159763.2 | 1 | 162005172 | 162016494 | 11323 | 1 | ENSMUST00 | 32779 | 2810442N19 | ENSMUSG0( | false | . | . | . | . | . | . | chr1 | 162036675 | 162039902 | 3.94508 | 3.47175 | 2.19039 |
| chr14 | 43939410 | 43939600 | 182 | Promoter (1-2kb) | 14 | 43940997 | 43941858 | 862 | 1 | ENSMUST00 | -1397 | Gm9550 | ENSMUSG0( | false | . | . | . | . | . | . | . | . | . | . | . | . |
| chr1 | 160044271 | 160044766 | 182 | Promoter (<=1kb) | 1 | 160044384 | 160071536 | 27153 | 1 | ENSMUST00 | 0 | 4930523C07 | ENSMUSG0( | false | chr1 | 160044086 | 160045572 | 25.53309 | 82.08177 | 78.89293 | chr1 | 160043756 | 160061722 | 5.00419 | 4.03176 | 3.02332 |
| chr18 | 78311395 | 78311645 | 182 | Distal Intergenic | 18 | 78349754 | 78350701 | 948 | 1 | ENSMUST00 | -38109 | Gm6133 | ENSMUSG0( | false | . | . | . | . | . | . | chr18 | 78311113 | 78313785 | 4.98111 | 4.09217 | 3.04672 |
| chr5 | 113844193 | 113844449 | 182 | Promoter (1-2kb) | 5 | 113842601 | 113843116 | 516 | 1 | ENSMUST00 | 1593 | Gm17122 | ENSMUSG0( | false | . | . | . | . | . | . | . | . | . | . | . | . |
| chr1 | 90953894 | 90954179 | 182 | Promoter (<=1kb) | 1 | 90953108 | 90954027 | 920 | 1 | ENSMUST00 | 787 | Prlh | ENSMUSG0( | false | . | . | . | . | . | . | . | . | . | . | . | . |
| chr10 | 99440985 | 99441358 | 182 | Promoter (2-3kb) | 10 | 99443730 | 99447007 | 3278 | 1 | ENSMUST00 | -2372 | Gad1-ps | ENSMUSG0( | false | . | . | . | . | . | . | chr10 | 99440376 | 99441474 | 3.51972 | 3.35549 | 1.83724 |
| chr14 | 45049366 | 45049545 | 182 | Distal Intergenic | 14 | 44972268 | 45006900 | 34633 | 2 | ENSMUST00 | -42467 | Gm17173 | ENSMUSG0( | false | . | . | . | . | . | . | . | . | . | . | . | . |
| chr17 | 34958909 | 34959523 | 182 | Promoter (<=1kb) | 17 | 34956436 | 34959238 | 2803 | 2 | ENSMUST00 | 0 | Hspa1b | ENSMUSG0( | false | chr17 | 34958204 | 34960170 | 16.96705 | 61.94942 | 59.07605 | . | . | . | . | . | . |
| chr14 | 24213361 | 24213615 | 182 | Intron (ENSMUST00000073687.7 | 14 | 24238172 | 24243391 | 5220 | 2 | ENSMUST00 | 29776 | Gm17105 | ENSMUSG0( | false | . | . | . | . | . | . | chr14 | 24191579 | 24244546 | 6.62149 | 4.76248 | 4.34728 |
| chr14 | 24222806 | 24223079 | 182 | Intron (ENSMUST00000073687.7 | 14 | 24238172 | 24243391 | 5220 | 2 | ENSMUST00 | 20312 | Gm17105 | ENSMUSG0( | false | . | . | . | . | . | . | chr14 | 24191579 | 24244546 | 6.62149 | 4.76248 | 4.34728 |
| chr10 | 111102323 | 111102593 | 182 | Distal Intergenic | 10 | 111125851 | 111126738 | 888 | 1 | ENSMUST00 | -23258 | Gm5428 | ENSMUSG0( | false | . | . | . | . | . | . | . | . | . | . | . | . |
| chr14 | 58253686 | 58253923 | 182 | Distal Intergenic | 14 | 58105772 | 58106129 | 358 | 2 | ENSMUST00 | -147558 | Gm17109 | ENSMUSG0( | false | . | . | . | . | . | . | . | . | . | . | . | . |
| chr4 | 117866097 | 117866426 | 182 | Promoter (2-3kb) | 4 | 117835299 | 117868558 | 33260 | 2 | ENSMUST00 | 2132 | Gm17114 | ENSMUSG0( | false | . | . | . | . | . | . | chr4 | 117865485 | 117868341 | 4.92715 | 4.04695 | 2.94373 |
| chr13 | 51229366 | 51229614 | 182 | Distal Intergenic | 13 | 51202688 | 51203065 | 378 | 2 | ENSMUST00 | -26302 | Hist1h2al | ENSMUSG0( | false | . | . | . | . | . | . | chr13 | 51229064 | 51229790 | 3.4725 | 3.30938 | 1.73824 |
| chr13 | 51295911 | 51296101 | 182 | Distal Intergenic | 13 | 51202688 | 51203065 | 378 | 2 | ENSMUST00 | -92847 | Hist1h2al | ENSMUSG0( | false | . | . | . | . | . | . | . | . | . | . | . | . |
| chr13 | 51231683 | 51231907 | 182 | Distal Intergenic | 13 | 51202688 | 51203065 | 378 | 2 | ENSMUST00 | -28619 | Hist1h2al | ENSMUSG0( | false | . | . | . | . | . | . | chr13 | 51231783 | 51233404 | 3.03724 | 3.04624 | 1.38837 |
| chr6 | 134186365 | 134186588 | 182 | Intron (ENSMUST00000081028.7 | 6 | 134168879 | 134172007 | 3129 | 2 | ENSMUST00 | -14359 | Gm17088 | ENSMUSG0( | false | . | . | . | . | . | . | chr6 | 134186494 | 134190488 | 5.7551 | 4.46071 | 3.63768 |
| chr1 | 177983310 | 177983691 | 182 | Distal Intergenic | 1 | 177991435 | 178048291 | 56857 | 1 | ENSMUST00 | -7744 | Gm16432 | ENSMUSG0( | false | chr1 | 177983280 | 177983856 | 36.31549 | 112.95179 | 109.03448 | . | . | . | . | . | . |
| chr7 | 98876121 | 98876346 | 182 | Distal Intergenic | 7 | 98854732 | 98855095 | 364 | 2 | ENSMUST00 | -21027 | Gm8398 | ENSMUSG0( | false | . | . | . | . | . | . | chr7 | 98875476 | 98876459 | 3.2608 | 3.21217 | 1.63855 |
| chr13 | 103226380 | 103226696 | 182 | Intron (ENSMUST00000166726.2 | 13 | 103159795 | 103164069 | 4275 | 2 | ENSMUST00 | -62312 | Gm6211 | ENSMUSG0( | false | . | . | . | . | . | . | . | . | . | . | . | . |
| chr9 | 32018145 | 32018386 | 182 | Distal Intergenic | 9 | 31936459 | 31936755 | 297 | 2 | ENSMUST00 | -81391 | Gm3756 | ENSMUSG0( | false | . | . | . | . | . | . | . | . | . | . | . | . |
| chr17 | 35331526 | 35331869 | 182 | Distal Intergenic | 17 | 35335909 | 35336292 | 384 | 1 | ENSMUST00 | -4040 | Gm8741 | ENSMUSG0( | false | . | . | . | . | . | . | . | . | . | . | . | . |
| chr8 | 69654434 | 69654984 | 182 | Distal Intergenic | 8 | 69659263 | 69664170 | 4908 | 1 | ENSMUST00 | -4279 | Zfp964 | ENSMUSG0( | false | chr8 | 69653962 | 69655366 | 24.82645 | 85.46523 | 82.21117 | chr8 | 69654874 | 69655781 | 3.23441 | 3.18606 | 1.53921 |
| chr5 | 15019483 | 15019816 | 182 | Distal Intergenic | 5 | 15028950 | 15032998 | 4049 | 2 | ENSMUST00 | 13182 | Gm17019 | ENSMUSG0( | false | . | . | . | . | . | . | . | . | . | . | . | . |
| chr17 | 34972121 | 34972515 | 182 | Promoter (<=1kb) | 17 | 34969190 | 34972156 | 2967 | 2 | ENSMUST00 | 0 | Hspa1a | ENSMUSG0( | false | chr17 | 34971678 | 34972891 | 28.56742 | 103.16996 | 99.51957 | chr17 | 34969123 | 34973097 | 4.29783 | 3.66937 | 2.43215 |
| chr1 | 171503652 | 171504036 | 182 | Promoter (<=1kb) | 1 | 171500568 | 171504111 | 3544 | 2 | ENSMUST00 | 75 | B930036N10 | ENSMUSG0( | false | chr1 | 171502954 | 171504470 | 21.18292 | 65.18933 | 62.27029 | . | . | . | . | . | . |
| chr13 | 98504725 | 98505086 | 182 | Distal Intergenic | 13 | 98489348 | 98492001 | 2654 | 2 | ENSMUST00 | -12725 | Gm10320 | ENSMUSG0( | false | chr13 | 98503874 | 98505195 | 20.9555 | 57.98476 | 55.16582 | chr13 | 98502445 | 98505712 | 4.6396 | 3.93808 | 2.71315 |
| chr12 | 84847599 | 84847828 | 182 | Intron (ENSMUST00000002073.7 | 12 | 84875769 | 84878132 | 2364 | 1 | ENSMUST00 | -27941 | D030025P21 | ENSMUSG0( | false | . | . | . | . | . | . | chr12 | 84846463 | 84847918 | 3.63957 | 3.46177 | 1.9225 |
| chr5 | 92745907 | 92746144 | 182 | Intron (ENSMUST00000113055.3 | 5 | 92736205 | 92737442 | 1238 | 1 | ENSMUST00 | 9703 | Gm20500 | ENSMUSG0( | false | . | . | . | . | . | . | chr5 | 92741029 | 92749681 | 4.65041 | 3.88261 | 2.72811 |
| chr16 | 50837835 | 50838064 | 182 | Distal Intergenic | 16 | 50783478 | 50784065 | 588 | 1 | ENSMUST00 | 54358 | 1700116B05 | ENSMUSG0( | false | . | . | . | . | . | . | chr16 | 50837680 | 50838127 | 2.8083 | 2.85598 | 1.26429 |
| chr16 | 50795611 | 50795885 | 182 | Distal Intergenic | 16 | 50783478 | 50784065 | 588 | 1 | ENSMUST00 | 12134 | 1700116B05 | ENSMUSG0( | false | . | . | . | . | . | . | chr16 | 50790830 | 50799832 | 4.14172 | 3.59695 | 2.3089 |
| chr19 | 5810713 | 5810910 | 182 | Distal Intergenic | 19 | 5795690 | 5802672 | 6983 | 2 | ENSMUST00 | -8042 | Malat1 | ENSMUSG0( | false | . | . | . | . | . | . | chr19 | 5803906 | 5840432 | 6.03605 | 4.50406 | 3.86334 |
| chr19 | 5802603 | 5802844 | 182 | Promoter (<=1kb) | 19 | 5795690 | 5802672 | 6983 | 2 | ENSMUST00 | 0 | Malat1 | ENSMUSG0( | false | chr19 | 5796007 | 5804263 | 15.48624 | 61.01235 | 58.15405 | . | . | . | . | . | . |
| chr3 | 52269077 | 52269268 | 182 | Promoter (<=1kb) | 3 | 52268435 | 52269435 | 1001 | 2 | ENSMUST00 | 167 | Gm20402 | ENSMUSG0( | false | chr3 | 52267261 | 52270933 | 33.7938 | 119.00099 | 114.88904 | . | . | . | . | . | . |
| chr3 | 52269666 | 52270366 | 182 | Promoter (<=1kb) | 3 | 52268435 | 52269435 | 1001 | 2 | ENSMUST00 | -232 | Gm20402 | ENSMUSG0( | false | chr3 | 52267261 | 52270933 | 33.7938 | 119.00099 | 114.88904 | chr3 | 52270355 | 52278010 | 3.39076 | 3.258 | 1.69225 |
| chr1 | 39313931 | 39314118 | 182 | Promoter (<=1kb) | 1 | 39314899 | 39315453 | 555 | 1 | ENSMUST00 | -781 | Gm3617 | ENSMUSG0( | false | . | . | . | . | . | . | chr1 | 39313740 | 39314414 | 3.10856 | 3.09984 | 1.49727 |
| chr18 | 75342610 | 75343055 | 182 | Distal Intergenic | 18 | 75365332 | 75367088 | 1757 | 2 | ENSMUST00 | 24033 | Gm20544 | ENSMUSG0( | false | chr18 | 75342214 | 75343517 | 28.96167 | 93.99567 | 90.56499 | chr18 | 75342076 | 75343896 | 5.24067 | 4.22706 | 3.22225 |
| chr4 | 43631339 | 43631790 | 182 | Promoter (<=1kb) | 4 | 43631425 | 43631557 | 133 | 1 | ENSMUST00 | 0 | Gm25262 | ENSMUSG0( | false | chr4 | 43631185 | 43632004 | 18.02468 | 55.19937 | 52.41551 | chr4 | 43629936 | 43632420 | 3.62371 | 3.32088 | 1.92539 |
| chr2 | 44976626 | 44976879 | 182 | Distal Intergenic | 2 | 45023100 | 45023177 | 78 | 2 | ENSMUST00 | 46298 | Mir5129 | ENSMUSG0( | false | . | . | . | . | . | . | chr2 | 44975552 | 44981626 | 5.83223 | 4.48649 | 3.69967 |
| chr18 | 55490773 | 55491031 | 182 | Distal Intergenic | 18 | 55622636 | 55622920 | 285 | 2 | ENSMUST00 | 131889 | Gm24514 | ENSMUSG0( | false | . | . | . | . | . | . | chr18 | 55489229 | 55492678 | 5.22289 | 4.18788 | 3.18833 |
| chr3 | 123508592 | 123508921 | 182 | Promoter (<=1kb) | 3 | 123507937 | 123508066 | 130 | 2 | ENSMUST00 | -527 | Snora24 | ENSMUSG0( | false | chr3 | 123507313 | 123509354 | 31.80867 | 99.00969 | 95.46461 | . | . | . | . | . | . |
| chr2 | 168058241 | 168058438 | 182 | Distal Intergenic | 2 | 168064820 | 168064897 | 78 | 2 | ENSMUST00 | 6459 | Gm24327 | ENSMUSG0( | false | . | . | . | . | . | . | chr2 | 168044068 | 168060701 | 5.7489 | 4.31854 | 3.64806 |
| chr17 | 65316847 | 65317138 | 182 | Intron (ENSMUST00000086722.4 | 17 | 65509368 | 65509492 | 125 | 1 | ENSMUST00 | -192230 | Gm25731 | ENSMUSG0( | false | . | . | . | . | . | . | . | . | . | . | . | . |
| chr6 | 48048368 | 48048667 | 182 | Promoter (<=1kb) | 6 | 48048703 | 48048795 | 93 | 1 | ENSMUST00 | -36 | Gm24563 | ENSMUSG0( | false | chr6 | 48048221 | 48049301 | 21.77823 | 66.26614 | 63.32996 | . | . | . | . | . | . |
| chr15 | 95906525 | 95906789 | 182 | Intron (ENSMUST00000071874.6 | 15 | 95879716 | 95879817 | 102 | 2 | ENSMUST00 | -26709 | Gm25070 | ENSMUSG0( | false | . | . | . | . | . | . | chr15 | 95905736 | 95907413 | 4.72366 | 3.92533 | 2.82298 |
| chr17 | 78322871 | 78323060 | 182 | Intron (ENSMUST00000112498.2 | 17 | 78302661 | 78302746 | 86 | 1 | ENSMUST00 | 20211 | Gm25069 | ENSMUSG0( | false | . | . | . | . | . | . | chr17 | 78315969 | 78336560 | 5.0014 | 4.08306 | 3.00503 |
| chr12 | 36724143 | 36724355 | 182 | Distal Intergenic | 12 | 36816205 | 36816278 | 74 | 1 | ENSMUST00 | -91850 | Mir5099 | ENSMUSG0( | false | . | . | . | . | . | . | chr12 | 36722792 | 36724339 | 3.57674 | 3.15869 | 1.87392 |
| chr7 | 112620395 | 112620646 | 182 | Promoter (2-3kb) | 7 | 112618096 | 112618225 | 130 | 2 | ENSMUST00 | -2171 | Gm24154 | ENSMUSG0( | false | chr7 | 112620319 | 112621241 | 9.76265 | 21.55577 | 19.19873 | chr7 | 112619110 | 112621979 | 3.76606 | 3.3977 | 1.99235 |
| chr7 | 112595326 | 112595637 | 182 | Distal Intergenic | 7 | 112618096 | 112618225 | 130 | 2 | ENSMUST00 | 22588 | Gm24154 | ENSMUSG0( | false | . | . | . | . | . | . | chr7 | 112592016 | 112605817 | 5.40057 | 4.28616 | 3.33023 |
| chr18 | 61369352 | 61369638 | 182 | Intron (ENSMUST00000075299.7 | 18 | 61397813 | 61397925 | 113 | 1 | ENSMUST00 | -28175 | Gm25301 | ENSMUSG0( | false | . | . | . | . | . | . | chr18 | 61366913 | 61371114 | 7.38415 | 5.03346 | 5.00481 |
| chr14 | 25533071 | 25533330 | 182 | Promoter (1-2kb) | 14 | 25534439 | 25534523 | 85 | 1 | ENSMUST00 | -1109 | Mir3075 | ENSMUSG0( | false | . | . | . | . | . | . | chr14 | 25530249 | 25537119 | 8.25302 | 5.48581 | 5.65915 |
| chr2 | 30743328 | 30743629 | 182 | Distal Intergenic | 2 | 30721210 | 30721293 | 84 | 1 | ENSMUST00 | 22119 | Mir3089 | ENSMUSG0( | false | . | . | . | . | . | . | chr2 | 30743149 | 30744848 | 8.39244 | 5.64319 | 5.7678 |
| chr7 | 35080173 | 35080444 | 182 | Promoter (1-2kb) | 7 | 35081910 | 35081979 | 70 | 1 | ENSMUST00 | -1466 | Gm25033 | ENSMUSG0( | false | . | . | . | . | . | . | chr7 | 35079794 | 35082018 | 4.40372 | 3.86609 | 2.50069 |
| chr2 | 172833266 | 172833519 | 182 | Distal Intergenic | 2 | 172864264 | 172864341 | 78 | 2 | ENSMUST00 | 30822 | Gm22773 | ENSMUSG0( | false | . | . | . | . | . | . | chr2 | 172831817 | 172833699 | 3.39898 | 3.26809 | 1.71382 |
| chr4 | 58413793 | 58414034 | 182 | Distal Intergenic | 4 | 58441012 | 58441098 | 87 | 2 | ENSMUST00 | 27064 | Mir3095 | ENSMUSG0( | false | . | . | . | . | . | . | chr4 | 58413260 | 58415406 | 4.80211 | 4.01267 | 2.83912 |
| chr12 | 54671356 | 54671619 | 182 | 3' UTR | 12 | 54666677 | 54666786 | 110 | 2 | ENSMUST00 | -4571 | Gm24971 | ENSMUSG0( | false | . | . | . | . | . | . | chr12 | 54670621 | 54672366 | 3.41531 | 3.08426 | 1.75368 |
| chr12 | 54666372 | 54666642 | 182 | Promoter (<=1kb) | 12 | 54666677 | 54666786 | 110 | 2 | ENSMUST00 | 144 | Gm24971 | ENSMUSG0( | false | . | . | . | . | . | . | chr12 | 54665007 | 54666824 | 5.39692 | 4.05991 | 3.36136 |
| chr11 | 51916099 | 51916434 | 182 | Distal Intergenic | 11 | 51964382 | 51964489 | 108 | 1 | ENSMUST00 | -47948 | Gm26061 | ENSMUSG0( | false | chr11 | 51915932 | 51916952 | 13.24688 | 29.26311 | 26.80765 | chr11 | 51915562 | 51918202 | 5.62534 | 4.46173 | 3.50587 |
| chr13 | 107831259 | 107831459 | 182 | Promoter (1-2kb) | 13 | 107829110 | 107829280 | 171 | 2 | ENSMUST00 | -1980 | Gm22118 | ENSMUSG0( | false | . | . | . | . | . | . | chr13 | 107830048 | 107858603 | 6.21396 | 4.54734 | 4.02799 |
| chr14 | 67344565 | 67344793 | 182 | Intron (ENSMUST00000022637.8 | 14 | 67324894 | 67335928 | 11035 | 2 | ENSMUST00 | -8638 | 1700120O09 | ENSMUSG0( | false | chr14 | 67344367 | 67345027 | 11.94783 | 25.26529 | 22.85895 | chr14 | 67344170 | 67345971 | 3.37633 | 3.08434 | 1.71634 |
| chr8 | 117136958 | 117137214 | 182 | Distal Intergenic | 8 | 117125335 | 117125738 | 404 | 1 | ENSMUST00 | 11624 | Gm20694 | ENSMUSG0( | false | . | . | . | . | . | . | chr8 | 117136664 | 117137793 | 5.04043 | 3.96925 | 3.07331 |
| chr17 | 66411936 | 66412147 | 182 | Promoter (1-2kb) | 17 | 66413631 | 66413984 | 354 | 2 | ENSMUST00 | 1837 | Gm4705 | ENSMUSG0( | false | . | . | . | . | . | . | chr17 | 66401889 | 66429482 | 6.16935 | 4.5489 | 3.97689 |
| chr8 | 108867978 | 108868151 | 182 | Intron (ENSMUST00000043896.9 | 8 | 108918264 | 108921874 | 3611 | 2 | ENSMUST00 | 53723 | Gm20686 | ENSMUSG0( | false | . | . | . | . | . | . | chr8 | 108867505 | 108870483 | 3.24169 | 3.06314 | 1.60321 |
| chr10 | 108454760 | 108455031 | 182 | Distal Intergenic | 10 | 108428067 | 108428160 | 94 | 1 | ENSMUST00 | 26694 | Gm25591 | ENSMUSG0( | false | . | . | . | . | . | . | chr10 | 108452094 | 108457073 | 4.88851 | 4.06098 | 2.92797 |
| chr10 | 108453606 | 108453859 | 182 | Distal Intergenic | 10 | 108428067 | 108428160 | 94 | 1 | ENSMUST00 | 25540 | Gm25591 | ENSMUSG0( | false | . | . | . | . | . | . | chr10 | 108452094 | 108457073 | 4.88851 | 4.06098 | 2.92797 |
| chr5 | 64112372 | 64112610 | 182 | Intron (ENSMUST00000087324.5 | 5 | 64107407 | 64107502 | 96 | 2 | ENSMUST00 | -4871 | Gm22395 | ENSMUSG0( | false | . | . | . | . | . | . | chr5 | 64112148 | 64113106 | 5.6858 | 4.38336 | 3.61191 |
| chr9 | 124423027 | 124423313 | 182 | Promoter (<=1kb) | 9 | 124422623 | 124423583 | 961 | 2 | ENSMUST00 | 270 | Ppp2r3d | ENSMUSG0( | false | chr9 | 124422515 | 124426377 | 33.91391 | 110.8507 | 106.99079 | . | . | . | . | . | . |
| chr15 | 88840098 | 88840389 | 182 | Distal Intergenic | 15 | 88835300 | 88835406 | 107 | 1 | ENSMUST00 | 4799 | Gm23144 | ENSMUSG0( | false | . | . | . | . | . | . | . | . | . | . | . | . |
| chr8 | 126945707 | 126946174 | 182 | Promoter (<=1kb) | 8 | 126930664 | 126945921 | 15258 | 2 | ENSMUST00 | 0 | Tomm20 | ENSMUSG0( | false | chr8 | 126943948 | 126946384 | 23.3092 | 84.4034 | 81.16875 | . | . | . | . | . | . |
| chr14 | 69447111 | 69447485 | 182 | Intron (ENSMUST00000111062.3 | 14 | 69419394 | 69426477 | 7084 | 2 | ENSMUST00 | -20635 | Gm28049 | ENSMUSG0( | false | . | . | . | . | . | . | chr14 | 69440416 | 69448720 | 4.10395 | 3.65347 | 2.25467 |
| chr1 | 72243792 | 72244331 | 182 | Promoter (<=1kb) | 1 | 72244213 | 72244403 | 191 | 1 | ENSMUST00 | 0 | Gm24497 | ENSMUSG0( | false | chr1 | 72243554 | 72244622 | 20.42851 | 61.25884 | 58.39448 | . | . | . | . | . | . |
| chrX | 123941190 | 123941400 | 182 | Distal Intergenic | 20 | 123957274 | 123957675 | 402 | 1 | ENSMUST00 | -15874 | Rps12-ps20 | ENSMUSG0( | false | . | . | . | . | . | . | . | . | . | . | . | . |
| chr10 | 23177273 | 23177552 | 182 | Intron (ENSMUST00000092665.5 | 10 | 23166865 | 23166971 | 107 | 2 | ENSMUST00 | -10303 | Gm24655 | ENSMUSG0( | false | . | . | . | . | . | . | chr10 | 23176188 | 23178770 | 4.21938 | 3.72221 | 2.39518 |
| chr1 | 72236478 | 72236988 | 182 | Promoter (<=1kb) | 1 | 72236946 | 72237136 | 191 | 1 | ENSMUST00 | 0 | Gm23472 | ENSMUSG0( | false | chr1 | 72236158 | 72237397 | 30.59503 | 95.89513 | 92.42037 | . | . | . | . | . | . |
| chr9 | 63091380 | 63091602 | 182 | Distal Intergenic | 9 | 63133283 | 63133389 | 107 | 1 | ENSMUST00 | -41681 | Gm25064 | ENSMUSG0( | false | . | . | . | . | . | . | chr9 | 63090225 | 63093748 | 4.55943 | 3.89487 | 2.64668 |
| chr9 | 49935870 | 49936072 | 182 | Distal Intergenic | 9 | 49825869 | 49825975 | 107 | 2 | ENSMUST00 | -109896 | Gm25633 | ENSMUSG0( | false | . | . | . | . | . | . | . | . | . | . | . | . |
| chr9 | 49988603 | 49988871 | 182 | Distal Intergenic | 9 | 49825869 | 49825975 | 107 | 2 | ENSMUST00 | -162629 | Gm25633 | ENSMUSG0( | false | . | . | . | . | . | . | chr9 | 49987558 | 49990811 | 5.24183 | 4.05424 | 3.24295 |
| chr13 | 29698737 | 29698976 | 182 | Intron (ENSMUST00000140278.2 | 13 | 29732942 | 29733036 | 95 | 1 | ENSMUST00 | -33966 | Gm23729 | ENSMUSG0( | false | . | . | . | . | . | . | chr13 | 29697735 | 29704646 | 4.65988 | 3.89142 | 2.72494 |
| chr4 | 32969560 | 32969777 | 182 | Distal Intergenic | 4 | 32964888 | 32965232 | 345 | 1 | ENSMUST00 | 4673 | Gm11942 | ENSMUSG0( | false | . | . | . | . | . | . | . | . | . | . | . | . |
| chr3 | 86745925 | 86746136 | 182 | Intron (ENSMUST00000107635.1 | 3 | 86672033 | 86672168 | 136 | 2 | ENSMUST00 | -73758 | Gm3788 | ENSMUSG0( | false | . | . | . | . | . | . | chr3 | 86744031 | 86751188 | 4.78929 | 3.88918 | 2.84191 |
| chr4 | 139602044 | 139602638 | 182 | Promoter (<=1kb) | 4 | 139603186 | 139662691 | 59506 | 1 | ENSMUST00 | -548 | Gm21969 | ENSMUSG0( | false | . | . | . | . | . | . | chr4 | 139576299 | 139606726 | 8.92638 | 5.72021 | 6.23508 |
| chr5 | 95906582 | 95906871 | 182 | Distal Intergenic | 5 | 95867334 | 95867440 | 107 | 1 | ENSMUST00 | 39249 | Gm25937 | ENSMUSG0( | false | . | . | . | . | . | . | . | . | . | . | . | . |
| chr3 | 89196510 | 89196708 | 182 | Distal Intergenic | 3 | 89201167 | 89201273 | 107 | 2 | ENSMUST00 | 4565 | Gm23269 | ENSMUSG0( | false | chr3 | 89196151 | 89196673 | 14.61479 | 34.85841 | 32.33254 | chr3 | 89195907 | 89197090 | 3.50439 | 3.28916 | 1.76821 |
| chr5 | 138843170 | 138843583 | 182 | Distal Intergenic | 5 | 138820080 | 138821619 | 1540 | 1 | ENSMUST00 | 23091 | Gm5294 | ENSMUSG0( | false | . | . | . | . | . | . | chr5 | 138842049 | 138844303 | 3.76869 | 3.45233 | 1.9923 |
| chr6 | 51809459 | 51809694 | 182 | Distal Intergenic | 6 | 51621632 | 51621708 | 77 | 1 | ENSMUST00 | 187828 | Gm22914 | ENSMUSG0( | false | . | . | . | . | . | . | chr6 | 51808893 | 51812949 | 5.35495 | 4.18951 | 3.30309 |
| chr13 | 97295534 | 97295845 | 182 | Distal Intergenic | 13 | 97292019 | 97292108 | 90 | 1 | ENSMUST00 | 3516 | Gm22007 | ENSMUSG0( | false | . | . | . | . | . | . | . | . | . | . | . | . |
| chr9 | 117956789 | 117957011 | 182 | Intron (ENSMUST00000111769.4 | 9 | 117999336 | 117999430 | 95 | 1 | ENSMUST00 | -42325 | Gm22008 | ENSMUSG0( | false | . | . | . | . | . | . | . | . | . | . | . | . |
| chr5 | 105859350 | 105859589 | 182 | Distal Intergenic | 5 | 105862701 | 105863702 | 1002 | 2 | ENSMUST00 | 4113 | Gm10359 | ENSMUSG0( | false | . | . | . | . | . | . | chr5 | 105859479 | 105860596 | 4.09797 | 3.65108 | 2.25411 |
| chr10 | 54403378 | 54403833 | 182 | Distal Intergenic | 10 | 54311225 | 54311331 | 107 | 2 | ENSMUST00 | -92048 | Gm26177 | ENSMUSG0( | false | . | . | . | . | . | . | . | . | . | . | . | . |
| chr6 | 47767630 | 47768104 | 182 | Distal Intergenic | 6 | 47773329 | 47773429 | 101 | 1 | ENSMUST00 | -5225 | Gm24975 | ENSMUSG0( | false | . | . | . | . | . | . | chr6 | 47767164 | 47768562 | 12.05524 | 4.54108 | 8.94963 |
| chr6 | 47744187 | 47744651 | 182 | Distal Intergenic | 6 | 47773329 | 47773429 | 101 | 1 | ENSMUST00 | -28678 | Gm24975 | ENSMUSG0( | false | . | . | . | . | . | . | . | . | . | . | . | . |
| chr6 | 47748520 | 47749046 | 182 | Distal Intergenic | 6 | 47773329 | 47773429 | 101 | 1 | ENSMUST00 | -24283 | Gm24975 | ENSMUSG0( | false | . | . | . | . | . | . | . | . | . | . | . | . |
| chr6 | 47752863 | 47753432 | 182 | Distal Intergenic | 6 | 47773329 | 47773429 | 101 | 1 | ENSMUST00 | -19897 | Gm24975 | ENSMUSG0( | false | . | . | . | . | . | . | . | . | . | . | . | . |
| chr6 | 47763297 | 47763741 | 182 | Distal Intergenic | 6 | 47773329 | 47773429 | 101 | 1 | ENSMUST00 | -9588 | Gm24975 | ENSMUSG0( | false | chr6 | 47763624 | 47764161 | 8.30651 | 102.0048 | 98.38202 | chr6 | 47763107 | 47765043 | 12.29093 | 4.35488 | 9.07091 |
| chr6 | 47739910 | 47740542 | 182 | Distal Intergenic | 6 | 47773329 | 47773429 | 101 | 1 | ENSMUST00 | -32787 | Gm24975 | ENSMUSG0( | false | chr6 | 47739814 | 47740114 | 6.03039 | 48.75505 | 46.05571 | chr6 | 47738986 | 47740437 | 8.06705 | 4.62677 | 5.53488 |
| chr13 | 119790656 | 119791336 | 182 | Promoter (<=1kb) | 13 | 119765187 | 119790805 | 25619 | 2 | ENSMUST00 | 0 | Zfp131 | ENSMUSG0( | false | chr13 | 119789745 | 119791716 | 32.87838 | 107.99121 | 104.21945 | . | . | . | . | . | . |
| chr17 | 48041285 | 48041519 | 182 | Promoter (2-3kb) | 17 | 48036745 | 48038293 | 1549 | 2 | ENSMUST00 | -2993 | 1700122O11 | ENSMUSG0( | false | . | . | . | . | . | . | . | . | . | . | . | . |
| chr9 | 13296598 | 13296849 | 182 | Distal Intergenic | 9 | 13275516 | 13275622 | 107 | 1 | ENSMUST00 | 21083 | Gm26189 | ENSMUSG0( | false | chr9 | 13296261 | 13298513 | 19.14982 | 55.63055 | 52.84086 | . | . | . | . | . | . |
| chr11 | 64496006 | 64496440 | 182 | Intron (ENSMUST00000058652.5 | 11 | 64435156 | 64436653 | 1498 | 2 | ENSMUST00 | -59354 | F930015N05 | ENSMUSG0( | false | . | . | . | . | . | . | chr11 | 64487471 | 64501485 | 4.71094 | 3.86043 | 2.79046 |
| chr17 | 32829918 | 32830190 | 182 | Distal Intergenic | 17 | 32815455 | 32822200 | 6746 | 2 | ENSMUST00 | -7719 | Zfp799 | ENSMUSG0( | false | chr17 | 32829461 | 32830595 | 37.30079 | 124.19571 | 119.91465 | chr17 | 32828376 | 32830709 | 3.79893 | 3.23935 | 2.05919 |
| chr17 | 8147631 | 8148079 | 182 | Promoter (<=1kb) | 17 | 8128610 | 8148097 | 19488 | 2 | ENSMUST00 | 18 | Rnaset2a | ENSMUSG0( | false | chr17 | 8147318 | 8148171 | 23.3092 | 84.4034 | 81.16875 | . | . | . | . | . | . |
| chr16 | 30722460 | 30722652 | 182 | Distal Intergenic | 16 | 30814162 | 30814268 | 107 | 1 | ENSMUST00 | -91510 | Gm26295 | ENSMUSG0( | false | . | . | . | . | . | . | . | . | . | . | . | . |
| chr9 | 60758038 | 60758327 | 182 | Intron (ENSMUST00000190053.1 | 9 | 60769602 | 60769955 | 354 | 1 | ENSMUST00 | -11275 | 1700036A12 | ENSMUSG0( | false | . | . | . | . | . | . | . | . | . | . | . | . |
| chr8 | 12395372 | 12395611 | 182 | Promoter (<=1kb) | 8 | 12395295 | 12400126 | 4832 | 1 | ENSMUST00 | 78 | Sox1 | ENSMUSG0( | false | chr8 | 12394922 | 12395972 | 16.90119 | 40.85703 | 38.25915 | chr8 | 12394738 | 12397650 | 5.69621 | 4.34814 | 3.59544 |
| chr11 | 20915177 | 20915391 | 182 | Distal Intergenic | 11 | 20937576 | 20937701 | 126 | 2 | ENSMUST00 | 22310 | Gm23681 | ENSMUSG0( | false | . | . | . | . | . | . | . | . | . | . | . | . |
| chr9 | 70847718 | 70848088 | 182 | Intron (ENSMUST00000034731.8 | 9 | 70852576 | 70853418 | 843 | 1 | ENSMUST00 | -4488 | Gm3436 | ENSMUSG0( | false | . | . | . | . | . | . | chr9 | 70847798 | 70849224 | 5.89747 | 4.5995 | 3.74662 |
| chr4 | 57866170 | 57866364 | 182 | Intron (ENSMUST00000098066.3 | 4 | 57877389 | 57877495 | 107 | 2 | ENSMUST00 | 11131 | Gm26138 | ENSMUSG0( | false | . | . | . | . | . | . | chr4 | 57846708 | 57868207 | 6.85521 | 4.86004 | 4.55185 |
| chr9 | 43858113 | 43858401 | 182 | Distal Intergenic | 9 | 43829963 | 43832516 | 2554 | 1 | ENSMUST00 | 28151 | Gm3898 | ENSMUSG0( | false | . | . | . | . | . | . | . | . | . | . | . | . |
| chr6 | 69814520 | 69814836 | 182 | Promoter (<=1kb) | 6 | 69814631 | 69815100 | 470 | 2 | ENSMUST00 | 264 | Igkv12-44 | ENSMUSG0( | false | . | . | . | . | . | . | chr6 | 69814390 | 69815610 | 3.70094 | 3.30429 | 1.97673 |
| chr18 | 80091205 | 80091412 | 182 | Promoter (1-2kb) | 18 | 80089297 | 80089962 | 666 | 2 | ENSMUST00 | -1244 | Gm21886 | ENSMUSG0( | false | . | . | . | . | . | . | . | . | . | . | . | . |
| chr18 | 80085694 | 80085920 | 182 | Intron (ENSMUST00000070219.7 | 18 | 80089297 | 80089962 | 666 | 2 | ENSMUST00 | 4042 | Gm21886 | ENSMUSG0( | false | . | . | . | . | . | . | . | . | . | . | . | . |
| chr1 | 134518793 | 134519001 | 182 | Promoter (<=1kb) | 1 | 134518081 | 134518563 | 483 | 1 | ENSMUST00 | 713 | Gm15454 | ENSMUSG0( | false | . | . | . | . | . | . | chr1 | 134515825 | 134519244 | 4.83117 | 3.99643 | 2.89892 |
| chr15 | 82213431 | 82213652 | 182 | Promoter (<=1kb) | 15 | 82206952 | 82212815 | 5864 | 2 | ENSMUST00 | -617 | AI848285 | ENSMUSG0( | false | . | . | . | . | . | . | chr15 | 82211519 | 82213435 | 5.33302 | 4.22274 | 3.2939 |
| chr1 | 195016858 | 195017100 | 182 | Promoter (<=1kb) | 1 | 195017399 | 195037908 | 20510 | 1 | ENSMUST00 | -299 | A330023F24 | ENSMUSG0( | false | chr1 | 195017069 | 195018317 | 20.9555 | 57.98476 | 55.16582 | . | . | . | . | . | . |
| chr8 | 10991244 | 10991434 | 182 | Intron (ENSMUST00000040514.6 | 8 | 11007850 | 11054536 | 46687 | 1 | ENSMUST00 | -16416 | 9530052E02 | ENSMUSG0( | false | . | . | . | . | . | . | chr8 | 10987277 | 10997404 | 4.95198 | 4.01815 | 2.9819 |
| chr18 | 54990465 | 54990757 | 182 | Promoter (<=1kb) | 18 | 54990310 | 55077337 | 87028 | 1 | ENSMUST00 | 156 | Gm4221 | ENSMUSG0( | false | chr18 | 54989542 | 54993168 | 25.04286 | 71.11688 | 68.10826 | . | . | . | . | . | . |
| chr18 | 54990973 | 54991407 | 182 | Promoter (<=1kb) | 18 | 54990310 | 55077337 | 87028 | 1 | ENSMUST00 | 664 | Gm4221 | ENSMUSG0( | false | chr18 | 54989542 | 54993168 | 25.04286 | 71.11688 | 68.10826 | . | . | . | . | . | . |
| chr16 | 25040075 | 25040254 | 182 | Distal Intergenic | 16 | 25059639 | 25069058 | 9420 | 1 | ENSMUST00 | -19385 | A230028O05 | ENSMUSG0( | false | . | . | . | . | . | . | . | . | . | . | . | . |
| chr15 | 10818200 | 10818585 | 182 | Distal Intergenic | 15 | 10717144 | 10722471 | 5328 | 1 | ENSMUST00 | 101057 | 4930556M19 | ENSMUSG0( | false | . | . | . | . | . | . | chr15 | 10818010 | 10818803 | 4.19607 | 3.54008 | 2.36086 |
| chr15 | 10731473 | 10731690 | 182 | Intron (ENSMUST00000180604.1 | 15 | 10717144 | 10722471 | 5328 | 1 | ENSMUST00 | 14330 | 4930556M19 | ENSMUSG0( | false | . | . | . | . | . | . | chr15 | 10731200 | 10731631 | 3.11911 | 3.07241 | 1.49923 |
| chr15 | 10780276 | 10780481 | 182 | Intron (ENSMUST00000180604.1 | 15 | 10717144 | 10722471 | 5328 | 1 | ENSMUST00 | 63133 | 4930556M19 | ENSMUSG0( | false | . | . | . | . | . | . | . | . | . | . | . | . |
| chr2 | 169943862 | 169944089 | 182 | Intron (ENSMUST00000123300.2 | 2 | 169633676 | 169888503 | 254828 | 2 | ENSMUST00 | -55360 | Gm26883 | ENSMUSG0( | false | . | . | . | . | . | . | chr2 | 169943381 | 169944411 | 3.29322 | 3.15239 | 1.62132 |
| chr8 | 72492643 | 72493028 | 182 | Promoter (<=1kb) | 8 | 72492934 | 72505499 | 12566 | 1 | ENSMUST00 | 0 | Gm17435 | ENSMUSG0( | false | chr8 | 72491599 | 72492739 | 36.19004 | 121.17046 | 116.98717 | . | . | . | . | . | . |
| chr15 | 103146750 | 103147179 | 182 | Promoter (<=1kb) | 15 | 103123070 | 103146828 | 23759 | 2 | ENSMUST00 | 0 | D930007P13 | ENSMUSG0( | false | chr15 | 103146571 | 103147229 | 18.38128 | 45.59164 | 42.93153 | chr15 | 103146557 | 103147497 | 3.2408 | 3.08937 | 1.60542 |
| chr13 | 69819203 | 69819421 | 182 | Distal Intergenic | 13 | 69809914 | 69816094 | 6181 | 2 | ENSMUST00 | -3110 | Gm26819 | ENSMUSG0( | false | . | . | . | . | . | . | . | . | . | . | . | . |
| chr15 | 62397126 | 62397401 | 182 | Distal Intergenic | 15 | 62222603 | 62242505 | 19903 | 1 | ENSMUST00 | 174524 | Pvt1 | ENSMUSG0( | false | . | . | . | . | . | . | . | . | . | . | . | . |
| chr6 | 86525169 | 86525424 | 182 | Promoter (<=1kb) | 6 | 86526271 | 86564442 | 38172 | 1 | ENSMUST00 | -847 | 1600020E01 | ENSMUSG0( | false | chr6 | 86523216 | 86527818 | 38.37027 | 120.70945 | 116.54665 | . | . | . | . | . | . |
| chr16 | 11024951 | 11025236 | 182 | Intron (ENSMUST00000162323.1 | 16 | 11008898 | 11015186 | 6289 | 1 | ENSMUST00 | 16054 | Gm4262 | ENSMUSG0( | false | . | . | . | . | . | . | chr16 | 11024489 | 11026954 | 5.58953 | 4.21217 | 3.53141 |
| chr13 | 48982507 | 48982771 | 182 | Distal Intergenic | 13 | 48968287 | 48972031 | 3745 | 1 | ENSMUST00 | 14221 | Fam120aos | ENSMUSG0( | false | . | . | . | . | . | . | chr13 | 48977668 | 48983230 | 5.0119 | 4.01873 | 3.0351 |
| chr10 | 54076296 | 54076682 | 182 | Promoter (<=1kb) | 10 | 54075922 | 54081081 | 5160 | 1 | ENSMUST00 | 375 | Gm16998 | ENSMUSG0( | false | chr10 | 54074002 | 54076827 | 34.34893 | 131.73137 | 127.17394 | chr10 | 54076481 | 54077206 | 4.79581 | 4.02213 | 2.82842 |
| chr12 | 111432088 | 111432419 | 182 | Promoter (1-2kb) | 12 | 111426874 | 111430946 | 4073 | 2 | ENSMUST00 | -1143 | Gm10425 | ENSMUSG0( | false | . | . | . | . | . | . | chr12 | 111425142 | 111437454 | 7.80275 | 5.329 | 5.29325 |
| chr8 | 121127407 | 121127797 | 182 | Promoter (<=1kb) | 8 | 121127940 | 121130644 | 2705 | 1 | ENSMUST00 | -143 | Foxl1 | ENSMUSG0( | false | chr8 | 121127717 | 121129604 | 19.85205 | 54.41815 | 51.64575 | chr8 | 121121895 | 121136533 | 5.26018 | 4.19829 | 3.22208 |
| chr1 | 192070360 | 192070576 | 182 | Exon (ENSMUST00000085573.6/ | 1 | 192092372 | 192099630 | 7259 | 1 | ENSMUST00 | -21796 | Gm26879 | ENSMUSG0( | false | . | . | . | . | . | . | chr1 | 192069811 | 192071176 | 3.73127 | 3.13638 | 2.00418 |
| chr8 | 121262730 | 121263064 | 182 | Intron (ENSMUST00000127664.1 | 8 | 121198714 | 121201835 | 3122 | 2 | ENSMUST00 | -60896 | 5033426O07 | ENSMUSG0( | false | . | . | . | . | . | . | chr8 | 121255503 | 121265207 | 5.96407 | 4.44635 | 3.80299 |
| chr8 | 121244768 | 121244972 | 182 | Intron (ENSMUST00000127664.1 | 8 | 121198714 | 121201835 | 3122 | 2 | ENSMUST00 | -42934 | 5033426O07 | ENSMUSG0( | false | . | . | . | . | . | . | chr8 | 121236124 | 121247254 | 7.60766 | 5.2125 | 5.14118 |
| chr13 | 80962796 | 80963141 | 182 | Promoter (<=1kb) | 13 | 80962888 | 81126424 | 163537 | 1 | ENSMUST00 | 0 | 9330111N05 | ENSMUSG0( | false | chr13 | 80962659 | 80963447 | 15.47449 | 37.71435 | 35.15295 | . | . | . | . | . | . |
| chr17 | 6827807 | 6828315 | 182 | Promoter (<=1kb) | 17 | 6798390 | 6827990 | 29601 | 2 | ENSMUST00 | 0 | Gm2885 | ENSMUSG0( | false | chr17 | 6827297 | 6828817 | 14.02077 | 44.3905 | 41.74443 | chr17 | 6826723 | 6829059 | 4.11359 | 3.62079 | 2.3073 |
| chr17 | 29492365 | 29493080 | 182 | Promoter (<=1kb) | 17 | 29461896 | 29491925 | 30030 | 2 | ENSMUST00 | -441 | Gm26885 | ENSMUSG0( | false | chr17 | 29490723 | 29492947 | 40.43883 | 128.57333 | 124.13428 | chr17 | 29491194 | 29493532 | 7.97827 | 5.47645 | 5.42471 |
| chr14 | 76513514 | 76513905 | 182 | Downstream (<1kb) | 14 | 76514558 | 76520855 | 6298 | 2 | ENSMUST00 | 6950 | 4930444M15 | ENSMUSG0( | false | . | . | . | . | . | . | chr14 | 76513610 | 76514220 | 3.4048 | 3.35217 | 1.67843 |
| chr13 | 91421665 | 91421968 | 182 | Distal Intergenic | 13 | 91368990 | 91388085 | 19096 | 2 | ENSMUST00 | -33581 | A830009L08 | ENSMUSG0( | false | chr13 | 91421549 | 91422152 | 7.14186 | 14.46535 | 12.21151 | chr13 | 91419905 | 91425156 | 4.40318 | 3.73853 | 2.51365 |
| chr3 | 145653327 | 145653646 | 182 | Distal Intergenic | 3 | 145650312 | 145652093 | 1782 | 1 | ENSMUST00 | 3016 | Gm17501 | ENSMUSG0( | false | chr3 | 145653019 | 145653876 | 25.36096 | 73.47037 | 70.42609 | chr3 | 145652582 | 145654049 | 6.46829 | 4.81326 | 4.21935 |
| chr3 | 145650571 | 145650866 | 182 | Promoter (<=1kb) | 3 | 145650312 | 145652093 | 1782 | 1 | ENSMUST00 | 260 | Gm17501 | ENSMUSG0( | false | chr3 | 145648777 | 145650997 | 33.55804 | 103.05054 | 99.40515 | chr3 | 145650372 | 145651310 | 3.96706 | 3.47274 | 2.18362 |
| chr4 | 130841051 | 130841395 | 182 | Distal Intergenic | 4 | 130824394 | 130826313 | 1920 | 2 | ENSMUST00 | -14739 | Gm26716 | ENSMUSG0( | false | . | . | . | . | . | . | . | . | . | . | . | . |
| chr9 | 117776038 | 117776271 | 182 | Distal Intergenic | 9 | 117873144 | 117881202 | 8059 | 1 | ENSMUST00 | -96873 | Gm26614 | ENSMUSG0( | false | . | . | . | . | . | . | . | . | . | . | . | . |
| chr14 | 70195976 | 70196198 | 182 | Intron (ENSMUST00000022682.5 | 14 | 70205324 | 70217659 | 12336 | 1 | ENSMUST00 | -9126 | Gm26908 | ENSMUSG0( | false | chr14 | 70195712 | 70196081 | 5.85266 | 9.15698 | 7.00004 | chr14 | 70190541 | 70199927 | 4.55845 | 3.79328 | 2.66728 |
| chr17 | 83862324 | 83862612 | 182 | Distal Intergenic | 17 | 83851675 | 83854393 | 2719 | 2 | ENSMUST00 | -7932 | C430042M11 | ENSMUSG0( | false | chr17 | 83861359 | 83863132 | 25.84672 | 90.2271 | 86.88143 | . | . | . | . | . | . |
| chr7 | 81523271 | 81523574 | 182 | Promoter (<=1kb) | 7 | 81523550 | 81529554 | 6005 | 1 | ENSMUST00 | 0 | 2900076A07 | ENSMUSG0( | false | chr7 | 81523006 | 81524204 | 25.98482 | 77.30067 | 74.19341 | chr7 | 81522733 | 81527118 | 4.61213 | 3.84195 | 2.70992 |
| chr10 | 14055662 | 14055906 | 182 | Intron (ENSMUST00000187083.1 | 10 | 13967024 | 14007405 | 40382 | 2 | ENSMUST00 | -48258 | Gm26835 | ENSMUSG0( | false | . | . | . | . | . | . | chr10 | 14054380 | 14062954 | 5.60088 | 4.29997 | 3.513 |
| chr16 | 29729246 | 29729519 | 182 | Distal Intergenic | 16 | 29811058 | 29837814 | 26757 | 1 | ENSMUST00 | -81539 | Gm26834 | ENSMUSG0( | false | . | . | . | . | . | . | chr16 | 29728710 | 29730319 | 3.9066 | 3.47994 | 2.15292 |
| chr1 | 194995245 | 194995540 | 182 | Intron (ENSMUST00000181226.2 | 1 | 194955759 | 194976959 | 21201 | 2 | ENSMUST00 | -18287 | Gm16897 | ENSMUSG0( | false | . | . | . | . | . | . | chr1 | 194990665 | 194996343 | 3.70869 | 3.40115 | 1.97407 |
| chr6 | 129220278 | 129220524 | 182 | Intron (ENSMUST00000181238.1 | 6 | 129213381 | 129233974 | 20594 | 2 | ENSMUST00 | 13450 | 2310001H17 | ENSMUSG0( | false | . | . | . | . | . | . | . | . | . | . | . | . |
| chr3 | 116183973 | 116184170 | 182 | Distal Intergenic | 3 | 116150479 | 116165821 | 15343 | 1 | ENSMUST00 | 33495 | Gm26544 | ENSMUSG0( | false | . | . | . | . | . | . | . | . | . | . | . | . |
| chr3 | 9500699 | 9500892 | 182 | Intron (ENSMUST00000041124.7 | 3 | 9403076 | 9413903 | 10828 | 1 | ENSMUST00 | 97624 | C030034L19 | ENSMUSG0( | false | . | . | . | . | . | . | chr3 | 9499389 | 9501374 | 3.41066 | 3.18997 | 1.73034 |
| chr12 | 80885391 | 80885638 | 182 | Promoter (2-3kb) | 12 | 80885549 | 80887674 | 2126 | 2 | ENSMUST00 | 2036 | Gm26545 | ENSMUSG0( | false | . | . | . | . | . | . | . | . | . | . | . | . |
| chr12 | 80846906 | 80847122 | 182 | Intron (ENSMUST00000068519.5 | 12 | 80885549 | 80887674 | 2126 | 2 | ENSMUST00 | 40552 | Gm26545 | ENSMUSG0( | false | . | . | . | . | . | . | chr12 | 80842883 | 80848124 | 5.84561 | 4.37417 | 3.7122 |
| chr8 | 121377802 | 121378071 | 182 | Intron (ENSMUST00000127664.1 | 8 | 121256747 | 121356836 | 100090 | 2 | ENSMUST00 | -20967 | Gm26815 | ENSMUSG0( | false | . | . | . | . | . | . | chr8 | 121372330 | 121385684 | 7.83977 | 5.33422 | 5.34287 |
| chr8 | 121334880 | 121335121 | 182 | Intron (ENSMUST00000127664.1 | 8 | 121256747 | 121356836 | 100090 | 2 | ENSMUST00 | 21715 | Gm26815 | ENSMUSG0( | false | . | . | . | . | . | . | chr8 | 121331338 | 121336444 | 7.7568 | 5.32028 | 5.25757 |
| chr17 | 46267697 | 46268065 | 182 | Intron (ENSMUST00000012440.7 | 17 | 46273479 | 46280060 | 6582 | 1 | ENSMUST00 | -5414 | Gm26785 | ENSMUSG0( | false | . | . | . | . | . | . | chr17 | 46263490 | 46282075 | 6.71144 | 4.73752 | 4.44242 |
| chr1 | 88757962 | 88758234 | 182 | Distal Intergenic | 1 | 88754954 | 88755709 | 756 | 1 | ENSMUST00 | 3009 | 2410088K16 | ENSMUSG0( | false | . | . | . | . | . | . | chr1 | 88755502 | 88758671 | 3.30363 | 3.17082 | 1.66115 |
| chr13 | 51962123 | 51962412 | 182 | Distal Intergenic | 13 | 51973214 | 51976583 | 3370 | 1 | ENSMUST00 | -10802 | Gm26651 | ENSMUSG0( | false | . | . | . | . | . | . | . | . | . | . | . | . |
| chr13 | 51970120 | 51970352 | 182 | Promoter (2-3kb) | 13 | 51973214 | 51976583 | 3370 | 1 | ENSMUST00 | -2862 | Gm26651 | ENSMUSG0( | false | . | . | . | . | . | . | chr13 | 51968572 | 51971263 | 3.5265 | 3.20761 | 1.84487 |
| chr13 | 51975941 | 51976298 | 182 | Promoter (2-3kb) | 13 | 51973214 | 51976583 | 3370 | 1 | ENSMUST00 | 2728 | Gm26651 | ENSMUSG0( | false | . | . | . | . | . | . | chr13 | 51973559 | 51977129 | 5.45013 | 4.33194 | 3.37717 |
| chr14 | 62845154 | 62845744 | 182 | Intron (ENSMUST00000014691.8 | 14 | 62836908 | 62838207 | 1300 | 2 | ENSMUST00 | -6948 | Gm26536 | ENSMUSG0( | false | . | . | . | . | . | . | chr14 | 62841867 | 62847132 | 6.79695 | 4.91293 | 4.47305 |
| chr9 | 119425280 | 119425529 | 182 | Intron (ENSMUST00000035093.9 | 9 | 119440278 | 119444830 | 4553 | 2 | ENSMUST00 | 19301 | 4930516B21 | ENSMUSG0( | false | . | . | . | . | . | . | chr9 | 119424302 | 119426929 | 6.24705 | 4.30112 | 4.06285 |
| chr6 | 72926987 | 72927225 | 182 | Distal Intergenic | 6 | 72953236 | 72957249 | 4014 | 1 | ENSMUST00 | -26011 | Gm26640 | ENSMUSG0( | false | . | . | . | . | . | . | chr6 | 72926728 | 72927867 | 5.31546 | 4.18471 | 3.28758 |
| chr15 | 89210423 | 89210706 | 182 | Downstream (<1kb) | 15 | 89211559 | 89215012 | 3454 | 2 | ENSMUST00 | 4306 | Gm26798 | ENSMUSG0( | false | chr15 | 89210046 | 89210714 | 12.8669 | 28.00426 | 25.56298 | . | . | . | . | . | . |
| chr11 | 119078757 | 119079086 | 182 | Promoter (1-2kb) | 11 | 119077573 | 119081865 | 4293 | 1 | ENSMUST00 | 1185 | Gm26508 | ENSMUSG0( | false | . | . | . | . | . | . | chr11 | 119077654 | 119084726 | 7.24094 | 5.11519 | 4.83739 |
| chr1 | 185489999 | 185490197 | 182 | Distal Intergenic | 1 | 185447116 | 185455568 | 8453 | 2 | ENSMUST00 | -34432 | Gm2061 | ENSMUSG0( | false | . | . | . | . | . | . | chr1 | 185488835 | 185491310 | 5.51555 | 4.39945 | 3.41913 |
| chr12 | 12904656 | 12904959 | 182 | Promoter (1-2kb) | 12 | 12902720 | 12907251 | 4532 | 1 | ENSMUST00 | 1937 | 4930519A11 | ENSMUSG0( | false | chr12 | 12904538 | 12904996 | 6.38327 | 11.47542 | 9.27082 | chr12 | 12904295 | 12905072 | 5.28304 | 3.97573 | 3.27 |
| chr3 | 41500558 | 41500834 | 182 | Distal Intergenic | 3 | 41488804 | 41493192 | 4389 | 2 | ENSMUST00 | -7367 | C430002E04 | ENSMUSG0( | false | . | . | . | . | . | . | chr3 | 41497652 | 41501687 | 4.01455 | 3.58945 | 2.20379 |
| chr1 | 55225389 | 55225593 | 182 | Promoter (<=1kb) | 1 | 55226251 | 55235622 | 9372 | 1 | ENSMUST00 | -658 | Gm10561 | ENSMUSG0( | false | . | . | . | . | . | . | chr1 | 55225009 | 55227552 | 3.6552 | 3.4054 | 1.91126 |
| chr19 | 45920715 | 45920971 | 182 | Intron (ENSMUST00000045396.7 | 19 | 45998137 | 46013386 | 15250 | 1 | ENSMUST00 | -77166 | 4930505N22 | ENSMUSG0( | false | . | . | . | . | . | . | chr19 | 45916903 | 45920732 | 5.3038 | 4.17016 | 3.28303 |
| chr10 | 52309214 | 52309458 | 182 | Intron (ENSMUST00000069004.6 | 10 | 52358767 | 52358853 | 87 | 1 | ENSMUST00 | -49309 | Gm26741 | ENSMUSG0( | false | . | . | . | . | . | . | . | . | . | . | . | . |
| chr18 | 75537635 | 75538340 | 182 | Intron (ENSMUST00000165559.1 | 18 | 75514645 | 75522806 | 8162 | 1 | ENSMUST00 | 22991 | Gm10532 | ENSMUSG0( | false | . | . | . | . | . | . | chr18 | 75537468 | 75538951 | 3.74726 | 3.5008 | 1.9639 |
| chr13 | 40722813 | 40723274 | 182 | Exon (ENSMUST00000021787.5/ | 13 | 40727808 | 40735162 | 7355 | 1 | ENSMUST00 | -4534 | Gm26688 | ENSMUSG0( | false | chr13 | 40722313 | 40723573 | 17.45709 | 49.39342 | 46.68804 | chr13 | 40722253 | 40723122 | 3.49287 | 3.33333 | 1.75947 |
| chr17 | 21474642 | 21475164 | 182 | Distal Intergenic | 17 | 21485177 | 21488875 | 3699 | 2 | ENSMUST00 | 13711 | Gm26753 | ENSMUSG0( | false | chr17 | 21474663 | 21475289 | 11.80365 | 26.21181 | 23.79474 | chr17 | 21475121 | 21476056 | 3.30323 | 3.13994 | 1.61116 |
| chr17 | 21535815 | 21536020 | 182 | Promoter (<=1kb) | 17 | 21528194 | 21535878 | 7685 | 2 | ENSMUST00 | 0 | 9330136K24 | ENSMUSG0( | false | chr17 | 21535333 | 21536358 | 20.56672 | 54.83671 | 52.06188 | chr17 | 21533109 | 21537002 | 4.61119 | 3.84394 | 2.71348 |
| chr10 | 77484222 | 77484587 | 182 | Promoter (2-3kb) | 10 | 77486664 | 77516430 | 29767 | 1 | ENSMUST00 | -2077 | 2810425M01 | ENSMUSG0( | false | . | . | . | . | . | . | chr10 | 77483144 | 77485075 | 3.67362 | 3.44493 | 1.94183 |
| chr6 | 18845723 | 18846219 | 182 | Promoter (1-2kb) | 6 | 18844220 | 18847360 | 3141 | 1 | ENSMUST00 | 1504 | Gm26809 | ENSMUSG0( | false | chr6 | 18845430 | 18846090 | 10.56924 | 21.28106 | 18.92771 | . | . | . | . | . | . |

| | | | | | | | | | | | | | | | | | | | | | | | | | | |
|---|---|---|---|---|---|---|---|---|---|---|---|---|---|---|---|---|---|---|---|---|---|---|---|---|---|---|
| chr9 | 114538197 | 114538379 | 182 | Intron (ENSMUST00000111816.2 | 9 | 114515697 | 114516424 | 728 | 1 | ENSMUST00 | 22501 | C130032M1C | ENSMUSG0C | false | . | . | . | . | . | . | . | . | . | . | . | . |
| chr12 | 110844686 | 110845302 | 182 | Promoter (<=1kb) | 12 | 110842276 | 110845216 | 2941 | 2 | ENSMUST00 | 0 | Gm26912 | ENSMUSG0C | false | chr12 | 110843843 | 110846184 | 16.16105 | 50.86513 | 48.14017 | . | . | . | . | . | . |
| chr2 | 79454222 | 79454509 | 182 | Promoter (<=1kb) | 2 | 79453985 | 79461107 | 7123 | 1 | ENSMUST00 | 238 | Gm26755 | ENSMUSG0C | false | chr2 | 79453846 | 79454353 | 11.00893 | 22.55865 | 20.19038 | . | . | . | . | . | . |
| chr2 | 52982519 | 52982823 | 182 | Intron (ENSMUST00000090952.5 | 2 | 52859723 | 52863516 | 3794 | 2 | ENSMUST00 | -119004 | Gm26603 | ENSMUSG0C | false | . | . | . | . | . | . | chr2 | 52976528 | 52990003 | 4.4322 | 3.76878 | 2.54508 |
| chr14 | 120366987 | 120367189 | 182 | Intron (ENSMUST00000088419.7 | 14 | 120384263 | 120388985 | 4723 | 2 | ENSMUST00 | 21796 | Gm26679 | ENSMUSG0C | false | . | . | . | . | . | . | chr14 | 120362275 | 120389547 | 5.43713 | 4.23022 | 3.37672 |
| chr14 | 120352185 | 120352378 | 182 | Intron (ENSMUST00000088419.7 | 14 | 120384263 | 120388985 | 4723 | 2 | ENSMUST00 | 36607 | Gm26679 | ENSMUSG0C | false | . | . | . | . | . | . | chr14 | 120347663 | 120357777 | 5.63093 | 4.35006 | 3.52524 |
| chr19 | 23044882 | 23045077 | 182 | Intron (ENSMUST00000181623.1 | 19 | 23037390 | 23075853 | 38464 | 2 | ENSMUST00 | 30776 | C330002G04 | ENSMUSG0C | false | . | . | . | . | . | . | chr19 | 23044667 | 23045224 | 3.55205 | 3.26387 | 1.8449 |
| chr13 | 97282097 | 97282440 | 182 | Promoter (<=1kb) | 13 | 97283182 | 97285934 | 2753 | 1 | ENSMUST00 | -742 | C430039J16F | ENSMUSG0C | false | . | . | . | . | . | . | . | . | . | . | . | . |
| chr8 | 120520907 | 120521291 | 182 | Promoter (<=1kb) | 8 | 120520615 | 120521686 | 1072 | 1 | ENSMUST00 | 293 | Gm26971 | ENSMUSG0C | false | . | . | . | . | . | . | chr8 | 120517809 | 120521647 | 5.68803 | 4.34228 | 3.58366 |
| chr7 | 70514747 | 70514967 | 182 | Distal Intergenic | 7 | 70490848 | 70491851 | 1004 | 1 | ENSMUST00 | 23900 | Gm7656 | ENSMUSG0C | false | . | . | . | . | . | . | chr7 | 70514295 | 70517035 | 3.77783 | 3.48235 | 2.03469 |
| chr1 | 135129842 | 135130033 | 182 | Promoter (1-2kb) | 1 | 135129472 | 135131163 | 1692 | 2 | ENSMUST00 | 1130 | Ptprv | ENSMUSG0C | false | . | . | . | . | . | . | chr1 | 135126075 | 135132880 | 5.21831 | 4.07933 | 3.22044 |
| chr1 | 135131128 | 135131308 | 182 | Promoter (<=1kb) | 1 | 135129472 | 135131163 | 1692 | 2 | ENSMUST00 | 0 | Ptprv | ENSMUSG0C | false | . | . | . | . | . | . | chr1 | 135126075 | 135132880 | 5.21831 | 4.07933 | 3.22044 |
| chr6 | 93224056 | 93224307 | 182 | Intron (ENSMUST00000181121.2 | 6 | 93137004 | 93137990 | 987 | 2 | ENSMUST00 | -86067 | Gm5313 | ENSMUSG0C | false | . | . | . | . | . | . | chr6 | 93222956 | 93224722 | 5.26625 | 4.24905 | 3.21239 |
| chr18 | 55877930 | 55878159 | 182 | Distal Intergenic | 18 | 55837972 | 55838949 | 978 | 2 | ENSMUST00 | -38982 | Gm8614 | ENSMUSG0C | false | . | . | . | . | . | . | . | . | . | . | . | . |
| chr7 | 30072646 | 30073004 | 182 | Promoter (<=1kb) | 7 | 30056037 | 30072900 | 16864 | 2 | ENSMUST00 | 0 | Zfp82 | ENSMUSG0C | false | chr7 | 30072094 | 30073231 | 30.03893 | 101.89226 | 98.2715 | chr7 | 30071915 | 30073163 | 4.21043 | 3.63143 | 2.42605 |
| chr7 | 136046582 | 136046891 | 182 | Distal Intergenic | 7 | 135952798 | 135953783 | 986 | 1 | ENSMUST00 | 93785 | Gm9341 | ENSMUSG0C | false | . | . | . | . | . | . | chr7 | 136046071 | 136049601 | 4.67747 | 3.75748 | 2.77723 |
| chr7 | 136048967 | 136049177 | 182 | Distal Intergenic | 7 | 135952798 | 135953783 | 986 | 1 | ENSMUST00 | 96170 | Gm9341 | ENSMUSG0C | false | . | . | . | . | . | . | chr7 | 136046071 | 136049601 | 4.67747 | 3.75748 | 2.77723 |
| chr18 | 61620542 | 61620815 | 182 | Distal Intergenic | 18 | 61639542 | 61647292 | 7751 | 1 | ENSMUST00 | -18727 | Gm20748 | ENSMUSG0C | false | . | . | . | . | . | . | chr18 | 61618885 | 61621535 | 4.86692 | 3.95755 | 2.9111 |
| chr13 | 44544797 | 44545081 | 182 | Distal Intergenic | 13 | 44400436 | 44455303 | 54868 | 2 | ENSMUST00 | -89495 | Gm27007 | ENSMUSG0C | false | . | . | . | . | . | . | . | . | . | . | . | . |
| chr13 | 44501556 | 44502160 | 182 | Distal Intergenic | 13 | 44400436 | 44455303 | 54868 | 2 | ENSMUST00 | -46254 | Gm27007 | ENSMUSG0C | false | . | . | . | . | . | . | chr13 | 44501171 | 44510788 | 6.13728 | 4.58773 | 3.94959 |
| chr7 | 112930697 | 112931019 | 182 | Distal Intergenic | 7 | 112953962 | 112957457 | 3496 | 1 | ENSMUST00 | -22943 | Rassf10 | ENSMUSG0C | false | . | . | . | . | . | . | chr7 | 112928775 | 112932203 | 4.42744 | 3.74158 | 2.54421 |
| chr13 | 45191949 | 45192128 | 182 | Distal Intergenic | 13 | 45186534 | 45187536 | 1003 | 2 | ENSMUST00 | -4414 | Gm8513 | ENSMUSG0C | false | . | . | . | . | . | . | chr13 | 45191595 | 45192721 | 3.83626 | 3.49529 | 2.08844 |
| chr10 | 122291205 | 122291449 | 182 | Distal Intergenic | 10 | 122169373 | 122170363 | 991 | 1 | ENSMUST00 | 121833 | Gm9081 | ENSMUSG0C | false | . | . | . | . | . | . | . | . | . | . | . | . |
| chr8 | 36016423 | 36016759 | 182 | Distal Intergenic | 8 | 36025753 | 36026133 | 381 | 1 | ENSMUST00 | -8994 | Rps12l2 | ENSMUSG0C | false | . | . | . | . | . | . | . | . | . | . | . | . |
| chr7 | 138919293 | 138919701 | 182 | Distal Intergenic | 7 | 138914388 | 138915314 | 927 | 1 | ENSMUST00 | 4906 | Gm9358 | ENSMUSG0C | false | chr7 | 138919098 | 138919991 | 10.54274 | 25.8059 | 23.39399 | chr7 | 138919025 | 138920078 | 4.66551 | 3.81163 | 2.75741 |
| chr1 | 7126192 | 7126409 | 182 | Intron (ENSMUST00000061280.1 | 1 | 7136199 | 7136586 | 388 | 2 | ENSMUST00 | 10177 | Gm26983 | ENSMUSG0C | false | . | . | . | . | . | . | chr1 | 7124522 | 7128133 | 5.67414 | 4.3411 | 3.5736 |
| chr1 | 7124941 | 7125122 | 182 | Intron (ENSMUST00000061280.1 | 1 | 7136199 | 7136586 | 388 | 2 | ENSMUST00 | 11464 | Gm26983 | ENSMUSG0C | false | . | . | . | . | . | . | chr1 | 7124522 | 7128133 | 5.67414 | 4.3411 | 3.5736 |
| chr18 | 76158247 | 76158437 | 182 | Distal Intergenic | 18 | 76170552 | 76170652 | 101 | 1 | ENSMUST00 | -12115 | Mir6358 | ENSMUSG0C | false | . | . | . | . | . | . | chr18 | 76157206 | 76161646 | 4.87318 | 4.02407 | 2.90771 |
| chr19 | 7422086 | 7422421 | 182 | Promoter (<=1kb) | 19 | 7422573 | 7422642 | 70 | 1 | ENSMUST00 | -152 | Mir6991 | ENSMUSG0C | false | chr19 | 7422035 | 7423817 | 11.15939 | 31.88065 | 29.39184 | chr19 | 7421090 | 7424070 | 5.83547 | 4.35906 | 3.72946 |
| chr4 | 33437978 | 33438185 | 182 | Promoter (<=1kb) | 4 | 33438081 | 33438209 | 129 | 1 | ENSMUST00 | 0 | Mir8118 | ENSMUSG0C | false | . | . | . | . | . | . | chr4 | 33437647 | 33438481 | 6.20534 | 4.75811 | 3.98753 |
| chr4 | 33405789 | 33406043 | 182 | Intron (ENSMUST00000108153.3 | 4 | 33438081 | 33438209 | 129 | 1 | ENSMUST00 | -32038 | Mir8118 | ENSMUSG0C | false | . | . | . | . | . | . | chr4 | 33405860 | 33406478 | 4.5069 | 3.76953 | 2.63976 |
| chr8 | 87439263 | 87439561 | 182 | Distal Intergenic | 8 | 87458444 | 87459435 | 992 | 1 | ENSMUST00 | -18883 | 4930535O05 | ENSMUSG0C | false | . | . | . | . | . | . | chr8 | 87438858 | 87439962 | 4.08986 | 3.38771 | 2.29562 |
| chr11 | 69503929 | 69504153 | 182 | Promoter (1-2kb) | 11 | 69505358 | 69505435 | 78 | 2 | ENSMUST00 | 1282 | Gm27493 | ENSMUSG0C | false | . | . | . | . | . | . | chr11 | 69501763 | 69504897 | 5.03474 | 4.15547 | 3.02535 |
| chr1 | 53851332 | 53851603 | 182 | Promoter (1-2kb) | 1 | 53849424 | 53849495 | 72 | 1 | ENSMUST00 | 1909 | Mir7681 | ENSMUSG0C | false | . | . | . | . | . | . | chr1 | 53849378 | 53851572 | 4.2838 | 3.69093 | 2.40793 |
| chr11 | 107648409 | 107648661 | 182 | Intron (ENSMUST00000100305.2 | 11 | 107637722 | 107637779 | 58 | 1 | ENSMUST00 | 10688 | Mir6932 | ENSMUSG0C | false | . | . | . | . | . | . | chr11 | 107647908 | 107653872 | 5.72702 | 4.35515 | 3.61075 |
| chr9 | 33077892 | 33078198 | 182 | Intron (ENSMUST00000184476.1 | 9 | 33031791 | 33090799 | 59009 | 1 | ENSMUST00 | 46102 | Gm27166 | ENSMUSG0C | false | . | . | . | . | . | . | . | . | . | . | . | . |
| chr9 | 102264983 | 102265220 | 182 | Intron (ENSMUST00000035129.8 | 9 | 102304924 | 102304975 | 52 | 2 | ENSMUST00 | 39755 | Mir7243 | ENSMUSG0C | false | . | . | . | . | . | . | chr9 | 102258607 | 102274961 | 4.8108 | 3.97089 | 2.85625 |
| chr16 | 85168517 | 85168703 | 182 | Promoter (<=1kb) | 16 | 85168849 | 85169001 | 153 | 1 | ENSMUST00 | -146 | Gm27295 | ENSMUSG0C | false | . | . | . | . | . | . | chr16 | 85167810 | 85172354 | 5.42256 | 4.17976 | 3.36286 |
| chr1 | 85751477 | 85751801 | 182 | Promoter (<=1kb) | 1 | 85751948 | 85752074 | 127 | 1 | ENSMUST00 | -147 | Mir8096 | ENSMUSG0C | false | . | . | . | . | . | . | chr1 | 85749089 | 85752462 | 4.06414 | 3.65882 | 2.22962 |
| chr15 | 89179019 | 89179416 | 182 | Downstream (2-3kb) | 15 | 89185465 | 89185537 | 73 | 2 | ENSMUST00 | 6121 | Mir6958 | ENSMUSG0C | false | chr15 | 89178489 | 89181434 | 28.29403 | 89.8816 | 86.54237 | chr15 | 89167407 | 89179959 | 7.78402 | 5.30321 | 5.27943 |
| chr6 | 108458134 | 108458375 | 182 | Promoter (1-2kb) | 6 | 108456183 | 108456536 | 354 | 1 | ENSMUST00 | 1952 | Gm27813 | ENSMUSG0C | false | . | . | . | . | . | . | chr6 | 108454833 | 108459631 | 5.66918 | 4.2365 | 3.5953 |
| chr19 | 47265528 | 47265725 | 182 | 3' UTR | 19 | 47273519 | 47273589 | 71 | 2 | ENSMUST00 | 7864 | Mir6995 | ENSMUSG0C | false | . | . | . | . | . | . | chr19 | 47264187 | 47266853 | 4.1213 | 3.37747 | 2.31459 |
| chr13 | 32680873 | 32681312 | 182 | Distal Intergenic | 13 | 32758202 | 32758310 | 109 | 1 | ENSMUST00 | -76890 | Gm27342 | ENSMUSG0C | false | . | . | . | . | . | . | chr13 | 32678441 | 32682534 | 5.54396 | 4.36904 | 3.43747 |
| chr13 | 32764635 | 32764952 | 182 | Intron (ENSMUST00000057428.5 | 13 | 32758202 | 32758310 | 109 | 1 | ENSMUST00 | 6434 | Gm27342 | ENSMUSG0C | false | . | . | . | . | . | . | . | . | . | . | . | . |
| chr1 | 169160876 | 169161063 | 182 | Distal Intergenic | 1 | 169018974 | 169019089 | 116 | 2 | ENSMUST00 | -141788 | Mir6354 | ENSMUSG0C | false | . | . | . | . | . | . | . | . | . | . | . | . |
| chr1 | 39128744 | 39128971 | 182 | Distal Intergenic | 1 | 39186896 | 39186992 | 97 | 2 | ENSMUST00 | 58021 | Mir6349 | ENSMUSG0C | false | . | . | . | . | . | . | . | . | . | . | . | . |
| chr14 | 67866492 | 67866700 | 182 | Intron (ENSMUST00000039135.4 | 14 | 67859636 | 67859745 | 110 | 2 | ENSMUST00 | -6748 | Mir6539 | ENSMUSG0C | false | . | . | . | . | . | . | chr14 | 67862320 | 67900048 | 6.11049 | 4.53842 | 3.92537 |
| chr5 | 36313820 | 36314056 | 182 | Intron (ENSMUST00000037370.8 | 5 | 36301570 | 36301628 | 59 | 1 | ENSMUST00 | 12251 | Mir7689 | ENSMUSG0C | false | . | . | . | . | . | . | chr5 | 36313194 | 36314360 | 3.84172 | 3.45333 | 2.11452 |
| chr8 | 8689702 | 8690034 | 182 | Promoter (<=1kb) | 8 | 8690093 | 8690162 | 70 | 2 | ENSMUST00 | 128 | Mir7654 | ENSMUSG0C | false | chr8 | 8688529 | 8690897 | 33.10969 | 106.34647 | 102.61562 | chr8 | 8678363 | 8689729 | 4.36174 | 3.73621 | 2.49372 |
| chr9 | 96493339 | 96493540 | 182 | Promoter (1-2kb) | 9 | 96495132 | 96495213 | 82 | 2 | ENSMUST00 | 1673 | Gm27289 | ENSMUSG0C | false | . | . | . | . | . | . | chr9 | 96491707 | 96493747 | 4.17095 | 3.58362 | 2.37404 |
| chr6 | 52160781 | 52161150 | 182 | Promoter (<=1kb) | 6 | 52161042 | 52161255 | 214 | 1 | ENSMUST00 | 0 | Gm27861 | ENSMUSG0C | false | chr6 | 52160172 | 52161353 | 10.10971 | 19.98882 | 17.65231 | chr6 | 52159328 | 52167512 | 5.47105 | 4.07894 | 3.42628 |
| chr5 | 34583755 | 34584038 | 182 | Intron (ENSMUST00000114338.3 | 5 | 34574151 | 34574213 | 63 | 1 | ENSMUST00 | 9605 | Mir7036b | ENSMUSG0C | false | . | . | . | . | . | . | chr5 | 34574694 | 34588910 | 5.00686 | 4.08485 | 3.01427 |
| chr7 | 143447428 | 143447689 | 182 | Promoter (<=1kb) | 7 | 143447605 | 143447665 | 61 | 1 | ENSMUST00 | 0 | Gm27901 | ENSMUSG0C | false | . | . | . | . | . | . | chr7 | 143437483 | 143448788 | 5.47047 | 4.33141 | 3.38873 |
| chr7 | 143442045 | 143442587 | 182 | Distal Intergenic | 7 | 143447605 | 143447665 | 61 | 1 | ENSMUST00 | -5018 | Gm27901 | ENSMUSG0C | false | . | . | . | . | . | . | chr7 | 143437483 | 143448788 | 5.47047 | 4.33141 | 3.38873 |
| chr18 | 38578351 | 38578557 | 182 | Distal Intergenic | 18 | 38594158 | 38594228 | 71 | 1 | ENSMUST00 | -15601 | Gm27369 | ENSMUSG0C | false | . | . | . | . | . | . | chr18 | 38576240 | 38578765 | 2.95177 | 2.9108 | 1.38494 |
| chr9 | 32764460 | 32764672 | 182 | Distal Intergenic | 9 | 32813734 | 32834104 | 20371 | 1 | ENSMUST00 | -49062 | Gm3331 | ENSMUSG0C | false | . | . | . | . | . | . | chr9 | 32764110 | 32765094 | 3.62647 | 3.28073 | 1.91462 |
| chr2 | 17583816 | 17584034 | 182 | Intron (ENSMUST00000028080.6 | 2 | 17563514 | 17563626 | 113 | 1 | ENSMUST00 | 20303 | Mir6419 | ENSMUSG0C | false | . | . | . | . | . | . | chr2 | 17582299 | 17588022 | 5.57183 | 4.36889 | 3.46548 |
| chr7 | 80057492 | 80057743 | 182 | Intron (ENSMUST00000164056.3 | 7 | 80026418 | 80026513 | 96 | 2 | ENSMUST00 | -30980 | Gm27983 | ENSMUSG0C | false | . | . | . | . | . | . | chr7 | 80043846 | 80073175 | 7.55169 | 5.20662 | 5.10629 |
| chr7 | 80055850 | 80056313 | 182 | Intron (ENSMUST00000164056.3 | 7 | 80026418 | 80026513 | 96 | 2 | ENSMUST00 | -29338 | Gm27983 | ENSMUSG0C | false | . | . | . | . | . | . | chr7 | 80043846 | 80073175 | 7.55169 | 5.20662 | 5.10629 |
| chrX | 169302700 | 169302962 | 182 | 3' UTR | 20 | 169295009 | 169295177 | 169 | 1 | ENSMUST00 | 7692 | Gm27980 | ENSMUSG0C | false | . | . | . | . | . | . | . | . | . | . | . | . |
| chr10 | 62208534 | 62208788 | 182 | Intron (ENSMUST00000047883.9 | 10 | 62194167 | 62194227 | 61 | 2 | ENSMUST00 | -14308 | Mir7662 | ENSMUSG0C | false | . | . | . | . | . | . | chr10 | 62185685 | 62229768 | 8.28034 | 5.46771 | 5.70297 |
| chr15 | 80004747 | 80005090 | 182 | Intron (ENSMUST00000000500.6 | 15 | 79998970 | 79999026 | 57 | 2 | ENSMUST00 | -5722 | Mir7213 | ENSMUSG0C | false | . | . | . | . | . | . | chr15 | 80003121 | 80008590 | 4.67086 | 3.88569 | 2.76659 |
| chr10 | 122718062 | 122718394 | 182 | Intron (ENSMUST00000162853.2 | 10 | 122679315 | 122679415 | 101 | 1 | ENSMUST00 | 38748 | Mir8104 | ENSMUSG0C | false | . | . | . | . | . | . | chr10 | 122701076 | 122720564 | 4.67114 | 3.91157 | 2.74683 |
| chr12 | 69361137 | 69361583 | 182 | Promoter (<=1kb) | 12 | 69361182 | 69361481 | 300 | 2 | ENSMUST00 | 0 | Gm26191 | ENSMUSG0C | false | chr12 | 69361102 | 69361383 | 13.18045 | 29.78312 | 27.32097 | . | . | . | . | . | . |
| chr19 | 3305423 | 3305634 | 182 | Distal Intergenic | 19 | 3288920 | 3288983 | 64 | 1 | ENSMUST00 | 16504 | Mir6984 | ENSMUSG0C | false | . | . | . | . | . | . | chr19 | 3305460 | 3305734 | 3.7679 | 3.42516 | 2.00531 |
| chr11 | 97138825 | 97139100 | 182 | 3' UTR | 11 | 97146364 | 97146419 | 56 | 2 | ENSMUST00 | 7319 | Mir7235 | ENSMUSG0C | false | chr11 | 97138628 | 97139531 | 12.76655 | 31.65917 | 29.17213 | chr11 | 97138554 | 97139744 | 3.89152 | 3.52379 | 2.13928 |
| chr1 | 88994923 | 88995343 | 182 | Distal Intergenic | 1 | 89016113 | 89022210 | 6098 | 1 | ENSMUST00 | -20770 | RP24-423I20 | ENSMUSG0C | false | . | . | . | . | . | . | chr1 | 88994776 | 88996817 | 3.87762 | 3.28546 | 2.12023 |
| chr1 | 138561389 | 138561710 | 182 | Exon (ENSMUST00000027642.3/ | 1 | 138515618 | 138520115 | 4498 | 1 | ENSMUST00 | 45772 | RP23-209N1 | ENSMUSG0C | false | . | . | . | . | . | . | chr1 | 138558996 | 138561915 | 4.87371 | 4.02638 | 2.9076 |
| chr11 | 109347554 | 109347922 | 182 | Promoter (<=1kb) | 11 | 109346839 | 109353651 | 6813 | 1 | ENSMUST00 | 716 | RP23-12N20 | ENSMUSG0C | false | . | . | . | . | . | . | chr11 | 109346151 | 109348426 | 5.72192 | 4.33546 | 3.63231 |
| chr7 | 110689109 | 110689673 | 182 | Distal Intergenic | 7 | 110626281 | 110639358 | 13078 | 2 | ENSMUST00 | -49752 | RP23-25I3.4 | ENSMUSG0C | false | . | . | . | . | . | . | chr7 | 110688531 | 110690876 | 3.35224 | 3.25066 | 1.65377 |
| chr7 | 71705443 | 71705811 | 182 | Distal Intergenic | 7 | 71370333 | 71379813 | 9481 | 1 | ENSMUST00 | 335111 | RP24-173C1 | ENSMUSG0C | false | chr7 | 71705310 | 71705898 | 16.11243 | 40.23772 | 37.64439 | chr7 | 71705224 | 71706119 | 3.79122 | 3.32563 | 2.0431 |
| chr8 | 8109920 | 8110408 | 182 | Distal Intergenic | 8 | 7855847 | 7857912 | 2066 | 1 | ENSMUST00 | 254074 | RP24-87L12. | ENSMUSG0C | false | . | . | . | . | . | . | chr8 | 8109619 | 8111477 | 5.39551 | 3.95271 | 3.37026 |
| chr14 | 20640841 | 20641100 | 182 | Promoter (<=1kb) | 14 | 20641009 | 20644048 | 3040 | 1 | ENSMUST00 | 0 | RP24-155O1 | ENSMUSG0C | false | chr14 | 20640227 | 20641954 | 34.54686 | 107.82351 | 104.0565 | . | . | . | . | . | . |
| chr19 | 53084652 | 53085093 | 182 | Distal Intergenic | 19 | 53076252 | 53084392 | 8141 | 1 | ENSMUST00 | 8401 | RP23-76E22. | ENSMUSG0C | false | . | . | . | . | . | . | chr19 | 53084059 | 53087368 | 3.42023 | 3.30519 | 1.69478 |
| chr13 | 37588226 | 37588507 | 182 | Distal Intergenic | 13 | 37508940 | 37512074 | 3135 | 1 | ENSMUST00 | 79287 | RP23-21N15 | ENSMUSG0C | false | . | . | . | . | . | . | . | . | . | . | . | . |
| chr5 | 100226891 | 100227072 | 182 | Intron (ENSMUST00000150506.1 | 5 | 100210708 | 100216741 | 6034 | 1 | ENSMUST00 | 16184 | RP23-52N17 | ENSMUSG0C | false | . | . | . | . | . | . | chr5 | 100226249 | 100228195 | 5.14112 | 3.9512 | 3.15502 |
| chr1 | 41426465 | 41426677 | 182 | Distal Intergenic | 1 | 41526931 | 41529543 | 2613 | 2 | ENSMUST00 | 102866 | RP23-59E7.1 | ENSMUSG0C | false | . | . | . | . | . | . | chr1 | 41426007 | 41428591 | 3.92994 | 3.56129 | 2.1546 |
| chr4 | 95162378 | 95162586 | 182 | Intron (ENSMUST00000125438.2 | 4 | 95151740 | 95157503 | 5764 | 1 | ENSMUST00 | 10639 | RP23-77I17. | ENSMUSG0C | false | . | . | . | . | . | . | chr4 | 95159243 | 95162809 | 4.78211 | 3.95901 | 2.84821 |
| chr4 | 95156909 | 95157129 | 182 | Exon (ENSMUST00000191391.1/ | 4 | 95151740 | 95157503 | 5764 | 1 | ENSMUST00 | 5170 | RP23-77I17. | ENSMUSG0C | false | . | . | . | . | . | . | chr4 | 95152754 | 95157985 | 3.5377 | 3.22179 | 1.82132 |
| chr18 | 35834836 | 35835078 | 182 | Intron (ENSMUST00000060722.6 | 18 | 35830389 | 35831093 | 705 | 2 | ENSMUST00 | -3744 | RP24-326K1 | ENSMUSG0C | false | . | . | . | . | . | . | chr18 | 35831800 | 35857612 | 7.70883 | 5.15861 | 5.24645 |
| chr18 | 35838753 | 35838942 | 182 | Intron (ENSMUST00000060722.6 | 18 | 35830389 | 35831093 | 705 | 2 | ENSMUST00 | -7661 | RP24-326K1 | ENSMUSG0C | false | . | . | . | . | . | . | chr18 | 35831800 | 35857612 | 7.70883 | 5.15861 | 5.24645 |
| chr8 | 26687567 | 26687814 | 182 | Distal Intergenic | 8 | 26814593 | 26824423 | 9831 | 2 | ENSMUST00 | 136609 | RP24-502D8. | ENSMUSG0C | false | . | . | . | . | . | . | chr8 | 26687046 | 26690598 | 2.93255 | 2.902 | 1.3714 |
| chr8 | 26780563 | 26780902 | 182 | Distal Intergenic | 8 | 26814593 | 26824423 | 9831 | 2 | ENSMUST00 | 43521 | RP24-502D8. | ENSMUSG0C | false | chr8 | 26780573 | 26780855 | 9.19064 | 17.46287 | 15.16242 | chr8 | 26778443 | 26781090 | 6.95771 | 5.07118 | 4.60599 |
| chr1 | 88639519 | 88639709 | 182 | Distal Intergenic | 1 | 88656308 | 88658100 | 1793 | 1 | ENSMUST00 | -16599 | RP24-108I6. | ENSMUSG0C | false | chr1 | 88639579 | 88640560 | 12.76655 | 31.65917 | 29.17213 | chr1 | 88638372 | 88641897 | 5.83065 | 4.49863 | 3.69173 |
| chr1 | 88640118 | 88640372 | 182 | Distal Intergenic | 1 | 88656308 | 88658100 | 1793 | 1 | ENSMUST00 | -15936 | RP24-108I6. | ENSMUSG0C | false | chr1 | 88639579 | 88640560 | 12.76655 | 31.65917 | 29.17213 | chr1 | 88638372 | 88641897 | 5.83065 | 4.49863 | 3.69173 |
| chr1 | 186705154 | 186705638 | 182 | Promoter (<=1kb) | 1 | 186705695 | 186707033 | 1339 | 1 | ENSMUST00 | -57 | RP24-276N2 | ENSMUSG0C | false | chr1 | 186703443 | 186705953 | 25.29353 | 74.26213 | 71.20484 | chr1 | 186705320 | 186705939 | 3.55975 | 3.36263 | 1.80966 |
| chr19 | 45330124 | 45330361 | 182 | Promoter (<=1kb) | 19 | 45329649 | 45330394 | 746 | 2 | ENSMUST00 | 33 | RP23-188J8. | ENSMUSG0C | false | . | . | . | . | . | . | chr19 | 45328946 | 45330740 | 4.83863 | 3.92205 | 2.9079 |
| chr1 | 62911741 | 62912291 | 182 | Distal Intergenic | 1 | 62821355 | 62832370 | 11016 | 2 | ENSMUST00 | -79372 | RP23-149A5. | ENSMUSG0C | false | . | . | . | . | . | . | chr1 | 62911161 | 62912391 | 3.61533 | 3.41656 | 1.86802 |
| chr8 | 34525393 | 34525636 | 182 | Distal Intergenic | 8 | 34359950 | 34381248 | 21299 | 1 | ENSMUST00 | 165444 | RP24-336D1 | ENSMUSG0C | false | . | . | . | . | . | . | . | . | . | . | . | . |
| chr5 | 118870781 | 118871007 | 182 | Distal Intergenic | 5 | 119108234 | 119114543 | 6310 | 1 | ENSMUST00 | -237227 | RP24-389L1C | ENSMUSG0C | false | . | . | . | . | . | . | . | . | . | . | . | . |
| chr15 | 95282920 | 95283146 | 182 | Intron (ENSMUST00000166170.2 | 15 | 95097542 | 95162394 | 64853 | 2 | ENSMUST00 | -120527 | RP24-303N1 | ENSMUSG0C | false | . | . | . | . | . | . | chr15 | 95280101 | 95284037 | 5.82739 | 4.36965 | 3.73075 |
| chr7 | 71223699 | 71223899 | 182 | Intron (ENSMUST00000186618.1 | 7 | 71217049 | 71224409 | 7361 | 1 | ENSMUST00 | 6651 | RP23-257M2 | ENSMUSG0C | false | . | . | . | . | . | . | chr7 | 71223586 | 71224831 | 2.85767 | 2.94726 | 1.29573 |
| chr7 | 71233155 | 71233592 | 182 | Distal Intergenic | 7 | 71217049 | 71224409 | 7361 | 1 | ENSMUST00 | 16107 | RP23-257M2 | ENSMUSG0C | false | . | . | . | . | . | . | . | . | . | . | . | . |
| chr17 | 86310942 | 86311228 | 182 | Intron (ENSMUST00000097275.3 | 17 | 86285164 | 86287178 | 2015 | 2 | ENSMUST00 | -23765 | RP23-458O2 | ENSMUSG0C | false | . | . | . | . | . | . | chr17 | 86309303 | 86315004 | 5.61066 | 4.27176 | 3.52062 |
| chr1 | 90973074 | 90973405 | 182 | Promoter (1-2kb) | 1 | 90974459 | 90975260 | 802 | 2 | ENSMUST00 | 1855 | RP23-448B1 | ENSMUSG0C | false | . | . | . | . | . | . | chr1 | 90972989 | 90975458 | 5.65816 | 4.37626 | 3.5539 |
| chr1 | 62729791 | 62730017 | 182 | Intron (ENSMUST00000126344.1 | 1 | 62714723 | 62718073 | 3351 | 2 | ENSMUST00 | -11719 | RP23-20G7.7 | ENSMUSG0C | false | . | . | . | . | . | . | . | . | . | . | . | . |
| chr9 | 25207073 | 25207264 | 182 | Distal Intergenic | 9 | 25175151 | 25175597 | 447 | 1 | ENSMUST00 | 31923 | RP23-308G1 | ENSMUSG0C | false | . | . | . | . | . | . | chr9 | 25206122 | 25207203 | 3.421 | 3.28092 | 1.74942 |
| chr9 | 107612499 | 107612983 | 182 | Promoter (1-2kb) | 9 | 107610287 | 107611005 | 719 | 2 | ENSMUST00 | -1495 | RP23-456B9. | ENSMUSG0C | false | chr9 | 107612415 | 107612844 | 11.18601 | 23.41478 | 21.03445 | chr9 | 107604044 | 107615326 | 5.94291 | 4.4466 | 3.78781 |
| chr17 | 87531557 | 87531843 | 182 | Distal Intergenic | 17 | 87490748 | 87502616 | 11869 | 1 | ENSMUST00 | 40810 | RP23-243O1 | ENSMUSG0C | false | chr17 | 87530230 | 87532496 | 15.30398 | 43.94659 | 41.30965 | chr17 | 87529319 | 87532887 | 3.88643 | 3.47761 | 2.12937 |
| chr17 | 87532316 | 87532606 | 182 | Distal Intergenic | 17 | 87490748 | 87502616 | 11869 | 1 | ENSMUST00 | 41569 | RP23-243O1 | ENSMUSG0C | false | chr17 | 87530230 | 87532496 | 15.30398 | 43.94659 | 41.30965 | chr17 | 87529319 | 87532887 | 3.88643 | 3.47761 | 2.12937 |
| chr1 | 16238951 | 16239212 | 182 | Intron (ENSMUST00000186343.1 | 1 | 16228506 | 16244062 | 15557 | 1 | ENSMUST00 | 10446 | RP24-73D23. | ENSMUSG0C | false | . | . | . | . | . | . | chr1 | 16237236 | 16239484 | 3.46005 | 3.24385 | 1.78548 |
| chr1 | 16228286 | 16228505 | 182 | Promoter (<=1kb) | 1 | 16228506 | 16244062 | 15557 | 1 | ENSMUST00 | -1 | RP24-73D23. | ENSMUSG0C | false | . | . | . | . | . | . | chr1 | 16227912 | 16230061 | 3.6683 | 3.41945 | 1.94861 |
| chr1 | 59943961 | 59944172 | 182 | Intron (ENSMUST00000036540.6 | 1 | 59950403 | 59952464 | 2062 | 2 | ENSMUST00 | 8292 | RP23-27C10. | ENSMUSG0C | false | . | . | . | . | . | . | chr1 | 59929477 | 59947471 | 4.4142 | 3.75043 | 2.54049 |
| chr4 | 114609386 | 114609594 | 182 | Intron (ENSMUST00000094894.3 | 4 | 114672697 | 114673340 | 644 | 2 | ENSMUST00 | 63746 | RP23-454F1 | ENSMUSG0C | false | . | . | . | . | . | . | chr4 | 114605726 | 114614447 | 5.56033 | 4.27375 | 3.4804 |
| chr14 | 76787539 | 76787829 | 182 | Distal Intergenic | 14 | 76681754 | 76686887 | 5134 | 1 | ENSMUST00 | 105786 | RP23-421C2 | ENSMUSG0C | false | . | . | . | . | . | . | . | . | . | . | . | . |
| chr1 | 90273411 | 90273642 | 182 | Distal Intergenic | 1 | 90280458 | 90283912 | 3455 | 1 | ENSMUST00 | -6816 | RP24-548K2. | ENSMUSG0C | false | . | . | . | . | . | . | chr1 | 90272952 | 90273968 | 4.11055 | 3.40856 | 2.30472 |
| chr1 | 23187707 | 23188056 | 182 | Intron (ENSMUST00000191045.1 | 1 | 23048295 | 23235673 | 187379 | 2 | ENSMUST00 | 47617 | RP24-495G2 | ENSMUSG0C | false | . | . | . | . | . | . | . | . | . | . | . | . |
| chr18 | 6489995 | 6490553 | 182 | Promoter (<=1kb) | 18 | 6490143 | 6496962 | 6820 | 1 | ENSMUST00 | 0 | RP23-258L8. | ENSMUSG0C | false | chr18 | 6489051 | 6491620 | 35.33424 | 109.18479 | 105.37405 | . | . | . | . | . | . |
| chr15 | 103101642 | 103101930 | 182 | Promoter (<=1kb) | 15 | 103096151 | 103102111 | 5961 | 2 | ENSMUST00 | 181 | RP24-89G21 | ENSMUSG0C | false | chr15 | 103101488 | 103102633 | 17.18351 | 44.2994 | 41.65401 | chr15 | 103100132 | 103102756 | 4.94116 | 4.10799 | 2.94652 |
| chr7 | 143376058 | 143376326 | 182 | Intron (ENSMUST00000009689.5 | 7 | 143322775 | 143330650 | 7876 | 2 | ENSMUST00 | -45409 | RP23-124B2. | ENSMUSG0C | false | . | . | . | . | . | . | chr7 | 143373469 | 143376609 | 3.42726 | 3.08852 | 1.74978 |
| chr9 | 99226347 | 99226537 | 182 | Downstream (2-3kb) | 9 | 99220048 | 99220567 | 520 | 2 | ENSMUST00 | -5781 | RP23-114N1 | ENSMUSG0C | false | . | . | . | . | . | . | . | . | . | . | . | . |
| chrY | 6045122 | 6045388 | 182 | Distal Intergenic | 21 | 6018394 | 6019091 | 698 | 1 | ENSMUST00 | 26729 | RP24-475E1 | ENSMUSG0C | false | . | . | . | . | . | . | . | . | . | . | . | . |
| chr12 | 79708339 | 79708545 | 182 | Distal Intergenic | 12 | 79963568 | 79963872 | 305 | 1 | ENSMUST00 | -255023 | RP23-179D3. | ENSMUSG0C | false | . | . | . | . | . | . | chr12 | 79707523 | 79709631 | 5.97923 | 4.61212 | 3.80673 |
| chr12 | 79936764 | 79937047 | 182 | Distal Intergenic | 12 | 79963568 | 79963872 | 305 | 1 | ENSMUST00 | -26521 | RP23-179D3. | ENSMUSG0C | false | . | . | . | . | . | . | chr12 | 79935640 | 79937951 | 3.33306 | 3.08397 | 1.66964 |
| chr12 | 79796309 | 79796552 | 182 | Distal Intergenic | 12 | 79963568 | 79963872 | 305 | 1 | ENSMUST00 | -167016 | RP23-179D3. | ENSMUSG0C | false | . | . | . | . | . | . | . | . | . | . | . | . |
| chr12 | 79880947 | 79881206 | 182 | Distal Intergenic | 12 | 79963568 | 79963872 | 305 | 1 | ENSMUST00 | -82362 | RP23-179D3. | ENSMUSG0C | false | . | . | . | . | . | . | chr12 | 79879029 | 79881714 | 6.24528 | 4.53851 | 4.09018 |
| chr12 | 80001760 | 80002085 | 182 | Distal Intergenic | 12 | 79963568 | 79963872 | 305 | 1 | ENSMUST00 | 38193 | RP23-179D3. | ENSMUSG0C | false | . | . | . | . | . | . | . | . | . | . | . | . |
| chr12 | 79966653 | 79966879 | 182 | Distal Intergenic | 12 | 79963568 | 79963872 | 305 | 1 | ENSMUST00 | 3086 | RP23-179D3. | ENSMUSG0C | false | . | . | . | . | . | . | chr12 | 79963528 | 79966903 | 5.11632 | 4.0702 | 3.14196 |
| chr12 | 79683515 | 79683713 | 182 | Distal Intergenic | 12 | 79963568 | 79963872 | 305 | 1 | ENSMUST00 | -279855 | RP23-179D3. | ENSMUSG0C | false | . | . | . | . | . | . | chr12 | 79683466 | 79685131 | 3.65929 | 3.19782 | 1.94233 |
| chr12 | 79712267 | 79712559 | 182 | Distal Intergenic | 12 | 79963568 | 79963872 | 305 | 1 | ENSMUST00 | -251009 | RP23-179D3. | ENSMUSG0C | false | . | . | . | . | . | . | chr12 | 79711141 | 79712960 | 4.72581 | 3.84176 | 2.80659 |
| chr1 | 44781677 | 44782039 | 182 | Intron (ENSMUST00000074525.4 | 1 | 44739676 | 44744949 | 5274 | 2 | ENSMUST00 | -36729 | RP24-541P1 | ENSMUSG0C | false | . | . | . | . | . | . | chr1 | 44780922 | 44782316 | 3.66648 | 3.39069 | 1.9124 |
| chr1 | 157075083 | 157075320 | 182 | Distal Intergenic | 1 | 156972760 | 157054034 | 81275 | 2 | ENSMUST00 | -21050 | RP24-329K2 | ENSMUSG0C | false | . | . | . | . | . | . | . | . | . | . | . | . |
| chr18 | 82797910 | 82798115 | 182 | Distal Intergenic | 18 | 82754505 | 82759041 | 4537 | 2 | ENSMUST00 | -38870 | RP23-321L1 | ENSMUSG0C | false | . | . | . | . | . | . | chr18 | 82797987 | 82798838 | 3.67815 | 3.31175 | 1.96343 |
| chr15 | 85634429 | 85634821 | 182 | Distal Intergenic | 15 | 85639747 | 85645934 | 6188 | 1 | ENSMUST00 | -4926 | RP23-64J7.2 | ENSMUSG0C | false | chr15 | 85634793 | 85635329 | 18.26114 | 53.21968 | 50.46473 | chr15 | 85633840 | 85635497 | 4.68718 | 3.9568 | 2.77934 |
| chr1 | 64301811 | 64302233 | 182 | Promoter (<=1kb) | 1 | 64185991 | 64303043 | 117053 | 2 | ENSMUST00 | 810 | RP23-45M23 | ENSMUSG0C | false | . | . | . | . | . | . | . | . | . | . | . | . |
| chr10 | 17101337 | 17101565 | 182 | Distal Intergenic | 10 | 16951294 | 17095782 | 144489 | 2 | ENSMUST00 | -5556 | Gm20125 | ENSMUSG0C | false | . | . | . | . | . | . | chr10 | 17100485 | 17103034 | 4.096 | 3.67824 | 2.25337 |
| chr10 | 16959259 | 16959573 | 182 | Exon (ENSMUST00000185722.1/ | 10 | 16951294 | 17095782 | 144489 | 2 | ENSMUST00 | 136209 | Gm20125 | ENSMUSG0C | false | . | . | . | . | . | . | chr10 | 16956885 | 16959912 | 3.51067 | 3.29589 | 1.78679 |
| chr13 | 107619526 | 107619766 | 182 | Distal Intergenic | 13 | 107687397 | 107692162 | 4766 | 1 | ENSMUST00 | -67631 | RP23-125L8. | ENSMUSG0C | false | . | . | . | . | . | . | chr13 | 107619129 | 107620120 | 3.39542 | 3.26988 | 1.66833 |
| chr9 | 79116807 | 79117003 | 182 | Distal Intergenic | 9 | 79238620 | 79239058 | 439 | 1 | ENSMUST00 | -121617 | RP23-261O3 | ENSMUSG0C | false | . | . | . | . | . | . | . | . | . | . | . | . |
| chr15 | 67245151 | 67245407 | 182 | Intron (ENSMUST00000187233.1 | 15 | 67226769 | 67377094 | 150326 | 1 | ENSMUST00 | 18383 | RP23-163N4 | ENSMUSG0C | false | . | . | . | . | . | . | chr15 | 67245026 | 67245971 | 3.99033 | 3.61286 | 2.17004 |
| chr1 | 89191151 | 89191422 | 182 | Distal Intergenic | 1 | 89256182 | 89256981 | 800 | 2 | ENSMUST00 | 65559 | RP24-217I2. | ENSMUSG0C | false | . | . | . | . | . | . | chr1 | 89189289 | 89192366 | 5.66554 | 4.42415 | 3.54749 |
| chr15 | 67245151 | 67245407 | 182 | Intron (ENSMUST00000187233.1 | 15 | 67226769 | 67377094 | 150326 | 1 | ENSMUST00 | 18383 | RP23-163N4 | ENSMUSG00000102069.1 | | | | | | | | | | | | | |
| chr1 | 89191151 | 89191422 | 182 | Distal Intergenic | 1 | 89256182 | 89256981 | 800 | 2 | ENSMUST00 | 65559 | RP24-217I2. | ENSMUSG00000102083.1 | | | | | | | | | | | | | |

| Ontology | # Term Name | Binom Ran | Binom Raw | Binom FDR | Binom Fold | Binom Obs | Binom Reg | Hyper Rank | Hyper FDR | Hyper Fold | Hyper Obs | Hyper Tota | Hyper Gene Set Coverage |
|---|---|---|---|---|---|---|---|---|---|---|---|---|---|
| GO Biological Process | cell junction organization | 94 | 1.14428e-5 | 1.59346e-4 | 2.1023 | 511 | 2.99% | 349 | 1.50354e-1 | 1.6630 | 116 | 147 | 1.14% |
| | cell-cell junction organization | 102 | 1.63499e-5 | 2.09823e-4 | 2.1646 | 467 | 2.73% | 353 | 2.50278e-1 | 1.6924 | 106 | 132 | 1.04% |
| | positive regulation of supramolecular fiber organization | 113 | 5.76727e-4 | 6.68085e-4 | 2.0991 | 455 | 2.66% | 476 | 2.61474e-9 | 1.5279 | 116 | 160 | 1.14% |
| | cell-substrate adhesion | 123 | 2.88692e-4 | 3.07234e-4 | 2.0434 | 456 | 2.67% | 601 | 1.99050e-7 | 1.4780 | 108 | 154 | 1.06% |
| | negative regulation of translation | 139 | 4.59033e-3 | 4.32284e-3 | 2.2410 | 332 | 1.94% | 463 | 1.21627e-9 | 1.5927 | 99 | 131 | 0.98% |
| | positive regulation of actin filament bundle assembly | 140 | 7.12985e-3 | 6.66641e-3 | 2.7725 | 221 | 1.29% | 778 | 1.19282e-5 | 1.6860 | 44 | 55 | 0.43% |
| | regulation of actin filament bundle assembly | 152 | 1.68395e-3 | 1.45020e-3 | 2.1338 | 345 | 2.02% | 685 | 1.94477e-6 | 1.5748 | 68 | 91 | 0.67% |
| | negative regulation of cellular amide metabolic process | 154 | 1.59850e-3 | 1.35873e-3 | 2.1080 | 345 | 2.02% | 503 | 1.09724e-8 | 1.5367 | 105 | 144 | 1.03% |
| | cell junction assembly | 164 | 6.38972e-3 | 5.10009e-3 | 2.0081 | 370 | 2.16% | 572 | 9.54584e-8 | 1.5959 | 78 | 103 | 0.77% |
| | negative regulation of erythrocyte differentiation | 165 | 2.94829e-3 | 2.33897e-3 | 5.6574 | 79 | 0.46% | 1244 | 2.88081e-3 | 2.1075 | 11 | 11 | 0.11% |
| | adherens junction organization | 166 | 3.55880e-3 | 2.80631e-3 | 2.7477 | 190 | 1.11% | 706 | 2.82628e-6 | 1.8134 | 37 | 43 | 0.36% |
| | positive regulation of nuclear-transcribed mRNA catabolic proc | 172 | 2.33448e-3 | 1.77665e-3 | 4.0467 | 108 | 0.63% | 1379 | 5.52819e-3 | 1.8595 | 15 | 17 | 0.15% |
| | positive regulation of mRNA catabolic process | 179 | 3.72130e-3 | 2.72133e-2 | 3.5027 | 124 | 0.73% | 954 | 2.05518e-4 | 1.8643 | 23 | 26 | 0.23% |
| | regulation of stress fiber assembly | 183 | 7.01087e-3 | 5.01488e-2 | 2.1018 | 300 | 1.75% | 656 | 8.22224e-7 | 1.6422 | 60 | 77 | 0.59% |
| | positive regulation of stress fiber assembly | 196 | 4.32757e-2 | 2.89020e-2 | 2.5999 | 182 | 1.06% | 790 | 1.48927e-5 | 1.7410 | 38 | 46 | 0.37% |
| | positive regulation of adherens junction organization | 199 | 5.12440e-2 | 3.37077e-2 | 3.1694 | 132 | 0.77% | 1446 | 8.04846e-3 | 1.6860 | 20 | 25 | 0.20% |
| | positive regulation of cell junction assembly | 206 | 9.15426e-2 | 5.81696e-2 | 2.8833 | 151 | 0.88% | 1480 | 9.42544e-3 | 1.6157 | 23 | 30 | 0.23% |
| | positive regulation of focal adhesion assembly | 209 | 1.62567e-2 | 1.01818e-2 | 3.1792 | 129 | 0.75% | 1367 | 5.46938e-3 | 1.7410 | 19 | 23 | 0.19% |
| | regulation of transcription from RNA polymerase II promoter in | 219 | 2.91150e-2 | 1.74025e-2 | 2.6338 | 166 | 0.97% | 1261 | 3.11779e-3 | 1.5367 | 35 | 48 | 0.34% |
| | p38MAPK cascade | 222 | 5.61615e-2 | 3.31151e-2 | 5.6849 | 63 | 0.37% | 1976 | 3.58390e-2 | 2.1075 | 7 | 7 | 0.07% |

The test set of 17,097 genomic regions picked 10,152 genes (47%) of all 21,395 genes.
*GO Biological Process* has 13,090 ontology terms were tested (100%) using an annotation count range of [1, 1000].
GREAT version 4.0.4
Species assembly: mm10
Association rule: Basal+extension: 5000 bp upstream, 1000 bp downstream, 1000000 bp max extension, curated regulatory domains included

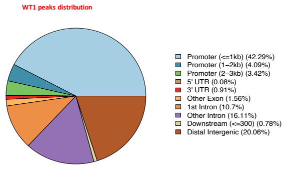

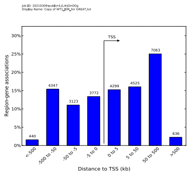

| Ontology | # Term Name | Binom Ran | Binom Raw | Binom FDR | Binom Fold | Binom Obs | Binom Reg | Hyper Rank | Hyper FDR | Hyper Fold | Hyper Obs | Hyper Tota | Hyper Gene Set Coverage |
|---|---|---|---|---|---|---|---|---|---|---|---|---|---|
| GO Cellular Component | ruffle | 20 | 2.41450e-7 | 2.04508e-7 | 2.5068 | 548 | 3.21% | 67 | 4.63813e-1 | 1.6162 | 125 | 163 | 1.23% |
| | ruffle membrane | 40 | 4.24824e-4 | 1.79913e-4 | 2.7117 | 280 | 1.64% | 124 | 2.44121e-6 | 1.5935 | 62 | 82 | 0.61% |
| | ribonucleoprotein granule | 41 | 1.80317e-4 | 7.45015e-4 | 2.1173 | 421 | 2.46% | 99 | 1.99685e-8 | 1.4523 | 133 | 193 | 1.31% |
| | cytoplasmic ribonucleoprotein granule | 43 | 5.11441e-4 | 2.01484e-4 | 2.1317 | 410 | 2.40% | 97 | 1.36478e-8 | 1.4706 | 127 | 182 | 1.25% |
| | cortical cytoskeleton | 46 | 2.55462e-3 | 9.40765e-4 | 2.3977 | 260 | 1.52% | 143 | 2.26157e-5 | 1.5119 | 66 | 92 | 0.65% |
| | actin filament bundle | 49 | 3.96890e-3 | 1.37211e-3 | 2.3982 | 243 | 1.42% | 134 | 1.35778e-5 | 1.5878 | 55 | 73 | 0.54% |
| | P-body | 51 | 6.14911e-3 | 2.04247e-2 | 2.4320 | 220 | 1.29% | 131 | 9.47426e-6 | 1.5739 | 59 | 79 | 0.58% |
| | stress fiber | 52 | 1.95882e-3 | 6.38123e-2 | 2.4754 | 209 | 1.22% | 135 | 1.41478e-5 | 1.6211 | 50 | 65 | 0.49% |
| | actomyosin | 55 | 1.12590e-2 | 3.46779e-2 | 2.2586 | 236 | 1.38% | 111 | 2.58752e-7 | 1.6638 | 60 | 76 | 0.59% |
| | PcG protein complex | 57 | 1.64892e-2 | 4.90047e-2 | 2.7051 | 160 | 0.94% | 153 | 4.82677e-5 | 1.6951 | 37 | 46 | 0.36% |
| | cortical actin cytoskeleton | 58 | 1.95833e-2 | 5.71967e-2 | 2.4430 | 192 | 1.12% | 219 | 3.55290e-3 | 1.4489 | 44 | 64 | 0.43% |
| | clathrin-coated pit | 67 | 3.76337e-2 | 9.51514e-2 | 2.2084 | 188 | 1.10% | 98 | 1.56396e-8 | 1.7782 | 54 | 64 | 0.53% |
| | pseudopodium | 75 | 1.09649e-1 | 2.47661e-1 | 5.1669 | 49 | 0.29% | 282 | 4.11595e-2 | 1.7832 | 11 | 13 | 0.11% |
| | cytoplasmic stress granule | 90 | 7.07921e-1 | 1.33246e-1 | 2.3143 | 118 | 0.69% | 180 | 4.81978e-4 | 1.6285 | 34 | 44 | 0.33% |
| | histone methyltransferase complex | 94 | 1.63184e-1 | 2.94079e-1 | 2.0195 | 157 | 0.92% | 173 | 3.12503e-4 | 1.5012 | 52 | 73 | 0.51% |
| | podosome | 103 | 2.47718e-1 | 4.07413e-1 | 2.4483 | 94 | 0.55% | 202 | 1.29446e-3 | 1.6996 | 25 | 31 | 0.25% |
| | ESC/E(Z) complex | 109 | 1.87539e-1 | 2.91460e-1 | 2.6629 | 75 | 0.44% | 253 | 1.30637e-2 | 1.7562 | 15 | 18 | 0.15% |
| | lamellipodium membrane | 113 | 4.64531e-1 | 6.96386e-1 | 2.7409 | 69 | 0.40% | 195 | 8.45828e-4 | 1.8326 | 20 | 23 | 0.20% |
| | cleavage furrow | 120 | 4.02729e-1 | 5.68519e-1 | 2.0377 | 116 | 0.68% | 157 | 6.20343e-5 | 1.6529 | 40 | 51 | 0.39% |
| | cytoplasmic microtubule | 121 | 7.81772e-1 | 1.09448e-1 | 2.0437 | 112 | 0.66% | 233 | 6.27345e-3 | 1.4401 | 41 | 60 | 0.40% |

The test set of 17,097 genomic regions picked 10,152 genes (47%) of all 21,395 genes.
*GO Cellular Component* ha 1,694 ontology terms were tested (100%) using an annotation count range of [1, 1000].
GREAT version 4.0.4
Species assembly: mm10
Association rule: Basal+extension: 5000 bp upstream, 1000 bp downstream, 1000000 bp max extension, curated regulatory domains included

| Ontology | # Term Name | Binom Ran | Binom Raw | Binom FDR | Binom Fold | Binom Obs | Binom Reg | Hyper Rank | Hyper FDR | Hyper Fold | Hyper Obs | Hyper Tota | Hyper Gene Set Coverage |
|---|---|---|---|---|---|---|---|---|---|---|---|---|---|
| GO Molecular Function | actin filament binding | 13 | 7.32144e-5 | 2.32653e-5 | 2.1670 | 488 | 2.85% | 102 | 1.24987e-7 | 1.4778 | 115 | 164 | 1.13% |
| | primary miRNA binding | 32 | 1.26126e-3 | 1.62821e-3 | 5.3757 | 89 | 0.52% | 279 | 3.79958e-2 | 2.1075 | 8 | 8 | 0.08% |
| | type I transforming growth factor beta receptor binding | 41 | 1.99758e-2 | 2.01269e-2 | 3.7893 | 98 | 0.57% | 230 | 1.03683e-2 | 2.1075 | 10 | 10 | 0.10% |
| | mRNA 3'-UTR binding | 58 | 1.39273e-2 | 9.91960e-2 | 2.2933 | 177 | 1.04% | 198 | 3.07567e-3 | 1.4956 | 44 | 62 | 0.43% |
| | R-SMAD binding | 64 | 4.14325e-2 | 2.67434e-1 | 2.5583 | 126 | 0.74% | 179 | 1.21014e-3 | 1.8440 | 21 | 24 | 0.21% |
| | protein phosphorylated amino acid binding | 71 | 8.20091e-1 | 4.77154e-1 | 2.1237 | 172 | 1.01% | 152 | 6.89889e-5 | 1.7039 | 38 | 47 | 0.37% |
| | mRNA 3'-UTR AU-rich region binding | 72 | 1.25694e-1 | 7.21168e-1 | 2.8363 | 97 | 0.57% | 267 | 2.87335e-2 | 1.9318 | 11 | 12 | 0.11% |
| | signaling adaptor activity | 73 | 1.42543e-1 | 8.06637e-1 | 2.0056 | 195 | 1.14% | 173 | 7.10296e-4 | 1.5905 | 40 | 53 | 0.39% |
| | 14-3-3 protein binding | 75 | 2.47777e-1 | 1.36475e-1 | 2.3684 | 132 | 0.77% | 191 | 2.16764e-3 | 1.7832 | 22 | 26 | 0.22% |
| | disordered domain specific binding | 91 | 3.86867e-1 | 1.75620e-1 | 2.3254 | 119 | 0.70% | 118 | 1.10589e-6 | 1.8913 | 35 | 39 | 0.34% |
| | translation regulator activity | 101 | 8.82058e-1 | 3.60770e-1 | 2.3672 | 104 | 0.61% | 223 | 8.18302e-3 | 1.5948 | 28 | 37 | 0.28% |
| | histone acetyltransferase binding | 104 | 1.10395e-1 | 4.38501e-1 | 2.1281 | 130 | 0.76% | 265 | 2.72254e-2 | 1.5988 | 22 | 29 | 0.22% |
| | unfolded protein binding | 120 | 2.53879e-1 | 8.73978e-1 | 2.1729 | 111 | 0.65% | 201 | 3.22594e-3 | 1.4310 | 55 | 81 | 0.54% |
| | cyclin-dependent protein serine/threonine kinase regulator acti | 122 | 4.29512e-1 | 1.45436e-1 | 2.4922 | 82 | 0.48% | 199 | 3.20427e-3 | 1.6996 | 25 | 31 | 0.25% |
| | glucocorticoid receptor binding | 128 | 1.14086e-1 | 3.68196e-1 | 2.3850 | 86 | 0.50% | 181 | 1.40518e-3 | 2.1075 | 13 | 13 | 0.13% |
| | histone acetyltransferase activity | 145 | 1.49840e-1 | 4.26888e-1 | 2.0430 | 109 | 0.64% | 178 | 1.16932e-3 | 1.5529 | 42 | 57 | 0.41% |
| | peptide-lysine-N-acetyltransferase activity | 149 | 2.71374e-1 | 7.52379e-1 | 2.0071 | 111 | 0.65% | 182 | 1.44470e-3 | 1.5359 | 43 | 59 | 0.42% |
| | histone-lysine N-methyltransferase activity | 156 | 8.28067e-1 | 2.19278e-9 | 2.0758 | 97 | 0.57% | 233 | 1.04095e-2 | 1.5327 | 32 | 44 | 0.32% |
| | poly(A) binding | 163 | 1.93213e-1 | 4.89669e-9 | 2.1927 | 82 | 0.48% | 235 | 1.05677e-2 | 1.7913 | 17 | 20 | 0.17% |
| | proline-rich region binding | 164 | 2.05873e-1 | 5.18574e-9 | 2.5329 | 61 | 0.36% | 202 | 3.32546e-3 | 1.8856 | 17 | 19 | 0.17% |

The test set of 17,097 genomic regions picked 10,152 genes (47%) of all 21,395 genes.
*GO Molecular Function* ha 4,131 ontology terms were tested (100%) using an annotation count range of [1, 1000].
GREAT version 4.0.4
Species assembly: mm10
Association rule: Basal+extension: 5000 bp upstream, 1000 bp downstream, 1000000 bp max extension, curated regulatory domains included

| Ontology | # Term Name | Binom Ran | Binom Raw | Binom FDR | Binom Fold | Binom Obs | Binom Reg | Hyper Rank | Hyper FDR | Hyper Fold | Hyper Obs | Hyper Tota | Hyper Gene Set Coverage |
|---|---|---|---|---|---|---|---|---|---|---|---|---|---|
| Human Phenotype | Hypoalbuminemia | 11 | 1.22760e-2 | 7.36447e-2 | 3.7054 | 82 | 0.48% | 491 | 4.32777e-3 | 1.7562 | 20 | 24 | 0.20% |
| | Focal segmental glomerulosclerosis | 19 | 2.54740e-2 | 8.84753e-1 | 2.7840 | 110 | 0.64% | 486 | 4.03941e-3 | 1.7172 | 22 | 27 | 0.22% |
| | Glomerulosclerosis | 22 | 1.33768e-1 | 4.01242e-1 | 2.6772 | 113 | 0.66% | 548 | 7.76257e-3 | 1.6316 | 24 | 31 | 0.24% |
| | Bruising susceptibility | 25 | 3.66334e-1 | 9.66976e-1 | 2.0771 | 185 | 1.08% | 462 | 3.18983e-3 | 1.4856 | 43 | 61 | 0.42% |
| | Cerebellar cyst | 43 | 1.95226e-1 | 2.99603e-1 | 2.0363 | 164 | 0.96% | 482 | 4.04629e-3 | 1.5110 | 38 | 53 | 0.37% |
| | Choanal atresia | 74 | 1.28797e-1 | 1.14855e-1 | 2.1097 | 132 | 0.77% | 392 | 1.34601e-3 | 1.6860 | 28 | 35 | 0.28% |
| | Stage 5 chronic kidney disease | 108 | 1.12668e-1 | 6.88421e-1 | 2.1808 | 104 | 0.61% | 685 | 2.86488e-2 | 1.4518 | 31 | 45 | 0.31% |
| | Tubular atrophy | 199 | 2.58006e-9 | 8.55569e-8 | 2.6275 | 50 | 0.29% | 524 | 6.07542e-3 | 1.9569 | 13 | 14 | 0.13% |
| | Diastasis recti | 224 | 1.01274e-8 | 2.98352e-7 | 2.6016 | 47 | 0.27% | 575 | 1.08914e-2 | 1.9454 | 12 | 13 | 0.12% |
| | Overgrowth | 242 | 1.89565e-8 | 5.16918e-7 | 2.7532 | 41 | 0.24% | 586 | 1.22463e-2 | 1.7747 | 16 | 19 | 0.16% |
| | Spinal muscular atrophy | 317 | 2.45802e-7 | 5.11687e-6 | 2.3266 | 47 | 0.27% | 586 | 1.22463e-2 | 1.7747 | 16 | 19 | 0.16% |
| | Vertebral compression fractures | 437 | 8.25240e-6 | 1.24617e-4 | 2.3514 | 34 | 0.20% | 711 | 3.21925e-2 | 1.7356 | 14 | 17 | 0.14% |
| | Renal corticomedullary cysts | 569 | 3.32084e-5 | 3.85136e-4 | 2.2532 | 32 | 0.19% | 766 | 4.66071e-2 | 2.1075 | 7 | 7 | 0.07% |

The test set of 17,097 genomic regions picked 10,152 genes (47%) of all 21,395 genes.
*Human Phenotype* has 6,5 6,599 ontology terms were tested (100%) using an annotation count range of [1, 1000].
GREAT version 4.0.4
Species assembly: mm10
Association rule: Basal+extension: 5000 bp upstream, 1000 bp downstream, 1000000 bp max extension, curated regulatory domains included

| Ontology | # Term Name | Binom Ran | Binom Raw | Binom FDR | Binom Fold | Binom Obs | Binom Reg | Hyper Rank | Hyper FDR | Hyper Fold | Hyper Obs | Hyper Tota | Hyper Gene Set Coverage |
|---|---|---|---|---|---|---|---|---|---|---|---|---|---|
| Mouse Phenotype Singl | abnormal placenta vasculature | 69 | 1.18337e-3 | 1.57269e-3 | 2.1270 | 359 | 2.10% | 480 | 7.41251e-7 | 1.5379 | 81 | 111 | 0.80% |
| | abnormal renal glomerulus basement membrane thickness | 81 | 1.22970e-3 | 1.39215e-3 | 3.4047 | 144 | 0.84% | 1108 | 2.66500e-3 | 1.7562 | 20 | 24 | 0.20% |
| | abnormal renal glomerulus basement membrane morphology | 86 | 7.23469e-3 | 7.71420e-3 | 2.9084 | 177 | 1.04% | 973 | 1.10912e-3 | 1.6518 | 29 | 37 | 0.29% |
| | pale kidney | 100 | 1.57719e-3 | 1.44629e-2 | 3.0052 | 156 | 0.91% | 805 | 1.94737e-4 | 1.7782 | 27 | 32 | 0.27% |
| | abnormal mesangial cell morphology | 102 | 1.91589e-3 | 1.72242e-2 | 3.1806 | 143 | 0.84% | 1379 | 1.20939e-2 | 1.5496 | 25 | 34 | 0.25% |
| | myocardial trabeculae hypoplasia | 105 | 6.58897e-3 | 5.75437e-2 | 2.2207 | 265 | 1.55% | 608 | 1.06925e-5 | 1.6860 | 44 | 55 | 0.43% |
| | extramedullary hematopoiesis | 110 | 8.78161e-3 | 7.32067e-2 | 2.1396 | 277 | 1.62% | 449 | 4.00250e-7 | 1.5519 | 81 | 110 | 0.80% |
| | abnormal podocyte morphology | 117 | 1.12739e-2 | 8.83607e-2 | 2.2958 | 228 | 1.33% | 559 | 4.31353e-6 | 1.7172 | 44 | 54 | 0.43% |
| | glomerulosclerosis | 120 | 5.51882e-2 | 4.21730e-2 | 2.2918 | 223 | 1.30% | 894 | 5.16809e-4 | 1.5529 | 42 | 57 | 0.41% |
| | trabecula carnea hypoplasia | 126 | 3.61244e-2 | 2.62906e-2 | 2.2440 | 226 | 1.32% | 705 | 5.67042e-5 | 1.6951 | 37 | 46 | 0.36% |
| | abnormal glomerular mesangium morphology | 128 | 6.78920e-2 | 4.86382e-2 | 2.1533 | 245 | 1.43% | 1106 | 2.57396e-3 | 1.4590 | 45 | 65 | 0.44% |
| | enhanced wound healing | 130 | 1.80219e-2 | 1.27123e-2 | 3.8691 | 92 | 0.54% | 885 | 4.60529e-4 | 1.9068 | 19 | 21 | 0.19% |
| | abnormal fibroblast migration | 136 | 1.16463e-2 | 7.85272e-2 | 2.4006 | 185 | 1.08% | 522 | 2.40053e-6 | 1.8440 | 35 | 40 | 0.34% |
| | abnormal placental labyrinth vasculature morphology | 139 | 2.73353e-2 | 1.80334e-2 | 2.2262 | 213 | 1.25% | 582 | 6.51293e-6 | 1.6017 | 57 | 75 | 0.56% |
| | cervical vertebral transformation | 142 | 4.13529e-2 | 2.67047e-2 | 3.0122 | 122 | 0.71% | 973 | 1.10912e-3 | 1.6518 | 29 | 37 | 0.29% |
| | abnormal glomerular capillary morphology | 157 | 5.28101e-2 | 3.08451e-2 | 2.3413 | 173 | 1.01% | 909 | 6.08061e-4 | 1.6449 | 32 | 41 | 0.32% |
| | lumbar vertebral transformation | 160 | 7.49165e-2 | 4.29365e-2 | 3.2180 | 100 | 0.58% | 1047 | 1.72109e-3 | 1.6391 | 28 | 36 | 0.28% |
| | abnormal kidney capillary morphology | 162 | 1.29276e-2 | 7.31764e-2 | 2.2950 | 177 | 1.04% | 935 | 8.26543e-4 | 1.6174 | 33 | 43 | 0.33% |
| | thoracic vertebral transformation | 163 | 1.37904e-2 | 7.75815e-2 | 2.5981 | 139 | 0.81% | 1150 | 3.72963e-3 | 1.5246 | 34 | 47 | 0.33% |
| | abnormal cell adhesion | 167 | 3.73278e-2 | 2.04967e-2 | 2.1000 | 211 | 1.23% | 1281 | 7.89821e-3 | 1.4420 | 39 | 57 | 0.38% |

The test set of 17,097 genomic regions picked 10,152 genes (47%) of all 21,395 genes.
*Mouse Phenotype Single K* 9,170 ontology terms were tested (100%) using an annotation count range of [1, 1000].
GREAT version 4.0.4
Species assembly: mm10
Association rule: Basal+extension: 5000 bp upstream, 1000 bp downstream, 1000000 bp max extension, curated regulatory domains included

| Ontology | # Term Name | Binom Ran | Binom Raw | Binom FDR | Binom Fold | Binom Obs | Binom Reg | Hyper Rank | Hyper FDR | Hyper Fold | Hyper Obs | Hyper Tota | Hyper Gene Set Coverage |
|---|---|---|---|---|---|---|---|---|---|---|---|---|---|
| Mouse Phenotype | abnormal podocyte morphology | 84 | 2.29489e-4 | 2.61591e-4 | 2.3637 | 347 | 2.03% | 525 | 1.58042e-9 | 1.7266 | 68 | 83 | 0.67% |
| | abnormal renal glomerulus basement membrane morphology | 90 | 8.88719e-4 | 9.45498e-4 | 2.8143 | 244 | 1.43% | 1107 | 9.79619e-5 | 1.6093 | 42 | 55 | 0.41% |
| | enlarged liver sinusoidal spaces | 100 | 2.74362e-4 | 2.62701e-3 | 3.1499 | 193 | 1.13% | 873 | 5.54137e-6 | 1.7991 | 35 | 41 | 0.34% |
| | abnormal renal glomerulus basement membrane thickness | 103 | 4.18100e-4 | 3.88671e-3 | 2.9467 | 207 | 1.21% | 1234 | 2.79686e-4 | 1.6559 | 33 | 42 | 0.33% |
| | glomerulosclerosis | 106 | 1.25195e-3 | 1.13089e-3 | 2.2998 | 319 | 1.87% | 1183 | 1.95611e-4 | 1.4830 | 57 | 81 | 0.56% |
| | pale kidney | 123 | 2.07285e-3 | 1.61362e-3 | 2.6842 | 218 | 1.28% | 774 | 1.10296e-6 | 1.7936 | 40 | 47 | 0.39% |
| | abnormal liver vasculature morphology | 127 | 3.66712e-3 | 2.76478e-3 | 2.0022 | 387 | 2.26% | 432 | 1.52212e-1 | 1.6935 | 90 | 112 | 0.89% |
| | abnormal podocyte foot process morphology | 128 | 4.21667e-3 | 3.15427e-3 | 2.3202 | 276 | 1.61% | 691 | 2.03791e-7 | 1.7184 | 53 | 65 | 0.52% |
| | abnormal liver sinusoid morphology | 129 | 7.61164e-3 | 5.64973e-3 | 2.3626 | 264 | 1.54% | 702 | 2.31257e-7 | 1.6638 | 60 | 76 | 0.59% |
| | myocardial trabeculae hypoplasia | 132 | 1.68692e-3 | 1.22365e-3 | 2.1318 | 326 | 1.91% | 856 | 4.62376e-6 | 1.6017 | 57 | 75 | 0.56% |
| | absent somites | 138 | 4.28461e-3 | 2.97283e-3 | 2.8805 | 181 | 1.06% | 2130 | 1.74582e-2 | 1.4193 | 33 | 49 | 0.33% |
| | trabecula carnea hypoplasia | 150 | 3.37101e-3 | 2.15183e-3 | 2.1944 | 284 | 1.66% | 935 | 1.35743e-5 | 1.6316 | 48 | 62 | 0.47% |
| | abnormal mesangial cell morphology | 157 | 2.73269e-3 | 1.66659e-2 | 2.7352 | 180 | 1.05% | 1567 | 2.27325e-3 | 1.5174 | 36 | 50 | 0.35% |
| | enhanced wound healing | 174 | 5.72364e-2 | 3.14964e-2 | 3.1993 | 130 | 0.76% | 1210 | 2.40856e-4 | 1.8064 | 24 | 28 | 0.24% |
| | abnormal glomerular mesangium morphology | 176 | 7.71850e-2 | 4.19913e-2 | 2.0773 | 287 | 1.68% | 1331 | 6.08996e-4 | 1.4380 | 58 | 85 | 0.57% |
| | colon polyps | 181 | 5.97033e-2 | 3.15834e-2 | 4.3150 | 86 | 0.50% | 2490 | 4.08275e-2 | 1.6860 | 12 | 15 | 0.12% |
| | abnormal neural plate morphology | 188 | 5.10793e-2 | 2.60151e-2 | 2.3912 | 197 | 1.15% | 1259 | 3.54833e-4 | 1.6977 | 29 | 36 | 0.29% |
| | renal cast | 192 | 1.13723e-2 | 5.67132e-2 | 2.3285 | 205 | 1.20% | 1312 | 5.12684e-4 | 1.6035 | 35 | 46 | 0.34% |
| | abnormal anterior definitive endoderm morphology | 194 | 1.89421e-2 | 9.34902e-2 | 4.8306 | 72 | 0.42% | 2340 | 2.85452e-2 | 1.8967 | 9 | 10 | 0.09% |
| | abnormal cell adhesion | 196 | 3.13774e-2 | 1.53285e-2 | 2.0595 | 265 | 1.55% | 1066 | 6.36677e-5 | 1.5379 | 54 | 74 | 0.53% |

The test set of 17,097 genomic regions picked 10,152 genes (47%) of all 21,395 genes.
*Mouse Phenotype* has 9,5 9,575 ontology terms were tested (100%) using an annotation count range of [1, 1000].
GREAT version 4.0.4
Species assembly: mm10
Association rule: Basal+extension: 5000 bp upstream, 1000 bp downstream, 1000000 bp max extension, curated regulatory domains included

| WT1 Peaks | | | | MAFB Peaks | |
|---|---|---|---|---|---|
| Chrom | Start | End | WT1 peakIDR score | Chrom | Start |
| chr1 | 93478131 | 93479270 | 1000 | chr1 | 93478233 |
| chr1 | 143503701 | 143504111 | 1000 | chr1 | 143503859 |
| chr1 | 152799911 | 152800393 | 1000 | chr1 | 152799996 |
| chr1 | 156309145 | 156309691 | 1000 | chr1 | 156309319 |
| chr1 | 127306400 | 127306861 | 1000 | chr1 | 127306458 |
| chr1 | 23263863 | 23264740 | 1000 | chr1 | 23264325 |
| chr1 | 93211355 | 93211932 | 1000 | chr1 | 93211537 |
| chr1 | 36352355 | 36352761 | 1000 | chr1 | 36352475 |
| chr1 | 90442551 | 90443219 | 1000 | chr1 | 90442898 |
| chr10 | 43540562 | 43541201 | 1000 | chr10 | 43540914 |
| chr10 | 86339936 | 86340518 | 1000 | chr10 | 86340102 |
| chr10 | 62216544 | 62217199 | 1000 | chr10 | 62216777 |
| chr10 | 22868297 | 22868932 | 1000 | chr10 | 22868403 |
| chr10 | 81401790 | 81402539 | 1000 | chr10 | 81402067 |
| chr10 | 119065414 | 119065910 | 1000 | chr10 | 119065504 |
| chr11 | 100619068 | 100619801 | 1000 | chr11 | 100619593 |
| chr11 | 8493988 | 8494521 | 1000 | chr11 | 8494100 |
| chr11 | 82870992 | 82871448 | 1000 | chr11 | 82871267 |
| chr11 | 32383770 | 32384358 | 1000 | chr11 | 32383789 |
| chr11 | 49203211 | 49203946 | 1000 | chr11 | 49203385 |
| chr11 | 69880837 | 69881570 | 1000 | chr11 | 69880992 |
| chr11 | 97437657 | 97438334 | 1000 | chr11 | 97437928 |
| chr11 | 121794576 | 121795090 | 1000 | chr11 | 121794597 |
| chr11 | 23067718 | 23068219 | 1000 | chr11 | 23067735 |
| chr11 | 31827994 | 31828381 | 1000 | chr11 | 31828057 |
| chr12 | 111516750 | 111517207 | 1000 | chr12 | 111516901 |
| chr12 | 85608000 | 85608396 | 1000 | chr12 | 85608219 |
| chr13 | 42242086 | 42242587 | 1000 | chr13 | 42242122 |
| chr13 | 119428710 | 119429621 | 1000 | chr13 | 119428834 |
| chr13 | 37531673 | 37532391 | 1000 | chr13 | 37531848 |
| chr14 | 60265083 | 60265571 | 1000 | chr14 | 60264972 |
| chr14 | 45506948 | 45507454 | 1000 | chr14 | 45507174 |
| chr14 | 70160632 | 70161121 | 1000 | chr14 | 70160831 |
| chr15 | 99702025 | 99702671 | 1000 | chr15 | 99701985 |
| chr15 | 84695997 | 84696622 | 1000 | chr15 | 84696178 |
| chr15 | 5617705 | 5618239 | 1000 | chr15 | 5617871 |
| chr15 | 38470131 | 38471083 | 1000 | chr15 | 38470642 |
| chr16 | 52000101 | 52000544 | 1000 | chr16 | 52000169 |
| chr16 | 58514614 | 58515115 | 1000 | chr16 | 58514787 |
| chr16 | 32877309 | 32878253 | 1000 | chr16 | 32877560 |
| chr16 | 33380577 | 33381582 | 1000 | chr16 | 33380600 |
| chr16 | 33871925 | 33872475 | 1000 | chr16 | 33872134 |
| chr16 | 87454653 | 87455249 | 1000 | chr16 | 87454878 |
| chr16 | 26049166 | 26050069 | 1000 | chr16 | 26049406 |
| chr16 | 50786043 | 50786723 | 1000 | chr16 | 50786294 |
| chr17 | 46051930 | 46052506 | 1000 | chr17 | 46052201 |
| chr17 | 28517495 | 28518459 | 1000 | chr17 | 28518117 |
| chr17 | 3114515 | 3115758 | 1000 | chr17 | 3114769 |
| chr17 | 49477271 | 49477791 | 1000 | chr17 | 49477431 |
| chr17 | 5189621 | 5190365 | 1000 | chr17 | 5189837 |
| chr17 | 66419458 | 66419852 | 1000 | chr17 | 66419507 |
| chr17 | 43554448 | 43555011 | 1000 | chr17 | 43554623 |
| chr17 | 4626079 | 4626671 | 1000 | chr17 | 4626163 |
| chr18 | 61894618 | 61894997 | 1000 | chr18 | 61894653 |
| chr18 | 33463727 | 33464497 | 1000 | chr18 | 33463794 |
| chr18 | 76087465 | 76088530 | 1000 | chr18 | 76088011 |
| chr18 | 35815452 | 35815931 | 1000 | chr18 | 35815583 |
| chr18 | 53744340 | 53745098 | 1000 | chr18 | 53744488 |
| chr18 | 5441156 | 5441554 | 1000 | chr18 | 5441180 |
| chr18 | 11973679 | 11974147 | 1000 | chr18 | 11973814 |
| chr19 | 47280763 | 47281346 | 1000 | chr19 | 47280914 |
| chr19 | 45304420 | 45305040 | 1000 | chr19 | 45304688 |
| chr2 | 25452632 | 25453208 | 1000 | chr2 | 25452963 |
| chr2 | 27758371 | 27759014 | 1000 | chr2 | 27758427 |
| chr2 | 32906947 | 32907675 | 1000 | chr2 | 32907336 |
| chr2 | 32616975 | 32617472 | 1000 | chr2 | 32617181 |
| chr2 | 122575383 | 122575807 | 1000 | chr2 | 122575492 |
| chr2 | 69080533 | 69081236 | 1000 | chr2 | 69080874 |
| chr2 | 129624840 | 129625790 | 1000 | chr2 | 129625322 |
| chr2 | 33605081 | 33606316 | 1000 | chr2 | 33606011 |
| chr2 | 33165609 | 33166278 | 1000 | chr2 | 33166087 |
| chr2 | 29696216 | 29696821 | 1000 | chr2 | 29696451 |
| chr2 | 153241259 | 153241725 | 1000 | chr2 | 153241358 |
| chr2 | 159270194 | 159271091 | 1000 | chr2 | 159270560 |
| chr3 | 75930714 | 75931247 | 1000 | chr3 | 75930732 |
| chr3 | 89858934 | 89859319 | 1000 | chr3 | 89859020 |
| chr3 | 51916079 | 51916782 | 1000 | chr3 | 51916199 |
| chr4 | 34687060 | 34687591 | 1000 | chr4 | 34687161 |
| chr4 | 66450072 | 66450460 | 1000 | chr4 | 66450124 |
| chr4 | 136602174 | 136602896 | 1000 | chr4 | 136602688 |
| chr4 | 132211840 | 132212463 | 1000 | chr4 | 132212035 |
| chr4 | 149860304 | 149860665 | 1000 | chr4 | 149860376 |
| chr4 | 116732309 | 116732745 | 1000 | chr4 | 116732409 |
| chr5 | 81519330 | 81519942 | 1000 | chr5 | 81519487 |
| chr5 | 34170782 | 34171111 | 1000 | chr5 | 34170814 |
| chr5 | 35233227 | 35233598 | 1000 | chr5 | 35233245 |
| chr5 | 140777048 | 140777488 | 1000 | chr5 | 140777154 |
| chr5 | 119325757 | 119326206 | 1000 | chr5 | 119325825 |
| chr5 | 103606238 | 103606750 | 1000 | chr5 | 103606423 |
| chr5 | 92834486 | 92834902 | 1000 | chr5 | 92834538 |
| chr5 | 19180792 | 19181379 | 1000 | chr5 | 19180868 |
| chr6 | 18543809 | 18544254 | 1000 | chr6 | 18543915 |
| chr6 | 52233382 | 52234275 | 1000 | chr6 | 52233552 |
| chr6 | 22355772 | 22356389 | 1000 | chr6 | 22356026 |
| chr6 | 116006706 | 116007185 | 1000 | chr6 | 116006861 |
| chr6 | 140433089 | 140433545 | 1000 | chr6 | 140433286 |
| chr6 | 114921632 | 114922233 | 1000 | chr6 | 114921773 |
| chr6 | 72743683 | 72744402 | 1000 | chr6 | 72743833 |
| chr6 | 70073781 | 70074409 | 1000 | chr6 | 70073928 |
| chr6 | 70116852 | 70117284 | 1000 | chr6 | 70116960 |
| chr6 | 70386409 | 70386889 | 1000 | chr6 | 70386500 |
| chr6 | 70215734 | 70216150 | 1000 | chr6 | 70215864 |
| chr6 | 70240204 | 70240610 | 1000 | chr6 | 70240248 |
| chr7 | 30459355 | 30459800 | 1000 | chr7 | 30459561 |
| chr7 | 110613978 | 110615726 | 1000 | chr7 | 110615259 |
| chr7 | 25316637 | 25317146 | 1000 | chr7 | 25316829 |
| chr7 | 112678900 | 112680254 | 1000 | chr7 | 112680121 |
| chr7 | 40192358 | 40192750 | 1000 | chr7 | 40192452 |
| chr7 | 16618438 | 16618982 | 1000 | chr7 | 16618636 |
| chr8 | 70418823 | 70419275 | 1000 | chr8 | 70418957 |
| chr8 | 61008386 | 61009136 | 1000 | chr8 | 61008825 |
| chr8 | 104886708 | 104887221 | 1000 | chr8 | 104886805 |
| chr8 | 87933550 | 87934511 | 1000 | chr8 | 87933878 |
| chr8 | 82939856 | 82940389 | 1000 | chr8 | 82939866 |
| chr8 | 91522206 | 91522591 | 1000 | chr8 | 91522387 |
| chr8 | 70609379 | 70611412 | 1000 | chr8 | 70610242 |
| chr8 | 121241670 | 121242435 | 1000 | chr8 | 121241958 |
| chr8 | 121280463 | 121281037 | 1000 | chr8 | 121280477 |
| chr9 | 57831347 | 57831930 | 1000 | chr9 | 57831471 |
| chr9 | 32646622 | 32647268 | 1000 | chr9 | 32646915 |
| chr9 | 32671858 | 32672986 | 1000 | chr9 | 32672456 |
| chr9 | 71674434 | 71674867 | 1000 | chr9 | 71674521 |
| chr9 | 63610824 | 63611500 | 1000 | chr9 | 63611068 |
| chr9 | 60637042 | 60637459 | 1000 | chr9 | 60637104 |
| chr19 | 38836383 | 38837069 | 991 | chr19 | 38836434 |
| chr2 | 157048462 | 157048881 | 982 | chr2 | 157048528 |
| chr9 | 58541860 | 58542160 | 960 | chr9 | 58541951 |
| chr4 | 62699276 | 62699759 | 954 | chr4 | 62699353 |
| chr6 | 132463605 | 132463906 | 901 | chr6 | 132463691 |
| chr12 | 94280920 | 94281302 | 882 | chr12 | 94281002 |
| chr17 | 83288075 | 83288521 | 871 | chr17 | 83288188 |
| chr17 | 5849397 | 5849838 | 851 | chr17 | 5849457 |
| chr9 | 14641789 | 14642122 | 846 | chr9 | 14641847 |
| chr8 | 104867376 | 104867853 | 841 | chr8 | 104867314 |
| chr11 | 59506135 | 59506554 | 836 | chr11 | 59506166 |
| chr13 | 21362662 | 21363031 | 828 | chr13 | 21362679 |
| chr7 | 105736374 | 105736893 | 817 | chr7 | 105736409 |
| chr19 | 10952042 | 10952505 | 816 | chr19 | 10952070 |
| chr5 | 64336868 | 64337476 | 807 | chr5 | 64337025 |
| chr11 | 76763499 | 76763743 | 794 | chr11 | 76763533 |
| chr4 | 150784639 | 150785198 | 783 | chr4 | 150784817 |
| chrX | 99216855 | 99217321 | 780 | chrX | 99216819 |
| chr6 | 47835525 | 47836083 | 752 | chr6 | 47835601 |
| chr18 | 53793534 | 53793802 | 751 | chr18 | 53793661 |
| chr9 | 65032249 | 65032986 | 737 | chr9 | 65032281 |
| chr6 | 114883233 | 114883677 | 733 | chr6 | 114883288 |
| chr6 | 106799921 | 106800274 | 730 | chr6 | 106800069 |
| chr16 | 52012986 | 52013474 | 726 | chr16 | 52013175 |
| chr11 | 96064932 | 96065574 | 682 | chr11 | 96065249 |
| chr1 | 179546245 | 179546853 | 652 | chr1 | 179546272 |
| chr17 | 45563725 | 45564432 | 638 | chr17 | 45563839 |
| chr11 | 58103852 | 58104403 | 622 | chr11 | 58103937 |
| chr9 | 32656591 | 32657525 | 611 | chr9 | 32657244 |
| chr4 | 123282555 | 123282850 | 598 | chr4 | 123282580 |
| chr7 | 122101691 | 122102006 | 597 | chr7 | 122101808 |
| chr5 | 32863427 | 32863772 | 595 | chr5 | 32863614 |
| chr13 | 45897633 | 45897975 | 586 | chr13 | 45897687 |
| chr13 | 21506211 | 21506610 | 585 | chr13 | 21506323 |
| chr2 | 120093700 | 120094085 | 564 | chr2 | 120093774 |
| chr17 | 56036473 | 56036838 | 563 | chr17 | 56036707 |
| chr6 | 83775658 | 83776217 | 560 | chr6 | 83775754 |
| chr10 | 122029835 | 122030100 | 551 | chr10 | 122029860 |
| chr11 | 76945407 | 76945839 | 528 | chr11 | 76945517 |
| chr7 | 46033572 | 46033857 | 489 | chr7 | 46033564 |
| chr14 | 30951216 | 30951557 | 487 | chr14 | 30951184 |
| chr18 | 34776834 | 34777883 | 475 | chr18 | 34776841 |
| chr5 | 24724859 | 24725277 | 455 | chr5 | 24724939 |
| chr10 | 22731019 | 22731591 | 451 | chr10 | 22731264 |
| chr2 | 26132678 | 26133360 | 437 | chr2 | 26132995 |
| chr14 | 69732403 | 69732877 | 435 | chr14 | 69732573 |
| chr7 | 127637002 | 127637326 | 432 | chr7 | 127636984 |
| chr1 | 163994494 | 163994953 | 431 | chr1 | 163994634 |
| chr6 | 121146874 | 121147200 | 412 | chr6 | 121146971 |
| chr9 | 66945906 | 66946270 | 410 | chr9 | 66945885 |
| chr7 | 127345363 | 127346412 | 402 | chr7 | 127345551 |
| chr15 | 80133045 | 80133542 | 399 | chr15 | 80132973 |
| chr5 | 143112466 | 143113162 | 398 | chr5 | 143112967 |
| chr5 | 136888503 | 136888915 | 381 | chr5 | 136888530 |
| chr5 | 114707701 | 114707881 | 376 | chr5 | 114707683 |
| chr15 | 102470338 | 102470604 | 374 | chr15 | 102470199 |
| chr1 | 39762073 | 39762299 | 371 | chr1 | 39762143 |
| chr5 | 103629497 | 103629775 | 367 | chr5 | 103629555 |
| chr1 | 133025053 | 133025489 | 364 | chr1 | 133025356 |
| chr11 | 48816967 | 48817318 | 356 | chr11 | 48816999 |
| chr4 | 115708753 | 115709283 | 356 | chr4 | 115709082 |
| chr11 | 54744248 | 54744609 | 353 | chr11 | 54744293 |

| | | | | | |
|---|---|---|---|---|---|
| chr19 | 10605299 | 10605848 | 346 | chr19 | 10605523 |
| chr18 | 69344189 | 69344633 | 343 | chr18 | 69344284 |
| chr2 | 50110152 | 50110565 | 333 | chr2 | 50110219 |
| chr17 | 45931103 | 45931562 | 326 | chr17 | 45931243 |
| chr12 | 84996080 | 84996874 | 323 | chr12 | 84996103 |
| chr14 | 57056374 | 57056798 | 317 | chr14 | 57056404 |
| chr19 | 8880507 | 8881032 | 313 | chr19 | 8880841 |
| chr4 | 106650872 | 106651124 | 311 | chr4 | 106650851 |
| chr3 | 96727444 | 96727973 | 301 | chr3 | 96727405 |
| chr1 | 133609822 | 133610342 | 299 | chr1 | 133610190 |
| chr1 | 143777006 | 143777411 | 182 | chr1 | 143777101 |
| chr1 | 13589770 | 13590112 | 182 | chr1 | 13589819 |
| chr1 | 74587674 | 74588490 | 182 | chr1 | 74587986 |
| chr1 | 52844724 | 52845010 | 182 | chr1 | 52844877 |
| chr1 | 153543095 | 153543310 | 182 | chr1 | 153543060 |
| chr1 | 44584879 | 44585073 | 182 | chr1 | 44584883 |
| chr1 | 118388829 | 118389438 | 182 | chr1 | 118388998 |
| chr11 | 100622764 | 100623120 | 182 | chr11 | 100622806 |
| chr12 | 83632018 | 83632484 | 182 | chr12 | 83631963 |
| chr12 | 69361137 | 69361583 | 182 | chr12 | 69361226 |
| chr14 | 50785341 | 50785642 | 182 | chr14 | 50785440 |
| chr15 | 102326045 | 102326904 | 182 | chr15 | 102325998 |
| chr15 | 27525377 | 27525650 | 182 | chr15 | 27525344 |
| chr15 | 75616371 | 75616731 | 182 | chr15 | 75616398 |
| chr16 | 25549627 | 25549908 | 182 | chr16 | 25549707 |
| chr16 | 74093169 | 74093369 | 182 | chr16 | 74093141 |
| chr17 | 56199898 | 56200186 | 182 | chr17 | 56199988 |
| chr17 | 36958185 | 36958665 | 182 | chr17 | 36958376 |
| chr17 | 35331526 | 35331869 | 182 | chr17 | 35331725 |
| chr18 | 9957794 | 9958294 | 182 | chr18 | 9958043 |
| chr18 | 38277883 | 38278168 | 182 | chr18 | 38277860 |
| chr2 | 106974365 | 106974599 | 182 | chr2 | 106974307 |
| chr2 | 167421492 | 167422036 | 182 | chr2 | 167421572 |
| chr2 | 164460841 | 164461347 | 182 | chr2 | 164460907 |
| chr3 | 132835122 | 132835435 | 182 | chr3 | 132835187 |
| chr7 | 120634949 | 120635247 | 182 | chr7 | 120635114 |
| chr8 | 88361830 | 88362259 | 182 | chr8 | 88362120 |
| chr8 | 126853015 | 126853352 | 182 | chr8 | 126853071 |
| chr8 | 128544475 | 128544830 | 182 | chr8 | 128544599 |
| chr9 | 57765001 | 57765645 | 182 | chr9 | 57765142 |
| chr9 | 58253063 | 58253269 | 182 | chr9 | 58253058 |
| chr9 | 120577128 | 120577462 | 182 | chr9 | 120577249 |

| End | MAFB peakIDR score | merged regions | | | annotation | geneChrom | geneStart | geneEnd | transcriptId | distanceToTSS | geneName | geneId | Podocytes specific | H3K4me3 Chrom | Start | End | -LOG10(pval | fold | -LOG10(q-value) | H3K4me1 Chrom | Start | End | -LOG10(pval | fold | -LOG10(q-value) |
|---|---|---|---|---|---|---|---|---|---|---|---|---|---|---|---|---|---|---|---|---|---|---|---|---|---|
| 93478572 | 642 | chr1 | 93478131 | 93479270 | Promoter (<=1kb) | 1 | 93478964 | 93510260 | ENSMUST00 | 0 | Sept2 | ENSMUSG0C | false | chr1 | 93477516 | 93480357 | 31.43325 | 99.35611 | 95.79983 | . | . | . | . | . | . |
| 143504022 | 608 | chr1 | 143503701 | 143504111 | Distal Intergenic | 1 | 143633851 | 143636577 | ENSMUST00 | 132466 | Cdc73 | ENSMUSG0C | false | chr1 | 143503168 | 143503845 | 4.29207 | 7.17162 | 5.05983 | . | . | . | . | . | . |
| 152800343 | 593 | chr1 | 152799911 | 152800393 | Promoter (<=1kb) | 1 | 152800194 | 152836991 | ENSMUST00 | 0 | Ncf2 | ENSMUSG0C | true | chr1 | 152800193 | 152801982 | 14.44046 | 33.88361 | 31.37022 | chr1 | 152788034 | 152800060 | 6.38067 | 4.71623 | 4.13727 |
| 156309613 | 591 | chr1 | 156309145 | 156309691 | Promoter (1-2kb) | 1 | 156310727 | 156328035 | ENSMUST00 | -1036 | Nphs2 | ENSMUSG0C | true | . | . | . | . | . | . | chr1 | 156307023 | 156309151 | 5.03822 | 4.14188 | 3.02784 |
| 127306781 | 636 | chr1 | 127306400 | 127306861 | Promoter (<=1kb) | 1 | 127306762 | 127483704 | ENSMUST00 | 0 | Mgat5 | ENSMUSG0C | true | chr1 | 127306652 | 127307946 | 25.3802 | 72.59858 | 69.5683 | . | . | . | . | . | . |
| 23264602 | 434 | chr1 | 23263863 | 23264740 | Distal Intergenic | 1 | 23272264 | 23272345 | ENSMUST00 | -7524 | Mir30a | ENSMUSG0C | false | . | . | . | . | . | . | chr1 | 23258890 | 23271405 | 4.63528 | 3.84316 | 2.71286 |
| 93211806 | 630 | chr1 | 93211355 | 93211932 | Distal Intergenic | 1 | 93215899 | 93231072 | ENSMUST00 | -3967 | E030010N08 | ENSMUSG0C | false | . | . | . | . | . | . | chr1 | 93210088 | 93211973 | 5.50157 | 4.34556 | 3.43814 |
| 36352590 | 599 | chr1 | 36352355 | 36352761 | Intron (ENSMUST0C | 1 | 36348682 | 36348758 | ENSMUST00 | -3598 | Mir6898 | ENSMUSG0C | false | . | . | . | . | . | . | chr1 | 36350640 | 36368234 | 5.46208 | 4.18784 | 3.39932 |
| 90443107 | 574 | chr1 | 90442551 | 90443219 | Distal Intergenic | 1 | 90453394 | 90454501 | ENSMUST00 | 11282 | RP24-154L11 | ENSMUSG0C | false | . | . | . | . | . | . | chr1 | 90438990 | 90445431 | 5.52456 | 4.20032 | 3.45273 |
| 43541106 | 604 | chr10 | 43540562 | 43541201 | Promoter (<=1kb) | 10 | 43525121 | 43540994 | ENSMUST00 | 0 | 1700021F05I | ENSMUSG0C | false | chr10 | 43540096 | 43541068 | 37.09717 | 122.76616 | 118.52905 | chr10 | 43536760 | 43541464 | 4.73951 | 3.86652 | 2.82674 |
| 86340227 | 619 | chr10 | 86339936 | 86340518 | Intron (ENSMUST0C | 10 | 86302854 | 86343981 | ENSMUST00 | 37083 | Timp3 | ENSMUSG0C | false | . | . | . | . | . | . | chr10 | 86303145 | 86347283 | 7.80266 | 5.33584 | 5.30789 |
| 62216860 | 605 | chr10 | 62216544 | 62217199 | Intron (ENSMUST0C | 10 | 62185396 | 62231218 | ENSMUST00 | 14019 | Tspan15 | ENSMUSG0C | false | . | . | . | . | . | . | chr10 | 62185685 | 62229768 | 8.28034 | 5.46771 | 5.70297 |
| 22868624 | 588 | chr10 | 22868297 | 22868932 | Distal Intergenic | 10 | 22817279 | 22820128 | ENSMUST00 | -48170 | Tcf21 | ENSMUSG0C | true | . | . | . | . | . | . | chr10 | 22866220 | 22874539 | 4.6225 | 3.87067 | 2.69769 |
| 81402224 | 387 | chr10 | 81401790 | 81402539 | Promoter (1-2kb) | 10 | 81399196 | 81400554 | ENSMUST00 | -1237 | Nfic | ENSMUSG0C | false | chr10 | 81401906 | 81402504 | 5.97392 | 9.35129 | 7.18948 | chr10 | 81392649 | 81426205 | 8.72751 | 5.63067 | 6.0679 |
| 119065739 | 464 | chr10 | 119065414 | 119065910 | Distal Intergenic | 10 | 119092905 | 119103107 | ENSMUST00 | -26995 | RP23-360L4.: | ENSMUSG0C | false | . | . | . | . | . | . | chr10 | 119061401 | 119066309 | 4.92666 | 3.97999 | 2.96336 |
| 100619867 | 586 | chr11 | 100619068 | 100619801 | Promoter (<=1kb) | 11 | 100590887 | 100619576 | ENSMUST00 | 0 | Dnajc7 | ENSMUSG0C | false | chr11 | 100618769 | 100620432 | 29.28796 | 98.53321 | 94.99628 | . | . | . | . | . | . |
| 8494421 | 628 | chr11 | 8493988 | 8494521 | Intron (ENSMUST0C | 11 | 8451070 | 8468675 | ENSMUST00 | -25314 | Tns3 | ENSMUSG0C | false | chr11 | 8494386 | 8495188 | 5.6323 | 9.47728 | 7.31216 | chr11 | 8476638 | 8500780 | 7.03559 | 4.95227 | 4.67872 |
| 82871429 | 603 | chr11 | 82870992 | 82871448 | Promoter (<=1kb) | 11 | 82803822 | 82871210 | ENSMUST00 | 0 | Rffl | ENSMUSG0C | false | chr11 | 82870035 | 82871276 | 36.78915 | 128.67926 | 124.23671 | chr11 | 82869837 | 82871608 | 3.13383 | 3.0708 | 1.52114 |
| 32384143 | 662 | chr11 | 32383770 | 32384358 | Intron (ENSMUST0C | 11 | 32347840 | 32390300 | ENSMUST00 | 35931 | Sh3pxd2b | ENSMUSG0C | false | . | . | . | . | . | . | chr11 | 32383288 | 32386252 | 5.8032 | 4.27592 | 3.68113 |
| 49203688 | 578 | chr11 | 49203211 | 49203946 | Promoter (<=1kb) | 11 | 49203292 | 49218553 | ENSMUST00 | 0 | Zfp62 | ENSMUSG0C | false | chr11 | 49202015 | 49204776 | 21.75525 | 76.8157 | 73.7174 | . | . | . | . | . | . |
| 69881443 | 399 | chr11 | 69880837 | 69881570 | Promoter (<=1kb) | 11 | 69878261 | 69881406 | ENSMUST00 | 0 | Kctd11 | ENSMUSG0C | false | chr11 | 69879287 | 69881992 | 16.78262 | 53.63071 | 50.86883 | . | . | . | . | . | . |
| 97438272 | 580 | chr11 | 97437657 | 97438334 | Promoter (1-2kb) | 11 | 97436285 | 97444228 | ENSMUST00 | 1373 | Arhgap23 | ENSMUSG0C | true | . | . | . | . | . | . | chr11 | 97431804 | 97440135 | 9.81823 | 6.18842 | 6.94298 |
| 121794898 | 579 | chr11 | 121794576 | 121795090 | Distal Intergenic | 11 | 121803107 | 121810319 | ENSMUST00 | -8017 | Gm12589 | ENSMUSG0C | false | . | . | . | . | . | . | chr11 | 121787526 | 121808717 | 6.67832 | 4.85129 | 4.38811 |
| 23067981 | 501 | chr11 | 23067718 | 23068219 | Promoter (<=1kb) | 11 | 23067998 | 23075574 | ENSMUST00 | 0 | Gm12059 | ENSMUSG0C | false | chr11 | 23067253 | 23068481 | 27.34985 | 78.73917 | 75.61149 | chr11 | 23067141 | 23068590 | 4.89236 | 3.95874 | 2.96174 |
| 31828264 | 452 | chr11 | 31827994 | 31828381 | Distal Intergenic | 11 | 31800688 | 31824524 | ENSMUST00 | -3471 | D630024D03 | ENSMUSG0C | false | . | . | . | . | . | . | chr11 | 31821641 | 31832598 | 5.80216 | 4.3489 | 3.6737 |
| 111517035 | 589 | chr12 | 111516750 | 111517207 | Distal Intergenic | 12 | 111538101 | 111546753 | ENSMUST00 | -20894 | Eif5 | ENSMUSG0C | false | . | . | . | . | . | . | chr12 | 111513836 | 111518852 | 5.17158 | 4.07062 | 3.1695 |
| 85608403 | 114 | chr12 | 85608000 | 85608396 | 5' UTR | 12 | 85600018 | 85639878 | ENSMUST00 | 7983 | Jdp2 | ENSMUSG0C | true | . | . | . | . | . | . | chr12 | 85599922 | 85615323 | 5.04005 | 3.93431 | 3.0761 |
| 42242453 | 613 | chr13 | 42242086 | 42242587 | Distal Intergenic | 13 | 42301270 | 42307989 | ENSMUST00 | -58683 | Edn1 | ENSMUSG0C | false | . | . | . | . | . | . | chr13 | 42235582 | 42245410 | 5.3203 | 4.22032 | 3.27886 |
| 119428999 | 593 | chr13 | 119428710 | 119429621 | Promoter (<=1kb) | 13 | 119428922 | 119458218 | ENSMUST00 | 0 | Paip1 | ENSMUSG0C | false | chr13 | 119428264 | 119430336 | 31.16539 | 106.97112 | 103.21663 | . | . | . | . | . | . |
| 37532120 | 620 | chr13 | 37531673 | 37532391 | Distal Intergenic | 13 | 37508940 | 37512074 | ENSMUST00 | 22734 | RP23-21N15. | ENSMUSG0C | false | . | . | . | . | . | . | chr13 | 37526947 | 37535595 | 4.83754 | 3.94 | 2.87988 |
| 60265292 | 468 | chr14 | 60265083 | 60265571 | Promoter (<=1kb) | 14 | 60265205 | 60302370 | ENSMUST00 | 0 | Mtmr6 | ENSMUSG0C | false | chr14 | 60265179 | 60266233 | 37.7199 | 126.00829 | 121.65485 | chr14 | 60264849 | 60267973 | 4.58611 | 3.73389 | 2.70745 |
| 45507316 | 584 | chr14 | 45506948 | 45507454 | Intron (ENSMUST0C | 14 | 45504672 | 45529786 | ENSMUST00 | 22332 | Fermt2 | ENSMUSG0C | true | . | . | . | . | . | . | chr14 | 45489964 | 45528839 | 6.12493 | 4.54024 | 3.93691 |
| 70161028 | 596 | chr14 | 70160632 | 70161121 | Promoter (1-2kb) | 14 | 70154405 | 70159502 | ENSMUST00 | -1131 | 9930012K11I | ENSMUSG0C | false | . | . | . | . | . | . | chr14 | 70156516 | 70175803 | 7.12683 | 5 | 4.75261 |
| 99702359 | 594 | chr15 | 99702025 | 99702671 | Promoter (<=1kb) | 15 | 99702287 | 99713995 | ENSMUST00 | 0 | Smarcd1 | ENSMUSG0C | false | chr15 | 99701792 | 99702839 | 37.30079 | 124.19571 | 119.91465 | chr15 | 99700422 | 99705143 | 3.90355 | 3.54268 | 2.10055 |
| 84696412 | 543 | chr15 | 84695997 | 84696622 | Intron (ENSMUST0C | 15 | 84681002 | 84703672 | ENSMUST00 | 14996 | Prr5 | ENSMUSG0C | false | . | . | . | . | . | . | chr15 | 84682178 | 84700461 | 6.48581 | 4.70787 | 4.24565 |
| 5618057 | 646 | chr15 | 5617705 | 5618239 | Distal Intergenic | 15 | 5864306 | 5864419 | ENSMUST00 | -246067 | Gm27760 | ENSMUSG0C | false | . | . | . | . | . | . | chr15 | 5617091 | 5619625 | 4.3034 | 3.71772 | 2.48127 |
| 38470826 | 436 | chr15 | 38470131 | 38471083 | Distal Intergenic | 15 | 38490473 | 38490547 | ENSMUST00 | 19464 | Mir6951 | ENSMUSG0C | false | chr15 | 38470059 | 38471389 | 13.87847 | 36.33116 | 33.78668 | chr15 | 38468227 | 38474123 | 4.55881 | 3.82285 | 2.65383 |
| 52000446 | 615 | chr16 | 52000101 | 52000544 | Distal Intergenic | 16 | 52031549 | 52208047 | ENSMUST00 | -31005 | Cblb | ENSMUSG0C | false | . | . | . | . | . | . | chr16 | 51999695 | 52002797 | 4.10448 | 3.61196 | 2.26807 |
| 58515018 | 538 | chr16 | 58514614 | 58515115 | Intron (ENSMUST0C | 16 | 58502861 | 58507898 | ENSMUST00 | -6717 | St3gal6 | ENSMUSG0C | true | chr16 | 58512380 | 58514683 | 5.72276 | 11.47427 | 9.27082 | chr16 | 58496830 | 58520680 | 5.70067 | 4.33546 | 3.59278 |
| 32877843 | 607 | chr16 | 32877309 | 32878253 | Promoter (<=1kb) | 16 | 32877500 | 32891409 | ENSMUST00 | 0 | Fyttd1 | ENSMUSG0C | false | chr16 | 32877139 | 32879191 | 27.85705 | 92.98368 | 89.57738 | . | . | . | . | . | . |
| 33380874 | 556 | chr16 | 33380577 | 33381582 | Promoter (<=1kb) | 16 | 33380775 | 33503903 | ENSMUST00 | 0 | Zfp148 | ENSMUSG0C | false | chr16 | 33380162 | 33382282 | 22.99841 | 82.87405 | 79.66897 | . | . | . | . | . | . |
| 33872340 | 593 | chr16 | 33871925 | 33872475 | Intron (ENSMUST0C | 16 | 33885026 | 33910585 | ENSMUST00 | -12551 | Itgb5 | ENSMUSG0C | true | . | . | . | . | . | . | chr16 | 33868993 | 33898226 | 6.80171 | 4.94523 | 4.47886 |
| 87455063 | 114 | chr16 | 87454653 | 87455249 | Promoter (<=1kb) | 16 | 87454703 | 87483517 | ENSMUST00 | 0 | Usp16 | ENSMUSG0C | false | chr16 | 87454436 | 87456031 | 38.65522 | 135.28891 | 130.59163 | . | . | . | . | . | . |
| 26049978 | 472 | chr16 | 26049166 | 26050069 | Intron (ENSMUST0C | 16 | 26031810 | 26040870 | ENSMUST00 | -8297 | AU015336 | ENSMUSG0C | false | . | . | . | . | . | . | chr16 | 26035252 | 26056185 | 6.03158 | 4.46161 | 3.87334 |
| 50786471 | 612 | chr16 | 50786043 | 50786723 | Promoter (2-3kb) | 16 | 50783478 | 50784065 | ENSMUST00 | 2566 | 1700116B05I | ENSMUSG0C | false | . | . | . | . | . | . | chr16 | 50781927 | 50788895 | 5.4458 | 4.28026 | 3.36614 |
| 46052365 | 577 | chr17 | 46051930 | 46052506 | Distal Intergenic | 17 | 46016996 | 46032377 | ENSMUST00 | -19554 | Vegfa | ENSMUSG0C | true | . | . | . | . | . | . | chr17 | 46048792 | 46077235 | 6.11576 | 4.54762 | 3.92959 |
| 28518304 | 503 | chr17 | 28517495 | 28518459 | Promoter (<=1kb) | 17 | 28399097 | 28517524 | ENSMUST00 | 0 | Fkbp5 | ENSMUSG0C | false | chr17 | 28517555 | 28518704 | 14.29631 | 42.77194 | 40.14703 | chr17 | 28514373 | 28519487 | 5.73469 | 4.2575 | 3.64316 |
| 3115224 | 619 | chr17 | 3114515 | 3115758 | Promoter (<=1kb) | 17 | 3114972 | 3198855 | ENSMUST00 | 0 | Scaf8 | ENSMUSG0C | false | chr17 | 3113719 | 3116459 | 36.98618 | 117.25745 | 113.20857 | chr17 | 3115708 | 3120462 | 4.42595 | 3.77349 | 2.5382 |
| 49477568 | 114 | chr17 | 49477271 | 49477791 | Intron (ENSMUST0C | 17 | 49452284 | 49455435 | ENSMUST00 | 24988 | Mocs1 | ENSMUSG0C | false | . | . | . | . | . | . | chr17 | 49468167 | 49480126 | 6.43614 | 4.71108 | 4.19031 |
| 5190082 | 582 | chr17 | 5189621 | 5190365 | Intron (ENSMUST0C | 17 | 5110781 | 5112119 | ENSMUST00 | -77503 | Gm15599 | ENSMUSG0C | false | chr17 | 5189499 | 5190676 | 15.70458 | 43.19576 | 40.56621 | chr17 | 5188721 | 5199127 | 4.36014 | 3.6534 | 2.51291 |
| 66419717 | 548 | chr17 | 66419458 | 66419852 | Intron (ENSMUST0C | 17 | 66413631 | 66413984 | ENSMUST00 | -5475 | Gm4705 | ENSMUSG0C | false | . | . | . | . | . | . | chr17 | 66401889 | 66429482 | 6.16935 | 4.5489 | 3.97689 |
| 43554811 | 114 | chr17 | 43554448 | 43555011 | Distal Intergenic | 17 | 43534174 | 43543639 | ENSMUST00 | -10810 | Ankrd66 | ENSMUSG0C | false | . | . | . | . | . | . | chr17 | 43554410 | 43555332 | 4.40255 | 3.73011 | 2.56571 |
| 4626599 | 719 | chr17 | 4626079 | 4626671 | Distal Intergenic | 17 | 4678020 | 4678126 | ENSMUST00 | -51349 | Gm23186 | ENSMUSG0C | false | chr17 | 4626080 | 4626682 | 79.22768 | 354.44785 | 346.18814 | . | . | . | . | . | . |
| 61894888 | 611 | chr18 | 61894618 | 61894997 | Intron (ENSMUST0C | 18 | 61799395 | 61911824 | ENSMUST00 | 16827 | Ablim3 | ENSMUSG0C | false | . | . | . | . | . | . | chr18 | 61877692 | 61898809 | 5.45644 | 4.15279 | 3.39554 |
| 33464200 | 548 | chr18 | 33463727 | 33464497 | Promoter (<=1kb) | 18 | 33437019 | 33463747 | ENSMUST00 | 0 | Nrep | ENSMUSG0C | false | chr18 | 33462104 | 33464772 | 29.3377 | 90.74815 | 87.38457 | . | . | . | . | . | . |
| 76088375 | 583 | chr18 | 76087465 | 76088530 | Intron (ENSMUST0C | 18 | 76059458 | 76146443 | ENSMUST00 | 28008 | Zbtb7c | ENSMUSG0C | true | . | . | . | . | . | . | chr18 | 76061445 | 76132293 | 6.70015 | 4.79833 | 4.4158 |
| 35815702 | 523 | chr18 | 35815452 | 35815931 | Distal Intergenic | 18 | 35829818 | 35861687 | ENSMUST00 | -13887 | Cxxc5 | ENSMUSG0C | false | chr18 | 35814542 | 35815564 | 10.17627 | 21.16558 | 18.81479 | chr18 | 35812838 | 35818671 | 7.77207 | 5.31683 | 5.31473 |
| 53744814 | 544 | chr18 | 53744340 | 53745098 | Promoter (<=1kb) | 18 | 53681724 | 53744547 | ENSMUST00 | 0 | Cep120 | ENSMUSG0C | false | chr18 | 53743447 | 53745668 | 31.14379 | 104.89771 | 101.20105 | . | . | . | . | . | . |
| 5441328 | 114 | chr18 | 5441156 | 5441554 | Distal Intergenic | 18 | 5518415 | 5521109 | ENSMUST00 | 79555 | Gm10125 | ENSMUSG0C | false | . | . | . | . | . | . | . | . | . | . | . | . |
| 11974028 | 114 | chr18 | 11973679 | 11974147 | Distal Intergenic | 18 | 11840361 | 11840438 | ENSMUST00 | -133242 | Mir1901 | ENSMUSG0C | false | . | . | . | . | . | . | chr18 | 11971839 | 11979509 | 5.50442 | 4.29031 | 3.42052 |
| 47281270 | 497 | chr19 | 47280763 | 47281346 | Intron (ENSMUST0C | 19 | 47273519 | 47273589 | ENSMUST00 | -7175 | Mir6995 | ENSMUSG0C | false | . | . | . | . | . | . | chr19 | 47274861 | 47283196 | 8.38768 | 5.49381 | 5.82214 |
| 45304861 | 600 | chr19 | 45304420 | 45305040 | Promoter (1-2kb) | 19 | 45300463 | 45303034 | ENSMUST00 | -1387 | RP23-188J8.: | ENSMUSG0C | false | . | . | . | . | . | . | chr19 | 45300529 | 45315943 | 6.02178 | 4.57633 | 3.84298 |
| 25453154 | 587 | chr2 | 25452632 | 25453208 | Downstream (1-2kb | 2 | 25456838 | 25458676 | ENSMUST00 | -3630 | Clic3 | ENSMUSG0C | true | . | . | . | . | . | . | chr2 | 25446553 | 25456355 | 6.09426 | 4.46882 | 3.92693 |
| 27758649 | 114 | chr2 | 27758371 | 27759014 | Intron (ENSMUST0C | 2 | 27740201 | 27748671 | ENSMUST00 | 18171 | Rxra | ENSMUSG0C | false | . | . | . | . | . | . | chr2 | 27756216 | 27760543 | 7.96442 | 5.34041 | 5.43403 |
| 32907628 | 506 | chr2 | 32906947 | 32907675 | Intron (ENSMUST0C | 2 | 32910908 | 32912587 | ENSMUST00 | -3233 | Fam129b | ENSMUSG0C | false | . | . | . | . | . | . | chr2 | 32877160 | 32918190 | 7.77045 | 5.19056 | 5.2936 |
| 32617356 | 574 | chr2 | 32616975 | 32617472 | Intron (ENSMUST0C | 2 | 32621758 | 32635058 | ENSMUST00 | -4286 | Ak1 | ENSMUSG0C | false | . | . | . | . | . | . | chr2 | 32605009 | 32639275 | 6.86759 | 4.92928 | 4.53618 |
| 122575750 | 561 | chr2 | 122575383 | 122575807 | Distal Intergenic | 2 | 122595489 | 122598534 | ENSMUST00 | 22727 | Gatm | ENSMUSG0C | false | . | . | . | . | . | . | chr2 | 122574487 | 122577662 | 5.6028 | 4.4306 | 3.49482 |
| 69081057 | 114 | chr2 | 69080533 | 69081236 | Intron (ENSMUST0C | 2 | 69135800 | 69189330 | ENSMUST00 | -54564 | Nostrin | ENSMUSG0C | false | . | . | . | . | . | . | chr2 | 69076857 | 69090376 | 5.44132 | 4.24372 | 3.37539 |
| 129625446 | 528 | chr2 | 129624840 | 129625790 | Intron (ENSMUST0C | 2 | 129620685 | 129628044 | ENSMUST00 | 4156 | Sirpa | ENSMUSG0C | true | . | . | . | . | . | . | chr2 | 129621436 | 129635375 | 5.74226 | 4.37432 | 3.64032 |
| 33606128 | 570 | chr2 | 33605081 | 33606316 | Intron (ENSMUST0C | 2 | 33567156 | 33632283 | ENSMUST00 | 25967 | Lmx1b | ENSMUSG0C | true | . | . | . | . | . | . | chr2 | 33565260 | 33638297 | 6.78481 | 4.81457 | 4.48235 |
| 33166290 | 114 | chr2 | 33165609 | 33166278 | Promoter (1-2kb) | 2 | 33137636 | 33168222 | ENSMUST00 | 1944 | Ralgps1 | ENSMUSG0C | false | . | . | . | . | . | . | chr2 | 33154669 | 33171007 | 7.40171 | 5.13174 | 4.97591 |
| 29696681 | 114 | chr2 | 29696216 | 29696821 | Intron (ENSMUST0C | 2 | 29702482 | 29740363 | ENSMUST00 | -5661 | Rapgef1 | ENSMUSG0C | false | . | . | . | . | . | . | chr2 | 29690471 | 29698739 | 5.92345 | 4.4721 | 3.78221 |
| 153241613 | 585 | chr2 | 153241259 | 153241725 | Promoter (<=1kb) | 2 | 153241533 | 153270247 | ENSMUST00 | 0 | Pofut1 | ENSMUSG0C | true | chr2 | 153239613 | 153242107 | 24.32174 | 95.77222 | 92.30251 | . | . | . | . | . | . |
| 159270753 | 114 | chr2 | 159270194 | 159271091 | Distal Intergenic | 2 | 159139361 | 159139492 | ENSMUST00 | 130834 | Gm25090 | ENSMUSG0C | false | . | . | . | . | . | . | chr2 | 159269480 | 159284862 | 4.11414 | 3.61003 | 2.27932 |
| 75931033 | 462 | chr3 | 75930714 | 75931247 | Intron (ENSMUST0C | 3 | 75909856 | 75956650 | ENSMUST00 | 25403 | Golim4 | ENSMUSG0C | true | . | . | . | . | . | . | chr3 | 75908736 | 75932827 | 4.90077 | 3.96047 | 2.93621 |
| 89859208 | 412 | chr3 | 89858934 | 89859319 | Distal Intergenic | 3 | 89831370 | 89858834 | ENSMUST00 | 27565 | She | ENSMUSG0C | false | . | . | . | . | . | . | chr3 | 89858315 | 89886122 | 6.15651 | 4.57192 | 3.96514 |
| 51916588 | 427 | chr3 | 51916079 | 51916782 | Intron (ENSMUST0C | 3 | 51828912 | 51829028 | ENSMUST00 | 87168 | n-R5s196 | ENSMUSG0C | false | . | . | . | . | . | . | chr3 | 51904229 | 51919287 | 6.09838 | 4.57081 | 3.90639 |
| 34687402 | 498 | chr4 | 34687060 | 34687591 | Promoter (<=1kb) | 4 | 34663257 | 34687438 | ENSMUST00 | 0 | Slc35a1 | ENSMUSG0C | false | chr4 | 34686600 | 34687906 | 28.9509 | 90.07051 | 86.72839 | . | . | . | . | . | . |
| 66450325 | 583 | chr4 | 66450072 | 66450460 | Distal Intergenic | 4 | 65380803 | 66404611 | ENSMUST00 | -45462 | Astn2 | ENSMUSG0C | true | . | . | . | . | . | . | chr4 | 66449477 | 66451280 | 4.51968 | 3.80583 | 2.65848 |
| 136602954 | 572 | chr4 | 136602174 | 136602896 | Promoter (<=1kb) | 4 | 136550540 | 136602723 | ENSMUST00 | 0 | Kdm1a | ENSMUSG0C | false | chr4 | 136601489 | 136602829 | 37.62274 | 127.9987 | 123.57949 | . | . | . | . | . | . |
| 132212466 | 579 | chr4 | 132211840 | 132212463 | Promoter (<=1kb) | 4 | 132184912 | 132212303 | ENSMUST00 | 0 | Ythdf2 | ENSMUSG0C | false | chr4 | 132210818 | 132212608 | 36.04346 | 118.78738 | 114.68494 | . | . | . | . | . | . |
| 149860597 | 588 | chr4 | 149860304 | 149860665 | Distal Intergenic | 4 | 149817732 | 149817840 | ENSMUST00 | 42573 | n-R5s193 | ENSMUSG0C | false | . | . | . | . | . | . | chr4 | 149859847 | 149861376 | 5.80362 | 4.28726 | 3.6996 |
| 116732670 | 542 | chr4 | 116732309 | 116732745 | Intron (ENSMUST0C | 4 | 116725899 | 116725989 | ENSMUST00 | 6411 | Gm27688 | ENSMUSG0C | false | . | . | . | . | . | . | chr4 | 116731176 | 116741002 | 4.98596 | 4.04488 | 2.98621 |
| 81519765 | 614 | chr5 | 81519330 | 81519942 | Intron (ENSMUST0C | 5 | 81645989 | 81688522 | ENSMUST00 | -126047 | Lphn3 | ENSMUSG0C | false | chr5 | 81519672 | 81520026 | 5.15049 | 9.68248 | 7.51107 | chr5 | 81518468 | 81520895 | 3.83839 | 3.51106 | 2.04204 |
| 34171018 | 532 | chr5 | 34170782 | 34171111 | Promoter (1-2kb) | 5 | 34007198 | 34169448 | ENSMUST00 | -1335 | Poln | ENSMUSG0C | false | . | . | . | . | . | . | chr5 | 34168693 | 34173571 | 5.15082 | 4.13032 | 3.13292 |
| 35233420 | 114 | chr5 | 35233227 | 35233598 | Distal Intergenic | 5 | 35278566 | 35281761 | ENSMUST00 | -44968 | Adra2c | ENSMUSG0C | false | . | . | . | . | . | . | chr5 | 35229105 | 35239186 | 4.86991 | 4.0349 | 2.9021 |
| 140777360 | 573 | chr5 | 140777048 | 140777488 | Intron (ENSMUST0C | 5 | 140749283 | 140761439 | ENSMUST00 | 27766 | Amz1 | ENSMUSG0C | false | . | . | . | . | . | . | chr5 | 140761728 | 140779292 | 6.36667 | 4.61767 | 4.14055 |
| 119326141 | 114 | chr5 | 119325757 | 119326206 | Distal Intergenic | 5 | 119297245 | 119297361 | ENSMUST00 | 28513 | n-R5s175 | ENSMUSG0C | false | . | . | . | . | . | . | . | . | . | . | . | . |
| 103606665 | 567 | chr5 | 103606238 | 103606750 | 3' UTR | 5 | 103602587 | 103602879 | ENSMUST00 | 3652 | Gm23085 | ENSMUSG0C | false | . | . | . | . | . | . | chr5 | 103602894 | 103607755 | 4.47885 | 3.74726 | 2.61032 |
| 92834783 | 448 | chr5 | 92834486 | 92834902 | Intron (ENSMUST0C | 5 | 92804220 | 92809488 | ENSMUST00 | -24999 | 4932430I15R | ENSMUSG0C | false | . | . | . | . | . | . | chr5 | 92813516 | 92845987 | 5.7053 | 4.3919 | 3.59815 |
| 19181188 | 510 | chr5 | 19180792 | 19181379 | Distal Intergenic | 5 | 19202368 | 19226555 | ENSMUST00 | 45176 | 4921504A21I | ENSMUSG0C | true | . | . | . | . | . | . | chr5 | 19177836 | 19184145 | 4.31014 | 3.73966 | 2.43836 |
| 18544306 | 615 | chr6 | 18543809 | 18544254 | Promoter (<=1kb) | 6 | 52231199 | 52233324 | ENSMUST00 | -59 | Hoxa10 | ENSMUSG0C | false | . | . | . | . | . | . | chr6 | 18539839 | 18548637 | 4.5475 | 3.89202 | 2.64481 |
| 52233733 | 114 | chr6 | 52233382 | 52234275 | Promoter (<=1kb) | 6 | 106778249 | 106800069 | ENSMUST00 | 0 | Crbn | ENSMUSG0C | false | chr6 | 52227601 | 52235632 | 41.69514 | 140.65352 | 135.74945 | . | . | . | . | . | . |
| 22356271 | 535 | chr6 | 22355772 | 22356389 | Promoter (<=1kb) | 6 | 85451505 | 85468477 | ENSMUST00 | 0 | Cct7 | ENSMUSG0C | false | chr6 | 22355441 | 22356405 | 28.45235 | 93.08366 | 89.67602 | chr6 | 22353838 | 22356671 | 3.43704 | 3.28121 | 1.74509 |
| 116007131 | 566 | chr6 | 116006706 | 116007185 | Distal Intergenic | 6 | 132361041 | 132364134 | ENSMUST00 | -99472 | Gm8882 | ENSMUSG0C | false | . | . | . | . | . | . | chr6 | 116004192 | 116008178 | 6.32655 | 4.46684 | 4.10455 |
| 140433494 | 114 | chr6 | 140433089 | 140433545 | Intron (ENSMUST0C | 6 | 114860628 | 114875107 | ENSMUST00 | -8127 | Vgll4 | ENSMUSG0C | true | . | . | . | . | . | . | chr6 | 140425415 | 140441290 | 5.09666 | 4.0883 | 3.09824 |
| 114922125 | 600 | chr6 | 114921632 | 114922233 | Intron (ENSMUST0C | 6 | 37327255 | 37442146 | ENSMUST00 | 10361 | Creb3l2 | ENSMUSG0C | true | chr6 | 114917117 | 114921811 | 29.60582 | 98.11335 | 94.5914 | chr6 | 114922002 | 114923292 | 3.84635 | 3.53185 | 2.04366 |
| 72744169 | 566 | chr6 | 72743683 | 72744402 | Promoter (<=1kb) | 6 | 47835661 | 47843647 | ENSMUST00 | 0 | Zfp398 | ENSMUSG0C | false | . | . | . | . | . | . | chr6 | 72739919 | 72749221 | 7.65916 | 5.17359 | 5.21352 |
| 70074050 | 587 | chr6 | 70073781 | 70074409 | Promoter (<=1kb) | 6 | 70117061 | 70117617 | ENSMUST00 | 333 | Igkv8-30 | ENSMUSG0C | false | . | . | . | . | . | . | chr6 | 70072263 | 70075068 | 3.76337 | 3.46811 | 1.98668 |
| 70117160 | 651 | chr6 | 70116852 | 70117284 | Promoter (<=1kb) | 6 | 70386672 | 70387238 | ENSMUST00 | 349 | Igkv8-16 | ENSMUSG0C | false | . | . | . | . | . | . | chr6 | 70115225 | 70118243 | 3.61941 | 3.39411 | 1.85895 |
| 70386896 | 114 | chr6 | 70386409 | 70386889 | Distal Intergenic | 6 | 12857397 | 12858390 | ENSMUST00 | 35570 | Gm15529 | ENSMUSG0C | false | . | . | . | . | . | . | chr6 | 70385956 | 70388311 | 6.31575 | 4.65551 | 4.07275 |
| 70216062 | 597 | chr6 | 70215734 | 70216150 | Distal Intergenic | 6 | 70260409 | 70260933 | ENSMUST00 | 20323 | Igkv6-23 | ENSMUSG0C | false | . | . | . | . | . | . | chr6 | 70213474 | 70219351 | 5.14256 | 4.19463 | 3.11118 |
| 70240516 | 591 | chr6 | 70240204 | 70240610 | Promoter (<=1kb) | 7 | 30459725 | 30487223 | ENSMUST00 | 0 | Nphs1 | ENSMUSG0C | true | . | . | . | . | . | . | chr6 | 70239615 | 70241403 | 4.75318 | 3.82127 | 2.83041 |
| 30459728 | 555 | chr7 | 30459355 | 30459800 | Promoter (<=1kb) | 7 | 46033698 | 46084212 | ENSMUST00 | 0 | Nomo1 | ENSMUSG0C | false | chr7 | 30459332 | 30463950 | 22.44583 | 74.56482 | 71.50171 | . | . | . | . | . | . |
| 110615685 | 576 | chr7 | 110613978 | 110615726 | Promoter (<=1kb) | 7 | 25317164 | 25365917 | ENSMUST00 | -18 | Megf8 | ENSMUSG0C | false | chr7 | 110613377 | 110615752 | 31.19317 | 93.97161 | 90.54182 | . | . | . | . | . | . |
| 25316988 | 570 | chr7 | 25316637 | 25317146 | Distal Intergenic | 7 | 127608756 | 127615797 | ENSMUST00 | -21206 | Zfp629 | ENSMUSG0C | false | chr7 | 25316780 | 25318001 | 32.44658 | 100.11784 | 96.54536 | chr7 | 25314793 | 25322508 | 5.81353 | 4.33708 | 3.6977 |
| 112680275 | 558 | chr7 | 112678900 | 112680254 | Promoter (1-2kb) | 7 | 40190935 | 40191324 | ENSMUST00 | 1424 | Gm6851 | ENSMUSG0C | false | chr7 | 112678205 | 112681471 | 36.76257 | 113.72284 | 109.77938 | chr7 | 112677714 | 112678974 | 5.79943 | 4.50423 | 3.65278 |
| 40192673 | 618 | chr7 | 40192358 | 40192750 | Promoter (<=1kb) | 7 | 16618627 | 16619832 | ENSMUST00 | 850 | RP24-490A2: | ENSMUSG0C | false | . | . | . | . | . | . | . | . | . | . | . | . |
| 16618859 | 538 | chr7 | 16618438 | 16618982 | Intron (ENSMUST0C | 8 | 70406127 | 70431285 | ENSMUST00 | 12010 | Crtc1 | ENSMUSG0C | false | . | . | . | . | . | . | chr7 | 16615607 | 16621870 | 5.3619 | 4.10589 | 3.33445 |
| 70419091 | 590 | chr8 | 70418823 | 70419275 | Intron (ENSMUST0C | 8 | 60993241 | 61030600 | ENSMUST00 | 15146 | Nek1 | ENSMUSG0C | true | chr8 | 70419225 | 70419465 | 5.08786 | 7.92609 | 5.7945 | chr8 | 70415799 | 70434143 | 6.2581 | 4.55457 | 4.06384 |
| 61009076 | 534 | chr8 | 61008386 | 61009136 | Promoter (<=1kb) | 8 | 88331059 | 88362194 | ENSMUST00 | 0 | Brd7 | ENSMUSG0C | false | . | . | . | . | . | . | chr8 | 61007494 | 61010706 | 4.67783 | 3.84507 | 2.76552 |
| 104887096 | 529 | chr8 | 104886708 | 104887221 | Intron (ENSMUST0C | 8 | 87783316 | 87944052 | ENSMUST00 | 9541 | Zfp423 | ENSMUSG0C | true | . | . | . | . | . | . | chr8 | 104884771 | 104889614 | 5.64618 | 4.36566 | 3.53177 |
| 87934027 | 499 | chr8 | 87933550 | 87934511 | Intron (ENSMUST0C | 8 | 82863356 | 83091271 | ENSMUST00 | 76501 | Rnf150 | ENSMUSG0C | true | . | . | . | . | . | . | chr8 | 87909819 | 87941142 | 6.06882 | 4.48607 | 3.90276 |
| 82940258 | 473 | chr8 | 82939856 | 82940389 | Intron (ENSMUST0C | 8 | 91467685 | 91666946 | ENSMUST00 | 54522 | Fto | ENSMUSG0C | false | . | . | . | . | . | . | chr8 | 82936463 | 82941511 | 3.87884 | 3.42569 | 2.13119 |
| 91522476 | 573 | chr8 | 91522206 | 91522591 | Distal Intergenic | 8 | 126944847 | 126944980 | ENSMUST00 | 91628 | Gm26397 | ENSMUSG0C | false | . | . | . | . | . | . | chr8 | 91521033 | 91528017 | 5.83853 | 4.42897 | 3.70928 |
| 70610426 | 576 | chr8 | 70609379 | 70611412 | Distal Intergenic | 8 | 128650997 | 128651103 | ENSMUST00 | -106167 | Gm26002 | ENSMUSG0C | false | chr8 | 70610311 | 70610820 | 10.04446 | 20.40655 | 18.06574 | chr8 | 70604127 | 70612475 | 6.23684 | 4.68532 | 4.01553 |
| 121242162 | 114 | chr8 | 121241670 | 121242435 | Intron (ENSMUST0C | 8 | 121256747 | 121356836 | ENSMUST00 | 75799 | Gm26815 | ENSMUSG0C | false | . | . | . | . | . | . | chr8 | 121236124 | 121247254 | 7.60766 | 5.2125 | 5.14118 |
| 121280768 | 424 | chr8 | 121280463 | 121281037 | Promoter (2-3kb) | 9 | 57790513 | 57834236 | ENSMUST00 | 2306 | Arid3b | ENSMUSG0C | false | . | . | . | . | . | . | chr8 | 121277448 | 121284034 | 5.99549 | 4.54477 | 3.8074 |
| 57831807 | 641 | chr9 | 57831347 | 57831930 | Distal Intergenic | 9 | 14593444 | 14615483 | ENSMUST00 | -26307 | Amotl1 | ENSMUSG0C | false | . | . | . | . | . | . | chr9 | 57827202 | 57836152 | 5.62883 | 4.39478 | 3.50932 |
| 32647076 | 114 | chr9 | 32646622 | 32647268 | Intron (ENSMUST0C | 9 | 32636239 | 32651476 | ENSMUST00 | 20353 | Ets1 | ENSMUSG0C | true | . | . | . | . | . | . | chr9 | 32645640 | 32647917 | 6.06489 | 4.58743 | 3.88733 |
| 32672608 | 603 | chr9 | 32671858 | 32672986 | Intron (ENSMUST0C | 9 | 71632633 | 71648350 | ENSMUST00 | -26085 | Cgnl1 | ENSMUSG0C | false | chr9 | 32671954 | 32672786 | 8.53631 | 15.91019 | 13.6344 | chr9 | 32666505 | 32679281 | 4.67115 | 3.90561 | 2.75084 |
| 71674693 | 114 | chr9 | 71674434 | 71674867 | Promoter (<=1kb) | 9 | 57750711 | 57765860 | ENSMUST00 | 215 | Clk3 | ENSMUSG0C | false | . | . | . | . | . | . | chr9 | 71672716 | 71681830 | 5.44209 | 4.19703 | 3.39991 |
| 63611246 | 571 | chr9 | 63610824 | 63611500 | Intron (ENSMUST0C | 9 | 60610270 | 60649710 | ENSMUST00 | 12251 | Lrrc49 | ENSMUSG0C | true | chr9 | 63611389 | 63611663 | 4.77198 | 8.97201 | 6.81752 | chr9 | 63603357 | 63616669 | 6.44347 | 4.69808 | 4.19638 |
| 60637360 | 550 | chr9 | 60637042 | 60637459 | Promoter (<=1kb) | 9 | 120577346 | 120578072 | ENSMUST00 | 0 | 5830454E08I | ENSMUSG0C | false | . | . | . | . | . | . | chr9 | 60623926 | 60640440 | 4.81608 | 3.88284 | 2.88797 |
| 38836633 | 627 | chr19 | 38836383 | 38837069 | Promoter (<=1kb) | 19 | 38836579 | 38919923 | ENSMUST00 | 0 | Tbc1d12 | ENSMUSG0C | false | chr19 | 38836126 | 38837723 | 33.26678 | 112.81776 | 108.90401 | . | . | . | . | . | . |
| 157048715 | 596 | chr2 | 157048462 | 157048881 | Intron (ENSMUST0C | 2 | 157037050 | 157040239 | ENSMUST00 | -8224 | Soga1 | ENSMUSG0C | false | . | . | . | . | . | . | chr2 | 157047505 | 157049845 | 3.42377 | 3.07497 | 1.74981 |
| 58542185 | 579 | chr9 | 58541860 | 58542160 | Promoter (<=1kb) | 9 | 66946086 | 66949512 | ENSMUST00 | 0 | Rps27l | ENSMUSG0C | false | . | . | . | . | . | . | chr9 | 58541298 | 58542496 | 5.43317 | 4.11368 | 3.40583 |
| 62699588 | 114 | chr4 | 62699276 | 62699759 | Promoter (<=1kb) | 4 | 62699466 | 62703018 | ENSMUST00 | 0 | Rgs3 | ENSMUSG0C | false | chr4 | 62699665 | 62702487 | 24.69334 | 83.34509 | 80.13418 | chr4 | 62694793 | 62699426 | 7.42731 | 5.01341 | 4.9919 |
| 132463814 | 805 | chr6 | 132463605 | 132463906 | Intron (ENSMUST0C | 6 | 140424099 | 140594906 | ENSMUST00 | 8991 | Plekha5 | ENSMUSG0C | true | chr6 | 132463619 | 132463896 | 29.41006 | 85.09721 | 81.85307 | . | . | . | . | . | . |
| 94281214 | 613 | chr12 | 94280920 | 94281302 | Distal Intergenic | 12 | 93927004 | 93929102 | ENSMUST00 | -351819 | Gm9726 | ENSMUSG0C | false | . | . | . | . | . | . | chr12 | 94279611 | 94284656 | 3.79808 | 3.5006 | 2.0053 |
| 83288417 | 604 | chr17 | 83288075 | 83288521 | Promoter (<=1kb) | 17 | 83288546 | 83289028 | ENSMUST00 | -25 | Rpl21-ps7 | ENSMUSG0C | false | . | . | . | . | . | . | chr17 | 83287263 | 83289778 | 7.57114 | 5.04054 | 5.14387 |
| 5849782 | 114 | chr17 | 5849397 | 5849838 | Intron (ENSMUST0C | 17 | 5841328 | 5931033 | ENSMUST00 | 8070 | Snx9 | ENSMUSG0C | false | chr17 | 5849210 | 5849793 | 9.38717 | 20.36066 | 18.02005 | chr17 | 5842208 | 5850409 | 5.53242 | 4.3591 | 3.4424 |
| 14642063 | 114 | chr9 | 14641789 | 14642122 | Intron (ENSMUST0C | 9 | 32636239 | 32651476 | ENSMUST00 | 10384 | Ets1 | ENSMUSG0C | true | . | . | . | . | . | . | . | . | . | . | . | . |
| 104867666 | 519 | chr8 | 104867376 | 104867853 | Distal Intergenic | 8 | 104867488 | 104874080 | ENSMUST00 | 19221 | Ces2d-ps | ENSMUSG0C | false | . | . | . | . | . | . | chr8 | 104864625 | 104870535 | 4.84736 | 4.01234 | 2.88046 |
| 59506345 | 508 | chr11 | 59506135 | 59506554 | Promoter (<=1kb) | 11 | 59499544 | 59506220 | ENSMUST00 | 0 | Zkscan17 | ENSMUSG0C | false | chr11 | 59505222 | 59507156 | 33.04283 | 115.53907 | 111.53828 | . | . | . | . | . | . |
| 21362925 | 532 | chr13 | 21362662 | 21363031 | Promoter (<=1kb) | 13 | 21362827 | 21369824 | ENSMUST00 | 0 | Zscan12 | ENSMUSG0C | false | chr13 | 21362385 | 21363516 | 25.17393 | 93.69764 | 90.27328 | chr13 | 21362419 | 21364810 | 3.63587 | 3.27994 | 1.90693 |
| 105736735 | 551 | chr7 | 105736374 | 105736893 | Promoter (<=1kb) | 7 | 110308013 | 110614922 | ENSMUST00 | 0 | Sbf2 | ENSMUSG0C | true | chr7 | 105735728 | 105738289 | 44.26889 | 153.06256 | 147.66058 | . | . | . | . | . | . |
| 10952268 | 456 | chr19 | 10952042 | 10952505 | Promoter (2-3kb) | 19 | 10941481 | 10949266 | ENSMUST00 | -2777 | Ccdc86 | ENSMUSG0C | false | . | . | . | . | . | . | chr19 | 10946315 | 10954190 | 5.98357 | 4.50647 | 3.81648 |
| 64337412 | 114 | chr5 | 64336868 | 64337476 | Intron (ENSMUST0C | 5 | 64349945 | 64350589 | ENSMUST00 | -12469 | Tbc1d1 | ENSMUSG0C | false | . | . | . | . | . | . | chr5 | 64333951 | 64340633 | 6.27533 | 4.60721 | 4.05113 |
| 76763713 | 559 | chr11 | 76763499 | 76763743 | Promoter (<=1kb) | 11 | 76726602 | 76763579 | ENSMUST00 | 0 | Gosr1 | ENSMUSG0C | false | chr11 | 76762717 | 76763684 | 29.3377 | 90.74815 | 87.38457 | . | . | . | . | . | . |
| 150785023 | 584 | chr4 | 150784639 | 150785198 | Distal Intergenic | 4 | 150825733 | 150839575 | ENSMUST00 | -40535 | Gm13049 | ENSMUSG0C | false | . | . | . | . | . | . | chr4 | 150784510 | 150786637 | 4.42467 | 3.8393 | 2.5188 |
| 99217086 | 564 | chrX | 99216855 | 99217321 | | | | | | | | | false | chrX | 99216303 | 99217267 | 11.94783 | 25.26529 | 22.85895 | . | . | . | . | . | . |
| 47835731 | 114 | chr6 | 47835525 | 47836083 | Distal Intergenic | 6 | 121183667 | 121196185 | ENSMUST00 | -36467 | Pex26 | ENSMUSG0C | false | chr6 | 47835563 | 47836836 | 41.84113 | 139.79433 | 134.92708 | chr6 | 47835331 | 47838254 | 3.87408 | 3.48275 | 2.12306 |
| 53793792 | 495 | chr18 | 53793534 | 53793802 | Distal Intergenic | 18 | 53681724 | 53744547 | ENSMUST00 | -48988 | Cep120 | ENSMUSG0C | false | . | . | . | . | . | . | chr18 | 53792269 | 53794132 | 4.9428 | 4.11433 | 2.94935 |
| 65032426 | 114 | chr9 | 65032249 | 65032986 | Promoter (<=1kb) | 9 | 58524300 | 58540941 | ENSMUST00 | -920 | Cd276 | ENSMUSG0C | false | chr9 | 65032063 | 65033438 | 35.98183 | 119.73972 | 115.60696 | chr9 | 65031928 | 65034604 | 4.72792 | 3.93879 | 2.82826 |
| 114883427 | 493 | chr6 | 114883233 | 114883677 | Promoter (<=1kb) | 6 | 114861424 | 114921821 | ENSMUST00 | 0 | Vgll4 | ENSMUSG0C | false | . | . | . | . | . | . | chr6 | 114853844 | 114919520 | 8.05355 | 5.36011 | 5.51688 |
| 106800186 | 525 | chr6 | 106799921 | 106800274 | Promoter (<=1kb) | 6 | 83768885 | 83775813 | ENSMUST00 | 0 | Tex261 | ENSMUSG0C | false | chr6 | 106799187 | 106800432 | 34.3275 | 112.61823 | 108.70669 | chr6 | 106798218 | 106800458 | 4.81515 | 3.86877 | 2.89302 |
| 52013380 | 114 | chr16 | 52012986 | 52013474 | Distal Intergenic | 16 | 52031549 | 52208047 | ENSMUST00 | -18075 | Cblb | ENSMUSG0C | false | . | . | . | . | . | . | chr16 | 52009607 | 52016621 | 4.28782 | 3.76463 | 2.41076 |
| 96065582 | 494 | chr11 | 96064932 | 96065574 | Promoter (<=1kb) | 11 | 96047432 | 96065388 | ENSMUST00 | 0 | Ube2z | ENSMUSG0C | false | chr11 | 96064220 | 96065900 | 22.47347 | 97.37944 | 93.87287 | . | . | . | . | . | . |
| 179546468 | 568 | chr1 | 179546245 | 179546853 | Promoter (<=1kb) | 1 | 179528055 | 179546267 | ENSMUST00 | 0 | Tfb2m | ENSMUSG0C | false | chr1 | 179545202 | 179547773 | 26.61085 | 108.21047 | 104.42454 | . | . | . | . | . | . |
| 45563943 | 114 | chr17 | 45563725 | 45564432 | Promoter (<=1kb) | 17 | 45563964 | 45567669 | ENSMUST00 | 0 | Slc35b2 | ENSMUSG0C | false | chr17 | 45563840 | 45564964 | 39.39634 | 133.30592 | 128.68834 | chr17 | 45563577 | 45571950 | 4.52176 | 3.76389 | 2.64295 |
| 58104177 | 563 | chr11 | 58103852 | 58104403 | Promoter (<=1kb) | 11 | 58104153 | 58118594 | ENSMUST00 | 0 | Cnot8 | ENSMUSG0C | false | chr11 | 58103618 | 58103927 | 9.78363 | 19.44713 | 17.12045 | . | . | . | . | . | . |
| 32657445 | 517 | chr9 | 32656591 | 32657525 | Intron (ENSMUST0C | 9 | 32696022 | 32757820 | ENSMUST00 | -23036 | Ets1 | ENSMUSG0C | true | . | . | . | . | . | . | chr9 | 32656242 | 32658110 | 6.87778 | 4.79195 | 4.58763 |
| 123282748 | 541 | chr4 | 123282555 | 123282850 | Promoter (<=1kb) | 4 | 123282825 | 123298925 | ENSMUST00 | 0 | Pabpc4 | ENSMUSG0C | false | chr4 | 123282579 | 123284220 | 24.32174 | 95.77222 | 92.30251 | . | . | . | . | . | . |
| 122102039 | 643 | chr7 | 122101691 | 122102006 | Promoter (<=1kb) | 7 | 120635176 | 120659478 | ENSMUST00 | 0 | Uqcrc2 | ENSMUSG0C | false | chr7 | 122100946 | 122102271 | 34.36326 | 108.53439 | 104.74065 | chr7 | 122098290 | 122102367 | 5.71206 | 4.36131 | 3.60933 |
| 32863801 | 569 | chr5 | 32863427 | 32863772 | Promoter (<=1kb) | 5 | 32863701 | 32887048 | ENSMUST00 | 0 | Depdc5 | ENSMUSG0C | false | chr5 | 32863170 | 32864374 | 20.32388 | 57.71489 | 54.90119 | . | . | . | . | . | . |
| 45897917 | 561 | chr13 | 45897633 | 45897975 | Intron (ENSMUST0C | 13 | 45549758 | 45964991 | ENSMUST00 | 67016 | Atxn1 | ENSMUSG0C | false | . | . | . | . | . | . | chr13 | 45897011 | 45898862 | 3.09204 | 2.93398 | 1.47583 |
| 21506671 | 477 | chr13 | 21506211 | 21506610 | Promoter (1-2kb) | 13 | 21504382 | 21505056 | ENSMUST00 | -1156 | Zfp389 | ENSMUSG0C | false | chr13 | 21505925 | 21506836 | 28.36989 | 85.2065 | 81.96005 | chr13 | 21505803 | 21506932 | 3.87999 | 3.3988 | 2.12773 |
| 120093938 | 598 | chr2 | 120093700 | 120094085 | Intron (ENSMUST0C | 2 | 120076214 | 120085772 | ENSMUST00 | -7929 | Sptbn5 | ENSMUSG0C | false | . | . | . | . | . | . | chr2 | 120090578 | 120107323 | 5.14183 | 4.07311 | 3.14279 |
| 56036892 | 568 | chr17 | 56036473 | 56036838 | Promoter (<=1kb) | 17 | 56016753 | 56036637 | ENSMUST00 | 0 | Sh3gl1 | ENSMUSG0C | false | chr17 | 56035704 | 56036685 | 31.62822 | 118.12274 | 114.03909 | . | . | . | . | . | . |
| 83775969 | 547 | chr6 | 83775658 | 83776217 | Promoter (<=1kb) | 6 | 22306520 | 22356166 | ENSMUST00 | 0 | Fam3c | ENSMUSG0C | false | chr6 | 83774759 | 83776337 | 32.94367 | 134.34938 | 129.69266 | chr6 | 83774182 | 83776885 | 4.59104 | 3.72868 | 2.70779 |
| 122030078 | 574 | chr10 | 122029835 | 122030100 | Intron (ENSMUST0C | 10 | 121780991 | 122047308 | ENSMUST00 | 17208 | Srgap1 | ENSMUSG0C | true | . | . | . | . | . | . | chr10 | 121945243 | 122036389 | 5.74397 | 4.3951 | 3.62974 |
| 76945801 | 114 | chr11 | 76945407 | 76945839 | Promoter (<=1kb) | 11 | 76945504 | 76987379 | ENSMUST00 | 0 | Blmh | ENSMUSG0C | false | chr11 | 76945142 | 76946667 | 29.48955 | 94.93548 | 91.48817 | chr11 | 76944160 | 76947151 | 4.10471 | 3.58068 | 2.28813 |
| 46033740 | 592 | chr7 | 46033572 | 46033857 | Promoter (<=1kb) | 7 | 122085403 | 122101886 | ENSMUST00 | 0 | Ndufab1 | ENSMUSG0C | false | chr7 | 46033517 | 46034659 | 31.66066 | 102.77801 | 99.14294 | . | . | . | . | . | . |
| 30951474 | 408 | chr14 | 30951216 | 30951557 | Promoter (<=1kb) | 14 | 30951486 | 30987439 | ENSMUST00 | 0 | Nek4 | ENSMUSG0C | false | chr14 | 30950835 | 30952082 | 29.63113 | 94.2009 | 90.76292 | chr14 | 30950690 | 30952434 | 5.14147 | 4.20193 | 3.1205 |
| 34776979 | 500 | chr18 | 34776834 | 34777883 | Promoter (<=1kb) | 18 | 34777008 | 34839370 | ENSMUST00 | 0 | Kdm3b | ENSMUSG0C | false | chr18 | 34776612 | 34778537 | 35.62435 | 116.99458 | 112.94973 | . | . | . | . | . | . |
| 24725267 | 589 | chr5 | 24724859 | 24725277 | Intron (ENSMUST0C | 5 | 24712265 | 24730680 | ENSMUST00 | 5403 | Wdr86 | ENSMUSG0C | false | . | . | . | . | . | . | chr5 | 24723425 | 24734625 | 5.82193 | 4.37932 | 3.69553 |
| 22731554 | 684 | chr10 | 22731019 | 22731591 | Promoter (<=1kb) | 10 | 22703879 | 22731447 | ENSMUST00 | 0 | Tbpl1 | ENSMUSG0C | false | chr10 | 22730228 | 22732192 | 39.29051 | 134.21745 | 129.56854 | . | . | . | . | . | . |
| 26133189 | 556 | chr2 | 26132678 | 26133360 | Distal Intergenic | 2 | 26138546 | 26139543 | ENSMUST00 | -5186 | 1810012K08I | ENSMUSG0C | false | chr2 | 26132583 | 26133600 | 9.51244 | 19.51414 | 17.18665 | chr2 | 26129136 | 26135084 | 6.01114 | 4.43692 | 3.85388 |
| 69732865 | 458 | chr14 | 69732403 | 69732877 | Promoter (<=1kb) | 14 | 69716979 | 69732570 | ENSMUST00 | 0 | Chmp7 | ENSMUSG0C | false | chr14 | 69731620 | 69733869 | 41.18432 | 138.64781 | 133.82259 | . | . | . | . | . | . |
| 127637190 | 550 | chr7 | 127637002 | 127637326 | Promoter (<=1kb) | 7 | 127342797 | 127345314 | ENSMUST00 | -50 | Zfp768 | ENSMUSG0C | false | chr7 | 127636649 | 127637757 | 14.64398 | 38.39655 | 35.82735 | chr7 | 127636481 | 127637911 | 5.47361 | 4.28826 | 3.42456 |
| 163994981 | 660 | chr1 | 163994494 | 163994953 | Promoter (<=1kb) | 1 | 163994889 | 163997243 | ENSMUST00 | 0 | Mettl18 | ENSMUSG0C | false | chr1 | 163994206 | 163995669 | 25.99992 | 84.80917 | 81.57088 | chr1 | 163991574 | 163996008 | 5.65475 | 4.33948 | 3.57649 |
| 121147218 | 582 | chr6 | 121146874 | 121147200 | Promoter (<=1kb) | 6 | 70074024 | 70074584 | ENSMUST00 | 175 | Igkv6-32 | ENSMUSG0C | false | . | . | . | . | . | . | chr6 | 121143778 | 121148146 | 5.47647 | 4.19044 | 3.41087 |
| 66946285 | 654 | chr9 | 66945906 | 66946270 | Intron (ENSMUST0C | 9 | 63602655 | 63641889 | ENSMUST00 | 8170 | Aagab | ENSMUSG0C | false | chr9 | 66945369 | 66946939 | 33.73597 | 126.30199 | 121.9427 | . | . | . | . | . | . |
| 127345781 | 714 | chr7 | 127345363 | 127346412 | Promoter (<=1kb) | 7 | 112679318 | 112899459 | ENSMUST00 | 0 | Tead1 | ENSMUSG0C | false | chr7 | 127344643 | 127346973 | 28.22734 | 101.53345 | 97.92473 | . | . | . | . | . | . |
| 80133203 | 560 | chr15 | 80133045 | 80133542 | Promoter (<=1kb) | 15 | 80133154 | 80161701 | ENSMUST00 | 0 | Tab1 | ENSMUSG0C | false | chr15 | 80132695 | 80134176 | 24.78212 | 78.86476 | 75.73294 | chr15 | 80132452 | 80135076 | 5.60141 | 4.37527 | 3.50624 |
| 143113208 | 557 | chr5 | 143112466 | 143113162 | Promoter (<=1kb) | 5 | 142990893 | 143113020 | ENSMUST00 | 0 | Rnf216 | ENSMUSG0C | false | chr5 | 143111903 | 143113458 | 29.34276 | 99.62727 | 96.06711 | chr5 | 143100232 | 143113412 | 4.82716 | 3.92416 | 2.88218 |
| 136888697 | 445 | chr5 | 136888503 | 136888915 | Distal Intergenic | 5 | 136757489 | 136883209 | ENSMUST00 | -5295 | Col26a1 | ENSMUSG0C | false | . | . | . | . | . | . | chr5 | 136885321 | 136889075 | 4.822 | 3.84455 | 2.90361 |
| 114707852 | 554 | chr5 | 114707701 | 114707881 | Promoter (<=1kb) | 5 | 114707760 | 114722327 | ENSMUST00 | 0 | Tchp | ENSMUSG0C | false | chr5 | 114707703 | 114708580 | 28.53699 | 95.19626 | 91.7378 | chr5 | 114707363 | 114708929 | 7.03659 | 5.04842 | 4.64939 |
| 102470541 | 403 | chr15 | 102470338 | 102470604 | Promoter (<=1kb) | 15 | 102470632 | 102500059 | ENSMUST00 | -28 | Pcbp2 | ENSMUSG0C | false | chr15 | 102470489 | 102471870 | 22.78592 | 76.10319 | 73.01487 | . | . | . | . | . | . |
| 39762248 | 601 | chr1 | 39762073 | 39762299 | Distal Intergenic | 1 | 39665306 | 39720997 | ENSMUST00 | -41077 | Rfx8 | ENSMUSG0C | false | chr1 | 39761731 | 39762300 | 12.68821 | 30.86953 | 28.39186 | chr1 | 39759438 | 39762551 | 4.33134 | 3.78958 | 2.45114 |
| 103629691 | 578 | chr5 | 103629497 | 103629775 | Promoter (<=1kb) | 5 | 103605711 | 103629403 | ENSMUST00 | -95 | Slc10a6 | ENSMUSG0C | false | . | . | . | . | . | . | chr5 | 103627004 | 103630959 | 5.1666 | 4.08182 | 3.15645 |
| 133025518 | 622 | chr1 | 133025053 | 133025489 | Promoter (<=1kb) | 1 | 132959484 | 133025338 | ENSMUST00 | 0 | Mdm4 | ENSMUSG0C | false | chr1 | 133024460 | 133026209 | 28.90751 | 104.81161 | 101.11721 | . | . | . | . | . | . |
| 48817429 | 558 | chr11 | 48816967 | 48817318 | Promoter (<=1kb) | 11 | 48809097 | 48816953 | ENSMUST00 | -15 | Trim41 | ENSMUSG0C | false | chr11 | 48815920 | 48818558 | 26.65955 | 86.95457 | 83.67183 | . | . | . | . | . | . |
| 115709344 | 575 | chr4 | 115708753 | 115709283 | Promoter (1-2kb) | 4 | 115707251 | 115708303 | ENSMUST00 | 1503 | Gm12849 | ENSMUSG0C | false | . | . | . | . | . | . | chr4 | 115707353 | 115710474 | 3.80958 | 3.35728 | 2.063 |
| 54744501 | 545 | chr11 | 54744248 | 54744609 | Promoter (2-3kb) | 11 | 54746357 | 54746824 | ENSMUST00 | 2215 | Gm12229 | ENSMUSG0C | false | . | . | . | . | . | . | chr11 | 54722256 | 54750513 | 5.8032 | 4.39 | 3.67574 |

| | | | | | | | | | | | | | | | | | | | | | | | | | |
|---|---|---|---|---|---|---|---|---|---|---|---|---|---|---|---|---|---|---|---|---|---|---|---|---|---|
| 10605623 | 114 | chr19 | 10605299 | 10605848 | Promoter (<=1kb) | 19 | 10605625 | 10629813 | ENSMUST00 | 0 | Ddb1 | ENSMUSG0C | false | chr19 | 10603045 | 10606602 | 41.22328 | 133.19821 | 128.59248 | . | . | . | . | . | . |
| 69344560 | 482 | chr18 | 69344189 | 69344633 | Promoter (<=1kb) | 18 | 69344492 | 69565506 | ENSMUST00 | 0 | Tcf4 | ENSMUSG0C | false | chr18 | 69343931 | 69345637 | 23.09106 | 69.49222 | 66.50756 | chr18 | 69343638 | 69345358 | 3.2489 | 3.19933 | 1.55274 |
| 50110384 | 607 | chr2 | 50110152 | 50110565 | Intron (ENSMUST0C | 2 | 50128369 | 50193563 | ENSMUST00 | -17804 | Lypd6 | ENSMUSG0C | false | . | . | . | . | . | . | chr2 | 50109251 | 50111092 | 3.6132 | 3.35253 | 1.86513 |
| 45931494 | 114 | chr17 | 45931103 | 45931562 | Distal Intergenic | 17 | 46018712 | 46021372 | ENSMUST00 | 89810 | Vegfa | ENSMUSG0C | true | . | . | . | . | . | . | chr17 | 45928605 | 45934413 | 4.63767 | 3.72884 | 2.75421 |
| 84996337 | 475 | chr12 | 84996080 | 84996874 | Promoter (<=1kb) | 12 | 84996321 | 85070492 | ENSMUST00 | 0 | Ylpm1 | ENSMUSG0C | false | chr12 | 84996250 | 84997380 | 39.12038 | 124.14072 | 119.86815 | . | . | . | . | . | . |
| 57056672 | 588 | chr14 | 57056374 | 57056798 | Promoter (1-2kb) | 14 | 57035608 | 57058030 | ENSMUST00 | 1232 | Gja3 | ENSMUSG0C | false | chr14 | 57056396 | 57058454 | 32.01914 | 97.37792 | 93.87154 | chr14 | 57053149 | 57056474 | 4.05021 | 3.5021 | 2.24184 |
| 8881036 | 629 | chr19 | 8880507 | 8881032 | Promoter (<=1kb) | 19 | 8880016 | 8880933 | ENSMUST00 | 0 | AI462493 | ENSMUSG0C | false | chr19 | 8879994 | 8881359 | 28.70854 | 85.79402 | 82.53136 | chr19 | 8878949 | 8881392 | 4.78404 | 4.03421 | 2.82388 |
| 106650998 | 114 | chr4 | 106650872 | 106651124 | Promoter (<=1kb) | 4 | 106651069 | 106653539 | ENSMUST00 | 0 | Pars2 | ENSMUSG0C | false | chr4 | 106651039 | 106651672 | 21.87981 | 58.7423 | 55.91244 | chr4 | 106649637 | 106652500 | 5.3602 | 4.00044 | 3.33642 |
| 96727638 | 560 | chr3 | 96727444 | 96727973 | Promoter (<=1kb) | 3 | 96727611 | 96791155 | ENSMUST00 | 0 | Rnf115 | ENSMUSG0C | false | chr3 | 96726884 | 96728769 | 34.74108 | 114.39095 | 110.42815 | . | . | . | . | . | . |
| 133610410 | 114 | chr1 | 133609822 | 133610342 | Promoter (<=1kb) | 1 | 133603871 | 133610291 | ENSMUST00 | 0 | Snrpe | ENSMUSG0C | false | chr1 | 133609373 | 133610411 | 25.61729 | 89.79346 | 86.4557 | . | . | . | . | . | . |
| 143777309 | 639 | chr1 | 143777006 | 143777411 | Promoter (<=1kb) | 1 | 143777272 | 143807305 | ENSMUST00 | 0 | Uchl5 | ENSMUSG0C | false | chr1 | 143776236 | 143778292 | 27.97104 | 107.99458 | 104.2211 | . | . | . | . | . | . |
| 13590108 | 521 | chr1 | 13589770 | 13590112 | Promoter (<=1kb) | 1 | 13564698 | 13589910 | ENSMUST00 | 0 | Tram1 | ENSMUSG0C | false | chr1 | 13588480 | 13590440 | 31.85236 | 101.09525 | 97.496 | . | . | . | . | . | . |
| 74588370 | 565 | chr1 | 74587674 | 74588490 | Promoter (<=1kb) | 1 | 74565127 | 74588028 | ENSMUST00 | 0 | Zfp142 | ENSMUSG0C | false | chr1 | 74587078 | 74588189 | 33.1634 | 104.39787 | 100.72088 | . | . | . | . | . | . |
| 52845014 | 649 | chr1 | 52844724 | 52845010 | Promoter (<=1kb) | 1 | 52844929 | 52848693 | ENSMUST00 | 0 | Hibch | ENSMUSG0C | false | chr1 | 52844968 | 52845904 | 18.77433 | 54.14365 | 51.37372 | chr1 | 52844108 | 52846226 | 4.34574 | 3.73717 | 2.50184 |
| 153543319 | 114 | chr1 | 153543095 | 153543310 | Intron (ENSMUST0C | 1 | 153548422 | 153549711 | ENSMUST00 | 6401 | Npl | ENSMUSG0C | false | . | . | . | . | . | . | chr1 | 153539148 | 153550339 | 5.26804 | 4.15937 | 3.23842 |
| 44585045 | 114 | chr1 | 44584879 | 44585073 | Intron (ENSMUST0C | 1 | 44552597 | 44708632 | ENSMUST00 | 32283 | Gulp1 | ENSMUSG0C | true | . | . | . | . | . | . | chr1 | 44583660 | 44586527 | 5.26874 | 4.13444 | 3.26857 |
| 118389476 | 546 | chr1 | 118388829 | 118389438 | Promoter (<=1kb) | 1 | 118389058 | 118609432 | ENSMUST00 | 0 | Clasp1 | ENSMUSG0C | false | chr1 | 118387964 | 118390405 | 29.66345 | 100.21 | 96.63225 | . | . | . | . | . | . |
| 100622968 | 536 | chr11 | 100622764 | 100623120 | Promoter (<=1kb) | 11 | 100622876 | 100627607 | ENSMUST00 | 0 | Nkiras2 | ENSMUSG0C | false | chr11 | 100622779 | 100623594 | 18.2599 | 77.14165 | 74.04132 | . | . | . | . | . | . |
| 83632324 | 681 | chr12 | 83632018 | 83632484 | Promoter (<=1kb) | 12 | 83632252 | 83683123 | ENSMUST00 | 0 | Rbm25 | ENSMUSG0C | false | chr12 | 83632162 | 83633479 | 32.16725 | 95.65015 | 92.18271 | . | . | . | . | . | . |
| 69361491 | 505 | chr12 | 69361137 | 69361583 | Promoter (<=1kb) | 12 | 69361182 | 69361481 | ENSMUST00 | 0 | Gm26191 | ENSMUSG0C | false | chr12 | 69361102 | 69361383 | 13.18045 | 29.78312 | 27.32097 | . | . | . | . | . | . |
| 50785674 | 576 | chr14 | 50785341 | 50785642 | Promoter (<=1kb) | 14 | 50765415 | 50785518 | ENSMUST00 | 0 | Ttc5 | ENSMUSG0C | false | chr14 | 50784823 | 50785681 | 31.62822 | 118.12274 | 114.03909 | chr14 | 50783693 | 50785856 | 3.59414 | 3.28458 | 1.88913 |
| 102326225 | 653 | chr15 | 102326045 | 102326904 | Promoter (<=1kb) | 15 | 102326116 | 102331873 | ENSMUST00 | 0 | Pfdn5 | ENSMUSG0C | false | chr15 | 102325753 | 102327357 | 28.48738 | 97.07422 | 93.57343 | . | . | . | . | . | . |
| 27525649 | 483 | chr15 | 27525377 | 27525650 | Intron (ENSMUST0C | 15 | 27571449 | 27593163 | ENSMUST00 | -45799 | Ank | ENSMUSG0C | false | . | . | . | . | . | . | . | . | . | . | . | . |
| 75616734 | 656 | chr15 | 75616371 | 75616731 | Promoter (<=1kb) | 15 | 75616679 | 75625300 | ENSMUST00 | 0 | Zfp41 | ENSMUSG0C | false | chr15 | 75615707 | 75617294 | 17.71499 | 57.83842 | 55.02271 | chr15 | 75615615 | 75617342 | 5.21478 | 4.18872 | 3.17278 |
| 25550005 | 599 | chr16 | 25549627 | 25549908 | Distal Intergenic | 16 | 25683763 | 25821318 | ENSMUST00 | -133855 | Trp63 | ENSMUSG0C | false | . | . | . | . | . | . | . | . | . | . | . | . |
| 74093367 | 595 | chr16 | 74093169 | 74093369 | Intron (ENSMUST0C | 16 | 74058899 | 74059033 | ENSMUST00 | -34137 | Gm22163 | ENSMUSG0C | false | . | . | . | . | . | . | chr16 | 74090760 | 74094201 | 4.77226 | 3.99214 | 2.80493 |
| 56200099 | 726 | chr17 | 56199898 | 56200186 | Intron (ENSMUST0C | 17 | 56175744 | 56183920 | ENSMUST00 | -15979 | D17Wsu104e | ENSMUSG0C | false | chr17 | 56199916 | 56200182 | 21.65864 | 58.36299 | 55.53803 | . | . | . | . | . | . |
| 36958657 | 626 | chr17 | 36958185 | 36958665 | Promoter (<=1kb) | 17 | 36958592 | 36965622 | ENSMUST00 | 0 | Znrd1as | ENSMUSG0C | false | chr17 | 36957679 | 36959243 | 19.17126 | 69.02763 | 66.05415 | chr17 | 36956323 | 36959716 | 5.61048 | 4.33555 | 3.51521 |
| 35331848 | 114 | chr17 | 35331526 | 35331869 | Distal Intergenic | 17 | 35335909 | 35336292 | ENSMUST00 | -4040 | Gm8741 | ENSMUSG0C | false | . | . | . | . | . | . | . | . | . | . | . | . |
| 9958217 | 580 | chr18 | 9957794 | 9958294 | Promoter (<=1kb) | 18 | 9958180 | 9995484 | ENSMUST00 | 0 | Thoc1 | ENSMUSG0C | false | chr18 | 9957507 | 9958955 | 23.46333 | 91.19009 | 87.82124 | . | . | . | . | . | . |
| 38278048 | 114 | chr18 | 38277883 | 38278168 | Exon (ENSMUST00C | 18 | 38267092 | 38284401 | ENSMUST00 | 6233 | Pcdh12 | ENSMUSG0C | false | . | . | . | . | . | . | chr18 | 38274249 | 38279359 | 5.63879 | 4.38394 | 3.54879 |
| 106974523 | 597 | chr2 | 106974365 | 106974599 | Promoter (<=1kb) | 2 | 106962527 | 106974424 | ENSMUST00 | 0 | Arl14ep | ENSMUSG0C | false | chr2 | 106973606 | 106974761 | 26.6505 | 78.45867 | 75.33303 | chr2 | 106973065 | 106974784 | 4.27265 | 3.80927 | 2.39771 |
| 167421945 | 114 | chr2 | 167421492 | 167422036 | Promoter (<=1kb) | 2 | 167421712 | 167477000 | ENSMUST00 | 0 | Slc9a8 | ENSMUSG0C | false | chr2 | 167420983 | 167422465 | 24.86315 | 92.1349 | 88.74321 | chr2 | 167420660 | 167422853 | 4.22416 | 3.64759 | 2.38613 |
| 164461079 | 114 | chr2 | 164460841 | 164461347 | Promoter (<=1kb) | 2 | 164460971 | 164465508 | ENSMUST00 | 0 | Sys1 | ENSMUSG0C | false | chr2 | 164460919 | 164462087 | 31.43241 | 100.35837 | 96.77719 | . | . | . | . | . | . |
| 132835388 | 592 | chr3 | 132835122 | 132835435 | Intron (ENSMUST0C | 3 | 132787225 | 132787335 | ENSMUST00 | -47788 | Gm22322 | ENSMUSG0C | false | . | . | . | . | . | . | . | . | . | . | . | . |
| 120635253 | 562 | chr7 | 120634949 | 120635247 | Promoter (<=1kb) | 7 | 105732207 | 105736781 | ENSMUST00 | 0 | Rrp8 | ENSMUSG0C | false | chr7 | 120634318 | 120636330 | 39.37288 | 136.56555 | 131.82602 | . | . | . | . | . | . |
| 88362280 | 509 | chr8 | 88361830 | 88362259 | Promoter (<=1kb) | 8 | 104867488 | 104874080 | ENSMUST00 | 0 | Ces2d-ps | ENSMUSG0C | false | chr8 | 88360901 | 88363027 | 43.4858 | 141.53311 | 136.59532 | . | . | . | . | . | . |
| 126853291 | 114 | chr8 | 126853015 | 126853352 | Promoter (1-2kb) | 8 | 70597491 | 70608314 | ENSMUST00 | -1066 | Ssbp4 | ENSMUSG0C | false | . | . | . | . | . | . | chr8 | 126847776 | 126853210 | 4.5877 | 3.85968 | 2.679 |
| 128544764 | 584 | chr8 | 128544475 | 128544830 | Distal Intergenic | 8 | 128650997 | 128651103 | ENSMUST00 | -114567 | Gm26002 | ENSMUSG0C | false | . | . | . | . | . | . | chr8 | 128519464 | 128548431 | 5.17648 | 4.1349 | 3.15487 |
| 57765292 | 114 | chr9 | 57765001 | 57765645 | Promoter (<=1kb) | 9 | 58253164 | 58262523 | ENSMUST00 | 0 | Stoml1 | ENSMUSG0C | false | chr9 | 57763942 | 57766341 | 37.17318 | 134.81126 | 130.13722 | . | . | . | . | . | . |
| 58253243 | 571 | chr9 | 58253063 | 58253269 | Promoter (<=1kb) | 9 | 65032458 | 65082650 | ENSMUST00 | 0 | Dpp8 | ENSMUSG0C | false | chr9 | 58253191 | 58253958 | 42.22279 | 143.81401 | 138.77701 | chr9 | 58252556 | 58255789 | 4.55946 | 3.87216 | 2.64575 |
| 120577403 | 443 | chr9 | 120577128 | 120577462 | Distal Intergenic | 20 | 99138131 | 99147230 | ENSMUST00 | 78725 | Efnb1 | ENSMUSG0C | false | chr9 | 120576838 | 120577979 | 22.37683 | 79.83291 | 76.68538 | chr9 | 120576533 | 120578356 | 5.20142 | 4.23175 | 3.16252 |

| Ontology | # Term Name | Binom Ran | Binom Raw P-\ | Binom FDR | Binom Fold | Binom Obs | Binom Reg | Hyper Ran | Hyper FDR Q-V | Hyper Fold | Hyper Obs | Hyper Tota | Hyper Gene Set Coverage |
|---|---|---|---|---|---|---|---|---|---|---|---|---|---|
| GO Biological Process | **mitochondrial respiratory chain complex III assembly** | 40 | 4.84716E-05 | 0.01586232 | 44.33615 | 3 | 1.32% | 152 | 0.028215581 | 20.61175 | 3 | 9 | 0.87% |
| | **mitochondrial respiratory chain complex III biogenesis** | 41 | 5.38827E-05 | 0.01720304 | 42.77334 | 3 | 1.32% | 159 | 0.038072191 | 18.55058 | 3 | 10 | 0.87% |
| | **DNA damage response, signal transduction by p53 class mediator** | 43 | 5.79165E-05 | 0.01763086 | 9.207579 | 6 | 2.63% | 140 | 0.01901402 | 9.093421 | 5 | 34 | 1.45% |
| | **cell-matrix adhesion** | 45 | 6.09594E-05 | 0.0177324 | 4.763289 | 10 | 4.39% | 81 | 0.001521453 | 5.778996 | 10 | 107 | 2.89% |
| | **lung vasculature development** | 47 | 8.3009E-05 | 0.02311888 | 18.00782 | 4 | 1.75% | 152 | 0.028215581 | 20.61175 | 3 | 9 | 0.87% |
| | **signal transduction by p53 class mediator** | 62 | 0.000220644 | 0.04658442 | 5.06532 | 8 | 3.51% | 154 | 0.028860789 | 5.343788 | 7 | 81 | 2.02% |

The test set of 228 genomic regions picked 387 genes (2%) of all 21,395 genes.
*GO Biological Process has 13,090 terms covering 17,925 (8* 13,090 ontology terms were tested (100%) using an annotation count range of [1, 1000].
GREAT version 4.0.4
Species assembly: mm10
Association rule: Basal+extension: 5000 bp upstream, 1000 bp downstream, 1000000 bp max extension, curated regulatory domains included

| Ontology | # Term Name | Binom Ran | Binom Raw P-\ | Binom FDR | Binom Fold | Binom Obs | Binom Reg | Hyper Ran | Hyper FDR Q-V | Hyper Fold | Hyper Obs | Hyper Tota | Hyper Gene Set Coverage |
|---|---|---|---|---|---|---|---|---|---|---|---|---|---|
| Mouse Phenotype | **dilated capillary** | 1 | 1.02E-07 | 0.00098112 | 19.50036 | 7 | 3.07% | 80 | 0.012555697 | 15.45882 | 4 | 16 | 0.01156069 |
| | **abnormal podocyte slit diaphragm morphology** | 2 | 1.94E-07 | 0.0009298 | 17.72073 | 7 | 3.07% | 3 | 0.000147201 | 26.50083 | 6 | 14 | 0.01734104 |
| | **abnormal mesangial matrix morphology** | 3 | 2.10E-07 | 0.00067124 | 9.160441 | 10 | 4.39% | 7 | 0.000186075 | 10.70226 | 9 | 52 | 0.02601156 |
| | **podocyte foot process effacement** | 4 | 2.79E-07 | 0.00066824 | 8.877908 | 10 | 4.39% | 4 | 0.000129494 | 11.84079 | 9 | 47 | 0.02601156 |
| | **abnormal podocyte foot process morphology** | 6 | 7.89E-07 | 0.00125858 | 6.934259 | 11 | 4.82% | 5 | 0.000167391 | 9.513117 | 10 | 65 | 0.02890173 |
| | **abnormal podocyte slit junction morphology** | 8 | 9.93E-07 | 0.00118828 | 13.85075 | 7 | 3.07% | 6 | 0.000190967 | 23.18822 | 6 | 16 | 0.01734104 |
| | **expanded mesangial matrix** | 9 | 1.53542E-06 | 0.00163351 | 8.528007 | 9 | 3.95% | 15 | 0.000818277 | 9.893642 | 8 | 50 | 0.02312139 |
| | **abnormal capillary morphology** | 10 | 1.6647E-06 | 0.00159395 | 4.85877 | 14 | 6.14% | 16 | 0.000861381 | 6.298036 | 11 | 108 | 0.03179191 |
| | **abnormal renal glomerulus basement membrane morphology** | 11 | 3.20156E-06 | 0.00278681 | 7.784058 | 9 | 3.95% | 17 | 0.001526289 | 8.99422 | 8 | 55 | 0.02312139 |
| | **abnormal glomerular endothelium fenestra morphology** | 12 | 4.00047E-06 | 0.00319204 | 14.87577 | 6 | 2.63% | 11 | 0.00039901 | 28.10694 | 5 | 11 | 0.01445087 |
| | **abnormal podocyte morphology** | 13 | 5.77868E-06 | 0.00425622 | 5.618764 | 11 | 4.82% | 13 | 0.00067617 | 7.450031 | 10 | 83 | 0.02890173 |
| | **abnormal choroid plexus morphology** | 14 | 9.20478E-06 | 0.00629541 | 5.968795 | 10 | 4.39% | 28 | 0.002634424 | 7.852097 | 8 | 63 | 0.02312139 |
| | **abnormal glomerular mesangium morphology** | 15 | 2.06166E-05 | 0.01316026 | 5.427581 | 10 | 4.39% | 32 | 0.002852215 | 6.547263 | 9 | 85 | 0.02601156 |
| | **dilated vasculature** | 16 | 2.30259E-05 | 0.01377956 | 7.042241 | 8 | 3.51% | 89 | 0.014339595 | 7.571665 | 6 | 49 | 0.01734104 |
| | **abnormal glomerular capillary morphology** | 17 | 3.34888E-05 | 0.0188621 | 5.770668 | 9 | 3.95% | 28 | 0.002634424 | 7.852097 | 8 | 63 | 0.02312139 |
| | **abnormal mesangial cell morphology** | 18 | 3.45488E-05 | 0.01837806 | 7.976363 | 7 | 3.07% | 93 | 0.01538408 | 7.420231 | 6 | 50 | 0.01734104 |
| | **abnormal fourth ventricle morphology** | 19 | 3.77671E-05 | 0.01903264 | 7.863635 | 7 | 3.07% | 10 | 0.000385396 | 14.42823 | 7 | 30 | 0.02023121 |
| | **abnormal kidney capillary morphology** | 20 | 4.12489E-05 | 0.01974793 | 5.615493 | 9 | 3.95% | 33 | 0.002832526 | 7.610494 | 8 | 65 | 0.02312139 |
| | **absent glomerular endothelium fenestra** | 21 | 4.29159E-05 | 0.01956761 | 13.37878 | 5 | 2.19% | 20 | 0.002140334 | 30.91763 | 4 | 8 | 0.01156069 |
| | **abnormal glomerular capillary endothelium morphology** | 22 | 4.81465E-05 | 0.02095465 | 9.523873 | 6 | 2.63% | 46 | 0.005922885 | 13.44245 | 5 | 23 | 0.01445087 |

The test set of 255 genomic regions picked 387 genes (2%) of all 21,395 genes.
*Mouse Phenotype has 9,575 terms covering 9,654 (45* 9,575 ontology terms were tested (100%) using an annotation count range of [1, 1000].
GREAT version 4.0.4
Species assembly: mm10
Association rule: Basal+extension: 5000 bp upstream, 1000 bp downstream, 1000000 bp max extension, curated regulatory domains included

MAFB-WT1 common peaks distribution

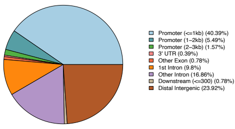

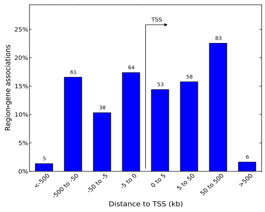
