## Supplementary Table 3 for "MAFB drives differentiation by permitting WT1 binding to podocyte specific promoters"

### RNA-Seq analysis

| | |
|---|---|
| **Worksheet 0** | **Summary** |
| **Worksheet 1** | **Differential expression analysis; p<0.05; downregulated genes** |
| **Worksheet 2** | **Differential expression analysis; p<0.05; upregulated genes** |

| gene_id | KO1 | KO2 | KO3 | KO4 | ctrl1 | ctrl2 | ctrl3 | ctrl4 | log2FoldChar | pvalue | padj | gene_name | Fold Change down | gene_chr | gene_start | gene_end | gene_strand | gene_length | gene_biotype | gene_descrip | tf_family | KO1_readcou | KO2_readcou | KO3_readcou | KO4_readcou | ctrl1_readco | ctrl2_readco | ctrl3_readco | ctrl4_readco | KO1_fpkm | KO2_fpkm | KO3_fpkm | KO4_fpkm | ctrl1_fpkm | ctrl2_fpkm | ctrl3_fpkm | ctrl4_fpkm |
|---|---|---|---|---|---|---|---|---|---|---|---|---|---|---|---|---|---|---|---|---|---|---|---|---|---|---|---|---|---|---|---|---|---|---|---|---|---|
| ENSMUSG0C | 884.716276 | 796.987798 | 626.327308 | 901.448792 | 19668.2368 | 19079.1008 | 17212.6882 | 21704.7578 | -4.5968113 | 0 | 0 | Clic3 | 24.20 | 2 | 25456838 | 25458776 | + | 1012 | protein_codi | Chloride intra | - | 934 | 774 | 666 | 974 | 20147 | 17622 | 15978 | 21854 | 25.7555682 | 23.2016457 | 18.2334338 | 26.2426796 | 572.575217 | 555.424485 | 501.090095 | 631.861743 |
| ENSMUSG0C | 1289.18507 | 877.304398 | 962.061316 | 1359.57729 | 33690.898 | 34879.8271 | 37862.9585 | 29790.1624 | -4.923364 | 6.34E-264 | 4.72E-260 | Mafb | 30.34 | 2 | 160363703 | 160367065 | - | 3363 | protein_codi | Transcription | bZIP_Maf | 1361 | 852 | 1023 | 1469 | 34511 | 32216 | 35147 | 29995 | 11.2936943 | 7.68548128 | 8.42798036 | 11.9103538 | 295.143586 | 305.559004 | 331.692237 | 260.971832 |
| ENSMUSG0C | 2164.42901 | 2211.79559 | 1915.65875 | 2044.45624 | 13281.6976 | 14307.7016 | 12140.8809 | 14252.003 | -2.6951674 | 4.44E-236 | 2.20E-232 | Paxx | 6.48 | 2 | 25455141 | 25461094 | - | 2066 | protein_codi | Protein PAXX | - | 2285 | 2148 | 2037 | 2209 | 13605 | 13215 | 11270 | 14350 | 30.8646004 | 31.5400445 | 27.3171546 | 29.1537975 | 189.396043 | 204.026785 | 173.12808 | 203.232528 |
| ENSMUSG0C | 13037.7247 | 11903.332 | 12672.313 | 12248.2272 | 30756.331 | 31118.5743 | 31230.1808 | 31949.3161 | -1.3265191 | 1.38E-138 | 5.13E-135 | Abca2 | 2.51 | 2 | 25428703 | 25448540 | + | 8346 | protein_codi | ATP-binding | ABC_tran | 13764 | 11560 | 13475 | 13234 | 31505 | 28742 | 28990 | 32169 | 46.0225899 | 42.018234 | 44.7327031 | 43.235699 | 108.568485 | 109.847187 | 110.241154 | 112.779669 |
| ENSMUSG0C | 1556.30497 | 1487.9165 | 1748.26196 | 1603.91248 | 5887.68235 | 6736.47516 | 6922.56441 | 7871.8729 | -2.09967 | 5.09E-96 | 1.51E-92 | Slc27a3 | 4.29 | 3 | 90385239 | 90389938 | - | 2569 | protein_codi | solute carrier | AMP-binding | 1643 | 1445 | 1859 | 1733 | 6031 | 6222 | 6426 | 7926 | 17.8475345 | 17.063263 | 20.0488762 | 18.3934922 | 67.5192947 | 77.2531575 | 79.3872084 | 90.273774 |
| ENSMUSG0C | 56089.4963 | 56288.5503 | 56782.3071 | 47202.043 | 146539.933 | 151161.436 | 154049.142 | 151660.183 | -1.4796965 | 1.21E-85 | 3.01E-82 | Plat | 2.79 | 8 | 22757727 | 22782844 | + | 6241 | protein_codi | Tissue-type p | Trypsin | 59214 | 54665 | 60379 | 51001 | 150107 | 139617 | 142999 | 152703 | 264.773787 | 265.713434 | 268.044242 | 222.820038 | 691.750426 | 713.566505 | 727.198094 | 715.920868 |
| ENSMUSG0C | 1184.04213 | 1107.9572 | 967.703904 | 1155.03911 | 3219.62799 | 3225.32297 | 3166.10905 | 3390.6856 | -1.5583607 | 6.00E-82 | 1.27E-78 | Prr5 | 2.95 | 15 | 84669620 | 84703673 | + | 1917 | protein_codi | Proline-rich p | - | 1250 | 1076 | 1029 | 1248 | 3298 | 2979 | 2939 | 3414 | 18.1967015 | 17.0274063 | 14.8719532 | 17.7509748 | 49.4801732 | 49.5676953 | 48.6576788 | 52.109036 |
| ENSMUSG0C | 52941.8387 | 69524.3141 | 66557.151 | 63519.7469 | 222998.776 | 197443.034 | 207305.273 | 204556.467 | -1.720587 | 5.63E-75 | 1.05E-71 | Nphs1 | 3.30 | 7 | 30458315 | 30487223 | + | 6620 | protein_codi | Nephrin [Sou | fn3 | 55891 | 67519 | 70773 | 68632 | 228427 | 182364 | 192435 | 205963 | 235.607256 | 309.404306 | 296.19953 | 282.682159 | 992.412257 | 878.681447 | 922.571403 | 910.338384 |
| ENSMUSG0C | 15320.5579 | 15837.8157 | 13391.743 | 17094.2086 | 46161.3431 | 51848.772 | 46722.462 | 53853.7012 | -1.687728 | 3.07E-73 | 5.08E-70 | Ddn | 3.22 | 15 | 98803777 | 98809490 | - | 3781 | protein_codi | Dendrin [Sou | - | 16174 | 15381 | 14240 | 18470 | 47285 | 47889 | 43371 | 54224 | 119.37559 | 123.405989 | 104.346542 | 133.195623 | 359.682567 | 403.998197 | 364.054725 | 419.620319 |
| ENSMUSG0C | 6648.63334 | 6209.09098 | 6153.24261 | 6727.54751 | 13438.8717 | 13890.8673 | 14265.2658 | 14870.7485 | -1.1333942 | 1.78E-71 | 2.65E-68 | Optn | 2.19 | 2 | 5020642 | 5064051 | - | 3753 | protein_codi | Mus musculu | - | 7019 | 6030 | 6543 | 7269 | 13766 | 12830 | 13242 | 14973 | 52.1917007 | 48.7412979 | 48.3028888 | 52.811176 | 105.494999 | 109.04316 | 111.982184 | 116.735217 |
| ENSMUSG0C | 1157.51958 | 927.759697 | 799.366685 | 966.23464 | 4268.10599 | 4129.36617 | 4503.00641 | 5509.11922 | -2.2572032 | 2.59E-67 | 3.51E-64 | Asb15 | 4.78 | 6 | 24528144 | 24573164 | + | 4497 | protein_codi | Ankyrin repe | SOCS_box | 1222 | 901 | 850 | 1044 | 4372 | 3814 | 4180 | 5547 | 7.58321011 | 6.07799369 | 5.23685786 | 6.33005299 | 27.9614661 | 27.0525456 | 29.5003595 | 36.0916647 |
| ENSMUSG0C | 1279.71273 | 857.740098 | 1185.88399 | 1239.26071 | 5497.18775 | 4785.47416 | 5117.05274 | 6038.47933 | -2.231886 | 8.93E-66 | 1.11E-62 | B3galt2 | 4.70 | 1 | 143640664 | 143654614 | + | 9288 | protein_codi | Beta-1,3-gal | Galactosyl_T | 1351 | 833 | 1261 | 1339 | 5631 | 4420 | 4750 | 6080 | 4.05917629 | 2.72070301 | 3.76155684 | 3.93086478 | 17.4367682 | 15.179253 | 16.2310015 | 19.1537144 |
| ENSMUSG0C | 14499.3063 | 14377.7011 | 13163.2182 | 13680.92 | 29509.677 | 29645.038 | 32183.5686 | 28528.8353 | -1.1051443 | 3.96E-64 | 4.54E-61 | Mpp5 | 2.15 | 12 | 78748907 | 78840714 | + | 8777 | protein_codi | membrane p | SH3_2 | 15307 | 13963 | 13997 | 14782 | 30228 | 27381 | 29875 | 28725 | 48.6685889 | 48.2604071 | 44.1838558 | 45.9215812 | 99.0526247 | 99.5069794 | 108.027849 | 95.7603168 |
| ENSMUSG0C | 9981.0015 | 8888.37038 | 8593.66208 | 8480.46744 | 23781.1212 | 30434.3164 | 29439.751 | 26959.6253 | -1.6217199 | 9.78E-64 | 1.04E-60 | Magi2 | 3.08 | 5 | 19227036 | 20704792 | + | 11348 | protein_codi | Membrane-a | PDZ | 10537 | 8632 | 9138 | 9163 | 24360 | 28110 | 27328 | 27145 | 25.9120888 | 23.0754642 | 22.3103598 | 22.0164906 | 61.7391477 | 79.0117816 | 76.4297494 | 69.9909928 |
| ENSMUSG0C | 5982.72805 | 5831.19108 | 5890.86225 | 5591.01864 | 11814.4142 | 13921.1825 | 13233.237 | 12990.6759 | -1.1572718 | 7.65E-63 | 7.59E-60 | Mtcl1 | 2.23 | 17 | 66336982 | 66449750 | - | 9268 | protein_codi | Microtubule | - | 6316 | 5663 | 6264 | 6041 | 12102 | 12858 | 12284 | 13080 | 19.0178259 | 18.536122 | 18.725804 | 17.7726646 | 37.5555214 | 44.252492 | 42.065659 | 41.2946086 |
| ENSMUSG0C | 653.591253 | 682.691098 | 705.323545 | 773.728121 | 2087.19364 | 1902.28011 | 2376.46702 | 2277.34097 | -1.6180621 | 1.22E-60 | 1.13E-57 | Bambi | 3.07 | 18 | 3507957 | 3516404 | + | 5040 | protein_codi | BMP and act | - | 690 | 663 | 750 | 836 | 2138 | 1757 | 2206 | 2293 | 3.82052746 | 3.99062881 | 4.12292539 | 4.52278013 | 12.200562 | 11.1196614 | 13.8914918 | 13.3120565 |
| ENSMUSG0C | 3190.2831 | 3254.88169 | 2843.86454 | 3021.79703 | 6834.63176 | 7342.77958 | 8159.27526 | 7783.48068 | -1.2908566 | 4.17E-57 | 3.65E-54 | Cttnbp2 | 2.45 | 6 | 18366478 | 18514843 | - | 8829 | protein_codi | Cortactin-bin | - | 3368 | 3161 | 3024 | 3265 | 7001 | 6782 | 7574 | 7837 | 10.6454822 | 10.8610377 | 9.48953689 | 10.0832702 | 22.806111 | 24.5017218 | 27.2262418 | 25.9722734 |
| ENSMUSG0C | 14854.5189 | 15998.4489 | 15082.6387 | 15686.5047 | 32733.21 | 29767.3816 | 30204.6157 | 34122.3741 | -1.0413897 | 4.77E-53 | 3.94E-50 | Calm2 | 2.06 | 17 | 87433412 | 87446935 | - | 1467 | protein_codi | Calmodulin-1 | - | 15682 | 15537 | 16038 | 16949 | 33530 | 27494 | 28038 | 34357 | 298.315697 | 321.288657 | 302.896911 | 315.024042 | 657.364296 | 597.803083 | 606.583833 | 685.262168 |
| ENSMUSG0C | 33288.6339 | 31036.1876 | 29842.7094 | 33631.2589 | 62982.8742 | 61723.9552 | 60440.4724 | 65288.0783 | -0.9705618 | 2.03E-52 | 1.59E-49 | Arhgap24 | 1.96 | 5 | 102481391 | 102897937 | + | 8792 | protein_codi | Rho GTPase | RhoGAP | 35143 | 30141 | 31733 | 36338 | 64516 | 57010 | 56105 | 65737 | 111.546494 | 103.998798 | 99.9995857 | 112.694592 | 211.048576 | 206.830079 | 202.529272 | 218.773057 |
| ENSMUSG0C | 23949.8569 | 22831.538 | 20068.8059 | 27464.5718 | 79878.5994 | 68799.3112 | 66111.2442 | 77493.156 | -1.6318064 | 1.26E-51 | 9.37E-49 | Angptl2 | 3.10 | 2 | 33216069 | 33247717 | + | 6286 | protein_codi | Angiopoietin | Fibrinogen_C | 25284 | 22173 | 21340 | 29675 | 81823 | 63545 | 61369 | 78026 | 112.24737 | 107.006072 | 94.0577936 | 128.720016 | 374.372289 | 322.44626 | 309.847919 | 363.192274 |
| ENSMUSG0C | 632.752112 | 560.156798 | 748.58339 | 358.173185 | 3386.56443 | 3564.20383 | 3298.61379 | 3672.74615 | -2.5979747 | 6.34E-50 | 4.49E-47 | Creb3l1 | 6.05 | 2 | 91982328 | 92024502 | - | 2603 | protein_codi | cAMP respon | bZIP_1 | 668 | 544 | 796 | 387 | 3469 | 3292 | 3062 | 3698 | 7.16155061 | 6.33990971 | 8.4725404 | 4.05383932 | 38.3294692 | 40.3400094 | 37.3340352 | 41.568502 |
| [illegible] | | | | | | | | | | | | | | | | | | | | | | | | | | | | | | | | | | | | | |

| | | | | | | | | | | | | | | | |
|---|---|---|---|---|---|---|---|---|---|---|---|---|---|---|---|
| ENSMUSG0( | 39905.0613 | 39382.9358 | 53354.4347 | 44518.9834 | 68250.6464 | 70767.6352 | 69629.6219 | 60381.8138 | -0.60269 | 2.58E-08 | 1.09E-06 | Mgat5 | 1.52 | | 1 |
| ENSMUSG0( | 107.984642 | 70.0195998 | 91.2218452 | 62.9348233 | 186.461172 | 192.71819 | 198.218464 | 149.968812 | -1.1293023 | 2.68E-08 | 1.13E-06 | Gm26815 | 2.19 | | 8 |
| ENSMUSG0( | 2956.31638 | 3083.95149 | 3077.09152 | 3221.70764 | 4656.64812 | 4314.50555 | 3746.75988 | 4882.42836 | -0.5125094 | 2.82E-08 | 1.18E-06 | Mtfr1l | 1.43 | | 4 |
| ENSMUSG0( | 15.1557392 | 9.26729997 | 1.88086279 | 17.58473 | 82.9801028 | 51.9689502 | 64.6364557 | 47.6722053 | -2.4918184 | 3.11E-08 | 1.29E-06 | Ces2d-ps | 5.62 | | 8 |
| ENSMUSG0( | 4386.63927 | 4375.19529 | 4212.19221 | 3789.97208 | 5417.13636 | 5805.36481 | 5467.16688 | 5347.23236 | -0.3945944 | 3.26E-08 | 1.34E-06 | Lyrm9 | 1.31 | | 11 |
| ENSMUSG0( | 6055.66505 | 6153.48718 | 5747.91668 | 6478.58475 | 7322.75001 | 8716.7087 | 8553.55764 | 8333.69738 | -0.430189 | 3.31E-08 | 1.36E-06 | Flrt2 | 1.35 | | 12 |
| ENSMUSG0( | 7962.44649 | 7680.53228 | 8500.55937 | 7131.9963 | 10105.024 | 10394.8727 | 10252.4191 | 11538.66 | -0.4353331 | 3.50E-08 | 1.43E-06 | Inppl1 | 1.35 | | 7 |
| ENSMUSG0( | 2344.40341 | 2229.30049 | 2507.1901 | 2284.16388 | 2939.44811 | 3009.86836 | 3048.68616 | 2938.79282 | -0.3498144 | 3.76E-08 | 1.53E-06 | Derl2 | 1.27 | | 11 |
| ENSMUSG0( | 1194.4617 | 983.363497 | 2009.70189 | 1267.95159 | 2767.63049 | 2358.09111 | 2868.78136 | 3216.88068 | -1.0390433 | 4.00E-08 | 1.62E-06 | Ube2ql1 | 2.05 | | 13 |
| ENSMUSG0( | 2674.04074 | 2784.30879 | 2786.49822 | 2395.22533 | 3492.97421 | 3837.04082 | 3979.45112 | 3389.69243 | -0.4661689 | 4.34E-08 | 1.74E-06 | Atg13 | 1.38 | | 2 | 91674618 |
| ENSMUSG0( | 4829.94464 | 5244.26209 | 4742.59552 | 4522.05215 | 6779.96252 | 6212.45492 | 6548.75024 | 5992.79347 | -0.4010498 | 5.49E-08 | 2.19E-06 | Il13ra1 | 1.32 | X | | 36112110 |
| ENSMUSG0( | 4642.39237 | 4962.12429 | 4881.77937 | 4288.8231 | 6435.35103 | 6660.68711 | 6318.21354 | 5732.58268 | -0.4215849 | 6.15E-08 | 2.43E-06 | Usp12 | 1.34 | | 5 | 146734809 |
| ENSMUSG0( | 107.037408 | 83.4056998 | 172.098945 | 75.8919928 | 309.466972 | 435.239958 | 184.213899 | 402.234232 | -1.6017323 | 6.65E-08 | 2.61E-06 | Pcp4l1 | 3.04 | | 1 | 171173262 |
| ENSMUSG0( | 8114.00388 | 7943.10578 | 11769.4989 | 8449.92554 | 13607.7606 | 15393.6361 | 16194.664 | 13909.3591 | -0.7042081 | 6.70E-08 | 2.62E-06 | Mlec | 1.63 | | 5 | 115142981 |
| ENSMUSG0( | 3059.56485 | 2467.16119 | 2541.98606 | 2483.14898 | 3574.97807 | 3768.83157 | 3553.92779 | 3703.53445 | -0.4685111 | 6.84E-08 | 2.67E-06 | Btrc | 1.38 | | 19 | 45363734 |
| ENSMUSG0( | 7862.03971 | 7880.29408 | 9925.31293 | 7570.68903 | 13093.284 | 11864.0782 | 12305.7039 | 11178.139 | -0.5433762 | 7.79E-08 | 3.02E-06 | Gpc1 | 1.46 | | 1 | 92831645 |
| ENSMUSG0( | 8337.55103 | 8781.28158 | 8944.44299 | 8624.84732 | 10255.3645 | 10942.7121 | 10952.6474 | 10424.3222 | -0.2955148 | 8.21E-08 | 3.17E-06 | Nbr1 | 1.23 | | 11 | 101552149 |
| ENSMUSG0( | 883.769042 | 867.007398 | 1040.11712 | 796.865924 | 1357.94498 | 1421.56732 | 1484.48393 | 1172.93488 | -0.5993982 | 8.43E-08 | 3.23E-06 | Rxra | 1.52 | | 2 | 27676440 |
| ENSMUSG0( | 59017.3957 | 54954.0591 | 61504.2132 | 58353.5383 | 76405.1499 | 71958.5903 | 70995.6057 | 71966.1597 | -0.3171732 | 8.57E-08 | 3.28E-06 | Golim4 | 1.25 | | 3 | 75875084 |
| ENSMUSG0( | 12568.844 | 11916.7181 | 12481.4055 | 13344.9591 | 15001.8263 | 16023.7596 | 16050.3092 | 15709.978 | -0.3194624 | 8.71E-08 | 3.32E-06 | Dynll2 | 1.25 | | 11 | 87979525 |
| ENSMUSG0( | 3872.29137 | 4114.68119 | 4228.17955 | 4049.11547 | 5205.29304 | 5429.67261 | 4807.87503 | 5420.72701 | -0.3593425 | 8.78E-08 | 3.34E-06 | Trim35 | 1.28 | | 14 | 66297031 |
| ENSMUSG0( | 359.948806 | 291.405099 | 401.564205 | 383.162012 | 644.316092 | 535.929799 | 533.25076 | 709.124053 | -0.7542117 | 9.26E-08 | 3.49E-06 | B3gnt9 | 1.69 | | 8 | 105252638 |
| ENSMUSG0( | 4551.45793 | 4867.39189 | 5023.78451 | 4097.2421 | 6615.95478 | 6251.43163 | 5914.2357 | 6363.24623 | -0.439761 | 9.27E-08 | 3.49E-06 | Fuca2 | 1.36 | | 10 | 13499540 |
| ENSMUSG0( | 11307.1287 | 9659.61567 | 9864.18489 | 9075.57172 | 13100.1176 | 14987.6287 | 14715.5664 | 12978.7579 | -0.4831345 | 9.67E-08 | 3.63E-06 | Chd3 | 1.40 | | 11 | 69343273 |
| ENSMUSG0( | 216.916517 | 234.771599 | 153.290317 | 167.517691 | 666.769532 | 358.369219 | 313.48681 | 499.564984 | -1.2522562 | 1.05E-07 | 3.91E-06 | Wnt4 | 2.38 | | 4 | 137277489 |
| ENSMUSG0( | 132.612718 | 74.1383998 | 169.277651 | 99.9553076 | 262.607619 | 265.258183 | 248.850354 | 353.568856 | -1.2464037 | 1.10E-07 | 4.06E-06 | Lrrc4 | 2.37 | | 6 | 28661831 |
| ENSMUSG0( | 232.072257 | 355.246499 | 110.970904 | 282.281193 | 692.151681 | 1476.78433 | 1464.01572 | 943.512396 | -2.2227576 | 1.12E-07 | 4.11E-06 | Nectin4 | 4.67 | | 1 | 171370099 |
| ENSMUSG0( | 11968.2978 | 13595.1291 | 11519.3441 | 10131.581 | 15736.9324 | 17127.0171 | 17047.8652 | 16649.5177 | -0.4955015 | 1.21E-07 | 4.46E-06 | Hipk3 | 1.41 | | 2 | 104426481 |
| ENSMUSG0( | 3626.95784 | 3537.01949 | 3803.10456 | 3634.48604 | 4555.11952 | 4393.54166 | 4388.81534 | 4735.43906 | -0.3077463 | 1.55E-07 | 5.64E-06 | Snx6 | 1.24 | | 12 | 54746349 |
| ENSMUSG0( | 31163.9887 | 28969.5797 | 33366.5059 | 33236.0653 | 42446.7632 | 39376.2239 | 39353.9061 | 41995.2402 | -0.3645588 | 1.58E-07 | 5.72E-06 | Htra1 | 1.29 | | 7 | 130936111 |
| ENSMUSG0( | 1755.22405 | 1574.4113 | 1718.16816 | 1877.86407 | 2384.94578 | 2298.54336 | 2310.75329 | 2172.06485 | -0.4040466 | 1.62E-07 | 5.84E-06 | Ube2e3 | 1.32 | | 2 | 78868124 |
| ENSMUSG0( | 1192.56723 | 1269.6201 | 1190.58614 | 1031.946 | 1520.00024 | 1720.38879 | 1639.61143 | 1571.19643 | -0.4617329 | 1.68E-07 | 6.03E-06 | Vps37c | 1.38 | | 19 | 10688815 |
| ENSMUSG0( | 9227.00348 | 8414.70838 | 9277.3557 | 7852.97023 | 12684.2409 | 11112.6938 | 11155.175 | 13145.6106 | -0.4680893 | 1.75E-07 | 6.25E-06 | Peg13 | 1.38 | | 15 | 72805600 |
| ENSMUSG0( | 8340.39273 | 6901.04938 | 8734.72679 | 7895.54378 | 10809.8668 | 11079.1306 | 11743.3667 | 10270.3807 | -0.461935 | 1.76E-07 | 6.25E-06 | St3gal3 | 1.38 | | 4 | 117932154 |
| ENSMUSG0( | 2100.96435 | 2163.39969 | 2049.20001 | 2090.73185 | 2533.33373 | 2541.06513 | 2592.99915 | 2594.1625 | -0.2881951 | 1.77E-07 | 6.27E-06 | Tbc1d20 | 1.22 | | 2 | 152293828 |
| ENSMUSG0( | 1190.67276 | 1315.9566 | 1247.01203 | 1217.04842 | 1623.4813 | 1593.71447 | 1569.5886 | 1539.41496 | -0.3482613 | 2.12E-07 | 7.39E-06 | Zfp180 | 1.27 | | 7 | 24081924 |
| ENSMUSG0( | 4679.33448 | 4638.79849 | 5000.27372 | 4439.68158 | 6105.38309 | 5575.83528 | 6038.12224 | 5938.16907 | -0.334841 | 2.58E-07 | 8.93E-06 | Rnf11 | 1.26 | | 4 | 109451098 |
| ENSMUSG0( | 1852.78912 | 1794.7671 | 1792.46224 | 1904.70392 | 2189.69848 | 2620.10124 | 2370.00338 | 2490.87273 | -0.3966267 | 2.59E-07 | 8.94E-06 | Cdk5rap2 | 1.32 | | 4 | 70216856 |
| ENSMUSG0( | 2.8417011 | 1.0297 | 5.64258836 | 7.40409686 | 16.5960206 | 38.9767126 | 58.1728101 | 28.8019574 | -3.0565679 | 2.73E-07 | 9.38E-06 | 0610012D04I | 8.32 | | 17 | 87362239 |
| ENSMUSG0( | 315.428822 | 374.810799 | 213.477926 | 245.260708 | 511.547928 | 567.327706 | 504.164354 | 497.578643 | -0.8578665 | 3.10E-07 | 1.05E-05 | Gm18588 | 1.81 | | 3 | 68902267 |
| ENSMUSG0( | 8822.53469 | 8792.60828 | 8139.43371 | 8168.56986 | 11247.2208 | 12683.6719 | 13685.6922 | 10282.2988 | -0.4976805 | 3.17E-07 | 1.07E-05 | Cflar | 1.41 | | 1 | 58711508 |
| ENSMUSG0( | 8009.80817 | 6834.11888 | 6367.66097 | 6743.28121 | 8173.05201 | 13248.8342 | 12656.8953 | 12887.3862 | -0.7484866 | 3.19E-07 | 1.08E-05 | Mcc | 1.68 | | 18 | 44425060 |
| ENSMUSG0( | 7132.66976 | 7442.67158 | 6484.27446 | 6760.86594 | 8417.11113 | 8893.1866 | 9023.24922 | 8786.58333 | -0.3361944 | 3.24E-07 | 1.09E-05 | Eif4b | 1.26 | | 15 | 102073773 |
| ENSMUSG0( | 675.377628 | 558.097398 | 525.701149 | 609.912478 | 974.28403 | 978.748561 | 905.987654 | 755.803088 | -0.6092103 | 4.09E-07 | 1.35E-05 | Aff3 | 1.53 | | 1 | 38177326 |
| ENSMUSG0( | 67167.3945 | 70339.8365 | 92561.0195 | 63273.5607 | 110173.171 | 143599.953 | 139403.599 | 99307.1626 | -0.7474896 | 4.44E-07 | 1.46E-05 | Rhpn1 | 1.68 | | 15 | 75704280 |
| ENSMUSG0( | 1136.68044 | 910.254797 | 1123.81552 | 1113.39106 | 1445.80626 | 1508.18224 | 1745.1843 | 1463.93397 | -0.5239809 | 4.45E-07 | 1.46E-05 | C430049B03I | 1.44 | X | | 53055046 |
| ENSMUSG0( | 2696.77434 | 2749.29899 | 2430.07472 | 2620.12477 | 3193.2696 | 3451.60444 | 3507.605 | 3251.64167 | -0.3527583 | 4.80E-07 | 1.57E-05 | Map2k7 | 1.28 | | 8 | 4238740 |
| ENSMUSG0( | 3271.7452 | 2887.27879 | 3182.41984 | 3357.75792 | 4678.12532 | 4537.53896 | 4763.70678 | 3741.27494 | -0.4805115 | 5.28E-07 | 1.71E-05 | Mocs2 | 1.40 | | 13 | 114818236 |
| ENSMUSG0( | 2979.04999 | 2472.30969 | 2886.18395 | 3539.1583 | 3904.94601 | 4644.72492 | 4679.67939 | 4456.35802 | -0.5742059 | 5.57E-07 | 1.79E-05 | Spats2l | 1.49 | | 1 | 57774162 |
| ENSMUSG0( | 608.124036 | 481.899599 | 601.876092 | 548.828679 | 773.179311 | 847.7435 | 980.319578 | 772.686994 | -0.5894158 | 6.06E-07 | 1.94E-05 | Ctnnbip1 | 1.50 | | 4 | 149518236 |
| ENSMUSG0( | 1875.52273 | 2020.27139 | 1743.5598 | 1840.84358 | 2361.5161 | 2314.78366 | 2396.93523 | 2532.58591 | -0.3612051 | 6.37E-07 | 2.02E-05 | Snta1 | 1.28 | | 2 | 154376313 |
| ENSMUSG0( | 4403.68947 | 4454.48219 | 4583.66261 | 4521.12664 | 5242.39002 | 5502.2126 | 5279.72116 | 5626.31339 | -0.2693585 | 6.54E-07 | 2.07E-05 | Ube2z | 1.21 | | 11 | 96047432 |
| ENSMUSG0( | 7682.06531 | 7991.50168 | 8093.35258 | 7846.49164 | 9383.58527 | 9782.07218 | 10044.5052 | 9161.00878 | -0.2794242 | 6.82E-07 | 2.14E-05 | Bcl2l2 | 1.21 | | 14 | 54883377 |
| ENSMUSG0( | 60.6229568 | 33.9800999 | 58.3067464 | 114.763501 | 203.057193 | 295.573404 | 296.250422 | 116.201 | -1.7640515 | 7.53E-07 | 2.35E-05 | 6030419C18I | 3.40 | | 9 | 58488603 |
| ENSMUSG0( | 3453.61407 | 3356.82199 | 2893.7074 | 3533.60522 | 4373.53953 | 4381.63211 | 4720.61581 | 4172.31113 | -0.4147983 | 7.61E-07 | 2.38E-05 | Col4a3bp | 1.33 | | 13 | 96542618 |
| ENSMUSG0( | 1646.29217 | 1720.6287 | 2009.70189 | 1655.74116 | 2411.30416 | 2315.86634 | 2324.75786 | 2294.22488 | -0.4103597 | 9.74E-07 | 2.98E-05 | Desi1 | 1.33 | | 15 | 81992523 |
| ENSMUSG0( | 4691.64852 | 4925.05509 | 5248.54761 | 4759.90877 | 6131.74148 | 6227.61253 | 6446.40918 | 5787.20709 | -0.3254771 | 1.02E-06 | 3.10E-05 | Tbc1d9b | 1.25 | | 11 | 50131396 |
| ENSMUSG0( | 29.3642447 | 27.8018999 | 57.366315 | 20.3612664 | 98.5998868 | 90.9456628 | 90.491038 | 91.3717268 | -1.4602795 | 1.19E-06 | 3.58E-05 | Mmp15 | 2.75 | | 8 | 95352268 |
| ENSMUSG0( | 12.3140381 | 4.11879999 | 9.40431394 | 4.62756053 | 26.3583856 | 44.3901449 | 56.0182616 | 26.8156155 | -2.3254272 | 1.22E-06 | 3.65E-05 | Igkv8-31 | 5.01 | | 6 | 70105860 |
| ENSMUSG0( | 6052.82335 | 5631.42928 | 5927.53908 | 5801.10989 | 6787.77241 | 7192.28617 | 6999.05088 | 7048.53418 | -0.259427 | 1.25E-06 | 3.72E-05 | Golga2 | 1.20 | | 2 | 32287384 |
| ENSMUSG0( | 480.247486 | 457.186799 | 398.742911 | 394.268158 | 588.670611 | 585.733376 | 672.219139 | 625.697694 | -0.5150618 | 1.29E-06 | 3.83E-05 | Slc39a7 | 1.43 | | 17 | 34028267 |
| ENSMUSG0( | 1546.83263 | 1785.4998 | 1873.33934 | 1729.78213 | 2224.84299 | 2302.8741 | 2438.94893 | 2900.05915 | -0.5086908 | 1.40E-06 | 4.12E-05 | Epha2 | 1.42 | | 4 | 141301240 |
| ENSMUSG0( | 776.731634 | 839.205498 | 730.715193 | 765.398512 | 941.091989 | 1317.62942 | 1139.75617 | 1135.19439 | -0.5426117 | 1.46E-06 | 4.25E-05 | Zfp641 | 1.46 | | 15 | 98285585 |
| ENSMUSG0( | 3039.67294 | 2914.05099 | 3055.4616 | 3078.25327 | 3525.19001 | 3693.04352 | 3636.87791 | 3868.40082 | -0.2844926 | 1.52E-06 | 4.40E-05 | Szrd1 | 1.22 | | 4 | 141113001 |
| ENSMUSG0( | 10894.1348 | 10525.5934 | 14428.0984 | 14112.2086 | 18048.6605 | 20836.301 | 22682.0096 | 16654.4835 | -0.6467402 | 1.56E-06 | 4.49E-05 | Pdia6 | 1.57 | | 12 | 17266545 |
| ENSMUSG0( | 352.370937 | 336.711899 | 269.90381 | 298.940411 | 556.454807 | 559.748901 | 580.650827 | 389.32301 | -0.7297329 | 1.74E-06 | 5.00E-05 | Mycbpap | 1.66 | | 11 | 94501347 |
| ENSMUSG0( | 569.287454 | 552.948898 | 615.042132 | 679.325887 | 880.565326 | 959.260205 | 1102.05157 | 760.768942 | -0.6147144 | 1.79E-06 | 5.14E-05 | Csrp2 | 1.53 | | 10 | 110919789 |
| ENSMUSG0( | 5675.82433 | 5163.94549 | 5304.97349 | 5466.0745 | 6313.32147 | 6946.51634 | 7005.51452 | 6531.09212 | -0.310118 | 1.83E-06 | 5.23E-05 | Myo10 | 1.24 | | 15 | 25622525 |
| ENSMUSG0( | 2404.07913 | 1972.90519 | 3176.77725 | 2693.24023 | 3656.0057 | 4195.41004 | 3842.63729 | 4243.81944 | -0.6370644 | 1.98E-06 | 5.64E-05 | Tubb2b | 1.56 | | 13 | 34126748 |
| ENSMUSG0( | 924.500092 | 827.878798 | 963.001747 | 851.471138 | 1090.45617 | 1216.93958 | 1228.09266 | 1192.7983 | -0.4060391 | 2.00E-06 | 5.70E-05 | Micall1 | 1.33 | | 15 | 79108898 |
| ENSMUSG0( | 26510.2296 | 22600.8852 | 24621.4343 | 22073.4638 | 30491.7709 | 30558.8254 | 30221.8521 | 30464.5255 | -0.3455707 | 2.12E-06 | 6.00E-05 | Lmx1b | 1.27 | | 2 | 33560965 |
| ENSMUSG0( | 311.639887 | 261.543799 | 391.21946 | 273.951584 | 474.450941 | 559.748901 | 453.532464 | 514.462549 | -0.6923674 | 2.18E-06 | 6.18E-05 | Ehbp1l1 | 1.62 | | 19 | 5707376 |
| ENSMUSG0( | 44.5199839 | 55.6037998 | 61.1280406 | 26.8398511 | 97.6236503 | 188.387444 | 126.041089 | 97.3307524 | -1.4375348 | 2.31E-06 | 6.47E-05 | Iigp1 | 2.71 | | 18 | 60376029 |
| ENSMUSG0( | 66.306359 | 93.7026997 | 94.0431394 | 58.3072627 | 163.031496 | 198.131622 | 186.368447 | 117.194171 | -1.0900417 | 2.57E-06 | 7.14E-05 | Syna | 2.13 | | 5 | 134558146 |
| ENSMUSG0( | 3988.80111 | 3735.75159 | 4000.59515 | 3447.5326 | 4561.95318 | 4892.66012 | 4801.41138 | 5082.05572 | -0.3499547 | 2.59E-06 | 7.17E-05 | Rab22a | 1.27 | | 2 | 173659760 |
| ENSMUSG0( | 5044.96669 | 4334.00729 | 3678.96761 | 4435.97953 | 5474.73431 | 6522.10325 | 6228.79978 | 6575.78481 | -0.5035324 | 2.63E-06 | 7.25E-05 | Crtc1 | 1.42 | | 8 | 70382355 |
| ENSMUSG0( | 6318.04878 | 5533.60778 | 7419.06327 | 6411.02236 | 9679.38493 | 9156.27941 | 10270.7328 | 7856.97533 | -0.5252705 | 2.73E-06 | 7.53E-05 | Abca9 | 1.44 | | 11 | 110100749 |
| ENSMUSG0( | 57.7812557 | 32.9503999 | 38.5576872 | 39.7970206 | 96.6474138 | 220.868038 | 198.218464 | 61.5765985 | -1.7689965 | 2.74E-06 | 7.54E-05 | Hoxb9 | 3.41 | | 11 | 96271457 |
| ENSMUSG0( | 3123.02951 | 3174.56509 | 2904.05214 | 3051.41342 | 3820.01344 | 4004.85722 | 4018.233 | 3531.71587 | -0.3273587 | 2.81E-06 | 7.70E-05 | Arhgap12 | 1.25 | | 18 | 6024427 |
| ENSMUSG0( | 1166.04469 | 1177.9768 | 1198.1096 | 1095.80633 | 1405.78056 | 1410.74046 | 1500.64305 | 1479.82471 | -0.3219117 | 3.03E-06 | 8.27E-05 | Dym | 1.25 | | 18 | 75018772 |
| ENSMUSG0( | 9360.56343 | 9288.92367 | 9624.37489 | 8965.43578 | 10626.3343 | 11505.709 | 11101.3113 | 11178.139 | -0.2540673 | 3.14E-06 | 8.55E-05 | Jak1 | 1.19 | | 4 | 101152367 |
| ENSMUSG0( | 69.1480601 | 50.4552999 | 48.9024325 | 37.0204843 | 117.14838 | 140.74924 | 136.813831 | 80.4468464 | -1.2080868 | 3.18E-06 | 8.64E-05 | Rsad2 | 2.31 | | 12 | 26442746 |
| ENSMUSG0( | 5591.52053 | 5776.61698 | 5774.24876 | 5357.78959 | 6495.87769 | 6719.15218 | 7145.56018 | 7077.33614 | -0.2862517 | 3.39E-06 | 9.12E-05 | Fxr2 | 1.22 | | 11 | 69632990 |
| ENSMUSG0( | 5764.8643 | 5575.82548 | 5969.85849 | 5841.83242 | 7132.38389 | 6634.70264 | 7214.50573 | 6852.87951 | -0.2656693 | 3.56E-06 | 9.48E-05 | Bzw1 | 1.20 | | 1 | 58392898 |
| ENSMUSG0( | 2046.02479 | 2084.11279 | 2099.04287 | 2007.43576 | 2455.23481 | 2600.61288 | 2449.72167 | 2435.25515 | -0.2712479 | 3.65E-06 | 9.71E-05 | Eapp | 1.21 | | 12 | 54670340 |
| ENSMUSG0( | 3423.30259 | 3554.52439 | 3367.68482 | 3320.73744 | 4262.24857 | 3997.27842 | 4053.78305 | 4064.0555 | -0.2613296 | 3.66E-06 | 9.71E-05 | Shc1 | 1.20 | | 3 | 89418443 |
| ENSMUSG0( | 995.542619 | 1019.403 | 932.907943 | 971.787712 | 1186.12735 | 1267.82585 | 1208.70172 | 1245.43636 | -0.3245867 | 3.84E-06 | 0.00010116 | Cyb561d1 | 1.25 | | 3 | 108195687 |
| ENSMUSG0( | 753.998025 | 588.988398 | 639.493348 | 724.67598 | 934.258334 | 1173.63212 | 990.015046 | 890.874336 | -0.5583037 | 3.95E-06 | 0.00010353 | Tceanc2 | 1.47 | | 4 | 107134155 |
| ENSMUSG0( | 11466.2639 | 13859.762 | 9785.18865 | 10408.3092 | 15797.4591 | 16259.7853 | 15458.8857 | 16811.4046 | -0.4990134 | 3.96E-06 | 0.00010383 | Epn3 | 1.41 | | 11 | 94489599 |
| ENSMUSG0( | 1447.37309 | 1208.8678 | 1152.02846 | 1556.71136 | 2012.02343 | 1889.28788 | 1896.0027 | 1800.61892 | -0.5018123 | 4.06E-06 | 0.00010572 | Marveld2 | 1.42 | | 13 | 100595957 |
| ENSMUSG0( | 9908.06451 | 10365.9899 | 9345.06676 | 9481.87154 | 11811.4855 | 13560.6479 | 13218.1552 | 11525.7488 | -0.3580405 | 4.31E-06 | 0.00011172 | Hivep1 | 1.28 | | 13 | 42052021 |
| ENSMUSG0( | 705.689107 | 686.809898 | 703.442683 | 777.43017 | 908.876185 | 1016.64259 | 893.060363 | 949.471422 | -0.3905165 | 4.48E-06 | 0.00011568 | Taf5l | 1.31 | | 8 | 123996318 |
| ENSMUSG0( | 1759.01298 | 1851.40059 | 1888.38624 | 1571.51956 | 2296.10826 | 2297.46067 | 2296.74873 | 2182.98973 | -0.3599991 | 4.55E-06 | 0.0001171 | Zbtb7b | 1.28 | | 3 | 89377644 |
| ENSMUSG0( | 10444.1988 | 9533.99227 | 10089.8884 | 9778.03541 | 13107.9275 | 11289.1717 | 12697.8317 | 13595.517 | -0.3473064 | 4.59E-06 | 0.00011785 | Micu1 | 1.27 | | 10 | 59702477 |
| ENSMUSG0( | 2514.90547 | 2501.14129 | 2674.58688 | 2559.96649 | 2998.99854 | 4091.47214 | 3562.54598 | 3378.76755 | -0.4527513 | 4.63E-06 | 0.00011881 | Ets1 | 1.37 | | 9 | 32636221 |
| ENSMUSG0( | 11614.9796 | 11372.0068 | 11876.7081 | 11361.5866 | 13029.8286 | 13616.9476 | 14213.5566 | 14118.9181 | -0.2501661 | 4.73E-06 | 0.0001205 | Spop | 1.19 | | 11 | 95414080 |
| ENSMUSG0( | 644.118916 | 729.027598 | 748.58339 | 678.400374 | 935.23457 | 902.960509 | 931.842236 | 875.976772 | -0.3810619 | 4.88E-06 | 0.00012386 | Bet1l | 1.30 | | 7 | 140853394 |
| ENSMUSG0( | 5238.20236 | 5822.95348 | 7679.56276 | 6026.00933 | 7739.603 | 10925.3891 | 10954.802 | 9177.89269 | -0.6475002 | 5.03E-06 | 0.00012722 | Mfge8 | 1.57 | | 7 | 79133768 |
| ENSMUSG0( | 99.4595385 | 90.6135997 | 106.268748 | 98.1042833 | 164.983969 | 147.245359 | 157.282042 | 165.859548 | -0.6875868 | 5.13E-06 | 0.00012914 | Rbm38 | 1.61 | | 2 | 173020498 |
| ENSMUSG0( | 1936.14568 | 1980.11309 | 1907.19487 | 1784.38734 | 2423.99524 | 2612.52243 | 2622.08555 | 2196.89413 | -0.3733266 | 5.51E-06 | 0.00013817 | Stk24 | 1.30 | | 14 | 121286343 |
| ENSMUSG0( | 29339.6166 | 32815.5092 | 25014.5346 | 21666.2384 | 40934.5728 | 43545.6495 | 43462.6301 | 35404.5578 | -0.5858011 | 5.67E-06 | 0.00014204 | Npr1 | 1.50 | | 3 | 90450591 |
| ENSMUSG0( | 309.74542 | 276.989299 | 293.414595 | 278.579144 | 472.498468 | 358.369219 | 411.518768 | 449.906437 | -0.5473234 | 5.73E-06 | 0.00014298 | Gm14494 | 1.46 | | 2 | 153826171 |
| ENSMUSG0( | 3659.16378 | 3225.02039 | 3536.02204 | 3404.03353 | 4113.86062 | 5168.74517 | 4782.02045 | 4307.38238 | -0.4100739 | 5.82E-06 | 0.000145 | Map3k20 | 1.33 | | 2 | 72285637 |
| ENSMUSG0( | 2877.69598 | 2586.60639 | 2739.47665 | 3179.13409 | 3718.48484 | 3676.80322 | 3772.61446 | 3486.03001 | -0.364179 | 6.08E-06 | 0.00015084 | Tle4 | 1.29 | | 19 | 14448072 |
| ENSMUSG0( | 985.123048 | 985.422897 | 941.371825 | 944.022349 | 1188.07982 | 1289.47958 | 1514.64761 | 1214.64806 | -0.4329745 | 6.32E-06 | 0.00015579 | Prkaa2 | 1.35 | | 4 | 105029874 |
| ENSMUSG0( | 15662.5092 | 17573.8899 | 17812.711 | 13917.8511 | 21881.365 | 22340.1525 | 23062.2874 | 20027.2921 | -0.4264696 | 6.37E-06 | 0.00015656 | Kremen1 | 1.34 | | 11 | 5191552 |
| ENSMUSG0( | 6021.56463 | 6039.19048 | 6521.89172 | 6350.86408 | 8456.16059 | 10801.9628 | 10670.4016 | 7178.63958 | -0.5735273 | 6.52E-06 | 0.00016008 | Smarca2 | 1.49 | | 19 | 26605050 |
| ENSMUSG0( | 2390.81786 | 2333.30019 | 2383.05315 | 2334.14153 | 2778.36909 | 2748.94093 | 2770.7494 | 2888.1411 | -0.2447202 | 6.59E-06 | 0.00016148 | Stxbp2 | 1.18 | | 8 | 3630955 |
| ENSMUSG0( | 3209.22778 | 3716.18729 | 2644.49308 | 2723.78213 | 5090.09713 | 4195.41004 | 4046.24213 | 4946.98447 | -0.5725003 | 6.64E-06 | 0.00016256 | Astn2 | 1.49 | | 4 | 65380803 |
| ENSMUSG0( | 15706.082 | 16021.1023 | 15574.4843 | 15768.8753 | 17595.6867 | 20088.1646 | 19132.3909 | 19119.5338 | -0.2677831 | 6.73E-06 | 0.00016435 | Hnrnpul2 | 1.20 | | 19 | 8819401 |
| ENSMUSG0( | 13716.8912 | 20527.0694 | 10163.2421 | 12622.1341 | 21666.593 | 27809.8845 | 25415.0544 | 24400.2237 | -0.7999966 | 6.86E-06 | 0.00016706 | Fam129b | 1.74 | | 2 | 32876114 |
| ENSMUSG0( | 2170.11241 | 2206.64709 | 2214.71593 | 2122.19926 | 2576.28813 | 2547.56124 | 2619.931 | 2573.30591 | -0.2437579 | 7.06E-06 | 0.00017133 | Thap12 | 1.18 | | 7 | 98703103 |
| ENSMUSG0( | 8111.16218 | 8427.06478 | 9690.20508 | 8371.25701 | 10862.5836 | 10921.0583 | 11090.5385 | 10541.5164 | -0.327424 | 7.08E-06 | 0.00017162 | Srpr | 1.25 | | 9 | 35200175 |
| ENSMUSG0( | 679.166563 | 633.265498 | 820.056175 | 560.860337 | 821.014899 | 1106.50556 | 1179.61532 | 1225.57294 | -0.6856004 | 7.11E-06 | 0.00017169 | Mafa | 1.61 | | 15 | 75745490 |
| ENSMUSG0( | 5020.33861 | 4787.07529 | 4797.14054 | 3852.9069 | 7828.44052 | 6675.84472 | 7213.42846 | 5462.44019 | -0.5583795 | 7.17E-06 | 0.00017281 | Ablim3 | 1.47 | | 18 | 61799395 |
| ENSMUSG0( | 2131.27583 | 1885.38069 | 2360.4828 | 2613.64619 | 3182.531 | 3136.54268 | 3311.54108 | 2899.06598 | -0.4785314 | 7.37E-06 | 0.00017711 | Rnf145 | 1.39 | | 11 | 44518964 |
| ENSMUSG0( | 0 | 7.20789998 | 1.88086279 | 1.85102421 | 16.5960206 | 27.0671615 | 18.3136624 | 26.8156155 | -3.0519316 | 7.52E-06 | 0.00018028 | Slc7a9 | 8.29 | | 7 | 35448796 |
| ENSMUSG0( | 7947.29075 | 10355.6929 | 10805.5567 | 7076.46557 | 15305.4359 | 13688.4049 | 14622.9208 | 12135.5558 | -0.6236715 | 7.56E-06 | 0.00018095 | Rasl11a | 1.54 | | 5 | 146845071 |
| ENSMUSG0( | 1064.69068 | 1072.9474 | 1234.78642 | 1424.36313 | 1740.62969 | 1634.85656 | 1607.2932 | 1689.38377 | -0.4758428 | 7.58E-06 | 0.00018108 | H2-DMb1 | 1.39 | | 17 | 34153072 |
| ENSMUSG0( | 1076.05748 | 1151.2046 | 921.622766 | 1000.47859 | 1405.78056 | 1276.48734 | 1509.26124 | 1392.42566 | -0.4288353 | 7.86E-06 | 0.00018601 | Acads | 1.35 | | 5 | 115110299 |
| ENSMUSG0( | 870.507771 | 911.284497 | 883.065079 | 944.947861 | 1081.67005 | 1168.21869 | 1202.23808 | 1108.37877 | -0.3368162 | 7.94E-06 | 0.00018753 | Nfu1 | 1.26 | | 6 | 87009236 |
| ENSMUSG0( | 89.9872015 | 112.2373 | 250.154751 | 87.9236502 | 305.562026 | 302.069523 | 392.127831 | 339.664463 | -1.3092984 | 8.32E-06 | 0.0001952 | Hist1h4i | 2.48 | | 13 | 22040636 |
| ENSMUSG0( | 1341.28292 | 1281.9765 | 1230.08426 | 1337.36499 | 1656.67335 | 1743.1252 | 1790.42982 | 1498.69495 | -0.3654541 | 8.45E-06 | 0.00019741 | Ubl7 | 1.29 | | 9 | 57910979 |
| ENSMUSG0( | 4447.26222 | 4256.77979 | 4244.16688 | 4136.11361 | 4906.56467 | 5504.37797 | 5649.22623 | 5099.93279 | -0.3085251 | 8.44E-06 | 0.00019741 | Rapgef1 | 1.24 | | 2 | 29619720 |
| ENSMUSG0( | 830815.837 | 806254.068 | 852378.802 | 787248.928 | 974038.019 | 967285.077 | 926855.534 | 983773.56 | -0.2333476 | 8.44E-06 | 0.00019741 | Podxl | 1.18 | | 6 | 31519488 |
| ENSMUSG0( | 197.02461 | 167.8411 | 138.243415 | 148.081937 | 238.201707 | 279.333107 | 272.550388 | 244.320052 | -0.6673108 | 8.55E-06 | 0.00019952 | Kcnu1 | 1.59 | | 8 | 25849623 |
| ENSMUSG0( | 7520.08835 | 6508.73368 | 5628.48189 | 6780.3017 | 7991.47202 | 10061.4053 | 10381.6921 | 9024.94436 | -0.5026797 | 8.68E-06 | 0.00020094 | C130021I20R | 1.42 | | 2 | 33641193 |
| ENSMUSG0( | 17550.346 | 16274.4085 | 15699.5617 | 15481.9665 | 18530.9213 | 22503.6381 | 21264.3167 | 20936.0435 | -0.3565839 | 8.64E-06 | 0.00020094 | Kif1c | 1.28 | | 11 | 70700548 |
| ENSMUSG0( | 1156.57235 | 1151.2046 | 1329.76999 | 1168.92179 | 1550.26357 | 1544.99358 | 1564.20223 | 1454.99543 | -0.3469566 | 8.72E-06 | 0.00020157 | Ccdc25 | 1.27 | | 14 | 65837302 |
| ENSMUSG0( | 1300.55187 | 1150.1749 | 1484.94117 | 1253.14339 | 1568.81206 | 1874.13027 | 1838.90716 | 1819.48917 | -0.4522181 | 9.06E-06 | 0.00020826 | Blmh | 1.37 | | 11 | 76924809 |
| ENSMUSG0( | 2172.00687 | 2622.64589 | 4319.40139 | 2571.99815 | 4838.22811 | 4631.73268 | 5943.3221 | 5096.95328 | -0.8115095 | 9.40E-06 | 0.0002142 | Slc3a2 | 1.76 | | 19 | 8706882 |
| ENSMUSG0( | 4178.24785 | 4583.19469 | 4180.21755 | 4459.11733 | 5234.58013 | 5299.75023 | 5253.86657 | 5082.05572 | -0.2623575 | 9.39E-06 | 0.0002142 | Htatsf1 | 1.20 | X | | 57053583 |
| ENSMUSG0( | 11037.1671 | 10611.0585 | 10687.0624 | 9867.81008 | 12308.3898 | 13184.9557 | 12864.8092 | 12424.5685 | -0.2669649 | 9.43E-06 | 0.00021459 | Dennd5a | 1.20 | | 7 | 109893780 |
| ENSMUSG0( | 8705.07771 | 8533.12388 | 8804.31811 | 8012.15831 | 10081.5944 | 11838.0938 | 12096.7127 | 10121.4051 | -0.3740928 | 9.47E-06 | 0.00021514 | Golga4 | 1.30 | | 9 | 118506267 |
| ENSMUSG0( | 8248.51106 | 7616.69088 | 9120.30366 | 10436.0745 | 12085.8079 | 11900.8896 | 11777.8395 | 12070.9996 | -0.4333924 | 1.10E-05 | 0.00024739 | Abr | 1.35 | | 11 | 76416734 |
| ENSMUSG0( | 2432.49614 | 2349.77539 | 2683.9912 | 2517.39293 | 3186.43595 | 3472.17548 | 2888.1723 | 3241.70996 | -0.3570881 | 1.12E-05 | 0.00025049 | Pik3r1 | 1.28 | | 13 | 101680563 |
| ENSMUSG0( | 1884.99506 | 1795.7968 | 1799.98569 | 1873.2365 | 2501.11792 | 2158.8768 | 2178.24856 | 2414.39856 | -0.3316069 | 1.14E-05 | 0.00025392 | Lamtor1 | 1.26 | | 7 | 101905904 |
| ENSMUSG0( | 2070.65287 | 1453.9364 | 1420.0514 | 1448.42645 | 2124.29063 | 2640.67228 | 2330.14423 | 2552.44932 | -0.5935308 | 1.15E-05 | 0.00025614 | Sptb | 1.51 | | 12 | 76580488 |
| ENSMUSG0( | 5160.5292 | 4506.99689 | 4987.10768 | 4538.71137 | 5690.48258 | 6046.80389 | 5986.41307 | 5969.95054 | -0.3037826 | 1.16E-05 | 0.0002579 | Stx16 | 1.23 | | 2 | 174076308 |
| ENSMUSG0( | 3757.67609 | 3832.54339 | 3824.73448 | 3388.29982 | 4364.75341 | 4600.33478 | 4514.85643 | 4566.6 | -0.2858695 | 1.18E-05 | 0.00026137 | Larp1 | 1.22 | | 11 | 58009064 |
| ENSMUSG0( | 5447.54101 | 6306.91248 | 5992.42884 | 5746.50467 | 7282.72431 | 7056.95036 | 7011.97817 | 7703.03384 | -0.3066573 | 1.22E-05 | 0.00027064 | Arpc5 | 1.24 | | 1 | 152766542 |
| ENSMUSG0( | 6947.95919 | 10812.8797 | 6354.49493 | 4007.46742 | 17014.826 | 16350.7309 | 16349.7915 | 18730.2108 | -1.2832527 | 1.22E-05 | 0.00027065 | Gadd45a | 2.43 | | 6 | 67035096 |
| ENSMUSG0( | 2047.91926 | 1888.46979 | 1774.59404 | 1899.15084 | 2442.54373 | 2247.65709 | 2392.62614 | 2354.80831 | -0.3106154 | 1.22E-05 | 0.00027065 | Eml3 | 1.24 | | 19 | 8929552 |
| ENSMUSG0( | 1907.72867 | 2043.95449 | 2634.14833 | 2157.36872 | 3150.3152 | 2914.59195 | 3275.99103 | 2839.47573 | -0.478213 | 1.27E-05 | 0.0002785 | Selenos | 1.39 | | 7 | 66079649 |
| ENSMUSG0( | 15776.1773 | 14020.3952 | 15244.3929 | 15258.9181 | 19390.0094 | 17463.7326 | 17931.2301 | 18644.7981 | -0.2842091 | 1.30E-05 | 0.00028434 | Eif3m | 1.22 | | 2 | 104999656 |
| ENSMUSG0( | 1000.27879 | 696.077198 | 774.915469 | 925.512107 | 1848.0157 | 1227.76645 | 1109.59249 | 1470.88617 | -0.7357595 | 1.31E-05 | 0.00028745 | Mylk3 | 1.67 | | 8 | 85324303 |
| ENSMUSG0( | 761.575895 | 467.483799 | 576.484444 | 1054.15829 | 1846.06323 | 2798.7445 | 3090.69986 | 1229.54563 | -1.6482433 | 1.32E-05 | 0.00028858 | R3hdml | 3.13 | | 2 | 163492318 |
| ENSMUSG0( | 600.546166 | 557.067698 | 605.637818 | 643.230914 | 830.777264 | 839.082008 | 880.133072 | 705.15137 | -0.4348919 | 1.34E-05 | 0.00029345 | Casp3 | 1.35 | | 8 | 46617291 |
| ENSMUSG0( | 4594.08345 | 3804.74149 | 4700.27611 | 3744.62198 | 6561.28554 | 5241.28516 | 5442.38957 | 6769.45315 | -0.511743 | 1.39E-05 | 0.00030101 | Npepl1 | 1.43 | | 2 | 174110349 |
| ENSMUSG0( | 3177.96906 | 3475.23749 | 2980.22709 | 3123.60336 | 3708.72248 | 4358.89569 | 3988.06932 | 4215.01748 | -0.3510431 | 1.42E-05 | 0.00030743 | Rreb1 | 1.28 | | 13 | 37778400 |
| ENSMUSG0( | 4623.44769 | 4473.01679 | 4653.25454 | 4711.78214 | 5361.49088 | 5717.6672 | 5273.25751 | 5874.60613 | -0.2677495 | 1.43E-05 | 0.00030805 | Ptov1 | 1.20 | | 7 | 44863067 |
| ENSMUSG0( | 1390.53907 | 1688.708 | 1419.11097 | 1205.94228 | 1745.51087 | 2204.34964 | 2396.93523 | 2032.02775 | -0.5547781 | 1.43E-05 | 0.00030889 | Srrm4 | 1.47 | | 5 | 116439275 |
| ENSMUSG0( | 1940.88185 | 2220.03319 | 2204.37119 | 2136.08194 | 2670.00684 | 2513.99796 | 2771.82668 | 2614.02592 | -0.3143101 | 1.46E-05 | 0.00031307 | Pfdn5 | 1.24 | | 15 | 102326116 |
| ENSMUSG0( | 9333.09365 | 9809.95187 | 9661.99214 | 9631.8045 | 11414.1572 | 11357.381 | 10991.4293 | 10840.4608 | -0.2146899 | 1.48E-05 | 0.00031706 | Ncl | 1.16 | | 1 | 86344719 |
| ENSMUSG0( | 52.0978535 | 54.5740998 | 29.1533732 | 69.413408 | 81.0276298 | 162.402969 | 121.731992 | 128.119052 | -1.263876 | 1.49E-05 | 0.00031855 | AC157583.2 | 2.40 | | 15 | 101395427 |
| ENSMUSG0( | 11489.9448 | 11990.8565 | 11697.0857 | 10832.1937 | 13246.5531 | 15198.7525 | 14925.6349 | 13623.3258 | -0.3088388 | 1.49E-05 | 0.00031855 | Eif3a | 1.24 | | 19 | 60761116 |
| ENSMUSG0( | 2849.27897 | 2471.27999 | 2964.23975 | 2904.25699 | 3923.49451 | 3582.6095 | 3400.95484 | 4559.6478 | -0.4670952 | 1.51E-05 | 0.00032085 | Slc9a3r1 | 1.38 | | 11 | 115163341 |
| ENSMUSG0( | 5672.0354 | 5961.96298 | 8626.57718 | 7915.90505 | 9602.26225 | 12400.008 | 12255.072 | 9802.59721 | -0.6449007 | 1.53E-05 | 0.00032497 | Pdia4 | 1.56 | | 6 | 47796141 |
| ENSMUSG0( | 37.889348 | 42.2176999 | 42.3194127 | 46.2756053 | 72.2415012 | 83.3668576 | 81.8728439 | 81.4400173 | -0.9174167 | 1.62E-05 | 0.0003422 | Gm7308 | 1.89 | | 6 | 127282063 |
| ENSMUSG0( | 9774.50455 | 8093.44198 | 11445.0501 | 9151.46371 | 13115.7374 | 15365.4863 | 13050.1004 | 12552.6876 | -0.4916155 | 1.65E-05 | 0.00034867 | Degs1 | 1.41 | | 1 | 182275772 |
| ENSMUSG0( | 5291.24745 | 6355.30838 | 5393.37404 | 6796.96091 | 7635.14569 | 8998.20718 | 9220.39041 | 7439.84354 | -0.482 | 1.71E-05 | 0.00035963 | Slk | 1.40 | | 19 | 47579678 |
| ENSMUSG0( | 4421.68691 | 4203.23539 | 4199.96661 | 4522.05215 | 5000.28337 | 5503.29528 | 5907.77205 | 5197.26355 | -0.3167351 | 1.74E-05 | 0.00036322 | Tpd52 | 1.25 | | 3 | 8925593 |
| ENSMUSG0( | 5524.26694 | 5320.45989 | 5372.68455 | 5399.43763 | 5992.13966 | 6904.29157 | 6732.96414 | 6572.8053 | -0.2773931 | 1.79E-05 | 0.00037188 | Dhx9 | 1.21 | | 1 | 153455758 |
| ENSMUSG0( | 39.7838154 | 56.6334998 | 31.9746674 | 32.3929237 | 69.3127917 | 132.087748 | 157.282042 | 70.515137 | -1.4173683 | 1.89E-05 | 0.0003915 | Gm29040 | 2.67 | | 1 | 43250156 |
| ENSMUSG0( | 151.557392 | 145.1877 | 261.439928 | 72.1899443 | 310.443208 | 319.392506 | 429.83243 | 460.831318 | -1.2702881 | 1.94E-05 | 0.00039998 | Kitl | 2.41 | | 10 | 100015630 |
| ENSMUSG0( | 2126.53966 | 2244.74599 | 2135.7197 | 2120.34824 | 2526.50007 | 2546.47856 | 2564.99002 | 2529.60639 | -0.2371469 | 2.01E-05 | 0.00041439 | Golga5 | 1.18 | | 12 | 102469135 |
| ENSMUSG0( | 12230.6815 | 10603.8506 | 12590.4955 | 10531.4023 | 15355.224 | 14300.1228 | 13925.9244 | 15112.0891 | -0.3529334 | 2.07E-05 | 0.00042362 | Vapb | 1.28 | | 2 | 173737511 |
| ENSMUSG0( | 733.158884 | 759.918598 | 799.366685 | 860.726259 | 936.210807 | 1028.55214 | 1136.52435 | 1060.70657 | -0.3998578 | 2.24E-05 | 0.00045588 | Trappc4 | 1.32 | | 9 | 44403698 |
| ENSMUSG0( | 2607.73438 | 2466.13149 | 2548.56908 | 2643.26258 | 3312.37046 | 3005.53762 | 2970.04514 | 3140.40652 | -0.2758957 | 2.25E-05 | 0.00045814 | Eif4a3 | 1.21 | | 11 | 119288363 |
| ENSMUSG0( | 823.146086 | 648.710998 | 707.204408 | 735.782125 | 925.472205 | 1474.61896 | 1238.8654 | 955.430447 | -0.6557775 | 2.34E-05 | 0.00047305 | Armcx4 | 1.58 | X | | 134686519 |
| ENSMUSG0( | 1144.25831 | 1084.2741 | 1069.27049 | 1019.91434 | 1291.56089 | 1396.66554 | 1335.82008 | 1312.97199 | -0.30561 | 2.35E-05 | 0.00047519 | Hmgxb3 | 1.24 | | 18 | 61131279 |
| ENSMUSG0( | 2494.06633 | 2095.43949 | 1975.84636 | 2098.13595 | 2933.59069 | 3368.23758 | 3446.20036 | 2520.66785 | -0.5017716 | 2.40E-05 | 0.00048255 | Foxn3 | 1.42 | | 12 | 99190078 |
| ENSMUSG0( | 3551.17914 | 3490.68299 | 2998.09528 | 3202.27189 | 4261.27234 | 4970.61355 | 5201.08014 | 3843.57155 | -0.4647615 | 2.57E-05 | 0.00051357 | Fbxl20 | 1.38 | | 11 | 98082556 |
| ENSMUSG0( | 2385.13446 | 2236.50839 | 2376.47013 | 2473.89386 | 2754.93941 | 2891.85554 | 2787.98579 | 2917.93623 | -0.2610451 | 2.70E-05 | 0.00053525 | Rbm8a | 1.20 | | 3 | 96629933 |
| ENSMUSG0( | 7946.34351 | 7339.70158 | 7832.85308 | 8036.22162 | 8888.63336 | 9614.25578 | 9231.16315 | 9181.86537 | -0.2446439 | 2.74E-05 | 0.00054246 | Mxd4 | 1.18 | | 5 | 34173883 |
| ENSMUSG0( | 53.0450872 | 27.8018999 | 43.2598441 | 35.1694601 | 76.1464473 | 109.351333 | 92.6455865 | 69.521966 | -1.1214222 | 2.77E-05 | 0.0005472 | Dnm1 | 2.18 | | 2 | 32308471 |
| ENSMUSG0( | 4439.68435 | 4359.74979 | 4091.81699 | 4057.44508 | 4697.65005 | 5610.48124 | 5487.63509 | 5554.80509 | -0.3330499 | 2.90E-05 | 0.00056892 | Pkp4 | 1.26 | | 2 | 59160850 |
| ENSMUSG0( | 6616.4274 | 6170.99208 | 5931.3008 | 6112.08195 | 7318.84506 | 8413.55649 | 8361.80282 | 7190.55763 | -0.3332274 | 2.96E-05 | 0.00058032 | Baz2a | 1.26 | | 10 | 128091577 |
| ENSMUSG0( | 18.944674 | 11.3267 | 12.2256081 | 16.6592179 | 29.2870951 | 55.2170095 | 54.9409873 | 34.760983 | -1.5511324 | 3.00E-05 | 0.00058747 | F2rl2 | 2.93 | | 13 | 95696853 |
| ENSMUSG0( | 57.7812557 | 48.3958999 | 39.4981185 | 72.1899443 | 90.7899948 | 103.9379 | 107.727426 | 122.160026 | -0.9608654 | 3.07E-05 | 0.00060115 | 4930551O13I | 1.95 | | 2 | 5037280 |
| ENSMUSG0( | 5159.58197 | 4902.40169 | 5109.36376 | 4758.98325 | 5901.34966 | 5782.62839 | 5776.34459 | 5824.94758 | -0.2244714 | 3.24E-05 | 0.00062802 | Dnajc5 | 1.17 | | 2 | 181520485 |
| ENSMUSG0( | 9476.12594 | 9786.26877 | 10825.3058 | 9222.72815 | 11776.3409 | 14047.8568 | 14250.1839 | 11646.9157 | -0.3957907 | 3.32E-05 | 0.00064097 | Tpcn1 | 1.32 | | 5 | 120534153 |
| ENSMUSG0( | 1859.41975 | 2318.88439 | 2625.68445 | 2057.41341 | 3061.47767 | 3173.35402 | 2945.26783 | 4448.41265 | -0.6211302 | 3.44E-05 | 0.0006615 | Slc26a7 | 1.54 | | 4 | 14502430 |
| ENSMUSG0( | 939.655831 | 1041.0267 | 979.929512 | 917.182498 | 1164.65015 | 1181.21093 | 1258.25634 | 1212.66172 | -0.3131626 | 3.66E-05 | 0.00069973 | Gmppa | 1.24 | | 1 | 75435930 |
| ENSMUSG0( | 1097.84386 | 992.630797 | 1137.92199 | 1031.946 | 1279.84606 | 1323.04286 | 1349.82465 | 1321.91053 | -0.30762 | 3.72E-05 | 0.00071033 | Arap1 | 1.24 | | 7 | 101348067 |
| ENSMUSG0( | 86.1982667 | 85.4650998 | 60.1876092 | 45.3500932 | 157.174077 | 153.741478 | 120.654717 | 115.207829 | -0.9825656 | 3.82E-05 | 0.00072398 | Gm867 | 1.98 | | 10 | 75937723 |
| ENSMUSG0( | 13345.5756 | 11940.4012 | 11993.3216 | 12323.1937 | 14640.6188 | 17457.2365 | 17627.4387 | 14462.5553 | -0.3718271 | 3.87E-05 | 0.00073084 | Rmnd5a | 1.29 | | 6 | 71388634 |
| ENSMUSG0( | 1958.87929 | 2032.62779 | 1913.77789 | 1893.59777 | 2279.51224 | 2302.8741 | 2391.54886 | 2275.35463 | -0.2461755 | 3.88E-05 | 0.00073195 | Pias2 | 1.19 | | 18 | 77065208 |
| ENSMUSG0( | 525.714704 | 445.860099 | 491.845619 | 471.085662 | 571.098354 | 657.190682 | 670.064591 | 722.035276 | -0.437189 | 3.91E-05 | 0.0007377 | Pex10 | 1.35 | | 4 | 155067016 |
| ENSMUSG0( | 1288.23783 | 1372.5901 | 1297.79532 | 1313.30168 | 1561.00217 | 1650.01417 | 1620.22049 | 1527.49691 | -0.2703492 | 3.93E-05 | 0.00073955 | Dhx8 | 1.21 | | 11 | 101732919 |
| ENSMUSG0( | 15137.7418 | 14011.1279 | 15701.4426 | 14064.082 | 19317.7679 | 29336.4724 | 27899.2488 | 16756.7802 | -0.6633863 | 3.98E-05 | 0.00074656 | Top1mt | 1.58 | | 15 | 75657035 |
| ENSMUSG0( | 18731.5464 | 18911.4701 | 22809.223 | 19713.4079 | 24915.508 | 27453.6806 | 28900.0366 | 23666.2704 | -0.3884141 | 3.99E-05 | 0.0007473 | Sema3e | 1.31 | | 5 | 14025276 |
| ENSMUSG0( | 1489.05138 | 1571.3222 | 1599.6738 | 1513.21229 | 1872.42161 | 1845.98042 | 1817.36168 | 1814.52331 | -0.2519205 | 4.02E-05 | 0.00075065 | Egln2 | 1.19 | | 7 | 27158658 |
| ENSMUSG0( | 1395.27524 | 1348.907 | 1382.43415 | 1324.40783 | 1864.61172 | 1660.84103 | 1546.96584 | 1876.09991 | -0.3505952 | 4.07E-05 | 0.0007582 | Mpp1 | 1.28 | X | | 75109733 |
| ENSMUSG0( | 496.350459 | 436.592799 | 562.377974 | 580.296091 | 653.102221 | 1076.19034 | 687.300979 | 1096.46072 | -0.7587681 | 4.09E-05 | 0.00076103 | Bend4 | 1.69 | | 5 | 67392147 |
| ENSMUSG0( | 13.2612718 | 18.5345999 | 10.3447453 | 12.9571695 | 42.9544061 | 38.9767126 | 44.1682447 | 26.8156155 | -1.4770463 | 4.14E-05 | 0.00076962 | Gbp4 | 2.78 | | 5 | 105115767 |
| ENSMUSG0( | 133.559952 | 65.9007998 | 86.5196882 | 37.0204843 | 192.318591 | 178.643266 | 246.695806 | 164.866377 | -1.2761299 | 4.28E-05 | 0.00079472 | Pcsk2 | 2.42 | | 2 | 143546156 |
| ENSMUSG0( | 1807.3219 | 1720.6287 | 1579.92474 | 1829.73744 | 2396.66062 | 2001.88727 | 2213.79861 | 2211.79169 | -0.3472035 | 4.34E-05 | 0.00080237 | Kctd2 | 1.27 | | 11 | 115420128 |
| ENSMUSG0( | 2243.99664 | 2159.28089 | 2099.9833 | 2265.65364 | 2524.5476 | 2636.34153 | 2919.41325 | 2655.7391 | -0.2917485 | 4.38E-05 | 0.00080834 | Fam184a | 1.22 | | 10 | 53633145 |
| ENSMUSG0( | 831.671189 | 939.086397 | 872.720334 | 788.536315 | 1047.50177 | 1066.44616 | 1128.98343 | 1096.46072 | -0.3388983 | 4.48E-05 | 0.00082167 | Tom1 | 1.26 | | 8 | 75033705 |
| ENSMUSG0( | 41.6782828 | 21.6236999 | 37.6172558 | 31.4674116 | 60.5266632 | 93.1110357 | 65.71373 | 71.5083079 | -1.1303536 | 4.51E-05 | 0.00082294 | Ces2e | 2.19 | | 8 | 104926260 |
| ENSMUSG0( | 2532.90291 | 2744.15049 | 3132.57697 | 2849.65178 | 4085.54977 | 3339.00505 | 3481.75041 | 3901.17546 | -0.3953356 | 4.56E-05 | 0.00083233 | Rer1 | 1.32 | | 4 | 155074110 |
| ENSMUSG0( | 997.437086 | 596.196298 | 974.286924 | 1012.51025 | 1085.57499 | 1876.29564 | 1754.87977 | 1483.79739 | -0.7916948 | 4.59E-05 | 0.00083521 | Cacna2d2 | 1.73 | | 9 | 107399612 |
| ENSMUSG0( | 472.669616 | 397.464199 | 390.279028 | 435.916202 | 501.785563 | 649.611877 | 646.364557 | 583.984515 | -0.4884861 | 4.69E-05 | 0.00084972 | Ftx | 1.40 | X | | 103560907 |
| ENSMUSG0( | 4065.52704 | 3965.37469 | 4380.52943 | 4521.12664 | 5767.60526 | 6345.62535 | 6534.74567 | 4618.24489 | -0.4582944 | 4.97E-05 | 0.0008943 | Snd1 | 1.37 | | 6 | 28475139 |
| ENSMUSG0( | 923.552858 | 914.373597 | 1139.80285 | 956.054006 | 1408.70927 | 1194.20317 | 1441.39296 | 1253.38173 | -0.4292973 | 4.98E-05 | 0.00089574 | Tmem50b | 1.35 | | 16 | 91574503 |
| ENSMUSG0( | 3532.23447 | 3810.91969 | 3596.20965 | 3664.10243 | 4154.86256 | 4300.43063 | 4350.03347 | 4242.82627 | -0.2233402 | 5.10E-05 | 0.00091218 | Naa50 | 1.17 | | 16 | 44139830 |
| ENSMUSG0( | 9790.60753 | 10183.733 | 9159.80178 | 9820.60897 | 11679.6935 | 12867.7286 | 13428.2237 | 10989.4365 | -0.3299148 | 5.23E-05 | 0.00093312 | Zfp91 | 1.26 | | 19 | 12763660 |
| ENSMUSG0( | 1413.27268 | 1287.125 | 1441.68133 | 1375.31099 | 1599.07539 | 1778.85386 | 1676.23875 | 1710.24036 | -0.2934795 | 5.24E-05 | 0.00093312 | Insig2 | 1.23 | | 1 | 121304353 |
| ENSMUSG0( | 3915.86412 | 6204.97218 | 4070.18707 | 2985.70206 | 7271.00948 | 6791.69217 | 7055.06914 | 6927.36733 | -0.7074251 | 5.37E-05 | 0.00095186 | Slc41a2 | 1.63 | | 10 | 83230848 |
| ENSMUSG0( | 1973.0878 | 1718.5693 | 2177.09868 | 1783.46183 | 2464.02093 | 2442.54066 | 2366.77155 | 2761.01522 | -0.3909436 | 5.49E-05 | 0.00097193 | Amigo1 | 1.31 | | 3 | 108186335 |
| ENSMUSG0( | 478.353019 | 440.711599 | 453.287932 | 410.927375 | 612.100288 | 592.229495 | 561.25989 | 551.209873 | -0.377782 | 5.60E-05 | 0.00098883 | Yrdc | 1.30 | | 4 | 124850679 |
| ENSMUSG0( | 10356.106 | 10971.4535 | 13075.7581 | 11070.9758 | 14491.2547 | 16726.4231 | 17834.2754 | 13261.8116 | -0.4544534 | 5.70E-05 | 0.00100438 | Dnajc3 | 1.37 | | 14 | 118937976 |
| ENSMUSG0( | 2938.31894 | 3074.68419 | 2783.67693 | 3073.62571 | 3436.35249 | 3641.07457 | 3759.68717 | 3459.2144 | -0.2682026 | 5.77E-05 | 0.00101408 | Ufm1 | 1.20 | | 3 | 53853376 |
| ENSMUSG0( | 2344.40341 | 2632.94289 | 2321.92511 | 2383.19368 | 2850.61059 | 2942.7418 | 2945.26783 | 2902.0455 | -0.2659328 | 5.79E-05 | 0.0010162 | Aaed1 | 1.20 | | 13 | 64275280 |
| ENSMUSG0( | 2589.73694 | 2498.05219 | 2368.94668 | 2204.56984 | 2881.85016 | 3934.4826 | 3713.36438 | 2915.94989 | -0.476692 | 5.98E-05 | 0.00104364 | Zfp609 | 1.39 | | 9 | 65692391 |
| ENSMUSG0( | 3633.58847 | 3894.32539 | 3907.49244 | 3902.88456 | 4542.42845 | 4512.63717 | 4325.25616 | 4653.99904 | -0.2337143 | 6.14E-05 | 0.00107032 | Anp32b | 1.18 | | 4 | 46450902 |
| ENSMUSG0( | 1616.92793 | 1550.7282 | 1874.27977 | 1735.3352 | 2576.28813 | 2313.70097 | 2312.90784 | 1917.81309 | -0.4282791 | 6.30E-05 | 0.00109385 | Slc2a12 | 1.35 | | 10 | 22645011 |
| ENSMUSG0( | 52.0978535 | 64.8710998 | 41.3789813 | 74.9664807 | 128.863218 | 85.5322305 | 106.650152 | 148.975641 | -1.012691 | 6.49E-05 | 0.00112101 | Gm9791 | 2.02 | | 3 | 34005017 |
| ENSMUSG0( | 1023.95963 | 906.135997 | 983.691238 | 892.193671 | 1188.07982 | 1188.78973 | 1235.63358 | 1133.20805 | -0.3180025 | 6.49E-05 | 0.00112101 | Ebag9 | 1.25 | | 15 | 44619175 |
| ENSMUSG0( | 13230.9603 | 13478.773 | 13163.2182 | 12159.3781 | 15226.3607 | 15701.1191 | 15369.4719 | 14759.5134 | -0.2307496 | 6.50E-05 | 0.00112101 | Brd2 | 1.17 | | 17 | 34112023 |
| ENSMUSG0( | 3336.15709 | 3135.43649 | 3141.04086 | 3239.29237 | 3575.95431 | 3816.46978 | 4015.00117 | 3959.77255 | -0.2577103 | 6.55E-05 | 0.00112735 | Ipo9 | 1.20 | | 1 | 135382312 |
| ENSMUSG0( | 35.9948806 | 15.4455 | 32.9150988 | 17.58473 | 67.3603187 | 57.3823825 | 71.1001013 | 52.63806 | -1.2816954 | 6.79E-05 | 0.00116402 | Ces2g | 2.43 | | 8 | 104961718 |
| ENSMUSG0( | 39.7838154 | 53.5443999 | 66.770629 | 24.9888269 | 133.744401 | 128.839689 | 101.263781 | 73.4946498 | -1.2418976 | 6.80E-05 | 0.00116433 | Cd74 | 2.37 | | 18 | 60803848 |
| ENSMUSG0( | 1992.97971 | 1821.53929 | 1561.11611 | 1723.30354 | 2265.84492 | 2360.25649 | 2585.45823 | 2125.38582 | -0.3952517 | 6.91E-05 | 0.0011803 | Stxbp5 | 1.32 | | 10 | 9755547 |
| ENSMUSG0( | 110.826343 | 121.5046 | 105.328316 | 131.422719 | 150.340422 | 205.710428 | 203.604835 | 179.763941 | -0.654941 | 7.04E-05 | 0.00119646 | Gm31166 | 1.57 | | 19 | 5512528 |
| ENSMUSG0( | 1779.85212 | 1940.98449 | 1740.73851 | 1933.39479 | 2207.27073 | 2267.14545 | 2232.11227 | 2185.96925 | -0.2661165 | 7.21E-05 | 0.00122313 | Slf2 | 1.20 | | 19 | 44931119 |
| ENSMUSG0( | 1904.88697 | 1975.99429 | 1644.81451 | 1839.91807 | 2289.2746 | 2450.11946 | 2322.60331 | 2165.11266 | -0.3251239 | 7.46E-05 | 0.00126203 | Ranbp10 | 1.25 | | 8 | 105768308 |
| ENSMUSG0( | 119.351446 | 93.7026997 | 140.124278 | 103.657356 | 175.722571 | 281.49848 | 277.93676 | 146.9893 | -0.9470112 | 7.53E-05 | 0.00127271 | Gm27028 | 1.93 | | 1 | 23291537 |
| ENSMUSG0( | 9048.92354 | 8657.71758 | 8011.53504 | 7896.4693 | 10201.6715 | 10848.5183 | 11173.4886 | 9532.45471 | -0.3128711 | 7.57E-05 | 0.00127507 | Mar-06 | 1.24 | | 15 | 31455891 |
| ENSMUSG0( | 4920.87907 | 5024.93599 | 4777.39148 | 4718.26072 | 5686.57763 | 6434.40564 | 6296.66806 | 5451.51531 | -0.2959196 | 7.58E-05 | 0.00127635 | Trim56 | 1.23 | | 5 | 137105644 |
| ENSMUSG0( | 49.2561524 | 36.0394999 | 29.1533732 | 37.0204843 | 51.7405347 | 131.005062 | 116.34562 | 72.5014789 | -1.2918631 | 7.68E-05 | 0.00128769 | Gm28722 | 2.45 | | 1 | 90453394 |
| ENSMUSG0( | 3558.75701 | 3565.85109 | 3416.58725 | 3421.61826 | 4133.38535 | 4068.73572 | 4331.71981 | 3908.12766 | -0.2357733 | 7.73E-05 | 0.00129302 | Ndfip2 | 1.18 | | 14 | 105258573 |
| ENSMUSG0( | 77.6731634 | 36.0394999 | 31.9746674 | 74.0409686 | 135.696874 | 114.764765 | 118.500169 | 117.194171 | -1.1435267 | 7.82E-05 | 0.00130644 | Gm13528 | 2.21 | | 2 | 33177902 |
| ENSMUSG0( | 4198.13976 | 4102.32479 | 3608.43526 | 3247.62198 | 4634.19468 | 5053.9804 | 5008.24804 | 4903.28495 | -0.3709593 | 7.99E-05 | 0.00133186 | Dlgap4 | 1.29 | | 2 | 156613705 |
| ENSMUSG0( | 4193.40359 | 3913.88969 | 3737.27436 | 3702.97394 | 4587.33533 | 5058.31115 | 5175.22555 | 4465.29656 | -0.3107659 | 8.07E-05 | 0.00134185 | Clcn3 | 1.24 | | 8 | 60910389 |
| ENSMUSG0( | 10316.3222 | 9947.93167 | 13454.752 | 12984.9349 | 14927.6324 | 19368.1781 | 21370.9668 | 14409.9172 | -0.5853382 | 8.13E-05 | 0.00134864 | Hyou1 | 1.50 | | 9 | 44379490 |
| ENSMUSG0( | 1094.05492 | 918.492397 | 920.682335 | 865.35382 | 1159.76897 | 1398.83091 | 1452.1657 | 1173.92805 | -0.448352 | 8.15E-05 | 0.00135037 | Ago4 | 1.36 | | 4 | 126489541 |
| ENSMUSG0( | 2666.46287 | 2507.31949 | 2352.01892 | 2616.42273 | 2890.63629 | 3623.75159 | 3242.59553 | 3140.40652 | -0.3464827 | 8.19E-05 | 0.00135451 | Zfp146 | 1.27 | | 7 | 30161269 |
| ENSMUSG0( | 1775.11595 | 1663.9952 | 1747.32153 | 1812.15271 | 1998.35612 | 2065.76577 | 2142.69851 | 2124.39265 | -0.2511069 | 8.25E-05 | 0.00136114 | Pggt1b | 1.19 | | 18 | 46239949 |
| ENSMUSG0( | 3149.55205 | 2784.30879 | 3137.27913 | 3114.34824 | 3502.73657 | 3871.68679 | 3669.19614 | 3706.51396 | -0.2752675 | 8.42E-05 | 0.00138384 | Bmi1 | 1.21 | | 2 | 18677018 |
| ENSMUSG0( | 170.502066 | 155.4847 | 239.810005 | 201.761639 | 279.20364 | 295.573404 | 294.095873 | 291.992257 | -0.595144 | 8.49E-05 | 0.00139092 | Spc25 | 1.51 | | 2 | 69193895 |
| ENSMUSG0( | 510.558964 | 541.622198 | 527.582012 | 456.277469 | 734.12985 | 721.069184 | 576.34173 | 731.966985 | -0.4416357 | 8.56E-05 | 0.00139996 | Cdc42ep4 | 1.36 | | 11 | 113726850 |
| ENSMUSG0( | 53.0450872 | 37.0691999 | 36.6768244 | 102.731844 | 234.296761 | 111.516706 | 147.586574 | 112.228317 | -1.3983443 | 8.74E-05 | 0.00142246 | Pld6 | 2.64 | | 11 | 59783897 |
| ENSMUSG0( | 2380.39829 | 2365.22089 | 2713.14457 | 2398.00187 | 2816.44231 | 3464.59668 | 3236.13188 | 3033.14406 | -0.3482628 | 8.73E-05 | 0.00142246 | Erc1 | 1.27 | | 6 | 119570796 |

| | | | | | | | | | | | | | | |
|---|---|---|---|---|---|---|---|---|---|---|---|---|---|---|
| ENSMUSG0( | 3952.80623 | 3495.83149 | 4471.75128 | 3969.52143 | 4739.62822 | 5814.0263 | 6494.88652 | 4907.25763 | -0.4663083 | 8.93E-05 | 0.00145121 | Tgfbr2 | 1.38 | |
| ENSMUSG0( | 28.417011 | 25.7424999 | 19.7490593 | 12.0316574 | 41.0019331 | 61.7131283 | 84.0273924 | 43.6995215 | -1.4233983 | 9.07E-05 | 0.00146849 | Gm29107 | 2.68 | |
| ENSMUSG0( | 21.7863751 | 14.4158 | 10.3447453 | 23.1378027 | 32.2158046 | 54.1343231 | 49.554616 | 46.6790343 | -1.384668 | 9.08E-05 | 0.00146849 | AC131178.1 | 2.61 | |
| ENSMUSG0( | 3396.78005 | 3463.91079 | 3266.11823 | 3313.33334 | 3909.8272 | 3815.38709 | 3926.66468 | 3842.57838 | -0.2053376 | 9.73E-05 | 0.00155811 | Pcif1 | 1.15 | |
| ENSMUSG0( | 1067.53238 | 1311.8378 | 1249.83332 | 1105.06146 | 1441.90132 | 1624.02969 | 1492.02485 | 1509.61983 | -0.3581794 | 9.90E-05 | 0.00157951 | Zfp710 | 1.28 | |
| ENSMUSG0( | 15293.0881 | 17471.9496 | 18545.3071 | 15291.311 | 20983.2274 | 20287.3789 | 20222.5924 | 21457.4582 | -0.3167242 | 9.97E-05 | 0.00158869 | Arhgdia | 1.25 | |
| ENSMUSG0( | 2217.47409 | 2250.92419 | 2616.28014 | 2358.20485 | 2932.61446 | 2809.57137 | 2832.15403 | 3001.36259 | -0.2937804 | 9.98E-05 | 0.0015888 | Sptlc1 | 1.23 | |
| ENSMUSG0( | 5054.43903 | 4614.08569 | 7151.04032 | 7049.62572 | 8820.29681 | 8336.68576 | 8483.53481 | 9149.09073 | -0.5434435 | 0.00010653 | 0.00168274 | Megf9 | 1.46 | |
| ENSMUSG0( | 656.432954 | 650.770398 | 686.514918 | 681.176911 | 887.398981 | 900.795136 | 919.992219 | 741.898695 | -0.3665301 | 0.00010701 | 0.0016868 | Phf5a | 1.29 | |
| ENSMUSG0( | 9767.87392 | 10342.3068 | 10492.3931 | 11281.0671 | 11855.4161 | 13100.5062 | 12883.1229 | 12474.227 | -0.2644886 | 0.00011095 | 0.00174155 | Copa | 1.20 | |
| ENSMUSG0( | 379.840714 | 543.681598 | 681.812761 | 441.469275 | 1142.19671 | 791.443804 | 1042.80149 | 627.684036 | -0.8163925 | 0.00011239 | 0.00176086 | Prr15 | 1.76 | |
| ENSMUSG0( | 32.2059458 | 19.5642999 | 11.2851767 | 22.2122906 | 38.0732236 | 75.7880523 | 67.8682785 | 47.6722053 | -1.424626 | 0.00011648 | 0.00180924 | Gm15594 | 2.68 | |
| ENSMUSG0( | 1618.82239 | 1760.787 | 1542.30749 | 1613.1676 | 2006.16601 | 1922.85116 | 1952.02096 | 1953.56725 | -0.2620985 | 0.00011639 | 0.00180924 | Prune1 | 1.20 | |
| ENSMUSG0( | 4148.88361 | 4011.71119 | 4664.53971 | 4168.50653 | 5241.41379 | 5085.37831 | 4984.54801 | 5067.15815 | -0.262 | 0.00011624 | 0.00180924 | Cct2 | 1.20 | |
| ENSMUSG0( | 2569.84503 | 2749.29899 | 2703.74026 | 2559.04098 | 3233.2953 | 3056.42388 | 3097.1635 | 3034.13723 | -0.2314082 | 0.00011593 | 0.00180924 | Enah | 1.17 | |
| ENSMUSG0( | 2089.59754 | 2039.83569 | 2338.85288 | 1764.95159 | 2586.0505 | 2735.94869 | 2694.26293 | 2550.46298 | -0.3599411 | 0.000118 | 0.00182708 | Abtb2 | 1.28 | |
| ENSMUSG0( | 15935.3125 | 20232.5752 | 12861.3397 | 13188.5475 | 23757.6915 | 29722.9914 | 31583.5268 | 17398.3686 | -0.7197041 | 0.0001199 | 0.00185072 | Adcy1 | 1.65 | |
| ENSMUSG0( | 1249.40125 | 1236.6697 | 1220.67995 | 1069.892 | 1573.69324 | 1398.83091 | 1443.54751 | 1573.18277 | -0.3269383 | 0.00012033 | 0.0018554 | Dock11 | 1.25 | X |
| ENSMUSG0( | 16685.5216 | 16757.3378 | 20018.0227 | 20115.0801 | 21336.625 | 27019.5233 | 31044.8897 | 23335.5445 | -0.4816074 | 0.00012194 | 0.00187825 | Lpin2 | 1.40 | |
| ENSMUSG0( | 2548.05865 | 2445.53749 | 2464.87068 | 2559.04098 | 2832.0621 | 2920.00539 | 2876.32228 | 3002.35576 | -0.2153151 | 0.00012779 | 0.00195629 | Ubap1 | 1.16 | |
| ENSMUSG0( | 2102.85881 | 1965.69729 | 2185.56256 | 2242.51584 | 2416.18535 | 2596.28214 | 2839.69495 | 2620.97812 | -0.3013387 | 0.00012974 | 0.00198412 | Med22 | 1.23 | |
| ENSMUSG0( | 78409.164 | 51731.0982 | 69538.3186 | 55190.138 | 93175.9168 | 97353.0013 | 94163.4659 | 78881.609 | -0.5124884 | 0.00013097 | 0.00199679 | Thsd7a | 1.43 | |
| ENSMUSG0( | 831.671189 | 865.977698 | 834.162646 | 705.240226 | 1039.69188 | 965.756324 | 1034.18329 | 1130.22853 | -0.3660222 | 0.00013128 | 0.00199943 | Arl10 | 1.29 | |
| ENSMUSG0( | 2786.76155 | 2798.72459 | 2561.73512 | 2694.16574 | 3074.16875 | 3303.27639 | 3148.87267 | 3257.60069 | -0.2378204 | 0.0001346 | 0.00203953 | Mta2 | 1.18 | |
| ENSMUSG0( | 2731.82199 | 2741.06139 | 2769.57046 | 2645.1136 | 3035.11929 | 3799.14679 | 3590.55511 | 3221.84654 | -0.3256379 | 0.00013544 | 0.00204812 | Smg7 | 1.25 | |
| ENSMUSG0( | 66.306359 | 65.9007998 | 36.6768244 | 54.6052143 | 93.7187043 | 125.59163 | 187.445722 | 84.4195302 | -1.1354621 | 0.00013693 | 0.00206647 | Gm45809 | 2.20 | |
| ENSMUSG0( | 16436.3992 | 14654.6904 | 17206.1328 | 13389.3837 | 18325.9116 | 19987.4748 | 19812.1509 | 20365.9634 | -0.347565 | 0.00013762 | 0.00207472 | Lasp1 | 1.27 | |
| ENSMUSG0( | 3788.9348 | 4010.68149 | 4253.57119 | 4014.87152 | 4609.78877 | 4822.2855 | 4824.03414 | 4625.19708 | -0.2326605 | 0.00013896 | 0.00209291 | Nars | 1.17 | |
| ENSMUSG0( | 9717.67053 | 9922.18917 | 9569.82986 | 9636.43206 | 10865.5123 | 11174.407 | 11433.1117 | 10760.014 | -0.1873294 | 0.00014089 | 0.00211126 | Dync1li2 | 1.14 | |
| ENSMUSG0( | 8979.77548 | 8695.81648 | 9456.03767 | 9091.30543 | 10391.0613 | 10240.0486 | 10387.0784 | 10408.4315 | -0.1936077 | 0.00014196 | 0.00212292 | Eif4h | 1.14 | |
| ENSMUSG0( | 3573.91275 | 3518.48489 | 3425.99157 | 3418.84172 | 3898.11236 | 4264.70197 | 4228.30148 | 3954.8067 | -0.2298341 | 0.00014339 | 0.00213999 | Appbp2 | 1.17 | |
| ENSMUSG0( | 754.945259 | 668.275298 | 682.753192 | 676.54935 | 828.824791 | 927.862298 | 886.596717 | 837.243105 | -0.3222245 | 0.00014978 | 0.00222283 | Efcab2 | 1.25 | |
| ENSMUSG0( | 1228.56211 | 1274.7686 | 1287.45058 | 1215.1974 | 1449.71121 | 1458.37866 | 1576.05224 | 1488.76324 | -0.2547368 | 0.00014998 | 0.00222283 | Atg12 | 1.19 | |
| ENSMUSG0( | 8212.51618 | 6768.21808 | 7398.37378 | 7830.75794 | 9022.37776 | 9535.21967 | 9192.38127 | 9029.91022 | -0.2837897 | 0.00015297 | 0.00226255 | Rdh1 | 1.22 | |
| ENSMUSG0( | 1178.35872 | 1203.7193 | 1200.93089 | 1180.02794 | 1602.0041 | 1417.23658 | 1333.66554 | 1521.53788 | -0.3031181 | 0.00015584 | 0.00230282 | Inafm2 | 1.23 | |
| ENSMUSG0( | 1892.57293 | 1901.85589 | 1926.00349 | 1887.11919 | 2192.62719 | 2233.58217 | 2193.3304 | 2160.1468 | -0.2066547 | 0.00015923 | 0.00234826 | Brap | 1.15 | |
| ENSMUSG0( | 94.72337 | 129.7422 | 127.89867 | 113.837989 | 205.985902 | 176.477893 | 216.532127 | 147.982471 | -0.6805351 | 0.00016239 | 0.00238533 | Mrnip | 1.60 | |
| ENSMUSG0( | 19190.9548 | 14456.988 | 16494.2262 | 16478.7431 | 22226.9527 | 22070.5635 | 23997.3615 | 19116.5543 | -0.3918108 | 0.00016421 | 0.00240842 | Syne1 | 1.31 | |
| ENSMUSG0( | 3778.51523 | 3618.36579 | 4515.95155 | 3628.00746 | 5044.21401 | 4710.76879 | 4754.01132 | 5078.08303 | -0.3338347 | 0.00017029 | 0.0024915 | Lrrc59 | 1.26 | |
| ENSMUSG0( | 5345.23977 | 5238.08389 | 4807.48529 | 5060.7002 | 5867.18138 | 6098.77284 | 5934.70391 | 5905.39443 | -0.2191368 | 0.00017407 | 0.00254433 | Cbx5 | 1.16 | |
| ENSMUSG0( | 5509.1112 | 5579.94428 | 5872.05362 | 5881.62944 | 6493.92522 | 6410.58654 | 6698.49136 | 6862.81122 | -0.2123639 | 0.0001745 | 0.00254816 | Eef1g | 1.16 | |
| ENSMUSG0( | 2236.41877 | 2447.59689 | 2367.06582 | 2238.81379 | 2782.27403 | 2845.30002 | 2688.87656 | 2656.73227 | -0.2403992 | 0.00017528 | 0.00255452 | Trappc6b | 1.18 | |
| ENSMUSG0( | 3089.87633 | 2662.80419 | 3758.90428 | 3246.69647 | 3943.01924 | 4189.99661 | 4869.27966 | 4182.24284 | -0.4294718 | 0.00017783 | 0.00258665 | Tmem64 | 1.35 | |
| ENSMUSG0( | 16734.7778 | 17922.9582 | 22370.0416 | 24961.0615 | 27114.9689 | 30550.1639 | 32534.76 | 25840.3216 | -0.5010978 | 0.00017908 | 0.00259968 | Calr | 1.42 | |
| ENSMUSG0( | 5974.20295 | 6154.51688 | 6378.00571 | 5954.7449 | 7490.66269 | 7079.68677 | 6734.04141 | 7456.72744 | -0.2337208 | 0.00018292 | 0.00264258 | Eif4a1 | 1.18 | |
| ENSMUSG0( | 88.0927341 | 83.4056998 | 125.077375 | 32.3929237 | 166.936442 | 189.470131 | 176.672979 | 188.702479 | -1.134041 | 0.00018342 | 0.00264728 | Ifi203 | 2.19 | |
| ENSMUSG0( | 3521.8149 | 3658.52409 | 3828.4962 | 3349.42832 | 4318.87029 | 4296.09988 | 4515.9337 | 4061.07599 | -0.2598351 | 0.00018398 | 0.0026528 | Pnpla8 | 1.20 | |
| ENSMUSG0( | 2046.97203 | 1738.1336 | 2344.49547 | 1826.03539 | 2476.71201 | 2716.46033 | 2686.72201 | 2523.64737 | -0.3867511 | 0.0001859 | 0.0026778 | Cbfa2t3 | 1.31 | |
| ENSMUSG0( | 6432.66406 | 5764.26058 | 6430.66987 | 5367.97022 | 7993.42449 | 12653.3567 | 12614.8816 | 6310.60817 | -0.7216677 | 0.00018916 | 0.00271608 | 2810039B14l | 1.65 | |
| ENSMUSG0( | 3001.7836 | 2938.76379 | 3189.94329 | 2707.12291 | 3366.06346 | 3608.59398 | 3869.56915 | 3638.97834 | -0.290904 | 0.00018929 | 0.00271608 | Cep120 | 1.22 | |
| ENSMUSG0( | 4526.82985 | 3665.73199 | 3998.71429 | 4703.45253 | 5069.59616 | 8405.97769 | 8213.13897 | 4969.8274 | -0.6579032 | 0.00019965 | 0.00284835 | Tet3 | 1.58 | |
| ENSMUSG0( | 4334.54141 | 4392.70019 | 6216.25151 | 7964.03168 | 8997.97185 | 9623.99996 | 10610.0742 | 7537.17429 | -0.6826012 | 0.00020478 | 0.00290198 | Ackr3 | 1.61 | |
| ENSMUSG0( | 2203.26559 | 2313.73589 | 2301.23562 | 2267.50466 | 2567.502 | 2572.46303 | 2683.49019 | 2724.2679 | -0.215338 | 0.00020654 | 0.00292134 | Ap3b1 | 1.16 | |
| ENSMUSG0( | 1930.46228 | 2014.09319 | 1928.82479 | 1928.76723 | 2204.34202 | 2339.68544 | 2234.26682 | 2286.27951 | -0.2164287 | 0.00020939 | 0.00295613 | Arfgap1 | 1.16 | |
| ENSMUSG0( | 2820.86196 | 2721.49709 | 2998.09528 | 2773.75978 | 3410.97034 | 3384.47788 | 3151.02722 | 3372.80852 | -0.2353759 | 0.00021187 | 0.00298266 | Ddx21 | 1.18 | |
| ENSMUSG0( | 9.472337 | 12.3564 | 3.76172558 | 3.70204843 | 46.8593522 | 36.8113397 | 31.2409536 | 10.9248804 | -2.1122513 | 0.00021418 | 0.00300694 | Best3 | 4.32 | |
| ENSMUSG0( | 3446.98344 | 3469.05929 | 3646.99295 | 3238.36686 | 3791.70258 | 4488.81807 | 4445.91088 | 4168.33845 | -0.2916171 | 0.0002142 | 0.00300694 | Rnpepl1 | 1.22 | |
| ENSMUSG0( | 4193.40359 | 4126.00789 | 4327.86527 | 4212.93111 | 4882.15875 | 4939.21564 | 4785.25227 | 4667.90343 | -0.1929702 | 0.0002172 | 0.00304039 | Ik | 1.14 | |
| ENSMUSG0( | 3095.55973 | 2993.33789 | 3090.25756 | 3056.04098 | 3532.02367 | 3354.16266 | 3689.66435 | 3669.76664 | -0.2194794 | 0.00021865 | 0.00305234 | Peli2 | 1.16 | |
| ENSMUSG0( | 2951.58021 | 2948.03109 | 3294.33117 | 3130.08195 | 3466.61582 | 3674.63785 | 3734.90987 | 3757.16568 | -0.2475969 | 0.00022164 | 0.00308795 | Snx1 | 1.19 | |
| ENSMUSG0( | 2092.43924 | 2011.00409 | 1813.15173 | 2008.36127 | 2373.23094 | 2485.84812 | 2578.99458 | 2243.57316 | -0.2886859 | 0.00022552 | 0.00312709 | Tbc1d17 | 1.22 | |
| ENSMUSG0( | 473.61685 | 419.087899 | 339.495733 | 511.808195 | 579.884483 | 689.671276 | 716.387384 | 556.175728 | -0.5428636 | 0.00022857 | 0.00316091 | Asap3 | 1.46 | |
| ENSMUSG0( | 37403.4171 | 39007.0953 | 39507.5229 | 36153.2794 | 43014.9328 | 43604.1146 | 44899.714 | 42936.7662 | -0.1981133 | 0.00023191 | 0.00320177 | Eef2 | 1.15 | |
| ENSMUSG0( | 19698.672 | 20715.5045 | 12106.1733 | 24928.6686 | 28211.2825 | 28651.1318 | 37135.7983 | 29039.3252 | -0.6677764 | 0.0002324 | 0.00320491 | Card10 | 1.59 | |
| ENSMUSG0( | 231.125023 | 211.088499 | 233.226986 | 319.301677 | 258.702673 | 462.307119 | 480.464321 | 526.3806 | -0.7955869 | 0.00023835 | 0.00327783 | Gm8822 | 1.74 | X |
| ENSMUSG0( | 429.096866 | 301.702099 | 137.302984 | 111.061453 | 609.171578 | 1192.03779 | 1036.33784 | 646.554284 | -1.8312799 | 0.00024218 | 0.00332748 | Adamts16 | 3.56 | |
| ENSMUSG0( | 8259.87787 | 8441.48058 | 7828.15092 | 7842.78959 | 8937.44519 | 9933.64829 | 10186.7054 | 9337.79321 | -0.2461445 | 0.0002441 | 0.00335078 | Mib1 | 1.19 | |
| ENSMUSG0( | 2088.65031 | 2145.89479 | 2334.15072 | 2213.82496 | 2559.69211 | 2546.47856 | 2553.14 | 2703.41131 | -0.2386842 | 0.00024578 | 0.00336089 | Ncln | 1.18 | |
| ENSMUSG0( | 41776.7951 | 39135.8078 | 45436.0024 | 40353.2534 | 49614.2916 | 49155.048 | 50671.7494 | 47558.9838 | -0.2409119 | 0.00024656 | 0.00336591 | Parva | 1.18 | |
| ENSMUSG0( | 1927.62058 | 2086.17219 | 1858.29243 | 1918.5866 | 2189.69848 | 2331.02395 | 2528.36269 | 2368.7127 | -0.2736774 | 0.00024788 | 0.00337969 | Ppp4r1 | 1.21 | |
| ENSMUSG0( | 379.840714 | 442.770999 | 371.470401 | 385.938549 | 550.597388 | 468.803238 | 498.777983 | 562.134754 | -0.39853 | 0.00024873 | 0.00338616 | Immp1l | 1.32 | |
| ENSMUSG0( | 73.8842286 | 66.9304998 | 71.4727859 | 74.9664807 | 109.338488 | 164.568342 | 165.900236 | 87.399043 | -0.8729969 | 0.00025195 | 0.00342687 | Gm5176 | 1.83 | |
| ENSMUSG0( | 6408.03598 | 5716.89438 | 8340.68603 | 6384.18251 | 9735.03041 | 11695.1792 | 14925.6349 | 7194.53031 | -0.6977167 | 0.00025264 | 0.0034331 | Arl5a | 1.62 | |
| ENSMUSG0( | 1718.28193 | 1707.2426 | 1831.01992 | 1843.62012 | 1995.42741 | 2086.33681 | 2247.19411 | 2154.18778 | -0.2564026 | 0.00025601 | 0.00346631 | Gle1 | 1.19 | |
| ENSMUSG0( | 2053.60266 | 1924.50929 | 1866.75632 | 1756.62198 | 2245.34396 | 2311.5356 | 2208.41224 | 2376.65807 | -0.2663707 | 0.00025882 | 0.00349475 | Rfwd3 | 1.20 | |
| ENSMUSG0( | 16955.4832 | 15933.5778 | 17243.75 | 16753.6202 | 18312.2443 | 19911.6867 | 20138.565 | 19223.8168 | -0.2140248 | 0.00026352 | 0.00354217 | Ctnnb1 | 1.16 | |
| ENSMUSG0( | 837.354591 | 789.779898 | 796.545391 | 890.342647 | 970.379084 | 1057.78467 | 1007.25143 | 1007.07534 | -0.2860047 | 0.00026703 | 0.00358289 | Mis12 | 1.22 | |
| ENSMUSG0( | 2846.43727 | 2867.71449 | 2525.99872 | 2749.69647 | 3269.41605 | 3358.4934 | 3289.9956 | 3120.5431 | -0.2466364 | 0.00027556 | 0.00367087 | Frs2 | 1.19 | |
| ENSMUSG0( | 762.523129 | 750.651298 | 710.025702 | 772.802609 | 905.947475 | 936.523789 | 891.983089 | 879.949456 | -0.2705427 | 0.00028045 | 0.00372264 | Ccdc174 | 1.21 | |
| ENSMUSG0( | 11.3668044 | 0 | 9.40431394 | 5.55307264 | 25.3821491 | 25.9844751 | 25.8545823 | 29.7951283 | -2.0112119 | 0.00028882 | 0.00382012 | Stpg4 | 4.03 | |
| ENSMUSG0( | 2102.85881 | 1951.28149 | 1928.82479 | 1893.59777 | 2337.11019 | 2572.46303 | 2256.88958 | 2306.14293 | -0.2660858 | 0.0002961 | 0.00389219 | Eloa | 1.20 | |
| ENSMUSG0( | 876.191173 | 812.433298 | 855.792568 | 880.162014 | 1014.30973 | 974.417816 | 1083.73791 | 1129.23536 | -0.2949094 | 0.00029709 | 0.00389834 | Imp3 | 1.23 | |
| ENSMUSG0( | 1656.71174 | 1632.0745 | 1661.74227 | 1753.84544 | 1941.7344 | 1900.11474 | 1999.42103 | 1976.41018 | -0.2214765 | 0.0003002 | 0.00393564 | Rpl7l1 | 1.17 | |
| ENSMUSG0( | 18320.447 | 17704.6618 | 24127.7078 | 23408.9777 | 26555.5854 | 30023.9783 | 33005.5288 | 25360.62 | -0.460005 | 0.00030219 | 0.00395478 | Hsp90b1 | 1.38 | |
| ENSMUSG0( | 5971.36125 | 5214.40079 | 5094.31686 | 5824.24769 | 6573.97661 | 7566.89568 | 8154.96616 | 6292.7311 | -0.3709835 | 0.00030215 | 0.00395478 | Tmem164 | 1.29 | X |
| ENSMUSG0( | 88.0927341 | 72.0789998 | 87.4601196 | 74.9664807 | 118.124617 | 123.426257 | 136.813831 | 118.187342 | -0.6201037 | 0.00030285 | 0.00395777 | Ly96 | 1.54 | |
| ENSMUSG0( | 3670.53059 | 3238.40649 | 3297.15247 | 3292.97208 | 3872.73021 | 4144.52378 | 4162.58775 | 3876.34619 | -0.25008 | 0.0003057 | 0.00398666 | Slc25a51 | 1.19 | |
| ENSMUSG0( | 2298.93619 | 1940.98449 | 1949.51428 | 2571.99815 | 2596.7891 | 3071.58149 | 2806.29945 | 3544.6271 | -0.4559909 | 0.00030774 | 0.00400012 | Plekhg5 | 1.37 | |
| ENSMUSG0( | 5020.33861 | 5091.86649 | 5157.32576 | 5444.78773 | 5950.16149 | 5754.47854 | 6100.60414 | 6445.67942 | -0.2274243 | 0.00031215 | 0.00404603 | Matr3 | 1.17 | |
| ENSMUSG0( | 713.266976 | 700.195998 | 818.175313 | 797.791436 | 983.070159 | 981.996621 | 859.664861 | 1018.00022 | -0.3429757 | 0.0003145 | 0.00407283 | Tsr1 | 1.27 | |
| ENSMUSG0( | 46.4144513 | 26.7721999 | 31.9746674 | 59.2327748 | 76.1464473 | 121.260884 | 105.572878 | 60.5834275 | -1.1399074 | 0.00032517 | 0.00419651 | Gm21057 | 2.20 | |
| ENSMUSG0( | 1858.47252 | 1640.3121 | 1811.27086 | 1693.68716 | 1985.66505 | 2650.41646 | 2424.94436 | 2076.72044 | -0.3833087 | 0.00032674 | 0.0042131 | Ankrd50 | 1.30 | |
| ENSMUSG0( | 5649.30179 | 5589.21158 | 5901.207 | 5626.1881 | 6327.96501 | 6473.38235 | 6443.17736 | 6812.1595 | -0.1947649 | 0.00033079 | 0.00426162 | Dnm2 | 1.14 | |
| ENSMUSG0( | 2623.83735 | 2502.17099 | 2797.7834 | 2604.39107 | 2959.94908 | 3120.30238 | 3076.69529 | 3198.01044 | -0.2306492 | 0.00033213 | 0.00427515 | Ctcf | 1.17 | |
| ENSMUSG0( | 238.702892 | 248.157699 | 266.142084 | 258.217878 | 318.2531 | 368.113397 | 433.064253 | 306.889821 | -0.4947303 | 0.00033459 | 0.00430312 | Ggct | 1.41 | |
| ENSMUSG0( | 3118.29334 | 3199.27789 | 3649.81424 | 3233.7393 | 3950.82913 | 4015.68409 | 4175.51504 | 3763.1247 | -0.2686892 | 0.0003376 | 0.00433803 | Gosr2 | 1.20 | |
| ENSMUSG0( | 2528.16675 | 2247.83509 | 2711.26371 | 2919.06519 | 3074.16875 | 4233.30407 | 4313.40614 | 3060.95285 | -0.4962982 | 0.00034347 | 0.00440583 | Pigt | 1.41 | |
| ENSMUSG0( | 3475.40045 | 2965.53599 | 3416.58725 | 3055.11547 | 3749.72441 | 4155.35064 | 4161.51047 | 3769.08373 | -0.2941566 | 0.00034411 | 0.00440653 | Hif1an | 1.23 | |
| ENSMUSG0( | 7.5778696 | 10.297 | 8.46388255 | 5.55307264 | 21.4772031 | 22.7364157 | 18.3136624 | 35.754154 | -1.6317015 | 0.00035716 | 0.00456187 | Bhlha15 | 3.10 | |
| ENSMUSG0( | 10755.8387 | 10486.4648 | 13179.2056 | 10355.555 | 13239.7195 | 14921.5848 | 14936.4076 | 13626.3053 | -0.3411525 | 0.00035833 | 0.0045729 | Hdlbp | 1.27 | |
| ENSMUSG0( | 3868.50243 | 3630.72219 | 3668.62287 | 3782.56798 | 4090.43095 | 4513.71986 | 4336.0289 | 4418.61753 | -0.2153063 | 0.00036382 | 0.00463097 | Sart1 | 1.16 | |
| ENSMUSG0( | 1049.53494 | 939.086397 | 972.406061 | 1076.37058 | 1201.74714 | 1189.87242 | 1381.0656 | 1231.53197 | -0.3091795 | 0.00037135 | 0.00471877 | Inip | 1.24 | |
| ENSMUSG0( | 1772.27425 | 2185.02339 | 1942.93126 | 1523.39293 | 2155.5302 | 2805.24062 | 2893.55867 | 2420.35759 | -0.4689279 | 0.00037232 | 0.00472712 | Mafg | 1.38 | |
| ENSMUSG0( | 697.164003 | 646.651598 | 750.464252 | 707.09125 | 976.236503 | 800.105295 | 917.837671 | 852.140669 | -0.3401372 | 0.00038374 | 0.00486787 | Ccdc107 | 1.27 | |
| ENSMUSG0( | 13168.4429 | 13367.5654 | 12519.9631 | 12483.3073 | 14376.0587 | 15789.8994 | 15592.4677 | 14350.327 | -0.2218783 | 0.0003915 | 0.00494729 | Ptpn11 | 1.17 | |
| ENSMUSG0( | 2279.99152 | 2375.51789 | 2800.60469 | 2588.65736 | 3063.43015 | 2934.08031 | 3109.01352 | 3235.75093 | -0.2971018 | 0.00039874 | 0.00502594 | Kdelr2 | 1.23 | |
| ENSMUSG0( | 2812.33686 | 2603.08159 | 3071.44893 | 2808.92924 | 3468.5683 | 3388.80863 | 3210.2773 | 3415.51487 | -0.2552841 | 0.00040051 | 0.00504197 | Sec22b | 1.19 | |
| ENSMUSG0( | 2419.23487 | 2407.43859 | 2434.77688 | 2296.19554 | 2643.64845 | 2765.18122 | 2796.60398 | 2830.53719 | -0.2074654 | 0.00040152 | 0.0050504 | Gsk3a | 1.15 | |
| ENSMUSG0( | 935.866896 | 891.720198 | 981.810375 | 987.521418 | 1166.60262 | 1122.74586 | 1088.047 | 1189.81879 | -0.2664076 | 0.00040318 | 0.00506696 | Gpr180 | 1.20 | |
| ENSMUSG0( | 947.2337 | 1064.7098 | 1054.22359 | 1079.14712 | 1206.62832 | 1289.47958 | 1241.01995 | 1248.41588 | -0.2661705 | 0.00040547 | 0.00508723 | Prcc | 1.20 | |
| ENSMUSG0( | 1790.27169 | 1870.96489 | 1851.70941 | 1616.86965 | 2220.93804 | 2118.81741 | 2053.28474 | 2528.61322 | -0.3238154 | 0.00040815 | 0.00511645 | Tshz2 | 1.25 | |
| ENSMUSG0( | 1229.50934 | 1342.7288 | 1320.36568 | 1167.07077 | 1543.42991 | 1583.97029 | 1502.79759 | 1464.92714 | -0.2688458 | 0.00041216 | 0.00516245 | Tatdn2 | 1.20 | |
| ENSMUSG0( | 5083.80327 | 4979.62919 | 4730.36991 | 4692.34638 | 5276.5583 | 6086.86329 | 6266.50438 | 5767.34367 | -0.2638004 | 0.00042511 | 0.00531115 | Marf1 | 1.20 | |
| ENSMUSG0( | 1339.38845 | 816.552098 | 931.967511 | 1107.83799 | 1571.74077 | 2458.78095 | 1829.2117 | 1154.06464 | -0.7409169 | 0.00042724 | 0.00532444 | Plxnd1 | 1.67 | |
| ENSMUSG0( | 1777.95766 | 1506.4511 | 2099.9833 | 1854.72626 | 2226.79546 | 2472.85588 | 2450.79895 | 2240.59365 | -0.3750588 | 0.00042862 | 0.00533708 | Lrrc8d | 1.30 | |
| ENSMUSG0( | 4488.94051 | 4636.73909 | 4830.99607 | 4864.49163 | 5415.18388 | 5468.64932 | 5263.56204 | 5383.97968 | -0.1940413 | 0.00043175 | 0.00537161 | Nono | 1.14 | X |
| ENSMUSG0( | 1640.60877 | 1606.332 | 1671.14659 | 1451.20298 | 2215.08063 | 1773.44042 | 1947.71187 | 2012.16433 | -0.3199009 | 0.00043609 | 0.00540581 | Btg3 | 1.25 | |
| ENSMUSG0( | 1128.15534 | 1191.3629 | 1299.67619 | 1173.54935 | 1515.11905 | 1383.6733 | 1395.07017 | 1508.62666 | -0.2761329 | 0.00043632 | 0.00540581 | Mydgf | 1.21 | |
| ENSMUSG0( | 3477.29491 | 3691.47449 | 3518.15384 | 3270.75979 | 3963.5202 | 4508.30643 | 4487.92457 | 3932.95693 | -0.2752678 | 0.00043774 | 0.00541895 | Cggbp1 | 1.21 | |
| ENSMUSG0( | 1407.58928 | 1298.4517 | 1301.55705 | 1259.62198 | 1548.31109 | 1551.4897 | 1666.54328 | 1514.58569 | -0.2537305 | 0.00043987 | 0.00543631 | Aptx | 1.19 | |
| ENSMUSG0( | 178.079936 | 148.2768 | 126.017807 | 161.039107 | 198.17601 | 239.273708 | 249.927629 | 210.55224 | -0.548251 | 0.00044332 | 0.00546493 | Rab43 | 1.46 | |
| ENSMUSG0( | 14035.1617 | 11590.3032 | 13220.5845 | 15445.8716 | 16841.0559 | 18229.192 | 16926.1332 | 16319.7849 | -0.3314298 | 0.00044366 | 0.00546493 | Nfic | 1.26 | |
| ENSMUSG0( | 3834.40202 | 4007.59239 | 4073.00837 | 3478.0745 | 4375.49201 | 4840.69117 | 5008.24804 | 4482.18047 | -0.2812162 | 0.00044769 | 0.00550552 | Golga3 | 1.22 | |
| ENSMUSG0( | 4375.27246 | 4575.98679 | 4116.26821 | 4158.3259 | 4861.65779 | 5202.30845 | 5547.96245 | 4916.19617 | -0.2530041 | 0.00044863 | 0.0055085 | Senp6 | 1.19 | |
| ENSMUSG0( | 3088.9291 | 3185.89179 | 2980.22709 | 3120.82682 | 3373.87336 | 3714.69725 | 3587.32329 | 3737.30226 | -0.2198442 | 0.00044868 | 0.0055085 | Ubap2 | 1.16 | |
| ENSMUSG0( | 3876.0803 | 3728.54369 | 3700.59754 | 3583.58288 | 4360.84846 | 4361.06107 | 4406.05173 | 4071.0077 | -0.2080451 | 0.00045185 | 0.00553837 | Nucks1 | 1.16 | |
| ENSMUSG0( | 1012.59283 | 1085.3038 | 1255.47591 | 1197.61267 | 1490.71314 | 1404.24434 | 1341.20646 | 1437.11835 | -0.3179857 | 0.0004566 | 0.00558279 | Srprb | 1.25 | |
| ENSMUSG0( | 256.700333 | 265.662599 | 242.6313 | 327.631286 | 536.930077 | 334.550117 | 339.341392 | 490.626446 | -0.6398176 | 0.00046174 | 0.00563639 | Tmem37 | 1.56 | |
| ENSMUSG0( | 6990.58471 | 7637.28488 | 7530.9746 | 7043.14713 | 8745.1266 | 8138.55413 | 8286.39362 | 8991.17655 | -0.2264223 | 0.00046347 | 0.0056452 | Rps3a1 | 1.17 | |
| ENSMUSG0( | 54.9395546 | 41.1879999 | 49.8428639 | 20.3612664 | 81.0276298 | 131.005062 | 54.9409873 | 124.146368 | -1.2342838 | 0.00048278 | 0.00586431 | Tmem182 | 2.35 | |
| ENSMUSG0( | 292.695213 | 256.395299 | 277.427261 | 161.964619 | 354.373851 | 475.299357 | 330.723198 | 415.145454 | -0.6727775 | 0.00048914 | 0.0059174 | Itpkb | 1.59 | |
| ENSMUSG0( | 972.80901 | 951.442797 | 968.644336 | 940.320301 | 1281.79853 | 1120.58049 | 1182.84714 | 1077.59047 | -0.2828258 | 0.00048899 | 0.0059174 | Atg101 | 1.22 | |
| ENSMUSG0( | 661.169123 | 359.365299 | 792.783665 | 477.564247 | 960.616719 | 1245.08943 | 733.623772 | 1006.08217 | -0.7838731 | 0.00049056 | 0.00592973 | Ptn | 1.72 | |
| ENSMUSG0( | 4308.01887 | 3665.73199 | 4244.16688 | 4259.20672 | 5209.19798 | 5223.96218 | 5165.53008 | 4536.80487 | -0.2890902 | 0.00049307 | 0.00594647 | Fez2 | 1.22 | |
| ENSMUSG0( | 1605.56112 | 1546.6094 | 1823.49647 | 1809.37617 | 2109.64708 | 1946.67026 | 2287.05326 | 2060.82971 | -0.3084759 | 0.00049648 | 0.00598191 | Mcrip1 | 1.24 | |
| ENSMUSG0( | 22.7336088 | 24.7127999 | 8.46388255 | 12.9571695 | 46.8593522 | 43.3074585 | 57.0955359 | 29.7951283 | -1.3666425 | 0.00051345 | 0.00613747 | Gm15691 | 2.58 | |
| ENSMUSG0( | 5936.3136 | 6164.81388 | 6009.35661 | 5740.02609 | 6557.38059 | 7183.62467 | 7485.97884 | 6741.64436 | -0.2297243 | 0.00051357 | 0.00613747 | AW549877 | 1.17 | |
| ENSMUSG0( | 826.93502 | 786.690798 | 757.987703 | 822.780263 | 937.187043 | 1005.81572 | 929.687688 | 959.403131 | -0.2623035 | 0.00052162 | 0.00621434 | Reps1 | 1.20 | |
| ENSMUSG0( | 1662.39514 | 1997.61799 | 1352.34034 | 1362.35382 | 1670.34066 | 2922.17076 | 2519.7445 | 2597.14202 | -0.6070797 | 0.00052373 | 0.00623454 | Etv6 | 1.52 | |
| ENSMUSG0( | 95.6706037 | 94.7323997 | 95.9240022 | 91.6256986 | 396.35202 | 206.793114 | 233.768515 | 71.5083079 | -1.2654102 | 0.00052807 | 0.00627365 | Fhod3 | 2.40 | |
| ENSMUSG0( | 1811.11084 | 1810.21259 | 1922.24177 | 1798.27002 | 2074.50257 | 2072.26189 | 2181.48038 | 2278.33414 | -0.2293132 | 0.00054577 | 0.00644014 | Fam114a2 | 1.17 | |
| ENSMUSG0( | 326.795627 | 309.939699 | 323.5084 | 363.726258 | 410.995568 | 459.05906 | 398.591477 | 436.002044 | -0.3637165 | 0.00055114 | 0.00648406 | Armc7 | 1.29 | |
| ENSMUSG0( | 1112.05236 | 958.650697 | 1118.17293 | 1061.56239 | 1341.34896 | 1274.32197 | 1305.65641 | 1230.5388 | -0.2770526 | 0.00055123 | 0.00648406 | Fitm2 | 1.21 | |
| ENSMUSG0( | 867.66607 | 904.076597 | 807.830567 | 841.290505 | 859.088123 | 1347.94464 | 1379.98833 | 1223.5866 | -0.4915401 | 0.00055597 | 0.00653466 | Plcd3 | 1.41 | |
| ENSMUSG0( | 2311.25023 | 2532.03229 | 2299.35476 | 1964.8622 | 2857.44425 | 2909.17852 | 2776.13577 | 2703.41131 | -0.3045323 | 0.00055801 | 0.00655338 | Bsdc1 | 1.24 | |
| ENSMUSG0( | 981.334114 | 917.462697 | 1038.23626 | 1111.54004 | 1167.57886 | 1530.91866 | 1358.44284 | 1213.65489 | -0.3797981 | 0.00055979 | 0.00656394 | Tacstd2 | 1.30 | |
| ENSMUSG0( | 28.417011 | 14.4158 | 10.3447453 | 21.2867785 | 27.3346221 | 85.5322305 | 105.572878 | 28.8019574 | -1.7272088 | 0.00056085 | 0.00656891 | Gm21009 | 3.31 | |
| ENSMUSG0( | 720.844846 | 794.928398 | 789.021939 | 770.951585 | 886.422745 | 993.906172 | 901.678557 | 1118.31048 | -0.3428985 | 0.00056317 | 0.0065829 | Trim27 | 1.27 | |
| ENSMUSG0( | 1856.57805 | 1836.98479 | 1890.2671 | 1878.78958 | 2065.71644 | 2210.84575 | 2126.53939 | 2180.01022 | -0.2015972 | 0.00056409 | 0.00658839 | Hmg20b | 1.15 | |
| ENSMUSG0( | 1091.21322 | 1171.7986 | 1085.25783 | 1085.6257 | 1291.56089 | 1249.42018 | 1343.361 | 1406.33006 | -0.255367 | 0.00056511 | 0.00659519 | Vps72 | 1.19 | |
| ENSMUSG0( | 107.037408 | 102.97 | 106.268748 | 102.731844 | 161.079023 | 138.583867 | 149.741122 | 147.982471 | -0.5124343 | 0.00057263 | 0.00667765 | Gm14584 | 1.43 | X |
| ENSMUSG0( | 11.3668044 | 5.14849999 | 10.3447453 | 4.62756053 | 14.6435475 | 48.7208908 | 21.5454852 | 26.8156155 | -1.8208883 | 0.00059387 | 0.00688764 | Cd40 | 3.53 | |
| ENSMUSG0( | 10589.1255 | 10328.9207 | 11214.6444 | 9852.07638 | 11929.6101 | 13406.9065 | 13082.4186 | 11699.5537 | -0.2554157 | 0.00059825 | 0.00692049 | Trim44 | 1.19 | |
| ENSMUSG0( | 309.74542 | 226.533999 | 312.223223 | 320.227189 | 473.474704 | 530.516366 | 456.764287 | 316.821531 | -0.6035989 | 0.00059959 | 0.00692162 | Kcnk1 | 1.52 | |
| ENSMUSG0( | 61.5701905 | 29.8612999 | 67.7110604 | 16.6592179 | 91.7662313 | 188.387444 | 75.4091983 | 112.228317 | -1.4105687 | 0.00063174 | 0.00727581 | Art3 | 2.66 | |
| ENSMUSG0( | 2625.73182 | 2568.07179 | 2620.04186 | 2317.48232 | 2922.85209 | 2929.74957 | 3546.38687 | 3080.81627 | -0.300714 | 0.00063431 | 0.00728849 | Dmwd | 1.23 | |
| ENSMUSG0( | 4409.37288 | 5391.50919 | 4129.43425 | 5204.15458 | 5921.85063 | 6331.55043 | 5893.76749 | 9018.98533 | -0.5057083 | 0.00063603 | 0.00729691 | Kcnf1 | 1.42 | |
| ENSMUSG0( | 53.0450872 | 43.2473999 | 42.3194127 | 27.7653632 | 61.5028997 | 105.020587 | 112.036523 | 61.5765985 | -1.0299081 | 0.00063847 | 0.00731923 | Ifi44 | 2.04 | |
| ENSMUSG0( | 1055.21834 | 956.591297 | 1057.98532 | 1079.14712 | 1160.7452 | 1390.16942 | 1261.48816 | 1268.27929 | -0.2915343 | 0.00064532 | 0.00738647 | Zkscan17 | 1.22 | |
| ENSMUSG0( | 4399.90054 | 4338.12609 | 4474.57257 | 4646.99629 | 5077.40605 | 4978.19235 | 5002.86167 | 5201.23623 | -0.1818858 | 0.00065415 | 0.00747028 | Ube2q1 | 1.13 | |
| ENSMUSG0( | 1640.60877 | 2000.70709 | 1745.44067 | 1498.4041 | 2012.02343 | 2402.48126 | 2269.81687 | 2141.27655 | -0.3583361 | 0.00065613 | 0.00748717 | Ppard | 1.28 | |
| ENSMUSG0( | 4418.84521 | 4167.19589 | 4201.84747 | 4152.77282 | 4568.78684 | 5337.64426 | 5252.7893 | 4899.31226 | -0.2435459 | 0.00065697 | 0.00749093 | Sp1 | 1.18 | |
| ENSMUSG0( | 965.231141 | 979.244697 | 1030.71281 | 1104.13594 | 1197.84219 | 1194.20317 | 1276.57 | 1221.60026 | -0.2611451 | 0.00066556 | 0.0075715 | Surf6 | 1.20 | |
| ENSMUSG0( | 4919.93184 | 5116.57929 | 5554.18781 | 5180.09126 | 6123.93158 | 5946.11405 | 5876.5311 | 5997.75933 | -0.2051397 | 0.00066738 | 0.00757485 | Rab2a | 1.15 | |
| ENSMUSG0( | 3478.24215 | 3746.04859 | 3712.82314 | 3655.77282 | 4181.22094 | 4184.58317 | 4108.72403 | 4099.80965 | -0.1837568 | 0.00066707 | 0.00757485 | Kpna4 | 1.14 | |
| ENSMUSG0( | 463.197279 | 388.196899 | 364.887381 | 470.16015 | 461.759866 | 734.061421 | 726.082852 | 565.114267 | -0.5593897 | 0.0006765 | 0.00766665 | Arid3b | 1.47 | |
| ENSMUSG0( | 2774.44751 | 3221.93129 | 3003.73787 | 2690.46369 | 3230.36659 | 3859.77724 | 3824.32363 | 3552.57246 | -0.3074183 | 0.00067868 | 0.0076855 | Uck2 | 1.24 | |
| ENSMUSG0( | 363.737741 | 433.503699 | 376.172558 | 315.599628 | 453.949974 | 644.198445 | 734.701046 | 442.954241 | -0.6117145 | 0.00068347 | 0.00771626 | Gm43595 | 1.53 | |
| ENSMUSG0( | 1148.04724 | 913.343897 | 1220.67995 | 880.162014 | 1364.77863 | 1667.33715 | 1493.10213 | 1221.60026 | -0.4649975 | 0.00070403 | 0.0078826 | Tbc1d16 | 1.38 | |
| ENSMUSG0( | 2309.35576 | 2542.32929 | 2217.53723 | 2393.37431 | 2706.12759 | 2990.38001 | 2870.93591 | 2684.54106 | -0.2498572 | 0.00071115 | 0.00793843 | Foxo4 | 1.19 | X |
| ENSMUSG0( | 2028.02735 | 2195.32039 | 2188.38385 | 1875.08753 | 2442.54373 | 2735.94869 | 2611.31281 | 2336.93123 | -0.2892286 | 0.00071808 | 0.00800376 | Elf2 | 1.22 | |
| ENSMUSG0( | 15958.0461 | 14119.2464 | 14698.0023 | 13224.6425 | 17815.3399 | 17674.8565 | 17727.6252 | 16173.7888 | -0.2586971 | 0.00072361 | 0.00804731 | Ddx3x | 1.20 | X |
| ENSMUSG0( | 7525.77175 | 7138.91008 | 5949.169 | 6132.44322 | 7990.49578 | 9007.95136 | 8250.84357 | 8089.37733 | -0.3178729 | 0.00073523 | 0.00813399 | Hnrnpa1 | 1.25 | |
| ENSMUSG0( | 1899.20357 | 1364.3525 | 937.6101 | 1203.16574 | 1794.32269 | 2469.60782 | 2123.30757 | 1957.53993 | -0.6266936 | 0.0007429 | 0.00818239 | Vtcn1 | 1.54 | |
| ENSMUSG0( | 248.175229 | 188.435099 | 151.409454 | 203.612664 | 299.704607 | 286.911912 | 256.391274 | 314.835189 | -0.5488835 | 0.00075707 | 0.00832817 | 2010300C02l | 1.46 | |
| ENSMUSG0( | 2382.29276 | 2395.08219 | 2394.33833 | 2234.18623 | 2823.27597 | 2774.9254 | 2762.13121 | 2563.3742 | -0.2158596 | 0.00076407 | 0.00837699 | Cers5 | 1.16 | |
| ENSMUSG0( | 6091.65993 | 6509.76338 | 5931.3008 | 6282.37618 | 6925.42175 | 7217.18795 | 7240.36031 | 6943.25806 | -0.1909313 | 0.00076451 | 0.00837699 | Bnip2 | 1.14 | |
| ENSMUSG0( | 174.291001 | 160.6332 | 193.728867 | 203.612664 | 243.082889 | 305.317582 | 241.309435 | 237.367855 | -0.4859667 | 0.00077636 | 0.008488 | Bcl2l11 | 1.40 | |
| ENSMUSG0( | 5255.25257 | 5014.63899 | 5273.93926 | 4845.05588 | 6066.33363 | 5948.27942 | 5748.33546 | 5654.12218 | -0.1997802 | 0.00078013 | 0.00851679 | Cul4a | 1.15 | |
| ENSMUSG0( | 4045.63513 | 3991.11719 | 5435.69346 | 5216.18623 | 5888.65859 | 6665.01786 | 7214.50573 | 5509.11922 | -0.4355433 | 0.00078136 | 0.00852394 | Dnajb11 | 1.35 | |
| ENSMUSG0( | 326.795627 | 540.592499 | 568.960993 | 285.057729 | 985.998868 | 593.312181 | 816.57389 | 592.923053 | -0.7965302 | 0.00078624 | 0.00857092 | Gm8116 | 1.74 | |
| ENSMUSG0( | 1718.28193 | 1843.16299 | 1646.69537 | 1579.84917 | 2040.33429 | 2355.92574 | 2022.04379 | 1937.67651 | -0.2998604 | 0.00079522 | 0.00865604 | Zfp646 | 1.23 | |
| ENSMUSG0( | 3056.72315 | 2872.86299 | 3524.73686 | 3034.7542 | 3618.90872 | 3908.49813 | 3882.49644 | 3676.71883 | -0.2723822 | 0.00080658 | 0.0087669 | Srp72 | 1.21 | |
| ENSMUSG0( | 7238.75994 | 7371.62228 | 8086.76956 | 7581.79518 | 8860.3225 | 8567.29797 | 8440.44384 | 9124.26145 | -0.2087453 | 0.00081158 | 0.00881477 | Pcbp1 | 1.16 | |
| ENSMUSG0( | 610.018503 | 634.295198 | 671.468015 | 630.273745 | 700.937809 | 837.999321 | 874.7467 | 775.666507 | -0.3242192 | 0.00081457 | 0.00884084 | Ccdc167 | 1.25 | |
| ENSMUSG0( | 4869.72845 | 4896.22349 | 5310.61608 | 4501.69089 | 5955.04267 | 6098.77284 | 6430.25007 | 5315.45089 | -0.2816123 | 0.00081879 | 0.00887365 | Atf6 | 1.22 | |
| ENSMUSG0( | 1477.68457 | 1590.8865 | 1697.47867 | 1592.80634 | 1867.54043 | 1840.56698 | 1836.75262 | 1918.80626 | -0.2312029 | 0.00083182 | 0.00897565 | Triobp | 1.17 | |
| ENSMUSG0( | 3541.70681 | 2542.32929 | 4547.92622 | 2764.50466 | 4790.39252 | 6105.26896 | 4832.65234 | 4314.33458 | -0.5811095 | 0.00084104 | 0.00905547 | Epas1 | 1.50 | |
| ENSMUSG0( | 156.293561 | 147.2471 | 144.826435 | 167.517691 | 185.484936 | 247.9352 | 218.686675 | 208.565898 | -0.4807123 | 0.0008432 | 0.00906559 | Tmem86b | 1.40 | |
| ENSMUSG0( | 284.17011 | 281.108099 | 249.214319 | 267.472999 | 329.967938 | 331.302057 | 353.345958 | 402.234232 | -0.3898077 | 0.00085506 | 0.00917984 | Pmf1 | 1.31 | |
| ENSMUSG0( | 705.689107 | 695.047498 | 714.727859 | 681.176911 | 865.921778 | 754.632464 | 918.914945 | 899.812875 | -0.2986979 | 0.00085724 | 0.00919666 | Thoc3 | 1.23 | |
| ENSMUSG0( | 4408.42564 | 3835.63249 | 5266.41581 | 3534.53074 | 6521.25984 | 5232.62367 | 5623.37165 | 5541.89386 | -0.4272184 | 0.0008626 | 0.0092408 | Tgfa | 1.34 | |
| ENSMUSG0( | 811.779281 | 736.235498 | 817.234881 | 881.087526 | 980.141449 | 1009.06378 | 1028.79692 | 935.567028 | -0.2834546 | 0.00087207 | 0.00932218 | Vhl | 1.22 | |
| ENSMUSG0( | 2663.62117 | 2638.09139 | 2438.5386 | 2438.7244 | 3052.69155 | 3732.02023 | 3608.86878 | 2712.34985 | -0.3644671 | 0.00087579 | 0.00935518 | Kdm7a | 1.29 | |
| ENSMUSG0( | 5139.69006 | 5088.77739 | 5077.3891 | 5050.51957 | 5463.01947 | 5854.0857 | 5768.80367 | 5928.23736 | -0.1769837 | 0.00087974 | 0.00939062 | Supt5 | 1.13 | |
| ENSMUSG0( | 769.153765 | 817.581798 | 795.604959 | 680.251399 | 1147.07789 | 937.606476 | 890.905814 | 953.444105 | -0.3602254 | 0.00088762 | 0.00945443 | Snap29 | 1.28 | |
| ENSMUSG0( | 139.243354 | 167.8411 | 186.205416 | 145.305401 | 1109.9809 | 256.596691 | 273.627662 | 436.995215 | -1.7020017 | 0.00089189 | 0.00948624 | Klhl30 | 3.25 | |
| ENSMUSG0( | 979.439646 | 1119.2839 | 1035.41496 | 1017.13781 | 1193.93724 | 1284.06614 | 1339.05191 | 1188.82562 | -0.2700305 | 0.0008959 | 0.00951538 | Esco1 | 1.21 | |
| ENSMUSG0( | 1447.37309 | 1482.768 | 1665.504 | 1522.46742 | 1734.77227 | 1860.05534 | 1830.28897 | 1808.56429 | -0.241337 | 0.00091999 | 0.00974337 | Cacfd1 | 1.18 | |
| ENSMUSG0( | 7706.69339 | 7655.81948 | 8892.71926 | 7888.13969 | 9604.21472 | 9220.15791 | 9162.2176 | 9881.05771 | -0.2364471 | 0.00092143 | 0.00975174 | Arcn1 | 1.18 | |
| ENSMUSG0( | 47.361685 | 53.5443999 | 63.9493348 | 56.4562385 | 68.3365552 | 150.493418 | 101.263781 | 89.3853849 | -0.8847902 | 0.00092462 | 0.00976464 | Plk2 | 1.85 | |
| ENSMUSG0( | 3976.48707 | 4339.15579 | 4524.41544 | 4199.97394 | 4622.47984 | 5074.55145 | 5467.16688 | 5162.50256 | -0.2543713 | 0.00092448 | 0.00976464 | Stk11 | 1.19 | |
| ENSMUSG0( | 5068.64753 | 5006.40139 | 5430.9913 | 5409.61827 | 6131.74148 | 5825.93585 | 5868.99018 | 6107.00813 | -0.1944447 | 0.00092918 | 0.00980587 | Cdc37 | 1.14 | |
| ENSMUSG0( | 19.8919077 | 14.4158 | 19.7490593 | 10.1806332 | 32.2158046 | 36.8113397 | 35.5500506 | 36.7473249 | -1.1366324 | 0.00093478 | 0.00985791 | Asb10 | 2.20 | |
| ENSMUSG0( | 4828.9974 | 4896.22349 | 5415.9444 | 5103.27376 | 5712.93602 | 5876.82211 | 5722.48088 | 6192.42083 | -0.2153888 | 0.00093988 | 0.00988373 | Set | 1.16 | |
| ENSMUSG0( | 1152.78341 | 1358.1743 | 1598.73337 | 1525.24395 | 1943.68688 | 1757.20013 | 2031.73926 | 1658.59548 | -0.3912615 | 0.00094921 | 0.00996773 | Fkbp2 | 1.31 | |
| ENSMUSG0( | 3438.45833 | 3719.27639 | 3986.48868 | 3732.59033 | 4205.62686 | 4318.8363 | 4314.48342 | 4600.36781 | -0.2292859 | 0.00096078 | 0.01007635 | Retreg2 | 1.17 | |
| ENSMUSG0( | 1915.30654 | 1786.5295 | 1666.44443 | 1646.48604 | 1993.47494 | 2590.8687 | 2206.25769 | 2070.76142 | -0.3369294 | 0.00096642 | 0.01009858 | Zbtb10 | 1.26 | |
| ENSMUSG0( | 9515.90975 | 9183.89427 | 9719.35846 | 9399.50096 | 12070.1881 | 10035.4208 | 10941.8747 | 12167.3372 | -0.2577573 | 0.00096604 | 0.01009858 | Cnbp | 1.20 | |
| ENSMUSG0( | 96.6178374 | 89.5838998 | 142.945572 | 112.912477 | 183.532463 | 254.431319 | 130.350186 | 180.757112 | -0.7594799 | 0.00097428 | 0.01015924 | Wdr86 | 1.69 | |
| ENSMUSG0( | 5793.28131 | 6017.56678 | 5341.65032 | 5156.95346 | 6111.24051 | 7170.63244 | 6787.90512 | 6741.64436 | -0.2652039 | 0.00098099 | 0.01022213 | Ptpn21 | 1.20 | |
| ENSMUSG0( | 399.732622 | 452.038299 | 324.448831 | 359.098698 | 485.189542 | 489.374281 | 514.937097 | 533.332796 | -0.3989883 | 0.00098724 | 0.01026563 | Fbxo44 | 1.32 | |
| ENSMUSG0( | 500.139394 | 507.642099 | 389.338597 | 481.266296 | 611.124051 | 674.513666 | 646.364557 | 540.284993 | -0.3962286 | 0.00099315 | 0.01029827 | Pdik1l | 1.32 | |
| ENSMUSG0( | 1383.90844 | 1269.6201 | 1227.26297 | 1384.56611 | 2308.79933 | 1419.40195 | 1574.97497 | 2070.76142 | -0.4862048 | 0.0009964 | 0.01031765 | Egln3 | 1.40 | |
| ENSMUSG0( | 220.705452 | 276.989299 | 314.104086 | 260.068902 | 341.682776 | 380.022948 | 373.814169 | 348.603001 | -0.4297085 | 0.00100767 | 0.01039814 | Mfhas1 | 1.35 | |
| ENSMUSG0( | 95.6706037 | 88.5541998 | 58.3067464 | 75.8919928 | 102.504833 | 118.012824 | 150.818397 | 155.927838 | -0.7282821 | 0.00101745 | 0.01047726 | Rasip1 | 1.66 | |
| ENSMUSG0( | 4134.6751 | 4127.03759 | 4423.78928 | 4038.00932 | 4925.11316 | 4956.53862 | 4898.36607 | 4525.87999 | -0.2070828 | 0.00101696 | 0.01047726 | Rab5a | 1.15 | |
| ENSMUSG0( | 1872.68103 | 1773.1434 | 1834.78165 | 1869.53446 | 2020.80956 | 2216.25919 | 2093.14389 | 2162.13314 | -0.2080694 | 0.00101838 | 0.01047963 | Gid8 | 1.16 | |
| ENSMUSG0( | 3918.70582 | 4079.67139 | 4234.76257 | 4237.91994 | 4850.91918 | 4924.05803 | 4557.9474 | 4594.40878 | -0.2004965 | 0.00102138 | 0.01050323 | Zbtb7a | 1.15 | |
| ENSMUSG0( | 1598.93049 | 1639.2824 | 1595.91208 | 1430.84172 | 1824.58602 | 1941.25683 | 1925.08911 | 1754.93306 | -0.2491157 | 0.00102765 | 0.01054583 | Osbpl11 | 1.19 | |
| ENSMUSG0( | 6416.56109 | 5511.98408 | 6241.64316 | 6191.676 | 6757.50908 | 8373.49709 | 8551.40309 | 6931.34001 | -0.3294057 | 0.00104327 | 0.010684 | 6530402F18l | 1.26 | |
| ENSMUSG0( | 2911.79639 | 2335.35959 | 2565.49684 | 2092.58287 | 3537.88109 | 3279.45729 | 3408.49576 | 2760.02205 | -0.3906322 | 0.00105091 | 0.01074749 | Gramd4 | 1.31 | |
| ENSMUSG0( | 6085.02929 | 6504.61488 | 6694.93109 | 5974.18065 | 7481.87656 | 7242.08974 | 7336.23772 | 7046.54784 | -0.2046224 | 0.00105985 | 0.01081658 | Rab10 | 1.15 | |
| ENSMUSG0( | 92540.9436 | 85102.6454 | 108349.922 | 98385.639 | 123052.659 | 114386.907 | 115515.042 | 112745.759 | -0.2768675 | 0.00106252 | 0.01082893 | Npnt | 1.21 | |
| ENSMUSG0( | 2037.49969 | 1974.96459 | 2261.7375 | 2313.78027 | 2404.47051 | 2609.27437 | 2622.08555 | 2655.7391 | -0.2607428 | 0.00106195 | 0.01082893 | Arf2 | 1.20 | |
| ENSMUSG0( | 1305.28804 | 1187.2441 | 1317.54438 | 1317.92924 | 1454.59239 | 1486.52851 | 1531.884 | 1536.43545 | -0.2283603 | 0.00107819 | 0.01095868 | Pgs1 | 1.17 | |
| ENSMUSG0( | 163.87143 | 146.2174 | 136.362552 | 139.752328 | 210.867085 | 242.521767 | 218.686675 | 166.852718 | -0.5159792 | 0.00109399 | 0.01110411 | Asic1 | 1.43 | |
| ENSMUSG0( | 7153.50891 | 7414.86968 | 8404.63537 | 7570.68903 | 9367.96549 | 8588.9517 | 8883.20356 | 9034.87607 | -0.2321201 | 0.00110102 | 0.01116782 | Sar1a | 1.17 | |
| ENSMUSG0( | 3722.62844 | 3754.28619 | 4909.05188 | 3760.35569 | 5099.85949 | 5196.89502 | 4835.88416 | 5213.15428 | -0.3334899 | 0.00110709 | 0.01122171 | Uso1 | 1.26 | |
| ENSMUSG0( | 7536.19132 | 6980.33628 | 7424.70585 | 7504.05216 | 8766.6038 | 8421.1353 | 8868.12172 | 7920.53827 | -0.2064097 | 0.00111379 | 0.01127426 | Nab1 | 1.15 | |
| ENSMUSG0( | 4469.0486 | 4149.69099 | 4809.36615 | 4758.98325 | 5164.2911 | 5325.73471 | 5470.3987 | 5286.64893 | -0.2241519 | 0.0011237 | 0.01135918 | Ttc3 | 1.17 | |
| ENSMUSG0( | 3704.631 | 3677.05869 | 3435.39588 | 3342.94973 | 4032.83299 | 4175.92168 | 4007.46025 | 4056.11013 | -0.2006925 | 0.00114068 | 0.01150732 | Gak | 1.15 | |
| ENSMUSG0( | 6536.85977 | 6427.38738 | 6705.27584 | 6760.86594 | 7224.15012 | 9278.62298 | 9690.08198 | 7404.08938 | -0.3460207 | 0.00114486 | 0.0115417 | Purb | 1.27 | |
| ENSMUSG0( | 458.461111 | 530.295499 | 645.135936 | 302.642459 | 746.820925 | 1198.53391 | 565.568987 | 826.318225 | -0.7852534 | 0.00116213 | 0.01169199 | Tinagl1 | 1.72 | |
| ENSMUSG0( | 14.2085055 | 5.14849999 | 7.52345115 | 7.40409686 | 24.4059126 | 31.3979074 | 33.3955021 | 13.9043932 | -1.5806714 | 0.00117218 | 0.01178517 | Gm43820 | 2.99 | |
| ENSMUSG0( | 157.240794 | 151.3659 | 99.6857278 | 80.5195533 | 196.223537 | 253.348632 | 433.064253 | 136.064419 | -1.0593011 | 0.00117435 | 0.01179899 | A430050A11l | 2.08 | |
| ENSMUSG0( | 1689.86492 | 1545.5797 | 1657.98055 | 1584.47673 | 1726.96237 | 1972.65473 | 2002.65285 | 2088.63849 | -0.265991 | 0.00119708 | 0.01196694 | Unc119b | 1.20 | |
| ENSMUSG0( | 2650.35989 | 2619.55679 | 3013.14219 | 2629.3799 | 3286.01207 | 3150.6176 | 3197.35001 | 3149.34506 | -0.2282419 | 0.00119806 | 0.01196694 | Ppp1r7 | 1.17 | |
| ENSMUSG0( | 4951.19055 | 4769.57039 | 4937.26482 | 4957.96836 | 5242.39002 | 5805.36481 | 5611.52163 | 5678.95145 | -0.1873259 | 0.0011983 | 0.01196694 | Ppp6r1 | 1.14 | |
| ENSMUSG0( | 955.758804 | 999.838697 | 1219.73952 | 1868.60894 | 2283.41718 | 1890.37056 | 1545.88857 | 2196.89413 | -0.6502401 | 0.00120725 | 0.0120321 | Paqr7 | 1.57 | |
| ENSMUSG0( | 13124.8702 | 13133.8235 | 12681.7173 | 12229.717 | 14366.2964 | 14398.6473 | 14415.0069 | 14090.1162 | -0.1625062 | 0.00121837 | 0.01211858 | Csde1 | 1.12 | |
| ENSMUSG0( | 1828.16104 | 2061.45939 | 2002.17844 | 1821.40783 | 2238.5103 | 2703.46809 | 2645.78559 | 2110.48825 | -0.3302301 | 0.00122122 | 0.01213072 | Retreg3 | 1.26 | |
| ENSMUSG0( | 2386.08169 | 2245.77569 | 2393.3979 | 2346.17319 | 2976.5451 | 2511.83259 | 2773.98122 | 2775.91279 | -0.2362946 | 0.00122535 | 0.01215546 | Snx2 | 1.18 | |
| ENSMUSG0( | 15989.3049 | 16324.8638 | 19700.1568 | 18074.3259 | 21247.7875 | 23125.1001 | 25174.8222 | 19154.2948 | -0.3397476 | 0.00122801 | 0.01217372 | Canx | 1.27 | |
| ENSMUSG0( | 42823.4884 | 37465.6344 | 40941.6807 | 42227.4154 | 44731.1566 | 46895.4814 | 53721.5129 | 59275.4214 | -0.3240444 | 0.0012341 | 0.01222598 | Sep-11 | 1.25 | |

[illegible]

[illegible]

| | | | | | | | | | | |
|---|---|---|---|---|---|---|---|---|---|---|
| ENSMUSG0C | 1437.90076 | 1354.0555 | 1222.56081 | 1259.62198 | 1512.19034 | 1705.23118 | 1605.13865 | 1534.44911 | -0.2692823 | 0.00123761 |
| ENSMUSG0C | 9492.22891 | 9337.31957 | 8795.85483 | 9474.46744 | 10329.5584 | 12773.5349 | 13236.4689 | 10184.968 | -0.3265088 | 0.00124593 |
| ENSMUSG0C | 779.573335 | 748.591898 | 722.251311 | 779.281194 | 922.543496 | 908.373941 | 1024.48782 | 840.222618 | -0.2861659 | 0.00124667 |
| ENSMUSG0C | 31.2587121 | 37.0691999 | 44.2002755 | 47.2011175 | 95.6711773 | 84.449544 | 88.3364895 | 42.7063506 | -0.9601114 | 0.00124883 |
| ENSMUSG0C | 966.178374 | 883.482598 | 821.937038 | 954.202982 | 1016.2622 | 1160.63989 | 1283.03365 | 1086.52901 | -0.3258668 | 0.0012509 |
| ENSMUSG0C | 791.887373 | 605.463598 | 1019.42763 | 811.674118 | 1095.33736 | 1211.52615 | 1204.39262 | 988.205088 | -0.4782348 | 0.0012584 |
| ENSMUSG0C | 2365.24255 | 2503.20069 | 2514.71355 | 2381.34265 | 2681.72167 | 2892.93823 | 2847.23587 | 2763.00156 | -0.1958401 | 0.00128711 |
| ENSMUSG0C | 15.1557392 | 5.14849999 | 23.5107848 | 10.1806332 | 41.0019331 | 32.4805939 | 62.4819072 | 22.8429317 | -1.5495365 | 0.00129065 |
| ENSMUSG0C | 18892.5762 | 18527.392 | 18920.5392 | 20116.9312 | 20861.1978 | 22855.5112 | 23045.051 | 21152.5547 | -0.2013861 | 0.00129163 |
| ENSMUSG0C | 60.6229568 | 47.3661999 | 54.5450208 | 41.6480448 | 71.2652647 | 93.1110357 | 100.186506 | 75.4809917 | -0.7331669 | 0.00129451 |
| ENSMUSG0C | 4479.46817 | 3735.75159 | 3962.03746 | 3998.2123 | 5276.5583 | 5247.78128 | 5600.74889 | 4221.96968 | -0.3308864 | 0.0012993 |
| ENSMUSG0C | 5337.6619 | 5411.07348 | 5999.01186 | 5726.14341 | 6579.83403 | 6536.17817 | 6438.86826 | 6258.96328 | -0.1998182 | 0.00131734 |
| ENSMUSG0C | 955.758804 | 1042.0564 | 1028.83194 | 1043.05214 | 1209.55703 | 1328.45629 | 1105.28339 | 1269.27247 | -0.2716613 | 0.00131924 |
| ENSMUSG0C | 920.711157 | 857.740098 | 870.839471 | 858.875235 | 1134.38682 | 1042.62706 | 1030.95147 | 979.26655 | -0.2553685 | 0.00132715 |
| ENSMUSG0C | 1801.6385 | 1893.61829 | 1965.50161 | 1821.40783 | 2315.63299 | 2023.541 | 2167.47581 | 2341.89708 | -0.2423305 | 0.00132655 |

[illegible]

| | | | | | | | | | | | | | | | | | | | | | | | | | | | | | | | | | | | | | |
|---|---|---|---|---|---|---|---|---|---|---|---|---|---|---|---|---|---|---|---|---|---|---|---|---|---|---|---|---|---|---|---|---|---|---|---|---|---|
| ENSMUSG0C | 48.3089187 | 55.6037998 | 96.8644336 | 31.4674116 | 86.8850488 | 149.410732 | 87.2592152 | 121.166855 | -0.936845 | 0.00445957 | 0.03312948 | Specc1 | 1.91 | 11 | 61956763 | 62223013 | + | 14840 | protein_codi | sperm antige | CH | 51 | 54 | 103 | 34 | 89 | 138 | 81 | 122 | 0.09590496 | 0.11038707 | 0.19229946 | 0.06247047 | 0.17248796 | 0.29661665 | 0.17323077 | 0.24054568 |
| ENSMUSG0C | 505.822796 | 413.939399 | 373.351263 | 492.372441 | 611.124051 | 542.425917 | 522.478017 | 611.793301 | -0.357833 | 0.00448745 | 0.03330334 | Ppcs | 1.28 | 4 | 119418530 | 119422420 | - | 1605 | protein_codi | Mus musculu | DFP | 534 | 402 | 397 | 532 | 626 | 501 | 485 | 616 | 9.28476661 | 7.59817616 | 6.8531497 | 9.03787499 | 11.2176522 | 9.95664506 | 9.590486 | 11.2299368 |
| ENSMUSG0C | 3067.14272 | 2740.03169 | 2755.46398 | 3046.78586 | 3241.10519 | 3565.28652 | 3716.5962 | 3152.32457 | -0.2359856 | 0.00450072 | 0.03335194 | Mpzl1 | 1.18 | 1 | 165592240 | 165634538 | - | 3550 | protein_codi | myelin prote | V-set | 3238 | 2661 | 2930 | 3292 | 3320 | 3293 | 3450 | 3174 | 25.4538372 | 22.7391833 | 22.867254 | 25.2848981 | 26.8975302 | 29.5878708 | 30.8435711 | 26.1607508 |
| ENSMUSG0C | 1928.56781 | 1848.31149 | 2060.48518 | 1737.18622 | 2156.50644 | 2224.92068 | 2123.30757 | 2291.24537 | -0.2157027 | 0.00453191 | 0.03350257 | Ddx54 | 1.16 | 5 | 120612739 | 120628592 | + | 6128 | protein_codi | ATP-depende | DEAD | 2036 | 1795 | 2191 | 1877 | 2209 | 2055 | 1971 | 2307 | 9.2717943 | 8.88595353 | 9.90600105 | 8.35170701 | 10.3676334 | 10.6965421 | 10.2080263 | 11.0154051 |
| ENSMUSG0C | 877.138407 | 821.700598 | 942.312257 | 915.331474 | 969.402848 | 1088.09989 | 1081.58336 | 1046.80217 | -0.2342317 | 0.00454307 | 0.03356549 | Kctd5 | 1.18 | 17 | 24047734 | 24073485 | - | 2751 | protein_codi | potassium ch | K_tetra | 926 | 798 | 1002 | 989 | 993 | 1005 | 1004 | 1054 | 9.39345085 | 8.79975625 | 10.0914107 | 9.80246805 | 10.3815292 | 11.6526797 | 11.5828928 | 11.2104141 |
| ENSMUSG0C | 2746.0305 | 2799.75429 | 2620.04186 | 2589.58288 | 3086.85982 | 3243.72864 | 3255.52282 | 2821.59865 | -0.2060982 | 0.00457874 | 0.03379544 | Zeb1 | 1.15 | 18 | 5591860 | 5775467 | + | 10266 | protein_codi | zinc finger E- | - | 2899 | 2719 | 2786 | 2798 | 3162 | 2996 | 3022 | 2841 | 7.88046286 | 8.03463753 | 7.51890505 | 7.43149491 | 8.85856301 | 9.30873968 | 9.34258621 | 8.09732571 |
| ENSMUSG0C | 1441.68969 | 1509.5402 | 1342.93603 | 1279.98324 | 1585.40808 | 1803.75565 | 1789.35255 | 1514.58569 | -0.2638412 | 0.00458756 | 0.03382704 | Tlk2 | 1.20 | 11 | 105178807 | 105283959 | + | 8070 | protein_codi | Serine/threo | Pkinase | 1522 | 1466 | 1428 | 1383 | 1624 | 1666 | 1661 | 1525 | 5.26315171 | 5.51085237 | 4.90263342 | 4.67281277 | 5.78782196 | 6.58493964 | 6.53235851 | 5.52927187 |
| ENSMUSG0C | 62.5174242 | 43.2473999 | 41.3789813 | 71.2644322 | 82.0038663 | 80.1187982 | 117.422895 | 81.4400173 | -0.7215624 | 0.00459009 | 0.03382889 | Gm12709 | 1.65 | 4 | 102967266 | 102989755 | - | 1912 | antisense | predicted ge | - | 66 | 42 | 44 | 77 | 84 | 74 | 109 | 82 | 0.96329835 | 0.66637661 | 0.63758712 | 1.09807644 | 1.26355477 | 1.23450874 | 1.80930808 | 1.25486671 |
| ENSMUSG0C | 1350.75526 | 1459.0849 | 1418.17054 | 1297.56797 | 1708.41388 | 1511.4303 | 1564.20223 | 1628.80035 | -0.2154453 | 0.00460946 | 0.033938 | Kat14 | 1.16 | 2 | 144368983 | 144407676 | + | 4699 | protein_codi | lysine acetylt | Acetyltransf_ | 1426 | 1417 | 1508 | 1402 | 1750 | 1396 | 1452 | 1640 | 8.46874101 | 9.14792819 | 8.89141017 | 8.13527621 | 10.7111297 | 9.47611472 | 9.80697538 | 10.2119819 |
| ENSMUSG0C | 25793.1737 | 22173.5597 | 25070.9605 | 23132.2496 | 25062.9197 | 30890.1274 | 30065.6474 | 28293.4538 | -0.2492857 | 0.00462055 | 0.03400287 | Fryl | 1.19 | 5 | 73019987 | 73256619 | - | 15192 | protein_codi | FRY like tran | - | 27230 | 21534 | 26659 | 24994 | 25673 | 28531 | 27909 | 28488 | 50.0192828 | 42.9999647 | 48.6187346 | 44.8590992 | 48.6031416 | 59.9035249 | 58.3046561 | 54.8679386 |
| ENSMUSG0C | 5014.65521 | 5160.85639 | 5631.30319 | 4955.19182 | 5962.85256 | 6070.62299 | 5651.38078 | 5938.16907 | -0.1862658 | 0.00466911 | 0.03427546 | Eif3b | 1.14 | 5 | 140419328 | 140443360 | + | 3346 | protein_codi | Eukaryotic tr | RRM_1 | 5294 | 5012 | 5988 | 5354 | 6108 | 5607 | 5246 | 5979 | 44.1532597 | 45.4405383 | 49.5827492 | 43.6296938 | 52.50199 | 53.4508918 | 49.7595293 | 52.2846556 |
| ENSMUSG0C | 3130.60738 | 3033.49619 | 2815.65159 | 3078.25327 | 3258.67745 | 3765.58351 | 4522.39735 | 3386.71292 | -0.3083782 | 0.00468588 | 0.03436131 | Rnf213 | 1.24 | 11 | 119393100 | 119487418 | + | 16888 | protein_codi | ring finger pr | - | 3305 | 2946 | 2994 | 3326 | 3338 | 3478 | 4198 | 3410 | 5.46132491 | 5.29191505 | 4.91188652 | 5.36999349 | 5.68474234 | 6.56903678 | 7.88929377 | 5.90809941 |
| ENSMUSG0C | 537.081508 | 613.701198 | 540.748051 | 563.636873 | 624.791362 | 737.30948 | 660.369122 | 691.246976 | -0.2671586 | 0.00468626 | 0.03436131 | Bud13 | 1.20 | 9 | 46283008 | 46299110 | + | 4315 | protein_codi | BUD13 homo | - | 567 | 596 | 575 | 609 | 640 | 681 | 613 | 696 | 3.66696724 | 4.19009434 | 3.69200086 | 3.84827613 | 4.26581333 | 5.03403984 | 4.50872335 | 4.71954438 |
| ENSMUSG0C | 4493.67667 | 4426.68029 | 4639.14807 | 4226.81379 | 4945.61413 | 5107.03204 | 5196.77104 | 4783.11126 | -0.1714811 | 0.00470075 | 0.03442283 | Ppm1b | 1.13 | 17 | 84956741 | 85023991 | + | 4971 | protein_codi | protein phos | PP2C | 4744 | 4299 | 4933 | 4567 | 5066 | 4717 | 4824 | 4816 | 26.6321169 | 26.2350577 | 27.4942642 | 25.0505337 | 29.3105586 | 30.2672141 | 30.7990591 | 28.347473 |
| ENSMUSG0C | 1794.06063 | 1812.27199 | 1792.46224 | 1740.88827 | 1978.83139 | 2227.08605 | 2066.21203 | 1928.73797 | -0.1996605 | 0.00470769 | 0.03445668 | Ddi2 | 1.15 | 4 | 141677549 | 141723419 | - | 9553 | protein_codi | 2 [Source:U | ubiquitin | 1894 | 1760 | 1906 | 1881 | 2027 | 2057 | 1918 | 1942 | 5.53279995 | 5.58896296 | 5.52787059 | 5.36881887 | 6.10262444 | 6.86823032 | 6.37210229 | 5.94813865 |
| ENSMUSG0C | 220.705452 | 212.118199 | 331.972282 | 155.486034 | 354.373851 | 534.847112 | 240.23216 | 395.282035 | -0.728261 | 0.00471785 | 0.03451405 | Fhl2 | 1.66 | 1 | 43123074 | 43196984 | - | 1897 | protein_codi | Four and a h | LIM | 233 | 206 | 353 | 168 | 363 | 494 | 223 | 398 | 3.42762547 | 3.29426272 | 5.15563452 | 2.41474728 | 5.50353797 | 8.30634479 | 3.73088142 | 6.13885501 |
| ENSMUSG0C | 3772.83183 | 4000.38449 | 4106.8639 | 3941.75606 | 4462.37706 | 4298.26525 | 4564.41105 | 4369.95215 | -0.1614472 | 0.0047606 | 0.03477551 | Ddx50 | 1.12 | 10 | 62615895 | 62651218 | - | 10129 | protein_codi | ATP-depende | DEAD | 3983 | 3885 | 4367 | 4259 | 4571 | 3970 | 4237 | 4400 | 10.9735843 | 11.6354395 | 11.9451434 | 11.4649141 | 12.979182 | 12.5018497 | 13.2759561 | 12.7103568 |
| ENSMUSG0C | 3187.4414 | 3398.00999 | 3581.16275 | 3436.42645 | 3692.12646 | 3806.7256 | 3935.28288 | 4002.4789 | -0.1823823 | 0.00477744 | 0.0348643 | Nudt3 | 1.13 | 17 | 27579382 | 27623465 | - | 2985 | protein_codi | Diphosphoino | NUDIX | 3365 | 3300 | 3808 | 3713 | 3782 | 3516 | 3653 | 4030 | 31.4590428 | 33.5372884 | 35.3449485 | 33.9164468 | 36.4401253 | 37.5711828 | 38.8400026 | 39.5032062 |
| ENSMUSG0C | 4847.94208 | 4915.78779 | 5869.23233 | 4982.95718 | 6154.19492 | 5752.31317 | 5918.54479 | 6443.69308 | -0.2353605 | 0.00481552 | 0.03512494 | Surf4 | 1.18 | 2 | 26920040 | 26933928 | - | 3281 | protein_codi | Surfeit locus | SURF4 | 5118 | 4774 | 6241 | 5384 | 6304 | 5313 | 5494 | 6488 | 43.5310179 | 44.1402233 | 52.7014666 | 44.743356 | 55.2602247 | 51.651617 | 53.1442569 | 57.8597091 |
| ENSMUSG0C | 30.3114784 | 19.5642999 | 23.5107848 | 43.499069 | 50.7642982 | 54.1343231 | 47.4000675 | 59.5902566 | -0.8540476 | 0.00485728 | 0.03537751 | Kcna6 | 1.81 | 6 | 126708329 | 126740674 | - | 5945 | protein_codi | Potassium vo | Ion_trans | 32 | 19 | 25 | 47 | 52 | 50 | 44 | 60 | 0.1502114 | 0.09695274 | 0.11650992 | 0.21556375 | 0.25156728 | 0.26826776 | 0.23489552 | 0.29530515 |
| ENSMUSG0C | 2537.63908 | 2348.74569 | 2762.047 | 2616.42273 | 2861.34919 | 3005.53762 | 3064.84527 | 2891.12062 | -0.2035186 | 0.00486527 | 0.03541837 | Pigq | 1.15 | 17 | 25926421 | 25944510 | - | 5992 | protein_codi | Phosphatidyl | - | 2679 | 2281 | 2937 | 2827 | 2931 | 2776 | 2845 | 2911 | 12.4768709 | 11.5481342 | 13.5802227 | 12.8642283 | 14.0684641 | 14.7773988 | 15.0689982 | 14.214842 |
| ENSMUSG0C | 2042.23586 | 2016.15259 | 2161.11134 | 2142.56053 | 2433.7576 | 2319.1144 | 2267.66232 | 2441.21418 | -0.1783107 | 0.00487588 | 0.03547828 | BC005624 | 1.13 | 2 | 30972177 | 30982201 | - | 3280 | protein_codi | Telomere ler | - | 2156 | 1958 | 2298 | 2315 | 2493 | 2142 | 2105 | 2458 | 18.3433935 | 18.1091133 | 19.411135 | 19.2445113 | 21.8600478 | 20.8303208 | 20.3681775 | 21.9270229 |
| ENSMUSG0C | 1005.01496 | 976.155597 | 1075.85351 | 1015.28678 | 1218.34316 | 1155.22645 | 1105.28339 | 1266.29295 | -0.2208266 | 0.00488267 | 0.03551032 | Golt1b | 1.17 | 6 | 142387221 | 142403858 | + | 2818 | protein_codi | golgi transpo | - | 1061 | 948 | 1144 | 1097 | 1248 | 1067 | 1026 | 1275 | 10.5070105 | 10.205298 | 11.247598 | 10.6143981 | 12.7372675 | 12.0774088 | 11.555275 | 13.2385625 |
| ENSMUSG0C | 4546.72176 | 4455.51189 | 4750.11897 | 4774.71696 | 4893.87359 | 5621.30811 | 5615.83073 | 5147.605 | -0.1995497 | 0.00490886 | 0.03564849 | Itpripl2 | 1.15 | 7 | 118485111 | 118491975 | - | 6865 | protein_codi | Inositol 1,4,5 | - | 4800 | 4327 | 5051 | 5159 | 5013 | 5192 | 5213 | 5183 | 19.5121653 | 19.12074 | 20.3850404 | 20.4906021 | 21.0019604 | 24.1237311 | 24.100225 | 22.0908437 |
| ENSMUSG0C | 3190.2831 | 3062.32779 | 2986.81011 | 3150.44321 | 3278.20218 | 3601.01517 | 3498.9868 | 3861.44863 | -0.2007089 | 0.00492242 | 0.03572955 | Iffo2 | 1.15 | 4 | 139530548 | 139620382 | + | 6216 | protein_codi | Intermediate | Filament | 3368 | 2974 | 3176 | 3404 | 3358 | 3326 | 3248 | 3888 | 15.1204894 | 14.5140395 | 14.15612 | 14.9316665 | 15.5371858 | 17.0671724 | 16.5836044 | 18.3015084 |
| ENSMUSG0C | 67.2535927 | 46.3364999 | 44.2002755 | 62.9348233 | 78.0989203 | 92.0283492 | 76.4864726 | 96.3375815 | -0.63398 | 0.00492752 | 0.03574917 | Gm12138 | 1.55 | 11 | 40672906 | 40673856 | - | 951 | processed_ps | predicted ge | Aldo_ket_red | 71 | 45 | 47 | 68 | 80 | 85 | 71 | 97 | 2.0834477 | 1.4354575 | 1.36927945 | 1.94965663 | 2.41942488 | 2.85094438 | 2.3694729 | 2.98444 |
| ENSMUSG0C | 1091.21322 | 1377.7386 | 834.162646 | 1217.97393 | 1544.40615 | 1380.42524 | 1416.61565 | 1625.82083 | -0.4008383 | 0.00493498 | 0.03578579 | Rpl17-ps10 | 1.32 | 7 | 72648757 | 72649311 | - | 555 | processed_ps | ribosomal pr | - | 1152 | 1338 | 887 | 1316 | 1582 | 1275 | 1315 | 1637 | 57.9247632 | 73.1343611 | 44.2797729 | 64.6535894 | 81.9815582 | 73.2769759 | 75.1980681 | 86.3032859 |
| ENSMUSG0C | 9.472337 | 5.14849999 | 8.46388255 | 3.70204843 | 15.6197841 | 21.6537292 | 19.3909367 | 14.8975641 | -1.4128874 | 0.00494094 | 0.03579415 | Gm47065 | 2.66 | 13 | 34087137 | 34089027 | - | 451 | transcribed_ | predicted ge | - | 10 | 5 | 9 | 4 | 16 | 20 | 18 | 15 | 0.61876855 | 0.33631931 | 0.55289252 | 0.24183167 | 1.02034282 | 1.41450273 | 1.26668864 | 0.97316471 |
| ENSMUSG0C | 20850.5082 | 21483.6607 | 21105.1613 | 20426.0522 | 22176.1884 | 24192.629 | 22876.9962 | 24132.0676 | -0.1550077 | 0.00493938 | 0.03579415 | Hspa12a | 1.11 | 19 | 58795751 | 58860984 | - | 5733 | protein_codi | Heat shock 7 | - | 22012 | 20864 | 22442 | 22070 | 22716 | 22345 | 21236 | 24298 | 107.147572 | 110.401246 | 108.456195 | 104.966357 | 113.960039 | 124.322219 | 117.561384 | 124.011003 |
| ENSMUSG0C | 3729.25908 | 3698.68239 | 3814.38973 | 3834.39666 | 4086.526 | 4173.75631 | 4276.77882 | 4119.67307 | -0.1436101 | 0.00500646 | 0.03616317 | Fgfr1op2 | 1.10 | 6 | 146577203 | 146599198 | + | 4619 | protein_codi | Mus musculu | - | 3937 | 3592 | 4056 | 4143 | 4186 | 3855 | 3970 | 4148 | 23.7860443 | 23.5910194 | 24.3290266 | 24.4566352 | 26.0647723 | 26.6211466 | 27.2782471 | 26.2761916 |
| ENSMUSG0C | 10573.0226 | 11067.2156 | 13608.9827 | 10224.1322 | 13654.62 | 14097.6604 | 14075.6655 | 13131.7062 | -0.273334 | 0.00501745 | 0.03620738 | Dusp3 | 1.21 | 11 | 101971143 | 101987013 | - | 4682 | protein_codi | Dual specific | DSPc | 11162 | 10748 | 14471 | 11047 | 13987 | 13021 | 13066 | 13222 | 66.5296716 | 69.6393309 | 85.6331427 | 64.3343147 | 85.9203106 | 88.7080976 | 88.5696969 | 82.6299288 |
| ENSMUSG0C | 11253.1364 | 10758.3056 | 13341.9002 | 11485.6052 | 14909.0839 | 14273.0556 | 12522.236 | 14138.7815 | -0.2536693 | 0.00503661 | 0.03632807 | Jup | 1.19 | 11 | 100368958 | 100397763 | - | 6436 | protein_codi | Junction plak | Arm | 11880 | 10448 | 14187 | 12410 | 15272 | 13183 | 11624 | 14236 | 51.5116162 | 49.2465113 | 61.073004 | 52.5757505 | 68.2468409 | 65.3353998 | 57.3209632 | 64.7207556 |
| ENSMUSG0C | 3807.87948 | 3785.17719 | 4841.34082 | 3815.88642 | 4794.29747 | 4797.38371 | 5313.11666 | 4767.22053 | -0.2755871 | 0.00505076 | 0.03641249 | Serp1 | 1.21 | 3 | 58519817 | 58525892 | - | 4611 | protein_codi | stress-associ | - | 4020 | 3676 | 5148 | 4123 | 4911 | 4431 | 4932 | 4800 | 24.3296411 | 24.1845896 | 30.9327239 | 24.3807997 | 30.6321503 | 30.6518692 | 33.9470359 | 30.4591479 |
| ENSMUSG0C | 83.3565656 | 66.9304998 | 125.077375 | 125.869647 | 192.318591 | 110.434019 | 162.668414 | 191.681992 | -0.710931 | 0.00507387 | 0.03656137 | Gm10824 | 1.64 | 10 | 22815749 | 22823330 | + | 2198 | antisense | - | - | 88 | 65 | 133 | 136 | 197 | 102 | 151 | 193 | 1.11727416 | 0.89710651 | 1.67648125 | 1.6871005 | 2.57775247 | 1.48020825 | 2.18033473 | 2.56921978 |
| ENSMUSG0C | 490.667057 | 418.058199 | 505.95209 | 598.806333 | 679.460606 | 623.627402 | 820.882987 | 561.141583 | -0.4143643 | 0.0050827 | 0.03660726 | Gm5641 | 1.33 | 3 | 54481438 | 54482925 | + | 1488 | transcribed_ | predicted ge | RRM_1 | 518 | 406 | 538 | 647 | 696 | 576 | 762 | 565 | 9.71474934 | 8.27716179 | 10.0173787 | 11.855806 | 13.4526852 | 12.34724 | 16.2527163 | 11.1100791 |
| ENSMUSG0C | 990.806451 | 1023.5218 | 1170.83709 | 1142.08194 | 1280.82229 | 1208.27809 | 1267.95181 | 1370.5759 | -0.2447951 | 0.00510788 | 0.03673528 | Pole4 | 1.18 | 6 | 82618992 | 82705365 | - | 2509 | protein_codi | polymerase ( | Histone | 1046 | 994 | 1245 | 1234 | 1312 | 1116 | 1177 | 1380 | 11.6341804 | 12.0183283 | 13.7481239 | 13.4104772 | 15.0395848 | 14.1877612 | 14.8884578 | 16.0934835 |
| ENSMUSG0C | 7902.77076 | 6668.33718 | 6530.3556 | 7055.17879 | 7648.813 | 8865.03675 | 9192.38127 | 7961.25828 | -0.2577824 | 0.00511759 | 0.03676954 | Gabpb2 | 1.20 | 3 | 95181766 | 95217916 | - | 15465 | protein_codi | GA-binding p | - | 8343 | 6476 | 6944 | 7623 | 7835 | 8188 | 8533 | 8016 | 15.0548728 | 12.7032621 | 12.4404055 | 13.4402 | 14.5710802 | 16.888001 | 17.5115962 | 15.1662922 |
| ENSMUSG0C | 370.368377 | 411.879999 | 365.827812 | 402.597767 | 499.83309 | 472.051297 | 424.446059 | 493.605959 | -0.2866021 | 0.00512675 | 0.03681761 | Zfp874a | 1.22 | 13 | 67424549 | 67451624 | - | 6733 | protein_codi | zinc finger pr | KRAB | 391 | 400 | 389 | 435 | 512 | 436 | 394 | 497 | 1.6205891 | 1.80222794 | 1.60072134 | 1.76161247 | 2.18707673 | 2.06551432 | 1.85721217 | 2.15982921 |
| ENSMUSG0C | 266.17267 | 266.692299 | 228.524829 | 266.547487 | 314.348154 | 276.085048 | 364.1187 | 423.090822 | -0.4231923 | 0.00516695 | 0.03708835 | Gm19426 | 1.34 | 2 | 84742169 | 84743655 | - | 726 | protein_codi | - | - | 281 | 259 | 243 | 288 | 322 | 255 | 338 | 426 | 10.8012612 | 10.8223478 | 9.27351548 | 10.8164713 | 12.7562177 | 11.2035046 | 14.7759017 | 17.1689847 |
| ENSMUSG0C | 2732.76923 | 2688.54669 | 2793.08124 | 2789.49349 | 3042.92918 | 3064.00269 | 2977.58606 | 3119.54993 | -0.1493092 | 0.00519181 | 0.03719505 | Psen1 | 1.11 | 12 | 83688152 | 83735199 | + | 4557 | protein_codi | presenilin 1 [ | - | 2885 | 2611 | 2970 | 3014 | 3117 | 2830 | 2764 | 3141 | 17.667356 | 17.3814573 | 18.0572732 | 18.0340784 | 19.6725404 | 19.8087807 | 19.2500971 | 20.1678937 |
| ENSMUSG0C | 1597.03602 | 1541.4609 | 1708.76384 | 1834.365 | 2007.14225 | 1931.51265 | 1823.82532 | 2161.13997 | -0.245935 | 0.00521145 | 0.03728744 | Dpp3 | 1.19 | 19 | 4907229 | 4928287 | - | 2683 | protein_codi | Dipeptidyl pe | - | 1686 | 1497 | 1817 | 1982 | 2056 | 1784 | 1693 | 2176 | 17.5364495 | 16.9262 | 18.7632905 | 20.1424694 | 22.0396712 | 21.209211 | 20.0267372 | 23.730662 |
| ENSMUSG0C | 5713.71368 | 5574.79578 | 6097.75716 | 5777.97208 | 6760.43779 | 6171.31283 | 6745.89143 | 6432.7682 | -0.1727089 | 0.00522133 | 0.03729879 | Ociad1 | 1.13 | 5 | 73292784 | 73314069 | + | 5515 | protein_codi | OCIA domain | - | 6032 | 5414 | 6484 | 6243 | 6925 | 5700 | 6262 | 6477 | 30.5225341 | 29.7804377 | 32.574086 | 30.865801 | 36.1141114 | 32.9670188 | 36.0364051 | 34.3637076 |
| ENSMUSG0C | 2156.85114 | 1927.59839 | 2065.18734 | 1928.76723 | 2329.3003 | 2238.9956 | 2283.82143 | 2341.89708 | -0.1866172 | 0.00532216 | 0.03788956 | Sipa1l3 | 1.14 | 7 | 29320372 | 29518641 | - | 16555 | protein_codi | signal-induce | - | 2277 | 1872 | 2196 | 2084 | 2386 | 2068 | 2120 | 2358 | 3.83829735 | 3.43032289 | 3.67517395 | 3.43240293 | 4.14518509 | 3.98448032 | 4.06425164 | 4.16760213 |
| ENSMUSG0C | 2664.5684 | 2514.52739 | 2532.58174 | 2665.47487 | 2780.32156 | 2947.07255 | 3000.20882 | 2926.87477 | -0.1672014 | 0.00538241 | 0.03823895 | Rnf40 | 1.12 | 7 | 127588767 | 127604799 | + | 5621 | protein_codi | ring finger pr | - | 2813 | 2442 | 2693 | 2880 | 2848 | 2722 | 2785 | 2947 | 13.9656424 | 13.1792415 | 13.2738687 | 13.9703934 | 14.5723325 | 15.4463145 | 15.7248144 | 15.340453 |
| ENSMUSG0C | 1977.82397 | 1861.69759 | 1880.86279 | 1763.10056 | 2136.98171 | 2136.14039 | 2102.83936 | 2085.65898 | -0.1772554 | 0.00539383 | 0.03827378 | Sfmbt1 | 1.13 | 14 | 30714849 | 30822721 | + | 8637 | protein_codi | Scm-like wit | MBT | 2088 | 1808 | 2000 | 1905 | 2189 | 1973 | 1952 | 2100 | 6.74640401 | 6.35029423 | 6.41566716 | 6.01397744 | 7.2892948 | 7.28642505 | 7.17283445 | 7.11423178 |
| ENSMUSG0C | 159.135262 | 119.4452 | 170.218082 | 139.752328 | 278.227403 | 180.808639 | 201.450287 | 185.722966 | -0.5239198 | 0.00541192 | 0.03836561 | Folr1 | 1.44 | 7 | 101858331 | 101870788 | - | 2297 | protein_codi | Folate recept | - | 168 | 116 | 181 | 151 | 285 | 167 | 187 | 187 | 2.04104724 | 1.5319879 | 2.183194 | 1.79244437 | 3.56850687 | 2.31902704 | 2.58377401 | 2.38205753 |
| ENSMUSG0C | 5159.58197 | 4934.32239 | 5524.09401 | 4508.16947 | 5777.36763 | 5896.31047 | 5696.6263 | 5922.27833 | -0.2108046 | 0.00541916 | 0.0383986 | Trim8 | 1.16 | 19 | 46501648 | 46516444 | + | 3230 | protein_codi | Probable E3 | - | 5447 | 4792 | 5874 | 4871 | 5918 | 5446 | 5288 | 5963 | 47.0608345 | 45.0062293 | 50.3855691 | 41.1192649 | 52.6956919 | 53.7805762 | 51.9592457 | 54.0174305 |
| ENSMUSG0C | 83.3565656 | 101.9403 | 90.2814138 | 100.88082 | 124.958272 | 201.379682 | 118.500169 | 124.146368 | -0.5948522 | 0.00543948 | 0.03852428 | Ptprm | 1.51 | 17 | 66666947 | 67354457 | - | 8828 | protein_codi | protein tyros | Y_phosphata | 88 | 99 | 96 | 109 | 128 | 186 | 110 | 125 | 0.2781795 | 0.34019757 | 0.30128926 | 0.33666185 | 0.4170137 | 0.67204903 | 0.39546156 | 0.41430419 |
| ENSMUSG0C | 500.139394 | 465.424399 | 521.939424 | 478.489759 | 549.621151 | 685.34053 | 656.060025 | 544.257677 | -0.3075826 | 0.00545649 | 0.03862637 | Slbp | 1.24 | 5 | 33634952 | 33652574 | - | 6076 | protein_codi | stem-loop bi | - | 528 | 452 | 555 | 517 | 563 | 633 | 609 | 548 | 2.42505129 | 2.25672693 | 2.5307542 | 2.32007761 | 2.664976 | 3.32304545 | 3.18107158 | 2.63896985 |
| ENSMUSG0C | 3708.41994 | 4315.47269 | 4432.25316 | 4794.15271 | 5471.8056 | 5138.42995 | 6068.28592 | 4600.36781 | -0.3027335 | 0.00547029 | 0.0387056 | Fads1 | 1.23 | 19 | 10182888 | 10196870 | + | 3456 | protein_codi | Fatty acid de | - | 3915 | 4191 | 4713 | 5180 | 5605 | 4746 | 5633 | 4632 | 31.6127883 | 36.7876689 | 37.7831754 | 40.8682235 | 46.6449418 | 43.8030485 | 51.7296966 | 39.2162852 |
| ENSMUSG0C | 11017.2752 | 10607.9694 | 10985.1791 | 8913.6071 | 12449.9441 | 12246.2666 | 12055.7763 | 11840.584 | -0.2268151 | 0.00548066 | 0.03874216 | Ddx6 | 1.17 | 9 | 44604892 | 44640731 | + | 8035 | protein_codi | Mus musculu | DEAD | 11631 | 10302 | 11681 | 9631 | 12753 | 11311 | 11191 | 11922 | 40.3957751 | 38.8950206 | 40.2780922 | 32.6824975 | 45.6487775 | 44.9019764 | 44.2035275 | 43.4145068 |
| ENSMUSG0C | 80.5148645 | 76.1977998 | 114.73263 | 31.4674116 | 111.290961 | 179.725953 | 108.8047 | 175.791257 | -0.9264854 | 0.00550038 | 0.03884464 | Ebf1 | 1.90 | 11 | 44617317 | 45008091 | + | 9756 | protein_codi | early B cell f | TIG | 85 | 74 | 122 | 34 | 114 | 166 | 101 | 177 | 0.24313748 | 0.23010087 | 0.34646773 | 0.09502478 | 0.33607463 | 0.5427335 | 0.32856666 | 0.53085157 |
| ENSMUSG0C | 6642.94994 | 6882.51478 | 5503.40452 | 5889.95905 | 7085.52454 | 7662.17209 | 7909.34763 | 7010.79369 | -0.251684 | 0.00556428 | 0.03920289 | Erbin | 1.19 | 13 | 103818787 | 103920514 | - | 7939 | protein_codi | Erbin [Sourc | PDZ | 7013 | 6684 | 5852 | 6364 | 7258 | 7077 | 7342 | 7059 | 24.6514692 | 25.5404756 | 20.422705 | 21.8571787 | 26.293829 | 28.4337231 | 29.3509723 | 26.0165087 |
| ENSMUSG0C | 576.865324 | 596.196298 | 534.165032 | 606.21043 | 618.933943 | 815.262906 | 696.996447 | 711.110395 | -0.2964922 | 0.00556904 | 0.03921618 | Zfp830 | 1.23 | 11 | 82764345 | 82765935 | + | 1591 | protein_codi | zinc finger pr | - | 609 | 579 | 568 | 655 | 634 | 753 | 647 | 716 | 10.681983 | 11.0399403 | 9.89128931 | 11.2253749 | 11.4609799 | 15.0964604 | 12.906486 | 13.1678381 |
| ENSMUSG0C | 412.993893 | 389.226599 | 435.419735 | 434.99069 | 451.997501 | 533.764426 | 498.777983 | 575.045976 | -0.2995468 | 0.0056253 | 0.03951154 | Snrnp35 | 1.23 | 5 | 124483134 | 124491124 | + | 1347 | protein_codi | U11/U12 sm | RRM_1 | 436 | 378 | 463 | 470 | 463 | 493 | 463 | 579 | 9.03282642 | 8.51299828 | 9.52331487 | 9.51393098 | 9.8858967 | 11.674268 | 10.9090595 | 12.5771605 |
| ENSMUSG0C | 862.929901 | 918.492397 | 878.362922 | 788.536315 | 979.165213 | 1150.89571 | 988.937772 | 991.184601 | -0.2533066 | 0.00563867 | 0.03957709 | Spata2 | 1.19 | 2 | 167481133 | 167492887 | - | 4510 | protein_codi | spermatoger | - | 911 | 892 | 934 | 852 | 1003 | 1063 | 918 | 998 | 5.63698145 | 5.9999365 | 5.73779573 | 5.15101468 | 6.39627406 | 7.51808201 | 6.46011208 | 6.47478921 |
| ENSMUSG0C | 27533.242 | 21160.3349 | 22762.2015 | 25031.4004 | 29458.9127 | 28019.9256 | 27823.8396 | 29196.2462 | -0.2469121 | 0.0056423 | 0.0395839 | Crb2 | 1.19 | 2 | 37776249 | 37799103 | + | 6698 | protein_codi | 2 [Source:U | EGF | 29067 | 20550 | 24204 | 27046 | 30176 | 25880 | 25828 | 29397 | 121.104377 | 93.0732812 | 100.119057 | 110.10008 | 129.574398 | 123.245044 | 122.382564 | 128.419065 |
| ENSMUSG0C | 5793.28131 | 5303.98469 | 5859.82802 | 6137.99629 | 6616.93102 | 6224.36447 | 6717.8823 | 6766.47364 | -0.1888084 | 0.0056535 | 0.03964375 | Nod1 | 1.14 | 6 | 54923949 | 54972612 | - | 5079 | protein_codi | Nucleotide-b | NACHT | 6116 | 5151 | 6231 | 6632 | 6778 | 5749 | 6236 | 6813 | 33.6042367 | 30.7660455 | 33.9902444 | 35.6037743 | 38.3818607 | 36.1047575 | 38.9674339 | 39.2492906 |
| ENSMUSG0C | 3254.69499 | 3593.65299 | 3627.24389 | 3393.8529 | 3982.0687 | 3698.45695 | 3998.84206 | 4141.52283 | -0.1900932 | 0.00567957 | 0.03980784 | Nop53 | 1.14 | 7 | 15936183 | 15946074 | - | 3847 | protein_codi | Ribosome bi | - | 3436 | 3490 | 3857 | 3667 | 4079 | 3416 | 3712 | 4170 | 24.9250329 | 27.5208335 | 27.7780785 | 25.9907288 | 30.49539 | 28.3234408 | 30.6238434 | 31.7165181 |
| ENSMUSG0C | 2756.45007 | 2840.94229 | 3189.00286 | 2578.47673 | 3338.72884 | 3235.06715 | 3362.17297 | 3291.36851 | -0.2189786 | 0.00572016 | 0.04003572 | Pdcl3 | 1.16 | 1 | 38985608 | 38997238 | + | 4219 | protein_codi | Phosducin-lik | - | 2910 | 2759 | 3391 | 2786 | 3420 | 2988 | 3121 | 3314 | 19.2481163 | 19.8381202 | 22.2686051 | 18.0053398 | 23.3141321 | 22.5902691 | 23.477841 | 22.9834179 |
| ENSMUSG0C | 792.834607 | 838.175798 | 868.958608 | 1042.12663 | 1156.84026 | 1004.73304 | 1019.10145 | 1232.52514 | -0.3173577 | 0.00573157 | 0.04007787 | Cdc26 | 1.25 | 4 | 62383654 | 62408642 | - | 7180 | protein_codi | Anaphase-pr | - | 837 | 814 | 924 | 1126 | 1185 | 928 | 946 | 1241 | 3.2531627 | 3.43920689 | 3.56551506 | 4.27605892 | 4.7467524 | 4.12262534 | 4.18158189 | 5.05730298 |
| ENSMUSG0C | 4135.62234 | 3102.48609 | 4287.42672 | 4150.9218 | 4645.90952 | 4944.62907 | 4550.40648 | 5217.12696 | -0.3042008 | 0.00576798 | 0.04029463 | Gpcpd1 | 1.23 | 2 | 132529082 | 132587729 | - | 7289 | protein_codi | glycerophosp | - | 4366 | 3013 | 4559 | 4485 | 4759 | 4567 | 4224 | 5253 | 16.7155454 | 12.5397686 | 17.3291153 | 16.7773835 | 18.7780472 | 19.9854254 | 18.3920387 | 21.0868198 |
| ENSMUSG0C | 9555.69357 | 8805.99438 | 8729.0842 | 10029.7747 | 9899.03814 | 11528.4454 | 10879.3928 | 10723.2667 | -0.2130409 | 0.00577205 | 0.04030414 | Lrch1 | 1.16 | 14 | 74754675 | 74947876 | - | 7734 | protein_codi | Leucine-rich | CH | 10088 | 8552 | 9282 | 10837 | 10140 | 10648 | 10099 | 10797 | 36.4003598 | 33.544542 | 33.2515692 | 38.20627 | 37.7082571 | 43.9151339 | 41.4427073 | 40.8479785 |
| ENSMUSG0C | 401.627089 | 449.978899 | 366.768244 | 452.57542 | 587.694375 | 455.811 | 504.164354 | 558.16207 | -0.334902 | 0.00577722 | 0.04032129 | Mrpl46 | 1.26 | 7 | 78775236 | 78783531 | - | 1415 | protein_codi | 39S ribosom | - | 424 | 437 | 390 | 489 | 602 | 421 | 468 | 562 | 8.36207749 | 9.36878643 | 7.63629884 | 9.42284731 | 12.2360917 | 9.49021371 | 10.4969548 | 11.6212143 |
| ENSMUSG0C | 31.2587121 | 31.9206999 | 39.4981185 | 12.9571695 | 58.5741902 | 73.6226794 | 43.0909705 | 49.6585472 | -0.9614001 | 0.00583475 | 0.04063583 | Fli1 | 1.95 | 9 | 32422204 | 32542861 | - | 3486 | protein_codi | Friend leuke | Ets | 33 | 31 | 42 | 14 | 60 | 68 | 40 | 50 | 0.26417476 | 0.26976938 | 0.33380794 | 0.1095041 | 0.49502433 | 0.62220266 | 0.36417198 | 0.41967612 |
| ENSMUSG0C | 4227.504 | 4417.41299 | 4351.37606 | 4809.88642 | 4893.87359 | 5068.05533 | 5223.70289 | 4969.8274 | -0.1786754 | 0.00585137 | 0.04072422 | Atxn7l3b | 1.13 | 10 | 112925430 | 112929001 | - | 3572 | protein_codi | ataxin 7-like | - | 4463 | 4290 | 4627 | 5197 | 5013 | 4681 | 4849 | 5004 | 34.8674517 | 36.433776 | 35.889119 | 39.6708039 | 40.3635102 | 41.8001199 | 43.0838641 | 40.9899592 |
| ENSMUSG0C | 1633.0309 | 1770.0543 | 1664.56357 | 1557.63688 | 1935.87699 | 2300.70873 | 2274.12597 | 1683.42475 | -0.3064204 | 0.00588133 | 0.04087858 | Ccdc117 | 1.24 | 11 | 5528887 | 5542187 | - | 3372 | protein_codi | Coiled-coil d | - | 1724 | 1719 | 1770 | 1683 | 1983 | 2125 | 2111 | 1695 | 14.2677163 | 15.4648835 | 14.5432158 | 13.6090022 | 16.9136688 | 20.101187 | 19.8689346 | 14.7080051 |
| ENSMUSG0C | 1044.79877 | 1184.155 | 965.823042 | 1155.96462 | 1315.96681 | 1235.34525 | 1250.71542 | 1370.5759 | -0.2501548 | 0.00594702 | 0.04123565 | Pfdn2 | 1.19 | 1 | 171345670 | 171359254 | + | 1937 | protein_codi | Prefoldin sub | - | 1103 | 1150 | 1027 | 1249 | 1348 | 1141 | 1161 | 1380 | 15.8909793 | 18.0105328 | 14.6897895 | 17.5817683 | 20.0153387 | 18.7891165 | 19.0228907 | 20.8459216 |
| ENSMUSG0C | 1321.39101 | 1231.5212 | 1396.54062 | 1257.77095 | 1470.21217 | 1533.08403 | 1433.85204 | 1564.24424 | -0.2046315 | 0.00595505 | 0.0412721 | 5031439G07 | 1.15 | 15 | 84943936 | 84988551 | - | 9604 | protein_codi | Uncharacteri | - | 1395 | 1196 | 1485 | 1359 | 1506 | 1416 | 1331 | 1575 | 4.05346873 | 3.77778615 | 4.28399594 | 3.8583093 | 4.50998911 | 4.70285337 | 4.39845161 | 4.7984397 |
| ENSMUSG0C | 2482.69953 | 2536.15109 | 2201.54989 | 2319.33334 | 2569.45448 | 3104.06209 | 3003.44064 | 2648.78691 | -0.2474569 | 0.00599169 | 0.04146864 | Rest | 1.19 | 5 | 77265491 | 77286432 | + | 7222 | protein_codi | RE1-silencing | - | 2621 | 2463 | 2341 | 2506 | 2632 | 2867 | 2788 | 2667 | 10.1277811 | 10.3458284 | 8.98087555 | 9.46135454 | 10.4816843 | 12.6625317 | 12.2520624 | 10.8053084 |
| ENSMUSG0C | 46.4144513 | 38.0988999 | 41.3789813 | 43.499069 | 44.9068792 | 97.4417816 | 128.195637 | 53.6312309 | -0.9326291 | 0.00599685 | 0.04148447 | Gm44669 | 1.91 | 7 | 67679335 | 67681303 | - | 1969 | TEC | predicted ge | - | 49 | 37 | 44 | 47 | 46 | 90 | 119 | 54 | 0.69447263 | 0.57005184 | 0.6191298 | 0.65085145 | 0.6719157 | 1.4579651 | 1.91811729 | 0.80245314 |
| ENSMUSG0C | 843.985227 | 903.046897 | 881.184216 | 901.448792 | 972.331557 | 979.831248 | 1053.57423 | 1071.63145 | -0.2082033 | 0.00600202 | 0.04150095 | Casp9 | 1.16 | 4 | 141793612 | 141815976 | + | 5197 | protein_codi | Mus musculu | Peptidase_C | 891 | 877 | 937 | 974 | 996 | 905 | 978 | 1079 | 4.78442508 | 5.11923679 | 4.99530054 | 5.11017737 | 5.51200108 | 5.5545157 | 5.97255354 | 6.0749172 |
| ENSMUSG0C | 14571.296 | 15103.6396 | 17898.2903 | 18302.0019 | 18983.895 | 19638.8497 | 21880.5175 | 18793.7738 | -0.2674903 | 0.00612435 | 0.04217044 | Pdia3 | 1.20 | 2 | 121413775 | 121438687 | + | 3139 | protein_codi | Protein disul | - | 15383 | 14668 | 19032 | 19775 | 19446 | 18139 | 20311 | 18923 | 136.758552 | 141.75485 | 167.983978 | 171.773004 | 178.172896 | 184.319957 | 205.359077 | 176.388517 |
| ENSMUSG0C | 2056.44436 | 2140.74629 | 1940.10997 | 2190.68716 | 2462.06846 | 2381.91022 | 2251.50321 | 2402.48051 | -0.1898667 | 0.00614933 | 0.04230328 | Axin1 | 1.14 | 17 | 26138688 | 26195811 | + | 3828 | protein_codi | axin 1 [Sourc | DIX | 2171 | 2079 | 2063 | 2367 | 2522 | 2200 | 2090 | 2419 | 15.8267836 | 16.4755872 | 14.9314522 | 16.8599415 | 18.9485432 | 18.3316302 | 17.3279933 | 18.4899431 |
| ENSMUSG0C | 17.9974403 | 9.26729997 | 12.2256081 | 3.70204843 | 30.2633316 | 34.6459668 | 23.7000338 | 18.8702479 | -1.3145657 | 0.00618522 | 0.04251011 | Gm29254 | 2.49 | 5 | 20702277 | 20706052 | + | 883 | 3prime_over | predicted ge | - | 19 | 9 | 13 | 4 | 31 | 32 | 22 | 19 | 0.60047878 | 0.30920047 | 0.40790347 | 0.12351765 | 1.00972629 | 1.1559515 | 0.79074397 | 0.62959973 |
| ENSMUSG0C | 20015.9953 | 14438.4534 | 21889.4811 | 17992.8809 | 26464.7954 | 21517.3107 | 22244.6362 | 24382.3467 | -0.3478933 | 0.00618796 | 0.04251011 | Sirpa | 1.27 | 2 | 129592835 | 129632228 | + | 4911 | protein_codi | signal-regula | V-set | 21131 | 14022 | 23276 | 19441 | 27109 | 19874 | 20649 | 24550 | 120.075634 | 86.6160492 | 131.314645 | 107.939003 | 158.761881 | 129.082001 | 133.445214 | 146.269305 |
| ENSMUSG0C | 4832.78634 | 4889.01559 | 4839.45995 | 4336.94973 | 5428.85119 | 6716.98681 | 6781.44148 | 4761.2615 | -0.3258501 | 0.0062131 | 0.0426434 | Slc38a2 | 1.25 | 15 | 96687392 | 96699730 | - | 8838 | protein_codi | Sodium-coup | Aa_trans | 5102 | 4748 | 5146 | 4686 | 5561 | 6204 | 6295 | 4794 | 16.10984 | 16.2972773 | 16.1320862 | 14.4569947 | 18.096791 | 22.3907236 | 22.6055798 | 15.8714157 |
| ENSMUSG0C | 1013.54006 | 979.244697 | 1094.66214 | 1018.98883 | 1203.69961 | 1182.29362 | 1286.26547 | 1109.37194 | -0.2191256 | 0.00622659 | 0.04271634 | Cln8 | 1.16 | 8 | 14881335 | 14901720 | + | 7606 | protein_codi | Protein CLN8 | LAG1 | 1070 | 951 | 1164 | 1101 | 1233 | 1092 | 1194 | 1117 | 3.92583667 | 3.79299733 | 4.2400542 | 3.94694189 | 4.66239891 | 4.57948513 | 4.98220874 | 4.297031 |
| ENSMUSG0C | 150.610158 | 140.0392 | 147.647729 | 134.199256 | 225.510632 | 168.899088 | 185.291173 | 178.77077 | -0.4071344 | 0.00626999 | 0.04295461 | Gm48488 | 1.33 | 13 | 60176682 | 60182862 | + | 5345 | lincRNA | predicted ge | - | 159 | 136 | 157 | 145 | 231 | 156 | 172 | 180 | 0.83014544 | 0.77187956 | 0.81381688 | 0.73969048 | 1.24298802 | 0.93095186 | 1.0213031 | 0.98536341 |
| ENSMUSG0C | 2196.63495 | 1861.69759 | 2825.99634 | 2236.03725 | 2799.84629 | 2746.77555 | 3151.02722 | 2804.71474 | -0.3345491 | 0.00631853 | 0.04326722 | Tubb2a | 1.26 | 13 | 34074274 | 34078007 | - | 1610 | protein_codi | tubulin, beta | Tubulin | 2319 | 1808 | 3005 | 2416 | 2868 | 2537 | 2925 | 2824 | 40.1957044 | 34.0667648 | 51.7122398 | 40.9167178 | 51.2337263 | 50.2625974 | 57.6599031 | 51.3228129 |
| ENSMUSG0C | 143.979522 | 131.8016 | 84.6388255 | 143.454377 | 153.269131 | 204.627741 | 177.750253 | 195.654676 | -0.537021 | 0.00633897 | 0.04337179 | B130046B21 | 1.45 | 15 | 98807024 | 98816077 | + | 2359 | antisense | RIKEN cDNA | CBS | 152 | 128 | 90 | 155 | 157 | 189 | 165 | 197 | 1.79812723 | 1.64603995 | 1.05703488 | 1.7915688 | 1.91414302 | 2.55554892 | 2.21988214 | 2.44348637 |
| ENSMUSG0C | 1365.911 | 902.017197 | 1281.80799 | 1100.4339 | 1566.85959 | 1931.51265 | 1276.57 | 1462.9408 | -0.4234726 | 0.00635272 | 0.04337808 | Amotl1 | 1.34 | 9 | 14541966 | 14645056 | - | 10064 | protein_codi | Angiomotin-l | - | 1442 | 876 | 1363 | 1189 | 1605 | 1784 | 1185 | 1473 | 3.9985212 | 2.64053434 | 3.75232093 | 3.2213726 | 4.58677125 | 5.65424416 | 3.73698742 | 4.28256293 |
| ENSMUSG0C | 3321.00135 | 3239.43619 | 3201.22846 | 3270.75979 | 3622.81366 | 3790.4853 | 3465.5913 | 4282.55311 | -0.2183936 | 0.00634453 | 0.04337808 | Zfand5 | 1.16 | 19 | 21272278 | 21282289 | + | 6193 | protein_codi | zinc finger, A | zf-A20 | 3506 | 3146 | 3404 | 3534 | 3711 | 3501 | 3217 | 4312 | 15.7984908 | 15.4104734 | 15.2287136 | 15.5594843 | 17.2342563 | 18.0318949 | 16.4863264 | 20.3727337 |
| ENSMUSG0C | 17689.5894 | 16707.9122 | 20734.6314 | 18180.7598 | 21627.5435 | 20371.8285 | 21785.7174 | 21374.0319 | -0.2160741 | 0.00635802 | 0.04337808 | Mtch1 | 1.16 | 17 | 29332072 | 29347904 | - | 2878 | protein_codi | Mus musculu | Mito_carr | 18675 | 16226 | 22048 | 19644 | 22154 | 18816 | 20223 | 21521 | 181.081712 | 171.03265 | 212.252669 | 186.109641 | 221.393076 | 208.538791 | 223.012243 | 218.797972 |
| ENSMUSG0C | 2130.32859 | 2097.49889 | 2080.23424 | 1970.41528 | 2200.43708 | 2485.84812 | 2516.51268 | 2295.21805 | -0.1980183 | 0.00638081 | 0.04351353 | Pcnt | 1.15 | 10 | 76351263 | 76442786 | - | 12154 | protein_codi | pericentrin [S | - | 2249 | 2037 | 2212 | 2129 | 2254 | 2296 | 2336 | 2311 | 5.16386636 | 5.08428795 | 5.04243884 | 4.77624025 | 5.33380768 | 6.02563732 | 6.09996749 | 5.56355452 |
| ENSMUSG0C | 162.924196 | 63.8413998 | 70.5323545 | 62.0093112 | 146.435475 | 236.025649 | 135.736557 | 144.009787 | -0.8809062 | 0.00640733 | 0.0436144 | Rn7sk | 1.84 | 9 | 78175303 | 78175633 | - | 331 | misc_RNA | RNA, 7SK, nu | - | 172 | 62 | 75 | 67 | 150 | 218 | 126 | 145 | 14.5012428 | 5.68227224 | 6.27780784 | 5.51920523 | 13.0336465 | 21.0077159 | 12.0813778 | 12.8177455 |
| ENSMUSG0C | 4082.57725 | 4026.12699 | 3867.99432 | 4651.62385 | 5034.45165 | 4942.4637 | 4369.42441 | 5585.59338 | -0.2614852 | 0.00640528 | 0.0436144 | Cdc42ep1 | 1.20 | 15 | 78842624 | 78850897 | + | 2560 | protein_codi | CDC42 effect | PBD | 4310 | 3910 | 4113 | 5026 | 5157 | 4565 | 4056 | 5624 | 46.983144 | 46.3335028 | 44.5136793 | 53.5318501 | 57.9375117 | 56.8788953 | 50.2842405 | 64.2801648 |
| ENSMUSG0C | 700.952938 | 688.869298 | 560.497111 | 633.975793 | 761.464473 | 822.841711 | 802.569325 | 732.960156 | -0.2717105 | 0.00649598 | 0.0441169 | Zfp790 | 1.21 | 7 | 29816114 | 29831005 | + | 3658 | protein_codi | zinc finger pr | KRAB | 740 | 669 | 596 | 685 | 780 | 760 | 745 | 738 | 5.64537492 | 5.54805501 | 4.51416373 | 5.10595055 | 6.13272618 | 6.6270497 | 6.46377879 | 5.90315648 |
| ENSMUSG0C | 1510.83775 | 1548.6688 | 1694.65737 | 1732.55866 | 2002.26107 | 1900.11474 | 1762.42069 | 1870.14089 | -0.2160861 | 0.00650336 | 0.04414176 | Fut11 | 1.16 | 14 | 20694968 | 20701016 | + | 4252 | protein_codi | Alpha-(1,3)-f | - | 1595 | 1504 | 1802 | 1872 | 2051 | 1755 | 1636 | 1883 | 10.4682046 | 10.7303261 | 11.7418433 | 12.0044515 | 13.8731498 | 13.1654043 | 12.2113578 | 12.9577232 |
| ENSMUSG0C | 1465.37053 | 1337.5803 | 1247.01203 | 681.176911 | 1746.4871 | 1795.09415 | 1697.78424 | 1533.45594 | -0.5177097 | 0.00652252 | 0.04421641 | Leo1 | 1.43 | 9 | 75441524 | 75466432 | + | 4846 | protein_codi | RNA polymer | - | 1547 | 1299 | 1326 | 736 | 1789 | 1658 | 1576 | 1544 | 8.90864545 | 8.13175121 | 7.58114604 | 4.1411803 | 10.6176793 | 10.9131834 | 10.3215927 | 9.32256717 |
| ENSMUSG0C | 3318.15965 | 3191.04029 | 3065.80634 | 3150.44321 | 3496.87915 | 3590.18831 | 3441.89127 | 3667.78029 | -0.1579072 | 0.00654445 | 0.04434492 | Zc3h14 | 1.12 | 12 | 98746964 | 98787753 | + | 7891 | protein_codi | zinc finger C | - | 3503 | 3099 | 3260 | 3404 | 3582 | 3316 | 3195 | 3693 | 12.3883328 | 11.9137333 | 11.446173 | 11.7621643 | 13.0555812 | 13.4039505 | 12.8502842 | 13.6936398 |
| ENSMUSG0C | 697.164003 | 749.621598 | 685.574486 | 708.942274 | 780.989203 | 834.751262 | 838.119376 | 832.27725 | -0.2099527 | 0.00657375 | 0.04444672 | Traf2 | 1.16 | 2 | 25517982 | 25546940 | - | 5101 | protein_codi | TNF receptor | zf-C3HC4 | 736 | 728 | 729 | 766 | 800 | 771 | 778 | 838 | 4.0264959 | 4.32946664 | 3.95956023 | 4.09452173 | 4.51063137 | 4.82113609 | 4.84058875 | 4.80684735 |
| ENSMUSG0C | 3745.36205 | 3534.96009 | 3819.09189 | 3433.64992 | 4121.67052 | 4015.68409 | 4044.08758 | 4105.76868 | -0.1644128 | 0.00656743 | 0.04444672 | Cep104 | 1.12 | 4 | 153975194 | 154008732 | + | 5279 | protein_codi | Centrosomal | - | 3954 | 3433 | 4061 | 3710 | 4222 | 3709 | 3754 | 4134 | 20.9020929 | 19.727883 | 21.3135639 | 19.1624918 | 23.0021928 | 22.4107044 | 22.5692184 | 22.9134479 |
| ENSMUSG0C | 242.491827 | 283.167499 | 233.226986 | 280.430168 | 306.538262 | 332.384744 | 327.491376 | 326.75324 | -0.3155352 | 0.00663722 | 0.04483064 | 2310030G06 | 1.24 | 9 | 50739691 | 50746536 | - | 1979 | protein_codi | Uncharacteri | - | 256 | 275 | 248 | 303 | 314 | 307 | 304 | 329 | 3.60993134 | 4.21546261 | 3.47200734 | 4.17471247 | 4.56337886 | 4.9481507 | 4.87530402 | 4.86431554 |
| ENSMUSG0C | 5139.69006 | 4870.48099 | 5378.32714 | 5795.55681 | 6071.21481 | 5756.64392 | 6184.63154 | 6385.09599 | -0.2036908 | 0.00665436 | 0.04490562 | Cxxc5 | 1.15 | 18 | 35829818 | 35861687 | + | 2271 | protein_codi | Mus musculu | zf-CXXC | 5426 | 4730 | 5719 | 6262 | 6219 | 5317 | 5741 | 6429 | 66.6756758 | 63.1833063 | 69.7714439 | 75.1840408 | 78.7600703 | 74.6792353 | 80.2313919 | 82.8319579 |
| ENSMUSG0C | 1901.09804 | 1854.48969 | 1851.70941 | 1890.82123 | 2138.93418 | 2117.73472 | 2088.83479 | 2014.15067 | -0.1568161 | 0.00668835 | 0.04511452 | Ngly1 | 1.11 | 14 | 16249280 | 16311926 | + | 7555 | protein_codi | Peptide-N(4) | Transglut_co | 2007 | 1801 | 1969 | 2043 | 2191 | 1956 | 1939 | 2028 | 7.41340411 | 7.23165311 | 7.2208113 | 7.37332933 | 8.3408552 | 8.2581871 | 8.14549074 | 7.85425717 |
| ENSMUSG0C | 955.758804 | 990.571397 | 1061.74704 | 953.27747 | 1144.14918 | 1095.6787 | 1148.37436 | 1157.04415 | -0.1985389 | 0.00671119 | 0.04522756 | Atg4d | 1.15 | 9 | 21265293 | 21277772 | + | 7264 | protein_codi | Cysteine prot | - | 1009 | 962 | 1129 | 1030 | 1172 | 1012 | 1066 | 1165 | 3.87632428 | 4.01751566 | 4.3061867 | 3.8662606 | 4.64038964 | 4.44380521 | 4.65752595 | 4.69268848 |
| ENSMUSG0C | 4418.84521 | 4584.22439 | 5167.67051 | 4728.44135 | 4945.61413 | 5814.0263 | 5774.19004 | 5522.03044 | -0.222695 | 0.0067186 | 0.04525453 | Nxn | 1.17 | 11 | 76257198 | 76399140 | - | 3239 | protein_codi | Nucleoredoxi | - | 4665 | 4452 | 5495 | 5109 | 5066 | 5370 | 5360 | 5560 | 40.1925417 | 41.696783 | 47.0036406 | 43.0085389 | 44.9838798 | 52.8827065 | 52.5203674 | 50.2267964 |
| ENSMUSG0C | 65.3591253 | 43.2473999 | 84.6388255 | 18.5102421 | 60.5266632 | 176.477893 | 96.9546835 | 131.098564 | -1.1340486 | 0.00672682 | 0.04527142 | Tril | 2.19 | 6 | 53815468 | 53820830 | - | 5363 | protein_codi | TLR4 interact | - | 69 | 42 | 90 | 20 | 62 | 163 | 90 | 132 | 0.35904267 | 0.23757451 | 0.46495344 | 0.10168384 | 0.33249611 | 0.96946056 | 0.53260915 | 0.72017455 |
| ENSMUSG0C | 2737.50539 | 3080.86239 | 3000.91658 | 3019.02049 | 3623.7899 | 3147.36954 | 3433.27307 | 3376.78121 | -0.1983561 | 0.00674489 | 0.0453579 | Syf2 | 1.15 | 4 | 134930898 | 134937548 | + | 2043 | protein_codi | Pre-mRNA-sp | - | 2890 | 2992 | 3191 | 3262 | 3712 | 2907 | 3187 | 3400 | 39.4761006 | 44.4274681 | 43.2746123 | 43.5356792 | 52.2567353 | 45.3865322 | 49.509394 | 48.6947551 |
| ENSMUSG0C | 662.116357 | 569.424098 | 601.876092 | 562.711361 | 799.537696 | 744.888286 | 654.982751 | 698.199173 | -0.2743635 | 0.00678666 | 0.04553001 | Aen | 1.21 | 7 | 78895854 | 78911209 | + | 5801 | protein_codi | Apoptosis-en | - | 699 | 553 | 640 | 608 | 819 | 688 | 608 | 703 | 3.36262998 | 2.89188226 | 3.05669324 | 2.85779089 | 4.06053921 | 3.78299624 | 3.32640119 | 3.5458805 |
| ENSMUSG0C | 190.393974 | 190.494499 | 244.512162 | 85.1471138 | 250.892781 | 411.420855 | 308.100439 | 232.402001 | -0.7591952 | 0.00683284 | 0.04577786 | Chst15 | 1.69 | 7 | 132235780 | 132317228 | - | 6754 | protein_codi | Carbohydrate | - | 201 | 185 | 260 | 92 | 257 | 380 | 286 | 234 | 0.83050026 | 0.83093875 | 1.06656429 | 0.37141249 | 1.09439661 | 1.79462154 | 1.34393694 | 1.01373966 |
| ENSMUSG0C | 940.603064 | 1507.4808 | 905.635432 | 655.262572 | 1933.92451 | 1410.74046 | 1156.99256 | 1578.14863 | -0.6013209 | 0.00685828 | 0.04592765 | Gabrb1 | 1.52 | 5 | 71658113 | 72149037 | + | 13838 | protein_codi | Gamma-ami | Neur_chan_L | 993 | 1464 | 963 | 708 | 1981 | 1303 | 1074 | 1589 | 2.00253766 | 3.20941658 | 1.92809181 | 1.39504965 | 4.11731242 | 3.00345705 | 2.46322945 | 3.35986793 |
| ENSMUSG0C | 7097.62212 | 7183.18718 | 8872.02977 | 7301.36501 | 8779.29487 | 9966.12888 | 9682.54106 | 8257.22322 | -0.2684772 | 0.00686426 | 0.04594505 | Rrbp1 | 1.20 | 2 | 143947395 | 144011263 | - | 8786 | protein_codi | Ribosome-bi | - | 7493 | 6976 | 9434 | 7889 | 8993 | 9205 | 8988 | 8314 | 23.7995806 | 24.0864954 | 29.7494829 | 24.4827666 | 29.4385264 | 33.4181905 | 32.4672704 | 27.687928 |
| ENSMUSG0C | 1005.01496 | 1113.1057 | 1029.77238 | 1047.67971 | 1221.27187 | 1172.54944 | 1220.55174 | 1170.94854 | -0.1901452 | 0.006913 | 0.04621091 | Fam104a | 1.14 | 11 | 113661319 | 113684155 | - | 2619 | protein_codi | family with s | - | 1061 | 1081 | 1095 | 1132 | 1251 | 1083 | 1133 | 1179 | 11.3053668 | 12.5212745 | 11.5838618 | 11.7853006 | 13.7380307 | 13.1899544 | 13.7299301 | 13.171946 |
| ENSMUSG0C | 8987.35335 | 9896.44667 | 9172.02738 | 9293.99258 | 10466.2316 | 10573.516 | 10035.887 | 10411.411 | -0.1516383 | 0.00693214 | 0.04631799 | Rab6a | 1.11 | 7 | 100607410 | 100641268 | + | 3728 | protein_codi | RAB6A, mem | Ras | 9488 | 9611 | 9753 | 10042 | 10721 | 9766 | 9316 | 10483 | 71.0237409 | 78.2079703 | 72.4831517 | 73.4469977 | 82.7107702 | 83.5585996 | 79.309916 | 82.2775436 |
| ENSMUSG0C | 4440.63159 | 4661.45189 | 4770.80846 | 4783.04657 | 5432.75614 | 5178.48935 | 4951.15251 | 5318.4304 | -0.1626411 | 0.00693974 | 0.04634799 | Ptp4a2 | 1.12 | 4 | 129811219 | 129850003 | + | 4377 | protein_codi | Protein tyros | Y_phosphata | 4688 | 4527 | 5073 | 5168 | 5565 | 4783 | 4596 | 5355 | 29.8893057 | 31.3756181 | 32.1116827 | 32.1940558 | 36.5671653 | 34.855729 | 33.3255547 | 35.7976538 |
| ENSMUSG0C | 5475.95802 | 5715.86468 | 7127.52953 | 5776.12106 | 6830.72681 | 7603.70702 | 7642.18361 | 6872.74293 | -0.264685 | 0.00699085 | 0.04660567 | Sec61a1 | 1.20 | 6 | 88503579 | 88518905 | - | 6732 | protein_codi | Protein trans | - | 5781 | 5551 | 7579 | 6241 | 6997 | 7023 | 7094 | 6920 | 23.9642385 | 25.0141334 | 31.1919516 | 25.2778312 | 29.8930645 | 33.27583 | 33.4442138 | 30.0769381 |
| ENSMUSG0C | 2378.50382 | 2407.43859 | 2550.44994 | 2546.08381 | 2978.49757 | 2647.1684 | 3044.37706 | 2692.48643 | -0.2012795 | 0.00700186 | 0.04665818 | Rnf5 | 1.15 | 17 | 34601091 | 34603690 | - | 1483 | protein_codi | E3 ubiquitin- | - | 2511 | 2338 | 2712 | 2751 | 3051 | 2445 | 2826 | 2711 | 47.2509269 | 47.8257399 | 50.6667774 | 50.5800406 | 59.1702943 | 52.5881689 | 60.4790444 | 53.4884486 |
| ENSMUSG0C | 4683.12341 | 4664.54099 | 5036.95055 | 4660.87897 | 5265.8197 | 5170.91054 | 5264.63932 | 5526.9963 | -0.1565657 | 0.00701231 | 0.04670103 | Tmem259 | 1.11 | 10 | 79974948 | 79984330 | - | 4663 | protein_codi | transmembr | - | 4944 | 4530 | 5356 | 5036 | 5394 | 4776 | 4887 | 5565 | 29.5881504 | 29.4707459 | 31.8236436 | 29.4476092 | 33.2696475 | 32.6700079 | 33.2621898 | 34.9197711 |
| ENSMUSG0C | 1791.21893 | 1578.5301 | 1747.32153 | 1851.02421 | 2209.22321 | 1848.14579 | 1996.18921 | 2160.1468 | -0.2373146 | 0.00704843 | 0.04686355 | Mpv17 | 1.18 | 5 | 31140654 | 31154251 | - | 3214 | protein_codi | Protein Mpv1 | Mpv17_PMP2 | 1891 | 1533 | 1858 | 2000 | 2263 | 1707 | 1853 | 2175 | 16.4191408 | 14.4695366 | 16.016757 | 16.9673437 | 20.2507613 | 16.9409588 | 18.2979932 | 19.8009043 |
| ENSMUSG0C | 176.185468 | 227.563699 | 155.17118 | 161.039107 | 318.2531 | 296.656091 | 465.382481 | 143.016616 | -0.7648136 | 0.00705992 | 0.04691896 | Col20a1 | 1.70 | 2 | 180986535 | 181018363 | + | 5142 | protein_codi | Collagen alph | VWA | 186 | 221 | 165 | 174 | 326 | 274 | 432 | 144 | 1.00945193 | 1.30382272 | 0.88905089 | 0.92267108 | 1.82342624 | 1.69968651 | 2.66640177 | 0.8194115 |
| ENSMUSG0C | 3250.90606 | 3764.58319 | 4185.86013 | 3294.8231 | 4404.7791 | 4742.1667 | 5360.51673 | 3767.09739 | -0.3340931 | 0.00709788 | 0.04710817 | Mef2d | 1.26 | 3 | 88142372 | 88172086 | + | 5985 | protein_codi | Myocyte-spe | SRF-TF | 3432 | 3656 | 4451 | 3560 | 4512 | 4380 | 4976 | 3793 | 16.0025022 | 18.5310648 | 20.6047898 | 16.2186827 | 21.6824128 | 23.3431947 | 26.3870069 | 18.5434408 |
| ENSMUSG0C | 3432.77493 | 3517.45519 | 3359.22094 | 3388.29982 | 3818.06096 | 4161.84676 | 3929.89651 | 3596.27199 | -0.178783 | 0.00709557 | 0.04710817 | Atxn1l | 1.13 | 8 | 109726451 | 109737739 | - | 7526 | protein_codi | Ataxin-1-like | - | 3624 | 3416 | 3572 | 3661 | 3911 | 3844 | 3648 | 3621 | 13.4378177 | 13.769304 | 13.1498859 | 13.2637171 | 14.9460446 | 16.2918162 | 15.383832 | 14.0778374 |
| ENSMUSG0C | 134.507185 | 133.861 | 125.077375 | 57.3817506 | 168.888915 | 273.919675 | 187.445722 | 137.05759 | -0.7673135 | 0.00716778 | 0.04750238 | Sp100 | 1.70 | 1 | 85649988 | 85709998 | + | 7556 | protein_codi | nuclear antig | Sp100 | 142 | 130 | 133 | 62 | 173 | 253 | 174 | 138 | 0.52444647 | 0.52192698 | 0.48767943 | 0.22373271 | 0.65850159 | 1.06801883 | 0.73085498 | 0.53439055 |
| ENSMUSG0C | 10.4195707 | 10.297 | 15.9873337 | 14.8081937 | 46.8593522 | 50.8862637 | 17.2363882 | 14.8975641 | -1.3317018 | 0.00724424 | 0.0479512 | Afp | 2.52 | 5 | 90490737 | 90508907 | + | 3368 | protein_codi | Alpha-fetopr | - | 11 | 10 | 17 | 16 | 48 | 47 | 16 | 15 | 0.09114343 | 0.09007126 | 0.1398465 | 0.12953217 | 0.40989425 | 0.44511898 | 0.15077239 | 0.13031392 |
| ENSMUSG0C | 1207.72297 | 1189.3035 | 1128.51767 | 1134.67784 | 1312.06186 | 1417.23658 | 1262.56543 | 1336.80809 | -0.1934449 | 0.00726233 | 0.04802818 | Fzr1 | 1.14 | 10 | 81366222 | 81378516 | - | 4488 | protein_codi | Fizzy-related | WD40 | 1275 | 1155 | 1200 | 1226 | 1344 | 1309 | 1172 | 1346 | 7.927972 | 7.80705929 | 7.40803706 | 7.44847486 | 8.61289382 | 9.30330233 | 8.28797967 | 8.77533787 |
| ENSMUSG0C | 1476.73734 | 1375.6792 | 1451.08564 | 1377.16202 | 1585.40808 | 1675.99864 | 1651.46144 | 1536.43545 | -0.1826721 | 0.00728515 | 0.04813635 | Gtf3c4 | 1.13 | 2 | 28822299 | 28840360 | - | 7230 | protein_codi | ral transcript | - | 1559 | 1336 | 1543 | 1488 | 1624 | 1548 | 1533 | 1547 | 6.01745136 | 5.60565677 | 5.91292509 | 5.611699 | 6.460266 | 6.829407 | 6.72942212 | 6.26071092 |
| ENSMUSG0C | 4106.25809 | 4181.61169 | 4226.29868 | 4082.4339 | 4473.11566 | 4727.00909 | 4937.14794 | 4441.46046 | -0.1626201 | 0.00730061 | 0.04818518 | Syncrip | 1.12 | 9 | 88447009 | 88482574 | - | 12297 | protein_codi | synaptotagm | RRM_1 | 4335 | 4061 | 4494 | 4411 | 4582 | 4366 | 4583 | 4472 | 9.83772548 | 10.0182568 | 10.1253173 | 9.78064777 | 10.7166386 | 11.3249135 | 11.8283617 | 10.6407994 |
| ENSMUSG0C | 3753.88715 | 4011.71119 | 3946.99056 | 3860.311 | 4186.10213 | 4276.61152 | 4523.47462 | 4332.21165 | -0.1532584 | 0.00730226 | 0.04818518 | Brox | 1.11 | 1 | 183276352 | 183297269 | - | 5010 | protein_codi | Mus musculu | - | 3963 | 3896 | 4197 | 4171 | 4288 | 3950 | 4199 | 4362 | 22.0745123 | 23.5906314 | 23.2100455 | 22.7003314 | 24.616127 | 25.1483622 | 26.6000261 | 25.4753155 |
| ENSMUSG0C | 2485.54123 | 2431.12169 | 2623.80359 | 2441.50094 | 2882.82639 | 2687.2278 | 2743.81754 | 2897.07964 | -0.16764 | 0.00735209 | 0.0484281 | Mtmr2 | 1.12 | 9 | 13748410 | 13806481 | + | 4913 | protein_codi | Myotubularin | GRAM | 2624 | 2361 | 2790 | 2638 | 2953 | 2482 | 2547 | 2917 | 14.9046519 | 14.5783229 | 15.7337479 | 14.6405625 | 17.286989 | 16.1140738 | 16.4534166 | 17.3724592 |
| ENSMUSG0C | 4937.92928 | 4663.51129 | 5254.1902 | 4593.31659 | 5419.08883 | 5537.94125 | 5328.1985 | 5881.55833 | -0.1887055 | 0.00736199 | 0.04847185 | Hnrnpab | 1.14 | 11 | 51600102 | 51606847 | - | 2782 | protein_codi | Heterogeneous nu | RRM_1 | 5213 | 4529 | 5587 | 4963 | 5551 | 5115 | 4946 | 5922 | 52.2920142 | 49.3859641 | 55.6411752 | 48.6426116 | 57.3874298 | 58.6460611 | 56.4249133 | 62.2849202 |
| ENSMUSG0C | 10754.8914 | 11117.6709 | 11802.414 | 10696.1434 | 13116.7137 | 11678.9389 | 12384.3449 | 13059.2047 | -0.1792514 | 0.00739109 | 0.04857749 | Rpl4 | 1.13 | 9 | 64173375 | 64178666 | + | 2007 | protein_codi | ribosomal pr | - | 11354 | 10797 | 12550 | 11557 | 13436 | 10787 | 11496 | 13149 | 157.872428 | 163.197714 | 173.249146 | 157.010058 | 192.541919 | 171.436639 | 181.791384 | 191.697738 |
| ENSMUSG0C | 2111.38392 | 2027.47929 | 2185.56256 | 2071.2961 | 2262.91621 | 2370.00066 | 2336.60787 | 2443.20052 | -0.1647975 | 0.00740996 | 0.04867998 | Sacm1l | 1.12 | 9 | 123529759 | 123592600 | + | 3738 | protein_codi | Mus musculu | - | 2229 | 1969 | 2324 | 2238 | 2318 | 2189 | 2169 | 2460 | 16.6408514 | 15.9795579 | 17.2254896 | 16.3248996 | 17.8351517 | 18.6791367 | 18.4159518 | 19.2560607 |
| ENSMUSG0C | 73.8842286 | 35.0097999 | 29.1533732 | 29.6163874 | 93.7187043 | 92.0283492 | 71.1001013 | 57.6039147 | -0.9067265 | 0.00741808 | 0.04871187 | Mcm10 | 1.87 | 2 | 4989714 | 5012791 | - | 7692 | protein_codi | minichromos | - | 78 | 34 | 31 | 32 | 96 | 85 | 66 | 58 | 0.28298284 | 0.13409049 | 0.11165988 | 0.11343327 | 0.35895056 | 0.35247635 | 0.2723194 | 0.22062786 |
| ENSMUSG0C | 1364.96376 | 1842.13329 | 835.103078 | 1751.06891 | 2696.36522 | 1679.2467 | 2712.57659 | 3667.78029 | -0.8927781 | 0.00742644 | 0.04874523 | Kcnj12 | 1.86 | 11 | 61022564 | 61071131 | + | 2750 | protein_codi | ATP-sensitive | - | 1441 | 1789 | 888 | 1892 | 2762 | 1551 | 2518 | 3693 | 14.6229858 | 19.7349481 | 8.9465382 | 18.7593667 | 28.8864155 | 17.9899286 | 29.0600894 | 39.2932769 |
| ENSMUSG0C | 8434.16887 | 8290.11468 | 7849.78084 | 8195.40971 | 9094.61926 | 9099.97971 | 9487.55442 | 8691.23892 | -0.1505008 | 0.00744954 | 0.04887526 | Zfp106 | 1.11 | 2 | 120506820 | 120563843 | - | 10429 | protein_codi | zinc finger pr | - | 8904 | 8051 | 8347 | 8855 | 9316 | 8405 | 8807 | 8751 | 23.825787 | 23.4188465 | 22.1749421 | 23.1513133 | 25.6915014 | 25.7066442 | 26.8015088 | 24.5519873 |
| ENSMUSG0C | 167.660365 | 156.5144 | 186.205416 | 82.3705775 | 219.653213 | 331.302057 | 176.672979 | 230.415659 | -0.6928085 | 0.00748645 | 0.04909578 | Tbx3 | 1.62 | 5 | 119670669 | 119684724 | + | 6624 | protein_codi | T-box 3 [Sou | T-box | 177 | 152 | 198 | 89 | 225 | 306 | 164 | 232 | 0.74568896 | 0.69611598 | 0.82817023 | 0.36635272 | 0.9769332 | 1.47350441 | 0.78577361 | 1.02480042 |
| ENSMUSG0C | 2923.1632 | 2930.52619 | 2783.67693 | 2525.72254 | 3024.38069 | 3446.19101 | 3531.30503 | 3078.82992 | -0.2286166 | 0.00750551 | 0.0491224 | Smurf2 | 1.17 | 11 | 106820066 | 106920715 | - | 6162 | protein_codi | E3 ubiquitin- | HECT | 3086 | 2846 | 2960 | 2729 | 3098 | 3183 | 3278 | 3100 | 13.9758747 | 14.0110778 | 13.3089798 | 12.0756794 | 14.4598035 | 16.4765121 | 16.883449 | 14.7201296 |
| ENSMUSG0C | 1021.11793 | 1153.264 | 1058.92575 | 985.670394 | 1250.55896 | 1275.40465 | 1242.09722 | 1143.13976 | -0.2194694 | 0.00749723 | 0.0491224 | Ttc33 | 1.16 | 15 | 5185525 | 5218336 | + | 2163 | protein_codi | tetratricopep | - | 1078 | 1120 | 1126 | 1065 | 1281 | 1178 | 1153 | 1151 | 13.9080746 | 15.7079621 | 14.4230337 | 13.4252636 | 17.0331622 | 17.3715715 | 16.9179097 | 15.5700655 |
| ENSMUSG0C | 1779.85212 | 1803.0047 | 1536.6649 | 1570.59405 | 1819.70484 | 2331.02395 | 2089.91207 | 1874.11357 | -0.2783858 | 0.0075146 | 0.04912884 | Wipf2 | 1.21 | 11 | 98863638 | 98905040 | + | 7819 | protein_codi | WAS/WASL i | - | 1879 | 1751 | 1634 | 1697 | 1864 | 2153 | 1940 | 1887 | 6.70625924 | 6.79349523 | 5.78996032 | 5.91780109 | 6.85641928 | 8.78300547 | 7.87452617 | 7.06142453 |
| ENSMUSG0C | 4776.89955 | 5114.51989 | 5360.45895 | 4680.31472 | 5808.60719 | 5399.35738 | 5862.52653 | 5587.57973 | -0.1849943 | 0.00766696 | 0.04988388 | Tmem248 | 1.14 | 5 | 130217081 | 130243765 | + | 4086 | protein_codi | Transmembr | - | 5043 | 4967 | 5700 | 5057 | 5950 | 4987 | 5442 | 5626 | 34.4425563 | 36.8768774 | 38.6501552 | 33.7461572 | 41.8814082 | 38.930622 | 42.2701792 | 40.2877488 |

| gene_id | KO1 | KO2 | KO3 | KO4 | ctrl1 | ctrl2 | ctrl3 | ctrl4 | log2FoldChar | pvalue | padj | gene_name | Fold Change up | gene_chr | gene_start | gene_end | gene_strand | gene_length | gene_biotype | gene_descrip | tf_family | KO1_readcou | KO2_readcou | KO3_readcou | KO4_readcou | ctrl1_readco | ctrl2_readco | ctrl3_readco | ctrl4_readco | KO1_fpkm | KO2_fpkm | KO3_fpkm | KO4_fpkm | ctrl1_fpkm | ctrl2_fpkm | ctrl3_fpkm | ctrl4_fpkm |
|---|---|---|---|---|---|---|---|---|---|---|---|---|---|---|---|---|---|---|---|---|---|---|---|---|---|---|---|---|---|---|---|---|---|---|---|---|---|
| ENSMUSG00 | 1180.25319 | 648.710998 | 863.31602 | 763.547488 | 53.6930077 | 75.7880523 | 56.0182616 | 132.091735 | 3.44247293 | 1.15E-38 | 5.91E-36 | Nptx1 | 10.8714535 | 11 | 119538719 | 119547753 | - | 5217 | protein_codi | neuronal pen | - | 1246 | 630 | 918 | 825 | 55 | 70 | 52 | 133 | 6.66502797 | 3.66334697 | 4.87524666 | 4.31184208 | 0.3032107 | 0.42798401 | 0.31634168 | 0.7459375 |
| ENSMUSG00 | 1958.87929 | 2107.79589 | 1931.64608 | 2123.12477 | 899.11382 | 822.841711 | 903.833106 | 663.43819 | 1.3042036 | 1.87E-33 | 7.52E-31 | Kirrel2 | 2.46947369 | 7 | 30447534 | 30457690 | - | 3939 | protein_codi | Kin of IRRE-li | I-set | 2068 | 2047 | 2054 | 2294 | 921 | 760 | 839 | 668 | 14.6510694 | 15.7648631 | 14.4473837 | 15.8795126 | 6.72475276 | 6.15428987 | 6.76004978 | 4.96206121 |
| ENSMUSG00 | 10304.9554 | 13272.833 | 13140.6479 | 12354.6611 | 3969.37762 | 2421.96961 | 1972.48917 | 3071.87773 | 2.101195 | 3.71E-31 | 1.38E-28 | Cyp2s1 | 4.29064636 | 7 | 25802475 | 25816913 | - | 4938 | protein_codi | Cytochrome I | p450 | 10879 | 12890 | 13973 | 13349 | 4066 | 2237 | 1831 | 3093 | 61.4812462 | 79.1881457 | 78.3995053 | 73.7101647 | 23.6820319 | 14.4499131 | 11.7682307 | 18.3273836 |
| ENSMUSG00 | 1823.42487 | 1799.9156 | 1827.2582 | 2257.32403 | 752.678344 | 673.430979 | 705.614641 | 898.819704 | 1.34636157 | 1.07E-29 | 3.55E-27 | Ccdc3 | 2.54270056 | 2 | 5137776 | 5230878 | + | 2670 | protein_codi | Coiled-coil dc | - | 1925 | 1748 | 1943 | 2439 | 771 | 622 | 655 | 905 | 20.1198271 | 19.8604237 | 20.1621243 | 24.9075077 | 8.30511766 | 7.43069541 | 7.78581271 | 9.917654 |
| ENSMUSG00 | 61478.3089 | 61223.9024 | 60863.7794 | 56889.3782 | 38934.2642 | 36185.5469 | 39270.9559 | 35823.6759 | 0.67874109 | 8.00E-29 | 2.38E-26 | St3gal1 | 1.60074232 | 15 | 67102875 | 67176830 | - | 6851 | protein_codi | ST3 beta-gal | Glyco_transf_ | 64903 | 59458 | 64719 | 61468 | 39882 | 33422 | 36454 | 36070 | 264.372072 | 263.278061 | 261.729442 | 244.638525 | 167.427054 | 155.606883 | 168.874912 | 154.050747 |
| ENSMUSG00 | 22316.826 | 23581.1596 | 24805.7589 | 20712.961 | 11314.5811 | 12227.8609 | 12670.8999 | 10315.0734 | 0.97438746 | 2.03E-27 | 5.50E-25 | Mast4 | 1.9648068 | 13 | 102732486 | 103334497 | - | 32326 | protein_codi | Microtubule- | Pkinase | 23560 | 22901 | 26377 | 22380 | 11590 | 11294 | 11762 | 10386 | 20.3389294 | 21.4912076 | 22.607273 | 18.8772118 | 10.3117919 | 11.1441295 | 11.5479028 | 9.40086865 |
| ENSMUSG00 | 672.535927 | 757.859198 | 726.013036 | 663.592181 | 314.348154 | 278.250421 | 256.391274 | 198.634189 | 1.42838429 | 8.91E-25 | 2.04E-22 | Wnk4 | 2.69145125 | 11 | 101260567 | 101277409 | + | 4148 | protein_codi | WNK lysine c | Pkinase | 710 | 736 | 772 | 717 | 322 | 257 | 238 | 200 | 4.77666047 | 5.3826657 | 5.15647957 | 4.71313785 | 2.23264563 | 1.97626287 | 1.82100913 | 1.41079166 |
| ENSMUSG00 | 7988.96903 | 8956.33058 | 8382.06501 | 8158.38922 | 5226.77024 | 4251.70974 | 4741.08403 | 4555.67512 | 0.83455976 | 1.35E-24 | 3.03E-22 | Serpinb6b | 1.78331278 | 13 | 32965209 | 32979067 | + | 2262 | protein_codi | serine (or cys | Serpin | 8434 | 8698 | 8913 | 8815 | 5354 | 3927 | 4401 | 4587 | 104.050882 | 116.650108 | 109.17069 | 106.257465 | 68.0751237 | 55.3756245 | 61.7493914 | 59.3345666 |
| ENSMUSG00 | 4163.09211 | 6827.94068 | 6271.73697 | 6597.97581 | 2020.80956 | 1967.2413 | 1681.62512 | 2400.49417 | 1.56382381 | 3.83E-23 | 8.15E-21 | Mmp17 | 2.95636379 | 5 | 129584169 | 129611099 | + | 6619 | protein_codi | Matrix metal | Hemopexin | 4395 | 6631 | 6669 | 7129 | 2070 | 1817 | 1561 | 2417 | 18.5298229 | 30.3909999 | 27.9153505 | 29.3674317 | 8.99457479 | 8.75614375 | 7.484873 | 10.6845419 |
| ENSMUSG00 | 114.615278 | 160.6332 | 129.779532 | 182.325885 | 34.1682776 | 20.5710428 | 29.0864051 | 21.8497607 | 2.47164573 | 1.18E-21 | 2.23E-19 | Cntnap2 | 5.54676165 | 6 | 45059357 | 47304213 | + | 10248 | protein_codi | Contactin-as: | F5_F8_type_ | 121 | 156 | 138 | 197 | 35 | 19 | 27 | 22 | 0.32949667 | 0.46178925 | 0.37309095 | 0.52415151 | 0.09822716 | 0.05913775 | 0.08361777 | 0.06281382 |
| ENSMUSG00 | 719.897612 | 884.512298 | 874.601196 | 905.150841 | 403.185676 | 263.09281 | 252.082177 | 291.992257 | 1.48180702 | 2.01E-21 | 3.73E-19 | Gm9885 | 2.79298343 | 7 | 78578830 | 78581038 | + | 2209 | antisense | - | - | 760 | 859 | 930 | 978 | 413 | 243 | 234 | 294 | 9.60113657 | 11.7965711 | 11.6643886 | 12.0718234 | 5.37721014 | 3.50881842 | 3.36197172 | 3.89424481 |
| ENSMUSG00 | 1060.90174 | 1337.5803 | 1218.79909 | 1259.62198 | 575.979537 | 500.201145 | 447.068819 | 624.704523 | 1.1819197 | 2.06E-21 | 3.78E-19 | Smarcd3 | 2.26878469 | 5 | 24590818 | 24624411 | - | 5120 | protein_codi | SWI/SNF rel: | - | 1120 | 1299 | 1296 | 1361 | 590 | 462 | 415 | 629 | 6.10453844 | 7.69657546 | 7.01309609 | 7.24799523 | 3.31424587 | 2.87820916 | 2.57248025 | 3.59461448 |
| ENSMUSG00 | 8653.92709 | 9320.84437 | 9425.00343 | 9134.8045 | 6453.89952 | 6178.89164 | 6441.02281 | 6451.63845 | 0.51726262 | 5.02E-21 | 8.79E-19 | Slc44a2 | 1.43123703 | 9 | 21320698 | 21355028 | + | 8677 | protein_codi | solute carrie | - | 9136 | 9052 | 10022 | 9870 | 6611 | 5707 | 5979 | 6496 | 29.3826705 | 31.6470541 | 32.0007052 | 31.0153931 | 21.912919 | 20.9791849 | 21.8691987 | 21.9052419 |
| ENSMUSG00 | 35149.0009 | 35458.7491 | 39793.414 | 41752.6277 | 19594.0429 | 14332.6034 | 15088.3033 | 11433.3839 | 1.33174772 | 1.83E-20 | 2.92E-18 | Ntrk3 | 2.51707416 | 7 | 78175959 | 78738012 | - | 30170 | protein_codi | NT-3 growth | Pkinase_Tyr | 37107 | 34436 | 42314 | 45113 | 20071 | 13238 | 14006 | 11512 | 34.3230051 | 34.6254743 | 38.8582752 | 40.7714477 | 19.133586 | 13.9957895 | 14.7337306 | 11.1647012 |
| ENSMUSG00 | 4691.64852 | 4362.83889 | 4829.11521 | 4799.70579 | 3249.89132 | 3069.41612 | 3105.7817 | 3233.76459 | 0.56159282 | 1.59E-19 | 2.39E-17 | Pitpnc1 | 1.47589779 | 11 | 107207892 | 107470699 | - | 6612 | protein_codi | Cytoplasmic | IP_trans | 4953 | 4237 | 5135 | 5186 | 3329 | 2835 | 2883 | 3256 | 20.9045226 | 19.4394494 | 21.5170313 | 21.3859921 | 14.4805022 | 13.676361 | 13.8383946 | 14.4086465 |
| ENSMUSG00 | 16441.1353 | 16194.0919 | 17198.6093 | 20952.6686 | 7193.88679 | 8631.17647 | 8649.43505 | 9846.29673 | 1.04442085 | 3.29E-19 | 4.79E-17 | Fam81a | 2.06253821 | 9 | 70088511 | 70142560 | - | 4397 | protein_codi | Protein FAM! | - | 17357 | 15727 | 18288 | 22639 | 7369 | 7972 | 8029 | 9914 | 110.159757 | 108.504503 | 115.235024 | 140.388168 | 48.2008574 | 57.8310611 | 57.9533981 | 65.9726735 |
| ENSMUSG00 | 141.137821 | 128.7125 | 103.447453 | 159.188082 | 32.2158046 | 32.4805939 | 29.0864051 | 34.760983 | 2.04961829 | 4.12E-19 | 5.95E-17 | Aard | 4.1399642 | 15 | 52040107 | 52045722 | + | 1348 | protein_codi | alanine and a | - | 149 | 125 | 110 | 172 | 33 | 30 | 27 | 35 | 3.08461629 | 2.81305655 | 2.26088015 | 3.47911103 | 0.70408764 | 0.7098747 | 0.63569352 | 0.7597134 |
| ENSMUSG00 | 1513.67945 | 1389.0653 | 1544.18835 | 1362.35382 | 974.28403 | 850.991559 | 852.123941 | 877.963114 | 0.70795894 | 7.95E-17 | 1.02E-14 | Cd59b | 1.63349149 | 2 | 104069849 | 104091187 | + | 1300 | protein_codi | CD59B glycop | - | 1598 | 1349 | 1642 | 1472 | 998 | 786 | 791 | 884 | 34.303481 | 31.4794357 | 34.9948833 | 30.8740919 | 22.0795319 | 19.285439 | 19.3111014 | 19.8966769 |
| ENSMUSG00 | 825.040553 | 1030.7297 | 896.231118 | 897.746744 | 440.282663 | 250.100573 | 277.93676 | 386.343497 | 1.42811571 | 8.09E-17 | 1.03E-14 | Gstt1 | 2.69095023 | 10 | 75783813 | 75798584 | - | 1528 | protein_codi | Glutathione : | GST_N | 871 | 1001 | 953 | 970 | 451 | 231 | 258 | 389 | 15.9074135 | 19.8732577 | 17.2800222 | 17.3092446 | 8.48898684 | 4.82213053 | 5.35883352 | 7.44899842 |
| ENSMUSG00 | 5401.12656 | 7291.30568 | 8140.37415 | 5719.66482 | 3507.61776 | 2559.4708 | 2406.6307 | 2387.58295 | 1.2895213 | 6.32E-16 | 7.52E-14 | Atp1b2 | 2.44446932 | 11 | 69599736 | 69605942 | - | 4085 | protein_codi | Sodium/pota | - | 5702 | 7081 | 8656 | 6180 | 3593 | 2364 | 2234 | 2404 | 38.9529114 | 52.5848785 | 58.7083582 | 41.2502086 | 25.2969306 | 18.4588971 | 17.3566147 | 17.2192423 |
| ENSMUSG00 | 150342.091 | 139525.379 | 153500.033 | 141495.067 | 106367.801 | 93716.2574 | 99633.8646 | 103647.32 | 0.53600587 | 7.19E-16 | 8.49E-14 | Dag1 | 1.44995274 | 9 | 108204634 | 108263958 | - | 6357 | protein_codi | Dystroglycan | - | 158717 | 135501 | 163223 | 152883 | 108957 | 86559 | 92487 | 104360 | 696.748441 | 646.619251 | 711.383672 | 655.747614 | 492.953096 | 434.320527 | 461.745206 | 480.345243 |
| ENSMUSG00 | 555.078948 | 380.988999 | 713.787428 | 675.623838 | 181.57999 | 178.643266 | 93.7228608 | 174.798086 | 1.88638114 | 7.55E-16 | 8.85E-14 | Fam189a2 | 3.69706687 | 19 | 23972751 | 24031019 | - | 2547 | protein_codi | family with s | - | 586 | 370 | 759 | 730 | 186 | 165 | 87 | 176 | 6.42056788 | 4.40687894 | 8.2563402 | 7.81490404 | 2.10032582 | 2.0663569 | 1.08408721 | 2.02187991 |
| ENSMUSG00 | 287.011811 | 540.592499 | 260.499496 | 401.672254 | 126.910745 | 84.449544 | 94.800135 | 81.4400173 | 1.9406626 | 2.75E-15 | 3.08E-13 | Dll1 | 3.83881916 | 17 | 15367354 | 15376872 | - | 4219 | protein_codi | Delta-like prc | EGF | 303 | 525 | 277 | 434 | 130 | 78 | 88 | 82 | 2.00418531 | 3.77492319 | 1.81905149 | 2.80485192 | 0.8862097 | 0.58970582 | 0.66198334 | 0.56869049 |
| ENSMUSG00 | 44054.8922 | 42862.2921 | 50629.0645 | 42439.3577 | 29992.9141 | 26732.6114 | 29782.3242 | 29689.8522 | 0.63129333 | 3.02E-15 | 3.35E-13 | Mme | 1.54895295 | 3 | 63241537 | 63386030 | + | 11542 | protein_codi | Neprilysin [S | - | 46509 | 41626 | 53836 | 45855 | 30723 | 24691 | 27646 | 29894 | 112.450322 | 109.406204 | 129.230928 | 108.326662 | 76.5570559 | 68.2351178 | 76.0195243 | 75.7834889 |
| ENSMUSG00 | 262.383735 | 229.623099 | 546.39064 | 394.268158 | 90.7899948 | 57.3823825 | 92.6455865 | 84.4195302 | 2.13870889 | 4.17E-15 | 4.53E-13 | Crispld1 | 4.40367771 | 1 | 17727045 | 17766344 | + | 4515 | protein_codi | Cysteine-rich | LCCL | 277 | 223 | 581 | 426 | 93 | 53 | 86 | 85 | 1.71209077 | 1.49832301 | 3.56527576 | 2.57265517 | 0.59241748 | 0.37442811 | 0.60452548 | 0.55084931 |
| ENSMUSG00 | 489.719823 | 487.048099 | 484.322168 | 472.011175 | 301.65708 | 237.108335 | 260.700371 | 222.470291 | 0.91877709 | 7.32E-15 | 7.78E-13 | Cyp46a1 | 1.8905121 | 12 | 108334381 | 108362234 | + | 3287 | protein_codi | Cholesterol 2 | p450 | 517 | 473 | 515 | 510 | 309 | 219 | 242 | 224 | 4.38930349 | 4.36535712 | 4.34092491 | 4.23058287 | 2.70371835 | 2.12517524 | 2.336628 | 1.9939761 |
| ENSMUSG00 | 76132.9614 | 82693.1474 | 92314.6265 | 77514.4155 | 56489.9253 | 51451.426 | 50485.381 | 54575.7365 | 0.62569478 | 1.44E-14 | 1.50E-12 | Crim1 | 1.54295371 | 17 | 78200248 | 78376592 | + | 5995 | protein_codi | Cysteine-rich | VWC | 80374 | 80308 | 98162 | 83753 | 57865 | 47522 | 46864 | 54951 | 374.137436 | 406.375919 | 453.658403 | 380.926266 | 277.60638 | 252.845867 | 248.098467 | 268.199552 |
| ENSMUSG00 | 1181.20042 | 1666.0546 | 1225.38211 | 1491.92552 | 800.513933 | 624.710088 | 619.4327 | 587.957198 | 1.07919559 | 2.96E-14 | 2.99E-12 | Ifitm1 | 2.11285767 | 7 | 140967221 | 140969825 | + | 1094 | protein_codi | Interferon-in | - | 1247 | 1618 | 1303 | 1612 | 820 | 577 | 575 | 592 | 31.8092846 | 44.8662257 | 32.9990808 | 40.1769949 | 21.5575401 | 16.8232085 | 16.6810904 | 15.833467 |
| ENSMUSG00 | 1175.51702 | 1273.7389 | 1361.74466 | 1324.40783 | 897.161346 | 769.790074 | 783.178388 | 759.775772 | 0.67758852 | 7.10E-14 | 6.90E-12 | Syt11 | 1.599464 | 3 | 88744700 | 88775164 | - | 5205 | protein_codi | Synaptotagm | C2 | 1241 | 1237 | 1448 | 1431 | 919 | 711 | 727 | 765 | 6.65358667 | 7.20953588 | 7.70766049 | 7.49632892 | 5.07805558 | 4.35711681 | 4.43289649 | 4.30043449 |
| ENSMUSG00 | 514.347899 | 474.691699 | 445.764481 | 422.033521 | 236.249234 | 216.537292 | 254.236726 | 279.081035 | 0.91208709 | 9.50E-14 | 9.12E-12 | Reep2 | 1.88176581 | 18 | 34840589 | 34847463 | + | 1926 | protein_codi | Receptor exp | TB2_DP1_HV | 543 | 461 | 474 | 456 | 242 | 200 | 236 | 281 | 7.86770953 | 7.26110947 | 6.81862502 | 6.45562499 | 3.61378038 | 3.31225717 | 3.88892559 | 4.26895594 |
| ENSMUSG00 | 27648.8045 | 20897.7614 | 37460.2037 | 25819.9368 | 11985.2556 | 12005.9102 | 11253.2069 | 14519.166 | 1.1681024 | 1.04E-13 | 9.93E-12 | Fgl2 | 2.2471593 | 5 | 21372642 | 21378374 | + | 3788 | protein_codi | Fibroleukin [ | Fibrinogen_C | 29189 | 20295 | 39833 | 27898 | 12277 | 11089 | 10446 | 14619 | 215.037408 | 162.531456 | 291.345151 | 200.813467 | 93.2148185 | 93.375459 | 87.5213413 | 112.9222 |
| ENSMUSG00 | 435.727502 | 447.919499 | 376.172558 | 406.299815 | 245.035362 | 206.793114 | 243.463983 | 223.463462 | 0.85764953 | 1.89E-13 | 1.75E-11 | Ralgps1 | 1.81208362 | 2 | 33133417 | 33371486 | - | 7657 | protein_codi | Ras-specific | PH | 460 | 435 | 400 | 439 | 251 | 191 | 226 | 225 | 1.67650154 | 1.72341136 | 1.44735842 | 1.56327581 | 0.94279603 | 0.79565548 | 0.93675001 | 0.85979617 |
| ENSMUSG00 | 12702.4039 | 11722.1048 | 13381.3983 | 11338.4488 | 7928.99288 | 8270.64188 | 7883.49305 | 8908.74336 | 0.5749079 | 2.48E-13 | 2.25E-11 | Pcsk5 | 1.48958236 | 19 | 17432832 | 17837632 | - | 9346 | protein_codi | Proprotein cc | Peptidase_S8 | 13410 | 11384 | 14229 | 12251 | 8122 | 7639 | 7318 | 8970 | 40.0412634 | 36.9511069 | 42.1816294 | 35.7417241 | 24.9942369 | 26.0712029 | 24.8508096 | 28.0826639 |
| ENSMUSG00 | 656.432954 | 898.928098 | 910.337589 | 711.71881 | 346.563959 | 313.979074 | 231.613966 | 409.186429 | 1.28664158 | 2.95E-13 | 2.66E-11 | Gnai1 | 2.43959486 | 5 | 18265135 | 18360355 | - | 3135 | protein_codi | guanine nucl | G-alpha | 693 | 873 | 968 | 769 | 355 | 290 | 215 | 412 | 6.16879674 | 8.44763304 | 8.55485318 | 6.68834288 | 3.25681793 | 2.95060306 | 2.17658097 | 3.84530955 |
| ENSMUSG00 | 322.059458 | 314.058499 | 214.418358 | 349.843576 | 113.243434 | 107.18596 | 114.191072 | 142.023445 | 1.33142678 | 4.86E-13 | 4.28E-11 | Ripor3 | 2.51651427 | 2 | 167980164 | 168010618 | - | 4244 | protein_codi | RIPOR family | - | 340 | 305 | 228 | 378 | 116 | 99 | 106 | 143 | 2.23567316 | 2.18013208 | 1.4884499 | 2.42854502 | 0.78611356 | 0.74406377 | 0.79269187 | 0.98589871 |
| ENSMUSG00 | 8044.85582 | 7528.13668 | 6910.28988 | 6492.46743 | 5013.95068 | 3923.65574 | 4351.11074 | 4125.6321 | 0.73444547 | 5.53E-13 | 4.78E-11 | Klhl5 | 1.66375784 | 5 | 65107539 | 65168188 | + | 7057 | protein_codi | Mus muscul | BACK | 8493 | 7311 | 7348 | 7015 | 5136 | 3624 | 4039 | 4154 | 33.5850329 | 31.4278743 | 28.848536 | 27.1042436 | 20.9318479 | 16.3801701 | 18.1646757 | 17.2233652 |
| ENSMUSG00 | 312.587121 | 335.682199 | 371.470401 | 482.191808 | 158.150314 | 176.477893 | 157.282042 | 178.77077 | 1.16376974 | 5.66E-13 | 4.87E-11 | Naaa | 2.24042082 | 5 | 92257659 | 92278170 | - | 3721 | protein_codi | N-acylethanc | - | 330 | 326 | 395 | 521 | 162 | 163 | 146 | 180 | 2.47490789 | 2.65776313 | 2.94111614 | 3.81775265 | 1.25215478 | 1.39726336 | 1.24528025 | 1.4154172 |
| ENSMUSG00 | 122.193147 | 103.9997 | 63.9493348 | 99.9553076 | 24.4059126 | 21.6537292 | 26.9318565 | 14.8975641 | 2.15214326 | 6.97E-13 | 5.92E-11 | Gm48822 | 4.44487626 | 9 | 75582739 | 75585109 | - | 2371 | TEC | predicted ge | - | 129 | 101 | 68 | 108 | 25 | 20 | 25 | 15 | 1.51831865 | 1.29225482 | 0.79460649 | 1.24200097 | 0.3032572 | 0.26905978 | 0.33464348 | 0.18511062 |
| ENSMUSG00 | 13948.0162 | 26263.5281 | 25244.9403 | 14213.0894 | 7356.91829 | 8582.45558 | 7517.2198 | 6915.44928 | 1.39128411 | 1.19E-12 | 9.80E-11 | F2r | 2.62312054 | 13 | 95601803 | 95618487 | - | 3336 | protein_codi | Proteinase-a | 7tm_1 | 14725 | 25506 | 26844 | 15357 | 7536 | 7927 | 6978 | 6963 | 123.178251 | 231.939467 | 222.944076 | 125.519175 | 64.9706962 | 75.7937077 | 66.386357 | 61.0719783 |
| ENSMUSG00 | 561.709584 | 1702.0941 | 1036.3554 | 1433.61825 | 219.653213 | 168.899088 | 196.063916 | 245.313223 | 2.51153036 | 1.24E-12 | 1.02E-10 | Bhlhe40 | 5.70224634 | 6 | 108660629 | 108666925 | + | 4705 | protein_codi | Class E basic | HLH | 593 | 1653 | 1102 | 1549 | 225 | 156 | 182 | 247 | 3.51722245 | 10.6578982 | 6.48928302 | 8.97679948 | 1.37538905 | 1.05758506 | 1.22768139 | 1.53606276 |
| ENSMUSG00 | 409.204959 | 572.513198 | 377.112989 | 448.873372 | 181.57999 | 156.989537 | 203.604835 | 234.388343 | 1.21755199 | 1.27E-12 | 1.04E-10 | Tppp3 | 2.32551781 | 8 | 105467493 | 105471526 | - | 1848 | protein_codi | Tubulin polyr | p25-alpha | 432 | 556 | 401 | 485 | 186 | 145 | 189 | 236 | 6.52358839 | 9.12706548 | 6.01197484 | 7.15598639 | 2.89476725 | 2.50274367 | 3.24588965 | 3.73664355 |
| ENSMUSG00 | 201.760778 | 271.840799 | 177.741533 | 197.134079 | 100.55236 | 68.2092471 | 72.1773755 | 61.5765985 | 1.48484702 | 1.48E-12 | 1.20E-10 | Acot11 | 2.79887493 | 4 | 106744555 | 106804998 | - | 6056 | protein_codi | acyl-CoA thic | START | 213 | 264 | 189 | 213 | 103 | 63 | 67 | 62 | 0.98151854 | 1.32244129 | 0.86467059 | 0.95901074 | 0.48916349 | 0.33182188 | 0.35112589 | 0.29955561 |
| ENSMUSG00 | 2898.53512 | 2084.11279 | 2650.13567 | 3130.08195 | 1110.95714 | 1295.97569 | 1354.13375 | 1596.02571 | 1.00670336 | 2.30E-12 | 1.83E-10 | Gm826 | 2.00931445 | 2 | 160311393 | 160334162 | - | 1493 | protein_codi | - | - | 3060 | 2024 | 2818 | 3382 | 1138 | 1197 | 1257 | 1607 | 57.1960964 | 41.1252965 | 52.2944897 | 61.7651542 | 21.9222501 | 25.5731767 | 26.7207955 | 31.4939914 |
| ENSMUSG00 | 382.682415 | 294.494199 | 236.988711 | 433.139666 | 83.9563393 | 58.4650689 | 49.554616 | 135.071248 | 2.03970359 | 2.51E-12 | 1.98E-10 | Ssc5d | 4.11161046 | 7 | 4925785 | 4944826 | + | 4590 | protein_codi | Soluble scav | SRCR | 404 | 286 | 252 | 468 | 86 | 54 | 46 | 136 | 2.45625499 | 1.89021711 | 1.52111694 | 2.7801159 | 0.5388755 | 0.37525925 | 0.31806733 | 0.8669576 |
| ENSMUSG00 | 806.095879 | 1717.5396 | 863.31602 | 1130.05028 | 477.37965 | 308.565642 | 318.873181 | 410.1796 | 1.57512232 | 5.54E-12 | 4.25E-10 | Hrh2 | 2.97960754 | 13 | 54192129 | 54236180 | + | 5935 | protein_codi | Histamine H: | 7tm_1 | 851 | 1668 | 918 | 1221 | 489 | 285 | 296 | 413 | 4.00141511 | 8.52577102 | 4.2854527 | 5.60950675 | 2.36968603 | 1.53170268 | 1.58286874 | 2.03610872 |
| ENSMUSG00 | 1047.64047 | 869.066798 | 1306.25921 | 1080.99814 | 565.240935 | 296.656091 | 286.554954 | 440.967899 | 1.4361733 | 5.52E-12 | 4.25E-10 | Lingo1 | 2.70602149 | 9 | 56618475 | 56796468 | - | 4715 | protein_codi | Leucine-rich | I-set | 1106 | 844 | 1389 | 1168 | 579 | 274 | 266 | 444 | 6.54603316 | 5.43024083 | 8.1619757 | 6.75446383 | 3.53182795 | 1.85361358 | 1.79049805 | 2.75532548 |
| ENSMUSG00 | 291.74798 | 253.306199 | 358.304361 | 376.683428 | 152.292895 | 141.831926 | 134.659283 | 165.859548 | 1.10603692 | 1.30E-11 | 9.40E-10 | Pcyox1l | 2.15253534 | 18 | 61696837 | 61707635 | - | 2645 | protein_codi | Prenylcystein | - | 308 | 246 | 381 | 407 | 156 | 131 | 125 | 167 | 3.24959929 | 2.82142021 | 3.99092943 | 4.19564242 | 1.69629583 | 1.57977761 | 1.499886 | 1.84740634 |
| ENSMUSG00 | 1493.78755 | 2622.64589 | 2502.48794 | 1470.63874 | 656.03093 | 344.294295 | 342.573215 | 497.578643 | 2.13553186 | 3.76E-11 | 2.62E-09 | Ajap1 | 4.39399083 | 4 | 153373221 | 153482811 | - | 3048 | protein_codi | Adherens jun | - | 1577 | 2547 | 2661 | 1589 | 672 | 318 | 318 | 501 | 14.4384809 | 25.3496708 | 24.1882618 | 14.2147318 | 6.34098876 | 3.32784043 | 3.31120501 | 4.80943875 |
| ENSMUSG00 | 399.732622 | 422.176999 | 590.590915 | 374.832403 | 241.130416 | 180.808639 | 151.895671 | 186.716137 | 1.23137353 | 3.98E-11 | 2.76E-09 | Ndrg4 | 2.34790418 | 8 | 95676980 | 95715119 | + | 4893 | protein_codi | Protein NDR( | Ndr | 422 | 410 | 628 | 405 | 247 | 167 | 141 | 188 | 2.4068111 | 2.54194988 | 3.55597891 | 2.25688558 | 1.4518589 | 1.08865831 | 0.91457181 | 1.12422767 |
| ENSMUSG00 | 1141.41661 | 2168.54819 | 1578.04388 | 1294.79144 | 758.535763 | 675.596352 | 577.419004 | 695.21966 | 1.19117128 | 5.00E-11 | 3.43E-09 | Adamtsl4 | 2.28338048 | 3 | 95676201 | 95687917 | - | 4198 | protein_codi | ADAMTS-like | TSP_1 | 1205 | 2106 | 1678 | 1399 | 777 | 624 | 536 | 700 | 8.01031111 | 15.218585 | 11.0745037 | 9.08667541 | 5.32330387 | 4.74124603 | 4.05225036 | 4.87895981 |
| ENSMUSG00 | 17397.8414 | 25409.9068 | 36058.0205 | 24746.3427 | 11852.4874 | 10283.356 | 12245.3765 | 12614.2641 | 1.14059183 | 7.42E-11 | 5.00E-09 | Chst1 | 2.20471447 | 2 | 92599707 | 92615250 | + | 2680 | protein_codi | Carbohydrate | Sulfotransfer | 18367 | 24677 | 38342 | 26738 | 12141 | 9498 | 11367 | 12701 | 191.252977 | 279.328924 | 396.382724 | 272.034421 | 130.293376 | 113.04405 | 134.612373 | 138.667523 |
| ENSMUSG00 | 2669.30457 | 2332.27049 | 2236.34585 | 2733.03725 | 1782.60785 | 1464.87478 | 1413.38383 | 1555.3057 | 0.68146779 | 7.89E-11 | 5.29E-09 | Tssc4 | 1.6037706 | 7 | 143069249 | 143071093 | + | 1845 | protein_codi | tumor-suppr | - | 2818 | 2265 | 2378 | 2953 | 1826 | 1353 | 1312 | 1566 | 42.6235275 | 37.2417583 | 35.7100311 | 43.6412128 | 28.4647304 | 23.3911602 | 22.5689512 | 24.8351634 |
| ENSMUSG00 | 2413.55147 | 2341.53779 | 2705.62112 | 2902.40597 | 1552.21604 | 1317.62942 | 1153.76073 | 1718.18573 | 0.85159705 | 7.95E-11 | 5.31E-09 | Loxl3 | 1.80449738 | 6 | 83034173 | 83052562 | + | 4603 | protein_codi | Lysyl oxidase | SRCR | 2548 | 2274 | 2877 | 3136 | 1590 | 1217 | 1071 | 1730 | 15.4476784 | 14.9867626 | 17.317039 | 18.5765394 | 9.93479299 | 8.43334641 | 7.38452236 | 10.9970643 |
| ENSMUSG00 | 6016.82846 | 6667.30748 | 6896.18341 | 6968.18065 | 4830.41822 | 4820.12013 | 4333.87435 | 4928.11422 | 0.48922126 | 1.40E-10 | 9.01E-09 | Grn | 1.40368698 | 11 | 102430315 | 102437048 | + | 3474 | protein_codi | granulin [Sou | - | 6352 | 6475 | 7333 | 7529 | 4948 | 4452 | 4023 | 4962 | 51.0252858 | 56.5416269 | 58.4825929 | 59.0931604 | 40.9640181 | 40.8766859 | 36.7531132 | 41.7925221 |
| ENSMUSG00 | 400.679855 | 402.612699 | 378.993852 | 457.202981 | 209.890848 | 132.087748 | 119.577443 | 220.483949 | 1.26275026 | 1.43E-10 | 9.14E-09 | Paqr8 | 2.39952736 | 1 | 20890606 | 20939650 | + | 5488 | protein_codi | Mus muscul | - | 423 | 391 | 403 | 494 | 215 | 122 | 111 | 222 | 2.15095357 | 2.16132958 | 2.03453747 | 2.45438439 | 1.12674861 | 0.70908135 | 0.64192278 | 1.18361513 |
| ENSMUSG00 | 831.671189 | 948.353697 | 1053.28316 | 994.925515 | 587.694375 | 645.281131 | 584.959924 | 644.567942 | 0.63669881 | 2.08E-10 | 1.28E-08 | Slc4a3 | 1.55476745 | 1 | 75546266 | 75562172 | + | 8599 | protein_codi | solute carrie | Band_3_cyto | 878 | 921 | 1120 | 1075 | 602 | 596 | 543 | 649 | 2.84938634 | 3.24915196 | 3.6086505 | 3.40871153 | 2.01349806 | 2.21079588 | 2.00412956 | 2.20835243 |
| ENSMUSG00 | 87.1455004 | 58.6928998 | 79.9366685 | 91.6256986 | 29.2870951 | 20.5710428 | 19.3909367 | 11.9180513 | 1.96723432 | 2.12E-10 | 1.29E-08 | Gm10535 | 3.91017809 | 1 | 135108961 | 135113378 | + | 915 | antisense | - | - | 92 | 57 | 85 | 99 | 30 | 19 | 18 | 12 | 2.80589558 | 1.88978372 | 2.57378687 | 2.95014821 | 0.94298076 | 0.66234284 | 0.62434599 | 0.38373533 |
| ENSMUSG00 | 2504.4859 | 2549.53719 | 3129.75568 | 2240.66481 | 1717.20001 | 1435.64225 | 1440.31569 | 1658.59548 | 0.73736842 | 2.88E-10 | 1.72E-08 | Lrrk2 | 1.66713209 | 15 | 91673175 | 91816120 | + | 11834 | protein_codi | Leucine-rich | Pkinase | 2644 | 2476 | 3328 | 2421 | 1759 | 1326 | 1337 | 1670 | 6.23497414 | 6.34713034 | 7.79159735 | 5.57818558 | 4.27500815 | 3.57406376 | 3.58569839 | 4.12911084 |
| ENSMUSG00 | 46.4144513 | 68.9898998 | 42.3194127 | 40.7225327 | 10.7386015 | 9.74417816 | 9.69546835 | 7.94536755 | 2.37733579 | 2.99E-10 | 1.78E-08 | Gm12216 | 5.1957636 | 11 | 53783418 | 53859256 | - | 2647 | protein_codi | - | - | 49 | 67 | 45 | 44 | 11 | 9 | 9 | 8 | 0.51659109 | 0.76785498 | 0.47101346 | 0.45324025 | 0.11952023 | 0.10845233 | 0.1079102 | 0.08843164 |
| ENSMUSG00 | 671.588694 | 789.779898 | 601.876092 | 596.955309 | 258.702673 | 303.152209 | 394.28238 | 383.363984 | 0.9898593 | 3.02E-10 | 1.79E-08 | Gm43128 | 1.98599129 | 2 | 35280390 | 35282010 | + | 1621 | TEC | predicted ge | - | 709 | 767 | 640 | 645 | 265 | 280 | 366 | 386 | 12.205849 | 14.3539256 | 10.938851 | 10.8494178 | 4.70181494 | 5.50966702 | 7.16592048 | 6.96748312 |
| ENSMUSG00 | 89443.4894 | 82546.93 | 89104.9341 | 90632.6241 | 70741.0257 | 60782.018 | 64481.3282 | 65554.2481 | 0.42732016 | 3.94E-10 | 2.32E-08 | Tmod3 | 1.34473338 | 9 | 75497707 | 75559657 | - | 11151 | protein_codi | tropomodulir | - | 94426 | 80166 | 94749 | 97927 | 72463 | 56140 | 59856 | 66005 | 236.310244 | 218.089492 | 235.415779 | 239.451945 | 186.898221 | 160.586462 | 170.360062 | 173.194723 |
| ENSMUSG00 | 1447.37309 | 1384.9465 | 1294.0336 | 1369.75792 | 987.951341 | 1004.73304 | 999.710515 | 1048.78852 | 0.443501 | 4.78E-10 | 2.79E-08 | Wdr19 | 1.35990041 | 5 | 65199696 | 65260415 | + | 11278 | protein_codi | WD repeat d | - | 1528 | 1345 | 1376 | 1480 | 1012 | 928 | 928 | 1056 | 3.78090735 | 3.61783317 | 3.38034551 | 3.57815673 | 2.58078066 | 2.62461872 | 2.61149861 | 2.73970286 |
| ENSMUSG00 | 747.36739 | 674.453498 | 884.945942 | 854.247675 | 512.524164 | 315.06176 | 353.345958 | 415.145454 | 0.98492963 | 5.05E-10 | 2.91E-08 | Ap3m2 | 1.97921677 | 8 | 22787354 | 22805622 | - | 4525 | protein_codi | AP-3 comple | Adap_comp_ | 789 | 655 | 941 | 923 | 525 | 291 | 328 | 418 | 4.86590013 | 4.39117816 | 5.76163561 | 5.56176778 | 3.33690154 | 2.05127904 | 2.30053674 | 2.70289599 |
| ENSMUSG00 | 1454.00373 | 1639.2824 | 1640.11235 | 1541.90317 | 1198.81843 | 1066.44616 | 1074.04244 | 1003.10265 | 0.53086628 | 5.27E-10 | 3.02E-08 | St3gal5 | 1.44479648 | 6 | 72097592 | 72154571 | + | 2618 | protein_codi | Lactosylcerar | Glyco_transf_ | 1535 | 1592 | 1744 | 1666 | 1228 | 985 | 997 | 1010 | 16.3622683 | 18.4472555 | 18.4565952 | 17.3514227 | 13.4906041 | 12.0009859 | 12.0864687 | 11.2881655 |
| ENSMUSG00 | 106.090174 | 155.4847 | 167.396788 | 121.242086 | 63.4553727 | 41.1420855 | 30.1636793 | 45.6858634 | 1.60316806 | 5.97E-10 | 3.41E-08 | Masp1 | 3.03809726 | 16 | 23449417 | 23520815 | - | 9833 | protein_codi | Mus muscul | Trypsin | 112 | 151 | 178 | 131 | 65 | 38 | 28 | 46 | 0.31786064 | 0.46585338 | 0.50154362 | 0.36325784 | 0.19012096 | 0.1232673 | 0.0903745 | 0.13688108 |
| ENSMUSG00 | 1007.85666 | 1027.6406 | 1140.74328 | 1069.892 | 674.579424 | 700.498141 | 751.937435 | 556.175728 | 0.6629667 | 6.55E-10 | 3.69E-08 | Pclo | 1.58333519 | 5 | 14514918 | 14863459 | + | 22457 | protein_codi | Protein picco | C2 | 1064 | 998 | 1213 | 1156 | 691 | 647 | 698 | 560 | 1.32219241 | 1.34814667 | 1.49652443 | 1.40357566 | 0.88497088 | 0.91897327 | 0.98645572 | 0.7296388 |
| ENSMUSG00 | 5370.81508 | 5217.48989 | 5184.59827 | 5717.8138 | 4143.14772 | 3563.12115 | 3310.46381 | 3878.33253 | 0.52870756 | 7.18E-10 | 4.03E-08 | Hk1 | 1.44263623 | 10 | 62268855 | 62379908 | - | 8532 | protein_codi | Hexokinase-1 | - | 5670 | 5067 | 5513 | 6178 | 4244 | 3291 | 3073 | 3905 | 18.5454332 | 18.0160011 | 17.9024263 | 19.7436204 | 14.3062958 | 12.3034631 | 11.4310369 | 13.3918886 |
| ENSMUSG00 | 5402.07379 | 6497.40698 | 6177.69383 | 5625.26259 | 4194.88825 | 4071.98378 | 3967.60111 | 4479.20095 | 0.50383639 | 8.44E-10 | 4.71E-08 | Esyt1 | 1.41797922 | 10 | 128509965 | 128525871 | - | 7267 | protein_codi | Extended syn | C2 | 5703 | 6310 | 6569 | 6078 | 4297 | 3761 | 3683 | 4510 | 21.9004472 | 26.3410171 | 25.044874 | 22.8052727 | 17.0064187 | 16.5081539 | 16.084978 | 18.1590455 |
| ENSMUSG00 | 41559.8786 | 44330.6443 | 40676.4791 | 40617.0243 | 30631.3728 | 19096.4238 | 19472.8096 | 23567.9465 | 0.84969732 | 9.68E-10 | 5.35E-08 | Tspan2 | 1.8021228 | 3 | 102734529 | 102801513 | + | 4755 | protein_codi | tetraspanin 2 | - | 43875 | 43052 | 43253 | 43886 | 31377 | 17638 | 18076 | 23730 | 257.496529 | 274.663628 | 252.023166 | 251.654796 | 189.785736 | 118.317546 | 120.649555 | 146.022188 |
| ENSMUSG00 | 715.161444 | 744.473098 | 690.276643 | 749.664807 | 544.739969 | 447.149509 | 480.464321 | 445.933753 | 0.59527673 | 1.13E-09 | 6.21E-08 | Caprin2 | 1.51076234 | 6 | 148842492 | 148896237 | - | 14140 | protein_codi | Caprin-2 [So | - | 755 | 723 | 734 | 810 | 558 | 413 | 446 | 449 | 1.49005505 | 1.55112654 | 1.438207 | 1.56194359 | 1.13497805 | 0.93164612 | 1.00105828 | 0.92911306 |

| | | | | | | | | | | | | | | | | | | | | | | | | | | | | | | | | | | | | | | |
|---|---|---|---|---|---|---|---|---|---|---|---|---|---|---|---|---|---|---|---|---|---|---|---|---|---|---|---|---|---|---|---|---|---|---|---|---|---|---|
| ENSMUSG0 | 340.056898 | 335.682199 | 233.226986 | 243.409684 | 183.532463 | 87.6976034 | 67.8682785 | 139.043932 | 1.26602534 | 4.49E-06 | 0.00011586 | Aox1 | 2.40498074 | | 1 | 58029931 | 58106413 | + | 8690 | protein_codi | aldehyde oxi | Fer2 | 359 | 326 | 248 | 263 | 188 | 81 | 63 | 140 | 1.15286762 | 1.13803643 | 0.79069074 | 0.82521233 | 0.62221539 | 0.29731415 | 0.23008838 | 0.47138949 |
| ENSMUSG0 | 7916.03203 | 12227.6875 | 10279.8556 | 6850.64062 | 5448.37592 | 5518.4529 | 5341.12579 | 6241.08621 | 0.72504695 | 4.64E-06 | 0.00011889 | Rusc2 | 1.65295443 | | 4 | 43381979 | 43427088 | + | 7923 | protein_codi | RUN and SH: | RUN | 8357 | 11875 | 10931 | 7402 | 5581 | 5097 | 4958 | 6284 | 29.4351001 | 45.4676286 | 38.2247793 | 25.4735316 | 20.2593282 | 20.5199035 | 19.8605275 | 23.2069548 |
| ENSMUSG0 | 4786.37189 | 4993.01529 | 5394.31448 | 5291.15272 | 3912.75591 | 3908.49813 | 4393.12444 | 4077.95989 | 0.32906382 | 4.69E-06 | 0.00011987 | 1700025G04 | 1.25619795 | | 1 | 151852403 | 152090125 | - | 11788 | protein_codi | RIKEN cDNA | - | 5053 | 4849 | 5736 | 5717 | 4008 | 3610 | 4078 | 4106 | 11.9622794 | 12.4787303 | 13.4816722 | 13.2238465 | 9.77890571 | 9.76826451 | 10.9794607 | 10.1917897 |
| ENSMUSG0 | 3321.00135 | 4086.87929 | 4122.85123 | 4075.95532 | 2751.03447 | 2373.24872 | 2630.70375 | 3197.01727 | 0.51078203 | 4.70E-06 | 0.00012004 | Tmem98 | 1.42482233 | | 11 | 80810175 | 80822033 | + | 1633 | protein_codi | Transmembr | - | 3506 | 3969 | 4384 | 4404 | 2818 | 2192 | 2442 | 3219 | 59.9143011 | 73.7315317 | 74.3805027 | 73.5344519 | 49.6315087 | 42.8158629 | 47.460618 | 57.6774999 |
| ENSMUSG0 | 8307.23955 | 7270.71168 | 7627.83904 | 8322.20487 | 6463.66189 | 5681.93855 | 5399.2986 | 6404.95941 | 0.39654544 | 4.83E-06 | 0.00012286 | Rcn2 | 1.31635211 | | 9 | 56041845 | 56061882 | + | 4745 | protein_codi | Reticulocalbi | - | 8770 | 7061 | 8111 | 8992 | 6621 | 5248 | 5012 | 6449 | 51.5784334 | 45.1427837 | 47.3601352 | 51.6713509 | 40.1319298 | 35.2783241 | 33.5234541 | 39.7674547 |
| ENSMUSG0 | 2231.6826 | 2417.73559 | 2676.46775 | 2323.9609 | 1974.92645 | 1856.80728 | 1857.22083 | 1739.04232 | 0.37749574 | 4.91E-06 | 0.00012437 | Bahcc1 | 1.29908492 | | 11 | 120232947 | 120292296 | + | 11118 | protein_codi | Mus musculu | BAH | 2356 | 2348 | 2846 | 2511 | 2023 | 1715 | 1724 | 1751 | 5.91361964 | 6.40663179 | 7.09223267 | 6.15814311 | 5.23325486 | 4.92025704 | 4.92135287 | 4.60819779 |
| ENSMUSG0 | 1714.493 | 1796.8265 | 1779.2962 | 1794.56798 | 1435.06766 | 1280.81808 | 1249.63814 | 1498.69495 | 0.37421591 | 5.06E-06 | 0.00012784 | Ghr | 1.29613492 | | 15 | 3317760 | 3583492 | - | 9053 | protein_codi | Growth horm | - | 1810 | 1745 | 1892 | 1939 | 1470 | 1183 | 1160 | 1509 | 5.57944275 | 5.84737912 | 5.7903306 | 5.84002926 | 4.67011405 | 4.16814252 | 4.06667422 | 4.87717518 |
| ENSMUSG0 | 66658.73 | 75347.2676 | 68670.3004 | 67623.4676 | 57277.7481 | 42390.423 | 44520.5134 | 54168.5364 | 0.48851916 | 5.11E-06 | 0.00012881 | Mylk | 1.40300404 | | 16 | 34745210 | 35002420 | + | 10159 | protein_codi | myosin light | Pkinase | 70372 | 73174 | 73020 | 73066 | 58672 | 39153 | 41327 | 54541 | 193.309726 | 218.506408 | 199.143262 | 196.107456 | 166.104962 | 122.931851 | 129.109094 | 157.088275 |
| ENSMUSG0 | 545.606611 | 566.334998 | 624.446446 | 565.487897 | 415.87675 | 429.826525 | 423.368785 | 452.88595 | 0.41878845 | 5.21E-06 | 0.00013089 | Gstcd | 1.33680446 | | 3 | 132981752 | 133092033 | - | 5401 | protein_codi | Mus musculu | - | 576 | 550 | 664 | 611 | 426 | 397 | 393 | 456 | 2.97613808 | 3.08920589 | 3.40618828 | 3.08458518 | 2.2684964 | 2.34458869 | 2.30936344 | 2.47037168 |
| ENSMUSG0 | 962.38944 | 1002.9278 | 962.061316 | 1006.95717 | 810.276298 | 718.903811 | 783.178388 | 783.611874 | 0.34514381 | 5.72E-06 | 0.00014298 | Osbpl1a | 1.2702776 | | 18 | 12755314 | 12941841 | - | 7392 | protein_codi | Oxysterol-bir | - | 1016 | 974 | 1023 | 1088 | 830 | 664 | 727 | 789 | 3.83562836 | 3.99719504 | 3.83432061 | 4.01325422 | 3.22937744 | 2.86521 | 3.12137801 | 3.12310568 |
| ENSMUSG0 | 26.5225436 | 39.1285999 | 17.8681965 | 38.8715085 | 2.92870951 | 5.41343231 | 4.30909705 | 10.9248804 | 2.37108864 | 5.92E-06 | 0.00014719 | Apob | 5.17331358 | | 12 | 7977648 | 8016835 | + | 13934 | protein_codi | Apolipoprote | Vitellogenin_ | 28 | 38 | 19 | 42 | 3 | 5 | 4 | 11 | 0.05607729 | 0.08273059 | 0.03777918 | 0.08218702 | 0.00619224 | 0.01144576 | 0.00911083 | 0.02309875 |
| ENSMUSG0 | 1047.64047 | 1304.6299 | 1498.10721 | 1286.46183 | 931.329624 | 780.616939 | 957.696819 | 767.721139 | 0.57957559 | 6.08E-06 | 0.00015084 | Jdp2 | 1.49440956 | | 12 | 85599027 | 85639878 | + | 2148 | protein_codi | Jun dimeriza | bZIP_1 | 1106 | 1267 | 1593 | 1390 | 954 | 721 | 889 | 773 | 14.3689694 | 17.8937217 | 20.5473703 | 17.6445366 | 12.7737017 | 10.7065937 | 13.1353424 | 10.5297207 |
| ENSMUSG0 | 2325.45873 | 4203.23539 | 1861.11373 | 3036.60522 | 1300.34702 | 1095.6787 | 918.914945 | 573.059634 | 1.55523058 | 6.19E-06 | 0.00015327 | Lrtm2 | 2.93880693 | | 6 | 119315133 | 119330766 | - | 4102 | protein_codi | Leucine-rich | - | 2455 | 4082 | 1979 | 3281 | 1332 | 1012 | 853 | 577 | 16.7016974 | 30.1880935 | 13.3667211 | 21.8092288 | 9.33923367 | 7.86928352 | 6.59974703 | 4.11577659 |
| ENSMUSG0 | 803.254178 | 695.047498 | 779.617626 | 893.119183 | 565.240935 | 543.508604 | 537.559857 | 627.684036 | 0.47979927 | 6.20E-06 | 0.00015336 | Stxbp4 | 1.39454962 | | 11 | 90476492 | 90638084 | - | 11834 | protein_codi | Syntaxin-binc | WW | 848 | 675 | 829 | 965 | 579 | 502 | 499 | 632 | 1.99971939 | 1.73033642 | 1.94087566 | 2.22344035 | 1.40718006 | 1.35307693 | 1.33826739 | 1.56263356 |
| ENSMUSG0 | 47.361685 | 100.9106 | 25.3916476 | 44.4245811 | 10.7386015 | 16.2402969 | 11.8500169 | 13.9043932 | 2.0469076 | 6.30E-06 | 0.00015551 | Gm26669 | 4.13219288 | | 12 | 80108034 | 80113000 | + | 1808 | lincRNA | predicted gei | - | 50 | 98 | 27 | 48 | 11 | 15 | 11 | 14 | 0.77174949 | 1.64431868 | 0.41375198 | 0.72388995 | 0.17498343 | 0.26463249 | 0.19309373 | 0.22656939 |
| ENSMUSG0 | 124.087615 | 132.8313 | 114.73263 | 154.560522 | 64.4316092 | 56.299696 | 68.9455527 | 89.3853849 | 0.91321617 | 6.86E-06 | 0.00016706 | Aph1c | 1.88323909 | | 9 | 66814994 | 66834726 | - | 5205 | protein_codi | Putative gam | - | 131 | 129 | 122 | 167 | 66 | 52 | 64 | 90 | 0.70235282 | 0.75184327 | 0.64940233 | 0.87483363 | 0.3646917 | 0.31866396 | 0.39024123 | 0.50593347 |
| ENSMUSG0 | 45.4672176 | 17.5049 | 26.332079 | 46.2756053 | 8.78612853 | 8.66149169 | 6.46364557 | 6.9521966 | 2.13724069 | 6.93E-06 | 0.00016857 | Pla2g5 | 4.39919846 | | 4 | 138799244 | 138863482 | - | 2884 | protein_codi | phospholipas | - | 48 | 17 | 28 | 50 | 9 | 8 | 6 | 7 | 0.4644626 | 0.17881832 | 0.26899086 | 0.47272055 | 0.0897532 | 0.08847999 | 0.06602827 | 0.07101898 |
| ENSMUSG0 | 8279.76977 | 10140.4856 | 6563.2707 | 7587.34825 | 5909.15955 | 4198.6581 | 3641.187 | 5548.84606 | 0.75500541 | 7.10E-06 | 0.00017169 | Tnfaip2 | 1.68763793 | | 12 | 111442469 | 111455018 | + | 4984 | protein_codi | tumor necros | - | 8741 | 9848 | 6979 | 8198 | 6053 | 3878 | 3380 | 5587 | 48.9426925 | 59.9416023 | 38.7962647 | 44.8497075 | 34.9297368 | 24.8187616 | 21.5234844 | 32.7998814 |
| ENSMUSG0 | 1292.974 | 2388.90399 | 1309.0805 | 1223.52701 | 905.947475 | 822.841711 | 888.751266 | 781.625532 | 0.87012926 | 7.33E-06 | 0.00017657 | Cyp2d22 | 1.82782667 | | 15 | 82370527 | 82380260 | - | 3909 | protein_codi | cytochrome I | p450 | 1365 | 2320 | 1392 | 1322 | 928 | 760 | 825 | 787 | 9.74477356 | 18.004483 | 9.86616362 | 9.22137151 | 6.82786583 | 6.20152156 | 6.69826294 | 5.89088707 |
| ENSMUSG0 | 15258.0404 | 23148.6856 | 22748.095 | 15397.7449 | 14301.8648 | 9546.04653 | 8316.5573 | 11099.6785 | 0.82323035 | 7.40E-06 | 0.00017769 | Cald1 | 1.76936335 | | 6 | 34598500 | 34775473 | + | 12041 | protein_codi | caldesmon 1 | - | 16108 | 22481 | 24189 | 16637 | 14650 | 8817 | 7720 | 11176 | 37.3322216 | 56.6384567 | 55.6583217 | 37.67404 | 34.9927232 | 23.3565461 | 20.3483246 | 27.1578554 |
| ENSMUSG0 | 323.953926 | 238.890399 | 153.290317 | 95.327747 | 48.8118252 | 87.6976034 | 87.2592152 | 51.644889 | 1.56071609 | 7.61E-06 | 0.00018146 | Pik3r2 | 2.95000232 | | 8 | 70768176 | 70776713 | - | 4176 | protein_codi | Phosphatidyl | SH2 | 342 | 232 | 163 | 103 | 50 | 81 | 81 | 52 | 2.28544296 | 1.68533343 | 1.08143859 | 0.67252196 | 0.34435959 | 0.61869252 | 0.61559976 | 0.36434641 |
| ENSMUSG0 | 264.278202 | 249.187399 | 213.477926 | 256.366854 | 127.886982 | 175.395207 | 154.050219 | 164.866377 | 0.66160863 | 7.70E-06 | 0.00018282 | Gm16364 | 1.58184543 | | 10 | 39556071 | 39558062 | - | 1313 | antisense | predicted gei | - | 279 | 242 | 227 | 277 | 131 | 162 | 143 | 166 | 5.92985738 | 5.59125091 | 4.79000404 | 5.75234304 | 2.86951991 | 3.93550642 | 3.45656896 | 3.6992612 |
| ENSMUSG0 | 641.277215 | 835.086698 | 656.421113 | 577.519555 | 491.046961 | 390.849813 | 363.041426 | 471.756198 | 0.65730425 | 7.70E-06 | 0.00018282 | Wfikkn2 | 1.57713291 | | 11 | 94235956 | 94246005 | - | 3827 | protein_codi | WAP, follista | Kunitz_BPTI | 677 | 811 | 698 | 624 | 503 | 361 | 337 | 475 | 4.93668001 | 6.42866409 | 5.05326075 | 4.44586081 | 3.78017752 | 3.00883988 | 2.79476536 | 3.63167338 |
| ENSMUSG0 | 3694.21143 | 3068.50599 | 3649.81424 | 3727.96277 | 2782.27403 | 2198.9362 | 2801.99035 | 2394.53514 | 0.47446433 | 7.69E-06 | 0.00018282 | Fa2h | 1.38940224 | | 8 | 111345135 | 111393824 | - | 3008 | protein_codi | Fatty acid 2-l | - | 3900 | 2980 | 3881 | 4028 | 2850 | 2031 | 2601 | 2411 | 36.1819147 | 30.0536188 | 35.7470789 | 36.5124827 | 27.2502004 | 21.5368621 | 27.4433064 | 23.4526009 |
| ENSMUSG0 | 25.5753099 | 33.9800999 | 35.736393 | 42.5735569 | 7.80989203 | 14.074924 | 1.07727426 | 5.95902566 | 2.25533406 | 7.80E-06 | 0.00018476 | D830026I12R | 4.77444837 | | 6 | 17197751 | 17210152 | + | 2112 | lincRNA | RIKEN cDNA | - | 27 | 33 | 38 | 46 | 8 | 13 | 1 | 6 | 0.35675874 | 0.47400003 | 0.49849916 | 0.5938731 | 0.10894285 | 0.19633593 | 0.01502727 | 0.08312449 |
| ENSMUSG0 | 518.136834 | 352.157399 | 456.109226 | 537.722534 | 331.920411 | 292.325345 | 239.154886 | 270.142497 | 0.71775125 | 7.98E-06 | 0.00018811 | Ttc22 | 1.64461655 | | 4 | 106622432 | 106640189 | + | 3318 | protein_codi | tetratricopep | - | 547 | 342 | 485 | 581 | 340 | 270 | 222 | 272 | 4.60061314 | 3.12685733 | 4.04986089 | 4.77451745 | 2.94717014 | 2.59559972 | 2.1234914 | 2.39863495 |
| ENSMUSG0 | 160.082495 | 131.8016 | 127.89867 | 248.037245 | 66.3840822 | 79.0361117 | 88.3364895 | 82.4331883 | 1.08070202 | 8.19E-06 | 0.00019284 | AC158986.1 | 2.11506503 | | 15 | 36499977 | 36503431 | - | 3455 | TEC | - | - | 169 | 128 | 136 | 268 | 68 | 73 | 82 | 83 | 1.36503386 | 1.12388082 | 1.09060028 | 2.11502973 | 0.56606139 | 0.67394607 | 0.753251 | 0.70291316 |
| ENSMUSG0 | 351.423703 | 280.078399 | 210.656632 | 211.942272 | 135.696874 | 175.395207 | 99.1092321 | 118.187342 | 0.99644715 | 8.24E-06 | 0.00019371 | 5330438I03R | 1.99508077 | | 1 | 166306741 | 166309585 | - | 2845 | antisense | RIKEN cDNA | - | 371 | 272 | 224 | 229 | 139 | 162 | 92 | 119 | 3.63912028 | 2.90031371 | 2.18142606 | 2.19473933 | 1.40519049 | 1.81628117 | 1.02631215 | 1.22387292 |
| ENSMUSG0 | 142.085055 | 136.9501 | 73.3536487 | 149.007449 | 58.5741902 | 28.149848 | 43.0909705 | 65.5492822 | 1.35564226 | 8.65E-06 | 0.00020094 | Tagln | 2.55911017 | | 9 | 45929619 | 45936058 | - | 1988 | protein_codi | Transgelin [! | CH | 150 | 133 | 78 | 161 | 60 | 26 | 40 | 66 | 2.10561832 | 2.02952125 | 1.08705864 | 2.20820419 | 0.86803561 | 0.41716446 | 0.63858325 | 0.97140244 |
| ENSMUSG0 | 1714.493 | 1295.3626 | 1268.64195 | 1714.04842 | 1014.30973 | 881.30678 | 1006.17416 | 1110.36511 | 0.57875393 | 8.68E-06 | 0.00020094 | BB014433 | 1.49355869 | | 8 | 15041444 | 15046078 | - | 1780 | protein_codi | expressed se | - | 1810 | 1258 | 1349 | 1852 | 1039 | 814 | 934 | 1118 | 28.376795 | 21.4397136 | 20.9974568 | 28.3694368 | 16.7879713 | 14.5866223 | 16.6533185 | 18.3777964 |
| ENSMUSG0 | 5556.47289 | 5990.79458 | 6611.2327 | 6400.84173 | 4463.35329 | 4286.3557 | 4203.52417 | 5286.64893 | 0.42909611 | 8.66E-06 | 0.00020094 | Sipa1l2 | 1.34638976 | | 8 | 125418063 | 125569808 | - | 9617 | protein_codi | Mus musculu | - | 5866 | 5818 | 7030 | 6916 | 4572 | 3959 | 3902 | 5323 | 17.0218678 | 18.3523821 | 20.2530511 | 19.608533 | 13.673172 | 13.1309522 | 12.8772036 | 16.1952809 |
| ENSMUSG0 | 2921.26873 | 3431.99009 | 2996.21442 | 3756.65364 | 2097.93225 | 2389.48902 | 2614.54463 | 2335.93806 | 0.47385825 | 8.75E-06 | 0.00020196 | Map2k3 | 1.38881867 | | 11 | 60932033 | 60952811 | + | 3314 | protein_codi | mitogen-acti | Pkinase | 3084 | 3333 | 3186 | 4059 | 2149 | 2207 | 2427 | 2352 | 25.9696683 | 30.5099258 | 26.6359393 | 33.3961407 | 18.6503327 | 21.2422329 | 23.2429467 | 20.7661721 |
| ENSMUSG0 | 10640.2762 | 8587.69798 | 10442.5502 | 13434.7337 | 7907.51568 | 6355.36953 | 7181.11023 | 7441.82988 | 0.57750787 | 8.99E-06 | 0.000207 | Snx31 | 1.49226925 | | 15 | 36504062 | 36555573 | - | 2935 | protein_codi | Mus musculu | PX | 11233 | 8340 | 11104 | 14516 | 8100 | 5870 | 6666 | 7493 | 106.805207 | 86.2017905 | 104.820468 | 134.855477 | 79.3742411 | 63.7940731 | 72.0827119 | 74.6997696 |
| ENSMUSG0 | 262.383735 | 282.137799 | 310.34236 | 332.258846 | 201.10472 | 103.9379 | 137.891105 | 192.675163 | 0.89891353 | 9.12E-06 | 0.00020939 | Mgst3 | 1.86466121 | | 1 | 167371966 | 167393841 | - | 1097 | protein_codi | Microsomal ; | - | 277 | 274 | 330 | 359 | 206 | 96 | 128 | 194 | 7.0465723 | 7.57708705 | 8.33454817 | 8.92313689 | 5.4008643 | 2.79135416 | 3.70320075 | 5.1744803 |
| ENSMUSG0 | 1557.2522 | 1607.3617 | 2219.41809 | 1752.91993 | 1026.02456 | 1277.57002 | 1306.73368 | 916.696781 | 0.65722682 | 9.23E-06 | 0.0002117 | Adamts9 | 1.57704827 | | 6 | 92772699 | 92943492 | - | 15048 | protein_codi | a disintegrin- | TSP_1 | 1644 | 1561 | 2360 | 1894 | 1051 | 1180 | 1213 | 923 | 3.04879204 | 3.14689652 | 4.34520331 | 3.43187078 | 2.00875331 | 2.50122961 | 2.55832628 | 1.79471112 |
| ENSMUSG0 | 6971.64003 | 9329.08197 | 7341.00746 | 7214.36687 | 5510.85506 | 5258.60814 | 5351.89853 | 6130.84423 | 0.47147805 | 9.31E-06 | 0.00021324 | Plekho1 | 1.38652925 | | 3 | 95988429 | 95996001 | - | 2172 | protein_codi | Pleckstrin ho | PH | 7360 | 9060 | 7806 | 7795 | 5645 | 4857 | 4968 | 6173 | 94.5633315 | 126.539676 | 99.5734316 | 97.8556786 | 74.7492429 | 71.3277655 | 72.5931564 | 83.1587765 |
| ENSMUSG0 | 1614.08623 | 1324.1942 | 1369.26811 | 1625.19926 | 1148.05413 | 1104.34019 | 1107.43794 | 1101.42658 | 0.41148307 | 9.33E-06 | 0.00021331 | Cc2d2a | 1.33005238 | | 5 | 43662346 | 43740972 | + | 6809 | protein_codi | Coiled-coil ar | - | 1704 | 1286 | 1456 | 1756 | 1176 | 1020 | 1028 | 1109 | 6.98378767 | 5.72949014 | 5.92451481 | 7.03187127 | 4.96737172 | 4.77823136 | 4.79163462 | 4.7656248 |
| ENSMUSG0 | 53968.6401 | 59030.6414 | 60869.422 | 57309.5607 | 48083.5527 | 41066.2975 | 40984.8993 | 48578.9703 | 0.37133542 | 9.80E-06 | 0.00022221 | Gsn | 1.29354964 | | 2 | 35256380 | 35307892 | + | 3157 | protein_codi | gelsolin [Sou | - | 56975 | 57328 | 64725 | 61922 | 49254 | 37930 | 38045 | 48913 | 503.633399 | 550.871812 | 568.031246 | 534.810749 | 448.713976 | 383.229204 | 382.469599 | 453.337196 |
| ENSMUSG0 | 58.7284894 | 56.6334998 | 58.3067464 | 48.1266296 | 28.3108586 | 18.4056698 | 5.38637131 | 16.883906 | 1.67642066 | 1.01E-05 | 0.0002284 | Zfp239 | 3.19633952 | | 6 | 117862100 | 117873291 | + | 4364 | protein_codi | zinc finger pr | zf-C2H2 | 62 | 55 | 62 | 52 | 29 | 17 | 5 | 17 | 0.39647127 | 0.38232816 | 0.39362411 | 0.32489897 | 0.19112431 | 0.12425518 | 0.03636296 | 0.11398188 |
| ENSMUSG0 | 520.031301 | 584.869598 | 584.007896 | 596.955309 | 404.161912 | 331.302057 | 347.959587 | 465.797172 | 0.55986893 | 1.02E-05 | 0.00022993 | Hexb | 1.47413529 | | 13 | 97176331 | 97198357 | - | 1906 | protein_codi | Beta-hexosa | Glyco_hydro_ | 549 | 568 | 621 | 645 | 414 | 306 | 323 | 469 | 8.03811507 | 9.04031953 | 9.02700021 | 9.22712816 | 6.24712387 | 5.12093032 | 5.37840547 | 7.19981905 |
| ENSMUSG0 | 1261.71529 | 1417.8969 | 1554.53309 | 1453.05401 | 1103.14725 | 1003.65035 | 1109.59249 | 1119.30365 | 0.39128871 | 1.07E-05 | 0.00024069 | Copz2 | 1.31156446 | | 11 | 96849870 | 96861203 | + | 2208 | protein_codi | Coatomer su | - | 1332 | 1377 | 1653 | 1570 | 1130 | 927 | 1030 | 1127 | 16.8348762 | 18.9187837 | 20.7418999 | 19.3878798 | 14.7191268 | 13.3915548 | 14.8051247 | 14.9346993 |
| ENSMUSG0 | 26985.7409 | 27749.3852 | 31686.8954 | 29083.2924 | 24590.4213 | 22990.847 | 22847.9098 | 22333.435 | 0.31634448 | 1.08E-05 | 0.0002429 | Fgfr1 | 1.24517152 | | 8 | 25513654 | 25575718 | + | 9875 | protein_codi | Fibroblast gr | Pkinase_Tyr | 28489 | 26949 | 33694 | 31424 | 25189 | 21235 | 21209 | 22487 | 80.5090815 | 82.7873329 | 94.5344748 | 86.766903 | 73.3629007 | 68.5907414 | 68.1643035 | 66.6294228 |
| ENSMUSG0 | 307.850953 | 291.405099 | 367.708675 | 320.227189 | 243.082889 | 190.552817 | 184.213899 | 136.064419 | 0.77172909 | 1.09E-05 | 0.00024464 | Mansc1 | 1.7073148 | | 6 | 134609207 | 134632488 | - | 2376 | protein_codi | MANSC dom | - | 325 | 283 | 391 | 346 | 249 | 176 | 171 | 137 | 3.8171717 | 3.61325274 | 4.55937244 | 3.97062979 | 3.01408561 | 2.36274345 | 2.28414457 | 1.68711919 |

| | | | | | | | | | | | | | | | | |
|---|---|---|---|---|---|---|---|---|---|---|---|---|---|---|---|---|
| ENSMUSG0C | 1926.67335 | 1944.07359 | 1974.90593 | 2262.8771 | 1561.97841 | 1710.64461 | 1410.15201 | 1678.45889 | 0.35019162 | 0.00016077 | 0.00236621 | Cnn3 | 1.27472993 | 3 | 121426497 | 121458207 + |
| ENSMUSG0C | 38690.7077 | 42248.5909 | 37066.163 | 40135.758 | 28752.1175 | 34005.0164 | 33771.4708 | 32451.8606 | 0.29407232 | 0.00016169 | 0.00237747 | Ptpn14 | 1.22609632 | 1 | 189728268 | 189876695 + |
| ENSMUSG0C | 504.875562 | 451.008599 | 558.616248 | 497.925514 | 435.40148 | 251.183259 | 289.786776 | 328.739582 | 0.62355198 | 0.00016428 | 0.00240842 | Car11 | 1.5406637 | 7 | 45699843 | 45704682 + |
| ENSMUSG0C | 59.6757231 | 78.2571998 | 116.613493 | 48.1266296 | 40.0256966 | 28.149848 | 31.2409536 | 12.9112223 | 1.42997567 | 0.0001669 | 0.00244438 | Dusp5 | 2.69442171 | 19 | 53529109 | 53542431 + |
| ENSMUSG0C | 849.668629 | 877.304398 | 745.762095 | 851.471138 | 722.415012 | 616.048597 | 629.128169 | 616.759156 | 0.36246373 | 0.0001748 | 0.00255002 | Tmem8b | 1.28561952 | 4 | 43668971 | 43692668 + |
| ENSMUSG0C | 1000.27879 | 1378.7683 | 778.677194 | 973.638737 | 753.654581 | 584.650689 | 648.519106 | 711.110395 | 0.61399257 | 0.0001776 | 0.00258582 | Pxmp2 | 1.53048888 | 5 | 110274282 | 110286186 - |
| ENSMUSG0C | 550.34278 | 507.642099 | 587.769621 | 607.135942 | 441.258899 | 334.550117 | 398.591477 | 458.844976 | 0.46334631 | 0.00017834 | 0.00259157 | A930004D18 | 1.37873607 | 2 | 18025193 | 18037737 - |
| ENSMUSG0C | 89.9872015 | 97.8214997 | 113.792199 | 104.582868 | 69.3127917 | 53.0516366 | 43.0909705 | 66.5424532 | 0.80425606 | 0.00018204 | 0.0026394 | Chtf18 | 1.74624509 | 17 | 25718926 | 25727419 - |
| ENSMUSG0C | 202.708012 | 244.038899 | 183.384122 | 216.569833 | 157.174077 | 134.253121 | 143.277477 | 144.009787 | 0.54686202 | 0.00018217 | 0.0026394 | Brca2 | 1.46090465 | 5 | 150522630 | 150570329 + |
| ENSMUSG0C | 35.9948806 | 60.7522998 | 77.1153743 | 22.2122906 | 19.5247301 | 11.9095511 | 14.0045654 | 17.877077 | 1.62549514 | 0.00018265 | 0.00264126 | AI839979 | 3.08548041 | 5 | 31569595 | 31571397 - |
| ENSMUSG0C | 272.803306 | 384.078099 | 408.147225 | 251.739293 | 231.368051 | 167.816402 | 212.22303 | 197.641018 | 0.70143368 | 0.00018254 | 0.00264126 | Plcd1 | 1.62611995 | 9 | 119071527 | 119093502 - |
| ENSMUSG0C | 3470.66428 | 4151.75039 | 3387.43388 | 3730.7393 | 2978.49757 | 2979.55314 | 2667.33107 | 3088.76163 | 0.33130211 | 0.00018679 | 0.00268804 | Fam78b | 1.25814841 | 1 | 167001417 | 167091302 + |
| ENSMUSG0C | 5206.94365 | 5392.53889 | 5575.81773 | 5210.63316 | 4773.7965 | 4139.11034 | 4197.06052 | 4638.1083 | 0.26879396 | 0.0001881 | 0.00270423 | Acadm | 1.20480024 | 3 | 153922357 | 153944632 - |
| ENSMUSG0C | 6160.80799 | 5765.29028 | 5729.10805 | 6226.84546 | 5005.16455 | 4760.57237 | 4957.61615 | 5382.98651 | 0.24822859 | 0.0001936 | 0.00277535 | Sppl2a | 1.18774784 | 2 | 126890391 | 126933235 - |
| ENSMUSG0C | 394.049219 | 371.721699 | 303.75934 | 458.128493 | 278.227403 | 218.702665 | 233.768515 | 288.019574 | 0.58355311 | 0.00019386 | 0.00277636 | Glrx | 1.49853534 | 13 | 75839868 | 75850154 + |
| ENSMUSG0C | 69.1480601 | 126.6531 | 42.3194127 | 77.743017 | 33.1920411 | 36.8113397 | 28.0091308 | 9.93170943 | 1.54979998 | 0.00019536 | 0.00279512 | Cacna2d4 | 2.92776546 | 6 | 119236526 | 119352407 + |
| ENSMUSG0C | 889.452445 | 1034.8485 | 731.655624 | 882.013038 | 620.886416 | 672.348293 | 700.22827 | 623.711352 | 0.43481129 | 0.00019584 | 0.00279926 | Traf3ip2 | 1.35173401 | 10 | 39612934 | 39655307 + |
| ENSMUSG0C | 77087.773 | 81800.3975 | 89555.4008 | 87124.0077 | 63896.6316 | 56144.8718 | 43207.3161 | 71875.7812 | 0.51316913 | 0.00019682 | 0.00281066 | Bcam | 1.42718181 | 7 | 19756131 | 19771016 - |
| ENSMUSG0C | 18999.6136 | 19815.5467 | 24122.0653 | 21791.1826 | 17677.6906 | 16094.1343 | 16460.7507 | 17297.0651 | 0.3273203 | 0.00020025 | 0.00285413 | Wls | 1.25468073 | 3 | 159839672 | 159938664 + |
| ENSMUSG0C | 1316.65484 | 1206.8084 | 1615.66113 | 1610.39107 | 1179.2937 | 980.913934 | 973.855933 | 1077.59047 | 0.44900509 | 0.00020074 | 0.00285841 | Igsf8 | 1.36509853 | 1 | 172261641 | 172319841 + |
| ENSMUSG0C | 1522.20456 | 1305.6596 | 1448.26435 | 1386.41714 | 1216.39068 | 1120.58049 | 1116.05614 | 1169.95537 | 0.29255641 | 0.00020256 | 0.0028815 | Cnpy4 | 1.22480868 | 5 | 138187485 | 138193918 + |
| ENSMUSG0C | 187.552273 | 327.444599 | 289.652869 | 247.111733 | 166.936442 | 166.733715 | 169.132059 | 103.289778 | 0.79541507 | 0.0002034 | 0.0028908 | Slc43a2 | 1.73557664 | 11 | 75531694 | 75577575 + |
| ENSMUSG0C | 1196.35616 | 1175.9174 | 1387.13631 | 1289.23836 | 1020.16715 | 912.704687 | 996.478692 | 1073.61779 | 0.33469009 | 0.00020417 | 0.00289609 | Gm5559 | 1.26110647 | 5 | 95861714 | 95862715 + |
| ENSMUSG0C | 26392.7726 | 30029.141 | 31676.5506 | 30110.6109 | 25470.0104 | 23473.7252 | 22892.0781 | 25103.3888 | 0.28616599 | 0.00020413 | 0.00289609 | Timp2 | 1.21939539 | 11 | 118301069 | 118355740 - |
| ENSMUSG0C | 259.542034 | 319.206999 | 332.912713 | 628.422721 | 248.940238 | 181.891326 | 192.832093 | 184.729795 | 0.92982143 | 0.0002058 | 0.00291371 | Parm1 | 1.90504019 | 5 | 91517615 | 91626894 + |
| ENSMUSG0C | 161.976963 | 371.721699 | 127.89867 | 140.67784 | 128.863218 | 67.1265606 | 52.7864388 | 62.5697694 | 1.36269347 | 0.0002069 | 0.00292368 | Perp | 2.57164851 | 10 | 18845020 | 18857073 + |
| ENSMUSG0C | 36.9421143 | 29.8612999 | 52.6641581 | 50.9031659 | 23.4296761 | 9.74417816 | 5.38637131 | 16.883906 | 1.61005283 | 0.00021045 | 0.00296746 | Cend1 | 3.0526302 | 7 | 141426446 | 141429491 - |
| ENSMUSG0C | 17430.0473 | 14347.8398 | 10923.1106 | 16075.2198 | 7814.77321 | 10921.0583 | 11340.4662 | 9534.44105 | 0.56938325 | 0.00021059 | 0.00296746 | Dock8 | 1.48388907 | 19 | 24999529 | 25202432 + |
| ENSMUSG0C | 51852.52 | 50581.953 | 49982.9882 | 52383.9853 | 37898.4773 | 44659.7339 | 45366.1737 | 42155.1407 | 0.26803842 | 0.00021329 | 0.00299984 | Agrn | 1.20416945 | 4 | 156165290 | 156197488 - |
| ENSMUSG0C | 13.2612718 | 93.7026997 | 27.2725104 | 53.6797022 | 3.90494601 | 14.074924 | 15.0818397 | 3.97268377 | 2.35122669 | 0.00021626 | 0.00303004 | 5830418P13F | 5.10257925 | 9 | 103423578 | 103461260 + |
| ENSMUSG0C | 1441.68969 | 1426.1345 | 1329.76999 | 1537.27561 | 1084.59876 | 1222.35302 | 1196.8517 | 1175.9144 | 0.29376283 | 0.00021623 | 0.00303004 | Metap1d | 1.22583332 | 2 | 71453276 | 71525194 + |
| ENSMUSG0C | 342.8986 | 244.038899 | 40.4385499 | 240.633148 | 41.9781696 | 59.5477554 | 46.3227933 | 43.6995215 | 2.1803965 | 0.00021835 | 0.00305234 | Gm9855 | 4.53278114 | 10 | 42054113 | 42055306 - |
| ENSMUSG0C | 2130.32859 | 2121.18199 | 2175.21781 | 1951.90503 | 1918.30473 | 1321.96017 | 1309.9655 | 1531.46959 | 0.46177819 | 0.00021867 | 0.00305234 | Tfpi | 1.37723829 | 2 | 84432855 | 84476775 - |
| ENSMUSG0C | 1563.88284 | 1425.1048 | 1846.06683 | 2186.98511 | 1414.56669 | 947.350654 | 1079.42881 | 1249.40905 | 0.58190291 | 0.00021995 | 0.00306734 | Il34 | 1.49682225 | 8 | 110741829 | 110805924 - |
| ENSMUSG0C | 167.660365 | 176.0787 | 189.967142 | 96.2532591 | 55.6454807 | 93.1110357 | 102.341055 | 72.5014789 | 0.96310783 | 0.00022231 | 0.00309448 | Matn4 | 1.94950497 | 2 | 164389393 | 164405160 - |
| ENSMUSG0C | 18377.281 | 22117.9559 | 21483.2148 | 19636.5904 | 16049.3281 | 15779.0725 | 15100.1533 | 18160.1307 | 0.32638572 | 0.00022382 | 0.0031126 | Myo1c | 1.25386821 | 11 | 75650504 | 75673910 + |
| ENSMUSG0C | 2770.65857 | 3201.33729 | 3139.15999 | 2901.48046 | 2688.55533 | 2232.49948 | 2294.59418 | 2398.50783 | 0.32097338 | 0.00022482 | 0.0031236 | Jade1 | 1.24917308 | 3 | 41555731 | 41616864 + |
| ENSMUSG0C | 357.107105 | 266.692299 | 322.567968 | 502.553074 | 259.67891 | 210.041174 | 157.282042 | 216.511266 | 0.78019856 | 0.00022519 | 0.0031258 | Olfml2a | 1.71736722 | 2 | 38931978 | 38963753 + |
| ENSMUSG0C | 119.351446 | 82.3759998 | 106.268748 | 99.9553076 | 66.3840822 | 60.6304419 | 59.2500844 | 58.5970856 | 0.73730145 | 0.00022571 | 0.00312709 | Hyal1 | 1.66705471 | 9 | 107576927 | 107581710 + |
| ENSMUSG0C | 149.662925 | 167.8411 | 173.979808 | 152.709498 | 117.14838 | 103.9379 | 107.727426 | 111.235146 | 0.54825893 | 0.00022854 | 0.00316091 | Mastl | 1.46231987 | 2 | 23115606 | 23156024 - |
| ENSMUSG0C | 579.707025 | 2214.88469 | 599.995229 | 559.934825 | 220.62945 | 193.800877 | 338.264118 | 244.320052 | 1.98772708 | 0.00023196 | 0.00320177 | Dlk2 | 3.96611656 | 17 | 46297421 | 46303271 + |
| ENSMUSG0C | 36240.2141 | 29620.3501 | 33025.1293 | 35361.0411 | 29265.6179 | 24886.631 | 27582.5302 | 24347.5857 | 0.33970972 | 0.00023327 | 0.00321397 | mt-Nd5 | 1.26550194 | MT | 11742 | 13565 + |
| ENSMUSG0C | 8541.20628 | 7952.37308 | 7167.02765 | 9049.65738 | 6687.22005 | 3952.88827 | 4740.00675 | 6468.52235 | 0.58209813 | 0.00023398 | 0.00322081 | Adcy5 | 1.49702481 | 16 | 35154877 | 35305737 + |
| ENSMUSG0C | 399.732622 | 576.631998 | 259.559065 | 165.666667 | 214.772031 | 135.335808 | 145.432025 | 129.112223 | 1.16465636 | 0.00024478 | 0.00335388 | Ikzf4 | 2.24179811 | 10 | 128630843 | 128645991 - |
| ENSMUSG0C | 227.336088 | 278.018999 | 156.111611 | 243.409684 | 178.65128 | 83.3668576 | 108.8047 | 69.521966 | 1.03729933 | 0.00024459 | 0.00335388 | Nkain4 | 2.05238207 | 2 | 180934772 | 180954699 - |
| ENSMUSG0C | 1408.53651 | 1234.6103 | 1448.26435 | 1615.94414 | 1195.88972 | 1088.09989 | 1050.34241 | 1104.40609 | 0.36279786 | 0.0002451 | 0.0033552 | Arsk | 1.2859173 | 13 | 76060422 | 76098660 - |
| ENSMUSG0C | 1000.27879 | 825.819398 | 933.848374 | 1175.40038 | 710.700174 | 661.521428 | 706.691916 | 803.475293 | 0.44936908 | 0.00024597 | 0.00336089 | Pcdh19 | 1.36544299 | X | 133582860 | 133688987 - |
| ENSMUSG0C | 160.082495 | 102.97 | 122.256081 | 248.962757 | 81.0276298 | 98.524468 | 66.7910042 | 53.6312309 | 1.08234599 | 0.00024803 | 0.00337969 | Prr33 | 2.11747655 | 7 | 142491074 | 142506771 - |
| ENSMUSG0C | 465.091747 | 2257.10239 | 520.998992 | 271.175047 | 158.150314 | 161.320283 | 109.881975 | 189.69565 | 2.50489542 | 0.00025359 | 0.00343971 | Ddit4 | 5.67608196 | 10 | 59949669 | 59951834 - |
| ENSMUSG0C | 34.1004132 | 47.3661999 | 11.2851767 | 29.6163874 | 11.714838 | 5.41343231 | 5.38637131 | 8.93853849 | 1.94844465 | 0.00025345 | 0.00343971 | Dhtkd1 | 3.85958211 | 2 | 5896115 | 5942792 - |
| ENSMUSG0C | 586.337661 | 515.879699 | 646.076368 | 663.592181 | 471.522231 | 370.27877 | 372.736895 | 493.605959 | 0.49710216 | 0.0002552 | 0.00345842 | Ttll1 | 1.41137578 | 15 | 83483769 | 83510893 - |
| ENSMUSG0C | 9924.16748 | 9061.35997 | 10150.076 | 9015.41343 | 8552.80801 | 7693.57 | 7914.734 | 7577.8943 | 0.26545162 | 0.00025756 | 0.00348408 | Bcar3 | 1.20201227 | 3 | 122294136 | 122530191 + |
| ENSMUSG0C | 97199.4389 | 128655.866 | 110969.024 | 102010.87 | 89035.6978 | 70851.0021 | 75735.6124 | 91691.5278 | 0.42299375 | 0.00025799 | 0.00348672 | Tmsb4x | 1.34070679 | X | 167207093 | 167209315 - |
| ENSMUSG0C | 1178.35872 | 1143.9967 | 1230.08426 | 1174.47486 | 1013.33349 | 928.944984 | 903.833106 | 1031.90461 | 0.28509229 | 0.00025983 | 0.00350523 | Pcbp4 | 1.21848821 | 9 | 106453291 | 106464012 + |
| ENSMUSG0C | 268.067137 | 316.117899 | 199.371456 | 388.715085 | 197.199774 | 148.328045 | 147.586574 | 194.661505 | 0.76797543 | 0.00026044 | 0.00351025 | Oaf | 1.70287842 | 9 | 43221235 | 43239911 - |
| ENSMUSG0C | 5992.20039 | 7268.65228 | 6534.11732 | 6481.36129 | 4825.53704 | 5835.68003 | 5326.04395 | 4919.17568 | 0.32986962 | 0.00026119 | 0.00351718 | Phldb2 | 1.25689978 | 16 | 45746243 | 45953598 - |
| ENSMUSG0C | 14594.9769 | 17034.3271 | 15573.5439 | 15189.5047 | 13328.557 | 12472.548 | 12009.4535 | 13703.7727 | 0.27631487 | 0.00026331 | 0.00354217 | Cds2 | 1.21109737 | 2 | 132263148 | 132312050 + |
| ENSMUSG0C | 547.501079 | 580.750798 | 621.625151 | 596.955309 | 496.90438 | 400.593991 | 413.673316 | 479.701566 | 0.38888263 | 0.00026663 | 0.00358079 | Tpgs2 | 1.3093789 | 18 | 25127223 | 25169007 - |
| ENSMUSG0C | 180.921637 | 192.553899 | 190.907573 | 228.60149 | 116.172144 | 165.651029 | 79.7182954 | 91.3717268 | 0.80930962 | 0.00026891 | 0.00360484 | Stk39 | 1.75237267 | 2 | 68210445 | 68472268 - |
| ENSMUSG0C | 655.485721 | 692.988098 | 696.859663 | 703.389201 | 624.791362 | 489.374281 | 490.159789 | 470.763027 | 0.40483023 | 0.00026959 | 0.00361072 | Tarsl2 | 1.32393311 | 7 | 65644898 | 65692091 + |
| ENSMUSG0C | 3257.5367 | 4159.98799 | 3651.6951 | 3477.14899 | 3006.80843 | 2885.35942 | 2942.03601 | 2490.87273 | 0.36105625 | 0.00027104 | 0.00362688 | Hivep2 | 1.28436589 | 10 | 13966075 | 14151374 + |
| ENSMUSG0C | 2607.73438 | 3299.15879 | 2714.085 | 2814.48232 | 2513.809 | 2076.59263 | 2036.04835 | 2176.03754 | 0.37714952 | 0.00027161 | 0.00363124 | Bicc1 | 1.2987732 | 10 | 70922832 | 71159700 - |
| ENSMUSG0C | 373.210078 | 354.216799 | 312.223223 | 333.184359 | 247.964072 | 283.663853 | 251.004903 | 241.340539 | 0.42353622 | 0.00027243 | 0.00363893 | Hoxa3 | 1.341211 | 6 | 52169062 | 52213336 - |
| ENSMUSG0C | 1115.8413 | 1239.7588 | 1055.16402 | 981.042833 | 920.591023 | 753.549777 | 733.623772 | 890.874336 | 0.41192238 | 0.00027326 | 0.00364674 | Acsf2 | 1.33045746 | 11 | 94557102 | 94601871 - |
| ENSMUSG0C | 3766.20119 | 3385.65359 | 3562.35412 | 4140.74117 | 2873.06403 | 3279.45729 | 2804.1449 | 2926.87477 | 0.3221843 | 0.00027397 | 0.00365295 | Nr2f2 | 1.250222 | 7 | 70351944 | 70366735 - |
| ENSMUSG0C | 1166.99192 | 2031.59809 | 1026.95108 | 1347.54563 | 740.963506 | 237.108335 | 239.154886 | 502.544497 | 1.69584965 | 0.00027582 | 0.00367098 | Nkd1 | 3.23967625 | 8 | 88521354 | 88594884 + |
| ENSMUSG0C | 418.677296 | 446.889799 | 499.36907 | 529.392925 | 409.043095 | 275.002361 | 311.332262 | 321.787386 | 0.52329317 | 0.00027738 | 0.0036885 | Hdac11 | 1.4372322 | 6 | 91156665 | 91174692 + |
| ENSMUSG0C | 295.536915 | 255.365599 | 334.793576 | 372.981379 | 253.821491 | 207.875801 | 189.60027 | 185.722966 | 0.58871304 | 0.00027928 | 0.00371043 | Gm28437 | 1.50390459 | 1 | 24613189 | 24613971 - |
| ENSMUSG0C | 82.4093319 | 93.7026997 | 87.4601196 | 119.391062 | 60.5266632 | 50.8862637 | 57.0955359 | 27.8087864 | 0.96571071 | 0.00028143 | 0.00373238 | Calr3 | 1.9530254 | 8 | 72424176 | 72443870 - |
| ENSMUSG0C | 646.013384 | 586.928998 | 603.756955 | 690.432032 | 517.405347 | 439.570703 | 483.696143 | 519.428403 | 0.36620561 | 0.00028832 | 0.00382012 | Ttc8 | 1.28895832 | 12 | 98920574 | 98983238 + |
| ENSMUSG0C | 2604.89268 | 2647.35869 | 2545.74778 | 2478.52142 | 1948.56806 | 2360.25649 | 1976.79827 | 2102.54289 | 0.29305598 | 0.00028876 | 0.00382012 | Bcl9l | 1.22523287 | 9 | 44482825 | 44511896 + |
| ENSMUSG0C | 76.7259297 | 65.9007998 | 93.102708 | 94.4022349 | 52.7167712 | 45.4728314 | 44.1682447 | 45.6858634 | 0.81258456 | 0.0002906 | 0.00384026 | Actr3b | 1.7563551 | 5 | 25759997 | 25850688 + |
| ENSMUSG0C | 316.376056 | 216.236999 | 266.142084 | 302.642459 | 227.463105 | 159.15491 | 161.591139 | 151.955154 | 0.65316952 | 0.00029167 | 0.00385057 | Rab40b | 1.57261936 | 11 | 121356125 | 121388251 - |
| ENSMUSG0C | 1497.57648 | 1621.7775 | 1419.11097 | 1348.47114 | 1252.51143 | 1150.89571 | 1148.37436 | 1214.64806 | 0.30391642 | 0.0002919 | 0.00385057 | Impact | 1.23449109 | 18 | 12972252 | 12992948 + |
| ENSMUSG0C | 647.907851 | 727.997898 | 902.814138 | 687.655495 | 529.120185 | 519.689502 | 516.014371 | 604.841104 | 0.45095114 | 0.0002933 | 0.00386558 | Limk1 | 1.36694116 | 5 | 134656039 | 134688598 - |
| ENSMUSG0C | 6453.5032 | 6313.09068 | 6849.16184 | 7268.04658 | 5559.66689 | 4274.44615 | 5552.27154 | 5478.33092 | 0.36562637 | 0.00029458 | 0.00387912 | Enpp5 | 1.2884409 | 17 | 44078813 | 44086567 + |
| ENSMUSG0C | 597.704465 | 505.582699 | 739.179076 | 638.603354 | 507.642982 | 365.948024 | 401.8233 | 444.940583 | 0.52787219 | 0.00029495 | 0.00388057 | Tmem97 | 1.44180114 | 11 | 78541817 | 78550777 - |
| ENSMUSG0C | 187.552273 | 171.9599 | 168.33722 | 193.43203 | 113.243434 | 124.508943 | 142.200203 | 87.399043 | 0.62888824 | 0.000297 | 0.00389834 | Abca4 | 1.54637288 | 3 | 122044443 | 122180123 + |
| ENSMUSG0C | 2872.01258 | 3301.21819 | 3007.4996 | 2252.69647 | 2244.36772 | 2158.8768 | 2064.05749 | 2128.36533 | 0.41135788 | 0.00030295 | 0.00395777 | Flrt1 | 1.32993329 | 19 | 7092014 | 7105729 - |
| ENSMUSG0C | 945.339233 | 968.947697 | 947.014414 | 1008.8082 | 846.397048 | 787.113058 | 681.914608 | 779.63919 | 0.32183852 | 0.00030413 | 0.0039697 | Extl2 | 1.2499224 | 3 | 116007462 | 116029017 + |
| ENSMUSG0C | 2110.43668 | 2015.12289 | 2673.64645 | 2460.93669 | 1735.7485 | 1587.21835 | 1655.77054 | 1978.39652 | 0.41251091 | 0.00030654 | 0.00399419 | Uxs1 | 1.33100031 | 1 | 43746966 | 43827800 - |
| ENSMUSG0C | 1496.62925 | 2451.71569 | 2802.48555 | 1983.37245 | 1706.46141 | 1273.23928 | 1159.14711 | 1396.39835 | 0.6577743 | 0.00030781 | 0.00400012 | Gpc4 | 1.57764685 | X | 52053021 | 52165252 - |
| ENSMUSG0C | 266.17267 | 247.127999 | 252.035614 | 221.197394 | 164.983969 | 210.041174 | 145.432025 | 155.927838 | 0.54531627 | 0.00030761 | 0.00400012 | Hspbap1 | 1.45934023 | 16 | 35770386 | 35828446 + |
| ENSMUSG0C | 54.9395546 | 37.0691999 | 25.3916476 | 37.0204843 | 7.80989203 | 18.4056698 | 8.61819409 | 18.8702479 | 1.52517449 | 0.00030852 | 0.00400586 | Susd1 | 2.87821526 | 4 | 59314683 | 59438633 - |
| ENSMUSG0C | 2621.94288 | 2736.94259 | 2999.03572 | 2550.71137 | 2335.15772 | 1988.89503 | 2006.96195 | 2344.8766 | 0.33006178 | 0.00031112 | 0.00403613 | Npc1 | 1.25706721 | 18 | 12189693 | 12236400 - |
| ENSMUSG0C | 1089.31876 | 1030.7297 | 890.58853 | 1096.73185 | 825.896082 | 847.7435 | 790.719308 | 679.328925 | 0.38603812 | 0.0003158 | 0.00408641 | Mkl2 | 1.30679978 | 16 | 13256481 | 13417529 + |
| ENSMUSG0C | 4446.31499 | 7511.66148 | 5785.53394 | 6265.71696 | 2611.43265 | 2729.45257 | 3438.65944 | 1140.16024 | 1.27525787 | 0.00031847 | 0.00411717 | Sulf2 | 2.42042078 | 2 | 166073089 | 166155663 - |
| ENSMUSG0C | 119.351446 | 135.9204 | 104.387885 | 126.795159 | 60.5266632 | 86.6149169 | 43.0909705 | 81.4400173 | 0.83944795 | 0.00032338 | 0.00417697 | Gm32856 | 1.78936531 | 8 | 129263641 | 129282844 + |
| ENSMUSG0C | 598.651699 | 547.800398 | 520.058561 | 660.815644 | 469.569758 | 451.480255 | 445.991544 | 407.200087 | 0.39193263 | 0.00034072 | 0.00437438 | Mccc2 | 1.31214998 | 13 | 99948530 | 100015639 - |
| ENSMUSG0C | 243.439061 | 805.225399 | 246.393025 | 338.737431 | 179.627517 | 103.9379 | 87.2592152 | 127.125881 | 1.71318357 | 0.00034386 | 0.00440653 | Ptgis | 3.2788356 | 2 | 167191805 | 167240604 - |
| ENSMUSG0C | 882.821809 | 587.958698 | 680.872329 | 611.763503 | 328.991702 | 279.333107 | 617.278152 | 345.623488 | 0.81511786 | 0.00034825 | 0.00445572 | Stmn1 | 1.75944188 | 4 | 134468320 | 134473843 + |
| ENSMUSG0C | 2615.31225 | 2410.52769 | 1594.03121 | 2370.23651 | 1643.00603 | 1574.22612 | 1642.84325 | 1504.65398 | 0.49819534 | 0.00035011 | 0.00447558 | Hoxc4 | 1.41244564 | 15 | 103018934 | 103036852 + |
| ENSMUSG0C | 671.588694 | 676.512898 | 697.800094 | 598.806333 | 508.619218 | 481.795475 | 482.618869 | 581.005002 | 0.36373298 | 0.00036156 | 0.00461006 | Rpgrip1l | 1.28675106 | 8 | 91217030 | 91313262 - |
| ENSMUSG0C | 228.283322 | 203.880599 | 248.273888 | 318.376165 | 193.249828 | 122.34357 | 110.959249 | 165.859548 | 0.75194541 | 0.00036349 | 0.00463082 | Slc7a4 | 1.68406219 | 16 | 17572018 | 17576998 - |
| ENSMUSG0C | 5001.39394 | 4972.42129 | 5324.72255 | 5258.75979 | 4527.7849 | 4469.32971 | 4274.62427 | 4572.55902 | 0.20417244 | 0.00037047 | 0.00471161 | Cyfip1 | 1.15202533 | 7 | 55841745 | 55932602 + |
| ENSMUSG0C | 2647.51819 | 2854.32839 | 2873.01791 | 2698.7933 | 2282.44094 | 2414.39081 | 2355.99881 | 2441.21418 | 0.22200059 | 0.00038676 | 0.00490206 | Ocrl | 1.16634985 | X | 47912387 | 47965868 + |
| ENSMUSG0C | 1545.8854 | 1888.46979 | 1732.27463 | 1576.14712 | 1410.66175 | 949.516027 | 1084.81518 | 1374.54859 | 0.48381028 | 0.00038903 | 0.00492662 | Acaa2 | 1.39843217 | 18 | 74779197 | 74806207 + |
| ENSMUSG0C | 92.8289026 | 97.8214997 | 62.068472 | 86.9981381 | 59.5504267 | 47.6382043 | 26.9318565 | 29.7951283 | 1.04848398 | 0.00039107 | 0.00494729 | 9530077C05F | 2.06835523 | 9 | 22411513 | 22444681 + |
| ENSMUSG0C | 481.19472 | 480.869899 | 459.870952 | 527.541901 | 403.185676 | 388.68444 | 383.509637 | 364.493736 | 0.34046546 | 0.00039166 | 0.00494729 | Ncapd2 | 1.26616503 | 6 | 125168007 | 125191701 - |
| ENSMUSG0C | 251.016931 | 2286.96369 | 222.88224 | 96.2532591 | 88.8375218 | 64.9611877 | 48.4773418 | 64.5561113 | 3.42024602 | 0.00039862 | 0.00502594 | Gadd45g | 10.7052458 | 13 | 51846678 | 51848468 + |
| ENSMUSG0C | 155.346327 | 88.5541998 | 126.958238 | 145.305401 | 89.8137583 | 76.8707388 | 67.8682785 | 53.6312309 | 0.84211687 | 0.0003989 | 0.00502594 | Ccdc136 | 1.79267861 | 6 | 29396309 | 29426994 + |
| ENSMUSG0C | 171.4493 | 193.583599 | 298.116752 | 235.080075 | 136.67311 | 128.839689 | 99.1092321 | 164.866377 | 0.7616183 | 0.00040458 | 0.00508031 | Pygl | 1.69539132 | 12 | 70190801 | 70231488 - |
| ENSMUSG0C | 1580.93305 | 2063.51879 | 1488.7029 | 1625.19926 | 1234.93918 | 1304.63719 | 1089.12428 | 1382.49395 | 0.43107887 | 0.00041666 | 0.00521438 | Akap12 | 1.34824144 | 10 | 4266380 | 4359470 + |
| ENSMUSG0C | 92.8289026 | 91.6432997 | 30.0938046 | 16.6592179 | 23.4296761 | 4.33074585 | 12.9272911 | 13.9043932 | 2.07550541 | 0.00041741 | 0.00521938 | Gm16425 | 4.21492046 | 8 | 19535443 | 19538583 + |
| ENSMUSG0C | 113333.671 | 103852.453 | 94415.5502 | 103715.663 | 87456.1471 | 87370.6321 | 83243.1367 | 90604.0056 | 0.25233207 | 0.00042548 | 0.00531141 | Myom2 | 1.19113098 | 8 | 15057653 | 15133541 + |
| ENSMUSG0C | 57.7812557 | 45.3067999 | 61.1280406 | 52.7541901 | 28.3108586 | 34.6459668 | 23.7000338 | 25.8224445 | 0.95052455 | 0.00042631 | 0.00531722 | F730043M19 | 1.9325752 | 12 | 33111711 | 33147597 - |
| ENSMUSG0C | 225.441621 | 180.1975 | 260.499496 | 195.283055 | 134.720637 | 168.899088 | 142.200203 | 120.173684 | 0.60828359 | 0.00043425 | 0.00539054 | Akap17b | 1.52444446 | X | 36608314 | 36645395 - |
| ENSMUSG0C | 145.87399 | 142.0986 | 128.839101 | 146.230913 | 97.6236503 | 74.7053659 | 107.727426 | 91.3717268 | 0.59947983 | 0.00043436 | 0.00539054 | Prox2 | 1.51517017 | 12 | 85086385 | 85110759 - |
| ENSMUSG0C | 815.568216 | 629.146698 | 872.720334 | 835.737433 | 592.575557 | 622.544715 | 591.42357 | 572.066463 | 0.40753352 | 0.00043404 | 0.00539054 | Soat1 | 1.32641619 | 1 | 156424525 | 156474331 - |
| ENSMUSG0C | 41.6782828 | 39.1285999 | 47.0215697 | 34.243948 | 18.5484936 | 17.3229834 | 22.6227595 | 17.877077 | 1.08695978 | 0.00043842 | 0.0054229 | Aox4 | 2.12425915 | 1 | 58210397 | 58268597 + |
| ENSMUSG0C | 37.889348 | 17.5049 | 47.0215697 | 19.4357542 | 12.6910745 | 2.16537292 | 4.30909705 | 10.9248804 | 2.00473004 | 0.00044285 | 0.00546493 | AC116487.3 | 4.01313598 | 15 | 76095575 | 76097668 + |
| ENSMUSG0C | 154.399093 | 227.563699 | 250.154751 | 241.55866 | 180.603753 | 83.3668576 | 96.9546835 | 113.221488 | 0.87928807 | 0.00044327 | 0.00546493 | Abhd3 | 1.83946735 | 18 | 10644411 | 10706771 - |
| ENSMUSG0C | 64.4118916 | 72.0789998 | 42.3194127 | 22.2122906 | 15.6197841 | 16.2402969 | 26.9318565 | 9.93170943 | 1.55086884 | 0.00044725 | 0.00550463 | 1700023H06I | 2.92993537 | 13 | 80884224 | 80885524 - |
| ENSMUSG0C | 503.928329 | 919.522097 | 807.830567 | 749.664807 | 444.187609 | 462.307119 | 327.491376 | 587.957198 | 0.70962584 | 0.00045104 | 0.005533 | Nradd | 1.63537993 | 9 | 110621135 | 110624452 - |
| ENSMUSG0C | 71.0425275 | 316.117899 | 45.1407069 | 13.8826816 | 12.6910745 | 12.9922375 | 3.23182278 | 13.9043932 | 3.37909054 | 0.00045311 | 0.00554919 | Prr15l | 10.4041741 | 11 | 96928104 | 96937830 + |
| ENSMUSG0C | 2325.45873 | 2500.11159 | 3184.3007 | 2717.30355 | 2142.83912 | 1737.71177 | 1894.92543 | 2233.64145 | 0.4213984 | 0.00045622 | 0.00558271 | Pon2 | 1.33922503 | 6 | 5264147 | 5298455 - |
| ENSMUSG0C | 8838.63766 | 6901.04938 | 7626.8986 | 7300.4395 | 4211.48428 | 5799.95138 | 3056.22708 | 6323.5194 | 0.6612726 | 0.00045944 | 0.00561283 | AC149090.1 | 1.58147702 | JH584304.1 | 52190 | 59690 - |
| ENSMUSG0C | 1598.93049 | 1226.3727 | 838.864803 | 799.64246 | 699.961573 | 255.514005 | 211.145755 | 310.862505 | 1.59472213 | 0.0004636 | 0.0056452 | Tmod2 | 3.02036337 | 9 | 75565621 | 75611325 - |
| ENSMUSG0C | 24.6280762 | 26.7721999 | 30.0938046 | 37.0204843 | 18.5484936 | 6.49611877 | 6.46364557 | 7.94536755 | 1.57612869 | 0.00046349 | 0.0056452 | Rdh12 | 2.98168673 | 12 | 79208914 | 79222665 + |
| ENSMUSG0C | 56.834022 | 43.2473999 | 101.566591 | 99.0297954 | 42.9544061 | 23.8191022 | 37.7045992 | 21.8497607 | 1.25189016 | 0.00047481 | 0.00577696 | 2900041M22 | 2.38153238 | 11 | 117611247 | 117613857 + |
| ENSMUSG0C | 203.655246 | 543.681598 | 196.550161 | 160.113595 | 115.195907 | 75.7880523 | 136.813831 | 143.016616 | 1.22803583 | 0.00047823 | 0.00581375 | Spp1 | 2.34247854 | 5 | 104435118 | 104441050 + |
| ENSMUSG0C | 35.0476469 | 27.8018999 | 25.3916476 | 32.3929237 | 16.5960206 | 10.8268646 | 10.7727426 | 9.93170943 | 1.32202679 | 0.00048341 | 0.00586718 | Mroh7 | 2.50017103 | 4 | 106680417 | 106730925 - |
| ENSMUSG0C | 1453.0565 | 1589.8568 | 1336.35301 | 1553.00932 | 1218.34316 | 1119.4978 | 1069.73334 | 1302.04711 | 0.33237846 | 0.00048735 | 0.00591012 | Ift122 | 1.25908742 | 6 | 115853470 | 115926699 + |
| ENSMUSG0C | 28.417011 | 111.2076 | 40.4385499 | 25.914339 | 14.6435475 | 5.41343231 | 20.468211 | 14.8975641 | 1.88968263 | 0.00048826 | 0.00591636 | Gm44421 | 3.70553701 | 6 | 90478466 | 90479686 + |
| ENSMUSG0C | 1262.66252 | 1274.7686 | 1115.35163 | 1320.70578 | 1002.59489 | 1023.13871 | 852.123941 | 1053.75437 | 0.33877832 | 0.00049314 | 0.00594647 | Enoph1 | 1.2646852 | 5 | 100039985 | 100068760 + |
| ENSMUSG0C | 6025.35357 | 5972.25998 | 6762.64215 | 6514.67972 | 5681.69645 | 5007.42489 | 5140.75278 | 5341.27333 | 0.25555932 | 0.00049311 | 0.00594647 | Tmx3 | 1.19379848 | 18 | 90510154 | 90543267 + |
| ENSMUSG0C | 738.842286 | 851.561898 | 857.673431 | 852.396651 | 613.076524 | 359.451905 | 463.227933 | 699.192344 | 0.62753268 | 0.0004973 | 0.0059869 | Cyba | 1.54492059 | 8 | 122424776 | 122432930 - |
| ENSMUSG0C | 11.3668044 | 91.6432997 | 7.52345115 | 0.92551211 | 0 | 0 | 0 | 0 | 7.19778214 | 0.00049807 | 0.00598796 | Serpina3n | 146.807528 | 12 | 104406729 | 104414329 + |
| ENSMUSG0C | 1543.99093 | 3231.19859 | 2805.30685 | 2697.86779 | 1762.10689 | 883.472153 | 968.469561 | 849.161156 | 1.20332128 | 0.00049819 | 0.00598796 | Pamr1 | 2.30269173 | 2 | 102550012 | 102643041 + |
| ENSMUSG0C | 860.0882 | 790.809598 | 851.090411 | 912.554937 | 728.272431 | 686.423217 | 638.823637 | 715.083079 | 0.30241486 | 0.00050049 | 0.00601069 | Bbs9 | 1.2332069 | 9 | 22475715 | 22888280 + |
| ENSMUSG0C | 48.3089187 | 50.4552999 | 63.0089034 | 30.5418995 | 3.90494601 | 23.8191022 | 23.7000338 | 9.93170943 | 1.65842597 | 0.00050147 | 0.00601759 | Ccdc81 | 3.15671928 | 7 | 89866148 | 89903629 - |
| ENSMUSG0C | 1081.74089 | 671.364398 | 542.628914 | 1143.93296 | 530.096421 | 544.59129 | 462.150658 | 531.346455 | 0.7340546 | 0.00050344 | 0.00603643 | Sox8 | 1.66330715 | 17 | 25565892 | 25570686 - |
| ENSMUSG0C | 297.431382 | 287.286299 | 344.19789 | 321.152701 | 249.916545 | 199.214309 | 189.60027 | 252.26542 | 0.48703939 | 0.00050601 | 0.0060623 | Selenom | 1.40156571 | 11 | 3514684 | 3517351 + |
| ENSMUSG0C | 916.922222 | 909.225097 | 1005.32116 | 953.27747 | 805.395115 | 529.43368 | 518.16892 | 767.721139 | 0.5293573 | 0.00050828 | 0.00608466 | Cryl1 | 1.44328609 | 14 | 57274993 | 57398529 - |
| ENSMUSG0C | 172.396533 | 254.335899 | 267.082516 | 215.644321 | 176.698807 | 127.757002 | 80.7955696 | 134.078077 | 0.80601545 | 0.00051393 | 0.00613747 | Cap2 | 1.74837596 | 13 | 46501848 | 46650281 + |
| ENSMUSG0C | 56.834022 | 29.8612999 | 59.2471778 | 39.7970206 | 25.3821491 | 17.3229834 | 16.1591139 | 21.8497607 | 1.19992043 | 0.00051556 | 0.00615198 | Tubb4a | 2.29727 | 17 | 57080066 | 57087782 - |
| ENSMUSG0C | 71.9897612 | 56.6334998 | 47.9620011 | 82.3705775 | 35.1445141 | 16.2402969 | 30.1636793 | 38.7336668 | 1.10230904 | 0.00051692 | 0.00616332 | Impa2 | 2.14698043 | 18 | 67289253 | 67319146 + |
| ENSMUSG0C | 1449.26756 | 1862.72729 | 2219.41809 | 1056.93483 | 1320.84799 | 828.255143 | 967.392287 | 674.36307 | 0.79720075 | 0.00052717 | 0.00627043 | Endou | 1.73772616 | 15 | 97711015 | 97731339 - |
| ENSMUSG0C | 2279.99152 | 2674.13089 | 2420.67041 | 2220.30354 | 1879.25527 | 1947.75294 | 2095.29844 | 1941.64919 | 0.28693912 | 0.00052871 | 0.00627365 | Cul4b | 1.22004902 | X | 38533274 | 38576196 - |
| ENSMUSG0C | 4943.61268 | 5006.40139 | 5063.28262 | 4771.01491 | 3996.71224 | 4602.50015 | 4031.16029 | 4161.38625 | 0.23669332 | 0.00052853 | 0.00627365 | Ctsa | 1.17828891 | 2 | 164832873 | 164841032 + |
| ENSMUSG0C | 68.2008264 | 53.5443999 | 173.979808 | 107.359404 | 65.4078457 | 19.4883563 | 18.3136624 | 7.94536755 | 1.8556113 | 0.00053454 | 0.0063378 | Spon2 | 3.61905064 | 5 | 33198184 | 33218455 - |
| ENSMUSG0C | 3261.32563 | 3445.37619 | 4022.22507 | 3350.35383 | 2921.87585 | 2358.09111 | 2652.24923 | 3021.22601 | 0.36200338 | 0.00053537 | 0.00634255 | Asah1 | 1.28520935 | 8 | 41340197 | 41374773 - |
| ENSMUSG0C | 1338.44122 | 1247.9964 | 1362.68509 | 1407.70391 | 1159.76897 | 893.216331 | 1118.21068 | 1054.74754 | 0.3419241 | 0.00053731 | 0.00636049 | Pcca | 1.26744584 | 14 | 122534324 | 122891100 + |
| ENSMUSG0C | 10045.4134 | 10226.9804 | 9421.2417 | 8759.97209 | 8322.41619 | 5921.21226 | 6253.57709 | 8149.96076 | 0.42460687 | 0.0005386 | 0.00637072 | Clstn1 | 1.3422067 | 4 | 149586468 | 149648899 + |
| ENSMUSG0C | 1496.62925 | 1542.4906 | 1630.70804 | 1666.8473 | 1346.23014 | 1180.12824 | 1166.68803 | 1410.30274 | 0.31185486 | 0.0005413 | 0.0063976 | Gbe1 | 1.24130261 | 16 | 70313949 | 70569716 + |
| ENSMUSG0C | 402.574323 | 394.375099 | 343.257459 | 440.543763 | 331.920411 | 262.010124 | 301.636793 | 262.197129 | 0.4488176 | 0.00054379 | 0.00642185 | Mfsd4b4 | 1.36492114 | 10 | 39890018 | 39899238 - |
| ENSMUSG0C | 927.341793 | 1002.9278 | 1260.17807 | 1101.35941 | 952.806827 | 594.394867 | 721.773755 | 718.062592 | 0.52241953 | 0.00054675 | 0.00644659 | Scg5 | 1.43636214 | 2 | 113776362 | 113829121 - |
| ENSMUSG0C | 121.245914 | 120.4749 | 153.290317 | 97.1787712 | 83.9563393 | 67.1265606 | 76.4864726 | 77.4673336 | 0.68872634 | 0.00054853 | 0.00646246 | Hspb6 | 1.61185989 | 7 | 30552178 | 30555443 + |
| ENSMUSG0C | 20.8391414 | 19.5642999 | 47.0215697 | 27.7653632 | 14.6435475 | 8.66149169 | 5.38637131 | 2.97951283 | 1.85977993 | 0.00055857 | 0.00655488 | Spx | 3.62952294 | 6 | 142413441 | 142419730 + |
| ENSMUSG0C | 3210.17501 | 4842.67909 | 3236.02443 | 2903.33148 | 2349.80126 | 2593.03408 | 2693.18565 | 2269.39561 | 0.51873019 | 0.00056109 | 0.00656891 | Lims2 | 1.43269369 | 18 | 31931325 | 31958619 + |
| ENSMUSG0C | 329.637328 | 263.603199 | 331.972282 | 401.672254 | 231.368051 | 221.950725 | 198.218464 | 262.197129 | 0.53832307 | 0.00056186 | 0.00657272 | Mdm1 | 1.45228346 | 10 | 118141811 | 118168997 + |
| ENSMUSG0C | 1436.00629 | 1496.1541 | 1598.73337 | 1650.18809 | 1358.92121 | 1023.13871 | 1199.00625 | 1259.34076 | 0.35234478 | 0.00058174 | 0.00677866 | mt-Rnr2 | 1.27663383 | MT | 1094 | 2675 + |
| ENSMUSG0C | 956.706037 | 813.462998 | 1116.29206 | 1011.58473 | 752.678344 | 660.438742 | 718.541933 | 815.393344 | 0.40348902 | 0.0005897 | 0.00686595 | Gm7336 | 1.32270288 | 7 | 51746647 | 51747878 + |
| ENSMUSG0C | 373.210078 | 313.028799 | 335.734008 | 317.450653 | 287.989768 | 245.769827 | 210.068481 | 186.716137 | 0.52519437 | 0.00059076 | 0.00687292 | Sfxn5 | 1.43912745 | 6 | 85213049 | 85333422 - |
| ENSMUSG0C | 2028.97459 | 1886.41039 | 2638.85049 | 2296.19554 | 1898.78 | 1438.89031 | 1516.80216 | 1677.46572 | 0.43811415 | 0.00059287 | 0.00688675 | Kdelc2 | 1.35483217 | 9 | 53384025 | 53401867 + |
| ENSMUSG0C | 25903.0528 | 30330.8431 | 31600.3757 | 28464.1248 | 22373.3882 | 25052.282 | 23990.8978 | 24651.496 | 0.2757074 | 0.00059249 | 0.00688675 | Tln1 | 1.21058754 | 4 | 43531519 | 43562691 - |
| ENSMUSG0C | 1177.41149 | 7076.09838 | 289.652869 | 437.767227 | 292.870951 | 202.462368 | 156.204768 | 220.483949 | 3.36432019 | 0.00059342 | 0.00688764 | Fkbp5 | 10.2981993 | 17 | 28399095 | 28517524 - |
| ENSMUSG0C | 5182.31557 | 7007.10848 | 6505.90438 | 5353.16203 | 4944.63789 | 3699.53964 | 3561.46871 | 4939.0391 | 0.48801108 | 0.00059486 | 0.00688838 | Ifitm2 | 1.40251003 | 7 | 140954837 | 140955987 - |
| ENSMUSG0C | 441.410904 | 469.543199 | 457.990089 | 429.437618 | 389.518365 | 303.152209 | 340.418667 | 353.568856 | 0.37344896 | 0.00059463 | 0.00688838 | Ispd | 1.29544608 | 12 | 36381450 | 36689503 + |
| ENSMUSG0C | 641.277215 | 692.988098 | 729.774762 | 715.420859 | 592.575557 | 545.637977 | 562.337165 | 566.107438 | 0.29389227 | 0.00059857 | 0.00692049 | Poc1b | 1.22594331 | 10 | 99107036 | 99198074 + |
| ENSMUSG0C | 134.507185 | 132.8313 | 95.9240022 | 205.463688 | 82.9801028 | 85.5322305 | 54.9409873 | 84.4195302 | 0.8848628 | 0.00059948 | 0.00692162 | Tspyl3 | 1.846589 | 2 | 153222370 | 153225441 - |
| ENSMUSG0C | 101.354006 | 72.0789998 | 52.6641581 | 66.6368717 | 41.9781696 | 27.0671615 | 43.0909705 | 36.7473249 | 0.97369864 | 0.0006061 | 0.00699129 | CT030170.3 | 1.96386892 | 12 | 17933164 | 17939628 + |
| ENSMUSG0C | 4095.83852 | 4220.74029 | 5008.7376 | 4330.47115 | 3828.79957 | 3301.11102 | 3492.52316 | 3700.55493 | 0.30171631 | 0.00061918 | 0.00713668 | Ermp1 | 1.23260992 | 19 | 29608214 | 29648415 - |
| ENSMUSG0C | 656.432954 | 513.820299 | 434.479304 | 590.476724 | 347.540195 | 356.203846 | 466.459755 | 279.081035 | 0.5999119 | 0.00063299 | 0.00727892 | Gdap10 | 1.51562401 | 12 | 32821872 | 32826904 + |
| ENSMUSG0C | 1049.53494 | 1176.9471 | 1025.07022 | 1170.77282 | 897.161346 | 981.996623 | 839.19665 | 803.475293 | 0.32858569 | 0.0006325 | 0.00727892 | Ror1 | 1.2557817 | 4 | 100095791 | 100444765 + |
| ENSMUSG0C | 485.930888 | 506.612399 | 536.045895 | 571.966482 | 404.161912 | 356.203846 | 356.577781 | 464.804001 | 0.40837904 | 0.00063553 | 0.00729684 | Dsel | 1.32719379 | 1 | 111858702 | 111864918 - |
| ENSMUSG0C | 39.7838154 | 84.4353998 | 33.8555302 | 110.135941 | 24.4059126 | 30.3152209 | 18.3136624 | 28.8019574 | 1.39624665 | 0.00064215 | 0.0073558 | Aldh1a1 | 2.63215901 | 19 | 20492715 | 20643462 + |
| ENSMUSG0C | 751.156324 | 662.097098 | 890.58853 | 924.586595 | 662.864586 | 542.425917 | 547.255325 | 619.738669 | 0.44448249 | 0.00065001 | 0.00743442 | Pdp1 | 1.36082589 | 4 | 11958184 | 11966452 - |
| ENSMUSG0C | 58.7284894 | 25.7424999 | 43.2598441 | 46.2756053 | 12.6910745 | 16.2402969 | 17.2363882 | 25.8224445 | 1.27478914 | 0.00065366 | 0.00747028 | Lgals12 | 2.41963451 | 19 | 7596660 | 7607193 - |
| ENSMUSG0C | 441.410904 | 382.018699 | 492.78605 | 519.212292 | 361.207506 | 245.769827 | 343.650489 | 333.705437 | 0.5148675 | 0.00066049 | 0.00752532 | Psph | 1.42886291 | 5 | 129765558 | 129787449 - |
| ENSMUSG0C | 933.972429 | 836.116398 | 1186.82442 | 1286.46183 | 802.466406 | 697.250081 | 728.237401 | 808.441148 | 0.48297561 | 0.00066217 | 0.00753876 | Mapk8ip1 | 1.39762334 | 2 | 92383676 | 92401263 - |
| ENSMUSG0C | 754.945259 | 711.522698 | 806.890136 | 919.033522 | 689.222971 | 525.102934 | 582.805376 | 611.793301 | 0.40594713 | 0.00066697 | 0.00757485 | Hhat | 1.32495846 | 1 | 192496711 | 192771223 - |
| ENSMUSG0C | 4154.56701 | 5149.52969 | 3299.03333 | 3652.07077 | 3341.65755 | 2827.97704 | 2647.94014 | 2947.73136 | 0.46614074 | 0.00066968 | 0.0075951 | Pbx1 | 1.3814092 | 1 | 168119364 | 168432270 - |
| ENSMUSG0C | 863.877135 | 907.165697 | 809.71143 | 910.703913 | 657.983403 | 612.800537 | 640.978186 | 802.482122 | 0.3625629 | 0.0006794 | 0.00768784 | Stk17b | 1.28570789 | 1 | 53755506 | 53785224 - |
| ENSMUSG0C | 197.02461 | 160.6332 | 142.945572 | 247.111733 | 95.6711773 | 115.847451 | 142.200203 | 95.3444105 | 0.7382107 | 0.00068171 | 0.00770225 | B130024G19 | 1.66810569 | 7 | 70364910 | 70489972 + |
| ENSMUSG0C | 492.561524 | 598.255698 | 426.015421 | 372.055867 | 319.229337 | 334.550117 | 291.941325 | 360.521052 | 0.53091684 | 0.0006813 | 0.00770225 | Hipk4 | 1.44484711 | 7 | 27523267 | 27531175 + |
| ENSMUSG0C | 179.027169 | 191.524199 | 201.252318 | 186.953446 | 149.364185 | 94.1937222 | 119.577443 | 144.009787 | 0.5782189 | 0.00068701 | 0.00775039 | Fam229b | 1.4930049 | 10 | 39118804 | 39133914 - |
| ENSMUSG0C | 548.448313 | 1149.1452 | 473.977423 | 708.942274 | 456.878684 | 463.389806 | 383.509637 | 341.650804 | 0.80744014 | 0.00069254 | 0.00780683 | Ier2 | 1.75010338 | 8 | 84661331 | 84662852 - |
| ENSMUSG0C | 789.045672 | 905.106297 | 866.137314 | 762.621976 | 731.201141 | 476.382043 | 526.787114 | 649.533797 | 0.47793111 | 0.0006945 | 0.00782298 | Fmo5 | 1.39274498 | 3 | 97628804 | 97655282 + |
| ENSMUSG0C | 53.0450872 | 98.8511997 | 74.2940801 | 54.6052143 | 42.9544061 | 23.8191022 | 20.468211 | 40.7200087 | 1.12605225 | 0.00069561 | 0.00782953 | Uchl1 | 2.1826068 | 5 | 66676091 | 66687234 + |
| ENSMUSG0C | 25.5753099 | 51.4849999 | 27.2725104 | 12.0316574 | 8.78612853 | 8.66149169 | 3.23182278 | 10.9248804 | 1.87085056 | 0.00069735 | 0.00784325 | Muc1 | 3.65748148 | 3 | 89229057 | 89233381 + |
| ENSMUSG0C | 2546.16419 | 2123.24139 | 2709.38285 | 2680.28306 | 2020.80956 | 1627.27775 | 1763.49797 | 2198.88047 | 0.40231746 | 0.00070188 | 0.00788226 | Asic2 | 1.3216292 | 11 | 80880169 | 81968457 - |
| ENSMUSG0C | 1341.28292 | 1260.3528 | 1532.90317 | 1355.87524 | 1087.52746 | 876.976034 | 1135.44707 | 1146.11927 | 0.37065923 | 0.0007018 | 0.00788226 | Fcgrt | 1.2929435 | 7 | 45092990 | 45103851 - |
| ENSMUSG0C | 46.4144513 | 152.3956 | 49.8428639 | 14.8081937 | 15.6197841 | 9.74417816 | 17.2363882 | 24.8292736 | 1.96167405 | 0.00070379 | 0.0078826 | Cxcl1 | 3.89513693 | 5 | 90891241 | 90893115 + |
| ENSMUSG0C | 130.718251 | 83.4056998 | 94.9835708 | 181.400373 | 65.4078457 | 53.0516366 | 80.7955696 | 31.7814702 | 1.08868925 | 0.00070272 | 0.0078826 | Hey1 | 2.12680719 | 3 | 8663359 | 8667256 - |
| ENSMUSG0C | 75.778696 | 71.0492998 | 96.8644336 | 66.6368717 | 27.3346221 | 42.224772 | 54.9409873 | 42.7063506 | 0.89651294 | 0.00070357 | 0.0078826 | Rgs11 | 1.86156107 | 17 | 26202951 | 26211324 + |
| ENSMUSG0C | 4913.3012 | 4865.33249 | 5377.38671 | 5489.21231 | 4731.81833 | 4005.93991 | 3858.79641 | 4255.73749 | 0.29276517 | 0.00070475 | 0.00788467 | Eml1 | 1.22498592 | 12 | 108370957 | 108539617 + |
| ENSMUSG0C | 757.78696 | 2562.92329 | 789.021939 | 807.972069 | 361.207506 | 356.203846 | 315.641359 | 546.244019 | 1.63847343 | 0.00071045 | 0.00793843 | Klf15 | 3.1133622 | 6 | 90462576 | 90475238 + |
| ENSMUSG0C | 605.282335 | 501.463899 | 618.803857 | 643.230914 | 479.332123 | 380.022948 | 416.905139 | 491.619617 | 0.42174036 | 0.00071113 | 0.00793843 | Rnpep | 1.3395425 | 1 | 135262712 | 135284084 - |
| ENSMUSG0C | 13.2612718 | 53.5443999 | 33.8555302 | 9.25512107 | 11.714838 | 3.24805939 | 1.07727426 | 1.98634189 | 2.59170567 | 0.00071672 | 0.0079946 | Gm16174 | 6.02810969 | 7 | 19816553 | 19818625 + |
| ENSMUSG0C | 35.0476469 | 67.9601998 | 35.736393 | 52.7541901 | 18.5484936 | 18.4056698 | 30.1636793 | 10.9248804 | 1.29785136 | 0.00072163 | 0.00803728 | Hsd17b1 | 2.45862441 | 11 | 101078411 | 101080527 + |
| ENSMUSG0C | 2952.52744 | 3174.56509 | 3233.20313 | 3049.56239 | 2814.48984 | 2026.78906 | 2320.44876 | 2553.44249 | 0.35285317 | 0.00072222 | 0.00803785 | Asph | 1.27708378 | 4 | 9448069 | 9669344 - |
| ENSMUSG0C | 107.984642 | 80.3165998 | 124.136944 | 117.540038 | 54.6692442 | 31.3979074 | 50.6318903 | 83.4263592 | 0.96349002 | 0.00072441 | 0.00805015 | Scml2 | 1.95002149 | X | 161117193 | 161258213 + |
| ENSMUSG0C | 1424.63949 | 1354.0555 | 1401.24278 | 1488.22347 | 1191.00853 | 1205.03003 | 1192.54261 | 1252.38856 | 0.22777193 | 0.00072793 | 0.0080833 | Mccc1 | 1.17102504 | 3 | 35959312 | 36000678 - |
| ENSMUSG0C | 189.44674 | 192.553899 | 195.60973 | 208.240224 | 139.60182 | 124.508943 | 118.500169 | 166.852768 | 0.51424497 | 0.00072878 | 0.00808673 | Abhd6 | 1.42824647 | 14 | 8002902 | 8056763 + |
| ENSMUSG0C | 281.328409 | 272.870499 | 242.6313 | 347.992552 | 233.320524 | 190.552817 | 184.213899 | 192.675163 | 0.51504466 | 0.00073049 | 0.00809958 | AA414768 | 1.42903837 | X | 12936872 | 12938128 + |
| ENSMUSG0C | 334.373496 | 1268.5904 | 293.414595 | 216.569833 | 105.433542 | 126.674316 | 170.209333 | 101.303436 | 2.06885378 | 0.00073161 | 0.00810596 | Slc16a9 | 4.19553208 | 10 | 70245100 | 70285968 + |
| ENSMUSG0C | 90.9344352 | 122.5343 | 109.090042 | 139.752328 | 87.8612853 | 61.7131283 | 49.554616 | 69.521966 | 0.77970092 | 0.00073348 | 0.00812066 | Gstm5 | 1.71677494 | 3 | 107895821 | 107898686 + |
| ENSMUSG0C | 97.5650711 | 156.5144 | 133.541258 | 85.1471138 | 64.4316092 | 61.7131283 | 31.2409536 | 78.4605045 | 0.99953106 | 0.00073691 | 0.00814267 | Tmem184a | 1.99935002 | 5 | 139802485 | 139819917 - |
| ENSMUSG0C | 1901.09804 | 1714.4505 | 1806.56871 | 1891.74675 | 1736.72474 | 1089.18258 | 1280.8791 | 1225.57294 | 0.45554491 | 0.00073758 | 0.00814267 | L3mbtl3 | 1.37130065 | 10 | 26274468 | 26375971 - |
| ENSMUSG0C | 848.721396 | 802.136298 | 855.792568 | 856.098699 | 689.222971 | 717.821124 | 729.314675 | 684.29478 | 0.25439985 | 0.00073766 | 0.00814267 | Dhx35 | 1.19283943 | 2 | 158794807 | 158858214 + |
| ENSMUSG0C | 454.672176 | 488.077799 | 398.742911 | 500.70205 | 329.967938 | 349.707727 | 356.577781 | 380.384471 | 0.37874568 | 0.00073907 | 0.0081522 | Dzip1l | 1.30021092 | 9 | 99629496 | 99669256 + |
| ENSMUSG0C | 3522.76213 | 3800.62269 | 3390.25518 | 3306.85476 | 3194.24584 | 2921.08807 | 2749.20392 | 2710.3635 | 0.27630407 | 0.00074072 | 0.00816433 | Rbm28 | 1.21108831 | 6 | 29123576 | 29165006 - |
| ENSMUSG0C | 36933.5892 | 44765.1777 | 40738.5476 | 37173.1938 | 33055.368 | 24915.8635 | 23186.1739 | 34876.1908 | 0.45998505 | 0.00075726 | 0.00832817 | Cavin1 | 1.37552757 | 11 | 100956733 | 100970887 - |
| ENSMUSG0C | 189.44674 | 197.702399 | 189.967142 | 209.165736 | 162.05526 | 113.682078 | 148.663848 | 109.248804 | 0.55849403 | 0.00075917 | 0.008343 | Zfp667 | 1.47273109 | 7 | 6286579 | 6307883 + |
| ENSMUSG0C | 897.977548 | 931.878497 | 973.346493 | 1047.67971 | 731.201141 | 813.097533 | 730.391949 | 838.236276 | 0.30713153 | 0.00076115 | 0.00835857 | Prkra | 1.23724527 | 2 | 76629898 | 76648015 - |
| ENSMUSG0C | 712.319743 | 620.909098 | 808.770999 | 590.476724 | 303.609553 | 438.488017 | 567.723536 | 484.66742 | 0.60738968 | 0.00076228 | 0.00836481 | Fah | 1.52350019 | 7 | 84585159 | 84606722 - |
| ENSMUSG0C | 2433.44338 | 1868.90549 | 1825.37734 | 2879.26816 | 1597.12292 | 1482.19777 | 1583.59316 | 1756.9194 | 0.48857208 | 0.00076534 | 0.0083799 | Gm45276 | 1.4030555 | 8 | 15044563 | 15045648 + |
| ENSMUSG0C | 934.919662 | 986.452597 | 820.996607 | 672.847302 | 790.751568 | 411.420855 | 476.155224 | 463.81083 | 0.67206545 | 0.00076796 | 0.00840237 | Tshb | 1.59335247 | 3 | 102775465 | 102782718 - |
| ENSMUSG0C | 6753.77628 | 7113.16758 | 6099.63802 | 6550.77469 | 5549.90452 | 5686.2693 | 5863.60381 | 5580.62753 | 0.22545554 | 0.00077853 | 0.00850548 | Pld1 | 1.16914635 | 3 | 27938695 | 28133362 + |
| ENSMUSG0C | 268.067137 | 230.652799 | 281.188987 | 311.89758 | 189.389882 | 200.296995 | 185.291173 | 218.497607 | 0.46130498 | 0.0007909 | 0.00861534 | Nek8 | 1.37678662 | 11 | 78166106 | 78176675 - |
| ENSMUSG0C | 2651.30713 | 2702.96249 | 2491.20276 | 2684.91062 | 2407.39922 | 2172.95173 | 2126.53939 | 2258.47072 | 0.23177283 | 0.00080618 | 0.0087669 | Nhlrc3 | 1.17427705 | 3 | 53448583 | 53463332 - |
| ENSMUSG0C | 1308.12974 | 1387.0059 | 1706.88298 | 1699.24023 | 1010.40478 | 1041.54438 | 999.710515 | 1385.47347 | 0.45940241 | 0.00081545 | 0.00884393 | Emp3 | 1.37497217 | 7 | 45918023 | 45921404 - |
| ENSMUSG0C | 42.6255165 | 58.6928998 | 56.4258836 | 61.0837991 | 38.0732236 | 27.0671615 | 22.6227595 | 24.8292736 | 0.95459064 | 0.00082007 | 0.00887641 | Tspoap1 | 1.93802965 | 11 | 87760541 | 87785928 + |
| ENSMUSG0C | 652.64402 | 651.800098 | 653.599819 | 670.996278 | 514.476637 | 533.764426 | 561.25989 | 567.100609 | 0.27274567 | 0.00082023 | 0.00887641 | Helb | 1.20810485 | 10 | 120083608 | 120112987 - |
| ENSMUSG0C | 266.17267 | 399.523599 | 306.580634 | 337.811919 | 266.515265 | 178.643266 | 196.063916 | 103.289778 | 0.81473404 | 0.00082247 | 0.00889416 | Lox | 1.75897386 | 18 | 52516069 | 52529867 - |
| ENSMUSG0C | 759.681428 | 1016.3139 | 703.442683 | 690.432032 | 639.43491 | 469.885924 | 373.814169 | 593.916224 | 0.60872331 | 0.00082523 | 0.00891751 | Plcb4 | 1.52490917 | 2 | 135659011 | 136014593 + |
| ENSMUSG0C | 2985.68062 | 3717.21699 | 3498.40479 | 3115.27375 | 2752.98694 | 2871.2845 | 2568.22184 | 2620.97812 | 0.30026155 | 0.00082874 | 0.00894897 | Sep-08 | 1.23136763 | 11 | 53519257 | 53549565 + |
| ENSMUSG0C | 299.325849 | 283.167499 | 290.593301 | 286.908753 | 223.558159 | 238.191022 | 227.304869 | 209.559069 | 0.36937235 | 0.00083478 | 0.0090011 | Fdxr | 1.29179071 | 11 | 115268024 | 115277050 - |
| ENSMUSG0C | 110.826343 | 181.227199 | 82.7579627 | 154.560522 | 61.5028997 | 37.8940262 | 38.7818734 | 100.310265 | 1.1464342 | 0.0008373 | 0.00902176 | Rbm11 | 2.21366084 | 16 | 75592844 | 75602829 + |
| ENSMUSG0C | 1748.59341 | 1979.08339 | 2047.31914 | 1766.80261 | 1618.60012 | 1241.84137 | 1299.19276 | 1625.82083 | 0.38194672 | 0.00084238 | 0.00906336 | Hcfc1r1 | 1.30309903 | 17 | 23673596 | 23675227 + |
| ENSMUSG0C | 677.272096 | 690.928698 | 619.744289 | 767.249537 | 588.670611 | 514.276069 | 531.096211 | 553.196215 | 0.33252213 | 0.00085414 | 0.00917657 | Tsen15 | 1.25921282 | 1 | 152370735 | 152386688 - |
| ENSMUSG0C | 127.87655 | 141.0689 | 187.145847 | 153.63501 | 85.9088123 | 98.524468 | 60.3273587 | 118.187342 | 0.74727585 | 0.00085934 | 0.00921255 | Atp10a | 1.67862021 | 7 | 58656166 | 58829420 + |
| ENSMUSG0C | 370.368377 | 519.998499 | 315.044517 | 306.344507 | 275.298694 | 182.974012 | 274.704937 | 217.504437 | 0.66819064 | 0.0008674 | 0.00928358 | Trpv4 | 1.58907877 | 5 | 114622152 | 114658421 - |
| ENSMUSG0C | 361.843274 | 301.702099 | 375.232126 | 362.800746 | 246.987835 | 208.958487 | 211.145755 | 311.855676 | 0.51680347 | 0.00086784 | 0.00928358 | Nap1l3 | 1.4307816 | X | 122394565 | 122397401 - |
| ENSMUSG0C | 294.589681 | 326.414899 | 352.661773 | 386.864061 | 275.298694 | 207.875801 | 228.382143 | 269.149326 | 0.47107193 | 0.00088123 | 0.00939976 | Gm10080 | 1.386139 | 9 | 44374744 | 44375181 - |
| ENSMUSG0C | 111.773577 | 74.1383998 | 62.068472 | 92.5512107 | 53.6930077 | 50.8862637 | 36.6273249 | 46.6790343 | 0.85719893 | 0.00088367 | 0.00941902 | Tubg2 | 1.81151774 | 11 | 101155907 | 101161787 + |
| ENSMUSG0C | 232.072257 | 235.801599 | 244.512162 | 244.335196 | 190.366118 | 153.741478 | 164.822962 | 196.647847 | 0.43733039 | 0.00089004 | 0.00947344 | Zcwpw1 | 1.35409635 | 5 | 137787798 | 137822621 + |
| ENSMUSG0C | 2097.17541 | 1863.75699 | 2333.21029 | 2250.84544 | 1843.13452 | 1650.01417 | 1633.14778 | 1783.73501 | 0.30639547 | 0.00089272 | 0.00948838 | Pfkm | 1.23661419 | 15 | 98092589 | 98132451 + |
| ENSMUSG0C | 7337.27224 | 6464.45658 | 7045.712 | 6870.07637 | 5499.14022 | 5951.52748 | 6335.44993 | 5616.38168 | 0.24430265 | 0.00090064 | 0.00955882 | Atp8a1 | 1.18452008 | 5 | 67618140 | 67847434 - |
| ENSMUSG0C | 4985.29096 | 4760.30309 | 5036.95055 | 5214.33521 | 4295.44061 | 4306.92674 | 4462.06999 | 3736.30909 | 0.2514248 | 0.00090191 | 0.00956554 | Tshz1 | 1.19038215 | 18 | 84011627 | 84086404 - |
| ENSMUSG0C | 263.330969 | 217.266699 | 314.104086 | 354.471137 | 201.10472 | 214.371919 | 168.054785 | 190.688821 | 0.57081809 | 0.00091308 | 0.00967711 | Bhlhb9 | 1.48536562 | X | 135797594 | 135891081 + |
| ENSMUSG0C | 1177.41149 | 1245.937 | 1445.44305 | 1093.0298 | 1125.60069 | 830.420516 | 844.583021 | 897.826533 | 0.42342849 | 0.00092247 | 0.00975578 | Aph1b | 1.34111085 | 9 | 66775202 | 66795490 - |
| ENSMUSG0C | 256.700333 | 261.543799 | 186.205416 | 199.910615 | 151.316658 | 160.237596 | 172.363882 | 135.071248 | 0.54704127 | 0.00093573 | 0.00986098 | Gm4924 | 1.46108618 | 10 | 82354255 | 82422882 + |
| ENSMUSG0C | 353.31817 | 407.761199 | 363.006518 | 365.577282 | 293.847187 | 289.077285 | 272.550388 | 309.869334 | 0.35278222 | 0.00093835 | 0.00988151 | Mettl8 | 1.27702097 | 2 | 70964561 | 71055583 - |
| ENSMUSG0C | 3217.75288 | 2976.86269 | 3484.29831 | 3661.3259 | 2809.60866 | 2402.48126 | 2435.71711 | 2946.73819 | 0.33236433 | 0.00093949 | 0.00988373 | Isoc1 | 1.25907509 | 18 | 58659482 | 58679570 + |
| ENSMUSG0C | 2510.16931 | 2467.16119 | 2599.35237 | 2520.16947 | 2207.27073 | 2221.67262 | 2093.14389 | 2280.32049 | 0.19786528 | 0.00094288 | 0.00990828 | Rbbp9 | 1.14699991 | 2 | 144542265 | 144550869 - |
| ENSMUSG0C | 23840.925 | 19419.1122 | 21813.3062 | 24577.8995 | 19967.9414 | 16475.2399 | 17601.5842 | 17944.6126 | 0.31653747 | 0.00096091 | 0.01007635 | mt-Nd1 | 1.2453381 | MT | 2751 | 3707 + |
| ENSMUSG0C | 2521.53611 | 2561.89359 | 2795.90253 | 2200.86779 | 2058.88279 | 2162.12486 | 2071.59841 | 1841.33893 | 0.30958577 | 0.00096191 | 0.0100797 | Lpp | 1.2393518 | 16 | 24393350 | 24992576 + |
| ENSMUSG0C | 1021.11793 | 818.611498 | 894.350256 | 947.724398 | 479.332123 | 624.710088 | 505.241629 | 862.072379 | 0.57503137 | 0.00096335 | 0.01008249 | Gm3716 | 1.48970985 | 5 | 64593857 | 64610700 - |
| ENSMUSG0C | 244.386295 | 272.870499 | 269.90381 | 306.344507 | 209.890848 | 190.552817 | 200.373013 | 218.497607 | 0.41578704 | 0.00096353 | 0.01008249 | Polr2h | 1.33402623 | 16 | 20717665 | 20722267 + |
| ENSMUSG0C | 145.87399 | 177.1084 | 73.3536487 | 170.294228 | 97.6236503 | 76.8707388 | 59.2500844 | 52.63806 | 0.98305793 | 0.0009716 | 0.0101455 | Tgfb2 | 1.97665067 | 1 | 186622792 | 186705989 - |
| ENSMUSG0C | 3944.28113 | 4064.22589 | 4241.34559 | 4522.97767 | 3859.0629 | 2979.55314 | 2756.74484 | 3420.48073 | 0.36566891 | 0.00097298 | 0.01015283 | Cyb5a | 1.2884789 | 18 | 84851338 | 84880401 + |
| ENSMUSG0C | 228.283322 | 201.821199 | 233.226986 | 247.111733 | 175.722571 | 137.501181 | 137.891105 | 187.709308 | 0.5095722 | 0.00098216 | 0.01022706 | Zranb3 | 1.42362799 | 1 | 127954184 | 128103047 - |
| ENSMUSG0C | 323.006692 | 386.137499 | 662.063701 | 459.979517 | 351.445141 | 268.506243 | 261.777646 | 241.340539 | 0.7052258 | 0.00098639 | 0.01026394 | Olfm1 | 1.63039983 | 2 | 28192992 | 28230736 + |

| | | | | | | | | | | | | |
|---|---|---|---|---|---|---|---|---|---|---|---|---|
| ENSMUSG0C | 987.964749 | 1023.5218 | 903.75457 | 993.074491 | 690.199208 | 865.066483 | 787.487485 | 806.454806 | 0.31195332 | 0.00098837 | 0.01027026 | Ccp110 |
| ENSMUSG0C | 179.027169 | 224.474599 | 181.503259 | 151.783986 | 148.387948 | 110.434019 | 95.8774093 | 123.153197 | 0.62105819 | 0.00099056 | 0.01028579 | Lurap1 |
| ENSMUSG0C | 1974.03503 | 1636.1933 | 1683.3722 | 1702.01676 | 1411.63798 | 1491.94194 | 1466.17027 | 1431.15933 | 0.27046514 | 0.0009929 | 0.01029827 | [illegible] |
| ENSMUSG0C | 306.903719 | 263.603199 | 245.452594 | 285.983241 | 182.556226 | 193.800877 | 237.000338 | 190.688821 | 0.45621731 | 0.00099526 | 0.01031302 | Ccdc62 |
| ENSMUSG0C | 32.2059458 | 26.7721999 | 23.5107848 | 56.4562385 | 4.88118252 | 9.74417816 | 19.3909367 | 12.9112223 | 1.57383307 | 0.0009994 | 0.01034145 | Ciart |
| ENSMUSG0C | 39.7838154 | 32.9503999 | 37.6172558 | 49.9776538 | 14.6435475 | 20.5710428 | 21.5454852 | 20.8565898 | 1.05181724 | 0.00100169 | 0.0103508 | [illegible] |
| ENSMUSG0C | 212.180349 | 179.1678 | 176.801102 | 160.113595 | 141.554293 | 108.268646 | 94.800135 | 138.050761 | 0.59054674 | 0.0010016 | 0.0103508 | Zfp982 |
| ENSMUSG0C | 3157.12992 | 4281.49259 | 2985.86968 | 2932.94787 | 2577.26437 | 2550.8093 | 2478.80808 | 2513.71566 | 0.40013317 | 0.0010061 | 0.01038916 | Ifit2 |
| ENSMUSG0C | 338.162431 | 316.117899 | 395.921617 | 397.970206 | 267.488802 | 273.919675 | 279.014034 | 284.04689 | 0.39194259 | 0.00101591 | 0.01047592 | Tm7sf2 |
| ENSMUSG0C | 620.438074 | 669.304998 | 711.906565 | 582.147115 | 596.480503 | 398.428618 | 434.141527 | 452.88595 | 0.45613473 | 0.00102464 | 0.01052943 | [illegible] |
| ENSMUSG0C | 265.225436 | 348.038599 | 295.295458 | 195.283055 | 172.793861 | 178.643266 | 175.595705 | 203.600043 | 0.59387869 | 0.00102637 | 0.01053992 | Naprt |
| ENSMUSG0C | 70.0952938 | 50.4552999 | 75.2345115 | 58.3072627 | 48.8118252 | 15.1576105 | 25.8545823 | 25.8224445 | 1.13020561 | 0.00103183 | 0.01058137 | Kcnn1 |
| ENSMUSG0C | 2376.60935 | 2188.11249 | 2220.35852 | 2131.45438 | 2095.97977 | 1628.36044 | 1432.77477 | 1704.28134 | 0.3776241 | 0.00103256 | 0.01058159 | Scrn1 |
| ENSMUSG0C | 2728.03306 | 2522.76499 | 2634.14833 | 2651.59219 | 2196.53213 | 2229.25142 | 2381.85339 | 2334.94489 | 0.20494356 | 0.00104806 | 0.01072569 | Hjurp |
| ENSMUSG0C | 127.87655 | 731.086998 | 100.626159 | 47.2011175 | 41.9781696 | 22.7364157 | 47.4000675 | 31.7814702 | 2.80616291 | 0.001052 | 0.01075123 | Bcl3 |
| ENSMUSG0C | 220.705452 | 367.602899 | 219.120515 | 289.685289 | 165.960206 | 195.96625 | 190.677544 | 161.886864 | 0.61850044 | 0.00105577 | 0.01078237 | Ccdc27 |
| ENSMUSG0C | 2388.92339 | 3317.69339 | 3994.01213 | 2820.03539 | 2158.45891 | 2217.34187 | 2238.57592 | 2455.11857 | 0.46516758 | 0.00106593 | 0.01085624 | [illegible] |
| ENSMUSG0C | 1192.56723 | 1397.3029 | 1346.69776 | 1178.17691 | 1186.12735 | 876.976034 | 940.46043 | 927.621661 | 0.37902807 | 0.00107584 | 0.01094975 | Gpr155 |
| ENSMUSG0C | 42.6255165 | 60.7522998 | 50.7832953 | 39.7970206 | 20.5009666 | 27.0671615 | 18.3136624 | 30.7882992 | 1.0012922 | 0.001077 | 0.01095409 | [illegible] |
| ENSMUSG0C | 1475.79011 | 1338.61 | 1207.51391 | 1336.43948 | 1028.95327 | 1125.99392 | 1130.0607 | 1123.27634 | 0.28178213 | 0.00108602 | 0.01103074 | [illegible] |
| ENSMUSG0C | 115.562511 | 137.9798 | 109.090042 | 198.985103 | 82.9801028 | 95.2764086 | 76.4864726 | 82.4331883 | 0.73692475 | 0.00111292 | 0.01127312 | Fbln5 |
| ENSMUSG0C | 277.539474 | 311.999099 | 266.142084 | 300.791435 | 246.987835 | 197.048936 | 219.763949 | 202.606872 | 0.41515561 | 0.00111845 | 0.01131374 | Thnsl1 |
| ENSMUSG0C | 1961.72099 | 1734.0148 | 1652.33796 | 2671.95345 | 1385.2796 | 1650.01417 | 1393.99289 | 1111.35829 | 0.53385024 | 0.00112913 | 0.01140626 | [illegible] |
| ENSMUSG0C | 847.774162 | 828.908498 | 899.052413 | 717.271883 | 689.222971 | 681.009784 | 647.441831 | 555.182557 | 0.35614651 | 0.0011364 | 0.01147189 | Bicd1 |
| ENSMUSG0C | 235.861191 | 232.712199 | 220.060946 | 200.836127 | 173.770098 | 162.402969 | 165.900236 | 164.866377 | 0.41427487 | 0.00115069 | 0.01159262 | Plppr2 |
| ENSMUSG0C | 521.925769 | 487.048099 | 513.475541 | 528.467413 | 436.377717 | 393.015186 | 389.973283 | 438.981557 | 0.3058918 | 0.00115432 | 0.01162131 | Bivm |
| ENSMUSG0C | 663.06359 | 931.878497 | 519.118129 | 753.366855 | 487.142015 | 495.870399 | 548.332599 | 443.947412 | 0.53746781 | 0.00117677 | 0.01181538 | Hes1 |
| ENSMUSG0C | 1051.42941 | 1073.9771 | 931.967511 | 1111.54004 | 893.2564 | 893.216331 | 775.637468 | 828.304567 | 0.2979658 | 0.001185 | 0.01188996 | [illegible] |
| ENSMUSG0C | 354.265404 | 340.830699 | 424.134559 | 422.959033 | 330.944175 | 230.612216 | 255.314 | 292.985428 | 0.47364305 | 0.00118664 | 0.01189711 | Triqk |
| ENSMUSG0C | 5467.43292 | 4456.54159 | 4630.68418 | 5248.57916 | 3550.57216 | 4230.05601 | 4286.47429 | 3857.47594 | 0.3146929 | 0.00118731 | 0.01189711 | [illegible] |
| ENSMUSG0C | 59.6757231 | 47.3661999 | 56.4258836 | 45.3500932 | 29.2870951 | 29.2325345 | 21.5454852 | 32.7746411 | 0.88581845 | 0.00119272 | 0.01194329 | [illegible] |
| ENSMUSG0C | 5167.15984 | 4345.33399 | 4611.87556 | 4966.29797 | 4084.57353 | 3691.96083 | 4017.15572 | 4174.29747 | 0.25768773 | 0.00119464 | 0.0119544 | Car13 |
| ENSMUSG0C | 48.3089187 | 60.7522998 | 70.5323545 | 32.3929237 | 35.1445141 | 11.9095511 | 14.0045654 | 24.8292736 | 1.29464094 | 0.001206 | 0.0120321 | Gng7 |
| ENSMUSG0C | 100.406772 | 87.5244998 | 75.2345115 | 80.5195533 | 49.7880617 | 67.1265606 | 24.777308 | 37.7404958 | 0.93788343 | 0.0012066 | 0.0120321 | [illegible] |
| ENSMUSG0C | 266.17267 | 122.5343 | 153.290317 | 346.141528 | 67.3603187 | 124.508943 | 110.959249 | 132.091735 | 1.0315369 | 0.00121306 | 0.0120819 | [illegible] |
| ENSMUSG0C | 44.5199839 | 43.2473999 | 72.4132173 | 72.1899443 | 32.2158046 | 31.3979074 | 23.7000338 | 31.7814702 | 0.96481545 | 0.00121819 | 0.01211858 | Sh3bgr |
| ENSMUSG0C | 9635.2612 | 10725.3552 | 9598.98324 | 9368.95906 | 8399.53887 | 8266.31114 | 8478.14844 | 8820.35115 | 0.21145456 | 0.00121994 | 0.0121261 | Lats2 |
| ENSMUSG0C | 843.037993 | 881.423198 | 1876.16063 | 888.491623 | 587.694375 | 648.529191 | 736.855595 | 706.144541 | 0.74480803 | 0.00122233 | 0.01213362 | Nmrk1 |
| ENSMUSG0C | 214.074816 | 583.839898 | 331.972282 | 176.772812 | 157.174077 | 62.7958148 | 74.3319241 | 137.05759 | 1.59756535 | 0.00124133 | 0.01228126 | Hrh1 |
| ENSMUSG0C | 84.3037993 | 128.7125 | 98.7452964 | 134.199256 | 80.0513933 | 66.0438742 | 67.8682785 | 57.6039147 | 0.71435969 | 0.00124797 | 0.01232236 | [illegible] |
| ENSMUSG0C | 1228.56211 | 1897.73709 | 1616.60157 | 1350.32216 | 1310.10939 | 882.389466 | 478.309772 | 907.758242 | 0.76737021 | 0.00126218 | 0.01242966 | Rcan2 |
| ENSMUSG0C | 825.987787 | 1020.4327 | 987.452964 | 821.854751 | 768.298128 | 593.312181 | 534.328034 | 762.755284 | 0.45841805 | 0.00126435 | 0.01244285 | [illegible] |
| ENSMUSG0C | 43.5727502 | 22.6533999 | 45.1407069 | 51.828678 | 17.5722571 | 18.4056698 | 12.9272911 | 22.8429317 | 1.18609467 | 0.0012946 | 0.01269856 | [illegible] |
| ENSMUSG0C | 37662.9592 | 36424.6077 | 36880.898 | 38902.9759 | 30355.0978 | 31558.145 | 32517.5236 | 35154.2787 | 0.2098308 | 0.00129947 | 0.01272953 | Nedd4 |
| ENSMUSG0C | 2468.49102 | 3616.30639 | 3457.0258 | 2582.17878 | 2454.25857 | 2280.13769 | 2009.1165 | 2219.73706 | 0.43556962 | 0.00130803 | 0.0128049 | [illegible] |
| ENSMUSG0C | 1454.00373 | 1472.471 | 1566.7587 | 1751.99442 | 1305.2282 | 1156.30914 | 1279.80182 | 1352.69882 | 0.29387175 | 0.00131554 | 0.01287001 | Dhrs7 |
| ENSMUSG0C | 1032.48473 | 1292.2735 | 1125.69638 | 903.299816 | 876.66038 | 778.451566 | 835.964827 | 861.079208 | 0.3764541 | 0.00131792 | 0.01287633 | [illegible] |
| ENSMUSG0C | 118.404213 | 180.1975 | 306.580634 | 165.666667 | 141.554293 | 92.0283492 | 80.7955696 | 44.6926924 | 1.101734 | 0.00132027 | 0.0128824 | Akap6 |
| ENSMUSG0C | 755.892493 | 761.977998 | 784.319783 | 822.780263 | 661.888349 | 539.177858 | 564.491713 | 699.192344 | 0.34151284 | 0.00132424 | 0.01291266 | Mras |
| ENSMUSG0C | 53.9923209 | 22.6533999 | 31.034236 | 32.3929237 | 14.6435475 | 8.66149169 | 17.2363882 | 15.8907351 | 1.3112646 | 0.00133638 | 0.01299697 | [illegible] |
| ENSMUSG0C | 771.048232 | 744.473098 | 806.890136 | 962.532591 | 639.43491 | 712.407692 | 622.664523 | 543.264506 | 0.38451882 | 0.00134832 | 0.01308741 | Tln2 |
| ENSMUSG0C | 1621.66409 | 1436.4315 | 1689.95521 | 1569.66853 | 1348.18261 | 1179.04556 | 1207.62445 | 1414.27542 | 0.2948362 | 0.00135099 | 0.01310481 | Wbp1 |
| ENSMUSG0C | 3610.85487 | 3360.94079 | 3999.65472 | 3777.94042 | 3302.60809 | 2951.40329 | 2921.5678 | 3163.24945 | 0.25740475 | 0.00135266 | 0.01311241 | Pigk |

[illegible]

| | | | | | | | | | | | | | | | | | | | | | | | | | | | | | | | | | | | | | |
|---|---|---|---|---|---|---|---|---|---|---|---|---|---|---|---|---|---|---|---|---|---|---|---|---|---|---|---|---|---|---|---|---|---|---|---|---|---|
| ENSMUSG0C | 5602.88734 | 5989.76488 | 4021.28464 | 5698.37804 | 3904.94601 | 4363.22644 | 3876.03279 | 4438.48095 | 0.3619632 | 0.00358469 | 0.0279561 | Plekhg2 | 1.28517356 | 7 | 28359604 | 28372599 | - | 6797 | protein_codi | pleckstrin ho | RhoGEF | 5915 | 5817 | 4276 | 6157 | 4000 | 4030 | 3598 | 4469 | 24.2852316 | 25.962119 | 17.4299112 | 24.6991278 | 16.9256515 | 18.9120285 | 16.8003296 | 19.2382126 |
| ENSMUSG0C | 1767.53808 | 1604.2726 | 1254.53548 | 2044.45624 | 1292.53713 | 882.389466 | 1023.41055 | 1463.93397 | 0.5165294 | 0.00364786 | 0.02835953 | Arsb | 1.43050982 | 13 | 93771630 | 93943016 | + | 6037 | protein_codi | arylsulfatase | Sulfatase | 1866 | 1558 | 1334 | 2209 | 1324 | 815 | 950 | 1474 | 8.62571758 | 7.82896982 | 6.12222663 | 9.97709883 | 6.30767752 | 4.30612636 | 4.99431975 | 7.14410687 |
| ENSMUSG0C | 41.6782828 | 203.880599 | 23.5107848 | 67.5623838 | 24.4059126 | 21.6537292 | 5.38637131 | 2.97951283 | 2.62772155 | 0.00365233 | 0.02836464 | Ankrd2 | 6.18049141 | 19 | 42036038 | 42045110 | + | 1100 | protein_codi | Ankyrin repe | - | 44 | 198 | 25 | 73 | 25 | 20 | 5 | 3 | 1.11625846 | 5.46048033 | 0.62968315 | 1.80950551 | 0.65365712 | 0.57994612 | 0.14426176 | 0.07979951 |
| ENSMUSG0C | 48.3089187 | 81.3462998 | 64.8897662 | 69.413408 | 49.7880617 | 25.9844751 | 26.9318565 | 41.7131796 | 0.86306746 | 0.00365636 | 0.02838113 | Bbof1 | 1.81890155 | 12 | 84409071 | 84442887 | + | 4257 | protein_codi | Basal body-o | - | 51 | 79 | 69 | 75 | 51 | 24 | 25 | 42 | 0.33432688 | 0.5629655 | 0.44907636 | 0.48038269 | 0.34456344 | 0.17982825 | 0.18638471 | 0.28868038 |
| ENSMUSG0C | 860.0882 | 955.561597 | 834.162646 | 833.886408 | 729.248668 | 701.580827 | 763.787452 | 746.864549 | 0.24345069 | 0.00366136 | 0.02840511 | Glb1l | 1.18382078 | 1 | 75198236 | 75210813 | - | 5905 | protein_codi | Beta-galacto | - | 908 | 928 | 887 | 901 | 747 | 648 | 709 | 752 | 4.29112057 | 4.76745295 | 4.16177375 | 4.16039555 | 3.63834077 | 3.50030139 | 3.8106604 | 3.72622928 |
| ENSMUSG0C | 669.694226 | 617.819998 | 819.115744 | 761.696464 | 614.052761 | 541.343231 | 565.568987 | 570.080121 | 0.32441666 | 0.00371698 | 0.0287499 | Siae | 1.25215804 | 9 | 37555698 | 37649655 | + | 10497 | protein_codi | Sialate O-ace | - | 707 | 600 | 871 | 823 | 629 | 500 | 525 | 574 | 1.87957209 | 1.73398124 | 2.29894037 | 2.13778671 | 1.72340807 | 1.5193406 | 1.58733291 | 1.59999391 |
| ENSMUSG0C | 5289.35298 | 5025.96569 | 6206.8472 | 5476.25514 | 4723.0322 | 4577.59836 | 4908.06154 | 4417.62436 | 0.24016879 | 0.00372383 | 0.02876908 | Erap1 | 1.18113084 | 13 | 74639568 | 74693201 | + | 7280 | protein_codi | Endoplasmic | - | 5584 | 4881 | 6600 | 5917 | 4838 | 4228 | 4556 | 4448 | 21.4051759 | 20.3392891 | 25.1181301 | 22.1615394 | 19.1133652 | 18.5248174 | 19.8621497 | 17.8774279 |
| ENSMUSG0C | 478.353019 | 372.751399 | 352.661773 | 512.733707 | 294.823424 | 232.777589 | 345.805038 | 343.637146 | 0.49613911 | 0.00375061 | 0.02890196 | Gm45274 | 1.41043396 | 8 | 15046212 | 15050770 | + | 647 | lincRNA | predicted ge | - | 505 | 362 | 375 | 554 | 302 | 215 | 321 | 346 | 21.7817048 | 16.9731571 | 16.0583802 | 23.3472223 | 13.4247231 | 10.5994789 | 15.7461602 | 15.6474457 |
| ENSMUSG0C | 9137.01627 | 10523.534 | 9711.835 | 9126.47489 | 8885.70465 | 7410.98883 | 8110.79792 | 8268.1481 | 0.23647721 | 0.00374842 | 0.02890196 | Agpat3 | 1.17811242 | 10 | 78269178 | 78352489 | - | 7790 | protein_codi | 1-acyl-sn-gly | Acyltransfera | 9646 | 10220 | 10327 | 9861 | 9102 | 6845 | 7529 | 8325 | 34.5552923 | 39.7989652 | 36.7291999 | 34.5154258 | 33.6048566 | 28.0276271 | 30.6742359 | 31.2693188 |
| ENSMUSG0C | 104.195707 | 185.345999 | 78.9962371 | 98.1042833 | 98.5998868 | 44.3901449 | 37.7045992 | 49.6585472 | 1.01384592 | 0.00378272 | 0.02913399 | Ifit1bl2 | 2.01928693 | 19 | 34617049 | 34640743 | - | 3367 | protein_codi | interferon in | - | 110 | 180 | 84 | 106 | 101 | 41 | 35 | 50 | 0.91170501 | 1.62176428 | 0.69121144 | 0.85840548 | 0.86274198 | 0.3884106 | 0.32991255 | 0.43450875 |
| ENSMUSG0C | 86.1982667 | 144.158 | 96.8644336 | 129.571695 | 68.3365552 | 77.9534252 | 78.6410211 | 70.515137 | 0.62838147 | 0.00378678 | 0.02915017 | Ccdc138 | 1.54582979 | 10 | 58497948 | 58576244 | + | 5102 | protein_codi | Coiled-coil d | - | 91 | 140 | 103 | 140 | 70 | 72 | 73 | 71 | 0.49774363 | 0.83242655 | 0.55933439 | 0.74819933 | 0.39460289 | 0.45013458 | 0.45410504 | 0.4071829 |
| ENSMUSG0C | 693.375069 | 710.492998 | 755.166409 | 704.314713 | 653.102221 | 511.02801 | 561.25989 | 613.779643 | 0.29070442 | 0.00380033 | 0.02922454 | Pus7 | 1.2232374 | 5 | 23740648 | 23783711 | - | 8289 | protein_codi | pseudouridyl | - | 732 | 690 | 803 | 761 | 669 | 472 | 521 | 618 | 2.46441426 | 2.52525531 | 2.68403487 | 2.50329627 | 2.3212753 | 1.81631093 | 1.99484657 | 2.18151383 |
| ENSMUSG0C | 298.378616 | 357.305899 | 323.5084 | 323.003725 | 239.177943 | 282.581167 | 261.777646 | 248.292736 | 0.33615965 | 0.00384046 | 0.02950232 | Fam161a | 1.26239173 | 11 | 23007531 | 23030788 | + | 4396 | protein_codi | - | - | 315 | 347 | 344 | 349 | 245 | 261 | 243 | 250 | 1.99966682 | 2.39458431 | 2.16808102 | 2.16469881 | 1.60291714 | 1.89379585 | 1.7543753 | 1.66400244 |
| ENSMUSG0C | 163.87143 | 119.4452 | 129.779532 | 138.826816 | 104.457306 | 81.2014846 | 101.263781 | 100.310265 | 0.51086416 | 0.00386158 | 0.02964929 | Kif9 | 1.42490344 | 9 | 110476958 | 110525179 | + | 4298 | protein_codi | Kinesin-like | Kinesin | 173 | 116 | 138 | 150 | 107 | 75 | 94 | 101 | 1.12327078 | 0.81874737 | 0.88958494 | 0.95160033 | 0.71601157 | 0.55660255 | 0.69412128 | 0.68758532 |
| ENSMUSG0C | 35.9948806 | 71.0492998 | 39.4981185 | 30.5418995 | 32.2158046 | 10.8268646 | 10.7727426 | 18.8702479 | 1.27387885 | 0.00386794 | 0.0296675 | Kcnh7 | 2.41810828 | 2 | 62693414 | 63184287 | - | 15226 | protein_codi | Potassium vo | Ion_trans | 38 | 69 | 42 | 33 | 33 | 10 | 10 | 19 | 0.06964702 | 0.13747433 | 0.07642549 | 0.05909597 | 0.06233483 | 0.02094906 | 0.02084434 | 0.03651232 |
| ENSMUSG0C | 12655.0422 | 15414.609 | 16899.5521 | 12292.6518 | 13088.4028 | 10669.8751 | 10044.5052 | 10850.3926 | 0.35876428 | 0.00386744 | 0.0296675 | Gns | 1.28232707 | 10 | 121365090 | 121397249 | + | 9442 | protein_codi | glucosamine | Sulfatase | 13360 | 14970 | 17970 | 13282 | 13407 | 9855 | 9324 | 10925 | 39.486372 | 48.0967959 | 52.7301285 | 38.3556383 | 40.8385474 | 33.2922363 | 31.3409519 | 33.8554885 |
| ENSMUSG0C | 51.1506198 | 71.0492998 | 123.196513 | 70.3389201 | 36.1207506 | 54.1343231 | 40.9364219 | 39.7268377 | 0.88793387 | 0.00387339 | 0.02969396 | Lrrc73 | 1.85052403 | 17 | 46254165 | 46257316 | + | 1384 | protein_codi | leucine rich r | - | 54 | 69 | 131 | 76 | 37 | 50 | 38 | 40 | 1.08883592 | 1.51241628 | 2.62246653 | 1.49729452 | 0.76889725 | 1.15234959 | 0.87140775 | 0.84565951 |
| ENSMUSG0C | 175.238235 | 161.6629 | 170.218082 | 161.964619 | 127.886982 | 123.426257 | 137.891105 | 114.214658 | 0.4115112 | 0.00388412 | 0.02974558 | Tacc3 | 1.33007832 | 5 | 33658128 | 33678995 | + | 3552 | protein_codi | transforming | - | 185 | 157 | 181 | 175 | 131 | 114 | 128 | 115 | 1.45346153 | 1.34086495 | 1.41182337 | 1.34336747 | 1.06072062 | 1.02372246 | 1.14369685 | 0.94731959 |
| ENSMUSG0C | 649.802318 | 1174.8877 | 817.234881 | 750.590319 | 640.411146 | 599.8083 | 537.559857 | 639.602087 | 0.48813477 | 0.00388752 | 0.02975633 | Mpi | 1.40263027 | 9 | 57544256 | 57552763 | - | 3142 | protein_codi | mannose pho | - | 686 | 1141 | 869 | 811 | 656 | 554 | 499 | 644 | 6.09288114 | 11.016352 | 7.66281506 | 7.03792133 | 6.00482467 | 5.62411148 | 5.04043802 | 5.99723852 |
| ENSMUSG0C | 321.112224 | 192.553899 | 326.329694 | 310.972068 | 163.031496 | 230.612216 | 218.686675 | 144.009787 | 0.60817098 | 0.00392025 | 0.02994524 | Pcdhb14 | 1.52432547 | 18 | 37447656 | 37456350 | + | 8695 | protein_codi | Protocadheri | Cadherin | 339 | 187 | 347 | 336 | 167 | 213 | 203 | 145 | 1.088015 | 0.65242465 | 1.1056932 | 1.05365741 | 0.55239477 | 0.78137651 | 0.74096956 | 0.48794408 |
| ENSMUSG0C | 46.4144513 | 57.6631998 | 35.736393 | 55.5307264 | 24.4059126 | 30.3152209 | 23.7000338 | 30.7882992 | 0.837631 | 0.00392711 | 0.02998222 | Tmem269 | 1.78711318 | 4 | 119205055 | 119218217 | - | 1504 | protein_codi | Transmembr | - | 49 | 56 | 38 | 60 | 25 | 28 | 22 | 31 | 0.90918658 | 1.12953198 | 0.7000201 | 1.08776016 | 0.47807369 | 0.59382781 | 0.46424663 | 0.60309467 |
| ENSMUSG0C | 43.5727502 | 42.2176999 | 33.8555302 | 14.8081937 | 7.80989203 | 7.57880523 | 24.777308 | 4.96585472 | 1.58119035 | 0.0039482 | 0.03011238 | Zfp459 | 2.99216627 | 13 | 67405709 | 67421418 | - | 3669 | protein_codi | zinc finger pr | KRAB | 46 | 41 | 36 | 16 | 8 | 7 | 23 | 5 | 0.34987659 | 0.33899593 | 0.27185013 | 0.11890552 | 0.06271118 | 0.06085562 | 0.19895462 | 0.03987438 |
| ENSMUSG0C | 177.132702 | 187.405399 | 183.384122 | 194.357542 | 141.554293 | 135.335808 | 136.813831 | 153.941496 | 0.38600842 | 0.00398162 | 0.03033614 | Lin9 | 1.30677288 | 1 | 180641150 | 180690694 | + | 7244 | protein_codi | lin-9 DREAM | DIRP | 187 | 182 | 195 | 210 | 145 | 125 | 127 | 155 | 0.72039043 | 0.76216901 | 0.74581466 | 0.79044305 | 0.5756947 | 0.55040441 | 0.55641547 | 0.62607288 |
| ENSMUSG0C | 28.417011 | 22.6533999 | 23.5107848 | 17.58473 | 11.714838 | 5.41343231 | 12.9272911 | 7.94536755 | 1.27574729 | 0.00398668 | 0.03035916 | 1810010D01I | 2.42124201 | 7 | 145205825 | 145208119 | - | 1278 | protein_codi | RIKEN cDNA | - | 30 | 22 | 25 | 19 | 12 | 5 | 12 | 8 | 0.65508125 | 0.52221599 | 0.5419808 | 0.40537082 | 0.27005552 | 0.12479279 | 0.29800552 | 0.18316006 |
| ENSMUSG0C | 321.112224 | 245.068599 | 265.201653 | 323.929237 | 227.463105 | 176.477893 | 132.504734 | 243.326881 | 0.56580471 | 0.00399282 | 0.03038271 | Ttll7 | 1.48021291 | 3 | 146852367 | 146984009 | + | 8266 | protein_codi | tubulin tyrosi | TTL | 339 | 238 | 282 | 350 | 233 | 163 | 123 | 245 | 1.14448227 | 0.87345372 | 0.94521032 | 1.15452244 | 0.81070564 | 0.62898826 | 0.47226268 | 0.86724603 |
| ENSMUSG0C | 411.099426 | 427.325499 | 363.006518 | 453.500932 | 322.158046 | 322.640566 | 330.723198 | 353.568856 | 0.31600084 | 0.00399386 | 0.03038271 | Arrb2 | 1.24487496 | 11 | 70432635 | 70440828 | + | 2845 | protein_codi | Beta-arrestir | - | 434 | 415 | 386 | 490 | 330 | 298 | 307 | 356 | 4.2570841 | 4.42511099 | 3.75906454 | 4.69616712 | 3.33606376 | 3.34106042 | 3.42475902 | 3.66133412 |
| ENSMUSG0C | 103.248473 | 113.267 | 91.2218452 | 83.2960896 | 69.3127917 | 72.5399929 | 65.71373 | 44.6926924 | 0.63271338 | 0.00399682 | 0.03038971 | Hoxaas2 | 1.55047835 | 6 | 52165674 | 52169564 | + | 3361 | processed_tr | - | - | 109 | 110 | 97 | 90 | 71 | 67 | 61 | 45 | 0.90502954 | 0.99284743 | 0.79960955 | 0.73013595 | 0.60756467 | 0.63585286 | 0.57601691 | 0.39175598 |
| ENSMUSG0C | 50.2033861 | 28.8315999 | 33.8555302 | 21.2867785 | 7.80989203 | 1.08268646 | 23.7000338 | 7.94536755 | 1.7303222 | 0.00403049 | 0.03056767 | 1700021N21 | 3.31801911 | 4 | 134448766 | 134450171 | - | 904 | antisense | RIKEN cDNA | - | 53 | 28 | 36 | 23 | 8 | 1 | 22 | 8 | 1.63610891 | 0.93961068 | 1.10333862 | 0.69372787 | 0.25452136 | 0.03528433 | 0.77237492 | 0.25893645 |
| ENSMUSG0C | 445.199839 | 725.938498 | 472.09656 | 397.044694 | 379.756 | 378.940262 | 321.02773 | 338.671292 | 0.52350608 | 0.00403998 | 0.03059279 | Acer2 | 1.43744433 | 4 | 86874396 | 86934822 | + | 7159 | protein_codi | alkaline cera | - | 470 | 705 | 502 | 429 | 389 | 350 | 298 | 341 | 1.83210461 | 2.98741183 | 1.94279109 | 1.63393458 | 1.56278744 | 1.55943048 | 1.32110646 | 1.39371397 |
| ENSMUSG0C | 263.330969 | 335.682199 | 329.150988 | 261.919926 | 218.676977 | 253.348632 | 225.150321 | 198.634189 | 0.41000447 | 0.00407004 | 0.03078911 | Irak2 | 1.32868993 | 6 | 113638467 | 113695026 | + | 4412 | protein_codi | Interleukin-1 | Pkinase | 278 | 326 | 350 | 283 | 224 | 234 | 209 | 200 | 1.75838538 | 2.24150875 | 2.19789676 | 1.74896318 | 1.46020956 | 1.69172859 | 1.50343514 | 1.32637439 |
| ENSMUSG0C | 1302.44634 | 1285.0656 | 1276.1654 | 1421.5866 | 1191.00853 | 1078.35572 | 1095.58792 | 1171.94171 | 0.22000049 | 0.00407429 | 0.03080561 | Gpx1 | 1.16473398 | 9 | 108338903 | 108340343 | + | 1441 | protein_codi | Mus muscul | GSHPx | 1375 | 1248 | 1357 | 1536 | 1220 | 996 | 1017 | 1180 | 26.6283029 | 26.2729565 | 26.0909934 | 29.064106 | 24.3499752 | 22.0468067 | 22.3991164 | 23.9601571 |
| ENSMUSG0C | 1055.21834 | 1502.3323 | 1310.96136 | 859.800747 | 956.711773 | 792.52649 | 921.069494 | 667.410874 | 0.50220126 | 0.00411309 | 0.03106741 | Tle3 | 1.41637301 | 9 | 61372366 | 61418497 | + | 11653 | protein_codi | transducin-lil | - | 1114 | 1459 | 1394 | 929 | 980 | 732 | 855 | 672 | 2.66779353 | 3.79818301 | 3.31436074 | 2.17374052 | 2.41875011 | 2.00365835 | 2.32863962 | 1.68734217 |
| ENSMUSG0C | 283.222876 | 267.721999 | 180.562828 | 371.130355 | 185.484936 | 202.462368 | 164.822962 | 183.736624 | 0.58244287 | 0.00414122 | 0.03124815 | Platr25 | 1.49738258 | 13 | 62671784 | 62760709 | - | 6180 | protein_codi | - | KRAB | 299 | 260 | 192 | 401 | 190 | 187 | 153 | 185 | 1.35016698 | 1.27627192 | 0.86077075 | 1.76923545 | 0.8842352 | 0.96516923 | 0.78573639 | 0.87590073 |
| ENSMUSG0C | 1717.3347 | 1397.3029 | 1776.4749 | 1713.12291 | 1455.56863 | 1256.99898 | 1417.69293 | 1280.19735 | 0.28786089 | 0.00416327 | 0.0313827 | Acad11 | 1.22082879 | 9 | 104063377 | 104127725 | + | 9798 | protein_codi | acyl-Coenzym | APH | 1813 | 1357 | 1889 | 1851 | 1491 | 1161 | 1316 | 1289 | 5.16374919 | 4.20146504 | 5.34157427 | 5.15108502 | 4.37666072 | 3.77959374 | 4.26277459 | 3.84934749 |
| ENSMUSG0C | 125.982082 | 139.0095 | 145.766866 | 136.975792 | 89.8137583 | 66.0438742 | 107.727426 | 110.241975 | 0.54973715 | 0.00417591 | 0.03144778 | Prr11 | 1.46381897 | 11 | 87089153 | 87108708 | - | 3824 | protein_codi | Proline-rich | - | 133 | 135 | 155 | 148 | 92 | 61 | 100 | 111 | 0.97059607 | 1.07096241 | 1.12302277 | 1.05529424 | 0.69194666 | 0.50881779 | 0.82995784 | 0.84933052 |
| ENSMUSG0C | 319.217757 | 245.068599 | 850.14998 | 552.530728 | 222.581923 | 239.273708 | 198.218464 | 225.449804 | 1.15158461 | 0.00419535 | 0.03156059 | Tnfrsf19 | 2.22157772 | 14 | 60963875 | 61046490 | - | 4738 | protein_codi | Tumor necro | - | 337 | 238 | 904 | 597 | 228 | 221 | 184 | 227 | 1.9849045 | 1.52384306 | 5.28625517 | 3.43565075 | 1.38402031 | 1.48781027 | 1.23252768 | 1.40185287 |
| ENSMUSG0C | 1657.65898 | 2794.60579 | 4602.47124 | 1903.7784 | 1335.49154 | 1319.7948 | 1019.10145 | 1565.23741 | 1.0644743 | 0.00423049 | 0.03179278 | Tmem176a | 2.09140765 | 6 | 48840919 | 48847071 | + | 3810 | protein_codi | Transmembr | - | 1750 | 2714 | 4894 | 2057 | 1368 | 1219 | 946 | 1576 | 12.8179285 | 21.6094249 | 35.5888322 | 14.7210589 | 10.3267531 | 10.2053773 | 7.88025143 | 12.1032742 |
| ENSMUSG0C | 236.808425 | 235.801299 | 229.46526 | 260.994414 | 158.150314 | 171.064461 | 175.595705 | 220.483949 | 0.40879683 | 0.004276 | 0.03208619 | Hspb2 | 1.32757819 | 9 | 50751078 | 50752354 | - | 834 | protein_codi | Heat shock p | - | 250 | 229 | 244 | 282 | 162 | 158 | 163 | 222 | 8.36524623 | 8.32966956 | 8.10584928 | 9.21961513 | 5.58665222 | 6.04284386 | 6.20290982 | 7.78858491 |
| ENSMUSG0C | 331.531795 | 583.839898 | 331.031851 | 338.737431 | 320.205573 | 126.674316 | 108.8047 | 128.119052 | 1.21189874 | 0.00428175 | 0.03211312 | Xkr4 | 2.31642302 | 1 | 3205901 | 3671498 | - | 6094 | protein_codi | X-linked Kx b | - | 350 | 567 | 352 | 366 | 328 | 117 | 101 | 129 | 1.6027669 | 2.8225325 | 1.60034996 | 1.63760204 | 1.54801109 | 0.61239798 | 0.52600859 | 0.61938245 |
| ENSMUSG0C | 484.983655 | 677.542598 | 637.612485 | 663.592181 | 479.332123 | 409.255483 | 367.350523 | 553.196215 | 0.44453125 | 0.00429212 | 0.03212611 | Ift81 | 1.36087188 | 5 | 122550204 | 122614518 | - | 5840 | protein_codi | Intraflagellar | - | 512 | 658 | 678 | 717 | 491 | 378 | 341 | 557 | 2.44659388 | 3.41799472 | 3.21655954 | 3.34761915 | 2.41808363 | 2.06456846 | 1.85317079 | 2.79070534 |
| ENSMUSG0C | 1664.28961 | 1706.2129 | 2031.33181 | 1671.47487 | 1609.81399 | 1331.70435 | 1422.00203 | 1461.94763 | 0.27976737 | 0.00428794 | 0.03212611 | Npdc1 | 1.21399912 | 2 | 25399351 | 25409494 | + | 3906 | protein_codi | Neural prolif | - | 1757 | 1657 | 2160 | 1806 | 1649 | 1230 | 1320 | 1472 | 12.5529065 | 12.8691129 | 15.3213227 | 12.6071013 | 12.1420241 | 10.0443817 | 10.7254521 | 11.0267418 |
| ENSMUSG0C | 1275.92379 | 1209.8975 | 1482.11988 | 1421.5866 | 1208.58079 | 1101.09213 | 1109.59249 | 1074.61096 | 0.2623802 | 0.00429865 | 0.03215881 | Xxylt1 | 1.19945597 | 16 | 30955627 | 31081432 | - | 3412 | protein_codi | xyloside xylos | Glyco_transf | 1347 | 1175 | 1576 | 1536 | 1238 | 1017 | 1030 | 1082 | 11.0169999 | 10.4468939 | 12.7974057 | 12.2747294 | 10.4355248 | 9.50741096 | 9.58080754 | 9.27875854 |
| ENSMUSG0C | 28.417011 | 60.7522998 | 35.736393 | 68.4878959 | 24.4059126 | 7.57880523 | 15.0818397 | 31.7814702 | 1.28705522 | 0.0043192 | 0.0322963 | Dpyd | 2.44029442 | 3 | 118562129 | 119432924 | + | 8884 | protein_codi | Dihydropyrim | DHO_dh | 30 | 59 | 38 | 74 | 25 | 7 | 14 | 32 | 0.09423614 | 0.20146602 | 0.11850858 | 0.22711871 | 0.08093458 | 0.02513274 | 0.05001421 | 0.10539331 |
| ENSMUSG0C | 2931.6883 | 3487.59389 | 3143.86215 | 3041.23278 | 2811.56113 | 2666.65676 | 2730.89025 | 2671.62984 | 0.21189094 | 0.00433571 | 0.03240353 | Arhgap21 | 1.15820525 | 2 | 20847919 | 20968881 | - | 9271 | protein_codi | Rho GTPase | RhoGAP | 3095 | 3387 | 3343 | 3286 | 2880 | 2463 | 2535 | 2690 | 9.31620085 | 11.0827352 | 9.99043835 | 9.66430689 | 8.93446557 | 8.47399429 | 8.67811291 | 8.48979755 |
| ENSMUSG0C | 3006.51976 | 1808.15319 | 2662.36128 | 3030.12664 | 1925.13838 | 1842.73236 | 2134.08031 | 1926.75163 | 0.42473836 | 0.00436493 | 0.0325891 | Cdh11 | 1.34232905 | 8 | 102632095 | 102785642 | - | 8904 | protein_codi | Cadherin-11 | Cadherin | 3174 | 1756 | 2831 | 3274 | 1972 | 1702 | 1981 | 1940 | 9.94778847 | 5.98270656 | 8.80905794 | 10.0258975 | 6.36977999 | 6.09711997 | 7.06111424 | 6.37511779 |
| ENSMUSG0C | 1824.37211 | 4177.49289 | 4359.83994 | 1553.93483 | 1613.71894 | 1423.7327 | 1500.64305 | 1106.39243 | 1.0779123 | 0.00440598 | 0.03281333 | Irs2 | 2.11097911 | 8 | 10984681 | 11008458 | - | 6323 | protein_codi | Insulin recep | - | 1926 | 4057 | 4636 | 1679 | 1653 | 1315 | 1393 | 1114 | 8.50037082 | 19.4643618 | 20.3139788 | 7.24031146 | 7.51886599 | 6.6336554 | 6.9920069 | 5.15505905 |
| ENSMUSG0C | 28.417011 | 56.6334998 | 61.1280406 | 44.4245811 | 30.2633316 | 20.5710428 | 25.8545823 | 21.8497607 | 0.94882419 | 0.00442422 | 0.03289972 | Trim46 | 1.93029881 | 3 | 89234177 | 89246309 | - | 3792 | protein_codi | tripartite mo | zf-B_box | 30 | 55 | 65 | 48 | 31 | 19 | 24 | 22 | 0.22077897 | 0.44000003 | 0.47491925 | 0.34514584 | 0.2351235 | 0.15982165 | 0.20087081 | 0.16975633 |
| ENSMUSG0C | 2082.96691 | 3028.34769 | 2162.99221 | 1725.15457 | 1902.68494 | 1660.84103 | 1386.45197 | 1579.1418 | 0.46261477 | 0.00442193 | 0.03289972 | Ceacam1 | 1.37803714 | 7 | 25461707 | 25477603 | - | 5806 | protein_codi | Carcinoembr | V-set | 2199 | 2941 | 2300 | 1864 | 1949 | 1534 | 1287 | 1590 | 10.5694641 | 15.3665486 | 10.9755313 | 8.75384013 | 9.65467103 | 8.42749812 | 7.03518349 | 8.012937 |
| ENSMUSG0C | 85.251033 | 113.267 | 68.6514918 | 56.4562385 | 74.1939742 | 19.4883563 | 20.468211 | 22.8429317 | 1.2340793 | 0.00446675 | 0.03316632 | Ifit1bl1 | 2.3523118 | 19 | 34592888 | 34601968 | - | 2275 | protein_codi | interferon in | - | 90 | 110 | 73 | 61 | 76 | 18 | 19 | 23 | 1.10399188 | 1.46679569 | 0.88902957 | 0.73110233 | 0.96080414 | 0.25237216 | 0.26506117 | 0.29581355 |
| ENSMUSG0C | 17.9974403 | 29.8612999 | 47.9620011 | 28.6908753 | 19.5247301 | 14.074924 | 9.69546835 | 6.9521966 | 1.30724162 | 0.00449472 | 0.03332411 | Gm29683 | 2.47467937 | 7 | 70178868 | 70337852 | - | 3344 | lincRNA | predicted ge | - | 19 | 29 | 51 | 31 | 20 | 13 | 9 | 7 | 0.15855944 | 0.26308136 | 0.42255054 | 0.25276979 | 0.17201503 | 0.12400164 | 0.08541815 | 0.06124962 |
| ENSMUSG0C | 27.4697773 | 20.5939999 | 28.2129418 | 31.4674116 | 8.78612853 | 12.9922375 | 14.0045654 | 14.8975641 | 1.09354571 | 0.00449332 | 0.03332411 | Tert | 2.13397859 | 13 | 73626911 | 73649843 | + | 5730 | protein_codi | telomerase r | RVT_1 | 29 | 20 | 30 | 34 | 9 | 12 | 13 | 15 | 0.14123689 | 0.10588482 | 0.1450579 | 0.16179088 | 0.04517421 | 0.06680008 | 0.072005 | 0.07659638 |
| ENSMUSG0C | 4524.93539 | 4047.75069 | 5269.2371 | 4560.92366 | 4282.74954 | 3280.53998 | 3604.55968 | 3701.54811 | 0.30752308 | 0.00451388 | 0.03343285 | Elovl5 | 1.23758111 | 9 | 77917364 | 77984519 | + | 3522 | protein_codi | Elongation of | ELO | 4777 | 3931 | 5603 | 4928 | 4387 | 3030 | 3346 | 3727 | 37.8504163 | 33.8588368 | 44.0763902 | 38.1514529 | 35.8245675 | 27.4412325 | 30.1516094 | 30.9629033 |
| ENSMUSG0C | 2454.28252 | 2036.74659 | 1026.95108 | 2086.10429 | 747.797161 | 757.880523 | 1457.55208 | 952.450935 | 0.95758248 | 0.00451844 | 0.03344997 | Nav2 | 1.94205288 | 7 | 48908716 | 49610090 | + | 32228 | protein_codi | neuron navig | CH | 2591 | 1978 | 1092 | 2254 | 766 | 700 | 1353 | 959 | 2.24356589 | 1.86187823 | 0.93878044 | 1.90699824 | 0.68359376 | 0.69281139 | 1.33241145 | 0.87067663 |
| ENSMUSG0C | 23704.5234 | 22833.5974 | 27203.8589 | 21816.1714 | 20467.7745 | 20247.3195 | 20926.0525 | 20063.0462 | 0.22597908 | 0.00452591 | 0.03348857 | Cpd | 1.1695707 | 11 | 76778424 | 76847018 | - | 8332 | protein_codi | Carboxypepti | - | 25025 | 22175 | 28927 | 23572 | 20966 | 18701 | 19425 | 20201 | 83.8165143 | 80.7370187 | 96.1897693 | 77.1395153 | 72.3717364 | 71.5922325 | 73.9921557 | 70.9406629 |
| ENSMUSG0C | 700.952938 | 743.443398 | 618.803857 | 619.1676 | 575.0033 | 555.418155 | 509.550726 | 562.134754 | 0.28357012 | 0.0045323 | 0.03350257 | Ehbp1 | 1.21720327 | 11 | 22005828 | 22342292 | - | 5825 | protein_codi | EH domain-b | CH | 740 | 722 | 658 | 669 | 589 | 513 | 473 | 566 | 3.54519853 | 3.76010186 | 3.12971443 | 3.13155413 | 2.90818505 | 2.80912957 | 2.57714661 | 2.84310001 |
| ENSMUSG0C | 765.36483 | 802.136298 | 704.383114 | 964.383615 | 645.292329 | 677.761725 | 652.828203 | 623.711352 | 0.3163949 | 0.0045518 | 0.03361327 | Abcd4 | 1.24521504 | 12 | 84601464 | 84617413 | - | 8006 | protein_codi | ATP-binding | - | 808 | 779 | 749 | 1042 | 661 | 626 | 606 | 628 | 2.81644027 | 2.95175436 | 2.59203571 | 3.5488028 | 2.37458952 | 2.49407256 | 2.40232053 | 2.29517441 |
| ENSMUSG0C | 487.825356 | 489.107499 | 387.457734 | 445.171323 | 321.18181 | 387.601753 | 361.964152 | 364.493736 | 0.33456885 | 0.00458458 | 0.03382177 | 2700097O09 | 1.2610005 | 12 | 55045661 | 55080110 | - | 5021 | protein_codi | RIKEN cDNA | - | 515 | 475 | 412 | 481 | 329 | 358 | 336 | 367 | 2.86234368 | 2.86986673 | 2.27343082 | 2.61206867 | 1.88455297 | 2.27427586 | 2.12384574 | 2.13868822 |
| ENSMUSG0C | 1804.4802 | 1766.9652 | 2401.86178 | 1985.22347 | 1499.49927 | 1641.35268 | 1614.83412 | 1671.5067 | 0.30855074 | 0.00460018 | 0.03388646 | Mllt3 | 1.23846298 | 4 | 87769925 | 88033364 | - | 12004 | protein_codi | myeloid/lym | - | 1905 | 1716 | 2554 | 2145 | 1536 | 1516 | 1499 | 1683 | 4.42867453 | 4.33660273 | 5.89480787 | 4.8722666 | 3.68017018 | 4.02831618 | 3.96323258 | 4.10232217 |
| ENSMUSG0C | 35.0476469 | 60.7522998 | 28.2129418 | 29.6163874 | 21.4772031 | 16.2402969 | 10.7727426 | 21.8497607 | 1.11789195 | 0.00462319 | 0.03400544 | Spry3 | 2.1702962 | GL456233.1 | 101352 | 106395 | - | 4349 | protein_codi | sprouty RTK | - | 37 | 59 | 30 | 32 | 22 | 15 | 10 | 22 | 0.23741988 | 0.41154843 | 0.1911202 | 0.20062743 | 0.14549094 | 0.11001507 | 0.07297675 | 0.14801472 |
| ENSMUSG0C | 2928.8466 | 3110.72369 | 3789.93852 | 3136.56053 | 2802.775 | 2231.4168 | 2223.49408 | 2930.84745 | 0.34759513 | 0.00463373 | 0.03406615 | Anxa6 | 1.27243779 | 11 | 54979108 | 55033445 | - | 2930 | protein_codi | Annexin A6 | - | 3092 | 3021 | 4030 | 3389 | 2871 | 2061 | 2064 | 2951 | 29.4494125 | 31.278178 | 38.1076506 | 31.5379662 | 28.1817686 | 22.4367892 | 22.3571266 | 29.469531 |
| ENSMUSG0C | 208.391414 | 220.355799 | 212.537495 | 217.495345 | 171.817625 | 149.410732 | 152.972945 | 186.716137 | 0.37572653 | 0.00464272 | 0.03411543 | Rpa3 | 1.2974928 | 6 | 8255936 | 8259173 | - | 1136 | protein_codi | Replication p | - | 220 | 214 | 226 | 235 | 176 | 138 | 142 | 188 | 5.40442035 | 5.71470457 | 5.51194476 | 5.64052158 | 4.45591614 | 3.87481606 | 3.96719846 | 4.84229399 |
| ENSMUSG0C | 2614.36501 | 2546.44809 | 2567.37771 | 2682.13409 | 2201.41331 | 2412.22544 | 2185.78948 | 2371.69221 | 0.18298175 | 0.00466097 | 0.03423257 | Srebf1 | 1.13522774 | 11 | 60199089 | 60222581 | - | 5634 | protein_codi | Mus muscul | HLH | 2760 | 2473 | 2730 | 2898 | 2255 | 2228 | 2029 | 2388 | 13.670897 | 13.3157495 | 13.4251935 | 14.0252714 | 11.5115122 | 12.6138796 | 11.4298128 | 12.4019254 |
| ENSMUSG0C | 754.945259 | 1652.6685 | 1697.47867 | 646.932963 | 610.147815 | 624.710088 | 554.796245 | 560.148412 | 1.01590754 | 0.00468773 | 0.03436131 | Csrnp1 | 2.02217456 | 9 | 119971163 | 119984658 | - | 3473 | protein_codi | Mus muscul | - | 797 | 1605 | 1805 | 699 | 625 | 577 | 515 | 564 | 6.40410301 | 14.0193731 | 14.399492 | 5.48784867 | 5.17580501 | 5.2993349 | 4.7062648 | 4.75166655 |
| ENSMUSG0C | 61.5701905 | 43.2473999 | 49.8428639 | 44.4245811 | 25.3821491 | 40.0593991 | 23.7000338 | 20.8565898 | 0.86074278 | 0.00469218 | 0.03437697 | Srl | 1.81597303 | 16 | 4480216 | 4541816 | - | 5325 | protein_codi | Sarcalumeni | - | 65 | 42 | 53 | 48 | 26 | 37 | 22 | 21 | 0.34064225 | 0.23926987 | 0.27575983 | 0.24578273 | 0.14042887 | 0.22163199 | 0.13112243 | 0.11539084 |
| ENSMUSG0C | 3313.42348 | 2672.07149 | 2773.33218 | 3422.54377 | 2720.77113 | 2115.56935 | 2346.30334 | 1325.88321 | 0.51775786 | 0.00473288 | 0.03460702 | Hist1h2bc | 1.43172842 | 13 | 23684199 | 23692908 | + | 1156 | protein_codi | Histone H2B | Histone | 3498 | 2595 | 2949 | 3698 | 2787 | 1954 | 2178 | 1335 | 84.4436004 | 68.0985508 | 70.6792101 | 87.224564 | 69.3396759 | 53.9159251 | 59.7962509 | 33.7905349 |
| ENSMUSG0C | 899.872015 | 840.235198 | 818.175313 | 852.396651 | 811.252534 | 672.348293 | 689.455527 | 612.786472 | 0.29164731 | 0.00473597 | 0.0346126 | Inpp1 | 1.22403712 | 1 | 52785427 | 52817688 | - | 6460 | protein_codi | Inositol polyp | - | 950 | 816 | 870 | 921 | 831 | 621 | 640 | 617 | 4.10389139 | 3.83191602 | 3.7313113 | 3.88737866 | 3.69973977 | 3.06626311 | 3.14428113 | 2.79462977 |
| ENSMUSG0C | 14788.2125 | 52590.8977 | 29309.4848 | 10508.2645 | 10022.0439 | 10031.0901 | 10112.3735 | 9223.57855 | 1.44438813 | 0.00477643 | 0.0348643 | Ctgf | 2.72147376 | 10 | 24595442 | 24598683 | + | 2861 | protein_codi | Connective ti | VWC | 15612 | 51074 | 31166 | 11354 | 10266 | 9265 | 9387 | 9287 | 152.280907 | 541.552239 | 301.81301 | 108.208348 | 103.201515 | 103.294668 | 104.13168 | 94.9793565 |
| ENSMUSG0C | 725.581014 | 781.542298 | 912.218452 | 850.545626 | 718.510066 | 556.500841 | 565.568987 | 718.062592 | 0.35315526 | 0.00482599 | 0.03518409 | Prcp | 1.27735122 | 7 | 92874470 | 92934583 | + | 5497 | protein_codi | Lysosomal Pr | Peptidase_S2 | 766 | 759 | 970 | 919 | 736 | 514 | 525 | 723 | 3.88873012 | 4.18865297 | 4.88900798 | 4.55847428 | 3.85083358 | 2.9825499 | 3.03115036 | 3.84843535 |
| ENSMUSG0C | 2907.06023 | 3311.51519 | 2456.4068 | 3153.21975 | 2495.2605 | 2408.97738 | 2054.36202 | 2543.51079 | 0.31567613 | 0.00484146 | 0.03527957 | Tacc2 | 1.2445948 | 7 | 130577438 | 130764785 | + | 15408 | protein_codi | Mus muscul | - | 3069 | 3216 | 2612 | 3407 | 2556 | 2225 | 1907 | 2561 | 5.55847158 | 6.33181346 | 4.6967955 | 6.02914319 | 4.77108608 | 4.60610763 | 3.92806203 | 4.86334349 |
| ENSMUSG0C | 552.237247 | 475.721399 | 448.585775 | 471.085662 | 358.278797 | 437.405331 | 354.423232 | 401.241061 | 0.32861339 | 0.00488764 | 0.03552914 | Zfp933 | 1.25580581 | 4 | 147822986 | 147848366 | - | 4270 | protein_codi | zinc finger pr | KRAB | 583 | 462 | 477 | 509 | 367 | 404 | 329 | 404 | 3.81017963 | 3.28225593 | 3.09503278 | 3.25027151 | 2.47195672 | 3.01789292 | 2.44535512 | 2.76837632 |
| ENSMUSG0C | 656.432954 | 663.126798 | 600.935661 | 745.037246 | 555.47857 | 555.418155 | 579.573553 | 477.715224 | 0.29851138 | 0.00489663 | 0.0355771 | Zfp932 | 1.22987474 | 5 | 109996521 | 110010411 | + | 3272 | protein_codi | zinc finger pr | KRAB | 693 | 644 | 639 | 805 | 569 | 513 | 538 | 481 | 5.91050665 | 5.97077786 | 5.41081035 | 6.70829758 | 5.00151579 | 5.00097181 | 5.21846644 | 4.30133647 |
| ENSMUSG0C | 506.77003 | 541.622198 | 694.9788 | 561.785849 | 346.563959 | 530.516366 | 442.759722 | 387.336668 | 0.43443093 | 0.00495128 | 0.03585157 | Wdr90 | 1.35137768 | 17 | 25844771 | 25861501 | - | 8466 | protein_codi | WD repeat d | - | 535 | 526 | 739 | 607 | 355 | 490 | 411 | 390 | 1.76351959 | 1.88480238 | 2.41847122 | 1.95497029 | 1.20601515 | 1.84615496 | 1.54076879 | 1.34790095 |
| ENSMUSG0C | 38.8365817 | 48.3958999 | 71.4727859 | 39.7970206 | 25.3821491 | 16.2402969 | 25.8545823 | 34.760983 | 0.95362775 | 0.00496058 | 0.03590147 | Xylb | 1.93673659 | 9 | 119357381 | 119393797 | + | 4106 | protein_codi | xylulokinase | - | 41 | 47 | 76 | 43 | 26 | 15 | 24 | 35 | 0.27865682 | 0.347246 | 0.51282524 | 0.28554808 | 0.18211976 | 0.11652595 | 0.18550952 | 0.24941395 |
| ENSMUSG0C | 1240.87615 | 1179.0065 | 1214.09693 | 1369.75792 | 1145.12542 | 972.252443 | 1025.5651 | 1077.59047 | 0.2453457 | 0.00499294 | 0.03611813 | Aga | 1.18537678 | 8 | 53511727 | 53523421 | + | 2321 | protein_codi | N(4)-(beta-N | Asparaginase | 1310 | 1145 | 1291 | 1480 | 1173 | 898 | 952 | 1085 | 15.7507387 | 14.9654124 | 15.4108236 | 17.3866659 | 14.5353517 | 12.3410335 | 13.0177439 | 13.6781144 |
| ENSMUSG0C | 1358.33313 | 1354.0555 | 1360.80423 | 1305.89758 | 1296.44208 | 1089.18258 | 1007.25143 | 1103.41292 | 0.25803875 | 0.00499638 | 0.03612542 | Socs5 | 1.19585192 | 17 | 87107679 | 87137583 | + | 4392 | protein_codi | Suppressor o | SOCS_box | 1434 | 1315 | 1447 | 1411 | 1328 | 1006 | 935 | 1111 | 9.1115359 | 9.08284207 | 9.12811175 | 8.75980456 | 8.69637815 | 7.30610628 | 6.75652197 | 7.40156168 |
| ENSMUSG0C | 28.417011 | 57.6631998 | 16.9277651 | 34.243948 | 23.4296761 | 5.41343231 | 10.7727426 | 9.93170943 | 1.45954201 | 0.0050008 | 0.03613982 | Sbk3 | 2.75021044 | 7 | 4965260 | 4971168 | - | 3197 | protein_codi | Uncharacteri | Pkinase | 30 | 56 | 18 | 37 | 24 | 5 | 10 | 10 | 0.26186858 | 0.5313782 | 0.15599282 | 0.315565 | 0.21590926 | 0.04988589 | 0.09927303 | 0.09152274 |
| ENSMUSG0C | 111.773577 | 84.4353998 | 117.553924 | 83.2960896 | 58.5741902 | 67.1265606 | 65.71373 | 73.4946498 | 0.58509284 | 0.00501137 | 0.03618109 | Tmem25 | 1.50013552 | 9 | 44793769 | 44799307 | - | 3212 | protein_codi | Mus muscul | - | 118 | 82 | 125 | 90 | 60 | 62 | 61 | 74 | 1.02520624 | 0.77445584 | 1.07822457 | 0.76400589 | 0.53725243 | 0.61569622 | 0.6027375 | 0.67410542 |
| ENSMUSG0C | 505.822796 | 719.760298 | 463.632677 | 458.128493 | 440.282663 | 333.46743 | 298.40497 | 433.022531 | 0.51103461 | 0.00509085 | 0.0366482 | Ankrd39 | 1.4250718 | 1 | 36537507 | 36547445 | - | 4654 | protein_codi | Ankyrin repe | - | 534 | 699 | 493 | 495 | 451 | 308 | 277 | 436 | 3.20198762 | 4.55626671 | 2.93491339 | 2.90007049 | 2.78710183 | 2.11093409 | 1.88897976 | 2.74114333 |
| ENSMUSG0C | 245.333528 | 220.355799 | 188.086279 | 223.048418 | 162.05526 | 192.71819 | 158.359316 | 137.05759 | 0.43269158 | 0.00509758 | 0.03667894 | Gm43268 | 1.34974941 | 5 | 53905164 | 53909218 | + | 4055 | TEC | predicted ge | - | 259 | 214 | 200 | 241 | 166 | 178 | 147 | 138 | 1.7824349 | 1.60096286 | 1.36651337 | 1.62052568 | 1.17738881 | 1.40016585 | 1.15053647 | 0.99577189 |
| ENSMUSG0C | 6097.34333 | 8199.50108 | 8445.07392 | 7096.82684 | 5497.18775 | 6557.8319 | 6303.1317 | 5376.03432 | 0.33026221 | 0.00511751 | 0.03676954 | Nbl1 | 1.25724186 | 4 | 139082284 | 139092993 | - | 2033 | protein_codi | Neuroblastor | - | 6437 | 7963 | 8980 | 7668 | 5631 | 6057 | 5851 | 5413 | 88.3590222 | 118.822225 | 122.380918 | 102.842934 | 79.6619295 | 95.0321449 | 91.3411833 | 77.906247 |
| ENSMUSG0C | 117.456979 | 153.4253 | 131.660395 | 69.413408 | 23.4296761 | 22.7364157 | 112.036523 | 35.754154 | 1.28494965 | 0.00516993 | 0.03709186 | F630042J09R | 2.43673549 | 13 | 67278577 | 67283361 | + | 4397 | lincRNA | - | - | 124 | 149 | 140 | 75 | 24 | 21 | 104 | 36 | 0.78699141 | 1.02798824 | 0.88215788 | 0.46508736 | 0.15698474 | 0.15233972 | 0.75067299 | 0.23956186 |
| ENSMUSG0C | 36.9421143 | 53.5443999 | 47.0215697 | 51.828678 | 33.1920411 | 24.9017886 | 20.468211 | 28.8019574 | 0.81301533 | 0.00517729 | 0.03712682 | Gm6180 | 1.7568796 | 8 | 42246911 | 42247408 | + | 498 | processed_p | predicted pse | Cofilin_ADF | 39 | 52 | 50 | 56 | 34 | 23 | 19 | 29 | 2.18544577 | 3.16761465 | 2.78173279 | 3.06611485 | 1.96359649 | 1.47315631 | 1.21087182 | 1.70388504 |
| ENSMUSG0C | 55.8867883 | 45.3067999 | 47.0215697 | 58.3072627 | 41.9781696 | 16.2402969 | 23.7000338 | 28.8019574 | 0.89335441 | 0.00518108 | 0.03713607 | Car7 | 1.85748996 | 8 | 104534689 | 104550343 | + | 3637 | protein_codi | Carbonic anh | - | 59 | 44 | 50 | 63 | 43 | 15 | 22 | 29 | 0.45270311 | 0.3670014 | 0.38089165 | 0.47230983 | 0.34003829 | 0.13155225 | 0.19197881 | 0.23330623 |
| ENSMUSG0C | 38.8365817 | 28.8315999 | 37.6172558 | 68.4878959 | 24.4059126 | 19.4883563 | 15.0818397 | 25.8224445 | 1.03384903 | 0.00519689 | 0.03721352 | Tmem35a | 2.04747953 | X | 134295225 | 134305969 | + | 1918 | protein_codi | Transmembr | - | 41 | 28 | 40 | 74 | 25 | 18 | 14 | 26 | 0.59654062 | 0.44286134 | 0.57781144 | 1.051993 | 0.37488156 | 0.29934654 | 0.23166122 | 0.39663988 |
| ENSMUSG0C | 580.654258 | 659.007998 | 659.242407 | 709.867786 | 495.928144 | 516.441442 | 484.773418 | 607.820617 | 0.30921513 | 0.00521222 | 0.03728744 | Btbd10 | 1.23903344 | 7 | 113315626 | 113369392 | - | 5801 | protein_codi | BTB/POZ dor | - | 613 | 640 | 701 | 767 | 508 | 477 | 450 | 612 | 2.94891585 | 3.34684385 | 3.34803432 | 3.60514082 | 2.51862506 | 2.62280408 | 2.46197457 | 3.08688316 |
| ENSMUSG0C | 36.9421143 | 10.297 | 11.2851767 | 18.5102421 | 3.90494601 | 6.49611877 | 9.69546835 | 1.98634189 | 1.81620674 | 0.00521847 | 0.03729627 | Gm38012 | 3.52154066 | 1 | 61677249 | 61680277 | + | 3029 | TEC | predicted ge | - | 39 | 10 | 12 | 20 | 4 | 6 | 9 | 2 | 0.35931066 | 0.10015187 | 0.10976319 | 0.18003646 | 0.03798074 | 0.0631833 | 0.09430118 | 0.01931979 |
| ENSMUSG0C | 211.233115 | 157.5441 | 226.643966 | 170.294228 | 139.60182 | 102.855214 | 151.895671 | 142.023445 | 0.51354388 | 0.00521691 | 0.03729627 | Rps6ka2 | 1.42755257 | 17 | 7170115 | 7303313 | + | 5404 | protein_codi | ribosomal pr | Pkinase | 223 | 153 | 241 | 184 | 143 | 95 | 141 | 143 | 1.15158048 | 0.85888384 | 1.23559588 | 0.92839377 | 0.76106784 | 0.56073621 | 0.82809028 | 0.77426982 |
| ENSMUSG0C | 81.4620982 | 54.5740998 | 59.2471778 | 55.5307264 | 43.9306426 | 43.3074585 | 32.3182278 | 30.7882992 | 0.73891205 | 0.00523238 | 0.03735982 | Gm15050 | 1.66891682 | 6 | 52172881 | 52174059 | + | 552 | antisense | predicted ge | - | 86 | 53 | 63 | 60 | 45 | 40 | 30 | 31 | 4.3477458 | 2.91269583 | 3.16210452 | 2.96375232 | 2.34463967 | 2.31137946 | 1.7248689 | 1.64321447 |
| ENSMUSG0C | 41.6782828 | 76.1977998 | 189.02671 | 31.4674116 | 28.3108586 | 36.8113397 | 42.0136962 | 22.8429317 | 1.38213112 | 0.00523754 | 0.03737867 | Gm8250 | 2.60653118 | 19 | 18581455 | 18582228 | + | 774 | processed_p | predicted ge | - | 44 | 74 | 201 | 34 | 29 | 34 | 39 | 23 | 1.58641383 | 2.90034126 | 7.19498421 | 1.19775417 | 1.07760528 | 1.40116181 | 1.59918078 | 0.86947783 |
| ENSMUSG0C | 268155.23 | 272593.51 | 319420.344 | 263706.165 | 252375.685 | 230606.803 | 249436.392 | 221875.382 | 0.23597609 | 0.00525843 | 0.03750979 | Itm2b | 1.17770327 | 14 | 73362226 | 73385289 | - | 2732 | protein_codi | Integral men | - | 283093 | 264731 | 339653 | 284930 | 258519 | 212995 | 231544 | 223401 | 2891.69981 | 2939.5608 | 3444.52633 | 2843.72252 | 2721.53827 | 2486.78964 | 2689.84188 | 2392.6328 |
| ENSMUSG0C | 1035.32643 | 1215.046 | 1023.18936 | 1118.01863 | 1006.49983 | 938.689162 | 791.796582 | 852.140669 | 0.29052832 | 0.0053115 | 0.03787018 | Adarb1 | 1.22308809 | 10 | 77290726 | 77418270 | - | 11272 | protein_codi | Double-stran | - | 1093 | 1180 | 1088 | 1208 | 1031 | 867 | 735 | 858 | 2.70597608 | 3.17569927 | 2.67425406 | 2.92210413 | 2.63063357 | 2.45340052 | 2.06947542 | 2.22719346 |
| ENSMUSG0C | 1484.31521 | 1542.4906 | 1913.77789 | 1621.49721 | 1444.83002 | 1048.04049 | 1249.63814 | 1413.28225 | 0.34764895 | 0.00531573 | 0.03788222 | Nagpa | 1.27248526 | 16 | 5195289 | 5204012 | - | 2370 | protein_codi | N-acetylgluc | - | 1567 | 1498 | 2035 | 1752 | 1480 | 968 | 1160 | 1423 | 18.4512342 | 19.1744011 | 23.7898014 | 20.156517 | 17.9604015 | 13.0279879 | 15.5340092 | 17.5682373 |
| ENSMUSG0C | 142.085055 | 149.3065 | 97.804865 | 147.156425 | 107.386015 | 89.8629763 | 77.5637468 | 89.3853849 | 0.5561536 | 0.0053244 | 0.03788956 | Pard6a | 1.47034388 | 8 | 105701148 | 105703496 | + | 1582 | protein_codi | Partitioning | PDZ | 150 | 145 | 104 | 159 | 110 | 83 | 72 | 90 | 2.64599824 | 2.78048057 | 1.82138438 | 2.74044051 | 1.99981066 | 1.67348548 | 1.44444141 | 1.66459147 |
| ENSMUSG0C | 1416.11438 | 1514.6887 | 1707.82341 | 1313.30168 | 1378.44594 | 1080.52109 | 1221.62901 | 1069.64511 | 0.32510655 | 0.00532069 | 0.03788956 | Ncald | 1.25275696 | 15 | 37366175 | 37792570 | - | 8581 | protein_codi | Neurocalcin- | efhand | 1495 | 1471 | 1816 | 1419 | 1412 | 998 | 1134 | 1077 | 4.86192283 | 5.20035645 | 5.86344277 | 4.50893763 | 4.73259638 | 3.70973575 | 4.19419933 | 3.67239541 |
| ENSMUSG0C | 1246.55955 | 1083.2444 | 1193.40744 | 991.223467 | 840.539629 | 679.927098 | 1097.74247 | 788.577729 | 0.40628304 | 0.00533057 | 0.03791536 | Mtrr | 1.325267 | 13 | 68560780 | 68582149 | - | 5198 | protein_codi | 5-methyltetr | NAD_binding | 1316 | 1052 | 1269 | 1071 | 861 | 628 | 1019 | 794 | 7.06519878 | 6.13956789 | 6.76394544 | 5.61801546 | 4.76397583 | 3.85366275 | 6.22173949 | 4.46946831 |
| ENSMUSG0C | 37672.4315 | 35187.938 | 40320.996 | 40709.5755 | 33914.4561 | 30075.9472 | 31160.158 | 35951.795 | 0.23120676 | 0.00537023 | 0.03817917 | Cldn5 | 1.17381639 | 16 | 18776847 | 18778260 | + | 1414 | protein_codi | Claudin-5 [S | - | 39771 | 34173 | 42875 | 43986 | 34740 | 27779 | 28925 | 36199 | 784.913633 | 733.148649 | 840.097074 | 848.193216 | 706.615366 | 626.639165 | 649.228942 | 749.063782 |
| ENSMUSG0C | 292.695213 | 250.217099 | 259.559065 | 281.355681 | 226.486869 | 213.289233 | 213.300304 | 211.545411 | 0.32633071 | 0.00538616 | 0.03823895 | Myo19 | 1.2538204 | 11 | 84880148 | 84911226 | + | 7113 | protein_codi | Unconventio | Myosin_head | 309 | 243 | 276 | 304 | 232 | 197 | 198 | 213 | 1.21230094 | 1.03636278 | 1.07505584 | 1.16533425 | 0.93807562 | 0.88341293 | 0.88345879 | 0.87619028 |
| ENSMUSG0C | 15828.2751 | 16607.0016 | 17654.7186 | 17034.0503 | 12944.896 | 14879.36 | 13961.4744 | 15808.3019 | 0.22092462 | 0.00538635 | 0.03823895 | Plxnb2 | 1.1654803 | 15 | 89155549 | 89180788 | - | 7720 | protein_codi | plexin B2 [S | Sema | 16710 | 16128 | 18773 | 18405 | 13260 | 13743 | 12960 | 15917 | 60.4037526 | 63.375523 | 67.3738132 | 65.0052235 | 49.4002215 | 56.7825095 | 53.2796809 | 60.3275309 |
| ENSMUSG0C | 1152.78341 | 1209.8975 | 1212.21607 | 1242.96276 | 1058.24037 | 1051.28855 | 1002.94234 | 1096.46072 | 0.19466878 | 0.00540804 | 0.03835635 | Vps18 | 1.14446138 | 2 | 119288740 | 119298453 | + | 4786 | protein_codi | VPS18 CORV | - | 1217 | 1175 | 1289 | 1343 | 1084 | 971 | 931 | 1104 | 7.09614784 | 7.44772297 | 7.46199531 | 7.65126163 | 6.51417259 | 6.47137955 | 6.17377646 | 6.74944425 |
| ENSMUSG0C | 1267.39869 | 1294.3329 | 1410.64709 | 1081.92365 | 1163.67391 | 813.097533 | 789.642034 | 1066.66559 | 0.39834075 | 0.0054755 | 0.03872404 | Adk | 1.31799121 | 14 | 21052574 | 21448569 | + | 8526 | protein_codi | Adenosine ki | PfkB | 1338 | 1257 | 1500 | 1169 | 1192 | 751 | 733 | 1074 | 4.3794095 | 4.4724788 | 4.87439455 | 3.73851319 | 4.02099562 | 2.80960292 | 2.72855404 | 3.68579001 |
| ENSMUSG0C | 700.952938 | 661.067398 | 783.379351 | 771.877097 | 583.789429 | 611.717851 | 644.210008 | 603.847933 | 0.25660061 | 0.00548946 | 0.0387859 | Tsga10 | 1.19466044 | 1 | 37754776 | 37866429 | - | 10662 | protein_codi | testis specifi | - | 740 | 642 | 833 | 834 | 598 | 565 | 598 | 608 | 1.93685814 | 1.82664727 | 2.16461703 | 2.13283425 | 1.61311444 | 1.69028566 | 1.78006729 | 1.66853967 |
| ENSMUSG0C | 106.090174 | 105.0294 | 119.434787 | 130.497207 | 107.386015 | 46.5555179 | 57.0955359 | 69.521966 | 0.71294983 | 0.00552665 | 0.03900351 | Cenpe | 1.63915222 | 3 | 135212537 | 135273611 | + | 8620 | protein_codi | centromere | Kinesin | 112 | 102 | 127 | 141 | 110 | 43 | 53 | 70 | 0.36258975 | 0.35896429 | 0.40819831 | 0.4460069 | 0.36701862 | 0.15911515 | 0.1951383 | 0.23760874 |
| ENSMUSG0C | 527.609171 | 642.532798 | 534.165032 | 487.74488 | 443.211372 | 426.578466 | 461.073384 | 440.967899 | 0.30608591 | 0.00552812 | 0.03900351 | Zmat1 | 1.23634888 | X | 134971372 | 135009209 | - | 5391 | protein_codi | zinc finger, m | - | 557 | 624 | 568 | 527 | 454 | 394 | 428 | 444 | 2.88330533 | 3.51134579 | 2.91913213 | 2.66545293 | 2.42208396 | 2.33118761 | 2.51969719 | 2.40982371 |
| ENSMUSG0C | 3477.29491 | 3205.45609 | 3836.96009 | 3727.03725 | 3126.88552 | 2702.38541 | 2839.69495 | 3273.49143 | 0.25445661 | 0.00555515 | 0.03917567 | Tmx4 | 1.19288636 | 2 | 134594185 | 134644145 | - | 10907 | protein_codi | thioredoxin-r | - | 3671 | 3113 | 4080 | 4027 | 3203 | 2496 | 2636 | 3296 | 9.39255706 | 8.65829042 | 10.3640524 | 10.0671387 | 8.44606265 | 7.29944103 | 7.67032925 | 8.84206137 |
| ENSMUSG0C | 30.3114784 | 23.6830999 | 19.7490593 | 16.6592179 | 16.5960206 | 5.41343231 | 6.46364557 | 4.96585472 | 1.42419346 | 0.00555999 | 0.03919122 | Ica1l | 2.6836443 | 1 | 59982490 | 60043184 | - | 5356 | protein_codi | Mus muscul | Arfaptin | 32 | 23 | 21 | 18 | 17 | 5 | 6 | 5 | 0.16673017 | 0.13027036 | 0.10863092 | 0.09163506 | 0.09128744 | 0.02977692 | 0.03555368 | 0.02731499 |
| ENSMUSG0C | 398.785388 | 579.721098 | 541.688483 | 540.49907 | 477.37965 | 316.144447 | 370.582346 | 344.630317 | 0.44877439 | 0.00557144 | 0.03921618 | Psmb10 | 1.36488026 | 8 | 105935735 | 105938444 | - | 1705 | protein_codi | Proteasome | Proteasome | 421 | 563 | 576 | 584 | 489 | 292 | 344 | 347 | 6.89068637 | 10.0171079 | 9.35993534 | 9.33938327 | 8.24873114 | 5.46271829 | 6.4033608 | 5.9549309 |
| ENSMUSG0C | 26.5225436 | 15.4455 | 17.8681965 | 13.8826816 | 3.90494601 | 10.8268646 | 7.54091983 | 0.99317094 | 1.68057002 | 0.0055796 | 0.03925505 | Hist1h2bf | 3.20554579 | 13 | 23573736 | 23574196 | - | 461 | protein_codi | Histone H2B | Histone | 28 | 15 | 19 | 15 | 4 | 10 | 7 | 1 | 1.69496946 | 0.98707164 | 1.14189829 | 0.88719701 | 0.24955239 | 0.69190969 | 0.48191565 | 0.06347032 |
| ENSMUSG0C | 60.6229568 | 60.7522998 | 62.068472 | 49.9776538 | 38.0732236 | 43.3074585 | 32.3182278 | 31.7814702 | 0.68244955 | 0.00559534 | 0.03934717 | Tnfrsf18 | 1.60486234 | 4 | 156026164 | 156028895 | + | 1524 | protein_codi | tumor necros | - | 64 | 59 | 66 | 54 | 39 | 40 | 30 | 32 | 1.17192489 | 1.17442527 | 1.19986868 | 0.96613658 | 0.73600762 | 0.83719256 | 0.62475566 | 0.6143794 |
| ENSMUSG0C | 25.5753099 | 22.6533999 | 27.2725104 | 38.8715085 | 12.6910745 | 11.9095511 | 15.0818397 | 15.8907351 | 1.04326768 | 0.00562668 | 0.03951154 | Nim1k | 2.06089025 | 13 | 119710094 | 119755882 | - | 8466 | protein_codi | Serine/threo | Pkinase | 27 | 22 | 29 | 42 | 13 | 11 | 14 | 16 | 0.08900005 | 0.07883204 | 0.09490618 | 0.13526977 | 0.04416393 | 0.0414443 | 0.05248361 | 0.0552985 |
| ENSMUSG0C | 62.5174242 | 54.5740998 | 67.7110604 | 43.499069 | 29.2870951 | 38.9767126 | 10.7727426 | 35.754154 | 0.98994119 | 0.00562347 | 0.03951154 | Gamt | 1.98610403 | 10 | 80258151 | 80261012 | - | 1444 | protein_codi | Mus muscul | - | 66 | 53 | 72 | 47 | 30 | 36 | 10 | 36 | 1.27550308 | 1.11344051 | 1.38146553 | 0.88748373 | 0.5975259 | 0.79521698 | 0.21978939 | 0.72946917 |
| ENSMUSG0C | 466.038981 | 231.682499 | 299.057183 | 317.450653 | 244.059126 | 153.741478 | 262.85492 | 183.736624 | 0.63852353 | 0.00568472 | 0.03982515 | Ccrl2 | 1.55673517 | 9 | 111054486 | 111057519 | - | 2598 | protein_codi | C-C chemokir | 7tm_1 | 492 | 225 | 318 | 343 | 250 | 142 | 244 | 185 | 5.28482641 | 2.62725189 | 3.39127276 | 3.59985251 | 2.76760136 | 1.74341001 | 2.98074342 | 2.08355139 |
| ENSMUSG0C | 122.193147 | 63.8413998 | 62.068472 | 63.8603354 | 52.7167712 | 28.149848 | 51.7091646 | 36.7473249 | 0.8807992 | 0.00570529 | 0.03995048 | Gm17235 | 1.84139509 | 11 | 78576237 | 78577355 | + | 964 | processed_p | predicted ge | - | 129 | 62 | 66 | 69 | 54 | 26 | 48 | 37 | 3.73437088 | 1.95107066 | 1.89688783 | 1.95164935 | 1.6110885 | 0.86029352 | 1.58029482 | 1.12304285 |
| ENSMUSG0C | 597.704465 | 583.839898 | 607.51868 | 644.156426 | 508.619218 | 515.358756 | 472.923401 | 550.216703 | 0.24913307 | 0.00572345 | 0.04003991 | Akr1b10 | 1.18849272 | 6 | 34384218 | 34396950 | + | 3535 | protein_codi | aldo-keto rec | Aldo_ket_red | 631 | 567 | 646 | 696 | 521 | 476 | 439 | 554 | 4.9813231 | 4.86577455 | 5.06311566 | 5.36845795 | 4.23887859 | 4.29504651 | 3.94138642 | 4.58555579 |
| ENSMUSG0C | 173.343767 | 151.3659 | 154.230749 | 158.26257 | 111.290961 | 137.501181 | 104.495603 | 121.166855 | 0.42653221 | 0.00574492 | 0.04015238 | Stard5 | 1.34399913 | 7 | 83631959 | 83653127 | + | 5460 | protein_codi | StAR-related | START | 183 | 147 | 164 | 171 | 114 | 127 | 97 | 122 | 0.93532645 | 0.81673851 | 0.83219663 | 0.85395149 | 0.60050259 | 0.74192741 | 0.56383626 | 0.65379083 |
| ENSMUSG0C | 384.576882 | 349.068299 | 442.943187 | 392.417133 | 303.609553 | 336.71549 | 312.409536 | 313.842018 | 0.31016529 | 0.00578652 | 0.04036729 | Tmem231 | 1.23984974 | 8 | 111912011 | 111933881 | - | 5372 | protein_codi | Mus muscul | - | 406 | 339 | 471 | 424 | 311 | 311 | 290 | 316 | 2.10908848 | 1.91435306 | 2.42917975 | 2.15208583 | 1.66504915 | 1.84660804 | 1.71330983 | 1.72116582 |
| ENSMUSG0C | 19232.6331 | 18953.6878 | 20070.6868 | 13287.5773 | 16553.0661 | 9489.74684 | 14230.793 | 8206.5715 | 0.56143417 | 0.00579018 | 0.0403739 | Fgfbp1 | 1.4757355 | 5 | 43978858 | 43981779 | - | 2263 | protein_codi | Fibroblast gr | - | 20304 | 18407 | 21342 | 14357 | 16956 | 8765 | 13210 | 8263 | 250.381261 | 246.749795 | 261.291517 | 172.985173 | 215.497148 | 123.542875 | 185.264487 | 106.837775 |
| ENSMUSG0C | 1792.16616 | 1691.7971 | 1956.0973 | 2460.01118 | 1395.04196 | 1308.96793 | 1125.7516 | 1909.86772 | 0.46078693 | 0.00580248 | 0.0404407 | Bmp1 | 1.37629233 | 14 | 70474558 | 70520234 | - | 8870 | protein_codi | Bone morph | Astacin | 1892 | 1643 | 2080 | 2658 | 1429 | 1209 | 1045 | 1923 | 5.95253946 | 5.61917147 | 6.49702389 | 8.17073436 | 4.63352256 | 4.34763441 | 3.73909573 | 6.34347598 |
| ENSMUSG0C | 102.30124 | 146.2174 | 180.562828 | 136.975792 | 107.386015 | 97.4417816 | 76.4864726 | 96.3375815 | 0.5824837 | 0.00583594 | 0.04063583 | Brsk1 | 1.49742495 | 7 | 4690604 | 4715997 | + | 4100 | protein_codi | Serine/threo | Pkinase | 108 | 142 | 192 | 148 | 110 | 90 | 71 | 97 | 0.73509703 | 1.05066153 | 1.29745445 | 0.98425492 | 0.77163426 | 0.70017885 | 0.54960213 | 0.6922445 |
| ENSMUSG0C | 6256.47859 | 6501.52578 | 7238.50044 | 5921.42646 | 5710.00731 | 5335.47888 | 5427.30773 | 5804.09099 | 0.21832279 | 0.00587323 | 0.04085726 | Ano6 | 1.16338031 | 15 | 95790843 | 95974751 | + | 12269 | protein_codi | Anoctamin-6 | - | 6605 | 6314 | 7697 | 6398 | 5849 | 4928 | 5038 | 5844 | 15.0234068 | 15.6118278 | 17.3814926 | 14.2188608 | 13.7111893 | 12.8118507 | 13.0323553 | 13.9371083 |
| ENSMUSG0C | 1084.58259 | 998.808997 | 1323.18697 | 1094.88082 | 896.18511 | 676.679039 | 673.296414 | 1070.63828 | 0.44013977 | 0.00588179 | 0.04087858 | Ust | 1.35673576 | 10 | 8204756 | 8518825 | - | 4626 | protein_codi | uronyl-2-sulf | - | 1145 | 970 | 1407 | 1183 | 918 | 625 | 625 | 1078 | 6.90724132 | 6.36098611 | 8.4268103 | 6.97282636 | 5.70741859 | 4.30947857 | 4.28793608 | 6.81843599 |
| ENSMUSG0C | 37.889348 | 29.8612999 | 32.9150988 | 49.9776538 | 15.6197841 | 22.7364157 | 23.7000338 | 17.877077 | 0.92137618 | 0.00588769 | 0.04090051 | 2610028D06I | 1.89392103 | 9 | 45797348 | 45799060 | - | 1713 | TEC | RIKEN cDNA | - | 40 | 29 | 35 | 54 | 16 | 21 | 22 | 18 | 0.6516395 | 0.51356921 | 0.56608993 | 0.85954008 | 0.26863667 | 0.39103197 | 0.40760475 | 0.30745869 |
| ENSMUSG0C | 1469.15947 | 1665.0249 | 1821.61561 | 1607.61453 | 1428.234 | 1161.72257 | 1039.56966 | 1501.67447 | 0.35467722 | 0.00591948 | 0.0411021 | Rab3d | 1.27869946 | 9 | 21907491 | 21918192 | - | 4196 | protein_codi | RAB3D, mem | Ras | 1551 | 1617 | 1937 | 1737 | 1463 | 1073 | 965 | 1512 | 10.3152816 | 11.6904945 | 12.7899513 | 11.2874041 | 10.0279352 | 8.15670168 | 7.29904007 | 10.5435763 |
| ENSMUSG0C | 1206.77573 | 1082.2147 | 1227.26297 | 1130.05028 | 1069.95521 | 965.756324 | 980.319578 | 971.321182 | 0.22061426 | 0.00592309 | 0.04110804 | Ctbs | 1.1652296 | 3 | 146449795 | 146465849 | + | 6052 | protein_codi | Di-N-acetylch | - | 1274 | 1051 | 1305 | 1221 | 1096 | 892 | 910 | 978 | 5.87455913 | 5.2681986 | 5.97429056 | 5.50106123 | 5.20851966 | 4.70128166 | 4.77217528 | 4.72837128 |
| ENSMUSG0C | 11856.5242 | 11852.8767 | 14058.5089 | 11642.0168 | 11220.8624 | 9858.94292 | 10168.3918 | 10803.7135 | 0.23259517 | 0.00593429 | 0.04116655 | Tmem123 | 1.17494658 | 9 | 7764041 | 7794333 | + | 3109 | protein_codi | transmembr | - | 12517 | 11511 | 14949 | 12579 | 11494 | 9106 | 9439 | 10878 | 112.352904 | 112.318339 | 133.219 | 110.320223 | 106.329346 | 93.4237424 | 96.3560921 | 102.376427 |
| ENSMUSG0C | 25.5753099 | 100.9106 | 121.31565 | 72.1899443 | 64.4316092 | 8.66149169 | 19.3909367 | 12.9112223 | 1.59761626 | 0.00595923 | 0.04128181 | Gdnf | 3.02642849 | 15 | 7810846 | 7837575 | + | 3674 | protein_codi | Mus muscul | - | 27 | 98 | 129 | 78 | 66 | 8 | 18 | 13 | 0.20508287 | 0.80918023 | 0.97280391 | 0.57887552 | 0.51666311 | 0.06945463 | 0.15549172 | 0.10353229 |
| ENSMUSG0C | 722.739313 | 814.492698 | 651.718956 | 804.270021 | 634.553727 | 597.642927 | 542.946228 | 656.485993 | 0.29891935 | 0.00599177 | 0.04146864 | Slx1b | 1.23022257 | 7 | 126689468 | 126695784 | - | 3521 | protein_codi | SLX1 structur | GIY-YIG | 763 | 791 | 693 | 869 | 650 | 552 | 504 | 661 | 6.04732464 | 6.81504613 | 5.45308112 | 6.72951067 | 5.30945573 | 5.00061465 | 4.5429549 | 5.49296801 |
| ENSMUSG0C | 15.1557392 | 21.6236999 | 21.6299221 | 12.0316574 | 8.78612853 | 7.57880523 | 5.38637131 | 4.96585472 | 1.39438227 | 0.00601055 | 0.04154051 | Celf4 | 2.6287597 | 18 | 25477632 | 25754157 | - | 8583 | protein_codi | CUGBP Elav- | RRM_1 | 16 | 21 | 23 | 13 | 9 | 7 | 5 | 5 | 0.05202183 | 0.074223 | 0.07424436 | 0.04129847 | 0.03015825 | 0.02601413 | 0.01848863 | 0.01704522 |
| ENSMUSG0C | 125.982082 | 85.4650998 | 124.136944 | 98.1042833 | 84.9325758 | 47.6382043 | 65.71373 | 78.4605045 | 0.64584691 | 0.00601398 | 0.04154051 | Tstd3 | 1.56465752 | 4 | 21757382 | 21767212 | - | 1588 | protein_codi | thiosulfate s | Rhodanese | 133 | 83 | 132 | 106 | 87 | 44 | 61 | 79 | 2.33725401 | 1.58557188 | 2.3030225 | 1.82005747 | 1.57569235 | 0.88379698 | 1.21913907 | 1.45562072 |
| ENSMUSG0C | 7063.5217 | [illegible] | 7323.43113 | 6293.48233 | 6008.73568 | 5563.00841 | 6037.51317 | 5690.8693 | 0.22143074 | 0.00601812 | 0.04154051 | Nlk | 1.16938924 | 11 | 78367168 | 78697373 | - | 5447 | protein_codi | Serine/threo | Pkinase | 7457 | 6128 | 8000 | 6800 | 6155 | 5140 | 5623 | 5730 | 38.2042377 | 34.1286982 | 40.691843 | 34.0393511 | 32.4992512 | 30.0992781 | 32.7630724 | 30.7800189 |
| ENSMUSG0C | 441.410904 | 419.087899 | 421.313264 | 468.309126 | 350.468905 | 306.400269 | 414.750591 | 312.848847 | 0.33882597 | 0.00603212 | 0.04163169 | Ap1s3 | 1.26472697 | 1 | 79606876 | 79671972 | - | 2878 | protein_codi | AP-1 comple | - | 466 | 407 | 448 | 506 | 359 | 283 | 385 | 315 | 4.51855838 | 4.29004612 | 4.31282636 | 4.79390542 | 3.58761914 | 3.13650499 | 4.24564672 | 3.20251667 |
| ENSMUSG0C | 30.3114784 | 24.7127999 | 36.6768244 | 23.1378027 | 9.76236503 | 14.074924 | 14.0045654 | 17.877077 | 1.04539385 | 0.00605695 | 0.04178368 | Gm43185 | 2.06392972 | 3 | 152250029 | 152252803 | - | 2775 | TEC | predicted ge | - | 32 | 24 | 39 | 25 | 10 | 13 | 13 | 18 | 0.32180424 | 0.26236542 | 0.38938245 | 0.24564434 | 0.10364293 | 0.14942756 | 0.14868059 | 0.18979342 |
| ENSMUSG0C | 691.480601 | 682.691098 | 710.025702 | 782.983242 | 545.716205 | 631.206207 | 648.519106 | 502.544497 | 0.30176413 | 0.00606058 | 0.04178937 | Tpk1 | 1.23265078 | 6 | 43345001 | 43666278 | - | 4603 | protein_codi | Thiamin pyro | - | 730 | 663 | 755 | 846 | 559 | 583 | 602 | 506 | 4.42574774 | 4.36949146 | 4.54444368 | 5.01140062 | 3.49279829 | 4.03996792 | 4.15077727 | 3.21648238 |
| ENSMUSG0C | 814.620982 | 813.462998 | 903.75457 | 834.81192 | 625.767599 | 714.573065 | 739.010143 | 749.844062 | 0.25150182 | 0.00606554 | 0.04180421 | Taf1a | 1.19044571 | 1 | 183388981 | 183410198 | + | 2409 | protein_codi | TATA box-bir | - | 860 | 790 | 961 | 902 | 641 | 660 | 686 | 755 | 9.96245613 | 9.94829449 | 11.0525207 | 10.2093824 | 7.65286236 | 8.73891413 | 9.03776885 | 9.17026291 |
| ENSMUSG0C | 264.278202 | 252.276499 | 206.894907 | 287.834265 | 231.368051 | 174.31252 | 143.277477 | 174.798086 | 0.48104346 | 0.00609912 | 0.04201616 | Clybl | 1.39575281 | 14 | 122181704 | 122402234 | + | 1444 | protein_codi | Citramalyl-C | - | 279 | 245 | 220 | 311 | 237 | 161 | 133 | 176 | 5.3918994 | 5.14703632 | 4.22114466 | 5.87249871 | 4.72045461 | 3.55638704 | 2.92319887 | 3.56629373 |
| ENSMUSG0C | 92.8289026 | 268.751699 | 179.622396 | 119.391062 | 82.0038663 | 94.1937222 | 120.654717 | 59.5902566 | 0.8902364 | 0.00612823 | 0.04217767 | Rasgef1b | 1.85347982 | 5 | 99217426 | 99729065 | - | 13805 | protein_codi | Ras-GEF don | - | 98 | 261 | 191 | 129 | 84 | 87 | 112 | 60 | 0.19810454 | 0.57353832 | 0.38332903 | 0.25479038 | 0.17500302 | 0.20101718 | 0.25748713 | 0.12717053 |
| ENSMUSG0C | 611.91297 | 663.126798 | 454.228363 | 565.487897 | 519.35782 | 359.451905 | 326.414101 | 454.872292 | 0.46568662 | 0.00617696 | 0.04247373 | Nr3c2 | 1.38097444 | 8 | 76899442 | 77245012 | + | 6474 | protein_codi | nuclear rece | Hormone_rec | 646 | 644 | 483 | 611 | 532 | 332 | 303 | 458 | 2.78461138 | 3.01766839 | 2.06704144 | 2.57334639 | 2.36342383 | 1.63574547 | 1.48540146 | 2.06997175 |
| ENSMUSG0C | 149.662925 | 128.7125 | 107.209179 | 161.039107 | 88.8375218 | 94.1937222 | 75.4091983 | 114.214658 | 0.55188117 | 0.00620201 | 0.04258691 | Bbc3 | 1.465996 | 7 | 16308393 | 16318205 | + | 2124 | protein_codi | Bcl-2-binding | - | 158 | 125 | 114 | 174 | 91 | 87 | 70 | 115 | 2.07590438 | 1.78531084 | 1.48704834 | 2.23369808 | 1.23222369 | 1.30651703 | 1.0459657 | 1.58421807 |
| ENSMUSG0C | 224.494387 | 364.513799 | 187.145847 | 246.18622 | 183.532463 | 182.974012 | 158.359316 | 171.818573 | 0.55169422 | 0.00625009 | 0.04285775 | Nfkbiz | 1.46580605 | 16 | 55811375 | 55838899 | - | 5018 | protein_codi | NF-kappa-B i | - | 237 | 354 | 199 | 266 | 188 | 169 | 147 | 173 | 1.31802139 | 2.14008462 | 1.09874565 | 1.44537558 | 1.07753123 | 1.07425253 | 0.92973802 | 1.00875821 |
| ENSMUSG0C | 11128.1015 | 13693.9803 | 14426.2176 | 10269.4823 | 9999.5905 | 10019.1805 | 9624.36825 | 10418.3632 | 0.30569291 | 0.00626196 | 0.04291939 | Tpm4 | 1.23601214 | 8 | 72134975 | 72153142 | + | 2203 | protein_codi | Tropomyosin | - | 11748 | 13299 | 15340 | 11096 | 10243 | 9254 | 8934 | 10490 | 148.817571 | 183.131406 | 192.923713 | 137.335143 | 133.725844 | 133.987824 | 128.707948 | 139.326147 |
| ENSMUSG0C | 343.845833 | 239.920099 | 374.291695 | 464.607078 | 254.797727 | 229.52953 | 232.691241 | 280.074206 | 0.51328007 | 0.00633963 | 0.04337179 | Arxes2 | 1.42729155 | X | 135993824 | 135995355 | + | 1532 | protein_codi | Adipocyte-re | - | 363 | 233 | 398 | 502 | 261 | 212 | 216 | 282 | 6.61230124 | 4.61376529 | 7.19778807 | 8.9345912 | 4.89986839 | 4.41395023 | 4.47475127 | 5.38594579 |
| ENSMUSG0C | 28.417011 | 36.0394999 | 15.9873337 | 52.7541901 | 6.83365552 | 5.41343231 | 3.23182278 | 25.8224445 | 1.68122601 | 0.00635653 | 0.04337808 | Fmo2 | 3.20700368 | 1 | 162874317 | 162898726 | - | 5609 | protein_codi | Dimethylanil | FMO-like | 30 | 35 | 17 | 57 | 7 | 5 | 3 | 26 | 0.14925902 | 0.18929579 | 0.08397272 | 0.27708891 | 0.03589346 | 0.0284338 | 0.016975 | 0.13563118 |
| ENSMUSG0C | 189.44674 | 192.553899 | 157.052043 | 164.741155 | 163.031496 | 109.351333 | 120.654717 | 107.262462 | 0.48994803 | 0.00635738 | 0.04337808 | Garnl3 | 1.40439428 | 2 | 32986224 | 33131654 | - | 12352 | protein_codi | Mus muscul | CNH | 200 | 187 | 167 | 178 | 167 | 101 | 112 | 108 | 0.45185333 | 0.45926428 | 0.37458807 | 0.39292753 | 0.38884978 | 0.26081612 | 0.28777605 | 0.25583391 |
| ENSMUSG0C | 2868.22364 | 3177.65419 | 3214.3945 | 2939.42645 | 2861.34919 | 2097.16368 | 2226.7259 | 2701.42497 | 0.30294969 | 0.00635804 | 0.04337808 | Lipa | 1.23366415 | 19 | 34492318 | 34527474 | - | 2969 | protein_codi | Lysosomal ac | Abhydrolase_ | 3028 | 3086 | 3418 | 3176 | 2931 | 1937 | 2067 | 2720 | 28.4610189 | 31.5314589 | 31.8960284 | 29.1675553 | 28.3928046 | 20.8098888 | 22.0955183 | 26.8058968 |
| ENSMUSG0C | 367.526676 | 465.424399 | 466.453971 | 395.19367 | 383.660946 | 336.71549 | 285.477679 | 228.429317 | 0.45688623 | 0.00638636 | 0.04353141 | Lgals9 | 1.37257619 | 11 | 78962974 | 78984946 | - | 2137 | protein_codi | Galectin-9 [ | - | 388 | 452 | 496 | 427 | 393 | 311 | 265 | 230 | 5.06677915 | 6.41641218 | 6.43060602 | 5.44820057 | 5.28920867 | 4.64201141 | 3.93563909 | 3.14916161 |
| ENSMUSG0C | 125.034848 | 124.5937 | 93.102708 | 108.284917 | 93.7187043 | 57.3823825 | 54.9409873 | 85.4127011 | 0.62507124 | 0.00640101 | 0.04361127 | Dglucy | 1.54228699 | 12 | 100779057 | 100896981 | + | 4599 | protein_codi | D-glutamate | - | 132 | 121 | 99 | 117 | 96 | 53 | 51 | 86 | 0.80096823 | 0.79814225 | 0.59641222 | 0.69366884 | 0.60035827 | 0.36758925 | 0.35194977 | 0.54715034 |
| ENSMUSG0C | 274.697773 | 255.365599 | 262.380359 | 325.780262 | 219.653213 | 211.12386 | 199.295738 | 240.347368 | 0.36130642 | 0.00641173 | 0.0436244 | Cep78 | 1.28458862 | 19 | 15955774 | 15984989 | - | 2548 | protein_codi | Centrosomal | - | 290 | 248 | 279 | 352 | 225 | 195 | 185 | 242 | 3.17616712 | 2.95264068 | 3.03374818 | 3.76680358 | 2.53971958 | 2.44109974 | 2.3043382 | 2.7789938 |
| ENSMUSG0C | 753.050792 | 692.988098 | 909.397158 | 1044.90317 | 690.199208 | 679.927098 | 668.987316 | 580.011831 | 0.3772034 | 0.00641838 | 0.04364971 | Akr1e1 | 1.29882171 | 13 | 4590750 | 4609174 | - | 3640 | protein_codi | 1,5-anhydro- | Aldo_ket_red | 795 | 673 | 967 | 1129 | 707 | 628 | 621 | 584 | 6.09495517 | 5.60882671 | 7.36037326 | 8.4571161 | 5.58625431 | 5.5031151 | 5.41457197 | 4.6944325 |
| ENSMUSG0C | 2651.30713 | 2806.96219 | 2637.91006 | 2344.32217 | 2325.39535 | 2315.86634 | 2248.27138 | 2063.80922 | 0.22158061 | 0.00643331 | 0.04373123 | Diaph2 | 1.16601036 | X | 129749742 | 130465834 | + | 11553 | protein_codi | 2 [Source:U | FH2 | 2799 | 2726 | 2805 | 2533 | 2382 | 2139 | 2087 | 2078 | 6.76103051 | 7.15796252 | 6.726867 | 5.97819602 | 5.92993123 | 5.90563154 | 5.73325937 | 5.26286712 |
| ENSMUSG0C | 384.576882 | 378.929599 | 355.483067 | 322.078213 | 279.20364 | 317.227133 | 258.545823 | 298.944454 | 0.32014252 | 0.00647447 | 0.04399092 | Kank3 | 1.24845387 | 17 | 33810520 | 33822918 | + | 3118 | protein_codi | Mus muscul | - | 406 | 368 | 378 | 348 | 286 | 293 | 240 | 301 | 3.63374706 | 3.58038764 | 3.35884867 | 3.04321662 | 2.63810815 | 2.99738028 | 2.44291887 | 2.82463294 |
| ENSMUSG0C | 3446.0362 | 3441.25739 | 3012.20175 | 3267.05774 | 2784.22651 | 2616.85318 | 2412.01707 | 3150.33823 | 0.26391448 | 0.00650557 | 0.04414176 | Syde2 | 1.20073224 | 3 | 145987870 | 146021720 | + | 5751 | protein_codi | synapse defe | RhoGAP | 3638 | 3342 | 3203 | 3530 | 2852 | 2417 | 2239 | 3172 | 17.6532267 | 17.628746 | 15.4307956 | 16.7363624 | 14.2629325 | 13.4055186 | 12.3561918 | 16.1384361 |
| ENSMUSG0C | 1006.90942 | 1064.7098 | 726.013036 | 821.854751 | 703.866519 | 756.797837 | 676.528236 | 697.206002 | 0.35236985 | 0.00651388 | 0.04417799 | Gba2 | 1.27665601 | 4 | 43566928 | 43578873 | - | 4481 | protein_codi | Non-lysosom | - | 1063 | 1034 | 772 | 888 | 721 | 699 | 628 | 702 | 6.62007777 | 7.00009504 | 4.77328213 | 5.40340794 | 4.62767652 | 4.97568145 | 4.44793686 | 4.58388595 |
| ENSMUSG0C | 178.079936 | 155.4847 | 194.669299 | 195.283055 | 155.221604 | 129.922375 | 129.272911 | 130.105394 | 0.41011397 | 0.00656006 | 0.04443046 | Ncs1 | 1.32879078 | 2 | 31245823 | 31295989 | + | 4653 | protein_codi | Neuronal cal | efhand | 188 | 151 | 207 | 211 | 159 | 120 | 120 | 131 | 1.1275338 | 0.98446943 | 1.23257127 | 1.23645734 | 0.98280361 | 0.82261861 | 0.81850645 | 0.82377741 |
| ENSMUSG0C | 130.718251 | 109.1482 | 126.017807 | 132.348231 | 96.6474138 | 85.5322305 | 76.4864726 | 100.310265 | 0.4718282 | 0.00657316 | 0.04444672 | Grtp1 | 1.38686581 | 8 | 13172022 | 13200620 | - | 4093 | protein_codi | Growth horm | TBC | 138 | 106 | 134 | 143 | 99 | 79 | 71 | 101 | 0.94089706 | 0.78563797 | 0.90706373 | 0.9526295 | 0.69565854 | 0.61565255 | 0.55054208 | 0.72202339 |
| ENSMUSG0C | 1636.81983 | 1727.8366 | 1730.39376 | 1877.86407 | 1551.2398 | 1151.9784 | 1131.13797 | 1617.87547 | 0.35445427 | 0.00657441 | 0.04444672 | Acat1 | 1.27850187 | 9 | 53580522 | 53610382 | - | 3758 | protein_codi | Acetyl-CoA a | Thiolase_N | 1728 | 1678 | 1840 | 2029 | 1589 | 1064 | 1050 | 1629 | 12.8319227 | 13.5454526 | 13.5654997 | 14.7215997 | 12.1610142 | 9.03098641 | 8.86760699 | 12.6834074 |
| ENSMUSG0C | 3004.6253 | 6061.84388 | 2313.46123 | 3573.40224 | 2047.16795 | 2062.51771 | 2129.77122 | 2020.1097 | 0.85627785 | 0.00659741 | 0.04458196 | Arhgef26 | 1.81036156 | 3 | 62338344 | 62462221 | + | 10116 | protein_codi | Rho guanine | RhoGEF | 3172 | 5887 | 2460 | 3861 | 2097 | 1905 | 1977 | 2034 | 8.75042464 | 17.6540177 | 6.73753501 | 10.406884 | 5.96200427 | 6.00670766 | 6.20257126 | 5.88320204 |
| ENSMUSG0C | 84.3037993 | 58.6928998 | 99.6857278 | 110.135941 | 66.3840822 | 59.5477554 | 46.3227933 | 44.6926924 | 0.70313468 | 0.00664719 | 0.04487757 | Scara5 | 1.62803835 | 14 | 65666403 | 65764826 | + | 6093 | protein_codi | Scavenger re | SRCR | 89 | 57 | 106 | 119 | 68 | 55 | 43 | 45 | 0.40762762 | 0.28379322 | 0.48200266 | 0.53253176 | 0.3209818 | 0.28792664 | 0.22398101 | 0.21609911 |
| ENSMUSG0C | 101.354006 | 59.7225998 | 62.068472 | 85.1471138 | 37.0969871 | 36.8113397 | 63.5591814 | 44.6926924 | 0.76225924 | 0.00670158 | 0.04518329 | 6430511E19I | 1.69614468 | 7 | 92639949 | 92643783 | + | 3835 | TEC | RIKEN cDNA | - | 107 | 58 | 66 | 92 | 38 | 34 | 59 | 45 | 0.77861574 | 0.45879742 | 0.47681874 | 0.65411212 | 0.28498428 | 0.28278989 | 0.48827058 | 0.34333556 |
| ENSMUSG0C | 1815.847 | 1830.80659 | 1887.44581 | 1886.19367 | 1727.93861 | 1566.64731 | 1650.38417 | 1639.72523 | 0.17216255 | 0.00672127 | 0.04525453 | Rhoq | 1.12674617 | 17 | 86963082 | 87000069 | + | [illegible] | [illegible] | [illegible] | [illegible] | [illegible] | [illegible] | [illegible] | [illegible] | [illegible] | [illegible] | [illegible] | [illegible] | [illegible] | [illegible] | [illegible] | [illegible] | [illegible] | [illegible] | [illegible] | [illegible] |
| ENSMUSG0C | 11048.5339 | 10796.4045 | 12499.2737 | 11609.6239 | 10633.168 | 8649.58214 | 9245.16771 | 10357.7798 | 0.24089363 | 0.00674577 | 0.0453579 | Anxa4 | 1.18172442 | 6 | 86736840 | 86793584 | - | [illegible] | [illegible] | [illegible] | [illegible] | [illegible] | [illegible] | [illegible] | [illegible] | [illegible] | [illegible] | [illegible] | [illegible] | [illegible] | [illegible] | [illegible] | [illegible] | [illegible] | [illegible] | [illegible] | [illegible] |
| ENSMUSG0C | 61.5701905 | 72.0789998 | 68.6514918 | 77.743017 | 39.0494601 | 49.8035772 | 49.554616 | 46.6790343 | 0.59979437 | 0.00676434 | 0.04546222 | 4732471J01R | 1.51550055 | 7 | 25376819 | 25566417 | + | [illegible] | [illegible] | [illegible] | [illegible] | [illegible] | [illegible] | [illegible] | [illegible] | [illegible] | [illegible] | [illegible] | [illegible] | [illegible] | [illegible] | [illegible] | [illegible] | [illegible] | [illegible] | [illegible] | [illegible] |
| ENSMUSG0C | 180.921637 | 226.533999 | 188.086279 | 203.612664 | 161.079023 | 108.268646 | 94.800135 | 174.798086 | 0.56501802 | 0.00677494 | 0.04551295 | Sh3yl1 | 1.47940599 | 12 | 30911668 | 30960162 | + | [illegible] | [illegible] | [illegible] | [illegible] | [illegible] | [illegible] | [illegible] | [illegible] | [illegible] | [illegible] | [illegible] | [illegible] | [illegible] | [illegible] | [illegible] | [illegible] | [illegible] | [illegible] | [illegible] | [illegible] |
| ENSMUSG0C | 400.679855 | 398.493899 | 511.594678 | 441.469275 | 420.757933 | 304.234896 | 305.94589 | 243.326881 | 0.45928142 | 0.00677896 | 0.04551934 | Elovl6 | 1.37485685 | 3 | 129532355 | 129638495 | + | [illegible] | [illegible] | [illegible] | [illegible] | [illegible] | [illegible] | [illegible] | [illegible] | [illegible] | [illegible] | [illegible] | [illegible] | [illegible] | [illegible] | [illegible] | [illegible] | [illegible] | [illegible] | [illegible] | [illegible] |
| ENSMUSG0C | 286.064578 | 283.167499 | 369.589538 | 380.385476 | 285.061059 | 250.100573 | 232.691241 | 233.395172 | 0.39815969 | 0.00678561 | 0.04553001 | Mrgpre | 1.31782581 | 7 | 143778363 | 143784500 | - | [illegible] | [illegible] | [illegible] | [illegible] | [illegible] | [illegible] | [illegible] | [illegible] | [illegible] | [illegible] | [illegible] | [illegible] | [illegible] | [illegible] | [illegible] | [illegible] | [illegible] | [illegible] | [illegible] | [illegible] |
| ENSMUSG0C | 114.615278 | 141.0689 | 124.136944 | 125.869647 | 90.7899948 | 102.855214 | 79.7182954 | 94.3512396 | 0.45855821 | 0.00678976 | 0.04553028 | Pcdhb18 | 1.37416783 | 18 | 37489465 | 37494505 | + | [illegible] | [illegible] | [illegible] | [illegible] | [illegible] | [illegible] | [illegible] | [illegible] | [illegible] | [illegible] | [illegible] | [illegible] | [illegible] | [illegible] | [illegible] | [illegible] | [illegible] | [illegible] | [illegible] | [illegible] |
| ENSMUSG0C | 55.8867883 | 39.1285999 | 70.5323545 | 49.9776538 | 30.2633316 | 32.4805939 | 38.7818734 | 20.8565898 | 0.82179761 | 0.00680768 | 0.04562987 | Gm44997 | 1.76760708 | 7 | 42561414 | 42578235 | - | [illegible] | [illegible] | [illegible] | [illegible] | [illegible] | [illegible] | [illegible] | [illegible] | [illegible] | [illegible] | [illegible] | [illegible] | [illegible] | [illegible] | [illegible] | [illegible] | [illegible] | [illegible] | [illegible] | [illegible] |
| ENSMUSG0C | 107.984642 | 75.1680998 | 101.566591 | 124.018622 | 65.4078457 | 55.2170095 | 63.5591814 | 83.4263592 | 0.61104184 | 0.00686706 | 0.04594505 | Cenpw | 1.52736179 | 10 | 30194569 | 30200560 | - | [illegible] | [illegible] | [illegible] | [illegible] | [illegible] | [illegible] | [illegible] | [illegible] | [illegible] | [illegible] | [illegible] | [illegible] | [illegible] | [illegible] | [illegible] | [illegible] | [illegible] | [illegible] | [illegible] | [illegible] |
| ENSMUSG0C | 1017.32899 | 1179.0065 | 675.229741 | 978.266297 | 636.5062 | 777.36888 | 673.296414 | 771.693823 | 0.42913709 | 0.00688502 | 0.04604453 | Igf2bp1 | 1.34642801 | 11 | 95957163 | 96005940 | - | [illegible] | [illegible] | [illegible] | [illegible] | [illegible] | [illegible] | [illegible] | [illegible] | [illegible] | [illegible] | [illegible] | [illegible] | [illegible] | [illegible] | [illegible] | [illegible] | [illegible] | [illegible] | [illegible] | [illegible] |
| ENSMUSG0C | 107.037408 | 115.3264 | 97.804865 | 113.837989 | 92.7424678 | 58.4650689 | 81.8728439 | 46.6790343 | 0.63266242 | 0.00695313 | 0.04641656 | Fez1 | 1.55042359 | 9 | 36821864 | 36878924 | + | [illegible] | [illegible] | [illegible] | [illegible] | [illegible] | [illegible] | [illegible] | [illegible] | [illegible] | [illegible] | [illegible] | [illegible] | [illegible] | [illegible] | [illegible] | [illegible] | [illegible] | [illegible] | [illegible] | [illegible] |
| ENSMUSG0C | 459.408345 | 390.256299 | 358.304361 | 513.659219 | 262.607619 | 350.790414 | 352.268684 | 332.712266 | 0.40835249 | 0.00696943 | 0.04650457 | Slc16a4 | 1.32716937 | 3 | 107291230 | 107312115 | + | [illegible] | [illegible] | [illegible] | [illegible] | [illegible] | [illegible] | [illegible] | [illegible] | [illegible] | [illegible] | [illegible] | [illegible] | [illegible] | [illegible] | [illegible] | [illegible] | [illegible] | [illegible] | [illegible] | [illegible] |
| ENSMUSG0C | 1917.20101 | 2107.79589 | 3846.3644 | 2149.96462 | 2004.21354 | 1539.58015 | 1600.82955 | 1790.68721 | 0.53098715 | 0.00698762 | 0.04660504 | Lgmn | 1.44491753 | 12 | 102394084 | 102439813 | - | [illegible] | [illegible] | [illegible] | [illegible] | [illegible] | [illegible] | [illegible] | [illegible] | [illegible] | [illegible] | [illegible] | [illegible] | [illegible] | [illegible] | [illegible] | [illegible] | [illegible] | [illegible] | [illegible] | [illegible] |
| ENSMUSG0C | 1285.39613 | 1510.5699 | 1395.60019 | 1401.22533 | 1352.08756 | 776.286193 | 785.332937 | 1151.08512 | 0.45974717 | 0.00701457 | 0.04670103 | Gulp1 | 1.37530078 | 1 | 44551511 | 44796838 | + | [illegible] | [illegible] | [illegible] | [illegible] | [illegible] | [illegible] | [illegible] | [illegible] | [illegible] | [illegible] | [illegible] | [illegible] | [illegible] | [illegible] | [illegible] | [illegible] | [illegible] | [illegible] | [illegible] | [illegible] |
| ENSMUSG0C | 88241.4498 | 123740.078 | 135343.124 | 87104.5719 | 83395.9795 | 75047.4948 | 89655.0732 | 83771.9827 | 0.3884977 | 0.00703192 | 0.04679564 | Itgb1 | 1.30902958 | 8 | 128685654 | 128733200 | + | 4081 | protein_codi | Integrin beta | - | 93157 | 120171 | 143916 | 94115 | 85426 | 69316 | 83224 | 84348 | 637.020884 | 893.28784 | 977.051 | 628.813687 | 602.041112 | 541.772847 | 647.225925 | 604.755504 |
| ENSMUSG0C | 421.518997 | 529.265799 | 533.2246 | 624.720672 | 476.403414 | 333.46743 | 370.582346 | 406.206916 | 0.40956656 | 0.00704188 | 0.04684092 | Epb41l4a | 1.32828669 | 18 | 33796327 | 34007206 | - | 3853 | protein_codi | Band 4.1-like | Band_41 | 445 | 514 | 567 | 675 | 488 | 308 | 344 | 409 | 3.22304057 | 4.0468998 | 4.0771698 | 4.77677184 | 3.64270067 | 2.54977609 | 2.8335661 | 3.10596054 |
| ENSMUSG0C | 173.343767 | 182.256899 | 254.856908 | 200.836127 | 149.364185 | 127.757002 | 141.122928 | 169.832231 | 0.46358972 | 0.00706967 | 0.04696283 | Morc4 | 1.37896871 | X | 139821632 | 139871677 | - | 6051 | protein_codi | MORC family | zf-CW | 183 | 177 | 271 | 217 | 153 | 118 | 131 | 171 | 0.8439733 | 0.88736941 | 1.24084315 | 0.97782765 | 0.7272219 | 0.62202121 | 0.68709701 | 0.82687639 |
| ENSMUSG0C | 107.037408 | 70.0195998 | 89.3409824 | 116.614525 | 70.2890282 | 72.5399929 | 49.554616 | 53.6312309 | 0.64029667 | 0.0071112 | 0.04716999 | Nmral1 | 1.55864964 | 16 | 4710059 | 4719356 | - | 3891 | protein_codi | NmrA-like fa | NmrA | 113 | 68 | 95 | 126 | 72 | 67 | 46 | 54 | 0.81044208 | 0.53015886 | 0.67645221 | 0.88295597 | 0.53219886 | 0.54924221 | 0.37520664 | 0.40607305 |
| ENSMUSG0C | 2651.30713 | 1826.68779 | 1738.85765 | 2117.5717 | 1986.64128 | 1355.52345 | 1453.24298 | 1389.44615 | 0.43021136 | 0.00711354 | 0.04716999 | Smyd1 | 1.34743097 | 6 | 71213940 | 71322233 | - | 4546 | protein_codi | SET and MYN | SET | 2799 | 1774 | 1849 | 2288 | 2035 | 1252 | 1349 | 1399 | 17.182179 | 11.8381142 | 11.2689183 | 13.7232294 | 12.8747159 | 8.78466559 | 9.41795112 | 9.00450655 |
| ENSMUSG0C | 148.715691 | 98.8511997 | 173.039376 | 248.962757 | 123.005799 | 87.6976034 | 81.8728439 | 112.228317 | 0.7258582 | 0.00717005 | 0.04750238 | Slc6a9 | 1.65388418 | 4 | 117834506 | 117875198 | + | 5141 | protein_codi | Sodium- and | SNF | 157 | 96 | 184 | 269 | 126 | 81 | 76 | 113 | 0.85223 | 0.56647659 | 0.99161929 | 1.42670575 | 0.70489692 | 0.50255981 | 0.46918044 | 0.64313549 |
| ENSMUSG0C | 5773.3894 | 5960.93328 | 6695.87152 | 5987.13782 | 5064.71498 | 5393.94395 | 5688.0081 | 5082.05572 | 0.20201489 | 0.00721017 | 0.04774693 | Pcyox1 | 1.15030376 | 6 | 86386006 | 86397154 | - | 4932 | protein_codi | Mus muscul | - | 6095 | 5789 | 7120 | 6469 | 5188 | 4982 | 5280 | 5117 | 34.4869997 | 35.6072819 | 39.997392 | 35.7638132 | 30.253775 | 32.2204047 | 33.9769795 | 30.3573588 |
| ENSMUSG0C | 65.3591253 | 64.8710998 | 67.7110604 | 81.4450654 | 43.9306426 | 42.224772 | 50.6318903 | 48.6653762 | 0.59243437 | 0.00726111 | 0.04802818 | Bard1 | 1.50778881 | 1 | 71027498 | 71103146 | - | 5659 | protein_codi | BRCA1-assoc | - | 69 | 63 | 72 | 88 | 45 | 39 | 47 | 49 | 0.34026256 | 0.33772188 | 0.35250684 | 0.42400669 | 0.22870491 | 0.21982407 | 0.26359191 | 0.25335415 |
| ENSMUSG0C | 96.6178374 | 165.7817 | 53.6045895 | 74.0409686 | 78.0989203 | 37.8940262 | 14.0045654 | 36.7473249 | 1.22142854 | 0.00727182 | 0.04806961 | Exph5 | 2.33177492 | 9 | 53301670 | 53377514 | + | 10519 | protein_codi | Exophilin-5 [ | - | 102 | 161 | 57 | 80 | 80 | 35 | 13 | 37 | 0.27060168 | 0.46431184 | 0.15013265 | 0.20736968 | 0.21873496 | 0.10613141 | 0.03922318 | 0.10291979 |
| ENSMUSG0C | 2333.0366 | 5087.74769 | 2480.85802 | 2752.47301 | 2132.10052 | 2026.78906 | 1525.42035 | 1674.48621 | 0.78196917 | 0.00729949 | 0.04818518 | Fosb | 1.71947623 | 7 | 19302696 | 19310051 | - | 6243 | protein_codi | Protein fosB | bZIP_1 | 2463 | 4941 | 2638 | 2974 | 2184 | 1872 | 1416 | 1686 | 11.0097092 | 24.00932 | 11.7072854 | 12.9890493 | 10.0614824 | 9.56451265 | 7.19853022 | 7.90197899 |
| ENSMUSG0C | 667.799759 | 750.651298 | 662.063701 | 750.590319 | 637.482437 | 578.154571 | 589.269021 | 587.957198 | 0.24195116 | 0.00733229 | 0.04836189 | Lmbr1 | 1.18259096 | 5 | 29229802 | 29378390 | - | 5850 | protein_codi | Limb region | - | 705 | 729 | 704 | 811 | 653 | 534 | 547 | 592 | 3.36308638 | 3.78033253 | 3.33419919 | 3.78002544 | 3.21040622 | 2.91162693 | 2.96760008 | 2.96099365 |
| ENSMUSG0C | 1844.26401 | 2154.13239 | 1570.52043 | 2066.66853 | 1560.02593 | 1507.09955 | 1018.02418 | 1642.70474 | 0.41443059 | 0.00734438 | 0.04842018 | Adgra2 | 1.33277254 | 8 | 27085583 | 27123436 | + | 5775 | protein_codi | Adhesion G | 7tm_2 | 1947 | 2092 | 1670 | 2233 | 1598 | 1392 | 945 | 1654 | 9.40846414 | 10.9892495 | 8.01196846 | 10.5430549 | 7.95843107 | 7.68842856 | 5.19342344 | 8.38021482 |
| ENSMUSG0C | 26.5225436 | 26.7721999 | 27.2725104 | 34.243948 | 19.5247301 | 12.9922375 | 14.0045654 | 11.9180513 | 0.97206019 | 0.00735054 | 0.0484281 | A930027P06I | 1.96163984 | 13 | 17951938 | 17954912 | + | 2975 | TEC | RIKEN cDNA | - | 28 | 26 | 29 | 37 | 20 | 12 | 13 | 12 | 0.26264905 | 0.26512136 | 0.27007587 | 0.33911304 | 0.19335068 | 0.12866032 | 0.13868526 | 0.1180228 |
| ENSMUSG0C | 32.2059458 | 19.5642999 | 31.9746674 | 27.7653632 | 19.5247301 | 12.9922375 | 6.46364557 | 11.9180513 | 1.12657695 | 0.00736715 | 0.04848439 | 1500026H17I | 2.18340075 | 10 | 89686371 | 89700866 | + | 3027 | bidirectional | RIKEN cDNA | - | 34 | 19 | 34 | 30 | 20 | 12 | 6 | 12 | 0.31345216 | 0.19041428 | 0.31120119 | 0.27023312 | 0.19002916 | 0.1264501 | 0.06290899 | 0.11599532 |
| ENSMUSG0C | 855.352031 | 790.809598 | 782.43892 | 767.249537 | 703.866519 | 692.919336 | 698.073722 | 665.424532 | 0.21145383 | 0.00737835 | 0.04853664 | Chic1 | 1.15785439 | X | 103356476 | 103396092 | + | 7375 | protein_codi | Cysteine-rich | - | 903 | 768 | 832 | 829 | 721 | 640 | 648 | 670 | 3.41688606 | 3.15905754 | 3.12561909 | 3.06494186 | 2.81174488 | 2.76801402 | 2.78860432 | 2.65818017 |
| ENSMUSG0C | 1269.29316 | 1171.7986 | 1217.85866 | 1115.24209 | 1088.5037 | 1046.95781 | 1014.79235 | 902.792387 | 0.23634392 | 0.00738896 | 0.04857749 | Inpp5f | 1.17800358 | 7 | 128611328 | 128696425 | + | 8323 | protein_codi | inositol polyp | - | 1340 | 1138 | 1295 | 1205 | 1115 | 967 | 942 | 909 | 4.49293023 | 4.14782771 | 4.31086698 | 3.94763405 | 3.8529879 | 3.70592747 | 3.59207108 | 3.19562363 |
| ENSMUSG0C | 87.1455004 | 78.2571998 | 92.1622766 | 107.359404 | 67.3603187 | 66.0438742 | 40.9364219 | 65.5492822 | 0.60430271 | 0.00750683 | 0.0491224 | 0610040J01R | 1.5202438 | 5 | 63812363 | 63899625 | + | 2403 | protein_codi | RIKEN cDNA | - | 92 | 76 | 98 | 116 | 69 | 61 | 38 | 66 | 1.06841217 | 0.95944076 | 1.12991833 | 1.31623657 | 0.82584395 | 0.80970422 | 0.50188445 | 0.80364047 |
| ENSMUSG0C | 219.758218 | 180.1975 | 266.142084 | 206.3892 | 145.459239 | 191.635504 | 147.586574 | 155.927838 | 0.44762679 | 0.00750701 | 0.0491224 | Pcdhb22 | 1.36379499 | 18 | 37517871 | 37524479 | + | 6609 | protein_codi | - | Cadherin | 232 | 175 | 283 | 223 | 149 | 177 | 137 | 157 | 0.97961856 | 0.80326832 | 1.18638441 | 0.92002334 | 0.6484152 | 0.85425563 | 0.65789825 | 0.69508119 |
| ENSMUSG0C | 1441.68969 | 1770.0543 | 1168.01579 | 1335.51397 | 1161.72144 | 1106.50556 | 1068.65607 | 1193.79147 | 0.33456704 | 0.00750627 | 0.0491224 | Mtmr12 | 1.26099892 | 15 | 12205028 | 12274496 | + | 7041 | protein_codi | Mus muscul | - | 1522 | 1719 | 1242 | 1443 | 1190 | 1022 | 992 | 1202 | 6.03232982 | 7.40627569 | 4.88722125 | 5.5880685 | 4.86088437 | 4.62984965 | 4.47147948 | 4.99507207 |
| ENSMUSG0C | 7624.28405 | 8759.65788 | 6088.35284 | 6767.34453 | 6691.12499 | 5311.65978 | 5290.49686 | 5890.20306 | 0.33146049 | 0.00751274 | 0.04912884 | Smim1 | 1.25828654 | 4 | 154020470 | 154026230 | - | 5761 | protein_codi | small integra | - | 8049 | 8507 | 6474 | 7312 | 6854 | 4906 | 4959 | 5931 | 38.9896039 | 44.795759 | 31.1350501 | 34.6073258 | 34.217549 | 27.1631421 | 27.3193365 | 30.1232401 |
| ENSMUSG0C | 166.713131 | 161.6629 | 187.145847 | 135.124768 | 114.219671 | 70.37462 | 43.0909705 | 142.023445 | 0.81244347 | 0.00755373 | 0.04936297 | Evc | 1.75618334 | 5 | 37289098 | 37336894 | - | 7701 | protein_codi | Ellis-van Cre | - | 176 | 157 | 199 | 146 | 117 | 65 | 40 | 143 | 0.63777915 | 0.61845894 | 0.71594672 | 0.51693444 | 0.43695973 | 0.26922573 | 0.16484918 | 0.54332608 |
| ENSMUSG0C | 713.266976 | 732.116698 | 735.41735 | 793.163876 | 616.005234 | 518.606815 | 617.278152 | 693.233318 | 0.28190705 | 0.00757793 | 0.04949941 | Ppcdc | 1.21580095 | 9 | 57385095 | 57440124 | - | 5803 | protein_codi | phosphopant | Flavoprotein | 753 | 711 | 782 | 857 | 631 | 479 | 573 | 698 | 3.62115552 | 3.71685288 | 3.73360983 | 4.02678077 | 3.12737141 | 2.63289342 | 3.13383584 | 3.51944747 |
| ENSMUSG0C | 373.210078 | 429.384899 | 316.92538 | 453.500932 | 293.847187 | 307.482955 | 291.941325 | 329.732753 | 0.36264725 | 0.00758621 | 0.04953176 | Ttc30b | 1.28578306 | 2 | 75935849 | 75938585 | - | 2737 | protein_codi | tetratricopep | - | 394 | 417 | 337 | 490 | 301 | 284 | 271 | 332 | 4.01722536 | 4.62188994 | 3.41137806 | 4.88147441 | 3.16296489 | 3.30974 | 3.14245022 | 3.54923636 |
| ENSMUSG0C | 4453.89286 | 5735.42898 | 4641.96936 | 4034.30727 | 4035.7617 | 3178.76745 | 3642.26428 | 4125.6321 | 0.33226728 | 0.00759267 | 0.04955217 | Ech1 | 1.2589904 | 7 | 28825217 | 28832247 | + | 2284 | protein_codi | Delta(3,5)-De | ECH | 4702 | 5570 | 4936 | 4359 | 4134 | 2936 | 3381 | 4154 | 57.4501671 | 73.9805298 | 59.8761407 | 52.0379889 | 52.0567494 | 41.0024953 | 46.9810788 | 53.2159755 |
| ENSMUSG0C | 2824.65089 | 2983.04089 | 2587.12676 | 2867.23651 | 2366.39728 | 2471.77319 | 2569.29911 | 2486.90004 | 0.18677278 | 0.00761543 | 0.04967894 | Zer1 | 1.13821475 | 2 | 30097283 | 30124585 | - | 9784 | protein_codi | zyg-11 relate | - | 2982 | 2897 | 2751 | 3098 | 2424 | 2283 | 2385 | 2504 | 8.50542395 | 8.98235867 | 7.79020515 | 8.63365527 | 7.12555742 | 7.44285921 | 7.73652357 | 7.48840829 |
| ENSMUSG0C | 14.2085055 | 27.8018999 | 34.7959616 | 12.0316574 | 11.714838 | 6.49611877 | 8.61819409 | 6.9521966 | 1.38934484 | 0.00762042 | 0.04968969 | Ncapg | 2.61959693 | 5 | 45669919 | 45700546 | + | 7066 | protein_codi | non-SMC con | - | 15 | 27 | 37 | 13 | 12 | 6 | 8 | 7 | 0.059241 | 0.11591736 | 0.14507843 | 0.05016484 | 0.04884389 | 0.02708494 | 0.03598119 | 0.02898652 |
| ENSMUSG0C | 183.763338 | 170.9302 | 151.409454 | 189.729982 | 131.791928 | 121.260884 | 86.1819409 | 150.961983 | 0.50330115 | 0.00762657 | 0.04970801 | Irak1bp1 | 1.41745325 | 9 | 82829540 | 82847687 | + | 2158 | protein_codi | Interleukin-1 | - | 194 | 166 | 161 | 205 | 135 | 112 | 80 | 152 | 2.50873657 | 2.3335386 | 2.06704144 | 2.59019318 | 1.79922303 | 1.65545324 | 1.17655561 | 2.06093257 |
| ENSMUSG0C | 415.835594 | 506.612399 | 447.645344 | 428.512106 | 389.518365 | 365.948024 | 325.336827 | 380.384471 | 0.29813916 | 0.00763437 | 0.04973704 | Cep126 | 1.22955746 | 9 | 8076461 | 8134294 | - | 4616 | protein_codi | Centrosomal | - | 439 | 492 | 476 | 463 | 399 | 338 | 302 | 383 | 2.65401572 | 3.23338668 | 2.85703724 | 2.73492188 | 2.48604948 | 2.3356149 | 2.0764193 | 2.42775361 |
| ENSMUSG0C | 1311.91867 | 1107.9572 | 1182.12226 | 1196.68715 | 1094.36112 | 1059.95005 | 977.087755 | 923.648977 | 0.24313902 | 0.00764633 | 0.04979316 | Dock9 | 1.18356507 | 14 | 121542046 | 121797837 | - | 13245 | protein_codi | dedicator of | Ded_cyto | 1385 | 1076 | 1257 | 1293 | 1121 | 979 | 907 | 930 | 2.9181162 | 2.46444228 | 2.62940851 | 2.66180537 | 2.43420036 | 2.3576594 | 2.1734783 | 2.05448333 |
| ENSMUSG0C | 162.924196 | 174.0193 | 226.643966 | 173.070764 | 149.364185 | 142.914613 | 119.577443 | 132.091735 | 0.43718788 | 0.00765426 | 0.04982298 | Fam57a | 1.35396259 | 11 | 76179671 | 76214827 | + | 6615 | protein_codi | Protein FAM | - | 172 | 169 | 241 | 187 | 153 | 132 | 111 | 133 | 0.72561018 | 0.77502408 | 1.00939486 | 0.77079962 | 0.6652184 | 0.63649415 | 0.53255816 | 0.58829266 |
