## Supplementary Table 4 for "MAFB drives differentiation by permitting WT1 binding to podocyte specific promoters"

primers figure 5 (embryo ChIP)

| | | |
|---|---|---|
| NPHS1_Prom | GTGCTGACAGGGGATTTCAT | GTTCACGAACCCAAAGCAGT |
| NPHS2_prom | TGAGTTCCTCCTCCGAATTG | TGCTCAGACCTTTGCTGAGA |
| Synpo_Prom | CTCTGTCAATCGGCCTCTTC | TGAGTCAGCTGTGGAGGAGA |
| NPHS2_neg | AGCTTAGTCTGGCCTAGGCT | CCTGCTCTGTTGGAGGACTG |
| NPHS1_neg | CCCTCTGAAGACCCTCCAGA | GGTTAGCAGACACGGACACA |

primers for qPCR

| | | |
|---|---|---|
| NPHS1 | TCTGGGTCCAAACCCTAAGATT | TCAATAAGCAGGTGGAACTCAC |
| NPHS2 | CCATCTGGTTCTGCATAAAGG | CCAGGACCTTTGGCTCTTC |
| MAFB | CATCACCATCATCACCAAGC | AGAAGCGGTCCTCCACACTA |
| PDXL | ACTACATTGCCCGTCTCCAC | AAATCCTCAGCTGGCTTGAA |
| SYNPO | CCTGCCCGTAACTTCCGTG | GAGCGGCGGTAGGGAAAAG |
